# Metabolic reprogramming during *Candida albicans* planktonic-biofilm transition is modulated by the *ZCF15* and *ZCF26* paralogs

**DOI:** 10.1101/2023.08.03.551767

**Authors:** Laxmi Shanker Rai, Murielle Chauvel, Hiram Sanchez, Lasse van Wijlick, Corinne Maufrais, Thomas Cokelaer, Natacha Sertour, Mélanie Legrand, David R Andes, Sophie Bachellier-Bassi, Christophe d’Enfert

## Abstract

*Candida albicans* is a commensal of the human microbiota that can form biofilms on implanted medical devices. These biofilms are tolerant to antifungals and to the host immune system. To identify novel genes modulating *C. albicans* biofilm formation, we performed a large-scale screen with 2454 *C. albicans* doxycycline-dependent overexpression strains and identified 16 genes whose overexpression significantly hampered biofilm formation. Among those, overexpression of the *ZCF15* and *ZCF26* paralogs that encode transcription factors and have orthologs only in biofilm-forming species of the *Candida* clade, caused impaired biofilm formation both *in vitro* and *in vivo*. Interestingly, overexpression of *ZCF15* specifically impeded biofilm formation without any defect in hyphal growth. Transcript profiling, transcription factor binding, and phenotypic microarray analyses conducted upon overexpression of *ZCF15* and *ZCF26* demonstrated their direct role in reprogramming cellular metabolism by regulating glycolytic cycle and tricarboxylic acid cycle genes. Taken together, this study has identified a new set of biofilm regulators, including *ZCF15* and *ZCF26,* that appear to control biofilm development through their specific role in metabolic remodeling.

## Introduction

*Candida albicans* is a commensal of the human microbiota that resides on the mucosal surfaces of the gastrointestinal and genital tracts. Under certain circumstances, such as if epithelial barriers are disturbed or the immune system is impaired, the fungus undergoes a transition from commensalism to pathogenicity [1]. This transition is well regulated by both the host immune system and fungal-specific virulence attributes.

*C. albicans* can form biofilms, which represent a major fungal virulence attribute [2,3]. Biofilms are microbial communities attached to surfaces and protected by self-produced extracellular substances [4]. Cells in a biofilm are more adherent and more tolerant to antimicrobials as compared to the free-floating planktonic cells and these properties make biofilm-associated infections a clinical challenge [2,5,6]. *C. albicans* biofilms are structured and composed of differentiated cell types encased in an extracellular matrix. Briefly, the *C. albicans* biofilm developmental process involves the attachment of yeast cells to a surface and their proliferation to establish a basal layer. Basal layer cells undergo cellular differentiation in multiple cell types including hyphae and pseudo-hyphae that become encased in a self-produced extracellular matrix, leading to a mature biofilm [4,7]. These biofilms can be the source of disseminated infections that can, in turn, lead to invasive systemic infections of tissues and organs [3,8,9].

Among the *Candida* clade, only a few species closely related to *C. albicans*, namely *Candida dubliniensis* and *Candida tropicalis*, can form a complex biofilm. The less closely related species, *Candida parapsilosis, Loderomyces elongisporus* and *Spathaspora passalidarum* are also able to form biofilms, but these are structurally different and of lesser biomass than those of *C. albicans* [10–14]. Transcript profiling, proteome analyses and metabolomic studies of *C. albicans* planktonic and biofilm cells have shown that cellular differentiation and metabolic reprogramming are two critical events that occur when *C. albicans* cells transition from the planktonic to the biofilm growth mode [13,15–21]. Studies on *C. albicans* transcription regulators have suggested that a well-coordinated crosstalk operates during biofilm development. For instance, transcription regulators, Ace2, Brg1, Efg1, Ndt80, Mss11, Tec1, Flo8, Rob1 and Ume6 are essential for *C. albicans* hyphal development and are also needed for *C. albicans* biofilm formation [13,22,23]. In parallel, Tye7 regulates the glycolytic flux, and the lack of this transcription factor leads to impaired biofilm formation (Bonhomme et al., 2011). In addition, amino acid metabolism is modulated during biofilm formation, and it has been shown that the Gcn4 regulator of the amino acid biosynthetic pathways is important for efficient biofilm formation [19]. Yet, it is notable that most modulators in the regulation of *C. albicans* biofilm formation identified so far are positive regulators. Only a few transcription regulators such as Nrg1, Rfx2, Gal4, Zcf32, and Upc2 have been shown to play a negative role during *C. albicans* biofilm formation [18,24,25]. This may be a consequence of the approach used to identify these modulators, as the biofilm growth conditions did not allow an increase in biofilm biomass to be observed when the target genes were inactivated, as expected for genes encoding negative regulators of biofilm formation [18]. In this study, we sought to identify additional negative regulators of biofilm formation and reasoned that their overexpression would result in reduced biofilm biomass in a biofilm formation assay.

Large collections of *C. albicans* overexpression strains are becoming available and have proven useful to identify genes with a role in *C. albicans* morphogenesis, genome plasticity, biofilm formation, antifungal tolerance, and intestinal colonization [26–30]. Using a novel collection of 2454 *C. albicans* doxycycline-dependent overexpression strains derived from the *C. albicans* ORFeome [27,28], we could identify 16 genes whose overexpression led to reduced biofilm formation. Among these genes, the *ZCF15* and *ZCF26* paralogs encode zinc cluster transcription factors whose overexpression leads to impaired biofilm growth *in vitro* and *in vivo*. Transcript profiling and ChIP-sequencing analyses demonstrated that both *ZCF15* and *ZCF26* directly regulate the expression of genes associated with cellular metabolism, including the genes of the glycolysis, glyoxylate cycle and tri-carboxylic acid (TCA) cycle, known to be differentially expressed when *C. albicans* proliferates as biofilms. Altogether, we discovered novel transcription regulators that recently appeared to regulate metabolic remodeling during planktonic to biofilm transition.

## Results

### A large-scale overexpression screen identifies *C. albicans* negative regulators of biofilm formation

In the frame of the *C. albicans* ORFeome project, 5099 ORFs representing approximately 83% of *C. albicans* predicted ORFs were cloned into a Gateway^TM^ donor vector [28]. A total of 2454 of these ORFs were then transferred in a tetracycline-dependent overexpression vector and introduced into a suitably engineered *C. albicans* strain [27] (**Figure 1A**). This unbiased *C. albicans* overexpression collection was used to uncover genes whose overexpression hampers *C. albicans* biofilm formation.

**Figure 1.**
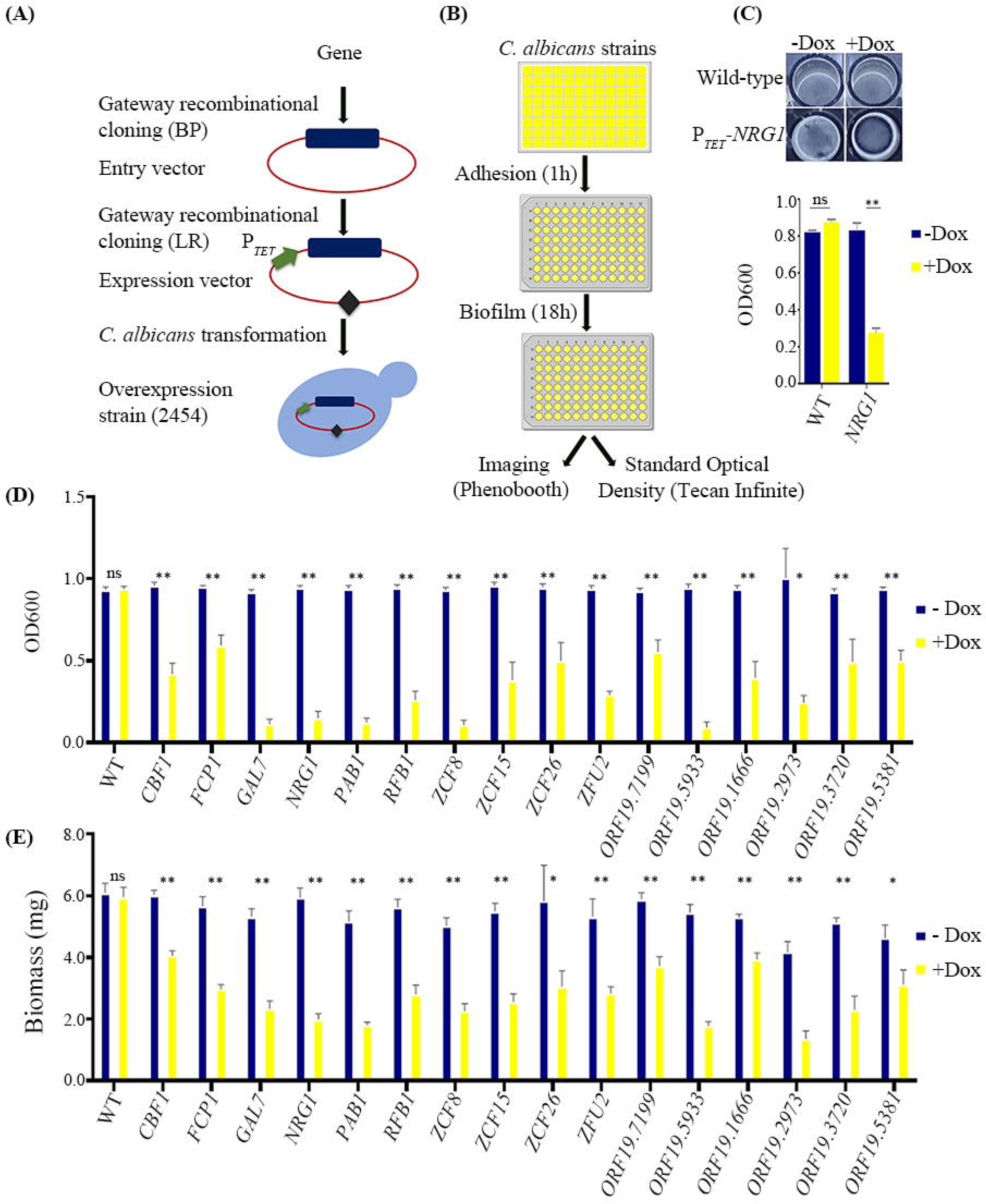
High-throughput screen for biofilm formation with *C. albicans* overexpression strains. (**A**) Schematic showing the construction of 2454 *C. albicans* P*_TET_* overexpression strains. (**B**) Overview of the *in vitro* screening strategy for the collection of *C. albicans* overexpression strain for biofilm formation. Cells were grown overnight in 96-deep-well plate in YPD with or without 25 µg/mL doxycycline. Then, 0.2 µL of culture was diluted in 200 µL of YPD medium with or without 25 µg/mL of doxycycline and transferred to FBS pre-coated 96-well polystyrene plates, that were incubated at 37°C for 1h for adhesion to occur. Then, the medium was aspirated, and the wells were washed with 1xPBS. A fresh aliquot of 200 µL of YPD medium with or without 25 µg/ml of doxycycline was added and biofilms were allowed to develop for 18h at 37°C at 110 rpm. After 18 h, the medium was discarded, the wells washed with 1xPBS and photographed. Quantification of biofilms was determined by measuring the standard optical density using Tecan infinite M200. (**C**) Biofilm formation by *C. albicans* wild-type and P*_TET_-NRG1* overexpression strains with or without doxycycline. (**D** and **E**) *C. albicans* overexpression strains identified in the screen and the WT control were grown overnight in YPD medium, with or without 25µg/mL doxycycline. Biofilms were allowed to develop in 96-well polystyrene plates (**D**) or in 12-well polystyrene plates (**E**) in YPD medium with or without 25µg/mL doxycycline at 37°C for 18 h. (**D**) Standard optical density was measured to quantify the extent of biofilm formation using a Tecan infinite M200. (**E**) Dry weight biomass of biofilms formed by the wild-type and the overexpression strains. Gene names are given below the bar. Statistical significance was determined using Holm-Sidak method by performing multiple *t*-test between uninduced and induced condition datasets.

We first set up the experimental conditions allowing the detection of genes whose overexpression would alter biofilm formation as compared to either the uninduced condition or the wild-type control. A strain overexpressing *NRG1*, a known negative transcription regulator of *C. albicans* morphogenesis and biofilm formation [25], was used to optimize the screening conditions. The wild-type control strain (CEC4665) and a P*_TET_-NRG1* overexpression strain (CEC6039) were induced to form biofilms in 96-well polystyrene plates at 37°C for 18h in YPD medium, with or without 25µg.mL^-1^ doxycycline (**Figure 1B**). The extent of biofilm formation was assessed by imaging and by quantifying standard optical density [31]. In these conditions, overexpression of *NRG1*, led to decreased biofilm formation as compared to either the uninduced condition or the wild-type control (**Figure 1C**). Then, the conditions optimized with the *NRG1* overexpression strain were individually applied to the 2454 doxycycline-dependent *C. albicans* overexpression strains. We identified 16 candidate genes that, when overexpressed, inhibited biofilm growth as compared to either wild-type or uninduced cells (**Figure S1A**). These genes encode transcription factors (*CBF1*, *NRG1*, *RBF1*, *ZFU2*, *ZCF8*, *ZCF15* and *ZCF26*), a protein phosphatase (*FCP1*), nucleic acid binding proteins (*PAB1*, *ORF19. 2973* and *ORF19. 5381*), or uncharacterized ORFs (*ORF19.1666, ORF19.3720, ORF19.5933, ORF19.7199* and *GAL7*). Of note, our large-scale screen identified Nrg1 as a negative regulator of biofilm formation.

To confirm that overexpression of the 16 identified genes genuinely hampered biofilm formation, independent overexpression strains for these genes were constructed and tested for their ability to form biofilms in the presence or absence of doxycycline. We could confirm the observed phenotypes for all candidate genes (**Figure 1D**). We also confirmed the reduction of biofilm formation upon overexpression of this set of genes by measuring the dry weight biomass produced on the surface of polystyrene plates (**Figure 1E**). To test whether the reduction in biofilm formation could be the result of a general growth defect upon overexpression, a growth assay was performed with the wild-type control and the 16 overexpression strains, with or without doxycycline. In total, 14 out of the 16 mutants showed no significant alteration in their doubling time upon induction (**Figure S1A**). Conversely, *PAB1* overexpression resulted in a ∼1.5-fold increase in doubling time as compared to both the wild-type strain or uninduced conditions, and the *ORF19.5381* overexpression strain grew poorly in the presence of doxycycline (∼1.9 fold increase in doubling time as compared to uninduced condition and ∼4 fold increase in doubling time as compared to the wild type control) (**Figure S1B**). Of note, although the *ORF19.5381* overexpression strain grew poorly in the absence of doxycycline (∼2.2 fold increase in doubling time as compared to the uninduced wild-type control), it could form robust biofilms in these conditions. Therefore, we did not investigate the *PAB1* and *ORF19.5381* genes further. The 14 remaining candidate genes did not cause any significant alteration of growth between uninduced and induced conditions, indicating a direct role in biofilm formation. We decided to focus on genes encoding transcription factors, namely *NRG1*, *RBF1*, *ZFU2*, *ZCF8*, *ZCF15* and *ZCF26*. *CBF1* was excluded from further studies as it is a characterized transcription factor that binds to the ribosomal protein gene promoters and whose knock-out mutant exhibits a slow growth phenotype [32].

To get further insight in the role of the six regulators in biofilm formation, we first determined the structure and thickness of biofilms formed upon their overexpression by performing confocal laser scanning microscopy (CLSM) with biofilms grown on silicone squares in 12-well polystyrene plates at 37°C for 18 h in YPD medium in the presence of doxycycline [22]. CLSM analysis revealed that biofilms formed upon overexpression of the six candidate genes were mostly composed of yeast cells (**Figure 2A**, top view) resulting in a reduction in the biofilm thickness as compared to the wild-type control strain (**Figure 2A**, side view). These results further confirmed that overexpression of *NRG1*, *RBF1*, *ZFU2*, *ZCF8*, *ZCF15* and *ZCF26* leads to impaired biofilm production.

**Figure 2.**
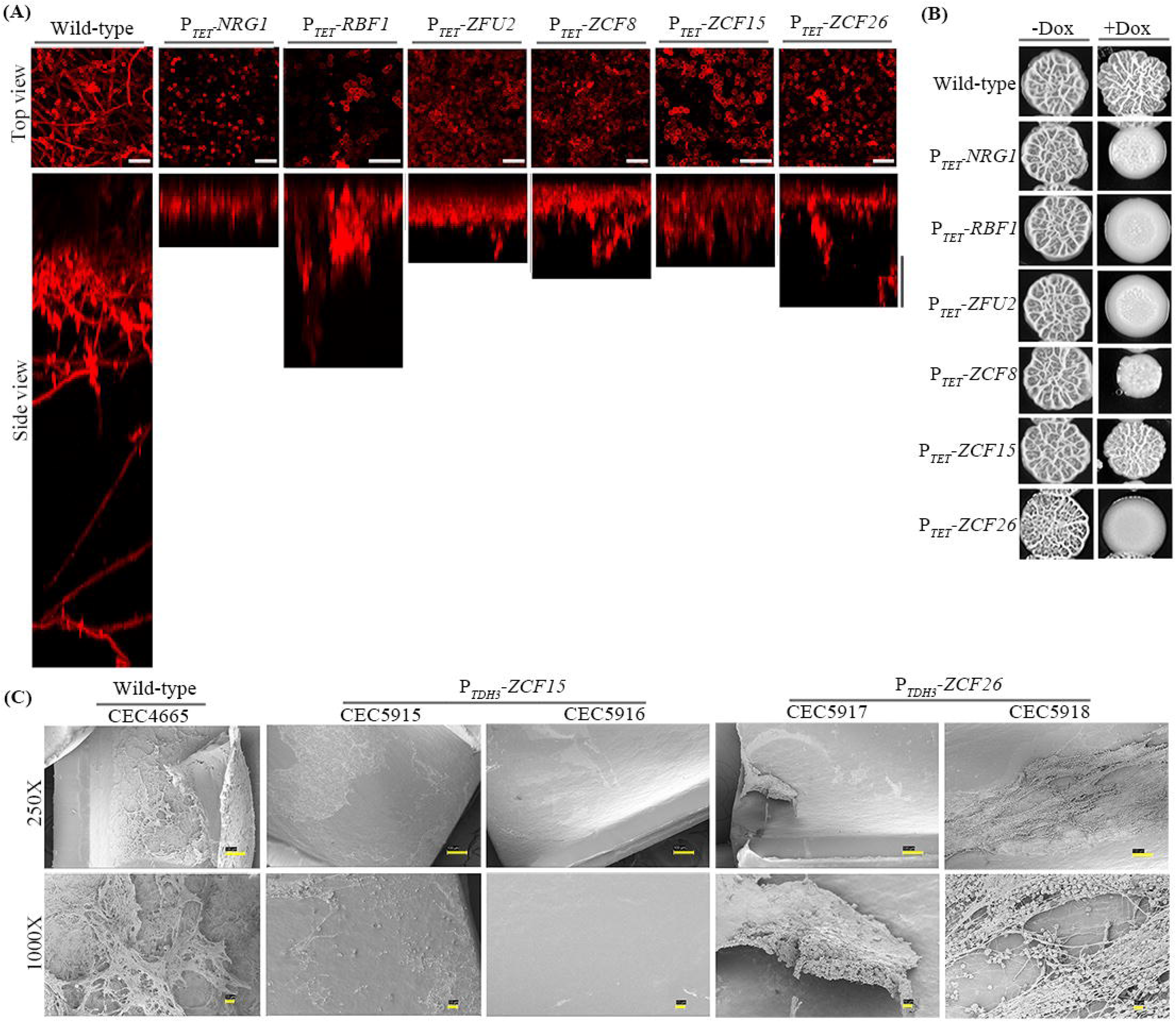
Overexpression of *ZCF15* and *ZCF26* leads to a rudimentary biofilm in rat catheter *in vivo* model. (**A**) Wild-type (CEC4665) and overexpression strains P*_TET_-NRG1* (CEC6039), P*_TET_-RBF1* (CEC6043), P*_TET_-ZFU2* (CEC6044), P*_TET_-ZCF8* (CEC6053), P*_TET_-ZCF15*(CEC6052), and P*_TET_-ZCF26* (CEC6051) were allowed to adhere to silicone squares in 12-well polystyrene plates in YPD medium supplemented with 25 µg/mL doxycycline at 37°C for 1h. Biofilms were allowed to grow for 18h at 110 rpm and stained with concanavalin A-Alexa Fluor^TM^ 594 conjugate for 2h. Biofilms were imaged by CLSM. Images are projections of the top and side views. Representative images of at least 3 replicates are shown. Scale bars for both top view and side view: 25 μm. (**B**) The extent of filamentation of wild-type, P*_TET_-NRG1*, P*_TET_-RBF1*, P*_TET_-ZFU2*, P*_TET_-ZCF8*, P*_TET_-ZCF15*, and P*_TET_-ZCF26* strains was estimated by spot assay on YPD agar containing 20% fetal bovine serum with or without 25 µg/mL doxycycline at 37°C and 3 days of incubation. (**C**) *In vivo* biofilm formation assay was performed using the rat catheter model. Wild-type (CEC4665), P*_TDH3_-ZCF15* (CEC5915 and CEC5916) and P*_TDH3_-ZCF26* (CEC5917 and CEC5918) strains were inoculated in a rat intravenous catheter and were allowed to form biofilms for 24 h. Then, biofilms were visualized using SEM. The images are 100 x and 1000 x magnification views of the catheter lumens. The scale bar for 250x magnification is 100µm and 10µm for 1000x magnification.

### Biofilm forming defect upon *ZCF15* overexpression is independent of hyphal growth

To test whether the defect in biofilm formation upon overexpression of the 6 transcription factor genes is merely the consequence of a defect in hyphal growth, their filamentation was tested by spot assays on solid YPD medium containing 20% fetal bovine serum (FBS) with or without 25µg.mL^-1^ doxycycline (**Figure 2B**). In these conditions, overexpression strains were forming smooth colonies (*RBF1*, *ZFU2* and *ZCF26*) or exhibited reduced wrinkling (*NRG1*, and *ZCF8*), as compared to the wild-type or uninduced conditions. Interestingly, cells overexpressing *ZCF15* still formed wrinkled colonies. We further examined the colony phenotype of the overexpression strains at the single colony level. *NRG1*, *RBF1*, *ZFU2* and *ZCF26* overexpression led to a defect in colony wrinkling. In contrast, *ZCF8* and *ZCF15*-overexpressing cells were able to form wrinkled colonies. (**Figure S2A**). We also inspected the extent of hyphal formation in liquid YPD medium containing 20% FBS with or without doxycycline. In these conditions, strains overexpressing *NRG1*, *RBF1*, and *ZCF26* were compromised for their ability to form hyphae as compared to wild-type control or the uninduced condition. However, overexpression of *ZCF8*, *ZFU2* and *ZCF15* did not prevent hyphal formation in liquid medium (**Figure S2B)**. In conclusion, these results indicate that *ZCF15* overexpression does not affect hyphal growth under hyphae inducing conditions. The phenotype observed is hence specific for biofilm formation.

### Evolutionary appearance of *C. albicans* transcription factors identified by overexpression approaches

In this study, we identified transcription regulators whose overexpression caused a reduction in biofilm formation. Therefore, we questioned the conservation of these novel biofilm regulators in different *Candida* species. To this aim, a search for orthologs of *NRG1*, *RBF1*, *ZCF8*, *ZCF15*, *ZCF26* and *ZFU2* was performed in the Saccharomycetes using PSI-BLAST and Hidden Markov models [33]. Further, the presence of orthologs of these transcription factor genes in closely related species was evaluated using Reciprocal Best Hits (RBH) analysis. The repressor of morphogenesis Nrg1 is present in most sequenced species of the Saccharomycetes, including *Saccharomyces cerevisiae*. Rbf1, another repressor of filamentation in *C. albicans* is present in other members of the CTG clade (ie species in which the CUG codon encodes serine instead of a universal leucine, **Figure S2C**). Zcf8, a regulator of vacuolar function [34], is present only in a few species of the CTG clade [35].

Interestingly, the three other regulators, namely Zfu2, Zcf15 and Zcf26 are restricted to CTG clade species able to form biofilms. Transcription factor Zcf15 is found in *C. albicans*, *C. dubliniensis*, *C. tropicalis*, and *C. parapsilosis*, Zcf26 in *C. albicans*, *C. dubliniensis*, *C. tropicalis*, *C. parapsilosis*, *L. elongisporus* and *S. passalidarum* and Zfu2 occurs in *C. albicans* and *C. dubliniensis* (**Figure S2C**). We also examined the phylogenetic relationship between the transcription factors identified in this study. Phylogenetic analyses suggested that *ZCF15* and *ZCF26* are paralogs and that *ZCF15* originates from a duplication of the *ZCF26* gene (**Figure S2D**). Our phylogenetic analysis confirms the published *ZCF15* and *ZCF26* phylogenetic relationship [36]. In conclusion, these analyses revealed a recent appearance of transcription factors Zcf15, Zcf26 and Zfu2 only in CTG clade species that form biofilms.

### Overexpression of *ZCF15* and *ZCF26* leads to impaired *in vivo* biofilm formation

We demonstrated that conditional overexpression of the *ZCF15* and *ZCF26* paralogs resulted in impaired biofilm formation and that overexpression of *ZCF15* specifically inhibited biofilm growth without any significant defect in hyphal development. To assess whether the results observed upon *in vitro* biofilm formation could be recapitulated *in vivo*, we placed *ZCF15* and *ZCF26* under the control of the constitutive *TDH3* promoter. Next, we examined the constitutive *ZCF15-* and *ZCF26-*overexpression strains for their ability to produce biofilms and filaments *in vitro*. Similar to conditional overexpression, constitutive overexpression of *ZCF15* and *ZCF26* resulted in impaired biofilm formation *in vitro* and overexpression of *ZCF26* alone resulted in impaired filamentation (**Figure S3A** and **S3B**). To investigate the impact of *ZCF15* and *ZCF26* overexpression on *C. albicans* biofilm formation *in vivo*, we used a well-established rat-catheter model [37]. Catheters were inoculated intraluminally with the wild-type control (CEC4665) and two independent clones of P*_TDH3_-ZCF15* (CEC5915 and CEC5916) and P*_TDH3-_ZCF26* (CEC5917 and CEC5918) overexpression strains. After 24 h of biofilm growth, the catheters were removed, and the luminal surfaces of the catheters were imaged by scanning electron microscopy (SEM). Overexpression of *ZCF15* failed to produce any biofilm on rat catheters, whereas overexpression of *ZCF26* resulted in less robust biofilm formation than the wild-type strain (**Figure 2C**). These *in vivo* results thus confirmed the role of transcription factors Zcf15 and Zcf26 in modulating *C. albicans* biofilm formation *in vitro* and *in vivo*.

### Transcriptome alterations upon Zcf15 and Zcf26 overexpression

To understand the mechanisms by which Zcf15 and Zcf26 inhibit *C. albicans* biofilm formation and to uncover the gene circuitry they orchestrate, we conducted a genome-wide transcript profiling with P*_TET_*-*ZCF15* (CEC6052), P*_TET_-ZCF26* (CEC6051) and the wild-type control strain (CEC4665) by RNA sequencing under conditions of biofilm formation in the presence of 25µg.mL^-1^ doxycycline. To rule out an effect of doxycycline on the overall gene expression profile, we also performed transcript profiling with the wild-type strain grown under biofilm conditions with or without doxycycline (**Datasheet A in S3 Table**). In the latter experiment, we considered as differentially expressed those genes that showed a change in expression level by Log2>1.2 or Log2<-1.2 and p<0.05 in response to doxycycline addition. Transcript profiling of *C. albicans* wild-type cells exposed to doxycycline revealed the upregulation of 1 gene and downregulation of 14 genes as compared with untreated wild-type cells (**Datasheets B and C in S3 Table**). Genes whose expression levels were altered by the presence of doxycycline in wild-type cells were excluded from transcriptome analysis of strains overexpressing *ZCF15* and *ZCF26*. RNA expression analysis with P*_TET_-ZCF15* and P*_TET_*-*ZCF26* overexpression strains displayed differential expression of 923 and 1239 genes, respectively, when Log2>1.2 or Log2<-1.2 and p<0.05 were used as the thresholds for differentially expressed genes as compared to the doxycycline-exposed wild-type strain (**Datasheets D and E in S3 Table**). Overexpression of *ZCF15* resulted in the upregulation of 406 coding genes and the downregulation of 517 genes as compared to the wild-type control (**Datasheets F and G in S3 Table**). Similarly, when *ZCF26* was overexpressed, 552 genes were upregulated, and 687 genes were downregulated (**Datasheets H and I in S3 Table**). Interestingly, comparison of differentially expressed genes upon overexpression of *ZCF15* or *ZCF26* showed a common set of 221 upregulated genes and 410 downregulated genes (**Figure S4A** and **S4B**).

To examine the altered pathways upon overexpression of the transcription factors Zcf15 and Zcf26, the differentially expressed genes were categorized into different functional classes using FungiFun 2 [38]. This analysis revealed that genes belonging to cellular metabolism (lipid, fatty acid and isoprenoid, amino acid, C-compound and carbohydrate, nitrogen, sulfur, and selenium metabolism), the tri-carboxylic acid pathway, NAD/NADP binding, and cellular transport were upregulated when *ZCF15* was overexpressed. Categories significantly downregulated included sugar, glucose, polyol and carboxylate metabolism, C-compound and carbohydrate metabolism, stress response, glycolysis, and gluconeogenesis (**Figure 3A**). Similarly, cellular metabolism (lipid, fatty acid and isoprenoid, amino acid, C-compound and carbohydrate and nitrogen, sulfur, and selenium metabolism), the tri-carboxylic acid pathway, protein synthesis (ribosomal proteins), NAD/NADP binding and electron transport were the categories significantly upregulated when *ZCF26* was overexpressed, while sugar, glucose, polyol and carboxylate metabolism, C-compound and carbohydrate metabolism, stress response, glycolysis and gluconeogenesis, filamentation and transcription control were the categories significantly downregulated (**Figure 3B**). Importantly, genes relevant to *C. albicans* morphogenesis including *ACE2*, *CPH2*, *EFG1*, *FKH2*, *ASH1*, *RAS1* etc. were downregulated when *ZCF26* was overexpressed, whereas no significant alterations in the expression of these genes were found when *ZCF15* was overexpressed, suggesting that Zcf26 plays a role in the regulation of both morphogenesis and biofilm formation.

**Figure 3.**
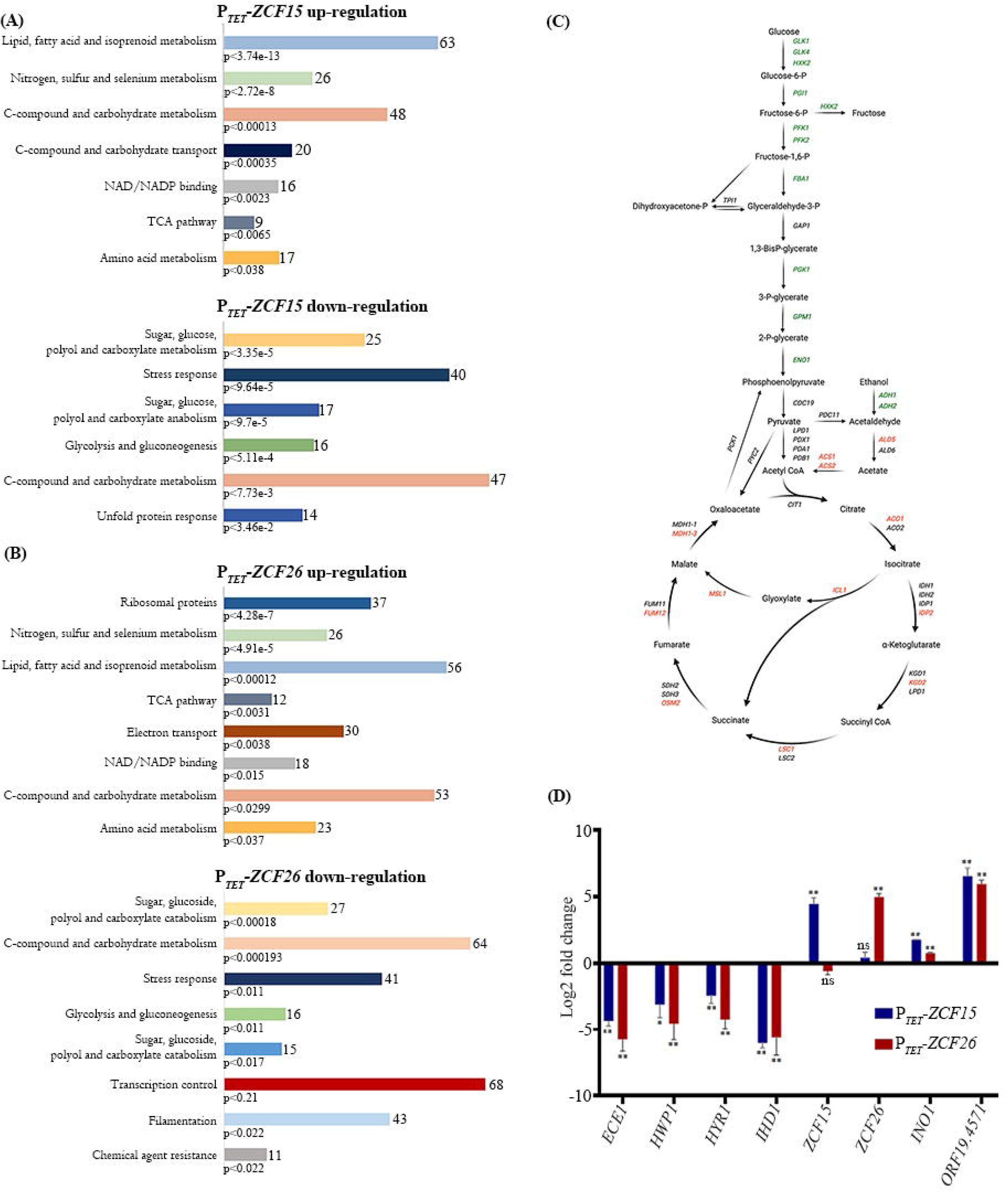
Transcript profiling of *C. albicans* transcription factors *ZCF15* and *ZCF26* during biofilm mode of growth. RNA expression profiling of P*_TET_-ZCF15* (**A**) and P*_TET_-ZCF26* (**B**) strains grown in biofilm condition with 25 µg/mL doxycycline was performed. Functional classification of genome-wide up and down regulated genes upon overexpression of *ZCF15* or *ZCF26* was determined using FungiFun2 and statistically significant altered categories are shown. (**C**) Central metabolic pathways of *C. albicans* is illustrated to show the genes of the glycolytic and tricarboxylic-acid pathways altered when *ZCF15* and *ZCF26* overexpressing strains are grown with 25µg/mL doxycycline in biofilm-forming conditions. Down-regulated genes are indicated in green, up-regulated genes in red and non-significantly altered genes in black. (**D**) qPCR analysis was performed to validate the altered expression of biofilm-related genes with wild-type parental cells, P*_TET_-ZCF15* and P*_TET_-ZCF26* overexpression strains grown in biofilm conditions in the presence of doxycycline. ΔC_T_ values were derived after normalizing the expression of genes of interest with that of *TEF3,* and ΔΔC_T_ values were calculated for the relative expression of the indicated genes. Statistical significance was determined using Holm-Sidak method by performing multiple *t*-tests.

Since transcript profiling pinpointed an alteration of central metabolic pathways of *C. albicans*, we specifically examined expression of glycolysis and TCA cycle genes. Interestingly, we noticed that overexpression of *ZCF15* and *ZCF26* severely hampered the expression of glycolysis genes known to be upregulated during *C. albicans* biofilm formation [15]. In contrast, genes of the glyoxylate shunt and TCA cycle were upregulated upon overexpression of *ZCF15* and *ZCF26* (**Figure 3C** and **S4C**). In addition, several critical biofilm-associated genes that are upregulated during *C. albicans* biofilm formation, such as *HWP1*, *ECE1*, *HYR1*, *HSP104*, and *IHD1* were downregulated upon overexpression of *ZCF15* and *ZCF26*. Similarly, biofilm-repressed genes such as *INO1* and *ORF19.4571*, were upregulated when *ZCF15* and *ZCF26* were overexpressed. These subsets of biofilm-critical genes were also validated by quantitative real-time PCR (**Figure 3D**).

In conclusion, global gene expression analyses with strains overexpressing the transcription factors Zcf15 or Zcf26 suggested a role in metabolic reprogramming during *C. albicans* biofilm development.

### Identification of directly bound targets of Zcf15 and Zcf26 by ChIP-sequencing

Genes directly regulated by Zcf15 and Zcf26 were identified by Chromatin Immunoprecipitation followed by high-throughput sequencing (ChIP-seq), which allowed us to map the binding sites of the regulators in the *C. albicans* genome. To implement the ChIP assay, we fused the N-terminus of the transcription factors Zcf15 or Zcf26 with a Tandem Affinity Purification (TAP) epitope tag. The functionality of TAP-Zcf15 and TAP-Zcf26 was verified by testing the impact of their overexpression on biofilm formation and filamentation. Overexpression of *TAP-ZCF15* or *TAP-ZCF26* phenocopied overexpression of *ZCF15* or *ZCF26*, respectively (**Figure S5A** and **S5B**). We then performed a ChIP assay followed by Illumina sequencing using an untagged *C. albicans* control strain and two independent clones of TAP-tagged Zcf15 and Zcf26 strains growing as biofilms. We detected the binding of Zcf15 and Zcf26 in 317 and 363 intergenic regions of the *C. albicans* genome, respectively. Among these regions, we then identified bona fide promoter regions and uncovered that Zcf15 binds to the promoters of 431 ORFs, whereas Zcf26 binds to the promoters of 494 ORFs (**Datasheets J and K in S3 Table)**. We then compared the results of transcript profiling and ChIP-seq to identify directly regulated genes. Zcf15 binds to the promoters of 89 upregulated and 43 downregulated genes. Similarly, Zcf26 binds to the promoters of 70 upregulated and 87 downregulated genes. A comparison of all genes directly bound either by Zcf15 or Zcf26 and differentially expressed upon their overexpression showed an overlap of 51 upregulated and 41 downregulated genes (**Figure 4A, Datasheet L in S3 Table**). Strikingly, both Zcf15 and Zcf26 bind to the promoter of the master regulator of glycolysis, *TYE7* (**Figure 4B**) as well as to the promoters of some of the TCA cycle genes, namely *IDP2*, *MDH1-3* and *OSM2*. The binding of Zcf15 and Zcf26 to the promoters of these subsets of genes was further verified by ChIP-quantitative PCR (ChIP-qPCR). Promoter region of *ORF19.4690* was used as a control since it is not bound either by Zcf15, Zcf26 or master regulators of biofilm formation (**Figure 4C**). Then, based on genome-wide binding events of Zcf15 and Zcf26, we determined their binding motif using MEME-ChIP [39]. Since Zcf15 and Zcf26 share many targets in which their binding area overlap, the motif identified here for Zcf15 and Zcf26 was very similar (WWWHTCCG) (**Figure 4D**) confirming their common evolutionary origin.

**Figure 4.**
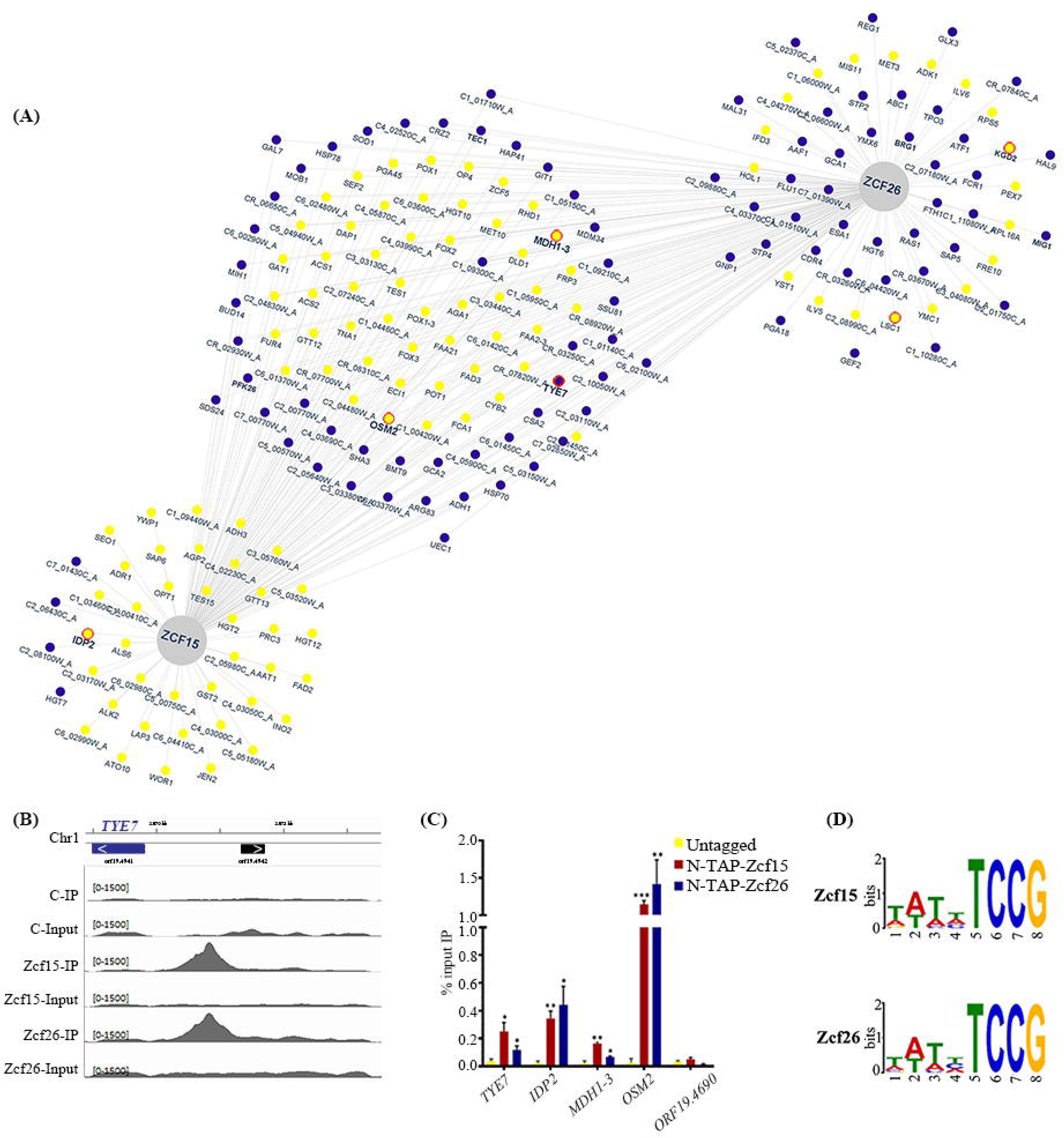
Binding of transcription factors Zcf15 and Zcf26 to the *C. albicans* genome. The DNA binding profile of Zcf15 and Zcf26 obtained by ChIP-sequencing was compared with gene expression data obtained from strains overexpressing *ZCF15* or *ZCF26*. (**A**) Network view for Zcf15 and Zcf26. Genes regulated and bound by Zcf15 (left), Zcf26 (right) or both (middle) are indicated in yellow (upregulation) or blue (downregulation). The interaction network was generated using Cytoscape [72]. Genes further analyzed in (**C**) are circle in red. (**B**) Binding of Zcf15 (middle lanes) and Zcf26 (bottom lanes) at the promoter of *TYE7*, a transcription factor that regulates the expression of genes of the glycolytic pathway. (**C**) ChIP assays were performed on wild-type untagged, *N-TAP-ZCF15* and *N-TAP-ZCF26* strains. Immunoprecipitated (IP) DNA fractions were analyzed by qPCR with primer pairs specific for *TYE7*, *IDP2*, *MDH1*-3 and *OSM2* promoter regions (see **S2 Table**); Zcf15 and Zcf26 unbound region of *ORF19.4690* was used as a negative control. Quantitative RT-PCR was performed on untagged strain samples to detect the background DNA elution in the ChIP assay. The enrichment of Zcf15 and Zcf26 to the promoters of indicated genes is represented as a percent input immunoprecipitated with standard error of mean (SEM). The values from three independent ChIP experiments were plotted. Statistical significance was determined using Holm-Sidak method by performing multiple *t*-test. (**D**) Genome-wide binding motifs of Zcf15 and Zcf26 were identified using MEME-ChIP.

### Metabolic profiling of *ZCF15* and *ZCF26* overexpression strains by phenotypic microarrays

Both transcript profiling and ChIP-sequencing highlighted the role of Zcf15 and Zcf26 in controlling metabolic remodeling during *C. albicans* biofilm formation. Therefore, we examined the metabolic profiles of *ZCF15* and *ZCF26* overexpression strains under different growth conditions. To this aim, we performed phenotypic microarrays (PM), a high-throughput tool to get the global metabolic profiles of microbial cells [40,41]. The growth of the wild-type and of the constitutive overexpression strains P*_TDH3_-ZCF15* and P*_TDH3_-ZCF26* were examined in PM plates (Biolog) coated with different nutrients and chemical substances: carbon sources (PM01 and PM02), nitrogen source (PM03), nutritional complements (PM05), and nitrogen peptides (PM06 and PM08). The global growth profile of the strains was monitored at 30°C for 96 h and is represented as heat-map (**Figure S6**). These PM-based results revealed an enhanced growth of strains overexpressing *ZCF15* and *ZCF26* when succinic acid, acetic acid, α-keto-glutaric acid and pyruvic acid were used as carbon sources. In addition, we noticed that strains overexpressing *ZCF15* and *ZCF26* showed a reduced growth when L-arginine was used either as a carbon source, a nitrogen source or provided as a nutrient supplement. Moreover, both *ZCF15* and *ZCF26* overexpression strains displayed a reduced growth when di-peptides containing Arg residues (Arg-Glu, Arg-Gln, Arg-Ile, Arg-Met, Ile-Arg, Arg-Lys, Arg-Asp, Arg-Leu, Arg-Ser, Arg-Val, Arg-Trp, Arg-Arg, Pro-Arg, Arg-Tyr, Leu-Arg, Arg-Ala) were used as nitrogen source (**Figure 5**). These results coincide with the previous reports on the role of arginine metabolism in *C. albicans* biofilm formation [42]. In conclusion, this PM-based growth analysis further establishes the role of the Zcf15 and Zcf26 transcription factors in controlling metabolic remodeling during *C. albicans* biofilm development.

**Figure 5.**
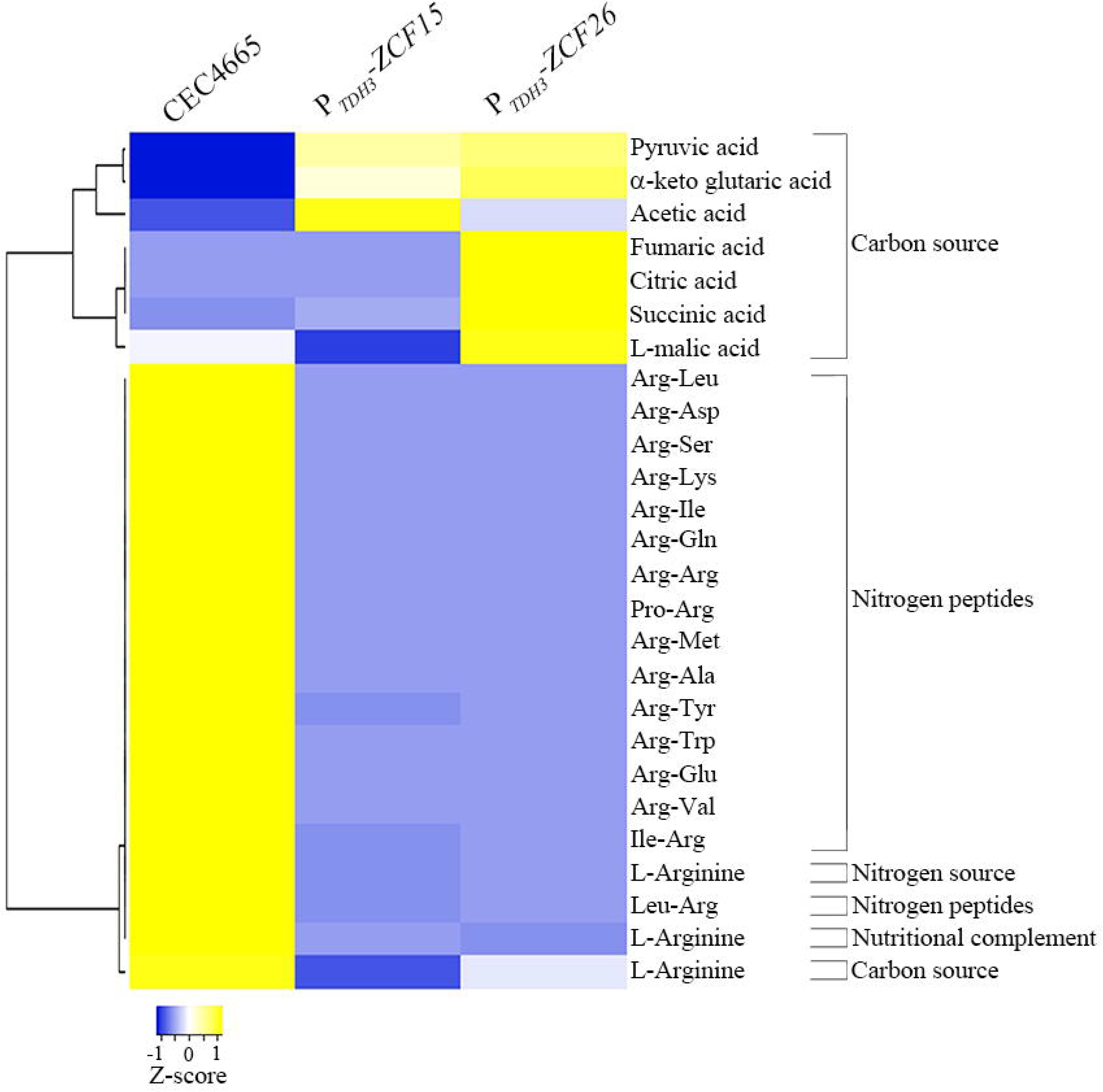
Metabolic activities profile of transcription factors *ZCF15* and *ZCF26*. Comparison of metabolic activities of parental reference strain and *ZCF15* and *ZCF26* overexpression strains is shown as a heat-map. Metabolic activities were monitored at 30°C for 96 h and were measured using the area under the curve (AUC). Metabolic activity in the indicated growth conditions is represented on a scale from -1 (minimum growth, blue) to +1 (maximum growth, yellow).

## Discussion

In their natural environment, many microbial species, including bacteria, archaea and fungi, alternate between planktonic and sessile states, alone or in association with other microbial species [6,43]. A radical shift in gene expression and cellular metabolism has been reported in bacteria and fungi during the transition from planktonic to community growth. Bacterial and fungal biofilms indeed show unique metabolic patterns, such as differential expression of glycolytic pathway genes, indicating significant metabolic reprogramming during microbial biofilm development [44–46].

Fungal biofilm formation is a complex developmental process that is associated to multiple traits, with each trait having a specific role during the transition from planktonic to biofilm growth. These traits are regulated by a different set of transcription regulators [47,48]. During *C. albicans* biofilm establishment, two major events occur: cell differentiation and metabolic reprogramming [19,21,42,49,50]. The regulators and their genetic networks modulating cell differentiation during *C. albicans* biofilm formation have been extensively studied. For instance, Ace2, Brg1, Efg1, Ndt80, Tec1, Flo8 and Ume6 regulate the expression of genes involved in *C. albicans* morphogenesis, which provides architectural stability to biofilms. In contrast, transcription regulators that modulate metabolic alterations during *C. albicans* biofilm formation have received less attention [48].

In this study, a large-scale overexpression approach identified a new set of transcription regulators involved in biofilm formation, associated with either morphogenesis (*NRG1*, *RBF1*, *ZFU2* and *ZCF8*), metabolic alteration (*ZCF15*) or both (*ZCF26*). We selected Zcf15 and Zcf26 for further study as they are paralogs whose occurrence is restricted to CTG clade species that form biofilms, and no prior information on their role in morphogenesis or biofilm formation was known.

Metabolic reprogramming is one of the major changes that occur during microbial biofilm formation [17,19–21]. Bonhomme et al. demonstrated the upregulation of glycolysis genes during *C. albicans* biofilm formation and highlighted the role of Tye7 in their regulation [15]. Furthermore, a comparative metabolomic study of *C. albicans* planktonic and biofilm cells revealed differential production of metabolites of the TCA cycle, lipid synthesis, amino-acid metabolism, glycolysis and oxidative stress [21]. These authors showed that the level of citrate decreased in all stages of biofilm formation, including early and intermediate biofilms, while other intermediates of the TCA cycle (succinate, fumarate, and malate) decreased only in mature biofilms. Moreover, comparison of transcript profiling of cells from planktonic cultures and biofilms also highlighted the role of the TCA cycle and mitochondrial activities during *C. albicans* biofilm formation [20]. These results suggest an inhibition of the TCA cycle during biofilm maturation and a reduction of the respiration rate in biofilm cells. Interestingly, transcript profiling of *ZCF15* and *ZCF26* overexpression strains demonstrated their role in the alteration of central metabolism, in particular the downregulation of genes of the glycolysis and upregulation of genes of the glyoxylate pathway and the TCA cycle. In a different study, Issi et al. also revealed the up-regulation of glucose metabolism in a *ZCF15* knockout strain [36], which further supports our results. In addition, *ZCF26* overexpression also impacted the expression level of genes associated with morphogenesis, which may be the cause of the defect in filamentous growth in the presence of doxycycline. For instance, *ACE2*, *BRG1, CPH2*, *EFG1*, *FKH2*, *ASH1*, or *RAS1,* which are involved in the yeast to hyphae transition, are downregulated when *ZCF26* is overexpressed. On the contrary, most genes whose expression is altered upon *ZCF15* overexpression are associated with metabolism, and no significant differences were observed for genes associated with morphogenesis (**Figure 3A**). These data were well supported by the genome-wide binding study and the locus-specific PCR (**Figure 4A** and **4C**). Chip-sequencing and ChIP-qPCR experiments revealed that both Zcf15 and Zcf26 bind to the regulatory region of *TYE7*, a key regulator of glycolysis. Furthermore, these two regulators bind to promoter regions of genes encoding enzymes of the TCA cycle, such as *IDP2*, *LSC1, KGD2*, *MDH1-3* and *OSM2,* demonstrating their role in modulating the expression levels of TCA cycle and glyoxylate cycle genes. In addition, Zcf15 and Zcf26 bind to the acetyl-CoA synthetase-encoding genes, *ACS1* and *ACS2*, which regulate the metabolism of nonfermentable carbon sources via gluconeogenesis, glyoxylate cycle and β-oxidation [51]. Phenotypic microarray results further highlighted the involvement of these two regulators in controlling metabolic remodeling; indeed, overexpression of *ZCF15* and *ZCF26* resulted in increased growth when precursors of the TCA cycle including succinic acid, α-keto-glutaric acid and pyruvic acid were used as a carbon source. Based on these results, we speculate that upon overexpression of *ZCF15* and *ZCF26*, an alteration in the rate of the glycolysis, TCA cycle, and glyoxylate cycle leads to the establishment of a non-fermentative environment, which favors the planktonic mode of growth and thus results in an impaired biofilm formation. Therefore, we posit that a higher occupancy of these two transcription factors at the promoters of critical genes of central metabolic pathways may prevent the regulators required for biofilm formation from accessing them.

Besides regulating genes of the central metabolism, Zcf15 and Zcf26 also directly regulate the expression of genes necessary for normal biofilm growth (*CRZ2, CSA2*, *RAS1*), involved in biofilm matrix formation (*GCA1*, *GCA2*), and of transcription regulators of biofilm gene networks (*BRG1*, *TEC1*) [6,52–54].

Apart from carbohydrate metabolism, amino-acid metabolism is also crucial for *C. albicans* biofilm formation. Garcia-Sanchez et al. observed that amino acid biosynthetic pathway genes are upregulated during biofilm formation under the aforementioned growth conditions, which led to the demonstration of a role in *C. albicans* biofilm formation of the *GCN4* gene encoding a master regulator of amino acid biosynthetic genes [19]. In addition, Rajendran et al. have shown that amino-acid biosynthetic pathway genes such as arginine and proline are upregulated in high biofilm forming *C. albicans* isolates [42]. Moreover, a recent study revealed the role of the amino acid permease Stp2 in *C. albicans* adherence and biofilm maturation [16]: *stp2* knock-out mutants are impaired for amino acid uptake and compensatory mechanisms in nutrient acquisition. We noticed a lower utilization of L-arginine by *ZCF15* and *ZCF26* overexpression strains when used as either carbon source, nitrogen source or provided as nutritional complement (**Figure 5**). In addition, overexpression of *ZCF15* and *ZCF26* resulted in slower growth when Arg-containing dipeptides were used as a nitrogen source. These results suggest that Zcf15 and Zcf26 may be involved in the regulation of L-arginine utilization. Strikingly, Zcf26 binds directly to the promoter region of *STP2,* which is downregulated during *ZCF26* overexpression. Therefore, this study establishes the role of arginine metabolism during *C. albicans* biofilm formation. Interestingly, the presence of the transcription regulators Zcf15 and Zcf26 is limited to species in the Candida clade that can form biofilms, suggesting their relatively recent acquisition in biofilm-forming species. Furthermore, the shared regulation of several genes by Zcf15 and Zcf26 argues for a common evolutionary origin of these regulators and may allow tight regulation of the set of regulated genes.

In summary, by using overexpression approaches, we discovered new biofilm regulators with either a role in architectural stability and/or a specific role in metabolic reprogramming. This study also identified several other regulators and genes whose further study will provide a better understanding of the mechanism of *C. albicans* biofilm formation. Altogether, this study highlights the role of metabolic reprogramming and its fine-tuned regulation during the shift from planktonic to biofilm growth. This could lead to the development of new antifungals designed to selectively disrupt the fungus central metabolism to treat biofilm-related infections.

## Materials and Methods

### Ethics Statement

All animal procedures were approved by the Institutional Animal Care and Use Committee at the University of Wisconsin according to the guidelines of the Animal Welfare Act, the Institute of Laboratory Animal Resources Guide for the Care and Use of Laboratory Animals, and Public Health Service Policy under protocol MV1947. Ketamine and xylazine were used for anesthesia. CO_2_ asphyxiation was used for euthanasia at the end of study.

### Data availability

Genome-wide RNA expression and ChIP-sequencing data are deposited to European Nucleotide Archive (ENA) under accession numbers E-MTAB-11383 and E-MTAB-11384, respectively.

### Media and growth conditions

*C. albicans* strains used in this study are listed in **S1 Table**. Cells were grown in YPD (1% yeast extract, 2% peptone and 2% dextrose) at 30°C for planktonic and at 37°C for biofilm growth. Solid media were obtained by adding 2% agar. Induction of P*_TET_* was achieved by adding 25 µg/mL doxycycline. Hyphal growth was induced by adding 20% fetal bovine serum to the medium.

### Biofilm measurement by standard optical density assay

To measure the extent of *C. albicans* biofilm formation, we performed 96-well standard optical density assays [31] for all *C. albicans* doxycycline-dependent P*_TET_* overexpression strains. Biofilms were allowed to grow at the bottom of 96-well polystyrene plate in YPD medium at 37°C for 18 h at 110 rpm with or without adding 25µg/mL doxycycline. Optical density was measured using Tecan I control infinite M200. We measured the optical density at nine independent locations per well; values from six independent wells were used to plot the graph and to estimate the statistical significance.

### *In vitro* biofilm formation and dry biomass measurement

To measure the dry biomass produced, biofilms were grown in 12-well polystyrene TPP plates (Cat. No. 92412) in 2 mL of YPD medium with or without 25µg/mL doxycycline. The plates were inoculated with cells at OD_600_=0.2 and incubated at 37°C for 60 min at 110 rpm agitation for initial adhesion of the cells. After 60 min, the plates were washed with 2 mL of 1x PBS, and 2 mL of fresh YPD medium with or without 25µg/mL doxycycline were added. Plates were then sealed with breathseal sealing membranes (Greiner bio-one) and incubated at 37°C for 18 h with shaking at 110 rpm. Then the medium was aspirated, and the wells were gently washed with 1x PBS. To estimate the dry biomass of biofilms produced, biofilms were scrapped, and the content of each well was transferred to pre-weighed nitrocellulose filters. Biofilm-containing filters were dried overnight at 60°C and weighed. The average total biomass for each strain was calculated from three independent samples after subtracting the mass of the empty filters [55].

### CLSM for biofilm imaging

Biofilms were grown on silicone squares in YPD medium with 25µg/mL doxycycline for 18 hours. The medium was discarded, and silicone squares were gently washed with 1x PBS and stained with 50 µg/mL of concanavalin A-Alexa Fluor 594 (Invitrogen) at 30°C for 2 h, with gentle shaking at 110 rpm. Silicone squares were then placed in a Petri dish and covered with 1x PBS. Biofilms were imaged as described previously [22]: CSLM was performed at the UtechS PBI facility of Institut Pasteur using an upright LSM700 microscope equipped with a Zeiss 40X/1.0 W plan-Apochromat immersion objective. Images were acquired and assembled into maximum intensity Z-stack projection using the ZEN software.

### RNA extraction and cDNA synthesis

RNAs were isolated using the RNeasy mini kit mirVana RNA isolation kit (Qiagen). Briefly, *C. albicans* strains were grown in YPD medium either in planktonic grown at 30°C or biofilm-growth conditions grown at 37°C in shaking mode for 18 h in polystyrene plates. Total RNA was isolated from four independent planktonic or biofilm cultures for each strain. Planktonic cells were grown in 50 mL YPD medium in flasks at 30°C till OD600=0.8 whereas biofilms were grown in 2 mL of YPD in 12-well polystyrene plates at 37°C for 18h. Cells were harvested by centrifugation at 4000 rpm both from planktonic and biofilms isolated cells and washed 3 times with 1xPBS and pelleted at 4000 rpm. Cells were resuspended in 700 µL of extraction buffer and lysed by adding 0.5mm of 500 µL of glass beads. Cells were broken in a bead-beater with 500 µL of 0.5mm of glass beads (six cycle of 2 min at 10). The RNeasy columns were used to isolate the total RNA. To remove the potential contaminating chromosomal DNA, RNA samples were treated on-column with DNAse for 15 min at room temperature (Cat. No. 79254, Qiagen). A total of 1µg of purified RNA was used to make cDNA by adding gDNA wipeout (2 μL), RT buffer 5x (4 μL) RT primer mix and Reverse transcriptase (1 μL) (Qiagen, Cat. No. 205311) added in a final volume of 20 μL. Reactions were carried out at 42°C for 15 min followed by heat inactivation at 95°C for 3 min.

### RNA sequencing and analysis

Libraries were built using a TruSeq Stranded mRNA library Preparation Kit (Illumina, USA) following the manufacturer’s protocol. Quality control was performed on a BioAnalyzer 2100 (Agilent Technologies). 75bp single-end RNA sequencing was performed on the Illumina NextSeq 500 platform.

The RNA-seq analysis was performed with Sequana [56]. In particular, we used RNA-seq pipeline (v0.9.16, https://github.com/sequana/sequana_rnaseq) built on top of Snakemake 5.8.1 [57]. Reads were trimmed from adapters using Cutadapt 2.10 [58] then mapped to the *C. albicans* (SC5314, version A22-s07-m01-r105) genome assembly and annotation from Candida Genome Database [59] using STAR 2.7.3a [60]. FeatureCounts 2.0.0 [61] was used to produce the count matrix, assigning reads to features with strand-specificity information. Quality control statistics were summarized using MultiQC 1.8 [62]. Statistical analysis on the count matrix was performed to identify differentially regulated genes, comparing biofilm and planktonic condition RNA expression. Clustering of transcriptomic profiles were assessed using a Principal Component Analysis (PCA). Differential expression testing was conducted using DESeq2 library 1.24.0 [63] scripts based on SARTools 1.7.0 [64] indicating the significance (Benjamini-Hochberg adjusted p-values, false discovery rate FDR < 0.05) and the effect size (fold-change) for each comparison. Functional categorization of up-and downregulated genes were achieved by using FungiFun2 [38].

### Quantitative PCR

*C. albicans* wild-type (CEC4665) and P*_TET_-ZCF15* and P*_TET_-ZCF26* strains were grown in biofilm forming condition in the presence of doxycycline as described earlier. RNAs were isolated as described above in RNA extraction section (Qiagen). The integrity of RNAs were examined on 1% agarose gel. cDNA was synthesized by reverse transcription using QuantiTech Reverse Transcription Kit. Primers designed for real time PCR reactions are listed in **S2 Table**. Analysis of melting curves were performed to ensure specific amplification without any secondary non-specific amplicons (melting curve temperatures used were 80°C (*TEF3*), 77°C (*ECE1*), 83°C (*HWP1*), 80°C (*HSP104*), 78°C (*HYR1*) 83°C (*ZCF15*), 80°C (*ZCF26*), 80°C (*INO1*), 80.5°C (*ORF19.4571*) and 81.5°C (*IHD1*). PCR was carried out in a final volume of 20 μL using SsoAdvanced^TM^ Universal SYBR Green supermix (BIO-RAD). The real time PCR analysis was achieved with an i-Cycler (BIO-RAD) using the following reaction conditions: 95°C for 2 min, then 40 cycles of 95°C for 30 s, 55°C for 30 s, 72°C for 30 s. Fold difference in expression of mRNA was calculated by the ΔΔC_T_ method (Real–time PCR applications guide BIO-RAD) [65] using *C. albicans* transcription elongation factor 3 (*TEF3*) transcript as normalization control.

### *In vivo* rat catheter biofilm formation

To perform *in vivo* biofilms, the rat central-venous catheter infection model was used, as described previously [13,37,66,67]. To achieve the *in vivo C. albicans* biofilm formation, specific pathogen free Sprague-Dawley rats weighing 400 g each were used. A heparinised (100 U/ml) polyethylene catheter with 0.76 mm inner and 1.52 mm outer diameters was inserted into the external jugular vein. The catheter was secured to the vein with the proximal end tunneled subcutaneously to the midscapular space and externalized through the skin. The catheters were inserted 24 h prior to infection to permit a conditioning period for a deposition of host protein on the catheter surface. Infection was achieved by intraluminal instillation of 500 μL *C. albicans* cells (10^6^ cells/ml). After a 4 h dwelling period, the catheter volume was withdrawn, and the catheter flushed with heparinized 0.15 M NaCl. Catheters were removed after 24 h of *C. albicans* infection to assay biofilm development on the intraluminal surface by scanning electron microscopy (SEM). Catheter segments were washed with 0.1 M phosphate buffer, pH 7.2, fixed in 1% glutaraldehyde/ 4% formaldehyde, washed again with phosphate buffer for 5 min, and placed in 1% osmium tetroxide for 30 min. The samples were dehydrated in a series of 10 min ethanol washes (30%, 50%, 85%, 95% and 100%), followed by critical point drying. Specimens were mounted on aluminum stubs, sputter coated with gold, and imaged using a Hitachi S-5700 or JEOL JSM-6100 scanning electron microscopy in the high-vacuum mode at 10kV. Images were processed using Adobe photoshop software.

### Chromatin immunoprecipitation (ChIP)

The ChIP assays were performed as described previously [68]. Briefly, each strain was grown in biofilm condition for 18 h and cells were cross-linked with 1% final concentration of formaldehyde for 25 min at 30°C. Chromatin was isolated and sonicated to yield an average fragment size of 300-500 bp. The DNA in 50 µL of water was immunoprecipitated with 20 µg/mL anti-protein A antibodies (Sigma Aldrich) and purified by phenol/chloroform extraction. The total, immunoprecipitated (IP) DNA, and beads only material were used to determine the binding of Zcf15 and Zcf26 across the genome by ChIP-sequencing, or to the promoters of a subset of biofilm-related genes by real time PCR (qPCR), as described before. The template used was as follows – 1 µL of a 1:50 dilution for input and 1 µL of a 1:3 dilution for immunoprecipitated DNA (IP) Zcf15-TAP, Zcf26-TAP, and an untagged control strain. The conditions used for qPCR were as follows: 95°C for 2 min; then 40 cycles of 95°C for 30 sec, 55°C for 30 sec, 72°C for 45 sec. The results were analyzed using CFX Manager Software. The graph was plotted according to the percent input method [69].

### Library preparation and ChIP-sequencing analysis and DNA binding motif identification

The ChIP DNA library was prepared using TruSeq ChIP sample preparation guidelines (Illumine) and sequencing was achieved by using Nextseq 500 run. The ChIP-seq analysis was performed with the ChIP-seq pipeline of the Sequana framework [56]. We checked the quality of the data by computing the ratio between data peak and so-called phantom peaks and found values >1.3, which indicates a good-quality ChIP-seq data according to best practices recommended by ENCODE [70]. We then mapped the data and identify narrow and broad peaks using Macs3 (https://github.com/macs3-project/MACS). Finally, we obtained the final list of peaks by computing IDR (Irreproducible Discovery Rate), which is the approach used in ChIP-sequencing analysis to provide stable thresholds based on reproducibility [71]. The DNA binding motif across the *C. albicans* genome was identified using Motif Analysis of Large Nucleotide Datasets (MEME-ChIP) [39]. The interaction network was generated with Cytoscape [72].

### Phenotype MicroArray and data analysis

Phenotypic Microarray (PM) plates and reagents (inoculating fluid IFY-0 base, redox dye mix D and E) were purchased from Biolog Inc. The composition of the PM plates can be found on the Biolog website (https://www.biolog.com/wp-content/uploads/2020/04/00A-042-Rev-C-Phenotype-MicroArrays-1-10-Plate-Maps.pdf). *C. albicans* strains were streaked to YPD plates and grown for 2 days at 30°C. A total of 2-6 colonies from each YPD plates were transferred to the 15 mL tubes in NS medium (nutrient supplement) and cell density was calculated using turbidimeter (Biolog). Turbidity of the suspension was measured by turbidimeter (Biolog) and transmittance was reached to 62%T (+/-1%). The PM panels represent 96-well plates containing different substrate in each well. In addition to the different substrate, PM wells were also containing the minimal components required for normal growth and prepared according to the manufacture’s guidelines. PM additives and dye were added according to the method provided by Biolog Inc. In summary, 0.5 mL of cell suspension were mixed to appropriate volume of PM inoculating fluids, A 100 µL of different cell suspension from the PM inoculating fluid was transferred to each well coated with different nutrients. Plates were sealed with PCR seal to keep wells from drying out and to avoid cross-well spreading of volatile chemicals. All PM plates were incubated in Omnilog at 30°C for 96h. The Omnilog software was used to analyze the data. Differential growth was considered when area under the curve (AUC) of mutants were differed by two times in both directions as compared to the reference strain. Differential growth was converted in the form of heat-map using Heatmapper [73]. Clustering was achieved by average linkage and distance was measured by using Pearson method.

### Statistical significance

Graphs were generated using GraphPad Prism. Statistical significance was determined by performing multiple *t*-test using Holm-Sidak method [74].

## Acknowledgments

This project was supported by a grant from the Fondation pour la Recherche Médicale (FRM, DBF20160635719) to CdE. We thank J. Fonseca, E. Turc, L. Lemee and E. Kornobis from the Biomics platform, C2R2, Institut Pasteur, Paris, France, supported by France Génomique (ANR-10-INBS-09-09) and IBISA, and Virginie Passet for her help during operating the Omnilog instrument. We also acknowledge the photonic bioimaging (UTechs PBI) facility of Institut Pasteur, Paris. Work in the laboratory of CdE is supported by the Agence Nationale de Recherche (ANR-10-LABX-62-IBEID).

