## Supplementary figures and images for "Metabolic reprogramming during *Candida albicans* planktonic-biofilm transition is modulated by the *ZCF15* and *ZCF26* paralogs"

### Supplementary figures

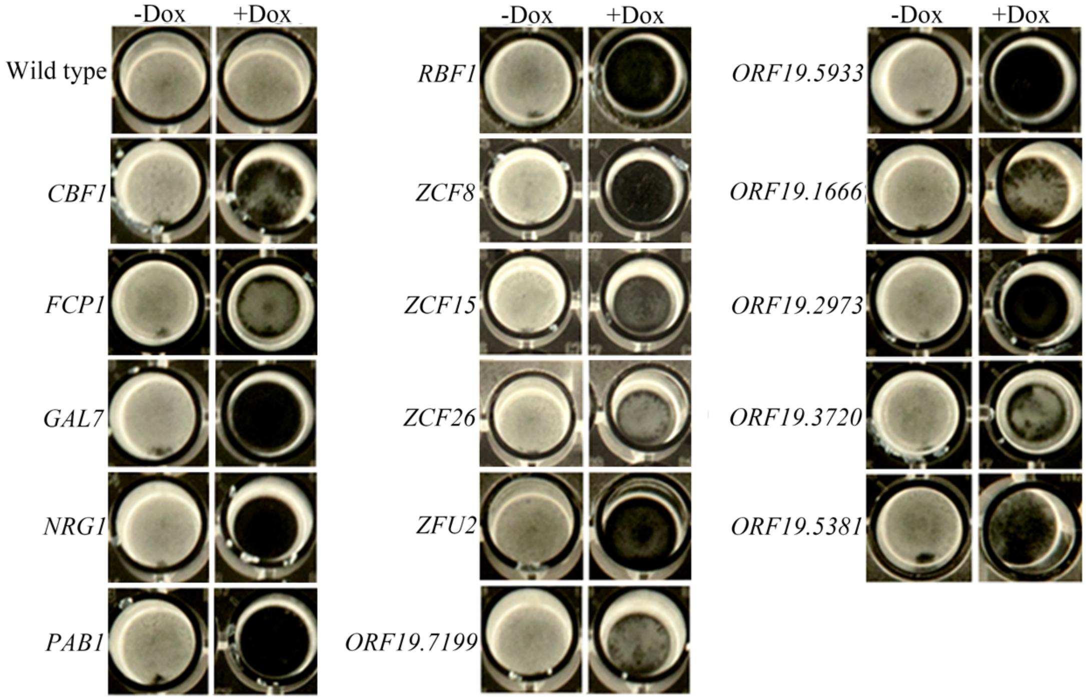

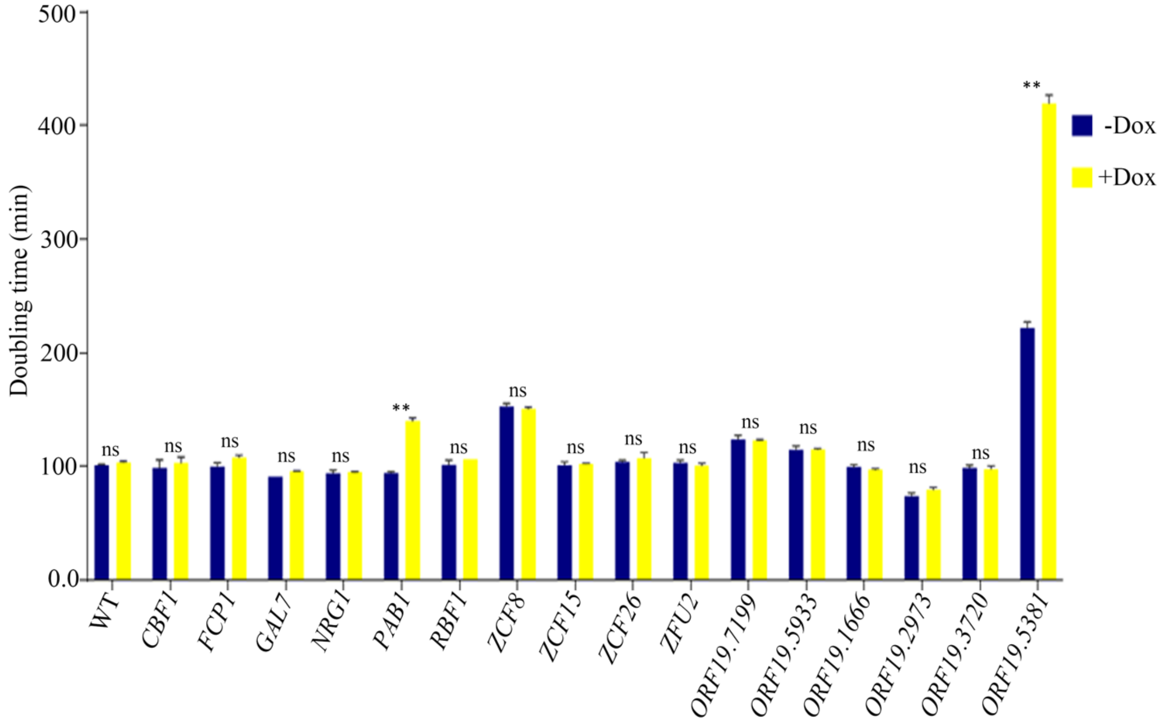

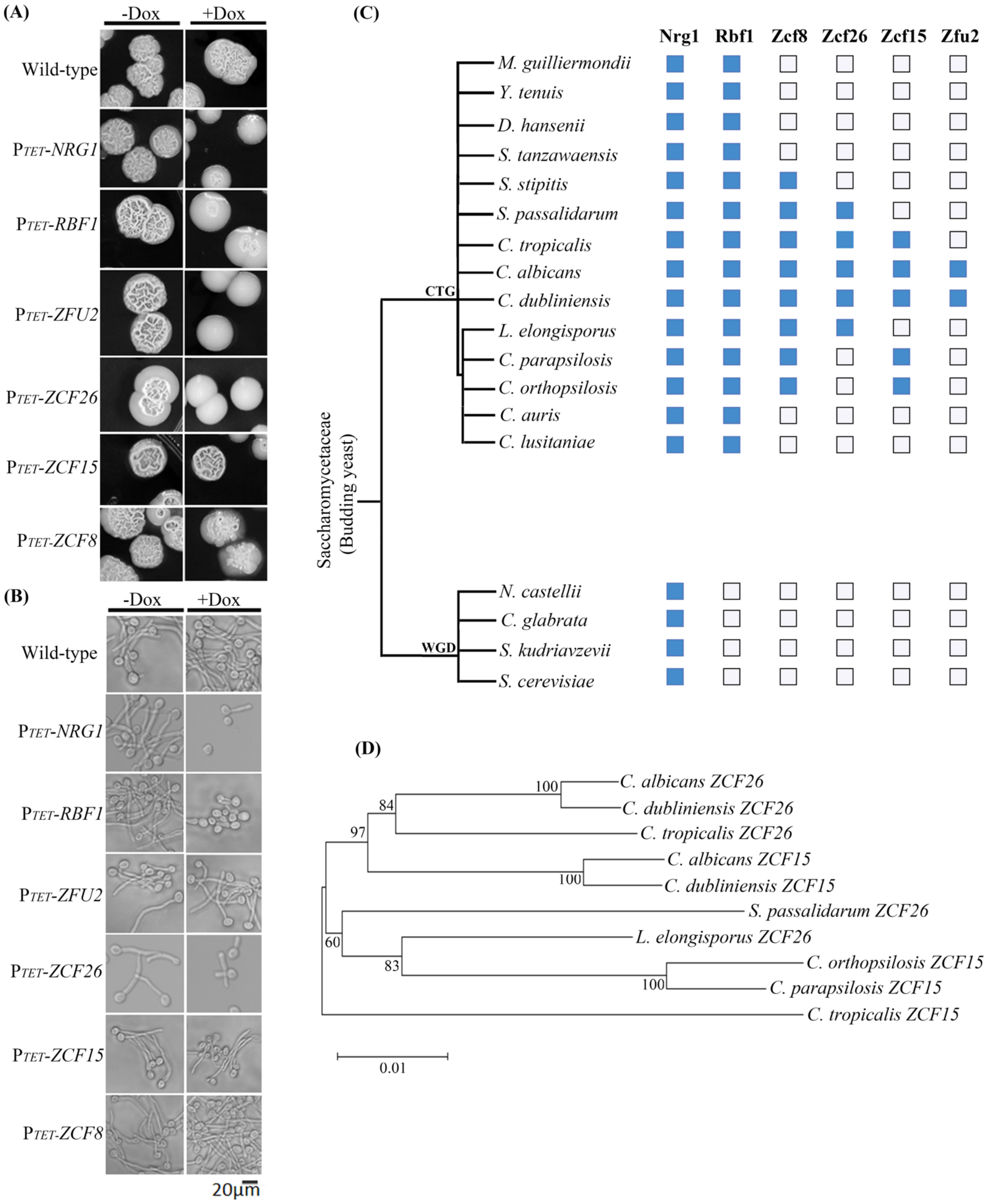

Figure S3

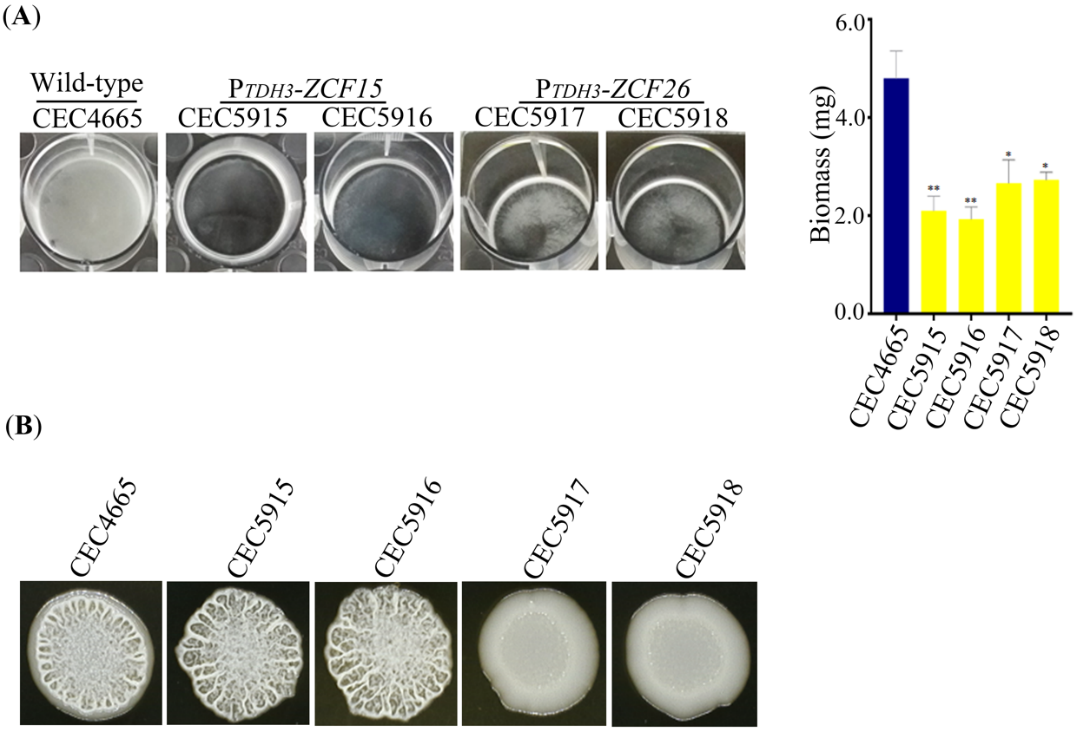


Figure S4

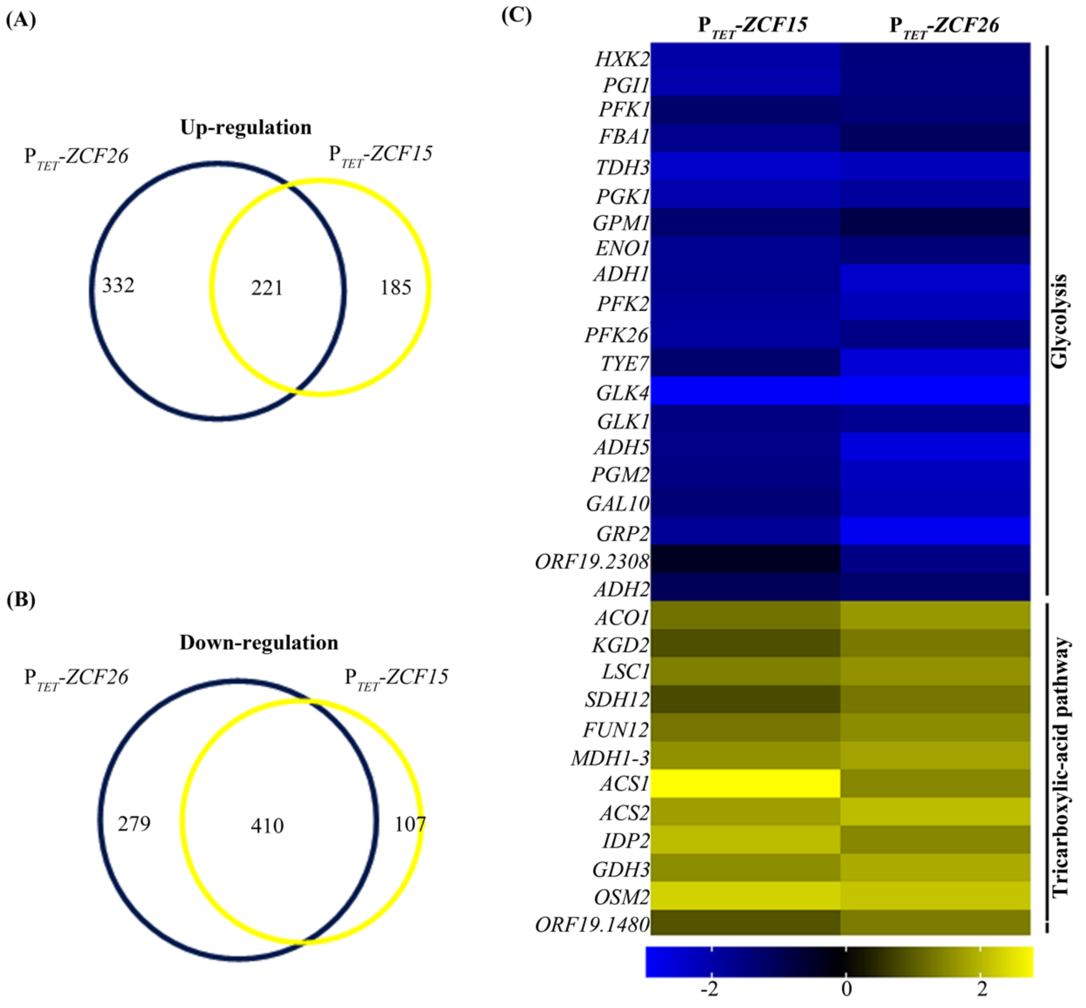

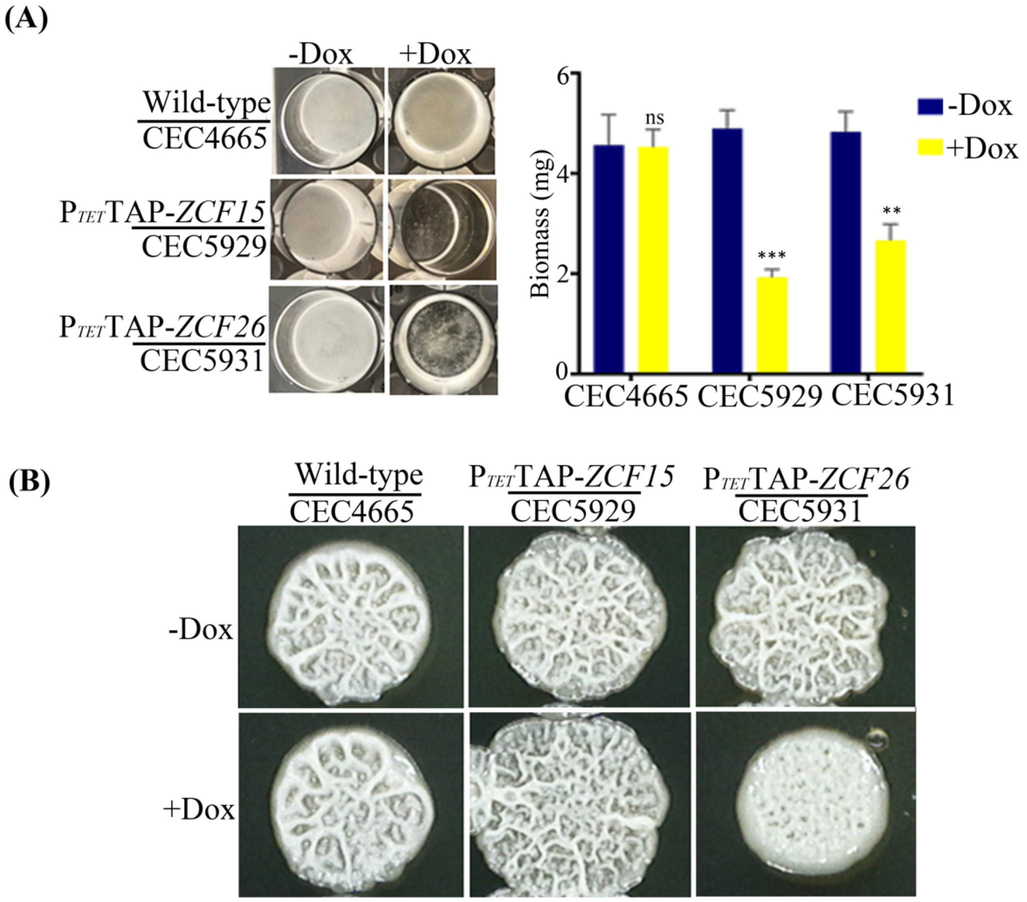

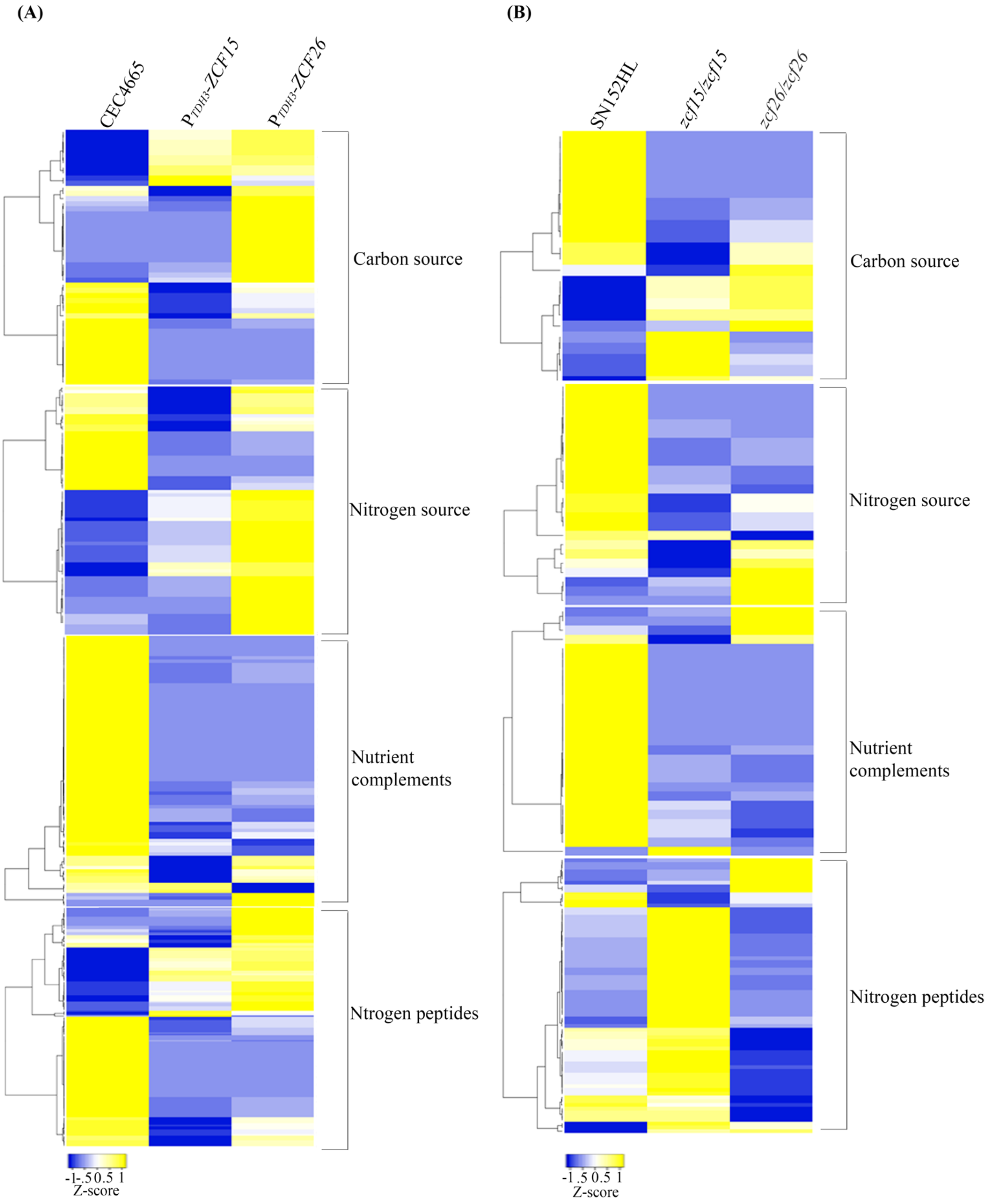
