## supplementary information for "Metabolic reprogramming during *Candida albicans* planktonic-biofilm transition is modulated by the *ZCF15* and *ZCF26* paralogs"

**Supplementary figures legend**

**Figure S1 Biofilm formation and growth measurements for candidate genes identified by overexpression approach. (A)** *C. albicans* wild-type and $P_{TET}$-overexpression strains were grown overnight in YPD medium with or without 25µg/mL doxycycline. Biofilm formation was allowed to develop in 96-well polystyrene plates in YPD medium with or without 25µg/mL doxycycline at 37°C for 18 h. (**B**) Wild-type (CEC4665) and $P_{TET}$-overexpression strains were grown in liquid YPD medium, with or without 25 µg/mL doxycycline until the stationary phase was reached. Optical density was measured using Tecan Sunrise.

**Figure S2 Transcription factors *ZCF15* and *ZCF26* are paralogous genes.** (**A**) The extent of filamentation of wild-type (CEC4665), $P_{TET}$-*NRG1* (CEC6039), $P_{TET}$-*RBF1* (CEC6043), $P_{TET}$-*ZCF8* (CEC6053), $P_{TET}$-*ZCF15* (CEC6052), $P_{TET}$-*ZCF26* (CEC6051) and $P_{TET}$-*ZFU2* (CEC6044) strains were examined at the single colony level on YPD plates containing 20% fetal bovine serum with or without 25µg/mL doxycycline and grown for five days at 37°C. (**B**) Similarly, filamentation assay was performed for 1h in YPD liquid medium with 10% FBS for the indicated strains in the absence or presence of 25µg/mL doxycycline at 37°C. Scale bars: 20 µm. (**C**) Orthologs of the indicated transcription factors in the budding yeasts of the Saccharomycetes class are shown. The presence (blue box) or absence (empty box) of the orthologs of the transcription factor indicated for each species was shown. This tree is illustrative as the branches are not drawn to the scale. (**D**) Phylogenetic analyses show that *ZCF15* and *ZCF26* transcription factors are paralogous genes. The yellow color mark indicates the branch duplication of the gene.

**Figure S3 Comparative analysis of transcriptome profile of $P_{TET}$-*ZCF15* and $P_{TET}$-*ZCF26*.** (**A**) Genome-wide expression data were compared for commonly up-regulated genes between $P_{TET}$-*ZCF15* and $P_{TET}$-*ZCF26* overexpression strains (blue and yellow circles, respectively) and represented as Venn diagrams. (**B**) Similarly, a Venn diagram was constructed for down-regulated genes in the two strains. A total of 221 up-regulated and 410 down-regulated genes are common between the two datasets. (**C**) A heat-map was illustrated to show the differentially expressed genes of the glycolytic and tricarboxylic-acid pathway when *ZCF15* and *ZCF26* were grown with 25µg/mL doxycycline in biofilm-forming condition.

**Figure S4 Constitutive expression of transcription factors *ZCF15* and *ZCF26* results in reduced biofilm formation.** (**A**) Biofilm formation assay of strains with constitutive expression of *ZCF15* or *ZCF26* placed under the control of $P_{TDH3}$, a constitutive promoter. The wild-type parental strain (CEC4665), two independent strains with $P_{TDH3}$-*ZCF15* (CEC5915 and CEC5916) or with $P_{TDH3}$-*ZCF26* (CEC5917 and CEC5918) were allowed to form biofilms in 12-well polystyrene microtiter plates in YPD medium at 37°C for 18 h before and dry weight biomass was estimated (**B**) The extent of filamentation of these strains was estimated by spot assay on YPD plates containing 20% fetal bovine serum.

**Figure S5 TAP-tagged *ZCF15* and *ZCF26* are functional upon overexpression.** (**A**) To examine the functionality of TAP-epitope tagged protein, the wild-type parental strain, *N-TAP-ZCF15* (CEC5929) and *N-TAP-ZCF26* (CEC5931) strains were allowed to form biofilms in 12-well polystyrene plates in YPD medium, with or without 25µg/mL doxycycline at 37°C for 18 h and dry weight biomass measured. (**B**) The extent of filamentation of these strains was estimated by spot assay on YPD plates containing 20% fetal bovine serum with or without 25µg/mL doxycycline.

**Figure S6 Metabolic activities of *ZCF15* and *ZCF26* overexpression and null mutant strains are illustrated as heat-map**.

(**A**) Comparison of metabolic activity profiles of the parental strain (CEC4665) and the overexpression strain of *ZCF15* and *ZCF26* on indicated PM plates is shown as a heat-map.

### Supporting Experimental Procedures

**Orthologues and phylogenetic analysis:** Orthologue searches were performed using Hidden Markov Model HMMER Web server [1]. Transcription factors orthologue was also confirmed by Best Reciprocal Hits (RBH). A phylogenetic tree was inferred using the Neighbor-Joining method. Evolutionary analyses were conducted using MEGA X.

#### Construction of *C. albicans* strains

**Construction of *C. albicans* overexpression strains:** Detailed methods of transferring *C. albicans* ORFs from pDONR207 into the expression plasmids and integrating the resulting expression plasmids at the *RPS1* locus have been described [2,3]. Briefly, ORFs were transferred from the entry clones into our collection of 2496 barcoded $P_{TET}$-driven overexpression plasmids using the Gateway™ LR clonase™ II Enzyme mix (Invitrogen).

After *E. coli* DH5α transformation, the ORF transfer into the expression plasmid was verified by EcoRV digestion. The *URA3*-bearing expression plasmids were digested by StuI and transformed into *C. albicans* strain CEC4642 (a SN76 derivative [4] according to Walther and Wendland [5], and adapted to the use of 96-well microplates. Transformants were selected for prototrophy and verified by PCR using primer pair CIpUL/CIpUR, which yields a 1 kb amplicon if the integration of the overexpression plasmid has occurred at the *RPS1* locus [6].

**Construction of CEC5929, CEC5930, CEC5931 and CEC5932:** To examine the genome-wide binding events of *ZCF15* and *ZCF26*, we constructed N-terminal TAP-tagged strains for both *ZCF15* and *ZCF26*. To generate the TAP-tagged strain of *ZCF15* and *ZCF26*, plasmids were isolated from *E. coli* strain ECC1983 and ECC1984 (see below plasmid construction) respectively. StuI digested DNA fragments were transformed to *C. albicans* strain CEC4642. Transformants were selected for prototrophy and proper integration at the *RPS1* locus was verified by PCR using primer pair CIpUL/CIpUR.

**Construction of CEC5915, CEC5916, CEC5917 and CEC5918:** Plasmids carrying *ZCF15* (ECC1985) or *ZCF26* (ECC1986) (see below plasmid construction) under the control of $P_{TDH3}$ were digested by StuI prior to transformation into *C. albicans* strain CEC4642. Transformants were selected for prototrophy and proper integration at the *RPS1* locus was verified by PCR using primers CIpUL and CIpUR.

**Plasmid construction**

**ECC1979:** 670 bp upstream *ZCF15* and 451 bp downstream *ZCF15*were PCR amplified with oligos ZCF15USSATF, ZCF15USSATR and ZCF15DSSATF, ZCF15DSSATR, respectively. The upstream fragment was cloned in pSFS2A digested with *Kpn*I and *Xho*I; the downstream fragment was inserted in the resulting plasmid digested with *Sac*I and *Sac*II.

**ECC1983 and ECC1984:** The Entry plasmids (BP clone) bearing the *ZCF15* or the *ZCF26* coding sequences, respectively, were used in a Gateway™ LR reaction together with the CIp-$P_{TET}$-TAP-GTW-SP vector (ECC1095). Recombination mixes were transformed into *E. coli* Top10. The resulting plasmids were digested with EcoRV and BamHI to verify the integration of the appropriate ORF.

**ECC1985:** The Entry plasmid (BP clone) bearing *ZCF15* coding sequences was used in a Gateway™ LR reaction together with the CIp-Ca$P_{TDH3}$-GTW (ECC843) vector. The recombination mixes were transformed into *E. coli* Top10. The resulting plasmids were digested with EcoRV and BamHI to verify the cloning of the appropriate ORF.

**ECC1986:** The Entry plasmid (BP clone) bearing *ZCF26* coding sequences were used in a Gateway™ LR reaction together with the ECC843 (CIp-CaP$_{TDH3}$-GTW) vector. The recombination mixes were transformed into *E. coli* Top10. Transformants were digested with EcoRV and BamHI to verify the cloning of the appropriate ORF.

**Table 1.** ***C. albicans*** **strains and** ***E. coli*** **plasmids used in this study**

| Yeast strain | Genotype | Reference |
|---|---|---|
| SN76 | *ura3Δ::λimm*$^{434}$*/ura3Δ::λimm*$^{434}$ *iro1Δ::λimm*$^{434}$*/iro1Δ::λimm*$^{434}$ *arg4Δ/arg4Δ his1Δ/his1Δ* | [1] |
| CEC4642 | SN76 *ADH1/adh1::*$P_{TDH3}$*-carTA::SAT1 arg4Δ/CaARG4 his1Δ::hisG/HIS1* | This study |
| CEC4665 | CEC4642 *RPS1/RPS1::*CIp10 | This study |
| CEC6038 | CEC4642 *RPS1/RPS1::*CIp10-$P_{TET}$-*ORF19.7199* | This study |
| CEC6039 | CEC4642 *RPS1/RPS1::*CIp10-$P_{TET}$-*NRG1* | This study |
| CEC6040 | CEC4642 *RPS1/RPS1::*CIp10-$P_{TET}$*CBF1* | This study |
| CEC6041 | CEC4642 *RPS1/RPS1::*CIp10-$P_{TET}$-*GAL7* | This study |
| CEC6042 | CEC4642 *RPS1/RPS1::*CIp10-$P_{TET}$-*ORF19.5933* | This study |
| CEC6043 | CEC4642 *RPS1/RPS1::*CIp10-$P_{TET}$-*RBF1* | This study |
| CEC6044 | CEC4642 *RPS1/RPS1::*CIp10-$P_{TET}$-*ZFU2* | This study |
| CEC6045 | CEC4642 *RPS1/RPS1::*CIp10-$P_{TET}$-*ORF19.1666* | This study |
| CEC6046 | CEC4642 *RPS1/RPS1::*CIp10-$P_{TET}$-*PAB1* | This study |
| CEC6047 | CEC4642 *RPS1/RPS1::*CIp10-$P_{TET}$-*ORF19.2973* | This study |
| CEC6048 | CEC4642 *RPS1/RPS1::*CIp10-$P_{TET}$-*3720* | This study |
| CEC6049 | CEC4642 *RPS1/RPS1::*CIp10-$P_{TET}$-*FCP1* | This study |
| CEC6050 | CEC4642 *RPS1/RPS1::*CIp10-$P_{TET}$-*ORF19.5381* | This study |
| CEC6051 | CEC4642 *RPS1/RPS1::*CIp10-$P_{TET}$-*ZCF26* | This study |
| CEC6052 | CEC4642 *RPS1/RPS1::*CIp10-$P_{TET}$-*ZCF15* | This study |
| CEC6053 | CEC4642 *RPS1/RPS1::*CIp10-$P_{TET}$-*ZCF8* | This study |
| CEC5915 | CEC4642 *RPS1/RPS1::*CIp10-$P_{TDH3}$-*ZCF15* | This study |
| CEC5916 | CEC4642 *RPS1/RPS1::*CIp10-$P_{TDH3}$-*ZCF15* | This study |
| CEC5917 | CEC4642 *RPS1/RPS1::*CIp10-$P_{TDH3}$-*ZCF26* | This study |
| CEC5918 | CEC4642 *RPS1/RPS1::*CIp10-$P_{TDH3}$-*ZCF26* | This study |
| CEC5929 | CEC4642 *RPS1/RPS1::*Clp10-$P_{TET}$-*TAP-ZCF15* | This study |
| CEC5930 | CEC4642 *RPS1/RPS1::*Clp10-$P_{TET}$-*TAP-ZCF15* | This study |
| CEC5931 | CEC4642 *RPS1/RPS1::*Clp10-$P_{TET}$-*TAP-ZCF26* | This study |

| | | |
|---|---|---|
| CEC5932 | CEC4642 *RPS1/RPS1::*Clp10-$P_{TET}$-*TAP-ZCF26* | This study |
| **Plasmid** | **Description** | **References** |
| ECC1979 | pSFS2A-5-UTR-3UTR-ZCF15 | This study |
| ECC1983 | CIp-pTET-NTAP-ZCF15 | This study |
| ECC1984 | CIp-pTET-NTAP-ZCF26 | This study |
| ECC1985 | CIp-pTDH3-ZCF15 | This study |
| ECC1986 | CIp-pTDH3-ZCF26 | This study |
| ECC843 | CIp-CaTDH3p-GTW | This study |
| ECC1095 | CIp-pTET-rTAP-GTW-SP | [2] |

1. Noble SM, Johnson AD. Strains and Strategies for Large-Scale Gene Deletion Studies of the Diploid Human Fungal Pathogen *Candida albicans.* Eukaryot Cell. 2005;4: 298–309. doi:10.1128/ec.4.2.298-309.2005

2. Legrand M, Bachellier-Bassi S, Lee KK, Chaudhari Y, Tournu H, Arbogast L, et al. Generating genomic platforms to study *Candida albicans* pathogenesis. Nucleic Acids Res. 2018;46: 6935–6949. doi:10.1093/nar/gky594

**Table 2. Oligonucleotide primers used in this study**

| Primer name | Sequence (5'-3') |
|---|---|
| TEC1qFP | GCAATTTCTGGGCAGATTG |
| TEC1qRP | TGGGAAATGTGCATTTAGG |
| Tye7qFP | GAATGACCTGGAAAAACCTAG |
| Tye7qRP | CAAATCTACTCTGAGAACAATC |
| IDP2qFP | CGTACACGGACACACACGAAAC |
| IDP2qRP | CATCGTCATTTTTTTCGCGTG |
| OSM2qFP | GGAGGGGGACACCGGAGAAAG |
| OSM2qRP | CGTTCCCGCGTCAAATTATCC |
| MDH1-3qFP | CGCAAAATGCTTGTTACACC |
| MDH1-3qRP | CGCCACCTTCGCAAACAAAC |
| orf19.4690-F | GAATATGCCGTTGGGTGTTT |
| orf19.4690-R | AACCCGAACCCAAAAACAAT |
| ZCF26FqRT | CATTGTTGTCCTTCCCGTTAC |
| ZCF26RqRT | GTCTTTTCTTTCATCGTCTGGTG |
| TEF3-F | GATCACAATTGGGTCCAAGG |
| TEF3-R | AGCAGCGGCAATCTTGTTAC |
| ZCF15FqRT | GAAAGGAGCGAGGAAGGAGAG |
| ZCFR15qRT | GTGGCTCTCGTTTCCCTGAATG |
| HWP2FqRT | CCCAGCATCTTCAACTACTAG |
| HWP2RqRT | CAGTGACAACAATAGCACC |
| HWP1RTFP | CACAACAGCCACAAGAACC |
| HWP1RTRP | AGGTTGAGGTGGATTGTCG |
| ECE1RTFP | CACCTACTGTTCCTGCACC |
| ECE1RTRP | ATTACTTGTGGAATGTTGCC |
| IHD1RTFP | GAATTGGCTCTGTGTGATTG |
| IHD1RTRP | ACTTGTAGATTGACCTTGAG |
| INO1RTFP | GCTGATGTTTTGCCAAATGTC |
| INO1RTRP | TTCCAAGATAGAAGCAACGGC |

| | |
|---|---|
| 4571RTFP | TCAGTAAAGTTCCCCCATTTG |
| 4571RTRP | ACAGAGTTCGAGCACTTTGAC |
| ZCF15USSATF | GGGGTACCGGTCCCTCGGATCAACAAG |
| ZCF15USSATR | CCGCTCGAGTTGAAATATTTGTGAATAG |
| ZCF15DSSATF | TTCCCGCGGTATAGACTCTTTTTTTACATATA |
| ZCF15DSSATR | CGAGCTCCCACCAATACTATTACTAC |
| CIpUL | ATACTACTGAAATTTCCTGACTTTC |
| CIpRL | ATTACTATTTACAATCAAAGGTGGTC |

**Supporting Experimental Procedures**

**Orthologues and phylogenetic analysis:** Orthologue searches were performed using Hidden Markov Model HMMER Web server [1]. Transcription factors orthologue was also confirmed by Best Reciprocal Hits (RBH). A phylogenetic tree was inferred using the Neighbor-Joining method. Evolutionary analyses were conducted using MEGA X.

**Construction of *C. albicans* strains**

**Construction of *C. albicans* overexpression strains:** Detailed methods of transferring *C. albicans* ORFs from pDONR207 into the expression plasmids and integrating the resulting expression plasmids at the *RPS1* locus have been described ([2,3]. Briefly, ORFs were transferred from the entry clones into our collection of 2496 barcoded $P_{TET}$-driven overexpression plasmids using the Gateway™ LR clonase™ II Enzyme mix (Invitrogen). After *E. coli* DH5α transformation, the ORF transfer into the expression plasmid was verified by EcoRV digestion. The *URA3*-bearing expression plasmids were digested by StuI and transformed into *C. albicans* strain CEC4642 (a SN76 derivative [4]) according to Walther and Wendland [5], and adapted to the use of 96-well microplates. Transformants were selected for prototrophy and verified by PCR using primer pair CIpUL/CIpUR, which yields a 1 kb amplicon if the integration of the overexpression plasmid has occurred at the *RPS1* locus [6].

**Construction of CEC5929, CEC5930, CEC5931 and CEC5932:** To examine the genome-wide binding events of *ZCF15* and *ZCF26*, we constructed N-terminal TAP-tagged strains for both *ZCF15* and *ZCF26*. To generate the TAP-tagged strain of *ZCF15* and *ZCF26*, plasmids were isolated from *E. coli* strain ECC1983 and ECC1984 (see below plasmid construction) respectively. StuI digested DNA fragments were transformed to *C. albicans* strain CEC4642. Transformants were selected for prototrophy and proper integration at the *RPS1* locus was verified by PCR using primer pair CIpUL/CIpUR.

**Construction of CEC5915, CEC5916, CEC5917 and CEC5918:** Plasmids carrying *ZCF15* (ECC1985) or *ZCF26* (ECC1986) (see below plasmid construction) under the control of $P_{TDH3}$ were digested by StuI prior to transformation into *C. albicans* strain CEC4642. Transformants were selected for prototrophy and proper integration at the *RPS1* locus was verified by PCR using primers CIpUL and CIpUR.

**Plasmid construction**

**ECC1979:** 670 bp upstream *ZCF15* and 451 bp downstream *ZCF15*were PCR amplified with oligos ZCF15USSATF, ZCF15USSATR and ZCF15DSSATF, ZCF15DSSATR, respectively (S2

Table). The upstream fragment was cloned in pSFS2A digested with *Kpn*I and *Xho*I; the downstream fragment was inserted in the resulting plasmid digested with *Sac*I and *Sac*II.

**ECC1983 and ECC1984:** The Entry plasmids (BP clone) bearing the *ZCF15* or the *ZCF26* coding sequences, respectively, were used in a Gateway™ LR reaction together with the CIp-P$_{TET}$-TAP-GTW-SP vector (ECC1095). Recombination mixes were transformed into *E. coli* Top10. The resulting plasmids were digested with EcoRV and BamHI to verify the integration of the appropriate ORF.

**ECC1985:** The Entry plasmid (BP clone) bearing *ZCF15* coding sequences was used in a Gateway™ LR reaction together with the CIp-CaP$_{TDH3}$-GTW (ECC843) vector. The recombination mixes were transformed into *E. coli* Top10. The resulting plasmids were digested with EcoRV and BamHI to verify the cloning of the appropriate ORF.

**ECC1986:** The Entry plasmid (BP clone) bearing *ZCF26* coding sequences were used in a Gateway™ LR reaction together with the ECC843 (CIp-CaP$_{TDH3}$-GTW) vector. The recombination mixes were transformed into *E. coli* Top10. Transformants were digested with EcoRV and BamHI to verify the cloning of the appropriate ORF.
