## supplementary data for "Metabolic reprogramming during *Candida albicans* planktonic-biofilm transition is modulated by the *ZCF15* and *ZCF26* paralogs"

### S3 Table description

**S3A**- Wild type (CEC4665) doxycycline-treated complete transcriptome data obtained on 4 independent samples. **Headers: A**- Gene ID (ORF19_ID, orf19 nomenclature at the *Candida albicans* genome database (CGD); **B-E**- Wild type without doxycycline treatment count; **F-I**- Wild type doxycycline treated count; **J-M**-Wild type without doxycycline treated normalized count; **N-Q**-Wild type doxycycline treated normalized count; **R**-Base mean over all samples; **S**-Wild type without doxycycline treated base mean; **T**-Wild type doxycycline treated mean; **U**-Fold change; **V**-log2 Fold Change; **X**-p-value; **Y**-Adjusted p value; **Z**-Dispersion parameter estimated from feature counts; **AA**- Dispersion parameter estimated from the model; **AB**-Dispersion parameter estimated from the Maximum A Posteriori model; **AC**-Final dispersion parameter used to perform the test; **AD**-Convergence of the coefficients of the model (TRUE or FALSE); **AE**-Maximum Cook's distance of the feature; **AF**-Indicates if the feature has been detected as a count outlier (it does not make sense if an infinite threshold is given for the Cook's distance).

**S3B**-Wild type (CEC4665) doxycycline-treated upregulated genes. **Header: A**- Gene ID (ORF19_ID, orf19 nomenclature at the *Candida albicans* genome database (CGD); **B-E-** Wild type without doxycycline trated count; **F-I**- Wild type doxycycline treated count; **J-M-**Wild type without doxycycline treated normalized count; **N-Q**-Wild type doxycycline treated normalized count; **R**-Base mean over all samples; **S**-Wild type without doxycycline treated base mean; **T**-Wild type doxycycline treated mean; **U**-Fold change; **V**-log2 FoldChange; **X**-p-value; **Y**-Adjusted p value; **Z**-Dispersion parameter estimated from feature counts; **AA**- Dispersion parameter estimated from the model; **AB**-dispersion parameter estimated from the Maximum A Posteriori model; **AC**-Final dispersion parameter used to perform the test; **AD**-Convergence of the coefficients of the model (TRUE or FALSE); **AE**-Maximum Cook's distance of the feature; **AF**-indicates if the feature has been detected as a count outlier (it does not make sense if an infinite threshold is given for the Cook's distance).

**S3C**- Wild type (CEC4665) doxycycline-treated downregulated genes. **Header: A**- Gene ID (ORF19_ID, orf19 nomenclature at the *Candida albicans* genome database (CGD); **B-E**- Wild type without doxycycline treated count; **F-I**- wild type doxycycline treated count; **J-M**-Wild type without doxycycline treated normalized count; **N-Q**-Wild type doxycycline treated normalized count; **R**-Base mean over all samples; **S**-Wild type without doxycycline treated base mean; **T**-Wild type doxycycline treated mean; **U**-Fold change; **V**-log2FoldChange; **X**-p-value; **Y**-Adjusted p value; Z-Dispersion parameter estimated from feature counts; **AA**- Dispersion parameter estimated from the model; **AB**-dispersion parameter estimated from the Maximum A Posteriori model; **AC**-final dispersion parameter used to perform the test; **AD**-Convergence of the coefficients of the model (TRUE or FALSE); **AE**-Maximum Cook's distance of the feature; **AF**-Indicates if the feature has been detected as a count outlier (it does not make sense if an infinite threshold is given for the Cook's distance).

**S3D**- $P_{TET}$-*ZCF15* doxycycline-treated complete transcriptome data. **Header: A**- Gene ID (ORF19_ID, orf19 nomenclature at the *Candida albicans* genome database (CGD); **B-E**- Wild type doxycycline treated count; **F-I**- ZCF15 overexpression count; **J-M**-Wild type normalized count; **N-Q**-*ZCF15* overexpression normalized count; **R**-Base mean over all samples; **S**-Wild type base mean; **T**- *ZCF15* overexpression mean; **U**-Fold change; **V**-log2 FoldChange; **X**-p-value; **Y**-Adjusted p value; **Z**-Dispersion parameter estimated from feature counts; **AA**- Dispersion parameter estimated from the model; **AB**-Dispersion parameter estimated from the Maximum A Posteriori model; **AC**-final dispersion parameter used to perform the test; **AD**-Convergence of the coefficients of the model (TRUE or FALSE); **AE**-Maximum Cook's distance of the feature; **AF**-indicates if the feature has been detected as a count outlier (it does not make sense if an infinite threshold is given for the Cook's distance).

**S3E**- $P_{TET}$-*ZCF26* doxycycline-treated complete transcriptome data. **Header: A**- Gene ID (ORF19_ID, orf19 nomenclature at the *Candida albicans* genome database (CGD); **B-E**- Wild type count; **F-I**- *ZCF26* overexpression count; **J-M**-wild type normalized count; **N-Q**-*ZCF26* overexpression normalized count; **R**-Base mean over all samples; **S**-Wild type base mean; **T**-*ZCF26* overexpression mean; **U**-Fold change; **V**-log2 FoldChange; **X**-p-value; **Y**-Adjusted p value; **Z**-Dispersion parameter estimated from feature counts; **AA**- Dispersion parameter estimated from the model; **AB**-Dispersion parameter estimated from the Maximum A Posteriori model; **AC**-Final dispersion parameter used to perform the test; **AD**-Convergence of the coefficients of the model (TRUE or FALSE); **AE**-Maximum Cook's distance of the feature; **AF**-indicates if the feature has been detected as a count outlier (it does not make sense if an infinite threshold is given for the Cook's distance).

**S3F**- $P_{TET}$-*ZCF15* doxycycline-treated upregulated genes. **Header: A**- Gene ID (ORF19_ID, orf19 nomenclature at the *Candida albicans* genome database (CGD); **B-E**- Wild type count; **F-I**- *ZCF15* overexpression count; **J-M**-Wild type normalized count; **N-Q**-*ZCF15* overexpression normalized count; **R**-Base mean over all samples; **S**-Wild type base mean; **T**-*ZCF15* overexpression mean; **U**-Fold change; **V**-log2FoldChange; **X**-p-value; **Y**-Adjusted p value; **Z**-Dispersion parameter estimated from feature counts; **AA**- Dispersion parameter estimated from the model; **AB**-dispersion parameter estimated from the Maximum A Posteriori model; **AC**-final dispersion parameter used to perform the test; **AD**-Convergence of the coefficients of the model (TRUE or FALSE); **AE**-Maximum Cook's distance of the feature; **AF**-Indicates if the feature has been detected as a count outlier (it does not make sense if an infinite threshold is given for the Cook's distance).

**S3G**- $P_{TET}$-*ZCF15* doxycycline-treated downregulated genes**. Header: A**- Gene ID (ORF19_ID, orf19 nomenclature at the *Candida albicans* genome database (CGD); **B-E**- Wild type count; **F-I**- *ZCF15* overexpression count; **J-M**-Wild type normalized count; **N-Q**-*ZCF15* overexpression normalized count; **R**-Base mean over all samples; **S**-Wild type base mean; **T**-*ZCF15* overexpression mean; **U**-Fold change; **V**-log2FoldChange; **X**-p-value; **Y**-Adjusted p value; **Z**-Dispersion parameter estimated from feature counts; **AA**- Dispersion parameter estimated from the model; **AB**-Dispersion parameter estimated from the Maximum A Posteriori model; **AC**-Final dispersion parameter used to perform the test; **AD**-Convergence of the coefficients of the model (TRUE or FALSE); **AE**-Maximum Cook's distance of the feature; **AF**-Indicates if the feature has been detected as a count outlier (it does not make sense if an infinite threshold is given for the Cook's distance).

**S3H**- $P_{TET}$-*ZCF26* doxycycline-treated upregulated genes. **Header: A**- Gene ID (ORF19_ID, orf19 nomenclature at the *Candida albicans* genome database (CGD); **B-E**- Wild type count; **F-I**- *ZCF26* overexpression count; **J-**M-Wild type normalized count; **N-Q**-*ZCF26* overexpression normalized count; **R**-Base mean over all samples; **S**-Wild type base mean; **T**-*ZCF15* overexpression mean; **U**-Fold change; **V**-log2FoldChange; **X**-p-value; **Y**-Adjusted p value; **Z**-Dispersion parameter estimated from feature counts; **AA**- Dispersion parameter estimated from the model; **AB**-Dispersion parameter estimated from the Maximum A Posteriori model; **AC**-Final dispersion parameter used to perform the test; **AD**-Convergence of the coefficients of the model (TRUE or FALSE); **AE**-Maximum Cook's distance of the feature; **AF**-Indicates if the feature has been detected as a count outlier (it does not make sense if an infinite threshold is given for the Cook's distance).

**S3I**- $P_{TET}$-*ZCF26* doxycycline-treated downregulated genes. **Header: A**- gene ID (ORF19_ID, orf19 nomenclature at the *Candida albicans* genome database (CGD); **B-E**- Wild type count; **F-I**- *ZCF15* overexpression count; **J-M**-Wild type normalized count; **N-Q**-*ZCF15* overexpression normalized count; **R**-Base mean over all samples; **S**-Wild type base mean; **T**-*ZCF15* overexpression mean; **U**-Fold change; **V**-log2 FoldChange; **X**-p-value; **Y**-Adjusted p value; **Z**-Dispersion parameter estimated from feature counts; **AA**- Dispersion parameter estimated from the model; **AB**-Dispersion parameter estimated from the Maximum A Posteriori model; **AC**-Final dispersion parameter used to perform the test; **AD**-Convergence of the coefficients of the model (TRUE or FALSE); **AE**-Maximum Cook's distance of the feature; **AF**-Indicates if the feature has been detected as a count outlier (it does not make sense if an infinite threshold is given for the Cook's distance).

**S3J**- TAP-Zcf15 ChIP-sequencing data. **Headers: A**- Chromosome number according to Assembly 22 of the *C. albicans* genome (CGD), start coordinate of the peak interval and, end coordinate of the peak; **B-**Score obtained; **C-**Signal Value; **D-**p-value; **E**-q-value; **F**- Left ORF to the binding region according to the CGD; **G-**Right ORF to the binding region according to the CGD.

**S3K**- TAP-Zcf26 ChIP sequencing data. **Headers: A-** Chromosome number according to Assembly 22 of the C. albicans genome (CGD), start coordinate of the peak interval and, end coordinate of the peak; **B-**Score obtained; **C**-Signal Value; **D-**p-value; **E-** q-value; **F**-Left ORF to the binding region according to the CGD; **G-**Right ORF to the binding region according to the CGD.

**S3L**- TAP-Zcf15 and TAP-Zcf26 bound and expression level altered genes. **Headers: A-** TAP-Zcf15 bound and up-regulated; **B**-TAP-Zcf15 bound and down-regulated; **C-**TAP-Zcf26 bound and up-regulated; **D**-TAP-Zcf26 bound and down-regulated.

Wild type doxycycline-treated complete data

| | WT | | | | WT dox | | | | norm. WT | | | | norm. WT dox | | | | | | | | | | | | | | | | | | |
|---|---|---|---|---|---|---|---|---|---|---|---|---|---|---|---|---|---|---|---|---|---|---|---|---|---|---|---|---|---|---|---|
| Gene ID | 1 | 2 | 3 | 4 | 1 | 2 | 3 | 4 | 1 | 2 | 3 | 4 | 1 | 2 | 3 | 4 | base Mean | WT | WT dox | FC | log2FC | stat | pvalue | padj | dispGeneEst | dispFit | dispMAP | dispersion | betaConv | maxCooks | outlier |
| CaalfMp02 | 2 | 0 | 0 | 4 | 1 | 3 | 4 | 3 | 2 | 0 | 0 | 3 | 1 | 3 | 4 | 4 | 2,08 | 1 | 3 | 2,029 | 1,021 | 0,963 | 0,33533848 | NA | 0,190633 | 0,7011328 | 0,5423307 | 0,5423307 | TRUE | 0,7924802 | No |
| CaalfMp03 | 4 | 6 | 3 | 8 | 8 | 6 | 7 | 5 | 4 | 8 | 3 | 7 | 7 | 6 | 6 | 7 | 5,84 | 6 | 6 | 1,248 | 0,32 | 0,569 | 0,56966195 | NA | 1,00E-08 | 0,2503538 | 0,1299654 | 0,1299654 | TRUE | 0,1478634 | No |
| CaalfMp04 | 0 | 0 | 0 | 0 | 0 | 0 | 0 | 0 | 0 | 0 | 0 | 0 | 0 | 0 | 0 | 0 | 0 | 0 | 0 | NA | NA | NA | NA | NA | NA | NA | NA | NA | NA | NA | NA |
| CaalfMp05 | 20 | 12 | 19 | 35 | 23 | 40 | 27 | 32 | 18 | 15 | 19 | 29 | 20 | 38 | 24 | 45 | 26,07 | 20 | 32 | 1,509 | 0,594 | 1,782 | 0,07478968 | 0,11918738 | 0,077248 | 0,0576581 | 0,0664841 | 0,0664841 | TRUE | 0,1597394 | No |
| CaalfMp06 | 5 | 11 | 11 | 19 | 5 | 19 | 13 | 13 | 4 | 14 | 11 | 16 | 4 | 18 | 12 | 18 | 12,22 | 11 | 13 | 1,135 | 0,183 | 0,361 | 0,7178705 | NA | 0,215241 | 0,1207858 | 0,162115 | 0,162115 | TRUE | 0,8547202 | No |
| CaalfMp07 | 0 | 0 | 1 | 0 | 0 | 0 | 1 | 0 | 0 | 0 | 1 | 0 | 0 | 0 | 1 | 0 | 0,24 | 0 | 0 | 1,016 | 0,023 | 0,008 | 0,99379373 | NA | 1,00E-08 | 6,0651585 | 6,4027826 | 6,4027826 | TRUE | 0,3731075 | No |
| CaalfMp08 | 20 | 25 | 22 | 26 | 21 | 21 | 20 | 17 | 18 | 32 | 23 | 21 | 18 | 20 | 18 | 24 | 21,69 | 24 | 20 | 0,854 | -0,227 | -0,753 | 0,45129743 | 0,53712261 | 1,00E-08 | 0,0689017 | 0,0403421 | 0,0403421 | TRUE | 0,2714542 | No |
| CaalfMp08.1 | 0 | 0 | 0 | 0 | 0 | 0 | 0 | 0 | 0 | 0 | 0 | 0 | 0 | 0 | 0 | 0 | 0 | 0 | 0 | NA | NA | NA | NA | NA | NA | NA | NA | NA | NA | NA | NA |
| CaalfMp08.2 | 0 | 0 | 0 | 0 | 0 | 0 | 0 | 0 | 0 | 0 | 0 | 0 | 0 | 0 | 0 | 0 | 0 | 0 | 0 | NA | NA | NA | NA | NA | NA | NA | NA | NA | NA | NA | NA |
| CaalfMp08.3 | 0 | 0 | 0 | 0 | 0 | 0 | 0 | 0 | 0 | 0 | 0 | 0 | 0 | 0 | 0 | 0 | 0 | 0 | 0 | NA | NA | NA | NA | NA | NA | NA | NA | NA | NA | NA | NA |
| CaalfMp08.4 | 0 | 0 | 0 | 0 | 0 | 0 | 0 | 0 | 0 | 0 | 0 | 0 | 0 | 0 | 0 | 0 | 0 | 0 | 0 | NA | NA | NA | NA | NA | NA | NA | NA | NA | NA | NA | NA |
| CaalfMp09 | 2 | 1 | 4 | 1 | 5 | 5 | 3 | 2 | 2 | 1 | 4 | 1 | 4 | 5 | 3 | 3 | 2,82 | 2 | 4 | 1,861 | 0,896 | 1,074 | 0,28298748 | NA | 1,00E-08 | 0,5174722 | 0,2829709 | 0,2829709 | TRUE | 0,4406206 | No |
| CaalfMp10 | 2 | 0 | 2 | 2 | 2 | 1 | 1 | 0 | 2 | 0 | 2 | 2 | 2 | 1 | 1 | 0 | 1,13 | 2 | 1 | 0,67 | -0,579 | -0,456 | 0,6483697 | NA | 1,00E-08 | 1,285734 | 0,6739356 | 0,6739356 | TRUE | 0,1914113 | No |
| CaalfMp11 | 72 | 71 | 78 | 112 | 80 | 76 | 97 | 82 | 65 | 90 | 80 | 92 | 68 | 73 | 87 | 116 | 83,79 | 82 | 86 | 1,037 | 0,052 | 0,271 | 0,78664439 | 0,83324392 | 0,028322 | 0,0193059 | 0,0233097 | 0,0233097 | TRUE | 0,4313831 | No |
| CaalfMp11.1 | 0 | 0 | 0 | 0 | 0 | 0 | 0 | 0 | 0 | 0 | 0 | 0 | 0 | 0 | 0 | 0 | 0 | 0 | 0 | NA | NA | NA | NA | NA | NA | NA | NA | NA | NA | NA | NA |
| CaalfMp12 | 0 | 0 | 0 | 0 | 0 | 0 | 0 | 0 | 0 | 0 | 0 | 0 | 0 | 0 | 0 | 0 | 0 | 0 | 0 | NA | NA | NA | NA | NA | NA | NA | NA | NA | NA | NA | NA |
| CaalfMp13 | 7 | 7 | 4 | 7 | 5 | 6 | 10 | 6 | 6 | 9 | 4 | 6 | 4 | 6 | 9 | 8 | 6,56 | 6 | 7 | 1,093 | 0,128 | 0,238 | 0,81185103 | NA | 1,00E-08 | 0,223431 | 0,1240075 | 0,1240075 | TRUE | 0,2058739 | No |
| CaalfMp14 | 9 | 7 | 3 | 7 | 3 | 3 | 5 | 3 | 8 | 9 | 3 | 6 | 3 | 3 | 4 | 4 | 4,99 | 6 | 4 | 0,544 | -0,878 | -1,386 | 0,16588484 | NA | 1,00E-08 | 0,292695 | 0,1649267 | 0,1649267 | TRUE | 0,2934463 | No |
| CaalfMp15 | 0 | 0 | 0 | 0 | 0 | 0 | 0 | 0 | 0 | 0 | 0 | 0 | 0 | 0 | 0 | 0 | 0 | 0 | 0 | NA | NA | NA | NA | NA | NA | NA | NA | NA | NA | NA | NA |
| CaalfMr16 | 2091 | 1662 | 1765 | 2753 | 1586 | 2242 | 2176 | 1673 | 1876 | 2116 | 1806 | 2270 | 1356 | 2139 | 1942 | 2360 | 1983,1 | 2017 | 1949 | 0,966 | -0,05 | -0,27 | 0,78694693 | 0,83341744 | 0,032333 | 0,0027116 | 0,0167063 | 0,0323329 | TRUE | 0,5094605 | No |
| CaalfMr17 | 549 | 448 | 479 | 766 | 399 | 590 | 654 | 470 | 493 | 570 | 490 | 632 | 341 | 563 | 584 | 663 | 541,91 | 546 | 538 | 0,983 | -0,025 | -0,112 | 0,91050714 | 0,93366221 | 0,044363 | 0,0046584 | 0,02326 | 0,044363 | TRUE | 0,7179476 | No |
| CaalfMt18 | 0 | 0 | 0 | 0 | 0 | 0 | 0 | 0 | 0 | 0 | 0 | 0 | 0 | 0 | 0 | 0 | 0 | 0 | 0 | NA | NA | NA | NA | NA | NA | NA | NA | NA | NA | NA | NA |
| CaalfMt19 | 0 | 0 | 0 | 0 | 0 | 0 | 0 | 0 | 0 | 0 | 0 | 0 | 0 | 0 | 0 | 0 | 0 | 0 | 0 | NA | NA | NA | NA | NA | NA | NA | NA | NA | NA | NA | NA |
| CaalfMt20 | 0 | 0 | 0 | 0 | 0 | 0 | 0 | 0 | 0 | 0 | 0 | 0 | 0 | 0 | 0 | 0 | 0 | 0 | 0 | NA | NA | NA | NA | NA | NA | NA | NA | NA | NA | NA | NA |
| CaalfMt21 | 0 | 0 | 0 | 0 | 0 | 0 | 0 | 0 | 0 | 0 | 0 | 0 | 0 | 0 | 0 | 0 | 0 | 0 | 0 | NA | NA | NA | NA | NA | NA | NA | NA | NA | NA | NA | NA |
| CaalfMt22 | 0 | 0 | 0 | 0 | 0 | 0 | 0 | 0 | 0 | 0 | 0 | 0 | 0 | 0 | 0 | 0 | 0 | 0 | 0 | NA | NA | NA | NA | NA | NA | NA | NA | NA | NA | NA | NA |
| CaalfMt23 | 0 | 0 | 0 | 0 | 0 | 0 | 0 | 0 | 0 | 0 | 0 | 0 | 0 | 0 | 0 | 0 | 0 | 0 | 0 | NA | NA | NA | NA | NA | NA | NA | NA | NA | NA | NA | NA |
| CaalfMt24 | 0 | 0 | 0 | 0 | 0 | 0 | 0 | 0 | 0 | 0 | 0 | 0 | 0 | 0 | 0 | 0 | 0 | 0 | 0 | NA | NA | NA | NA | NA | NA | NA | NA | NA | NA | NA | NA |
| CaalfMt25 | 0 | 0 | 0 | 0 | 0 | 0 | 0 | 0 | 0 | 0 | 0 | 0 | 0 | 0 | 0 | 0 | 0 | 0 | 0 | NA | NA | NA | NA | NA | NA | NA | NA | NA | NA | NA | NA |
| CaalfMt26 | 0 | 0 | 0 | 0 | 0 | 0 | 1 | 0 | 0 | 0 | 0 | 0 | 0 | 0 | 1 | 0 | 0,11 | 0 | 0 | 1,675 | 0,744 | 0,211 | 0,83317474 | NA | 1,00E-08 | 13,01685 | 10 | 10 | TRUE | 0,2773144 | No |
| CaalfMt27 | 1 | 0 | 0 | 0 | 2 | 0 | 1 | 0 | 1 | 0 | 0 | 0 | 2 | 0 | 1 | 0 | 0,44 | 0 | 1 | 2,286 | 1,193 | 0,551 | 0,58139296 | NA | 1,00E-08 | 3,3202472 | 2,7675089 | 2,7675089 | TRUE | 0,4198638 | No |
| CaalfMt28 | 0 | 0 | 0 | 0 | 0 | 0 | 1 | 0 | 0 | 0 | 0 | 0 | 0 | 0 | 1 | 0 | 0,11 | 0 | 0 | 1,675 | 0,744 | 0,211 | 0,83317474 | NA | 1,00E-08 | 13,01685 | 10 | 10 | TRUE | 0,2773144 | No |
| CaalfMt29 | 0 | 0 | 0 | 0 | 0 | 0 | 0 | 0 | 0 | 0 | 0 | 0 | 0 | 0 | 0 | 0 | 0 | 0 | 0 | NA | NA | NA | NA | NA | NA | NA | NA | NA | NA | NA | NA |
| CaalfMt30 | 1 | 0 | 0 | 0 | 0 | 0 | 0 | 0 | 1 | 0 | 0 | 0 | 0 | 0 | 0 | 0 | 0,11 | 0 | 0 | 0,616 | -0,698 | -0,198 | 0,84334499 | NA | 1,00E-08 | 12,947638 | 10 | 10 | TRUE | 0,289592 | No |
| CaalfMt31 | 0 | 0 | 0 | 0 | 0 | 0 | 0 | 0 | 0 | 0 | 0 | 0 | 0 | 0 | 0 | 0 | 0 | 0 | 0 | NA | NA | NA | NA | NA | NA | NA | NA | NA | NA | NA | NA |
| CaalfMt32 | 0 | 0 | 0 | 0 | 0 | 0 | 0 | 0 | 0 | 0 | 0 | 0 | 0 | 0 | 0 | 0 | 0 | 0 | 0 | NA | NA | NA | NA | NA | NA | NA | NA | NA | NA | NA | NA |
| CaalfMt33 | 0 | 0 | 0 | 0 | 0 | 0 | 0 | 0 | 0 | 0 | 0 | 0 | 0 | 0 | 0 | 0 | 0 | 0 | 0 | NA | NA | NA | NA | NA | NA | NA | NA | NA | NA | NA | NA |
| CaalfMt34 | 0 | 0 | 0 | 0 | 0 | 0 | 0 | 0 | 0 | 0 | 0 | 0 | 0 | 0 | 0 | 0 | 0 | 0 | 0 | NA | NA | NA | NA | NA | NA | NA | NA | NA | NA | NA | NA |
| CaalfMt35 | 0 | 0 | 0 | 0 | 0 | 0 | 0 | 0 | 0 | 0 | 0 | 0 | 0 | 0 | 0 | 0 | 0 | 0 | 0 | NA | NA | NA | NA | NA | NA | NA | NA | NA | NA | NA | NA |
| CaalfMt36 | 0 | 0 | 0 | 0 | 0 | 0 | 0 | 0 | 0 | 0 | 0 | 0 | 0 | 0 | 0 | 0 | 0 | 0 | 0 | NA | NA | NA | NA | NA | NA | NA | NA | NA | NA | NA | NA |
| CaalfMt37 | 0 | 0 | 0 | 0 | 0 | 0 | 0 | 0 | 0 | 0 | 0 | 0 | 0 | 0 | 0 | 0 | 0 | 0 | 0 | NA | NA | NA | NA | NA | NA | NA | NA | NA | NA | NA | NA |
| CaalfMt38 | 0 | 2 | 0 | 0 | 0 | 1 | 1 | 1 | 0 | 3 | 0 | 0 | 0 | 1 | 1 | 1 | 0,73 | 1 | 1 | 1,507 | 0,592 | 0,321 | 0,74826407 | NA | 1,00E-08 | 2,0032473 | 1,7107573 | 1,7107573 | TRUE | 1,2569317 | No |
| CaalfMt39 | 0 | 0 | 0 | 0 | 0 | 0 | 0 | 0 | 0 | 0 | 0 | 0 | 0 | 0 | 0 | 0 | 0 | 0 | 0 | NA | NA | NA | NA | NA | NA | NA | NA | NA | NA | NA | NA |
| CaalfMt40 | 0 | 0 | 0 | 0 | 0 | 0 | 0 | 0 | 0 | 0 | 0 | 0 | 0 | 0 | 0 | 0 | 0 | 0 | 0 | NA | NA | NA | NA | NA | NA | NA | NA | NA | NA | NA | NA |
| CaalfMt41 | 0 | 0 | 0 | 0 | 0 | 0 | 0 | 0 | 0 | 0 | 0 | 0 | 0 | 0 | 0 | 0 | 0 | 0 | 0 | NA | NA | NA | NA | NA | NA | NA | NA | NA | NA | NA | NA |
| CaalfMt42 | 0 | 0 | 0 | 0 | 0 | 0 | 0 | 0 | 0 | 0 | 0 | 0 | 0 | 0 | 0 | 0 | 0 | 0 | 0 | NA | NA | NA | NA | NA | NA | NA | NA | NA | NA | NA | NA |
| CaalfMt43 | 0 | 0 | 0 | 0 | 0 | 0 | 0 | 0 | 0 | 0 | 0 | 0 | 0 | 0 | 0 | 0 | 0 | 0 | 0 | NA | NA | NA | NA | NA | NA | NA | NA | NA | NA | NA | NA |
| CaalfMt44 | 0 | 0 | 0 | 0 | 0 | 0 | 0 | 0 | 0 | 0 | 0 | 0 | 0 | 0 | 0 | 0 | 0 | 0 | 0 | NA | NA | NA | NA | NA | NA | NA | NA | NA | NA | NA | NA |
| CaalfMt45 | 0 | 0 | 0 | 0 | 0 | 0 | 0 | 0 | 0 | 0 | 0 | 0 | 0 | 0 | 0 | 0 | 0 | 0 | 0 | NA | NA | NA | NA | NA | NA | NA | NA | NA | NA | NA | NA |
| CaalfMt46 | 0 | 0 | 0 | 0 | 0 | 0 | 0 | 0 | 0 | 0 | 0 | 0 | 0 | 0 | 0 | 0 | 0 | 0 | 0 | NA | NA | NA | NA | NA | NA | NA | NA | NA | NA | NA | NA |
| CaalfMt47 | 0 | 0 | 0 | 0 | 0 | 0 | 0 | 0 | 0 | 0 | 0 | 0 | 0 | 0 | 0 | 0 | 0 | 0 | 0 | NA | NA | NA | NA | NA | NA | NA | NA | NA | NA | NA | NA |
| CD39 | 0 | 0 | 0 | 0 | 0 | 0 | 0 | 0 | 0 | 0 | 0 | 0 | 0 | 0 | 0 | 0 | 0 | 0 | 0 | NA | NA | NA | NA | NA | NA | NA | NA | NA | NA | NA | NA |
| ITS1 | 171 | 144 | 142 | 213 | 192 | 224 | 222 | 233 | 153 | 183 | 145 | 176 | 164 | 214 | 198 | 329 | 195,29 | 164 | 226 | 1,366 | 0,449 | 2,461 | 0,01383903 | 0,02688972 | 0,045716 | 0,0094131 | 0,0267202 | 0,0267202 | TRUE | 0,9508392 | No |
| ITS2 | 297 | 193 | 242 | 286 | 444 | 314 | 315 | 222 | 266 | 246 | 248 | 236 | 380 | 300 | 281 | 313 | 283,64 | 249 | 318 | 1,281 | 0,357 | 3,324 | 0,0008874 | 0,0022553 | 0,007893 | 0,0070977 | 0,0075056 | 0,0075056 | TRUE | 0,2058077 | No |
| LSU-C2809 | 87 | 87 | 79 | 122 | 108 | 116 | 123 | 93 | 78 | 111 | 81 | 101 | 92 | 111 | 110 | 131 | 101,78 | 93 | 111 | 1,194 | 0,256 | 1,591 | 0,11156245 | 0,16732094 | 0,013232 | 0,0162432 | 0,0149578 | 0,0149578 | TRUE | 0,141135 | No |

| | | | | | | | | | | | | | | | | | | | | | | | | | | | | | | | |
|---|---|---|---|---|---|---|---|---|---|---|---|---|---|---|---|---|---|---|---|---|---|---|---|---|---|---|---|---|---|---|---|
| LSU-G1431 | 1 | 2 | 2 | 13 | 11 | 18 | 17 | 9 | 1 | 3 | 2 | 11 | 9 | 17 | 15 | 13 | 8,83 | 4 | 14 | 3,217 | 1,686 | 2,751 | 0,0059451 | NA | 0,285987 | 0,1663492 | 0,2066629 | 0,2066629 | TRUE | 2,6067912 | No |
| orf19.10 | 82 | 68 | 63 | 71 | 54 | 62 | 48 | 40 | 74 | 87 | 64 | 59 | 46 | 59 | 43 | 56 | 60,97 | 71 | 51 | 0,723 | -0,468 | -2,401 | 0,0163664 | 0,0311662 | 0,010672 | 0,0257913 | 0,0194746 | 0,0194746 | TRUE | 0,1738337 | No |
| orf19.100 | 2282 | 1637 | 2021 | 2498 | 2501 | 2105 | 2433 | 1417 | 2047 | 2084 | 2068 | 2060 | 2139 | 2009 | 2171 | 1999 | 2072 | 2065 | 2080 | 1,009 | 0,013 | 0,293 | 0,76954841 | 0,8201283 | 0,000443 | 0,0026802 | 0,0013042 | 0,0013042 | TRUE | 0,0105513 | No |
| orf19.1002 | 38 | 32 | 30 | 31 | 38 | 55 | 61 | 34 | 34 | 41 | 31 | 26 | 32 | 52 | 54 | 48 | 39,81 | 33 | 46 | 1,442 | 0,528 | 2,15 | 0,03156416 | 0,05612236 | 0,023001 | 0,0384487 | 0,0317675 | 0,0317675 | TRUE | 0,3924786 | No |
| orf19.1005 | 125 | 103 | 130 | 146 | 97 | 115 | 104 | 76 | 112 | 131 | 133 | 120 | 83 | 110 | 93 | 107 | 111,17 | 124 | 98 | 0,788 | -0,344 | -2,447 | 0,01441554 | 0,02790224 | 1,00E-08 | 0,0150381 | 0,0098486 | 0,0098486 | TRUE | 0,1156073 | No |
| orf19.1007 | 5391 | 3703 | 4701 | 5650 | 5607 | 5297 | 5561 | 3619 | 4836 | 4714 | 4810 | 4659 | 4795 | 5054 | 4962 | 5104 | 4866,9 | 4755 | 4979 | 1,046 | 0,066 | 1,9 | 0,05737945 | 0,09468161 | 0,00034 | 0,0022778 | 0,0009346 | 0,0009346 | TRUE | 0,0076453 | No |
| orf19.1008 | 63 | 51 | 43 | 65 | 57 | 54 | 43 | 41 | 57 | 65 | 44 | 54 | 49 | 52 | 38 | 58 | 51,94 | 55 | 49 | 0,892 | -0,165 | -0,813 | 0,41617859 | 0,50428611 | 0,004563 | 0,0299298 | 0,0202591 | 0,0202591 | TRUE | 0,1862089 | No |
| orf19.1009 | 127 | 73 | 88 | 135 | 100 | 63 | 92 | 75 | 114 | 93 | 90 | 111 | 86 | 60 | 82 | 106 | 92,72 | 102 | 84 | 0,805 | -0,314 | -1,8 | 0,07193166 | 0,11539797 | 0,019091 | 0,0176369 | 0,0182412 | 0,0182412 | TRUE | 0,3185501 | No |
| orf19.101 | 955 | 676 | 870 | 1134 | 1027 | 926 | 1006 | 703 | 857 | 861 | 890 | 935 | 878 | 884 | 898 | 992 | 899,21 | 886 | 913 | 1,026 | 0,037 | 0,623 | 0,53358454 | 0,6160259 | 0,001046 | 0,0035939 | 0,0023081 | 0,0023081 | TRUE | 0,035543 | No |
| orf19.1010 | 1835 | 1705 | 1767 | 2255 | 1853 | 1986 | 1805 | 1295 | 1646 | 2171 | 1808 | 1860 | 1585 | 1895 | 1611 | 1826 | 1800,1 | 1871 | 1729 | 0,924 | -0,113 | -1,303 | 0,19246574 | 0,26573622 | 0,010141 | 0,002786 | 0,0066993 | 0,0066993 | TRUE | 0,1375702 | No |
| orf19.1011 | 2422 | 1732 | 2294 | 2743 | 2041 | 1971 | 2098 | 1314 | 2173 | 2205 | 2347 | 2262 | 1746 | 1881 | 1872 | 1853 | 2042,3 | 2247 | 1838 | 0,818 | -0,291 | -6,403 | 1,52E-10 | 1,07E-09 | 0,000733 | 0,0026903 | 0,0014805 | 0,0014805 | TRUE | 0,0146511 | No |
| orf19.1012 | 670 | 475 | 531 | 734 | 523 | 442 | 470 | 241 | 601 | 605 | 543 | 605 | 447 | 422 | 419 | 340 | 497,83 | 588 | 407 | 0,696 | -0,522 | -6,174 | 6,64E-10 | 4,33E-09 | 0,00466 | 0,0048956 | 0,0047974 | 0,0047974 | TRUE | 0,1216013 | No |
| orf19.102 | 792 | 630 | 742 | 963 | 766 | 742 | 784 | 457 | 711 | 802 | 759 | 794 | 655 | 708 | 700 | 645 | 721,63 | 766 | 677 | 0,885 | -0,176 | -2,738 | 0,00618773 | 0,01309933 | 0,001251 | 0,0039912 | 0,0025548 | 0,0025548 | TRUE | 0,0296268 | No |
| orf19.1026 | 4778 | 3479 | 4262 | 5313 | 4451 | 4136 | 4317 | 2865 | 4286 | 4429 | 4360 | 4381 | 3807 | 3946 | 3852 | 4041 | 4137,9 | 4364 | 3912 | 0,896 | -0,159 | -4,68 | 2,87E-06 | 1,13E-05 | 0,000123 | 0,0023304 | 0,0008659 | 0,0008659 | TRUE | 0,0055696 | No |
| orf19.1026.1 | 340 | 277 | 293 | 392 | 353 | 299 | 296 | 186 | 305 | 353 | 300 | 323 | 302 | 285 | 264 | 262 | 299,29 | 320 | 278 | 0,875 | -0,193 | -2,149 | 0,03166801 | 0,05627371 | 0,001359 | 0,00683 | 0,0043934 | 0,0043934 | TRUE | 0,0461188 | No |
| orf19.1027 | 4014 | 2940 | 3542 | 4266 | 4527 | 4448 | 4524 | 2647 | 3601 | 3743 | 3624 | 3518 | 3872 | 4244 | 4037 | 3733 | 3796,5 | 3622 | 3972 | 1,098 | 0,135 | 2,875 | 0,0040361 | 0,00889 | 0,001513 | 0,0023619 | 0,0018484 | 0,0018484 | TRUE | 0,025775 | No |
| orf19.1028 | 1471 | 1250 | 1307 | 1712 | 2052 | 2048 | 2180 | 1512 | 1320 | 1591 | 1337 | 1412 | 1755 | 1954 | 1945 | 2133 | 1680,9 | 1415 | 1947 | 1,376 | 0,46 | 6,221 | 4,93E-10 | 3,29E-09 | 0,006177 | 0,0028432 | 0,0046419 | 0,0046419 | TRUE | 0,08151 | No |
| orf19.1029 | 4817 | 4333 | 4498 | 5876 | 4948 | 5425 | 5372 | 3297 | 4321 | 5516 | 4602 | 4845 | 4232 | 5176 | 4794 | 4650 | 4767,1 | 4821 | 4713 | 0,978 | -0,032 | -0,407 | 0,68423078 | 0,74756475 | 0,007721 | 0,002284 | 0,0058206 | 0,0058206 | TRUE | 0,1137145 | No |
| orf19.103 | 151 | 104 | 124 | 168 | 153 | 151 | 165 | 119 | 135 | 132 | 127 | 139 | 131 | 144 | 147 | 168 | 140,41 | 133 | 148 | 1,096 | 0,132 | 1,091 | 0,27506871 | 0,35941992 | 1,00E-08 | 0,0123186 | 0,0068338 | 0,0068338 | TRUE | 0,0725494 | No |
| orf19.1030 | 29590 | 20295 | 25021 | 30976 | 26080 | 23650 | 25185 | 14868 | 26545 | 25838 | 25599 | 25543 | 22304 | 22566 | 22473 | 20970 | 23980 | 25881 | 22078 | 0,853 | -0,229 | -6,779 | 1,21E-11 | 9,79E-11 | 0,000675 | 0,0020401 | 0,0010532 | 0,0010532 | TRUE | 0,0137538 | No |
| orf19.1031 | 1460 | 1098 | 1346 | 1634 | 1617 | 1427 | 1672 | 1007 | 1310 | 1398 | 1377 | 1347 | 1383 | 1362 | 1492 | 1420 | 1386,1 | 1358 | 1414 | 1,042 | 0,06 | 1,218 | 0,22308626 | 0,30207398 | 0,000587 | 0,0030268 | 0,0015949 | 0,0015949 | TRUE | 0,0176652 | No |
| orf19.1032 | 3715 | 2400 | 3426 | 4119 | 4068 | 3561 | 3902 | 2678 | 3333 | 3055 | 3505 | 3397 | 3479 | 3398 | 3482 | 3777 | 3428,2 | 3322 | 3534 | 1,062 | 0,087 | 1,644 | 0,10018453 | 0,15300169 | 0,002388 | 0,002403 | 0,0023824 | 0,0023824 | TRUE | 0,0343525 | No |
| orf19.1033 | 4421 | 3173 | 3894 | 4856 | 4691 | 4344 | 4561 | 2484 | 3966 | 4040 | 3984 | 4004 | 4012 | 4145 | 4070 | 3503 | 3965,5 | 3998 | 3932 | 0,985 | -0,022 | -0,418 | 0,67627388 | 0,74116262 | 0,002161 | 0,0023456 | 0,0024349 | 0,0024349 | TRUE | 0,0630446 | No |
| orf19.1034 | 1678 | 1219 | 1484 | 1802 | 2002 | 1872 | 1942 | 1091 | 1505 | 1552 | 1518 | 1486 | 1712 | 1786 | 1733 | 1539 | 1603,9 | 1515 | 1692 | 1,12 | 0,164 | 3,117 | 0,00182566 | 0,00434744 | 0,001337 | 0,0028846 | 0,0020232 | 0,0020232 | TRUE | 0,0409611 | No |
| orf19.1035 | 3536 | 2529 | 3233 | 3919 | 3933 | 3545 | 3802 | 2767 | 3172 | 3220 | 3308 | 3232 | 3364 | 3383 | 3393 | 3903 | 3371,6 | 3233 | 3511 | 1,084 | 0,117 | 2,199 | 0,02784973 | 0,0501412 | 0,002535 | 0,0024101 | 0,0024078 | 0,0024078 | TRUE | 0,0661767 | No |
| orf19.1036 | 1074 | 685 | 977 | 1202 | 1184 | 1100 | 1068 | 666 | 963 | 872 | 1000 | 991 | 1013 | 1050 | 953 | 939 | 972,61 | 956 | 989 | 1,033 | 0,047 | 0,751 | 0,45243048 | 0,53815132 | 0,001855 | 0,0034721 | 0,0026814 | 0,0026814 | TRUE | 0,0386959 | No |
| orf19.1037 | 79 | 62 | 74 | 102 | 93 | 97 | 93 | 52 | 71 | 79 | 76 | 84 | 80 | 93 | 83 | 73 | 79,76 | 78 | 82 | 1,066 | 0,092 | 0,592 | 0,5540637 | 0,63486787 | 1,00E-08 | 0,0201814 | 0,0106075 | 0,0106075 | TRUE | 0,0666414 | No |
| orf19.1039 | 219 | 231 | 217 | 274 | 259 | 390 | 313 | 181 | 196 | 294 | 222 | 226 | 222 | 372 | 279 | 255 | 258,34 | 234 | 282 | 1,207 | 0,272 | 1,682 | 0,09256681 | 0,14340933 | 0,036813 | 0,0075989 | 0,0211724 | 0,0211724 | TRUE | 0,52792 | No |
| orf19.104 | 233 | 167 | 223 | 264 | 252 | 258 | 310 | 182 | 209 | 213 | 228 | 218 | 216 | 246 | 277 | 257 | 232,81 | 217 | 249 | 1,145 | 0,195 | 1,905 | 0,05678905 | 0,09381045 | 0,002997 | 0,008215 | 0,0057359 | 0,0057359 | TRUE | 0,099355 | No |
| orf19.1040 | 241 | 222 | 215 | 290 | 343 | 305 | 333 | 189 | 216 | 283 | 220 | 239 | 293 | 291 | 297 | 267 | 263,25 | 240 | 287 | 1,209 | 0,274 | 2,734 | 0,00626456 | 0,01323864 | 0,003466 | 0,007494 | 0,0057976 | 0,0057976 | TRUE | 0,1426311 | No |
| orf19.1041 | 444 | 372 | 385 | 518 | 428 | 512 | 467 | 354 | 398 | 474 | 394 | 427 | 366 | 489 | 417 | 499 | 432,94 | 423 | 443 | 1,044 | 0,062 | 0,58 | 0,56213117 | 0,6421913 | 0,011614 | 0,0053326 | 0,0084871 | 0,0084871 | TRUE | 0,1618514 | No |
| orf19.1042 | 14649 | 9835 | 12437 | 15955 | 16601 | 14194 | 14985 | 10665 | 13142 | 12521 | 12724 | 13157 | 14198 | 13543 | 13372 | 15042 | 13462 | 12886 | 14039 | 1,089 | 0,123 | 2,79 | 0,00526557 | 0,01132267 | 0,001656 | 0,0020874 | 0,0017898 | 0,0017898 | TRUE | 0,0278983 | No |
| orf19.1043 | 649 | 600 | 599 | 750 | 690 | 794 | 858 | 434 | 582 | 764 | 613 | 618 | 590 | 758 | 766 | 612 | 662,85 | 644 | 682 | 1,061 | 0,086 | 0,79 | 0,42963392 | 0,51666145 | 0,014263 | 0,0041696 | 0,0097686 | 0,0097686 | TRUE | 0,176344 | No |
| orf19.1045 | 2343 | 1966 | 2025 | 2570 | 2228 | 2289 | 2250 | 1022 | 2102 | 2503 | 2072 | 2119 | 1905 | 2184 | 2008 | 1441 | 2041,8 | 2199 | 1884 | 0,859 | -0,22 | -2,045 | 0,04089076 | 0,07035272 | 0,017785 | 0,0026905 | 0,0106157 | 0,0106157 | TRUE | 0,2952319 | No |
| orf19.1047 | 8349 | 6648 | 7668 | 9292 | 8238 | 8502 | 8740 | 5130 | 7490 | 8464 | 7845 | 7662 | 7045 | 8112 | 7799 | 7235 | 7706,6 | 7865 | 7548 | 0,96 | -0,059 | -1,038 | 0,29910708 | 0,38460482 | 0,003412 | 0,0021679 | 0,0029416 | 0,0029416 | TRUE | 0,031716 | No |
| orf19.1048 | 736 | 536 | 626 | 796 | 564 | 522 | 546 | 326 | 660 | 682 | 640 | 656 | 482 | 498 | 487 | 460 | 570,87 | 660 | 482 | 0,733 | -0,449 | -7,209 | 5,63E-13 | 5,34E-12 | 1,00E-08 | 0,0045225 | 0,0019305 | 0,0019305 | TRUE | 0,0088484 | No |
| orf19.1049 | 62 | 60 | 54 | 79 | 71 | 101 | 91 | 59 | 56 | 76 | 55 | 65 | 61 | 96 | 81 | 83 | 71,74 | 63 | 80 | 1,277 | 0,352 | 1,868 | 0,06172043 | 0,10101235 | 0,01714 | 0,0222155 | 0,0199751 | 0,0199751 | TRUE | 0,2841043 | No |
| orf19.105 | 1026 | 674 | 941 | 1093 | 1075 | 747 | 801 | 537 | 920 | 858 | 963 | 901 | 919 | 713 | 715 | 757 | 843,35 | 910 | 776 | 0,853 | -0,23 | -2,7 | 0,00693138 | 0,01449667 | 0,007475 | 0,0037009 | 0,0057576 | 0,0057576 | TRUE | 0,1912387 | No |
| orf19.1050 | 142 | 160 | 176 | 211 | 176 | 207 | 215 | 110 | 127 | 204 | 180 | 174 | 151 | 198 | 192 | 155 | 172,52 | 171 | 174 | 1,022 | 0,031 | 0,206 | 0,8364358 | 0,87489706 | 0,022733 | 0,0103941 | 0,0162092 | 0,0162092 | TRUE | 0,327824 | No |
| orf19.1051 | 2990 | 1950 | 2349 | 2888 | 3102 | 2566 | 2657 | 2028 | 2682 | 2483 | 2403 | 2381 | 2653 | 2448 | 2371 | 2860 | 2535,3 | 2487 | 2583 | 1,037 | 0,053 | 0,803 | 0,42219825 | 0,50977463 | 0,004478 | 0,0025521 | 0,0037479 | 0,0037479 | TRUE | 0,0609491 | No |
| orf19.1052 | 1360 | 880 | 1090 | 1375 | 1519 | 1302 | 1421 | 1077 | 1220 | 1120 | 1115 | 1134 | 1299 | 1242 | 1268 | 1519 | 1239,7 | 1147 | 1332 | 1,157 | 0,211 | 3,061 | 0,00220804 | 0,0051719 | 0,004361 | 0,0031505 | 0,0037313 | 0,0037313 | TRUE | 0,1004332 | No |
| orf19.1053 | 1077 | 828 | 947 | 1237 | 1174 | 1122 | 1222 | 701 | 966 | 1054 | 969 | 1020 | 1004 | 1071 | 1090 | 989 | 1020,4 | 1002 | 1038 | 1,039 | 0,055 | 0,971 | 0,33140414 | 0,41811761 | 0,000963 | 0,0034022 | 0,0020748 | 0,0020748 | TRUE | 0,0136347 | No |
| orf19.1054 | 313 | 245 | 289 | 371 | 413 | 407 | 451 | 317 | 281 | 312 | 296 | 306 | 353 | 388 | 402 | 447 | 348,18 | 299 | 398 | 1,327 | 0,408 | 4,586 | 4,52E-06 | 1,73E-05 | 0,00305 | 0,006149 | 0,0046579 | 0,0046579 | TRUE | 0,0699639 | No |
| orf19.1055 | 1777 | 1195 | 1487 | 1796 | 2136 | 2050 | 2055 | 1365 | 1594 | 1521 | 1521 | 1481 | 1827 | 1956 | 1834 | 1925 | 1707,5 | 1529 | 1886 | 1,232 | 0,301 | 6,49 | 8,56E-11 | 6,29E-10 | 0,000588 | 0,0028297 | 0,0014652 | 0,0014652 | TRUE | 0,0103143 | No |
| orf19.1057 | 1522 | 956 | 1171 | 1433 | 1437 | 1302 | 1294 | 859 | 1365 | 1217 | 1198 | 1182 | 1229 | 1242 | 1155 | 1212 | 1225 | 1240 | 1210 | 0,974 | -0,038 | -0,633 | 0,52692372 | 0,60997258 | 0,002124 | 0,0031646 | 0,0025605 | 0,0025605 | TRUE | 0,0570044 | No |
| orf19.1058 | 4692 | 3314 | 3803 | 4956 | 3888 | 3456 | 3695 | 2232 | 4209 | 4219 | 3891 | 4087 | 3325 | 3298 | 3297 | 3148 | 3684,2 | 4102 | 3267 | 0,797 | -0,327 | -8,102 | 5,42E-16 | 6,83E-15 | 0,00063 | 0,0023736 | 0,0012847 | 0,0012847 | TRUE | 0,0143824 | No |
| orf19.1059 | 3789 | 2448 | 2836 | 3792 | 3262 | 2761 | 2890 | 1699 | 3399 | 3117 | 2902 | 3127 | 2790 | 2634 | 2579 | 2396 | 2867,9 | 3136 | 2600 | 0,83 | -0,269 | -4,43 | 9,41E-06 | 3,44E-05 | 0,003596 | 0,0024857 | 0,0031944 | 0,0031944 | TRUE | 0,0391813 | No |
| orf19.1060 | 85 | 84 | 94 | 134 | 108 | 88 | 98 | 56 | 76 | 107 | 96 | 110 | 92 | 84 | 87 | 79 | 91,58 | 97 | 86 | 0,886 | -0,175 | -1,114 | 0,2653364 | 0,3493625 | 2,84E-05 | 0,0178315 | 0,0128853 | 0,0128853 | TRUE | 0,2260941 | No |
| orf19.1061 | 1956 | 1305 | 1617 | 1968 | 2149 | 1731 | 1842 | 953 | 1755 | 1661 | 1654 | 1623 | 1838 | 1652 | 1644 | 1344 | 1646,3 | 1673 | 1620 | 0,97 | -0,044 | -0,549 | 0,58319397 | 0,6615992 | 0,007991 | 0,0028613 | 0,0056175 | 0,0056175 | TRUE | 0,1508468 | No |
| orf19.1062 | 89 | 49 | 83 | 94 | 92 | 93 | 78 | 47 | 80 | 62 | 85 | 78 | 79 | 89 | 70 | 66 | 76 | 76 | 76 | 0,995 | -0,007 | -0,04 | 0,96801446 | 0,9775232 | 1,00E-08 | 0,0210819 | 0,0135379 | 0,0135379 | TRUE | 0,118071 | No |
| orf19.1063 | 446 | 475 | 433 | 546 | 409 | 484 | 453 | 284 | 400 | 605 | 443 | 450 | 350 | 462 | 404 | 401 | 439,31 | 474 | 404 | 0,854 | -0,227 | -1,81 | 0,07021855 | 0,11308981 | 0,021188 | 0,005284 | 0,0128416 | 0,0128416 | TRUE | 0,379971 | No |
| orf19.1064 | 5625 | 3909 | 5001 | 5973 | 4444 | 4108 | 4375 | 2750 | 5046 | 4977 | 5117 | 4925 | 3801 | 3920 | 3904 | 3879 | 4446 | 5016 | 3876 | 0,773 | -0,372 | -11,802 | 3,82E-32 | 1,58E-30 | 1,00E-08 | 0,002306 | 0,0007252 | 0,0007252 | TRUE | 0,0022052 | No |
| orf19.1065 | 1447 | 938 | 1256 | 1552 | 1289 | 1045 | 1228 | 786 | 1298 | 1194 | 1285 | 1280 | 1102 | 997 | 1096 | 1109 | 1170,1 | 1264 | 1076 | 0,849 | -0,236 | -4,33 | 1,49E-05 | 5,29E-05 | 0,000963 | 0,0032202 | 0,0019797 | 0,0019797 | TRUE | 0,0296148 | No |
| orf19.1066 | 211 | 153 | 193 | 222 | 177 | 164 | 210 | 89 | 189 | 195 | 197 | 183 | 151 | 156 | 187 | 126 | 173,17 | 191 | 155 | 0,821 | -0,285 | -2,342 | 0,01915605 | 0,03577542 | 0,005657 | 0,0103625 | 0,0084172 | 0,0084172 | TRUE | 0,2042247 | No |
| orf19.1067 | 872 | 702 | 697 | 927 | 1172 | 1077 | 1095 | 633 | 782 | 894 | 713 | 764 | 1002 | 1028 | 977 | 893 | 881,67 | 788 | 975 | 1,242 | 0,312 | 4,19 | 2,79E-05 | 9,43E-05 | 0,004932 | 0,0036261 | 0,0041874 | 0,0041874 | TRUE | 0,0894045 | No |
| orf19.1069 | 40819 | 28196 | 35756 | 43916 | 44639 | 39590 | 43620 | 27657 | 36619 | 35897 | 36582 | 36214 | 38177 | 37776 | 38924 | 39008 | 37399 | 36328 | 38471 | 1,059 | 0,083 | 3,467 | 0,00052688 | 0,00141443 | 0,000128 | 0,0020183 | 0,0005174 | 0,0005174 | TRUE | 0,0018094 | No |
| orf19.107 | 6294 | 4370 | 5684 | 6908 | 7797 | 7215 | 7799 | 4674 | 5646 | 5563 | 5815 | 5696 | 6668 | 6884 | 6959 | 6592 | 6228,2 | 5680 | 6776 | 1,193 | 0,255 | 7,71 | 1,26E-14 | 1,39E-13 | 0,000285 | 0,0022126 | 0,000886 | 0,000886 | TRUE | 0,004045 | No |
| orf19.1070 | 264 | 219 | 230 | 322 | 338 | 308 | 366 | 198 | 237 | 279 | 235 | 266 | 289 | 294 | 327 | 279 | 275,66 | 254 | 297 | 1,176 | 0,233 | 2,474 | 0,01334344 | 0,02601149 | 0,002005 | 0,0072458 | 0,0048933 | 0,0048933 | TRUE | 0,0515095 | No |
| orf19.1075 | 127 | 73 | 82 | 146 | 116 | 114 | 110 | 86 | 114 | 93 | 84 | 120 | 99 | 109 | 98 | 121 | 104,82 | 103 | 107 | 1,025 | 0,035 | 0,228 | 0,81986139 | 0,86114112 | 0,00941 | 0,0158285 | 0,0130633 | 0,0130633 | TRUE | 0,1573961 | No |

| | | | | | | | | | | | | | | | | | | | | | | | | | | | | | | | |
|---|---|---|---|---|---|---|---|---|---|---|---|---|---|---|---|---|---|---|---|---|---|---|---|---|---|---|---|---|---|---|---|
| orf19.1077 | 2020 | 1563 | 1801 | 2270 | 2593 | 2494 | 2552 | 1720 | 1812 | 1990 | 1843 | 1872 | 2218 | 2380 | 2277 | 2426 | 2102,1 | 1879 | 2325 | 1,237 | 0,307 | 6,396 | 1,59E-10 | 1,12E-09 | 0,001046 | 0,0026701 | 0,0017298 | 0,0017298 | TRUE | 0,0178165 | No |
| orf19.1078 | 2473 | 1910 | 2100 | 2821 | 2823 | 2775 | 2704 | 1813 | 2219 | 2432 | 2149 | 2326 | 2414 | 2648 | 2413 | 2557 | 2394,6 | 2282 | 2508 | 1,099 | 0,137 | 2,585 | 0,00974709 | 0,01969082 | 0,001975 | 0,0025858 | 0,0022702 | 0,0022702 | TRUE | 0,0226217 | No |
| orf19.1080 | 18 | 10 | 26 | 21 | 18 | 18 | 13 | 16 | 16 | 13 | 27 | 17 | 15 | 17 | 12 | 23 | 17,44 | 18 | 17 | 0,893 | -0,163 | -0,462 | 0,64377013 | 0,7138743 | 0,022648 | 0,0852118 | 0,0609026 | 0,0609026 | TRUE | 0,4670465 | No |
| orf19.1082 | 922 | 775 | 760 | 1161 | 1029 | 1053 | 1009 | 606 | 827 | 987 | 778 | 957 | 880 | 1005 | 900 | 855 | 898,57 | 887 | 910 | 1,027 | 0,039 | 0,458 | 0,64696093 | 0,71652497 | 0,007821 | 0,0035951 | 0,0058355 | 0,0058355 | TRUE | 0,0810255 | No |
| orf19.1082.1 | 5977 | 3808 | 4661 | 5974 | 6626 | 5220 | 5190 | 4152 | 5362 | 4848 | 4769 | 4926 | 5667 | 4981 | 4631 | 5856 | 5130 | 4976 | 5284 | 1,061 | 0,086 | 1,16 | 0,24624555 | 0,32818174 | 0,00708 | 0,0022625 | 0,0050459 | 0,0050459 | TRUE | 0,0845115 | No |
| orf19.1083 | 782 | 556 | 629 | 772 | 682 | 600 | 640 | 415 | 702 | 708 | 644 | 637 | 583 | 573 | 571 | 585 | 625,21 | 673 | 578 | 0,861 | -0,216 | -3,415 | 0,00063817 | 0,0016743 | 1,00E-08 | 0,0043015 | 0,0022441 | 0,0022441 | TRUE | 0,0161412 | No |
| orf19.1084 | 6465 | 4614 | 5723 | 6732 | 8396 | 8020 | 8317 | 5363 | 5800 | 5874 | 5855 | 5551 | 7181 | 7652 | 7422 | 7564 | 6612,4 | 5770 | 7455 | 1,292 | 0,37 | 10,32 | 5,73E-25 | 1,44E-23 | 0,000511 | 0,0021991 | 0,0010778 | 0,0010778 | TRUE | 0,0081315 | No |
| orf19.1085 | 765 | 565 | 679 | 905 | 966 | 883 | 971 | 627 | 686 | 719 | 695 | 746 | 826 | 843 | 866 | 884 | 783,25 | 712 | 855 | 1,199 | 0,262 | 4,591 | 4,41E-06 | 1,69E-05 | 1,00E-08 | 0,003833 | 0,0018333 | 0,0018333 | TRUE | 0,0143291 | No |
| orf19.1086 | 2367 | 1703 | 2179 | 2661 | 2821 | 2880 | 3071 | 2000 | 2123 | 2168 | 2229 | 2194 | 2413 | 2748 | 2740 | 2821 | 2429,6 | 2178 | 2680 | 1,229 | 0,297 | 5,483 | 4,18E-08 | 2,14E-07 | 0,002303 | 0,002577 | 0,0024077 | 0,0024077 | TRUE | 0,0560607 | No |
| orf19.1087 | 1368 | 1123 | 1312 | 1586 | 1773 | 1951 | 2091 | 1119 | 1227 | 1430 | 1342 | 1308 | 1516 | 1862 | 1866 | 1578 | 1516,1 | 1327 | 1706 | 1,288 | 0,365 | 4,605 | 4,13E-06 | 1,59E-05 | 0,006952 | 0,002937 | 0,0053488 | 0,0053488 | TRUE | 0,070184 | No |
| orf19.1089 | 1316 | 1403 | 1318 | 1645 | 1271 | 1531 | 1466 | 870 | 1181 | 1786 | 1348 | 1356 | 1087 | 1461 | 1308 | 1227 | 1344,3 | 1418 | 1271 | 0,897 | -0,156 | -1,324 | 0,18534685 | 0,25776721 | 0,023184 | 0,0030594 | 0,0126127 | 0,0126127 | TRUE | 0,3622603 | No |
| orf19.109 | 1170 | 827 | 927 | 1230 | 1541 | 1287 | 1426 | 870 | 1050 | 1053 | 948 | 1014 | 1318 | 1228 | 1272 | 1227 | 1138,8 | 1016 | 1261 | 1,243 | 0,314 | 5,839 | 5,24E-09 | 3,04E-08 | 0,000775 | 0,0032543 | 0,0018868 | 0,0018868 | TRUE | 0,023543 | No |
| orf19.1090 | 198 | 148 | 184 | 231 | 257 | 213 | 220 | 170 | 178 | 188 | 188 | 190 | 220 | 203 | 196 | 240 | 200,49 | 186 | 215 | 1,147 | 0,198 | 1,932 | 0,05341869 | 0,08907563 | 1,00E-08 | 0,0092204 | 0,0051079 | 0,0051079 | TRUE | 0,0543884 | No |
| orf19.1091 | 6638 | 5133 | 5886 | 7330 | 6769 | 6749 | 7154 | 3907 | 5955 | 6535 | 6022 | 6044 | 5789 | 6440 | 6384 | 5510 | 6084,9 | 6139 | 6031 | 0,983 | -0,025 | -0,425 | 0,67117523 | 0,73772792 | 0,003142 | 0,0022181 | 0,0030466 | 0,0030466 | TRUE | 0,0402073 | No |
| orf19.1092 | 809 | 786 | 810 | 1049 | 910 | 1083 | 1107 | 656 | 726 | 1001 | 829 | 865 | 778 | 1033 | 988 | 925 | 893,1 | 855 | 931 | 1,09 | 0,125 | 1,187 | 0,23523732 | 0,31610508 | 0,014455 | 0,003605 | 0,0095184 | 0,0095184 | TRUE | 0,1512589 | No |
| orf19.1093 | 7829 | 5085 | 6754 | 8446 | 8049 | 6936 | 7919 | 5419 | 7023 | 6474 | 6910 | 6965 | 6884 | 6618 | 7066 | 7643 | 6947,9 | 6843 | 7053 | 1,03 | 0,042 | 0,84 | 0,40096889 | 0,48947984 | 0,002347 | 0,0021885 | 0,0022922 | 0,0022922 | TRUE | 0,0378757 | No |
| orf19.1095 | 2402 | 1639 | 2066 | 2631 | 2590 | 2383 | 2527 | 1772 | 2155 | 2087 | 2114 | 2170 | 2215 | 2274 | 2255 | 2499 | 2221 | 2132 | 2311 | 1,081 | 0,113 | 2,393 | 0,01673259 | 0,03177289 | 0,001092 | 0,0026332 | 0,0016855 | 0,0016855 | TRUE | 0,0339093 | No |
| orf19.1096 | 1036 | 929 | 1047 | 1210 | 1151 | 1313 | 1370 | 772 | 929 | 1183 | 1071 | 998 | 984 | 1253 | 1222 | 1089 | 1091,2 | 1045 | 1137 | 1,09 | 0,124 | 1,356 | 0,17509565 | 0,24630641 | 0,010125 | 0,0033099 | 0,0071401 | 0,0071401 | TRUE | 0,1018571 | No |
| orf19.1097 | 2757 | 1913 | 2536 | 2877 | 2298 | 1849 | 2067 | 1588 | 2473 | 2435 | 2595 | 2372 | 1965 | 1764 | 1844 | 2240 | 2211,2 | 2469 | 1953 | 0,79 | -0,34 | -4,872 | 1,10E-06 | 4,62E-06 | 0,005774 | 0,002636 | 0,0042247 | 0,0042247 | TRUE | 0,1126997 | No |
| orf19.11 | 378 | 225 | 298 | 351 | 480 | 348 | 412 | 230 | 339 | 286 | 305 | 289 | 411 | 332 | 368 | 324 | 331,81 | 305 | 359 | 1,179 | 0,238 | 2,416 | 0,01567759 | 0,0300258 | 0,006233 | 0,0063546 | 0,0062824 | 0,0062824 | TRUE | 0,1112749 | No |
| orf19.1105 | 48 | 65 | 63 | 66 | 44 | 47 | 64 | 34 | 43 | 83 | 64 | 54 | 38 | 45 | 57 | 48 | 54,03 | 61 | 47 | 0,775 | -0,369 | -1,599 | 0,10974617 | 0,16504506 | 0,03591 | 0,0288484 | 0,0320255 | 0,0320255 | TRUE | 0,2721422 | No |
| orf19.1105.2 | 60742 | 39769 | 52790 | 64073 | 69796 | 53337 | 58691 | 41323 | 54492 | 50630 | 54009 | 52835 | 59692 | 50892 | 52372 | 58282 | 54151 | 52992 | 55310 | 1,044 | 0,062 | 1,102 | 0,27044978 | 0,35485091 | 0,003564 | 0,0020063 | 0,0029922 | 0,0029922 | TRUE | 0,0354197 | No |
| orf19.1105.3 | 22 | 6 | 9 | 14 | 13 | 8 | 7 | 13 | 20 | 8 | 9 | 12 | 11 | 8 | 6 | 18 | 11,43 | 12 | 11 | 0,857 | -0,222 | -0,476 | 0,63391902 | NA | 0,109638 | 0,1289591 | 0,1207785 | 0,1207785 | TRUE | 0,7922111 | No |
| orf19.1106 | 174 | 149 | 179 | 242 | 277 | 302 | 300 | 195 | 156 | 190 | 183 | 200 | 237 | 288 | 268 | 275 | 224,53 | 182 | 267 | 1,464 | 0,55 | 5,047 | 4,50E-07 | 2,01E-06 | 0,004852 | 0,0084449 | 0,0067783 | 0,0067783 | TRUE | 0,1076143 | No |
| orf19.1107 | 4958 | 3207 | 4098 | 5332 | 4921 | 4095 | 4409 | 3251 | 4448 | 4083 | 4193 | 4397 | 4209 | 3907 | 3934 | 4585 | 4219,5 | 4280 | 4159 | 0,97 | -0,043 | -0,757 | 0,44913889 | 0,53561462 | 0,003094 | 0,0023236 | 0,00291 | 0,00291 | TRUE | 0,0562794 | No |
| orf19.1108 | 3045 | 2228 | 2507 | 3216 | 3164 | 2859 | 2984 | 2078 | 2732 | 2836 | 2565 | 2652 | 2706 | 2728 | 2663 | 2931 | 2726,6 | 2696 | 2757 | 1,022 | 0,031 | 0,654 | 0,51304457 | 0,59724313 | 0,001357 | 0,002512 | 0,001793 | 0,001793 | TRUE | 0,0205676 | No |
| orf19.1109 | 1838 | 1453 | 1598 | 2156 | 2217 | 2192 | 2289 | 1250 | 1649 | 1850 | 1635 | 1778 | 1896 | 2092 | 2043 | 1763 | 1838,1 | 1728 | 1948 | 1,13 | 0,176 | 2,707 | 0,0067895 | 0,01423284 | 0,003991 | 0,0027693 | 0,0035104 | 0,0035104 | TRUE | 0,0467502 | No |
| orf19.111 | 23662 | 16718 | 21817 | 26266 | 25665 | 23125 | 24607 | 15595 | 21227 | 21284 | 22321 | 21659 | 21950 | 22065 | 21958 | 21995 | 21807 | 21623 | 21992 | 1,017 | 0,024 | 0,91 | 0,36275978 | 0,45151456 | 0,000213 | 0,0020461 | 0,0006422 | 0,0006422 | TRUE | 0,0057594 | No |
| orf19.1110 | 451 | 327 | 368 | 481 | 426 | 377 | 409 | 270 | 405 | 416 | 377 | 397 | 364 | 360 | 365 | 381 | 382,98 | 399 | 368 | 0,921 | -0,118 | -1,609 | 0,10755307 | 0,16227495 | 1,00E-08 | 0,00577 | 0,0025826 | 0,0025826 | TRUE | 0,0149453 | No |
| orf19.1111 | 863 | 569 | 809 | 953 | 756 | 720 | 837 | 557 | 774 | 724 | 828 | 786 | 647 | 687 | 747 | 786 | 747,27 | 778 | 717 | 0,917 | -0,125 | -1,714 | 0,08652249 | 0,13530118 | 0,003425 | 0,0039222 | 0,0037692 | 0,0037692 | TRUE | 0,0521229 | No |
| orf19.1112 | 943 | 740 | 816 | 1064 | 1208 | 1156 | 1244 | 727 | 846 | 942 | 835 | 877 | 1033 | 1103 | 1110 | 1025 | 971,48 | 875 | 1068 | 1,224 | 0,292 | 5,01 | 5,45E-07 | 2,40E-06 | 0,001011 | 0,0034738 | 0,0022108 | 0,0022108 | TRUE | 0,0285132 | No |
| orf19.1113 | 409 | 323 | 414 | 517 | 535 | 444 | 511 | 279 | 367 | 411 | 424 | 426 | 458 | 424 | 456 | 394 | 419,84 | 407 | 433 | 1,068 | 0,095 | 1,168 | 0,24268881 | 0,32423268 | 0,002406 | 0,0054373 | 0,0039749 | 0,0039749 | TRUE | 0,0539474 | No |
| orf19.1114 | 4588 | 3119 | 3883 | 4808 | 5176 | 4457 | 4583 | 2909 | 4116 | 3971 | 3973 | 3965 | 4427 | 4253 | 4090 | 4103 | 4112 | 4006 | 4218 | 1,053 | 0,075 | 1,935 | 0,05303468 | 0,08853252 | 0,000541 | 0,0023326 | 0,0012001 | 0,0012001 | TRUE | 0,013806 | No |
| orf19.1115 | 2432 | 1623 | 1886 | 2448 | 2402 | 2249 | 2357 | 1372 | 2182 | 2066 | 1930 | 2019 | 2054 | 2146 | 2103 | 1935 | 2054,3 | 2049 | 2060 | 1,006 | 0,009 | 0,173 | 0,862981 | 0,89578709 | 0,001755 | 0,0026862 | 0,0021288 | 0,0021288 | TRUE | 0,0236839 | No |
| orf19.1116 | 151 | 102 | 128 | 171 | 192 | 165 | 175 | 110 | 135 | 130 | 131 | 141 | 164 | 157 | 156 | 155 | 146,28 | 134 | 158 | 1,177 | 0,235 | 2,088 | 0,03680212 | 0,06425665 | 1,00E-08 | 0,0119037 | 0,0052852 | 0,0052852 | TRUE | 0,0126047 | No |
| orf19.1117 | 208 | 138 | 217 | 205 | 299 | 264 | 271 | 152 | 187 | 176 | 222 | 169 | 256 | 252 | 242 | 214 | 214,65 | 188 | 241 | 1,287 | 0,363 | 3,273 | 0,00106255 | 0,00266325 | 0,005022 | 0,0087428 | 0,0071223 | 0,0071223 | TRUE | 0,1547745 | No |
| orf19.1119 | 1538 | 1303 | 1500 | 1807 | 2020 | 1955 | 2126 | 1329 | 1380 | 1659 | 1535 | 1490 | 1728 | 1865 | 1897 | 1874 | 1678,5 | 1516 | 1841 | 1,216 | 0,282 | 4,614 | 3,96E-06 | 1,53E-05 | 0,002971 | 0,0028444 | 0,0029886 | 0,0029886 | TRUE | 0,0465065 | No |
| orf19.1120 | 473 | 292 | 404 | 543 | 465 | 393 | 460 | 280 | 424 | 372 | 413 | 448 | 398 | 375 | 410 | 395 | 404,4 | 414 | 394 | 0,948 | -0,076 | -0,971 | 0,33130596 | 0,41811761 | 0,001102 | 0,0055693 | 0,0034752 | 0,0034752 | TRUE | 0,0491948 | No |
| orf19.1121 | 4397 | 3492 | 3911 | 5104 | 4245 | 4242 | 4295 | 2583 | 3945 | 4446 | 4001 | 4209 | 3630 | 4048 | 3833 | 3643 | 3969,3 | 4150 | 3788 | 0,914 | -0,13 | -2,455 | 0,0140946 | 0,02733388 | 0,002503 | 0,0023453 | 0,0024559 | 0,0024559 | TRUE | 0,0272569 | No |
| orf19.1122 | 74 | 55 | 65 | 86 | 60 | 77 | 79 | 53 | 66 | 70 | 67 | 71 | 51 | 73 | 70 | 75 | 67,98 | 68 | 67 | 0,978 | -0,032 | -0,182 | 0,85547008 | 0,88998515 | 1,00E-08 | 0,0233337 | 0,0145707 | 0,0145707 | TRUE | 0,2628392 | No |
| orf19.1123 | 2 | 0 | 0 | 4 | 4 | 0 | 3 | 1 | 2 | 0 | 0 | 3 | 3 | 0 | 3 | 1 | 1,58 | 1 | 2 | 1,406 | 0,492 | 0,392 | 0,69476813 | NA | 0,781528 | 0,9236216 | 0,8827149 | 0,8827149 | TRUE | 0,6187286 | No |
| orf19.1124 | 221 | 154 | 209 | 266 | 317 | 256 | 260 | 161 | 198 | 196 | 214 | 219 | 271 | 244 | 232 | 227 | 225,24 | 207 | 244 | 1,18 | 0,238 | 2,396 | 0,01657184 | 0,03149751 | 1,00E-08 | 0,0084245 | 0,0050558 | 0,0050558 | TRUE | 0,0676909 | No |
| orf19.1124.2 | 339 | 236 | 306 | 340 | 374 | 215 | 270 | 183 | 304 | 300 | 313 | 280 | 320 | 205 | 241 | 258 | 277,76 | 299 | 256 | 0,856 | -0,224 | -1,812 | 0,07004517 | 0,11288396 | 0,014891 | 0,0072061 | 0,0109883 | 0,0109883 | TRUE | 0,3385844 | No |
| orf19.113 | 2122 | 1565 | 1904 | 2397 | 2155 | 2036 | 2074 | 1309 | 1904 | 1992 | 1948 | 1977 | 1843 | 1943 | 1851 | 1846 | 1912,9 | 1955 | 1871 | 0,957 | -0,063 | -1,531 | 0,12578166 | 0,1858117 | 1,00E-08 | 0,0027384 | 0,0010886 | 0,0010886 | TRUE | 0,0082871 | No |
| orf19.1133 | 732 | 479 | 645 | 765 | 957 | 908 | 946 | 699 | 657 | 610 | 660 | 631 | 818 | 866 | 844 | 986 | 759,01 | 640 | 878 | 1,368 | 0,452 | 6,535 | 6,34E-11 | 4,76E-10 | 0,002432 | 0,0038921 | 0,0032349 | 0,0032349 | TRUE | 0,0728232 | No |
| orf19.1135 | 399 | 299 | 407 | 465 | 411 | 507 | 497 | 265 | 358 | 381 | 416 | 383 | 352 | 484 | 443 | 374 | 398,87 | 384 | 413 | 1,076 | 0,106 | 1,006 | 0,31448994 | 0,40107313 | 0,010688 | 0,0056191 | 0,0081918 | 0,0081918 | TRUE | 0,1537038 | No |
| orf19.1136 | 152 | 145 | 107 | 170 | 197 | 202 | 173 | 113 | 136 | 185 | 109 | 140 | 168 | 193 | 154 | 159 | 155,7 | 142 | 168 | 1,192 | 0,253 | 1,695 | 0,09016624 | 0,1403053 | 0,019413 | 0,0113033 | 0,0148996 | 0,0148996 | TRUE | 0,3599648 | No |
| orf19.1137 | 1895 | 1437 | 1605 | 2189 | 1646 | 1506 | 1529 | 1031 | 1700 | 1829 | 1642 | 1805 | 1408 | 1437 | 1364 | 1454 | 1580 | 1744 | 1416 | 0,811 | -0,302 | -5,966 | 2,43E-09 | 1,46E-08 | 0,00106 | 0,0028983 | 0,0018122 | 0,0018122 | TRUE | 0,0182989 | No |
| orf19.1139 | 6494 | 4324 | 5763 | 7098 | 7250 | 6628 | 7452 | 4614 | 5826 | 5505 | 5896 | 5853 | 6200 | 6324 | 6650 | 6508 | 6095,2 | 5770 | 6420 | 1,112 | 0,153 | 3,999 | 6,35E-05 | 0,00020117 | 0,000739 | 0,0022177 | 0,0012428 | 0,0012428 | TRUE | 0,0112607 | No |
| orf19.114 | 191 | 152 | 169 | 221 | 153 | 187 | 188 | 107 | 171 | 194 | 173 | 182 | 131 | 178 | 168 | 151 | 168,49 | 180 | 157 | 0,874 | -0,194 | -1,605 | 0,10839708 | 0,16326139 | 0,004847 | 0,0105952 | 0,0080648 | 0,0080648 | TRUE | 0,1528863 | No |
| orf19.1140 | 695 | 578 | 645 | 790 | 743 | 743 | 768 | 459 | 623 | 736 | 660 | 651 | 635 | 709 | 685 | 647 | 668,47 | 668 | 669 | 1,006 | 0,009 | 0,128 | 0,89795829 | 0,92271167 | 0,002146 | 0,0041512 | 0,0031218 | 0,0031218 | TRUE | 0,0511844 | No |
| orf19.1142 | 1296 | 993 | 1328 | 1581 | 1322 | 1274 | 1378 | 853 | 1163 | 1264 | 1359 | 1304 | 1131 | 1216 | 1230 | 1203 | 1233,5 | 1272 | 1195 | 0,939 | -0,091 | -1,526 | 0,12689534 | 0,1871348 | 0,002182 | 0,0031564 | 0,0025944 | 0,0025944 | TRUE | 0,0419685 | No |
| orf19.1143 | 321 | 241 | 283 | 308 | 429 | 394 | 422 | 221 | 288 | 307 | 290 | 254 | 367 | 376 | 377 | 312 | 321,18 | 285 | 358 | 1,269 | 0,343 | 3,721 | 0,00019879 | 0,00058006 | 0,003241 | 0,0064995 | 0,0050308 | 0,0050308 | TRUE | 0,072643 | No |
| orf19.1144 | 756 | 510 | 689 | 806 | 730 | 677 | 738 | 427 | 678 | 649 | 705 | 665 | 624 | 646 | 659 | 602 | 653,52 | 674 | 633 | 0,94 | -0,089 | -1,474 | 0,14042987 | 0,20399 | 1,00E-08 | 0,0042009 | 0,0019957 | 0,0019957 | TRUE | 0,010515 | No |
| orf19.1146 | 74 | 36 | 60 | 77 | 54 | 42 | 49 | 38 | 66 | 46 | 61 | 63 | 46 | 40 | 44 | 54 | 52,58 | 59 | 46 | 0,76 | -0,396 | -2,016 | 0,04382396 | 0,07490594 | 1,00E-08 | 0,0295865 | 0,017833 | 0,017833 | TRUE | 0,161726 | No |
| orf19.1148 | 7 | 6 | 4 | 10 | 4 | 3 | 7 | 6 | 6 | 8 | 4 | 8 | 3 | 3 | 6 | 8 | 5,91 | 6 | 5 | 0,769 | -0,38 | -0,656 | 0,51187625 | NA | 1,00E-08 | 0,2477747 | 0,1476124 | 0,1476124 | TRUE | 0,2274558 | No |
| orf19.1149 | 2907 | 1843 | 2431 | 3059 | 2892 | 2328 | 2534 | 1631 | 2608 | 2346 | 2487 | 2522 | 2473 | 2221 | 2261 | 2300 | 2402,5 | 2491 | 2314 | 0,928 | -0,107 | -2,118 | 0,03414502 | 0,06014169 | 0,001551 | 0,0025838 | 0,0020355 | 0,0020355 | TRUE | 0,0270856 | No |
| orf19.115 | 155 | 114 | 120 | 166 | 129 | 149 | 110 | 59 | 139 | 145 | 123 | 137 | 110 | 142 | 98 | 83 | 122,21 | 136 | 108 | 0,805 | -0,312 | -2,007 | 0,04478098 | 0,07638958 | 0,016596 | 0,0138579 | 0,0150142 | 0,0150142 | TRUE | 0,424839 | No |
| orf19.1150 | 5105 | 3490 | 4472 | 5683 | 5702 | 5076 | 5481 | 3883 | 4580 | 4443 | 4575 | 4686 | 4877 | 4843 | 4891 | 5477 | 4796,5 | 4571 | 5022 | 1,097 | 0,134 | 2,837 | 0,00455508 | 0,0099167 | 0,001789 | 0,0022822 | 0,0019233 | 0,0019233 | TRUE | 0,043743 | No |

| | | | | | | | | | | | | | | | | | | | | | | | | | | | | | | | |
|---|---|---|---|---|---|---|---|---|---|---|---|---|---|---|---|---|---|---|---|---|---|---|---|---|---|---|---|---|---|---|---|
| orf19.1150.1 | 303 | 214 | 248 | 329 | 200 | 184 | 194 | 149 | 272 | 272 | 254 | 271 | 171 | 176 | 173 | 210 | 224,9 | 267 | 182 | 0,676 | -0,565 | -5,733 | 9,86E-09 | 5,45E-08 | 1,00E-08 | 0,0084345 | 0,0046849 | 0,0046849 | TRUE | 0,0896054 | No |
| orf19.1152 | 39048 | 27680 | 34891 | 43113 | 55367 | 49905 | 53037 | 32479 | 35030 | 35240 | 35697 | 35552 | 47352 | 47618 | 47327 | 45809 | 41203 | 35380 | 47026 | 1,329 | 0,411 | 16,857 | 9,25E-64 | 1,74E-61 | 0,000156 | 0,0020148 | 0,0005453 | 0,0005453 | TRUE | 0,0036663 | No |
| orf19.1153 | 6250 | 4433 | 5511 | 6672 | 7748 | 6907 | 7682 | 4561 | 5607 | 5644 | 5638 | 5502 | 6626 | 6590 | 6855 | 6433 | 6111,9 | 5598 | 6626 | 1,185 | 0,245 | 7,616 | 2,62E-14 | 2,79E-13 | 0,000195 | 0,002217 | 0,0008241 | 0,0008241 | TRUE | 0,006592 | No |
| orf19.1154 | 2387 | 1520 | 1935 | 2377 | 2071 | 1698 | 1779 | 1223 | 2141 | 1935 | 1980 | 1960 | 1771 | 1620 | 1587 | 1725 | 1840 | 2004 | 1676 | 0,835 | -0,26 | -4,808 | 1,53E-06 | 6,27E-06 | 0,001684 | 0,0027685 | 0,0022474 | 0,0022474 | TRUE | 0,0261473 | No |
| orf19.1155 | 231 | 157 | 236 | 285 | 274 | 256 | 256 | 177 | 207 | 200 | 241 | 235 | 234 | 244 | 228 | 250 | 230,03 | 221 | 239 | 1,077 | 0,106 | 1,085 | 0,27793504 | 0,36198433 | 1,00E-08 | 0,0082904 | 0,004895 | 0,004895 | TRUE | 0,0390125 | No |
| orf19.1156 | 201 | 118 | 173 | 223 | 208 | 189 | 220 | 136 | 180 | 150 | 177 | 184 | 178 | 180 | 196 | 192 | 179,72 | 173 | 186 | 1,071 | 0,099 | 0,924 | 0,35531117 | 0,4428858 | 1,00E-08 | 0,010057 | 0,0055536 | 0,0055536 | TRUE | 0,0700391 | No |
| orf19.1158 | 392 | 271 | 325 | 452 | 458 | 423 | 417 | 335 | 352 | 345 | 333 | 373 | 392 | 404 | 372 | 472 | 380,23 | 351 | 410 | 1,161 | 0,215 | 2,446 | 0,01442585 | 0,02791321 | 0,003607 | 0,0057975 | 0,004755 | 0,004755 | TRUE | 0,1062943 | No |
| orf19.1159 | 4107 | 2655 | 3601 | 4310 | 3963 | 3282 | 3733 | 2139 | 3684 | 3380 | 3684 | 3554 | 3389 | 3132 | 3331 | 3017 | 3396,5 | 3576 | 3217 | 0,9 | -0,151 | -3,034 | 0,00241461 | 0,00559252 | 0,001869 | 0,0024069 | 0,0020965 | 0,0020965 | TRUE | 0,0203366 | No |
| orf19.1160 | 1053 | 751 | 986 | 1209 | 1247 | 1013 | 1183 | 776 | 945 | 956 | 1009 | 997 | 1066 | 967 | 1056 | 1094 | 1011,2 | 977 | 1046 | 1,069 | 0,096 | 1,709 | 0,08753183 | 0,13663062 | 0,000929 | 0,0034151 | 0,0020549 | 0,0020549 | TRUE | 0,0311491 | No |
| orf19.1161 | 3142 | 2080 | 2822 | 3256 | 3654 | 3059 | 3446 | 2090 | 2819 | 2648 | 2887 | 2685 | 3125 | 2919 | 3075 | 2948 | 2888,2 | 2760 | 3017 | 1,093 | 0,129 | 2,904 | 0,0036786 | 0,00817043 | 0,000904 | 0,0024822 | 0,0015426 | 0,0015426 | TRUE | 0,0114843 | No |
| orf19.1162 | 1673 | 1088 | 1334 | 1688 | 2002 | 1668 | 1764 | 971 | 1501 | 1385 | 1365 | 1392 | 1712 | 1592 | 1574 | 1370 | 1486,3 | 1411 | 1562 | 1,109 | 0,15 | 2,207 | 0,02733187 | 0,04946081 | 0,004184 | 0,0029563 | 0,0037376 | 0,0037376 | TRUE | 0,077762 | No |
| orf19.1162.1 | 538 | 366 | 499 | 624 | 502 | 447 | 444 | 278 | 483 | 466 | 511 | 515 | 429 | 427 | 396 | 392 | 452,23 | 494 | 411 | 0,833 | -0,263 | -3,654 | 0,00025787 | 0,00073636 | 1,00E-08 | 0,0051896 | 0,0027485 | 0,0027485 | TRUE | 0,0143848 | No |
| orf19.1163 | 1603 | 1105 | 1327 | 1795 | 1732 | 1526 | 1683 | 1209 | 1438 | 1407 | 1358 | 1480 | 1481 | 1456 | 1502 | 1705 | 1478,4 | 1421 | 1536 | 1,078 | 0,108 | 1,827 | 0,06767271 | 0,10958965 | 0,002526 | 0,0029615 | 0,0026766 | 0,0026766 | TRUE | 0,0614015 | No |
| orf19.1164 | 11158 | 6667 | 8999 | 11393 | 9610 | 7379 | 8223 | 5485 | 10010 | 8488 | 9207 | 9395 | 8219 | 7041 | 7338 | 7736 | 8429,1 | 9275 | 7584 | 0,817 | -0,291 | -4,722 | 2,33E-06 | 9,39E-06 | 0,00439 | 0,0021517 | 0,0035245 | 0,0035245 | TRUE | 0,039555 | No |
| orf19.1166 | 5310 | 3499 | 4930 | 5759 | 6185 | 5158 | 5779 | 3889 | 4764 | 4455 | 5044 | 4749 | 5290 | 4922 | 5157 | 5485 | 4983 | 4753 | 5214 | 1,096 | 0,132 | 2,68 | 0,0073689 | 0,01529269 | 0,002044 | 0,0022709 | 0,0021372 | 0,0021372 | TRUE | 0,0210883 | No |
| orf19.1167 | 1077 | 732 | 911 | 1133 | 898 | 786 | 808 | 550 | 966 | 932 | 932 | 934 | 768 | 750 | 721 | 776 | 847,39 | 941 | 754 | 0,799 | -0,323 | -6,053 | 1,42E-09 | 8,86E-09 | 1,00E-08 | 0,0036927 | 0,0015406 | 0,0015406 | TRUE | 0,0104292 | No |
| orf19.1168 | 8344 | 5477 | 6868 | 8921 | 9210 | 7383 | 8350 | 5658 | 7485 | 6973 | 7027 | 7356 | 7877 | 7045 | 7451 | 7980 | 7399,2 | 7210 | 7588 | 1,052 | 0,073 | 1,507 | 0,13179636 | 0,19350745 | 0,002015 | 0,0021757 | 0,0021092 | 0,0021092 | TRUE | 0,0283852 | No |
| orf19.1169 | 6005 | 4060 | 5029 | 6121 | 6683 | 5818 | 6486 | 3468 | 5387 | 5169 | 5145 | 5047 | 5716 | 5551 | 5788 | 4891 | 5336,8 | 5187 | 5486 | 1,059 | 0,082 | 1,493 | 0,13530953 | 0,19769933 | 0,002909 | 0,0022515 | 0,0027207 | 0,0027207 | TRUE | 0,0633459 | No |
| orf19.117 | 3774 | 2748 | 3264 | 4190 | 4006 | 3732 | 4081 | 2527 | 3386 | 3498 | 3339 | 3455 | 3426 | 3561 | 3642 | 3564 | 3483,9 | 3420 | 3548 | 1,038 | 0,053 | 1,467 | 0,14236689 | 0,20635471 | 0,00028 | 0,0023962 | 0,0009788 | 0,0009788 | TRUE | 0,0068782 | No |
| orf19.1170 | 1820 | 1388 | 1514 | 1922 | 1777 | 1564 | 1681 | 1061 | 1633 | 1767 | 1549 | 1585 | 1520 | 1492 | 1500 | 1496 | 1567,8 | 1634 | 1502 | 0,921 | -0,118 | -2,357 | 0,01844106 | 0,03457986 | 0,000889 | 0,0029055 | 0,0017772 | 0,0017772 | TRUE | 0,0340257 | No |
| orf19.1171 | 43 | 25 | 43 | 49 | 30 | 31 | 29 | 22 | 39 | 32 | 44 | 40 | 26 | 30 | 26 | 31 | 33,37 | 39 | 28 | 0,714 | -0,486 | -2,055 | 0,03992294 | 0,06894425 | 1,00E-08 | 0,0454851 | 0,0230697 | 0,0230697 | TRUE | 0,0814702 | No |
| orf19.1177 | 427 | 285 | 366 | 487 | 479 | 363 | 445 | 251 | 383 | 363 | 374 | 402 | 410 | 346 | 397 | 354 | 378,63 | 380 | 377 | 0,992 | -0,012 | -0,146 | 0,88416159 | 0,9122519 | 0,001881 | 0,0058136 | 0,0039244 | 0,0039244 | TRUE | 0,0405786 | No |
| orf19.1178 | 160 | 81 | 132 | 154 | 130 | 104 | 123 | 84 | 144 | 103 | 135 | 127 | 111 | 99 | 110 | 118 | 118,42 | 127 | 110 | 0,854 | -0,228 | -1,667 | 0,09560506 | 0,14731859 | 0,002427 | 0,0142387 | 0,0095657 | 0,0095657 | TRUE | 0,1385949 | No |
| orf19.1179 | 1607 | 975 | 1255 | 1556 | 1708 | 1222 | 1268 | 1048 | 1442 | 1241 | 1284 | 1283 | 1461 | 1166 | 1131 | 1478 | 1310,8 | 1312 | 1309 | 0,996 | -0,006 | -0,067 | 0,94638197 | 0,96142423 | 0,011547 | 0,003087 | 0,0075509 | 0,0075509 | TRUE | 0,100979 | No |
| orf19.118 | 22420 | 17390 | 20316 | 25865 | 18331 | 17865 | 18334 | 11007 | 20113 | 22139 | 20785 | 21329 | 15677 | 17046 | 16360 | 15524 | 18622 | 21092 | 16152 | 0,766 | -0,385 | -8,731 | 2,52E-18 | 4,04E-17 | 0,001687 | 0,0020575 | 0,0018091 | 0,0018091 | TRUE | 0,0169889 | No |
| orf19.1180 | 250 | 154 | 228 | 262 | 296 | 209 | 281 | 190 | 224 | 196 | 233 | 216 | 253 | 199 | 251 | 268 | 230,12 | 217 | 243 | 1,111 | 0,151 | 1,375 | 0,16907838 | 0,23895053 | 0,006056 | 0,008288 | 0,0072787 | 0,0072787 | TRUE | 0,1624264 | No |
| orf19.1181 | 100 | 59 | 82 | 100 | 93 | 69 | 91 | 50 | 90 | 75 | 84 | 82 | 80 | 66 | 81 | 71 | 78,53 | 83 | 74 | 0,898 | -0,156 | -0,996 | 0,3190197 | 0,40590223 | 1,00E-08 | 0,0204644 | 0,0107783 | 0,0107783 | TRUE | 0,0610908 | No |
| orf19.1182 | 589 | 383 | 474 | 636 | 707 | 608 | 642 | 405 | 528 | 488 | 485 | 524 | 605 | 580 | 573 | 571 | 544,28 | 506 | 582 | 1,148 | 0,199 | 3,064 | 0,00218338 | 0,00511614 | 1,00E-08 | 0,0046467 | 0,0022309 | 0,0022309 | TRUE | 0,0102931 | No |
| orf19.1183 | 2245 | 1447 | 1899 | 2296 | 2597 | 2139 | 2513 | 1473 | 2014 | 1842 | 1943 | 1893 | 2221 | 2041 | 2242 | 2078 | 2034,3 | 1923 | 2146 | 1,116 | 0,158 | 3,184 | 0,00145376 | 0,00353755 | 0,001177 | 0,0026931 | 0,0018797 | 0,0018797 | TRUE | 0,013847 | No |
| orf19.1185 | 4199 | 2629 | 3752 | 4604 | 4333 | 3438 | 3895 | 2614 | 3767 | 3347 | 3839 | 3797 | 3706 | 3280 | 3476 | 3687 | 3612,2 | 3688 | 3537 | 0,958 | -0,061 | -1,062 | 0,28826957 | 0,37269713 | 0,00316 | 0,0023814 | 0,0029317 | 0,0029317 | TRUE | 0,0457688 | No |
| orf19.1186 | 876 | 571 | 738 | 875 | 949 | 727 | 807 | 481 | 786 | 727 | 755 | 722 | 812 | 694 | 720 | 678 | 736,65 | 748 | 726 | 0,974 | -0,038 | -0,549 | 0,5828165 | 0,66129586 | 0,002799 | 0,0039502 | 0,003331 | 0,003331 | TRUE | 0,0770824 | No |
| orf19.1187 | 28617 | 20377 | 26600 | 31787 | 36280 | 33438 | 37018 | 23905 | 25672 | 25942 | 27214 | 26212 | 31028 | 31905 | 33032 | 33716 | 29340 | 26260 | 32420 | 1,234 | 0,304 | 8,236 | 1,79E-16 | 2,35E-15 | 0,000958 | 0,002029 | 0,001273 | 0,001273 | TRUE | 0,0102558 | No |
| orf19.1189 | 22553 | 14642 | 19618 | 24300 | 27688 | 22718 | 25701 | 17458 | 20232 | 18641 | 20071 | 20038 | 23680 | 21677 | 22934 | 24623 | 21487 | 19746 | 23228 | 1,176 | 0,234 | 4,964 | 6,92E-07 | 2,99E-06 | 0,002077 | 0,0020471 | 0,0020879 | 0,0020879 | TRUE | 0,0247301 | No |
| orf19.119 | 299 | 292 | 298 | 433 | 255 | 248 | 254 | 171 | 268 | 372 | 305 | 357 | 218 | 237 | 227 | 241 | 278,06 | 326 | 231 | 0,709 | -0,497 | -4,438 | 9,07E-06 | 3,32E-05 | 0,009395 | 0,0072004 | 0,0082979 | 0,0082979 | TRUE | 0,1645515 | No |
| orf19.1190 | 1199 | 934 | 1187 | 1451 | 1669 | 1433 | 1656 | 971 | 1076 | 1189 | 1214 | 1197 | 1427 | 1367 | 1478 | 1370 | 1289,7 | 1169 | 1410 | 1,209 | 0,273 | 4,942 | 7,73E-07 | 3,32E-06 | 0,001448 | 0,0031051 | 0,0021555 | 0,0021555 | TRUE | 0,0358698 | No |
| orf19.1191 | 3992 | 2632 | 3532 | 4199 | 4888 | 4380 | 4864 | 2929 | 3581 | 3351 | 3614 | 3463 | 4180 | 4179 | 4340 | 4131 | 3854,9 | 3502 | 4208 | 1,201 | 0,265 | 6,931 | 4,17E-12 | 3,55E-11 | 0,000473 | 0,0023561 | 0,0011371 | 0,0011371 | TRUE | 0,0096275 | No |
| orf19.1192 | 1193 | 772 | 1049 | 1206 | 745 | 680 | 680 | 471 | 1070 | 983 | 1073 | 994 | 637 | 649 | 607 | 664 | 834,73 | 1030 | 639 | 0,619 | -0,692 | -11,672 | 1,77E-31 | 7,00E-30 | 0,000709 | 0,0037186 | 0,0021073 | 0,0021073 | TRUE | 0,0139589 | No |
| orf19.1193 | 95974 | 62634 | 86423 | 1E+05 | 1E+05 | 1E+05 | 1E+05 | 73853 | 86099 | 79740 | 88419 | 85563 | 1E+05 | 99977 | 1E+05 | 1E+05 | 94311 | 84955 | 1E+05 | 1,22 | 0,287 | 7,21 | 5,62E-13 | 5,33E-12 | 0,001284 | 0,0019949 | 0,0015132 | 0,0015132 | TRUE | 0,0208811 | No |
| orf19.1195 | 1629 | 1093 | 1472 | 1683 | 1529 | 1366 | 1531 | 922 | 1461 | 1392 | 1506 | 1388 | 1308 | 1303 | 1366 | 1300 | 1378 | 1437 | 1319 | 0,919 | -0,122 | -2,535 | 0,01123355 | 0,02235548 | 0,000399 | 0,003033 | 0,0015038 | 0,0015038 | TRUE | 0,0124273 | No |
| orf19.1196 | 964 | 711 | 1006 | 1186 | 1441 | 1320 | 1443 | 915 | 865 | 905 | 1029 | 978 | 1232 | 1260 | 1288 | 1291 | 1105,9 | 944 | 1268 | 1,341 | 0,423 | 6,913 | 4,75E-12 | 4,02E-11 | 0,002027 | 0,0032922 | 0,0026737 | 0,0026737 | TRUE | 0,0427621 | No |
| orf19.1198 | 1394 | 962 | 1224 | 1510 | 1540 | 1364 | 1439 | 940 | 1251 | 1225 | 1252 | 1245 | 1317 | 1301 | 1284 | 1326 | 1275,1 | 1243 | 1307 | 1,05 | 0,071 | 1,615 | 0,10625615 | 0,16060049 | 1,00E-08 | 0,003118 | 0,0010574 | 0,0010574 | TRUE | 0,0017173 | No |
| orf19.1199 | 27392 | 18670 | 23961 | 29425 | 22316 | 19379 | 21051 | 14824 | 24573 | 23769 | 24514 | 24264 | 19085 | 18491 | 18785 | 20908 | 21799 | 24280 | 19317 | 0,795 | -0,33 | -7,688 | 1,50E-14 | 1,63E-13 | 0,001638 | 0,0020461 | 0,0017273 | 0,0017273 | TRUE | 0,0372031 | No |
| orf19.120 | 250 | 214 | 282 | 362 | 197 | 233 | 206 | 138 | 224 | 272 | 289 | 299 | 168 | 222 | 184 | 195 | 231,63 | 271 | 192 | 0,709 | -0,497 | -4,093 | 4,27E-05 | 0,00013951 | 0,011197 | 0,008247 | 0,0096892 | 0,0096892 | TRUE | 0,1592156 | No |
| orf19.1200 | 2496 | 1774 | 2156 | 2745 | 2386 | 2034 | 2188 | 1489 | 2239 | 2258 | 2206 | 2264 | 2041 | 1941 | 1952 | 2100 | 2125,1 | 2242 | 2008 | 0,895 | -0,161 | -3,887 | 0,00010139 | 0,00031015 | 0,000147 | 0,0026626 | 0,0011651 | 0,0011651 | TRUE | 0,010111 | No |
| orf19.1201 | 2292 | 1559 | 2075 | 2529 | 2793 | 2281 | 2559 | 1640 | 2056 | 1985 | 2123 | 2085 | 2389 | 2176 | 2283 | 2313 | 2176,4 | 2062 | 2290 | 1,11 | 0,15 | 3,376 | 0,00073524 | 0,00190565 | 0,000708 | 0,0026466 | 0,0014427 | 0,0014427 | TRUE | 0,0139523 | No |
| orf19.1202 | 149 | 96 | 113 | 163 | 127 | 142 | 146 | 113 | 134 | 122 | 116 | 134 | 109 | 135 | 130 | 159 | 129,96 | 126 | 133 | 1,041 | 0,058 | 0,421 | 0,67380141 | 0,73939705 | 0,006777 | 0,01315 | 0,0104087 | 0,0104087 | TRUE | 0,1686593 | No |
| orf19.1203 | 951 | 619 | 821 | 947 | 976 | 831 | 874 | 523 | 853 | 788 | 840 | 781 | 835 | 793 | 780 | 738 | 800,9 | 816 | 786 | 0,966 | -0,049 | -0,809 | 0,41867153 | 0,50647075 | 0,000992 | 0,0037921 | 0,0023145 | 0,0023145 | TRUE | 0,020485 | No |
| orf19.1203.1 | 397 | 299 | 366 | 438 | 380 | 314 | 313 | 242 | 356 | 381 | 374 | 361 | 325 | 300 | 279 | 341 | 339,71 | 368 | 311 | 0,843 | -0,246 | -2,919 | 0,00351144 | 0,00782809 | 0,000815 | 0,0062529 | 0,0038512 | 0,0038512 | TRUE | 0,0552875 | No |
| orf19.1204 | 804 | 641 | 799 | 941 | 1244 | 1075 | 1232 | 763 | 721 | 816 | 817 | 776 | 1064 | 1026 | 1099 | 1076 | 924,49 | 782 | 1066 | 1,365 | 0,449 | 7,708 | 1,28E-14 | 1,41E-13 | 0,000894 | 0,0035498 | 0,0021456 | 0,0021456 | TRUE | 0,0337302 | No |
| orf19.1207 | 18 | 12 | 5 | 8 | 9 | 3 | 9 | 4 | 16 | 15 | 5 | 7 | 8 | 3 | 8 | 6 | 8,42 | 11 | 6 | 0,573 | -0,803 | -1,464 | 0,14324877 | NA | 0,142088 | 0,1743655 | 0,1604423 | 0,1604423 | TRUE | 0,1093145 | No |
| orf19.1208 | 22 | 23 | 22 | 35 | 20 | 20 | 27 | 19 | 20 | 29 | 23 | 29 | 17 | 19 | 24 | 27 | 23,43 | 25 | 22 | 0,858 | -0,22 | -0,755 | 0,45016243 | 0,53623917 | 1,00E-08 | 0,0639299 | 0,0385007 | 0,0385007 | TRUE | 0,1407963 | No |
| orf19.121 | 249 | 240 | 227 | 279 | 339 | 379 | 347 | 174 | 223 | 306 | 232 | 230 | 290 | 362 | 310 | 245 | 274,73 | 248 | 302 | 1,228 | 0,296 | 2,251 | 0,02436726 | 0,04456575 | 0,02003 | 0,0072636 | 0,0129466 | 0,0129466 | TRUE | 0,253277 | No |
| orf19.1210 | 3557 | 2741 | 3348 | 4294 | 4505 | 4400 | 4617 | 2773 | 3191 | 3490 | 3425 | 3541 | 3853 | 4198 | 4120 | 3911 | 3716,1 | 3412 | 4020 | 1,179 | 0,238 | 4,984 | 6,23E-07 | 2,72E-06 | 0,001506 | 0,0023702 | 0,0019096 | 0,0019096 | TRUE | 0,0235014 | No |
| orf19.1212 | 621 | 507 | 631 | 770 | 804 | 711 | 753 | 557 | 557 | 645 | 646 | 635 | 688 | 678 | 672 | 786 | 663,33 | 621 | 706 | 1,135 | 0,182 | 2,444 | 0,01453405 | 0,02805931 | 0,003227 | 0,004168 | 0,0038078 | 0,0038078 | TRUE | 0,0613459 | No |
| orf19.1214 | 7571 | 5360 | 6556 | 8006 | 6763 | 6081 | 6547 | 3806 | 6792 | 6824 | 6707 | 6602 | 5784 | 5802 | 5842 | 5368 | 6215,2 | 6731 | 5699 | 0,848 | -0,239 | -6,471 | 9,71E-11 | 7,10E-10 | 0,000652 | 0,0022131 | 0,0011416 | 0,0011416 | TRUE | 0,0176828 | No |
| orf19.1215 | 11 | 4 | 8 | 13 | 11 | 14 | 3 | 12 | 10 | 5 | 8 | 11 | 9 | 13 | 3 | 17 | 9,53 | 8 | 10 | 1,194 | 0,256 | 0,488 | 0,62563113 | NA | 0,161097 | 0,1543228 | 0,1571487 | 0,1571487 | TRUE | 0,3908105 | No |
| orf19.1217 | 685 | 563 | 694 | 734 | 763 | 821 | 858 | 466 | 615 | 717 | 710 | 605 | 653 | 783 | 766 | 657 | 688,17 | 662 | 715 | 1,084 | 0,117 | 1,341 | 0,1800361 | 0,25195828 | 0,006886 | 0,004089 | 0,0058269 | 0,0058269 | TRUE | 0,0490328 | No |
| orf19.1219 | 325 | 231 | 272 | 361 | 309 | 313 | 320 | 157 | 292 | 294 | 278 | 298 | 264 | 299 | 286 | 221 | 278,94 | 290 | 268 | 0,927 | -0,109 | -1,115 | 0,2646652 | 0,34855514 | 0,00323 | 0,0071839 | 0,0055325 | 0,0055325 | TRUE | 0,1227312 | No |
| orf19.122 | 715 | 517 | 641 | 781 | 707 | 678 | 716 | 445 | 641 | 658 | 656 | 644 | 605 | 647 | 639 | 628 | 639,7 | 650 | 630 | 0,969 | -0,045 | -0,77 | 0,44115395 | 0,52787352 | 1,00E-08 | 0,0042489 | 0,0017376 | 0,0017376 | TRUE | 0,0094164 | No |

| | | | | | | | | | | | | | | | | | | | | | | | | | | | | | | | |
|---|---|---|---|---|---|---|---|---|---|---|---|---|---|---|---|---|---|---|---|---|---|---|---|---|---|---|---|---|---|---|---|
| orf19.1220 | 11049 | 7434 | 9526 | 11595 | 12106 | 10859 | 12070 | 7778 | 9912 | 9464 | 9746 | 9561 | 10353 | 10361 | 10770 | 10970 | 10142 | 9671 | 10614 | 1,097 | 0,133 | 3,995 | 6,48E-05 | 0,00020512 | 0,000498 | 0,0021227 | 0,000973 | 0,000973 | TRUE | 0,0060372 | No |
| orf19.1221 | 753 | 609 | 685 | 844 | 900 | 862 | 925 | 540 | 676 | 775 | 701 | 696 | 770 | 822 | 825 | 762 | 753,36 | 712 | 795 | 1,121 | 0,165 | 2,613 | 0,00897267 | 0,01829864 | 0,001013 | 0,0039065 | 0,0024932 | 0,0024932 | TRUE | 0,0386351 | No |
| orf19.1223 | 755 | 654 | 682 | 919 | 958 | 1086 | 1084 | 642 | 677 | 833 | 698 | 758 | 819 | 1036 | 967 | 905 | 836,73 | 742 | 932 | 1,259 | 0,333 | 3,826 | 0,00013021 | 0,00039019 | 0,007707 | 0,0037145 | 0,0060447 | 0,0060447 | TRUE | 0,0830359 | No |
| orf19.1224 | 7973 | 6110 | 7505 | 9584 | 5623 | 5785 | 5913 | 4173 | 7153 | 7779 | 7678 | 7903 | 4809 | 5520 | 5276 | 5886 | 6500,5 | 7628 | 5373 | 0,704 | -0,507 | -8,238 | 1,76E-16 | 2,32E-15 | 0,004258 | 0,0022028 | 0,003473 | 0,003473 | TRUE | 0,0610851 | No |
| orf19.1225 | 2012 | 1707 | 1732 | 2232 | 1597 | 1639 | 1637 | 937 | 1805 | 2173 | 1772 | 1841 | 1366 | 1564 | 1461 | 1322 | 1662,8 | 1898 | 1428 | 0,754 | -0,407 | -5,356 | 8,49E-08 | 4,20E-07 | 0,006814 | 0,0028525 | 0,0049329 | 0,0049329 | TRUE | 0,1123525 | No |
| orf19.1227 | 136 | 122 | 121 | 167 | 133 | 120 | 129 | 76 | 122 | 155 | 124 | 138 | 114 | 115 | 115 | 107 | 123,67 | 135 | 113 | 0,843 | -0,247 | -1,921 | 0,05478961 | 0,09098286 | 1,00E-08 | 0,0137178 | 0,0077303 | 0,0077303 | TRUE | 0,0917566 | No |
| orf19.1228 | 3268 | 2444 | 3091 | 3829 | 3775 | 3579 | 3831 | 2404 | 2932 | 3111 | 3162 | 3157 | 3228 | 3415 | 3419 | 3391 | 3227 | 3090 | 3363 | 1,088 | 0,122 | 2,95 | 0,00318001 | 0,00717982 | 0,000741 | 0,0024294 | 0,001322 | 0,001322 | TRUE | 0,0150017 | No |
| orf19.1229 | 1322 | 1320 | 1316 | 1720 | 1634 | 2169 | 2077 | 1214 | 1186 | 1681 | 1346 | 1418 | 1397 | 2070 | 1853 | 1712 | 1583 | 1408 | 1758 | 1,25 | 0,322 | 2,709 | 0,00675221 | 0,01418436 | 0,022713 | 0,0028966 | 0,0129118 | 0,0129118 | TRUE | 0,2337415 | No |
| orf19.123 | 328 | 336 | 351 | 448 | 414 | 456 | 492 | 228 | 294 | 428 | 359 | 369 | 354 | 435 | 439 | 322 | 375,04 | 362 | 388 | 1,075 | 0,104 | 0,817 | 0,41404608 | 0,50213744 | 0,020242 | 0,0058503 | 0,0128454 | 0,0128454 | TRUE | 0,1872377 | No |
| orf19.1232 | 3146 | 2836 | 3011 | 3841 | 3432 | 3901 | 3708 | 2047 | 2822 | 3611 | 3081 | 3167 | 2935 | 3722 | 3309 | 2887 | 3191,7 | 3170 | 3213 | 1,014 | 0,021 | 0,23 | 0,81788184 | 0,8596231 | 0,0106 | 0,0024344 | 0,007494 | 0,007494 | TRUE | 0,1381925 | No |
| orf19.1233 | 10506 | 8480 | 9709 | 11898 | 9158 | 9322 | 9884 | 5906 | 9425 | 10796 | 9933 | 9811 | 7832 | 8895 | 8820 | 8330 | 9230,3 | 9991 | 8469 | 0,848 | -0,238 | -4,3 | 1,71E-05 | 6,01E-05 | 0,003225 | 0,0021368 | 0,0028347 | 0,0028347 | TRUE | 0,0356188 | No |
| orf19.1234 | 62 | 45 | 38 | 43 | 39 | 42 | 53 | 33 | 56 | 57 | 39 | 35 | 33 | 40 | 47 | 47 | 44,31 | 47 | 42 | 0,895 | -0,161 | -0,689 | 0,49068697 | 0,57612807 | 0,022098 | 0,0347389 | 0,0292955 | 0,0292955 | TRUE | 0,1849757 | No |
| orf19.1235 | 2244 | 1607 | 2121 | 2522 | 2594 | 2448 | 2579 | 1626 | 2013 | 2046 | 2170 | 2080 | 2218 | 2336 | 2301 | 2293 | 2182,2 | 2077 | 2287 | 1,101 | 0,139 | 3,335 | 0,00085385 | 0,00218112 | 0,000325 | 0,0026448 | 0,0011991 | 0,0011991 | TRUE | 0,0108319 | No |
| orf19.1236 | 2097 | 1636 | 1837 | 2391 | 2785 | 2663 | 2654 | 1989 | 1881 | 2083 | 1879 | 1972 | 2382 | 2541 | 2368 | 2805 | 2238,9 | 1954 | 2524 | 1,291 | 0,368 | 5,878 | 4,17E-09 | 2,45E-08 | 0,003848 | 0,0026279 | 0,0033105 | 0,0033105 | TRUE | 0,0656242 | No |
| orf19.1237 | 900 | 702 | 835 | 1036 | 823 | 778 | 772 | 528 | 807 | 894 | 854 | 854 | 704 | 742 | 689 | 745 | 786,18 | 852 | 720 | 0,844 | -0,244 | -4,169 | 3,06E-05 | 0,0001028 | 1,00E-08 | 0,003826 | 0,0020098 | 0,0020098 | TRUE | 0,0151582 | No |
| orf19.1238 | 191 | 142 | 174 | 207 | 235 | 230 | 226 | 144 | 171 | 181 | 178 | 171 | 201 | 219 | 202 | 203 | 190,76 | 175 | 206 | 1,18 | 0,239 | 2,391 | 0,01682284 | 0,031914 | 1,00E-08 | 0,0095898 | 0,004349 | 0,004349 | TRUE | 0,0210456 | No |
| orf19.1239 | 5 | 7 | 9 | 8 | 10 | 3 | 6 | 4 | 4 | 9 | 9 | 7 | 9 | 3 | 5 | 6 | 6,45 | 7 | 6 | 0,785 | -0,349 | -0,631 | 0,52805627 | NA | 1,00E-08 | 0,2269949 | 0,1370193 | 0,1370193 | TRUE | 0,3574215 | No |
| orf19.124 | 8651 | 6873 | 7727 | 9934 | 8265 | 8415 | 8613 | 5433 | 7761 | 8750 | 7905 | 8192 | 7068 | 8029 | 7686 | 7663 | 7881,8 | 8152 | 7612 | 0,934 | -0,099 | -1,891 | 0,05863845 | 0,09654694 | 0,002642 | 0,0021637 | 0,0024916 | 0,0024916 | TRUE | 0,0293988 | No |
| orf19.1240 | 6807 | 4844 | 6356 | 7499 | 9978 | 9008 | 9729 | 6093 | 6107 | 6167 | 6503 | 6184 | 8533 | 8595 | 8682 | 8594 | 7420,5 | 6240 | 8601 | 1,378 | 0,463 | 14,733 | 3,93E-49 | 3,89E-47 | 0,000249 | 0,0021752 | 0,0008096 | 0,0008096 | TRUE | 0,0097854 | No |
| orf19.1244 | 2196 | 1537 | 1918 | 2294 | 2597 | 2492 | 2626 | 1542 | 1970 | 1957 | 1962 | 1892 | 2221 | 2378 | 2343 | 2175 | 2112,2 | 1945 | 2279 | 1,173 | 0,231 | 5,22 | 1,79E-07 | 8,45E-07 | 0,000615 | 0,0026668 | 0,0013984 | 0,0013984 | TRUE | 0,0103799 | No |
| orf19.1246 | 4012 | 2799 | 3726 | 4375 | 4766 | 4092 | 4683 | 3028 | 3599 | 3563 | 3812 | 3608 | 4076 | 3904 | 4179 | 4271 | 3876,6 | 3646 | 4108 | 1,126 | 0,171 | 4,093 | 4,25E-05 | 0,00013921 | 0,000862 | 0,002354 | 0,0014179 | 0,0014179 | TRUE | 0,0133943 | No |
| orf19.1247 | 319 | 226 | 249 | 341 | 266 | 232 | 249 | 144 | 286 | 288 | 255 | 281 | 227 | 221 | 222 | 203 | 248 | 278 | 218 | 0,791 | -0,338 | -3,698 | 0,0002176 | 0,00063003 | 1,00E-08 | 0,0078332 | 0,0039491 | 0,0039491 | TRUE | 0,0331316 | No |
| orf19.1248 | 3944 | 2389 | 3254 | 4139 | 4050 | 3417 | 3707 | 2565 | 3538 | 3041 | 3329 | 3413 | 3464 | 3260 | 3308 | 3618 | 3371,4 | 3330 | 3412 | 1,023 | 0,033 | 0,594 | 0,55247063 | 0,63344342 | 0,002536 | 0,0024101 | 0,002637 | 0,002637 | TRUE | 0,0403331 | No |
| orf19.1249 | 770 | 569 | 756 | 949 | 738 | 587 | 640 | 414 | 691 | 724 | 773 | 783 | 631 | 560 | 571 | 584 | 664,68 | 743 | 586 | 0,79 | -0,34 | -4,979 | 6,38E-07 | 2,77E-06 | 0,001917 | 0,0041636 | 0,002961 | 0,002961 | TRUE | 0,0334878 | No |
| orf19.125 | 8389 | 6482 | 7780 | 9550 | 9690 | 9205 | 9602 | 5680 | 7526 | 8252 | 7960 | 7875 | 8287 | 8783 | 8568 | 8011 | 8157,8 | 7903 | 8412 | 1,065 | 0,091 | 2,13 | 0,03317433 | 0,05858653 | 0,001349 | 0,0021575 | 0,0016315 | 0,0016315 | TRUE | 0,0126332 | No |
| orf19.1250 | 4644 | 2965 | 3924 | 4811 | 3919 | 3250 | 3574 | 1957 | 4166 | 3775 | 4015 | 3967 | 3352 | 3101 | 3189 | 2760 | 3540,6 | 3981 | 3100 | 0,78 | -0,359 | -5,986 | 2,15E-09 | 1,30E-08 | 0,003677 | 0,0023895 | 0,0031658 | 0,0031658 | TRUE | 0,0638449 | No |
| orf19.1251 | 1801 | 1304 | 1657 | 2018 | 1912 | 1739 | 1863 | 1230 | 1616 | 1660 | 1695 | 1664 | 1635 | 1659 | 1662 | 1735 | 1665,9 | 1659 | 1673 | 1,007 | 0,01 | 0,245 | 0,80623381 | 0,84979872 | 1,00E-08 | 0,002851 | 0,0011085 | 0,0011085 | TRUE | 0,0065745 | No |
| orf19.1252 | 17171 | 11829 | 15327 | 18719 | 20845 | 17709 | 19667 | 12803 | 15404 | 15060 | 15681 | 15436 | 17827 | 16897 | 17550 | 18057 | 16489 | 15395 | 17583 | 1,142 | 0,191 | 6,041 | 1,53E-09 | 9,50E-09 | 0,000464 | 0,0020676 | 0,0009022 | 0,0009022 | TRUE | 0,0084266 | No |
| orf19.1253 | 1512 | 966 | 1328 | 1553 | 2165 | 1835 | 2106 | 1387 | 1356 | 1230 | 1359 | 1281 | 1852 | 1751 | 1879 | 1956 | 1582,9 | 1306 | 1860 | 1,42 | 0,506 | 9,55 | 1,30E-21 | 2,64E-20 | 0,001321 | 0,0028966 | 0,0020455 | 0,0020455 | TRUE | 0,0184314 | No |
| orf19.1254 | 771 | 502 | 664 | 788 | 1002 | 837 | 839 | 532 | 692 | 639 | 679 | 650 | 857 | 799 | 749 | 750 | 726,81 | 665 | 789 | 1,187 | 0,248 | 3,803 | 0,00014303 | 0,00042541 | 0,001597 | 0,0039769 | 0,002694 | 0,002694 | TRUE | 0,0420059 | No |
| orf19.1255 | 47 | 39 | 58 | 64 | 61 | 42 | 58 | 43 | 42 | 50 | 59 | 53 | 52 | 40 | 52 | 61 | 51,07 | 51 | 51 | 0,996 | -0,006 | -0,028 | 0,97749406 | 0,98378316 | 1,00E-08 | 0,030404 | 0,0198739 | 0,0198739 | TRUE | 0,1739199 | No |
| orf19.1257 | 12 | 14 | 13 | 11 | 12 | 9 | 10 | 9 | 11 | 18 | 13 | 9 | 10 | 9 | 9 | 13 | 11,43 | 13 | 10 | 0,8 | -0,322 | -0,784 | 0,4327985 | NA | 1,00E-08 | 0,1290054 | 0,0719738 | 0,0719738 | TRUE | 0,2314778 | No |
| orf19.1258 | 99 | 69 | 76 | 99 | 92 | 91 | 70 | 52 | 89 | 88 | 78 | 82 | 79 | 87 | 62 | 73 | 79,67 | 84 | 75 | 0,898 | -0,155 | -0,971 | 0,33140388 | 0,41811761 | 1,00E-08 | 0,0202008 | 0,0119648 | 0,0119648 | TRUE | 0,1392621 | No |
| orf19.1259 | 3690 | 2476 | 3229 | 3913 | 3926 | 3647 | 3837 | 2424 | 3310 | 3152 | 3304 | 3227 | 3358 | 3480 | 3424 | 3419 | 3334,1 | 3248 | 3420 | 1,052 | 0,073 | 2,118 | 0,03413308 | 0,0601402 | 1,00E-08 | 0,0024149 | 0,0008535 | 0,0008535 | TRUE | 0,0044292 | No |
| orf19.1260 | 712 | 469 | 518 | 716 | 681 | 614 | 598 | 379 | 639 | 597 | 530 | 590 | 582 | 586 | 534 | 535 | 574,08 | 589 | 559 | 0,95 | -0,074 | -0,996 | 0,31931332 | 0,40610381 | 0,002627 | 0,0045083 | 0,0035418 | 0,0035418 | TRUE | 0,0534205 | No |
| orf19.1261 | 2954 | 1982 | 2482 | 3131 | 3116 | 2822 | 3221 | 1903 | 2650 | 2523 | 2539 | 2582 | 2665 | 2693 | 2874 | 2684 | 2651,3 | 2574 | 2729 | 1,06 | 0,084 | 2,049 | 0,04049488 | 0,06979153 | 0,00055 | 0,0025271 | 0,0012514 | 0,0012514 | TRUE | 0,0160914 | No |
| orf19.1263 | 2438 | 1693 | 2127 | 2552 | 2360 | 2188 | 2438 | 1560 | 2187 | 2155 | 2176 | 2104 | 2018 | 2088 | 2176 | 2200 | 2138,1 | 2156 | 2120 | 0,983 | -0,025 | -0,59 | 0,55493873 | 0,63540861 | 0,000481 | 0,0026585 | 0,0013037 | 0,0013037 | TRUE | 0,0130412 | No |
| orf19.1264 | 617 | 447 | 608 | 734 | 619 | 561 | 575 | 404 | 554 | 569 | 622 | 605 | 529 | 535 | 513 | 570 | 562,18 | 588 | 537 | 0,911 | -0,135 | -1,997 | 0,04581352 | 0,07790654 | 0,000578 | 0,0045618 | 0,0026083 | 0,0026083 | TRUE | 0,0194563 | No |
| orf19.1265 | 327 | 260 | 308 | 393 | 351 | 354 | 319 | 208 | 293 | 331 | 315 | 324 | 300 | 338 | 285 | 293 | 309,94 | 316 | 304 | 0,965 | -0,052 | -0,591 | 0,55475646 | 0,63540861 | 0,001219 | 0,0066633 | 0,0041624 | 0,0041624 | TRUE | 0,0650353 | No |
| orf19.1266 | 1779 | 1372 | 1500 | 1745 | 1542 | 1499 | 1509 | 802 | 1596 | 1747 | 1535 | 1439 | 1319 | 1430 | 1347 | 1131 | 1442,9 | 1579 | 1307 | 0,83 | -0,269 | -3,415 | 0,00063678 | 0,00167137 | 0,007078 | 0,0029856 | 0,0052389 | 0,0052389 | TRUE | 0,0924839 | No |
| orf19.1267 | 8289 | 5846 | 7233 | 8883 | 7180 | 6286 | 7137 | 4305 | 7436 | 7443 | 7400 | 7325 | 6141 | 5998 | 6369 | 6072 | 6772,8 | 7401 | 6145 | 0,831 | -0,268 | -8,628 | 6,22E-18 | 9,60E-17 | 0,000168 | 0,0021939 | 0,0007748 | 0,0007748 | TRUE | 0,0074892 | No |
| orf19.1267.1 | 1228 | 958 | 1017 | 1414 | 1501 | 1396 | 1383 | 871 | 1102 | 1220 | 1040 | 1166 | 1284 | 1332 | 1234 | 1228 | 1200,8 | 1132 | 1270 | 1,124 | 0,168 | 2,82 | 0,00479859 | 0,01040041 | 0,001973 | 0,0031885 | 0,0025705 | 0,0025705 | TRUE | 0,0343004 | No |
| orf19.1268 | 193 | 135 | 188 | 202 | 190 | 156 | 158 | 138 | 173 | 172 | 192 | 167 | 162 | 149 | 141 | 195 | 168,86 | 176 | 162 | 0,911 | -0,134 | -1,112 | 0,26615855 | 0,35029147 | 0,004841 | 0,0105765 | 0,0080864 | 0,0080864 | TRUE | 0,1628596 | No |
| orf19.1270 | 1294 | 887 | 1095 | 1346 | 1179 | 1091 | 1183 | 738 | 1161 | 1129 | 1120 | 1110 | 1008 | 1041 | 1056 | 1041 | 1083,3 | 1130 | 1036 | 0,917 | -0,125 | -2,619 | 0,00882775 | 0,0180337 | 1,00E-08 | 0,0033196 | 0,0012733 | 0,0012733 | TRUE | 0,0043873 | No |
| orf19.1272 | 1272 | 891 | 1075 | 1265 | 1009 | 930 | 922 | 621 | 1141 | 1134 | 1100 | 1043 | 863 | 887 | 823 | 876 | 983,42 | 1104 | 862 | 0,781 | -0,357 | -6,523 | 6,89E-11 | 5,13E-10 | 1,00E-08 | 0,0034557 | 0,0018422 | 0,0018422 | TRUE | 0,0179357 | No |
| orf19.1273 | 152 | 87 | 110 | 117 | 178 | 125 | 122 | 82 | 136 | 111 | 113 | 96 | 152 | 119 | 109 | 116 | 119,02 | 114 | 124 | 1,092 | 0,127 | 0,815 | 0,41528289 | 0,50343414 | 0,015403 | 0,0141766 | 0,0147567 | 0,0147567 | TRUE | 0,2592825 | No |
| orf19.1274 | 11 | 5 | 21 | 15 | 12 | 8 | 15 | 13 | 10 | 6 | 21 | 12 | 10 | 8 | 13 | 18 | 12,46 | 12 | 12 | 0,961 | -0,057 | -0,128 | 0,89784014 | NA | 0,097957 | 0,1184599 | 0,1100858 | 0,1100858 | TRUE | 0,7368875 | No |
| orf19.1275 | 423 | 272 | 309 | 395 | 282 | 239 | 258 | 185 | 379 | 346 | 316 | 326 | 241 | 228 | 230 | 261 | 291 | 342 | 240 | 0,699 | -0,516 | -5,568 | 2,58E-08 | 1,36E-07 | 0,002215 | 0,0069682 | 0,0047059 | 0,0047059 | TRUE | 0,0669098 | No |
| orf19.1276 | 145 | 152 | 150 | 197 | 201 | 223 | 225 | 136 | 130 | 194 | 153 | 162 | 172 | 213 | 201 | 192 | 177,1 | 160 | 194 | 1,223 | 0,29 | 2,235 | 0,02540891 | 0,04630171 | 0,010865 | 0,0101767 | 0,0104554 | 0,0104554 | TRUE | 0,187261 | No |
| orf19.1277 | 13657 | 8998 | 12234 | 14727 | 15782 | 13278 | 14984 | 10449 | 12252 | 11455 | 12517 | 12144 | 13497 | 12669 | 13371 | 14737 | 12830 | 12092 | 13568 | 1,122 | 0,166 | 3,256 | 0,00112885 | 0,00281184 | 0,002604 | 0,0020927 | 0,0024064 | 0,0024064 | TRUE | 0,040645 | No |
| orf19.1279 | 5748 | 4341 | 5411 | 6588 | 4956 | 4818 | 5094 | 3458 | 5157 | 5527 | 5536 | 5433 | 4239 | 4597 | 4546 | 4877 | 4988,8 | 5413 | 4565 | 0,843 | -0,247 | -5,078 | 3,82E-07 | 1,72E-06 | 0,00191 | 0,0022705 | 0,0020673 | 0,0020673 | TRUE | 0,028318 | No |
| orf19.1280 | 19072 | 15658 | 16576 | 21237 | 17903 | 17965 | 17399 | 9922 | 17110 | 19934 | 16959 | 17512 | 15311 | 17142 | 15526 | 13994 | 16686 | 17879 | 15493 | 0,867 | -0,206 | -2,993 | 0,00275949 | 0,00631822 | 0,006413 | 0,0020665 | 0,0044998 | 0,0044998 | TRUE | 0,0730689 | No |
| orf19.1281 | 4475 | 2964 | 3671 | 4716 | 4474 | 3926 | 4186 | 2621 | 4015 | 3773 | 3756 | 3889 | 3826 | 3746 | 3735 | 3697 | 3804,6 | 3858 | 3751 | 0,972 | -0,041 | -1,128 | 0,25938127 | 0,34279865 | 0,000337 | 0,0023611 | 0,0009982 | 0,0009982 | TRUE | 0,0091046 | No |
| orf19.1282 | 1203 | 849 | 965 | 1196 | 1174 | 1140 | 1195 | 669 | 1079 | 1081 | 987 | 986 | 1004 | 1088 | 1066 | 944 | 1029,4 | 1033 | 1026 | 0,995 | -0,007 | -0,107 | 0,91497709 | 0,93746461 | 0,002246 | 0,0033897 | 0,0027681 | 0,0027681 | TRUE | 0,0307202 | No |
| orf19.1283 | 597 | 465 | 542 | 673 | 669 | 661 | 667 | 467 | 536 | 592 | 555 | 555 | 572 | 631 | 595 | 659 | 586,72 | 560 | 614 | 1,096 | 0,133 | 1,954 | 0,05069908 | 0,085179 | 0,00096 | 0,0044538 | 0,0027213 | 0,0027213 | TRUE | 0,0251364 | No |
| orf19.1285 | 4827 | 3296 | 4378 | 5298 | 5777 | 5533 | 5973 | 3978 | 4330 | 4196 | 4479 | 4369 | 4941 | 5279 | 5330 | 5611 | 4816,9 | 4344 | 5290 | 1,217 | 0,283 | 6,225 | 4,83E-10 | 3,23E-09 | 0,001431 | 0,0022809 | 0,0017728 | 0,0017728 | TRUE | 0,0242614 | No |
| orf19.1286 | 112 | 89 | 101 | 128 | 144 | 88 | 116 | 67 | 100 | 113 | 103 | 106 | 123 | 84 | 104 | 94 | 103,47 | 106 | 101 | 0,967 | -0,049 | -0,329 | 0,74185534 | 0,79804241 | 0,005988 | 0,0160091 | 0,0116653 | 0,0116653 | TRUE | 0,2263643 | No |
| orf19.1287 | 7496 | 5528 | 6512 | 8215 | 10769 | 10359 | 10676 | 6717 | 6725 | 7038 | 6662 | 6774 | 9210 | 9884 | 9527 | 9474 | 8161,7 | 6800 | 9524 | 1,401 | 0,486 | 13,884 | 7,92E-44 | 6,10E-42 | 0,000551 | 0,0021574 | 0,0010523 | 0,0010523 | TRUE | 0,0080391 | No |
| orf19.1288 | 7506 | 4995 | 6733 | 7882 | 6296 | 5354 | 5631 | 3891 | 6734 | 6359 | 6889 | 6500 | 5385 | 5109 | 5025 | 5488 | 5935,8 | 6620 | 5252 | 0,793 | -0,335 | -7,753 | 8,99E-15 | 1,01E-13 | 0,001288 | 0,0022241 | 0,0016229 | 0,0016229 | TRUE | 0,0107345 | No |

| | | | | | | | | | | | | | | | | | | | | | | | | | | | | | | | |
|---|---|---|---|---|---|---|---|---|---|---|---|---|---|---|---|---|---|---|---|---|---|---|---|---|---|---|---|---|---|---|---|
| orf19.1289 | 1565 | 1154 | 1459 | 1636 | 2030 | 1898 | 2044 | 1165 | 1404 | 1469 | 1493 | 1349 | 1736 | 1811 | 1824 | 1643 | 1591,1 | 1429 | 1754 | 1,231 | 0,3 | 5,64 | 1,70E-08 | 9,14E-08 | 0,001314 | 0,0028919 | 0,002072 | 0,002072 | TRUE | 0,0197899 | No |
| orf19.1290 | 2935 | 2155 | 2660 | 3363 | 4791 | 4396 | 4762 | 2926 | 2633 | 2744 | 2721 | 2773 | 4097 | 4195 | 4249 | 4127 | 3442,4 | 2718 | 4167 | 1,534 | 0,617 | 17,627 | 1,54E-69 | 3,55E-67 | 1,00E-08 | 0,0024012 | 0,0008723 | 0,0008723 | TRUE | 0,0055878 | No |
| orf19.1291 | 1720 | 1207 | 1491 | 1891 | 2356 | 2071 | 2223 | 1355 | 1543 | 1537 | 1525 | 1559 | 2015 | 1976 | 1984 | 1911 | 1756,3 | 1541 | 1972 | 1,28 | 0,357 | 8,86 | 7,97E-19 | 1,32E-17 | 1,00E-08 | 0,0028061 | 0,0009785 | 0,0009785 | TRUE | 0,004516 | No |
| orf19.1295 | 5233 | 4105 | 4871 | 5816 | 5607 | 5834 | 5999 | 3443 | 4695 | 5226 | 4984 | 4796 | 4795 | 5567 | 5353 | 4856 | 5033,9 | 4925 | 5143 | 1,045 | 0,063 | 1,087 | 0,27695476 | 0,36102065 | 0,003551 | 0,0022679 | 0,0030707 | 0,0030707 | TRUE | 0,037523 | No |
| orf19.1296 | 1155 | 890 | 1001 | 1241 | 1436 | 1533 | 1586 | 901 | 1036 | 1133 | 1024 | 1023 | 1228 | 1463 | 1415 | 1271 | 1199,2 | 1054 | 1344 | 1,278 | 0,354 | 5,177 | 2,26E-07 | 1,05E-06 | 0,003634 | 0,0031901 | 0,003632 | 0,003632 | TRUE | 0,0434804 | No |
| orf19.1297 | 592 | 472 | 579 | 705 | 924 | 949 | 985 | 612 | 531 | 601 | 592 | 581 | 790 | 906 | 879 | 863 | 717,95 | 576 | 860 | 1,493 | 0,578 | 8,545 | 1,28E-17 | 1,91E-16 | 0,002024 | 0,0040015 | 0,0029386 | 0,0029386 | TRUE | 0,037482 | No |
| orf19.1298 | 2302 | 2043 | 2205 | 2871 | 3162 | 3341 | 3549 | 1888 | 2065 | 2601 | 2256 | 2367 | 2704 | 3188 | 3167 | 2663 | 2626,4 | 2322 | 2930 | 1,263 | 0,337 | 4,122 | 3,76E-05 | 0,00012411 | 0,008882 | 0,0025323 | 0,0060406 | 0,0060406 | TRUE | 0,0775053 | No |
| orf19.1299 | 1771 | 1642 | 1693 | 2277 | 1890 | 2400 | 2371 | 1527 | 1589 | 2090 | 1732 | 1878 | 1616 | 2290 | 2116 | 2154 | 1933,1 | 1822 | 2044 | 1,122 | 0,166 | 1,559 | 0,11895442 | 0,17694433 | 0,01661 | 0,0027305 | 0,0103385 | 0,0103385 | TRUE | 0,2428206 | No |
| orf19.130 | 426 | 361 | 408 | 500 | 406 | 444 | 465 | 277 | 382 | 460 | 417 | 412 | 347 | 424 | 415 | 391 | 406 | 418 | 394 | 0,946 | -0,081 | -0,915 | 0,36007685 | 0,44836097 | 0,004466 | 0,0055552 | 0,0050133 | 0,0050133 | TRUE | 0,0812019 | No |
| orf19.1300 | 2125 | 1668 | 1914 | 2400 | 2655 | 2469 | 2605 | 1538 | 1906 | 2124 | 1958 | 1979 | 2271 | 2356 | 2325 | 2169 | 2135,9 | 1992 | 2280 | 1,147 | 0,198 | 4,149 | 3,33E-05 | 0,00011129 | 0,001107 | 0,0026592 | 0,0017228 | 0,0017228 | TRUE | 0,0225608 | No |
| orf19.1301 | 574 | 438 | 400 | 574 | 433 | 482 | 448 | 319 | 515 | 558 | 409 | 473 | 370 | 460 | 400 | 450 | 454,38 | 489 | 420 | 0,858 | -0,22 | -2,103 | 0,03543664 | 0,06219787 | 0,011393 | 0,0051744 | 0,0082943 | 0,0082943 | TRUE | 0,1363706 | No |
| orf19.1302 | 36 | 28 | 34 | 53 | 44 | 34 | 27 | 23 | 32 | 36 | 35 | 44 | 38 | 32 | 24 | 32 | 34,13 | 37 | 32 | 0,861 | -0,217 | -0,893 | 0,37166395 | 0,46021372 | 1,00E-08 | 0,0445145 | 0,0271211 | 0,0271211 | TRUE | 0,2055393 | No |
| orf19.1303 | 1970 | 1436 | 1696 | 2178 | 1904 | 1746 | 1887 | 1149 | 1767 | 1828 | 1735 | 1796 | 1628 | 1666 | 1684 | 1621 | 1715,7 | 1782 | 1650 | 0,927 | -0,109 | -2,654 | 0,00795607 | 0,01641849 | 1,00E-08 | 0,0028257 | 0,0010482 | 0,0010482 | TRUE | 0,0035437 | No |
| orf19.1304 | 5031 | 3747 | 4345 | 5345 | 5115 | 4919 | 5232 | 2790 | 4513 | 4770 | 4445 | 4408 | 4375 | 4694 | 4669 | 3935 | 4476,1 | 4534 | 4418 | 0,976 | -0,036 | -0,609 | 0,54266789 | 0,62494704 | 0,003203 | 0,0023038 | 0,0030841 | 0,0030841 | TRUE | 0,0639833 | No |
| orf19.1305 | 1159 | 1074 | 1065 | 1362 | 1202 | 1419 | 1458 | 887 | 1040 | 1367 | 1090 | 1123 | 1028 | 1354 | 1301 | 1251 | 1194,2 | 1155 | 1234 | 1,069 | 0,096 | 0,967 | 0,33351699 | 0,42025379 | 0,012933 | 0,0031951 | 0,0086973 | 0,0086973 | TRUE | 0,1788383 | No |
| orf19.1306 | 2872 | 2084 | 2554 | 2996 | 2372 | 2332 | 2436 | 1514 | 2576 | 2653 | 2613 | 2471 | 2029 | 2225 | 2174 | 2135 | 2359,5 | 2578 | 2141 | 0,831 | -0,267 | -5,958 | 2,55E-09 | 1,52E-08 | 0,000863 | 0,0025948 | 0,001505 | 0,001505 | TRUE | 0,0159864 | No |
| orf19.1307 | 838 | 603 | 743 | 963 | 764 | 711 | 698 | 501 | 752 | 768 | 760 | 794 | 653 | 678 | 623 | 707 | 716,87 | 768 | 665 | 0,863 | -0,213 | -3,539 | 0,00040179 | 0,0011063 | 1,00E-08 | 0,0040046 | 0,0020795 | 0,0020795 | TRUE | 0,0216148 | No |
| orf19.1308 | 3350 | 2322 | 3049 | 3653 | 4137 | 3649 | 3881 | 2514 | 3005 | 2956 | 3119 | 3012 | 3538 | 3482 | 3463 | 3546 | 3265,3 | 3023 | 3507 | 1,159 | 0,213 | 6,223 | 4,88E-10 | 3,26E-09 | 1,00E-08 | 0,0024241 | 0,0008204 | 0,0008204 | TRUE | 0,0054075 | No |
| orf19.1309 | 573 | 345 | 447 | 552 | 747 | 542 | 604 | 380 | 514 | 439 | 457 | 455 | 639 | 517 | 539 | 536 | 512,09 | 466 | 558 | 1,196 | 0,258 | 2,996 | 0,0027337 | 0,00626872 | 0,005486 | 0,0048144 | 0,0051678 | 0,0051678 | TRUE | 0,1179328 | No |
| orf19.131 | 426 | 318 | 336 | 497 | 368 | 331 | 334 | 206 | 382 | 405 | 344 | 410 | 315 | 316 | 298 | 291 | 344,97 | 385 | 305 | 0,793 | -0,334 | -3,963 | 7,40E-05 | 0,00023106 | 1,00E-08 | 0,0061877 | 0,0039043 | 0,0039043 | TRUE | 0,0591024 | No |
| orf19.1310 | 768 | 559 | 675 | 900 | 745 | 673 | 819 | 493 | 689 | 712 | 691 | 742 | 637 | 642 | 731 | 695 | 692,36 | 708 | 676 | 0,953 | -0,069 | -1,042 | 0,2975121 | 0,38275218 | 0,001588 | 0,0040763 | 0,0027481 | 0,0027481 | TRUE | 0,0385525 | No |
| orf19.1311 | 18 | 7 | 10 | 11 | 14 | 13 | 12 | 1 | 16 | 9 | 10 | 9 | 12 | 12 | 11 | 1 | 10,11 | 11 | 9 | 0,852 | -0,23 | -0,478 | 0,63294457 | NA | 0,082882 | 0,1456106 | 0,1262901 | 0,1262901 | TRUE | 0,6555223 | No |
| orf19.131.2 | 2780 | 2048 | 2292 | 3156 | 3045 | 2581 | 2569 | 1887 | 2494 | 2607 | 2345 | 2602 | 2604 | 2463 | 2292 | 2661 | 2508,7 | 2512 | 2505 | 0,996 | -0,005 | -0,089 | 0,92879271 | 0,94901133 | 0,002999 | 0,0025582 | 0,002822 | 0,002822 | TRUE | 0,0400355 | No |
| orf19.1313 | 479 | 328 | 412 | 548 | 602 | 517 | 550 | 394 | 430 | 418 | 422 | 452 | 515 | 493 | 491 | 556 | 471,92 | 430 | 514 | 1,187 | 0,248 | 3,491 | 0,00048091 | 0,0013056 | 1,00E-08 | 0,0050557 | 0,002701 | 0,002701 | TRUE | 0,029887 | No |
| orf19.1314 | 3720 | 3216 | 3439 | 4261 | 4584 | 4462 | 4678 | 2902 | 3337 | 4094 | 3518 | 3514 | 3920 | 4257 | 4174 | 4093 | 3863,6 | 3616 | 4111 | 1,138 | 0,186 | 2,987 | 0,00282108 | 0,00644205 | 0,004428 | 0,0023553 | 0,0034824 | 0,0034824 | TRUE | 0,0947056 | No |
| orf19.1317 | 1280 | 919 | 1166 | 1418 | 1246 | 1141 | 1211 | 802 | 1148 | 1170 | 1193 | 1169 | 1066 | 1089 | 1081 | 1131 | 1130,8 | 1170 | 1092 | 0,931 | -0,103 | -2,18 | 0,0292492 | 0,05239391 | 1,00E-08 | 0,0032633 | 0,0012432 | 0,0012432 | TRUE | 0,0061235 | No |
| orf19.1318 | 1319 | 999 | 1200 | 1502 | 1137 | 1183 | 1195 | 668 | 1183 | 1272 | 1228 | 1239 | 972 | 1129 | 1066 | 942 | 1128,9 | 1230 | 1027 | 0,837 | -0,256 | -4,028 | 5,63E-05 | 0,00018065 | 0,002738 | 0,0032655 | 0,0029929 | 0,0029929 | TRUE | 0,0519015 | No |
| orf19.132 | 2539 | 1911 | 2277 | 2858 | 3268 | 3043 | 3222 | 2075 | 2278 | 2433 | 2330 | 2357 | 2795 | 2904 | 2875 | 2927 | 2612,1 | 2350 | 2875 | 1,224 | 0,292 | 7,596 | 3,05E-14 | 3,23E-13 | 0,000149 | 0,0025353 | 0,0010273 | 0,0010273 | TRUE | 0,0064203 | No |
| orf19.1321 | 6396 | 4113 | 5898 | 6766 | 13694 | 11416 | 13298 | 8965 | 5738 | 5236 | 6034 | 5579 | 11712 | 10893 | 11866 | 12644 | 8712,8 | 5647 | 11779 | 2,085 | 1,06 | 18,74 | 2,36E-78 | 6,15E-76 | 0,003415 | 0,0021461 | 0,0029418 | 0,0029418 | TRUE | 0,0312921 | No |
| orf19.1323 | 1946 | 1352 | 1743 | 2013 | 1728 | 1630 | 1735 | 1078 | 1746 | 1721 | 1783 | 1660 | 1478 | 1555 | 1548 | 1520 | 1626,5 | 1728 | 1525 | 0,883 | -0,179 | -4,027 | 5,65E-05 | 0,00018112 | 1,00E-08 | 0,0028721 | 0,0012774 | 0,0012774 | TRUE | 0,0090176 | No |
| orf19.1324 | 1504 | 993 | 1250 | 1545 | 1809 | 1568 | 1755 | 1084 | 1349 | 1264 | 1279 | 1274 | 1547 | 1496 | 1566 | 1529 | 1413,1 | 1292 | 1534 | 1,188 | 0,248 | 5,478 | 4,30E-08 | 2,20E-07 | 1,00E-08 | 0,0030069 | 0,0012564 | 0,0012564 | TRUE | 0,0112022 | No |
| orf19.1325 | 507 | 310 | 425 | 496 | 369 | 307 | 353 | 204 | 455 | 395 | 435 | 409 | 316 | 293 | 315 | 288 | 363,07 | 424 | 303 | 0,716 | -0,481 | -5,952 | 2,66E-09 | 1,58E-08 | 1,00E-08 | 0,0059779 | 0,0034557 | 0,0034557 | TRUE | 0,029663 | No |
| orf19.1326 | 321 | 244 | 310 | 333 | 267 | 212 | 204 | 136 | 288 | 311 | 317 | 275 | 228 | 202 | 182 | 192 | 249,36 | 298 | 201 | 0,68 | -0,557 | -5,492 | 3,97E-08 | 2,04E-07 | 0,003228 | 0,0078013 | 0,0057033 | 0,0057033 | TRUE | 0,0989244 | No |
| orf19.1327 | 127 | 97 | 159 | 164 | 159 | 138 | 150 | 109 | 114 | 123 | 163 | 135 | 136 | 132 | 134 | 154 | 136,32 | 134 | 139 | 1,032 | 0,046 | 0,338 | 0,73563298 | 0,79347988 | 0,006798 | 0,0126286 | 0,0101111 | 0,0101111 | TRUE | 0,2119982 | No |
| orf19.133 | 0 | 1 | 0 | 2 | 0 | 0 | 0 | 0 | 0 | 1 | 0 | 2 | 0 | 0 | 0 | 0 | 0,37 | 1 | 0 | 0,254 | -1,977 | -0,763 | 0,4454552 | NA | 1,00E-08 | 3,976069 | 4,763999 | 4,763999 | TRUE | 0,093742 | No |
| orf19.1330 | 169 | 110 | 147 | 179 | 204 | 164 | 209 | 120 | 152 | 140 | 150 | 148 | 174 | 156 | 186 | 169 | 159,54 | 148 | 171 | 1,164 | 0,219 | 1,971 | 0,04877204 | 0,08242544 | 1,00E-08 | 0,0110786 | 0,0056282 | 0,0056282 | TRUE | 0,041307 | No |
| orf19.1331 | 3629 | 3243 | 3347 | 4476 | 5106 | 5566 | 5505 | 3189 | 3256 | 4129 | 3424 | 3691 | 4367 | 5311 | 4912 | 4498 | 4198,4 | 3625 | 4772 | 1,317 | 0,398 | 4,922 | 8,56E-07 | 3,65E-06 | 0,009213 | 0,0023253 | 0,0060231 | 0,0060231 | TRUE | 0,1051377 | No |
| orf19.1332 | 975 | 690 | 867 | 1074 | 966 | 826 | 936 | 623 | 875 | 878 | 887 | 886 | 826 | 788 | 835 | 879 | 856,75 | 882 | 832 | 0,942 | -0,087 | -1,608 | 0,10774415 | 0,16244358 | 1,00E-08 | 0,003674 | 0,0016283 | 0,0016283 | TRUE | 0,0143081 | No |
| orf19.1333 | 1269 | 983 | 1188 | 1438 | 987 | 898 | 1009 | 651 | 1138 | 1251 | 1215 | 1186 | 844 | 857 | 900 | 918 | 1038,8 | 1198 | 880 | 0,734 | -0,446 | -8,196 | 2,48E-16 | 3,23E-15 | 0,000579 | 0,003377 | 0,0018496 | 0,0018496 | TRUE | 0,013334 | No |
| orf19.1334 | 731 | 547 | 648 | 880 | 783 | 731 | 882 | 484 | 656 | 696 | 663 | 726 | 670 | 697 | 787 | 683 | 697,2 | 685 | 709 | 1,036 | 0,051 | 0,727 | 0,46692094 | 0,55276079 | 0,002673 | 0,0040617 | 0,0033096 | 0,0033096 | TRUE | 0,0672993 | No |
| orf19.1335 | 7933 | 5954 | 7556 | 8944 | 7704 | 7521 | 8033 | 4862 | 7117 | 7580 | 7731 | 7375 | 6589 | 7176 | 7168 | 6857 | 7199,2 | 7451 | 6948 | 0,933 | -0,1 | -2,343 | 0,01911299 | 0,0357061 | 0,001273 | 0,0021812 | 0,0016245 | 0,0016245 | TRUE | 0,0151566 | No |
| orf19.1336 | 4726 | 3438 | 3871 | 5059 | 4453 | 3706 | 3812 | 2386 | 4240 | 4377 | 3960 | 4172 | 3808 | 3536 | 3402 | 3365 | 3857,5 | 4187 | 3528 | 0,843 | -0,246 | -4,813 | 1,49E-06 | 6,12E-06 | 0,002134 | 0,0023559 | 0,0022417 | 0,0022417 | TRUE | 0,0353274 | No |
| orf19.1336.2 | 1150 | 859 | 953 | 1181 | 1645 | 1464 | 1435 | 728 | 1032 | 1094 | 975 | 974 | 1407 | 1397 | 1280 | 1027 | 1148,2 | 1019 | 1278 | 1,258 | 0,332 | 3,631 | 0,00028194 | 0,00080078 | 0,009963 | 0,0032439 | 0,0071245 | 0,0071245 | TRUE | 0,1999548 | No |
| orf19.1338 | 1147 | 755 | 919 | 1143 | 1371 | 1212 | 1338 | 748 | 1029 | 961 | 940 | 943 | 1173 | 1156 | 1194 | 1055 | 1056,4 | 968 | 1144 | 1,185 | 0,245 | 4,244 | 2,19E-05 | 7,57E-05 | 0,0013 | 0,0033538 | 0,0022358 | 0,0022358 | TRUE | 0,0295682 | No |
| orf19.1339 | 13872 | 9386 | 11922 | 15282 | 10661 | 9007 | 9824 | 6833 | 12445 | 11949 | 12197 | 12602 | 9118 | 8594 | 8766 | 9637 | 10664 | 12298 | 9029 | 0,734 | -0,447 | -10,506 | 8,08E-26 | 2,23E-24 | 0,001443 | 0,0021157 | 0,0016401 | 0,0016401 | TRUE | 0,0245276 | No |
| orf19.134 | 0 | 0 | 2 | 2 | 0 | 1 | 0 | 0 | 0 | 0 | 2 | 2 | 0 | 1 | 0 | 0 | 0,58 | 1 | 0 | 0,324 | -1,627 | -0,813 | 0,41605507 | NA | 1,00E-08 | 2,4997496 | 2,3173122 | 2,3173122 | TRUE | 0,1949575 | No |
| orf19.1340 | 213 | 132 | 152 | 218 | 142 | 118 | 153 | 76 | 191 | 168 | 156 | 180 | 121 | 113 | 137 | 107 | 146,52 | 174 | 120 | 0,69 | -0,534 | -4,282 | 1,85E-05 | 6,46E-05 | 0,002427 | 0,0118874 | 0,0079257 | 0,0079257 | TRUE | 0,0953593 | No |
| orf19.1341 | 557 | 379 | 495 | 598 | 445 | 391 | 401 | 252 | 500 | 483 | 506 | 493 | 381 | 373 | 358 | 355 | 431,08 | 496 | 367 | 0,741 | -0,432 | -6,221 | 4,94E-10 | 3,29E-09 | 1,00E-08 | 0,0053471 | 0,002271 | 0,002271 | TRUE | 0,0077483 | No |
| orf19.1342 | 6401 | 4156 | 5408 | 6413 | 6082 | 5453 | 5850 | 3576 | 5742 | 5291 | 5533 | 5288 | 5202 | 5203 | 5220 | 5044 | 5315,4 | 5464 | 5167 | 0,946 | -0,08 | -2,092 | 0,0364642 | 0,06377786 | 0,000676 | 0,0022526 | 0,0012253 | 0,0012253 | TRUE | 0,0145803 | No |
| orf19.1343 | 556 | 445 | 488 | 628 | 809 | 813 | 822 | 473 | 499 | 567 | 499 | 518 | 692 | 776 | 733 | 667 | 618,84 | 521 | 717 | 1,384 | 0,468 | 6,572 | 4,96E-11 | 3,78E-10 | 0,002116 | 0,0043254 | 0,0032171 | 0,0032171 | TRUE | 0,0367843 | No |
| orf19.1344 | 2221 | 2420 | 2258 | 3213 | 1589 | 1830 | 1816 | 1063 | 1992 | 3081 | 2310 | 2649 | 1359 | 1746 | 1620 | 1499 | 2032,2 | 2508 | 1556 | 0,621 | -0,688 | -4,467 | 7,95E-06 | 2,94E-05 | 0,022276 | 0,0026939 | 0,012526 | 0,0222763 | TRUE | 0,1922579 | No |
| orf19.1345 | 88 | 54 | 84 | 88 | 47 | 28 | 38 | 15 | 79 | 69 | 86 | 73 | 40 | 27 | 34 | 21 | 53,52 | 77 | 30 | 0,407 | -1,296 | -6,115 | 9,67E-10 | 6,17E-09 | 0,002164 | 0,0291029 | 0,0206272 | 0,0206272 | TRUE | 0,318968 | No |
| orf19.1348 | 1 | 4 | 3 | 0 | 1 | 0 | 3 | 3 | 1 | 5 | 3 | 0 | 1 | 0 | 3 | 4 | 2,1 | 2 | 2 | 0,858 | -0,221 | -0,191 | 0,84880799 | NA | 0,947576 | 0,6923498 | 0,7835902 | 0,7835902 | TRUE | 0,2994121 | No |
| orf19.1349 | 119 | 79 | 131 | 135 | 95 | 60 | 88 | 62 | 107 | 101 | 134 | 111 | 81 | 57 | 79 | 87 | 94,64 | 113 | 76 | 0,668 | -0,582 | -3,563 | 0,00036674 | 0,00101915 | 0,010697 | 0,0173182 | 0,0146451 | 0,0146451 | TRUE | 0,2612768 | No |
| orf19.135 | 1103 | 873 | 986 | 1224 | 1168 | 1183 | 1288 | 664 | 990 | 1111 | 1009 | 1009 | 999 | 1129 | 1149 | 937 | 1041,6 | 1030 | 1054 | 1,027 | 0,038 | 0,513 | 0,60810597 | 0,6831527 | 0,00499 | 0,0033733 | 0,0042431 | 0,0042431 | TRUE | 0,0612301 | No |
| orf19.1350 | 7062 | 5636 | 6496 | 8210 | 7818 | 7918 | 8155 | 5370 | 6335 | 7175 | 6646 | 6770 | 6686 | 7555 | 7277 | 7574 | 7002,4 | 6732 | 7273 | 1,08 | 0,112 | 2,086 | 0,03693952 | 0,06444037 | 0,002764 | 0,0021868 | 0,0026018 | 0,0026018 | TRUE | 0,0363064 | No |
| orf19.1351 | 447 | 285 | 347 | 444 | 430 | 395 | 453 | 262 | 401 | 363 | 355 | 366 | 368 | 377 | 404 | 370 | 375,42 | 371 | 380 | 1,022 | 0,032 | 0,41 | 0,68199537 | 0,74593632 | 1,00E-08 | 0,0058464 | 0,003166 | 0,003166 | TRUE | 0,0349135 | No |
| orf19.1352 | 3741 | 2471 | 3272 | 4017 | 4902 | 4264 | 4794 | 2897 | 3356 | 3146 | 3348 | 3312 | 4192 | 4069 | 4278 | 4086 | 3723,3 | 3290 | 4156 | 1,263 | 0,336 | 8,986 | 2,57E-19 | 4,43E-18 | 0,00036 | 0,0023694 | 0,0010733 | 0,0010733 | TRUE | 0,009985 | No |
| orf19.1353 | 223 | 121 | 207 | 232 | 244 | 173 | 205 | 176 | 200 | 154 | 212 | 191 | 209 | 165 | 183 | 248 | 195,26 | 189 | 201 | 1,052 | 0,073 | 0,526 | 0,59901108 | 0,67533469 | 0,0175 | 0,0094141 | 0,0133748 | 0,0133748 | TRUE | 0,2344491 | No |

| | | | | | | | | | | | | | | | | | | | | | | | | | | | | | | | |
|---|---|---|---|---|---|---|---|---|---|---|---|---|---|---|---|---|---|---|---|---|---|---|---|---|---|---|---|---|---|---|---|
| orf19.1354 | 1E+05 | 91264 | 1E+05 | 1E+05 | 1E+05 | 1E+05 | 1E+05 | 82571 | 1E+05 | 1E+05 | 1E+05 | 1E+05 | 1E+05 | 1E+05 | 1E+05 | 1E+05 | 117537 | 1E+05 | 1E+05 | 1,196 | 0,259 | 4,314 | 1,61E-05 | 5,67E-05 | 0,004398 | 0,0019919 | 0,0034503 | 0,0034503 | TRUE | 0,0454581 | No |
| orf19.1355 | 3020 | 2301 | 2641 | 3511 | 3259 | 3300 | 3399 | 2180 | 2709 | 2929 | 2702 | 2895 | 2787 | 3149 | 3033 | 3075 | 2910 | 2809 | 3011 | 1,072 | 0,1 | 1,948 | 0,0514606 | 0,08619328 | 0,00181 | 0,0024784 | 0,0021761 | 0,0021761 | TRUE | 0,0311876 | No |
| orf19.1356 | 533 | 372 | 444 | 630 | 552 | 572 | 613 | 388 | 478 | 474 | 454 | 520 | 472 | 546 | 547 | 547 | 504,7 | 482 | 528 | 1,093 | 0,128 | 1,663 | 0,09639055 | 0,14807324 | 0,002635 | 0,0048559 | 0,0037064 | 0,0037064 | TRUE | 0,0634925 | No |
| orf19.1357 | 24163 | 16889 | 21966 | 26528 | 25327 | 23935 | 26421 | 15406 | 21677 | 21501 | 22473 | 21875 | 21660 | 22838 | 23576 | 21729 | 22166 | 21882 | 22451 | 1,026 | 0,037 | 0,996 | 0,3192964 | 0,40610381 | 0,000974 | 0,002045 | 0,0012933 | 0,0012933 | TRUE | 0,0139823 | No |
| orf19.1358 | 13615 | 9592 | 12105 | 14950 | 19307 | 16929 | 18143 | 12938 | 12214 | 12212 | 12385 | 12328 | 16512 | 16153 | 16190 | 18248 | 14530 | 12285 | 16776 | 1,365 | 0,449 | 10,207 | 1,85E-24 | 4,53E-23 | 0,001688 | 0,0020794 | 0,0017879 | 0,0017879 | TRUE | 0,0421987 | No |
| orf19.1359 | 1261 | 1065 | 1227 | 1459 | 1236 | 1324 | 1410 | 945 | 1131 | 1356 | 1255 | 1203 | 1057 | 1263 | 1258 | 1333 | 1232,1 | 1236 | 1228 | 0,993 | -0,011 | -0,132 | 0,8947378 | 0,92047683 | 0,006982 | 0,0031577 | 0,0052437 | 0,0052437 | TRUE | 0,107445 | No |
| orf19.136 | 3311 | 2373 | 3138 | 3859 | 3774 | 3525 | 3872 | 2522 | 2970 | 3021 | 3210 | 3182 | 3228 | 3363 | 3455 | 3557 | 3248,4 | 3096 | 3401 | 1,097 | 0,134 | 2,934 | 0,00334793 | 0,00751095 | 0,001164 | 0,0024264 | 0,0016873 | 0,0016873 | TRUE | 0,0144291 | No |
| orf19.1360 | 545 | 423 | 484 | 637 | 533 | 489 | 543 | 334 | 489 | 539 | 495 | 525 | 456 | 467 | 485 | 471 | 490,74 | 512 | 470 | 0,918 | -0,123 | -1,824 | 0,06819091 | 0,1102802 | 1,00E-08 | 0,0049377 | 0,0023569 | 0,0023569 | TRUE | 0,0122077 | No |
| orf19.1360.1 | 209 | 168 | 204 | 290 | 274 | 204 | 196 | 176 | 187 | 214 | 209 | 239 | 234 | 195 | 175 | 248 | 212,67 | 212 | 213 | 0,997 | -0,004 | -0,029 | 0,9771417 | 0,98378316 | 0,013409 | 0,0088057 | 0,0110056 | 0,0110056 | TRUE | 0,1632759 | No |
| orf19.1361 | 3839 | 2845 | 3457 | 4272 | 4349 | 4166 | 4281 | 2433 | 3444 | 3622 | 3537 | 3523 | 3719 | 3975 | 3820 | 3432 | 3634 | 3532 | 3736 | 1,06 | 0,083 | 1,72 | 0,08543211 | 0,13387484 | 0,001723 | 0,002379 | 0,0019848 | 0,0019848 | TRUE | 0,0348926 | No |
| orf19.1362 | 6860 | 5195 | 6223 | 7657 | 7737 | 7457 | 7946 | 4596 | 6154 | 6614 | 6367 | 6314 | 6617 | 7115 | 7090 | 6482 | 6594,2 | 6362 | 6826 | 1,074 | 0,103 | 2,355 | 0,01852238 | 0,03470845 | 0,001374 | 0,0021997 | 0,0016726 | 0,0016726 | TRUE | 0,0135632 | No |
| orf19.1363 | 29370 | 23082 | 26071 | 33827 | 34139 | 32582 | 33618 | 22204 | 26348 | 29386 | 26673 | 27894 | 29197 | 31089 | 29998 | 31317 | 28988 | 27575 | 30400 | 1,102 | 0,141 | 3,201 | 0,00136824 | 0,00334705 | 0,001731 | 0,0020296 | 0,0018217 | 0,0018217 | TRUE | 0,0237887 | No |
| orf19.1364 | 2591 | 1781 | 2333 | 2799 | 2693 | 2579 | 2780 | 1850 | 2324 | 2267 | 2387 | 2308 | 2303 | 2461 | 2481 | 2609 | 2392,6 | 2322 | 2464 | 1,059 | 0,083 | 1,785 | 0,07431935 | 0,11859512 | 0,000969 | 0,0025863 | 0,001653 | 0,001653 | TRUE | 0,0235554 | No |
| orf19.1365 | 1284 | 860 | 1198 | 1356 | 1250 | 1059 | 1103 | 704 | 1152 | 1095 | 1226 | 1118 | 1069 | 1010 | 984 | 993 | 1080,9 | 1148 | 1014 | 0,884 | -0,178 | -3,192 | 0,00141106 | 0,003442 | 0,001025 | 0,0033226 | 0,0020394 | 0,0020394 | TRUE | 0,0244525 | No |
| orf19.1366 | 93 | 58 | 82 | 96 | 94 | 64 | 81 | 50 | 83 | 74 | 84 | 79 | 80 | 61 | 72 | 71 | 75,57 | 80 | 71 | 0,887 | -0,172 | -1,08 | 0,28027499 | 0,36439994 | 1,00E-08 | 0,0211887 | 0,0112799 | 0,0112799 | TRUE | 0,089058 | No |
| orf19.1367 | 100 | 69 | 104 | 120 | 162 | 186 | 177 | 96 | 90 | 88 | 106 | 99 | 139 | 177 | 158 | 135 | 124,03 | 96 | 152 | 1,594 | 0,673 | 4,877 | 1,08E-06 | 4,53E-06 | 0,004866 | 0,0136835 | 0,0098133 | 0,0098133 | TRUE | 0,1288081 | No |
| orf19.1367.1 | 1166 | 880 | 1023 | 1329 | 1521 | 1200 | 1250 | 639 | 1046 | 1120 | 1047 | 1096 | 1301 | 1145 | 1115 | 901 | 1096,4 | 1077 | 1116 | 1,039 | 0,055 | 0,602 | 0,54708957 | 0,62847306 | 0,010621 | 0,0033036 | 0,0071002 | 0,0071002 | TRUE | 0,1898777 | No |
| orf19.1368 | 1707 | 1160 | 1483 | 1883 | 2545 | 2250 | 2518 | 1694 | 1531 | 1477 | 1517 | 1553 | 2177 | 2147 | 2247 | 2389 | 1879,7 | 1520 | 2240 | 1,47 | 0,556 | 11,792 | 4,28E-32 | 1,75E-30 | 0,000804 | 0,0027518 | 0,0015816 | 0,0015816 | TRUE | 0,0224231 | No |
| orf19.1369 | 503 | 372 | 440 | 568 | 457 | 422 | 505 | 245 | 451 | 474 | 450 | 468 | 391 | 403 | 451 | 346 | 429,13 | 461 | 398 | 0,868 | -0,205 | -2,47 | 0,0135037 | 0,02628126 | 0,003111 | 0,0053624 | 0,0042722 | 0,0042722 | TRUE | 0,0924613 | No |
| orf19.137 | 3441 | 3169 | 3358 | 4489 | 3162 | 3713 | 3615 | 2285 | 3087 | 4034 | 3436 | 3702 | 2704 | 3543 | 3226 | 3223 | 3369,3 | 3565 | 3174 | 0,891 | -0,167 | -1,836 | 0,06635209 | 0,10785805 | 0,011942 | 0,0024104 | 0,0076726 | 0,0076726 | TRUE | 0,1219225 | No |
| orf19.1370 | 11 | 11 | 8 | 10 | 5 | 14 | 5 | 4 | 10 | 14 | 8 | 8 | 4 | 13 | 4 | 6 | 8,51 | 10 | 7 | 0,698 | -0,518 | -1,007 | 0,31386325 | NA | 0,071169 | 0,1726648 | 0,1317824 | 0,1317824 | TRUE | 1,1204955 | No |
| orf19.1371 | 409 | 306 | 325 | 446 | 571 | 538 | 541 | 344 | 367 | 390 | 333 | 368 | 488 | 513 | 483 | 485 | 428,3 | 364 | 492 | 1,355 | 0,438 | 5,956 | 2,58E-09 | 1,54E-08 | 1,00E-08 | 0,005369 | 0,0028058 | 0,0028058 | TRUE | 0,0370759 | No |
| orf19.1372 | 391 | 225 | 321 | 410 | 431 | 442 | 420 | 281 | 351 | 286 | 328 | 338 | 369 | 422 | 375 | 396 | 358,15 | 326 | 390 | 1,191 | 0,252 | 2,914 | 0,0035706 | 0,00795408 | 0,002398 | 0,0060329 | 0,0043541 | 0,0043541 | TRUE | 0,0664132 | No |
| orf19.1373 | 133 | 96 | 119 | 170 | 170 | 141 | 163 | 106 | 119 | 122 | 122 | 140 | 145 | 135 | 145 | 150 | 134,79 | 126 | 144 | 1,136 | 0,184 | 1,529 | 0,12626731 | 0,18639165 | 1,00E-08 | 0,0127493 | 0,0065155 | 0,0065155 | TRUE | 0,0681704 | No |
| orf19.1374 | 429 | 316 | 343 | 480 | 381 | 377 | 337 | 232 | 385 | 402 | 351 | 396 | 326 | 360 | 301 | 327 | 355,92 | 384 | 328 | 0,856 | -0,224 | -2,648 | 0,00810568 | 0,0166976 | 0,001756 | 0,0060582 | 0,0040478 | 0,0040478 | TRUE | 0,0497163 | No |
| orf19.1375 | 22999 | 15404 | 20241 | 24661 | 27705 | 23957 | 25968 | 16330 | 20633 | 19611 | 20709 | 20336 | 23694 | 22859 | 23172 | 23032 | 21756 | 20322 | 23189 | 1,141 | 0,19 | 6,461 | 1,04E-10 | 7,55E-10 | 0,000362 | 0,0020463 | 0,0007867 | 0,0007867 | TRUE | 0,0066717 | No |
| orf19.1376 | 3209 | 2275 | 2897 | 3535 | 2669 | 2541 | 2555 | 1746 | 2879 | 2896 | 2964 | 2915 | 2283 | 2425 | 2280 | 2463 | 2638 | 2914 | 2363 | 0,81 | -0,305 | -7,483 | 7,25E-14 | 7,43E-13 | 0,000377 | 0,0025298 | 0,0012065 | 0,0012065 | TRUE | 0,0090463 | No |
| orf19.1377 | 2265 | 1687 | 2029 | 2615 | 2005 | 1824 | 1801 | 1227 | 2032 | 2148 | 2076 | 2156 | 1715 | 1740 | 1607 | 1731 | 1900,6 | 2103 | 1698 | 0,807 | -0,309 | -6,843 | 7,73E-12 | 6,39E-11 | 0,000583 | 0,0027433 | 0,0014249 | 0,0014249 | TRUE | 0,0163298 | No |
| orf19.1378 | 10516 | 6761 | 9167 | 11198 | 10508 | 8709 | 9980 | 6277 | 9434 | 8607 | 9379 | 9234 | 8987 | 8310 | 8905 | 8853 | 8963,7 | 9164 | 8764 | 0,956 | -0,065 | -1,521 | 0,12826043 | 0,18891607 | 0,001328 | 0,0021415 | 0,0016323 | 0,0016323 | TRUE | 0,0200211 | No |
| orf19.138 | 101 | 85 | 113 | 136 | 100 | 83 | 90 | 74 | 91 | 108 | 116 | 112 | 86 | 79 | 80 | 104 | 97 | 107 | 87 | 0,812 | -0,301 | -2,008 | 0,0446554 | 0,07621865 | 0,002722 | 0,0169459 | 0,0111683 | 0,0111683 | TRUE | 0,126367 | No |
| orf19.1381 | 2213 | 1361 | 1756 | 2260 | 2848 | 2370 | 2455 | 1564 | 1985 | 1733 | 1797 | 1864 | 2436 | 2261 | 2191 | 2206 | 2059 | 1845 | 2274 | 1,232 | 0,301 | 5,377 | 7,59E-08 | 3,79E-07 | 0,002281 | 0,0026846 | 0,0025142 | 0,0025142 | TRUE | 0,0318822 | No |
| orf19.1382 | 585 | 390 | 502 | 629 | 574 | 493 | 494 | 316 | 525 | 497 | 514 | 519 | 491 | 470 | 441 | 446 | 487,68 | 514 | 462 | 0,9 | -0,151 | -2,221 | 0,02636396 | 0,04795479 | 1,00E-08 | 0,0049563 | 0,0024089 | 0,0024089 | TRUE | 0,022029 | No |
| orf19.1383 | 756 | 518 | 666 | 779 | 748 | 722 | 752 | 474 | 678 | 659 | 681 | 642 | 640 | 689 | 671 | 669 | 666,2 | 665 | 667 | 1,002 | 0,003 | 0,056 | 0,95503765 | 0,96792615 | 1,00E-08 | 0,0041586 | 0,0018438 | 0,0018438 | TRUE | 0,0099319 | No |
| orf19.1386 | 2805 | 2005 | 2484 | 3016 | 2390 | 2177 | 2332 | 1439 | 2516 | 2553 | 2541 | 2487 | 2044 | 2077 | 2081 | 2030 | 2291,1 | 2524 | 2058 | 0,816 | -0,293 | -8,131 | 4,26E-16 | 5,43E-15 | 1,00E-08 | 0,0026131 | 0,0008071 | 0,0008071 | TRUE | 0,0012412 | No |
| orf19.1387 | 4788 | 3261 | 4183 | 5088 | 4628 | 3768 | 4093 | 2744 | 4295 | 4152 | 4280 | 4196 | 3958 | 3595 | 3652 | 3870 | 3999,8 | 4231 | 3769 | 0,89 | -0,167 | -4,017 | 5,90E-05 | 0,00018863 | 0,000849 | 0,0023425 | 0,0014155 | 0,0014155 | TRUE | 0,0147329 | No |
| orf19.1388 | 6111 | 4203 | 5129 | 6484 | 5760 | 5207 | 5575 | 3128 | 5482 | 5351 | 5247 | 5347 | 4926 | 4968 | 4975 | 4412 | 5088,5 | 5357 | 4820 | 0,901 | -0,151 | -3,323 | 0,00089153 | 0,00226386 | 0,001505 | 0,0022648 | 0,0017799 | 0,0017799 | TRUE | 0,0379821 | No |
| orf19.1389 | 3311 | 2273 | 2670 | 3313 | 3758 | 3363 | 3496 | 2040 | 2970 | 2894 | 2732 | 2732 | 3214 | 3209 | 3120 | 2877 | 2968,4 | 2832 | 3105 | 1,098 | 0,135 | 2,697 | 0,00700332 | 0,01461477 | 0,001736 | 0,0024686 | 0,0020616 | 0,0020616 | TRUE | 0,0279966 | No |
| orf19.139 | 845 | 679 | 804 | 921 | 1031 | 970 | 1069 | 682 | 758 | 864 | 823 | 759 | 882 | 926 | 954 | 962 | 865,95 | 801 | 931 | 1,164 | 0,219 | 3,521 | 0,0004304 | 0,00117698 | 0,001563 | 0,0036559 | 0,0025408 | 0,0025408 | TRUE | 0,0312937 | No |
| orf19.1390 | 600 | 372 | 517 | 647 | 586 | 507 | 539 | 382 | 538 | 474 | 529 | 534 | 501 | 484 | 481 | 539 | 509,87 | 519 | 501 | 0,96 | -0,059 | -0,826 | 0,40858821 | 0,49686764 | 0,000494 | 0,0048267 | 0,0028547 | 0,0028547 | TRUE | 0,0352741 | No |
| orf19.1391 | 131 | 89 | 114 | 150 | 115 | 100 | 86 | 70 | 118 | 113 | 117 | 124 | 98 | 95 | 77 | 99 | 105,05 | 118 | 92 | 0,779 | -0,361 | -2,612 | 0,00901275 | 0,01837415 | 1,00E-08 | 0,0157988 | 0,0087167 | 0,0087167 | TRUE | 0,1357692 | No |
| orf19.1392 | 542 | 388 | 435 | 559 | 430 | 390 | 441 | 259 | 486 | 494 | 445 | 461 | 368 | 372 | 394 | 365 | 423,11 | 472 | 375 | 0,797 | -0,328 | -4,515 | 6,33E-06 | 2,38E-05 | 1,00E-08 | 0,0054105 | 0,0026772 | 0,0026772 | TRUE | 0,0156034 | No |
| orf19.1393 | 2665 | 1764 | 2398 | 2816 | 3387 | 3010 | 3417 | 2147 | 2391 | 2246 | 2453 | 2322 | 2897 | 2872 | 3049 | 3028 | 2657,3 | 2353 | 2962 | 1,257 | 0,33 | 7,646 | 2,08E-14 | 2,23E-13 | 0,000675 | 0,0025258 | 0,0014091 | 0,0014091 | TRUE | 0,0105412 | No |
| orf19.1394 | 1324 | 933 | 1155 | 1365 | 1485 | 1325 | 1348 | 656 | 1188 | 1188 | 1182 | 1126 | 1270 | 1264 | 1203 | 925 | 1168,2 | 1171 | 1166 | 0,999 | -0,002 | -0,019 | 0,98458523 | 0,98820382 | 0,009686 | 0,0032223 | 0,0065534 | 0,0065534 | TRUE | 0,2182349 | No |
| orf19.1395 | 314 | 208 | 223 | 336 | 356 | 269 | 296 | 193 | 282 | 265 | 228 | 277 | 304 | 257 | 264 | 272 | 268,65 | 263 | 274 | 1,043 | 0,06 | 0,608 | 0,54321865 | 0,62534157 | 0,003981 | 0,0073832 | 0,0057779 | 0,0057779 | TRUE | 0,0890449 | No |
| orf19.1396 | 2017 | 1338 | 1713 | 2219 | 2146 | 1796 | 1962 | 1300 | 1809 | 1703 | 1753 | 1830 | 1835 | 1714 | 1751 | 1834 | 1778,6 | 1774 | 1784 | 1,003 | 0,005 | 0,107 | 0,91515145 | 0,93746461 | 0,000526 | 0,0027957 | 0,0014362 | 0,0014362 | TRUE | 0,0083132 | No |
| orf19.1397 | 777 | 603 | 736 | 955 | 1110 | 925 | 1019 | 688 | 697 | 768 | 753 | 788 | 949 | 883 | 909 | 970 | 839,6 | 752 | 928 | 1,234 | 0,303 | 5,003 | 5,64E-07 | 2,48E-06 | 0,001102 | 0,0037086 | 0,0023168 | 0,0023168 | TRUE | 0,0296122 | No |
| orf19.1400 | 1177 | 989 | 1109 | 1353 | 1463 | 1455 | 1542 | 825 | 1056 | 1259 | 1135 | 1116 | 1251 | 1388 | 1376 | 1164 | 1218,1 | 1142 | 1295 | 1,138 | 0,187 | 2,565 | 0,01032327 | 0,0207155 | 0,005125 | 0,0031713 | 0,0042661 | 0,0042661 | TRUE | 0,0552888 | No |
| orf19.1401 | 1054 | 650 | 861 | 971 | 1009 | 880 | 998 | 713 | 946 | 828 | 881 | 801 | 863 | 840 | 891 | 1006 | 881,68 | 864 | 900 | 1,038 | 0,054 | 0,728 | 0,46634385 | 0,55229525 | 0,004568 | 0,003626 | 0,0041809 | 0,0041809 | TRUE | 0,0687663 | No |
| orf19.1402 | 5808 | 3676 | 4993 | 6035 | 5678 | 4683 | 5075 | 3636 | 5210 | 4680 | 5108 | 4977 | 4856 | 4468 | 4529 | 5128 | 4869,6 | 4994 | 4745 | 0,949 | -0,075 | -1,378 | 0,16818576 | 0,23786724 | 0,002879 | 0,0022776 | 0,0026469 | 0,0026469 | TRUE | 0,0349065 | No |
| orf19.1403 | 4407 | 2966 | 3976 | 4820 | 5164 | 4503 | 4983 | 3275 | 3954 | 3776 | 4068 | 3975 | 4416 | 4297 | 4446 | 4619 | 4193,8 | 3943 | 4444 | 1,126 | 0,171 | 4,42 | 9,85E-06 | 3,59E-05 | 0,000607 | 0,0023257 | 0,0011966 | 0,0011966 | TRUE | 0,0093703 | No |
| orf19.1404 | 5757 | 3867 | 5069 | 6192 | 7218 | 5914 | 6527 | 4326 | 5165 | 4923 | 5186 | 5106 | 6173 | 5643 | 5824 | 6101 | 5515,2 | 5095 | 5935 | 1,164 | 0,219 | 5,433 | 5,55E-08 | 2,81E-07 | 0,000892 | 0,0022427 | 0,0013835 | 0,0013835 | TRUE | 0,0134699 | No |
| orf19.1405 | 665 | 504 | 653 | 735 | 722 | 655 | 633 | 435 | 597 | 642 | 668 | 606 | 617 | 625 | 565 | 614 | 616,65 | 628 | 605 | 0,965 | -0,052 | -0,784 | 0,43305323 | 0,52004474 | 0,00099 | 0,0043337 | 0,0026131 | 0,0026131 | TRUE | 0,0252424 | No |
| orf19.1406 | 930 | 750 | 787 | 1054 | 787 | 819 | 804 | 465 | 834 | 955 | 805 | 869 | 673 | 781 | 717 | 656 | 786,41 | 866 | 707 | 0,819 | -0,288 | -3,852 | 0,00011695 | 0,00035325 | 0,004231 | 0,0038255 | 0,0040702 | 0,0040702 | TRUE | 0,0591712 | No |
| orf19.1409 | 642 | 406 | 615 | 724 | 929 | 791 | 901 | 580 | 576 | 517 | 629 | 597 | 795 | 755 | 804 | 818 | 686,29 | 580 | 793 | 1,363 | 0,446 | 6,566 | 5,17E-11 | 3,93E-10 | 0,001671 | 0,0040948 | 0,0029438 | 0,0029438 | TRUE | 0,0552299 | No |
| orf19.1409.1 | 9157 | 6481 | 7399 | 9650 | 9708 | 8118 | 8666 | 5476 | 8215 | 8251 | 7570 | 7958 | 8303 | 7746 | 7733 | 7723 | 7937,3 | 7998 | 7876 | 0,985 | -0,022 | -0,515 | 0,60670101 | 0,68182934 | 0,001161 | 0,0021624 | 0,00158 | 0,00158 | TRUE | 0,0164275 | No |
| orf19.1409.2 | 461 | 370 | 447 | 524 | 327 | 302 | 301 | 209 | 414 | 471 | 457 | 432 | 280 | 288 | 269 | 295 | 363,15 | 444 | 283 | 0,638 | -0,649 | -8,124 | 4,50E-16 | 5,72E-15 | 1,00E-08 | 0,005977 | 0,0032273 | 0,0032273 | TRUE | 0,0237814 | No |
| orf19.1409.3 | 43 | 48 | 30 | 50 | 19 | 14 | 18 | 9 | 39 | 61 | 31 | 41 | 16 | 13 | 16 | 13 | 28,75 | 43 | 14 | 0,347 | -1,526 | -5,045 | 4,54E-07 | 2,02E-06 | 1,00E-08 | 0,0524797 | 0,0422115 | 0,0422115 | TRUE | 0,3966769 | No |
| orf19.1411 | 1244 | 848 | 1100 | 1319 | 1185 | 1100 | 1151 | 770 | 1116 | 1080 | 1125 | 1088 | 1013 | 1050 | 1027 | 1086 | 1073,1 | 1102 | 1044 | 0,945 | -0,082 | -1,671 | 0,09467699 | 0,14627012 | 1,00E-08 | 0,0033323 | 0,0013614 | 0,0013614 | TRUE | 0,0075594 | No |
| orf19.1412 | 2293 | 1579 | 1868 | 2400 | 2297 | 2042 | 2225 | 1329 | 2057 | 2010 | 1911 | 1979 | 1964 | 1948 | 1985 | 1874 | 1966,3 | 1989 | 1943 | 0,978 | -0,032 | -0,758 | 0,44825537 | 0,53477329 | 0,000229 | 0,0027178 | 0,0012153 | 0,0012153 | TRUE | 0,0083991 | No |

| | | | | | | | | | | | | | | | | | | | | | | | | | | | | | | | |
|---|---|---|---|---|---|---|---|---|---|---|---|---|---|---|---|---|---|---|---|---|---|---|---|---|---|---|---|---|---|---|---|
| orf19.1413 | 1056 | 650 | 857 | 1080 | 1225 | 1027 | 1099 | 702 | 947 | 828 | 877 | 891 | 1048 | 980 | 981 | 990 | 942,58 | 886 | 1000 | 1,127 | 0,173 | 2,992 | 0,00276774 | 0,0063347 | 0,000938 | 0,0035197 | 0,0021269 | 0,0021269 | TRUE | 0,026126 | No |
| orf19.1414 | 3439 | 2004 | 2813 | 3480 | 2293 | 1824 | 2049 | 1220 | 3085 | 2551 | 2878 | 2870 | 1961 | 1740 | 1828 | 1721 | 2329,3 | 2846 | 1812 | 0,637 | -0,651 | -9,957 | 2,34E-23 | 5,31E-22 | 0,004217 | 0,0026028 | 0,0036446 | 0,0036446 | TRUE | 0,0574737 | No |
| orf19.1415 | 19672 | 13803 | 17940 | 22171 | 16264 | 14770 | 15800 | 10398 | 17648 | 17573 | 18354 | 18282 | 13909 | 14093 | 14099 | 14665 | 16078 | 17964 | 14192 | 0,79 | -0,341 | -10,889 | 1,30E-27 | 4,20E-26 | 0,000445 | 0,0020698 | 0,0008767 | 0,0008767 | TRUE | 0,0060107 | No |
| orf19.1416 | 2053 | 1451 | 1727 | 2193 | 2652 | 2190 | 2213 | 1487 | 1842 | 1847 | 1767 | 1808 | 2268 | 2090 | 1975 | 2097 | 1961,8 | 1816 | 2108 | 1,161 | 0,215 | 4,243 | 2,21E-05 | 7,60E-05 | 0,001491 | 0,0027195 | 0,0019583 | 0,0019583 | TRUE | 0,0334671 | No |
| orf19.1417 | 754 | 511 | 666 | 767 | 1139 | 988 | 1012 | 600 | 676 | 651 | 681 | 632 | 974 | 943 | 903 | 846 | 788,37 | 660 | 916 | 1,393 | 0,478 | 7,666 | 1,78E-14 | 1,93E-13 | 0,001192 | 0,0038209 | 0,0024278 | 0,0024278 | TRUE | 0,0279449 | No |
| orf19.1418 | 509 | 416 | 489 | 548 | 684 | 644 | 685 | 425 | 457 | 530 | 500 | 452 | 585 | 614 | 611 | 599 | 543,57 | 485 | 602 | 1,249 | 0,321 | 4,567 | 4,95E-06 | 1,89E-05 | 0,000909 | 0,0046502 | 0,0028688 | 0,0028688 | TRUE | 0,0415696 | No |
| orf19.1420 | 116 | 79 | 91 | 113 | 124 | 109 | 121 | 78 | 104 | 101 | 93 | 93 | 106 | 104 | 108 | 110 | 102,37 | 98 | 107 | 1,094 | 0,13 | 0,975 | 0,32950159 | 0,41685525 | 1,00E-08 | 0,0161604 | 0,0073806 | 0,0073806 | TRUE | 0,0215983 | No |
| orf19.1421 | 203 | 122 | 144 | 184 | 162 | 120 | 143 | 109 | 182 | 155 | 147 | 152 | 139 | 115 | 128 | 154 | 146,36 | 159 | 134 | 0,833 | -0,264 | -2,056 | 0,0397736 | 0,06872584 | 0,004998 | 0,0118983 | 0,0089325 | 0,0089325 | TRUE | 0,1063274 | No |
| orf19.1422 | 7337 | 5262 | 6635 | 7837 | 8541 | 7897 | 8494 | 5312 | 6582 | 6699 | 6788 | 6462 | 7305 | 7535 | 7579 | 7492 | 7055,4 | 6633 | 7478 | 1,128 | 0,173 | 5,6 | 2,14E-08 | 1,14E-07 | 0,00024 | 0,0021853 | 0,0007768 | 0,0007768 | TRUE | 0,0037397 | No |
| orf19.1424 | 813 | 541 | 694 | 911 | 901 | 819 | 868 | 529 | 729 | 689 | 710 | 751 | 771 | 781 | 775 | 746 | 744 | 720 | 768 | 1,066 | 0,092 | 1,638 | 0,10135864 | 0,15444147 | 1,00E-08 | 0,0039307 | 0,0017112 | 0,0017112 | TRUE | 0,0104815 | No |
| orf19.1426 | 5866 | 3888 | 5284 | 6485 | 6851 | 6139 | 6695 | 4436 | 5262 | 4950 | 5406 | 5348 | 5859 | 5858 | 5974 | 6257 | 5614,2 | 5242 | 5987 | 1,141 | 0,19 | 4,665 | 3,09E-06 | 1,22E-05 | 0,001005 | 0,0022381 | 0,001417 | 0,001417 | TRUE | 0,0165175 | No |
| orf19.1427 | 285 | 190 | 260 | 290 | 296 | 269 | 295 | 210 | 256 | 242 | 266 | 239 | 253 | 257 | 263 | 296 | 258,99 | 251 | 267 | 1,06 | 0,084 | 0,921 | 0,35725328 | 0,44493733 | 1,00E-08 | 0,0075847 | 0,0041663 | 0,0041663 | TRUE | 0,0486713 | No |
| orf19.1428 | 173 | 108 | 154 | 168 | 168 | 156 | 168 | 87 | 155 | 137 | 158 | 139 | 144 | 149 | 150 | 123 | 144,24 | 147 | 142 | 0,966 | -0,05 | -0,422 | 0,67320796 | 0,73915084 | 1,00E-08 | 0,0120439 | 0,0067628 | 0,0067628 | TRUE | 0,0627458 | No |
| orf19.1429 | 62 | 65 | 69 | 117 | 85 | 76 | 85 | 46 | 56 | 83 | 71 | 96 | 73 | 73 | 76 | 65 | 73,92 | 76 | 72 | 0,94 | -0,089 | -0,476 | 0,63386555 | 0,70510354 | 0,018319 | 0,0216175 | 0,0201211 | 0,0201211 | TRUE | 0,2462661 | No |
| orf19.1430 | 77 | 63 | 83 | 97 | 76 | 69 | 93 | 53 | 69 | 80 | 85 | 80 | 65 | 66 | 83 | 75 | 75,35 | 78 | 72 | 0,919 | -0,122 | -0,75 | 0,4530118 | 0,53873612 | 1,00E-08 | 0,0212469 | 0,0123056 | 0,0123056 | TRUE | 0,1095153 | No |
| orf19.1433 | 1911 | 1332 | 1711 | 2130 | 2255 | 2007 | 2046 | 1300 | 1714 | 1696 | 1751 | 1756 | 1929 | 1915 | 1826 | 1834 | 1802,5 | 1729 | 1876 | 1,085 | 0,118 | 2,815 | 0,00487251 | 0,01053403 | 1,00E-08 | 0,0027849 | 0,0011215 | 0,0011215 | TRUE | 0,0044721 | No |
| orf19.1433.1 | 12 | 6 | 14 | 7 | 4 | 4 | 5 | 9 | 11 | 8 | 14 | 6 | 3 | 4 | 4 | 13 | 7,86 | 10 | 6 | 0,595 | -0,749 | -1,337 | 0,18125783 | NA | 0,124956 | 0,1866375 | 0,1618015 | 0,1618015 | TRUE | 0,7341943 | No |
| orf19.1434 | 1283 | 895 | 1155 | 1299 | 1169 | 1044 | 1184 | 733 | 1151 | 1139 | 1182 | 1071 | 1000 | 996 | 1057 | 1034 | 1078,7 | 1136 | 1022 | 0,9 | -0,152 | -2,888 | 0,00387356 | 0,00857809 | 1,72E-06 | 0,0033253 | 0,0017366 | 0,0017366 | TRUE | 0,0189491 | No |
| orf19.1435 | 11108 | 6220 | 9088 | 11181 | 11795 | 8193 | 9107 | 5622 | 9965 | 7919 | 9298 | 9220 | 10087 | 7818 | 8126 | 7929 | 8795,3 | 9100 | 8490 | 0,933 | -0,1 | -1,136 | 0,25583499 | 0,33840334 | 0,01169 | 0,0021446 | 0,0073449 | 0,0073449 | TRUE | 0,1965736 | No |
| orf19.1438 | 383 | 261 | 326 | 417 | 378 | 356 | 390 | 239 | 344 | 332 | 334 | 344 | 323 | 340 | 348 | 337 | 337,67 | 338 | 337 | 0,995 | -0,008 | -0,102 | 0,91905321 | 0,94033918 | 1,00E-08 | 0,0062787 | 0,0026843 | 0,0026843 | TRUE | 0,0098311 | No |
| orf19.1439 | 1846 | 1572 | 1694 | 2153 | 1628 | 1784 | 1863 | 1137 | 1656 | 2001 | 1733 | 1775 | 1392 | 1702 | 1662 | 1604 | 1690,8 | 1791 | 1590 | 0,888 | -0,171 | -2,247 | 0,02465898 | 0,04504444 | 0,006402 | 0,0028381 | 0,0049699 | 0,0049699 | TRUE | 0,0871189 | No |
| orf19.144 | 1971 | 1823 | 2011 | 2425 | 2494 | 2821 | 2959 | 1674 | 1768 | 2321 | 2057 | 2000 | 2133 | 2692 | 2640 | 2361 | 2246,5 | 2036 | 2456 | 1,207 | 0,272 | 3,016 | 0,0025584 | 0,00589826 | 0,011221 | 0,0026257 | 0,0073525 | 0,0073525 | TRUE | 0,1051382 | No |
| orf19.1440 | 1027 | 730 | 883 | 1149 | 960 | 1021 | 1009 | 632 | 921 | 929 | 903 | 947 | 821 | 974 | 900 | 891 | 911,07 | 925 | 896 | 0,968 | -0,046 | -0,753 | 0,45156757 | 0,53723127 | 0,001726 | 0,0035729 | 0,002549 | 0,002549 | TRUE | 0,04213 | No |
| orf19.1440.1 | 16 | 19 | 9 | 26 | 9 | 23 | 9 | 9 | 14 | 24 | 9 | 21 | 8 | 22 | 8 | 13 | 14,94 | 17 | 13 | 0,727 | -0,461 | -1,041 | 0,2980958 | NA | 0,144214 | 0,0991172 | 0,1190551 | 0,1190551 | TRUE | 0,5807646 | No |
| orf19.1440.2 | 374 | 288 | 333 | 367 | 240 | 212 | 213 | 101 | 336 | 367 | 341 | 303 | 205 | 202 | 190 | 142 | 260,69 | 337 | 185 | 0,557 | -0,843 | -7,494 | 6,71E-14 | 6,95E-13 | 0,008603 | 0,0075481 | 0,0079769 | 0,0079769 | TRUE | 0,2088221 | No |
| orf19.1441 | 5233 | 3829 | 4617 | 5895 | 4737 | 4366 | 4800 | 2814 | 4695 | 4875 | 4724 | 4861 | 4051 | 4166 | 4283 | 3969 | 4452,9 | 4789 | 4117 | 0,861 | -0,217 | -5,876 | 4,19E-09 | 2,46E-08 | 0,000439 | 0,0023055 | 0,001077 | 0,001077 | TRUE | 0,0089724 | No |
| orf19.1442 | 486 | 353 | 458 | 515 | 441 | 374 | 372 | 265 | 436 | 449 | 469 | 425 | 377 | 357 | 332 | 374 | 402,3 | 445 | 360 | 0,809 | -0,305 | -3,998 | 6,39E-05 | 0,00020239 | 1,00E-08 | 0,0055881 | 0,003087 | 0,003087 | TRUE | 0,0332852 | No |
| orf19.1444 | 2192 | 1590 | 2106 | 2539 | 2408 | 2138 | 2475 | 1734 | 1966 | 2024 | 2155 | 2094 | 2059 | 2040 | 2209 | 2446 | 2124,1 | 2060 | 2188 | 1,061 | 0,085 | 1,353 | 0,17610957 | 0,24750089 | 0,003825 | 0,002663 | 0,0033003 | 0,0033003 | TRUE | 0,0723615 | No |
| orf19.1445 | 68 | 48 | 62 | 86 | 85 | 74 | 78 | 44 | 61 | 61 | 63 | 71 | 73 | 71 | 70 | 62 | 66,43 | 64 | 69 | 1,075 | 0,104 | 0,626 | 0,53116226 | 0,61381475 | 1,00E-08 | 0,0238333 | 0,0115093 | 0,0115093 | TRUE | 0,0518144 | No |
| orf19.1446 | 3355 | 2422 | 3165 | 3998 | 3728 | 3390 | 3747 | 2246 | 3010 | 3083 | 3238 | 3297 | 3188 | 3235 | 3344 | 3168 | 3195,3 | 3157 | 3234 | 1,024 | 0,035 | 0,809 | 0,41868356 | 0,50647075 | 0,000767 | 0,0024338 | 0,0014433 | 0,0014433 | TRUE | 0,0125769 | No |
| orf19.1447 | 293 | 213 | 276 | 315 | 288 | 237 | 293 | 193 | 263 | 271 | 282 | 260 | 246 | 226 | 261 | 272 | 260,28 | 269 | 251 | 0,933 | -0,101 | -1,098 | 0,27210913 | 0,35656089 | 1,00E-08 | 0,0075569 | 0,0042133 | 0,0042133 | TRUE | 0,0503044 | No |
| orf19.1448 | 4689 | 3229 | 3719 | 4879 | 3386 | 2972 | 3084 | 1870 | 4207 | 4111 | 3805 | 4023 | 2896 | 2836 | 2752 | 2637 | 3408,3 | 4036 | 2780 | 0,69 | -0,536 | -11,543 | 7,98E-31 | 2,96E-29 | 0,001309 | 0,0024055 | 0,0017668 | 0,0017668 | TRUE | 0,0180803 | No |
| orf19.1448.1 | 619 | 394 | 463 | 599 | 542 | 536 | 495 | 380 | 555 | 502 | 474 | 494 | 464 | 511 | 442 | 536 | 497,15 | 506 | 488 | 0,96 | -0,059 | -0,715 | 0,47437044 | 0,56014496 | 0,004165 | 0,0048996 | 0,004529 | 0,004529 | TRUE | 0,0519862 | No |
| orf19.1449 | 186 | 133 | 172 | 216 | 221 | 238 | 260 | 138 | 167 | 169 | 176 | 178 | 189 | 227 | 232 | 195 | 191,63 | 172 | 211 | 1,223 | 0,291 | 2,678 | 0,00741708 | 0,01538205 | 1,00E-08 | 0,0095552 | 0,0060805 | 0,0060805 | TRUE | 0,063056 | No |
| orf19.145 | 1272 | 1023 | 1131 | 1428 | 1372 | 1362 | 1420 | 861 | 1141 | 1302 | 1157 | 1178 | 1173 | 1300 | 1267 | 1214 | 1216,6 | 1194 | 1238 | 1,039 | 0,055 | 0,944 | 0,34513234 | 0,4324287 | 0,001811 | 0,0031728 | 0,0024741 | 0,0024741 | TRUE | 0,0412993 | No |
| orf19.1450 | 6 | 3 | 12 | 8 | 13 | 12 | 19 | 9 | 5 | 4 | 12 | 7 | 11 | 11 | 17 | 13 | 10,04 | 7 | 13 | 1,848 | 0,886 | 1,925 | 0,05421921 | NA | 0,004068 | 0,1466215 | 0,0952871 | 0,0952871 | TRUE | 0,6291425 | No |
| orf19.1451 | 659 | 549 | 629 | 748 | 822 | 760 | 750 | 537 | 591 | 699 | 644 | 617 | 703 | 725 | 669 | 757 | 675,66 | 638 | 714 | 1,12 | 0,164 | 2,362 | 0,01819231 | 0,03416675 | 0,001989 | 0,0041281 | 0,0031265 | 0,0031265 | TRUE | 0,0454555 | No |
| orf19.1453 | 12598 | 8212 | 11188 | 13219 | 14603 | 11721 | 13429 | 7928 | 11302 | 10455 | 11446 | 10901 | 12489 | 11184 | 11983 | 11182 | 11368 | 11026 | 11710 | 1,062 | 0,087 | 1,789 | 0,07359239 | 0,11762253 | 0,002202 | 0,0021072 | 0,002181 | 0,002181 | TRUE | 0,0246313 | No |
| orf19.1456 | 12 | 8 | 12 | 15 | 20 | 6 | 2 | 4 | 11 | 10 | 12 | 12 | 17 | 6 | 2 | 6 | 9,48 | 11 | 8 | 0,674 | -0,568 | -1,023 | 0,30645791 | NA | 0,233665 | 0,1550875 | 0,187832 | 0,187832 | TRUE | 2,2652513 | No |
| orf19.1457 | 2094 | 1320 | 1844 | 2295 | 1954 | 1750 | 1922 | 1222 | 1879 | 1681 | 1887 | 1892 | 1671 | 1670 | 1715 | 1724 | 1764,7 | 1835 | 1695 | 0,922 | -0,117 | -2,373 | 0,01762999 | 0,03319367 | 0,000955 | 0,0028022 | 0,0017796 | 0,0017796 | TRUE | 0,0360523 | No |
| orf19.1458 | 506 | 386 | 453 | 595 | 446 | 437 | 417 | 277 | 454 | 491 | 463 | 491 | 381 | 417 | 372 | 391 | 432,58 | 475 | 390 | 0,822 | -0,283 | -3,88 | 0,00010434 | 0,00031838 | 1,00E-08 | 0,0053354 | 0,0027744 | 0,0027744 | TRUE | 0,0260976 | No |
| orf19.1459 | 658 | 401 | 589 | 690 | 554 | 513 | 595 | 387 | 590 | 511 | 603 | 569 | 474 | 489 | 531 | 546 | 539,06 | 568 | 510 | 0,893 | -0,163 | -2,148 | 0,03167739 | 0,05627373 | 0,002623 | 0,0046726 | 0,0036714 | 0,0036714 | TRUE | 0,0489047 | No |
| orf19.1460 | 3232 | 2263 | 2989 | 3662 | 2833 | 2316 | 2719 | 1620 | 2899 | 2881 | 3058 | 3020 | 2423 | 2210 | 2426 | 2285 | 2650,3 | 2964 | 2336 | 0,788 | -0,343 | -7,489 | 6,92E-14 | 7,12E-13 | 0,000971 | 0,0025273 | 0,0016304 | 0,0016304 | TRUE | 0,0168591 | No |
| orf19.1461 | 1463 | 983 | 1226 | 1535 | 1663 | 1352 | 1600 | 1061 | 1312 | 1251 | 1254 | 1266 | 1422 | 1290 | 1428 | 1496 | 1340,1 | 1271 | 1409 | 1,106 | 0,146 | 2,679 | 0,00737535 | 0,0153008 | 0,001373 | 0,0030628 | 0,0020973 | 0,0020973 | TRUE | 0,0387362 | No |
| orf19.1462 | 2936 | 2013 | 2622 | 3132 | 2175 | 1851 | 2029 | 1372 | 2634 | 2563 | 2683 | 2583 | 1860 | 1766 | 1811 | 1935 | 2229,2 | 2616 | 1843 | 0,703 | -0,508 | -12,003 | 3,42E-33 | 1,50E-31 | 0,000367 | 0,0026307 | 0,0012519 | 0,0012519 | TRUE | 0,0121042 | No |
| orf19.1464 | 4424 | 3115 | 4119 | 4911 | 5196 | 4681 | 5211 | 2968 | 3969 | 3966 | 4214 | 4050 | 4444 | 4466 | 4650 | 4186 | 4243,1 | 4050 | 4436 | 1,096 | 0,133 | 3,128 | 0,00176273 | 0,00421094 | 0,001023 | 0,0023217 | 0,0014956 | 0,0014956 | TRUE | 0,0166916 | No |
| orf19.1465 | 102 | 70 | 82 | 76 | 85 | 76 | 73 | 48 | 92 | 89 | 84 | 63 | 73 | 73 | 65 | 68 | 75,65 | 82 | 70 | 0,856 | -0,224 | -1,314 | 0,1889581 | 0,26182202 | 1,00E-08 | 0,021168 | 0,0145831 | 0,0145831 | TRUE | 0,2664578 | No |
| orf19.1466 | 1875 | 1340 | 1618 | 1957 | 1958 | 1709 | 1790 | 951 | 1682 | 1706 | 1655 | 1614 | 1675 | 1631 | 1597 | 1341 | 1612,6 | 1664 | 1561 | 0,941 | -0,088 | -1,332 | 0,18301672 | 0,25521338 | 0,00406 | 0,0028797 | 0,0035809 | 0,0035809 | TRUE | 0,1011289 | No |
| orf19.1467 | 7605 | 5029 | 6126 | 7806 | 8783 | 7302 | 7533 | 5809 | 6822 | 6402 | 6268 | 6437 | 7511 | 6967 | 6722 | 8193 | 6915,4 | 6482 | 7348 | 1,133 | 0,18 | 2,923 | 0,00346842 | 0,00774657 | 0,004425 | 0,0021894 | 0,0034995 | 0,0034995 | TRUE | 0,0720435 | No |
| orf19.1468 | 4078 | 2822 | 3493 | 4212 | 4421 | 3907 | 4124 | 2494 | 3658 | 3593 | 3574 | 3473 | 3781 | 3728 | 3680 | 3518 | 3625,6 | 3574 | 3677 | 1,03 | 0,042 | 1,132 | 0,25782818 | 0,34082105 | 0,000341 | 0,0023799 | 0,0010767 | 0,0010767 | TRUE | 0,0095391 | No |
| orf19.147 | 9112 | 7719 | 8712 | 11163 | 13321 | 13905 | 15000 | 9272 | 8174 | 9827 | 8913 | 9205 | 11393 | 13268 | 13385 | 13077 | 10905 | 9030 | 12781 | 1,416 | 0,501 | 7,521 | 5,42E-14 | 5,65E-13 | 0,00543 | 0,0021126 | 0,0041732 | 0,0041732 | TRUE | 0,06568 | No |
| orf19.1470 | 10858 | 6818 | 8864 | 11237 | 10651 | 8522 | 9081 | 8039 | 9741 | 8680 | 9069 | 9266 | 9109 | 8131 | 8103 | 11338 | 9179,7 | 9189 | 9170 | 0,998 | -0,004 | -0,038 | 0,96963846 | 0,97826833 | 0,012615 | 0,0021377 | 0,0082301 | 0,0082301 | TRUE | 0,3078931 | No |
| orf19.1471 | 5693 | 3729 | 4642 | 5675 | 5897 | 5129 | 5138 | 3238 | 5107 | 4747 | 4749 | 4680 | 5043 | 4894 | 4585 | 4567 | 4796,6 | 4821 | 4772 | 0,99 | -0,014 | -0,295 | 0,76832979 | 0,81918817 | 0,001544 | 0,0022822 | 0,0019733 | 0,0019733 | TRUE | 0,0197464 | No |
| orf19.1473 | 363 | 245 | 311 | 391 | 356 | 357 | 380 | 202 | 326 | 312 | 318 | 322 | 304 | 341 | 339 | 285 | 318,41 | 320 | 317 | 0,996 | -0,005 | -0,062 | 0,95055661 | 0,96468651 | 1,00E-08 | 0,0065388 | 0,0037772 | 0,0037772 | TRUE | 0,043118 | No |
| orf19.1474 | 4286 | 2799 | 3909 | 4773 | 5234 | 4465 | 4970 | 3282 | 3845 | 3563 | 3999 | 3936 | 4476 | 4260 | 4435 | 4629 | 4143 | 3836 | 4450 | 1,159 | 0,213 | 4,582 | 4,61E-06 | 1,76E-05 | 0,001514 | 0,0023299 | 0,0018229 | 0,0018229 | TRUE | 0,0267949 | No |
| orf19.1476 | 36 | 28 | 32 | 47 | 33 | 29 | 32 | 20 | 32 | 36 | 33 | 39 | 28 | 28 | 29 | 28 | 31,51 | 35 | 28 | 0,807 | -0,31 | -1,304 | 0,19235915 | 0,26570586 | 1,00E-08 | 0,0480479 | 0,02223 | 0,02223 | TRUE | 0,0502185 | No |
| orf19.1477 | 1342 | 1121 | 1201 | 1645 | 1899 | 1910 | 1981 | 1264 | 1204 | 1427 | 1229 | 1356 | 1624 | 1822 | 1768 | 1783 | 1526,7 | 1304 | 1749 | 1,342 | 0,425 | 6,525 | 6,80E-11 | 5,07E-10 | 0,003626 | 0,0029304 | 0,003392 | 0,003392 | TRUE | 0,0456927 | No |
| orf19.1478 | 958 | 608 | 861 | 1090 | 1181 | 1104 | 1173 | 738 | 859 | 774 | 881 | 899 | 1010 | 1053 | 1047 | 1041 | 945,53 | 853 | 1038 | 1,212 | 0,278 | 4,811 | 1,50E-06 | 6,18E-06 | 0,000842 | 0,0035149 | 0,002133 | 0,002133 | TRUE | 0,0419277 | No |
| orf19.1479 | 2827 | 1891 | 2459 | 3044 | 3460 | 2979 | 3222 | 2238 | 2536 | 2407 | 2516 | 2510 | 2959 | 2842 | 2875 | 3156 | 2725,3 | 2492 | 2958 | 1,185 | 0,244 | 5,504 | 3,70E-08 | 1,91E-07 | 0,000911 | 0,0025122 | 0,0015209 | 0,0015209 | TRUE | 0,0230168 | No |

| | | | | | | | | | | | | | | | | | | | | | | | | | | | | | | | |
|---|---|---|---|---|---|---|---|---|---|---|---|---|---|---|---|---|---|---|---|---|---|---|---|---|---|---|---|---|---|---|---|
| orf19.1480 | 1711 | 1152 | 1413 | 1794 | 1974 | 1675 | 1770 | 1287 | 1535 | 1467 | 1446 | 1479 | 1688 | 1598 | 1579 | 1815 | 1576 | 1482 | 1670 | 1,124 | 0,169 | 3,134 | 0,00172719 | 0,00413096 | 0,00156 | 0,0029007 | 0,002161 | 0,002161 | TRUE | 0,037798 | No |
| orf19.1481 | 4380 | 2943 | 3821 | 4805 | 7286 | 6788 | 7489 | 4549 | 3929 | 3747 | 3909 | 3962 | 6231 | 6477 | 6683 | 6416 | 5169,3 | 3887 | 6452 | 1,659 | 0,73 | 19,938 | 1,90E-88 | 7,14E-86 | 0,000446 | 0,0022604 | 0,001081 | 0,001081 | TRUE | 0,0072782 | No |
| orf19.1482 | 1392 | 904 | 1073 | 1458 | 1737 | 1310 | 1469 | 792 | 1249 | 1151 | 1098 | 1202 | 1486 | 1250 | 1311 | 1117 | 1232,9 | 1175 | 1291 | 1,101 | 0,138 | 1,694 | 0,09027773 | 0,14044241 | 0,007473 | 0,003157 | 0,0055752 | 0,0055752 | TRUE | 0,1242143 | No |
| orf19.1483 | 1195 | 888 | 977 | 1310 | 1290 | 1339 | 1284 | 737 | 1072 | 1131 | 1000 | 1080 | 1103 | 1278 | 1146 | 1039 | 1106,1 | 1071 | 1142 | 1,069 | 0,096 | 1,382 | 0,16688049 | 0,23641075 | 0,003954 | 0,003292 | 0,0037017 | 0,0037017 | TRUE | 0,0765236 | No |
| orf19.1484 | 923 | 944 | 919 | 1177 | 1039 | 1252 | 1224 | 773 | 828 | 1202 | 940 | 971 | 889 | 1195 | 1092 | 1090 | 1025,8 | 985 | 1066 | 1,084 | 0,116 | 1,033 | 0,30138121 | 0,38731515 | 0,017522 | 0,0033947 | 0,011108 | 0,011108 | TRUE | 0,2552639 | No |
| orf19.1485 | 888 | 773 | 753 | 981 | 923 | 822 | 823 | 465 | 797 | 984 | 770 | 809 | 789 | 784 | 734 | 656 | 790,5 | 840 | 741 | 0,886 | -0,174 | -1,985 | 0,04714316 | 0,07987481 | 0,008523 | 0,003816 | 0,0061112 | 0,0061112 | TRUE | 0,1510931 | No |
| orf19.1486 | 1485 | 1330 | 1399 | 1934 | 2013 | 2099 | 2271 | 1619 | 1332 | 1693 | 1431 | 1595 | 1722 | 2003 | 2026 | 2283 | 1760,7 | 1513 | 2008 | 1,327 | 0,408 | 4,445 | 8,79E-06 | 3,23E-05 | 0,011485 | 0,002804 | 0,0075153 | 0,0075153 | TRUE | 0,1126429 | No |
| orf19.1488 | 359 | 332 | 359 | 450 | 460 | 504 | 478 | 317 | 322 | 423 | 367 | 371 | 393 | 481 | 427 | 447 | 403,88 | 371 | 437 | 1,181 | 0,24 | 2,508 | 0,01215307 | 0,02395526 | 0,007057 | 0,0055739 | 0,0063089 | 0,0063089 | TRUE | 0,0932857 | No |
| orf19.149 | 13 | 8 | 13 | 19 | 17 | 12 | 11 | 15 | 12 | 10 | 13 | 16 | 15 | 11 | 10 | 21 | 13,47 | 13 | 14 | 1,079 | 0,11 | 0,286 | 0,77499137 | NA | 1,00E-08 | 0,1097365 | 0,0672573 | 0,0672573 | TRUE | 0,3211691 | No |
| orf19.1490 | 2860 | 1983 | 2633 | 3186 | 3578 | 3311 | 3619 | 2607 | 2566 | 2525 | 2694 | 2627 | 3060 | 3159 | 3229 | 3677 | 2942,1 | 2603 | 3281 | 1,259 | 0,332 | 5,642 | 1,68E-08 | 9,07E-08 | 0,003397 | 0,0024729 | 0,0029766 | 0,0029766 | TRUE | 0,0773539 | No |
| orf19.1490.1 | 44 | 31 | 25 | 34 | 43 | 36 | 30 | 15 | 39 | 39 | 26 | 28 | 37 | 34 | 27 | 21 | 31,45 | 33 | 30 | 0,917 | -0,124 | -0,466 | 0,64097421 | 0,71103638 | 0,019156 | 0,0481377 | 0,036687 | 0,036687 | TRUE | 0,1748579 | No |
| orf19.1491 | 1736 | 1377 | 1557 | 1930 | 2251 | 2271 | 2313 | 1388 | 1557 | 1753 | 1593 | 1592 | 1925 | 2167 | 2064 | 1958 | 1826,1 | 1624 | 2028 | 1,252 | 0,324 | 5,765 | 8,16E-09 | 4,58E-08 | 0,002185 | 0,0027745 | 0,0024705 | 0,0024705 | TRUE | 0,0328867 | No |
| orf19.1492 | 565 | 577 | 575 | 755 | 856 | 992 | 930 | 686 | 507 | 735 | 588 | 623 | 732 | 947 | 830 | 968 | 741,04 | 613 | 869 | 1,418 | 0,504 | 4,416 | 1,00E-05 | 3,66E-05 | 0,017952 | 0,0039385 | 0,0111101 | 0,0111101 | TRUE | 0,2010987 | No |
| orf19.1493 | 3762 | 2640 | 3318 | 4035 | 3457 | 3088 | 3362 | 1999 | 3375 | 3361 | 3395 | 3327 | 2957 | 2946 | 3000 | 2819 | 3147,5 | 3364 | 2930 | 0,872 | -0,197 | -5,642 | 1,68E-08 | 9,06E-08 | 1,00E-08 | 0,0024407 | 0,0008521 | 0,0008521 | TRUE | 0,0070939 | No |
| orf19.1494 | 9155 | 6395 | 7983 | 10116 | 9007 | 7641 | 8335 | 5984 | 8213 | 8142 | 8167 | 8342 | 7703 | 7291 | 7438 | 8440 | 7966,9 | 8216 | 7718 | 0,939 | -0,091 | -1,914 | 0,0556158 | 0,09212565 | 0,002081 | 0,0021617 | 0,0020578 | 0,0020578 | TRUE | 0,0473495 | No |
| orf19.1495 | 613 | 593 | 590 | 765 | 622 | 747 | 718 | 457 | 550 | 755 | 604 | 631 | 532 | 713 | 641 | 645 | 633,66 | 635 | 633 | 0,998 | -0,003 | -0,03 | 0,97570401 | 0,98257286 | 0,014635 | 0,0042705 | 0,0095495 | 0,0095495 | TRUE | 0,1829847 | No |
| orf19.1496 | 289 | 269 | 325 | 420 | 462 | 510 | 489 | 284 | 259 | 342 | 333 | 346 | 395 | 487 | 436 | 401 | 374,9 | 320 | 430 | 1,346 | 0,428 | 3,99 | 6,61E-05 | 0,00020902 | 0,010879 | 0,0058517 | 0,0083408 | 0,0083408 | TRUE | 0,1939363 | No |
| orf19.1497 | 414 | 342 | 353 | 431 | 489 | 462 | 509 | 294 | 371 | 435 | 361 | 355 | 418 | 441 | 454 | 415 | 406,41 | 380 | 432 | 1,141 | 0,191 | 2,316 | 0,0205453 | 0,03816827 | 0,002366 | 0,0055516 | 0,0040478 | 0,0040478 | TRUE | 0,0957714 | No |
| orf19.1499 | 2981 | 2222 | 2700 | 3404 | 2920 | 2793 | 3238 | 2021 | 2674 | 2829 | 2762 | 2807 | 2497 | 2665 | 2889 | 2850 | 2746,8 | 2768 | 2725 | 0,984 | -0,023 | -0,439 | 0,66056387 | 0,72873076 | 0,002215 | 0,002508 | 0,0023275 | 0,0023275 | TRUE | 0,0394303 | No |
| orf19.150 | 4362 | 3066 | 3746 | 4768 | 5012 | 4113 | 4425 | 2809 | 3913 | 3903 | 3833 | 3932 | 4286 | 3924 | 3949 | 3962 | 3962,8 | 3895 | 4030 | 1,035 | 0,05 | 1,248 | 0,2121331 | 0,28952651 | 0,00063 | 0,0023459 | 0,001277 | 0,001277 | TRUE | 0,0229527 | No |
| orf19.1500 | 699 | 534 | 644 | 823 | 945 | 925 | 987 | 613 | 627 | 680 | 659 | 679 | 808 | 883 | 881 | 865 | 760,07 | 661 | 859 | 1,3 | 0,378 | 6,343 | 2,25E-10 | 1,56E-09 | 1,00E-08 | 0,0038895 | 0,0020712 | 0,0020712 | TRUE | 0,0204967 | No |
| orf19.1502 | 339 | 340 | 310 | 446 | 483 | 561 | 566 | 346 | 304 | 433 | 317 | 368 | 413 | 535 | 505 | 488 | 420,42 | 356 | 485 | 1,369 | 0,453 | 3,873 | 0,00010758 | 0,00032792 | 0,016003 | 0,0054325 | 0,0106938 | 0,0106938 | TRUE | 0,2285562 | No |
| orf19.1504 | 2457 | 2001 | 2360 | 2840 | 2559 | 2611 | 2706 | 1516 | 2204 | 2547 | 2415 | 2342 | 2189 | 2491 | 2415 | 2138 | 2342,6 | 2377 | 2308 | 0,973 | -0,04 | -0,629 | 0,52951118 | 0,61226005 | 0,003709 | 0,0025992 | 0,0034648 | 0,0034648 | TRUE | 0,0342763 | No |
| orf19.1505 | 2451 | 2081 | 2318 | 2917 | 2297 | 2270 | 2269 | 1430 | 2199 | 2649 | 2372 | 2405 | 1964 | 2166 | 2025 | 2017 | 2224,6 | 2406 | 2043 | 0,85 | -0,235 | -3,873 | 0,00010768 | 0,00032808 | 0,003262 | 0,0026321 | 0,0030659 | 0,0030659 | TRUE | 0,054299 | No |
| orf19.1506 | 0 | 0 | 0 | 0 | 0 | 0 | 0 | 0 | 0 | 0 | 0 | 0 | 0 | 0 | 0 | 0 | 0 | 0 | 0 | NA | NA | NA | NA | NA | NA | NA | NA | NA | NA | NA | NA |
| orf19.1507 | 211 | 188 | 205 | 269 | 255 | 281 | 303 | 147 | 189 | 239 | 210 | 222 | 218 | 268 | 270 | 207 | 228,01 | 215 | 241 | 1,129 | 0,175 | 1,485 | 0,1375005 | 0,2002674 | 0,009547 | 0,0083463 | 0,0089045 | 0,0089045 | TRUE | 0,0823712 | No |
| orf19.1508 | 21 | 14 | 17 | 22 | 26 | 20 | 19 | 15 | 19 | 18 | 17 | 18 | 22 | 19 | 17 | 21 | 18,95 | 18 | 20 | 1,095 | 0,131 | 0,43 | 0,66685794 | 0,73405689 | 1,00E-08 | 0,0785729 | 0,036605 | 0,036605 | TRUE | 0,0602106 | No |
| orf19.1509 | 8150 | 6298 | 7433 | 9415 | 9211 | 9071 | 9649 | 5932 | 7311 | 8018 | 7605 | 7764 | 7878 | 8655 | 8610 | 8367 | 8025,9 | 7674 | 8378 | 1,092 | 0,127 | 2,862 | 0,00421451 | 0,00922432 | 0,001483 | 0,0021604 | 0,0017575 | 0,0017575 | TRUE | 0,0200503 | No |
| orf19.151 | 1035 | 822 | 1074 | 1216 | 1172 | 1129 | 1268 | 797 | 929 | 1046 | 1099 | 1003 | 1002 | 1077 | 1131 | 1124 | 1051,5 | 1019 | 1084 | 1,064 | 0,089 | 1,34 | 0,18028208 | 0,25201289 | 0,003189 | 0,0033602 | 0,0032647 | 0,0032647 | TRUE | 0,0436169 | No |
| orf19.1510 | 1244 | 920 | 1133 | 1504 | 1222 | 1151 | 1251 | 742 | 1116 | 1171 | 1159 | 1240 | 1045 | 1098 | 1116 | 1047 | 1124,1 | 1172 | 1076 | 0,919 | -0,122 | -2,267 | 0,02337475 | 0,04301241 | 0,000833 | 0,003271 | 0,0018674 | 0,0018674 | TRUE | 0,0201665 | No |
| orf19.1513 | 4918 | 3706 | 4628 | 5288 | 6521 | 6281 | 6545 | 3767 | 4412 | 4718 | 4735 | 4361 | 5577 | 5993 | 5840 | 5313 | 5118,6 | 4556 | 5681 | 1,248 | 0,32 | 6,434 | 1,24E-10 | 8,92E-10 | 0,002067 | 0,0022631 | 0,0021724 | 0,0021724 | TRUE | 0,0224292 | No |
| orf19.1514 | 2152 | 1522 | 1852 | 2408 | 2396 | 2069 | 2312 | 1342 | 1931 | 1938 | 1895 | 1986 | 2049 | 1974 | 2063 | 1893 | 1966 | 1938 | 1995 | 1,031 | 0,044 | 1,02 | 0,30756805 | 0,39358229 | 0,000382 | 0,0027179 | 0,0013112 | 0,0013112 | TRUE | 0,0129052 | No |
| orf19.1515 | 276 | 201 | 216 | 264 | 259 | 257 | 237 | 151 | 248 | 256 | 221 | 218 | 222 | 245 | 211 | 213 | 229,17 | 236 | 223 | 0,95 | -0,073 | -0,734 | 0,46298448 | 0,54885769 | 1,00E-08 | 0,0083141 | 0,0052632 | 0,0052632 | TRUE | 0,0512501 | No |
| orf19.1516 | 1624 | 1210 | 1460 | 1849 | 1185 | 1164 | 1198 | 653 | 1457 | 1540 | 1494 | 1525 | 1013 | 1111 | 1069 | 921 | 1266,2 | 1504 | 1028 | 0,686 | -0,543 | -9,118 | 7,69E-20 | 1,36E-18 | 0,002096 | 0,003126 | 0,0025875 | 0,0025875 | TRUE | 0,0523849 | No |
| orf19.1517 | 1710 | 1108 | 1358 | 1786 | 2073 | 1709 | 1802 | 1143 | 1534 | 1411 | 1389 | 1473 | 1773 | 1631 | 1608 | 1612 | 1553,8 | 1452 | 1656 | 1,141 | 0,19 | 3,546 | 0,00039096 | 0,00107994 | 0,001546 | 0,0029138 | 0,0021006 | 0,0021006 | TRUE | 0,0283041 | No |
| orf19.1519 | 933 | 668 | 829 | 1017 | 1148 | 1003 | 1108 | 713 | 837 | 850 | 848 | 839 | 982 | 957 | 989 | 1006 | 913,42 | 844 | 984 | 1,165 | 0,22 | 4,401 | 1,08E-05 | 3,90E-05 | 1,00E-08 | 0,0035688 | 0,0013101 | 0,0013101 | TRUE | 0,0037494 | No |
| orf19.1522 | 74 | 44 | 60 | 65 | 72 | 73 | 73 | 42 | 66 | 56 | 61 | 54 | 62 | 70 | 65 | 59 | 61,62 | 59 | 64 | 1,079 | 0,11 | 0,634 | 0,52598844 | 0,60935954 | 1,00E-08 | 0,0255367 | 0,0129293 | 0,0129293 | TRUE | 0,0610396 | No |
| orf19.1523 | 1423 | 971 | 1336 | 1501 | 1491 | 1198 | 1390 | 898 | 1277 | 1236 | 1367 | 1238 | 1275 | 1143 | 1240 | 1267 | 1255,3 | 1280 | 1231 | 0,962 | -0,056 | -0,995 | 0,31954936 | 0,40629615 | 0,001573 | 0,003136 | 0,0022458 | 0,0022458 | TRUE | 0,028353 | No |
| orf19.1524 | 10 | 4 | 6 | 4 | 5 | 6 | 5 | 4 | 9 | 5 | 6 | 3 | 4 | 6 | 4 | 6 | 5,45 | 6 | 5 | 0,846 | -0,241 | -0,411 | 0,6814285 | NA | 1,00E-08 | 0,2683162 | 0,1452303 | 0,1452303 | TRUE | 0,3486111 | No |
| orf19.1525 | 1618 | 1250 | 1469 | 1764 | 1543 | 1524 | 1580 | 1082 | 1452 | 1591 | 1503 | 1455 | 1320 | 1454 | 1410 | 1526 | 1463,8 | 1500 | 1428 | 0,951 | -0,073 | -1,284 | 0,19930704 | 0,27382499 | 0,001864 | 0,0029713 | 0,0024016 | 0,0024016 | TRUE | 0,0315163 | No |
| orf19.1526 | 3088 | 2228 | 3004 | 3428 | 3719 | 3285 | 3732 | 2354 | 2770 | 2836 | 3073 | 2827 | 3181 | 3134 | 3330 | 3320 | 3059 | 2876 | 3241 | 1,127 | 0,172 | 3,781 | 0,00015635 | 0,00046274 | 0,001116 | 0,0024541 | 0,0016544 | 0,0016544 | TRUE | 0,0257326 | No |
| orf19.1527 | 1260 | 911 | 1015 | 1316 | 1130 | 1019 | 1074 | 723 | 1130 | 1160 | 1038 | 1085 | 966 | 972 | 958 | 1020 | 1041,3 | 1103 | 979 | 0,887 | -0,173 | -3,228 | 0,00124795 | 0,00308293 | 0,000453 | 0,0033736 | 0,0018065 | 0,0018065 | TRUE | 0,0179661 | No |
| orf19.1528 | 3833 | 2894 | 3379 | 4031 | 3731 | 3727 | 3950 | 2253 | 3439 | 3684 | 3457 | 3324 | 3191 | 3556 | 3525 | 3178 | 3419,2 | 3476 | 3362 | 0,969 | -0,046 | -0,863 | 0,3881389 | 0,47726945 | 0,00241 | 0,0024041 | 0,0024533 | 0,0024533 | TRUE | 0,0193176 | No |
| orf19.1529 | 15 | 11 | 12 | 13 | 17 | 9 | 12 | 14 | 13 | 14 | 12 | 11 | 15 | 9 | 11 | 20 | 13 | 12 | 14 | 1,042 | 0,06 | 0,153 | 0,8781536 | NA | 1,00E-08 | 0,113608 | 0,0676496 | 0,0676496 | TRUE | 0,2884961 | No |
| orf19.1530 | 4 | 0 | 4 | 3 | 1 | 5 | 7 | 0 | 4 | 0 | 4 | 2 | 1 | 5 | 6 | 0 | 2,75 | 2 | 3 | 1,18 | 0,238 | 0,244 | 0,80748614 | NA | 0,643875 | 0,5292223 | 0,5657601 | 0,5657601 | TRUE | 0,1857398 | No |
| orf19.1531 | 2716 | 2582 | 2760 | 3465 | 3032 | 3448 | 3332 | 2018 | 2437 | 3287 | 2824 | 2857 | 2593 | 3290 | 2973 | 2846 | 2888,4 | 2851 | 2926 | 1,027 | 0,038 | 0,421 | 0,67375108 | 0,73939705 | 0,011974 | 0,0024821 | 0,0075221 | 0,0075221 | TRUE | 0,1271197 | No |
| orf19.1532 | 494 | 472 | 471 | 580 | 466 | 494 | 499 | 355 | 443 | 601 | 482 | 478 | 399 | 471 | 445 | 501 | 477,51 | 501 | 454 | 0,906 | -0,142 | -1,375 | 0,16898295 | 0,23893848 | 0,011189 | 0,0050197 | 0,0080811 | 0,0080811 | TRUE | 0,1982685 | No |
| orf19.1533 | 2404 | 1893 | 2104 | 2611 | 2622 | 2660 | 2720 | 1558 | 2157 | 2410 | 2153 | 2153 | 2242 | 2538 | 2427 | 2197 | 2284,7 | 2218 | 2351 | 1,062 | 0,087 | 1,437 | 0,15078905 | 0,21660258 | 0,003376 | 0,0026149 | 0,0030548 | 0,0030548 | TRUE | 0,0396901 | No |
| orf19.1534 | 580 | 467 | 524 | 629 | 776 | 818 | 882 | 470 | 520 | 595 | 536 | 519 | 664 | 781 | 787 | 663 | 632,97 | 542 | 724 | 1,34 | 0,422 | 5,187 | 2,14E-07 | 1,00E-06 | 0,00516 | 0,004273 | 0,0047372 | 0,0047372 | TRUE | 0,0454285 | No |
| orf19.1535 | 85 | 63 | 84 | 113 | 144 | 113 | 136 | 66 | 76 | 80 | 86 | 93 | 123 | 108 | 121 | 93 | 97,62 | 84 | 111 | 1,336 | 0,418 | 2,823 | 0,00475697 | 0,01031764 | 1,00E-08 | 0,0168497 | 0,0106857 | 0,0106857 | TRUE | 0,0954751 | No |
| orf19.1536 | 1309 | 1080 | 1278 | 1560 | 1257 | 1403 | 1423 | 1030 | 1174 | 1375 | 1308 | 1286 | 1075 | 1339 | 1270 | 1453 | 1284,9 | 1286 | 1284 | 0,997 | -0,004 | -0,044 | 0,96474613 | 0,97464179 | 0,009195 | 0,0031093 | 0,0063683 | 0,0063683 | TRUE | 0,1462128 | No |
| orf19.1538 | 172 | 132 | 130 | 214 | 247 | 223 | 226 | 143 | 154 | 168 | 133 | 176 | 211 | 213 | 202 | 202 | 182,4 | 158 | 207 | 1,31 | 0,389 | 3,488 | 0,00048608 | 0,00131784 | 0,001086 | 0,0099384 | 0,0063927 | 0,0063927 | TRUE | 0,1174674 | No |
| orf19.1539 | 206 | 178 | 211 | 254 | 277 | 318 | 302 | 212 | 185 | 227 | 216 | 209 | 237 | 303 | 269 | 299 | 243,19 | 209 | 277 | 1,324 | 0,405 | 3,71 | 0,00020736 | 0,00060272 | 0,006456 | 0,0079488 | 0,0072365 | 0,0072365 | TRUE | 0,113638 | No |
| orf19.154 | 2189 | 1838 | 1980 | 2612 | 2489 | 2664 | 2685 | 1616 | 1964 | 2340 | 2026 | 2154 | 2129 | 2542 | 2396 | 2279 | 2228,6 | 2121 | 2336 | 1,103 | 0,141 | 2,025 | 0,04281614 | 0,07342118 | 0,005284 | 0,0026309 | 0,0041964 | 0,0041964 | TRUE | 0,0568215 | No |
| orf19.1541 | 61 | 33 | 58 | 47 | 30 | 40 | 40 | 23 | 55 | 42 | 59 | 39 | 26 | 38 | 36 | 32 | 40,85 | 49 | 33 | 0,676 | -0,565 | -2,396 | 0,01657029 | 0,03149751 | 0,015234 | 0,0375181 | 0,0279829 | 0,0279829 | TRUE | 0,1650703 | No |
| orf19.1542 | 3776 | 2642 | 3366 | 4144 | 4469 | 3948 | 4337 | 2562 | 3387 | 3364 | 3444 | 3417 | 3822 | 3767 | 3870 | 3613 | 3585,6 | 3403 | 3768 | 1,108 | 0,148 | 4,208 | 2,58E-05 | 8,78E-05 | 0,000117 | 0,0023844 | 0,0009147 | 0,0009147 | TRUE | 0,0085262 | No |
| orf19.1543 | 1060 | 733 | 881 | 1137 | 1017 | 916 | 966 | 637 | 951 | 933 | 901 | 938 | 870 | 874 | 862 | 898 | 903,41 | 931 | 876 | 0,94 | -0,09 | -1,766 | 0,07745272 | 0,12310474 | 1,00E-08 | 0,0035864 | 0,0013844 | 0,0013844 | TRUE | 0,005416 | No |
| orf19.1544 | 413 | 259 | 300 | 425 | 312 | 271 | 294 | 219 | 371 | 330 | 307 | 350 | 267 | 259 | 262 | 309 | 306,78 | 340 | 274 | 0,801 | -0,32 | -3,439 | 0,00058339 | 0,00154882 | 0,002981 | 0,0067115 | 0,0050035 | 0,0050035 | TRUE | 0,0681124 | No |
| orf19.1545 | 1117 | 780 | 899 | 1155 | 1507 | 1230 | 1313 | 855 | 1002 | 993 | 920 | 952 | 1289 | 1174 | 1172 | 1206 | 1088,4 | 967 | 1210 | 1,253 | 0,325 | 5,868 | 4,42E-09 | 2,59E-08 | 0,001032 | 0,0033133 | 0,0020121 | 0,0020121 | TRUE | 0,0247301 | No |

| | | | | | | | | | | | | | | | | | | | | | | | | | | | | | | | |
|---|---|---|---|---|---|---|---|---|---|---|---|---|---|---|---|---|---|---|---|---|---|---|---|---|---|---|---|---|---|---|---|
| orf19.1546 | 1323 | 864 | 1091 | 1278 | 1567 | 1420 | 1459 | 895 | 1187 | 1100 | 1116 | 1054 | 1340 | 1355 | 1302 | 1262 | 1214,5 | 1114 | 1315 | 1,182 | 0,241 | 4,514 | 6,37E-06 | 2,39E-05 | 0,000934 | 0,0031748 | 0,0018999 | 0,0018999 | TRUE | 0,0244322 | No |
| orf19.1547 | 358 | 270 | 343 | 418 | 352 | 333 | 340 | 227 | 321 | 344 | 351 | 345 | 301 | 318 | 303 | 320 | 325,36 | 340 | 310 | 0,912 | -0,133 | -1,681 | 0,092822 | 0,14373057 | 1,00E-08 | 0,0064414 | 0,002944 | 0,002944 | TRUE | 0,0170778 | No |
| orf19.1548 | 925 | 676 | 807 | 1075 | 1059 | 814 | 898 | 519 | 830 | 861 | 826 | 886 | 906 | 777 | 801 | 732 | 827,28 | 851 | 804 | 0,948 | -0,077 | -1,094 | 0,27403932 | 0,35838665 | 0,003147 | 0,0037343 | 0,0035487 | 0,0035487 | TRUE | 0,0878112 | No |
| orf19.1549 | 2768 | 2045 | 2419 | 2778 | 2575 | 2496 | 2742 | 1400 | 2483 | 2604 | 2475 | 2291 | 2202 | 2382 | 2447 | 1975 | 2357,2 | 2463 | 2252 | 0,916 | -0,127 | -1,837 | 0,06615289 | 0,10759246 | 0,005218 | 0,0025954 | 0,0041488 | 0,0041488 | TRUE | 0,0793431 | No |
| orf19.155 | 1845 | 1685 | 1735 | 2298 | 2285 | 2371 | 2402 | 1551 | 1655 | 2145 | 1775 | 1895 | 1954 | 2262 | 2143 | 2188 | 2002,2 | 1868 | 2137 | 1,145 | 0,195 | 2,504 | 0,01226276 | 0,02414771 | 0,007502 | 0,0027046 | 0,005337 | 0,005337 | TRUE | 0,1178468 | No |
| orf19.1551 | 16 | 5 | 11 | 12 | 10 | 13 | 12 | 4 | 14 | 6 | 11 | 10 | 9 | 12 | 11 | 6 | 9,9 | 10 | 10 | 0,895 | -0,159 | -0,36 | 0,71873516 | NA | 1,00E-08 | 0,1486627 | 0,0886732 | 0,0886732 | TRUE | 0,2381982 | No |
| orf19.1552 | 3224 | 2106 | 2521 | 3299 | 3135 | 2407 | 2418 | 1645 | 2892 | 2681 | 2579 | 2720 | 2681 | 2297 | 2158 | 2320 | 2541,1 | 2718 | 2364 | 0,87 | -0,201 | -2,962 | 0,00305511 | 0,00692646 | 0,005085 | 0,0025508 | 0,0040374 | 0,0040374 | TRUE | 0,1011988 | No |
| orf19.1553 | 6587 | 4521 | 5745 | 7377 | 6783 | 5957 | 6498 | 4109 | 5909 | 5756 | 5878 | 6083 | 5801 | 5684 | 5798 | 5795 | 5838,1 | 5906 | 5770 | 0,976 | -0,035 | -1,122 | 0,26190798 | 0,3455299 | 0,00015 | 0,0022282 | 0,0007455 | 0,0007455 | TRUE | 0,0050354 | No |
| orf19.1555 | 2180 | 1545 | 1977 | 2300 | 2182 | 2111 | 2191 | 1361 | 1956 | 1967 | 2023 | 1897 | 1866 | 2014 | 1955 | 1920 | 1949,6 | 1961 | 1939 | 0,99 | -0,015 | -0,349 | 0,72692997 | 0,7859268 | 0,000436 | 0,0027241 | 0,0013054 | 0,0013054 | TRUE | 0,008541 | No |
| orf19.1557 | 189 | 148 | 173 | 244 | 149 | 106 | 141 | 95 | 170 | 188 | 177 | 201 | 127 | 101 | 126 | 134 | 153,07 | 184 | 122 | 0,66 | -0,598 | -4,842 | 1,29E-06 | 5,33E-06 | 0,002734 | 0,0114635 | 0,0078441 | 0,0078441 | TRUE | 0,1375951 | No |
| orf19.1559 | 5152 | 3326 | 4327 | 5539 | 5067 | 4228 | 4786 | 3069 | 4622 | 4234 | 4427 | 4568 | 4333 | 4034 | 4271 | 4329 | 4352,2 | 4463 | 4242 | 0,95 | -0,075 | -1,756 | 0,07908422 | 0,12530011 | 0,000999 | 0,0023131 | 0,0015006 | 0,0015006 | TRUE | 0,0139913 | No |
| orf19.156 | 2847 | 1972 | 2624 | 3156 | 3098 | 2892 | 3073 | 1769 | 2554 | 2511 | 2685 | 2602 | 2650 | 2759 | 2742 | 2495 | 2624,7 | 2588 | 2662 | 1,03 | 0,042 | 0,933 | 0,35090526 | 0,4384856 | 0,000942 | 0,0025326 | 0,0015606 | 0,0015606 | TRUE | 0,0198374 | No |
| orf19.1560 | 5899 | 3963 | 5201 | 6485 | 5786 | 5000 | 5449 | 3380 | 5292 | 5045 | 5321 | 5348 | 4948 | 4771 | 4862 | 4767 | 5044,4 | 5252 | 4837 | 0,921 | -0,119 | -3,502 | 0,00046156 | 0,00125706 | 0,000246 | 0,0022673 | 0,0009063 | 0,0009063 | TRUE | 0,0081245 | No |
| orf19.1562 | 26899 | 19289 | 23871 | 30259 | 32160 | 29870 | 31589 | 22185 | 24131 | 24557 | 24422 | 24952 | 27504 | 28501 | 28188 | 31290 | 26693 | 24516 | 28871 | 1,177 | 0,236 | 5,407 | 6,40E-08 | 3,22E-07 | 0,001711 | 0,0020339 | 0,001786 | 0,001786 | TRUE | 0,038701 | No |
| orf19.1563 | 3324 | 2246 | 2974 | 3639 | 2834 | 2517 | 2683 | 1762 | 2982 | 2859 | 3043 | 3001 | 2424 | 2402 | 2394 | 2485 | 2698,7 | 2971 | 2426 | 0,815 | -0,295 | -7,89 | 3,02E-15 | 3,54E-14 | 1,00E-08 | 0,0025175 | 0,0009629 | 0,0009629 | TRUE | 0,0072131 | No |
| orf19.1564 | 1427 | 1152 | 1412 | 1773 | 1483 | 1381 | 1438 | 1016 | 1280 | 1467 | 1445 | 1462 | 1268 | 1318 | 1283 | 1433 | 1369,5 | 1414 | 1326 | 0,937 | -0,095 | -1,534 | 0,12503976 | 0,18476117 | 0,002603 | 0,0030396 | 0,0029174 | 0,0029174 | TRUE | 0,0495746 | No |
| orf19.1565 | 854 | 589 | 770 | 984 | 984 | 875 | 921 | 589 | 766 | 750 | 788 | 811 | 842 | 835 | 822 | 831 | 805,53 | 779 | 832 | 1,067 | 0,094 | 1,73 | 0,08361145 | 0,13146757 | 1,00E-08 | 0,0037817 | 0,0015663 | 0,0015663 | TRUE | 0,009969 | No |
| orf19.1566 | 18607 | 13439 | 17034 | 20148 | 18399 | 16883 | 17981 | 11461 | 16692 | 17109 | 17427 | 16614 | 15735 | 16109 | 16045 | 16165 | 16487 | 16960 | 16014 | 0,944 | -0,083 | -2,939 | 0,00328968 | 0,00740239 | 0,000241 | 0,0020676 | 0,000702 | 0,000702 | TRUE | 0,004237 | No |
| orf19.1567 | 2473 | 1871 | 2225 | 2658 | 3284 | 3156 | 3346 | 1915 | 2219 | 2382 | 2276 | 2192 | 2809 | 3011 | 2986 | 2701 | 2571,9 | 2267 | 2877 | 1,271 | 0,346 | 7,021 | 2,20E-12 | 1,94E-11 | 0,001486 | 0,002544 | 0,0019408 | 0,0019408 | TRUE | 0,0192202 | No |
| orf19.1569 | 9463 | 7089 | 8815 | 10579 | 10173 | 9891 | 10562 | 6839 | 8489 | 9025 | 9019 | 8724 | 8700 | 9438 | 9425 | 9646 | 9058,1 | 8814 | 9302 | 1,055 | 0,078 | 1,837 | 0,06624334 | 0,10771042 | 0,001271 | 0,0021398 | 0,0016005 | 0,0016005 | TRUE | 0,0233866 | No |
| orf19.1570 | 790 | 559 | 703 | 896 | 1001 | 889 | 925 | 526 | 709 | 712 | 719 | 739 | 856 | 848 | 825 | 742 | 768,76 | 720 | 818 | 1,14 | 0,189 | 3,113 | 0,00184929 | 0,00440086 | 0,00063 | 0,0038679 | 0,0022478 | 0,0022478 | TRUE | 0,0401078 | No |
| orf19.1571 | 1 | 1 | 0 | 1 | 1 | 3 | 2 | 1 | 1 | 1 | 0 | 1 | 1 | 3 | 2 | 1 | 1,24 | 1 | 2 | 2,352 | 1,234 | 0,974 | 0,32996718 | NA | 1,00E-08 | 1,1741669 | 0,582708 | 0,582708 | TRUE | 0,1673246 | No |
| orf19.1573 | 1958 | 1312 | 1700 | 2140 | 2338 | 2099 | 2171 | 1376 | 1757 | 1670 | 1739 | 1765 | 2000 | 2003 | 1937 | 1941 | 1851,4 | 1733 | 1970 | 1,136 | 0,184 | 4,526 | 6,01E-06 | 2,26E-05 | 1,00E-08 | 0,0027636 | 0,0010489 | 0,0010489 | TRUE | 0,006467 | No |
| orf19.1574 | 1949 | 1171 | 1619 | 1989 | 2188 | 1819 | 2106 | 1194 | 1748 | 1491 | 1656 | 1640 | 1871 | 1736 | 1879 | 1684 | 1713,3 | 1634 | 1792 | 1,097 | 0,134 | 2,221 | 0,02634905 | 0,04794219 | 0,002842 | 0,0028269 | 0,0028915 | 0,0028915 | TRUE | 0,0398147 | No |
| orf19.1575 | 11415 | 7593 | 9926 | 12482 | 11725 | 10114 | 10546 | 6747 | 10240 | 9667 | 10155 | 10293 | 10028 | 9650 | 9411 | 9516 | 9870 | 10089 | 9651 | 0,956 | -0,064 | -1,805 | 0,0710361 | 0,11417466 | 0,000659 | 0,0021266 | 0,0011124 | 0,0011124 | TRUE | 0,0094911 | No |
| orf19.1576 | 1462 | 1123 | 1293 | 1565 | 1574 | 1379 | 1547 | 967 | 1312 | 1430 | 1323 | 1291 | 1346 | 1316 | 1380 | 1364 | 1345,1 | 1339 | 1352 | 1,012 | 0,017 | 0,336 | 0,7366309 | 0,79398609 | 0,000404 | 0,0030588 | 0,0015658 | 0,0015658 | TRUE | 0,023251 | No |
| orf19.1577 | 87 | 49 | 61 | 96 | 69 | 69 | 65 | 53 | 78 | 62 | 62 | 79 | 59 | 66 | 58 | 75 | 67,45 | 70 | 64 | 0,897 | -0,157 | -0,897 | 0,36967763 | 0,45822637 | 1,00E-08 | 0,023502 | 0,0144988 | 0,0144988 | TRUE | 0,0787067 | No |
| orf19.1578 | 13411 | 10868 | 12633 | 14847 | 12235 | 12656 | 12802 | 7283 | 12031 | 13836 | 12925 | 12243 | 10464 | 12076 | 11424 | 10272 | 11909 | 12759 | 11059 | 0,867 | -0,206 | -3,275 | 0,00105709 | 0,00265177 | 0,004836 | 0,0021014 | 0,0037078 | 0,0037078 | TRUE | 0,0468366 | No |
| orf19.158 | 5174 | 4882 | 4808 | 6060 | 5391 | 6152 | 6296 | 3163 | 4642 | 6215 | 4919 | 4997 | 4611 | 5870 | 5618 | 4461 | 5166,6 | 5193 | 5140 | 0,99 | -0,014 | -0,136 | 0,89198219 | 0,91842828 | 0,015703 | 0,0022605 | 0,0100707 | 0,0100707 | TRUE | 0,212911 | No |
| orf19.1580 | 344 | 258 | 332 | 427 | 293 | 308 | 303 | 201 | 309 | 328 | 340 | 352 | 251 | 294 | 270 | 283 | 303,4 | 332 | 274 | 0,824 | -0,28 | -3,175 | 0,00150086 | 0,00363889 | 1,00E-08 | 0,0067643 | 0,0041272 | 0,0041272 | TRUE | 0,0421184 | No |
| orf19.1581 | 20 | 23 | 20 | 26 | 21 | 12 | 18 | 7 | 18 | 29 | 20 | 21 | 18 | 11 | 16 | 10 | 18,06 | 22 | 14 | 0,643 | -0,638 | -1,898 | 0,05764351 | 0,09503903 | 1,00E-08 | 0,0823671 | 0,050617 | 0,050617 | TRUE | 0,2022175 | No |
| orf19.1582 | 88 | 56 | 86 | 73 | 89 | 55 | 84 | 51 | 79 | 71 | 88 | 60 | 76 | 52 | 75 | 72 | 71,74 | 74 | 69 | 0,925 | -0,112 | -0,605 | 0,54485497 | 0,62638513 | 0,014431 | 0,0222157 | 0,0188468 | 0,0188468 | TRUE | 0,2432088 | No |
| orf19.1584 | 13043 | 8834 | 11609 | 14064 | 17607 | 14400 | 15754 | 9476 | 11701 | 11247 | 11877 | 11597 | 15058 | 13740 | 14058 | 13365 | 12830 | 11606 | 14055 | 1,211 | 0,277 | 6,442 | 1,18E-10 | 8,49E-10 | 0,001491 | 0,0020927 | 0,0016913 | 0,0016913 | TRUE | 0,0283135 | No |
| orf19.1585 | 1509 | 997 | 1304 | 1647 | 1219 | 1034 | 1158 | 797 | 1354 | 1269 | 1334 | 1358 | 1043 | 987 | 1033 | 1124 | 1187,7 | 1329 | 1047 | 0,785 | -0,35 | -6,561 | 5,35E-11 | 4,06E-10 | 0,000829 | 0,0032018 | 0,0018753 | 0,0018753 | TRUE | 0,025924 | No |
| orf19.1586 | 123 | 79 | 84 | 140 | 149 | 117 | 111 | 73 | 110 | 101 | 86 | 115 | 127 | 112 | 99 | 103 | 106,67 | 103 | 110 | 1,069 | 0,096 | 0,654 | 0,51284958 | 0,59716911 | 0,006299 | 0,0155885 | 0,0115484 | 0,0115484 | TRUE | 0,1257957 | No |
| orf19.1587 | 178 | 96 | 165 | 160 | 102 | 110 | 99 | 79 | 160 | 122 | 169 | 132 | 87 | 105 | 88 | 111 | 121,83 | 146 | 98 | 0,665 | -0,588 | -3,965 | 7,34E-05 | 0,00022977 | 0,010744 | 0,0138957 | 0,0125062 | 0,0125062 | TRUE | 0,1130628 | No |
| orf19.1588 | 928 | 591 | 788 | 965 | 1124 | 940 | 1089 | 598 | 833 | 752 | 806 | 796 | 961 | 897 | 972 | 843 | 857,53 | 797 | 918 | 1,155 | 0,207 | 3,303 | 0,0009556 | 0,00241027 | 0,001656 | 0,0036724 | 0,0026112 | 0,0026112 | TRUE | 0,0319419 | No |
| orf19.1589 | 2142 | 1518 | 2012 | 2263 | 1835 | 1628 | 1789 | 1134 | 1922 | 1933 | 2058 | 1866 | 1569 | 1553 | 1596 | 1599 | 1762,2 | 1945 | 1579 | 0,812 | -0,3 | -6,603 | 4,03E-11 | 3,10E-10 | 0,000527 | 0,0028033 | 0,0014022 | 0,0014022 | TRUE | 0,0189381 | No |
| orf19.1589.1 | 275 | 163 | 217 | 262 | 172 | 139 | 155 | 75 | 247 | 208 | 222 | 216 | 147 | 133 | 138 | 106 | 177,01 | 223 | 131 | 0,593 | -0,755 | -6,376 | 1,82E-10 | 1,27E-09 | 0,003315 | 0,0101806 | 0,007451 | 0,007451 | TRUE | 0,1326955 | No |
| orf19.1590 | 644 | 618 | 596 | 724 | 695 | 744 | 813 | 452 | 578 | 787 | 610 | 597 | 594 | 710 | 725 | 638 | 654,82 | 643 | 667 | 1,041 | 0,058 | 0,561 | 0,57453682 | 0,65276423 | 0,013066 | 0,0041965 | 0,0087182 | 0,0087182 | TRUE | 0,2566691 | No |
| orf19.1591 | 1810 | 1169 | 1543 | 2004 | 1521 | 1327 | 1370 | 1009 | 1624 | 1488 | 1579 | 1653 | 1301 | 1266 | 1222 | 1423 | 1444,5 | 1586 | 1303 | 0,819 | -0,289 | -4,96 | 7,04E-07 | 3,04E-06 | 0,002278 | 0,0029845 | 0,0025488 | 0,0025488 | TRUE | 0,042175 | No |
| orf19.1592 | 1469 | 1088 | 1292 | 1677 | 1555 | 1505 | 1551 | 1165 | 1318 | 1385 | 1322 | 1383 | 1330 | 1436 | 1384 | 1643 | 1400,1 | 1352 | 1448 | 1,068 | 0,095 | 1,449 | 0,14734227 | 0,21238781 | 0,003894 | 0,0030164 | 0,0034419 | 0,0034419 | TRUE | 0,0929243 | No |
| orf19.1595 | 585 | 496 | 545 | 751 | 659 | 778 | 778 | 447 | 525 | 631 | 558 | 619 | 564 | 742 | 694 | 630 | 620,47 | 583 | 658 | 1,129 | 0,175 | 1,841 | 0,06566737 | 0,10686066 | 0,009528 | 0,0043192 | 0,0070489 | 0,0070489 | TRUE | 0,1160621 | No |
| orf19.1596 | 449 | 314 | 356 | 445 | 447 | 419 | 491 | 318 | 403 | 400 | 364 | 367 | 382 | 400 | 438 | 449 | 400,31 | 384 | 417 | 1,086 | 0,118 | 1,461 | 0,14405887 | 0,20838012 | 0,001876 | 0,005606 | 0,0038052 | 0,0038052 | TRUE | 0,0379712 | No |
| orf19.1597 | 1716 | 1260 | 1489 | 1861 | 1744 | 1683 | 1667 | 1141 | 1539 | 1604 | 1523 | 1535 | 1492 | 1606 | 1488 | 1609 | 1549,5 | 1550 | 1549 | 0,998 | -0,003 | -0,062 | 0,95046334 | 0,96468651 | 0,000559 | 0,0029164 | 0,0015387 | 0,0015387 | TRUE | 0,0084684 | No |
| orf19.1598 | 622 | 443 | 586 | 705 | 649 | 489 | 570 | 408 | 558 | 564 | 600 | 581 | 555 | 467 | 509 | 575 | 551,07 | 576 | 526 | 0,911 | -0,134 | -1,766 | 0,07740047 | 0,12305426 | 0,002718 | 0,0046138 | 0,003678 | 0,003678 | TRUE | 0,0677742 | No |
| orf19.1599 | 12 | 12 | 10 | 27 | 13 | 15 | 8 | 5 | 11 | 15 | 10 | 22 | 11 | 14 | 7 | 7 | 12,27 | 14 | 10 | 0,681 | -0,554 | -1,284 | 0,19898206 | NA | 0,061364 | 0,1202934 | 0,0946921 | 0,0946921 | TRUE | 0,6845079 | No |
| orf19.1600 | 1156 | 754 | 1003 | 1263 | 1483 | 1348 | 1386 | 983 | 1037 | 960 | 1026 | 1041 | 1268 | 1286 | 1237 | 1386 | 1155,3 | 1016 | 1294 | 1,269 | 0,343 | 6,339 | 2,31E-10 | 1,61E-09 | 0,000777 | 0,0032361 | 0,0019343 | 0,0019343 | TRUE | 0,0247207 | No |
| orf19.1601 | 27521 | 16599 | 22648 | 27974 | 27325 | 20869 | 22874 | 15894 | 24689 | 21132 | 23171 | 23068 | 23369 | 19913 | 20411 | 22417 | 22271 | 23015 | 21528 | 0,935 | -0,097 | -1,54 | 0,1236705 | 0,18309816 | 0,004836 | 0,0020447 | 0,0037361 | 0,0037361 | TRUE | 0,0408066 | No |
| orf19.1604 | 2402 | 1772 | 2185 | 2697 | 2610 | 2321 | 2521 | 1608 | 2155 | 2256 | 2235 | 2224 | 2232 | 2215 | 2250 | 2268 | 2229,3 | 2218 | 2241 | 1,011 | 0,015 | 0,416 | 0,67752403 | 0,7421266 | 1,00E-08 | 0,0026307 | 0,0008766 | 0,0008766 | TRUE | 0,004469 | No |
| orf19.1605 | 130 | 109 | 96 | 126 | 90 | 81 | 90 | 58 | 117 | 139 | 98 | 104 | 77 | 77 | 80 | 82 | 96,74 | 114 | 79 | 0,695 | -0,525 | -3,472 | 0,00051728 | 0,00139168 | 1,00E-08 | 0,0169865 | 0,0112355 | 0,0112355 | TRUE | 0,1767686 | No |
| orf19.1606 | 1810 | 1349 | 1558 | 1914 | 1299 | 1369 | 1365 | 817 | 1624 | 1717 | 1594 | 1578 | 1111 | 1306 | 1218 | 1152 | 1412,6 | 1628 | 1197 | 0,736 | -0,442 | -7,429 | 1,10E-13 | 1,11E-12 | 0,002299 | 0,0030072 | 0,0026704 | 0,0026704 | TRUE | 0,0460185 | No |
| orf19.1607 | 8815 | 6528 | 8179 | 9933 | 11431 | 11329 | 12154 | 6959 | 7908 | 8311 | 8368 | 8191 | 9776 | 10810 | 10845 | 9815 | 9253 | 8194 | 10312 | 1,259 | 0,332 | 7,121 | 1,07E-12 | 9,80E-12 | 0,001838 | 0,0021364 | 0,0019783 | 0,0019783 | TRUE | 0,0152771 | No |
| orf19.1608 | 726 | 549 | 699 | 828 | 750 | 712 | 749 | 434 | 651 | 699 | 715 | 683 | 641 | 679 | 668 | 612 | 668,68 | 687 | 650 | 0,95 | -0,074 | -1,193 | 0,23281478 | 0,31334032 | 1,00E-08 | 0,0041505 | 0,0021991 | 0,0021991 | TRUE | 0,0153875 | No |
| orf19.1609 | 11491 | 8118 | 9979 | 12552 | 10109 | 9374 | 10243 | 6100 | 10309 | 10335 | 10209 | 10351 | 8646 | 8944 | 9140 | 8604 | 9567,2 | 10301 | 8834 | 0,858 | -0,221 | -7,167 | 7,68E-13 | 7,13E-12 | 0,000294 | 0,0021313 | 0,0008093 | 0,0008093 | TRUE | 0,0067226 | No |
| orf19.1610 | 70 | 58 | 76 | 63 | 116 | 97 | 109 | 55 | 63 | 74 | 78 | 52 | 99 | 93 | 97 | 78 | 79,12 | 67 | 92 | 1,401 | 0,486 | 2,845 | 0,00443429 | 0,00968066 | 0,006919 | 0,0203281 | 0,0150824 | 0,0150824 | TRUE | 0,220979 | No |
| orf19.1611 | 270 | 241 | 199 | 376 | 328 | 358 | 337 | 215 | 242 | 307 | 204 | 310 | 281 | 342 | 301 | 303 | 286,09 | 266 | 307 | 1,154 | 0,207 | 1,58 | 0,11411449 | 0,17042005 | 0,019317 | 0,0070537 | 0,0129525 | 0,0129525 | TRUE | 0,2727279 | No |
| orf19.1613 | 5643 | 4041 | 5148 | 6241 | 5363 | 4999 | 5488 | 3636 | 5062 | 5145 | 5267 | 5146 | 4587 | 4770 | 4897 | 5128 | 5000,3 | 5155 | 4846 | 0,939 | -0,091 | -2,206 | 0,02737681 | 0,04952722 | 0,000969 | 0,0022698 | 0,0014199 | 0,0014199 | TRUE | 0,0179683 | No |
| orf19.1614 | 12763 | 9478 | 11548 | 14484 | 13144 | 12209 | 13313 | 7459 | 11450 | 12067 | 11815 | 11944 | 11241 | 11649 | 11880 | 10520 | 11571 | 11819 | 11322 | 0,959 | -0,061 | -1,404 | 0,16017798 | 0,22799084 | 0,001527 | 0,002105 | 0,0017336 | 0,0017336 | TRUE | 0,0272674 | No |

| | | | | | | | | | | | | | | | | | | | | | | | | | | | | | | | |
|---|---|---|---|---|---|---|---|---|---|---|---|---|---|---|---|---|---|---|---|---|---|---|---|---|---|---|---|---|---|---|---|
| orf19.1616 | 1879 | 1346 | 1679 | 2046 | 2455 | 2233 | 2377 | 1584 | 1686 | 1714 | 1718 | 1687 | 2100 | 2131 | 2121 | 2234 | 1923,7 | 1701 | 2146 | 1,26 | 0,334 | 8,32 | 8,80E-17 | 1,22E-15 | 1,00E-08 | 0,0027342 | 0,0010179 | 0,0010179 | TRUE | 0,0081712 | No |
| orf19.1617 | 543 | 449 | 532 | 672 | 497 | 527 | 521 | 296 | 487 | 572 | 544 | 554 | 425 | 503 | 465 | 417 | 495,93 | 539 | 452 | 0,842 | -0,248 | -3,015 | 0,00257374 | 0,00593134 | 0,003968 | 0,0049067 | 0,0044362 | 0,0044362 | TRUE | 0,0643195 | No |
| orf19.1618 | 9997 | 7600 | 9111 | 11062 | 8029 | 8034 | 8233 | 4951 | 8968 | 9676 | 9321 | 9122 | 6867 | 7666 | 7347 | 6983 | 8243,7 | 9272 | 7216 | 0,779 | -0,361 | -8,01 | 1,15E-15 | 1,40E-14 | 0,001646 | 0,0021556 | 0,0018279 | 0,0018279 | TRUE | 0,0216064 | No |
| orf19.1618.1 | 495 | 360 | 415 | 524 | 528 | 543 | 515 | 343 | 444 | 458 | 425 | 432 | 452 | 518 | 460 | 484 | 459,01 | 440 | 478 | 1,087 | 0,121 | 1,658 | 0,09731458 | 0,14937813 | 1,00E-08 | 0,0051422 | 0,0029219 | 0,0029219 | TRUE | 0,0396031 | No |
| orf19.1619 | 4297 | 2996 | 3634 | 4686 | 4620 | 4301 | 4516 | 2801 | 3855 | 3814 | 3718 | 3864 | 3951 | 4104 | 4030 | 3951 | 3910,8 | 3813 | 4009 | 1,051 | 0,072 | 2,179 | 0,029324 | 0,05248097 | 1,00E-08 | 0,0023507 | 0,0008032 | 0,0008032 | TRUE | 0,0034469 | No |
| orf19.1620 | 14 | 6 | 16 | 12 | 7 | 13 | 13 | 6 | 13 | 8 | 16 | 10 | 6 | 12 | 12 | 8 | 10,61 | 12 | 10 | 0,824 | -0,279 | -0,639 | 0,52309417 | NA | 1,00E-08 | 0,1387443 | 0,0886808 | 0,0886808 | TRUE | 0,2856593 | No |
| orf19.1621 | 712 | 537 | 619 | 765 | 745 | 689 | 725 | 483 | 639 | 684 | 633 | 631 | 637 | 657 | 647 | 681 | 651,16 | 647 | 656 | 1,014 | 0,02 | 0,336 | 0,73692804 | 0,79401475 | 1,00E-08 | 0,0042089 | 0,0018966 | 0,0018966 | TRUE | 0,0154868 | No |
| orf19.1622 | 2856 | 2120 | 2560 | 3090 | 2920 | 2744 | 3050 | 1790 | 2562 | 2699 | 2619 | 2548 | 2497 | 2618 | 2722 | 2525 | 2598,8 | 2607 | 2590 | 0,995 | -0,007 | -0,168 | 0,86635054 | 0,898039 | 0,000739 | 0,0025381 | 0,0013953 | 0,0013953 | TRUE | 0,0144497 | No |
| orf19.1623 | 9804 | 6651 | 8493 | 10540 | 10715 | 9628 | 10333 | 6261 | 8795 | 8467 | 8689 | 8691 | 9164 | 9187 | 9220 | 8831 | 8880,6 | 8660 | 9100 | 1,051 | 0,072 | 2,444 | 0,01451981 | 0,02804985 | 0,000175 | 0,002143 | 0,0007136 | 0,0007136 | TRUE | 0,0046204 | No |
| orf19.1624 | 1233 | 1065 | 1244 | 1501 | 1069 | 1218 | 1133 | 703 | 1106 | 1356 | 1273 | 1238 | 914 | 1162 | 1011 | 992 | 1131,4 | 1243 | 1020 | 0,821 | -0,284 | -3,427 | 0,00061081 | 0,00161025 | 0,007478 | 0,0032626 | 0,0056997 | 0,0056997 | TRUE | 0,107611 | No |
| orf19.1624.1 | 25 | 16 | 33 | 21 | 28 | 37 | 25 | 25 | 22 | 20 | 34 | 17 | 24 | 35 | 22 | 35 | 26,34 | 23 | 29 | 1,234 | 0,303 | 1,007 | 0,31372711 | 0,40027023 | 0,036329 | 0,0570998 | 0,0483229 | 0,0483229 | TRUE | 0,4635468 | No |
| orf19.1625 | 2397 | 1581 | 1894 | 2524 | 2435 | 2137 | 2200 | 1443 | 2150 | 2013 | 1938 | 2081 | 2082 | 2039 | 1963 | 2035 | 2037,8 | 2046 | 2030 | 0,992 | -0,012 | -0,265 | 0,79080659 | 0,83690924 | 0,000862 | 0,0026919 | 0,0015615 | 0,0015615 | TRUE | 0,0155405 | No |
| orf19.1626 | 2653 | 1763 | 2206 | 2872 | 2469 | 2107 | 2412 | 1416 | 2380 | 2244 | 2257 | 2368 | 2112 | 2010 | 2152 | 1997 | 2190,2 | 2312 | 2068 | 0,895 | -0,161 | -3,625 | 0,00028912 | 0,00081859 | 0,000678 | 0,0026424 | 0,0014306 | 0,0014306 | TRUE | 0,0090797 | No |
| orf19.1628 | 1283 | 871 | 1141 | 1370 | 1150 | 1005 | 1053 | 631 | 1151 | 1109 | 1167 | 1130 | 984 | 959 | 940 | 890 | 1041,1 | 1139 | 943 | 0,83 | -0,27 | -5,252 | 1,51E-07 | 7,20E-07 | 1,00E-08 | 0,0033739 | 0,0015608 | 0,0015608 | TRUE | 0,0145709 | No |
| orf19.163 | 1973 | 1433 | 1741 | 2208 | 2298 | 2164 | 2302 | 1443 | 1770 | 1824 | 1781 | 1821 | 1965 | 2065 | 2054 | 2035 | 1914,5 | 1799 | 2030 | 1,128 | 0,174 | 4,358 | 1,31E-05 | 4,70E-05 | 1,00E-08 | 0,0027378 | 0,001007 | 0,001007 | TRUE | 0,0059707 | No |
| orf19.1630 | 2848 | 1907 | 2417 | 3063 | 3281 | 2978 | 3368 | 2246 | 2555 | 2428 | 2473 | 2526 | 2806 | 2842 | 3005 | 3168 | 2725,3 | 2496 | 2955 | 1,182 | 0,241 | 5,041 | 4,62E-07 | 2,06E-06 | 0,001345 | 0,0025122 | 0,0018272 | 0,0018272 | TRUE | 0,0268648 | No |
| orf19.1631 | 2828 | 2081 | 2461 | 3096 | 2288 | 2192 | 2248 | 1314 | 2537 | 2649 | 2518 | 2553 | 1957 | 2092 | 2006 | 1853 | 2270,6 | 2564 | 1977 | 0,773 | -0,372 | -8,127 | 4,41E-16 | 5,61E-15 | 0,000928 | 0,0026189 | 0,0015651 | 0,0015651 | TRUE | 0,019281 | No |
| orf19.1632 | 485 | 416 | 453 | 573 | 404 | 419 | 435 | 241 | 435 | 530 | 463 | 473 | 346 | 400 | 388 | 340 | 421,76 | 475 | 368 | 0,779 | -0,359 | -4,16 | 3,18E-05 | 0,00010662 | 0,004039 | 0,0054216 | 0,0047522 | 0,0047522 | TRUE | 0,0632692 | No |
| orf19.1633 | 10855 | 8724 | 10329 | 12382 | 10624 | 10652 | 11253 | 6640 | 9738 | 11107 | 10568 | 10210 | 9086 | 10164 | 10041 | 9365 | 10035 | 10406 | 9664 | 0,929 | -0,106 | -1,997 | 0,04582585 | 0,07790654 | 0,002864 | 0,0021242 | 0,0026139 | 0,0026139 | TRUE | 0,0249356 | No |
| orf19.1634 | 3055 | 2160 | 2656 | 3237 | 3381 | 3035 | 3201 | 1969 | 2741 | 2750 | 2717 | 2669 | 2892 | 2896 | 2856 | 2777 | 2787,3 | 2719 | 2855 | 1,051 | 0,072 | 2,061 | 0,0393283 | 0,06805427 | 1,00E-08 | 0,0025004 | 0,0008233 | 0,0008233 | TRUE | 0,0036657 | No |
| orf19.1635 | 19013 | 12374 | 15166 | 19051 | 17038 | 14160 | 14512 | 10374 | 17057 | 15753 | 15516 | 15710 | 14571 | 13511 | 12950 | 14632 | 14962 | 16009 | 13916 | 0,869 | -0,202 | -3,96 | 7,51E-05 | 0,00023436 | 0,002638 | 0,0020765 | 0,0024441 | 0,0024441 | TRUE | 0,0268016 | No |
| orf19.1636 | 1928 | 1217 | 1628 | 1973 | 1930 | 1569 | 1694 | 1150 | 1730 | 1549 | 1666 | 1627 | 1651 | 1497 | 1512 | 1622 | 1606,6 | 1643 | 1570 | 0,954 | -0,068 | -1,257 | 0,20867102 | 0,28519006 | 0,001433 | 0,0028831 | 0,0021448 | 0,0021448 | TRUE | 0,0165657 | No |
| orf19.1637 | 603 | 370 | 478 | 630 | 627 | 475 | 532 | 317 | 541 | 471 | 489 | 520 | 536 | 453 | 475 | 447 | 491,48 | 505 | 478 | 0,947 | -0,079 | -0,983 | 0,32556073 | 0,41256462 | 0,00344 | 0,0049333 | 0,0041769 | 0,0041769 | TRUE | 0,0819453 | No |
| orf19.1638 | 672 | 450 | 560 | 690 | 729 | 664 | 761 | 493 | 603 | 573 | 573 | 569 | 623 | 634 | 679 | 695 | 618,64 | 580 | 658 | 1,132 | 0,179 | 2,806 | 0,00501394 | 0,0108242 | 1,00E-08 | 0,0043261 | 0,0022732 | 0,0022732 | TRUE | 0,0148527 | No |
| orf19.1639 | 76 | 56 | 72 | 112 | 104 | 66 | 82 | 46 | 68 | 71 | 74 | 92 | 89 | 63 | 73 | 65 | 74,43 | 76 | 72 | 0,952 | -0,072 | -0,402 | 0,68766339 | 0,75076898 | 0,011665 | 0,021483 | 0,0171046 | 0,0171046 | TRUE | 0,2298952 | No |
| orf19.164 | 1313 | 1096 | 1185 | 1451 | 1170 | 1198 | 1195 | 734 | 1178 | 1395 | 1212 | 1197 | 1001 | 1143 | 1066 | 1035 | 1153,4 | 1246 | 1061 | 0,854 | -0,228 | -3,347 | 0,00081615 | 0,0020919 | 0,003754 | 0,0032381 | 0,0035625 | 0,0035625 | TRUE | 0,0749824 | No |
| orf19.1641 | 89 | 41 | 78 | 86 | 38 | 37 | 50 | 32 | 80 | 52 | 80 | 71 | 32 | 35 | 45 | 45 | 55,04 | 71 | 39 | 0,548 | -0,867 | -4,17 | 3,05E-05 | 0,00010266 | 0,011405 | 0,0283555 | 0,0217443 | 0,0217443 | TRUE | 0,2311695 | No |
| orf19.1642 | 8578 | 5894 | 6903 | 9010 | 7872 | 6899 | 7351 | 4468 | 7695 | 7504 | 7062 | 7430 | 6732 | 6583 | 6560 | 6302 | 6983,5 | 7423 | 6544 | 0,882 | -0,181 | -4,697 | 2,64E-06 | 1,05E-05 | 0,000819 | 0,0021874 | 0,001283 | 0,001283 | TRUE | 0,0130319 | No |
| orf19.1643 | 1990 | 1477 | 1837 | 2292 | 2767 | 2427 | 2667 | 1841 | 1785 | 1880 | 1879 | 1890 | 2366 | 2316 | 2380 | 2597 | 2136,7 | 1858 | 2415 | 1,297 | 0,375 | 7,842 | 4,44E-15 | 5,13E-14 | 0,001043 | 0,0026589 | 0,001722 | 0,001722 | TRUE | 0,0289433 | No |
| orf19.1646 | 12092 | 8132 | 10696 | 13220 | 12545 | 10706 | 12504 | 7437 | 10848 | 10353 | 10943 | 10901 | 10729 | 10215 | 11158 | 10489 | 10705 | 10761 | 10648 | 0,989 | -0,015 | -0,404 | 0,68630334 | 0,74942029 | 0,000817 | 0,0021151 | 0,0012941 | 0,0012941 | TRUE | 0,0128091 | No |
| orf19.1647 | 55 | 30 | 47 | 45 | 31 | 33 | 27 | 21 | 49 | 38 | 48 | 37 | 27 | 31 | 24 | 30 | 35,55 | 43 | 28 | 0,642 | -0,639 | -2,708 | 0,00676307 | 0,01419601 | 1,00E-08 | 0,0428095 | 0,0240489 | 0,0240489 | TRUE | 0,0878928 | No |
| orf19.1648 | 359 | 277 | 329 | 381 | 443 | 420 | 453 | 268 | 322 | 353 | 337 | 314 | 379 | 401 | 404 | 378 | 360,92 | 332 | 390 | 1,184 | 0,244 | 3,157 | 0,00159375 | 0,00383777 | 1,00E-08 | 0,0060018 | 0,002948 | 0,002948 | TRUE | 0,0189179 | No |
| orf19.1649 | 3387 | 2357 | 2873 | 3699 | 3494 | 3137 | 3161 | 2089 | 3038 | 3001 | 2939 | 3050 | 2988 | 2993 | 2821 | 2946 | 2972,2 | 3007 | 2937 | 0,976 | -0,035 | -0,929 | 0,35263323 | 0,44020389 | 0,000233 | 0,002468 | 0,0009965 | 0,0009965 | TRUE | 0,0090846 | No |
| orf19.1650 | 1139 | 701 | 890 | 1074 | 995 | 738 | 710 | 482 | 1022 | 892 | 911 | 886 | 851 | 704 | 634 | 680 | 822,37 | 928 | 717 | 0,774 | -0,37 | -4,066 | 4,79E-05 | 0,00015543 | 0,00928 | 0,0037448 | 0,0067007 | 0,0067007 | TRUE | 0,1934801 | No |
| orf19.1651 | 26 | 12 | 20 | 25 | 14 | 22 | 16 | 9 | 23 | 15 | 20 | 21 | 12 | 21 | 14 | 13 | 17,45 | 20 | 15 | 0,748 | -0,418 | -1,251 | 0,21101556 | 0,28813215 | 1,00E-08 | 0,0851625 | 0,0496264 | 0,0496264 | TRUE | 0,3507571 | No |
| orf19.1652 | 15913 | 10561 | 14015 | 16933 | 11107 | 9726 | 10252 | 7484 | 14276 | 13445 | 14339 | 13963 | 9499 | 9280 | 9148 | 10556 | 11813 | 14006 | 9621 | 0,686 | -0,543 | -10,884 | 1,38E-27 | 4,42E-26 | 0,002519 | 0,0021024 | 0,0022998 | 0,0022998 | TRUE | 0,0514426 | No |
| orf19.1653 | 2089 | 1728 | 1796 | 2448 | 2277 | 2200 | 2223 | 1227 | 1874 | 2200 | 1837 | 2019 | 1947 | 2099 | 1984 | 1731 | 1961,4 | 1982 | 1940 | 0,981 | -0,028 | -0,391 | 0,6960121 | 0,75867659 | 0,005918 | 0,0027197 | 0,0044989 | 0,0044989 | TRUE | 0,0638368 | No |
| orf19.1654 | 2105 | 1452 | 1712 | 2252 | 1520 | 1294 | 1379 | 1024 | 1888 | 1849 | 1752 | 1857 | 1300 | 1235 | 1231 | 1444 | 1569,4 | 1836 | 1302 | 0,707 | -0,5 | -8,614 | 7,06E-18 | 1,09E-16 | 0,002579 | 0,0029045 | 0,0025759 | 0,0025759 | TRUE | 0,0590964 | No |
| orf19.1655 | 5852 | 3731 | 5021 | 6147 | 2404 | 2002 | 2213 | 1558 | 5250 | 4750 | 5137 | 5069 | 2056 | 1910 | 1975 | 2197 | 3543 | 5052 | 2034 | 0,402 | -1,315 | -25,179 | 6,74E-140 | 1,35E-136 | 0,002294 | 0,0023893 | 0,00227 | 0,00227 | TRUE | 0,0326763 | No |
| orf19.1655.3 | 3388 | 2519 | 2947 | 3885 | 3569 | 3178 | 3256 | 2046 | 3039 | 3207 | 3015 | 3204 | 3052 | 3032 | 2905 | 2886 | 3042,6 | 3116 | 2969 | 0,954 | -0,069 | -1,675 | 0,09397783 | 0,14544534 | 0,000526 | 0,0024566 | 0,0012777 | 0,0012777 | TRUE | 0,0056893 | No |
| orf19.1656 | 1178 | 912 | 1126 | 1351 | 1069 | 1042 | 1055 | 714 | 1057 | 1161 | 1152 | 1114 | 914 | 994 | 941 | 1007 | 1042,6 | 1121 | 964 | 0,859 | -0,219 | -3,898 | 9,68E-05 | 0,00029719 | 0,000977 | 0,0033719 | 0,0020482 | 0,0020482 | TRUE | 0,0180141 | No |
| orf19.1658 | 12093 | 7808 | 10356 | 12868 | 11833 | 10175 | 11442 | 7319 | 10849 | 9940 | 10595 | 10611 | 10120 | 9709 | 10210 | 10323 | 10295 | 10499 | 10090 | 0,961 | -0,058 | -1,517 | 0,12915034 | 0,18992454 | 0,000912 | 0,0021205 | 0,0013035 | 0,0013035 | TRUE | 0,0153824 | No |
| orf19.1659 | 668 | 622 | 656 | 902 | 661 | 751 | 795 | 477 | 599 | 792 | 671 | 744 | 565 | 717 | 709 | 673 | 683,77 | 702 | 666 | 0,95 | -0,074 | -0,738 | 0,46028513 | 0,54598087 | 0,011841 | 0,0041026 | 0,0081227 | 0,0081227 | TRUE | 0,1276863 | No |
| orf19.166 | 2269 | 1934 | 2244 | 2673 | 2737 | 3004 | 3160 | 1966 | 2036 | 2462 | 2296 | 2204 | 2341 | 2866 | 2820 | 2773 | 2474,7 | 2250 | 2700 | 1,201 | 0,264 | 3,49 | 0,0004829 | 0,00131041 | 0,006625 | 0,0025661 | 0,0050831 | 0,0050831 | TRUE | 0,0989457 | No |
| orf19.1661 | 4864 | 3355 | 4267 | 5250 | 5015 | 4533 | 4883 | 2904 | 4364 | 4271 | 4366 | 4329 | 4289 | 4325 | 4357 | 4096 | 4299,6 | 4332 | 4267 | 0,986 | -0,021 | -0,623 | 0,53345085 | 0,6160259 | 0,00012 | 0,0023172 | 0,0008407 | 0,0008407 | TRUE | 0,0081309 | No |
| orf19.1662 | 2711 | 2119 | 2400 | 2970 | 3147 | 2860 | 3085 | 1904 | 2432 | 2698 | 2455 | 2449 | 2691 | 2729 | 2753 | 2685 | 2611,6 | 2508 | 2714 | 1,084 | 0,116 | 2,654 | 0,00794341 | 0,0164 | 0,000829 | 0,0025354 | 0,0014574 | 0,0014574 | TRUE | 0,0295455 | No |
| orf19.1663 | 2034 | 1643 | 1913 | 2233 | 2194 | 2211 | 2411 | 1324 | 1825 | 2092 | 1957 | 1841 | 1876 | 2110 | 2151 | 1867 | 1965 | 1929 | 2001 | 1,04 | 0,056 | 0,855 | 0,39277491 | 0,48158455 | 0,004076 | 0,0027183 | 0,0036214 | 0,0036214 | TRUE | 0,0380059 | No |
| orf19.1664 | 292 | 225 | 238 | 303 | 363 | 326 | 298 | 238 | 262 | 286 | 243 | 250 | 310 | 311 | 266 | 336 | 283,11 | 260 | 306 | 1,173 | 0,23 | 2,374 | 0,01760129 | 0,03317082 | 0,003432 | 0,0071073 | 0,0054234 | 0,0054234 | TRUE | 0,0894969 | No |
| orf19.1665 | 10043 | 7458 | 9224 | 11135 | 9438 | 8913 | 9468 | 6098 | 9010 | 9495 | 9437 | 9182 | 8072 | 8505 | 8449 | 8601 | 8843,6 | 9281 | 8407 | 0,906 | -0,143 | -4,147 | 3,37E-05 | 0,0001124 | 0,000526 | 0,0021437 | 0,0010264 | 0,0010264 | TRUE | 0,0088892 | No |
| orf19.1666 | 1280 | 769 | 1119 | 1388 | 1296 | 1056 | 1256 | 859 | 1148 | 979 | 1145 | 1145 | 1108 | 1008 | 1121 | 1212 | 1108,1 | 1104 | 1112 | 1,004 | 0,005 | 0,072 | 0,94287617 | 0,95883548 | 0,004626 | 0,0032896 | 0,0040073 | 0,0040073 | TRUE | 0,0658177 | No |
| orf19.1667 | 813 | 586 | 777 | 933 | 1016 | 989 | 1066 | 598 | 729 | 746 | 795 | 769 | 869 | 944 | 951 | 843 | 830,87 | 760 | 902 | 1,189 | 0,25 | 4,062 | 4,87E-05 | 0,00015783 | 0,001273 | 0,0037267 | 0,0024177 | 0,0024177 | TRUE | 0,0196717 | No |
| orf19.1667.1 | 484 | 350 | 390 | 513 | 599 | 476 | 474 | 244 | 434 | 446 | 399 | 423 | 512 | 454 | 423 | 344 | 429,42 | 426 | 433 | 1,025 | 0,036 | 0,337 | 0,73592065 | 0,79364768 | 0,011507 | 0,0053601 | 0,0083443 | 0,0083443 | TRUE | 0,1993647 | No |
| orf19.1668 | 627 | 443 | 591 | 706 | 849 | 814 | 861 | 534 | 562 | 564 | 605 | 582 | 726 | 777 | 768 | 753 | 667,19 | 578 | 756 | 1,307 | 0,386 | 6,474 | 9,56E-11 | 7,00E-10 | 1,00E-08 | 0,0041554 | 0,0018886 | 0,0018886 | TRUE | 0,0105603 | No |
| orf19.1669 | 5172 | 3578 | 4528 | 5554 | 6266 | 5727 | 6336 | 3908 | 4640 | 4555 | 4633 | 4580 | 5359 | 5465 | 5654 | 5512 | 5049,6 | 4602 | 5498 | 1,194 | 0,256 | 8,114 | 4,89E-16 | 6,18E-15 | 0,000149 | 0,002267 | 0,0007566 | 0,0007566 | TRUE | 0,0047086 | No |
| orf19.1670 | 946 | 759 | 941 | 1104 | 1433 | 1482 | 1608 | 860 | 849 | 966 | 963 | 910 | 1226 | 1414 | 1435 | 1213 | 1121,9 | 922 | 1322 | 1,437 | 0,523 | 7,121 | 1,07E-12 | 9,82E-12 | 0,005055 | 0,0032734 | 0,004262 | 0,004262 | TRUE | 0,0399388 | No |
| orf19.1671 | 8071 | 5983 | 7482 | 9093 | 8775 | 8378 | 8615 | 5593 | 7241 | 7617 | 7655 | 7498 | 7505 | 7994 | 7687 | 7888 | 7635,6 | 7503 | 7768 | 1,035 | 0,05 | 1,428 | 0,15334688 | 0,21977769 | 0,000527 | 0,0021696 | 0,0010538 | 0,0010538 | TRUE | 0,0068066 | No |
| orf19.1671.1 | 7 | 6 | 7 | 10 | 10 | 7 | 4 | 2 | 6 | 8 | 7 | 8 | 9 | 7 | 4 | 3 | 6,37 | 7 | 6 | 0,761 | -0,393 | -0,722 | 0,4704201 | NA | 1,00E-08 | 0,2299305 | 0,12806 | 0,12806 | TRUE | 0,3800207 | No |
| orf19.1672 | 3161 | 2618 | 3014 | 3492 | 3731 | 3582 | 3875 | 2278 | 2836 | 3333 | 3084 | 2880 | 3191 | 3418 | 3458 | 3213 | 3176,4 | 3033 | 3320 | 1,096 | 0,132 | 2,279 | 0,02265467 | 0,04176413 | 0,003245 | 0,0024365 | 0,002922 | 0,002922 | TRUE | 0,0526689 | No |
| orf19.1673 | 2577 | 2188 | 2321 | 2886 | 2705 | 2787 | 2733 | 1535 | 2312 | 2786 | 2375 | 2380 | 2313 | 2659 | 2439 | 2165 | 2428,5 | 2463 | 2394 | 0,974 | -0,039 | -0,513 | 0,60827628 | 0,68321628 | 0,006422 | 0,0025773 | 0,0050135 | 0,0050135 | TRUE | 0,0923882 | No |

| | | | | | | | | | | | | | | | | | | | | | | | | | | | | | | | |
|---|---|---|---|---|---|---|---|---|---|---|---|---|---|---|---|---|---|---|---|---|---|---|---|---|---|---|---|---|---|---|---|
| orf19.1674 | 2244 | 1831 | 2067 | 2621 | 2799 | 2830 | 2917 | 1787 | 2013 | 2331 | 2115 | 2161 | 2394 | 2700 | 2603 | 2520 | 2354,7 | 2155 | 2554 | 1,187 | 0,247 | 4,344 | 1,40E-05 | 4,97E-05 | 0,002616 | 0,002596 | 0,002667 | 0,002667 | TRUE | 0,0351295 | No |
| orf19.1675 | 1957 | 1319 | 1673 | 2007 | 1765 | 1510 | 1707 | 989 | 1756 | 1679 | 1712 | 1655 | 1509 | 1441 | 1523 | 1395 | 1583,7 | 1700 | 1467 | 0,864 | -0,21 | -4,478 | 7,54E-06 | 2,80E-05 | 0,000467 | 0,0028962 | 0,001478 | 0,001478 | TRUE | 0,0117361 | No |
| orf19.1676 | 1444 | 1171 | 1336 | 1648 | 1555 | 1546 | 1587 | 953 | 1295 | 1491 | 1367 | 1359 | 1330 | 1475 | 1416 | 1344 | 1384,7 | 1378 | 1391 | 1,012 | 0,017 | 0,296 | 0,76691321 | 0,81840401 | 0,002095 | 0,0030279 | 0,0024951 | 0,0024951 | TRUE | 0,0344357 | No |
| orf19.1677 | 2217 | 1649 | 2158 | 2388 | 2522 | 2048 | 2404 | 1529 | 1989 | 2099 | 2208 | 1969 | 2157 | 1954 | 2145 | 2157 | 2084,8 | 2066 | 2103 | 1,018 | 0,026 | 0,483 | 0,62887393 | 0,70045065 | 0,002121 | 0,0026759 | 0,0023407 | 0,0023407 | TRUE | 0,0278858 | No |
| orf19.1678 | 1112 | 806 | 1000 | 1190 | 1147 | 972 | 1070 | 713 | 998 | 1026 | 1023 | 981 | 981 | 927 | 955 | 1006 | 987,11 | 1007 | 967 | 0,96 | -0,059 | -1,152 | 0,24940621 | 0,33152996 | 1,00E-08 | 0,0034502 | 0,0014951 | 0,0014951 | TRUE | 0,008838 | No |
| orf19.1679 | 44 | 45 | 52 | 65 | 64 | 39 | 51 | 37 | 39 | 57 | 53 | 54 | 55 | 37 | 46 | 52 | 49,15 | 51 | 48 | 0,934 | -0,099 | -0,471 | 0,6373714 | 0,70808568 | 1,00E-08 | 0,0315153 | 0,021831 | 0,021831 | TRUE | 0,2016712 | No |
| orf19.168 | 6211 | 4158 | 5060 | 6645 | 6503 | 5851 | 6045 | 3561 | 5572 | 5294 | 5177 | 5480 | 5562 | 5583 | 5394 | 5022 | 5385,4 | 5381 | 5390 | 1,002 | 0,003 | 0,074 | 0,94066527 | 0,95726727 | 0,001276 | 0,0022491 | 0,0017581 | 0,0017581 | TRUE | 0,0247361 | No |
| orf19.1680 | 1873 | 1258 | 1706 | 1924 | 2017 | 1821 | 1929 | 1198 | 1680 | 1602 | 1745 | 1587 | 1725 | 1738 | 1721 | 1690 | 1685,9 | 1654 | 1718 | 1,04 | 0,056 | 1,227 | 0,21973575 | 0,29854645 | 0,000451 | 0,0028406 | 0,0014228 | 0,0014228 | TRUE | 0,0166546 | No |
| orf19.1681 | 293 | 186 | 254 | 282 | 241 | 264 | 267 | 191 | 263 | 237 | 260 | 233 | 206 | 252 | 238 | 269 | 244,71 | 248 | 241 | 0,967 | -0,049 | -0,471 | 0,63780927 | 0,7082894 | 0,00418 | 0,0079118 | 0,0061986 | 0,0061986 | TRUE | 0,112693 | No |
| orf19.1682 | 5884 | 3799 | 4910 | 6196 | 5701 | 4614 | 5189 | 3426 | 5279 | 4837 | 5023 | 5109 | 4876 | 4403 | 4630 | 4832 | 4873,6 | 5062 | 4685 | 0,925 | -0,113 | -2,47 | 0,01349627 | 0,0262753 | 0,001455 | 0,0022774 | 0,00179 | 0,00179 | TRUE | 0,0201842 | No |
| orf19.1683 | 1938 | 1257 | 1716 | 2021 | 1890 | 1694 | 1851 | 1167 | 1739 | 1600 | 1756 | 1667 | 1616 | 1616 | 1652 | 1646 | 1661,4 | 1690 | 1632 | 0,965 | -0,052 | -1,152 | 0,24942001 | 0,33152996 | 0,00026 | 0,0028533 | 0,0013473 | 0,0013473 | TRUE | 0,0140102 | No |
| orf19.1684 | 452 | 351 | 458 | 563 | 701 | 669 | 754 | 486 | 405 | 447 | 469 | 464 | 600 | 638 | 673 | 685 | 547,67 | 446 | 649 | 1,452 | 0,538 | 7,242 | 4,44E-13 | 4,27E-12 | 0,002209 | 0,0046302 | 0,0033993 | 0,0033993 | TRUE | 0,0468902 | No |
| orf19.1685 | 343 | 291 | 313 | 418 | 433 | 383 | 435 | 301 | 308 | 370 | 320 | 345 | 370 | 365 | 388 | 425 | 361,44 | 336 | 387 | 1,152 | 0,204 | 2,368 | 0,01787278 | 0,03361917 | 0,002106 | 0,0059959 | 0,0043032 | 0,0043032 | TRUE | 0,0482635 | No |
| orf19.1686 | 677 | 412 | 592 | 709 | 680 | 586 | 735 | 387 | 607 | 525 | 606 | 585 | 582 | 559 | 656 | 546 | 583,07 | 581 | 586 | 1,009 | 0,013 | 0,168 | 0,86681886 | 0,89822733 | 0,003786 | 0,0044693 | 0,0041224 | 0,0041224 | TRUE | 0,0780225 | No |
| orf19.1687 | 15381 | 10654 | 13527 | 16755 | 13647 | 12223 | 13389 | 8458 | 13798 | 13564 | 13839 | 13816 | 11671 | 11663 | 11947 | 11929 | 12779 | 13754 | 11802 | 0,858 | -0,221 | -8,849 | 8,81E-19 | 1,45E-17 | 2,68E-05 | 0,0020931 | 0,000521 | 0,000521 | TRUE | 0,0010309 | No |
| orf19.169 | 1649 | 1554 | 1556 | 2068 | 1686 | 1865 | 1977 | 1049 | 1479 | 1978 | 1592 | 1705 | 1442 | 1780 | 1764 | 1480 | 1652,5 | 1688 | 1616 | 0,959 | -0,061 | -0,633 | 0,52645833 | 0,60955129 | 0,012229 | 0,002858 | 0,0083346 | 0,0083346 | TRUE | 0,157737 | No |
| orf19.1690 | 2921 | 1981 | 2598 | 3155 | 3664 | 3064 | 3341 | 2157 | 2620 | 2522 | 2658 | 2602 | 3134 | 2924 | 2981 | 3042 | 2810,4 | 2600 | 3020 | 1,161 | 0,215 | 5,502 | 3,77E-08 | 1,94E-07 | 0,000359 | 0,0024961 | 0,0011085 | 0,0011085 | TRUE | 0,0082887 | No |
| orf19.1691 | 290 | 203 | 224 | 294 | 406 | 373 | 376 | 268 | 260 | 258 | 229 | 242 | 347 | 356 | 336 | 378 | 300,86 | 247 | 354 | 1,427 | 0,513 | 5,995 | 2,04E-09 | 1,24E-08 | 1,00E-08 | 0,0068048 | 0,003604 | 0,003604 | TRUE | 0,026319 | No |
| orf19.1693 | 774 | 544 | 694 | 783 | 815 | 737 | 821 | 488 | 694 | 693 | 710 | 646 | 697 | 703 | 733 | 688 | 695,47 | 686 | 705 | 1,031 | 0,044 | 0,746 | 0,45580951 | 0,5415273 | 1,00E-08 | 0,0040669 | 0,0019591 | 0,0019591 | TRUE | 0,0199078 | No |
| orf19.1697 | 3086 | 2064 | 2528 | 3198 | 3724 | 3316 | 3435 | 2043 | 2768 | 2628 | 2586 | 2637 | 3185 | 3164 | 3065 | 2881 | 2864,4 | 2655 | 3074 | 1,159 | 0,213 | 4,732 | 2,22E-06 | 8,97E-06 | 0,000988 | 0,0024863 | 0,0015898 | 0,0015898 | TRUE | 0,0201383 | No |
| orf19.1698 | 720 | 586 | 703 | 832 | 764 | 671 | 776 | 508 | 646 | 746 | 719 | 686 | 653 | 640 | 692 | 716 | 687,48 | 699 | 675 | 0,966 | -0,049 | -0,738 | 0,46047067 | 0,54609313 | 0,001674 | 0,0040911 | 0,0028508 | 0,0028508 | TRUE | 0,0312771 | No |
| orf19.1699 | 696 | 505 | 618 | 811 | 870 | 882 | 891 | 563 | 624 | 643 | 632 | 669 | 744 | 842 | 795 | 794 | 717,89 | 642 | 794 | 1,235 | 0,304 | 4,941 | 7,78E-07 | 3,34E-06 | 1,00E-08 | 0,0040017 | 0,0022321 | 0,0022321 | TRUE | 0,0229127 | No |
| orf19.17 | 694 | 724 | 706 | 1008 | 738 | 850 | 827 | 469 | 623 | 922 | 722 | 831 | 631 | 811 | 738 | 661 | 742,44 | 774 | 710 | 0,919 | -0,122 | -1,044 | 0,29630493 | 0,38136247 | 0,018747 | 0,0039348 | 0,0116754 | 0,0116754 | TRUE | 0,2088493 | No |
| orf19.1700 | 10997 | 8408 | 9594 | 11924 | 10363 | 9784 | 9502 | 6754 | 9865 | 10704 | 9816 | 9833 | 8863 | 9336 | 8479 | 9526 | 9552,7 | 10054 | 9051 | 0,9 | -0,152 | -3,139 | 0,0016959 | 0,00406097 | 0,002152 | 0,0021315 | 0,0021433 | 0,0021433 | TRUE | 0,0228651 | No |
| orf19.1701 | 1485 | 1089 | 1275 | 1608 | 1628 | 1564 | 1584 | 1105 | 1332 | 1386 | 1304 | 1326 | 1392 | 1492 | 1413 | 1559 | 1400,7 | 1337 | 1464 | 1,093 | 0,128 | 2,516 | 0,01185396 | 0,02341946 | 0,000741 | 0,0030159 | 0,0017851 | 0,0017851 | TRUE | 0,0205678 | No |
| orf19.1702 | 1235 | 887 | 1031 | 1362 | 1072 | 969 | 942 | 687 | 1108 | 1129 | 1055 | 1123 | 917 | 925 | 841 | 969 | 1008,3 | 1104 | 913 | 0,825 | -0,278 | -4,857 | 1,19E-06 | 4,97E-06 | 0,001015 | 0,0034193 | 0,002134 | 0,002134 | TRUE | 0,0342212 | No |
| orf19.1704 | 9692 | 6747 | 8362 | 10334 | 6422 | 5352 | 5903 | 3504 | 8695 | 8590 | 8555 | 8522 | 5492 | 5107 | 5267 | 4942 | 6896,2 | 8590 | 5202 | 0,606 | -0,722 | -18,824 | 4,83E-79 | 1,32E-76 | 0,00085 | 0,00219 | 0,0012582 | 0,0012582 | TRUE | 0,0172395 | No |
| orf19.1706 | 357 | 391 | 348 | 386 | 324 | 414 | 398 | 245 | 320 | 498 | 356 | 318 | 277 | 395 | 355 | 346 | 358,15 | 373 | 343 | 0,923 | -0,116 | -0,781 | 0,43478678 | 0,52191787 | 0,031267 | 0,0060328 | 0,0181871 | 0,0181871 | TRUE | 0,5579049 | No |
| orf19.1707 | 12 | 39 | 25 | 29 | 20 | 48 | 34 | 12 | 11 | 50 | 26 | 24 | 17 | 46 | 30 | 17 | 27,51 | 28 | 28 | 1,024 | 0,035 | 0,079 | 0,93687222 | 0,95418347 | 0,249843 | 0,0547503 | 0,1456788 | 0,1456788 | TRUE | 0,562339 | No |
| orf19.1708 | 3003 | 2564 | 2720 | 3702 | 3571 | 3682 | 3816 | 2282 | 2694 | 3264 | 2783 | 3053 | 3054 | 3513 | 3405 | 3219 | 3123,1 | 2948 | 3298 | 1,119 | 0,162 | 2,379 | 0,01733673 | 0,03278535 | 0,005354 | 0,0024444 | 0,0041546 | 0,0041546 | TRUE | 0,0616875 | No |
| orf19.1709 | 2797 | 1790 | 2152 | 2726 | 1862 | 1714 | 1679 | 921 | 2509 | 2279 | 2202 | 2248 | 1592 | 1635 | 1498 | 1299 | 1907,9 | 2310 | 1506 | 0,653 | -0,614 | -8,46 | 2,67E-17 | 3,87E-16 | 0,005958 | 0,0027404 | 0,0045023 | 0,0045023 | TRUE | 0,0974377 | No |
| orf19.171 | 53864 | 37492 | 49232 | 60597 | 49630 | 46105 | 49665 | 30828 | 48322 | 47731 | 50369 | 49969 | 42445 | 43992 | 44318 | 43480 | 46328 | 49098 | 43559 | 0,887 | -0,173 | -5,682 | 1,33E-08 | 7,25E-08 | 0,000483 | 0,0020109 | 0,0008665 | 0,0008665 | TRUE | 0,0042712 | No |
| orf19.1710 | 1763 | 1489 | 1604 | 2120 | 1819 | 1910 | 1883 | 1190 | 1582 | 1896 | 1641 | 1748 | 1556 | 1822 | 1680 | 1678 | 1700,4 | 1717 | 1684 | 0,982 | -0,026 | -0,386 | 0,69935145 | 0,76145406 | 0,00466 | 0,0028333 | 0,0038735 | 0,0038735 | TRUE | 0,0572219 | No |
| orf19.1711 | 1208 | 1065 | 1202 | 1476 | 1458 | 1620 | 1653 | 803 | 1084 | 1356 | 1230 | 1217 | 1247 | 1546 | 1475 | 1133 | 1285,8 | 1222 | 1350 | 1,108 | 0,148 | 1,498 | 0,13424596 | 0,19648444 | 0,013335 | 0,0031085 | 0,0085409 | 0,0085409 | TRUE | 0,1356123 | No |
| orf19.1712 | 839 | 783 | 804 | 1044 | 929 | 1013 | 967 | 658 | 753 | 997 | 823 | 861 | 795 | 967 | 863 | 928 | 873,12 | 858 | 888 | 1,036 | 0,05 | 0,558 | 0,57665876 | 0,6548036 | 0,009397 | 0,0036422 | 0,006669 | 0,006669 | TRUE | 0,1346051 | No |
| orf19.1714 | 112 | 86 | 105 | 102 | 96 | 125 | 114 | 111 | 100 | 109 | 107 | 84 | 82 | 119 | 102 | 157 | 107,64 | 100 | 115 | 1,135 | 0,183 | 0,976 | 0,32904753 | 0,4164562 | 0,03422 | 0,0154656 | 0,0241741 | 0,0241741 | TRUE | 0,3680556 | No |
| orf19.1715 | 680 | 458 | 634 | 708 | 740 | 715 | 814 | 515 | 610 | 583 | 649 | 584 | 633 | 682 | 726 | 726 | 649,18 | 606 | 692 | 1,139 | 0,187 | 2,707 | 0,00679437 | 0,01423808 | 0,001976 | 0,0042157 | 0,0030403 | 0,0030403 | TRUE | 0,0409801 | No |
| orf19.1716 | 2358 | 1516 | 1906 | 2450 | 2608 | 2200 | 2285 | 1392 | 2115 | 1930 | 1950 | 2020 | 2230 | 2099 | 2039 | 1963 | 2043,5 | 2004 | 2083 | 1,04 | 0,057 | 1,078 | 0,28119715 | 0,3651248 | 0,001691 | 0,0026899 | 0,0021601 | 0,0021601 | TRUE | 0,0277922 | No |
| orf19.1717 | 122 | 110 | 139 | 179 | 166 | 177 | 184 | 134 | 109 | 140 | 142 | 148 | 142 | 169 | 164 | 189 | 150,42 | 135 | 166 | 1,225 | 0,293 | 2,185 | 0,02888031 | 0,05181036 | 0,009034 | 0,0116306 | 0,010468 | 0,010468 | TRUE | 0,1786236 | No |
| orf19.1718 | 778 | 726 | 813 | 988 | 898 | 982 | 1004 | 584 | 698 | 924 | 832 | 815 | 768 | 937 | 896 | 824 | 836,66 | 817 | 856 | 1,05 | 0,071 | 0,78 | 0,43547879 | 0,52243541 | 0,009281 | 0,0037146 | 0,0066484 | 0,0066484 | TRUE | 0,115548 | No |
| orf19.1719 | 2889 | 2349 | 2683 | 3375 | 2259 | 2591 | 2404 | 1616 | 2592 | 2991 | 2745 | 2783 | 1932 | 2472 | 2145 | 2279 | 2492,4 | 2778 | 2207 | 0,795 | -0,332 | -4,472 | 7,75E-06 | 2,88E-05 | 0,006512 | 0,002562 | 0,004879 | 0,004879 | TRUE | 0,0868664 | No |
| orf19.172 | 4919 | 3896 | 4339 | 5426 | 4746 | 4901 | 4703 | 3027 | 4413 | 4960 | 4439 | 4474 | 4059 | 4676 | 4197 | 4269 | 4436 | 4572 | 4300 | 0,941 | -0,088 | -1,549 | 0,12145362 | 0,18026022 | 0,003224 | 0,0023068 | 0,0028462 | 0,0028462 | TRUE | 0,0427066 | No |
| orf19.1720 | 4760 | 3956 | 4543 | 5743 | 4388 | 4671 | 4879 | 2981 | 4270 | 5036 | 4648 | 4736 | 3753 | 4457 | 4354 | 4204 | 4432,3 | 4672 | 4192 | 0,897 | -0,156 | -2,395 | 0,01660708 | 0,03155451 | 0,004725 | 0,0023071 | 0,0038606 | 0,0038606 | TRUE | 0,0615443 | No |
| orf19.1721 | 24123 | 19277 | 21562 | 26774 | 21670 | 22389 | 22725 | 13009 | 21641 | 24542 | 22060 | 22078 | 18533 | 21363 | 20278 | 18348 | 21105 | 22580 | 19630 | 0,87 | -0,202 | -3,361 | 0,00077717 | 0,00200138 | 0,004365 | 0,0020483 | 0,0034118 | 0,0034118 | TRUE | 0,043188 | No |
| orf19.1722 | 55 | 41 | 56 | 79 | 72 | 48 | 54 | 55 | 49 | 52 | 57 | 65 | 62 | 46 | 48 | 78 | 57,14 | 56 | 58 | 1,022 | 0,031 | 0,152 | 0,87946259 | 0,90849575 | 0,018326 | 0,027386 | 0,0236151 | 0,0236151 | TRUE | 0,3290954 | No |
| orf19.1723 | 1567 | 1042 | 1287 | 1695 | 1457 | 1343 | 1434 | 965 | 1406 | 1327 | 1317 | 1398 | 1246 | 1281 | 1280 | 1361 | 1326,9 | 1362 | 1292 | 0,946 | -0,081 | -1,633 | 0,10238896 | 0,15569548 | 0,000433 | 0,0030736 | 0,0015845 | 0,0015845 | TRUE | 0,0137624 | No |
| orf19.1724 | 4 | 1 | 1 | 5 | 0 | 1 | 1 | 1 | 4 | 1 | 1 | 4 | 0 | 1 | 1 | 1 | 1,66 | 2 | 1 | 0,294 | -1,765 | -1,479 | 0,13919345 | NA | 1,00E-08 | 0,8775136 | 0,5203984 | 0,5203984 | TRUE | 0,162504 | No |
| orf19.1725 | 1071 | 650 | 845 | 1029 | 1347 | 1075 | 1355 | 965 | 961 | 828 | 865 | 849 | 1152 | 1026 | 1209 | 1361 | 1031,2 | 876 | 1187 | 1,352 | 0,435 | 5,146 | 2,66E-07 | 1,23E-06 | 0,007807 | 0,0033874 | 0,0058723 | 0,0058723 | TRUE | 0,1108123 | No |
| orf19.1727 | 15912 | 11842 | 14909 | 17721 | 14135 | 13832 | 14788 | 9072 | 14275 | 15076 | 15253 | 14613 | 12089 | 13198 | 13196 | 12795 | 13812 | 14804 | 12820 | 0,866 | -0,208 | -5,128 | 2,93E-07 | 1,34E-06 | 0,001185 | 0,0020846 | 0,0015002 | 0,0015002 | TRUE | 0,018256 | No |
| orf19.1728 | 685 | 535 | 595 | 730 | 509 | 500 | 543 | 328 | 615 | 681 | 609 | 602 | 435 | 477 | 485 | 463 | 545,74 | 627 | 465 | 0,744 | -0,427 | -6,104 | 1,04E-09 | 6,58E-09 | 0,00095 | 0,0046396 | 0,0028222 | 0,0028222 | TRUE | 0,0366241 | No |
| orf19.1729 | 911 | 875 | 843 | 1136 | 1054 | 1274 | 1251 | 700 | 817 | 1114 | 862 | 937 | 901 | 1216 | 1116 | 987 | 993,89 | 932 | 1055 | 1,133 | 0,181 | 1,666 | 0,09572613 | 0,14746734 | 0,017093 | 0,0034402 | 0,0102888 | 0,0102888 | TRUE | 0,1983251 | No |
| orf19.173 | 18751 | 13466 | 17175 | 20812 | 24307 | 22932 | 25099 | 15373 | 16822 | 17144 | 17572 | 17162 | 20788 | 21881 | 22397 | 21682 | 19431 | 17175 | 21687 | 1,263 | 0,337 | 10,209 | 1,81E-24 | 4,47E-23 | 0,000569 | 0,0020542 | 0,0009916 | 0,0009916 | TRUE | 0,0096469 | No |
| orf19.1730 | 869 | 570 | 798 | 942 | 1097 | 920 | 1021 | 584 | 780 | 726 | 816 | 777 | 938 | 878 | 911 | 824 | 831,16 | 775 | 888 | 1,148 | 0,199 | 3,232 | 0,00122893 | 0,00304472 | 0,00109 | 0,0037261 | 0,0024278 | 0,0024278 | TRUE | 0,0247444 | No |
| orf19.1734 | 2301 | 1825 | 2221 | 2653 | 2399 | 2498 | 2449 | 1590 | 2064 | 2323 | 2272 | 2188 | 2052 | 2384 | 2185 | 2243 | 2213,8 | 2212 | 2216 | 1,002 | 0,003 | 0,051 | 0,95893877 | 0,97028049 | 0,002848 | 0,0026353 | 0,0027628 | 0,0027628 | TRUE | 0,0324386 | No |
| orf19.1735 | 448 | 346 | 374 | 489 | 485 | 479 | 473 | 269 | 402 | 440 | 383 | 403 | 415 | 457 | 422 | 379 | 412,7 | 407 | 418 | 1,034 | 0,048 | 0,599 | 0,54905689 | 0,63025101 | 0,001431 | 0,0054971 | 0,0036828 | 0,0036828 | TRUE | 0,0428268 | No |
| orf19.1736 | 153 | 97 | 126 | 140 | 156 | 107 | 126 | 93 | 137 | 123 | 129 | 115 | 133 | 102 | 112 | 131 | 123,03 | 126 | 120 | 0,946 | -0,081 | -0,601 | 0,54803256 | 0,62931568 | 1,00E-08 | 0,0137793 | 0,0092524 | 0,0092524 | TRUE | 0,101138 | No |
| orf19.1737 | 18 | 15 | 24 | 28 | 21 | 29 | 29 | 15 | 16 | 19 | 25 | 23 | 18 | 28 | 26 | 21 | 21,94 | 21 | 23 | 1,118 | 0,162 | 0,541 | 0,5885513 | 0,66666972 | 1,00E-08 | 0,0681341 | 0,0401948 | 0,0401948 | TRUE | 0,1700726 | No |
| orf19.1738 | 6344 | 4352 | 5418 | 6809 | 8865 | 7398 | 8135 | 4875 | 5691 | 5541 | 5543 | 5615 | 7582 | 7059 | 7259 | 6876 | 6395,7 | 5598 | 7194 | 1,286 | 0,363 | 9,459 | 3,12E-21 | 6,13E-20 | 0,000752 | 0,0022065 | 0,0012524 | 0,0012524 | TRUE | 0,0161448 | No |
| orf19.1738.1 | 391 | 227 | 251 | 368 | 374 | 318 | 299 | 180 | 351 | 289 | 257 | 303 | 320 | 303 | 267 | 254 | 293 | 300 | 286 | 0,955 | -0,067 | -0,59 | 0,5551592 | 0,63540861 | 0,01083 | 0,0069342 | 0,0088372 | 0,0088372 | TRUE | 0,1503236 | No |

| | | | | | | | | | | | | | | | | | | | | | | | | | | | | | | | |
|---|---|---|---|---|---|---|---|---|---|---|---|---|---|---|---|---|---|---|---|---|---|---|---|---|---|---|---|---|---|---|---|
| orf19.1741 | 3 | 12 | 4 | 6 | 3 | 4 | 8 | 1 | 3 | 15 | 4 | 5 | 3 | 4 | 7 | 1 | 5,24 | 7 | 4 | 0,592 | -0,758 | -1,046 | 0,29544786 | NA | 0,317829 | 0,2788889 | 0,2965595 | 0,2965595 | TRUE | 1,4754834 | No |
| orf19.1742 | 6585 | 5767 | 6392 | 8832 | 6501 | 7783 | 6691 | 4831 | 5907 | 7342 | 6540 | 7283 | 5560 | 7426 | 5971 | 6814 | 6605,3 | 6768 | 6443 | 0,952 | -0,071 | -0,768 | 0,44263865 | 0,52938385 | 0,011982 | 0,0021993 | 0,0081208 | 0,0081208 | TRUE | 0,1297351 | No |
| orf19.1743 | 4548 | 3148 | 4108 | 4887 | 3455 | 2959 | 3085 | 1981 | 4080 | 4008 | 4203 | 4030 | 2955 | 2823 | 2753 | 2794 | 3455,7 | 4080 | 2831 | 0,694 | -0,527 | -13,853 | 1,22E-43 | 9,14E-42 | 0,000407 | 0,0023996 | 0,0010863 | 0,0010863 | TRUE | 0,0109648 | No |
| orf19.1744 | 2276 | 2081 | 2116 | 2649 | 2570 | 2766 | 2846 | 1631 | 2042 | 2649 | 2165 | 2184 | 2198 | 2639 | 2540 | 2300 | 2339,7 | 2260 | 2419 | 1,072 | 0,1 | 1,182 | 0,23707125 | 0,31792948 | 0,009759 | 0,0026 | 0,0064343 | 0,0064343 | TRUE | 0,1600202 | No |
| orf19.1745 | 623 | 438 | 545 | 700 | 726 | 749 | 774 | 524 | 559 | 558 | 558 | 577 | 621 | 715 | 691 | 739 | 627,08 | 563 | 692 | 1,224 | 0,291 | 4,245 | 2,18E-05 | 7,54E-05 | 0,001658 | 0,0042945 | 0,0028992 | 0,0028992 | TRUE | 0,0577536 | No |
| orf19.1746 | 109 | 82 | 102 | 118 | 96 | 96 | 109 | 69 | 98 | 104 | 104 | 97 | 82 | 92 | 97 | 97 | 96,52 | 101 | 92 | 0,91 | -0,135 | -0,97 | 0,33213949 | 0,41895739 | 1,00E-08 | 0,0170206 | 0,0083209 | 0,0083209 | TRUE | 0,0571122 | No |
| orf19.1747 | 98 | 63 | 117 | 136 | 102 | 102 | 93 | 57 | 88 | 80 | 120 | 112 | 87 | 97 | 83 | 80 | 93,49 | 100 | 87 | 0,868 | -0,205 | -1,26 | 0,20760027 | 0,28385581 | 0,010746 | 0,0175076 | 0,0146139 | 0,0146139 | TRUE | 0,1406202 | No |
| orf19.1748 | 405 | 270 | 359 | 423 | 168 | 151 | 173 | 126 | 363 | 344 | 367 | 349 | 144 | 144 | 154 | 178 | 255,38 | 356 | 155 | 0,431 | -1,214 | -12,665 | 9,26E-37 | 5,15E-35 | 1,00E-08 | 0,0076641 | 0,0041713 | 0,0041713 | TRUE | 0,0794407 | No |
| orf19.175 | 137 | 122 | 135 | 186 | 167 | 166 | 203 | 105 | 123 | 155 | 138 | 153 | 143 | 158 | 181 | 148 | 150,02 | 142 | 158 | 1,112 | 0,153 | 1,202 | 0,22954829 | 0,3095882 | 0,005575 | 0,0116562 | 0,0089211 | 0,0089211 | TRUE | 0,1135394 | No |
| orf19.1750 | 18722 | 14091 | 17131 | 22424 | 23658 | 22443 | 24658 | 14735 | 16796 | 17939 | 17527 | 18491 | 20233 | 21414 | 22003 | 20782 | 19398 | 17688 | 21108 | 1,193 | 0,255 | 6,09 | 1,13E-09 | 7,12E-09 | 0,001407 | 0,0020544 | 0,0016331 | 0,0016331 | TRUE | 0,0142099 | No |
| orf19.1751 | 11990 | 8949 | 10856 | 13294 | 12327 | 12083 | 12800 | 8129 | 10756 | 11393 | 11107 | 10962 | 10542 | 11529 | 11422 | 11465 | 11147 | 11054 | 11240 | 1,017 | 0,024 | 0,605 | 0,54502617 | 0,62646206 | 0,000986 | 0,0021098 | 0,0014104 | 0,0014104 | TRUE | 0,0215649 | No |
| orf19.1753 | 7901 | 6132 | 7186 | 9027 | 8119 | 7681 | 8093 | 4812 | 7088 | 7807 | 7352 | 7444 | 6944 | 7329 | 7222 | 6787 | 7246,5 | 7423 | 7070 | 0,953 | -0,069 | -1,643 | 0,10029131 | 0,15312584 | 0,001238 | 0,0021799 | 0,0015603 | 0,0015603 | TRUE | 0,0145031 | No |
| orf19.1754 | 2186 | 1612 | 1983 | 2389 | 1930 | 1775 | 1772 | 1146 | 1961 | 2052 | 2029 | 1970 | 1651 | 1694 | 1581 | 1616 | 1819,2 | 2003 | 1636 | 0,817 | -0,291 | -6,822 | 8,98E-12 | 7,38E-11 | 1,00E-08 | 0,0027775 | 0,0011878 | 0,0011878 | TRUE | 0,0070435 | No |
| orf19.1755 | 2209 | 1723 | 1977 | 2572 | 3383 | 3287 | 3501 | 2094 | 1982 | 2194 | 2023 | 2121 | 2893 | 3136 | 3124 | 2953 | 2553,2 | 2080 | 3026 | 1,457 | 0,543 | 11,16 | 6,41E-29 | 2,19E-27 | 0,001333 | 0,0025481 | 0,0018655 | 0,0018655 | TRUE | 0,0154316 | No |
| orf19.1756 | 6708 | 5000 | 6087 | 7477 | 7614 | 7039 | 7851 | 4780 | 6018 | 6366 | 6228 | 6166 | 6512 | 6716 | 7006 | 6742 | 6469 | 6194 | 6744 | 1,089 | 0,123 | 3,442 | 0,00057685 | 0,00153215 | 0,000516 | 0,0022039 | 0,0010737 | 0,0010737 | TRUE | 0,0085823 | No |
| orf19.1757 | 3852 | 2813 | 3402 | 4315 | 3194 | 3068 | 3278 | 1939 | 3456 | 3581 | 3481 | 3558 | 2732 | 2927 | 2925 | 2735 | 3174,3 | 3519 | 2830 | 0,805 | -0,313 | -7,799 | 6,26E-15 | 7,17E-14 | 0,000497 | 0,0024368 | 0,0012278 | 0,0012278 | TRUE | 0,0073338 | No |
| orf19.1759 | 2452 | 1857 | 2194 | 2684 | 2705 | 2442 | 2502 | 1677 | 2200 | 2364 | 2245 | 2213 | 2313 | 2330 | 2233 | 2365 | 2282,9 | 2256 | 2310 | 1,025 | 0,035 | 0,852 | 0,3940765 | 0,48273084 | 0,000262 | 0,0026154 | 0,0012085 | 0,0012085 | TRUE | 0,0119293 | No |
| orf19.176 | 1754 | 1291 | 1613 | 2103 | 2920 | 2632 | 2665 | 1796 | 1574 | 1644 | 1650 | 1734 | 2497 | 2511 | 2378 | 2533 | 2065,2 | 1650 | 2480 | 1,501 | 0,586 | 12,864 | 7,18E-38 | 4,40E-36 | 0,000741 | 0,0026825 | 0,0014876 | 0,0014876 | TRUE | 0,0151616 | No |
| orf19.1760 | 9562 | 7082 | 8334 | 10332 | 11030 | 10671 | 11070 | 7002 | 8578 | 9016 | 8527 | 8520 | 9433 | 10182 | 9878 | 9876 | 9251,2 | 8660 | 9842 | 1,137 | 0,185 | 5,066 | 4,07E-07 | 1,83E-06 | 0,000726 | 0,0021364 | 0,0011703 | 0,0011703 | TRUE | 0,0097451 | No |
| orf19.1761 | 269 | 189 | 218 | 269 | 273 | 300 | 253 | 169 | 241 | 241 | 223 | 222 | 233 | 286 | 226 | 238 | 238,83 | 232 | 246 | 1,063 | 0,088 | 0,86 | 0,38957904 | 0,47845275 | 0,0035 | 0,0080579 | 0,0059129 | 0,0059129 | TRUE | 0,1408705 | No |
| orf19.1762 | 1458 | 1135 | 1270 | 1746 | 1517 | 1333 | 1568 | 882 | 1308 | 1445 | 1299 | 1440 | 1297 | 1272 | 1399 | 1244 | 1338,1 | 1373 | 1303 | 0,951 | -0,073 | -1,23 | 0,21887646 | 0,29758085 | 0,002056 | 0,0030644 | 0,0026158 | 0,0026158 | TRUE | 0,0299526 | No |
| orf19.1763 | 1328 | 1043 | 1294 | 1554 | 1395 | 1483 | 1516 | 938 | 1191 | 1328 | 1324 | 1281 | 1193 | 1415 | 1353 | 1323 | 1301,1 | 1281 | 1321 | 1,032 | 0,045 | 0,701 | 0,48339907 | 0,56879389 | 0,003185 | 0,0030953 | 0,003147 | 0,003147 | TRUE | 0,0536593 | No |
| orf19.1764 | 4492 | 3780 | 4113 | 5315 | 5674 | 5573 | 5979 | 3800 | 4030 | 4812 | 4208 | 4383 | 4853 | 5318 | 5335 | 5360 | 4787,2 | 4358 | 5216 | 1,197 | 0,26 | 4,375 | 1,21E-05 | 4,35E-05 | 0,003764 | 0,0022828 | 0,0031726 | 0,0031726 | TRUE | 0,0588142 | No |
| orf19.1765 | 1154 | 957 | 1090 | 1311 | 880 | 822 | 921 | 461 | 1035 | 1218 | 1115 | 1081 | 753 | 784 | 822 | 650 | 932,35 | 1112 | 752 | 0,68 | -0,557 | -7,2 | 6,01E-13 | 5,67E-12 | 0,005758 | 0,0035366 | 0,0046309 | 0,0046309 | TRUE | 0,089102 | No |
| orf19.1766 | 136 | 116 | 142 | 161 | 138 | 136 | 127 | 69 | 122 | 148 | 145 | 133 | 118 | 130 | 113 | 97 | 125,77 | 137 | 114 | 0,847 | -0,239 | -1,821 | 0,06853545 | 0,11077777 | 1,00E-08 | 0,013522 | 0,0086134 | 0,0086134 | TRUE | 0,0785612 | No |
| orf19.1767 | 426 | 416 | 382 | 515 | 485 | 622 | 652 | 350 | 382 | 530 | 391 | 425 | 415 | 593 | 582 | 494 | 476,38 | 432 | 521 | 1,21 | 0,275 | 2,133 | 0,03292099 | 0,05816155 | 0,022362 | 0,0050269 | 0,0138117 | 0,0138117 | TRUE | 0,2560779 | No |
| orf19.1768 | 268 | 236 | 264 | 318 | 293 | 334 | 368 | 219 | 240 | 300 | 270 | 262 | 251 | 319 | 328 | 309 | 284,97 | 268 | 302 | 1,127 | 0,172 | 1,591 | 0,11164336 | 0,16739538 | 0,008308 | 0,0070738 | 0,0076773 | 0,0076773 | TRUE | 0,1587657 | No |
| orf19.1769 | 907 | 744 | 749 | 1027 | 886 | 852 | 915 | 507 | 814 | 947 | 766 | 847 | 758 | 813 | 816 | 715 | 809,54 | 844 | 776 | 0,923 | -0,116 | -1,546 | 0,12202103 | 0,18096825 | 0,004429 | 0,0037728 | 0,0041264 | 0,0041264 | TRUE | 0,0757929 | No |
| orf19.177 | 451 | 353 | 453 | 508 | 496 | 532 | 509 | 372 | 405 | 449 | 463 | 419 | 424 | 508 | 454 | 525 | 455,88 | 434 | 478 | 1,098 | 0,135 | 1,567 | 0,11713645 | 0,17454261 | 0,004624 | 0,0051639 | 0,0048924 | 0,0048924 | TRUE | 0,0676063 | No |
| orf19.1770 | 7538 | 5286 | 6108 | 7975 | 8255 | 6835 | 6983 | 4786 | 6762 | 6730 | 6249 | 6576 | 7060 | 6522 | 6231 | 6750 | 6610,1 | 6579 | 6641 | 1,009 | 0,013 | 0,281 | 0,778767 | 0,82650276 | 0,001619 | 0,0021991 | 0,0020128 | 0,0020128 | TRUE | 0,0225688 | No |
| orf19.1771 | 1412 | 1202 | 1252 | 1579 | 1740 | 1803 | 1813 | 1138 | 1267 | 1530 | 1281 | 1302 | 1488 | 1720 | 1618 | 1605 | 1476,4 | 1345 | 1608 | 1,197 | 0,26 | 3,686 | 0,00022803 | 0,00065739 | 0,005113 | 0,0029628 | 0,0040847 | 0,0040847 | TRUE | 0,0991127 | No |
| orf19.1772 | 2984 | 2649 | 2791 | 3611 | 3054 | 3485 | 3620 | 2194 | 2677 | 3372 | 2855 | 2978 | 2612 | 3325 | 3230 | 3094 | 3018,1 | 2970 | 3065 | 1,032 | 0,046 | 0,539 | 0,58971875 | 0,66748517 | 0,010123 | 0,0024605 | 0,0066128 | 0,0066128 | TRUE | 0,1221632 | No |
| orf19.1773 | 2405 | 1800 | 2227 | 2839 | 2507 | 2391 | 2437 | 1586 | 2158 | 2292 | 2278 | 2341 | 2144 | 2281 | 2175 | 2237 | 2238,2 | 2267 | 2209 | 0,974 | -0,038 | -0,874 | 0,38188314 | 0,4709948 | 0,000628 | 0,0026281 | 0,0013638 | 0,0013638 | TRUE | 0,0133454 | No |
| orf19.1774 | 510 | 396 | 463 | 580 | 501 | 499 | 472 | 343 | 458 | 504 | 474 | 478 | 428 | 476 | 421 | 484 | 465,4 | 478 | 452 | 0,944 | -0,084 | -1,128 | 0,25950176 | 0,34288247 | 0,000999 | 0,0050988 | 0,003148 | 0,003148 | TRUE | 0,0245491 | No |
| orf19.1776 | 392 | 393 | 390 | 502 | 311 | 305 | 334 | 188 | 352 | 500 | 399 | 414 | 266 | 291 | 298 | 265 | 348,14 | 416 | 280 | 0,677 | -0,564 | -5,236 | 1,64E-07 | 7,81E-07 | 0,010103 | 0,0061493 | 0,0081308 | 0,0081308 | TRUE | 0,1995013 | No |
| orf19.1777 | 8203 | 6785 | 7748 | 9499 | 11727 | 12362 | 12818 | 7314 | 7359 | 8638 | 7927 | 7833 | 10029 | 11795 | 11438 | 10316 | 9416,9 | 7939 | 10894 | 1,373 | 0,457 | 7,057 | 1,71E-12 | 1,52E-11 | 0,005143 | 0,0021337 | 0,0039212 | 0,0039212 | TRUE | 0,0425509 | No |
| orf19.1778 | 124 | 98 | 103 | 132 | 93 | 97 | 84 | 58 | 111 | 125 | 105 | 109 | 80 | 93 | 75 | 82 | 97,39 | 112 | 82 | 0,733 | -0,449 | -3,151 | 0,00162819 | 0,00391286 | 1,00E-08 | 0,0168863 | 0,0089937 | 0,0089937 | TRUE | 0,0728858 | No |
| orf19.1779 | 17106 | 11834 | 15853 | 19831 | 20322 | 17443 | 19167 | 13593 | 15346 | 15066 | 16219 | 16353 | 17380 | 16644 | 17103 | 19172 | 16660 | 15746 | 17575 | 1,116 | 0,158 | 3,091 | 0,00199205 | 0,00470448 | 0,002721 | 0,0020667 | 0,0024495 | 0,0024495 | TRUE | 0,0454001 | No |
| orf19.178 | 950 | 815 | 833 | 1090 | 959 | 1077 | 1094 | 670 | 852 | 1038 | 852 | 899 | 820 | 1028 | 976 | 945 | 926,24 | 910 | 942 | 1,037 | 0,052 | 0,608 | 0,54304326 | 0,62525947 | 0,007874 | 0,0035468 | 0,0058577 | 0,0058577 | TRUE | 0,1008128 | No |
| orf19.1780 | 50 | 40 | 44 | 67 | 24 | 34 | 47 | 27 | 45 | 51 | 45 | 55 | 21 | 32 | 42 | 38 | 41,13 | 49 | 33 | 0,671 | -0,575 | -2,38 | 0,01732163 | 0,03277582 | 0,020565 | 0,0372757 | 0,0307788 | 0,0307788 | TRUE | 0,5444982 | No |
| orf19.1782 | 668 | 388 | 501 | 654 | 535 | 472 | 474 | 352 | 599 | 494 | 513 | 539 | 458 | 450 | 423 | 496 | 496,56 | 536 | 457 | 0,847 | -0,24 | -2,962 | 0,00305652 | 0,00692704 | 0,003642 | 0,0049031 | 0,0042694 | 0,0042694 | TRUE | 0,0736605 | No |
| orf19.1782.1 | 34 | 17 | 29 | 28 | 26 | 19 | 29 | 10 | 31 | 22 | 30 | 23 | 22 | 18 | 26 | 14 | 23,16 | 26 | 20 | 0,778 | -0,362 | -1,233 | 0,21774366 | 0,29629488 | 1,00E-08 | 0,0646709 | 0,0395939 | 0,0395939 | TRUE | 0,2048421 | No |
| orf19.1783 | 12391 | 9395 | 11549 | 13486 | 12123 | 12235 | 12792 | 7193 | 11116 | 11961 | 11816 | 11121 | 10368 | 11674 | 11415 | 10145 | 11202 | 11504 | 10900 | 0,948 | -0,077 | -1,428 | 0,15329218 | 0,21975171 | 0,00305 | 0,0021091 | 0,0027088 | 0,0027088 | TRUE | 0,027838 | No |
| orf19.1785 | 19602 | 13978 | 16982 | 21486 | 29607 | 27220 | 29944 | 20314 | 17585 | 17795 | 17374 | 17718 | 25321 | 25972 | 26720 | 28651 | 22142 | 17618 | 26666 | 1,513 | 0,598 | 14,281 | 2,86E-46 | 2,42E-44 | 0,001453 | 0,0020451 | 0,0016351 | 0,0016351 | TRUE | 0,0304981 | No |
| orf19.1789 | 202 | 201 | 193 | 237 | 273 | 394 | 338 | 182 | 181 | 256 | 197 | 195 | 233 | 376 | 302 | 257 | 249,72 | 207 | 292 | 1,415 | 0,501 | 3,252 | 0,00114591 | 0,00284961 | 0,030293 | 0,0077929 | 0,0186085 | 0,0186085 | TRUE | 0,4277063 | No |
| orf19.1789.1 | 1711 | 1192 | 1484 | 1882 | 1763 | 1697 | 1743 | 1186 | 1535 | 1518 | 1518 | 1552 | 1508 | 1619 | 1555 | 1673 | 1559,7 | 1531 | 1589 | 1,035 | 0,05 | 1,072 | 0,28377343 | 0,36799283 | 0,000414 | 0,0029103 | 0,0014485 | 0,0014485 | TRUE | 0,0142537 | No |
| orf19.1791 | 7050 | 5777 | 6324 | 8158 | 6886 | 6867 | 7214 | 4031 | 6325 | 7355 | 6470 | 6727 | 5889 | 6552 | 6437 | 5685 | 6430,1 | 6719 | 6141 | 0,915 | -0,129 | -2,086 | 0,03702099 | 0,06456375 | 0,004421 | 0,0022053 | 0,003509 | 0,003509 | TRUE | 0,0489372 | No |
| orf19.1792 | 427 | 356 | 374 | 487 | 488 | 513 | 583 | 320 | 383 | 453 | 383 | 402 | 417 | 489 | 520 | 451 | 437,36 | 405 | 469 | 1,163 | 0,218 | 2,427 | 0,01521942 | 0,02921351 | 0,005616 | 0,0052987 | 0,0054592 | 0,0054592 | TRUE | 0,0719638 | No |
| orf19.1793 | 1352 | 1024 | 1263 | 1634 | 1556 | 1416 | 1537 | 927 | 1213 | 1304 | 1292 | 1347 | 1331 | 1351 | 1372 | 1307 | 1314,6 | 1289 | 1340 | 1,041 | 0,057 | 1,161 | 0,24546999 | 0,3274206 | 0,000517 | 0,0030838 | 0,0015911 | 0,0015911 | TRUE | 0,0200951 | No |
| orf19.1794 | 111 | 113 | 116 | 139 | 183 | 200 | 175 | 83 | 100 | 144 | 119 | 115 | 157 | 191 | 156 | 117 | 137,16 | 120 | 155 | 1,319 | 0,4 | 2,522 | 0,01166998 | 0,02310155 | 0,021895 | 0,0125633 | 0,0167161 | 0,0167161 | TRUE | 0,1950941 | No |
| orf19.1795 | 1778 | 1370 | 1474 | 1975 | 2354 | 2365 | 2381 | 1364 | 1595 | 1744 | 1508 | 1629 | 2013 | 2257 | 2125 | 1924 | 1849,3 | 1619 | 2080 | 1,287 | 0,364 | 5,82 | 5,88E-09 | 3,37E-08 | 0,00348 | 0,0027645 | 0,0031996 | 0,0031996 | TRUE | 0,0393715 | No |
| orf19.1795.1 | 262 | 208 | 215 | 329 | 268 | 266 | 270 | 184 | 235 | 265 | 220 | 271 | 229 | 254 | 241 | 260 | 246,82 | 248 | 246 | 0,989 | -0,016 | -0,158 | 0,87472039 | 0,90421888 | 0,002511 | 0,0078611 | 0,0054319 | 0,0054319 | TRUE | 0,0619062 | No |
| orf19.1796 | 247 | 203 | 276 | 305 | 269 | 268 | 293 | 153 | 222 | 258 | 282 | 252 | 230 | 256 | 261 | 216 | 247,12 | 254 | 241 | 0,956 | -0,065 | -0,624 | 0,53286299 | 0,61554333 | 0,00483 | 0,0078541 | 0,0064623 | 0,0064623 | TRUE | 0,0835277 | No |
| orf19.1797 | 7515 | 6248 | 6816 | 8805 | 6377 | 6646 | 6891 | 4306 | 6742 | 7954 | 6973 | 7261 | 5454 | 6341 | 6149 | 6073 | 6618,5 | 7232 | 6004 | 0,83 | -0,268 | -4,29 | 1,78E-05 | 6,25E-05 | 0,004486 | 0,0021989 | 0,0036019 | 0,0036019 | TRUE | 0,0545483 | No |
| orf19.1798 | 590 | 463 | 578 | 668 | 724 | 759 | 839 | 463 | 529 | 589 | 591 | 551 | 619 | 724 | 749 | 653 | 625,75 | 565 | 686 | 1,218 | 0,284 | 3,658 | 0,00025464 | 0,00072816 | 0,004055 | 0,0042995 | 0,0041688 | 0,0041688 | TRUE | 0,0563486 | No |
| orf19.1799 | 13238 | 8916 | 11631 | 14386 | 15248 | 13446 | 14503 | 8524 | 11876 | 11351 | 11900 | 11863 | 13041 | 12830 | 12942 | 12022 | 12228 | 11748 | 12709 | 1,082 | 0,114 | 3,105 | 0,00190384 | 0,00451393 | 0,000815 | 0,0020982 | 0,0012079 | 0,0012079 | TRUE | 0,0157752 | No |
| orf19.18 | 33111 | 23368 | 28911 | 36072 | 32285 | 27614 | 28971 | 18080 | 29704 | 29750 | 29579 | 29745 | 27611 | 26348 | 25852 | 25500 | 28011 | 29694 | 26328 | 0,887 | -0,173 | -5,405 | 6,49E-08 | 3,26E-07 | 0,000571 | 0,0020313 | 0,0009522 | 0,0009522 | TRUE | 0,0132348 | No |
| orf19.1800 | 10817 | 7987 | 9821 | 12593 | 10424 | 9833 | 10849 | 6225 | 9704 | 10168 | 10048 | 10384 | 8915 | 9382 | 9681 | 8780 | 9632,8 | 10076 | 9190 | 0,912 | -0,132 | -3,16 | 0,0015757 | 0,00380345 | 0,00129 | 0,0021302 | 0,0015808 | 0,0015808 | TRUE | 0,0158717 | No |
| orf19.1801 | 1669 | 1291 | 1490 | 1860 | 1830 | 1796 | 1931 | 1080 | 1497 | 1644 | 1524 | 1534 | 1565 | 1714 | 1723 | 1523 | 1590,5 | 1550 | 1631 | 1,055 | 0,078 | 1,375 | 0,16910054 | 0,23895053 | 0,001984 | 0,0028922 | 0,0024212 | 0,0024212 | TRUE | 0,0219067 | No |

| | | | | | | | | | | | | | | | | | | | | | | | | | | | | | | | |
|---|---|---|---|---|---|---|---|---|---|---|---|---|---|---|---|---|---|---|---|---|---|---|---|---|---|---|---|---|---|---|---|
| orf19.1802 | 6740 | 5034 | 6030 | 7282 | 6960 | 6655 | 6950 | 3922 | 6046 | 6409 | 6169 | 6005 | 5952 | 6350 | 6202 | 5532 | 6083,2 | 6157 | 6009 | 0,977 | -0,034 | -0,698 | 0,48505915 | 0,57052376 | 0,001869 | 0,0022182 | 0,0020826 | 0,0020826 | TRUE | 0,0337872 | No |
| orf19.1803 | 153 | 119 | 150 | 192 | 163 | 192 | 163 | 130 | 137 | 151 | 153 | 158 | 139 | 183 | 145 | 183 | 156,49 | 150 | 162 | 1,077 | 0,107 | 0,837 | 0,40261217 | 0,49095622 | 0,006866 | 0,0112559 | 0,0092584 | 0,0092584 | TRUE | 0,1036334 | No |
| orf19.1804 | 4748 | 3472 | 4025 | 5105 | 4051 | 3892 | 4245 | 2461 | 4259 | 4420 | 4118 | 4210 | 3465 | 3714 | 3788 | 3471 | 3930,6 | 4252 | 3610 | 0,85 | -0,235 | -5,325 | 1,01E-07 | 4,96E-07 | 0,001162 | 0,0023489 | 0,0016127 | 0,0016127 | TRUE | 0,0136294 | No |
| orf19.1805 | 3148 | 2453 | 2820 | 3559 | 3067 | 2917 | 3158 | 2082 | 2824 | 3123 | 2885 | 2935 | 2623 | 2783 | 2818 | 2936 | 2866 | 2942 | 2790 | 0,948 | -0,077 | -1,57 | 0,11641598 | 0,17355514 | 0,00157 | 0,0024861 | 0,001946 | 0,001946 | TRUE | 0,0200387 | No |
| orf19.1806 | 34 | 22 | 29 | 54 | 45 | 38 | 43 | 22 | 31 | 28 | 30 | 45 | 38 | 36 | 38 | 31 | 34,61 | 34 | 36 | 1,082 | 0,113 | 0,468 | 0,64013631 | 0,71023804 | 1,00E-08 | 0,0439283 | 0,0276635 | 0,0276635 | TRUE | 0,4257908 | No |
| orf19.1808 | 270 | 273 | 250 | 355 | 349 | 389 | 434 | 248 | 242 | 348 | 256 | 293 | 298 | 371 | 387 | 350 | 318,12 | 285 | 352 | 1,24 | 0,311 | 2,545 | 0,01092913 | 0,02180021 | 0,015501 | 0,0065428 | 0,011104 | 0,011104 | TRUE | 0,2298323 | No |
| orf19.1813 | 3102 | 2155 | 2854 | 3464 | 3086 | 2757 | 2898 | 1770 | 2783 | 2744 | 2920 | 2856 | 2639 | 2631 | 2586 | 2496 | 2706,9 | 2826 | 2588 | 0,917 | -0,126 | -3,222 | 0,00127268 | 0,00313628 | 0,00021 | 0,0025158 | 0,0010914 | 0,0010914 | TRUE | 0,0062198 | No |
| orf19.1814 | 1886 | 1467 | 1810 | 2120 | 2185 | 2220 | 2239 | 1202 | 1692 | 1868 | 1852 | 1748 | 1869 | 2118 | 1998 | 1695 | 1855 | 1790 | 1920 | 1,075 | 0,104 | 1,518 | 0,1289169 | 0,18969697 | 0,004837 | 0,0027621 | 0,0039949 | 0,0039949 | TRUE | 0,0711115 | No |
| orf19.1815 | 13303 | 8899 | 11093 | 14215 | 12663 | 10467 | 10748 | 6838 | 11934 | 11329 | 11349 | 11722 | 10830 | 9987 | 9591 | 9644 | 10798 | 11584 | 10013 | 0,865 | -0,21 | -4,584 | 4,56E-06 | 1,75E-05 | 0,001833 | 0,002114 | 0,0019221 | 0,0019221 | TRUE | 0,0371368 | No |
| orf19.1816 | 6759 | 4569 | 6213 | 7428 | 7074 | 6242 | 6875 | 4597 | 6064 | 5817 | 6357 | 6125 | 6050 | 5956 | 6135 | 6484 | 6123,3 | 6091 | 6156 | 1,01 | 0,014 | 0,34 | 0,73390152 | 0,79203886 | 0,001035 | 0,0022166 | 0,0014956 | 0,0014956 | TRUE | 0,0150771 | No |
| orf19.182 | 2301 | 1654 | 2072 | 2586 | 2718 | 2414 | 2571 | 1495 | 2064 | 2106 | 2120 | 2132 | 2325 | 2303 | 2294 | 2109 | 2181,6 | 2106 | 2258 | 1,074 | 0,103 | 2,383 | 0,01719541 | 0,03256937 | 0,000472 | 0,0026449 | 0,0013501 | 0,0013501 | TRUE | 0,0216831 | No |
| orf19.1821 | 0 | 0 | 0 | 1 | 0 | 0 | 2 | 1 | 0 | 0 | 0 | 1 | 0 | 0 | 2 | 1 | 0,5 | 0 | 1 | 2,585 | 1,37 | 0,647 | 0,51795137 | NA | 1,00E-08 | 2,8911592 | 2,6636908 | 2,6636908 | TRUE | 0,1530619 | No |
| orf19.1822 | 51 | 49 | 51 | 71 | 77 | 65 | 63 | 47 | 46 | 62 | 52 | 59 | 66 | 62 | 56 | 66 | 58,66 | 55 | 62 | 1,146 | 0,197 | 1,082 | 0,27907119 | 0,36322556 | 1,00E-08 | 0,0267293 | 0,0146529 | 0,0146529 | TRUE | 0,109565 | No |
| orf19.1823 | 130 | 101 | 140 | 143 | 167 | 180 | 180 | 122 | 117 | 129 | 143 | 118 | 143 | 172 | 161 | 172 | 144,2 | 127 | 162 | 1,277 | 0,353 | 2,846 | 0,00442436 | 0,00966251 | 1,00E-08 | 0,0120466 | 0,0077448 | 0,0077448 | TRUE | 0,0825845 | No |
| orf19.1824 | 0 | 1 | 0 | 0 | 0 | 1 | 0 | 0 | 0 | 1 | 0 | 0 | 0 | 1 | 0 | 0 | 0,28 | 0 | 0 | 1,016 | 0,023 | 0,008 | 0,99349273 | NA | 1,00E-08 | 5,2162474 | 5,6438415 | 5,6438415 | TRUE | 0,544438 | No |
| orf19.1825 | 72 | 42 | 56 | 74 | 62 | 49 | 66 | 47 | 65 | 53 | 57 | 61 | 53 | 47 | 59 | 66 | 57,67 | 59 | 56 | 0,938 | -0,093 | -0,505 | 0,6132523 | 0,68687916 | 1,00E-08 | 0,0271532 | 0,0151727 | 0,0151727 | TRUE | 0,1049589 | No |
| orf19.1826 | 12658 | 10220 | 11652 | 14797 | 20168 | 20208 | 21038 | 13363 | 11356 | 13011 | 11921 | 12202 | 17248 | 19282 | 18773 | 18847 | 15330 | 12122 | 18538 | 1,529 | 0,613 | 11,907 | 1,09E-32 | 4,63E-31 | 0,002673 | 0,0020742 | 0,0024776 | 0,0024776 | TRUE | 0,0295296 | No |
| orf19.1827 | 12 | 23 | 27 | 24 | 17 | 18 | 17 | 8 | 11 | 29 | 28 | 20 | 15 | 17 | 15 | 11 | 18,2 | 22 | 14 | 0,683 | -0,549 | -1,502 | 0,13317639 | 0,19519974 | 0,057295 | 0,081728 | 0,071334 | 0,071334 | TRUE | 0,5062437 | No |
| orf19.1828 | 391 | 317 | 416 | 455 | 453 | 400 | 462 | 226 | 351 | 404 | 426 | 375 | 387 | 382 | 412 | 319 | 381,91 | 389 | 375 | 0,972 | -0,041 | -0,433 | 0,66491712 | 0,7328604 | 0,006113 | 0,0057807 | 0,0059378 | 0,0059378 | TRUE | 0,1009746 | No |
| orf19.183 | 1248 | 868 | 1041 | 1360 | 1367 | 1214 | 1175 | 803 | 1120 | 1105 | 1065 | 1121 | 1169 | 1158 | 1048 | 1133 | 1115 | 1103 | 1127 | 1,021 | 0,031 | 0,573 | 0,56687253 | 0,64650155 | 0,000752 | 0,0032815 | 0,0018413 | 0,0018413 | TRUE | 0,0282001 | No |
| orf19.1830 | 3527 | 2895 | 3150 | 3945 | 3333 | 3492 | 3543 | 2149 | 3164 | 3686 | 3223 | 3253 | 2850 | 3332 | 3162 | 3031 | 3212,6 | 3332 | 3094 | 0,929 | -0,106 | -1,668 | 0,09538545 | 0,14709338 | 0,004326 | 0,0024314 | 0,0035411 | 0,0035411 | TRUE | 0,0610096 | No |
| orf19.1831 | 3 | 1 | 3 | 7 | 1 | 3 | 0 | 2 | 3 | 1 | 3 | 6 | 1 | 3 | 0 | 3 | 2,42 | 3 | 2 | 0,467 | -1,1 | -1,134 | 0,25673328 | NA | 1,00E-08 | 0,6023347 | 0,4274121 | 0,4274121 | TRUE | 0,4894201 | No |
| orf19.1832 | 2639 | 1925 | 2415 | 3027 | 2551 | 2236 | 2561 | 1485 | 2367 | 2451 | 2471 | 2496 | 2182 | 2134 | 2285 | 2094 | 2310 | 2446 | 2174 | 0,89 | -0,169 | -3,951 | 7,78E-05 | 0,00024166 | 0,000534 | 0,002608 | 0,0013125 | 0,0013125 | TRUE | 0,0145281 | No |
| orf19.1833 | 20418 | 15372 | 18066 | 22916 | 18265 | 17982 | 18976 | 10723 | 18317 | 19570 | 18483 | 18897 | 15621 | 17158 | 16933 | 15124 | 17513 | 18817 | 16209 | 0,862 | -0,215 | -4,455 | 8,39E-06 | 3,10E-05 | 0,002234 | 0,0020624 | 0,0021764 | 0,0021764 | TRUE | 0,0245482 | No |
| orf19.1834 | 260 | 182 | 241 | 330 | 299 | 341 | 385 | 227 | 233 | 232 | 247 | 272 | 256 | 325 | 344 | 320 | 278,55 | 246 | 311 | 1,26 | 0,334 | 3,056 | 0,00224217 | 0,00524162 | 0,008408 | 0,0071911 | 0,007802 | 0,007802 | TRUE | 0,1761301 | No |
| orf19.1835 | 3922 | 3170 | 3647 | 4488 | 4228 | 4317 | 4578 | 2829 | 3518 | 4036 | 3731 | 3701 | 3616 | 4119 | 4085 | 3990 | 3849,6 | 3746 | 3952 | 1,055 | 0,078 | 1,372 | 0,17020411 | 0,2403258 | 0,002999 | 0,0023566 | 0,0028285 | 0,0028285 | TRUE | 0,040834 | No |
| orf19.1836 | 135 | 127 | 133 | 169 | 152 | 152 | 139 | 85 | 121 | 162 | 136 | 139 | 130 | 145 | 124 | 120 | 134,65 | 140 | 130 | 0,938 | -0,092 | -0,713 | 0,47577963 | 0,56124065 | 0,002722 | 0,0127611 | 0,0085507 | 0,0085507 | TRUE | 0,1017415 | No |
| orf19.1837 | 5772 | 3971 | 4978 | 6209 | 5851 | 5002 | 5690 | 3193 | 5178 | 5056 | 5093 | 5120 | 5004 | 4773 | 5077 | 4503 | 4975,5 | 5112 | 4839 | 0,948 | -0,078 | -1,797 | 0,07228193 | 0,115898 | 0,001213 | 0,0022713 | 0,0015884 | 0,0015884 | TRUE | 0,0254487 | No |
| orf19.1839 | 14780 | 10511 | 13679 | 16069 | 13989 | 13377 | 14033 | 7808 | 13259 | 13382 | 13995 | 13251 | 11964 | 12764 | 12522 | 11012 | 12769 | 13472 | 12066 | 0,896 | -0,158 | -3,212 | 0,00131769 | 0,00323129 | 0,002319 | 0,0020932 | 0,0022588 | 0,0022588 | TRUE | 0,041577 | No |
| orf19.184 | 472 | 431 | 512 | 660 | 567 | 579 | 661 | 340 | 423 | 549 | 524 | 544 | 485 | 552 | 590 | 480 | 518,37 | 510 | 527 | 1,036 | 0,051 | 0,509 | 0,61070668 | 0,68517754 | 0,010414 | 0,00478 | 0,007662 | 0,007662 | TRUE | 0,1569039 | No |
| orf19.1840 | 1354 | 985 | 1318 | 1494 | 2296 | 2068 | 2174 | 1340 | 1215 | 1254 | 1348 | 1232 | 1964 | 1973 | 1940 | 1890 | 1602 | 1262 | 1942 | 1,541 | 0,623 | 13,15 | 1,71E-39 | 1,13E-37 | 0,000409 | 0,0028857 | 0,0015042 | 0,0015042 | TRUE | 0,0255378 | No |
| orf19.1841 | 61 | 76 | 78 | 83 | 51 | 87 | 80 | 40 | 55 | 97 | 80 | 68 | 44 | 83 | 71 | 56 | 69,27 | 75 | 64 | 0,857 | -0,222 | -0,964 | 0,33518356 | 0,42196664 | 0,052304 | 0,0229368 | 0,0364619 | 0,0364619 | TRUE | 0,2976917 | No |
| orf19.1842 | 3280 | 2611 | 3137 | 3707 | 3217 | 3298 | 3462 | 1969 | 2943 | 3324 | 3209 | 3057 | 2751 | 3147 | 3089 | 2777 | 3037,2 | 3133 | 2941 | 0,94 | -0,09 | -1,498 | 0,13412165 | 0,19639359 | 0,003445 | 0,0024575 | 0,0031036 | 0,0031036 | TRUE | 0,0268366 | No |
| orf19.1843 | 259 | 196 | 237 | 297 | 209 | 265 | 252 | 174 | 232 | 250 | 242 | 245 | 179 | 253 | 225 | 245 | 233,89 | 242 | 226 | 0,926 | -0,11 | -0,969 | 0,33263608 | 0,41941681 | 0,008074 | 0,0081862 | 0,0081261 | 0,0081261 | TRUE | 0,2304226 | No |
| orf19.1844 | 15 | 7 | 17 | 19 | 11 | 17 | 16 | 13 | 13 | 9 | 17 | 16 | 9 | 16 | 14 | 18 | 14,21 | 14 | 14 | 1,018 | 0,026 | 0,069 | 0,94506561 | NA | 1,00E-08 | 0,1041493 | 0,0643937 | 0,0643937 | TRUE | 0,298703 | No |
| orf19.1847 | 531 | 408 | 477 | 596 | 1025 | 872 | 1033 | 685 | 476 | 519 | 488 | 491 | 877 | 832 | 922 | 966 | 696,48 | 494 | 899 | 1,82 | 0,864 | 12,985 | 1,49E-38 | 9,50E-37 | 0,00143 | 0,0040639 | 0,0026804 | 0,0026804 | TRUE | 0,0292731 | No |
| orf19.1848.1 | 298 | 189 | 241 | 346 | 503 | 388 | 342 | 267 | 267 | 241 | 247 | 285 | 430 | 370 | 305 | 377 | 315,25 | 260 | 370 | 1,421 | 0,507 | 4,538 | 5,69E-06 | 2,15E-05 | 0,010725 | 0,0065845 | 0,0086998 | 0,0086998 | TRUE | 0,1712776 | No |
| orf19.1849 | 4770 | 3718 | 4609 | 5598 | 5328 | 5114 | 5436 | 3209 | 4279 | 4733 | 4715 | 4616 | 4557 | 4880 | 4851 | 4526 | 4644,7 | 4586 | 4704 | 1,026 | 0,038 | 0,801 | 0,42285393 | 0,51044935 | 0,001548 | 0,0022921 | 0,0018887 | 0,0018887 | TRUE | 0,0249357 | No |
| orf19.185 | 2799 | 2001 | 2371 | 3000 | 3020 | 2755 | 2755 | 1614 | 2511 | 2547 | 2426 | 2474 | 2583 | 2629 | 2458 | 2276 | 2488,1 | 2490 | 2486 | 1,001 | 0,001 | 0,019 | 0,98446085 | 0,98820382 | 0,001497 | 0,002563 | 0,0020461 | 0,0020461 | TRUE | 0,0366197 | No |
| orf19.1850 | 852 | 572 | 704 | 990 | 1002 | 908 | 981 | 601 | 764 | 728 | 720 | 816 | 857 | 866 | 875 | 848 | 809,44 | 757 | 862 | 1,136 | 0,183 | 3,122 | 0,0017989 | 0,0042905 | 0,000553 | 0,003773 | 0,0020756 | 0,0020756 | TRUE | 0,0343952 | No |
| orf19.1852 | 1041 | 590 | 873 | 1030 | 1066 | 868 | 960 | 628 | 934 | 751 | 893 | 849 | 912 | 828 | 857 | 886 | 863,73 | 857 | 871 | 1,013 | 0,019 | 0,27 | 0,78710484 | 0,83343776 | 0,00377 | 0,0036603 | 0,0037182 | 0,0037182 | TRUE | 0,0759207 | No |
| orf19.1853 | 1175 | 751 | 876 | 1118 | 1338 | 1030 | 1032 | 648 | 1054 | 956 | 896 | 922 | 1144 | 983 | 921 | 914 | 973,78 | 957 | 990 | 1,036 | 0,052 | 0,625 | 0,53200576 | 0,61467127 | 0,007251 | 0,0034703 | 0,0055264 | 0,0055264 | TRUE | 0,1334504 | No |
| orf19.1854 | 3497 | 2398 | 2762 | 3447 | 3489 | 3062 | 3215 | 2048 | 3137 | 3053 | 2826 | 2842 | 2984 | 2922 | 2869 | 2889 | 2940,2 | 2964 | 2916 | 0,984 | -0,023 | -0,506 | 0,61274988 | 0,68671822 | 0,001045 | 0,0024733 | 0,0016661 | 0,0016661 | TRUE | 0,01948 | No |
| orf19.1855 | 344 | 252 | 310 | 410 | 335 | 345 | 362 | 234 | 309 | 321 | 317 | 338 | 287 | 329 | 323 | 330 | 319,18 | 321 | 317 | 0,984 | -0,023 | -0,277 | 0,78185411 | 0,82890056 | 1,00E-08 | 0,0065278 | 0,0036535 | 0,0036535 | TRUE | 0,0521558 | No |
| orf19.1856 | 2961 | 2067 | 2644 | 3213 | 2920 | 2761 | 2893 | 1716 | 2656 | 2632 | 2705 | 2649 | 2497 | 2634 | 2582 | 2420 | 2597 | 2660 | 2533 | 0,953 | -0,069 | -1,74 | 0,08180931 | 0,1290731 | 0,000248 | 0,0025385 | 0,0011271 | 0,0011271 | TRUE | 0,0098136 | No |
| orf19.1857 | 893 | 577 | 720 | 977 | 763 | 661 | 773 | 439 | 801 | 735 | 737 | 806 | 653 | 631 | 690 | 619 | 708,77 | 770 | 648 | 0,842 | -0,248 | -3,888 | 0,00010098 | 0,00030921 | 0,001054 | 0,0040277 | 0,0024793 | 0,0024793 | TRUE | 0,0223593 | No |
| orf19.1860 | 5041 | 3084 | 4369 | 5495 | 5006 | 3788 | 4333 | 3096 | 4522 | 3926 | 4470 | 4531 | 4281 | 3614 | 3866 | 4367 | 4197,3 | 4362 | 4032 | 0,923 | -0,115 | -1,651 | 0,09868734 | 0,15117632 | 0,005776 | 0,0023254 | 0,0044081 | 0,0044081 | TRUE | 0,0592133 | No |
| orf19.1860.1 | 178 | 122 | 174 | 194 | 233 | 242 | 235 | 137 | 160 | 155 | 178 | 160 | 199 | 231 | 210 | 193 | 185,76 | 163 | 208 | 1,279 | 0,355 | 3,319 | 0,00090395 | 0,00229249 | 1,00E-08 | 0,0097943 | 0,0055412 | 0,0055412 | TRUE | 0,0579464 | No |
| orf19.1861 | 11259 | 8090 | 9974 | 12288 | 15241 | 13677 | 14933 | 9177 | 10100 | 10299 | 10204 | 10133 | 13035 | 13050 | 13325 | 12943 | 11636 | 10184 | 13088 | 1,286 | 0,362 | 14,474 | 1,78E-47 | 1,58E-45 | 3,71E-07 | 0,0021043 | 0,0005145 | 0,0005145 | TRUE | 0,0018312 | No |
| orf19.1862 | 4784 | 3190 | 3940 | 5093 | 7477 | 6116 | 6479 | 3788 | 4292 | 4061 | 4031 | 4200 | 6395 | 5836 | 5781 | 5343 | 4992,3 | 4146 | 5839 | 1,409 | 0,495 | 8,996 | 2,34E-19 | 4,06E-18 | 0,002887 | 0,0022703 | 0,0026959 | 0,0026959 | TRUE | 0,0502365 | No |
| orf19.1863 | 24 | 13 | 24 | 35 | 22 | 21 | 17 | 12 | 22 | 17 | 25 | 29 | 19 | 20 | 15 | 17 | 20,31 | 23 | 18 | 0,766 | -0,384 | -1,248 | 0,2121918 | 0,28954084 | 1,00E-08 | 0,0734726 | 0,041356 | 0,041356 | TRUE | 0,204769 | No |
| orf19.1864 | 3284 | 2284 | 3008 | 3673 | 3094 | 2645 | 3026 | 2047 | 2946 | 2908 | 3077 | 3029 | 2646 | 2524 | 2700 | 2887 | 2839,7 | 2990 | 2689 | 0,898 | -0,155 | -3,262 | 0,00110718 | 0,00276817 | 0,001441 | 0,0024907 | 0,0018233 | 0,0018233 | TRUE | 0,0278263 | No |
| orf19.1865 | 925 | 626 | 815 | 1038 | 948 | 845 | 882 | 541 | 830 | 797 | 834 | 856 | 811 | 806 | 787 | 763 | 810,46 | 829 | 792 | 0,955 | -0,066 | -1,219 | 0,22278641 | 0,30173596 | 1,00E-08 | 0,0037707 | 0,0016083 | 0,0016083 | TRUE | 0,0072638 | No |
| orf19.1866 | 439 | 311 | 364 | 446 | 436 | 317 | 340 | 236 | 394 | 396 | 372 | 368 | 373 | 302 | 303 | 333 | 355,19 | 382 | 328 | 0,859 | -0,22 | -2,489 | 0,01281553 | 0,02509638 | 0,003204 | 0,0060666 | 0,0046704 | 0,0046704 | TRUE | 0,1085237 | No |
| orf19.1867 | 623 | 386 | 518 | 674 | 619 | 529 | 551 | 394 | 559 | 491 | 530 | 556 | 529 | 505 | 492 | 556 | 527,2 | 534 | 520 | 0,969 | -0,046 | -0,65 | 0,51589703 | 0,60005891 | 0,000933 | 0,0047331 | 0,0029242 | 0,0029242 | TRUE | 0,0302901 | No |
| orf19.1868 | 5860 | 4003 | 5035 | 6258 | 7428 | 6458 | 7075 | 4361 | 5257 | 5096 | 5151 | 5160 | 6353 | 6162 | 6313 | 6151 | 5705,5 | 5166 | 6245 | 1,209 | 0,274 | 9,19 | 3,91E-20 | 7,10E-19 | 1,00E-08 | 0,002234 | 0,0006742 | 0,0006742 | TRUE | 0,0017155 | No |
| orf19.1871 | 2571 | 1760 | 2290 | 2690 | 3420 | 2924 | 3239 | 2041 | 2306 | 2241 | 2343 | 2218 | 2925 | 2790 | 2890 | 2879 | 2574 | 2277 | 2871 | 1,261 | 0,334 | 8,726 | 2,63E-18 | 4,19E-17 | 1,00E-08 | 0,0025435 | 0,0010161 | 0,0010161 | TRUE | 0,0045322 | No |
| orf19.1872 | 3309 | 2169 | 2817 | 3525 | 3379 | 2693 | 2876 | 2111 | 2969 | 2761 | 2882 | 2907 | 2890 | 2570 | 2566 | 2977 | 2815,2 | 2880 | 2751 | 0,954 | -0,068 | -1,182 | 0,23717237 | 0,31799411 | 0,00301 | 0,0024952 | 0,0028327 | 0,0028327 | TRUE | 0,0355897 | No |
| orf19.1873 | 1322 | 950 | 1124 | 1513 | 1242 | 1056 | 1152 | 807 | 1186 | 1209 | 1150 | 1248 | 1062 | 1008 | 1028 | 1138 | 1128,6 | 1198 | 1059 | 0,881 | -0,183 | -3,331 | 0,00086457 | 0,00220287 | 0,000869 | 0,0032658 | 0,0019904 | 0,0019904 | TRUE | 0,0266664 | No |

| | | | | | | | | | | | | | | | | | | | | | | | | | | | | | | | |
|---|---|---|---|---|---|---|---|---|---|---|---|---|---|---|---|---|---|---|---|---|---|---|---|---|---|---|---|---|---|---|---|
| orf19.1874 | 2 | 7 | 6 | 4 | 4 | 2 | 3 | 0 | 2 | 9 | 6 | 3 | 3 | 2 | 3 | 0 | 3,52 | 5 | 2 | 0,44 | -1,183 | -1,442 | 0,14935145 | NA | 1,00E-08 | 0,4145519 | 0,3117246 | 0,3117246 | TRUE | 0,1670028 | No |
| orf19.1876 | 1749 | 1229 | 1551 | 1912 | 1881 | 1508 | 1691 | 1213 | 1569 | 1565 | 1587 | 1577 | 1609 | 1439 | 1509 | 1711 | 1570,6 | 1574 | 1567 | 0,993 | -0,01 | -0,175 | 0,86125753 | 0,89507615 | 0,00182 | 0,0029038 | 0,0024224 | 0,0024224 | TRUE | 0,0423348 | No |
| orf19.1877 | 535 | 365 | 453 | 626 | 632 | 583 | 644 | 402 | 480 | 465 | 463 | 516 | 541 | 556 | 575 | 567 | 520,34 | 481 | 560 | 1,159 | 0,213 | 3,175 | 0,00150039 | 0,00363889 | 1,00E-08 | 0,0047694 | 0,0024067 | 0,0024067 | TRUE | 0,030144 | No |
| orf19.1878 | 1240 | 823 | 1076 | 1310 | 1323 | 1072 | 1233 | 786 | 1112 | 1048 | 1101 | 1080 | 1131 | 1023 | 1100 | 1109 | 1088,1 | 1085 | 1091 | 1,004 | 0,006 | 0,109 | 0,91287161 | 0,93576738 | 0,000409 | 0,0033137 | 0,0017041 | 0,0017041 | TRUE | 0,021692 | No |
| orf19.188 | 1909 | 1372 | 1647 | 2192 | 2284 | 2117 | 2236 | 1384 | 1713 | 1747 | 1685 | 1808 | 1953 | 2020 | 1995 | 1952 | 1859,1 | 1738 | 1980 | 1,139 | 0,188 | 4,487 | 7,22E-06 | 2,69E-05 | 1,00E-08 | 0,0027604 | 0,0011466 | 0,0011466 | TRUE | 0,0094981 | No |
| orf19.1880 | 4514 | 3040 | 3926 | 4740 | 3800 | 3321 | 3640 | 2206 | 4050 | 3870 | 4017 | 3909 | 3250 | 3169 | 3248 | 3111 | 3577,9 | 3962 | 3194 | 0,807 | -0,31 | -8,819 | 1,16E-18 | 1,88E-17 | 0,000149 | 0,0023853 | 0,0008988 | 0,0008988 | TRUE | 0,0033987 | No |
| orf19.1881 | 1046 | 739 | 885 | 1130 | 1402 | 1276 | 1230 | 835 | 938 | 941 | 905 | 932 | 1199 | 1218 | 1098 | 1178 | 1051 | 929 | 1173 | 1,262 | 0,336 | 6,34 | 2,29E-10 | 1,59E-09 | 1,00E-08 | 0,0033607 | 0,0017308 | 0,0017308 | TRUE | 0,0239031 | No |
| orf19.1883 | 1771 | 1155 | 1552 | 1878 | 1956 | 1663 | 1853 | 1157 | 1589 | 1470 | 1588 | 1549 | 1673 | 1587 | 1653 | 1632 | 1592,6 | 1549 | 1636 | 1,055 | 0,078 | 1,725 | 0,08452006 | 0,13268788 | 0,000173 | 0,0028911 | 0,0013264 | 0,0013264 | TRUE | 0,0126822 | No |
| orf19.1885 | 2762 | 1690 | 2200 | 2849 | 3025 | 2545 | 2838 | 1638 | 2478 | 2152 | 2251 | 2349 | 2587 | 2428 | 2532 | 2310 | 2386 | 2308 | 2464 | 1,068 | 0,095 | 1,706 | 0,08805509 | 0,13734034 | 0,002493 | 0,002588 | 0,0025705 | 0,0025705 | TRUE | 0,0298444 | No |
| orf19.1886 | 7308 | 4980 | 6386 | 7552 | 6333 | 5347 | 5799 | 3325 | 6556 | 6340 | 6534 | 6227 | 5416 | 5102 | 5175 | 4690 | 5754,9 | 6414 | 5096 | 0,795 | -0,331 | -6,986 | 2,82E-12 | 2,45E-11 | 0,001826 | 0,0022318 | 0,0019738 | 0,0019738 | TRUE | 0,0338342 | No |
| orf19.1887 | 1491 | 988 | 1360 | 1554 | 1359 | 1146 | 1254 | 768 | 1338 | 1258 | 1391 | 1281 | 1162 | 1093 | 1119 | 1083 | 1215,8 | 1317 | 1114 | 0,847 | -0,24 | -4,603 | 4,17E-06 | 1,60E-05 | 0,000696 | 0,0031736 | 0,0017746 | 0,0017746 | TRUE | 0,0168129 | No |
| orf19.1888 | 1451 | 940 | 1146 | 1464 | 1198 | 1028 | 1091 | 703 | 1302 | 1197 | 1172 | 1207 | 1025 | 981 | 974 | 992 | 1106,1 | 1220 | 993 | 0,813 | -0,298 | -5,633 | 1,77E-08 | 9,47E-08 | 0,000644 | 0,003292 | 0,0017699 | 0,0017699 | TRUE | 0,0254146 | No |
| orf19.1889 | 628 | 386 | 487 | 574 | 528 | 476 | 442 | 316 | 563 | 491 | 498 | 473 | 452 | 454 | 394 | 446 | 471,53 | 506 | 436 | 0,86 | -0,217 | -2,671 | 0,00756549 | 0,01566281 | 0,003429 | 0,0050582 | 0,004217 | 0,004217 | TRUE | 0,0696886 | No |
| orf19.1890 | 3855 | 2501 | 3330 | 4108 | 4080 | 3460 | 3845 | 2680 | 3458 | 3184 | 3407 | 3388 | 3489 | 3301 | 3431 | 3780 | 3429,8 | 3359 | 3500 | 1,04 | 0,057 | 1,145 | 0,25230977 | 0,33463071 | 0,001923 | 0,0024028 | 0,0020881 | 0,0020881 | TRUE | 0,0335023 | No |
| orf19.1891 | 12177 | 7652 | 10477 | 12949 | 12297 | 9732 | 10461 | 6787 | 10924 | 9742 | 10719 | 10678 | 10517 | 9286 | 9335 | 9572 | 10097 | 10516 | 9678 | 0,92 | -0,12 | -2,26 | 0,0238372 | 0,04368941 | 0,002879 | 0,0021233 | 0,0026098 | 0,0026098 | TRUE | 0,0420377 | No |
| orf19.1893 | 1385 | 870 | 1138 | 1391 | 1426 | 1209 | 1309 | 803 | 1242 | 1108 | 1164 | 1147 | 1220 | 1154 | 1168 | 1133 | 1166,9 | 1165 | 1169 | 1,003 | 0,004 | 0,073 | 0,94174689 | 0,95784921 | 0,000813 | 0,0032236 | 0,0018657 | 0,0018657 | TRUE | 0,0241357 | No |
| orf19.1896 | 18600 | 12041 | 16661 | 19703 | 22154 | 18958 | 20649 | 13991 | 16686 | 15329 | 17046 | 16247 | 18947 | 18089 | 18426 | 19733 | 17563 | 16327 | 18799 | 1,151 | 0,203 | 4,603 | 4,17E-06 | 1,61E-05 | 0,001679 | 0,0020622 | 0,0018098 | 0,0018098 | TRUE | 0,020492 | No |
| orf19.1897 | 3136 | 2129 | 2713 | 3314 | 3151 | 2661 | 2890 | 1827 | 2813 | 2710 | 2776 | 2733 | 2695 | 2539 | 2579 | 2577 | 2677,7 | 2758 | 2598 | 0,942 | -0,086 | -2,298 | 0,02154011 | 0,03988072 | 1,00E-08 | 0,0025217 | 0,0009805 | 0,0009805 | TRUE | 0,0081608 | No |
| orf19.190 | 446 | 387 | 434 | 513 | 502 | 570 | 499 | 294 | 400 | 493 | 444 | 423 | 429 | 544 | 445 | 415 | 449,12 | 440 | 458 | 1,047 | 0,066 | 0,66 | 0,50908831 | 0,59321229 | 0,008772 | 0,0052118 | 0,0072467 | 0,0072467 | TRUE | 0,184389 | No |
| orf19.1900 | 3389 | 2081 | 2906 | 3496 | 3124 | 2732 | 2941 | 1864 | 3040 | 2649 | 2973 | 2883 | 2672 | 2607 | 2624 | 2629 | 2759,7 | 2886 | 2633 | 0,911 | -0,134 | -2,801 | 0,00508862 | 0,01097359 | 0,001319 | 0,0025056 | 0,0018305 | 0,0018305 | TRUE | 0,0353651 | No |
| orf19.1901 | 9182 | 6164 | 8278 | 9853 | 9793 | 8687 | 9435 | 6000 | 8237 | 7847 | 8469 | 8125 | 8375 | 8289 | 8419 | 8462 | 8278,1 | 8170 | 8386 | 1,026 | 0,037 | 1,143 | 0,2531488 | 0,33559531 | 0,000378 | 0,0021549 | 0,0008963 | 0,0008963 | TRUE | 0,0082786 | No |
| orf19.1902 | 9628 | 6947 | 8633 | 10428 | 10423 | 9557 | 10461 | 6067 | 8637 | 8844 | 8832 | 8599 | 8914 | 9119 | 9335 | 8557 | 8854,7 | 8728 | 8981 | 1,03 | 0,042 | 1,19 | 0,23424194 | 0,31511992 | 0,000636 | 0,0021435 | 0,0010962 | 0,0010962 | TRUE | 0,0119001 | No |
| orf19.1906 | 83 | 60 | 75 | 111 | 78 | 81 | 68 | 55 | 74 | 76 | 77 | 92 | 67 | 77 | 61 | 78 | 75,17 | 80 | 71 | 0,875 | -0,193 | -1,174 | 0,24058197 | 0,32191904 | 1,00E-08 | 0,0212918 | 0,0126664 | 0,0126664 | TRUE | 0,1044274 | No |
| orf19.1907 | 4172 | 2561 | 3852 | 4241 | 4181 | 3427 | 4085 | 2558 | 3743 | 3260 | 3941 | 3497 | 3576 | 3270 | 3645 | 3608 | 3567,5 | 3610 | 3525 | 0,976 | -0,035 | -0,564 | 0,57283751 | 0,65132623 | 0,00382 | 0,0023864 | 0,0035054 | 0,0035054 | TRUE | 0,0504482 | No |
| orf19.191 | 3658 | 2683 | 3449 | 4304 | 4539 | 4218 | 4559 | 2940 | 3282 | 3416 | 3529 | 3549 | 3882 | 4025 | 4068 | 4147 | 3737,1 | 3444 | 4030 | 1,169 | 0,226 | 5,553 | 2,81E-08 | 1,47E-07 | 0,000659 | 0,002368 | 0,001318 | 0,001318 | TRUE | 0,0126791 | No |
| orf19.1910 | 693 | 425 | 617 | 716 | 648 | 477 | 585 | 381 | 622 | 541 | 631 | 590 | 554 | 455 | 522 | 537 | 556,64 | 596 | 517 | 0,866 | -0,208 | -2,608 | 0,00911493 | 0,01853843 | 0,004047 | 0,0045875 | 0,0043164 | 0,0043164 | TRUE | 0,0781159 | No |
| orf19.1911 | 5122 | 3174 | 4475 | 5492 | 4663 | 3646 | 3960 | 2731 | 4595 | 4041 | 4578 | 4529 | 3988 | 3479 | 3534 | 3852 | 4074,4 | 4436 | 3713 | 0,836 | -0,258 | -4,301 | 1,70E-05 | 5,98E-05 | 0,00365 | 0,0023358 | 0,0031971 | 0,0031971 | TRUE | 0,0428947 | No |
| orf19.1913 | 737 | 441 | 617 | 813 | 875 | 582 | 681 | 570 | 661 | 561 | 631 | 670 | 748 | 555 | 608 | 804 | 654,94 | 631 | 679 | 1,071 | 0,1 | 0,897 | 0,36981538 | 0,45830256 | 0,015773 | 0,0041961 | 0,0103002 | 0,0103002 | TRUE | 0,168818 | No |
| orf19.1914 | 620 | 371 | 528 | 662 | 543 | 417 | 458 | 335 | 556 | 472 | 540 | 546 | 464 | 398 | 409 | 472 | 482,26 | 528 | 436 | 0,82 | -0,286 | -3,452 | 0,00055564 | 0,00148237 | 0,003992 | 0,0049897 | 0,0045051 | 0,0045051 | TRUE | 0,0548126 | No |
| orf19.1915 | 7467 | 4468 | 6388 | 7964 | 6484 | 4823 | 5370 | 3078 | 6699 | 5688 | 6536 | 6567 | 5545 | 4602 | 4792 | 4341 | 5596,3 | 6372 | 4820 | 0,757 | -0,402 | -5,236 | 1,64E-07 | 7,81E-07 | 0,008145 | 0,0022389 | 0,0054898 | 0,0054898 | TRUE | 0,125481 | No |
| orf19.1917 | 3174 | 1959 | 2829 | 3470 | 3139 | 2434 | 2834 | 1814 | 2847 | 2494 | 2894 | 2861 | 2685 | 2322 | 2529 | 2558 | 2648,9 | 2774 | 2524 | 0,909 | -0,138 | -2,24 | 0,02510114 | 0,04579668 | 0,003558 | 0,0025276 | 0,0032566 | 0,0032566 | TRUE | 0,0544546 | No |
| orf19.192 | 1768 | 1396 | 1636 | 1975 | 1765 | 1785 | 1882 | 1271 | 1586 | 1777 | 1674 | 1629 | 1509 | 1703 | 1679 | 1793 | 1668,8 | 1666 | 1671 | 1,002 | 0,003 | 0,053 | 0,95791041 | 0,96952927 | 0,003107 | 0,0028494 | 0,0029981 | 0,0029981 | TRUE | 0,0520908 | No |
| orf19.1920 | 23 | 17 | 31 | 23 | 32 | 12 | 18 | 10 | 21 | 22 | 32 | 19 | 27 | 11 | 16 | 14 | 20,24 | 24 | 17 | 0,756 | -0,403 | -1,159 | 0,24649033 | 0,3284351 | 0,056069 | 0,0736942 | 0,0660817 | 0,0660817 | TRUE | 0,8980045 | No |
| orf19.1923 | 2248 | 1307 | 1994 | 2323 | 2683 | 2023 | 2429 | 1775 | 2017 | 1664 | 2040 | 1916 | 2295 | 1930 | 2167 | 2503 | 2066,5 | 1909 | 2224 | 1,163 | 0,218 | 2,582 | 0,00983533 | 0,01984239 | 0,00923 | 0,002682 | 0,0063518 | 0,0063518 | TRUE | 0,0954532 | No |
| orf19.1925 | 25 | 22 | 26 | 34 | 37 | 37 | 32 | 28 | 22 | 28 | 27 | 28 | 32 | 35 | 29 | 39 | 30,01 | 26 | 34 | 1,274 | 0,349 | 1,401 | 0,16128671 | 0,22935161 | 1,00E-08 | 0,050356 | 0,0258792 | 0,0258792 | TRUE | 0,0771313 | No |
| orf19.1926 | 279 | 257 | 283 | 347 | 247 | 234 | 217 | 189 | 250 | 327 | 290 | 286 | 211 | 223 | 194 | 267 | 255,98 | 288 | 224 | 0,772 | -0,372 | -3,224 | 0,00126536 | 0,00311954 | 0,010299 | 0,0076506 | 0,0088077 | 0,0088077 | TRUE | 0,1583694 | No |
| orf19.1927 | 2016 | 1372 | 1756 | 2167 | 1726 | 1644 | 1713 | 1062 | 1809 | 1747 | 1797 | 1787 | 1476 | 1569 | 1529 | 1498 | 1651,3 | 1785 | 1518 | 0,85 | -0,234 | -5,559 | 2,71E-08 | 1,42E-07 | 1,00E-08 | 0,0028587 | 0,0010929 | 0,0010929 | TRUE | 0,0062604 | No |
| orf19.193 | 364 | 341 | 353 | 409 | 433 | 509 | 497 | 287 | 327 | 434 | 361 | 337 | 370 | 486 | 443 | 405 | 395,42 | 365 | 426 | 1,174 | 0,231 | 2,101 | 0,03560821 | 0,06244428 | 0,012237 | 0,0056508 | 0,0090473 | 0,0090473 | TRUE | 0,1758228 | No |
| orf19.1930 | 126 | 102 | 118 | 174 | 147 | 141 | 144 | 86 | 113 | 130 | 121 | 143 | 126 | 135 | 128 | 121 | 127,14 | 127 | 128 | 1,007 | 0,01 | 0,079 | 0,93716158 | 0,95431641 | 1,00E-08 | 0,0133973 | 0,0076481 | 0,0076481 | TRUE | 0,0946837 | No |
| orf19.1932 | 1125 | 706 | 933 | 1213 | 1051 | 867 | 910 | 655 | 1009 | 899 | 955 | 1000 | 899 | 827 | 812 | 924 | 915,6 | 966 | 866 | 0,893 | -0,164 | -2,581 | 0,00985146 | 0,01986827 | 0,001879 | 0,003565 | 0,0027729 | 0,0027729 | TRUE | 0,0241665 | No |
| orf19.1933 | 1631 | 1174 | 1455 | 1743 | 1685 | 1569 | 1660 | 997 | 1463 | 1495 | 1489 | 1437 | 1441 | 1497 | 1481 | 1406 | 1463,7 | 1471 | 1456 | 0,992 | -0,011 | -0,256 | 0,79801535 | 0,84335554 | 1,00E-08 | 0,0029713 | 0,0011751 | 0,0011751 | TRUE | 0,0055549 | No |
| orf19.1934 | 2625 | 1758 | 2273 | 2771 | 2698 | 2438 | 2704 | 1821 | 2355 | 2238 | 2326 | 2285 | 2307 | 2326 | 2413 | 2568 | 2352,3 | 2301 | 2404 | 1,042 | 0,06 | 1,317 | 0,18794784 | 0,2607023 | 0,000915 | 0,0025967 | 0,0015629 | 0,0015629 | TRUE | 0,0238159 | No |
| orf19.1935 | 14 | 8 | 11 | 22 | 11 | 13 | 10 | 7 | 13 | 10 | 11 | 18 | 9 | 12 | 9 | 10 | 11,59 | 13 | 10 | 0,766 | -0,385 | -0,957 | 0,33874557 | NA | 1,00E-08 | 0,1271966 | 0,0692311 | 0,0692311 | TRUE | 0,3459724 | No |
| orf19.1936 | 6421 | 4356 | 5841 | 7180 | 8020 | 6740 | 7596 | 4880 | 5760 | 5546 | 5976 | 5921 | 6859 | 6431 | 6778 | 6883 | 6269,2 | 5801 | 6738 | 1,161 | 0,215 | 5,52 | 3,38E-08 | 1,76E-07 | 0,000822 | 0,0022111 | 0,001298 | 0,001298 | TRUE | 0,0115131 | No |
| orf19.1938 | 1790 | 1115 | 1436 | 1686 | 1076 | 963 | 1037 | 687 | 1606 | 1420 | 1469 | 1390 | 920 | 919 | 925 | 969 | 1202,3 | 1471 | 933 | 0,633 | -0,659 | -11,266 | 1,92E-29 | 6,68E-28 | 0,001865 | 0,003187 | 0,0024065 | 0,0024065 | TRUE | 0,0470394 | No |
| orf19.1939 | 25 | 14 | 22 | 22 | 20 | 18 | 19 | 8 | 22 | 18 | 23 | 18 | 17 | 17 | 17 | 11 | 17,93 | 20 | 16 | 0,785 | -0,349 | -1,097 | 0,27267443 | 0,357068 | 1,00E-08 | 0,0829568 | 0,0415284 | 0,0415284 | TRUE | 0,10341 | No |
| orf19.194 | 1367 | 1412 | 1348 | 1798 | 1356 | 1648 | 1508 | 982 | 1226 | 1798 | 1379 | 1483 | 1160 | 1572 | 1346 | 1385 | 1418,6 | 1472 | 1366 | 0,929 | -0,107 | -0,944 | 0,34519442 | 0,4324287 | 0,019405 | 0,0030029 | 0,0115318 | 0,0115318 | TRUE | 0,2636522 | No |
| orf19.1940 | 2336 | 1456 | 2010 | 2542 | 2851 | 2350 | 2646 | 1669 | 2096 | 1854 | 2056 | 2096 | 2438 | 2242 | 2361 | 2354 | 2187,2 | 2026 | 2349 | 1,158 | 0,212 | 4,113 | 3,90E-05 | 0,00012829 | 0,001618 | 0,0026433 | 0,0020832 | 0,0020832 | TRUE | 0,0373778 | No |
| orf19.1941 | 841 | 600 | 746 | 897 | 935 | 794 | 849 | 493 | 754 | 764 | 763 | 740 | 800 | 758 | 758 | 695 | 753,93 | 755 | 753 | 1,001 | 0,001 | 0,018 | 0,98572267 | 0,98918019 | 1,00E-08 | 0,003905 | 0,0020334 | 0,0020334 | TRUE | 0,0264949 | No |
| orf19.1942 | 3312 | 2258 | 2930 | 3521 | 3049 | 2818 | 3009 | 1970 | 2971 | 2875 | 2998 | 2903 | 2608 | 2689 | 2685 | 2779 | 2813,4 | 2937 | 2690 | 0,915 | -0,129 | -3,451 | 0,00055753 | 0,00148674 | 0,000109 | 0,0024955 | 0,0009773 | 0,0009773 | TRUE | 0,0054003 | No |
| orf19.1943 | 381 | 343 | 418 | 519 | 431 | 446 | 430 | 296 | 342 | 437 | 428 | 428 | 369 | 426 | 384 | 417 | 403,68 | 409 | 399 | 0,976 | -0,035 | -0,371 | 0,71052665 | 0,77124555 | 0,006489 | 0,0055757 | 0,0060351 | 0,0060351 | TRUE | 0,1445969 | No |
| orf19.1944 | 14305 | 9899 | 12895 | 16136 | 20649 | 17719 | 20142 | 12906 | 12833 | 12602 | 13193 | 13306 | 17660 | 16907 | 17973 | 18203 | 15335 | 12984 | 17686 | 1,362 | 0,445 | 12,587 | 2,50E-36 | 1,36E-34 | 0,000733 | 0,0020742 | 0,0011358 | 0,0011358 | TRUE | 0,0107506 | No |
| orf19.1945 | 19091 | 12755 | 16656 | 20125 | 19365 | 16882 | 18378 | 11219 | 17127 | 16238 | 17041 | 16595 | 16562 | 16108 | 16399 | 15823 | 16487 | 16750 | 16223 | 0,969 | -0,046 | -1,483 | 0,13801075 | 0,20086448 | 0,000425 | 0,0020676 | 0,0008627 | 0,0008627 | TRUE | 0,0050741 | No |
| orf19.1946 | 2758 | 1900 | 2294 | 2845 | 2556 | 2191 | 2292 | 1374 | 2474 | 2419 | 2347 | 2346 | 2186 | 2091 | 2045 | 1938 | 2230,7 | 2396 | 2065 | 0,863 | -0,212 | -4,524 | 6,07E-06 | 2,29E-05 | 0,000978 | 0,0026303 | 0,0016584 | 0,0016584 | TRUE | 0,0189916 | No |
| orf19.1948 | 1093 | 703 | 977 | 1215 | 1151 | 939 | 1037 | 733 | 981 | 895 | 1000 | 1002 | 984 | 896 | 925 | 1034 | 964,56 | 970 | 960 | 0,986 | -0,02 | -0,314 | 0,75327066 | 0,80671125 | 0,001887 | 0,0034846 | 0,0027485 | 0,0027485 | TRUE | 0,02925 | No |
| orf19.1949 | 4379 | 2996 | 3890 | 4663 | 4098 | 3521 | 3831 | 2389 | 3928 | 3814 | 3980 | 3845 | 3505 | 3360 | 3419 | 3369 | 3652,5 | 3892 | 3413 | 0,877 | -0,189 | -5,511 | 3,56E-08 | 1,84E-07 | 0,000116 | 0,002377 | 0,0008522 | 0,0008522 | TRUE | 0,0041006 | No |
| orf19.1950 | 506 | 354 | 443 | 510 | 583 | 442 | 497 | 389 | 454 | 451 | 453 | 421 | 499 | 422 | 443 | 549 | 461,36 | 445 | 478 | 1,072 | 0,1 | 1,139 | 0,25479063 | 0,33739963 | 0,004885 | 0,0051261 | 0,0051836 | 0,0051836 | TRUE | 0,1011824 | No |
| orf19.1953 | 2519 | 1743 | 2223 | 2791 | 3206 | 2721 | 2961 | 1953 | 2260 | 2219 | 2274 | 2301 | 2742 | 2596 | 2642 | 2755 | 2473,7 | 2264 | 2684 | 1,184 | 0,244 | 6,313 | 2,74E-10 | 1,89E-09 | 1,00E-08 | 0,0025664 | 0,0010239 | 0,0010239 | TRUE | 0,0057774 | No |
| orf19.1954 | 4926 | 3174 | 4239 | 5144 | 4939 | 4418 | 4921 | 2826 | 4419 | 4041 | 4337 | 4242 | 4224 | 4216 | 4391 | 3986 | 4231,9 | 4260 | 4204 | 0,987 | -0,018 | -0,421 | 0,67379748 | 0,73939705 | 0,001107 | 0,0023226 | 0,0016133 | 0,0016133 | TRUE | 0,0141351 | No |

| | | | | | | | | | | | | | | | | | | | | | | | | | | | | | | | |
|---|---|---|---|---|---|---|---|---|---|---|---|---|---|---|---|---|---|---|---|---|---|---|---|---|---|---|---|---|---|---|---|
| orf19.1955 | 961 | 633 | 785 | 881 | 887 | 766 | 802 | 541 | 862 | 806 | 803 | 726 | 759 | 731 | 716 | 763 | 770,72 | 799 | 742 | 0,928 | -0,108 | -1,64 | 0,10107194 | 0,15418076 | 0,001991 | 0,0038631 | 0,0028302 | 0,0028302 | TRUE | 0,0485675 | No |
| orf19.1956 | 3326 | 2473 | 3069 | 3602 | 3764 | 3379 | 3537 | 2316 | 2984 | 3148 | 3140 | 2970 | 3219 | 3224 | 3156 | 3267 | 3138,5 | 3060 | 3216 | 1,051 | 0,072 | 1,935 | 0,05294067 | 0,08840121 | 0,000275 | 0,0024421 | 0,0010244 | 0,0010244 | TRUE | 0,0048419 | No |
| orf19.1957 | 9032 | 5803 | 7701 | 9452 | 10981 | 8991 | 9929 | 5810 | 8103 | 7388 | 7879 | 7794 | 9391 | 8579 | 8860 | 8194 | 8273,5 | 7791 | 8756 | 1,124 | 0,169 | 3,422 | 0,00062051 | 0,00163153 | 0,002227 | 0,002155 | 0,0022148 | 0,0022148 | TRUE | 0,0291761 | No |
| orf19.1958 | 21 | 14 | 22 | 19 | 13 | 13 | 17 | 13 | 19 | 18 | 23 | 16 | 11 | 12 | 15 | 18 | 16,48 | 19 | 14 | 0,75 | -0,415 | -1,226 | 0,22030498 | 0,29898177 | 1,00E-08 | 0,090051 | 0,0480365 | 0,0480365 | TRUE | 0,108217 | No |
| orf19.1959 | 1815 | 1274 | 1630 | 1890 | 1320 | 1259 | 1266 | 894 | 1628 | 1622 | 1668 | 1559 | 1129 | 1201 | 1130 | 1261 | 1399,6 | 1619 | 1180 | 0,727 | -0,46 | -8,899 | 5,61E-19 | 9,39E-18 | 0,000865 | 0,0030167 | 0,0018235 | 0,0018235 | TRUE | 0,0227111 | No |
| orf19.1960 | 4892 | 3718 | 4588 | 5469 | 6339 | 5973 | 6402 | 4019 | 4389 | 4733 | 4694 | 4510 | 5421 | 5699 | 5713 | 5668 | 5103,4 | 4582 | 5625 | 1,228 | 0,297 | 7,711 | 1,25E-14 | 1,38E-13 | 0,000654 | 0,002264 | 0,0012233 | 0,0012233 | TRUE | 0,0097703 | No |
| orf19.1961 | 8360 | 6161 | 7564 | 9403 | 9144 | 8859 | 9615 | 5557 | 7500 | 7844 | 7739 | 7754 | 7820 | 8453 | 8580 | 7838 | 7940,8 | 7709 | 8173 | 1,061 | 0,085 | 2,029 | 0,04240924 | 0,0728193 | 0,001213 | 0,0021623 | 0,0015537 | 0,0015537 | TRUE | 0,0137907 | No |
| orf19.1963 | 5017 | 3488 | 4531 | 5254 | 5059 | 4545 | 5046 | 2765 | 4501 | 4441 | 4636 | 4333 | 4327 | 4337 | 4503 | 3900 | 4371,9 | 4478 | 4267 | 0,954 | -0,068 | -1,384 | 0,16621186 | 0,23563022 | 0,001914 | 0,0023116 | 0,0020812 | 0,0020812 | TRUE | 0,0390874 | No |
| orf19.1964 | 3317 | 2365 | 3017 | 3634 | 2680 | 2592 | 2703 | 1789 | 2976 | 3011 | 3087 | 2997 | 2292 | 2473 | 2412 | 2523 | 2721,3 | 3018 | 2425 | 0,803 | -0,317 | -7,646 | 2,08E-14 | 2,23E-13 | 0,000478 | 0,002513 | 0,0012778 | 0,0012778 | TRUE | 0,0168897 | No |
| orf19.1966 | 4642 | 3058 | 3896 | 4931 | 4567 | 3888 | 4219 | 2598 | 4164 | 3893 | 3986 | 4066 | 3906 | 3710 | 3765 | 3664 | 3894,3 | 4027 | 3761 | 0,934 | -0,099 | -2,584 | 0,0097695 | 0,01972609 | 0,000457 | 0,0023523 | 0,0011389 | 0,0011389 | TRUE | 0,0082884 | No |
| orf19.1967 | 1519 | 1047 | 1302 | 1584 | 1565 | 1301 | 1449 | 789 | 1363 | 1333 | 1332 | 1306 | 1338 | 1241 | 1293 | 1113 | 1289,9 | 1334 | 1246 | 0,938 | -0,093 | -1,563 | 0,11810783 | 0,17588853 | 0,002164 | 0,0031049 | 0,0025943 | 0,0025943 | TRUE | 0,0567003 | No |
| orf19.1968 | 123 | 107 | 158 | 149 | 106 | 95 | 117 | 78 | 110 | 136 | 162 | 123 | 91 | 91 | 104 | 110 | 115,85 | 133 | 99 | 0,744 | -0,427 | -2,83 | 0,00465759 | 0,01012768 | 0,011118 | 0,0145104 | 0,0129546 | 0,0129546 | TRUE | 0,2262112 | No |
| orf19.1968.1 | 275 | 207 | 214 | 278 | 245 | 233 | 210 | 137 | 247 | 264 | 219 | 229 | 210 | 222 | 187 | 193 | 221,36 | 240 | 203 | 0,852 | -0,231 | -2,256 | 0,02409614 | 0,04411017 | 0,001533 | 0,0085376 | 0,0055403 | 0,0055403 | TRUE | 0,0442051 | No |
| orf19.1969 | 68 | 41 | 61 | 58 | 50 | 31 | 40 | 34 | 61 | 52 | 62 | 48 | 43 | 30 | 36 | 48 | 47,43 | 56 | 39 | 0,691 | -0,534 | -2,502 | 0,01236601 | 0,02431118 | 9,86E-05 | 0,032588 | 0,0218993 | 0,0218993 | TRUE | 0,1948891 | No |
| orf19.1970 | 2299 | 1621 | 1997 | 2540 | 2499 | 2112 | 2370 | 1632 | 2062 | 2064 | 2043 | 2095 | 2137 | 2015 | 2115 | 2302 | 2104,1 | 2066 | 2142 | 1,035 | 0,049 | 1,053 | 0,29224125 | 0,37686797 | 0,000958 | 0,0026695 | 0,0016415 | 0,0016415 | TRUE | 0,0278635 | No |
| orf19.1971 | 13220 | 9023 | 11254 | 13976 | 12433 | 10958 | 11911 | 7586 | 11860 | 11487 | 11514 | 11525 | 10633 | 10456 | 10629 | 10699 | 11100 | 11596 | 10604 | 0,914 | -0,129 | -4,954 | 7,27E-07 | 3,14E-06 | 8,20E-05 | 0,0021103 | 0,0005644 | 0,0005644 | TRUE | 0,0028913 | No |
| orf19.1972 | 4205 | 2783 | 3676 | 4454 | 4432 | 3654 | 3903 | 2581 | 3772 | 3543 | 3761 | 3673 | 3790 | 3487 | 3483 | 3640 | 3643,6 | 3687 | 3600 | 0,976 | -0,035 | -0,829 | 0,40707856 | 0,49524111 | 0,000876 | 0,0023779 | 0,0014645 | 0,0014645 | TRUE | 0,0161914 | No |
| orf19.1973 | 1137 | 658 | 849 | 1145 | 1248 | 950 | 1152 | 692 | 1020 | 838 | 869 | 944 | 1067 | 906 | 1028 | 976 | 956,03 | 918 | 994 | 1,082 | 0,114 | 1,502 | 0,13314239 | 0,19519753 | 0,005004 | 0,003498 | 0,0045089 | 0,0045089 | TRUE | 0,067088 | No |
| orf19.1974 | 3930 | 2734 | 3386 | 4448 | 4953 | 4094 | 4594 | 3278 | 3526 | 3481 | 3464 | 3668 | 4236 | 3906 | 4099 | 4623 | 3875,4 | 3535 | 4216 | 1,191 | 0,253 | 4,716 | 2,41E-06 | 9,66E-06 | 0,002582 | 0,0023541 | 0,0024946 | 0,0024946 | TRUE | 0,049696 | No |
| orf19.1975 | 662 | 460 | 569 | 718 | 588 | 558 | 534 | 298 | 594 | 586 | 582 | 592 | 503 | 532 | 477 | 420 | 535,73 | 588 | 483 | 0,825 | -0,278 | -3,732 | 0,00019005 | 0,00055591 | 0,001893 | 0,0046893 | 0,0034323 | 0,0034323 | TRUE | 0,0748002 | No |
| orf19.1976 | 512 | 364 | 451 | 595 | 451 | 432 | 469 | 270 | 459 | 463 | 461 | 491 | 386 | 412 | 419 | 381 | 434 | 468 | 400 | 0,853 | -0,23 | -3,218 | 0,00128972 | 0,00317308 | 1,00E-08 | 0,0053244 | 0,0025788 | 0,0025788 | TRUE | 0,0126955 | No |
| orf19.1977 | 1197 | 874 | 1147 | 1313 | 1387 | 1203 | 1265 | 925 | 1074 | 1113 | 1173 | 1083 | 1186 | 1148 | 1129 | 1305 | 1151,3 | 1111 | 1192 | 1,071 | 0,099 | 1,668 | 0,09525016 | 0,14692245 | 0,001852 | 0,0032405 | 0,0024901 | 0,0024901 | TRUE | 0,044094 | No |
| orf19.1978 | 1068 | 704 | 889 | 1127 | 955 | 873 | 937 | 575 | 958 | 896 | 910 | 929 | 817 | 833 | 836 | 811 | 873,76 | 923 | 824 | 0,892 | -0,165 | -3,159 | 0,00158436 | 0,00381822 | 1,00E-08 | 0,003641 | 0,0014656 | 0,0014656 | TRUE | 0,007658 | No |
| orf19.1979 | 15042 | 10432 | 13631 | 16654 | 22144 | 20492 | 21615 | 12990 | 13494 | 13281 | 13946 | 13733 | 18938 | 19553 | 19288 | 18321 | 16319 | 13614 | 19025 | 1,398 | 0,483 | 14,816 | 1,15E-49 | 1,17E-47 | 0,000532 | 0,0020685 | 0,0009574 | 0,0009574 | TRUE | 0,0074282 | No |
| orf19.198 | 5296 | 3831 | 4818 | 5921 | 5538 | 4894 | 5401 | 3618 | 4751 | 4877 | 4929 | 4883 | 4736 | 4670 | 4819 | 5103 | 4846,1 | 4860 | 4832 | 0,993 | -0,01 | -0,254 | 0,79933431 | 0,84445236 | 0,000569 | 0,0022791 | 0,0011663 | 0,0011663 | TRUE | 0,0163743 | No |
| orf19.1980 | 100 | 83 | 101 | 116 | 111 | 107 | 110 | 54 | 90 | 106 | 103 | 96 | 95 | 102 | 98 | 76 | 95,71 | 99 | 93 | 0,955 | -0,066 | -0,458 | 0,64699533 | 0,71652497 | 1,00E-08 | 0,0171466 | 0,009817 | 0,009817 | TRUE | 0,1048853 | No |
| orf19.1981 | 1756 | 1162 | 1400 | 1802 | 2014 | 1736 | 1842 | 1073 | 1575 | 1479 | 1432 | 1486 | 1722 | 1656 | 1644 | 1513 | 1563,6 | 1493 | 1634 | 1,096 | 0,132 | 2,476 | 0,0132794 | 0,02592029 | 0,001379 | 0,002908 | 0,0020967 | 0,0020967 | TRUE | 0,027188 | No |
| orf19.1985 | 578 | 483 | 577 | 712 | 643 | 584 | 680 | 414 | 519 | 615 | 590 | 587 | 550 | 557 | 607 | 584 | 576,09 | 578 | 574 | 0,996 | -0,006 | -0,084 | 0,93310455 | 0,95179832 | 0,002062 | 0,0044994 | 0,0032482 | 0,0032482 | TRUE | 0,0573132 | No |
| orf19.1986 | 2507 | 1572 | 2120 | 2576 | 2656 | 2283 | 2489 | 1519 | 2249 | 2001 | 2169 | 2124 | 2271 | 2178 | 2221 | 2142 | 2169,6 | 2136 | 2203 | 1,031 | 0,044 | 0,954 | 0,34013036 | 0,42720132 | 0,000879 | 0,0026486 | 0,0016192 | 0,0016192 | TRUE | 0,0203705 | No |
| orf19.1988 | 1330 | 863 | 1132 | 1360 | 1378 | 1139 | 1245 | 858 | 1193 | 1099 | 1158 | 1121 | 1179 | 1087 | 1111 | 1210 | 1144,7 | 1143 | 1147 | 1,001 | 0,001 | 0,02 | 0,98397818 | 0,98801577 | 0,000942 | 0,0032477 | 0,0019704 | 0,0019704 | TRUE | 0,0146469 | No |
| orf19.1989 | 4213 | 2893 | 3793 | 4627 | 4210 | 3733 | 4028 | 2580 | 3780 | 3683 | 3881 | 3815 | 3601 | 3562 | 3594 | 3639 | 3694,3 | 3790 | 3599 | 0,949 | -0,076 | -2,302 | 0,0213116 | 0,03949416 | 1,00E-08 | 0,0023725 | 0,0007652 | 0,0007652 | TRUE | 0,0040277 | No |
| orf19.199 | 2506 | 2223 | 2533 | 3018 | 2665 | 2790 | 2955 | 1650 | 2248 | 2830 | 2592 | 2489 | 2279 | 2662 | 2637 | 2327 | 2508 | 2540 | 2476 | 0,976 | -0,035 | -0,451 | 0,65207502 | 0,72108719 | 0,007423 | 0,0025584 | 0,0052685 | 0,0052685 | TRUE | 0,0725168 | No |
| orf19.1990 | 2784 | 2348 | 2681 | 3203 | 3063 | 3345 | 3462 | 2059 | 2498 | 2989 | 2743 | 2641 | 2620 | 3192 | 3089 | 2904 | 2834,4 | 2718 | 2951 | 1,087 | 0,12 | 1,664 | 0,09605097 | 0,14771653 | 0,006052 | 0,0024917 | 0,0046297 | 0,0046297 | TRUE | 0,0709636 | No |
| orf19.1991 | 3310 | 2515 | 2805 | 3656 | 3881 | 3731 | 3837 | 2448 | 2969 | 3202 | 2870 | 3015 | 3319 | 3560 | 3424 | 3453 | 3226,5 | 3014 | 3439 | 1,142 | 0,191 | 4,302 | 1,69E-05 | 5,96E-05 | 0,001065 | 0,0024295 | 0,0015806 | 0,0015806 | TRUE | 0,0202859 | No |
| orf19.1992 | 3845 | 3081 | 3466 | 4352 | 3360 | 3456 | 3692 | 2316 | 3449 | 3922 | 3546 | 3589 | 2874 | 3298 | 3294 | 3267 | 3404,9 | 3626 | 3183 | 0,878 | -0,188 | -3,186 | 0,00144347 | 0,00351678 | 0,003324 | 0,0024059 | 0,0030482 | 0,0030482 | TRUE | 0,0531174 | No |
| orf19.1993 | 1216 | 1538 | 1217 | 1796 | 1265 | 1651 | 1531 | 935 | 1091 | 1958 | 1245 | 1481 | 1082 | 1575 | 1366 | 1319 | 1389,6 | 1444 | 1336 | 0,925 | -0,112 | -0,531 | 0,59551201 | 0,67277851 | 0,041906 | 0,0030242 | 0,0215632 | 0,0419057 | TRUE | 0,660375 | No |
| orf19.1994 | 1514 | 1186 | 1383 | 1743 | 1492 | 1642 | 1744 | 1157 | 1358 | 1510 | 1415 | 1437 | 1276 | 1567 | 1556 | 1632 | 1468,9 | 1430 | 1508 | 1,053 | 0,075 | 0,995 | 0,3197547 | 0,40640708 | 0,006181 | 0,0029678 | 0,0047593 | 0,0047593 | TRUE | 0,1316745 | No |
| orf19.1995 | 552 | 515 | 524 | 695 | 544 | 543 | 525 | 302 | 495 | 656 | 536 | 573 | 465 | 518 | 468 | 426 | 517,23 | 565 | 469 | 0,835 | -0,26 | -2,773 | 0,00555314 | 0,01188151 | 0,00794 | 0,0047862 | 0,0064835 | 0,0064835 | TRUE | 0,1280463 | No |
| orf19.1996 | 7085 | 5069 | 6284 | 7620 | 7192 | 6588 | 7097 | 4291 | 6356 | 6453 | 6429 | 6284 | 6151 | 6286 | 6333 | 6052 | 6293 | 6380 | 6206 | 0,973 | -0,039 | -1,296 | 0,19499858 | 0,26858126 | 7,36E-05 | 0,0022102 | 0,0007093 | 0,0007093 | TRUE | 0,0031339 | No |
| orf19.1998 | 344 | 220 | 322 | 368 | 367 | 319 | 337 | 221 | 309 | 280 | 329 | 303 | 314 | 304 | 301 | 312 | 306,53 | 305 | 308 | 1,005 | 0,007 | 0,081 | 0,93554538 | 0,95380225 | 1,00E-08 | 0,0067154 | 0,0033373 | 0,0033373 | TRUE | 0,0293652 | No |
| orf19.1999 | 2076 | 1335 | 1635 | 2145 | 1911 | 1601 | 1723 | 1055 | 1862 | 1700 | 1673 | 1769 | 1634 | 1528 | 1537 | 1488 | 1648,9 | 1751 | 1547 | 0,884 | -0,178 | -3,431 | 0,00060244 | 0,00159237 | 0,001384 | 0,0028599 | 0,0019804 | 0,0019804 | TRUE | 0,0223827 | No |
| orf19.20 | 1987 | 1414 | 1807 | 2152 | 2552 | 2444 | 2486 | 1629 | 1783 | 1800 | 1849 | 1775 | 2183 | 2332 | 2218 | 2298 | 2029,6 | 1802 | 2258 | 1,253 | 0,325 | 7,816 | 5,46E-15 | 6,27E-14 | 1,00E-08 | 0,0026948 | 0,0011591 | 0,0011591 | TRUE | 0,0064622 | No |
| orf19.200 | 1328 | 824 | 1071 | 1373 | 1072 | 998 | 1118 | 599 | 1191 | 1049 | 1096 | 1132 | 917 | 952 | 998 | 845 | 1022,5 | 1117 | 928 | 0,832 | -0,266 | -4,138 | 3,50E-05 | 0,00011631 | 0,002547 | 0,0033993 | 0,0029759 | 0,0029759 | TRUE | 0,038538 | No |
| orf19.2000 | 3775 | 3011 | 3408 | 4054 | 3654 | 3792 | 4004 | 2245 | 3387 | 3833 | 3487 | 3343 | 3125 | 3618 | 3573 | 3166 | 3441,5 | 3512 | 3370 | 0,961 | -0,058 | -0,897 | 0,36967518 | 0,45822637 | 0,004441 | 0,0024013 | 0,0037043 | 0,0037043 | TRUE | 0,0452251 | No |
| orf19.2001 | 445 | 408 | 405 | 560 | 607 | 678 | 702 | 388 | 399 | 519 | 414 | 462 | 519 | 647 | 626 | 547 | 516,81 | 448 | 585 | 1,308 | 0,388 | 3,866 | 0,00011084 | 0,00033685 | 0,010174 | 0,0047885 | 0,0076731 | 0,0076731 | TRUE | 0,1211814 | No |
| orf19.2002 | 2785 | 2361 | 2734 | 3289 | 3030 | 3061 | 3031 | 2009 | 2498 | 3006 | 2797 | 2712 | 2591 | 2921 | 2705 | 2834 | 2758 | 2753 | 2763 | 1,004 | 0,005 | 0,088 | 0,9301837 | 0,94946375 | 0,00392 | 0,0025059 | 0,0033416 | 0,0033416 | TRUE | 0,0472151 | No |
| orf19.2003 | 9040 | 6514 | 8185 | 10105 | 8360 | 7676 | 8230 | 4854 | 8110 | 8293 | 8374 | 8333 | 7150 | 7324 | 7344 | 6846 | 7721,7 | 8278 | 7166 | 0,866 | -0,207 | -6,156 | 7,45E-10 | 4,82E-09 | 0,000442 | 0,0021675 | 0,0009545 | 0,0009545 | TRUE | 0,0104657 | No |
| orf19.2005 | 14516 | 9869 | 12843 | 15978 | 18724 | 16446 | 17945 | 11023 | 13022 | 12564 | 13140 | 13176 | 16013 | 15692 | 16013 | 15547 | 14396 | 12976 | 15816 | 1,219 | 0,286 | 9,965 | 2,18E-23 | 4,96E-22 | 0,000253 | 0,0020804 | 0,000718 | 0,000718 | TRUE | 0,0054211 | No |
| orf19.2006 | 1095 | 861 | 1006 | 1237 | 908 | 867 | 935 | 566 | 982 | 1096 | 1029 | 1020 | 777 | 827 | 834 | 798 | 920,52 | 1032 | 809 | 0,786 | -0,348 | -6,184 | 6,24E-10 | 4,08E-09 | 0,000532 | 0,0035566 | 0,0019314 | 0,0019314 | TRUE | 0,0192371 | No |
| orf19.2006.1 | 1531 | 1284 | 1324 | 1837 | 2255 | 1972 | 1955 | 1188 | 1373 | 1635 | 1355 | 1515 | 1929 | 1882 | 1745 | 1676 | 1638,5 | 1470 | 1808 | 1,233 | 0,302 | 4,257 | 2,08E-05 | 7,19E-05 | 0,005291 | 0,0028655 | 0,0042081 | 0,0042081 | TRUE | 0,0660013 | No |
| orf19.2007 | 460 | 341 | 424 | 509 | 536 | 526 | 510 | 317 | 413 | 434 | 434 | 420 | 458 | 502 | 455 | 447 | 445,35 | 425 | 466 | 1,098 | 0,135 | 1,908 | 0,05640893 | 0,09320815 | 1,00E-08 | 0,0052392 | 0,00259 | 0,00259 | TRUE | 0,0321232 | No |
| orf19.2008 | 1586 | 1216 | 1361 | 1723 | 1670 | 1670 | 1786 | 1042 | 1423 | 1548 | 1392 | 1421 | 1428 | 1593 | 1594 | 1470 | 1483,7 | 1446 | 1521 | 1,054 | 0,076 | 1,34 | 0,18035409 | 0,25201289 | 0,001822 | 0,002958 | 0,0023982 | 0,0023982 | TRUE | 0,0254816 | No |
| orf19.2009 | 1697 | 1407 | 1647 | 1941 | 1934 | 1752 | 1864 | 1136 | 1522 | 1791 | 1685 | 1601 | 1654 | 1672 | 1663 | 1602 | 1648,8 | 1650 | 1648 | 1,001 | 0,002 | 0,035 | 0,97226191 | 0,98075043 | 0,001925 | 0,00286 | 0,0023157 | 0,0023157 | TRUE | 0,0383358 | No |
| orf19.201 | 290 | 179 | 246 | 292 | 262 | 213 | 254 | 130 | 260 | 228 | 252 | 241 | 224 | 203 | 227 | 183 | 227,23 | 245 | 209 | 0,858 | -0,22 | -2,201 | 0,02771868 | 0,04993518 | 0,000727 | 0,0083682 | 0,0052184 | 0,0052184 | TRUE | 0,061273 | No |
| orf19.2010 | 24 | 9 | 12 | 23 | 14 | 20 | 23 | 9 | 22 | 11 | 12 | 19 | 12 | 19 | 21 | 13 | 16,06 | 16 | 16 | 0,992 | -0,012 | -0,032 | 0,97463608 | 0,98199121 | 0,024114 | 0,0923538 | 0,065211 | 0,065211 | TRUE | 0,2541481 | No |
| orf19.2012 | 3304 | 2351 | 2932 | 3602 | 3759 | 3403 | 3540 | 2238 | 2964 | 2993 | 3000 | 2970 | 3215 | 3247 | 3159 | 3156 | 3088 | 2982 | 3194 | 1,072 | 0,1 | 3,058 | 0,00223116 | 0,00521995 | 1,00E-08 | 0,0024496 | 0,0007053 | 0,0007053 | TRUE | 0,0014798 | No |
| orf19.2013 | 8393 | 5574 | 7367 | 8694 | 10555 | 8738 | 9585 | 6144 | 7529 | 7096 | 7537 | 7169 | 9027 | 8338 | 8553 | 8666 | 7989,4 | 7333 | 8646 | 1,179 | 0,237 | 6,115 | 9,68E-10 | 6,18E-09 | 0,000893 | 0,0021612 | 0,0013215 | 0,0013215 | TRUE | 0,0109935 | No |
| orf19.2014 | 6593 | 4361 | 5675 | 6809 | 6808 | 5818 | 6304 | 4158 | 5915 | 5552 | 5806 | 5615 | 5822 | 5551 | 5625 | 5864 | 5718,9 | 5722 | 5716 | 0,998 | -0,002 | -0,066 | 0,94766331 | 0,96239402 | 0,000492 | 0,0022334 | 0,0011082 | 0,0011082 | TRUE | 0,0063275 | No |
| orf19.2016 | 236 | 214 | 212 | 313 | 219 | 247 | 207 | 138 | 212 | 272 | 217 | 258 | 187 | 236 | 185 | 195 | 220,19 | 240 | 201 | 0,838 | -0,254 | -2,11 | 0,03484116 | 0,06126007 | 0,010194 | 0,0085726 | 0,0093419 | 0,0093419 | TRUE | 0,1573105 | No |

| | | | | | | | | | | | | | | | | | | | | | | | | | | | | | | | |
|---|---|---|---|---|---|---|---|---|---|---|---|---|---|---|---|---|---|---|---|---|---|---|---|---|---|---|---|---|---|---|---|
| orf19.2017 | 6818 | 4840 | 5983 | 7194 | 6121 | 5487 | 5798 | 3879 | 6116 | 6162 | 6121 | 5932 | 5235 | 5236 | 5174 | 5471 | 5680,9 | 6083 | 5279 | 0,867 | -0,205 | -6,31 | 2,79E-10 | 1,92E-09 | 0,000216 | 0,0022351 | 0,0008384 | 0,0008384 | TRUE | 0,0068383 | No |
| orf19.2018 | 1684 | 1181 | 1462 | 1764 | 1651 | 1361 | 1460 | 1108 | 1511 | 1504 | 1496 | 1455 | 1412 | 1299 | 1303 | 1563 | 1442,6 | 1492 | 1394 | 0,933 | -0,101 | -1,62 | 0,10521084 | 0,15914066 | 0,003179 | 0,0029858 | 0,0030057 | 0,0030057 | TRUE | 0,0739527 | No |
| orf19.2018.1 | 136 | 109 | 110 | 149 | 116 | 106 | 122 | 83 | 122 | 139 | 113 | 123 | 99 | 101 | 109 | 117 | 115,31 | 124 | 106 | 0,857 | -0,222 | -1,693 | 0,09048683 | 0,14071567 | 1,00E-08 | 0,0145693 | 0,0078212 | 0,0078212 | TRUE | 0,0537315 | No |
| orf19.2018.2 | 10 | 18 | 9 | 20 | 18 | 12 | 9 | 7 | 9 | 23 | 9 | 16 | 15 | 11 | 8 | 10 | 12,79 | 14 | 11 | 0,796 | -0,329 | -0,768 | 0,44226086 | NA | 0,071304 | 0,1154655 | 0,096904 | 0,096904 | TRUE | 0,409077 | No |
| orf19.2019 | 2292 | 1538 | 1884 | 2363 | 2861 | 2363 | 2529 | 1850 | 2056 | 1958 | 1928 | 1949 | 2447 | 2255 | 2257 | 2609 | 2182,2 | 1973 | 2392 | 1,211 | 0,276 | 4,948 | 7,51E-07 | 3,23E-06 | 0,002348 | 0,0026448 | 0,0025162 | 0,0025162 | TRUE | 0,0428503 | No |
| orf19.202 | 5640 | 3878 | 5036 | 6071 | 5247 | 4506 | 4771 | 3194 | 5060 | 4937 | 5152 | 5006 | 4487 | 4299 | 4257 | 4505 | 4713,1 | 5039 | 4387 | 0,87 | -0,201 | -5,736 | 9,69E-09 | 5,37E-08 | 0,000298 | 0,0022875 | 0,0009619 | 0,0009619 | TRUE | 0,0048785 | No |
| orf19.2020 | 8584 | 5722 | 7688 | 9525 | 11044 | 9528 | 10493 | 6820 | 7701 | 7285 | 7866 | 7854 | 9445 | 9091 | 9363 | 9619 | 8528 | 7676 | 9380 | 1,221 | 0,288 | 7,828 | 4,97E-15 | 5,73E-14 | 0,000743 | 0,0021497 | 0,0011825 | 0,0011825 | TRUE | 0,0140054 | No |
| orf19.2021 | 29902 | 20045 | 26373 | 33700 | 30152 | 25220 | 27159 | 16389 | 26825 | 25519 | 26982 | 27789 | 25787 | 24064 | 24235 | 23115 | 25540 | 26779 | 24300 | 0,907 | -0,14 | -3,252 | 0,00114438 | 0,00284696 | 0,001601 | 0,0020364 | 0,0017418 | 0,0017418 | TRUE | 0,0207957 | No |
| orf19.2023 | 31297 | 20271 | 28198 | 35671 | 27155 | 21870 | 24432 | 15567 | 28077 | 25807 | 28849 | 29415 | 23224 | 20868 | 21801 | 21956 | 25000 | 28037 | 21962 | 0,783 | -0,352 | -7,021 | 2,21E-12 | 1,95E-11 | 0,002516 | 0,0020376 | 0,0023801 | 0,0023801 | TRUE | 0,0349586 | No |
| orf19.2024 | 28 | 23 | 35 | 39 | 39 | 32 | 26 | 17 | 25 | 29 | 36 | 32 | 33 | 31 | 23 | 24 | 29,18 | 30 | 28 | 0,916 | -0,127 | -0,491 | 0,62355914 | 0,6964758 | 1,00E-08 | 0,0517307 | 0,0300757 | 0,0300757 | TRUE | 0,1326585 | No |
| orf19.2025 | 5 | 4 | 3 | 11 | 1 | 4 | 3 | 0 | 4 | 5 | 3 | 9 | 1 | 4 | 3 | 0 | 3,63 | 5 | 2 | 0,353 | -1,504 | -1,851 | 0,06417957 | NA | 0,089731 | 0,4015272 | 0,2862564 | 0,2862564 | TRUE | 0,5538669 | No |
| orf19.2026 | 595 | 375 | 510 | 579 | 569 | 540 | 540 | 329 | 534 | 477 | 522 | 477 | 487 | 515 | 482 | 464 | 494,77 | 502 | 487 | 0,97 | -0,044 | -0,622 | 0,53416117 | 0,61633192 | 1,00E-08 | 0,0049136 | 0,0028457 | 0,0028457 | TRUE | 0,0212382 | No |
| orf19.2028 | 4303 | 2821 | 3588 | 4495 | 4196 | 3596 | 3849 | 2568 | 3860 | 3591 | 3671 | 3707 | 3589 | 3431 | 3435 | 3622 | 3613,2 | 3707 | 3519 | 0,948 | -0,077 | -1,964 | 0,04947636 | 0,08345143 | 0,000501 | 0,0023813 | 0,0011911 | 0,0011911 | TRUE | 0,0094195 | No |
| orf19.2029 | 547 | 427 | 562 | 654 | 517 | 522 | 549 | 330 | 491 | 544 | 575 | 539 | 442 | 498 | 490 | 465 | 505,52 | 537 | 474 | 0,883 | -0,179 | -2,397 | 0,01652379 | 0,03142696 | 0,001899 | 0,0048512 | 0,0033524 | 0,0033524 | TRUE | 0,0412665 | No |
| orf19.203 | 2554 | 1682 | 2324 | 2729 | 3379 | 2955 | 3321 | 2300 | 2291 | 2141 | 2378 | 2250 | 2890 | 2820 | 2963 | 3244 | 2622,2 | 2265 | 2979 | 1,313 | 0,393 | 7,256 | 3,98E-13 | 3,87E-12 | 0,002369 | 0,0025331 | 0,0024221 | 0,0024221 | TRUE | 0,0414705 | No |
| orf19.2030 | 10825 | 6809 | 8920 | 10855 | 11158 | 8570 | 9276 | 6447 | 9711 | 8669 | 9126 | 8951 | 9543 | 8177 | 8277 | 9093 | 8943,4 | 9114 | 8772 | 0,962 | -0,055 | -0,948 | 0,34327258 | 0,43029036 | 0,003656 | 0,0021418 | 0,0031786 | 0,0031786 | TRUE | 0,0432432 | No |
| orf19.2031 | 1964 | 1303 | 1636 | 2108 | 2101 | 1922 | 2084 | 1208 | 1762 | 1659 | 1674 | 1738 | 1797 | 1834 | 1860 | 1704 | 1753,4 | 1708 | 1799 | 1,054 | 0,075 | 1,649 | 0,09912183 | 0,15176451 | 0,000382 | 0,0028075 | 0,0014283 | 0,0014283 | TRUE | 0,0135488 | No |
| orf19.2032 | 3 | 6 | 2 | 4 | 8 | 4 | 9 | 3 | 3 | 8 | 2 | 3 | 7 | 4 | 8 | 4 | 4,82 | 4 | 6 | 1,551 | 0,633 | 0,973 | 0,33074676 | NA | 1,00E-08 | 0,3028864 | 0,1894306 | 0,1894306 | TRUE | 0,4870664 | No |
| orf19.2033 | 8 | 5 | 5 | 9 | 6 | 2 | 11 | 8 | 7 | 6 | 5 | 7 | 5 | 2 | 10 | 11 | 6,78 | 6 | 7 | 1,042 | 0,06 | 0,105 | 0,91636806 | NA | 0,071772 | 0,2161814 | 0,1632634 | 0,1632634 | TRUE | 0,3034136 | No |
| orf19.2034 | 13 | 14 | 14 | 8 | 16 | 14 | 17 | 5 | 12 | 18 | 14 | 7 | 14 | 13 | 15 | 7 | 12,46 | 13 | 12 | 1,025 | 0,036 | 0,086 | 0,93138403 | NA | 0,032131 | 0,1185002 | 0,0864264 | 0,0864264 | TRUE | 0,4016566 | No |
| orf19.2035 | 110 | 89 | 119 | 138 | 141 | 128 | 111 | 89 | 99 | 113 | 122 | 114 | 121 | 122 | 99 | 126 | 114,35 | 112 | 117 | 1,042 | 0,059 | 0,43 | 0,66730691 | 0,73428204 | 1,00E-08 | 0,0146743 | 0,0091415 | 0,0091415 | TRUE | 0,1131109 | No |
| orf19.2036 | 460 | 322 | 401 | 517 | 487 | 421 | 459 | 309 | 413 | 410 | 410 | 426 | 416 | 402 | 410 | 436 | 415,35 | 415 | 416 | 0,999 | -0,001 | -0,017 | 0,98664705 | 0,98961194 | 1,00E-08 | 0,0054747 | 0,0023284 | 0,0023284 | TRUE | 0,0096943 | No |
| orf19.2037 | 554 | 410 | 503 | 632 | 545 | 464 | 474 | 306 | 497 | 522 | 515 | 521 | 466 | 443 | 423 | 432 | 477,27 | 514 | 441 | 0,86 | -0,218 | -3,212 | 0,00131768 | 0,00323129 | 1,00E-08 | 0,0050212 | 0,0023103 | 0,0023103 | TRUE | 0,0187924 | No |
| orf19.2038 | 704 | 574 | 710 | 802 | 576 | 615 | 609 | 427 | 632 | 731 | 726 | 661 | 493 | 587 | 543 | 602 | 621,9 | 688 | 556 | 0,808 | -0,308 | -3,816 | 0,00013554 | 0,00040454 | 0,004868 | 0,0043138 | 0,0046009 | 0,0046009 | TRUE | 0,0717835 | No |
| orf19.2039 | 2232 | 1472 | 1816 | 2395 | 2359 | 1901 | 2235 | 1324 | 2002 | 1874 | 1858 | 1975 | 2017 | 1814 | 1994 | 1867 | 1925,3 | 1927 | 1923 | 0,998 | -0,003 | -0,054 | 0,95680527 | 0,96917621 | 0,00153 | 0,0027335 | 0,0020051 | 0,0020051 | TRUE | 0,0187724 | No |
| orf19.204 | 1084 | 787 | 963 | 1223 | 880 | 746 | 862 | 532 | 972 | 1002 | 985 | 1009 | 753 | 712 | 769 | 750 | 869,01 | 992 | 746 | 0,752 | -0,411 | -7,75 | 9,19E-15 | 1,03E-13 | 1,00E-08 | 0,00365 | 0,0015258 | 0,0015258 | TRUE | 0,0118273 | No |
| orf19.2040 | 2731 | 1892 | 2398 | 2863 | 2568 | 2253 | 2474 | 1488 | 2450 | 2409 | 2453 | 2361 | 2196 | 2150 | 2208 | 2099 | 2290,7 | 2418 | 2163 | 0,896 | -0,159 | -4,15 | 3,33E-05 | 0,00011118 | 1,00E-08 | 0,0026133 | 0,0009683 | 0,0009683 | TRUE | 0,0043126 | No |
| orf19.2041 | 850 | 560 | 742 | 908 | 1001 | 985 | 1039 | 582 | 763 | 713 | 759 | 749 | 856 | 940 | 927 | 821 | 815,91 | 746 | 886 | 1,189 | 0,25 | 4,058 | 4,95E-05 | 0,00016038 | 0,001193 | 0,0037588 | 0,0024127 | 0,0024127 | TRUE | 0,0252731 | No |
| orf19.2042 | 1568 | 973 | 1240 | 1581 | 1669 | 1369 | 1496 | 876 | 1407 | 1239 | 1269 | 1304 | 1427 | 1306 | 1335 | 1236 | 1315,2 | 1305 | 1326 | 1,017 | 0,025 | 0,411 | 0,68140529 | 0,74556237 | 0,00251 | 0,0030833 | 0,002758 | 0,002758 | TRUE | 0,0336943 | No |
| orf19.2043 | 839 | 594 | 715 | 895 | 899 | 777 | 857 | 567 | 753 | 756 | 732 | 738 | 769 | 741 | 765 | 800 | 756,64 | 745 | 769 | 1,031 | 0,044 | 0,798 | 0,4246908 | 0,51210612 | 1,00E-08 | 0,0038981 | 0,0015789 | 0,0015789 | TRUE | 0,0072758 | No |
| orf19.2044 | 404 | 340 | 371 | 468 | 488 | 489 | 471 | 321 | 362 | 433 | 380 | 386 | 417 | 467 | 420 | 453 | 414,72 | 390 | 439 | 1,128 | 0,173 | 2,168 | 0,03014082 | 0,05381458 | 0,001722 | 0,00548 | 0,0037089 | 0,0037089 | TRUE | 0,0553747 | No |
| orf19.2045 | 6481 | 4557 | 5704 | 7089 | 5803 | 5492 | 5705 | 3795 | 5814 | 5802 | 5836 | 5846 | 4963 | 5240 | 5091 | 5353 | 5493 | 5824 | 5162 | 0,885 | -0,176 | -5,175 | 2,28E-07 | 1,06E-06 | 0,000307 | 0,0022438 | 0,0009221 | 0,0009221 | TRUE | 0,0082239 | No |
| orf19.2046 | 779 | 523 | 658 | 792 | 798 | 645 | 749 | 438 | 699 | 666 | 673 | 653 | 682 | 615 | 668 | 618 | 659,38 | 673 | 646 | 0,963 | -0,055 | -0,876 | 0,38126067 | 0,47054522 | 1,00E-08 | 0,0041812 | 0,0022587 | 0,0022587 | TRUE | 0,0172923 | No |
| orf19.2047 | 1371 | 892 | 1153 | 1434 | 1643 | 1426 | 1521 | 931 | 1230 | 1136 | 1180 | 1182 | 1405 | 1361 | 1357 | 1313 | 1270,5 | 1182 | 1359 | 1,15 | 0,202 | 4,214 | 2,51E-05 | 8,55E-05 | 1,00E-08 | 0,0031222 | 0,0014088 | 0,0014088 | TRUE | 0,0089379 | No |
| orf19.2048 | 8219 | 5586 | 6873 | 8860 | 15558 | 12582 | 13136 | 9718 | 7373 | 7112 | 7032 | 7306 | 13306 | 12005 | 11722 | 13706 | 9945,2 | 7206 | 12685 | 1,76 | 0,815 | 14,979 | 1,01E-50 | 1,10E-48 | 0,003022 | 0,0021255 | 0,0027371 | 0,0027371 | TRUE | 0,0355431 | No |
| orf19.2049 | 817 | 708 | 819 | 1025 | 681 | 658 | 623 | 377 | 733 | 901 | 838 | 845 | 582 | 628 | 556 | 532 | 701,92 | 829 | 574 | 0,695 | -0,524 | -6,689 | 2,24E-11 | 1,77E-10 | 0,004751 | 0,0040477 | 0,0044173 | 0,0044173 | TRUE | 0,0739303 | No |
| orf19.2050 | 5051 | 3483 | 4179 | 5401 | 7142 | 6226 | 6871 | 4275 | 4531 | 4434 | 4276 | 4454 | 6108 | 5941 | 6131 | 6030 | 5238 | 4424 | 6052 | 1,368 | 0,452 | 13,893 | 6,95E-44 | 5,42E-42 | 0,000135 | 0,0022567 | 0,0008206 | 0,0008206 | TRUE | 0,0062174 | No |
| orf19.2051 | 509 | 341 | 484 | 570 | 406 | 387 | 437 | 292 | 457 | 434 | 495 | 470 | 347 | 369 | 390 | 412 | 421,78 | 464 | 380 | 0,813 | -0,298 | -3,793 | 0,00014871 | 0,00044099 | 0,001319 | 0,0054214 | 0,003533 | 0,003533 | TRUE | 0,0388641 | No |
| orf19.2054 | 1885 | 1219 | 1608 | 1955 | 2150 | 1905 | 2224 | 1356 | 1691 | 1552 | 1645 | 1612 | 1839 | 1818 | 1985 | 1913 | 1756,7 | 1625 | 1889 | 1,161 | 0,215 | 4,45 | 8,60E-06 | 3,17E-05 | 0,000912 | 0,0028059 | 0,0016651 | 0,0016651 | TRUE | 0,0151682 | No |
| orf19.2055 | 2847 | 1873 | 2334 | 2943 | 2806 | 2307 | 2498 | 1493 | 2554 | 2385 | 2388 | 2427 | 2400 | 2201 | 2229 | 2106 | 2336,2 | 2438 | 2234 | 0,917 | -0,125 | -2,503 | 0,01230898 | 0,02421491 | 0,001463 | 0,0026009 | 0,0019553 | 0,0019553 | TRUE | 0,0305694 | No |
| orf19.2057 | 6032 | 4136 | 5424 | 6569 | 6606 | 5900 | 6590 | 4203 | 5411 | 5266 | 5549 | 5417 | 5650 | 5630 | 5880 | 5928 | 5591,3 | 5411 | 5772 | 1,066 | 0,092 | 2,68 | 0,00735327 | 0,01526554 | 0,000348 | 0,0022392 | 0,0009543 | 0,0009543 | TRUE | 0,003806 | No |
| orf19.2059 | 1823 | 1233 | 1606 | 2019 | 1442 | 1292 | 1403 | 957 | 1635 | 1570 | 1643 | 1665 | 1233 | 1233 | 1252 | 1350 | 1447,6 | 1628 | 1267 | 0,775 | -0,367 | -7,701 | 1,35E-14 | 1,48E-13 | 0,00023 | 0,0029823 | 0,0014792 | 0,0014792 | TRUE | 0,020395 | No |
| orf19.206 | 1935 | 1306 | 1728 | 2046 | 1439 | 1228 | 1302 | 945 | 1736 | 1663 | 1768 | 1687 | 1231 | 1172 | 1162 | 1333 | 1468,8 | 1714 | 1224 | 0,712 | -0,489 | -9,152 | 5,60E-20 | 1,01E-18 | 0,001275 | 0,0029679 | 0,0020421 | 0,0020421 | TRUE | 0,0382454 | No |
| orf19.2060 | 70 | 33 | 54 | 58 | 54 | 36 | 41 | 33 | 63 | 42 | 55 | 48 | 46 | 34 | 37 | 47 | 46,44 | 52 | 41 | 0,778 | -0,362 | -1,692 | 0,09073698 | 0,14097434 | 1,00E-08 | 0,033237 | 0,0222805 | 0,0222805 | TRUE | 0,1695528 | No |
| orf19.2061 | 7 | 5 | 2 | 7 | 7 | 4 | 1 | 1 | 6 | 6 | 2 | 6 | 6 | 4 | 1 | 1 | 4,07 | 5 | 3 | 0,613 | -0,706 | -0,958 | 0,33787011 | NA | 0,095633 | 0,3585556 | 0,265091 | 0,265091 | TRUE | 0,6783944 | No |
| orf19.2062 | 511 | 346 | 433 | 602 | 340 | 282 | 279 | 250 | 458 | 440 | 443 | 496 | 291 | 269 | 249 | 353 | 374,97 | 459 | 290 | 0,626 | -0,676 | -6,699 | 2,09E-11 | 1,65E-10 | 0,008301 | 0,005851 | 0,0069542 | 0,0069542 | TRUE | 0,2038006 | No |
| orf19.2063 | 3191 | 2116 | 2956 | 3388 | 3845 | 3511 | 3841 | 2453 | 2863 | 2694 | 3024 | 2794 | 3288 | 3350 | 3427 | 3460 | 3112,5 | 2844 | 3381 | 1,188 | 0,249 | 5,618 | 1,93E-08 | 1,03E-07 | 0,001023 | 0,0024459 | 0,001554 | 0,001554 | TRUE | 0,0217218 | No |
| orf19.2064 | 2088 | 1376 | 1840 | 2261 | 2696 | 2179 | 2351 | 1439 | 1873 | 1752 | 1883 | 1864 | 2306 | 2079 | 2098 | 2030 | 1985,5 | 1843 | 2128 | 1,155 | 0,208 | 4,008 | 6,13E-05 | 0,00019517 | 0,001517 | 0,0027107 | 0,0020816 | 0,0020816 | TRUE | 0,0390256 | No |
| orf19.2065 | 148 | 84 | 139 | 174 | 111 | 88 | 134 | 54 | 133 | 107 | 142 | 143 | 95 | 84 | 120 | 76 | 112,5 | 131 | 94 | 0,716 | -0,482 | -2,996 | 0,00273869 | 0,00627538 | 0,016534 | 0,014883 | 0,0157325 | 0,0157325 | TRUE | 0,3415876 | No |
| orf19.2066 | 1215 | 821 | 1036 | 1297 | 1060 | 944 | 987 | 654 | 1090 | 1045 | 1060 | 1070 | 907 | 901 | 881 | 922 | 984,38 | 1066 | 903 | 0,845 | -0,243 | -4,952 | 7,36E-07 | 3,17E-06 | 1,00E-08 | 0,0034543 | 0,0012919 | 0,0012919 | TRUE | 0,0031804 | No |
| orf19.2066.1 | 2254 | 1523 | 1928 | 2488 | 2408 | 1878 | 1897 | 1279 | 2022 | 1939 | 1973 | 2052 | 2059 | 1792 | 1693 | 1804 | 1916,7 | 1996 | 1837 | 0,92 | -0,12 | -1,943 | 0,05203042 | 0,08705062 | 0,003245 | 0,0027369 | 0,0031294 | 0,0031294 | TRUE | 0,0830345 | No |
| orf19.2067 | 4015 | 2653 | 3320 | 4249 | 4618 | 4025 | 4257 | 2472 | 3602 | 3378 | 3397 | 3504 | 3949 | 3841 | 3799 | 3487 | 3619,4 | 3470 | 3769 | 1,087 | 0,12 | 2,565 | 0,01031327 | 0,02070235 | 0,001469 | 0,0023806 | 0,0018328 | 0,0018328 | TRUE | 0,029393 | No |
| orf19.2068 | 506 | 418 | 489 | 537 | 544 | 494 | 531 | 320 | 454 | 532 | 500 | 443 | 465 | 471 | 474 | 451 | 473,87 | 482 | 465 | 0,971 | -0,043 | -0,565 | 0,57223715 | 0,65101322 | 0,001755 | 0,005043 | 0,0034334 | 0,0034334 | TRUE | 0,0520024 | No |
| orf19.2069 | 2594 | 1762 | 2298 | 2799 | 2614 | 2210 | 2411 | 1584 | 2327 | 2243 | 2351 | 2308 | 2236 | 2109 | 2151 | 2234 | 2244,9 | 2307 | 2182 | 0,945 | -0,082 | -2,051 | 0,0402477 | 0,06944519 | 0,000123 | 0,0026262 | 0,0010856 | 0,0010856 | TRUE | 0,0062421 | No |
| orf19.207 | 2554 | 1732 | 2412 | 2900 | 2516 | 2078 | 2292 | 1647 | 2291 | 2205 | 2468 | 2391 | 2152 | 1983 | 2045 | 2323 | 2232,3 | 2339 | 2126 | 0,907 | -0,141 | -2,399 | 0,01645603 | 0,03131702 | 0,003059 | 0,0026299 | 0,0028533 | 0,0028533 | TRUE | 0,044611 | No |
| orf19.2070 | 2973 | 2000 | 2601 | 3232 | 3312 | 3040 | 3241 | 2039 | 2667 | 2546 | 2661 | 2665 | 2833 | 2901 | 2892 | 2876 | 2755,1 | 2635 | 2876 | 1,09 | 0,125 | 3,51 | 0,00044741 | 0,00122129 | 1,00E-08 | 0,0025064 | 0,0008439 | 0,0008439 | TRUE | 0,0057181 | No |
| orf19.2071 | 0 | 1 | 0 | 1 | 1 | 1 | 3 | 1 | 0 | 1 | 0 | 1 | 1 | 1 | 3 | 1 | 1 | 0 | 2 | 3,01 | 1,59 | 1,086 | 0,27745273 | NA | 1,00E-08 | 1,4546737 | 0,7833508 | 0,7833508 | TRUE | 0,2487985 | No |
| orf19.2072 | 8 | 10 | 17 | 14 | 6 | 9 | 10 | 17 | 7 | 13 | 17 | 12 | 5 | 9 | 9 | 24 | 11,93 | 12 | 12 | 0,916 | -0,127 | -0,256 | 0,79817329 | NA | 0,185787 | 0,1236334 | 0,1509746 | 0,1509746 | TRUE | 1,1574198 | No |
| orf19.2073 | 8990 | 5669 | 7852 | 9383 | 10479 | 8744 | 9716 | 6082 | 8065 | 7217 | 8033 | 7737 | 8962 | 8343 | 8670 | 8578 | 8200,8 | 7763 | 8638 | 1,112 | 0,154 | 3,45 | 0,00056012 | 0,00149233 | 0,001556 | 0,0021565 | 0,0017837 | 0,0017837 | TRUE | 0,0268148 | No |

| | | | | | | | | | | | | | | | | | | | | | | | | | | | | | | | |
|---|---|---|---|---|---|---|---|---|---|---|---|---|---|---|---|---|---|---|---|---|---|---|---|---|---|---|---|---|---|---|---|
| orf19.2074 | 171 | 132 | 172 | 190 | 254 | 194 | 204 | 122 | 153 | 168 | 176 | 157 | 217 | 185 | 182 | 172 | 176,32 | 164 | 189 | 1,166 | 0,221 | 1,951 | 0,05106128 | 0,08566775 | 1,00E-08 | 0,0102129 | 0,006694 | 0,006694 | TRUE | 0,1141 | No |
| orf19.2075 | 3720 | 2481 | 3301 | 4044 | 3332 | 2879 | 3019 | 1780 | 3337 | 3159 | 3377 | 3335 | 2850 | 2747 | 2694 | 2511 | 3001,1 | 3302 | 2700 | 0,819 | -0,289 | -6,149 | 7,79E-10 | 5,03E-09 | 0,001371 | 0,0024632 | 0,0017781 | 0,0017781 | TRUE | 0,0253534 | No |
| orf19.2076 | 174 | 97 | 149 | 177 | 184 | 199 | 193 | 119 | 156 | 123 | 152 | 146 | 157 | 190 | 172 | 168 | 158,16 | 144 | 172 | 1,183 | 0,242 | 2,048 | 0,04051971 | 0,06981428 | 1,00E-08 | 0,0111582 | 0,0070803 | 0,0070803 | TRUE | 0,0830291 | No |
| orf19.2077 | 3014 | 2043 | 2749 | 3175 | 3291 | 2852 | 3063 | 2069 | 2704 | 2601 | 2812 | 2618 | 2815 | 2721 | 2733 | 2918 | 2740,3 | 2684 | 2797 | 1,041 | 0,058 | 1,35 | 0,17707838 | 0,24874606 | 0,000692 | 0,0025093 | 0,0013936 | 0,0013936 | TRUE | 0,0124398 | No |
| orf19.2078 | 1195 | 871 | 1019 | 1294 | 1352 | 1238 | 1309 | 740 | 1072 | 1109 | 1043 | 1067 | 1156 | 1181 | 1168 | 1044 | 1105 | 1073 | 1137 | 1,064 | 0,09 | 1,665 | 0,09582198 | 0,14750009 | 0,000664 | 0,0032933 | 0,0018867 | 0,0018867 | TRUE | 0,0321859 | No |
| orf19.2079 | 3752 | 2322 | 2859 | 3757 | 4523 | 3582 | 3630 | 2955 | 3366 | 2956 | 2925 | 3098 | 3868 | 3418 | 3239 | 4168 | 3379,8 | 3086 | 3673 | 1,189 | 0,25 | 3,16 | 0,00157806 | 0,0038074 | 0,008348 | 0,002409 | 0,0056984 | 0,0056984 | TRUE | 0,0980691 | No |
| orf19.2081 | 2093 | 1379 | 1925 | 2290 | 2518 | 2298 | 2486 | 1523 | 1878 | 1756 | 1969 | 1888 | 2153 | 2193 | 2218 | 2148 | 2025,5 | 1873 | 2178 | 1,162 | 0,217 | 4,848 | 1,25E-06 | 5,18E-06 | 0,000565 | 0,0026963 | 0,0014228 | 0,0014228 | TRUE | 0,0196266 | No |
| orf19.2082 | 7 | 2 | 3 | 3 | 4 | 4 | 1 | 3 | 6 | 3 | 3 | 2 | 3 | 4 | 1 | 4 | 3,34 | 4 | 3 | 0,829 | -0,271 | -0,353 | 0,72411441 | NA | 1,00E-08 | 0,436454 | 0,2618925 | 0,2618925 | TRUE | 0,4454829 | No |
| orf19.2084 | 2282 | 1604 | 2180 | 2654 | 2196 | 1800 | 2087 | 1323 | 2047 | 2042 | 2230 | 2189 | 1878 | 1718 | 1862 | 1866 | 1979 | 2127 | 1831 | 0,86 | -0,217 | -4,311 | 1,63E-05 | 5,74E-05 | 0,001419 | 0,0027131 | 0,0019296 | 0,0019296 | TRUE | 0,0214042 | No |
| orf19.2087 | 756 | 540 | 582 | 793 | 902 | 761 | 833 | 570 | 678 | 687 | 595 | 654 | 771 | 726 | 743 | 804 | 707,48 | 654 | 761 | 1,163 | 0,218 | 3,334 | 0,00085569 | 0,00218488 | 0,001299 | 0,0040315 | 0,0026666 | 0,0026666 | TRUE | 0,0412877 | No |
| orf19.2088 | 693 | 414 | 513 | 649 | 695 | 516 | 620 | 382 | 622 | 527 | 525 | 535 | 594 | 492 | 553 | 539 | 548,44 | 552 | 544 | 0,986 | -0,02 | -0,244 | 0,80687169 | 0,85017277 | 0,004879 | 0,0046265 | 0,0047587 | 0,0047587 | TRUE | 0,0864152 | No |
| orf19.2089 | 3766 | 2710 | 3143 | 4182 | 3928 | 3486 | 3597 | 2257 | 3378 | 3450 | 3216 | 3449 | 3359 | 3326 | 3210 | 3183 | 3321,4 | 3373 | 3270 | 0,97 | -0,044 | -1,106 | 0,26857366 | 0,3527745 | 0,00048 | 0,0024166 | 0,0012158 | 0,0012158 | TRUE | 0,0119248 | No |
| orf19.209 | 276 | 197 | 250 | 318 | 350 | 304 | 269 | 218 | 248 | 251 | 256 | 262 | 299 | 290 | 240 | 307 | 269,16 | 254 | 284 | 1,114 | 0,155 | 1,595 | 0,11076553 | 0,16636982 | 0,003137 | 0,0073729 | 0,0053795 | 0,0053795 | TRUE | 0,1295011 | No |
| orf19.2090 | 9689 | 6451 | 8786 | 10108 | 8730 | 7576 | 8205 | 4715 | 8692 | 8213 | 8989 | 8335 | 7466 | 7229 | 7322 | 6650 | 7862 | 8557 | 7167 | 0,838 | -0,255 | -5,391 | 7,02E-08 | 3,51E-07 | 0,001924 | 0,0021642 | 0,0020217 | 0,0020217 | TRUE | 0,028013 | No |
| orf19.2091 | 3661 | 2570 | 2986 | 3899 | 3481 | 3031 | 3331 | 2161 | 3284 | 3272 | 3055 | 3215 | 2977 | 2892 | 2972 | 3048 | 3089,5 | 3206 | 2972 | 0,926 | -0,11 | -2,818 | 0,00482959 | 0,01045251 | 0,000338 | 0,0024494 | 0,0011427 | 0,0011427 | TRUE | 0,012183 | No |
| orf19.2092 | 2855 | 1799 | 2364 | 2854 | 2851 | 2409 | 2570 | 1741 | 2561 | 2290 | 2419 | 2353 | 2438 | 2299 | 2293 | 2456 | 2388,7 | 2406 | 2372 | 0,984 | -0,023 | -0,468 | 0,63968889 | 0,70987271 | 0,00122 | 0,0025873 | 0,0018552 | 0,0018552 | TRUE | 0,0231418 | No |
| orf19.2093 | 3824 | 2500 | 3317 | 4069 | 3738 | 3094 | 3453 | 2388 | 3431 | 3183 | 3394 | 3355 | 3197 | 2952 | 3081 | 3368 | 3245,1 | 3341 | 3150 | 0,941 | -0,087 | -1,786 | 0,07402387 | 0,11823634 | 0,001699 | 0,0024269 | 0,0019762 | 0,0019762 | TRUE | 0,0250413 | No |
| orf19.2094 | 4575 | 3262 | 4046 | 4871 | 6578 | 6162 | 6594 | 4214 | 4104 | 4153 | 4139 | 4017 | 5626 | 5880 | 5884 | 5943 | 4968,3 | 4103 | 5833 | 1,421 | 0,507 | 15,249 | 1,66E-52 | 2,00E-50 | 0,000259 | 0,0022717 | 0,0008546 | 0,0008546 | TRUE | 0,0071528 | No |
| orf19.2095 | 10121 | 6951 | 8987 | 10880 | 13608 | 11724 | 12599 | 8669 | 9080 | 8849 | 9195 | 8972 | 11638 | 11187 | 11243 | 12227 | 10299 | 9024 | 11574 | 1,282 | 0,358 | 9,568 | 1,09E-21 | 2,23E-20 | 0,000861 | 0,0021205 | 0,0012476 | 0,0012476 | TRUE | 0,0172178 | No |
| orf19.2097 | 724 | 475 | 633 | 747 | 701 | 615 | 703 | 402 | 650 | 605 | 648 | 616 | 600 | 587 | 627 | 567 | 612,31 | 630 | 595 | 0,947 | -0,078 | -1,252 | 0,21039445 | 0,28734936 | 1,00E-08 | 0,0043504 | 0,0021378 | 0,0021378 | TRUE | 0,0154252 | No |
| orf19.2098 | 1745 | 1174 | 1582 | 2005 | 1986 | 1579 | 1793 | 1247 | 1565 | 1495 | 1619 | 1653 | 1698 | 1507 | 1600 | 1759 | 1612 | 1583 | 1641 | 1,034 | 0,049 | 0,826 | 0,40862566 | 0,49686764 | 0,002411 | 0,0028801 | 0,0026893 | 0,0026893 | TRUE | 0,0363316 | No |
| orf19.2099 | 1495 | 1027 | 1406 | 1646 | 1887 | 1552 | 1750 | 1128 | 1341 | 1307 | 1438 | 1357 | 1614 | 1481 | 1562 | 1591 | 1461,5 | 1361 | 1562 | 1,147 | 0,197 | 3,942 | 8,09E-05 | 0,00025063 | 0,000805 | 0,0029728 | 0,0017165 | 0,0017165 | TRUE | 0,0170667 | No |
| orf19.21 | 582 | 391 | 475 | 592 | 630 | 532 | 573 | 428 | 522 | 498 | 486 | 488 | 539 | 508 | 511 | 604 | 519,43 | 498 | 540 | 1,079 | 0,109 | 1,494 | 0,1352518 | 0,19766305 | 0,001362 | 0,0047743 | 0,0031934 | 0,0031934 | TRUE | 0,0627616 | No |
| orf19.210 | 120 | 81 | 138 | 154 | 146 | 152 | 128 | 97 | 108 | 103 | 141 | 127 | 125 | 145 | 114 | 137 | 124,98 | 120 | 130 | 1,08 | 0,112 | 0,792 | 0,42846272 | 0,51566586 | 0,00778 | 0,0135945 | 0,0110871 | 0,0110871 | TRUE | 0,1377527 | No |
| orf19.2101 | 984 | 632 | 755 | 984 | 1135 | 920 | 1009 | 631 | 883 | 805 | 772 | 811 | 971 | 878 | 900 | 890 | 863,76 | 818 | 910 | 1,113 | 0,154 | 2,479 | 0,01315742 | 0,02569889 | 0,001616 | 0,0036602 | 0,0025369 | 0,0025369 | TRUE | 0,0350492 | No |
| orf19.2102 | 4900 | 3292 | 4269 | 5309 | 4914 | 4087 | 4480 | 2726 | 4396 | 4191 | 4368 | 4378 | 4203 | 3900 | 3998 | 3845 | 4159,6 | 4333 | 3986 | 0,92 | -0,12 | -3,004 | 0,00266547 | 0,00612864 | 0,000702 | 0,0023285 | 0,001285 | 0,001285 | TRUE | 0,0164866 | No |
| orf19.2103 | 41 | 22 | 39 | 41 | 25 | 23 | 32 | 16 | 37 | 28 | 40 | 34 | 21 | 22 | 29 | 23 | 29,12 | 35 | 24 | 0,68 | -0,556 | -2,167 | 0,03022262 | 0,05392857 | 1,00E-08 | 0,0518346 | 0,0278643 | 0,0278643 | TRUE | 0,1358439 | No |
| orf19.2104 | 546 | 350 | 463 | 591 | 593 | 507 | 492 | 348 | 490 | 446 | 474 | 487 | 507 | 484 | 439 | 491 | 477,15 | 474 | 480 | 1,009 | 0,014 | 0,187 | 0,85150117 | 0,88661042 | 1,00E-08 | 0,005022 | 0,0029935 | 0,0029935 | TRUE | 0,0416009 | No |
| orf19.2105 | 1409 | 907 | 1235 | 1420 | 1134 | 966 | 1052 | 779 | 1264 | 1155 | 1264 | 1171 | 970 | 922 | 939 | 1099 | 1097,8 | 1214 | 982 | 0,806 | -0,31 | -4,755 | 1,99E-06 | 8,03E-06 | 0,00327 | 0,0033019 | 0,003161 | 0,003161 | TRUE | 0,0690749 | No |
| orf19.2106 | 1614 | 1105 | 1430 | 1745 | 1498 | 1128 | 1307 | 805 | 1448 | 1407 | 1463 | 1439 | 1281 | 1076 | 1166 | 1135 | 1302 | 1439 | 1164 | 0,81 | -0,304 | -5,199 | 2,01E-07 | 9,43E-07 | 0,001832 | 0,0030945 | 0,0024974 | 0,0024974 | TRUE | 0,056703 | No |
| orf19.2107 | 1053 | 690 | 850 | 1113 | 869 | 683 | 739 | 520 | 945 | 878 | 870 | 918 | 743 | 652 | 659 | 733 | 799,78 | 903 | 697 | 0,77 | -0,377 | -5,839 | 5,26E-09 | 3,05E-08 | 0,001767 | 0,0037946 | 0,0027266 | 0,0027266 | TRUE | 0,0273912 | No |
| orf19.2107.1 | 78725 | 46798 | 58582 | 75143 | 1E+05 | 78158 | 83372 | 44334 | 70625 | 59579 | 59935 | 61964 | 94630 | 74576 | 74396 | 62529 | 69779 | 63026 | 76533 | 1,214 | 0,28 | 2,019 | 0,04349513 | 0,07438621 | 0,018489 | 0,0020003 | 0,009992 | 0,0184887 | TRUE | 0,3105718 | No |
| orf19.2108 | 3371 | 2177 | 2741 | 3569 | 4061 | 3383 | 3667 | 2304 | 3024 | 2772 | 2804 | 2943 | 3473 | 3228 | 3272 | 3250 | 3095,7 | 2886 | 3306 | 1,145 | 0,195 | 4,379 | 1,19E-05 | 4,29E-05 | 0,000995 | 0,0024485 | 0,0015866 | 0,0015866 | TRUE | 0,0145116 | No |
| orf19.211 | 1457 | 919 | 1221 | 1528 | 1916 | 1627 | 1802 | 1232 | 1307 | 1170 | 1249 | 1260 | 1639 | 1552 | 1608 | 1738 | 1440,4 | 1246 | 1634 | 1,307 | 0,386 | 7,239 | 4,51E-13 | 4,32E-12 | 0,001228 | 0,0029874 | 0,0020206 | 0,0020206 | TRUE | 0,0199389 | No |
| orf19.2110 | 283 | 170 | 241 | 278 | 250 | 200 | 213 | 118 | 254 | 216 | 247 | 229 | 214 | 191 | 190 | 166 | 213,41 | 236 | 190 | 0,809 | -0,307 | -2,913 | 0,00358175 | 0,00797595 | 0,002014 | 0,008782 | 0,0059255 | 0,0059255 | TRUE | 0,0754998 | No |
| orf19.2111 | 3762 | 2285 | 2996 | 3871 | 3763 | 2907 | 3351 | 2192 | 3375 | 2909 | 3065 | 3192 | 3218 | 2774 | 2990 | 3092 | 3076,9 | 3135 | 3018 | 0,962 | -0,056 | -0,931 | 0,35210814 | 0,43980576 | 0,003442 | 0,0024513 | 0,0031485 | 0,0031485 | TRUE | 0,0365321 | No |
| orf19.2111.2 | 6820 | 4434 | 5101 | 6849 | 6678 | 5169 | 5233 | 3337 | 6118 | 5645 | 5219 | 5648 | 5711 | 4932 | 4670 | 4707 | 5331,2 | 5658 | 5005 | 0,885 | -0,176 | -2,488 | 0,01285256 | 0,02514432 | 0,006553 | 0,0022518 | 0,0046406 | 0,0046406 | TRUE | 0,1108197 | No |
| orf19.2112 | 807 | 494 | 688 | 962 | 1015 | 782 | 957 | 671 | 724 | 629 | 704 | 793 | 868 | 746 | 854 | 946 | 783,08 | 712 | 854 | 1,193 | 0,254 | 2,94 | 0,00328546 | 0,00739997 | 0,007444 | 0,0038334 | 0,0058956 | 0,0058956 | TRUE | 0,0850006 | No |
| orf19.2113 | 2197 | 1531 | 1932 | 2355 | 2533 | 2180 | 2340 | 1552 | 1971 | 1949 | 1977 | 1942 | 2166 | 2080 | 2088 | 2189 | 2045,3 | 1960 | 2131 | 1,086 | 0,12 | 3,065 | 0,00217972 | 0,00510955 | 1,00E-08 | 0,0026893 | 0,0009706 | 0,0009706 | TRUE | 0,003564 | No |
| orf19.2114 | 228 | 145 | 197 | 255 | 201 | 157 | 183 | 154 | 205 | 185 | 202 | 210 | 172 | 150 | 163 | 217 | 187,9 | 200 | 176 | 0,865 | -0,209 | -1,74 | 0,08191864 | 0,12914767 | 0,006725 | 0,0097056 | 0,0084735 | 0,0084735 | TRUE | 0,227248 | No |
| orf19.2115 | 3600 | 2220 | 3124 | 3715 | 2784 | 2207 | 2510 | 1548 | 3230 | 2826 | 3196 | 3063 | 2381 | 2106 | 2240 | 2183 | 2653,2 | 3079 | 2228 | 0,723 | -0,467 | -8,278 | 1,26E-16 | 1,69E-15 | 0,00261 | 0,0025267 | 0,0026687 | 0,0026687 | TRUE | 0,0358071 | No |
| orf19.2116 | 2530 | 1607 | 2146 | 2546 | 2491 | 1970 | 2162 | 1317 | 2270 | 2046 | 2196 | 2099 | 2130 | 1880 | 1929 | 1858 | 2050,9 | 2153 | 1949 | 0,906 | -0,142 | -2,501 | 0,01237765 | 0,02432611 | 0,002457 | 0,0026873 | 0,0026145 | 0,0026145 | TRUE | 0,0483167 | No |
| orf19.2117 | 3148 | 1970 | 2583 | 3246 | 3896 | 3145 | 3511 | 2333 | 2824 | 2508 | 2643 | 2677 | 3332 | 3001 | 3133 | 3290 | 2926 | 2663 | 3189 | 1,196 | 0,258 | 5,056 | 4,27E-07 | 1,91E-06 | 0,001864 | 0,0024757 | 0,0021631 | 0,0021631 | TRUE | 0,0201161 | No |
| orf19.2119 | 7164 | 4353 | 6688 | 7938 | 8972 | 7230 | 8601 | 6332 | 6427 | 5542 | 6842 | 6546 | 7673 | 6899 | 7675 | 8931 | 7066,8 | 6339 | 7794 | 1,229 | 0,297 | 3,652 | 0,00025999 | 0,00074204 | 0,009529 | 0,0021849 | 0,006221 | 0,006221 | TRUE | 0,1166871 | No |
| orf19.212 | 1695 | 1128 | 1389 | 1789 | 1767 | 1562 | 1724 | 1180 | 1521 | 1436 | 1421 | 1475 | 1511 | 1490 | 1538 | 1664 | 1507,2 | 1463 | 1551 | 1,057 | 0,08 | 1,587 | 0,11257717 | 0,16850116 | 0,000842 | 0,0029427 | 0,0017564 | 0,0017564 | TRUE | 0,0262484 | No |
| orf19.2123 | 1 | 0 | 0 | 0 | 1 | 0 | 2 | 0 | 1 | 0 | 0 | 0 | 1 | 0 | 2 | 0 | 0,44 | 0 | 1 | 2,306 | 1,205 | 0,558 | 0,576871 | NA | 1,00E-08 | 3,2854393 | 2,7564786 | 2,7564786 | TRUE | 0,4448116 | No |
| orf19.2124 | 2540 | 1767 | 2189 | 2726 | 2947 | 2637 | 2731 | 1792 | 2279 | 2250 | 2240 | 2248 | 2520 | 2516 | 2437 | 2527 | 2377,1 | 2254 | 2500 | 1,109 | 0,149 | 4,135 | 3,54E-05 | 0,00011765 | 1,00E-08 | 0,0025902 | 0,0008214 | 0,0008214 | TRUE | 0,0036484 | No |
| orf19.2125 | 3834 | 2413 | 3096 | 3873 | 5095 | 3726 | 3907 | 3136 | 3439 | 3072 | 3168 | 3194 | 4357 | 3555 | 3486 | 4423 | 3586,9 | 3218 | 3955 | 1,228 | 0,297 | 3,652 | 0,00026039 | 0,00074283 | 0,008908 | 0,0023843 | 0,0060515 | 0,0060515 | TRUE | 0,077829 | No |
| orf19.2127 | 394 | 261 | 313 | 372 | 449 | 376 | 422 | 266 | 353 | 332 | 320 | 307 | 384 | 359 | 377 | 375 | 350,9 | 328 | 374 | 1,14 | 0,189 | 2,376 | 0,01748783 | 0,03298804 | 1,00E-08 | 0,0061166 | 0,0031922 | 0,0031922 | TRUE | 0,0341636 | No |
| orf19.2128 | 738 | 464 | 590 | 807 | 755 | 563 | 595 | 444 | 662 | 591 | 604 | 665 | 646 | 537 | 531 | 626 | 607,74 | 630 | 585 | 0,925 | -0,113 | -1,366 | 0,17199547 | 0,24239944 | 0,005446 | 0,0043682 | 0,0049297 | 0,0049297 | TRUE | 0,0640321 | No |
| orf19.2129 | 865 | 536 | 779 | 905 | 706 | 667 | 642 | 477 | 776 | 682 | 797 | 746 | 604 | 636 | 573 | 673 | 685,94 | 750 | 622 | 0,824 | -0,279 | -3,904 | 9,46E-05 | 0,00029094 | 0,002635 | 0,0040959 | 0,0034366 | 0,0034366 | TRUE | 0,0401038 | No |
| orf19.213 | 2806 | 2297 | 2503 | 3432 | 2749 | 2736 | 2776 | 1906 | 2517 | 2924 | 2561 | 2830 | 2351 | 2611 | 2477 | 2688 | 2619,9 | 2708 | 2532 | 0,934 | -0,098 | -1,554 | 0,12017687 | 0,17849756 | 0,003937 | 0,0025336 | 0,00342 | 0,00342 | TRUE | 0,0339449 | No |
| orf19.2131 | 2093 | 1435 | 1959 | 2296 | 3005 | 2622 | 2809 | 1697 | 1878 | 1827 | 2004 | 1893 | 2570 | 2502 | 2507 | 2393 | 2196,7 | 1900 | 2493 | 1,313 | 0,392 | 8,877 | 6,84E-19 | 1,14E-17 | 0,000586 | 0,0026404 | 0,0014118 | 0,0014118 | TRUE | 0,0158569 | No |
| orf19.2132 | 1311 | 862 | 1113 | 1349 | 1510 | 1166 | 1277 | 787 | 1176 | 1097 | 1139 | 1112 | 1291 | 1113 | 1140 | 1110 | 1147,3 | 1131 | 1164 | 1,03 | 0,042 | 0,688 | 0,4915569 | 0,57669866 | 0,002384 | 0,0032449 | 0,0027523 | 0,0027523 | TRUE | 0,0680971 | No |
| orf19.2133 | 56 | 37 | 50 | 64 | 57 | 49 | 56 | 41 | 50 | 47 | 51 | 53 | 49 | 47 | 50 | 58 | 50,57 | 50 | 51 | 0,998 | -0,003 | -0,014 | 0,98883543 | 0,99114502 | 1,00E-08 | 0,0306854 | 0,0144879 | 0,0144879 | TRUE | 0,0508247 | No |
| orf19.2135 | 1623 | 1070 | 1481 | 1678 | 1967 | 1721 | 1805 | 1156 | 1456 | 1362 | 1515 | 1384 | 1682 | 1642 | 1611 | 1630 | 1535,3 | 1429 | 1641 | 1,148 | 0,199 | 4,123 | 3,74E-05 | 0,00012373 | 0,000606 | 0,0029251 | 0,0015883 | 0,0015883 | TRUE | 0,0191155 | No |
| orf19.2137 | 689 | 401 | 595 | 734 | 814 | 671 | 708 | 498 | 618 | 511 | 609 | 605 | 696 | 640 | 632 | 702 | 626,65 | 586 | 668 | 1,134 | 0,182 | 2,432 | 0,01501994 | 0,02891375 | 0,003081 | 0,0042961 | 0,0037757 | 0,0037757 | TRUE | 0,0792991 | No |
| orf19.2138 | 11832 | 7933 | 10727 | 12492 | 13730 | 11723 | 12719 | 8135 | 10615 | 10100 | 10975 | 10301 | 11742 | 11186 | 11350 | 11474 | 10968 | 10498 | 11438 | 1,089 | 0,124 | 3,417 | 0,00063342 | 0,00166327 | 0,000752 | 0,0021119 | 0,0011649 | 0,0011649 | TRUE | 0,0113584 | No |
| orf19.2143 | 4217 | 2852 | 3872 | 4601 | 4455 | 3651 | 4245 | 2821 | 3783 | 3631 | 3961 | 3794 | 3810 | 3484 | 3788 | 3979 | 3778,7 | 3792 | 3765 | 0,992 | -0,012 | -0,241 | 0,80947447 | 0,85201868 | 0,001568 | 0,0023637 | 0,0020322 | 0,0020322 | TRUE | 0,0307368 | No |

| | | | | | | | | | | | | | | | | | | | | | | | | | | | | | | | |
|---|---|---|---|---|---|---|---|---|---|---|---|---|---|---|---|---|---|---|---|---|---|---|---|---|---|---|---|---|---|---|---|
| orf19.2146 | 664 | 396 | 600 | 744 | 943 | 758 | 834 | 570 | 596 | 504 | 614 | 614 | 806 | 723 | 744 | 804 | 675,64 | 582 | 769 | 1,317 | 0,397 | 5,298 | 1,17E-07 | 5,68E-07 | 0,003553 | 0,0041282 | 0,0038798 | 0,0038798 | TRUE | 0,0858416 | No |
| orf19.2149 | 866 | 534 | 789 | 943 | 854 | 646 | 750 | 490 | 777 | 680 | 807 | 778 | 730 | 616 | 669 | 691 | 718,58 | 760 | 676 | 0,888 | -0,171 | -2,314 | 0,02066663 | 0,03836994 | 0,003713 | 0,0039997 | 0,0038503 | 0,0038503 | TRUE | 0,0554102 | No |
| orf19.215 | 8469 | 5881 | 7603 | 9293 | 10232 | 8792 | 9481 | 6411 | 7598 | 7487 | 7779 | 7663 | 8751 | 8389 | 8460 | 9042 | 8146,1 | 7632 | 8660 | 1,134 | 0,182 | 5,21 | 1,89E-07 | 8,91E-07 | 0,000558 | 0,0021577 | 0,0010423 | 0,0010423 | TRUE | 0,0103309 | No |
| orf19.2150 | 1223 | 713 | 1012 | 1254 | 1079 | 842 | 907 | 639 | 1097 | 908 | 1035 | 1034 | 923 | 803 | 809 | 901 | 938,89 | 1018 | 859 | 0,841 | -0,25 | -3,406 | 0,00065865 | 0,00172275 | 0,004196 | 0,0035257 | 0,0040752 | 0,0040752 | TRUE | 0,0604152 | No |
| orf19.2151 | 9341 | 5834 | 8180 | 10068 | 12561 | 10014 | 11528 | 7117 | 8380 | 7427 | 8369 | 8302 | 10743 | 9555 | 10287 | 10038 | 9137,6 | 8120 | 10156 | 1,25 | 0,322 | 6,159 | 7,34E-10 | 4,75E-09 | 0,002666 | 0,0021384 | 0,002522 | 0,002522 | TRUE | 0,0396925 | No |
| orf19.2154 | 1960 | 1212 | 1680 | 2075 | 2297 | 1716 | 2033 | 1220 | 1758 | 1543 | 1719 | 1711 | 1964 | 1637 | 1814 | 1721 | 1733,5 | 1683 | 1784 | 1,06 | 0,084 | 1,28 | 0,20048304 | 0,27493771 | 0,003928 | 0,002817 | 0,0035738 | 0,0035738 | TRUE | 0,0570974 | No |
| orf19.2156 | 1724 | 1026 | 1442 | 1772 | 2096 | 1576 | 1804 | 1201 | 1547 | 1306 | 1475 | 1461 | 1793 | 1504 | 1610 | 1694 | 1548,7 | 1447 | 1650 | 1,138 | 0,187 | 2,724 | 0,00644526 | 0,01359184 | 0,004352 | 0,0029169 | 0,0038732 | 0,0038732 | TRUE | 0,049508 | No |
| orf19.2157 | 3952 | 2357 | 3438 | 4243 | 5435 | 4302 | 4715 | 3257 | 3545 | 3001 | 3517 | 3499 | 4648 | 4105 | 4207 | 4594 | 3889,6 | 3390 | 4388 | 1,293 | 0,371 | 5,756 | 8,60E-09 | 4,80E-08 | 0,004534 | 0,0023528 | 0,0037221 | 0,0037221 | TRUE | 0,0712399 | No |
| orf19.2158 | 308 | 180 | 234 | 319 | 272 | 212 | 237 | 166 | 276 | 229 | 239 | 263 | 233 | 202 | 211 | 234 | 236,06 | 252 | 220 | 0,868 | -0,205 | -2,045 | 0,04086846 | 0,07033449 | 0,001945 | 0,0081294 | 0,0053917 | 0,0053917 | TRUE | 0,046023 | No |
| orf19.216 | 6510 | 4736 | 5669 | 7214 | 5434 | 4831 | 5353 | 3051 | 5840 | 6029 | 5800 | 5949 | 4647 | 4610 | 4777 | 4303 | 5244,4 | 5904 | 4584 | 0,777 | -0,363 | -9,184 | 4,17E-20 | 7,54E-19 | 0,000855 | 0,0022563 | 0,0013084 | 0,0013084 | TRUE | 0,0196034 | No |
| orf19.2160 | 55 | 32 | 51 | 54 | 44 | 14 | 27 | 21 | 49 | 41 | 52 | 45 | 38 | 13 | 24 | 30 | 36,44 | 47 | 26 | 0,559 | -0,84 | -3,038 | 0,00238539 | 0,00553124 | 0,046636 | 0,0418221 | 0,0435673 | 0,0435673 | TRUE | 0,5792057 | No |
| orf19.216.1 | 2065 | 1302 | 1530 | 2148 | 2472 | 1853 | 1767 | 1275 | 1853 | 1658 | 1565 | 1771 | 2114 | 1768 | 1577 | 1798 | 1763 | 1712 | 1814 | 1,06 | 0,084 | 0,973 | 0,33078012 | 0,41785681 | 0,009421 | 0,002803 | 0,0065272 | 0,0065272 | TRUE | 0,1527847 | No |
| orf19.2163 | 1070 | 692 | 1051 | 1241 | 1074 | 945 | 1083 | 605 | 960 | 881 | 1075 | 1023 | 919 | 902 | 966 | 853 | 947,43 | 985 | 910 | 0,924 | -0,114 | -1,632 | 0,10263166 | 0,15602506 | 0,003526 | 0,0035118 | 0,0036089 | 0,0036089 | TRUE | 0,0558964 | No |
| orf19.2164.1 | 0 | 0 | 0 | 2 | 0 | 0 | 1 | 2 | 0 | 0 | 0 | 2 | 0 | 0 | 1 | 3 | 0,67 | 0 | 1 | 1,818 | 0,862 | 0,421 | 0,67351524 | NA | 3,568365 | 2,1677359 | 2,478998 | 2,478998 | TRUE | 0,8241687 | No |
| orf19.2165 | 1528 | 1363 | 1471 | 2032 | 1430 | 1650 | 1664 | 938 | 1371 | 1735 | 1505 | 1676 | 1223 | 1574 | 1485 | 1323 | 1486,5 | 1572 | 1401 | 0,892 | -0,164 | -1,785 | 0,07431861 | 0,11859512 | 0,011012 | 0,0029561 | 0,0074527 | 0,0074527 | TRUE | 0,0912528 | No |
| orf19.2166 | 1357 | 1301 | 1352 | 1721 | 1339 | 1573 | 1587 | 980 | 1217 | 1656 | 1383 | 1419 | 1145 | 1501 | 1416 | 1382 | 1390,1 | 1419 | 1361 | 0,96 | -0,059 | -0,597 | 0,55063354 | 0,63145759 | 0,013922 | 0,0030239 | 0,0086915 | 0,0086915 | TRUE | 0,1492376 | No |
| orf19.2167 | 11868 | 8539 | 9829 | 12922 | 9837 | 9265 | 9602 | 5517 | 10647 | 10871 | 10056 | 10656 | 8413 | 8840 | 8568 | 7781 | 9479 | 10558 | 8400 | 0,796 | -0,329 | -7,105 | 1,20E-12 | 1,09E-11 | 0,001836 | 0,0021327 | 0,0019505 | 0,0019505 | TRUE | 0,0293708 | No |
| orf19.2168 | 554 | 428 | 503 | 639 | 459 | 435 | 466 | 270 | 497 | 545 | 515 | 527 | 393 | 415 | 416 | 381 | 460,96 | 521 | 401 | 0,773 | -0,371 | -5,315 | 1,06E-07 | 5,21E-07 | 1,00E-08 | 0,0051288 | 0,0024851 | 0,0024851 | TRUE | 0,0112751 | No |
| orf19.2168.3 | 2828 | 2180 | 2483 | 3215 | 2647 | 2487 | 2522 | 1765 | 2537 | 2775 | 2540 | 2651 | 2264 | 2373 | 2250 | 2489 | 2485,1 | 2626 | 2344 | 0,892 | -0,165 | -3,348 | 0,0008129 | 0,00208447 | 0,001533 | 0,0025637 | 0,0019243 | 0,0019243 | TRUE | 0,0197461 | No |
| orf19.2169 | 16 | 13 | 22 | 20 | 12 | 7 | 13 | 14 | 14 | 17 | 23 | 16 | 10 | 7 | 12 | 20 | 14,77 | 18 | 12 | 0,668 | -0,583 | -1,511 | 0,13085141 | NA | 0,019031 | 0,1002396 | 0,0713274 | 0,0713274 | TRUE | 0,486992 | No |
| orf19.217 | 141 | 98 | 126 | 178 | 138 | 131 | 121 | 67 | 126 | 125 | 129 | 147 | 118 | 125 | 108 | 94 | 121,55 | 132 | 111 | 0,849 | -0,236 | -1,782 | 0,07476154 | 0,11917414 | 1,00E-08 | 0,0139223 | 0,0086113 | 0,0086113 | TRUE | 0,0786926 | No |
| orf19.2170 | 6086 | 4375 | 5608 | 6680 | 6274 | 5988 | 6388 | 4084 | 5460 | 5570 | 5738 | 5508 | 5366 | 5714 | 5700 | 5760 | 5601,9 | 5569 | 5635 | 1,012 | 0,017 | 0,452 | 0,65142484 | 0,72063348 | 0,000507 | 0,0022387 | 0,0011093 | 0,0011093 | TRUE | 0,0128812 | No |
| orf19.2172 | 7990 | 5990 | 6724 | 8694 | 11777 | 11357 | 11125 | 6229 | 7168 | 7626 | 6879 | 7169 | 10072 | 10836 | 9927 | 8785 | 8557,9 | 7210 | 9905 | 1,374 | 0,459 | 7,416 | 1,20E-13 | 1,21E-12 | 0,004408 | 0,0021492 | 0,0035567 | 0,0035567 | TRUE | 0,0699294 | No |
| orf19.2173 | 10121 | 6904 | 8669 | 10845 | 13142 | 11478 | 12638 | 7630 | 9080 | 8790 | 8869 | 8943 | 11239 | 10952 | 11277 | 10761 | 9988,9 | 8920 | 11057 | 1,24 | 0,31 | 10,528 | 6,44E-26 | 1,80E-24 | 0,000206 | 0,0021249 | 0,0007316 | 0,0007316 | TRUE | 0,0038173 | No |
| orf19.2174 | 771 | 469 | 652 | 735 | 575 | 484 | 516 | 290 | 692 | 597 | 667 | 606 | 492 | 462 | 460 | 409 | 548,12 | 640 | 456 | 0,714 | -0,486 | -6,227 | 4,75E-10 | 3,18E-09 | 0,003306 | 0,004628 | 0,003968 | 0,003968 | TRUE | 0,0473771 | No |
| orf19.2175 | 18359 | 12227 | 15719 | 19212 | 21385 | 18393 | 20029 | 12354 | 16470 | 15566 | 16082 | 15842 | 18289 | 17550 | 17873 | 17424 | 16887 | 15990 | 17784 | 1,112 | 0,153 | 4,912 | 9,01E-07 | 3,82E-06 | 0,000443 | 0,0020655 | 0,0008776 | 0,0008776 | TRUE | 0,0050044 | No |
| orf19.2176 | 1517 | 922 | 1309 | 1538 | 1796 | 1396 | 1636 | 980 | 1361 | 1174 | 1339 | 1268 | 1536 | 1332 | 1460 | 1382 | 1356,5 | 1286 | 1428 | 1,11 | 0,151 | 2,348 | 0,0188618 | 0,03525878 | 0,003129 | 0,0030497 | 0,0032193 | 0,0032193 | TRUE | 0,0385874 | No |
| orf19.2177 | 810 | 514 | 689 | 913 | 520 | 436 | 495 | 388 | 727 | 654 | 705 | 753 | 445 | 416 | 442 | 547 | 586,06 | 710 | 462 | 0,646 | -0,629 | -7,333 | 2,25E-13 | 2,22E-12 | 0,006169 | 0,0044566 | 0,0052724 | 0,0052724 | TRUE | 0,1571504 | No |
| orf19.2178 | 3824 | 2621 | 3395 | 4262 | 3490 | 3007 | 3359 | 2511 | 3431 | 3337 | 3473 | 3514 | 2985 | 2869 | 2997 | 3542 | 3268,5 | 3439 | 3098 | 0,899 | -0,153 | -2,432 | 0,01503507 | 0,02892139 | 0,00456 | 0,0024237 | 0,0034889 | 0,0034889 | TRUE | 0,1091637 | No |
| orf19.2178.1 | 894 | 532 | 716 | 874 | 836 | 666 | 660 | 399 | 802 | 677 | 733 | 721 | 715 | 635 | 589 | 563 | 679,34 | 733 | 626 | 0,855 | -0,226 | -2,693 | 0,00707463 | 0,01473799 | 0,006209 | 0,0041165 | 0,0053047 | 0,0053047 | TRUE | 0,1113815 | No |
| orf19.2179 | 14953 | 9638 | 13124 | 16307 | 11681 | 9583 | 10435 | 6393 | 13414 | 12270 | 13427 | 13447 | 9990 | 9144 | 9312 | 9017 | 11253 | 13140 | 9366 | 0,713 | -0,488 | -10,458 | 1,35E-25 | 3,60E-24 | 0,001923 | 0,0021085 | 0,0020028 | 0,0020028 | TRUE | 0,0247683 | No |
| orf19.2179.2 | 15016 | 9935 | 11944 | 15050 | 15850 | 12553 | 12815 | 8071 | 13471 | 12648 | 12220 | 12410 | 13555 | 11978 | 11435 | 11383 | 12388 | 12687 | 12088 | 0,953 | -0,07 | -1,156 | 0,24759559 | 0,32954238 | 0,00431 | 0,0020967 | 0,0033959 | 0,0033959 | TRUE | 0,0819752 | No |
| orf19.218 | 66 | 53 | 55 | 58 | 28 | 20 | 24 | 15 | 59 | 67 | 56 | 48 | 24 | 19 | 21 | 21 | 39,55 | 58 | 21 | 0,375 | -1,415 | -5,974 | 2,32E-09 | 1,40E-08 | 1,00E-08 | 0,0386869 | 0,0219898 | 0,0219898 | TRUE | 0,1261572 | No |
| orf19.2180 | 2386 | 1600 | 2223 | 2677 | 3070 | 2811 | 3147 | 2115 | 2140 | 2037 | 2274 | 2207 | 2626 | 2682 | 2808 | 2983 | 2469,8 | 2164 | 2775 | 1,279 | 0,355 | 6,624 | 3,49E-11 | 2,71E-10 | 0,002176 | 0,0025673 | 0,0023493 | 0,0023493 | TRUE | 0,0294828 | No |
| orf19.2181 | 33 | 29 | 37 | 30 | 23 | 27 | 19 | 12 | 30 | 37 | 38 | 25 | 20 | 26 | 17 | 17 | 26,05 | 32 | 20 | 0,625 | -0,678 | -2,389 | 0,01690848 | 0,03205621 | 1,00E-08 | 0,0576991 | 0,0368618 | 0,0368618 | TRUE | 0,2247365 | No |
| orf19.2182 | 6769 | 4768 | 6255 | 7542 | 7587 | 7146 | 7412 | 4787 | 6073 | 6070 | 6399 | 6219 | 6489 | 6818 | 6614 | 6752 | 6429,3 | 6190 | 6668 | 1,077 | 0,107 | 3,161 | 0,00157042 | 0,00379376 | 0,000352 | 0,0022053 | 0,0009367 | 0,0009367 | TRUE | 0,006235 | No |
| orf19.2183 | 26463 | 17722 | 23196 | 28008 | 24152 | 20980 | 22497 | 15119 | 23740 | 22562 | 23732 | 23096 | 20656 | 20018 | 20075 | 21324 | 21900 | 23282 | 20518 | 0,881 | -0,183 | -5,405 | 6,49E-08 | 3,26E-07 | 0,000677 | 0,0020458 | 0,0010521 | 0,0010521 | TRUE | 0,008407 | No |
| orf19.2184 | 244 | 145 | 165 | 227 | 234 | 182 | 201 | 135 | 219 | 185 | 169 | 187 | 200 | 174 | 179 | 190 | 187,88 | 190 | 186 | 0,976 | -0,035 | -0,309 | 0,75719617 | 0,81004889 | 0,003399 | 0,0097063 | 0,006849 | 0,006849 | TRUE | 0,1200583 | No |
| orf19.2185 | 5648 | 3760 | 4686 | 5916 | 4931 | 4107 | 4640 | 2764 | 5067 | 4787 | 4794 | 4878 | 4217 | 3919 | 4140 | 3898 | 4462,6 | 4882 | 4044 | 0,829 | -0,271 | -6,654 | 2,86E-11 | 2,23E-10 | 0,000831 | 0,0023048 | 0,0013646 | 0,0013646 | TRUE | 0,010196 | No |
| orf19.2186 | 882 | 568 | 814 | 944 | 1090 | 917 | 1026 | 668 | 791 | 723 | 833 | 778 | 932 | 875 | 916 | 942 | 848,81 | 781 | 916 | 1,17 | 0,226 | 3,834 | 0,00012597 | 0,0003786 | 0,000618 | 0,0036898 | 0,0021526 | 0,0021526 | TRUE | 0,0263131 | No |
| orf19.2187 | 901 | 635 | 860 | 1030 | 835 | 707 | 859 | 562 | 808 | 808 | 880 | 849 | 714 | 675 | 767 | 793 | 786,73 | 836 | 737 | 0,878 | -0,187 | -2,856 | 0,00429241 | 0,00938798 | 0,001736 | 0,0038248 | 0,0028416 | 0,0028416 | TRUE | 0,0376205 | No |
| orf19.2190 | 709 | 402 | 682 | 789 | 1007 | 797 | 951 | 612 | 636 | 512 | 698 | 651 | 861 | 760 | 849 | 863 | 728,71 | 624 | 833 | 1,331 | 0,413 | 4,679 | 2,88E-06 | 1,14E-05 | 0,007944 | 0,0039717 | 0,0060626 | 0,0060626 | TRUE | 0,1601126 | No |
| orf19.2191 | 1716 | 1135 | 1439 | 1852 | 1661 | 1359 | 1447 | 944 | 1539 | 1445 | 1472 | 1527 | 1421 | 1297 | 1291 | 1331 | 1415,5 | 1496 | 1335 | 0,892 | -0,165 | -3,286 | 0,00101588 | 0,00255265 | 0,000814 | 0,0030051 | 0,0017174 | 0,0017174 | TRUE | 0,0240606 | No |
| orf19.2191.1 | 221 | 150 | 204 | 231 | 251 | 199 | 194 | 172 | 198 | 191 | 209 | 190 | 215 | 190 | 173 | 243 | 201,08 | 197 | 205 | 1,032 | 0,046 | 0,398 | 0,69056963 | 0,75339429 | 0,00605 | 0,0091989 | 0,0078014 | 0,0078014 | TRUE | 0,1388164 | No |
| orf19.2192 | 73577 | 51713 | 67481 | 80279 | 81189 | 72951 | 78116 | 50388 | 66006 | 65836 | 69040 | 66199 | 69435 | 69608 | 69706 | 71068 | 68362 | 66770 | 69954 | 1,048 | 0,067 | 2,513 | 0,01198604 | 0,023663 | 0,000297 | 0,0020008 | 0,0006712 | 0,0006712 | TRUE | 0,0064101 | No |
| orf19.2193 | 8501 | 5502 | 7324 | 9077 | 9165 | 7676 | 8212 | 5366 | 7626 | 7005 | 7493 | 7485 | 7838 | 7324 | 7328 | 7568 | 7458,5 | 7402 | 7514 | 1,015 | 0,021 | 0,524 | 0,60026055 | 0,6763626 | 0,000961 | 0,0021742 | 0,0014253 | 0,0014253 | TRUE | 0,015587 | No |
| orf19.2194 | 830 | 621 | 694 | 895 | 984 | 903 | 928 | 583 | 745 | 791 | 710 | 738 | 842 | 862 | 828 | 822 | 792,1 | 746 | 838 | 1,127 | 0,172 | 3,07 | 0,00214068 | 0,00502979 | 1,00E-08 | 0,0038123 | 0,0017585 | 0,0017585 | TRUE | 0,0169183 | No |
| orf19.2195 | 12 | 6 | 6 | 13 | 7 | 9 | 3 | 5 | 11 | 8 | 6 | 11 | 6 | 9 | 3 | 7 | 7,45 | 9 | 6 | 0,671 | -0,576 | -1,108 | 0,2676811 | NA | 1,00E-08 | 0,1969511 | 0,1203866 | 0,1203866 | TRUE | 0,402083 | No |
| orf19.2196 | 639 | 468 | 588 | 752 | 983 | 939 | 1045 | 589 | 573 | 596 | 602 | 620 | 841 | 896 | 932 | 831 | 736,33 | 598 | 875 | 1,466 | 0,552 | 8,836 | 9,90E-19 | 1,62E-17 | 0,000957 | 0,0039511 | 0,0023392 | 0,0023392 | TRUE | 0,0236844 | No |
| orf19.2197 | 72 | 46 | 68 | 78 | 65 | 38 | 45 | 41 | 65 | 59 | 70 | 64 | 56 | 36 | 40 | 58 | 55,86 | 64 | 48 | 0,73 | -0,455 | -2,292 | 0,02192619 | 0,04055804 | 0,004109 | 0,027968 | 0,0194696 | 0,0194696 | TRUE | 0,2009982 | No |
| orf19.2198 | 4248 | 3011 | 3854 | 4744 | 3973 | 3634 | 3934 | 2495 | 3811 | 3833 | 3943 | 3912 | 3398 | 3467 | 3510 | 3519 | 3674,2 | 3875 | 3474 | 0,896 | -0,159 | -4,809 | 1,51E-06 | 6,22E-06 | 1,00E-08 | 0,0023746 | 0,0007697 | 0,0007697 | TRUE | 0,002716 | No |
| orf19.2199 | 4774 | 3445 | 4229 | 5208 | 3640 | 3281 | 3538 | 2438 | 4283 | 4386 | 4327 | 4295 | 3113 | 3131 | 3157 | 3439 | 3766,2 | 4323 | 3210 | 0,742 | -0,431 | -10,632 | 2,11E-26 | 6,26E-25 | 0,00084 | 0,002365 | 0,0013076 | 0,0013076 | TRUE | 0,0260505 | No |
| orf19.22 | 4342 | 3025 | 3759 | 4712 | 3427 | 3108 | 3220 | 2059 | 3895 | 3851 | 3846 | 3886 | 2931 | 2966 | 2873 | 2904 | 3393,9 | 3870 | 2918 | 0,754 | -0,407 | -12,721 | 4,53E-37 | 2,64E-35 | 1,00E-08 | 0,0024073 | 0,0006815 | 0,0006815 | TRUE | 0,0014613 | No |
| orf19.220 | 17645 | 12107 | 15556 | 19838 | 14471 | 12613 | 13578 | 9559 | 15829 | 15414 | 15915 | 16359 | 12376 | 12035 | 12116 | 13482 | 14191 | 15879 | 12502 | 0,787 | -0,346 | -7,954 | 1,80E-15 | 2,17E-14 | 0,001635 | 0,0020818 | 0,0017426 | 0,0017426 | TRUE | 0,0334776 | No |
| orf19.2200 | 1587 | 1090 | 1378 | 1681 | 1503 | 1372 | 1509 | 924 | 1424 | 1388 | 1410 | 1386 | 1285 | 1309 | 1347 | 1303 | 1356,5 | 1402 | 1311 | 0,935 | -0,097 | -2,21 | 0,02711978 | 0,04912138 | 1,00E-08 | 0,0030497 | 0,0010952 | 0,0010952 | TRUE | 0,0042719 | No |
| orf19.2201 | 1050 | 730 | 899 | 1165 | 1142 | 1015 | 999 | 620 | 942 | 929 | 920 | 961 | 977 | 968 | 891 | 874 | 932,85 | 938 | 928 | 0,991 | -0,013 | -0,237 | 0,81273846 | 0,85530438 | 0,000651 | 0,0035357 | 0,001985 | 0,001985 | TRUE | 0,0154262 | No |
| orf19.2202 | 149 | 89 | 131 | 164 | 128 | 122 | 125 | 86 | 134 | 113 | 134 | 135 | 109 | 116 | 112 | 121 | 121,87 | 129 | 114 | 0,88 | -0,184 | -1,465 | 0,14290314 | 0,20693228 | 1,00E-08 | 0,0138915 | 0,0069553 | 0,0069553 | TRUE | 0,0573003 | No |
| orf19.2203 | 10 | 4 | 4 | 5 | 3 | 5 | 5 | 3 | 9 | 5 | 4 | 4 | 3 | 5 | 4 | 4 | 4,79 | 6 | 4 | 0,71 | -0,495 | -0,785 | 0,43229202 | NA | 1,00E-08 | 0,3051407 | 0,1666415 | 0,1666415 | TRUE | 0,4372483 | No |
| orf19.2203.1 | 44 | 29 | 38 | 40 | 59 | 56 | 55 | 43 | 39 | 37 | 39 | 33 | 50 | 53 | 49 | 61 | 45,23 | 37 | 53 | 1,432 | 0,518 | 2,549 | 0,01079532 | 0,02156321 | 1,00E-08 | 0,0340726 | 0,0168323 | 0,0168323 | TRUE | 0,0517319 | No |

| | | | | | | | | | | | | | | | | | | | | | | | | | | | | | | | |
|---|---|---|---|---|---|---|---|---|---|---|---|---|---|---|---|---|---|---|---|---|---|---|---|---|---|---|---|---|---|---|---|
| orf19.2204 | 7164 | 4684 | 6383 | 7775 | 6978 | 5840 | 6374 | 4202 | 6427 | 5963 | 6530 | 6411 | 5968 | 5572 | 5688 | 5927 | 6060,8 | 6333 | 5789 | 0,913 | -0,131 | -3,126 | 0,00177033 | 0,00422739 | 0,001109 | 0,002219 | 0,0015098 | 0,0015098 | TRUE | 0,0183246 | No |
| orf19.2204.2 | 1277 | 865 | 1083 | 1376 | 1005 | 815 | 890 | 645 | 1146 | 1101 | 1108 | 1135 | 860 | 778 | 794 | 910 | 978,82 | 1122 | 836 | 0,741 | -0,432 | -7,284 | 3,23E-13 | 3,16E-12 | 0,001227 | 0,0034626 | 0,002323 | 0,002323 | TRUE | 0,0372008 | No |
| orf19.2208 | 5993 | 3923 | 5259 | 6401 | 7226 | 6154 | 6634 | 4384 | 5376 | 4994 | 5380 | 5278 | 6180 | 5872 | 5920 | 6183 | 5648,1 | 5257 | 6039 | 1,148 | 0,199 | 5,166 | 2,39E-07 | 1,11E-06 | 0,000738 | 0,0022365 | 0,0012423 | 0,0012423 | TRUE | 0,0132907 | No |
| orf19.2209 | 1318 | 1026 | 1277 | 1498 | 1356 | 1086 | 1289 | 743 | 1182 | 1306 | 1306 | 1235 | 1160 | 1036 | 1150 | 1048 | 1178,1 | 1257 | 1098 | 0,876 | -0,191 | -3,19 | 0,00142456 | 0,0034721 | 0,002086 | 0,0032118 | 0,002577 | 0,002577 | TRUE | 0,0195621 | No |
| orf19.2210 | 1768 | 1130 | 1583 | 1936 | 2050 | 1824 | 1985 | 1351 | 1586 | 1439 | 1620 | 1596 | 1753 | 1740 | 1771 | 1905 | 1676,4 | 1560 | 1792 | 1,145 | 0,196 | 3,729 | 0,00019207 | 0,00056127 | 0,001496 | 0,0028455 | 0,0020474 | 0,0020474 | TRUE | 0,0308462 | No |
| orf19.2211 | 7 | 9 | 11 | 16 | 11 | 13 | 9 | 9 | 6 | 11 | 11 | 13 | 9 | 12 | 8 | 13 | 10,59 | 10 | 10 | 0,996 | -0,005 | -0,012 | 0,99046173 | NA | 1,00E-08 | 0,1390585 | 0,079854 | 0,079854 | TRUE | 0,3147261 | No |
| orf19.2212 | 30 | 21 | 23 | 45 | 35 | 23 | 42 | 15 | 27 | 27 | 24 | 37 | 30 | 22 | 37 | 21 | 28,1 | 29 | 28 | 0,973 | -0,039 | -0,139 | 0,88960947 | 0,9164564 | 0,025057 | 0,0536414 | 0,0415899 | 0,0415899 | TRUE | 0,3856974 | No |
| orf19.2213 | 181 | 133 | 142 | 222 | 125 | 129 | 152 | 88 | 162 | 169 | 145 | 183 | 107 | 123 | 136 | 124 | 143,72 | 165 | 122 | 0,74 | -0,435 | -3,432 | 0,00059897 | 0,00158414 | 1,00E-08 | 0,0120802 | 0,0083183 | 0,0083183 | TRUE | 0,0850603 | No |
| orf19.2214 | 2096 | 1449 | 1755 | 2342 | 2492 | 2131 | 2307 | 1546 | 1880 | 1845 | 1796 | 1931 | 2131 | 2033 | 2059 | 2180 | 1981,9 | 1863 | 2101 | 1,126 | 0,171 | 3,889 | 0,00010062 | 0,00030828 | 0,000468 | 0,002712 | 0,0013443 | 0,0013443 | TRUE | 0,0076824 | No |
| orf19.2215 | 4442 | 3130 | 4011 | 4992 | 5108 | 4387 | 4811 | 3150 | 3985 | 3985 | 4104 | 4116 | 4369 | 4186 | 4293 | 4443 | 4185 | 4048 | 4323 | 1,067 | 0,094 | 2,713 | 0,00666096 | 0,01401719 | 0,000163 | 0,0023264 | 0,0009034 | 0,0009034 | TRUE | 0,0054468 | No |
| orf19.2216 | 212 | 159 | 181 | 210 | 238 | 229 | 225 | 145 | 190 | 202 | 185 | 173 | 204 | 219 | 201 | 205 | 197,29 | 188 | 207 | 1,106 | 0,146 | 1,451 | 0,14679134 | 0,21174475 | 1,00E-08 | 0,0093379 | 0,0046504 | 0,0046504 | TRUE | 0,0318478 | No |
| orf19.2219 | 3088 | 2166 | 2719 | 3450 | 3019 | 2689 | 3026 | 1971 | 2770 | 2758 | 2782 | 2845 | 2582 | 2566 | 2700 | 2780 | 2722,8 | 2789 | 2657 | 0,951 | -0,072 | -1,803 | 0,07136134 | 0,11458053 | 0,000337 | 0,0025127 | 0,0011687 | 0,0011687 | TRUE | 0,0107263 | No |
| orf19.2220 | 363 | 255 | 301 | 415 | 285 | 266 | 226 | 161 | 326 | 325 | 308 | 342 | 244 | 254 | 202 | 227 | 278,34 | 325 | 232 | 0,712 | -0,49 | -5,181 | 2,21E-07 | 1,03E-06 | 0,002024 | 0,007195 | 0,0049053 | 0,0049053 | TRUE | 0,0934626 | No |
| orf19.2222 | 2861 | 1935 | 2553 | 3021 | 2727 | 2514 | 2842 | 1696 | 2567 | 2463 | 2612 | 2491 | 2332 | 2399 | 2536 | 2392 | 2474 | 2533 | 2415 | 0,953 | -0,069 | -1,625 | 0,10410545 | 0,15773775 | 0,000627 | 0,0025663 | 0,0013351 | 0,0013351 | TRUE | 0,0144592 | No |
| orf19.2227 | 162 | 114 | 144 | 174 | 146 | 158 | 131 | 88 | 145 | 145 | 147 | 143 | 125 | 151 | 117 | 124 | 137,24 | 145 | 129 | 0,89 | -0,169 | -1,375 | 0,1691108 | 0,23895053 | 1,00E-08 | 0,0125575 | 0,0071646 | 0,0071646 | TRUE | 0,1359524 | No |
| orf19.2228 | 131 | 149 | 140 | 149 | 195 | 226 | 203 | 146 | 118 | 190 | 143 | 123 | 167 | 216 | 181 | 206 | 167,85 | 144 | 192 | 1,351 | 0,434 | 2,824 | 0,00473837 | 0,01028102 | 0,023546 | 0,0106284 | 0,0165529 | 0,0165529 | TRUE | 0,4503401 | No |
| orf19.2229 | 1334 | 882 | 1160 | 1491 | 1259 | 1083 | 1175 | 727 | 1197 | 1123 | 1187 | 1229 | 1077 | 1033 | 1048 | 1025 | 1115 | 1184 | 1046 | 0,883 | -0,18 | -3,597 | 0,00032131 | 0,00090461 | 1,00E-08 | 0,0032815 | 0,0015021 | 0,0015021 | TRUE | 0,0132037 | No |
| orf19.223 | 8485 | 6190 | 7690 | 9285 | 10082 | 9243 | 9935 | 6231 | 7612 | 7881 | 7868 | 7657 | 8622 | 8819 | 8865 | 8788 | 8264 | 7754 | 8774 | 1,132 | 0,179 | 6,291 | 3,15E-10 | 2,15E-09 | 0,000121 | 0,0021552 | 0,0006523 | 0,0006523 | TRUE | 0,0018439 | No |
| orf19.2230 | 542 | 469 | 519 | 634 | 550 | 490 | 534 | 290 | 486 | 597 | 531 | 523 | 470 | 468 | 477 | 409 | 495,07 | 534 | 456 | 0,86 | -0,218 | -2,698 | 0,0069676 | 0,01455034 | 0,003404 | 0,0049118 | 0,0042499 | 0,0042499 | TRUE | 0,0674549 | No |
| orf19.2231 | 2024 | 1611 | 1820 | 2436 | 2191 | 2044 | 2094 | 1567 | 1816 | 2051 | 1862 | 2009 | 1874 | 1950 | 1869 | 2210 | 1955 | 1934 | 1976 | 1,02 | 0,028 | 0,435 | 0,66329367 | 0,7313394 | 0,003939 | 0,0027221 | 0,0035831 | 0,0035831 | TRUE | 0,0736724 | No |
| orf19.2232 | 13181 | 8606 | 10347 | 13037 | 12829 | 10735 | 10913 | 6754 | 11825 | 10956 | 10586 | 10750 | 10972 | 10243 | 9738 | 9526 | 10575 | 11029 | 10120 | 0,918 | -0,124 | -2,279 | 0,0226514 | 0,04176413 | 0,003141 | 0,0021168 | 0,0027409 | 0,0027409 | TRUE | 0,0394095 | No |
| orf19.2233 | 4845 | 3469 | 4217 | 5354 | 4739 | 4026 | 4291 | 2560 | 4346 | 4416 | 4314 | 4415 | 4053 | 3841 | 3829 | 3611 | 4103,3 | 4373 | 3834 | 0,878 | -0,188 | -4,587 | 4,50E-06 | 1,73E-05 | 0,000852 | 0,0023333 | 0,0013693 | 0,0013693 | TRUE | 0,018075 | No |
| orf19.2236 | 3430 | 2391 | 3018 | 3766 | 4447 | 4008 | 4436 | 2735 | 3077 | 3044 | 3088 | 3105 | 3803 | 3824 | 3958 | 3857 | 3469,7 | 3078 | 3860 | 1,254 | 0,326 | 9,928 | 3,14E-23 | 7,09E-22 | 1,00E-08 | 0,0023979 | 0,0007442 | 0,0007442 | TRUE | 0,0037142 | No |
| orf19.2237 | 64 | 57 | 70 | 84 | 76 | 56 | 67 | 41 | 57 | 73 | 72 | 69 | 65 | 53 | 60 | 58 | 63,36 | 68 | 59 | 0,877 | -0,189 | -1,084 | 0,27833332 | 0,36242448 | 1,00E-08 | 0,0248901 | 0,0133114 | 0,0133114 | TRUE | 0,1004209 | No |
| orf19.2237.1 | 2654 | 2105 | 2228 | 3040 | 2362 | 2450 | 2452 | 1556 | 2381 | 2680 | 2279 | 2507 | 2020 | 2338 | 2188 | 2195 | 2323,4 | 2462 | 2185 | 0,888 | -0,172 | -2,754 | 0,00589124 | 0,01253348 | 0,003649 | 0,0026043 | 0,0032917 | 0,0032917 | TRUE | 0,04162 | No |
| orf19.2238 | 1077 | 912 | 957 | 1311 | 1040 | 1095 | 1113 | 658 | 966 | 1161 | 979 | 1081 | 889 | 1045 | 993 | 928 | 1005,4 | 1047 | 964 | 0,922 | -0,117 | -1,547 | 0,12193852 | 0,18089053 | 0,005075 | 0,0034235 | 0,0044571 | 0,0044571 | TRUE | 0,0609414 | No |
| orf19.2241 | 3174 | 2123 | 2754 | 3206 | 5701 | 4703 | 5235 | 3228 | 2847 | 2703 | 2818 | 2644 | 4876 | 4487 | 4671 | 4553 | 3699,9 | 2753 | 4647 | 1,689 | 0,756 | 17,49 | 1,71E-68 | 3,79E-66 | 0,000893 | 0,0023719 | 0,0015035 | 0,0015035 | TRUE | 0,0136326 | No |
| orf19.2242 | 9942 | 6870 | 8630 | 10680 | 10748 | 9808 | 10517 | 6796 | 8919 | 8746 | 8829 | 8807 | 9192 | 9358 | 9385 | 9585 | 9102,7 | 8825 | 9380 | 1,062 | 0,087 | 3,229 | 0,00124191 | 0,0030718 | 3,39E-05 | 0,002139 | 0,0005885 | 0,0005885 | TRUE | 0,0025057 | No |
| orf19.2244 | 1338 | 964 | 1204 | 1398 | 805 | 711 | 788 | 549 | 1200 | 1227 | 1232 | 1153 | 688 | 678 | 703 | 774 | 957,07 | 1203 | 711 | 0,589 | -0,763 | -13,32 | 1,76E-40 | 1,22E-38 | 0,000596 | 0,0034963 | 0,0020262 | 0,0020262 | TRUE | 0,0361612 | No |
| orf19.2245 | 788 | 581 | 685 | 825 | 642 | 597 | 603 | 365 | 707 | 740 | 701 | 680 | 549 | 570 | 538 | 515 | 624,91 | 707 | 543 | 0,771 | -0,375 | -6,04 | 1,55E-09 | 9,56E-09 | 1,00E-08 | 0,0043026 | 0,0020691 | 0,0020691 | TRUE | 0,0121006 | No |
| orf19.2246 | 1295 | 877 | 1045 | 1346 | 1003 | 876 | 896 | 505 | 1162 | 1117 | 1069 | 1110 | 858 | 836 | 800 | 712 | 957,85 | 1114 | 802 | 0,722 | -0,47 | -7,397 | 1,39E-13 | 1,38E-12 | 0,002346 | 0,0034951 | 0,0028039 | 0,0028039 | TRUE | 0,0582805 | No |
| orf19.2247 | 130 | 98 | 140 | 137 | 102 | 96 | 90 | 81 | 117 | 125 | 143 | 113 | 87 | 92 | 80 | 114 | 108,87 | 124 | 93 | 0,743 | -0,428 | -2,895 | 0,00379721 | 0,00841832 | 0,006261 | 0,0153135 | 0,0115816 | 0,0115816 | TRUE | 0,173467 | No |
| orf19.2248 | 1242 | 929 | 1106 | 1419 | 1205 | 1128 | 1244 | 804 | 1114 | 1183 | 1132 | 1170 | 1031 | 1076 | 1110 | 1134 | 1118,7 | 1150 | 1088 | 0,945 | -0,082 | -1,589 | 0,1121033 | 0,16791738 | 0,000345 | 0,0032772 | 0,0016496 | 0,0016496 | TRUE | 0,0154523 | No |
| orf19.2249 | 390 | 271 | 305 | 394 | 361 | 273 | 328 | 202 | 350 | 345 | 312 | 325 | 309 | 260 | 293 | 285 | 309,83 | 333 | 287 | 0,863 | -0,213 | -2,445 | 0,01446701 | 0,02796582 | 1,00E-08 | 0,006665 | 0,0040251 | 0,0040251 | TRUE | 0,0459833 | No |
| orf19.225 | 3157 | 2028 | 2772 | 3367 | 3602 | 3276 | 3568 | 2043 | 2832 | 2582 | 2836 | 2776 | 3081 | 3126 | 3184 | 2881 | 2912,3 | 2756 | 3068 | 1,113 | 0,155 | 3,257 | 0,00112594 | 0,00280575 | 0,001399 | 0,002478 | 0,001822 | 0,001822 | TRUE | 0,0210276 | No |
| orf19.2250 | 4817 | 3210 | 3957 | 4971 | 4273 | 3530 | 3599 | 2541 | 4321 | 4087 | 4048 | 4099 | 3654 | 3368 | 3212 | 3584 | 3796,7 | 4139 | 3454 | 0,834 | -0,262 | -5,295 | 1,19E-07 | 5,77E-07 | 0,001812 | 0,0023619 | 0,0020779 | 0,0020779 | TRUE | 0,0277109 | No |
| orf19.2251 | 9039 | 6440 | 7767 | 9797 | 9027 | 7569 | 7943 | 5390 | 8109 | 8199 | 7946 | 8079 | 7720 | 7222 | 7088 | 7602 | 7745,6 | 8083 | 7408 | 0,916 | -0,126 | -3,38 | 0,00072574 | 0,00188346 | 0,000738 | 0,0021669 | 0,0012065 | 0,0012065 | TRUE | 0,0104643 | No |
| orf19.2252 | 3 | 5 | 7 | 6 | 5 | 8 | 3 | 3 | 3 | 6 | 7 | 5 | 4 | 8 | 3 | 4 | 5 | 5 | 5 | 0,903 | -0,147 | -0,234 | 0,81488775 | NA | 1,00E-08 | 0,2924363 | 0,1750034 | 0,1750034 | TRUE | 0,3717637 | No |
| orf19.2253 | 313 | 217 | 336 | 347 | 273 | 201 | 231 | 152 | 281 | 276 | 344 | 286 | 233 | 192 | 206 | 214 | 254,09 | 297 | 211 | 0,713 | -0,488 | -4,645 | 3,40E-06 | 1,33E-05 | 0,005323 | 0,0076928 | 0,0065652 | 0,0065652 | TRUE | 0,1262202 | No |
| orf19.2256 | 4452 | 2774 | 3578 | 4398 | 4175 | 3294 | 3645 | 2261 | 3994 | 3532 | 3661 | 3627 | 3571 | 3143 | 3253 | 3189 | 3496 | 3704 | 3289 | 0,888 | -0,171 | -3,074 | 0,00210857 | 0,00496017 | 0,00282 | 0,0023948 | 0,0026818 | 0,0026818 | TRUE | 0,0411402 | No |
| orf19.2257 | 1739 | 1224 | 1580 | 1945 | 1604 | 1403 | 1507 | 898 | 1560 | 1558 | 1616 | 1604 | 1372 | 1339 | 1345 | 1267 | 1457,6 | 1584 | 1331 | 0,841 | -0,25 | -5,554 | 2,79E-08 | 1,46E-07 | 1,00E-08 | 0,0029755 | 0,001248 | 0,001248 | TRUE | 0,0108661 | No |
| orf19.2258 | 1165 | 806 | 1039 | 1194 | 1115 | 1015 | 1138 | 602 | 1045 | 1026 | 1063 | 985 | 954 | 968 | 1015 | 849 | 988,18 | 1030 | 946 | 0,923 | -0,116 | -1,871 | 0,06129957 | 0,10040563 | 0,001932 | 0,0034486 | 0,0026601 | 0,0026601 | TRUE | 0,0508305 | No |
| orf19.2259 | 1877 | 1218 | 1470 | 1898 | 1951 | 1698 | 1774 | 1044 | 1684 | 1551 | 1504 | 1565 | 1669 | 1620 | 1583 | 1472 | 1581 | 1576 | 1586 | 1,008 | 0,011 | 0,202 | 0,83956968 | 0,87739339 | 0,001847 | 0,0028978 | 0,0022886 | 0,0022886 | TRUE | 0,0261881 | No |
| orf19.2260 | 2569 | 1706 | 2159 | 2690 | 2216 | 1976 | 2275 | 1293 | 2305 | 2172 | 2209 | 2218 | 1895 | 1885 | 2030 | 1824 | 2067,3 | 2226 | 1908 | 0,858 | -0,221 | -4,805 | 1,55E-06 | 6,34E-06 | 0,000707 | 0,0026818 | 0,0015363 | 0,0015363 | TRUE | 0,0223822 | No |
| orf19.2261 | 2004 | 1461 | 1767 | 2126 | 1930 | 1745 | 1848 | 1167 | 1798 | 1860 | 1808 | 1753 | 1651 | 1665 | 1649 | 1646 | 1728,7 | 1805 | 1653 | 0,917 | -0,125 | -3,083 | 0,00204591 | 0,00482032 | 1,00E-08 | 0,0028193 | 0,0009989 | 0,0009989 | TRUE | 0,0047562 | No |
| orf19.2262 | 2418 | 1732 | 2047 | 2554 | 2957 | 2610 | 2879 | 1698 | 2169 | 2205 | 2094 | 2106 | 2529 | 2490 | 2569 | 2395 | 2319,7 | 2144 | 2496 | 1,167 | 0,222 | 5,479 | 4,28E-08 | 2,19E-07 | 0,000168 | 0,0026053 | 0,0011452 | 0,0011452 | TRUE | 0,008135 | No |
| orf19.2263 | 864 | 592 | 734 | 985 | 875 | 789 | 838 | 539 | 775 | 754 | 751 | 812 | 748 | 753 | 748 | 760 | 762,64 | 773 | 752 | 0,971 | -0,043 | -0,767 | 0,44308959 | 0,52976762 | 1,00E-08 | 0,003883 | 0,0016562 | 0,0016562 | TRUE | 0,0147207 | No |
| orf19.2264 | 5847 | 3928 | 4985 | 5928 | 5565 | 4724 | 5115 | 3406 | 5245 | 5001 | 5100 | 4888 | 4759 | 4507 | 4564 | 4804 | 4858,7 | 5058 | 4658 | 0,92 | -0,12 | -3,082 | 0,00205402 | 0,00483754 | 0,000658 | 0,0022783 | 0,0012361 | 0,0012361 | TRUE | 0,0077099 | No |
| orf19.2265 | 2308 | 1572 | 1964 | 2525 | 2284 | 2001 | 2199 | 1487 | 2071 | 2001 | 2009 | 2082 | 1953 | 1909 | 1962 | 2097 | 2010,7 | 2041 | 1980 | 0,968 | -0,047 | -1,08 | 0,28013274 | 0,36430166 | 0,000383 | 0,0027015 | 0,0013161 | 0,0013161 | TRUE | 0,0171079 | No |
| orf19.2266 | 1524 | 1037 | 1394 | 1689 | 1425 | 1305 | 1294 | 866 | 1367 | 1320 | 1426 | 1393 | 1219 | 1245 | 1155 | 1221 | 1293,3 | 1376 | 1210 | 0,878 | -0,188 | -3,869 | 0,00010912 | 0,00033229 | 4,36E-06 | 0,003102 | 0,0014928 | 0,0014928 | TRUE | 0,0119896 | No |
| orf19.2267 | 2705 | 1826 | 2219 | 2873 | 2681 | 2058 | 2194 | 1545 | 2427 | 2325 | 2270 | 2369 | 2293 | 1964 | 1958 | 2179 | 2223 | 2348 | 2098 | 0,893 | -0,163 | -2,76 | 0,00578342 | 0,01232593 | 0,002912 | 0,0026326 | 0,0028938 | 0,0028938 | TRUE | 0,0496504 | No |
| orf19.2268 | 7186 | 4847 | 6436 | 7876 | 7054 | 6247 | 6832 | 4505 | 6447 | 6171 | 6585 | 6495 | 6033 | 5961 | 6096 | 6354 | 6267,6 | 6424 | 6111 | 0,95 | -0,074 | -2,047 | 0,04061597 | 0,06996008 | 0,000565 | 0,0022111 | 0,0010788 | 0,0010788 | TRUE | 0,0083063 | No |
| orf19.2269 | 731 | 528 | 587 | 831 | 689 | 699 | 654 | 409 | 656 | 672 | 601 | 685 | 589 | 667 | 584 | 577 | 628,81 | 654 | 604 | 0,925 | -0,112 | -1,563 | 0,11812753 | 0,17588853 | 0,002445 | 0,0042882 | 0,0033333 | 0,0033333 | TRUE | 0,0582415 | No |
| orf19.227 | 2296 | 2003 | 2046 | 2980 | 2901 | 2558 | 2673 | 1582 | 2060 | 2550 | 2093 | 2457 | 2481 | 2441 | 2385 | 2231 | 2337,3 | 2290 | 2384 | 1,042 | 0,06 | 0,813 | 0,41623749 | 0,50428611 | 0,005871 | 0,0026006 | 0,004807 | 0,004807 | TRUE | 0,0687467 | No |
| orf19.2270 | 8309 | 5981 | 7502 | 9567 | 8753 | 7895 | 8530 | 5228 | 7454 | 7614 | 7675 | 7889 | 7486 | 7533 | 7612 | 7374 | 7579,6 | 7658 | 7501 | 0,98 | -0,03 | -0,966 | 0,33417911 | 0,42099881 | 0,000188 | 0,002171 | 0,0007753 | 0,0007753 | TRUE | 0,0052036 | No |
| orf19.2272 | 2813 | 1868 | 2602 | 3137 | 2942 | 2668 | 3017 | 1954 | 2524 | 2378 | 2662 | 2587 | 2516 | 2546 | 2692 | 2756 | 2582,6 | 2538 | 2628 | 1,034 | 0,048 | 0,961 | 0,33673812 | 0,4236906 | 0,001571 | 0,0025416 | 0,001978 | 0,001978 | TRUE | 0,0206021 | No |
| orf19.2274 | 46 | 26 | 49 | 75 | 49 | 53 | 57 | 36 | 41 | 33 | 50 | 62 | 42 | 51 | 51 | 51 | 47,56 | 46 | 49 | 1,025 | 0,035 | 0,159 | 0,87359354 | 0,9035204 | 0,017887 | 0,0325047 | 0,026377 | 0,026377 | TRUE | 0,3500179 | No |
| orf19.2275 | 1327 | 911 | 1007 | 1325 | 1519 | 1271 | 1361 | 931 | 1190 | 1160 | 1030 | 1093 | 1299 | 1213 | 1214 | 1313 | 1189,1 | 1118 | 1260 | 1,126 | 0,171 | 2,866 | 0,00415254 | 0,00911192 | 0,002076 | 0,0032004 | 0,0025698 | 0,0025698 | TRUE | 0,0330244 | No |
| orf19.2276 | 1244 | 779 | 993 | 1280 | 1020 | 824 | 846 | 475 | 1116 | 992 | 1016 | 1056 | 872 | 786 | 755 | 670 | 907,83 | 1045 | 771 | 0,74 | -0,435 | -5,604 | 2,10E-08 | 1,12E-07 | 0,005728 | 0,0035786 | 0,004649 | 0,004649 | TRUE | 0,0933207 | No |

| | | | | | | | | | | | | | | | | | | | | | | | | | | | | | | | |
|---|---|---|---|---|---|---|---|---|---|---|---|---|---|---|---|---|---|---|---|---|---|---|---|---|---|---|---|---|---|---|---|
| orf19.2277 | 1757 | 1166 | 1548 | 1900 | 2187 | 1956 | 2155 | 1416 | 1576 | 1484 | 1584 | 1567 | 1870 | 1866 | 1923 | 1997 | 1733,5 | 1553 | 1914 | 1,23 | 0,298 | 6,723 | 1,78E-11 | 1,41E-10 | 0,000171 | 0,002817 | 0,0013043 | 0,0013043 | TRUE | 0,0095986 | No |
| orf19.2278 | 2381 | 1645 | 2147 | 2626 | 2482 | 2255 | 2357 | 1588 | 2136 | 2094 | 2197 | 2165 | 2123 | 2152 | 2103 | 2240 | 2151,2 | 2148 | 2154 | 1,001 | 0,001 | 0,033 | 0,97372017 | 0,98139755 | 1,00E-08 | 0,0026544 | 0,0010591 | 0,0010591 | TRUE | 0,0077143 | No |
| orf19.2280 | 873 | 620 | 892 | 921 | 727 | 653 | 732 | 458 | 783 | 789 | 913 | 759 | 622 | 623 | 653 | 646 | 723,57 | 811 | 636 | 0,784 | -0,351 | -5,004 | 5,61E-07 | 2,47E-06 | 0,002742 | 0,0039858 | 0,003306 | 0,003306 | TRUE | 0,084436 | No |
| orf19.2281 | 99 | 63 | 83 | 98 | 125 | 99 | 107 | 60 | 89 | 80 | 85 | 81 | 107 | 94 | 95 | 85 | 89,53 | 84 | 95 | 1,147 | 0,197 | 1,354 | 0,17572442 | 0,24703302 | 1,00E-08 | 0,0181947 | 0,009285 | 0,009285 | TRUE | 0,0662998 | No |
| orf19.2282 | 949 | 615 | 816 | 924 | 1179 | 1030 | 1047 | 618 | 851 | 783 | 835 | 762 | 1008 | 983 | 934 | 872 | 878,52 | 808 | 949 | 1,178 | 0,236 | 3,662 | 0,00025009 | 0,00071585 | 0,002188 | 0,003632 | 0,0028556 | 0,0028556 | TRUE | 0,0322659 | No |
| orf19.2283 | 1768 | 1149 | 1468 | 1808 | 1933 | 1515 | 1650 | 1115 | 1586 | 1463 | 1502 | 1491 | 1653 | 1446 | 1472 | 1573 | 1523,2 | 1510 | 1536 | 1,016 | 0,023 | 0,408 | 0,68325864 | 0,74677422 | 0,001968 | 0,0029326 | 0,0023624 | 0,0023624 | TRUE | 0,0341522 | No |
| orf19.2284 | 1495 | 943 | 1243 | 1619 | 1792 | 1466 | 1613 | 1136 | 1341 | 1201 | 1272 | 1335 | 1533 | 1399 | 1439 | 1602 | 1390,2 | 1287 | 1493 | 1,156 | 0,21 | 3,543 | 0,00039557 | 0,00109167 | 0,002226 | 0,0030238 | 0,0026372 | 0,0026372 | TRUE | 0,026712 | No |
| orf19.2285 | 116 | 87 | 110 | 121 | 111 | 108 | 112 | 70 | 104 | 111 | 113 | 100 | 95 | 103 | 100 | 99 | 102,97 | 107 | 99 | 0,932 | -0,102 | -0,765 | 0,44428616 | 0,53098692 | 1,00E-08 | 0,0160772 | 0,0074619 | 0,0074619 | TRUE | 0,0213196 | No |
| orf19.2286 | 5425 | 3597 | 4636 | 5845 | 5580 | 4521 | 4802 | 3309 | 4867 | 4579 | 4743 | 4820 | 4772 | 4314 | 4285 | 4667 | 4630,9 | 4752 | 4510 | 0,948 | -0,077 | -1,645 | 0,09988374 | 0,15270371 | 0,00156 | 0,002293 | 0,0018676 | 0,0018676 | TRUE | 0,01957 | No |
| orf19.2287 | 1961 | 1469 | 1680 | 2057 | 1802 | 1401 | 1536 | 1042 | 1759 | 1870 | 1719 | 1696 | 1541 | 1337 | 1371 | 1470 | 1595,3 | 1761 | 1430 | 0,812 | -0,3 | -5,154 | 2,55E-07 | 1,18E-06 | 0,002248 | 0,0028895 | 0,0026151 | 0,0026151 | TRUE | 0,0356959 | No |
| orf19.2288 | 4974 | 3141 | 4315 | 5082 | 5162 | 4275 | 4932 | 2885 | 4462 | 3999 | 4415 | 4191 | 4415 | 4079 | 4401 | 4069 | 4253,8 | 4267 | 4241 | 0,994 | -0,008 | -0,17 | 0,86528166 | 0,89755045 | 0,001739 | 0,0023208 | 0,0021442 | 0,0021442 | TRUE | 0,0210596 | No |
| orf19.2289 | 1107 | 719 | 929 | 1265 | 1312 | 1199 | 1189 | 842 | 993 | 915 | 950 | 1043 | 1122 | 1144 | 1061 | 1188 | 1052,1 | 975 | 1129 | 1,152 | 0,205 | 3,447 | 0,00056787 | 0,00151232 | 0,001662 | 0,0033593 | 0,0024243 | 0,0024243 | TRUE | 0,026583 | No |
| orf19.229 | 1925 | 1463 | 1719 | 2213 | 1953 | 1910 | 1896 | 1086 | 1727 | 1863 | 1759 | 1825 | 1670 | 1822 | 1692 | 1532 | 1736,2 | 1794 | 1679 | 0,938 | -0,092 | -1,609 | 0,10758612 | 0,16228407 | 0,002335 | 0,0028157 | 0,0025377 | 0,0025377 | TRUE | 0,038957 | No |
| orf19.2290 | 4146 | 3001 | 3867 | 4618 | 6307 | 5542 | 6064 | 3802 | 3719 | 3821 | 3956 | 3808 | 5394 | 5288 | 5411 | 5362 | 4595 | 3826 | 5364 | 1,402 | 0,488 | 14,843 | 7,76E-50 | 8,04E-48 | 7,97E-05 | 0,0022955 | 0,0008127 | 0,0008127 | TRUE | 0,006427 | No |
| orf19.2296 | 83635 | 54658 | 74020 | 88951 | 1E+05 | 82301 | 92018 | 62502 | 75029 | 69585 | 75730 | 73350 | 86670 | 78529 | 82111 | 88153 | 78645 | 73424 | 83866 | 1,142 | 0,192 | 4,137 | 3,52E-05 | 0,00011688 | 0,002053 | 0,001998 | 0,0020514 | 0,0020514 | TRUE | 0,0224859 | No |
| orf19.2297 | 216 | 142 | 196 | 218 | 228 | 198 | 233 | 145 | 194 | 181 | 201 | 180 | 195 | 189 | 208 | 205 | 193,9 | 189 | 199 | 1,053 | 0,075 | 0,744 | 0,45693719 | 0,54254518 | 1,00E-08 | 0,0094664 | 0,0045939 | 0,0045939 | TRUE | 0,0184247 | No |
| orf19.2298 | 1226 | 773 | 1041 | 1351 | 1555 | 1439 | 1412 | 1060 | 1100 | 984 | 1065 | 1114 | 1330 | 1373 | 1260 | 1495 | 1215,1 | 1066 | 1364 | 1,275 | 0,351 | 5,448 | 5,08E-08 | 2,58E-07 | 0,002994 | 0,0031742 | 0,003136 | 0,003136 | TRUE | 0,0464668 | No |
| orf19.2299 | 1279 | 842 | 1018 | 1301 | 905 | 682 | 779 | 492 | 1147 | 1072 | 1042 | 1073 | 774 | 651 | 695 | 694 | 893,43 | 1084 | 704 | 0,65 | -0,622 | -9,528 | 1,60E-21 | 3,22E-20 | 0,002327 | 0,0036044 | 0,002915 | 0,002915 | TRUE | 0,0576573 | No |
| orf19.23 | 3719 | 2492 | 3349 | 4085 | 3901 | 3437 | 3770 | 2275 | 3336 | 3173 | 3426 | 3369 | 3336 | 3279 | 3364 | 3209 | 3311,5 | 3326 | 3297 | 0,991 | -0,013 | -0,328 | 0,74256236 | 0,79851703 | 0,00035 | 0,0024179 | 0,0010974 | 0,0010974 | TRUE | 0,0109798 | No |
| orf19.230 | 511 | 477 | 527 | 626 | 670 | 811 | 691 | 405 | 458 | 607 | 539 | 516 | 573 | 774 | 617 | 571 | 581,97 | 530 | 634 | 1,199 | 0,262 | 2,355 | 0,01852696 | 0,03470845 | 0,015851 | 0,004474 | 0,0101245 | 0,0101245 | TRUE | 0,2628824 | No |
| orf19.2301 | 2163 | 1448 | 1786 | 2245 | 2001 | 1714 | 1891 | 1101 | 1940 | 1843 | 1827 | 1851 | 1711 | 1635 | 1687 | 1553 | 1756,2 | 1865 | 1646 | 0,884 | -0,178 | -3,817 | 0,00013485 | 0,00040268 | 0,000517 | 0,0028061 | 0,0015046 | 0,0015046 | TRUE | 0,0159156 | No |
| orf19.2302 | 1182 | 820 | 1100 | 1209 | 945 | 861 | 1002 | 616 | 1060 | 1044 | 1125 | 997 | 808 | 822 | 894 | 869 | 952,42 | 1056 | 848 | 0,802 | -0,318 | -5,3 | 1,16E-07 | 5,64E-07 | 0,001493 | 0,0035037 | 0,0023816 | 0,0023816 | TRUE | 0,0231159 | No |
| orf19.2303 | 306 | 187 | 250 | 306 | 233 | 225 | 211 | 153 | 275 | 238 | 256 | 252 | 199 | 215 | 188 | 216 | 229,84 | 255 | 204 | 0,796 | -0,329 | -3,407 | 0,00065768 | 0,00172098 | 1,00E-08 | 0,0082956 | 0,0045328 | 0,0045328 | TRUE | 0,0318646 | No |
| orf19.2304 | 656 | 494 | 540 | 700 | 778 | 709 | 710 | 544 | 589 | 629 | 552 | 577 | 665 | 677 | 634 | 767 | 636,23 | 587 | 686 | 1,165 | 0,221 | 3,019 | 0,00253782 | 0,0058598 | 0,002865 | 0,0042613 | 0,0035537 | 0,0035537 | TRUE | 0,0677502 | No |
| orf19.2305 | 437 | 314 | 367 | 447 | 499 | 398 | 399 | 239 | 392 | 400 | 375 | 369 | 427 | 380 | 356 | 337 | 379,44 | 384 | 375 | 0,982 | -0,026 | -0,295 | 0,7678895 | 0,81915485 | 0,003247 | 0,0058054 | 0,0045625 | 0,0045625 | TRUE | 0,1025061 | No |
| orf19.2306 | 649 | 478 | 554 | 691 | 702 | 625 | 666 | 341 | 582 | 609 | 567 | 570 | 600 | 596 | 594 | 481 | 574,92 | 582 | 568 | 0,983 | -0,025 | -0,327 | 0,74371025 | 0,79917925 | 0,003241 | 0,0045046 | 0,0039361 | 0,0039361 | TRUE | 0,1083866 | No |
| orf19.2307 | 1676 | 1168 | 1462 | 1800 | 2298 | 1945 | 2264 | 1328 | 1504 | 1487 | 1496 | 1484 | 1965 | 1856 | 2020 | 1873 | 1710,6 | 1493 | 1928 | 1,294 | 0,371 | 8,286 | 1,17E-16 | 1,58E-15 | 0,000396 | 0,0028282 | 0,0013335 | 0,0013335 | TRUE | 0,0124623 | No |
| orf19.2308 | 28512 | 20799 | 24690 | 32197 | 30929 | 28640 | 30843 | 19118 | 25578 | 26479 | 25260 | 26550 | 26451 | 27327 | 27522 | 26964 | 26517 | 25967 | 27066 | 1,042 | 0,06 | 1,998 | 0,04566897 | 0,07770579 | 0,000411 | 0,0020343 | 0,000824 | 0,000824 | TRUE | 0,0040971 | No |
| orf19.2309 | 5875 | 4132 | 5303 | 6189 | 5332 | 4940 | 5377 | 3548 | 5270 | 5260 | 5425 | 5104 | 4560 | 4714 | 4798 | 5004 | 5017 | 5265 | 4769 | 0,905 | -0,144 | -3,626 | 0,0002873 | 0,00081495 | 0,000799 | 0,0022689 | 0,0013078 | 0,0013078 | TRUE | 0,0127731 | No |
| orf19.2309.2 | 28800 | 18529 | 24199 | 30270 | 30325 | 24404 | 25995 | 17536 | 25837 | 23589 | 24758 | 24961 | 25935 | 23286 | 23196 | 24733 | 24537 | 24786 | 24288 | 0,98 | -0,029 | -0,629 | 0,5294413 | 0,61226005 | 0,001898 | 0,0020387 | 0,002072 | 0,002072 | TRUE | 0,025701 | No |
| orf19.231 | 562 | 525 | 535 | 670 | 725 | 739 | 773 | 418 | 504 | 668 | 547 | 552 | 620 | 705 | 690 | 590 | 609,61 | 568 | 651 | 1,152 | 0,204 | 2,199 | 0,02790055 | 0,05021765 | 0,009058 | 0,0043609 | 0,0066209 | 0,0066209 | TRUE | 0,1559817 | No |
| orf19.2310 | 1000 | 698 | 907 | 1079 | 1102 | 1055 | 1135 | 720 | 897 | 889 | 928 | 890 | 942 | 1007 | 1013 | 1015 | 947,61 | 901 | 994 | 1,102 | 0,141 | 2,69 | 0,00715344 | 0,01489185 | 1,00E-08 | 0,0035115 | 0,001568 | 0,001568 | TRUE | 0,015711 | No |
| orf19.2310.1 | 9669 | 5778 | 7574 | 9652 | 9479 | 7163 | 7316 | 5824 | 8674 | 7356 | 7749 | 7959 | 8107 | 6835 | 6528 | 8214 | 7677,8 | 7934 | 7421 | 0,935 | -0,097 | -1,218 | 0,22326667 | 0,30218206 | 0,008796 | 0,0021686 | 0,00598 | 0,00598 | TRUE | 0,0802434 | No |
| orf19.2311 | 15816 | 10431 | 13785 | 16776 | 14631 | 12221 | 13426 | 9909 | 14189 | 13280 | 14103 | 13834 | 12513 | 11661 | 11980 | 13976 | 13192 | 13852 | 12532 | 0,904 | -0,145 | -2,563 | 0,01038684 | 0,02082221 | 0,003677 | 0,0020896 | 0,0029982 | 0,0029982 | TRUE | 0,0727727 | No |
| orf19.2312 | 79 | 48 | 76 | 73 | 69 | 46 | 65 | 47 | 71 | 61 | 78 | 60 | 59 | 44 | 58 | 66 | 62,14 | 68 | 57 | 0,835 | -0,26 | -1,399 | 0,16168184 | 0,22985908 | 1,00E-08 | 0,025341 | 0,0169819 | 0,0169819 | TRUE | 0,1999992 | No |
| orf19.2313 | 1326 | 942 | 1172 | 1476 | 1326 | 1173 | 1215 | 884 | 1190 | 1199 | 1199 | 1217 | 1134 | 1119 | 1084 | 1247 | 1173,7 | 1201 | 1146 | 0,951 | -0,073 | -1,368 | 0,17124535 | 0,24145554 | 0,000792 | 0,0032164 | 0,0018606 | 0,0018606 | TRUE | 0,0371784 | No |
| orf19.2314 | 8144 | 5526 | 6573 | 8289 | 7229 | 6271 | 6724 | 4081 | 7306 | 7035 | 6725 | 6835 | 6182 | 5984 | 6000 | 5756 | 6477,9 | 6975 | 5980 | 0,858 | -0,221 | -5,599 | 2,16E-08 | 1,15E-07 | 0,000889 | 0,0022036 | 0,0013426 | 0,0013426 | TRUE | 0,0126621 | No |
| orf19.2315 | 2516 | 1685 | 2224 | 2721 | 3416 | 2831 | 3249 | 2193 | 2257 | 2145 | 2275 | 2244 | 2921 | 2701 | 2899 | 3093 | 2567,1 | 2230 | 2904 | 1,3 | 0,378 | 7,788 | 6,79E-15 | 7,75E-14 | 0,001338 | 0,002545 | 0,0018667 | 0,0018667 | TRUE | 0,0264817 | No |
| orf19.2317 | 94 | 50 | 76 | 105 | 101 | 77 | 67 | 54 | 84 | 64 | 78 | 87 | 86 | 73 | 60 | 76 | 76,02 | 78 | 74 | 0,939 | -0,091 | -0,53 | 0,59593708 | 0,67281569 | 0,007582 | 0,0210771 | 0,015483 | 0,015483 | TRUE | 0,1707135 | No |
| orf19.2318.1 | 1614 | 1175 | 1374 | 1780 | 1527 | 1269 | 1357 | 922 | 1448 | 1496 | 1406 | 1468 | 1306 | 1211 | 1211 | 1300 | 1355,7 | 1454 | 1257 | 0,864 | -0,211 | -4,305 | 1,67E-05 | 5,89E-05 | 0,00044 | 0,0030504 | 0,0015694 | 0,0015694 | TRUE | 0,0095796 | No |
| orf19.2319 | 6666 | 4422 | 5880 | 6914 | 7428 | 6436 | 7092 | 3844 | 5980 | 5630 | 6016 | 5701 | 6353 | 6141 | 6328 | 5422 | 5946,3 | 5832 | 6061 | 1,04 | 0,057 | 1,032 | 0,30195629 | 0,38788825 | 0,002927 | 0,0022237 | 0,0027173 | 0,0027173 | TRUE | 0,0600687 | No |
| orf19.232 | 277 | 211 | 249 | 316 | 384 | 395 | 418 | 277 | 248 | 269 | 255 | 261 | 328 | 377 | 373 | 391 | 312,68 | 258 | 367 | 1,42 | 0,506 | 5,836 | 5,34E-09 | 3,09E-08 | 1,00E-08 | 0,0066223 | 0,0039078 | 0,0039078 | TRUE | 0,0615418 | No |
| orf19.2320 | 10585 | 7046 | 8869 | 11166 | 10101 | 8433 | 9526 | 4994 | 9496 | 8970 | 9074 | 9208 | 8639 | 8047 | 8500 | 7044 | 8622,1 | 9187 | 8058 | 0,877 | -0,189 | -3,098 | 0,00195126 | 0,00461725 | 0,004162 | 0,0021479 | 0,0034424 | 0,0034424 | TRUE | 0,0863445 | No |
| orf19.2322.3 | 1173 | 816 | 1022 | 1248 | 1122 | 930 | 1025 | 613 | 1052 | 1039 | 1046 | 1029 | 960 | 887 | 915 | 865 | 974,01 | 1042 | 907 | 0,873 | -0,196 | -3,752 | 0,00017555 | 0,00051605 | 1,00E-08 | 0,00347 | 0,0015997 | 0,0015997 | TRUE | 0,018837 | No |
| orf19.2324 | 2005 | 1398 | 1814 | 2226 | 1697 | 1393 | 1507 | 1115 | 1799 | 1780 | 1856 | 1836 | 1451 | 1329 | 1345 | 1573 | 1621 | 1818 | 1424 | 0,782 | -0,355 | -6,192 | 5,95E-10 | 3,91E-09 | 0,002476 | 0,0028751 | 0,0025347 | 0,0025347 | TRUE | 0,0540827 | No |
| orf19.2325 | 832 | 568 | 726 | 911 | 952 | 795 | 927 | 607 | 746 | 723 | 743 | 751 | 814 | 759 | 827 | 856 | 777,45 | 741 | 814 | 1,096 | 0,132 | 2,28 | 0,02260784 | 0,04171619 | 1,00E-08 | 0,0038468 | 0,0019129 | 0,0019129 | TRUE | 0,0246354 | No |
| orf19.2326 | 2602 | 1614 | 2168 | 2733 | 2812 | 2444 | 2731 | 1548 | 2334 | 2055 | 2218 | 2254 | 2405 | 2332 | 2437 | 2183 | 2277,3 | 2215 | 2339 | 1,056 | 0,079 | 1,486 | 0,1372375 | 0,19993281 | 0,001984 | 0,002617 | 0,0022683 | 0,0022683 | TRUE | 0,0274844 | No |
| orf19.2328 | 1924 | 1398 | 1661 | 2141 | 2428 | 2099 | 2363 | 1393 | 1726 | 1780 | 1699 | 1765 | 2077 | 2003 | 2109 | 1965 | 1890,4 | 1742 | 2038 | 1,171 | 0,228 | 5,363 | 8,17E-08 | 4,05E-07 | 0,000194 | 0,0027475 | 0,0012075 | 0,0012075 | TRUE | 0,0064061 | No |
| orf19.2329.1 | 17099 | 11789 | 13137 | 17264 | 15691 | 13976 | 13706 | 8625 | 15340 | 15009 | 13440 | 14236 | 13419 | 13335 | 12230 | 12165 | 13647 | 14506 | 12787 | 0,882 | -0,182 | -3,373 | 0,00074341 | 0,00192434 | 0,003034 | 0,0020859 | 0,0027119 | 0,0027119 | TRUE | 0,0298868 | No |
| orf19.2330 | 6296 | 4151 | 5388 | 6571 | 5895 | 5040 | 5533 | 3286 | 5648 | 5285 | 5512 | 5419 | 5042 | 4809 | 4937 | 4635 | 5160,8 | 5466 | 4856 | 0,889 | -0,17 | -4,324 | 1,53E-05 | 5,44E-05 | 0,000786 | 0,0022608 | 0,0012898 | 0,0012898 | TRUE | 0,0108693 | No |
| orf19.2331 | 1280 | 889 | 1120 | 1395 | 1727 | 1573 | 1774 | 1026 | 1148 | 1132 | 1146 | 1150 | 1477 | 1501 | 1583 | 1447 | 1323 | 1144 | 1502 | 1,314 | 0,394 | 8,333 | 7,91E-17 | 1,10E-15 | 1,00E-08 | 0,0030768 | 0,0013786 | 0,0013786 | TRUE | 0,0161366 | No |
| orf19.233.1 | 1995 | 1438 | 1719 | 2079 | 2106 | 1818 | 1766 | 1177 | 1790 | 1831 | 1759 | 1714 | 1801 | 1735 | 1576 | 1660 | 1733,2 | 1774 | 1693 | 0,956 | -0,065 | -1,244 | 0,21332755 | 0,29076041 | 0,00154 | 0,0028171 | 0,0020484 | 0,0020484 | TRUE | 0,0279281 | No |
| orf19.2332 | 0 | 1 | 0 | 1 | 1 | 2 | 0 | 1 | 0 | 1 | 0 | 1 | 1 | 2 | 0 | 1 | 0,78 | 0 | 1 | 2,07 | 1,05 | 0,642 | 0,52090404 | NA | 1,00E-08 | 1,8537282 | 1,1394303 | 1,1394303 | TRUE | 0,2655038 | No |
| orf19.2333 | 8195 | 5258 | 7130 | 8684 | 9513 | 8095 | 9249 | 5565 | 7352 | 6694 | 7295 | 7161 | 8136 | 7724 | 8253 | 7849 | 7557,9 | 7126 | 7990 | 1,121 | 0,165 | 3,973 | 7,09E-05 | 0,00022282 | 0,001173 | 0,0021716 | 0,0015235 | 0,0015235 | TRUE | 0,0197787 | No |
| orf19.2334 | 768 | 571 | 651 | 891 | 943 | 959 | 883 | 526 | 689 | 727 | 666 | 735 | 806 | 915 | 788 | 742 | 758,5 | 704 | 813 | 1,157 | 0,21 | 2,883 | 0,00393392 | 0,00869253 | 0,003667 | 0,0038934 | 0,003773 | 0,003773 | TRUE | 0,084342 | No |
| orf19.2335 | 1089 | 792 | 969 | 1200 | 1251 | 1210 | 1298 | 769 | 977 | 1008 | 991 | 990 | 1070 | 1155 | 1158 | 1085 | 1054,2 | 992 | 1117 | 1,128 | 0,174 | 3,397 | 0,00068113 | 0,00177613 | 1,00E-08 | 0,0033566 | 0,0015612 | 0,0015612 | TRUE | 0,0110541 | No |
| orf19.2336 | 250 | 168 | 202 | 263 | 229 | 195 | 190 | 136 | 224 | 214 | 207 | 217 | 196 | 186 | 170 | 192 | 200,62 | 216 | 186 | 0,86 | -0,217 | -2,178 | 0,02939819 | 0,05259807 | 1,00E-08 | 0,0092156 | 0,0045542 | 0,0045542 | TRUE | 0,0408269 | No |
| orf19.2337 | 14287 | 9555 | 12822 | 15334 | 14599 | 12810 | 13661 | 7770 | 12817 | 12165 | 13118 | 12645 | 12486 | 12223 | 12190 | 10959 | 12325 | 12686 | 11964 | 0,943 | -0,084 | -1,774 | 0,07606679 | 0,12112623 | 0,002017 | 0,0020973 | 0,002074 | 0,002074 | TRUE | 0,038525 | No |
| orf19.234 | 572 | 476 | 542 | 662 | 727 | 653 | 719 | 469 | 513 | 606 | 555 | 546 | 622 | 623 | 642 | 661 | 595,93 | 555 | 637 | 1,15 | 0,202 | 3,008 | 0,00263358 | 0,00606005 | 0,000583 | 0,0044155 | 0,0026253 | 0,0026253 | TRUE | 0,0404991 | No |

| | | | | | | | | | | | | | | | | | | | | | | | | | | | | | | | |
|---|---|---|---|---|---|---|---|---|---|---|---|---|---|---|---|---|---|---|---|---|---|---|---|---|---|---|---|---|---|---|---|
| orf19.2340 | 21446 | 14038 | 18606 | 23039 | 21992 | 17998 | 19671 | 12859 | 19239 | 17872 | 19036 | 18998 | 18808 | 17173 | 17553 | 18136 | 18352 | 18786 | 17918 | 0,954 | -0,069 | -1,689 | 0,09116769 | 0,14153372 | 0,001263 | 0,0020586 | 0,0015259 | 0,0015259 | TRUE | 0,0138549 | No |
| orf19.2341 | 4283 | 3133 | 3718 | 4766 | 5624 | 4982 | 5407 | 3320 | 3842 | 3989 | 3804 | 3930 | 4810 | 4754 | 4825 | 4683 | 4329,5 | 3891 | 4768 | 1,226 | 0,294 | 9,094 | 9,58E-20 | 1,68E-18 | 1,18E-05 | 0,0023148 | 0,0007714 | 0,0007714 | TRUE | 0,0031817 | No |
| orf19.2342 | 1829 | 1251 | 1535 | 2023 | 1949 | 1623 | 1750 | 992 | 1641 | 1593 | 1570 | 1668 | 1667 | 1549 | 1562 | 1399 | 1581 | 1618 | 1544 | 0,956 | -0,065 | -1,14 | 0,25433434 | 0,33694392 | 0,00208 | 0,0028977 | 0,0024597 | 0,0024597 | TRUE | 0,0442034 | No |
| orf19.2343 | 216 | 172 | 207 | 263 | 270 | 267 | 253 | 195 | 194 | 219 | 212 | 217 | 231 | 255 | 226 | 275 | 228,48 | 210 | 247 | 1,167 | 0,222 | 2,217 | 0,02659921 | 0,04832418 | 1,00E-08 | 0,0083332 | 0,0052523 | 0,0052523 | TRUE | 0,0561506 | No |
| orf19.2343.1 | 429 | 298 | 365 | 490 | 369 | 312 | 345 | 241 | 385 | 379 | 373 | 404 | 316 | 298 | 308 | 340 | 350,35 | 385 | 316 | 0,813 | -0,298 | -3,801 | 0,00014407 | 0,00042806 | 1,00E-08 | 0,0061231 | 0,0030232 | 0,0030232 | TRUE | 0,0252487 | No |
| orf19.2344 | 1677 | 1073 | 1487 | 1791 | 2352 | 1820 | 1963 | 1502 | 1504 | 1366 | 1521 | 1477 | 2012 | 1737 | 1752 | 2118 | 1685,9 | 1467 | 1905 | 1,295 | 0,373 | 5,205 | 1,94E-07 | 9,15E-07 | 0,005328 | 0,0028406 | 0,0043379 | 0,0043379 | TRUE | 0,0659963 | No |
| orf19.2346 | 1601 | 1082 | 1364 | 1560 | 989 | 891 | 962 | 692 | 1436 | 1378 | 1396 | 1286 | 846 | 850 | 858 | 976 | 1128,3 | 1374 | 882 | 0,64 | -0,643 | -10,447 | 1,51E-25 | 4,01E-24 | 0,002327 | 0,0032662 | 0,0027023 | 0,0027023 | TRUE | 0,0539688 | No |
| orf19.2347 | 3241 | 2261 | 2924 | 3479 | 2887 | 2579 | 2664 | 1745 | 2908 | 2878 | 2992 | 2869 | 2469 | 2461 | 2377 | 2461 | 2676,8 | 2912 | 2442 | 0,839 | -0,254 | -6,986 | 2,84E-12 | 2,46E-11 | 1,00E-08 | 0,0025218 | 0,0008926 | 0,0008926 | TRUE | 0,0041145 | No |
| orf19.2350 | 506 | 339 | 442 | 558 | 528 | 413 | 437 | 277 | 454 | 432 | 452 | 460 | 452 | 394 | 390 | 391 | 428,02 | 450 | 407 | 0,906 | -0,143 | -1,882 | 0,05981746 | 0,09824585 | 1,00E-08 | 0,0053712 | 0,0032137 | 0,0032137 | TRUE | 0,0687826 | No |
| orf19.2351 | 848 | 541 | 711 | 852 | 1290 | 1159 | 1164 | 827 | 761 | 689 | 727 | 703 | 1103 | 1106 | 1039 | 1166 | 911,71 | 720 | 1104 | 1,528 | 0,612 | 10,436 | 1,70E-25 | 4,49E-24 | 0,00092 | 0,0035718 | 0,0021519 | 0,0021519 | TRUE | 0,0187333 | No |
| orf19.2352 | 453 | 293 | 438 | 503 | 437 | 357 | 383 | 208 | 406 | 373 | 448 | 415 | 374 | 341 | 342 | 293 | 373,98 | 410 | 338 | 0,825 | -0,277 | -3,089 | 0,00200898 | 0,00474261 | 0,004039 | 0,0058613 | 0,005038 | 0,005038 | TRUE | 0,0742834 | No |
| orf19.2356 | 24541 | 17259 | 21982 | 27864 | 28074 | 25541 | 28171 | 17897 | 22016 | 21973 | 22490 | 22977 | 24010 | 24370 | 25138 | 25242 | 23527 | 22364 | 24690 | 1,104 | 0,142 | 4,616 | 3,91E-06 | 1,51E-05 | 0,000458 | 0,0020412 | 0,0008717 | 0,0008717 | TRUE | 0,0042289 | No |
| orf19.236 | 12020 | 7994 | 9188 | 11854 | 10652 | 9144 | 9387 | 5525 | 10783 | 10177 | 9400 | 9775 | 9110 | 8725 | 8376 | 7793 | 9267,4 | 10034 | 8501 | 0,848 | -0,239 | -4,098 | 4,16E-05 | 0,00013637 | 0,003759 | 0,0021362 | 0,0031473 | 0,0031473 | TRUE | 0,0378809 | No |
| orf19.2360 | 10077 | 7361 | 9717 | 11219 | 8826 | 8153 | 8552 | 5580 | 9040 | 9371 | 9941 | 9251 | 7548 | 7779 | 7631 | 7870 | 8554,2 | 9401 | 7707 | 0,82 | -0,287 | -7,503 | 6,24E-14 | 6,47E-13 | 0,000859 | 0,0021492 | 0,0012843 | 0,0012843 | TRUE | 0,0183519 | No |
| orf19.2361 | 2929 | 2108 | 2597 | 3098 | 3823 | 3544 | 3918 | 2484 | 2628 | 2684 | 2657 | 2555 | 3270 | 3382 | 3496 | 3503 | 3021,7 | 2631 | 3413 | 1,297 | 0,375 | 9,507 | 1,97E-21 | 3,92E-20 | 0,000482 | 0,0024599 | 0,0011591 | 0,0011591 | TRUE | 0,0099413 | No |
| orf19.2362 | 3403 | 2465 | 2892 | 3646 | 3635 | 3188 | 3316 | 2188 | 3053 | 3138 | 2959 | 3007 | 3109 | 3042 | 2959 | 3086 | 3044 | 3039 | 3049 | 1,003 | 0,005 | 0,136 | 0,89218506 | 0,91847975 | 0,000203 | 0,0024564 | 0,0009922 | 0,0009922 | TRUE | 0,0054363 | No |
| orf19.2363 | 4116 | 2749 | 3408 | 4432 | 3989 | 3693 | 4079 | 2430 | 3692 | 3500 | 3487 | 3655 | 3412 | 3524 | 3640 | 3427 | 3542 | 3584 | 3501 | 0,977 | -0,034 | -0,86 | 0,38955558 | 0,47845275 | 0,000508 | 0,0023894 | 0,0012183 | 0,0012183 | TRUE | 0,0088961 | No |
| orf19.2364 | 1617 | 1028 | 1448 | 1706 | 1588 | 1419 | 1502 | 1014 | 1451 | 1309 | 1481 | 1407 | 1358 | 1354 | 1340 | 1430 | 1391,3 | 1412 | 1370 | 0,968 | -0,047 | -0,906 | 0,36501518 | 0,45385166 | 0,000897 | 0,003023 | 0,0018475 | 0,0018475 | TRUE | 0,0265942 | No |
| orf19.2365 | 2374 | 1709 | 2139 | 2542 | 2473 | 2329 | 2400 | 1404 | 2130 | 2176 | 2188 | 2096 | 2115 | 2222 | 2142 | 1980 | 2131,1 | 2148 | 2115 | 0,987 | -0,019 | -0,42 | 0,67454747 | 0,73994545 | 0,000702 | 0,0026607 | 0,0014945 | 0,0014945 | TRUE | 0,0200495 | No |
| orf19.2366 | 4 | 1 | 2 | 1 | 1 | 3 | 2 | 2 | 4 | 1 | 2 | 1 | 1 | 3 | 2 | 3 | 2,01 | 2 | 2 | 1,036 | 0,051 | 0,053 | 0,95792953 | NA | 1,00E-08 | 0,7253183 | 0,3946768 | 0,3946768 | TRUE | 0,3436761 | No |
| orf19.2367 | 1155 | 1059 | 1107 | 1318 | 1423 | 1439 | 1467 | 672 | 1036 | 1348 | 1133 | 1087 | 1217 | 1373 | 1309 | 948 | 1181,3 | 1151 | 1212 | 1,056 | 0,079 | 0,714 | 0,47526291 | 0,56086812 | 0,018456 | 0,0032084 | 0,0107969 | 0,0107969 | TRUE | 0,2481598 | No |
| orf19.2368 | 2101 | 1558 | 1788 | 2174 | 2116 | 2133 | 2214 | 1125 | 1885 | 1984 | 1829 | 1793 | 1810 | 2035 | 1976 | 1587 | 1862,2 | 1873 | 1852 | 0,991 | -0,013 | -0,174 | 0,861809 | 0,89542891 | 0,006299 | 0,0027591 | 0,0047078 | 0,0047078 | TRUE | 0,1068819 | No |
| orf19.2369 | 621 | 508 | 595 | 783 | 621 | 631 | 642 | 410 | 557 | 647 | 609 | 646 | 531 | 602 | 573 | 578 | 592,82 | 615 | 571 | 0,929 | -0,106 | -1,462 | 0,14381789 | 0,20815655 | 0,002346 | 0,0044283 | 0,0033377 | 0,0033377 | TRUE | 0,0484592 | No |
| orf19.2369.1 | 1459 | 1077 | 1163 | 1470 | 1469 | 1332 | 1377 | 702 | 1309 | 1371 | 1190 | 1212 | 1256 | 1271 | 1229 | 990 | 1228,5 | 1270 | 1186 | 0,937 | -0,094 | -1,155 | 0,24801921 | 0,32996001 | 0,007333 | 0,0031612 | 0,005499 | 0,005499 | TRUE | 0,1398463 | No |
| orf19.2370 | 271 | 244 | 276 | 316 | 342 | 346 | 367 | 204 | 243 | 311 | 282 | 261 | 292 | 330 | 327 | 288 | 291,82 | 274 | 309 | 1,137 | 0,186 | 1,899 | 0,05759797 | 0,09499001 | 0,004333 | 0,0069542 | 0,0057492 | 0,0057492 | TRUE | 0,0813277 | No |
| orf19.2371 | 1706 | 1218 | 1509 | 1871 | 2275 | 1898 | 1992 | 1259 | 1530 | 1551 | 1544 | 1543 | 1946 | 1811 | 1778 | 1776 | 1684,7 | 1542 | 1828 | 1,187 | 0,247 | 5,377 | 7,59E-08 | 3,79E-07 | 0,000552 | 0,0028412 | 0,0014278 | 0,0014278 | TRUE | 0,0238397 | No |
| orf19.2372 | 2491 | 1891 | 2293 | 2729 | 2856 | 2500 | 2645 | 1648 | 2235 | 2407 | 2346 | 2250 | 2443 | 2385 | 2360 | 2324 | 2343,9 | 2310 | 2378 | 1,032 | 0,045 | 1,09 | 0,27578251 | 0,36015751 | 0,000379 | 0,0025989 | 0,0012161 | 0,0012161 | TRUE | 0,009322 | No |
| orf19.2376 | 94 | 55 | 58 | 91 | 65 | 67 | 69 | 45 | 84 | 70 | 59 | 75 | 56 | 64 | 62 | 63 | 66,66 | 72 | 61 | 0,84 | -0,251 | -1,439 | 0,15016396 | 0,21591023 | 1,00E-08 | 0,0237569 | 0,0141153 | 0,0141153 | TRUE | 0,1319846 | No |
| orf19.2378 | 541 | 351 | 446 | 564 | 468 | 417 | 430 | 305 | 485 | 447 | 456 | 465 | 400 | 398 | 384 | 430 | 433,2 | 463 | 403 | 0,865 | -0,209 | -2,931 | 0,00337547 | 0,00756711 | 1,00E-08 | 0,0053306 | 0,0025746 | 0,0025746 | TRUE | 0,0195212 | No |
| orf19.2379 | 4197 | 2682 | 3705 | 4651 | 4079 | 3420 | 3826 | 2559 | 3765 | 3414 | 3791 | 3835 | 3488 | 3263 | 3414 | 3609 | 3572,6 | 3701 | 3444 | 0,929 | -0,106 | -2,128 | 0,03333702 | 0,05885655 | 0,001875 | 0,0023859 | 0,0021059 | 0,0021059 | TRUE | 0,0320732 | No |
| orf19.238 | 4846 | 3380 | 4192 | 5115 | 7210 | 6274 | 6632 | 3910 | 4347 | 4303 | 4289 | 4218 | 6166 | 5986 | 5918 | 5515 | 5092,8 | 4289 | 5896 | 1,376 | 0,46 | 11,248 | 2,37E-29 | 8,18E-28 | 0,000899 | 0,0022646 | 0,0014073 | 0,0014073 | TRUE | 0,0221957 | No |
| orf19.2381 | 491 | 381 | 452 | 532 | 802 | 635 | 690 | 500 | 440 | 485 | 462 | 439 | 686 | 606 | 616 | 705 | 554,92 | 456 | 653 | 1,431 | 0,517 | 6,962 | 3,36E-12 | 2,89E-11 | 0,002303 | 0,0045956 | 0,0034227 | 0,0034227 | TRUE | 0,0296003 | No |
| orf19.2382 | 482 | 349 | 436 | 522 | 453 | 438 | 455 | 278 | 432 | 444 | 446 | 430 | 387 | 418 | 406 | 392 | 419,59 | 438 | 401 | 0,916 | -0,126 | -1,801 | 0,07162527 | 0,11493714 | 1,00E-08 | 0,0054394 | 0,0023358 | 0,0023358 | TRUE | 0,0095852 | No |
| orf19.2383 | 187 | 110 | 169 | 205 | 133 | 130 | 127 | 112 | 168 | 140 | 173 | 169 | 114 | 124 | 113 | 158 | 144,85 | 162 | 127 | 0,77 | -0,376 | -2,824 | 0,00473756 | 0,01028102 | 0,006985 | 0,0120014 | 0,0100019 | 0,0100019 | TRUE | 0,2207466 | No |
| orf19.2384 | 2639 | 1733 | 2248 | 2787 | 2831 | 2343 | 2477 | 1600 | 2367 | 2206 | 2300 | 2298 | 2421 | 2236 | 2210 | 2257 | 2287 | 2293 | 2281 | 0,995 | -0,008 | -0,176 | 0,86067668 | 0,89462725 | 0,00089 | 0,0026143 | 0,0015332 | 0,0015332 | TRUE | 0,0217475 | No |
| orf19.2385 | 2822 | 1838 | 2342 | 2829 | 2872 | 2557 | 2624 | 1650 | 2532 | 2340 | 2396 | 2333 | 2456 | 2440 | 2341 | 2327 | 2395,7 | 2400 | 2391 | 0,997 | -0,005 | -0,113 | 0,91023837 | 0,93354594 | 0,00073 | 0,0025855 | 0,0014137 | 0,0014137 | TRUE | 0,0169113 | No |
| orf19.2386 | 5140 | 3691 | 4389 | 5723 | 5500 | 4865 | 5232 | 3364 | 4611 | 4699 | 4490 | 4719 | 4704 | 4642 | 4669 | 4745 | 4659,9 | 4630 | 4690 | 1,013 | 0,018 | 0,57 | 0,56894 | 0,64812126 | 8,21E-05 | 0,0022911 | 0,0007579 | 0,0007579 | TRUE | 0,0049737 | No |
| orf19.2387 | 4569 | 3062 | 4132 | 4898 | 4580 | 4030 | 4499 | 2747 | 4099 | 3898 | 4227 | 4039 | 3917 | 3845 | 4015 | 3874 | 3989,4 | 4066 | 3913 | 0,962 | -0,056 | -1,487 | 0,13713433 | 0,19983097 | 0,000409 | 0,0023434 | 0,0010902 | 0,0010902 | TRUE | 0,008817 | No |
| orf19.2389 | 1042 | 639 | 895 | 1050 | 1166 | 1001 | 1010 | 640 | 935 | 814 | 916 | 866 | 997 | 955 | 901 | 903 | 910,76 | 883 | 939 | 1,064 | 0,089 | 1,426 | 0,15398727 | 0,22053767 | 0,001909 | 0,0035735 | 0,0026783 | 0,0026783 | TRUE | 0,0298591 | No |
| orf19.239 | 3791 | 2657 | 3410 | 4059 | 4165 | 3995 | 4302 | 2513 | 3401 | 3383 | 3489 | 3347 | 3562 | 3812 | 3839 | 3544 | 3547,1 | 3405 | 3689 | 1,084 | 0,117 | 2,837 | 0,00455561 | 0,0099167 | 0,000693 | 0,0023888 | 0,0013452 | 0,0013452 | TRUE | 0,0090858 | No |
| orf19.2391 | 1008 | 630 | 774 | 1001 | 899 | 727 | 828 | 601 | 904 | 802 | 792 | 825 | 769 | 694 | 739 | 848 | 796,59 | 831 | 762 | 0,914 | -0,13 | -1,791 | 0,07335222 | 0,11736356 | 0,003809 | 0,0038019 | 0,0038287 | 0,0038287 | TRUE | 0,0605732 | No |
| orf19.2392 | 0 | 0 | 0 | 0 | 0 | 0 | 0 | 0 | 0 | 0 | 0 | 0 | 0 | 0 | 0 | 0 | 0 | 0 | 0 | NA | NA | NA | NA | NA | NA | NA | NA | NA | NA | NA | NA |
| orf19.2393 | 7006 | 4578 | 5929 | 7428 | 6773 | 5616 | 6314 | 3769 | 6285 | 5828 | 6066 | 6125 | 5792 | 5359 | 5634 | 5316 | 5800,7 | 6076 | 5525 | 0,91 | -0,137 | -3,274 | 0,00106001 | 0,00265798 | 0,001101 | 0,0022298 | 0,0015024 | 0,0015024 | TRUE | 0,0129894 | No |
| orf19.2394 | 46 | 41 | 39 | 43 | 28 | 31 | 35 | 32 | 41 | 52 | 40 | 35 | 24 | 30 | 31 | 45 | 37,34 | 42 | 32 | 0,759 | -0,398 | -1,611 | 0,10721789 | 0,16185052 | 0,014921 | 0,0408582 | 0,030879 | 0,030879 | TRUE | 0,3498964 | No |
| orf19.2395 | 21 | 12 | 24 | 10 | 14 | 13 | 17 | 12 | 19 | 15 | 25 | 8 | 12 | 12 | 15 | 17 | 15,42 | 17 | 14 | 0,841 | -0,249 | -0,642 | 0,5210726 | 0,60413048 | 0,055537 | 0,0961014 | 0,078729 | 0,078729 | TRUE | 0,295118 | No |
| orf19.2396 | 503 | 344 | 448 | 541 | 643 | 568 | 628 | 396 | 451 | 438 | 458 | 446 | 550 | 542 | 560 | 559 | 500,56 | 448 | 553 | 1,231 | 0,3 | 4,687 | 2,78E-06 | 1,10E-05 | 1,00E-08 | 0,0048797 | 0,001926 | 0,001926 | TRUE | 0,0023684 | No |
| orf19.2397 | 215 | 158 | 195 | 219 | 229 | 208 | 176 | 138 | 193 | 201 | 200 | 181 | 196 | 198 | 157 | 195 | 190,02 | 194 | 186 | 0,964 | -0,053 | -0,483 | 0,62911376 | 0,70046617 | 1,00E-08 | 0,0096194 | 0,0064199 | 0,0064199 | TRUE | 0,1318303 | No |
| orf19.2397.3 | 6105 | 4648 | 5277 | 6513 | 6566 | 6653 | 6954 | 3814 | 5477 | 5917 | 5399 | 5371 | 5615 | 6348 | 6205 | 5379 | 5714 | 5541 | 5887 | 1,063 | 0,089 | 1,48 | 0,13888233 | 0,20188847 | 0,003929 | 0,0022336 | 0,0032612 | 0,0032612 | TRUE | 0,0402054 | No |
| orf19.2398 | 13587 | 9647 | 12043 | 15300 | 19972 | 17786 | 19466 | 12883 | 12189 | 12282 | 12321 | 12617 | 17081 | 16971 | 17370 | 18170 | 14875 | 12352 | 17398 | 1,408 | 0,494 | 15,235 | 2,07E-52 | 2,44E-50 | 0,000504 | 0,0020771 | 0,0009386 | 0,0009386 | TRUE | 0,0106915 | No |
| orf19.2399 | 441 | 363 | 391 | 455 | 463 | 489 | 520 | 253 | 396 | 462 | 400 | 375 | 396 | 467 | 464 | 357 | 414,55 | 408 | 421 | 1,038 | 0,054 | 0,536 | 0,59196069 | 0,66952181 | 0,009371 | 0,0054814 | 0,0073777 | 0,0073777 | TRUE | 0,1068116 | No |
| orf19.24 | 6532 | 4465 | 5634 | 7017 | 4698 | 4031 | 4267 | 2773 | 5860 | 5684 | 5764 | 5786 | 4018 | 3846 | 3808 | 3911 | 4834,7 | 5774 | 3896 | 0,675 | -0,568 | -17,466 | 2,58E-68 | 5,55E-66 | 0,000136 | 0,0022798 | 0,0007991 | 0,0007991 | TRUE | 0,005809 | No |
| orf19.240 | 1459 | 1092 | 1300 | 1623 | 1857 | 1820 | 1766 | 908 | 1309 | 1390 | 1330 | 1338 | 1588 | 1737 | 1576 | 1281 | 1443,6 | 1342 | 1546 | 1,155 | 0,207 | 2,608 | 0,00910715 | 0,01852886 | 0,007039 | 0,0029851 | 0,0053662 | 0,0053662 | TRUE | 0,1520776 | No |
| orf19.2400 | 4440 | 3169 | 3933 | 4824 | 4499 | 4240 | 4511 | 2486 | 3983 | 4034 | 4024 | 3978 | 3848 | 4046 | 4025 | 3506 | 3930,5 | 4005 | 3856 | 0,964 | -0,053 | -1,086 | 0,27752328 | 0,36152642 | 0,001782 | 0,0023489 | 0,0020013 | 0,0020013 | TRUE | 0,0432446 | No |
| orf19.2401 | 336 | 280 | 334 | 364 | 306 | 330 | 336 | 205 | 301 | 356 | 342 | 300 | 262 | 315 | 300 | 289 | 308,16 | 325 | 292 | 0,9 | -0,152 | -1,602 | 0,10911977 | 0,16414411 | 0,003921 | 0,0066903 | 0,0053614 | 0,0053614 | TRUE | 0,0590767 | No |
| orf19.2402 | 3350 | 2275 | 2904 | 3591 | 3380 | 2821 | 2988 | 2034 | 3005 | 2896 | 2971 | 2961 | 2891 | 2692 | 2666 | 2869 | 2868,9 | 2958 | 2780 | 0,938 | -0,092 | -2,204 | 0,02751466 | 0,04969792 | 0,000535 | 0,0024855 | 0,0013066 | 0,0013066 | TRUE | 0,0097124 | No |
| orf19.2404 | 3590 | 2544 | 3129 | 3817 | 3935 | 3729 | 4122 | 2436 | 3221 | 3239 | 3201 | 3148 | 3365 | 3558 | 3678 | 3436 | 3355,7 | 3202 | 3509 | 1,097 | 0,133 | 3,349 | 0,00081223 | 0,00208445 | 0,000615 | 0,0024121 | 0,001218 | 0,001218 | TRUE | 0,0130958 | No |
| orf19.2406 | 2062 | 1511 | 1939 | 2335 | 2487 | 2323 | 2451 | 1420 | 1850 | 1924 | 1984 | 1925 | 2127 | 2217 | 2187 | 2003 | 2027 | 1921 | 2134 | 1,113 | 0,154 | 3,327 | 0,00087798 | 0,00223418 | 0,000734 | 0,0026957 | 0,0015655 | 0,0015655 | TRUE | 0,0186472 | No |
| orf19.2407 | 4012 | 2701 | 3651 | 4187 | 4061 | 3657 | 3741 | 2374 | 3599 | 3439 | 3735 | 3453 | 3473 | 3489 | 3338 | 3348 | 3484,4 | 3556 | 3412 | 0,96 | -0,059 | -1,451 | 0,14686091 | 0,21174475 | 0,000663 | 0,0023962 | 0,0013199 | 0,0013199 | TRUE | 0,01377 | No |
| orf19.2408 | 1194 | 853 | 1024 | 1282 | 1691 | 1617 | 1604 | 1030 | 1071 | 1086 | 1048 | 1057 | 1446 | 1543 | 1431 | 1453 | 1266,9 | 1066 | 1468 | 1,379 | 0,464 | 9,73 | 2,25E-22 | 4,84E-21 | 1,00E-08 | 0,0031254 | 0,0013698 | 0,0013698 | TRUE | 0,0145323 | No |

| | | | | | | | | | | | | | | | | | | | | | | | | | | | | | | | |
|---|---|---|---|---|---|---|---|---|---|---|---|---|---|---|---|---|---|---|---|---|---|---|---|---|---|---|---|---|---|---|---|
| orf19.2410 | 2320 | 1716 | 1951 | 2463 | 2360 | 2205 | 2397 | 1452 | 2081 | 2185 | 1996 | 2031 | 2018 | 2104 | 2139 | 2048 | 2075,3 | 2073 | 2077 | 1,003 | 0,005 | 0,107 | 0,91467081 | 0,93745176 | 0,000475 | 0,002679 | 0,0014006 | 0,0014006 | TRUE | 0,0147033 | No |
| orf19.2411 | 1505 | 1087 | 1297 | 1599 | 1476 | 1352 | 1429 | 866 | 1350 | 1384 | 1327 | 1319 | 1262 | 1290 | 1275 | 1221 | 1303,6 | 1345 | 1262 | 0,941 | -0,088 | -1,951 | 0,05100485 | 0,08564481 | 1,00E-08 | 0,0030932 | 0,0011909 | 0,0011909 | TRUE | 0,0048276 | No |
| orf19.2412 | 354 | 257 | 285 | 389 | 383 | 345 | 369 | 189 | 318 | 327 | 292 | 321 | 328 | 329 | 329 | 267 | 313,71 | 314 | 313 | 1,004 | 0,005 | 0,059 | 0,95323769 | 0,96659106 | 0,001909 | 0,006607 | 0,004569 | 0,004569 | TRUE | 0,0936685 | No |
| orf19.2414 | 6549 | 4373 | 5382 | 6649 | 9629 | 8265 | 8602 | 5697 | 5875 | 5567 | 5506 | 5483 | 8235 | 7886 | 7676 | 8035 | 6783 | 5608 | 7958 | 1,419 | 0,505 | 13,178 | 1,17E-39 | 7,80E-38 | 0,000763 | 0,0021935 | 0,0012559 | 0,0012559 | TRUE | 0,0127839 | No |
| orf19.2415 | 728 | 487 | 655 | 727 | 602 | 492 | 570 | 331 | 653 | 620 | 670 | 599 | 515 | 469 | 509 | 467 | 562,81 | 636 | 490 | 0,773 | -0,371 | -5,397 | 6,79E-08 | 3,41E-07 | 0,00095 | 0,0045589 | 0,0027374 | 0,0027374 | TRUE | 0,0192712 | No |
| orf19.2416.1 | 291 | 190 | 234 | 275 | 401 | 308 | 322 | 243 | 261 | 242 | 239 | 227 | 343 | 294 | 287 | 343 | 279,5 | 242 | 317 | 1,304 | 0,383 | 3,955 | 7,67E-05 | 0,00023851 | 0,003406 | 0,0071734 | 0,0053613 | 0,0053613 | TRUE | 0,045569 | No |
| orf19.2417 | 985 | 689 | 880 | 1038 | 811 | 720 | 810 | 504 | 884 | 877 | 900 | 856 | 694 | 687 | 723 | 711 | 791,42 | 879 | 704 | 0,8 | -0,322 | -5,941 | 2,82E-09 | 1,68E-08 | 1,00E-08 | 0,0038138 | 0,001535 | 0,001535 | TRUE | 0,0045788 | No |
| orf19.2418 | 766 | 488 | 638 | 756 | 903 | 656 | 723 | 495 | 687 | 621 | 653 | 623 | 772 | 626 | 645 | 698 | 665,77 | 646 | 685 | 1,06 | 0,084 | 1,082 | 0,2791299 | 0,36322556 | 0,004438 | 0,00416 | 0,0043102 | 0,0043102 | TRUE | 0,0926856 | No |
| orf19.2419 | 1365 | 936 | 1181 | 1471 | 1732 | 1461 | 1542 | 985 | 1225 | 1192 | 1208 | 1213 | 1481 | 1394 | 1376 | 1389 | 1309,8 | 1210 | 1410 | 1,166 | 0,222 | 4,763 | 1,90E-06 | 7,71E-06 | 1,00E-08 | 0,0030879 | 0,0013188 | 0,0013188 | TRUE | 0,0147699 | No |
| orf19.242 | 116 | 74 | 112 | 148 | 156 | 154 | 165 | 99 | 104 | 94 | 115 | 122 | 133 | 147 | 147 | 140 | 125,27 | 109 | 142 | 1,296 | 0,374 | 2,924 | 0,00345826 | 0,00772675 | 1,00E-08 | 0,0135685 | 0,0076491 | 0,0076491 | TRUE | 0,0726882 | No |
| orf19.2422 | 3254 | 2133 | 2765 | 3408 | 3373 | 2930 | 3210 | 1947 | 2919 | 2716 | 2829 | 2810 | 2885 | 2796 | 2864 | 2746 | 2820,6 | 2818 | 2823 | 1,002 | 0,002 | 0,061 | 0,95136179 | 0,96517758 | 0,000221 | 0,0024942 | 0,0010887 | 0,0010887 | TRUE | 0,0070165 | No |
| orf19.242.2 | 125 | 96 | 90 | 131 | 153 | 147 | 153 | 68 | 112 | 122 | 92 | 108 | 131 | 140 | 137 | 96 | 117,25 | 108 | 126 | 1,174 | 0,232 | 1,575 | 0,11524342 | 0,17193505 | 0,008893 | 0,0143607 | 0,0122155 | 0,0122155 | TRUE | 0,2083925 | No |
| orf19.2423 | 2286 | 1482 | 1968 | 2363 | 2322 | 1941 | 2221 | 1539 | 2051 | 1887 | 2013 | 1949 | 1986 | 1852 | 1982 | 2171 | 1986,2 | 1975 | 1998 | 1,009 | 0,013 | 0,242 | 0,80867423 | 0,85177328 | 0,00196 | 0,0027104 | 0,0023781 | 0,0023781 | TRUE | 0,0383445 | No |
| orf19.2425 | 206 | 139 | 186 | 238 | 263 | 238 | 252 | 155 | 185 | 177 | 190 | 196 | 225 | 227 | 225 | 219 | 205,48 | 187 | 224 | 1,195 | 0,257 | 2,673 | 0,0075149 | 0,01557954 | 1,00E-08 | 0,0090445 | 0,0039848 | 0,0039848 | TRUE | 0,012553 | No |
| orf19.2426 | 412 | 292 | 387 | 482 | 494 | 434 | 440 | 316 | 370 | 372 | 396 | 397 | 422 | 414 | 393 | 446 | 401,21 | 384 | 419 | 1,087 | 0,12 | 1,603 | 0,10900849 | 0,16405887 | 1,00E-08 | 0,0055978 | 0,0028645 | 0,0028645 | TRUE | 0,0202942 | No |
| orf19.2429 | 53 | 40 | 33 | 57 | 35 | 29 | 32 | 17 | 48 | 51 | 34 | 47 | 30 | 28 | 29 | 24 | 36,17 | 45 | 28 | 0,621 | -0,687 | -2,925 | 0,00344254 | 0,00769735 | 1,00E-08 | 0,0421134 | 0,0239692 | 0,0239692 | TRUE | 0,2076927 | No |
| orf19.243 | 810 | 553 | 668 | 800 | 1195 | 974 | 1087 | 678 | 727 | 704 | 683 | 660 | 1022 | 929 | 970 | 956 | 831,42 | 694 | 969 | 1,4 | 0,486 | 8,273 | 1,30E-16 | 1,75E-15 | 0,000727 | 0,0037256 | 0,002074 | 0,002074 | TRUE | 0,0168882 | No |
| orf19.2430 | 10 | 19 | 8 | 16 | 9 | 8 | 4 | 1 | 9 | 24 | 8 | 13 | 8 | 8 | 4 | 1 | 9,36 | 14 | 5 | 0,395 | -1,339 | -2,423 | 0,01539301 | NA | 0,168194 | 0,1571404 | 0,1613972 | 0,1613972 | TRUE | 0,9328264 | No |
| orf19.2431 | 34 | 32 | 39 | 53 | 32 | 30 | 36 | 24 | 31 | 41 | 40 | 44 | 27 | 29 | 32 | 34 | 34,6 | 39 | 30 | 0,783 | -0,353 | -1,496 | 0,13464994 | 0,1970231 | 1,00E-08 | 0,0439343 | 0,0240108 | 0,0240108 | TRUE | 0,1701123 | No |
| orf19.2432 | 7865 | 6199 | 7301 | 8859 | 6562 | 6629 | 7026 | 4325 | 7056 | 7892 | 7470 | 7305 | 5612 | 6325 | 6270 | 6100 | 6753,7 | 7431 | 6077 | 0,818 | -0,29 | -5,687 | 1,29E-08 | 7,05E-08 | 0,002343 | 0,0021945 | 0,0023447 | 0,0023447 | TRUE | 0,0328836 | No |
| orf19.2433 | 186 | 163 | 154 | 191 | 150 | 181 | 174 | 94 | 167 | 208 | 158 | 158 | 128 | 173 | 155 | 133 | 159,78 | 173 | 147 | 0,861 | -0,215 | -1,584 | 0,11310958 | 0,16912951 | 0,011865 | 0,0110649 | 0,0113904 | 0,0113904 | TRUE | 0,1800848 | No |
| orf19.2434 | 1831 | 1528 | 1783 | 2321 | 1819 | 1857 | 1904 | 1177 | 1643 | 1945 | 1824 | 1914 | 1556 | 1772 | 1699 | 1660 | 1751,6 | 1832 | 1672 | 0,913 | -0,131 | -2,035 | 0,04180232 | 0,07181822 | 0,003448 | 0,0028083 | 0,0033996 | 0,0033996 | TRUE | 0,0591319 | No |
| orf19.2435 | 11326 | 8670 | 10089 | 12759 | 10574 | 10403 | 10638 | 6488 | 10161 | 11038 | 10322 | 10521 | 9043 | 9926 | 9493 | 9151 | 9956,8 | 10510 | 9403 | 0,895 | -0,16 | -3,744 | 0,00018076 | 0,00053053 | 0,001419 | 0,0021253 | 0,0016541 | 0,0016541 | TRUE | 0,0171797 | No |
| orf19.2436 | 6989 | 5000 | 6211 | 7878 | 8296 | 7833 | 8404 | 5187 | 6270 | 6366 | 6354 | 6496 | 7095 | 7474 | 7499 | 7316 | 6858,8 | 6372 | 7346 | 1,153 | 0,205 | 6,375 | 1,84E-10 | 1,28E-09 | 0,000329 | 0,0021912 | 0,0008475 | 0,0008475 | TRUE | 0,0067184 | No |
| orf19.2437 | 693 | 621 | 631 | 874 | 742 | 870 | 841 | 596 | 622 | 791 | 646 | 721 | 635 | 830 | 750 | 841 | 729,29 | 695 | 764 | 1,099 | 0,136 | 1,344 | 0,17897568 | 0,25070783 | 0,012013 | 0,0039701 | 0,0084812 | 0,0084812 | TRUE | 0,1576237 | No |
| orf19.2438 | 2093 | 1485 | 1752 | 2362 | 2572 | 2316 | 2340 | 1327 | 1878 | 1891 | 1792 | 1948 | 2200 | 2210 | 2088 | 1872 | 1984,7 | 1877 | 2092 | 1,116 | 0,159 | 2,711 | 0,00671126 | 0,01410327 | 0,002729 | 0,002711 | 0,0027923 | 0,0027923 | TRUE | 0,0572818 | No |
| orf19.2439 | 1661 | 1164 | 1239 | 1652 | 1668 | 1518 | 1481 | 900 | 1490 | 1482 | 1268 | 1362 | 1427 | 1448 | 1322 | 1269 | 1383,5 | 1400 | 1366 | 0,978 | -0,033 | -0,488 | 0,62564675 | 0,69774954 | 0,00406 | 0,0030288 | 0,0036079 | 0,0036079 | TRUE | 0,0483482 | No |
| orf19.2439.1 | 1965 | 1284 | 1597 | 2137 | 2202 | 1831 | 1829 | 1080 | 1763 | 1635 | 1634 | 1762 | 1883 | 1747 | 1632 | 1523 | 1697,4 | 1698 | 1696 | 1 | 0 | -0,002 | 0,99853943 | 0,99870566 | 0,00441 | 0,0028348 | 0,0037282 | 0,0037282 | TRUE | 0,0666298 | No |
| orf19.244 | 85 | 48 | 62 | 63 | 79 | 66 | 58 | 42 | 76 | 61 | 63 | 52 | 68 | 63 | 52 | 59 | 61,78 | 63 | 60 | 0,957 | -0,063 | -0,337 | 0,73621593 | 0,79382363 | 1,00E-08 | 0,0254757 | 0,0173559 | 0,0173559 | TRUE | 0,1905479 | No |
| orf19.2440 | 722 | 599 | 736 | 867 | 965 | 1037 | 1029 | 615 | 648 | 763 | 753 | 715 | 825 | 989 | 918 | 867 | 809,83 | 720 | 900 | 1,254 | 0,326 | 4,314 | 1,61E-05 | 5,67E-05 | 0,004621 | 0,0037721 | 0,0042314 | 0,0042314 | TRUE | 0,0542892 | No |
| orf19.2442 | 306 | 263 | 239 | 335 | 408 | 474 | 492 | 296 | 275 | 335 | 245 | 276 | 349 | 452 | 439 | 417 | 348,48 | 283 | 414 | 1,471 | 0,557 | 5,003 | 5,65E-07 | 2,48E-06 | 0,011802 | 0,0061454 | 0,0089024 | 0,0089024 | TRUE | 0,158179 | No |
| orf19.2443 | 1053 | 929 | 1018 | 1230 | 1099 | 1112 | 1115 | 677 | 945 | 1183 | 1042 | 1014 | 940 | 1061 | 995 | 955 | 1016,7 | 1046 | 988 | 0,947 | -0,078 | -1,067 | 0,28579238 | 0,37013162 | 0,00476 | 0,0034073 | 0,0041983 | 0,0041983 | TRUE | 0,0885978 | No |
| orf19.2444 | 3636 | 2773 | 3241 | 4237 | 3217 | 3034 | 3106 | 1945 | 3262 | 3530 | 3316 | 3494 | 2751 | 2895 | 2772 | 2743 | 3095,4 | 3400 | 2790 | 0,821 | -0,284 | -6,783 | 1,17E-11 | 9,49E-11 | 0,000768 | 0,0024485 | 0,0013595 | 0,0013595 | TRUE | 0,0092863 | No |
| orf19.2445 | 120 | 80 | 111 | 131 | 92 | 79 | 84 | 68 | 108 | 102 | 114 | 108 | 79 | 75 | 75 | 96 | 94,5 | 108 | 81 | 0,744 | -0,426 | -2,968 | 0,00299596 | 0,00680519 | 1,00E-08 | 0,0173412 | 0,0089981 | 0,0089981 | TRUE | 0,1039562 | No |
| orf19.2446 | 567 | 464 | 502 | 635 | 678 | 715 | 682 | 473 | 509 | 591 | 514 | 524 | 580 | 682 | 609 | 667 | 584,3 | 534 | 634 | 1,188 | 0,249 | 3,219 | 0,00128476 | 0,00316345 | 0,003566 | 0,004464 | 0,0040043 | 0,0040043 | TRUE | 0,0541739 | No |
| orf19.2447 | 1581 | 1257 | 1405 | 1717 | 1484 | 1404 | 1483 | 907 | 1418 | 1600 | 1437 | 1416 | 1269 | 1340 | 1323 | 1279 | 1385,4 | 1468 | 1303 | 0,89 | -0,169 | -3,141 | 0,0016838 | 0,00403521 | 0,001201 | 0,0030274 | 0,0020399 | 0,0020399 | TRUE | 0,0415751 | No |
| orf19.2449 | 9 | 4 | 13 | 13 | 10 | 7 | 12 | 3 | 8 | 5 | 13 | 11 | 9 | 7 | 11 | 4 | 8,42 | 9 | 8 | 0,823 | -0,281 | -0,577 | 0,56397467 | NA | 1,00E-08 | 0,1743974 | 0,1107384 | 0,1107384 | TRUE | 0,1720401 | No |
| orf19.245 | 471 | 432 | 480 | 579 | 541 | 562 | 600 | 377 | 423 | 550 | 491 | 477 | 463 | 536 | 535 | 532 | 500,89 | 485 | 516 | 1,067 | 0,093 | 1,046 | 0,29545829 | 0,38035428 | 0,006274 | 0,0048778 | 0,0055854 | 0,0055854 | TRUE | 0,0886253 | No |
| orf19.2451 | 879 | 720 | 849 | 1021 | 568 | 590 | 571 | 409 | 789 | 917 | 869 | 842 | 486 | 563 | 510 | 577 | 693,86 | 854 | 534 | 0,624 | -0,681 | -9,147 | 5,88E-20 | 1,05E-18 | 0,003515 | 0,0040717 | 0,003786 | 0,003786 | TRUE | 0,0437278 | No |
| orf19.2452 | 1267 | 1077 | 1195 | 1472 | 974 | 1041 | 1057 | 629 | 1137 | 1371 | 1223 | 1214 | 833 | 993 | 943 | 887 | 1075,1 | 1236 | 914 | 0,741 | -0,433 | -5,869 | 4,37E-09 | 2,56E-08 | 0,00475 | 0,0033298 | 0,0042557 | 0,0042557 | TRUE | 0,0622608 | No |
| orf19.2454 | 10799 | 8432 | 10043 | 11920 | 12521 | 12228 | 12522 | 8083 | 9688 | 10735 | 10275 | 9829 | 10708 | 11668 | 11174 | 11400 | 10685 | 10132 | 11238 | 1,109 | 0,15 | 3,372 | 0,00074645 | 0,00193139 | 0,001621 | 0,0021154 | 0,0017996 | 0,0017996 | TRUE | 0,0194058 | No |
| orf19.2455 | 3245 | 2594 | 3087 | 3721 | 3630 | 3562 | 3736 | 2227 | 2911 | 3302 | 3158 | 3068 | 3104 | 3399 | 3334 | 3141 | 3177,3 | 3110 | 3244 | 1,044 | 0,063 | 1,227 | 0,21973376 | 0,29854645 | 0,001953 | 0,0024364 | 0,0021933 | 0,0021933 | TRUE | 0,0225284 | No |
| orf19.2457 | 804 | 676 | 811 | 952 | 1179 | 1160 | 1173 | 742 | 721 | 861 | 830 | 785 | 1008 | 1107 | 1047 | 1047 | 925,63 | 799 | 1052 | 1,32 | 0,4 | 6,172 | 6,75E-10 | 4,39E-09 | 0,002396 | 0,0035479 | 0,0029313 | 0,0029313 | TRUE | 0,0514839 | No |
| orf19.2458 | 16060 | 11954 | 14679 | 19026 | 21292 | 19620 | 21289 | 14171 | 14407 | 15219 | 15018 | 15689 | 18210 | 18721 | 18997 | 19987 | 17031 | 15083 | 18979 | 1,258 | 0,331 | 8,064 | 7,41E-16 | 9,25E-15 | 0,001316 | 0,0020648 | 0,0015596 | 0,0015596 | TRUE | 0,015473 | No |
| orf19.2459 | 5233 | 3473 | 4616 | 5903 | 5436 | 4999 | 5332 | 3531 | 4695 | 4421 | 4723 | 4868 | 4649 | 4770 | 4758 | 4980 | 4732,9 | 4677 | 4789 | 1,023 | 0,032 | 0,787 | 0,43109232 | 0,51820782 | 0,000929 | 0,0022862 | 0,0014068 | 0,0014068 | TRUE | 0,015878 | No |
| orf19.246 | 440 | 284 | 393 | 463 | 481 | 428 | 457 | 287 | 395 | 362 | 402 | 382 | 411 | 408 | 408 | 405 | 396,56 | 385 | 408 | 1,058 | 0,082 | 1,136 | 0,25587546 | 0,33840334 | 1,00E-08 | 0,0056402 | 0,0024734 | 0,0024734 | TRUE | 0,016229 | No |
| orf19.2460 | 6 | 3 | 5 | 16 | 13 | 9 | 7 | 6 | 5 | 4 | 5 | 13 | 11 | 9 | 6 | 8 | 7,74 | 7 | 8 | 1,214 | 0,279 | 0,528 | 0,59737278 | NA | 0,07456 | 0,1895209 | 0,140499 | 0,140499 | TRUE | 1,2133451 | No |
| orf19.2461 | 1456 | 995 | 1233 | 1542 | 1161 | 1029 | 1132 | 678 | 1306 | 1267 | 1261 | 1272 | 993 | 982 | 1010 | 956 | 1130,9 | 1276 | 985 | 0,773 | -0,372 | -7,912 | 2,53E-15 | 2,99E-14 | 1,00E-08 | 0,0032632 | 0,0012222 | 0,0012222 | TRUE | 0,0039497 | No |
| orf19.2462 | 3618 | 2517 | 3187 | 3933 | 2895 | 2573 | 2811 | 1655 | 3246 | 3204 | 3261 | 3243 | 2476 | 2455 | 2508 | 2334 | 2840,9 | 3238 | 2443 | 0,756 | -0,404 | -11,04 | 2,44E-28 | 8,20E-27 | 1,00E-08 | 0,0024905 | 0,0009278 | 0,0009278 | TRUE | 0,0097402 | No |
| orf19.2463 | 1150 | 705 | 948 | 1121 | 939 | 942 | 979 | 523 | 1032 | 898 | 970 | 924 | 803 | 899 | 874 | 738 | 892,08 | 956 | 828 | 0,868 | -0,204 | -2,796 | 0,00517215 | 0,01114571 | 0,004241 | 0,0036068 | 0,0039875 | 0,0039875 | TRUE | 0,0578395 | No |
| orf19.2465 | 4484 | 2985 | 3798 | 4748 | 3805 | 3248 | 3540 | 2109 | 4023 | 3800 | 3886 | 3915 | 3254 | 3099 | 3159 | 2975 | 3513,8 | 3906 | 3122 | 0,8 | -0,322 | -8,066 | 7,28E-16 | 9,11E-15 | 0,000595 | 0,0023927 | 0,0012421 | 0,0012421 | TRUE | 0,0113293 | No |
| orf19.2467 | 3723 | 2543 | 3186 | 4017 | 3007 | 2443 | 2694 | 1692 | 3340 | 3238 | 3260 | 3312 | 2572 | 2331 | 2404 | 2386 | 2855,3 | 3288 | 2423 | 0,737 | -0,439 | -10,545 | 5,37E-26 | 1,52E-24 | 0,00054 | 0,0024879 | 0,001307 | 0,001307 | TRUE | 0,021513 | No |
| orf19.2468 | 22 | 20 | 13 | 23 | 8 | 8 | 3 | 3 | 20 | 25 | 13 | 19 | 7 | 8 | 3 | 4 | 12,36 | 19 | 6 | 0,282 | -1,828 | -4,028 | 5,61E-05 | NA | 1,00E-08 | 0,1194688 | 0,079884 | 0,079884 | TRUE | 0,2104432 | No |
| orf19.2469 | 198 | 156 | 201 | 246 | 161 | 146 | 149 | 94 | 178 | 199 | 206 | 203 | 138 | 139 | 133 | 133 | 165,91 | 196 | 136 | 0,693 | -0,529 | -4,826 | 1,39E-06 | 5,75E-06 | 1,00E-08 | 0,0107295 | 0,005337 | 0,005337 | TRUE | 0,0474184 | No |
| orf19.247 | 165 | 127 | 133 | 203 | 202 | 203 | 188 | 98 | 148 | 162 | 136 | 167 | 173 | 194 | 168 | 138 | 160,7 | 153 | 168 | 1,104 | 0,143 | 1,139 | 0,25459787 | 0,33721869 | 0,006183 | 0,0110131 | 0,0089693 | 0,0089693 | TRUE | 0,1226716 | No |
| orf19.2471 | 979 | 644 | 833 | 1024 | 898 | 804 | 853 | 549 | 878 | 820 | 852 | 844 | 768 | 767 | 761 | 774 | 808,18 | 848 | 768 | 0,903 | -0,147 | -2,768 | 0,00563976 | 0,01204539 | 1,00E-08 | 0,0037758 | 0,0014773 | 0,0014773 | TRUE | 0,0065673 | No |
| orf19.2472 | 777 | 560 | 703 | 808 | 814 | 695 | 734 | 502 | 697 | 713 | 719 | 666 | 696 | 663 | 655 | 708 | 689,73 | 699 | 680 | 0,974 | -0,039 | -0,646 | 0,51842365 | 0,60187233 | 1,00E-08 | 0,0040843 | 0,0019678 | 0,0019678 | TRUE | 0,0124747 | No |
| orf19.2472.1 | 1387 | 922 | 976 | 1454 | 1720 | 1359 | 1361 | 911 | 1244 | 1174 | 999 | 1199 | 1471 | 1297 | 1214 | 1285 | 1235,3 | 1154 | 1317 | 1,141 | 0,191 | 2,378 | 0,01739616 | 0,03283572 | 0,006693 | 0,0031547 | 0,0053489 | 0,0053489 | TRUE | 0,0986164 | No |
| orf19.2473 | 2880 | 1972 | 2663 | 3207 | 3305 | 3002 | 3155 | 2074 | 2584 | 2511 | 2725 | 2645 | 2827 | 2864 | 2815 | 2925 | 2736,8 | 2616 | 2858 | 1,091 | 0,126 | 3,201 | 0,0013712 | 0,00335294 | 0,000265 | 0,00251 | 0,0011139 | 0,0011139 | TRUE | 0,0090969 | No |

| | | | | | | | | | | | | | | | | | | | | | | | | | | | | | | | |
|---|---|---|---|---|---|---|---|---|---|---|---|---|---|---|---|---|---|---|---|---|---|---|---|---|---|---|---|---|---|---|---|
| orf19.2474 | 1347 | 864 | 1159 | 1318 | 1256 | 1113 | 1202 | 813 | 1208 | 1100 | 1186 | 1087 | 1074 | 1062 | 1073 | 1147 | 1117,1 | 1145 | 1089 | 0,949 | -0,076 | -1,369 | 0,17091493 | 0,24104622 | 0,001119 | 0,0032791 | 0,0020703 | 0,0020703 | TRUE | 0,0171477 | No |
| orf19.2475 | 5 | 6 | 11 | 6 | 5 | 6 | 8 | 2 | 4 | 8 | 11 | 5 | 4 | 6 | 7 | 3 | 6,04 | 7 | 5 | 0,734 | -0,446 | -0,773 | 0,43965983 | NA | 1,00E-08 | 0,2424937 | 0,1510587 | 0,1510587 | TRUE | 0,438001 | No |
| orf19.2476 | 1276 | 1001 | 1210 | 1373 | 1644 | 1575 | 1583 | 1012 | 1145 | 1274 | 1238 | 1132 | 1406 | 1503 | 1413 | 1427 | 1317,2 | 1197 | 1437 | 1,203 | 0,267 | 4,887 | 1,02E-06 | 4,32E-06 | 0,001339 | 0,0030816 | 0,0020907 | 0,0020907 | TRUE | 0,0213148 | No |
| orf19.2478 | 52 | 40 | 37 | 53 | 94 | 75 | 77 | 34 | 47 | 51 | 38 | 44 | 80 | 72 | 69 | 48 | 55,97 | 45 | 67 | 1,527 | 0,61 | 3,025 | 0,0024863 | 0,00574747 | 0,009232 | 0,0279172 | 0,0206198 | 0,0206198 | TRUE | 0,2505254 | No |
| orf19.2478.1 | 23826 | 15555 | 19610 | 24492 | 22386 | 18853 | 19644 | 14457 | 21374 | 19803 | 20063 | 20196 | 19145 | 17989 | 17529 | 20390 | 19561 | 20359 | 18763 | 0,921 | -0,118 | -2,278 | 0,02272052 | 0,04184698 | 0,002844 | 0,0020537 | 0,0025327 | 0,0025327 | TRUE | 0,0413683 | No |
| orf19.2479 | 186 | 128 | 159 | 167 | 161 | 155 | 163 | 98 | 167 | 163 | 163 | 138 | 138 | 148 | 145 | 138 | 149,93 | 158 | 142 | 0,907 | -0,14 | -1,203 | 0,22890825 | 0,30884365 | 1,00E-08 | 0,0116619 | 0,006368 | 0,006368 | TRUE | 0,086216 | No |
| orf19.248 | 1049 | 779 | 935 | 1191 | 1031 | 1087 | 1111 | 678 | 941 | 992 | 957 | 982 | 882 | 1037 | 991 | 956 | 967,26 | 968 | 966 | 0,999 | -0,002 | -0,025 | 0,98000084 | 0,98508366 | 0,001774 | 0,0034804 | 0,0025242 | 0,0025242 | TRUE | 0,0451923 | No |
| orf19.2480.1 | 13817 | 9160 | 11999 | 14843 | 17088 | 14277 | 15811 | 9393 | 12395 | 11662 | 12276 | 12240 | 14614 | 13623 | 14109 | 13248 | 13021 | 12143 | 13898 | 1,145 | 0,195 | 4,854 | 1,21E-06 | 5,04E-06 | 0,001172 | 0,002091 | 0,0014718 | 0,0014718 | TRUE | 0,0147194 | No |
| orf19.2481 | 571 | 418 | 448 | 620 | 541 | 462 | 499 | 379 | 512 | 532 | 458 | 511 | 463 | 441 | 445 | 535 | 487,17 | 503 | 471 | 0,931 | -0,103 | -1,282 | 0,19994165 | 0,27450855 | 0,0035 | 0,0049594 | 0,0041766 | 0,0041766 | TRUE | 0,0840558 | No |
| orf19.2483 | 1847 | 1173 | 1425 | 1816 | 2182 | 1678 | 1694 | 1469 | 1657 | 1493 | 1458 | 1497 | 1866 | 1601 | 1512 | 2072 | 1644,6 | 1526 | 1763 | 1,153 | 0,205 | 2,251 | 0,02441497 | 0,04463942 | 0,011362 | 0,0028623 | 0,0073652 | 0,0073652 | TRUE | 0,1633643 | No |
| orf19.2484 | 2318 | 1556 | 2057 | 2508 | 2262 | 1830 | 1975 | 1572 | 2079 | 1981 | 2105 | 2068 | 1935 | 1746 | 1762 | 2217 | 1986,7 | 2058 | 1915 | 0,928 | -0,107 | -1,489 | 0,1365159 | 0,19926811 | 0,00619 | 0,0027102 | 0,0044737 | 0,0044737 | TRUE | 0,1307878 | No |
| orf19.2485 | 3153 | 2433 | 3032 | 3455 | 3296 | 3290 | 3556 | 2191 | 2829 | 3097 | 3102 | 2849 | 2819 | 3139 | 3173 | 3090 | 3012,3 | 2969 | 3055 | 1,029 | 0,042 | 0,777 | 0,43686808 | 0,52377325 | 0,0022 | 0,0024614 | 0,0024491 | 0,0024491 | TRUE | 0,0339252 | No |
| orf19.2487 | 2707 | 1906 | 2249 | 2839 | 2634 | 2332 | 2555 | 1545 | 2428 | 2427 | 2301 | 2341 | 2253 | 2225 | 2280 | 2179 | 2304,2 | 2374 | 2234 | 0,942 | -0,086 | -2,191 | 0,0284681 | 0,05117784 | 1,00E-08 | 0,0026095 | 0,001039 | 0,001039 | TRUE | 0,0050562 | No |
| orf19.2488 | 5040 | 3447 | 4349 | 5413 | 4821 | 4260 | 4675 | 2668 | 4521 | 4388 | 4449 | 4464 | 4123 | 4065 | 4172 | 3763 | 4243,2 | 4456 | 4031 | 0,906 | -0,143 | -3,556 | 0,00037604 | 0,00104256 | 0,000799 | 0,0023216 | 0,0013164 | 0,0013164 | TRUE | 0,0228388 | No |
| orf19.2489 | 19412 | 13817 | 17689 | 21223 | 17737 | 16166 | 17228 | 11717 | 17415 | 17591 | 18098 | 17501 | 15169 | 15425 | 15373 | 16526 | 16637 | 17651 | 15623 | 0,885 | -0,177 | -4,92 | 8,64E-07 | 3,67E-06 | 0,000837 | 0,0020668 | 0,0011759 | 0,0011759 | TRUE | 0,0181559 | No |
| orf19.2492 | 41 | 19 | 35 | 33 | 38 | 22 | 24 | 23 | 37 | 24 | 36 | 27 | 32 | 21 | 21 | 32 | 28,92 | 31 | 26 | 0,853 | -0,229 | -0,835 | 0,40359535 | 0,4919458 | 0,013582 | 0,0521817 | 0,0372314 | 0,0372314 | TRUE | 0,173434 | No |
| orf19.2494 | 542 | 393 | 476 | 585 | 534 | 428 | 521 | 317 | 486 | 500 | 487 | 482 | 457 | 408 | 465 | 447 | 466,63 | 489 | 444 | 0,91 | -0,136 | -1,95 | 0,05114622 | 0,0857145 | 1,00E-08 | 0,0050906 | 0,0025482 | 0,0025482 | TRUE | 0,0362321 | No |
| orf19.2495 | 477 | 392 | 457 | 587 | 663 | 564 | 657 | 368 | 428 | 499 | 468 | 484 | 567 | 538 | 586 | 519 | 511,13 | 470 | 552 | 1,182 | 0,242 | 3,287 | 0,00101189 | 0,00254369 | 0,001531 | 0,0048197 | 0,0032148 | 0,0032148 | TRUE | 0,042986 | No |
| orf19.2496 | 4072 | 2714 | 3668 | 4608 | 2143 | 1778 | 1878 | 1371 | 3653 | 3455 | 3753 | 3800 | 1833 | 1697 | 1676 | 1934 | 2724,9 | 3665 | 1785 | 0,486 | -1,041 | -18,837 | 3,79E-79 | 1,08E-76 | 0,002623 | 0,0025123 | 0,0025144 | 0,0025144 | TRUE | 0,0354306 | No |
| orf19.2498 | 10012 | 6666 | 9115 | 11167 | 11072 | 9143 | 10296 | 6861 | 8982 | 8487 | 9326 | 9208 | 9469 | 8724 | 9187 | 9677 | 9132,5 | 9001 | 9264 | 1,029 | 0,041 | 0,901 | 0,36733495 | 0,45616957 | 0,001695 | 0,0021385 | 0,0018707 | 0,0018707 | TRUE | 0,0188235 | No |
| orf19.2499 | 512 | 419 | 508 | 595 | 688 | 569 | 625 | 357 | 459 | 533 | 520 | 491 | 588 | 543 | 558 | 504 | 524,46 | 501 | 548 | 1,102 | 0,14 | 1,865 | 0,06211488 | 0,10151964 | 0,002127 | 0,0047475 | 0,0034585 | 0,0034585 | TRUE | 0,0364458 | No |
| orf19.25 | 2469 | 1687 | 2045 | 2665 | 3180 | 2748 | 2978 | 1896 | 2215 | 2148 | 2092 | 2198 | 2720 | 2622 | 2657 | 2674 | 2415,7 | 2163 | 2668 | 1,233 | 0,302 | 7,924 | 2,30E-15 | 2,74E-14 | 1,00E-08 | 0,0025805 | 0,0009752 | 0,0009752 | TRUE | 0,0060344 | No |
| orf19.250 | 3949 | 3274 | 3537 | 4607 | 3790 | 4159 | 4322 | 2654 | 3543 | 4168 | 3619 | 3799 | 3241 | 3968 | 3857 | 3743 | 3742,3 | 3782 | 3702 | 0,979 | -0,03 | -0,426 | 0,67034326 | 0,73694827 | 0,006324 | 0,0023674 | 0,0046303 | 0,0046303 | TRUE | 0,0867725 | No |
| orf19.2500 | 409 | 284 | 385 | 447 | 524 | 495 | 494 | 324 | 367 | 362 | 394 | 369 | 448 | 472 | 441 | 457 | 413,65 | 373 | 454 | 1,218 | 0,285 | 3,98 | 6,90E-05 | 0,00021727 | 1,00E-08 | 0,005489 | 0,0024897 | 0,0024897 | TRUE | 0,0161128 | No |
| orf19.2501 | 8295 | 6387 | 7906 | 9702 | 9940 | 9948 | 10472 | 6720 | 7441 | 8131 | 8089 | 8000 | 8501 | 9492 | 9345 | 9478 | 8559,7 | 7915 | 9204 | 1,163 | 0,217 | 4,502 | 6,72E-06 | 2,52E-05 | 0,001986 | 0,0021491 | 0,0021226 | 0,0021226 | TRUE | 0,0325849 | No |
| orf19.2503 | 13 | 15 | 19 | 23 | 16 | 24 | 14 | 9 | 12 | 19 | 19 | 19 | 14 | 23 | 12 | 13 | 16,37 | 17 | 16 | 0,903 | -0,147 | -0,412 | 0,67999661 | 0,74429216 | 1,00E-08 | 0,0906777 | 0,0612675 | 0,0612675 | TRUE | 0,5161103 | No |
| orf19.2504 | 16389 | 12005 | 14730 | 17875 | 14899 | 14026 | 14711 | 8875 | 14703 | 15284 | 15070 | 14740 | 12742 | 13383 | 13127 | 12517 | 13946 | 14949 | 12942 | 0,866 | -0,207 | -6,335 | 2,38E-10 | 1,65E-09 | 0,000523 | 0,0020836 | 0,0009562 | 0,0009562 | TRUE | 0,0063846 | No |
| orf19.2505 | 12 | 8 | 18 | 15 | 21 | 13 | 17 | 11 | 11 | 10 | 18 | 12 | 18 | 12 | 15 | 16 | 14,1 | 13 | 15 | 1,18 | 0,238 | 0,65 | 0,51591363 | NA | 1,00E-08 | 0,1049523 | 0,0584014 | 0,0584014 | TRUE | 0,3207572 | No |
| orf19.2506 | 1028 | 709 | 931 | 1121 | 782 | 710 | 778 | 478 | 922 | 903 | 953 | 924 | 669 | 677 | 694 | 674 | 802,05 | 926 | 678 | 0,733 | -0,448 | -8,372 | 5,69E-17 | 8,04E-16 | 1,00E-08 | 0,0037895 | 0,0014713 | 0,0014713 | TRUE | 0,004368 | No |
| orf19.2507 | 1373 | 1197 | 1321 | 1652 | 1462 | 1582 | 1610 | 943 | 1232 | 1524 | 1352 | 1362 | 1250 | 1509 | 1437 | 1330 | 1374,5 | 1368 | 1382 | 1,012 | 0,017 | 0,225 | 0,82200764 | 0,86279209 | 0,006513 | 0,0030357 | 0,0049546 | 0,0049546 | TRUE | 0,0689727 | No |
| orf19.2508 | 137 | 122 | 115 | 147 | 154 | 173 | 159 | 106 | 123 | 155 | 118 | 121 | 132 | 165 | 142 | 150 | 138,16 | 129 | 147 | 1,143 | 0,193 | 1,468 | 0,14211282 | 0,20608588 | 0,005077 | 0,0124871 | 0,0093422 | 0,0093422 | TRUE | 0,1618882 | No |
| orf19.2509 | 0 | 0 | 0 | 0 | 0 | 0 | 0 | 0 | 0 | 0 | 0 | 0 | 0 | 0 | 0 | 0 | 0 | 0 | 0 | NA | NA | NA | NA | NA | NA | NA | NA | NA | NA | NA | NA |
| orf19.2509.1 | 208 | 157 | 147 | 181 | 88 | 89 | 71 | 55 | 187 | 200 | 150 | 149 | 75 | 85 | 63 | 78 | 123,4 | 172 | 75 | 0,439 | -1,186 | -7,775 | 7,55E-15 | 8,58E-14 | 0,011576 | 0,0137433 | 0,0127567 | 0,0127567 | TRUE | 0,1182859 | No |
| orf19.251 | 10057 | 6880 | 8489 | 10693 | 11165 | 9856 | 9932 | 5974 | 9022 | 8759 | 8685 | 8818 | 9549 | 9404 | 8863 | 8426 | 8940,7 | 8821 | 9060 | 1,028 | 0,039 | 0,879 | 0,37946968 | 0,46900923 | 0,001607 | 0,0021419 | 0,001805 | 0,001805 | TRUE | 0,0265324 | No |
| orf19.2510 | 107 | 113 | 121 | 133 | 113 | 172 | 192 | 83 | 96 | 144 | 124 | 110 | 97 | 164 | 171 | 117 | 127,81 | 118 | 137 | 1,169 | 0,225 | 1,165 | 0,24391379 | 0,32565201 | 0,044164 | 0,0133379 | 0,0280452 | 0,0280452 | TRUE | 0,249347 | No |
| orf19.2511.1 | 1494 | 999 | 1170 | 1591 | 1647 | 1315 | 1294 | 748 | 1340 | 1272 | 1197 | 1312 | 1409 | 1255 | 1155 | 1055 | 1249,3 | 1280 | 1218 | 0,953 | -0,069 | -0,84 | 0,40100196 | 0,48947984 | 0,0072 | 0,0031416 | 0,0056698 | 0,0056698 | TRUE | 0,133542 | No |
| orf19.2511.2 | 718 | 698 | 729 | 848 | 911 | 1060 | 1015 | 559 | 644 | 889 | 746 | 699 | 779 | 1011 | 906 | 788 | 807,82 | 744 | 871 | 1,174 | 0,231 | 2,146 | 0,03187877 | 0,05656458 | 0,016178 | 0,0037766 | 0,009903 | 0,009903 | TRUE | 0,1948403 | No |
| orf19.2512 | 137 | 119 | 148 | 194 | 170 | 174 | 196 | 128 | 123 | 151 | 151 | 160 | 145 | 166 | 175 | 181 | 156,58 | 146 | 167 | 1,134 | 0,182 | 1,457 | 0,14508528 | 0,20963741 | 0,00516 | 0,0112508 | 0,0085362 | 0,0085362 | TRUE | 0,1325747 | No |
| orf19.2513 | 358 | 392 | 382 | 500 | 431 | 611 | 587 | 365 | 321 | 499 | 391 | 412 | 369 | 583 | 524 | 515 | 451,69 | 406 | 498 | 1,228 | 0,296 | 2,016 | 0,04375541 | 0,07481005 | 0,032594 | 0,0051934 | 0,0184332 | 0,0184332 | TRUE | 0,3658746 | No |
| orf19.2514 | 818 | 664 | 803 | 996 | 1130 | 1128 | 1199 | 666 | 734 | 845 | 822 | 821 | 966 | 1076 | 1070 | 939 | 909,25 | 806 | 1013 | 1,261 | 0,335 | 4,898 | 9,68E-07 | 4,09E-06 | 0,003198 | 0,0035761 | 0,0033711 | 0,0033711 | TRUE | 0,0436445 | No |
| orf19.2515 | 8191 | 5974 | 7306 | 9332 | 11265 | 11190 | 11752 | 6926 | 7348 | 7606 | 7475 | 7695 | 9634 | 10677 | 10487 | 9769 | 8836,3 | 7531 | 10142 | 1,347 | 0,43 | 10,006 | 1,44E-23 | 3,32E-22 | 0,001349 | 0,0021438 | 0,001655 | 0,001655 | TRUE | 0,0154388 | No |
| orf19.2516 | 432 | 366 | 430 | 586 | 497 | 538 | 540 | 316 | 388 | 466 | 440 | 483 | 425 | 513 | 482 | 446 | 455,33 | 444 | 466 | 1,052 | 0,073 | 0,804 | 0,42163508 | 0,50942997 | 0,006174 | 0,0051678 | 0,005684 | 0,005684 | TRUE | 0,089328 | No |
| orf19.2517 | 236 | 160 | 228 | 262 | 281 | 264 | 256 | 174 | 212 | 204 | 233 | 216 | 240 | 252 | 228 | 245 | 228,85 | 216 | 241 | 1,114 | 0,156 | 1,664 | 0,09613996 | 0,14776384 | 1,00E-08 | 0,008323 | 0,0040924 | 0,0040924 | TRUE | 0,0291059 | No |
| orf19.2518 | 3594 | 2491 | 3091 | 3866 | 3974 | 3679 | 3902 | 2350 | 3224 | 3171 | 3162 | 3188 | 3399 | 3510 | 3482 | 3314 | 3306,4 | 3186 | 3426 | 1,076 | 0,106 | 3,052 | 0,00227543 | 0,00530904 | 1,00E-08 | 0,0024186 | 0,000849 | 0,000849 | TRUE | 0,0052868 | No |
| orf19.2519 | 420 | 269 | 275 | 420 | 522 | 446 | 466 | 328 | 377 | 342 | 281 | 346 | 446 | 426 | 416 | 463 | 387,17 | 336 | 438 | 1,295 | 0,373 | 4,045 | 5,24E-05 | 0,00016889 | 0,005339 | 0,005729 | 0,0055449 | 0,0055449 | TRUE | 0,1395483 | No |
| orf19.252 | 416 | 355 | 384 | 519 | 445 | 515 | 518 | 236 | 373 | 452 | 393 | 428 | 381 | 491 | 462 | 333 | 414,13 | 412 | 417 | 1,018 | 0,026 | 0,219 | 0,82651958 | 0,86677075 | 0,016309 | 0,0054849 | 0,0107583 | 0,0107583 | TRUE | 0,1913515 | No |
| orf19.2520 | 2567 | 1707 | 2180 | 2730 | 2829 | 2561 | 2757 | 1755 | 2303 | 2173 | 2230 | 2251 | 2419 | 2444 | 2460 | 2475 | 2344,5 | 2239 | 2450 | 1,092 | 0,127 | 3,425 | 0,00061529 | 0,00162064 | 1,00E-08 | 0,0025987 | 0,0009024 | 0,0009024 | TRUE | 0,0044103 | No |
| orf19.2521 | 597 | 408 | 535 | 610 | 430 | 392 | 445 | 257 | 536 | 519 | 547 | 503 | 368 | 374 | 397 | 362 | 450,84 | 526 | 375 | 0,715 | -0,484 | -6,858 | 6,99E-12 | 5,80E-11 | 1,00E-08 | 0,0051995 | 0,0025141 | 0,0025141 | TRUE | 0,0181491 | No |
| orf19.2524 | 5949 | 3739 | 4829 | 6050 | 7296 | 5970 | 6383 | 4267 | 5337 | 4760 | 4941 | 4989 | 6240 | 5696 | 5696 | 6018 | 5459,6 | 5007 | 5912 | 1,18 | 0,239 | 4,937 | 7,95E-07 | 3,41E-06 | 0,001925 | 0,0022454 | 0,0020693 | 0,0020693 | TRUE | 0,0241012 | No |
| orf19.2525 | 1802 | 1225 | 1664 | 1945 | 2070 | 1662 | 1791 | 1222 | 1617 | 1560 | 1702 | 1604 | 1770 | 1586 | 1598 | 1724 | 1645 | 1621 | 1670 | 1,029 | 0,042 | 0,78 | 0,435419 | 0,52243541 | 0,001592 | 0,002862 | 0,0021089 | 0,0021089 | TRUE | 0,0217152 | No |
| orf19.2527 | 3925 | 2602 | 3458 | 4210 | 4143 | 3418 | 3829 | 2335 | 3521 | 3313 | 3538 | 3472 | 3543 | 3261 | 3417 | 3293 | 3419,7 | 3461 | 3378 | 0,976 | -0,034 | -0,82 | 0,41249382 | 0,50058759 | 0,000774 | 0,002404 | 0,0014002 | 0,0014002 | TRUE | 0,0133265 | No |
| orf19.2528 | 3259 | 2001 | 2809 | 3546 | 3618 | 2957 | 3330 | 2146 | 2924 | 2547 | 2874 | 2924 | 3094 | 2821 | 2971 | 3027 | 2897,9 | 2817 | 2978 | 1,056 | 0,079 | 1,446 | 0,14828883 | 0,21354729 | 0,002372 | 0,0024805 | 0,0024868 | 0,0024868 | TRUE | 0,0486775 | No |
| orf19.2529 | 1393 | 993 | 1164 | 1493 | 1302 | 1190 | 1246 | 786 | 1250 | 1264 | 1191 | 1231 | 1114 | 1135 | 1112 | 1109 | 1175,7 | 1234 | 1118 | 0,906 | -0,142 | -3,068 | 0,00215528 | 0,00505621 | 1,00E-08 | 0,0032143 | 0,0012114 | 0,0012114 | TRUE | 0,0063522 | No |
| orf19.2529.1 | 15810 | 9921 | 12019 | 15186 | 17392 | 12949 | 13002 | 7510 | 14183 | 12630 | 12297 | 12523 | 14874 | 12356 | 11602 | 10592 | 12632 | 12908 | 12356 | 0,957 | -0,063 | -0,701 | 0,48315523 | 0,56861834 | 0,011429 | 0,0020944 | 0,0076296 | 0,0076296 | TRUE | 0,2304673 | No |
| orf19.2530 | 8 | 3 | 7 | 8 | 8 | 9 | 8 | 4 | 7 | 4 | 7 | 7 | 7 | 9 | 7 | 6 | 6,62 | 6 | 7 | 1,132 | 0,179 | 0,347 | 0,72864878 | NA | 1,00E-08 | 0,2212514 | 0,1060609 | 0,1060609 | TRUE | 0,1100904 | No |
| orf19.2531 | 8738 | 6332 | 7336 | 9278 | 11271 | 10210 | 10723 | 7177 | 7839 | 8061 | 7505 | 7651 | 9639 | 9742 | 9568 | 10123 | 8766,1 | 7764 | 9768 | 1,258 | 0,331 | 9,334 | 1,02E-20 | 1,96E-19 | 0,000645 | 0,0021451 | 0,0010922 | 0,0010922 | TRUE | 0,0078595 | No |
| orf19.2532 | 1001 | 705 | 888 | 1102 | 1132 | 994 | 1138 | 679 | 898 | 898 | 909 | 909 | 968 | 948 | 1015 | 958 | 937,81 | 904 | 972 | 1,077 | 0,108 | 2,116 | 0,0343154 | 0,06041817 | 1,00E-08 | 0,0035275 | 0,0014135 | 0,0014135 | TRUE | 0,0111023 | No |
| orf19.2533.1 | 1329 | 911 | 1099 | 1386 | 1152 | 997 | 1061 | 706 | 1192 | 1160 | 1124 | 1143 | 985 | 951 | 947 | 996 | 1062,3 | 1155 | 970 | 0,839 | -0,254 | -5,15 | 2,61E-07 | 1,20E-06 | 1,00E-08 | 0,0033461 | 0,0013822 | 0,0013822 | TRUE | 0,0060584 | No |
| orf19.2534 | 3247 | 2323 | 3087 | 3774 | 3945 | 3523 | 3920 | 2587 | 2913 | 2957 | 3158 | 3112 | 3374 | 3362 | 3498 | 3649 | 3252,9 | 3035 | 3471 | 1,142 | 0,192 | 4,282 | 1,85E-05 | 6,46E-05 | 0,001093 | 0,0024258 | 0,001613 | 0,001613 | TRUE | 0,0137096 | No |

| | | | | | | | | | | | | | | | | | | | | | | | | | | | | | | | |
|---|---|---|---|---|---|---|---|---|---|---|---|---|---|---|---|---|---|---|---|---|---|---|---|---|---|---|---|---|---|---|---|
| orf19.2537 | 38 | 22 | 33 | 35 | 22 | 26 | 30 | 19 | 34 | 28 | 34 | 29 | 19 | 25 | 27 | 27 | 27,74 | 31 | 24 | 0,771 | -0,376 | -1,444 | 0,14871733 | 0,21411304 | 1,00E-08 | 0,0543135 | 0,0283503 | 0,0283503 | TRUE | 0,1650634 | No |
| orf19.2538 | 24643 | 18195 | 22367 | 27718 | 22341 | 21422 | 23143 | 13111 | 22107 | 23164 | 22884 | 22857 | 19107 | 20440 | 20651 | 18492 | 21213 | 22753 | 19672 | 0,865 | -0,21 | -4,899 | 9,62E-07 | 4,07E-06 | 0,001539 | 0,002048 | 0,0017093 | 0,0017093 | TRUE | 0,0197329 | No |
| orf19.2539 | 0 | 0 | 2 | 2 | 1 | 1 | 2 | 3 | 0 | 0 | 2 | 2 | 1 | 1 | 2 | 4 | 1,44 | 1 | 2 | 1,923 | 0,943 | 0,759 | 0,44763383 | NA | 1,00E-08 | 1,0100422 | 0,6943315 | 0,6943315 | TRUE | 0,3575151 | No |
| orf19.254 | 9092 | 6248 | 8480 | 10053 | 7588 | 6946 | 7546 | 5258 | 8156 | 7954 | 8676 | 8290 | 6489 | 6628 | 6734 | 7416 | 7542,9 | 8269 | 6817 | 0,824 | -0,28 | -5,656 | 1,55E-08 | 8,42E-08 | 0,002323 | 0,002172 | 0,0022166 | 0,0022166 | TRUE | 0,0417896 | No |
| orf19.2540 | 1039 | 755 | 874 | 1162 | 1289 | 1168 | 1259 | 675 | 932 | 961 | 894 | 958 | 1102 | 1114 | 1123 | 952 | 1004,8 | 936 | 1073 | 1,15 | 0,202 | 3,218 | 0,00129082 | 0,00317365 | 0,002077 | 0,0034244 | 0,0027652 | 0,0027652 | TRUE | 0,0618646 | No |
| orf19.2541 | 606 | 546 | 583 | 718 | 726 | 769 | 790 | 431 | 544 | 695 | 596 | 592 | 621 | 734 | 705 | 608 | 636,85 | 607 | 667 | 1,104 | 0,143 | 1,598 | 0,11000813 | 0,16535622 | 0,007699 | 0,004259 | 0,006067 | 0,006067 | TRUE | 0,1062848 | No |
| orf19.2542 | 468 | 435 | 440 | 570 | 526 | 535 | 538 | 339 | 420 | 554 | 450 | 470 | 450 | 510 | 480 | 478 | 476,55 | 474 | 480 | 1,017 | 0,024 | 0,266 | 0,78988031 | 0,83622923 | 0,005991 | 0,0050258 | 0,0055063 | 0,0055063 | TRUE | 0,1399 | No |
| orf19.2544 | 403 | 265 | 331 | 422 | 396 | 358 | 351 | 234 | 362 | 337 | 339 | 348 | 339 | 342 | 313 | 330 | 338,63 | 346 | 331 | 0,954 | -0,068 | -0,877 | 0,3805641 | 0,4698022 | 1,00E-08 | 0,0062665 | 0,0028835 | 0,0028835 | TRUE | 0,0164121 | No |
| orf19.2545 | 5760 | 3755 | 5071 | 6166 | 5566 | 4620 | 5235 | 3245 | 5167 | 4781 | 5188 | 5085 | 4760 | 4408 | 4671 | 4577 | 4829,6 | 5055 | 4604 | 0,911 | -0,135 | -3,258 | 0,00112336 | 0,00280165 | 0,000953 | 0,0022801 | 0,0014459 | 0,0014459 | TRUE | 0,0157263 | No |
| orf19.2546 | 1906 | 1348 | 1673 | 2139 | 2366 | 2240 | 2432 | 1491 | 1710 | 1716 | 1712 | 1764 | 2023 | 2137 | 2170 | 2103 | 1916,9 | 1726 | 2108 | 1,221 | 0,289 | 6,914 | 4,72E-12 | 3,99E-11 | 1,00E-08 | 0,0027368 | 0,0011453 | 0,0011453 | TRUE | 0,0096422 | No |
| orf19.2547 | 3064 | 2409 | 2931 | 3470 | 3365 | 3428 | 3570 | 2131 | 2749 | 3067 | 2999 | 2861 | 2878 | 3271 | 3186 | 3006 | 3002 | 2919 | 3085 | 1,058 | 0,081 | 1,491 | 0,13598903 | 0,19859555 | 0,002384 | 0,0024631 | 0,0025074 | 0,0025074 | TRUE | 0,0258238 | No |
| orf19.2549 | 3554 | 2428 | 3043 | 3636 | 3335 | 2969 | 3134 | 1878 | 3188 | 3091 | 3113 | 2998 | 2852 | 2833 | 2797 | 2649 | 2940,2 | 3098 | 2783 | 0,9 | -0,153 | -3,811 | 0,00013838 | 0,00041239 | 0,000413 | 0,0024733 | 0,0011983 | 0,0011983 | TRUE | 0,0116198 | No |
| orf19.255 | 14644 | 11260 | 13961 | 16782 | 11443 | 11114 | 11718 | 7165 | 13137 | 14335 | 14283 | 13839 | 9786 | 10605 | 10456 | 10106 | 12068 | 13898 | 10238 | 0,737 | -0,441 | -10,487 | 9,95E-26 | 2,72E-24 | 0,001324 | 0,0020998 | 0,0016107 | 0,0016107 | TRUE | 0,016678 | No |
| orf19.2551 | 11153 | 7465 | 9800 | 11837 | 7293 | 6391 | 6606 | 4724 | 10005 | 9504 | 10026 | 9761 | 6237 | 6098 | 5895 | 6663 | 8023,7 | 9824 | 6223 | 0,633 | -0,66 | -15,161 | 6,38E-52 | 7,37E-50 | 0,001501 | 0,0021605 | 0,0016872 | 0,0016872 | TRUE | 0,0266937 | No |
| orf19.2552 | 616 | 435 | 563 | 691 | 943 | 906 | 930 | 601 | 553 | 554 | 576 | 570 | 806 | 864 | 830 | 848 | 700,09 | 563 | 837 | 1,485 | 0,57 | 9,859 | 6,26E-23 | 1,38E-21 | 1,00E-08 | 0,0040531 | 0,0017299 | 0,0017299 | TRUE | 0,0076903 | No |
| orf19.2553 | 220 | 189 | 224 | 299 | 329 | 348 | 357 | 200 | 197 | 241 | 229 | 247 | 281 | 332 | 319 | 282 | 265,97 | 228 | 304 | 1,333 | 0,415 | 4,059 | 4,92E-05 | 0,00015949 | 0,004799 | 0,0074376 | 0,0062009 | 0,0062009 | TRUE | 0,0985061 | No |
| orf19.2555 | 1872 | 1391 | 1701 | 2157 | 1849 | 1680 | 1703 | 1148 | 1679 | 1771 | 1740 | 1779 | 1581 | 1603 | 1520 | 1619 | 1661,6 | 1742 | 1581 | 0,907 | -0,141 | -3,22 | 0,0012837 | 0,00316213 | 1,00E-08 | 0,0028532 | 0,0012474 | 0,0012474 | TRUE | 0,0083314 | No |
| orf19.2557 | 2057 | 1487 | 1767 | 2211 | 2302 | 1944 | 2108 | 1303 | 1845 | 1893 | 1808 | 1823 | 1969 | 1855 | 1881 | 1838 | 1864 | 1842 | 1886 | 1,025 | 0,036 | 0,854 | 0,39285052 | 0,48158455 | 1,00E-08 | 0,0027583 | 0,0011749 | 0,0011749 | TRUE | 0,0109935 | No |
| orf19.2558 | 950 | 785 | 805 | 1093 | 881 | 821 | 880 | 560 | 852 | 999 | 824 | 901 | 753 | 783 | 785 | 790 | 836,06 | 894 | 778 | 0,871 | -0,199 | -2,97 | 0,00297549 | 0,00676381 | 0,002348 | 0,0037159 | 0,0030882 | 0,0030882 | TRUE | 0,0688564 | No |
| orf19.2559 | 2936 | 2098 | 2746 | 3164 | 4400 | 3795 | 4130 | 2542 | 2634 | 2671 | 2809 | 2609 | 3763 | 3621 | 3685 | 3585 | 3172,3 | 2681 | 3664 | 1,368 | 0,452 | 11,586 | 4,86E-31 | 1,84E-29 | 0,000353 | 0,0024371 | 0,001138 | 0,001138 | TRUE | 0,0127625 | No |
| orf19.2560 | 7859 | 5367 | 7160 | 8394 | 7977 | 6872 | 7414 | 4871 | 7050 | 6833 | 7325 | 6922 | 6822 | 6557 | 6616 | 6870 | 6874,4 | 7032 | 6716 | 0,955 | -0,067 | -1,894 | 0,05821778 | 0,09590688 | 0,000532 | 0,0021907 | 0,0010544 | 0,0010544 | TRUE | 0,0094979 | No |
| orf19.2563 | 1508 | 1057 | 1290 | 1656 | 1618 | 1406 | 1619 | 1016 | 1353 | 1346 | 1320 | 1366 | 1384 | 1342 | 1445 | 1433 | 1373,4 | 1346 | 1401 | 1,04 | 0,056 | 1,22 | 0,22260106 | 0,30162093 | 1,00E-08 | 0,0030366 | 0,001292 | 0,001292 | TRUE | 0,0097489 | No |
| orf19.2564 | 1732 | 1250 | 1414 | 1842 | 2040 | 1739 | 1959 | 1200 | 1554 | 1591 | 1447 | 1519 | 1745 | 1659 | 1748 | 1692 | 1619,4 | 1528 | 1711 | 1,121 | 0,165 | 3,557 | 0,00037554 | 0,00104167 | 0,000432 | 0,002876 | 0,0014551 | 0,0014551 | TRUE | 0,0148014 | No |
| orf19.2565 | 1981 | 1365 | 1851 | 2135 | 2550 | 2343 | 2507 | 1446 | 1777 | 1738 | 1894 | 1761 | 2181 | 2236 | 2237 | 2039 | 1982,8 | 1792 | 2173 | 1,214 | 0,28 | 5,819 | 5,91E-09 | 3,39E-08 | 0,000988 | 0,0027117 | 0,0017123 | 0,0017123 | TRUE | 0,0190424 | No |
| orf19.2568 | 7271 | 4263 | 5936 | 7220 | 6834 | 5240 | 6051 | 4055 | 6523 | 5427 | 6073 | 5954 | 5845 | 5000 | 5400 | 5719 | 5742,5 | 5994 | 5491 | 0,916 | -0,127 | -1,953 | 0,05079324 | 0,08531334 | 0,00495 | 0,0022323 | 0,003902 | 0,003902 | TRUE | 0,0488623 | No |
| orf19.257 | 2 | 2 | 0 | 4 | 4 | 3 | 4 | 1 | 2 | 3 | 0 | 3 | 3 | 3 | 4 | 1 | 2,36 | 2 | 3 | 1,5 | 0,585 | 0,647 | 0,51766303 | NA | 1,00E-08 | 0,6163895 | 0,355189 | 0,355189 | TRUE | 0,3771028 | No |
| orf19.2570 | 2457 | 1629 | 2042 | 2416 | 1982 | 1728 | 1912 | 1149 | 2204 | 2074 | 2089 | 1992 | 1695 | 1649 | 1706 | 1621 | 1878,8 | 2090 | 1668 | 0,799 | -0,324 | -7 | 2,55E-12 | 2,23E-11 | 0,000764 | 0,0027522 | 0,0015159 | 0,0015159 | TRUE | 0,0171645 | No |
| orf19.2571 | 4184 | 3175 | 3941 | 4794 | 4179 | 3852 | 4319 | 2484 | 3753 | 4042 | 4032 | 3953 | 3574 | 3675 | 3854 | 3503 | 3798,5 | 3945 | 3652 | 0,927 | -0,11 | -2,5 | 0,01241831 | 0,02439007 | 0,001093 | 0,0023617 | 0,0015916 | 0,0015916 | TRUE | 0,0170268 | No |
| orf19.2573 | 6295 | 4505 | 5638 | 7037 | 6129 | 5396 | 5837 | 3713 | 5647 | 5735 | 5768 | 5803 | 5242 | 5149 | 5209 | 5237 | 5473,7 | 5738 | 5209 | 0,908 | -0,14 | -4,935 | 8,03E-07 | 3,44E-06 | 1,00E-08 | 0,0022447 | 0,0005871 | 0,0005871 | TRUE | 0,0014558 | No |
| orf19.2574 | 3484 | 2263 | 3120 | 3878 | 3906 | 3302 | 3693 | 2232 | 3126 | 2881 | 3192 | 3198 | 3341 | 3151 | 3295 | 3148 | 3166,4 | 3099 | 3234 | 1,043 | 0,061 | 1,327 | 0,18434845 | 0,25649965 | 0,001205 | 0,002438 | 0,0016968 | 0,0016968 | TRUE | 0,0260349 | No |
| orf19.2575 | 4841 | 3297 | 4121 | 4988 | 4112 | 3627 | 3907 | 2502 | 4343 | 4197 | 4216 | 4113 | 3517 | 3461 | 3486 | 3529 | 3857,8 | 4217 | 3498 | 0,829 | -0,27 | -8,118 | 4,73E-16 | 5,99E-15 | 1,00E-08 | 0,0023558 | 0,0007998 | 0,0007998 | TRUE | 0,0050892 | No |
| orf19.2579 | 816 | 512 | 714 | 792 | 1081 | 948 | 1129 | 679 | 732 | 652 | 730 | 653 | 925 | 905 | 1007 | 958 | 820,2 | 692 | 949 | 1,37 | 0,454 | 6,964 | 3,32E-12 | 2,86E-11 | 0,002027 | 0,0037495 | 0,0028284 | 0,0028284 | TRUE | 0,0222929 | No |
| orf19.258 | 696 | 570 | 647 | 858 | 428 | 397 | 410 | 283 | 624 | 726 | 662 | 708 | 366 | 379 | 366 | 399 | 528,67 | 680 | 378 | 0,554 | -0,852 | -11,585 | 4,91E-31 | 1,85E-29 | 0,001618 | 0,0047255 | 0,00312 | 0,00312 | TRUE | 0,0366366 | No |
| orf19.2580 | 736 | 470 | 608 | 753 | 827 | 744 | 843 | 449 | 660 | 598 | 622 | 621 | 707 | 710 | 752 | 633 | 663,04 | 625 | 700 | 1,123 | 0,167 | 2,484 | 0,0129966 | 0,02541783 | 0,001525 | 0,004169 | 0,0028491 | 0,0028491 | TRUE | 0,0429471 | No |
| orf19.2581 | 69 | 45 | 53 | 62 | 95 | 80 | 100 | 62 | 62 | 57 | 54 | 51 | 81 | 76 | 89 | 87 | 69,85 | 56 | 83 | 1,488 | 0,573 | 3,464 | 0,00053261 | 0,00142727 | 1,00E-08 | 0,0227627 | 0,0114727 | 0,0114727 | TRUE | 0,047662 | No |
| orf19.2582 | 1641 | 1145 | 1404 | 1754 | 1939 | 1850 | 1916 | 1106 | 1472 | 1458 | 1436 | 1446 | 1658 | 1765 | 1710 | 1560 | 1563,2 | 1453 | 1673 | 1,154 | 0,206 | 4,253 | 2,11E-05 | 7,30E-05 | 0,000559 | 0,0029082 | 0,0016151 | 0,0016151 | TRUE | 0,0223487 | No |
| orf19.2583 | 5 | 2 | 2 | 3 | 9 | 7 | 13 | 4 | 4 | 3 | 2 | 2 | 8 | 7 | 12 | 6 | 5,4 | 3 | 8 | 2,749 | 1,459 | 2,317 | 0,02051162 | NA | 1,00E-08 | 0,2709998 | 0,149701 | 0,149701 | TRUE | 0,3180294 | No |
| orf19.2585 | 9 | 15 | 11 | 13 | 9 | 12 | 17 | 19 | 8 | 19 | 11 | 11 | 8 | 11 | 15 | 27 | 13,78 | 12 | 15 | 1,227 | 0,296 | 0,663 | 0,50739904 | NA | 0,126164 | 0,1073096 | 0,1155037 | 0,1155037 | TRUE | 0,6084371 | No |
| orf19.2587 | 10 | 3 | 11 | 8 | 6 | 7 | 5 | 6 | 9 | 4 | 11 | 7 | 5 | 7 | 4 | 8 | 6,92 | 8 | 6 | 0,777 | -0,364 | -0,685 | 0,49305603 | NA | 1,00E-08 | 0,2117021 | 0,1245765 | 0,1245765 | TRUE | 0,1766134 | No |
| orf19.259 | 1393 | 958 | 1207 | 1603 | 1562 | 1439 | 1487 | 905 | 1250 | 1220 | 1235 | 1322 | 1336 | 1373 | 1327 | 1276 | 1292,3 | 1257 | 1328 | 1,057 | 0,08 | 1,634 | 0,10216878 | 0,15543936 | 0,000307 | 0,0031029 | 0,0015137 | 0,0015137 | TRUE | 0,015435 | No |
| orf19.2590 | 6 | 4 | 10 | 13 | 13 | 4 | 4 | 10 | 5 | 5 | 10 | 11 | 11 | 4 | 4 | 14 | 8 | 8 | 8 | 0,995 | -0,008 | -0,013 | 0,98937137 | NA | 0,172414 | 0,1833455 | 0,1789401 | 0,1789401 | TRUE | 0,2283395 | No |
| orf19.2591 | 59 | 46 | 62 | 76 | 51 | 33 | 47 | 29 | 53 | 59 | 63 | 63 | 44 | 31 | 42 | 41 | 49,44 | 60 | 40 | 0,665 | -0,589 | -2,958 | 0,00309641 | 0,00700687 | 1,00E-08 | 0,0313409 | 0,0170311 | 0,0170311 | TRUE | 0,1496801 | No |
| orf19.2593 | 124 | 75 | 100 | 113 | 94 | 84 | 93 | 69 | 111 | 95 | 102 | 93 | 80 | 80 | 83 | 97 | 92,88 | 100 | 85 | 0,839 | -0,253 | -1,74 | 0,08187959 | 0,12914767 | 1,00E-08 | 0,0176089 | 0,0094664 | 0,0094664 | TRUE | 0,065643 | No |
| orf19.2594 | 5896 | 3937 | 4743 | 6308 | 4970 | 4331 | 4753 | 2970 | 5289 | 5012 | 4853 | 5202 | 4250 | 4133 | 4241 | 4189 | 4646,1 | 5089 | 4203 | 0,826 | -0,276 | -7,247 | 4,25E-13 | 4,12E-12 | 0,000657 | 0,002292 | 0,0011773 | 0,0011773 | TRUE | 0,0120924 | No |
| orf19.2597 | 662 | 458 | 562 | 760 | 708 | 627 | 721 | 431 | 594 | 583 | 575 | 627 | 606 | 598 | 643 | 608 | 604,21 | 595 | 614 | 1,031 | 0,044 | 0,703 | 0,48233975 | 0,56778278 | 1,00E-08 | 0,0043822 | 0,0021122 | 0,0021122 | TRUE | 0,01661 | No |
| orf19.2598 | 1977 | 1395 | 1711 | 2128 | 2272 | 2080 | 2118 | 1285 | 1774 | 1776 | 1751 | 1755 | 1943 | 1985 | 1890 | 1812 | 1835,6 | 1764 | 1908 | 1,083 | 0,116 | 2,7 | 0,00693734 | 0,01450227 | 0,000138 | 0,0027704 | 0,0012167 | 0,0012167 | TRUE | 0,01204 | No |
| orf19.2599 | 2790 | 1857 | 2376 | 2922 | 2339 | 1889 | 2144 | 1456 | 2503 | 2364 | 2431 | 2410 | 2000 | 1802 | 1913 | 2054 | 2184,6 | 2427 | 1942 | 0,799 | -0,324 | -6,606 | 3,96E-11 | 3,05E-10 | 0,001264 | 0,002644 | 0,001838 | 0,001838 | TRUE | 0,028261 | No |
| orf19.26 | 1574 | 1076 | 1387 | 1801 | 2122 | 1983 | 2055 | 1365 | 1412 | 1370 | 1419 | 1485 | 1815 | 1892 | 1834 | 1925 | 1644 | 1422 | 1866 | 1,31 | 0,389 | 8,591 | 8,59E-18 | 1,31E-16 | 0,000296 | 0,0028626 | 0,0013489 | 0,0013489 | TRUE | 0,0113454 | No |
| orf19.260 | 6085 | 4376 | 5526 | 6561 | 6578 | 6064 | 6574 | 3882 | 5459 | 5571 | 5654 | 5410 | 5626 | 5786 | 5866 | 5475 | 5605,9 | 5524 | 5688 | 1,031 | 0,044 | 1,24 | 0,21496207 | 0,29285535 | 0,000426 | 0,0022385 | 0,001016 | 0,001016 | TRUE | 0,0073023 | No |
| orf19.2600 | 1355 | 1068 | 1241 | 1551 | 1203 | 1164 | 1234 | 807 | 1216 | 1360 | 1270 | 1279 | 1029 | 1111 | 1101 | 1138 | 1187,8 | 1281 | 1095 | 0,854 | -0,227 | -4,152 | 3,29E-05 | 0,00010997 | 0,001095 | 0,0032017 | 0,0020204 | 0,0020204 | TRUE | 0,0203257 | No |
| orf19.2601 | 10525 | 7266 | 9628 | 11430 | 10697 | 9158 | 9865 | 6861 | 9442 | 9250 | 9850 | 9425 | 9148 | 8738 | 8803 | 9677 | 9291,8 | 9492 | 9092 | 0,957 | -0,063 | -1,503 | 0,13288524 | 0,19486808 | 0,001332 | 0,0021358 | 0,0015787 | 0,0015787 | TRUE | 0,0223676 | No |
| orf19.2602 | 1036 | 708 | 1068 | 1264 | 994 | 801 | 915 | 609 | 929 | 901 | 1093 | 1042 | 850 | 764 | 816 | 859 | 906,94 | 991 | 822 | 0,828 | -0,273 | -3,712 | 0,00020582 | 0,00059854 | 0,004467 | 0,0035802 | 0,0040611 | 0,0040611 | TRUE | 0,0545904 | No |
| orf19.2604 | 5240 | 3442 | 4473 | 5592 | 6324 | 5295 | 5911 | 3611 | 4701 | 4382 | 4576 | 4611 | 5408 | 5052 | 5275 | 5093 | 4887,4 | 4568 | 5207 | 1,14 | 0,189 | 4,884 | 1,04E-06 | 4,38E-06 | 0,000663 | 0,0022766 | 0,0012301 | 0,0012301 | TRUE | 0,0087549 | No |
| orf19.2605 | 2631 | 1847 | 2430 | 2863 | 2641 | 2409 | 2564 | 1618 | 2360 | 2351 | 2486 | 2361 | 2259 | 2299 | 2288 | 2282 | 2335,7 | 2390 | 2282 | 0,955 | -0,067 | -1,761 | 0,07829138 | 0,12410939 | 1,00E-08 | 0,002601 | 0,0009483 | 0,0009483 | TRUE | 0,0088469 | No |
| orf19.2606 | 2468 | 1766 | 2129 | 2611 | 2151 | 1989 | 2107 | 1415 | 2214 | 2248 | 2178 | 2153 | 1840 | 1898 | 1880 | 1996 | 2050,9 | 2198 | 1904 | 0,865 | -0,21 | -5,042 | 4,61E-07 | 2,05E-06 | 0,000132 | 0,0026874 | 0,00117 | 0,00117 | TRUE | 0,0115499 | No |
| orf19.2607 | 806 | 583 | 694 | 967 | 939 | 773 | 895 | 633 | 723 | 742 | 710 | 797 | 803 | 738 | 799 | 893 | 775,6 | 743 | 808 | 1,083 | 0,115 | 1,648 | 0,09937255 | 0,15207974 | 0,002749 | 0,0038512 | 0,0033848 | 0,0033848 | TRUE | 0,0530197 | No |
| orf19.2608 | 1E+05 | 70472 | 97786 | 1E+05 | 2E+05 | 1E+05 | 1E+05 | 82223 | 99498 | 89718 | 1E+05 | 98486 | 1E+05 | 1E+05 | 1E+05 | 1E+05 | 108436 | 96937 | 1E+05 | 1,237 | 0,307 | 6,191 | 5,96E-10 | 3,92E-09 | 0,002534 | 0,0019929 | 0,0023553 | 0,0023553 | TRUE | 0,0307562 | No |
| orf19.2609 | 9164 | 5928 | 7809 | 9529 | 9664 | 8023 | 9319 | 5670 | 8221 | 7547 | 7989 | 7858 | 8265 | 7655 | 8316 | 7997 | 7981 | 7904 | 8058 | 1,019 | 0,028 | 0,668 | 0,50417438 | 0,58874241 | 0,001129 | 0,0021614 | 0,0015284 | 0,0015284 | TRUE | 0,0140518 | No |
| orf19.261 | 1881 | 1540 | 1769 | 2127 | 1738 | 1831 | 1810 | 1080 | 1687 | 1961 | 1810 | 1754 | 1486 | 1747 | 1615 | 1523 | 1698 | 1803 | 1593 | 0,885 | -0,176 | -2,692 | 0,00710678 | 0,01479983 | 0,003953 | 0,0028345 | 0,0035212 | 0,0035212 | TRUE | 0,0515292 | No |

| | | | | | | | | | | | | | | | | | | | | | | | | | | | | | | | |
|---|---|---|---|---|---|---|---|---|---|---|---|---|---|---|---|---|---|---|---|---|---|---|---|---|---|---|---|---|---|---|---|
| orf19.2610 | 1448 | 982 | 1186 | 1483 | 1320 | 1044 | 1093 | 766 | 1299 | 1250 | 1213 | 1223 | 1129 | 996 | 975 | 1080 | 1145,8 | 1246 | 1045 | 0,838 | -0,255 | -4,267 | 1,98E-05 | 6,88E-05 | 0,002009 | 0,0032465 | 0,0025466 | 0,0025466 | TRUE | 0,0383488 | No |
| orf19.2611 | 2490 | 1637 | 2189 | 2574 | 2317 | 1938 | 2134 | 1281 | 2234 | 2084 | 2240 | 2123 | 1982 | 1849 | 1904 | 1807 | 2027,7 | 2170 | 1886 | 0,87 | -0,201 | -4,293 | 1,76E-05 | 6,18E-05 | 0,000797 | 0,0026955 | 0,0016156 | 0,0016156 | TRUE | 0,0143907 | No |
| orf19.2612 | 2267 | 1559 | 1934 | 2548 | 2433 | 1736 | 2067 | 1535 | 2034 | 1985 | 1979 | 2101 | 2081 | 1656 | 1844 | 2165 | 1980,6 | 2025 | 1936 | 0,955 | -0,066 | -0,862 | 0,38846957 | 0,4775783 | 0,007069 | 0,0027125 | 0,0051218 | 0,0051218 | TRUE | 0,1146302 | No |
| orf19.2613 | 14190 | 9428 | 12153 | 15047 | 16506 | 13696 | 15090 | 9402 | 12730 | 12003 | 12434 | 12408 | 14116 | 13068 | 13465 | 13261 | 12936 | 12394 | 13478 | 1,087 | 0,121 | 3,367 | 0,00076013 | 0,0019617 | 0,000759 | 0,0020917 | 0,0011619 | 0,0011619 | TRUE | 0,0125744 | No |
| orf19.2614 | 2142 | 1479 | 1932 | 2424 | 2263 | 1910 | 2120 | 1425 | 1922 | 1883 | 1977 | 1999 | 1935 | 1822 | 1892 | 2010 | 1929,9 | 1945 | 1915 | 0,982 | -0,026 | -0,565 | 0,57222936 | 0,65101322 | 0,000516 | 0,0027317 | 0,0014494 | 0,0014494 | TRUE | 0,0124383 | No |
| orf19.2615 | 457 | 351 | 463 | 535 | 743 | 626 | 743 | 405 | 410 | 447 | 474 | 441 | 635 | 597 | 663 | 571 | 529,83 | 443 | 616 | 1,399 | 0,484 | 6,511 | 7,46E-11 | 5,52E-10 | 0,002006 | 0,0047194 | 0,0033658 | 0,0033658 | TRUE | 0,0303032 | No |
| orf19.2616 | 4629 | 3048 | 4109 | 4932 | 4883 | 4392 | 4796 | 2839 | 4153 | 3880 | 4204 | 4067 | 4176 | 4191 | 4280 | 4004 | 4119,3 | 4076 | 4163 | 1,021 | 0,031 | 0,777 | 0,43694266 | 0,52377325 | 0,000667 | 0,0023319 | 0,0012471 | 0,0012471 | TRUE | 0,0120644 | No |
| orf19.2618 | 7319 | 4816 | 6120 | 7605 | 9361 | 7726 | 8456 | 5278 | 6566 | 6131 | 6261 | 6271 | 8006 | 7372 | 7546 | 7444 | 6949,7 | 6307 | 7592 | 1,204 | 0,267 | 6,738 | 1,61E-11 | 1,28E-10 | 0,000935 | 0,0021884 | 0,001367 | 0,001367 | TRUE | 0,0166902 | No |
| orf19.2619 | 375 | 259 | 347 | 416 | 423 | 416 | 429 | 310 | 336 | 330 | 355 | 343 | 362 | 397 | 383 | 437 | 367,87 | 341 | 395 | 1,15 | 0,201 | 2,488 | 0,01283462 | 0,0251174 | 0,000815 | 0,0059258 | 0,0035604 | 0,0035604 | TRUE | 0,0517917 | No |
| orf19.262 | 96 | 82 | 93 | 103 | 95 | 93 | 121 | 66 | 86 | 104 | 95 | 85 | 81 | 89 | 108 | 93 | 92,71 | 92 | 93 | 1,007 | 0,011 | 0,071 | 0,94374779 | 0,95939708 | 1,00E-08 | 0,0176388 | 0,0112998 | 0,0112998 | TRUE | 0,1335601 | No |
| orf19.2620 | 190 | 131 | 194 | 264 | 262 | 203 | 255 | 184 | 170 | 167 | 198 | 218 | 224 | 194 | 228 | 260 | 207,28 | 188 | 226 | 1,19 | 0,251 | 2,058 | 0,03956076 | 0,06843682 | 0,009952 | 0,0089831 | 0,0094286 | 0,0094286 | TRUE | 0,1271716 | No |
| orf19.2621 | 805 | 646 | 708 | 960 | 795 | 833 | 787 | 502 | 722 | 822 | 724 | 792 | 680 | 795 | 702 | 708 | 743,2 | 765 | 721 | 0,944 | -0,084 | -1,171 | 0,24177094 | 0,32322203 | 0,003193 | 0,0039328 | 0,003538 | 0,003538 | TRUE | 0,0575881 | No |
| orf19.2622 | 2566 | 1818 | 2135 | 2671 | 2251 | 2154 | 2175 | 1297 | 2302 | 2315 | 2184 | 2203 | 1925 | 2055 | 1941 | 1829 | 2094,2 | 2251 | 1938 | 0,862 | -0,214 | -4,531 | 5,87E-06 | 2,22E-05 | 0,000944 | 0,0026727 | 0,0016509 | 0,0016509 | TRUE | 0,0197652 | No |
| orf19.2623 | 13221 | 9070 | 12003 | 14779 | 15021 | 13342 | 14429 | 9492 | 11861 | 11547 | 12280 | 12187 | 12846 | 12731 | 12875 | 13388 | 12464 | 11969 | 12960 | 1,082 | 0,114 | 3,444 | 0,00057241 | 0,00152305 | 0,000535 | 0,002096 | 0,0009723 | 0,0009723 | TRUE | 0,0067194 | No |
| orf19.2624 | 13 | 8 | 12 | 10 | 5 | 14 | 6 | 3 | 12 | 10 | 12 | 8 | 4 | 13 | 5 | 4 | 8,7 | 10 | 6 | 0,652 | -0,617 | -1,228 | 0,21954262 | NA | 0,049395 | 0,168865 | 0,1227991 | 0,1227991 | TRUE | 1,150498 | No |
| orf19.2625 | 1 | 0 | 0 | 0 | 1 | 2 | 2 | 0 | 1 | 0 | 0 | 0 | 1 | 2 | 2 | 0 | 0,68 | 0 | 1 | 3,931 | 1,975 | 1,148 | 0,25080659 | NA | 1,00E-08 | 2,1347404 | 1,4142346 | 1,4142346 | TRUE | 0,2895963 | No |
| orf19.2629 | 1124 | 835 | 985 | 1176 | 1181 | 1149 | 1264 | 736 | 1008 | 1063 | 1008 | 970 | 1010 | 1096 | 1128 | 1038 | 1040,2 | 1012 | 1068 | 1,058 | 0,081 | 1,432 | 0,15200574 | 0,21806363 | 0,001103 | 0,0033752 | 0,0021016 | 0,0021016 | TRUE | 0,0179554 | No |
| orf19.2630 | 555 | 405 | 469 | 614 | 649 | 655 | 590 | 372 | 498 | 516 | 480 | 506 | 555 | 625 | 526 | 525 | 528,85 | 500 | 558 | 1,118 | 0,161 | 2,169 | 0,03011129 | 0,05377784 | 0,00214 | 0,0047245 | 0,0034085 | 0,0034085 | TRUE | 0,0774143 | No |
| orf19.2631 | 965 | 788 | 944 | 1082 | 1311 | 1287 | 1251 | 815 | 866 | 1003 | 966 | 892 | 1121 | 1228 | 1116 | 1149 | 1042,8 | 932 | 1154 | 1,241 | 0,312 | 4,992 | 5,98E-07 | 2,61E-06 | 0,002048 | 0,0033717 | 0,0027682 | 0,0027682 | TRUE | 0,0299719 | No |
| orf19.263.1 | 8866 | 5947 | 7428 | 9434 | 7503 | 5863 | 6582 | 3739 | 7954 | 7571 | 7600 | 7779 | 6417 | 5594 | 5873 | 5274 | 6757,7 | 7726 | 5790 | 0,75 | -0,415 | -7,256 | 3,99E-13 | 3,88E-12 | 0,003568 | 0,0021943 | 0,0029963 | 0,0029963 | TRUE | 0,0650564 | No |
| orf19.2633 | 3279 | 2180 | 2998 | 3786 | 3499 | 2988 | 3282 | 2188 | 2942 | 2775 | 3067 | 3122 | 2992 | 2851 | 2929 | 3086 | 2970,5 | 2976 | 2964 | 0,994 | -0,008 | -0,17 | 0,8649284 | 0,89733894 | 0,001294 | 0,0024682 | 0,0018697 | 0,0018697 | TRUE | 0,0240901 | No |
| orf19.2633.1 | 210 | 126 | 155 | 191 | 236 | 217 | 230 | 140 | 188 | 160 | 159 | 158 | 202 | 207 | 205 | 197 | 184,56 | 166 | 203 | 1,22 | 0,287 | 2,716 | 0,00659837 | 0,01389521 | 1,00E-08 | 0,0098453 | 0,0052668 | 0,0052668 | TRUE | 0,0920214 | No |
| orf19.2637 | 928 | 672 | 819 | 1070 | 1217 | 1136 | 1164 | 694 | 833 | 856 | 838 | 882 | 1041 | 1084 | 1039 | 979 | 943,82 | 852 | 1036 | 1,217 | 0,284 | 5,278 | 1,30E-07 | 6,30E-07 | 1,00E-08 | 0,0035176 | 0,0017084 | 0,0017084 | TRUE | 0,0139436 | No |
| orf19.2638 | 591 | 401 | 536 | 665 | 467 | 443 | 460 | 300 | 530 | 511 | 548 | 548 | 399 | 423 | 410 | 423 | 474,14 | 534 | 414 | 0,772 | -0,373 | -5,539 | 3,05E-08 | 1,59E-07 | 1,00E-08 | 0,0050413 | 0,0022214 | 0,0022214 | TRUE | 0,0091953 | No |
| orf19.2638.1 | 3370 | 2377 | 2760 | 3448 | 2999 | 2620 | 2603 | 1476 | 3023 | 3026 | 2824 | 2843 | 2565 | 2500 | 2323 | 2082 | 2648,2 | 2929 | 2368 | 0,81 | -0,305 | -4,724 | 2,31E-06 | 9,30E-06 | 0,004372 | 0,0025277 | 0,0036086 | 0,0036086 | TRUE | 0,0763314 | No |
| orf19.2639 | 1283 | 910 | 1167 | 1408 | 1774 | 1532 | 1725 | 1026 | 1151 | 1159 | 1194 | 1161 | 1517 | 1462 | 1539 | 1447 | 1328,7 | 1166 | 1491 | 1,281 | 0,357 | 7,802 | 6,10E-15 | 6,99E-14 | 1,00E-08 | 0,0030721 | 0,0012479 | 0,0012479 | TRUE | 0,0055447 | No |
| orf19.2639.1 | 1434 | 1024 | 1188 | 1439 | 1255 | 1147 | 1199 | 625 | 1286 | 1304 | 1215 | 1187 | 1073 | 1094 | 1070 | 882 | 1138,9 | 1248 | 1030 | 0,829 | -0,271 | -3,843 | 0,00012173 | 0,0003664 | 0,004563 | 0,0032542 | 0,0038995 | 0,0038995 | TRUE | 0,1025863 | No |
| orf19.264 | 937 | 755 | 859 | 1137 | 881 | 940 | 1004 | 603 | 841 | 961 | 879 | 938 | 753 | 897 | 896 | 850 | 876,87 | 905 | 849 | 0,939 | -0,091 | -1,248 | 0,21202075 | 0,28943892 | 0,004132 | 0,0036351 | 0,003908 | 0,003908 | TRUE | 0,0728383 | No |
| orf19.2640 | 1793 | 1234 | 1547 | 1817 | 1693 | 1574 | 1560 | 913 | 1609 | 1571 | 1583 | 1498 | 1448 | 1502 | 1392 | 1288 | 1486,3 | 1565 | 1408 | 0,902 | -0,15 | -2,708 | 0,00675884 | 0,01419333 | 0,00167 | 0,0029563 | 0,0022483 | 0,0022483 | TRUE | 0,0355391 | No |
| orf19.2641 | 117 | 98 | 123 | 137 | 135 | 114 | 125 | 70 | 105 | 125 | 126 | 113 | 115 | 109 | 112 | 99 | 112,88 | 117 | 109 | 0,937 | -0,094 | -0,711 | 0,47729717 | 0,56266491 | 1,00E-08 | 0,0148401 | 0,0078612 | 0,0078612 | TRUE | 0,0504849 | No |
| orf19.2642 | 70 | 40 | 63 | 55 | 35 | 49 | 51 | 28 | 63 | 51 | 64 | 45 | 30 | 47 | 46 | 39 | 48,15 | 56 | 40 | 0,723 | -0,469 | -2,135 | 0,03278663 | 0,05797001 | 0,015103 | 0,0321276 | 0,0249599 | 0,0249599 | TRUE | 0,2770209 | No |
| orf19.2643 | 5559 | 3682 | 4780 | 5872 | 5342 | 4441 | 5065 | 3075 | 4987 | 4688 | 4890 | 4842 | 4569 | 4237 | 4520 | 4337 | 4633,7 | 4852 | 4416 | 0,91 | -0,136 | -3,477 | 0,00050639 | 0,00136613 | 0,00066 | 0,0022928 | 0,0012457 | 0,0012457 | TRUE | 0,0093731 | No |
| orf19.2644 | 2728 | 1948 | 2533 | 3014 | 2837 | 2685 | 2665 | 1956 | 2447 | 2480 | 2592 | 2485 | 2426 | 2562 | 2378 | 2759 | 2516,2 | 2501 | 2531 | 1,01 | 0,015 | 0,284 | 0,77663606 | 0,82491479 | 0,00187 | 0,0025565 | 0,0022429 | 0,0022429 | TRUE | 0,0420763 | No |
| orf19.2645 | 42 | 30 | 41 | 45 | 40 | 50 | 35 | 27 | 38 | 38 | 42 | 37 | 34 | 48 | 31 | 38 | 38,27 | 39 | 38 | 0,974 | -0,038 | -0,168 | 0,86683123 | 0,89822733 | 1,00E-08 | 0,0399131 | 0,0228332 | 0,0228332 | TRUE | 0,2522658 | No |
| orf19.2646 | 1910 | 1483 | 1836 | 2154 | 1871 | 2001 | 2141 | 1261 | 1713 | 1888 | 1878 | 1776 | 1600 | 1909 | 1910 | 1779 | 1806,8 | 1814 | 1800 | 0,993 | -0,011 | -0,163 | 0,87085844 | 0,90111898 | 0,00405 | 0,002783 | 0,0035199 | 0,0035199 | TRUE | 0,0700026 | No |
| orf19.2647 | 114 | 107 | 128 | 140 | 131 | 141 | 135 | 82 | 102 | 136 | 131 | 115 | 112 | 135 | 120 | 116 | 120,95 | 121 | 121 | 1,003 | 0,005 | 0,033 | 0,9734928 | 0,98139755 | 0,003627 | 0,0139822 | 0,0095651 | 0,0095651 | TRUE | 0,1147197 | No |
| orf19.2649 | 194 | 158 | 168 | 229 | 267 | 273 | 334 | 169 | 174 | 201 | 172 | 189 | 228 | 260 | 298 | 238 | 220,14 | 184 | 256 | 1,4 | 0,485 | 4,324 | 1,54E-05 | 5,45E-05 | 0,006196 | 0,0085739 | 0,0074255 | 0,0074255 | TRUE | 0,1405507 | No |
| orf19.265 | 3696 | 3109 | 3477 | 4363 | 5820 | 5882 | 5786 | 3606 | 3316 | 3958 | 3557 | 3598 | 4977 | 5612 | 5163 | 5086 | 4408,5 | 3607 | 5210 | 1,445 | 0,531 | 8,901 | 5,54E-19 | 9,29E-18 | 0,00382 | 0,0023088 | 0,0031879 | 0,0031879 | TRUE | 0,0510692 | No |
| orf19.2650 | 1179 | 1034 | 1137 | 1331 | 957 | 1128 | 1099 | 679 | 1058 | 1316 | 1163 | 1098 | 818 | 1076 | 981 | 958 | 1058,5 | 1159 | 958 | 0,828 | -0,272 | -3,008 | 0,00263362 | 0,00606005 | 0,009717 | 0,003351 | 0,0069196 | 0,0069196 | TRUE | 0,1194337 | No |
| orf19.2650.1 | 396 | 226 | 318 | 433 | 494 | 369 | 465 | 206 | 355 | 288 | 325 | 357 | 422 | 352 | 415 | 291 | 350,68 | 331 | 370 | 1,12 | 0,163 | 1,366 | 0,1719174 | 0,24234625 | 0,015917 | 0,0061192 | 0,0108138 | 0,0108138 | TRUE | 0,2157718 | No |
| orf19.2651 | 3771 | 2910 | 3317 | 4271 | 3548 | 3566 | 3520 | 2213 | 3383 | 3705 | 3394 | 3522 | 3034 | 3403 | 3141 | 3121 | 3337,8 | 3501 | 3175 | 0,907 | -0,14 | -2,838 | 0,00453526 | 0,00988315 | 0,001767 | 0,0024144 | 0,0020394 | 0,0020394 | TRUE | 0,0287504 | No |
| orf19.2652 | 148 | 121 | 125 | 150 | 95 | 106 | 118 | 62 | 133 | 154 | 128 | 124 | 81 | 101 | 105 | 87 | 114,19 | 135 | 94 | 0,702 | -0,51 | -3,631 | 0,00028203 | 0,00080078 | 0,003053 | 0,0146925 | 0,009878 | 0,009878 | TRUE | 0,0958831 | No |
| orf19.2653 | 2360 | 1788 | 2178 | 2751 | 2485 | 2546 | 2559 | 1619 | 2117 | 2276 | 2228 | 2269 | 2125 | 2429 | 2283 | 2283 | 2251,5 | 2222 | 2280 | 1,026 | 0,037 | 0,728 | 0,46667847 | 0,55258263 | 0,001682 | 0,0026243 | 0,0020418 | 0,0020418 | TRUE | 0,0265169 | No |
| orf19.2654 | 3205 | 2554 | 2992 | 3571 | 3420 | 3528 | 3554 | 2110 | 2875 | 3252 | 3061 | 2945 | 2925 | 3366 | 3171 | 2976 | 3071,4 | 3033 | 3110 | 1,026 | 0,038 | 0,648 | 0,51667029 | 0,60041684 | 0,002919 | 0,0024522 | 0,0028926 | 0,0028926 | TRUE | 0,037646 | No |
| orf19.2655 | 18199 | 13371 | 17100 | 20815 | 24787 | 23324 | 24963 | 14754 | 16326 | 17023 | 17495 | 17164 | 21199 | 22255 | 22275 | 20809 | 19318 | 17002 | 21634 | 1,273 | 0,348 | 9,322 | 1,14E-20 | 2,17E-19 | 0,000942 | 0,0020547 | 0,001285 | 0,001285 | TRUE | 0,0088154 | No |
| orf19.2657 | 30 | 16 | 23 | 25 | 22 | 32 | 28 | 20 | 27 | 20 | 24 | 21 | 19 | 31 | 25 | 28 | 24,25 | 23 | 26 | 1,107 | 0,147 | 0,52 | 0,60310343 | 0,67870822 | 1,00E-08 | 0,0618523 | 0,0356991 | 0,0356991 | TRUE | 0,233057 | No |
| orf19.2660 | 2742 | 2254 | 2613 | 3258 | 3036 | 3255 | 3287 | 2209 | 2460 | 2870 | 2673 | 2687 | 2596 | 3106 | 2933 | 3116 | 2805,1 | 2672 | 2938 | 1,099 | 0,136 | 2,012 | 0,04421437 | 0,07555174 | 0,004896 | 0,0024971 | 0,0040505 | 0,0040505 | TRUE | 0,0752186 | No |
| orf19.2661 | 1167 | 661 | 979 | 1327 | 1402 | 1053 | 1151 | 699 | 1047 | 842 | 1002 | 1094 | 1199 | 1005 | 1027 | 986 | 1025,1 | 996 | 1054 | 1,057 | 0,081 | 0,91 | 0,36296094 | 0,45167136 | 0,008758 | 0,0033956 | 0,0065564 | 0,0065564 | TRUE | 0,1250733 | No |
| orf19.2662 | 228 | 110 | 168 | 230 | 212 | 180 | 194 | 114 | 205 | 140 | 172 | 190 | 181 | 172 | 173 | 161 | 174,14 | 177 | 172 | 0,968 | -0,047 | -0,377 | 0,706115 | 0,76701119 | 0,007113 | 0,0103162 | 0,0089376 | 0,0089376 | TRUE | 0,1809168 | No |
| orf19.2663 | 709 | 482 | 634 | 771 | 794 | 668 | 815 | 465 | 636 | 614 | 649 | 636 | 679 | 637 | 727 | 656 | 654,2 | 634 | 675 | 1,066 | 0,092 | 1,461 | 0,1440764 | 0,20838012 | 1,00E-08 | 0,0041986 | 0,0023169 | 0,0023169 | TRUE | 0,0334484 | No |
| orf19.2664 | 1837 | 1478 | 1740 | 2025 | 2568 | 2423 | 2558 | 1618 | 1648 | 1882 | 1780 | 1670 | 2196 | 2312 | 2283 | 2282 | 2006,6 | 1745 | 2268 | 1,302 | 0,381 | 7,475 | 7,71E-14 | 7,88E-13 | 0,001481 | 0,002703 | 0,0019826 | 0,0019826 | TRUE | 0,0319983 | No |
| orf19.2665 | 25075 | 17657 | 22993 | 27412 | 39545 | 35601 | 38783 | 23638 | 22495 | 22479 | 23524 | 22604 | 33820 | 33969 | 34607 | 33339 | 28355 | 22776 | 33934 | 1,49 | 0,575 | 20,483 | 3,05E-93 | 1,67E-90 | 0,000313 | 0,0020307 | 0,0007211 | 0,0007211 | TRUE | 0,0059808 | No |
| orf19.2667 | 13427 | 8789 | 11369 | 14156 | 11398 | 9230 | 10163 | 5968 | 12045 | 11189 | 11632 | 11673 | 9748 | 8807 | 9069 | 8417 | 10323 | 11635 | 9010 | 0,775 | -0,368 | -7,54 | 4,71E-14 | 4,93E-13 | 0,002253 | 0,0021202 | 0,0021935 | 0,0021935 | TRUE | 0,0372083 | No |
| orf19.2668 | 1321 | 955 | 1121 | 1402 | 1099 | 946 | 996 | 707 | 1185 | 1216 | 1147 | 1156 | 940 | 903 | 889 | 997 | 1054 | 1176 | 932 | 0,791 | -0,338 | -6,317 | 2,67E-10 | 1,84E-09 | 0,000419 | 0,0033568 | 0,0017871 | 0,0017871 | TRUE | 0,0226944 | No |
| orf19.2669 | 1915 | 1421 | 1886 | 2040 | 1653 | 1511 | 1691 | 1034 | 1718 | 1809 | 1930 | 1682 | 1414 | 1442 | 1509 | 1458 | 1620,2 | 1785 | 1456 | 0,816 | -0,293 | -5,364 | 8,16E-08 | 4,05E-07 | 0,001804 | 0,0028755 | 0,0022316 | 0,0022316 | TRUE | 0,0367454 | No |
| orf19.267 | 3006 | 2006 | 2688 | 3271 | 3106 | 2788 | 2964 | 1951 | 2697 | 2554 | 2750 | 2697 | 2656 | 2660 | 2645 | 2752 | 2676,4 | 2674 | 2678 | 1 | 0 | -0,011 | 0,99094898 | 0,9929322 | 0,000168 | 0,0025219 | 0,0010577 | 0,0010577 | TRUE | 0,0103359 | No |
| orf19.2670 | 1481 | 1046 | 1297 | 1608 | 1383 | 1240 | 1299 | 815 | 1329 | 1332 | 1327 | 1326 | 1183 | 1183 | 1159 | 1149 | 1248,5 | 1328 | 1168 | 0,881 | -0,183 | -4,177 | 2,96E-05 | 9,98E-05 | 1,00E-08 | 0,0031423 | 0,001049 | 0,001049 | TRUE | 0,0012235 | No |
| orf19.2671 | 574 | 389 | 518 | 598 | 545 | 441 | 457 | 363 | 515 | 495 | 530 | 493 | 466 | 421 | 408 | 512 | 479,99 | 508 | 452 | 0,884 | -0,178 | -2,204 | 0,02752907 | 0,04969792 | 0,003296 | 0,005004 | 0,0041525 | 0,0041525 | TRUE | 0,0810239 | No |
| orf19.2672 | 2611 | 1842 | 2443 | 2940 | 2730 | 2450 | 2582 | 1696 | 2342 | 2345 | 2499 | 2424 | 2335 | 2338 | 2304 | 2392 | 2372,5 | 2402 | 2342 | 0,974 | -0,038 | -0,974 | 0,33021456 | 0,41730435 | 0,000154 | 0,0025914 | 0,0010678 | 0,0010678 | TRUE | 0,0086497 | No |

| | | | | | | | | | | | | | | | | | | | | | | | | | | | | | | | |
|---|---|---|---|---|---|---|---|---|---|---|---|---|---|---|---|---|---|---|---|---|---|---|---|---|---|---|---|---|---|---|---|
| orf19.2673 | 663 | 499 | 516 | 725 | 474 | 463 | 457 | 248 | 595 | 635 | 528 | 598 | 405 | 442 | 408 | 350 | 495,07 | 589 | 401 | 0,685 | -0,546 | -6,464 | 1,02E-10 | 7,45E-10 | 0,004588 | 0,0049118 | 0,0047502 | 0,0047502 | TRUE | 0,0726569 | No |
| orf19.2674 | 381 | 255 | 375 | 434 | 413 | 292 | 342 | 240 | 342 | 325 | 384 | 358 | 353 | 279 | 305 | 338 | 335,44 | 352 | 319 | 0,903 | -0,147 | -1,542 | 0,12305198 | 0,18231721 | 0,005037 | 0,0063073 | 0,0057009 | 0,0057009 | TRUE | 0,0835665 | No |
| orf19.2675 | 1954 | 1426 | 1796 | 2227 | 1887 | 1698 | 1844 | 1167 | 1753 | 1815 | 1837 | 1836 | 1614 | 1620 | 1645 | 1646 | 1721 | 1810 | 1631 | 0,901 | -0,151 | -3,704 | 0,00021218 | 0,00061523 | 1,00E-08 | 0,0028231 | 0,0010065 | 0,0010065 | TRUE | 0,0058438 | No |
| orf19.2676 | 1872 | 1403 | 1610 | 2124 | 1979 | 1749 | 1845 | 1267 | 1679 | 1786 | 1647 | 1751 | 1693 | 1669 | 1646 | 1787 | 1707,4 | 1716 | 1699 | 0,989 | -0,016 | -0,346 | 0,7289959 | 0,78787685 | 0,000575 | 0,0028298 | 0,0015388 | 0,0015388 | TRUE | 0,0132902 | No |
| orf19.2677 | 316 | 213 | 281 | 361 | 363 | 327 | 376 | 262 | 283 | 271 | 287 | 298 | 310 | 312 | 336 | 370 | 308,42 | 285 | 332 | 1,156 | 0,209 | 2,406 | 0,01614885 | 0,03079094 | 1,00E-08 | 0,0066865 | 0,0039609 | 0,0039609 | TRUE | 0,0556305 | No |
| orf19.2678 | 514 | 344 | 449 | 526 | 726 | 573 | 659 | 438 | 461 | 438 | 459 | 434 | 621 | 547 | 588 | 618 | 520,7 | 448 | 594 | 1,322 | 0,403 | 5,797 | 6,77E-09 | 3,84E-08 | 1,00E-08 | 0,0047675 | 0,0026912 | 0,0026912 | TRUE | 0,0332681 | No |
| orf19.268 | 553 | 320 | 477 | 575 | 566 | 461 | 530 | 333 | 496 | 407 | 488 | 474 | 484 | 440 | 473 | 470 | 466,52 | 466 | 467 | 0,997 | -0,004 | -0,056 | 0,95528282 | 0,96801133 | 0,002019 | 0,0050913 | 0,0036575 | 0,0036575 | TRUE | 0,0746847 | No |
| orf19.2680 | 1914 | 1287 | 1711 | 2011 | 1992 | 1714 | 1821 | 1211 | 1717 | 1638 | 1751 | 1658 | 1704 | 1635 | 1625 | 1708 | 1679,6 | 1691 | 1668 | 0,985 | -0,021 | -0,486 | 0,6268957 | 0,69864392 | 0,000194 | 0,0028439 | 0,0012776 | 0,0012776 | TRUE | 0,006507 | No |
| orf19.2681 | 113 | 74 | 107 | 145 | 106 | 86 | 106 | 63 | 101 | 94 | 109 | 120 | 91 | 82 | 95 | 89 | 97,6 | 106 | 89 | 0,835 | -0,261 | -1,827 | 0,06774105 | 0,10967078 | 1,00E-08 | 0,0168539 | 0,0092711 | 0,0092711 | TRUE | 0,0777956 | No |
| orf19.2682 | 3079 | 2117 | 2605 | 3250 | 2891 | 2683 | 2910 | 1583 | 2762 | 2695 | 2665 | 2680 | 2472 | 2560 | 2597 | 2233 | 2583,1 | 2700 | 2466 | 0,915 | -0,129 | -2,536 | 0,01121866 | 0,02233324 | 0,001813 | 0,0025415 | 0,0020924 | 0,0020924 | TRUE | 0,0457106 | No |
| orf19.2684 | 1456 | 1176 | 1336 | 1560 | 1866 | 1854 | 1941 | 1128 | 1306 | 1497 | 1367 | 1286 | 1596 | 1769 | 1732 | 1591 | 1518,1 | 1364 | 1672 | 1,229 | 0,297 | 4,796 | 1,62E-06 | 6,62E-06 | 0,003054 | 0,0029358 | 0,003025 | 0,003025 | TRUE | 0,0494907 | No |
| orf19.2685 | 286 | 213 | 265 | 330 | 321 | 307 | 299 | 238 | 257 | 271 | 271 | 272 | 275 | 293 | 267 | 336 | 280,12 | 268 | 293 | 1,085 | 0,117 | 1,262 | 0,20694067 | 0,28314724 | 0,001609 | 0,007162 | 0,0047211 | 0,0047211 | TRUE | 0,0943357 | No |
| orf19.2686 | 2325 | 1657 | 2140 | 2548 | 2340 | 2121 | 2202 | 1516 | 2086 | 2110 | 2189 | 2101 | 2001 | 2024 | 1965 | 2138 | 2076,8 | 2122 | 2032 | 0,956 | -0,064 | -1,517 | 0,12916639 | 0,18992454 | 0,000274 | 0,0026785 | 0,0012486 | 0,0012486 | TRUE | 0,0134819 | No |
| orf19.2687.1 | 5232 | 3539 | 4254 | 5522 | 4727 | 4066 | 4203 | 2461 | 4694 | 4506 | 4352 | 4554 | 4043 | 3880 | 3750 | 3471 | 4156,1 | 4526 | 3786 | 0,837 | -0,256 | -5,022 | 5,10E-07 | 2,26E-06 | 0,002197 | 0,0023288 | 0,002256 | 0,002256 | TRUE | 0,0365644 | No |
| orf19.2688 | 15274 | 11701 | 14177 | 17543 | 15612 | 15136 | 16085 | 10202 | 13702 | 14897 | 14504 | 14466 | 13352 | 14442 | 14353 | 14389 | 14263 | 14392 | 14134 | 0,982 | -0,026 | -0,645 | 0,51918607 | 0,60240825 | 0,001124 | 0,0020813 | 0,0015062 | 0,0015062 | TRUE | 0,0171257 | No |
| orf19.269 | 4806 | 3145 | 4067 | 4994 | 4359 | 3994 | 4151 | 2791 | 4311 | 4004 | 4161 | 4118 | 3728 | 3811 | 3704 | 3936 | 3971,7 | 4148 | 3795 | 0,913 | -0,131 | -3,404 | 0,00066327 | 0,00173334 | 0,000516 | 0,002345 | 0,0011596 | 0,0011596 | TRUE | 0,0085396 | No |
| orf19.2690 | 9454 | 6786 | 8615 | 10218 | 9652 | 8819 | 9507 | 6244 | 8481 | 8639 | 8814 | 8426 | 8255 | 8415 | 8483 | 8807 | 8540 | 8590 | 8490 | 0,988 | -0,017 | -0,528 | 0,59755238 | 0,67432282 | 0,000415 | 0,0021495 | 0,0009298 | 0,0009298 | TRUE | 0,0073956 | No |
| orf19.2691 | 979 | 741 | 858 | 1048 | 1018 | 972 | 979 | 731 | 878 | 943 | 878 | 864 | 871 | 927 | 874 | 1031 | 908,29 | 891 | 926 | 1,037 | 0,052 | 0,79 | 0,42925113 | 0,51630446 | 0,002605 | 0,0035778 | 0,0030233 | 0,0030233 | TRUE | 0,0635039 | No |
| orf19.2693 | 2568 | 1681 | 2023 | 2611 | 2146 | 1846 | 1969 | 1151 | 2304 | 2140 | 2070 | 2153 | 1835 | 1761 | 1757 | 1623 | 1955,5 | 2167 | 1744 | 0,806 | -0,311 | -5,926 | 3,11E-09 | 1,84E-08 | 0,001622 | 0,0027219 | 0,002121 | 0,002121 | TRUE | 0,0241351 | No |
| orf19.2694 | 5977 | 4060 | 5208 | 6357 | 6547 | 5720 | 6138 | 3697 | 5362 | 5169 | 5328 | 5242 | 5599 | 5458 | 5477 | 5214 | 5356,2 | 5275 | 5437 | 1,031 | 0,045 | 1,298 | 0,1943414 | 0,26787477 | 0,000327 | 0,0022506 | 0,0009413 | 0,0009413 | TRUE | 0,0087237 | No |
| orf19.2695 | 3217 | 2282 | 2840 | 3426 | 3680 | 3381 | 3602 | 2171 | 2886 | 2905 | 2906 | 2825 | 3147 | 3226 | 3214 | 3062 | 3021,4 | 2880 | 3162 | 1,099 | 0,137 | 3,861 | 0,00011297 | 0,00034228 | 1,00E-08 | 0,00246 | 0,0008696 | 0,0008696 | TRUE | 0,0049755 | No |
| orf19.2697 | 960 | 721 | 895 | 1034 | 1148 | 1087 | 1141 | 687 | 861 | 918 | 916 | 853 | 982 | 1037 | 1018 | 969 | 944,19 | 887 | 1002 | 1,133 | 0,18 | 3,329 | 0,00087002 | 0,00221487 | 1,00E-08 | 0,003517 | 0,0017371 | 0,0017371 | TRUE | 0,0082265 | No |
| orf19.2698 | 483 | 321 | 400 | 528 | 539 | 487 | 534 | 304 | 433 | 409 | 409 | 435 | 461 | 465 | 477 | 429 | 439,69 | 422 | 458 | 1,087 | 0,12 | 1,704 | 0,08838759 | 0,1377874 | 1,00E-08 | 0,0052812 | 0,00251 | 0,00251 | TRUE | 0,0172857 | No |
| orf19.2699 | 3428 | 2092 | 3090 | 3629 | 3394 | 2739 | 3107 | 1907 | 3075 | 2663 | 3161 | 2993 | 2903 | 2613 | 2772 | 2690 | 2858,9 | 2973 | 2744 | 0,923 | -0,116 | -1,958 | 0,05023008 | 0,08453287 | 0,003232 | 0,0024873 | 0,003014 | 0,003014 | TRUE | 0,0578462 | No |
| orf19.27 | 1626 | 1111 | 1448 | 1787 | 1896 | 1604 | 1809 | 1189 | 1459 | 1414 | 1481 | 1474 | 1622 | 1530 | 1614 | 1677 | 1533,9 | 1457 | 1611 | 1,103 | 0,142 | 3,106 | 0,00189757 | 0,0045026 | 0,000173 | 0,0029259 | 0,0013468 | 0,0013468 | TRUE | 0,0133748 | No |
| orf19.270 | 611 | 367 | 476 | 629 | 413 | 316 | 317 | 230 | 548 | 467 | 487 | 519 | 353 | 302 | 283 | 324 | 410,38 | 505 | 316 | 0,623 | -0,684 | -7,667 | 1,76E-14 | 1,92E-13 | 0,004567 | 0,005517 | 0,0050524 | 0,0050524 | TRUE | 0,0833092 | No |
| orf19.2701 | 2891 | 2155 | 2430 | 3111 | 2932 | 2679 | 2535 | 1733 | 2594 | 2744 | 2486 | 2565 | 2508 | 2556 | 2262 | 2444 | 2519,8 | 2597 | 2442 | 0,941 | -0,088 | -1,708 | 0,08756858 | 0,13665248 | 0,001719 | 0,0025556 | 0,0021366 | 0,0021366 | TRUE | 0,0310389 | No |
| orf19.2703 | 4773 | 3179 | 4116 | 5042 | 5572 | 4691 | 5270 | 3273 | 4282 | 4047 | 4211 | 4158 | 4765 | 4476 | 4703 | 4616 | 4407,3 | 4174 | 4640 | 1,111 | 0,152 | 4,204 | 2,62E-05 | 8,92E-05 | 0,000342 | 0,0023089 | 0,001031 | 0,001031 | TRUE | 0,0071389 | No |
| orf19.2704 | 4 | 4 | 3 | 7 | 2 | 4 | 3 | 2 | 4 | 5 | 3 | 6 | 2 | 4 | 3 | 3 | 3,57 | 4 | 3 | 0,623 | -0,683 | -0,949 | 0,34248255 | NA | 1,00E-08 | 0,4087981 | 0,201213 | 0,201213 | TRUE | 0,1139676 | No |
| orf19.2706 | 682 | 434 | 564 | 779 | 608 | 553 | 570 | 400 | 612 | 553 | 577 | 642 | 520 | 528 | 509 | 564 | 563,02 | 596 | 530 | 0,884 | -0,178 | -2,535 | 0,01124467 | 0,02236279 | 0,00134 | 0,0045579 | 0,0029458 | 0,0029458 | TRUE | 0,0331088 | No |
| orf19.2707.1 | 1931 | 1205 | 1494 | 1871 | 2067 | 1418 | 1569 | 1108 | 1732 | 1534 | 1529 | 1543 | 1768 | 1353 | 1400 | 1563 | 1552,7 | 1584 | 1521 | 0,96 | -0,06 | -0,711 | 0,47734738 | 0,56266491 | 0,008735 | 0,0029145 | 0,0060991 | 0,0060991 | TRUE | 0,1484128 | No |
| orf19.2708 | 2393 | 1605 | 1995 | 2581 | 2343 | 2066 | 2185 | 1443 | 2147 | 2043 | 2041 | 2128 | 2004 | 1971 | 1950 | 2035 | 2040 | 2090 | 1990 | 0,951 | -0,073 | -1,818 | 0,06913334 | 0,11153062 | 1,00E-08 | 0,0026912 | 0,0010646 | 0,0010646 | TRUE | 0,003986 | No |
| orf19.2709 | 13082 | 8458 | 10828 | 13371 | 12457 | 10702 | 11615 | 7308 | 11736 | 10768 | 11078 | 11026 | 10654 | 10212 | 10364 | 10307 | 10768 | 11152 | 10384 | 0,931 | -0,103 | -2,862 | 0,00420987 | 0,00921752 | 0,000727 | 0,0021143 | 0,0011522 | 0,0011522 | TRUE | 0,0152547 | No |
| orf19.271 | 105 | 68 | 99 | 128 | 71 | 63 | 84 | 49 | 94 | 87 | 101 | 106 | 61 | 60 | 75 | 69 | 81,56 | 97 | 66 | 0,679 | -0,559 | -3,556 | 0,0003763 | 0,00104281 | 1,00E-08 | 0,019778 | 0,011089 | 0,011089 | TRUE | 0,0841347 | No |
| orf19.2710 | 4402 | 3052 | 3630 | 4583 | 4553 | 4036 | 4442 | 2258 | 3949 | 3886 | 3714 | 3779 | 3894 | 3851 | 3964 | 3185 | 3777,6 | 3832 | 3724 | 0,973 | -0,04 | -0,604 | 0,54582911 | 0,62714501 | 0,005004 | 0,0023638 | 0,0038845 | 0,0038845 | TRUE | 0,1121337 | No |
| orf19.2711 | 7693 | 4989 | 6646 | 8028 | 7602 | 6535 | 7165 | 4133 | 6901 | 6352 | 6800 | 6620 | 6501 | 6235 | 6394 | 5829 | 6454 | 6668 | 6240 | 0,936 | -0,095 | -2,108 | 0,03504603 | 0,06153026 | 0,001562 | 0,0022045 | 0,0017997 | 0,0017997 | TRUE | 0,0231999 | No |
| orf19.2711.1 | 211 | 149 | 217 | 226 | 197 | 179 | 177 | 116 | 189 | 190 | 222 | 186 | 168 | 171 | 158 | 164 | 181,02 | 197 | 165 | 0,84 | -0,251 | -2,339 | 0,01933991 | 0,03607973 | 1,00E-08 | 0,009999 | 0,0054804 | 0,0054804 | TRUE | 0,0798001 | No |
| orf19.2712 | 9628 | 6443 | 8365 | 10215 | 10876 | 9470 | 10241 | 6073 | 8637 | 8203 | 8558 | 8423 | 9301 | 9036 | 9138 | 8565 | 8732,9 | 8455 | 9010 | 1,066 | 0,092 | 2,531 | 0,01138107 | 0,02258918 | 0,000716 | 0,0021458 | 0,0011598 | 0,0011598 | TRUE | 0,0130541 | No |
| orf19.2713 | 74 | 43 | 68 | 91 | 59 | 55 | 58 | 49 | 66 | 55 | 70 | 75 | 50 | 52 | 52 | 69 | 61,19 | 66 | 56 | 0,824 | -0,28 | -1,52 | 0,12860064 | 0,18937075 | 1,00E-08 | 0,0257029 | 0,0160422 | 0,0160422 | TRUE | 0,1591223 | No |
| orf19.2715 | 5092 | 3450 | 4463 | 5560 | 5595 | 4991 | 5479 | 3214 | 4568 | 4392 | 4566 | 4585 | 4785 | 4762 | 4889 | 4533 | 4635,1 | 4528 | 4742 | 1,048 | 0,067 | 1,866 | 0,06211041 | 0,10151964 | 0,000409 | 0,0022927 | 0,0010382 | 0,0010382 | TRUE | 0,0100694 | No |
| orf19.2717 | 15667 | 10930 | 13225 | 17004 | 16771 | 15058 | 15637 | 8737 | 14055 | 13915 | 13530 | 14022 | 14343 | 14368 | 13953 | 12323 | 13814 | 13880 | 13747 | 0,991 | -0,013 | -0,263 | 0,79222147 | 0,83782197 | 0,002189 | 0,0020846 | 0,002434 | 0,002434 | TRUE | 0,058762 | No |
| orf19.272 | 3394 | 2499 | 3087 | 3868 | 2394 | 2254 | 2424 | 1579 | 3045 | 3181 | 3158 | 3190 | 2047 | 2151 | 2163 | 2227 | 2645,3 | 3144 | 2147 | 0,682 | -0,552 | -13,596 | 4,20E-42 | 2,97E-40 | 0,0004 | 0,0025283 | 0,001187 | 0,001187 | TRUE | 0,0120677 | No |
| orf19.2720 | 6011 | 3931 | 5086 | 6231 | 6055 | 5245 | 5753 | 4136 | 5392 | 5005 | 5203 | 5138 | 5178 | 5005 | 5134 | 5833 | 5236,1 | 5184 | 5288 | 1,019 | 0,027 | 0,509 | 0,61108689 | 0,6854761 | 0,002479 | 0,0022568 | 0,0024643 | 0,0024643 | TRUE | 0,0573271 | No |
| orf19.2721 | 3761 | 2673 | 3262 | 4111 | 3544 | 3235 | 3305 | 2169 | 3374 | 3403 | 3337 | 3390 | 3031 | 3087 | 2949 | 3059 | 3203,8 | 3376 | 3032 | 0,898 | -0,156 | -4,63 | 3,66E-06 | 1,42E-05 | 1,00E-08 | 0,0024326 | 0,0007735 | 0,0007735 | TRUE | 0,0042019 | No |
| orf19.2722 | 555 | 435 | 496 | 577 | 723 | 668 | 696 | 398 | 498 | 554 | 507 | 476 | 618 | 637 | 621 | 561 | 559,13 | 509 | 609 | 1,206 | 0,27 | 3,828 | 0,00012915 | 0,00038719 | 0,001163 | 0,0045759 | 0,0029879 | 0,0029879 | TRUE | 0,0376807 | No |
| orf19.2723 | 2913 | 2334 | 2616 | 3330 | 2769 | 2725 | 2840 | 1980 | 2613 | 2971 | 2676 | 2746 | 2368 | 2600 | 2534 | 2793 | 2662,8 | 2752 | 2574 | 0,935 | -0,097 | -1,628 | 0,10347664 | 0,15699184 | 0,003434 | 0,0025247 | 0,003055 | 0,003055 | TRUE | 0,0381721 | No |
| orf19.2724 | 38805 | 32078 | 36574 | 47415 | 58125 | 58917 | 61853 | 41939 | 34812 | 40839 | 37419 | 39099 | 49710 | 56217 | 55194 | 59151 | 46555 | 38042 | 55068 | 1,448 | 0,534 | 8,512 | 1,70E-17 | 2,51E-16 | 0,004813 | 0,0020107 | 0,0037532 | 0,0037532 | TRUE | 0,052588 | No |
| orf19.2725 | 9 | 6 | 7 | 14 | 6 | 12 | 5 | 10 | 8 | 8 | 7 | 12 | 5 | 11 | 4 | 14 | 8,7 | 9 | 8 | 0,972 | -0,041 | -0,082 | 0,9350366 | NA | 0,062879 | 0,1689232 | 0,1264354 | 0,1264354 | TRUE | 0,2107827 | No |
| orf19.2726 | 4556 | 3345 | 4138 | 5290 | 4309 | 3895 | 4249 | 2632 | 4087 | 4259 | 4234 | 4362 | 3685 | 3716 | 3792 | 3712 | 3980,9 | 4236 | 3726 | 0,88 | -0,185 | -5,315 | 1,07E-07 | 5,21E-07 | 0,000262 | 0,0023442 | 0,0009051 | 0,0009051 | TRUE | 0,0070162 | No |
| orf19.2727 | 4546 | 3124 | 3814 | 4915 | 4548 | 4225 | 4265 | 2855 | 4078 | 3977 | 3902 | 4053 | 3890 | 4031 | 3806 | 4027 | 3970,5 | 4002 | 3938 | 0,983 | -0,025 | -0,686 | 0,49249699 | 0,57757601 | 0,000348 | 0,0023451 | 0,0009973 | 0,0009973 | TRUE | 0,0062908 | No |
| orf19.2728 | 1000 | 720 | 924 | 1117 | 881 | 889 | 918 | 506 | 897 | 917 | 945 | 921 | 753 | 848 | 819 | 714 | 851,84 | 920 | 784 | 0,854 | -0,227 | -3,59 | 0,00033018 | 0,00092742 | 0,001709 | 0,0036837 | 0,0026602 | 0,0026602 | TRUE | 0,0369591 | No |
| orf19.273 | 1269 | 1020 | 1166 | 1461 | 1362 | 1316 | 1394 | 939 | 1138 | 1299 | 1193 | 1205 | 1165 | 1256 | 1244 | 1324 | 1227,9 | 1209 | 1247 | 1,031 | 0,045 | 0,763 | 0,44558382 | 0,53190295 | 0,001803 | 0,0031618 | 0,0024697 | 0,0024697 | TRUE | 0,028084 | No |
| orf19.2730 | 1566 | 912 | 1387 | 1625 | 2418 | 1715 | 2096 | 1483 | 1405 | 1161 | 1419 | 1340 | 2068 | 1636 | 1870 | 2092 | 1623,9 | 1331 | 1916 | 1,437 | 0,523 | 6,046 | 1,49E-09 | 9,23E-09 | 0,009205 | 0,0028735 | 0,0065583 | 0,0065583 | TRUE | 0,1174151 | No |
| orf19.2731 | 179 | 122 | 137 | 135 | 147 | 122 | 117 | 57 | 161 | 155 | 140 | 111 | 126 | 116 | 104 | 80 | 124,29 | 142 | 106 | 0,763 | -0,39 | -2,407 | 0,0160921 | 0,03070224 | 0,021203 | 0,0136596 | 0,0170407 | 0,0170407 | TRUE | 0,2385589 | No |
| orf19.2733 | 718 | 639 | 629 | 810 | 829 | 765 | 795 | 491 | 644 | 814 | 644 | 668 | 709 | 730 | 709 | 693 | 701,24 | 692 | 710 | 1,03 | 0,043 | 0,545 | 0,58554341 | 0,66388844 | 0,004749 | 0,0040497 | 0,0044846 | 0,0044846 | TRUE | 0,1533198 | No |
| orf19.2734 | 176 | 147 | 145 | 194 | 114 | 103 | 100 | 68 | 158 | 187 | 148 | 160 | 97 | 98 | 89 | 96 | 129,28 | 163 | 95 | 0,585 | -0,773 | -6,031 | 1,63E-09 | 1,00E-08 | 1,00E-08 | 0,0132082 | 0,0074538 | 0,0074538 | TRUE | 0,0872922 | No |
| orf19.2735 | 11714 | 8551 | 10436 | 12946 | 11850 | 11019 | 11883 | 7323 | 10509 | 10886 | 10677 | 10675 | 10134 | 10514 | 10604 | 10328 | 10541 | 10687 | 10395 | 0,973 | -0,04 | -1,373 | 0,16975912 | 0,23975384 | 0,000179 | 0,0021172 | 0,0007024 | 0,0007024 | TRUE | 0,0036187 | No |
| orf19.2736 | 1138 | 792 | 992 | 1164 | 903 | 801 | 882 | 557 | 1021 | 1008 | 1015 | 960 | 772 | 764 | 787 | 786 | 889,15 | 1001 | 777 | 0,777 | -0,364 | -6,982 | 2,92E-12 | 2,53E-11 | 1,00E-08 | 0,0036122 | 0,0014737 | 0,0014737 | TRUE | 0,0099971 | No |
| orf19.2737 | 6447 | 4168 | 5621 | 6921 | 8395 | 6930 | 7605 | 5080 | 5784 | 5306 | 5751 | 5707 | 7180 | 6612 | 6786 | 7165 | 6286,4 | 5637 | 6936 | 1,23 | 0,298 | 6,765 | 1,33E-11 | 1,07E-10 | 0,001394 | 0,0022105 | 0,0017032 | 0,0017032 | TRUE | 0,0185094 | No |

| | | | | | | | | | | | | | | | | | | | | | | | | | | | | | | | |
|---|---|---|---|---|---|---|---|---|---|---|---|---|---|---|---|---|---|---|---|---|---|---|---|---|---|---|---|---|---|---|---|
| orf19.2738 | 4483 | 3089 | 4166 | 5189 | 3229 | 2865 | 3144 | 2100 | 4022 | 3933 | 4262 | 4279 | 2762 | 2734 | 2805 | 2962 | 3469,8 | 4124 | 2816 | 0,682 | -0,553 | -12,346 | 5,14E-35 | 2,51E-33 | 0,00114 | 0,0023979 | 0,0016244 | 0,0016244 | TRUE | 0,0138429 | No |
| orf19.2739 | 344 | 216 | 312 | 392 | 307 | 289 | 311 | 181 | 309 | 275 | 319 | 323 | 263 | 276 | 278 | 255 | 287,15 | 306 | 268 | 0,872 | -0,197 | -2,264 | 0,02358342 | 0,04330354 | 1,00E-08 | 0,0070352 | 0,0038216 | 0,0038216 | TRUE | 0,046828 | No |
| orf19.274 | 296 | 244 | 266 | 349 | 499 | 452 | 526 | 269 | 266 | 311 | 272 | 288 | 427 | 431 | 469 | 379 | 355,37 | 284 | 426 | 1,513 | 0,598 | 6,742 | 1,56E-11 | 1,25E-10 | 0,00294 | 0,0060646 | 0,0046158 | 0,0046158 | TRUE | 0,0552131 | No |
| orf19.2740 | 772 | 561 | 725 | 870 | 659 | 605 | 668 | 416 | 693 | 714 | 742 | 717 | 564 | 577 | 596 | 587 | 648,7 | 716 | 581 | 0,811 | -0,303 | -5,129 | 2,92E-07 | 1,34E-06 | 1,00E-08 | 0,0042174 | 0,0017949 | 0,0017949 | TRUE | 0,0066633 | No |
| orf19.2742 | 143 | 88 | 105 | 136 | 135 | 94 | 108 | 49 | 128 | 112 | 107 | 112 | 115 | 90 | 96 | 69 | 103,82 | 115 | 92 | 0,816 | -0,293 | -1,865 | 0,06218288 | 0,10157838 | 0,011462 | 0,015963 | 0,0140983 | 0,0140983 | TRUE | 0,26606 | No |
| orf19.2743 | 813 | 471 | 675 | 803 | 1197 | 922 | 1006 | 773 | 729 | 600 | 691 | 662 | 1024 | 880 | 898 | 1090 | 821,64 | 670 | 973 | 1,446 | 0,532 | 6,274 | 3,52E-10 | 2,39E-09 | 0,007046 | 0,0037464 | 0,0056321 | 0,0056321 | TRUE | 0,0736113 | No |
| orf19.2745 | 14 | 15 | 10 | 18 | 20 | 17 | 21 | 11 | 13 | 19 | 10 | 15 | 17 | 16 | 19 | 16 | 15,54 | 14 | 17 | 1,212 | 0,277 | 0,803 | 0,42221014 | 0,50977463 | 1,00E-08 | 0,0954048 | 0,0496303 | 0,0496303 | TRUE | 0,1734528 | No |
| orf19.2746 | 2155 | 1845 | 2104 | 2583 | 2523 | 2607 | 2658 | 1418 | 1933 | 2349 | 2153 | 2130 | 2158 | 2488 | 2372 | 2000 | 2197,7 | 2141 | 2254 | 1,055 | 0,077 | 0,998 | 0,31832598 | 0,40527707 | 0,007062 | 0,0026401 | 0,0052206 | 0,0052206 | TRUE | 0,0672772 | No |
| orf19.2747 | 885 | 749 | 885 | 1062 | 923 | 1046 | 1125 | 739 | 794 | 954 | 905 | 876 | 789 | 998 | 1004 | 1042 | 920,29 | 882 | 958 | 1,086 | 0,119 | 1,311 | 0,18973439 | 0,26277645 | 0,009627 | 0,003557 | 0,0067784 | 0,0067784 | TRUE | 0,1725843 | No |
| orf19.2748 | 1138 | 1048 | 1082 | 1375 | 1392 | 1582 | 1567 | 949 | 1021 | 1334 | 1107 | 1134 | 1190 | 1509 | 1398 | 1338 | 1254,1 | 1149 | 1359 | 1,185 | 0,244 | 2,712 | 0,00668851 | 0,0140653 | 0,010366 | 0,0031371 | 0,0069958 | 0,0069958 | TRUE | 0,1367586 | No |
| orf19.2749 | 5188 | 4240 | 4838 | 6011 | 5668 | 6031 | 6434 | 3841 | 4654 | 5398 | 4950 | 4957 | 4847 | 5755 | 5741 | 5417 | 5214,9 | 4990 | 5440 | 1,091 | 0,125 | 1,943 | 0,05207502 | 0,08710098 | 0,00468 | 0,0022579 | 0,0037945 | 0,0037945 | TRUE | 0,0664934 | No |
| orf19.275 | 575 | 413 | 437 | 552 | 463 | 362 | 381 | 227 | 516 | 526 | 447 | 455 | 396 | 345 | 340 | 320 | 418,18 | 486 | 350 | 0,725 | -0,464 | -5,163 | 2,43E-07 | 1,13E-06 | 0,00511 | 0,005451 | 0,005282 | 0,005282 | TRUE | 0,0907041 | No |
| orf19.2751 | 1053 | 888 | 1043 | 1258 | 1022 | 1118 | 1151 | 637 | 945 | 1131 | 1067 | 1037 | 874 | 1067 | 1027 | 898 | 1005,7 | 1045 | 966 | 0,927 | -0,109 | -1,366 | 0,17207135 | 0,2424495 | 0,006208 | 0,0034229 | 0,0051194 | 0,0051194 | TRUE | 0,0580366 | No |
| orf19.2752 | 2053 | 1559 | 1982 | 2276 | 2150 | 2164 | 2305 | 1391 | 1842 | 1985 | 2028 | 1877 | 1839 | 2065 | 2057 | 1962 | 1956,7 | 1933 | 1981 | 1,026 | 0,036 | 0,671 | 0,50252362 | 0,58727134 | 0,002048 | 0,0027214 | 0,0023141 | 0,0023141 | TRUE | 0,0295977 | No |
| orf19.2753 | 1931 | 1465 | 1802 | 2177 | 2106 | 2040 | 2206 | 1375 | 1732 | 1865 | 1844 | 1795 | 1801 | 1947 | 1968 | 1939 | 1861,5 | 1809 | 1914 | 1,058 | 0,082 | 1,725 | 0,0844642 | 0,13263484 | 0,000885 | 0,0027594 | 0,0016122 | 0,0016122 | TRUE | 0,0200053 | No |
| orf19.2754 | 509 | 371 | 427 | 549 | 527 | 494 | 473 | 375 | 457 | 472 | 437 | 453 | 451 | 471 | 422 | 529 | 461,45 | 455 | 468 | 1,025 | 0,036 | 0,45 | 0,65238258 | 0,72129455 | 0,002281 | 0,0051255 | 0,0038256 | 0,0038256 | TRUE | 0,0769725 | No |
| orf19.2755 | 5863 | 4430 | 5068 | 6506 | 5445 | 5081 | 5411 | 3402 | 5260 | 5640 | 5185 | 5365 | 4657 | 4848 | 4828 | 4798 | 5072,6 | 5362 | 4783 | 0,892 | -0,164 | -4,377 | 1,20E-05 | 4,32E-05 | 0,00059 | 0,0022657 | 0,0011561 | 0,0011561 | TRUE | 0,014198 | No |
| orf19.2756 | 165 | 213 | 171 | 266 | 193 | 310 | 271 | 129 | 148 | 271 | 175 | 219 | 165 | 296 | 242 | 182 | 212,26 | 203 | 221 | 1,093 | 0,128 | 0,627 | 0,53055216 | 0,61322766 | 0,065051 | 0,0088187 | 0,0350798 | 0,0350798 | TRUE | 0,3994912 | No |
| orf19.2757 | 715 | 690 | 680 | 906 | 670 | 696 | 678 | 518 | 641 | 878 | 696 | 747 | 573 | 664 | 605 | 731 | 691,92 | 740 | 643 | 0,868 | -0,205 | -1,993 | 0,04624016 | 0,07847765 | 0,013275 | 0,0040776 | 0,0086683 | 0,0086683 | TRUE | 0,1781442 | No |
| orf19.2758 | 267 | 199 | 248 | 288 | 414 | 335 | 360 | 269 | 240 | 253 | 254 | 237 | 354 | 320 | 321 | 379 | 294,81 | 246 | 344 | 1,393 | 0,479 | 5,363 | 8,19E-08 | 4,05E-07 | 1,00E-08 | 0,0069038 | 0,00416 | 0,00416 | TRUE | 0,0469397 | No |
| orf19.276 | 723 | 593 | 707 | 867 | 897 | 858 | 862 | 538 | 649 | 755 | 723 | 715 | 767 | 819 | 769 | 759 | 744,45 | 710 | 778 | 1,099 | 0,136 | 2,143 | 0,03207738 | 0,0568666 | 0,0012 | 0,0039295 | 0,0025148 | 0,0025148 | TRUE | 0,041392 | No |
| orf19.2760 | 1955 | 1685 | 1892 | 2176 | 1950 | 2103 | 2125 | 1189 | 1754 | 2145 | 1936 | 1794 | 1668 | 2007 | 1896 | 1677 | 1859,6 | 1907 | 1812 | 0,952 | -0,071 | -0,898 | 0,3689888 | 0,45756135 | 0,007547 | 0,0027602 | 0,005537 | 0,005537 | TRUE | 0,0835075 | No |
| orf19.2761 | 328 | 230 | 260 | 334 | 253 | 291 | 290 | 185 | 294 | 293 | 266 | 275 | 216 | 278 | 259 | 261 | 267,78 | 282 | 254 | 0,897 | -0,158 | -1,604 | 0,10860674 | 0,16353617 | 0,003323 | 0,0074008 | 0,0055053 | 0,0055053 | TRUE | 0,1186922 | No |
| orf19.2762 | 13730 | 9615 | 11388 | 14740 | 15534 | 13772 | 14189 | 9355 | 12317 | 12241 | 11651 | 12155 | 13285 | 13141 | 12661 | 13194 | 12581 | 12091 | 13070 | 1,081 | 0,112 | 3,502 | 0,00046201 | 0,00125769 | 0,000431 | 0,0020949 | 0,0009064 | 0,0009064 | TRUE | 0,007322 | No |
| orf19.2763 | 151 | 157 | 164 | 234 | 139 | 148 | 131 | 59 | 135 | 200 | 168 | 193 | 119 | 141 | 117 | 83 | 144,54 | 174 | 115 | 0,669 | -0,58 | -3,506 | 0,00045407 | 0,00123777 | 0,027902 | 0,0120234 | 0,0190818 | 0,0190818 | TRUE | 0,2835959 | No |
| orf19.2765 | 85072 | 53677 | 75284 | 94318 | 92176 | 76457 | 85080 | 71014 | 76318 | 68337 | 77023 | 77776 | 78832 | 72953 | 75920 | 1E+05 | 78415 | 74864 | 81966 | 1,095 | 0,131 | 1,481 | 0,1387023 | 0,20167556 | 0,01308 | 0,001998 | 0,0074717 | 0,0074717 | TRUE | 0,2734318 | No |
| orf19.2766 | 4 | 11 | 18 | 10 | 9 | 12 | 12 | 4 | 4 | 14 | 18 | 8 | 8 | 11 | 11 | 6 | 9,97 | 11 | 9 | 0,83 | -0,269 | -0,523 | 0,60110552 | NA | 0,158252 | 0,1476024 | 0,1521722 | 0,1521722 | TRUE | 0,1902461 | No |
| orf19.2767 | 23644 | 14938 | 20624 | 25757 | 27267 | 22772 | 24868 | 18897 | 21211 | 19018 | 21100 | 21240 | 23320 | 21728 | 22191 | 26653 | 22057 | 20642 | 23473 | 1,137 | 0,185 | 2,792 | 0,00524157 | 0,0112751 | 0,005803 | 0,0020453 | 0,0041747 | 0,0041747 | TRUE | 0,1014275 | No |
| orf19.2768 | 31373 | 25194 | 29277 | 36360 | 40960 | 40990 | 42868 | 23944 | 28145 | 32075 | 29953 | 29983 | 35030 | 39111 | 38253 | 33771 | 33290 | 30039 | 36541 | 1,217 | 0,283 | 4,922 | 8,55E-07 | 3,64E-06 | 0,003823 | 0,0020231 | 0,003143 | 0,003143 | TRUE | 0,0317835 | No |
| orf19.2769 | 13517 | 9306 | 11097 | 14124 | 16305 | 13916 | 14030 | 8185 | 12126 | 11848 | 11353 | 11647 | 13945 | 13278 | 12519 | 11544 | 12283 | 11744 | 12822 | 1,092 | 0,127 | 2,243 | 0,02487803 | 0,04543076 | 0,003513 | 0,0020977 | 0,0030064 | 0,0030064 | TRUE | 0,0544411 | No |
| orf19.277 | 964 | 742 | 926 | 1113 | 955 | 846 | 867 | 624 | 865 | 945 | 947 | 918 | 817 | 807 | 774 | 880 | 869,04 | 919 | 820 | 0,89 | -0,168 | -2,812 | 0,00492076 | 0,01063451 | 0,000981 | 0,00365 | 0,0022506 | 0,0022506 | TRUE | 0,0257512 | No |
| orf19.2770 | 4037 | 2558 | 3194 | 4288 | 5078 | 4615 | 4754 | 3049 | 3622 | 3257 | 3268 | 3536 | 4343 | 4403 | 4242 | 4300 | 3871,4 | 3421 | 4322 | 1,262 | 0,336 | 7,489 | 6,92E-14 | 7,12E-13 | 0,001256 | 0,0023545 | 0,0016726 | 0,0016726 | TRUE | 0,018913 | No |
| orf19.2770.1 | 5487 | 3695 | 4624 | 5833 | 7699 | 6124 | 6291 | 4266 | 4922 | 4704 | 4731 | 4810 | 6584 | 5843 | 5614 | 6017 | 5403,2 | 4792 | 6014 | 1,255 | 0,328 | 6,367 | 1,93E-10 | 1,34E-09 | 0,002331 | 0,0022482 | 0,0023568 | 0,0023568 | TRUE | 0,0505377 | No |
| orf19.2771 | 1370 | 1017 | 1292 | 1554 | 1508 | 1440 | 1549 | 919 | 1229 | 1295 | 1322 | 1281 | 1290 | 1374 | 1382 | 1296 | 1308,6 | 1282 | 1336 | 1,044 | 0,061 | 1,247 | 0,21247218 | 0,28972602 | 0,000446 | 0,0030888 | 0,001566 | 0,001566 | TRUE | 0,0094844 | No |
| orf19.2772 | 5193 | 3751 | 4744 | 5732 | 6301 | 5902 | 6326 | 3990 | 4659 | 4775 | 4854 | 4727 | 5389 | 5632 | 5645 | 5628 | 5163,4 | 4754 | 5574 | 1,172 | 0,229 | 6,999 | 2,58E-12 | 2,25E-11 | 0,000241 | 0,0022607 | 0,0008358 | 0,0008358 | TRUE | 0,0062527 | No |
| orf19.2774 | 5307 | 3340 | 4488 | 5422 | 6012 | 5104 | 5308 | 3173 | 4761 | 4252 | 4592 | 4471 | 5142 | 4870 | 4737 | 4475 | 4662,4 | 4519 | 4806 | 1,064 | 0,089 | 1,696 | 0,08982517 | 0,13981079 | 0,002501 | 0,0022909 | 0,0024441 | 0,0024441 | TRUE | 0,0270043 | No |
| orf19.2775 | 4816 | 3524 | 4082 | 5333 | 4236 | 3852 | 4125 | 2520 | 4320 | 4486 | 4176 | 4398 | 3623 | 3675 | 3681 | 3554 | 3989,3 | 4345 | 3633 | 0,837 | -0,257 | -7,153 | 8,47E-13 | 7,83E-12 | 0,000252 | 0,0023434 | 0,0009849 | 0,0009849 | TRUE | 0,0081688 | No |
| orf19.2778 | 798 | 871 | 830 | 1111 | 822 | 1053 | 924 | 618 | 716 | 1109 | 849 | 916 | 703 | 1005 | 825 | 872 | 874,25 | 898 | 851 | 0,949 | -0,076 | -0,59 | 0,55497784 | 0,63540861 | 0,026045 | 0,00364 | 0,0146058 | 0,0146058 | TRUE | 0,2925464 | No |
| orf19.2779 | 1 | 4 | 1 | 4 | 2 | 1 | 2 | 1 | 1 | 5 | 1 | 3 | 2 | 1 | 2 | 1 | 2,02 | 2 | 2 | 0,59 | -0,762 | -0,764 | 0,4447142 | NA | 1,00E-08 | 0,7201639 | 0,4187962 | 0,4187962 | TRUE | 0,2871242 | No |
| orf19.278 | 2028 | 1412 | 1877 | 2296 | 1349 | 1157 | 1295 | 908 | 1819 | 1798 | 1920 | 1893 | 1154 | 1104 | 1156 | 1281 | 1515,6 | 1858 | 1174 | 0,63 | -0,668 | -12,43 | 1,80E-35 | 9,30E-34 | 0,001344 | 0,0029374 | 0,00207 | 0,00207 | TRUE | 0,0404279 | No |
| orf19.2781 | 1551 | 1529 | 1614 | 1956 | 1796 | 1995 | 2022 | 1200 | 1391 | 1947 | 1651 | 1613 | 1536 | 1904 | 1804 | 1692 | 1692,3 | 1650 | 1734 | 1,052 | 0,073 | 0,764 | 0,44496837 | 0,53146187 | 0,013028 | 0,0028373 | 0,0081715 | 0,0081715 | TRUE | 0,1726695 | No |
| orf19.2782 | 2103 | 1581 | 1795 | 2217 | 2141 | 2019 | 2099 | 1433 | 1887 | 2013 | 1836 | 1828 | 1831 | 1926 | 1873 | 2021 | 1902 | 1891 | 1913 | 1,011 | 0,016 | 0,322 | 0,74769449 | 0,80202615 | 0,001161 | 0,0027428 | 0,0018557 | 0,0018557 | TRUE | 0,0214051 | No |
| orf19.2783 | 6 | 5 | 5 | 6 | 1 | 3 | 6 | 1 | 5 | 6 | 5 | 5 | 1 | 3 | 5 | 1 | 4,04 | 5 | 2 | 0,497 | -1,009 | -1,415 | 0,15700261 | NA | 1,00E-08 | 0,3616066 | 0,2111974 | 0,2111974 | TRUE | 0,6617128 | No |
| orf19.2784 | 24 | 23 | 20 | 29 | 22 | 31 | 26 | 33 | 22 | 29 | 20 | 24 | 19 | 30 | 23 | 47 | 26,67 | 24 | 30 | 1,215 | 0,281 | 0,901 | 0,3677238 | 0,45646376 | 0,052823 | 0,0564201 | 0,0549476 | 0,0549476 | TRUE | 0,6797569 | No |
| orf19.2785 | 2164 | 1771 | 1882 | 2332 | 2304 | 2291 | 2085 | 1316 | 1941 | 2255 | 1925 | 1923 | 1970 | 2186 | 1861 | 1856 | 1989,7 | 2011 | 1968 | 0,98 | -0,029 | -0,405 | 0,68532977 | 0,74862932 | 0,005067 | 0,0027091 | 0,0042927 | 0,0042927 | TRUE | 0,0781782 | No |
| orf19.2786 | 508 | 436 | 482 | 617 | 625 | 683 | 759 | 416 | 456 | 555 | 493 | 509 | 535 | 652 | 677 | 587 | 557,87 | 503 | 613 | 1,221 | 0,288 | 3,181 | 0,00146956 | 0,00357166 | 0,007369 | 0,0045817 | 0,0060373 | 0,0060373 | TRUE | 0,0932044 | No |
| orf19.2787 | 9 | 8 | 7 | 10 | 9 | 11 | 11 | 7 | 8 | 10 | 7 | 8 | 8 | 10 | 10 | 10 | 8,94 | 8 | 10 | 1,129 | 0,175 | 0,394 | 0,69343211 | NA | 1,00E-08 | 0,1642984 | 0,0766285 | 0,0766285 | TRUE | 0,0664301 | No |
| orf19.2788 | 574 | 404 | 491 | 581 | 601 | 557 | 627 | 398 | 515 | 514 | 502 | 479 | 514 | 531 | 559 | 561 | 522,13 | 502 | 541 | 1,077 | 0,107 | 1,603 | 0,10899233 | 0,16405887 | 1,00E-08 | 0,0047599 | 0,0023357 | 0,0023357 | TRUE | 0,0144831 | No |
| orf19.2789 | 1647 | 1191 | 1383 | 1758 | 1925 | 1914 | 1891 | 1257 | 1478 | 1516 | 1415 | 1450 | 1646 | 1826 | 1687 | 1773 | 1598,9 | 1465 | 1733 | 1,183 | 0,242 | 4,873 | 1,10E-06 | 4,61E-06 | 0,000976 | 0,0028875 | 0,0017438 | 0,0017438 | TRUE | 0,0168754 | No |
| orf19.279 | 66 | 39 | 57 | 51 | 45 | 26 | 28 | 19 | 59 | 50 | 58 | 42 | 38 | 25 | 25 | 27 | 40,54 | 52 | 29 | 0,555 | -0,85 | -3,564 | 0,00036479 | 0,00101467 | 0,013643 | 0,0377899 | 0,0277891 | 0,0277891 | TRUE | 0,3973922 | No |
| orf19.2790 | 1536 | 1021 | 1344 | 1698 | 1718 | 1429 | 1545 | 965 | 1378 | 1300 | 1375 | 1400 | 1469 | 1364 | 1379 | 1361 | 1378,2 | 1363 | 1393 | 1,022 | 0,031 | 0,634 | 0,52628199 | 0,60955129 | 0,000478 | 0,0030329 | 0,001555 | 0,001555 | TRUE | 0,0170455 | No |
| orf19.2791 | 3966 | 2532 | 3565 | 4228 | 5422 | 4782 | 5346 | 3286 | 3558 | 3224 | 3647 | 3486 | 4637 | 4563 | 4770 | 4635 | 4065 | 3479 | 4651 | 1,336 | 0,418 | 9,501 | 2,08E-21 | 4,12E-20 | 0,001214 | 0,0023366 | 0,0016063 | 0,0016063 | TRUE | 0,0282886 | No |
| orf19.2792 | 636 | 410 | 540 | 675 | 978 | 818 | 955 | 599 | 571 | 522 | 552 | 557 | 836 | 781 | 852 | 845 | 689,44 | 550 | 828 | 1,501 | 0,586 | 9,632 | 5,83E-22 | 1,21E-20 | 1,00E-08 | 0,0040851 | 0,0020493 | 0,0020493 | TRUE | 0,0184784 | No |
| orf19.2794 | 1032 | 869 | 909 | 1149 | 1123 | 1018 | 1037 | 624 | 926 | 1106 | 930 | 947 | 960 | 971 | 925 | 880 | 955,85 | 977 | 934 | 0,96 | -0,059 | -0,865 | 0,38676496 | 0,47587219 | 0,003486 | 0,0034983 | 0,0034282 | 0,0034282 | TRUE | 0,0883497 | No |
| orf19.2795 | 5158 | 4239 | 4626 | 6105 | 4382 | 4566 | 4507 | 2704 | 4627 | 5397 | 4733 | 5034 | 3748 | 4357 | 4022 | 3814 | 4466,4 | 4948 | 3985 | 0,806 | -0,311 | -4,938 | 7,87E-07 | 3,38E-06 | 0,004498 | 0,0023046 | 0,0035899 | 0,0035899 | TRUE | 0,0481824 | No |
| orf19.2796 | 500 | 356 | 441 | 561 | 522 | 422 | 482 | 310 | 449 | 453 | 451 | 463 | 446 | 403 | 430 | 437 | 441,5 | 454 | 429 | 0,945 | -0,082 | -1,18 | 0,23795247 | 0,31882659 | 1,00E-08 | 0,0052676 | 0,0023772 | 0,0023772 | TRUE | 0,0206629 | No |
| orf19.2797 | 108 | 46 | 58 | 82 | 89 | 82 | 73 | 54 | 97 | 59 | 59 | 68 | 76 | 78 | 65 | 76 | 72,26 | 71 | 74 | 1,037 | 0,053 | 0,276 | 0,78262726 | 0,82942752 | 0,020234 | 0,0220699 | 0,021246 | 0,021246 | TRUE | 0,5906668 | No |
| orf19.2798 | 95 | 71 | 99 | 137 | 187 | 155 | 168 | 116 | 85 | 90 | 101 | 113 | 160 | 148 | 150 | 164 | 126,4 | 97 | 156 | 1,583 | 0,663 | 5,057 | 4,25E-07 | 1,90E-06 | 1,00E-08 | 0,0134643 | 0,0081857 | 0,0081857 | TRUE | 0,1279145 | No |
| orf19.2799 | 1473 | 1075 | 1251 | 1483 | 1657 | 1367 | 1576 | 971 | 1321 | 1369 | 1280 | 1223 | 1417 | 1304 | 1406 | 1370 | 1336,3 | 1298 | 1374 | 1,061 | 0,085 | 1,606 | 0,10834443 | 0,163223 | 0,001129 | 0,0030659 | 0,0019449 | 0,0019449 | TRUE | 0,0190117 | No |

| | | | | | | | | | | | | | | | | | | | | | | | | | | | | | | | |
|---|---|---|---|---|---|---|---|---|---|---|---|---|---|---|---|---|---|---|---|---|---|---|---|---|---|---|---|---|---|---|---|
| orf19.28 | 7554 | 5111 | 6640 | 7978 | 8061 | 7261 | 7685 | 4231 | 6777 | 6507 | 6793 | 6579 | 6894 | 6928 | 6858 | 5967 | 6662,9 | 6664 | 6662 | 1 | 0,001 | 0,01 | 0,99185296 | 0,99367226 | 0,002122 | 0,0021974 | 0,0024097 | 0,0024097 | TRUE | 0,0586209 | No |
| orf19.280 | 32 | 27 | 37 | 33 | 35 | 30 | 43 | 25 | 29 | 34 | 38 | 27 | 30 | 29 | 38 | 35 | 32,54 | 32 | 33 | 1,037 | 0,052 | 0,213 | 0,83108474 | 0,8701912 | 1,00E-08 | 0,0465895 | 0,0263919 | 0,0263919 | TRUE | 0,1095987 | No |
| orf19.2800 | 0 | 0 | 0 | 1 | 0 | 0 | 0 | 0 | 0 | 0 | 0 | 1 | 0 | 0 | 0 | 0 | 0,1 | 0 | 0 | 0,616 | -0,698 | -0,198 | 0,84334499 | NA | 1,00E-08 | 14,085693 | 10 | 10 | TRUE | 0,2428614 | No |
| orf19.2803 | 2247 | 1465 | 1938 | 2456 | 2875 | 2482 | 2689 | 1703 | 2016 | 1865 | 1983 | 2025 | 2459 | 2368 | 2399 | 2402 | 2189,7 | 1972 | 2407 | 1,219 | 0,286 | 6,883 | 5,88E-12 | 4,90E-11 | 0,000206 | 0,0026425 | 0,0011904 | 0,0011904 | TRUE | 0,0149448 | No |
| orf19.2804 | 1 | 3 | 2 | 3 | 5 | 1 | 6 | 1 | 1 | 4 | 2 | 2 | 4 | 1 | 5 | 1 | 2,65 | 2 | 3 | 1,385 | 0,47 | 0,534 | 0,59303195 | NA | 1,00E-08 | 0,5489863 | 0,3594897 | 0,3594897 | TRUE | 0,1955497 | No |
| orf19.2805 | 253 | 193 | 249 | 298 | 288 | 291 | 321 | 194 | 227 | 246 | 255 | 246 | 246 | 278 | 286 | 274 | 257,15 | 244 | 271 | 1,114 | 0,156 | 1,69 | 0,09105586 | 0,14139664 | 1,00E-08 | 0,0076249 | 0,0042422 | 0,0042422 | TRUE | 0,0473315 | No |
| orf19.2808 | 699 | 546 | 716 | 830 | 754 | 832 | 853 | 493 | 627 | 695 | 733 | 684 | 645 | 794 | 761 | 695 | 704,3 | 685 | 724 | 1,058 | 0,082 | 1,03 | 0,30293805 | 0,38889996 | 0,005159 | 0,0040407 | 0,0046414 | 0,0046414 | TRUE | 0,0689771 | No |
| orf19.2809 | 1693 | 1186 | 1509 | 1804 | 1565 | 1381 | 1406 | 988 | 1519 | 1510 | 1544 | 1488 | 1338 | 1318 | 1255 | 1393 | 1420,6 | 1515 | 1326 | 0,874 | -0,195 | -4,128 | 3,65E-05 | 0,00012107 | 0,000197 | 0,0030015 | 0,0014285 | 0,0014285 | TRUE | 0,0158416 | No |
| orf19.281 | 192 | 132 | 163 | 196 | 205 | 183 | 176 | 134 | 172 | 168 | 167 | 162 | 175 | 175 | 157 | 189 | 170,58 | 167 | 174 | 1,036 | 0,051 | 0,476 | 0,63416608 | 0,70517672 | 1,00E-08 | 0,0104898 | 0,0052425 | 0,0052425 | TRUE | 0,0469262 | No |
| orf19.2810 | 641 | 462 | 551 | 735 | 387 | 320 | 361 | 241 | 575 | 588 | 564 | 606 | 331 | 305 | 322 | 340 | 453,92 | 583 | 324 | 0,555 | -0,849 | -11,95 | 6,53E-33 | 2,82E-31 | 1,00E-08 | 0,0051776 | 0,0024557 | 0,0024557 | TRUE | 0,0177244 | No |
| orf19.2812 | 435 | 281 | 398 | 462 | 357 | 297 | 351 | 205 | 390 | 358 | 407 | 381 | 305 | 283 | 313 | 289 | 340,9 | 384 | 298 | 0,776 | -0,366 | -4,55 | 5,35E-06 | 2,03E-05 | 1,00E-08 | 0,006238 | 0,0032542 | 0,0032542 | TRUE | 0,0208126 | No |
| orf19.2813 | 3 | 1 | 2 | 1 | 3 | 2 | 0 | 0 | 3 | 1 | 2 | 1 | 3 | 2 | 0 | 0 | 1,41 | 2 | 1 | 0,695 | -0,526 | -0,439 | 0,66079028 | NA | 1,00E-08 | 1,0288909 | 0,6680193 | 0,6680193 | TRUE | 0,4009054 | No |
| orf19.2818 | 1326 | 916 | 1220 | 1442 | 1591 | 1498 | 1656 | 1037 | 1190 | 1166 | 1248 | 1189 | 1361 | 1429 | 1478 | 1463 | 1315,4 | 1198 | 1433 | 1,194 | 0,256 | 5,224 | 1,75E-07 | 8,31E-07 | 0,000417 | 0,0030831 | 0,0015371 | 0,0015371 | TRUE | 0,0144968 | No |
| orf19.2819 | 148 | 133 | 135 | 165 | 138 | 181 | 180 | 107 | 133 | 169 | 138 | 136 | 118 | 173 | 161 | 151 | 147,32 | 144 | 151 | 1,049 | 0,069 | 0,488 | 0,62582853 | 0,69774954 | 0,013233 | 0,0118338 | 0,0124788 | 0,0124788 | TRUE | 0,2466695 | No |
| orf19.2820 | 4438 | 3137 | 3934 | 4679 | 4432 | 3959 | 4294 | 2769 | 3981 | 3994 | 4025 | 3858 | 3790 | 3778 | 3832 | 3905 | 3895,4 | 3964 | 3826 | 0,965 | -0,052 | -1,571 | 0,11627887 | 0,17339375 | 1,00E-08 | 0,0023522 | 0,0007749 | 0,0007749 | TRUE | 0,0041419 | No |
| orf19.2821 | 2093 | 1661 | 1840 | 2471 | 2194 | 2140 | 2187 | 1407 | 1878 | 2115 | 1883 | 2038 | 1876 | 2042 | 1952 | 1984 | 1970,8 | 1978 | 1964 | 0,993 | -0,01 | -0,191 | 0,84832824 | 0,88415642 | 0,001848 | 0,0027161 | 0,002196 | 0,002196 | TRUE | 0,0245742 | No |
| orf19.2822 | 305 | 269 | 258 | 319 | 255 | 258 | 239 | 154 | 274 | 342 | 264 | 263 | 218 | 246 | 213 | 217 | 254,73 | 286 | 224 | 0,788 | -0,345 | -3,2 | 0,0013736 | 0,00335611 | 0,006511 | 0,0076785 | 0,0071317 | 0,0071317 | TRUE | 0,1820388 | No |
| orf19.2823 | 3202 | 2368 | 3081 | 3746 | 3512 | 3236 | 3505 | 2598 | 2873 | 3015 | 3152 | 3089 | 3004 | 3088 | 3128 | 3664 | 3126,4 | 3032 | 3221 | 1,061 | 0,085 | 1,325 | 0,18521687 | 0,25764829 | 0,004702 | 0,0024439 | 0,0036456 | 0,0036456 | TRUE | 0,1014258 | No |
| orf19.2825 | 6610 | 5393 | 5671 | 7616 | 7112 | 6561 | 6923 | 4081 | 5930 | 6866 | 5802 | 6280 | 6082 | 6260 | 6178 | 5756 | 6144,3 | 6220 | 6069 | 0,977 | -0,034 | -0,608 | 0,54345262 | 0,62537126 | 0,003133 | 0,0022158 | 0,0028952 | 0,0028952 | TRUE | 0,0587983 | No |
| orf19.2826 | 1565 | 1022 | 1338 | 1664 | 2118 | 1840 | 2048 | 1246 | 1404 | 1301 | 1369 | 1372 | 1811 | 1756 | 1827 | 1757 | 1574,8 | 1362 | 1788 | 1,312 | 0,392 | 8,893 | 5,97E-19 | 9,96E-18 | 1,00E-08 | 0,0029014 | 0,0012204 | 0,0012204 | TRUE | 0,0096727 | No |
| orf19.2827 | 256 | 233 | 268 | 328 | 296 | 348 | 374 | 194 | 230 | 297 | 274 | 270 | 253 | 332 | 334 | 274 | 282,94 | 268 | 298 | 1,119 | 0,162 | 1,388 | 0,16528616 | 0,23462569 | 0,011974 | 0,0071103 | 0,0095035 | 0,0095035 | TRUE | 0,1293018 | No |
| orf19.2828 | 336 | 318 | 332 | 474 | 451 | 427 | 460 | 241 | 301 | 405 | 340 | 391 | 386 | 407 | 410 | 340 | 372,54 | 359 | 386 | 1,08 | 0,111 | 1,07 | 0,2845947 | 0,36881901 | 0,009615 | 0,0058763 | 0,0076956 | 0,0076956 | TRUE | 0,1384018 | No |
| orf19.2829 | 1817 | 1528 | 1508 | 2034 | 1627 | 1535 | 1548 | 907 | 1630 | 1945 | 1543 | 1677 | 1391 | 1465 | 1381 | 1279 | 1539 | 1699 | 1379 | 0,814 | -0,297 | -4,058 | 4,95E-05 | 0,00016037 | 0,005867 | 0,0029228 | 0,0044945 | 0,0044945 | TRUE | 0,1111672 | No |
| orf19.2830 | 11983 | 9152 | 10555 | 13253 | 10887 | 10294 | 10555 | 6740 | 10750 | 11651 | 10799 | 10929 | 9311 | 9822 | 9419 | 9506 | 10273 | 11032 | 9514 | 0,863 | -0,213 | -5,697 | 1,22E-08 | 6,70E-08 | 0,000861 | 0,0021208 | 0,0012458 | 0,0012458 | TRUE | 0,0171398 | No |
| orf19.2831 | 3203 | 2237 | 2673 | 3404 | 2771 | 2448 | 2656 | 1440 | 2873 | 2848 | 2735 | 2807 | 2370 | 2336 | 2370 | 2031 | 2546,2 | 2816 | 2277 | 0,81 | -0,304 | -5,64 | 1,70E-08 | 9,16E-08 | 0,002314 | 0,0025497 | 0,0023827 | 0,0023827 | TRUE | 0,0598673 | No |
| orf19.2832 | 4807 | 3462 | 4308 | 5197 | 4022 | 3903 | 4244 | 2561 | 4312 | 4407 | 4408 | 4286 | 3440 | 3724 | 3787 | 3612 | 3997 | 4353 | 3641 | 0,837 | -0,257 | -6,413 | 1,43E-10 | 1,01E-09 | 0,000646 | 0,0023427 | 0,0012934 | 0,0012934 | TRUE | 0,017687 | No |
| orf19.2833 | 78 | 56 | 84 | 104 | 88 | 84 | 100 | 75 | 70 | 71 | 86 | 86 | 75 | 80 | 89 | 106 | 82,92 | 78 | 88 | 1,104 | 0,143 | 0,873 | 0,3827774 | 0,47193241 | 0,004643 | 0,0194859 | 0,0135055 | 0,0135055 | TRUE | 0,146925 | No |
| orf19.2834 | 2700 | 1848 | 2305 | 3010 | 3465 | 3230 | 3535 | 2148 | 2422 | 2353 | 2358 | 2482 | 2963 | 3082 | 3154 | 3030 | 2730,6 | 2404 | 3057 | 1,271 | 0,346 | 8,787 | 1,53E-18 | 2,49E-17 | 0,000369 | 0,0025112 | 0,0011157 | 0,0011157 | TRUE | 0,0059946 | No |
| orf19.2835 | 605 | 518 | 567 | 748 | 591 | 618 | 639 | 359 | 543 | 659 | 580 | 617 | 505 | 590 | 570 | 506 | 571,35 | 600 | 543 | 0,908 | -0,138 | -1,692 | 0,09073024 | 0,14097434 | 0,004818 | 0,0045204 | 0,0046731 | 0,0046731 | TRUE | 0,0487852 | No |
| orf19.2836 | 237 | 148 | 197 | 234 | 262 | 178 | 205 | 143 | 213 | 188 | 202 | 193 | 224 | 170 | 183 | 202 | 196,76 | 199 | 195 | 0,977 | -0,034 | -0,299 | 0,76465713 | 0,81628643 | 0,004369 | 0,0093576 | 0,00706 | 0,00706 | TRUE | 0,1286826 | No |
| orf19.2837 | 201 | 169 | 166 | 226 | 204 | 174 | 176 | 112 | 180 | 215 | 170 | 186 | 174 | 166 | 157 | 158 | 175,9 | 188 | 164 | 0,878 | -0,188 | -1,692 | 0,09065821 | 0,14092484 | 1,00E-08 | 0,0102326 | 0,0061088 | 0,0061088 | TRUE | 0,0876089 | No |
| orf19.2838 | 1457 | 1128 | 1376 | 1682 | 2453 | 2390 | 2564 | 1499 | 1307 | 1436 | 1408 | 1387 | 2098 | 2280 | 2288 | 2114 | 1789,8 | 1384 | 2195 | 1,588 | 0,667 | 12,969 | 1,84E-38 | 1,16E-36 | 0,001396 | 0,0027906 | 0,0019525 | 0,0019525 | TRUE | 0,0172644 | No |
| orf19.2839 | 2969 | 1911 | 2446 | 2953 | 4738 | 4177 | 4483 | 2617 | 2664 | 2433 | 2502 | 2435 | 4052 | 3986 | 4000 | 3691 | 3220,4 | 2508 | 3932 | 1,568 | 0,649 | 13,785 | 3,12E-43 | 2,29E-41 | 0,001359 | 0,0024303 | 0,0018029 | 0,0018029 | TRUE | 0,021388 | No |
| orf19.284 | 120 | 69 | 91 | 109 | 117 | 84 | 82 | 72 | 108 | 88 | 93 | 90 | 100 | 80 | 73 | 102 | 91,68 | 95 | 89 | 0,929 | -0,106 | -0,667 | 0,50483158 | 0,58928077 | 0,00667 | 0,0178145 | 0,0131192 | 0,0131192 | TRUE | 0,140755 | No |
| orf19.2841 | 7844 | 6130 | 7162 | 8800 | 8829 | 9009 | 9147 | 5089 | 7037 | 7804 | 7327 | 7257 | 7551 | 8596 | 8162 | 7178 | 7614 | 7356 | 7872 | 1,071 | 0,099 | 1,653 | 0,09828492 | 0,15059826 | 0,004001 | 0,0021702 | 0,0032805 | 0,0032805 | TRUE | 0,0467266 | No |
| orf19.2842 | 7931 | 5659 | 7249 | 9187 | 9774 | 8734 | 9780 | 6401 | 7115 | 7205 | 7416 | 7576 | 8359 | 8334 | 8727 | 9028 | 7969,9 | 7328 | 8612 | 1,174 | 0,232 | 5,883 | 4,02E-09 | 2,37E-08 | 0,000971 | 0,0021617 | 0,0013668 | 0,0013668 | TRUE | 0,0125435 | No |
| orf19.2843 | 5209 | 3397 | 4444 | 5713 | 4508 | 3923 | 3958 | 2346 | 4673 | 4325 | 4547 | 4711 | 3855 | 3743 | 3532 | 3309 | 4086,8 | 4564 | 3610 | 0,791 | -0,337 | -6,276 | 3,48E-10 | 2,37E-09 | 0,002633 | 0,0023347 | 0,0025272 | 0,0025272 | TRUE | 0,036791 | No |
| orf19.2844 | 599 | 513 | 555 | 698 | 724 | 754 | 789 | 436 | 537 | 653 | 568 | 576 | 619 | 719 | 704 | 615 | 623,94 | 584 | 664 | 1,144 | 0,194 | 2,391 | 0,01680159 | 0,03188376 | 0,004852 | 0,0043062 | 0,0046923 | 0,0046923 | TRUE | 0,0702062 | No |
| orf19.2846 | 278 | 198 | 235 | 315 | 342 | 291 | 361 | 215 | 249 | 252 | 240 | 260 | 292 | 278 | 322 | 303 | 274,65 | 250 | 299 | 1,192 | 0,254 | 2,88 | 0,00397402 | 0,00877468 | 1,00E-08 | 0,0072652 | 0,0037919 | 0,0037919 | TRUE | 0,0346671 | No |
| orf19.2847 | 3672 | 3254 | 3559 | 4459 | 3999 | 4431 | 4452 | 2800 | 3294 | 4143 | 3641 | 3677 | 3420 | 4228 | 3973 | 3949 | 3790,6 | 3689 | 3892 | 1,056 | 0,078 | 1,002 | 0,31612483 | 0,40290157 | 0,007611 | 0,0023625 | 0,0055523 | 0,0055523 | TRUE | 0,0824732 | No |
| orf19.2848 | 2602 | 1752 | 2319 | 2935 | 2954 | 2498 | 2742 | 1786 | 2334 | 2230 | 2373 | 2420 | 2526 | 2384 | 2447 | 2519 | 2404,2 | 2339 | 2469 | 1,054 | 0,076 | 1,79 | 0,07342519 | 0,11743112 | 0,000514 | 0,0025834 | 0,001294 | 0,001294 | TRUE | 0,0111411 | No |
| orf19.2849 | 26967 | 16930 | 23963 | 29337 | 24746 | 19196 | 20827 | 12641 | 24192 | 21554 | 24517 | 24192 | 21164 | 18316 | 18585 | 17829 | 21293 | 23614 | 18974 | 0,804 | -0,316 | -5,092 | 3,54E-07 | 1,60E-06 | 0,004746 | 0,0020477 | 0,0036418 | 0,0036418 | TRUE | 0,0740584 | No |
| orf19.285 | 5135 | 3588 | 4251 | 5316 | 6130 | 5272 | 5735 | 3081 | 4607 | 4568 | 4349 | 4384 | 5243 | 5030 | 5118 | 4345 | 4705,4 | 4477 | 4934 | 1,103 | 0,142 | 2,421 | 0,01548609 | 0,02968744 | 0,00345 | 0,002288 | 0,0030804 | 0,0030804 | TRUE | 0,0765553 | No |
| orf19.2850 | 6239 | 4274 | 5406 | 6435 | 6961 | 6192 | 6785 | 4059 | 5597 | 5441 | 5531 | 5306 | 5953 | 5908 | 6054 | 5725 | 5689,6 | 5469 | 5910 | 1,081 | 0,113 | 3,333 | 0,00085974 | 0,00219242 | 0,000302 | 0,0022347 | 0,0009261 | 0,0009261 | TRUE | 0,0051248 | No |
| orf19.2851 | 918 | 668 | 816 | 978 | 857 | 884 | 908 | 602 | 824 | 850 | 835 | 806 | 733 | 843 | 810 | 849 | 818,88 | 829 | 809 | 0,975 | -0,037 | -0,6 | 0,54831723 | 0,62952225 | 0,001365 | 0,0037523 | 0,0024663 | 0,0024663 | TRUE | 0,0498051 | No |
| orf19.2852 | 3434 | 2507 | 2928 | 3774 | 3546 | 3193 | 3264 | 2106 | 3081 | 3192 | 2996 | 3112 | 3033 | 3047 | 2913 | 2970 | 3042,8 | 3095 | 2991 | 0,967 | -0,049 | -1,31 | 0,19023235 | 0,26328403 | 0,000207 | 0,0024566 | 0,0009972 | 0,0009972 | TRUE | 0,0054836 | No |
| orf19.2853 | 1264 | 936 | 1144 | 1371 | 1457 | 1446 | 1509 | 1029 | 1134 | 1192 | 1170 | 1131 | 1246 | 1380 | 1347 | 1451 | 1256,3 | 1157 | 1356 | 1,171 | 0,227 | 4,045 | 5,22E-05 | 0,00016852 | 0,001549 | 0,0031351 | 0,0022265 | 0,0022265 | TRUE | 0,0363775 | No |
| orf19.2857 | 9121 | 6273 | 8050 | 9590 | 8865 | 7774 | 8404 | 5119 | 8182 | 7986 | 8236 | 7908 | 7582 | 7418 | 7499 | 7220 | 7753,9 | 8078 | 7430 | 0,92 | -0,12 | -3,864 | 0,00011161 | 0,00033867 | 0,000232 | 0,0021667 | 0,0007957 | 0,0007957 | TRUE | 0,0041784 | No |
| orf19.2859 | 7956 | 4983 | 6430 | 8206 | 7255 | 6151 | 6816 | 4134 | 7137 | 6344 | 6579 | 6767 | 6205 | 5869 | 6082 | 5831 | 6351,6 | 6707 | 5997 | 0,894 | -0,162 | -3,622 | 0,00029219 | 0,00082687 | 0,001476 | 0,0022081 | 0,0017518 | 0,0017518 | TRUE | 0,0228096 | No |
| orf19.286 | 99 | 55 | 64 | 107 | 80 | 67 | 74 | 55 | 89 | 70 | 65 | 88 | 68 | 64 | 66 | 78 | 73,56 | 78 | 69 | 0,87 | -0,201 | -1,183 | 0,23677416 | 0,31767288 | 1,00E-08 | 0,0217138 | 0,0140585 | 0,0140585 | TRUE | 0,1164267 | No |
| orf19.2862 | 3946 | 2514 | 3408 | 4125 | 4231 | 3440 | 3887 | 2436 | 3540 | 3201 | 3487 | 3402 | 3618 | 3282 | 3469 | 3436 | 3429,2 | 3408 | 3451 | 1,013 | 0,018 | 0,384 | 0,70126746 | 0,76298713 | 0,001254 | 0,0024029 | 0,0018053 | 0,0018053 | TRUE | 0,0194257 | No |
| orf19.2863 | 291 | 203 | 262 | 369 | 447 | 365 | 409 | 250 | 261 | 258 | 268 | 304 | 382 | 348 | 365 | 353 | 317,49 | 273 | 362 | 1,324 | 0,405 | 4,69 | 2,73E-06 | 1,08E-05 | 1,00E-08 | 0,0065519 | 0,0039527 | 0,0039527 | TRUE | 0,0730444 | No |
| orf19.2863.1 | 512 | 307 | 400 | 463 | 593 | 458 | 545 | 315 | 459 | 391 | 409 | 382 | 507 | 437 | 486 | 444 | 439,49 | 410 | 468 | 1,144 | 0,194 | 2,286 | 0,02223408 | 0,04107699 | 0,00402 | 0,0052826 | 0,0046431 | 0,0046431 | TRUE | 0,0781615 | No |
| orf19.2864.1 | 19323 | 12278 | 14882 | 18743 | 20529 | 16209 | 17035 | 10274 | 17335 | 15631 | 15226 | 15456 | 17557 | 15466 | 15201 | 14491 | 15795 | 15912 | 15679 | 0,985 | -0,021 | -0,329 | 0,74212461 | 0,79818916 | 0,004576 | 0,0020714 | 0,0038866 | 0,0038866 | TRUE | 0,0797347 | No |
| orf19.2866 | 78 | 43 | 74 | 96 | 109 | 74 | 80 | 59 | 70 | 55 | 76 | 79 | 93 | 71 | 71 | 83 | 74,75 | 70 | 80 | 1,128 | 0,174 | 1 | 0,31745123 | 0,40433474 | 0,007452 | 0,0213996 | 0,0156605 | 0,0156605 | TRUE | 0,156375 | No |
| orf19.2867 | 3679 | 2618 | 3307 | 3988 | 4169 | 3762 | 4061 | 2608 | 3300 | 3333 | 3383 | 3289 | 3565 | 3590 | 3624 | 3678 | 3470,3 | 3326 | 3614 | 1,086 | 0,119 | 3,69 | 0,00022462 | 0,00064851 | 1,00E-08 | 0,0023978 | 0,0007146 | 0,0007146 | TRUE | 0,0016541 | No |
| orf19.2868 | 7 | 7 | 3 | 3 | 3 | 4 | 4 | 2 | 6 | 9 | 3 | 2 | 3 | 4 | 4 | 3 | 4,19 | 5 | 4 | 0,637 | -0,65 | -0,934 | 0,35055592 | NA | 1,00E-08 | 0,3485806 | 0,2109655 | 0,2109655 | TRUE | 0,263201 | No |
| orf19.2869 | 2 | 1 | 2 | 1 | 5 | 2 | 5 | 0 | 2 | 1 | 2 | 1 | 4 | 2 | 4 | 0 | 2,07 | 2 | 2 | 1,906 | 0,93 | 0,935 | 0,34984141 | NA | 1,00E-08 | 0,7022573 | 0,4377885 | 0,4377885 | TRUE | 0,2342587 | No |
| orf19.287 | 4593 | 2837 | 3569 | 4483 | 4030 | 3267 | 3560 | 2283 | 4120 | 3612 | 3651 | 3697 | 3447 | 3117 | 3177 | 3220 | 3505,1 | 3770 | 3240 | 0,859 | -0,219 | -4,032 | 5,54E-05 | 0,00017778 | 0,002565 | 0,0023937 | 0,002541 | 0,002541 | TRUE | 0,047993 | No |

| | | | | | | | | | | | | | | | | | | | | | | | | | | | | | | | |
|---|---|---|---|---|---|---|---|---|---|---|---|---|---|---|---|---|---|---|---|---|---|---|---|---|---|---|---|---|---|---|---|
| orf19.2870 | 2473 | 1747 | 1948 | 2501 | 2828 | 2341 | 2453 | 1299 | 2219 | 2224 | 1993 | 2062 | 2419 | 2234 | 2189 | 1832 | 2146,4 | 2124 | 2168 | 1,023 | 0,032 | 0,414 | 0,6789416 | 0,7432728 | 0,007521 | 0,0026559 | 0,0053191 | 0,0053191 | TRUE | 0,1272248 | No |
| orf19.2871 | 6664 | 4353 | 5964 | 7014 | 7784 | 6696 | 7059 | 4763 | 5978 | 5542 | 6102 | 5784 | 6657 | 6389 | 6299 | 6718 | 6183,6 | 5852 | 6516 | 1,113 | 0,154 | 3,688 | 0,00022636 | 0,0006529 | 0,001142 | 0,0022143 | 0,0015162 | 0,0015162 | TRUE | 0,0149679 | No |
| orf19.2873 | 2944 | 2067 | 2598 | 3142 | 2513 | 2349 | 2581 | 1551 | 2641 | 2632 | 2658 | 2591 | 2149 | 2241 | 2303 | 2188 | 2425,3 | 2630 | 2220 | 0,845 | -0,244 | -6,295 | 3,08E-10 | 2,11E-09 | 1,00E-08 | 0,0025781 | 0,0010224 | 0,0010224 | TRUE | 0,0079763 | No |
| orf19.2875 | 1983 | 1528 | 1593 | 2054 | 1788 | 1654 | 1710 | 961 | 1779 | 1945 | 1630 | 1694 | 1529 | 1578 | 1526 | 1355 | 1629,6 | 1762 | 1497 | 0,852 | -0,231 | -3,455 | 0,00055055 | 0,0014714 | 0,004304 | 0,0028704 | 0,0036695 | 0,0036695 | TRUE | 0,0569554 | No |
| orf19.2876 | 2735 | 1747 | 2158 | 2868 | 3371 | 2850 | 3096 | 1994 | 2454 | 2224 | 2208 | 2365 | 2883 | 2719 | 2763 | 2812 | 2553,5 | 2313 | 2794 | 1,207 | 0,271 | 5,808 | 6,30E-09 | 3,60E-08 | 0,00108 | 0,002548 | 0,0016996 | 0,0016996 | TRUE | 0,0203708 | No |
| orf19.2877 | 21945 | 14839 | 19427 | 23789 | 28896 | 24558 | 26727 | 18340 | 19687 | 18892 | 19876 | 19617 | 24713 | 23432 | 23849 | 25867 | 21992 | 19518 | 24465 | 1,253 | 0,326 | 8,332 | 7,92E-17 | 1,10E-15 | 0,00115 | 0,0020455 | 0,0014202 | 0,0014202 | TRUE | 0,018008 | No |
| orf19.2878 | 85 | 53 | 90 | 90 | 81 | 72 | 95 | 48 | 76 | 67 | 92 | 74 | 69 | 69 | 85 | 68 | 75,06 | 77 | 73 | 0,939 | -0,091 | -0,547 | 0,58409094 | 0,66236662 | 1,00E-08 | 0,0213205 | 0,0133843 | 0,0133843 | TRUE | 0,1402096 | No |
| orf19.2879 | 244 | 156 | 210 | 260 | 230 | 193 | 211 | 154 | 219 | 199 | 215 | 214 | 197 | 184 | 188 | 217 | 204,14 | 212 | 196 | 0,921 | -0,119 | -1,193 | 0,23278653 | 0,31334032 | 1,00E-08 | 0,0090909 | 0,0046108 | 0,0046108 | TRUE | 0,0428283 | No |
| orf19.288 | 2291 | 1607 | 2050 | 2576 | 2035 | 1854 | 2040 | 1341 | 2055 | 2046 | 2097 | 2124 | 1740 | 1769 | 1820 | 1891 | 1943 | 2080 | 1805 | 0,866 | -0,208 | -4,896 | 9,78E-07 | 4,13E-06 | 0,000138 | 0,0027267 | 0,0012152 | 0,0012152 | TRUE | 0,01108 | No |
| orf19.2881 | 7067 | 6011 | 7111 | 8357 | 8320 | 8986 | 8936 | 5199 | 6340 | 7653 | 7275 | 6891 | 7116 | 8574 | 7974 | 7333 | 7394,4 | 7040 | 7749 | 1,101 | 0,139 | 1,951 | 0,0510499 | 0,08566775 | 0,006616 | 0,0021758 | 0,0047447 | 0,0047447 | TRUE | 0,0628355 | No |
| orf19.2882 | 2653 | 1801 | 2300 | 2926 | 2708 | 2496 | 2670 | 1610 | 2380 | 2293 | 2353 | 2413 | 2316 | 2382 | 2383 | 2271 | 2348,7 | 2360 | 2338 | 0,991 | -0,014 | -0,351 | 0,72582174 | 0,7852939 | 1,00E-08 | 0,0025976 | 0,001 | 0,001 | TRUE | 0,0041101 | No |
| orf19.2883 | 1336 | 979 | 1175 | 1497 | 1281 | 1229 | 1243 | 886 | 1199 | 1246 | 1202 | 1234 | 1096 | 1173 | 1109 | 1250 | 1188,6 | 1220 | 1157 | 0,945 | -0,081 | -1,503 | 0,13281666 | 0,19481506 | 0,00092 | 0,0032009 | 0,0019639 | 0,0019639 | TRUE | 0,0314068 | No |
| orf19.2884 | 3681 | 3002 | 3516 | 4262 | 4127 | 4181 | 4262 | 2427 | 3302 | 3822 | 3597 | 3514 | 3530 | 3989 | 3803 | 3423 | 3622,6 | 3559 | 3686 | 1,037 | 0,052 | 0,857 | 0,39149084 | 0,48040788 | 0,00375 | 0,0023803 | 0,0033073 | 0,0033073 | TRUE | 0,0371004 | No |
| orf19.2885 | 30 | 37 | 52 | 40 | 54 | 63 | 52 | 33 | 27 | 47 | 53 | 33 | 46 | 60 | 46 | 47 | 44,93 | 40 | 50 | 1,261 | 0,335 | 1,385 | 0,16610038 | 0,23552775 | 0,032504 | 0,0342895 | 0,0335671 | 0,0335671 | TRUE | 0,2448595 | No |
| orf19.2886 | 941 | 836 | 955 | 1177 | 964 | 1134 | 1136 | 773 | 844 | 1064 | 977 | 971 | 824 | 1082 | 1014 | 1090 | 983,32 | 964 | 1002 | 1,04 | 0,056 | 0,585 | 0,55871384 | 0,639138 | 0,011701 | 0,0034559 | 0,0077798 | 0,0077798 | TRUE | 0,1747632 | No |
| orf19.2887 | 5201 | 3645 | 4329 | 5470 | 4514 | 3934 | 4215 | 2808 | 4666 | 4640 | 4429 | 4511 | 3861 | 3754 | 3761 | 3960 | 4197,7 | 4562 | 3834 | 0,84 | -0,252 | -7,009 | 2,41E-12 | 2,11E-11 | 0,000301 | 0,0023254 | 0,0009952 | 0,0009952 | TRUE | 0,0055262 | No |
| orf19.2888 | 214 | 171 | 193 | 275 | 170 | 223 | 197 | 98 | 192 | 218 | 197 | 227 | 145 | 213 | 176 | 138 | 188,26 | 208 | 168 | 0,81 | -0,304 | -2,211 | 0,02703898 | 0,04900459 | 0,016398 | 0,0096906 | 0,0127476 | 0,0127476 | TRUE | 0,3397435 | No |
| orf19.2889 | 818 | 607 | 700 | 875 | 861 | 785 | 831 | 502 | 734 | 773 | 716 | 722 | 736 | 749 | 742 | 708 | 734,91 | 736 | 734 | 1 | 0 | 0,005 | 0,99584529 | 0,99717303 | 1,00E-08 | 0,0039549 | 0,0016954 | 0,0016954 | TRUE | 0,0117439 | No |
| orf19.2890 | 927 | 753 | 872 | 1171 | 1012 | 986 | 919 | 623 | 832 | 959 | 892 | 966 | 865 | 941 | 820 | 879 | 894,13 | 912 | 876 | 0,961 | -0,057 | -0,843 | 0,39908506 | 0,48783378 | 0,003071 | 0,0036031 | 0,0033091 | 0,0033091 | TRUE | 0,0433316 | No |
| orf19.2891 | 787 | 520 | 677 | 931 | 1006 | 1074 | 1112 | 722 | 706 | 662 | 693 | 768 | 860 | 1025 | 992 | 1018 | 840,51 | 707 | 974 | 1,372 | 0,457 | 6,193 | 5,91E-10 | 3,89E-09 | 0,004236 | 0,0037067 | 0,0039974 | 0,0039974 | TRUE | 0,0764162 | No |
| orf19.2892 | 3342 | 2746 | 3150 | 3836 | 3694 | 4010 | 4122 | 2655 | 2998 | 3496 | 3223 | 3163 | 3159 | 3826 | 3678 | 3745 | 3411 | 3220 | 3602 | 1,119 | 0,162 | 2,375 | 0,0175574 | 0,03309848 | 0,005175 | 0,0024051 | 0,0041704 | 0,0041704 | TRUE | 0,08443 | No |
| orf19.2893 | 876 | 893 | 918 | 1138 | 903 | 1121 | 1176 | 642 | 786 | 1137 | 939 | 938 | 772 | 1070 | 1049 | 905 | 949,64 | 950 | 949 | 1,001 | 0,001 | 0,008 | 0,99388561 | 0,99554264 | 0,021459 | 0,0035082 | 0,0124513 | 0,0124513 | TRUE | 0,2041675 | No |
| orf19.2895 | 665 | 453 | 534 | 689 | 696 | 590 | 645 | 387 | 597 | 577 | 546 | 568 | 595 | 563 | 576 | 546 | 570,92 | 572 | 570 | 0,999 | -0,002 | -0,031 | 0,97519454 | 0,98222445 | 1,00E-08 | 0,0045223 | 0,0021474 | 0,0021474 | TRUE | 0,0105942 | No |
| orf19.2896 | 1369 | 927 | 1094 | 1533 | 1669 | 1318 | 1522 | 956 | 1228 | 1180 | 1119 | 1264 | 1427 | 1258 | 1358 | 1348 | 1272,9 | 1198 | 1348 | 1,124 | 0,169 | 2,885 | 0,00391583 | 0,00865892 | 0,002057 | 0,00312 | 0,0025062 | 0,0025062 | TRUE | 0,0253437 | No |
| orf19.2897 | 77 | 56 | 60 | 67 | 112 | 80 | 105 | 69 | 69 | 71 | 61 | 55 | 96 | 76 | 94 | 97 | 77,52 | 64 | 91 | 1,418 | 0,504 | 3,069 | 0,00214965 | 0,00504496 | 1,00E-08 | 0,020707 | 0,0126204 | 0,0126204 | TRUE | 0,1133074 | No |
| orf19.2898 | 1573 | 1072 | 1405 | 1719 | 1419 | 1360 | 1449 | 910 | 1411 | 1365 | 1437 | 1418 | 1214 | 1298 | 1293 | 1283 | 1339,8 | 1408 | 1272 | 0,902 | -0,148 | -3,197 | 0,00138871 | 0,00338885 | 1,00E-08 | 0,003063 | 0,0013209 | 0,0013209 | TRUE | 0,0123562 | No |
| orf19.2899 | 119 | 87 | 101 | 102 | 112 | 90 | 121 | 75 | 107 | 111 | 103 | 84 | 96 | 86 | 108 | 106 | 100,05 | 101 | 99 | 0,979 | -0,03 | -0,204 | 0,83853978 | 0,87649723 | 1,00E-08 | 0,0164898 | 0,0110779 | 0,0110779 | TRUE | 0,1432821 | No |
| orf19.29 | 4112 | 3512 | 3793 | 4861 | 5432 | 5613 | 5711 | 3754 | 3689 | 4471 | 3881 | 4008 | 4646 | 5356 | 5096 | 5295 | 4555,2 | 4012 | 5098 | 1,271 | 0,346 | 5,243 | 1,58E-07 | 7,55E-07 | 0,005132 | 0,0022982 | 0,0039574 | 0,0039574 | TRUE | 0,0710008 | No |
| orf19.290 | 907 | 626 | 847 | 925 | 1005 | 913 | 1046 | 578 | 814 | 797 | 867 | 763 | 860 | 871 | 933 | 815 | 839,9 | 810 | 870 | 1,077 | 0,107 | 1,686 | 0,09184023 | 0,14243993 | 0,001773 | 0,0037079 | 0,002667 | 0,002667 | TRUE | 0,0288302 | No |
| orf19.2901 | 2403 | 1956 | 2206 | 2776 | 2995 | 2899 | 3018 | 2073 | 2156 | 2490 | 2257 | 2289 | 2561 | 2766 | 2693 | 2924 | 2517,1 | 2298 | 2736 | 1,19 | 0,252 | 4,403 | 1,07E-05 | 3,87E-05 | 0,002851 | 0,0025563 | 0,0027325 | 0,0027325 | TRUE | 0,0370007 | No |
| orf19.2903 | 192 | 199 | 186 | 256 | 195 | 270 | 237 | 138 | 172 | 253 | 190 | 211 | 167 | 258 | 211 | 195 | 207,19 | 206 | 208 | 1,009 | 0,012 | 0,082 | 0,93466904 | 0,95307053 | 0,025482 | 0,0089862 | 0,0169414 | 0,0169414 | TRUE | 0,2945993 | No |
| orf19.2905 | 13 | 21 | 17 | 17 | 16 | 18 | 15 | 4 | 12 | 27 | 17 | 14 | 14 | 17 | 13 | 6 | 14,96 | 18 | 12 | 0,75 | -0,415 | -1,041 | 0,29769272 | NA | 0,060938 | 0,099007 | 0,0844995 | 0,0844995 | TRUE | 0,5182708 | No |
| orf19.2906 | 159 | 92 | 135 | 197 | 161 | 123 | 158 | 111 | 143 | 117 | 138 | 162 | 138 | 117 | 141 | 157 | 139,12 | 140 | 138 | 0,974 | -0,038 | -0,284 | 0,77637889 | 0,82484803 | 0,007666 | 0,0124147 | 0,0103757 | 0,0103757 | TRUE | 0,1250253 | No |
| orf19.2907 | 6 | 5 | 12 | 12 | 4 | 6 | 6 | 7 | 5 | 6 | 12 | 10 | 3 | 6 | 5 | 10 | 7,29 | 8 | 6 | 0,685 | -0,545 | -1,018 | 0,30874133 | NA | 1,00E-08 | 0,201206 | 0,1315474 | 0,1315474 | TRUE | 0,2622376 | No |
| orf19.2907.1 | 233 | 210 | 182 | 291 | 353 | 258 | 294 | 150 | 209 | 267 | 186 | 240 | 302 | 246 | 262 | 212 | 240,57 | 226 | 256 | 1,141 | 0,191 | 1,437 | 0,15080702 | 0,21660258 | 0,017565 | 0,0080141 | 0,0127535 | 0,0127535 | TRUE | 0,1679631 | No |
| orf19.2908 | 320 | 225 | 256 | 324 | 353 | 318 | 333 | 244 | 287 | 286 | 262 | 267 | 302 | 303 | 297 | 344 | 293,65 | 276 | 312 | 1,126 | 0,171 | 1,974 | 0,04840326 | 0,08184825 | 1,00E-08 | 0,0069231 | 0,0037773 | 0,0037773 | TRUE | 0,0462882 | No |
| orf19.2909 | 861 | 616 | 792 | 963 | 942 | 877 | 919 | 582 | 772 | 784 | 810 | 794 | 806 | 837 | 820 | 821 | 805,55 | 790 | 821 | 1,038 | 0,054 | 1,03 | 0,30322699 | 0,38910461 | 1,00E-08 | 0,0037817 | 0,0014481 | 0,0014481 | TRUE | 0,0033834 | No |
| orf19.291 | 1667 | 1047 | 1387 | 1724 | 1536 | 1329 | 1452 | 908 | 1495 | 1333 | 1419 | 1422 | 1314 | 1268 | 1296 | 1281 | 1353,4 | 1417 | 1290 | 0,909 | -0,138 | -2,816 | 0,00486837 | 0,01052886 | 0,000451 | 0,0030522 | 0,0015673 | 0,0015673 | TRUE | 0,0177016 | No |
| orf19.2910 | 100 | 69 | 99 | 99 | 130 | 98 | 107 | 81 | 90 | 88 | 101 | 82 | 111 | 94 | 95 | 114 | 96,86 | 90 | 104 | 1,147 | 0,197 | 1,357 | 0,17471378 | 0,2458268 | 1,00E-08 | 0,016967 | 0,0099541 | 0,0099541 | TRUE | 0,0669274 | No |
| orf19.2911 | 1419 | 942 | 1316 | 1533 | 2162 | 1930 | 2137 | 1251 | 1273 | 1199 | 1346 | 1264 | 1849 | 1842 | 1907 | 1764 | 1555,6 | 1270 | 1840 | 1,449 | 0,535 | 10,84 | 2,23E-27 | 7,07E-26 | 0,000742 | 0,0029127 | 0,0016724 | 0,0016724 | TRUE | 0,0184055 | No |
| orf19.2914 | 458 | 313 | 414 | 517 | 688 | 542 | 637 | 376 | 411 | 398 | 424 | 426 | 588 | 517 | 568 | 530 | 482,94 | 415 | 551 | 1,33 | 0,411 | 5,747 | 9,09E-09 | 5,05E-08 | 1,00E-08 | 0,0049855 | 0,0028074 | 0,0028074 | TRUE | 0,0252606 | No |
| orf19.2915 | 2755 | 1954 | 2474 | 2986 | 3275 | 3220 | 3381 | 2047 | 2472 | 2488 | 2531 | 2462 | 2801 | 3072 | 3017 | 2887 | 2716,3 | 2488 | 2944 | 1,184 | 0,243 | 5,792 | 6,95E-09 | 3,94E-08 | 0,000684 | 0,002514 | 0,0013245 | 0,0013245 | TRUE | 0,0139932 | No |
| orf19.2916 | 66 | 37 | 63 | 67 | 63 | 79 | 85 | 32 | 59 | 47 | 64 | 55 | 54 | 75 | 76 | 45 | 59,53 | 56 | 62 | 1,115 | 0,157 | 0,763 | 0,44533508 | 0,53171168 | 0,020273 | 0,0263646 | 0,0238586 | 0,0238586 | TRUE | 0,208957 | No |
| orf19.2917 | 18417 | 12386 | 15710 | 19628 | 15512 | 13669 | 14351 | 8884 | 16522 | 15769 | 16073 | 16185 | 13266 | 13043 | 12806 | 12530 | 14524 | 16137 | 12911 | 0,8 | -0,322 | -10,425 | 1,91E-25 | 5,01E-24 | 0,000393 | 0,0020795 | 0,0008434 | 0,0008434 | TRUE | 0,0046774 | No |
| orf19.2917.1 | 559 | 353 | 421 | 579 | 715 | 550 | 585 | 455 | 501 | 449 | 431 | 477 | 611 | 525 | 522 | 642 | 519,89 | 464 | 575 | 1,231 | 0,3 | 3,444 | 0,00057416 | 0,00152704 | 0,005911 | 0,0047719 | 0,0053553 | 0,0053553 | TRUE | 0,0641899 | No |
| orf19.2919 | 563 | 391 | 470 | 614 | 599 | 558 | 585 | 355 | 505 | 498 | 481 | 506 | 512 | 532 | 522 | 501 | 507,18 | 498 | 517 | 1,04 | 0,056 | 0,872 | 0,38312049 | 0,47216162 | 1,00E-08 | 0,0048418 | 0,0020125 | 0,0020125 | TRUE | 0,0059987 | No |
| orf19.292 | 110 | 64 | 85 | 103 | 106 | 77 | 70 | 60 | 99 | 81 | 87 | 85 | 91 | 73 | 62 | 85 | 82,91 | 88 | 78 | 0,878 | -0,187 | -1,143 | 0,25284151 | 0,33526193 | 0,005741 | 0,0194891 | 0,0136172 | 0,0136172 | TRUE | 0,1793257 | No |
| orf19.2920 | 2521 | 1681 | 2048 | 2715 | 2723 | 2173 | 2371 | 1479 | 2262 | 2140 | 2095 | 2239 | 2329 | 2073 | 2116 | 2086 | 2167,5 | 2184 | 2151 | 0,985 | -0,022 | -0,416 | 0,67727113 | 0,74198486 | 0,001771 | 0,0026493 | 0,0021102 | 0,0021102 | TRUE | 0,0385852 | No |
| orf19.2921 | 156 | 113 | 151 | 153 | 105 | 121 | 114 | 80 | 140 | 144 | 154 | 126 | 90 | 115 | 102 | 113 | 123,03 | 141 | 105 | 0,742 | -0,431 | -3,217 | 0,00129654 | 0,00318464 | 1,00E-08 | 0,0137787 | 0,0089431 | 0,0089431 | TRUE | 0,1024279 | No |
| orf19.2922 | 23 | 20 | 15 | 23 | 31 | 14 | 17 | 12 | 21 | 25 | 15 | 19 | 27 | 13 | 15 | 17 | 19,05 | 20 | 18 | 0,909 | -0,137 | -0,409 | 0,68283598 | 0,7465839 | 0,022411 | 0,0781977 | 0,0555219 | 0,0555219 | TRUE | 0,6210786 | No |
| orf19.2923 | 204 | 170 | 167 | 234 | 288 | 260 | 245 | 170 | 183 | 216 | 171 | 193 | 246 | 248 | 219 | 240 | 214,5 | 191 | 238 | 1,253 | 0,325 | 3,145 | 0,00166278 | 0,00398802 | 0,001026 | 0,0087472 | 0,0055352 | 0,0055352 | TRUE | 0,0742972 | No |
| orf19.2925 | 307 | 289 | 265 | 366 | 289 | 316 | 315 | 174 | 275 | 368 | 271 | 302 | 247 | 302 | 281 | 245 | 286,43 | 304 | 269 | 0,89 | -0,169 | -1,472 | 0,14092628 | 0,20461216 | 0,011284 | 0,0070478 | 0,0090843 | 0,0090843 | TRUE | 0,2069827 | No |
| orf19.2926 | 1845 | 1431 | 1720 | 2121 | 2136 | 2073 | 2309 | 1310 | 1655 | 1822 | 1760 | 1749 | 1827 | 1978 | 2060 | 1848 | 1837,3 | 1746 | 1928 | 1,106 | 0,145 | 2,687 | 0,00720272 | 0,01497887 | 0,001695 | 0,0027696 | 0,002255 | 0,002255 | TRUE | 0,0262151 | No |
| orf19.2927 | 1024 | 1125 | 1061 | 1316 | 1507 | 1968 | 1961 | 1166 | 919 | 1432 | 1086 | 1085 | 1289 | 1878 | 1750 | 1645 | 1385,3 | 1130 | 1640 | 1,452 | 0,538 | 3,049 | 0,00229478 | 0,00534796 | 0,029158 | 0,0030274 | 0,0155788 | 0,0291577 | TRUE | 0,3828586 | No |
| orf19.2928 | 715 | 698 | 667 | 863 | 778 | 857 | 909 | 521 | 641 | 889 | 682 | 712 | 665 | 818 | 811 | 735 | 744,14 | 731 | 757 | 1,039 | 0,055 | 0,53 | 0,59583748 | 0,67281569 | 0,013522 | 0,0039304 | 0,0088373 | 0,0088373 | TRUE | 0,2396531 | No |
| orf19.2929 | 10438 | 8115 | 10407 | 13143 | 9400 | 9250 | 9451 | 6113 | 9364 | 10331 | 10647 | 10838 | 8039 | 8826 | 8433 | 8622 | 9387,6 | 10295 | 8480 | 0,824 | -0,28 | -5,315 | 1,07E-07 | 5,21E-07 | 0,00266 | 0,0021342 | 0,0025574 | 0,0025574 | TRUE | 0,0456289 | No |
| orf19.2930 | 4674 | 3367 | 4245 | 5126 | 4404 | 4052 | 4384 | 2995 | 4193 | 4287 | 4343 | 4227 | 3766 | 3866 | 3912 | 4224 | 4102,3 | 4262 | 3942 | 0,924 | -0,114 | -2,7 | 0,00694018 | 0,01450316 | 0,001051 | 0,0023334 | 0,001476 | 0,001476 | TRUE | 0,0267475 | No |
| orf19.2933 | 2185 | 1490 | 1782 | 2350 | 1767 | 1719 | 1840 | 1082 | 1960 | 1897 | 1823 | 1938 | 1511 | 1640 | 1642 | 1526 | 1742,2 | 1904 | 1580 | 0,83 | -0,269 | -5,53 | 3,21E-08 | 1,67E-07 | 0,000939 | 0,0028128 | 0,0016937 | 0,0016937 | TRUE | 0,0115636 | No |
| orf19.2934 | 6184 | 4998 | 5473 | 7288 | 6255 | 6512 | 6561 | 4012 | 5548 | 6363 | 5599 | 6010 | 5349 | 6214 | 5855 | 5659 | 5824,5 | 5880 | 5769 | 0,981 | -0,027 | -0,454 | 0,649868 | 0,71930858 | 0,003434 | 0,0022288 | 0,0032302 | 0,0032302 | TRUE | 0,036742 | No |

| | | | | | | | | | | | | | | | | | | | | | | | | | | | | | | | |
|---|---|---|---|---|---|---|---|---|---|---|---|---|---|---|---|---|---|---|---|---|---|---|---|---|---|---|---|---|---|---|---|
| orf19.2935 | 28277 | 18282 | 23440 | 29562 | 26374 | 21971 | 23633 | 14688 | 25367 | 23275 | 23981 | 24377 | 22556 | 20964 | 21089 | 20716 | 22791 | 24250 | 21331 | 0,88 | -0,185 | -4,526 | 6,01E-06 | 2,26E-05 | 0,001349 | 0,0020432 | 0,0015608 | 0,0015608 | TRUE | 0,01836 | No |
| orf19.2936 | 350 | 330 | 340 | 442 | 311 | 324 | 340 | 177 | 314 | 420 | 348 | 364 | 266 | 309 | 303 | 250 | 321,83 | 362 | 282 | 0,786 | -0,348 | -3,269 | 0,00108043 | 0,00270355 | 0,009086 | 0,0064903 | 0,0077354 | 0,0077354 | TRUE | 0,1251676 | No |
| orf19.2937 | 1395 | 1347 | 1269 | 1717 | 1402 | 1385 | 1392 | 783 | 1251 | 1715 | 1298 | 1416 | 1199 | 1322 | 1242 | 1104 | 1318,4 | 1420 | 1217 | 0,859 | -0,22 | -2,328 | 0,01991001 | 0,0370568 | 0,012507 | 0,0030806 | 0,0077754 | 0,0077754 | TRUE | 0,2292192 | No |
| orf19.2938 | 1595 | 1293 | 1440 | 1753 | 1498 | 1581 | 1616 | 993 | 1431 | 1646 | 1473 | 1446 | 1281 | 1509 | 1442 | 1401 | 1453,5 | 1499 | 1408 | 0,941 | -0,088 | -1,348 | 0,17760583 | 0,24931212 | 0,003597 | 0,0029783 | 0,0034393 | 0,0034393 | TRUE | 0,0504488 | No |
| orf19.2939 | 362 | 278 | 328 | 429 | 457 | 375 | 407 | 230 | 325 | 354 | 336 | 354 | 391 | 358 | 363 | 324 | 350,53 | 342 | 359 | 1,056 | 0,079 | 0,958 | 0,33781211 | 0,42495292 | 1,00E-08 | 0,006121 | 0,003656 | 0,003656 | TRUE | 0,0404135 | No |
| orf19.2940 | 388 | 319 | 323 | 456 | 523 | 485 | 556 | 315 | 348 | 406 | 330 | 376 | 447 | 463 | 496 | 444 | 413,89 | 365 | 462 | 1,272 | 0,347 | 4,228 | 2,36E-05 | 8,10E-05 | 0,002167 | 0,0054869 | 0,0040194 | 0,0040194 | TRUE | 0,0566643 | No |
| orf19.2941 | 5 | 1 | 3 | 5 | 3 | 7 | 5 | 3 | 4 | 1 | 3 | 4 | 3 | 7 | 4 | 4 | 3,86 | 3 | 4 | 1,334 | 0,415 | 0,596 | 0,55129378 | NA | 1,00E-08 | 0,3779612 | 0,2070652 | 0,2070652 | TRUE | 0,2241168 | No |
| orf19.2942 | 46227 | 32276 | 42180 | 50632 | 43201 | 38206 | 41933 | 25897 | 41470 | 41091 | 43154 | 41752 | 36947 | 36455 | 37418 | 36525 | 39352 | 41867 | 36836 | 0,88 | -0,185 | -6,887 | 5,68E-12 | 4,74E-11 | 0,00027 | 0,0020164 | 0,0006647 | 0,0006647 | TRUE | 0,0052358 | No |
| orf19.2943.5 | 2 | 2 | 1 | 4 | 2 | 2 | 1 | 3 | 2 | 3 | 1 | 3 | 2 | 2 | 1 | 4 | 2,18 | 2 | 2 | 0,941 | -0,087 | -0,094 | 0,92541896 | NA | 1,00E-08 | 0,669262 | 0,3605252 | 0,3605252 | TRUE | 0,246511 | No |
| orf19.2945 | 80 | 44 | 56 | 72 | 56 | 54 | 52 | 17 | 72 | 56 | 57 | 59 | 48 | 52 | 46 | 24 | 51,78 | 61 | 42 | 0,708 | -0,499 | -2,286 | 0,02223193 | 0,04107699 | 0,017952 | 0,030015 | 0,0260005 | 0,0260005 | TRUE | 0,4658055 | No |
| orf19.2946 | 362 | 208 | 283 | 382 | 245 | 257 | 250 | 166 | 325 | 265 | 290 | 315 | 210 | 245 | 223 | 234 | 263,26 | 299 | 228 | 0,758 | -0,399 | -4,079 | 4,52E-05 | 0,00014763 | 0,002746 | 0,0074939 | 0,0053208 | 0,0053208 | TRUE | 0,0582708 | No |
| orf19.2947 | 3848 | 2522 | 3204 | 4007 | 4150 | 3605 | 3792 | 2438 | 3452 | 3211 | 3278 | 3304 | 3549 | 3440 | 3384 | 3439 | 3382,1 | 3311 | 3453 | 1,042 | 0,06 | 1,581 | 0,11391095 | 0,17024304 | 0,000392 | 0,0024088 | 0,0010719 | 0,0010719 | TRUE | 0,0100239 | No |
| orf19.2948 | 1555 | 1073 | 1327 | 1728 | 1707 | 1436 | 1615 | 1078 | 1395 | 1366 | 1358 | 1425 | 1460 | 1370 | 1441 | 1520 | 1416,9 | 1386 | 1448 | 1,042 | 0,06 | 1,244 | 0,21354017 | 0,2909842 | 0,00037 | 0,0030041 | 0,0015031 | 0,0015031 | TRUE | 0,0153554 | No |
| orf19.2949 | 1092 | 789 | 1017 | 1249 | 1273 | 1268 | 1302 | 805 | 980 | 1004 | 1040 | 1030 | 1089 | 1210 | 1162 | 1135 | 1081,3 | 1014 | 1149 | 1,133 | 0,181 | 3,403 | 0,00066654 | 0,00174113 | 0,000604 | 0,0033221 | 0,0017762 | 0,0017762 | TRUE | 0,0163996 | No |
| orf19.295 | 138 | 97 | 125 | 160 | 158 | 119 | 141 | 88 | 124 | 123 | 128 | 132 | 135 | 114 | 126 | 124 | 125,72 | 127 | 125 | 0,983 | -0,024 | -0,198 | 0,84310863 | 0,88047917 | 1,00E-08 | 0,013527 | 0,0066045 | 0,0066045 | TRUE | 0,0400522 | No |
| orf19.2951 | 746 | 516 | 598 | 775 | 699 | 637 | 691 | 468 | 669 | 657 | 612 | 639 | 598 | 608 | 617 | 660 | 632,42 | 644 | 621 | 0,96 | -0,058 | -0,937 | 0,34898827 | 0,4365278 | 1,00E-08 | 0,004275 | 0,0021612 | 0,0021612 | TRUE | 0,0183383 | No |
| orf19.2952 | 717 | 439 | 629 | 724 | 568 | 493 | 566 | 375 | 643 | 559 | 644 | 597 | 486 | 470 | 505 | 529 | 554,1 | 611 | 498 | 0,811 | -0,302 | -4,234 | 2,30E-05 | 7,90E-05 | 0,001438 | 0,0045994 | 0,0030618 | 0,0030618 | TRUE | 0,033967 | No |
| orf19.2953 | 2509 | 1642 | 2145 | 2602 | 2016 | 1753 | 1780 | 1150 | 2251 | 2090 | 2195 | 2146 | 1724 | 1673 | 1588 | 1622 | 1911,1 | 2170 | 1652 | 0,761 | -0,394 | -8,642 | 5,53E-18 | 8,61E-17 | 0,000634 | 0,0027391 | 0,0014593 | 0,0014593 | TRUE | 0,0110284 | No |
| orf19.2954 | 4650 | 2978 | 3762 | 4638 | 4402 | 3651 | 3861 | 2482 | 4172 | 3791 | 3849 | 3825 | 3765 | 3484 | 3445 | 3501 | 3728,8 | 3909 | 3549 | 0,908 | -0,14 | -2,952 | 0,00315282 | 0,00712379 | 0,001507 | 0,0023688 | 0,0018762 | 0,0018762 | TRUE | 0,0251529 | No |
| orf19.2956 | 1363 | 906 | 1106 | 1393 | 1360 | 1200 | 1286 | 876 | 1223 | 1153 | 1132 | 1149 | 1163 | 1145 | 1148 | 1236 | 1168,5 | 1164 | 1173 | 1,005 | 0,007 | 0,138 | 0,89042513 | 0,91713941 | 0,000307 | 0,0032219 | 0,001606 | 0,001606 | TRUE | 0,0143573 | No |
| orf19.2957 | 22 | 7 | 22 | 22 | 19 | 29 | 25 | 15 | 20 | 9 | 23 | 18 | 16 | 28 | 22 | 21 | 19,59 | 18 | 22 | 1,238 | 0,308 | 0,924 | 0,35530092 | 0,4428858 | 0,018954 | 0,0761015 | 0,0556276 | 0,0556276 | TRUE | 0,3893911 | No |
| orf19.2958 | 270 | 186 | 258 | 290 | 399 | 328 | 386 | 240 | 242 | 237 | 264 | 239 | 341 | 313 | 344 | 338 | 289,91 | 246 | 334 | 1,361 | 0,445 | 5,214 | 1,85E-07 | 8,74E-07 | 1,00E-08 | 0,006987 | 0,0034787 | 0,0034787 | TRUE | 0,0274931 | No |
| orf19.2959 | 18 | 9 | 15 | 23 | 7 | 7 | 8 | 3 | 16 | 11 | 15 | 19 | 6 | 7 | 7 | 4 | 10,74 | 15 | 6 | 0,391 | -1,354 | -3,095 | 0,00197067 | NA | 1,00E-08 | 0,1370936 | 0,070922 | 0,070922 | TRUE | 0,1212047 | No |
| orf19.2959.1 | 2735 | 2083 | 2388 | 3269 | 2842 | 2460 | 2334 | 1436 | 2454 | 2652 | 2443 | 2696 | 2431 | 2347 | 2083 | 2025 | 2391,3 | 2561 | 2222 | 0,868 | -0,203 | -3,046 | 0,00231721 | 0,00539395 | 0,004717 | 0,0025866 | 0,0038605 | 0,0038605 | TRUE | 0,0488672 | No |
| orf19.296 | 41 | 31 | 43 | 43 | 53 | 48 | 53 | 39 | 37 | 39 | 44 | 35 | 45 | 46 | 47 | 55 | 43,64 | 39 | 48 | 1,237 | 0,307 | 1,494 | 0,13512899 | 0,1975797 | 1,00E-08 | 0,0352443 | 0,0174354 | 0,0174354 | TRUE | 0,0591077 | No |
| orf19.2960 | 3931 | 2608 | 3296 | 4126 | 3469 | 3054 | 3227 | 2007 | 3527 | 3320 | 3372 | 3402 | 2967 | 2914 | 2880 | 2831 | 3151,5 | 3405 | 2898 | 0,851 | -0,233 | -6,334 | 2,39E-10 | 1,65E-09 | 0,00021 | 0,0024402 | 0,0009741 | 0,0009741 | TRUE | 0,007026 | No |
| orf19.2961 | 832 | 639 | 762 | 929 | 720 | 663 | 719 | 420 | 746 | 814 | 780 | 766 | 616 | 633 | 642 | 592 | 698,49 | 776 | 621 | 0,803 | -0,317 | -5,372 | 7,81E-08 | 3,89E-07 | 1,00E-08 | 0,0040579 | 0,0019016 | 0,0019016 | TRUE | 0,0111053 | No |
| orf19.2962 | 1089 | 813 | 970 | 1183 | 916 | 899 | 972 | 572 | 977 | 1035 | 992 | 976 | 783 | 858 | 867 | 807 | 911,9 | 995 | 829 | 0,835 | -0,261 | -4,647 | 3,36E-06 | 1,32E-05 | 0,000459 | 0,0035715 | 0,0019107 | 0,0019107 | TRUE | 0,0185457 | No |
| orf19.2963 | 145 | 108 | 123 | 147 | 156 | 140 | 156 | 73 | 130 | 137 | 126 | 121 | 133 | 134 | 139 | 103 | 127,97 | 128 | 127 | 1,003 | 0,004 | 0,033 | 0,97333538 | 0,98139755 | 1,00E-08 | 0,0133232 | 0,0080825 | 0,0080825 | TRUE | 0,1303217 | No |
| orf19.2964 | 4447 | 2939 | 3983 | 4699 | 5263 | 4679 | 4934 | 3174 | 3989 | 3742 | 4075 | 3875 | 4501 | 4465 | 4403 | 4477 | 4190,8 | 3920 | 4462 | 1,137 | 0,186 | 5,052 | 4,38E-07 | 1,95E-06 | 0,000404 | 0,0023259 | 0,0010565 | 0,0010565 | TRUE | 0,0107279 | No |
| orf19.2965 | 2394 | 1721 | 2144 | 2604 | 2193 | 2016 | 2040 | 1416 | 2148 | 2191 | 2194 | 2147 | 1876 | 1924 | 1820 | 1997 | 2037 | 2170 | 1904 | 0,876 | -0,191 | -4,519 | 6,22E-06 | 2,34E-05 | 0,000189 | 0,0026922 | 0,0012128 | 0,0012128 | TRUE | 0,0117208 | No |
| orf19.2965.1 | 381 | 275 | 333 | 412 | 443 | 410 | 434 | 249 | 342 | 350 | 341 | 340 | 379 | 391 | 387 | 351 | 360,11 | 343 | 377 | 1,104 | 0,143 | 1,899 | 0,05756165 | 0,09495617 | 1,00E-08 | 0,0060108 | 0,002678 | 0,002678 | TRUE | 0,0197756 | No |
| orf19.2966 | 245 | 167 | 223 | 258 | 268 | 265 | 241 | 135 | 220 | 213 | 228 | 213 | 229 | 253 | 215 | 190 | 220,1 | 218 | 222 | 1,023 | 0,032 | 0,311 | 0,75598307 | 0,80947181 | 0,001886 | 0,0085751 | 0,0057679 | 0,0057679 | TRUE | 0,0916789 | No |
| orf19.2967 | 9018 | 5902 | 7684 | 9418 | 8542 | 7044 | 7680 | 4656 | 8090 | 7514 | 7861 | 7766 | 7305 | 6721 | 6853 | 6567 | 7334,8 | 7808 | 6862 | 0,879 | -0,186 | -4,321 | 1,55E-05 | 5,50E-05 | 0,001359 | 0,0021774 | 0,001641 | 0,001641 | TRUE | 0,0233017 | No |
| orf19.2968 | 385 | 266 | 296 | 382 | 352 | 318 | 331 | 191 | 345 | 339 | 303 | 315 | 301 | 303 | 295 | 269 | 308,89 | 326 | 292 | 0,903 | -0,148 | -1,742 | 0,08153826 | 0,12867923 | 1,00E-08 | 0,0066793 | 0,0036768 | 0,0036768 | TRUE | 0,0252825 | No |
| orf19.2969 | 19429 | 12886 | 17097 | 20765 | 22894 | 19524 | 21950 | 13394 | 17430 | 16405 | 17492 | 17123 | 19580 | 18629 | 19587 | 18891 | 18142 | 17112 | 19172 | 1,12 | 0,164 | 4,786 | 1,70E-06 | 6,95E-06 | 0,000673 | 0,0020595 | 0,0010703 | 0,0010703 | TRUE | 0,009327 | No |
| orf19.297 | 873 | 471 | 710 | 877 | 900 | 618 | 696 | 432 | 783 | 600 | 726 | 723 | 770 | 590 | 621 | 609 | 677,77 | 708 | 648 | 0,914 | -0,13 | -1,281 | 0,20021351 | 0,27475764 | 0,012126 | 0,0041214 | 0,0083559 | 0,0083559 | TRUE | 0,196258 | No |
| orf19.2970 | 2657 | 1728 | 2421 | 2818 | 2833 | 2542 | 2639 | 1669 | 2384 | 2200 | 2477 | 2324 | 2423 | 2425 | 2355 | 2354 | 2367,7 | 2346 | 2389 | 1,018 | 0,026 | 0,572 | 0,5676528 | 0,64690024 | 0,000802 | 0,0025927 | 0,0015211 | 0,0015211 | TRUE | 0,0197911 | No |
| orf19.2971 | 745 | 532 | 598 | 831 | 739 | 781 | 815 | 511 | 668 | 677 | 612 | 685 | 632 | 745 | 727 | 721 | 683,49 | 660 | 706 | 1,067 | 0,094 | 1,324 | 0,18538817 | 0,25776721 | 0,002742 | 0,0041035 | 0,0033756 | 0,0033756 | TRUE | 0,0632507 | No |
| orf19.2972 | 8609 | 6503 | 7831 | 9824 | 11936 | 11120 | 11897 | 7501 | 7723 | 8279 | 8012 | 8101 | 10208 | 10610 | 10616 | 10579 | 9266,1 | 8029 | 10503 | 1,308 | 0,388 | 11,659 | 2,07E-31 | 8,14E-30 | 0,000448 | 0,0021362 | 0,0009525 | 0,0009525 | TRUE | 0,0080643 | No |
| orf19.2973 | 4983 | 3583 | 4723 | 5519 | 7081 | 6572 | 7268 | 4692 | 4470 | 4562 | 4832 | 4551 | 6056 | 6271 | 6485 | 6618 | 5480,6 | 4604 | 6358 | 1,38 | 0,465 | 11,04 | 2,46E-28 | 8,20E-27 | 0,001089 | 0,0022444 | 0,0015161 | 0,0015161 | TRUE | 0,013599 | No |
| orf19.2974 | 1422 | 1058 | 1273 | 1486 | 1466 | 1459 | 1523 | 850 | 1276 | 1347 | 1302 | 1225 | 1254 | 1392 | 1359 | 1199 | 1294,3 | 1288 | 1301 | 1,013 | 0,019 | 0,323 | 0,74639536 | 0,80106007 | 0,002284 | 0,0031012 | 0,0026418 | 0,0026418 | TRUE | 0,0302804 | No |
| orf19.2975 | 977 | 683 | 875 | 1076 | 1002 | 909 | 932 | 565 | 876 | 870 | 895 | 887 | 857 | 867 | 832 | 797 | 860,16 | 882 | 838 | 0,952 | -0,071 | -1,337 | 0,1811494 | 0,25292717 | 1,00E-08 | 0,0036672 | 0,0015351 | 0,0015351 | TRUE | 0,0109329 | No |
| orf19.2977 | 5996 | 4076 | 5045 | 6471 | 5616 | 4918 | 5560 | 3655 | 5379 | 5189 | 5162 | 5336 | 4803 | 4693 | 4961 | 5155 | 5084,7 | 5266 | 4903 | 0,93 | -0,105 | -2,668 | 0,00762215 | 0,01577467 | 0,000785 | 0,002265 | 0,0012818 | 0,0012818 | TRUE | 0,0138345 | No |
| orf19.2978 | 2038 | 1333 | 1722 | 2142 | 1714 | 1373 | 1509 | 937 | 1828 | 1697 | 1762 | 1766 | 1466 | 1310 | 1347 | 1322 | 1562,2 | 1763 | 1361 | 0,772 | -0,373 | -7,256 | 3,97E-13 | 3,87E-12 | 0,001157 | 0,0029088 | 0,0018817 | 0,0018817 | TRUE | 0,0337459 | No |
| orf19.298 | 54 | 38 | 40 | 50 | 41 | 31 | 41 | 25 | 48 | 48 | 41 | 41 | 35 | 30 | 37 | 35 | 39,43 | 44 | 34 | 0,765 | -0,387 | -1,79 | 0,07345341 | 0,11743163 | 1,00E-08 | 0,0387937 | 0,0192236 | 0,0192236 | TRUE | 0,0603241 | No |
| orf19.2980 | 1688 | 1153 | 1563 | 1913 | 1748 | 1491 | 1590 | 1055 | 1514 | 1468 | 1599 | 1577 | 1495 | 1423 | 1419 | 1488 | 1497,9 | 1540 | 1456 | 0,944 | -0,083 | -1,746 | 0,08086683 | 0,12772028 | 0,00044 | 0,0029487 | 0,0014967 | 0,0014967 | TRUE | 0,0108218 | No |
| orf19.2982 | 343 | 226 | 274 | 359 | 300 | 305 | 303 | 180 | 308 | 288 | 280 | 296 | 257 | 291 | 270 | 254 | 280,45 | 293 | 268 | 0,915 | -0,128 | -1,468 | 0,14207594 | 0,20608588 | 1,00E-08 | 0,0071558 | 0,0037265 | 0,0037265 | TRUE | 0,0374762 | No |
| orf19.2983 | 1318 | 873 | 1138 | 1460 | 1205 | 1038 | 1141 | 802 | 1182 | 1111 | 1164 | 1204 | 1031 | 990 | 1018 | 1131 | 1104 | 1165 | 1042 | 0,891 | -0,167 | -3,001 | 0,00268686 | 0,00617546 | 0,001097 | 0,0032944 | 0,002067 | 0,002067 | TRUE | 0,0345365 | No |
| orf19.2984 | 1274 | 907 | 1196 | 1436 | 1743 | 1468 | 1625 | 991 | 1143 | 1155 | 1224 | 1184 | 1491 | 1401 | 1450 | 1398 | 1305,6 | 1176 | 1435 | 1,221 | 0,288 | 5,996 | 2,03E-09 | 1,23E-08 | 1,00E-08 | 0,0030915 | 0,0014434 | 0,0014434 | TRUE | 0,0085521 | No |
| orf19.2985 | 1246 | 837 | 1098 | 1261 | 1329 | 1076 | 1191 | 775 | 1118 | 1066 | 1123 | 1040 | 1137 | 1027 | 1063 | 1093 | 1083,2 | 1087 | 1080 | 0,994 | -0,009 | -0,167 | 0,86737672 | 0,89863758 | 0,000865 | 0,0033197 | 0,0019264 | 0,0019264 | TRUE | 0,0165098 | No |
| orf19.2986 | 422 | 267 | 340 | 463 | 547 | 411 | 447 | 335 | 379 | 340 | 348 | 382 | 468 | 392 | 399 | 472 | 397,44 | 362 | 433 | 1,189 | 0,25 | 2,82 | 0,00480155 | 0,01040308 | 0,004347 | 0,0056322 | 0,0049942 | 0,0049942 | TRUE | 0,0454897 | No |
| orf19.2987 | 3828 | 2593 | 3276 | 4076 | 4280 | 3590 | 3913 | 2598 | 3434 | 3301 | 3352 | 3361 | 3660 | 3425 | 3492 | 3664 | 3461,2 | 3362 | 3560 | 1,058 | 0,081 | 2,145 | 0,03191362 | 0,0566097 | 0,000422 | 0,0023989 | 0,0010917 | 0,0010917 | TRUE | 0,0078836 | No |
| orf19.2988 | 1715 | 1129 | 1598 | 1908 | 2356 | 2005 | 2120 | 1461 | 1539 | 1437 | 1635 | 1573 | 2015 | 1913 | 1892 | 2061 | 1758,1 | 1546 | 1970 | 1,272 | 0,347 | 6,608 | 3,90E-11 | 3,00E-10 | 0,001502 | 0,0028053 | 0,0020627 | 0,0020627 | TRUE | 0,0249898 | No |
| orf19.2989 | 3087 | 2039 | 2686 | 3239 | 4404 | 3676 | 4081 | 2654 | 2769 | 2596 | 2748 | 2671 | 3766 | 3508 | 3642 | 3743 | 3180,4 | 2696 | 3665 | 1,358 | 0,442 | 10,86 | 1,78E-27 | 5,66E-26 | 0,000519 | 0,002436 | 0,0012644 | 0,0012644 | TRUE | 0,0102146 | No |
| orf19.299 | 2704 | 1855 | 2376 | 2957 | 2983 | 2681 | 2891 | 2089 | 2426 | 2362 | 2431 | 2438 | 2551 | 2558 | 2580 | 2946 | 2536,5 | 2414 | 2659 | 1,099 | 0,136 | 2,585 | 0,00973159 | 0,01966613 | 0,00219 | 0,0025518 | 0,0022639 | 0,0022639 | TRUE | 0,060976 | No |
| orf19.2990 | 2606 | 1903 | 2511 | 2800 | 2176 | 1913 | 2027 | 1418 | 2338 | 2423 | 2569 | 2309 | 1861 | 1825 | 1809 | 2000 | 2141,7 | 2410 | 1874 | 0,777 | -0,364 | -7,085 | 1,39E-12 | 1,25E-11 | 0,001616 | 0,0026574 | 0,0020624 | 0,0020624 | TRUE | 0,0243341 | No |
| orf19.2991 | 122 | 65 | 97 | 99 | 123 | 93 | 104 | 90 | 109 | 83 | 99 | 82 | 105 | 89 | 93 | 127 | 98,34 | 93 | 104 | 1,096 | 0,132 | 0,817 | 0,41391084 | 0,50213744 | 0,012297 | 0,0167411 | 0,014836 | 0,014836 | TRUE | 0,1868343 | No |
| orf19.2992 | 7391 | 4892 | 6203 | 8009 | 6796 | 5684 | 5906 | 4283 | 6630 | 6228 | 6346 | 6604 | 5812 | 5423 | 5270 | 6041 | 6044,5 | 6452 | 5636 | 0,873 | -0,196 | -3,94 | 8,15E-05 | 0,0002525 | 0,002206 | 0,0022197 | 0,0022156 | 0,0022156 | TRUE | 0,0275862 | No |

| | | | | | | | | | | | | | | | | | | | | | | | | | | | | | | | |
|---|---|---|---|---|---|---|---|---|---|---|---|---|---|---|---|---|---|---|---|---|---|---|---|---|---|---|---|---|---|---|---|
| orf19.2994 | 24238 | 14687 | 19368 | 24018 | 21751 | 17635 | 19237 | 12741 | 21744 | 18698 | 19815 | 19806 | 18602 | 16827 | 17166 | 17970 | 18828 | 20016 | 17641 | 0,881 | -0,182 | -3,459 | 0,00054286 | 0,00145277 | 0,002958 | 0,0020566 | 0,0026181 | 0,0026181 | TRUE | 0,041388 | No |
| orf19.2994.1 | 12613 | 8248 | 10234 | 12820 | 12694 | 10396 | 10736 | 7298 | 11315 | 10501 | 10470 | 10572 | 10856 | 9920 | 9580 | 10293 | 10438 | 10714 | 10162 | 0,948 | -0,077 | -1,61 | 0,107391 | 0,16207111 | 0,002026 | 0,0021186 | 0,002073 | 0,002073 | TRUE | 0,026218 | No |
| orf19.2995 | 1070 | 685 | 915 | 1028 | 1175 | 947 | 1102 | 722 | 960 | 872 | 936 | 848 | 1005 | 904 | 983 | 1018 | 940,75 | 904 | 978 | 1,08 | 0,111 | 1,763 | 0,07785923 | 0,12365272 | 0,002123 | 0,0035227 | 0,0027508 | 0,0027508 | TRUE | 0,031166 | No |
| orf19.2996 | 1782 | 1294 | 1724 | 1960 | 1800 | 1702 | 1918 | 1109 | 1599 | 1647 | 1764 | 1616 | 1539 | 1624 | 1711 | 1564 | 1633,1 | 1656 | 1610 | 0,973 | -0,04 | -0,748 | 0,45431288 | 0,54017648 | 0,001559 | 0,0028684 | 0,0020875 | 0,0020875 | TRUE | 0,0230853 | No |
| orf19.2998 | 2798 | 1762 | 2336 | 2875 | 2590 | 2077 | 2170 | 1361 | 2510 | 2243 | 2390 | 2371 | 2215 | 1982 | 1936 | 1920 | 2195,9 | 2378 | 2013 | 0,847 | -0,24 | -4,146 | 3,39E-05 | 0,00011275 | 0,002763 | 0,0026406 | 0,0027563 | 0,0027563 | TRUE | 0,056336 | No |
| orf19.30 | 2387 | 1841 | 2302 | 2794 | 2565 | 2389 | 2494 | 1575 | 2141 | 2344 | 2355 | 2304 | 2194 | 2280 | 2225 | 2221 | 2258,1 | 2286 | 2230 | 0,976 | -0,035 | -0,791 | 0,4292351 | 0,51630446 | 0,000695 | 0,0026224 | 0,001414 | 0,001414 | TRUE | 0,0224521 | No |
| orf19.300 | 1617 | 1021 | 1402 | 1761 | 1720 | 1360 | 1463 | 998 | 1451 | 1300 | 1434 | 1452 | 1471 | 1298 | 1305 | 1408 | 1389,8 | 1409 | 1370 | 0,971 | -0,043 | -0,716 | 0,47372635 | 0,5597054 | 0,00224 | 0,003024 | 0,0027243 | 0,0027243 | TRUE | 0,0315733 | No |
| orf19.3000 | 656 | 473 | 564 | 677 | 715 | 605 | 672 | 469 | 589 | 602 | 577 | 558 | 611 | 577 | 600 | 661 | 596,98 | 582 | 612 | 1,051 | 0,072 | 1,11 | 0,26701863 | 0,35130558 | 1,00E-08 | 0,0044113 | 0,0023817 | 0,0023817 | TRUE | 0,0289051 | No |
| orf19.3001 | 194 | 101 | 129 | 178 | 216 | 199 | 247 | 140 | 174 | 129 | 132 | 147 | 185 | 190 | 220 | 197 | 171,73 | 146 | 198 | 1,355 | 0,439 | 3,547 | 0,00038919 | 0,00107641 | 0,006653 | 0,0104328 | 0,0087498 | 0,0087498 | TRUE | 0,1878044 | No |
| orf19.3002 | 14303 | 9337 | 11665 | 14714 | 14698 | 12055 | 12543 | 8012 | 12831 | 11887 | 11934 | 12133 | 12570 | 11502 | 11193 | 11300 | 11919 | 12196 | 11641 | 0,955 | -0,067 | -1,431 | 0,15232795 | 0,2184737 | 0,002004 | 0,0021013 | 0,0020265 | 0,0020265 | TRUE | 0,0355305 | No |
| orf19.3003 | 1779 | 1224 | 1765 | 2020 | 2850 | 2656 | 2765 | 1658 | 1596 | 1558 | 1806 | 1666 | 2437 | 2534 | 2467 | 2338 | 2050,4 | 1656 | 2444 | 1,476 | 0,562 | 10,447 | 1,51E-25 | 4,01E-24 | 0,002046 | 0,0026875 | 0,0022674 | 0,0022674 | TRUE | 0,0435101 | No |
| orf19.3003.1 | 14497 | 9513 | 11804 | 14988 | 14572 | 12058 | 12889 | 9273 | 13005 | 12111 | 12077 | 12359 | 12462 | 11505 | 11501 | 13079 | 12263 | 12388 | 12137 | 0,979 | -0,03 | -0,597 | 0,55036142 | 0,63126602 | 0,002271 | 0,0020979 | 0,0023606 | 0,0023606 | TRUE | 0,0329218 | No |
| orf19.3004 | 6390 | 4444 | 5553 | 6505 | 6796 | 6407 | 6991 | 4045 | 5732 | 5658 | 5681 | 5364 | 5812 | 6113 | 6238 | 5705 | 5788,1 | 5609 | 5967 | 1,065 | 0,09 | 2,157 | 0,03104128 | 0,05525956 | 0,001086 | 0,0022303 | 0,0015121 | 0,0015121 | TRUE | 0,0114138 | No |
| orf19.3006 | 2980 | 1947 | 2600 | 3234 | 3130 | 2872 | 3002 | 1994 | 2673 | 2479 | 2660 | 2667 | 2677 | 2740 | 2679 | 2812 | 2673,4 | 2620 | 2727 | 1,039 | 0,055 | 1,347 | 0,17811065 | 0,24972901 | 0,000423 | 0,0025225 | 0,0012244 | 0,0012244 | TRUE | 0,0149018 | No |
| orf19.3007 | 792 | 572 | 696 | 888 | 759 | 605 | 664 | 423 | 711 | 728 | 712 | 732 | 649 | 577 | 593 | 597 | 662,32 | 721 | 604 | 0,839 | -0,253 | -4,133 | 3,58E-05 | 0,00011888 | 1,00E-08 | 0,0041714 | 0,0020869 | 0,0020869 | TRUE | 0,032675 | No |
| orf19.3007.2 | 1100 | 716 | 942 | 1173 | 912 | 802 | 859 | 441 | 987 | 912 | 964 | 967 | 780 | 765 | 767 | 622 | 845,39 | 958 | 734 | 0,769 | -0,379 | -5,265 | 1,40E-07 | 6,73E-07 | 0,004131 | 0,0036967 | 0,0037629 | 0,0037629 | TRUE | 0,1098772 | No |
| orf19.3008 | 1304 | 903 | 1138 | 1361 | 1449 | 1221 | 1343 | 805 | 1170 | 1150 | 1164 | 1122 | 1239 | 1165 | 1198 | 1135 | 1168 | 1152 | 1184 | 1,031 | 0,044 | 0,9 | 0,36811814 | 0,45683831 | 1,00E-08 | 0,0032224 | 0,0014535 | 0,0014535 | TRUE | 0,0116906 | No |
| orf19.3009 | 481 | 378 | 459 | 564 | 655 | 672 | 730 | 470 | 432 | 481 | 470 | 465 | 560 | 641 | 651 | 663 | 545,39 | 462 | 629 | 1,361 | 0,444 | 5,919 | 3,24E-09 | 1,92E-08 | 0,00251 | 0,0046413 | 0,0035253 | 0,0035253 | TRUE | 0,0671168 | No |
| orf19.301 | 1622 | 979 | 1463 | 1610 | 2355 | 1843 | 2078 | 1646 | 1455 | 1246 | 1497 | 1328 | 2014 | 1759 | 1854 | 2322 | 1684,3 | 1382 | 1987 | 1,436 | 0,522 | 5,943 | 2,80E-09 | 1,67E-08 | 0,010366 | 0,0028414 | 0,0067851 | 0,0067851 | TRUE | 0,1508193 | No |
| orf19.3010 | 436 | 321 | 368 | 449 | 602 | 468 | 500 | 277 | 391 | 409 | 377 | 370 | 515 | 447 | 446 | 391 | 418,1 | 387 | 450 | 1,171 | 0,227 | 2,567 | 0,01024714 | 0,02058336 | 0,004806 | 0,0054517 | 0,0051317 | 0,0051317 | TRUE | 0,1117902 | No |
| orf19.3010.1 | 9211 | 6493 | 8605 | 10298 | 9256 | 7967 | 8283 | 6021 | 8263 | 8266 | 8804 | 8492 | 7916 | 7602 | 7391 | 8492 | 8153,3 | 8456 | 7850 | 0,928 | -0,108 | -2,232 | 0,02560947 | 0,04665304 | 0,002154 | 0,0021576 | 0,0021311 | 0,0021311 | TRUE | 0,0362003 | No |
| orf19.3012 | 1296 | 859 | 1161 | 1390 | 1251 | 1127 | 1248 | 866 | 1163 | 1094 | 1188 | 1146 | 1070 | 1075 | 1114 | 1221 | 1133,8 | 1148 | 1120 | 0,972 | -0,041 | -0,717 | 0,4731577 | 0,55926254 | 0,001397 | 0,0032599 | 0,0022039 | 0,0022039 | TRUE | 0,0398226 | No |
| orf19.3013 | 992 | 686 | 842 | 1118 | 1270 | 1113 | 1134 | 715 | 890 | 873 | 861 | 922 | 1086 | 1062 | 1012 | 1008 | 964,39 | 886 | 1042 | 1,176 | 0,233 | 4,378 | 1,20E-05 | 4,31E-05 | 1,00E-08 | 0,0034848 | 0,0016862 | 0,0016862 | TRUE | 0,0099802 | No |
| orf19.3014 | 8939 | 6045 | 7557 | 9582 | 9551 | 8316 | 8868 | 6199 | 8019 | 7696 | 7732 | 7901 | 8168 | 7935 | 7913 | 8743 | 8013,5 | 7837 | 8190 | 1,044 | 0,062 | 1,54 | 0,12354738 | 0,18296097 | 0,001133 | 0,0021607 | 0,0014492 | 0,0014492 | TRUE | 0,0245218 | No |
| orf19.3015 | 12993 | 8738 | 11763 | 14410 | 11540 | 9842 | 10909 | 7214 | 11656 | 11124 | 12035 | 11883 | 9869 | 9391 | 9734 | 10175 | 10733 | 11674 | 9792 | 0,838 | -0,254 | -6,534 | 6,42E-11 | 4,81E-10 | 0,001007 | 0,0021148 | 0,0013613 | 0,0013613 | TRUE | 0,0120873 | No |
| orf19.3016 | 1422 | 919 | 1170 | 1450 | 1434 | 1138 | 1319 | 717 | 1276 | 1170 | 1197 | 1196 | 1226 | 1086 | 1177 | 1011 | 1167,4 | 1210 | 1125 | 0,932 | -0,101 | -1,543 | 0,12275106 | 0,1819611 | 0,003187 | 0,0032231 | 0,0032585 | 0,0032585 | TRUE | 0,0509602 | No |
| orf19.3018 | 1330 | 859 | 1148 | 1396 | 1072 | 961 | 1032 | 645 | 1193 | 1094 | 1175 | 1151 | 917 | 917 | 921 | 910 | 1034,6 | 1153 | 916 | 0,794 | -0,334 | -6,649 | 2,94E-11 | 2,30E-10 | 1,00E-08 | 0,0033827 | 0,0014379 | 0,0014379 | TRUE | 0,0129124 | No |
| orf19.3019 | 512 | 321 | 459 | 594 | 414 | 359 | 385 | 223 | 459 | 409 | 470 | 490 | 354 | 343 | 344 | 315 | 397,76 | 457 | 339 | 0,741 | -0,432 | -5,365 | 8,11E-08 | 4,02E-07 | 0,00139 | 0,0056292 | 0,0036555 | 0,0036555 | TRUE | 0,0522864 | No |
| orf19.302 | 266 | 211 | 286 | 308 | 344 | 306 | 314 | 185 | 239 | 269 | 293 | 254 | 294 | 292 | 280 | 261 | 272,64 | 264 | 282 | 1,077 | 0,106 | 1,14 | 0,25408373 | 0,33668617 | 0,001367 | 0,0073041 | 0,0046862 | 0,0046862 | TRUE | 0,0639008 | No |
| orf19.3021 | 2103 | 1416 | 1787 | 2235 | 2105 | 1788 | 1903 | 1197 | 1887 | 1803 | 1828 | 1843 | 1800 | 1706 | 1698 | 1688 | 1781,7 | 1840 | 1723 | 0,937 | -0,094 | -2,217 | 0,02662292 | 0,04833802 | 1,00E-08 | 0,0027943 | 0,001173 | 0,001173 | TRUE | 0,011465 | No |
| orf19.3022 | 3627 | 2303 | 2994 | 3680 | 3917 | 3129 | 3530 | 2247 | 3254 | 2932 | 3063 | 3035 | 3350 | 2986 | 3150 | 3169 | 3117,3 | 3071 | 3164 | 1,03 | 0,042 | 0,859 | 0,39044278 | 0,47931757 | 0,001638 | 0,0024452 | 0,0020164 | 0,0020164 | TRUE | 0,0197538 | No |
| orf19.3023 | 1575 | 1138 | 1327 | 1661 | 1908 | 1522 | 1653 | 988 | 1413 | 1449 | 1358 | 1370 | 1632 | 1452 | 1475 | 1393 | 1442,7 | 1398 | 1488 | 1,068 | 0,095 | 1,657 | 0,09761143 | 0,14971904 | 0,001858 | 0,0029858 | 0,0024385 | 0,0024385 | TRUE | 0,0517922 | No |
| orf19.3026 | 2822 | 1795 | 2386 | 3040 | 3352 | 2780 | 3176 | 2058 | 2532 | 2285 | 2441 | 2507 | 2867 | 2653 | 2834 | 2903 | 2627,6 | 2441 | 2814 | 1,151 | 0,203 | 4,222 | 2,42E-05 | 8,29E-05 | 0,0013 | 0,002532 | 0,0018288 | 0,0018288 | TRUE | 0,0213645 | No |
| orf19.3027 | 4664 | 3297 | 4105 | 5106 | 5752 | 5138 | 5548 | 3635 | 4184 | 4197 | 4200 | 4210 | 4919 | 4903 | 4951 | 5127 | 4586,4 | 4198 | 4975 | 1,184 | 0,244 | 7,919 | 2,39E-15 | 2,83E-14 | 1,00E-08 | 0,002296 | 0,0006893 | 0,0006893 | TRUE | 0,0047357 | No |
| orf19.3029 | 3475 | 2425 | 2978 | 3719 | 3221 | 3017 | 3105 | 2089 | 3117 | 3087 | 3047 | 3067 | 2755 | 2879 | 2771 | 2946 | 2958,6 | 3080 | 2838 | 0,92 | -0,12 | -3,226 | 0,00125536 | 0,00309869 | 0,000111 | 0,0024702 | 0,0009887 | 0,0009887 | TRUE | 0,0073806 | No |
| orf19.3030 | 1998 | 1316 | 1716 | 1999 | 2078 | 1852 | 2059 | 1314 | 1792 | 1675 | 1756 | 1648 | 1777 | 1767 | 1837 | 1853 | 1763,3 | 1718 | 1808 | 1,052 | 0,073 | 1,599 | 0,10986167 | 0,1651774 | 0,000524 | 0,0028028 | 0,0014244 | 0,0014244 | TRUE | 0,0107088 | No |
| orf19.3031 | 1953 | 1213 | 1594 | 1994 | 1859 | 1452 | 1532 | 934 | 1752 | 1544 | 1631 | 1644 | 1590 | 1385 | 1367 | 1317 | 1528,9 | 1643 | 1415 | 0,862 | -0,214 | -3,215 | 0,00130286 | 0,00319884 | 0,004073 | 0,002929 | 0,0036042 | 0,0036042 | TRUE | 0,0849904 | No |
| orf19.3034 | 12099 | 8194 | 10891 | 12882 | 10893 | 9671 | 10661 | 6499 | 10854 | 10432 | 11143 | 10623 | 9316 | 9228 | 9513 | 9166 | 10034 | 10763 | 9306 | 0,865 | -0,21 | -6,5 | 8,03E-11 | 5,92E-10 | 0,000404 | 0,0021242 | 0,0008989 | 0,0008989 | TRUE | 0,0068223 | No |
| orf19.3035 | 4352 | 2894 | 3932 | 4428 | 5029 | 3987 | 4634 | 2806 | 3904 | 3684 | 4023 | 3651 | 4301 | 3804 | 4135 | 3958 | 3932,6 | 3816 | 4050 | 1,062 | 0,086 | 1,674 | 0,09414783 | 0,14563341 | 0,002162 | 0,0023487 | 0,0022972 | 0,0022972 | TRUE | 0,0215877 | No |
| orf19.3037 | 28281 | 17948 | 24753 | 30083 | 25733 | 21208 | 23228 | 15940 | 25371 | 22850 | 25325 | 24807 | 22008 | 20236 | 20727 | 22482 | 22976 | 24588 | 21363 | 0,869 | -0,203 | -4,155 | 3,26E-05 | 0,00010898 | 0,002327 | 0,0020427 | 0,0022528 | 0,0022528 | TRUE | 0,027589 | No |
| orf19.3038 | 54323 | 36996 | 49374 | 58046 | 70134 | 61043 | 66769 | 41592 | 48733 | 47100 | 50514 | 47865 | 59981 | 58245 | 59580 | 58662 | 53835 | 48553 | 59117 | 1,218 | 0,284 | 9,223 | 2,88E-20 | 5,36E-19 | 0,000524 | 0,0020065 | 0,0008921 | 0,0008921 | TRUE | 0,0090393 | No |
| orf19.304 | 1911 | 1374 | 1699 | 2139 | 1662 | 1502 | 1561 | 959 | 1714 | 1749 | 1738 | 1764 | 1421 | 1433 | 1393 | 1353 | 1570,7 | 1741 | 1400 | 0,805 | -0,313 | -7,424 | 1,13E-13 | 1,14E-12 | 1,00E-08 | 0,0029037 | 0,0010581 | 0,0010581 | TRUE | 0,0053241 | No |
| orf19.3040 | 6745 | 4544 | 5706 | 6978 | 6462 | 5643 | 6012 | 3867 | 6051 | 5785 | 5838 | 5754 | 5527 | 5384 | 5365 | 5454 | 5644,7 | 5857 | 5432 | 0,927 | -0,109 | -3,426 | 0,00061278 | 0,00161472 | 0,000212 | 0,0022367 | 0,0007908 | 0,0007908 | TRUE | 0,006199 | No |
| orf19.3041 | 7198 | 4981 | 6557 | 7856 | 7375 | 6559 | 7263 | 4379 | 6457 | 6341 | 6708 | 6478 | 6307 | 6258 | 6481 | 6176 | 6401 | 6496 | 6306 | 0,971 | -0,043 | -1,303 | 0,1926348 | 0,26581301 | 0,00028 | 0,0022063 | 0,0008697 | 0,0008697 | TRUE | 0,0058058 | No |
| orf19.3042 | 176 | 155 | 150 | 207 | 207 | 206 | 203 | 164 | 158 | 197 | 153 | 171 | 177 | 197 | 181 | 231 | 183,18 | 170 | 196 | 1,153 | 0,205 | 1,686 | 0,09186851 | 0,14243993 | 0,00731 | 0,0099046 | 0,0086954 | 0,0086954 | TRUE | 0,1313803 | No |
| orf19.3043 | 344 | 219 | 291 | 382 | 415 | 374 | 385 | 289 | 309 | 279 | 298 | 315 | 355 | 357 | 344 | 408 | 332,88 | 300 | 366 | 1,208 | 0,273 | 3,218 | 0,00129192 | 0,00317458 | 0,000727 | 0,0063405 | 0,0038704 | 0,0038704 | TRUE | 0,0576289 | No |
| orf19.3045 | 2400 | 1494 | 2018 | 2418 | 2070 | 1896 | 2045 | 1304 | 2153 | 1902 | 2065 | 1994 | 1770 | 1809 | 1825 | 1839 | 1919,6 | 2028 | 1811 | 0,891 | -0,166 | -3,466 | 0,00052797 | 0,00141672 | 0,000986 | 0,0027358 | 0,0016767 | 0,0016767 | TRUE | 0,02078 | No |
| orf19.3047 | 3904 | 2839 | 3571 | 4252 | 5448 | 5084 | 5647 | 3470 | 3502 | 3614 | 3653 | 3506 | 4659 | 4851 | 5039 | 4894 | 4215 | 3569 | 4861 | 1,362 | 0,446 | 11,765 | 5,92E-32 | 2,40E-30 | 0,000582 | 0,0023239 | 0,0011361 | 0,0011361 | TRUE | 0,0098422 | No |
| orf19.3048 | 164 | 93 | 144 | 209 | 161 | 133 | 169 | 117 | 147 | 118 | 147 | 172 | 138 | 127 | 151 | 165 | 145,7 | 146 | 145 | 0,979 | -0,031 | -0,226 | 0,82125205 | 0,86245103 | 0,010055 | 0,011943 | 0,0111168 | 0,0111168 | TRUE | 0,1559545 | No |
| orf19.3049 | 57 | 36 | 49 | 75 | 66 | 50 | 44 | 40 | 51 | 46 | 50 | 62 | 56 | 48 | 39 | 56 | 51,1 | 52 | 50 | 0,943 | -0,085 | -0,418 | 0,67620998 | 0,74116262 | 1,00E-08 | 0,0303901 | 0,0198742 | 0,0198742 | TRUE | 0,1850579 | No |
| orf19.305 | 370 | 297 | 345 | 447 | 419 | 403 | 433 | 256 | 332 | 378 | 353 | 369 | 358 | 385 | 386 | 361 | 365,24 | 358 | 372 | 1,044 | 0,062 | 0,797 | 0,42532452 | 0,51260777 | 1,00E-08 | 0,0059542 | 0,0031381 | 0,0031381 | TRUE | 0,0283704 | No |
| orf19.3050 | 4282 | 3046 | 3785 | 4665 | 3777 | 3618 | 3968 | 2156 | 3841 | 3878 | 3872 | 3847 | 3230 | 3452 | 3541 | 3041 | 3587,8 | 3860 | 3316 | 0,86 | -0,217 | -4,359 | 1,31E-05 | 4,68E-05 | 0,001943 | 0,0023841 | 0,0021024 | 0,0021024 | TRUE | 0,0359351 | No |
| orf19.3051 | 1846 | 1171 | 1494 | 1973 | 1054 | 947 | 975 | 623 | 1656 | 1491 | 1529 | 1627 | 901 | 904 | 870 | 879 | 1232 | 1576 | 888 | 0,563 | -0,828 | -15,623 | 5,08E-55 | 6,63E-53 | 0,000872 | 0,0031578 | 0,0018167 | 0,0018167 | TRUE | 0,0149184 | No |
| orf19.3052 | 9743 | 6589 | 8153 | 10429 | 8718 | 7438 | 7966 | 4471 | 8740 | 8388 | 8341 | 8600 | 7456 | 7097 | 7108 | 6306 | 7754,7 | 8517 | 6992 | 0,821 | -0,284 | -5,539 | 3,04E-08 | 1,59E-07 | 0,002488 | 0,0021667 | 0,0023918 | 0,0023918 | TRUE | 0,0520237 | No |
| orf19.3053 | 1390 | 958 | 1251 | 1539 | 1732 | 1474 | 1508 | 1156 | 1247 | 1220 | 1280 | 1269 | 1481 | 1406 | 1346 | 1630 | 1359,9 | 1254 | 1466 | 1,166 | 0,221 | 3,583 | 0,00034002 | 0,00095016 | 0,002753 | 0,003047 | 0,0029163 | 0,0029163 | TRUE | 0,0637966 | No |
| orf19.3054 | 8611 | 6036 | 7633 | 9384 | 7752 | 6709 | 7329 | 4716 | 7725 | 7684 | 7809 | 7738 | 6630 | 6402 | 6540 | 6651 | 7147,5 | 7739 | 6556 | 0,847 | -0,24 | -8,612 | 7,19E-18 | 1,10E-16 | 1,00E-08 | 0,0021826 | 0,0006035 | 0,0006035 | TRUE | 0,0030599 | No |
| orf19.3055 | 1661 | 1165 | 1509 | 1835 | 1848 | 1666 | 1752 | 1145 | 1490 | 1483 | 1544 | 1513 | 1580 | 1590 | 1563 | 1615 | 1547,3 | 1508 | 1587 | 1,052 | 0,073 | 1,749 | 0,08031612 | 0,12698401 | 1,00E-08 | 0,0029177 | 0,0010068 | 0,0010068 | TRUE | 0,0030284 | No |
| orf19.3057 | 603 | 412 | 545 | 626 | 742 | 578 | 690 | 366 | 541 | 525 | 558 | 516 | 635 | 552 | 616 | 516 | 557,16 | 535 | 580 | 1,089 | 0,122 | 1,591 | 0,11156586 | 0,16732094 | 0,003211 | 0,0045851 | 0,0038896 | 0,0038896 | TRUE | 0,0558323 | No |
| orf19.3058 | 1208 | 784 | 1080 | 1286 | 1142 | 970 | 1024 | 702 | 1084 | 998 | 1105 | 1060 | 977 | 926 | 914 | 990 | 1006,7 | 1062 | 952 | 0,894 | -0,162 | -2,954 | 0,00313976 | 0,00709695 | 0,000546 | 0,0034216 | 0,001898 | 0,001898 | TRUE | 0,0175137 | No |

| | | | | | | | | | | | | | | | | | | | | | | | | | | | | | | | |
|---|---|---|---|---|---|---|---|---|---|---|---|---|---|---|---|---|---|---|---|---|---|---|---|---|---|---|---|---|---|---|---|
| orf19.3059 | 27978 | 17911 | 24558 | 29976 | 32410 | 26084 | 28912 | 18704 | 25099 | 22803 | 25125 | 24719 | 27718 | 24889 | 25799 | 26380 | 25316 | 24436 | 26196 | 1,072 | 0,1 | 2,159 | 0,03087212 | 0,05498953 | 0,001994 | 0,0020369 | 0,0020299 | 0,0020299 | TRUE | 0,0246596 | No |
| orf19.3059.1 | 6 | 1 | 4 | 4 | 0 | 4 | 1 | 2 | 5 | 1 | 4 | 3 | 0 | 4 | 1 | 3 | 2,7 | 3 | 2 | 0,5 | -1 | -1,08 | 0,28026156 | NA | 0,007234 | 0,5402345 | 0,4060358 | 0,4060358 | TRUE | 0,4134054 | No |
| orf19.306 | 150 | 122 | 143 | 188 | 211 | 191 | 198 | 133 | 135 | 155 | 146 | 155 | 180 | 182 | 177 | 188 | 164,77 | 148 | 182 | 1,229 | 0,298 | 2,748 | 0,00599716 | 0,01273845 | 1,00E-08 | 0,0107898 | 0,0051671 | 0,0051671 | TRUE | 0,0409188 | No |
| orf19.3060 | 1308 | 915 | 1128 | 1447 | 963 | 829 | 850 | 519 | 1173 | 1165 | 1154 | 1193 | 824 | 791 | 758 | 732 | 973,83 | 1171 | 776 | 0,664 | -0,59 | -10,993 | 4,12E-28 | 1,36E-26 | 1,00E-08 | 0,0034702 | 0,001695 | 0,001695 | TRUE | 0,0202798 | No |
| orf19.3061 | 359 | 234 | 256 | 392 | 402 | 300 | 331 | 203 | 322 | 298 | 262 | 323 | 344 | 286 | 295 | 286 | 302,11 | 301 | 303 | 1,006 | 0,009 | 0,089 | 0,9289953 | 0,94901598 | 0,005742 | 0,0067848 | 0,0062761 | 0,0062761 | TRUE | 0,0985614 | No |
| orf19.3061.1 | 6573 | 4627 | 5633 | 7117 | 7120 | 6400 | 6402 | 4724 | 5897 | 5891 | 5763 | 5869 | 6089 | 6107 | 5713 | 6663 | 5998,8 | 5855 | 6143 | 1,048 | 0,068 | 1,439 | 0,15008178 | 0,21587055 | 0,001864 | 0,0022215 | 0,0019833 | 0,0019833 | TRUE | 0,0385202 | No |
| orf19.3062 | 1036 | 812 | 998 | 1148 | 996 | 1070 | 1073 | 689 | 929 | 1034 | 1021 | 947 | 852 | 1021 | 957 | 972 | 966,61 | 983 | 950 | 0,968 | -0,048 | -0,697 | 0,4858977 | 0,57128638 | 0,00339 | 0,0034814 | 0,0034311 | 0,0034311 | TRUE | 0,0612614 | No |
| orf19.3063 | 809 | 497 | 661 | 782 | 786 | 626 | 662 | 451 | 726 | 633 | 676 | 645 | 672 | 597 | 591 | 636 | 646,99 | 670 | 624 | 0,93 | -0,104 | -1,477 | 0,13974631 | 0,20306957 | 0,002377 | 0,0042233 | 0,0032453 | 0,0032453 | TRUE | 0,0381597 | No |
| orf19.3064 | 3001 | 2014 | 2453 | 2995 | 3040 | 2479 | 2638 | 1653 | 2692 | 2564 | 2510 | 2470 | 2600 | 2365 | 2354 | 2331 | 2485,8 | 2559 | 2412 | 0,944 | -0,084 | -1,663 | 0,09634613 | 0,14804285 | 0,001536 | 0,0025635 | 0,0020199 | 0,0020199 | TRUE | 0,0339264 | No |
| orf19.3064.1 | 411 | 335 | 377 | 443 | 465 | 369 | 398 | 292 | 369 | 426 | 386 | 365 | 398 | 352 | 355 | 412 | 382,87 | 386 | 379 | 0,982 | -0,027 | -0,315 | 0,75246736 | 0,80613835 | 0,00272 | 0,0057711 | 0,0043356 | 0,0043356 | TRUE | 0,0504737 | No |
| orf19.3065 | 1260 | 818 | 1037 | 1384 | 1035 | 935 | 1024 | 648 | 1130 | 1041 | 1061 | 1141 | 885 | 892 | 914 | 914 | 997,37 | 1093 | 901 | 0,822 | -0,283 | -5,32 | 1,04E-07 | 5,09E-07 | 1,00E-08 | 0,003435 | 0,0017038 | 0,0017038 | TRUE | 0,0114418 | No |
| orf19.3066 | 1999 | 1300 | 1799 | 2103 | 2177 | 1998 | 2210 | 1464 | 1793 | 1655 | 1841 | 1734 | 1862 | 1906 | 1972 | 2065 | 1853,5 | 1756 | 1951 | 1,109 | 0,149 | 2,926 | 0,00343173 | 0,00767604 | 0,001339 | 0,0027627 | 0,0019477 | 0,0019477 | TRUE | 0,0171616 | No |
| orf19.307 | 2123 | 1519 | 1883 | 2402 | 2587 | 2168 | 2440 | 1674 | 1905 | 1934 | 1926 | 1981 | 2212 | 2069 | 2177 | 2361 | 2070,6 | 1936 | 2205 | 1,137 | 0,185 | 3,864 | 0,00011144 | 0,00033833 | 0,000991 | 0,0026806 | 0,0017074 | 0,0017074 | TRUE | 0,0253016 | No |
| orf19.3071 | 5856 | 4051 | 5643 | 6628 | 7979 | 6977 | 7846 | 5231 | 5253 | 5157 | 5773 | 5466 | 6824 | 6657 | 7001 | 7378 | 6188,7 | 5412 | 6965 | 1,286 | 0,363 | 7,492 | 6,79E-14 | 7,01E-13 | 0,002036 | 0,0022141 | 0,0020888 | 0,0020888 | TRUE | 0,0243638 | No |
| orf19.3073 | 0 | 0 | 0 | 0 | 0 | 0 | 0 | 0 | 0 | 0 | 0 | 0 | 0 | 0 | 0 | 0 | 0 | 0 | 0 | NA | NA | NA | NA | NA | NA | NA | NA | NA | NA | NA | NA |
| orf19.3074 | 126 | 63 | 111 | 131 | 131 | 105 | 135 | 84 | 113 | 80 | 114 | 108 | 112 | 100 | 120 | 118 | 108,25 | 104 | 112 | 1,077 | 0,107 | 0,739 | 0,4598429 | 0,54556401 | 0,003199 | 0,0153903 | 0,0108066 | 0,0108066 | TRUE | 0,1889372 | No |
| orf19.3076 | 784 | 487 | 678 | 896 | 859 | 680 | 728 | 410 | 703 | 620 | 694 | 739 | 735 | 649 | 650 | 578 | 670,9 | 689 | 653 | 0,949 | -0,076 | -0,928 | 0,35321217 | 0,44063512 | 0,005437 | 0,0041433 | 0,0049164 | 0,0049164 | TRUE | 0,0844579 | No |
| orf19.3077 | 1583 | 1007 | 1468 | 1816 | 1978 | 1515 | 1807 | 1255 | 1420 | 1282 | 1502 | 1497 | 1692 | 1446 | 1612 | 1770 | 1527,7 | 1425 | 1630 | 1,141 | 0,19 | 2,619 | 0,00882256 | 0,01802924 | 0,005341 | 0,0029298 | 0,0044072 | 0,0044072 | TRUE | 0,0698637 | No |
| orf19.308 | 1811 | 1290 | 1563 | 1983 | 1505 | 1358 | 1386 | 999 | 1625 | 1642 | 1599 | 1635 | 1287 | 1296 | 1237 | 1409 | 1466,2 | 1625 | 1307 | 0,802 | -0,318 | -6,481 | 9,14E-11 | 6,71E-10 | 0,000497 | 0,0029696 | 0,0016251 | 0,0016251 | TRUE | 0,0293495 | No |
| orf19.3080 | 1112 | 727 | 1083 | 1230 | 1504 | 1116 | 1390 | 976 | 998 | 926 | 1108 | 1014 | 1286 | 1065 | 1240 | 1377 | 1126,7 | 1012 | 1242 | 1,225 | 0,293 | 3,583 | 0,00033936 | 0,00094919 | 0,007117 | 0,003268 | 0,0055331 | 0,0055331 | TRUE | 0,1106237 | No |
| orf19.3083 | 1508 | 927 | 1324 | 1599 | 2265 | 1788 | 2188 | 1480 | 1353 | 1180 | 1355 | 1319 | 1937 | 1706 | 1952 | 2087 | 1611,1 | 1302 | 1920 | 1,472 | 0,558 | 8,05 | 8,26E-16 | 1,03E-14 | 0,004549 | 0,0028806 | 0,0039697 | 0,0039697 | TRUE | 0,0683167 | No |
| orf19.3084 | 1482 | 953 | 1241 | 1583 | 2281 | 1752 | 1942 | 1114 | 1330 | 1213 | 1270 | 1305 | 1951 | 1672 | 1733 | 1571 | 1505,6 | 1280 | 1732 | 1,355 | 0,438 | 6,434 | 1,24E-10 | 8,92E-10 | 0,004173 | 0,0029438 | 0,0037652 | 0,0037652 | TRUE | 0,0883779 | No |
| orf19.3086 | 2005 | 1332 | 1710 | 2085 | 2221 | 1752 | 1873 | 1160 | 1799 | 1696 | 1749 | 1719 | 1899 | 1672 | 1671 | 1636 | 1730,2 | 1741 | 1720 | 0,989 | -0,016 | -0,287 | 0,77387301 | 0,82334497 | 0,002197 | 0,0028185 | 0,0024479 | 0,0024479 | TRUE | 0,0614025 | No |
| orf19.3087 | 20020 | 12405 | 16533 | 20500 | 17946 | 13740 | 14644 | 10516 | 17960 | 15793 | 16915 | 16905 | 15348 | 13110 | 13067 | 14832 | 15491 | 16893 | 14089 | 0,834 | -0,262 | -4,19 | 2,79E-05 | 9,44E-05 | 0,00476 | 0,0020732 | 0,0036938 | 0,0036938 | TRUE | 0,0446063 | No |
| orf19.3087.1 | 169 | 102 | 123 | 171 | 195 | 125 | 132 | 90 | 152 | 130 | 126 | 141 | 167 | 119 | 118 | 127 | 134,89 | 137 | 133 | 0,969 | -0,046 | -0,321 | 0,74806917 | 0,80214164 | 0,01176 | 0,012742 | 0,01233 | 0,01233 | TRUE | 0,3377776 | No |
| orf19.3087.2 | 450 | 285 | 342 | 462 | 417 | 348 | 341 | 281 | 404 | 363 | 350 | 381 | 357 | 332 | 304 | 396 | 360,84 | 374 | 347 | 0,921 | -0,118 | -1,276 | 0,20201932 | 0,2769813 | 0,004908 | 0,0060027 | 0,005485 | 0,005485 | TRUE | 0,0889225 | No |
| orf19.3088 | 3047 | 1925 | 2389 | 3223 | 2957 | 2450 | 2577 | 1644 | 2733 | 2451 | 2444 | 2658 | 2529 | 2338 | 2300 | 2319 | 2471,4 | 2572 | 2372 | 0,922 | -0,118 | -2,187 | 0,02874561 | 0,05161495 | 0,002089 | 0,0025669 | 0,0023659 | 0,0023659 | TRUE | 0,0249626 | No |
| orf19.3089 | 7413 | 4943 | 6154 | 7799 | 7642 | 6313 | 6747 | 3775 | 6650 | 6293 | 6296 | 6431 | 6536 | 6024 | 6021 | 5324 | 6196,8 | 6418 | 5976 | 0,932 | -0,102 | -1,755 | 0,07979765 | 0,12560514 | 0,003591 | 0,0022138 | 0,003078 | 0,003078 | TRUE | 0,0643784 | No |
| orf19.309 | 280 | 196 | 240 | 321 | 375 | 315 | 362 | 242 | 251 | 250 | 246 | 265 | 321 | 301 | 323 | 341 | 287,07 | 253 | 322 | 1,266 | 0,34 | 4,01 | 6,07E-05 | 0,00019352 | 1,00E-08 | 0,0070364 | 0,0033906 | 0,0033906 | TRUE | 0,0208333 | No |
| orf19.3091 | 5241 | 3471 | 4748 | 5555 | 4879 | 4260 | 4653 | 2808 | 4702 | 4419 | 4858 | 4581 | 4173 | 4065 | 4152 | 3960 | 4363,6 | 4640 | 4088 | 0,881 | -0,182 | -4,535 | 5,76E-06 | 2,18E-05 | 0,000775 | 0,0023122 | 0,0013225 | 0,0013225 | TRUE | 0,0119237 | No |
| orf19.3093 | 2645 | 1713 | 2312 | 2821 | 2796 | 2320 | 2559 | 1712 | 2373 | 2181 | 2365 | 2326 | 2391 | 2214 | 2283 | 2415 | 2318,5 | 2311 | 2326 | 1,005 | 0,007 | 0,14 | 0,88843668 | 0,91556219 | 0,000915 | 0,0026057 | 0,0016643 | 0,0016643 | TRUE | 0,0164567 | No |
| orf19.3097 | 13516 | 8312 | 11675 | 13797 | 14516 | 11428 | 12973 | 7962 | 12125 | 10582 | 11945 | 11377 | 12415 | 10904 | 11576 | 11230 | 11519 | 11507 | 11531 | 1,002 | 0,003 | 0,053 | 0,95771404 | 0,96952927 | 0,002833 | 0,0021055 | 0,0028661 | 0,0028661 | TRUE | 0,0354949 | No |
| orf19.3098 | 1851 | 1214 | 1576 | 1905 | 1577 | 1434 | 1503 | 972 | 1661 | 1546 | 1612 | 1571 | 1349 | 1368 | 1341 | 1371 | 1477,3 | 1598 | 1357 | 0,849 | -0,237 | -5,35 | 8,79E-08 | 4,34E-07 | 1,00E-08 | 0,0029622 | 0,0011994 | 0,0011994 | TRUE | 0,0087182 | No |
| orf19.3099 | 815 | 604 | 817 | 904 | 1047 | 869 | 972 | 652 | 731 | 769 | 836 | 745 | 895 | 829 | 867 | 920 | 824,12 | 770 | 878 | 1,139 | 0,188 | 3,001 | 0,00269001 | 0,00618034 | 0,001493 | 0,003741 | 0,0025479 | 0,0025479 | TRUE | 0,0393592 | No |
| orf19.31 | 207 | 151 | 191 | 263 | 210 | 157 | 153 | 112 | 186 | 192 | 195 | 217 | 180 | 150 | 137 | 158 | 176,77 | 198 | 156 | 0,788 | -0,343 | -2,93 | 0,00338743 | 0,00758543 | 0,004139 | 0,0101921 | 0,0074431 | 0,0074431 | TRUE | 0,1261593 | No |
| orf19.310 | 50 | 25 | 39 | 66 | 41 | 25 | 34 | 21 | 45 | 32 | 40 | 54 | 35 | 24 | 30 | 30 | 36,24 | 43 | 30 | 0,688 | -0,539 | -2,189 | 0,02857654 | 0,05134205 | 1,00E-08 | 0,0420424 | 0,0299614 | 0,0299614 | TRUE | 0,2230501 | No |
| orf19.3100 | 245 | 167 | 212 | 274 | 265 | 190 | 241 | 150 | 220 | 213 | 217 | 226 | 227 | 181 | 215 | 212 | 213,72 | 219 | 209 | 0,953 | -0,07 | -0,692 | 0,48909988 | 0,57452496 | 1,00E-08 | 0,008772 | 0,00516 | 0,00516 | TRUE | 0,0902442 | No |
| orf19.3102 | 1447 | 933 | 1214 | 1452 | 1733 | 1469 | 1581 | 989 | 1298 | 1188 | 1242 | 1197 | 1482 | 1402 | 1411 | 1395 | 1326,9 | 1231 | 1422 | 1,155 | 0,208 | 4,203 | 2,64E-05 | 8,96E-05 | 0,000522 | 0,0030736 | 0,0015998 | 0,0015998 | TRUE | 0,0164737 | No |
| orf19.3103 | 5881 | 4106 | 5400 | 6476 | 6955 | 5874 | 6537 | 3873 | 5276 | 5227 | 5525 | 5340 | 5948 | 5605 | 5833 | 5463 | 5527,1 | 5342 | 5712 | 1,07 | 0,098 | 2,493 | 0,01267576 | 0,02483887 | 0,000792 | 0,0022422 | 0,0012884 | 0,0012884 | TRUE | 0,0101069 | No |
| orf19.3104 | 1195 | 860 | 1107 | 1383 | 1316 | 1161 | 1297 | 706 | 1072 | 1095 | 1133 | 1140 | 1125 | 1108 | 1157 | 996 | 1103,3 | 1110 | 1096 | 0,991 | -0,013 | -0,233 | 0,81575421 | 0,85802719 | 0,001225 | 0,0032953 | 0,0022282 | 0,0022282 | TRUE | 0,0406674 | No |
| orf19.3105 | 1438 | 921 | 1266 | 1565 | 950 | 787 | 890 | 595 | 1290 | 1173 | 1295 | 1291 | 812 | 751 | 794 | 839 | 1030,6 | 1262 | 799 | 0,631 | -0,664 | -11,593 | 4,47E-31 | 1,70E-29 | 0,001004 | 0,0033881 | 0,0021248 | 0,0021248 | TRUE | 0,0252769 | No |
| orf19.3106 | 1495 | 1043 | 1369 | 1624 | 1358 | 1124 | 1185 | 878 | 1341 | 1328 | 1401 | 1339 | 1161 | 1072 | 1057 | 1238 | 1242,3 | 1352 | 1132 | 0,835 | -0,26 | -4,451 | 8,55E-06 | 3,15E-05 | 0,001978 | 0,0031481 | 0,0024579 | 0,0024579 | TRUE | 0,0427109 | No |
| orf19.3107 | 176 | 168 | 177 | 234 | 97 | 102 | 94 | 45 | 158 | 214 | 181 | 193 | 83 | 97 | 84 | 63 | 134,18 | 186 | 82 | 0,446 | -1,165 | -7,94 | 2,03E-15 | 2,43E-14 | 0,01062 | 0,0127985 | 0,011947 | 0,011947 | TRUE | 0,1588014 | No |
| orf19.3108 | 619 | 376 | 455 | 592 | 407 | 371 | 367 | 198 | 555 | 479 | 466 | 488 | 348 | 354 | 327 | 279 | 412,06 | 497 | 327 | 0,661 | -0,596 | -6,552 | 5,67E-11 | 4,30E-10 | 0,005238 | 0,0055025 | 0,0054171 | 0,0054171 | TRUE | 0,0930001 | No |
| orf19.3109 | 25 | 28 | 17 | 34 | 27 | 26 | 42 | 20 | 22 | 36 | 17 | 28 | 23 | 25 | 37 | 28 | 27,14 | 26 | 28 | 1,108 | 0,148 | 0,506 | 0,61274087 | 0,68671822 | 0,031214 | 0,0554763 | 0,0455298 | 0,0455298 | TRUE | 0,3011236 | No |
| orf19.3110 | 3568 | 2463 | 3112 | 3888 | 3812 | 3332 | 3537 | 2156 | 3201 | 3136 | 3184 | 3206 | 3260 | 3179 | 3156 | 3041 | 3170,4 | 3182 | 3159 | 0,994 | -0,009 | -0,252 | 0,8010511 | 0,84552267 | 8,21E-05 | 0,0024374 | 0,0009116 | 0,0009116 | TRUE | 0,0070036 | No |
| orf19.3111 | 55 | 39 | 66 | 74 | 51 | 38 | 50 | 37 | 49 | 50 | 68 | 61 | 44 | 36 | 45 | 52 | 50,53 | 57 | 44 | 0,767 | -0,383 | -1,884 | 0,0595993 | 0,09791431 | 1,00E-08 | 0,0307108 | 0,0195847 | 0,0195847 | TRUE | 0,1258982 | No |
| orf19.3112 | 585 | 410 | 548 | 737 | 513 | 486 | 535 | 348 | 525 | 522 | 561 | 608 | 439 | 464 | 477 | 491 | 510,73 | 554 | 468 | 0,841 | -0,251 | -3,345 | 0,00082143 | 0,00210454 | 0,0021 | 0,0048219 | 0,0034159 | 0,0034159 | TRUE | 0,0534599 | No |
| orf19.3113 | 418 | 210 | 323 | 385 | 501 | 413 | 471 | 304 | 375 | 267 | 330 | 317 | 428 | 394 | 420 | 429 | 370,23 | 322 | 418 | 1,289 | 0,366 | 3,804 | 0,00014242 | 0,00042382 | 0,006531 | 0,0059006 | 0,0061431 | 0,0061431 | TRUE | 0,1355324 | No |
| orf19.3114 | 707 | 512 | 647 | 735 | 884 | 757 | 796 | 497 | 634 | 652 | 662 | 606 | 756 | 722 | 710 | 701 | 680,47 | 638 | 722 | 1,135 | 0,183 | 3,04 | 0,00236187 | 0,00548092 | 1,00E-08 | 0,0041129 | 0,0020025 | 0,0020025 | TRUE | 0,0147894 | No |
| orf19.3115 | 757 | 539 | 623 | 803 | 838 | 778 | 800 | 496 | 679 | 686 | 637 | 662 | 717 | 742 | 714 | 700 | 692,17 | 666 | 718 | 1,08 | 0,111 | 1,918 | 0,0550585 | 0,09130723 | 1,00E-08 | 0,0040769 | 0,0017504 | 0,0017504 | TRUE | 0,0095137 | No |
| orf19.3116 | 1374 | 905 | 1144 | 1354 | 1793 | 1444 | 1593 | 787 | 1233 | 1152 | 1170 | 1117 | 1533 | 1378 | 1421 | 1110 | 1264,3 | 1168 | 1360 | 1,168 | 0,224 | 2,609 | 0,00909062 | 0,01851406 | 0,008507 | 0,0031277 | 0,0062791 | 0,0062791 | TRUE | 0,1760684 | No |
| orf19.3117 | 308 | 172 | 267 | 279 | 318 | 240 | 215 | 165 | 276 | 219 | 273 | 230 | 272 | 229 | 192 | 233 | 240,51 | 250 | 232 | 0,926 | -0,112 | -0,899 | 0,36866189 | 0,45725034 | 0,013339 | 0,0080155 | 0,0106079 | 0,0106079 | TRUE | 0,1685424 | No |
| orf19.3120 | 62 | 38 | 56 | 67 | 40 | 29 | 41 | 26 | 56 | 48 | 57 | 55 | 34 | 28 | 37 | 37 | 43,96 | 54 | 34 | 0,619 | -0,692 | -3,325 | 0,00088505 | 0,00225027 | 1,00E-08 | 0,035003 | 0,0175896 | 0,0175896 | TRUE | 0,107998 | No |
| orf19.3121 | 35 | 27 | 34 | 34 | 40 | 35 | 40 | 25 | 31 | 34 | 35 | 28 | 34 | 33 | 36 | 35 | 33,39 | 32 | 34 | 1,084 | 0,116 | 0,503 | 0,61476656 | 0,68806213 | 1,00E-08 | 0,0454512 | 0,0212474 | 0,0212474 | TRUE | 0,0599493 | No |
| orf19.3122 | 1449 | 966 | 1247 | 1607 | 1464 | 1229 | 1344 | 875 | 1300 | 1230 | 1276 | 1325 | 1252 | 1173 | 1199 | 1234 | 1248,6 | 1283 | 1214 | 0,945 | -0,082 | -1,683 | 0,09228936 | 0,14305327 | 1,00E-08 | 0,0031422 | 0,0014508 | 0,0014508 | TRUE | 0,0084992 | No |
| orf19.3122.2 | 2247 | 1332 | 1569 | 2035 | 2489 | 1882 | 1914 | 1233 | 2016 | 1696 | 1605 | 1678 | 2129 | 1796 | 1708 | 1739 | 1795,8 | 1749 | 1843 | 1,054 | 0,076 | 0,877 | 0,3803329 | 0,4697862 | 0,009637 | 0,0027879 | 0,0066623 | 0,0066623 | TRUE | 0,1336093 | No |
| orf19.3123 | 7624 | 5140 | 6574 | 8067 | 9063 | 7261 | 7977 | 5172 | 6840 | 6544 | 6726 | 6652 | 7751 | 6928 | 7118 | 7295 | 6981,7 | 6690 | 7273 | 1,087 | 0,12 | 2,881 | 0,00396513 | 0,00875826 | 0,001153 | 0,0021875 | 0,0015321 | 0,0015321 | TRUE | 0,0244246 | No |
| orf19.3123.2 | 333 | 191 | 252 | 333 | 308 | 242 | 253 | 182 | 299 | 243 | 258 | 275 | 263 | 231 | 226 | 257 | 256,39 | 269 | 244 | 0,904 | -0,146 | -1,47 | 0,14163209 | 0,20558724 | 0,003061 | 0,0076417 | 0,0055373 | 0,0055373 | TRUE | 0,0636958 | No |

| | | | | | | | | | | | | | | | | | | | | | | | | | | | | | | |
|---|---|---|---|---|---|---|---|---|---|---|---|---|---|---|---|---|---|---|---|---|---|---|---|---|---|---|---|---|---|---|
| orf19.3124 | 7086 | 4630 | 6079 | 7804 | 6407 | 5283 | 5760 | 3975 | 6357 | 5894 | 6219 | 6435 | 5479 | 5041 | 5140 | 5606 | 5771,6 | 6226 | 5316 | 0,853 | -0,229 | -4,859 | 1,18E-06 | 4,92E-06 | 0,001786 | 0,002231 | 0,0019619 | 0,0019619 | TRUE | 0,0158214 | No |
| orf19.3125 | 491 | 399 | 448 | 595 | 540 | 491 | 508 | 315 | 440 | 508 | 458 | 491 | 462 | 468 | 453 | 444 | 465,67 | 474 | 457 | 0,966 | -0,049 | -0,689 | 0,49097901 | 0,57626595 | 1,00E-08 | 0,005097 | 0,0027644 | 0,0027644 | TRUE | 0,0276326 | No |
| orf19.3126 | 5036 | 3373 | 4330 | 5348 | 5261 | 4631 | 5049 | 3184 | 4518 | 4294 | 4430 | 4410 | 4499 | 4419 | 4505 | 4491 | 4445,8 | 4413 | 4478 | 1,014 | 0,02 | 0,649 | 0,5164165 | 0,60035417 | 1,00E-08 | 0,0023061 | 0,0007292 | 0,0007292 | TRUE | 0,0037483 | No |
| orf19.3127 | 1497 | 1004 | 1405 | 1706 | 2096 | 1679 | 2029 | 1234 | 1343 | 1278 | 1437 | 1407 | 1793 | 1602 | 1811 | 1740 | 1551,4 | 1366 | 1736 | 1,27 | 0,345 | 6,011 | 1,85E-09 | 1,13E-08 | 0,002199 | 0,0029153 | 0,0024996 | 0,0024996 | TRUE | 0,0336741 | No |
| orf19.3128 | 952 | 713 | 913 | 1023 | 1328 | 1207 | 1334 | 866 | 854 | 908 | 934 | 844 | 1136 | 1152 | 1190 | 1221 | 1029,8 | 885 | 1175 | 1,328 | 0,409 | 7,439 | 1,02E-13 | 1,03E-12 | 0,000671 | 0,0033892 | 0,001917 | 0,001917 | TRUE | 0,0176029 | No |
| orf19.3129 | 4723 | 2995 | 3955 | 4887 | 5692 | 4711 | 5196 | 3219 | 4237 | 3813 | 4046 | 4030 | 4868 | 4495 | 4637 | 4540 | 4333,2 | 4032 | 4635 | 1,149 | 0,201 | 4,519 | 6,21E-06 | 2,34E-05 | 0,001259 | 0,0023145 | 0,0016646 | 0,0016646 | TRUE | 0,0156417 | No |
| orf19.313 | 9 | 4 | 5 | 7 | 7 | 8 | 4 | 5 | 8 | 5 | 5 | 6 | 6 | 8 | 4 | 7 | 6,04 | 6 | 6 | 0,985 | -0,022 | -0,04 | 0,96826993 | NA | 1,00E-08 | 0,2424493 | 0,1248701 | 0,1248701 | TRUE | 0,2134016 | No |
| orf19.3130 | 6597 | 4431 | 5414 | 6894 | 6433 | 5721 | 5947 | 3831 | 5918 | 5641 | 5539 | 5685 | 5502 | 5459 | 5307 | 5403 | 5556,7 | 5696 | 5418 | 0,951 | -0,073 | -2,141 | 0,03225995 | 0,0571342 | 0,000364 | 0,0022408 | 0,0009214 | 0,0009214 | TRUE | 0,0085463 | No |
| orf19.3131 | 1824 | 1189 | 1629 | 1981 | 2124 | 1851 | 1877 | 1326 | 1636 | 1514 | 1667 | 1634 | 1817 | 1766 | 1675 | 1870 | 1697,3 | 1613 | 1782 | 1,102 | 0,141 | 2,745 | 0,00606004 | 0,01285618 | 0,001176 | 0,0028348 | 0,0019267 | 0,0019267 | TRUE | 0,0199385 | No |
| orf19.3132 | 2558 | 1726 | 2232 | 2879 | 2856 | 2593 | 2639 | 1758 | 2295 | 2197 | 2284 | 2374 | 2443 | 2474 | 2355 | 2480 | 2362,6 | 2288 | 2438 | 1,064 | 0,089 | 2,179 | 0,0293032 | 0,05245936 | 0,000351 | 0,002594 | 0,0011868 | 0,0011868 | TRUE | 0,0080151 | No |
| orf19.3133 | 22457 | 15120 | 20006 | 24737 | 20477 | 17515 | 18699 | 12784 | 20146 | 19249 | 20468 | 20398 | 17513 | 16712 | 16686 | 18031 | 18650 | 20065 | 17236 | 0,859 | -0,22 | -5,755 | 8,65E-09 | 4,82E-08 | 0,001041 | 0,0020574 | 0,0013473 | 0,0013473 | TRUE | 0,0116168 | No |
| orf19.3134 | 4 | 12 | 7 | 3 | 4 | 4 | 5 | 5 | 4 | 15 | 7 | 2 | 3 | 4 | 4 | 7 | 5,91 | 7 | 4 | 0,669 | -0,58 | -0,873 | 0,38251824 | NA | 0,231248 | 0,2477569 | 0,2408616 | 0,2408616 | TRUE | 0,8892101 | No |
| orf19.3135 | 2284 | 1635 | 1973 | 2371 | 2420 | 2263 | 2402 | 1513 | 2049 | 2082 | 2019 | 1955 | 2070 | 2159 | 2143 | 2134 | 2076,3 | 2026 | 2126 | 1,05 | 0,071 | 1,759 | 0,07863519 | 0,12462152 | 1,00E-08 | 0,0026787 | 0,0010735 | 0,0010735 | TRUE | 0,007015 | No |
| orf19.3136 | 5240 | 3640 | 4658 | 5533 | 6056 | 5488 | 5859 | 3512 | 4701 | 4634 | 4766 | 4563 | 5179 | 5236 | 5228 | 4953 | 4907,6 | 4666 | 5149 | 1,105 | 0,144 | 4,227 | 2,37E-05 | 8,13E-05 | 0,000236 | 0,0022753 | 0,0009021 | 0,0009021 | TRUE | 0,0074549 | No |
| orf19.3138 | 26230 | 15294 | 22232 | 27285 | 24277 | 17839 | 20810 | 13803 | 23531 | 19471 | 22746 | 22500 | 20762 | 17021 | 18569 | 19468 | 20509 | 22062 | 18955 | 0,859 | -0,219 | -3,084 | 0,00204473 | 0,00481943 | 0,006639 | 0,0020503 | 0,0048013 | 0,0048013 | TRUE | 0,0763007 | No |
| orf19.3139 | 3202 | 2025 | 2705 | 3247 | 3333 | 2802 | 2991 | 1875 | 2873 | 2578 | 2767 | 2678 | 2850 | 2674 | 2669 | 2645 | 2716,6 | 2724 | 2710 | 0,995 | -0,008 | -0,166 | 0,86823866 | 0,89922045 | 0,001293 | 0,0025139 | 0,001749 | 0,001749 | TRUE | 0,016451 | No |
| orf19.314 | 698 | 483 | 670 | 703 | 922 | 850 | 954 | 592 | 626 | 615 | 685 | 580 | 789 | 811 | 851 | 835 | 724,01 | 626 | 822 | 1,312 | 0,391 | 5,96 | 2,52E-09 | 1,51E-08 | 0,001576 | 0,0039846 | 0,00273 | 0,00273 | TRUE | 0,0475159 | No |
| orf19.3140 | 24 | 13 | 14 | 27 | 23 | 19 | 18 | 18 | 22 | 17 | 14 | 22 | 20 | 18 | 16 | 25 | 19,24 | 19 | 20 | 1,032 | 0,045 | 0,143 | 0,88605449 | 0,91373419 | 1,00E-08 | 0,0774329 | 0,0441862 | 0,0441862 | TRUE | 0,1335948 | No |
| orf19.3140.1 | 4797 | 2891 | 3668 | 4705 | 4355 | 3321 | 3793 | 2510 | 4303 | 3681 | 3753 | 3880 | 3725 | 3169 | 3385 | 3540 | 3679,3 | 3904 | 3455 | 0,884 | -0,177 | -2,766 | 0,005674 | 0,0121056 | 0,004512 | 0,0023741 | 0,0036685 | 0,0036685 | TRUE | 0,0579208 | No |
| orf19.3141 | 3683 | 2477 | 3344 | 3963 | 5012 | 4268 | 4808 | 3249 | 3304 | 3153 | 3421 | 3268 | 4286 | 4072 | 4290 | 4582 | 3797,3 | 3286 | 4308 | 1,309 | 0,389 | 8,461 | 2,65E-17 | 3,87E-16 | 0,001387 | 0,0023618 | 0,0017567 | 0,0017567 | TRUE | 0,0213938 | No |
| orf19.3142 | 503 | 394 | 473 | 605 | 646 | 610 | 614 | 442 | 451 | 502 | 484 | 499 | 552 | 582 | 548 | 623 | 530,19 | 484 | 576 | 1,188 | 0,248 | 3,522 | 0,00042861 | 0,00117316 | 0,000937 | 0,0047176 | 0,0028761 | 0,0028761 | TRUE | 0,0306236 | No |
| orf19.3147 | 491 | 452 | 479 | 640 | 571 | 795 | 858 | 425 | 440 | 575 | 490 | 528 | 488 | 759 | 766 | 599 | 580,71 | 508 | 653 | 1,286 | 0,363 | 2,695 | 0,00703268 | 0,01467095 | 0,027231 | 0,0044794 | 0,0156934 | 0,0156934 | TRUE | 0,3517528 | No |
| orf19.3148 | 568 | 442 | 526 | 650 | 731 | 682 | 788 | 429 | 510 | 563 | 538 | 536 | 625 | 651 | 703 | 605 | 591,32 | 537 | 646 | 1,209 | 0,274 | 3,977 | 6,98E-05 | 0,00021971 | 0,00133 | 0,0044345 | 0,0028478 | 0,0028478 | TRUE | 0,0424276 | No |
| orf19.3149 | 7901 | 5722 | 6921 | 8338 | 7794 | 7369 | 7374 | 4847 | 7088 | 7285 | 7081 | 6876 | 6666 | 7031 | 6580 | 6836 | 6930,3 | 7082 | 6778 | 0,957 | -0,063 | -1,783 | 0,07460676 | 0,11895898 | 0,000519 | 0,002189 | 0,0010565 | 0,0010565 | TRUE | 0,0079149 | No |
| orf19.315 | 87 | 56 | 77 | 93 | 93 | 66 | 66 | 59 | 78 | 71 | 79 | 77 | 80 | 63 | 59 | 83 | 73,68 | 76 | 71 | 0,924 | -0,114 | -0,679 | 0,49690992 | 0,58184269 | 1,00E-08 | 0,0216827 | 0,0134853 | 0,0134853 | TRUE | 0,1278032 | No |
| orf19.3150 | 2320 | 1772 | 2095 | 2483 | 2578 | 2455 | 2396 | 1612 | 2081 | 2256 | 2143 | 2048 | 2205 | 2342 | 2138 | 2274 | 2185,9 | 2132 | 2240 | 1,051 | 0,072 | 1,498 | 0,13424911 | 0,19648444 | 0,001087 | 0,0026437 | 0,0017698 | 0,0017698 | TRUE | 0,017675 | No |
| orf19.3151 | 447 | 416 | 438 | 558 | 421 | 450 | 402 | 259 | 401 | 530 | 448 | 460 | 360 | 429 | 359 | 365 | 419,04 | 460 | 378 | 0,826 | -0,276 | -2,821 | 0,0047935 | 0,01039313 | 0,008067 | 0,0054439 | 0,0067604 | 0,0067604 | TRUE | 0,113159 | No |
| orf19.3152 | 2076 | 1566 | 1879 | 2392 | 1455 | 1354 | 1331 | 860 | 1862 | 1994 | 1922 | 1972 | 1244 | 1292 | 1188 | 1213 | 1586 | 1938 | 1234 | 0,638 | -0,649 | -13,746 | 5,42E-43 | 3,92E-41 | 0,000482 | 0,0028949 | 0,0014771 | 0,0014771 | TRUE | 0,0120683 | No |
| orf19.3153 | 6630 | 4867 | 5911 | 7403 | 6349 | 6114 | 6861 | 3927 | 5948 | 6196 | 6048 | 6105 | 5430 | 5834 | 6122 | 5539 | 5902,6 | 6074 | 5731 | 0,944 | -0,083 | -1,877 | 0,06057547 | 0,0993279 | 0,001399 | 0,0022255 | 0,0017154 | 0,0017154 | TRUE | 0,0260438 | No |
| orf19.3154 | 1536 | 1232 | 1363 | 1751 | 1949 | 1996 | 2060 | 1221 | 1378 | 1568 | 1394 | 1444 | 1667 | 1905 | 1838 | 1722 | 1614,6 | 1446 | 1783 | 1,235 | 0,304 | 5,029 | 4,94E-07 | 2,19E-06 | 0,002777 | 0,0028787 | 0,0028894 | 0,0028894 | TRUE | 0,0368668 | No |
| orf19.3156 | 1911 | 1268 | 1718 | 2117 | 1810 | 1628 | 1779 | 1091 | 1714 | 1614 | 1758 | 1746 | 1548 | 1553 | 1587 | 1539 | 1632,5 | 1708 | 1557 | 0,911 | -0,135 | -3,042 | 0,0023481 | 0,0054574 | 0,000103 | 0,0028688 | 0,0012756 | 0,0012756 | TRUE | 0,0150128 | No |
| orf19.3158 | 1860 | 1259 | 1601 | 1957 | 2542 | 2333 | 2520 | 1614 | 1669 | 1603 | 1638 | 1614 | 2174 | 2226 | 2249 | 2276 | 1931 | 1631 | 2231 | 1,366 | 0,45 | 11,339 | 8,37E-30 | 2,98E-28 | 1,00E-08 | 0,0027313 | 0,0009846 | 0,0009846 | TRUE | 0,0037082 | No |
| orf19.3159 | 4345 | 3791 | 4332 | 5284 | 3380 | 3547 | 3504 | 2001 | 3898 | 4826 | 4432 | 4357 | 2891 | 3384 | 3127 | 2822 | 3717,2 | 4378 | 3056 | 0,699 | -0,517 | -7,014 | 2,32E-12 | 2,04E-11 | 0,006887 | 0,0023701 | 0,0049475 | 0,0049475 | TRUE | 0,0666037 | No |
| orf19.3159.4 | 2 | 1 | 4 | 1 | 0 | 2 | 1 | 0 | 2 | 1 | 4 | 1 | 0 | 2 | 1 | 0 | 1,35 | 2 | 1 | 0,369 | -1,438 | -1,104 | 0,26966818 | NA | 1,00E-08 | 1,07881 | 0,6985236 | 0,6985236 | TRUE | 0,4495898 | No |
| orf19.316 | 1015 | 649 | 854 | 1023 | 1212 | 939 | 1062 | 724 | 911 | 826 | 874 | 844 | 1037 | 896 | 948 | 1021 | 919,43 | 864 | 976 | 1,127 | 0,173 | 2,713 | 0,00666689 | 0,01402474 | 0,002193 | 0,0035584 | 0,0028062 | 0,0028062 | TRUE | 0,0360819 | No |
| orf19.3160 | 1424 | 894 | 1130 | 1428 | 2415 | 1873 | 1853 | 1233 | 1277 | 1138 | 1156 | 1178 | 2065 | 1787 | 1653 | 1739 | 1499,3 | 1187 | 1811 | 1,526 | 0,609 | 8,325 | 8,41E-17 | 1,17E-15 | 0,005283 | 0,0029478 | 0,0044441 | 0,0044441 | TRUE | 0,1101912 | No |
| orf19.3161 | 6031 | 4011 | 4990 | 6430 | 6650 | 5573 | 5989 | 3475 | 5410 | 5106 | 5105 | 5302 | 5687 | 5318 | 5344 | 4901 | 5271,8 | 5231 | 5312 | 1,016 | 0,023 | 0,473 | 0,63654879 | 0,7073026 | 0,001825 | 0,0022549 | 0,0021125 | 0,0021125 | TRUE | 0,0320816 | No |
| orf19.3163 | 854 | 601 | 748 | 858 | 831 | 704 | 790 | 484 | 766 | 765 | 765 | 708 | 711 | 672 | 705 | 683 | 721,76 | 751 | 693 | 0,924 | -0,113 | -1,937 | 0,05268787 | 0,08800354 | 1,00E-08 | 0,0039909 | 0,0019061 | 0,0019061 | TRUE | 0,0195633 | No |
| orf19.3166 | 680 | 484 | 602 | 748 | 619 | 578 | 599 | 385 | 610 | 616 | 616 | 617 | 529 | 552 | 535 | 543 | 577,17 | 615 | 540 | 0,877 | -0,189 | -3,15 | 0,00163125 | 0,0039171 | 1,00E-08 | 0,0044947 | 0,0017164 | 0,0017164 | TRUE | 0,0028798 | No |
| orf19.3167 | 3692 | 2368 | 3154 | 3960 | 3675 | 2737 | 3090 | 2226 | 3312 | 3015 | 3227 | 3265 | 3143 | 2612 | 2757 | 3140 | 3058,8 | 3205 | 2913 | 0,908 | -0,139 | -2,115 | 0,0344358 | 0,06060056 | 0,004712 | 0,0024541 | 0,003836 | 0,003836 | TRUE | 0,0588911 | No |
| orf19.3168 | 7241 | 4758 | 6270 | 7419 | 6312 | 5632 | 6024 | 3595 | 6496 | 6057 | 6415 | 6118 | 5398 | 5374 | 5375 | 5070 | 5788 | 6272 | 5304 | 0,846 | -0,241 | -6,18 | 6,40E-10 | 4,18E-09 | 0,000798 | 0,0022303 | 0,0012839 | 0,0012839 | TRUE | 0,0102266 | No |
| orf19.3169 | 7805 | 5265 | 6484 | 8345 | 7882 | 6592 | 7370 | 4604 | 7002 | 6703 | 6634 | 6881 | 6741 | 6290 | 6576 | 6494 | 6665,1 | 6805 | 6525 | 0,959 | -0,061 | -1,703 | 0,08847864 | 0,13789356 | 0,000517 | 0,0021973 | 0,0010689 | 0,0010689 | TRUE | 0,0073762 | No |
| orf19.317 | 876 | 522 | 716 | 836 | 1105 | 914 | 1027 | 634 | 786 | 665 | 733 | 689 | 945 | 872 | 916 | 894 | 812,51 | 718 | 907 | 1,262 | 0,335 | 5,172 | 2,32E-07 | 1,08E-06 | 0,001918 | 0,0037662 | 0,0027924 | 0,0027924 | TRUE | 0,0487447 | No |
| orf19.3170 | 3100 | 2221 | 2846 | 3277 | 2908 | 2698 | 2999 | 1721 | 2781 | 2828 | 2912 | 2702 | 2487 | 2574 | 2676 | 2427 | 2673,4 | 2806 | 2541 | 0,907 | -0,141 | -3,128 | 0,00175896 | 0,00420359 | 0,000896 | 0,0025225 | 0,0015642 | 0,0015642 | TRUE | 0,0155543 | No |
| orf19.3171 | 6861 | 4751 | 6153 | 7241 | 9767 | 8394 | 8929 | 6676 | 6155 | 6049 | 6295 | 5971 | 8353 | 8009 | 7968 | 9416 | 7277 | 6118 | 8436 | 1,378 | 0,463 | 8,357 | 6,44E-17 | 9,06E-16 | 0,00328 | 0,002179 | 0,0028056 | 0,0028056 | TRUE | 0,0734835 | No |
| orf19.3172 | 16 | 7 | 15 | 12 | 13 | 7 | 16 | 14 | 14 | 9 | 15 | 10 | 11 | 7 | 14 | 20 | 12,54 | 12 | 13 | 1,036 | 0,051 | 0,121 | 0,90332336 | NA | 0,036356 | 0,1177365 | 0,0871093 | 0,0871093 | TRUE | 0,228812 | No |
| orf19.3173 | 722 | 435 | 530 | 643 | 613 | 494 | 531 | 295 | 648 | 554 | 542 | 530 | 524 | 471 | 474 | 416 | 519,94 | 568 | 471 | 0,832 | -0,265 | -2,96 | 0,00307378 | 0,0069609 | 0,006267 | 0,0047716 | 0,0057626 | 0,0057626 | TRUE | 0,1069729 | No |
| orf19.3174 | 1034 | 738 | 965 | 1110 | 1202 | 1102 | 1231 | 811 | 928 | 940 | 987 | 915 | 1028 | 1051 | 1098 | 1144 | 1011,4 | 942 | 1080 | 1,145 | 0,195 | 3,54 | 0,00039945 | 0,00110037 | 0,00066 | 0,0034148 | 0,0019107 | 0,0019107 | TRUE | 0,0165066 | No |
| orf19.3175 | 4602 | 3027 | 4010 | 4739 | 6107 | 5247 | 5637 | 3617 | 4128 | 3854 | 4103 | 3908 | 5223 | 5007 | 5030 | 5101 | 4544,2 | 3998 | 5090 | 1,273 | 0,348 | 9,343 | 9,40E-21 | 1,80E-19 | 0,000478 | 0,002299 | 0,0011074 | 0,0011074 | TRUE | 0,0067691 | No |
| orf19.3176 | 1670 | 1132 | 1488 | 1775 | 2033 | 1734 | 1920 | 1278 | 1498 | 1441 | 1522 | 1464 | 1739 | 1655 | 1713 | 1803 | 1604,3 | 1481 | 1728 | 1,164 | 0,219 | 4,846 | 1,26E-06 | 5,24E-06 | 0,000194 | 0,0028844 | 0,0013331 | 0,0013331 | TRUE | 0,0094076 | No |
| orf19.3177 | 623 | 424 | 546 | 585 | 509 | 444 | 477 | 258 | 559 | 540 | 559 | 482 | 435 | 424 | 426 | 364 | 473,52 | 535 | 412 | 0,775 | -0,367 | -4,548 | 5,41E-06 | 2,05E-05 | 0,003124 | 0,0050453 | 0,0041029 | 0,0041029 | TRUE | 0,0606434 | No |
| orf19.3178 | 2560 | 1892 | 2216 | 2694 | 2878 | 2583 | 2843 | 1554 | 2297 | 2409 | 2267 | 2222 | 2461 | 2465 | 2537 | 2192 | 2356,1 | 2299 | 2414 | 1,053 | 0,074 | 1,399 | 0,16174946 | 0,22990082 | 0,002017 | 0,0025957 | 0,0022668 | 0,0022668 | TRUE | 0,0435127 | No |
| orf19.318 | 981 | 678 | 856 | 1098 | 1006 | 908 | 944 | 576 | 880 | 863 | 876 | 905 | 860 | 866 | 842 | 812 | 863,24 | 881 | 845 | 0,96 | -0,059 | -1,117 | 0,26383474 | 0,34753763 | 1,00E-08 | 0,0036612 | 0,0014833 | 0,0014833 | TRUE | 0,0068452 | No |
| orf19.3180 | 1006 | 755 | 873 | 1094 | 1346 | 1152 | 1249 | 778 | 902 | 961 | 893 | 902 | 1151 | 1099 | 1115 | 1097 | 1015,1 | 914 | 1116 | 1,223 | 0,29 | 5,738 | 9,57E-09 | 5,31E-08 | 1,00E-08 | 0,0034096 | 0,0014604 | 0,0014604 | TRUE | 0,0123394 | No |
| orf19.3181.1 | 420 | 316 | 370 | 432 | 424 | 318 | 357 | 209 | 377 | 402 | 379 | 356 | 363 | 303 | 319 | 295 | 349,16 | 378 | 320 | 0,851 | -0,232 | -2,658 | 0,00786275 | 0,01625023 | 0,002599 | 0,0061373 | 0,0044609 | 0,0044609 | TRUE | 0,0960205 | No |
| orf19.3182 | 13184 | 8520 | 11155 | 13699 | 13072 | 10295 | 11098 | 7595 | 11827 | 10847 | 11413 | 11296 | 11180 | 9823 | 9903 | 10712 | 10875 | 11346 | 10404 | 0,917 | -0,125 | -2,476 | 0,01329836 | 0,02594391 | 0,002487 | 0,002113 | 0,0023672 | 0,0023672 | TRUE | 0,0311919 | No |
| orf19.3183 | 1194 | 879 | 995 | 1343 | 1268 | 1043 | 1181 | 903 | 1071 | 1119 | 1018 | 1107 | 1084 | 995 | 1054 | 1274 | 1090,3 | 1079 | 1102 | 1,018 | 0,026 | 0,35 | 0,72630378 | 0,78553251 | 0,005163 | 0,0033109 | 0,004413 | 0,004413 | TRUE | 0,123228 | No |
| orf19.3184 | 2211 | 1557 | 1938 | 2410 | 2536 | 2111 | 2241 | 1405 | 1983 | 1982 | 1983 | 1987 | 2169 | 2014 | 2000 | 1982 | 2012,5 | 1984 | 2041 | 1,03 | 0,043 | 0,984 | 0,3251916 | 0,41218378 | 0,000469 | 0,0027009 | 0,0013053 | 0,0013053 | TRUE | 0,0222901 | No |
| orf19.3185 | 4628 | 3253 | 4156 | 5159 | 4943 | 4496 | 4768 | 3135 | 4152 | 4141 | 4252 | 4254 | 4227 | 4290 | 4255 | 4422 | 4249,1 | 4200 | 4298 | 1,022 | 0,032 | 0,987 | 0,3237024 | 0,41072728 | 1,00E-08 | 0,0023212 | 0,0007647 | 0,0007647 | TRUE | 0,0041689 | No |

| | | | | | | | | | | | | | | | | | | | | | | | | | | | | | | | |
|---|---|---|---|---|---|---|---|---|---|---|---|---|---|---|---|---|---|---|---|---|---|---|---|---|---|---|---|---|---|---|---|
| orf19.3186 | 0 | 0 | 1 | 0 | 0 | 1 | 3 | 0 | 0 | 0 | 1 | 0 | 0 | 1 | 3 | 0 | 0,58 | 0 | 1 | 3,121 | 1,642 | 0,807 | 0,41990578 | NA | 1,00E-08 | 2,4972394 | 2,4418387 | 2,4418387 | TRUE | 0,8211518 | No |
| orf19.3187 | 3920 | 2910 | 3694 | 4451 | 3880 | 3734 | 3933 | 2465 | 3517 | 3705 | 3779 | 3670 | 3318 | 3563 | 3510 | 3477 | 3567,3 | 3668 | 3467 | 0,945 | -0,081 | -2,023 | 0,04308778 | 0,07383668 | 0,000691 | 0,0023865 | 0,001262 | 0,001262 | TRUE | 0,0105696 | No |
| orf19.3188 | 1807 | 1347 | 1785 | 2088 | 1955 | 1880 | 1967 | 1265 | 1621 | 1715 | 1826 | 1722 | 1672 | 1794 | 1755 | 1784 | 1736,2 | 1721 | 1751 | 1,017 | 0,025 | 0,493 | 0,62195697 | 0,69507394 | 0,001174 | 0,0028157 | 0,0018274 | 0,0018274 | TRUE | 0,0205405 | No |
| orf19.319 | 228 | 172 | 210 | 248 | 192 | 158 | 181 | 99 | 205 | 219 | 215 | 205 | 164 | 151 | 162 | 140 | 182,37 | 211 | 154 | 0,737 | -0,439 | -4,21 | 2,56E-05 | 8,72E-05 | 1,00E-08 | 0,0099396 | 0,0048755 | 0,0048755 | TRUE | 0,0321506 | No |
| orf19.3190 | 5834 | 4239 | 5250 | 6589 | 7022 | 6486 | 6792 | 4490 | 5234 | 5397 | 5371 | 5433 | 6005 | 6189 | 6061 | 6333 | 5752,8 | 5359 | 6147 | 1,146 | 0,197 | 6,107 | 1,02E-09 | 6,48E-09 | 0,000181 | 0,0022319 | 0,0008249 | 0,0008249 | TRUE | 0,0047708 | No |
| orf19.3192 | 4611 | 3344 | 4059 | 4964 | 6510 | 6062 | 6419 | 4334 | 4137 | 4257 | 4153 | 4093 | 5568 | 5784 | 5728 | 6113 | 4979 | 4160 | 5798 | 1,393 | 0,478 | 12,46 | 1,23E-35 | 6,43E-34 | 0,000644 | 0,0022711 | 0,0012074 | 0,0012074 | TRUE | 0,0155691 | No |
| orf19.3193 | 2076 | 1605 | 2027 | 2372 | 1620 | 1598 | 1642 | 1104 | 1862 | 2043 | 2074 | 1956 | 1385 | 1525 | 1465 | 1557 | 1733,5 | 1984 | 1483 | 0,747 | -0,421 | -7,713 | 1,23E-14 | 1,36E-13 | 0,001884 | 0,002817 | 0,0022655 | 0,0022655 | TRUE | 0,0241324 | No |
| orf19.3195 | 9418 | 6530 | 8692 | 10322 | 10526 | 9494 | 10692 | 6442 | 8449 | 8313 | 8893 | 8512 | 9002 | 9059 | 9541 | 9086 | 8856,8 | 8542 | 9172 | 1,074 | 0,103 | 2,887 | 0,00388853 | 0,00860806 | 0,000638 | 0,0021434 | 0,0011017 | 0,0011017 | TRUE | 0,0092759 | No |
| orf19.3196 | 208 | 159 | 195 | 268 | 259 | 278 | 265 | 205 | 187 | 202 | 200 | 221 | 222 | 265 | 236 | 289 | 227,74 | 202 | 253 | 1,241 | 0,311 | 2,85 | 0,00437312 | 0,00956102 | 0,005473 | 0,008354 | 0,0070042 | 0,0070042 | TRUE | 0,0885823 | No |
| orf19.3197 | 1322 | 928 | 1119 | 1310 | 1435 | 1273 | 1349 | 825 | 1186 | 1181 | 1145 | 1080 | 1227 | 1215 | 1204 | 1164 | 1175,2 | 1148 | 1202 | 1,05 | 0,07 | 1,384 | 0,16638838 | 0,23582481 | 0,000409 | 0,0032148 | 0,0016231 | 0,0016231 | TRUE | 0,0203146 | No |
| orf19.3198 | 2230 | 1412 | 1835 | 2204 | 2695 | 2190 | 2390 | 1464 | 2001 | 1798 | 1877 | 1817 | 2305 | 2090 | 2133 | 2065 | 2010,6 | 1873 | 2148 | 1,147 | 0,198 | 3,7 | 0,00021566 | 0,00062501 | 0,001794 | 0,0027015 | 0,0022503 | 0,0022503 | TRUE | 0,0298584 | No |
| orf19.3199 | 1353 | 882 | 1194 | 1352 | 1411 | 1256 | 1344 | 784 | 1214 | 1123 | 1222 | 1115 | 1207 | 1198 | 1199 | 1106 | 1172,9 | 1168 | 1178 | 1,01 | 0,014 | 0,256 | 0,7978296 | 0,84330757 | 0,001062 | 0,0032172 | 0,0020245 | 0,0020245 | TRUE | 0,0178997 | No |
| orf19.320 | 2203 | 1352 | 1803 | 2152 | 2226 | 1882 | 2062 | 1541 | 1976 | 1721 | 1845 | 1775 | 1904 | 1796 | 1840 | 2173 | 1878,7 | 1829 | 1928 | 1,052 | 0,073 | 1,068 | 0,28549191 | 0,36982221 | 0,004967 | 0,0027522 | 0,0039154 | 0,0039154 | TRUE | 0,0845911 | No |
| orf19.3200 | 101 | 102 | 120 | 147 | 103 | 124 | 106 | 92 | 91 | 130 | 123 | 121 | 88 | 118 | 95 | 130 | 111,9 | 116 | 108 | 0,922 | -0,117 | -0,713 | 0,47585884 | 0,56124065 | 0,019467 | 0,0149526 | 0,0169802 | 0,0169802 | TRUE | 0,2310655 | No |
| orf19.3201 | 1136 | 786 | 888 | 1153 | 1140 | 967 | 1064 | 704 | 1019 | 1001 | 909 | 951 | 975 | 923 | 949 | 993 | 964,88 | 970 | 960 | 0,989 | -0,016 | -0,277 | 0,78200411 | 0,82891332 | 0,000781 | 0,0034841 | 0,0020221 | 0,0020221 | TRUE | 0,0210063 | No |
| orf19.3202 | 574 | 518 | 552 | 651 | 870 | 1042 | 978 | 628 | 515 | 659 | 565 | 537 | 744 | 994 | 873 | 886 | 721,59 | 569 | 874 | 1,54 | 0,623 | 6,299 | 3,00E-10 | 2,05E-09 | 0,011193 | 0,0039913 | 0,0079225 | 0,0079225 | TRUE | 0,1278365 | No |
| orf19.3203 | 329 | 289 | 297 | 389 | 464 | 541 | 587 | 320 | 295 | 368 | 304 | 321 | 397 | 516 | 524 | 451 | 396,98 | 322 | 472 | 1,471 | 0,557 | 5,225 | 1,74E-07 | 8,25E-07 | 0,010885 | 0,0056363 | 0,0082966 | 0,0082966 | TRUE | 0,1434961 | No |
| orf19.3204 | 1579 | 1496 | 1473 | 1818 | 1935 | 2394 | 2341 | 1262 | 1417 | 1905 | 1507 | 1499 | 1655 | 2284 | 2089 | 1780 | 1766,9 | 1582 | 1952 | 1,236 | 0,305 | 2,784 | 0,00537644 | 0,01154042 | 0,01963 | 0,0028011 | 0,0109789 | 0,0109789 | TRUE | 0,223925 | No |
| orf19.3205 | 3817 | 2640 | 3289 | 4142 | 4323 | 3781 | 4093 | 2300 | 3424 | 3361 | 3365 | 3416 | 3697 | 3608 | 3652 | 3244 | 3470,9 | 3392 | 3550 | 1,048 | 0,068 | 1,467 | 0,14236545 | 0,20635471 | 0,001371 | 0,0023978 | 0,0017701 | 0,0017701 | TRUE | 0,0388117 | No |
| orf19.3206 | 5782 | 4374 | 5175 | 6547 | 5872 | 5819 | 6254 | 3661 | 5187 | 5569 | 5295 | 5399 | 5022 | 5552 | 5581 | 5164 | 5345,9 | 5362 | 5330 | 0,994 | -0,008 | -0,174 | 0,86192023 | 0,89542891 | 0,00139 | 0,0022511 | 0,0018676 | 0,0018676 | TRUE | 0,0190789 | No |
| orf19.3207 | 133 | 143 | 127 | 179 | 135 | 137 | 160 | 90 | 119 | 182 | 130 | 148 | 115 | 131 | 143 | 127 | 136,85 | 145 | 129 | 0,897 | -0,156 | -1,072 | 0,283533 | 0,36783983 | 0,013696 | 0,0125875 | 0,0130325 | 0,0130325 | TRUE | 0,2775925 | No |
| orf19.3208 | 786 | 639 | 740 | 957 | 843 | 844 | 973 | 515 | 705 | 814 | 757 | 789 | 721 | 805 | 868 | 726 | 773,22 | 766 | 780 | 1,021 | 0,03 | 0,394 | 0,69352559 | 0,75642174 | 0,004651 | 0,003857 | 0,0042838 | 0,0042838 | TRUE | 0,0698627 | No |
| orf19.3209 | 26 | 12 | 13 | 30 | 23 | 18 | 22 | 15 | 23 | 15 | 13 | 25 | 20 | 17 | 20 | 21 | 19,28 | 19 | 20 | 0,991 | -0,013 | -0,039 | 0,9685383 | 0,97781517 | 1,00E-08 | 0,0772589 | 0,0472179 | 0,0472179 | TRUE | 0,1896021 | No |
| orf19.321 | 2063 | 1348 | 1868 | 2219 | 1413 | 1290 | 1354 | 985 | 1851 | 1716 | 1911 | 1830 | 1208 | 1231 | 1208 | 1389 | 1543,1 | 1827 | 1259 | 0,687 | -0,542 | -9,346 | 9,06E-21 | 1,75E-19 | 0,002508 | 0,0029203 | 0,0025567 | 0,0025567 | TRUE | 0,0531122 | No |
| orf19.3210 | 30 | 17 | 20 | 25 | 23 | 35 | 36 | 18 | 27 | 22 | 20 | 21 | 20 | 33 | 32 | 25 | 25,03 | 22 | 28 | 1,233 | 0,302 | 1,057 | 0,29043814 | 0,37501663 | 1,00E-08 | 0,0599864 | 0,0380655 | 0,0380655 | TRUE | 0,2989458 | No |
| orf19.3211 | 1098 | 862 | 1063 | 1327 | 1266 | 1187 | 1365 | 808 | 985 | 1097 | 1088 | 1094 | 1083 | 1133 | 1218 | 1140 | 1104,7 | 1066 | 1144 | 1,073 | 0,102 | 1,73 | 0,08371093 | 0,13158955 | 0,001776 | 0,0032937 | 0,0024272 | 0,0024272 | TRUE | 0,03235 | No |
| orf19.3212 | 1387 | 1114 | 1228 | 1525 | 1980 | 2007 | 2057 | 1261 | 1244 | 1418 | 1256 | 1258 | 1693 | 1915 | 1836 | 1779 | 1549,9 | 1294 | 1806 | 1,398 | 0,483 | 8,152 | 3,58E-16 | 4,61E-15 | 0,00251 | 0,0029162 | 0,0027083 | 0,0027083 | TRUE | 0,0471922 | No |
| orf19.3213 | 2105 | 1604 | 2004 | 2473 | 2120 | 2136 | 2317 | 1444 | 1888 | 2042 | 2050 | 2039 | 1813 | 2038 | 2068 | 2037 | 1996,9 | 2005 | 1989 | 0,992 | -0,012 | -0,222 | 0,8239912 | 0,8642701 | 0,002189 | 0,0027065 | 0,0023941 | 0,0023941 | TRUE | 0,0443751 | No |
| orf19.3214 | 436 | 451 | 460 | 589 | 424 | 477 | 504 | 297 | 391 | 574 | 471 | 486 | 363 | 455 | 450 | 419 | 451 | 480 | 422 | 0,88 | -0,185 | -1,614 | 0,10657507 | 0,16100151 | 0,015433 | 0,0051984 | 0,0103445 | 0,0103445 | TRUE | 0,1908136 | No |
| orf19.3215 | 2017 | 1448 | 1734 | 2174 | 2122 | 2040 | 2101 | 1237 | 1809 | 1843 | 1774 | 1793 | 1815 | 1947 | 1875 | 1745 | 1825,1 | 1805 | 1846 | 1,024 | 0,035 | 0,756 | 0,44959074 | 0,53583439 | 0,000488 | 0,002775 | 0,0014595 | 0,0014595 | TRUE | 0,0160417 | No |
| orf19.3216 | 1032 | 886 | 1025 | 1262 | 1142 | 1277 | 1381 | 894 | 926 | 1128 | 1049 | 1041 | 977 | 1218 | 1232 | 1261 | 1103,9 | 1036 | 1172 | 1,131 | 0,178 | 2,028 | 0,04258836 | 0,07308507 | 0,00924 | 0,0032945 | 0,0064832 | 0,0064832 | TRUE | 0,154588 | No |
| orf19.3218 | 706 | 606 | 650 | 832 | 766 | 781 | 804 | 466 | 633 | 772 | 665 | 686 | 655 | 745 | 717 | 657 | 691,37 | 689 | 694 | 1,011 | 0,015 | 0,204 | 0,83856626 | 0,87649723 | 0,004005 | 0,0040793 | 0,0040408 | 0,0040408 | TRUE | 0,0711684 | No |
| orf19.3219 | 277 | 272 | 271 | 330 | 250 | 372 | 366 | 200 | 248 | 346 | 277 | 272 | 214 | 355 | 327 | 282 | 290,2 | 286 | 294 | 1,032 | 0,046 | 0,302 | 0,76250887 | 0,81471692 | 0,030499 | 0,0069819 | 0,0184218 | 0,0184218 | TRUE | 0,4048954 | No |
| orf19.322 | 11 | 7 | 4 | 14 | 9 | 6 | 6 | 9 | 10 | 9 | 4 | 12 | 8 | 6 | 5 | 13 | 8,24 | 9 | 8 | 0,874 | -0,193 | -0,383 | 0,70167064 | NA | 1,00E-08 | 0,178246 | 0,1220555 | 0,1220555 | TRUE | 0,3845964 | No |
| orf19.3220 | 8292 | 6127 | 7055 | 9033 | 7007 | 6462 | 6648 | 3854 | 7439 | 7800 | 7218 | 7449 | 5993 | 6166 | 5932 | 5436 | 6679 | 7476 | 5882 | 0,787 | -0,345 | -7,472 | 7,89E-14 | 8,05E-13 | 0,001735 | 0,0021969 | 0,0018919 | 0,0018919 | TRUE | 0,0306866 | No |
| orf19.3221 | 10972 | 7890 | 9968 | 11838 | 11925 | 11127 | 12024 | 7275 | 9843 | 10045 | 10198 | 9762 | 10199 | 10617 | 10729 | 10261 | 10207 | 9962 | 10452 | 1,05 | 0,07 | 2,181 | 0,02915268 | 0,05225219 | 0,000378 | 0,0021218 | 0,0008829 | 0,0008829 | TRUE | 0,0039516 | No |
| orf19.3222 | 771 | 588 | 680 | 849 | 811 | 824 | 930 | 521 | 692 | 749 | 696 | 700 | 694 | 786 | 830 | 735 | 735,07 | 709 | 761 | 1,076 | 0,106 | 1,527 | 0,12672108 | 0,1869237 | 0,002592 | 0,0039544 | 0,0032173 | 0,0032173 | TRUE | 0,046816 | No |
| orf19.3223 | 5308 | 3662 | 4431 | 5522 | 5584 | 5052 | 5083 | 3465 | 4762 | 4662 | 4533 | 4554 | 4776 | 4820 | 4536 | 4887 | 4691,2 | 4628 | 4755 | 1,027 | 0,038 | 1,024 | 0,30588735 | 0,39175282 | 0,000495 | 0,002289 | 0,0011304 | 0,0011304 | TRUE | 0,0118493 | No |
| orf19.3223.1 | 1858 | 1371 | 1522 | 1960 | 1620 | 1469 | 1384 | 934 | 1667 | 1745 | 1557 | 1616 | 1385 | 1402 | 1235 | 1317 | 1490,6 | 1646 | 1335 | 0,812 | -0,301 | -5,249 | 1,53E-07 | 7,30E-07 | 0,002129 | 0,0029534 | 0,0024771 | 0,0024771 | TRUE | 0,0323478 | No |
| orf19.3225 | 661 | 550 | 708 | 843 | 815 | 814 | 888 | 500 | 593 | 700 | 724 | 695 | 697 | 777 | 792 | 705 | 710,5 | 678 | 743 | 1,098 | 0,135 | 1,734 | 0,08297162 | 0,13052985 | 0,004695 | 0,0040227 | 0,0043795 | 0,0043795 | TRUE | 0,0871416 | No |
| orf19.3226 | 1093 | 828 | 916 | 1261 | 965 | 988 | 996 | 647 | 981 | 1054 | 937 | 1040 | 825 | 943 | 889 | 913 | 947,62 | 1003 | 892 | 0,889 | -0,17 | -2,708 | 0,00676877 | 0,01419601 | 0,002052 | 0,0035115 | 0,0027081 | 0,0027081 | TRUE | 0,0321317 | No |
| orf19.3227 | 2191 | 1858 | 2090 | 2528 | 2375 | 2526 | 2660 | 1402 | 1966 | 2365 | 2138 | 2085 | 2031 | 2410 | 2374 | 1977 | 2168,3 | 2138 | 2198 | 1,03 | 0,042 | 0,535 | 0,5927063 | 0,6701128 | 0,006939 | 0,002649 | 0,0054614 | 0,0054614 | TRUE | 0,0605506 | No |
| orf19.3228 | 1344 | 1258 | 1340 | 1754 | 1208 | 1410 | 1465 | 853 | 1206 | 1602 | 1371 | 1446 | 1033 | 1345 | 1307 | 1203 | 1314,2 | 1406 | 1222 | 0,87 | -0,201 | -2,07 | 0,03845706 | 0,06681608 | 0,012357 | 0,0030842 | 0,0082918 | 0,0082918 | TRUE | 0,1339198 | No |
| orf19.3229 | 4 | 5 | 6 | 6 | 4 | 8 | 2 | 0 | 4 | 6 | 6 | 5 | 3 | 8 | 2 | 0 | 4,23 | 5 | 3 | 0,646 | -0,63 | -0,863 | 0,38793405 | NA | 0,106668 | 0,3447746 | 0,2669255 | 0,2669255 | TRUE | 0,8646409 | No |
| orf19.323 | 1475 | 967 | 1270 | 1428 | 2120 | 1786 | 1879 | 1158 | 1323 | 1231 | 1299 | 1178 | 1813 | 1704 | 1677 | 1633 | 1482,3 | 1258 | 1707 | 1,359 | 0,442 | 7,976 | 1,51E-15 | 1,83E-14 | 0,001766 | 0,0029589 | 0,0022602 | 0,0022602 | TRUE | 0,0239755 | No |
| orf19.3230 | 4776 | 3429 | 4362 | 5317 | 6073 | 5393 | 6068 | 3870 | 4285 | 4365 | 4463 | 4384 | 5194 | 5146 | 5415 | 5458 | 4838,7 | 4374 | 5303 | 1,212 | 0,277 | 7,909 | 2,60E-15 | 3,06E-14 | 0,000296 | 0,0022795 | 0,0009693 | 0,0009693 | TRUE | 0,0047803 | No |
| orf19.3231 | 443 | 327 | 423 | 548 | 476 | 584 | 547 | 331 | 397 | 416 | 433 | 452 | 407 | 557 | 488 | 467 | 452,21 | 424 | 480 | 1,129 | 0,175 | 1,81 | 0,07029853 | 0,11317083 | 0,008228 | 0,0051898 | 0,0067737 | 0,0067737 | TRUE | 0,1416155 | No |
| orf19.3232 | 775 | 532 | 710 | 840 | 438 | 422 | 411 | 290 | 695 | 677 | 726 | 693 | 375 | 403 | 367 | 409 | 543,08 | 698 | 388 | 0,554 | -0,852 | -12,712 | 5,07E-37 | 2,93E-35 | 1,00E-08 | 0,0046526 | 0,0023064 | 0,0023064 | TRUE | 0,0156672 | No |
| orf19.3233 | 2831 | 1877 | 2387 | 2927 | 2864 | 2531 | 2750 | 1820 | 2540 | 2390 | 2442 | 2414 | 2449 | 2415 | 2454 | 2567 | 2458,8 | 2446 | 2471 | 1,009 | 0,012 | 0,308 | 0,75810975 | 0,81088185 | 0,000178 | 0,0025699 | 0,0011184 | 0,0011184 | TRUE | 0,0081755 | No |
| orf19.3234 | 381 | 313 | 353 | 452 | 255 | 243 | 248 | 151 | 342 | 398 | 361 | 373 | 218 | 232 | 221 | 213 | 294,8 | 368 | 221 | 0,602 | -0,731 | -8,38 | 5,28E-17 | 7,48E-16 | 1,00E-08 | 0,0069039 | 0,0036957 | 0,0036957 | TRUE | 0,0301447 | No |
| orf19.3234.1 | 530 | 436 | 489 | 604 | 749 | 644 | 629 | 363 | 475 | 555 | 500 | 498 | 641 | 614 | 561 | 512 | 544,65 | 507 | 582 | 1,155 | 0,208 | 2,504 | 0,01226867 | 0,02415143 | 0,004592 | 0,0046449 | 0,0047569 | 0,0047569 | TRUE | 0,0683997 | No |
| orf19.3235 | 647 | 553 | 555 | 757 | 673 | 728 | 743 | 462 | 580 | 704 | 568 | 624 | 576 | 695 | 663 | 652 | 632,67 | 619 | 646 | 1,045 | 0,064 | 0,755 | 0,45023387 | 0,53623917 | 0,006236 | 0,0042741 | 0,0052921 | 0,0052921 | TRUE | 0,0930515 | No |
| orf19.3237 | 2064 | 1516 | 1805 | 2276 | 2265 | 2207 | 2163 | 1261 | 1852 | 1930 | 1847 | 1877 | 1937 | 2106 | 1930 | 1779 | 1907,1 | 1876 | 1938 | 1,035 | 0,05 | 0,921 | 0,35710842 | 0,44484914 | 0,001889 | 0,0027407 | 0,0022516 | 0,0022516 | TRUE | 0,040369 | No |
| orf19.3238 | 4 | 5 | 2 | 6 | 5 | 6 | 6 | 5 | 4 | 6 | 2 | 5 | 4 | 6 | 5 | 7 | 4,92 | 4 | 6 | 1,317 | 0,397 | 0,646 | 0,51824189 | NA | 1,00E-08 | 0,2970779 | 0,1538972 | 0,1538972 | TRUE | 0,1962101 | No |
| orf19.3239 | 896 | 605 | 773 | 1005 | 952 | 761 | 878 | 512 | 804 | 770 | 791 | 829 | 814 | 726 | 783 | 722 | 779,94 | 798 | 761 | 0,955 | -0,067 | -1,088 | 0,27638322 | 0,36035382 | 0,000923 | 0,0038408 | 0,0023033 | 0,0023033 | TRUE | 0,0262041 | No |
| orf19.3240 | 538 | 380 | 505 | 615 | 562 | 476 | 609 | 408 | 483 | 484 | 517 | 507 | 481 | 454 | 543 | 575 | 505,49 | 498 | 513 | 1,027 | 0,039 | 0,471 | 0,63798453 | 0,7082894 | 0,004261 | 0,0048514 | 0,0045571 | 0,0045571 | TRUE | 0,0684588 | No |
| orf19.3241 | 1283 | 922 | 1215 | 1494 | 1411 | 1265 | 1350 | 889 | 1151 | 1174 | 1243 | 1232 | 1207 | 1207 | 1205 | 1254 | 1209 | 1200 | 1218 | 1,013 | 0,019 | 0,394 | 0,69359644 | 0,75642174 | 1,00E-08 | 0,0031803 | 0,0014353 | 0,0014353 | TRUE | 0,0098752 | No |
| orf19.3242 | 1345 | 849 | 1197 | 1427 | 1619 | 1311 | 1354 | 904 | 1207 | 1081 | 1225 | 1177 | 1385 | 1251 | 1208 | 1275 | 1226 | 1172 | 1280 | 1,091 | 0,125 | 2,058 | 0,03962007 | 0,06851968 | 0,0024 | 0,0031637 | 0,0027311 | 0,0027311 | TRUE | 0,0387037 | No |
| orf19.3243 | 3917 | 2656 | 3550 | 4173 | 3528 | 3020 | 3361 | 2058 | 3514 | 3381 | 3632 | 3441 | 3017 | 2882 | 2999 | 2903 | 3221,1 | 3492 | 2950 | 0,845 | -0,243 | -6,287 | 3,24E-10 | 2,21E-09 | 0,000421 | 0,0024302 | 0,0011186 | 0,0011186 | TRUE | 0,0086553 | No |

| | | | | | | | | | | | | | | | | | | | | | | | | | | | | | | | |
|---|---|---|---|---|---|---|---|---|---|---|---|---|---|---|---|---|---|---|---|---|---|---|---|---|---|---|---|---|---|---|---|
| orf19.3244 | 347 | 258 | 327 | 352 | 389 | 336 | 339 | 214 | 311 | 328 | 335 | 290 | 333 | 321 | 303 | 302 | 315,27 | 316 | 315 | 1 | 0 | -0,001 | 0,99933385 | 0,99933385 | 1,00E-08 | 0,0065841 | 0,003717 | 0,003717 | TRUE | 0,0372234 | No |
| orf19.3245 | 1456 | 1011 | 1326 | 1567 | 1839 | 1731 | 1774 | 1123 | 1306 | 1287 | 1357 | 1292 | 1573 | 1652 | 1583 | 1584 | 1454,2 | 1310 | 1598 | 1,219 | 0,286 | 6,455 | 1,09E-10 | 7,86E-10 | 1,00E-08 | 0,0029778 | 0,0011915 | 0,0011915 | TRUE | 0,0065938 | No |
| orf19.3247 | 356 | 362 | 335 | 472 | 547 | 667 | 626 | 396 | 319 | 461 | 343 | 389 | 468 | 636 | 559 | 559 | 466,69 | 378 | 556 | 1,473 | 0,558 | 4,64 | 3,49E-06 | 1,36E-05 | 0,018354 | 0,0050901 | 0,0116684 | 0,0116684 | TRUE | 0,2343842 | No |
| orf19.3248 | 321 | 251 | 307 | 398 | 394 | 473 | 401 | 277 | 288 | 320 | 314 | 328 | 337 | 451 | 358 | 391 | 348,32 | 312 | 384 | 1,228 | 0,296 | 2,92 | 0,00349848 | 0,0078021 | 0,007771 | 0,0061472 | 0,0069803 | 0,0069803 | TRUE | 0,1678613 | No |
| orf19.3249 | 1886 | 1523 | 1888 | 2245 | 1883 | 1982 | 1977 | 1148 | 1692 | 1939 | 1932 | 1851 | 1610 | 1891 | 1764 | 1619 | 1787,3 | 1854 | 1721 | 0,93 | -0,105 | -1,572 | 0,11590305 | 0,17287625 | 0,004175 | 0,0027917 | 0,0036965 | 0,0036965 | TRUE | 0,0533928 | No |
| orf19.325 | 2924 | 1953 | 2654 | 3034 | 3062 | 2807 | 3029 | 1921 | 2623 | 2486 | 2715 | 2502 | 2619 | 2678 | 2703 | 2709 | 2629,5 | 2582 | 2677 | 1,036 | 0,052 | 1,232 | 0,21797424 | 0,2964884 | 0,000488 | 0,0025316 | 0,001302 | 0,001302 | TRUE | 0,0144513 | No |
| orf19.3250 | 1626 | 1328 | 1417 | 1785 | 1316 | 1340 | 1391 | 783 | 1459 | 1691 | 1450 | 1472 | 1125 | 1279 | 1241 | 1104 | 1352,6 | 1518 | 1187 | 0,784 | -0,35 | -5,064 | 4,10E-07 | 1,84E-06 | 0,004496 | 0,0030528 | 0,0038389 | 0,0038389 | TRUE | 0,0680446 | No |
| orf19.3251 | 832 | 735 | 724 | 1013 | 964 | 965 | 958 | 641 | 746 | 936 | 741 | 835 | 824 | 921 | 855 | 904 | 845,29 | 814 | 876 | 1,077 | 0,106 | 1,335 | 0,18179539 | 0,2537112 | 0,006035 | 0,0036969 | 0,0049139 | 0,0049139 | TRUE | 0,111817 | No |
| orf19.3252 | 1848 | 1472 | 1857 | 2155 | 2365 | 2300 | 2501 | 1574 | 1658 | 1874 | 1900 | 1777 | 2023 | 2195 | 2232 | 2220 | 1984,7 | 1802 | 2168 | 1,203 | 0,267 | 4,721 | 2,35E-06 | 9,43E-06 | 0,002194 | 0,002711 | 0,0025533 | 0,0025533 | TRUE | 0,0354716 | No |
| orf19.3254 | 677 | 499 | 499 | 734 | 573 | 536 | 593 | 355 | 607 | 635 | 511 | 605 | 490 | 511 | 529 | 501 | 548,72 | 590 | 508 | 0,862 | -0,214 | -2,766 | 0,00567008 | 0,01210155 | 0,003202 | 0,0046251 | 0,0039036 | 0,0039036 | TRUE | 0,0937072 | No |
| orf19.3255 | 520 | 513 | 503 | 636 | 495 | 570 | 525 | 262 | 466 | 653 | 515 | 524 | 423 | 544 | 468 | 370 | 495,49 | 540 | 451 | 0,841 | -0,251 | -2,019 | 0,043437 | 0,07430794 | 0,021573 | 0,0049094 | 0,0127398 | 0,0127398 | TRUE | 0,2250421 | No |
| orf19.3256 | 5947 | 4377 | 5376 | 6570 | 7104 | 6884 | 7177 | 4526 | 5335 | 5572 | 5500 | 5418 | 6076 | 6568 | 6404 | 6384 | 5907,2 | 5456 | 6358 | 1,165 | 0,221 | 6,14 | 8,24E-10 | 5,32E-09 | 0,000473 | 0,0022253 | 0,0010711 | 0,0010711 | TRUE | 0,011234 | No |
| orf19.3259 | 397 | 281 | 363 | 478 | 492 | 438 | 478 | 265 | 356 | 358 | 371 | 394 | 421 | 418 | 427 | 374 | 389,8 | 370 | 410 | 1,111 | 0,151 | 1,962 | 0,04970622 | 0,08372161 | 1,00E-08 | 0,0057037 | 0,0031541 | 0,0031541 | TRUE | 0,0333668 | No |
| orf19.326 | 320 | 183 | 260 | 309 | 372 | 302 | 304 | 205 | 287 | 233 | 266 | 255 | 318 | 288 | 271 | 289 | 275,95 | 260 | 292 | 1,118 | 0,161 | 1,697 | 0,08975085 | 0,13973131 | 0,002401 | 0,0072403 | 0,0050134 | 0,0050134 | TRUE | 0,0542751 | No |
| orf19.3260 | 3962 | 2518 | 3594 | 4410 | 3857 | 3425 | 3825 | 2324 | 3554 | 3206 | 3677 | 3637 | 3299 | 3268 | 3413 | 3278 | 3416,4 | 3518 | 3314 | 0,941 | -0,087 | -1,783 | 0,07454807 | 0,11889695 | 0,001675 | 0,0024044 | 0,0020012 | 0,0020012 | TRUE | 0,0419441 | No |
| orf19.3261 | 144 | 91 | 104 | 151 | 102 | 70 | 86 | 50 | 129 | 116 | 106 | 125 | 87 | 67 | 77 | 71 | 97,16 | 119 | 76 | 0,635 | -0,655 | -4,427 | 9,56E-06 | 3,49E-05 | 1,00E-08 | 0,0169216 | 0,0102784 | 0,0102784 | TRUE | 0,1148609 | No |
| orf19.3263 | 29 | 34 | 39 | 44 | 27 | 26 | 31 | 19 | 26 | 43 | 40 | 36 | 23 | 25 | 28 | 27 | 30,98 | 36 | 26 | 0,708 | -0,498 | -1,947 | 0,05157496 | 0,08634119 | 1,00E-08 | 0,0488378 | 0,0294996 | 0,0294996 | TRUE | 0,2859501 | No |
| orf19.3264 | 306 | 303 | 342 | 431 | 340 | 381 | 377 | 237 | 275 | 386 | 350 | 355 | 291 | 364 | 336 | 334 | 336,32 | 342 | 331 | 0,972 | -0,041 | -0,365 | 0,715292 | 0,77557739 | 0,011735 | 0,0062959 | 0,0089974 | 0,0089974 | TRUE | 0,203678 | No |
| orf19.3264.1 | 6919 | 4023 | 5560 | 6303 | 6478 | 4812 | 5025 | 2697 | 6207 | 5122 | 5688 | 5198 | 5540 | 4591 | 4484 | 3804 | 5079,3 | 5554 | 4605 | 0,83 | -0,27 | -2,737 | 0,00619558 | 0,01311132 | 0,015869 | 0,0022653 | 0,009121 | 0,009121 | TRUE | 0,2281278 | No |
| orf19.3265 | 1593 | 1142 | 1428 | 1740 | 1512 | 1421 | 1519 | 878 | 1429 | 1454 | 1461 | 1435 | 1293 | 1356 | 1355 | 1238 | 1377,7 | 1445 | 1310 | 0,909 | -0,137 | -2,921 | 0,00348833 | 0,00778814 | 2,19E-05 | 0,0030332 | 0,0013823 | 0,0013823 | TRUE | 0,0142861 | No |
| orf19.3265.1 | 345 | 197 | 269 | 320 | 293 | 249 | 271 | 175 | 310 | 251 | 275 | 264 | 251 | 238 | 242 | 247 | 259,53 | 275 | 244 | 0,886 | -0,175 | -1,868 | 0,06179522 | 0,10110721 | 1,00E-08 | 0,0075732 | 0,0045941 | 0,0045941 | TRUE | 0,0831827 | No |
| orf19.3266 | 201 | 142 | 172 | 227 | 188 | 157 | 202 | 164 | 180 | 181 | 176 | 187 | 161 | 150 | 180 | 231 | 180,8 | 181 | 180 | 0,986 | -0,021 | -0,159 | 0,873925 | 0,90359019 | 0,011759 | 0,0100088 | 0,0108025 | 0,0108025 | TRUE | 0,32944 | No |
| orf19.3267 | 118 | 79 | 78 | 120 | 83 | 74 | 89 | 65 | 106 | 101 | 80 | 99 | 71 | 71 | 79 | 92 | 87,23 | 96 | 78 | 0,803 | -0,317 | -2,011 | 0,04430157 | 0,07567922 | 1,00E-08 | 0,0186209 | 0,0121607 | 0,0121607 | TRUE | 0,1270633 | No |
| orf19.3268 | 12601 | 7750 | 10144 | 12855 | 11881 | 9287 | 9628 | 7778 | 11304 | 9867 | 10378 | 10600 | 10161 | 8861 | 8591 | 10970 | 10092 | 10537 | 9646 | 0,915 | -0,128 | -1,682 | 0,09261072 | 0,14344037 | 0,008181 | 0,0021234 | 0,0054749 | 0,0054749 | TRUE | 0,1035982 | No |
| orf19.3269 | 333 | 243 | 300 | 350 | 445 | 371 | 431 | 262 | 299 | 309 | 307 | 289 | 381 | 354 | 385 | 370 | 336,54 | 301 | 372 | 1,241 | 0,311 | 3,978 | 6,95E-05 | 0,00021893 | 1,00E-08 | 0,0062931 | 0,0028743 | 0,0028743 | TRUE | 0,0134627 | No |
| orf19.327 | 500 | 330 | 377 | 476 | 544 | 496 | 526 | 342 | 449 | 420 | 386 | 393 | 465 | 473 | 469 | 482 | 442,14 | 412 | 472 | 1,147 | 0,198 | 2,688 | 0,00719613 | 0,01497033 | 1,00E-08 | 0,0052629 | 0,0029252 | 0,0029252 | TRUE | 0,0457511 | No |
| orf19.3271 | 3 | 2 | 4 | 2 | 2 | 5 | 2 | 1 | 3 | 3 | 4 | 2 | 2 | 5 | 2 | 1 | 2,58 | 3 | 2 | 0,904 | -0,146 | -0,172 | 0,86372092 | NA | 1,00E-08 | 0,5642313 | 0,305082 | 0,305082 | TRUE | 0,4828628 | No |
| orf19.3272 | 976 | 745 | 861 | 1084 | 1023 | 891 | 950 | 714 | 876 | 948 | 881 | 894 | 875 | 850 | 848 | 1007 | 897,33 | 900 | 895 | 0,992 | -0,012 | -0,177 | 0,85974758 | 0,89381614 | 0,002521 | 0,0035973 | 0,0030321 | 0,0030321 | TRUE | 0,0762611 | No |
| orf19.3273 | 254 | 154 | 184 | 283 | 317 | 272 | 260 | 190 | 228 | 196 | 188 | 233 | 271 | 260 | 232 | 268 | 234,52 | 211 | 258 | 1,211 | 0,276 | 2,631 | 0,00850167 | 0,01741493 | 0,00396 | 0,0081696 | 0,0062529 | 0,0062529 | TRUE | 0,0630857 | No |
| orf19.3275 | 2112 | 1518 | 1715 | 2263 | 2624 | 2125 | 2246 | 1380 | 1895 | 1933 | 1755 | 1866 | 2244 | 2028 | 2004 | 1946 | 1958,8 | 1862 | 2056 | 1,106 | 0,145 | 2,596 | 0,00943886 | 0,01913249 | 0,002171 | 0,0027206 | 0,0024786 | 0,0024786 | TRUE | 0,0469961 | No |
| orf19.3276 | 9939 | 6998 | 9013 | 10915 | 8846 | 7907 | 8591 | 5343 | 8916 | 8909 | 9221 | 9001 | 7565 | 7545 | 7666 | 7536 | 8294,9 | 9012 | 7578 | 0,841 | -0,25 | -9,205 | 3,43E-20 | 6,26E-19 | 1,00E-08 | 0,0021545 | 0,0005854 | 0,0005854 | TRUE | 0,0029597 | No |
| orf19.3277 | 34 | 27 | 26 | 49 | 26 | 17 | 24 | 20 | 31 | 34 | 27 | 40 | 22 | 16 | 21 | 28 | 27,5 | 33 | 22 | 0,655 | -0,61 | -2,216 | 0,02666836 | 0,04840588 | 1,00E-08 | 0,0547775 | 0,0345655 | 0,0345655 | TRUE | 0,1927339 | No |
| orf19.3278 | 28598 | 19325 | 25657 | 31519 | 26585 | 22972 | 24811 | 16191 | 25655 | 24603 | 26250 | 25991 | 22736 | 21919 | 22140 | 22836 | 24016 | 25625 | 22408 | 0,874 | -0,194 | -6,049 | 1,46E-09 | 9,07E-09 | 0,000543 | 0,00204 | 0,0009444 | 0,0009444 | TRUE | 0,0087232 | No |
| orf19.3279 | 2049 | 1368 | 1944 | 2137 | 2485 | 2175 | 2412 | 1536 | 1838 | 1742 | 1989 | 1762 | 2125 | 2075 | 2152 | 2166 | 1981,3 | 1833 | 2130 | 1,161 | 0,216 | 4,283 | 1,85E-05 | 6,45E-05 | 0,001371 | 0,0027122 | 0,0019311 | 0,0019311 | TRUE | 0,0392026 | No |
| orf19.328 | 400 | 323 | 420 | 492 | 533 | 518 | 559 | 313 | 359 | 411 | 430 | 406 | 456 | 494 | 499 | 441 | 436,98 | 402 | 472 | 1,182 | 0,241 | 3,011 | 0,00260614 | 0,00600143 | 0,002348 | 0,0053376 | 0,0038729 | 0,0038729 | TRUE | 0,0612698 | No |
| orf19.3280 | 2 | 2 | 1 | 1 | 2 | 1 | 2 | 1 | 2 | 3 | 1 | 1 | 2 | 1 | 2 | 1 | 1,51 | 2 | 2 | 0,986 | -0,021 | -0,019 | 0,98471911 | NA | 1,00E-08 | 0,965937 | 0,4419158 | 0,4419158 | TRUE | 0,0950392 | No |
| orf19.3281 | 518 | 333 | 452 | 549 | 829 | 745 | 752 | 543 | 465 | 424 | 462 | 453 | 709 | 711 | 671 | 766 | 582,57 | 451 | 714 | 1,576 | 0,656 | 9,782 | 1,35E-22 | 2,95E-21 | 1,00E-08 | 0,0044714 | 0,002513 | 0,002513 | TRUE | 0,024438 | No |
| orf19.3282 | 2347 | 1712 | 2113 | 2552 | 1894 | 1646 | 1888 | 1054 | 2106 | 2180 | 2162 | 2104 | 1620 | 1571 | 1685 | 1487 | 1864,1 | 2138 | 1591 | 0,746 | -0,423 | -8,948 | 3,63E-19 | 6,20E-18 | 0,000812 | 0,0027583 | 0,0015915 | 0,0015915 | TRUE | 0,0208448 | No |
| orf19.3283 | 112 | 80 | 94 | 118 | 95 | 93 | 104 | 57 | 100 | 102 | 96 | 97 | 81 | 89 | 93 | 80 | 92,37 | 99 | 86 | 0,871 | -0,199 | -1,415 | 0,15717697 | 0,22427679 | 1,00E-08 | 0,0176952 | 0,0082449 | 0,0082449 | TRUE | 0,0300385 | No |
| orf19.3284 | 0 | 1 | 2 | 1 | 0 | 1 | 0 | 1 | 0 | 1 | 2 | 1 | 0 | 1 | 0 | 1 | 0,81 | 1 | 0 | 0,51 | -0,972 | -0,596 | 0,55088006 | NA | 1,00E-08 | 1,7863565 | 1,1580578 | 1,1580578 | TRUE | 0,2124588 | No |
| orf19.3285 | 104 | 75 | 88 | 135 | 151 | 124 | 110 | 77 | 93 | 95 | 90 | 111 | 129 | 118 | 98 | 109 | 105,54 | 97 | 114 | 1,163 | 0,218 | 1,505 | 0,1322375 | 0,19410771 | 1,00E-08 | 0,015734 | 0,0106675 | 0,0106675 | TRUE | 0,1000874 | No |
| orf19.3286 | 618 | 423 | 498 | 694 | 845 | 771 | 787 | 490 | 554 | 539 | 510 | 572 | 723 | 736 | 702 | 691 | 628,3 | 544 | 713 | 1,311 | 0,39 | 6,227 | 4,77E-10 | 3,19E-09 | 1,00E-08 | 0,00429 | 0,0021553 | 0,0021553 | TRUE | 0,0215989 | No |
| orf19.3287 | 8491 | 5912 | 7243 | 9274 | 8810 | 7443 | 8325 | 4770 | 7617 | 7527 | 7410 | 7647 | 7535 | 7102 | 7429 | 6728 | 7374,3 | 7550 | 7198 | 0,954 | -0,068 | -1,631 | 0,10284768 | 0,15627437 | 0,001196 | 0,0021764 | 0,0015314 | 0,0015314 | TRUE | 0,0229018 | No |
| orf19.3288 | 10733 | 7164 | 9432 | 11472 | 9764 | 8266 | 9233 | 5676 | 9629 | 9121 | 9650 | 9460 | 8350 | 7887 | 8239 | 8005 | 8792,6 | 9465 | 8120 | 0,858 | -0,221 | -6,435 | 1,24E-10 | 8,90E-10 | 0,000522 | 0,0021446 | 0,0010173 | 0,0010173 | TRUE | 0,00712 | No |
| orf19.3289 | 523 | 337 | 421 | 568 | 620 | 531 | 566 | 322 | 469 | 429 | 431 | 468 | 530 | 507 | 505 | 454 | 474,18 | 449 | 499 | 1,112 | 0,154 | 2,09 | 0,03657548 | 0,06393526 | 0,000878 | 0,005041 | 0,0030904 | 0,0030904 | TRUE | 0,0361659 | No |
| orf19.329 | 2916 | 1783 | 2580 | 3016 | 3430 | 2897 | 3175 | 2006 | 2616 | 2270 | 2640 | 2487 | 2933 | 2764 | 2833 | 2829 | 2671,6 | 2503 | 2840 | 1,134 | 0,181 | 3,436 | 0,00059134 | 0,00156716 | 0,002089 | 0,0025229 | 0,0022868 | 0,0022868 | TRUE | 0,0454524 | No |
| orf19.3290 | 2513 | 1578 | 2120 | 2723 | 2577 | 2078 | 2241 | 1441 | 2254 | 2009 | 2169 | 2245 | 2204 | 1983 | 2000 | 2032 | 2112,1 | 2169 | 2055 | 0,946 | -0,08 | -1,469 | 0,14183787 | 0,20583621 | 0,001963 | 0,0026669 | 0,0023371 | 0,0023371 | TRUE | 0,0300307 | No |
| orf19.3291 | 11772 | 7924 | 9937 | 12394 | 10237 | 8696 | 9281 | 5296 | 10561 | 10088 | 10167 | 10220 | 8755 | 8297 | 8282 | 7470 | 9229,9 | 10259 | 8201 | 0,8 | -0,322 | -6,601 | 4,08E-11 | 3,14E-10 | 0,002186 | 0,0021368 | 0,0021783 | 0,0021783 | TRUE | 0,0430996 | No |
| orf19.3292 | 2109 | 1419 | 1770 | 2258 | 2618 | 2151 | 2270 | 1634 | 1892 | 1807 | 1811 | 1862 | 2239 | 2052 | 2026 | 2305 | 1999,1 | 1843 | 2156 | 1,167 | 0,223 | 4,249 | 2,15E-05 | 7,42E-05 | 0,001615 | 0,0027057 | 0,0021485 | 0,0021485 | TRUE | 0,0244383 | No |
| orf19.3293 | 928 | 668 | 853 | 1060 | 875 | 783 | 847 | 485 | 833 | 850 | 873 | 874 | 748 | 747 | 756 | 684 | 795,63 | 858 | 734 | 0,859 | -0,22 | -3,909 | 9,28E-05 | 0,00028589 | 1,00E-08 | 0,0038041 | 0,0017819 | 0,0017819 | TRUE | 0,020665 | No |
| orf19.3294 | 14185 | 8942 | 11598 | 14614 | 15847 | 12436 | 13316 | 8048 | 12725 | 11384 | 11866 | 12051 | 13553 | 11866 | 11882 | 11351 | 12085 | 12006 | 12163 | 1,013 | 0,019 | 0,32 | 0,74885099 | 0,80276166 | 0,003567 | 0,0020996 | 0,0032456 | 0,0032456 | TRUE | 0,0725708 | No |
| orf19.3295 | 965 | 692 | 915 | 1049 | 1248 | 1127 | 1172 | 731 | 866 | 881 | 936 | 865 | 1067 | 1075 | 1046 | 1031 | 970,92 | 887 | 1055 | 1,191 | 0,253 | 4,876 | 1,08E-06 | 4,55E-06 | 1,00E-08 | 0,0034747 | 0,0015388 | 0,0015388 | TRUE | 0,0166991 | No |
| orf19.3296 | 150 | 88 | 124 | 169 | 197 | 164 | 171 | 100 | 135 | 112 | 127 | 139 | 168 | 156 | 153 | 141 | 141,43 | 128 | 154 | 1,204 | 0,268 | 2,214 | 0,02684904 | 0,04870442 | 1,00E-08 | 0,0122442 | 0,0070228 | 0,0070228 | TRUE | 0,0621653 | No |
| orf19.3297 | 2295 | 1543 | 1817 | 2327 | 2134 | 1984 | 2038 | 1426 | 2059 | 1964 | 1859 | 1919 | 1825 | 1893 | 1819 | 2011 | 1918,6 | 1950 | 1887 | 0,966 | -0,05 | -0,989 | 0,32271434 | 0,40956226 | 0,001362 | 0,0027362 | 0,001964 | 0,001964 | TRUE | 0,0219798 | No |
| orf19.3298 | 2312 | 1769 | 2105 | 2648 | 2271 | 2093 | 2338 | 1349 | 2074 | 2252 | 2154 | 2184 | 1942 | 1997 | 2086 | 1903 | 2074 | 2166 | 1982 | 0,917 | -0,125 | -2,708 | 0,00676485 | 0,01419601 | 0,000741 | 0,0026795 | 0,0015694 | 0,0015694 | TRUE | 0,0152913 | No |
| orf19.33 | 2952 | 2186 | 2446 | 3038 | 4209 | 3677 | 3783 | 2480 | 2648 | 2783 | 2502 | 2505 | 3600 | 3508 | 3376 | 3498 | 3052,6 | 2610 | 3496 | 1,341 | 0,423 | 9,211 | 3,22E-20 | 5,93E-19 | 0,00122 | 0,0024551 | 0,0016892 | 0,0016892 | TRUE | 0,0230424 | No |
| orf19.3300 | 18204 | 12766 | 16073 | 19296 | 17731 | 15674 | 16858 | 10315 | 16331 | 16252 | 16444 | 15912 | 15164 | 14956 | 15043 | 14548 | 15581 | 16235 | 14928 | 0,92 | -0,121 | -4,455 | 8,40E-06 | 3,10E-05 | 0,000174 | 0,0020727 | 0,0006383 | 0,0006383 | TRUE | 0,0034556 | No |
| orf19.3301 | 29912 | 20732 | 26853 | 33792 | 29617 | 26467 | 29114 | 17610 | 26834 | 26394 | 27473 | 27865 | 25329 | 25254 | 25979 | 24837 | 26246 | 27142 | 25350 | 0,934 | -0,099 | -3,29 | 0,00100288 | 0,00252211 | 0,000414 | 0,0020348 | 0,000823 | 0,000823 | TRUE | 0,0041645 | No |
| orf19.330.1 | 419 | 241 | 338 | 388 | 444 | 340 | 381 | 269 | 376 | 307 | 346 | 320 | 380 | 324 | 340 | 379 | 346,5 | 337 | 356 | 1,051 | 0,072 | 0,783 | 0,43364272 | 0,52064857 | 0,004271 | 0,0061692 | 0,0052538 | 0,0052538 | TRUE | 0,0700957 | No |

| | | | | | | | | | | | | | | | | | | | | | | | | | | | | | | | |
|---|---|---|---|---|---|---|---|---|---|---|---|---|---|---|---|---|---|---|---|---|---|---|---|---|---|---|---|---|---|---|---|
| orf19.3302 | 16153 | 11687 | 14385 | 17775 | 23393 | 22188 | 23519 | 14726 | 14491 | 14879 | 14717 | 14657 | 20006 | 21171 | 20987 | 20770 | 17710 | 14686 | 20734 | 1,412 | 0,498 | 17,146 | 6,69E-66 | 1,34E-63 | 0,000287 | 0,0020615 | 0,0007504 | 0,0007504 | TRUE | 0,0069229 | No |
| orf19.3303 | 3063 | 2170 | 2754 | 3309 | 3710 | 3263 | 3506 | 2095 | 2748 | 2763 | 2818 | 2729 | 3173 | 3113 | 3129 | 2955 | 2928,3 | 2764 | 3092 | 1,121 | 0,164 | 4,393 | 1,12E-05 | 4,05E-05 | 0,000112 | 0,0024753 | 0,0009975 | 0,0009975 | TRUE | 0,009924 | No |
| orf19.3304 | 5341 | 3691 | 4627 | 5794 | 6525 | 5849 | 6098 | 3329 | 4791 | 4699 | 4734 | 4778 | 5580 | 5581 | 5441 | 4695 | 5037,5 | 4750 | 5324 | 1,122 | 0,166 | 2,978 | 0,00289685 | 0,00660254 | 0,002954 | 0,0022677 | 0,0027751 | 0,0027751 | TRUE | 0,0751027 | No |
| orf19.3305 | 2455 | 1772 | 2314 | 2658 | 2598 | 2352 | 2493 | 1549 | 2202 | 2256 | 2367 | 2192 | 2222 | 2244 | 2225 | 2185 | 2236,6 | 2254 | 2219 | 0,985 | -0,021 | -0,523 | 0,60092805 | 0,67698776 | 0,000275 | 0,0026286 | 0,0011527 | 0,0011527 | TRUE | 0,0140006 | No |
| orf19.3306 | 21 | 13 | 21 | 29 | 20 | 20 | 21 | 14 | 19 | 17 | 21 | 24 | 17 | 19 | 19 | 20 | 19,43 | 20 | 19 | 0,911 | -0,134 | -0,443 | 0,65793507 | 0,72649768 | 1,00E-08 | 0,0766838 | 0,0372022 | 0,0372022 | TRUE | 0,0973258 | No |
| orf19.3307 | 202 | 156 | 199 | 238 | 261 | 242 | 276 | 177 | 181 | 199 | 204 | 196 | 223 | 231 | 246 | 250 | 216,22 | 195 | 238 | 1,217 | 0,283 | 2,921 | 0,00348989 | 0,00778872 | 1,00E-08 | 0,0086937 | 0,0043638 | 0,0043638 | TRUE | 0,0257436 | No |
| orf19.3308 | 503 | 366 | 451 | 519 | 506 | 470 | 528 | 321 | 451 | 466 | 461 | 428 | 433 | 448 | 471 | 453 | 451,46 | 452 | 451 | 1,001 | 0,001 | 0,021 | 0,98355194 | 0,98782682 | 1,00E-08 | 0,0051951 | 0,0024711 | 0,0024711 | TRUE | 0,0155167 | No |
| orf19.3309 | 3435 | 2377 | 3033 | 3720 | 3325 | 2736 | 2885 | 1842 | 3082 | 3026 | 3103 | 3068 | 2844 | 2611 | 2574 | 2598 | 2863,1 | 3070 | 2657 | 0,866 | -0,208 | -4,833 | 1,35E-06 | 5,56E-06 | 0,000705 | 0,0024866 | 0,0014184 | 0,0014184 | TRUE | 0,0282673 | No |
| orf19.331 | 1219 | 793 | 1003 | 1280 | 1157 | 922 | 1006 | 710 | 1094 | 1010 | 1026 | 1056 | 990 | 880 | 898 | 1001 | 994,14 | 1046 | 942 | 0,898 | -0,155 | -2,575 | 0,01002686 | 0,02018137 | 0,001683 | 0,0034398 | 0,0024846 | 0,0024846 | TRUE | 0,0232344 | No |
| orf19.3310 | 690 | 450 | 583 | 728 | 942 | 785 | 920 | 599 | 619 | 573 | 596 | 600 | 806 | 749 | 821 | 845 | 701,14 | 597 | 805 | 1,344 | 0,426 | 6,924 | 4,39E-12 | 3,73E-11 | 1,00E-08 | 0,00405 | 0,0021833 | 0,0021833 | TRUE | 0,0259047 | No |
| orf19.3311 | 167 | 133 | 133 | 167 | 189 | 161 | 164 | 92 | 150 | 169 | 136 | 138 | 162 | 154 | 146 | 130 | 148,04 | 148 | 148 | 1,009 | 0,014 | 0,111 | 0,91152864 | 0,93455019 | 0,001219 | 0,011786 | 0,0075675 | 0,0075675 | TRUE | 0,0821407 | No |
| orf19.3312 | 470 | 323 | 448 | 503 | 545 | 517 | 524 | 403 | 422 | 411 | 458 | 415 | 466 | 493 | 468 | 568 | 462,67 | 426 | 499 | 1,163 | 0,218 | 2,682 | 0,00731743 | 0,01519637 | 0,003024 | 0,0051172 | 0,0041556 | 0,0041556 | TRUE | 0,0906002 | No |
| orf19.3314 | 206 | 163 | 195 | 279 | 244 | 217 | 229 | 173 | 185 | 208 | 200 | 230 | 209 | 207 | 204 | 244 | 210,75 | 206 | 216 | 1,044 | 0,062 | 0,574 | 0,56567973 | 0,64538621 | 0,003085 | 0,0088679 | 0,0062763 | 0,0062763 | TRUE | 0,0806876 | No |
| orf19.3315 | 1732 | 1214 | 1533 | 1935 | 2353 | 2155 | 2460 | 1559 | 1554 | 1546 | 1568 | 1596 | 2012 | 2056 | 2195 | 2199 | 1840,7 | 1566 | 2116 | 1,349 | 0,432 | 9,333 | 1,03E-20 | 1,96E-19 | 0,000719 | 0,0027682 | 0,0014988 | 0,0014988 | TRUE | 0,0133826 | No |
| orf19.3317 | 304 | 277 | 249 | 339 | 289 | 318 | 286 | 217 | 273 | 353 | 255 | 280 | 247 | 303 | 255 | 306 | 283,94 | 290 | 278 | 0,959 | -0,06 | -0,514 | 0,60739649 | 0,68248329 | 0,011953 | 0,0070922 | 0,0096921 | 0,0096921 | TRUE | 0,2190945 | No |
| orf19.3318 | 603 | 411 | 584 | 724 | 313 | 316 | 327 | 187 | 541 | 523 | 597 | 597 | 268 | 302 | 292 | 264 | 422,93 | 564 | 282 | 0,498 | -1,005 | -12,277 | 1,21E-34 | 5,79E-33 | 0,002014 | 0,005412 | 0,0037719 | 0,0037719 | TRUE | 0,025843 | No |
| orf19.3319 | 1776 | 1236 | 1466 | 1904 | 1453 | 1316 | 1316 | 967 | 1593 | 1574 | 1500 | 1570 | 1243 | 1256 | 1174 | 1364 | 1409,2 | 1559 | 1259 | 0,805 | -0,313 | -5,838 | 5,29E-09 | 3,06E-08 | 0,001325 | 0,0030097 | 0,002032 | 0,002032 | TRUE | 0,033908 | No |
| orf19.332 | 4179 | 2859 | 3562 | 4163 | 4369 | 3789 | 4280 | 2607 | 3749 | 3640 | 3644 | 3433 | 3737 | 3615 | 3819 | 3677 | 3664,2 | 3616 | 3712 | 1,027 | 0,038 | 0,954 | 0,34016829 | 0,42720132 | 0,000743 | 0,0023757 | 0,001288 | 0,001288 | TRUE | 0,0148347 | No |
| orf19.3320 | 32 | 33 | 32 | 40 | 25 | 11 | 20 | 20 | 29 | 42 | 33 | 33 | 21 | 10 | 18 | 28 | 26,8 | 34 | 19 | 0,563 | -0,829 | -2,771 | 0,00558237 | 0,01193553 | 0,022583 | 0,0561538 | 0,0450267 | 0,0450267 | TRUE | 0,4168056 | No |
| orf19.3321 | 6507 | 4286 | 5620 | 7063 | 5751 | 4933 | 5504 | 3324 | 5837 | 5457 | 5750 | 5824 | 4918 | 4707 | 4911 | 4688 | 5261,6 | 5717 | 4806 | 0,841 | -0,25 | -6,713 | 1,90E-11 | 1,51E-10 | 0,000571 | 0,0022554 | 0,0011431 | 0,0011431 | TRUE | 0,0110795 | No |
| orf19.332.1 | 1487 | 1021 | 1148 | 1566 | 1695 | 1366 | 1391 | 963 | 1334 | 1300 | 1175 | 1291 | 1450 | 1303 | 1241 | 1358 | 1306,5 | 1275 | 1338 | 1,049 | 0,069 | 1,098 | 0,27233881 | 0,35670625 | 0,003039 | 0,0030906 | 0,0030607 | 0,0030607 | TRUE | 0,040224 | No |
| orf19.3322 | 506 | 350 | 442 | 524 | 579 | 389 | 469 | 355 | 454 | 446 | 452 | 432 | 495 | 371 | 419 | 501 | 446,17 | 446 | 446 | 0,998 | -0,002 | -0,023 | 0,98136891 | 0,98612886 | 0,008017 | 0,0052332 | 0,0066295 | 0,0066295 | TRUE | 0,1495331 | No |
| orf19.3323 | 917 | 605 | 773 | 988 | 704 | 725 | 703 | 482 | 823 | 770 | 791 | 815 | 602 | 692 | 627 | 680 | 724,93 | 800 | 650 | 0,81 | -0,304 | -4,768 | 1,86E-06 | 7,54E-06 | 0,001111 | 0,0039821 | 0,0025094 | 0,0025094 | TRUE | 0,0305136 | No |
| orf19.3324 | 21375 | 14289 | 18831 | 22610 | 21151 | 18138 | 19788 | 11826 | 19176 | 18191 | 19266 | 18644 | 18089 | 17307 | 17657 | 16680 | 18126 | 18819 | 17433 | 0,927 | -0,11 | -3,025 | 0,00248408 | 0,00574456 | 0,000864 | 0,0020596 | 0,0012172 | 0,0012172 | TRUE | 0,0101924 | No |
| orf19.3325 | 1816 | 1194 | 1739 | 2005 | 1907 | 1657 | 1718 | 1132 | 1629 | 1520 | 1779 | 1653 | 1631 | 1581 | 1533 | 1597 | 1615,4 | 1645 | 1586 | 0,962 | -0,055 | -1,019 | 0,30807488 | 0,39397912 | 0,001591 | 0,0028782 | 0,0021915 | 0,0021915 | TRUE | 0,0349116 | No |
| orf19.3325.3 | 8111 | 5006 | 6347 | 8149 | 8301 | 6301 | 6284 | 3776 | 7276 | 6373 | 6494 | 6720 | 7099 | 6012 | 5607 | 5326 | 6363,5 | 6716 | 6011 | 0,895 | -0,159 | -1,954 | 0,05065196 | 0,08514744 | 0,009948 | 0,0022077 | 0,0062387 | 0,0062387 | TRUE | 0,1817385 | No |
| orf19.3327 | 6049 | 4122 | 5220 | 6335 | 5071 | 4450 | 4798 | 3124 | 5427 | 5248 | 5341 | 5224 | 4337 | 4246 | 4281 | 4406 | 4813,7 | 5310 | 4318 | 0,813 | -0,299 | -9,503 | 2,05E-21 | 4,07E-20 | 8,21E-05 | 0,0022811 | 0,0007415 | 0,0007415 | TRUE | 0,0027462 | No |
| orf19.3328 | 2631 | 1683 | 2437 | 2751 | 2513 | 2197 | 2410 | 1584 | 2360 | 2143 | 2493 | 2269 | 2149 | 2096 | 2151 | 2234 | 2236,9 | 2316 | 2158 | 0,93 | -0,104 | -2,001 | 0,04534248 | 0,07721588 | 0,001793 | 0,0026285 | 0,0021596 | 0,0021596 | TRUE | 0,0313239 | No |
| orf19.3329 | 960 | 641 | 804 | 1094 | 861 | 640 | 724 | 491 | 861 | 816 | 823 | 902 | 736 | 611 | 646 | 693 | 760,95 | 850 | 672 | 0,788 | -0,344 | -4,854 | 1,21E-06 | 5,04E-06 | 0,003126 | 0,0038873 | 0,0034806 | 0,0034806 | TRUE | 0,0552021 | No |
| orf19.333 | 717 | 493 | 638 | 672 | 560 | 489 | 514 | 302 | 643 | 628 | 653 | 554 | 479 | 467 | 459 | 426 | 538,48 | 620 | 458 | 0,742 | -0,43 | -5,732 | 9,92E-09 | 5,48E-08 | 0,002444 | 0,0046754 | 0,003508 | 0,003508 | TRUE | 0,063876 | No |
| orf19.3330 | 2599 | 1709 | 2285 | 2880 | 2407 | 1909 | 2264 | 1529 | 2332 | 2176 | 2338 | 2375 | 2059 | 1822 | 2020 | 2157 | 2159,6 | 2305 | 2014 | 0,872 | -0,197 | -3,46 | 0,00054028 | 0,00144718 | 0,002553 | 0,0026517 | 0,0026518 | 0,0026518 | TRUE | 0,0500698 | No |
| orf19.3330.3 | 149 | 112 | 116 | 145 | 155 | 160 | 131 | 108 | 134 | 143 | 119 | 120 | 133 | 153 | 117 | 152 | 133,62 | 129 | 139 | 1,075 | 0,105 | 0,792 | 0,42830803 | 0,51558298 | 0,00457 | 0,012844 | 0,0092925 | 0,0092925 | TRUE | 0,120022 | No |
| orf19.3331 | 21339 | 14288 | 19236 | 22612 | 23800 | 21332 | 23257 | 15506 | 19143 | 18190 | 19680 | 18646 | 20355 | 20354 | 20753 | 21870 | 19874 | 18915 | 20833 | 1,101 | 0,139 | 3,6 | 0,00031793 | 0,00089592 | 0,0011 | 0,0020526 | 0,0013796 | 0,0013796 | TRUE | 0,0135696 | No |
| orf19.3332 | 1691 | 1220 | 1618 | 1919 | 2421 | 2041 | 2272 | 1462 | 1517 | 1553 | 1655 | 1582 | 2071 | 1947 | 2027 | 2062 | 1801,9 | 1577 | 2027 | 1,285 | 0,362 | 7,958 | 1,75E-15 | 2,11E-14 | 0,000503 | 0,0027852 | 0,0014182 | 0,0014182 | TRUE | 0,0133471 | No |
| orf19.3333 | 8489 | 5618 | 7335 | 8954 | 7684 | 6479 | 7065 | 5050 | 7616 | 7152 | 7504 | 7384 | 6572 | 6182 | 6304 | 7123 | 6979,6 | 7414 | 6545 | 0,882 | -0,181 | -3,698 | 0,00021708 | 0,00062884 | 0,002205 | 0,0021875 | 0,0021569 | 0,0021569 | TRUE | 0,0419519 | No |
| orf19.3334 | 19962 | 12268 | 16094 | 20264 | 20239 | 15571 | 16619 | 10620 | 17908 | 15618 | 16466 | 16710 | 17309 | 14857 | 14830 | 14979 | 16085 | 16676 | 15494 | 0,929 | -0,106 | -1,738 | 0,08214924 | 0,12940551 | 0,004584 | 0,0020698 | 0,0035132 | 0,0035132 | TRUE | 0,0764012 | No |
| orf19.3335 | 12679 | 8605 | 10499 | 13531 | 13244 | 11188 | 11997 | 8482 | 11374 | 10955 | 10742 | 11158 | 11327 | 10675 | 10705 | 11963 | 11112 | 11057 | 11168 | 1,009 | 0,014 | 0,307 | 0,75911346 | 0,81166643 | 0,001481 | 0,0021102 | 0,0018082 | 0,0018082 | TRUE | 0,0275983 | No |
| orf19.3336 | 8 | 6 | 9 | 13 | 6 | 7 | 6 | 2 | 7 | 8 | 9 | 11 | 5 | 7 | 5 | 3 | 6,84 | 9 | 5 | 0,587 | -0,769 | -1,475 | 0,14020177 | NA | 1,00E-08 | 0,2141827 | 0,1069194 | 0,1069194 | TRUE | 0,0990051 | No |
| orf19.3337 | 15530 | 10732 | 13597 | 17329 | 28155 | 24424 | 27176 | 18157 | 13932 | 13663 | 13911 | 14290 | 24079 | 23305 | 24250 | 25609 | 19130 | 13949 | 24311 | 1,742 | 0,801 | 21,922 | 1,61E-106 | 1,93E-103 | 0,000873 | 0,0020554 | 0,0012256 | 0,0012256 | TRUE | 0,0156224 | No |
| orf19.334 | 4912 | 3952 | 4636 | 5980 | 6384 | 6225 | 6353 | 4352 | 4407 | 5031 | 4743 | 4931 | 5460 | 5940 | 5669 | 6138 | 5289,8 | 4778 | 5802 | 1,214 | 0,28 | 5,19 | 2,10E-07 | 9,87E-07 | 0,002722 | 0,002254 | 0,0025984 | 0,0025984 | TRUE | 0,0336189 | No |
| orf19.3340 | 2961 | 1914 | 2426 | 3043 | 2726 | 2290 | 2416 | 1834 | 2656 | 2437 | 2482 | 2509 | 2331 | 2185 | 2156 | 2587 | 2417,9 | 2521 | 2315 | 0,916 | -0,126 | -2,057 | 0,03970915 | 0,06865398 | 0,003775 | 0,0025799 | 0,0031939 | 0,0031939 | TRUE | 0,07233 | No |
| orf19.3341 | 9951 | 6929 | 8743 | 10771 | 9456 | 8392 | 8889 | 5851 | 8927 | 8821 | 8945 | 8882 | 8087 | 8007 | 7932 | 8252 | 8481,8 | 8894 | 8070 | 0,907 | -0,141 | -5,254 | 1,49E-07 | 7,12E-07 | 6,25E-06 | 0,0021507 | 0,0005736 | 0,0005736 | TRUE | 0,0026483 | No |
| orf19.3342 | 2687 | 1895 | 2393 | 3059 | 2859 | 2643 | 2751 | 1700 | 2411 | 2413 | 2448 | 2522 | 2445 | 2522 | 2455 | 2398 | 2451,7 | 2448 | 2455 | 1,003 | 0,004 | 0,104 | 0,91717282 | 0,93905492 | 1,00E-08 | 0,0025717 | 0,0009705 | 0,0009705 | TRUE | 0,0052914 | No |
| orf19.3344 | 2222 | 1478 | 1925 | 2283 | 2486 | 2210 | 2375 | 1503 | 1993 | 1882 | 1969 | 1883 | 2126 | 2109 | 2119 | 2120 | 2025,1 | 1932 | 2118 | 1,096 | 0,133 | 3,33 | 0,00086741 | 0,00220916 | 1,00E-08 | 0,0026964 | 0,0010258 | 0,0010258 | TRUE | 0,0057578 | No |
| orf19.3345 | 2373 | 1600 | 2111 | 2517 | 3220 | 2805 | 3003 | 1943 | 2129 | 2037 | 2160 | 2076 | 2754 | 2676 | 2680 | 2740 | 2406,4 | 2100 | 2712 | 1,291 | 0,368 | 9,702 | 2,97E-22 | 6,32E-21 | 1,00E-08 | 0,0025828 | 0,0009587 | 0,0009587 | TRUE | 0,004448 | No |
| orf19.3347 | 1058 | 755 | 900 | 1166 | 1078 | 983 | 1075 | 640 | 949 | 961 | 921 | 961 | 922 | 938 | 959 | 903 | 939,3 | 948 | 930 | 0,983 | -0,025 | -0,497 | 0,61923244 | 0,6924155 | 1,00E-08 | 0,003525 | 0,0014036 | 0,0014036 | TRUE | 0,0051579 | No |
| orf19.3348 | 1706 | 1195 | 1420 | 1750 | 2046 | 1669 | 1664 | 1056 | 1530 | 1521 | 1453 | 1443 | 1750 | 1593 | 1485 | 1489 | 1533 | 1487 | 1579 | 1,064 | 0,089 | 1,463 | 0,14334164 | 0,20751725 | 0,002686 | 0,0029265 | 0,0029245 | 0,0029245 | TRUE | 0,0653538 | No |
| orf19.3349 | 5609 | 3828 | 5025 | 5889 | 6569 | 5799 | 6459 | 3756 | 5032 | 4873 | 5141 | 4856 | 5618 | 5533 | 5764 | 5298 | 5264,4 | 4976 | 5553 | 1,117 | 0,159 | 4,106 | 4,03E-05 | 0,00013227 | 0,000735 | 0,0022553 | 0,0012559 | 0,0012559 | TRUE | 0,0111518 | No |
| orf19.335 | 627 | 650 | 625 | 894 | 1305 | 1536 | 1503 | 999 | 562 | 828 | 639 | 737 | 1116 | 1466 | 1341 | 1409 | 1012,3 | 692 | 1333 | 1,929 | 0,948 | 8,304 | 1,01E-16 | 1,38E-15 | 0,019064 | 0,0034136 | 0,0114135 | 0,0114135 | TRUE | 0,1988382 | No |
| orf19.3350 | 3149 | 2382 | 2763 | 3392 | 3404 | 3188 | 3288 | 2186 | 2825 | 3033 | 2827 | 2797 | 2911 | 3042 | 2934 | 3083 | 2931,5 | 2870 | 2992 | 1,043 | 0,06 | 1,447 | 0,1479325 | 0,21313632 | 0,000651 | 0,0024747 | 0,0013334 | 0,0013334 | TRUE | 0,0166616 | No |
| orf19.3351 | 1676 | 1336 | 1525 | 1937 | 1945 | 2047 | 2105 | 1246 | 1504 | 1701 | 1560 | 1597 | 1663 | 1953 | 1878 | 1757 | 1701,8 | 1590 | 1813 | 1,141 | 0,191 | 3,069 | 0,00214483 | 0,00503562 | 0,003081 | 0,0028326 | 0,003115 | 0,003115 | TRUE | 0,0392589 | No |
| orf19.3352 | 727 | 525 | 648 | 759 | 749 | 723 | 754 | 409 | 652 | 668 | 663 | 626 | 641 | 690 | 673 | 577 | 648,69 | 652 | 645 | 0,994 | -0,009 | -0,13 | 0,89636181 | 0,92167409 | 0,001348 | 0,0042174 | 0,002838 | 0,002838 | TRUE | 0,0508953 | No |
| orf19.3353 | 3855 | 2784 | 3365 | 4145 | 3046 | 2741 | 2832 | 1647 | 3458 | 3544 | 3443 | 3418 | 2605 | 2615 | 2527 | 2323 | 2991,7 | 3466 | 2518 | 0,728 | -0,458 | -10,248 | 1,20E-24 | 2,99E-23 | 0,001119 | 0,0024648 | 0,0015729 | 0,0015729 | TRUE | 0,0302384 | No |
| orf19.3354 | 9860 | 6129 | 7855 | 9477 | 9353 | 7345 | 7884 | 4287 | 8845 | 7803 | 8036 | 7815 | 7999 | 7008 | 7035 | 6046 | 7573,6 | 8125 | 7022 | 0,865 | -0,21 | -2,745 | 0,00604217 | 0,0128228 | 0,008102 | 0,0021712 | 0,0054777 | 0,0054777 | TRUE | 0,1070801 | No |
| orf19.3355 | 2306 | 1696 | 2152 | 2535 | 2775 | 2693 | 2808 | 1810 | 2069 | 2159 | 2202 | 2090 | 2373 | 2570 | 2506 | 2553 | 2315,2 | 2130 | 2500 | 1,174 | 0,231 | 5,29 | 1,22E-07 | 5,92E-07 | 0,000691 | 0,0026066 | 0,0013962 | 0,0013962 | TRUE | 0,0146948 | No |
| orf19.3356 | 398 | 343 | 360 | 455 | 468 | 590 | 518 | 293 | 357 | 437 | 368 | 375 | 400 | 563 | 462 | 413 | 421,99 | 384 | 460 | 1,201 | 0,264 | 2,34 | 0,01926618 | 0,03595874 | 0,014036 | 0,0054197 | 0,009801 | 0,009801 | TRUE | 0,2679435 | No |
| orf19.3357 | 974 | 703 | 813 | 1082 | 1014 | 885 | 990 | 556 | 874 | 895 | 832 | 892 | 867 | 844 | 883 | 784 | 859 | 873 | 844 | 0,971 | -0,043 | -0,746 | 0,45594381 | 0,54157975 | 0,000329 | 0,0036695 | 0,0020184 | 0,0020184 | TRUE | 0,0238989 | No |
| orf19.3358 | 2323 | 1454 | 2007 | 2502 | 2489 | 1927 | 2081 | 1468 | 2084 | 1851 | 2053 | 2063 | 2129 | 1839 | 1857 | 2070 | 1993,3 | 2013 | 1974 | 0,979 | -0,031 | -0,484 | 0,62805771 | 0,69980911 | 0,003755 | 0,0027078 | 0,0033152 | 0,0033152 | TRUE | 0,0357257 | No |
| orf19.3359 | 1305 | 1009 | 1215 | 1381 | 1665 | 1692 | 1727 | 938 | 1171 | 1285 | 1243 | 1139 | 1424 | 1614 | 1541 | 1323 | 1342,5 | 1210 | 1476 | 1,224 | 0,291 | 4,207 | 2,58E-05 | 8,80E-05 | 0,0043 | 0,0030609 | 0,0038535 | 0,0038535 | TRUE | 0,05445 | No |

| | | | | | | | | | | | | | | | | | | | | | | | | | | | | | | | |
|---|---|---|---|---|---|---|---|---|---|---|---|---|---|---|---|---|---|---|---|---|---|---|---|---|---|---|---|---|---|---|---|
| orf19.336 | 2181 | 1476 | 1822 | 2208 | 2640 | 2257 | 2385 | 1426 | 1957 | 1879 | 1864 | 1821 | 2258 | 2154 | 2128 | 2011 | 2008,9 | 1880 | 2138 | 1,139 | 0,188 | 3,913 | 9,10E-05 | 0,00028094 | 0,001067 | 0,0027022 | 0,0017077 | 0,0017077 | TRUE | 0,0176571 | No |
| orf19.3360 | 6292 | 4979 | 5794 | 7010 | 9339 | 9626 | 9941 | 6252 | 5645 | 6339 | 5928 | 5781 | 7987 | 9185 | 8871 | 8818 | 7319 | 5923 | 8715 | 1,472 | 0,558 | 10,371 | 3,36E-25 | 8,69E-24 | 0,002821 | 0,0021779 | 0,0026343 | 0,0026343 | TRUE | 0,0390328 | No |
| orf19.3362 | 287 | 166 | 248 | 325 | 253 | 229 | 242 | 161 | 257 | 211 | 254 | 268 | 216 | 219 | 216 | 227 | 233,55 | 248 | 220 | 0,879 | -0,186 | -1,887 | 0,0591142 | 0,09722368 | 1,00E-08 | 0,0081952 | 0,0050136 | 0,0050136 | TRUE | 0,0937147 | No |
| orf19.3363 | 23733 | 16752 | 21088 | 25417 | 30621 | 27690 | 29468 | 17871 | 21291 | 21327 | 21575 | 20959 | 26188 | 26421 | 26295 | 25205 | 23658 | 21288 | 26027 | 1,223 | 0,29 | 10,67 | 1,41E-26 | 4,25E-25 | 0,000242 | 0,0020409 | 0,0006685 | 0,0006685 | TRUE | 0,0054159 | No |
| orf19.3364 | 636 | 494 | 551 | 741 | 834 | 790 | 864 | 488 | 571 | 629 | 564 | 611 | 713 | 754 | 771 | 688 | 662,57 | 594 | 732 | 1,237 | 0,306 | 4,684 | 2,82E-06 | 1,12E-05 | 0,001007 | 0,0041705 | 0,0025859 | 0,0025859 | TRUE | 0,0164554 | No |
| orf19.3365 | 3157 | 2095 | 2622 | 3155 | 3069 | 2566 | 2773 | 1751 | 2832 | 2667 | 2683 | 2602 | 2625 | 2448 | 2474 | 2470 | 2600,1 | 2696 | 2504 | 0,929 | -0,106 | -2,408 | 0,01605851 | 0,0306631 | 0,000866 | 0,0025379 | 0,0014641 | 0,0014641 | TRUE | 0,0145213 | No |
| orf19.3366 | 4443 | 2926 | 3871 | 4757 | 3792 | 3066 | 3231 | 2094 | 3986 | 3725 | 3960 | 3923 | 3243 | 2925 | 2883 | 2953 | 3449,9 | 3898 | 3001 | 0,77 | -0,377 | -7,872 | 3,48E-15 | 4,05E-14 | 0,001557 | 0,0024003 | 0,001909 | 0,001909 | TRUE | 0,0368529 | No |
| orf19.3366.1 | 1741 | 1117 | 1440 | 1677 | 1480 | 1317 | 1366 | 896 | 1562 | 1422 | 1473 | 1383 | 1266 | 1257 | 1219 | 1264 | 1355,6 | 1460 | 1252 | 0,856 | -0,223 | -4,304 | 1,68E-05 | 5,91E-05 | 0,001031 | 0,0030504 | 0,0018448 | 0,0018448 | TRUE | 0,0276429 | No |
| orf19.3367 | 4352 | 3324 | 3748 | 4816 | 4334 | 4123 | 4139 | 2605 | 3904 | 4232 | 3835 | 3971 | 3707 | 3934 | 3693 | 3674 | 3868,8 | 3986 | 3752 | 0,942 | -0,086 | -1,963 | 0,04965595 | 0,0837011 | 0,001131 | 0,0023548 | 0,0015755 | 0,0015755 | TRUE | 0,0202415 | No |
| orf19.3368 | 320 | 272 | 261 | 372 | 319 | 333 | 307 | 241 | 287 | 346 | 267 | 307 | 273 | 318 | 274 | 340 | 301,44 | 302 | 301 | 0,995 | -0,007 | -0,065 | 0,94808707 | 0,96250543 | 0,008208 | 0,0067953 | 0,0074589 | 0,0074589 | TRUE | 0,0997888 | No |
| orf19.3369 | 16390 | 15184 | 15796 | 20567 | 22127 | 23586 | 24885 | 15033 | 14704 | 19331 | 16161 | 16960 | 18924 | 22505 | 22206 | 21203 | 18999 | 16789 | 21210 | 1,263 | 0,337 | 4,198 | 2,69E-05 | 9,14E-05 | 0,009497 | 0,0020559 | 0,0061509 | 0,0061509 | TRUE | 0,1268229 | No |
| orf19.337 | 2740 | 1995 | 2358 | 2950 | 3055 | 2932 | 2842 | 1826 | 2458 | 2540 | 2412 | 2433 | 2613 | 2798 | 2536 | 2575 | 2545,6 | 2461 | 2630 | 1,07 | 0,097 | 2,215 | 0,02676079 | 0,048559 | 0,000703 | 0,0025498 | 0,001463 | 0,001463 | TRUE | 0,0225055 | No |
| orf19.3370 | 12597 | 9398 | 11707 | 14309 | 12294 | 11725 | 12505 | 7878 | 11301 | 11965 | 11977 | 11799 | 10514 | 11188 | 11159 | 11111 | 11377 | 11760 | 10993 | 0,935 | -0,097 | -2,74 | 0,00614187 | 0,01301601 | 0,000672 | 0,0021071 | 0,0011246 | 0,0011246 | TRUE | 0,0106682 | No |
| orf19.3371 | 1732 | 1306 | 1581 | 1831 | 2231 | 2097 | 2287 | 1256 | 1554 | 1663 | 1618 | 1510 | 1908 | 2001 | 2041 | 1771 | 1758,1 | 1586 | 1930 | 1,221 | 0,288 | 5,128 | 2,92E-07 | 1,34E-06 | 0,002062 | 0,0028052 | 0,0024434 | 0,0024434 | TRUE | 0,0343908 | No |
| orf19.3373 | 10718 | 7067 | 9330 | 11526 | 14416 | 12283 | 13125 | 9620 | 9615 | 8997 | 9546 | 9504 | 12329 | 11720 | 11712 | 13568 | 10874 | 9416 | 12332 | 1,309 | 0,389 | 7,429 | 1,09E-13 | 1,11E-12 | 0,002832 | 0,002113 | 0,0025337 | 0,0025337 | TRUE | 0,0550031 | No |
| orf19.3374 | 11080 | 6949 | 9590 | 11043 | 16501 | 13556 | 14242 | 10402 | 9940 | 8847 | 9812 | 9106 | 14112 | 12935 | 12709 | 14671 | 11516 | 9426 | 13607 | 1,443 | 0,529 | 9,054 | 1,37E-19 | 2,41E-18 | 0,003858 | 0,0021056 | 0,0031904 | 0,0031904 | TRUE | 0,0336164 | No |
| orf19.3375 | 1 | 2 | 2 | 3 | 2 | 3 | 1 | 0 | 1 | 3 | 2 | 2 | 2 | 3 | 1 | 0 | 1,68 | 2 | 2 | 0,733 | -0,448 | -0,426 | 0,67041551 | NA | 1,00E-08 | 0,8668174 | 0,4665904 | 0,4665904 | TRUE | 0,3211827 | No |
| orf19.3376 | 146 | 113 | 120 | 179 | 135 | 149 | 134 | 89 | 131 | 144 | 123 | 148 | 115 | 142 | 120 | 126 | 130,99 | 136 | 126 | 0,921 | -0,119 | -0,935 | 0,34991573 | 0,43743106 | 1,00E-08 | 0,0130618 | 0,0079946 | 0,0079946 | TRUE | 0,0860872 | No |
| orf19.3378 | 84 | 45 | 57 | 91 | 94 | 70 | 68 | 44 | 75 | 57 | 58 | 75 | 80 | 67 | 61 | 62 | 66,99 | 66 | 68 | 1,011 | 0,016 | 0,089 | 0,92911318 | 0,94901598 | 0,00644 | 0,0236498 | 0,0164721 | 0,0164721 | TRUE | 0,1639409 | No |
| orf19.338 | 2929 | 2017 | 2673 | 3098 | 3947 | 3291 | 3542 | 2355 | 2628 | 2568 | 2735 | 2555 | 3376 | 3140 | 3161 | 3322 | 2935,4 | 2622 | 3250 | 1,239 | 0,31 | 7,242 | 4,42E-13 | 4,26E-12 | 0,000819 | 0,0024741 | 0,0014077 | 0,0014077 | TRUE | 0,0102334 | No |
| orf19.3380 | 10 | 8 | 14 | 10 | 23 | 18 | 13 | 11 | 9 | 10 | 14 | 8 | 20 | 17 | 12 | 16 | 13,21 | 10 | 16 | 1,55 | 0,632 | 1,634 | 0,10218935 | NA | 1,00E-08 | 0,1118678 | 0,0646492 | 0,0646492 | TRUE | 0,2293719 | No |
| orf19.339 | 8315 | 5783 | 7115 | 8845 | 8216 | 7279 | 7632 | 5111 | 7459 | 7362 | 7279 | 7294 | 7027 | 6945 | 6810 | 7209 | 7173,2 | 7348 | 6998 | 0,952 | -0,071 | -2,352 | 0,018653 | 0,03491191 | 0,00016 | 0,0021819 | 0,0007451 | 0,0007451 | TRUE | 0,0047253 | No |
| orf19.3391 | 1124 | 826 | 1031 | 1258 | 1260 | 1086 | 1165 | 810 | 1008 | 1052 | 1055 | 1037 | 1078 | 1036 | 1040 | 1142 | 1056 | 1038 | 1074 | 1,033 | 0,046 | 0,895 | 0,37079285 | 0,45922973 | 9,20E-05 | 0,0033543 | 0,0016279 | 0,0016279 | TRUE | 0,0190964 | No |
| orf19.3392 | 856 | 584 | 682 | 931 | 670 | 584 | 628 | 409 | 768 | 743 | 698 | 768 | 573 | 557 | 560 | 577 | 655,55 | 744 | 567 | 0,76 | -0,395 | -6,499 | 8,11E-11 | 5,98E-10 | 1,00E-08 | 0,004194 | 0,0020017 | 0,0020017 | TRUE | 0,0212147 | No |
| orf19.3393 | 11681 | 8487 | 10518 | 12953 | 10852 | 10150 | 10835 | 6735 | 10479 | 10805 | 10761 | 10681 | 9281 | 9685 | 9668 | 9499 | 10107 | 10682 | 9533 | 0,893 | -0,164 | -5,714 | 1,11E-08 | 6,08E-08 | 0,000153 | 0,0021231 | 0,0006898 | 0,0006898 | TRUE | 0,004029 | No |
| orf19.3394 | 483 | 350 | 414 | 550 | 483 | 445 | 462 | 350 | 433 | 446 | 424 | 454 | 413 | 425 | 412 | 494 | 437,45 | 439 | 436 | 0,987 | -0,02 | -0,253 | 0,80045556 | 0,84519104 | 0,00139 | 0,0052981 | 0,0034488 | 0,0034488 | TRUE | 0,0788891 | No |
| orf19.3395 | 2426 | 1667 | 2165 | 2702 | 2124 | 1850 | 1972 | 1211 | 2176 | 2122 | 2215 | 2228 | 1817 | 1765 | 1760 | 1708 | 1973,9 | 2185 | 1762 | 0,807 | -0,31 | -7,621 | 2,52E-14 | 2,69E-13 | 1,00E-08 | 0,002715 | 0,0010709 | 0,0010709 | TRUE | 0,0051876 | No |
| orf19.3396 | 896 | 688 | 835 | 915 | 1314 | 1025 | 1258 | 711 | 804 | 876 | 854 | 755 | 1124 | 978 | 1123 | 1003 | 939,46 | 822 | 1057 | 1,29 | 0,368 | 5,194 | 2,05E-07 | 9,64E-07 | 0,003891 | 0,0035248 | 0,0037266 | 0,0037266 | TRUE | 0,037561 | No |
| orf19.3399 | 1691 | 1126 | 1379 | 1745 | 1922 | 1540 | 1770 | 1095 | 1517 | 1434 | 1411 | 1439 | 1644 | 1469 | 1579 | 1544 | 1504,7 | 1450 | 1559 | 1,076 | 0,105 | 2,065 | 0,03888362 | 0,06742072 | 0,00105 | 0,0029443 | 0,001815 | 0,001815 | TRUE | 0,0190798 | No |
| orf19.34 | 286 | 189 | 270 | 337 | 383 | 334 | 354 | 206 | 257 | 241 | 276 | 278 | 328 | 319 | 316 | 291 | 288 | 263 | 314 | 1,192 | 0,253 | 2,908 | 0,0036328 | 0,00807645 | 1,00E-08 | 0,0070202 | 0,0038165 | 0,0038165 | TRUE | 0,0307619 | No |
| orf19.340 | 426 | 295 | 348 | 436 | 581 | 551 | 588 | 324 | 382 | 376 | 356 | 360 | 497 | 526 | 525 | 457 | 434,7 | 368 | 501 | 1,366 | 0,45 | 6,054 | 1,41E-09 | 8,82E-09 | 1,00E-08 | 0,0053191 | 0,0029641 | 0,0029641 | TRUE | 0,0341912 | No |
| orf19.3400 | 1546 | 1049 | 1321 | 1603 | 1551 | 1301 | 1400 | 750 | 1387 | 1335 | 1352 | 1322 | 1326 | 1241 | 1249 | 1058 | 1283,8 | 1349 | 1218 | 0,906 | -0,142 | -2,163 | 0,03056192 | 0,05450163 | 0,003627 | 0,0031103 | 0,0033453 | 0,0033453 | TRUE | 0,0870777 | No |
| orf19.3401 | 2560 | 1705 | 2165 | 2712 | 2327 | 1948 | 2170 | 1365 | 2297 | 2171 | 2215 | 2236 | 1990 | 1859 | 1936 | 1925 | 2078,6 | 2230 | 1928 | 0,864 | -0,211 | -5,097 | 3,45E-07 | 1,56E-06 | 0,000218 | 0,0026779 | 0,0011542 | 0,0011542 | TRUE | 0,0073577 | No |
| orf19.3402 | 2390 | 1507 | 2049 | 2520 | 2872 | 2334 | 2584 | 1850 | 2144 | 1919 | 2096 | 2078 | 2456 | 2227 | 2306 | 2609 | 2229,4 | 2059 | 2400 | 1,163 | 0,218 | 3,694 | 0,00022036 | 0,00063772 | 0,003009 | 0,0026307 | 0,0028804 | 0,0028804 | TRUE | 0,039815 | No |
| orf19.3404 | 17 | 9 | 9 | 14 | 7 | 15 | 12 | 4 | 15 | 11 | 9 | 12 | 6 | 14 | 11 | 6 | 10,51 | 12 | 9 | 0,781 | -0,356 | -0,806 | 0,42028747 | NA | 1,00E-08 | 0,140056 | 0,0924237 | 0,0924237 | TRUE | 0,4712282 | No |
| orf19.3405 | 711 | 440 | 587 | 749 | 470 | 424 | 477 | 343 | 638 | 560 | 601 | 618 | 402 | 405 | 426 | 484 | 516,52 | 604 | 429 | 0,705 | -0,505 | -6,576 | 4,85E-11 | 3,70E-10 | 0,00237 | 0,0047901 | 0,0036653 | 0,0036653 | TRUE | 0,0732572 | No |
| orf19.3406 | 931 | 622 | 831 | 1061 | 1231 | 1091 | 1167 | 754 | 835 | 792 | 850 | 875 | 1053 | 1041 | 1041 | 1063 | 943,85 | 838 | 1050 | 1,249 | 0,321 | 6,164 | 7,10E-10 | 4,61E-09 | 1,00E-08 | 0,0035176 | 0,0015293 | 0,0015293 | TRUE | 0,014543 | No |
| orf19.3407 | 1077 | 698 | 901 | 1107 | 1384 | 1128 | 1301 | 874 | 966 | 889 | 922 | 913 | 1184 | 1076 | 1161 | 1233 | 1042,9 | 922 | 1164 | 1,258 | 0,331 | 5,771 | 7,87E-09 | 4,42E-08 | 0,001222 | 0,0033715 | 0,0021869 | 0,0021869 | TRUE | 0,0301439 | No |
| orf19.3409 | 1311 | 919 | 1194 | 1382 | 1570 | 1330 | 1476 | 948 | 1176 | 1170 | 1222 | 1140 | 1343 | 1269 | 1317 | 1337 | 1246,7 | 1177 | 1316 | 1,119 | 0,162 | 3,407 | 0,00065596 | 0,00171721 | 1,00E-08 | 0,003144 | 0,0013652 | 0,0013652 | TRUE | 0,0079413 | No |
| orf19.341 | 236 | 179 | 198 | 292 | 376 | 291 | 346 | 213 | 212 | 228 | 203 | 241 | 322 | 278 | 309 | 300 | 261,42 | 221 | 302 | 1,369 | 0,453 | 4,773 | 1,82E-06 | 7,37E-06 | 1,00E-08 | 0,0075327 | 0,0047441 | 0,0047441 | TRUE | 0,0474613 | No |
| orf19.3411 | 301 | 243 | 303 | 394 | 204 | 166 | 179 | 102 | 270 | 309 | 310 | 325 | 174 | 158 | 160 | 144 | 231,34 | 304 | 159 | 0,528 | -0,922 | -8,91 | 5,12E-19 | 8,62E-18 | 1,00E-08 | 0,0082546 | 0,005523 | 0,005523 | TRUE | 0,0668312 | No |
| orf19.3412 | 4012 | 2669 | 3558 | 4401 | 5005 | 4056 | 4652 | 2881 | 3599 | 3398 | 3640 | 3629 | 4280 | 3870 | 4151 | 4063 | 3828,9 | 3566 | 4091 | 1,147 | 0,198 | 4,513 | 6,40E-06 | 2,40E-05 | 0,001046 | 0,0023587 | 0,0015763 | 0,0015763 | TRUE | 0,0165521 | No |
| orf19.3413 | 7 | 2 | 1 | 6 | 3 | 7 | 0 | 1 | 6 | 3 | 1 | 5 | 3 | 7 | 0 | 1 | 3,18 | 4 | 3 | 0,715 | -0,485 | -0,526 | 0,5991529 | NA | 0,561871 | 0,458274 | 0,4977959 | 0,4977959 | TRUE | 0,6619576 | No |
| orf19.3414 | 10648 | 7511 | 9219 | 11608 | 9053 | 8538 | 9260 | 5360 | 9552 | 9562 | 9432 | 9572 | 7742 | 8147 | 8263 | 7560 | 8728,8 | 9530 | 7928 | 0,832 | -0,265 | -7,224 | 5,05E-13 | 4,81E-12 | 0,000745 | 0,0021458 | 0,0011736 | 0,0011736 | TRUE | 0,0114597 | No |
| orf19.3415 | 11974 | 7597 | 10580 | 12360 | 11953 | 10233 | 11804 | 7584 | 10742 | 9672 | 10824 | 10192 | 10223 | 9764 | 10533 | 10697 | 10331 | 10358 | 10304 | 0,994 | -0,008 | -0,169 | 0,86577353 | 0,89777419 | 0,001741 | 0,00212 | 0,0020691 | 0,0020691 | TRUE | 0,0239366 | No |
| orf19.3415.1 | 14939 | 9978 | 12547 | 15843 | 15953 | 13458 | 13657 | 8936 | 13402 | 12703 | 12837 | 13064 | 13644 | 12841 | 12187 | 12603 | 12910 | 13002 | 12819 | 0,986 | -0,02 | -0,492 | 0,6227833 | 0,69573857 | 0,001185 | 0,002092 | 0,0015777 | 0,0015777 | TRUE | 0,0231758 | No |
| orf19.3417 | 102 | 65 | 91 | 104 | 114 | 114 | 128 | 86 | 92 | 83 | 93 | 86 | 97 | 109 | 114 | 121 | 99,36 | 88 | 110 | 1,241 | 0,312 | 2,227 | 0,02597466 | 0,04730396 | 1,00E-08 | 0,0165896 | 0,0086652 | 0,0086652 | TRUE | 0,0664996 | No |
| orf19.3418 | 127 | 99 | 119 | 144 | 132 | 120 | 133 | 57 | 114 | 126 | 122 | 119 | 113 | 115 | 119 | 80 | 113,37 | 120 | 107 | 0,901 | -0,15 | -1,073 | 0,28306892 | 0,36731708 | 0,000648 | 0,014785 | 0,0098717 | 0,0098717 | TRUE | 0,2064492 | No |
| orf19.3419 | 736 | 546 | 692 | 855 | 913 | 845 | 859 | 585 | 660 | 695 | 708 | 705 | 781 | 806 | 767 | 825 | 743,39 | 692 | 795 | 1,146 | 0,197 | 3,435 | 0,00059197 | 0,00156746 | 1,00E-08 | 0,0039323 | 0,0018054 | 0,0018054 | TRUE | 0,0120799 | No |
| orf19.342 | 31 | 23 | 25 | 48 | 35 | 33 | 24 | 23 | 28 | 29 | 26 | 40 | 30 | 31 | 21 | 32 | 29,69 | 31 | 28 | 0,929 | -0,107 | -0,405 | 0,68574308 | 0,74894464 | 1,00E-08 | 0,0508734 | 0,0333721 | 0,0333721 | TRUE | 0,3141104 | No |
| orf19.3420 | 0 | 1 | 1 | 2 | 0 | 0 | 0 | 0 | 0 | 1 | 1 | 2 | 0 | 0 | 0 | 0 | 0,49 | 1 | 0 | 0,189 | -2,407 | -1,179 | 0,23822303 | NA | 1,00E-08 | 2,9455389 | 2,4961694 | 2,4961694 | TRUE | 0,2448382 | No |
| orf19.3421.1 | 1598 | 1129 | 1245 | 1587 | 1902 | 1763 | 1771 | 1216 | 1434 | 1437 | 1274 | 1309 | 1627 | 1682 | 1580 | 1715 | 1507,2 | 1364 | 1651 | 1,211 | 0,276 | 4,965 | 6,86E-07 | 2,97E-06 | 0,00168 | 0,0029427 | 0,0022928 | 0,0022928 | TRUE | 0,0227287 | No |
| orf19.3422 | 971 | 691 | 938 | 1007 | 824 | 771 | 868 | 502 | 871 | 880 | 960 | 830 | 705 | 736 | 775 | 708 | 807,98 | 885 | 731 | 0,827 | -0,274 | -4,274 | 1,92E-05 | 6,68E-05 | 0,001781 | 0,0037762 | 0,0026846 | 0,0026846 | TRUE | 0,0386706 | No |
| orf19.3423 | 11106 | 7469 | 9760 | 11877 | 10311 | 8900 | 10229 | 6156 | 9963 | 9509 | 9985 | 9794 | 8818 | 8492 | 9128 | 8682 | 9296,5 | 9813 | 8780 | 0,895 | -0,16 | -4,622 | 3,81E-06 | 1,48E-05 | 0,000572 | 0,0021357 | 0,0010487 | 0,0010487 | TRUE | 0,0087822 | No |
| orf19.3425 | 2792 | 2049 | 2562 | 3154 | 2533 | 2377 | 2564 | 1634 | 2505 | 2609 | 2621 | 2601 | 2166 | 2268 | 2288 | 2305 | 2420,3 | 2584 | 2257 | 0,873 | -0,196 | -4,966 | 6,83E-07 | 2,96E-06 | 0,000222 | 0,0025793 | 0,0010802 | 0,0010802 | TRUE | 0,00917 | No |
| orf19.3426 | 13467 | 9477 | 11530 | 14898 | 13165 | 11876 | 12008 | 9136 | 12081 | 12065 | 11796 | 12285 | 11259 | 11332 | 10715 | 12886 | 11802 | 12057 | 11548 | 0,957 | -0,063 | -1,155 | 0,24823853 | 0,3301056 | 0,003249 | 0,0021025 | 0,0027684 | 0,0027684 | TRUE | 0,0735333 | No |
| orf19.3427 | 168 | 132 | 163 | 203 | 224 | 213 | 243 | 188 | 151 | 168 | 167 | 167 | 192 | 203 | 217 | 265 | 191,22 | 163 | 219 | 1,332 | 0,414 | 3,494 | 0,00047551 | 0,00129211 | 0,006371 | 0,0095715 | 0,0081211 | 0,0081211 | TRUE | 0,1873602 | No |
| orf19.3428 | 7544 | 5256 | 6618 | 8062 | 9425 | 8804 | 9674 | 5936 | 6768 | 6691 | 6771 | 6648 | 8061 | 8400 | 8632 | 8372 | 7543 | 6720 | 8366 | 1,245 | 0,316 | 9,869 | 5,67E-23 | 1,26E-21 | 0,000355 | 0,002172 | 0,0008507 | 0,0008507 | TRUE | 0,0076429 | No |
| orf19.3429 | 359 | 278 | 310 | 443 | 298 | 318 | 317 | 216 | 322 | 354 | 317 | 365 | 255 | 303 | 283 | 305 | 313,03 | 340 | 286 | 0,84 | -0,251 | -2,743 | 0,00608955 | 0,01291423 | 0,002866 | 0,0066171 | 0,0048365 | 0,0048365 | TRUE | 0,0659044 | No |

| | | | | | | | | | | | | | | | | | | | | | | | | | | | | | | | |
|---|---|---|---|---|---|---|---|---|---|---|---|---|---|---|---|---|---|---|---|---|---|---|---|---|---|---|---|---|---|---|---|
| orf19.3430 | 120 | 92 | 91 | 122 | 142 | 137 | 145 | 84 | 108 | 117 | 93 | 101 | 121 | 131 | 129 | 118 | 114,81 | 105 | 125 | 1,203 | 0,266 | 2,044 | 0,04095772 | 0,07042758 | 1,00E-08 | 0,0146235 | 0,0075742 | 0,0075742 | TRUE | 0,051768 | No |
| orf19.3431 | 1337 | 1048 | 1244 | 1597 | 1615 | 1689 | 1692 | 977 | 1199 | 1334 | 1273 | 1317 | 1381 | 1612 | 1510 | 1378 | 1375,5 | 1281 | 1470 | 1,15 | 0,201 | 3,144 | 0,00166452 | 0,0039906 | 0,003057 | 0,0030349 | 0,0031969 | 0,0031969 | TRUE | 0,0504356 | No |
| orf19.3432 | 284 | 202 | 238 | 296 | 354 | 335 | 350 | 192 | 255 | 257 | 243 | 244 | 303 | 320 | 312 | 271 | 275,63 | 250 | 302 | 1,213 | 0,279 | 3,183 | 0,00145731 | 0,00354474 | 1,00E-08 | 0,0072464 | 0,0037445 | 0,0037445 | TRUE | 0,0424033 | No |
| orf19.3432.1 | 25 | 26 | 21 | 46 | 28 | 32 | 28 | 31 | 22 | 33 | 21 | 38 | 24 | 31 | 25 | 44 | 29,77 | 28 | 31 | 1,049 | 0,069 | 0,235 | 0,81419846 | 0,85654077 | 0,044698 | 0,0507488 | 0,0481614 | 0,0481614 | TRUE | 0,4069315 | No |
| orf19.3433 | 470 | 338 | 383 | 497 | 568 | 570 | 554 | 365 | 422 | 430 | 392 | 410 | 486 | 544 | 494 | 515 | 461,55 | 414 | 510 | 1,233 | 0,302 | 4,218 | 2,47E-05 | 8,44E-05 | 1,00E-08 | 0,0051248 | 0,0027286 | 0,0027286 | TRUE | 0,025728 | No |
| orf19.3434 | 671 | 567 | 679 | 845 | 602 | 651 | 679 | 425 | 602 | 722 | 695 | 697 | 515 | 621 | 606 | 599 | 632,08 | 679 | 585 | 0,862 | -0,214 | -2,618 | 0,00884844 | 0,0180649 | 0,005263 | 0,0042762 | 0,0047974 | 0,0047974 | TRUE | 0,081918 | No |
| orf19.3435 | 215 | 133 | 194 | 241 | 210 | 168 | 214 | 154 | 193 | 169 | 198 | 199 | 180 | 160 | 191 | 217 | 188,43 | 190 | 187 | 0,974 | -0,037 | -0,327 | 0,74353031 | 0,79917925 | 0,003704 | 0,0096835 | 0,00728 | 0,00728 | TRUE | 0,1046117 | No |
| orf19.3436 | 48 | 28 | 49 | 44 | 69 | 67 | 76 | 30 | 43 | 36 | 50 | 36 | 59 | 64 | 68 | 42 | 49,77 | 41 | 58 | 1,428 | 0,514 | 2,43 | 0,01508022 | 0,02898335 | 0,00806 | 0,0311453 | 0,022481 | 0,022481 | TRUE | 0,2162447 | No |
| orf19.3437 | 106 | 90 | 96 | 121 | 88 | 89 | 89 | 69 | 95 | 115 | 98 | 100 | 75 | 85 | 79 | 97 | 93,07 | 102 | 84 | 0,822 | -0,283 | -1,923 | 0,05446916 | 0,09055083 | 1,00E-08 | 0,0175769 | 0,009857 | 0,009857 | TRUE | 0,0795717 | No |
| orf19.3438 | 1003 | 693 | 869 | 1149 | 1087 | 975 | 1120 | 571 | 900 | 882 | 889 | 947 | 930 | 930 | 999 | 805 | 910,41 | 904 | 916 | 1,015 | 0,022 | 0,327 | 0,74370432 | 0,79917925 | 0,003045 | 0,0035741 | 0,0032968 | 0,0032968 | TRUE | 0,0708832 | No |
| orf19.3439 | 2355 | 2031 | 2233 | 3062 | 3312 | 3561 | 3734 | 2208 | 2113 | 2586 | 2285 | 2525 | 2833 | 3398 | 3332 | 3114 | 2773,1 | 2377 | 3169 | 1,334 | 0,416 | 5,466 | 4,61E-08 | 2,35E-07 | 0,00699 | 0,002503 | 0,0051826 | 0,0051826 | TRUE | 0,0684722 | No |
| orf19.344 | 1005 | 745 | 937 | 1168 | 939 | 864 | 852 | 630 | 902 | 948 | 959 | 963 | 803 | 824 | 760 | 889 | 881,02 | 943 | 819 | 0,866 | -0,208 | -3,458 | 0,00054462 | 0,0014562 | 0,001057 | 0,0036273 | 0,0023312 | 0,0023312 | TRUE | 0,0340905 | No |
| orf19.3440 | 138 | 106 | 137 | 171 | 118 | 133 | 119 | 81 | 124 | 135 | 140 | 141 | 101 | 127 | 106 | 114 | 123,52 | 135 | 112 | 0,828 | -0,272 | -2,109 | 0,03494047 | 0,06139876 | 1,00E-08 | 0,0137321 | 0,0078681 | 0,0078681 | TRUE | 0,0883172 | No |
| orf19.3441 | 3760 | 2767 | 3450 | 4307 | 3189 | 2984 | 3111 | 2135 | 3373 | 3523 | 3530 | 3552 | 2727 | 2847 | 2776 | 3011 | 3167,4 | 3494 | 2840 | 0,812 | -0,301 | -7,111 | 1,15E-12 | 1,04E-11 | 0,000801 | 0,0024379 | 0,0013984 | 0,0013984 | TRUE | 0,0185114 | No |
| orf19.3442 | 4024 | 3033 | 3558 | 4547 | 3612 | 3387 | 3625 | 2371 | 3610 | 3861 | 3640 | 3750 | 3089 | 3232 | 3235 | 3344 | 3470,1 | 3715 | 3225 | 0,868 | -0,205 | -5,075 | 3,88E-07 | 1,75E-06 | 0,000613 | 0,0023979 | 0,0012738 | 0,0012738 | TRUE | 0,0098827 | No |
| orf19.3443 | 2659 | 1878 | 2342 | 2848 | 2896 | 2805 | 2867 | 1845 | 2385 | 2391 | 2396 | 2348 | 2477 | 2676 | 2558 | 2602 | 2479,3 | 2380 | 2578 | 1,083 | 0,115 | 2,931 | 0,00338261 | 0,00757745 | 0,000257 | 0,002565 | 0,0010716 | 0,0010716 | TRUE | 0,0089672 | No |
| orf19.3444 | 410 | 253 | 305 | 425 | 509 | 349 | 373 | 289 | 368 | 322 | 312 | 350 | 435 | 333 | 333 | 408 | 357,65 | 338 | 377 | 1,112 | 0,153 | 1,433 | 0,15198682 | 0,21806363 | 0,010106 | 0,0060385 | 0,008096 | 0,008096 | TRUE | 0,1367988 | No |
| orf19.3445 | 941 | 757 | 863 | 1048 | 807 | 855 | 853 | 581 | 844 | 964 | 883 | 864 | 690 | 816 | 761 | 819 | 830,2 | 889 | 772 | 0,868 | -0,205 | -2,88 | 0,00398232 | 0,00878978 | 0,003554 | 0,0037281 | 0,0036351 | 0,0036351 | TRUE | 0,0620836 | No |
| orf19.3446 | 500 | 458 | 462 | 637 | 667 | 692 | 703 | 348 | 449 | 583 | 473 | 525 | 570 | 660 | 627 | 491 | 547,31 | 508 | 587 | 1,163 | 0,218 | 2,084 | 0,03714879 | 0,06475948 | 0,012473 | 0,004632 | 0,0086541 | 0,0086541 | TRUE | 0,131491 | No |
| orf19.3447 | 367 | 334 | 337 | 426 | 304 | 383 | 331 | 244 | 329 | 425 | 345 | 351 | 260 | 365 | 295 | 344 | 339,43 | 362 | 316 | 0,872 | -0,198 | -1,667 | 0,09548266 | 0,1472055 | 0,01433 | 0,0062564 | 0,0105733 | 0,0105733 | TRUE | 0,1698384 | No |
| orf19.3448 | 1813 | 2144 | 1787 | 2480 | 1776 | 2920 | 2753 | 1405 | 1626 | 2730 | 1828 | 2045 | 1519 | 2786 | 2457 | 1982 | 2121,6 | 2057 | 2186 | 1,063 | 0,088 | 0,355 | 0,72292472 | 0,7828644 | 0,058427 | 0,0026638 | 0,0271361 | 0,0584272 | TRUE | 0,3026619 | No |
| orf19.3449 | 2239 | 1688 | 1862 | 2618 | 2699 | 2425 | 2747 | 1853 | 2009 | 2149 | 1905 | 2159 | 2308 | 2314 | 2451 | 2613 | 2238,5 | 2056 | 2422 | 1,177 | 0,235 | 4,018 | 5,87E-05 | 0,00018763 | 0,002913 | 0,002628 | 0,0028336 | 0,0028336 | TRUE | 0,0327955 | No |
| orf19.3449.2 | 850 | 596 | 718 | 868 | 909 | 962 | 936 | 629 | 763 | 759 | 735 | 716 | 777 | 918 | 835 | 887 | 798,67 | 743 | 854 | 1,149 | 0,2 | 3,05 | 0,00229089 | 0,00534096 | 0,002108 | 0,0037972 | 0,0028668 | 0,0028668 | TRUE | 0,0458242 | No |
| orf19.345 | 359 | 266 | 320 | 344 | 203 | 194 | 208 | 132 | 322 | 339 | 327 | 284 | 174 | 185 | 186 | 186 | 250,28 | 318 | 183 | 0,576 | -0,797 | -8,281 | 1,23E-16 | 1,65E-15 | 1,00E-08 | 0,0077798 | 0,0045794 | 0,0045794 | TRUE | 0,0643808 | No |
| orf19.3450.1 | 293 | 207 | 228 | 293 | 356 | 337 | 368 | 221 | 263 | 264 | 233 | 242 | 304 | 322 | 328 | 312 | 283,42 | 250 | 316 | 1,267 | 0,342 | 3,966 | 7,30E-05 | 0,00022889 | 1,00E-08 | 0,0071016 | 0,0035592 | 0,0035592 | TRUE | 0,0217121 | No |
| orf19.3453 | 151 | 106 | 123 | 162 | 164 | 166 | 169 | 112 | 135 | 135 | 126 | 134 | 140 | 158 | 151 | 158 | 142,16 | 132 | 152 | 1,143 | 0,192 | 1,666 | 0,09576999 | 0,14749708 | 1,00E-08 | 0,0121914 | 0,0057511 | 0,0057511 | TRUE | 0,0301766 | No |
| orf19.3455 | 1595 | 1469 | 1559 | 1914 | 2098 | 2156 | 2258 | 1292 | 1431 | 1870 | 1595 | 1578 | 1794 | 2057 | 2015 | 1822 | 1770,4 | 1618 | 1922 | 1,19 | 0,251 | 3,129 | 0,00175563 | 0,00419731 | 0,008047 | 0,0027995 | 0,0055905 | 0,0055905 | TRUE | 0,1286513 | No |
| orf19.3456 | 888 | 675 | 746 | 1009 | 866 | 870 | 900 | 591 | 797 | 859 | 763 | 832 | 741 | 830 | 803 | 834 | 807,33 | 813 | 802 | 0,985 | -0,021 | -0,335 | 0,73731829 | 0,79401475 | 0,001538 | 0,0037777 | 0,0025876 | 0,0025876 | TRUE | 0,032846 | No |
| orf19.3457 | 517 | 386 | 499 | 589 | 542 | 539 | 525 | 314 | 464 | 491 | 511 | 486 | 464 | 514 | 468 | 443 | 480,08 | 488 | 472 | 0,971 | -0,042 | -0,584 | 0,55900285 | 0,63924007 | 0,000578 | 0,0050034 | 0,0028942 | 0,0028942 | TRUE | 0,0410419 | No |
| orf19.3458 | 1369 | 1163 | 1286 | 1669 | 1614 | 1531 | 1625 | 843 | 1228 | 1481 | 1316 | 1376 | 1380 | 1461 | 1450 | 1189 | 1360,1 | 1350 | 1370 | 1,018 | 0,025 | 0,331 | 0,74033636 | 0,79669369 | 0,006502 | 0,0030469 | 0,0049121 | 0,0049121 | TRUE | 0,0896205 | No |
| orf19.3459 | 199 | 183 | 185 | 232 | 303 | 307 | 315 | 205 | 179 | 233 | 189 | 191 | 259 | 293 | 281 | 289 | 239,3 | 198 | 280 | 1,424 | 0,51 | 4,971 | 6,65E-07 | 2,88E-06 | 0,002646 | 0,0080461 | 0,0057805 | 0,0057805 | TRUE | 0,1333998 | No |
| orf19.346 | 3058 | 2074 | 2615 | 3209 | 3617 | 3121 | 3398 | 2043 | 2743 | 2640 | 2675 | 2646 | 3093 | 2978 | 3032 | 2881 | 2836,3 | 2676 | 2996 | 1,121 | 0,164 | 4,315 | 1,60E-05 | 5,65E-05 | 0,000231 | 0,0024914 | 0,0010375 | 0,0010375 | TRUE | 0,0073822 | No |
| orf19.3460 | 467 | 382 | 418 | 517 | 416 | 461 | 474 | 271 | 419 | 486 | 428 | 426 | 356 | 440 | 423 | 382 | 420,01 | 440 | 400 | 0,913 | -0,131 | -1,487 | 0,13706328 | 0,19977588 | 0,004624 | 0,0054359 | 0,0050341 | 0,0050341 | TRUE | 0,0721359 | No |
| orf19.3461 | 21 | 11 | 20 | 12 | 11 | 10 | 18 | 11 | 19 | 14 | 20 | 10 | 9 | 10 | 16 | 16 | 14,22 | 16 | 13 | 0,792 | -0,336 | -0,863 | 0,38825901 | NA | 1,00E-08 | 0,1040983 | 0,0743304 | 0,0743304 | TRUE | 0,3809593 | No |
| orf19.3462 | 1359 | 1003 | 1119 | 1511 | 1679 | 1582 | 1494 | 993 | 1219 | 1277 | 1145 | 1246 | 1436 | 1509 | 1333 | 1401 | 1320,8 | 1222 | 1420 | 1,163 | 0,218 | 3,868 | 0,00010993 | 0,00033424 | 0,001683 | 0,0030787 | 0,0022761 | 0,0022761 | TRUE | 0,022235 | No |
| orf19.3463 | 14418 | 10740 | 13384 | 15792 | 15181 | 14783 | 15533 | 9084 | 12934 | 13673 | 13693 | 13022 | 12983 | 14105 | 13861 | 12812 | 13386 | 13330 | 13440 | 1,009 | 0,012 | 0,287 | 0,77427734 | 0,8236293 | 0,001317 | 0,002088 | 0,0017018 | 0,0017018 | TRUE | 0,0135297 | No |
| orf19.3465 | 22926 | 15567 | 18511 | 23000 | 20221 | 17666 | 17905 | 11984 | 20567 | 19818 | 18939 | 18966 | 17294 | 16856 | 15977 | 16902 | 18165 | 19572 | 16757 | 0,856 | -0,224 | -5,507 | 3,65E-08 | 1,89E-07 | 0,001268 | 0,0020594 | 0,0015342 | 0,0015342 | TRUE | 0,0144224 | No |
| orf19.3466 | 1231 | 982 | 1204 | 1359 | 1273 | 1332 | 1372 | 821 | 1104 | 1250 | 1232 | 1121 | 1089 | 1271 | 1224 | 1158 | 1181,1 | 1177 | 1186 | 1,009 | 0,013 | 0,2 | 0,84168153 | 0,8792945 | 0,003519 | 0,0032086 | 0,0033835 | 0,0033835 | TRUE | 0,0385885 | No |
| orf19.3467 | 2725 | 2110 | 2434 | 2959 | 3079 | 3107 | 3430 | 1839 | 2445 | 2686 | 2490 | 2440 | 2633 | 2965 | 3061 | 2594 | 2664,2 | 2515 | 2813 | 1,12 | 0,164 | 2,589 | 0,0096263 | 0,019473 | 0,004035 | 0,0025244 | 0,0034528 | 0,0034528 | TRUE | 0,0427579 | No |
| orf19.3468 | 3613 | 3091 | 3447 | 4385 | 3958 | 4238 | 4200 | 2689 | 3241 | 3935 | 3527 | 3616 | 3385 | 4044 | 3748 | 3793 | 3661 | 3580 | 3742 | 1,046 | 0,065 | 0,941 | 0,34660879 | 0,43392907 | 0,005254 | 0,0023761 | 0,004236 | 0,004236 | TRUE | 0,0534355 | No |
| orf19.3469 | 3543 | 2598 | 3389 | 4281 | 3772 | 3481 | 3908 | 2989 | 3178 | 3308 | 3467 | 3530 | 3226 | 3321 | 3487 | 4216 | 3466,7 | 3371 | 3562 | 1,056 | 0,078 | 0,996 | 0,31901882 | 0,40590223 | 0,008654 | 0,0023983 | 0,0056073 | 0,0056073 | TRUE | 0,1819841 | No |
| orf19.347 | 3430 | 2427 | 3052 | 3952 | 4986 | 4248 | 4594 | 3217 | 3077 | 3090 | 3122 | 3259 | 4264 | 4053 | 4099 | 4537 | 3687,8 | 3137 | 4238 | 1,349 | 0,432 | 9,446 | 3,53E-21 | 6,91E-20 | 0,001316 | 0,0023732 | 0,0017322 | 0,0017322 | TRUE | 0,026155 | No |
| orf19.3470 | 4827 | 3461 | 4408 | 5487 | 5746 | 5429 | 5881 | 3561 | 4330 | 4406 | 4510 | 4525 | 4914 | 5180 | 5248 | 5022 | 4767 | 4443 | 5091 | 1,146 | 0,197 | 5,43 | 5,62E-08 | 2,84E-07 | 0,000491 | 0,0022841 | 0,001047 | 0,001047 | TRUE | 0,0070542 | No |
| orf19.3471 | 170 | 107 | 138 | 173 | 211 | 166 | 190 | 105 | 153 | 136 | 141 | 143 | 180 | 158 | 170 | 148 | 153,63 | 143 | 164 | 1,151 | 0,203 | 1,771 | 0,07658712 | 0,12179339 | 1,00E-08 | 0,0114287 | 0,006124 | 0,006124 | TRUE | 0,0481261 | No |
| orf19.3473 | 538 | 324 | 476 | 558 | 498 | 461 | 508 | 366 | 483 | 412 | 487 | 460 | 426 | 440 | 453 | 516 | 459,69 | 460 | 459 | 0,989 | -0,016 | -0,196 | 0,84446344 | 0,88158755 | 0,00338 | 0,0051375 | 0,0043691 | 0,0043691 | TRUE | 0,0720843 | No |
| orf19.3474 | 1665 | 1207 | 1448 | 1913 | 1928 | 1942 | 2126 | 1201 | 1494 | 1537 | 1481 | 1577 | 1649 | 1853 | 1897 | 1694 | 1647,8 | 1522 | 1773 | 1,166 | 0,221 | 3,888 | 0,00010124 | 0,00030985 | 0,002239 | 0,0028605 | 0,0024899 | 0,0024899 | TRUE | 0,0289889 | No |
| orf19.3475 | 1730 | 1250 | 1576 | 1907 | 1546 | 1442 | 1516 | 1079 | 1552 | 1591 | 1612 | 1573 | 1322 | 1376 | 1353 | 1522 | 1487,6 | 1582 | 1393 | 0,878 | -0,188 | -3,606 | 0,00031082 | 0,00087755 | 0,001253 | 0,0029554 | 0,001917 | 0,001917 | TRUE | 0,042104 | No |
| orf19.3476 | 11056 | 7816 | 9592 | 12429 | 12550 | 11587 | 11980 | 7398 | 9918 | 9951 | 9814 | 10249 | 10733 | 11056 | 10690 | 10434 | 10356 | 9983 | 10728 | 1,075 | 0,104 | 3,348 | 0,00081255 | 0,00208445 | 0,000329 | 0,0021197 | 0,0008313 | 0,0008313 | TRUE | 0,0051231 | No |
| orf19.3477 | 9354 | 7428 | 8690 | 10579 | 8942 | 8989 | 9557 | 5114 | 8392 | 9457 | 8891 | 8724 | 7647 | 8577 | 8528 | 7213 | 8428,5 | 8866 | 7991 | 0,902 | -0,149 | -2,377 | 0,01744346 | 0,03291467 | 0,00464 | 0,0021518 | 0,0036528 | 0,0036528 | TRUE | 0,0517495 | No |
| orf19.3478 | 7881 | 5514 | 6740 | 8381 | 8103 | 7344 | 7716 | 4105 | 7070 | 7020 | 6896 | 6911 | 6930 | 7007 | 6885 | 5790 | 6813,6 | 6974 | 6653 | 0,955 | -0,067 | -1,137 | 0,25539031 | 0,33804472 | 0,003633 | 0,0021926 | 0,003178 | 0,003178 | TRUE | 0,0913295 | No |
| orf19.348 | 1176 | 774 | 1060 | 1298 | 1110 | 934 | 1107 | 631 | 1055 | 985 | 1084 | 1070 | 949 | 891 | 988 | 890 | 989,19 | 1048 | 930 | 0,887 | -0,173 | -3,017 | 0,0025557 | 0,00589429 | 0,001037 | 0,0034471 | 0,0021549 | 0,0021549 | TRUE | 0,0213096 | No |
| orf19.3480 | 3057 | 2537 | 2684 | 3571 | 3799 | 3650 | 3723 | 2551 | 2742 | 3230 | 2746 | 2945 | 3249 | 3483 | 3322 | 3598 | 3164,4 | 2916 | 3413 | 1,171 | 0,227 | 3,776 | 0,00015906 | 0,0004703 | 0,003742 | 0,0024383 | 0,0031603 | 0,0031603 | TRUE | 0,0620515 | No |
| orf19.3481 | 7168 | 5365 | 6367 | 7925 | 6129 | 5967 | 6303 | 4220 | 6430 | 6830 | 6514 | 6535 | 5242 | 5694 | 5624 | 5952 | 6102,7 | 6577 | 5628 | 0,855 | -0,226 | -5,038 | 4,71E-07 | 2,09E-06 | 0,001492 | 0,0022174 | 0,0017602 | 0,0017602 | TRUE | 0,0262071 | No |
| orf19.3482 | 3081 | 2602 | 2848 | 3582 | 3365 | 3394 | 3530 | 2048 | 2764 | 3313 | 2914 | 2954 | 2878 | 3238 | 3150 | 2889 | 3012,4 | 2986 | 3039 | 1,019 | 0,027 | 0,422 | 0,67276281 | 0,73893217 | 0,003889 | 0,0024614 | 0,0035824 | 0,0035824 | TRUE | 0,0643607 | No |
| orf19.3483 | 8969 | 8123 | 8364 | 10663 | 10181 | 11277 | 11144 | 6422 | 8046 | 10341 | 8557 | 8793 | 8707 | 10760 | 9944 | 9058 | 9275,8 | 8934 | 9617 | 1,077 | 0,107 | 1,281 | 0,20026027 | 0,27475764 | 0,010547 | 0,002136 | 0,0065586 | 0,0065586 | TRUE | 0,136734 | No |
| orf19.3487 | 662 | 505 | 549 | 742 | 614 | 489 | 585 | 350 | 594 | 643 | 562 | 612 | 525 | 467 | 522 | 494 | 552,21 | 603 | 502 | 0,835 | -0,26 | -3,691 | 0,00022312 | 0,00064478 | 0,001212 | 0,0046084 | 0,0029398 | 0,0029398 | TRUE | 0,0282204 | No |
| orf19.349 | 1274 | 1000 | 1186 | 1512 | 1427 | 1304 | 1405 | 1001 | 1143 | 1273 | 1213 | 1247 | 1220 | 1244 | 1254 | 1412 | 1250,8 | 1219 | 1282 | 1,05 | 0,07 | 1,169 | 0,24250612 | 0,32413276 | 0,002318 | 0,0031401 | 0,0026788 | 0,0026788 | TRUE | 0,0507886 | No |
| orf19.3490 | 31 | 24 | 25 | 28 | 39 | 28 | 29 | 18 | 28 | 31 | 26 | 23 | 33 | 27 | 26 | 25 | 27,3 | 27 | 28 | 1,056 | 0,078 | 0,302 | 0,76288974 | 0,81497894 | 1,00E-08 | 0,0551634 | 0,0282805 | 0,0282805 | TRUE | 0,1316878 | No |
| orf19.3494 | 53 | 39 | 62 | 75 | 75 | 69 | 63 | 26 | 48 | 50 | 63 | 62 | 64 | 66 | 56 | 37 | 55,67 | 56 | 56 | 1,015 | 0,021 | 0,102 | 0,91882435 | 0,94026515 | 0,017782 | 0,0280574 | 0,0241231 | 0,0241231 | TRUE | 0,3308357 | No |

| | | | | | | | | | | | | | | | | | | | | | | | | | | | | | | | |
|---|---|---|---|---|---|---|---|---|---|---|---|---|---|---|---|---|---|---|---|---|---|---|---|---|---|---|---|---|---|---|---|
| orf19.3496 | 2468 | 1706 | 2282 | 2552 | 2746 | 2782 | 2770 | 1811 | 2214 | 2172 | 2335 | 2104 | 2348 | 2654 | 2472 | 2554 | 2356,8 | 2206 | 2507 | 1,136 | 0,184 | 3,522 | 0,00042884 | 0,00117325 | 0,00173 | 0,0025955 | 0,0021916 | 0,0021916 | TRUE | 0,0226609 | No |
| orf19.3498 | 305 | 204 | 280 | 367 | 339 | 323 | 312 | 221 | 274 | 260 | 286 | 303 | 290 | 308 | 278 | 312 | 288,83 | 281 | 297 | 1,052 | 0,073 | 0,829 | 0,40712314 | 0,49524111 | 1,00E-08 | 0,0070056 | 0,0039389 | 0,0039389 | TRUE | 0,0332641 | No |
| orf19.3499 | 2471 | 1768 | 2057 | 2737 | 2073 | 1863 | 1968 | 1327 | 2217 | 2251 | 2105 | 2257 | 1773 | 1778 | 1756 | 1872 | 2000,9 | 2208 | 1795 | 0,812 | -0,301 | -6,952 | 3,60E-12 | 3,09E-11 | 0,000364 | 0,002705 | 0,0012915 | 0,0012915 | TRUE | 0,0117892 | No |
| orf19.35 | 12431 | 10646 | 11889 | 14474 | 18923 | 20636 | 21636 | 11880 | 11152 | 13553 | 12164 | 11935 | 16184 | 19690 | 19307 | 16756 | 15093 | 12201 | 17984 | 1,474 | 0,56 | 7,385 | 1,52E-13 | 1,51E-12 | 0,008118 | 0,0020757 | 0,005456 | 0,005456 | TRUE | 0,0678416 | No |
| orf19.350 | 4210 | 3008 | 3668 | 4567 | 4484 | 3876 | 4153 | 2598 | 3777 | 3830 | 3753 | 3766 | 3835 | 3698 | 3706 | 3664 | 3753,6 | 3782 | 3726 | 0,986 | -0,02 | -0,615 | 0,53838939 | 0,62061464 | 1,00E-08 | 0,0023663 | 0,0007492 | 0,0007492 | TRUE | 0,0048618 | No |
| orf19.3501 | 1519 | 1089 | 1505 | 1662 | 1665 | 1511 | 1653 | 1052 | 1363 | 1386 | 1540 | 1371 | 1424 | 1442 | 1475 | 1484 | 1435,5 | 1415 | 1456 | 1,029 | 0,041 | 0,785 | 0,43245882 | 0,51964251 | 0,001202 | 0,0029908 | 0,0019495 | 0,0019495 | TRUE | 0,0425034 | No |
| orf19.3503 | 148 | 76 | 83 | 164 | 244 | 146 | 149 | 96 | 133 | 97 | 85 | 135 | 209 | 139 | 133 | 135 | 133,25 | 112 | 154 | 1,368 | 0,452 | 2,348 | 0,0188527 | 0,03525273 | 0,043319 | 0,0128739 | 0,0278562 | 0,0278562 | TRUE | 0,6273426 | No |
| orf19.3504 | 17079 | 11396 | 13611 | 17552 | 18030 | 14785 | 15592 | 9821 | 15322 | 14508 | 13925 | 14474 | 15420 | 14107 | 13913 | 13852 | 14440 | 14557 | 14323 | 0,984 | -0,023 | -0,497 | 0,61941862 | 0,6924948 | 0,001808 | 0,0020801 | 0,0020312 | 0,0020312 | TRUE | 0,0326806 | No |
| orf19.3505 | 6275 | 4392 | 5556 | 6685 | 8757 | 7587 | 8261 | 5476 | 5629 | 5591 | 5684 | 5513 | 7489 | 7239 | 7372 | 7723 | 6530,2 | 5604 | 7456 | 1,33 | 0,411 | 12,666 | 9,15E-37 | 5,14E-35 | 0,000254 | 0,0022018 | 0,0008549 | 0,0008549 | TRUE | 0,0067489 | No |
| orf19.3506 | 215 | 162 | 190 | 244 | 242 | 216 | 199 | 119 | 193 | 206 | 194 | 201 | 207 | 206 | 178 | 168 | 194,15 | 198 | 190 | 0,961 | -0,057 | -0,538 | 0,59058491 | 0,66821735 | 1,00E-08 | 0,0094568 | 0,005632 | 0,005632 | TRUE | 0,0510862 | No |
| orf19.3507 | 3166 | 1998 | 2528 | 3204 | 2570 | 2105 | 2141 | 1403 | 2840 | 2544 | 2586 | 2642 | 2198 | 2009 | 1910 | 1979 | 2338,5 | 2653 | 2024 | 0,763 | -0,39 | -6,928 | 4,27E-12 | 3,63E-11 | 0,002488 | 0,0026003 | 0,002612 | 0,002612 | TRUE | 0,041852 | No |
| orf19.3508 | 503 | 330 | 404 | 508 | 482 | 405 | 414 | 306 | 451 | 420 | 413 | 419 | 412 | 386 | 369 | 432 | 412,91 | 426 | 400 | 0,935 | -0,097 | -1,268 | 0,2047125 | 0,28029004 | 1,00E-08 | 0,0054953 | 0,0032105 | 0,0032105 | TRUE | 0,030514 | No |
| orf19.351 | 207 | 136 | 182 | 197 | 268 | 206 | 218 | 123 | 186 | 173 | 186 | 162 | 229 | 197 | 195 | 173 | 187,66 | 177 | 198 | 1,13 | 0,177 | 1,56 | 0,11887498 | 0,17686995 | 0,003748 | 0,0097154 | 0,0070185 | 0,0070185 | TRUE | 0,1215961 | No |
| orf19.35.1 | 17 | 8 | 11 | 16 | 15 | 5 | 8 | 4 | 15 | 10 | 11 | 13 | 13 | 5 | 7 | 6 | 10,03 | 12 | 8 | 0,618 | -0,694 | -1,532 | 0,12562109 | NA | 1,00E-08 | 0,1466737 | 0,0933587 | 0,0933587 | TRUE | 0,7261609 | No |
| orf19.3512 | 417 | 273 | 380 | 438 | 350 | 306 | 334 | 197 | 374 | 348 | 389 | 361 | 299 | 292 | 298 | 278 | 329,85 | 368 | 292 | 0,795 | -0,332 | -4,172 | 3,02E-05 | 0,00010156 | 1,00E-08 | 0,0063806 | 0,003014 | 0,003014 | TRUE | 0,0155777 | No |
| orf19.3515 | 1384 | 889 | 1149 | 1437 | 1651 | 1467 | 1567 | 904 | 1242 | 1132 | 1176 | 1185 | 1412 | 1400 | 1398 | 1275 | 1277,4 | 1184 | 1371 | 1,16 | 0,214 | 4,073 | 4,64E-05 | 0,00015131 | 0,000783 | 0,003116 | 0,0018637 | 0,0018637 | TRUE | 0,0239368 | No |
| orf19.3516 | 1674 | 1065 | 1419 | 1769 | 1382 | 1080 | 1140 | 681 | 1502 | 1356 | 1452 | 1459 | 1182 | 1031 | 1017 | 960 | 1244,8 | 1442 | 1048 | 0,728 | -0,459 | -6,808 | 9,91E-12 | 8,12E-11 | 0,00389 | 0,0031457 | 0,0035335 | 0,0035335 | TRUE | 0,0909485 | No |
| orf19.3517 | 1199 | 773 | 1000 | 1266 | 1195 | 1111 | 1155 | 613 | 1076 | 984 | 1023 | 1044 | 1022 | 1060 | 1031 | 865 | 1013 | 1032 | 994 | 0,966 | -0,05 | -0,74 | 0,45899922 | 0,54467061 | 0,003236 | 0,0034126 | 0,0033399 | 0,0033399 | TRUE | 0,0830588 | No |
| orf19.3518 | 3181 | 1998 | 2628 | 3308 | 2873 | 2570 | 2657 | 1717 | 2854 | 2544 | 2689 | 2728 | 2457 | 2452 | 2371 | 2422 | 2564,5 | 2704 | 2426 | 0,896 | -0,158 | -3,579 | 0,00034473 | 0,00096199 | 0,000727 | 0,0025456 | 0,0014818 | 0,0014818 | TRUE | 0,0182204 | No |
| orf19.3519 | 523 | 336 | 433 | 523 | 534 | 482 | 539 | 362 | 469 | 428 | 443 | 431 | 457 | 460 | 481 | 511 | 459,92 | 443 | 477 | 1,072 | 0,101 | 1,414 | 0,1573294 | 0,22436321 | 1,00E-08 | 0,0051359 | 0,0027017 | 0,0027017 | TRUE | 0,0218958 | No |
| orf19.352 | 1596 | 1292 | 1504 | 1827 | 1831 | 1739 | 1804 | 1169 | 1432 | 1645 | 1539 | 1507 | 1566 | 1659 | 1610 | 1649 | 1575,7 | 1531 | 1621 | 1,06 | 0,084 | 1,625 | 0,10420012 | 0,15781052 | 0,001165 | 0,0029008 | 0,0019577 | 0,0019577 | TRUE | 0,0286766 | No |
| orf19.3521 | 451 | 313 | 409 | 531 | 451 | 390 | 401 | 286 | 405 | 398 | 418 | 438 | 386 | 372 | 358 | 403 | 397,3 | 415 | 380 | 0,911 | -0,135 | -1,796 | 0,07243865 | 0,11608733 | 1,00E-08 | 0,0056334 | 0,0028946 | 0,0028946 | TRUE | 0,0173024 | No |
| orf19.3522 | 4 | 1 | 3 | 2 | 2 | 1 | 2 | 1 | 4 | 1 | 3 | 2 | 2 | 1 | 2 | 1 | 1,93 | 2 | 2 | 0,608 | -0,717 | -0,74 | 0,45929958 | NA | 1,00E-08 | 0,7541682 | 0,35919 | 0,35919 | TRUE | 0,1355494 | No |
| orf19.3523 | 2293 | 1452 | 1923 | 2301 | 2441 | 2073 | 2270 | 1392 | 2057 | 1849 | 1967 | 1897 | 2088 | 1978 | 2026 | 1963 | 1978,1 | 1942 | 2014 | 1,037 | 0,052 | 1,096 | 0,27307062 | 0,35735314 | 0,000945 | 0,0027134 | 0,0016391 | 0,0016391 | TRUE | 0,0192614 | No |
| orf19.3526 | 6031 | 4270 | 5495 | 6810 | 8574 | 7359 | 8023 | 5125 | 5410 | 5436 | 5622 | 5616 | 7333 | 7022 | 7159 | 7228 | 6353,3 | 5521 | 7186 | 1,301 | 0,38 | 12,043 | 2,11E-33 | 9,46E-32 | 0,00023 | 0,002208 | 0,0007942 | 0,0007942 | TRUE | 0,0029182 | No |
| orf19.3527 | 3993 | 2564 | 3507 | 4236 | 4427 | 3683 | 3989 | 2647 | 3582 | 3264 | 3588 | 3493 | 3786 | 3514 | 3560 | 3733 | 3565,1 | 3482 | 3648 | 1,047 | 0,066 | 1,457 | 0,14521593 | 0,20977574 | 0,001224 | 0,0023867 | 0,0016865 | 0,0016865 | TRUE | 0,0205948 | No |
| orf19.3528 | 880 | 671 | 715 | 890 | 746 | 695 | 738 | 402 | 789 | 854 | 732 | 734 | 638 | 663 | 659 | 567 | 704,48 | 777 | 632 | 0,817 | -0,291 | -3,982 | 6,83E-05 | 0,00021536 | 0,003339 | 0,0040402 | 0,0036981 | 0,0036981 | TRUE | 0,0489553 | No |
| orf19.3529 | 256 | 195 | 241 | 254 | 231 | 221 | 249 | 152 | 230 | 248 | 247 | 209 | 198 | 211 | 222 | 214 | 222,37 | 234 | 211 | 0,908 | -0,14 | -1,406 | 0,15963042 | 0,22731917 | 1,00E-08 | 0,0085079 | 0,0049587 | 0,0049587 | TRUE | 0,0573809 | No |
| orf19.353 | 1602 | 1075 | 1257 | 1695 | 1340 | 1123 | 1271 | 736 | 1437 | 1369 | 1286 | 1398 | 1146 | 1072 | 1134 | 1038 | 1234,9 | 1372 | 1098 | 0,801 | -0,32 | -5,772 | 7,81E-09 | 4,40E-08 | 0,001352 | 0,0031551 | 0,0021353 | 0,0021353 | TRUE | 0,0217809 | No |
| orf19.3530 | 3185 | 2603 | 2894 | 3753 | 3933 | 4027 | 4170 | 2384 | 2857 | 3314 | 2961 | 3095 | 3364 | 3842 | 3721 | 3362 | 3314,5 | 3057 | 3572 | 1,17 | 0,226 | 3,639 | 0,00027343 | 0,00077745 | 0,00406 | 0,0024175 | 0,0034105 | 0,0034105 | TRUE | 0,0380012 | No |
| orf19.3531 | 588 | 476 | 535 | 683 | 538 | 499 | 568 | 339 | 527 | 606 | 547 | 563 | 460 | 476 | 507 | 478 | 520,66 | 561 | 480 | 0,858 | -0,22 | -3,151 | 0,0016289 | 0,00391302 | 1,09E-05 | 0,0047677 | 0,0027658 | 0,0027658 | TRUE | 0,0304367 | No |
| orf19.3532 | 2177 | 1529 | 1855 | 2268 | 2157 | 1954 | 2084 | 1352 | 1953 | 1947 | 1898 | 1870 | 1845 | 1864 | 1860 | 1907 | 1892,9 | 1917 | 1869 | 0,975 | -0,037 | -0,939 | 0,34788132 | 0,4352501 | 1,00E-08 | 0,0027464 | 0,0009689 | 0,0009689 | TRUE | 0,0034643 | No |
| orf19.3533 | 1174 | 785 | 919 | 1129 | 940 | 753 | 846 | 540 | 1053 | 999 | 940 | 931 | 804 | 718 | 755 | 762 | 870,34 | 981 | 760 | 0,775 | -0,368 | -5,828 | 5,60E-09 | 3,23E-08 | 0,001852 | 0,0036475 | 0,0026476 | 0,0026476 | TRUE | 0,0312258 | No |
| orf19.3534 | 612 | 465 | 541 | 736 | 703 | 623 | 686 | 421 | 549 | 592 | 553 | 607 | 601 | 594 | 612 | 594 | 587,88 | 575 | 600 | 1,044 | 0,062 | 0,987 | 0,3237796 | 0,41072728 | 1,00E-08 | 0,0044489 | 0,0021518 | 0,0021518 | TRUE | 0,0184802 | No |
| orf19.3535 | 3724 | 2223 | 3150 | 3828 | 4274 | 3367 | 3892 | 2412 | 3341 | 2830 | 3223 | 3157 | 3655 | 3213 | 3473 | 3402 | 3286,6 | 3138 | 3436 | 1,094 | 0,13 | 2,184 | 0,02897871 | 0,05197137 | 0,003441 | 0,0024212 | 0,0031069 | 0,0031069 | TRUE | 0,0514032 | No |
| orf19.3536 | 1309 | 989 | 1200 | 1523 | 1091 | 1031 | 1148 | 714 | 1174 | 1259 | 1228 | 1256 | 933 | 984 | 1024 | 1007 | 1108,2 | 1229 | 987 | 0,803 | -0,317 | -6,034 | 1,60E-09 | 9,86E-09 | 0,000534 | 0,0032895 | 0,0017395 | 0,0017395 | TRUE | 0,017329 | No |
| orf19.3537 | 2716 | 1670 | 2095 | 2757 | 2762 | 2208 | 2140 | 1683 | 2437 | 2126 | 2143 | 2273 | 2362 | 2107 | 1910 | 2374 | 2216,5 | 2245 | 2188 | 0,973 | -0,039 | -0,516 | 0,60615573 | 0,68159903 | 0,006875 | 0,0026345 | 0,0050085 | 0,0050085 | TRUE | 0,0897599 | No |
| orf19.3538 | 5603 | 3743 | 4809 | 6046 | 5906 | 5193 | 5902 | 3730 | 5026 | 4765 | 4920 | 4986 | 5051 | 4955 | 5267 | 5261 | 5028,9 | 4924 | 5134 | 1,041 | 0,059 | 1,608 | 0,10774595 | 0,16244358 | 0,000465 | 0,0022682 | 0,001076 | 0,001076 | TRUE | 0,0066351 | No |
| orf19.3539 | 2671 | 1769 | 2321 | 2810 | 2769 | 2420 | 2535 | 1680 | 2396 | 2252 | 2375 | 2317 | 2368 | 2309 | 2262 | 2369 | 2331,1 | 2335 | 2327 | 0,995 | -0,007 | -0,167 | 0,86771297 | 0,89883095 | 0,000166 | 0,0026023 | 0,0010883 | 0,0010883 | TRUE | 0,006343 | No |
| orf19.354 | 949 | 636 | 881 | 1093 | 1130 | 966 | 1085 | 695 | 851 | 810 | 901 | 901 | 966 | 922 | 968 | 980 | 912,53 | 866 | 959 | 1,105 | 0,144 | 2,568 | 0,01023345 | 0,02056272 | 0,000354 | 0,0035704 | 0,0019084 | 0,0019084 | TRUE | 0,020308 | No |
| orf19.3540 | 8951 | 5754 | 7471 | 9185 | 7946 | 6403 | 6933 | 4422 | 8030 | 7325 | 7644 | 7574 | 6796 | 6110 | 6187 | 6237 | 6987,7 | 7643 | 6332 | 0,828 | -0,271 | -5,851 | 4,90E-09 | 2,86E-08 | 0,001761 | 0,0021873 | 0,0019227 | 0,0019227 | TRUE | 0,030103 | No |
| orf19.3541 | 11266 | 7774 | 9706 | 12194 | 11370 | 9837 | 10769 | 6355 | 10107 | 9897 | 9930 | 10055 | 9724 | 9386 | 9610 | 8963 | 9709 | 9997 | 9421 | 0,943 | -0,085 | -2,495 | 0,01261161 | 0,02473737 | 0,000578 | 0,002129 | 0,0010109 | 0,0010109 | TRUE | 0,0126014 | No |
| orf19.3542 | 5953 | 3969 | 5216 | 6210 | 6455 | 5412 | 5729 | 3709 | 5340 | 5053 | 5336 | 5121 | 5521 | 5164 | 5112 | 5231 | 5234,8 | 5212 | 5257 | 1,008 | 0,012 | 0,309 | 0,75714221 | 0,81004889 | 0,000688 | 0,0022568 | 0,0012865 | 0,0012865 | TRUE | 0,0143134 | No |
| orf19.3543 | 0 | 3 | 0 | 1 | 2 | 0 | 1 | 2 | 0 | 4 | 0 | 1 | 2 | 0 | 1 | 3 | 1,26 | 1 | 2 | 1,213 | 0,279 | 0,195 | 0,84518967 | NA | 0,994192 | 1,155548 | 1,1057566 | 1,1057566 | TRUE | 0,8118057 | No |
| orf19.3544 | 335 | 209 | 296 | 317 | 292 | 317 | 319 | 196 | 301 | 266 | 303 | 261 | 250 | 302 | 285 | 276 | 280,52 | 283 | 278 | 0,983 | -0,024 | -0,254 | 0,79932679 | 0,84445236 | 0,003129 | 0,0071546 | 0,0052481 | 0,0052481 | TRUE | 0,0600298 | No |
| orf19.3545 | 1274 | 905 | 1302 | 1365 | 1592 | 1480 | 1473 | 975 | 1143 | 1152 | 1332 | 1126 | 1362 | 1412 | 1314 | 1375 | 1277 | 1188 | 1366 | 1,15 | 0,201 | 3,235 | 0,00121836 | 0,00301976 | 0,00269 | 0,0031163 | 0,002928 | 0,002928 | TRUE | 0,0797246 | No |
| orf19.3546 | 2245 | 1491 | 1905 | 2449 | 2610 | 2213 | 2320 | 1375 | 2014 | 1898 | 1949 | 2019 | 2232 | 2112 | 2070 | 1939 | 2029,2 | 1970 | 2088 | 1,061 | 0,085 | 1,676 | 0,09370984 | 0,14506795 | 0,001443 | 0,0026949 | 0,001997 | 0,001997 | TRUE | 0,0260256 | No |
| orf19.3547 | 16104 | 11205 | 14063 | 16948 | 14660 | 13512 | 14167 | 8585 | 14447 | 14265 | 14388 | 13976 | 12538 | 12893 | 12642 | 12108 | 13407 | 14269 | 12545 | 0,88 | -0,185 | -6,094 | 1,10E-09 | 6,97E-09 | 0,000348 | 0,0020878 | 0,0008112 | 0,0008112 | TRUE | 0,0064738 | No |
| orf19.3548 | 2867 | 1898 | 2406 | 2959 | 2531 | 2099 | 2335 | 1413 | 2572 | 2416 | 2462 | 2440 | 2165 | 2003 | 2084 | 1993 | 2266,7 | 2472 | 2061 | 0,834 | -0,261 | -5,929 | 3,04E-09 | 1,81E-08 | 0,000692 | 0,00262 | 0,0014203 | 0,0014203 | TRUE | 0,0141344 | No |
| orf19.3548.1 | 70943 | 44461 | 50950 | 69735 | 82538 | 59849 | 60697 | 46945 | 63643 | 56604 | 52127 | 57504 | 70589 | 57106 | 54162 | 66212 | 59743 | 57470 | 62017 | 1,079 | 0,11 | 1,298 | 0,19439647 | 0,26787477 | 0,011045 | 0,0020038 | 0,0068667 | 0,0068667 | TRUE | 0,1061949 | No |
| orf19.3549 | 1629 | 1007 | 1421 | 1771 | 1433 | 1191 | 1257 | 790 | 1461 | 1282 | 1454 | 1460 | 1226 | 1136 | 1122 | 1114 | 1281,9 | 1414 | 1150 | 0,812 | -0,3 | -5,076 | 3,86E-07 | 1,74E-06 | 0,002142 | 0,003112 | 0,0025662 | 0,0025662 | TRUE | 0,044804 | No |
| orf19.355 | 81 | 58 | 78 | 79 | 99 | 88 | 81 | 62 | 73 | 74 | 80 | 65 | 85 | 84 | 72 | 87 | 77,48 | 73 | 82 | 1,126 | 0,172 | 1,09 | 0,27581303 | 0,36015751 | 1,00E-08 | 0,0207169 | 0,0108689 | 0,0108689 | TRUE | 0,0651952 | No |
| orf19.3550 | 953 | 701 | 875 | 1010 | 1176 | 1035 | 1117 | 694 | 855 | 892 | 895 | 833 | 1006 | 988 | 997 | 979 | 930,54 | 869 | 992 | 1,145 | 0,195 | 3,783 | 0,00015524 | 0,0004599 | 1,00E-08 | 0,0035396 | 0,0014795 | 0,0014795 | TRUE | 0,0097271 | No |
| orf19.3551 | 210 | 141 | 147 | 204 | 311 | 243 | 257 | 140 | 188 | 180 | 150 | 168 | 266 | 232 | 229 | 197 | 201,39 | 172 | 231 | 1,356 | 0,439 | 3,753 | 0,00017472 | 0,00051429 | 0,006767 | 0,0091878 | 0,00809 | 0,00809 | TRUE | 0,1146669 | No |
| orf19.3552 | 2582 | 1960 | 2312 | 2798 | 2964 | 2904 | 3115 | 1866 | 2316 | 2495 | 2365 | 2307 | 2535 | 2771 | 2780 | 2632 | 2525,2 | 2371 | 2680 | 1,131 | 0,178 | 3,773 | 0,00016142 | 0,00047703 | 0,001103 | 0,0025544 | 0,0017397 | 0,0017397 | TRUE | 0,0169372 | No |
| orf19.3553 | 11997 | 8943 | 9866 | 12377 | 9901 | 9452 | 9631 | 5028 | 10763 | 11385 | 10094 | 10206 | 8468 | 9019 | 8594 | 7092 | 9452,5 | 10612 | 8293 | 0,782 | -0,355 | -4,993 | 5,96E-07 | 2,61E-06 | 0,006485 | 0,0021331 | 0,0047476 | 0,0047476 | TRUE | 0,1148896 | No |
| orf19.3554 | 3441 | 2695 | 3192 | 4008 | 3454 | 3385 | 3386 | 2089 | 3087 | 3431 | 3266 | 3305 | 2954 | 3230 | 3021 | 2946 | 3155 | 3272 | 3038 | 0,929 | -0,106 | -2,209 | 0,02714418 | 0,04915075 | 0,00147 | 0,0024396 | 0,0018823 | 0,0018823 | TRUE | 0,0221 | No |
| orf19.3555 | 6434 | 4667 | 5816 | 7169 | 10230 | 8880 | 9872 | 5584 | 5772 | 5942 | 5950 | 5912 | 8749 | 8473 | 8809 | 7876 | 7185,3 | 5894 | 8477 | 1,439 | 0,525 | 12,49 | 8,44E-36 | 4,45E-34 | 0,001173 | 0,0021816 | 0,0015548 | 0,0015548 | TRUE | 0,027064 | No |
| orf19.3556 | 3082 | 2671 | 3092 | 3647 | 3688 | 3937 | 4117 | 2720 | 2765 | 3400 | 3163 | 3007 | 3154 | 3757 | 3674 | 3836 | 3344,6 | 3084 | 3605 | 1,169 | 0,225 | 2,99 | 0,002794 | 0,00638508 | 0,007 | 0,0024136 | 0,0051593 | 0,0051593 | TRUE | 0,0871026 | No |

| | | | | | | | | | | | | | | | | | | | | | | | | | | | | | | | |
|---|---|---|---|---|---|---|---|---|---|---|---|---|---|---|---|---|---|---|---|---|---|---|---|---|---|---|---|---|---|---|---|
| orf19.3557 | 167 | 102 | 143 | 186 | 281 | 246 | 254 | 137 | 150 | 130 | 146 | 153 | 240 | 235 | 227 | 193 | 184,28 | 145 | 224 | 1,547 | 0,629 | 5,686 | 1,30E-08 | 7,11E-08 | 1,00E-08 | 0,009857 | 0,0061349 | 0,0061349 | TRUE | 0,075566 | No |
| orf19.3558 | 544 | 515 | 508 | 675 | 749 | 848 | 788 | 481 | 488 | 656 | 520 | 557 | 641 | 809 | 703 | 678 | 631,41 | 555 | 708 | 1,279 | 0,355 | 3,563 | 0,00036613 | 0,00101791 | 0,011654 | 0,0042787 | 0,0079235 | 0,0079235 | TRUE | 0,1648427 | No |
| orf19.3559 | 5722 | 4043 | 4858 | 6081 | 6055 | 5602 | 5562 | 3670 | 5133 | 5147 | 4970 | 5014 | 5178 | 5345 | 4963 | 5176 | 5116 | 5066 | 5166 | 1,02 | 0,028 | 0,797 | 0,42550991 | 0,51272834 | 0,000348 | 0,0022633 | 0,0009994 | 0,0009994 | TRUE | 0,0087578 | No |
| orf19.356 | 221 | 145 | 183 | 209 | 208 | 195 | 193 | 105 | 198 | 185 | 187 | 172 | 178 | 186 | 172 | 148 | 178,34 | 186 | 171 | 0,929 | -0,107 | -0,975 | 0,32957118 | 0,41685551 | 1,00E-08 | 0,0101197 | 0,005905 | 0,005905 | TRUE | 0,0679815 | No |
| orf19.3561 | 2890 | 2164 | 2801 | 3309 | 2966 | 2920 | 3228 | 1978 | 2593 | 2755 | 2866 | 2729 | 2537 | 2786 | 2880 | 2790 | 2741,9 | 2736 | 2748 | 1,005 | 0,007 | 0,127 | 0,89881664 | 0,92340807 | 0,002032 | 0,002509 | 0,0022122 | 0,0022122 | TRUE | 0,033689 | No |
| orf19.3563 | 2216 | 1522 | 1924 | 2541 | 3398 | 3019 | 3348 | 2390 | 1988 | 1938 | 1968 | 2095 | 2906 | 2881 | 2988 | 3371 | 2516,8 | 1997 | 3036 | 1,517 | 0,601 | 10,485 | 1,01E-25 | 2,74E-24 | 0,002847 | 0,0025563 | 0,0027402 | 0,0027402 | TRUE | 0,0641966 | No |
| orf19.3564 | 8138 | 5770 | 6945 | 8582 | 8036 | 7118 | 7665 | 4263 | 7301 | 7346 | 7105 | 7077 | 6873 | 6792 | 6840 | 6013 | 6918,2 | 7207 | 6630 | 0,921 | -0,119 | -2,5 | 0,01242509 | 0,0243954 | 0,001919 | 0,0021894 | 0,002041 | 0,002041 | TRUE | 0,0465114 | No |
| orf19.3565 | 345 | 265 | 312 | 349 | 480 | 427 | 457 | 302 | 310 | 337 | 319 | 288 | 411 | 407 | 408 | 426 | 363,19 | 314 | 413 | 1,321 | 0,402 | 5,151 | 2,59E-07 | 1,20E-06 | 1,00E-08 | 0,0059766 | 0,0030417 | 0,0030417 | TRUE | 0,0366437 | No |
| orf19.3567 | 1471 | 1064 | 1286 | 1549 | 2209 | 2099 | 2057 | 1385 | 1320 | 1355 | 1316 | 1277 | 1889 | 2003 | 1836 | 1953 | 1618,5 | 1317 | 1920 | 1,459 | 0,545 | 11,65 | 2,28E-31 | 8,79E-30 | 0,000542 | 0,0028765 | 0,0014573 | 0,0014573 | TRUE | 0,0109182 | No |
| orf19.3568 | 1284 | 1010 | 1259 | 1628 | 1758 | 1674 | 1869 | 1115 | 1152 | 1286 | 1288 | 1342 | 1503 | 1597 | 1668 | 1573 | 1426,2 | 1267 | 1585 | 1,251 | 0,324 | 5,457 | 4,83E-08 | 2,46E-07 | 0,002423 | 0,0029974 | 0,0026631 | 0,0026631 | TRUE | 0,0467594 | No |
| orf19.3569 | 2469 | 1745 | 2079 | 2550 | 2642 | 2606 | 2672 | 1556 | 2215 | 2222 | 2127 | 2103 | 2260 | 2487 | 2384 | 2195 | 2248,9 | 2167 | 2332 | 1,078 | 0,108 | 2,175 | 0,02959698 | 0,05292222 | 0,001353 | 0,002625 | 0,0019138 | 0,0019138 | TRUE | 0,0240541 | No |
| orf19.357 | 1787 | 1121 | 1517 | 1945 | 1917 | 1532 | 1748 | 1066 | 1603 | 1427 | 1552 | 1604 | 1639 | 1462 | 1560 | 1503 | 1543,8 | 1546 | 1541 | 0,996 | -0,006 | -0,107 | 0,91500914 | 0,93746461 | 0,002024 | 0,0029198 | 0,0024007 | 0,0024007 | TRUE | 0,0307139 | No |
| orf19.3571 | 153 | 136 | 139 | 203 | 255 | 224 | 222 | 180 | 137 | 173 | 142 | 167 | 218 | 214 | 198 | 254 | 187,97 | 155 | 221 | 1,42 | 0,506 | 4,265 | 2,00E-05 | 6,94E-05 | 0,005832 | 0,0097024 | 0,0080269 | 0,0080269 | TRUE | 0,0940659 | No |
| orf19.3572 | 229 | 186 | 204 | 285 | 322 | 376 | 347 | 205 | 205 | 237 | 209 | 235 | 275 | 359 | 310 | 289 | 264,86 | 222 | 308 | 1,395 | 0,48 | 4,502 | 6,75E-06 | 2,53E-05 | 0,00661 | 0,0074605 | 0,007043 | 0,007043 | TRUE | 0,1407409 | No |
| orf19.3572.3 | 11598 | 8413 | 9439 | 12178 | 11419 | 9967 | 10117 | 6395 | 10405 | 10711 | 9657 | 10042 | 9766 | 9510 | 9028 | 9020 | 9767,2 | 10204 | 9331 | 0,915 | -0,128 | -2,867 | 0,00414963 | 0,00910888 | 0,001645 | 0,0021282 | 0,0018241 | 0,0018241 | TRUE | 0,0157822 | No |
| orf19.3573 | 1472 | 1066 | 1381 | 1719 | 1537 | 1469 | 1540 | 916 | 1321 | 1357 | 1413 | 1418 | 1314 | 1402 | 1374 | 1292 | 1361,3 | 1377 | 1346 | 0,978 | -0,032 | -0,647 | 0,51734407 | 0,60096736 | 0,000565 | 0,0030459 | 0,0016166 | 0,0016166 | TRUE | 0,0098357 | No |
| orf19.3574 | 774 | 524 | 687 | 811 | 628 | 577 | 589 | 411 | 694 | 667 | 703 | 669 | 537 | 551 | 526 | 580 | 615,75 | 683 | 548 | 0,799 | -0,323 | -5,253 | 1,49E-07 | 7,14E-07 | 1,00E-08 | 0,0043371 | 0,0019825 | 0,0019825 | TRUE | 0,0145853 | No |
| orf19.3575 | 3579 | 2410 | 3252 | 3829 | 4633 | 3803 | 4074 | 2877 | 3211 | 3068 | 3327 | 3157 | 3962 | 3629 | 3635 | 4058 | 3506 | 3191 | 3821 | 1,196 | 0,259 | 5,171 | 2,33E-07 | 1,08E-06 | 0,001875 | 0,0023936 | 0,0021121 | 0,0021121 | TRUE | 0,0202206 | No |
| orf19.3577 | 1748 | 1164 | 1464 | 1865 | 1807 | 1458 | 1520 | 1020 | 1568 | 1482 | 1498 | 1538 | 1545 | 1391 | 1356 | 1439 | 1477,2 | 1522 | 1433 | 0,941 | -0,087 | -1,643 | 0,10037443 | 0,15321381 | 0,001393 | 0,0029623 | 0,002031 | 0,002031 | TRUE | 0,0360229 | No |
| orf19.3577.1 | 116 | 64 | 111 | 126 | 97 | 86 | 84 | 69 | 104 | 81 | 114 | 104 | 83 | 82 | 75 | 97 | 92,54 | 101 | 84 | 0,825 | -0,278 | -1,818 | 0,06911302 | 0,11153062 | 0,000942 | 0,0176673 | 0,0116063 | 0,0116063 | TRUE | 0,1335887 | No |
| orf19.3578 | 698 | 443 | 563 | 723 | 721 | 579 | 633 | 502 | 626 | 564 | 576 | 596 | 617 | 552 | 565 | 708 | 600,54 | 590 | 610 | 1,028 | 0,039 | 0,478 | 0,63296015 | 0,70422678 | 0,005442 | 0,0043968 | 0,0048911 | 0,0048911 | TRUE | 0,1225849 | No |
| orf19.3579 | 4484 | 3071 | 3886 | 4762 | 5120 | 4222 | 4582 | 3379 | 4023 | 3910 | 3976 | 3927 | 4379 | 4028 | 4089 | 4766 | 4137,1 | 3959 | 4316 | 1,089 | 0,123 | 2,261 | 0,02374068 | 0,04356567 | 0,002808 | 0,0023304 | 0,0025947 | 0,0025947 | TRUE | 0,0581973 | No |
| orf19.3581 | 611 | 440 | 472 | 610 | 466 | 455 | 487 | 309 | 548 | 560 | 483 | 503 | 399 | 434 | 435 | 436 | 474,66 | 524 | 426 | 0,813 | -0,298 | -3,977 | 6,97E-05 | 0,00021932 | 0,001434 | 0,0050379 | 0,0032615 | 0,0032615 | TRUE | 0,0301477 | No |
| orf19.3582 | 1511 | 1053 | 1303 | 1560 | 1188 | 1011 | 1047 | 599 | 1356 | 1341 | 1333 | 1286 | 1016 | 965 | 934 | 845 | 1134,4 | 1329 | 940 | 0,71 | -0,494 | -8,319 | 8,86E-17 | 1,22E-15 | 0,00196 | 0,0032592 | 0,0024707 | 0,0024707 | TRUE | 0,0488269 | No |
| orf19.3583 | 3811 | 2649 | 3218 | 4129 | 3846 | 3414 | 3707 | 2183 | 3419 | 3372 | 3292 | 3405 | 3289 | 3258 | 3308 | 3079 | 3302,8 | 3372 | 3234 | 0,96 | -0,059 | -1,578 | 0,11461331 | 0,17112246 | 0,000265 | 0,0024191 | 0,0010371 | 0,0010371 | TRUE | 0,0114927 | No |
| orf19.3585 | 1108 | 865 | 890 | 1213 | 988 | 913 | 889 | 531 | 994 | 1101 | 911 | 1000 | 845 | 871 | 793 | 749 | 908,05 | 1002 | 814 | 0,816 | -0,293 | -4,119 | 3,81E-05 | 0,00012587 | 0,003785 | 0,0035782 | 0,0037402 | 0,0037402 | TRUE | 0,0500778 | No |
| orf19.3586 | 2872 | 1900 | 2682 | 3253 | 2153 | 1827 | 2092 | 1295 | 2576 | 2419 | 2744 | 2682 | 1841 | 1743 | 1867 | 1826 | 2212,5 | 2605 | 1819 | 0,698 | -0,519 | -10,352 | 4,11E-25 | 1,05E-23 | 0,001505 | 0,0026357 | 0,001946 | 0,001946 | TRUE | 0,0267766 | No |
| orf19.3589 | 127 | 95 | 123 | 138 | 99 | 78 | 90 | 59 | 114 | 121 | 126 | 114 | 85 | 74 | 80 | 83 | 99,64 | 119 | 80 | 0,681 | -0,555 | -4,026 | 5,67E-05 | 0,00018158 | 1,00E-08 | 0,0165488 | 0,0078397 | 0,0078397 | TRUE | 0,0262181 | No |
| orf19.359 | 40 | 28 | 30 | 34 | 28 | 18 | 31 | 22 | 36 | 36 | 31 | 28 | 24 | 17 | 28 | 31 | 28,76 | 33 | 25 | 0,76 | -0,396 | -1,494 | 0,13523742 | 0,19766305 | 1,00E-08 | 0,0524573 | 0,0318281 | 0,0318281 | TRUE | 0,2715076 | No |
| orf19.3590 | 2356 | 1549 | 1973 | 2361 | 2096 | 1812 | 1956 | 1446 | 2114 | 1972 | 2019 | 1947 | 1793 | 1729 | 1745 | 2039 | 1919,7 | 2013 | 1826 | 0,905 | -0,144 | -2,424 | 0,01534446 | 0,02942531 | 0,003097 | 0,0027357 | 0,0028389 | 0,0028389 | TRUE | 0,0698576 | No |
| orf19.3591 | 2300 | 1682 | 2025 | 2571 | 1787 | 1663 | 1750 | 1123 | 2063 | 2141 | 2072 | 2120 | 1528 | 1587 | 1562 | 1584 | 1832,1 | 2099 | 1565 | 0,745 | -0,424 | -10,588 | 3,39E-26 | 9,78E-25 | 1,00E-08 | 0,0027719 | 0,000981 | 0,000981 | TRUE | 0,0031739 | No |
| orf19.3592 | 1157 | 876 | 1092 | 1267 | 1408 | 1407 | 1444 | 928 | 1038 | 1115 | 1117 | 1045 | 1204 | 1343 | 1289 | 1309 | 1182,4 | 1079 | 1286 | 1,193 | 0,254 | 4,642 | 3,44E-06 | 1,34E-05 | 0,00115 | 0,0032073 | 0,0020253 | 0,0020253 | TRUE | 0,0232157 | No |
| orf19.3593 | 6251 | 4073 | 5181 | 6478 | 6056 | 5226 | 5400 | 3536 | 5608 | 5185 | 5301 | 5342 | 5179 | 4986 | 4819 | 4987 | 5175,9 | 5359 | 4993 | 0,931 | -0,103 | -2,618 | 0,0088494 | 0,0180649 | 0,000734 | 0,00226 | 0,0012794 | 0,0012794 | TRUE | 0,0119735 | No |
| orf19.3594 | 601 | 414 | 479 | 638 | 632 | 565 | 551 | 363 | 539 | 527 | 490 | 526 | 541 | 539 | 492 | 512 | 520,71 | 520 | 521 | 1,001 | 0,001 | 0,016 | 0,98744246 | 0,98991395 | 1,00E-08 | 0,0047675 | 0,0025004 | 0,0025004 | TRUE | 0,018674 | No |
| orf19.3599 | 16331 | 11056 | 14410 | 17457 | 15362 | 13806 | 15036 | 9075 | 14651 | 14075 | 14743 | 14395 | 13138 | 13173 | 13417 | 12799 | 13799 | 14466 | 13132 | 0,908 | -0,139 | -4,706 | 2,52E-06 | 1,01E-05 | 0,000304 | 0,0020847 | 0,0007693 | 0,0007693 | TRUE | 0,0039377 | No |
| orf19.36 | 321 | 259 | 311 | 347 | 276 | 326 | 253 | 158 | 288 | 330 | 318 | 286 | 236 | 311 | 226 | 223 | 277,22 | 306 | 249 | 0,819 | -0,288 | -2,483 | 0,01301545 | 0,02544641 | 0,011797 | 0,0072162 | 0,0092887 | 0,0092887 | TRUE | 0,3179059 | No |
| orf19.360 | 3 | 0 | 3 | 3 | 3 | 3 | 0 | 5 | 3 | 0 | 3 | 2 | 3 | 3 | 0 | 7 | 2,59 | 2 | 3 | 1,379 | 0,464 | 0,47 | 0,63814913 | NA | 0,458007 | 0,5626253 | 0,5305425 | 0,5305425 | TRUE | 0,3081351 | No |
| orf19.3600 | 96 | 81 | 117 | 125 | 121 | 136 | 141 | 94 | 86 | 103 | 120 | 103 | 103 | 130 | 126 | 133 | 112,96 | 103 | 123 | 1,19 | 0,251 | 1,729 | 0,08383631 | 0,13175217 | 0,00682 | 0,0148311 | 0,0113668 | 0,0113668 | TRUE | 0,1288055 | No |
| orf19.3601 | 1120 | 998 | 1098 | 1391 | 1260 | 1346 | 1335 | 788 | 1005 | 1271 | 1123 | 1147 | 1078 | 1284 | 1191 | 1111 | 1151,3 | 1136 | 1166 | 1,028 | 0,04 | 0,506 | 0,61276571 | 0,68671822 | 0,00677 | 0,0032405 | 0,005161 | 0,005161 | TRUE | 0,0733045 | No |
| orf19.3603 | 1111 | 776 | 1051 | 1325 | 1338 | 1250 | 1356 | 858 | 997 | 988 | 1075 | 1093 | 1144 | 1193 | 1210 | 1210 | 1113,7 | 1038 | 1189 | 1,143 | 0,193 | 3,583 | 0,00033988 | 0,00095016 | 0,000768 | 0,003283 | 0,0018817 | 0,0018817 | TRUE | 0,0156673 | No |
| orf19.3604 | 929 | 669 | 836 | 1036 | 881 | 875 | 898 | 478 | 833 | 852 | 855 | 854 | 753 | 835 | 801 | 674 | 807,32 | 848 | 766 | 0,906 | -0,142 | -2,118 | 0,03413417 | 0,0601402 | 0,00256 | 0,0037777 | 0,0030915 | 0,0030915 | TRUE | 0,0673413 | No |
| orf19.3605 | 1614 | 1427 | 1439 | 1809 | 1349 | 1561 | 1492 | 820 | 1448 | 1817 | 1472 | 1492 | 1154 | 1489 | 1331 | 1157 | 1420 | 1557 | 1283 | 0,825 | -0,277 | -2,898 | 0,00375259 | 0,00832246 | 0,013161 | 0,0030019 | 0,0080543 | 0,0080543 | TRUE | 0,1486012 | No |
| orf19.3606 | 5910 | 3915 | 5237 | 6360 | 5398 | 4769 | 5342 | 3079 | 5302 | 4984 | 5358 | 5245 | 4617 | 4550 | 4767 | 4343 | 4895,6 | 5222 | 4569 | 0,875 | -0,192 | -4,664 | 3,10E-06 | 1,22E-05 | 0,000969 | 0,0022761 | 0,0014225 | 0,0014225 | TRUE | 0,0128744 | No |
| orf19.3607 | 351 | 360 | 329 | 460 | 538 | 587 | 578 | 318 | 315 | 458 | 337 | 379 | 460 | 560 | 516 | 449 | 434,2 | 372 | 496 | 1,339 | 0,422 | 3,63 | 0,00028297 | 0,00080305 | 0,016765 | 0,0053229 | 0,0105843 | 0,0105843 | TRUE | 0,2596196 | No |
| orf19.3608 | 737 | 697 | 728 | 899 | 611 | 727 | 782 | 413 | 661 | 887 | 745 | 741 | 523 | 694 | 698 | 582 | 691,4 | 758 | 624 | 0,825 | -0,278 | -2,542 | 0,01102538 | 0,02197032 | 0,015293 | 0,0040792 | 0,0100161 | 0,0100161 | TRUE | 0,1493497 | No |
| orf19.3609 | 8737 | 6870 | 8028 | 9866 | 8219 | 7901 | 8259 | 5478 | 7838 | 8746 | 8213 | 8136 | 7029 | 7539 | 7370 | 7726 | 7824,7 | 8233 | 7416 | 0,901 | -0,151 | -3,316 | 0,00091344 | 0,00231168 | 0,001682 | 0,0021651 | 0,0018599 | 0,0018599 | TRUE | 0,0211603 | No |
| orf19.36.1 | 249 | 131 | 204 | 255 | 196 | 118 | 120 | 91 | 223 | 167 | 209 | 210 | 168 | 113 | 107 | 128 | 165,6 | 202 | 129 | 0,636 | -0,653 | -4,268 | 1,98E-05 | 6,86E-05 | 0,02229 | 0,0107459 | 0,0160942 | 0,0160942 | TRUE | 0,4623215 | No |
| orf19.3610 | 1520 | 1140 | 1313 | 1638 | 1407 | 1484 | 1533 | 878 | 1364 | 1451 | 1343 | 1351 | 1203 | 1416 | 1368 | 1238 | 1341,8 | 1377 | 1306 | 0,95 | -0,074 | -1,183 | 0,23700534 | 0,31791205 | 0,0027 | 0,0030614 | 0,0029768 | 0,0029768 | TRUE | 0,0383687 | No |
| orf19.3611 | 2903 | 2062 | 2395 | 3133 | 3032 | 2681 | 2717 | 1710 | 2604 | 2625 | 2450 | 2584 | 2593 | 2558 | 2424 | 2412 | 2531,3 | 2566 | 2497 | 0,974 | -0,038 | -0,87 | 0,38427101 | 0,47319128 | 0,000627 | 0,002553 | 0,0014177 | 0,0014177 | TRUE | 0,0109946 | No |
| orf19.3612 | 1146 | 748 | 954 | 1160 | 896 | 862 | 847 | 598 | 1028 | 952 | 976 | 957 | 766 | 822 | 756 | 843 | 887,62 | 978 | 797 | 0,812 | -0,301 | -5,151 | 2,59E-07 | 1,20E-06 | 0,000821 | 0,003615 | 0,0021303 | 0,0021303 | TRUE | 0,0160183 | No |
| orf19.3613 | 3010 | 2357 | 2628 | 3358 | 2823 | 2887 | 3141 | 1666 | 2700 | 3001 | 2689 | 2769 | 2414 | 2755 | 2803 | 2350 | 2685 | 2790 | 2580 | 0,926 | -0,111 | -1,659 | 0,09707032 | 0,14904126 | 0,004876 | 0,0025202 | 0,0038858 | 0,0038858 | TRUE | 0,0421097 | No |
| orf19.3615 | 1435 | 1237 | 1297 | 1590 | 1254 | 1325 | 1316 | 844 | 1287 | 1575 | 1327 | 1311 | 1072 | 1264 | 1174 | 1190 | 1275,2 | 1375 | 1175 | 0,856 | -0,225 | -2,972 | 0,00295708 | 0,00672645 | 0,005967 | 0,0031179 | 0,0046914 | 0,0046914 | TRUE | 0,1110165 | No |
| orf19.3616 | 1578 | 1339 | 1496 | 1816 | 1593 | 1687 | 1678 | 937 | 1416 | 1705 | 1531 | 1497 | 1362 | 1610 | 1497 | 1322 | 1492,4 | 1537 | 1448 | 0,944 | -0,083 | -1,098 | 0,27228598 | 0,35670625 | 0,006368 | 0,0029522 | 0,0048231 | 0,0048231 | TRUE | 0,0678949 | No |
| orf19.3617 | 771 | 863 | 783 | 1073 | 823 | 1036 | 1074 | 595 | 692 | 1099 | 801 | 885 | 704 | 989 | 958 | 839 | 870,77 | 869 | 872 | 1,005 | 0,007 | 0,039 | 0,96908073 | 0,97799346 | 0,029389 | 0,0036467 | 0,016088 | 0,0293895 | TRUE | 0,3708172 | No |
| orf19.3618 | 2005 | 1359 | 1914 | 2242 | 2010 | 1675 | 1842 | 1420 | 1799 | 1730 | 1958 | 1849 | 1719 | 1598 | 1644 | 2003 | 1787,4 | 1834 | 1741 | 0,947 | -0,079 | -1,09 | 0,27572451 | 0,36015751 | 0,005864 | 0,0027917 | 0,0044246 | 0,0044246 | TRUE | 0,1182621 | No |
| orf19.362 | 79 | 52 | 50 | 58 | 70 | 55 | 57 | 44 | 71 | 66 | 51 | 48 | 60 | 52 | 51 | 62 | 57,67 | 59 | 56 | 0,954 | -0,067 | -0,346 | 0,72939696 | 0,78816851 | 1,00E-08 | 0,0271542 | 0,0190874 | 0,0190874 | TRUE | 0,1855737 | No |
| orf19.3621 | 510 | 397 | 485 | 624 | 747 | 688 | 755 | 504 | 458 | 505 | 496 | 515 | 639 | 656 | 674 | 711 | 581,7 | 494 | 670 | 1,355 | 0,438 | 6,533 | 6,45E-11 | 4,83E-10 | 1,00E-08 | 0,0044751 | 0,002562 | 0,002562 | TRUE | 0,0297673 | No |
| orf19.3622 | 872 | 662 | 844 | 969 | 1240 | 1275 | 1280 | 757 | 782 | 843 | 863 | 799 | 1060 | 1217 | 1142 | 1068 | 971,82 | 822 | 1122 | 1,368 | 0,452 | 7,14 | 9,33E-13 | 8,58E-12 | 0,002251 | 0,0034733 | 0,0027916 | 0,0027916 | TRUE | 0,0389122 | No |
| orf19.3623 | 1126 | 1028 | 1079 | 1334 | 1211 | 1152 | 1326 | 809 | 1010 | 1309 | 1104 | 1100 | 1036 | 1099 | 1183 | 1141 | 1122,8 | 1131 | 1115 | 0,987 | -0,018 | -0,23 | 0,81810805 | 0,8596231 | 0,006627 | 0,0032725 | 0,0050876 | 0,0050876 | TRUE | 0,1294003 | No |

| | | | | | | | | | | | | | | | | | | | | | | | | | | | | | | | |
|---|---|---|---|---|---|---|---|---|---|---|---|---|---|---|---|---|---|---|---|---|---|---|---|---|---|---|---|---|---|---|---|
| orf19.3624 | 3644 | 3214 | 3528 | 4509 | 4243 | 4595 | 4628 | 2674 | 3269 | 4092 | 3609 | 3718 | 3629 | 4384 | 4130 | 3771 | 3825,4 | 3672 | 3978 | 1,084 | 0,117 | 1,527 | 0,1266995 | 0,1869237 | 0,007625 | 0,002359 | 0,0053361 | 0,0053361 | TRUE | 0,0710504 | No |
| orf19.3625 | 419 | 339 | 337 | 433 | 492 | 493 | 521 | 265 | 376 | 432 | 345 | 357 | 421 | 470 | 465 | 374 | 404,89 | 378 | 432 | 1,154 | 0,207 | 2,157 | 0,03104211 | 0,05525956 | 0,007274 | 0,0055649 | 0,0063698 | 0,0063698 | TRUE | 0,0981115 | No |
| orf19.3626 | 1110 | 1079 | 1138 | 1455 | 1586 | 1742 | 1679 | 1029 | 996 | 1374 | 1164 | 1200 | 1356 | 1662 | 1498 | 1451 | 1337,7 | 1184 | 1492 | 1,263 | 0,337 | 3,642 | 0,00027011 | 0,0007691 | 0,011235 | 0,0030647 | 0,0074352 | 0,0074352 | TRUE | 0,1373754 | No |
| orf19.3626.1 | 39 | 31 | 30 | 49 | 46 | 61 | 51 | 24 | 35 | 39 | 31 | 40 | 39 | 58 | 46 | 34 | 40,31 | 36 | 44 | 1,227 | 0,295 | 1,271 | 0,20367744 | 0,27906363 | 0,010795 | 0,0379956 | 0,0267964 | 0,0267964 | TRUE | 0,3483485 | No |
| orf19.3627 | 1202 | 981 | 1044 | 1369 | 1186 | 1125 | 1240 | 647 | 1078 | 1249 | 1068 | 1129 | 1014 | 1073 | 1106 | 913 | 1078,9 | 1131 | 1026 | 0,911 | -0,134 | -1,846 | 0,06485156 | 0,10564755 | 0,004887 | 0,0033251 | 0,0041374 | 0,0041374 | TRUE | 0,0613961 | No |
| orf19.3628 | 5098 | 3755 | 4649 | 5617 | 6245 | 5797 | 6014 | 3660 | 4573 | 4781 | 4756 | 4632 | 5341 | 5531 | 5366 | 5162 | 5017,9 | 4686 | 5350 | 1,143 | 0,193 | 5,484 | 4,15E-08 | 2,13E-07 | 0,000356 | 0,0022688 | 0,0009869 | 0,0009869 | TRUE | 0,0063897 | No |
| orf19.3629 | 1402 | 1203 | 1356 | 1752 | 1706 | 1740 | 1741 | 1081 | 1258 | 1532 | 1387 | 1445 | 1459 | 1660 | 1554 | 1525 | 1477,4 | 1406 | 1550 | 1,104 | 0,143 | 2,126 | 0,03349045 | 0,05911005 | 0,0039 | 0,0029622 | 0,0036342 | 0,0036342 | TRUE | 0,0607952 | No |
| orf19.3630 | 8252 | 6585 | 7285 | 9366 | 7191 | 7372 | 7583 | 4734 | 7403 | 8383 | 7453 | 7723 | 6150 | 7034 | 6767 | 6677 | 7198,8 | 7740 | 6657 | 0,86 | -0,217 | -3,964 | 7,37E-05 | 0,00023047 | 0,003059 | 0,0021812 | 0,0027466 | 0,0027466 | TRUE | 0,0376734 | No |
| orf19.3631 | 215 | 192 | 213 | 248 | 229 | 247 | 267 | 165 | 193 | 244 | 218 | 205 | 196 | 236 | 238 | 233 | 220,28 | 215 | 226 | 1,053 | 0,075 | 0,684 | 0,49403713 | 0,57904312 | 0,004902 | 0,0085698 | 0,0068886 | 0,0068886 | TRUE | 0,0953299 | No |
| orf19.3633 | 337 | 254 | 280 | 384 | 360 | 366 | 352 | 228 | 302 | 323 | 286 | 317 | 308 | 349 | 314 | 322 | 315,2 | 307 | 323 | 1,052 | 0,074 | 0,875 | 0,38145153 | 0,47068408 | 1,00E-08 | 0,0065852 | 0,0036155 | 0,0036155 | TRUE | 0,0355335 | No |
| orf19.3635 | 1192 | 838 | 1143 | 1321 | 1546 | 1394 | 1479 | 931 | 1069 | 1067 | 1169 | 1089 | 1322 | 1330 | 1320 | 1313 | 1210 | 1098 | 1321 | 1,203 | 0,266 | 5,475 | 4,37E-08 | 2,23E-07 | 1,00E-08 | 0,0031793 | 0,0014371 | 0,0014371 | TRUE | 0,0219543 | No |
| orf19.3637 | 0 | 2 | 1 | 2 | 0 | 1 | 0 | 1 | 0 | 3 | 1 | 2 | 0 | 1 | 0 | 1 | 0,95 | 2 | 0 | 0,406 | -1,3 | -0,833 | 0,40492402 | NA | 1,00E-08 | 1,5334883 | 1,0455352 | 1,0455352 | TRUE | 0,1876382 | No |
| orf19.3638 | 20 | 17 | 26 | 25 | 41 | 42 | 42 | 23 | 18 | 22 | 27 | 21 | 35 | 40 | 37 | 32 | 28,98 | 22 | 36 | 1,688 | 0,756 | 2,942 | 0,00325752 | 0,00734103 | 1,00E-08 | 0,0520689 | 0,0267165 | 0,0267165 | TRUE | 0,132146 | No |
| orf19.3639 | 829 | 611 | 636 | 875 | 879 | 825 | 943 | 561 | 744 | 778 | 651 | 722 | 752 | 787 | 841 | 791 | 758,18 | 724 | 793 | 1,097 | 0,134 | 1,962 | 0,04974607 | 0,08376525 | 0,002449 | 0,0038942 | 0,0031374 | 0,0031374 | TRUE | 0,0524931 | No |
| orf19.364 | 1662 | 1132 | 1543 | 1853 | 1595 | 1317 | 1392 | 913 | 1491 | 1441 | 1579 | 1528 | 1364 | 1257 | 1242 | 1288 | 1398,7 | 1510 | 1288 | 0,853 | -0,23 | -4,486 | 7,26E-06 | 2,71E-05 | 0,000933 | 0,0030174 | 0,001807 | 0,001807 | TRUE | 0,0206972 | No |
| orf19.3642 | 8392 | 5679 | 7663 | 9362 | 9544 | 8061 | 8767 | 5478 | 7529 | 7230 | 7840 | 7720 | 8162 | 7692 | 7823 | 7726 | 7715,2 | 7580 | 7851 | 1,036 | 0,051 | 1,331 | 0,18334713 | 0,25552066 | 0,000818 | 0,0021677 | 0,0012579 | 0,0012579 | TRUE | 0,0114444 | No |
| orf19.3643 | 474 | 298 | 400 | 576 | 557 | 460 | 535 | 362 | 425 | 379 | 409 | 475 | 476 | 439 | 477 | 511 | 449,01 | 422 | 476 | 1,12 | 0,163 | 1,938 | 0,05266078 | 0,08798275 | 0,003866 | 0,0052126 | 0,0045504 | 0,0045504 | TRUE | 0,0847371 | No |
| orf19.3644 | 3816 | 2599 | 3508 | 4279 | 4364 | 3774 | 4106 | 2778 | 3423 | 3309 | 3589 | 3529 | 3732 | 3601 | 3664 | 3918 | 3595,6 | 3462 | 3729 | 1,075 | 0,105 | 2,466 | 0,01367798 | 0,02659461 | 0,000937 | 0,0023833 | 0,0014571 | 0,0014571 | TRUE | 0,0133623 | No |
| orf19.3646 | 43808 | 28841 | 36299 | 46586 | 41323 | 33937 | 35671 | 23957 | 39300 | 36718 | 37137 | 38415 | 35341 | 32382 | 31830 | 33789 | 35614 | 37892 | 33336 | 0,88 | -0,185 | -4,353 | 1,34E-05 | 4,79E-05 | 0,001561 | 0,0020203 | 0,0017061 | 0,0017061 | TRUE | 0,0201984 | No |
| orf19.3647 | 446 | 358 | 430 | 500 | 537 | 490 | 546 | 335 | 400 | 456 | 440 | 412 | 459 | 468 | 487 | 472 | 449,33 | 427 | 472 | 1,108 | 0,148 | 2,071 | 0,03840033 | 0,06675991 | 1,00E-08 | 0,0052104 | 0,0026879 | 0,0026879 | TRUE | 0,0213855 | No |
| orf19.3648 | 311 | 239 | 290 | 358 | 425 | 335 | 360 | 291 | 279 | 304 | 297 | 295 | 363 | 320 | 321 | 410 | 323,75 | 294 | 354 | 1,198 | 0,261 | 2,753 | 0,0059037 | 0,01255556 | 0,004441 | 0,0064636 | 0,0055112 | 0,0055112 | TRUE | 0,1150906 | No |
| orf19.3649 | 2215 | 1696 | 1944 | 2426 | 2262 | 2236 | 2263 | 1504 | 1987 | 2159 | 1989 | 2001 | 1935 | 2134 | 2019 | 2121 | 2043 | 2034 | 2052 | 1,009 | 0,013 | 0,259 | 0,79577485 | 0,84143177 | 0,001345 | 0,0026901 | 0,0018773 | 0,0018773 | TRUE | 0,0196539 | No |
| orf19.3651 | 13345 | 8497 | 11100 | 13659 | 20804 | 16377 | 17502 | 11579 | 11972 | 10818 | 11356 | 11263 | 17792 | 15626 | 15618 | 16331 | 13847 | 11352 | 16342 | 1,439 | 0,525 | 10,166 | 2,80E-24 | 6,73E-23 | 0,002737 | 0,0020844 | 0,0024911 | 0,0024911 | TRUE | 0,0439642 | No |
| orf19.3653 | 6093 | 4217 | 5442 | 6786 | 5454 | 4726 | 5023 | 3349 | 5466 | 5369 | 5568 | 5596 | 4664 | 4509 | 4482 | 4723 | 5047,2 | 5500 | 4594 | 0,835 | -0,261 | -7,716 | 1,20E-14 | 1,33E-13 | 0,000238 | 0,0022671 | 0,0008947 | 0,0008947 | TRUE | 0,0039951 | No |
| orf19.3655 | 47 | 31 | 46 | 51 | 41 | 44 | 39 | 29 | 42 | 39 | 47 | 42 | 35 | 42 | 35 | 41 | 40,44 | 42 | 38 | 0,888 | -0,172 | -0,811 | 0,41716872 | 0,50510877 | 1,00E-08 | 0,0378795 | 0,0183669 | 0,0183669 | TRUE | 0,0404189 | No |
| orf19.3656 | 3587 | 2417 | 3274 | 3675 | 4086 | 3935 | 4167 | 2923 | 3218 | 3077 | 3350 | 3030 | 3494 | 3755 | 3718 | 4123 | 3470,7 | 3169 | 3772 | 1,189 | 0,25 | 4,411 | 1,03E-05 | 3,74E-05 | 0,003032 | 0,0023978 | 0,0027944 | 0,0027944 | TRUE | 0,0460363 | No |
| orf19.3658 | 225 | 124 | 193 | 263 | 237 | 218 | 204 | 143 | 202 | 158 | 197 | 217 | 203 | 208 | 182 | 202 | 196,06 | 194 | 199 | 1,018 | 0,025 | 0,224 | 0,8229004 | 0,86342746 | 0,004448 | 0,0093839 | 0,0072707 | 0,0072707 | TRUE | 0,1446232 | No |
| orf19.3659 | 1385 | 934 | 1202 | 1472 | 1756 | 1592 | 1751 | 1030 | 1242 | 1189 | 1230 | 1214 | 1502 | 1519 | 1562 | 1453 | 1363,9 | 1219 | 1509 | 1,239 | 0,309 | 6,798 | 1,06E-11 | 8,66E-11 | 1,00E-08 | 0,0030439 | 0,0012466 | 0,0012466 | TRUE | 0,00676 | No |
| orf19.366 | 1613 | 1010 | 1301 | 1664 | 1549 | 1091 | 1257 | 650 | 1447 | 1286 | 1331 | 1372 | 1325 | 1041 | 1122 | 917 | 1230 | 1359 | 1101 | 0,812 | -0,301 | -3,184 | 0,00145083 | 0,00353184 | 0,012275 | 0,0031597 | 0,0077434 | 0,0077434 | TRUE | 0,2249482 | No |
| orf19.3660 | 2114 | 1477 | 1902 | 2352 | 2320 | 2008 | 2273 | 1338 | 1896 | 1880 | 1946 | 1939 | 1984 | 1916 | 2028 | 1887 | 1934,7 | 1915 | 1954 | 1,021 | 0,03 | 0,717 | 0,47338804 | 0,55942473 | 1,00E-08 | 0,0027299 | 0,0011588 | 0,0011588 | TRUE | 0,0078546 | No |
| orf19.3661 | 4580 | 3467 | 4475 | 5348 | 4994 | 4921 | 5164 | 3355 | 4109 | 4414 | 4578 | 4410 | 4271 | 4695 | 4608 | 4732 | 4477,2 | 4378 | 4576 | 1,045 | 0,064 | 1,31 | 0,1902951 | 0,26331022 | 0,001757 | 0,0023038 | 0,0020431 | 0,0020431 | TRUE | 0,0249732 | No |
| orf19.3663 | 8383 | 6076 | 7702 | 9654 | 7744 | 7396 | 7846 | 4857 | 7520 | 7735 | 7880 | 7961 | 6623 | 7057 | 7001 | 6850 | 7328,5 | 7774 | 6883 | 0,885 | -0,176 | -4,963 | 6,96E-07 | 3,01E-06 | 0,000523 | 0,0021776 | 0,0010668 | 0,0010668 | TRUE | 0,0081646 | No |
| orf19.3663.1 | 676 | 552 | 615 | 818 | 873 | 754 | 784 | 504 | 606 | 703 | 629 | 675 | 747 | 719 | 700 | 711 | 686,18 | 653 | 719 | 1,104 | 0,142 | 2,204 | 0,02749599 | 0,04968297 | 0,00102 | 0,0040952 | 0,0025319 | 0,0025319 | TRUE | 0,0279634 | No |
| orf19.3664 | 11522 | 8070 | 10274 | 12417 | 10653 | 9513 | 9835 | 6309 | 10336 | 10274 | 10511 | 10239 | 9111 | 9077 | 8776 | 8898 | 9652,9 | 10340 | 8966 | 0,867 | -0,206 | -7,431 | 1,08E-13 | 1,09E-12 | 8,29E-05 | 0,0021299 | 0,0006302 | 0,0006302 | TRUE | 0,002588 | No |
| orf19.3665 | 2017 | 1350 | 1786 | 2178 | 1607 | 1467 | 1502 | 959 | 1809 | 1719 | 1827 | 1796 | 1374 | 1400 | 1340 | 1353 | 1577,3 | 1788 | 1367 | 0,764 | -0,388 | -8,98 | 2,71E-19 | 4,67E-18 | 1,00E-08 | 0,0028999 | 0,0011494 | 0,0011494 | TRUE | 0,0074161 | No |
| orf19.3666 | 885 | 646 | 841 | 965 | 635 | 626 | 661 | 385 | 794 | 822 | 860 | 796 | 543 | 597 | 590 | 543 | 693,22 | 818 | 568 | 0,696 | -0,522 | -8,331 | 8,02E-17 | 1,12E-15 | 0,000515 | 0,0040537 | 0,002277 | 0,002277 | TRUE | 0,0147053 | No |
| orf19.3667 | 54 | 32 | 45 | 49 | 56 | 49 | 46 | 33 | 48 | 41 | 46 | 40 | 48 | 47 | 41 | 47 | 44,73 | 44 | 46 | 1,035 | 0,049 | 0,243 | 0,80778115 | 0,85098179 | 1,00E-08 | 0,0344319 | 0,0167046 | 0,0167046 | TRUE | 0,0415568 | No |
| orf19.3668 | 6560 | 4353 | 5867 | 7220 | 4258 | 3577 | 3770 | 2602 | 5885 | 5542 | 6003 | 5954 | 3642 | 3413 | 3364 | 3670 | 4684 | 5846 | 3522 | 0,602 | -0,732 | -16,417 | 1,43E-60 | 2,39E-58 | 0,001312 | 0,0022895 | 0,0016818 | 0,0016818 | TRUE | 0,0145455 | No |
| orf19.3669 | 11131 | 8024 | 10105 | 12502 | 17247 | 15782 | 17267 | 11387 | 9986 | 10215 | 10338 | 10309 | 14750 | 15059 | 15408 | 16060 | 12766 | 10212 | 15319 | 1,5 | 0,585 | 16,585 | 9,01E-62 | 1,55E-59 | 0,000688 | 0,0020932 | 0,0011113 | 0,0011113 | TRUE | 0,0127137 | No |
| orf19.367 | 4043 | 2565 | 3597 | 4487 | 4440 | 3473 | 4148 | 2873 | 3627 | 3266 | 3680 | 3700 | 3797 | 3314 | 3701 | 4052 | 3642,2 | 3568 | 3716 | 1,04 | 0,057 | 0,874 | 0,38193852 | 0,4709948 | 0,00457 | 0,0023781 | 0,0037997 | 0,0037997 | TRUE | 0,0645241 | No |
| orf19.3670 | 4604 | 3411 | 4080 | 5030 | 6097 | 5715 | 6019 | 3658 | 4130 | 4343 | 4174 | 4148 | 5214 | 5453 | 5371 | 5159 | 4749,1 | 4199 | 5299 | 1,263 | 0,337 | 9,612 | 7,13E-22 | 1,47E-20 | 0,000315 | 0,0022852 | 0,0009688 | 0,0009688 | TRUE | 0,0061602 | No |
| orf19.3671 | 0 | 7 | 1 | 5 | 1 | 1 | 0 | 2 | 0 | 9 | 1 | 4 | 1 | 1 | 0 | 3 | 2,34 | 4 | 1 | 0,31 | -1,69 | -1,376 | 0,16870184 | NA | 1,157388 | 0,6234226 | 0,827169 | 0,827169 | TRUE | 0,3475483 | No |
| orf19.3672 | 5376 | 3748 | 4759 | 5856 | 6920 | 6218 | 6519 | 4217 | 4823 | 4772 | 4869 | 4829 | 5918 | 5933 | 5817 | 5948 | 5363,5 | 4823 | 5904 | 1,224 | 0,291 | 10,337 | 4,80E-25 | 1,22E-23 | 1,00E-08 | 0,0022502 | 0,0005729 | 0,0005729 | TRUE | 0,0012216 | No |
| orf19.3673 | 126 | 76 | 93 | 127 | 182 | 139 | 163 | 104 | 113 | 97 | 95 | 105 | 156 | 133 | 145 | 147 | 123,76 | 102 | 145 | 1,411 | 0,497 | 3,9 | 9,63E-05 | 0,00029587 | 1,00E-08 | 0,0137095 | 0,0073413 | 0,0073413 | TRUE | 0,0489342 | No |
| orf19.3674 | 1972 | 1528 | 1830 | 2153 | 2682 | 2505 | 2592 | 1716 | 1769 | 1945 | 1872 | 1775 | 2294 | 2390 | 2313 | 2420 | 2097,4 | 1840 | 2354 | 1,28 | 0,356 | 7,77 | 7,83E-15 | 8,86E-14 | 0,000736 | 0,0026717 | 0,0015334 | 0,0015334 | TRUE | 0,0167858 | No |
| orf19.3675 | 5500 | 4250 | 4813 | 6266 | 7745 | 7553 | 7704 | 5487 | 4934 | 5411 | 4924 | 5167 | 6624 | 7207 | 6875 | 7739 | 6110 | 5109 | 7111 | 1,391 | 0,477 | 8,641 | 5,55E-18 | 8,62E-17 | 0,003089 | 0,0022171 | 0,0027522 | 0,0027522 | TRUE | 0,0424099 | No |
| orf19.3676 | 3675 | 2924 | 3230 | 3998 | 3237 | 3287 | 3270 | 2101 | 3297 | 3723 | 3305 | 3297 | 2768 | 3136 | 2918 | 2963 | 3175,9 | 3406 | 2946 | 0,866 | -0,208 | -3,718 | 0,00020047 | 0,00058467 | 0,002831 | 0,0024366 | 0,0026843 | 0,0026843 | TRUE | 0,0466419 | No |
| orf19.3677 | 334 | 255 | 315 | 399 | 302 | 270 | 355 | 202 | 300 | 325 | 322 | 329 | 258 | 258 | 317 | 285 | 299,14 | 319 | 280 | 0,876 | -0,191 | -2,056 | 0,03973807 | 0,06868421 | 0,002828 | 0,0068324 | 0,004933 | 0,004933 | TRUE | 0,101518 | No |
| orf19.3678 | 2246 | 1661 | 2167 | 2678 | 2926 | 2855 | 3033 | 1925 | 2015 | 2115 | 2217 | 2208 | 2502 | 2724 | 2706 | 2715 | 2400,4 | 2139 | 2662 | 1,244 | 0,315 | 6,422 | 1,35E-10 | 9,60E-10 | 0,00146 | 0,0025843 | 0,0018839 | 0,0018839 | TRUE | 0,0203863 | No |
| orf19.3679 | 1380 | 1140 | 1339 | 1626 | 1802 | 1721 | 1785 | 1095 | 1238 | 1451 | 1370 | 1341 | 1541 | 1642 | 1593 | 1544 | 1465,1 | 1350 | 1580 | 1,173 | 0,23 | 4,127 | 3,68E-05 | 0,00012175 | 0,001669 | 0,0029704 | 0,0022988 | 0,0022988 | TRUE | 0,0376749 | No |
| orf19.3680 | 376 | 310 | 390 | 411 | 388 | 428 | 425 | 261 | 337 | 395 | 399 | 339 | 332 | 408 | 379 | 368 | 369,68 | 368 | 372 | 1,015 | 0,022 | 0,231 | 0,8174621 | 0,85952263 | 0,005558 | 0,0059064 | 0,0057327 | 0,0057327 | TRUE | 0,0660016 | No |
| orf19.3681 | 2380 | 1765 | 2187 | 2679 | 2377 | 2202 | 2398 | 1691 | 2135 | 2247 | 2238 | 2209 | 2033 | 2101 | 2140 | 2385 | 2186 | 2207 | 2165 | 0,979 | -0,031 | -0,571 | 0,56828796 | 0,64750125 | 0,001983 | 0,0026436 | 0,002307 | 0,002307 | TRUE | 0,0534464 | No |
| orf19.3682 | 1487 | 1421 | 1509 | 1874 | 1634 | 1943 | 2008 | 1208 | 1334 | 1809 | 1544 | 1545 | 1397 | 1854 | 1792 | 1704 | 1622,4 | 1558 | 1687 | 1,083 | 0,116 | 1,151 | 0,24980642 | 0,33182334 | 0,01401 | 0,0028743 | 0,0090549 | 0,0090549 | TRUE | 0,1637937 | No |
| orf19.3683 | 1888 | 1439 | 1681 | 2184 | 1814 | 1819 | 1977 | 1047 | 1694 | 1832 | 1720 | 1801 | 1551 | 1736 | 1764 | 1477 | 1696,8 | 1762 | 1632 | 0,928 | -0,107 | -1,686 | 0,09172613 | 0,14232711 | 0,003584 | 0,0028351 | 0,0033047 | 0,0033047 | TRUE | 0,0460671 | No |
| orf19.3684 | 3511 | 2384 | 3090 | 3847 | 2405 | 2012 | 2137 | 1482 | 3150 | 3035 | 3161 | 3172 | 2057 | 1920 | 1907 | 2090 | 2561,5 | 3130 | 1994 | 0,636 | -0,653 | -14,863 | 5,71E-50 | 6,02E-48 | 0,000763 | 0,0025463 | 0,0014416 | 0,0014416 | TRUE | 0,0116146 | No |
| orf19.3685 | 436 | 376 | 407 | 540 | 620 | 650 | 696 | 392 | 391 | 479 | 416 | 445 | 530 | 620 | 621 | 553 | 506,99 | 433 | 581 | 1,347 | 0,43 | 5,055 | 4,30E-07 | 1,92E-06 | 0,004966 | 0,0048429 | 0,0049208 | 0,0049208 | TRUE | 0,053044 | No |
| orf19.3686 | 618 | 516 | 602 | 766 | 780 | 778 | 730 | 487 | 554 | 657 | 616 | 632 | 667 | 742 | 651 | 687 | 650,82 | 615 | 687 | 1,119 | 0,162 | 2,248 | 0,02455981 | 0,04487693 | 0,002755 | 0,0042101 | 0,0034484 | 0,0034484 | TRUE | 0,0526545 | No |
| orf19.3687 | 242 | 174 | 201 | 283 | 238 | 225 | 227 | 144 | 217 | 222 | 206 | 233 | 204 | 215 | 203 | 203 | 212,69 | 220 | 206 | 0,938 | -0,093 | -0,964 | 0,33505883 | 0,42193114 | 1,00E-08 | 0,008805 | 0,0041587 | 0,0041587 | TRUE | 0,0229539 | No |
| orf19.3688 | 262 | 189 | 241 | 307 | 296 | 278 | 303 | 135 | 235 | 241 | 247 | 253 | 253 | 265 | 270 | 190 | 244,32 | 244 | 244 | 1,012 | 0,018 | 0,163 | 0,87085285 | 0,90111898 | 0,006095 | 0,0079213 | 0,0073084 | 0,0073084 | TRUE | 0,2061696 | No |

| | | | | | | | | | | | | | | | | | | | | | | | | | | | | | | | |
|---|---|---|---|---|---|---|---|---|---|---|---|---|---|---|---|---|---|---|---|---|---|---|---|---|---|---|---|---|---|---|---|
| orf19.3689 | 3416 | 2591 | 3113 | 3869 | 3880 | 3766 | 4007 | 2278 | 3065 | 3299 | 3185 | 3190 | 3318 | 3593 | 3576 | 3213 | 3304,8 | 3185 | 3425 | 1,077 | 0,107 | 2,217 | 0,02659549 | 0,04832418 | 0,001573 | 0,0024188 | 0,0019282 | 0,0019282 | TRUE | 0,0199576 | No |
| orf19.3690.2 | 6136 | 5274 | 5134 | 7158 | 6008 | 5918 | 5344 | 3462 | 5505 | 6714 | 5253 | 5903 | 5138 | 5647 | 4769 | 4883 | 5476,3 | 5844 | 5109 | 0,875 | -0,193 | -2,472 | 0,01343848 | 0,02617977 | 0,008657 | 0,0022446 | 0,0056789 | 0,0056789 | TRUE | 0,1216947 | No |
| orf19.3691 | 792 | 618 | 707 | 927 | 941 | 966 | 1014 | 496 | 711 | 787 | 723 | 764 | 805 | 922 | 905 | 700 | 789,49 | 746 | 833 | 1,12 | 0,164 | 1,925 | 0,05425764 | 0,09024914 | 0,007039 | 0,0038183 | 0,0056608 | 0,0056608 | TRUE | 0,1263824 | No |
| orf19.3693 | 38 | 29 | 26 | 21 | 33 | 32 | 27 | 13 | 34 | 37 | 27 | 17 | 28 | 31 | 24 | 18 | 27,01 | 29 | 25 | 0,904 | -0,146 | -0,489 | 0,62477115 | 0,697311 | 0,038628 | 0,0557184 | 0,0486052 | 0,0486052 | TRUE | 0,5528305 | No |
| orf19.3694 | 388 | 326 | 373 | 445 | 411 | 408 | 451 | 227 | 348 | 415 | 382 | 367 | 352 | 389 | 402 | 320 | 371,89 | 378 | 366 | 0,976 | -0,036 | -0,39 | 0,69616911 | 0,75867659 | 0,004369 | 0,0058831 | 0,0053264 | 0,0053264 | TRUE | 0,0686645 | No |
| orf19.3695 | 1965 | 1776 | 1912 | 2451 | 2220 | 2456 | 2459 | 1161 | 1763 | 2261 | 1956 | 2021 | 1899 | 2343 | 2194 | 1637 | 2009,4 | 2000 | 2018 | 1,011 | 0,015 | 0,146 | 0,8838282 | 0,91220357 | 0,01706 | 0,002702 | 0,0099678 | 0,0099678 | TRUE | 0,1904015 | No |
| orf19.3696 | 1590 | 1288 | 1306 | 1719 | 1753 | 1579 | 1540 | 890 | 1426 | 1640 | 1336 | 1418 | 1499 | 1507 | 1374 | 1255 | 1431,9 | 1455 | 1409 | 0,971 | -0,042 | -0,554 | 0,57960281 | 0,65802224 | 0,006067 | 0,0029934 | 0,0049218 | 0,0049218 | TRUE | 0,0848206 | No |
| orf19.3697 | 231 | 220 | 209 | 302 | 243 | 314 | 333 | 191 | 207 | 280 | 214 | 249 | 208 | 300 | 297 | 269 | 253,02 | 238 | 268 | 1,132 | 0,179 | 1,321 | 0,18650045 | 0,25913384 | 0,01999 | 0,0077171 | 0,0137182 | 0,0137182 | TRUE | 0,2769531 | No |
| orf19.3698 | 170 | 192 | 178 | 225 | 338 | 436 | 437 | 232 | 153 | 244 | 182 | 186 | 289 | 416 | 390 | 327 | 273,36 | 191 | 356 | 1,872 | 0,905 | 6,086 | 1,16E-09 | 7,31E-09 | 0,028182 | 0,0072902 | 0,0171772 | 0,0171772 | TRUE | 0,2913536 | No |
| orf19.3699 | 410 | 405 | 401 | 528 | 522 | 615 | 657 | 359 | 368 | 516 | 410 | 435 | 446 | 587 | 586 | 506 | 481,87 | 432 | 531 | 1,233 | 0,303 | 2,622 | 0,0087507 | 0,01788847 | 0,015763 | 0,0049922 | 0,0106903 | 0,0106903 | TRUE | 0,184072 | No |
| orf19.3700 | 2806 | 2437 | 2763 | 3394 | 3232 | 3644 | 3679 | 2364 | 2517 | 3103 | 2827 | 2799 | 2764 | 3477 | 3283 | 3334 | 3013 | 2812 | 3214 | 1,144 | 0,194 | 2,446 | 0,01445508 | 0,02795175 | 0,007865 | 0,0024613 | 0,0056859 | 0,0056859 | TRUE | 0,1094665 | No |
| orf19.3701 | 987 | 764 | 927 | 1209 | 1029 | 1056 | 1074 | 624 | 885 | 973 | 948 | 997 | 880 | 1008 | 958 | 880 | 941,2 | 951 | 932 | 0,981 | -0,027 | -0,423 | 0,67262531 | 0,73891623 | 0,002541 | 0,0035219 | 0,0029803 | 0,0029803 | TRUE | 0,0358975 | No |
| orf19.3704 | 7919 | 5814 | 7181 | 8639 | 7712 | 7132 | 7592 | 4801 | 7104 | 7402 | 7347 | 7124 | 6596 | 6805 | 6775 | 6771 | 6990,4 | 7244 | 6737 | 0,93 | -0,104 | -3,439 | 0,00058463 | 0,00155075 | 0,00019 | 0,0021872 | 0,000741 | 0,000741 | TRUE | 0,0025454 | No |
| orf19.3704.1 | 1065 | 646 | 858 | 1106 | 1256 | 928 | 949 | 588 | 955 | 822 | 878 | 912 | 1074 | 885 | 847 | 829 | 900,43 | 892 | 909 | 1,02 | 0,028 | 0,328 | 0,74289594 | 0,79873279 | 0,008189 | 0,0035917 | 0,0060581 | 0,0060581 | TRUE | 0,1824053 | No |
| orf19.3705 | 3001 | 1838 | 2544 | 3298 | 2870 | 2492 | 2680 | 1966 | 2692 | 2340 | 2603 | 2720 | 2455 | 2378 | 2391 | 2773 | 2543,9 | 2589 | 2499 | 0,963 | -0,054 | -0,835 | 0,40370407 | 0,49197851 | 0,004218 | 0,0025502 | 0,0036189 | 0,0036189 | TRUE | 0,0633987 | No |
| orf19.3706 | 4371 | 3238 | 3937 | 4657 | 3454 | 3379 | 3557 | 1954 | 3921 | 4122 | 4028 | 3840 | 2954 | 3224 | 3174 | 2756 | 3502,5 | 3978 | 3027 | 0,762 | -0,392 | -7,239 | 4,50E-13 | 4,32E-12 | 0,002625 | 0,002394 | 0,0025209 | 0,0025209 | TRUE | 0,0419409 | No |
| orf19.3707 | 40601 | 28141 | 34888 | 42807 | 29758 | 26245 | 28293 | 17328 | 36423 | 35826 | 35694 | 35299 | 25450 | 25042 | 25247 | 24440 | 30428 | 35810 | 25045 | 0,7 | -0,516 | -20,096 | 8,06E-90 | 3,72E-87 | 0,000191 | 0,0020272 | 0,0005981 | 0,0005981 | TRUE | 0,0031637 | No |
| orf19.3708 | 13 | 11 | 9 | 15 | 15 | 13 | 17 | 5 | 12 | 14 | 9 | 12 | 13 | 12 | 15 | 7 | 11,84 | 12 | 12 | 1,034 | 0,049 | 0,123 | 0,90201254 | NA | 1,00E-08 | 0,1246179 | 0,0667485 | 0,0667485 | TRUE | 0,1826327 | No |
| orf19.371 | 1058 | 655 | 907 | 1129 | 1115 | 911 | 947 | 695 | 949 | 834 | 928 | 931 | 954 | 869 | 845 | 980 | 911,26 | 910 | 912 | 0,998 | -0,003 | -0,047 | 0,96239861 | 0,97258153 | 0,002911 | 0,0035726 | 0,0032196 | 0,0032196 | TRUE | 0,0354153 | No |
| orf19.3710 | 3275 | 2995 | 3232 | 4273 | 3704 | 3854 | 3906 | 2143 | 2938 | 3813 | 3307 | 3524 | 3168 | 3677 | 3485 | 3023 | 3366,8 | 3396 | 3338 | 0,984 | -0,023 | -0,279 | 0,78061461 | 0,82817104 | 0,009619 | 0,0024107 | 0,0063328 | 0,0063328 | TRUE | 0,1003295 | No |
| orf19.3711 | 69 | 52 | 53 | 78 | 59 | 50 | 61 | 29 | 62 | 66 | 54 | 64 | 50 | 48 | 54 | 41 | 55,02 | 62 | 48 | 0,793 | -0,334 | -1,803 | 0,0713649 | 0,11458053 | 1,00E-08 | 0,0283653 | 0,0146659 | 0,0146659 | TRUE | 0,0614269 | No |
| orf19.3712 | 13090 | 11299 | 12642 | 15674 | 15372 | 16687 | 16874 | 9718 | 11743 | 14385 | 12934 | 12925 | 13147 | 15922 | 15057 | 13706 | 13727 | 12997 | 14458 | 1,113 | 0,154 | 2,123 | 0,03371912 | 0,05949618 | 0,007181 | 0,0020853 | 0,0049823 | 0,0049823 | TRUE | 0,0629868 | No |
| orf19.3713 | 3709 | 4108 | 3515 | 4870 | 4025 | 5711 | 4514 | 3380 | 3327 | 5230 | 3596 | 4016 | 3442 | 5449 | 4028 | 4767 | 4232 | 4042 | 4422 | 1,094 | 0,13 | 0,656 | 0,51199585 | 0,59648459 | 0,037238 | 0,0023226 | 0,0192601 | 0,0372376 | TRUE | 0,3510117 | No |
| orf19.3714 | 1122 | 757 | 973 | 1184 | 868 | 818 | 897 | 547 | 1007 | 964 | 995 | 976 | 742 | 781 | 800 | 771 | 879,61 | 986 | 774 | 0,785 | -0,35 | -6,622 | 3,54E-11 | 2,75E-10 | 1,00E-08 | 0,0036299 | 0,001531 | 0,001531 | TRUE | 0,0099805 | No |
| orf19.3715 | 1023 | 678 | 884 | 1181 | 1176 | 921 | 997 | 582 | 918 | 863 | 904 | 974 | 1006 | 879 | 890 | 821 | 906,78 | 915 | 899 | 0,984 | -0,024 | -0,336 | 0,73652472 | 0,79398609 | 0,003713 | 0,0035805 | 0,0036519 | 0,0036519 | TRUE | 0,0777471 | No |
| orf19.3719 | 17 | 9 | 8 | 16 | 14 | 9 | 12 | 8 | 15 | 11 | 8 | 13 | 12 | 9 | 11 | 11 | 11,33 | 12 | 11 | 0,876 | -0,191 | -0,474 | 0,63522779 | NA | 1,00E-08 | 0,1301094 | 0,0681953 | 0,0681953 | TRUE | 0,1834322 | No |
| orf19.372 | 509 | 316 | 420 | 567 | 717 | 524 | 617 | 394 | 457 | 402 | 430 | 468 | 613 | 500 | 551 | 556 | 496,96 | 439 | 555 | 1,261 | 0,334 | 4,077 | 4,56E-05 | 0,00014876 | 0,003954 | 0,0049007 | 0,0044129 | 0,0044129 | TRUE | 0,0630615 | No |
| orf19.3720 | 3389 | 2361 | 3070 | 3810 | 4107 | 3550 | 3899 | 2297 | 3040 | 3006 | 3141 | 3142 | 3512 | 3387 | 3479 | 3240 | 3243,4 | 3082 | 3404 | 1,106 | 0,145 | 3,64 | 0,00027232 | 0,00077467 | 0,000486 | 0,0024271 | 0,0012075 | 0,0012075 | TRUE | 0,0120089 | No |
| orf19.3721 | 291 | 185 | 228 | 289 | 251 | 222 | 239 | 182 | 261 | 236 | 233 | 238 | 215 | 212 | 213 | 257 | 233,08 | 242 | 224 | 0,917 | -0,124 | -1,263 | 0,20675615 | 0,28295921 | 0,000389 | 0,008208 | 0,004994 | 0,004994 | TRUE | 0,0863632 | No |
| orf19.3722 | 2744 | 1948 | 2419 | 3021 | 3306 | 2793 | 2978 | 2003 | 2462 | 2480 | 2475 | 2491 | 2827 | 2665 | 2657 | 2825 | 2610,3 | 2477 | 2744 | 1,107 | 0,147 | 3,752 | 0,00017557 | 0,00051605 | 0,00028 | 0,0025357 | 0,0010793 | 0,0010793 | TRUE | 0,0057679 | No |
| orf19.3724 | 3326 | 2141 | 2652 | 3533 | 3361 | 3008 | 3138 | 2012 | 2984 | 2726 | 2713 | 2913 | 2874 | 2870 | 2800 | 2838 | 2839,8 | 2834 | 2846 | 1,003 | 0,004 | 0,103 | 0,91817447 | 0,93976017 | 0,00087 | 0,0024907 | 0,0014565 | 0,0014565 | TRUE | 0,015257 | No |
| orf19.3726 | 970 | 679 | 855 | 959 | 895 | 816 | 917 | 523 | 870 | 864 | 875 | 791 | 765 | 779 | 818 | 738 | 812,52 | 850 | 775 | 0,915 | -0,129 | -2,135 | 0,03277083 | 0,05795913 | 0,000956 | 0,0037662 | 0,0022492 | 0,0022492 | TRUE | 0,0281635 | No |
| orf19.3727 | 117 | 74 | 85 | 105 | 115 | 112 | 108 | 71 | 105 | 94 | 87 | 87 | 98 | 107 | 96 | 100 | 96,81 | 93 | 100 | 1,077 | 0,108 | 0,764 | 0,44503739 | 0,53146187 | 1,00E-08 | 0,0169755 | 0,0087601 | 0,0087601 | TRUE | 0,0784184 | No |
| orf19.3728 | 2122 | 1457 | 1909 | 2329 | 2122 | 2011 | 2183 | 1454 | 1904 | 1855 | 1953 | 1921 | 1815 | 1919 | 1948 | 2051 | 1920,6 | 1908 | 1933 | 1,011 | 0,015 | 0,325 | 0,74500045 | 0,79999333 | 0,000796 | 0,0027354 | 0,0016294 | 0,0016294 | TRUE | 0,0207239 | No |
| orf19.3730 | 508 | 462 | 457 | 651 | 413 | 432 | 411 | 277 | 456 | 588 | 468 | 537 | 353 | 412 | 367 | 391 | 446,39 | 512 | 381 | 0,744 | -0,427 | -4,481 | 7,42E-06 | 2,76E-05 | 0,007518 | 0,0052316 | 0,006395 | 0,006395 | TRUE | 0,1079683 | No |
| orf19.3732 | 5465 | 4180 | 5009 | 6198 | 4912 | 4720 | 4657 | 3050 | 4903 | 5322 | 5125 | 5111 | 4201 | 4504 | 4156 | 4302 | 4702,7 | 5115 | 4291 | 0,839 | -0,253 | -6,118 | 9,45E-10 | 6,05E-09 | 0,000924 | 0,0022882 | 0,0014286 | 0,0014286 | TRUE | 0,0139664 | No |
| orf19.3733 | 1679 | 1242 | 1541 | 1950 | 1623 | 1452 | 1525 | 996 | 1506 | 1581 | 1577 | 1608 | 1388 | 1385 | 1361 | 1405 | 1476,4 | 1568 | 1385 | 0,883 | -0,18 | -4,1 | 4,13E-05 | 0,00013554 | 1,00E-08 | 0,0029628 | 0,0011697 | 0,0011697 | TRUE | 0,008952 | No |
| orf19.3734 | 3339 | 2495 | 3125 | 3798 | 3928 | 3761 | 3996 | 2289 | 2995 | 3176 | 3197 | 3132 | 3359 | 3589 | 3566 | 3228 | 3280,4 | 3125 | 3436 | 1,101 | 0,138 | 2,996 | 0,00273764 | 0,00627537 | 0,001272 | 0,0024221 | 0,0017432 | 0,0017432 | TRUE | 0,0188296 | No |
| orf19.3735 | 36 | 42 | 27 | 39 | 12 | 11 | 9 | 18 | 32 | 53 | 28 | 32 | 10 | 10 | 8 | 25 | 24,97 | 36 | 13 | 0,357 | -1,485 | -4,037 | 5,42E-05 | 0,00017426 | 0,104615 | 0,060127 | 0,0770807 | 0,0770807 | TRUE | 1,046267 | No |
| orf19.3736 | 6 | 8 | 7 | 8 | 6 | 9 | 11 | 3 | 5 | 10 | 7 | 7 | 5 | 9 | 10 | 4 | 7,14 | 7 | 7 | 0,985 | -0,022 | -0,043 | 0,96575547 | NA | 1,00E-08 | 0,2053992 | 0,1167635 | 0,1167635 | TRUE | 0,2210876 | No |
| orf19.3737 | 4143 | 3008 | 3889 | 4525 | 4730 | 4397 | 4693 | 3076 | 3717 | 3830 | 3979 | 3731 | 4045 | 4195 | 4188 | 4338 | 4002,9 | 3814 | 4192 | 1,099 | 0,136 | 3,471 | 0,00051771 | 0,00139168 | 0,000575 | 0,0023422 | 0,0012142 | 0,0012142 | TRUE | 0,0104125 | No |
| orf19.3738 | 27 | 18 | 31 | 29 | 34 | 37 | 29 | 18 | 24 | 23 | 32 | 24 | 29 | 35 | 26 | 25 | 27,3 | 26 | 29 | 1,131 | 0,178 | 0,676 | 0,49927706 | 0,58404529 | 1,00E-08 | 0,0551518 | 0,0301662 | 0,0301662 | TRUE | 0,1481707 | No |
| orf19.374 | 6156 | 3638 | 5391 | 6450 | 6878 | 5462 | 6148 | 3701 | 5523 | 4632 | 5516 | 5319 | 5882 | 5212 | 5486 | 5220 | 5348,5 | 5248 | 5450 | 1,039 | 0,055 | 0,852 | 0,39408605 | 0,48273084 | 0,004365 | 0,0022509 | 0,0037434 | 0,0037434 | TRUE | 0,0748912 | No |
| orf19.3740 | 350 | 226 | 330 | 420 | 370 | 281 | 289 | 262 | 314 | 288 | 338 | 346 | 316 | 268 | 258 | 370 | 312,2 | 322 | 303 | 0,935 | -0,097 | -0,839 | 0,40137727 | 0,48983844 | 0,013108 | 0,0066294 | 0,0097073 | 0,0097073 | TRUE | 0,2198015 | No |
| orf19.3742 | 665 | 433 | 525 | 661 | 808 | 655 | 702 | 485 | 597 | 551 | 537 | 545 | 691 | 625 | 626 | 684 | 607,06 | 558 | 656 | 1,175 | 0,233 | 3,453 | 0,00055367 | 0,00147843 | 0,001178 | 0,0043709 | 0,0027207 | 0,0027207 | TRUE | 0,0274307 | No |
| orf19.3743 | 440 | 284 | 390 | 483 | 510 | 414 | 514 | 370 | 395 | 362 | 399 | 398 | 436 | 395 | 459 | 522 | 420,66 | 388 | 453 | 1,158 | 0,212 | 2,379 | 0,01738232 | 0,03282211 | 0,004722 | 0,0054305 | 0,005221 | 0,005221 | TRUE | 0,1072524 | No |
| orf19.3744 | 465 | 390 | 417 | 532 | 485 | 473 | 524 | 315 | 417 | 497 | 427 | 439 | 415 | 451 | 468 | 444 | 444,62 | 445 | 444 | 1,003 | 0,004 | 0,048 | 0,9613684 | 0,97188311 | 0,001903 | 0,0052446 | 0,003641 | 0,003641 | TRUE | 0,0636397 | No |
| orf19.3746 | 3611 | 2648 | 3476 | 3975 | 3145 | 3152 | 3323 | 2078 | 3239 | 3371 | 3556 | 3278 | 2690 | 3008 | 2965 | 2931 | 3129,8 | 3361 | 2898 | 0,862 | -0,214 | -4,288 | 1,80E-05 | 6,31E-05 | 0,001666 | 0,0024434 | 0,0020646 | 0,0020646 | TRUE | 0,0294511 | No |
| orf19.3748 | 77 | 54 | 63 | 106 | 59 | 51 | 76 | 34 | 69 | 69 | 64 | 87 | 50 | 49 | 68 | 48 | 63,07 | 72 | 54 | 0,743 | -0,429 | -2,255 | 0,02413551 | 0,0441688 | 0,010354 | 0,0249958 | 0,0186485 | 0,0186485 | TRUE | 0,2802466 | No |
| orf19.3749 | 18484 | 13138 | 17024 | 20283 | 15278 | 14480 | 15053 | 9386 | 16582 | 16726 | 17417 | 16726 | 13066 | 13816 | 13432 | 13238 | 15126 | 16863 | 13388 | 0,794 | -0,333 | -10,517 | 7,19E-26 | 2,00E-24 | 0,000449 | 0,0020755 | 0,0008939 | 0,0008939 | TRUE | 0,0059952 | No |
| orf19.3751 | 2649 | 1870 | 2413 | 2886 | 2854 | 2557 | 2802 | 1745 | 2376 | 2381 | 2469 | 2380 | 2441 | 2440 | 2500 | 2461 | 2431 | 2402 | 2460 | 1,025 | 0,035 | 0,972 | 0,33089506 | 0,41791413 | 1,00E-08 | 0,0025767 | 0,0008491 | 0,0008491 | TRUE | 0,0042859 | No |
| orf19.3752 | 7571 | 4872 | 6178 | 7893 | 6264 | 5149 | 5566 | 3660 | 6792 | 6203 | 6321 | 6509 | 5357 | 4913 | 4967 | 5162 | 5777,9 | 6456 | 5100 | 0,79 | -0,341 | -7,738 | 1,01E-14 | 1,13E-13 | 0,001345 | 0,0022308 | 0,0016861 | 0,0016861 | TRUE | 0,0149899 | No |
| orf19.3753 | 8293 | 5791 | 7741 | 9455 | 9456 | 8391 | 9476 | 6216 | 7440 | 7373 | 7920 | 7797 | 8087 | 8006 | 8456 | 8767 | 7980,6 | 7632 | 8329 | 1,091 | 0,125 | 2,938 | 0,00330113 | 0,0074226 | 0,001336 | 0,0021614 | 0,0016167 | 0,0016167 | TRUE | 0,0149074 | No |
| orf19.3755 | 1823 | 1321 | 1535 | 1794 | 2450 | 2307 | 2254 | 1392 | 1635 | 1682 | 1570 | 1479 | 2095 | 2201 | 2011 | 1963 | 1829,8 | 1592 | 2068 | 1,302 | 0,381 | 6,806 | 1,00E-11 | 8,20E-11 | 0,002052 | 0,0027729 | 0,0024436 | 0,0024436 | TRUE | 0,0281238 | No |
| orf19.3756 | 11839 | 8908 | 10739 | 13314 | 13268 | 12851 | 13770 | 8438 | 10621 | 11341 | 10987 | 10979 | 11347 | 12262 | 12287 | 11901 | 11466 | 10982 | 11949 | 1,088 | 0,122 | 3,2 | 0,00137361 | 0,00335611 | 0,000915 | 0,0021061 | 0,0013096 | 0,0013096 | TRUE | 0,0143163 | No |
| orf19.3757 | 1804 | 1317 | 1505 | 1886 | 1652 | 1673 | 1621 | 1262 | 1618 | 1677 | 1540 | 1555 | 1413 | 1596 | 1446 | 1780 | 1578,2 | 1598 | 1559 | 0,974 | -0,038 | -0,527 | 0,59812143 | 0,67483822 | 0,005037 | 0,0028994 | 0,0043817 | 0,0043817 | TRUE | 0,1048637 | No |
| orf19.3758 | 4268 | 3385 | 3768 | 4988 | 4932 | 4944 | 5002 | 3324 | 3829 | 4309 | 3855 | 4113 | 4218 | 4717 | 4463 | 4688 | 4274,2 | 4026 | 4522 | 1,123 | 0,167 | 3,111 | 0,00186285 | 0,00442898 | 0,002609 | 0,0023192 | 0,0025385 | 0,0025385 | TRUE | 0,0264798 | No |
| orf19.3759 | 5287 | 4178 | 5098 | 6239 | 6728 | 6236 | 6900 | 4342 | 4743 | 5319 | 5216 | 5145 | 5754 | 5950 | 6157 | 6124 | 5551 | 5106 | 5996 | 1,175 | 0,232 | 5,128 | 2,93E-07 | 1,34E-06 | 0,001455 | 0,002241 | 0,0017874 | 0,0017874 | TRUE | 0,0280709 | No |
| orf19.376 | 2064 | 1180 | 1777 | 2113 | 1601 | 1269 | 1512 | 882 | 1852 | 1502 | 1818 | 1742 | 1369 | 1211 | 1349 | 1244 | 1511 | 1728 | 1293 | 0,748 | -0,419 | -5,797 | 6,74E-09 | 3,83E-08 | 0,005063 | 0,0029403 | 0,0043343 | 0,0043343 | TRUE | 0,0902413 | No |

| | | | | | | | | | | | | | | | | | | | | | | | | | | | | | | | |
|---|---|---|---|---|---|---|---|---|---|---|---|---|---|---|---|---|---|---|---|---|---|---|---|---|---|---|---|---|---|---|---|
| orf19.3760 | 100 | 67 | 99 | 106 | 111 | 95 | 88 | 52 | 90 | 85 | 101 | 87 | 95 | 91 | 79 | 73 | 87,64 | 91 | 84 | 0,935 | -0,097 | -0,645 | 0,51896234 | 0,60226497 | 1,00E-08 | 0,0185432 | 0,0106024 | 0,0106024 | TRUE | 0,0693368 | No |
| orf19.3761 | 3134 | 2300 | 2729 | 3428 | 3014 | 2889 | 3052 | 1760 | 2812 | 2928 | 2792 | 2827 | 2578 | 2757 | 2723 | 2482 | 2737,3 | 2840 | 2635 | 0,929 | -0,105 | -2,373 | 0,01762255 | 0,03319006 | 0,000922 | 0,0025099 | 0,0015289 | 0,0015289 | TRUE | 0,0169437 | No |
| orf19.3762 | 588 | 460 | 544 | 753 | 713 | 732 | 654 | 473 | 527 | 586 | 557 | 621 | 610 | 698 | 584 | 667 | 606,2 | 573 | 640 | 1,115 | 0,157 | 1,973 | 0,04844644 | 0,08189821 | 0,004439 | 0,0043743 | 0,0044084 | 0,0044084 | TRUE | 0,0484413 | No |
| orf19.3763 | 894 | 731 | 778 | 1052 | 961 | 985 | 1041 | 598 | 802 | 931 | 796 | 867 | 822 | 940 | 929 | 843 | 866,27 | 849 | 884 | 1,043 | 0,061 | 0,855 | 0,39275926 | 0,48158455 | 0,003513 | 0,0036553 | 0,003717 | 0,003717 | TRUE | 0,0460742 | No |
| orf19.3764 | 490 | 456 | 463 | 545 | 656 | 780 | 790 | 422 | 440 | 581 | 474 | 449 | 561 | 744 | 705 | 595 | 568,58 | 486 | 651 | 1,346 | 0,429 | 3,868 | 0,00010981 | 0,00033403 | 0,015451 | 0,0045327 | 0,0100056 | 0,0100056 | TRUE | 0,1910375 | No |
| orf19.3765 | 968 | 869 | 963 | 1146 | 1131 | 1151 | 1179 | 682 | 868 | 1106 | 985 | 945 | 967 | 1098 | 1052 | 962 | 998,06 | 976 | 1020 | 1,048 | 0,068 | 0,866 | 0,38648107 | 0,4756203 | 0,005683 | 0,0034341 | 0,0048519 | 0,0048519 | TRUE | 0,0919327 | No |
| orf19.3767 | 599 | 438 | 596 | 700 | 740 | 657 | 659 | 384 | 537 | 558 | 610 | 577 | 633 | 627 | 588 | 542 | 583,92 | 570 | 598 | 1,051 | 0,072 | 0,999 | 0,31790713 | 0,4048296 | 0,001977 | 0,0044656 | 0,0032328 | 0,0032328 | TRUE | 0,039977 | No |
| orf19.3769 | 810 | 614 | 723 | 949 | 1209 | 1129 | 1135 | 715 | 727 | 782 | 740 | 783 | 1034 | 1077 | 1013 | 1008 | 895,38 | 758 | 1033 | 1,365 | 0,449 | 8,211 | 2,19E-16 | 2,86E-15 | 1,00E-08 | 0,0036008 | 0,0017328 | 0,0017328 | TRUE | 0,0098847 | No |
| orf19.377 | 174 | 95 | 131 | 139 | 143 | 125 | 91 | 80 | 156 | 121 | 134 | 115 | 122 | 119 | 81 | 113 | 120,16 | 132 | 109 | 0,826 | -0,275 | -1,72 | 0,0854891 | 0,13392921 | 0,018707 | 0,0140608 | 0,0162072 | 0,0162072 | TRUE | 0,3180398 | No |
| orf19.3770 | 5996 | 4625 | 5291 | 6478 | 7402 | 7204 | 7481 | 4199 | 5379 | 5888 | 5413 | 5342 | 6330 | 6874 | 6676 | 5922 | 5978 | 5506 | 6450 | 1,173 | 0,23 | 4,219 | 2,46E-05 | 8,40E-05 | 0,002958 | 0,0022224 | 0,0026817 | 0,0026817 | TRUE | 0,036202 | No |
| orf19.3773 | 2428 | 1737 | 2200 | 2695 | 2334 | 2196 | 2490 | 1560 | 2178 | 2211 | 2251 | 2222 | 1996 | 2095 | 2222 | 2200 | 2172 | 2216 | 2128 | 0,96 | -0,059 | -1,297 | 0,1946456 | 0,26815656 | 0,000883 | 0,0026479 | 0,0015504 | 0,0015504 | TRUE | 0,021956 | No |
| orf19.3774 | 1567 | 1137 | 1484 | 1726 | 1724 | 1535 | 1621 | 981 | 1406 | 1448 | 1518 | 1423 | 1474 | 1465 | 1446 | 1384 | 1445,5 | 1449 | 1442 | 0,998 | -0,003 | -0,07 | 0,94447973 | 0,95997872 | 0,000218 | 0,0029838 | 0,0014046 | 0,0014046 | TRUE | 0,0127262 | No |
| orf19.3774.1 | 112 | 94 | 88 | 142 | 93 | 87 | 108 | 68 | 100 | 120 | 90 | 117 | 80 | 83 | 96 | 96 | 97,76 | 107 | 89 | 0,828 | -0,272 | -1,81 | 0,07022938 | 0,11308981 | 1,00E-08 | 0,0168287 | 0,0114203 | 0,0114203 | TRUE | 0,1063279 | No |
| orf19.3775 | 4288 | 2841 | 3758 | 4368 | 4353 | 3920 | 4215 | 2584 | 3847 | 3617 | 3845 | 3602 | 3723 | 3740 | 3761 | 3645 | 3722,4 | 3728 | 3717 | 0,997 | -0,004 | -0,096 | 0,9232747 | 0,94385475 | 0,000499 | 0,0023695 | 0,0011261 | 0,0011261 | TRUE | 0,0067304 | No |
| orf19.3777 | 188 | 320 | 226 | 314 | 293 | 503 | 488 | 193 | 169 | 407 | 231 | 259 | 251 | 480 | 435 | 272 | 313,05 | 266 | 360 | 1,352 | 0,435 | 1,218 | 0,22316966 | 0,30211882 | 0,118983 | 0,0066168 | 0,0540171 | 0,1189832 | TRUE | 0,3717305 | No |
| orf19.3778 | 8748 | 6036 | 7729 | 9489 | 8931 | 7877 | 8274 | 5299 | 7848 | 7684 | 7908 | 7825 | 7638 | 7516 | 7383 | 7474 | 7659,5 | 7816 | 7503 | 0,96 | -0,059 | -2,141 | 0,03223464 | 0,05711168 | 1,00E-08 | 0,0021691 | 0,0006019 | 0,0006019 | TRUE | 0,0018716 | No |
| orf19.3780 | 369 | 269 | 282 | 411 | 354 | 361 | 349 | 235 | 331 | 342 | 289 | 339 | 303 | 344 | 311 | 331 | 323,88 | 325 | 322 | 0,99 | -0,015 | -0,172 | 0,86363523 | 0,89630686 | 0,001689 | 0,0064618 | 0,0042672 | 0,0042672 | TRUE | 0,064228 | No |
| orf19.3781 | 1411 | 1048 | 1299 | 1648 | 1433 | 1333 | 1449 | 978 | 1266 | 1334 | 1329 | 1359 | 1226 | 1272 | 1293 | 1379 | 1307,2 | 1322 | 1292 | 0,975 | -0,036 | -0,695 | 0,48676949 | 0,5721994 | 0,000826 | 0,00309 | 0,0018075 | 0,0018075 | TRUE | 0,0220549 | No |
| orf19.3782 | 493 | 360 | 448 | 519 | 549 | 490 | 559 | 369 | 442 | 458 | 458 | 428 | 470 | 468 | 499 | 520 | 467,9 | 446 | 489 | 1,093 | 0,129 | 1,84 | 0,06569945 | 0,10688391 | 1,00E-08 | 0,0050821 | 0,0025603 | 0,0025603 | TRUE | 0,01815 | No |
| orf19.3782.2 | 1268 | 762 | 963 | 1187 | 1261 | 1009 | 1120 | 559 | 1138 | 970 | 985 | 979 | 1078 | 963 | 999 | 788 | 987,59 | 1018 | 957 | 0,943 | -0,085 | -0,935 | 0,34985823 | 0,43743106 | 0,009451 | 0,0034495 | 0,0069425 | 0,0069425 | TRUE | 0,1580714 | No |
| orf19.3783 | 141 | 132 | 132 | 182 | 149 | 138 | 175 | 89 | 126 | 168 | 135 | 150 | 127 | 132 | 156 | 126 | 140,06 | 145 | 135 | 0,941 | -0,088 | -0,67 | 0,50281437 | 0,58749684 | 0,005902 | 0,0123446 | 0,0095392 | 0,0095392 | TRUE | 0,1186878 | No |
| orf19.3784 | 297 | 270 | 282 | 316 | 280 | 296 | 325 | 164 | 266 | 344 | 289 | 261 | 239 | 282 | 290 | 231 | 275,31 | 290 | 260 | 0,908 | -0,14 | -1,225 | 0,22053255 | 0,29922302 | 0,010181 | 0,0072525 | 0,0088239 | 0,0088239 | TRUE | 0,1635924 | No |
| orf19.3785 | 832 | 567 | 689 | 963 | 1005 | 921 | 918 | 636 | 746 | 722 | 705 | 794 | 860 | 879 | 819 | 897 | 802,72 | 742 | 864 | 1,16 | 0,214 | 3,535 | 0,00040719 | 0,00112014 | 0,000932 | 0,003788 | 0,0022658 | 0,0022658 | TRUE | 0,0281999 | No |
| orf19.3787 | 1521 | 908 | 1313 | 1631 | 1233 | 977 | 1036 | 727 | 1364 | 1156 | 1343 | 1345 | 1055 | 932 | 924 | 1025 | 1143,2 | 1302 | 984 | 0,754 | -0,408 | -5,842 | 5,15E-09 | 3,00E-08 | 0,003861 | 0,0032494 | 0,0037745 | 0,0037745 | TRUE | 0,0651509 | No |
| orf19.3788 | 878 | 658 | 749 | 980 | 893 | 733 | 808 | 480 | 788 | 838 | 766 | 808 | 764 | 699 | 721 | 677 | 757,61 | 800 | 715 | 0,898 | -0,156 | -2,576 | 0,01000308 | 0,02014024 | 0,000561 | 0,0038957 | 0,0021946 | 0,0021946 | TRUE | 0,025074 | No |
| orf19.3788.1 | 17863 | 12073 | 14279 | 18692 | 19120 | 15439 | 15834 | 11064 | 16025 | 15370 | 14609 | 15414 | 16352 | 14731 | 14129 | 15605 | 15279 | 15354 | 15204 | 0,99 | -0,014 | -0,277 | 0,78143411 | 0,82874778 | 0,002327 | 0,0020745 | 0,0024884 | 0,0024884 | TRUE | 0,0318948 | No |
| orf19.3789 | 8841 | 5978 | 7136 | 8765 | 7622 | 6689 | 6565 | 4638 | 7931 | 7611 | 7301 | 7228 | 6519 | 6382 | 5858 | 6541 | 6921,4 | 7518 | 6325 | 0,841 | -0,25 | -5,132 | 2,86E-07 | 1,31E-06 | 0,001995 | 0,0021893 | 0,0021234 | 0,0021234 | TRUE | 0,0304209 | No |
| orf19.3791 | 475 | 333 | 382 | 569 | 565 | 463 | 522 | 327 | 426 | 424 | 391 | 469 | 483 | 442 | 466 | 461 | 445,26 | 428 | 463 | 1,081 | 0,113 | 1,468 | 0,14207985 | 0,20608588 | 0,001555 | 0,0052399 | 0,0034134 | 0,0034134 | TRUE | 0,0540317 | No |
| orf19.3792 | 4008 | 2905 | 3649 | 4360 | 4616 | 4119 | 4475 | 2872 | 3596 | 3698 | 3733 | 3595 | 3948 | 3930 | 3993 | 4051 | 3818,1 | 3656 | 3980 | 1,089 | 0,123 | 3,745 | 0,00018026 | 0,00052933 | 1,00E-08 | 0,0023597 | 0,0007701 | 0,0007701 | TRUE | 0,0025902 | No |
| orf19.3793 | 1804 | 1367 | 1656 | 2037 | 1609 | 1591 | 1677 | 1057 | 1618 | 1740 | 1694 | 1680 | 1376 | 1518 | 1496 | 1491 | 1576,8 | 1683 | 1470 | 0,874 | -0,195 | -3,973 | 7,10E-05 | 0,00022294 | 0,000832 | 0,0029002 | 0,0016721 | 0,0016721 | TRUE | 0,0237924 | No |
| orf19.3794 | 706 | 526 | 619 | 737 | 446 | 447 | 484 | 317 | 633 | 670 | 633 | 608 | 381 | 427 | 432 | 447 | 528,87 | 636 | 422 | 0,662 | -0,595 | -8,357 | 6,44E-17 | 9,06E-16 | 0,00087 | 0,0047244 | 0,0028924 | 0,0028924 | TRUE | 0,0507501 | No |
| orf19.3795 | 4268 | 2936 | 3882 | 4632 | 3281 | 2867 | 3182 | 2048 | 3829 | 3738 | 3972 | 3820 | 2806 | 2736 | 2839 | 2889 | 3328,4 | 3840 | 2818 | 0,733 | -0,448 | -12,172 | 4,39E-34 | 2,03E-32 | 0,000169 | 0,0024157 | 0,0009909 | 0,0009909 | TRUE | 0,0063882 | No |
| orf19.3796 | 2016 | 1376 | 1774 | 2159 | 1726 | 1643 | 1716 | 1167 | 1809 | 1752 | 1815 | 1780 | 1476 | 1568 | 1531 | 1646 | 1672,1 | 1789 | 1555 | 0,867 | -0,206 | -4,469 | 7,85E-06 | 2,91E-05 | 0,000378 | 0,0028477 | 0,0014339 | 0,0014339 | TRUE | 0,0166642 | No |
| orf19.3797 | 1753 | 1312 | 1556 | 1865 | 1969 | 1877 | 1854 | 1155 | 1573 | 1670 | 1592 | 1538 | 1684 | 1791 | 1654 | 1629 | 1641,4 | 1593 | 1690 | 1,063 | 0,088 | 1,807 | 0,07070565 | 0,11370437 | 0,000827 | 0,002864 | 0,0016636 | 0,0016636 | TRUE | 0,0196512 | No |
| orf19.3798 | 5205 | 3507 | 4396 | 5558 | 5657 | 4686 | 5077 | 2970 | 4669 | 4465 | 4498 | 4583 | 4838 | 4471 | 4530 | 4189 | 4530,4 | 4554 | 4507 | 0,991 | -0,014 | -0,292 | 0,77046212 | 0,82073341 | 0,001462 | 0,0023 | 0,0019019 | 0,0019019 | TRUE | 0,0297932 | No |
| orf19.3799 | 3937 | 2826 | 3558 | 4519 | 3643 | 3440 | 3721 | 2526 | 3532 | 3598 | 3640 | 3726 | 3116 | 3282 | 3320 | 3563 | 3472,2 | 3624 | 3320 | 0,915 | -0,129 | -2,768 | 0,00564488 | 0,01205205 | 0,001367 | 0,0023976 | 0,0017797 | 0,0017797 | TRUE | 0,0278866 | No |
| orf19.3801 | 100 | 80 | 93 | 116 | 92 | 66 | 98 | 42 | 90 | 102 | 95 | 96 | 79 | 63 | 87 | 59 | 83,84 | 96 | 72 | 0,765 | -0,386 | -2,394 | 0,01666667 | 0,03165771 | 0,002303 | 0,0192951 | 0,0129328 | 0,0129328 | TRUE | 0,1859345 | No |
| orf19.3802 | 598 | 433 | 598 | 697 | 601 | 519 | 646 | 382 | 536 | 551 | 612 | 575 | 514 | 495 | 576 | 539 | 549,84 | 568 | 531 | 0,934 | -0,099 | -1,347 | 0,17797478 | 0,24959675 | 0,002177 | 0,0046197 | 0,0033681 | 0,0033681 | TRUE | 0,0421689 | No |
| orf19.3803 | 2091 | 1536 | 1951 | 2253 | 1660 | 1484 | 1654 | 1036 | 1876 | 1955 | 1996 | 1858 | 1420 | 1416 | 1476 | 1461 | 1682,3 | 1921 | 1443 | 0,752 | -0,412 | -9,216 | 3,08E-20 | 5,69E-19 | 1,00E-08 | 0,0028425 | 0,0013104 | 0,0013104 | TRUE | 0,008637 | No |
| orf19.3804 | 1797 | 1417 | 1614 | 2024 | 1827 | 1733 | 1759 | 988 | 1612 | 1804 | 1651 | 1669 | 1563 | 1654 | 1570 | 1393 | 1614,5 | 1684 | 1545 | 0,92 | -0,12 | -1,994 | 0,04610309 | 0,07833353 | 0,002889 | 0,0028787 | 0,002859 | 0,002859 | TRUE | 0,0482774 | No |
| orf19.3806 | 2002 | 1363 | 1720 | 2133 | 2227 | 1906 | 2095 | 1285 | 1796 | 1735 | 1760 | 1759 | 1905 | 1819 | 1869 | 1812 | 1806,9 | 1762 | 1851 | 1,051 | 0,072 | 1,771 | 0,07658151 | 0,12179339 | 1,00E-08 | 0,002783 | 0,0010216 | 0,0010216 | TRUE | 0,0046362 | No |
| orf19.3809 | 3615 | 2254 | 3014 | 3701 | 3798 | 3160 | 3604 | 2272 | 3243 | 2870 | 3084 | 3052 | 3248 | 3015 | 3216 | 3204 | 3116,5 | 3062 | 3171 | 1,035 | 0,049 | 1,038 | 0,2991442 | 0,38460482 | 0,001376 | 0,0024453 | 0,0018303 | 0,0018303 | TRUE | 0,0208221 | No |
| orf19.3810 | 1642 | 1067 | 1397 | 1724 | 1757 | 1552 | 1686 | 1048 | 1473 | 1358 | 1429 | 1422 | 1503 | 1481 | 1504 | 1478 | 1456,1 | 1420 | 1492 | 1,049 | 0,069 | 1,559 | 0,11899189 | 0,17695626 | 1,00E-08 | 0,0029765 | 0,0011925 | 0,0011925 | TRUE | 0,0094071 | No |
| orf19.3811 | 862 | 544 | 743 | 963 | 978 | 858 | 931 | 543 | 773 | 693 | 760 | 794 | 836 | 819 | 831 | 766 | 783,98 | 755 | 813 | 1,076 | 0,106 | 1,726 | 0,08433114 | 0,13246052 | 0,000776 | 0,0038312 | 0,0023346 | 0,0023346 | TRUE | 0,0332753 | No |
| orf19.3812 | 14353 | 9391 | 12457 | 15193 | 13382 | 11210 | 12271 | 8501 | 12876 | 11956 | 12745 | 12528 | 11445 | 10696 | 10950 | 11990 | 11898 | 12526 | 11270 | 0,899 | -0,153 | -3,429 | 0,00060635 | 0,0016013 | 0,001701 | 0,0021015 | 0,0018315 | 0,0018315 | TRUE | 0,0221817 | No |
| orf19.3813 | 1128 | 791 | 925 | 1213 | 1342 | 1058 | 1230 | 782 | 1012 | 1007 | 946 | 1000 | 1148 | 1010 | 1098 | 1103 | 1040,4 | 991 | 1090 | 1,099 | 0,136 | 2,413 | 0,01582709 | 0,03029282 | 0,001085 | 0,0033748 | 0,0020876 | 0,0020876 | TRUE | 0,0302322 | No |
| orf19.3815 | 6736 | 4488 | 6032 | 7412 | 7932 | 6947 | 7849 | 5236 | 6043 | 5714 | 6171 | 6112 | 6784 | 6629 | 7004 | 7385 | 6480,1 | 6010 | 6950 | 1,155 | 0,208 | 4,698 | 2,63E-06 | 1,05E-05 | 0,001486 | 0,0022035 | 0,0017337 | 0,0017337 | TRUE | 0,0209995 | No |
| orf19.3817 | 898 | 568 | 769 | 945 | 717 | 568 | 652 | 432 | 806 | 723 | 787 | 779 | 613 | 542 | 582 | 609 | 680,13 | 774 | 586 | 0,756 | -0,404 | -6,222 | 4,92E-10 | 3,28E-09 | 0,000953 | 0,004114 | 0,0025421 | 0,0025421 | TRUE | 0,0311807 | No |
| orf19.3818 | 399 | 291 | 365 | 434 | 295 | 245 | 306 | 191 | 358 | 370 | 373 | 358 | 252 | 234 | 273 | 269 | 311,03 | 365 | 257 | 0,704 | -0,507 | -6,054 | 1,41E-09 | 8,84E-09 | 1,00E-08 | 0,0066469 | 0,0034126 | 0,0034126 | TRUE | 0,0420135 | No |
| orf19.3819 | 130 | 81 | 105 | 127 | 98 | 77 | 93 | 61 | 117 | 103 | 107 | 105 | 84 | 73 | 83 | 86 | 94,78 | 108 | 82 | 0,752 | -0,411 | -2,909 | 0,00362792 | 0,00806981 | 1,00E-08 | 0,0172968 | 0,0084076 | 0,0084076 | TRUE | 0,0425819 | No |
| orf19.382 | 10422 | 6084 | 8579 | 10442 | 10247 | 7690 | 8246 | 6829 | 9350 | 7746 | 8777 | 8611 | 8764 | 7338 | 7358 | 9632 | 8446,7 | 8621 | 8273 | 0,959 | -0,06 | -0,68 | 0,49646415 | 0,5815474 | 0,010617 | 0,0021514 | 0,0073803 | 0,0073803 | TRUE | 0,1483913 | No |
| orf19.3820 | 50 | 23 | 39 | 41 | 43 | 34 | 38 | 23 | 45 | 29 | 40 | 34 | 37 | 32 | 34 | 32 | 35,43 | 37 | 34 | 0,914 | -0,13 | -0,562 | 0,57430679 | 0,65262629 | 1,00E-08 | 0,0429574 | 0,0236706 | 0,0236706 | TRUE | 0,1585831 | No |
| orf19.3821 | 699 | 451 | 556 | 718 | 550 | 476 | 496 | 300 | 627 | 574 | 569 | 592 | 470 | 454 | 443 | 423 | 519,06 | 590 | 448 | 0,759 | -0,398 | -5,836 | 5,34E-09 | 3,09E-08 | 1,00E-08 | 0,0047763 | 0,0024988 | 0,0024988 | TRUE | 0,0208008 | No |
| orf19.3822 | 16169 | 10460 | 13836 | 17094 | 17957 | 14290 | 15634 | 10440 | 14505 | 13317 | 14156 | 14096 | 15357 | 13635 | 13951 | 14725 | 14218 | 14018 | 14417 | 1,028 | 0,04 | 0,856 | 0,39177957 | 0,48066401 | 0,001964 | 0,0020816 | 0,0020428 | 0,0020428 | TRUE | 0,0238594 | No |
| orf19.3823 | 3567 | 2294 | 3113 | 3731 | 3874 | 3202 | 3586 | 2208 | 3200 | 2921 | 3185 | 3077 | 3313 | 3055 | 3200 | 3114 | 3133,1 | 3096 | 3170 | 1,024 | 0,034 | 0,764 | 0,44496396 | 0,53146187 | 0,001067 | 0,0024429 | 0,0016223 | 0,0016223 | TRUE | 0,0167117 | No |
| orf19.3825 | 1459 | 961 | 1249 | 1548 | 1059 | 948 | 1043 | 692 | 1309 | 1223 | 1278 | 1277 | 906 | 905 | 931 | 976 | 1100,5 | 1272 | 930 | 0,728 | -0,458 | -9,109 | 8,29E-20 | 1,47E-18 | 1,00E-08 | 0,0032987 | 0,0014907 | 0,0014907 | TRUE | 0,0116499 | No |
| orf19.3826 | 5348 | 3619 | 4916 | 5910 | 5473 | 4596 | 4910 | 3452 | 4798 | 4607 | 5030 | 4873 | 4681 | 4385 | 4381 | 4869 | 4703 | 4827 | 4579 | 0,948 | -0,078 | -1,639 | 0,10124082 | 0,15430108 | 0,001734 | 0,0022882 | 0,0019364 | 0,0019364 | TRUE | 0,0212156 | No |
| orf19.3827 | 2611 | 1803 | 2323 | 2810 | 2671 | 2191 | 2503 | 1599 | 2342 | 2295 | 2377 | 2317 | 2284 | 2091 | 2234 | 2255 | 2274,4 | 2333 | 2216 | 0,949 | -0,075 | -1,776 | 0,07580816 | 0,12074639 | 0,000447 | 0,0026178 | 0,0012637 | 0,0012637 | TRUE | 0,0179228 | No |
| orf19.3828 | 23 | 13 | 16 | 18 | 10 | 12 | 12 | 7 | 21 | 17 | 16 | 15 | 9 | 11 | 11 | 10 | 13,62 | 17 | 10 | 0,593 | -0,755 | -2,042 | 0,04116885 | NA | 1,00E-08 | 0,1085463 | 0,0531305 | 0,0531305 | TRUE | 0,1108487 | No |

| | | | | | | | | | | | | | | | | | | | | | | | | | | | | | | | |
|---|---|---|---|---|---|---|---|---|---|---|---|---|---|---|---|---|---|---|---|---|---|---|---|---|---|---|---|---|---|---|---|
| orf19.3829 | 3432 | 2228 | 3059 | 3761 | 3002 | 2551 | 2704 | 1736 | 3079 | 2836 | 3130 | 3101 | 2567 | 2434 | 2413 | 2448 | 2751,2 | 3036 | 2466 | 0,811 | -0,301 | -6,71 | 1,95E-11 | 1,54E-10 | 0,000894 | 0,0025072 | 0,0015679 | 0,0015679 | TRUE | 0,0227177 | No |
| orf19.3831 | 7707 | 4828 | 6007 | 7721 | 7764 | 6118 | 6643 | 4026 | 6914 | 6147 | 6146 | 6367 | 6640 | 5838 | 5928 | 5678 | 6207,1 | 6394 | 6021 | 0,942 | -0,086 | -1,46 | 0,14427416 | 0,20861593 | 0,003888 | 0,0022134 | 0,0031912 | 0,0031912 | TRUE | 0,0588137 | No |
| orf19.3833 | 4629 | 3134 | 4257 | 4922 | 4956 | 4323 | 4882 | 3053 | 4153 | 3990 | 4355 | 4059 | 4239 | 4125 | 4356 | 4306 | 4197,8 | 4139 | 4256 | 1,028 | 0,04 | 1 | 0,31718297 | 0,40407873 | 0,000695 | 0,0023253 | 0,00128 | 0,00128 | TRUE | 0,0148452 | No |
| orf19.3836 | 2719 | 1718 | 2140 | 2682 | 2599 | 2209 | 2409 | 1592 | 2439 | 2187 | 2189 | 2212 | 2223 | 2108 | 2150 | 2245 | 2219,1 | 2257 | 2182 | 0,965 | -0,051 | -1,027 | 0,30437196 | 0,39024045 | 0,00129 | 0,0026337 | 0,0018972 | 0,0018972 | TRUE | 0,0364751 | No |
| orf19.3838 | 14733 | 9652 | 12428 | 15410 | 14581 | 12035 | 12630 | 8657 | 13217 | 12288 | 12715 | 12707 | 12470 | 11483 | 11270 | 12210 | 12295 | 12732 | 11858 | 0,931 | -0,103 | -2,379 | 0,01735334 | 0,03278949 | 0,001504 | 0,0020976 | 0,0017158 | 0,0017158 | TRUE | 0,0150361 | No |
| orf19.3839 | 2294 | 1473 | 1888 | 2472 | 2254 | 2082 | 2253 | 1386 | 2058 | 1875 | 1932 | 2038 | 1928 | 1987 | 2010 | 1955 | 1972,9 | 1976 | 1970 | 0,996 | -0,006 | -0,141 | 0,88792704 | 0,91519397 | 0,000619 | 0,0027154 | 0,0014399 | 0,0014399 | TRUE | 0,0133733 | No |
| orf19.384 | 2032 | 1380 | 1849 | 2364 | 1991 | 1665 | 1842 | 1240 | 1823 | 1757 | 1892 | 1949 | 1703 | 1589 | 1644 | 1749 | 1763,1 | 1855 | 1671 | 0,899 | -0,154 | -3,047 | 0,00231284 | 0,00538587 | 0,001126 | 0,0028029 | 0,0018864 | 0,0018864 | TRUE | 0,0144977 | No |
| orf19.3840 | 1234 | 779 | 1140 | 1265 | 1750 | 1486 | 1655 | 1033 | 1107 | 992 | 1166 | 1043 | 1497 | 1418 | 1477 | 1457 | 1269,6 | 1077 | 1462 | 1,356 | 0,44 | 7,737 | 1,02E-14 | 1,14E-13 | 0,001674 | 0,003123 | 0,0022942 | 0,0022942 | TRUE | 0,0357598 | No |
| orf19.3841 | 5960 | 3998 | 5368 | 6478 | 7339 | 6369 | 7080 | 4522 | 5347 | 5090 | 5492 | 5342 | 6277 | 6077 | 6318 | 6378 | 5790 | 5318 | 6262 | 1,177 | 0,235 | 6,61 | 3,85E-11 | 2,97E-10 | 0,000478 | 0,0022303 | 0,0010381 | 0,0010381 | TRUE | 0,00966 | No |
| orf19.3843 | 1846 | 1307 | 1523 | 1916 | 1542 | 1382 | 1494 | 843 | 1656 | 1664 | 1558 | 1580 | 1319 | 1319 | 1333 | 1189 | 1452,2 | 1614 | 1290 | 0,802 | -0,319 | -6,204 | 5,49E-10 | 3,62E-09 | 0,00099 | 0,0029792 | 0,001842 | 0,001842 | TRUE | 0,0294873 | No |
| orf19.3844 | 852 | 572 | 708 | 809 | 1038 | 968 | 977 | 651 | 764 | 728 | 724 | 667 | 888 | 924 | 872 | 918 | 810,67 | 721 | 900 | 1,249 | 0,32 | 5,339 | 9,37E-08 | 4,62E-07 | 0,000854 | 0,0037703 | 0,0022099 | 0,0022099 | TRUE | 0,0330287 | No |
| orf19.3845 | 1974 | 1268 | 1673 | 2080 | 2399 | 1990 | 2398 | 1534 | 1771 | 1614 | 1712 | 1715 | 2052 | 1899 | 2140 | 2164 | 1883,2 | 1703 | 2064 | 1,209 | 0,274 | 5,094 | 3,51E-07 | 1,59E-06 | 0,001703 | 0,0027504 | 0,0022478 | 0,0022478 | TRUE | 0,0349706 | No |
| orf19.3846 | 2033 | 1399 | 1782 | 2163 | 1735 | 1571 | 1630 | 1012 | 1824 | 1781 | 1823 | 1784 | 1484 | 1499 | 1455 | 1427 | 1634,5 | 1803 | 1466 | 0,814 | -0,297 | -7,207 | 5,73E-13 | 5,42E-12 | 1,00E-08 | 0,0028677 | 0,0010111 | 0,0010111 | TRUE | 0,0032532 | No |
| orf19.3848 | 482 | 315 | 413 | 547 | 573 | 517 | 528 | 309 | 432 | 401 | 423 | 451 | 490 | 493 | 471 | 436 | 449,67 | 427 | 472 | 1,108 | 0,148 | 2,029 | 0,04245448 | 0,07287615 | 1,00E-08 | 0,0052079 | 0,0028939 | 0,0028939 | TRUE | 0,0265365 | No |
| orf19.385 | 6796 | 4578 | 6268 | 7430 | 7046 | 6168 | 6720 | 4617 | 6097 | 5828 | 6413 | 6127 | 6026 | 5885 | 5996 | 6512 | 6110,5 | 6116 | 6105 | 0,997 | -0,004 | -0,088 | 0,92958134 | 0,94902236 | 0,001377 | 0,0022171 | 0,0017809 | 0,0017809 | TRUE | 0,0237866 | No |
| orf19.3851 | 748 | 540 | 660 | 795 | 735 | 627 | 702 | 460 | 671 | 687 | 675 | 656 | 629 | 598 | 626 | 649 | 648,92 | 672 | 626 | 0,93 | -0,105 | -1,783 | 0,0745403 | 0,11889695 | 1,00E-08 | 0,0042166 | 0,0017729 | 0,0017729 | TRUE | 0,0098412 | No |
| orf19.3852 | 154 | 79 | 124 | 151 | 181 | 186 | 200 | 122 | 138 | 101 | 127 | 125 | 155 | 177 | 178 | 172 | 146,61 | 123 | 170 | 1,381 | 0,466 | 3,753 | 0,0001748 | 0,00051429 | 0,001556 | 0,011881 | 0,0078494 | 0,0078494 | TRUE | 0,1219919 | No |
| orf19.3854 | 8228 | 5610 | 7281 | 9223 | 8583 | 7519 | 8436 | 5254 | 7381 | 7142 | 7449 | 7605 | 7340 | 7174 | 7528 | 7410 | 7378,9 | 7394 | 7363 | 0,995 | -0,007 | -0,207 | 0,8363237 | 0,87489706 | 0,000331 | 0,0021763 | 0,0009157 | 0,0009157 | TRUE | 0,0062419 | No |
| orf19.3856 | 654 | 513 | 643 | 786 | 654 | 646 | 647 | 338 | 587 | 653 | 658 | 648 | 559 | 616 | 577 | 477 | 596,95 | 636 | 557 | 0,88 | -0,184 | -2,276 | 0,0228269 | 0,04201716 | 0,004707 | 0,0044114 | 0,0045691 | 0,0045691 | TRUE | 0,0973061 | No |
| orf19.3858 | 2876 | 1934 | 2388 | 3164 | 2774 | 2401 | 2560 | 1454 | 2580 | 2462 | 2443 | 2609 | 2372 | 2291 | 2284 | 2051 | 2386,6 | 2524 | 2250 | 0,893 | -0,164 | -3,154 | 0,00161231 | 0,00387781 | 0,001877 | 0,0025878 | 0,0021726 | 0,0021726 | TRUE | 0,0400176 | No |
| orf19.3859 | 1761 | 1197 | 1501 | 1863 | 1562 | 1419 | 1511 | 1010 | 1580 | 1524 | 1536 | 1536 | 1336 | 1354 | 1348 | 1425 | 1454,8 | 1544 | 1366 | 0,882 | -0,18 | -4,116 | 3,85E-05 | 0,00012705 | 1,00E-08 | 0,0029774 | 0,0011551 | 0,0011551 | TRUE | 0,0087688 | No |
| orf19.386 | 1821 | 1271 | 1615 | 1954 | 1943 | 1654 | 1725 | 1236 | 1634 | 1618 | 1652 | 1611 | 1662 | 1578 | 1539 | 1743 | 1629,7 | 1629 | 1630 | 0,999 | -0,001 | -0,027 | 0,97856725 | 0,98436381 | 0,000761 | 0,0028703 | 0,0017013 | 0,0017013 | TRUE | 0,0234985 | No |
| orf19.3861 | 5978 | 4020 | 5240 | 6312 | 7648 | 6665 | 7311 | 4313 | 5363 | 5118 | 5361 | 5205 | 6541 | 6360 | 6524 | 6083 | 5819,3 | 5262 | 6377 | 1,213 | 0,278 | 7,55 | 4,34E-14 | 4,55E-13 | 0,000587 | 0,002229 | 0,0011277 | 0,0011277 | TRUE | 0,0111991 | No |
| orf19.3862 | 2677 | 1833 | 2342 | 2977 | 3247 | 2623 | 2902 | 1923 | 2402 | 2334 | 2396 | 2455 | 2777 | 2503 | 2590 | 2712 | 2521 | 2397 | 2646 | 1,103 | 0,141 | 3,141 | 0,0016849 | 0,00403624 | 0,000954 | 0,0025554 | 0,0015398 | 0,0015398 | TRUE | 0,0161433 | No |
| orf19.3863 | 112 | 90 | 110 | 117 | 97 | 107 | 119 | 68 | 100 | 115 | 113 | 96 | 83 | 102 | 106 | 96 | 101,4 | 106 | 97 | 0,917 | -0,125 | -0,871 | 0,38389319 | 0,47291988 | 1,00E-08 | 0,0162957 | 0,0098217 | 0,0098217 | TRUE | 0,1055266 | No |
| orf19.3865 | 2063 | 1450 | 1864 | 2318 | 2295 | 2101 | 2282 | 1473 | 1851 | 1846 | 1907 | 1911 | 1963 | 2005 | 2036 | 2078 | 1949,6 | 1879 | 2020 | 1,074 | 0,102 | 2,547 | 0,01085204 | 0,02166801 | 1,00E-08 | 0,0027241 | 0,0010379 | 0,0010379 | TRUE | 0,0045028 | No |
| orf19.3867 | 9897 | 7789 | 8680 | 10973 | 8785 | 8491 | 9122 | 5490 | 8879 | 9916 | 8880 | 9048 | 7513 | 8102 | 8140 | 7743 | 8527,7 | 9181 | 7874 | 0,858 | -0,221 | -4,632 | 3,63E-06 | 1,41E-05 | 0,002031 | 0,0021498 | 0,0020654 | 0,0020654 | TRUE | 0,0350403 | No |
| orf19.3868 | 0 | 1 | 0 | 2 | 1 | 1 | 1 | 1 | 0 | 1 | 0 | 2 | 1 | 1 | 1 | 1 | 0,88 | 1 | 1 | 1,378 | 0,463 | 0,316 | 0,75185508 | NA | 1,00E-08 | 1,6529353 | 0,881185 | 0,881185 | TRUE | 0,3622976 | No |
| orf19.3869 | 13744 | 9960 | 12946 | 15254 | 16240 | 15531 | 16956 | 11323 | 12330 | 12680 | 13245 | 12579 | 13889 | 14819 | 15130 | 15970 | 13830 | 12708 | 14952 | 1,176 | 0,234 | 4,973 | 6,59E-07 | 2,86E-06 | 0,002018 | 0,0020845 | 0,0020576 | 0,0020576 | TRUE | 0,0280884 | No |
| orf19.387 | 2258 | 1677 | 2054 | 2704 | 2184 | 1978 | 2263 | 1419 | 2026 | 2135 | 2101 | 2230 | 1868 | 1887 | 2019 | 2001 | 2033,5 | 2123 | 1944 | 0,915 | -0,128 | -2,652 | 0,0080004 | 0,016495 | 0,00118 | 0,0026934 | 0,0017575 | 0,0017575 | TRUE | 0,0147261 | No |
| orf19.3870 | 6181 | 4424 | 5474 | 6622 | 6592 | 6038 | 6279 | 4048 | 5545 | 5632 | 5600 | 5461 | 5638 | 5761 | 5603 | 5709 | 5618,7 | 5560 | 5678 | 1,021 | 0,031 | 1,05 | 0,29358791 | 0,37827067 | 1,00E-08 | 0,0022379 | 0,0006371 | 0,0006371 | TRUE | 0,001794 | No |
| orf19.3871 | 25 | 24 | 19 | 38 | 31 | 29 | 43 | 21 | 22 | 31 | 19 | 31 | 27 | 28 | 38 | 30 | 28,24 | 26 | 31 | 1,181 | 0,24 | 0,885 | 0,3763668 | 0,46555729 | 1,00E-08 | 0,0533835 | 0,0353821 | 0,0353821 | TRUE | 0,2281551 | No |
| orf19.3872 | 160 | 108 | 135 | 166 | 211 | 184 | 217 | 125 | 144 | 137 | 138 | 137 | 180 | 176 | 194 | 176 | 160,25 | 139 | 182 | 1,308 | 0,387 | 3,561 | 0,00036965 | 0,00102626 | 1,00E-08 | 0,0110385 | 0,0050228 | 0,0050228 | TRUE | 0,0230631 | No |
| orf19.3873 | 1135 | 700 | 940 | 1228 | 1406 | 1184 | 1287 | 822 | 1018 | 891 | 962 | 1013 | 1202 | 1130 | 1148 | 1159 | 1065,5 | 971 | 1160 | 1,192 | 0,253 | 4,486 | 7,27E-06 | 2,71E-05 | 0,000934 | 0,003342 | 0,0021093 | 0,0021093 | TRUE | 0,0334711 | No |
| orf19.3874 | 4033 | 3028 | 3731 | 4645 | 4173 | 3962 | 4226 | 2632 | 3618 | 3855 | 3817 | 3830 | 3569 | 3780 | 3771 | 3712 | 3744,1 | 3780 | 3708 | 0,981 | -0,027 | -0,714 | 0,47505577 | 0,56082038 | 0,000553 | 0,0023637 | 0,0011509 | 0,0011509 | TRUE | 0,0103264 | No |
| orf19.3876 | 63 | 59 | 67 | 93 | 79 | 63 | 91 | 48 | 57 | 75 | 69 | 77 | 68 | 60 | 81 | 68 | 69,18 | 70 | 69 | 1,003 | 0,005 | 0,028 | 0,97772457 | 0,98378316 | 1,00E-08 | 0,0229637 | 0,0153637 | 0,0153637 | TRUE | 0,1495874 | No |
| orf19.3877 | 473 | 343 | 476 | 580 | 486 | 513 | 573 | 324 | 424 | 437 | 487 | 478 | 416 | 489 | 511 | 457 | 462,46 | 456 | 468 | 1,025 | 0,036 | 0,423 | 0,67254442 | 0,73891623 | 0,004504 | 0,0051186 | 0,0048034 | 0,0048034 | TRUE | 0,0735707 | No |
| orf19.3878 | 775 | 567 | 707 | 856 | 1015 | 962 | 1011 | 575 | 695 | 722 | 723 | 706 | 868 | 918 | 902 | 811 | 793,18 | 712 | 875 | 1,234 | 0,303 | 5,2 | 1,99E-07 | 9,38E-07 | 1,00E-08 | 0,0038098 | 0,0019903 | 0,0019903 | TRUE | 0,0244979 | No |
| orf19.3879 | 78 | 49 | 69 | 81 | 69 | 69 | 72 | 64 | 70 | 62 | 71 | 67 | 59 | 66 | 64 | 90 | 68,64 | 68 | 70 | 1,016 | 0,023 | 0,13 | 0,89684997 | 0,92201824 | 0,002 | 0,0231295 | 0,0151123 | 0,0151123 | TRUE | 0,272245 | No |
| orf19.388 | 1431 | 970 | 1335 | 1546 | 2042 | 1646 | 1855 | 1246 | 1284 | 1235 | 1366 | 1275 | 1746 | 1571 | 1655 | 1757 | 1486,1 | 1290 | 1682 | 1,303 | 0,381 | 7,024 | 2,16E-12 | 1,91E-11 | 0,001558 | 0,0029564 | 0,0021448 | 0,0021448 | TRUE | 0,0241715 | No |
| orf19.3881 | 1137 | 747 | 907 | 1283 | 1512 | 1140 | 1272 | 845 | 1020 | 951 | 928 | 1058 | 1293 | 1088 | 1135 | 1192 | 1083,1 | 989 | 1177 | 1,188 | 0,249 | 3,604 | 0,00031345 | 0,00088414 | 0,003885 | 0,0033199 | 0,0036381 | 0,0036381 | TRUE | 0,0559079 | No |
| orf19.3884 | 429 | 284 | 453 | 494 | 338 | 270 | 299 | 202 | 385 | 362 | 463 | 407 | 289 | 258 | 267 | 285 | 339,46 | 404 | 275 | 0,677 | -0,562 | -5,984 | 2,18E-09 | 1,32E-08 | 0,004515 | 0,0062561 | 0,0054054 | 0,0054054 | TRUE | 0,1076469 | No |
| orf19.3885 | 166 | 114 | 135 | 193 | 173 | 182 | 156 | 118 | 149 | 145 | 138 | 159 | 148 | 174 | 139 | 166 | 152,32 | 148 | 157 | 1,053 | 0,075 | 0,628 | 0,53005882 | 0,61277533 | 1,00E-08 | 0,0115101 | 0,0071015 | 0,0071015 | TRUE | 0,0635275 | No |
| orf19.3886 | 2720 | 1951 | 2428 | 2964 | 2852 | 2453 | 2654 | 1779 | 2440 | 2484 | 2484 | 2444 | 2439 | 2341 | 2368 | 2509 | 2438,7 | 2463 | 2414 | 0,979 | -0,03 | -0,784 | 0,43301787 | 0,52004474 | 1,00E-08 | 0,0025748 | 0,0010038 | 0,0010038 | TRUE | 0,0075401 | No |
| orf19.3887 | 2862 | 2051 | 2345 | 2972 | 2821 | 2414 | 2691 | 1398 | 2568 | 2611 | 2399 | 2451 | 2413 | 2303 | 2401 | 1972 | 2389,7 | 2507 | 2272 | 0,908 | -0,139 | -2,133 | 0,03292401 | 0,05816155 | 0,004337 | 0,002587 | 0,003649 | 0,003649 | TRUE | 0,0916221 | No |
| orf19.3888 | 2414 | 1514 | 2070 | 2512 | 4151 | 3525 | 3802 | 2528 | 2166 | 1927 | 2118 | 2071 | 3550 | 3363 | 3393 | 3566 | 2769,3 | 2070 | 3468 | 1,672 | 0,742 | 15,994 | 1,40E-57 | 2,00E-55 | 0,001175 | 0,0025037 | 0,0016781 | 0,0016781 | TRUE | 0,0244942 | No |
| orf19.3888.2 | 3 | 1 | 5 | 2 | 1 | 1 | 3 | 0 | 3 | 1 | 5 | 2 | 1 | 1 | 3 | 0 | 1,9 | 3 | 1 | 0,446 | -1,165 | -1,103 | 0,27000313 | NA | 1,00E-08 | 0,7652524 | 0,4712505 | 0,4712505 | TRUE | 0,4847177 | No |
| orf19.389 | 134 | 81 | 94 | 143 | 168 | 147 | 136 | 104 | 120 | 103 | 96 | 118 | 144 | 140 | 121 | 147 | 123,68 | 109 | 138 | 1,251 | 0,323 | 2,446 | 0,01443068 | 0,02791357 | 1,00E-08 | 0,0137175 | 0,0085969 | 0,0085969 | TRUE | 0,0725703 | No |
| orf19.3893 | 113 | 85 | 95 | 108 | 156 | 129 | 162 | 112 | 101 | 108 | 97 | 89 | 133 | 123 | 145 | 158 | 119,36 | 99 | 140 | 1,411 | 0,497 | 3,704 | 0,00021181 | 0,00061447 | 1,00E-08 | 0,0141421 | 0,0086158 | 0,0086158 | TRUE | 0,0640226 | No |
| orf19.3894 | 2909 | 1966 | 2324 | 2999 | 3210 | 2833 | 2999 | 1850 | 2610 | 2503 | 2378 | 2473 | 2745 | 2703 | 2676 | 2609 | 2587,1 | 2491 | 2683 | 1,078 | 0,108 | 2,589 | 0,0096319 | 0,01947777 | 0,000566 | 0,0025406 | 0,0012918 | 0,0012918 | TRUE | 0,0129355 | No |
| orf19.3895 | 843 | 567 | 767 | 966 | 825 | 659 | 728 | 566 | 756 | 722 | 785 | 797 | 706 | 629 | 650 | 798 | 730,21 | 765 | 696 | 0,905 | -0,144 | -1,865 | 0,0621947 | 0,10157838 | 0,00505 | 0,0039676 | 0,0043815 | 0,0043815 | TRUE | 0,1054538 | No |
| orf19.3897 | 32 | 25 | 35 | 46 | 24 | 15 | 31 | 10 | 29 | 32 | 36 | 38 | 21 | 14 | 28 | 14 | 26,36 | 34 | 19 | 0,58 | -0,785 | -2,719 | 0,0065396 | 0,01377626 | 0,009088 | 0,0570516 | 0,0396026 | 0,0396026 | TRUE | 0,50255 | No |
| orf19.3898 | 224 | 151 | 207 | 262 | 298 | 267 | 258 | 158 | 201 | 192 | 212 | 216 | 255 | 255 | 230 | 223 | 222,96 | 205 | 241 | 1,174 | 0,231 | 2,378 | 0,01738348 | 0,03282211 | 1,00E-08 | 0,0084905 | 0,0045816 | 0,0045816 | TRUE | 0,0221104 | No |
| orf19.3899 | 304 | 223 | 275 | 349 | 324 | 289 | 287 | 164 | 273 | 284 | 281 | 288 | 277 | 276 | 256 | 231 | 270,75 | 282 | 260 | 0,93 | -0,105 | -1,18 | 0,23818177 | 0,31906267 | 1,00E-08 | 0,0073412 | 0,003938 | 0,003938 | TRUE | 0,0493113 | No |
| orf19.390 | 1776 | 1202 | 1534 | 1909 | 1875 | 1591 | 1858 | 1088 | 1593 | 1530 | 1569 | 1574 | 1604 | 1518 | 1658 | 1535 | 1572,7 | 1566 | 1579 | 1,008 | 0,012 | 0,253 | 0,80029754 | 0,84517273 | 0,000365 | 0,0029026 | 0,001403 | 0,001403 | TRUE | 0,0140151 | No |
| orf19.3900 | 8206 | 5598 | 7193 | 9030 | 9397 | 7973 | 9040 | 5511 | 7362 | 7127 | 7359 | 7446 | 8037 | 7608 | 8067 | 7773 | 7597,2 | 7324 | 7871 | 1,075 | 0,104 | 3,111 | 0,00186726 | 0,00443594 | 0,000389 | 0,0021706 | 0,0009392 | 0,0009392 | TRUE | 0,0063839 | No |
| orf19.3901 | 610 | 407 | 549 | 660 | 897 | 741 | 822 | 569 | 547 | 518 | 562 | 544 | 767 | 707 | 733 | 803 | 647,69 | 543 | 752 | 1,381 | 0,466 | 7,302 | 2,83E-13 | 2,76E-12 | 1,00E-08 | 0,0042209 | 0,002326 | 0,002326 | TRUE | 0,0203127 | No |
| orf19.3902 | 53 | 52 | 55 | 63 | 36 | 48 | 44 | 38 | 48 | 66 | 56 | 52 | 31 | 46 | 39 | 54 | 48,93 | 56 | 42 | 0,756 | -0,403 | -1,859 | 0,06296156 | 0,10273575 | 0,012014 | 0,0316503 | 0,0240266 | 0,0240266 | TRUE | 0,2844264 | No |
| orf19.3903 | 1 | 1 | 0 | 0 | 1 | 1 | 1 | 0 | 1 | 1 | 0 | 0 | 1 | 1 | 1 | 0 | 0,61 | 0 | 1 | 1,431 | 0,517 | 0,298 | 0,76564036 | NA | 1,00E-08 | 2,3857522 | 1,2633306 | 1,2633306 | TRUE | 0,1915996 | No |

| | | | | | | | | | | | | | | | | | | | | | | | | | | | | | | | |
|---|---|---|---|---|---|---|---|---|---|---|---|---|---|---|---|---|---|---|---|---|---|---|---|---|---|---|---|---|---|---|---|
| orf19.3904 | 18 | 7 | 18 | 26 | 10 | 19 | 20 | 6 | 16 | 9 | 18 | 21 | 9 | 18 | 18 | 8 | 14,74 | 16 | 13 | 0,814 | -0,296 | -0,737 | 0,4612482 | NA | 0,068329 | 0,1004784 | 0,0879593 | 0,0879593 | TRUE | 0,1994582 | No |
| orf19.3905 | 8 | 7 | 5 | 8 | 2 | 7 | 8 | 2 | 7 | 9 | 5 | 7 | 2 | 7 | 7 | 3 | 5,77 | 7 | 5 | 0,676 | -0,564 | -0,946 | 0,343921 | NA | 1,00E-08 | 0,2536281 | 0,1622438 | 0,1622438 | TRUE | 0,3783525 | No |
| orf19.3906 | 13 | 5 | 5 | 12 | 5 | 9 | 15 | 9 | 12 | 6 | 5 | 10 | 4 | 9 | 13 | 13 | 9 | 8 | 10 | 1,138 | 0,187 | 0,373 | 0,70886475 | NA | 0,077696 | 0,163322 | 0,1284957 | 0,1284957 | TRUE | 0,5921232 | No |
| orf19.3907 | 1 | 1 | 1 | 0 | 1 | 0 | 0 | 0 | 1 | 1 | 1 | 0 | 1 | 0 | 0 | 0 | 0,51 | 1 | 0 | 0,396 | -1,338 | -0,7 | 0,48405801 | NA | 1,00E-08 | 2,8705706 | 1,8671604 | 1,8671604 | TRUE | 0,2537767 | No |
| orf19.3908 | 55 | 32 | 37 | 44 | 23 | 20 | 26 | 19 | 49 | 41 | 38 | 36 | 20 | 19 | 23 | 27 | 31,62 | 41 | 22 | 0,533 | -0,908 | -3,58 | 0,00034393 | 0,00096018 | 1,00E-08 | 0,0478886 | 0,0269151 | 0,0269151 | TRUE | 0,1574746 | No |
| orf19.391 | 1191 | 814 | 1077 | 1355 | 1017 | 937 | 1019 | 718 | 1068 | 1036 | 1102 | 1117 | 870 | 894 | 909 | 1013 | 1001,2 | 1081 | 922 | 0,848 | -0,237 | -4,006 | 6,16E-05 | 0,00019591 | 0,001552 | 0,0034295 | 0,0023595 | 0,0023595 | TRUE | 0,0470552 | No |
| orf19.3910 | 374 | 252 | 326 | 378 | 328 | 274 | 310 | 206 | 336 | 321 | 334 | 312 | 281 | 261 | 277 | 291 | 301,34 | 326 | 278 | 0,85 | -0,234 | -2,842 | 0,00448465 | 0,00977996 | 1,00E-08 | 0,006797 | 0,0031662 | 0,0031662 | TRUE | 0,0161646 | No |
| orf19.3911 | 9260 | 5947 | 7898 | 9705 | 7965 | 6358 | 6457 | 4885 | 8307 | 7571 | 8080 | 8003 | 6812 | 6067 | 5762 | 6890 | 7186,5 | 7990 | 6383 | 0,798 | -0,325 | -5,267 | 1,39E-07 | 6,69E-07 | 0,004347 | 0,0021815 | 0,0035138 | 0,0035138 | TRUE | 0,0525008 | No |
| orf19.3912 | 3204 | 2194 | 2895 | 3703 | 3820 | 3473 | 3842 | 2564 | 2874 | 2793 | 2962 | 3054 | 3267 | 3314 | 3428 | 3616 | 3163,6 | 2921 | 3406 | 1,164 | 0,219 | 4,689 | 2,75E-06 | 1,09E-05 | 0,001356 | 0,0024384 | 0,0017811 | 0,0017811 | TRUE | 0,0198474 | No |
| orf19.3914 | 7688 | 5817 | 6881 | 8627 | 8283 | 7991 | 8517 | 5103 | 6897 | 7406 | 7040 | 7114 | 7084 | 7625 | 7600 | 7197 | 7245,3 | 7114 | 7376 | 1,037 | 0,053 | 1,331 | 0,18332409 | 0,25552066 | 0,000972 | 0,0021799 | 0,0013841 | 0,0013841 | TRUE | 0,0090616 | No |
| orf19.3915 | 4297 | 2994 | 3634 | 4672 | 4783 | 4081 | 4368 | 2734 | 3855 | 3812 | 3718 | 3853 | 4091 | 3894 | 3898 | 3856 | 3871,9 | 3810 | 3935 | 1,033 | 0,047 | 1,339 | 0,18046828 | 0,25209333 | 0,000273 | 0,0023545 | 0,0009393 | 0,0009393 | TRUE | 0,0089087 | No |
| orf19.3916 | 1086 | 895 | 940 | 1164 | 1058 | 1069 | 1104 | 700 | 974 | 1139 | 962 | 960 | 905 | 1020 | 985 | 987 | 991,56 | 1009 | 974 | 0,967 | -0,048 | -0,688 | 0,49134119 | 0,57655818 | 0,003665 | 0,0034436 | 0,0035949 | 0,0035949 | TRUE | 0,08562 | No |
| orf19.3919 | 1166 | 855 | 993 | 1248 | 1483 | 1374 | 1464 | 960 | 1046 | 1089 | 1016 | 1029 | 1268 | 1311 | 1306 | 1354 | 1177,4 | 1045 | 1310 | 1,253 | 0,326 | 6,746 | 1,52E-11 | 1,22E-10 | 1,00E-08 | 0,0032125 | 0,0013779 | 0,0013779 | TRUE | 0,0083743 | No |
| orf19.3919.1 | 764 | 497 | 532 | 701 | 792 | 635 | 662 | 457 | 685 | 633 | 544 | 578 | 677 | 606 | 591 | 645 | 619,87 | 610 | 630 | 1,032 | 0,045 | 0,544 | 0,58627668 | 0,66459439 | 0,005618 | 0,0043215 | 0,005007 | 0,005007 | TRUE | 0,0854144 | No |
| orf19.392 | 836 | 577 | 734 | 980 | 731 | 588 | 714 | 433 | 750 | 735 | 751 | 808 | 625 | 561 | 637 | 611 | 684,71 | 761 | 608 | 0,799 | -0,324 | -5,012 | 5,39E-07 | 2,38E-06 | 0,001106 | 0,0040997 | 0,0025361 | 0,0025361 | TRUE | 0,0341194 | No |
| orf19.3920 | 797 | 509 | 650 | 825 | 639 | 529 | 570 | 368 | 715 | 648 | 665 | 680 | 546 | 505 | 509 | 519 | 598,4 | 677 | 520 | 0,767 | -0,383 | -5,995 | 2,04E-09 | 1,24E-08 | 1,00E-08 | 0,0044055 | 0,0022288 | 0,0022288 | TRUE | 0,0168264 | No |
| orf19.3921 | 1433 | 882 | 1169 | 1445 | 1590 | 1209 | 1418 | 895 | 1286 | 1123 | 1196 | 1192 | 1360 | 1154 | 1265 | 1262 | 1229,6 | 1199 | 1260 | 1,05 | 0,071 | 1,108 | 0,26789751 | 0,35204794 | 0,003045 | 0,0031601 | 0,0030945 | 0,0030945 | TRUE | 0,0402728 | No |
| orf19.3922 | 3971 | 2742 | 3327 | 4144 | 4759 | 3962 | 4227 | 2873 | 3562 | 3491 | 3404 | 3417 | 4070 | 3780 | 3772 | 4052 | 3693,6 | 3468 | 3918 | 1,129 | 0,175 | 4,225 | 2,38E-05 | 8,19E-05 | 0,000744 | 0,0023726 | 0,00138 | 0,00138 | TRUE | 0,0088605 | No |
| orf19.3923 | 450 | 290 | 374 | 443 | 420 | 374 | 371 | 297 | 404 | 369 | 383 | 365 | 359 | 357 | 331 | 419 | 373,36 | 380 | 366 | 0,957 | -0,063 | -0,734 | 0,46274265 | 0,54867926 | 0,002612 | 0,0058678 | 0,0043856 | 0,0043856 | TRUE | 0,0909263 | No |
| orf19.3924 | 50 | 32 | 44 | 54 | 26 | 27 | 36 | 12 | 45 | 41 | 45 | 45 | 22 | 26 | 32 | 17 | 34,02 | 44 | 24 | 0,563 | -0,828 | -3,393 | 0,00069096 | 0,00179787 | 1,00E-08 | 0,0446472 | 0,0256547 | 0,0256547 | TRUE | 0,2935929 | No |
| orf19.3925 | 8 | 8 | 14 | 12 | 8 | 12 | 10 | 5 | 7 | 10 | 14 | 10 | 7 | 11 | 9 | 7 | 9,48 | 10 | 8 | 0,835 | -0,26 | -0,581 | 0,56100011 | NA | 1,00E-08 | 0,155097 | 0,0867061 | 0,0867061 | TRUE | 0,2318442 | No |
| orf19.3926 | 43 | 41 | 39 | 46 | 51 | 33 | 29 | 29 | 39 | 52 | 40 | 38 | 44 | 31 | 26 | 41 | 38,81 | 42 | 36 | 0,844 | -0,244 | -1,022 | 0,30691431 | 0,3928294 | 0,012953 | 0,0393837 | 0,028858 | 0,028858 | TRUE | 0,2648389 | No |
| orf19.3928 | 469 | 348 | 428 | 513 | 652 | 516 | 602 | 356 | 421 | 443 | 438 | 423 | 558 | 492 | 537 | 502 | 476,74 | 431 | 522 | 1,216 | 0,283 | 3,992 | 6,56E-05 | 0,00020755 | 1,00E-08 | 0,0050246 | 0,0026981 | 0,0026981 | TRUE | 0,0251089 | No |
| orf19.3929 | 955 | 642 | 880 | 1059 | 778 | 667 | 694 | 480 | 857 | 817 | 900 | 873 | 665 | 636 | 619 | 677 | 755,72 | 862 | 649 | 0,751 | -0,413 | -6,975 | 3,06E-12 | 2,65E-11 | 1,00E-08 | 0,0039005 | 0,0020098 | 0,0020098 | TRUE | 0,012801 | No |
| orf19.393 | 984 | 726 | 821 | 1087 | 1059 | 863 | 908 | 651 | 883 | 924 | 840 | 896 | 906 | 823 | 810 | 918 | 875,11 | 886 | 864 | 0,974 | -0,037 | -0,601 | 0,54789301 | 0,6292757 | 0,001592 | 0,0036384 | 0,0025464 | 0,0025464 | TRUE | 0,0213833 | No |
| orf19.3930 | 3111 | 2072 | 2635 | 3405 | 3271 | 2821 | 3061 | 2032 | 2791 | 2638 | 2696 | 2808 | 2797 | 2692 | 2731 | 2866 | 2752,4 | 2733 | 2772 | 1,012 | 0,018 | 0,44 | 0,66025371 | 0,72865619 | 0,000329 | 0,002507 | 0,0011673 | 0,0011673 | TRUE | 0,0063379 | No |
| orf19.3931 | 2778 | 1746 | 2337 | 2888 | 1858 | 1574 | 1686 | 1158 | 2492 | 2223 | 2391 | 2381 | 1589 | 1502 | 1504 | 1633 | 1964,5 | 2372 | 1557 | 0,655 | -0,61 | -12,074 | 1,45E-33 | 6,55E-32 | 0,00126 | 0,0027185 | 0,0019188 | 0,0019188 | TRUE | 0,0205931 | No |
| orf19.3931.2 | 17 | 20 | 14 | 22 | 12 | 9 | 15 | 16 | 15 | 25 | 14 | 18 | 10 | 9 | 13 | 23 | 16 | 18 | 14 | 0,731 | -0,452 | -1,186 | 0,23553144 | 0,31641017 | 0,043166 | 0,0927253 | 0,0740737 | 0,0740737 | TRUE | 0,5643321 | No |
| orf19.3932 | 9780 | 6306 | 7832 | 10181 | 11765 | 9706 | 10734 | 6896 | 8774 | 8028 | 8013 | 8395 | 10062 | 9261 | 9578 | 9726 | 8979,7 | 8302 | 9657 | 1,163 | 0,218 | 5,072 | 3,94E-07 | 1,77E-06 | 0,001386 | 0,0021412 | 0,0016549 | 0,0016549 | TRUE | 0,0178571 | No |
| orf19.3932.1 | 1456 | 944 | 1217 | 1517 | 1946 | 1361 | 1406 | 1212 | 1306 | 1202 | 1245 | 1251 | 1664 | 1299 | 1255 | 1709 | 1366,4 | 1251 | 1482 | 1,183 | 0,242 | 2,503 | 0,01233166 | 0,02425158 | 0,012528 | 0,003042 | 0,0082331 | 0,0082331 | TRUE | 0,1242267 | No |
| orf19.3934 | 16588 | 10907 | 14266 | 17380 | 13578 | 11100 | 12103 | 7436 | 14881 | 13886 | 14596 | 14332 | 11612 | 10591 | 10800 | 10488 | 12648 | 14424 | 10873 | 0,754 | -0,407 | -9,639 | 5,47E-22 | 1,14E-20 | 0,001422 | 0,0020943 | 0,001635 | 0,001635 | TRUE | 0,0257995 | No |
| orf19.3935 | 4 | 2 | 4 | 6 | 4 | 3 | 2 | 2 | 4 | 3 | 4 | 5 | 3 | 3 | 2 | 3 | 3,26 | 4 | 3 | 0,704 | -0,506 | -0,687 | 0,49224265 | NA | 1,00E-08 | 0,4475664 | 0,2097291 | 0,2097291 | TRUE | 0,0818001 | No |
| orf19.3936 | 2948 | 2179 | 2701 | 3148 | 2512 | 2332 | 2419 | 1497 | 2645 | 2774 | 2763 | 2596 | 2148 | 2225 | 2159 | 2111 | 2427,7 | 2694 | 2161 | 0,803 | -0,316 | -7,718 | 1,18E-14 | 1,31E-13 | 0,000398 | 0,0025775 | 0,0011915 | 0,0011915 | TRUE | 0,0075176 | No |
| orf19.3937 | 526 | 386 | 510 | 602 | 621 | 604 | 634 | 406 | 472 | 491 | 522 | 496 | 531 | 576 | 566 | 573 | 528,41 | 495 | 562 | 1,132 | 0,179 | 2,694 | 0,00705888 | 0,01471538 | 1,00E-08 | 0,0047268 | 0,0023522 | 0,0023522 | TRUE | 0,0166874 | No |
| orf19.3938 | 1795 | 1252 | 1489 | 1940 | 2352 | 2134 | 2281 | 1782 | 1610 | 1594 | 1523 | 1600 | 2012 | 2036 | 2035 | 2513 | 1865,5 | 1582 | 2149 | 1,356 | 0,439 | 6,105 | 1,03E-09 | 6,54E-09 | 0,005978 | 0,0027577 | 0,0044156 | 0,0044156 | TRUE | 0,1520492 | No |
| orf19.3939 | 690 | 514 | 603 | 737 | 652 | 613 | 676 | 435 | 619 | 654 | 617 | 608 | 558 | 585 | 603 | 614 | 607,16 | 624 | 590 | 0,944 | -0,083 | -1,328 | 0,18406805 | 0,25622819 | 1,00E-08 | 0,0043705 | 0,002101 | 0,002101 | TRUE | 0,0166265 | No |
| orf19.394 | 1534 | 1045 | 1280 | 1596 | 1906 | 1538 | 1738 | 1173 | 1376 | 1330 | 1310 | 1316 | 1630 | 1468 | 1551 | 1654 | 1454,4 | 1333 | 1576 | 1,18 | 0,239 | 4,642 | 3,45E-06 | 1,35E-05 | 0,00107 | 0,0029777 | 0,001859 | 0,001859 | TRUE | 0,0256652 | No |
| orf19.3940 | 119 | 78 | 108 | 144 | 80 | 74 | 70 | 53 | 107 | 99 | 110 | 119 | 68 | 71 | 62 | 75 | 88,94 | 109 | 69 | 0,628 | -0,671 | -4,524 | 6,08E-06 | 2,29E-05 | 1,00E-08 | 0,0183013 | 0,0093132 | 0,0093132 | TRUE | 0,0403521 | No |
| orf19.3940.1 | 61567 | 33145 | 45302 | 59023 | 74206 | 42907 | 49512 | 40372 | 55232 | 42197 | 46348 | 48671 | 63463 | 40941 | 44181 | 56941 | 49747 | 48112 | 51382 | 1,068 | 0,095 | 0,578 | 0,56335391 | 0,64322127 | 0,025853 | 0,0020087 | 0,01414 | 0,0258534 | TRUE | 0,2454343 | No |
| orf19.3941 | 18021 | 12069 | 15940 | 19477 | 18056 | 15293 | 16580 | 10269 | 16167 | 15365 | 16308 | 16061 | 15442 | 14592 | 14795 | 14484 | 15402 | 15975 | 14828 | 0,928 | -0,107 | -3,112 | 0,00185663 | 0,00441592 | 0,000676 | 0,0020738 | 0,0010791 | 0,0010791 | TRUE | 0,0095673 | No |
| orf19.3942 | 408 | 280 | 332 | 419 | 386 | 357 | 361 | 206 | 366 | 356 | 340 | 346 | 330 | 341 | 322 | 291 | 336,39 | 352 | 321 | 0,916 | -0,126 | -1,565 | 0,11769638 | 0,17533346 | 1,00E-08 | 0,0062951 | 0,0032616 | 0,0032616 | TRUE | 0,0368102 | No |
| orf19.3942.1 | 15060 | 10115 | 11755 | 15416 | 15321 | 12228 | 12211 | 7621 | 13510 | 12877 | 12027 | 12712 | 13103 | 11668 | 10896 | 10749 | 12193 | 12782 | 11604 | 0,908 | -0,139 | -2,148 | 0,03172964 | 0,05634988 | 0,00534 | 0,0020986 | 0,0039508 | 0,0039508 | TRUE | 0,0927515 | No |
| orf19.3944 | 7870 | 5504 | 7199 | 8515 | 10661 | 9403 | 10300 | 6634 | 7060 | 7007 | 7365 | 7022 | 9118 | 8972 | 9191 | 9357 | 8136,5 | 7114 | 9160 | 1,287 | 0,364 | 11,658 | 2,09E-31 | 8,14E-30 | 0,000269 | 0,0021579 | 0,000812 | 0,000812 | TRUE | 0,0068929 | No |
| orf19.3945 | 4023 | 3134 | 3709 | 4529 | 4851 | 4673 | 4942 | 3022 | 3609 | 3990 | 3795 | 3735 | 4149 | 4459 | 4410 | 4262 | 4051 | 3782 | 4320 | 1,143 | 0,193 | 4,445 | 8,81E-06 | 3,24E-05 | 0,001086 | 0,0023379 | 0,0015632 | 0,0015632 | TRUE | 0,0160785 | No |
| orf19.3946 | 405 | 278 | 358 | 444 | 461 | 417 | 420 | 312 | 363 | 354 | 366 | 366 | 394 | 398 | 375 | 440 | 382,08 | 362 | 402 | 1,102 | 0,14 | 1,834 | 0,06671305 | 0,10832757 | 1,00E-08 | 0,005779 | 0,0029799 | 0,0029799 | TRUE | 0,0397046 | No |
| orf19.3947 | 1107 | 787 | 949 | 1190 | 1341 | 1088 | 1106 | 688 | 993 | 1002 | 971 | 981 | 1147 | 1038 | 987 | 970 | 1011,2 | 987 | 1036 | 1,052 | 0,073 | 1,185 | 0,23582805 | 0,31661563 | 0,0021 | 0,0034152 | 0,0026754 | 0,0026754 | TRUE | 0,0644844 | No |
| orf19.3949 | 3652 | 2527 | 3246 | 3859 | 4641 | 3976 | 4353 | 2756 | 3276 | 3217 | 3321 | 3182 | 3969 | 3794 | 3884 | 3887 | 3566,4 | 3249 | 3884 | 1,195 | 0,258 | 7,512 | 5,84E-14 | 6,07E-13 | 1,00E-08 | 0,0023866 | 0,0008456 | 0,0008456 | TRUE | 0,0030551 | No |
| orf19.395 | 9088 | 5549 | 7716 | 9223 | 14978 | 11641 | 12454 | 8741 | 8153 | 7064 | 7894 | 7605 | 12810 | 11107 | 11113 | 12328 | 9759,5 | 7679 | 11840 | 1,541 | 0,624 | 10,165 | 2,84E-24 | 6,81E-23 | 0,004373 | 0,0021283 | 0,0035141 | 0,0035141 | TRUE | 0,037557 | No |
| orf19.3950 | 301 | 194 | 251 | 305 | 337 | 301 | 336 | 207 | 270 | 247 | 257 | 252 | 288 | 287 | 300 | 292 | 274,06 | 256 | 292 | 1,137 | 0,185 | 2,191 | 0,02848203 | 0,05118756 | 1,00E-08 | 0,0072765 | 0,0031972 | 0,0031972 | TRUE | 0,0150317 | No |
| orf19.3951 | 1453 | 991 | 1221 | 1559 | 1364 | 1142 | 1326 | 807 | 1303 | 1262 | 1249 | 1286 | 1167 | 1090 | 1183 | 1138 | 1209,7 | 1275 | 1144 | 0,898 | -0,156 | -3,218 | 0,00129101 | 0,00317365 | 1,00E-08 | 0,0031796 | 0,0014236 | 0,0014236 | TRUE | 0,0132346 | No |
| orf19.3952 | 2863 | 2019 | 2622 | 3264 | 2684 | 2275 | 2537 | 1576 | 2568 | 2570 | 2683 | 2692 | 2295 | 2171 | 2264 | 2223 | 2433,2 | 2628 | 2238 | 0,852 | -0,232 | -5,833 | 5,43E-09 | 3,13E-08 | 0,000262 | 0,0025761 | 0,0011005 | 0,0011005 | TRUE | 0,0052959 | No |
| orf19.3954 | 1073 | 722 | 929 | 1050 | 1285 | 1301 | 1324 | 804 | 963 | 919 | 950 | 866 | 1099 | 1241 | 1181 | 1134 | 1044,2 | 924 | 1164 | 1,26 | 0,334 | 5,589 | 2,28E-08 | 1,21E-07 | 0,001745 | 0,0033697 | 0,0024463 | 0,0024463 | TRUE | 0,0244468 | No |
| orf19.3954.1 | 41 | 32 | 25 | 45 | 39 | 38 | 52 | 32 | 37 | 41 | 26 | 37 | 33 | 36 | 46 | 45 | 37,67 | 35 | 40 | 1,144 | 0,194 | 0,825 | 0,40936826 | 0,49736795 | 1,00E-08 | 0,0405178 | 0,0264221 | 0,0264221 | TRUE | 0,2253526 | No |
| orf19.3955 | 4460 | 2999 | 3780 | 4597 | 4608 | 4169 | 4278 | 3020 | 4001 | 3818 | 3867 | 3791 | 3941 | 3978 | 3817 | 4259 | 3934,1 | 3869 | 3999 | 1,032 | 0,046 | 1,066 | 0,28660597 | 0,37110531 | 0,001043 | 0,0023485 | 0,0015087 | 0,0015087 | TRUE | 0,0222192 | No |
| orf19.3956 | 1110 | 717 | 910 | 1117 | 1275 | 1107 | 1163 | 810 | 996 | 913 | 931 | 921 | 1090 | 1056 | 1038 | 1142 | 1011 | 940 | 1082 | 1,147 | 0,198 | 3,604 | 0,00031276 | 0,0008826 | 0,00063 | 0,0034155 | 0,0019051 | 0,0019051 | TRUE | 0,019514 | No |
| orf19.3957 | 2062 | 1388 | 1798 | 2269 | 2248 | 1887 | 2067 | 1230 | 1850 | 1767 | 1840 | 1871 | 1923 | 1801 | 1844 | 1735 | 1828,7 | 1832 | 1826 | 0,997 | -0,004 | -0,08 | 0,93618994 | 0,95395767 | 0,000505 | 0,0027734 | 0,0014838 | 0,0014838 | TRUE | 0,0155083 | No |
| orf19.3959 | 4168 | 2727 | 3435 | 4312 | 4464 | 3652 | 3919 | 2647 | 3739 | 3472 | 3514 | 3556 | 3818 | 3485 | 3497 | 3733 | 3601,7 | 3570 | 3633 | 1,017 | 0,024 | 0,53 | 0,59620336 | 0,67292689 | 0,001196 | 0,0023826 | 0,0017173 | 0,0017173 | TRUE | 0,0150114 | No |
| orf19.396 | 385 | 280 | 327 | 397 | 314 | 286 | 310 | 185 | 345 | 356 | 335 | 327 | 269 | 273 | 277 | 261 | 305,35 | 341 | 270 | 0,794 | -0,332 | -4,106 | 4,03E-05 | 0,00013227 | 1,00E-08 | 0,0067338 | 0,0029774 | 0,0029774 | TRUE | 0,0093482 | No |
| orf19.3960 | 3049 | 1975 | 2800 | 3223 | 2927 | 2767 | 2918 | 1773 | 2735 | 2514 | 2865 | 2658 | 2503 | 2640 | 2604 | 2501 | 2627,5 | 2693 | 2562 | 0,951 | -0,072 | -1,501 | 0,13327281 | 0,19529343 | 0,001355 | 0,002532 | 0,0018446 | 0,0018446 | TRUE | 0,0228417 | No |

| | | | | | | | | | | | | | | | | | | | | | | | | | | | | | | | |
|---|---|---|---|---|---|---|---|---|---|---|---|---|---|---|---|---|---|---|---|---|---|---|---|---|---|---|---|---|---|---|---|
| orf19.3962 | 16232 | 11316 | 14342 | 17531 | 16540 | 14753 | 16086 | 10102 | 14562 | 14406 | 14673 | 14456 | 14146 | 14077 | 14354 | 14248 | 14365 | 14524 | 14206 | 0,978 | -0,032 | -1,379 | 0,16784482 | 0,23749687 | 1,00E-08 | 0,0020806 | 0,0004498 | 0,0004498 | TRUE | 0,0006241 | No |
| orf19.3963 | 1242 | 920 | 1072 | 1352 | 1465 | 1343 | 1417 | 1012 | 1114 | 1171 | 1097 | 1115 | 1253 | 1281 | 1264 | 1427 | 1215,4 | 1124 | 1306 | 1,16 | 0,214 | 3,868 | 0,00010951 | 0,0003333 | 0,001253 | 0,0031739 | 0,0021001 | 0,0021001 | TRUE | 0,0421593 | No |
| orf19.3964 | 687 | 427 | 579 | 672 | 716 | 580 | 638 | 397 | 616 | 544 | 592 | 554 | 612 | 553 | 569 | 560 | 575,18 | 576 | 574 | 0,995 | -0,007 | -0,098 | 0,92157265 | 0,94259593 | 0,001173 | 0,0045034 | 0,0028067 | 0,0028067 | TRUE | 0,0259412 | No |
| orf19.3965 | 405 | 300 | 376 | 470 | 406 | 381 | 370 | 252 | 363 | 382 | 385 | 388 | 347 | 364 | 330 | 355 | 364,23 | 380 | 349 | 0,919 | -0,122 | -1,608 | 0,10781402 | 0,16250543 | 1,00E-08 | 0,0059652 | 0,002755 | 0,002755 | TRUE | 0,0161253 | No |
| orf19.3966 | 22 | 23 | 17 | 25 | 25 | 32 | 26 | 13 | 20 | 29 | 17 | 21 | 21 | 31 | 23 | 18 | 22,56 | 22 | 23 | 1,098 | 0,135 | 0,451 | 0,65179463 | 0,72090982 | 1,00E-08 | 0,0663294 | 0,0414797 | 0,0414797 | TRUE | 0,2509905 | No |
| orf19.3967 | 26208 | 17798 | 23451 | 27398 | 33248 | 29380 | 31722 | 19913 | 23511 | 22659 | 23993 | 22593 | 28435 | 28033 | 28307 | 28086 | 25702 | 23189 | 28215 | 1,217 | 0,283 | 9,524 | 1,67E-21 | 3,35E-20 | 0,0004 | 0,002036 | 0,0008089 | 0,0008089 | TRUE | 0,0066455 | No |
| orf19.3968 | 3 | 6 | 5 | 9 | 6 | 14 | 9 | 8 | 3 | 8 | 5 | 7 | 5 | 13 | 8 | 11 | 7,58 | 6 | 9 | 1,646 | 0,719 | 1,338 | 0,18093755 | NA | 1,00E-08 | 0,193399 | 0,1360111 | 0,1360111 | TRUE | 0,2296295 | No |
| orf19.3969 | 4275 | 2940 | 3959 | 4686 | 6451 | 5738 | 6325 | 3964 | 3835 | 3743 | 4050 | 3864 | 5517 | 5475 | 5644 | 5591 | 4715 | 3873 | 5557 | 1,434 | 0,52 | 14,697 | 6,78E-49 | 6,57E-47 | 0,000354 | 0,0022874 | 0,0009829 | 0,0009829 | TRUE | 0,0112985 | No |
| orf19.397 | 1113 | 715 | 811 | 1028 | 1138 | 984 | 1031 | 796 | 998 | 910 | 830 | 848 | 973 | 939 | 920 | 1123 | 942,63 | 896 | 989 | 1,1 | 0,137 | 1,713 | 0,0866417 | 0,13545234 | 0,006525 | 0,0035196 | 0,0050971 | 0,0050971 | TRUE | 0,0929327 | No |
| orf19.3970 | 1637 | 1105 | 1341 | 1798 | 1983 | 1706 | 1635 | 1011 | 1469 | 1407 | 1372 | 1483 | 1696 | 1628 | 1459 | 1426 | 1492,3 | 1433 | 1552 | 1,085 | 0,117 | 1,827 | 0,06767077 | 0,10958965 | 0,003295 | 0,0029523 | 0,0032789 | 0,0032789 | TRUE | 0,0473031 | No |
| orf19.3971 | 263 | 219 | 230 | 305 | 367 | 329 | 355 | 232 | 236 | 279 | 235 | 252 | 314 | 314 | 317 | 327 | 284,17 | 250 | 318 | 1,273 | 0,348 | 4,006 | 6,17E-05 | 0,00019593 | 1,00E-08 | 0,0070881 | 0,0036888 | 0,0036888 | TRUE | 0,0549306 | No |
| orf19.3972 | 713 | 788 | 765 | 1056 | 687 | 781 | 836 | 567 | 640 | 1003 | 783 | 871 | 588 | 745 | 746 | 800 | 771,84 | 824 | 720 | 0,873 | -0,196 | -1,538 | 0,12403652 | 0,18350441 | 0,023558 | 0,0038603 | 0,0142318 | 0,0142318 | TRUE | 0,217314 | No |
| orf19.3973 | 1252 | 915 | 1127 | 1462 | 1606 | 1351 | 1375 | 966 | 1123 | 1165 | 1153 | 1206 | 1374 | 1289 | 1227 | 1362 | 1237,3 | 1162 | 1313 | 1,129 | 0,175 | 3,235 | 0,00121621 | 0,00301568 | 0,001155 | 0,0031528 | 0,001991 | 0,001991 | TRUE | 0,0241203 | No |
| orf19.3974 | 30678 | 21466 | 27814 | 33637 | 34189 | 31460 | 33836 | 22477 | 27521 | 27329 | 28456 | 27737 | 29240 | 30018 | 30193 | 31702 | 29025 | 27761 | 30288 | 1,091 | 0,125 | 3,734 | 0,0001888 | 0,00055278 | 0,000689 | 0,0020295 | 0,0010488 | 0,0010488 | TRUE | 0,0119578 | No |
| orf19.3976 | 342 | 254 | 284 | 384 | 423 | 386 | 443 | 240 | 307 | 323 | 291 | 317 | 362 | 368 | 395 | 338 | 337,66 | 310 | 366 | 1,188 | 0,249 | 3,041 | 0,00235837 | 0,00547704 | 1,00E-08 | 0,0062788 | 0,0034606 | 0,0034606 | TRUE | 0,0331785 | No |
| orf19.3977 | 3475 | 2313 | 2948 | 3558 | 3682 | 3088 | 3297 | 1965 | 3117 | 2945 | 3016 | 2934 | 3149 | 2946 | 2942 | 2771 | 2977,6 | 3003 | 2952 | 0,984 | -0,023 | -0,489 | 0,62475578 | 0,697311 | 0,00124 | 0,0024671 | 0,001785 | 0,001785 | TRUE | 0,0245431 | No |
| orf19.3978 | 6268 | 4854 | 5266 | 6562 | 5622 | 5602 | 5736 | 2606 | 5623 | 6180 | 5388 | 5411 | 4808 | 5345 | 5118 | 3676 | 5193,6 | 5650 | 4737 | 0,839 | -0,253 | -2,608 | 0,00909575 | 0,01851822 | 0,014102 | 0,002259 | 0,0088634 | 0,0088634 | TRUE | 0,2739597 | No |
| orf19.398 | 127 | 87 | 109 | 123 | 124 | 113 | 127 | 96 | 114 | 111 | 112 | 101 | 106 | 108 | 113 | 135 | 112,53 | 110 | 116 | 1,048 | 0,068 | 0,509 | 0,61041391 | 0,68510495 | 1,00E-08 | 0,0148802 | 0,0083176 | 0,0083176 | TRUE | 0,1047566 | No |
| orf19.3980 | 1060 | 805 | 954 | 1199 | 985 | 924 | 1079 | 635 | 951 | 1025 | 976 | 989 | 842 | 882 | 963 | 896 | 940,38 | 985 | 896 | 0,91 | -0,136 | -2,339 | 0,01934301 | 0,03607973 | 0,001095 | 0,0035233 | 0,0021833 | 0,0021833 | TRUE | 0,032833 | No |
| orf19.3981 | 436 | 333 | 410 | 531 | 412 | 344 | 390 | 260 | 391 | 424 | 419 | 438 | 352 | 328 | 348 | 367 | 383,47 | 418 | 349 | 0,832 | -0,264 | -3,459 | 0,00054128 | 0,00144921 | 1,00E-08 | 0,0057653 | 0,0029846 | 0,0029846 | TRUE | 0,0234901 | No |
| orf19.3982 | 266 | 217 | 259 | 280 | 234 | 195 | 219 | 124 | 239 | 276 | 265 | 231 | 200 | 186 | 195 | 175 | 220,91 | 253 | 189 | 0,756 | -0,404 | -3,986 | 6,72E-05 | 0,00021254 | 1,00E-08 | 0,008551 | 0,0052536 | 0,0052536 | TRUE | 0,0399462 | No |
| orf19.3983 | 1195 | 990 | 1107 | 1342 | 992 | 951 | 974 | 642 | 1072 | 1260 | 1133 | 1107 | 848 | 907 | 869 | 905 | 1012,8 | 1143 | 882 | 0,773 | -0,371 | -6,001 | 1,96E-09 | 1,20E-08 | 0,002057 | 0,0034129 | 0,0026629 | 0,0026629 | TRUE | 0,0538936 | No |
| orf19.3984 | 9141 | 7493 | 8390 | 10763 | 11600 | 12418 | 12609 | 7504 | 8200 | 9539 | 8584 | 8875 | 9921 | 11849 | 11251 | 10584 | 9850,5 | 8800 | 10901 | 1,239 | 0,309 | 4,873 | 1,10E-06 | 4,61E-06 | 0,004848 | 0,0021269 | 0,003767 | 0,003767 | TRUE | 0,0451964 | No |
| orf19.3986 | 845 | 673 | 850 | 1021 | 1116 | 1270 | 1269 | 737 | 758 | 857 | 870 | 842 | 954 | 1212 | 1132 | 1039 | 958,06 | 832 | 1084 | 1,306 | 0,385 | 4,793 | 1,64E-06 | 6,71E-06 | 0,006451 | 0,0034948 | 0,0051173 | 0,0051173 | TRUE | 0,0821989 | No |
| orf19.3988 | 6050 | 4146 | 5356 | 6569 | 9452 | 8183 | 8926 | 5710 | 5427 | 5278 | 5480 | 5417 | 8084 | 7808 | 7965 | 8053 | 6689,1 | 5400 | 7978 | 1,476 | 0,562 | 19,11 | 2,10E-81 | 6,31E-79 | 8,52E-05 | 0,0021965 | 0,0006752 | 0,0006752 | TRUE | 0,0026564 | No |
| orf19.399 | 7459 | 4986 | 6767 | 8059 | 7580 | 6709 | 7297 | 4881 | 6692 | 6348 | 6923 | 6646 | 6483 | 6402 | 6511 | 6884 | 6611 | 6652 | 6570 | 0,987 | -0,019 | -0,483 | 0,62898324 | 0,70045065 | 0,00089 | 0,0021991 | 0,0013592 | 0,0013592 | TRUE | 0,0121759 | No |
| orf19.3990 | 596 | 548 | 545 | 725 | 668 | 743 | 719 | 424 | 535 | 698 | 558 | 598 | 571 | 709 | 642 | 598 | 613,45 | 597 | 630 | 1,059 | 0,082 | 0,864 | 0,38760368 | 0,47670889 | 0,009196 | 0,004346 | 0,0070572 | 0,0070572 | TRUE | 0,1428919 | No |
| orf19.3991 | 2335 | 1692 | 2135 | 2448 | 2627 | 2320 | 2628 | 1517 | 2095 | 2154 | 2184 | 2019 | 2247 | 2214 | 2345 | 2140 | 2174,6 | 2113 | 2236 | 1,061 | 0,085 | 1,856 | 0,06348087 | 0,1035269 | 0,000861 | 0,0026471 | 0,0015453 | 0,0015453 | TRUE | 0,0129427 | No |
| orf19.3992 | 67 | 43 | 57 | 62 | 67 | 40 | 46 | 41 | 60 | 55 | 58 | 51 | 57 | 38 | 41 | 58 | 52,33 | 56 | 48 | 0,86 | -0,218 | -1,076 | 0,28206673 | 0,36609568 | 1,00E-08 | 0,0297212 | 0,0201558 | 0,0201558 | TRUE | 0,1679906 | No |
| orf19.3994 | 493 | 340 | 429 | 529 | 464 | 421 | 426 | 307 | 442 | 433 | 439 | 436 | 397 | 402 | 380 | 433 | 420,24 | 438 | 403 | 0,917 | -0,126 | -1,763 | 0,07794449 | 0,12369004 | 1,00E-08 | 0,005434 | 0,0025051 | 0,0025051 | TRUE | 0,0239823 | No |
| orf19.3995 | 340 | 257 | 327 | 322 | 382 | 379 | 363 | 197 | 305 | 327 | 335 | 266 | 327 | 362 | 324 | 278 | 315,3 | 308 | 323 | 1,055 | 0,078 | 0,754 | 0,45098287 | 0,53696098 | 0,007499 | 0,0065838 | 0,0070181 | 0,0070181 | TRUE | 0,104385 | No |
| orf19.3996 | 89 | 74 | 73 | 89 | 103 | 83 | 102 | 58 | 80 | 94 | 75 | 73 | 88 | 79 | 91 | 82 | 82,78 | 80 | 85 | 1,067 | 0,094 | 0,611 | 0,54096069 | 0,62321991 | 1,00E-08 | 0,0195165 | 0,0105685 | 0,0105685 | TRUE | 0,1008961 | No |
| orf19.3997 | 55114 | 34420 | 48266 | 57595 | 75794 | 57465 | 64327 | 42099 | 49443 | 43820 | 49381 | 47494 | 64821 | 54831 | 57401 | 59377 | 53321 | 47534 | 59108 | 1,243 | 0,314 | 5,36 | 8,31E-08 | 4,11E-07 | 0,004085 | 0,0020067 | 0,0032851 | 0,0032851 | TRUE | 0,0519625 | No |
| orf19.3998 | 155 | 129 | 159 | 193 | 158 | 173 | 160 | 122 | 139 | 164 | 163 | 159 | 135 | 165 | 143 | 172 | 155,02 | 156 | 154 | 0,979 | -0,031 | -0,251 | 0,80164322 | 0,84555346 | 0,00302 | 0,0113442 | 0,0077843 | 0,0077843 | TRUE | 0,0723611 | No |
| orf19.3999 | 745 | 551 | 674 | 817 | 966 | 909 | 976 | 612 | 668 | 701 | 690 | 674 | 826 | 867 | 871 | 863 | 770,09 | 683 | 857 | 1,255 | 0,328 | 5,973 | 2,33E-09 | 1,41E-08 | 1,00E-08 | 0,0038646 | 0,0015789 | 0,0015789 | TRUE | 0,0075682 | No |
| orf19.400 | 3238 | 2149 | 2819 | 3436 | 2649 | 2312 | 2435 | 1652 | 2905 | 2736 | 2884 | 2833 | 2266 | 2206 | 2173 | 2330 | 2541,6 | 2840 | 2244 | 0,789 | -0,342 | -8,461 | 2,66E-17 | 3,87E-16 | 0,000296 | 0,0025507 | 0,0011723 | 0,0011723 | TRUE | 0,0073058 | No |
| orf19.4000 | 13190 | 9475 | 12288 | 15016 | 15040 | 13746 | 15383 | 9817 | 11833 | 12063 | 12572 | 12382 | 12863 | 13116 | 13727 | 13846 | 12800 | 12212 | 13388 | 1,096 | 0,132 | 3,533 | 0,00041135 | 0,00113054 | 0,000886 | 0,0020929 | 0,0012659 | 0,0012659 | TRUE | 0,0085679 | No |
| orf19.4001 | 269 | 206 | 249 | 323 | 377 | 366 | 393 | 233 | 241 | 262 | 255 | 266 | 322 | 349 | 351 | 329 | 296,96 | 256 | 338 | 1,32 | 0,401 | 4,775 | 1,80E-06 | 7,32E-06 | 1,00E-08 | 0,0068681 | 0,0033556 | 0,0033556 | TRUE | 0,018384 | No |
| orf19.4002 | 436 | 320 | 404 | 491 | 462 | 395 | 481 | 247 | 391 | 407 | 413 | 405 | 395 | 377 | 429 | 348 | 395,79 | 404 | 387 | 0,964 | -0,053 | -0,665 | 0,5062588 | 0,59054776 | 0,001127 | 0,0056474 | 0,0035688 | 0,0035688 | TRUE | 0,0597514 | No |
| orf19.4003 | 537 | 421 | 507 | 549 | 778 | 748 | 744 | 483 | 482 | 536 | 519 | 453 | 665 | 714 | 664 | 681 | 589,17 | 498 | 681 | 1,375 | 0,459 | 6,549 | 5,80E-11 | 4,40E-10 | 0,001471 | 0,0044435 | 0,0029755 | 0,0029755 | TRUE | 0,0443995 | No |
| orf19.4004 | 5338 | 3598 | 4687 | 5779 | 5834 | 4850 | 5512 | 3351 | 4789 | 4581 | 4795 | 4765 | 4989 | 4628 | 4919 | 4726 | 4774 | 4732 | 4816 | 1,018 | 0,025 | 0,666 | 0,50552531 | 0,58997592 | 0,000558 | 0,0022836 | 0,0011651 | 0,0011651 | TRUE | 0,0087583 | No |
| orf19.4005 | 432 | 269 | 358 | 460 | 407 | 378 | 409 | 282 | 388 | 342 | 366 | 379 | 348 | 361 | 365 | 398 | 368,38 | 369 | 368 | 0,99 | -0,015 | -0,191 | 0,84854292 | 0,88415642 | 1,00E-08 | 0,0059203 | 0,0032405 | 0,0032405 | TRUE | 0,0286075 | No |
| orf19.4006 | 1022 | 707 | 849 | 1103 | 1278 | 1143 | 1126 | 802 | 917 | 900 | 869 | 910 | 1093 | 1091 | 1005 | 1131 | 989,33 | 899 | 1080 | 1,199 | 0,261 | 4,755 | 1,98E-06 | 8,01E-06 | 0,000615 | 0,0034469 | 0,0018831 | 0,0018831 | TRUE | 0,0267903 | No |
| orf19.4007 | 223 | 188 | 240 | 269 | 259 | 200 | 231 | 164 | 200 | 239 | 246 | 222 | 222 | 191 | 206 | 231 | 219,57 | 227 | 212 | 0,937 | -0,094 | -0,89 | 0,37346024 | 0,46215224 | 0,00301 | 0,0085912 | 0,0061142 | 0,0061142 | TRUE | 0,0710247 | No |
| orf19.4009 | 1063 | 792 | 966 | 1178 | 1113 | 983 | 984 | 618 | 954 | 1008 | 988 | 971 | 952 | 938 | 878 | 872 | 945,14 | 980 | 910 | 0,931 | -0,103 | -1,914 | 0,05568077 | 0,09220784 | 1,00E-08 | 0,0035155 | 0,0017443 | 0,0017443 | TRUE | 0,0117626 | No |
| orf19.401 | 3572 | 2433 | 3161 | 3825 | 3427 | 2824 | 3035 | 2541 | 3204 | 3097 | 3234 | 3154 | 2931 | 2695 | 2708 | 3584 | 3076 | 3172 | 2980 | 0,938 | -0,093 | -1,145 | 0,25203548 | 0,33441457 | 0,008846 | 0,0024515 | 0,0059523 | 0,0059523 | TRUE | 0,221924 | No |
| orf19.4010 | 1473 | 1060 | 1335 | 1667 | 1985 | 1807 | 1893 | 1426 | 1321 | 1349 | 1366 | 1375 | 1698 | 1724 | 1689 | 2011 | 1566,7 | 1353 | 1780 | 1,313 | 0,393 | 6,395 | 1,61E-10 | 1,13E-09 | 0,002883 | 0,0029061 | 0,002965 | 0,002965 | TRUE | 0,0865792 | No |
| orf19.4011 | 51 | 49 | 64 | 64 | 46 | 34 | 53 | 29 | 46 | 62 | 65 | 53 | 39 | 32 | 47 | 41 | 48,3 | 56 | 40 | 0,712 | -0,49 | -2,32 | 0,02034129 | 0,03782436 | 1,00E-08 | 0,0320381 | 0,0215679 | 0,0215679 | TRUE | 0,1478451 | No |
| orf19.4012 | 9215 | 7589 | 8361 | 10819 | 9852 | 10704 | 11112 | 6808 | 8267 | 9662 | 8554 | 8921 | 8426 | 10213 | 9916 | 9602 | 9195,1 | 8851 | 9539 | 1,078 | 0,108 | 1,612 | 0,10698376 | 0,16153768 | 0,005547 | 0,0021374 | 0,0042191 | 0,0042191 | TRUE | 0,0759668 | No |
| orf19.4013 | 639 | 480 | 572 | 665 | 522 | 460 | 504 | 302 | 573 | 611 | 585 | 548 | 446 | 439 | 450 | 426 | 509,87 | 579 | 440 | 0,763 | -0,39 | -5,837 | 5,32E-09 | 3,08E-08 | 1,00E-08 | 0,0048267 | 0,0022856 | 0,0022856 | TRUE | 0,0159565 | No |
| orf19.4014 | 1335 | 849 | 1072 | 1302 | 1087 | 980 | 1063 | 699 | 1198 | 1081 | 1097 | 1074 | 930 | 935 | 949 | 986 | 1031 | 1112 | 950 | 0,852 | -0,231 | -4,177 | 2,96E-05 | 9,98E-05 | 0,000866 | 0,0033876 | 0,0019597 | 0,0019597 | TRUE | 0,0331812 | No |
| orf19.4015 | 989 | 734 | 845 | 1066 | 918 | 826 | 917 | 590 | 887 | 934 | 865 | 879 | 785 | 788 | 818 | 832 | 848,61 | 891 | 806 | 0,904 | -0,145 | -2,684 | 0,0072658 | 0,01509436 | 1,00E-08 | 0,0036902 | 0,0016382 | 0,0016382 | TRUE | 0,0112219 | No |
| orf19.4016 | 6311 | 4030 | 5438 | 6403 | 6671 | 5375 | 5913 | 4318 | 5662 | 5131 | 5564 | 5280 | 5705 | 5129 | 5276 | 6090 | 5479,5 | 5409 | 5550 | 1,025 | 0,036 | 0,606 | 0,54440971 | 0,6261563 | 0,003573 | 0,0022445 | 0,003214 | 0,003214 | TRUE | 0,0512122 | No |
| orf19.4017 | 463 | 338 | 426 | 470 | 417 | 338 | 376 | 267 | 415 | 430 | 436 | 388 | 357 | 323 | 336 | 377 | 382,54 | 417 | 348 | 0,832 | -0,265 | -3,307 | 0,00094385 | 0,00238364 | 0,000915 | 0,0057744 | 0,0035068 | 0,0035068 | TRUE | 0,0290288 | No |
| orf19.4018 | 1853 | 1248 | 1619 | 1979 | 1836 | 1551 | 1788 | 1125 | 1662 | 1589 | 1656 | 1632 | 1570 | 1480 | 1595 | 1587 | 1596,5 | 1635 | 1558 | 0,952 | -0,071 | -1,589 | 0,11201726 | 0,16783034 | 1,00E-08 | 0,0028888 | 0,0012872 | 0,0012872 | TRUE | 0,0140023 | No |
| orf19.4019 | 53 | 46 | 42 | 56 | 53 | 48 | 60 | 30 | 48 | 59 | 43 | 46 | 45 | 46 | 54 | 42 | 47,78 | 49 | 47 | 0,971 | -0,043 | -0,214 | 0,83067672 | 0,86991559 | 1,00E-08 | 0,0323627 | 0,017494 | 0,017494 | TRUE | 0,1164329 | No |
| orf19.4021 | 1321 | 748 | 1128 | 1246 | 1228 | 907 | 1086 | 651 | 1185 | 952 | 1154 | 1027 | 1050 | 865 | 969 | 918 | 1015,2 | 1080 | 950 | 0,88 | -0,184 | -2,208 | 0,02721877 | 0,04927098 | 0,007195 | 0,0034095 | 0,0056437 | 0,0056437 | TRUE | 0,0717991 | No |
| orf19.4022 | 1189 | 745 | 1014 | 1252 | 1077 | 845 | 930 | 606 | 1067 | 948 | 1037 | 1032 | 921 | 806 | 830 | 855 | 937,11 | 1021 | 853 | 0,834 | -0,261 | -4,216 | 2,49E-05 | 8,50E-05 | 0,001851 | 0,0035286 | 0,0026154 | 0,0026154 | TRUE | 0,0371391 | No |
| orf19.4023 | 1209 | 763 | 1062 | 1315 | 1273 | 1005 | 1033 | 756 | 1085 | 971 | 1087 | 1084 | 1089 | 959 | 922 | 1066 | 1032,8 | 1057 | 1009 | 0,952 | -0,071 | -1,026 | 0,30472422 | 0,39052541 | 0,003391 | 0,0033851 | 0,0035644 | 0,0035644 | TRUE | 0,041425 | No |

| | | | | | | | | | | | | | | | | | | | | | | | | | | | | | | | |
|---|---|---|---|---|---|---|---|---|---|---|---|---|---|---|---|---|---|---|---|---|---|---|---|---|---|---|---|---|---|---|---|
| orf19.4024 | 1045 | 654 | 858 | 1116 | 1087 | 967 | 1055 | 647 | 937 | 833 | 878 | 920 | 930 | 923 | 941 | 913 | 909,3 | 892 | 927 | 1,037 | 0,052 | 0,941 | 0,34658384 | 0,43392907 | 0,000218 | 0,003576 | 0,0018244 | 0,0018244 | TRUE | 0,0217474 | No |
| orf19.4025 | 5254 | 3288 | 4182 | 5223 | 4929 | 3940 | 4086 | 2573 | 4713 | 4186 | 4279 | 4307 | 4215 | 3759 | 3646 | 3629 | 4091,9 | 4371 | 3812 | 0,872 | -0,197 | -3,316 | 0,00091257 | 0,00231045 | 0,003702 | 0,0023343 | 0,0031394 | 0,0031394 | TRUE | 0,0623396 | No |
| orf19.4026 | 1107 | 742 | 890 | 1215 | 1179 | 863 | 981 | 521 | 993 | 945 | 911 | 1002 | 1008 | 823 | 875 | 735 | 911,52 | 963 | 860 | 0,896 | -0,158 | -1,826 | 0,06784963 | 0,10978739 | 0,008355 | 0,0035721 | 0,0060715 | 0,0060715 | TRUE | 0,1599585 | No |
| orf19.4028 | 3050 | 1871 | 2549 | 3117 | 2811 | 2373 | 2570 | 1619 | 2736 | 2382 | 2608 | 2570 | 2404 | 2264 | 2293 | 2283 | 2442,7 | 2574 | 2311 | 0,897 | -0,156 | -3,147 | 0,00164753 | 0,00395461 | 0,001443 | 0,0025738 | 0,0019562 | 0,0019562 | TRUE | 0,029164 | No |
| orf19.4029 | 17982 | 11764 | 15498 | 19393 | 17983 | 14420 | 16288 | 10090 | 16132 | 14977 | 15856 | 15992 | 15380 | 13759 | 14534 | 14231 | 15108 | 15739 | 14476 | 0,92 | -0,121 | -2,778 | 0,00546388 | 0,01171975 | 0,001567 | 0,0020756 | 0,0017469 | 0,0017469 | TRUE | 0,021751 | No |
| orf19.403 | 81 | 63 | 62 | 90 | 26 | 23 | 19 | 17 | 73 | 80 | 63 | 74 | 22 | 22 | 17 | 24 | 46,95 | 72 | 21 | 0,291 | -1,783 | -7,948 | 1,90E-15 | 2,28E-14 | 1,00E-08 | 0,0328971 | 0,0178259 | 0,0178259 | TRUE | 0,117728 | No |
| orf19.4030 | 822 | 505 | 722 | 898 | 813 | 672 | 774 | 466 | 737 | 643 | 739 | 741 | 695 | 641 | 691 | 657 | 692,99 | 715 | 671 | 0,937 | -0,094 | -1,421 | 0,15527547 | 0,2221005 | 0,001365 | 0,0040744 | 0,0027176 | 0,0027176 | TRUE | 0,0489776 | No |
| orf19.4031 | 3976 | 2516 | 3524 | 4279 | 4283 | 3484 | 3914 | 2647 | 3567 | 3203 | 3605 | 3529 | 3663 | 3324 | 3493 | 3733 | 3514,7 | 3476 | 3553 | 1,021 | 0,03 | 0,565 | 0,5717924 | 0,65087699 | 0,002194 | 0,0023926 | 0,0024179 | 0,0024179 | TRUE | 0,0328749 | No |
| orf19.4032 | 5062 | 3588 | 4383 | 5392 | 5117 | 4407 | 4531 | 2936 | 4541 | 4568 | 4484 | 4446 | 4376 | 4205 | 4043 | 4141 | 4350,6 | 4510 | 4191 | 0,93 | -0,105 | -2,894 | 0,00379876 | 0,00841864 | 0,000435 | 0,0023132 | 0,0010287 | 0,0010287 | TRUE | 0,0111465 | No |
| orf19.4033 | 719 | 525 | 705 | 795 | 587 | 459 | 619 | 335 | 645 | 668 | 721 | 656 | 502 | 438 | 552 | 472 | 581,88 | 672 | 491 | 0,732 | -0,45 | -5,634 | 1,77E-08 | 9,47E-08 | 0,004223 | 0,0044743 | 0,0043467 | 0,0043467 | TRUE | 0,0847046 | No |
| orf19.4035 | 6714 | 4066 | 5817 | 6994 | 9465 | 7588 | 8482 | 5375 | 6023 | 5176 | 5951 | 5767 | 8095 | 7240 | 7569 | 7581 | 6675,4 | 5729 | 7621 | 1,33 | 0,411 | 7,401 | 1,35E-13 | 1,35E-12 | 0,003117 | 0,002197 | 0,0028096 | 0,0028096 | TRUE | 0,0506016 | No |
| orf19.4036 | 873 | 643 | 814 | 1013 | 762 | 643 | 732 | 470 | 783 | 819 | 833 | 835 | 652 | 614 | 653 | 663 | 731,4 | 818 | 646 | 0,789 | -0,342 | -5,935 | 2,93E-09 | 1,74E-08 | 1,00E-08 | 0,0039643 | 0,00181 | 0,00181 | TRUE | 0,0130295 | No |
| orf19.4040 | 3128 | 1970 | 2815 | 3325 | 3025 | 2327 | 2603 | 1809 | 2806 | 2508 | 2880 | 2742 | 2587 | 2220 | 2323 | 2551 | 2577,2 | 2734 | 2420 | 0,884 | -0,177 | -2,817 | 0,00485323 | 0,0104999 | 0,003901 | 0,0025428 | 0,0034158 | 0,0034158 | TRUE | 0,0376086 | No |
| orf19.4041 | 931 | 593 | 806 | 1051 | 793 | 618 | 691 | 460 | 835 | 755 | 825 | 867 | 678 | 590 | 617 | 649 | 726,84 | 820 | 634 | 0,77 | -0,377 | -5,569 | 2,56E-08 | 1,35E-07 | 0,002117 | 0,0039768 | 0,0030026 | 0,0030026 | TRUE | 0,0316225 | No |
| orf19.4043 | 2922 | 1891 | 2593 | 3228 | 3422 | 2750 | 3215 | 2187 | 2621 | 2407 | 2653 | 2662 | 2927 | 2624 | 2869 | 3085 | 2730,9 | 2586 | 2876 | 1,11 | 0,151 | 2,653 | 0,00798953 | 0,01647824 | 0,0028 | 0,0025111 | 0,0027467 | 0,0027467 | TRUE | 0,042105 | No |
| orf19.4044 | 586 | 364 | 529 | 607 | 535 | 430 | 456 | 364 | 526 | 463 | 541 | 501 | 458 | 410 | 407 | 513 | 477,38 | 508 | 447 | 0,875 | -0,193 | -2,205 | 0,0274315 | 0,04959628 | 0,005666 | 0,0050205 | 0,0052268 | 0,0052268 | TRUE | 0,1018719 | No |
| orf19.4045 | 5915 | 3837 | 5451 | 6508 | 6662 | 5359 | 5995 | 3656 | 5306 | 4885 | 5577 | 5367 | 5698 | 5113 | 5350 | 5156 | 5306,5 | 5284 | 5329 | 1,009 | 0,012 | 0,236 | 0,81306576 | 0,85549897 | 0,002188 | 0,0022531 | 0,0024357 | 0,0024357 | TRUE | 0,0307641 | No |
| orf19.4046 | 825 | 547 | 690 | 908 | 1070 | 804 | 824 | 575 | 740 | 696 | 706 | 749 | 915 | 767 | 735 | 811 | 764,96 | 723 | 807 | 1,116 | 0,158 | 2,102 | 0,03552457 | 0,06232104 | 0,004296 | 0,0038773 | 0,0041086 | 0,0041086 | TRUE | 0,1026354 | No |
| orf19.4048 | 9360 | 6014 | 8196 | 9994 | 11704 | 9201 | 10285 | 6809 | 8397 | 7656 | 8385 | 8241 | 10010 | 8779 | 9178 | 9603 | 8781,2 | 8170 | 9392 | 1,149 | 0,201 | 3,983 | 6,81E-05 | 0,00021476 | 0,002364 | 0,0021448 | 0,0023246 | 0,0023246 | TRUE | 0,0242718 | No |
| orf19.405 | 869 | 567 | 746 | 920 | 974 | 818 | 911 | 595 | 780 | 722 | 763 | 759 | 833 | 781 | 813 | 839 | 786,12 | 756 | 816 | 1,077 | 0,108 | 1,922 | 0,0546681 | 0,09080617 | 1,00E-08 | 0,0038262 | 0,0017356 | 0,0017356 | TRUE | 0,0103983 | No |
| orf19.4051 | 4185 | 2548 | 3674 | 4488 | 4386 | 3332 | 3890 | 2249 | 3754 | 3244 | 3759 | 3701 | 3751 | 3179 | 3471 | 3172 | 3503,9 | 3614 | 3393 | 0,939 | -0,091 | -1,341 | 0,18001773 | 0,25195828 | 0,005294 | 0,0023938 | 0,004125 | 0,004125 | TRUE | 0,0616115 | No |
| orf19.4054 | 478 | 263 | 372 | 546 | 560 | 399 | 521 | 310 | 429 | 335 | 381 | 450 | 479 | 381 | 465 | 437 | 419,53 | 399 | 440 | 1,101 | 0,139 | 1,321 | 0,18662865 | 0,25925201 | 0,011073 | 0,0054398 | 0,0082443 | 0,0082443 | TRUE | 0,1244398 | No |
| orf19.4055 | 225 | 134 | 186 | 207 | 243 | 172 | 178 | 117 | 202 | 171 | 190 | 171 | 208 | 164 | 159 | 165 | 178,65 | 184 | 174 | 0,951 | -0,073 | -0,598 | 0,54960897 | 0,63052333 | 0,007001 | 0,0101053 | 0,0086453 | 0,0086453 | TRUE | 0,1999879 | No |
| orf19.4056 | 21468 | 14614 | 19410 | 24430 | 26344 | 22994 | 25417 | 17877 | 19259 | 18605 | 19858 | 20145 | 22530 | 21940 | 22680 | 25214 | 21279 | 19467 | 23091 | 1,186 | 0,246 | 4,953 | 7,32E-07 | 3,16E-06 | 0,002526 | 0,0020477 | 0,0023204 | 0,0023204 | TRUE | 0,046563 | No |
| orf19.4057 | 18 | 19 | 19 | 14 | 14 | 11 | 10 | 7 | 16 | 24 | 19 | 12 | 12 | 10 | 9 | 10 | 14,07 | 18 | 10 | 0,592 | -0,756 | -1,973 | 0,04850656 | NA | 1,00E-08 | 0,1051331 | 0,0642163 | 0,0642163 | TRUE | 0,1713059 | No |
| orf19.4059 | 2308 | 1615 | 2063 | 2397 | 2198 | 1932 | 2172 | 1262 | 2071 | 2056 | 2111 | 1977 | 1880 | 1843 | 1938 | 1780 | 1956,9 | 2054 | 1860 | 0,908 | -0,14 | -3,163 | 0,00155975 | 0,00376949 | 0,000477 | 0,0027214 | 0,0013563 | 0,0013563 | TRUE | 0,0094235 | No |
| orf19.406 | 2705 | 1817 | 2494 | 2916 | 2738 | 2393 | 2712 | 1778 | 2427 | 2313 | 2552 | 2405 | 2342 | 2283 | 2420 | 2508 | 2406,1 | 2424 | 2388 | 0,984 | -0,024 | -0,507 | 0,61204379 | 0,68642134 | 0,001057 | 0,0025829 | 0,0016915 | 0,0016915 | TRUE | 0,0148372 | No |
| orf19.4060 | 6333 | 4129 | 5424 | 6511 | 6784 | 5642 | 5990 | 4353 | 5681 | 5257 | 5549 | 5369 | 5802 | 5383 | 5345 | 6140 | 5565,8 | 5464 | 5668 | 1,036 | 0,052 | 0,987 | 0,3238374 | 0,41072728 | 0,002563 | 0,0022403 | 0,0024379 | 0,0024379 | TRUE | 0,0373285 | No |
| orf19.4061 | 634 | 401 | 564 | 629 | 612 | 591 | 574 | 425 | 569 | 511 | 577 | 519 | 523 | 564 | 512 | 599 | 546,74 | 544 | 550 | 1,006 | 0,009 | 0,118 | 0,90591012 | 0,92974171 | 0,002487 | 0,0046347 | 0,0035663 | 0,0035663 | TRUE | 0,0381957 | No |
| orf19.4062 | 2837 | 1973 | 2558 | 3064 | 3227 | 2851 | 2980 | 1732 | 2545 | 2512 | 2617 | 2527 | 2760 | 2720 | 2659 | 2443 | 2597,9 | 2550 | 2646 | 1,039 | 0,055 | 1,201 | 0,22956316 | 0,3095882 | 0,001081 | 0,0025383 | 0,0016547 | 0,0016547 | TRUE | 0,029929 | No |
| orf19.4063 | 340 | 209 | 245 | 339 | 243 | 196 | 228 | 150 | 305 | 266 | 251 | 280 | 208 | 187 | 203 | 212 | 238,89 | 276 | 202 | 0,732 | -0,45 | -4,566 | 4,96E-06 | 1,89E-05 | 1,00E-08 | 0,0080563 | 0,005034 | 0,005034 | TRUE | 0,060683 | No |
| orf19.4064 | 744 | 624 | 712 | 816 | 715 | 670 | 714 | 415 | 667 | 794 | 728 | 673 | 611 | 639 | 637 | 585 | 667,05 | 716 | 618 | 0,868 | -0,204 | -2,863 | 0,00419097 | 0,0091862 | 0,002585 | 0,0041558 | 0,0033429 | 0,0033429 | TRUE | 0,0604029 | No |
| orf19.4066 | 1513 | 1439 | 1615 | 2077 | 1124 | 1439 | 1401 | 873 | 1357 | 1832 | 1652 | 1713 | 961 | 1373 | 1250 | 1231 | 1421,3 | 1638 | 1204 | 0,735 | -0,445 | -4,054 | 5,03E-05 | 0,00016244 | 0,017016 | 0,0030009 | 0,0108229 | 0,0108229 | TRUE | 0,2254052 | No |
| orf19.4067 | 4 | 2 | 8 | 6 | 3 | 6 | 3 | 0 | 4 | 3 | 8 | 5 | 3 | 6 | 3 | 0 | 3,78 | 5 | 3 | 0,594 | -0,752 | -0,989 | 0,3228001 | NA | 0,075136 | 0,3860935 | 0,2793703 | 0,2793703 | TRUE | 0,439903 | No |
| orf19.4068 | 647 | 474 | 637 | 745 | 561 | 524 | 563 | 362 | 580 | 603 | 652 | 614 | 480 | 500 | 502 | 511 | 555,33 | 612 | 498 | 0,812 | -0,3 | -4,575 | 4,77E-06 | 1,82E-05 | 1,00E-08 | 0,0045936 | 0,0023034 | 0,0023034 | TRUE | 0,0216446 | No |
| orf19.4069 | 105 | 76 | 83 | 105 | 55 | 67 | 54 | 46 | 94 | 97 | 85 | 87 | 47 | 64 | 48 | 65 | 73,31 | 91 | 56 | 0,612 | -0,709 | -4,155 | 3,25E-05 | 0,00010871 | 1,00E-08 | 0,0217816 | 0,013432 | 0,013432 | TRUE | 0,1041558 | No |
| orf19.407 | 4058 | 2650 | 3483 | 4207 | 4134 | 3390 | 3744 | 2269 | 3640 | 3374 | 3563 | 3469 | 3536 | 3235 | 3341 | 3200 | 3419,8 | 3512 | 3328 | 0,948 | -0,077 | -1,702 | 0,08867926 | 0,13817038 | 0,001179 | 0,002404 | 0,0016662 | 0,0016662 | TRUE | 0,0217725 | No |
| orf19.4070 | 295 | 229 | 270 | 293 | 258 | 234 | 222 | 146 | 265 | 292 | 276 | 242 | 221 | 223 | 198 | 206 | 240,25 | 269 | 212 | 0,794 | -0,332 | -3,408 | 0,0006534 | 0,00171126 | 1,00E-08 | 0,0080221 | 0,004915 | 0,004915 | TRUE | 0,0541322 | No |
| orf19.4072 | 2752 | 1784 | 2528 | 3125 | 3314 | 2573 | 2971 | 2006 | 2469 | 2271 | 2586 | 2577 | 2834 | 2455 | 2651 | 2829 | 2584,1 | 2476 | 2692 | 1,086 | 0,119 | 1,954 | 0,05066734 | 0,08514947 | 0,00346 | 0,0025413 | 0,0031814 | 0,0031814 | TRUE | 0,0428919 | No |
| orf19.4076 | 9080 | 6168 | 8008 | 9321 | 8342 | 7612 | 8011 | 5007 | 8146 | 7853 | 8193 | 7686 | 7134 | 7263 | 7148 | 7062 | 7560,7 | 7970 | 7152 | 0,898 | -0,156 | -4,714 | 2,43E-06 | 9,72E-06 | 0,00035 | 0,0021715 | 0,0009158 | 0,0009158 | TRUE | 0,0072412 | No |
| orf19.4077 | 10459 | 7973 | 9603 | 12023 | 12385 | 11968 | 12832 | 7787 | 9383 | 10150 | 9825 | 9914 | 10592 | 11419 | 11450 | 10983 | 10465 | 9818 | 11111 | 1,132 | 0,179 | 4,474 | 7,69E-06 | 2,86E-05 | 0,001077 | 0,0021182 | 0,0014384 | 0,0014384 | TRUE | 0,0123279 | No |
| orf19.4078 | 3687 | 2822 | 3255 | 4138 | 3819 | 3672 | 3739 | 2374 | 3308 | 3593 | 3330 | 3412 | 3266 | 3504 | 3336 | 3348 | 3387,2 | 3411 | 3364 | 0,987 | -0,019 | -0,456 | 0,64818192 | 0,71770678 | 0,000755 | 0,0024081 | 0,0013852 | 0,0013852 | TRUE | 0,0149301 | No |
| orf19.4079 | 1482 | 1081 | 1350 | 1724 | 1517 | 1427 | 1537 | 1010 | 1330 | 1376 | 1381 | 1422 | 1297 | 1362 | 1372 | 1425 | 1370,5 | 1377 | 1364 | 0,988 | -0,017 | -0,353 | 0,72409485 | 0,78384898 | 0,000417 | 0,0030388 | 0,0015181 | 0,0015181 | TRUE | 0,0131355 | No |
| orf19.4080 | 794 | 676 | 775 | 1032 | 781 | 859 | 993 | 523 | 712 | 861 | 793 | 851 | 668 | 820 | 886 | 738 | 791,02 | 804 | 778 | 0,969 | -0,046 | -0,492 | 0,62259745 | 0,6956603 | 0,010163 | 0,0038148 | 0,0071727 | 0,0071727 | TRUE | 0,1134916 | No |
| orf19.4081 | 0 | 1 | 1 | 0 | 3 | 0 | 0 | 0 | 0 | 1 | 1 | 0 | 3 | 0 | 0 | 0 | 0,61 | 0 | 1 | 1,307 | 0,387 | 0,184 | 0,85363788 | NA | 1,00E-08 | 2,3906774 | 2,5821321 | 2,5821321 | TRUE | 1,4355494 | No |
| orf19.4082 | 6243 | 3955 | 5042 | 6356 | 7723 | 6095 | 6396 | 4586 | 5601 | 5035 | 5158 | 5241 | 6605 | 5816 | 5707 | 6468 | 5704 | 5259 | 6149 | 1,169 | 0,225 | 3,865 | 0,00011116 | 0,00033764 | 0,003548 | 0,002234 | 0,003074 | 0,003074 | TRUE | 0,031105 | No |
| orf19.4084 | 3605 | 2306 | 3183 | 4006 | 4276 | 3680 | 4179 | 3006 | 3234 | 2936 | 3257 | 3303 | 3657 | 3511 | 3729 | 4240 | 3483,4 | 3182 | 3784 | 1,187 | 0,248 | 3,903 | 9,51E-05 | 0,00029233 | 0,004495 | 0,0023963 | 0,0035747 | 0,0035747 | TRUE | 0,0777656 | No |
| orf19.4085 | 277 | 263 | 251 | 379 | 345 | 379 | 342 | 186 | 248 | 335 | 257 | 313 | 295 | 362 | 305 | 262 | 297,11 | 288 | 306 | 1,068 | 0,095 | 0,772 | 0,44030814 | 0,52707139 | 0,015659 | 0,0068657 | 0,0111039 | 0,0111039 | TRUE | 0,1669829 | No |
| orf19.4086 | 678 | 450 | 572 | 616 | 716 | 613 | 557 | 411 | 608 | 573 | 585 | 508 | 612 | 585 | 497 | 580 | 568,53 | 568 | 568 | 1 | 0 | 0,004 | 0,99641654 | 0,99757883 | 0,005598 | 0,0045329 | 0,0050974 | 0,0050974 | TRUE | 0,0884924 | No |
| orf19.4088 | 933 | 654 | 854 | 990 | 852 | 768 | 777 | 548 | 837 | 833 | 874 | 816 | 729 | 733 | 693 | 773 | 785,93 | 840 | 732 | 0,869 | -0,202 | -3,523 | 0,00042664 | 0,00116895 | 1,00E-08 | 0,0038266 | 0,0018745 | 0,0018745 | TRUE | 0,0147213 | No |
| orf19.4089 | 4439 | 2966 | 3789 | 4666 | 4572 | 3903 | 4305 | 2609 | 3982 | 3776 | 3877 | 3848 | 3910 | 3724 | 3841 | 3680 | 3829,7 | 3871 | 3789 | 0,979 | -0,03 | -0,825 | 0,40913205 | 0,49728209 | 0,000286 | 0,0023586 | 0,0010207 | 0,0010207 | TRUE | 0,0056673 | No |
| orf19.409 | 47471 | 30329 | 39109 | 49427 | 53612 | 43511 | 47000 | 30771 | 42586 | 38612 | 40012 | 40758 | 45851 | 41517 | 41940 | 43400 | 41834 | 40492 | 43177 | 1,066 | 0,093 | 2,076 | 0,03787597 | 0,06592086 | 0,001831 | 0,0020142 | 0,0018841 | 0,0018841 | TRUE | 0,0213668 | No |
| orf19.4090 | 120 | 84 | 106 | 130 | 132 | 120 | 135 | 102 | 108 | 107 | 108 | 107 | 113 | 115 | 120 | 144 | 115,24 | 108 | 123 | 1,132 | 0,178 | 1,36 | 0,17377066 | 0,24455706 | 1,00E-08 | 0,0145762 | 0,0077991 | 0,0077991 | TRUE | 0,1055626 | No |
| orf19.4090.1 | 538 | 316 | 412 | 585 | 523 | 385 | 422 | 270 | 483 | 402 | 422 | 482 | 447 | 367 | 377 | 381 | 420,11 | 447 | 393 | 0,877 | -0,189 | -2,032 | 0,04219943 | 0,07247975 | 0,00633 | 0,0054351 | 0,0058881 | 0,0058881 | TRUE | 0,1058602 | No |
| orf19.4091 | 736 | 494 | 654 | 831 | 767 | 631 | 754 | 427 | 660 | 629 | 669 | 685 | 656 | 602 | 673 | 602 | 647,08 | 661 | 633 | 0,959 | -0,06 | -0,934 | 0,35043675 | 0,43799126 | 0,000695 | 0,004223 | 0,0024486 | 0,0024486 | TRUE | 0,0206236 | No |
| orf19.4092 | 951 | 681 | 782 | 994 | 846 | 754 | 743 | 482 | 853 | 867 | 800 | 820 | 724 | 719 | 663 | 680 | 765,71 | 835 | 696 | 0,836 | -0,259 | -4,387 | 1,15E-05 | 4,14E-05 | 1,00E-08 | 0,0038754 | 0,002025 | 0,002025 | TRUE | 0,0140873 | No |
| orf19.4093 | 22810 | 15663 | 19862 | 24578 | 18677 | 16417 | 18002 | 10855 | 20463 | 19941 | 20321 | 20267 | 15973 | 15665 | 16064 | 15310 | 18000 | 20248 | 15753 | 0,778 | -0,362 | -13,234 | 5,57E-40 | 3,76E-38 | 0,000219 | 0,0020602 | 0,000661 | 0,000661 | TRUE | 0,0042426 | No |
| orf19.4094 | 65 | 56 | 81 | 81 | 60 | 65 | 59 | 55 | 58 | 71 | 83 | 67 | 51 | 62 | 53 | 78 | 65,35 | 70 | 61 | 0,862 | -0,215 | -1,127 | 0,25958628 | 0,34291873 | 0,012197 | 0,0241927 | 0,0192599 | 0,0192599 | TRUE | 0,2302517 | No |
| orf19.4095 | 13 | 11 | 11 | 8 | 6 | 7 | 1 | 7 | 12 | 14 | 11 | 7 | 5 | 7 | 1 | 10 | 8,26 | 11 | 6 | 0,5 | -0,999 | -1,802 | 0,07154759 | NA | 0,107965 | 0,1776949 | 0,1557939 | 0,1557939 | TRUE | 0,6419556 | No |

| | | | | | | | | | | | | | | | | | | | | | | | | | | | | | | | |
|---|---|---|---|---|---|---|---|---|---|---|---|---|---|---|---|---|---|---|---|---|---|---|---|---|---|---|---|---|---|---|---|
| orf19.4096 | 2012 | 1220 | 1680 | 2109 | 1983 | 1653 | 1886 | 1175 | 1805 | 1553 | 1719 | 1739 | 1696 | 1577 | 1683 | 1657 | 1678,7 | 1704 | 1653 | 0,969 | -0,046 | -0,838 | 0,40218789 | 0,49072803 | 0,001698 | 0,0028443 | 0,0022676 | 0,0022676 | TRUE | 0,0404051 | No |
| orf19.4097 | 1122 | 806 | 1016 | 1204 | 939 | 961 | 942 | 579 | 1007 | 1026 | 1039 | 993 | 803 | 917 | 841 | 817 | 930,28 | 1016 | 844 | 0,832 | -0,265 | -4,609 | 4,04E-06 | 1,56E-05 | 0,000853 | 0,00354 | 0,0020979 | 0,0020979 | TRUE | 0,0406937 | No |
| orf19.4099 | 15561 | 10637 | 13821 | 16405 | 14727 | 12743 | 13838 | 8735 | 13960 | 13542 | 14140 | 13528 | 12595 | 12159 | 12348 | 12320 | 13074 | 13792 | 12356 | 0,896 | -0,159 | -5,448 | 5,10E-08 | 2,59E-07 | 0,000251 | 0,0020906 | 0,0007375 | 0,0007375 | TRUE | 0,003501 | No |
| orf19.4101 | 227 | 143 | 182 | 247 | 265 | 169 | 207 | 134 | 204 | 182 | 186 | 204 | 227 | 161 | 185 | 189 | 192,15 | 194 | 190 | 0,981 | -0,027 | -0,23 | 0,81787023 | 0,8596231 | 0,007319 | 0,0095347 | 0,0084902 | 0,0084902 | TRUE | 0,1955555 | No |
| orf19.4102 | 5497 | 3689 | 4704 | 5874 | 5541 | 4865 | 5162 | 3595 | 4931 | 4696 | 4813 | 4844 | 4739 | 4642 | 4606 | 5070 | 4792,7 | 4821 | 4764 | 0,987 | -0,019 | -0,471 | 0,63753857 | 0,70814045 | 0,000839 | 0,0022824 | 0,0013571 | 0,0013571 | TRUE | 0,0216557 | No |
| orf19.410.3 | 1574 | 1111 | 1299 | 1666 | 1153 | 931 | 1045 | 717 | 1412 | 1414 | 1329 | 1374 | 986 | 888 | 932 | 1011 | 1168,4 | 1382 | 954 | 0,689 | -0,537 | -9,806 | 1,06E-22 | 2,32E-21 | 0,000902 | 0,003222 | 0,0019855 | 0,0019855 | TRUE | 0,0255983 | No |
| orf19.4104 | 179 | 108 | 127 | 174 | 148 | 131 | 137 | 74 | 161 | 137 | 130 | 143 | 127 | 125 | 122 | 104 | 131,21 | 143 | 120 | 0,84 | -0,251 | -1,987 | 0,04693513 | 0,07958969 | 1,00E-08 | 0,0130434 | 0,0076839 | 0,0076839 | TRUE | 0,0754806 | No |
| orf19.4105 | 600 | 383 | 578 | 702 | 769 | 666 | 698 | 442 | 538 | 488 | 591 | 579 | 658 | 635 | 623 | 623 | 591,94 | 549 | 635 | 1,154 | 0,206 | 2,908 | 0,00363494 | 0,00807645 | 0,001699 | 0,004432 | 0,0031336 | 0,0031336 | TRUE | 0,0588967 | No |
| orf19.4106 | 1390 | 1023 | 1270 | 1553 | 1269 | 1164 | 1303 | 794 | 1247 | 1302 | 1299 | 1281 | 1085 | 1111 | 1163 | 1120 | 1201 | 1282 | 1120 | 0,874 | -0,195 | -4,107 | 4,01E-05 | 0,00013187 | 1,00E-08 | 0,0031883 | 0,0013248 | 0,0013248 | TRUE | 0,0087897 | No |
| orf19.4107 | 1846 | 1244 | 1619 | 1973 | 1744 | 1442 | 1621 | 952 | 1656 | 1584 | 1656 | 1627 | 1492 | 1376 | 1446 | 1343 | 1522,5 | 1631 | 1414 | 0,869 | -0,203 | -4,214 | 2,51E-05 | 8,55E-05 | 0,000474 | 0,002933 | 0,0015737 | 0,0015737 | TRUE | 0,0163686 | No |
| orf19.4109 | 785 | 617 | 760 | 896 | 1223 | 1071 | 1108 | 754 | 704 | 786 | 778 | 739 | 1046 | 1022 | 989 | 1063 | 890,77 | 752 | 1030 | 1,372 | 0,456 | 7,997 | 1,28E-15 | 1,56E-14 | 0,000464 | 0,0036092 | 0,0019731 | 0,0019731 | TRUE | 0,0213776 | No |
| orf19.411 | 20561 | 14657 | 17326 | 21752 | 19940 | 17605 | 18706 | 10985 | 18445 | 18660 | 17726 | 17937 | 17053 | 16798 | 16692 | 15493 | 17351 | 18192 | 16509 | 0,908 | -0,139 | -3,583 | 0,00034031 | 0,00095053 | 0,001091 | 0,0020632 | 0,0013977 | 0,0013977 | TRUE | 0,020651 | No |
| orf19.4110 | 1600 | 1066 | 1447 | 1702 | 1805 | 1613 | 1806 | 1209 | 1435 | 1357 | 1480 | 1403 | 1544 | 1539 | 1612 | 1705 | 1509,5 | 1419 | 1600 | 1,124 | 0,169 | 3,318 | 0,00090682 | 0,00229882 | 0,000962 | 0,0029412 | 0,0018302 | 0,0018302 | TRUE | 0,021471 | No |
| orf19.4112 | 1719 | 1149 | 1474 | 1898 | 1806 | 1475 | 1580 | 1034 | 1542 | 1463 | 1508 | 1565 | 1545 | 1407 | 1410 | 1458 | 1487,3 | 1520 | 1455 | 0,957 | -0,064 | -1,29 | 0,19697256 | 0,2709895 | 0,000805 | 0,0029556 | 0,0016793 | 0,0016793 | TRUE | 0,0222345 | No |
| orf19.4114 | 2721 | 1756 | 2380 | 2850 | 2387 | 2152 | 2370 | 1423 | 2441 | 2236 | 2435 | 2350 | 2041 | 2053 | 2115 | 2007 | 2209,8 | 2366 | 2054 | 0,868 | -0,204 | -4,684 | 2,81E-06 | 1,11E-05 | 0,000457 | 0,0026365 | 0,0013621 | 0,0013621 | TRUE | 0,0153713 | No |
| orf19.4115 | 17 | 7 | 14 | 9 | 16 | 8 | 8 | 4 | 15 | 9 | 14 | 7 | 14 | 8 | 7 | 6 | 10 | 11 | 9 | 0,759 | -0,397 | -0,857 | 0,39137646 | NA | 0,045343 | 0,147141 | 0,1057556 | 0,1057556 | TRUE | 0,4348824 | No |
| orf19.4116 | 679 | 468 | 573 | 723 | 716 | 589 | 641 | 421 | 609 | 596 | 586 | 596 | 612 | 562 | 572 | 594 | 590,94 | 597 | 585 | 0,98 | -0,03 | -0,482 | 0,6299248 | 0,70110934 | 1,00E-08 | 0,0044361 | 0,0019591 | 0,0019591 | TRUE | 0,0133927 | No |
| orf19.4117 | 1480 | 1098 | 1357 | 1633 | 1557 | 1450 | 1581 | 898 | 1328 | 1398 | 1388 | 1347 | 1332 | 1384 | 1411 | 1267 | 1356,6 | 1365 | 1348 | 0,991 | -0,014 | -0,277 | 0,78163111 | 0,82881039 | 0,000428 | 0,0030496 | 0,00163 | 0,00163 | TRUE | 0,017496 | No |
| orf19.4118 | 26667 | 19861 | 25374 | 30588 | 28153 | 26406 | 27868 | 17511 | 23923 | 25285 | 25960 | 25223 | 24077 | 25196 | 24868 | 24698 | 24904 | 25098 | 24710 | 0,985 | -0,022 | -0,655 | 0,51258661 | 0,59705707 | 0,000682 | 0,0020378 | 0,0010872 | 0,0010872 | TRUE | 0,0122089 | No |
| orf19.4119 | 4743 | 3308 | 4184 | 4893 | 5369 | 5083 | 5394 | 3191 | 4255 | 4211 | 4281 | 4035 | 4592 | 4850 | 4813 | 4501 | 4442,2 | 4196 | 4689 | 1,119 | 0,162 | 4,067 | 4,76E-05 | 0,00015473 | 0,000702 | 0,0023063 | 0,0012905 | 0,0012905 | TRUE | 0,0083795 | No |
| orf19.412 | 2150 | 2128 | 2169 | 2941 | 2540 | 2879 | 2684 | 1470 | 1929 | 2709 | 2219 | 2425 | 2172 | 2747 | 2395 | 2073 | 2333,7 | 2320 | 2347 | 1,012 | 0,018 | 0,169 | 0,86584958 | 0,89777419 | 0,017547 | 0,0026016 | 0,0101776 | 0,0101776 | TRUE | 0,1596133 | No |
| orf19.4120 | 5736 | 3865 | 4815 | 6122 | 6174 | 5406 | 5653 | 3595 | 5146 | 4921 | 4926 | 5048 | 5280 | 5158 | 5044 | 5070 | 5074,3 | 5010 | 5138 | 1,025 | 0,036 | 1,089 | 0,27609577 | 0,36025364 | 0,000265 | 0,0022656 | 0,0008676 | 0,0008676 | TRUE | 0,0043297 | No |
| orf19.4121 | 1923 | 1200 | 1600 | 1960 | 1907 | 1599 | 1766 | 1102 | 1725 | 1528 | 1637 | 1616 | 1631 | 1526 | 1576 | 1554 | 1599,1 | 1626 | 1572 | 0,966 | -0,051 | -1,011 | 0,31212513 | 0,3984327 | 0,000994 | 0,0028873 | 0,0017732 | 0,0017732 | TRUE | 0,0200856 | No |
| orf19.4122 | 1988 | 1378 | 1870 | 2260 | 1979 | 1839 | 1890 | 1267 | 1783 | 1754 | 1913 | 1864 | 1693 | 1755 | 1687 | 1787 | 1779,4 | 1828 | 1730 | 0,944 | -0,083 | -1,795 | 0,07265365 | 0,11636981 | 0,000566 | 0,0027954 | 0,0014708 | 0,0014708 | TRUE | 0,0112159 | No |
| orf19.4123 | 3137 | 2263 | 2825 | 3310 | 3394 | 2973 | 3285 | 2003 | 2814 | 2881 | 2890 | 2729 | 2903 | 2837 | 2931 | 2825 | 2851,3 | 2828 | 2874 | 1,017 | 0,025 | 0,667 | 0,50466221 | 0,58919754 | 0,000188 | 0,0024887 | 0,0009874 | 0,0009874 | TRUE | 0,0070273 | No |
| orf19.4125 | 3470 | 2615 | 2959 | 3853 | 3310 | 3100 | 3241 | 2025 | 3113 | 3329 | 3027 | 3177 | 2831 | 2958 | 2892 | 2856 | 3023 | 3162 | 2884 | 0,913 | -0,131 | -3,206 | 0,00134372 | 0,00328842 | 0,000553 | 0,0024597 | 0,0012782 | 0,0012782 | TRUE | 0,0145269 | No |
| orf19.4127 | 9182 | 6138 | 8108 | 9904 | 10413 | 8902 | 9841 | 5774 | 8237 | 7814 | 8295 | 8167 | 8906 | 8494 | 8781 | 8144 | 8354,8 | 8128 | 8581 | 1,056 | 0,079 | 2,004 | 0,04512047 | 0,0768814 | 0,000972 | 0,0021533 | 0,0013587 | 0,0013587 | TRUE | 0,0139659 | No |
| orf19.4128 | 1433 | 1076 | 1258 | 1564 | 1045 | 955 | 1083 | 594 | 1286 | 1370 | 1287 | 1290 | 894 | 911 | 966 | 838 | 1105,2 | 1308 | 902 | 0,693 | -0,53 | -9,555 | 1,23E-21 | 2,51E-20 | 0,000973 | 0,0032931 | 0,0020158 | 0,0020158 | TRUE | 0,0269317 | No |
| orf19.413 | 4167 | 3066 | 3716 | 4648 | 4678 | 4447 | 4784 | 2895 | 3738 | 3903 | 3802 | 3833 | 4001 | 4243 | 4269 | 4083 | 3984 | 3819 | 4149 | 1,087 | 0,12 | 3,277 | 0,00104981 | 0,00263462 | 0,000425 | 0,0023439 | 0,0010438 | 0,0010438 | TRUE | 0,0075275 | No |
| orf19.4131 | 842 | 645 | 814 | 1034 | 975 | 1049 | 1024 | 602 | 755 | 821 | 833 | 853 | 834 | 1001 | 914 | 849 | 857,44 | 816 | 900 | 1,104 | 0,143 | 1,99 | 0,04657804 | 0,07902876 | 0,003858 | 0,0036726 | 0,0037739 | 0,0037739 | TRUE | 0,0692946 | No |
| orf19.413.1 | 4796 | 4430 | 4341 | 5562 | 4665 | 4937 | 4999 | 2167 | 4303 | 5640 | 4441 | 4586 | 3990 | 4711 | 4461 | 3056 | 4398,5 | 4742 | 4054 | 0,855 | -0,226 | -1,416 | 0,15665653 | 0,22377376 | 0,024133 | 0,0023096 | 0,0132206 | 0,0241333 | TRUE | 0,3321907 | No |
| orf19.4132 | 2221 | 1531 | 1893 | 2351 | 2179 | 1969 | 1975 | 1178 | 1992 | 1949 | 1937 | 1939 | 1864 | 1879 | 1762 | 1661 | 1872,9 | 1954 | 1792 | 0,919 | -0,123 | -2,538 | 0,01114974 | 0,02221075 | 0,000963 | 0,0027546 | 0,001702 | 0,001702 | TRUE | 0,0260372 | No |
| orf19.4133 | 2487 | 1706 | 2340 | 2727 | 2890 | 2478 | 2771 | 1673 | 2231 | 2172 | 2394 | 2249 | 2472 | 2364 | 2473 | 2360 | 2339,3 | 2262 | 2417 | 1,069 | 0,097 | 2,191 | 0,02845445 | 0,05116862 | 0,000645 | 0,0026001 | 0,0014371 | 0,0014371 | TRUE | 0,0184504 | No |
| orf19.4134 | 967 | 723 | 936 | 1089 | 912 | 880 | 937 | 587 | 868 | 920 | 958 | 898 | 780 | 840 | 836 | 828 | 865,91 | 911 | 821 | 0,902 | -0,15 | -2,637 | 0,00836096 | 0,01715596 | 1,00E-08 | 0,003656 | 0,0019299 | 0,0019299 | TRUE | 0,0145793 | No |
| orf19.4135 | 941 | 584 | 866 | 1011 | 946 | 863 | 814 | 562 | 844 | 743 | 886 | 834 | 809 | 823 | 726 | 793 | 807,36 | 827 | 788 | 0,951 | -0,073 | -1,062 | 0,28812592 | 0,37259159 | 0,00281 | 0,0037776 | 0,0032687 | 0,0032687 | TRUE | 0,0499465 | No |
| orf19.4136 | 381 | 226 | 297 | 367 | 385 | 397 | 415 | 231 | 342 | 288 | 304 | 303 | 329 | 379 | 370 | 326 | 330,03 | 309 | 351 | 1,136 | 0,184 | 2,054 | 0,03995877 | 0,06896647 | 0,002752 | 0,0063783 | 0,0046664 | 0,0046664 | TRUE | 0,059193 | No |
| orf19.414 | 5411 | 3817 | 4858 | 5772 | 6693 | 5998 | 6511 | 3678 | 4854 | 4859 | 4970 | 4760 | 5724 | 5723 | 5810 | 5187 | 5236 | 4861 | 5611 | 1,156 | 0,209 | 4,842 | 1,29E-06 | 5,33E-06 | 0,001194 | 0,0022568 | 0,0015898 | 0,0015898 | TRUE | 0,0302412 | No |
| orf19.4142 | 579 | 455 | 489 | 627 | 681 | 597 | 578 | 403 | 519 | 579 | 500 | 517 | 582 | 570 | 516 | 568 | 544,03 | 529 | 559 | 1,059 | 0,083 | 1,151 | 0,24960155 | 0,33169788 | 0,001511 | 0,004648 | 0,0031016 | 0,0031016 | TRUE | 0,0426831 | No |
| orf19.4143 | 1220 | 832 | 1070 | 1332 | 1748 | 1455 | 1615 | 981 | 1094 | 1059 | 1095 | 1098 | 1495 | 1388 | 1441 | 1384 | 1256,9 | 1086 | 1427 | 1,314 | 0,394 | 8,187 | 2,68E-16 | 3,47E-15 | 1,00E-08 | 0,0031346 | 0,0014138 | 0,0014138 | TRUE | 0,0127002 | No |
| orf19.4144 | 2330 | 1572 | 2047 | 2645 | 3381 | 2954 | 3185 | 2139 | 2090 | 2001 | 2094 | 2181 | 2892 | 2819 | 2842 | 3017 | 2492 | 2092 | 2892 | 1,38 | 0,464 | 10,883 | 1,38E-27 | 4,42E-26 | 0,000534 | 0,0025621 | 0,0013343 | 0,0013343 | TRUE | 0,0102165 | No |
| orf19.4145 | 2645 | 1956 | 2325 | 2807 | 2624 | 2462 | 2615 | 1556 | 2373 | 2490 | 2379 | 2315 | 2244 | 2349 | 2333 | 2195 | 2334,7 | 2389 | 2280 | 0,956 | -0,064 | -1,517 | 0,12932988 | 0,19002542 | 0,000419 | 0,0026013 | 0,0012947 | 0,0012947 | TRUE | 0,0092821 | No |
| orf19.4146 | 1289 | 913 | 1018 | 1294 | 1416 | 1204 | 1192 | 617 | 1156 | 1162 | 1042 | 1067 | 1211 | 1149 | 1064 | 870 | 1090,1 | 1107 | 1074 | 0,973 | -0,04 | -0,437 | 0,66180892 | 0,72997026 | 0,010328 | 0,0033112 | 0,0069329 | 0,0069329 | TRUE | 0,1834236 | No |
| orf19.4147 | 2655 | 1912 | 2362 | 2830 | 2664 | 2430 | 2554 | 1686 | 2382 | 2434 | 2417 | 2334 | 2278 | 2319 | 2279 | 2378 | 2352,5 | 2392 | 2314 | 0,967 | -0,048 | -1,288 | 0,1975845 | 0,27176916 | 1,00E-08 | 0,0025966 | 0,0009288 | 0,0009288 | TRUE | 0,0038039 | No |
| orf19.4148 | 485 | 352 | 405 | 521 | 557 | 522 | 558 | 327 | 435 | 448 | 414 | 430 | 476 | 498 | 498 | 461 | 457,6 | 432 | 483 | 1,123 | 0,167 | 2,444 | 0,01452784 | 0,02805634 | 1,00E-08 | 0,005152 | 0,0023225 | 0,0023225 | TRUE | 0,0090348 | No |
| orf19.4149 | 31 | 14 | 17 | 9 | 17 | 16 | 11 | 14 | 28 | 18 | 17 | 7 | 15 | 15 | 10 | 20 | 16,23 | 18 | 15 | 0,833 | -0,263 | -0,633 | 0,52644897 | 0,60955129 | 0,117421 | 0,0914423 | 0,1031776 | 0,1031776 | TRUE | 0,2334488 | No |
| orf19.4149.1 | 13951 | 9414 | 11928 | 14770 | 14018 | 11936 | 12490 | 8860 | 12516 | 11985 | 12204 | 12180 | 11989 | 11389 | 11145 | 12496 | 11988 | 12221 | 11755 | 0,961 | -0,057 | -1,349 | 0,17732331 | 0,24903189 | 0,001389 | 0,0021006 | 0,0016164 | 0,0016164 | TRUE | 0,0216204 | No |
| orf19.415 | 1263 | 824 | 1046 | 1288 | 1405 | 1155 | 1247 | 853 | 1133 | 1049 | 1070 | 1062 | 1202 | 1102 | 1113 | 1203 | 1116,7 | 1078 | 1155 | 1,069 | 0,096 | 1,765 | 0,07754822 | 0,12322394 | 0,000877 | 0,0032795 | 0,0019275 | 0,0019275 | TRUE | 0,0141826 | No |
| orf19.4150 | 7720 | 5497 | 6641 | 8683 | 7482 | 6617 | 6985 | 4566 | 6926 | 6998 | 6794 | 7160 | 6399 | 6314 | 6233 | 6440 | 6658 | 6970 | 6346 | 0,91 | -0,136 | -4,39 | 1,13E-05 | 4,08E-05 | 0,000215 | 0,0021976 | 0,0007668 | 0,0007668 | TRUE | 0,0042896 | No |
| orf19.4151 | 178 | 122 | 144 | 162 | 111 | 107 | 103 | 66 | 160 | 155 | 147 | 134 | 95 | 102 | 92 | 93 | 122,24 | 149 | 96 | 0,643 | -0,636 | -4,949 | 7,44E-07 | 3,20E-06 | 1,00E-08 | 0,0138551 | 0,0073025 | 0,0073025 | TRUE | 0,0573428 | No |
| orf19.4152 | 61084 | 40723 | 56007 | 64559 | 59685 | 52446 | 56865 | 37633 | 54799 | 51845 | 57301 | 53236 | 51044 | 50042 | 50743 | 53078 | 52761 | 54295 | 51227 | 0,943 | -0,084 | -2,147 | 0,03182312 | 0,056495 | 0,001226 | 0,002007 | 0,0014536 | 0,0014536 | TRUE | 0,0169923 | No |
| orf19.4153 | 792 | 797 | 806 | 985 | 898 | 1074 | 1080 | 581 | 711 | 1015 | 825 | 812 | 768 | 1025 | 964 | 819 | 867,25 | 841 | 894 | 1,066 | 0,093 | 0,813 | 0,4159721 | 0,50416792 | 0,018804 | 0,0036534 | 0,0112675 | 0,0112675 | TRUE | 0,2250067 | No |
| orf19.4157 | 405 | 285 | 317 | 432 | 403 | 331 | 348 | 237 | 363 | 363 | 324 | 356 | 345 | 316 | 311 | 334 | 339 | 352 | 326 | 0,927 | -0,109 | -1,347 | 0,1779131 | 0,24959675 | 1,00E-08 | 0,0062618 | 0,0033513 | 0,0033513 | TRUE | 0,0306117 | No |
| orf19.4159 | 1720 | 1199 | 1503 | 1867 | 1746 | 1500 | 1655 | 1075 | 1543 | 1526 | 1538 | 1540 | 1493 | 1431 | 1477 | 1516 | 1508 | 1537 | 1479 | 0,962 | -0,056 | -1,34 | 0,18024149 | 0,25201289 | 1,00E-08 | 0,0029422 | 0,0010412 | 0,0010412 | TRUE | 0,005688 | No |
| orf19.416 | 163 | 77 | 115 | 158 | 126 | 104 | 134 | 80 | 146 | 98 | 118 | 130 | 108 | 99 | 120 | 113 | 116,45 | 123 | 110 | 0,885 | -0,177 | -1,216 | 0,22409755 | 0,30310178 | 0,008118 | 0,0144458 | 0,0117326 | 0,0117326 | TRUE | 0,16351 | No |
| orf19.4160 | 2406 | 1665 | 2085 | 2617 | 2536 | 2238 | 2407 | 1712 | 2158 | 2120 | 2133 | 2158 | 2169 | 2135 | 2148 | 2415 | 2179,5 | 2142 | 2217 | 1,032 | 0,046 | 0,955 | 0,33949479 | 0,42653381 | 0,001219 | 0,0026456 | 0,001736 | 0,001736 | TRUE | 0,0404842 | No |
| orf19.4161 | 565 | 430 | 469 | 626 | 727 | 698 | 682 | 460 | 507 | 547 | 480 | 516 | 622 | 666 | 609 | 649 | 574,43 | 512 | 636 | 1,242 | 0,312 | 4,693 | 2,70E-06 | 1,07E-05 | 1,00E-08 | 0,0045067 | 0,002494 | 0,002494 | TRUE | 0,0210631 | No |
| orf19.4162 | 208 | 189 | 220 | 278 | 295 | 307 | 342 | 185 | 187 | 241 | 225 | 229 | 252 | 293 | 305 | 261 | 249,11 | 220 | 278 | 1,267 | 0,341 | 3,173 | 0,00151103 | 0,00366059 | 0,006178 | 0,0078071 | 0,0070432 | 0,0070432 | TRUE | 0,1221051 | No |
| orf19.4163 | 1228 | 1010 | 1144 | 1470 | 1234 | 1208 | 1325 | 830 | 1102 | 1286 | 1170 | 1212 | 1055 | 1153 | 1182 | 1171 | 1166,4 | 1192 | 1140 | 0,957 | -0,064 | -1,029 | 0,30342852 | 0,38928007 | 0,002574 | 0,0032241 | 0,0028577 | 0,0028577 | TRUE | 0,0318115 | No |
| orf19.4164 | 1488 | 1240 | 1281 | 1620 | 1547 | 1559 | 1556 | 922 | 1335 | 1579 | 1311 | 1336 | 1323 | 1488 | 1388 | 1300 | 1382,4 | 1390 | 1375 | 0,992 | -0,012 | -0,173 | 0,8625682 | 0,89566363 | 0,004455 | 0,0030296 | 0,004041 | 0,004041 | TRUE | 0,096295 | No |

| | | | | | | | | | | | | | | | | | | | | | | | | | | | | | | | |
|---|---|---|---|---|---|---|---|---|---|---|---|---|---|---|---|---|---|---|---|---|---|---|---|---|---|---|---|---|---|---|---|
| orf19.4166 | 4224 | 3033 | 3889 | 4695 | 5157 | 4319 | 4909 | 3594 | 3789 | 3861 | 3979 | 3872 | 4410 | 4121 | 4380 | 5069 | 4185,3 | 3875 | 4495 | 1,159 | 0,213 | 3,55 | 0,00038576 | 0,00106754 | 0,003921 | 0,0023264 | 0,0032081 | 0,0032081 | TRUE | 0,0877133 | No |
| orf19.4167 | 867 | 561 | 719 | 883 | 800 | 731 | 781 | 443 | 778 | 714 | 736 | 728 | 684 | 697 | 697 | 625 | 707,39 | 739 | 676 | 0,917 | -0,125 | -2,054 | 0,03994596 | 0,06896417 | 0,000218 | 0,0040317 | 0,0021641 | 0,0021641 | TRUE | 0,0256846 | No |
| orf19.4168 | 1468 | 937 | 1208 | 1562 | 1753 | 1343 | 1471 | 895 | 1317 | 1193 | 1236 | 1288 | 1499 | 1281 | 1313 | 1262 | 1298,7 | 1258 | 1339 | 1,064 | 0,09 | 1,385 | 0,16592124 | 0,23532928 | 0,003236 | 0,0030974 | 0,0032929 | 0,0032929 | TRUE | 0,08003 | No |
| orf19.4169 | 64 | 42 | 64 | 76 | 87 | 76 | 71 | 52 | 57 | 53 | 65 | 63 | 74 | 73 | 63 | 73 | 65,33 | 60 | 71 | 1,179 | 0,238 | 1,408 | 0,15903846 | 0,22663736 | 1,00E-08 | 0,0242 | 0,0121234 | 0,0121234 | TRUE | 0,0514125 | No |
| orf19.417 | 3524 | 2453 | 3235 | 3971 | 3621 | 3201 | 3499 | 2264 | 3161 | 3123 | 3310 | 3275 | 3097 | 3054 | 3122 | 3193 | 3166,9 | 3217 | 3116 | 0,968 | -0,047 | -1,291 | 0,19684932 | 0,27088197 | 0,000137 | 0,0024379 | 0,0009823 | 0,0009823 | TRUE | 0,0043885 | No |
| orf19.4170 | 877 | 636 | 822 | 952 | 1398 | 1276 | 1297 | 824 | 787 | 810 | 841 | 785 | 1196 | 1218 | 1157 | 1162 | 994,39 | 806 | 1183 | 1,471 | 0,557 | 10,82 | 2,78E-27 | 8,69E-26 | 1,00E-08 | 0,0034394 | 0,0015027 | 0,0015027 | TRUE | 0,0106689 | No |
| orf19.4171 | 413 | 269 | 306 | 412 | 354 | 340 | 403 | 188 | 371 | 342 | 313 | 340 | 303 | 324 | 360 | 265 | 327,21 | 342 | 313 | 0,921 | -0,118 | -1,186 | 0,23556969 | 0,31641017 | 0,006273 | 0,0064161 | 0,0065202 | 0,0065202 | TRUE | 0,1127555 | No |
| orf19.4172 | 1357 | 976 | 1144 | 1526 | 1343 | 1353 | 1282 | 731 | 1217 | 1243 | 1170 | 1258 | 1149 | 1291 | 1144 | 1031 | 1187,9 | 1222 | 1154 | 0,946 | -0,08 | -1,201 | 0,22961603 | 0,30959002 | 0,003631 | 0,0032016 | 0,0034284 | 0,0034284 | TRUE | 0,0755839 | No |
| orf19.4173 | 1120 | 664 | 954 | 1140 | 1257 | 929 | 1087 | 609 | 1005 | 845 | 976 | 940 | 1075 | 886 | 970 | 859 | 944,57 | 942 | 948 | 1,007 | 0,011 | 0,131 | 0,89591358 | 0,9213709 | 0,006756 | 0,0035164 | 0,0052454 | 0,0052454 | TRUE | 0,098978 | No |
| orf19.4174 | 2261 | 1455 | 2066 | 2387 | 2658 | 2222 | 2304 | 1676 | 2028 | 1852 | 2114 | 1968 | 2273 | 2120 | 2056 | 2364 | 2097 | 1990 | 2203 | 1,105 | 0,144 | 2,441 | 0,01464461 | 0,02825459 | 0,00292 | 0,0026718 | 0,002854 | 0,002854 | TRUE | 0,0275539 | No |
| orf19.4175 | 4413 | 3099 | 4120 | 4944 | 4573 | 3997 | 4360 | 2816 | 3959 | 3945 | 4215 | 4077 | 3911 | 3814 | 3891 | 3972 | 3972,9 | 4049 | 3897 | 0,962 | -0,056 | -1,554 | 0,12022854 | 0,17853017 | 0,000284 | 0,0023449 | 0,0010042 | 0,0010042 | TRUE | 0,0091088 | No |
| orf19.4176 | 1332 | 897 | 1067 | 1317 | 1335 | 1201 | 1245 | 743 | 1195 | 1142 | 1092 | 1086 | 1142 | 1146 | 1111 | 1048 | 1120,2 | 1129 | 1112 | 0,987 | -0,019 | -0,341 | 0,73336792 | 0,79160519 | 0,000912 | 0,0032755 | 0,0019595 | 0,0019595 | TRUE | 0,0199751 | No |
| orf19.4177 | 1869 | 1259 | 1671 | 2043 | 2146 | 1951 | 2048 | 1394 | 1677 | 1603 | 1710 | 1685 | 1835 | 1862 | 1827 | 1966 | 1770,5 | 1669 | 1872 | 1,119 | 0,163 | 3,689 | 0,00022547 | 0,00065063 | 0,000207 | 0,0027994 | 0,0012993 | 0,0012993 | TRUE | 0,0122591 | No |
| orf19.4179 | 54 | 33 | 61 | 65 | 56 | 40 | 47 | 31 | 48 | 42 | 62 | 54 | 48 | 38 | 42 | 44 | 47,27 | 52 | 43 | 0,828 | -0,272 | -1,329 | 0,18395017 | 0,2561382 | 1,00E-08 | 0,0326882 | 0,0191243 | 0,0191243 | TRUE | 0,1492422 | No |
| orf19.418 | 2553 | 1799 | 2410 | 2732 | 3209 | 3003 | 3226 | 1865 | 2290 | 2290 | 2466 | 2253 | 2744 | 2865 | 2879 | 2630 | 2552,3 | 2325 | 2780 | 1,197 | 0,26 | 5,483 | 4,18E-08 | 2,14E-07 | 0,001195 | 0,0025483 | 0,0017569 | 0,0017569 | TRUE | 0,0201846 | No |
| orf19.4180 | 1196 | 804 | 936 | 1210 | 1146 | 1014 | 1015 | 686 | 1073 | 1024 | 958 | 998 | 980 | 968 | 906 | 968 | 984,1 | 1013 | 956 | 0,942 | -0,086 | -1,521 | 0,1282536 | 0,18891607 | 0,000889 | 0,0034547 | 0,0020274 | 0,0020274 | TRUE | 0,0199933 | No |
| orf19.4181 | 517 | 350 | 448 | 543 | 569 | 421 | 452 | 309 | 464 | 446 | 458 | 448 | 487 | 402 | 403 | 436 | 442,87 | 454 | 432 | 0,952 | -0,071 | -0,896 | 0,37042004 | 0,45894872 | 0,002498 | 0,0052574 | 0,0038571 | 0,0038571 | TRUE | 0,0933268 | No |
| orf19.4182 | 1419 | 1120 | 1349 | 1768 | 1777 | 1580 | 1576 | 1069 | 1273 | 1426 | 1380 | 1458 | 1520 | 1508 | 1406 | 1508 | 1434,8 | 1384 | 1486 | 1,073 | 0,102 | 1,822 | 0,06846189 | 0,11068866 | 0,001794 | 0,0029913 | 0,0022881 | 0,0022881 | TRUE | 0,0364661 | No |
| orf19.4183 | 3453 | 2226 | 3080 | 3569 | 4016 | 3370 | 3730 | 2399 | 3098 | 2834 | 3151 | 2943 | 3435 | 3216 | 3328 | 3384 | 3173,5 | 3006 | 3341 | 1,11 | 0,151 | 3,335 | 0,00085386 | 0,00218112 | 0,001137 | 0,002437 | 0,0016527 | 0,0016527 | TRUE | 0,0171226 | No |
| orf19.4184 | 2076 | 1450 | 2032 | 2340 | 2389 | 2104 | 2475 | 1717 | 1862 | 1846 | 2079 | 1930 | 2043 | 2008 | 2209 | 2422 | 2049,7 | 1929 | 2170 | 1,123 | 0,167 | 2,509 | 0,01209633 | 0,0238591 | 0,004658 | 0,0026878 | 0,0037794 | 0,0037794 | TRUE | 0,0705835 | No |
| orf19.4185 | 240 | 192 | 227 | 268 | 230 | 237 | 222 | 133 | 215 | 244 | 232 | 221 | 197 | 226 | 198 | 188 | 215,19 | 228 | 202 | 0,891 | -0,166 | -1,655 | 0,09788654 | 0,15006438 | 1,00E-08 | 0,0087257 | 0,0050095 | 0,0050095 | TRUE | 0,0692555 | No |
| orf19.4186 | 1221 | 838 | 1041 | 1322 | 1179 | 1043 | 1211 | 728 | 1095 | 1067 | 1065 | 1090 | 1008 | 995 | 1081 | 1027 | 1053,5 | 1079 | 1028 | 0,952 | -0,071 | -1,422 | 0,15515049 | 0,22199193 | 1,00E-08 | 0,0033574 | 0,0014713 | 0,0014713 | TRUE | 0,0155 | No |
| orf19.4187 | 1226 | 838 | 1036 | 1321 | 1380 | 1159 | 1379 | 803 | 1100 | 1067 | 1060 | 1089 | 1180 | 1106 | 1231 | 1133 | 1120,6 | 1079 | 1162 | 1,078 | 0,108 | 2,083 | 0,03722116 | 0,06483756 | 0,000499 | 0,0032749 | 0,0017025 | 0,0017025 | TRUE | 0,0191718 | No |
| orf19.4188 | 6651 | 5243 | 6256 | 7405 | 6954 | 6801 | 6959 | 4124 | 5967 | 6675 | 6401 | 6106 | 5947 | 6489 | 6210 | 5817 | 6201,4 | 6287 | 6116 | 0,974 | -0,039 | -0,775 | 0,43812334 | 0,52497906 | 0,002153 | 0,0022136 | 0,0022326 | 0,0022326 | TRUE | 0,020741 | No |
| orf19.4189 | 12 | 15 | 16 | 26 | 22 | 15 | 15 | 11 | 11 | 19 | 16 | 21 | 19 | 14 | 13 | 16 | 16,21 | 17 | 16 | 0,92 | -0,12 | -0,342 | 0,7323364 | 0,79091804 | 1,00E-08 | 0,0915226 | 0,0557978 | 0,0557978 | TRUE | 0,3402082 | No |
| orf19.419 | 5817 | 3875 | 4863 | 6041 | 7407 | 6750 | 6900 | 4020 | 5218 | 4933 | 4975 | 4981 | 6335 | 6441 | 6157 | 5670 | 5588,9 | 5027 | 6151 | 1,224 | 0,292 | 6,299 | 3,00E-10 | 2,05E-09 | 0,001628 | 0,0022393 | 0,0018795 | 0,0018795 | TRUE | 0,0326718 | No |
| orf19.4190 | 3217 | 2170 | 2539 | 3233 | 3479 | 2861 | 3195 | 2078 | 2886 | 2763 | 2598 | 2666 | 2975 | 2730 | 2851 | 2931 | 2799,9 | 2728 | 2872 | 1,052 | 0,074 | 1,546 | 0,12215344 | 0,18111991 | 0,001279 | 0,002498 | 0,0018264 | 0,0018264 | TRUE | 0,0190157 | No |
| orf19.4191 | 14071 | 9338 | 11406 | 14431 | 11630 | 10606 | 10870 | 6369 | 12623 | 11888 | 11669 | 11900 | 9946 | 10120 | 9700 | 8983 | 10854 | 12020 | 9687 | 0,806 | -0,311 | -6,786 | 1,15E-11 | 9,34E-11 | 0,001807 | 0,0021133 | 0,0019203 | 0,0019203 | TRUE | 0,0286596 | No |
| orf19.4191.1 | 825 | 742 | 824 | 1019 | 1438 | 1479 | 1591 | 929 | 740 | 945 | 843 | 840 | 1230 | 1411 | 1420 | 1310 | 1092,4 | 842 | 1343 | 1,599 | 0,677 | 8,736 | 2,42E-18 | 3,89E-17 | 0,005887 | 0,0033085 | 0,0047919 | 0,0047919 | TRUE | 0,0793895 | No |
| orf19.4192 | 2895 | 1891 | 2459 | 3110 | 3207 | 2828 | 3000 | 2033 | 2597 | 2407 | 2516 | 2565 | 2743 | 2698 | 2677 | 2867 | 2633,8 | 2521 | 2746 | 1,087 | 0,12 | 2,886 | 0,00390032 | 0,00862781 | 0,000514 | 0,0025307 | 0,0012848 | 0,0012848 | TRUE | 0,0105343 | No |
| orf19.4192.1 | 807 | 580 | 683 | 840 | 864 | 655 | 734 | 434 | 724 | 738 | 699 | 693 | 739 | 625 | 655 | 612 | 685,6 | 714 | 658 | 0,926 | -0,111 | -1,576 | 0,11512509 | 0,1718366 | 0,002643 | 0,0040969 | 0,0033309 | 0,0033309 | TRUE | 0,0841081 | No |
| orf19.4193 | 538 | 360 | 420 | 543 | 552 | 501 | 512 | 371 | 483 | 458 | 430 | 448 | 472 | 478 | 457 | 523 | 468,59 | 455 | 482 | 1,057 | 0,08 | 1,097 | 0,27275893 | 0,35710082 | 0,000515 | 0,0050776 | 0,002935 | 0,002935 | TRUE | 0,0319109 | No |
| orf19.4193.1 | 18132 | 11956 | 14612 | 18505 | 18375 | 14749 | 15131 | 10531 | 16266 | 15221 | 14950 | 15259 | 15715 | 14073 | 13502 | 14853 | 14980 | 15424 | 14536 | 0,942 | -0,086 | -1,644 | 0,10009683 | 0,15292889 | 0,002802 | 0,0020764 | 0,0025433 | 0,0025433 | TRUE | 0,036823 | No |
| orf19.4194 | 525 | 385 | 471 | 634 | 634 | 567 | 610 | 393 | 471 | 490 | 482 | 523 | 542 | 541 | 544 | 554 | 518,46 | 492 | 545 | 1,108 | 0,148 | 2,254 | 0,02421381 | 0,04429859 | 1,00E-08 | 0,0047796 | 0,002197 | 0,002197 | TRUE | 0,0241647 | No |
| orf19.4195 | 236 | 183 | 181 | 269 | 264 | 221 | 241 | 137 | 212 | 233 | 185 | 222 | 226 | 211 | 215 | 193 | 212,08 | 213 | 211 | 0,998 | -0,003 | -0,025 | 0,98043797 | 0,9853582 | 0,001219 | 0,0088236 | 0,0056596 | 0,0056596 | TRUE | 0,0806855 | No |
| orf19.4195.1 | 2771 | 1863 | 2229 | 2883 | 2467 | 2025 | 2161 | 1220 | 2486 | 2372 | 2280 | 2377 | 2110 | 1932 | 1928 | 1721 | 2150,8 | 2379 | 1923 | 0,81 | -0,304 | -4,997 | 5,83E-07 | 2,56E-06 | 0,003421 | 0,0026545 | 0,0030924 | 0,0030924 | TRUE | 0,0570088 | No |
| orf19.4196 | 34 | 29 | 41 | 49 | 46 | 23 | 38 | 26 | 31 | 37 | 42 | 40 | 39 | 22 | 34 | 37 | 35,21 | 38 | 33 | 0,879 | -0,187 | -0,753 | 0,45141531 | 0,53715651 | 1,00E-08 | 0,0432151 | 0,0305281 | 0,0305281 | TRUE | 0,372106 | No |
| orf19.4197 | 2239 | 1411 | 1838 | 2397 | 2265 | 1711 | 1969 | 1269 | 2009 | 1796 | 1880 | 1977 | 1937 | 1633 | 1757 | 1790 | 1847,3 | 1916 | 1779 | 0,928 | -0,108 | -1,762 | 0,07809603 | 0,12386508 | 0,003018 | 0,0027654 | 0,0030426 | 0,0030426 | TRUE | 0,0449688 | No |
| orf19.4199 | 9918 | 7267 | 9020 | 11545 | 10509 | 9793 | 10586 | 7065 | 8897 | 9252 | 9228 | 9520 | 8988 | 9344 | 9446 | 9965 | 9330 | 9224 | 9436 | 1,022 | 0,032 | 0,785 | 0,43254653 | 0,51964398 | 0,001155 | 0,0021351 | 0,0014806 | 0,0014806 | TRUE | 0,0169877 | No |
| orf19.420 | 22 | 15 | 11 | 20 | 32 | 23 | 12 | 13 | 20 | 19 | 11 | 16 | 27 | 22 | 11 | 18 | 18,12 | 16 | 20 | 1,183 | 0,242 | 0,672 | 0,50183423 | 0,58669392 | 0,05297 | 0,0821088 | 0,0697524 | 0,0697524 | TRUE | 0,3406608 | No |
| orf19.4200 | 52 | 29 | 45 | 62 | 22 | 26 | 35 | 36 | 47 | 37 | 46 | 51 | 19 | 25 | 31 | 51 | 38,3 | 45 | 32 | 0,671 | -0,577 | -2,069 | 0,03857375 | 0,06698009 | 0,058237 | 0,0398873 | 0,0471444 | 0,0471444 | TRUE | 0,8960327 | No |
| orf19.4201 | 3054 | 2073 | 2828 | 3360 | 2689 | 2275 | 2637 | 1770 | 2740 | 2639 | 2893 | 2771 | 2300 | 2171 | 2353 | 2496 | 2545,4 | 2761 | 2330 | 0,843 | -0,247 | -4,82 | 1,44E-06 | 5,93E-06 | 0,001865 | 0,0025499 | 0,002127 | 0,002127 | TRUE | 0,0261812 | No |
| orf19.4203 | 2717 | 2253 | 2663 | 3238 | 3341 | 3449 | 3659 | 2388 | 2437 | 2868 | 2725 | 2670 | 2857 | 3291 | 3265 | 3368 | 2935,2 | 2675 | 3195 | 1,195 | 0,256 | 3,931 | 8,47E-05 | 0,00026184 | 0,004365 | 0,0024741 | 0,0037415 | 0,0037415 | TRUE | 0,0624382 | No |
| orf19.4204 | 2869 | 1997 | 2500 | 3058 | 3618 | 3217 | 3350 | 2150 | 2574 | 2542 | 2558 | 2522 | 3094 | 3070 | 2989 | 3032 | 2797,6 | 2549 | 3046 | 1,195 | 0,258 | 7,519 | 5,50E-14 | 5,73E-13 | 1,00E-08 | 0,0024984 | 0,0007654 | 0,0007654 | TRUE | 0,0020971 | No |
| orf19.4205.1 | 806 | 519 | 553 | 763 | 886 | 684 | 759 | 359 | 723 | 661 | 566 | 629 | 758 | 653 | 677 | 506 | 646,6 | 645 | 648 | 1,009 | 0,014 | 0,121 | 0,90375082 | 0,92768408 | 0,017034 | 0,0042247 | 0,0105645 | 0,0105645 | TRUE | 0,2401632 | No |
| orf19.4206 | 540 | 430 | 548 | 715 | 603 | 572 | 623 | 364 | 484 | 547 | 561 | 590 | 516 | 546 | 556 | 513 | 539,12 | 546 | 533 | 0,978 | -0,033 | -0,433 | 0,66515974 | 0,73299334 | 0,002717 | 0,0046723 | 0,0036534 | 0,0036534 | TRUE | 0,0709388 | No |
| orf19.4207 | 237 | 197 | 252 | 325 | 235 | 195 | 220 | 160 | 213 | 251 | 258 | 268 | 201 | 186 | 196 | 226 | 224,78 | 248 | 202 | 0,814 | -0,297 | -2,782 | 0,00540753 | 0,01160302 | 0,004139 | 0,0084378 | 0,0064562 | 0,0064562 | TRUE | 0,1064018 | No |
| orf19.4208 | 1106 | 669 | 978 | 1228 | 1178 | 960 | 1082 | 686 | 992 | 852 | 1001 | 1013 | 1007 | 916 | 966 | 968 | 964,2 | 964 | 964 | 0,997 | -0,004 | -0,056 | 0,95498837 | 0,96792615 | 0,002669 | 0,0034851 | 0,0030499 | 0,0030499 | TRUE | 0,0682319 | No |
| orf19.4209 | 1287 | 952 | 1142 | 1384 | 1391 | 1410 | 1495 | 852 | 1155 | 1212 | 1168 | 1141 | 1190 | 1345 | 1334 | 1202 | 1218,4 | 1169 | 1268 | 1,087 | 0,12 | 2,087 | 0,03692539 | 0,06443443 | 0,001736 | 0,003171 | 0,0023508 | 0,0023508 | TRUE | 0,023003 | No |
| orf19.4210 | 1064 | 745 | 929 | 1108 | 1055 | 974 | 1005 | 649 | 955 | 948 | 950 | 914 | 902 | 929 | 897 | 915 | 926,36 | 942 | 911 | 0,967 | -0,048 | -0,948 | 0,34324798 | 0,43029036 | 1,00E-08 | 0,0035466 | 0,0013525 | 0,0013525 | TRUE | 0,0052799 | No |
| orf19.4211 | 2474 | 1719 | 2224 | 2737 | 1934 | 1835 | 1865 | 1212 | 2219 | 2188 | 2275 | 2257 | 1654 | 1751 | 1664 | 1709 | 1964,9 | 2235 | 1694 | 0,757 | -0,401 | -9,896 | 4,32E-23 | 9,68E-22 | 1,00E-08 | 0,0027184 | 0,0010556 | 0,0010556 | TRUE | 0,0065095 | No |
| orf19.4212 | 421 | 285 | 368 | 418 | 370 | 318 | 344 | 206 | 378 | 363 | 377 | 345 | 316 | 303 | 307 | 291 | 334,88 | 366 | 304 | 0,836 | -0,259 | -3,273 | 0,00106364 | 0,00266486 | 1,00E-08 | 0,0063145 | 0,0029942 | 0,0029942 | TRUE | 0,0190447 | No |
| orf19.4213 | 893 | 648 | 816 | 987 | 975 | 818 | 1000 | 585 | 801 | 825 | 835 | 814 | 834 | 781 | 892 | 825 | 825,83 | 819 | 833 | 1,019 | 0,027 | 0,458 | 0,64693791 | 0,71652497 | 0,000648 | 0,0037374 | 0,0020719 | 0,0020719 | TRUE | 0,0289463 | No |
| orf19.4214 | 49 | 35 | 47 | 51 | 42 | 38 | 44 | 26 | 44 | 45 | 48 | 42 | 36 | 36 | 39 | 37 | 40,85 | 45 | 37 | 0,831 | -0,267 | -1,28 | 0,20046725 | 0,27493771 | 1,00E-08 | 0,0375202 | 0,0170719 | 0,0170719 | TRUE | 0,0212118 | No |
| orf19.4215 | 18524 | 12601 | 16477 | 20247 | 16848 | 14225 | 15404 | 9726 | 16618 | 16042 | 16858 | 16696 | 14409 | 13573 | 13746 | 13718 | 15207 | 16554 | 13862 | 0,837 | -0,256 | -7,913 | 2,52E-15 | 2,98E-14 | 0,000503 | 0,002075 | 0,0009392 | 0,0009392 | TRUE | 0,0087695 | No |
| orf19.4216 | 1490 | 929 | 1082 | 1454 | 2318 | 1924 | 1917 | 1335 | 1337 | 1183 | 1107 | 1199 | 1982 | 1836 | 1711 | 1883 | 1529,6 | 1206 | 1853 | 1,535 | 0,618 | 9,124 | 7,24E-20 | 1,29E-18 | 0,004089 | 0,0029286 | 0,0037162 | 0,0037162 | TRUE | 0,0646573 | No |
| orf19.4217 | 4 | 1 | 8 | 7 | 7 | 3 | 3 | 0 | 4 | 1 | 8 | 6 | 6 | 3 | 3 | 0 | 3,79 | 5 | 3 | 0,64 | -0,645 | -0,817 | 0,41379871 | NA | 0,223161 | 0,384702 | 0,3301704 | 0,3301704 | TRUE | 0,2746687 | No |
| orf19.4219 | 7565 | 4917 | 6586 | 7966 | 6798 | 5688 | 6356 | 3968 | 6787 | 6260 | 6738 | 6569 | 5814 | 5427 | 5672 | 5597 | 6107,9 | 6588 | 5628 | 0,854 | -0,228 | -5,784 | 7,31E-09 | 4,13E-08 | 0,000842 | 0,0022172 | 0,0013235 | 0,0013235 | TRUE | 0,0133305 | No |
| orf19.422 | 756 | 518 | 693 | 868 | 947 | 797 | 899 | 652 | 678 | 659 | 709 | 716 | 810 | 760 | 802 | 920 | 756,83 | 690 | 823 | 1,186 | 0,246 | 3,607 | 0,00030982 | 0,00087514 | 0,002362 | 0,0038977 | 0,0031323 | 0,0031323 | TRUE | 0,0662382 | No |

| | | | | | | | | | | | | | | | | | | | | | | | | | | | | | | | |
|---|---|---|---|---|---|---|---|---|---|---|---|---|---|---|---|---|---|---|---|---|---|---|---|---|---|---|---|---|---|---|---|
| orf19.4220 | 1208 | 870 | 1001 | 1356 | 1520 | 1306 | 1424 | 959 | 1084 | 1108 | 1024 | 1118 | 1300 | 1246 | 1271 | 1353 | 1187,9 | 1084 | 1292 | 1,191 | 0,252 | 4,915 | 8,87E-07 | 3,76E-06 | 0,00041 | 0,0032016 | 0,0016732 | 0,0016732 | TRUE | 0,0160618 | No |
| orf19.4221 | 2352 | 1690 | 2132 | 2641 | 2397 | 2052 | 2278 | 1491 | 2110 | 2152 | 2181 | 2178 | 2050 | 1958 | 2033 | 2103 | 2095,5 | 2155 | 2036 | 0,944 | -0,084 | -2,089 | 0,0367271 | 0,06414431 | 1,00E-08 | 0,0026723 | 0,0010584 | 0,0010584 | TRUE | 0,0078855 | No |
| orf19.4222 | 1805 | 1307 | 1583 | 1939 | 2136 | 2065 | 2138 | 1438 | 1619 | 1664 | 1620 | 1599 | 1827 | 1970 | 1908 | 2028 | 1779,4 | 1626 | 1933 | 1,188 | 0,249 | 5,394 | 6,88E-08 | 3,45E-07 | 0,000609 | 0,0027954 | 0,0014722 | 0,0014722 | TRUE | 0,0168435 | No |
| orf19.4223 | 12543 | 8033 | 11271 | 13538 | 13098 | 10504 | 11788 | 7254 | 11252 | 10227 | 11531 | 11164 | 11202 | 10023 | 10519 | 10231 | 10769 | 11044 | 10494 | 0,95 | -0,074 | -1,473 | 0,14084179 | 0,20453891 | 0,002392 | 0,0021143 | 0,0023154 | 0,0023154 | TRUE | 0,0299512 | No |
| orf19.4225 | 5168 | 3229 | 4505 | 5374 | 5405 | 4861 | 5354 | 3318 | 4636 | 4111 | 4609 | 4431 | 4623 | 4638 | 4778 | 4680 | 4563,2 | 4447 | 4680 | 1,052 | 0,073 | 1,634 | 0,10224224 | 0,15551174 | 0,001296 | 0,0022977 | 0,0016738 | 0,0016738 | TRUE | 0,0303915 | No |
| orf19.4225.1 | 3671 | 2459 | 3053 | 3937 | 4465 | 3724 | 4043 | 2785 | 3293 | 3131 | 3124 | 3246 | 3819 | 3553 | 3608 | 3928 | 3462,7 | 3198 | 3727 | 1,164 | 0,219 | 4,925 | 8,43E-07 | 3,60E-06 | 0,001081 | 0,0023988 | 0,0016014 | 0,0016014 | TRUE | 0,015132 | No |
| orf19.4227 | 98 | 68 | 98 | 114 | 94 | 76 | 98 | 67 | 88 | 87 | 100 | 94 | 80 | 73 | 87 | 94 | 87,95 | 92 | 84 | 0,9 | -0,151 | -1,013 | 0,31120244 | 0,39747115 | 1,00E-08 | 0,0184852 | 0,0100344 | 0,0100344 | TRUE | 0,0737588 | No |
| orf19.4228 | 1368 | 966 | 1143 | 1410 | 1165 | 982 | 936 | 665 | 1227 | 1230 | 1169 | 1163 | 996 | 937 | 835 | 938 | 1062 | 1197 | 926 | 0,774 | -0,369 | -5,97 | 2,38E-09 | 1,43E-08 | 0,002207 | 0,0033465 | 0,0027132 | 0,0027132 | TRUE | 0,0557506 | No |
| orf19.4229 | 1042 | 663 | 863 | 1107 | 850 | 731 | 812 | 495 | 935 | 844 | 883 | 913 | 727 | 697 | 725 | 698 | 802,73 | 894 | 712 | 0,796 | -0,33 | -5,806 | 6,38E-09 | 3,64E-08 | 1,00E-08 | 0,003788 | 0,0018349 | 0,0018349 | TRUE | 0,0150796 | No |
| orf19.423 | 1229 | 885 | 1046 | 1253 | 1197 | 990 | 1053 | 873 | 1103 | 1127 | 1070 | 1033 | 1024 | 945 | 940 | 1231 | 1059 | 1083 | 1035 | 0,953 | -0,07 | -0,847 | 0,39725896 | 0,48599711 | 0,007388 | 0,0033504 | 0,0055765 | 0,0055765 | TRUE | 0,1840866 | No |
| orf19.4230 | 6169 | 4136 | 5199 | 6520 | 6338 | 5515 | 5827 | 3761 | 5534 | 5266 | 5319 | 5376 | 5420 | 5262 | 5200 | 5305 | 5335,3 | 5374 | 5297 | 0,985 | -0,021 | -0,661 | 0,50884925 | 0,59304875 | 0,00022 | 0,0022516 | 0,0008187 | 0,0008187 | TRUE | 0,0049486 | No |
| orf19.4231 | 761 | 405 | 601 | 663 | 608 | 540 | 599 | 420 | 683 | 516 | 615 | 547 | 520 | 515 | 535 | 592 | 565,25 | 590 | 540 | 0,912 | -0,133 | -1,441 | 0,14954008 | 0,21519444 | 0,007754 | 0,0045478 | 0,006349 | 0,006349 | TRUE | 0,1334105 | No |
| orf19.4232 | 1890 | 1340 | 1756 | 2038 | 2001 | 1896 | 1994 | 1188 | 1696 | 1706 | 1797 | 1681 | 1711 | 1809 | 1779 | 1676 | 1731,7 | 1720 | 1744 | 1,016 | 0,022 | 0,488 | 0,62533596 | 0,69768216 | 0,000457 | 0,0028178 | 0,0014154 | 0,0014154 | TRUE | 0,0109814 | No |
| orf19.4233 | 4244 | 2809 | 3707 | 4461 | 4160 | 3747 | 3954 | 2433 | 3807 | 3576 | 3793 | 3679 | 3558 | 3575 | 3528 | 3432 | 3618,4 | 3714 | 3523 | 0,949 | -0,076 | -2,09 | 0,03665062 | 0,06404797 | 0,000222 | 0,0023807 | 0,0009841 | 0,0009841 | TRUE | 0,0070674 | No |
| orf19.4234 | 4983 | 3694 | 4447 | 5702 | 5399 | 4839 | 5320 | 3373 | 4470 | 4703 | 4550 | 4702 | 4617 | 4617 | 4747 | 4757 | 4645,5 | 4606 | 4684 | 1,017 | 0,024 | 0,716 | 0,47381186 | 0,5597054 | 0,00025 | 0,002292 | 0,000881 | 0,000881 | TRUE | 0,0049089 | No |
| orf19.4235 | 1900 | 1393 | 1652 | 2240 | 2049 | 1933 | 2172 | 1449 | 1704 | 1773 | 1690 | 1847 | 1752 | 1844 | 1938 | 2044 | 1824,2 | 1754 | 1894 | 1,078 | 0,109 | 1,902 | 0,05718609 | 0,09438847 | 0,002288 | 0,0027753 | 0,0025832 | 0,0025832 | TRUE | 0,0318234 | No |
| orf19.4236 | 1183 | 837 | 1090 | 1326 | 1352 | 1245 | 1246 | 771 | 1061 | 1066 | 1115 | 1093 | 1156 | 1188 | 1112 | 1087 | 1109,9 | 1084 | 1136 | 1,049 | 0,07 | 1,379 | 0,16804689 | 0,2377268 | 1,00E-08 | 0,0032875 | 0,0015426 | 0,0015426 | TRUE | 0,010986 | No |
| orf19.4239 | 23 | 12 | 16 | 14 | 11 | 16 | 19 | 15 | 21 | 15 | 16 | 12 | 9 | 15 | 17 | 21 | 15,83 | 16 | 16 | 0,964 | -0,053 | -0,146 | 0,88396292 | 0,91220357 | 1,00E-08 | 0,0937073 | 0,064746 | 0,064746 | TRUE | 0,3980451 | No |
| orf19.424 | 1814 | 1103 | 1411 | 1769 | 1760 | 1415 | 1473 | 1008 | 1627 | 1404 | 1444 | 1459 | 1505 | 1350 | 1314 | 1422 | 1440,7 | 1484 | 1398 | 0,941 | -0,087 | -1,371 | 0,17030247 | 0,24040819 | 0,003103 | 0,0029872 | 0,0031966 | 0,0031966 | TRUE | 0,0519224 | No |
| orf19.4240 | 551 | 380 | 455 | 578 | 709 | 622 | 663 | 453 | 494 | 484 | 466 | 477 | 606 | 593 | 592 | 639 | 543,83 | 480 | 608 | 1,262 | 0,336 | 5,276 | 1,32E-07 | 6,37E-07 | 1,00E-08 | 0,0046489 | 0,0020382 | 0,0020382 | TRUE | 0,0118369 | No |
| orf19.4241 | 618 | 446 | 571 | 707 | 780 | 707 | 744 | 545 | 554 | 568 | 584 | 583 | 667 | 675 | 664 | 769 | 632,96 | 572 | 694 | 1,206 | 0,27 | 4,068 | 4,73E-05 | 0,00015389 | 0,000976 | 0,004273 | 0,0026435 | 0,0026435 | TRUE | 0,0549066 | No |
| orf19.4242 | 1915 | 1345 | 1859 | 1999 | 1901 | 1708 | 1824 | 1217 | 1718 | 1712 | 1902 | 1648 | 1626 | 1630 | 1628 | 1716 | 1697,5 | 1745 | 1650 | 0,945 | -0,082 | -1,529 | 0,12630117 | 0,18639583 | 0,001533 | 0,0028347 | 0,0021589 | 0,0021589 | TRUE | 0,0443148 | No |
| orf19.4243 | 168 | 108 | 131 | 154 | 215 | 198 | 181 | 105 | 151 | 137 | 134 | 127 | 184 | 189 | 162 | 148 | 153,95 | 137 | 171 | 1,251 | 0,323 | 2,655 | 0,00792602 | 0,01636973 | 0,002722 | 0,011409 | 0,0076504 | 0,0076504 | TRUE | 0,067258 | No |
| orf19.4244 | 64 | 56 | 59 | 79 | 83 | 113 | 115 | 58 | 57 | 71 | 60 | 65 | 71 | 108 | 103 | 82 | 77,18 | 63 | 91 | 1,438 | 0,524 | 2,918 | 0,00351911 | 0,0078423 | 0,013804 | 0,0207887 | 0,0176072 | 0,0176072 | TRUE | 0,2325733 | No |
| orf19.4245 | 1006 | 748 | 931 | 1151 | 1241 | 1045 | 1139 | 769 | 902 | 952 | 953 | 949 | 1061 | 997 | 1016 | 1085 | 989,48 | 939 | 1040 | 1,106 | 0,146 | 2,775 | 0,0055285 | 0,01183899 | 1,00E-08 | 0,0034467 | 0,0016316 | 0,0016316 | TRUE | 0,0088364 | No |
| orf19.4246 | 13399 | 9192 | 11974 | 14529 | 18310 | 15829 | 17288 | 10706 | 12020 | 11702 | 12251 | 11981 | 15659 | 15104 | 15427 | 15100 | 13655 | 11988 | 15322 | 1,278 | 0,354 | 12,384 | 3,17E-35 | 1,62E-33 | 0,000234 | 0,0020858 | 0,0007101 | 0,0007101 | TRUE | 0,0030514 | No |
| orf19.4247 | 551 | 396 | 476 | 652 | 726 | 585 | 654 | 382 | 494 | 504 | 487 | 538 | 621 | 558 | 584 | 539 | 540,57 | 506 | 576 | 1,14 | 0,189 | 2,685 | 0,0072572 | 0,01508173 | 0,001208 | 0,004665 | 0,0029094 | 0,0029094 | TRUE | 0,0338925 | No |
| orf19.4248 | 1368 | 845 | 1068 | 1256 | 1263 | 1090 | 1164 | 714 | 1227 | 1076 | 1093 | 1036 | 1080 | 1040 | 1039 | 1007 | 1074,7 | 1108 | 1042 | 0,941 | -0,088 | -1,411 | 0,15814791 | 0,22542174 | 0,002422 | 0,0033304 | 0,0028109 | 0,0028109 | TRUE | 0,0653322 | No |
| orf19.425 | 362 | 256 | 280 | 342 | 326 | 274 | 313 | 201 | 325 | 326 | 286 | 282 | 279 | 261 | 279 | 283 | 290,27 | 305 | 276 | 0,906 | -0,142 | -1,607 | 0,1081564 | 0,16298061 | 1,00E-08 | 0,0069807 | 0,0040763 | 0,0040763 | TRUE | 0,0311219 | No |
| orf19.4250 | 121 | 87 | 109 | 146 | 156 | 121 | 113 | 89 | 109 | 111 | 112 | 120 | 133 | 115 | 101 | 126 | 115,81 | 113 | 119 | 1,049 | 0,069 | 0,512 | 0,60874645 | 0,68361658 | 1,00E-08 | 0,0145151 | 0,0088734 | 0,0088734 | TRUE | 0,1148515 | No |
| orf19.4251 | 303 | 236 | 303 | 345 | 193 | 189 | 198 | 149 | 272 | 300 | 310 | 284 | 165 | 180 | 177 | 210 | 237,37 | 292 | 183 | 0,623 | -0,682 | -6,788 | 1,14E-11 | 9,27E-11 | 0,001255 | 0,0080952 | 0,0052305 | 0,0052305 | TRUE | 0,0865694 | No |
| orf19.4252 | 2328 | 1532 | 1951 | 2541 | 3116 | 2755 | 2905 | 1929 | 2088 | 1950 | 1996 | 2095 | 2665 | 2629 | 2592 | 2721 | 2342,1 | 2032 | 2652 | 1,302 | 0,381 | 9,257 | 2,10E-20 | 3,93E-19 | 0,000245 | 0,0025994 | 0,0011869 | 0,0011869 | TRUE | 0,0084036 | No |
| orf19.4253 | 580 | 461 | 520 | 676 | 524 | 555 | 555 | 307 | 520 | 587 | 532 | 557 | 448 | 530 | 495 | 433 | 512,83 | 549 | 476 | 0,871 | -0,199 | -2,501 | 0,01238848 | 0,02433943 | 0,003139 | 0,0048103 | 0,0040902 | 0,0040902 | TRUE | 0,0640974 | No |
| orf19.4255 | 347 | 211 | 270 | 378 | 434 | 339 | 381 | 292 | 311 | 269 | 276 | 312 | 371 | 323 | 340 | 412 | 326,79 | 292 | 362 | 1,228 | 0,297 | 3,067 | 0,00216441 | 0,00507563 | 0,005269 | 0,0064218 | 0,0058794 | 0,0058794 | TRUE | 0,0867752 | No |
| orf19.4257 | 3008 | 2128 | 2735 | 3303 | 3042 | 2815 | 3159 | 1848 | 2698 | 2709 | 2798 | 2724 | 2602 | 2686 | 2819 | 2606 | 2705,3 | 2732 | 2678 | 0,981 | -0,028 | -0,685 | 0,49356519 | 0,57860286 | 0,000501 | 0,0025161 | 0,0012196 | 0,0012196 | TRUE | 0,0155322 | No |
| orf19.4258 | 3012 | 1910 | 2591 | 3108 | 3397 | 2832 | 3266 | 2035 | 2702 | 2432 | 2651 | 2563 | 2905 | 2702 | 2914 | 2870 | 2717,4 | 2587 | 2848 | 1,1 | 0,138 | 2,939 | 0,00329412 | 0,00740961 | 0,00116 | 0,0025137 | 0,0017334 | 0,0017334 | TRUE | 0,0187121 | No |
| orf19.426 | 1066 | 766 | 884 | 1089 | 1107 | 1018 | 1119 | 666 | 956 | 975 | 904 | 898 | 947 | 971 | 999 | 939 | 948,73 | 933 | 964 | 1,035 | 0,05 | 0,93 | 0,35256842 | 0,44020389 | 1,00E-08 | 0,0035097 | 0,0017129 | 0,0017129 | TRUE | 0,0097019 | No |
| orf19.4261 | 3996 | 3095 | 3570 | 4409 | 4059 | 4250 | 4304 | 2625 | 3585 | 3940 | 3652 | 3636 | 3471 | 4055 | 3841 | 3702 | 3735,4 | 3703 | 3767 | 1,018 | 0,026 | 0,465 | 0,64158791 | 0,71158578 | 0,002447 | 0,0023682 | 0,0026261 | 0,0026261 | TRUE | 0,0350173 | No |
| orf19.4262 | 907 | 619 | 841 | 961 | 986 | 840 | 953 | 684 | 814 | 788 | 860 | 792 | 843 | 802 | 850 | 965 | 839,31 | 814 | 865 | 1,059 | 0,083 | 1,247 | 0,212454 | 0,28972602 | 0,002505 | 0,0037092 | 0,003045 | 0,003045 | TRUE | 0,0642815 | No |
| orf19.4263 | 609 | 483 | 606 | 729 | 592 | 610 | 724 | 418 | 546 | 615 | 620 | 601 | 506 | 582 | 646 | 590 | 588,29 | 596 | 581 | 0,976 | -0,035 | -0,428 | 0,66834477 | 0,73502021 | 0,005273 | 0,0044472 | 0,004884 | 0,004884 | TRUE | 0,0941076 | No |
| orf19.4264 | 450 | 392 | 413 | 562 | 473 | 491 | 495 | 322 | 404 | 499 | 423 | 463 | 405 | 468 | 442 | 454 | 444,7 | 447 | 442 | 0,99 | -0,015 | -0,174 | 0,86204407 | 0,89542891 | 0,004175 | 0,005244 | 0,0047126 | 0,0047126 | TRUE | 0,0636675 | No |
| orf19.4265 | 3981 | 3482 | 3834 | 4727 | 4080 | 4514 | 4450 | 2716 | 3571 | 4433 | 3923 | 3898 | 3489 | 4307 | 3971 | 3831 | 3927,9 | 3956 | 3900 | 0,986 | -0,02 | -0,266 | 0,79053821 | 0,83677829 | 0,006506 | 0,0023491 | 0,0052474 | 0,0052474 | TRUE | 0,0791654 | No |
| orf19.4266 | 790 | 717 | 764 | 967 | 860 | 943 | 896 | 592 | 709 | 913 | 782 | 797 | 735 | 900 | 800 | 835 | 808,79 | 800 | 818 | 1,023 | 0,033 | 0,38 | 0,70430943 | 0,76574214 | 0,007723 | 0,0037744 | 0,0058632 | 0,0058632 | TRUE | 0,1014882 | No |
| orf19.4268 | 13072 | 9745 | 11715 | 14234 | 12445 | 11899 | 12805 | 6819 | 11727 | 12406 | 11986 | 11738 | 10643 | 11354 | 11426 | 9618 | 11362 | 11964 | 10760 | 0,9 | -0,152 | -2,739 | 0,0061566 | 0,01304261 | 0,003281 | 0,0021073 | 0,0028797 | 0,0028797 | TRUE | 0,0616279 | No |
| orf19.4269 | 1482 | 1169 | 1515 | 1719 | 2088 | 2070 | 2149 | 1339 | 1330 | 1488 | 1550 | 1418 | 1786 | 1975 | 1918 | 1889 | 1669 | 1446 | 1892 | 1,309 | 0,389 | 6,699 | 2,10E-11 | 1,65E-10 | 0,002215 | 0,0028493 | 0,0026196 | 0,0026196 | TRUE | 0,0360586 | No |
| orf19.427 | 2438 | 1803 | 2221 | 2590 | 2706 | 2368 | 2671 | 1513 | 2187 | 2295 | 2272 | 2136 | 2314 | 2259 | 2383 | 2134 | 2247,7 | 2222 | 2272 | 1,025 | 0,036 | 0,756 | 0,44987167 | 0,53606287 | 0,001044 | 0,0026254 | 0,0017085 | 0,0017085 | TRUE | 0,0188905 | No |
| orf19.4270 | 820 | 722 | 836 | 995 | 1055 | 1112 | 1134 | 710 | 736 | 919 | 855 | 820 | 902 | 1061 | 1012 | 1001 | 913,4 | 832 | 994 | 1,196 | 0,258 | 3,353 | 0,000798 | 0,0020515 | 0,005446 | 0,0035689 | 0,0045819 | 0,0045819 | TRUE | 0,0735997 | No |
| orf19.4271 | 1624 | 1274 | 1398 | 1738 | 1913 | 1795 | 1869 | 864 | 1457 | 1622 | 1430 | 1433 | 1636 | 1713 | 1668 | 1219 | 1522,2 | 1486 | 1559 | 1,052 | 0,073 | 0,771 | 0,44046737 | 0,52715696 | 0,012753 | 0,0029332 | 0,0079759 | 0,0079759 | TRUE | 0,2504673 | No |
| orf19.4273 | 3509 | 3268 | 3347 | 4290 | 5223 | 6084 | 5950 | 3470 | 3148 | 4161 | 3424 | 3538 | 4467 | 5805 | 5309 | 4894 | 4343,2 | 3568 | 5119 | 1,436 | 0,522 | 5,669 | 1,44E-08 | 7,82E-08 | 0,013125 | 0,0023138 | 0,0078973 | 0,0078973 | TRUE | 0,1508285 | No |
| orf19.4274 | 32874 | 28973 | 31924 | 40483 | 36042 | 40494 | 41408 | 26188 | 29491 | 36886 | 32661 | 33383 | 30824 | 38638 | 36950 | 36936 | 34471 | 33105 | 35837 | 1,083 | 0,114 | 1,455 | 0,14576111 | 0,21046209 | 0,008759 | 0,0020216 | 0,0059152 | 0,0059152 | TRUE | 0,1087374 | No |
| orf19.4275 | 657 | 558 | 653 | 779 | 835 | 776 | 851 | 496 | 589 | 710 | 668 | 642 | 714 | 740 | 759 | 700 | 690,47 | 652 | 728 | 1,121 | 0,165 | 2,41 | 0,01593334 | 0,03045736 | 0,002028 | 0,004082 | 0,0030336 | 0,0030336 | TRUE | 0,0502728 | No |
| orf19.4278 | 4654 | 3202 | 3809 | 4917 | 4571 | 4079 | 4295 | 2659 | 4175 | 4076 | 3897 | 4055 | 3909 | 3892 | 3833 | 3750 | 3948,4 | 4051 | 3846 | 0,95 | -0,074 | -2,062 | 0,03917277 | 0,0678401 | 0,000321 | 0,0023472 | 0,0009854 | 0,0009854 | TRUE | 0,0079491 | No |
| orf19.4279 | 21173 | 15435 | 19112 | 23224 | 21197 | 19906 | 21241 | 12835 | 18994 | 19650 | 19553 | 19151 | 18128 | 18994 | 18954 | 18103 | 18941 | 19337 | 18545 | 0,959 | -0,06 | -1,957 | 0,0503517 | 0,08467798 | 0,00042 | 0,0020562 | 0,0008497 | 0,0008497 | TRUE | 0,0032353 | No |
| orf19.428 | 1172 | 809 | 981 | 1292 | 1758 | 1659 | 1708 | 1038 | 1051 | 1030 | 1004 | 1065 | 1503 | 1583 | 1524 | 1464 | 1278,1 | 1038 | 1518 | 1,464 | 0,55 | 11,423 | 3,20E-30 | 1,16E-28 | 1,00E-08 | 0,0031153 | 0,0014146 | 0,0014146 | TRUE | 0,0095831 | No |
| orf19.4280 | 8266 | 5808 | 7176 | 8925 | 8023 | 7385 | 7803 | 4774 | 7415 | 7394 | 7342 | 7360 | 6862 | 7047 | 6963 | 6733 | 7139,4 | 7378 | 6901 | 0,936 | -0,096 | -3,391 | 0,00069626 | 0,00181087 | 1,46E-05 | 0,0021829 | 0,0006228 | 0,0006228 | TRUE | 0,0030237 | No |
| orf19.4282 | 4569 | 3174 | 4182 | 5061 | 4752 | 4237 | 4647 | 2752 | 4099 | 4041 | 4279 | 4173 | 4064 | 4043 | 4147 | 3881 | 4090,8 | 4148 | 4034 | 0,973 | -0,039 | -1,078 | 0,28097916 | 0,36492062 | 0,000359 | 0,0023344 | 0,0010342 | 0,0010342 | TRUE | 0,007296 | No |
| orf19.4283 | 5493 | 3924 | 4731 | 5828 | 5800 | 5222 | 5522 | 3320 | 4928 | 4996 | 4840 | 4806 | 4960 | 4983 | 4927 | 4683 | 4890,3 | 4892 | 4888 | 1 | 0,001 | 0,017 | 0,98630171 | 0,98959597 | 0,000271 | 0,0022764 | 0,0009278 | 0,0009278 | TRUE | 0,0091026 | No |
| orf19.4284 | 555 | 446 | 524 | 619 | 545 | 515 | 559 | 303 | 498 | 568 | 536 | 510 | 466 | 491 | 499 | 427 | 499,49 | 528 | 471 | 0,898 | -0,155 | -2,091 | 0,03651244 | 0,06384363 | 0,001378 | 0,0048859 | 0,0032983 | 0,0032983 | TRUE | 0,0377894 | No |
| orf19.4286 | 4127 | 3023 | 3641 | 4683 | 4476 | 4169 | 4432 | 2616 | 3702 | 3849 | 3725 | 3862 | 3828 | 3978 | 3955 | 3690 | 3823,5 | 3784 | 3863 | 1,022 | 0,031 | 0,808 | 0,41888368 | 0,50647075 | 0,000479 | 0,0023592 | 0,0011482 | 0,0011482 | TRUE | 0,0102437 | No |

| | | | | | | | | | | | | | | | | | | | | | | | | | | | | | | | |
|---|---|---|---|---|---|---|---|---|---|---|---|---|---|---|---|---|---|---|---|---|---|---|---|---|---|---|---|---|---|---|---|
| orf19.4287 | 1147 | 800 | 923 | 1217 | 808 | 629 | 729 | 432 | 1029 | 1018 | 944 | 1004 | 691 | 600 | 651 | 609 | 818,29 | 999 | 638 | 0,64 | -0,643 | -10,186 | 2,30E-24 | 5,57E-23 | 0,001426 | 0,0037536 | 0,0025451 | 0,0025451 | TRUE | 0,0387407 | No |
| orf19.4288 | 2967 | 1976 | 2559 | 3292 | 2319 | 2049 | 2223 | 1550 | 2662 | 2516 | 2618 | 2715 | 1983 | 1955 | 1984 | 2186 | 2327,3 | 2628 | 2027 | 0,769 | -0,378 | -7,885 | 3,15E-15 | 3,69E-14 | 0,001324 | 0,0026033 | 0,0017718 | 0,0017718 | TRUE | 0,0310842 | No |
| orf19.429 | 3324 | 2160 | 2883 | 3563 | 3300 | 3023 | 3207 | 1923 | 2982 | 2750 | 2950 | 2938 | 2822 | 2884 | 2862 | 2712 | 2862,5 | 2905 | 2820 | 0,971 | -0,043 | -1,037 | 0,29987151 | 0,38545744 | 0,000578 | 0,0024867 | 0,0012822 | 0,0012822 | TRUE | 0,0147498 | No |
| orf19.4290 | 5566 | 3579 | 4599 | 5723 | 4759 | 3993 | 4653 | 2749 | 4993 | 4556 | 4705 | 4719 | 4070 | 3810 | 4152 | 3877 | 4360,4 | 4743 | 3977 | 0,839 | -0,254 | -5,72 | 1,06E-08 | 5,86E-08 | 0,001241 | 0,0023124 | 0,0016611 | 0,0016611 | TRUE | 0,0153642 | No |
| orf19.4292 | 1434 | 1011 | 1193 | 1499 | 1419 | 1210 | 1231 | 832 | 1286 | 1287 | 1221 | 1236 | 1214 | 1155 | 1098 | 1173 | 1208,8 | 1258 | 1160 | 0,923 | -0,116 | -2,286 | 0,02225838 | 0,04110924 | 0,000501 | 0,0031805 | 0,0016502 | 0,0016502 | TRUE | 0,0163003 | No |
| orf19.4293 | 2953 | 2038 | 2540 | 3191 | 3879 | 3097 | 3442 | 2222 | 2649 | 2595 | 2599 | 2631 | 3317 | 2955 | 3071 | 3134 | 2869 | 2618 | 3119 | 1,191 | 0,252 | 5,73 | 1,00E-08 | 5,53E-08 | 0,000988 | 0,0024855 | 0,0015112 | 0,0015112 | TRUE | 0,0232178 | No |
| orf19.4294 | 2356 | 1858 | 2140 | 2601 | 2734 | 2541 | 2822 | 1642 | 2114 | 2365 | 2189 | 2145 | 2338 | 2425 | 2518 | 2316 | 2301,3 | 2203 | 2399 | 1,091 | 0,126 | 2,533 | 0,0112952 | 0,0224447 | 0,001459 | 0,0026104 | 0,0019299 | 0,0019299 | TRUE | 0,028341 | No |
| orf19.4295 | 1106 | 866 | 1009 | 1215 | 1366 | 1213 | 1382 | 778 | 992 | 1103 | 1032 | 1002 | 1168 | 1157 | 1233 | 1097 | 1098,1 | 1032 | 1164 | 1,132 | 0,179 | 3,172 | 0,00151255 | 0,00366281 | 0,001092 | 0,0033015 | 0,0021499 | 0,0021499 | TRUE | 0,0232695 | No |
| orf19.4297 | 2585 | 1920 | 2322 | 2737 | 2319 | 2366 | 2443 | 1542 | 2319 | 2444 | 2376 | 2257 | 1983 | 2258 | 2180 | 2175 | 2249 | 2349 | 2149 | 0,915 | -0,128 | -2,509 | 0,01209634 | 0,0238591 | 0,001693 | 0,002625 | 0,0020521 | 0,0020521 | TRUE | 0,0339474 | No |
| orf19.4299 | 1484 | 972 | 1220 | 1606 | 1788 | 1584 | 1745 | 1145 | 1331 | 1237 | 1248 | 1324 | 1529 | 1511 | 1557 | 1615 | 1419,2 | 1285 | 1553 | 1,205 | 0,269 | 5,609 | 2,04E-08 | 1,08E-07 | 0,000328 | 0,0030024 | 0,0014925 | 0,0014925 | TRUE | 0,0076762 | No |
| orf19.430 | 1348 | 944 | 1215 | 1495 | 2152 | 1795 | 1993 | 1257 | 1209 | 1202 | 1243 | 1233 | 1840 | 1713 | 1778 | 1773 | 1498,9 | 1222 | 1776 | 1,454 | 0,54 | 12,087 | 1,24E-33 | 5,65E-32 | 1,00E-08 | 0,002948 | 0,0012244 | 0,0012244 | TRUE | 0,0076853 | No |
| orf19.4301 | 337 | 216 | 305 | 360 | 435 | 367 | 414 | 248 | 302 | 275 | 312 | 297 | 372 | 350 | 369 | 350 | 328,45 | 296 | 360 | 1,215 | 0,28 | 3,515 | 0,00043915 | 0,00119983 | 1,00E-08 | 0,0063993 | 0,0030584 | 0,0030584 | TRUE | 0,0225383 | No |
| orf19.4304 | 29382 | 21013 | 26419 | 32537 | 36040 | 32669 | 34838 | 21350 | 26359 | 26752 | 27029 | 26830 | 30823 | 31172 | 31087 | 30112 | 28770 | 26742 | 30798 | 1,152 | 0,204 | 8,355 | 6,56E-17 | 9,21E-16 | 0,000131 | 0,00203 | 0,0005373 | 0,0005373 | TRUE | 0,0026926 | No |
| orf19.4305 | 133 | 104 | 114 | 148 | 154 | 144 | 149 | 94 | 119 | 132 | 117 | 122 | 132 | 137 | 133 | 133 | 128,13 | 122 | 134 | 1,094 | 0,129 | 1,079 | 0,28058566 | 0,36464603 | 1,00E-08 | 0,0133095 | 0,0060041 | 0,0060041 | TRUE | 0,0239465 | No |
| orf19.4305.1 | 1217 | 831 | 1015 | 1260 | 1232 | 1018 | 1050 | 615 | 1092 | 1058 | 1038 | 1039 | 1054 | 971 | 937 | 867 | 1007,1 | 1057 | 957 | 0,909 | -0,138 | -2,187 | 0,02874236 | 0,05161495 | 0,002222 | 0,003421 | 0,0028229 | 0,0028229 | TRUE | 0,0550979 | No |
| orf19.4306 | 1498 | 1034 | 1264 | 1594 | 1329 | 1154 | 1247 | 846 | 1344 | 1316 | 1293 | 1314 | 1137 | 1101 | 1113 | 1193 | 1226,4 | 1317 | 1136 | 0,86 | -0,217 | -4,591 | 4,42E-06 | 1,70E-05 | 1,00E-08 | 0,0031632 | 0,00132 | 0,00132 | TRUE | 0,0117954 | No |
| orf19.4307 | 3396 | 2337 | 2740 | 3564 | 3301 | 2955 | 2888 | 1883 | 3047 | 2975 | 2803 | 2939 | 2823 | 2820 | 2577 | 2656 | 2830 | 2941 | 2719 | 0,925 | -0,113 | -2,412 | 0,01587767 | 0,03037996 | 0,001148 | 0,0024925 | 0,0017419 | 0,0017419 | TRUE | 0,0158295 | No |
| orf19.4308 | 246 | 211 | 232 | 287 | 397 | 353 | 373 | 261 | 221 | 269 | 237 | 237 | 340 | 337 | 333 | 368 | 292,58 | 241 | 344 | 1,432 | 0,518 | 5,861 | 4,61E-09 | 2,70E-08 | 1,00E-08 | 0,0069412 | 0,0039785 | 0,0039785 | TRUE | 0,0577616 | No |
| orf19.4309 | 3110 | 2228 | 2572 | 3169 | 3683 | 3306 | 3350 | 2226 | 2790 | 2836 | 2631 | 2613 | 3150 | 3154 | 2989 | 3140 | 2913 | 2718 | 3108 | 1,144 | 0,194 | 4,528 | 5,95E-06 | 2,24E-05 | 0,000719 | 0,0024779 | 0,0014238 | 0,0014238 | TRUE | 0,0099238 | No |
| orf19.431 | 5751 | 3798 | 5277 | 6352 | 7604 | 6593 | 7260 | 4616 | 5159 | 4835 | 5399 | 5238 | 6503 | 6291 | 6478 | 6510 | 5801,8 | 5158 | 6446 | 1,249 | 0,321 | 7,977 | 1,50E-15 | 1,83E-14 | 0,000954 | 0,0022297 | 0,0013764 | 0,0013764 | TRUE | 0,0207961 | No |
| orf19.4310 | 0 | 0 | 1 | 3 | 1 | 0 | 1 | 1 | 0 | 0 | 1 | 2 | 1 | 0 | 1 | 1 | 0,83 | 1 | 1 | 0,823 | -0,282 | -0,179 | 0,85829379 | NA | 1,00E-08 | 1,7470978 | 1,194914 | 1,194914 | TRUE | 0,7725611 | No |
| orf19.4311 | 2635 | 1916 | 2118 | 2726 | 2458 | 2068 | 2212 | 1327 | 2364 | 2439 | 2167 | 2248 | 2102 | 1973 | 1974 | 1872 | 2142,4 | 2304 | 1980 | 0,861 | -0,216 | -4,058 | 4,94E-05 | 0,00016017 | 0,001886 | 0,0026571 | 0,0022459 | 0,0022459 | TRUE | 0,0209237 | No |
| orf19.4312 | 839 | 571 | 732 | 836 | 913 | 771 | 847 | 487 | 753 | 727 | 749 | 689 | 781 | 736 | 756 | 687 | 734,63 | 730 | 740 | 1,018 | 0,026 | 0,417 | 0,67647907 | 0,74125228 | 0,000844 | 0,0039556 | 0,0023445 | 0,0023445 | TRUE | 0,0237336 | No |
| orf19.4315 | 1875 | 1637 | 1837 | 2259 | 2033 | 2201 | 2308 | 1282 | 1682 | 2084 | 1879 | 1863 | 1739 | 2100 | 2060 | 1808 | 1901,9 | 1877 | 1927 | 1,028 | 0,04 | 0,501 | 0,6162053 | 0,68954394 | 0,007669 | 0,0027428 | 0,0054599 | 0,0054599 | TRUE | 0,0650459 | No |
| orf19.4316 | 945 | 705 | 822 | 1020 | 700 | 694 | 722 | 461 | 848 | 898 | 841 | 841 | 599 | 662 | 644 | 650 | 747,84 | 857 | 639 | 0,746 | -0,424 | -7,184 | 6,78E-13 | 6,35E-12 | 1,00E-08 | 0,0039207 | 0,0019698 | 0,0019698 | TRUE | 0,0228933 | No |
| orf19.4317 | 1412 | 1225 | 1258 | 1568 | 2151 | 2085 | 2080 | 1282 | 1267 | 1560 | 1287 | 1293 | 1840 | 1989 | 1856 | 1808 | 1612,5 | 1352 | 1873 | 1,389 | 0,475 | 6,825 | 8,81E-12 | 7,25E-11 | 0,005358 | 0,0028798 | 0,0040017 | 0,0040017 | TRUE | 0,1237587 | No |
| orf19.4318 | 11084 | 8276 | 10382 | 12870 | 17510 | 16362 | 18446 | 11314 | 9944 | 10536 | 10622 | 10613 | 14975 | 15612 | 16460 | 15957 | 13090 | 10429 | 15751 | 1,51 | 0,595 | 14,69 | 7,47E-49 | 7,12E-47 | 0,001165 | 0,0020904 | 0,0014952 | 0,0014952 | TRUE | 0,0135823 | No |
| orf19.432 | 3012 | 2051 | 2696 | 3177 | 2978 | 2591 | 2827 | 1748 | 2702 | 2611 | 2758 | 2620 | 2547 | 2472 | 2523 | 2465 | 2587,3 | 2673 | 2502 | 0,936 | -0,095 | -2,518 | 0,01181868 | 0,02336514 | 1,00E-08 | 0,0025406 | 0,0009729 | 0,0009729 | TRUE | 0,0054913 | No |
| orf19.4320 | 5 | 9 | 3 | 9 | 11 | 5 | 6 | 3 | 4 | 11 | 3 | 7 | 9 | 5 | 5 | 4 | 6,27 | 6 | 6 | 0,938 | -0,093 | -0,159 | 0,87338325 | NA | 0,061406 | 0,2333359 | 0,1657825 | 0,1657825 | TRUE | 0,3774281 | No |
| orf19.4321 | 22 | 16 | 25 | 40 | 34 | 10 | 20 | 20 | 20 | 20 | 26 | 33 | 29 | 10 | 18 | 28 | 22,92 | 25 | 21 | 0,844 | -0,245 | -0,69 | 0,49001091 | 0,57544675 | 0,093612 | 0,0653233 | 0,0773477 | 0,0773477 | TRUE | 0,5985922 | No |
| orf19.4322 | 6784 | 5158 | 6158 | 7602 | 7371 | 7081 | 7496 | 4323 | 6086 | 6567 | 6300 | 6269 | 6304 | 6756 | 6689 | 6097 | 6383,5 | 6306 | 6462 | 1,026 | 0,037 | 0,823 | 0,41068758 | 0,49876914 | 0,001464 | 0,0022069 | 0,0017346 | 0,0017346 | TRUE | 0,0169792 | No |
| orf19.4323 | 1006 | 860 | 910 | 1176 | 1469 | 1444 | 1402 | 781 | 902 | 1095 | 931 | 970 | 1256 | 1378 | 1251 | 1102 | 1110,6 | 974 | 1247 | 1,284 | 0,36 | 4,563 | 5,04E-06 | 1,92E-05 | 0,006685 | 0,0032866 | 0,0050676 | 0,0050676 | TRUE | 0,0785107 | No |
| orf19.4324 | 427 | 329 | 359 | 448 | 498 | 438 | 438 | 291 | 383 | 419 | 367 | 369 | 426 | 418 | 391 | 410 | 397,97 | 384 | 411 | 1,073 | 0,102 | 1,34 | 0,18028907 | 0,25201289 | 1,00E-08 | 0,0056273 | 0,0029982 | 0,0029982 | TRUE | 0,0359529 | No |
| orf19.4325 | 634 | 486 | 601 | 747 | 746 | 814 | 838 | 496 | 569 | 619 | 615 | 616 | 638 | 777 | 748 | 700 | 660,05 | 605 | 716 | 1,184 | 0,244 | 3,296 | 0,00098103 | 0,00247129 | 0,003363 | 0,0041789 | 0,0037371 | 0,0037371 | TRUE | 0,0682831 | No |
| orf19.4326 | 1798 | 1431 | 1605 | 1977 | 1519 | 1563 | 1636 | 957 | 1613 | 1822 | 1642 | 1630 | 1299 | 1491 | 1460 | 1350 | 1538,4 | 1677 | 1400 | 0,836 | -0,258 | -4,179 | 2,93E-05 | 9,90E-05 | 0,002909 | 0,0029232 | 0,0029905 | 0,0029905 | TRUE | 0,0391476 | No |
| orf19.4328 | 9746 | 7720 | 9067 | 11037 | 9486 | 9638 | 10104 | 6236 | 8743 | 9828 | 9276 | 9101 | 8113 | 9196 | 9016 | 8795 | 9008,7 | 9237 | 8780 | 0,951 | -0,073 | -1,421 | 0,1553003 | 0,2221005 | 0,002509 | 0,0021407 | 0,0024194 | 0,0024194 | TRUE | 0,0323752 | No |
| orf19.433 | 217 | 164 | 235 | 253 | 212 | 169 | 199 | 120 | 195 | 209 | 240 | 209 | 181 | 161 | 178 | 169 | 192,74 | 213 | 172 | 0,811 | -0,302 | -2,82 | 0,00480347 | 0,01040346 | 1,00E-08 | 0,0095115 | 0,0057726 | 0,0057726 | TRUE | 0,0812986 | No |
| orf19.4330 | 394 | 266 | 359 | 433 | 485 | 362 | 418 | 238 | 353 | 339 | 367 | 357 | 415 | 345 | 373 | 336 | 360,67 | 354 | 367 | 1,041 | 0,058 | 0,673 | 0,50107122 | 0,58602996 | 0,00246 | 0,0060046 | 0,004297 | 0,004297 | TRUE | 0,0898768 | No |
| orf19.4332 | 754 | 508 | 604 | 774 | 822 | 713 | 782 | 494 | 676 | 647 | 618 | 638 | 703 | 680 | 698 | 697 | 669,65 | 645 | 694 | 1,077 | 0,106 | 1,818 | 0,06910896 | 0,11153062 | 1,00E-08 | 0,0041474 | 0,0017999 | 0,0017999 | TRUE | 0,0136439 | No |
| orf19.4334 | 89 | 49 | 87 | 116 | 94 | 71 | 87 | 68 | 80 | 62 | 89 | 96 | 80 | 68 | 78 | 96 | 81,07 | 82 | 80 | 0,967 | -0,049 | -0,281 | 0,77870744 | 0,82650276 | 0,01115 | 0,0198861 | 0,0163877 | 0,0163877 | TRUE | 0,197964 | No |
| orf19.4335 | 1590 | 1145 | 1479 | 1815 | 278 | 296 | 240 | 175 | 1426 | 1458 | 1513 | 1497 | 238 | 282 | 214 | 247 | 859,39 | 1474 | 245 | 0,166 | -2,59 | -34,682 | 1,46E-263 | 8,75E-260 | 0,001437 | 0,0036687 | 0,0029702 | 0,0029702 | TRUE | 0,1256677 | No |
| orf19.4336 | 12769 | 8593 | 10064 | 13207 | 12170 | 10109 | 10141 | 7408 | 11455 | 10940 | 10296 | 10891 | 10408 | 9646 | 9049 | 10448 | 10392 | 10896 | 9888 | 0,907 | -0,14 | -2,563 | 0,01037127 | 0,02079793 | 0,003114 | 0,0021192 | 0,0027848 | 0,0027848 | TRUE | 0,039998 | No |
| orf19.4337 | 3225 | 2294 | 2961 | 3572 | 3537 | 3275 | 3482 | 2133 | 2893 | 2921 | 3029 | 2946 | 3025 | 3125 | 3107 | 3008 | 3006,7 | 2947 | 3066 | 1,041 | 0,058 | 1,629 | 0,10338907 | 0,15693621 | 1,00E-08 | 0,0024623 | 0,0008821 | 0,0008821 | TRUE | 0,0042462 | No |
| orf19.4339 | 3807 | 2457 | 3306 | 4009 | 3345 | 2920 | 3231 | 2192 | 3415 | 3128 | 3382 | 3306 | 2861 | 2786 | 2883 | 3092 | 3106,7 | 3308 | 2906 | 0,877 | -0,19 | -4,094 | 4,24E-05 | 0,00013887 | 0,001336 | 0,0024468 | 0,0017395 | 0,0017395 | TRUE | 0,021194 | No |
| orf19.434 | 537 | 357 | 454 | 551 | 546 | 500 | 559 | 326 | 482 | 454 | 464 | 454 | 467 | 477 | 499 | 460 | 469,72 | 464 | 476 | 1,027 | 0,038 | 0,564 | 0,57304441 | 0,65143819 | 1,00E-08 | 0,0050701 | 0,0022686 | 0,0022686 | TRUE | 0,0128282 | No |
| orf19.4340 | 829 | 628 | 708 | 917 | 1014 | 929 | 925 | 551 | 744 | 800 | 724 | 756 | 867 | 886 | 825 | 777 | 797,49 | 756 | 839 | 1,114 | 0,156 | 2,546 | 0,01088749 | 0,02173157 | 0,000882 | 0,0037999 | 0,0023635 | 0,0023635 | TRUE | 0,0253953 | No |
| orf19.4340.1 | 1199 | 925 | 988 | 1375 | 1132 | 1056 | 1083 | 566 | 1076 | 1178 | 1011 | 1134 | 968 | 1008 | 966 | 798 | 1017,3 | 1100 | 935 | 0,854 | -0,228 | -2,963 | 0,00304581 | 0,00690797 | 0,005811 | 0,0034065 | 0,0047052 | 0,0047052 | TRUE | 0,1060844 | No |
| orf19.4341 | 364 | 307 | 375 | 417 | 369 | 413 | 403 | 261 | 327 | 391 | 384 | 344 | 316 | 394 | 360 | 368 | 360,29 | 362 | 360 | 0,996 | -0,005 | -0,057 | 0,95429155 | 0,9674964 | 0,005446 | 0,0060088 | 0,0057303 | 0,0057303 | TRUE | 0,0827455 | No |
| orf19.4342 | 1761 | 1363 | 1540 | 1962 | 2662 | 2589 | 2835 | 1674 | 1580 | 1735 | 1576 | 1618 | 2277 | 2470 | 2530 | 2361 | 2018,3 | 1627 | 2410 | 1,483 | 0,568 | 11,02 | 3,07E-28 | 1,02E-26 | 0,001459 | 0,0026988 | 0,0020385 | 0,0020385 | TRUE | 0,0225709 | No |
| orf19.4346 | 1415 | 992 | 1340 | 1544 | 1880 | 1623 | 1763 | 1198 | 1269 | 1263 | 1371 | 1273 | 1608 | 1549 | 1573 | 1690 | 1449,5 | 1294 | 1605 | 1,238 | 0,308 | 6,207 | 5,40E-10 | 3,58E-09 | 0,000594 | 0,0029811 | 0,0016677 | 0,0016677 | TRUE | 0,0189452 | No |
| orf19.4347 | 6547 | 4661 | 6137 | 7432 | 8411 | 7558 | 8546 | 5481 | 5873 | 5934 | 6279 | 6129 | 7193 | 7212 | 7626 | 7730 | 6747 | 6054 | 7440 | 1,228 | 0,297 | 7,406 | 1,31E-13 | 1,31E-12 | 0,000967 | 0,0021947 | 0,0013922 | 0,0013922 | TRUE | 0,0081352 | No |
| orf19.4348 | 634 | 624 | 659 | 780 | 843 | 992 | 1054 | 584 | 569 | 794 | 674 | 643 | 721 | 947 | 941 | 824 | 764,04 | 670 | 858 | 1,283 | 0,36 | 3,282 | 0,00102957 | 0,00258598 | 0,015886 | 0,0038796 | 0,0102124 | 0,0102124 | TRUE | 0,1774578 | No |
| orf19.4349 | 784 | 538 | 748 | 963 | 732 | 687 | 786 | 407 | 703 | 685 | 765 | 794 | 626 | 656 | 701 | 574 | 688,07 | 737 | 639 | 0,869 | -0,203 | -2,654 | 0,00795783 | 0,01641849 | 0,003966 | 0,0040893 | 0,0041494 | 0,0041494 | TRUE | 0,0499295 | No |
| orf19.4353 | 1245 | 1044 | 1170 | 1396 | 2075 | 2111 | 2270 | 1345 | 1117 | 1329 | 1197 | 1151 | 1775 | 2014 | 2026 | 1897 | 1563,2 | 1198 | 1928 | 1,612 | 0,689 | 10,381 | 3,02E-25 | 7,86E-24 | 0,003969 | 0,0029082 | 0,003547 | 0,003547 | TRUE | 0,0615335 | No |
| orf19.4354 | 3701 | 2717 | 3469 | 4103 | 3915 | 3629 | 3917 | 2299 | 3320 | 3459 | 3549 | 3383 | 3348 | 3463 | 3495 | 3243 | 3407,6 | 3428 | 3387 | 0,989 | -0,015 | -0,38 | 0,7040477 | 0,76567505 | 0,000592 | 0,0024055 | 0,001276 | 0,001276 | TRUE | 0,0092956 | No |
| orf19.4355 | 1609 | 1230 | 1507 | 1829 | 1715 | 1749 | 1803 | 1096 | 1443 | 1566 | 1542 | 1508 | 1467 | 1669 | 1609 | 1546 | 1543,7 | 1515 | 1573 | 1,039 | 0,055 | 1,028 | 0,30383767 | 0,38963857 | 0,001592 | 0,0029199 | 0,0021315 | 0,0021315 | TRUE | 0,0265306 | No |
| orf19.4356 | 195 | 136 | 135 | 193 | 188 | 219 | 196 | 136 | 175 | 173 | 138 | 159 | 161 | 209 | 175 | 192 | 172,73 | 161 | 184 | 1,138 | 0,187 | 1,519 | 0,12879753 | 0,18956775 | 0,006778 | 0,0103842 | 0,0087441 | 0,0087441 | TRUE | 0,0980269 | No |
| orf19.4357 | 1833 | 1376 | 1602 | 2032 | 1706 | 1619 | 1652 | 1019 | 1644 | 1752 | 1639 | 1676 | 1459 | 1545 | 1474 | 1437 | 1578,3 | 1678 | 1479 | 0,883 | -0,179 | -3,936 | 8,27E-05 | 0,00025579 | 0,000211 | 0,0028993 | 0,0013578 | 0,0013578 | TRUE | 0,0108279 | No |
| orf19.4358 | 1084 | 720 | 948 | 1242 | 1458 | 1217 | 1269 | 766 | 972 | 917 | 970 | 1024 | 1247 | 1161 | 1132 | 1080 | 1063 | 971 | 1155 | 1,19 | 0,252 | 4,186 | 2,84E-05 | 9,62E-05 | 0,001861 | 0,0033452 | 0,0025189 | 0,0025189 | TRUE | 0,0347282 | No |

| | | | | | | | | | | | | | | | | | | | | | | | | | | | | | | | |
|---|---|---|---|---|---|---|---|---|---|---|---|---|---|---|---|---|---|---|---|---|---|---|---|---|---|---|---|---|---|---|---|
| orf19.4361 | 31 | 19 | 24 | 35 | 75 | 49 | 50 | 34 | 28 | 24 | 25 | 29 | 64 | 47 | 45 | 48 | 38,61 | 26 | 51 | 1,926 | 0,946 | 4,04 | 5,34E-05 | 0,00017172 | 1,00E-08 | 0,0395783 | 0,0241682 | 0,0241682 | TRUE | 0,2871169 | No |
| orf19.4362 | 1891 | 1360 | 1619 | 2021 | 2260 | 2206 | 2436 | 1335 | 1696 | 1731 | 1656 | 1667 | 1933 | 2105 | 2174 | 1883 | 1855,6 | 1688 | 2024 | 1,201 | 0,264 | 4,85 | 1,24E-06 | 5,14E-06 | 0,001803 | 0,0027618 | 0,0023076 | 0,0023076 | TRUE | 0,0302272 | No |
| orf19.4363 | 7656 | 5259 | 6658 | 8119 | 8027 | 7439 | 8204 | 4695 | 6868 | 6695 | 6812 | 6695 | 6865 | 7098 | 7321 | 6622 | 6872 | 6768 | 6976 | 1,031 | 0,045 | 1,162 | 0,2450673 | 0,3269741 | 0,000818 | 0,0021908 | 0,0012692 | 0,0012692 | TRUE | 0,013702 | No |
| orf19.4365 | 1283 | 900 | 1170 | 1367 | 1836 | 1874 | 1941 | 1259 | 1151 | 1146 | 1197 | 1127 | 1570 | 1788 | 1732 | 1776 | 1435,9 | 1155 | 1716 | 1,484 | 0,57 | 10,354 | 4,03E-25 | 1,03E-23 | 0,001643 | 0,0029905 | 0,0021829 | 0,0021829 | TRUE | 0,0411771 | No |
| orf19.4366 | 49 | 36 | 42 | 63 | 44 | 33 | 42 | 34 | 44 | 46 | 43 | 52 | 38 | 31 | 37 | 48 | 42,41 | 46 | 38 | 0,823 | -0,281 | -1,315 | 0,18834846 | 0,26109773 | 1,00E-08 | 0,0362117 | 0,0200649 | 0,0200649 | TRUE | 0,1387361 | No |
| orf19.4367 | 756 | 480 | 641 | 736 | 617 | 594 | 603 | 395 | 678 | 611 | 656 | 607 | 528 | 567 | 538 | 557 | 592,71 | 638 | 548 | 0,856 | -0,224 | -3,397 | 0,00068223 | 0,00177823 | 1,00E-08 | 0,0044288 | 0,0024717 | 0,0024717 | TRUE | 0,0221891 | No |
| orf19.4368 | 1455 | 1050 | 1385 | 1582 | 1220 | 1074 | 1216 | 826 | 1305 | 1337 | 1417 | 1305 | 1043 | 1025 | 1085 | 1165 | 1210,2 | 1341 | 1080 | 0,803 | -0,316 | -5,65 | 1,60E-08 | 8,66E-08 | 0,001367 | 0,0031791 | 0,0021604 | 0,0021604 | TRUE | 0,0302122 | No |
| orf19.4369 | 1725 | 1240 | 1577 | 1863 | 2076 | 1987 | 2095 | 1332 | 1548 | 1579 | 1613 | 1536 | 1775 | 1896 | 1869 | 1879 | 1711,9 | 1569 | 1855 | 1,182 | 0,241 | 5,547 | 2,91E-08 | 1,52E-07 | 1,00E-08 | 0,0028275 | 0,0012296 | 0,0012296 | TRUE | 0,010599 | No |
| orf19.437 | 5141 | 3560 | 4864 | 5673 | 4873 | 4353 | 4774 | 2780 | 4612 | 4532 | 4976 | 4678 | 4168 | 4153 | 4260 | 3921 | 4412,6 | 4700 | 4126 | 0,878 | -0,187 | -4,319 | 1,57E-05 | 5,56E-05 | 0,001159 | 0,0023085 | 0,0015719 | 0,0015719 | TRUE | 0,0189605 | No |
| orf19.4370 | 2509 | 1682 | 2130 | 2555 | 2691 | 2490 | 2641 | 1615 | 2251 | 2141 | 2179 | 2107 | 2301 | 2376 | 2357 | 2278 | 2248,8 | 2170 | 2328 | 1,074 | 0,102 | 2,555 | 0,01062548 | 0,02124389 | 0,000194 | 0,0026251 | 0,001094 | 0,001094 | TRUE | 0,0080507 | No |
| orf19.4371 | 3262 | 2126 | 2810 | 3426 | 2963 | 2517 | 2659 | 1830 | 2926 | 2707 | 2875 | 2825 | 2534 | 2402 | 2373 | 2581 | 2652,8 | 2833 | 2472 | 0,871 | -0,199 | -4,453 | 8,47E-06 | 3,13E-05 | 0,000878 | 0,0025268 | 0,0015356 | 0,0015356 | TRUE | 0,0103973 | No |
| orf19.4372 | 4353 | 3075 | 4072 | 4797 | 5333 | 5067 | 5410 | 3454 | 3905 | 3915 | 4166 | 3956 | 4561 | 4835 | 4828 | 4872 | 4379,6 | 3986 | 4774 | 1,197 | 0,26 | 6,652 | 2,88E-11 | 2,25E-10 | 0,000614 | 0,002311 | 0,0012321 | 0,0012321 | TRUE | 0,0112647 | No |
| orf19.4373 | 1823 | 1366 | 1527 | 2013 | 1735 | 1617 | 1577 | 1130 | 1635 | 1739 | 1562 | 1660 | 1484 | 1543 | 1407 | 1594 | 1578,1 | 1649 | 1507 | 0,913 | -0,132 | -2,433 | 0,01495834 | 0,02880439 | 0,001547 | 0,0028995 | 0,0021741 | 0,0021741 | TRUE | 0,0239441 | No |
| orf19.4374 | 689 | 495 | 632 | 762 | 598 | 599 | 632 | 354 | 618 | 630 | 647 | 628 | 511 | 572 | 564 | 499 | 583,68 | 631 | 536 | 0,853 | -0,229 | -3,443 | 0,00057617 | 0,00153103 | 0,000409 | 0,0044667 | 0,0025292 | 0,0025292 | TRUE | 0,0214717 | No |
| orf19.4375 | 2460 | 1676 | 2122 | 2580 | 2839 | 2354 | 2562 | 1656 | 2207 | 2134 | 2171 | 2128 | 2428 | 2246 | 2286 | 2336 | 2241,9 | 2160 | 2324 | 1,076 | 0,106 | 2,559 | 0,01050438 | 0,02102275 | 0,00037 | 0,0026271 | 0,0011877 | 0,0011877 | TRUE | 0,0118107 | No |
| orf19.4375.1 | 10571 | 6810 | 8075 | 10749 | 10056 | 7792 | 8021 | 6341 | 9483 | 8670 | 8262 | 8864 | 8600 | 7435 | 7157 | 8943 | 8426,8 | 8820 | 8034 | 0,911 | -0,135 | -1,83 | 0,06729187 | 0,10914944 | 0,007379 | 0,0021518 | 0,0051219 | 0,0051219 | TRUE | 0,0702344 | No |
| orf19.4376 | 457 | 334 | 367 | 454 | 384 | 371 | 379 | 256 | 410 | 425 | 375 | 374 | 328 | 354 | 338 | 361 | 370,84 | 396 | 345 | 0,871 | -0,199 | -2,506 | 0,01222337 | 0,02408592 | 1,00E-08 | 0,0058942 | 0,0033196 | 0,0033196 | TRUE | 0,0245185 | No |
| orf19.4377 | 2187 | 1547 | 1919 | 2369 | 2574 | 2458 | 2626 | 1636 | 1962 | 1969 | 1963 | 1954 | 2201 | 2345 | 2343 | 2307 | 2130,7 | 1962 | 2299 | 1,172 | 0,229 | 5,757 | 8,57E-09 | 4,78E-08 | 1,00E-08 | 0,0026608 | 0,0010403 | 0,0010403 | TRUE | 0,0106694 | No |
| orf19.4378 | 703 | 468 | 572 | 690 | 715 | 561 | 600 | 357 | 631 | 596 | 585 | 569 | 611 | 535 | 535 | 504 | 570,8 | 595 | 546 | 0,922 | -0,118 | -1,581 | 0,11396273 | 0,17027806 | 0,002694 | 0,0045228 | 0,0035779 | 0,0035779 | TRUE | 0,0775121 | No |
| orf19.4379 | 1139 | 737 | 985 | 1198 | 1701 | 1283 | 1587 | 1019 | 1022 | 938 | 1008 | 988 | 1455 | 1224 | 1416 | 1437 | 1186 | 989 | 1383 | 1,397 | 0,482 | 7,433 | 1,06E-13 | 1,08E-12 | 0,003138 | 0,0032036 | 0,0031662 | 0,0031662 | TRUE | 0,0730159 | No |
| orf19.438 | 856 | 583 | 796 | 950 | 720 | 765 | 741 | 541 | 768 | 742 | 814 | 783 | 616 | 730 | 661 | 763 | 734,73 | 777 | 692 | 0,887 | -0,173 | -2,338 | 0,01940197 | 0,03616725 | 0,003536 | 0,0039553 | 0,0038644 | 0,0038644 | TRUE | 0,0668975 | No |
| orf19.4380.1 | 1226 | 766 | 1114 | 1236 | 1803 | 1455 | 1626 | 1059 | 1100 | 975 | 1140 | 1019 | 1542 | 1388 | 1451 | 1494 | 1263,6 | 1058 | 1469 | 1,386 | 0,471 | 7,695 | 1,41E-14 | 1,55E-13 | 0,0024 | 0,0031284 | 0,0027806 | 0,0027806 | TRUE | 0,0307303 | No |
| orf19.4381 | 2201 | 1715 | 2087 | 2409 | 2444 | 2317 | 2488 | 1717 | 1975 | 2183 | 2135 | 1986 | 2090 | 2211 | 2220 | 2422 | 2152,8 | 2070 | 2236 | 1,08 | 0,11 | 1,947 | 0,05157765 | 0,08634119 | 0,002559 | 0,0026539 | 0,0026208 | 0,0026208 | TRUE | 0,0360059 | No |
| orf19.4382 | 1148 | 905 | 1014 | 1323 | 1278 | 1228 | 1202 | 720 | 1030 | 1152 | 1037 | 1091 | 1093 | 1172 | 1073 | 1015 | 1082,9 | 1078 | 1088 | 1,013 | 0,019 | 0,306 | 0,75939803 | 0,81182622 | 0,001788 | 0,0033201 | 0,0025882 | 0,0025882 | TRUE | 0,0312697 | No |
| orf19.4383 | 170 | 142 | 169 | 189 | 151 | 149 | 162 | 128 | 153 | 181 | 173 | 156 | 129 | 142 | 145 | 181 | 157,31 | 166 | 149 | 0,895 | -0,16 | -1,269 | 0,20437555 | 0,27989248 | 0,005757 | 0,0112081 | 0,0088506 | 0,0088506 | TRUE | 0,1742433 | No |
| orf19.4384 | 28 | 9 | 13 | 17 | 28 | 11 | 28 | 18 | 25 | 11 | 13 | 14 | 24 | 10 | 25 | 25 | 18,59 | 16 | 21 | 1,305 | 0,384 | 1,017 | 0,30907826 | 0,39509408 | 0,085873 | 0,0800749 | 0,0825471 | 0,0825471 | TRUE | 0,7664786 | No |
| orf19.4384.1 | 22 | 11 | 30 | 26 | 33 | 20 | 22 | 12 | 20 | 14 | 31 | 21 | 28 | 19 | 20 | 17 | 21,22 | 22 | 21 | 0,982 | -0,027 | -0,083 | 0,93382929 | 0,9523759 | 0,025852 | 0,0704013 | 0,0527321 | 0,0527321 | TRUE | 0,4171583 | No |
| orf19.4388 | 1912 | 1335 | 1588 | 2099 | 1920 | 1551 | 1835 | 1084 | 1715 | 1700 | 1625 | 1731 | 1642 | 1480 | 1637 | 1529 | 1632,3 | 1693 | 1572 | 0,93 | -0,105 | -2,083 | 0,03726316 | 0,06489191 | 0,001136 | 0,0028689 | 0,0018389 | 0,0018389 | TRUE | 0,0200711 | No |
| orf19.439 | 2360 | 1527 | 1961 | 2403 | 2395 | 1919 | 2177 | 1469 | 2117 | 1944 | 2006 | 1982 | 2048 | 1831 | 1943 | 2072 | 1992,9 | 2012 | 1974 | 0,979 | -0,03 | -0,577 | 0,56416275 | 0,64402238 | 0,001732 | 0,002708 | 0,0021229 | 0,0021229 | TRUE | 0,028524 | No |
| orf19.4390 | 627 | 421 | 518 | 627 | 624 | 592 | 606 | 377 | 562 | 536 | 530 | 517 | 534 | 565 | 541 | 532 | 539,56 | 536 | 543 | 1,013 | 0,018 | 0,284 | 0,77638409 | 0,82484803 | 1,00E-08 | 0,0046701 | 0,0021233 | 0,0021233 | TRUE | 0,0137316 | No |
| orf19.4391 | 68 | 52 | 59 | 69 | 65 | 52 | 50 | 39 | 61 | 66 | 60 | 57 | 56 | 50 | 45 | 55 | 56,16 | 61 | 52 | 0,838 | -0,255 | -1,399 | 0,16184282 | 0,22997911 | 1,00E-08 | 0,027828 | 0,0138519 | 0,0138519 | TRUE | 0,0678584 | No |
| orf19.4392 | 505 | 326 | 446 | 570 | 407 | 365 | 381 | 225 | 453 | 415 | 456 | 470 | 348 | 348 | 340 | 317 | 393,51 | 448 | 338 | 0,755 | -0,406 | -5,37 | 7,87E-08 | 3,91E-07 | 1,00E-08 | 0,0056686 | 0,0029081 | 0,0029081 | TRUE | 0,0259444 | No |
| orf19.4393 | 13893 | 9191 | 12234 | 14905 | 12760 | 10656 | 11509 | 8198 | 12463 | 11701 | 12517 | 12291 | 10913 | 10168 | 10270 | 11563 | 11486 | 12243 | 10728 | 0,876 | -0,191 | -3,982 | 6,83E-05 | 0,00021536 | 0,002166 | 0,0021059 | 0,0021284 | 0,0021284 | TRUE | 0,0329569 | No |
| orf19.4394 | 35 | 22 | 37 | 28 | 17 | 11 | 18 | 19 | 31 | 28 | 38 | 23 | 15 | 10 | 16 | 27 | 23,53 | 30 | 17 | 0,547 | -0,87 | -2,7 | 0,00693225 | 0,01449667 | 0,033804 | 0,0636735 | 0,0525237 | 0,0525237 | TRUE | 0,5155517 | No |
| orf19.4395 | 1319 | 929 | 1274 | 1396 | 1295 | 1164 | 1246 | 823 | 1183 | 1183 | 1303 | 1151 | 1108 | 1111 | 1112 | 1161 | 1163,9 | 1205 | 1123 | 0,931 | -0,103 | -1,911 | 0,0560207 | 0,0926176 | 0,00096 | 0,0032268 | 0,0019405 | 0,0019405 | TRUE | 0,0362223 | No |
| orf19.4396 | 3139 | 2054 | 2745 | 3360 | 3583 | 3083 | 3534 | 2146 | 2816 | 2615 | 2808 | 2771 | 3064 | 2942 | 3154 | 3027 | 2899,5 | 2752 | 3047 | 1,106 | 0,146 | 3,523 | 0,00042668 | 0,00116895 | 0,000535 | 0,0024802 | 0,0012924 | 0,0012924 | TRUE | 0,0128795 | No |
| orf19.4398 | 2321 | 1558 | 2000 | 2458 | 2151 | 1853 | 1975 | 1378 | 2082 | 1984 | 2046 | 2027 | 1840 | 1768 | 1762 | 1944 | 1931,5 | 2035 | 1828 | 0,896 | -0,158 | -3,488 | 0,00048731 | 0,00132 | 0,000578 | 0,0027311 | 0,0014428 | 0,0014428 | TRUE | 0,0195027 | No |
| orf19.4399 | 9792 | 6599 | 8679 | 10546 | 9811 | 8410 | 9100 | 5659 | 8784 | 8401 | 8879 | 8696 | 8391 | 8025 | 8120 | 7982 | 8409,8 | 8690 | 8130 | 0,935 | -0,096 | -2,948 | 0,00319372 | 0,00720808 | 0,00038 | 0,0021521 | 0,0009026 | 0,0009026 | TRUE | 0,0059206 | No |
| orf19.440 | 465 | 295 | 428 | 550 | 499 | 430 | 457 | 341 | 417 | 376 | 438 | 454 | 427 | 410 | 408 | 481 | 426,24 | 421 | 432 | 1,017 | 0,024 | 0,284 | 0,77604801 | 0,82478267 | 0,003205 | 0,0053853 | 0,0043776 | 0,0043776 | TRUE | 0,0597763 | No |
| orf19.4400 | 444 | 275 | 346 | 460 | 471 | 358 | 422 | 287 | 398 | 350 | 354 | 379 | 403 | 342 | 377 | 405 | 375,94 | 370 | 382 | 1,025 | 0,036 | 0,424 | 0,6715581 | 0,73801373 | 0,002253 | 0,0058411 | 0,0041352 | 0,0041352 | TRUE | 0,057292 | No |
| orf19.4401 | 3844 | 2569 | 3287 | 4152 | 3461 | 2937 | 3128 | 2080 | 3448 | 3271 | 3363 | 3424 | 2960 | 2802 | 2791 | 2934 | 3124,1 | 3376 | 2872 | 0,85 | -0,235 | -6,069 | 1,29E-09 | 8,09E-09 | 0,000403 | 0,0024442 | 0,0011182 | 0,0011182 | TRUE | 0,0057202 | No |
| orf19.4403 | 770 | 654 | 741 | 857 | 1007 | 1012 | 1077 | 615 | 691 | 833 | 758 | 707 | 861 | 966 | 961 | 867 | 830,43 | 747 | 914 | 1,228 | 0,297 | 4,043 | 5,28E-05 | 0,00016992 | 0,003996 | 0,0037276 | 0,0039435 | 0,0039435 | TRUE | 0,066285 | No |
| orf19.4404 | 34 | 17 | 32 | 26 | 15 | 15 | 16 | 8 | 31 | 22 | 33 | 21 | 13 | 14 | 14 | 11 | 19,88 | 27 | 13 | 0,499 | -1,003 | -3,12 | 0,00180589 | 0,00430375 | 1,00E-08 | 0,0750096 | 0,0429864 | 0,0429864 | TRUE | 0,1341666 | No |
| orf19.4405 | 40 | 44 | 32 | 50 | 36 | 33 | 30 | 31 | 36 | 56 | 33 | 41 | 31 | 31 | 27 | 44 | 37,33 | 42 | 33 | 0,795 | -0,332 | -1,33 | 0,18360104 | 0,25575587 | 0,019095 | 0,0408679 | 0,0322944 | 0,0322944 | TRUE | 0,3463884 | No |
| orf19.4406 | 468 | 304 | 408 | 456 | 426 | 397 | 366 | 332 | 420 | 387 | 417 | 376 | 364 | 379 | 327 | 468 | 392,29 | 400 | 384 | 0,954 | -0,067 | -0,65 | 0,51538071 | 0,59969468 | 0,009844 | 0,0056801 | 0,0076947 | 0,0076947 | TRUE | 0,2228506 | No |
| orf19.4407 | 3282 | 2216 | 2830 | 3517 | 3110 | 2827 | 3012 | 1886 | 2944 | 2821 | 2895 | 2900 | 2660 | 2697 | 2688 | 2660 | 2783,3 | 2890 | 2676 | 0,925 | -0,112 | -3,253 | 0,00114099 | 0,00283971 | 1,00E-08 | 0,0025011 | 0,0007715 | 0,0007715 | TRUE | 0,0028927 | No |
| orf19.4409 | 2084 | 1362 | 1802 | 2124 | 2378 | 2019 | 2249 | 1525 | 1870 | 1734 | 1844 | 1751 | 2034 | 1926 | 2007 | 2151 | 1914,6 | 1800 | 2030 | 1,126 | 0,171 | 3,489 | 0,00048447 | 0,00131409 | 0,001041 | 0,0027378 | 0,0017707 | 0,0017707 | TRUE | 0,0179612 | No |
| orf19.441 | 9019 | 5957 | 7669 | 9400 | 9876 | 7984 | 8973 | 5326 | 8091 | 7584 | 7846 | 7751 | 8446 | 7618 | 8007 | 7512 | 7856,9 | 7818 | 7896 | 1,01 | 0,015 | 0,326 | 0,74454899 | 0,79979444 | 0,001459 | 0,0021643 | 0,0018313 | 0,0018313 | TRUE | 0,027031 | No |
| orf19.4410 | 1052 | 764 | 975 | 1231 | 1201 | 1088 | 1140 | 704 | 944 | 973 | 998 | 1015 | 1027 | 1038 | 1017 | 993 | 1000,6 | 982 | 1019 | 1,038 | 0,054 | 1,065 | 0,28678266 | 0,37125408 | 1,00E-08 | 0,0034304 | 0,0014481 | 0,0014481 | TRUE | 0,009123 | No |
| orf19.4411 | 1266 | 882 | 1041 | 1349 | 1246 | 1099 | 1242 | 755 | 1136 | 1123 | 1065 | 1112 | 1066 | 1049 | 1108 | 1065 | 1090,4 | 1109 | 1072 | 0,967 | -0,049 | -0,992 | 0,32103078 | 0,40768399 | 1,00E-08 | 0,0033108 | 0,0013962 | 0,0013962 | TRUE | 0,0084127 | No |
| orf19.4412 | 975 | 746 | 940 | 1092 | 1220 | 1032 | 1192 | 679 | 875 | 950 | 962 | 900 | 1043 | 985 | 1064 | 958 | 967 | 922 | 1012 | 1,103 | 0,141 | 2,427 | 0,01521772 | 0,02921351 | 0,001116 | 0,0034808 | 0,0021946 | 0,0021946 | TRUE | 0,0140638 | No |
| orf19.4413 | 8440 | 5731 | 7040 | 8687 | 9297 | 7643 | 8344 | 5335 | 7572 | 7296 | 7203 | 7163 | 7951 | 7293 | 7446 | 7525 | 7431 | 7308 | 7554 | 1,034 | 0,048 | 1,239 | 0,21537922 | 0,29329064 | 0,000834 | 0,0021749 | 0,0012912 | 0,0012912 | TRUE | 0,0156694 | No |
| orf19.4414 | 970 | 670 | 829 | 1097 | 929 | 792 | 849 | 594 | 870 | 853 | 848 | 905 | 795 | 756 | 758 | 838 | 827,69 | 869 | 787 | 0,901 | -0,15 | -2,618 | 0,00885204 | 0,0180649 | 0,000218 | 0,0037334 | 0,0019298 | 0,0019298 | TRUE | 0,0195595 | No |
| orf19.4415 | 1203 | 829 | 1057 | 1335 | 1534 | 1327 | 1496 | 1008 | 1079 | 1055 | 1081 | 1101 | 1312 | 1266 | 1335 | 1422 | 1206,5 | 1079 | 1334 | 1,232 | 0,301 | 5,919 | 3,23E-09 | 1,91E-08 | 0,000393 | 0,0031828 | 0,0016428 | 0,0016428 | TRUE | 0,0209621 | No |
| orf19.4416 | 885 | 614 | 855 | 933 | 1284 | 1103 | 1271 | 769 | 794 | 782 | 875 | 769 | 1098 | 1052 | 1134 | 1085 | 948,63 | 805 | 1092 | 1,358 | 0,442 | 7,589 | 3,23E-14 | 3,42E-13 | 0,001012 | 0,0035098 | 0,0021734 | 0,0021734 | TRUE | 0,0402211 | No |
| orf19.4418 | 959 | 659 | 850 | 1102 | 1114 | 963 | 1123 | 704 | 860 | 839 | 870 | 909 | 953 | 919 | 1002 | 993 | 918,03 | 870 | 967 | 1,109 | 0,149 | 2,711 | 0,00670188 | 0,01408849 | 1,00E-08 | 0,0035608 | 0,0018233 | 0,0018233 | TRUE | 0,0132419 | No |
| orf19.4418.1 | 217 | 190 | 204 | 264 | 326 | 246 | 281 | 237 | 195 | 242 | 209 | 218 | 279 | 235 | 251 | 334 | 245,19 | 216 | 275 | 1,269 | 0,343 | 2,865 | 0,0041661 | 0,009135 | 0,011388 | 0,0079003 | 0,0096177 | 0,0096177 | TRUE | 0,2109042 | No |
| orf19.442 | 64 | 38 | 45 | 85 | 85 | 52 | 58 | 45 | 57 | 48 | 46 | 70 | 73 | 50 | 52 | 63 | 57,43 | 55 | 60 | 1,059 | 0,083 | 0,398 | 0,69038806 | 0,75333299 | 0,020252 | 0,0272562 | 0,0241398 | 0,0241398 | TRUE | 0,2916376 | No |
| orf19.4420 | 2190 | 1564 | 1985 | 2329 | 1769 | 1612 | 1751 | 1121 | 1965 | 1991 | 2031 | 1921 | 1513 | 1538 | 1562 | 1581 | 1762,7 | 1977 | 1548 | 0,783 | -0,353 | -8,477 | 2,30E-17 | 3,37E-16 | 1,00E-08 | 0,0028031 | 0,0010835 | 0,0010835 | TRUE | 0,0046776 | No |

| | | | | | | | | | | | | | | | | | | | | | | | | | | | | | | | |
|---|---|---|---|---|---|---|---|---|---|---|---|---|---|---|---|---|---|---|---|---|---|---|---|---|---|---|---|---|---|---|---|
| orf19.4423 | 1442 | 956 | 1236 | 1473 | 1507 | 1412 | 1548 | 925 | 1294 | 1217 | 1265 | 1215 | 1289 | 1347 | 1381 | 1305 | 1289 | 1248 | 1330 | 1,067 | 0,093 | 1,923 | 0,0545093 | 0,0905925 | 1,00E-08 | 0,0031057 | 0,0014806 | 0,0014806 | TRUE | 0,0082649 | No |
| orf19.4424 | 67 | 43 | 69 | 64 | 43 | 42 | 63 | 37 | 60 | 55 | 71 | 53 | 37 | 40 | 56 | 52 | 52,93 | 60 | 46 | 0,772 | -0,372 | -1,803 | 0,07131807 | 0,11458053 | 0,010573 | 0,0294043 | 0,0216977 | 0,0216977 | TRUE | 0,2046866 | No |
| orf19.4426 | 1958 | 1567 | 1773 | 2313 | 1976 | 1999 | 2118 | 1219 | 1757 | 1995 | 1814 | 1907 | 1690 | 1907 | 1890 | 1719 | 1834,9 | 1868 | 1802 | 0,966 | -0,05 | -0,84 | 0,40086151 | 0,48947984 | 0,002813 | 0,0027707 | 0,0028892 | 0,0028892 | TRUE | 0,0240439 | No |
| orf19.4427 | 6192 | 4632 | 5006 | 6743 | 6930 | 5924 | 6384 | 3598 | 5555 | 5897 | 5122 | 5560 | 5927 | 5652 | 5697 | 5075 | 5560,6 | 5534 | 5588 | 1,011 | 0,015 | 0,264 | 0,79195622 | 0,83782197 | 0,003066 | 0,0022406 | 0,0030569 | 0,0030569 | TRUE | 0,0455338 | No |
| orf19.4428 | 1281 | 999 | 1176 | 1531 | 1415 | 1355 | 1442 | 872 | 1149 | 1272 | 1203 | 1262 | 1210 | 1293 | 1287 | 1230 | 1238,3 | 1222 | 1255 | 1,028 | 0,041 | 0,769 | 0,4421052 | 0,52890642 | 0,000896 | 0,0031519 | 0,0018588 | 0,0018588 | TRUE | 0,0197249 | No |
| orf19.443 | 1217 | 789 | 925 | 1203 | 863 | 769 | 804 | 493 | 1092 | 1004 | 946 | 992 | 738 | 734 | 717 | 695 | 864,9 | 1008 | 721 | 0,716 | -0,483 | -7,958 | 1,75E-15 | 2,11E-14 | 0,001302 | 0,003658 | 0,0023419 | 0,0023419 | TRUE | 0,0381924 | No |
| orf19.4430 | 5285 | 3789 | 4764 | 6092 | 4595 | 4147 | 4472 | 2718 | 4741 | 4824 | 4874 | 5024 | 3930 | 3957 | 3991 | 3833 | 4396,7 | 4866 | 3928 | 0,808 | -0,308 | -9,044 | 1,51E-19 | 2,64E-18 | 0,000254 | 0,0023097 | 0,0008854 | 0,0008854 | TRUE | 0,0060634 | No |
| orf19.4432 | 12640 | 8695 | 11484 | 13852 | 18400 | 16196 | 17266 | 10464 | 11339 | 11070 | 11749 | 11423 | 15736 | 15454 | 15407 | 14759 | 13367 | 11395 | 15339 | 1,346 | 0,429 | 12,838 | 1,01E-37 | 6,05E-36 | 0,000563 | 0,0020881 | 0,0009957 | 0,0009957 | TRUE | 0,0077517 | No |
| orf19.4433 | 323 | 196 | 295 | 349 | 314 | 248 | 282 | 208 | 290 | 250 | 302 | 288 | 269 | 237 | 252 | 293 | 272,38 | 282 | 263 | 0,923 | -0,116 | -1,2 | 0,23025476 | 0,31031193 | 0,002617 | 0,0073091 | 0,0053281 | 0,0053281 | TRUE | 0,0593369 | No |
| orf19.4435 | 3700 | 2565 | 3353 | 4127 | 4059 | 3940 | 4004 | 2733 | 3319 | 3266 | 3430 | 3403 | 3471 | 3759 | 3573 | 3855 | 3509,6 | 3354 | 3664 | 1,091 | 0,125 | 2,87 | 0,0041101 | 0,00904192 | 0,001003 | 0,0023932 | 0,001548 | 0,001548 | TRUE | 0,015382 | No |
| orf19.4436 | 375 | 240 | 324 | 364 | 412 | 392 | 394 | 271 | 336 | 306 | 331 | 300 | 352 | 374 | 352 | 382 | 341,72 | 318 | 365 | 1,143 | 0,192 | 2,391 | 0,01678657 | 0,03186531 | 1,00E-08 | 0,0062277 | 0,0032857 | 0,0032857 | TRUE | 0,0201178 | No |
| orf19.4437 | 2709 | 1886 | 2399 | 2876 | 2715 | 2615 | 2826 | 1727 | 2430 | 2401 | 2454 | 2372 | 2322 | 2495 | 2522 | 2436 | 2429 | 2414 | 2444 | 1,012 | 0,017 | 0,422 | 0,6729776 | 0,73903298 | 0,000462 | 0,0025772 | 0,0012175 | 0,0012175 | TRUE | 0,0145794 | No |
| orf19.4438 | 835 | 541 | 777 | 995 | 911 | 811 | 931 | 654 | 749 | 689 | 795 | 820 | 779 | 774 | 831 | 922 | 794,92 | 763 | 826 | 1,077 | 0,107 | 1,415 | 0,15705928 | 0,22418916 | 0,004663 | 0,0038057 | 0,0042621 | 0,0042621 | TRUE | 0,0663177 | No |
| orf19.4439 | 16 | 13 | 23 | 23 | 17 | 11 | 11 | 5 | 14 | 17 | 24 | 19 | 15 | 10 | 10 | 7 | 14,41 | 18 | 10 | 0,585 | -0,774 | -2,069 | 0,03852394 | NA | 1,00E-08 | 0,1027012 | 0,0606997 | 0,0606997 | TRUE | 0,2737655 | No |
| orf19.4439.1 | 380 | 262 | 313 | 416 | 425 | 345 | 358 | 287 | 341 | 334 | 320 | 343 | 363 | 329 | 319 | 405 | 344,33 | 334 | 354 | 1,052 | 0,073 | 0,826 | 0,40892272 | 0,49712823 | 0,002881 | 0,0061956 | 0,0046688 | 0,0046688 | TRUE | 0,0899539 | No |
| orf19.444 | 452 | 317 | 374 | 491 | 468 | 394 | 419 | 265 | 405 | 404 | 383 | 405 | 400 | 376 | 374 | 374 | 390,05 | 399 | 381 | 0,956 | -0,066 | -0,902 | 0,36687997 | 0,45569875 | 1,00E-08 | 0,0057013 | 0,0025271 | 0,0025271 | TRUE | 0,0145079 | No |
| orf19.4440 | 838 | 575 | 748 | 858 | 893 | 739 | 832 | 509 | 752 | 732 | 765 | 708 | 764 | 705 | 742 | 718 | 735,72 | 739 | 732 | 0,993 | -0,011 | -0,184 | 0,85406576 | 0,88898598 | 1,00E-08 | 0,0039527 | 0,0019191 | 0,0019191 | TRUE | 0,010954 | No |
| orf19.4441 | 490 | 332 | 432 | 515 | 417 | 376 | 436 | 264 | 440 | 423 | 442 | 425 | 357 | 359 | 389 | 372 | 400,71 | 432 | 369 | 0,853 | -0,229 | -3,159 | 0,00158537 | 0,00381912 | 1,00E-08 | 0,0056023 | 0,0025246 | 0,0025246 | TRUE | 0,0170692 | No |
| orf19.4442 | 771 | 522 | 730 | 859 | 1075 | 955 | 1039 | 701 | 692 | 665 | 747 | 708 | 919 | 911 | 927 | 989 | 819,73 | 703 | 936 | 1,328 | 0,409 | 7,032 | 2,04E-12 | 1,81E-11 | 0,000252 | 0,0037505 | 0,0020085 | 0,0020085 | TRUE | 0,0197467 | No |
| orf19.4443 | 1959 | 1304 | 1650 | 2010 | 1474 | 1230 | 1377 | 787 | 1757 | 1660 | 1688 | 1657 | 1261 | 1174 | 1229 | 1110 | 1442 | 1690 | 1194 | 0,708 | -0,499 | -9,66 | 4,45E-22 | 9,36E-21 | 0,000961 | 0,0029862 | 0,0018471 | 0,0018471 | TRUE | 0,0234541 | No |
| orf19.4444 | 1494 | 973 | 1238 | 1588 | 982 | 913 | 924 | 593 | 1340 | 1239 | 1267 | 1309 | 840 | 871 | 825 | 836 | 1065,9 | 1289 | 843 | 0,653 | -0,614 | -12,01 | 3,17E-33 | 1,40E-31 | 1,00E-08 | 0,0033415 | 0,001531 | 0,001531 | TRUE | 0,0085413 | No |
| orf19.4445 | 18068 | 13732 | 16176 | 20249 | 23556 | 21849 | 22926 | 15876 | 16209 | 17482 | 16550 | 16698 | 20146 | 20848 | 20458 | 22392 | 18848 | 16735 | 20961 | 1,252 | 0,325 | 7,571 | 3,70E-14 | 3,90E-13 | 0,001576 | 0,0020565 | 0,0017127 | 0,0017127 | TRUE | 0,0256048 | No |
| orf19.4446 | 4327 | 2847 | 3741 | 4740 | 4959 | 4347 | 4732 | 2975 | 3882 | 3625 | 3827 | 3909 | 4241 | 4148 | 4223 | 4196 | 4006,2 | 3811 | 4202 | 1,102 | 0,14 | 3,885 | 0,0001022 | 0,00031247 | 0,000282 | 0,0023419 | 0,00099 | 0,00099 | TRUE | 0,0124549 | No |
| orf19.4447 | 2110 | 1424 | 1808 | 2289 | 1600 | 1246 | 1444 | 880 | 1893 | 1813 | 1850 | 1888 | 1368 | 1189 | 1289 | 1241 | 1566,3 | 1861 | 1272 | 0,684 | -0,548 | -10,515 | 7,37E-26 | 2,04E-24 | 0,001113 | 0,0029064 | 0,0019436 | 0,0019436 | TRUE | 0,0328416 | No |
| orf19.4448 | 2196 | 1473 | 1953 | 2369 | 2370 | 2010 | 2175 | 1492 | 1970 | 1875 | 1998 | 1954 | 2027 | 1918 | 1941 | 2104 | 1973,4 | 1949 | 1998 | 1,023 | 0,032 | 0,709 | 0,47845867 | 0,56375361 | 0,000601 | 0,0027152 | 0,0014729 | 0,0014729 | TRUE | 0,014156 | No |
| orf19.4449 | 2982 | 1938 | 2502 | 3221 | 3557 | 2996 | 3294 | 2121 | 2675 | 2467 | 2560 | 2656 | 3042 | 2859 | 2939 | 2991 | 2773,7 | 2590 | 2958 | 1,141 | 0,19 | 4,529 | 5,92E-06 | 2,23E-05 | 0,000553 | 0,0025029 | 0,0013264 | 0,0013264 | TRUE | 0,0116147 | No |
| orf19.445 | 3293 | 2118 | 2949 | 3480 | 3621 | 3191 | 3452 | 2040 | 2954 | 2696 | 3017 | 2870 | 3097 | 3045 | 3080 | 2877 | 2954,6 | 2884 | 3025 | 1,049 | 0,069 | 1,485 | 0,13754599 | 0,2002851 | 0,001265 | 0,0024709 | 0,0017304 | 0,0017304 | TRUE | 0,0221266 | No |
| orf19.4450 | 536 | 346 | 502 | 616 | 493 | 397 | 446 | 295 | 481 | 440 | 514 | 508 | 422 | 379 | 398 | 416 | 444,67 | 486 | 404 | 0,828 | -0,272 | -3,577 | 0,00034779 | 0,00096961 | 0,001116 | 0,0052442 | 0,0033045 | 0,0033045 | TRUE | 0,0404288 | No |
| orf19.4450.1 | 2745 | 1826 | 2172 | 2691 | 4231 | 3527 | 3514 | 2520 | 2463 | 2325 | 2222 | 2219 | 3618 | 3365 | 3136 | 3554 | 2862,8 | 2307 | 3418 | 1,481 | 0,567 | 9,88 | 5,09E-23 | 1,14E-21 | 0,002766 | 0,0024866 | 0,0027951 | 0,0027951 | TRUE | 0,0382228 | No |
| orf19.4450.2 | 547 | 386 | 445 | 591 | 580 | 458 | 436 | 344 | 491 | 491 | 455 | 487 | 496 | 437 | 389 | 485 | 466,51 | 481 | 452 | 0,937 | -0,094 | -1,101 | 0,27095881 | 0,35536357 | 0,00466 | 0,0050914 | 0,0048744 | 0,0048744 | TRUE | 0,1059253 | No |
| orf19.4451 | 6210 | 4165 | 5516 | 6597 | 6857 | 6024 | 6743 | 3823 | 5571 | 5302 | 5643 | 5440 | 5864 | 5748 | 6017 | 5392 | 5622,3 | 5489 | 5755 | 1,049 | 0,069 | 1,62 | 0,10528371 | 0,15921081 | 0,001206 | 0,0022377 | 0,0015713 | 0,0015713 | TRUE | 0,0211767 | No |
| orf19.4455 | 3576 | 2270 | 2954 | 3765 | 3509 | 2887 | 3357 | 2192 | 3208 | 2890 | 3022 | 3105 | 3001 | 2755 | 2996 | 3092 | 3008,5 | 3056 | 2961 | 0,967 | -0,048 | -0,958 | 0,33804979 | 0,4251629 | 0,001711 | 0,0024621 | 0,0020554 | 0,0020554 | TRUE | 0,0266785 | No |
| orf19.4456 | 36188 | 23881 | 32641 | 40363 | 37945 | 31914 | 34616 | 21680 | 32464 | 30403 | 33395 | 33284 | 32452 | 30451 | 30889 | 30578 | 31740 | 32386 | 31092 | 0,96 | -0,059 | -1,453 | 0,14630086 | 0,21113994 | 0,001316 | 0,0020253 | 0,0015474 | 0,0015474 | TRUE | 0,020702 | No |
| orf19.4457 | 2314 | 1667 | 2137 | 2433 | 2268 | 2179 | 2362 | 1324 | 2076 | 2122 | 2186 | 2006 | 1940 | 2079 | 2108 | 1867 | 2048,1 | 2098 | 1998 | 0,955 | -0,067 | -1,286 | 0,19842272 | 0,27273478 | 0,001634 | 0,0026883 | 0,0021223 | 0,0021223 | TRUE | 0,0217053 | No |
| orf19.4459 | 2344 | 2047 | 2279 | 2824 | 2428 | 2452 | 2565 | 1583 | 2103 | 2606 | 2332 | 2329 | 2077 | 2340 | 2289 | 2233 | 2288,4 | 2342 | 2235 | 0,955 | -0,067 | -1 | 0,31714281 | 0,40407873 | 0,004583 | 0,0026139 | 0,0038329 | 0,0038329 | TRUE | 0,0678475 | No |
| orf19.446 | 892 | 657 | 845 | 1016 | 868 | 851 | 868 | 543 | 800 | 836 | 865 | 838 | 742 | 812 | 775 | 766 | 804,21 | 835 | 774 | 0,927 | -0,109 | -1,92 | 0,05480746 | 0,09098734 | 1,00E-08 | 0,0037846 | 0,0018426 | 0,0018426 | TRUE | 0,01375 | No |
| orf19.4461 | 0 | 1 | 0 | 1 | 0 | 0 | 1 | 0 | 0 | 1 | 0 | 1 | 0 | 0 | 1 | 0 | 0,37 | 0 | 0 | 0,623 | -0,682 | -0,299 | 0,76478835 | NA | 1,00E-08 | 3,8860606 | 3,0564468 | 3,0564468 | TRUE | 0,2773135 | No |
| orf19.446.1 | 592 | 443 | 471 | 614 | 533 | 467 | 462 | 294 | 531 | 564 | 482 | 506 | 456 | 446 | 412 | 415 | 476,45 | 521 | 432 | 0,833 | -0,264 | -3,527 | 0,0004197 | 0,0011514 | 0,001325 | 0,0050264 | 0,0032469 | 0,0032469 | TRUE | 0,0322791 | No |
| orf19.446.2 | 460 | 382 | 412 | 523 | 470 | 431 | 397 | 312 | 413 | 486 | 422 | 431 | 402 | 411 | 354 | 440 | 419,91 | 438 | 402 | 0,917 | -0,126 | -1,458 | 0,14487747 | 0,20943788 | 0,003954 | 0,0054367 | 0,0047267 | 0,0047267 | TRUE | 0,0746422 | No |
| orf19.4463 | 9 | 5 | 8 | 9 | 9 | 4 | 8 | 3 | 8 | 6 | 8 | 7 | 8 | 4 | 7 | 4 | 6,62 | 7 | 6 | 0,775 | -0,367 | -0,704 | 0,48152341 | NA | 1,00E-08 | 0,2213964 | 0,1108285 | 0,1108285 | TRUE | 0,1392917 | No |
| orf19.4464 | 704 | 603 | 610 | 888 | 804 | 740 | 726 | 439 | 632 | 768 | 624 | 732 | 688 | 706 | 648 | 619 | 677,04 | 689 | 665 | 0,969 | -0,046 | -0,565 | 0,57236764 | 0,65103839 | 0,005135 | 0,0041237 | 0,0048592 | 0,0048592 | TRUE | 0,0645281 | No |
| orf19.4465 | 1356 | 1104 | 1188 | 1547 | 1520 | 1582 | 1634 | 1039 | 1216 | 1406 | 1215 | 1276 | 1300 | 1509 | 1458 | 1465 | 1355,8 | 1278 | 1433 | 1,122 | 0,166 | 2,521 | 0,01170874 | 0,02317066 | 0,003505 | 0,0030503 | 0,0034194 | 0,0034194 | TRUE | 0,0510284 | No |
| orf19.4466 | 328 | 256 | 314 | 382 | 371 | 373 | 339 | 249 | 294 | 326 | 321 | 315 | 317 | 356 | 303 | 351 | 322,91 | 314 | 332 | 1,054 | 0,076 | 0,882 | 0,37803581 | 0,46733316 | 1,00E-08 | 0,0064752 | 0,0040379 | 0,0040379 | TRUE | 0,0407629 | No |
| orf19.4467 | 533 | 418 | 497 | 601 | 648 | 709 | 653 | 433 | 478 | 532 | 508 | 496 | 554 | 677 | 583 | 611 | 554,81 | 504 | 606 | 1,204 | 0,268 | 3,479 | 0,00050408 | 0,00136053 | 0,003255 | 0,0045961 | 0,0038888 | 0,0038888 | TRUE | 0,0760843 | No |
| orf19.4468 | 2407 | 1850 | 2107 | 2816 | 2860 | 2540 | 2581 | 1404 | 2159 | 2355 | 2156 | 2322 | 2446 | 2424 | 2303 | 1980 | 2268,2 | 2248 | 2288 | 1,02 | 0,028 | 0,414 | 0,67857803 | 0,74314561 | 0,005097 | 0,0026196 | 0,0040174 | 0,0040174 | TRUE | 0,0952061 | No |
| orf19.4470 | 246 | 177 | 205 | 248 | 298 | 330 | 299 | 171 | 221 | 225 | 210 | 205 | 255 | 315 | 267 | 241 | 242,25 | 215 | 270 | 1,259 | 0,333 | 3,207 | 0,00134168 | 0,00328478 | 0,004121 | 0,0079721 | 0,0061613 | 0,0061613 | TRUE | 0,1436307 | No |
| orf19.4471 | 800 | 661 | 785 | 936 | 907 | 899 | 974 | 571 | 718 | 842 | 803 | 772 | 776 | 858 | 869 | 805 | 805,27 | 784 | 827 | 1,058 | 0,082 | 1,221 | 0,22195028 | 0,30094274 | 0,002462 | 0,0037823 | 0,0030729 | 0,0030729 | TRUE | 0,0381352 | No |
| orf19.4472 | 50 | 35 | 35 | 49 | 41 | 34 | 34 | 29 | 45 | 45 | 36 | 40 | 35 | 32 | 30 | 41 | 38,05 | 42 | 34 | 0,83 | -0,269 | -1,215 | 0,22428971 | 0,3032934 | 1,00E-08 | 0,040135 | 0,0206385 | 0,0206385 | TRUE | 0,0722349 | No |
| orf19.4473 | 359 | 376 | 356 | 473 | 396 | 469 | 389 | 260 | 322 | 479 | 364 | 390 | 339 | 448 | 347 | 367 | 381,88 | 389 | 375 | 0,968 | -0,048 | -0,378 | 0,70532976 | 0,76629678 | 0,019528 | 0,005781 | 0,0125746 | 0,0125746 | TRUE | 0,2638305 | No |
| orf19.4474 | 2394 | 2042 | 2121 | 2720 | 2004 | 2266 | 2273 | 1275 | 2148 | 2600 | 2170 | 2243 | 1714 | 2162 | 2028 | 1798 | 2107,9 | 2290 | 1926 | 0,842 | -0,248 | -2,975 | 0,00293216 | 0,00667541 | 0,009314 | 0,0026682 | 0,006214 | 0,006214 | TRUE | 0,098485 | No |
| orf19.4475 | 2228 | 2038 | 2113 | 2764 | 2123 | 2460 | 2509 | 1427 | 1999 | 2595 | 2162 | 2279 | 1816 | 2347 | 2239 | 2013 | 2181,1 | 2259 | 2104 | 0,932 | -0,101 | -1,103 | 0,26986488 | 0,35423819 | 0,011759 | 0,0026451 | 0,0076465 | 0,0076465 | TRUE | 0,1195564 | No |
| orf19.4476 | 732 | 449 | 582 | 813 | 510 | 431 | 444 | 352 | 657 | 572 | 595 | 670 | 436 | 411 | 396 | 496 | 529,28 | 624 | 435 | 0,692 | -0,53 | -6,222 | 4,91E-10 | 3,28E-09 | 0,005456 | 0,0047223 | 0,0050118 | 0,0050118 | TRUE | 0,0918 | No |
| orf19.4477 | 821 | 527 | 650 | 810 | 956 | 825 | 795 | 503 | 737 | 671 | 665 | 668 | 818 | 787 | 709 | 709 | 720,5 | 685 | 756 | 1,105 | 0,144 | 2,067 | 0,03874827 | 0,06722483 | 0,002641 | 0,0039944 | 0,0032679 | 0,0032679 | TRUE | 0,0367466 | No |
| orf19.4478 | 299 | 203 | 266 | 342 | 282 | 233 | 277 | 176 | 268 | 258 | 272 | 282 | 241 | 222 | 247 | 248 | 254,97 | 270 | 240 | 0,884 | -0,178 | -2,004 | 0,04510123 | 0,07687077 | 1,00E-08 | 0,0076732 | 0,0036567 | 0,0036567 | TRUE | 0,0268516 | No |
| orf19.4479 | 14935 | 11138 | 13317 | 16445 | 14988 | 13470 | 14449 | 8285 | 13398 | 14180 | 13625 | 13561 | 12818 | 12853 | 12893 | 11685 | 13127 | 13691 | 12562 | 0,918 | -0,123 | -2,989 | 0,00280067 | 0,00639787 | 0,001292 | 0,0020901 | 0,0015573 | 0,0015573 | TRUE | 0,0264882 | No |
| orf19.448 | 2877 | 1862 | 2589 | 3085 | 2735 | 2327 | 2446 | 1532 | 2581 | 2371 | 2649 | 2544 | 2339 | 2220 | 2183 | 2161 | 2380,9 | 2536 | 2226 | 0,878 | -0,188 | -3,936 | 8,29E-05 | 0,00025619 | 0,001169 | 0,0025893 | 0,0017745 | 0,0017745 | TRUE | 0,022099 | No |
| orf19.4481 | 17 | 12 | 11 | 18 | 10 | 8 | 10 | 6 | 15 | 15 | 11 | 15 | 9 | 8 | 9 | 8 | 11,27 | 14 | 8 | 0,593 | -0,754 | -1,873 | 0,06102848 | NA | 1,00E-08 | 0,1307383 | 0,0612353 | 0,0612353 | TRUE | 0,0810947 | No |
| orf19.4482 | 321 | 246 | 266 | 373 | 358 | 357 | 352 | 203 | 288 | 313 | 272 | 308 | 306 | 341 | 314 | 286 | 303,51 | 295 | 312 | 1,061 | 0,085 | 0,964 | 0,33522749 | 0,42196664 | 0,000815 | 0,0067625 | 0,004172 | 0,004172 | TRUE | 0,0423722 | No |
| orf19.4488 | 10031 | 7327 | 8678 | 11021 | 11215 | 10307 | 11046 | 6873 | 8999 | 9328 | 8878 | 9088 | 9591 | 9835 | 9857 | 9694 | 9408,7 | 9073 | 9744 | 1,074 | 0,103 | 3,577 | 0,00034692 | 0,00096763 | 0,00016 | 0,0021338 | 0,0006941 | 0,0006941 | TRUE | 0,0041834 | No |

| | | | | | | | | | | | | | | | | | | | | | | | | | | | | | | | |
|---|---|---|---|---|---|---|---|---|---|---|---|---|---|---|---|---|---|---|---|---|---|---|---|---|---|---|---|---|---|---|---|
| orf19.449 | 24474 | 17284 | 21782 | 26625 | 38067 | 35063 | 37919 | 24321 | 21956 | 22004 | 22285 | 21955 | 32556 | 33456 | 33836 | 34303 | 27794 | 22050 | 33538 | 1,521 | 0,605 | 22,723 | 2,65E-114 | 3,98E-111 | 0,000219 | 0,0020318 | 0,0006427 | 0,0006427 | TRUE | 0,0047729 | No |
| orf19.4490 | 15883 | 10634 | 13027 | 16761 | 14942 | 12863 | 12830 | 9295 | 14249 | 13538 | 13328 | 13821 | 12779 | 12273 | 11449 | 13110 | 13068 | 13734 | 12403 | 0,903 | -0,148 | -3,121 | 0,00180517 | 0,00430374 | 0,002017 | 0,0020906 | 0,0020716 | 0,0020716 | TRUE | 0,0328857 | No |
| orf19.4490.2 | 2866 | 1896 | 2345 | 3223 | 2873 | 2472 | 2541 | 1731 | 2571 | 2414 | 2399 | 2658 | 2457 | 2359 | 2267 | 2441 | 2445,8 | 2510 | 2381 | 0,947 | -0,078 | -1,587 | 0,1126057 | 0,16850188 | 0,001371 | 0,0025731 | 0,0019366 | 0,0019366 | TRUE | 0,0192193 | No |
| orf19.4491 | 3632 | 3299 | 3283 | 4363 | 3871 | 4162 | 4377 | 2259 | 3258 | 4200 | 3359 | 3598 | 3311 | 3971 | 3906 | 3186 | 3598,6 | 3604 | 3594 | 0,998 | -0,003 | -0,031 | 0,97511254 | 0,98222445 | 0,010984 | 0,0023829 | 0,0077269 | 0,0077269 | TRUE | 0,1494114 | No |
| orf19.4492 | 15013 | 10883 | 13201 | 16751 | 14470 | 12945 | 13404 | 8915 | 13468 | 13855 | 13506 | 13813 | 12375 | 12352 | 11961 | 12574 | 12988 | 13660 | 12316 | 0,901 | -0,15 | -5,239 | 1,62E-07 | 7,71E-07 | 0,000223 | 0,0020913 | 0,0007081 | 0,0007081 | TRUE | 0,0046153 | No |
| orf19.4494 | 1103 | 824 | 921 | 1267 | 888 | 918 | 917 | 597 | 990 | 1049 | 942 | 1045 | 759 | 876 | 818 | 842 | 915,16 | 1006 | 824 | 0,818 | -0,29 | -4,59 | 4,43E-06 | 1,70E-05 | 0,002037 | 0,0035658 | 0,00273 | 0,00273 | TRUE | 0,0349121 | No |
| orf19.4495 | 5236 | 3604 | 4533 | 5689 | 5836 | 5236 | 5610 | 3384 | 4697 | 4588 | 4638 | 4691 | 4991 | 4996 | 5006 | 4773 | 4797,6 | 4654 | 4942 | 1,062 | 0,087 | 2,757 | 0,0058353 | 0,01242326 | 4,18E-05 | 0,0022821 | 0,0007546 | 0,0007546 | TRUE | 0,0059254 | No |
| orf19.4496 | 2060 | 1487 | 1870 | 2290 | 2247 | 2178 | 2312 | 1421 | 1848 | 1893 | 1913 | 1888 | 1922 | 2078 | 2063 | 2004 | 1951,2 | 1886 | 2017 | 1,07 | 0,097 | 2,278 | 0,02270782 | 0,04183643 | 0,000339 | 0,0027235 | 0,0012344 | 0,0012344 | TRUE | 0,0131853 | No |
| orf19.4497 | 947 | 806 | 800 | 1079 | 1167 | 1199 | 1086 | 607 | 850 | 1026 | 818 | 890 | 998 | 1144 | 969 | 856 | 943,9 | 896 | 992 | 1,111 | 0,151 | 1,627 | 0,10382012 | 0,15739371 | 0,010943 | 0,0035175 | 0,0072392 | 0,0072392 | TRUE | 0,1260982 | No |
| orf19.4498 | 512 | 399 | 396 | 505 | 388 | 463 | 480 | 247 | 459 | 508 | 405 | 416 | 332 | 442 | 428 | 348 | 417,4 | 447 | 388 | 0,87 | -0,201 | -1,821 | 0,06858945 | 0,11083524 | 0,012507 | 0,0054575 | 0,0092476 | 0,0092476 | TRUE | 0,1183377 | No |
| orf19.4499 | 2729 | 1854 | 2278 | 2767 | 2846 | 2643 | 2772 | 1721 | 2448 | 2360 | 2331 | 2282 | 2434 | 2522 | 2474 | 2427 | 2409,7 | 2355 | 2464 | 1,047 | 0,066 | 1,667 | 0,09553658 | 0,14725085 | 0,000246 | 0,002582 | 0,0010867 | 0,0010867 | TRUE | 0,0090589 | No |
| orf19.450 | 62 | 44 | 59 | 74 | 51 | 51 | 59 | 58 | 56 | 56 | 60 | 61 | 44 | 49 | 53 | 82 | 57,47 | 58 | 57 | 0,95 | -0,073 | -0,352 | 0,72491887 | 0,78445832 | 0,019765 | 0,0272401 | 0,0242128 | 0,0242128 | TRUE | 0,5966124 | No |
| orf19.4502 | 880 | 628 | 876 | 959 | 1347 | 1212 | 1311 | 722 | 789 | 800 | 896 | 791 | 1152 | 1156 | 1170 | 1018 | 971,58 | 819 | 1124 | 1,377 | 0,461 | 7,039 | 1,93E-12 | 1,71E-11 | 0,002577 | 0,0034737 | 0,0030646 | 0,0030646 | TRUE | 0,0478038 | No |
| orf19.4503 | 1189 | 699 | 948 | 1189 | 967 | 720 | 864 | 581 | 1067 | 890 | 970 | 980 | 827 | 687 | 771 | 819 | 876,42 | 977 | 776 | 0,793 | -0,335 | -4,41 | 1,03E-05 | 3,76E-05 | 0,00464 | 0,0036359 | 0,0043921 | 0,0043921 | TRUE | 0,0716822 | No |
| orf19.4504 | 525 | 396 | 464 | 545 | 510 | 443 | 489 | 273 | 471 | 504 | 475 | 449 | 436 | 423 | 436 | 385 | 447,44 | 475 | 420 | 0,891 | -0,166 | -2,278 | 0,02274924 | 0,04188705 | 1,00E-08 | 0,005224 | 0,0028829 | 0,0028829 | TRUE | 0,0300548 | No |
| orf19.4505 | 124 | 98 | 104 | 145 | 162 | 138 | 129 | 99 | 111 | 125 | 106 | 120 | 139 | 132 | 115 | 140 | 123,37 | 116 | 132 | 1,135 | 0,182 | 1,422 | 0,15501251 | 0,22184734 | 1,00E-08 | 0,0137468 | 0,0076525 | 0,0076525 | TRUE | 0,0753534 | No |
| orf19.4506 | 2529 | 1865 | 2199 | 2829 | 3545 | 3177 | 3357 | 1939 | 2269 | 2374 | 2250 | 2333 | 3032 | 3031 | 2996 | 2735 | 2627,4 | 2306 | 2948 | 1,281 | 0,357 | 7,826 | 5,04E-15 | 5,80E-14 | 0,000979 | 0,002532 | 0,0016101 | 0,0016101 | TRUE | 0,0269027 | No |
| orf19.4507 | 62 | 52 | 72 | 77 | 79 | 91 | 70 | 48 | 56 | 66 | 74 | 63 | 68 | 87 | 62 | 68 | 67,94 | 65 | 71 | 1,103 | 0,142 | 0,797 | 0,42528288 | 0,51260777 | 1,00E-08 | 0,0233463 | 0,0156073 | 0,0156073 | TRUE | 0,2097217 | No |
| orf19.4508 | 35 | 18 | 36 | 36 | 31 | 43 | 29 | 26 | 31 | 23 | 37 | 30 | 27 | 41 | 26 | 37 | 31,37 | 30 | 33 | 1,059 | 0,083 | 0,315 | 0,75293048 | 0,8064907 | 0,013582 | 0,0482633 | 0,0341696 | 0,0341696 | TRUE | 0,250486 | No |
| orf19.4509 | 8 | 11 | 4 | 11 | 8 | 9 | 10 | 4 | 7 | 14 | 4 | 9 | 7 | 9 | 9 | 6 | 8,04 | 8 | 8 | 0,901 | -0,15 | -0,299 | 0,76463829 | NA | 1,00E-08 | 0,1824875 | 0,1152503 | 0,1152503 | TRUE | 0,2866406 | No |
| orf19.451 | 13359 | 10148 | 12611 | 15371 | 18265 | 17169 | 18307 | 12512 | 11984 | 12919 | 12902 | 12675 | 15621 | 16382 | 16336 | 17647 | 14558 | 12620 | 16496 | 1,307 | 0,386 | 8,505 | 1,81E-17 | 2,67E-16 | 0,001814 | 0,0020792 | 0,0019099 | 0,0019099 | TRUE | 0,026681 | No |
| orf19.4510 | 369 | 248 | 329 | 346 | 375 | 364 | 385 | 215 | 331 | 316 | 337 | 285 | 321 | 347 | 344 | 303 | 322,94 | 317 | 329 | 1,041 | 0,058 | 0,668 | 0,50415798 | 0,58874241 | 0,001726 | 0,0064748 | 0,0042618 | 0,0042618 | TRUE | 0,0580613 | No |
| orf19.4511 | 524 | 391 | 479 | 494 | 684 | 604 | 754 | 444 | 470 | 498 | 490 | 407 | 585 | 576 | 673 | 626 | 540,7 | 466 | 615 | 1,323 | 0,403 | 4,834 | 1,34E-06 | 5,55E-06 | 0,004917 | 0,0046644 | 0,0047956 | 0,0047956 | TRUE | 0,0892413 | No |
| orf19.4513 | 4931 | 3931 | 4559 | 5910 | 7353 | 7474 | 8078 | 4696 | 4424 | 5005 | 4664 | 4873 | 6289 | 7131 | 7208 | 6623 | 5777,2 | 4742 | 6813 | 1,437 | 0,523 | 9,206 | 3,38E-20 | 6,19E-19 | 0,003239 | 0,0022308 | 0,0029257 | 0,0029257 | TRUE | 0,0333384 | No |
| orf19.4515 | 213 | 151 | 190 | 216 | 202 | 168 | 207 | 114 | 191 | 192 | 194 | 178 | 173 | 160 | 185 | 161 | 179,3 | 189 | 170 | 0,903 | -0,147 | -1,401 | 0,16124025 | 0,22933982 | 1,00E-08 | 0,0100761 | 0,0050736 | 0,0050736 | TRUE | 0,0392087 | No |
| orf19.4516 | 1075 | 826 | 971 | 1182 | 1258 | 1102 | 1265 | 691 | 964 | 1052 | 993 | 975 | 1076 | 1051 | 1129 | 975 | 1026,9 | 996 | 1058 | 1,067 | 0,093 | 1,589 | 0,11196683 | 0,16779664 | 0,001302 | 0,0033933 | 0,0023139 | 0,0023139 | TRUE | 0,0297144 | No |
| orf19.4517 | 2541 | 2842 | 2567 | 3564 | 2900 | 3679 | 3368 | 2042 | 2280 | 3618 | 2626 | 2939 | 2480 | 3510 | 3005 | 2880 | 2917,4 | 2866 | 2969 | 1,036 | 0,051 | 0,295 | 0,76819244 | 0,81918711 | 0,028916 | 0,0024771 | 0,0150909 | 0,0289157 | TRUE | 0,3777081 | No |
| orf19.4518 | 977 | 724 | 869 | 1079 | 1121 | 982 | 1107 | 670 | 876 | 922 | 889 | 890 | 959 | 937 | 988 | 945 | 925,69 | 894 | 957 | 1,072 | 0,101 | 1,981 | 0,04758954 | 0,08054027 | 1,00E-08 | 0,0035478 | 0,0014064 | 0,0014064 | TRUE | 0,0057789 | No |
| orf19.4519 | 5595 | 4419 | 5172 | 6152 | 6786 | 6707 | 6934 | 4591 | 5019 | 5626 | 5291 | 5073 | 5804 | 6400 | 6187 | 6475 | 5734,4 | 5252 | 6216 | 1,184 | 0,243 | 4,785 | 1,71E-06 | 6,98E-06 | 0,002312 | 0,0022327 | 0,0023075 | 0,0023075 | TRUE | 0,0274508 | No |
| orf19.4520 | 223 | 134 | 161 | 197 | 165 | 163 | 169 | 84 | 200 | 171 | 165 | 162 | 141 | 156 | 151 | 118 | 157,97 | 174 | 142 | 0,817 | -0,292 | -2,365 | 0,01804487 | 0,03392164 | 0,004536 | 0,0111694 | 0,0083191 | 0,0083191 | TRUE | 0,1117132 | No |
| orf19.4521 | 387 | 291 | 400 | 459 | 458 | 436 | 486 | 276 | 347 | 370 | 409 | 378 | 392 | 416 | 434 | 389 | 392,01 | 376 | 408 | 1,086 | 0,119 | 1,49 | 0,13622158 | 0,19888681 | 0,00117 | 0,0056828 | 0,003554 | 0,003554 | TRUE | 0,0394468 | No |
| orf19.4522 | 884 | 569 | 687 | 915 | 962 | 757 | 809 | 596 | 793 | 724 | 703 | 755 | 823 | 722 | 722 | 841 | 760,3 | 744 | 777 | 1,041 | 0,058 | 0,81 | 0,41783501 | 0,50581357 | 0,003397 | 0,0038889 | 0,0036266 | 0,0036266 | TRUE | 0,0322322 | No |
| orf19.4523 | 258 | 209 | 210 | 296 | 322 | 292 | 304 | 203 | 231 | 266 | 215 | 244 | 275 | 279 | 271 | 286 | 258,51 | 239 | 278 | 1,164 | 0,219 | 2,382 | 0,0172132 | 0,03258251 | 1,00E-08 | 0,0075952 | 0,0042111 | 0,0042111 | TRUE | 0,0533848 | No |
| orf19.4524 | 917 | 688 | 823 | 1023 | 918 | 886 | 986 | 580 | 823 | 876 | 842 | 844 | 785 | 845 | 880 | 818 | 839,06 | 846 | 832 | 0,985 | -0,022 | -0,378 | 0,70519518 | 0,76628914 | 1,00E-08 | 0,0037097 | 0,0019672 | 0,0019672 | TRUE | 0,0194176 | No |
| orf19.4525 | 2656 | 2026 | 2378 | 2970 | 2829 | 2566 | 2690 | 1725 | 2383 | 2579 | 2433 | 2449 | 2419 | 2448 | 2400 | 2433 | 2443,2 | 2461 | 2425 | 0,986 | -0,02 | -0,504 | 0,61402669 | 0,68736209 | 6,90E-05 | 0,0025737 | 0,0010528 | 0,0010528 | TRUE | 0,0117829 | No |
| orf19.4526 | 42280 | 30913 | 37260 | 46355 | 39009 | 35560 | 36958 | 23720 | 37930 | 39356 | 38121 | 38225 | 33362 | 33930 | 32979 | 33455 | 35920 | 38408 | 33432 | 0,871 | -0,2 | -8,042 | 8,84E-16 | 1,10E-14 | 0,000172 | 0,0020199 | 0,0005664 | 0,0005664 | TRUE | 0,0033414 | No |
| orf19.4527 | 8867 | 5831 | 7727 | 9503 | 9545 | 7787 | 8528 | 5721 | 7955 | 7423 | 7905 | 7836 | 8163 | 7430 | 7610 | 8069 | 7799 | 7780 | 7818 | 1,004 | 0,006 | 0,147 | 0,88281251 | 0,91146189 | 0,001174 | 0,0021657 | 0,0016432 | 0,0016432 | TRUE | 0,0136422 | No |
| orf19.4528 | 1699 | 1217 | 1461 | 1835 | 1355 | 1300 | 1306 | 892 | 1524 | 1549 | 1495 | 1513 | 1159 | 1240 | 1165 | 1258 | 1363 | 1520 | 1206 | 0,792 | -0,337 | -7,118 | 1,10E-12 | 9,99E-12 | 1,23E-05 | 0,0030446 | 0,0014087 | 0,0014087 | TRUE | 0,0090657 | No |
| orf19.4528.1 | 4 | 1 | 4 | 4 | 3 | 2 | 2 | 4 | 4 | 1 | 4 | 3 | 3 | 2 | 2 | 6 | 3,02 | 3 | 3 | 0,898 | -0,156 | -0,197 | 0,84363531 | NA | 1,00E-08 | 0,4828203 | 0,2628845 | 0,2628845 | TRUE | 0,2981381 | No |
| orf19.4529 | 835 | 602 | 666 | 932 | 910 | 819 | 828 | 580 | 749 | 766 | 681 | 769 | 778 | 781 | 739 | 818 | 760,25 | 741 | 779 | 1,049 | 0,069 | 1,103 | 0,27002884 | 0,35437599 | 0,000973 | 0,003889 | 0,0023901 | 0,0023901 | TRUE | 0,0346441 | No |
| orf19.4530.1 | 6131 | 4130 | 5455 | 6575 | 8342 | 6990 | 7902 | 5093 | 5500 | 5258 | 5581 | 5422 | 7134 | 6670 | 7051 | 7183 | 6224,9 | 5440 | 7010 | 1,288 | 0,365 | 9,738 | 2,08E-22 | 4,49E-21 | 0,000656 | 0,0022127 | 0,0011843 | 0,0011843 | TRUE | 0,0130645 | No |
| orf19.4531 | 29378 | 20761 | 26603 | 31437 | 26415 | 24471 | 26708 | 15803 | 26355 | 26431 | 27217 | 25923 | 22591 | 23349 | 23832 | 22289 | 24749 | 26482 | 23015 | 0,869 | -0,202 | -6,125 | 9,09E-10 | 5,83E-09 | 0,000621 | 0,0020382 | 0,0010053 | 0,0010053 | TRUE | 0,0069979 | No |
| orf19.4532 | 3025 | 2113 | 2642 | 3343 | 2466 | 2173 | 2272 | 1497 | 2714 | 2690 | 2703 | 2757 | 2109 | 2073 | 2027 | 2111 | 2398,1 | 2716 | 2080 | 0,765 | -0,386 | -10,599 | 3,02E-26 | 8,82E-25 | 1,00E-08 | 0,0025849 | 0,0008481 | 0,0008481 | TRUE | 0,0036646 | No |
| orf19.4533 | 1096 | 692 | 985 | 1133 | 961 | 853 | 956 | 644 | 983 | 881 | 1008 | 934 | 822 | 814 | 853 | 908 | 900,43 | 952 | 849 | 0,889 | -0,17 | -2,746 | 0,00603342 | 0,01280877 | 0,001523 | 0,0035918 | 0,0025414 | 0,0025414 | TRUE | 0,0268775 | No |
| orf19.4534 | 9675 | 6477 | 8652 | 10535 | 11102 | 9086 | 10092 | 6261 | 8679 | 8246 | 8852 | 8687 | 9495 | 8670 | 9005 | 8831 | 8808,1 | 8616 | 9000 | 1,045 | 0,063 | 1,561 | 0,11848914 | 0,17638323 | 0,001089 | 0,0021443 | 0,0014452 | 0,0014452 | TRUE | 0,0169247 | No |
| orf19.4535 | 358 | 251 | 292 | 353 | 520 | 436 | 475 | 270 | 321 | 320 | 299 | 291 | 445 | 416 | 424 | 381 | 361,99 | 308 | 416 | 1,362 | 0,445 | 5,513 | 3,54E-08 | 1,83E-07 | 1,00E-08 | 0,0059898 | 0,0034507 | 0,0034507 | TRUE | 0,0320728 | No |
| orf19.4536 | 9140 | 6153 | 7748 | 9567 | 7762 | 6685 | 7329 | 4656 | 8200 | 7833 | 7927 | 7889 | 6638 | 6379 | 6540 | 6567 | 7246,6 | 7962 | 6531 | 0,82 | -0,286 | -9,321 | 1,15E-20 | 2,18E-19 | 0,00023 | 0,0021799 | 0,0007651 | 0,0007651 | TRUE | 0,0049713 | No |
| orf19.4537 | 3472 | 2155 | 2875 | 3487 | 3559 | 2940 | 3216 | 2012 | 3115 | 2744 | 2941 | 2875 | 3044 | 2805 | 2870 | 2838 | 2904 | 2919 | 2889 | 0,99 | -0,015 | -0,305 | 0,76006674 | 0,81239654 | 0,001714 | 0,0024794 | 0,0020021 | 0,0020021 | TRUE | 0,0249059 | No |
| orf19.4538 | 3793 | 2475 | 3080 | 3889 | 3732 | 2967 | 3206 | 1869 | 3403 | 3151 | 3151 | 3207 | 3192 | 2831 | 2861 | 2636 | 3053,9 | 3228 | 2880 | 0,893 | -0,163 | -2,755 | 0,00587451 | 0,01250231 | 0,003451 | 0,0024549 | 0,0030401 | 0,0030401 | TRUE | 0,0648921 | No |
| orf19.4539 | 1767 | 1130 | 1467 | 1796 | 1829 | 1469 | 1699 | 959 | 1585 | 1439 | 1501 | 1481 | 1564 | 1402 | 1516 | 1353 | 1480 | 1502 | 1459 | 0,973 | -0,04 | -0,678 | 0,49755754 | 0,58226056 | 0,002181 | 0,0029604 | 0,0026075 | 0,0026075 | TRUE | 0,0285094 | No |
| orf19.454 | 6740 | 4553 | 5989 | 7359 | 7286 | 6316 | 7187 | 4852 | 6046 | 5796 | 6127 | 6068 | 6231 | 6027 | 6413 | 6843 | 6194,1 | 6009 | 6378 | 1,06 | 0,085 | 1,88 | 0,06010469 | 0,09866366 | 0,001567 | 0,0022139 | 0,001782 | 0,001782 | TRUE | 0,0284516 | No |
| orf19.4540 | 6192 | 4375 | 5448 | 6856 | 6306 | 5452 | 5987 | 3765 | 5555 | 5570 | 5574 | 5654 | 5393 | 5202 | 5342 | 5310 | 5450 | 5588 | 5312 | 0,951 | -0,073 | -2,518 | 0,01179229 | 0,02332064 | 1,00E-08 | 0,0022459 | 0,0006249 | 0,0006249 | TRUE | 0,0024507 | No |
| orf19.4542 | 956 | 731 | 929 | 1094 | 894 | 859 | 923 | 568 | 858 | 931 | 950 | 902 | 765 | 820 | 824 | 801 | 856,23 | 910 | 802 | 0,882 | -0,181 | -3,145 | 0,00166261 | 0,00398802 | 1,00E-08 | 0,003675 | 0,001996 | 0,001996 | TRUE | 0,0184072 | No |
| orf19.4543 | 1753 | 1268 | 1617 | 1889 | 2325 | 1920 | 2186 | 1403 | 1573 | 1614 | 1654 | 1558 | 1988 | 1832 | 1951 | 1979 | 1768,6 | 1600 | 1938 | 1,212 | 0,277 | 6,031 | 1,63E-09 | 1,00E-08 | 0,000579 | 0,0028003 | 0,0014511 | 0,0014511 | TRUE | 0,0164905 | No |
| orf19.4544 | 1062 | 728 | 837 | 1109 | 1125 | 1002 | 1010 | 639 | 953 | 927 | 856 | 914 | 962 | 956 | 901 | 901 | 921,39 | 912 | 930 | 1,02 | 0,029 | 0,515 | 0,60638354 | 0,68160006 | 0,000589 | 0,0035551 | 0,0019601 | 0,0019601 | TRUE | 0,0203364 | No |
| orf19.4545 | 2245 | 1522 | 2015 | 2438 | 2386 | 2102 | 2341 | 1504 | 2014 | 1938 | 2062 | 2010 | 2041 | 2006 | 2089 | 2121 | 2035 | 2006 | 2064 | 1,027 | 0,039 | 0,957 | 0,33881295 | 0,42600097 | 1,00E-08 | 0,0026929 | 0,0011019 | 0,0011019 | TRUE | 0,0057435 | No |
| orf19.4546 | 8182 | 5603 | 7324 | 9040 | 6584 | 5836 | 6460 | 3871 | 7340 | 7133 | 7493 | 7454 | 5631 | 5569 | 5764 | 5460 | 6480,6 | 7355 | 5606 | 0,762 | -0,391 | -11,932 | 8,03E-33 | 3,45E-31 | 0,000289 | 0,0022035 | 0,0008754 | 0,0008754 | TRUE | 0,0048504 | No |
| orf19.4548 | 291 | 203 | 245 | 289 | 321 | 293 | 339 | 206 | 261 | 258 | 251 | 238 | 275 | 280 | 303 | 291 | 269,45 | 252 | 287 | 1,139 | 0,187 | 2,15 | 0,03155222 | 0,05611775 | 1,00E-08 | 0,0073671 | 0,0035604 | 0,0035604 | TRUE | 0,0175675 | No |
| orf19.4549 | 413 | 267 | 331 | 426 | 525 | 461 | 513 | 310 | 371 | 340 | 339 | 351 | 449 | 440 | 458 | 437 | 398,03 | 350 | 446 | 1,273 | 0,348 | 4,803 | 1,57E-06 | 6,41E-06 | 1,00E-08 | 0,0056268 | 0,0025111 | 0,0025111 | TRUE | 0,0180135 | No |

| | | | | | | | | | | | | | | | | | | | | | | | | | | | | | | | |
|---|---|---|---|---|---|---|---|---|---|---|---|---|---|---|---|---|---|---|---|---|---|---|---|---|---|---|---|---|---|---|---|
| orf19.4550 | 924 | 689 | 871 | 1067 | 1022 | 926 | 974 | 634 | 829 | 877 | 891 | 880 | 874 | 884 | 869 | 894 | 874,75 | 869 | 880 | 1,012 | 0,018 | 0,338 | 0,7354504 | 0,79342539 | 1,00E-08 | 0,0036391 | 0,0014962 | 0,0014962 | TRUE | 0,0122274 | No |
| orf19.4551 | 2876 | 2272 | 2712 | 3322 | 2484 | 2417 | 2567 | 1702 | 2580 | 2892 | 2775 | 2739 | 2124 | 2306 | 2291 | 2401 | 2513,5 | 2746 | 2280 | 0,83 | -0,269 | -5,172 | 2,32E-07 | 1,08E-06 | 0,001834 | 0,0025571 | 0,0021944 | 0,0021944 | TRUE | 0,0261048 | No |
| orf19.4553 | 4970 | 3370 | 4651 | 5524 | 6165 | 5595 | 6243 | 4119 | 4459 | 4290 | 4758 | 4555 | 5273 | 5339 | 5571 | 5809 | 5006,8 | 4516 | 5498 | 1,216 | 0,283 | 6,074 | 1,24E-09 | 7,82E-09 | 0,001652 | 0,0022695 | 0,0018752 | 0,0018752 | TRUE | 0,0171745 | No |
| orf19.4555 | 5768 | 4005 | 5540 | 6515 | 4196 | 3375 | 3868 | 2814 | 5174 | 5099 | 5668 | 5372 | 3589 | 3220 | 3452 | 3969 | 4442,9 | 5328 | 3558 | 0,667 | -0,584 | -9,204 | 3,45E-20 | 6,29E-19 | 0,004711 | 0,0023063 | 0,0036361 | 0,0036361 | TRUE | 0,0716688 | No |
| orf19.4557 | 381 | 302 | 329 | 447 | 576 | 539 | 535 | 320 | 342 | 384 | 337 | 369 | 493 | 514 | 477 | 451 | 420,89 | 358 | 484 | 1,358 | 0,442 | 5,758 | 8,53E-09 | 4,78E-08 | 0,000515 | 0,0054286 | 0,003231 | 0,003231 | TRUE | 0,0243468 | No |
| orf19.4560 | 5689 | 3845 | 4903 | 6007 | 4915 | 4294 | 4683 | 2990 | 5104 | 4895 | 5016 | 4953 | 4203 | 4097 | 4179 | 4217 | 4583,1 | 4992 | 4174 | 0,836 | -0,259 | -8,271 | 1,33E-16 | 1,78E-15 | 1,00E-08 | 0,0022963 | 0,0007202 | 0,0007202 | TRUE | 0,0027826 | No |
| orf19.4563 | 6061 | 4177 | 5022 | 6341 | 5851 | 4982 | 5502 | 3469 | 5437 | 5318 | 5138 | 5229 | 5004 | 4754 | 4910 | 4893 | 5085,2 | 5280 | 4890 | 0,926 | -0,111 | -3,232 | 0,00123037 | 0,003047 | 0,000347 | 0,002265 | 0,0009284 | 0,0009284 | TRUE | 0,0050248 | No |
| orf19.4565 | 6467 | 4263 | 5499 | 6780 | 5431 | 4383 | 4582 | 3334 | 5802 | 5427 | 5626 | 5591 | 4645 | 4182 | 4089 | 4702 | 5008 | 5612 | 4404 | 0,784 | -0,351 | -6,588 | 4,46E-11 | 3,42E-10 | 0,002644 | 0,0022694 | 0,002515 | 0,002515 | TRUE | 0,0284909 | No |
| orf19.4566 | 21 | 18 | 33 | 30 | 27 | 23 | 26 | 13 | 19 | 23 | 34 | 25 | 23 | 22 | 23 | 18 | 23,35 | 25 | 22 | 0,874 | -0,195 | -0,671 | 0,50240382 | 0,58724555 | 1,00E-08 | 0,0641413 | 0,0382343 | 0,0382343 | TRUE | 0,3271624 | No |
| orf19.4567 | 60 | 32 | 44 | 50 | 41 | 41 | 36 | 23 | 54 | 41 | 45 | 41 | 35 | 39 | 32 | 32 | 39,95 | 45 | 34 | 0,767 | -0,383 | -1,754 | 0,07935007 | 0,12565505 | 1,00E-08 | 0,0383219 | 0,0204128 | 0,0204128 | TRUE | 0,1407258 | No |
| orf19.4568 | 239 | 176 | 185 | 263 | 274 | 233 | 261 | 164 | 214 | 224 | 189 | 217 | 234 | 222 | 233 | 231 | 220,69 | 211 | 230 | 1,091 | 0,126 | 1,314 | 0,18883865 | 0,26171686 | 1,00E-08 | 0,0085577 | 0,0042838 | 0,0042838 | TRUE | 0,0510819 | No |
| orf19.4569 | 58 | 36 | 53 | 66 | 66 | 48 | 54 | 36 | 52 | 46 | 54 | 54 | 56 | 46 | 48 | 51 | 50,96 | 52 | 50 | 0,97 | -0,043 | -0,229 | 0,81858222 | 0,85994789 | 1,00E-08 | 0,0304639 | 0,0150511 | 0,0150511 | TRUE | 0,0663375 | No |
| orf19.457 | 2593 | 1942 | 2440 | 2768 | 3486 | 3493 | 3749 | 2089 | 2326 | 2472 | 2496 | 2283 | 2981 | 3333 | 3345 | 2946 | 2772,9 | 2394 | 3151 | 1,318 | 0,398 | 6,916 | 4,63E-12 | 3,93E-11 | 0,002863 | 0,002503 | 0,0028163 | 0,0028163 | TRUE | 0,0222844 | No |
| orf19.4570 | 143 | 84 | 106 | 143 | 140 | 129 | 135 | 91 | 128 | 107 | 108 | 118 | 120 | 123 | 120 | 128 | 119,15 | 115 | 123 | 1,057 | 0,08 | 0,631 | 0,52818288 | 0,61119468 | 1,00E-08 | 0,014163 | 0,0070643 | 0,0070643 | TRUE | 0,0575292 | No |
| orf19.4571 | 70 | 60 | 73 | 73 | 86 | 65 | 82 | 42 | 63 | 76 | 75 | 60 | 74 | 62 | 73 | 59 | 67,76 | 68 | 67 | 0,994 | -0,009 | -0,055 | 0,95652245 | 0,96910403 | 1,00E-08 | 0,023405 | 0,0137938 | 0,0137938 | TRUE | 0,0650144 | No |
| orf19.4573 | 232 | 149 | 203 | 225 | 239 | 168 | 220 | 165 | 208 | 190 | 208 | 186 | 204 | 160 | 196 | 233 | 198,1 | 198 | 198 | 0,997 | -0,005 | -0,039 | 0,96898853 | 0,97799346 | 0,00746 | 0,0093077 | 0,0084904 | 0,0084904 | TRUE | 0,1797402 | No |
| orf19.4574 | 1194 | 836 | 1072 | 1304 | 1416 | 1270 | 1372 | 857 | 1071 | 1064 | 1097 | 1075 | 1211 | 1212 | 1224 | 1209 | 1145,4 | 1077 | 1214 | 1,127 | 0,173 | 3,814 | 0,00013662 | 0,00040756 | 1,00E-08 | 0,0032469 | 0,0010989 | 0,0010989 | TRUE | 0,0017713 | No |
| orf19.4575 | 983 | 701 | 943 | 1110 | 999 | 952 | 1030 | 683 | 882 | 892 | 965 | 915 | 854 | 908 | 919 | 963 | 912,45 | 914 | 911 | 0,995 | -0,007 | -0,128 | 0,89798521 | 0,92271167 | 0,000861 | 0,0035705 | 0,0021181 | 0,0021181 | TRUE | 0,0215126 | No |
| orf19.4577 | 313 | 226 | 310 | 326 | 381 | 326 | 342 | 195 | 281 | 288 | 317 | 269 | 326 | 311 | 305 | 275 | 296,45 | 289 | 304 | 1,061 | 0,086 | 0,956 | 0,3391679 | 0,42630141 | 0,00135 | 0,0068765 | 0,0044101 | 0,0044101 | TRUE | 0,050869 | No |
| orf19.4577.3 | 1469 | 1039 | 1131 | 1615 | 1602 | 1355 | 1379 | 922 | 1318 | 1323 | 1157 | 1332 | 1370 | 1293 | 1231 | 1300 | 1290,4 | 1282 | 1298 | 1,012 | 0,018 | 0,295 | 0,76817721 | 0,81918711 | 0,00245 | 0,0031045 | 0,0027305 | 0,0027305 | TRUE | 0,0516973 | No |
| orf19.4578 | 1472 | 993 | 1229 | 1568 | 1655 | 1451 | 1495 | 887 | 1321 | 1264 | 1257 | 1293 | 1415 | 1384 | 1334 | 1251 | 1315 | 1284 | 1346 | 1,051 | 0,071 | 1,381 | 0,16719167 | 0,23676361 | 0,000662 | 0,0030835 | 0,0017894 | 0,0017894 | TRUE | 0,0241951 | No |
| orf19.4579 | 365 | 268 | 319 | 419 | 476 | 417 | 452 | 290 | 327 | 341 | 326 | 346 | 407 | 398 | 403 | 409 | 369,73 | 335 | 404 | 1,206 | 0,27 | 3,682 | 0,00023111 | 0,00066436 | 1,00E-08 | 0,0059059 | 0,0024514 | 0,0024514 | TRUE | 0,0059252 | No |
| orf19.458 | 1825 | 1217 | 1558 | 1908 | 1700 | 1432 | 1565 | 970 | 1637 | 1549 | 1594 | 1573 | 1454 | 1366 | 1397 | 1368 | 1492,4 | 1588 | 1396 | 0,879 | -0,185 | -4,125 | 3,71E-05 | 0,00012264 | 1,00E-08 | 0,0029523 | 0,0012663 | 0,0012663 | TRUE | 0,0097179 | No |
| orf19.4580 | 1251 | 870 | 991 | 1190 | 1319 | 1236 | 1238 | 705 | 1122 | 1108 | 1014 | 981 | 1128 | 1179 | 1105 | 994 | 1078,9 | 1056 | 1102 | 1,046 | 0,065 | 0,962 | 0,33605061 | 0,42291414 | 0,003424 | 0,003325 | 0,003494 | 0,003494 | TRUE | 0,0469021 | No |
| orf19.4581 | 194 | 160 | 182 | 224 | 194 | 202 | 205 | 118 | 174 | 204 | 186 | 185 | 166 | 193 | 183 | 166 | 182,08 | 187 | 177 | 0,952 | -0,072 | -0,669 | 0,50362742 | 0,5883324 | 1,00E-08 | 0,0099523 | 0,0055218 | 0,0055218 | TRUE | 0,0381824 | No |
| orf19.458.1 | 264 | 175 | 186 | 270 | 294 | 201 | 233 | 141 | 237 | 223 | 190 | 223 | 251 | 192 | 208 | 199 | 215,32 | 218 | 212 | 0,977 | -0,033 | -0,288 | 0,7730684 | 0,82263459 | 0,007727 | 0,0087215 | 0,0082479 | 0,0082479 | TRUE | 0,1775365 | No |
| orf19.4582 | 1887 | 1357 | 1639 | 2012 | 1948 | 1685 | 1812 | 1136 | 1693 | 1728 | 1677 | 1659 | 1666 | 1608 | 1617 | 1602 | 1656,2 | 1689 | 1623 | 0,962 | -0,055 | -1,344 | 0,17883029 | 0,25056259 | 1,00E-08 | 0,0028561 | 0,0010142 | 0,0010142 | TRUE | 0,0039781 | No |
| orf19.4583 | 616 | 346 | 492 | 625 | 167 | 159 | 161 | 98 | 553 | 440 | 503 | 515 | 143 | 152 | 144 | 138 | 323,54 | 503 | 144 | 0,286 | -1,805 | -18,172 | 8,53E-74 | 2,05E-71 | 1,00E-08 | 0,0064665 | 0,0050209 | 0,0050209 | TRUE | 0,0745945 | No |
| orf19.4584 | 408 | 276 | 405 | 493 | 383 | 359 | 357 | 182 | 366 | 351 | 414 | 407 | 328 | 343 | 319 | 257 | 347,96 | 384 | 312 | 0,813 | -0,298 | -3,041 | 0,00236097 | 0,00548092 | 0,006649 | 0,0061516 | 0,0063167 | 0,0063167 | TRUE | 0,1336367 | No |
| orf19.4585 | 10431 | 6696 | 8906 | 11078 | 10863 | 9315 | 10334 | 6040 | 9358 | 8525 | 9112 | 9135 | 9290 | 8888 | 9221 | 8519 | 9006 | 9032 | 8980 | 0,994 | -0,008 | -0,192 | 0,84778859 | 0,8841371 | 0,001229 | 0,0021407 | 0,0016665 | 0,0016665 | TRUE | 0,0171889 | No |
| orf19.4587 | 4926 | 3562 | 4278 | 5388 | 4524 | 4038 | 4209 | 2528 | 4419 | 4535 | 4377 | 4443 | 3869 | 3853 | 3756 | 3566 | 4102,1 | 4444 | 3761 | 0,848 | -0,239 | -6,381 | 1,76E-10 | 1,23E-09 | 0,000473 | 0,0023334 | 0,0010948 | 0,0010948 | TRUE | 0,0137609 | No |
| orf19.4589 | 1337 | 861 | 1050 | 1408 | 1129 | 1007 | 1102 | 652 | 1199 | 1096 | 1074 | 1161 | 966 | 961 | 983 | 920 | 1045 | 1132 | 958 | 0,845 | -0,243 | -4,413 | 1,02E-05 | 3,71E-05 | 0,000798 | 0,0033687 | 0,0019375 | 0,0019375 | TRUE | 0,0188047 | No |
| orf19.459 | 2911 | 2078 | 2683 | 3176 | 2550 | 2306 | 2492 | 1436 | 2611 | 2646 | 2745 | 2619 | 2181 | 2200 | 2224 | 2025 | 2406,4 | 2655 | 2158 | 0,814 | -0,296 | -6,911 | 4,81E-12 | 4,06E-11 | 0,000576 | 0,0025828 | 0,0013452 | 0,0013452 | TRUE | 0,0185672 | No |
| orf19.4590 | 424 | 332 | 403 | 453 | 418 | 424 | 455 | 327 | 380 | 423 | 412 | 374 | 357 | 405 | 406 | 461 | 402,27 | 397 | 407 | 1,022 | 0,032 | 0,362 | 0,71698471 | 0,77727249 | 0,004424 | 0,0055883 | 0,0050232 | 0,0050232 | TRUE | 0,0805529 | No |
| orf19.4591 | 3732 | 2611 | 3324 | 4224 | 2955 | 2653 | 2862 | 1929 | 3348 | 3324 | 3401 | 3483 | 2527 | 2531 | 2554 | 2721 | 2986,2 | 3389 | 2583 | 0,761 | -0,395 | -10,034 | 1,08E-23 | 2,51E-22 | 0,0004 | 0,0024657 | 0,0011418 | 0,0011418 | TRUE | 0,0141449 | No |
| orf19.4592 | 2204 | 1464 | 1929 | 2298 | 2153 | 2011 | 2093 | 1331 | 1977 | 1864 | 1974 | 1895 | 1841 | 1919 | 1868 | 1877 | 1901,8 | 1928 | 1876 | 0,973 | -0,04 | -0,965 | 0,33468109 | 0,42154381 | 1,00E-08 | 0,0027428 | 0,0011169 | 0,0011169 | TRUE | 0,0053577 | No |
| orf19.4593 | 7011 | 4639 | 6298 | 7543 | 10418 | 9199 | 10176 | 6298 | 6290 | 5906 | 6443 | 6220 | 8910 | 8777 | 9080 | 8883 | 7563,7 | 6215 | 8912 | 1,434 | 0,52 | 14,733 | 3,95E-49 | 3,89E-47 | 0,000579 | 0,0021715 | 0,0010572 | 0,0010572 | TRUE | 0,0130979 | No |
| orf19.4593.1 | 405 | 260 | 304 | 399 | 334 | 306 | 310 | 193 | 363 | 331 | 311 | 329 | 286 | 292 | 277 | 272 | 307,6 | 334 | 282 | 0,845 | -0,244 | -2,886 | 0,00389565 | 0,00862066 | 1,00E-08 | 0,0066989 | 0,0035806 | 0,0035806 | TRUE | 0,0432837 | No |
| orf19.4594 | 982 | 701 | 809 | 983 | 929 | 818 | 924 | 514 | 881 | 892 | 828 | 811 | 795 | 781 | 825 | 725 | 817,02 | 853 | 782 | 0,92 | -0,121 | -1,967 | 0,0492288 | 0,08312737 | 0,001109 | 0,0037563 | 0,0023781 | 0,0023781 | TRUE | 0,0240754 | No |
| orf19.4595 | 1693 | 1460 | 1604 | 1921 | 2087 | 2132 | 2130 | 1564 | 1519 | 1859 | 1641 | 1584 | 1785 | 2034 | 1901 | 2206 | 1816,1 | 1651 | 1982 | 1,2 | 0,263 | 3,379 | 0,0007266 | 0,00188489 | 0,007423 | 0,0027789 | 0,0052722 | 0,0052722 | TRUE | 0,0846065 | No |
| orf19.4596 | 1464 | 1001 | 1326 | 1653 | 1822 | 1512 | 1664 | 1097 | 1313 | 1274 | 1357 | 1363 | 1558 | 1443 | 1485 | 1547 | 1417,6 | 1327 | 1508 | 1,135 | 0,182 | 3,788 | 0,00015176 | 0,00044982 | 0,000432 | 0,0030036 | 0,0015158 | 0,0015158 | TRUE | 0,0103601 | No |
| orf19.4597 | 2022 | 1330 | 1676 | 2004 | 2418 | 1982 | 2207 | 1448 | 1814 | 1693 | 1715 | 1653 | 2068 | 1891 | 1969 | 2042 | 1855,7 | 1719 | 1992 | 1,159 | 0,213 | 4,335 | 1,46E-05 | 5,18E-05 | 0,001132 | 0,0027618 | 0,0017669 | 0,0017669 | TRUE | 0,0174972 | No |
| orf19.4599 | 258 | 168 | 227 | 288 | 238 | 242 | 234 | 179 | 231 | 214 | 232 | 237 | 204 | 231 | 209 | 252 | 226,35 | 228 | 224 | 0,969 | -0,046 | -0,456 | 0,64844371 | 0,71786435 | 0,000815 | 0,0083931 | 0,0051839 | 0,0051839 | TRUE | 0,0673993 | No |
| orf19.460 | 1678 | 1172 | 1512 | 1934 | 1593 | 1358 | 1528 | 1067 | 1505 | 1492 | 1547 | 1595 | 1362 | 1296 | 1363 | 1505 | 1458,2 | 1535 | 1382 | 0,897 | -0,156 | -2,878 | 0,00400479 | 0,00883289 | 0,001571 | 0,0029751 | 0,0021415 | 0,0021415 | TRUE | 0,0392758 | No |
| orf19.4600 | 2169 | 1278 | 1629 | 2095 | 2641 | 2135 | 2400 | 1407 | 1946 | 1627 | 1667 | 1728 | 2259 | 2037 | 2142 | 1984 | 1923,6 | 1742 | 2106 | 1,209 | 0,274 | 4,155 | 3,26E-05 | 0,00010901 | 0,00429 | 0,0027342 | 0,0036368 | 0,0036368 | TRUE | 0,0756462 | No |
| orf19.4600.1 | 588 | 426 | 474 | 604 | 662 | 530 | 533 | 372 | 527 | 542 | 485 | 498 | 566 | 506 | 476 | 525 | 515,63 | 513 | 518 | 1,011 | 0,015 | 0,204 | 0,83823558 | 0,87645655 | 0,002301 | 0,0047949 | 0,0035175 | 0,0035175 | TRUE | 0,0507654 | No |
| orf19.4601 | 503 | 342 | 430 | 517 | 552 | 480 | 558 | 366 | 451 | 435 | 440 | 426 | 472 | 458 | 498 | 516 | 462,14 | 438 | 486 | 1,106 | 0,145 | 2,078 | 0,03766511 | 0,06557288 | 1,00E-08 | 0,0051208 | 0,0025258 | 0,0025258 | TRUE | 0,0168625 | No |
| orf19.4602 | 2065 | 1266 | 1786 | 2240 | 2671 | 2047 | 2192 | 1783 | 1853 | 1612 | 1827 | 1847 | 2284 | 1953 | 1956 | 2515 | 1980,9 | 1785 | 2177 | 1,218 | 0,284 | 3,371 | 0,00074839 | 0,00193475 | 0,009321 | 0,0027124 | 0,006301 | 0,006301 | TRUE | 0,1282926 | No |
| orf19.4603 | 2348 | 1697 | 2099 | 2528 | 2547 | 2344 | 2541 | 1482 | 2106 | 2160 | 2147 | 2085 | 2178 | 2237 | 2267 | 2090 | 2158,9 | 2124 | 2193 | 1,034 | 0,049 | 1,181 | 0,23759267 | 0,31848656 | 0,000151 | 0,0026519 | 0,0011709 | 0,0011709 | TRUE | 0,0107725 | No |
| orf19.4605 | 1362 | 868 | 1124 | 1419 | 1126 | 1013 | 1121 | 708 | 1222 | 1105 | 1150 | 1170 | 963 | 967 | 1000 | 999 | 1071,9 | 1162 | 982 | 0,843 | -0,246 | -4,781 | 1,75E-06 | 7,10E-06 | 1,00E-08 | 0,0033338 | 0,0015986 | 0,0015986 | TRUE | 0,0144314 | No |
| orf19.4606 | 1083 | 693 | 948 | 1153 | 1023 | 942 | 1024 | 623 | 972 | 882 | 970 | 951 | 875 | 899 | 914 | 879 | 917,58 | 944 | 892 | 0,944 | -0,084 | -1,558 | 0,11913752 | 0,17712899 | 1,00E-08 | 0,0035616 | 0,0016936 | 0,0016936 | TRUE | 0,0203269 | No |
| orf19.4607 | 294 | 225 | 301 | 371 | 263 | 251 | 303 | 133 | 264 | 286 | 308 | 306 | 225 | 239 | 270 | 188 | 260,81 | 291 | 230 | 0,798 | -0,325 | -2,934 | 0,00334683 | 0,00751095 | 0,008432 | 0,0075457 | 0,0079229 | 0,0079229 | TRUE | 0,1462731 | No |
| orf19.4608 | 280 | 241 | 267 | 314 | 244 | 240 | 262 | 158 | 251 | 307 | 273 | 259 | 209 | 229 | 234 | 223 | 248,05 | 272 | 224 | 0,824 | -0,279 | -2,877 | 0,00401051 | 0,0088419 | 1,00E-08 | 0,0078319 | 0,0049254 | 0,0049254 | TRUE | 0,071743 | No |
| orf19.4609 | 1568 | 1008 | 1288 | 1568 | 1641 | 1387 | 1465 | 1077 | 1407 | 1283 | 1318 | 1293 | 1403 | 1323 | 1307 | 1519 | 1356,7 | 1325 | 1388 | 1,045 | 0,063 | 1,057 | 0,29061463 | 0,37516388 | 0,002346 | 0,0030495 | 0,0026746 | 0,0026746 | TRUE | 0,0444232 | No |
| orf19.4610 | 287 | 210 | 251 | 319 | 344 | 334 | 341 | 183 | 257 | 267 | 257 | 263 | 294 | 319 | 304 | 258 | 277,49 | 261 | 294 | 1,132 | 0,179 | 2,006 | 0,0448785 | 0,07653421 | 1,00E-08 | 0,007211 | 0,0040623 | 0,0040623 | TRUE | 0,0608233 | No |
| orf19.4611 | 4588 | 2981 | 3733 | 4741 | 4824 | 3868 | 4228 | 2772 | 4116 | 3795 | 3819 | 3909 | 4126 | 3691 | 3773 | 3910 | 3892,3 | 3910 | 3875 | 0,991 | -0,014 | -0,287 | 0,77441703 | 0,82363206 | 0,001658 | 0,0023525 | 0,0019063 | 0,0019063 | TRUE | 0,023943 | No |
| orf19.4612 | 59 | 45 | 59 | 81 | 66 | 47 | 65 | 39 | 53 | 57 | 60 | 67 | 56 | 45 | 58 | 55 | 56,46 | 59 | 54 | 0,901 | -0,151 | -0,816 | 0,41445195 | 0,50252821 | 1,00E-08 | 0,0276919 | 0,0152605 | 0,0152605 | TRUE | 0,1071075 | No |
| orf19.4614 | 1189 | 927 | 1095 | 1311 | 1481 | 1438 | 1441 | 862 | 1067 | 1180 | 1120 | 1081 | 1267 | 1372 | 1286 | 1216 | 1198,6 | 1112 | 1285 | 1,159 | 0,213 | 3,819 | 0,0001338 | 0,00040012 | 0,001207 | 0,0031907 | 0,0021544 | 0,0021544 | TRUE | 0,0245833 | No |
| orf19.4615 | 2103 | 1627 | 1933 | 2248 | 2359 | 2136 | 2192 | 1213 | 1887 | 2071 | 1978 | 1854 | 2017 | 2038 | 1956 | 1711 | 1939 | 1948 | 1930 | 0,994 | -0,008 | -0,132 | 0,8946703 | 0,92047683 | 0,003692 | 0,0027282 | 0,0032755 | 0,0032755 | TRUE | 0,066871 | No |

| | | | | | | | | | | | | | | | | | | | | | | | | | | | | | | | |
|---|---|---|---|---|---|---|---|---|---|---|---|---|---|---|---|---|---|---|---|---|---|---|---|---|---|---|---|---|---|---|---|
| orf19.4616 | 700 | 523 | 636 | 706 | 489 | 443 | 543 | 353 | 628 | 666 | 651 | 582 | 418 | 423 | 485 | 498 | 543,75 | 632 | 456 | 0,72 | -0,473 | -5,961 | 2,51E-09 | 1,51E-08 | 0,003643 | 0,0046493 | 0,0041463 | 0,0041463 | TRUE | 0,0387977 | No |
| orf19.4617 | 1089 | 862 | 945 | 1192 | 1383 | 1431 | 1331 | 836 | 977 | 1097 | 967 | 983 | 1183 | 1365 | 1188 | 1179 | 1117,4 | 1006 | 1229 | 1,224 | 0,292 | 4,358 | 1,31E-05 | 4,68E-05 | 0,003403 | 0,0032787 | 0,0033939 | 0,0033939 | TRUE | 0,0680964 | No |
| orf19.4618 | 7206 | 4759 | 6531 | 7569 | 10564 | 8608 | 9382 | 6692 | 6465 | 6059 | 6682 | 6241 | 9035 | 8213 | 8372 | 9438 | 7563,1 | 6362 | 8764 | 1,377 | 0,462 | 8,623 | 6,49E-18 | 1,00E-16 | 0,002859 | 0,0021715 | 0,0026155 | 0,0026155 | TRUE | 0,0321742 | No |
| orf19.4620 | 418 | 323 | 347 | 472 | 388 | 396 | 361 | 236 | 375 | 411 | 355 | 389 | 332 | 378 | 322 | 333 | 361,89 | 382 | 341 | 0,893 | -0,163 | -1,941 | 0,05221035 | 0,08727873 | 0,00164 | 0,005991 | 0,0039694 | 0,0039694 | TRUE | 0,0620272 | No |
| orf19.4621 | 1160 | 949 | 1060 | 1289 | 998 | 1071 | 1093 | 670 | 1041 | 1208 | 1084 | 1063 | 854 | 1022 | 975 | 945 | 1024 | 1099 | 949 | 0,865 | -0,21 | -2,967 | 0,00300364 | 0,00682006 | 0,003817 | 0,0033972 | 0,0038139 | 0,0038139 | TRUE | 0,0579405 | No |
| orf19.4622 | 8309 | 5555 | 7271 | 8774 | 8890 | 7445 | 8149 | 5254 | 7454 | 7072 | 7439 | 7235 | 7603 | 7104 | 7272 | 7410 | 7323,6 | 7300 | 7347 | 1,006 | 0,009 | 0,251 | 0,80147733 | 0,84555346 | 0,000495 | 0,0021777 | 0,0010654 | 0,0010654 | TRUE | 0,0069448 | No |
| orf19.4623 | 3115 | 1923 | 2798 | 3530 | 3540 | 2850 | 3449 | 2585 | 2794 | 2448 | 2863 | 2911 | 3028 | 2719 | 3078 | 3646 | 2935,8 | 2754 | 3118 | 1,13 | 0,177 | 2,089 | 0,03671381 | 0,06413975 | 0,010107 | 0,002474 | 0,0065346 | 0,0065346 | TRUE | 0,1551256 | No |
| orf19.4623.3 | 9289 | 6393 | 7490 | 9045 | 8847 | 8470 | 8249 | 4941 | 8333 | 8139 | 7663 | 7459 | 7566 | 8082 | 7361 | 6969 | 7696,4 | 7898 | 7494 | 0,949 | -0,075 | -1,373 | 0,16964188 | 0,23964458 | 0,003063 | 0,0021681 | 0,0027368 | 0,0027368 | TRUE | 0,0338645 | No |
| orf19.4624 | 326 | 266 | 305 | 387 | 422 | 378 | 351 | 207 | 292 | 339 | 312 | 319 | 361 | 361 | 313 | 292 | 323,63 | 316 | 332 | 1,059 | 0,082 | 0,882 | 0,37752378 | 0,46679623 | 0,003826 | 0,0064652 | 0,0052393 | 0,0052393 | TRUE | 0,0622573 | No |
| orf19.4625 | 3565 | 2855 | 3102 | 4253 | 3173 | 3345 | 3356 | 1820 | 3198 | 3635 | 3174 | 3507 | 2714 | 3192 | 2995 | 2567 | 3122,6 | 3378 | 2867 | 0,849 | -0,235 | -3,224 | 0,0012648 | 0,00311942 | 0,006629 | 0,0024444 | 0,0047998 | 0,0047998 | TRUE | 0,0703196 | No |
| orf19.4626 | 2235 | 1753 | 2044 | 2513 | 2090 | 2037 | 2088 | 1100 | 2005 | 2232 | 2091 | 2072 | 1787 | 1944 | 1863 | 1551 | 1943,3 | 2100 | 1786 | 0,853 | -0,23 | -3,384 | 0,00071484 | 0,001856 | 0,00482 | 0,0027266 | 0,0039039 | 0,0039039 | TRUE | 0,0893767 | No |
| orf19.4627 | 1412 | 1232 | 1371 | 1751 | 1438 | 1575 | 1697 | 980 | 1267 | 1568 | 1403 | 1444 | 1230 | 1503 | 1514 | 1382 | 1413,9 | 1420 | 1407 | 0,992 | -0,012 | -0,148 | 0,8826489 | 0,91146189 | 0,007798 | 0,0030063 | 0,0056308 | 0,0056308 | TRUE | 0,089811 | No |
| orf19.4628 | 1524 | 983 | 1323 | 1599 | 1657 | 1523 | 1697 | 970 | 1367 | 1251 | 1354 | 1319 | 1417 | 1453 | 1514 | 1368 | 1380,4 | 1323 | 1438 | 1,088 | 0,121 | 2,374 | 0,01762019 | 0,03319006 | 0,000713 | 0,0030311 | 0,0017806 | 0,0017806 | TRUE | 0,0152607 | No |
| orf19.4629 | 2606 | 2163 | 2526 | 3132 | 2561 | 2841 | 2799 | 1800 | 2338 | 2754 | 2584 | 2583 | 2190 | 2711 | 2498 | 2539 | 2524,5 | 2565 | 2484 | 0,969 | -0,046 | -0,645 | 0,51884944 | 0,60225027 | 0,005824 | 0,0025546 | 0,0044249 | 0,0044249 | TRUE | 0,0785214 | No |
| orf19.4630 | 10495 | 7495 | 9069 | 11711 | 11614 | 10251 | 10895 | 7145 | 9415 | 9542 | 9278 | 9657 | 9933 | 9781 | 9722 | 10077 | 9675,7 | 9473 | 9878 | 1,042 | 0,06 | 2,101 | 0,03567505 | 0,06252501 | 0,000141 | 0,0021296 | 0,0006773 | 0,0006773 | TRUE | 0,0023383 | No |
| orf19.4631 | 2764 | 2270 | 2611 | 3386 | 3360 | 3463 | 3468 | 2520 | 2480 | 2890 | 2671 | 2792 | 2874 | 3304 | 3095 | 3554 | 2957,5 | 2708 | 3207 | 1,183 | 0,243 | 3,443 | 0,00057577 | 0,00153063 | 0,005789 | 0,0024704 | 0,0044362 | 0,0044362 | TRUE | 0,0632133 | No |
| orf19.4632 | 11722 | 8393 | 9646 | 12129 | 11607 | 10271 | 10175 | 7688 | 10516 | 10685 | 9869 | 10002 | 9927 | 9800 | 9079 | 10843 | 10090 | 10268 | 9912 | 0,965 | -0,051 | -0,929 | 0,35306282 | 0,44054027 | 0,003115 | 0,0021234 | 0,0028287 | 0,0028287 | TRUE | 0,0482179 | No |
| orf19.4633 | 1544 | 1052 | 1283 | 1666 | 1580 | 1408 | 1566 | 960 | 1385 | 1339 | 1313 | 1374 | 1351 | 1343 | 1397 | 1354 | 1357,1 | 1353 | 1361 | 1,006 | 0,009 | 0,193 | 0,84734527 | 0,88395321 | 1,00E-08 | 0,0030492 | 0,0011848 | 0,0011848 | TRUE | 0,0049566 | No |
| orf19.4634 | 4387 | 3172 | 3988 | 4814 | 4596 | 4248 | 4624 | 2807 | 3936 | 4038 | 4080 | 3970 | 3931 | 4053 | 4126 | 3959 | 4011,6 | 4006 | 4017 | 1,003 | 0,005 | 0,149 | 0,88149409 | 0,91043776 | 0,000159 | 0,0023414 | 0,0008471 | 0,0008471 | TRUE | 0,0041552 | No |
| orf19.4635 | 8724 | 7434 | 8142 | 10267 | 7645 | 7972 | 7982 | 4619 | 7826 | 9464 | 8330 | 8466 | 6538 | 7607 | 7123 | 6515 | 7733,6 | 8522 | 6946 | 0,815 | -0,294 | -4,341 | 1,42E-05 | 5,04E-05 | 0,005899 | 0,0021672 | 0,0042868 | 0,0042868 | TRUE | 0,0672573 | No |
| orf19.4636 | 2 | 1 | 1 | 5 | 1 | 5 | 2 | 0 | 2 | 1 | 1 | 4 | 1 | 5 | 2 | 0 | 1,95 | 2 | 2 | 0,909 | -0,138 | -0,132 | 0,89486952 | NA | 0,26871 | 0,7452876 | 0,5521715 | 0,5521715 | TRUE | 0,9446606 | No |
| orf19.4637 | 353 | 296 | 309 | 414 | 445 | 463 | 489 | 285 | 317 | 377 | 316 | 341 | 381 | 442 | 436 | 402 | 376,47 | 338 | 415 | 1,234 | 0,303 | 3,498 | 0,00046828 | 0,00127363 | 0,003083 | 0,0058357 | 0,0045229 | 0,0045229 | TRUE | 0,0607907 | No |
| orf19.4639 | 645 | 557 | 618 | 797 | 738 | 697 | 753 | 435 | 579 | 709 | 632 | 657 | 631 | 665 | 672 | 614 | 644,86 | 644 | 646 | 1,006 | 0,008 | 0,115 | 0,90818582 | 0,93175895 | 0,002756 | 0,0042307 | 0,0034763 | 0,0034763 | TRUE | 0,0562286 | No |
| orf19.4640 | 10819 | 7642 | 9583 | 12070 | 11180 | 9625 | 10689 | 6320 | 9706 | 9729 | 9804 | 9953 | 9561 | 9184 | 9538 | 8914 | 9548,7 | 9798 | 9299 | 0,95 | -0,075 | -2,24 | 0,02510422 | 0,04579668 | 0,00048 | 0,0021316 | 0,000962 | 0,000962 | TRUE | 0,0091708 | No |
| orf19.4641 | 2622 | 1916 | 2274 | 2807 | 2845 | 3015 | 2946 | 1904 | 2352 | 2439 | 2327 | 2315 | 2433 | 2877 | 2629 | 2685 | 2507,1 | 2358 | 2656 | 1,126 | 0,172 | 3,161 | 0,00157392 | 0,00380069 | 0,002115 | 0,0025586 | 0,0024275 | 0,0024275 | TRUE | 0,0399001 | No |
| orf19.4642 | 781 | 776 | 642 | 947 | 891 | 1006 | 992 | 504 | 701 | 988 | 657 | 781 | 762 | 960 | 885 | 711 | 805,53 | 782 | 830 | 1,064 | 0,09 | 0,718 | 0,4724757 | 0,55889624 | 0,024308 | 0,0037817 | 0,0137992 | 0,0137992 | TRUE | 0,3640098 | No |
| orf19.4643 | 1598 | 1157 | 1419 | 1762 | 1842 | 1777 | 1910 | 1030 | 1434 | 1473 | 1452 | 1453 | 1575 | 1696 | 1704 | 1453 | 1529,9 | 1453 | 1607 | 1,109 | 0,149 | 2,64 | 0,00829338 | 0,01704057 | 0,001906 | 0,0029284 | 0,0024039 | 0,0024039 | TRUE | 0,0460114 | No |
| orf19.4645 | 4011 | 3282 | 3987 | 4866 | 5027 | 5704 | 5826 | 3310 | 3598 | 4178 | 4079 | 4013 | 4299 | 5443 | 5199 | 4668 | 4434,7 | 3967 | 4902 | 1,236 | 0,306 | 4,043 | 5,28E-05 | 0,00016992 | 0,007299 | 0,0023069 | 0,0052705 | 0,0052705 | TRUE | 0,0847359 | No |
| orf19.4646 | 310 | 200 | 269 | 313 | 315 | 251 | 295 | 260 | 278 | 255 | 275 | 258 | 269 | 239 | 263 | 367 | 275,61 | 266 | 284 | 1,059 | 0,082 | 0,679 | 0,49738923 | 0,58217699 | 0,014161 | 0,0072468 | 0,0104992 | 0,0104992 | TRUE | 0,3780705 | No |
| orf19.4647 | 334 | 216 | 277 | 342 | 323 | 236 | 289 | 176 | 300 | 275 | 283 | 282 | 276 | 225 | 258 | 248 | 268,45 | 285 | 252 | 0,884 | -0,177 | -1,922 | 0,05456331 | 0,09065718 | 1,00E-08 | 0,0073873 | 0,0044279 | 0,0044279 | TRUE | 0,0616267 | No |
| orf19.4649 | 760 | 631 | 729 | 916 | 740 | 887 | 801 | 563 | 682 | 803 | 746 | 755 | 633 | 846 | 715 | 794 | 746,79 | 746 | 747 | 1 | 0 | -0,003 | 0,99726661 | 0,99812386 | 0,009021 | 0,0039234 | 0,0066173 | 0,0066173 | TRUE | 0,1293618 | No |
| orf19.465 | 24 | 15 | 17 | 31 | 36 | 18 | 28 | 32 | 22 | 19 | 17 | 26 | 31 | 17 | 25 | 45 | 25,21 | 21 | 30 | 1,373 | 0,457 | 1,426 | 0,15390474 | 0,22047202 | 0,055142 | 0,0595682 | 0,05776 | 0,05776 | TRUE | 0,295277 | No |
| orf19.4650 | 1318 | 906 | 1038 | 1493 | 1129 | 969 | 1088 | 649 | 1182 | 1153 | 1062 | 1231 | 966 | 925 | 971 | 915 | 1050,7 | 1157 | 944 | 0,816 | -0,293 | -4,989 | 6,06E-07 | 2,65E-06 | 0,001591 | 0,0033612 | 0,0023572 | 0,0023572 | TRUE | 0,0373209 | No |
| orf19.4651 | 1893 | 1430 | 1752 | 2233 | 1459 | 1492 | 1505 | 992 | 1698 | 1821 | 1792 | 1841 | 1248 | 1424 | 1343 | 1399 | 1570,8 | 1788 | 1354 | 0,756 | -0,404 | -7,404 | 1,32E-13 | 1,32E-12 | 0,001705 | 0,0029037 | 0,0022006 | 0,0022006 | TRUE | 0,0344808 | No |
| orf19.4652 | 3 | 6 | 6 | 3 | 1 | 0 | 1 | 0 | 3 | 8 | 6 | 2 | 1 | 0 | 1 | 0 | 2,59 | 5 | 0 | 0,108 | -3,212 | -2,652 | 0,00800376 | NA | 1,00E-08 | 0,563296 | 0,3753487 | 0,3753487 | TRUE | 0,0992406 | No |
| orf19.4653 | 14 | 10 | 12 | 15 | 12 | 8 | 14 | 14 | 13 | 13 | 12 | 12 | 10 | 8 | 12 | 20 | 12,51 | 12 | 12 | 0,972 | -0,041 | -0,103 | 0,91806467 | NA | 1,00E-08 | 0,1180327 | 0,0706702 | 0,0706702 | TRUE | 0,410309 | No |
| orf19.4654 | 24 | 16 | 17 | 35 | 28 | 15 | 23 | 14 | 22 | 20 | 17 | 29 | 24 | 14 | 21 | 20 | 20,84 | 22 | 20 | 0,885 | -0,176 | -0,568 | 0,57034425 | 0,64935157 | 1,00E-08 | 0,0716544 | 0,0449855 | 0,0449855 | TRUE | 0,2945461 | No |
| orf19.4655 | 415 | 363 | 385 | 447 | 412 | 470 | 473 | 306 | 372 | 462 | 394 | 369 | 352 | 448 | 422 | 432 | 406,43 | 399 | 414 | 1,038 | 0,054 | 0,539 | 0,58965939 | 0,66748517 | 0,008647 | 0,0055514 | 0,0071012 | 0,0071012 | TRUE | 0,1213645 | No |
| orf19.4656 | 184 | 209 | 165 | 244 | 195 | 250 | 236 | 136 | 165 | 266 | 169 | 201 | 167 | 239 | 211 | 192 | 201,11 | 200 | 202 | 1,014 | 0,019 | 0,122 | 0,90312531 | 0,92751742 | 0,031094 | 0,009198 | 0,0196212 | 0,0196212 | TRUE | 0,4922098 | No |
| orf19.4657 | 9322 | 6782 | 8437 | 10536 | 11544 | 11204 | 12027 | 6889 | 8363 | 8634 | 8632 | 8688 | 9873 | 10691 | 10732 | 9716 | 9416,1 | 8579 | 10253 | 1,196 | 0,258 | 6,039 | 1,55E-09 | 9,58E-09 | 0,001357 | 0,0021337 | 0,0016403 | 0,0016403 | TRUE | 0,0148072 | No |
| orf19.4658 | 3543 | 2756 | 3177 | 4070 | 3247 | 3301 | 3505 | 1850 | 3178 | 3509 | 3250 | 3356 | 2777 | 3150 | 3128 | 2609 | 3119,7 | 3323 | 2916 | 0,879 | -0,187 | -2,868 | 0,0041324 | 0,009081 | 0,004675 | 0,0024449 | 0,0037538 | 0,0037538 | TRUE | 0,0585508 | No |
| orf19.4659 | 2960 | 2230 | 2727 | 3495 | 3284 | 3059 | 3296 | 2038 | 2655 | 2839 | 2790 | 2882 | 2809 | 2919 | 2941 | 2874 | 2838,7 | 2792 | 2886 | 1,034 | 0,048 | 1,187 | 0,23521736 | 0,31610508 | 0,000548 | 0,0024909 | 0,0012252 | 0,0012252 | TRUE | 0,0135275 | No |
| orf19.4660 | 20286 | 13994 | 16751 | 21331 | 20674 | 17816 | 18599 | 11675 | 18199 | 17816 | 17138 | 17590 | 17681 | 16999 | 16597 | 16467 | 17311 | 17686 | 16936 | 0,958 | -0,062 | -1,763 | 0,07788187 | 0,12365599 | 0,000757 | 0,0020634 | 0,0011382 | 0,0011382 | TRUE | 0,0107946 | No |
| orf19.4662 | 1245 | 893 | 1264 | 1412 | 1378 | 1111 | 1245 | 909 | 1117 | 1137 | 1293 | 1164 | 1179 | 1060 | 1111 | 1282 | 1167,9 | 1178 | 1158 | 0,981 | -0,028 | -0,39 | 0,69639325 | 0,75878321 | 0,00455 | 0,0032226 | 0,0039486 | 0,0039486 | TRUE | 0,0579265 | No |
| orf19.4664 | 2860 | 2183 | 2517 | 3122 | 3626 | 3316 | 3423 | 1968 | 2566 | 2779 | 2575 | 2574 | 3101 | 3164 | 3054 | 2776 | 2823,7 | 2624 | 3024 | 1,155 | 0,208 | 4,07 | 4,71E-05 | 0,00015315 | 0,001915 | 0,0024936 | 0,0021462 | 0,0021462 | TRUE | 0,0350311 | No |
| orf19.4665 | 2000 | 1348 | 1804 | 2205 | 2799 | 2139 | 2429 | 1335 | 1794 | 1716 | 1846 | 1818 | 2394 | 2041 | 2167 | 1883 | 1957,4 | 1794 | 2121 | 1,184 | 0,244 | 3,53 | 0,00041486 | 0,00113916 | 0,00497 | 0,0027212 | 0,0040669 | 0,0040669 | TRUE | 0,0907746 | No |
| orf19.4666 | 3167 | 2237 | 2612 | 3413 | 3481 | 3208 | 3108 | 1984 | 2841 | 2848 | 2672 | 2814 | 2977 | 3061 | 2773 | 2798 | 2848,2 | 2794 | 2902 | 1,04 | 0,056 | 1,21 | 0,22646984 | 0,30589256 | 0,001118 | 0,0024892 | 0,0017006 | 0,0017006 | TRUE | 0,0163394 | No |
| orf19.4668 | 1401 | 889 | 1245 | 1472 | 1407 | 1270 | 1440 | 919 | 1257 | 1132 | 1274 | 1214 | 1203 | 1212 | 1285 | 1296 | 1234,1 | 1219 | 1249 | 1,022 | 0,031 | 0,574 | 0,56602756 | 0,64566045 | 0,001019 | 0,0031559 | 0,0020339 | 0,0020339 | TRUE | 0,0253818 | No |
| orf19.4669 | 417 | 360 | 404 | 500 | 485 | 582 | 534 | 334 | 374 | 458 | 413 | 412 | 415 | 555 | 477 | 471 | 446,97 | 414 | 480 | 1,159 | 0,213 | 2,169 | 0,03010974 | 0,05377784 | 0,008657 | 0,0052274 | 0,0069893 | 0,0069893 | TRUE | 0,1367182 | No |
| orf19.467 | 9156 | 5858 | 8364 | 10261 | 11616 | 10141 | 11279 | 7755 | 8214 | 7458 | 8557 | 8461 | 9934 | 9676 | 10065 | 10938 | 9162,9 | 8172 | 10153 | 1,242 | 0,312 | 5,69 | 1,27E-08 | 6,96E-08 | 0,003162 | 0,002138 | 0,0027808 | 0,0027808 | TRUE | 0,0418044 | No |
| orf19.4670 | 5085 | 3599 | 4617 | 5825 | 5949 | 5515 | 6067 | 3678 | 4562 | 4582 | 4724 | 4803 | 5088 | 5262 | 5414 | 5187 | 4952,8 | 4668 | 5238 | 1,122 | 0,166 | 4,62 | 3,84E-06 | 1,49E-05 | 0,000484 | 0,0022726 | 0,0010361 | 0,0010361 | TRUE | 0,0064055 | No |
| orf19.4672 | 7 | 2 | 3 | 12 | 3 | 4 | 6 | 4 | 6 | 3 | 3 | 10 | 3 | 4 | 5 | 6 | 4,9 | 6 | 4 | 0,755 | -0,405 | -0,619 | 0,53579162 | NA | 1,00E-08 | 0,2984825 | 0,2059266 | 0,2059266 | TRUE | 0,6654511 | No |
| orf19.4673 | 1316 | 979 | 1135 | 1468 | 1537 | 1419 | 1564 | 941 | 1181 | 1246 | 1161 | 1211 | 1314 | 1354 | 1396 | 1327 | 1273,8 | 1200 | 1348 | 1,125 | 0,17 | 3,576 | 0,00034842 | 0,00097093 | 1,00E-08 | 0,0031192 | 0,0013796 | 0,0013796 | TRUE | 0,0072817 | No |
| orf19.4674.1 | 322 | 208 | 212 | 317 | 310 | 276 | 255 | 149 | 289 | 265 | 217 | 261 | 265 | 263 | 228 | 210 | 249,77 | 258 | 242 | 0,94 | -0,089 | -0,787 | 0,43155121 | 0,51865567 | 0,00907 | 0,0077917 | 0,008388 | 0,008388 | TRUE | 0,1277138 | No |
| orf19.4675 | 650 | 468 | 589 | 682 | 709 | 600 | 712 | 390 | 583 | 596 | 603 | 562 | 606 | 573 | 635 | 550 | 588,52 | 586 | 591 | 1,014 | 0,019 | 0,29 | 0,77174909 | 0,82174247 | 0,000687 | 0,0044462 | 0,0025841 | 0,0025841 | TRUE | 0,0293634 | No |
| orf19.4676 | 359 | 317 | 322 | 397 | 370 | 364 | 370 | 178 | 322 | 404 | 329 | 327 | 316 | 347 | 330 | 251 | 328,43 | 346 | 311 | 0,909 | -0,137 | -1,247 | 0,21225347 | 0,28955923 | 0,011354 | 0,0063997 | 0,008575 | 0,008575 | TRUE | 0,1661639 | No |
| orf19.4677 | 1328 | 946 | 1031 | 1325 | 1119 | 1133 | 1149 | 709 | 1191 | 1204 | 1055 | 1093 | 957 | 1081 | 1025 | 1000 | 1075,8 | 1136 | 1016 | 0,895 | -0,16 | -2,533 | 0,01131916 | 0,02248115 | 0,00252 | 0,0033289 | 0,0028758 | 0,0028758 | TRUE | 0,026692 | No |
| orf19.4678 | 1152 | 869 | 1033 | 1262 | 1182 | 1196 | 1231 | 703 | 1033 | 1106 | 1057 | 1041 | 1011 | 1141 | 1098 | 992 | 1059,9 | 1059 | 1060 | 1,004 | 0,006 | 0,098 | 0,92184439 | 0,94271337 | 0,001656 | 0,0033492 | 0,0024196 | 0,0024196 | TRUE | 0,0308074 | No |
| orf19.4679 | 21521 | 14626 | 19618 | 23977 | 23209 | 19566 | 21758 | 14503 | 19307 | 18620 | 20071 | 19772 | 19849 | 18669 | 19415 | 20455 | 19520 | 19442 | 19597 | 1,008 | 0,011 | 0,278 | 0,78107028 | 0,82850817 | 0,00105 | 0,0020539 | 0,0014663 | 0,0014663 | TRUE | 0,0124117 | No |

| | | | | | | | | | | | | | | | | | | | | | | | | | | | | | | | |
|---|---|---|---|---|---|---|---|---|---|---|---|---|---|---|---|---|---|---|---|---|---|---|---|---|---|---|---|---|---|---|---|
| orf19.4680 | 392 | 277 | 338 | 409 | 435 | 408 | 437 | 279 | 352 | 353 | 346 | 337 | 372 | 389 | 390 | 394 | 366,52 | 347 | 386 | 1,113 | 0,155 | 2,096 | 0,03607721 | 0,06315614 | 1,00E-08 | 0,0059403 | 0,0024924 | 0,0024924 | TRUE | 0,0075422 | No |
| orf19.4681 | 7356 | 5110 | 6588 | 7769 | 7015 | 6617 | 7222 | 4585 | 6599 | 6506 | 6740 | 6406 | 5999 | 6314 | 6444 | 6467 | 6434,5 | 6563 | 6306 | 0,961 | -0,058 | -1,575 | 0,11514882 | 0,1718366 | 0,00061 | 0,0022051 | 0,0011507 | 0,0011507 | TRUE | 0,013311 | No |
| orf19.4682 | 325 | 231 | 317 | 352 | 273 | 288 | 317 | 200 | 292 | 294 | 324 | 290 | 233 | 275 | 283 | 282 | 284,18 | 300 | 268 | 0,892 | -0,165 | -1,771 | 0,07658038 | 0,12179339 | 0,002203 | 0,0070879 | 0,0048308 | 0,0048308 | TRUE | 0,0931038 | No |
| orf19.4683 | 3626 | 2744 | 3249 | 3937 | 5045 | 4886 | 5143 | 3165 | 3253 | 3493 | 3324 | 3246 | 4315 | 4662 | 4589 | 4464 | 3918,4 | 3329 | 4508 | 1,355 | 0,438 | 10,547 | 5,25E-26 | 1,49E-24 | 0,00082 | 0,00235 | 0,0013971 | 0,0013971 | TRUE | 0,0128458 | No |
| orf19.4684.2 | 4947 | 3151 | 3896 | 4934 | 4860 | 3794 | 3905 | 2553 | 4438 | 4012 | 3986 | 4069 | 4156 | 3620 | 3485 | 3601 | 3920,8 | 4126 | 3716 | 0,901 | -0,151 | -2,431 | 0,01504412 | 0,02892322 | 0,004224 | 0,0023498 | 0,0034455 | 0,0034455 | TRUE | 0,0788323 | No |
| orf19.4685 | 25 | 19 | 20 | 22 | 11 | 10 | 18 | 6 | 22 | 24 | 20 | 18 | 9 | 10 | 16 | 8 | 16,09 | 21 | 11 | 0,522 | -0,939 | -2,648 | 0,00810701 | 0,0166976 | 1,00E-08 | 0,0922217 | 0,0521186 | 0,0521186 | TRUE | 0,4293641 | No |
| orf19.4686 | 3036 | 2439 | 2964 | 3652 | 3317 | 3273 | 3465 | 2046 | 2724 | 3105 | 3032 | 3011 | 2837 | 3123 | 3092 | 2886 | 2976,3 | 2968 | 2984 | 1,006 | 0,009 | 0,169 | 0,86594564 | 0,89777419 | 0,002499 | 0,0024673 | 0,0024869 | 0,0024869 | TRUE | 0,0377483 | No |
| orf19.4688 | 2155 | 1522 | 1870 | 2326 | 1842 | 1626 | 1834 | 1243 | 1933 | 1938 | 1913 | 1918 | 1575 | 1551 | 1637 | 1753 | 1777,3 | 1926 | 1629 | 0,844 | -0,245 | -5,179 | 2,23E-07 | 1,04E-06 | 0,000634 | 0,0027963 | 0,0015712 | 0,0015712 | TRUE | 0,0284483 | No |
| orf19.4689 | 843 | 575 | 702 | 894 | 936 | 781 | 897 | 590 | 756 | 732 | 718 | 737 | 800 | 745 | 800 | 832 | 765,25 | 736 | 794 | 1,077 | 0,107 | 1,866 | 0,06209552 | 0,10151964 | 1,00E-08 | 0,0038766 | 0,0018652 | 0,0018652 | TRUE | 0,0204713 | No |
| orf19.469 | 3001 | 1954 | 2589 | 3291 | 3398 | 2984 | 3327 | 2128 | 2692 | 2488 | 2649 | 2714 | 2906 | 2847 | 2969 | 3001 | 2783,2 | 2636 | 2931 | 1,11 | 0,151 | 3,625 | 0,00028898 | 0,00081857 | 0,000545 | 0,0025011 | 0,001296 | 0,001296 | TRUE | 0,0163306 | No |
| orf19.4690 | 195 | 165 | 170 | 219 | 181 | 158 | 171 | 96 | 175 | 210 | 174 | 181 | 155 | 151 | 153 | 135 | 166,63 | 185 | 148 | 0,811 | -0,302 | -2,698 | 0,00698146 | 0,01457422 | 1,00E-08 | 0,0106915 | 0,0060124 | 0,0060124 | TRUE | 0,0777672 | No |
| orf19.4691 | 223 | 162 | 211 | 238 | 183 | 170 | 192 | 107 | 200 | 206 | 216 | 196 | 157 | 162 | 171 | 151 | 182,42 | 204 | 160 | 0,787 | -0,345 | -3,335 | 0,00085365 | 0,00218112 | 1,00E-08 | 0,0099374 | 0,0047533 | 0,0047533 | TRUE | 0,0233926 | No |
| orf19.4697 | 1550 | 1275 | 1484 | 1772 | 982 | 930 | 1015 | 564 | 1391 | 1623 | 1518 | 1461 | 840 | 887 | 906 | 795 | 1177,7 | 1498 | 857 | 0,574 | -0,8 | -12,528 | 5,24E-36 | 2,79E-34 | 0,002603 | 0,0032122 | 0,0029971 | 0,0029971 | TRUE | 0,0363153 | No |
| orf19.4698 | 1933 | 1604 | 1818 | 2307 | 2517 | 2627 | 2843 | 1456 | 1734 | 2042 | 1860 | 1902 | 2153 | 2507 | 2537 | 2054 | 2098,5 | 1884 | 2313 | 1,229 | 0,297 | 3,833 | 0,00012676 | 0,0003804 | 0,007258 | 0,0026713 | 0,005298 | 0,005298 | TRUE | 0,0663617 | No |
| orf19.4699 | 2708 | 2325 | 2597 | 3397 | 2994 | 3125 | 3198 | 1923 | 2429 | 2960 | 2657 | 2801 | 2561 | 2982 | 2854 | 2712 | 2744,5 | 2712 | 2777 | 1,025 | 0,035 | 0,516 | 0,60589327 | 0,68155902 | 0,005292 | 0,0025085 | 0,0041151 | 0,0041151 | TRUE | 0,0600552 | No |
| orf19.470 | 1781 | 1107 | 1594 | 1863 | 2629 | 2213 | 2484 | 1560 | 1598 | 1409 | 1631 | 1536 | 2248 | 2112 | 2217 | 2200 | 1868,9 | 1544 | 2194 | 1,42 | 0,506 | 9,67 | 4,03E-22 | 8,51E-21 | 0,001623 | 0,0027563 | 0,0020711 | 0,0020711 | TRUE | 0,0381932 | No |
| orf19.4701 | 314 | 270 | 285 | 404 | 344 | 445 | 403 | 266 | 282 | 344 | 292 | 333 | 294 | 425 | 360 | 375 | 337,97 | 313 | 364 | 1,162 | 0,217 | 1,865 | 0,06220154 | 0,10157838 | 0,013637 | 0,0062749 | 0,0099857 | 0,0099857 | TRUE | 0,1988236 | No |
| orf19.4702 | 118 | 84 | 93 | 127 | 117 | 93 | 116 | 78 | 106 | 107 | 95 | 105 | 100 | 89 | 104 | 110 | 101,87 | 103 | 101 | 0,97 | -0,043 | -0,316 | 0,7523431 | 0,80613835 | 1,00E-08 | 0,0162294 | 0,0082454 | 0,0082454 | TRUE | 0,0606053 | No |
| orf19.4703 | 351 | 228 | 307 | 368 | 410 | 367 | 379 | 224 | 315 | 290 | 314 | 303 | 351 | 350 | 338 | 316 | 322,21 | 306 | 339 | 1,11 | 0,151 | 1,885 | 0,05937534 | 0,09759974 | 1,00E-08 | 0,006485 | 0,0030658 | 0,0030658 | TRUE | 0,0189038 | No |
| orf19.4704 | 5721 | 3923 | 5348 | 5898 | 7364 | 6644 | 7070 | 4433 | 5132 | 4994 | 5472 | 4864 | 6298 | 6339 | 6309 | 6252 | 5707,6 | 5116 | 6300 | 1,232 | 0,301 | 7,232 | 4,74E-13 | 4,54E-12 | 0,001087 | 0,0022339 | 0,0014818 | 0,0014818 | TRUE | 0,0267611 | No |
| orf19.4705 | 1612 | 1174 | 1394 | 1782 | 1562 | 1414 | 1510 | 1013 | 1446 | 1495 | 1426 | 1469 | 1336 | 1349 | 1347 | 1429 | 1412,2 | 1459 | 1365 | 0,934 | -0,098 | -2,185 | 0,02885994 | 0,05180065 | 1,00E-08 | 0,0030075 | 0,0012265 | 0,0012265 | TRUE | 0,0102371 | No |
| orf19.4706 | 2057 | 1489 | 1873 | 2192 | 1293 | 1163 | 1224 | 815 | 1845 | 1896 | 1916 | 1808 | 1106 | 1110 | 1092 | 1149 | 1490,3 | 1866 | 1114 | 0,597 | -0,745 | -16,599 | 7,14E-62 | 1,26E-59 | 1,00E-08 | 0,0029537 | 0,001215 | 0,001215 | TRUE | 0,005625 | No |
| orf19.4707 | 203 | 156 | 204 | 224 | 181 | 132 | 167 | 105 | 182 | 199 | 209 | 185 | 155 | 126 | 149 | 148 | 169 | 194 | 144 | 0,748 | -0,418 | -3,694 | 0,00022096 | 0,00063916 | 1,00E-08 | 0,0105695 | 0,0062602 | 0,0062602 | TRUE | 0,0819925 | No |
| orf19.4711 | 2797 | 1815 | 2494 | 2926 | 3544 | 3118 | 3431 | 2066 | 2509 | 2311 | 2552 | 2413 | 3031 | 2975 | 3062 | 2914 | 2720,7 | 2446 | 2996 | 1,224 | 0,292 | 6,821 | 9,01E-12 | 7,40E-11 | 0,000703 | 0,0025131 | 0,0013889 | 0,0013889 | TRUE | 0,0156095 | No |
| orf19.4712 | 20 | 10 | 12 | 25 | 25 | 17 | 15 | 15 | 18 | 13 | 12 | 21 | 21 | 16 | 13 | 21 | 16,96 | 16 | 18 | 1,107 | 0,147 | 0,43 | 0,66752602 | 0,73438863 | 1,00E-08 | 0,0875572 | 0,0542272 | 0,0542272 | TRUE | 0,2182621 | No |
| orf19.4713 | 336 | 252 | 311 | 349 | 392 | 412 | 410 | 297 | 301 | 321 | 318 | 288 | 335 | 393 | 366 | 419 | 342,67 | 307 | 378 | 1,229 | 0,297 | 3,324 | 0,00088778 | 0,00225531 | 0,003111 | 0,006216 | 0,0047165 | 0,0047165 | TRUE | 0,0690051 | No |
| orf19.4715 | 1114 | 773 | 1020 | 1253 | 1818 | 1685 | 1831 | 1117 | 999 | 984 | 1044 | 1033 | 1555 | 1608 | 1634 | 1575 | 1304 | 1015 | 1593 | 1,568 | 0,649 | 13,971 | 2,34E-44 | 1,85E-42 | 1,00E-08 | 0,0030928 | 0,0012688 | 0,0012688 | TRUE | 0,0044017 | No |
| orf19.4716 | 23986 | 15377 | 20088 | 24920 | 16048 | 13178 | 14286 | 9270 | 21518 | 19577 | 20552 | 20549 | 13725 | 12574 | 12748 | 13075 | 16790 | 20549 | 13030 | 0,634 | -0,657 | -15,63 | 4,55E-55 | 6,07E-53 | 0,001418 | 0,002066 | 0,0016358 | 0,0016358 | TRUE | 0,0159401 | No |
| orf19.4718 | 9447 | 6337 | 8473 | 10220 | 10656 | 9119 | 9850 | 6451 | 8475 | 8068 | 8669 | 8428 | 9113 | 8701 | 8789 | 9099 | 8667,7 | 8410 | 8926 | 1,061 | 0,085 | 2,445 | 0,0144721 | 0,02796666 | 0,000571 | 0,002147 | 0,0010503 | 0,0010503 | TRUE | 0,008871 | No |
| orf19.4719 | 760 | 526 | 682 | 787 | 798 | 758 | 784 | 431 | 682 | 670 | 698 | 649 | 682 | 723 | 700 | 608 | 676,42 | 675 | 678 | 1,01 | 0,014 | 0,217 | 0,82815952 | 0,86788459 | 0,001192 | 0,0041257 | 0,0027089 | 0,0027089 | TRUE | 0,0493031 | No |
| orf19.4720 | 3958 | 2631 | 3277 | 4042 | 3542 | 3073 | 3343 | 2143 | 3551 | 3350 | 3353 | 3333 | 3029 | 2932 | 2983 | 3023 | 3194,1 | 3397 | 2992 | 0,88 | -0,184 | -4,921 | 8,62E-07 | 3,67E-06 | 0,000322 | 0,002434 | 0,0010235 | 0,0010235 | TRUE | 0,011665 | No |
| orf19.4721 | 5149 | 3512 | 4223 | 5370 | 4739 | 4327 | 4485 | 2515 | 4619 | 4471 | 4321 | 4428 | 4053 | 4129 | 4002 | 3547 | 4196,3 | 4460 | 3933 | 0,883 | -0,18 | -3,483 | 0,00049578 | 0,00133933 | 0,002289 | 0,0023255 | 0,0023191 | 0,0023191 | TRUE | 0,0508286 | No |
| orf19.4722 | 2424 | 1647 | 2031 | 2584 | 2758 | 2324 | 2631 | 1528 | 2175 | 2097 | 2078 | 2131 | 2359 | 2217 | 2348 | 2155 | 2194,9 | 2120 | 2270 | 1,072 | 0,1 | 2,257 | 0,02399242 | 0,0439605 | 0,00068 | 0,0026409 | 0,0014339 | 0,0014339 | TRUE | 0,012842 | No |
| orf19.4723 | 1252 | 773 | 1031 | 1458 | 1040 | 864 | 932 | 593 | 1123 | 984 | 1055 | 1202 | 889 | 824 | 832 | 836 | 968,28 | 1091 | 845 | 0,773 | -0,371 | -5,411 | 6,28E-08 | 3,16E-07 | 0,003404 | 0,0034788 | 0,0034545 | 0,0034545 | TRUE | 0,0570709 | No |
| orf19.4724 | 1782 | 1216 | 1487 | 1819 | 1179 | 1017 | 1145 | 657 | 1599 | 1548 | 1521 | 1500 | 1008 | 970 | 1022 | 927 | 1261,9 | 1542 | 982 | 0,639 | -0,647 | -12,875 | 6,20E-38 | 3,84E-36 | 0,000363 | 0,0031299 | 0,0015919 | 0,0015919 | TRUE | 0,0146724 | No |
| orf19.4725 | 1236 | 809 | 1044 | 1340 | 1147 | 1030 | 1033 | 753 | 1109 | 1030 | 1068 | 1105 | 981 | 983 | 922 | 1062 | 1032,4 | 1078 | 987 | 0,912 | -0,134 | -2,349 | 0,01881506 | 0,0351933 | 0,000993 | 0,0033856 | 0,0021342 | 0,0021342 | TRUE | 0,0277752 | No |
| orf19.4726 | 954 | 596 | 834 | 976 | 1102 | 1020 | 1109 | 609 | 856 | 759 | 853 | 805 | 942 | 973 | 990 | 859 | 879,62 | 818 | 941 | 1,152 | 0,204 | 3,18 | 0,00147132 | 0,00357449 | 0,001967 | 0,0036299 | 0,0027978 | 0,0027978 | TRUE | 0,0365246 | No |
| orf19.4727 | 1011 | 713 | 843 | 1030 | 967 | 855 | 848 | 540 | 907 | 908 | 862 | 849 | 827 | 816 | 757 | 762 | 835,96 | 882 | 790 | 0,899 | -0,154 | -2,666 | 0,00768383 | 0,01589137 | 0,000515 | 0,0037161 | 0,0020076 | 0,0020076 | TRUE | 0,0120386 | No |
| orf19.4728 | 3888 | 2545 | 3352 | 4007 | 4659 | 4068 | 4648 | 2625 | 3488 | 3240 | 3429 | 3304 | 3985 | 3882 | 4148 | 3702 | 3647,2 | 3365 | 3929 | 1,168 | 0,224 | 4,888 | 1,02E-06 | 4,30E-06 | 0,001328 | 0,0023776 | 0,0017438 | 0,0017438 | TRUE | 0,0174649 | No |
| orf19.473 | 1470 | 934 | 1279 | 1481 | 1877 | 1494 | 1689 | 1054 | 1319 | 1189 | 1309 | 1221 | 1605 | 1426 | 1507 | 1487 | 1382,8 | 1260 | 1506 | 1,196 | 0,258 | 4,568 | 4,92E-06 | 1,88E-05 | 0,001823 | 0,0030294 | 0,0023302 | 0,0023302 | TRUE | 0,0246811 | No |
| orf19.4730 | 2172 | 1587 | 1821 | 2371 | 2151 | 1831 | 2053 | 1225 | 1949 | 2020 | 1863 | 1955 | 1840 | 1747 | 1832 | 1728 | 1866,7 | 1947 | 1787 | 0,919 | -0,121 | -2,708 | 0,00676957 | 0,01419601 | 0,000453 | 0,0027572 | 0,0013855 | 0,0013855 | TRUE | 0,009791 | No |
| orf19.4731 | 1164 | 888 | 1080 | 1242 | 1040 | 934 | 1113 | 619 | 1044 | 1131 | 1105 | 1024 | 889 | 891 | 993 | 873 | 993,84 | 1076 | 912 | 0,85 | -0,234 | -3,839 | 0,00012375 | 0,00037213 | 0,001849 | 0,0034402 | 0,0025602 | 0,0025602 | TRUE | 0,0443431 | No |
| orf19.4732 | 1538 | 1066 | 1452 | 1692 | 2160 | 1889 | 2080 | 1402 | 1380 | 1357 | 1486 | 1395 | 1847 | 1802 | 1856 | 1977 | 1637,6 | 1404 | 1870 | 1,33 | 0,411 | 8,396 | 4,62E-17 | 6,60E-16 | 0,000773 | 0,002866 | 0,0016758 | 0,0016758 | TRUE | 0,0177631 | No |
| orf19.4733 | 1588 | 1017 | 1304 | 1629 | 1265 | 1132 | 1201 | 621 | 1425 | 1295 | 1334 | 1343 | 1082 | 1080 | 1072 | 876 | 1188,3 | 1349 | 1028 | 0,764 | -0,389 | -5,558 | 2,73E-08 | 1,44E-07 | 0,004516 | 0,0032012 | 0,0038402 | 0,0038402 | TRUE | 0,1075426 | No |
| orf19.4734 | 940 | 663 | 759 | 1002 | 768 | 639 | 717 | 489 | 843 | 844 | 777 | 826 | 657 | 610 | 640 | 690 | 735,77 | 822 | 649 | 0,787 | -0,345 | -5,662 | 1,50E-08 | 8,14E-08 | 0,000376 | 0,0039526 | 0,0021904 | 0,0021904 | TRUE | 0,0188254 | No |
| orf19.4735 | 1638 | 1117 | 1523 | 1800 | 1643 | 1432 | 1570 | 1052 | 1469 | 1422 | 1558 | 1484 | 1405 | 1366 | 1401 | 1484 | 1448,8 | 1483 | 1414 | 0,951 | -0,073 | -1,499 | 0,13380375 | 0,19602364 | 0,000418 | 0,0029815 | 0,0015535 | 0,0015535 | TRUE | 0,0132944 | No |
| orf19.4736 | 1577 | 1020 | 1382 | 1694 | 1177 | 1053 | 1164 | 729 | 1415 | 1299 | 1414 | 1397 | 1007 | 1005 | 1039 | 1028 | 1200,3 | 1381 | 1020 | 0,737 | -0,441 | -8,943 | 3,80E-19 | 6,47E-18 | 1,00E-08 | 0,003189 | 0,0014771 | 0,0014771 | TRUE | 0,017628 | No |
| orf19.4737 | 1825 | 1297 | 1614 | 1982 | 1674 | 1540 | 1589 | 1033 | 1637 | 1651 | 1651 | 1634 | 1432 | 1469 | 1418 | 1457 | 1543,8 | 1643 | 1444 | 0,878 | -0,187 | -4,566 | 4,97E-06 | 1,89E-05 | 1,00E-08 | 0,0029199 | 0,0009633 | 0,0009633 | TRUE | 0,0018861 | No |
| orf19.4738 | 971 | 672 | 831 | 1047 | 1356 | 1173 | 1314 | 811 | 871 | 856 | 850 | 863 | 1160 | 1119 | 1173 | 1144 | 1004,4 | 860 | 1149 | 1,336 | 0,418 | 8,566 | 1,08E-17 | 1,62E-16 | 1,00E-08 | 0,0034248 | 0,0012693 | 0,0012693 | TRUE | 0,0038697 | No |
| orf19.4739 | 25357 | 17385 | 22219 | 27479 | 21910 | 18915 | 20381 | 13354 | 22748 | 22133 | 22732 | 22660 | 18738 | 18048 | 18187 | 18835 | 20510 | 22568 | 18452 | 0,817 | -0,291 | -10,567 | 4,22E-26 | 1,21E-24 | 0,000244 | 0,0020503 | 0,0006782 | 0,0006782 | TRUE | 0,0026422 | No |
| orf19.474 | 1509 | 1060 | 1258 | 1601 | 1350 | 1332 | 1308 | 829 | 1354 | 1349 | 1287 | 1320 | 1155 | 1271 | 1167 | 1169 | 1259,1 | 1328 | 1190 | 0,897 | -0,157 | -3,11 | 0,00187175 | 0,00444484 | 0,000502 | 0,0031325 | 0,0016391 | 0,0016391 | TRUE | 0,0255365 | No |
| orf19.4740 | 575 | 404 | 502 | 623 | 577 | 485 | 557 | 354 | 516 | 514 | 514 | 514 | 493 | 463 | 497 | 499 | 501,26 | 514 | 488 | 0,948 | -0,077 | -1,185 | 0,23601591 | 0,31679705 | 1,00E-08 | 0,0048756 | 0,0020331 | 0,0020331 | TRUE | 0,0145088 | No |
| orf19.4741 | 26 | 16 | 21 | 25 | 18 | 13 | 16 | 7 | 23 | 20 | 21 | 21 | 15 | 12 | 14 | 10 | 17,22 | 21 | 13 | 0,616 | -0,699 | -2,145 | 0,03193214 | 0,05662582 | 1,00E-08 | 0,0862929 | 0,0413229 | 0,0413229 | TRUE | 0,0674889 | No |
| orf19.4742 | 1137 | 833 | 1020 | 1254 | 1149 | 973 | 1053 | 690 | 1020 | 1060 | 1044 | 1034 | 983 | 928 | 940 | 973 | 997,75 | 1040 | 956 | 0,92 | -0,121 | -2,428 | 0,01517423 | 0,02914539 | 1,00E-08 | 0,0034345 | 0,0013615 | 0,0013615 | TRUE | 0,0049629 | No |
| orf19.4743 | 383 | 280 | 380 | 434 | 443 | 456 | 488 | 289 | 344 | 356 | 389 | 358 | 379 | 435 | 435 | 408 | 387,97 | 362 | 414 | 1,146 | 0,197 | 2,447 | 0,01438904 | 0,02785995 | 0,001367 | 0,0057213 | 0,0036203 | 0,0036203 | TRUE | 0,0430119 | No |
| orf19.4744 | 2431 | 1800 | 2215 | 2810 | 2931 | 2607 | 2733 | 1804 | 2181 | 2292 | 2266 | 2317 | 2507 | 2488 | 2439 | 2544 | 2379,1 | 2264 | 2494 | 1,101 | 0,139 | 3,592 | 0,00032784 | 0,00092255 | 1,00E-08 | 0,0025897 | 0,0010184 | 0,0010184 | TRUE | 0,0077154 | No |
| orf19.4746 | 9650 | 6662 | 8387 | 10560 | 9692 | 8253 | 9329 | 5456 | 8657 | 8481 | 8581 | 8708 | 8289 | 7875 | 8325 | 7695 | 8326,3 | 8607 | 8046 | 0,935 | -0,097 | -2,693 | 0,00707153 | 0,01473664 | 0,000649 | 0,0021539 | 0,0011135 | 0,0011135 | TRUE | 0,0101347 | No |
| orf19.4747 | 314 | 226 | 284 | 358 | 339 | 290 | 343 | 171 | 282 | 288 | 291 | 295 | 290 | 277 | 306 | 241 | 283,63 | 289 | 278 | 0,971 | -0,043 | -0,47 | 0,63802633 | 0,7082894 | 0,000404 | 0,0070978 | 0,0043457 | 0,0043457 | TRUE | 0,074268 | No |
| orf19.4748 | 421 | 363 | 347 | 457 | 341 | 391 | 355 | 268 | 378 | 462 | 355 | 377 | 292 | 373 | 317 | 378 | 366,4 | 393 | 340 | 0,864 | -0,211 | -1,926 | 0,05407084 | 0,08998826 | 0,011409 | 0,0059416 | 0,0087373 | 0,0087373 | TRUE | 0,1504489 | No |

| | | | | | | | | | | | | | | | | | | | | | | | | | | | | | | | |
|---|---|---|---|---|---|---|---|---|---|---|---|---|---|---|---|---|---|---|---|---|---|---|---|---|---|---|---|---|---|---|---|
| orf19.4749 | 1227 | 905 | 1123 | 1367 | 969 | 862 | 982 | 656 | 1101 | 1152 | 1149 | 1127 | 829 | 822 | 876 | 925 | 997,73 | 1132 | 863 | 0,76 | -0,395 | -7,245 | 4,33E-13 | 4,19E-12 | 0,000394 | 0,0034345 | 0,0018295 | 0,0018295 | TRUE | 0,0239792 | No |
| orf19.475 | 4972 | 3570 | 4214 | 5192 | 4586 | 4108 | 4425 | 2932 | 4460 | 4545 | 4311 | 4281 | 3922 | 3920 | 3949 | 4135 | 4190,5 | 4399 | 3982 | 0,905 | -0,145 | -3,916 | 9,01E-05 | 0,00027805 | 0,00042 | 0,002326 | 0,0010687 | 0,0010687 | TRUE | 0,0076706 | No |
| orf19.4750 | 519 | 480 | 492 | 648 | 839 | 883 | 954 | 545 | 466 | 611 | 503 | 534 | 718 | 843 | 851 | 769 | 661,8 | 528 | 795 | 1,509 | 0,594 | 6,621 | 3,56E-11 | 2,76E-10 | 0,007923 | 0,0041731 | 0,006134 | 0,006134 | TRUE | 0,1201116 | No |
| orf19.4751 | 2219 | 1780 | 1917 | 2596 | 2558 | 2528 | 2564 | 1548 | 1991 | 2266 | 1961 | 2141 | 2188 | 2412 | 2288 | 2183 | 2178,7 | 2090 | 2268 | 1,087 | 0,12 | 2,068 | 0,03863012 | 0,06705858 | 0,002806 | 0,0026458 | 0,0027584 | 0,0027584 | TRUE | 0,0372879 | No |
| orf19.4751.1 | 1031 | 701 | 793 | 996 | 1268 | 1008 | 996 | 702 | 925 | 892 | 811 | 821 | 1084 | 962 | 889 | 990 | 921,89 | 862 | 981 | 1,138 | 0,187 | 2,536 | 0,01120449 | 0,02231242 | 0,004574 | 0,0035542 | 0,0041199 | 0,0041199 | TRUE | 0,063431 | No |
| orf19.4752 | 1812 | 1230 | 1662 | 2067 | 1643 | 1398 | 1548 | 979 | 1626 | 1566 | 1700 | 1704 | 1405 | 1334 | 1381 | 1381 | 1512,2 | 1649 | 1375 | 0,833 | -0,264 | -5,607 | 2,06E-08 | 1,09E-07 | 0,000398 | 0,0029395 | 0,0014535 | 0,0014535 | TRUE | 0,0127661 | No |
| orf19.4753 | 7672 | 5434 | 6864 | 8188 | 8483 | 7877 | 8677 | 4862 | 6883 | 6918 | 7023 | 6752 | 7255 | 7516 | 7743 | 6857 | 7118,3 | 6894 | 7343 | 1,066 | 0,092 | 2,161 | 0,03071486 | 0,05475813 | 0,001283 | 0,0021835 | 0,0016026 | 0,0016026 | TRUE | 0,0234022 | No |
| orf19.4754 | 1596 | 1397 | 1546 | 1893 | 1540 | 1661 | 1687 | 978 | 1432 | 1779 | 1582 | 1561 | 1317 | 1585 | 1505 | 1379 | 1517,5 | 1588 | 1446 | 0,912 | -0,132 | -1,712 | 0,08690973 | 0,13573009 | 0,006675 | 0,0029362 | 0,005071 | 0,005071 | TRUE | 0,0761769 | No |
| orf19.4755 | 3598 | 2535 | 3293 | 4033 | 3667 | 3409 | 3733 | 2299 | 3228 | 3227 | 3369 | 3326 | 3136 | 3253 | 3331 | 3243 | 3264 | 3288 | 3241 | 0,985 | -0,021 | -0,576 | 0,56464845 | 0,64445439 | 0,000277 | 0,0024243 | 0,0009956 | 0,0009956 | TRUE | 0,0060873 | No |
| orf19.4756 | 1153 | 950 | 1060 | 1282 | 1106 | 1097 | 1072 | 689 | 1034 | 1209 | 1084 | 1057 | 946 | 1047 | 957 | 972 | 1038,3 | 1096 | 980 | 0,896 | -0,158 | -2,496 | 0,01254305 | 0,02461093 | 0,002332 | 0,0033777 | 0,0028898 | 0,0028898 | TRUE | 0,0543149 | No |
| orf19.4757 | 7008 | 4979 | 6182 | 7674 | 7216 | 6665 | 7218 | 4237 | 6287 | 6339 | 6325 | 6328 | 6171 | 6360 | 6441 | 5976 | 6278,3 | 6320 | 6237 | 0,988 | -0,018 | -0,534 | 0,59312014 | 0,67045453 | 0,000347 | 0,0022107 | 0,0009207 | 0,0009207 | TRUE | 0,0091145 | No |
| orf19.4758 | 2750 | 2255 | 2522 | 3183 | 3194 | 3180 | 3266 | 2118 | 2467 | 2871 | 2580 | 2625 | 2732 | 3034 | 2914 | 2987 | 2776,3 | 2636 | 2917 | 1,107 | 0,147 | 2,637 | 0,0083753 | 0,01717952 | 0,002644 | 0,0025024 | 0,0026177 | 0,0026177 | TRUE | 0,0422865 | No |
| orf19.4759 | 3967 | 2594 | 3149 | 4122 | 4186 | 3699 | 3632 | 2978 | 3559 | 3302 | 3222 | 3399 | 3580 | 3529 | 3241 | 4200 | 3504,1 | 3370 | 3638 | 1,078 | 0,108 | 1,498 | 0,13411053 | 0,19639359 | 0,006701 | 0,0023938 | 0,0047234 | 0,0047234 | TRUE | 0,1291331 | No |
| orf19.4760 | 4464 | 4151 | 4399 | 5658 | 4783 | 5144 | 5289 | 2714 | 4005 | 5285 | 4501 | 4666 | 4091 | 4908 | 4720 | 3828 | 4500,2 | 4614 | 4387 | 0,951 | -0,072 | -0,778 | 0,43632206 | 0,52323811 | 0,011927 | 0,0023021 | 0,0079628 | 0,0079628 | TRUE | 0,1158487 | No |
| orf19.4761 | 1586 | 1251 | 1438 | 1811 | 1440 | 1461 | 1529 | 915 | 1423 | 1593 | 1471 | 1493 | 1232 | 1394 | 1364 | 1291 | 1407,6 | 1495 | 1320 | 0,884 | -0,177 | -3,108 | 0,00188316 | 0,00447018 | 0,001971 | 0,0030109 | 0,0024106 | 0,0024106 | TRUE | 0,0265028 | No |
| orf19.4763 | 134 | 118 | 130 | 183 | 203 | 183 | 217 | 143 | 120 | 150 | 133 | 151 | 174 | 175 | 194 | 202 | 162,24 | 138 | 186 | 1,339 | 0,421 | 3,547 | 0,00038932 | 0,00107641 | 0,002164 | 0,0109276 | 0,0072113 | 0,0072113 | TRUE | 0,0887074 | No |
| orf19.4764 | 951 | 885 | 892 | 1148 | 1214 | 1403 | 1364 | 841 | 853 | 1127 | 913 | 947 | 1038 | 1339 | 1217 | 1186 | 1077,4 | 960 | 1195 | 1,247 | 0,319 | 3,366 | 0,00076197 | 0,00196562 | 0,011623 | 0,0033269 | 0,0076544 | 0,0076544 | TRUE | 0,1579381 | No |
| orf19.4765 | 118 | 94 | 93 | 129 | 159 | 146 | 132 | 116 | 106 | 120 | 95 | 106 | 136 | 139 | 118 | 164 | 122,97 | 107 | 139 | 1,297 | 0,375 | 2,724 | 0,00645636 | 0,01361046 | 0,004338 | 0,0137851 | 0,0098773 | 0,0098773 | TRUE | 0,1190651 | No |
| orf19.4766 | 2412 | 1910 | 2290 | 2797 | 3187 | 3210 | 3423 | 2020 | 2164 | 2432 | 2343 | 2306 | 2726 | 3063 | 3054 | 2849 | 2617,1 | 2311 | 2923 | 1,266 | 0,34 | 6,236 | 4,49E-10 | 3,02E-09 | 0,002231 | 0,0025342 | 0,0024693 | 0,0024693 | TRUE | 0,0261793 | No |
| orf19.4767 | 570 | 477 | 556 | 637 | 511 | 450 | 526 | 319 | 511 | 607 | 569 | 525 | 437 | 429 | 469 | 450 | 499,8 | 553 | 446 | 0,81 | -0,304 | -4,033 | 5,50E-05 | 0,00017662 | 0,00201 | 0,0048841 | 0,0034372 | 0,0034372 | TRUE | 0,0468743 | No |
| orf19.4768 | 681 | 648 | 630 | 858 | 659 | 808 | 803 | 432 | 611 | 825 | 645 | 708 | 564 | 771 | 717 | 609 | 681,05 | 697 | 665 | 0,957 | -0,064 | -0,564 | 0,57272007 | 0,65131595 | 0,01757 | 0,0041111 | 0,010894 | 0,010894 | TRUE | 0,1737817 | No |
| orf19.4769 | 2401 | 1819 | 2197 | 2687 | 3245 | 3134 | 3270 | 2001 | 2154 | 2316 | 2248 | 2216 | 2775 | 2990 | 2918 | 2822 | 2554,9 | 2234 | 2876 | 1,29 | 0,367 | 8,63 | 6,14E-18 | 9,50E-17 | 0,000638 | 0,0025477 | 0,0013358 | 0,0013358 | TRUE | 0,0087103 | No |
| orf19.477 | 1121 | 827 | 1019 | 1225 | 1123 | 1069 | 1134 | 685 | 1006 | 1053 | 1043 | 1010 | 960 | 1020 | 1012 | 966 | 1008,7 | 1028 | 990 | 0,965 | -0,052 | -1,024 | 0,30594278 | 0,39175282 | 1,00E-08 | 0,0034187 | 0,0014626 | 0,0014626 | TRUE | 0,0055629 | No |
| orf19.4771 | 5987 | 4792 | 5715 | 7084 | 5948 | 5715 | 6135 | 3878 | 5371 | 6101 | 5847 | 5842 | 5087 | 5453 | 5474 | 5470 | 5580,5 | 5790 | 5371 | 0,928 | -0,108 | -2,267 | 0,02339533 | 0,0430371 | 0,001766 | 0,0022397 | 0,0020062 | 0,0020062 | TRUE | 0,0290969 | No |
| orf19.4772 | 1023 | 739 | 930 | 1129 | 1200 | 1128 | 1179 | 813 | 918 | 941 | 951 | 931 | 1026 | 1076 | 1052 | 1147 | 1005,3 | 935 | 1075 | 1,147 | 0,198 | 3,74 | 0,00018418 | 0,00054004 | 1,00E-08 | 0,0034236 | 0,0016826 | 0,0016826 | TRUE | 0,0209925 | No |
| orf19.4773 | 10048 | 7082 | 8635 | 11152 | 7716 | 6976 | 7287 | 4438 | 9014 | 9016 | 8834 | 9196 | 6599 | 6656 | 6502 | 6259 | 7759,7 | 9015 | 6504 | 0,722 | -0,47 | -14,645 | 1,45E-48 | 1,36E-46 | 0,00032 | 0,0021666 | 0,0008564 | 0,0008564 | TRUE | 0,0073758 | No |
| orf19.4774 | 728 | 494 | 620 | 819 | 1072 | 938 | 992 | 700 | 653 | 629 | 634 | 675 | 917 | 895 | 885 | 987 | 784,5 | 648 | 921 | 1,415 | 0,501 | 8,455 | 2,79E-17 | 4,04E-16 | 0,000365 | 0,00383 | 0,002057 | 0,002057 | TRUE | 0,0243868 | No |
| orf19.4775 | 7638 | 5195 | 6868 | 8473 | 10028 | 8998 | 9702 | 6360 | 6852 | 6614 | 7027 | 6987 | 8576 | 8586 | 8657 | 8970 | 7783,6 | 6870 | 8697 | 1,265 | 0,339 | 10,178 | 2,50E-24 | 6,03E-23 | 0,000422 | 0,002166 | 0,0009351 | 0,0009351 | TRUE | 0,0073927 | No |
| orf19.4776 | 1760 | 1274 | 1594 | 1901 | 1721 | 1652 | 1711 | 1129 | 1579 | 1622 | 1631 | 1568 | 1472 | 1576 | 1527 | 1592 | 1570,8 | 1600 | 1542 | 0,963 | -0,054 | -1,209 | 0,22651731 | 0,30589256 | 1,00E-08 | 0,0029037 | 0,0013092 | 0,0013092 | TRUE | 0,0115498 | No |
| orf19.4777 | 2918 | 2060 | 2544 | 3119 | 3414 | 3062 | 3315 | 1916 | 2618 | 2623 | 2603 | 2572 | 2920 | 2922 | 2958 | 2702 | 2739,6 | 2604 | 2876 | 1,106 | 0,146 | 3,626 | 0,0002879 | 0,00081627 | 0,000375 | 0,0025094 | 0,0011868 | 0,0011868 | TRUE | 0,0182168 | No |
| orf19.4778 | 340 | 228 | 263 | 375 | 422 | 367 | 423 | 237 | 305 | 290 | 269 | 309 | 361 | 350 | 377 | 334 | 324,55 | 293 | 356 | 1,213 | 0,279 | 3,341 | 0,00083341 | 0,00213341 | 1,00E-08 | 0,0064525 | 0,003608 | 0,003608 | TRUE | 0,0359967 | No |
| orf19.4779 | 7780 | 5663 | 7151 | 8655 | 7221 | 6279 | 6822 | 4356 | 6979 | 7210 | 7316 | 7137 | 6176 | 5991 | 6088 | 6144 | 6630,1 | 7160 | 6100 | 0,852 | -0,231 | -7,682 | 1,57E-14 | 1,71E-13 | 0,000154 | 0,0021985 | 0,0007169 | 0,0007169 | TRUE | 0,0035809 | No |
| orf19.478 | 1149 | 882 | 1070 | 1283 | 1565 | 1347 | 1466 | 986 | 1031 | 1123 | 1095 | 1058 | 1338 | 1285 | 1308 | 1391 | 1203,6 | 1077 | 1330 | 1,236 | 0,306 | 6,058 | 1,38E-09 | 8,64E-09 | 0,000289 | 0,0031856 | 0,0016071 | 0,0016071 | TRUE | 0,0097421 | No |
| orf19.4780 | 2251 | 1591 | 1980 | 2527 | 1804 | 1482 | 1721 | 1090 | 2019 | 2026 | 2026 | 2084 | 1543 | 1414 | 1536 | 1537 | 1773,1 | 2039 | 1508 | 0,739 | -0,436 | -9,738 | 2,08E-22 | 4,49E-21 | 0,000331 | 0,0027983 | 0,0013495 | 0,0013495 | TRUE | 0,0214583 | No |
| orf19.4781 | 489 | 304 | 405 | 512 | 433 | 336 | 407 | 314 | 439 | 387 | 414 | 422 | 370 | 321 | 363 | 443 | 394,9 | 416 | 374 | 0,894 | -0,162 | -1,707 | 0,08777213 | 0,13693455 | 0,007153 | 0,0056556 | 0,0061405 | 0,0061405 | TRUE | 0,1532655 | No |
| orf19.4783 | 149 | 103 | 124 | 136 | 115 | 103 | 134 | 77 | 134 | 131 | 127 | 112 | 98 | 98 | 120 | 109 | 116,08 | 126 | 106 | 0,845 | -0,243 | -1,818 | 0,06911756 | 0,11153062 | 1,00E-08 | 0,0144859 | 0,0085508 | 0,0085508 | TRUE | 0,0822476 | No |
| orf19.4784 | 1417 | 1014 | 1341 | 1512 | 1763 | 1507 | 1579 | 1077 | 1271 | 1291 | 1372 | 1247 | 1508 | 1438 | 1409 | 1519 | 1381,8 | 1295 | 1468 | 1,133 | 0,181 | 3,554 | 0,00038011 | 0,00105287 | 0,00083 | 0,0030301 | 0,0017533 | 0,0017533 | TRUE | 0,0193533 | No |
| orf19.4785 | 1419 | 874 | 1224 | 1516 | 1951 | 1572 | 1798 | 1090 | 1273 | 1113 | 1252 | 1250 | 1669 | 1500 | 1604 | 1537 | 1399,8 | 1222 | 1578 | 1,29 | 0,367 | 6,336 | 2,36E-10 | 1,63E-09 | 0,00196 | 0,0030166 | 0,002494 | 0,002494 | TRUE | 0,0404275 | No |
| orf19.4786 | 458 | 338 | 405 | 548 | 588 | 575 | 589 | 371 | 411 | 430 | 414 | 452 | 503 | 549 | 526 | 523 | 475,97 | 427 | 525 | 1,229 | 0,297 | 4,28 | 1,87E-05 | 6,51E-05 | 1,00E-08 | 0,0050295 | 0,0025163 | 0,0025163 | TRUE | 0,0204998 | No |
| orf19.4787 | 1226 | 859 | 1161 | 1375 | 1885 | 1624 | 1875 | 1191 | 1100 | 1094 | 1188 | 1134 | 1612 | 1550 | 1673 | 1680 | 1378,7 | 1129 | 1629 | 1,441 | 0,527 | 10,493 | 9,27E-26 | 2,54E-24 | 0,000655 | 0,0030325 | 0,0016724 | 0,0016724 | TRUE | 0,0143855 | No |
| orf19.4788 | 13643 | 9249 | 12071 | 14649 | 13751 | 11957 | 13040 | 8093 | 12239 | 11775 | 12350 | 12080 | 11760 | 11409 | 11636 | 11414 | 11833 | 12111 | 11555 | 0,954 | -0,068 | -2,359 | 0,01834063 | 0,03441302 | 0,000212 | 0,0021022 | 0,0007081 | 0,0007081 | TRUE | 0,0041278 | No |
| orf19.4789 | 60 | 33 | 38 | 53 | 62 | 60 | 76 | 50 | 54 | 42 | 39 | 44 | 53 | 57 | 68 | 71 | 53,38 | 45 | 62 | 1,376 | 0,461 | 2,336 | 0,01947823 | 0,03629814 | 1,00E-08 | 0,0291755 | 0,0182105 | 0,0182105 | TRUE | 0,1633206 | No |
| orf19.4791 | 1366 | 1148 | 1358 | 1612 | 1776 | 1962 | 2039 | 1151 | 1225 | 1462 | 1389 | 1329 | 1519 | 1872 | 1819 | 1623 | 1529,9 | 1351 | 1708 | 1,266 | 0,34 | 4,377 | 1,21E-05 | 4,33E-05 | 0,006545 | 0,0029284 | 0,0051415 | 0,0051415 | TRUE | 0,069954 | No |
| orf19.4792 | 15732 | 10879 | 14219 | 18047 | 17765 | 15331 | 17092 | 11488 | 14113 | 13850 | 14547 | 14882 | 15193 | 14628 | 15252 | 16203 | 14834 | 14348 | 15319 | 1,067 | 0,094 | 2,279 | 0,02264989 | 0,04176413 | 0,001326 | 0,0020774 | 0,0015608 | 0,0015608 | TRUE | 0,0181648 | No |
| orf19.479.2 | 346 | 283 | 318 | 357 | 449 | 384 | 417 | 256 | 310 | 360 | 325 | 294 | 384 | 366 | 372 | 361 | 346,75 | 322 | 371 | 1,158 | 0,212 | 2,545 | 0,01091876 | 0,02178675 | 0,000727 | 0,0061661 | 0,003732 | 0,003732 | TRUE | 0,0629875 | No |
| orf19.4793 | 5263 | 3941 | 4716 | 5707 | 5251 | 4956 | 5108 | 3095 | 4721 | 5017 | 4825 | 4706 | 4491 | 4729 | 4558 | 4365 | 4676,6 | 4817 | 4536 | 0,943 | -0,085 | -2,18 | 0,02929036 | 0,05245201 | 0,000715 | 0,0022899 | 0,0012522 | 0,0012522 | TRUE | 0,0098985 | No |
| orf19.4795 | 1638 | 1159 | 1514 | 1796 | 2047 | 2018 | 2077 | 1296 | 1469 | 1476 | 1549 | 1481 | 1751 | 1926 | 1853 | 1828 | 1666,6 | 1494 | 1840 | 1,231 | 0,3 | 6,432 | 1,26E-10 | 9,04E-10 | 0,000607 | 0,0028506 | 0,001484 | 0,001484 | TRUE | 0,0136784 | No |
| orf19.4796 | 2727 | 1824 | 2258 | 2831 | 2698 | 2322 | 2497 | 1495 | 2446 | 2322 | 2310 | 2334 | 2307 | 2216 | 2228 | 2109 | 2284,1 | 2353 | 2215 | 0,942 | -0,086 | -1,994 | 0,04611775 | 0,07833628 | 0,000453 | 0,0026151 | 0,0013345 | 0,0013345 | TRUE | 0,0114974 | No |
| orf19.4798 | 3725 | 2699 | 3428 | 4121 | 4024 | 3799 | 3874 | 2680 | 3342 | 3436 | 3507 | 3398 | 3441 | 3625 | 3457 | 3780 | 3498,3 | 3421 | 3576 | 1,044 | 0,062 | 1,488 | 0,13682933 | 0,1996286 | 0,000819 | 0,0023945 | 0,0014025 | 0,0014025 | TRUE | 0,0169461 | No |
| orf19.4799 | 5698 | 3906 | 5140 | 6072 | 5708 | 5207 | 5770 | 3627 | 5112 | 4973 | 5259 | 5007 | 4882 | 4968 | 5149 | 5116 | 5058,1 | 5088 | 5029 | 0,988 | -0,018 | -0,495 | 0,62076377 | 0,69386953 | 0,000367 | 0,0022665 | 0,0010048 | 0,0010048 | TRUE | 0,0061746 | No |
| orf19.48 | 3055 | 2832 | 2945 | 3612 | 4019 | 4635 | 4748 | 3005 | 2741 | 3605 | 3013 | 2979 | 3437 | 4423 | 4237 | 4238 | 3584,1 | 3084 | 4084 | 1,325 | 0,405 | 4,412 | 1,02E-05 | 3,71E-05 | 0,012531 | 0,0023846 | 0,007825 | 0,007825 | TRUE | 0,1554855 | No |
| orf19.480 | 4000 | 2663 | 3573 | 4502 | 4638 | 3914 | 4286 | 2741 | 3588 | 3390 | 3656 | 3712 | 3967 | 3735 | 3825 | 3866 | 3717,3 | 3586 | 3848 | 1,072 | 0,1 | 2,451 | 0,01426366 | 0,02764389 | 0,00072 | 0,00237 | 0,0013354 | 0,0013354 | TRUE | 0,0157644 | No |
| orf19.4800 | 1806 | 1354 | 1717 | 2033 | 2528 | 2253 | 2426 | 1470 | 1620 | 1724 | 1757 | 1676 | 2162 | 2150 | 2165 | 2073 | 1915,9 | 1694 | 2138 | 1,264 | 0,338 | 7,883 | 3,19E-15 | 3,72E-14 | 0,000204 | 0,0027372 | 0,0012359 | 0,0012359 | TRUE | 0,0106139 | No |
| orf19.4801 | 584 | 401 | 490 | 592 | 634 | 598 | 605 | 363 | 524 | 511 | 501 | 488 | 542 | 571 | 540 | 512 | 523,57 | 506 | 541 | 1,073 | 0,101 | 1,537 | 0,1242922 | 0,18383741 | 1,00E-08 | 0,0047522 | 0,0022607 | 0,0022607 | TRUE | 0,014911 | No |
| orf19.4802 | 5249 | 3863 | 4794 | 5975 | 5052 | 4436 | 4921 | 2801 | 4709 | 4918 | 4905 | 4927 | 4321 | 4233 | 4391 | 3951 | 4544,2 | 4865 | 4224 | 0,869 | -0,202 | -4,849 | 1,24E-06 | 5,15E-06 | 0,000982 | 0,002299 | 0,001443 | 0,001443 | TRUE | 0,0218365 | No |
| orf19.4803 | 27 | 22 | 22 | 23 | 14 | 11 | 14 | 9 | 24 | 28 | 23 | 19 | 12 | 10 | 12 | 13 | 17,67 | 24 | 12 | 0,512 | -0,966 | -2,911 | 0,00360329 | 0,00801798 | 1,00E-08 | 0,0841358 | 0,0423624 | 0,0423624 | TRUE | 0,113541 | No |
| orf19.4804 | 101 | 51 | 102 | 102 | 69 | 68 | 82 | 34 | 91 | 65 | 104 | 84 | 59 | 65 | 73 | 48 | 73,63 | 86 | 61 | 0,716 | -0,482 | -2,58 | 0,00988839 | 0,01993606 | 0,016575 | 0,0216963 | 0,0196349 | 0,0196349 | TRUE | 0,2145086 | No |
| orf19.4805 | 4479 | 3149 | 4154 | 4855 | 5039 | 4495 | 4949 | 3108 | 4018 | 4009 | 4250 | 4003 | 4310 | 4289 | 4416 | 4384 | 4209,9 | 4070 | 4350 | 1,069 | 0,096 | 2,733 | 0,00627513 | 0,01325632 | 0,000213 | 0,0023244 | 0,0009374 | 0,0009374 | TRUE | 0,0107217 | No |
| orf19.4807 | 1137 | 824 | 927 | 1246 | 1352 | 1273 | 1380 | 860 | 1020 | 1049 | 948 | 1027 | 1156 | 1215 | 1231 | 1213 | 1107,5 | 1011 | 1204 | 1,19 | 0,251 | 4,836 | 1,33E-06 | 5,49E-06 | 0,000365 | 0,0032903 | 0,0016832 | 0,0016832 | TRUE | 0,0203186 | No |

| | | | | | | | | | | | | | | | | | | | | | | | | | | | | | | | |
|---|---|---|---|---|---|---|---|---|---|---|---|---|---|---|---|---|---|---|---|---|---|---|---|---|---|---|---|---|---|---|---|
| orf19.4808 | 3788 | 2721 | 3403 | 4138 | 4085 | 4000 | 3968 | 2509 | 3398 | 3464 | 3482 | 3412 | 3494 | 3817 | 3541 | 3539 | 3518,3 | 3439 | 3598 | 1,046 | 0,065 | 1,645 | 0,10004868 | 0,15291082 | 0,00065 | 0,0023921 | 0,0012371 | 0,0012371 | TRUE | 0,0208084 | No |
| orf19.4809 | 1039 | 757 | 1012 | 1255 | 1293 | 1139 | 1123 | 736 | 932 | 964 | 1035 | 1035 | 1106 | 1087 | 1002 | 1038 | 1024,9 | 992 | 1058 | 1,067 | 0,094 | 1,596 | 0,11043818 | 0,16596113 | 0,001535 | 0,003396 | 0,0023503 | 0,0023503 | TRUE | 0,0209121 | No |
| orf19.481 | 4015 | 2726 | 3446 | 4214 | 3700 | 2997 | 3298 | 2215 | 3602 | 3470 | 3526 | 3475 | 3164 | 2860 | 2943 | 3124 | 3270,5 | 3518 | 3023 | 0,859 | -0,22 | -5,067 | 4,04E-07 | 1,82E-06 | 0,000895 | 0,0024234 | 0,0014996 | 0,0014996 | TRUE | 0,0160772 | No |
| orf19.4811 | 2055 | 1306 | 1818 | 2244 | 2435 | 2068 | 2112 | 1599 | 1844 | 1663 | 1860 | 1850 | 2082 | 1973 | 1885 | 2255 | 1926,5 | 1804 | 2049 | 1,133 | 0,18 | 2,845 | 0,00443784 | 0,00968491 | 0,003713 | 0,0027331 | 0,0033162 | 0,0033162 | TRUE | 0,052862 | No |
| orf19.4812 | 518 | 323 | 426 | 546 | 476 | 386 | 382 | 341 | 465 | 411 | 436 | 450 | 407 | 368 | 341 | 481 | 419,9 | 440 | 399 | 0,9 | -0,153 | -1,518 | 0,12900786 | 0,18978434 | 0,009761 | 0,0054368 | 0,0073006 | 0,0073006 | TRUE | 0,1956316 | No |
| orf19.4813 | 23689 | 15436 | 20647 | 25474 | 22379 | 18635 | 19951 | 12995 | 21252 | 19652 | 21124 | 21006 | 19139 | 17781 | 17803 | 18328 | 19511 | 20758 | 18263 | 0,88 | -0,185 | -4,656 | 3,22E-06 | 1,27E-05 | 0,001182 | 0,0020539 | 0,0014642 | 0,0014642 | TRUE | 0,0156244 | No |
| orf19.4814 | 107 | 77 | 101 | 114 | 82 | 70 | 65 | 39 | 96 | 98 | 103 | 94 | 70 | 67 | 58 | 55 | 80,16 | 98 | 62 | 0,645 | -0,633 | -4,072 | 4,66E-05 | 0,00015186 | 1,00E-08 | 0,0200894 | 0,0102138 | 0,0102138 | TRUE | 0,0624349 | No |
| orf19.4815 | 9916 | 6628 | 8595 | 10722 | 10958 | 9181 | 10036 | 6101 | 8896 | 8438 | 8794 | 8841 | 9372 | 8760 | 8955 | 8605 | 8832,6 | 8742 | 8923 | 1,021 | 0,03 | 0,793 | 0,42785086 | 0,51513587 | 0,00082 | 0,0021439 | 0,0012403 | 0,0012403 | TRUE | 0,0140787 | No |
| orf19.4816 | 1900 | 1324 | 1704 | 2038 | 2145 | 1845 | 1983 | 1301 | 1704 | 1686 | 1743 | 1681 | 1834 | 1760 | 1769 | 1835 | 1751,7 | 1704 | 1800 | 1,056 | 0,079 | 1,918 | 0,05512969 | 0,09139602 | 1,00E-08 | 0,0028083 | 0,001038 | 0,001038 | TRUE | 0,002954 | No |
| orf19.4817 | 658 | 534 | 560 | 745 | 562 | 551 | 560 | 384 | 590 | 680 | 573 | 614 | 481 | 526 | 500 | 542 | 563,14 | 614 | 512 | 0,833 | -0,264 | -3,591 | 0,00032957 | 0,00092656 | 0,002191 | 0,0045574 | 0,0033862 | 0,0033862 | TRUE | 0,0547274 | No |
| orf19.4818 | 3612 | 2585 | 3086 | 3921 | 3642 | 3003 | 3281 | 2218 | 3240 | 3291 | 3157 | 3233 | 3115 | 2865 | 2928 | 3128 | 3119,8 | 3230 | 3009 | 0,931 | -0,103 | -2,463 | 0,01377205 | 0,02676885 | 0,000687 | 0,0024448 | 0,0013644 | 0,0013644 | TRUE | 0,0124599 | No |
| orf19.4819 | 256 | 189 | 215 | 237 | 211 | 257 | 196 | 126 | 230 | 241 | 220 | 195 | 180 | 245 | 175 | 178 | 207,99 | 222 | 194 | 0,883 | -0,18 | -1,403 | 0,16062149 | 0,22851383 | 0,01309 | 0,008959 | 0,0109341 | 0,0109341 | TRUE | 0,3405801 | No |
| orf19.482 | 5201 | 3512 | 4469 | 5490 | 5045 | 4295 | 4796 | 2717 | 4666 | 4471 | 4572 | 4527 | 4315 | 4098 | 4280 | 3832 | 4345,1 | 4559 | 4131 | 0,907 | -0,141 | -3,175 | 0,00149919 | 0,00363779 | 0,001284 | 0,0023136 | 0,001654 | 0,001654 | TRUE | 0,0275055 | No |
| orf19.4820 | 2208 | 1580 | 2060 | 2585 | 2598 | 2283 | 2503 | 1544 | 1981 | 2012 | 2108 | 2132 | 2222 | 2178 | 2234 | 2178 | 2130,4 | 2058 | 2203 | 1,07 | 0,098 | 2,364 | 0,01807528 | 0,03396818 | 0,000286 | 0,002661 | 0,001181 | 0,001181 | TRUE | 0,0083363 | No |
| orf19.4821 | 9 | 5 | 5 | 6 | 15 | 19 | 14 | 8 | 8 | 6 | 5 | 5 | 13 | 18 | 12 | 11 | 9,9 | 6 | 14 | 2,252 | 1,171 | 2,593 | 0,00951693 | NA | 1,00E-08 | 0,1485489 | 0,0788892 | 0,0788892 | TRUE | 0,2120503 | No |
| orf19.4822 | 4 | 1 | 1 | 4 | 5 | 5 | 7 | 3 | 4 | 1 | 1 | 3 | 4 | 5 | 6 | 4 | 3,59 | 2 | 5 | 2,052 | 1,037 | 1,412 | 0,15795005 | NA | 1,00E-08 | 0,4065272 | 0,2098556 | 0,2098556 | TRUE | 0,150028 | No |
| orf19.4823 | 224 | 152 | 229 | 265 | 282 | 248 | 260 | 178 | 201 | 194 | 234 | 219 | 241 | 237 | 232 | 251 | 226,02 | 212 | 240 | 1,129 | 0,175 | 1,803 | 0,07136297 | 0,11458053 | 1,00E-08 | 0,0084025 | 0,0045902 | 0,0045902 | TRUE | 0,0516482 | No |
| orf19.4824 | 1525 | 1087 | 1428 | 1635 | 1610 | 1504 | 1591 | 1027 | 1368 | 1384 | 1461 | 1348 | 1377 | 1435 | 1420 | 1448 | 1405,2 | 1390 | 1420 | 1,021 | 0,03 | 0,644 | 0,51974193 | 0,60293677 | 1,00E-08 | 0,0030126 | 0,0013807 | 0,0013807 | TRUE | 0,0142422 | No |
| orf19.4825 | 1433 | 1054 | 1101 | 1518 | 1407 | 1184 | 1251 | 853 | 1286 | 1342 | 1126 | 1252 | 1203 | 1130 | 1116 | 1203 | 1207,3 | 1252 | 1163 | 0,929 | -0,106 | -1,701 | 0,08887913 | 0,1384459 | 0,002354 | 0,003182 | 0,0028672 | 0,0028672 | TRUE | 0,0530997 | No |
| orf19.4826 | 15281 | 9870 | 13191 | 16135 | 14758 | 11921 | 12683 | 8864 | 13709 | 12566 | 13496 | 13305 | 12622 | 11375 | 11317 | 12502 | 12611 | 13269 | 11954 | 0,901 | -0,151 | -3,056 | 0,00224526 | 0,00524477 | 0,002329 | 0,0020946 | 0,0022657 | 0,0022657 | TRUE | 0,0175842 | No |
| orf19.4827 | 12144 | 7579 | 10471 | 12890 | 11444 | 9282 | 9855 | 6333 | 10894 | 9649 | 10713 | 10629 | 9787 | 8857 | 8794 | 8932 | 9781,9 | 10471 | 9092 | 0,868 | -0,204 | -3,949 | 7,84E-05 | 0,00024329 | 0,002607 | 0,0021279 | 0,0024563 | 0,0024563 | TRUE | 0,0338089 | No |
| orf19.4828 | 8512 | 5623 | 7544 | 9208 | 12077 | 10530 | 11767 | 6704 | 7636 | 7159 | 7718 | 7593 | 10329 | 10047 | 10500 | 9455 | 8804,7 | 7526 | 10083 | 1,34 | 0,422 | 9,727 | 2,32E-22 | 4,96E-21 | 0,001432 | 0,0021444 | 0,0016922 | 0,0016922 | TRUE | 0,0209906 | No |
| orf19.4829 | 1551 | 1039 | 1443 | 1705 | 1904 | 1548 | 1683 | 1150 | 1391 | 1323 | 1476 | 1406 | 1628 | 1477 | 1502 | 1622 | 1478,2 | 1399 | 1557 | 1,111 | 0,152 | 2,8 | 0,00510758 | 0,01101053 | 0,001397 | 0,0029616 | 0,0021523 | 0,0021523 | TRUE | 0,0159375 | No |
| orf19.4830 | 361 | 271 | 293 | 449 | 510 | 406 | 381 | 289 | 324 | 345 | 300 | 370 | 436 | 387 | 340 | 408 | 363,75 | 335 | 393 | 1,171 | 0,228 | 2,308 | 0,02098075 | 0,03892908 | 0,00725 | 0,0059704 | 0,006618 | 0,006618 | TRUE | 0,0995512 | No |
| orf19.4831 | 3242 | 2152 | 2791 | 3358 | 3253 | 2724 | 2989 | 1867 | 2908 | 2740 | 2855 | 2769 | 2782 | 2599 | 2667 | 2633 | 2744,3 | 2818 | 2670 | 0,948 | -0,077 | -1,914 | 0,05557959 | 0,09209105 | 0,000487 | 0,0025085 | 0,0012037 | 0,0012037 | TRUE | 0,0099828 | No |
| orf19.4833 | 15000 | 9637 | 12731 | 15710 | 11923 | 9778 | 10533 | 7209 | 13457 | 12269 | 13025 | 12955 | 10197 | 9330 | 9399 | 10168 | 11350 | 12926 | 9774 | 0,756 | -0,404 | -8,866 | 7,58E-19 | 1,25E-17 | 0,001774 | 0,0021074 | 0,0019033 | 0,0019033 | TRUE | 0,0141792 | No |
| orf19.4834 | 29 | 26 | 18 | 37 | 26 | 31 | 24 | 16 | 26 | 33 | 18 | 31 | 22 | 30 | 21 | 23 | 25,48 | 27 | 24 | 0,89 | -0,168 | -0,593 | 0,5533662 | 0,6343492 | 1,00E-08 | 0,0589534 | 0,0373809 | 0,0373809 | TRUE | 0,2769892 | No |
| orf19.4835 | 13341 | 8910 | 11678 | 13922 | 13975 | 11804 | 12981 | 8113 | 11968 | 11343 | 11948 | 11480 | 11952 | 11263 | 11583 | 11443 | 11623 | 11685 | 11560 | 0,989 | -0,015 | -0,452 | 0,65094043 | 0,72023022 | 0,000542 | 0,0021044 | 0,0010315 | 0,0010315 | TRUE | 0,006444 | No |
| orf19.4836 | 2228 | 1393 | 1861 | 2392 | 2272 | 1798 | 1930 | 1347 | 1999 | 1773 | 1904 | 1972 | 1943 | 1716 | 1722 | 1900 | 1866,2 | 1912 | 1820 | 0,95 | -0,074 | -1,235 | 0,21684309 | 0,29515027 | 0,002785 | 0,0027574 | 0,0028764 | 0,0028764 | TRUE | 0,0276619 | No |
| orf19.4837 | 390 | 251 | 362 | 432 | 514 | 383 | 466 | 349 | 350 | 320 | 370 | 356 | 440 | 365 | 416 | 492 | 388,64 | 349 | 428 | 1,22 | 0,286 | 3,004 | 0,0026644 | 0,00612853 | 0,006556 | 0,0057149 | 0,0061211 | 0,0061211 | TRUE | 0,1107785 | No |
| orf19.4837.1 | 529 | 403 | 423 | 550 | 666 | 577 | 612 | 319 | 475 | 513 | 433 | 454 | 570 | 551 | 546 | 450 | 498,76 | 469 | 529 | 1,137 | 0,185 | 2,156 | 0,03106899 | 0,05529103 | 0,005148 | 0,0048901 | 0,0050769 | 0,0050769 | TRUE | 0,1044577 | No |
| orf19.4839 | 1276 | 829 | 1128 | 1486 | 1240 | 933 | 1082 | 738 | 1145 | 1055 | 1154 | 1225 | 1060 | 890 | 966 | 1041 | 1067,1 | 1145 | 989 | 0,862 | -0,214 | -3,076 | 0,00209478 | 0,00492966 | 0,00368 | 0,00334 | 0,0037104 | 0,0037104 | TRUE | 0,0547601 | No |
| orf19.484 | 6152 | 4148 | 5296 | 6498 | 6909 | 5819 | 6356 | 3942 | 5519 | 5281 | 5418 | 5358 | 5909 | 5552 | 5672 | 5560 | 5533,6 | 5394 | 5673 | 1,052 | 0,073 | 2,1 | 0,03569519 | 0,06254205 | 0,000362 | 0,0022419 | 0,000981 | 0,000981 | TRUE | 0,0097216 | No |
| orf19.4841 | 2044 | 1295 | 1741 | 2213 | 3360 | 2828 | 3135 | 1863 | 1834 | 1649 | 1781 | 1825 | 2874 | 2698 | 2797 | 2628 | 2260,7 | 1772 | 2749 | 1,551 | 0,633 | 12,847 | 8,95E-38 | 5,43E-36 | 0,001319 | 0,0026217 | 0,0018649 | 0,0018649 | TRUE | 0,0249268 | No |
| orf19.4842 | 769 | 530 | 637 | 901 | 1438 | 1104 | 1183 | 940 | 690 | 675 | 652 | 743 | 1230 | 1053 | 1056 | 1326 | 927,99 | 690 | 1166 | 1,685 | 0,753 | 9,004 | 2,18E-19 | 3,79E-18 | 0,006715 | 0,0035439 | 0,0055589 | 0,0055589 | TRUE | 0,0961046 | No |
| orf19.4843 | 6138 | 4268 | 5333 | 6662 | 6186 | 5757 | 6168 | 4012 | 5506 | 5434 | 5456 | 5494 | 5290 | 5493 | 5504 | 5659 | 5479,5 | 5472 | 5486 | 1,002 | 0,002 | 0,074 | 0,94137359 | 0,95779382 | 0,000154 | 0,0022445 | 0,0008134 | 0,0008134 | TRUE | 0,007129 | No |
| orf19.4844 | 3444 | 2199 | 2938 | 3722 | 3863 | 3033 | 3467 | 2089 | 3090 | 2800 | 3006 | 3069 | 3304 | 2894 | 3094 | 2946 | 3025,3 | 2991 | 3060 | 1,023 | 0,033 | 0,606 | 0,54444749 | 0,6261563 | 0,00221 | 0,0024594 | 0,0024454 | 0,0024454 | TRUE | 0,0356461 | No |
| orf19.4845 | 633 | 501 | 603 | 772 | 499 | 497 | 512 | 270 | 568 | 638 | 617 | 637 | 427 | 474 | 457 | 381 | 524,74 | 615 | 435 | 0,711 | -0,492 | -6,251 | 4,07E-10 | 2,75E-09 | 0,003227 | 0,0047461 | 0,0039904 | 0,0039904 | TRUE | 0,0677367 | No |
| orf19.4846 | 1637 | 1131 | 1432 | 1687 | 1795 | 1459 | 1644 | 1030 | 1469 | 1440 | 1465 | 1391 | 1535 | 1392 | 1467 | 1453 | 1451,5 | 1441 | 1462 | 1,015 | 0,022 | 0,453 | 0,6504987 | 0,71987404 | 0,000574 | 0,0029797 | 0,0015462 | 0,0015462 | TRUE | 0,0146248 | No |
| orf19.4847 | 37 | 30 | 43 | 35 | 50 | 31 | 35 | 22 | 33 | 38 | 44 | 29 | 43 | 30 | 31 | 31 | 34,86 | 36 | 34 | 0,947 | -0,078 | -0,319 | 0,75003062 | 0,80381448 | 1,00E-08 | 0,0436289 | 0,0290614 | 0,0290614 | TRUE | 0,2673494 | No |
| orf19.4848 | 1908 | 1394 | 1775 | 2107 | 2511 | 2226 | 2367 | 1497 | 1712 | 1775 | 1816 | 1737 | 2147 | 2124 | 2112 | 2111 | 1941,9 | 1760 | 2124 | 1,208 | 0,273 | 6,888 | 5,67E-12 | 4,74E-11 | 1,00E-08 | 0,0027271 | 0,0009832 | 0,0009832 | TRUE | 0,0057217 | No |
| orf19.4849 | 6368 | 4897 | 5601 | 7241 | 7368 | 6896 | 7254 | 4868 | 5713 | 6234 | 5730 | 5971 | 6301 | 6580 | 6473 | 6866 | 6233,6 | 5912 | 6555 | 1,109 | 0,149 | 3,466 | 0,00052886 | 0,00141785 | 0,001297 | 0,0022124 | 0,001605 | 0,001605 | TRUE | 0,0159647 | No |
| orf19.485 | 984 | 695 | 797 | 1007 | 893 | 794 | 891 | 530 | 883 | 885 | 815 | 830 | 764 | 758 | 795 | 748 | 809,66 | 853 | 766 | 0,9 | -0,153 | -2,702 | 0,00689426 | 0,01443734 | 1,00E-08 | 0,0037725 | 0,0018289 | 0,0018289 | TRUE | 0,0100439 | No |
| orf19.4850 | 9753 | 7546 | 8791 | 10870 | 10183 | 9749 | 10635 | 6082 | 8749 | 9607 | 8994 | 8964 | 8709 | 9302 | 9490 | 8578 | 9049,1 | 9078 | 9020 | 0,994 | -0,009 | -0,183 | 0,8547883 | 0,88950275 | 0,001645 | 0,0021399 | 0,0019778 | 0,0019778 | TRUE | 0,0185067 | No |
| orf19.4851 | 2297 | 1475 | 1974 | 2498 | 1782 | 1578 | 1693 | 1194 | 2061 | 1878 | 2020 | 2060 | 1524 | 1506 | 1511 | 1684 | 1780,3 | 2005 | 1556 | 0,774 | -0,37 | -7,014 | 2,32E-12 | 2,04E-11 | 0,001686 | 0,0027949 | 0,0021014 | 0,0021014 | TRUE | 0,0337424 | No |
| orf19.4853 | 248 | 162 | 229 | 272 | 169 | 145 | 167 | 118 | 222 | 206 | 234 | 224 | 145 | 138 | 149 | 166 | 185,71 | 222 | 150 | 0,668 | -0,581 | -5,507 | 3,64E-08 | 1,88E-07 | 1,00E-08 | 0,0097967 | 0,0050945 | 0,0050945 | TRUE | 0,0465625 | No |
| orf19.4855 | 885 | 633 | 733 | 889 | 930 | 730 | 829 | 518 | 794 | 806 | 750 | 733 | 795 | 697 | 740 | 731 | 755,63 | 771 | 741 | 0,963 | -0,054 | -0,848 | 0,3965047 | 0,485272 | 0,001387 | 0,0039007 | 0,002551 | 0,002551 | TRUE | 0,0314155 | No |
| orf19.4856 | 18 | 10 | 15 | 24 | 21 | 9 | 21 | 8 | 16 | 13 | 15 | 20 | 18 | 9 | 19 | 11 | 15,07 | 16 | 14 | 0,888 | -0,171 | -0,463 | 0,64319555 | NA | 1,00E-08 | 0,0982894 | 0,0657712 | 0,0657712 | TRUE | 0,3276599 | No |
| orf19.4857 | 2911 | 1977 | 2595 | 3134 | 2683 | 2392 | 2679 | 1914 | 2611 | 2517 | 2655 | 2584 | 2295 | 2282 | 2391 | 2700 | 2504,3 | 2592 | 2417 | 0,93 | -0,104 | -1,836 | 0,0664166 | 0,10793371 | 0,002958 | 0,0025592 | 0,0026815 | 0,0026815 | TRUE | 0,0715046 | No |
| orf19.4858 | 745 | 527 | 593 | 770 | 634 | 564 | 614 | 366 | 668 | 671 | 607 | 635 | 542 | 538 | 548 | 516 | 590,67 | 645 | 536 | 0,833 | -0,263 | -4,147 | 3,36E-05 | 0,00011217 | 1,00E-08 | 0,0044372 | 0,0021638 | 0,0021638 | TRUE | 0,0182026 | No |
| orf19.486 | 407 | 331 | 380 | 468 | 592 | 508 | 553 | 320 | 365 | 421 | 389 | 386 | 506 | 485 | 493 | 451 | 437,13 | 390 | 484 | 1,248 | 0,32 | 4,315 | 1,60E-05 | 5,66E-05 | 1,00E-08 | 0,0053005 | 0,0029684 | 0,0029684 | TRUE | 0,0291414 | No |
| orf19.4861 | 34 | 28 | 40 | 28 | 15 | 27 | 32 | 26 | 31 | 36 | 41 | 23 | 13 | 26 | 29 | 37 | 29,25 | 33 | 26 | 0,787 | -0,346 | -1,089 | 0,2762147 | 0,36025364 | 0,074984 | 0,051615 | 0,0615579 | 0,0615579 | TRUE | 0,6058161 | No |
| orf19.4861.1 | 14 | 9 | 16 | 11 | 8 | 10 | 15 | 1 | 13 | 11 | 16 | 9 | 7 | 10 | 13 | 1 | 10,08 | 12 | 8 | 0,661 | -0,598 | -1,233 | 0,21747073 | NA | 0,077894 | 0,1460034 | 0,1238903 | 0,1238903 | TRUE | 0,372134 | No |
| orf19.4862 | 1178 | 904 | 1043 | 1281 | 1066 | 1006 | 1144 | 749 | 1057 | 1151 | 1067 | 1056 | 912 | 960 | 1021 | 1056 | 1035 | 1083 | 987 | 0,911 | -0,135 | -2,223 | 0,02623443 | 0,04775937 | 0,001714 | 0,0033822 | 0,0025543 | 0,0025543 | TRUE | 0,0323681 | No |
| orf19.4862.2 | 2047 | 1385 | 1594 | 2052 | 2195 | 1999 | 1972 | 1292 | 1836 | 1763 | 1631 | 1692 | 1877 | 1907 | 1760 | 1822 | 1786,1 | 1730 | 1842 | 1,064 | 0,09 | 1,752 | 0,07981107 | 0,12625194 | 0,001434 | 0,0027923 | 0,0019751 | 0,0019751 | TRUE | 0,0213299 | No |
| orf19.4863 | 1408 | 1062 | 1300 | 1620 | 1858 | 1699 | 1870 | 1156 | 1263 | 1352 | 1330 | 1336 | 1589 | 1621 | 1669 | 1630 | 1473,8 | 1320 | 1627 | 1,233 | 0,303 | 6,743 | 1,55E-11 | 1,24E-10 | 1,00E-08 | 0,0029645 | 0,0012474 | 0,0012474 | TRUE | 0,0105126 | No |
| orf19.4864 | 830 | 692 | 810 | 1016 | 993 | 914 | 956 | 562 | 745 | 881 | 829 | 838 | 849 | 872 | 853 | 793 | 832,4 | 823 | 842 | 1,027 | 0,038 | 0,59 | 0,55524221 | 0,63540861 | 0,001933 | 0,0037235 | 0,0027792 | 0,0027792 | TRUE | 0,0498672 | No |
| orf19.4865 | 982 | 831 | 893 | 1111 | 1019 | 1125 | 1138 | 646 | 881 | 1058 | 914 | 916 | 871 | 1073 | 1015 | 911 | 955,03 | 942 | 968 | 1,03 | 0,043 | 0,523 | 0,60108685 | 0,67703971 | 0,00685 | 0,0034996 | 0,0052978 | 0,0052978 | TRUE | 0,0779033 | No |
| orf19.4866 | 4812 | 3503 | 4509 | 5479 | 4864 | 4501 | 5017 | 3522 | 4317 | 4460 | 4613 | 4518 | 4160 | 4295 | 4477 | 4967 | 4475,8 | 4477 | 4475 | 0,998 | -0,002 | -0,039 | 0,96920324 | 0,97799346 | 0,002772 | 0,0023039 | 0,0028008 | 0,0028008 | TRUE | 0,0651459 | No |

| | | | | | | | | | | | | | | | | | | | | | | | | | | | | | | | |
|---|---|---|---|---|---|---|---|---|---|---|---|---|---|---|---|---|---|---|---|---|---|---|---|---|---|---|---|---|---|---|---|
| orf19.4867 | 7846 | 5346 | 6883 | 8564 | 11634 | 10837 | 11881 | 7471 | 7039 | 6806 | 7042 | 7062 | 9950 | 10340 | 10602 | 10537 | 8672,2 | 6987 | 10357 | 1,482 | 0,567 | 17,112 | 1,22E-65 | 2,36E-63 | 0,000392 | 0,0021469 | 0,0009343 | 0,0009343 | TRUE | 0,008711 | No |
| orf19.4869 | 4173 | 2642 | 3614 | 4590 | 3752 | 3035 | 3591 | 2069 | 3744 | 3364 | 3697 | 3785 | 3209 | 2896 | 3204 | 2918 | 3352,1 | 3648 | 3057 | 0,838 | -0,255 | -4,644 | 3,42E-06 | 1,34E-05 | 0,002598 | 0,0024126 | 0,0025871 | 0,0025871 | TRUE | 0,0325138 | No |
| orf19.487 | 1370 | 865 | 1236 | 1501 | 1847 | 1627 | 1827 | 1228 | 1229 | 1101 | 1265 | 1238 | 1580 | 1552 | 1630 | 1732 | 1415,9 | 1208 | 1624 | 1,339 | 0,422 | 7,337 | 2,18E-13 | 2,15E-12 | 0,002044 | 0,0030048 | 0,0024465 | 0,0024465 | TRUE | 0,0393416 | No |
| orf19.4870 | 14345 | 10300 | 12859 | 15621 | 13486 | 12435 | 13531 | 8673 | 12869 | 13113 | 13156 | 12881 | 11534 | 11865 | 12074 | 12232 | 12466 | 13005 | 11926 | 0,917 | -0,125 | -4,216 | 2,49E-05 | 8,50E-05 | 0,000281 | 0,002096 | 0,0007663 | 0,0007663 | TRUE | 0,0060589 | No |
| orf19.4871 | 18776 | 14102 | 16977 | 20983 | 18055 | 17150 | 18083 | 11063 | 16844 | 17953 | 17369 | 17303 | 15441 | 16364 | 16136 | 15603 | 16627 | 17367 | 15886 | 0,915 | -0,128 | -3,779 | 0,00015735 | 0,00046547 | 0,000639 | 0,0020668 | 0,0010453 | 0,0010453 | TRUE | 0,006232 | No |
| orf19.4872 | 159 | 154 | 178 | 220 | 139 | 154 | 163 | 112 | 143 | 196 | 182 | 181 | 119 | 147 | 145 | 158 | 158,93 | 176 | 142 | 0,809 | -0,306 | -2,294 | 0,0217987 | 0,04033465 | 0,010198 | 0,0111136 | 0,0106965 | 0,0106965 | TRUE | 0,1774803 | No |
| orf19.4873 | 8485 | 6323 | 7483 | 9331 | 12611 | 11711 | 11972 | 7825 | 7612 | 8050 | 7656 | 7694 | 10785 | 11174 | 10683 | 11036 | 9336,4 | 7753 | 10920 | 1,409 | 0,494 | 15,255 | 1,52E-52 | 1,87E-50 | 0,000401 | 0,002135 | 0,0008981 | 0,0008981 | TRUE | 0,0078437 | No |
| orf19.4874 | 4551 | 3285 | 4149 | 5024 | 5444 | 5082 | 5469 | 3197 | 4083 | 4182 | 4245 | 4143 | 4656 | 4849 | 4880 | 4509 | 4443,4 | 4163 | 4724 | 1,136 | 0,184 | 4,87 | 1,12E-06 | 4,67E-06 | 0,000511 | 0,0023062 | 0,0011382 | 0,0011382 | TRUE | 0,010665 | No |
| orf19.4875 | 829 | 582 | 668 | 797 | 923 | 757 | 857 | 453 | 744 | 741 | 683 | 657 | 789 | 722 | 765 | 639 | 717,58 | 706 | 729 | 1,037 | 0,053 | 0,689 | 0,49099623 | 0,57626595 | 0,004395 | 0,0040026 | 0,0042055 | 0,0042055 | TRUE | 0,0733627 | No |
| orf19.4878 | 811 | 528 | 734 | 908 | 1139 | 921 | 1094 | 658 | 728 | 672 | 751 | 749 | 974 | 879 | 976 | 928 | 832,08 | 725 | 939 | 1,294 | 0,372 | 6,053 | 1,42E-09 | 8,87E-09 | 0,001237 | 0,0037242 | 0,0024062 | 0,0024062 | TRUE | 0,0253401 | No |
| orf19.4879.2 | 3158 | 2351 | 2672 | 3412 | 3664 | 3277 | 3239 | 2201 | 2833 | 2993 | 2734 | 2814 | 3134 | 3127 | 2890 | 3104 | 2953,6 | 2844 | 3064 | 1,078 | 0,108 | 2,406 | 0,01613285 | 0,03077021 | 0,000999 | 0,002471 | 0,001603 | 0,001603 | TRUE | 0,0181585 | No |
| orf19.488 | 4519 | 3068 | 4180 | 4855 | 4631 | 4238 | 4526 | 2871 | 4054 | 3906 | 4277 | 4003 | 3961 | 4044 | 4039 | 4049 | 4041,5 | 4060 | 4023 | 0,99 | -0,014 | -0,366 | 0,71471034 | 0,77508659 | 0,000465 | 0,0023387 | 0,001126 | 0,001126 | TRUE | 0,0154325 | No |
| orf19.4880 | 266 | 176 | 246 | 289 | 263 | 207 | 252 | 126 | 239 | 224 | 252 | 238 | 225 | 198 | 225 | 178 | 222,21 | 238 | 206 | 0,872 | -0,198 | -1,931 | 0,05344897 | 0,08907667 | 0,001305 | 0,0085124 | 0,0055962 | 0,0055962 | TRUE | 0,0764297 | No |
| orf19.4882 | 5328 | 3431 | 4622 | 5731 | 5917 | 4942 | 5415 | 2990 | 4780 | 4368 | 4729 | 4726 | 5060 | 4716 | 4832 | 4217 | 4678,4 | 4651 | 4706 | 1,013 | 0,018 | 0,309 | 0,75697326 | 0,81004889 | 0,003014 | 0,0022898 | 0,0030197 | 0,0030197 | TRUE | 0,0580539 | No |
| orf19.4883 | 3411 | 2503 | 3133 | 3823 | 3483 | 3258 | 3473 | 2209 | 3060 | 3187 | 3205 | 3152 | 2979 | 3109 | 3099 | 3116 | 3113,3 | 3151 | 3076 | 0,976 | -0,035 | -0,976 | 0,32897436 | 0,41645132 | 0,000154 | 0,0024458 | 0,0009278 | 0,0009278 | TRUE | 0,005674 | No |
| orf19.4884 | 166 | 106 | 139 | 151 | 111 | 120 | 118 | 89 | 149 | 135 | 142 | 125 | 95 | 115 | 105 | 126 | 123,86 | 138 | 110 | 0,793 | -0,335 | -2,526 | 0,01154695 | 0,02288062 | 1,00E-08 | 0,0137004 | 0,0087134 | 0,0087134 | TRUE | 0,089137 | No |
| orf19.4885 | 15610 | 10251 | 13559 | 16531 | 15225 | 12508 | 13116 | 9831 | 14004 | 13051 | 13872 | 13632 | 13021 | 11935 | 11704 | 13866 | 13135 | 13640 | 12632 | 0,926 | -0,111 | -1,979 | 0,04784899 | 0,08095656 | 0,003532 | 0,00209 | 0,0029652 | 0,0029652 | TRUE | 0,0523949 | No |
| orf19.4886 | 93 | 70 | 83 | 90 | 86 | 82 | 80 | 42 | 83 | 89 | 85 | 74 | 74 | 78 | 71 | 59 | 76,76 | 83 | 70 | 0,864 | -0,212 | -1,332 | 0,18292681 | 0,25517165 | 1,00E-08 | 0,0208912 | 0,0111942 | 0,0111942 | TRUE | 0,0766212 | No |
| orf19.4887 | 10268 | 7606 | 9439 | 11324 | 13227 | 12112 | 12871 | 8651 | 9211 | 9683 | 9657 | 9338 | 11312 | 11557 | 11485 | 12201 | 10556 | 9472 | 11639 | 1,228 | 0,297 | 8,243 | 1,68E-16 | 2,22E-15 | 0,000736 | 0,0021171 | 0,0011495 | 0,0011495 | TRUE | 0,0126209 | No |
| orf19.4888 | 3251 | 2374 | 2745 | 3395 | 3555 | 3353 | 3399 | 2097 | 2916 | 3022 | 2808 | 2800 | 3040 | 3199 | 3033 | 2958 | 2972,1 | 2886 | 3058 | 1,061 | 0,085 | 1,989 | 0,0467348 | 0,07927235 | 0,000764 | 0,002468 | 0,0014149 | 0,0014149 | TRUE | 0,0117622 | No |
| orf19.4889 | 5175 | 3943 | 5092 | 6023 | 5898 | 5561 | 5937 | 3608 | 4643 | 5020 | 5210 | 4967 | 5044 | 5306 | 5298 | 5089 | 5071,9 | 4960 | 5184 | 1,046 | 0,065 | 1,477 | 0,13976243 | 0,20306957 | 0,001217 | 0,0022657 | 0,0016374 | 0,0016374 | TRUE | 0,0228534 | No |
| orf19.489 | 846 | 633 | 703 | 931 | 553 | 498 | 535 | 345 | 759 | 806 | 719 | 768 | 473 | 475 | 477 | 487 | 620,49 | 763 | 478 | 0,627 | -0,674 | -10,79 | 3,85E-27 | 1,19E-25 | 1,00E-08 | 0,0043191 | 0,002048 | 0,002048 | TRUE | 0,0166484 | No |
| orf19.4890 | 1668 | 1192 | 1493 | 1807 | 1905 | 1875 | 1924 | 1301 | 1496 | 1518 | 1527 | 1490 | 1629 | 1789 | 1717 | 1835 | 1625,2 | 1508 | 1742 | 1,154 | 0,206 | 4,233 | 2,31E-05 | 7,94E-05 | 0,000852 | 0,0028728 | 0,0016621 | 0,0016621 | TRUE | 0,0235433 | No |
| orf19.4892 | 12680 | 9643 | 11353 | 14128 | 12078 | 11737 | 12118 | 7205 | 11375 | 12277 | 11615 | 11650 | 10329 | 11199 | 10813 | 10162 | 11178 | 11729 | 10626 | 0,906 | -0,142 | -3,352 | 0,000802 | 0,00206004 | 0,001403 | 0,0021094 | 0,0016303 | 0,0016303 | TRUE | 0,0160753 | No |
| orf19.4893 | 2223 | 1588 | 2043 | 2373 | 2906 | 2713 | 2949 | 1773 | 1994 | 2022 | 2090 | 1957 | 2485 | 2589 | 2631 | 2501 | 2283,6 | 2016 | 2552 | 1,267 | 0,342 | 8,249 | 1,60E-16 | 2,13E-15 | 0,000376 | 0,0026152 | 0,0012016 | 0,0012016 | TRUE | 0,0076355 | No |
| orf19.4894 | 533 | 334 | 428 | 515 | 530 | 460 | 456 | 365 | 478 | 425 | 438 | 425 | 453 | 439 | 407 | 515 | 447,48 | 442 | 454 | 1,021 | 0,03 | 0,356 | 0,72184236 | 0,78197419 | 0,003626 | 0,0052237 | 0,0045583 | 0,0045583 | TRUE | 0,0841986 | No |
| orf19.4895 | 1493 | 966 | 1245 | 1535 | 1604 | 1318 | 1542 | 893 | 1339 | 1230 | 1274 | 1266 | 1372 | 1258 | 1376 | 1259 | 1296,7 | 1277 | 1316 | 1,031 | 0,045 | 0,834 | 0,40449939 | 0,49273601 | 0,001147 | 0,0030991 | 0,0019799 | 0,0019799 | TRUE | 0,0130518 | No |
| orf19.4896 | 4848 | 3181 | 4019 | 5037 | 4036 | 3590 | 3811 | 2489 | 4349 | 4050 | 4112 | 4154 | 3452 | 3425 | 3401 | 3511 | 3806,6 | 4166 | 3447 | 0,827 | -0,275 | -7,595 | 3,08E-14 | 3,27E-13 | 0,000333 | 0,0023609 | 0,0009908 | 0,0009908 | TRUE | 0,010742 | No |
| orf19.4897 | 860 | 594 | 725 | 899 | 682 | 627 | 707 | 410 | 772 | 756 | 742 | 741 | 583 | 598 | 631 | 578 | 675,19 | 753 | 598 | 0,795 | -0,33 | -5,604 | 2,10E-08 | 1,12E-07 | 1,00E-08 | 0,0041296 | 0,0018363 | 0,0018363 | TRUE | 0,0170634 | No |
| orf19.4898 | 1825 | 1088 | 1405 | 1705 | 2257 | 1904 | 2154 | 1244 | 1637 | 1385 | 1437 | 1406 | 1930 | 1817 | 1922 | 1755 | 1661,2 | 1466 | 1856 | 1,266 | 0,34 | 5,42 | 5,97E-08 | 3,01E-07 | 0,003358 | 0,0028534 | 0,0031694 | 0,0031694 | TRUE | 0,0752454 | No |
| orf19.4899 | 658 | 446 | 585 | 772 | 1082 | 863 | 936 | 660 | 590 | 568 | 599 | 637 | 925 | 823 | 835 | 931 | 738,51 | 598 | 878 | 1,463 | 0,549 | 8,174 | 2,98E-16 | 3,85E-15 | 0,002054 | 0,0039452 | 0,0029308 | 0,0029308 | TRUE | 0,0228363 | No |
| orf19.490 | 1365 | 895 | 1171 | 1480 | 1169 | 1014 | 1066 | 675 | 1225 | 1139 | 1198 | 1220 | 1000 | 968 | 951 | 952 | 1081,6 | 1196 | 968 | 0,809 | -0,306 | -6,137 | 8,39E-10 | 5,41E-09 | 1,00E-08 | 0,0033217 | 0,0014565 | 0,0014565 | TRUE | 0,0108985 | No |
| orf19.4900 | 1302 | 1050 | 1170 | 1551 | 1156 | 1224 | 1241 | 755 | 1168 | 1337 | 1197 | 1279 | 989 | 1168 | 1107 | 1065 | 1163,7 | 1245 | 1082 | 0,87 | -0,201 | -3,018 | 0,00254533 | 0,00587263 | 0,003526 | 0,003227 | 0,0033946 | 0,0033946 | TRUE | 0,0431711 | No |
| orf19.4901 | 659 | 506 | 592 | 801 | 700 | 636 | 649 | 455 | 591 | 644 | 606 | 661 | 599 | 607 | 579 | 642 | 615,99 | 626 | 607 | 0,967 | -0,048 | -0,729 | 0,4657987 | 0,55186651 | 0,000727 | 0,0043362 | 0,0024874 | 0,0024874 | TRUE | 0,0191962 | No |
| orf19.4903 | 933 | 948 | 942 | 1286 | 847 | 994 | 936 | 589 | 837 | 1207 | 964 | 1060 | 724 | 948 | 835 | 831 | 925,86 | 1017 | 834 | 0,822 | -0,284 | -2,6 | 0,00932669 | 0,01892426 | 0,016349 | 0,0035475 | 0,0103274 | 0,0103274 | TRUE | 0,1836291 | No |
| orf19.4904 | 804 | 647 | 718 | 953 | 979 | 967 | 1003 | 668 | 721 | 824 | 735 | 786 | 837 | 923 | 895 | 942 | 832,82 | 766 | 899 | 1,173 | 0,23 | 3,586 | 0,00033529 | 0,00093826 | 0,001877 | 0,0037226 | 0,0027418 | 0,0027418 | TRUE | 0,0271394 | No |
| orf19.4905 | 920 | 657 | 832 | 1057 | 1129 | 1021 | 1181 | 679 | 825 | 836 | 851 | 872 | 966 | 974 | 1054 | 958 | 916,98 | 846 | 988 | 1,168 | 0,224 | 4,07 | 4,70E-05 | 0,00015299 | 1,00E-08 | 0,0035626 | 0,0018221 | 0,0018221 | TRUE | 0,0250569 | No |
| orf19.4906 | 6112 | 4075 | 5613 | 6926 | 7828 | 6904 | 7642 | 4890 | 5483 | 5188 | 5743 | 5711 | 6695 | 6588 | 6819 | 6897 | 6140,4 | 5531 | 6750 | 1,219 | 0,286 | 6,973 | 3,10E-12 | 2,68E-11 | 0,00106 | 0,0022159 | 0,0014508 | 0,0014508 | TRUE | 0,0206117 | No |
| orf19.4907 | 10625 | 7891 | 9550 | 12144 | 12038 | 11315 | 11786 | 7531 | 9532 | 10046 | 9771 | 10014 | 10295 | 10796 | 10517 | 10622 | 10199 | 9841 | 10558 | 1,073 | 0,101 | 3,191 | 0,00141783 | 0,00345712 | 0,000363 | 0,0021219 | 0,0008725 | 0,0008725 | TRUE | 0,0055132 | No |
| orf19.4908 | 151 | 92 | 96 | 137 | 117 | 89 | 109 | 74 | 135 | 117 | 98 | 113 | 100 | 85 | 97 | 104 | 106,3 | 116 | 96 | 0,83 | -0,27 | -1,857 | 0,06324841 | 0,10317579 | 1,00E-08 | 0,0156362 | 0,010745 | 0,010745 | TRUE | 0,1388612 | No |
| orf19.4909 | 2248 | 1518 | 1951 | 2400 | 2528 | 2258 | 2474 | 1572 | 2017 | 1933 | 1996 | 1979 | 2162 | 2155 | 2208 | 2217 | 2083,2 | 1981 | 2186 | 1,102 | 0,14 | 3,684 | 0,00022949 | 0,00066128 | 1,00E-08 | 0,0026764 | 0,0009007 | 0,0009007 | TRUE | 0,0029635 | No |
| orf19.4909.1 | 11410 | 7203 | 8342 | 10971 | 10814 | 8262 | 8737 | 5351 | 10236 | 9170 | 8535 | 9047 | 9248 | 7883 | 7796 | 7547 | 8682,9 | 9247 | 8118 | 0,878 | -0,187 | -2,583 | 0,009781 | 0,01973942 | 0,007287 | 0,0021467 | 0,0049392 | 0,0049392 | TRUE | 0,1075614 | No |
| orf19.491 | 2896 | 1973 | 2565 | 3136 | 3241 | 2838 | 3188 | 1882 | 2598 | 2512 | 2624 | 2586 | 2772 | 2708 | 2845 | 2654 | 2662,4 | 2580 | 2745 | 1,064 | 0,09 | 2,338 | 0,01938648 | 0,03614958 | 0,000195 | 0,0025248 | 0,0010504 | 0,0010504 | TRUE | 0,0072502 | No |
| orf19.4910 | 124 | 98 | 100 | 137 | 178 | 164 | 153 | 101 | 111 | 125 | 102 | 113 | 152 | 156 | 137 | 142 | 129,87 | 113 | 147 | 1,308 | 0,388 | 3,135 | 0,00171872 | 0,00411234 | 1,00E-08 | 0,0131574 | 0,0068936 | 0,0068936 | TRUE | 0,040934 | No |
| orf19.4911 | 389 | 258 | 345 | 391 | 478 | 426 | 472 | 301 | 349 | 328 | 353 | 322 | 409 | 406 | 421 | 425 | 376,73 | 338 | 415 | 1,226 | 0,294 | 3,955 | 7,66E-05 | 0,00023849 | 1,00E-08 | 0,005833 | 0,0026468 | 0,0026468 | TRUE | 0,0133649 | No |
| orf19.4912 | 1783 | 1155 | 1528 | 1894 | 2198 | 1832 | 2055 | 1327 | 1600 | 1470 | 1563 | 1562 | 1880 | 1748 | 1834 | 1872 | 1691 | 1549 | 1834 | 1,182 | 0,241 | 5,183 | 2,19E-07 | 1,02E-06 | 0,000522 | 0,002838 | 0,0014762 | 0,0014762 | TRUE | 0,0128194 | No |
| orf19.4913 | 1131 | 1013 | 1144 | 1448 | 1526 | 1596 | 1654 | 979 | 1015 | 1290 | 1170 | 1194 | 1305 | 1523 | 1476 | 1381 | 1294,2 | 1167 | 1421 | 1,22 | 0,286 | 3,708 | 0,0002089 | 0,00060661 | 0,006079 | 0,0031012 | 0,0049405 | 0,0049405 | TRUE | 0,0938113 | No |
| orf19.4914 | 227 | 195 | 193 | 260 | 206 | 171 | 176 | 111 | 204 | 248 | 197 | 214 | 176 | 163 | 157 | 157 | 189,59 | 216 | 163 | 0,761 | -0,393 | -3,588 | 0,00033261 | 0,00093206 | 1,00E-08 | 0,0096367 | 0,0061536 | 0,0061536 | TRUE | 0,0964459 | No |
| orf19.4914.1 | 32755 | 23651 | 27793 | 33747 | 42186 | 39346 | 39854 | 21332 | 29385 | 30110 | 28435 | 27828 | 36079 | 37543 | 35563 | 30087 | 31879 | 28940 | 34818 | 1,203 | 0,267 | 4,164 | 3,13E-05 | 0,00010504 | 0,004959 | 0,0020251 | 0,0039193 | 0,0039193 | TRUE | 0,102152 | No |
| orf19.4916 | 663 | 440 | 583 | 734 | 843 | 774 | 866 | 535 | 595 | 560 | 596 | 605 | 721 | 739 | 773 | 755 | 667,94 | 589 | 747 | 1,264 | 0,338 | 5,694 | 1,24E-08 | 6,80E-08 | 1,00E-08 | 0,0041529 | 0,0018695 | 0,0018695 | TRUE | 0,0112866 | No |
| orf19.4917 | 89 | 56 | 84 | 113 | 93 | 83 | 103 | 51 | 80 | 71 | 86 | 93 | 80 | 79 | 92 | 72 | 81,6 | 82 | 81 | 0,976 | -0,034 | -0,219 | 0,82677324 | 0,86684949 | 1,00E-08 | 0,0197691 | 0,0116084 | 0,0116084 | TRUE | 0,0846254 | No |
| orf19.4918 | 356 | 240 | 280 | 391 | 253 | 238 | 301 | 209 | 319 | 306 | 286 | 322 | 216 | 227 | 269 | 295 | 280,08 | 308 | 252 | 0,809 | -0,305 | -2,877 | 0,00401182 | 0,0088419 | 0,00711 | 0,0071627 | 0,0071936 | 0,0071936 | TRUE | 0,1267146 | No |
| orf19.4919 | 432 | 282 | 388 | 417 | 376 | 295 | 320 | 226 | 388 | 359 | 397 | 344 | 322 | 281 | 286 | 319 | 336,84 | 372 | 302 | 0,81 | -0,304 | -3,494 | 0,00047586 | 0,00129249 | 0,001983 | 0,0062893 | 0,0042418 | 0,0042418 | TRUE | 0,0332496 | No |
| orf19.492 | 6342 | 4099 | 5659 | 6722 | 7265 | 6248 | 6879 | 4822 | 5689 | 5218 | 5790 | 5543 | 6213 | 5962 | 6138 | 6801 | 5919,4 | 5560 | 6278 | 1,128 | 0,174 | 3,396 | 0,00068474 | 0,00178324 | 0,002459 | 0,0022248 | 0,00235 | 0,00235 | TRUE | 0,0373552 | No |
| orf19.4921 | 978 | 693 | 884 | 1038 | 1125 | 1002 | 1019 | 657 | 877 | 882 | 904 | 856 | 962 | 956 | 909 | 927 | 909,27 | 880 | 938 | 1,068 | 0,095 | 1,826 | 0,06781037 | 0,10975342 | 1,00E-08 | 0,0035761 | 0,0014886 | 0,0014886 | TRUE | 0,0059458 | No |
| orf19.4921.1 | 1138 | 1028 | 1020 | 1455 | 977 | 919 | 865 | 471 | 1021 | 1309 | 1044 | 1200 | 836 | 877 | 772 | 664 | 965,21 | 1144 | 787 | 0,691 | -0,534 | -5,426 | 5,75E-08 | 2,90E-07 | 0,012565 | 0,0034836 | 0,0082045 | 0,0082045 | TRUE | 0,1223589 | No |
| orf19.4922 | 942 | 698 | 861 | 1058 | 764 | 760 | 799 | 461 | 845 | 889 | 881 | 872 | 653 | 725 | 713 | 650 | 778,6 | 872 | 685 | 0,788 | -0,343 | -5,781 | 7,44E-09 | 4,20E-08 | 0,000345 | 0,003844 | 0,0020805 | 0,0020805 | TRUE | 0,0174944 | No |
| orf19.4923 | 175 | 133 | 137 | 177 | 239 | 195 | 229 | 139 | 157 | 169 | 140 | 146 | 204 | 186 | 204 | 196 | 175,41 | 153 | 198 | 1,298 | 0,376 | 3,487 | 0,00048935 | 0,00132432 | 1,00E-08 | 0,0102555 | 0,005405 | 0,005405 | TRUE | 0,0443223 | No |
| orf19.4923.1 | 26 | 23 | 26 | 23 | 33 | 28 | 22 | 15 | 23 | 29 | 27 | 19 | 28 | 27 | 20 | 21 | 24,24 | 24 | 24 | 0,995 | -0,008 | -0,027 | 0,97862933 | 0,98436381 | 1,00E-08 | 0,0618745 | 0,036041 | 0,036041 | TRUE | 0,1658421 | No |

| | | | | | | | | | | | | | | | | | | | | | | | | | | | | | | | |
|---|---|---|---|---|---|---|---|---|---|---|---|---|---|---|---|---|---|---|---|---|---|---|---|---|---|---|---|---|---|---|---|
| orf19.4924 | 1792 | 1359 | 1666 | 1988 | 2550 | 2534 | 2645 | 1596 | 1608 | 1730 | 1704 | 1639 | 2181 | 2418 | 2360 | 2251 | 1986,4 | 1670 | 2302 | 1,38 | 0,465 | 9,467 | 2,88E-21 | 5,70E-20 | 0,001226 | 0,0027103 | 0,0017977 | 0,0017977 | TRUE | 0,0163815 | No |
| orf19.4927 | 3028 | 2044 | 2718 | 3133 | 3915 | 3605 | 3949 | 2350 | 2716 | 2602 | 2781 | 2584 | 3348 | 3440 | 3524 | 3314 | 3038,7 | 2671 | 3406 | 1,276 | 0,352 | 8,535 | 1,40E-17 | 2,09E-16 | 0,000558 | 0,0024573 | 0,0012959 | 0,0012959 | TRUE | 0,0092166 | No |
| orf19.4928 | 4695 | 3509 | 4361 | 5278 | 4362 | 4309 | 4628 | 2668 | 4212 | 4467 | 4462 | 4352 | 3731 | 4112 | 4130 | 3763 | 4153,5 | 4373 | 3934 | 0,9 | -0,152 | -3,219 | 0,00128561 | 0,00316425 | 0,001567 | 0,002329 | 0,0018895 | 0,0018895 | TRUE | 0,0155219 | No |
| orf19.4929 | 2541 | 1828 | 2358 | 2615 | 3113 | 2998 | 3193 | 1845 | 2280 | 2327 | 2412 | 2156 | 2662 | 2861 | 2849 | 2602 | 2518,8 | 2294 | 2744 | 1,198 | 0,26 | 5,084 | 3,69E-07 | 1,67E-06 | 0,001692 | 0,0025559 | 0,0021172 | 0,0021172 | TRUE | 0,0203731 | No |
| orf19.493 | 25460 | 16005 | 20499 | 25526 | 25663 | 20052 | 20963 | 12759 | 22840 | 20376 | 20972 | 21049 | 21948 | 19133 | 18706 | 17995 | 20378 | 21309 | 19446 | 0,913 | -0,132 | -2,084 | 0,03715478 | 0,06475948 | 0,005152 | 0,0020508 | 0,0038002 | 0,0038002 | TRUE | 0,0920049 | No |
| orf19.4930 | 455 | 285 | 345 | 466 | 502 | 464 | 481 | 342 | 408 | 363 | 353 | 384 | 429 | 443 | 429 | 482 | 411,49 | 377 | 446 | 1,175 | 0,233 | 2,985 | 0,00283722 | 0,00647643 | 0,000891 | 0,0055075 | 0,003387 | 0,003387 | TRUE | 0,0358879 | No |
| orf19.4931 | 4430 | 3157 | 4033 | 4793 | 4585 | 4225 | 4546 | 2897 | 3974 | 4019 | 4126 | 3952 | 3921 | 4031 | 4057 | 4086 | 4020,9 | 4018 | 4024 | 1,001 | 0,002 | 0,053 | 0,95789715 | 0,96952927 | 0,000123 | 0,0023406 | 0,0008183 | 0,0008183 | TRUE | 0,0040486 | No |
| orf19.4931.1 | 11373 | 7566 | 8943 | 11319 | 11232 | 9633 | 9728 | 6462 | 10203 | 9632 | 9150 | 9334 | 9606 | 9192 | 8681 | 9114 | 9363,8 | 9580 | 9148 | 0,955 | -0,066 | -1,422 | 0,15491783 | 0,22176467 | 0,001872 | 0,0021346 | 0,0019901 | 0,0019901 | TRUE | 0,0236449 | No |
| orf19.4932 | 3898 | 2691 | 3533 | 4248 | 4548 | 3951 | 4094 | 2596 | 3497 | 3426 | 3615 | 3503 | 3890 | 3770 | 3653 | 3661 | 3626,8 | 3510 | 3744 | 1,067 | 0,093 | 2,494 | 0,01263594 | 0,02476892 | 0,000413 | 0,0023798 | 0,0010692 | 0,0010692 | TRUE | 0,0086112 | No |
| orf19.4933 | 8302 | 5767 | 7355 | 9413 | 5888 | 5212 | 5547 | 3822 | 7448 | 7342 | 7525 | 7762 | 5036 | 4973 | 4950 | 5391 | 6303,2 | 7519 | 5088 | 0,676 | -0,565 | -14,567 | 4,56E-48 | 4,15E-46 | 0,000877 | 0,0022098 | 0,0012807 | 0,0012807 | TRUE | 0,0186749 | No |
| orf19.4934 | 15 | 9 | 9 | 21 | 25 | 21 | 12 | 14 | 13 | 11 | 9 | 17 | 21 | 20 | 11 | 20 | 15,41 | 12 | 18 | 1,371 | 0,455 | 1,224 | 0,22090554 | NA | 1,00E-08 | 0,0961607 | 0,0667136 | 0,0667136 | TRUE | 0,4352009 | No |
| orf19.4936 | 95 | 64 | 75 | 124 | 96 | 87 | 101 | 59 | 85 | 81 | 77 | 102 | 82 | 83 | 90 | 83 | 85,52 | 86 | 84 | 0,974 | -0,038 | -0,25 | 0,80278134 | 0,8464567 | 1,00E-08 | 0,0189549 | 0,0109437 | 0,0109437 | TRUE | 0,1614025 | No |
| orf19.4936.1 | 11908 | 8910 | 10284 | 13204 | 17231 | 15923 | 16570 | 9585 | 10683 | 11343 | 10522 | 10888 | 14736 | 15193 | 14786 | 13519 | 12709 | 10859 | 14558 | 1,341 | 0,424 | 9,509 | 1,93E-21 | 3,86E-20 | 0,001663 | 0,0020938 | 0,001826 | 0,001826 | TRUE | 0,0278556 | No |
| orf19.4937 | 7474 | 5410 | 6792 | 8218 | 8210 | 7445 | 7933 | 4811 | 6705 | 6888 | 6949 | 6777 | 7021 | 7104 | 7079 | 6785 | 6913,5 | 6830 | 6997 | 1,025 | 0,036 | 1,188 | 0,23496963 | 0,31588667 | 0,000151 | 0,0021895 | 0,0007471 | 0,0007471 | TRUE | 0,0047595 | No |
| orf19.4939 | 241 | 249 | 291 | 313 | 382 | 436 | 419 | 275 | 216 | 317 | 298 | 258 | 327 | 416 | 374 | 388 | 324,19 | 272 | 376 | 1,386 | 0,471 | 3,901 | 9,60E-05 | 0,00029489 | 0,015492 | 0,0064575 | 0,0108258 | 0,0108258 | TRUE | 0,2187802 | No |
| orf19.494 | 7870 | 5075 | 6677 | 8584 | 7920 | 6339 | 7163 | 4856 | 7060 | 6461 | 6831 | 7078 | 6773 | 6048 | 6392 | 6849 | 6686,7 | 6858 | 6516 | 0,949 | -0,075 | -1,482 | 0,13846734 | 0,20143142 | 0,002293 | 0,0021966 | 0,0022976 | 0,0022976 | TRUE | 0,0284366 | No |
| orf19.4940 | 9423 | 6228 | 8261 | 10162 | 11721 | 9842 | 10929 | 7392 | 8453 | 7929 | 8452 | 8380 | 10024 | 9391 | 9752 | 10426 | 9100,9 | 8304 | 9898 | 1,191 | 0,253 | 6,023 | 1,72E-09 | 1,05E-08 | 0,001287 | 0,002139 | 0,0015778 | 0,0015778 | TRUE | 0,0153469 | No |
| orf19.4941 | 24263 | 16747 | 20201 | 25802 | 28783 | 25073 | 28101 | 17479 | 21766 | 21321 | 20668 | 21277 | 24616 | 23924 | 25075 | 24653 | 22912 | 21258 | 24567 | 1,156 | 0,209 | 7,11 | 1,16E-12 | 1,06E-11 | 0,000355 | 0,0020429 | 0,0007834 | 0,0007834 | TRUE | 0,0042802 | No |
| orf19.4942 | 54 | 35 | 39 | 75 | 41 | 46 | 40 | 34 | 48 | 45 | 40 | 62 | 35 | 44 | 36 | 48 | 44,67 | 49 | 41 | 0,818 | -0,29 | -1,308 | 0,19098034 | 0,26413671 | 0,010795 | 0,0344785 | 0,0244946 | 0,0244946 | TRUE | 0,3015619 | No |
| orf19.4943 | 9302 | 6875 | 8316 | 10362 | 8732 | 7984 | 8528 | 5467 | 8345 | 8753 | 8508 | 8545 | 7468 | 7618 | 7610 | 7711 | 8069,6 | 8538 | 7602 | 0,89 | -0,168 | -5,757 | 8,56E-09 | 4,78E-08 | 0,000114 | 0,0021594 | 0,0006878 | 0,0006878 | TRUE | 0,0033794 | No |
| orf19.4945 | 515 | 338 | 432 | 541 | 547 | 516 | 525 | 276 | 462 | 430 | 442 | 446 | 468 | 492 | 468 | 389 | 449,79 | 445 | 454 | 1,026 | 0,037 | 0,46 | 0,64571898 | 0,71577114 | 0,002359 | 0,005207 | 0,0038681 | 0,0038681 | TRUE | 0,0914472 | No |
| orf19.4946 | 14 | 7 | 8 | 11 | 6 | 5 | 5 | 5 | 13 | 9 | 8 | 9 | 5 | 5 | 4 | 7 | 7,52 | 10 | 5 | 0,538 | -0,895 | -1,782 | 0,07471141 | NA | 1,00E-08 | 0,1950804 | 0,0959421 | 0,0959421 | TRUE | 0,1539401 | No |
| orf19.4947 | 5284 | 4055 | 4576 | 6052 | 7612 | 6886 | 6472 | 4453 | 4740 | 5162 | 4682 | 4991 | 6510 | 6570 | 5775 | 6281 | 5588,9 | 4894 | 6284 | 1,284 | 0,361 | 6,896 | 5,33E-12 | 4,48E-11 | 0,002457 | 0,0022393 | 0,0024504 | 0,0024504 | TRUE | 0,0367796 | No |
| orf19.4948 | 840 | 578 | 789 | 897 | 609 | 614 | 638 | 323 | 754 | 736 | 807 | 740 | 521 | 586 | 569 | 456 | 645,99 | 759 | 533 | 0,705 | -0,504 | -6,416 | 1,40E-10 | 9,95E-10 | 0,004422 | 0,0042268 | 0,0043202 | 0,0043202 | TRUE | 0,0965603 | No |
| orf19.4949 | 8 | 6 | 7 | 6 | 1 | 4 | 1 | 5 | 7 | 8 | 7 | 5 | 1 | 4 | 1 | 7 | 4,94 | 7 | 3 | 0,429 | -1,221 | -1,751 | 0,07995856 | NA | 1,19E-08 | 0,2956891 | 0,2178571 | 0,2178571 | TRUE | 0,6254954 | No |
| orf19.4950 | 7575 | 5876 | 7109 | 8781 | 6524 | 6414 | 6792 | 4498 | 6796 | 7481 | 7273 | 7241 | 5580 | 6120 | 6061 | 6344 | 6611,9 | 7198 | 6026 | 0,837 | -0,257 | -5,255 | 1,48E-07 | 7,07E-07 | 0,002024 | 0,0021991 | 0,0021387 | 0,0021387 | TRUE | 0,0306022 | No |
| orf19.4951 | 2620 | 2091 | 2348 | 3082 | 2724 | 2847 | 2993 | 1878 | 2350 | 2662 | 2402 | 2541 | 2330 | 2717 | 2671 | 2649 | 2540,2 | 2489 | 2592 | 1,041 | 0,058 | 0,951 | 0,34146496 | 0,42838202 | 0,003401 | 0,002551 | 0,0032032 | 0,0032032 | TRUE | 0,0574584 | No |
| orf19.4952 | 5230 | 3889 | 4638 | 5781 | 6275 | 5791 | 6262 | 3664 | 4692 | 4951 | 4745 | 4767 | 5367 | 5526 | 5588 | 5168 | 5100,4 | 4789 | 5412 | 1,131 | 0,178 | 4,727 | 2,27E-06 | 9,17E-06 | 0,000615 | 0,0022641 | 0,0011657 | 0,0011657 | TRUE | 0,0106352 | No |
| orf19.4952.1 | 1315 | 859 | 1078 | 1362 | 1304 | 1078 | 1146 | 632 | 1180 | 1094 | 1103 | 1123 | 1115 | 1029 | 1023 | 891 | 1069,6 | 1125 | 1014 | 0,904 | -0,145 | -2,151 | 0,03149291 | 0,05602884 | 0,00348 | 0,0033367 | 0,0034198 | 0,0034198 | TRUE | 0,072442 | No |
| orf19.4953 | 1634 | 1382 | 1443 | 1941 | 1588 | 1808 | 1646 | 1046 | 1466 | 1759 | 1476 | 1601 | 1358 | 1725 | 1469 | 1475 | 1541,2 | 1576 | 1507 | 0,957 | -0,063 | -0,772 | 0,4398384 | 0,52671897 | 0,007639 | 0,0029215 | 0,0057773 | 0,0057773 | TRUE | 0,116274 | No |
| orf19.4954 | 917 | 757 | 839 | 1029 | 835 | 788 | 820 | 536 | 823 | 964 | 858 | 849 | 714 | 752 | 732 | 756 | 805,87 | 874 | 738 | 0,847 | -0,24 | -3,801 | 0,00014426 | 0,00042844 | 0,001472 | 0,0037809 | 0,0025622 | 0,0025622 | TRUE | 0,0533461 | No |
| orf19.4955 | 75 | 69 | 57 | 100 | 46 | 66 | 69 | 47 | 67 | 88 | 58 | 82 | 39 | 63 | 62 | 66 | 65,76 | 74 | 58 | 0,773 | -0,372 | -1,799 | 0,07196753 | 0,1154247 | 0,027321 | 0,0240552 | 0,0254653 | 0,0254653 | TRUE | 0,4333744 | No |
| orf19.4956 | 8225 | 5984 | 7408 | 9016 | 8033 | 7704 | 7989 | 4956 | 7379 | 7618 | 7579 | 7435 | 6870 | 7351 | 7129 | 6990 | 7293,8 | 7503 | 7085 | 0,945 | -0,082 | -2,477 | 0,01326176 | 0,02589426 | 0,000358 | 0,0021786 | 0,0009184 | 0,0009184 | TRUE | 0,0078743 | No |
| orf19.4958 | 728 | 760 | 688 | 953 | 995 | 1088 | 1181 | 666 | 653 | 968 | 704 | 786 | 851 | 1038 | 1054 | 939 | 874,08 | 778 | 970 | 1,251 | 0,323 | 2,869 | 0,00411547 | 0,00904711 | 0,01893 | 0,0036403 | 0,0110019 | 0,0110019 | TRUE | 0,3103596 | No |
| orf19.4959 | 5310 | 3597 | 4830 | 5874 | 6207 | 5421 | 5885 | 3898 | 4764 | 4579 | 4942 | 4844 | 5308 | 5173 | 5251 | 5498 | 5044,8 | 4782 | 5308 | 1,109 | 0,149 | 3,984 | 6,77E-05 | 0,00021377 | 0,000591 | 0,0022673 | 0,0011411 | 0,0011411 | TRUE | 0,0094626 | No |
| orf19.496 | 603 | 385 | 502 | 570 | 551 | 498 | 587 | 360 | 541 | 490 | 514 | 470 | 471 | 475 | 524 | 508 | 499,09 | 504 | 494 | 0,98 | -0,028 | -0,387 | 0,69871071 | 0,76103226 | 0,001547 | 0,0048882 | 0,0031852 | 0,0031852 | TRUE | 0,0309116 | No |
| orf19.4960 | 11629 | 8731 | 10247 | 13206 | 11514 | 10913 | 11334 | 7025 | 10432 | 11115 | 10484 | 10890 | 9847 | 10413 | 10114 | 9908 | 10400 | 10730 | 10070 | 0,939 | -0,091 | -2,58 | 0,00989416 | 0,01994101 | 0,000661 | 0,0021191 | 0,0011024 | 0,0011024 | TRUE | 0,0069651 | No |
| orf19.4961 | 27309 | 19492 | 24715 | 31001 | 39259 | 35803 | 38734 | 25804 | 24499 | 24815 | 25286 | 25564 | 33576 | 34162 | 34564 | 36394 | 29857 | 25041 | 34674 | 1,384 | 0,469 | 13,649 | 2,04E-42 | 1,46E-40 | 0,00075 | 0,0020281 | 0,001101 | 0,001101 | TRUE | 0,0135365 | No |
| orf19.4962 | 1069 | 924 | 990 | 1270 | 1159 | 1266 | 1347 | 814 | 959 | 1176 | 1013 | 1047 | 991 | 1208 | 1202 | 1148 | 1093,1 | 1049 | 1137 | 1,085 | 0,118 | 1,466 | 0,14257253 | 0,20655311 | 0,006608 | 0,0033076 | 0,0053286 | 0,0053286 | TRUE | 0,0930234 | No |
| orf19.4963 | 1825 | 1966 | 1908 | 2330 | 1926 | 2286 | 2214 | 1145 | 1637 | 2503 | 1952 | 1921 | 1647 | 2181 | 1976 | 1615 | 1929,1 | 2003 | 1855 | 0,926 | -0,11 | -0,679 | 0,49715768 | 0,58201935 | 0,024771 | 0,0027321 | 0,013818 | 0,0247711 | TRUE | 0,338572 | No |
| orf19.4964 | 1570 | 1230 | 1373 | 1770 | 1876 | 1723 | 1846 | 1046 | 1408 | 1566 | 1405 | 1460 | 1604 | 1644 | 1647 | 1475 | 1526,2 | 1460 | 1592 | 1,095 | 0,13 | 2,376 | 0,01749836 | 0,03299754 | 0,001733 | 0,0029307 | 0,0022335 | 0,0022335 | TRUE | 0,0270781 | No |
| orf19.4965 | 1020 | 868 | 973 | 1182 | 1215 | 1400 | 1350 | 769 | 915 | 1105 | 995 | 975 | 1039 | 1336 | 1205 | 1085 | 1081,8 | 998 | 1166 | 1,171 | 0,228 | 2,657 | 0,00789474 | 0,01631072 | 0,008429 | 0,0033214 | 0,0061489 | 0,0061489 | TRUE | 0,1158292 | No |
| orf19.4966 | 4788 | 4393 | 4710 | 6073 | 5207 | 5879 | 5875 | 3746 | 4295 | 5593 | 4819 | 5008 | 4453 | 5610 | 5242 | 5283 | 5037,9 | 4929 | 5147 | 1,045 | 0,063 | 0,741 | 0,45888193 | 0,54463901 | 0,009248 | 0,0022677 | 0,0067095 | 0,0067095 | TRUE | 0,1011614 | No |
| orf19.4967 | 2458 | 1751 | 2102 | 2719 | 2609 | 2352 | 2310 | 1396 | 2205 | 2229 | 2151 | 2242 | 2231 | 2244 | 2061 | 1969 | 2166,6 | 2207 | 2126 | 0,965 | -0,051 | -1,007 | 0,31404819 | 0,4005948 | 0,00154 | 0,0026496 | 0,0020157 | 0,0020157 | TRUE | 0,0277976 | No |
| orf19.4969 | 2330 | 1870 | 2304 | 2702 | 2959 | 3033 | 3107 | 1813 | 2090 | 2381 | 2357 | 2228 | 2531 | 2894 | 2772 | 2557 | 2476,3 | 2264 | 2688 | 1,189 | 0,25 | 4,128 | 3,65E-05 | 0,00012107 | 0,003316 | 0,0025658 | 0,0031047 | 0,0031047 | TRUE | 0,0325082 | No |
| orf19.497 | 2114 | 1378 | 1739 | 2185 | 2315 | 1901 | 2228 | 1353 | 1896 | 1754 | 1779 | 1802 | 1980 | 1814 | 1988 | 1908 | 1865,2 | 1808 | 1922 | 1,063 | 0,088 | 1,833 | 0,06680977 | 0,10842601 | 0,001003 | 0,0027578 | 0,0016883 | 0,0016883 | TRUE | 0,0182688 | No |
| orf19.4970 | 2226 | 1648 | 2053 | 2486 | 2740 | 2574 | 2731 | 1583 | 1997 | 2098 | 2100 | 2050 | 2343 | 2456 | 2437 | 2233 | 2214,3 | 2061 | 2367 | 1,151 | 0,202 | 4,545 | 5,50E-06 | 2,08E-05 | 0,00061 | 0,0026351 | 0,0014508 | 0,0014508 | TRUE | 0,0161381 | No |
| orf19.4972 | 413 | 280 | 358 | 431 | 258 | 263 | 275 | 168 | 371 | 356 | 366 | 355 | 221 | 251 | 245 | 237 | 300,32 | 362 | 238 | 0,658 | -0,604 | -7,227 | 4,93E-13 | 4,71E-12 | 1,00E-08 | 0,0068133 | 0,0032363 | 0,0032363 | TRUE | 0,0326437 | No |
| orf19.4975 | 2413 | 1648 | 2306 | 2677 | 5466 | 4689 | 5034 | 3614 | 2165 | 2098 | 2359 | 2207 | 4675 | 4474 | 4492 | 5097 | 3446 | 2207 | 4684 | 2,12 | 1,084 | 19,599 | 1,57E-85 | 5,55E-83 | 0,002728 | 0,0024008 | 0,002603 | 0,002603 | TRUE | 0,0416154 | No |
| orf19.4976 | 2 | 6 | 5 | 7 | 2 | 5 | 7 | 0 | 2 | 8 | 5 | 6 | 2 | 5 | 6 | 0 | 4,13 | 5 | 3 | 0,668 | -0,581 | -0,761 | 0,44688861 | NA | 0,229277 | 0,353393 | 0,3114394 | 0,3114394 | TRUE | 0,2876667 | No |
| orf19.4979 | 369 | 296 | 327 | 470 | 526 | 511 | 534 | 249 | 331 | 377 | 335 | 388 | 450 | 488 | 477 | 351 | 399,39 | 358 | 442 | 1,241 | 0,312 | 2,961 | 0,00306629 | 0,00694655 | 0,010766 | 0,0056143 | 0,0080959 | 0,0080959 | TRUE | 0,1952196 | No |
| orf19.498 | 2944 | 2041 | 2778 | 3404 | 3763 | 3372 | 3682 | 2370 | 2641 | 2598 | 2842 | 2807 | 3218 | 3217 | 3286 | 3343 | 2994,1 | 2722 | 3266 | 1,198 | 0,261 | 6,193 | 5,92E-10 | 3,89E-09 | 0,000699 | 0,0024644 | 0,0013677 | 0,0013677 | TRUE | 0,0106493 | No |
| orf19.4980 | 50178 | 34776 | 44848 | 53187 | 61165 | 56134 | 59582 | 36289 | 45015 | 44274 | 45884 | 43859 | 52310 | 53561 | 53167 | 51182 | 48657 | 44758 | 52555 | 1,174 | 0,232 | 8,141 | 3,93E-16 | 5,03E-15 | 0,000371 | 0,0020094 | 0,0007581 | 0,0007581 | TRUE | 0,0037506 | No |
| orf19.4981 | 797 | 662 | 763 | 1021 | 688 | 804 | 798 | 480 | 715 | 843 | 781 | 842 | 588 | 767 | 712 | 677 | 740,62 | 795 | 686 | 0,863 | -0,212 | -2,41 | 0,01594235 | 0,03046491 | 0,007951 | 0,0039396 | 0,0060689 | 0,0060689 | TRUE | 0,1149789 | No |
| orf19.4982 | 823 | 789 | 814 | 1011 | 1212 | 1257 | 1341 | 732 | 738 | 1004 | 833 | 834 | 1037 | 1199 | 1197 | 1032 | 984,28 | 852 | 1116 | 1,314 | 0,394 | 4,278 | 1,89E-05 | 6,58E-05 | 0,01066 | 0,0034544 | 0,0070945 | 0,0070945 | TRUE | 0,1651752 | No |
| orf19.4983 | 1562 | 1156 | 1287 | 1671 | 1643 | 1579 | 1663 | 988 | 1401 | 1472 | 1317 | 1378 | 1405 | 1507 | 1484 | 1393 | 1419,6 | 1392 | 1447 | 1,042 | 0,059 | 1,146 | 0,25189569 | 0,33430292 | 0,000878 | 0,0030021 | 0,0018512 | 0,0018512 | TRUE | 0,016397 | No |
| orf19.4984 | 2605 | 2010 | 2447 | 2824 | 2986 | 2802 | 3027 | 1676 | 2337 | 2559 | 2504 | 2329 | 2554 | 2674 | 2701 | 2364 | 2502,6 | 2432 | 2573 | 1,06 | 0,084 | 1,537 | 0,12440289 | 0,18395583 | 0,002398 | 0,0025596 | 0,0025011 | 0,0025011 | TRUE | 0,0342931 | No |
| orf19.4985 | 427 | 350 | 386 | 561 | 594 | 657 | 644 | 317 | 383 | 446 | 395 | 463 | 508 | 627 | 575 | 447 | 480,35 | 422 | 539 | 1,283 | 0,36 | 3,37 | 0,00075165 | 0,00194151 | 0,01191 | 0,0050017 | 0,0088328 | 0,0088328 | TRUE | 0,1408003 | No |
| orf19.4987 | 3450 | 2430 | 3143 | 3891 | 3956 | 3551 | 3934 | 2449 | 3095 | 3094 | 3216 | 3209 | 3383 | 3388 | 3510 | 3454 | 3293,6 | 3154 | 3434 | 1,088 | 0,122 | 3,483 | 0,00049566 | 0,00133933 | 1,00E-08 | 0,0024203 | 0,0008789 | 0,0008789 | TRUE | 0,0029232 | No |

| | | | | | | | | | | | | | | | | | | | | | | | | | | | | | | | |
|---|---|---|---|---|---|---|---|---|---|---|---|---|---|---|---|---|---|---|---|---|---|---|---|---|---|---|---|---|---|---|---|
| orf19.4988 | 1226 | 978 | 1156 | 1350 | 1659 | 1646 | 1677 | 1023 | 1100 | 1245 | 1183 | 1113 | 1419 | 1571 | 1496 | 1443 | 1321,2 | 1160 | 1482 | 1,281 | 0,357 | 6,234 | 4,53E-10 | 3,04E-09 | 0,001685 | 0,0030783 | 0,0023769 | 0,0023769 | TRUE | 0,0274009 | No |
| orf19.499 | 1963 | 1229 | 1561 | 1920 | 1866 | 1576 | 1729 | 1004 | 1761 | 1565 | 1597 | 1583 | 1596 | 1504 | 1543 | 1416 | 1570,6 | 1626 | 1515 | 0,932 | -0,101 | -1,788 | 0,07376325 | 0,11786425 | 0,001955 | 0,0029038 | 0,0024346 | 0,0024346 | TRUE | 0,0380903 | No |
| orf19.4991 | 1276 | 1078 | 1262 | 1601 | 974 | 937 | 1036 | 692 | 1145 | 1372 | 1291 | 1320 | 833 | 894 | 924 | 976 | 1094,5 | 1282 | 907 | 0,706 | -0,501 | -7,106 | 1,19E-12 | 1,08E-11 | 0,004252 | 0,0033059 | 0,0038287 | 0,0038287 | TRUE | 0,0634759 | No |
| orf19.4993 | 2419 | 1843 | 2115 | 2690 | 2522 | 2523 | 2644 | 1485 | 2170 | 2346 | 2164 | 2218 | 2157 | 2407 | 2359 | 2094 | 2239,6 | 2224 | 2254 | 1,015 | 0,022 | 0,391 | 0,69608846 | 0,75867659 | 0,00219 | 0,0026277 | 0,0025279 | 0,0025279 | TRUE | 0,0258583 | No |
| orf19.4995 | 37 | 33 | 48 | 47 | 53 | 49 | 49 | 30 | 33 | 42 | 49 | 39 | 45 | 47 | 44 | 42 | 42,65 | 41 | 44 | 1,102 | 0,14 | 0,661 | 0,50837322 | 0,5926089 | 1,00E-08 | 0,0360182 | 0,0193574 | 0,0193574 | TRUE | 0,1478384 | No |
| orf19.4996 | 27 | 17 | 31 | 22 | 30 | 42 | 39 | 26 | 24 | 22 | 32 | 18 | 26 | 40 | 35 | 37 | 29,12 | 24 | 34 | 1,431 | 0,517 | 1,888 | 0,05896238 | 0,09700054 | 1,00E-08 | 0,0518392 | 0,0363436 | 0,0363436 | TRUE | 0,2844957 | No |
| orf19.4997 | 1799 | 1341 | 1629 | 2071 | 2097 | 1907 | 2072 | 1305 | 1614 | 1707 | 1667 | 1708 | 1793 | 1820 | 1849 | 1841 | 1749,8 | 1674 | 1826 | 1,091 | 0,125 | 3,024 | 0,00249404 | 0,00576316 | 1,00E-08 | 0,0028092 | 0,0010749 | 0,0010749 | TRUE | 0,0073 | No |
| orf19.4998 | 3387 | 2584 | 3116 | 3872 | 3675 | 3554 | 3875 | 2293 | 3038 | 3290 | 3188 | 3193 | 3143 | 3391 | 3458 | 3234 | 3241,9 | 3177 | 3306 | 1,042 | 0,059 | 1,301 | 0,19322727 | 0,26650814 | 0,001067 | 0,0024273 | 0,0016428 | 0,0016428 | TRUE | 0,014255 | No |
| orf19.50 | 1484 | 1158 | 1295 | 1663 | 1863 | 1929 | 1919 | 1129 | 1331 | 1474 | 1325 | 1371 | 1593 | 1841 | 1712 | 1592 | 1530,1 | 1375 | 1684 | 1,227 | 0,295 | 4,815 | 1,47E-06 | 6,06E-06 | 0,002841 | 0,0029283 | 0,0029469 | 0,0029469 | TRUE | 0,0469093 | No |
| orf19.500 | 6817 | 4803 | 5870 | 7320 | 6365 | 5734 | 6214 | 3698 | 6116 | 6115 | 6006 | 6036 | 5444 | 5471 | 5545 | 5216 | 5743,4 | 6068 | 5419 | 0,894 | -0,162 | -5,127 | 2,94E-07 | 1,35E-06 | 0,000153 | 0,0022323 | 0,0007816 | 0,0007816 | TRUE | 0,0072058 | No |
| orf19.5000 | 3366 | 2370 | 3064 | 3694 | 4355 | 3924 | 4197 | 2534 | 3020 | 3017 | 3135 | 3046 | 3725 | 3744 | 3745 | 3574 | 3375,7 | 3054 | 3697 | 1,211 | 0,277 | 7,927 | 2,25E-15 | 2,68E-14 | 1,34E-05 | 0,0024096 | 0,0008701 | 0,0008701 | TRUE | 0,0055559 | No |
| orf19.5001 | 2352 | 1706 | 2118 | 2638 | 2765 | 2430 | 2654 | 1619 | 2110 | 2172 | 2167 | 2175 | 2365 | 2319 | 2368 | 2283 | 2244,9 | 2156 | 2334 | 1,084 | 0,116 | 3,123 | 0,00179171 | 0,00427675 | 1,00E-08 | 0,0026262 | 0,0008765 | 0,0008765 | TRUE | 0,0026043 | No |
| orf19.5003 | 1553 | 1254 | 1455 | 1797 | 1762 | 1798 | 1819 | 988 | 1393 | 1596 | 1489 | 1482 | 1507 | 1716 | 1623 | 1393 | 1524,9 | 1490 | 1560 | 1,049 | 0,07 | 1,004 | 0,31522204 | 0,4018362 | 0,004709 | 0,0029315 | 0,0039627 | 0,0039627 | TRUE | 0,05816 | No |
| orf19.5004 | 460 | 332 | 375 | 494 | 373 | 350 | 368 | 194 | 413 | 423 | 384 | 407 | 319 | 334 | 328 | 274 | 360,16 | 407 | 314 | 0,777 | -0,364 | -4,432 | 9,34E-06 | 3,41E-05 | 0,000154 | 0,0060102 | 0,0036594 | 0,0036594 | TRUE | 0,0677138 | No |
| orf19.5005 | 2780 | 1854 | 2404 | 2874 | 3667 | 3324 | 3587 | 2231 | 2494 | 2360 | 2460 | 2370 | 3136 | 3172 | 3201 | 3147 | 2792,4 | 2421 | 3164 | 1,306 | 0,386 | 10,579 | 3,73E-26 | 1,07E-24 | 1,00E-08 | 0,0024994 | 0,0009112 | 0,0009112 | TRUE | 0,005131 | No |
| orf19.5006 | 2426 | 1788 | 2112 | 2726 | 2633 | 2420 | 2428 | 1608 | 2176 | 2276 | 2161 | 2248 | 2252 | 2309 | 2167 | 2268 | 2232,1 | 2215 | 2249 | 1,015 | 0,022 | 0,533 | 0,59432581 | 0,67156469 | 0,00024 | 0,0026299 | 0,001137 | 0,001137 | TRUE | 0,0076425 | No |
| orf19.5006.1 | 4802 | 3508 | 4220 | 5330 | 4834 | 4694 | 4879 | 2610 | 4308 | 4466 | 4317 | 4395 | 4134 | 4479 | 4354 | 3681 | 4266,8 | 4372 | 4162 | 0,953 | -0,069 | -1,194 | 0,23230844 | 0,31286911 | 0,003382 | 0,0023198 | 0,002999 | 0,002999 | TRUE | 0,0712462 | No |
| orf19.5007 | 4944 | 3323 | 4095 | 5175 | 5516 | 4594 | 4905 | 3441 | 4435 | 4231 | 4190 | 4267 | 4717 | 4383 | 4377 | 4853 | 4431,7 | 4281 | 4582 | 1,07 | 0,097 | 2,141 | 0,03229459 | 0,05716733 | 0,001392 | 0,0023071 | 0,0017462 | 0,0017462 | TRUE | 0,01838 | No |
| orf19.5008.1 | 265 | 174 | 223 | 274 | 245 | 218 | 238 | 159 | 238 | 222 | 228 | 226 | 210 | 208 | 212 | 224 | 220,94 | 228 | 214 | 0,931 | -0,103 | -1,108 | 0,26778218 | 0,35204274 | 1,00E-08 | 0,0085501 | 0,0037516 | 0,0037516 | TRUE | 0,0098185 | No |
| orf19.5009 | 1687 | 1291 | 1501 | 1863 | 1782 | 1729 | 1799 | 1030 | 1513 | 1644 | 1536 | 1536 | 1524 | 1650 | 1605 | 1453 | 1557,6 | 1557 | 1558 | 1,003 | 0,005 | 0,089 | 0,92930041 | 0,94902236 | 0,001301 | 0,0029115 | 0,0020733 | 0,0020733 | TRUE | 0,0224536 | No |
| orf19.501 | 18575 | 12235 | 16068 | 19930 | 16976 | 14359 | 15808 | 9554 | 16664 | 15576 | 16439 | 16435 | 14518 | 13701 | 14106 | 13475 | 15114 | 16278 | 13950 | 0,857 | -0,223 | -6,01 | 1,86E-09 | 1,14E-08 | 0,00089 | 0,0020756 | 0,0012503 | 0,0012503 | TRUE | 0,0101805 | No |
| orf19.5010 | 3222 | 2376 | 2794 | 3531 | 3305 | 3009 | 3145 | 1814 | 2890 | 3025 | 2859 | 2912 | 2827 | 2871 | 2806 | 2558 | 2843,5 | 2922 | 2766 | 0,949 | -0,076 | -1,673 | 0,09434821 | 0,14583074 | 0,001132 | 0,0024901 | 0,0016329 | 0,0016329 | TRUE | 0,0286219 | No |
| orf19.5011 | 678 | 550 | 635 | 822 | 922 | 891 | 928 | 574 | 608 | 700 | 650 | 678 | 789 | 850 | 828 | 810 | 739,04 | 659 | 819 | 1,245 | 0,316 | 5,07 | 3,98E-07 | 1,79E-06 | 0,00088 | 0,0039438 | 0,0023713 | 0,0023713 | TRUE | 0,0325818 | No |
| orf19.5012 | 217 | 142 | 158 | 225 | 185 | 193 | 160 | 98 | 195 | 181 | 162 | 186 | 158 | 184 | 143 | 138 | 168,25 | 181 | 156 | 0,865 | -0,209 | -1,72 | 0,08540028 | 0,13385987 | 0,005107 | 0,0106077 | 0,0082232 | 0,0082232 | TRUE | 0,1561954 | No |
| orf19.5013 | 201 | 202 | 198 | 273 | 277 | 286 | 300 | 248 | 180 | 257 | 203 | 225 | 237 | 273 | 268 | 350 | 249,06 | 216 | 282 | 1,299 | 0,378 | 2,771 | 0,00558034 | 0,01193545 | 0,020576 | 0,0078083 | 0,0137251 | 0,0137251 | TRUE | 0,269639 | No |
| orf19.5014 | 219 | 159 | 166 | 232 | 203 | 172 | 179 | 118 | 196 | 202 | 170 | 191 | 174 | 164 | 160 | 166 | 177,99 | 190 | 166 | 0,875 | -0,193 | -1,82 | 0,06880301 | 0,11112055 | 1,00E-08 | 0,0101356 | 0,0052137 | 0,0052137 | TRUE | 0,0527883 | No |
| orf19.5015 | 980 | 774 | 857 | 973 | 1362 | 1312 | 1292 | 798 | 879 | 985 | 877 | 802 | 1165 | 1252 | 1153 | 1126 | 1029,9 | 886 | 1174 | 1,33 | 0,411 | 6,085 | 1,16E-09 | 7,34E-09 | 0,003373 | 0,0033891 | 0,0033908 | 0,0033908 | TRUE | 0,0641431 | No |
| orf19.5016 | 815 | 643 | 680 | 942 | 1137 | 1023 | 1016 | 765 | 731 | 819 | 696 | 777 | 972 | 976 | 907 | 1079 | 869,54 | 756 | 984 | 1,3 | 0,378 | 5,342 | 9,19E-08 | 4,54E-07 | 0,003571 | 0,003649 | 0,0036414 | 0,0036414 | TRUE | 0,0468023 | No |
| orf19.5017 | 55 | 43 | 54 | 69 | 70 | 63 | 61 | 42 | 49 | 55 | 55 | 57 | 60 | 60 | 54 | 59 | 56,23 | 54 | 58 | 1,08 | 0,111 | 0,621 | 0,5345582 | 0,6165532 | 1,00E-08 | 0,0277945 | 0,01288 | 0,01288 | TRUE | 0,0325646 | No |
| orf19.5019 | 2788 | 1853 | 2342 | 3054 | 2799 | 2295 | 2572 | 1660 | 2501 | 2359 | 2396 | 2518 | 2394 | 2190 | 2295 | 2341 | 2374,3 | 2444 | 2305 | 0,942 | -0,086 | -1,926 | 0,0540497 | 0,08997801 | 0,000825 | 0,0025909 | 0,0014848 | 0,0014848 | TRUE | 0,0139736 | No |
| orf19.5020 | 6457 | 4289 | 5683 | 7092 | 6710 | 5853 | 6368 | 4469 | 5793 | 5460 | 5814 | 5848 | 5739 | 5585 | 5682 | 6303 | 5778 | 5729 | 5827 | 1,016 | 0,023 | 0,49 | 0,6243442 | 0,69709347 | 0,001669 | 0,0022308 | 0,0019236 | 0,0019236 | TRUE | 0,0357377 | No |
| orf19.5021 | 5644 | 3703 | 4899 | 6065 | 5121 | 4428 | 4828 | 3118 | 5063 | 4714 | 5012 | 5001 | 4380 | 4225 | 4308 | 4398 | 4637,7 | 4948 | 4328 | 0,874 | -0,195 | -5,386 | 7,20E-08 | 3,60E-07 | 0,000385 | 0,0022925 | 0,001035 | 0,001035 | TRUE | 0,0117465 | No |
| orf19.5022 | 3273 | 2101 | 2882 | 3502 | 4509 | 3939 | 4295 | 2809 | 2936 | 2675 | 2949 | 2888 | 3856 | 3758 | 3833 | 3962 | 3357,1 | 2862 | 3852 | 1,344 | 0,427 | 10,164 | 2,86E-24 | 6,82E-23 | 0,000823 | 0,002412 | 0,0013855 | 0,0013855 | TRUE | 0,022128 | No |
| orf19.5023 | 198 | 127 | 166 | 204 | 205 | 168 | 167 | 117 | 178 | 162 | 170 | 168 | 175 | 160 | 149 | 165 | 165,88 | 170 | 162 | 0,957 | -0,064 | -0,584 | 0,55923828 | 0,63937271 | 1,00E-08 | 0,0107311 | 0,0053828 | 0,0053828 | TRUE | 0,0365611 | No |
| orf19.5024 | 5567 | 3710 | 4863 | 6033 | 4833 | 4186 | 4469 | 2966 | 4994 | 4723 | 4975 | 4975 | 4133 | 3994 | 3988 | 4183 | 4495,8 | 4917 | 4074 | 0,828 | -0,273 | -7,62 | 2,53E-14 | 2,70E-13 | 0,000341 | 0,0023024 | 0,001004 | 0,001004 | TRUE | 0,0081793 | No |
| orf19.5025 | 19828 | 13744 | 17069 | 20630 | 14259 | 13157 | 13528 | 8291 | 17788 | 17498 | 17463 | 17012 | 12195 | 12554 | 12071 | 11694 | 14784 | 17440 | 12128 | 0,696 | -0,523 | -16,176 | 7,43E-59 | 1,14E-56 | 0,000504 | 0,0020777 | 0,0009353 | 0,0009353 | TRUE | 0,0068852 | No |
| orf19.5026 | 15530 | 11088 | 14554 | 17446 | 19789 | 18176 | 19551 | 12735 | 13932 | 14116 | 14890 | 14386 | 16924 | 17343 | 17446 | 17962 | 15875 | 14331 | 17419 | 1,215 | 0,281 | 8,281 | 1,22E-16 | 1,65E-15 | 0,000638 | 0,002071 | 0,0010432 | 0,0010432 | TRUE | 0,0084138 | No |
| orf19.5027 | 3435 | 2307 | 3013 | 3588 | 3227 | 3039 | 3103 | 1829 | 3082 | 2937 | 3083 | 2959 | 2760 | 2900 | 2769 | 2580 | 2883,5 | 3015 | 2752 | 0,914 | -0,13 | -2,913 | 0,00358353 | 0,00797698 | 0,001013 | 0,002483 | 0,0015669 | 0,0015669 | TRUE | 0,0199185 | No |
| orf19.5029 | 2600 | 1892 | 2418 | 2962 | 2356 | 2207 | 2459 | 1489 | 2332 | 2409 | 2474 | 2443 | 2015 | 2106 | 2194 | 2100 | 2259,1 | 2414 | 2104 | 0,871 | -0,199 | -4,633 | 3,61E-06 | 1,40E-05 | 0,000559 | 0,0026221 | 0,0013185 | 0,0013185 | TRUE | 0,0107784 | No |
| orf19.5030 | 1605 | 1071 | 1383 | 1645 | 1773 | 1568 | 1688 | 949 | 1440 | 1363 | 1415 | 1356 | 1516 | 1496 | 1506 | 1338 | 1429 | 1394 | 1464 | 1,053 | 0,075 | 1,415 | 0,15700353 | 0,22416983 | 0,001142 | 0,0029954 | 0,0019788 | 0,0019788 | TRUE | 0,03624 | No |
| orf19.5031 | 8897 | 5932 | 7820 | 9660 | 10198 | 8895 | 9985 | 6676 | 7982 | 7552 | 8001 | 7966 | 8722 | 8487 | 8910 | 9416 | 8379,4 | 7875 | 8884 | 1,127 | 0,173 | 4,193 | 2,75E-05 | 9,32E-05 | 0,001202 | 0,0021528 | 0,0015102 | 0,0015102 | TRUE | 0,0193431 | No |
| orf19.5032 | 35002 | 23235 | 30945 | 37038 | 33778 | 27708 | 29922 | 20112 | 31400 | 29581 | 31660 | 30542 | 28888 | 26438 | 26700 | 28366 | 29197 | 30796 | 27598 | 0,896 | -0,158 | -3,846 | 0,00012007 | 0,00036197 | 0,001384 | 0,0020292 | 0,0015933 | 0,0015933 | TRUE | 0,0122334 | No |
| orf19.5033 | 177 | 135 | 163 | 189 | 189 | 169 | 159 | 100 | 159 | 172 | 167 | 156 | 162 | 161 | 142 | 141 | 157,39 | 164 | 152 | 0,934 | -0,099 | -0,878 | 0,3799722 | 0,46953373 | 1,00E-08 | 0,0112033 | 0,0057547 | 0,0057547 | TRUE | 0,0240617 | No |
| orf19.5034 | 1059 | 776 | 948 | 1160 | 1087 | 1093 | 1144 | 787 | 950 | 988 | 970 | 957 | 930 | 1043 | 1021 | 1110 | 995,97 | 966 | 1026 | 1,059 | 0,083 | 1,369 | 0,1708715 | 0,24104155 | 0,001777 | 0,0034371 | 0,0025241 | 0,0025241 | TRUE | 0,0486234 | No |
| orf19.5035 | 1593 | 1026 | 1357 | 1741 | 1539 | 1335 | 1496 | 913 | 1429 | 1306 | 1388 | 1436 | 1316 | 1274 | 1335 | 1288 | 1346,5 | 1390 | 1303 | 0,936 | -0,095 | -1,951 | 0,05102589 | 0,0856562 | 0,000337 | 0,0030577 | 0,0015212 | 0,0015212 | TRUE | 0,0181552 | No |
| orf19.5037 | 5579 | 5004 | 5313 | 6639 | 6654 | 7434 | 7287 | 4734 | 5005 | 6371 | 5436 | 5475 | 5691 | 7093 | 6502 | 6677 | 6031,2 | 5572 | 6491 | 1,165 | 0,221 | 2,742 | 0,00611294 | 0,01295925 | 0,009185 | 0,0022202 | 0,0060551 | 0,0060551 | TRUE | 0,1128678 | No |
| orf19.5038 | 3684 | 3520 | 3780 | 4777 | 4898 | 5528 | 5744 | 3301 | 3305 | 4481 | 3867 | 3939 | 4189 | 5275 | 5126 | 4656 | 4354,7 | 3898 | 4812 | 1,235 | 0,304 | 3,316 | 0,00091235 | 0,00231045 | 0,012529 | 0,0023129 | 0,0078539 | 0,0078539 | TRUE | 0,1280442 | No |
| orf19.5039 | 2733 | 2117 | 2358 | 3026 | 3020 | 2804 | 2676 | 1682 | 2452 | 2695 | 2412 | 2495 | 2583 | 2675 | 2388 | 2372 | 2509,2 | 2514 | 2504 | 0,998 | -0,003 | -0,054 | 0,95691632 | 0,96917621 | 0,002505 | 0,0025581 | 0,0025269 | 0,0025269 | TRUE | 0,0275324 | No |
| orf19.504 | 3493 | 2298 | 2976 | 3869 | 3426 | 3010 | 3318 | 2183 | 3134 | 2926 | 3045 | 3190 | 2930 | 2872 | 2961 | 3079 | 3017 | 3074 | 2960 | 0,961 | -0,057 | -1,352 | 0,176307 | 0,24772041 | 0,000696 | 0,0024607 | 0,0013675 | 0,0013675 | TRUE | 0,0122021 | No |
| orf19.5040 | 3529 | 2864 | 3370 | 4097 | 4089 | 4330 | 4573 | 2453 | 3166 | 3646 | 3448 | 3378 | 3497 | 4132 | 4081 | 3460 | 3600,9 | 3410 | 3792 | 1,113 | 0,155 | 2,204 | 0,02752238 | 0,04969792 | 0,005907 | 0,0023827 | 0,0044583 | 0,0044583 | TRUE | 0,0439458 | No |
| orf19.5041 | 145 | 89 | 125 | 162 | 225 | 202 | 176 | 125 | 130 | 113 | 128 | 134 | 192 | 193 | 157 | 176 | 152,92 | 126 | 180 | 1,417 | 0,503 | 4,185 | 2,85E-05 | 9,64E-05 | 1,00E-08 | 0,0114726 | 0,0071716 | 0,0071716 | TRUE | 0,0861138 | No |
| orf19.5042 | 50 | 44 | 46 | 60 | 59 | 87 | 77 | 45 | 45 | 56 | 47 | 49 | 50 | 83 | 69 | 63 | 57,88 | 49 | 66 | 1,352 | 0,435 | 2,171 | 0,02993774 | 0,0535156 | 0,013072 | 0,0270596 | 0,020834 | 0,020834 | TRUE | 0,2188811 | No |
| orf19.5043 | 169 | 149 | 168 | 213 | 203 | 195 | 209 | 112 | 152 | 190 | 172 | 176 | 174 | 186 | 186 | 158 | 174,12 | 172 | 176 | 1,031 | 0,045 | 0,399 | 0,69001398 | 0,75306158 | 1,00E-08 | 0,0103169 | 0,0063556 | 0,0063556 | TRUE | 0,0715573 | No |
| orf19.5045 | 5462 | 4863 | 5386 | 6862 | 8709 | 9723 | 10201 | 5931 | 4900 | 6191 | 5510 | 5658 | 7448 | 9277 | 9103 | 8365 | 7056,7 | 5565 | 8548 | 1,537 | 0,62 | 7,661 | 1,84E-14 | 1,99E-13 | 0,009105 | 0,0021852 | 0,0061359 | 0,0061359 | TRUE | 0,0922863 | No |
| orf19.5046 | 238 | 207 | 207 | 257 | 270 | 284 | 293 | 180 | 214 | 264 | 212 | 212 | 231 | 271 | 261 | 254 | 239,75 | 226 | 254 | 1,135 | 0,183 | 1,772 | 0,07639586 | 0,12158579 | 0,003575 | 0,0080347 | 0,0060579 | 0,0060579 | TRUE | 0,127495 | No |
| orf19.5049 | 4844 | 3434 | 4015 | 4902 | 5125 | 4417 | 4635 | 2984 | 4346 | 4372 | 4108 | 4042 | 4383 | 4215 | 4136 | 4209 | 4226,2 | 4217 | 4236 | 1,005 | 0,007 | 0,181 | 0,85655596 | 0,89096056 | 0,000869 | 0,002323 | 0,0013545 | 0,0013545 | TRUE | 0,0096833 | No |
| orf19.505 | 2834 | 1881 | 2398 | 2884 | 3397 | 2905 | 3202 | 2081 | 2542 | 2395 | 2453 | 2378 | 2905 | 2772 | 2857 | 2935 | 2654,8 | 2442 | 2867 | 1,173 | 0,23 | 5,737 | 9,63E-09 | 5,34E-08 | 0,000404 | 0,0025264 | 0,0011684 | 0,0011684 | TRUE | 0,0095514 | No |
| orf19.5050 | 1921 | 1252 | 1694 | 2004 | 2426 | 2217 | 2349 | 1467 | 1723 | 1594 | 1733 | 1653 | 2075 | 2115 | 2096 | 2069 | 1882,3 | 1676 | 2089 | 1,246 | 0,317 | 7,445 | 9,71E-14 | 9,87E-13 | 9,36E-05 | 0,0027508 | 0,001201 | 0,001201 | TRUE | 0,0116856 | No |
| orf19.5051 | 2281 | 1528 | 2067 | 2574 | 2815 | 2485 | 2680 | 1768 | 2046 | 1945 | 2115 | 2123 | 2407 | 2371 | 2391 | 2494 | 2236,6 | 2057 | 2416 | 1,172 | 0,229 | 5,342 | 9,21E-08 | 4,54E-07 | 0,000445 | 0,0026286 | 0,0013105 | 0,0013105 | TRUE | 0,0150212 | No |

| | | | | | | | | | | | | | | | | | | | | | | | | | | | | | | | |
|---|---|---|---|---|---|---|---|---|---|---|---|---|---|---|---|---|---|---|---|---|---|---|---|---|---|---|---|---|---|---|---|
| orf19.5052 | 500 | 367 | 452 | 548 | 646 | 559 | 551 | 410 | 449 | 467 | 462 | 452 | 552 | 533 | 492 | 578 | 498,24 | 458 | 539 | 1,175 | 0,233 | 3,29 | 0,00100139 | 0,00251965 | 1,00E-08 | 0,0048932 | 0,0028012 | 0,0028012 | TRUE | 0,0415524 | No |
| orf19.5053 | 975 | 808 | 864 | 1142 | 1292 | 1213 | 1340 | 844 | 875 | 1029 | 884 | 942 | 1105 | 1157 | 1196 | 1190 | 1047,2 | 932 | 1162 | 1,248 | 0,32 | 5,142 | 2,72E-07 | 1,25E-06 | 0,002238 | 0,0033658 | 0,0027437 | 0,0027437 | TRUE | 0,0531215 | No |
| orf19.5054 | 2092 | 1487 | 1783 | 2234 | 2484 | 2270 | 2239 | 1322 | 1877 | 1893 | 1824 | 1842 | 2124 | 2166 | 1998 | 1865 | 1948,6 | 1859 | 2038 | 1,099 | 0,136 | 2,572 | 0,01009719 | 0,02030833 | 0,001685 | 0,0027245 | 0,002157 | 0,002157 | TRUE | 0,0367748 | No |
| orf19.5056 | 110 | 111 | 109 | 157 | 162 | 131 | 131 | 91 | 99 | 141 | 112 | 129 | 139 | 125 | 117 | 128 | 123,72 | 120 | 127 | 1,063 | 0,088 | 0,632 | 0,527237 | 0,61021767 | 0,006656 | 0,0137132 | 0,010689 | 0,010689 | TRUE | 0,1543453 | No |
| orf19.5057 | 7 | 10 | 6 | 18 | 18 | 22 | 14 | 8 | 6 | 13 | 6 | 15 | 15 | 21 | 12 | 11 | 12,52 | 10 | 15 | 1,519 | 0,603 | 1,415 | 0,15700589 | NA | 0,05145 | 0,1179368 | 0,0919021 | 0,0919021 | TRUE | 0,4641415 | No |
| orf19.5058 | 17451 | 12149 | 15100 | 18867 | 21958 | 19774 | 21530 | 12778 | 15655 | 15467 | 15449 | 15558 | 18779 | 18868 | 19212 | 18022 | 17126 | 15532 | 18720 | 1,206 | 0,27 | 9,3 | 1,41E-20 | 2,65E-19 | 0,000299 | 0,0020643 | 0,0007486 | 0,0007486 | TRUE | 0,0075121 | No |
| orf19.5059 | 9473 | 7740 | 9137 | 10956 | 9309 | 9485 | 10118 | 5908 | 8498 | 9854 | 9348 | 9034 | 7961 | 9050 | 9029 | 8333 | 8888,5 | 9184 | 8593 | 0,936 | -0,095 | -1,637 | 0,10166634 | 0,15477633 | 0,003717 | 0,0021428 | 0,003148 | 0,003148 | TRUE | 0,0308543 | No |
| orf19.506 | 9685 | 6414 | 8536 | 10131 | 9252 | 7847 | 8248 | 5234 | 8688 | 8166 | 8733 | 8354 | 7913 | 7487 | 7360 | 7382 | 8010,4 | 8485 | 7536 | 0,888 | -0,171 | -4,381 | 1,18E-05 | 4,26E-05 | 0,000931 | 0,0021607 | 0,0013396 | 0,0013396 | TRUE | 0,0140631 | No |
| orf19.5061 | 9275 | 6442 | 8385 | 10135 | 8545 | 7739 | 8296 | 5017 | 8321 | 8201 | 8579 | 8357 | 7308 | 7384 | 7403 | 7076 | 7828,7 | 8364 | 7293 | 0,872 | -0,197 | -6,437 | 1,22E-10 | 8,76E-10 | 0,000218 | 0,002165 | 0,0007723 | 0,0007723 | TRUE | 0,0046033 | No |
| orf19.5063 | 4115 | 3740 | 3643 | 5244 | 4013 | 3818 | 3650 | 2698 | 3692 | 4761 | 3727 | 4324 | 3432 | 3643 | 3257 | 3805 | 3830,2 | 4126 | 3534 | 0,857 | -0,223 | -2,699 | 0,00696112 | 0,01454186 | 0,009695 | 0,0023585 | 0,00632 | 0,00632 | TRUE | 0,1293121 | No |
| orf19.5064 | 4528 | 3680 | 3916 | 5109 | 4917 | 4810 | 4772 | 2908 | 4062 | 4685 | 4006 | 4213 | 4205 | 4590 | 4258 | 4101 | 4265,1 | 4242 | 4288 | 1,012 | 0,017 | 0,296 | 0,76712193 | 0,81848137 | 0,003082 | 0,0023199 | 0,0029973 | 0,0029973 | TRUE | 0,0590637 | No |
| orf19.5064.1 | 710 | 442 | 582 | 670 | 725 | 577 | 568 | 340 | 637 | 563 | 595 | 552 | 620 | 551 | 507 | 480 | 563,07 | 587 | 540 | 0,922 | -0,118 | -1,343 | 0,17930422 | 0,25110951 | 0,006191 | 0,0045577 | 0,0055888 | 0,0055888 | TRUE | 0,1214817 | No |
| orf19.5065 | 431 | 449 | 471 | 571 | 457 | 596 | 544 | 318 | 387 | 572 | 482 | 471 | 391 | 569 | 485 | 449 | 475,56 | 478 | 474 | 0,993 | -0,009 | -0,073 | 0,94162103 | 0,95784921 | 0,022833 | 0,0050321 | 0,0138425 | 0,0138425 | TRUE | 0,217101 | No |
| orf19.5066 | 4572 | 3397 | 3892 | 5020 | 4123 | 3803 | 3849 | 2223 | 4102 | 4325 | 3982 | 4140 | 3526 | 3629 | 3435 | 3135 | 3784,1 | 4137 | 3431 | 0,831 | -0,268 | -5,258 | 1,46E-07 | 6,99E-07 | 0,002153 | 0,0023632 | 0,0022216 | 0,0022216 | TRUE | 0,038988 | No |
| orf19.5067 | 2876 | 2197 | 2457 | 3143 | 3528 | 3413 | 3382 | 1884 | 2580 | 2797 | 2514 | 2592 | 3017 | 3257 | 3018 | 2657 | 2803,9 | 2621 | 2987 | 1,142 | 0,191 | 3,048 | 0,00230512 | 0,00536998 | 0,004038 | 0,0024973 | 0,0034213 | 0,0034213 | TRUE | 0,0645306 | No |
| orf19.5068 | 2428 | 1852 | 2269 | 2773 | 3256 | 3356 | 3443 | 2040 | 2178 | 2358 | 2321 | 2287 | 2785 | 3202 | 3072 | 2877 | 2635,1 | 2286 | 2984 | 1,306 | 0,386 | 7,198 | 6,13E-13 | 5,77E-12 | 0,002054 | 0,0025304 | 0,0023694 | 0,0023694 | TRUE | 0,0295666 | No |
| orf19.5069 | 8527 | 6113 | 7452 | 9531 | 10880 | 10053 | 10907 | 6945 | 7650 | 7783 | 7624 | 7859 | 9305 | 9592 | 9733 | 9795 | 8667,6 | 7729 | 9606 | 1,242 | 0,313 | 10,317 | 5,91E-25 | 1,48E-23 | 0,000206 | 0,002147 | 0,0007677 | 0,0007677 | TRUE | 0,0055289 | No |
| orf19.5070 | 2192 | 1573 | 2059 | 2570 | 2148 | 1979 | 2207 | 1497 | 1966 | 2003 | 2107 | 2119 | 1837 | 1888 | 1969 | 2111 | 2000,1 | 2049 | 1951 | 0,95 | -0,073 | -1,37 | 0,17081839 | 0,24102322 | 0,001897 | 0,0027053 | 0,0022476 | 0,0022476 | TRUE | 0,034304 | No |
| orf19.5071 | 204 | 138 | 213 | 243 | 218 | 190 | 241 | 161 | 183 | 176 | 218 | 200 | 186 | 181 | 215 | 227 | 198,36 | 194 | 202 | 1,035 | 0,049 | 0,431 | 0,66657193 | 0,73387652 | 0,004949 | 0,0092981 | 0,0073877 | 0,0073877 | TRUE | 0,0687228 | No |
| orf19.5073 | 547 | 375 | 450 | 623 | 733 | 629 | 674 | 383 | 491 | 477 | 460 | 514 | 627 | 600 | 601 | 540 | 538,87 | 486 | 592 | 1,222 | 0,289 | 4,114 | 3,89E-05 | 0,00012805 | 0,000982 | 0,0046735 | 0,0028844 | 0,0028844 | TRUE | 0,0351326 | No |
| orf19.5074 | 904 | 640 | 813 | 980 | 984 | 913 | 896 | 590 | 811 | 815 | 832 | 808 | 842 | 871 | 800 | 832 | 826,26 | 816 | 836 | 1,024 | 0,035 | 0,642 | 0,52105945 | 0,60413048 | 1,00E-08 | 0,0037365 | 0,0016005 | 0,0016005 | TRUE | 0,0112612 | No |
| orf19.5076 | 1440 | 903 | 1263 | 1482 | 1550 | 1368 | 1384 | 924 | 1292 | 1150 | 1292 | 1222 | 1326 | 1305 | 1235 | 1303 | 1265,6 | 1239 | 1292 | 1,041 | 0,058 | 1,087 | 0,27714173 | 0,36118601 | 0,001088 | 0,0031266 | 0,0019856 | 0,0019856 | TRUE | 0,0257156 | No |
| orf19.5076.1 | 927 | 615 | 814 | 930 | 902 | 832 | 884 | 713 | 832 | 783 | 833 | 767 | 771 | 794 | 789 | 1006 | 821,75 | 804 | 840 | 1,041 | 0,058 | 0,682 | 0,49523904 | 0,58033863 | 0,007847 | 0,0037461 | 0,0056277 | 0,0056277 | TRUE | 0,195966 | No |
| orf19.5077 | 3246 | 2201 | 2729 | 3314 | 3045 | 2617 | 2909 | 1806 | 2912 | 2802 | 2792 | 2733 | 2604 | 2497 | 2596 | 2547 | 2685,4 | 2810 | 2561 | 0,912 | -0,133 | -3,473 | 0,00051488 | 0,00138656 | 0,000255 | 0,0025201 | 0,0010374 | 0,0010374 | TRUE | 0,007637 | No |
| orf19.5078 | 695 | 514 | 625 | 786 | 792 | 657 | 713 | 417 | 623 | 654 | 639 | 648 | 677 | 627 | 636 | 588 | 636,76 | 641 | 632 | 0,99 | -0,015 | -0,233 | 0,81599993 | 0,8581354 | 1,00E-08 | 0,0042594 | 0,0022435 | 0,0022435 | TRUE | 0,0273918 | No |
| orf19.5079 | 55895 | 37902 | 51225 | 61228 | 81762 | 70731 | 78184 | 51925 | 50144 | 48253 | 52408 | 50489 | 69925 | 67489 | 69766 | 73236 | 60214 | 50324 | 70104 | 1,393 | 0,478 | 12,56 | 3,52E-36 | 1,91E-34 | 0,001119 | 0,0020036 | 0,0013753 | 0,0013753 | TRUE | 0,0110347 | No |
| orf19.5079.1 | 2598 | 1778 | 2097 | 2660 | 2261 | 1808 | 2037 | 1200 | 2331 | 2264 | 2145 | 2193 | 1934 | 1725 | 1818 | 1692 | 2012,8 | 2233 | 1792 | 0,804 | -0,315 | -5,902 | 3,59E-09 | 2,12E-08 | 0,001809 | 0,0027008 | 0,0022287 | 0,0022287 | TRUE | 0,0345426 | No |
| orf19.508 | 2018 | 1424 | 1894 | 2300 | 2538 | 2303 | 2476 | 1642 | 1810 | 1813 | 1938 | 1897 | 2171 | 2197 | 2209 | 2316 | 2043,9 | 1864 | 2223 | 1,19 | 0,251 | 5,806 | 6,41E-09 | 3,65E-08 | 0,000329 | 0,0026898 | 0,001307 | 0,001307 | TRUE | 0,0086475 | No |
| orf19.5081 | 13096 | 8579 | 11850 | 14366 | 10991 | 9044 | 9989 | 6339 | 11748 | 10922 | 12124 | 11846 | 9400 | 8629 | 8914 | 8941 | 10316 | 11660 | 8971 | 0,769 | -0,378 | -8,688 | 3,68E-18 | 5,82E-17 | 0,001483 | 0,0021203 | 0,0017233 | 0,0017233 | TRUE | 0,0218912 | No |
| orf19.5083 | 3350 | 2160 | 2901 | 3467 | 3301 | 2642 | 3015 | 1933 | 3005 | 2750 | 2968 | 2859 | 2823 | 2521 | 2690 | 2726 | 2792,9 | 2896 | 2690 | 0,929 | -0,107 | -2,201 | 0,02770561 | 0,04992661 | 0,001411 | 0,0024993 | 0,0019061 | 0,0019061 | TRUE | 0,0221795 | No |
| orf19.5085 | 4414 | 2911 | 4054 | 4787 | 4489 | 4023 | 4373 | 2870 | 3960 | 3706 | 4148 | 3947 | 3839 | 3839 | 3902 | 4048 | 3923,6 | 3940 | 3907 | 0,99 | -0,014 | -0,326 | 0,74474216 | 0,79985894 | 0,000944 | 0,0023495 | 0,0015056 | 0,0015056 | TRUE | 0,0185904 | No |
| orf19.5087 | 2493 | 1544 | 2138 | 2556 | 3086 | 2535 | 2895 | 1682 | 2236 | 1966 | 2187 | 2108 | 2639 | 2419 | 2583 | 2372 | 2313,9 | 2124 | 2503 | 1,179 | 0,237 | 4,317 | 1,58E-05 | 5,59E-05 | 0,002279 | 0,0026069 | 0,0024592 | 0,0024592 | TRUE | 0,0290921 | No |
| orf19.5089 | 402 | 289 | 353 | 413 | 427 | 438 | 476 | 309 | 361 | 368 | 361 | 341 | 365 | 418 | 425 | 436 | 384,24 | 358 | 411 | 1,147 | 0,198 | 2,45 | 0,01427849 | 0,02766371 | 1,00E-08 | 0,0057576 | 0,003659 | 0,003659 | TRUE | 0,0689441 | No |
| orf19.5090 | 478 | 287 | 405 | 455 | 566 | 471 | 515 | 362 | 429 | 365 | 414 | 375 | 484 | 449 | 460 | 511 | 435,92 | 396 | 476 | 1,197 | 0,259 | 3,269 | 0,00107831 | 0,00269938 | 0,001926 | 0,0053097 | 0,0037131 | 0,0037131 | TRUE | 0,0370632 | No |
| orf19.5092 | 11 | 15 | 10 | 10 | 19 | 19 | 16 | 9 | 10 | 19 | 10 | 8 | 16 | 18 | 14 | 13 | 13,6 | 12 | 15 | 1,345 | 0,428 | 1,107 | 0,26812971 | NA | 1,00E-08 | 0,1087306 | 0,0678503 | 0,0678503 | TRUE | 0,5069812 | No |
| orf19.5093 | 3049 | 2120 | 2708 | 3341 | 3272 | 2830 | 3066 | 2003 | 2735 | 2699 | 2771 | 2755 | 2798 | 2700 | 2736 | 2825 | 2752,4 | 2740 | 2765 | 1,008 | 0,012 | 0,331 | 0,74063255 | 0,79686969 | 1,00E-08 | 0,0025069 | 0,0008287 | 0,0008287 | TRUE | 0,0029363 | No |
| orf19.5094 | 1475 | 997 | 1287 | 1578 | 1565 | 1401 | 1487 | 890 | 1323 | 1269 | 1317 | 1301 | 1338 | 1337 | 1327 | 1255 | 1308,5 | 1302 | 1314 | 1,01 | 0,015 | 0,323 | 0,74666052 | 0,80106007 | 1,00E-08 | 0,003089 | 0,001233 | 0,001233 | TRUE | 0,0094558 | No |
| orf19.5095 | 4468 | 3085 | 4104 | 4813 | 5115 | 4358 | 4944 | 2903 | 4008 | 3928 | 4199 | 3969 | 4375 | 4158 | 4412 | 4094 | 4142,8 | 4026 | 4260 | 1,059 | 0,082 | 2,022 | 0,04319348 | 0,0739756 | 0,0008 | 0,0023299 | 0,0013556 | 0,0013556 | TRUE | 0,0100803 | No |
| orf19.5097 | 870 | 610 | 810 | 979 | 740 | 683 | 731 | 424 | 780 | 777 | 829 | 807 | 633 | 652 | 652 | 598 | 715,99 | 798 | 634 | 0,796 | -0,33 | -5,651 | 1,60E-08 | 8,64E-08 | 1,00E-08 | 0,0040071 | 0,0018601 | 0,0018601 | TRUE | 0,0139552 | No |
| orf19.5098 | 2136 | 1399 | 1772 | 2349 | 1832 | 1465 | 1633 | 1049 | 1916 | 1781 | 1813 | 1937 | 1567 | 1398 | 1457 | 1480 | 1668,6 | 1862 | 1476 | 0,792 | -0,337 | -6,472 | 9,67E-11 | 7,08E-10 | 0,001411 | 0,0028495 | 0,0019924 | 0,0019924 | TRUE | 0,0223463 | No |
| orf19.51 | 1890 | 1687 | 1849 | 2339 | 2375 | 2738 | 2647 | 1457 | 1696 | 2148 | 1892 | 1929 | 2031 | 2613 | 2362 | 2055 | 2090,6 | 1916 | 2265 | 1,184 | 0,243 | 2,694 | 0,00705882 | 0,01471538 | 0,011474 | 0,0026739 | 0,0073482 | 0,0073482 | TRUE | 0,1292081 | No |
| orf19.510 | 38 | 27 | 30 | 54 | 44 | 41 | 42 | 19 | 34 | 34 | 31 | 45 | 38 | 39 | 37 | 27 | 35,59 | 36 | 35 | 0,988 | -0,018 | -0,074 | 0,94069673 | 0,95726727 | 1,00E-08 | 0,0427701 | 0,025635 | 0,025635 | TRUE | 0,2203246 | No |
| orf19.5100 | 4174 | 2969 | 3925 | 4288 | 3541 | 3111 | 3497 | 2178 | 3745 | 3780 | 4016 | 3536 | 3028 | 2968 | 3120 | 3072 | 3408,1 | 3769 | 3047 | 0,809 | -0,306 | -6,648 | 2,96E-11 | 2,31E-10 | 0,001189 | 0,0024055 | 0,001737 | 0,001737 | TRUE | 0,0238773 | No |
| orf19.5101 | 3961 | 2684 | 3452 | 4169 | 4760 | 4154 | 4725 | 2954 | 3553 | 3417 | 3532 | 3438 | 4071 | 3964 | 4216 | 4166 | 3794,6 | 3485 | 4104 | 1,177 | 0,235 | 6,519 | 7,09E-11 | 5,26E-10 | 0,00032 | 0,0023621 | 0,0009839 | 0,0009839 | TRUE | 0,0065266 | No |
| orf19.5102 | 1844 | 1090 | 1633 | 1825 | 1683 | 1473 | 1590 | 971 | 1654 | 1388 | 1671 | 1505 | 1439 | 1405 | 1419 | 1370 | 1481,3 | 1554 | 1408 | 0,906 | -0,143 | -2,279 | 0,02266509 | 0,04177051 | 0,003022 | 0,0029595 | 0,0031065 | 0,0031065 | TRUE | 0,0592039 | No |
| orf19.5103 | 506 | 354 | 491 | 526 | 569 | 470 | 517 | 356 | 454 | 451 | 502 | 434 | 487 | 448 | 461 | 502 | 467,4 | 460 | 474 | 1,03 | 0,043 | 0,574 | 0,56563639 | 0,64538621 | 0,001142 | 0,0050854 | 0,0031791 | 0,0031791 | TRUE | 0,0442858 | No |
| orf19.5104 | 1872 | 1244 | 1531 | 1981 | 1753 | 1438 | 1462 | 1260 | 1679 | 1584 | 1566 | 1634 | 1499 | 1372 | 1305 | 1777 | 1552 | 1616 | 1488 | 0,919 | -0,122 | -1,453 | 0,14627907 | 0,21113994 | 0,009481 | 0,0029149 | 0,006154 | 0,006154 | TRUE | 0,1976578 | No |
| orf19.5105 | 3030 | 2056 | 2903 | 3336 | 3309 | 2854 | 3186 | 2153 | 2718 | 2618 | 2970 | 2751 | 2830 | 2723 | 2843 | 3037 | 2811,2 | 2764 | 2858 | 1,033 | 0,046 | 0,9 | 0,36824526 | 0,45683831 | 0,002012 | 0,0024959 | 0,0021888 | 0,0021888 | TRUE | 0,0300758 | No |
| orf19.5106 | 14629 | 10197 | 13212 | 15485 | 16350 | 14711 | 16180 | 9220 | 13124 | 12982 | 13517 | 12769 | 13983 | 14037 | 14438 | 13004 | 13482 | 13098 | 13866 | 1,059 | 0,083 | 2,069 | 0,03853246 | 0,06692772 | 0,001155 | 0,0020872 | 0,0014607 | 0,0014607 | TRUE | 0,0209958 | No |
| orf19.5107 | 5404 | 3809 | 4971 | 5828 | 5907 | 4916 | 5484 | 3731 | 4848 | 4849 | 5086 | 4806 | 5052 | 4691 | 4894 | 5262 | 4935,9 | 4897 | 4975 | 1,015 | 0,022 | 0,498 | 0,61824116 | 0,69143576 | 0,001181 | 0,0022736 | 0,001622 | 0,001622 | TRUE | 0,0178247 | No |
| orf19.511 | 2517 | 1551 | 2079 | 2506 | 2728 | 2347 | 2506 | 1613 | 2258 | 1975 | 2127 | 2066 | 2333 | 2239 | 2236 | 2275 | 2188,7 | 2106 | 2271 | 1,077 | 0,107 | 2,201 | 0,02775478 | 0,04998523 | 0,001178 | 0,0026428 | 0,0018099 | 0,0018099 | TRUE | 0,028555 | No |
| orf19.5110 | 7679 | 4968 | 6940 | 8249 | 6078 | 5183 | 5939 | 3857 | 6889 | 6325 | 7100 | 6802 | 5198 | 4945 | 5300 | 5440 | 5999,9 | 6779 | 5221 | 0,77 | -0,378 | -8,057 | 7,82E-16 | 9,75E-15 | 0,001741 | 0,0022215 | 0,0019419 | 0,0019419 | TRUE | 0,0242887 | No |
| orf19.5112 | 5001 | 3469 | 4535 | 5394 | 5727 | 4746 | 5123 | 3670 | 4486 | 4416 | 4640 | 4448 | 4898 | 4528 | 4571 | 5176 | 4645,6 | 4498 | 4793 | 1,065 | 0,091 | 1,853 | 0,06389244 | 0,10414156 | 0,001963 | 0,002292 | 0,0020824 | 0,0020824 | TRUE | 0,0339437 | No |
| orf19.5113 | 4366 | 2499 | 3618 | 4395 | 4356 | 3074 | 3294 | 2418 | 3917 | 3181 | 3702 | 3624 | 3725 | 2933 | 2939 | 3410 | 3429 | 3606 | 3252 | 0,901 | -0,15 | -1,75 | 0,08011597 | 0,1267009 | 0,010349 | 0,0024029 | 0,0067478 | 0,0067478 | TRUE | 0,1186591 | No |
| orf19.5114 | 906 | 599 | 710 | 927 | 1064 | 894 | 926 | 509 | 813 | 763 | 726 | 764 | 910 | 853 | 826 | 718 | 796,67 | 766 | 827 | 1,082 | 0,114 | 1,523 | 0,12783207 | 0,1883775 | 0,004139 | 0,0038017 | 0,0041073 | 0,0041073 | TRUE | 0,0844894 | No |
| orf19.5114.1 | 14841 | 9312 | 11854 | 15057 | 22816 | 16304 | 17523 | 9896 | 13314 | 11855 | 12128 | 12416 | 19513 | 15557 | 15636 | 13957 | 14297 | 12428 | 16166 | 1,301 | 0,379 | 4,406 | 1,05E-05 | 3,81E-05 | 0,011874 | 0,0020811 | 0,0070549 | 0,0070549 | TRUE | 0,2379501 | No |
| orf19.5117 | 1E+05 | 87955 | 1E+05 | 1E+05 | 2E+05 | 1E+05 | 1E+05 | 95503 | 1E+05 | 1E+05 | 1E+05 | 1E+05 | 1E+05 | 1E+05 | 1E+05 | 1E+05 | 124436 | 1E+05 | 1E+05 | 1,096 | 0,133 | 3,208 | 0,00133728 | 0,00327667 | 0,00145 | 0,0019912 | 0,0016329 | 0,0016329 | TRUE | 0,0178946 | No |
| orf19.5118 | 36127 | 23684 | 32031 | 39287 | 48886 | 42179 | 46387 | 31412 | 32410 | 30152 | 32771 | 32397 | 41809 | 40246 | 41393 | 44304 | 36935 | 31932 | 41938 | 1,313 | 0,393 | 9,358 | 8,14E-21 | 1,57E-19 | 0,0015 | 0,0020188 | 0,0016666 | 0,0016666 | TRUE | 0,017572 | No |
| orf19.512 | 10799 | 7411 | 9459 | 11307 | 9834 | 8762 | 9776 | 6272 | 9688 | 9435 | 9677 | 9324 | 8410 | 8360 | 8723 | 8846 | 9058,1 | 9531 | 8585 | 0,9 | -0,151 | -4,62 | 3,84E-06 | 1,49E-05 | 0,000413 | 0,0021398 | 0,0009199 | 0,0009199 | TRUE | 0,0048996 | No |

| | | | | | | | | | | | | | | | | | | | | | | | | | | | | | | | |
|---|---|---|---|---|---|---|---|---|---|---|---|---|---|---|---|---|---|---|---|---|---|---|---|---|---|---|---|---|---|---|---|
| orf19.5121 | 68 | 45 | 60 | 68 | 54 | 68 | 69 | 24 | 61 | 57 | 61 | 56 | 46 | 65 | 62 | 34 | 55,28 | 59 | 52 | 0,889 | -0,169 | -0,817 | 0,41399955 | 0,50213744 | 0,014508 | 0,0282404 | 0,0231474 | 0,0231474 | TRUE | 0,3194192 | No |
| orf19.5124 | 1276 | 755 | 1226 | 1445 | 1514 | 1207 | 1404 | 1028 | 1145 | 961 | 1254 | 1192 | 1295 | 1152 | 1253 | 1450 | 1212,6 | 1138 | 1288 | 1,128 | 0,174 | 1,966 | 0,04931059 | 0,08320357 | 0,00967 | 0,0031767 | 0,0066718 | 0,0066718 | TRUE | 0,1257277 | No |
| orf19.5125 | 5586 | 4564 | 5158 | 6729 | 6690 | 6414 | 6442 | 4394 | 5011 | 5810 | 5277 | 5549 | 5721 | 6120 | 5748 | 6197 | 5679,4 | 5412 | 5946 | 1,099 | 0,136 | 2,566 | 0,01030109 | 0,0206848 | 0,002654 | 0,0022351 | 0,0025203 | 0,0025203 | TRUE | 0,0304031 | No |
| orf19.5126 | 2270 | 1608 | 2133 | 2632 | 2656 | 2442 | 2515 | 1882 | 2036 | 2047 | 2182 | 2170 | 2271 | 2330 | 2244 | 2654 | 2242,1 | 2109 | 2375 | 1,124 | 0,168 | 2,828 | 0,00468568 | 0,01018508 | 0,003274 | 0,002627 | 0,0029487 | 0,0029487 | TRUE | 0,0725507 | No |
| orf19.5129 | 464 | 326 | 419 | 585 | 423 | 408 | 448 | 260 | 416 | 415 | 429 | 482 | 362 | 389 | 400 | 367 | 407,49 | 436 | 380 | 0,87 | -0,201 | -2,514 | 0,01192983 | 0,02356161 | 0,001878 | 0,0055421 | 0,0036943 | 0,0036943 | TRUE | 0,0649951 | No |
| orf19.513 | 1 | 2 | 4 | 5 | 0 | 2 | 4 | 1 | 1 | 3 | 4 | 4 | 0 | 2 | 4 | 1 | 2,32 | 3 | 2 | 0,592 | -0,757 | -0,777 | 0,43696872 | NA | 1,00E-08 | 0,6281561 | 0,4513809 | 0,4513809 | TRUE | 0,4905934 | No |
| orf19.5130 | 3968 | 2703 | 3410 | 4348 | 3746 | 3192 | 3354 | 2272 | 3560 | 3441 | 3489 | 3585 | 3204 | 3046 | 2993 | 3204 | 3315,2 | 3519 | 3112 | 0,883 | -0,179 | -4,61 | 4,02E-06 | 1,55E-05 | 0,000383 | 0,0024174 | 0,0011444 | 0,0011444 | TRUE | 0,0081456 | No |
| orf19.5131 | 4806 | 3449 | 4189 | 5212 | 5690 | 5102 | 5669 | 3449 | 4311 | 4391 | 4286 | 4298 | 4866 | 4868 | 5059 | 4865 | 4618 | 4322 | 4914 | 1,138 | 0,186 | 5,936 | 2,92E-09 | 1,74E-08 | 1,00E-08 | 0,0022939 | 0,0007279 | 0,0007279 | TRUE | 0,0048926 | No |
| orf19.5132 | 116 | 99 | 125 | 146 | 112 | 126 | 116 | 72 | 104 | 126 | 128 | 120 | 96 | 120 | 104 | 102 | 112,43 | 120 | 106 | 0,883 | -0,179 | -1,304 | 0,19222656 | 0,26561573 | 1,00E-08 | 0,0148914 | 0,0091526 | 0,0091526 | TRUE | 0,0950758 | No |
| orf19.5133 | 3756 | 2655 | 3569 | 3979 | 4115 | 3688 | 4043 | 2614 | 3370 | 3380 | 3651 | 3281 | 3519 | 3519 | 3608 | 3687 | 3501,9 | 3420 | 3583 | 1,047 | 0,067 | 1,538 | 0,12400291 | 0,18349988 | 0,000969 | 0,0023941 | 0,0015163 | 0,0015163 | TRUE | 0,0252049 | No |
| orf19.5134 | 2203 | 1850 | 2093 | 2584 | 2455 | 2342 | 2499 | 1475 | 1976 | 2355 | 2141 | 2131 | 2100 | 2235 | 2230 | 2080 | 2156 | 2151 | 2161 | 1,007 | 0,009 | 0,164 | 0,8696274 | 0,90050352 | 0,002786 | 0,0026528 | 0,0027324 | 0,0027324 | TRUE | 0,0476967 | No |
| orf19.5136 | 1720 | 1151 | 1518 | 1831 | 1439 | 1312 | 1395 | 824 | 1543 | 1465 | 1553 | 1510 | 1231 | 1252 | 1245 | 1162 | 1370,1 | 1518 | 1222 | 0,806 | -0,31 | -6,676 | 2,45E-11 | 1,92E-10 | 1,00E-08 | 0,0030391 | 0,0013367 | 0,0013367 | TRUE | 0,0112971 | No |
| orf19.5137 | 27 | 16 | 25 | 27 | 12 | 19 | 14 | 9 | 24 | 20 | 26 | 22 | 10 | 18 | 12 | 13 | 18,25 | 23 | 13 | 0,577 | -0,794 | -2,434 | 0,01494131 | 0,02878083 | 1,00E-08 | 0,0815183 | 0,0433891 | 0,0433891 | TRUE | 0,2658307 | No |
| orf19.5137.1 | 1862 | 1237 | 1500 | 1961 | 1841 | 1524 | 1549 | 1120 | 1670 | 1575 | 1535 | 1617 | 1574 | 1454 | 1382 | 1580 | 1548,4 | 1599 | 1498 | 0,935 | -0,097 | -1,715 | 0,08632448 | 0,13506185 | 0,001868 | 0,0029171 | 0,0024423 | 0,0024423 | TRUE | 0,033028 | No |
| orf19.5138 | 231 | 171 | 265 | 295 | 336 | 350 | 342 | 206 | 207 | 218 | 271 | 243 | 287 | 334 | 305 | 291 | 269,54 | 235 | 304 | 1,296 | 0,374 | 3,613 | 0,00030315 | 0,00085709 | 0,005537 | 0,0073653 | 0,0065237 | 0,0065237 | TRUE | 0,1158965 | No |
| orf19.5139 | 2247 | 1604 | 2057 | 2460 | 2678 | 2458 | 2651 | 1496 | 2016 | 2042 | 2105 | 2029 | 2290 | 2345 | 2366 | 2110 | 2162,8 | 2048 | 2278 | 1,115 | 0,157 | 3,372 | 0,00074721 | 0,00193252 | 0,00087 | 0,0026507 | 0,0016048 | 0,0016048 | TRUE | 0,0272599 | No |
| orf19.514 | 848 | 603 | 703 | 974 | 861 | 806 | 796 | 514 | 761 | 768 | 719 | 803 | 736 | 769 | 710 | 725 | 748,94 | 763 | 735 | 0,963 | -0,054 | -0,903 | 0,366742 | 0,45562158 | 1,00E-08 | 0,0039179 | 0,002085 | 0,002085 | TRUE | 0,0176535 | No |
| orf19.5140 | 1475 | 1000 | 1251 | 1530 | 1597 | 1337 | 1486 | 903 | 1323 | 1273 | 1280 | 1262 | 1366 | 1276 | 1326 | 1274 | 1297,4 | 1284 | 1310 | 1,021 | 0,031 | 0,646 | 0,51800244 | 0,60149954 | 1,00E-08 | 0,0030985 | 0,0013755 | 0,0013755 | TRUE | 0,0100373 | No |
| orf19.5141 | 24 | 18 | 22 | 24 | 15 | 25 | 20 | 18 | 22 | 23 | 23 | 20 | 13 | 24 | 18 | 25 | 20,83 | 22 | 20 | 0,906 | -0,142 | -0,458 | 0,64690542 | 0,71652497 | 1,00E-08 | 0,0716631 | 0,0436798 | 0,0436798 | TRUE | 0,3636678 | No |
| orf19.5142 | 79 | 50 | 67 | 98 | 81 | 86 | 82 | 52 | 71 | 64 | 69 | 81 | 69 | 82 | 73 | 73 | 72,72 | 71 | 74 | 1,041 | 0,058 | 0,361 | 0,71829309 | 0,77855041 | 1,00E-08 | 0,0219434 | 0,0115897 | 0,0115897 | TRUE | 0,0866105 | No |
| orf19.5143 | 3746 | 2464 | 3231 | 4061 | 3716 | 2939 | 3394 | 2010 | 3361 | 3137 | 3306 | 3349 | 3178 | 2804 | 3029 | 2835 | 3124,7 | 3288 | 2962 | 0,901 | -0,15 | -2,991 | 0,0027827 | 0,00636652 | 0,001817 | 0,0024441 | 0,002103 | 0,002103 | TRUE | 0,0298199 | No |
| orf19.5144 | 188 | 114 | 176 | 201 | 182 | 142 | 166 | 112 | 169 | 145 | 180 | 166 | 156 | 135 | 148 | 158 | 157,1 | 165 | 149 | 0,9 | -0,152 | -1,319 | 0,18713661 | 0,25977743 | 1,00E-08 | 0,0112198 | 0,0063824 | 0,0063824 | TRUE | 0,0573159 | No |
| orf19.5145 | 482 | 335 | 448 | 532 | 482 | 429 | 468 | 289 | 432 | 426 | 458 | 439 | 412 | 409 | 418 | 408 | 425,34 | 439 | 412 | 0,938 | -0,092 | -1,338 | 0,18087447 | 0,252602 | 1,00E-08 | 0,0053926 | 0,0022348 | 0,0022348 | TRUE | 0,0097019 | No |
| orf19.5147 | 1376 | 1072 | 1202 | 1493 | 1647 | 1644 | 1602 | 1001 | 1234 | 1365 | 1230 | 1231 | 1409 | 1569 | 1430 | 1412 | 1359,8 | 1265 | 1455 | 1,152 | 0,205 | 3,591 | 0,00032906 | 0,00092555 | 0,001783 | 0,0030471 | 0,0023739 | 0,0023739 | TRUE | 0,0339616 | No |
| orf19.5148 | 2864 | 1964 | 2680 | 3099 | 4093 | 3575 | 3891 | 2390 | 2569 | 2500 | 2742 | 2555 | 3500 | 3411 | 3472 | 3371 | 3015,2 | 2592 | 3438 | 1,327 | 0,408 | 10,127 | 4,19E-24 | 9,87E-23 | 0,000506 | 0,002461 | 0,0012214 | 0,0012214 | TRUE | 0,0181058 | No |
| orf19.515 | 458 | 313 | 357 | 458 | 489 | 393 | 450 | 282 | 411 | 398 | 365 | 378 | 418 | 375 | 402 | 398 | 393,09 | 388 | 398 | 1,027 | 0,039 | 0,505 | 0,61366709 | 0,68708757 | 1,00E-08 | 0,0056725 | 0,0030686 | 0,0030686 | TRUE | 0,0198629 | No |
| orf19.5155 | 233 | 187 | 250 | 296 | 340 | 282 | 292 | 203 | 209 | 238 | 256 | 244 | 291 | 269 | 261 | 286 | 256,71 | 237 | 277 | 1,169 | 0,225 | 2,36 | 0,01827845 | 0,0343178 | 1,00E-08 | 0,0076345 | 0,0047907 | 0,0047907 | TRUE | 0,0737716 | No |
| orf19.5156 | 771 | 580 | 731 | 844 | 708 | 652 | 604 | 442 | 692 | 738 | 748 | 696 | 606 | 622 | 539 | 623 | 657,99 | 718 | 598 | 0,831 | -0,267 | -3,953 | 7,72E-05 | 0,00023974 | 0,001563 | 0,0041858 | 0,0028403 | 0,0028403 | TRUE | 0,0533106 | No |
| orf19.5157 | 365 | 292 | 331 | 445 | 260 | 266 | 262 | 161 | 327 | 372 | 339 | 367 | 222 | 254 | 234 | 227 | 292,73 | 351 | 234 | 0,668 | -0,581 | -6,544 | 6,00E-11 | 4,53E-10 | 1,00E-08 | 0,0069387 | 0,0040203 | 0,0040203 | TRUE | 0,0359742 | No |
| orf19.5158 | 1019 | 783 | 930 | 1091 | 1254 | 1225 | 1186 | 810 | 914 | 997 | 951 | 900 | 1072 | 1169 | 1058 | 1142 | 1025,5 | 940 | 1110 | 1,182 | 0,241 | 4,176 | 2,97E-05 | 0,00010005 | 0,001257 | 0,0033951 | 0,0022081 | 0,0022081 | TRUE | 0,018018 | No |
| orf19.5159 | 3326 | 2461 | 2883 | 3729 | 3966 | 3631 | 3816 | 2338 | 2984 | 3133 | 2950 | 3075 | 3392 | 3465 | 3405 | 3298 | 3212,6 | 3036 | 3390 | 1,118 | 0,161 | 4,364 | 1,28E-05 | 4,58E-05 | 0,000174 | 0,0024314 | 0,0009923 | 0,0009923 | TRUE | 0,0052568 | No |
| orf19.516 | 310 | 204 | 277 | 303 | 299 | 300 | 339 | 178 | 278 | 260 | 283 | 250 | 256 | 286 | 303 | 251 | 270,82 | 268 | 274 | 1,027 | 0,038 | 0,399 | 0,68988161 | 0,75305391 | 0,002261 | 0,0073398 | 0,0049994 | 0,0049994 | TRUE | 0,0567034 | No |
| orf19.5160 | 2305 | 1588 | 2038 | 2500 | 2369 | 2014 | 2272 | 1312 | 2068 | 2022 | 2085 | 2062 | 2026 | 1922 | 2027 | 1850 | 2007,7 | 2059 | 1956 | 0,952 | -0,071 | -1,629 | 0,10341386 | 0,15693621 | 0,000392 | 0,0027026 | 0,0013466 | 0,0013466 | TRUE | 0,0144905 | No |
| orf19.5161 | 1378 | 929 | 1130 | 1366 | 1797 | 1459 | 1456 | 1054 | 1236 | 1183 | 1156 | 1126 | 1537 | 1392 | 1299 | 1487 | 1302 | 1175 | 1429 | 1,215 | 0,281 | 4,469 | 7,87E-06 | 2,92E-05 | 0,002955 | 0,0030945 | 0,0030132 | 0,0030132 | TRUE | 0,0460899 | No |
| orf19.5162 | 927 | 607 | 817 | 923 | 1214 | 1168 | 1243 | 754 | 832 | 773 | 836 | 761 | 1038 | 1114 | 1109 | 1063 | 940,84 | 800 | 1081 | 1,351 | 0,434 | 7,652 | 1,97E-14 | 2,13E-13 | 0,000727 | 0,0035225 | 0,0020055 | 0,0020055 | TRUE | 0,0147196 | No |
| orf19.5163 | 1246 | 881 | 1198 | 1475 | 1342 | 1263 | 1325 | 825 | 1118 | 1122 | 1226 | 1216 | 1148 | 1205 | 1182 | 1164 | 1172,5 | 1170 | 1175 | 1,003 | 0,004 | 0,08 | 0,93633295 | 0,95395767 | 0,000659 | 0,0032176 | 0,0017742 | 0,0017742 | TRUE | 0,0120882 | No |
| orf19.5164 | 376 | 273 | 371 | 417 | 358 | 341 | 351 | 271 | 337 | 348 | 380 | 344 | 306 | 325 | 313 | 382 | 341,91 | 352 | 332 | 0,936 | -0,095 | -1,069 | 0,28512734 | 0,3694296 | 0,002955 | 0,0062254 | 0,0046778 | 0,0046778 | TRUE | 0,1022886 | No |
| orf19.5165 | 1211 | 738 | 981 | 1167 | 1155 | 1057 | 1172 | 650 | 1086 | 940 | 1004 | 962 | 988 | 1009 | 1046 | 917 | 993,86 | 998 | 990 | 0,993 | -0,01 | -0,163 | 0,87041045 | 0,90111898 | 0,002453 | 0,0034402 | 0,0029017 | 0,0029017 | TRUE | 0,0433227 | No |
| orf19.5166 | 1677 | 1105 | 1372 | 1773 | 1661 | 1379 | 1626 | 1060 | 1504 | 1407 | 1404 | 1462 | 1421 | 1316 | 1451 | 1495 | 1432,4 | 1444 | 1421 | 0,981 | -0,027 | -0,511 | 0,60940998 | 0,68410598 | 0,001253 | 0,002993 | 0,0019936 | 0,0019936 | TRUE | 0,0295161 | No |
| orf19.5167 | 5091 | 3333 | 4435 | 5437 | 5764 | 4837 | 5481 | 3292 | 4567 | 4243 | 4537 | 4483 | 4930 | 4615 | 4891 | 4643 | 4613,8 | 4458 | 4770 | 1,07 | 0,098 | 2,401 | 0,01633563 | 0,03111746 | 0,000843 | 0,0022942 | 0,0013666 | 0,0013666 | TRUE | 0,012274 | No |
| orf19.5168 | 3756 | 2442 | 3093 | 3968 | 4005 | 3439 | 3767 | 2363 | 3370 | 3109 | 3164 | 3272 | 3425 | 3281 | 3361 | 3333 | 3289,5 | 3229 | 3350 | 1,037 | 0,052 | 1,34 | 0,18013716 | 0,25198231 | 0,000397 | 0,0024208 | 0,0011583 | 0,0011583 | TRUE | 0,0104141 | No |
| orf19.5169 | 413 | 307 | 363 | 448 | 201 | 179 | 175 | 131 | 371 | 391 | 371 | 369 | 172 | 171 | 156 | 185 | 273,22 | 376 | 171 | 0,453 | -1,142 | -12,68 | 7,68E-37 | 4,35E-35 | 1,00E-08 | 0,0072928 | 0,0035207 | 0,0035207 | TRUE | 0,0364756 | No |
| orf19.517 | 1336 | 919 | 1196 | 1498 | 1288 | 1097 | 1225 | 870 | 1199 | 1170 | 1224 | 1235 | 1102 | 1047 | 1093 | 1227 | 1162 | 1207 | 1117 | 0,922 | -0,117 | -2,065 | 0,03896133 | 0,06753598 | 0,001524 | 0,0032289 | 0,0022234 | 0,0022234 | TRUE | 0,0469699 | No |
| orf19.5170 | 7542 | 5461 | 6831 | 8260 | 6910 | 6510 | 6886 | 4192 | 6766 | 6952 | 6989 | 6811 | 5910 | 6212 | 6145 | 5912 | 6462,1 | 6880 | 6045 | 0,879 | -0,186 | -5,736 | 9,72E-09 | 5,38E-08 | 0,000251 | 0,0022042 | 0,0008519 | 0,0008519 | TRUE | 0,0041971 | No |
| orf19.5171 | 1277 | 921 | 1030 | 1370 | 1216 | 1195 | 1325 | 876 | 1146 | 1173 | 1054 | 1130 | 1040 | 1140 | 1182 | 1236 | 1137,5 | 1126 | 1150 | 1,02 | 0,028 | 0,44 | 0,66002604 | 0,72853876 | 0,002745 | 0,0032558 | 0,0029712 | 0,0029712 | TRUE | 0,0505106 | No |
| orf19.5172 | 11 | 3 | 7 | 10 | 13 | 7 | 8 | 4 | 10 | 4 | 7 | 8 | 11 | 7 | 7 | 6 | 7,46 | 7 | 8 | 1,046 | 0,065 | 0,129 | 0,89699702 | NA | 1,00E-08 | 0,1966011 | 0,1130716 | 0,1130716 | TRUE | 0,3109843 | No |
| orf19.5174 | 855 | 556 | 622 | 874 | 925 | 672 | 694 | 452 | 767 | 708 | 636 | 721 | 791 | 641 | 619 | 638 | 690,13 | 708 | 672 | 0,951 | -0,073 | -0,811 | 0,4171572 | 0,50510877 | 0,008368 | 0,004083 | 0,0063631 | 0,0063631 | TRUE | 0,1733983 | No |
| orf19.5175 | 3341 | 2420 | 3346 | 3817 | 3734 | 3341 | 3649 | 2526 | 2997 | 3081 | 3423 | 3148 | 3193 | 3188 | 3256 | 3563 | 3231,1 | 3162 | 3300 | 1,043 | 0,06 | 1,093 | 0,27451599 | 0,35880679 | 0,002742 | 0,0024288 | 0,0025937 | 0,0025937 | TRUE | 0,037443 | No |
| orf19.5177 | 2127 | 1472 | 1882 | 2296 | 2857 | 2534 | 2720 | 1662 | 1908 | 1874 | 1925 | 1893 | 2443 | 2418 | 2427 | 2344 | 2154,2 | 1900 | 2408 | 1,268 | 0,343 | 9,174 | 4,58E-20 | 8,26E-19 | 1,00E-08 | 0,0026534 | 0,0008706 | 0,0008706 | TRUE | 0,0034362 | No |
| orf19.5178 | 3293 | 2157 | 2777 | 3580 | 3075 | 2641 | 2897 | 1801 | 2954 | 2746 | 2841 | 2952 | 2630 | 2520 | 2585 | 2540 | 2721,1 | 2873 | 2569 | 0,893 | -0,163 | -4,055 | 5,01E-05 | 0,00016204 | 0,00042 | 0,002513 | 0,001174 | 0,001174 | TRUE | 0,0102329 | No |
| orf19.5179 | 45 | 26 | 29 | 45 | 23 | 19 | 29 | 25 | 40 | 33 | 30 | 37 | 20 | 18 | 26 | 35 | 29,9 | 35 | 25 | 0,685 | -0,547 | -2,021 | 0,04324148 | 0,0740367 | 0,008368 | 0,0505344 | 0,0353523 | 0,0353523 | TRUE | 0,3513708 | No |
| orf19.518 | 8926 | 6128 | 7741 | 9463 | 7857 | 7052 | 7651 | 4481 | 8008 | 7802 | 7920 | 7803 | 6720 | 6729 | 6827 | 6320 | 7266 | 7883 | 6649 | 0,844 | -0,245 | -7,175 | 7,23E-13 | 6,74E-12 | 0,000466 | 0,0021793 | 0,0009762 | 0,0009762 | TRUE | 0,0128347 | No |
| orf19.5180 | 1143 | 701 | 971 | 1119 | 1165 | 862 | 1007 | 680 | 1025 | 892 | 993 | 923 | 996 | 822 | 899 | 959 | 938,81 | 958 | 919 | 0,957 | -0,063 | -0,858 | 0,39085486 | 0,47972544 | 0,00449 | 0,0035258 | 0,0040517 | 0,0040517 | TRUE | 0,0606063 | No |
| orf19.5181 | 8887 | 5887 | 8083 | 9317 | 9089 | 7678 | 8503 | 4947 | 7973 | 7495 | 8270 | 7683 | 7773 | 7326 | 7588 | 6977 | 7635,5 | 7855 | 7416 | 0,944 | -0,082 | -1,763 | 0,07794382 | 0,12369004 | 0,001841 | 0,0021696 | 0,0019668 | 0,0019668 | TRUE | 0,0188968 | No |
| orf19.5182 | 3296 | 2223 | 2917 | 3385 | 3613 | 3139 | 3584 | 2115 | 2957 | 2830 | 2984 | 2791 | 3090 | 2995 | 3198 | 2983 | 2978,6 | 2890 | 3066 | 1,062 | 0,086 | 2,055 | 0,03990473 | 0,06893261 | 0,000741 | 0,0024669 | 0,0013532 | 0,0013532 | TRUE | 0,0102077 | No |
| orf19.5184 | 4280 | 2718 | 3826 | 4582 | 4994 | 3980 | 4524 | 2704 | 3840 | 3460 | 3914 | 3778 | 4271 | 3798 | 4037 | 3814 | 3864 | 3748 | 3980 | 1,062 | 0,087 | 1,594 | 0,11082691 | 0,16642042 | 0,002642 | 0,0023552 | 0,0025714 | 0,0025714 | TRUE | 0,0315796 | No |
| orf19.5188 | 191 | 113 | 182 | 224 | 217 | 212 | 214 | 137 | 171 | 144 | 186 | 185 | 186 | 202 | 191 | 193 | 182,27 | 172 | 193 | 1,118 | 0,16 | 1,461 | 0,14389787 | 0,20822216 | 1,00E-08 | 0,009944 | 0,0060856 | 0,0060856 | TRUE | 0,1055652 | No |
| orf19.519 | 1126 | 786 | 992 | 1243 | 1112 | 1039 | 1080 | 610 | 1010 | 1001 | 1015 | 1025 | 951 | 991 | 964 | 860 | 977,15 | 1013 | 942 | 0,933 | -0,101 | -1,828 | 0,06751558 | 0,10942368 | 0,000305 | 0,0034652 | 0,001889 | 0,001889 | TRUE | 0,0345166 | No |
| orf19.5190 | 0 | 3 | 1 | 0 | 1 | 2 | 1 | 1 | 0 | 4 | 1 | 0 | 1 | 2 | 1 | 1 | 1,24 | 1 | 1 | 1,153 | 0,206 | 0,153 | 0,87858052 | NA | 0,362835 | 1,1740377 | 0,8820981 | 0,8820981 | TRUE | 0,9344686 | No |

| | | | | | | | | | | | | | | | | | | | | | | | | | | | | | | | |
|---|---|---|---|---|---|---|---|---|---|---|---|---|---|---|---|---|---|---|---|---|---|---|---|---|---|---|---|---|---|---|---|
| orf19.5191 | 25 | 23 | 29 | 36 | 41 | 32 | 25 | 23 | 22 | 29 | 30 | 30 | 35 | 31 | 22 | 32 | 28,93 | 28 | 30 | 1,083 | 0,115 | 0,439 | 0,66048178 | 0,72873076 | 1,00E-08 | 0,0521657 | 0,0306948 | 0,0306948 | TRUE | 0,2306288 | No |
| orf19.5194.1 | 664 | 424 | 608 | 715 | 876 | 703 | 791 | 491 | 596 | 540 | 622 | 590 | 749 | 671 | 706 | 693 | 645,68 | 587 | 705 | 1,2 | 0,263 | 3,97 | 7,19E-05 | 0,00022543 | 0,001075 | 0,0042279 | 0,0026466 | 0,0026466 | TRUE | 0,029888 | No |
| orf19.5195 | 2333 | 1599 | 2032 | 2439 | 2321 | 1915 | 2105 | 1367 | 2093 | 2036 | 2079 | 2011 | 1985 | 1827 | 1878 | 1928 | 1979,7 | 2055 | 1904 | 0,927 | -0,11 | -2,556 | 0,0105744 | 0,02114879 | 0,000361 | 0,0027128 | 0,0012566 | 0,0012566 | TRUE | 0,0106561 | No |
| orf19.5196 | 2194 | 1361 | 1788 | 2256 | 2115 | 1632 | 1841 | 1094 | 1968 | 1733 | 1829 | 1860 | 1809 | 1557 | 1643 | 1543 | 1742,8 | 1848 | 1638 | 0,887 | -0,173 | -2,757 | 0,00583415 | 0,01242326 | 0,003392 | 0,0028125 | 0,0032025 | 0,0032025 | TRUE | 0,0605292 | No |
| orf19.5197 | 8924 | 5949 | 7839 | 9185 | 9023 | 7635 | 8235 | 5573 | 8006 | 7574 | 8020 | 7574 | 7717 | 7285 | 7348 | 7860 | 7673 | 7794 | 7552 | 0,969 | -0,046 | -1,143 | 0,25322696 | 0,33562487 | 0,001042 | 0,0021687 | 0,0014227 | 0,0014227 | TRUE | 0,0088631 | No |
| orf19.5198 | 27861 | 18549 | 23568 | 29095 | 24411 | 21059 | 22778 | 13004 | 24994 | 23615 | 24112 | 23992 | 20877 | 20094 | 20326 | 18341 | 22044 | 24178 | 19910 | 0,824 | -0,28 | -6,278 | 3,44E-10 | 2,34E-09 | 0,001746 | 0,0020454 | 0,0018643 | 0,0018643 | TRUE | 0,0340645 | No |
| orf19.52 | 2391 | 1683 | 2154 | 2688 | 2521 | 2346 | 2439 | 1556 | 2145 | 2143 | 2204 | 2217 | 2156 | 2238 | 2176 | 2195 | 2184,2 | 2177 | 2191 | 1,006 | 0,009 | 0,225 | 0,82164144 | 0,86270915 | 1,00E-08 | 0,0026442 | 0,0009119 | 0,0009119 | TRUE | 0,0026814 | No |
| orf19.520 | 1363 | 964 | 1185 | 1486 | 1483 | 1169 | 1360 | 947 | 1223 | 1227 | 1212 | 1225 | 1268 | 1115 | 1214 | 1336 | 1227,6 | 1222 | 1233 | 1,007 | 0,01 | 0,177 | 0,8595363 | 0,89375114 | 0,001925 | 0,0031621 | 0,0024739 | 0,0024739 | TRUE | 0,0490882 | No |
| orf19.5200 | 3284 | 2412 | 2957 | 3781 | 3239 | 2810 | 3164 | 1967 | 2946 | 3071 | 3025 | 3118 | 2770 | 2681 | 2823 | 2774 | 2901,1 | 3040 | 2762 | 0,909 | -0,138 | -3,681 | 0,00023231 | 0,00066716 | 0,000242 | 0,0024799 | 0,0010056 | 0,0010056 | TRUE | 0,0054663 | No |
| orf19.5201 | 2476 | 1687 | 1979 | 2655 | 2906 | 2511 | 2676 | 1745 | 2221 | 2148 | 2025 | 2189 | 2485 | 2396 | 2388 | 2461 | 2289,2 | 2146 | 2432 | 1,133 | 0,18 | 4,174 | 3,00E-05 | 0,00010096 | 0,000572 | 0,0026137 | 0,0013434 | 0,0013434 | TRUE | 0,0175958 | No |
| orf19.5201.1 | 7128 | 5023 | 5414 | 7221 | 7393 | 6652 | 6500 | 3999 | 6395 | 6395 | 5539 | 5955 | 6323 | 6347 | 5800 | 5640 | 6049,2 | 6071 | 6028 | 0,993 | -0,01 | -0,16 | 0,87276333 | 0,90281716 | 0,003349 | 0,0022195 | 0,0032656 | 0,0032656 | TRUE | 0,0422588 | No |
| orf19.5203 | 663 | 426 | 563 | 692 | 507 | 418 | 497 | 298 | 595 | 542 | 576 | 571 | 434 | 399 | 443 | 420 | 497,5 | 571 | 424 | 0,742 | -0,43 | -6,27 | 3,61E-10 | 2,45E-09 | 1,00E-08 | 0,0048975 | 0,0024651 | 0,0024651 | TRUE | 0,0201015 | No |
| orf19.5204 | 2283 | 1365 | 2007 | 2334 | 2145 | 1739 | 1910 | 1462 | 2048 | 1738 | 2053 | 1925 | 1834 | 1659 | 1704 | 2062 | 1878 | 1941 | 1815 | 0,933 | -0,1 | -1,304 | 0,19214225 | 0,26556031 | 0,007146 | 0,0027525 | 0,0051093 | 0,0051093 | TRUE | 0,0973831 | No |
| orf19.5205 | 682 | 524 | 630 | 839 | 727 | 696 | 758 | 551 | 612 | 667 | 645 | 692 | 622 | 664 | 676 | 777 | 669,34 | 654 | 685 | 1,043 | 0,061 | 0,791 | 0,42874478 | 0,51590199 | 0,00398 | 0,0041484 | 0,0041301 | 0,0041301 | TRUE | 0,0879766 | No |
| orf19.5206 | 1732 | 1310 | 1607 | 1990 | 1856 | 1730 | 1828 | 1255 | 1554 | 1668 | 1644 | 1641 | 1587 | 1651 | 1631 | 1770 | 1643,2 | 1627 | 1660 | 1,019 | 0,027 | 0,549 | 0,58272866 | 0,66129586 | 0,00077 | 0,002863 | 0,0016843 | 0,0016843 | TRUE | 0,0219214 | No |
| orf19.5207 | 3963 | 2696 | 3243 | 4046 | 4564 | 3931 | 4190 | 2407 | 3555 | 3432 | 3318 | 3336 | 3903 | 3751 | 3739 | 3395 | 3553,7 | 3410 | 3697 | 1,085 | 0,118 | 2,385 | 0,01707766 | 0,03236674 | 0,001822 | 0,002388 | 0,0020735 | 0,0020735 | TRUE | 0,0351395 | No |
| orf19.5209 | 250 | 216 | 249 | 273 | 364 | 406 | 409 | 233 | 224 | 275 | 255 | 225 | 311 | 387 | 365 | 329 | 296,43 | 245 | 348 | 1,431 | 0,517 | 5,016 | 5,29E-07 | 2,34E-06 | 0,006549 | 0,0068768 | 0,0067182 | 0,0067182 | TRUE | 0,0697628 | No |
| orf19.5210 | 428 | 281 | 383 | 500 | 395 | 315 | 393 | 291 | 384 | 358 | 392 | 412 | 338 | 301 | 351 | 410 | 368,17 | 386 | 350 | 0,898 | -0,155 | -1,619 | 0,10549277 | 0,1594868 | 0,006577 | 0,0059226 | 0,0061083 | 0,0061083 | TRUE | 0,1349729 | No |
| orf19.5210.1 | 48 | 42 | 45 | 69 | 59 | 59 | 71 | 65 | 43 | 53 | 46 | 57 | 50 | 56 | 63 | 92 | 57,66 | 50 | 65 | 1,29 | 0,368 | 1,705 | 0,08813373 | 0,13742732 | 0,026469 | 0,0271577 | 0,0268343 | 0,0268343 | TRUE | 0,5217188 | No |
| orf19.5211 | 888 | 552 | 836 | 947 | 628 | 489 | 505 | 377 | 797 | 703 | 855 | 781 | 537 | 467 | 451 | 532 | 640,2 | 784 | 497 | 0,631 | -0,664 | -8,07 | 7,04E-16 | 8,84E-15 | 0,005408 | 0,0042471 | 0,0048387 | 0,0048387 | TRUE | 0,0534259 | No |
| orf19.5212 | 30 | 21 | 25 | 28 | 11 | 17 | 17 | 13 | 27 | 27 | 26 | 23 | 9 | 16 | 15 | 18 | 20,18 | 26 | 14 | 0,568 | -0,816 | -2,605 | 0,00918043 | 0,018659 | 1,00E-08 | 0,0739134 | 0,0403625 | 0,0403625 | TRUE | 0,3249904 | No |
| orf19.5213 | 3110 | 2137 | 2805 | 3353 | 2701 | 2517 | 2825 | 1812 | 2790 | 2721 | 2870 | 2765 | 2310 | 2402 | 2521 | 2556 | 2616,7 | 2786 | 2447 | 0,877 | -0,189 | -4,267 | 1,98E-05 | 6,87E-05 | 0,000801 | 0,0025343 | 0,001505 | 0,001505 | TRUE | 0,0175876 | No |
| orf19.5213.1 | 6819 | 4469 | 5317 | 6910 | 7334 | 5872 | 6094 | 3668 | 6117 | 5690 | 5440 | 5698 | 6272 | 5603 | 5438 | 5173 | 5678,9 | 5736 | 5622 | 0,98 | -0,028 | -0,459 | 0,64620154 | 0,71617392 | 0,003903 | 0,0022352 | 0,0035201 | 0,0035201 | TRUE | 0,0743812 | No |
| orf19.5213.2 | 3750 | 2512 | 2872 | 3860 | 4602 | 3548 | 3554 | 2991 | 3364 | 3198 | 2938 | 3183 | 3936 | 3385 | 3171 | 4219 | 3424,3 | 3171 | 3678 | 1,159 | 0,213 | 2,535 | 0,01123935 | 0,0223596 | 0,009776 | 0,0024035 | 0,0064759 | 0,0064759 | TRUE | 0,1172162 | No |
| orf19.5214 | 820 | 549 | 761 | 918 | 979 | 892 | 1021 | 611 | 736 | 699 | 779 | 757 | 837 | 851 | 911 | 862 | 803,92 | 743 | 865 | 1,164 | 0,219 | 3,737 | 0,00018588 | 0,00054476 | 0,000459 | 0,0037853 | 0,0020375 | 0,0020375 | TRUE | 0,0162688 | No |
| orf19.5215 | 2037 | 1302 | 1712 | 2271 | 2342 | 1940 | 2077 | 1465 | 1827 | 1658 | 1752 | 1873 | 2003 | 1851 | 1853 | 2066 | 1860,4 | 1778 | 1943 | 1,091 | 0,125 | 2,22 | 0,02644411 | 0,04807791 | 0,002242 | 0,0027599 | 0,0025154 | 0,0025154 | TRUE | 0,0237442 | No |
| orf19.5216 | 566 | 365 | 492 | 596 | 677 | 661 | 633 | 427 | 508 | 465 | 503 | 491 | 579 | 631 | 565 | 602 | 543,01 | 492 | 594 | 1,205 | 0,269 | 4,001 | 6,31E-05 | 0,00020012 | 1,00E-08 | 0,004653 | 0,0024709 | 0,0024709 | TRUE | 0,0217922 | No |
| orf19.5217 | 627 | 423 | 567 | 651 | 494 | 443 | 447 | 301 | 562 | 539 | 580 | 537 | 422 | 423 | 399 | 425 | 485,81 | 554 | 417 | 0,751 | -0,412 | -6,104 | 1,03E-09 | 6,57E-09 | 1,00E-08 | 0,0049677 | 0,0022843 | 0,0022843 | TRUE | 0,011 | No |
| orf19.5219 | 3794 | 2857 | 3349 | 3999 | 4002 | 3935 | 4095 | 2418 | 3404 | 3637 | 3426 | 3298 | 3423 | 3755 | 3654 | 3410 | 3500,8 | 3441 | 3560 | 1,036 | 0,051 | 1,057 | 0,29038475 | 0,37501663 | 0,00163 | 0,0023942 | 0,001945 | 0,001945 | TRUE | 0,0173514 | No |
| orf19.522 | 5839 | 3979 | 5273 | 6159 | 6315 | 5637 | 5977 | 3673 | 5238 | 5066 | 5395 | 5079 | 5401 | 5379 | 5333 | 5180 | 5258,9 | 5194 | 5323 | 1,025 | 0,036 | 1,031 | 0,30262139 | 0,38857647 | 0,000365 | 0,0022556 | 0,0009802 | 0,0009802 | TRUE | 0,0081346 | No |
| orf19.5220 | 3471 | 2578 | 2944 | 3788 | 3641 | 3213 | 3570 | 2191 | 3114 | 3282 | 3012 | 3124 | 3114 | 3066 | 3186 | 3090 | 3123,4 | 3133 | 3114 | 0,995 | -0,007 | -0,191 | 0,84845059 | 0,88415642 | 0,000345 | 0,0024443 | 0,0011183 | 0,0011183 | TRUE | 0,0116632 | No |
| orf19.5221 | 1212 | 866 | 1113 | 1361 | 1654 | 1556 | 1734 | 1039 | 1087 | 1103 | 1139 | 1122 | 1415 | 1485 | 1547 | 1465 | 1295,4 | 1113 | 1478 | 1,328 | 0,41 | 8,432 | 3,39E-17 | 4,87E-16 | 1,00E-08 | 0,0031002 | 0,0014788 | 0,0014788 | TRUE | 0,0127558 | No |
| orf19.5223 | 26 | 14 | 22 | 25 | 29 | 28 | 29 | 15 | 23 | 18 | 23 | 21 | 25 | 27 | 26 | 21 | 22,85 | 21 | 25 | 1,172 | 0,229 | 0,82 | 0,41200962 | 0,50017252 | 1,00E-08 | 0,065503 | 0,0310756 | 0,0310756 | TRUE | 0,0436403 | No |
| orf19.5224 | 2705 | 2025 | 2487 | 3088 | 2674 | 2587 | 2788 | 1771 | 2427 | 2578 | 2544 | 2546 | 2287 | 2468 | 2488 | 2498 | 2479,6 | 2524 | 2435 | 0,964 | -0,052 | -1,177 | 0,23899676 | 0,3200117 | 0,000853 | 0,002565 | 0,0014815 | 0,0014815 | TRUE | 0,021136 | No |
| orf19.5225 | 501 | 347 | 451 | 508 | 556 | 561 | 519 | 346 | 449 | 442 | 461 | 419 | 476 | 535 | 463 | 488 | 466,68 | 443 | 490 | 1,108 | 0,148 | 1,999 | 0,0456366 | 0,07767271 | 0,001086 | 0,0050902 | 0,0031109 | 0,0031109 | TRUE | 0,0464382 | No |
| orf19.5225.2 | 13722 | 9296 | 10795 | 14296 | 14223 | 11965 | 12432 | 9059 | 12310 | 11835 | 11044 | 11789 | 12164 | 11417 | 11093 | 12777 | 11804 | 11744 | 11863 | 1,01 | 0,014 | 0,264 | 0,79211844 | 0,83782197 | 0,002544 | 0,0021025 | 0,0026225 | 0,0026225 | TRUE | 0,0325064 | No |
| orf19.5226 | 1902 | 1525 | 1718 | 2153 | 2211 | 2020 | 2267 | 1319 | 1706 | 1941 | 1758 | 1775 | 1891 | 1927 | 2023 | 1860 | 1860,3 | 1795 | 1925 | 1,075 | 0,104 | 2,002 | 0,04523496 | 0,07705462 | 0,001563 | 0,0027599 | 0,0020631 | 0,0020631 | TRUE | 0,0343195 | No |
| orf19.5227 | 1529 | 1167 | 1384 | 1813 | 1789 | 1614 | 1781 | 993 | 1372 | 1486 | 1416 | 1495 | 1530 | 1540 | 1589 | 1401 | 1478,5 | 1442 | 1515 | 1,053 | 0,075 | 1,389 | 0,16496961 | 0,23431145 | 0,00134 | 0,0029614 | 0,0021018 | 0,0021018 | TRUE | 0,0281975 | No |
| orf19.5228 | 6049 | 4480 | 5316 | 6705 | 6173 | 5548 | 6109 | 3945 | 5427 | 5704 | 5439 | 5529 | 5279 | 5294 | 5451 | 5564 | 5460,8 | 5525 | 5397 | 0,977 | -0,034 | -0,991 | 0,32157667 | 0,40820465 | 0,00034 | 0,0022454 | 0,0009435 | 0,0009435 | TRUE | 0,0055411 | No |
| orf19.5229 | 9079 | 7058 | 8398 | 10232 | 9449 | 9124 | 9410 | 5797 | 8145 | 8986 | 8592 | 8437 | 8081 | 8706 | 8397 | 8176 | 8440 | 8540 | 8340 | 0,977 | -0,033 | -0,804 | 0,42167031 | 0,50942997 | 0,001236 | 0,0021515 | 0,0015494 | 0,0015494 | TRUE | 0,0148266 | No |
| orf19.5230 | 4825 | 3460 | 4193 | 5197 | 5030 | 4569 | 4987 | 2798 | 4329 | 4405 | 4290 | 4286 | 4302 | 4360 | 4450 | 3946 | 4295,8 | 4328 | 4264 | 0,987 | -0,019 | -0,444 | 0,65735147 | 0,72598669 | 0,001026 | 0,0023175 | 0,0015386 | 0,0015386 | TRUE | 0,0290882 | No |
| orf19.5231 | 76 | 58 | 61 | 53 | 38 | 49 | 39 | 18 | 68 | 74 | 62 | 44 | 32 | 47 | 35 | 25 | 48,45 | 62 | 35 | 0,573 | -0,804 | -3,373 | 0,00074318 | 0,00192434 | 0,032456 | 0,0319443 | 0,0321761 | 0,0321761 | TRUE | 0,3867315 | No |
| orf19.5231.2 | 1426 | 1004 | 1082 | 1412 | 1311 | 1117 | 1193 | 685 | 1279 | 1278 | 1107 | 1164 | 1121 | 1066 | 1065 | 966 | 1130,8 | 1207 | 1054 | 0,876 | -0,191 | -2,871 | 0,00409095 | 0,0090031 | 0,003216 | 0,0032633 | 0,003351 | 0,003351 | TRUE | 0,0366408 | No |
| orf19.5232 | 4403 | 3947 | 4160 | 5657 | 5019 | 5436 | 5175 | 3463 | 3950 | 5025 | 4256 | 4665 | 4292 | 5187 | 4618 | 4884 | 4609,7 | 4474 | 4745 | 1,061 | 0,085 | 1,08 | 0,28013876 | 0,36430166 | 0,00815 | 0,0022944 | 0,0057519 | 0,0057519 | TRUE | 0,0828002 | No |
| orf19.5233 | 0 | 1 | 0 | 2 | 0 | 0 | 1 | 1 | 0 | 1 | 0 | 2 | 0 | 0 | 1 | 1 | 0,65 | 1 | 0 | 0,699 | -0,516 | -0,28 | 0,7792861 | NA | 1,00E-08 | 2,2246473 | 1,6670821 | 1,6670821 | TRUE | 0,2969363 | No |
| orf19.5234 | 1612 | 1267 | 1566 | 1842 | 1540 | 1373 | 1507 | 894 | 1446 | 1613 | 1602 | 1519 | 1317 | 1310 | 1345 | 1261 | 1426,6 | 1545 | 1308 | 0,849 | -0,236 | -4,603 | 4,17E-06 | 1,60E-05 | 0,000977 | 0,0029971 | 0,0018188 | 0,0018188 | TRUE | 0,0224994 | No |
| orf19.5235 | 729 | 909 | 706 | 992 | 797 | 1044 | 1025 | 537 | 654 | 1157 | 722 | 818 | 682 | 996 | 915 | 757 | 837,67 | 838 | 838 | 1 | 0,001 | 0,003 | 0,99749976 | 0,99816432 | 0,045595 | 0,0037125 | 0,023059 | 0,0455952 | TRUE | 0,7732405 | No |
| orf19.5236 | 322 | 266 | 325 | 363 | 484 | 528 | 537 | 330 | 289 | 339 | 333 | 299 | 414 | 504 | 479 | 465 | 390,21 | 315 | 466 | 1,483 | 0,569 | 6,386 | 1,70E-10 | 1,19E-09 | 0,004201 | 0,0056998 | 0,0049488 | 0,0049488 | TRUE | 0,0701999 | No |
| orf19.5237 | 489 | 358 | 397 | 498 | 679 | 592 | 645 | 378 | 439 | 456 | 406 | 411 | 581 | 565 | 576 | 533 | 495,69 | 428 | 564 | 1,324 | 0,405 | 5,847 | 5,01E-09 | 2,92E-08 | 1,00E-08 | 0,0049082 | 0,0025519 | 0,0025519 | TRUE | 0,019372 | No |
| orf19.5238 | 2821 | 2006 | 2506 | 3057 | 3024 | 2814 | 2967 | 1979 | 2531 | 2554 | 2564 | 2521 | 2586 | 2685 | 2648 | 2791 | 2609,9 | 2542 | 2678 | 1,052 | 0,073 | 1,918 | 0,05505096 | 0,09130723 | 0,000116 | 0,0025357 | 0,0010024 | 0,0010024 | TRUE | 0,0090124 | No |
| orf19.5239 | 325 | 280 | 301 | 338 | 535 | 513 | 513 | 327 | 292 | 356 | 308 | 279 | 458 | 489 | 458 | 461 | 387,59 | 309 | 466 | 1,522 | 0,606 | 7,099 | 1,26E-12 | 1,13E-11 | 0,002447 | 0,005725 | 0,00429 | 0,00429 | TRUE | 0,1105302 | No |
| orf19.524 | 991 | 698 | 919 | 1061 | 1362 | 1123 | 1324 | 757 | 889 | 889 | 940 | 875 | 1165 | 1072 | 1181 | 1068 | 1009,8 | 898 | 1122 | 1,251 | 0,324 | 5,692 | 1,25E-08 | 6,85E-08 | 0,001058 | 0,0034172 | 0,0021006 | 0,0021006 | TRUE | 0,0153315 | No |
| orf19.5241 | 6489 | 4154 | 5534 | 6918 | 7932 | 6839 | 7505 | 4650 | 5821 | 5288 | 5662 | 5705 | 6784 | 6526 | 6697 | 6558 | 6130,1 | 5619 | 6641 | 1,181 | 0,24 | 6,24 | 4,37E-10 | 2,94E-09 | 0,000787 | 0,0022163 | 0,0012605 | 0,0012605 | TRUE | 0,0184971 | No |
| orf19.5242 | 109 | 92 | 85 | 106 | 75 | 97 | 81 | 42 | 98 | 117 | 87 | 87 | 64 | 93 | 72 | 59 | 84,69 | 97 | 72 | 0,75 | -0,415 | -2,366 | 0,01798165 | 0,03381338 | 0,015196 | 0,0191215 | 0,0175386 | 0,0175386 | TRUE | 0,3322238 | No |
| orf19.5243 | 1033 | 851 | 949 | 1190 | 1473 | 1538 | 1596 | 854 | 927 | 1083 | 971 | 981 | 1260 | 1468 | 1424 | 1204 | 1164,8 | 990 | 1339 | 1,356 | 0,439 | 5,817 | 6,01E-09 | 3,44E-08 | 0,005541 | 0,0032259 | 0,0045877 | 0,0045877 | TRUE | 0,051385 | No |
| orf19.5244 | 373 | 363 | 411 | 470 | 450 | 517 | 552 | 323 | 335 | 462 | 420 | 388 | 385 | 493 | 493 | 456 | 428,89 | 401 | 457 | 1,141 | 0,191 | 1,719 | 0,08556383 | 0,13397641 | 0,013483 | 0,0053643 | 0,0094505 | 0,0094505 | TRUE | 0,1451733 | No |
| orf19.5245 | 1343 | 1040 | 1304 | 1541 | 1781 | 1677 | 1871 | 1222 | 1205 | 1324 | 1334 | 1271 | 1523 | 1600 | 1670 | 1724 | 1456,3 | 1284 | 1629 | 1,269 | 0,343 | 6,136 | 8,47E-10 | 5,45E-09 | 0,001824 | 0,0029764 | 0,0023076 | 0,0023076 | TRUE | 0,0234818 | No |
| orf19.5246 | 262 | 183 | 226 | 297 | 327 | 285 | 328 | 220 | 235 | 233 | 231 | 245 | 280 | 272 | 293 | 310 | 262,34 | 236 | 289 | 1,216 | 0,283 | 3,208 | 0,00133471 | 0,0032717 | 1,00E-08 | 0,0075132 | 0,0036143 | 0,0036143 | TRUE | 0,0232819 | No |
| orf19.5247 | 1586 | 1129 | 1388 | 1728 | 1863 | 1666 | 1678 | 1053 | 1423 | 1437 | 1420 | 1425 | 1593 | 1590 | 1497 | 1485 | 1483,8 | 1426 | 1541 | 1,082 | 0,114 | 2,533 | 0,0113233 | 0,02248196 | 1,00E-08 | 0,0029579 | 0,0012787 | 0,0012787 | TRUE | 0,0063429 | No |

| | | | | | | | | | | | | | | | | | | | | | | | | | | | | | | | |
|---|---|---|---|---|---|---|---|---|---|---|---|---|---|---|---|---|---|---|---|---|---|---|---|---|---|---|---|---|---|---|---|
| orf19.5248 | 3498 | 2167 | 2865 | 3784 | 4183 | 3305 | 3756 | 2379 | 3138 | 2759 | 2931 | 3120 | 3577 | 3154 | 3352 | 3355 | 3173,3 | 2987 | 3360 | 1,124 | 0,168 | 3,021 | 0,00252357 | 0,00582915 | 0,002697 | 0,002437 | 0,0026665 | 0,0026665 | TRUE | 0,03126 | No |
| orf19.5249 | 599 | 434 | 508 | 635 | 778 | 612 | 677 | 432 | 537 | 553 | 520 | 524 | 665 | 584 | 604 | 609 | 574,5 | 534 | 616 | 1,157 | 0,211 | 3,164 | 0,00155412 | 0,00375741 | 1,00E-08 | 0,0045064 | 0,0025017 | 0,0025017 | TRUE | 0,0377144 | No |
| orf19.5250 | 1088 | 733 | 941 | 1126 | 1216 | 1095 | 1135 | 751 | 976 | 933 | 963 | 929 | 1040 | 1045 | 1013 | 1059 | 994,66 | 950 | 1039 | 1,092 | 0,127 | 2,573 | 0,01007635 | 0,02027417 | 1,00E-08 | 0,003439 | 0,00135 | 0,00135 | TRUE | 0,0042493 | No |
| orf19.5251 | 4795 | 3286 | 4354 | 5320 | 5538 | 4921 | 5500 | 3773 | 4302 | 4183 | 4455 | 4387 | 4736 | 4695 | 4908 | 5321 | 4623,5 | 4332 | 4915 | 1,133 | 0,181 | 3,811 | 0,00013854 | 0,00041267 | 0,001768 | 0,0022935 | 0,0019382 | 0,0019382 | TRUE | 0,0363942 | No |
| orf19.5253 | 948 | 674 | 822 | 1091 | 869 | 815 | 905 | 529 | 850 | 858 | 841 | 900 | 743 | 778 | 808 | 746 | 815,46 | 862 | 769 | 0,892 | -0,165 | -2,924 | 0,00345433 | 0,00772083 | 1,00E-08 | 0,0037597 | 0,0018411 | 0,0018411 | TRUE | 0,0143276 | No |
| orf19.5254 | 494 | 387 | 432 | 582 | 429 | 423 | 439 | 312 | 443 | 493 | 442 | 480 | 367 | 404 | 392 | 440 | 432,51 | 464 | 401 | 0,859 | -0,219 | -2,786 | 0,00532816 | 0,01145316 | 0,001642 | 0,005336 | 0,0036049 | 0,0036049 | TRUE | 0,0434395 | No |
| orf19.5255 | 3547 | 2389 | 3080 | 3846 | 3542 | 3163 | 3491 | 2224 | 3182 | 3041 | 3151 | 3171 | 3029 | 3018 | 3115 | 3137 | 3105,7 | 3136 | 3075 | 0,979 | -0,031 | -0,865 | 0,38719508 | 0,47630386 | 1,00E-08 | 0,002447 | 0,000882 | 0,000882 | TRUE | 0,0047163 | No |
| orf19.5256 | 68 | 56 | 83 | 101 | 72 | 57 | 78 | 43 | 61 | 71 | 85 | 83 | 62 | 54 | 70 | 61 | 68,34 | 75 | 62 | 0,82 | -0,286 | -1,615 | 0,10629382 | 0,16061701 | 1,00E-08 | 0,0232221 | 0,0154717 | 0,0154717 | TRUE | 0,1646999 | No |
| orf19.5257 | 719 | 451 | 653 | 778 | 979 | 834 | 905 | 607 | 645 | 574 | 668 | 642 | 837 | 796 | 808 | 856 | 728,19 | 632 | 824 | 1,299 | 0,377 | 6,032 | 1,62E-09 | 9,95E-09 | 0,000548 | 0,0039731 | 0,0023522 | 0,0023522 | TRUE | 0,03946 | No |
| orf19.5258 | 1046 | 820 | 915 | 1235 | 1249 | 1257 | 1330 | 791 | 938 | 1044 | 936 | 1018 | 1068 | 1199 | 1187 | 1116 | 1063,4 | 984 | 1142 | 1,162 | 0,217 | 3,534 | 0,00040872 | 0,00112385 | 0,002136 | 0,0033447 | 0,0026685 | 0,0026685 | TRUE | 0,0249509 | No |
| orf19.5259 | 455 | 321 | 420 | 553 | 664 | 619 | 667 | 433 | 408 | 409 | 430 | 456 | 568 | 591 | 595 | 611 | 508,37 | 426 | 591 | 1,383 | 0,468 | 6,846 | 7,59E-12 | 6,29E-11 | 1,00E-08 | 0,0048351 | 0,0024735 | 0,0024735 | TRUE | 0,0291083 | No |
| orf19.526 | 13047 | 9404 | 10684 | 13629 | 10843 | 9646 | 9999 | 5503 | 11705 | 11972 | 10931 | 11239 | 9273 | 9204 | 8922 | 7761 | 10126 | 11462 | 8790 | 0,767 | -0,382 | -6,454 | 1,09E-10 | 7,90E-10 | 0,003885 | 0,0021229 | 0,0032644 | 0,0032644 | TRUE | 0,0747547 | No |
| orf19.5260 | 6555 | 4583 | 6004 | 7046 | 6356 | 5629 | 6119 | 4134 | 5881 | 5835 | 6143 | 5810 | 5436 | 5371 | 5460 | 5831 | 5720,7 | 5917 | 5524 | 0,933 | -0,1 | -2,59 | 0,00958394 | 0,01940038 | 0,000808 | 0,0022333 | 0,0012607 | 0,0012607 | TRUE | 0,016188 | No |
| orf19.5262 | 25 | 18 | 27 | 26 | 43 | 21 | 24 | 14 | 22 | 23 | 28 | 21 | 37 | 20 | 21 | 20 | 24,05 | 24 | 24 | 1,055 | 0,077 | 0,257 | 0,79716682 | 0,84275528 | 0,024619 | 0,0623472 | 0,0458344 | 0,0458344 | TRUE | 0,7803444 | No |
| orf19.5263 | 8588 | 5862 | 7458 | 9377 | 7556 | 6613 | 7013 | 4695 | 7704 | 7463 | 7630 | 7732 | 6462 | 6310 | 6258 | 6622 | 7022,7 | 7632 | 6413 | 0,839 | -0,252 | -7,939 | 2,03E-15 | 2,43E-14 | 0,000261 | 0,0021862 | 0,0008266 | 0,0008266 | TRUE | 0,0055152 | No |
| orf19.5264 | 4 | 4 | 15 | 8 | 3 | 11 | 2 | 7 | 4 | 5 | 15 | 7 | 3 | 10 | 2 | 10 | 6,92 | 8 | 6 | 0,783 | -0,354 | -0,521 | 0,60224684 | NA | 0,388419 | 0,2118262 | 0,2925579 | 0,2925579 | TRUE | 0,36836 | No |
| orf19.5265 | 3010 | 2102 | 2711 | 3114 | 3359 | 3061 | 3214 | 1910 | 2700 | 2676 | 2774 | 2568 | 2873 | 2921 | 2868 | 2694 | 2759,1 | 2680 | 2839 | 1,061 | 0,086 | 2,011 | 0,04432282 | 0,07569401 | 0,000636 | 0,0025057 | 0,0013773 | 0,0013773 | TRUE | 0,0131941 | No |
| orf19.5266 | 33 | 24 | 26 | 30 | 64 | 45 | 47 | 22 | 30 | 31 | 27 | 25 | 55 | 43 | 42 | 31 | 35,27 | 28 | 43 | 1,562 | 0,644 | 2,629 | 0,00855108 | 0,01751018 | 1,00E-08 | 0,0431422 | 0,0280884 | 0,0280884 | TRUE | 0,1958227 | No |
| orf19.5267 | 127 | 80 | 99 | 147 | 138 | 120 | 122 | 95 | 114 | 102 | 101 | 121 | 118 | 115 | 109 | 134 | 114,21 | 110 | 119 | 1,071 | 0,099 | 0,741 | 0,45856946 | 0,54437568 | 1,00E-08 | 0,0146906 | 0,0083668 | 0,0083668 | TRUE | 0,0584438 | No |
| orf19.5268 | 117 | 110 | 122 | 175 | 169 | 139 | 163 | 122 | 105 | 140 | 125 | 144 | 145 | 133 | 145 | 172 | 138,6 | 128 | 149 | 1,151 | 0,203 | 1,487 | 0,13696975 | 0,19977588 | 0,008285 | 0,0124534 | 0,010644 | 0,010644 | TRUE | 0,1676875 | No |
| orf19.5269 | 854 | 591 | 758 | 911 | 658 | 545 | 599 | 353 | 766 | 752 | 776 | 751 | 563 | 520 | 535 | 498 | 645,05 | 761 | 529 | 0,697 | -0,521 | -8,521 | 1,58E-17 | 2,34E-16 | 1,00E-08 | 0,00423 | 0,0019855 | 0,0019855 | TRUE | 0,0220883 | No |
| orf19.5269.1 | 327 | 250 | 311 | 392 | 374 | 382 | 320 | 215 | 293 | 318 | 318 | 323 | 320 | 364 | 286 | 303 | 315,77 | 313 | 318 | 1,018 | 0,025 | 0,272 | 0,78556282 | 0,83224501 | 0,003701 | 0,0065768 | 0,0051856 | 0,0051856 | TRUE | 0,1108039 | No |
| orf19.527 | 1683 | 1166 | 1437 | 1829 | 1662 | 1487 | 1582 | 1111 | 1510 | 1484 | 1470 | 1508 | 1421 | 1419 | 1412 | 1567 | 1473,9 | 1493 | 1455 | 0,971 | -0,042 | -0,877 | 0,38047981 | 0,4698022 | 0,000505 | 0,0029644 | 0,0015424 | 0,0015424 | TRUE | 0,0289387 | No |
| orf19.5270 | 265 | 202 | 260 | 309 | 182 | 170 | 196 | 113 | 238 | 257 | 266 | 255 | 156 | 162 | 175 | 159 | 208,48 | 254 | 163 | 0,644 | -0,636 | -6,409 | 1,47E-10 | 1,04E-09 | 1,00E-08 | 0,0089428 | 0,0044292 | 0,0044292 | TRUE | 0,027927 | No |
| orf19.5271 | 196 | 153 | 182 | 199 | 213 | 232 | 253 | 141 | 176 | 195 | 186 | 164 | 182 | 221 | 226 | 199 | 193,63 | 180 | 207 | 1,155 | 0,208 | 1,871 | 0,06140764 | 0,10055521 | 0,003367 | 0,0094766 | 0,006678 | 0,006678 | TRUE | 0,0824813 | No |
| orf19.5274 | 2729 | 1787 | 2457 | 2779 | 3597 | 3374 | 3837 | 2271 | 2448 | 2275 | 2514 | 2292 | 3076 | 3219 | 3424 | 3203 | 2806,4 | 2382 | 3230 | 1,356 | 0,439 | 8,682 | 3,90E-18 | 6,12E-17 | 0,001719 | 0,0024968 | 0,0020923 | 0,0020923 | TRUE | 0,0202423 | No |
| orf19.5275 | 348 | 267 | 280 | 361 | 488 | 517 | 510 | 333 | 312 | 340 | 286 | 298 | 417 | 493 | 455 | 470 | 383,96 | 309 | 459 | 1,487 | 0,572 | 6,746 | 1,52E-11 | 1,22E-10 | 0,00252 | 0,0057604 | 0,004197 | 0,004197 | TRUE | 0,046046 | No |
| orf19.5276 | 496 | 534 | 479 | 718 | 805 | 866 | 867 | 538 | 445 | 680 | 490 | 592 | 688 | 826 | 774 | 759 | 656,77 | 552 | 762 | 1,384 | 0,468 | 4,036 | 5,43E-05 | 0,00017446 | 0,018899 | 0,0041899 | 0,0113622 | 0,0113622 | TRUE | 0,2728516 | No |
| orf19.5277 | 1405 | 1056 | 1247 | 1612 | 1814 | 1676 | 1873 | 1058 | 1260 | 1344 | 1276 | 1329 | 1551 | 1599 | 1671 | 1492 | 1440,5 | 1302 | 1578 | 1,215 | 0,28 | 5,512 | 3,55E-08 | 1,84E-07 | 0,000912 | 0,0029873 | 0,0017842 | 0,0017842 | TRUE | 0,0188526 | No |
| orf19.5278 | 2208 | 1647 | 1961 | 2489 | 1940 | 1784 | 1824 | 1071 | 1981 | 2097 | 2006 | 2052 | 1659 | 1702 | 1628 | 1511 | 1829,5 | 2034 | 1625 | 0,801 | -0,32 | -6,723 | 1,78E-11 | 1,42E-10 | 0,000857 | 0,002773 | 0,0016224 | 0,0016224 | TRUE | 0,0241186 | No |
| orf19.5279 | 2705 | 1807 | 2097 | 2570 | 2962 | 2606 | 2608 | 1482 | 2427 | 2301 | 2145 | 2119 | 2533 | 2487 | 2327 | 2090 | 2303,6 | 2248 | 2359 | 1,051 | 0,072 | 1,048 | 0,29457262 | 0,37929539 | 0,005047 | 0,0026097 | 0,0040847 | 0,0040847 | TRUE | 0,068532 | No |
| orf19.528 | 2867 | 2018 | 2661 | 3175 | 3213 | 3089 | 3182 | 1957 | 2572 | 2569 | 2722 | 2618 | 2748 | 2947 | 2839 | 2760 | 2722,1 | 2620 | 2824 | 1,078 | 0,108 | 2,638 | 0,00833099 | 0,01710031 | 0,00053 | 0,0025128 | 0,0012452 | 0,0012452 | TRUE | 0,0106241 | No |
| orf19.5280 | 76433 | 49698 | 67412 | 83411 | 68553 | 57925 | 63100 | 41383 | 68568 | 63271 | 68969 | 68782 | 58629 | 55270 | 56306 | 58367 | 62270 | 67398 | 57143 | 0,848 | -0,238 | -6,052 | 1,43E-09 | 8,88E-09 | 0,001218 | 0,0020028 | 0,0014723 | 0,0014723 | TRUE | 0,0207593 | No |
| orf19.5281 | 1592 | 992 | 1366 | 1574 | 1603 | 1477 | 1600 | 1002 | 1428 | 1263 | 1398 | 1298 | 1371 | 1409 | 1428 | 1413 | 1376 | 1347 | 1405 | 1,042 | 0,06 | 1,136 | 0,25588655 | 0,33840334 | 0,001087 | 0,0030346 | 0,0019178 | 0,0019178 | TRUE | 0,0202616 | No |
| orf19.5282 | 8649 | 5929 | 7506 | 9363 | 9071 | 7863 | 8643 | 5760 | 7759 | 7548 | 7679 | 7721 | 7758 | 7503 | 7712 | 8124 | 7725,6 | 7677 | 7774 | 1,012 | 0,017 | 0,506 | 0,61290228 | 0,68674317 | 0,000451 | 0,0021674 | 0,000956 | 0,000956 | TRUE | 0,0106908 | No |
| orf19.5284 | 94 | 78 | 96 | 114 | 63 | 58 | 60 | 51 | 84 | 99 | 98 | 94 | 54 | 55 | 54 | 72 | 76,32 | 94 | 59 | 0,617 | -0,697 | -4,251 | 2,13E-05 | 7,37E-05 | 1,00E-08 | 0,0210012 | 0,0118498 | 0,0118498 | TRUE | 0,1464725 | No |
| orf19.5285 | 1738 | 1179 | 1476 | 1894 | 1366 | 1089 | 1071 | 848 | 1559 | 1501 | 1510 | 1562 | 1168 | 1039 | 956 | 1196 | 1311,4 | 1533 | 1090 | 0,709 | -0,496 | -7,098 | 1,26E-12 | 1,14E-11 | 0,004503 | 0,0030865 | 0,0038941 | 0,0038941 | TRUE | 0,083389 | No |
| orf19.5286 | 4876 | 3013 | 4108 | 5084 | 3661 | 2881 | 3100 | 2443 | 4374 | 3836 | 4203 | 4192 | 3131 | 2749 | 2766 | 3446 | 3587,2 | 4151 | 3023 | 0,727 | -0,46 | -6,214 | 5,16E-10 | 3,43E-09 | 0,007147 | 0,0023842 | 0,0049641 | 0,0049641 | TRUE | 0,1049096 | No |
| orf19.5287 | 164 | 74 | 107 | 153 | 128 | 103 | 116 | 98 | 147 | 94 | 109 | 126 | 109 | 98 | 104 | 138 | 115,81 | 119 | 112 | 0,927 | -0,109 | -0,672 | 0,50132077 | 0,5862077 | 0,018874 | 0,0145151 | 0,0166953 | 0,0166953 | TRUE | 0,2498532 | No |
| orf19.5288 | 1E+05 | 94938 | 1E+05 | 2E+05 | 2E+05 | 1E+05 | 2E+05 | 1E+05 | 1E+05 | 1E+05 | 1E+05 | 1E+05 | 1E+05 | 1E+05 | 1E+05 | 1E+05 | 133881 | 1E+05 | 1E+05 | 1,15 | 0,202 | 5,302 | 1,14E-07 | 5,57E-07 | 0,001136 | 0,0019904 | 0,0013864 | 0,0013864 | TRUE | 0,0114412 | No |
| orf19.5288.1 | 716 | 532 | 596 | 691 | 796 | 765 | 759 | 441 | 642 | 677 | 610 | 570 | 681 | 730 | 677 | 622 | 651,15 | 625 | 678 | 1,09 | 0,124 | 1,693 | 0,09050023 | 0,14071567 | 0,003064 | 0,004209 | 0,003624 | 0,003624 | TRUE | 0,0432116 | No |
| orf19.5289 | 5 | 5 | 4 | 6 | 5 | 3 | 11 | 4 | 4 | 6 | 4 | 5 | 4 | 3 | 10 | 6 | 5,31 | 5 | 6 | 1,151 | 0,203 | 0,334 | 0,73835674 | NA | 1,00E-08 | 0,2753239 | 0,1656951 | 0,1656951 | TRUE | 0,6679855 | No |
| orf19.529 | 1670 | 1119 | 1596 | 1806 | 2659 | 2208 | 2530 | 1680 | 1498 | 1425 | 1633 | 1489 | 2274 | 2107 | 2258 | 2369 | 1881,6 | 1511 | 2252 | 1,488 | 0,574 | 10,354 | 4,02E-25 | 1,03E-23 | 0,002101 | 0,002751 | 0,0023916 | 0,0023916 | TRUE | 0,0346069 | No |
| orf19.5290 | 0 | 0 | 0 | 0 | 0 | 0 | 0 | 0 | 0 | 0 | 0 | 0 | 0 | 0 | 0 | 0 | 0 | 0 | 0 | NA | NA | NA | NA | NA | NA | NA | NA | NA | NA | NA | NA |
| orf19.5291 | 961 | 782 | 876 | 1127 | 919 | 1049 | 1032 | 703 | 862 | 996 | 896 | 929 | 786 | 1001 | 921 | 992 | 922,82 | 921 | 925 | 1,003 | 0,005 | 0,061 | 0,95152698 | 0,96518219 | 0,006888 | 0,0035526 | 0,0053492 | 0,0053492 | TRUE | 0,1254703 | No |
| orf19.529.1 | 62 | 53 | 73 | 69 | 64 | 77 | 97 | 58 | 56 | 67 | 75 | 57 | 55 | 73 | 87 | 82 | 68,91 | 64 | 74 | 1,164 | 0,219 | 1,146 | 0,25175915 | 0,33419553 | 0,01743 | 0,0230475 | 0,0205441 | 0,0205441 | TRUE | 0,3177301 | No |
| orf19.5292 | 755 | 584 | 764 | 898 | 914 | 945 | 873 | 538 | 677 | 743 | 782 | 741 | 782 | 902 | 779 | 759 | 770,52 | 736 | 806 | 1,097 | 0,133 | 1,833 | 0,06680968 | 0,10842601 | 0,00367 | 0,0038636 | 0,0037597 | 0,0037597 | TRUE | 0,0775972 | No |
| orf19.5293 | 2667 | 2261 | 2518 | 3239 | 2404 | 2684 | 2742 | 1651 | 2393 | 2878 | 2576 | 2671 | 2056 | 2561 | 2447 | 2329 | 2488,8 | 2630 | 2348 | 0,893 | -0,163 | -2,161 | 0,03072891 | 0,05476691 | 0,006675 | 0,0025628 | 0,0050405 | 0,0050405 | TRUE | 0,0869597 | No |
| orf19.5294 | 9343 | 6493 | 8045 | 9944 | 9527 | 8395 | 8787 | 6011 | 8382 | 8266 | 8231 | 8200 | 8148 | 8010 | 7841 | 8478 | 8194,5 | 8270 | 8119 | 0,981 | -0,027 | -0,823 | 0,41042698 | 0,49855344 | 0,000437 | 0,0021567 | 0,0009395 | 0,0009395 | TRUE | 0,0103394 | No |
| orf19.5295 | 934 | 735 | 802 | 1054 | 1104 | 1055 | 1023 | 670 | 838 | 936 | 821 | 869 | 944 | 1007 | 913 | 945 | 908,99 | 866 | 952 | 1,102 | 0,14 | 2,33 | 0,01980513 | 0,03689588 | 0,001099 | 0,0035766 | 0,0023449 | 0,0023449 | TRUE | 0,0317408 | No |
| orf19.5296 | 1337 | 934 | 1089 | 1387 | 1687 | 1473 | 1589 | 915 | 1199 | 1189 | 1114 | 1144 | 1443 | 1405 | 1418 | 1291 | 1275,4 | 1162 | 1389 | 1,199 | 0,262 | 4,996 | 5,85E-07 | 2,57E-06 | 0,000717 | 0,0031178 | 0,0018523 | 0,0018523 | TRUE | 0,0245615 | No |
| orf19.5297 | 842 | 734 | 829 | 1047 | 1103 | 1141 | 1159 | 712 | 755 | 934 | 848 | 863 | 943 | 1089 | 1034 | 1004 | 933,97 | 850 | 1018 | 1,199 | 0,262 | 3,586 | 0,00033529 | 0,00093826 | 0,004087 | 0,0035338 | 0,004033 | 0,004033 | TRUE | 0,0680307 | No |
| orf19.5299 | 4681 | 3282 | 3734 | 4587 | 3697 | 3624 | 3606 | 1957 | 4199 | 4178 | 3820 | 3782 | 3162 | 3458 | 3218 | 2760 | 3572,3 | 3995 | 3150 | 0,789 | -0,341 | -5,014 | 5,34E-07 | 2,36E-06 | 0,005425 | 0,0023859 | 0,0041672 | 0,0041672 | TRUE | 0,0814144 | No |
| orf19.5299.1 | 1258 | 912 | 1136 | 1390 | 1261 | 1184 | 1348 | 588 | 1129 | 1161 | 1162 | 1146 | 1078 | 1130 | 1203 | 829 | 1104,8 | 1150 | 1060 | 0,925 | -0,112 | -1,212 | 0,22559352 | 0,30478207 | 0,011259 | 0,0032935 | 0,0073368 | 0,0073368 | TRUE | 0,2425235 | No |
| orf19.53 | 940 | 668 | 861 | 1018 | 942 | 815 | 957 | 532 | 843 | 850 | 881 | 839 | 806 | 778 | 854 | 750 | 825,2 | 853 | 797 | 0,937 | -0,094 | -1,626 | 0,10387814 | 0,15744194 | 0,000421 | 0,0037387 | 0,0020238 | 0,0020238 | TRUE | 0,0274989 | No |
| orf19.530 | 209 | 192 | 200 | 253 | 284 | 314 | 326 | 190 | 187 | 244 | 205 | 209 | 243 | 300 | 291 | 268 | 243,32 | 211 | 276 | 1,31 | 0,39 | 3,589 | 0,00033256 | 0,00093206 | 0,0062 | 0,0079458 | 0,007136 | 0,007136 | TRUE | 0,1077819 | No |
| orf19.5300 | 2203 | 1636 | 1979 | 2464 | 2077 | 1980 | 2055 | 1318 | 1976 | 2083 | 2025 | 2032 | 1776 | 1889 | 1834 | 1859 | 1934,2 | 2029 | 1840 | 0,907 | -0,141 | -3,428 | 0,00060833 | 0,0016044 | 1,00E-08 | 0,0027301 | 0,0011119 | 0,0011119 | TRUE | 0,0068341 | No |
| orf19.5302 | 299 | 207 | 250 | 290 | 127 | 104 | 111 | 80 | 268 | 264 | 256 | 239 | 109 | 99 | 99 | 113 | 180,8 | 257 | 105 | 0,408 | -1,295 | -11,647 | 2,38E-31 | 9,11E-30 | 1,00E-08 | 0,0100088 | 0,0051505 | 0,0051505 | TRUE | 0,0261205 | No |
| orf19.5303 | 2 | 2 | 0 | 1 | 1 | 4 | 1 | 0 | 2 | 3 | 0 | 1 | 1 | 4 | 1 | 0 | 1,34 | 2 | 2 | 1,152 | 0,205 | 0,162 | 0,87144771 | NA | 0,284638 | 1,0844021 | 0,7942552 | 0,7942552 | TRUE | 0,755983 | No |

| | | | | | | | | | | | | | | | | | | | | | | | | | | | | | | | |
|---|---|---|---|---|---|---|---|---|---|---|---|---|---|---|---|---|---|---|---|---|---|---|---|---|---|---|---|---|---|---|---|
| orf19.5304 | 0 | 1 | 0 | 0 | 0 | 0 | 0 | 0 | 0 | 1 | 0 | 0 | 0 | 0 | 0 | 0 | 0,16 | 0 | 0 | 0,616 | -0,698 | -0,198 | 0,84334499 | NA | 1,00E-08 | 9,1242388 | 10 | 10 | TRUE | 0,5444342 | No |
| orf19.5305 | 3290 | 2116 | 2765 | 3517 | 3624 | 2964 | 3130 | 2256 | 2951 | 2694 | 2829 | 2900 | 3099 | 2828 | 2793 | 3182 | 2909,6 | 2844 | 2976 | 1,045 | 0,063 | 1,161 | 0,24551095 | 0,3274206 | 0,00244 | 0,0024785 | 0,0024997 | 0,0024997 | TRUE | 0,0251855 | No |
| orf19.5306 | 3 | 2 | 3 | 3 | 0 | 0 | 0 | 0 | 3 | 3 | 3 | 2 | 0 | 0 | 0 | 0 | 1,35 | 3 | 0 | 0,069 | -3,863 | -2,806 | 0,00501884 | NA | 1,00E-08 | 1,0792427 | 0,6366978 | 0,6366978 | TRUE | 0,0477013 | No |
| orf19.5307 | 1622 | 1065 | 1506 | 1822 | 1372 | 1154 | 1221 | 781 | 1455 | 1356 | 1541 | 1502 | 1173 | 1101 | 1090 | 1102 | 1290 | 1464 | 1116 | 0,762 | -0,392 | -7,159 | 8,15E-13 | 7,55E-12 | 0,001303 | 0,0031049 | 0,0020807 | 0,0020807 | TRUE | 0,0273315 | No |
| orf19.5308 | 191 | 123 | 142 | 152 | 92 | 81 | 82 | 48 | 171 | 157 | 145 | 125 | 79 | 77 | 73 | 68 | 111,93 | 150 | 74 | 0,5 | -1 | -6,798 | 1,06E-11 | 8,67E-11 | 1,00E-08 | 0,0149498 | 0,0107568 | 0,0107568 | TRUE | 0,1406241 | No |
| orf19.5311 | 2087 | 1409 | 1940 | 2265 | 2132 | 1880 | 2105 | 1252 | 1872 | 1794 | 1985 | 1868 | 1823 | 1794 | 1878 | 1766 | 1847,5 | 1880 | 1815 | 0,966 | -0,05 | -1,084 | 0,27847254 | 0,3625272 | 0,000508 | 0,0027653 | 0,0014747 | 0,0014747 | TRUE | 0,016589 | No |
| orf19.5312 | 2238 | 1575 | 2022 | 2531 | 2645 | 2323 | 2580 | 1684 | 2008 | 2005 | 2069 | 2087 | 2262 | 2217 | 2302 | 2375 | 2165,6 | 2042 | 2289 | 1,119 | 0,162 | 4,013 | 6,00E-05 | 0,00019158 | 0,000123 | 0,0026499 | 0,0011044 | 0,0011044 | TRUE | 0,0069531 | No |
| orf19.5316 | 148 | 112 | 146 | 178 | 165 | 147 | 128 | 88 | 133 | 143 | 149 | 147 | 141 | 140 | 114 | 124 | 136,4 | 143 | 130 | 0,912 | -0,133 | -1,079 | 0,28078724 | 0,36475021 | 1,00E-08 | 0,0126223 | 0,0073077 | 0,0073077 | TRUE | 0,0793737 | No |
| orf19.5318 | 1076 | 760 | 959 | 1105 | 1453 | 1342 | 1474 | 937 | 965 | 968 | 981 | 911 | 1243 | 1280 | 1315 | 1322 | 1123,2 | 956 | 1290 | 1,349 | 0,432 | 8,584 | 9,19E-18 | 1,39E-16 | 1,00E-08 | 0,0032721 | 0,0015209 | 0,0015209 | TRUE | 0,0130244 | No |
| orf19.532 | 1362 | 892 | 1202 | 1494 | 723 | 605 | 558 | 386 | 1222 | 1136 | 1230 | 1232 | 618 | 577 | 498 | 544 | 882,14 | 1205 | 559 | 0,465 | -1,106 | -15,741 | 7,87E-56 | 1,10E-53 | 0,003146 | 0,0036252 | 0,0034274 | 0,0034274 | TRUE | 0,0708551 | No |
| orf19.5320 | 206 | 147 | 161 | 235 | 271 | 227 | 274 | 184 | 185 | 187 | 165 | 194 | 232 | 217 | 244 | 260 | 210,35 | 183 | 238 | 1,297 | 0,375 | 3,656 | 0,0002558 | 0,0007308 | 1,00E-08 | 0,0088808 | 0,0052813 | 0,0052813 | TRUE | 0,046527 | No |
| orf19.5321 | 2404 | 1603 | 2183 | 2601 | 2304 | 1929 | 2211 | 1375 | 2157 | 2041 | 2233 | 2145 | 1970 | 1841 | 1973 | 1939 | 2037,4 | 2144 | 1931 | 0,9 | -0,152 | -3,36 | 0,00077804 | 0,00200277 | 0,000673 | 0,0026921 | 0,0014682 | 0,0014682 | TRUE | 0,0123704 | No |
| orf19.5322 | 741 | 517 | 620 | 802 | 776 | 604 | 696 | 388 | 665 | 658 | 634 | 661 | 664 | 576 | 621 | 547 | 628,36 | 654 | 602 | 0,923 | -0,115 | -1,641 | 0,10083799 | 0,1538823 | 0,001961 | 0,0042898 | 0,0031146 | 0,0031146 | TRUE | 0,05609 | No |
| orf19.5323 | 918 | 568 | 825 | 980 | 1022 | 784 | 868 | 609 | 824 | 723 | 844 | 808 | 874 | 748 | 775 | 859 | 806,81 | 800 | 814 | 1,015 | 0,021 | 0,293 | 0,76962107 | 0,8201283 | 0,003786 | 0,0037788 | 0,0037808 | 0,0037808 | TRUE | 0,0454332 | No |
| orf19.5325 | 2541 | 1609 | 2220 | 2728 | 3061 | 2351 | 2700 | 1573 | 2280 | 2048 | 2271 | 2250 | 2618 | 2243 | 2409 | 2219 | 2292,2 | 2212 | 2372 | 1,073 | 0,101 | 1,627 | 0,10378415 | 0,15737889 | 0,003671 | 0,0026128 | 0,0032856 | 0,0032856 | TRUE | 0,0594838 | No |
| orf19.5326 | 2243 | 1276 | 1967 | 2353 | 2953 | 2411 | 2639 | 1916 | 2012 | 1624 | 2012 | 1940 | 2525 | 2300 | 2355 | 2702 | 2184,1 | 1897 | 2470 | 1,3 | 0,378 | 4,99 | 6,03E-07 | 2,64E-06 | 0,006839 | 0,0026442 | 0,0050496 | 0,0050496 | TRUE | 0,1098231 | No |
| orf19.5328 | 1867 | 1243 | 1744 | 2005 | 1885 | 1605 | 1722 | 1045 | 1675 | 1582 | 1784 | 1653 | 1612 | 1531 | 1537 | 1474 | 1606,1 | 1674 | 1538 | 0,92 | -0,12 | -2,346 | 0,01895089 | 0,0354143 | 0,001095 | 0,0028834 | 0,0019011 | 0,0019011 | TRUE | 0,0231588 | No |
| orf19.533 | 1 | 2 | 0 | 0 | 0 | 0 | 0 | 1 | 1 | 3 | 0 | 0 | 0 | 0 | 0 | 1 | 0,61 | 1 | 0 | 0,375 | -1,414 | -0,692 | 0,48866075 | NA | 1,00E-08 | 2,3946987 | 2,3962181 | 2,3962181 | TRUE | 0,6248744 | No |
| orf19.5334 | 2435 | 1602 | 1993 | 2472 | 3949 | 3043 | 3312 | 2275 | 2184 | 2040 | 2039 | 2038 | 3377 | 2904 | 2955 | 3209 | 2593,3 | 2075 | 3111 | 1,498 | 0,583 | 10,187 | 2,26E-24 | 5,50E-23 | 0,002622 | 0,0025393 | 0,0027463 | 0,0027463 | TRUE | 0,0417978 | No |
| orf19.5335 | 2189 | 1391 | 1829 | 2293 | 2468 | 2020 | 2303 | 1477 | 1964 | 1771 | 1871 | 1891 | 2111 | 1927 | 2055 | 2083 | 1959,1 | 1874 | 2044 | 1,089 | 0,123 | 2,519 | 0,01178465 | 0,0233132 | 0,001004 | 0,0027205 | 0,0017815 | 0,0017815 | TRUE | 0,0181625 | No |
| orf19.5337 | 401 | 257 | 330 | 443 | 416 | 316 | 335 | 300 | 360 | 327 | 338 | 365 | 356 | 302 | 299 | 423 | 346,15 | 348 | 345 | 0,984 | -0,023 | -0,206 | 0,83645285 | 0,87489706 | 0,011407 | 0,0061734 | 0,0086821 | 0,0086821 | TRUE | 0,2383376 | No |
| orf19.5338 | 2442 | 1564 | 2153 | 2669 | 1709 | 1466 | 1610 | 1124 | 2191 | 1991 | 2203 | 2201 | 1462 | 1399 | 1437 | 1585 | 1808,5 | 2146 | 1471 | 0,683 | -0,55 | -10,136 | 3,81E-24 | 9,01E-23 | 0,001908 | 0,0027822 | 0,0022499 | 0,0022499 | TRUE | 0,0302794 | No |
| orf19.5339 | 2587 | 1615 | 2028 | 2624 | 2825 | 1961 | 2175 | 1511 | 2321 | 2056 | 2075 | 2164 | 2416 | 1871 | 1941 | 2131 | 2121,8 | 2154 | 2090 | 0,97 | -0,044 | -0,56 | 0,57560488 | 0,65385406 | 0,007746 | 0,0026637 | 0,0054933 | 0,0054933 | TRUE | 0,1370169 | No |
| orf19.5340 | 5069 | 3420 | 4397 | 5425 | 5155 | 4336 | 4802 | 2916 | 4547 | 4354 | 4499 | 4474 | 4409 | 4137 | 4285 | 4113 | 4352,2 | 4468 | 4236 | 0,948 | -0,076 | -2,087 | 0,03688251 | 0,0643783 | 0,000394 | 0,0023131 | 0,0010579 | 0,0010579 | TRUE | 0,0092891 | No |
| orf19.5341 | 7491 | 4515 | 6085 | 7601 | 7571 | 5535 | 5844 | 4185 | 6720 | 5748 | 6226 | 6268 | 6475 | 5281 | 5215 | 5903 | 5979,4 | 6240 | 5718 | 0,916 | -0,126 | -1,722 | 0,08502805 | 0,13335765 | 0,00716 | 0,0022223 | 0,00501 | 0,00501 | TRUE | 0,0979144 | No |
| orf19.5342 | 1131 | 773 | 1067 | 1323 | 1211 | 1098 | 1182 | 841 | 1015 | 984 | 1092 | 1091 | 1036 | 1048 | 1055 | 1186 | 1063,2 | 1046 | 1081 | 1,03 | 0,042 | 0,686 | 0,49241515 | 0,57757601 | 0,002232 | 0,0033449 | 0,0027235 | 0,0027235 | TRUE | 0,0465288 | No |
| orf19.5342.1 | 1447 | 924 | 1120 | 1532 | 1495 | 1066 | 1269 | 800 | 1298 | 1176 | 1146 | 1263 | 1279 | 1017 | 1132 | 1128 | 1180 | 1221 | 1139 | 0,933 | -0,1 | -1,356 | 0,17519603 | 0,24638993 | 0,005017 | 0,0032098 | 0,0044167 | 0,0044167 | TRUE | 0,0845732 | No |
| orf19.5342.2 | 236 | 139 | 141 | 193 | 190 | 123 | 157 | 101 | 212 | 177 | 144 | 159 | 162 | 117 | 140 | 142 | 156,81 | 173 | 140 | 0,813 | -0,299 | -2,04 | 0,04139254 | 0,07113455 | 0,017287 | 0,0112372 | 0,0141417 | 0,0141417 | TRUE | 0,2569547 | No |
| orf19.5343 | 317 | 209 | 292 | 367 | 323 | 288 | 345 | 231 | 284 | 266 | 299 | 303 | 276 | 275 | 308 | 326 | 292,07 | 288 | 296 | 1,021 | 0,03 | 0,326 | 0,74410289 | 0,79945818 | 0,001257 | 0,00695 | 0,0044515 | 0,0044515 | TRUE | 0,0426387 | No |
| orf19.5345 | 2757 | 1728 | 2349 | 3011 | 3409 | 2671 | 2985 | 2179 | 2473 | 2200 | 2403 | 2483 | 2915 | 2549 | 2664 | 3073 | 2595,1 | 2390 | 2800 | 1,17 | 0,226 | 3,428 | 0,00060712 | 0,00160244 | 0,004607 | 0,0025389 | 0,0037901 | 0,0037901 | TRUE | 0,0504342 | No |
| orf19.5346 | 1456 | 879 | 1193 | 1443 | 1446 | 1213 | 1423 | 821 | 1306 | 1119 | 1221 | 1190 | 1237 | 1157 | 1270 | 1158 | 1207,2 | 1209 | 1206 | 0,997 | -0,004 | -0,071 | 0,94329314 | 0,95909717 | 0,002269 | 0,0031821 | 0,0026686 | 0,0026686 | TRUE | 0,035346 | No |
| orf19.5348 | 7463 | 4872 | 6872 | 8059 | 11880 | 9871 | 11245 | 7228 | 6695 | 6203 | 7031 | 6646 | 10160 | 9419 | 10034 | 10194 | 8297,7 | 6644 | 9952 | 1,497 | 0,582 | 12,555 | 3,71E-36 | 1,99E-34 | 0,001805 | 0,0021545 | 0,0019409 | 0,0019409 | TRUE | 0,023794 | No |
| orf19.535 | 87 | 60 | 69 | 84 | 58 | 38 | 39 | 18 | 78 | 76 | 71 | 69 | 50 | 36 | 35 | 25 | 55,04 | 74 | 36 | 0,508 | -0,978 | -4,781 | 1,75E-06 | 7,10E-06 | 0,002427 | 0,0283533 | 0,0199895 | 0,0199895 | TRUE | 0,445941 | No |
| orf19.5350 | 895 | 627 | 820 | 980 | 907 | 749 | 869 | 549 | 803 | 798 | 839 | 808 | 776 | 715 | 775 | 774 | 786,04 | 812 | 760 | 0,935 | -0,096 | -1,712 | 0,08693949 | 0,13574129 | 1,00E-08 | 0,0038264 | 0,0017663 | 0,0017663 | TRUE | 0,0196958 | No |
| orf19.5351 | 6145 | 4006 | 5020 | 6439 | 5765 | 4410 | 4902 | 2748 | 5513 | 5100 | 5136 | 5310 | 4930 | 4208 | 4374 | 3876 | 4805,8 | 5265 | 4347 | 0,826 | -0,275 | -4,11 | 3,95E-05 | 0,00012997 | 0,005502 | 0,0022816 | 0,0040983 | 0,0040983 | TRUE | 0,0996575 | No |
| orf19.5352 | 2489 | 1649 | 2217 | 2730 | 2701 | 2219 | 2469 | 1659 | 2233 | 2099 | 2268 | 2251 | 2310 | 2117 | 2203 | 2340 | 2227,8 | 2213 | 2242 | 1,012 | 0,017 | 0,352 | 0,72490526 | 0,78445832 | 0,001007 | 0,0026312 | 0,0017182 | 0,0017182 | TRUE | 0,0170386 | No |
| orf19.5353 | 25 | 17 | 17 | 34 | 13 | 11 | 14 | 6 | 22 | 22 | 17 | 28 | 11 | 10 | 12 | 8 | 16,51 | 22 | 10 | 0,479 | -1,061 | -3,037 | 0,00238648 | 0,00553163 | 1,00E-08 | 0,0899155 | 0,0489339 | 0,0489339 | TRUE | 0,1989292 | No |
| orf19.5356 | 6871 | 4534 | 6111 | 7241 | 6200 | 5202 | 5729 | 3921 | 6164 | 5772 | 6252 | 5971 | 5302 | 4964 | 5112 | 5530 | 5633,5 | 6040 | 5227 | 0,865 | -0,21 | -4,681 | 2,86E-06 | 1,13E-05 | 0,001486 | 0,0022372 | 0,001749 | 0,001749 | TRUE | 0,0178492 | No |
| orf19.5357 | 2503 | 1563 | 2159 | 2684 | 2319 | 1975 | 2249 | 1587 | 2245 | 1990 | 2209 | 2213 | 1983 | 1884 | 2007 | 2238 | 2096,3 | 2164 | 2028 | 0,935 | -0,097 | -1,585 | 0,11297488 | 0,16897014 | 0,003562 | 0,002672 | 0,0031455 | 0,0031455 | TRUE | 0,0557382 | No |
| orf19.5358 | 1554 | 974 | 1378 | 1593 | 1585 | 1302 | 1351 | 1011 | 1394 | 1240 | 1410 | 1314 | 1356 | 1242 | 1206 | 1426 | 1323,4 | 1340 | 1308 | 0,973 | -0,039 | -0,583 | 0,5601758 | 0,64020091 | 0,003821 | 0,0030765 | 0,0034893 | 0,0034893 | TRUE | 0,0415884 | No |
| orf19.536 | 2460 | 1641 | 2155 | 2668 | 2944 | 2502 | 2835 | 1698 | 2207 | 2089 | 2205 | 2200 | 2518 | 2387 | 2530 | 2395 | 2316,3 | 2175 | 2458 | 1,13 | 0,176 | 4,244 | 2,20E-05 | 7,58E-05 | 0,000372 | 0,0026062 | 0,0012182 | 0,0012182 | TRUE | 0,0079466 | No |
| orf19.5360 | 1083 | 605 | 865 | 1053 | 1095 | 799 | 843 | 612 | 972 | 770 | 885 | 868 | 936 | 762 | 752 | 863 | 851,17 | 874 | 828 | 0,946 | -0,079 | -0,893 | 0,37201561 | 0,46055425 | 0,008875 | 0,0036851 | 0,0064321 | 0,0064321 | TRUE | 0,0967212 | No |
| orf19.5362 | 293 | 180 | 235 | 352 | 261 | 180 | 225 | 159 | 263 | 229 | 240 | 290 | 223 | 172 | 201 | 224 | 230,34 | 256 | 205 | 0,796 | -0,329 | -2,85 | 0,00437783 | 0,00956784 | 0,00853 | 0,008282 | 0,008397 | 0,008397 | TRUE | 0,1317575 | No |
| orf19.5363 | 1067 | 660 | 977 | 1080 | 1149 | 881 | 967 | 642 | 957 | 840 | 1000 | 891 | 983 | 841 | 863 | 905 | 909,91 | 922 | 898 | 0,973 | -0,039 | -0,542 | 0,588162 | 0,66635438 | 0,004226 | 0,003575 | 0,003929 | 0,003929 | TRUE | 0,0511472 | No |
| orf19.5364 | 1972 | 1222 | 1650 | 2151 | 1712 | 1361 | 1529 | 997 | 1769 | 1556 | 1688 | 1774 | 1464 | 1299 | 1364 | 1406 | 1540 | 1697 | 1383 | 0,814 | -0,297 | -5,059 | 4,22E-07 | 1,89E-06 | 0,002198 | 0,0029222 | 0,0026493 | 0,0026493 | TRUE | 0,035945 | No |
| orf19.5365 | 1242 | 794 | 1087 | 1242 | 841 | 782 | 861 | 562 | 1114 | 1011 | 1112 | 1024 | 719 | 746 | 768 | 793 | 910,96 | 1065 | 756 | 0,708 | -0,498 | -8,384 | 5,14E-17 | 7,30E-16 | 0,001147 | 0,0035731 | 0,0022522 | 0,0022522 | TRUE | 0,0133501 | No |
| orf19.5365.1 | 766 | 507 | 597 | 835 | 743 | 559 | 556 | 424 | 687 | 645 | 611 | 689 | 635 | 533 | 496 | 598 | 611,87 | 658 | 566 | 0,858 | -0,221 | -2,59 | 0,00958749 | 0,01940103 | 0,00626 | 0,0043521 | 0,005351 | 0,005351 | TRUE | 0,0889003 | No |
| orf19.5367 | 624 | 370 | 542 | 654 | 559 | 493 | 517 | 341 | 560 | 471 | 555 | 539 | 478 | 470 | 461 | 481 | 501,93 | 531 | 472 | 0,886 | -0,174 | -2,419 | 0,01557899 | 0,02984648 | 0,000808 | 0,0048718 | 0,0029892 | 0,0029892 | TRUE | 0,0599314 | No |
| orf19.5368 | 726 | 430 | 653 | 766 | 828 | 657 | 732 | 617 | 651 | 547 | 668 | 632 | 708 | 627 | 653 | 870 | 669,61 | 624 | 714 | 1,138 | 0,187 | 1,829 | 0,06746694 | 0,10937437 | 0,013337 | 0,0041475 | 0,008521 | 0,008521 | TRUE | 0,2392356 | No |
| orf19.5369 | 1825 | 1065 | 1519 | 1806 | 1293 | 1064 | 1173 | 901 | 1637 | 1356 | 1554 | 1489 | 1106 | 1015 | 1047 | 1271 | 1309,4 | 1509 | 1110 | 0,733 | -0,449 | -5,67 | 1,43E-08 | 7,76E-08 | 0,007313 | 0,0030882 | 0,0052195 | 0,0052195 | TRUE | 0,1077115 | No |
| orf19.537 | 271 | 148 | 243 | 256 | 260 | 190 | 239 | 147 | 243 | 188 | 249 | 211 | 222 | 181 | 213 | 207 | 214,44 | 223 | 206 | 0,923 | -0,116 | -1,014 | 0,31079942 | 0,39704081 | 0,007008 | 0,0087493 | 0,0079598 | 0,0079598 | TRUE | 0,1026533 | No |
| orf19.5370 | 692 | 454 | 575 | 768 | 746 | 603 | 637 | 429 | 621 | 578 | 588 | 633 | 638 | 575 | 568 | 605 | 600,9 | 605 | 596 | 0,984 | -0,024 | -0,353 | 0,72390407 | 0,78378368 | 0,000914 | 0,0043954 | 0,0026045 | 0,0026045 | TRUE | 0,0286609 | No |
| orf19.5376 | 513 | 383 | 534 | 605 | 648 | 605 | 674 | 404 | 460 | 488 | 546 | 499 | 554 | 577 | 601 | 570 | 536,97 | 498 | 576 | 1,156 | 0,209 | 2,939 | 0,0032876 | 0,00740048 | 0,001297 | 0,004683 | 0,0030042 | 0,0030042 | TRUE | 0,0487671 | No |
| orf19.5377 | 1901 | 1273 | 1600 | 2086 | 2526 | 2198 | 2396 | 1439 | 1705 | 1621 | 1637 | 1720 | 2160 | 2097 | 2138 | 2030 | 1888,5 | 1671 | 2106 | 1,261 | 0,334 | 7,785 | 6,95E-15 | 7,90E-14 | 0,000208 | 0,0027482 | 0,0012327 | 0,0012327 | TRUE | 0,0065559 | No |
| orf19.5378 | 3152 | 2051 | 2579 | 3169 | 2644 | 2321 | 2545 | 1377 | 2828 | 2611 | 2639 | 2613 | 2261 | 2215 | 2271 | 1942 | 2422,5 | 2673 | 2172 | 0,814 | -0,297 | -5,216 | 1,83E-07 | 8,66E-07 | 0,00276 | 0,0025788 | 0,002687 | 0,002687 | TRUE | 0,0578804 | No |
| orf19.5379 | 1855 | 1339 | 1654 | 2095 | 1985 | 1766 | 1751 | 1284 | 1664 | 1705 | 1692 | 1728 | 1698 | 1685 | 1562 | 1811 | 1693,1 | 1697 | 1689 | 0,993 | -0,01 | -0,194 | 0,84608594 | 0,88297452 | 0,001248 | 0,0028369 | 0,0019061 | 0,0019061 | TRUE | 0,030738 | No |
| orf19.538 | 323 | 227 | 259 | 310 | 314 | 279 | 297 | 234 | 290 | 289 | 265 | 256 | 269 | 266 | 265 | 330 | 278,65 | 275 | 282 | 1,022 | 0,031 | 0,32 | 0,74891463 | 0,80276166 | 0,003496 | 0,0071893 | 0,0055274 | 0,0055274 | TRUE | 0,1229125 | No |
| orf19.5380 | 856 | 593 | 753 | 931 | 761 | 734 | 767 | 432 | 768 | 755 | 770 | 768 | 651 | 700 | 684 | 609 | 713,24 | 765 | 661 | 0,867 | -0,207 | -3,433 | 0,00059695 | 0,00157995 | 2,00E-05 | 0,0040149 | 0,0020677 | 0,0020677 | TRUE | 0,0273791 | No |
| orf19.5381 | 1041 | 736 | 914 | 1105 | 1352 | 1333 | 1412 | 971 | 934 | 937 | 935 | 911 | 1156 | 1272 | 1260 | 1370 | 1096,9 | 929 | 1264 | 1,357 | 0,441 | 7,493 | 6,73E-14 | 6,96E-13 | 0,001694 | 0,003303 | 0,0023877 | 0,0023877 | TRUE | 0,0403228 | No |

| | | | | | | | | | | | | | | | | | | | | | | | | | | | | | | | |
|---|---|---|---|---|---|---|---|---|---|---|---|---|---|---|---|---|---|---|---|---|---|---|---|---|---|---|---|---|---|---|---|
| orf19.5382 | 0 | 0 | 0 | 0 | 0 | 0 | 0 | 0 | 0 | 0 | 0 | 0 | 0 | 0 | 0 | 0 | 0 | 0 | 0 | NA | NA | NA | NA | NA | NA | NA | NA | NA | NA | NA | NA |
| orf19.5383 | 3944 | 2677 | 3742 | 4441 | 4131 | 3571 | 3894 | 2657 | 3538 | 3408 | 3828 | 3662 | 3533 | 3407 | 3475 | 3747 | 3574,9 | 3609 | 3540 | 0,98 | -0,03 | -0,61 | 0,54168449 | 0,62393412 | 0,001624 | 0,0023856 | 0,0019653 | 0,0019653 | TRUE | 0,0199582 | No |
| orf19.5384 | 1249 | 887 | 1036 | 1318 | 969 | 910 | 962 | 626 | 1120 | 1129 | 1060 | 1087 | 829 | 868 | 858 | 883 | 979,36 | 1099 | 860 | 0,781 | -0,356 | -6,937 | 4,02E-12 | 3,44E-11 | 1,00E-08 | 0,0034618 | 0,0014933 | 0,0014933 | TRUE | 0,0071955 | No |
| orf19.5387 | 863 | 676 | 848 | 904 | 777 | 834 | 806 | 527 | 774 | 861 | 868 | 745 | 665 | 796 | 719 | 743 | 771,33 | 812 | 731 | 0,901 | -0,151 | -1,985 | 0,04719881 | 0,07993934 | 0,004539 | 0,0038616 | 0,0042287 | 0,0042287 | TRUE | 0,0463525 | No |
| orf19.5388 | 1242 | 865 | 1077 | 1421 | 1108 | 958 | 1065 | 669 | 1114 | 1101 | 1102 | 1172 | 948 | 914 | 950 | 944 | 1030,6 | 1122 | 939 | 0,836 | -0,259 | -5,187 | 2,13E-07 | 9,99E-07 | 1,00E-08 | 0,0033881 | 0,0014188 | 0,0014188 | TRUE | 0,0112333 | No |
| orf19.5389 | 1084 | 796 | 1093 | 1246 | 1226 | 1116 | 1192 | 866 | 972 | 1013 | 1118 | 1027 | 1049 | 1065 | 1064 | 1221 | 1066,3 | 1032 | 1100 | 1,062 | 0,086 | 1,304 | 0,19238024 | 0,26570586 | 0,00327 | 0,003341 | 0,0032739 | 0,0032739 | TRUE | 0,0612581 | No |
| orf19.539 | 679 | 506 | 612 | 658 | 831 | 739 | 780 | 491 | 609 | 644 | 626 | 543 | 711 | 705 | 696 | 693 | 653,3 | 606 | 701 | 1,162 | 0,217 | 3,259 | 0,00111849 | 0,00279298 | 0,0013 | 0,0042016 | 0,0027164 | 0,0027164 | TRUE | 0,0612164 | No |
| orf19.5390 | 0 | 0 | 0 | 0 | 0 | 0 | 0 | 0 | 0 | 0 | 0 | 0 | 0 | 0 | 0 | 0 | 0 | 0 | 0 | NA | NA | NA | NA | NA | NA | NA | NA | NA | NA | NA | NA |
| orf19.5391 | 1692 | 1373 | 1587 | 1928 | 1707 | 1737 | 1818 | 1133 | 1518 | 1748 | 1624 | 1590 | 1460 | 1657 | 1622 | 1598 | 1602,1 | 1620 | 1584 | 0,979 | -0,031 | -0,516 | 0,60608599 | 0,68159903 | 0,002659 | 0,0028856 | 0,0027537 | 0,0027537 | TRUE | 0,0352565 | No |
| orf19.5392 | 41 | 39 | 44 | 60 | 54 | 34 | 35 | 22 | 37 | 50 | 45 | 49 | 46 | 32 | 31 | 31 | 40,23 | 45 | 35 | 0,788 | -0,343 | -1,495 | 0,13501228 | 0,1974571 | 1,00E-08 | 0,0380679 | 0,0254466 | 0,0254466 | TRUE | 0,3561369 | No |
| orf19.5393 | 6424 | 4600 | 5668 | 7147 | 5202 | 4989 | 5375 | 3109 | 5763 | 5856 | 5799 | 5894 | 4449 | 4760 | 4796 | 4385 | 5212,8 | 5828 | 4598 | 0,789 | -0,341 | -8,588 | 8,84E-18 | 1,34E-16 | 0,000841 | 0,002258 | 0,0013188 | 0,0013188 | TRUE | 0,0111144 | No |
| orf19.5394.1 | 1192 | 843 | 992 | 1282 | 1696 | 1395 | 1450 | 998 | 1069 | 1073 | 1015 | 1057 | 1450 | 1331 | 1294 | 1408 | 1212,2 | 1054 | 1371 | 1,3 | 0,379 | 7,04 | 1,92E-12 | 1,71E-11 | 0,001062 | 0,0031771 | 0,0019401 | 0,0019401 | TRUE | 0,0203829 | No |
| orf19.5395 | 2024 | 1467 | 1843 | 2120 | 2957 | 2706 | 2877 | 1838 | 1816 | 1868 | 1886 | 1748 | 2529 | 2582 | 2567 | 2592 | 2198,5 | 1830 | 2568 | 1,405 | 0,49 | 12,136 | 6,83E-34 | 3,13E-32 | 2,30E-06 | 0,0026399 | 0,001099 | 0,001099 | TRUE | 0,0112909 | No |
| orf19.5397 | 293 | 233 | 259 | 349 | 324 | 312 | 327 | 202 | 263 | 297 | 265 | 288 | 277 | 298 | 292 | 285 | 282,97 | 278 | 288 | 1,037 | 0,053 | 0,615 | 0,53871613 | 0,62087215 | 1,00E-08 | 0,0071098 | 0,0035376 | 0,0035376 | TRUE | 0,0189481 | No |
| orf19.5399 | 39 | 23 | 37 | 33 | 31 | 31 | 15 | 18 | 35 | 29 | 38 | 27 | 27 | 30 | 13 | 25 | 28,02 | 32 | 24 | 0,73 | -0,455 | -1,585 | 0,11293973 | 0,16895964 | 0,024018 | 0,0537799 | 0,0424214 | 0,0424214 | TRUE | 0,5895201 | No |
| orf19.54 | 429 | 344 | 396 | 479 | 302 | 250 | 282 | 203 | 385 | 438 | 405 | 395 | 258 | 239 | 252 | 286 | 332,21 | 406 | 259 | 0,636 | -0,653 | -7,65 | 2,01E-14 | 2,16E-13 | 0,000409 | 0,0063493 | 0,0038177 | 0,0038177 | TRUE | 0,046341 | No |
| orf19.540 | 481 | 293 | 373 | 514 | 393 | 303 | 385 | 215 | 432 | 373 | 382 | 424 | 336 | 289 | 344 | 303 | 360,25 | 403 | 318 | 0,79 | -0,34 | -3,844 | 0,00012105 | 0,00036473 | 0,003302 | 0,0060092 | 0,0047121 | 0,0047121 | TRUE | 0,0469856 | No |
| orf19.5402 | 0 | 0 | 0 | 0 | 0 | 0 | 0 | 0 | 0 | 0 | 0 | 0 | 0 | 0 | 0 | 0 | 0 | 0 | 0 | NA | NA | NA | NA | NA | NA | NA | NA | NA | NA | NA | NA |
| orf19.5404.1 | 1113 | 716 | 1000 | 1173 | 1129 | 993 | 1111 | 744 | 998 | 912 | 1023 | 967 | 966 | 947 | 991 | 1049 | 981,77 | 975 | 988 | 1,01 | 0,014 | 0,253 | 0,79993903 | 0,84494263 | 0,000915 | 0,0034582 | 0,0021081 | 0,0021081 | TRUE | 0,0205927 | No |
| orf19.5406 | 4260 | 2979 | 3932 | 4744 | 4036 | 3538 | 3864 | 2249 | 3822 | 3793 | 4023 | 3912 | 3452 | 3376 | 3448 | 3172 | 3624,6 | 3888 | 3362 | 0,866 | -0,208 | -5,08 | 3,78E-07 | 1,71E-06 | 0,000749 | 0,00238 | 0,0013293 | 0,0013293 | TRUE | 0,0163617 | No |
| orf19.5407 | 13370 | 8988 | 11660 | 14218 | 14021 | 11945 | 12862 | 7879 | 11994 | 11443 | 11929 | 11724 | 11991 | 11398 | 11477 | 11113 | 11634 | 11772 | 11495 | 0,977 | -0,034 | -0,993 | 0,3205008 | 0,40718308 | 0,000615 | 0,0021043 | 0,0010505 | 0,0010505 | TRUE | 0,0103907 | No |
| orf19.5408 | 6241 | 4076 | 5479 | 6661 | 7561 | 6412 | 7223 | 4628 | 5599 | 5189 | 5606 | 5493 | 6466 | 6118 | 6445 | 6527 | 5930,4 | 5472 | 6389 | 1,167 | 0,223 | 5,683 | 1,32E-08 | 7,19E-08 | 0,000825 | 0,0022243 | 0,0013037 | 0,0013037 | TRUE | 0,0141904 | No |
| orf19.5409 | 469 | 289 | 355 | 451 | 501 | 418 | 490 | 308 | 421 | 368 | 363 | 372 | 428 | 399 | 437 | 434 | 402,84 | 381 | 424 | 1,112 | 0,154 | 1,964 | 0,04950132 | 0,08347009 | 1,28E-05 | 0,0055832 | 0,0033834 | 0,0033834 | TRUE | 0,0592795 | No |
| orf19.541 | 3103 | 2143 | 2674 | 3245 | 2366 | 2025 | 2264 | 1499 | 2784 | 2728 | 2736 | 2676 | 2023 | 1932 | 2020 | 2114 | 2376,7 | 2731 | 2022 | 0,74 | -0,435 | -10,711 | 9,08E-27 | 2,77E-25 | 0,000232 | 0,0025903 | 0,0011522 | 0,0011522 | TRUE | 0,0106196 | No |
| orf19.5410 | 914 | 582 | 836 | 1025 | 1293 | 1044 | 1253 | 759 | 820 | 741 | 855 | 845 | 1106 | 996 | 1118 | 1071 | 944 | 815 | 1073 | 1,313 | 0,393 | 6,205 | 5,46E-10 | 3,61E-09 | 0,002123 | 0,0035173 | 0,0027724 | 0,0027724 | TRUE | 0,040737 | No |
| orf19.5411 | 1075 | 717 | 902 | 1120 | 967 | 799 | 963 | 639 | 964 | 913 | 923 | 924 | 827 | 762 | 859 | 901 | 884,2 | 931 | 837 | 0,897 | -0,157 | -2,572 | 0,01010008 | 0,02030833 | 0,001235 | 0,0036214 | 0,0024404 | 0,0024404 | TRUE | 0,042595 | No |
| orf19.5412 | 1372 | 944 | 1197 | 1591 | 1549 | 1305 | 1470 | 941 | 1231 | 1202 | 1225 | 1312 | 1325 | 1245 | 1312 | 1327 | 1272,3 | 1242 | 1302 | 1,046 | 0,065 | 1,306 | 0,19146705 | 0,26474891 | 0,000483 | 0,0031206 | 0,0016 | 0,0016 | TRUE | 0,0179208 | No |
| orf19.5413 | 1328 | 809 | 1164 | 1344 | 1467 | 1196 | 1324 | 859 | 1191 | 1030 | 1191 | 1108 | 1255 | 1141 | 1181 | 1212 | 1163,7 | 1130 | 1197 | 1,058 | 0,081 | 1,34 | 0,18036875 | 0,25201289 | 0,001997 | 0,0032271 | 0,0026467 | 0,0026467 | TRUE | 0,039219 | No |
| orf19.5416 | 7326 | 4786 | 6486 | 7696 | 7673 | 6941 | 7685 | 4787 | 6572 | 6093 | 6636 | 6346 | 6562 | 6623 | 6858 | 6752 | 6555,2 | 6412 | 6699 | 1,044 | 0,062 | 1,655 | 0,09802425 | 0,15023716 | 0,000728 | 0,002201 | 0,0012132 | 0,0012132 | TRUE | 0,0131759 | No |
| orf19.5417 | 14665 | 8975 | 11897 | 14870 | 13564 | 10673 | 11939 | 7925 | 13156 | 11426 | 12172 | 12262 | 11600 | 10184 | 10654 | 11178 | 11579 | 12254 | 10904 | 0,89 | -0,169 | -3,089 | 0,00201001 | 0,00474319 | 0,003155 | 0,0021049 | 0,0027805 | 0,0027805 | TRUE | 0,0300382 | No |
| orf19.5418 | 1664 | 1051 | 1416 | 1679 | 1564 | 1341 | 1413 | 937 | 1493 | 1338 | 1449 | 1385 | 1338 | 1280 | 1261 | 1322 | 1358 | 1416 | 1300 | 0,917 | -0,126 | -2,461 | 0,01384319 | 0,02688972 | 0,00082 | 0,0030486 | 0,0017619 | 0,0017619 | TRUE | 0,0161752 | No |
| orf19.5419 | 4970 | 3246 | 4051 | 5164 | 4880 | 4043 | 4373 | 2986 | 4459 | 4133 | 4145 | 4258 | 4174 | 3858 | 3902 | 4211 | 4142,4 | 4249 | 4036 | 0,949 | -0,075 | -1,665 | 0,0958138 | 0,14750009 | 0,001327 | 0,00233 | 0,0017211 | 0,0017211 | TRUE | 0,0135701 | No |
| orf19.542 | 4033 | 2694 | 3509 | 4207 | 4983 | 4098 | 4554 | 2720 | 3618 | 3430 | 3590 | 3469 | 4262 | 3910 | 4064 | 3836 | 3772,4 | 3527 | 4018 | 1,14 | 0,189 | 4,308 | 1,65E-05 | 5,82E-05 | 0,001076 | 0,0023643 | 0,0015824 | 0,0015824 | TRUE | 0,0204535 | No |
| orf19.5420 | 5065 | 2997 | 4216 | 5091 | 5162 | 4070 | 4609 | 3058 | 4544 | 3816 | 4313 | 4198 | 4415 | 3883 | 4113 | 4313 | 4199,4 | 4218 | 4181 | 0,991 | -0,014 | -0,224 | 0,82277342 | 0,86342746 | 0,003978 | 0,0023252 | 0,0033118 | 0,0033118 | TRUE | 0,0491735 | No |
| orf19.5422 | 41 | 21 | 43 | 54 | 56 | 43 | 66 | 46 | 37 | 27 | 44 | 45 | 48 | 41 | 59 | 65 | 45,59 | 38 | 53 | 1,37 | 0,454 | 1,984 | 0,04720785 | 0,07993934 | 0,01861 | 0,0338208 | 0,027847 | 0,027847 | TRUE | 0,2004389 | No |
| orf19.542.2 | 918 | 652 | 793 | 987 | 1253 | 1019 | 1130 | 741 | 824 | 830 | 811 | 814 | 1072 | 972 | 1008 | 1045 | 922,02 | 820 | 1024 | 1,25 | 0,322 | 5,988 | 2,13E-09 | 1,29E-08 | 1,00E-08 | 0,003554 | 0,0016707 | 0,0016707 | TRUE | 0,0143046 | No |
| orf19.5423 | 54 | 30 | 48 | 65 | 42 | 49 | 35 | 32 | 48 | 38 | 49 | 54 | 36 | 47 | 31 | 45 | 43,55 | 47 | 40 | 0,824 | -0,279 | -1,274 | 0,20250224 | 0,27758007 | 1,00E-08 | 0,0353152 | 0,0228871 | 0,0228871 | TRUE | 0,1675323 | No |
| orf19.5424 | 4 | 3 | 9 | 2 | 3 | 0 | 3 | 0 | 4 | 4 | 9 | 2 | 3 | 0 | 3 | 0 | 2,94 | 5 | 2 | 0,314 | -1,67 | -1,708 | 0,08769063 | NA | 0,381751 | 0,496017 | 0,4557905 | 0,4557905 | TRUE | 0,6881341 | No |
| orf19.5425 | 6391 | 4291 | 5678 | 6925 | 6491 | 5568 | 6298 | 3975 | 5733 | 5463 | 5809 | 5710 | 5551 | 5313 | 5620 | 5606 | 5600,8 | 5679 | 5522 | 0,972 | -0,041 | -1,157 | 0,24746539 | 0,32951288 | 0,000446 | 0,0022387 | 0,0010358 | 0,0010358 | TRUE | 0,0080412 | No |
| orf19.5426 | 2070 | 1366 | 1926 | 2194 | 1874 | 1631 | 1751 | 1118 | 1857 | 1739 | 1970 | 1809 | 1603 | 1556 | 1562 | 1577 | 1709,3 | 1844 | 1574 | 0,853 | -0,229 | -4,72 | 2,36E-06 | 9,48E-06 | 0,000867 | 0,0028288 | 0,0016628 | 0,0016628 | TRUE | 0,0250416 | No |
| orf19.5428 | 125 | 82 | 91 | 116 | 134 | 150 | 122 | 89 | 112 | 104 | 93 | 96 | 115 | 143 | 109 | 126 | 112,18 | 101 | 123 | 1,212 | 0,277 | 1,966 | 0,04931557 | 0,08320357 | 1,00E-08 | 0,0149208 | 0,0100982 | 0,0100982 | TRUE | 0,1340665 | No |
| orf19.5429 | 1486 | 1049 | 1367 | 1703 | 2080 | 1836 | 2032 | 1303 | 1333 | 1335 | 1399 | 1404 | 1779 | 1752 | 1813 | 1838 | 1581,7 | 1368 | 1796 | 1,311 | 0,39 | 8,93 | 4,26E-19 | 7,22E-18 | 1,00E-08 | 0,0028974 | 0,0011892 | 0,0011892 | TRUE | 0,0041123 | No |
| orf19.543 | 660 | 433 | 592 | 678 | 774 | 689 | 713 | 465 | 592 | 551 | 606 | 559 | 662 | 657 | 636 | 656 | 614,94 | 577 | 653 | 1,13 | 0,177 | 2,879 | 0,00398664 | 0,00879609 | 1,00E-08 | 0,0043402 | 0,0019892 | 0,0019892 | TRUE | 0,0123898 | No |
| orf19.5430 | 5458 | 3354 | 4450 | 5871 | 5163 | 4137 | 4540 | 2727 | 4896 | 4270 | 4553 | 4841 | 4416 | 3947 | 4051 | 3846 | 4352,6 | 4640 | 4065 | 0,876 | -0,191 | -3,261 | 0,00111046 | 0,00277522 | 0,003497 | 0,002313 | 0,0030573 | 0,0030573 | TRUE | 0,0413227 | No |
| orf19.5431 | 1015 | 699 | 954 | 1156 | 851 | 781 | 854 | 629 | 911 | 890 | 976 | 953 | 728 | 745 | 762 | 887 | 856,49 | 932 | 780 | 0,833 | -0,264 | -3,802 | 0,00014353 | 0,00042669 | 0,003225 | 0,0036745 | 0,0034453 | 0,0034453 | TRUE | 0,090417 | No |
| orf19.5432 | 1208 | 754 | 972 | 1217 | 1224 | 1102 | 1183 | 780 | 1084 | 960 | 994 | 1004 | 1047 | 1051 | 1056 | 1100 | 1037 | 1010 | 1064 | 1,049 | 0,069 | 1,286 | 0,1985762 | 0,27288331 | 0,000547 | 0,0033795 | 0,0018395 | 0,0018395 | TRUE | 0,0288346 | No |
| orf19.5433 | 447 | 297 | 332 | 458 | 556 | 474 | 475 | 300 | 401 | 378 | 340 | 378 | 476 | 452 | 424 | 423 | 408,9 | 374 | 444 | 1,187 | 0,248 | 3,113 | 0,00184956 | 0,00440086 | 0,001573 | 0,0055298 | 0,0036316 | 0,0036316 | TRUE | 0,0439698 | No |
| orf19.5436 | 11522 | 7875 | 9912 | 12207 | 12961 | 11348 | 12340 | 7960 | 10336 | 10026 | 10141 | 10066 | 11085 | 10828 | 11011 | 11227 | 10590 | 10142 | 11038 | 1,088 | 0,122 | 4,504 | 6,67E-06 | 2,50E-05 | 7,95E-05 | 0,0021166 | 0,000604 | 0,000604 | TRUE | 0,0020531 | No |
| orf19.5437 | 82184 | 52276 | 68841 | 85669 | 86361 | 67816 | 73595 | 49806 | 73728 | 66553 | 70431 | 70644 | 73859 | 64708 | 65671 | 70247 | 69480 | 70339 | 68621 | 0,976 | -0,036 | -0,699 | 0,48425708 | 0,56969189 | 0,002528 | 0,0020004 | 0,0024915 | 0,0024915 | TRUE | 0,0324152 | No |
| orf19.5438 | 2114 | 1507 | 1779 | 2215 | 1702 | 1341 | 1437 | 933 | 1896 | 1919 | 1820 | 1827 | 1456 | 1280 | 1282 | 1316 | 1599,4 | 1866 | 1334 | 0,716 | -0,483 | -8,966 | 3,06E-19 | 5,26E-18 | 0,001441 | 0,0028872 | 0,0021357 | 0,0021357 | TRUE | 0,0484369 | No |
| orf19.5439 | 659 | 410 | 598 | 678 | 462 | 454 | 461 | 314 | 591 | 522 | 612 | 559 | 395 | 433 | 411 | 443 | 495,83 | 571 | 420 | 0,733 | -0,448 | -6,008 | 1,88E-09 | 1,15E-08 | 0,001516 | 0,0049074 | 0,003267 | 0,003267 | TRUE | 0,0346028 | No |
| orf19.544 | 1017 | 769 | 905 | 1146 | 888 | 793 | 802 | 546 | 912 | 979 | 926 | 945 | 759 | 757 | 716 | 770 | 845,52 | 940 | 750 | 0,798 | -0,326 | -5,965 | 2,45E-09 | 1,47E-08 | 1,00E-08 | 0,0036965 | 0,001669 | 0,001669 | TRUE | 0,0119924 | No |
| orf19.5440 | 16900 | 10523 | 14132 | 17626 | 17491 | 13971 | 15724 | 9461 | 15161 | 13397 | 14458 | 14535 | 14959 | 13331 | 14031 | 13344 | 14152 | 14388 | 13916 | 0,967 | -0,048 | -0,929 | 0,35264669 | 0,44020389 | 0,002654 | 0,0020821 | 0,0024983 | 0,0024983 | TRUE | 0,0312271 | No |
| orf19.5441 | 1543 | 1064 | 1398 | 1602 | 1554 | 1406 | 1593 | 986 | 1384 | 1355 | 1430 | 1321 | 1329 | 1342 | 1421 | 1391 | 1371,6 | 1372 | 1371 | 0,998 | -0,002 | -0,048 | 0,96207257 | 0,97243136 | 0,000438 | 0,0030379 | 0,001516 | 0,001516 | TRUE | 0,0096584 | No |
| orf19.544.1 | 7260 | 4753 | 5862 | 7472 | 7242 | 5679 | 6078 | 3804 | 6513 | 6051 | 5997 | 6162 | 6194 | 5419 | 5424 | 5365 | 5890,5 | 6181 | 5600 | 0,906 | -0,142 | -2,608 | 0,00910475 | 0,01852886 | 0,002982 | 0,002226 | 0,0026717 | 0,0026717 | TRUE | 0,0626701 | No |
| orf19.5442 | 2164 | 1379 | 1843 | 2205 | 2757 | 2331 | 2642 | 1564 | 1941 | 1756 | 1886 | 1818 | 2358 | 2224 | 2358 | 2206 | 2068,3 | 1850 | 2286 | 1,236 | 0,306 | 6,4 | 1,55E-10 | 1,09E-09 | 0,000953 | 0,0026814 | 0,0016989 | 0,0016989 | TRUE | 0,0133603 | No |
| orf19.5443 | 4357 | 2804 | 3794 | 4734 | 6086 | 5043 | 5542 | 3193 | 3909 | 3570 | 3882 | 3904 | 5205 | 4812 | 4945 | 4503 | 4341,2 | 3816 | 4866 | 1,276 | 0,351 | 6,683 | 2,34E-11 | 1,84E-10 | 0,002416 | 0,0023139 | 0,0024165 | 0,0024165 | TRUE | 0,0297956 | No |
| orf19.5444 | 4043 | 2508 | 3365 | 4094 | 3661 | 3043 | 3342 | 2071 | 3627 | 3193 | 3443 | 3376 | 3131 | 2904 | 2982 | 2921 | 3197 | 3410 | 2984 | 0,875 | -0,192 | -3,963 | 7,39E-05 | 0,00023088 | 0,001563 | 0,0024336 | 0,0019475 | 0,0019475 | TRUE | 0,0224094 | No |
| orf19.5445 | 2623 | 1594 | 2255 | 2650 | 2890 | 2208 | 2669 | 1741 | 2353 | 2029 | 2307 | 2185 | 2472 | 2107 | 2382 | 2456 | 2286,3 | 2218 | 2354 | 1,06 | 0,084 | 1,287 | 0,19822009 | 0,27258099 | 0,004132 | 0,0026145 | 0,0036372 | 0,0036372 | TRUE | 0,0609749 | No |

| | | | | | | | | | | | | | | | | | | | | | | | | | | | | | | | |
|---|---|---|---|---|---|---|---|---|---|---|---|---|---|---|---|---|---|---|---|---|---|---|---|---|---|---|---|---|---|---|---|
| orf19.5446 | 2 | 0 | 1 | 2 | 0 | 2 | 2 | 4 | 2 | 0 | 1 | 2 | 0 | 2 | 2 | 6 | 1,73 | 1 | 2 | 1,842 | 0,881 | 0,75 | 0,45305844 | NA | 1,00E-08 | 0,8434735 | 0,6724933 | 0,6724933 | TRUE | 0,3929969 | No |
| orf19.5447 | 2021 | 1281 | 1844 | 2177 | 3376 | 2733 | 2974 | 2074 | 1813 | 1631 | 1887 | 1795 | 2887 | 2608 | 2654 | 2925 | 2275 | 1782 | 2768 | 1,552 | 0,634 | 10,819 | 2,79E-27 | 8,69E-26 | 0,002908 | 0,0026176 | 0,0028314 | 0,0028314 | TRUE | 0,0370358 | No |
| orf19.5449 | 99 | 55 | 77 | 78 | 60 | 51 | 54 | 44 | 89 | 70 | 79 | 64 | 51 | 49 | 48 | 62 | 64,02 | 76 | 52 | 0,688 | -0,539 | -2,951 | 0,00316875 | 0,00715708 | 1,00E-08 | 0,0246556 | 0,0158334 | 0,0158334 | TRUE | 0,1423396 | No |
| orf19.5450 | 1933 | 1219 | 1661 | 1933 | 2658 | 1985 | 2180 | 1502 | 1734 | 1552 | 1699 | 1594 | 2273 | 1894 | 1945 | 2118 | 1851,3 | 1645 | 2058 | 1,25 | 0,322 | 4,832 | 1,35E-06 | 5,58E-06 | 0,004161 | 0,0027637 | 0,0037184 | 0,0037184 | TRUE | 0,0625046 | No |
| orf19.5451 | 444 | 283 | 418 | 469 | 681 | 529 | 652 | 574 | 398 | 360 | 428 | 387 | 582 | 505 | 582 | 810 | 506,44 | 393 | 620 | 1,568 | 0,649 | 5,171 | 2,33E-07 | 1,08E-06 | 0,022361 | 0,004846 | 0,0130416 | 0,0130416 | TRUE | 0,4767899 | No |
| orf19.5454 | 503 | 364 | 483 | 552 | 603 | 469 | 520 | 363 | 451 | 463 | 494 | 455 | 516 | 448 | 464 | 512 | 475,4 | 466 | 485 | 1,039 | 0,056 | 0,745 | 0,45614864 | 0,54171596 | 0,001381 | 0,0050332 | 0,0032333 | 0,0032333 | TRUE | 0,031118 | No |
| orf19.5455 | 2276 | 1498 | 1896 | 2507 | 2806 | 2170 | 2244 | 1369 | 2042 | 1907 | 1940 | 2067 | 2400 | 2071 | 2002 | 1931 | 2045 | 1989 | 2101 | 1,057 | 0,08 | 1,168 | 0,24274771 | 0,32423927 | 0,004974 | 0,0026894 | 0,004001 | 0,004001 | TRUE | 0,1124139 | No |
| orf19.5457 | 2050 | 1346 | 1892 | 2286 | 2483 | 1887 | 2130 | 1363 | 1839 | 1714 | 1936 | 1885 | 2124 | 1801 | 1901 | 1922 | 1890,1 | 1844 | 1937 | 1,05 | 0,071 | 1,161 | 0,24565344 | 0,32753792 | 0,003066 | 0,0027476 | 0,0030382 | 0,0030382 | TRUE | 0,0526992 | No |
| orf19.5459 | 5389 | 3410 | 4766 | 5997 | 6804 | 4955 | 5986 | 3946 | 4834 | 4341 | 4876 | 4945 | 5819 | 4728 | 5342 | 5565 | 5056,4 | 4749 | 5364 | 1,129 | 0,175 | 2,604 | 0,00922213 | 0,01873742 | 0,005203 | 0,0022666 | 0,0041283 | 0,0041283 | TRUE | 0,0779597 | No |
| orf19.5461 | 38 | 16 | 27 | 27 | 51 | 23 | 28 | 24 | 34 | 20 | 28 | 22 | 44 | 22 | 25 | 34 | 28,59 | 26 | 31 | 1,184 | 0,244 | 0,819 | 0,41251817 | 0,50058759 | 0,04653 | 0,0527503 | 0,0499671 | 0,0499671 | TRUE | 0,5123542 | No |
| orf19.5462 | 6 | 1 | 2 | 6 | 5 | 1 | 2 | 0 | 5 | 1 | 2 | 5 | 4 | 1 | 2 | 0 | 2,58 | 3 | 2 | 0,536 | -0,9 | -0,968 | 0,33324558 | NA | 0,174241 | 0,5639845 | 0,4233982 | 0,4233982 | TRUE | 0,483325 | No |
| orf19.5463 | 1092 | 722 | 1005 | 1223 | 1533 | 1236 | 1466 | 816 | 980 | 919 | 1028 | 1009 | 1311 | 1179 | 1308 | 1151 | 1110,6 | 984 | 1237 | 1,259 | 0,332 | 5,297 | 1,18E-07 | 5,73E-07 | 0,002267 | 0,0032866 | 0,0028585 | 0,0028585 | TRUE | 0,0244075 | No |
| orf19.5464 | 94 | 66 | 107 | 105 | 103 | 76 | 103 | 65 | 84 | 84 | 109 | 87 | 88 | 73 | 92 | 92 | 88,58 | 91 | 86 | 0,943 | -0,085 | -0,543 | 0,58740353 | 0,66574616 | 1,00E-08 | 0,018369 | 0,012278 | 0,012278 | TRUE | 0,1728328 | No |
| orf19.5465 | 1223 | 693 | 1094 | 1323 | 1283 | 1053 | 1248 | 782 | 1097 | 882 | 1119 | 1091 | 1097 | 1005 | 1114 | 1103 | 1063,5 | 1047 | 1080 | 1,028 | 0,04 | 0,526 | 0,59861422 | 0,67514065 | 0,005816 | 0,0033445 | 0,0046556 | 0,0046556 | TRUE | 0,1273429 | No |
| orf19.5466 | 13339 | 7979 | 10745 | 13241 | 12074 | 9100 | 9664 | 6308 | 11966 | 10158 | 10993 | 10919 | 10326 | 8683 | 8624 | 8897 | 10071 | 11009 | 9132 | 0,83 | -0,27 | -3,984 | 6,77E-05 | 0,00021377 | 0,006014 | 0,0021237 | 0,0043002 | 0,0043002 | TRUE | 0,0951599 | No |
| orf19.5467 | 10 | 6 | 8 | 14 | 12 | 12 | 17 | 9 | 9 | 8 | 8 | 12 | 10 | 11 | 15 | 13 | 10,74 | 9 | 12 | 1,343 | 0,426 | 1,034 | 0,30114196 | NA | 1,00E-08 | 0,1371547 | 0,0686478 | 0,0686478 | TRUE | 0,1321383 | No |
| orf19.5468 | 127 | 79 | 111 | 126 | 94 | 80 | 77 | 44 | 114 | 101 | 114 | 104 | 80 | 76 | 69 | 62 | 89,93 | 108 | 72 | 0,67 | -0,577 | -3,889 | 0,0001005 | 0,00030821 | 1,00E-08 | 0,0181215 | 0,0096426 | 0,0096426 | TRUE | 0,0588939 | No |
| orf19.5469 | 316 | 223 | 336 | 365 | 363 | 321 | 300 | 197 | 283 | 284 | 344 | 301 | 310 | 306 | 268 | 278 | 296,8 | 303 | 290 | 0,96 | -0,058 | -0,613 | 0,53958097 | 0,62174961 | 0,003605 | 0,0068707 | 0,0053323 | 0,0053323 | TRUE | 0,0902092 | No |
| orf19.547 | 954 | 646 | 855 | 1079 | 1083 | 887 | 995 | 602 | 856 | 822 | 875 | 890 | 926 | 846 | 888 | 849 | 869,03 | 861 | 877 | 1,02 | 0,028 | 0,499 | 0,61771393 | 0,69097473 | 1,00E-08 | 0,00365 | 0,0019016 | 0,0019016 | TRUE | 0,017319 | No |
| orf19.5472 | 0 | 0 | 0 | 1 | 2 | 3 | 4 | 2 | 0 | 0 | 0 | 1 | 2 | 3 | 4 | 3 | 1,47 | 0 | 3 | 9,093 | 3,185 | 2,367 | 0,0179328 | NA | 1,00E-08 | 0,9872075 | 0,5546477 | 0,5546477 | TRUE | 0,2428654 | No |
| orf19.5474 | 244 | 158 | 232 | 241 | 359 | 275 | 274 | 141 | 219 | 201 | 237 | 199 | 307 | 262 | 244 | 199 | 233,62 | 214 | 253 | 1,191 | 0,252 | 1,986 | 0,04698498 | 0,07965173 | 0,014553 | 0,0081936 | 0,011203 | 0,011203 | TRUE | 0,2012886 | No |
| orf19.5475 | 18 | 9 | 14 | 18 | 15 | 14 | 22 | 20 | 16 | 11 | 14 | 15 | 13 | 13 | 20 | 28 | 16,35 | 14 | 18 | 1,257 | 0,329 | 0,912 | 0,3619221 | 0,45056526 | 0,021499 | 0,0907695 | 0,0629492 | 0,0629492 | TRUE | 0,458803 | No |
| orf19.5479 | 4 | 3 | 4 | 6 | 4 | 3 | 1 | 0 | 4 | 4 | 4 | 5 | 3 | 3 | 1 | 0 | 2,95 | 4 | 2 | 0,465 | -1,104 | -1,336 | 0,1814002 | NA | 1,00E-08 | 0,4935905 | 0,2755278 | 0,2755278 | TRUE | 0,3301769 | No |
| orf19.548 | 453 | 331 | 377 | 528 | 579 | 553 | 587 | 364 | 406 | 421 | 386 | 435 | 495 | 528 | 524 | 513 | 463,61 | 412 | 515 | 1,248 | 0,32 | 4,557 | 5,18E-06 | 1,97E-05 | 1,00E-08 | 0,0051108 | 0,0025558 | 0,0025558 | TRUE | 0,0214892 | No |
| orf19.5480 | 1805 | 1234 | 1622 | 1906 | 1810 | 1497 | 1718 | 1057 | 1619 | 1571 | 1659 | 1572 | 1548 | 1428 | 1533 | 1491 | 1552,7 | 1605 | 1500 | 0,935 | -0,097 | -2,093 | 0,03631691 | 0,06355724 | 0,000394 | 0,0029145 | 0,0014253 | 0,0014253 | TRUE | 0,0131841 | No |
| orf19.5483 | 1289 | 913 | 1108 | 1409 | 1524 | 1363 | 1598 | 922 | 1156 | 1162 | 1134 | 1162 | 1303 | 1301 | 1426 | 1300 | 1243,1 | 1154 | 1332 | 1,156 | 0,21 | 4,168 | 3,07E-05 | 0,00010335 | 0,000534 | 0,0031474 | 0,0016169 | 0,0016169 | TRUE | 0,0278177 | No |
| orf19.5484 | 2092 | 1280 | 1736 | 2114 | 2515 | 2300 | 2363 | 1492 | 1877 | 1630 | 1776 | 1743 | 2151 | 2195 | 2109 | 2104 | 1948 | 1756 | 2140 | 1,217 | 0,283 | 5,728 | 1,02E-08 | 5,60E-08 | 0,001186 | 0,0027247 | 0,0018262 | 0,0018262 | TRUE | 0,0265984 | No |
| orf19.5485 | 850 | 567 | 676 | 852 | 605 | 527 | 551 | 316 | 763 | 722 | 692 | 703 | 517 | 503 | 492 | 446 | 604,53 | 720 | 490 | 0,683 | -0,55 | -8,145 | 3,78E-16 | 4,86E-15 | 0,000861 | 0,0043809 | 0,0026638 | 0,0026638 | TRUE | 0,0350397 | No |
| orf19.5486 | 3809 | 2549 | 3484 | 3881 | 4142 | 3734 | 4158 | 2319 | 3417 | 3245 | 3564 | 3200 | 3542 | 3563 | 3710 | 3271 | 3439,2 | 3356 | 3522 | 1,05 | 0,071 | 1,354 | 0,17573554 | 0,24703302 | 0,002203 | 0,0024016 | 0,0023115 | 0,0023115 | TRUE | 0,0267242 | No |
| orf19.5486.1 | 1897 | 1382 | 1554 | 1987 | 2202 | 1891 | 1791 | 1056 | 1702 | 1759 | 1590 | 1639 | 1883 | 1804 | 1598 | 1489 | 1683,1 | 1672 | 1694 | 1,015 | 0,021 | 0,29 | 0,7718197 | 0,82174247 | 0,006059 | 0,002842 | 0,0046265 | 0,0046265 | TRUE | 0,0757078 | No |
| orf19.5487 | 5002 | 3352 | 4463 | 5321 | 4707 | 4159 | 4413 | 2724 | 4487 | 4267 | 4566 | 4388 | 4026 | 3968 | 3938 | 3842 | 4185,3 | 4427 | 3944 | 0,891 | -0,167 | -4,65 | 3,31E-06 | 1,30E-05 | 0,000295 | 0,0023264 | 0,0009904 | 0,0009904 | TRUE | 0,0067548 | No |
| orf19.5488 | 471 | 309 | 336 | 413 | 321 | 276 | 298 | 200 | 423 | 393 | 344 | 341 | 275 | 263 | 266 | 282 | 323,27 | 375 | 272 | 0,723 | -0,467 | -4,982 | 6,28E-07 | 2,73E-06 | 0,00407 | 0,0064703 | 0,0052512 | 0,0052512 | TRUE | 0,090731 | No |
| orf19.5489 | 246 | 163 | 226 | 264 | 137 | 150 | 151 | 98 | 221 | 208 | 231 | 218 | 117 | 143 | 135 | 138 | 176,3 | 220 | 133 | 0,605 | -0,725 | -6,62 | 3,60E-11 | 2,78E-10 | 1,00E-08 | 0,0102139 | 0,0055019 | 0,0055019 | TRUE | 0,0772081 | No |
| orf19.549 | 3205 | 2124 | 2673 | 3386 | 2955 | 2689 | 2748 | 2061 | 2875 | 2704 | 2735 | 2792 | 2527 | 2566 | 2452 | 2907 | 2694,8 | 2776 | 2613 | 0,939 | -0,091 | -1,624 | 0,10437016 | 0,15798839 | 0,002846 | 0,0025182 | 0,0026216 | 0,0026216 | TRUE | 0,0663414 | No |
| orf19.5491 | 50 | 49 | 65 | 64 | 66 | 107 | 84 | 64 | 45 | 62 | 67 | 53 | 56 | 102 | 75 | 90 | 68,78 | 57 | 81 | 1,429 | 0,515 | 2,411 | 0,01591498 | 0,03043195 | 0,035157 | 0,0230845 | 0,0287111 | 0,0287111 | TRUE | 0,2609368 | No |
| orf19.5491.1 | 3026 | 2060 | 2404 | 3134 | 3179 | 2682 | 2762 | 2170 | 2715 | 2623 | 2460 | 2584 | 2719 | 2559 | 2465 | 3061 | 2648 | 2596 | 2701 | 1,039 | 0,055 | 0,83 | 0,40648052 | 0,49465971 | 0,004942 | 0,0025277 | 0,0038973 | 0,0038973 | TRUE | 0,0943693 | No |
| orf19.5492 | 2493 | 1734 | 2052 | 2643 | 2253 | 1846 | 1926 | 1120 | 2236 | 2208 | 2099 | 2179 | 1927 | 1761 | 1719 | 1580 | 1963,7 | 2180 | 1747 | 0,803 | -0,317 | -5,333 | 9,65E-08 | 4,75E-07 | 0,003031 | 0,0027188 | 0,0028785 | 0,0028785 | TRUE | 0,0584757 | No |
| orf19.5493 | 10044 | 6871 | 8535 | 10486 | 9408 | 8328 | 8640 | 5398 | 9011 | 8748 | 8732 | 8647 | 8046 | 7946 | 7710 | 7613 | 8306,6 | 8784 | 7829 | 0,892 | -0,166 | -5,159 | 2,48E-07 | 1,15E-06 | 0,00033 | 0,0021543 | 0,0008669 | 0,0008669 | TRUE | 0,0042744 | No |
| orf19.5495 | 1801 | 1187 | 1563 | 1828 | 2099 | 1920 | 2177 | 1340 | 1616 | 1511 | 1599 | 1507 | 1795 | 1832 | 1943 | 1890 | 1711,6 | 1558 | 1865 | 1,196 | 0,258 | 5,428 | 5,68E-08 | 2,87E-07 | 0,000762 | 0,0028277 | 0,0015779 | 0,0015779 | TRUE | 0,010269 | No |
| orf19.5496 | 2351 | 1654 | 2148 | 2566 | 1897 | 1673 | 1959 | 1312 | 2109 | 2106 | 2198 | 2116 | 1622 | 1596 | 1748 | 1850 | 1918,2 | 2132 | 1704 | 0,798 | -0,326 | -6,117 | 9,54E-10 | 6,10E-09 | 0,001898 | 0,0027363 | 0,0021976 | 0,0021976 | TRUE | 0,0370633 | No |
| orf19.5498 | 317 | 235 | 321 | 383 | 282 | 260 | 299 | 173 | 284 | 299 | 328 | 316 | 241 | 248 | 267 | 244 | 278,48 | 307 | 250 | 0,815 | -0,295 | -3,319 | 0,00090272 | 0,00229035 | 1,00E-08 | 0,0071924 | 0,0039411 | 0,0039411 | TRUE | 0,030328 | No |
| orf19.5499 | 192 | 160 | 196 | 197 | 160 | 182 | 179 | 98 | 172 | 204 | 201 | 162 | 137 | 174 | 160 | 138 | 168,42 | 185 | 152 | 0,83 | -0,268 | -2,141 | 0,03226637 | 0,0571342 | 0,007206 | 0,010599 | 0,0090621 | 0,0090621 | TRUE | 0,0954244 | No |
| orf19.55 | 3492 | 3338 | 3373 | 4383 | 3222 | 3467 | 3505 | 2029 | 3133 | 4250 | 3451 | 3614 | 2756 | 3308 | 3128 | 2862 | 3312,6 | 3612 | 3014 | 0,835 | -0,26 | -2,947 | 0,00321023 | 0,00723989 | 0,011531 | 0,0024178 | 0,0071916 | 0,0071916 | TRUE | 0,1701382 | No |
| orf19.550 | 20151 | 12695 | 16945 | 21161 | 23973 | 20045 | 22250 | 14692 | 18078 | 16162 | 17336 | 17450 | 20502 | 19126 | 19854 | 20722 | 18654 | 17256 | 20051 | 1,162 | 0,216 | 4,925 | 8,44E-07 | 3,60E-06 | 0,001642 | 0,0020573 | 0,0017963 | 0,0017963 | TRUE | 0,0221138 | No |
| orf19.5500 | 20281 | 13662 | 17307 | 21185 | 17716 | 15895 | 17628 | 9014 | 18194 | 17393 | 17707 | 17469 | 15151 | 15167 | 15730 | 12713 | 16191 | 17691 | 14690 | 0,831 | -0,268 | -4,364 | 1,28E-05 | 4,58E-05 | 0,004318 | 0,0020692 | 0,0035532 | 0,0035532 | TRUE | 0,0995861 | No |
| orf19.5501 | 930 | 710 | 749 | 1018 | 1088 | 1050 | 1074 | 704 | 834 | 904 | 766 | 839 | 930 | 1002 | 958 | 993 | 903,45 | 836 | 971 | 1,162 | 0,216 | 3,546 | 0,00039088 | 0,00107994 | 0,00135 | 0,0035864 | 0,0024596 | 0,0024596 | TRUE | 0,0357195 | No |
| orf19.5502 | 356 | 296 | 372 | 446 | 336 | 397 | 420 | 334 | 319 | 377 | 381 | 368 | 287 | 379 | 375 | 471 | 369,57 | 361 | 378 | 1,042 | 0,059 | 0,462 | 0,64389322 | 0,71387903 | 0,020312 | 0,0059076 | 0,012964 | 0,012964 | TRUE | 0,23916 | No |
| orf19.5503 | 46 | 27 | 41 | 64 | 51 | 43 | 45 | 27 | 41 | 34 | 42 | 53 | 44 | 41 | 40 | 38 | 41,66 | 42 | 41 | 0,949 | -0,075 | -0,35 | 0,72610491 | 0,78545882 | 1,00E-08 | 0,0368296 | 0,0207472 | 0,0207472 | TRUE | 0,2221317 | No |
| orf19.5504 | 1720 | 1430 | 1678 | 2195 | 2270 | 2688 | 2726 | 1554 | 1543 | 1821 | 1717 | 1810 | 1941 | 2565 | 2433 | 2192 | 2002,6 | 1723 | 2283 | 1,326 | 0,407 | 4,715 | 2,42E-06 | 9,70E-06 | 0,009357 | 0,0027044 | 0,0066364 | 0,0066364 | TRUE | 0,1253483 | No |
| orf19.5505 | 1455 | 1154 | 1373 | 1675 | 2317 | 2231 | 2531 | 1360 | 1305 | 1469 | 1405 | 1381 | 1982 | 2129 | 2258 | 1918 | 1730,9 | 1390 | 2072 | 1,494 | 0,579 | 9,31 | 1,27E-20 | 2,41E-19 | 0,003096 | 0,0028182 | 0,0031079 | 0,0031079 | TRUE | 0,0447416 | No |
| orf19.5506 | 940 | 798 | 883 | 1059 | 1084 | 1194 | 1105 | 713 | 843 | 1016 | 903 | 873 | 927 | 1139 | 986 | 1006 | 961,73 | 909 | 1014 | 1,118 | 0,161 | 2,046 | 0,04071395 | 0,07010874 | 0,005955 | 0,003489 | 0,0048973 | 0,0048973 | TRUE | 0,0840813 | No |
| orf19.5507 | 7777 | 5735 | 7011 | 8575 | 9368 | 9190 | 9617 | 5654 | 6977 | 7301 | 7173 | 7071 | 8012 | 8769 | 8582 | 7974 | 7732,3 | 7130 | 8334 | 1,169 | 0,226 | 5,432 | 5,57E-08 | 2,82E-07 | 0,001177 | 0,0021673 | 0,0015285 | 0,0015285 | TRUE | 0,0150054 | No |
| orf19.5508 | 437 | 307 | 374 | 477 | 527 | 497 | 506 | 349 | 392 | 391 | 383 | 393 | 451 | 474 | 452 | 492 | 428,44 | 390 | 467 | 1,195 | 0,256 | 3,658 | 0,00025435 | 0,00072769 | 1,00E-08 | 0,0053678 | 0,0023732 | 0,0023732 | TRUE | 0,0127158 | No |
| orf19.551 | 17281 | 11797 | 15385 | 18522 | 18926 | 17083 | 18210 | 11867 | 15503 | 15019 | 15740 | 15273 | 16186 | 16300 | 16249 | 16737 | 15876 | 15384 | 16368 | 1,064 | 0,089 | 3,181 | 0,00146886 | 0,0035714 | 0,000235 | 0,002071 | 0,000688 | 0,000688 | TRUE | 0,0030329 | No |
| orf19.5510 | 2711 | 2040 | 2478 | 2931 | 2157 | 2075 | 2314 | 1261 | 2432 | 2597 | 2535 | 2417 | 1845 | 1980 | 2065 | 1779 | 2206,2 | 2495 | 1917 | 0,77 | -0,377 | -6,891 | 5,55E-12 | 4,65E-11 | 0,002164 | 0,0026375 | 0,0024151 | 0,0024151 | TRUE | 0,032604 | No |
| orf19.5513 | 1695 | 1260 | 1457 | 1786 | 2189 | 1966 | 2118 | 1531 | 1521 | 1604 | 1491 | 1473 | 1872 | 1876 | 1890 | 2159 | 1735,7 | 1522 | 1949 | 1,279 | 0,355 | 6,15 | 7,77E-10 | 5,02E-09 | 0,002538 | 0,0028159 | 0,0026164 | 0,0026164 | TRUE | 0,0597928 | No |
| orf19.5514 | 958 | 651 | 803 | 1021 | 793 | 748 | 765 | 488 | 859 | 829 | 822 | 842 | 678 | 714 | 683 | 688 | 764,32 | 838 | 691 | 0,824 | -0,28 | -5,126 | 2,95E-07 | 1,35E-06 | 1,00E-08 | 0,0038789 | 0,001547 | 0,001547 | TRUE | 0,0062861 | No |
| orf19.5515 | 2707 | 2034 | 2391 | 3013 | 2828 | 2756 | 2798 | 1789 | 2428 | 2589 | 2446 | 2485 | 2419 | 2630 | 2497 | 2523 | 2502,1 | 2487 | 2517 | 1,012 | 0,018 | 0,419 | 0,67485517 | 0,74014784 | 0,000621 | 0,0025597 | 0,0013345 | 0,0013345 | TRUE | 0,0114595 | No |
| orf19.5516 | 1349 | 1125 | 1277 | 1555 | 1274 | 1387 | 1474 | 884 | 1210 | 1432 | 1306 | 1282 | 1090 | 1323 | 1315 | 1247 | 1275,8 | 1308 | 1244 | 0,952 | -0,071 | -0,953 | 0,34060178 | 0,42765632 | 0,005253 | 0,0031174 | 0,0045521 | 0,0045521 | TRUE | 0,0871024 | No |

| | | | | | | | | | | | | | | | | | | | | | | | | | | | | | | | |
|---|---|---|---|---|---|---|---|---|---|---|---|---|---|---|---|---|---|---|---|---|---|---|---|---|---|---|---|---|---|---|---|
| orf19.5517 | 2350 | 1732 | 2084 | 2672 | 2123 | 1805 | 1901 | 1279 | 2108 | 2205 | 2132 | 2203 | 1816 | 1722 | 1696 | 1804 | 1960,9 | 2162 | 1760 | 0,813 | -0,298 | -6,932 | 4,16E-12 | 3,55E-11 | 0,000324 | 0,0027199 | 0,0012568 | 0,0012568 | TRUE | 0,0072608 | No |
| orf19.5518 | 2298 | 1827 | 1984 | 2720 | 2492 | 2431 | 2613 | 1700 | 2062 | 2326 | 2030 | 2243 | 2131 | 2320 | 2332 | 2398 | 2230,1 | 2165 | 2295 | 1,06 | 0,084 | 1,435 | 0,15129488 | 0,21720093 | 0,002813 | 0,0026305 | 0,0028159 | 0,0028159 | TRUE | 0,0288951 | No |
| orf19.5519 | 1740 | 1163 | 1426 | 1808 | 1282 | 1161 | 1245 | 725 | 1561 | 1481 | 1459 | 1491 | 1096 | 1108 | 1111 | 1023 | 1291,1 | 1498 | 1084 | 0,725 | -0,463 | -9,462 | 3,02E-21 | 5,94E-20 | 0,000213 | 0,0031039 | 0,0015058 | 0,0015058 | TRUE | 0,0151243 | No |
| orf19.552 | 0 | 3 | 1 | 3 | 4 | 2 | 1 | 2 | 0 | 4 | 1 | 2 | 3 | 2 | 1 | 3 | 2,04 | 2 | 2 | 1,273 | 0,349 | 0,344 | 0,73118048 | NA | 1,00E-08 | 0,7119154 | 0,4835151 | 0,4835151 | TRUE | 0,2963115 | No |
| orf19.5520 | 0 | 0 | 2 | 2 | 3 | 2 | 2 | 2 | 0 | 0 | 2 | 2 | 3 | 2 | 2 | 3 | 1,6 | 1 | 2 | 2,333 | 1,222 | 1,074 | 0,28285499 | NA | 1,00E-08 | 0,9110724 | 0,5076935 | 0,5076935 | TRUE | 0,2539724 | No |
| orf19.5521 | 3964 | 2728 | 3377 | 4326 | 4509 | 3933 | 4260 | 2598 | 3556 | 3473 | 3455 | 3567 | 3856 | 3753 | 3801 | 3664 | 3640,8 | 3513 | 3768 | 1,073 | 0,102 | 2,997 | 0,00273077 | 0,0062644 | 1,00E-08 | 0,0023783 | 0,0008366 | 0,0008366 | TRUE | 0,0038759 | No |
| orf19.5522 | 540 | 430 | 509 | 605 | 505 | 527 | 508 | 257 | 484 | 547 | 521 | 499 | 432 | 503 | 453 | 362 | 475,25 | 513 | 438 | 0,859 | -0,219 | -2,372 | 0,01770483 | 0,03331369 | 0,007468 | 0,0050341 | 0,0060876 | 0,0060876 | TRUE | 0,1347683 | No |
| orf19.5523 | 439 | 319 | 410 | 457 | 443 | 499 | 416 | 320 | 394 | 406 | 419 | 377 | 379 | 476 | 371 | 451 | 409,23 | 399 | 419 | 1,049 | 0,068 | 0,725 | 0,46866805 | 0,55471978 | 0,006674 | 0,005527 | 0,0061101 | 0,0061101 | TRUE | 0,1048909 | No |
| orf19.5524 | 42 | 25 | 47 | 40 | 49 | 58 | 56 | 37 | 38 | 32 | 48 | 33 | 42 | 55 | 50 | 52 | 43,75 | 38 | 50 | 1,317 | 0,398 | 1,832 | 0,06701211 | 0,10872502 | 1,00E-08 | 0,0351631 | 0,0220546 | 0,0220546 | TRUE | 0,2449788 | No |
| orf19.5525 | 1124 | 664 | 900 | 1072 | 1285 | 1065 | 1175 | 796 | 1008 | 845 | 921 | 884 | 1099 | 1016 | 1048 | 1123 | 993,1 | 914 | 1072 | 1,168 | 0,225 | 3,451 | 0,00055919 | 0,0014905 | 0,002561 | 0,0034413 | 0,0030536 | 0,0030536 | TRUE | 0,0575725 | No |
| orf19.5526 | 84 | 56 | 59 | 79 | 130 | 115 | 119 | 90 | 75 | 71 | 60 | 65 | 111 | 110 | 106 | 127 | 90,77 | 68 | 114 | 1,659 | 0,73 | 4,897 | 9,72E-07 | 4,11E-06 | 1,00E-08 | 0,017972 | 0,0096417 | 0,0096417 | TRUE | 0,0540532 | No |
| orf19.5527 | 235 | 174 | 225 | 269 | 335 | 327 | 326 | 238 | 211 | 222 | 230 | 222 | 287 | 312 | 291 | 336 | 263,68 | 221 | 306 | 1,379 | 0,464 | 5,083 | 3,71E-07 | 1,68E-06 | 1,00E-08 | 0,007485 | 0,0040944 | 0,0040944 | TRUE | 0,039782 | No |
| orf19.5528 | 701 | 505 | 620 | 762 | 1002 | 886 | 972 | 624 | 629 | 643 | 634 | 628 | 857 | 845 | 867 | 880 | 748,03 | 634 | 862 | 1,361 | 0,445 | 8,194 | 2,53E-16 | 3,28E-15 | 1,00E-08 | 0,0039202 | 0,0014619 | 0,0014619 | TRUE | 0,002002 | No |
| orf19.553 | 13 | 10 | 13 | 14 | 11 | 16 | 18 | 9 | 12 | 13 | 13 | 12 | 9 | 15 | 16 | 13 | 12,83 | 12 | 13 | 1,089 | 0,124 | 0,328 | 0,7430376 | NA | 1,00E-08 | 0,1150974 | 0,0588136 | 0,0588136 | TRUE | 0,2165441 | No |
| orf19.5530 | 5823 | 3823 | 5277 | 6205 | 6582 | 5652 | 6443 | 4001 | 5224 | 4867 | 5399 | 5117 | 5629 | 5393 | 5749 | 5643 | 5377,6 | 5152 | 5604 | 1,087 | 0,121 | 2,904 | 0,00368864 | 0,0081897 | 0,001044 | 0,0022495 | 0,0014723 | 0,0014723 | TRUE | 0,016219 | No |
| orf19.5531 | 1592 | 1048 | 1374 | 1718 | 2360 | 2050 | 2154 | 1444 | 1428 | 1334 | 1406 | 1417 | 2018 | 1956 | 1922 | 2037 | 1689,7 | 1396 | 1983 | 1,417 | 0,503 | 11,392 | 4,59E-30 | 1,64E-28 | 0,000154 | 0,0028386 | 0,0012626 | 0,0012626 | TRUE | 0,0097368 | No |
| orf19.5532 | 147 | 139 | 121 | 174 | 182 | 155 | 143 | 94 | 132 | 177 | 124 | 143 | 156 | 148 | 128 | 133 | 142,48 | 144 | 141 | 0,987 | -0,018 | -0,134 | 0,8930799 | 0,91924346 | 0,00885 | 0,0121683 | 0,0107308 | 0,0107308 | TRUE | 0,214088 | No |
| orf19.5533 | 1166 | 798 | 1039 | 1370 | 1586 | 1421 | 1579 | 961 | 1046 | 1016 | 1063 | 1130 | 1356 | 1356 | 1409 | 1355 | 1216,4 | 1064 | 1369 | 1,285 | 0,362 | 7,204 | 5,85E-13 | 5,53E-12 | 0,000343 | 0,0031729 | 0,0015902 | 0,0015902 | TRUE | 0,0219361 | No |
| orf19.5534 | 1595 | 1046 | 1346 | 1671 | 1593 | 1341 | 1370 | 935 | 1431 | 1332 | 1377 | 1378 | 1362 | 1280 | 1222 | 1319 | 1337,6 | 1380 | 1296 | 0,938 | -0,092 | -1,803 | 0,07139157 | 0,11459272 | 0,000815 | 0,0030648 | 0,0017571 | 0,0017571 | TRUE | 0,0183525 | No |
| orf19.5535 | 1021 | 687 | 966 | 1090 | 1008 | 930 | 1012 | 699 | 916 | 875 | 988 | 899 | 862 | 887 | 903 | 986 | 914,51 | 920 | 910 | 0,986 | -0,021 | -0,332 | 0,74018209 | 0,79667037 | 0,001621 | 0,0035669 | 0,0025889 | 0,0025889 | TRUE | 0,0339236 | No |
| orf19.5536 | 399 | 253 | 368 | 469 | 521 | 451 | 504 | 334 | 358 | 322 | 377 | 387 | 446 | 430 | 450 | 471 | 405 | 361 | 449 | 1,237 | 0,307 | 3,938 | 8,22E-05 | 0,00025439 | 0,000474 | 0,005564 | 0,003342 | 0,003342 | TRUE | 0,0519306 | No |
| orf19.5537 | 7536 | 4655 | 6643 | 8187 | 8943 | 7389 | 8145 | 5499 | 6761 | 5926 | 6796 | 6751 | 7648 | 7050 | 7268 | 7756 | 6994,6 | 6558 | 7430 | 1,132 | 0,179 | 3,315 | 0,00091558 | 0,00231614 | 0,002857 | 0,0021871 | 0,0026598 | 0,0026598 | TRUE | 0,0505885 | No |
| orf19.5539 | 647 | 495 | 633 | 744 | 644 | 622 | 694 | 420 | 580 | 630 | 648 | 614 | 551 | 593 | 619 | 592 | 603,46 | 618 | 589 | 0,954 | -0,068 | -1,026 | 0,30467461 | 0,39052541 | 0,000915 | 0,0043852 | 0,0025741 | 0,0025741 | TRUE | 0,0246958 | No |
| orf19.554 | 415 | 314 | 435 | 420 | 652 | 580 | 610 | 408 | 372 | 400 | 445 | 346 | 558 | 553 | 544 | 575 | 474,28 | 391 | 558 | 1,43 | 0,516 | 6,29 | 3,17E-10 | 2,17E-09 | 0,003202 | 0,0050404 | 0,0042813 | 0,0042813 | TRUE | 0,1038293 | No |
| orf19.5541 | 4159 | 2873 | 3722 | 4536 | 4160 | 3858 | 4315 | 2692 | 3731 | 3658 | 3808 | 3740 | 3558 | 3681 | 3850 | 3797 | 3727,9 | 3734 | 3722 | 0,996 | -0,006 | -0,159 | 0,87396182 | 0,90359019 | 0,000506 | 0,0023689 | 0,001124 | 0,001124 | TRUE | 0,0110312 | No |
| orf19.5542 | 58 | 45 | 55 | 77 | 67 | 55 | 82 | 58 | 52 | 57 | 56 | 63 | 57 | 52 | 73 | 82 | 61,73 | 57 | 66 | 1,141 | 0,19 | 0,998 | 0,31851073 | 0,40542638 | 0,008951 | 0,0254964 | 0,0186567 | 0,0186567 | TRUE | 0,1731399 | No |
| orf19.5543 | 1407 | 1029 | 1262 | 1565 | 1582 | 1472 | 1642 | 932 | 1262 | 1310 | 1291 | 1291 | 1353 | 1405 | 1465 | 1315 | 1336,4 | 1288 | 1384 | 1,077 | 0,107 | 2,169 | 0,03007941 | 0,05375285 | 0,000464 | 0,0030658 | 0,0015748 | 0,0015748 | TRUE | 0,0186965 | No |
| orf19.5544 | 6826 | 4824 | 6139 | 7441 | 7138 | 6360 | 6793 | 3988 | 6124 | 6141 | 6281 | 6136 | 6105 | 6069 | 6062 | 5625 | 6067,7 | 6170 | 5965 | 0,968 | -0,047 | -1,309 | 0,19049976 | 0,26353271 | 0,00058 | 0,0022188 | 0,001098 | 0,001098 | TRUE | 0,0170576 | No |
| orf19.5547 | 2317 | 1618 | 1997 | 2499 | 2470 | 2066 | 2220 | 1274 | 2079 | 2060 | 2043 | 2061 | 2112 | 1971 | 1981 | 1797 | 2013 | 2061 | 1965 | 0,956 | -0,065 | -1,271 | 0,20378529 | 0,27914775 | 0,001532 | 0,0027007 | 0,0020181 | 0,0020181 | TRUE | 0,0371904 | No |
| orf19.5548 | 626 | 461 | 563 | 726 | 720 | 690 | 710 | 457 | 562 | 587 | 576 | 599 | 616 | 658 | 634 | 645 | 609,43 | 581 | 638 | 1,098 | 0,134 | 2,213 | 0,02691493 | 0,04880921 | 1,00E-08 | 0,0043616 | 0,0018979 | 0,0018979 | TRUE | 0,0070315 | No |
| orf19.5549 | 37 | 22 | 33 | 39 | 28 | 32 | 38 | 25 | 33 | 28 | 34 | 32 | 24 | 31 | 34 | 35 | 31,35 | 32 | 31 | 0,958 | -0,061 | -0,251 | 0,80191252 | 0,84568903 | 1,00E-08 | 0,0482909 | 0,0255295 | 0,0255295 | TRUE | 0,179297 | No |
| orf19.555 | 2 | 3 | 1 | 4 | 2 | 1 | 2 | 3 | 2 | 4 | 1 | 3 | 2 | 1 | 2 | 4 | 2,33 | 2 | 2 | 0,832 | -0,265 | -0,291 | 0,77102853 | NA | 1,00E-08 | 0,6258436 | 0,3484094 | 0,3484094 | TRUE | 0,2484447 | No |
| orf19.5550 | 10947 | 8047 | 9157 | 11385 | 9470 | 9208 | 9334 | 4899 | 9821 | 10245 | 9369 | 9388 | 8099 | 8786 | 8329 | 6910 | 8868,2 | 9706 | 8031 | 0,828 | -0,272 | -4,007 | 6,15E-05 | 0,00019572 | 0,005729 | 0,0021432 | 0,0043256 | 0,0043256 | TRUE | 0,1065435 | No |
| orf19.5551 | 5244 | 3629 | 4805 | 5805 | 6529 | 5747 | 6293 | 3782 | 4704 | 4620 | 4916 | 4787 | 5584 | 5484 | 5615 | 5334 | 5130,6 | 4757 | 5504 | 1,158 | 0,211 | 6,048 | 1,46E-09 | 9,10E-09 | 0,000354 | 0,0022625 | 0,0009721 | 0,0009721 | TRUE | 0,0060293 | No |
| orf19.5552 | 3398 | 2350 | 2982 | 3446 | 4808 | 4416 | 4769 | 2861 | 3048 | 2992 | 3051 | 2842 | 4112 | 4214 | 4256 | 4035 | 3568,6 | 2983 | 4154 | 1,394 | 0,479 | 12,234 | 2,04E-34 | 9,60E-33 | 0,000559 | 0,0023863 | 0,0011834 | 0,0011834 | TRUE | 0,0130445 | No |
| orf19.5553 | 968 | 673 | 820 | 1037 | 1033 | 969 | 1052 | 603 | 868 | 857 | 839 | 855 | 883 | 925 | 939 | 850 | 877,06 | 855 | 899 | 1,054 | 0,076 | 1,407 | 0,15940374 | 0,22705018 | 1,00E-08 | 0,0036347 | 0,0016797 | 0,0016797 | TRUE | 0,0134451 | No |
| orf19.5555 | 2044 | 1427 | 1841 | 2157 | 2266 | 2032 | 2197 | 1396 | 1834 | 1817 | 1884 | 1779 | 1938 | 1939 | 1960 | 1969 | 1889,9 | 1828 | 1952 | 1,067 | 0,094 | 2,382 | 0,01721215 | 0,03258251 | 1,00E-08 | 0,0027477 | 0,0009721 | 0,0009721 | TRUE | 0,0050584 | No |
| orf19.5556 | 0 | 3 | 1 | 2 | 1 | 2 | 1 | 0 | 0 | 4 | 1 | 2 | 1 | 2 | 1 | 0 | 1,27 | 2 | 1 | 0,623 | -0,682 | -0,527 | 0,59848267 | NA | 1,00E-08 | 1,1464542 | 0,7692074 | 0,7692074 | TRUE | 0,352388 | No |
| orf19.5557 | 831 | 670 | 827 | 990 | 630 | 685 | 640 | 444 | 745 | 853 | 846 | 816 | 539 | 654 | 571 | 626 | 706,33 | 815 | 598 | 0,732 | -0,45 | -5,894 | 3,77E-09 | 2,22E-08 | 0,004235 | 0,0040348 | 0,0041401 | 0,0041401 | TRUE | 0,0534734 | No |
| orf19.5558 | 1239 | 811 | 1058 | 1350 | 1625 | 1262 | 1508 | 1082 | 1112 | 1032 | 1082 | 1113 | 1390 | 1204 | 1346 | 1526 | 1225,7 | 1085 | 1366 | 1,256 | 0,329 | 4,687 | 2,78E-06 | 1,10E-05 | 0,004214 | 0,0031639 | 0,0038889 | 0,0038889 | TRUE | 0,0760614 | No |
| orf19.5559 | 656 | 516 | 615 | 714 | 621 | 555 | 559 | 350 | 589 | 657 | 629 | 589 | 531 | 530 | 499 | 494 | 564,57 | 616 | 514 | 0,837 | -0,256 | -3,819 | 0,0001342 | 0,00040113 | 1,00E-08 | 0,0045509 | 0,0025266 | 0,0025266 | TRUE | 0,0217459 | No |
| orf19.556 | 1317 | 1077 | 1128 | 1653 | 995 | 1094 | 991 | 651 | 1181 | 1371 | 1154 | 1363 | 851 | 1044 | 884 | 918 | 1095,9 | 1267 | 924 | 0,729 | -0,455 | -5,557 | 2,74E-08 | 1,44E-07 | 0,007075 | 0,0033042 | 0,0055041 | 0,0055041 | TRUE | 0,092767 | No |
| orf19.5561 | 4448 | 3442 | 4313 | 5191 | 3914 | 3929 | 4145 | 2547 | 3990 | 4382 | 4413 | 4281 | 3347 | 3749 | 3699 | 3592 | 3931,6 | 4266 | 3597 | 0,843 | -0,246 | -4,88 | 1,06E-06 | 4,46E-06 | 0,00191 | 0,0023488 | 0,0021819 | 0,0021819 | TRUE | 0,0273289 | No |
| orf19.5563 | 1995 | 1363 | 1763 | 2157 | 1649 | 1344 | 1471 | 896 | 1790 | 1735 | 1804 | 1779 | 1410 | 1282 | 1313 | 1264 | 1547,1 | 1777 | 1317 | 0,742 | -0,43 | -8,912 | 5,00E-19 | 8,44E-18 | 0,000482 | 0,0029179 | 0,0015685 | 0,0015685 | TRUE | 0,0280641 | No |
| orf19.5564 | 557 | 408 | 507 | 662 | 437 | 422 | 427 | 267 | 500 | 519 | 519 | 546 | 374 | 403 | 381 | 377 | 452,22 | 521 | 384 | 0,736 | -0,442 | -6,314 | 2,73E-10 | 1,88E-09 | 1,00E-08 | 0,0051897 | 0,0024482 | 0,0024482 | TRUE | 0,01362 | No |
| orf19.5565 | 488 | 278 | 400 | 447 | 298 | 302 | 264 | 199 | 438 | 354 | 409 | 369 | 255 | 288 | 236 | 281 | 328,6 | 392 | 265 | 0,671 | -0,575 | -5,816 | 6,03E-09 | 3,45E-08 | 0,005998 | 0,0063973 | 0,0062024 | 0,0062024 | TRUE | 0,0713548 | No |
| orf19.5566 | 1583 | 1083 | 1328 | 1627 | 1986 | 1748 | 1947 | 1198 | 1420 | 1379 | 1359 | 1342 | 1698 | 1668 | 1737 | 1690 | 1536,6 | 1375 | 1698 | 1,236 | 0,306 | 7,081 | 1,43E-12 | 1,29E-11 | 1,00E-08 | 0,0029243 | 0,0011302 | 0,0011302 | TRUE | 0,0063938 | No |
| orf19.5567 | 2042 | 1359 | 1661 | 2091 | 2333 | 2020 | 2251 | 1339 | 1832 | 1730 | 1699 | 1724 | 1995 | 1927 | 2009 | 1889 | 1850,7 | 1746 | 1955 | 1,12 | 0,164 | 3,683 | 0,00023049 | 0,00066328 | 0,000441 | 0,0027639 | 0,0013587 | 0,0013587 | TRUE | 0,0135589 | No |
| orf19.5568 | 334 | 238 | 307 | 383 | 363 | 348 | 435 | 243 | 300 | 303 | 314 | 316 | 310 | 332 | 388 | 343 | 325,74 | 308 | 343 | 1,114 | 0,155 | 1,739 | 0,08195694 | 0,12917032 | 0,002676 | 0,0064361 | 0,0045642 | 0,0045642 | TRUE | 0,0963494 | No |
| orf19.5569 | 9768 | 6820 | 9019 | 10838 | 10795 | 9605 | 10421 | 6263 | 8763 | 8683 | 9227 | 8937 | 9232 | 9165 | 9299 | 8833 | 9017,4 | 8902 | 9132 | 1,026 | 0,037 | 1,112 | 0,26595923 | 0,35010584 | 0,000475 | 0,0021405 | 0,000963 | 0,000963 | TRUE | 0,0072954 | No |
| orf19.557 | 48 | 38 | 61 | 78 | 51 | 58 | 49 | 28 | 43 | 48 | 62 | 64 | 44 | 55 | 44 | 39 | 50,04 | 54 | 46 | 0,836 | -0,258 | -1,221 | 0,22214502 | 0,30113883 | 0,011666 | 0,0309887 | 0,0228555 | 0,0228555 | TRUE | 0,19343 | No |
| orf19.5572 | 459 | 322 | 359 | 466 | 412 | 378 | 398 | 228 | 412 | 410 | 367 | 384 | 352 | 361 | 355 | 322 | 370,38 | 393 | 348 | 0,888 | -0,172 | -2,203 | 0,02757661 | 0,04972397 | 1,00E-08 | 0,005899 | 0,0031593 | 0,0031593 | TRUE | 0,0231977 | No |
| orf19.5573 | 289 | 174 | 233 | 289 | 266 | 264 | 294 | 167 | 259 | 222 | 238 | 238 | 227 | 252 | 262 | 236 | 241,84 | 239 | 244 | 1,019 | 0,027 | 0,284 | 0,77672153 | 0,82491479 | 1,00E-08 | 0,0079821 | 0,0045884 | 0,0045884 | TRUE | 0,0349253 | No |
| orf19.5574 | 1169 | 868 | 1084 | 1361 | 1238 | 1139 | 1203 | 731 | 1049 | 1105 | 1109 | 1122 | 1059 | 1087 | 1073 | 1031 | 1079,4 | 1096 | 1062 | 0,971 | -0,043 | -0,872 | 0,38305718 | 0,47216162 | 1,00E-08 | 0,0033244 | 0,0014103 | 0,0014103 | TRUE | 0,0108233 | No |
| orf19.5575 | 1469 | 1210 | 1374 | 1746 | 1616 | 1734 | 1692 | 1163 | 1318 | 1540 | 1406 | 1440 | 1382 | 1655 | 1510 | 1640 | 1486,3 | 1426 | 1547 | 1,084 | 0,117 | 1,664 | 0,09608459 | 0,14771653 | 0,004587 | 0,0029562 | 0,0040482 | 0,0040482 | TRUE | 0,0630922 | No |
| orf19.5576 | 5498 | 4270 | 4936 | 6031 | 4774 | 4731 | 4807 | 3119 | 4932 | 5436 | 5050 | 4973 | 4083 | 4514 | 4289 | 4399 | 4709,7 | 5098 | 4321 | 0,848 | -0,238 | -5,071 | 3,96E-07 | 1,78E-06 | 0,001641 | 0,0022878 | 0,0019019 | 0,0019019 | TRUE | 0,0237607 | No |
| orf19.5578 | 551 | 402 | 481 | 619 | 464 | 430 | 477 | 321 | 494 | 512 | 492 | 510 | 397 | 410 | 426 | 453 | 461,77 | 502 | 422 | 0,836 | -0,259 | -3,712 | 0,00020569 | 0,00059844 | 1,00E-08 | 0,0051233 | 0,0024815 | 0,0024815 | TRUE | 0,0240167 | No |
| orf19.5579 | 272 | 251 | 238 | 291 | 306 | 330 | 322 | 222 | 244 | 320 | 243 | 240 | 262 | 315 | 287 | 313 | 278,01 | 262 | 294 | 1,128 | 0,173 | 1,561 | 0,1186022 | 0,17650781 | 0,00968 | 0,0072014 | 0,0082267 | 0,0082267 | TRUE | 0,2246113 | No |
| orf19.558 | 5761 | 4031 | 5184 | 6329 | 6477 | 5626 | 6252 | 3885 | 5168 | 5132 | 5304 | 5219 | 5539 | 5368 | 5579 | 5479 | 5348,6 | 5206 | 5491 | 1,055 | 0,077 | 2,535 | 0,01125722 | 0,02238034 | 1,00E-08 | 0,0022509 | 0,0007001 | 0,0007001 | TRUE | 0,0028878 | No |

| | | | | | | | | | | | | | | | | | | | | | | | | | | | | | | | |
|---|---|---|---|---|---|---|---|---|---|---|---|---|---|---|---|---|---|---|---|---|---|---|---|---|---|---|---|---|---|---|---|
| orf19.5580 | 610 | 415 | 568 | 694 | 707 | 626 | 696 | 365 | 547 | 528 | 581 | 572 | 605 | 597 | 621 | 515 | 570,85 | 557 | 584 | 1,052 | 0,074 | 1,02 | 0,30766647 | 0,3936244 | 0,001929 | 0,0045226 | 0,0032773 | 0,0032773 | TRUE | 0,0650144 | No |
| orf19.5584 | 563 | 429 | 496 | 618 | 611 | 615 | 629 | 371 | 505 | 546 | 507 | 510 | 523 | 587 | 561 | 523 | 532,78 | 517 | 548 | 1,064 | 0,09 | 1,318 | 0,18738193 | 0,26005789 | 1,00E-08 | 0,0047043 | 0,0026072 | 0,0026072 | TRUE | 0,0257906 | No |
| orf19.5585 | 1163 | 777 | 1016 | 1220 | 920 | 792 | 833 | 591 | 1043 | 989 | 1039 | 1006 | 787 | 756 | 743 | 834 | 899,68 | 1019 | 780 | 0,762 | -0,392 | -7,043 | 1,88E-12 | 1,68E-11 | 0,000152 | 0,0035931 | 0,0018384 | 0,0018384 | TRUE | 0,0216656 | No |
| orf19.5586 | 511 | 371 | 460 | 512 | 801 | 656 | 727 | 497 | 458 | 472 | 471 | 422 | 685 | 626 | 649 | 701 | 560,53 | 456 | 665 | 1,46 | 0,546 | 7,888 | 3,07E-15 | 3,59E-14 | 0,000903 | 0,0045694 | 0,0027526 | 0,0027526 | TRUE | 0,0310423 | No |
| orf19.5587 | 775 | 641 | 712 | 961 | 931 | 894 | 889 | 580 | 695 | 816 | 728 | 792 | 796 | 853 | 793 | 818 | 786,6 | 758 | 815 | 1,076 | 0,106 | 1,622 | 0,10484639 | 0,1586693 | 0,001981 | 0,0038251 | 0,0028463 | 0,0028463 | TRUE | 0,0377861 | No |
| orf19.5588 | 33 | 22 | 34 | 35 | 86 | 72 | 63 | 43 | 30 | 28 | 35 | 29 | 74 | 69 | 56 | 61 | 47,55 | 30 | 65 | 2,144 | 1,1 | 5,258 | 1,45E-07 | 6,98E-07 | 1,00E-08 | 0,0325116 | 0,018039 | 0,018039 | TRUE | 0,0829387 | No |
| orf19.5589 | 241 | 170 | 201 | 251 | 217 | 211 | 243 | 136 | 216 | 216 | 206 | 207 | 186 | 201 | 217 | 192 | 205,1 | 211 | 199 | 0,944 | -0,084 | -0,847 | 0,39717134 | 0,48598889 | 1,00E-08 | 0,0090575 | 0,004552 | 0,004552 | TRUE | 0,0433252 | No |
| orf19.559 | 2394 | 1831 | 2216 | 2577 | 2942 | 2859 | 3148 | 1886 | 2148 | 2331 | 2267 | 2125 | 2516 | 2728 | 2809 | 2660 | 2448 | 2218 | 2678 | 1,209 | 0,274 | 5,431 | 5,60E-08 | 2,83E-07 | 0,0015 | 0,0025725 | 0,0020268 | 0,0020268 | TRUE | 0,0211113 | No |
| orf19.5591 | 2684 | 2009 | 2430 | 3046 | 2676 | 2392 | 2503 | 1782 | 2408 | 2558 | 2486 | 2512 | 2289 | 2282 | 2234 | 2513 | 2410,2 | 2491 | 2330 | 0,934 | -0,099 | -2,101 | 0,03567245 | 0,06252501 | 0,001204 | 0,0025818 | 0,0017062 | 0,0017062 | TRUE | 0,0317424 | No |
| orf19.5592 | 22 | 24 | 16 | 27 | 20 | 19 | 23 | 8 | 20 | 31 | 16 | 22 | 17 | 18 | 21 | 11 | 19,5 | 22 | 17 | 0,778 | -0,363 | -1,12 | 0,26254203 | 0,34621434 | 1,00E-08 | 0,0764421 | 0,0489555 | 0,0489555 | TRUE | 0,2814065 | No |
| orf19.5595 | 535 | 372 | 492 | 583 | 750 | 667 | 729 | 486 | 480 | 474 | 503 | 481 | 641 | 636 | 651 | 685 | 568,94 | 484 | 653 | 1,345 | 0,428 | 6,841 | 7,89E-12 | 6,51E-11 | 1,00E-08 | 0,0045311 | 0,0019671 | 0,0019671 | TRUE | 0,01078 | No |
| orf19.5596 | 1588 | 1191 | 1413 | 1706 | 1744 | 1591 | 1619 | 1175 | 1425 | 1516 | 1446 | 1407 | 1492 | 1518 | 1445 | 1657 | 1488,1 | 1448 | 1528 | 1,054 | 0,075 | 1,404 | 0,16041957 | 0,22828062 | 0,001442 | 0,0029551 | 0,0020829 | 0,0020829 | TRUE | 0,0358157 | No |
| orf19.5597 | 3777 | 3059 | 3530 | 4265 | 3895 | 3773 | 4036 | 2318 | 3388 | 3894 | 3612 | 3517 | 3331 | 3600 | 3601 | 3269 | 3526,7 | 3603 | 3450 | 0,959 | -0,06 | -1,104 | 0,26951332 | 0,35385402 | 0,002691 | 0,0023912 | 0,0025869 | 0,0025869 | TRUE | 0,0352784 | No |
| orf19.5597.1 | 1245 | 909 | 1002 | 1363 | 1238 | 1097 | 1089 | 900 | 1117 | 1157 | 1025 | 1124 | 1059 | 1047 | 972 | 1269 | 1096,2 | 1106 | 1087 | 0,98 | -0,029 | -0,371 | 0,71050091 | 0,77124555 | 0,006689 | 0,0033038 | 0,0051069 | 0,0051069 | TRUE | 0,1444892 | No |
| orf19.5599 | 2717 | 1937 | 2549 | 3183 | 3638 | 3169 | 3542 | 2221 | 2437 | 2466 | 2608 | 2625 | 3111 | 3024 | 3161 | 3133 | 2820,5 | 2534 | 3107 | 1,225 | 0,293 | 7,179 | 7,03E-13 | 6,56E-12 | 0,000538 | 0,0024942 | 0,0012422 | 0,0012422 | TRUE | 0,0085589 | No |
| orf19.56 | 1509 | 1144 | 1380 | 1752 | 2318 | 2141 | 2291 | 1440 | 1354 | 1456 | 1412 | 1445 | 1982 | 2043 | 2044 | 2031 | 1720,9 | 1417 | 2025 | 1,43 | 0,516 | 12,038 | 2,24E-33 | 9,98E-32 | 1,00E-08 | 0,0028231 | 0,0011659 | 0,0011659 | TRUE | 0,0111359 | No |
| orf19.5601 | 1124 | 774 | 991 | 1275 | 1288 | 1141 | 1272 | 879 | 1008 | 985 | 1014 | 1051 | 1102 | 1089 | 1135 | 1240 | 1078 | 1014 | 1142 | 1,12 | 0,164 | 2,932 | 0,00336303 | 0,00754203 | 0,000924 | 0,0033262 | 0,0020573 | 0,0020573 | TRUE | 0,0360538 | No |
| orf19.5602 | 3086 | 2046 | 2738 | 3297 | 2277 | 2055 | 2197 | 1428 | 2768 | 2605 | 2801 | 2719 | 1947 | 1961 | 1960 | 2014 | 2347 | 2723 | 1970 | 0,722 | -0,469 | -11,8 | 3,92E-32 | 1,61E-30 | 0,000154 | 0,0025981 | 0,0010777 | 0,0010777 | TRUE | 0,0096219 | No |
| orf19.5604 | 2523 | 1786 | 2385 | 2907 | 1404 | 1236 | 1370 | 847 | 2263 | 2274 | 2440 | 2397 | 1201 | 1179 | 1222 | 1195 | 1771,5 | 2344 | 1199 | 0,512 | -0,967 | -21,229 | 5,20E-100 | 4,46E-97 | 1,00E-08 | 0,002799 | 0,0013588 | 0,0013588 | TRUE | 0,0090211 | No |
| orf19.5605 | 10157 | 6803 | 9301 | 11206 | 14609 | 12284 | 14029 | 9124 | 9112 | 8661 | 9516 | 9241 | 12494 | 11721 | 12519 | 12869 | 10766 | 9132 | 12401 | 1,357 | 0,441 | 10,334 | 4,94E-25 | 1,25E-23 | 0,001398 | 0,0021144 | 0,0016517 | 0,0016517 | TRUE | 0,0166395 | No |
| orf19.5606 | 79 | 55 | 61 | 87 | 55 | 53 | 55 | 43 | 71 | 70 | 62 | 72 | 47 | 51 | 49 | 61 | 60,3 | 69 | 52 | 0,744 | -0,426 | -2,411 | 0,01590298 | 0,03041869 | 1,00E-08 | 0,0260553 | 0,0130847 | 0,0130847 | TRUE | 0,0787862 | No |
| orf19.5607 | 8184 | 5553 | 6922 | 8442 | 10441 | 9627 | 10041 | 6245 | 7342 | 7070 | 7082 | 6961 | 8929 | 9186 | 8960 | 8808 | 8042,2 | 7114 | 8971 | 1,261 | 0,335 | 10,748 | 6,03E-27 | 1,86E-25 | 0,000244 | 0,00216 | 0,0008056 | 0,0008056 | TRUE | 0,0058321 | No |
| orf19.5608 | 2974 | 2409 | 2698 | 3318 | 3616 | 3465 | 3508 | 1902 | 2668 | 3067 | 2760 | 2736 | 3093 | 3306 | 3130 | 2683 | 2930,4 | 2808 | 3053 | 1,089 | 0,123 | 1,812 | 0,06999344 | 0,11283086 | 0,005268 | 0,0024749 | 0,0040982 | 0,0040982 | TRUE | 0,0783135 | No |
| orf19.5609 | 2284 | 1659 | 1994 | 2561 | 2229 | 1848 | 2033 | 1299 | 2049 | 2112 | 2040 | 2112 | 1906 | 1763 | 1814 | 1832 | 1953,6 | 2078 | 1829 | 0,88 | -0,184 | -4,354 | 1,34E-05 | 4,77E-05 | 0,000244 | 0,0027226 | 0,0011945 | 0,0011945 | TRUE | 0,0105679 | No |
| orf19.5610 | 2744 | 2223 | 2435 | 3191 | 3599 | 3539 | 3692 | 2163 | 2462 | 2830 | 2491 | 2631 | 3078 | 3377 | 3294 | 3051 | 2901,8 | 2604 | 3200 | 1,231 | 0,299 | 5,3 | 1,16E-07 | 5,64E-07 | 0,002881 | 0,0024798 | 0,0027148 | 0,0027148 | TRUE | 0,0401194 | No |
| orf19.5611 | 167 | 133 | 152 | 163 | 236 | 236 | 240 | 118 | 150 | 169 | 156 | 134 | 202 | 225 | 214 | 166 | 177,08 | 152 | 202 | 1,343 | 0,425 | 3,502 | 0,00046104 | 0,00125622 | 0,00601 | 0,0101773 | 0,0084014 | 0,0084014 | TRUE | 0,1249134 | No |
| orf19.5612 | 1109 | 864 | 1042 | 1316 | 1036 | 1070 | 1138 | 602 | 995 | 1100 | 1066 | 1085 | 886 | 1021 | 1015 | 849 | 1002,2 | 1062 | 943 | 0,891 | -0,167 | -2,351 | 0,01870443 | 0,03499726 | 0,004013 | 0,003428 | 0,0038346 | 0,0038346 | TRUE | 0,0483153 | No |
| orf19.5614 | 6117 | 3796 | 5134 | 6328 | 7951 | 6518 | 7279 | 4557 | 5488 | 4833 | 5253 | 5218 | 6800 | 6219 | 6495 | 6427 | 5841,6 | 5198 | 6485 | 1,247 | 0,319 | 6,716 | 1,86E-11 | 1,48E-10 | 0,001817 | 0,002228 | 0,001989 | 0,001989 | TRUE | 0,026574 | No |
| orf19.5615 | 1750 | 1274 | 1537 | 1951 | 1827 | 1525 | 1615 | 1061 | 1570 | 1622 | 1573 | 1609 | 1563 | 1455 | 1441 | 1496 | 1541,1 | 1594 | 1489 | 0,935 | -0,097 | -2,143 | 0,0320962 | 0,05688318 | 1,00E-08 | 0,0029215 | 0,0013274 | 0,0013274 | TRUE | 0,0146574 | No |
| orf19.5616 | 66 | 40 | 58 | 49 | 42 | 40 | 39 | 35 | 59 | 51 | 59 | 40 | 36 | 38 | 35 | 49 | 46,02 | 52 | 40 | 0,745 | -0,425 | -1,95 | 0,05114134 | 0,0857145 | 1,00E-08 | 0,033527 | 0,0232251 | 0,0232251 | TRUE | 0,2362775 | No |
| orf19.5617 | 4088 | 2790 | 3480 | 4349 | 4949 | 4286 | 4572 | 2695 | 3667 | 3552 | 3560 | 3586 | 4233 | 4090 | 4080 | 3801 | 3821,1 | 3591 | 4051 | 1,129 | 0,175 | 4,283 | 1,84E-05 | 6,45E-05 | 0,00075 | 0,0023594 | 0,0013406 | 0,0013406 | TRUE | 0,0197239 | No |
| orf19.5618 | 1320 | 1074 | 1192 | 1453 | 1455 | 1489 | 1539 | 980 | 1184 | 1367 | 1220 | 1198 | 1244 | 1421 | 1373 | 1382 | 1298,7 | 1242 | 1355 | 1,092 | 0,127 | 2,008 | 0,04468347 | 0,0762449 | 0,002891 | 0,0030973 | 0,0030722 | 0,0030722 | TRUE | 0,0522035 | No |
| orf19.5619 | 1865 | 1293 | 1663 | 2003 | 1628 | 1463 | 1536 | 966 | 1673 | 1646 | 1701 | 1652 | 1392 | 1396 | 1371 | 1362 | 1524,2 | 1668 | 1380 | 0,828 | -0,272 | -6,578 | 4,76E-11 | 3,64E-10 | 1,00E-08 | 0,002932 | 0,0009856 | 0,0009856 | TRUE | 0,0021184 | No |
| orf19.562 | 718 | 513 | 611 | 807 | 698 | 617 | 730 | 452 | 644 | 653 | 625 | 665 | 597 | 589 | 651 | 638 | 632,8 | 647 | 619 | 0,955 | -0,067 | -1,07 | 0,2844001 | 0,36867082 | 1,00E-08 | 0,0042736 | 0,0021754 | 0,0021754 | TRUE | 0,017255 | No |
| orf19.5620 | 572 | 403 | 492 | 573 | 795 | 777 | 751 | 485 | 513 | 513 | 503 | 473 | 680 | 741 | 670 | 684 | 597,2 | 500 | 694 | 1,389 | 0,474 | 7,245 | 4,33E-13 | 4,19E-12 | 1,00E-08 | 0,0044104 | 0,0023882 | 0,0023882 | TRUE | 0,026011 | No |
| orf19.5621 | 553 | 553 | 568 | 772 | 681 | 730 | 763 | 434 | 496 | 704 | 581 | 637 | 582 | 697 | 681 | 612 | 623,72 | 604 | 643 | 1,067 | 0,093 | 0,9 | 0,36825365 | 0,45683831 | 0,012773 | 0,004307 | 0,0086296 | 0,0086296 | TRUE | 0,1736824 | No |
| orf19.5622 | 3483 | 2607 | 3095 | 3906 | 3019 | 2963 | 3198 | 1763 | 3125 | 3319 | 3166 | 3221 | 2582 | 2827 | 2854 | 2487 | 2947,6 | 3208 | 2688 | 0,839 | -0,253 | -4,838 | 1,31E-06 | 5,42E-06 | 0,002164 | 0,002472 | 0,0022873 | 0,0022873 | TRUE | 0,0288616 | No |
| orf19.5623 | 1783 | 1333 | 1506 | 1976 | 1560 | 1568 | 1648 | 999 | 1600 | 1697 | 1541 | 1629 | 1334 | 1496 | 1471 | 1409 | 1522,1 | 1617 | 1428 | 0,884 | -0,179 | -3,35 | 0,00080715 | 0,00207238 | 0,00145 | 0,0029333 | 0,0020659 | 0,0020659 | TRUE | 0,0251026 | No |
| orf19.5625 | 739 | 621 | 754 | 937 | 703 | 763 | 767 | 438 | 663 | 791 | 771 | 773 | 601 | 728 | 684 | 618 | 703,63 | 750 | 658 | 0,88 | -0,185 | -2,26 | 0,02381966 | 0,04368237 | 0,005811 | 0,0040427 | 0,0049796 | 0,0049796 | TRUE | 0,0731503 | No |
| orf19.5626 | 18174 | 13652 | 16323 | 21157 | 27896 | 26829 | 28764 | 19222 | 16304 | 17380 | 16700 | 17446 | 23858 | 25599 | 25667 | 27111 | 21258 | 16958 | 25559 | 1,507 | 0,592 | 13,075 | 4,59E-39 | 2,96E-37 | 0,001819 | 0,0020478 | 0,0019182 | 0,0019182 | TRUE | 0,0246161 | No |
| orf19.5627 | 10748 | 7849 | 9674 | 12337 | 10552 | 10280 | 11084 | 6798 | 9642 | 9993 | 9897 | 10173 | 9024 | 9809 | 9891 | 9588 | 9752,2 | 9926 | 9578 | 0,965 | -0,052 | -1,324 | 0,18556601 | 0,25795479 | 0,000943 | 0,0021284 | 0,0013586 | 0,0013586 | TRUE | 0,0188489 | No |
| orf19.5628 | 1860 | 1288 | 1510 | 1947 | 1875 | 1727 | 1822 | 1156 | 1669 | 1640 | 1545 | 1606 | 1604 | 1648 | 1626 | 1630 | 1620,8 | 1615 | 1627 | 1,007 | 0,011 | 0,245 | 0,80638344 | 0,84980735 | 1,00E-08 | 0,0028752 | 0,0011873 | 0,0011873 | TRUE | 0,0100466 | No |
| orf19.5629 | 2517 | 1845 | 2006 | 2775 | 2815 | 2579 | 2543 | 1579 | 2258 | 2349 | 2052 | 2288 | 2407 | 2461 | 2269 | 2227 | 2289 | 2237 | 2341 | 1,048 | 0,067 | 1,229 | 0,21919371 | 0,29794476 | 0,002195 | 0,0026137 | 0,0024367 | 0,0024367 | TRUE | 0,0369702 | No |
| orf19.563 | 9106 | 5968 | 7350 | 9151 | 8292 | 6797 | 7622 | 4362 | 8169 | 7598 | 7520 | 7546 | 7092 | 6485 | 6801 | 6152 | 7170,4 | 7708 | 6632 | 0,861 | -0,216 | -4,221 | 2,43E-05 | 8,32E-05 | 0,002479 | 0,002182 | 0,0023747 | 0,0023747 | TRUE | 0,0283645 | No |
| orf19.5630 | 1541 | 1176 | 1420 | 1892 | 1680 | 1761 | 1795 | 1096 | 1382 | 1497 | 1453 | 1560 | 1437 | 1680 | 1602 | 1546 | 1519,7 | 1473 | 1566 | 1,063 | 0,088 | 1,439 | 0,15005701 | 0,21587055 | 0,002946 | 0,0029348 | 0,0029448 | 0,0029448 | TRUE | 0,0392469 | No |
| orf19.5633 | 5657 | 3755 | 5231 | 6102 | 6258 | 5481 | 5717 | 3714 | 5075 | 4781 | 5352 | 5032 | 5352 | 5230 | 5101 | 5238 | 5145,1 | 5060 | 5230 | 1,033 | 0,047 | 1,145 | 0,25213756 | 0,33447614 | 0,001002 | 0,0022617 | 0,0014433 | 0,0014433 | TRUE | 0,018083 | No |
| orf19.5634 | 4826 | 3180 | 4190 | 5162 | 4214 | 3498 | 3830 | 2331 | 4329 | 4048 | 4287 | 4257 | 3604 | 3338 | 3418 | 3288 | 3821 | 4230 | 3412 | 0,807 | -0,31 | -7,329 | 2,32E-13 | 2,28E-12 | 0,000911 | 0,0023594 | 0,0014483 | 0,0014483 | TRUE | 0,0177486 | No |
| orf19.5635 | 2775 | 1708 | 2312 | 2798 | 3366 | 2570 | 2830 | 2270 | 2489 | 2174 | 2365 | 2307 | 2879 | 2452 | 2525 | 3202 | 2549,3 | 2334 | 2764 | 1,183 | 0,242 | 2,975 | 0,00292739 | 0,00666961 | 0,00887 | 0,002549 | 0,00596 | 0,00596 | TRUE | 0,1343236 | No |
| orf19.5636 | 17862 | 11010 | 15847 | 19376 | 23644 | 18043 | 19957 | 15914 | 16024 | 14017 | 16213 | 15978 | 20221 | 17216 | 17808 | 22445 | 17490 | 15558 | 19422 | 1,248 | 0,32 | 3,946 | 7,96E-05 | 0,00024685 | 0,009745 | 0,0020625 | 0,0062525 | 0,0062525 | TRUE | 0,1338963 | No |
| orf19.5639 | 4343 | 2884 | 3964 | 4785 | 5998 | 5118 | 5728 | 3708 | 3896 | 3672 | 4056 | 3946 | 5130 | 4883 | 5111 | 5230 | 4490,4 | 3892 | 5088 | 1,306 | 0,385 | 9,241 | 2,44E-20 | 4,55E-19 | 0,000972 | 0,0023028 | 0,0014412 | 0,0014412 | TRUE | 0,0169092 | No |
| orf19.564 | 306 | 206 | 262 | 314 | 435 | 401 | 492 | 268 | 275 | 262 | 268 | 259 | 372 | 383 | 439 | 378 | 329,43 | 266 | 393 | 1,48 | 0,565 | 6,573 | 4,93E-11 | 3,76E-10 | 1,00E-08 | 0,0063863 | 0,0039605 | 0,0039605 | TRUE | 0,0758928 | No |
| orf19.5640 | 12495 | 7967 | 10946 | 13283 | 13546 | 10993 | 12318 | 8335 | 11209 | 10143 | 11199 | 10953 | 11585 | 10489 | 10992 | 11756 | 11041 | 10876 | 11206 | 1,03 | 0,042 | 0,859 | 0,39034551 | 0,47929611 | 0,00223 | 0,002111 | 0,0022552 | 0,0022552 | TRUE | 0,0248886 | No |
| orf19.5641 | 16480 | 10417 | 14349 | 17622 | 17672 | 13823 | 15433 | 9742 | 14784 | 13262 | 14680 | 14531 | 15114 | 13189 | 13771 | 13740 | 14134 | 14314 | 13954 | 0,975 | -0,037 | -0,703 | 0,48234366 | 0,56778278 | 0,00263 | 0,0020822 | 0,0025838 | 0,0025838 | TRUE | 0,0385334 | No |
| orf19.5642 | 7044 | 4347 | 5635 | 7152 | 7754 | 6023 | 6412 | 4436 | 6319 | 5534 | 5765 | 5898 | 6631 | 5747 | 5722 | 6257 | 5984,1 | 5879 | 6089 | 1,035 | 0,05 | 0,837 | 0,40250696 | 0,49095622 | 0,003774 | 0,0022221 | 0,0032719 | 0,0032719 | TRUE | 0,0446082 | No |
| orf19.5643 | 986 | 727 | 956 | 1082 | 980 | 835 | 943 | 600 | 885 | 926 | 978 | 892 | 838 | 797 | 841 | 846 | 875,37 | 920 | 830 | 0,903 | -0,147 | -2,602 | 0,00926562 | 0,01881942 | 1,00E-08 | 0,0036379 | 0,0019046 | 0,0019046 | TRUE | 0,0219242 | No |
| orf19.5644 | 270 | 242 | 264 | 339 | 274 | 251 | 275 | 187 | 242 | 308 | 270 | 280 | 234 | 239 | 245 | 264 | 260,36 | 275 | 246 | 0,894 | -0,162 | -1,671 | 0,09468117 | 0,14627012 | 0,002153 | 0,0075552 | 0,0051434 | 0,0051434 | TRUE | 0,0735158 | No |
| orf19.5645 | 4969 | 3191 | 4326 | 5333 | 3024 | 2384 | 2672 | 1892 | 4458 | 4062 | 4426 | 4398 | 2586 | 2275 | 2384 | 2668 | 3407,2 | 4336 | 2478 | 0,571 | -0,809 | -14,117 | 2,97E-45 | 2,41E-43 | 0,003139 | 0,0024056 | 0,0028331 | 0,0028331 | TRUE | 0,0368766 | No |
| orf19.5646 | 596 | 400 | 570 | 670 | 407 | 311 | 342 | 233 | 535 | 509 | 583 | 552 | 348 | 297 | 305 | 329 | 432,28 | 545 | 320 | 0,586 | -0,772 | -9,679 | 3,70E-22 | 7,84E-21 | 0,001675 | 0,0053378 | 0,0036252 | 0,0036252 | TRUE | 0,0471758 | No |
| orf19.5647 | 5484 | 3691 | 4898 | 5805 | 5038 | 4306 | 4677 | 2995 | 4920 | 4699 | 5011 | 4787 | 4309 | 4109 | 4173 | 4224 | 4529 | 4854 | 4204 | 0,866 | -0,208 | -5,834 | 5,42E-09 | 3,13E-08 | 0,000313 | 0,0023001 | 0,0009969 | 0,0009969 | TRUE | 0,005637 | No |

| | | | | | | | | | | | | | | | | | | | | | | | | | | | | | | | |
|---|---|---|---|---|---|---|---|---|---|---|---|---|---|---|---|---|---|---|---|---|---|---|---|---|---|---|---|---|---|---|---|
| orf19.5648 | 787 | 496 | 611 | 817 | 531 | 454 | 444 | 279 | 706 | 631 | 625 | 674 | 454 | 433 | 396 | 394 | 539,17 | 659 | 419 | 0,637 | -0,651 | -8,705 | 3,18E-18 | 5,04E-17 | 0,002161 | 0,004672 | 0,0034166 | 0,0034166 | TRUE | 0,037668 | No |
| orf19.5649 | 63 | 48 | 59 | 91 | 45 | 42 | 30 | 27 | 57 | 61 | 60 | 75 | 38 | 40 | 27 | 38 | 49,56 | 63 | 36 | 0,562 | -0,831 | -3,956 | 7,61E-05 | 0,0002375 | 1,00E-08 | 0,0312742 | 0,0205378 | 0,0205378 | TRUE | 0,2385574 | No |
| orf19.5650 | 7412 | 4828 | 6286 | 7898 | 6629 | 5221 | 5722 | 3631 | 6649 | 6147 | 6431 | 6513 | 5669 | 4982 | 5106 | 5121 | 5827,3 | 6435 | 5220 | 0,811 | -0,302 | -6,185 | 6,21E-10 | 4,07E-09 | 0,002035 | 0,0022286 | 0,0021146 | 0,0021146 | TRUE | 0,0417217 | No |
| orf19.5651 | 6220 | 4185 | 5708 | 6920 | 7528 | 6128 | 6905 | 4362 | 5580 | 5328 | 5840 | 5706 | 6438 | 5847 | 6162 | 6152 | 5881,7 | 5614 | 6150 | 1,095 | 0,131 | 3,014 | 0,00258193 | 0,00594794 | 0,001297 | 0,0022263 | 0,0016504 | 0,0016504 | TRUE | 0,0138791 | No |
| orf19.5653 | 12347 | 7591 | 10480 | 12835 | 13174 | 10753 | 11642 | 7883 | 11077 | 9664 | 10722 | 10584 | 11267 | 10260 | 10389 | 11118 | 10635 | 10512 | 10758 | 1,023 | 0,033 | 0,637 | 0,52436457 | 0,60759543 | 0,002407 | 0,002116 | 0,0024772 | 0,0024772 | TRUE | 0,0356266 | No |
| orf19.5654 | 2941 | 2113 | 2559 | 3293 | 2799 | 2492 | 2610 | 1758 | 2638 | 2690 | 2618 | 2715 | 2394 | 2378 | 2329 | 2480 | 2530,3 | 2665 | 2395 | 0,897 | -0,156 | -4,141 | 3,45E-05 | 0,0001149 | 1,00E-08 | 0,0025533 | 0,0009682 | 0,0009682 | TRUE | 0,006099 | No |
| orf19.5655 | 202 | 153 | 214 | 265 | 135 | 166 | 123 | 74 | 181 | 195 | 219 | 219 | 115 | 158 | 110 | 104 | 162,68 | 204 | 122 | 0,602 | -0,732 | -5,181 | 2,21E-07 | 1,03E-06 | 0,014777 | 0,0109032 | 0,0126092 | 0,0126092 | TRUE | 0,4131202 | No |
| orf19.5657 | 2805 | 1876 | 2529 | 3215 | 3759 | 3125 | 3593 | 2251 | 2516 | 2388 | 2587 | 2651 | 3215 | 2982 | 3206 | 3175 | 2840,1 | 2536 | 3144 | 1,239 | 0,309 | 6,674 | 2,48E-11 | 1,94E-10 | 0,001121 | 0,0024907 | 0,0016977 | 0,0016977 | TRUE | 0,0176613 | No |
| orf19.5658 | 1152 | 733 | 900 | 1167 | 1039 | 805 | 850 | 584 | 1033 | 933 | 921 | 962 | 889 | 768 | 758 | 824 | 886,08 | 962 | 810 | 0,84 | -0,251 | -3,683 | 0,00023052 | 0,00066328 | 0,003062 | 0,0036179 | 0,0033116 | 0,0033116 | TRUE | 0,0554321 | No |
| orf19.5660 | 1204 | 814 | 1029 | 1300 | 828 | 695 | 715 | 495 | 1080 | 1036 | 1053 | 1072 | 708 | 663 | 638 | 698 | 868,58 | 1060 | 677 | 0,637 | -0,651 | -11,679 | 1,64E-31 | 6,51E-30 | 1,00E-08 | 0,0036509 | 0,0017681 | 0,0017681 | TRUE | 0,018594 | No |
| orf19.5660.1 | 2846 | 1842 | 2386 | 2905 | 2592 | 2101 | 2156 | 1572 | 2553 | 2345 | 2441 | 2395 | 2217 | 2005 | 1924 | 2217 | 2262,2 | 2434 | 2091 | 0,858 | -0,221 | -3,862 | 0,00011234 | 0,00034066 | 0,002632 | 0,0026213 | 0,0027006 | 0,0027006 | TRUE | 0,035345 | No |
| orf19.5661 | 2223 | 1413 | 1913 | 2266 | 2061 | 1649 | 1765 | 1210 | 1994 | 1799 | 1957 | 1869 | 1763 | 1573 | 1575 | 1707 | 1779,6 | 1905 | 1654 | 0,868 | -0,205 | -3,672 | 0,00024084 | 0,00069035 | 0,001979 | 0,0027953 | 0,002424 | 0,002424 | TRUE | 0,0252087 | No |
| orf19.5662 | 1196 | 791 | 1029 | 1344 | 1436 | 1225 | 1328 | 834 | 1073 | 1007 | 1053 | 1108 | 1228 | 1169 | 1185 | 1176 | 1124,9 | 1060 | 1190 | 1,12 | 0,164 | 3,259 | 0,00111936 | 0,00279399 | 1,00E-08 | 0,00327 | 0,00154 | 0,00154 | TRUE | 0,0124795 | No |
| orf19.5663 | 1588 | 1030 | 1319 | 1730 | 1534 | 1349 | 1391 | 856 | 1425 | 1311 | 1349 | 1427 | 1312 | 1287 | 1241 | 1207 | 1320 | 1378 | 1262 | 0,916 | -0,127 | -2,488 | 0,01283032 | 0,02511716 | 0,000722 | 0,0030793 | 0,0017413 | 0,0017413 | TRUE | 0,011939 | No |
| orf19.5664 | 219 | 139 | 198 | 231 | 144 | 159 | 170 | 86 | 196 | 177 | 203 | 190 | 123 | 152 | 152 | 121 | 164,29 | 192 | 137 | 0,717 | -0,48 | -4,073 | 4,65E-05 | 0,0001514 | 0,001254 | 0,0108156 | 0,007108 | 0,007108 | TRUE | 0,0611287 | No |
| orf19.5665 | 629 | 363 | 503 | 651 | 546 | 452 | 498 | 322 | 564 | 462 | 515 | 537 | 467 | 431 | 444 | 454 | 484,33 | 520 | 449 | 0,861 | -0,215 | -2,826 | 0,00470874 | 0,01023069 | 0,001952 | 0,0049769 | 0,0034979 | 0,0034979 | TRUE | 0,0581045 | No |
| orf19.5666 | 890 | 579 | 727 | 955 | 899 | 729 | 768 | 436 | 798 | 737 | 744 | 788 | 769 | 696 | 685 | 615 | 728,94 | 767 | 691 | 0,904 | -0,146 | -2,025 | 0,04282087 | 0,07342118 | 0,003122 | 0,003971 | 0,0035891 | 0,0035891 | TRUE | 0,0678275 | No |
| orf19.5667 | 5534 | 3601 | 4781 | 5906 | 6904 | 5628 | 6325 | 3781 | 4965 | 4584 | 4891 | 4870 | 5905 | 5370 | 5644 | 5333 | 5195,3 | 4828 | 5563 | 1,152 | 0,205 | 4,501 | 6,77E-06 | 2,53E-05 | 0,001491 | 0,002259 | 0,0017915 | 0,0017915 | TRUE | 0,0209987 | No |
| orf19.5669 | 834 | 537 | 698 | 873 | 730 | 592 | 649 | 391 | 748 | 684 | 714 | 720 | 624 | 565 | 579 | 551 | 648,2 | 716 | 580 | 0,811 | -0,303 | -4,743 | 2,11E-06 | 8,52E-06 | 0,000515 | 0,0042191 | 0,0023634 | 0,0023634 | TRUE | 0,0323161 | No |
| orf19.567 | 887 | 781 | 837 | 1021 | 927 | 1030 | 1070 | 670 | 796 | 994 | 856 | 842 | 793 | 983 | 955 | 945 | 895,46 | 872 | 919 | 1,055 | 0,077 | 0,896 | 0,37048956 | 0,45894872 | 0,007461 | 0,0036007 | 0,0059815 | 0,0059815 | TRUE | 0,1054358 | No |
| orf19.5671 | 3860 | 2547 | 3309 | 3956 | 5360 | 4573 | 4938 | 3464 | 3463 | 3243 | 3385 | 3262 | 4584 | 4363 | 4406 | 4886 | 3949,1 | 3338 | 4560 | 1,364 | 0,448 | 9,614 | 6,98E-22 | 1,44E-20 | 0,001497 | 0,0023471 | 0,0018269 | 0,0018269 | TRUE | 0,0270236 | No |
| orf19.5672 | 76 | 37 | 57 | 68 | 42 | 51 | 43 | 40 | 68 | 47 | 58 | 56 | 36 | 49 | 38 | 56 | 51,13 | 57 | 45 | 0,763 | -0,39 | -1,855 | 0,06354311 | 0,10360027 | 0,010161 | 0,0303716 | 0,0224586 | 0,0224586 | TRUE | 0,1794776 | No |
| orf19.5673 | 3306 | 2208 | 3088 | 3595 | 2127 | 1841 | 2087 | 1499 | 2966 | 2811 | 3159 | 2964 | 1819 | 1757 | 1862 | 2114 | 2431,6 | 2975 | 1888 | 0,633 | -0,659 | -10,648 | 1,77E-26 | 5,30E-25 | 0,004025 | 0,0025765 | 0,0032451 | 0,0032451 | TRUE | 0,0744087 | No |
| orf19.5674 | 471 | 281 | 361 | 493 | 460 | 332 | 410 | 308 | 423 | 358 | 369 | 407 | 393 | 317 | 366 | 434 | 383,33 | 389 | 378 | 0,964 | -0,053 | -0,521 | 0,60206428 | 0,67775946 | 0,008638 | 0,0057666 | 0,0071579 | 0,0071579 | TRUE | 0,133498 | No |
| orf19.5675 | 4769 | 3230 | 4334 | 5048 | 5733 | 4972 | 5471 | 3400 | 4278 | 4112 | 4434 | 4163 | 4903 | 4744 | 4882 | 4795 | 4539 | 4247 | 4831 | 1,138 | 0,186 | 5,139 | 2,77E-07 | 1,27E-06 | 0,00039 | 0,0022994 | 0,001036 | 0,001036 | TRUE | 0,0105617 | No |
| orf19.5676 | 2628 | 1786 | 2325 | 2837 | 2752 | 2123 | 2554 | 1503 | 2358 | 2274 | 2379 | 2339 | 2354 | 2026 | 2279 | 2120 | 2266 | 2338 | 2195 | 0,94 | -0,09 | -1,686 | 0,09186996 | 0,14243993 | 0,001901 | 0,0026202 | 0,0022897 | 0,0022897 | TRUE | 0,0339473 | No |
| orf19.5677 | 94 | 73 | 94 | 104 | 80 | 81 | 86 | 47 | 84 | 93 | 96 | 86 | 68 | 77 | 77 | 66 | 80,99 | 90 | 72 | 0,81 | -0,304 | -1,997 | 0,04582118 | 0,07790654 | 1,00E-08 | 0,0199036 | 0,0098209 | 0,0098209 | TRUE | 0,023325 | No |
| orf19.5678 | 1356 | 892 | 1149 | 1470 | 1741 | 1309 | 1566 | 981 | 1216 | 1136 | 1176 | 1212 | 1489 | 1249 | 1397 | 1384 | 1282,4 | 1185 | 1380 | 1,164 | 0,219 | 3,598 | 0,00032033 | 0,00090228 | 0,002501 | 0,0031116 | 0,0027586 | 0,0027586 | TRUE | 0,0505295 | No |
| orf19.568 | 3059 | 2411 | 2794 | 3503 | 3187 | 3160 | 3311 | 1847 | 2744 | 3069 | 2859 | 2889 | 2726 | 3015 | 2955 | 2605 | 2857,7 | 2890 | 2825 | 0,979 | -0,031 | -0,537 | 0,59134827 | 0,66895507 | 0,002691 | 0,0024875 | 0,0027787 | 0,0027787 | TRUE | 0,0317269 | No |
| orf19.5680 | 2186 | 1378 | 1880 | 2373 | 1929 | 1591 | 1725 | 1075 | 1961 | 1754 | 1923 | 1957 | 1650 | 1518 | 1539 | 1516 | 1727,4 | 1899 | 1556 | 0,819 | -0,288 | -5,473 | 4,43E-08 | 2,26E-07 | 0,00138 | 0,0028199 | 0,0020752 | 0,0020752 | TRUE | 0,029998 | No |
| orf19.5681 | 1263 | 751 | 1044 | 1288 | 1139 | 965 | 969 | 642 | 1133 | 956 | 1068 | 1062 | 974 | 921 | 865 | 905 | 985,55 | 1055 | 916 | 0,867 | -0,205 | -3,17 | 0,00152428 | 0,00368973 | 0,002634 | 0,0034525 | 0,0030032 | 0,0030032 | TRUE | 0,0441813 | No |
| orf19.5682 | 3042 | 1936 | 2618 | 3362 | 2944 | 2365 | 2582 | 1812 | 2729 | 2465 | 2678 | 2772 | 2518 | 2257 | 2304 | 2556 | 2534,8 | 2661 | 2409 | 0,904 | -0,146 | -2,572 | 0,01012067 | 0,02034292 | 0,002729 | 0,0025522 | 0,002713 | 0,002713 | TRUE | 0,0289791 | No |
| orf19.5683 | 1515 | 968 | 1403 | 1731 | 1761 | 1460 | 1616 | 923 | 1359 | 1232 | 1435 | 1427 | 1506 | 1393 | 1442 | 1302 | 1387,2 | 1363 | 1411 | 1,035 | 0,05 | 0,774 | 0,43879676 | 0,52565895 | 0,003214 | 0,0030261 | 0,0032215 | 0,0032215 | TRUE | 0,047629 | No |
| orf19.5684 | 1678 | 1056 | 1314 | 1613 | 2145 | 1556 | 1731 | 1068 | 1505 | 1344 | 1344 | 1330 | 1834 | 1485 | 1545 | 1506 | 1486,8 | 1381 | 1592 | 1,154 | 0,207 | 2,739 | 0,0061624 | 0,01305029 | 0,006161 | 0,0029559 | 0,0047852 | 0,0047852 | TRUE | 0,1286078 | No |
| orf19.5684.1 | 105 | 64 | 82 | 106 | 110 | 95 | 90 | 78 | 94 | 81 | 84 | 87 | 94 | 91 | 80 | 110 | 90,25 | 86 | 94 | 1,067 | 0,093 | 0,622 | 0,53369088 | 0,6160259 | 1,00E-08 | 0,0180644 | 0,0104841 | 0,0104841 | TRUE | 0,1022674 | No |
| orf19.5685 | 4477 | 2849 | 3864 | 4541 | 4357 | 3552 | 3982 | 2430 | 4016 | 3627 | 3953 | 3745 | 3726 | 3389 | 3553 | 3427 | 3679,7 | 3835 | 3524 | 0,919 | -0,122 | -2,512 | 0,01198905 | 0,023663 | 0,00168 | 0,002374 | 0,0019866 | 0,0019866 | TRUE | 0,0184423 | No |
| orf19.5686 | 3613 | 2143 | 3002 | 3591 | 5857 | 4399 | 4676 | 2874 | 3241 | 2728 | 3071 | 2961 | 5009 | 4197 | 4173 | 4054 | 3679,3 | 3000 | 4358 | 1,452 | 0,539 | 7,224 | 5,03E-13 | 4,80E-12 | 0,007253 | 0,0023741 | 0,0050546 | 0,0050546 | TRUE | 0,1239691 | No |
| orf19.5688 | 2031 | 1243 | 1677 | 2230 | 2757 | 2157 | 2306 | 1372 | 1822 | 1582 | 1716 | 1839 | 2358 | 2058 | 2058 | 1935 | 1921 | 1740 | 2102 | 1,208 | 0,273 | 3,884 | 0,0001029 | 0,00031431 | 0,005274 | 0,0027352 | 0,0042171 | 0,0042171 | TRUE | 0,0819229 | No |
| orf19.5689 | 1026 | 591 | 893 | 1000 | 1096 | 845 | 957 | 619 | 920 | 752 | 914 | 825 | 937 | 806 | 854 | 873 | 860,21 | 853 | 868 | 1,016 | 0,023 | 0,3 | 0,76396795 | 0,81569566 | 0,005144 | 0,0036671 | 0,0044546 | 0,0044546 | TRUE | 0,0693621 | No |
| orf19.5691 | 562 | 366 | 465 | 602 | 742 | 597 | 670 | 367 | 504 | 466 | 476 | 496 | 635 | 570 | 598 | 518 | 532,75 | 486 | 580 | 1,198 | 0,26 | 3,476 | 0,00050966 | 0,00137434 | 0,002283 | 0,0047044 | 0,0034994 | 0,0034994 | TRUE | 0,0534681 | No |
| orf19.5692 | 448 | 287 | 372 | 461 | 484 | 398 | 413 | 320 | 402 | 365 | 381 | 380 | 414 | 380 | 369 | 451 | 392,7 | 382 | 404 | 1,049 | 0,069 | 0,841 | 0,40055087 | 0,48942641 | 0,002014 | 0,0056763 | 0,0039728 | 0,0039728 | TRUE | 0,0627666 | No |
| orf19.5693 | 922 | 575 | 885 | 1067 | 1036 | 792 | 922 | 560 | 827 | 732 | 905 | 880 | 886 | 756 | 823 | 790 | 824,84 | 836 | 814 | 0,972 | -0,041 | -0,524 | 0,60013992 | 0,67635352 | 0,00527 | 0,0037395 | 0,0045491 | 0,0045491 | TRUE | 0,0778574 | No |
| orf19.5694 | 180 | 114 | 153 | 185 | 153 | 129 | 171 | 118 | 161 | 145 | 157 | 153 | 131 | 123 | 153 | 166 | 148,58 | 154 | 143 | 0,921 | -0,118 | -0,955 | 0,3394939 | 0,42653381 | 0,002311 | 0,0117499 | 0,0079582 | 0,0079582 | TRUE | 0,099139 | No |
| orf19.5698 | 1904 | 1147 | 1566 | 1939 | 2260 | 1775 | 2001 | 1481 | 1708 | 1460 | 1602 | 1599 | 1933 | 1694 | 1786 | 2089 | 1733,8 | 1592 | 1876 | 1,175 | 0,233 | 3,24 | 0,0011962 | 0,00296974 | 0,005526 | 0,0028168 | 0,0043641 | 0,0043641 | TRUE | 0,0678735 | No |
| orf19.57 | 636 | 454 | 538 | 672 | 693 | 625 | 649 | 405 | 571 | 578 | 550 | 554 | 593 | 596 | 579 | 571 | 574,06 | 563 | 585 | 1,04 | 0,057 | 0,942 | 0,34613906 | 0,43352168 | 1,00E-08 | 0,0045084 | 0,0018102 | 0,0018102 | TRUE | 0,0031159 | No |
| orf19.570 | 193 | 152 | 165 | 205 | 194 | 166 | 190 | 111 | 173 | 194 | 169 | 169 | 166 | 158 | 170 | 157 | 169,36 | 176 | 163 | 0,93 | -0,105 | -0,984 | 0,32517196 | 0,41218378 | 1,00E-08 | 0,010551 | 0,0051187 | 0,0051187 | TRUE | 0,0401218 | No |
| orf19.5700 | 66 | 54 | 50 | 82 | 57 | 49 | 59 | 33 | 59 | 69 | 51 | 68 | 49 | 47 | 53 | 47 | 55,18 | 62 | 49 | 0,793 | -0,335 | -1,794 | 0,07282369 | 0,11661107 | 1,00E-08 | 0,028289 | 0,0152786 | 0,0152786 | TRUE | 0,1099084 | No |
| orf19.5701 | 16 | 7 | 14 | 23 | 26 | 25 | 30 | 13 | 14 | 9 | 14 | 19 | 22 | 24 | 27 | 18 | 18,47 | 14 | 23 | 1,593 | 0,671 | 2,051 | 0,04029273 | 0,06950294 | 1,00E-08 | 0,0805824 | 0,0469662 | 0,0469662 | TRUE | 0,2650863 | No |
| orf19.5702 | 1436 | 1071 | 1314 | 1608 | 1671 | 1657 | 1690 | 1033 | 1288 | 1363 | 1344 | 1326 | 1429 | 1581 | 1508 | 1457 | 1412,2 | 1330 | 1494 | 1,124 | 0,169 | 3,419 | 0,00062883 | 0,00165195 | 0,000667 | 0,0030075 | 0,0016297 | 0,0016297 | TRUE | 0,0189064 | No |
| orf19.5704 | 2869 | 2026 | 2616 | 3145 | 4242 | 3745 | 4335 | 2622 | 2574 | 2579 | 2676 | 2593 | 3628 | 3573 | 3868 | 3698 | 3148,8 | 2606 | 3692 | 1,417 | 0,503 | 12,69 | 6,72E-37 | 3,84E-35 | 0,000544 | 0,0024406 | 0,0011786 | 0,0011786 | TRUE | 0,0131022 | No |
| orf19.5705 | 816 | 563 | 756 | 817 | 938 | 889 | 960 | 583 | 732 | 717 | 773 | 674 | 802 | 848 | 857 | 822 | 778,17 | 724 | 832 | 1,151 | 0,203 | 3,326 | 0,00088113 | 0,00224124 | 0,000918 | 0,0038451 | 0,0022868 | 0,0022868 | TRUE | 0,0283493 | No |
| orf19.5710 | 4174 | 2802 | 3858 | 4586 | 5921 | 5328 | 5784 | 3667 | 3745 | 3567 | 3947 | 3782 | 5064 | 5084 | 5161 | 5172 | 4440,2 | 3760 | 5120 | 1,361 | 0,444 | 11,655 | 2,16E-31 | 8,35E-30 | 0,000608 | 0,0023065 | 0,0011638 | 0,0011638 | TRUE | 0,0136548 | No |
| orf19.5711 | 1273 | 892 | 1170 | 1359 | 958 | 929 | 1002 | 663 | 1142 | 1136 | 1197 | 1121 | 819 | 886 | 894 | 935 | 1016,3 | 1149 | 884 | 0,767 | -0,382 | -6,843 | 7,73E-12 | 6,39E-11 | 0,000768 | 0,003408 | 0,0019886 | 0,0019886 | TRUE | 0,0295998 | No |
| orf19.5713 | 7026 | 4921 | 6225 | 7486 | 6403 | 5970 | 6462 | 4157 | 6303 | 6265 | 6369 | 6173 | 5476 | 5696 | 5766 | 5863 | 5989 | 6278 | 5700 | 0,908 | -0,14 | -4,204 | 2,62E-05 | 8,91E-05 | 0,000278 | 0,0022219 | 0,0008948 | 0,0008948 | TRUE | 0,0087005 | No |
| orf19.5714 | 946 | 566 | 764 | 966 | 842 | 713 | 777 | 476 | 849 | 721 | 782 | 797 | 720 | 680 | 693 | 671 | 739,07 | 787 | 691 | 0,878 | -0,189 | -2,944 | 0,00323658 | 0,00729657 | 0,001322 | 0,0039437 | 0,0025777 | 0,0025777 | TRUE | 0,0347438 | No |
| orf19.5716 | 91 | 86 | 79 | 105 | 74 | 80 | 60 | 51 | 82 | 109 | 81 | 87 | 63 | 76 | 54 | 72 | 77,95 | 90 | 66 | 0,74 | -0,434 | -2,505 | 0,01223357 | 0,02409813 | 0,008489 | 0,0206023 | 0,0156771 | 0,0156771 | TRUE | 0,1811945 | No |
| orf19.5717 | 5 | 1 | 8 | 3 | 2 | 11 | 5 | 4 | 4 | 1 | 8 | 2 | 2 | 10 | 4 | 6 | 4,84 | 4 | 6 | 1,335 | 0,417 | 0,574 | 0,56615272 | NA | 0,288985 | 0,3018645 | 0,2965947 | 0,2965947 | TRUE | 0,4397968 | No |
| orf19.5718 | 485 | 361 | 463 | 568 | 541 | 547 | 553 | 375 | 435 | 460 | 474 | 468 | 463 | 522 | 493 | 529 | 480,47 | 459 | 502 | 1,09 | 0,124 | 1,719 | 0,08552367 | 0,13394844 | 1,00E-08 | 0,005001 | 0,0028817 | 0,0028817 | TRUE | 0,0333843 | No |
| orf19.5719 | 2206 | 1457 | 1718 | 2221 | 1905 | 1592 | 1728 | 1074 | 1979 | 1855 | 1758 | 1831 | 1629 | 1519 | 1542 | 1515 | 1703,5 | 1856 | 1551 | 0,836 | -0,258 | -5,039 | 4,67E-07 | 2,08E-06 | 0,00132 | 0,0028317 | 0,0019173 | 0,0019173 | TRUE | 0,0248763 | No |

| | | | | | | | | | | | | | | | | | | | | | | | | | | | | | | | |
|---|---|---|---|---|---|---|---|---|---|---|---|---|---|---|---|---|---|---|---|---|---|---|---|---|---|---|---|---|---|---|---|
| orf19.5720 | 3633 | 2656 | 3347 | 4234 | 4265 | 3693 | 3969 | 2277 | 3259 | 3381 | 3424 | 3491 | 3648 | 3524 | 3542 | 3212 | 3435,1 | 3389 | 3482 | 1,028 | 0,04 | 0,855 | 0,39258087 | 0,48154877 | 0,001484 | 0,0024021 | 0,0018611 | 0,0018611 | TRUE | 0,03133 | No |
| orf19.5722 | 1185 | 720 | 1014 | 1259 | 1100 | 872 | 938 | 611 | 1063 | 917 | 1037 | 1038 | 941 | 832 | 837 | 862 | 940,86 | 1014 | 868 | 0,855 | -0,226 | -3,439 | 0,00058407 | 0,00154995 | 0,002715 | 0,0035225 | 0,0030829 | 0,0030829 | TRUE | 0,046251 | No |
| orf19.5723 | 4219 | 2877 | 3530 | 4478 | 3726 | 3173 | 3397 | 2376 | 3785 | 3663 | 3612 | 3693 | 3187 | 3028 | 3031 | 3351 | 3418,5 | 3688 | 3149 | 0,853 | -0,23 | -5,333 | 9,68E-08 | 4,77E-07 | 0,000993 | 0,0024042 | 0,0014896 | 0,0014896 | TRUE | 0,0211402 | No |
| orf19.5724 | 9 | 13 | 6 | 4 | 8 | 5 | 11 | 4 | 8 | 17 | 6 | 3 | 7 | 5 | 10 | 6 | 7,64 | 8 | 7 | 0,833 | -0,263 | -0,465 | 0,64157668 | NA | 0,148114 | 0,1919574 | 0,1728456 | 0,1728456 | TRUE | 0,7103455 | No |
| orf19.5725 | 496 | 364 | 454 | 521 | 379 | 391 | 477 | 294 | 445 | 463 | 464 | 430 | 324 | 373 | 426 | 415 | 417,5 | 450 | 384 | 0,851 | -0,232 | -2,542 | 0,01100872 | 0,02194439 | 0,005743 | 0,0054567 | 0,0055979 | 0,0055979 | TRUE | 0,1359601 | No |
| orf19.5727 | 1032 | 720 | 921 | 1182 | 1374 | 1123 | 1189 | 705 | 926 | 917 | 942 | 975 | 1175 | 1072 | 1061 | 994 | 1007,7 | 940 | 1076 | 1,146 | 0,197 | 3,259 | 0,00111651 | 0,00278918 | 0,001797 | 0,0034202 | 0,002516 | 0,002516 | TRUE | 0,0471519 | No |
| orf19.5728 | 487 | 332 | 422 | 489 | 269 | 262 | 243 | 173 | 437 | 423 | 432 | 403 | 230 | 250 | 217 | 244 | 329,43 | 424 | 235 | 0,554 | -0,852 | -10,251 | 1,17E-24 | 2,93E-23 | 1,00E-08 | 0,0063862 | 0,0033322 | 0,0033322 | TRUE | 0,0320908 | No |
| orf19.5729 | 378 | 296 | 364 | 444 | 418 | 383 | 404 | 277 | 339 | 377 | 372 | 366 | 357 | 365 | 361 | 391 | 366,08 | 364 | 368 | 1,012 | 0,017 | 0,223 | 0,82351904 | 0,86392568 | 1,00E-08 | 0,0059451 | 0,0029121 | 0,0029121 | TRUE | 0,0244391 | No |
| orf19.5730 | 2005 | 1438 | 1912 | 2182 | 2136 | 1966 | 2092 | 1379 | 1799 | 1831 | 1956 | 1799 | 1827 | 1876 | 1867 | 1945 | 1862,4 | 1846 | 1879 | 1,017 | 0,024 | 0,527 | 0,59827792 | 0,67488805 | 0,000589 | 0,002759 | 0,0014496 | 0,0014496 | TRUE | 0,0195117 | No |
| orf19.5731 | 2080 | 1482 | 1792 | 2309 | 2012 | 1847 | 1969 | 1175 | 1866 | 1887 | 1833 | 1904 | 1721 | 1762 | 1757 | 1657 | 1798,4 | 1872 | 1724 | 0,922 | -0,117 | -2,838 | 0,00454213 | 0,00989454 | 1,00E-08 | 0,0027867 | 0,0010744 | 0,0010744 | TRUE | 0,0071786 | No |
| orf19.5732 | 27759 | 18936 | 23953 | 29285 | 29322 | 26514 | 28239 | 16621 | 24903 | 24108 | 24506 | 24149 | 25077 | 25299 | 25199 | 23442 | 24585 | 24416 | 24754 | 1,014 | 0,02 | 0,592 | 0,55398781 | 0,63486787 | 0,000658 | 0,0020386 | 0,0010703 | 0,0010703 | TRUE | 0,0153553 | No |
| orf19.5734 | 2480 | 1791 | 2339 | 2866 | 3364 | 2896 | 3246 | 2088 | 2225 | 2280 | 2393 | 2363 | 2877 | 2763 | 2897 | 2945 | 2592,9 | 2315 | 2870 | 1,239 | 0,309 | 7,449 | 9,43E-14 | 9,60E-13 | 0,000517 | 0,0025394 | 0,0012609 | 0,0012609 | TRUE | 0,0088589 | No |
| orf19.5735 | 2126 | 1481 | 1809 | 2348 | 1926 | 1627 | 1746 | 982 | 1907 | 1885 | 1851 | 1936 | 1647 | 1552 | 1558 | 1385 | 1715,3 | 1895 | 1536 | 0,812 | -0,3 | -5,489 | 4,05E-08 | 2,08E-07 | 0,001932 | 0,0028259 | 0,0022712 | 0,0022712 | TRUE | 0,047968 | No |
| orf19.5735.3 | 2680 | 1827 | 2257 | 2772 | 3054 | 2483 | 2673 | 1687 | 2404 | 2326 | 2309 | 2286 | 2612 | 2369 | 2385 | 2379 | 2383,9 | 2331 | 2436 | 1,046 | 0,065 | 1,408 | 0,15917773 | 0,22678203 | 0,001046 | 0,0025885 | 0,0016105 | 0,0016105 | TRUE | 0,0294928 | No |
| orf19.5736 | 4361 | 2899 | 4098 | 4684 | 4827 | 4138 | 4711 | 3143 | 3912 | 3691 | 4193 | 3862 | 4128 | 3948 | 4204 | 4433 | 4046,4 | 3914 | 4178 | 1,066 | 0,093 | 1,81 | 0,07036299 | 0,11324427 | 0,00224 | 0,0023383 | 0,0022723 | 0,0022723 | TRUE | 0,0275261 | No |
| orf19.5741 | 63068 | 43513 | 59137 | 70644 | 51254 | 43385 | 48418 | 33794 | 56579 | 55397 | 60503 | 58254 | 43834 | 41397 | 43205 | 47663 | 50854 | 57683 | 44025 | 0,763 | -0,39 | -7,932 | 2,16E-15 | 2,58E-14 | 0,002504 | 0,0020081 | 0,0023027 | 0,0023027 | TRUE | 0,0377794 | No |
| orf19.5742 | 1394 | 966 | 1294 | 1482 | 1742 | 1472 | 1669 | 1051 | 1251 | 1230 | 1324 | 1222 | 1490 | 1405 | 1489 | 1482 | 1361,5 | 1257 | 1466 | 1,167 | 0,223 | 4,615 | 3,93E-06 | 1,52E-05 | 0,000346 | 0,0030457 | 0,0014995 | 0,0014995 | TRUE | 0,0154477 | No |
| orf19.5746 | 8405 | 6053 | 7571 | 8792 | 8669 | 8205 | 8664 | 5168 | 7540 | 7706 | 7746 | 7250 | 7414 | 7829 | 7731 | 7289 | 7563,2 | 7560 | 7566 | 1,001 | 0,002 | 0,051 | 0,95913712 | 0,97028049 | 0,000737 | 0,0021715 | 0,0012863 | 0,0012863 | TRUE | 0,0094729 | No |
| orf19.5747 | 2269 | 1799 | 2085 | 2677 | 2620 | 2536 | 2663 | 1764 | 2036 | 2290 | 2133 | 2207 | 2241 | 2420 | 2376 | 2488 | 2273,9 | 2166 | 2381 | 1,099 | 0,136 | 2,643 | 0,00822332 | 0,01690819 | 0,001624 | 0,0026179 | 0,0020929 | 0,0020929 | TRUE | 0,020264 | No |
| orf19.5748 | 14 | 10 | 12 | 13 | 17 | 18 | 20 | 12 | 13 | 13 | 12 | 11 | 15 | 17 | 18 | 17 | 14,35 | 12 | 17 | 1,381 | 0,465 | 1,323 | 0,18577438 | NA | 1,00E-08 | 0,1031669 | 0,0474363 | 0,0474363 | TRUE | 0,0427992 | No |
| orf19.5749 | 3453 | 2472 | 3077 | 3989 | 4462 | 4053 | 4311 | 2718 | 3098 | 3147 | 3148 | 3289 | 3816 | 3867 | 3847 | 3833 | 3505,7 | 3170 | 3841 | 1,211 | 0,276 | 8,02 | 1,06E-15 | 1,30E-14 | 1,00E-08 | 0,0023936 | 0,0008492 | 0,0008492 | TRUE | 0,0081988 | No |
| orf19.575 | 8203 | 5265 | 7435 | 9169 | 9303 | 7467 | 8420 | 5451 | 7359 | 6703 | 7607 | 7561 | 7956 | 7125 | 7513 | 7688 | 7439 | 7308 | 7570 | 1,036 | 0,05 | 0,975 | 0,3294743 | 0,41685525 | 0,00251 | 0,0021747 | 0,0024387 | 0,0024387 | TRUE | 0,0372717 | No |
| orf19.5750 | 4918 | 3503 | 4466 | 5372 | 4209 | 3815 | 4109 | 2438 | 4412 | 4460 | 4569 | 4430 | 3600 | 3640 | 3667 | 3439 | 4027 | 4468 | 3586 | 0,804 | -0,315 | -9,039 | 1,58E-19 | 2,74E-18 | 0,000196 | 0,00234 | 0,000915 | 0,000915 | TRUE | 0,0085161 | No |
| orf19.5751 | 4353 | 3321 | 3880 | 4967 | 4615 | 4492 | 4752 | 3051 | 3905 | 4228 | 3970 | 4096 | 3947 | 4286 | 4240 | 4303 | 4121,9 | 4050 | 4194 | 1,036 | 0,05 | 1,155 | 0,24819707 | 0,3301056 | 0,001091 | 0,0023317 | 0,0015864 | 0,0015864 | TRUE | 0,0195829 | No |
| orf19.5752 | 1918 | 1362 | 1526 | 2013 | 2129 | 2086 | 2203 | 1253 | 1721 | 1734 | 1561 | 1660 | 1821 | 1990 | 1966 | 1767 | 1777,5 | 1669 | 1886 | 1,132 | 0,179 | 3,198 | 0,00138615 | 0,00338536 | 0,002028 | 0,0027962 | 0,0024279 | 0,0024279 | TRUE | 0,0224015 | No |
| orf19.5753 | 5466 | 3952 | 5052 | 6142 | 5390 | 4940 | 5312 | 3376 | 4904 | 5031 | 5169 | 5065 | 4610 | 4714 | 4740 | 4762 | 4874,2 | 5042 | 4706 | 0,933 | -0,1 | -3,096 | 0,00196281 | 0,00463908 | 0,000159 | 0,0022774 | 0,0007915 | 0,0007915 | TRUE | 0,0042888 | No |
| orf19.5754 | 8369 | 6690 | 7652 | 9597 | 10340 | 10691 | 11027 | 6505 | 7508 | 8517 | 7829 | 7914 | 8843 | 10201 | 9840 | 9175 | 8728,3 | 7942 | 9515 | 1,198 | 0,261 | 4,654 | 3,26E-06 | 1,28E-05 | 0,003359 | 0,0021458 | 0,0029096 | 0,0029096 | TRUE | 0,0288754 | No |
| orf19.5755 | 2348 | 1686 | 2089 | 2535 | 3292 | 3064 | 3368 | 1981 | 2106 | 2146 | 2137 | 2090 | 2815 | 2924 | 3005 | 2794 | 2502,4 | 2120 | 2884 | 1,362 | 0,446 | 11,15 | 7,15E-29 | 2,43E-27 | 0,000343 | 0,0025597 | 0,0011267 | 0,0011267 | TRUE | 0,0097626 | No |
| orf19.5757 | 1787 | 1444 | 1656 | 2143 | 1916 | 1884 | 2030 | 1243 | 1603 | 1838 | 1694 | 1767 | 1639 | 1798 | 1811 | 1753 | 1738 | 1726 | 1750 | 1,015 | 0,021 | 0,38 | 0,70374929 | 0,76554874 | 0,002193 | 0,0028148 | 0,0024461 | 0,0024461 | TRUE | 0,0279081 | No |
| orf19.5758 | 6451 | 4898 | 5980 | 7212 | 7575 | 7146 | 7461 | 4425 | 5787 | 6236 | 6118 | 5947 | 6478 | 6818 | 6658 | 6241 | 6285,5 | 6022 | 6549 | 1,088 | 0,122 | 2,997 | 0,00272852 | 0,00626164 | 0,001035 | 0,0022105 | 0,0014396 | 0,0014396 | TRUE | 0,0117315 | No |
| orf19.5759 | 4696 | 3468 | 4482 | 5378 | 4075 | 3883 | 4187 | 2456 | 4213 | 4415 | 4586 | 4435 | 3485 | 3705 | 3736 | 3464 | 4004,8 | 4412 | 3598 | 0,816 | -0,293 | -6,752 | 1,46E-11 | 1,17E-10 | 0,001055 | 0,002342 | 0,001559 | 0,001559 | TRUE | 0,011451 | No |
| orf19.576 | 307 | 233 | 258 | 296 | 270 | 223 | 240 | 165 | 275 | 297 | 264 | 244 | 231 | 213 | 214 | 233 | 246,33 | 270 | 223 | 0,827 | -0,275 | -2,873 | 0,00407108 | 0,00896265 | 1,00E-08 | 0,0078728 | 0,0046896 | 0,0046896 | TRUE | 0,0499148 | No |
| orf19.5760 | 5673 | 3544 | 4944 | 6034 | 5180 | 3905 | 4443 | 3473 | 5089 | 4512 | 5058 | 4976 | 4430 | 3726 | 3965 | 4898 | 4581,8 | 4909 | 4255 | 0,866 | -0,208 | -2,629 | 0,00855553 | 0,01751333 | 0,008653 | 0,0022964 | 0,0057662 | 0,0057662 | TRUE | 0,1242312 | No |
| orf19.5762 | 114 | 89 | 117 | 130 | 106 | 63 | 71 | 59 | 102 | 113 | 120 | 107 | 91 | 60 | 63 | 83 | 92,48 | 110 | 74 | 0,671 | -0,576 | -3,53 | 0,00041562 | 0,00114072 | 0,009297 | 0,0176775 | 0,0143029 | 0,0143029 | TRUE | 0,2470492 | No |
| orf19.5763 | 158 | 105 | 143 | 152 | 195 | 136 | 143 | 98 | 142 | 134 | 146 | 125 | 167 | 130 | 128 | 138 | 138,68 | 137 | 141 | 1,031 | 0,044 | 0,342 | 0,73217783 | 0,79088896 | 0,004815 | 0,0124477 | 0,008959 | 0,008959 | TRUE | 0,1828802 | No |
| orf19.5764 | 1138 | 723 | 1078 | 1237 | 1259 | 1007 | 1085 | 669 | 1021 | 920 | 1103 | 1020 | 1077 | 961 | 968 | 944 | 1001,7 | 1016 | 988 | 0,972 | -0,041 | -0,605 | 0,54519649 | 0,62653797 | 0,003164 | 0,0034288 | 0,0034249 | 0,0034249 | TRUE | 0,0457578 | No |
| orf19.5765 | 1419 | 974 | 1205 | 1510 | 1644 | 1503 | 1666 | 1005 | 1273 | 1240 | 1233 | 1245 | 1406 | 1434 | 1487 | 1417 | 1341,9 | 1248 | 1436 | 1,151 | 0,203 | 4,538 | 5,67E-06 | 2,15E-05 | 1,00E-08 | 0,0030613 | 0,0011723 | 0,0011723 | TRUE | 0,0071102 | No |
| orf19.5767 | 890 | 546 | 710 | 867 | 730 | 677 | 703 | 410 | 798 | 695 | 726 | 715 | 624 | 646 | 627 | 578 | 676,34 | 734 | 619 | 0,845 | -0,244 | -3,684 | 0,00022982 | 0,00066191 | 0,001393 | 0,0041259 | 0,0027145 | 0,0027145 | TRUE | 0,0427146 | No |
| orf19.5768 | 7194 | 5172 | 6278 | 7993 | 7483 | 6481 | 6947 | 4334 | 6454 | 6585 | 6423 | 6591 | 6400 | 6184 | 6199 | 6113 | 6368,5 | 6513 | 6224 | 0,956 | -0,065 | -2,149 | 0,0316316 | 0,05622564 | 0,000133 | 0,0022075 | 0,0007143 | 0,0007143 | TRUE | 0,0044926 | No |
| orf19.577 | 4228 | 2945 | 3726 | 4383 | 4089 | 3534 | 3854 | 2133 | 3793 | 3749 | 3812 | 3614 | 3497 | 3372 | 3439 | 3008 | 3535,6 | 3742 | 3329 | 0,891 | -0,166 | -3,258 | 0,00112192 | 0,00279921 | 0,002086 | 0,0023901 | 0,0022188 | 0,0022188 | TRUE | 0,0486884 | No |
| orf19.5770 | 6967 | 4840 | 6367 | 7668 | 7355 | 6628 | 7127 | 4140 | 6250 | 6162 | 6514 | 6323 | 6290 | 6324 | 6360 | 5839 | 6257,8 | 6312 | 6203 | 0,983 | -0,024 | -0,621 | 0,53439382 | 0,61648197 | 0,000824 | 0,0022115 | 0,0012921 | 0,0012921 | TRUE | 0,0182265 | No |
| orf19.5771 | 8637 | 5739 | 7486 | 9125 | 8586 | 7005 | 7871 | 5502 | 7748 | 7306 | 7659 | 7525 | 7343 | 6684 | 7024 | 7760 | 7381,1 | 7560 | 7203 | 0,952 | -0,071 | -1,443 | 0,14909286 | 0,21460228 | 0,002171 | 0,0021762 | 0,0021719 | 0,0021719 | TRUE | 0,0323403 | No |
| orf19.5772 | 1441 | 930 | 1191 | 1525 | 1302 | 1125 | 1213 | 759 | 1293 | 1184 | 1219 | 1258 | 1114 | 1073 | 1082 | 1071 | 1161,6 | 1238 | 1085 | 0,875 | -0,192 | -3,894 | 9,87E-05 | 0,00030273 | 1,00E-08 | 0,0032293 | 0,0014671 | 0,0014671 | TRUE | 0,0103621 | No |
| orf19.5773 | 1957 | 1373 | 1769 | 2203 | 1921 | 1831 | 1916 | 1247 | 1756 | 1748 | 1810 | 1817 | 1643 | 1747 | 1710 | 1759 | 1748,6 | 1783 | 1715 | 0,96 | -0,059 | -1,365 | 0,17240191 | 0,24285831 | 1,00E-08 | 0,0028097 | 0,0011957 | 0,0011957 | TRUE | 0,0099075 | No |
| orf19.5775 | 49 | 27 | 38 | 48 | 47 | 57 | 50 | 22 | 44 | 34 | 39 | 40 | 40 | 54 | 45 | 31 | 40,88 | 39 | 42 | 1,093 | 0,128 | 0,572 | 0,56751517 | 0,64690024 | 1,00E-08 | 0,0374929 | 0,0242742 | 0,0242742 | TRUE | 0,2201033 | No |
| orf19.5776 | 3451 | 2533 | 3208 | 3743 | 4457 | 4037 | 4286 | 2571 | 3096 | 3225 | 3282 | 3087 | 3812 | 3852 | 3825 | 3626 | 3475,5 | 3172 | 3779 | 1,193 | 0,255 | 6,542 | 6,09E-11 | 4,59E-10 | 0,000449 | 0,0023972 | 0,0011628 | 0,0011628 | TRUE | 0,0083306 | No |
| orf19.5777 | 261 | 174 | 195 | 302 | 286 | 213 | 300 | 188 | 234 | 222 | 200 | 249 | 245 | 203 | 268 | 265 | 235,61 | 226 | 245 | 1,079 | 0,11 | 0,968 | 0,33324513 | 0,41999932 | 0,008155 | 0,0081409 | 0,0081483 | 0,0081483 | TRUE | 0,1496415 | No |
| orf19.5779 | 4677 | 3191 | 4165 | 4953 | 4492 | 3859 | 4266 | 2823 | 4196 | 4062 | 4261 | 4084 | 3842 | 3682 | 3807 | 3982 | 3989,5 | 4151 | 3828 | 0,921 | -0,118 | -3,152 | 0,00161988 | 0,00389446 | 0,000438 | 0,0023434 | 0,0010982 | 0,0010982 | TRUE | 0,0082103 | No |
| orf19.578 | 3525 | 2446 | 2965 | 3799 | 3785 | 3431 | 3620 | 2075 | 3162 | 3114 | 3033 | 3133 | 3237 | 3274 | 3230 | 2927 | 3138,8 | 3110 | 3167 | 1,02 | 0,028 | 0,63 | 0,52891344 | 0,6119222 | 0,00103 | 0,002442 | 0,0015606 | 0,0015606 | TRUE | 0,029584 | No |
| orf19.5780 | 1200 | 819 | 1053 | 1321 | 1378 | 1218 | 1336 | 896 | 1077 | 1043 | 1077 | 1089 | 1179 | 1162 | 1192 | 1264 | 1135,3 | 1072 | 1199 | 1,116 | 0,158 | 3,227 | 0,00125268 | 0,00309333 | 1,00E-08 | 0,0032582 | 0,0014113 | 0,0014113 | TRUE | 0,0136585 | No |
| orf19.5782 | 7221 | 4934 | 6330 | 7535 | 8804 | 7776 | 8659 | 4841 | 6478 | 6282 | 6476 | 6213 | 7529 | 7420 | 7727 | 6828 | 6869,1 | 6362 | 7376 | 1,16 | 0,214 | 4,88 | 1,06E-06 | 4,45E-06 | 0,001407 | 0,0021909 | 0,001704 | 0,001704 | TRUE | 0,0296623 | No |
| orf19.5783 | 1075 | 690 | 860 | 1119 | 1112 | 909 | 1003 | 652 | 964 | 878 | 880 | 923 | 951 | 867 | 895 | 920 | 909,8 | 911 | 908 | 0,995 | -0,007 | -0,129 | 0,8974436 | 0,92247068 | 0,000775 | 0,0035752 | 0,0020471 | 0,0020471 | TRUE | 0,0183494 | No |
| orf19.5784 | 1345 | 902 | 1179 | 1461 | 786 | 710 | 707 | 463 | 1207 | 1148 | 1206 | 1205 | 672 | 677 | 631 | 653 | 924,94 | 1192 | 658 | 0,552 | -0,857 | -16,112 | 2,11E-58 | 3,10E-56 | 1,00E-08 | 0,003549 | 0,0015368 | 0,0015368 | TRUE | 0,0105099 | No |
| orf19.5785 | 2521 | 1871 | 2316 | 2917 | 1876 | 1758 | 1884 | 1175 | 2262 | 2382 | 2369 | 2405 | 1604 | 1677 | 1681 | 1657 | 2004,8 | 2354 | 1655 | 0,703 | -0,509 | -12,2 | 3,10E-34 | 1,44E-32 | 1,00E-08 | 0,0027036 | 0,0011525 | 0,0011525 | TRUE | 0,0088932 | No |
| orf19.5788 | 40825 | 25818 | 36355 | 42738 | 41699 | 33997 | 37247 | 24690 | 36624 | 32869 | 37195 | 35242 | 35662 | 32439 | 33237 | 34823 | 34761 | 35482 | 34040 | 0,959 | -0,06 | -1,227 | 0,21980162 | 0,29856843 | 0,002362 | 0,0020213 | 0,002268 | 0,002268 | TRUE | 0,0299975 | No |
| orf19.5789 | 3683 | 2367 | 3180 | 3903 | 3918 | 3191 | 3447 | 2275 | 3304 | 3013 | 3253 | 3218 | 3351 | 3045 | 3076 | 3209 | 3183,7 | 3197 | 3170 | 0,991 | -0,013 | -0,286 | 0,77476608 | 0,82367931 | 0,001455 | 0,0024355 | 0,0018151 | 0,0018151 | TRUE | 0,0187408 | No |
| orf19.579 | 1217 | 915 | 1101 | 1309 | 1371 | 1244 | 1277 | 834 | 1092 | 1165 | 1126 | 1079 | 1173 | 1187 | 1140 | 1176 | 1142,2 | 1116 | 1169 | 1,049 | 0,069 | 1,43 | 0,15275743 | 0,21903738 | 1,00E-08 | 0,0032505 | 0,0013869 | 0,0013869 | TRUE | 0,0097125 | No |
| orf19.5791 | 7691 | 5098 | 6883 | 8725 | 7603 | 6005 | 6648 | 4725 | 6900 | 6490 | 7042 | 7195 | 6502 | 5730 | 5932 | 6664 | 6556,9 | 6907 | 6207 | 0,898 | -0,155 | -2,736 | 0,0062154 | 0,013144 | 0,003349 | 0,0022009 | 0,0029298 | 0,0029298 | TRUE | 0,0326154 | No |
| orf19.5792 | 559 | 391 | 470 | 607 | 477 | 457 | 497 | 335 | 501 | 498 | 481 | 501 | 408 | 436 | 443 | 472 | 467,58 | 495 | 440 | 0,885 | -0,177 | -2,527 | 0,0115027 | 0,02280799 | 1,00E-08 | 0,0050842 | 0,0025661 | 0,0025661 | TRUE | 0,0284787 | No |

| | | | | | | | | | | | | | | | | | | | | | | | | | | | | | | | |
|---|---|---|---|---|---|---|---|---|---|---|---|---|---|---|---|---|---|---|---|---|---|---|---|---|---|---|---|---|---|---|---|
| orf19.5793 | 10088 | 6559 | 8395 | 10557 | 9273 | 7706 | 8210 | 4965 | 9050 | 8350 | 8589 | 8705 | 7931 | 7353 | 7326 | 7003 | 8038,4 | 8674 | 7403 | 0,854 | -0,228 | -4,973 | 6,59E-07 | 2,86E-06 | 0,001756 | 0,0021601 | 0,0018946 | 0,0018946 | TRUE | 0,0282289 | No |
| orf19.5796 | 1369 | 879 | 1176 | 1308 | 1222 | 1010 | 1139 | 654 | 1228 | 1119 | 1203 | 1079 | 1045 | 964 | 1016 | 922 | 1072,1 | 1157 | 987 | 0,855 | -0,227 | -3,603 | 0,00031511 | 0,00088839 | 0,002477 | 0,0033336 | 0,0028541 | 0,0028541 | TRUE | 0,0270867 | No |
| orf19.5797 | 2839 | 1890 | 2401 | 2898 | 2083 | 1866 | 1854 | 1217 | 2547 | 2406 | 2456 | 2390 | 1781 | 1780 | 1654 | 1716 | 2091,5 | 2450 | 1733 | 0,708 | -0,499 | -11,267 | 1,91E-29 | 6,68E-28 | 0,000606 | 0,0026736 | 0,0013899 | 0,0013899 | TRUE | 0,0122029 | No |
| orf19.5798 | 2458 | 1709 | 2134 | 2471 | 2431 | 2190 | 2350 | 1408 | 2205 | 2176 | 2183 | 2038 | 2079 | 2090 | 2097 | 1986 | 2106,7 | 2150 | 2063 | 0,961 | -0,057 | -1,32 | 0,18667733 | 0,25925968 | 0,000522 | 0,0026686 | 0,0013329 | 0,0013329 | TRUE | 0,0159677 | No |
| orf19.5799 | 3616 | 2278 | 3135 | 3591 | 2656 | 2254 | 2404 | 1746 | 3244 | 2900 | 3207 | 2961 | 2271 | 2151 | 2145 | 2463 | 2667,8 | 3078 | 2258 | 0,732 | -0,45 | -7,669 | 1,74E-14 | 1,89E-13 | 0,003231 | 0,0025237 | 0,0029171 | 0,0029171 | TRUE | 0,0429937 | No |
| orf19.58 | 15009 | 10411 | 13312 | 16369 | 14747 | 13186 | 14392 | 9359 | 13465 | 13254 | 13619 | 13498 | 12612 | 12582 | 12842 | 13200 | 13134 | 13459 | 12809 | 0,951 | -0,072 | -2,564 | 0,0103472 | 0,02075659 | 0,00021 | 0,0020901 | 0,0006832 | 0,0006832 | TRUE | 0,0049662 | No |
| orf19.580 | 729 | 562 | 629 | 792 | 1155 | 1018 | 1056 | 607 | 654 | 715 | 644 | 653 | 988 | 971 | 942 | 856 | 802,96 | 666 | 939 | 1,417 | 0,502 | 7,846 | 4,30E-15 | 4,98E-14 | 0,001461 | 0,0037875 | 0,0026549 | 0,0026549 | TRUE | 0,0376297 | No |
| orf19.5801 | 11042 | 7716 | 9421 | 11501 | 10033 | 8863 | 9426 | 5789 | 9906 | 9823 | 9639 | 9484 | 8581 | 8457 | 8411 | 8165 | 9058,1 | 9713 | 8404 | 0,866 | -0,208 | -6,786 | 1,15E-11 | 9,36E-11 | 0,000257 | 0,0021398 | 0,0007909 | 0,0007909 | TRUE | 0,0042425 | No |
| orf19.5802 | 1731 | 1115 | 1453 | 1837 | 1579 | 1217 | 1411 | 841 | 1553 | 1420 | 1487 | 1515 | 1350 | 1161 | 1259 | 1186 | 1366,3 | 1494 | 1239 | 0,83 | -0,268 | -4,549 | 5,39E-06 | 2,04E-05 | 0,002076 | 0,003042 | 0,0025976 | 0,0025976 | TRUE | 0,0450495 | No |
| orf19.5804 | 1230 | 801 | 1130 | 1311 | 1701 | 1502 | 1696 | 1081 | 1103 | 1020 | 1156 | 1081 | 1455 | 1433 | 1513 | 1525 | 1285,8 | 1090 | 1482 | 1,356 | 0,44 | 8,45 | 2,91E-17 | 4,20E-16 | 0,00073 | 0,0031086 | 0,0018017 | 0,0018017 | TRUE | 0,0201081 | No |
| orf19.5805 | 639 | 419 | 543 | 647 | 773 | 610 | 726 | 475 | 573 | 533 | 556 | 534 | 661 | 582 | 648 | 670 | 594,58 | 549 | 640 | 1,164 | 0,219 | 3,236 | 0,00121305 | 0,00300909 | 0,00106 | 0,0044211 | 0,0027122 | 0,0027122 | TRUE | 0,0446135 | No |
| orf19.5806 | 82731 | 52036 | 70747 | 87076 | 60096 | 48214 | 53462 | 36525 | 74218 | 66247 | 72381 | 71804 | 51396 | 46004 | 47706 | 51515 | 60159 | 71162 | 49155 | 0,691 | -0,534 | -10,47 | 1,19E-25 | 3,18E-24 | 0,002702 | 0,0020037 | 0,0024809 | 0,0024809 | TRUE | 0,0264511 | No |
| orf19.5808 | 3657 | 2491 | 3237 | 4089 | 3724 | 3232 | 3539 | 2332 | 3281 | 3171 | 3312 | 3372 | 3185 | 3084 | 3158 | 3289 | 3231,4 | 3284 | 3179 | 0,966 | -0,049 | -1,303 | 0,19249068 | 0,26573622 | 0,000272 | 0,0024288 | 0,0010563 | 0,0010563 | TRUE | 0,0061153 | No |
| orf19.5809 | 123 | 76 | 113 | 124 | 66 | 55 | 67 | 57 | 110 | 97 | 116 | 102 | 56 | 52 | 60 | 80 | 84,26 | 106 | 62 | 0,575 | -0,798 | -4,883 | 1,04E-06 | 4,39E-06 | 0,000138 | 0,0192087 | 0,0128245 | 0,0128245 | TRUE | 0,250735 | No |
| orf19.581 | 14486 | 9525 | 13101 | 15930 | 14233 | 12148 | 13839 | 9161 | 12995 | 12126 | 13404 | 13136 | 12173 | 11591 | 12349 | 12921 | 12587 | 12915 | 12258 | 0,949 | -0,076 | -1,665 | 0,0958456 | 0,14750009 | 0,001807 | 0,0020949 | 0,0019181 | 0,0019181 | TRUE | 0,0204463 | No |
| orf19.5811 | 5520 | 3290 | 4883 | 5978 | 6038 | 4433 | 5083 | 3639 | 4952 | 4189 | 4996 | 4930 | 5164 | 4230 | 4536 | 5132 | 4766 | 4767 | 4766 | 0,999 | -0,001 | -0,016 | 0,9869941 | 0,9896901 | 0,007938 | 0,0022841 | 0,0054374 | 0,0054374 | TRUE | 0,0802526 | No |
| orf19.5812 | 5668 | 3725 | 4755 | 5822 | 5939 | 5284 | 5749 | 3719 | 5085 | 4742 | 4865 | 4801 | 5079 | 5042 | 5130 | 5245 | 4998,7 | 4873 | 5124 | 1,051 | 0,071 | 2,024 | 0,0429995 | 0,07370642 | 0,000354 | 0,0022699 | 0,0009879 | 0,0009879 | TRUE | 0,0105426 | No |
| orf19.5813 | 13699 | 9412 | 12696 | 15747 | 15116 | 13131 | 14448 | 9098 | 12289 | 11982 | 12989 | 12985 | 12928 | 12529 | 12892 | 12832 | 12678 | 12561 | 12795 | 1,018 | 0,026 | 0,718 | 0,47269681 | 0,55898623 | 0,000791 | 0,002094 | 0,0012049 | 0,0012049 | TRUE | 0,0115703 | No |
| orf19.5814 | 992 | 840 | 800 | 1097 | 963 | 863 | 872 | 573 | 890 | 1069 | 818 | 905 | 824 | 823 | 778 | 808 | 864,47 | 920 | 808 | 0,88 | -0,184 | -2,379 | 0,01734828 | 0,03278949 | 0,005741 | 0,0036588 | 0,0046039 | 0,0046039 | TRUE | 0,1331056 | No |
| orf19.5814.1 | 706 | 535 | 615 | 803 | 574 | 493 | 535 | 310 | 633 | 681 | 629 | 662 | 491 | 470 | 477 | 437 | 560,22 | 651 | 469 | 0,723 | -0,467 | -7,121 | 1,07E-12 | 9,82E-12 | 1,00E-08 | 0,0045708 | 0,0023007 | 0,0023007 | TRUE | 0,0198606 | No |
| orf19.5815 | 1237 | 917 | 1133 | 1321 | 1175 | 1092 | 1127 | 722 | 1110 | 1167 | 1159 | 1089 | 1005 | 1042 | 1006 | 1018 | 1074,6 | 1131 | 1018 | 0,901 | -0,151 | -3,039 | 0,00237501 | 0,00550928 | 1,00E-08 | 0,0033305 | 0,0014284 | 0,0014284 | TRUE | 0,007748 | No |
| orf19.5816 | 1295 | 845 | 1101 | 1326 | 1102 | 961 | 1098 | 715 | 1162 | 1076 | 1126 | 1093 | 942 | 917 | 980 | 1008 | 1038,1 | 1114 | 962 | 0,861 | -0,216 | -4,053 | 5,06E-05 | 0,00016334 | 0,00037 | 0,0033779 | 0,0017597 | 0,0017597 | TRUE | 0,0113342 | No |
| orf19.5817 | 1085 | 779 | 936 | 1230 | 989 | 950 | 988 | 593 | 973 | 992 | 958 | 1014 | 846 | 906 | 882 | 836 | 925,91 | 984 | 868 | 0,882 | -0,181 | -3,4 | 0,00067276 | 0,0017566 | 1,00E-08 | 0,0035474 | 0,0016248 | 0,0016248 | TRUE | 0,0106103 | No |
| orf19.5818 | 50160 | 33822 | 44526 | 54978 | 49848 | 43607 | 47803 | 28523 | 44999 | 43059 | 45554 | 45336 | 42632 | 41608 | 42656 | 40229 | 43259 | 44737 | 41781 | 0,934 | -0,099 | -2,972 | 0,00295794 | 0,00672645 | 0,00067 | 0,0020131 | 0,0010328 | 0,0010328 | TRUE | 0,007763 | No |
| orf19.5820 | 33 | 19 | 21 | 25 | 21 | 8 | 25 | 16 | 30 | 24 | 21 | 21 | 18 | 8 | 22 | 23 | 20,8 | 24 | 18 | 0,727 | -0,46 | -1,347 | 0,17793738 | 0,24959675 | 0,045998 | 0,0717888 | 0,0624078 | 0,0624078 | TRUE | 0,7730336 | No |
| orf19.5821 | 744 | 590 | 742 | 932 | 1290 | 1225 | 1298 | 773 | 667 | 751 | 759 | 769 | 1103 | 1169 | 1158 | 1090 | 933,36 | 736 | 1130 | 1,537 | 0,62 | 10,195 | 2,10E-24 | 5,12E-23 | 0,001451 | 0,0035349 | 0,002429 | 0,002429 | TRUE | 0,049752 | No |
| orf19.5823 | 4868 | 3128 | 4181 | 5088 | 5255 | 4252 | 4555 | 3078 | 4367 | 3982 | 4278 | 4196 | 4494 | 4057 | 4065 | 4341 | 4222,5 | 4206 | 4239 | 1,007 | 0,011 | 0,219 | 0,82688323 | 0,86684949 | 0,001543 | 0,0023233 | 0,0020036 | 0,0020036 | TRUE | 0,0207317 | No |
| orf19.5824 | 7278 | 4863 | 6138 | 7812 | 7128 | 6190 | 6573 | 4555 | 6529 | 6191 | 6280 | 6442 | 6096 | 5906 | 5865 | 6424 | 6216,8 | 6360 | 6073 | 0,954 | -0,068 | -1,714 | 0,08649992 | 0,1353011 | 0,000956 | 0,002213 | 0,00136 | 0,00136 | TRUE | 0,0177533 | No |
| orf19.5825 | 2202 | 1490 | 1835 | 2345 | 2571 | 2102 | 2238 | 1429 | 1975 | 1897 | 1877 | 1934 | 2199 | 2006 | 1997 | 2015 | 1987,6 | 1921 | 2054 | 1,07 | 0,097 | 2,071 | 0,0383452 | 0,06669889 | 0,000949 | 0,0027099 | 0,0016131 | 0,0016131 | TRUE | 0,0283438 | No |
| orf19.5825.1 | 755 | 476 | 582 | 792 | 899 | 732 | 737 | 509 | 677 | 606 | 595 | 653 | 769 | 698 | 658 | 718 | 671,84 | 633 | 711 | 1,121 | 0,164 | 2,304 | 0,02123268 | 0,03936006 | 0,002723 | 0,0041403 | 0,0033823 | 0,0033823 | TRUE | 0,0391517 | No |
| orf19.5826 | 30 | 18 | 20 | 20 | 16 | 16 | 17 | 10 | 27 | 23 | 20 | 16 | 14 | 15 | 15 | 14 | 18,13 | 22 | 14 | 0,675 | -0,568 | -1,752 | 0,07969974 | 0,12610904 | 1,00E-08 | 0,0820693 | 0,0436991 | 0,0436991 | TRUE | 0,1855913 | No |
| orf19.5827 | 134 | 76 | 108 | 138 | 153 | 154 | 159 | 126 | 120 | 97 | 110 | 114 | 131 | 147 | 142 | 178 | 129,83 | 110 | 150 | 1,333 | 0,415 | 3,096 | 0,00195854 | 0,00463203 | 0,004083 | 0,013161 | 0,0093876 | 0,0093876 | TRUE | 0,1421905 | No |
| orf19.5828 | 8549 | 5585 | 7611 | 9039 | 8705 | 7485 | 8333 | 5177 | 7669 | 7110 | 7787 | 7454 | 7445 | 7142 | 7436 | 7302 | 7418,1 | 7505 | 7331 | 0,977 | -0,034 | -0,901 | 0,36751553 | 0,4562995 | 0,000792 | 0,0021752 | 0,0012459 | 0,0012459 | TRUE | 0,0148432 | No |
| orf19.583 | 11259 | 7349 | 9831 | 12398 | 12120 | 10405 | 11307 | 7465 | 10100 | 9356 | 10058 | 10224 | 10365 | 9928 | 10090 | 10529 | 10081 | 9934 | 10228 | 1,029 | 0,041 | 1,054 | 0,29178105 | 0,37642701 | 0,000994 | 0,0021235 | 0,0013626 | 0,0013626 | TRUE | 0,0184374 | No |
| orf19.5830 | 1099 | 778 | 918 | 1144 | 1376 | 1189 | 1405 | 833 | 986 | 990 | 939 | 943 | 1177 | 1135 | 1254 | 1175 | 1074,9 | 964 | 1185 | 1,23 | 0,298 | 5,653 | 1,58E-08 | 8,53E-08 | 0,000515 | 0,0033301 | 0,001736 | 0,001736 | TRUE | 0,0192571 | No |
| orf19.5831 | 93 | 64 | 83 | 84 | 37 | 41 | 42 | 27 | 83 | 81 | 85 | 69 | 32 | 39 | 37 | 38 | 58,18 | 80 | 36 | 0,458 | -1,127 | -5,959 | 2,53E-09 | 1,52E-08 | 1,00E-08 | 0,0269326 | 0,0144055 | 0,0144055 | TRUE | 0,0843142 | No |
| orf19.5832 | 11159 | 7403 | 9382 | 11618 | 10298 | 8643 | 9182 | 5574 | 10011 | 9425 | 9599 | 9580 | 8807 | 8247 | 8193 | 7862 | 8965,5 | 9654 | 8277 | 0,858 | -0,221 | -5,269 | 1,37E-07 | 6,61E-07 | 0,00131 | 0,0021414 | 0,0015828 | 0,0015828 | TRUE | 0,0227899 | No |
| orf19.5833 | 5550 | 3543 | 4586 | 5923 | 5304 | 4432 | 4877 | 3085 | 4979 | 4511 | 4692 | 4884 | 4536 | 4229 | 4352 | 4351 | 4566,7 | 4766 | 4367 | 0,916 | -0,127 | -2,959 | 0,0030876 | 0,00698957 | 0,001082 | 0,0022974 | 0,0015491 | 0,0015491 | TRUE | 0,0154545 | No |
| orf19.5834 | 6532 | 4454 | 5816 | 6975 | 6435 | 5757 | 6201 | 4021 | 5860 | 5670 | 5950 | 5752 | 5503 | 5493 | 5533 | 5671 | 5679,2 | 5808 | 5550 | 0,955 | -0,067 | -2,141 | 0,03230454 | 0,05716809 | 9,80E-05 | 0,0022351 | 0,0007507 | 0,0007507 | TRUE | 0,0032329 | No |
| orf19.5835 | 9288 | 5991 | 7717 | 9577 | 8778 | 7343 | 8215 | 5251 | 8332 | 7627 | 7895 | 7897 | 7507 | 7006 | 7331 | 7406 | 7625,3 | 7938 | 7312 | 0,921 | -0,119 | -3,018 | 0,00254168 | 0,00586646 | 0,000939 | 0,0021699 | 0,0013592 | 0,0013592 | TRUE | 0,0136904 | No |
| orf19.5837 | 1244 | 768 | 1030 | 1237 | 1129 | 936 | 1090 | 643 | 1116 | 978 | 1054 | 1020 | 966 | 893 | 973 | 907 | 988,22 | 1042 | 935 | 0,897 | -0,157 | -2,638 | 0,0083291 | 0,01710031 | 0,0015 | 0,0034485 | 0,0023739 | 0,0023739 | TRUE | 0,027711 | No |
| orf19.5838 | 868 | 590 | 829 | 981 | 924 | 791 | 839 | 547 | 779 | 751 | 848 | 809 | 790 | 755 | 749 | 771 | 781,51 | 797 | 766 | 0,961 | -0,058 | -0,992 | 0,32114144 | 0,40773832 | 1,00E-08 | 0,0038371 | 0,0020179 | 0,0020179 | TRUE | 0,0211078 | No |
| orf19.5839 | 617 | 418 | 539 | 666 | 634 | 493 | 565 | 393 | 554 | 532 | 551 | 549 | 542 | 470 | 504 | 554 | 532,17 | 546 | 518 | 0,944 | -0,083 | -1,186 | 0,23562349 | 0,31641169 | 0,000746 | 0,0047074 | 0,0028189 | 0,0028189 | TRUE | 0,0438704 | No |
| orf19.584 | 4 | 4 | 5 | 5 | 7 | 10 | 11 | 7 | 4 | 5 | 5 | 4 | 6 | 10 | 10 | 10 | 6,64 | 4 | 9 | 1,97 | 0,979 | 1,818 | 0,06902143 | NA | 1,00E-08 | 0,2205423 | 0,1092348 | 0,1092348 | TRUE | 0,1812181 | No |
| orf19.5840 | 44 | 32 | 55 | 53 | 47 | 42 | 36 | 44 | 39 | 41 | 56 | 44 | 40 | 40 | 32 | 62 | 44,33 | 45 | 44 | 0,948 | -0,077 | -0,323 | 0,74653846 | 0,80106007 | 0,026357 | 0,0347271 | 0,0313192 | 0,0313192 | TRUE | 0,4849735 | No |
| orf19.5841 | 516 | 339 | 431 | 438 | 534 | 435 | 473 | 332 | 463 | 432 | 441 | 361 | 457 | 415 | 422 | 468 | 432,34 | 424 | 440 | 1,038 | 0,054 | 0,608 | 0,54332544 | 0,62534469 | 0,005447 | 0,0053373 | 0,0053933 | 0,0053933 | TRUE | 0,1253015 | No |
| orf19.5842 | 713 | 501 | 637 | 809 | 655 | 612 | 682 | 409 | 640 | 638 | 652 | 667 | 560 | 584 | 609 | 577 | 615,73 | 649 | 582 | 0,897 | -0,157 | -2,596 | 0,00944491 | 0,01913828 | 1,00E-08 | 0,0043372 | 0,0018978 | 0,0018978 | TRUE | 0,011717 | No |
| orf19.5843 | 2430 | 1789 | 2210 | 2741 | 2802 | 2790 | 2926 | 1856 | 2180 | 2278 | 2261 | 2260 | 2396 | 2662 | 2611 | 2618 | 2408,3 | 2245 | 2572 | 1,145 | 0,196 | 4,302 | 1,69E-05 | 5,95E-05 | 0,000987 | 0,0025823 | 0,001568 | 0,001568 | TRUE | 0,026595 | No |
| orf19.5844 | 8 | 3 | 8 | 8 | 6 | 3 | 4 | 4 | 7 | 4 | 8 | 7 | 5 | 3 | 4 | 6 | 5,37 | 6 | 4 | 0,647 | -0,628 | -1,067 | 0,28615443 | NA | 1,00E-08 | 0,272172 | 0,1388968 | 0,1388968 | TRUE | 0,1306809 | No |
| orf19.5845 | 4781 | 3274 | 4306 | 5269 | 4843 | 4288 | 4752 | 2993 | 4289 | 4168 | 4405 | 4345 | 4142 | 4091 | 4240 | 4221 | 4237,8 | 4302 | 4174 | 0,97 | -0,045 | -1,329 | 0,18396077 | 0,2561382 | 9,97E-05 | 0,0023221 | 0,0008487 | 0,0008487 | TRUE | 0,0049973 | No |
| orf19.5846 | 3401 | 2216 | 2828 | 3447 | 3361 | 2888 | 3216 | 2093 | 3051 | 2821 | 2893 | 2842 | 2874 | 2756 | 2870 | 2952 | 2882,5 | 2902 | 2863 | 0,985 | -0,021 | -0,506 | 0,61253167 | 0,68671822 | 0,000577 | 0,0024832 | 0,0013279 | 0,0013279 | TRUE | 0,0147848 | No |
| orf19.5847 | 10898 | 7233 | 9565 | 11819 | 10411 | 8906 | 9811 | 6183 | 9777 | 9208 | 9786 | 9746 | 8904 | 8498 | 8755 | 8721 | 9174,2 | 9629 | 8720 | 0,905 | -0,144 | -4,29 | 1,78E-05 | 6,25E-05 | 0,000465 | 0,0021378 | 0,0009668 | 0,0009668 | TRUE | 0,0102955 | No |
| orf19.5848 | 4868 | 3134 | 4248 | 5015 | 6486 | 5569 | 5956 | 4021 | 4367 | 3990 | 4346 | 4135 | 5547 | 5314 | 5315 | 5671 | 4835,7 | 4210 | 5462 | 1,296 | 0,374 | 8,712 | 2,99E-18 | 4,75E-17 | 0,001163 | 0,0022797 | 0,0015618 | 0,0015618 | TRUE | 0,0144057 | No |
| orf19.5849 | 2788 | 1795 | 2419 | 3118 | 3485 | 2757 | 3144 | 2030 | 2501 | 2285 | 2475 | 2571 | 2980 | 2631 | 2805 | 2863 | 2639 | 2458 | 2820 | 1,146 | 0,197 | 3,693 | 0,0002219 | 0,00064157 | 0,002063 | 0,0025296 | 0,0023379 | 0,0023379 | TRUE | 0,0261203 | No |
| orf19.585 | 3569 | 2501 | 3005 | 3645 | 3872 | 3501 | 3660 | 2382 | 3202 | 3184 | 3074 | 3006 | 3311 | 3341 | 3266 | 3360 | 3217,9 | 3116 | 3320 | 1,065 | 0,091 | 2,489 | 0,01280792 | 0,02508966 | 0,000239 | 0,0024307 | 0,0009779 | 0,0009779 | TRUE | 0,007284 | No |
| orf19.5850 | 18398 | 12733 | 16320 | 19935 | 16022 | 14594 | 16116 | 10022 | 16505 | 16210 | 16697 | 16439 | 13703 | 13925 | 14381 | 14135 | 15249 | 16463 | 14036 | 0,852 | -0,23 | -8,319 | 8,84E-17 | 1,22E-15 | 0,0002 | 0,0020747 | 0,0006697 | 0,0006697 | TRUE | 0,003449 | No |
| orf19.5851 | 956 | 695 | 909 | 1070 | 771 | 703 | 747 | 513 | 858 | 885 | 930 | 882 | 659 | 671 | 667 | 724 | 784,38 | 889 | 680 | 0,763 | -0,39 | -6,764 | 1,34E-11 | 1,08E-10 | 1,00E-08 | 0,0038303 | 0,0018918 | 0,0018918 | TRUE | 0,0185307 | No |
| orf19.5852 | 344 | 267 | 352 | 409 | 259 | 281 | 271 | 168 | 309 | 340 | 360 | 337 | 222 | 268 | 242 | 237 | 289,29 | 336 | 242 | 0,72 | -0,473 | -5,113 | 3,18E-07 | 1,45E-06 | 0,001961 | 0,0069977 | 0,0046778 | 0,0046778 | TRUE | 0,0610832 | No |
| orf19.5854 | 4813 | 3473 | 4199 | 5090 | 4547 | 4216 | 4487 | 2757 | 4318 | 4421 | 4296 | 4197 | 3889 | 4023 | 4004 | 3889 | 4129,6 | 4308 | 3951 | 0,918 | -0,123 | -3,666 | 0,00024682 | 0,00070682 | 0,000156 | 0,0023311 | 0,0008461 | 0,0008461 | TRUE | 0,0037278 | No |

| | | | | | | | | | | | | | | | | | | | | | | | | | | | | | | | |
|---|---|---|---|---|---|---|---|---|---|---|---|---|---|---|---|---|---|---|---|---|---|---|---|---|---|---|---|---|---|---|---|
| orf19.5854.1 | 145 | 111 | 134 | 146 | 126 | 122 | 134 | 90 | 130 | 141 | 137 | 120 | 108 | 116 | 120 | 127 | 124,95 | 132 | 118 | 0,889 | -0,169 | -1,348 | 0,17760399 | 0,24931212 | 1,00E-08 | 0,0135982 | 0,0070555 | 0,0070555 | TRUE | 0,0406483 | No |
| orf19.5855 | 3287 | 2352 | 2975 | 3606 | 4107 | 3523 | 4013 | 2631 | 2949 | 2994 | 3044 | 2974 | 3512 | 3362 | 3581 | 3711 | 3265,8 | 2990 | 3542 | 1,183 | 0,243 | 6,037 | 1,57E-09 | 9,70E-09 | 0,000523 | 0,002424 | 0,001246 | 0,001246 | TRUE | 0,0140141 | No |
| orf19.5856 | 2526 | 1850 | 2238 | 2803 | 2142 | 2040 | 2143 | 1280 | 2266 | 2355 | 2290 | 2311 | 1832 | 1947 | 1912 | 1805 | 2089,8 | 2306 | 1874 | 0,814 | -0,297 | -7,159 | 8,15E-13 | 7,55E-12 | 0,000117 | 0,0026742 | 0,0011652 | 0,0011652 | TRUE | 0,0079792 | No |
| orf19.5857 | 1104 | 788 | 1013 | 1252 | 1231 | 1070 | 1207 | 761 | 990 | 1003 | 1036 | 1032 | 1053 | 1021 | 1077 | 1073 | 1035,8 | 1015 | 1056 | 1,039 | 0,055 | 1,118 | 0,26356874 | 0,34731961 | 1,00E-08 | 0,003381 | 0,0013618 | 0,0013618 | TRUE | 0,0060209 | No |
| orf19.5858 | 2498 | 1637 | 2008 | 2458 | 2808 | 2122 | 2282 | 1372 | 2241 | 2084 | 2054 | 2027 | 2401 | 2025 | 2036 | 1935 | 2100,5 | 2102 | 2099 | 1 | 0 | 0,002 | 0,99832711 | 0,99865955 | 0,005188 | 0,0026706 | 0,0041164 | 0,0041164 | TRUE | 0,1149871 | No |
| orf19.5859 | 947 | 647 | 830 | 1078 | 759 | 589 | 695 | 430 | 850 | 824 | 849 | 889 | 649 | 562 | 620 | 606 | 731,14 | 853 | 609 | 0,714 | -0,485 | -7,734 | 1,04E-14 | 1,16E-13 | 0,000855 | 0,003965 | 0,0023722 | 0,0023722 | TRUE | 0,0344859 | No |
| orf19.586 | 1482 | 1092 | 1275 | 1610 | 1429 | 1282 | 1308 | 772 | 1330 | 1390 | 1304 | 1328 | 1222 | 1223 | 1167 | 1089 | 1256,7 | 1338 | 1175 | 0,882 | -0,182 | -3,484 | 0,00049352 | 0,00133442 | 0,000762 | 0,0031347 | 0,0018147 | 0,0018147 | TRUE | 0,0257887 | No |
| orf19.5860 | 185 | 132 | 135 | 188 | 134 | 128 | 128 | 77 | 166 | 168 | 138 | 155 | 115 | 122 | 114 | 109 | 135,84 | 157 | 115 | 0,736 | -0,443 | -3,625 | 0,00028887 | 0,00081857 | 1,00E-08 | 0,0126664 | 0,006811 | 0,006811 | TRUE | 0,0647894 | No |
| orf19.5861 | 5346 | 3624 | 4533 | 5977 | 4917 | 4308 | 4513 | 3276 | 4796 | 4614 | 4638 | 4929 | 4205 | 4111 | 4027 | 4621 | 4492,4 | 4744 | 4241 | 0,893 | -0,164 | -3,305 | 0,00094961 | 0,00239617 | 0,002112 | 0,0023027 | 0,0021351 | 0,0021351 | TRUE | 0,0425972 | No |
| orf19.5861.1 | 565 | 389 | 453 | 627 | 451 | 397 | 460 | 261 | 507 | 495 | 463 | 517 | 386 | 379 | 410 | 368 | 440,71 | 496 | 386 | 0,779 | -0,36 | -4,934 | 8,05E-07 | 3,44E-06 | 1,00E-08 | 0,0052735 | 0,0028013 | 0,0028013 | TRUE | 0,0226274 | No |
| orf19.5862 | 736 | 520 | 684 | 875 | 744 | 635 | 710 | 470 | 660 | 662 | 700 | 722 | 636 | 606 | 634 | 663 | 660,28 | 686 | 635 | 0,922 | -0,117 | -1,891 | 0,05867101 | 0,09656821 | 1,00E-08 | 0,0041781 | 0,0021435 | 0,0021435 | TRUE | 0,0154996 | No |
| orf19.5863 | 87 | 76 | 87 | 89 | 71 | 79 | 73 | 32 | 78 | 97 | 89 | 73 | 61 | 75 | 65 | 45 | 72,95 | 84 | 62 | 0,745 | -0,424 | -2,356 | 0,01849701 | 0,03467396 | 0,009448 | 0,0218802 | 0,0171205 | 0,0171205 | TRUE | 0,2044619 | No |
| orf19.5864 | 5465 | 4277 | 5018 | 6099 | 5885 | 5804 | 6192 | 3443 | 4903 | 5445 | 5134 | 5029 | 5033 | 5538 | 5525 | 4856 | 5182,9 | 5128 | 5238 | 1,022 | 0,032 | 0,584 | 0,55901587 | 0,63924007 | 0,002707 | 0,0022596 | 0,0027009 | 0,0027009 | TRUE | 0,0285432 | No |
| orf19.5865 | 2324 | 1651 | 2147 | 2652 | 2181 | 2142 | 2415 | 1499 | 2085 | 2102 | 2197 | 2187 | 1865 | 2044 | 2155 | 2114 | 2093,6 | 2143 | 2044 | 0,953 | -0,069 | -1,3 | 0,19353951 | 0,26687753 | 0,001957 | 0,0026729 | 0,0022373 | 0,0022373 | TRUE | 0,0436244 | No |
| orf19.5866 | 20 | 20 | 16 | 22 | 27 | 24 | 20 | 14 | 18 | 25 | 16 | 18 | 23 | 23 | 18 | 20 | 20,19 | 19 | 21 | 1,09 | 0,125 | 0,411 | 0,68121741 | 0,74549257 | 1,00E-08 | 0,0738909 | 0,0387825 | 0,0387825 | TRUE | 0,1724665 | No |
| orf19.5867 | 3675 | 2461 | 3280 | 4084 | 4279 | 3874 | 4088 | 2645 | 3297 | 3133 | 3356 | 3368 | 3660 | 3696 | 3648 | 3731 | 3486 | 3288 | 3684 | 1,119 | 0,162 | 4,454 | 8,43E-06 | 3,11E-05 | 0,000208 | 0,002396 | 0,0009787 | 0,0009787 | TRUE | 0,0114919 | No |
| orf19.5869 | 4940 | 3309 | 4302 | 5247 | 4730 | 4341 | 4722 | 3001 | 4432 | 4213 | 4401 | 4327 | 4045 | 4142 | 4214 | 4233 | 4250,8 | 4343 | 4158 | 0,957 | -0,064 | -1,875 | 0,06085379 | 0,09972984 | 0,000147 | 0,002321 | 0,0008866 | 0,0008866 | TRUE | 0,0046789 | No |
| orf19.587 | 331 | 262 | 294 | 355 | 357 | 310 | 304 | 195 | 297 | 334 | 301 | 293 | 305 | 296 | 271 | 275 | 296,43 | 306 | 287 | 0,943 | -0,084 | -0,975 | 0,32967759 | 0,41690233 | 1,00E-08 | 0,0068768 | 0,0038336 | 0,0038336 | TRUE | 0,036594 | No |
| orf19.5870 | 490 | 317 | 398 | 518 | 484 | 449 | 450 | 266 | 440 | 404 | 407 | 427 | 414 | 428 | 402 | 375 | 412,07 | 420 | 405 | 0,967 | -0,049 | -0,662 | 0,50792509 | 0,59231637 | 1,00E-08 | 0,0055025 | 0,0028556 | 0,0028556 | TRUE | 0,0227254 | No |
| orf19.5871 | 396 | 292 | 354 | 442 | 640 | 539 | 597 | 367 | 355 | 372 | 362 | 364 | 547 | 514 | 533 | 518 | 445,71 | 363 | 528 | 1,456 | 0,542 | 7,899 | 2,81E-15 | 3,31E-14 | 1,00E-08 | 0,0052366 | 0,0022157 | 0,0022157 | TRUE | 0,0074759 | No |
| orf19.5873 | 663 | 483 | 615 | 663 | 539 | 527 | 518 | 328 | 595 | 615 | 629 | 547 | 461 | 503 | 462 | 463 | 534,28 | 596 | 472 | 0,794 | -0,333 | -4,684 | 2,81E-06 | 1,11E-05 | 0,001323 | 0,0046966 | 0,0029569 | 0,0029569 | TRUE | 0,0394783 | No |
| orf19.5874 | 5 | 3 | 2 | 4 | 2 | 1 | 1 | 1 | 4 | 4 | 2 | 3 | 2 | 1 | 1 | 1 | 2,33 | 3 | 1 | 0,362 | -1,466 | -1,562 | 0,11829267 | NA | 1,00E-08 | 0,6258014 | 0,2985208 | 0,2985208 | TRUE | 0,102477 | No |
| orf19.5875 | 407 | 282 | 308 | 452 | 254 | 198 | 222 | 151 | 365 | 359 | 315 | 373 | 217 | 189 | 198 | 213 | 278,65 | 353 | 204 | 0,577 | -0,793 | -8,41 | 4,10E-17 | 5,87E-16 | 1,00E-08 | 0,0071893 | 0,0046672 | 0,0046672 | TRUE | 0,0597214 | No |
| orf19.5876 | 20763 | 13156 | 16852 | 21323 | 21643 | 17430 | 18729 | 11093 | 18627 | 16749 | 17241 | 17583 | 18510 | 16631 | 16713 | 15646 | 17212 | 17550 | 16875 | 0,962 | -0,056 | -1,017 | 0,30894782 | 0,39501139 | 0,003354 | 0,0020639 | 0,0028974 | 0,0028974 | TRUE | 0,0521382 | No |
| orf19.5877 | 10143 | 6918 | 9014 | 10724 | 13681 | 12497 | 13790 | 8256 | 9099 | 8807 | 9222 | 8843 | 11700 | 11924 | 12305 | 11644 | 10443 | 8993 | 11893 | 1,323 | 0,403 | 12,363 | 4,17E-35 | 2,05E-33 | 0,000437 | 0,0021185 | 0,0009248 | 0,0009248 | TRUE | 0,0066854 | No |
| orf19.5879 | 165 | 128 | 148 | 179 | 150 | 110 | 112 | 66 | 148 | 163 | 151 | 148 | 128 | 105 | 100 | 93 | 129,53 | 152 | 106 | 0,707 | -0,5 | -3,821 | 0,00013289 | 0,00039782 | 0,001533 | 0,0131866 | 0,0085453 | 0,0085453 | TRUE | 0,1989377 | No |
| orf19.588 | 674 | 482 | 548 | 752 | 600 | 556 | 527 | 372 | 605 | 614 | 561 | 620 | 513 | 531 | 470 | 525 | 554,71 | 600 | 510 | 0,848 | -0,238 | -3,472 | 0,00051728 | 0,00139168 | 8,56E-06 | 0,0045966 | 0,0026875 | 0,0026875 | TRUE | 0,0332165 | No |
| orf19.5880 | 506 | 340 | 441 | 540 | 552 | 437 | 524 | 341 | 454 | 433 | 451 | 445 | 472 | 417 | 468 | 481 | 452,61 | 446 | 460 | 1,028 | 0,04 | 0,551 | 0,5818742 | 0,66047614 | 1,00E-08 | 0,0051869 | 0,0027666 | 0,0027666 | TRUE | 0,0454431 | No |
| orf19.5884 | 2283 | 1696 | 2115 | 2514 | 1836 | 1650 | 1800 | 1050 | 2048 | 2159 | 2164 | 2073 | 1570 | 1574 | 1606 | 1481 | 1834,5 | 2111 | 1558 | 0,74 | -0,434 | -9,813 | 9,90E-23 | 2,18E-21 | 0,000245 | 0,0027709 | 0,0013221 | 0,0013221 | TRUE | 0,0117096 | No |
| orf19.5885 | 17759 | 11926 | 14456 | 18581 | 15771 | 13780 | 14068 | 9321 | 15932 | 15183 | 14790 | 15322 | 13488 | 13148 | 12553 | 13146 | 14195 | 15307 | 13084 | 0,855 | -0,227 | -6,185 | 6,23E-10 | 4,08E-09 | 0,000825 | 0,0020818 | 0,0012172 | 0,0012172 | TRUE | 0,0092877 | No |
| orf19.5886 | 4474 | 2980 | 3798 | 4821 | 5306 | 4320 | 4492 | 3243 | 4014 | 3794 | 3886 | 3975 | 4538 | 4122 | 4008 | 4574 | 4113,9 | 3917 | 4310 | 1,099 | 0,137 | 2,647 | 0,00812928 | 0,016732 | 0,002213 | 0,0023324 | 0,0023171 | 0,0023171 | TRUE | 0,0272766 | No |
| orf19.5887 | 3189 | 2271 | 2891 | 3371 | 3704 | 3473 | 3611 | 2287 | 2861 | 2891 | 2958 | 2780 | 3168 | 3314 | 3222 | 3226 | 3052,4 | 2872 | 3232 | 1,126 | 0,171 | 4,629 | 3,67E-06 | 1,43E-05 | 0,000223 | 0,0024551 | 0,0009807 | 0,0009807 | TRUE | 0,0059933 | No |
| orf19.589 | 19364 | 12584 | 17107 | 21182 | 20209 | 16518 | 18226 | 11740 | 17372 | 16021 | 17502 | 17467 | 17283 | 15761 | 16264 | 16558 | 16778 | 17090 | 16466 | 0,963 | -0,054 | -1,246 | 0,21272544 | 0,29000555 | 0,001543 | 0,002066 | 0,0017361 | 0,0017361 | TRUE | 0,0215203 | No |
| orf19.5890 | 3205 | 2197 | 2848 | 3462 | 4013 | 3530 | 3759 | 2387 | 2875 | 2797 | 2914 | 2855 | 3432 | 3368 | 3354 | 3367 | 3120,3 | 2860 | 3380 | 1,182 | 0,241 | 7,197 | 6,16E-13 | 5,79E-12 | 1,00E-08 | 0,0024448 | 0,0007508 | 0,0007508 | TRUE | 0,0024245 | No |
| orf19.5892 | 219 | 153 | 202 | 234 | 249 | 230 | 257 | 175 | 196 | 195 | 207 | 193 | 213 | 219 | 229 | 247 | 212,43 | 198 | 227 | 1,143 | 0,193 | 1,994 | 0,04617789 | 0,07841628 | 1,00E-08 | 0,0088133 | 0,0043129 | 0,0043129 | TRUE | 0,0303541 | No |
| orf19.5893 | 6607 | 4336 | 5501 | 7124 | 6832 | 5490 | 5945 | 3696 | 5927 | 5520 | 5628 | 5875 | 5843 | 5238 | 5305 | 5213 | 5568,6 | 5738 | 5400 | 0,941 | -0,087 | -1,829 | 0,06736694 | 0,10924172 | 0,00186 | 0,0022402 | 0,0020095 | 0,0020095 | TRUE | 0,0376979 | No |
| orf19.5894 | 24 | 23 | 20 | 17 | 30 | 25 | 27 | 22 | 22 | 29 | 20 | 14 | 26 | 24 | 24 | 31 | 23,74 | 21 | 26 | 1,24 | 0,311 | 1,053 | 0,29233805 | 0,3769028 | 1,00E-08 | 0,0631277 | 0,0405739 | 0,0405739 | TRUE | 0,3548809 | No |
| orf19.5896 | 88 | 62 | 81 | 90 | 121 | 108 | 102 | 75 | 79 | 79 | 83 | 74 | 103 | 103 | 91 | 106 | 89,79 | 79 | 101 | 1,279 | 0,355 | 2,461 | 0,01385194 | 0,02689802 | 1,00E-08 | 0,0181478 | 0,0087267 | 0,0087267 | TRUE | 0,0451831 | No |
| orf19.5897 | 452 | 289 | 434 | 449 | 550 | 492 | 579 | 380 | 405 | 368 | 444 | 370 | 470 | 469 | 517 | 536 | 447,52 | 397 | 498 | 1,251 | 0,323 | 3,856 | 0,00011506 | 0,00034809 | 0,003675 | 0,0052234 | 0,0044701 | 0,0044701 | TRUE | 0,0716153 | No |
| orf19.59 | 12093 | 8370 | 10180 | 12849 | 11987 | 10379 | 11078 | 7019 | 10849 | 10656 | 10415 | 10595 | 10252 | 9903 | 9885 | 9900 | 10307 | 10629 | 9985 | 0,94 | -0,09 | -3,122 | 0,00179771 | 0,00428937 | 0,000171 | 0,0021204 | 0,0006994 | 0,0006994 | TRUE | 0,0040392 | No |
| orf19.590 | 1931 | 1189 | 1591 | 1970 | 1739 | 1356 | 1516 | 1012 | 1732 | 1514 | 1628 | 1624 | 1487 | 1294 | 1353 | 1427 | 1507,4 | 1624 | 1390 | 0,855 | -0,227 | -3,769 | 0,00016417 | 0,00048492 | 0,002472 | 0,0029425 | 0,0028032 | 0,0028032 | TRUE | 0,0284169 | No |
| orf19.5901 | 3373 | 2289 | 2955 | 3646 | 4463 | 3720 | 4137 | 2726 | 3026 | 2914 | 3023 | 3007 | 3817 | 3550 | 3692 | 3845 | 3359,1 | 2992 | 3726 | 1,244 | 0,315 | 8,022 | 1,04E-15 | 1,29E-14 | 0,000529 | 0,0024117 | 0,0011754 | 0,0011754 | TRUE | 0,0123208 | No |
| orf19.5902 | 4288 | 3097 | 3864 | 4999 | 5768 | 5316 | 5793 | 3986 | 3847 | 3943 | 3953 | 4122 | 4933 | 5072 | 5169 | 5622 | 4582,7 | 3966 | 5199 | 1,309 | 0,389 | 8,139 | 4,00E-16 | 5,11E-15 | 0,001764 | 0,0022963 | 0,0019696 | 0,0019696 | TRUE | 0,0353602 | No |
| orf19.5903 | 3505 | 2609 | 3466 | 4029 | 3698 | 3340 | 3896 | 2379 | 3144 | 3322 | 3546 | 3322 | 3163 | 3187 | 3477 | 3355 | 3314,5 | 3334 | 3296 | 0,988 | -0,017 | -0,332 | 0,73972868 | 0,79646772 | 0,001952 | 0,0024175 | 0,0021245 | 0,0021245 | TRUE | 0,0224594 | No |
| orf19.5904 | 23980 | 15749 | 19443 | 24433 | 23427 | 19785 | 20655 | 13114 | 21513 | 20050 | 19892 | 20148 | 20036 | 18878 | 18431 | 18496 | 19680 | 20401 | 18960 | 0,929 | -0,106 | -2,56 | 0,01047841 | 0,02097776 | 0,001374 | 0,0020533 | 0,0015825 | 0,0015825 | TRUE | 0,017932 | No |
| orf19.5905 | 5226 | 3565 | 4161 | 5410 | 4868 | 4107 | 4387 | 2411 | 4688 | 4539 | 4257 | 4461 | 4163 | 3919 | 3915 | 3400 | 4167,8 | 4486 | 3849 | 0,859 | -0,219 | -3,59 | 0,0003301 | 0,00092742 | 0,003956 | 0,0023278 | 0,0033426 | 0,0033426 | TRUE | 0,0727209 | No |
| orf19.5906 | 9529 | 6434 | 8538 | 10382 | 9784 | 8326 | 8706 | 6151 | 8549 | 8191 | 8735 | 8561 | 8368 | 7944 | 7769 | 8675 | 8349 | 8509 | 8189 | 0,962 | -0,056 | -1,294 | 0,19567759 | 0,26939298 | 0,001456 | 0,0021534 | 0,0016927 | 0,0016927 | TRUE | 0,0190178 | No |
| orf19.5908 | 8315 | 5539 | 7535 | 9176 | 10566 | 9139 | 10326 | 6845 | 7459 | 7052 | 7709 | 7567 | 9036 | 8720 | 9214 | 9654 | 8301,5 | 7447 | 9156 | 1,229 | 0,297 | 6,82 | 9,13E-12 | 7,49E-11 | 0,001463 | 0,0021544 | 0,0017014 | 0,0017014 | TRUE | 0,0159943 | No |
| orf19.5910 | 855 | 624 | 735 | 894 | 834 | 684 | 757 | 495 | 767 | 794 | 752 | 737 | 713 | 653 | 675 | 698 | 723,77 | 762 | 685 | 0,899 | -0,154 | -2,617 | 0,0088801 | 0,01811601 | 1,00E-08 | 0,0039853 | 0,0019188 | 0,0019188 | TRUE | 0,012157 | No |
| orf19.5911 | 2353 | 1442 | 2022 | 2521 | 1957 | 1728 | 1852 | 1318 | 2111 | 1836 | 2069 | 2079 | 1674 | 1649 | 1653 | 1859 | 1866 | 2024 | 1709 | 0,842 | -0,249 | -4,168 | 3,08E-05 | 0,0001034 | 0,002976 | 0,0027575 | 0,0028705 | 0,0028705 | TRUE | 0,0451895 | No |
| orf19.5912 | 26861 | 17922 | 23801 | 28377 | 24050 | 21002 | 23064 | 13614 | 24097 | 22817 | 24351 | 23400 | 20568 | 20039 | 20581 | 19201 | 21882 | 23666 | 20097 | 0,849 | -0,236 | -6,461 | 1,04E-10 | 7,57E-10 | 0,00089 | 0,0020459 | 0,001231 | 0,001231 | TRUE | 0,0108661 | No |
| orf19.5915 | 281 | 224 | 268 | 276 | 280 | 220 | 259 | 174 | 252 | 285 | 274 | 228 | 239 | 210 | 231 | 245 | 245,62 | 260 | 231 | 0,894 | -0,162 | -1,594 | 0,11101634 | 0,16666321 | 0,003412 | 0,0078899 | 0,0058204 | 0,0058204 | TRUE | 0,082451 | No |
| orf19.5917 | 3882 | 2717 | 3399 | 4342 | 2916 | 2522 | 2873 | 1721 | 3483 | 3459 | 3478 | 3580 | 2494 | 2406 | 2564 | 2427 | 2986,4 | 3500 | 2473 | 0,707 | -0,501 | -13,447 | 3,21E-41 | 2,24E-39 | 0,000177 | 0,0024656 | 0,000984 | 0,000984 | TRUE | 0,0075489 | No |
| orf19.5917.3 | 7036 | 5124 | 6263 | 7991 | 7939 | 7433 | 7900 | 5492 | 6312 | 6523 | 6408 | 6589 | 6790 | 7092 | 7049 | 7746 | 6813,8 | 6458 | 7169 | 1,109 | 0,15 | 3,333 | 0,00086023 | 0,00219272 | 0,001597 | 0,0021926 | 0,0017887 | 0,0017887 | TRUE | 0,034875 | No |
| orf19.5918 | 14 | 6 | 13 | 9 | 9 | 10 | 7 | 2 | 13 | 8 | 13 | 7 | 8 | 10 | 6 | 3 | 8,4 | 10 | 7 | 0,662 | -0,595 | -1,211 | 0,22572254 | NA | 1,00E-08 | 0,1747347 | 0,1093604 | 0,1093604 | TRUE | 0,2076795 | No |
| orf19.5919 | 491 | 490 | 523 | 693 | 464 | 553 | 567 | 376 | 440 | 624 | 535 | 571 | 397 | 528 | 506 | 530 | 516,45 | 542 | 490 | 0,903 | -0,147 | -1,253 | 0,21010857 | 0,28708944 | 0,017762 | 0,0047905 | 0,0113043 | 0,0113043 | TRUE | 0,1980906 | No |
| orf19.592 | 1971 | 1197 | 1740 | 2189 | 2635 | 2036 | 2442 | 1536 | 1768 | 1524 | 1780 | 1805 | 2254 | 1943 | 2179 | 2166 | 1927,4 | 1719 | 2136 | 1,24 | 0,311 | 4,68 | 2,88E-06 | 1,14E-05 | 0,004183 | 0,0027327 | 0,0037055 | 0,0037055 | TRUE | 0,0678275 | No |
| orf19.5920 | 1893 | 1331 | 1516 | 2096 | 2437 | 2071 | 2191 | 1059 | 1698 | 1695 | 1551 | 1728 | 2084 | 1976 | 1955 | 1494 | 1772,6 | 1668 | 1877 | 1,127 | 0,173 | 1,935 | 0,05304878 | 0,08853252 | 0,010114 | 0,0027985 | 0,0071069 | 0,0071069 | TRUE | 0,2210809 | No |
| orf19.5921 | 967 | 704 | 884 | 1123 | 1046 | 1063 | 993 | 732 | 868 | 896 | 904 | 926 | 895 | 1014 | 886 | 1032 | 927,7 | 898 | 957 | 1,062 | 0,086 | 1,32 | 0,18672772 | 0,25926973 | 0,002581 | 0,0035444 | 0,0030149 | 0,0030149 | TRUE | 0,030873 | No |

| | | | | | | | | | | | | | | | | | | | | | | | | | | | | | | | |
|---|---|---|---|---|---|---|---|---|---|---|---|---|---|---|---|---|---|---|---|---|---|---|---|---|---|---|---|---|---|---|---|
| orf19.5924 | 834 | 585 | 748 | 862 | 802 | 810 | 906 | 500 | 748 | 745 | 765 | 711 | 686 | 773 | 808 | 705 | 742,68 | 742 | 743 | 1,003 | 0,005 | 0,069 | 0,94512122 | 0,96046825 | 0,002277 | 0,0039342 | 0,0030403 | 0,0030403 | TRUE | 0,0428943 | No |
| orf19.5925 | 600 | 444 | 458 | 654 | 712 | 700 | 730 | 417 | 538 | 565 | 469 | 539 | 609 | 668 | 651 | 588 | 578,47 | 528 | 629 | 1,195 | 0,257 | 3,432 | 0,00059906 | 0,00158414 | 0,002774 | 0,0044891 | 0,0036443 | 0,0036443 | TRUE | 0,0649499 | No |
| orf19.5926 | 920 | 627 | 807 | 1014 | 1050 | 963 | 1052 | 640 | 825 | 798 | 826 | 836 | 898 | 919 | 939 | 903 | 867,95 | 821 | 915 | 1,113 | 0,154 | 2,983 | 0,00285809 | 0,00651666 | 1,00E-08 | 0,0036521 | 0,0014113 | 0,0014113 | TRUE | 0,0039634 | No |
| orf19.5927 | 13352 | 8720 | 10283 | 13122 | 13795 | 11278 | 11473 | 8004 | 11978 | 11101 | 10521 | 10821 | 11798 | 10761 | 10238 | 11289 | 11063 | 11105 | 11022 | 0,992 | -0,011 | -0,199 | 0,84252181 | 0,8800193 | 0,002891 | 0,0021107 | 0,0028902 | 0,0028902 | TRUE | 0,0344098 | No |
| orf19.5928 | 6927 | 4295 | 5758 | 7372 | 7115 | 5470 | 5372 | 4482 | 6214 | 5468 | 5891 | 6079 | 6085 | 5219 | 4794 | 6321 | 5759 | 5913 | 5605 | 0,947 | -0,078 | -0,951 | 0,3415721 | 0,42839515 | 0,008687 | 0,0022316 | 0,0063038 | 0,0063038 | TRUE | 0,1160846 | No |
| orf19.5929 | 613 | 463 | 579 | 686 | 860 | 769 | 803 | 567 | 550 | 589 | 592 | 566 | 735 | 734 | 717 | 800 | 660,37 | 574 | 746 | 1,297 | 0,376 | 6,041 | 1,53E-09 | 9,50E-09 | 1,00E-08 | 0,0041778 | 0,0021731 | 0,0021731 | TRUE | 0,0236553 | No |
| orf19.593 | 2380 | 1526 | 2221 | 2551 | 2883 | 2439 | 2722 | 1724 | 2135 | 1943 | 2272 | 2104 | 2466 | 2327 | 2429 | 2432 | 2263,4 | 2114 | 2414 | 1,141 | 0,19 | 3,681 | 0,00023214 | 0,00066701 | 0,001765 | 0,0026209 | 0,0021153 | 0,0021153 | TRUE | 0,0336116 | No |
| orf19.5930 | 1550 | 979 | 1299 | 1717 | 1929 | 1481 | 1650 | 1180 | 1391 | 1246 | 1329 | 1416 | 1650 | 1413 | 1472 | 1664 | 1447,7 | 1346 | 1550 | 1,149 | 0,201 | 2,973 | 0,00295163 | 0,0067172 | 0,003961 | 0,0029823 | 0,0036733 | 0,0036733 | TRUE | 0,0423578 | No |
| orf19.5931 | 150 | 157 | 178 | 193 | 166 | 183 | 165 | 74 | 135 | 200 | 182 | 159 | 142 | 175 | 147 | 104 | 155,49 | 169 | 142 | 0,852 | -0,232 | -1,435 | 0,15129592 | 0,21720093 | 0,027423 | 0,011316 | 0,0185717 | 0,0185717 | TRUE | 0,2770596 | No |
| orf19.5932 | 1775 | 1315 | 1674 | 2008 | 2542 | 2238 | 2471 | 1486 | 1592 | 1674 | 1713 | 1656 | 2174 | 2135 | 2205 | 2096 | 1905,7 | 1659 | 2152 | 1,3 | 0,378 | 8,955 | 3,39E-19 | 5,80E-18 | 1,00E-08 | 0,0027413 | 0,0011789 | 0,0011789 | TRUE | 0,0090061 | No |
| orf19.5933 | 66 | 56 | 60 | 77 | 72 | 71 | 86 | 45 | 59 | 71 | 61 | 63 | 62 | 68 | 77 | 63 | 65,61 | 64 | 68 | 1,064 | 0,089 | 0,525 | 0,5999201 | 0,67623263 | 1,00E-08 | 0,0241042 | 0,0125997 | 0,0125997 | TRUE | 0,0851991 | No |
| orf19.5934 | 652 | 395 | 540 | 588 | 281 | 275 | 281 | 156 | 585 | 503 | 552 | 485 | 240 | 262 | 251 | 220 | 387,33 | 531 | 243 | 0,46 | -1,121 | -12,369 | 3,87E-35 | 1,92E-33 | 0,00406 | 0,0057275 | 0,0048908 | 0,0048908 | TRUE | 0,0563136 | No |
| orf19.5935 | 1147 | 791 | 1052 | 1255 | 1090 | 970 | 1076 | 708 | 1029 | 1007 | 1076 | 1035 | 932 | 926 | 960 | 999 | 995,46 | 1037 | 954 | 0,918 | -0,124 | -2,402 | 0,01629077 | 0,0310517 | 1,00E-08 | 0,0034378 | 0,0015358 | 0,0015358 | TRUE | 0,0100222 | No |
| orf19.5938 | 2337 | 1512 | 2022 | 2454 | 2342 | 2185 | 2308 | 1478 | 2097 | 1925 | 2069 | 2024 | 2003 | 2085 | 2060 | 2085 | 2043,2 | 2029 | 2058 | 1,013 | 0,018 | 0,433 | 0,66475068 | 0,73281139 | 0,000297 | 0,00269 | 0,0012476 | 0,0012476 | TRUE | 0,0131111 | No |
| orf19.5940 | 1220 | 799 | 1141 | 1294 | 1169 | 996 | 1083 | 749 | 1094 | 1017 | 1167 | 1067 | 1000 | 950 | 966 | 1056 | 1039,9 | 1086 | 993 | 0,912 | -0,134 | -2,26 | 0,02382609 | 0,04368237 | 0,001492 | 0,0033756 | 0,0023909 | 0,0023909 | TRUE | 0,0289728 | No |
| orf19.5941 | 644 | 442 | 534 | 704 | 642 | 546 | 572 | 333 | 578 | 563 | 546 | 581 | 549 | 521 | 510 | 470 | 539,68 | 567 | 512 | 0,907 | -0,14 | -2,068 | 0,03867071 | 0,06710965 | 1,00E-08 | 0,0046695 | 0,0025714 | 0,0025714 | TRUE | 0,0309455 | No |
| orf19.5942 | 2632 | 1798 | 2385 | 2841 | 2792 | 2452 | 2670 | 1548 | 2361 | 2289 | 2440 | 2343 | 2388 | 2340 | 2383 | 2183 | 2340,8 | 2358 | 2324 | 0,987 | -0,019 | -0,446 | 0,6556169 | 0,72433732 | 0,000632 | 0,0025997 | 0,0014027 | 0,0014027 | TRUE | 0,0181758 | No |
| orf19.5943 | 1482 | 965 | 1296 | 1576 | 1312 | 1289 | 1359 | 768 | 1330 | 1229 | 1326 | 1300 | 1122 | 1230 | 1213 | 1083 | 1228,9 | 1296 | 1162 | 0,897 | -0,156 | -2,77 | 0,00559808 | 0,01196062 | 0,001415 | 0,0031608 | 0,0022307 | 0,0022307 | TRUE | 0,022048 | No |
| orf19.5943.1 | 7749 | 5173 | 6511 | 8130 | 7773 | 6414 | 6865 | 4467 | 6952 | 6586 | 6661 | 6704 | 6648 | 6120 | 6126 | 6300 | 6512,1 | 6726 | 6298 | 0,936 | -0,095 | -2,426 | 0,01525181 | 0,02926632 | 0,000844 | 0,0022024 | 0,0013131 | 0,0013131 | TRUE | 0,0174811 | No |
| orf19.5947 | 2481 | 1749 | 2238 | 2575 | 2604 | 2335 | 2558 | 1658 | 2226 | 2227 | 2290 | 2123 | 2227 | 2228 | 2283 | 2338 | 2242,7 | 2216 | 2269 | 1,023 | 0,033 | 0,808 | 0,4188826 | 0,50647075 | 0,000335 | 0,0026268 | 0,0011902 | 0,0011902 | TRUE | 0,0102135 | No |
| orf19.5949 | 10537 | 7091 | 9620 | 10840 | 6066 | 5619 | 6144 | 3856 | 9453 | 9028 | 9842 | 8939 | 5188 | 5361 | 5482 | 5439 | 7341,5 | 9316 | 5368 | 0,576 | -0,796 | -19,157 | 8,44E-82 | 2,67E-79 | 0,001098 | 0,0021773 | 0,0015086 | 0,0015086 | TRUE | 0,0176677 | No |
| orf19.5952 | 481 | 328 | 419 | 491 | 272 | 249 | 295 | 159 | 432 | 418 | 429 | 405 | 233 | 238 | 263 | 224 | 330,04 | 421 | 240 | 0,572 | -0,807 | -9,754 | 1,77E-22 | 3,85E-21 | 1,00E-08 | 0,006378 | 0,0033097 | 0,0033097 | TRUE | 0,0513718 | No |
| orf19.5953 | 7783 | 4856 | 7153 | 8506 | 7255 | 5963 | 6990 | 4468 | 6982 | 6182 | 7318 | 7014 | 6205 | 5690 | 6237 | 6302 | 6491,3 | 6874 | 6108 | 0,888 | -0,171 | -2,99 | 0,00278972 | 0,00637772 | 0,003354 | 0,0022032 | 0,0029906 | 0,0029906 | TRUE | 0,0552417 | No |
| orf19.5954 | 66 | 53 | 83 | 83 | 100 | 71 | 63 | 41 | 59 | 67 | 85 | 68 | 86 | 68 | 56 | 58 | 68,42 | 70 | 67 | 0,963 | -0,054 | -0,28 | 0,7797278 | 0,82737631 | 0,017839 | 0,0231972 | 0,0208161 | 0,0208161 | TRUE | 0,3439783 | No |
| orf19.5956 | 12759 | 7841 | 10577 | 13757 | 12060 | 9884 | 10840 | 7587 | 11446 | 9982 | 10821 | 11344 | 10314 | 9431 | 9673 | 10701 | 10464 | 10898 | 10030 | 0,92 | -0,121 | -2,135 | 0,03276663 | 0,05795913 | 0,003443 | 0,0021183 | 0,0029682 | 0,0029682 | TRUE | 0,0388416 | No |
| orf19.5958 | 3621 | 2526 | 3289 | 3910 | 3771 | 3399 | 3634 | 2015 | 3248 | 3216 | 3365 | 3224 | 3225 | 3243 | 3243 | 2842 | 3200,8 | 3263 | 3138 | 0,963 | -0,054 | -1,093 | 0,27425141 | 0,35858595 | 0,00172 | 0,0024331 | 0,0020445 | 0,0020445 | TRUE | 0,0463838 | No |
| orf19.5959 | 14051 | 10726 | 12807 | 15745 | 13492 | 13161 | 13982 | 7954 | 12605 | 13655 | 13103 | 12984 | 11539 | 12558 | 12477 | 11218 | 12517 | 13087 | 11948 | 0,913 | -0,131 | -2,774 | 0,00553642 | 0,01184996 | 0,002022 | 0,0020955 | 0,0020534 | 0,0020534 | TRUE | 0,0203075 | No |
| orf19.5960 | 31606 | 23314 | 29894 | 37030 | 38765 | 35479 | 38516 | 28931 | 28354 | 29681 | 30585 | 30535 | 33153 | 33853 | 34369 | 40805 | 32667 | 29789 | 35545 | 1,193 | 0,255 | 3,954 | 7,70E-05 | 0,00023931 | 0,005535 | 0,002024 | 0,0039559 | 0,0039559 | TRUE | 0,1212298 | No |
| orf19.5961 | 3601 | 2519 | 3281 | 4084 | 4536 | 3928 | 4210 | 2650 | 3230 | 3207 | 3357 | 3368 | 3879 | 3748 | 3757 | 3738 | 3535,4 | 3290 | 3780 | 1,149 | 0,2 | 5,637 | 1,74E-08 | 9,32E-08 | 0,000197 | 0,0023901 | 0,0009245 | 0,0009245 | TRUE | 0,0038996 | No |
| orf19.596.1 | 3478 | 2090 | 2664 | 3358 | 2835 | 2267 | 2588 | 1696 | 3120 | 2661 | 2726 | 2769 | 2425 | 2163 | 2309 | 2392 | 2570,6 | 2819 | 2322 | 0,823 | -0,281 | -4,643 | 3,44E-06 | 1,34E-05 | 0,003401 | 0,0025443 | 0,0031188 | 0,0031188 | TRUE | 0,0632908 | No |
| orf19.5962 | 2638 | 1957 | 2518 | 2873 | 2440 | 2397 | 2366 | 1416 | 2367 | 2491 | 2576 | 2369 | 2087 | 2287 | 2111 | 1997 | 2285,7 | 2451 | 2120 | 0,867 | -0,207 | -3,917 | 8,98E-05 | 0,00027727 | 0,001916 | 0,0026146 | 0,0022269 | 0,0022269 | TRUE | 0,0335994 | No |
| orf19.596.2 | 541 | 373 | 399 | 600 | 549 | 430 | 494 | 276 | 485 | 475 | 408 | 495 | 470 | 410 | 441 | 389 | 446,64 | 466 | 428 | 0,92 | -0,12 | -1,379 | 0,16778423 | 0,23746706 | 0,004749 | 0,0052298 | 0,0049909 | 0,0049909 | TRUE | 0,0805774 | No |
| orf19.5963 | 7057 | 4977 | 6120 | 7655 | 7248 | 6400 | 6908 | 3731 | 6331 | 6336 | 6261 | 6312 | 6199 | 6107 | 6164 | 5262 | 6121,6 | 6310 | 5933 | 0,941 | -0,088 | -1,649 | 0,09904827 | 0,15169055 | 0,002657 | 0,0022167 | 0,0025469 | 0,0025469 | TRUE | 0,0688307 | No |
| orf19.5964 | 4169 | 3110 | 3600 | 4498 | 4698 | 4202 | 4580 | 2603 | 3740 | 3959 | 3683 | 3709 | 4018 | 4009 | 4087 | 3671 | 3859,6 | 3773 | 3946 | 1,048 | 0,067 | 1,487 | 0,13705863 | 0,19977588 | 0,001334 | 0,0023556 | 0,0017042 | 0,0017042 | TRUE | 0,025425 | No |
| orf19.5964.2 | 9352 | 7209 | 7915 | 10383 | 10104 | 9677 | 9414 | 5548 | 8390 | 9178 | 8098 | 8562 | 8641 | 9233 | 8400 | 7825 | 8540,9 | 8557 | 8525 | 0,997 | -0,005 | -0,08 | 0,93602391 | 0,95395767 | 0,003107 | 0,0021495 | 0,0030323 | 0,0030323 | TRUE | 0,0380834 | No |
| orf19.5965 | 1405 | 1095 | 1156 | 1520 | 1689 | 1691 | 1772 | 1060 | 1260 | 1394 | 1183 | 1253 | 1444 | 1613 | 1581 | 1495 | 1403,1 | 1272 | 1533 | 1,207 | 0,272 | 4,433 | 9,29E-06 | 3,40E-05 | 0,002725 | 0,0030142 | 0,002881 | 0,002881 | TRUE | 0,0463853 | No |
| orf19.5966 | 3 | 10 | 4 | 14 | 12 | 7 | 9 | 8 | 3 | 13 | 4 | 12 | 10 | 7 | 8 | 11 | 8,41 | 8 | 9 | 1,173 | 0,23 | 0,427 | 0,66961761 | NA | 0,130496 | 0,1745043 | 0,1573224 | 0,1573224 | TRUE | 0,2023141 | No |
| orf19.5967 | 1208 | 801 | 1051 | 1328 | 1233 | 1009 | 1199 | 777 | 1084 | 1020 | 1075 | 1095 | 1055 | 963 | 1070 | 1096 | 1057,1 | 1068 | 1046 | 0,976 | -0,035 | -0,622 | 0,53363116 | 0,6160259 | 0,000991 | 0,0033528 | 0,0020708 | 0,0020708 | TRUE | 0,0342025 | No |
| orf19.5968 | 2910 | 2111 | 2430 | 2903 | 3161 | 2990 | 2907 | 1975 | 2611 | 2688 | 2486 | 2394 | 2703 | 2853 | 2594 | 2786 | 2639,3 | 2545 | 2734 | 1,075 | 0,104 | 2,063 | 0,03908969 | 0,0677194 | 0,001638 | 0,0025296 | 0,0020676 | 0,0020676 | TRUE | 0,0197006 | No |
| orf19.597 | 3573 | 2160 | 3045 | 3668 | 3467 | 2676 | 3094 | 2125 | 3205 | 2750 | 3115 | 3025 | 2965 | 2553 | 2761 | 2997 | 2921,5 | 3024 | 2819 | 0,931 | -0,103 | -1,593 | 0,11106099 | 0,1666886 | 0,004328 | 0,0024764 | 0,0036515 | 0,0036515 | TRUE | 0,0489579 | No |
| orf19.5970 | 576 | 370 | 509 | 614 | 696 | 628 | 718 | 373 | 517 | 471 | 521 | 506 | 595 | 599 | 641 | 526 | 547,01 | 504 | 590 | 1,175 | 0,232 | 3,16 | 0,0015786 | 0,0038074 | 0,001791 | 0,0046334 | 0,0033615 | 0,0033615 | TRUE | 0,0543749 | No |
| orf19.5971 | 788 | 598 | 763 | 954 | 1174 | 1046 | 1066 | 681 | 707 | 761 | 781 | 787 | 1004 | 998 | 951 | 960 | 868,67 | 759 | 978 | 1,291 | 0,368 | 6,466 | 1,01E-10 | 7,34E-10 | 1,00E-08 | 0,0036507 | 0,00194 | 0,00194 | TRUE | 0,0269165 | No |
| orf19.5973 | 3546 | 2582 | 3024 | 3736 | 3931 | 3704 | 3813 | 2424 | 3181 | 3287 | 3094 | 3081 | 3362 | 3534 | 3402 | 3419 | 3295 | 3161 | 3429 | 1,086 | 0,119 | 3,139 | 0,00169739 | 0,00406291 | 0,000282 | 0,0024201 | 0,0010685 | 0,0010685 | TRUE | 0,0082915 | No |
| orf19.5974 | 1428 | 1214 | 1381 | 1696 | 1379 | 1557 | 1646 | 946 | 1281 | 1546 | 1413 | 1399 | 1179 | 1486 | 1469 | 1334 | 1388,3 | 1410 | 1367 | 0,971 | -0,043 | -0,522 | 0,60161732 | 0,67751019 | 0,007889 | 0,0030252 | 0,0056861 | 0,0056861 | TRUE | 0,1064853 | No |
| orf19.5975 | 2283 | 1478 | 1967 | 2477 | 1678 | 1391 | 1493 | 898 | 2048 | 1882 | 2012 | 2043 | 1435 | 1327 | 1332 | 1267 | 1668,2 | 1996 | 1340 | 0,672 | -0,573 | -10,991 | 4,20E-28 | 1,38E-26 | 0,001291 | 0,0028497 | 0,0019866 | 0,0019866 | TRUE | 0,0276936 | No |
| orf19.5976 | 1292 | 1002 | 1124 | 1487 | 1505 | 1421 | 1548 | 969 | 1159 | 1276 | 1150 | 1226 | 1287 | 1356 | 1381 | 1367 | 1275,2 | 1203 | 1348 | 1,121 | 0,165 | 3,159 | 0,00158195 | 0,00381394 | 0,000833 | 0,0031179 | 0,0018288 | 0,0018288 | TRUE | 0,0180683 | No |
| orf19.5977 | 1523 | 1039 | 1359 | 1616 | 1607 | 1323 | 1451 | 912 | 1366 | 1323 | 1390 | 1333 | 1374 | 1262 | 1295 | 1286 | 1328,7 | 1353 | 1304 | 0,965 | -0,052 | -1,079 | 0,2807478 | 0,36475021 | 1,00E-08 | 0,0030721 | 0,0014575 | 0,0014575 | TRUE | 0,0166124 | No |
| orf19.5978 | 278 | 199 | 264 | 302 | 264 | 248 | 281 | 209 | 249 | 253 | 270 | 249 | 226 | 237 | 251 | 295 | 253,73 | 255 | 252 | 0,979 | -0,031 | -0,31 | 0,75627551 | 0,80964064 | 0,002734 | 0,007701 | 0,0055371 | 0,0055371 | TRUE | 0,1225046 | No |
| orf19.598 | 1 | 0 | 3 | 2 | 5 | 6 | 1 | 1 | 1 | 0 | 3 | 2 | 4 | 6 | 1 | 1 | 2,24 | 2 | 3 | 2,173 | 1,12 | 1,102 | 0,27025344 | NA | 0,213619 | 0,6500771 | 0,4880798 | 0,4880798 | TRUE | 0,2012814 | No |
| orf19.5980 | 1203 | 897 | 1164 | 1310 | 1288 | 1224 | 1287 | 806 | 1079 | 1142 | 1191 | 1080 | 1102 | 1168 | 1148 | 1137 | 1130,9 | 1123 | 1139 | 1,015 | 0,022 | 0,41 | 0,68197047 | 0,74593632 | 0,000663 | 0,0032632 | 0,0017911 | 0,0017911 | TRUE | 0,0204677 | No |
| orf19.5982 | 17464 | 10791 | 14255 | 17626 | 17187 | 13356 | 14325 | 8991 | 15667 | 13738 | 14584 | 14535 | 14699 | 12744 | 12783 | 12681 | 13929 | 14631 | 13227 | 0,904 | -0,146 | -2,467 | 0,01360619 | 0,02646358 | 0,004128 | 0,0020837 | 0,0032693 | 0,0032693 | TRUE | 0,0689535 | No |
| orf19.5984 | 819 | 556 | 682 | 898 | 979 | 839 | 898 | 606 | 735 | 708 | 698 | 741 | 837 | 801 | 801 | 855 | 771,83 | 720 | 824 | 1,14 | 0,189 | 3,369 | 0,00075408 | 0,00194694 | 1,00E-08 | 0,0038604 | 0,0017235 | 0,0017235 | TRUE | 0,0065273 | No |
| orf19.5985 | 1644 | 1072 | 1284 | 1627 | 1665 | 1386 | 1477 | 944 | 1475 | 1365 | 1314 | 1342 | 1424 | 1322 | 1318 | 1331 | 1361,3 | 1374 | 1349 | 0,982 | -0,026 | -0,479 | 0,63186013 | 0,70313311 | 0,001386 | 0,0030459 | 0,0020685 | 0,0020685 | TRUE | 0,0305951 | No |
| orf19.5986 | 630 | 466 | 579 | 707 | 764 | 631 | 656 | 545 | 565 | 593 | 592 | 583 | 653 | 602 | 585 | 769 | 617,92 | 583 | 652 | 1,113 | 0,155 | 1,844 | 0,06523901 | 0,10619234 | 0,006131 | 0,0043289 | 0,005155 | 0,005155 | TRUE | 0,1547158 | No |
| orf19.5987 | 4091 | 2752 | 3608 | 4304 | 3805 | 3187 | 3415 | 2167 | 3670 | 3504 | 3691 | 3549 | 3254 | 3041 | 3047 | 3056 | 3351,6 | 3604 | 3100 | 0,86 | -0,217 | -5,461 | 4,72E-08 | 2,41E-07 | 0,000588 | 0,0024127 | 0,0012132 | 0,0012132 | TRUE | 0,0142039 | No |
| orf19.5989 | 5993 | 3979 | 5332 | 6781 | 5557 | 4555 | 5195 | 3959 | 5376 | 5066 | 5455 | 5592 | 4753 | 4346 | 4636 | 5584 | 5100,9 | 5372 | 4830 | 0,898 | -0,155 | -2,194 | 0,02823682 | 0,05080767 | 0,006644 | 0,0022641 | 0,0046023 | 0,0046023 | TRUE | 0,1324626 | No |
| orf19.599 | 0 | 0 | 0 | 1 | 1 | 0 | 0 | 0 | 0 | 0 | 0 | 1 | 1 | 0 | 0 | 0 | 0,21 | 0 | 0 | 1,016 | 0,023 | 0,007 | 0,9940485 | NA | 1,00E-08 | 6,9154834 | 7,1370638 | 7,1370638 | TRUE | 0,2537766 | No |
| orf19.5991 | 20773 | 14323 | 18702 | 22385 | 19515 | 17617 | 19336 | 11538 | 18636 | 18235 | 19134 | 18459 | 16690 | 16810 | 17254 | 16273 | 17686 | 18616 | 16757 | 0,9 | -0,152 | -4,919 | 8,68E-07 | 3,69E-06 | 0,000425 | 0,0020616 | 0,0008541 | 0,0008541 | TRUE | 0,0048805 | No |
| orf19.5992 | 2859 | 2050 | 2675 | 3468 | 3279 | 2860 | 3159 | 2280 | 2565 | 2610 | 2737 | 2860 | 2804 | 2729 | 2819 | 3216 | 2792,4 | 2693 | 2892 | 1,072 | 0,1 | 1,671 | 0,09476407 | 0,14629026 | 0,00362 | 0,0024994 | 0,00311 | 0,00311 | TRUE | 0,0663684 | No |

| | | | | | | | | | | | | | | | | | | | | | | | | | | | | | | | |
|---|---|---|---|---|---|---|---|---|---|---|---|---|---|---|---|---|---|---|---|---|---|---|---|---|---|---|---|---|---|---|---|
| orf19.5994 | 2950 | 2189 | 2510 | 3307 | 2362 | 2254 | 2221 | 1393 | 2646 | 2787 | 2568 | 2727 | 2020 | 2151 | 1982 | 1965 | 2355,7 | 2682 | 2030 | 0,757 | -0,401 | -8,682 | 3,88E-18 | 6,10E-17 | 0,000915 | 0,0025958 | 0,0016116 | 0,0016116 | TRUE | 0,0196547 | No |
| orf19.5995 | 3434 | 2287 | 3161 | 3805 | 3682 | 3173 | 3311 | 2256 | 3081 | 2912 | 3234 | 3138 | 3149 | 3028 | 2955 | 3182 | 3084,6 | 3091 | 3078 | 0,995 | -0,008 | -0,173 | 0,8629853 | 0,89578709 | 0,001005 | 0,0024502 | 0,0016435 | 0,0016435 | TRUE | 0,0176076 | No |
| orf19.5996.1 | 17731 | 11474 | 14613 | 17738 | 17196 | 13051 | 13830 | 8931 | 15907 | 14608 | 14951 | 14627 | 14707 | 12453 | 12341 | 12596 | 14024 | 15023 | 13024 | 0,867 | -0,206 | -3,41 | 0,00064964 | 0,00170291 | 0,004468 | 0,002083 | 0,0034285 | 0,0034285 | TRUE | 0,092911 | No |
| orf19.5999 | 1206 | 870 | 1107 | 1337 | 1264 | 1134 | 1196 | 712 | 1082 | 1108 | 1133 | 1103 | 1081 | 1082 | 1067 | 1004 | 1082,4 | 1106 | 1058 | 0,96 | -0,059 | -1,208 | 0,22687222 | 0,30630299 | 1,00E-08 | 0,0033207 | 0,0013982 | 0,0013982 | TRUE | 0,0121724 | No |
| orf19.600 | 3688 | 2334 | 3311 | 3677 | 3736 | 3053 | 3602 | 2016 | 3309 | 2971 | 3387 | 3032 | 3195 | 2913 | 3214 | 2843 | 3108,2 | 3175 | 3041 | 0,959 | -0,061 | -1,011 | 0,31187678 | 0,39824776 | 0,003602 | 0,0024466 | 0,0031779 | 0,0031779 | TRUE | 0,024423 | No |
| orf19.6000 | 77734 | 56586 | 73652 | 85652 | 75534 | 71159 | 76660 | 44869 | 69736 | 72040 | 75353 | 70630 | 64599 | 67898 | 68406 | 63284 | 68993 | 71940 | 66047 | 0,918 | -0,123 | -3,106 | 0,00189921 | 0,00450472 | 0,001283 | 0,0020006 | 0,0014976 | 0,0014976 | TRUE | 0,0125109 | No |
| orf19.6001 | 983 | 743 | 888 | 1144 | 612 | 631 | 672 | 378 | 882 | 946 | 909 | 943 | 523 | 602 | 600 | 533 | 742,24 | 920 | 564 | 0,615 | -0,701 | -10,611 | 2,66E-26 | 7,80E-25 | 0,001429 | 0,0039354 | 0,0027597 | 0,0027597 | TRUE | 0,0331057 | No |
| orf19.6002 | 11378 | 8177 | 9730 | 11690 | 10673 | 10072 | 10401 | 7337 | 10207 | 10410 | 9955 | 9640 | 9128 | 9610 | 9281 | 10348 | 9822,4 | 10053 | 9592 | 0,954 | -0,068 | -1,447 | 0,14800996 | 0,2131968 | 0,002013 | 0,0021273 | 0,0020339 | 0,0020339 | TRUE | 0,0338387 | No |
| orf19.6003 | 2753 | 2090 | 2555 | 3207 | 3127 | 3122 | 3481 | 2140 | 2470 | 2661 | 2614 | 2645 | 2674 | 2979 | 3106 | 3018 | 2770,9 | 2598 | 2944 | 1,133 | 0,181 | 3,342 | 0,0008315 | 0,00212943 | 0,002114 | 0,0025034 | 0,0024417 | 0,0024417 | TRUE | 0,0475568 | No |
| orf19.6005 | 266 | 216 | 261 | 311 | 348 | 350 | 361 | 229 | 239 | 275 | 267 | 256 | 298 | 334 | 322 | 323 | 289,22 | 259 | 319 | 1,233 | 0,303 | 3,475 | 0,00051072 | 0,00137659 | 1,00E-08 | 0,0069988 | 0,0037938 | 0,0037938 | TRUE | 0,032573 | No |
| orf19.6007 | 3304 | 3001 | 3055 | 3987 | 2445 | 2961 | 2822 | 1801 | 2964 | 3821 | 3126 | 3288 | 2091 | 2825 | 2518 | 2540 | 2896,6 | 3300 | 2494 | 0,756 | -0,404 | -4,281 | 1,86E-05 | 6,50E-05 | 0,013242 | 0,0024807 | 0,0081886 | 0,0081886 | TRUE | 0,1451005 | No |
| orf19.6008 | 320 | 273 | 318 | 415 | 312 | 392 | 401 | 249 | 287 | 348 | 325 | 342 | 267 | 374 | 358 | 351 | 331,51 | 326 | 338 | 1,036 | 0,051 | 0,447 | 0,6550283 | 0,72382012 | 0,012244 | 0,0063586 | 0,0093107 | 0,0093107 | TRUE | 0,2407872 | No |
| orf19.6008.3 | 329 | 229 | 290 | 369 | 295 | 265 | 292 | 168 | 295 | 292 | 297 | 304 | 252 | 253 | 261 | 237 | 273,79 | 297 | 251 | 0,846 | -0,241 | -2,862 | 0,00420453 | 0,00921254 | 1,00E-08 | 0,0072817 | 0,0031505 | 0,0031505 | TRUE | 0,0118224 | No |
| orf19.6008.4 | 2039 | 1217 | 1368 | 1885 | 2460 | 1903 | 1999 | 1306 | 1829 | 1549 | 1400 | 1554 | 2104 | 1816 | 1784 | 1842 | 1734,8 | 1583 | 1886 | 1,191 | 0,253 | 3,056 | 0,00224187 | 0,00524162 | 0,008657 | 0,0028164 | 0,0059783 | 0,0059783 | TRUE | 0,1335536 | No |
| orf19.6009 | 1201 | 865 | 1137 | 1250 | 1345 | 1278 | 1342 | 846 | 1077 | 1101 | 1163 | 1031 | 1150 | 1219 | 1198 | 1193 | 1141,6 | 1093 | 1190 | 1,09 | 0,124 | 2,323 | 0,02018899 | 0,03756441 | 0,00079 | 0,0032511 | 0,0018542 | 0,0018542 | TRUE | 0,0229203 | No |
| orf19.6010 | 452 | 344 | 429 | 527 | 557 | 531 | 549 | 339 | 405 | 438 | 439 | 435 | 476 | 507 | 490 | 478 | 458,5 | 429 | 488 | 1,138 | 0,187 | 2,725 | 0,00642961 | 0,01356359 | 1,00E-08 | 0,0051457 | 0,0023364 | 0,0023364 | TRUE | 0,0168502 | No |
| orf19.6010.1 | 1320 | 1022 | 1150 | 1468 | 1383 | 1257 | 1241 | 833 | 1184 | 1301 | 1177 | 1211 | 1183 | 1199 | 1107 | 1175 | 1192,1 | 1218 | 1166 | 0,958 | -0,061 | -1,157 | 0,24742401 | 0,32951288 | 0,000768 | 0,0031973 | 0,0018468 | 0,0018468 | TRUE | 0,0230972 | No |
| orf19.6011 | 5450 | 3934 | 5194 | 5860 | 5898 | 5257 | 5762 | 3514 | 4889 | 5008 | 5314 | 4832 | 5044 | 5016 | 5142 | 4956 | 5025,2 | 5011 | 5040 | 1,006 | 0,009 | 0,23 | 0,81812997 | 0,8596231 | 0,000692 | 0,0022684 | 0,0013022 | 0,0013022 | TRUE | 0,0203519 | No |
| orf19.6012 | 1111 | 867 | 1045 | 1186 | 1435 | 1402 | 1498 | 902 | 997 | 1104 | 1069 | 978 | 1227 | 1338 | 1337 | 1272 | 1165,2 | 1037 | 1294 | 1,251 | 0,323 | 5,588 | 2,29E-08 | 1,21E-07 | 0,001635 | 0,0032254 | 0,0023291 | 0,0023291 | TRUE | 0,0212801 | No |
| orf19.6013 | 1275 | 935 | 1171 | 1416 | 1262 | 1068 | 1205 | 743 | 1144 | 1190 | 1198 | 1168 | 1079 | 1019 | 1075 | 1048 | 1115,2 | 1175 | 1055 | 0,9 | -0,153 | -3,166 | 0,00154456 | 0,00373729 | 1,00E-08 | 0,0032813 | 0,0013368 | 0,0013368 | TRUE | 0,0069492 | No |
| orf19.6014 | 8495 | 5747 | 6837 | 8795 | 7947 | 6759 | 7524 | 3949 | 7621 | 7317 | 6995 | 7252 | 6797 | 6449 | 6714 | 5570 | 6839,3 | 7296 | 6382 | 0,875 | -0,192 | -3,1 | 0,0019329 | 0,0045774 | 0,004298 | 0,0021918 | 0,0035395 | 0,0035395 | TRUE | 0,0880531 | No |
| orf19.6016 | 1260 | 961 | 1158 | 1427 | 1869 | 1828 | 1954 | 1200 | 1130 | 1223 | 1185 | 1177 | 1598 | 1744 | 1744 | 1692 | 1436,8 | 1179 | 1694 | 1,439 | 0,525 | 10,485 | 1,01E-25 | 2,74E-24 | 0,000803 | 0,0029899 | 0,0016874 | 0,0016874 | TRUE | 0,018868 | No |
| orf19.6017 | 376 | 263 | 344 | 443 | 506 | 451 | 494 | 313 | 337 | 335 | 352 | 365 | 433 | 430 | 441 | 441 | 391,84 | 347 | 436 | 1,253 | 0,325 | 4,485 | 7,30E-06 | 2,72E-05 | 1,00E-08 | 0,0056843 | 0,0024804 | 0,0024804 | TRUE | 0,0151207 | No |
| orf19.6018 | 1537 | 1024 | 1270 | 1646 | 1476 | 1357 | 1524 | 906 | 1379 | 1304 | 1299 | 1357 | 1262 | 1295 | 1360 | 1278 | 1316,8 | 1335 | 1299 | 0,972 | -0,04 | -0,835 | 0,40379663 | 0,49199152 | 1,00E-08 | 0,003082 | 0,0014756 | 0,0014756 | TRUE | 0,0126849 | No |
| orf19.6020 | 1157 | 1048 | 1120 | 1403 | 1226 | 1367 | 1357 | 736 | 1038 | 1334 | 1146 | 1157 | 1049 | 1304 | 1211 | 1038 | 1159,6 | 1169 | 1150 | 0,987 | -0,019 | -0,212 | 0,83242104 | 0,87143851 | 0,010683 | 0,0032314 | 0,0071575 | 0,0071575 | TRUE | 0,1056639 | No |
| orf19.6021 | 228 | 147 | 205 | 253 | 283 | 247 | 251 | 169 | 205 | 187 | 210 | 209 | 242 | 236 | 224 | 238 | 218,76 | 203 | 235 | 1,156 | 0,209 | 2,21 | 0,0271008 | 0,0491018 | 1,00E-08 | 0,0086155 | 0,004015 | 0,004015 | TRUE | 0,0236596 | No |
| orf19.6022 | 44 | 48 | 48 | 61 | 96 | 83 | 110 | 70 | 39 | 61 | 49 | 50 | 82 | 79 | 98 | 99 | 69,77 | 50 | 90 | 1,802 | 0,849 | 4,704 | 2,55E-06 | 1,02E-05 | 0,004563 | 0,0227859 | 0,0156647 | 0,0156647 | TRUE | 0,1711804 | No |
| orf19.6023 | 2010 | 1472 | 1845 | 2304 | 1500 | 1423 | 1564 | 922 | 1803 | 1874 | 1888 | 1900 | 1283 | 1358 | 1396 | 1300 | 1600,2 | 1866 | 1334 | 0,716 | -0,483 | -10,4 | 2,48E-25 | 6,47E-24 | 0,000409 | 0,0028867 | 0,0014251 | 0,0014251 | TRUE | 0,0119971 | No |
| orf19.6024 | 188 | 234 | 208 | 283 | 282 | 308 | 323 | 183 | 169 | 298 | 213 | 233 | 241 | 294 | 288 | 258 | 249,27 | 228 | 270 | 1,192 | 0,253 | 1,715 | 0,08626624 | 0,13500588 | 0,026634 | 0,0078034 | 0,0168169 | 0,0168169 | TRUE | 0,4166916 | No |
| orf19.6025 | 55 | 35 | 36 | 56 | 36 | 50 | 51 | 27 | 49 | 45 | 37 | 46 | 31 | 48 | 46 | 38 | 42,37 | 44 | 41 | 0,914 | -0,129 | -0,583 | 0,55984272 | 0,63994198 | 1,00E-08 | 0,0362383 | 0,0236811 | 0,0236811 | TRUE | 0,2413569 | No |
| orf19.6026 | 174 | 131 | 162 | 218 | 166 | 188 | 183 | 112 | 156 | 167 | 166 | 180 | 142 | 179 | 163 | 158 | 163,87 | 167 | 160 | 0,96 | -0,059 | -0,511 | 0,60908297 | 0,68386665 | 1,00E-08 | 0,0108381 | 0,0066737 | 0,0066737 | TRUE | 0,0747351 | No |
| orf19.6027 | 99 | 67 | 68 | 107 | 146 | 140 | 152 | 80 | 89 | 85 | 70 | 88 | 125 | 134 | 136 | 113 | 104,85 | 83 | 127 | 1,532 | 0,615 | 4,365 | 1,27E-05 | 4,56E-05 | 1,00E-08 | 0,0158245 | 0,0091816 | 0,0091816 | TRUE | 0,1101354 | No |
| orf19.6028 | 199 | 177 | 177 | 226 | 332 | 342 | 333 | 214 | 179 | 225 | 181 | 186 | 284 | 326 | 297 | 302 | 247,57 | 193 | 302 | 1,577 | 0,657 | 6,534 | 6,39E-11 | 4,80E-10 | 0,002188 | 0,0078433 | 0,0054613 | 0,0054613 | TRUE | 0,1201006 | No |
| orf19.6029 | 593 | 477 | 507 | 693 | 558 | 570 | 574 | 364 | 532 | 607 | 519 | 571 | 477 | 544 | 512 | 513 | 534,51 | 557 | 512 | 0,919 | -0,122 | -1,645 | 0,09988775 | 0,15270371 | 0,002084 | 0,0046955 | 0,0033904 | 0,0033904 | TRUE | 0,037888 | No |
| orf19.603 | 12362 | 8207 | 10313 | 13070 | 12142 | 10075 | 10608 | 7014 | 11090 | 10448 | 10551 | 10778 | 10384 | 9613 | 9466 | 9893 | 10278 | 10717 | 9839 | 0,918 | -0,124 | -3,111 | 0,00186652 | 0,00443594 | 0,001066 | 0,0021208 | 0,0014167 | 0,0014167 | TRUE | 0,0173422 | No |
| orf19.6030 | 2309 | 1427 | 1887 | 2330 | 2492 | 2098 | 2276 | 1358 | 2071 | 1817 | 1931 | 1921 | 2131 | 2002 | 2031 | 1915 | 1977,4 | 1935 | 2020 | 1,044 | 0,062 | 1,173 | 0,24076277 | 0,32207828 | 0,001757 | 0,0027137 | 0,0021964 | 0,0021964 | TRUE | 0,0273084 | No |
| orf19.6031 | 4596 | 3508 | 4153 | 5197 | 5005 | 4838 | 5076 | 3124 | 4123 | 4466 | 4249 | 4286 | 4280 | 4616 | 4529 | 4406 | 4369,5 | 4281 | 4458 | 1,042 | 0,059 | 1,466 | 0,1425164 | 0,20652159 | 0,000782 | 0,0023118 | 0,0013454 | 0,0013454 | TRUE | 0,0099222 | No |
| orf19.6032 | 4199 | 3518 | 4062 | 5107 | 4484 | 4487 | 4773 | 3010 | 3767 | 4479 | 4156 | 4211 | 3835 | 4281 | 4259 | 4245 | 4154,2 | 4153 | 4155 | 1,001 | 0,001 | 0,017 | 0,98663981 | 0,98961194 | 0,003678 | 0,002329 | 0,0031439 | 0,0031439 | TRUE | 0,0479293 | No |
| orf19.6033 | 1810 | 1259 | 1553 | 1940 | 1539 | 1396 | 1539 | 1004 | 1624 | 1603 | 1589 | 1600 | 1316 | 1332 | 1373 | 1416 | 1481,6 | 1604 | 1359 | 0,846 | -0,241 | -5,506 | 3,66E-08 | 1,89E-07 | 1,00E-08 | 0,0029593 | 0,0011646 | 0,0011646 | TRUE | 0,0081912 | No |
| orf19.6034 | 1662 | 1215 | 1483 | 1769 | 2073 | 1876 | 2014 | 1274 | 1491 | 1547 | 1517 | 1459 | 1773 | 1790 | 1797 | 1797 | 1646,3 | 1504 | 1789 | 1,191 | 0,253 | 6,129 | 8,84E-10 | 5,68E-09 | 1,00E-08 | 0,0028613 | 0,0010201 | 0,0010201 | TRUE | 0,0049777 | No |
| orf19.6035 | 883 | 716 | 675 | 1013 | 971 | 933 | 919 | 508 | 792 | 912 | 691 | 835 | 830 | 890 | 820 | 716 | 810,85 | 808 | 814 | 1,012 | 0,017 | 0,188 | 0,85059129 | 0,88598344 | 0,009072 | 0,0037699 | 0,0064945 | 0,0064945 | TRUE | 0,1106292 | No |
| orf19.6036 | 126 | 127 | 122 | 147 | 177 | 213 | 215 | 126 | 113 | 162 | 125 | 121 | 151 | 203 | 192 | 178 | 155,62 | 130 | 181 | 1,402 | 0,487 | 3,469 | 0,00052331 | 0,00140612 | 0,013499 | 0,0113082 | 0,0123 | 0,0123 | TRUE | 0,2400863 | No |
| orf19.6037 | 818 | 614 | 792 | 922 | 930 | 796 | 900 | 624 | 734 | 782 | 810 | 760 | 795 | 760 | 803 | 880 | 790,52 | 772 | 810 | 1,047 | 0,066 | 1,057 | 0,29069092 | 0,37518175 | 0,001159 | 0,0038159 | 0,0025131 | 0,0025131 | TRUE | 0,0358362 | No |
| orf19.6038 | 631 | 454 | 592 | 730 | 518 | 395 | 434 | 301 | 566 | 578 | 606 | 602 | 443 | 377 | 387 | 425 | 497,93 | 588 | 408 | 0,693 | -0,529 | -7,195 | 6,23E-13 | 5,85E-12 | 0,001169 | 0,004895 | 0,0031198 | 0,0031198 | TRUE | 0,0451864 | No |
| orf19.6039 | 1172 | 779 | 993 | 1146 | 1325 | 1182 | 1298 | 851 | 1051 | 992 | 1016 | 945 | 1133 | 1128 | 1158 | 1200 | 1078 | 1001 | 1155 | 1,152 | 0,205 | 3,854 | 0,00011638 | 0,00035171 | 0,000528 | 0,0033262 | 0,0017712 | 0,0017712 | TRUE | 0,0187989 | No |
| orf19.604 | 32 | 35 | 47 | 52 | 32 | 33 | 36 | 36 | 29 | 45 | 48 | 43 | 27 | 31 | 32 | 51 | 38,25 | 41 | 35 | 0,848 | -0,238 | -0,924 | 0,35542066 | 0,44293038 | 0,032518 | 0,0399343 | 0,0368704 | 0,0368704 | TRUE | 0,4599622 | No |
| orf19.6040 | 5358 | 3814 | 4658 | 5491 | 5253 | 4686 | 5237 | 3037 | 4807 | 4856 | 4766 | 4528 | 4493 | 4471 | 4673 | 4283 | 4609,5 | 4739 | 4480 | 0,947 | -0,079 | -1,961 | 0,04993717 | 0,08406347 | 0,000814 | 0,0022945 | 0,0013517 | 0,0013517 | TRUE | 0,0112029 | No |
| orf19.6041 | 9976 | 6696 | 9032 | 10335 | 11197 | 10288 | 10978 | 7332 | 8950 | 8525 | 9241 | 8522 | 9576 | 9816 | 9796 | 10341 | 9345,9 | 8810 | 9882 | 1,121 | 0,165 | 4,044 | 5,25E-05 | 0,00016919 | 0,001189 | 0,0021349 | 0,0014944 | 0,0014944 | TRUE | 0,0132132 | No |
| orf19.6043 | 434 | 302 | 394 | 492 | 525 | 426 | 481 | 326 | 389 | 384 | 403 | 406 | 449 | 406 | 429 | 460 | 415,89 | 396 | 436 | 1,099 | 0,136 | 1,86 | 0,06285335 | 0,10258705 | 1,00E-08 | 0,0054701 | 0,0026993 | 0,0026993 | TRUE | 0,0238502 | No |
| orf19.6044 | 2279 | 1498 | 2043 | 2359 | 2775 | 2381 | 2564 | 1874 | 2045 | 1907 | 2090 | 1945 | 2373 | 2272 | 2288 | 2643 | 2195,4 | 1997 | 2394 | 1,197 | 0,259 | 4,481 | 7,43E-06 | 2,77E-05 | 0,002872 | 0,0026408 | 0,0027453 | 0,0027453 | TRUE | 0,0564751 | No |
| orf19.6045 | 4594 | 3046 | 4102 | 4947 | 5504 | 4507 | 5259 | 3024 | 4121 | 3878 | 4197 | 4079 | 4707 | 4300 | 4693 | 4265 | 4280,1 | 4069 | 4491 | 1,104 | 0,143 | 2,993 | 0,00275864 | 0,00631822 | 0,001693 | 0,0023187 | 0,0019592 | 0,0019592 | TRUE | 0,0135084 | No |
| orf19.6046 | 685 | 441 | 608 | 705 | 984 | 800 | 816 | 565 | 615 | 561 | 622 | 581 | 842 | 763 | 728 | 797 | 688,66 | 595 | 782 | 1,314 | 0,393 | 5,842 | 5,17E-09 | 3,01E-08 | 0,001828 | 0,0040875 | 0,002877 | 0,002877 | TRUE | 0,0338301 | No |
| orf19.6047 | 8085 | 5256 | 7051 | 8561 | 8705 | 7055 | 8010 | 5150 | 7253 | 6691 | 7214 | 7060 | 7445 | 6732 | 7148 | 7264 | 7100,7 | 7054 | 7147 | 1,013 | 0,018 | 0,418 | 0,67562399 | 0,74072061 | 0,001241 | 0,002184 | 0,0016831 | 0,0016831 | TRUE | 0,0188256 | No |
| orf19.6048 | 1243 | 823 | 1013 | 1223 | 1440 | 1248 | 1314 | 871 | 1115 | 1048 | 1036 | 1009 | 1232 | 1191 | 1173 | 1228 | 1128,9 | 1052 | 1206 | 1,146 | 0,196 | 3,8 | 0,00014459 | 0,00042921 | 0,000459 | 0,0032655 | 0,0016661 | 0,0016661 | TRUE | 0,020834 | No |
| orf19.6049 | 2092 | 1350 | 1769 | 2148 | 2375 | 1921 | 2124 | 1461 | 1877 | 1719 | 1810 | 1771 | 2031 | 1833 | 1895 | 2061 | 1874,6 | 1794 | 1955 | 1,088 | 0,121 | 2,313 | 0,02071078 | 0,03844003 | 0,00152 | 0,0027539 | 0,0021081 | 0,0021081 | TRUE | 0,021181 | No |
| orf19.605 | 105 | 91 | 99 | 128 | 85 | 79 | 85 | 76 | 94 | 116 | 101 | 106 | 73 | 75 | 76 | 107 | 93,5 | 104 | 83 | 0,784 | -0,351 | -2,22 | 0,02644765 | 0,04807791 | 0,005735 | 0,0175058 | 0,0130263 | 0,0130263 | TRUE | 0,2918499 | No |
| orf19.6052 | 3124 | 2077 | 2714 | 3355 | 3722 | 3277 | 3596 | 2358 | 2803 | 2644 | 2777 | 2767 | 3183 | 3127 | 3209 | 3326 | 2979,3 | 2748 | 3211 | 1,167 | 0,222 | 5,857 | 4,72E-09 | 2,75E-08 | 0,000219 | 0,0024668 | 0,0010463 | 0,0010463 | TRUE | 0,0072269 | No |
| orf19.6053 | 3566 | 2348 | 3159 | 3875 | 2990 | 2660 | 3051 | 1801 | 3199 | 2989 | 3232 | 3195 | 2557 | 2538 | 2723 | 2540 | 2871,7 | 3154 | 2590 | 0,821 | -0,285 | -6,598 | 4,16E-11 | 3,19E-10 | 0,000743 | 0,002485 | 0,0014355 | 0,0014355 | TRUE | 0,0148789 | No |
| orf19.6054 | 10271 | 7071 | 8989 | 11442 | 10249 | 8847 | 9859 | 6110 | 9214 | 9002 | 9197 | 9435 | 8765 | 8442 | 8798 | 8618 | 8933,8 | 9212 | 8656 | 0,939 | -0,09 | -2,99 | 0,00278588 | 0,00637137 | 0,000212 | 0,002142 | 0,0007604 | 0,0007604 | TRUE | 0,003474 | No |

| | | | | | | | | | | | | | | | | | | | | | | | | | | | | | | | |
|---|---|---|---|---|---|---|---|---|---|---|---|---|---|---|---|---|---|---|---|---|---|---|---|---|---|---|---|---|---|---|---|
| orf19.6055 | 2252 | 1478 | 2005 | 2396 | 2592 | 2210 | 2410 | 1590 | 2020 | 1882 | 2051 | 1976 | 2217 | 2109 | 2151 | 2243 | 2081 | 1982 | 2180 | 1,098 | 0,135 | 3,07 | 0,00214201 | 0,00503097 | 0,000446 | 0,0026771 | 0,0013632 | 0,0013632 | TRUE | 0,0129498 | No |
| orf19.6056 | 2397 | 1511 | 1863 | 2358 | 2443 | 2053 | 2243 | 1248 | 2150 | 1924 | 1906 | 1944 | 2089 | 1959 | 2002 | 1760 | 1966,8 | 1981 | 1952 | 0,987 | -0,019 | -0,307 | 0,75884109 | 0,81151963 | 0,003598 | 0,0027176 | 0,0032244 | 0,0032244 | TRUE | 0,0501994 | No |
| orf19.6057 | 1004 | 594 | 767 | 993 | 876 | 638 | 783 | 499 | 901 | 756 | 785 | 819 | 749 | 609 | 699 | 704 | 752,61 | 815 | 690 | 0,845 | -0,242 | -3,06 | 0,00221144 | 0,00517783 | 0,005358 | 0,0039084 | 0,0046815 | 0,0046815 | TRUE | 0,0765892 | No |
| orf19.6058 | 1083 | 700 | 883 | 1145 | 1234 | 956 | 1007 | 591 | 972 | 891 | 903 | 944 | 1055 | 912 | 899 | 834 | 926,25 | 928 | 925 | 0,999 | -0,001 | -0,02 | 0,98406896 | 0,98801577 | 0,004625 | 0,0035468 | 0,0041322 | 0,0041322 | TRUE | 0,1092149 | No |
| orf19.6059 | 6149 | 4071 | 4820 | 6418 | 7134 | 5375 | 5420 | 4169 | 5516 | 5183 | 4931 | 5292 | 6101 | 5129 | 4836 | 5880 | 5358,6 | 5230 | 5486 | 1,049 | 0,068 | 0,936 | 0,34904786 | 0,4365278 | 0,006733 | 0,0022504 | 0,0049344 | 0,0049344 | TRUE | 0,0780162 | No |
| orf19.606 | 5868 | 4037 | 5341 | 6356 | 5343 | 4866 | 5310 | 3504 | 5264 | 5140 | 5464 | 5241 | 4570 | 4643 | 4738 | 4942 | 5000,3 | 5277 | 4723 | 0,894 | -0,162 | -4,255 | 2,09E-05 | 7,25E-05 | 0,000649 | 0,0022698 | 0,0011823 | 0,0011823 | TRUE | 0,0112103 | No |
| orf19.6060 | 2939 | 1905 | 2532 | 3016 | 2821 | 2402 | 2661 | 1613 | 2637 | 2425 | 2590 | 2487 | 2413 | 2292 | 2375 | 2275 | 2436,7 | 2535 | 2339 | 0,923 | -0,116 | -2,672 | 0,00753873 | 0,01561817 | 0,000567 | 0,0025753 | 0,0013934 | 0,0013934 | TRUE | 0,0095691 | No |
| orf19.6061 | 1961 | 1300 | 1679 | 2151 | 1810 | 1587 | 1735 | 1030 | 1759 | 1655 | 1718 | 1774 | 1548 | 1514 | 1548 | 1453 | 1621,1 | 1726 | 1516 | 0,878 | -0,187 | -4,166 | 3,09E-05 | 0,00010393 | 0,000173 | 0,002875 | 0,0013181 | 0,0013181 | TRUE | 0,0086891 | No |
| orf19.6062 | 4961 | 3053 | 3854 | 4724 | 5416 | 4374 | 4552 | 2898 | 4451 | 3887 | 3943 | 3895 | 4632 | 4174 | 4062 | 4087 | 4141,3 | 4044 | 4239 | 1,048 | 0,068 | 1,125 | 0,26039862 | 0,34376508 | 0,003919 | 0,0023301 | 0,0032653 | 0,0032653 | TRUE | 0,0562222 | No |
| orf19.6062.3 | 1290 | 818 | 1054 | 1351 | 1198 | 929 | 1033 | 725 | 1157 | 1041 | 1078 | 1114 | 1025 | 886 | 922 | 1023 | 1030,8 | 1098 | 964 | 0,876 | -0,192 | -2,985 | 0,00283829 | 0,00647643 | 0,002392 | 0,0033879 | 0,0029784 | 0,0029784 | TRUE | 0,0347679 | No |
| orf19.6063 | 8153 | 5754 | 7190 | 8775 | 7959 | 7195 | 7891 | 4519 | 7314 | 7325 | 7356 | 7236 | 6807 | 6865 | 7041 | 6374 | 7039,8 | 7308 | 6772 | 0,927 | -0,109 | -2,935 | 0,00333016 | 0,00747948 | 0,000711 | 0,0021857 | 0,0011728 | 0,0011728 | TRUE | 0,018279 | No |
| orf19.6064 | 1835 | 1219 | 1607 | 2052 | 1716 | 1499 | 1623 | 1091 | 1646 | 1552 | 1644 | 1692 | 1468 | 1430 | 1448 | 1539 | 1552,4 | 1634 | 1471 | 0,898 | -0,155 | -3,32 | 0,00090137 | 0,0022879 | 0,000308 | 0,0029147 | 0,0014579 | 0,0014579 | TRUE | 0,0126536 | No |
| orf19.6065 | 1513 | 1055 | 1293 | 1556 | 1117 | 1020 | 1061 | 615 | 1357 | 1343 | 1323 | 1283 | 955 | 973 | 947 | 867 | 1131,1 | 1326 | 936 | 0,708 | -0,498 | -9,585 | 9,22E-22 | 1,88E-20 | 0,000334 | 0,0032629 | 0,001684 | 0,001684 | TRUE | 0,0244697 | No |
| orf19.6066 | 1587 | 1119 | 1372 | 1646 | 1586 | 1433 | 1488 | 980 | 1424 | 1425 | 1404 | 1357 | 1356 | 1367 | 1328 | 1382 | 1380,4 | 1402 | 1358 | 0,969 | -0,046 | -1,041 | 0,29797612 | 0,38326708 | 1,00E-08 | 0,0030312 | 0,0011468 | 0,0011468 | TRUE | 0,0060229 | No |
| orf19.6068 | 10076 | 6574 | 8749 | 10655 | 9829 | 8271 | 8910 | 5979 | 9039 | 8369 | 8951 | 8786 | 8406 | 7892 | 7951 | 8433 | 8478,4 | 8786 | 8170 | 0,929 | -0,106 | -2,646 | 0,00813354 | 0,01673503 | 0,001039 | 0,0021507 | 0,0014122 | 0,0014122 | TRUE | 0,0122049 | No |
| orf19.607 | 4968 | 3228 | 4459 | 5164 | 4544 | 3904 | 4436 | 2690 | 4457 | 4110 | 4562 | 4258 | 3886 | 3725 | 3958 | 3794 | 4093,8 | 4347 | 3841 | 0,884 | -0,179 | -4,089 | 4,33E-05 | 0,00014146 | 0,001109 | 0,0023341 | 0,0015849 | 0,0015849 | TRUE | 0,0157236 | No |
| orf19.6070 | 11201 | 7492 | 10206 | 12153 | 9478 | 8431 | 9024 | 6144 | 10048 | 9538 | 10442 | 10022 | 8106 | 8045 | 8052 | 8666 | 9114,8 | 10012 | 8217 | 0,82 | -0,286 | -6,998 | 2,59E-12 | 2,26E-11 | 0,001202 | 0,0021388 | 0,0014936 | 0,0014936 | TRUE | 0,0160233 | No |
| orf19.6071 | 2739 | 2151 | 2621 | 3174 | 2626 | 2530 | 2726 | 1726 | 2457 | 2738 | 2682 | 2617 | 2246 | 2414 | 2433 | 2434 | 2502,7 | 2624 | 2382 | 0,908 | -0,139 | -2,857 | 0,00427997 | 0,00936419 | 0,00147 | 0,0025596 | 0,0018813 | 0,0018813 | TRUE | 0,0222916 | No |
| orf19.6072 | 719 | 498 | 645 | 793 | 370 | 333 | 346 | 232 | 645 | 634 | 660 | 654 | 316 | 318 | 309 | 327 | 482,87 | 648 | 318 | 0,489 | -1,034 | -15,314 | 6,15E-53 | 7,70E-51 | 1,00E-08 | 0,0049859 | 0,0020306 | 0,0020306 | TRUE | 0,0038341 | No |
| orf19.6072.1 | 734 | 599 | 667 | 898 | 852 | 823 | 833 | 627 | 658 | 763 | 682 | 741 | 729 | 785 | 743 | 884 | 748,19 | 711 | 785 | 1,102 | 0,14 | 1,827 | 0,06763998 | 0,10958965 | 0,00455 | 0,0039198 | 0,0042742 | 0,0042742 | TRUE | 0,0782199 | No |
| orf19.6073 | 38316 | 24992 | 32918 | 41018 | 25710 | 21227 | 23025 | 14898 | 34373 | 31817 | 33678 | 33824 | 21988 | 20254 | 20546 | 21012 | 27187 | 33423 | 20950 | 0,627 | -0,674 | -17,224 | 1,75E-66 | 3,62E-64 | 0,001158 | 0,0020329 | 0,0014319 | 0,0014319 | TRUE | 0,0137365 | No |
| orf19.6074 | 5336 | 3513 | 4290 | 5317 | 5195 | 4332 | 4447 | 2619 | 4787 | 4472 | 4389 | 4384 | 4443 | 4133 | 3968 | 3694 | 4283,9 | 4508 | 4060 | 0,901 | -0,15 | -2,557 | 0,01056351 | 0,02113406 | 0,003556 | 0,0023184 | 0,0030715 | 0,0030715 | TRUE | 0,049284 | No |
| orf19.6075 | 1258 | 784 | 1111 | 1342 | 1355 | 1126 | 1171 | 928 | 1129 | 998 | 1137 | 1107 | 1159 | 1074 | 1045 | 1309 | 1119,6 | 1093 | 1147 | 1,046 | 0,065 | 0,852 | 0,39410698 | 0,48273084 | 0,005861 | 0,0032761 | 0,0046023 | 0,0046023 | TRUE | 0,1017541 | No |
| orf19.6076 | 448 | 295 | 395 | 503 | 569 | 469 | 532 | 399 | 402 | 376 | 404 | 415 | 487 | 448 | 475 | 563 | 446 | 399 | 493 | 1,226 | 0,294 | 3,589 | 0,0003317 | 0,00093036 | 0,002863 | 0,0052345 | 0,0041815 | 0,0041815 | TRUE | 0,0926299 | No |
| orf19.6077 | 2124 | 1419 | 1982 | 2314 | 2331 | 1919 | 2112 | 1586 | 1905 | 1807 | 2028 | 1908 | 1994 | 1831 | 1885 | 2237 | 1949,3 | 1912 | 1987 | 1,037 | 0,052 | 0,788 | 0,43056819 | 0,51768134 | 0,004479 | 0,0027243 | 0,0036775 | 0,0036775 | TRUE | 0,0829795 | No |
| orf19.6078 | 2106 | 1462 | 1969 | 2296 | 3114 | 2668 | 2897 | 1646 | 1889 | 1861 | 2014 | 1893 | 2663 | 2546 | 2585 | 2322 | 2221,7 | 1914 | 2529 | 1,323 | 0,404 | 7,765 | 8,15E-15 | 9,21E-14 | 0,001701 | 0,0026329 | 0,0021344 | 0,0021344 | TRUE | 0,0346894 | No |
| orf19.6078.1 | 0 | 0 | 0 | 0 | 0 | 4 | 3 | 3 | 0 | 0 | 0 | 0 | 0 | 4 | 3 | 4 | 1,34 | 0 | 3 | 14,406 | 3,849 | 2,433 | 0,01495619 | NA | 1,89238 | 1,0848429 | 1,2098828 | 1,2098828 | TRUE | 0,6491409 | No |
| orf19.6079 | 281 | 183 | 234 | 305 | 468 | 366 | 354 | 245 | 252 | 233 | 239 | 252 | 400 | 349 | 316 | 346 | 298,36 | 244 | 353 | 1,444 | 0,53 | 5,582 | 2,37E-08 | 1,25E-07 | 0,003636 | 0,0068451 | 0,0052142 | 0,0052142 | TRUE | 0,1021923 | No |
| orf19.6080 | 2712 | 1776 | 2318 | 2905 | 3012 | 2491 | 2717 | 1655 | 2433 | 2261 | 2372 | 2395 | 2576 | 2377 | 2424 | 2334 | 2396,6 | 2365 | 2428 | 1,027 | 0,038 | 0,833 | 0,4045717 | 0,49273601 | 0,000874 | 0,0025853 | 0,0015891 | 0,0015891 | TRUE | 0,0208234 | No |
| orf19.6081 | 14465 | 9518 | 12996 | 15946 | 13887 | 11419 | 12710 | 8623 | 12977 | 12117 | 13296 | 13149 | 11877 | 10896 | 11342 | 12162 | 12227 | 12885 | 11569 | 0,898 | -0,156 | -3,352 | 0,00080198 | 0,00206004 | 0,001915 | 0,0020983 | 0,0019981 | 0,0019981 | TRUE | 0,0194481 | No |
| orf19.6082 | 4576 | 2940 | 3979 | 4917 | 4258 | 3623 | 3947 | 2541 | 4105 | 3743 | 4071 | 4055 | 3642 | 3457 | 3522 | 3584 | 3772,3 | 3994 | 3551 | 0,888 | -0,171 | -4,117 | 3,84E-05 | 0,00012659 | 0,000784 | 0,0023644 | 0,0013824 | 0,0013824 | TRUE | 0,0207713 | No |
| orf19.6083 | 689 | 442 | 498 | 649 | 613 | 539 | 564 | 357 | 618 | 563 | 510 | 535 | 524 | 514 | 503 | 504 | 533,86 | 556 | 511 | 0,919 | -0,121 | -1,639 | 0,10113019 | 0,15421071 | 0,002143 | 0,0046988 | 0,0033717 | 0,0033717 | TRUE | 0,0692215 | No |
| orf19.6084 | 9602 | 6586 | 7921 | 10054 | 11794 | 10222 | 10864 | 7148 | 8614 | 8385 | 8104 | 8291 | 10087 | 9754 | 9694 | 10082 | 9126,2 | 8348 | 9904 | 1,186 | 0,246 | 7,533 | 4,95E-14 | 5,17E-13 | 0,000399 | 0,0021386 | 0,0009146 | 0,0009146 | TRUE | 0,005709 | No |
| orf19.6085 | 21124 | 13593 | 17264 | 21616 | 18708 | 15845 | 16961 | 10695 | 18950 | 17305 | 17663 | 17825 | 16000 | 15119 | 15135 | 15084 | 16635 | 17936 | 15334 | 0,855 | -0,226 | -5,772 | 7,85E-09 | 4,42E-08 | 0,001123 | 0,0020668 | 0,0014132 | 0,0014132 | TRUE | 0,0177829 | No |
| orf19.6086 | 974 | 556 | 754 | 992 | 486 | 386 | 406 | 278 | 874 | 708 | 771 | 818 | 416 | 368 | 362 | 392 | 588,67 | 793 | 384 | 0,484 | -1,048 | -12,791 | 1,85E-37 | 1,10E-35 | 0,004553 | 0,0044457 | 0,0045018 | 0,0045018 | TRUE | 0,0579802 | No |
| orf19.609 | 37 | 22 | 27 | 44 | 48 | 45 | 45 | 37 | 33 | 28 | 28 | 36 | 41 | 43 | 40 | 52 | 37,68 | 31 | 44 | 1,381 | 0,465 | 2,066 | 0,03886908 | 0,06741496 | 1,00E-08 | 0,040507 | 0,0216206 | 0,0216206 | TRUE | 0,0890339 | No |
| orf19.6090 | 37847 | 24315 | 32237 | 39113 | 31343 | 25417 | 27699 | 18183 | 33953 | 30956 | 32982 | 32253 | 26806 | 24252 | 24717 | 25645 | 28945 | 32536 | 25355 | 0,779 | -0,36 | -8,215 | 2,13E-16 | 2,79E-15 | 0,001695 | 0,0020297 | 0,0018084 | 0,0018084 | TRUE | 0,0182832 | No |
| orf19.6091 | 4013 | 2484 | 3624 | 4441 | 4651 | 3889 | 4425 | 2969 | 3600 | 3162 | 3708 | 3662 | 3978 | 3711 | 3949 | 4188 | 3744,6 | 3533 | 3956 | 1,118 | 0,161 | 2,753 | 0,00591065 | 0,01256589 | 0,003384 | 0,0023672 | 0,003031 | 0,003031 | TRUE | 0,0591059 | No |
| orf19.6092 | 1624 | 948 | 1411 | 1651 | 1430 | 1173 | 1358 | 933 | 1457 | 1207 | 1444 | 1361 | 1223 | 1119 | 1212 | 1316 | 1292,3 | 1367 | 1218 | 0,888 | -0,171 | -2,415 | 0,01574517 | 0,03014563 | 0,004678 | 0,0031028 | 0,0040565 | 0,0040565 | TRUE | 0,0712106 | No |
| orf19.6094 | 1015 | 667 | 913 | 1071 | 1082 | 863 | 927 | 545 | 911 | 849 | 934 | 883 | 925 | 823 | 827 | 769 | 865,21 | 894 | 836 | 0,937 | -0,094 | -1,413 | 0,15752346 | 0,22458495 | 0,002333 | 0,0036574 | 0,003044 | 0,003044 | TRUE | 0,0623877 | No |
| orf19.6096 | 787 | 510 | 655 | 857 | 768 | 604 | 663 | 410 | 706 | 649 | 670 | 707 | 657 | 576 | 592 | 578 | 641,89 | 683 | 601 | 0,88 | -0,185 | -2,76 | 0,00577342 | 0,01231158 | 0,001335 | 0,0042411 | 0,0027385 | 0,0027385 | TRUE | 0,0490348 | No |
| orf19.6099 | 4630 | 2822 | 4056 | 4732 | 5230 | 3867 | 4519 | 3373 | 4154 | 3593 | 4150 | 3902 | 4473 | 3690 | 4032 | 4757 | 4093,8 | 3950 | 4238 | 1,072 | 0,101 | 1,286 | 0,19839217 | 0,27273478 | 0,008127 | 0,0023341 | 0,005627 | 0,005627 | TRUE | 0,0922624 | No |
| orf19.610 | 4462 | 2848 | 4155 | 5091 | 7007 | 5678 | 6610 | 4667 | 4003 | 3626 | 4251 | 4198 | 5993 | 5418 | 5898 | 6582 | 4996,1 | 4020 | 5973 | 1,485 | 0,57 | 8,482 | 2,22E-17 | 3,25E-16 | 0,005409 | 0,0022701 | 0,0041303 | 0,0041303 | TRUE | 0,0566829 | No |
| orf19.6100 | 557 | 364 | 537 | 623 | 689 | 486 | 517 | 380 | 500 | 463 | 549 | 514 | 589 | 464 | 461 | 536 | 509,56 | 506 | 512 | 1,01 | 0,014 | 0,154 | 0,87755552 | 0,90683756 | 0,007816 | 0,0048284 | 0,0063725 | 0,0063725 | TRUE | 0,1285406 | No |
| orf19.6102 | 1537 | 864 | 1277 | 1559 | 1897 | 1533 | 1858 | 1059 | 1379 | 1100 | 1306 | 1286 | 1622 | 1463 | 1658 | 1494 | 1413,5 | 1268 | 1559 | 1,229 | 0,297 | 4,083 | 4,45E-05 | 0,00014534 | 0,00531 | 0,0030066 | 0,0043725 | 0,0043725 | TRUE | 0,090764 | No |
| orf19.6103 | 285 | 233 | 268 | 343 | 353 | 300 | 332 | 223 | 256 | 297 | 274 | 283 | 302 | 286 | 296 | 315 | 288,53 | 278 | 300 | 1,081 | 0,112 | 1,312 | 0,1894522 | 0,26244612 | 1,00E-08 | 0,0070108 | 0,0035877 | 0,0035877 | TRUE | 0,0319215 | No |
| orf19.6105 | 1419 | 820 | 1183 | 1442 | 1705 | 1139 | 1382 | 1107 | 1273 | 1044 | 1210 | 1189 | 1458 | 1087 | 1233 | 1561 | 1257 | 1179 | 1335 | 1,13 | 0,176 | 1,691 | 0,09090571 | 0,14119998 | 0,015252 | 0,0031344 | 0,0096062 | 0,0096062 | TRUE | 0,1879729 | No |
| orf19.6109 | 5043 | 2982 | 4372 | 5127 | 6045 | 4454 | 5209 | 3291 | 4524 | 3796 | 4473 | 4228 | 5170 | 4250 | 4648 | 4642 | 4466,4 | 4255 | 4678 | 1,099 | 0,136 | 1,939 | 0,05253906 | 0,0878038 | 0,006004 | 0,0023046 | 0,0045057 | 0,0045057 | TRUE | 0,063163 | No |
| orf19.6110 | 17 | 4 | 9 | 12 | 10 | 5 | 5 | 4 | 15 | 5 | 9 | 10 | 9 | 5 | 4 | 6 | 7,86 | 10 | 6 | 0,586 | -0,77 | -1,488 | 0,13680342 | NA | 1,00E-08 | 0,1866951 | 0,123048 | 0,123048 | TRUE | 0,2422617 | No |
| orf19.6112 | 281 | 153 | 228 | 260 | 322 | 290 | 304 | 180 | 252 | 195 | 233 | 214 | 275 | 277 | 271 | 254 | 246,47 | 224 | 269 | 1,202 | 0,266 | 2,702 | 0,00689897 | 0,01444217 | 0,001375 | 0,0078695 | 0,0052382 | 0,0052382 | TRUE | 0,0818275 | No |
| orf19.6113 | 73 | 51 | 52 | 69 | 90 | 64 | 77 | 37 | 65 | 65 | 53 | 57 | 77 | 61 | 69 | 52 | 62,43 | 60 | 65 | 1,092 | 0,127 | 0,698 | 0,48526133 | 0,57064985 | 1,00E-08 | 0,0252324 | 0,0159747 | 0,0159747 | TRUE | 0,1455334 | No |
| orf19.6114 | 2400 | 1470 | 2014 | 2539 | 2421 | 1762 | 2145 | 1200 | 2153 | 1871 | 2061 | 2094 | 2071 | 1681 | 1914 | 1692 | 1942,1 | 2045 | 1840 | 0,9 | -0,152 | -2,046 | 0,04079722 | 0,070232 | 0,006282 | 0,002727 | 0,0047635 | 0,0047635 | TRUE | 0,0870424 | No |
| orf19.6115 | 4 | 1 | 2 | 1 | 4 | 0 | 2 | 0 | 4 | 1 | 2 | 1 | 3 | 0 | 2 | 0 | 1,62 | 2 | 1 | 0,719 | -0,476 | -0,409 | 0,68217719 | NA | 0,282685 | 0,8996201 | 0,683848 | 0,683848 | TRUE | 0,8153808 | No |
| orf19.6116 | 1346 | 888 | 1116 | 1348 | 2015 | 1700 | 1939 | 1119 | 1208 | 1131 | 1142 | 1112 | 1723 | 1622 | 1730 | 1578 | 1405,7 | 1148 | 1663 | 1,451 | 0,537 | 10,355 | 3,97E-25 | 1,02E-23 | 0,001015 | 0,0030123 | 0,0018487 | 0,0018487 | TRUE | 0,0153559 | No |
| orf19.6117 | 7388 | 4831 | 6447 | 7904 | 8545 | 6833 | 7600 | 5667 | 6628 | 6150 | 6596 | 6518 | 7308 | 6520 | 6782 | 7993 | 6811,8 | 6473 | 7151 | 1,104 | 0,143 | 2,299 | 0,02150395 | 0,03982606 | 0,004514 | 0,0021926 | 0,0035499 | 0,0035499 | TRUE | 0,0756053 | No |
| orf19.6118 | 379 | 269 | 338 | 393 | 350 | 301 | 279 | 176 | 340 | 342 | 346 | 324 | 299 | 287 | 249 | 248 | 304,51 | 338 | 271 | 0,806 | -0,311 | -3,481 | 0,00049873 | 0,00134669 | 0,001396 | 0,0067469 | 0,0043693 | 0,0043693 | TRUE | 0,0581176 | No |
| orf19.6119 | 197 | 162 | 213 | 239 | 243 | 191 | 239 | 164 | 177 | 206 | 218 | 197 | 208 | 182 | 213 | 231 | 204,08 | 200 | 208 | 1,044 | 0,062 | 0,568 | 0,5701224 | 0,64928276 | 0,003464 | 0,009093 | 0,0066324 | 0,0066324 | TRUE | 0,0780122 | No |
| orf19.612 | 527 | 317 | 413 | 530 | 532 | 501 | 447 | 325 | 473 | 404 | 423 | 437 | 455 | 478 | 399 | 458 | 440,78 | 434 | 448 | 1,027 | 0,039 | 0,474 | 0,63578115 | 0,70658031 | 0,003129 | 0,005273 | 0,004204 | 0,004204 | TRUE | 0,0658217 | No |
| orf19.6121 | 5842 | 3873 | 5235 | 6170 | 8452 | 7340 | 8294 | 5299 | 5241 | 4931 | 5356 | 5088 | 7228 | 7004 | 7401 | 7474 | 6215,3 | 5154 | 7277 | 1,411 | 0,497 | 12,732 | 3,92E-37 | 2,31E-35 | 0,000835 | 0,0022131 | 0,0012954 | 0,0012954 | TRUE | 0,0099093 | No |

| | | | | | | | | | | | | | | | | | | | | | | | | | | | | | | | |
|---|---|---|---|---|---|---|---|---|---|---|---|---|---|---|---|---|---|---|---|---|---|---|---|---|---|---|---|---|---|---|---|
| orf19.6124 | 1805 | 1220 | 1750 | 1994 | 1746 | 1630 | 1700 | 1077 | 1619 | 1553 | 1790 | 1644 | 1493 | 1555 | 1517 | 1519 | 1586,5 | 1652 | 1521 | 0,92 | -0,12 | -2,314 | 0,02065724 | 0,03836435 | 0,001136 | 0,0028946 | 0,0019445 | 0,0019445 | TRUE | 0,0375443 | No |
| orf19.6126 | 6416 | 3949 | 5704 | 6732 | 5981 | 5076 | 5667 | 3410 | 5756 | 5027 | 5836 | 5551 | 5115 | 4843 | 5057 | 4809 | 5249,4 | 5542 | 4956 | 0,894 | -0,162 | -3,119 | 0,00181611 | 0,00432641 | 0,002371 | 0,0022561 | 0,0023841 | 0,0023841 | TRUE | 0,0466838 | No |
| orf19.6127 | 19480 | 12323 | 16803 | 20387 | 19612 | 15662 | 17550 | 12195 | 17476 | 15688 | 17191 | 16811 | 16773 | 14944 | 15660 | 17200 | 16468 | 16792 | 16144 | 0,961 | -0,057 | -1,063 | 0,28780878 | 0,37229078 | 0,003029 | 0,0020677 | 0,0027117 | 0,0027117 | TRUE | 0,0306221 | No |
| orf19.6129 | 2196 | 1350 | 1792 | 2128 | 2108 | 1831 | 1970 | 1176 | 1970 | 1719 | 1833 | 1755 | 1803 | 1747 | 1758 | 1659 | 1780,4 | 1819 | 1742 | 0,958 | -0,062 | -1,151 | 0,24978712 | 0,33182334 | 0,001725 | 0,0027949 | 0,0022462 | 0,0022462 | TRUE | 0,0381985 | No |
| orf19.6131 | 820 | 515 | 668 | 853 | 781 | 596 | 715 | 513 | 736 | 656 | 683 | 703 | 668 | 569 | 638 | 724 | 672,03 | 694 | 650 | 0,931 | -0,103 | -1,337 | 0,18128781 | 0,25306162 | 0,004121 | 0,0041397 | 0,0042209 | 0,0042209 | TRUE | 0,0817099 | No |
| orf19.6132 | 1619 | 1065 | 1255 | 1675 | 1463 | 1224 | 1291 | 744 | 1452 | 1356 | 1284 | 1381 | 1251 | 1168 | 1152 | 1049 | 1261,7 | 1368 | 1155 | 0,846 | -0,241 | -3,833 | 0,00012659 | 0,00038028 | 0,002799 | 0,0031301 | 0,0030095 | 0,0030095 | TRUE | 0,0415201 | No |
| orf19.6133 | 4376 | 2956 | 4010 | 4774 | 4407 | 3780 | 4345 | 2519 | 3926 | 3763 | 4103 | 3937 | 3769 | 3607 | 3877 | 3553 | 3816,8 | 3932 | 3702 | 0,942 | -0,087 | -1,986 | 0,04699947 | 0,07965382 | 0,001081 | 0,0023599 | 0,0015591 | 0,0015591 | TRUE | 0,0123918 | No |
| orf19.6134 | 412 | 289 | 387 | 434 | 805 | 720 | 793 | 506 | 370 | 368 | 396 | 358 | 688 | 687 | 708 | 714 | 536,01 | 373 | 699 | 1,875 | 0,907 | 13,822 | 1,88E-43 | 1,39E-41 | 1,00E-08 | 0,0046879 | 0,0020892 | 0,0020892 | TRUE | 0,0199922 | No |
| orf19.6135 | 839 | 576 | 773 | 937 | 1003 | 840 | 941 | 598 | 753 | 733 | 791 | 773 | 858 | 802 | 840 | 843 | 798,99 | 762 | 836 | 1,095 | 0,131 | 2,359 | 0,01832103 | 0,03438699 | 1,00E-08 | 0,0037964 | 0,0016965 | 0,0016965 | TRUE | 0,0093248 | No |
| orf19.6135.1 | 895 | 590 | 646 | 866 | 981 | 836 | 792 | 465 | 803 | 751 | 661 | 714 | 839 | 798 | 707 | 656 | 741,04 | 732 | 750 | 1,027 | 0,038 | 0,431 | 0,66613308 | 0,73352778 | 0,008129 | 0,0039385 | 0,0061266 | 0,0061266 | TRUE | 0,0775855 | No |
| orf19.6136 | 2879 | 1920 | 2390 | 3158 | 4031 | 3178 | 3607 | 2111 | 2583 | 2444 | 2445 | 2604 | 3447 | 3032 | 3219 | 2977 | 2844 | 2519 | 3169 | 1,259 | 0,332 | 6,094 | 1,10E-09 | 6,98E-09 | 0,002358 | 0,00249 | 0,0024877 | 0,0024877 | TRUE | 0,0428833 | No |
| orf19.6137 | 3135 | 2132 | 2826 | 3347 | 2891 | 2544 | 2778 | 1702 | 2812 | 2714 | 2891 | 2760 | 2472 | 2427 | 2479 | 2401 | 2619,7 | 2794 | 2445 | 0,875 | -0,192 | -5,113 | 3,18E-07 | 1,45E-06 | 1,00E-08 | 0,0025337 | 0,0009739 | 0,0009739 | TRUE | 0,0063998 | No |
| orf19.6138 | 55 | 31 | 51 | 63 | 44 | 37 | 67 | 27 | 49 | 39 | 52 | 52 | 38 | 35 | 60 | 38 | 45,47 | 48 | 43 | 0,885 | -0,176 | -0,772 | 0,44014019 | 0,52697535 | 0,019535 | 0,0339081 | 0,0278072 | 0,0278072 | TRUE | 0,6129889 | No |
| orf19.6139 | 2265 | 1463 | 2051 | 2570 | 2189 | 1950 | 2103 | 1322 | 2032 | 1863 | 2098 | 2119 | 1872 | 1861 | 1877 | 1865 | 1948,3 | 2028 | 1869 | 0,92 | -0,12 | -2,48 | 0,0131452 | 0,02568337 | 0,00097 | 0,0027247 | 0,0017465 | 0,0017465 | TRUE | 0,0342508 | No |
| orf19.6140 | 1011 | 728 | 974 | 1213 | 889 | 857 | 955 | 620 | 907 | 927 | 996 | 1000 | 760 | 818 | 852 | 874 | 891,9 | 958 | 826 | 0,86 | -0,217 | -3,43 | 0,00060321 | 0,00159369 | 0,001989 | 0,0036072 | 0,0027256 | 0,0027256 | TRUE | 0,035435 | No |
| orf19.6141 | 995 | 650 | 936 | 1075 | 867 | 764 | 847 | 549 | 893 | 828 | 958 | 886 | 741 | 729 | 756 | 774 | 820,6 | 891 | 750 | 0,84 | -0,252 | -4,224 | 2,41E-05 | 8,25E-05 | 0,00073 | 0,0037486 | 0,0021909 | 0,0021909 | TRUE | 0,0285743 | No |
| orf19.6142 | 10 | 7 | 14 | 12 | 13 | 6 | 5 | 4 | 9 | 9 | 14 | 10 | 11 | 6 | 4 | 6 | 8,63 | 10 | 7 | 0,65 | -0,621 | -1,282 | 0,1997863 | NA | 1,00E-08 | 0,170176 | 0,105021 | 0,105021 | TRUE | 0,6117658 | No |
| orf19.6143 | 5175 | 3466 | 4682 | 5646 | 4771 | 3909 | 4233 | 2864 | 4643 | 4413 | 4790 | 4656 | 4080 | 3730 | 3777 | 4039 | 4266 | 4626 | 3906 | 0,844 | -0,245 | -5,448 | 5,10E-08 | 2,59E-07 | 0,001305 | 0,0023198 | 0,0016996 | 0,0016996 | TRUE | 0,0115377 | No |
| orf19.6146 | 9270 | 5968 | 8242 | 9762 | 12158 | 9752 | 10461 | 7078 | 8316 | 7598 | 8432 | 8050 | 10398 | 9305 | 9335 | 9983 | 8927,1 | 8099 | 9755 | 1,204 | 0,268 | 5,327 | 9,97E-08 | 4,90E-07 | 0,002375 | 0,0021421 | 0,0023171 | 0,0023171 | TRUE | 0,0244152 | No |
| orf19.6147 | 933 | 617 | 620 | 886 | 729 | 632 | 621 | 317 | 837 | 786 | 634 | 731 | 623 | 603 | 554 | 447 | 651,9 | 747 | 557 | 0,749 | -0,418 | -3,847 | 0,00011962 | 0,00036079 | 0,014654 | 0,0042064 | 0,0097528 | 0,0097528 | TRUE | 0,1903403 | No |
| orf19.6148 | 397 | 307 | 362 | 397 | 352 | 341 | 403 | 245 | 356 | 391 | 370 | 327 | 301 | 325 | 360 | 346 | 347,04 | 361 | 333 | 0,923 | -0,115 | -1,295 | 0,19548775 | 0,26919331 | 0,003163 | 0,0061626 | 0,0047028 | 0,0047028 | TRUE | 0,0511316 | No |
| orf19.6151 | 1679 | 1118 | 1381 | 1896 | 2333 | 1833 | 1963 | 1225 | 1506 | 1423 | 1413 | 1563 | 1995 | 1749 | 1752 | 1728 | 1641,2 | 1476 | 1806 | 1,223 | 0,291 | 4,756 | 1,98E-06 | 8,00E-06 | 0,002811 | 0,0028641 | 0,002967 | 0,002967 | TRUE | 0,0616434 | No |
| orf19.6152 | 2143 | 1508 | 1948 | 2356 | 2487 | 2149 | 2511 | 1445 | 1922 | 1920 | 1993 | 1943 | 2127 | 2051 | 2241 | 2038 | 2029,3 | 1944 | 2114 | 1,088 | 0,122 | 2,728 | 0,00636847 | 0,01344877 | 0,00065 | 0,0026949 | 0,0014318 | 0,0014318 | TRUE | 0,019894 | No |
| orf19.6155 | 5064 | 3769 | 4470 | 5555 | 4935 | 4521 | 4708 | 2965 | 4543 | 4798 | 4573 | 4581 | 4221 | 4314 | 4201 | 4182 | 4426,6 | 4624 | 4230 | 0,916 | -0,127 | -3,82 | 0,00013363 | 0,00039982 | 0,000117 | 0,0023075 | 0,00084 | 0,00084 | TRUE | 0,0074514 | No |
| orf19.6156 | 1263 | 781 | 1047 | 1271 | 1496 | 1119 | 1200 | 708 | 1133 | 994 | 1071 | 1048 | 1279 | 1068 | 1071 | 999 | 1082,9 | 1062 | 1104 | 1,041 | 0,058 | 0,746 | 0,45556411 | 0,54144989 | 0,006106 | 0,0033201 | 0,0049697 | 0,0049697 | TRUE | 0,1389911 | No |
| orf19.6160 | 8499 | 5485 | 7359 | 8682 | 8309 | 7308 | 7834 | 5127 | 7624 | 6983 | 7529 | 7159 | 7106 | 6973 | 6991 | 7231 | 7199,6 | 7324 | 7075 | 0,966 | -0,051 | -1,324 | 0,18562296 | 0,25797426 | 0,000814 | 0,0021812 | 0,0012652 | 0,0012652 | TRUE | 0,0116187 | No |
| orf19.6163 | 1445 | 917 | 1252 | 1513 | 1205 | 975 | 1071 | 601 | 1296 | 1167 | 1281 | 1248 | 1031 | 930 | 956 | 848 | 1094,6 | 1248 | 941 | 0,756 | -0,404 | -6,206 | 5,44E-10 | 3,60E-09 | 0,003029 | 0,0033058 | 0,0031355 | 0,0031355 | TRUE | 0,0487442 | No |
| orf19.6164 | 12 | 12 | 11 | 10 | 17 | 10 | 6 | 2 | 11 | 15 | 11 | 8 | 15 | 10 | 5 | 3 | 9,72 | 11 | 8 | 0,746 | -0,422 | -0,858 | 0,3906535 | NA | 0,090054 | 0,1512582 | 0,1279627 | 0,1279627 | TRUE | 0,5414396 | No |
| orf19.6165 | 15650 | 10200 | 13735 | 16521 | 15448 | 12889 | 14048 | 9354 | 14040 | 12986 | 14052 | 13623 | 13212 | 12298 | 12536 | 13193 | 13242 | 13675 | 12810 | 0,936 | -0,095 | -2,329 | 0,01983411 | 0,03693841 | 0,001226 | 0,0020891 | 0,0015147 | 0,0015147 | TRUE | 0,0138973 | No |
| orf19.6166 | 816 | 574 | 769 | 940 | 1062 | 1014 | 1037 | 624 | 732 | 731 | 787 | 775 | 908 | 968 | 925 | 880 | 838,24 | 756 | 920 | 1,218 | 0,284 | 4,995 | 5,88E-07 | 2,58E-06 | 1,00E-08 | 0,0037114 | 0,0019081 | 0,0019081 | TRUE | 0,0138657 | No |
| orf19.6167 | 800 | 524 | 719 | 868 | 1000 | 810 | 977 | 656 | 718 | 667 | 736 | 716 | 855 | 773 | 872 | 925 | 782,66 | 709 | 856 | 1,203 | 0,267 | 4,007 | 6,14E-05 | 0,00019539 | 0,002179 | 0,0038343 | 0,0029685 | 0,0029685 | TRUE | 0,0505604 | No |
| orf19.6168 | 664 | 463 | 581 | 706 | 771 | 733 | 777 | 440 | 596 | 589 | 594 | 582 | 659 | 699 | 693 | 621 | 629,3 | 590 | 668 | 1,135 | 0,183 | 2,954 | 0,00313889 | 0,00709695 | 1,00E-08 | 0,0042864 | 0,0020995 | 0,0020995 | TRUE | 0,0226359 | No |
| orf19.6169 | 1316 | 844 | 1168 | 1426 | 450 | 443 | 490 | 339 | 1181 | 1075 | 1195 | 1176 | 385 | 423 | 437 | 478 | 793,61 | 1157 | 431 | 0,37 | -1,435 | -20,221 | 6,46E-91 | 3,23E-88 | 0,002565 | 0,0038088 | 0,0032293 | 0,0032293 | TRUE | 0,0608939 | No |
| orf19.6170 | 854 | 507 | 664 | 900 | 907 | 756 | 797 | 494 | 766 | 645 | 679 | 742 | 776 | 721 | 711 | 697 | 717,26 | 708 | 726 | 1,024 | 0,034 | 0,487 | 0,62623007 | 0,69803159 | 0,002725 | 0,0040035 | 0,0033332 | 0,0033332 | TRUE | 0,039059 | No |
| orf19.6171 | 1350 | 856 | 1220 | 1422 | 1320 | 1222 | 1212 | 863 | 1211 | 1090 | 1248 | 1173 | 1129 | 1166 | 1082 | 1217 | 1164,4 | 1180 | 1148 | 0,97 | -0,044 | -0,754 | 0,45110844 | 0,53700407 | 0,001757 | 0,0032263 | 0,0024676 | 0,0024676 | TRUE | 0,0293995 | No |
| orf19.6173 | 2463 | 1593 | 2178 | 2749 | 2791 | 2320 | 2548 | 2083 | 2210 | 2028 | 2228 | 2267 | 2387 | 2214 | 2274 | 2938 | 2318,1 | 2183 | 2453 | 1,121 | 0,165 | 1,996 | 0,04592531 | 0,07805354 | 0,009715 | 0,0026058 | 0,0061524 | 0,0061524 | TRUE | 0,2094548 | No |
| orf19.6175 | 3006 | 1895 | 2360 | 3064 | 2810 | 2401 | 2628 | 1719 | 2697 | 2413 | 2415 | 2527 | 2403 | 2291 | 2345 | 2424 | 2439,3 | 2513 | 2366 | 0,94 | -0,089 | -1,848 | 0,06455032 | 0,10518534 | 0,001195 | 0,0025747 | 0,0018001 | 0,0018001 | TRUE | 0,0297029 | No |
| orf19.6176 | 1850 | 1071 | 1485 | 1789 | 1976 | 1580 | 1684 | 1158 | 1660 | 1363 | 1519 | 1475 | 1690 | 1508 | 1503 | 1633 | 1543,9 | 1504 | 1584 | 1,051 | 0,072 | 1,059 | 0,28948282 | 0,37410471 | 0,004187 | 0,0029198 | 0,0037286 | 0,0037286 | TRUE | 0,0583105 | No |
| orf19.6177 | 645 | 391 | 514 | 642 | 599 | 472 | 494 | 371 | 579 | 498 | 526 | 529 | 512 | 450 | 441 | 523 | 507,3 | 533 | 482 | 0,9 | -0,153 | -1,896 | 0,05800843 | 0,09561444 | 0,003647 | 0,0048411 | 0,0042438 | 0,0042438 | TRUE | 0,0393472 | No |
| orf19.6178 | 3101 | 1896 | 2747 | 3254 | 3497 | 2647 | 2813 | 2131 | 2782 | 2414 | 2810 | 2683 | 2991 | 2526 | 2510 | 3006 | 2715,2 | 2672 | 2758 | 1,031 | 0,044 | 0,585 | 0,55868947 | 0,639138 | 0,006284 | 0,0025142 | 0,005037 | 0,005037 | TRUE | 0,0503173 | No |
| orf19.6179 | 8 | 4 | 6 | 8 | 7 | 4 | 6 | 6 | 7 | 5 | 6 | 7 | 6 | 4 | 5 | 8 | 6,08 | 6 | 6 | 0,912 | -0,133 | -0,246 | 0,80593702 | NA | 1,00E-08 | 0,2408222 | 0,1178589 | 0,1178589 | TRUE | 0,1288256 | No |
| orf19.6180 | 792 | 536 | 684 | 907 | 1063 | 817 | 905 | 571 | 711 | 682 | 700 | 748 | 909 | 780 | 808 | 805 | 767,77 | 710 | 826 | 1,162 | 0,217 | 3,27 | 0,0010757 | 0,00269395 | 0,002118 | 0,0038703 | 0,0029045 | 0,0029045 | TRUE | 0,0585408 | No |
| orf19.6181 | 15 | 12 | 19 | 23 | 23 | 17 | 22 | 18 | 13 | 15 | 19 | 19 | 20 | 16 | 20 | 25 | 18,51 | 16 | 20 | 1,185 | 0,245 | 0,768 | 0,44268051 | 0,52938385 | 1,00E-08 | 0,080424 | 0,0430435 | 0,0430435 | TRUE | 0,1091831 | No |
| orf19.6182 | 1581 | 977 | 1401 | 1708 | 1685 | 1325 | 1459 | 1056 | 1418 | 1244 | 1433 | 1408 | 1441 | 1264 | 1302 | 1489 | 1375,1 | 1376 | 1374 | 0,996 | -0,006 | -0,085 | 0,93256211 | 0,95140655 | 0,004343 | 0,0030352 | 0,0037605 | 0,0037605 | TRUE | 0,0477766 | No |
| orf19.6183 | 342 | 229 | 278 | 367 | 450 | 319 | 354 | 270 | 307 | 292 | 284 | 303 | 385 | 304 | 316 | 381 | 321,42 | 296 | 346 | 1,165 | 0,22 | 2,257 | 0,02402155 | 0,04400045 | 0,005425 | 0,0064961 | 0,0059678 | 0,0059678 | TRUE | 0,076039 | No |
| orf19.6184 | 49 | 31 | 43 | 54 | 52 | 38 | 60 | 24 | 44 | 39 | 44 | 45 | 44 | 36 | 54 | 34 | 42,51 | 43 | 42 | 0,987 | -0,019 | -0,088 | 0,9295933 | 0,94902236 | 1,00E-08 | 0,0361305 | 0,0219453 | 0,0219453 | TRUE | 0,2677665 | No |
| orf19.6185 | 0 | 0 | 0 | 0 | 0 | 0 | 0 | 0 | 0 | 0 | 0 | 0 | 0 | 0 | 0 | 0 | 0 | 0 | 0 | NA | NA | NA | NA | NA | NA | NA | NA | NA | NA | NA | NA |
| orf19.6186 | 447 | 254 | 397 | 466 | 571 | 458 | 565 | 385 | 401 | 323 | 406 | 384 | 488 | 437 | 504 | 543 | 435,92 | 378 | 493 | 1,294 | 0,372 | 4,063 | 4,85E-05 | 0,00015742 | 0,0059 | 0,0053097 | 0,0057104 | 0,0057104 | TRUE | 0,0993229 | No |
| orf19.6187 | 1141 | 695 | 1047 | 1217 | 1285 | 1022 | 1133 | 737 | 1024 | 885 | 1071 | 1004 | 1099 | 975 | 1011 | 1039 | 1013,5 | 996 | 1031 | 1,034 | 0,048 | 0,711 | 0,47699837 | 0,56247423 | 0,003107 | 0,0034119 | 0,0033745 | 0,0033745 | TRUE | 0,0622231 | No |
| orf19.6188 | 10 | 2 | 7 | 4 | 9 | 3 | 3 | 1 | 9 | 3 | 7 | 3 | 8 | 3 | 3 | 1 | 4,58 | 6 | 4 | 0,687 | -0,542 | -0,764 | 0,4446909 | NA | 0,184418 | 0,3190798 | 0,2625935 | 0,2625935 | TRUE | 0,5338886 | No |
| orf19.6189 | 1827 | 1056 | 1551 | 1829 | 2163 | 1688 | 1866 | 1145 | 1639 | 1344 | 1587 | 1508 | 1850 | 1611 | 1665 | 1615 | 1602,4 | 1520 | 1685 | 1,108 | 0,149 | 2,11 | 0,0348639 | 0,06128212 | 0,004977 | 0,0028855 | 0,0041349 | 0,0041349 | TRUE | 0,0693927 | No |
| orf19.6190 | 5276 | 3137 | 4412 | 5325 | 5399 | 4050 | 4371 | 2805 | 4733 | 3994 | 4514 | 4391 | 4617 | 3864 | 3900 | 3956 | 4246,3 | 4408 | 4084 | 0,927 | -0,11 | -1,582 | 0,11354627 | 0,16974024 | 0,005987 | 0,0023214 | 0,0044053 | 0,0044053 | TRUE | 0,0949301 | No |
| orf19.6191 | 32 | 16 | 16 | 31 | 22 | 14 | 18 | 9 | 29 | 20 | 16 | 26 | 19 | 13 | 16 | 13 | 18,99 | 23 | 15 | 0,672 | -0,574 | -1,762 | 0,07812975 | 0,12388586 | 1,00E-08 | 0,0784156 | 0,0479344 | 0,0479344 | TRUE | 0,1902274 | No |
| orf19.6192 | 287 | 169 | 286 | 320 | 394 | 267 | 288 | 279 | 257 | 215 | 293 | 264 | 337 | 255 | 257 | 394 | 283,92 | 257 | 311 | 1,199 | 0,262 | 1,808 | 0,07055931 | 0,11349942 | 0,026415 | 0,0070927 | 0,0165125 | 0,0165125 | TRUE | 0,3364234 | No |
| orf19.6193 | 3321 | 2156 | 2794 | 3454 | 3691 | 3229 | 3538 | 2219 | 2979 | 2745 | 2859 | 2848 | 3157 | 3081 | 3157 | 3130 | 2994,4 | 2858 | 3131 | 1,095 | 0,131 | 3,455 | 0,00055102 | 0,00147201 | 0,000183 | 0,0024643 | 0,0010382 | 0,0010382 | TRUE | 0,0099703 | No |
| orf19.6194 | 132 | 123 | 134 | 192 | 178 | 188 | 175 | 108 | 118 | 157 | 137 | 158 | 152 | 179 | 156 | 152 | 151,32 | 142 | 160 | 1,126 | 0,171 | 1,34 | 0,18011589 | 0,25198231 | 0,005863 | 0,0115734 | 0,0090508 | 0,0090508 | TRUE | 0,1451664 | No |
| orf19.6195 | 6124 | 4194 | 5612 | 6692 | 6493 | 5972 | 6584 | 3848 | 5494 | 5339 | 5742 | 5518 | 5553 | 5698 | 5875 | 5427 | 5580,9 | 5523 | 5638 | 1,021 | 0,03 | 0,774 | 0,43886573 | 0,52565895 | 0,000796 | 0,0022396 | 0,0012821 | 0,0012821 | TRUE | 0,0097517 | No |
| orf19.6196 | 1234 | 782 | 1050 | 1317 | 1090 | 939 | 992 | 725 | 1107 | 996 | 1074 | 1086 | 932 | 896 | 885 | 1023 | 999,85 | 1066 | 934 | 0,872 | -0,197 | -3,225 | 0,00125809 | 0,00310415 | 0,001935 | 0,0034314 | 0,0025815 | 0,0025815 | TRUE | 0,0432969 | No |
| orf19.6197 | 425 | 317 | 406 | 529 | 591 | 517 | 549 | 359 | 381 | 404 | 415 | 436 | 505 | 493 | 490 | 506 | 453,93 | 409 | 498 | 1,217 | 0,283 | 4,005 | 6,21E-05 | 0,00019715 | 1,00E-08 | 0,0051776 | 0,0025929 | 0,0025929 | TRUE | 0,0271238 | No |

| | | | | | | | | | | | | | | | | | | | | | | | | | | | | | | | |
|---|---|---|---|---|---|---|---|---|---|---|---|---|---|---|---|---|---|---|---|---|---|---|---|---|---|---|---|---|---|---|---|
| orf19.6198.1 | 1964 | 1479 | 1655 | 2210 | 1742 | 1516 | 1605 | 899 | 1762 | 1883 | 1693 | 1822 | 1490 | 1447 | 1432 | 1268 | 1599,6 | 1790 | 1409 | 0,79 | -0,341 | -5,794 | 6,86E-09 | 3,89E-08 | 0,002567 | 0,0028871 | 0,0026856 | 0,0026856 | TRUE | 0,0501066 | No |
| orf19.6199 | 799 | 593 | 731 | 831 | 731 | 604 | 716 | 398 | 717 | 755 | 748 | 685 | 625 | 576 | 639 | 561 | 663,33 | 726 | 600 | 0,831 | -0,267 | -4,024 | 5,72E-05 | 0,00018307 | 0,00129 | 0,004168 | 0,0027007 | 0,0027007 | TRUE | 0,0212287 | No |
| orf19.6200 | 33 | 11 | 19 | 28 | 27 | 17 | 22 | 16 | 30 | 14 | 19 | 23 | 23 | 16 | 20 | 23 | 20,96 | 22 | 20 | 0,927 | -0,11 | -0,349 | 0,7267156 | 0,78583646 | 1,00E-08 | 0,0712538 | 0,0473091 | 0,0473091 | TRUE | 0,2657638 | No |
| orf19.6202 | 1285 | 849 | 1191 | 1371 | 1446 | 1319 | 1317 | 1027 | 1153 | 1081 | 1219 | 1131 | 1237 | 1259 | 1175 | 1448 | 1212,7 | 1146 | 1280 | 1,113 | 0,155 | 2,236 | 0,02536183 | 0,04622994 | 0,004384 | 0,0031766 | 0,0037701 | 0,0037701 | TRUE | 0,0886917 | No |
| orf19.6203 | 1677 | 1140 | 1606 | 1871 | 2545 | 2226 | 2445 | 1549 | 1504 | 1451 | 1643 | 1543 | 2177 | 2124 | 2182 | 2185 | 1851,1 | 1535 | 2167 | 1,41 | 0,496 | 10,593 | 3,20E-26 | 9,29E-25 | 0,000737 | 0,0027638 | 0,0015435 | 0,0015435 | TRUE | 0,0258601 | No |
| orf19.6205 | 239 | 161 | 196 | 280 | 323 | 286 | 319 | 222 | 214 | 205 | 201 | 231 | 276 | 273 | 285 | 313 | 249,71 | 213 | 287 | 1,337 | 0,419 | 4,474 | 7,68E-06 | 2,85E-05 | 1,00E-08 | 0,007793 | 0,0043505 | 0,0043505 | TRUE | 0,0394657 | No |
| orf19.6208 | 2271 | 1481 | 1982 | 2432 | 2809 | 2225 | 2468 | 1794 | 2037 | 1885 | 2028 | 2005 | 2402 | 2123 | 2202 | 2530 | 2151,7 | 1989 | 2314 | 1,161 | 0,216 | 3,571 | 0,00035545 | 0,00099006 | 0,003223 | 0,0026542 | 0,0030417 | 0,0030417 | TRUE | 0,0453172 | No |
| orf19.6209 | 1341 | 879 | 1277 | 1440 | 1278 | 1045 | 1179 | 731 | 1203 | 1119 | 1306 | 1187 | 1093 | 997 | 1052 | 1031 | 1123,7 | 1204 | 1043 | 0,866 | -0,207 | -3,485 | 0,00049185 | 0,00133049 | 0,001863 | 0,0032715 | 0,0024855 | 0,0024855 | TRUE | 0,0381704 | No |
| orf19.6211 | 4331 | 2894 | 3864 | 4704 | 5670 | 4992 | 5503 | 3288 | 3885 | 3684 | 3953 | 3879 | 4849 | 4763 | 4911 | 4637 | 4320,3 | 3850 | 4790 | 1,244 | 0,315 | 8,521 | 1,58E-17 | 2,34E-16 | 0,000445 | 0,0023155 | 0,0010777 | 0,0010777 | TRUE | 0,00968 | No |
| orf19.6213 | 7281 | 4605 | 6135 | 7759 | 7472 | 5913 | 6336 | 4333 | 6532 | 5863 | 6277 | 6398 | 6390 | 5642 | 5654 | 6111 | 6108,4 | 6268 | 5949 | 0,949 | -0,076 | -1,418 | 0,15628878 | 0,2234078 | 0,002741 | 0,0022172 | 0,0025828 | 0,0025828 | TRUE | 0,0310486 | No |
| orf19.6214 | 1923 | 1338 | 1831 | 2127 | 1834 | 1572 | 1705 | 1163 | 1725 | 1703 | 1873 | 1754 | 1568 | 1500 | 1521 | 1640 | 1660,8 | 1764 | 1557 | 0,881 | -0,182 | -3,67 | 0,00024264 | 0,00069516 | 0,000913 | 0,0028536 | 0,0017568 | 0,0017568 | TRUE | 0,0206528 | No |
| orf19.6217 | 917 | 591 | 790 | 987 | 1285 | 1010 | 1223 | 832 | 823 | 752 | 808 | 814 | 1099 | 964 | 1091 | 1173 | 940,58 | 799 | 1082 | 1,349 | 0,432 | 6,421 | 1,36E-10 | 9,68E-10 | 0,00303 | 0,0035229 | 0,0032547 | 0,0032547 | TRUE | 0,0643717 | No |
| orf19.6219 | 134 | 121 | 152 | 170 | 134 | 128 | 93 | 75 | 120 | 154 | 156 | 140 | 115 | 122 | 83 | 106 | 124,43 | 142 | 106 | 0,749 | -0,417 | -2,794 | 0,00519997 | 0,01118963 | 0,012525 | 0,0136462 | 0,0131484 | 0,0131484 | TRUE | 0,2407579 | No |
| orf19.6220 | 2607 | 1726 | 2331 | 2842 | 3015 | 2781 | 2881 | 1967 | 2339 | 2197 | 2385 | 2344 | 2579 | 2654 | 2571 | 2774 | 2480,2 | 2316 | 2644 | 1,139 | 0,188 | 4,292 | 1,77E-05 | 6,22E-05 | 0,000721 | 0,0025648 | 0,001435 | 0,001435 | TRUE | 0,0134593 | No |
| orf19.6220.3 | 1000 | 656 | 750 | 995 | 1528 | 1173 | 1331 | 969 | 897 | 835 | 767 | 820 | 1307 | 1119 | 1188 | 1367 | 1037,6 | 830 | 1245 | 1,497 | 0,582 | 7,772 | 7,72E-15 | 8,75E-14 | 0,004785 | 0,0033787 | 0,0043821 | 0,0043821 | TRUE | 0,0551092 | No |
| orf19.6220.4 | 14748 | 9405 | 11749 | 14795 | 14906 | 11699 | 12207 | 8082 | 13230 | 11974 | 12020 | 12200 | 12748 | 11163 | 10893 | 11399 | 11953 | 12356 | 11551 | 0,935 | -0,097 | -1,724 | 0,08472999 | 0,1329827 | 0,003581 | 0,002101 | 0,0029737 | 0,0029737 | TRUE | 0,0599666 | No |
| orf19.6222 | 133 | 89 | 116 | 134 | 106 | 103 | 97 | 67 | 119 | 113 | 119 | 110 | 91 | 98 | 87 | 94 | 103,97 | 115 | 92 | 0,8 | -0,323 | -2,419 | 0,01554849 | 0,02979755 | 1,00E-08 | 0,0159418 | 0,0073879 | 0,0073879 | TRUE | 0,0194929 | No |
| orf19.6222.1 | 2336 | 1448 | 1771 | 2289 | 3104 | 2416 | 2458 | 1473 | 2096 | 1843 | 1812 | 1888 | 2655 | 2305 | 2193 | 2078 | 2108,7 | 1910 | 2308 | 1,209 | 0,274 | 3,538 | 0,0004036 | 0,00111077 | 0,007436 | 0,002668 | 0,0052843 | 0,0052843 | TRUE | 0,1250546 | No |
| orf19.6224 | 197 | 142 | 188 | 262 | 155 | 132 | 173 | 84 | 177 | 181 | 192 | 216 | 133 | 126 | 154 | 118 | 162,16 | 192 | 133 | 0,696 | -0,523 | -4,25 | 2,14E-05 | 7,40E-05 | 0,004772 | 0,0109319 | 0,0081738 | 0,0081738 | TRUE | 0,1254529 | No |
| orf19.6225 | 1569 | 1121 | 1416 | 1741 | 1549 | 1433 | 1644 | 1067 | 1408 | 1427 | 1449 | 1436 | 1325 | 1367 | 1467 | 1505 | 1422,9 | 1430 | 1416 | 0,988 | -0,017 | -0,325 | 0,74545015 | 0,80033318 | 0,001088 | 0,0029998 | 0,0018956 | 0,0018956 | TRUE | 0,0230267 | No |
| orf19.6225.1 | 1137 | 727 | 990 | 1254 | 1282 | 1076 | 1107 | 687 | 1020 | 926 | 1013 | 1034 | 1096 | 1027 | 988 | 969 | 1009 | 998 | 1020 | 1,022 | 0,031 | 0,52 | 0,60313302 | 0,67870822 | 0,001653 | 0,0034182 | 0,002454 | 0,002454 | TRUE | 0,0312826 | No |
| orf19.6227 | 4415 | 2781 | 3826 | 4887 | 4893 | 4025 | 4614 | 2789 | 3961 | 3541 | 3914 | 4030 | 4185 | 3841 | 4117 | 3934 | 3940,2 | 3862 | 4019 | 1,04 | 0,057 | 1,123 | 0,26130377 | 0,34488424 | 0,002058 | 0,002348 | 0,0022228 | 0,0022228 | TRUE | 0,037005 | No |
| orf19.6229 | 7404 | 4651 | 6506 | 7891 | 7493 | 6494 | 7055 | 5035 | 6642 | 5921 | 6656 | 6507 | 6408 | 6196 | 6295 | 7101 | 6466 | 6432 | 6500 | 1,01 | 0,014 | 0,252 | 0,80135773 | 0,84555346 | 0,002831 | 0,002204 | 0,0028178 | 0,0028178 | TRUE | 0,0463526 | No |
| orf19.6230 | 4331 | 3332 | 3855 | 4808 | 4745 | 4696 | 5005 | 2758 | 3885 | 4242 | 3944 | 3965 | 4058 | 4481 | 4466 | 3890 | 4116,4 | 4009 | 4224 | 1,055 | 0,077 | 1,386 | 0,16578392 | 0,23519003 | 0,002925 | 0,0023322 | 0,0027112 | 0,0027112 | TRUE | 0,0332561 | No |
| orf19.6231 | 1903 | 1315 | 1603 | 1957 | 2310 | 1747 | 1922 | 1478 | 1707 | 1674 | 1640 | 1614 | 1976 | 1667 | 1715 | 2085 | 1759,7 | 1659 | 1861 | 1,12 | 0,164 | 2,284 | 0,02234289 | 0,04125263 | 0,005418 | 0,0028045 | 0,0043587 | 0,0043587 | TRUE | 0,0757797 | No |
| orf19.6232 | 4406 | 2961 | 3944 | 4814 | 5510 | 4758 | 5183 | 3157 | 3953 | 3770 | 4035 | 3970 | 4712 | 4540 | 4625 | 4453 | 4257,1 | 3932 | 4582 | 1,166 | 0,221 | 6,027 | 1,67E-09 | 1,02E-08 | 0,0004 | 0,0023205 | 0,0010545 | 0,0010545 | TRUE | 0,0088442 | No |
| orf19.6233 | 2934 | 2022 | 2668 | 3307 | 3894 | 3135 | 3293 | 2039 | 2632 | 2574 | 2730 | 2727 | 3330 | 2991 | 2938 | 2876 | 2849,9 | 2666 | 3034 | 1,139 | 0,187 | 3,53 | 0,00041486 | 0,00113916 | 0,002177 | 0,0024889 | 0,0023537 | 0,0023537 | TRUE | 0,0533209 | No |
| orf19.6234 | 9459 | 6494 | 8675 | 10456 | 9983 | 8670 | 9419 | 5888 | 8486 | 8268 | 8875 | 8622 | 8538 | 8273 | 8405 | 8304 | 8471,3 | 8563 | 8380 | 0,979 | -0,031 | -0,956 | 0,33885797 | 0,42600097 | 0,000386 | 0,0021509 | 0,0009047 | 0,0009047 | TRUE | 0,0072532 | No |
| orf19.6235 | 893 | 549 | 750 | 912 | 923 | 714 | 816 | 600 | 801 | 699 | 767 | 752 | 789 | 681 | 728 | 846 | 758,06 | 755 | 761 | 1,004 | 0,006 | 0,08 | 0,93585338 | 0,95395438 | 0,004437 | 0,0038945 | 0,0041763 | 0,0041763 | TRUE | 0,0603511 | No |
| orf19.6236 | 7386 | 4894 | 6025 | 7610 | 6354 | 5751 | 5984 | 3488 | 6626 | 6231 | 6164 | 6275 | 5434 | 5487 | 5340 | 4920 | 5809,6 | 6324 | 5295 | 0,838 | -0,255 | -5,697 | 1,22E-08 | 6,68E-08 | 0,001487 | 0,0022294 | 0,001752 | 0,001752 | TRUE | 0,0267174 | No |
| orf19.6237 | 2646 | 1889 | 2317 | 2899 | 2774 | 2491 | 2682 | 1722 | 2374 | 2405 | 2371 | 2391 | 2372 | 2377 | 2393 | 2429 | 2388,9 | 2385 | 2393 | 1,003 | 0,004 | 0,118 | 0,90629007 | 0,92997279 | 1,00E-08 | 0,0025872 | 0,0007533 | 0,0007533 | TRUE | 0,0010826 | No |
| orf19.6238 | 328 | 239 | 329 | 390 | 337 | 330 | 337 | 202 | 294 | 304 | 337 | 322 | 288 | 315 | 301 | 285 | 305,68 | 314 | 297 | 0,947 | -0,079 | -0,929 | 0,35291707 | 0,44044989 | 1,00E-08 | 0,0067286 | 0,003596 | 0,003596 | TRUE | 0,025198 | No |
| orf19.6239 | 1587 | 1009 | 1278 | 1622 | 1614 | 1377 | 1477 | 1020 | 1424 | 1285 | 1308 | 1338 | 1380 | 1314 | 1318 | 1439 | 1350,5 | 1339 | 1363 | 1,015 | 0,022 | 0,414 | 0,67880732 | 0,74326123 | 0,000987 | 0,0030544 | 0,0019538 | 0,0019538 | TRUE | 0,022383 | No |
| orf19.6240 | 679 | 428 | 614 | 666 | 665 | 581 | 600 | 394 | 609 | 545 | 628 | 549 | 569 | 554 | 535 | 556 | 568,2 | 583 | 554 | 0,949 | -0,076 | -1,089 | 0,27619466 | 0,36025364 | 0,001242 | 0,0045344 | 0,0028718 | 0,0028718 | TRUE | 0,0309438 | No |
| orf19.6242 | 302 | 215 | 285 | 324 | 270 | 242 | 275 | 153 | 271 | 274 | 292 | 267 | 231 | 231 | 245 | 216 | 253,3 | 276 | 231 | 0,841 | -0,251 | -2,805 | 0,00503344 | 0,0108585 | 1,00E-08 | 0,0077107 | 0,0037022 | 0,0037022 | TRUE | 0,0197827 | No |
| orf19.6243 | 579 | 452 | 539 | 654 | 814 | 775 | 871 | 500 | 519 | 575 | 551 | 539 | 696 | 739 | 777 | 705 | 637,96 | 546 | 729 | 1,34 | 0,422 | 6,49 | 8,57E-11 | 6,29E-10 | 0,000815 | 0,0042551 | 0,002457 | 0,002457 | TRUE | 0,0238798 | No |
| orf19.6244 | 1251 | 888 | 1131 | 1385 | 1352 | 1173 | 1324 | 905 | 1122 | 1131 | 1157 | 1142 | 1156 | 1119 | 1181 | 1276 | 1160,7 | 1138 | 1183 | 1,037 | 0,052 | 0,995 | 0,3195998 | 0,40629615 | 0,000547 | 0,0032303 | 0,0017828 | 0,0017828 | TRUE | 0,029652 | No |
| orf19.6245 | 9625 | 6949 | 8642 | 10311 | 14100 | 12871 | 13791 | 8222 | 8635 | 8847 | 8842 | 8503 | 12059 | 12281 | 12306 | 11596 | 10384 | 8707 | 12060 | 1,386 | 0,471 | 14,473 | 1,79E-47 | 1,58E-45 | 0,000426 | 0,0021193 | 0,0009181 | 0,0009181 | TRUE | 0,0079394 | No |
| orf19.6246 | 1322 | 876 | 1043 | 1394 | 1461 | 1196 | 1269 | 759 | 1186 | 1115 | 1067 | 1150 | 1249 | 1141 | 1132 | 1071 | 1138,9 | 1130 | 1148 | 1,018 | 0,026 | 0,43 | 0,66725377 | 0,73428204 | 0,002187 | 0,0032542 | 0,0026539 | 0,0026539 | TRUE | 0,0427585 | No |
| orf19.6247 | 45 | 34 | 42 | 42 | 80 | 63 | 81 | 49 | 40 | 43 | 43 | 35 | 68 | 60 | 72 | 69 | 53,9 | 40 | 67 | 1,685 | 0,753 | 3,972 | 7,14E-05 | 0,00022416 | 1,00E-08 | 0,028914 | 0,0147569 | 0,0147569 | TRUE | 0,080172 | No |
| orf19.6247.1 | 3996 | 2642 | 3287 | 3955 | 4590 | 3806 | 3990 | 1710 | 3585 | 3364 | 3363 | 3261 | 3926 | 3632 | 3560 | 2412 | 3387,7 | 3393 | 3382 | 0,997 | -0,004 | -0,025 | 0,9796641 | 0,98508366 | 0,022454 | 0,002408 | 0,0121967 | 0,0224543 | TRUE | 0,4500555 | No |
| orf19.6248 | 71 | 68 | 61 | 90 | 45 | 66 | 60 | 32 | 64 | 87 | 62 | 74 | 38 | 63 | 54 | 45 | 60,88 | 72 | 50 | 0,703 | -0,509 | -2,529 | 0,01145168 | 0,02271433 | 0,016318 | 0,0258256 | 0,0219217 | 0,0219217 | TRUE | 0,2574129 | No |
| orf19.6249 | 1960 | 1517 | 1819 | 2239 | 2885 | 2568 | 2819 | 1622 | 1758 | 1931 | 1861 | 1846 | 2467 | 2450 | 2515 | 2288 | 2139,7 | 1849 | 2430 | 1,317 | 0,398 | 8,431 | 3,42E-17 | 4,90E-16 | 0,000932 | 0,002658 | 0,0016589 | 0,0016589 | TRUE | 0,0174455 | No |
| orf19.6250 | 5216 | 3118 | 4434 | 5549 | 6215 | 4632 | 5435 | 3645 | 4679 | 3970 | 4536 | 4576 | 5315 | 4420 | 4850 | 5141 | 4685,9 | 4440 | 4932 | 1,11 | 0,15 | 2,205 | 0,02746489 | 0,04964172 | 0,005455 | 0,0022893 | 0,0042575 | 0,0042575 | TRUE | 0,0611207 | No |
| orf19.6252 | 3557 | 2236 | 3013 | 3553 | 3544 | 2996 | 3344 | 2096 | 3191 | 2847 | 3083 | 2930 | 3031 | 2859 | 2984 | 2956 | 2985 | 3013 | 2958 | 0,981 | -0,027 | -0,592 | 0,55413567 | 0,63486787 | 0,001151 | 0,0024659 | 0,0017093 | 0,0017093 | TRUE | 0,0194739 | No |
| orf19.6253 | 17782 | 11268 | 14391 | 17774 | 16846 | 13493 | 14563 | 9070 | 15952 | 14345 | 14723 | 14657 | 14407 | 12875 | 12995 | 12792 | 14093 | 14919 | 13267 | 0,889 | -0,169 | -3,311 | 0,00093062 | 0,00235121 | 0,002691 | 0,0020825 | 0,0024376 | 0,0024376 | TRUE | 0,0411146 | No |
| orf19.6254 | 5491 | 3764 | 4900 | 6106 | 3570 | 3022 | 3291 | 2158 | 4926 | 4792 | 5013 | 5035 | 3053 | 2883 | 2937 | 3044 | 3960,4 | 4942 | 2979 | 0,602 | -0,731 | -19,99 | 6,78E-89 | 2,72E-86 | 0,000277 | 0,0023461 | 0,0010144 | 0,0010144 | TRUE | 0,0056819 | No |
| orf19.6255 | 1091 | 755 | 944 | 1273 | 1307 | 1219 | 1289 | 751 | 979 | 961 | 966 | 1050 | 1118 | 1163 | 1150 | 1059 | 1055,7 | 989 | 1122 | 1,136 | 0,184 | 3,365 | 0,0007655 | 0,00197303 | 0,00071 | 0,0033546 | 0,0019061 | 0,0019061 | TRUE | 0,0216532 | No |
| orf19.6257 | 4339 | 2976 | 4063 | 4728 | 4608 | 4220 | 4502 | 2739 | 3893 | 3789 | 4157 | 3899 | 3941 | 4027 | 4017 | 3863 | 3948,1 | 3934 | 3962 | 1,007 | 0,01 | 0,265 | 0,79094053 | 0,83690924 | 0,000592 | 0,0023472 | 0,0012435 | 0,0012435 | TRUE | 0,0174012 | No |
| orf19.6259 | 4640 | 3394 | 3987 | 5196 | 4470 | 4158 | 4464 | 2610 | 4163 | 4321 | 4079 | 4285 | 3823 | 3967 | 3983 | 3681 | 4037,8 | 4212 | 3864 | 0,918 | -0,123 | -3,087 | 0,00202042 | 0,00476588 | 0,00069 | 0,0023391 | 0,0012728 | 0,0012728 | TRUE | 0,0114758 | No |
| orf19.6260 | 3287 | 2284 | 2968 | 3631 | 3616 | 3299 | 3566 | 2277 | 2949 | 2908 | 3037 | 2994 | 3093 | 3148 | 3182 | 3212 | 3065,2 | 2972 | 3159 | 1,062 | 0,086 | 2,494 | 0,01263175 | 0,02476879 | 1,00E-08 | 0,0024531 | 0,0008258 | 0,0008258 | TRUE | 0,0024846 | No |
| orf19.6261 | 1416 | 1177 | 1403 | 1624 | 2125 | 2013 | 2199 | 1245 | 1270 | 1498 | 1435 | 1339 | 1817 | 1921 | 1962 | 1756 | 1625 | 1386 | 1864 | 1,349 | 0,432 | 6,968 | 3,22E-12 | 2,77E-11 | 0,003104 | 0,0028729 | 0,003053 | 0,003053 | TRUE | 0,0377625 | No |
| orf19.6263 | 4385 | 2959 | 3835 | 4829 | 4770 | 3944 | 4101 | 2890 | 3934 | 3767 | 3924 | 3982 | 4079 | 3763 | 3659 | 4076 | 3898,1 | 3902 | 3894 | 0,997 | -0,004 | -0,087 | 0,9307665 | 0,94989725 | 0,001566 | 0,0023519 | 0,0018485 | 0,0018485 | TRUE | 0,0202935 | No |
| orf19.6264 | 477 | 301 | 407 | 462 | 425 | 362 | 401 | 278 | 428 | 383 | 416 | 381 | 363 | 345 | 358 | 392 | 383,41 | 402 | 364 | 0,903 | -0,147 | -1,875 | 0,06085338 | 0,09972984 | 1,00E-08 | 0,0057658 | 0,0033076 | 0,0033076 | TRUE | 0,0241471 | No |
| orf19.6264.3 | 5809 | 4667 | 5080 | 6620 | 5554 | 5495 | 5505 | 2746 | 5211 | 5942 | 5197 | 5459 | 4750 | 5243 | 4912 | 3873 | 5073,4 | 5452 | 4694 | 0,862 | -0,215 | -2,591 | 0,00957607 | 0,01939098 | 0,00953 | 0,0022657 | 0,0063927 | 0,0063927 | TRUE | 0,166527 | No |
| orf19.6264.4 | 6697 | 4247 | 5392 | 6707 | 7119 | 5447 | 5871 | 3676 | 6008 | 5407 | 5517 | 5531 | 6088 | 5197 | 5239 | 5185 | 5521,4 | 5616 | 5427 | 0,967 | -0,049 | -0,805 | 0,42100207 | 0,50885846 | 0,003891 | 0,0022424 | 0,0033622 | 0,0033622 | TRUE | 0,0827266 | No |
| orf19.6265 | 8406 | 5386 | 6297 | 8369 | 8804 | 6569 | 6532 | 4459 | 7541 | 6857 | 6442 | 6901 | 7529 | 6268 | 5829 | 6289 | 6707,1 | 6935 | 6479 | 0,934 | -0,098 | -1,281 | 0,20025036 | 0,27475764 | 0,008236 | 0,002196 | 0,0054908 | 0,0054908 | TRUE | 0,1464411 | No |
| orf19.6265.1 | 15490 | 9940 | 12410 | 15827 | 16128 | 12924 | 13608 | 8399 | 13896 | 12655 | 12697 | 13051 | 13793 | 12332 | 12143 | 11846 | 12802 | 13075 | 12528 | 0,958 | -0,061 | -1,126 | 0,26024758 | 0,34364121 | 0,003257 | 0,0020929 | 0,0027776 | 0,0027776 | TRUE | 0,0566888 | No |
| orf19.6266 | 2624 | 2216 | 2525 | 2918 | 2490 | 2635 | 2700 | 1599 | 2354 | 2821 | 2583 | 2406 | 2130 | 2514 | 2409 | 2255 | 2434,1 | 2541 | 2327 | 0,917 | -0,125 | -1,786 | 0,07405142 | 0,11823634 | 0,005541 | 0,0025759 | 0,0043129 | 0,0043129 | TRUE | 0,0654682 | No |

| | | | | | | | | | | | | | | | | | | | | | | | | | | | | | | | |
|---|---|---|---|---|---|---|---|---|---|---|---|---|---|---|---|---|---|---|---|---|---|---|---|---|---|---|---|---|---|---|---|
| orf19.6267 | 437 | 366 | 421 | 519 | 537 | 500 | 534 | 309 | 392 | 466 | 431 | 428 | 459 | 477 | 477 | 436 | 445,67 | 429 | 462 | 1,083 | 0,115 | 1,536 | 0,12456851 | 0,18415541 | 0,000644 | 0,0052369 | 0,0031501 | 0,0031501 | TRUE | 0,0395096 | No |
| orf19.6268 | 1470 | 1044 | 1356 | 1640 | 1758 | 1647 | 1757 | 1080 | 1319 | 1329 | 1387 | 1352 | 1503 | 1572 | 1568 | 1523 | 1444,2 | 1347 | 1542 | 1,145 | 0,195 | 4,423 | 9,72E-06 | 3,54E-05 | 1,00E-08 | 0,0029847 | 0,0011705 | 0,0011705 | TRUE | 0,0047861 | No |
| orf19.6269 | 502 | 432 | 465 | 559 | 686 | 766 | 804 | 465 | 450 | 550 | 476 | 461 | 587 | 731 | 717 | 656 | 578,49 | 484 | 673 | 1,394 | 0,479 | 5,323 | 1,02E-07 | 5,00E-07 | 0,007382 | 0,004489 | 0,0059913 | 0,0059913 | TRUE | 0,0931707 | No |
| orf19.6271 | 3395 | 2497 | 2968 | 3834 | 3313 | 3188 | 3529 | 2311 | 3046 | 3179 | 3037 | 3162 | 2833 | 3042 | 3149 | 3259 | 3088,3 | 3106 | 3071 | 0,988 | -0,018 | -0,36 | 0,71906147 | 0,77924266 | 0,001719 | 0,0024496 | 0,0019934 | 0,0019934 | TRUE | 0,0334008 | No |
| orf19.6272 | 880 | 1027 | 899 | 1338 | 879 | 1258 | 1200 | 705 | 789 | 1307 | 920 | 1103 | 752 | 1200 | 1071 | 994 | 1017,2 | 1030 | 1004 | 0,975 | -0,036 | -0,17 | 0,86468375 | 0,89724006 | 0,041681 | 0,0034067 | 0,0212077 | 0,0416814 | TRUE | 0,3073717 | No |
| orf19.6273 | 681 | 469 | 571 | 703 | 696 | 532 | 570 | 361 | 611 | 597 | 584 | 580 | 595 | 508 | 509 | 509 | 561,57 | 593 | 530 | 0,896 | -0,158 | -2,185 | 0,02886628 | 0,05180065 | 0,00196 | 0,0045646 | 0,0032158 | 0,0032158 | TRUE | 0,0850224 | No |
| orf19.6274 | 290 | 198 | 232 | 320 | 201 | 164 | 156 | 131 | 260 | 252 | 237 | 264 | 172 | 156 | 139 | 185 | 208,23 | 253 | 163 | 0,638 | -0,648 | -6,031 | 1,63E-09 | 1,00E-08 | 0,001636 | 0,0089512 | 0,0060183 | 0,0060183 | TRUE | 0,1063821 | No |
| orf19.6275 | 1023 | 815 | 893 | 1167 | 1265 | 1314 | 1339 | 721 | 918 | 1038 | 914 | 962 | 1082 | 1254 | 1195 | 1017 | 1047,3 | 958 | 1137 | 1,191 | 0,252 | 3,396 | 0,0006835 | 0,00178076 | 0,004965 | 0,0033656 | 0,0043036 | 0,0043036 | TRUE | 0,0563465 | No |
| orf19.6276 | 1205 | 959 | 1097 | 1358 | 1667 | 1613 | 1804 | 1009 | 1081 | 1221 | 1122 | 1120 | 1426 | 1539 | 1610 | 1423 | 1317,7 | 1136 | 1500 | 1,323 | 0,404 | 6,732 | 1,68E-11 | 1,34E-10 | 0,002202 | 0,0030812 | 0,002687 | 0,002687 | TRUE | 0,0299952 | No |
| orf19.6277 | 31 | 12 | 20 | 26 | 50 | 46 | 31 | 26 | 28 | 15 | 20 | 21 | 43 | 44 | 28 | 37 | 29,5 | 21 | 38 | 1,753 | 0,81 | 2,937 | 0,00330907 | 0,00743768 | 1,00E-08 | 0,0511948 | 0,0367264 | 0,0367264 | TRUE | 0,2725116 | No |
| orf19.6280 | 5 | 3 | 3 | 1 | 5 | 4 | 3 | 2 | 4 | 4 | 3 | 1 | 4 | 4 | 3 | 3 | 3,22 | 3 | 4 | 1,151 | 0,203 | 0,265 | 0,79085688 | NA | 1,00E-08 | 0,4523043 | 0,2486258 | 0,2486258 | TRUE | 0,4363902 | No |
| orf19.6282 | 518 | 348 | 456 | 587 | 798 | 773 | 761 | 529 | 465 | 443 | 467 | 484 | 682 | 738 | 679 | 746 | 587,94 | 465 | 711 | 1,524 | 0,608 | 9,222 | 2,92E-20 | 5,42E-19 | 1,00E-08 | 0,0044486 | 0,0023952 | 0,0023952 | TRUE | 0,0112806 | No |
| orf19.6283 | 2913 | 2175 | 2420 | 3142 | 3410 | 3145 | 3277 | 2005 | 2613 | 2769 | 2476 | 2591 | 2916 | 3001 | 2924 | 2828 | 2764,8 | 2612 | 2917 | 1,118 | 0,161 | 3,677 | 0,00023585 | 0,00067669 | 0,000876 | 0,0025046 | 0,0014853 | 0,0014853 | TRUE | 0,0185769 | No |
| orf19.6284 | 3406 | 2586 | 2965 | 3877 | 3713 | 3568 | 3822 | 2065 | 3056 | 3292 | 3033 | 3197 | 3175 | 3404 | 3410 | 2912 | 3185,2 | 3144 | 3225 | 1,027 | 0,039 | 0,68 | 0,49644455 | 0,5815474 | 0,002845 | 0,0024353 | 0,0028093 | 0,0028093 | TRUE | 0,0497131 | No |
| orf19.6285 | 6044 | 4271 | 5360 | 6517 | 6181 | 5510 | 6224 | 3629 | 5422 | 5437 | 5484 | 5374 | 5286 | 5257 | 5554 | 5118 | 5366,7 | 5429 | 5304 | 0,978 | -0,033 | -0,931 | 0,35175856 | 0,43946047 | 0,000375 | 0,00225 | 0,0009839 | 0,0009839 | TRUE | 0,012315 | No |
| orf19.6286 | 551 | 365 | 500 | 586 | 474 | 369 | 431 | 242 | 494 | 465 | 512 | 483 | 405 | 352 | 385 | 341 | 429,64 | 488 | 371 | 0,762 | -0,392 | -5,059 | 4,21E-07 | 1,89E-06 | 0,001041 | 0,0053584 | 0,0033925 | 0,0033925 | TRUE | 0,0456285 | No |
| orf19.6286.2 | 9584 | 6725 | 7838 | 10754 | 9817 | 8388 | 8621 | 7149 | 8598 | 8562 | 8019 | 8868 | 8396 | 8004 | 7693 | 10083 | 8527,7 | 8512 | 8544 | 1,003 | 0,005 | 0,062 | 0,95074876 | 0,96471855 | 0,007324 | 0,0021498 | 0,0054612 | 0,0054612 | TRUE | 0,178288 | No |
| orf19.6287 | 5289 | 3595 | 4595 | 5860 | 4746 | 4065 | 4354 | 3002 | 4745 | 4577 | 4701 | 4832 | 4059 | 3879 | 3885 | 4234 | 4364 | 4714 | 4014 | 0,85 | -0,234 | -5,794 | 6,88E-09 | 3,90E-08 | 0,000829 | 0,0023122 | 0,0013313 | 0,0013313 | TRUE | 0,0155447 | No |
| orf19.6288 | 148 | 92 | 106 | 150 | 158 | 127 | 130 | 92 | 133 | 117 | 108 | 124 | 135 | 121 | 116 | 130 | 123,01 | 120 | 126 | 1,037 | 0,052 | 0,408 | 0,68301696 | 0,74664591 | 1,00E-08 | 0,0137807 | 0,0076207 | 0,0076207 | TRUE | 0,0491394 | No |
| orf19.6291 | 4289 | 3136 | 4008 | 4855 | 3531 | 3142 | 3291 | 2054 | 3848 | 3992 | 4101 | 4003 | 3020 | 2998 | 2937 | 2897 | 3474,5 | 3986 | 2963 | 0,744 | -0,427 | -11,741 | 7,88E-32 | 3,18E-30 | 0,000272 | 0,0023973 | 0,0009725 | 0,0009725 | TRUE | 0,0068195 | No |
| orf19.6293 | 664 | 484 | 584 | 712 | 832 | 690 | 785 | 445 | 596 | 616 | 597 | 587 | 712 | 658 | 700 | 628 | 636,81 | 599 | 674 | 1,131 | 0,178 | 2,82 | 0,00480519 | 0,01040346 | 1,00E-08 | 0,0042592 | 0,00225 | 0,00225 | TRUE | 0,0221321 | No |
| orf19.6294 | 296 | 228 | 283 | 308 | 310 | 387 | 370 | 229 | 266 | 290 | 290 | 254 | 265 | 369 | 330 | 323 | 298,36 | 275 | 322 | 1,173 | 0,23 | 2,122 | 0,03384508 | 0,05970089 | 0,008911 | 0,0068452 | 0,0078843 | 0,0078843 | TRUE | 0,1729579 | No |
| orf19.6295 | 2654 | 1934 | 2413 | 3001 | 3489 | 3209 | 3505 | 2096 | 2381 | 2462 | 2469 | 2475 | 2984 | 3062 | 3128 | 2956 | 2739,5 | 2447 | 3032 | 1,241 | 0,311 | 8,272 | 1,32E-16 | 1,76E-15 | 1,00E-08 | 0,0025094 | 0,0009865 | 0,0009865 | TRUE | 0,0054771 | No |
| orf19.6296 | 254 | 212 | 245 | 255 | 325 | 355 | 342 | 189 | 228 | 270 | 251 | 210 | 278 | 339 | 305 | 267 | 268,39 | 240 | 297 | 1,25 | 0,322 | 2,966 | 0,00301771 | 0,00684941 | 0,007691 | 0,0073884 | 0,0075305 | 0,0075305 | TRUE | 0,0993175 | No |
| orf19.6297 | 3076 | 2330 | 2676 | 3286 | 3003 | 2718 | 2874 | 1740 | 2759 | 2966 | 2738 | 2710 | 2568 | 2593 | 2565 | 2454 | 2669,2 | 2793 | 2545 | 0,913 | -0,131 | -3,07 | 0,00213737 | 0,00502398 | 0,000675 | 0,0025234 | 0,001378 | 0,001378 | TRUE | 0,0200136 | No |
| orf19.6298 | 2855 | 2208 | 2585 | 3088 | 3120 | 3013 | 3036 | 1780 | 2561 | 2811 | 2645 | 2546 | 2668 | 2875 | 2709 | 2511 | 2665,8 | 2641 | 2691 | 1,021 | 0,03 | 0,568 | 0,57017574 | 0,64928276 | 0,00196 | 0,0025241 | 0,0022715 | 0,0022715 | TRUE | 0,0253951 | No |
| orf19.6302 | 23 | 17 | 22 | 28 | 31 | 23 | 11 | 7 | 21 | 22 | 23 | 23 | 27 | 22 | 10 | 10 | 19,5 | 22 | 17 | 0,792 | -0,336 | -0,941 | 0,34681094 | 0,43400127 | 0,062801 | 0,0764161 | 0,0710659 | 0,0710659 | TRUE | 0,3847421 | No |
| orf19.6304 | 66 | 98 | 73 | 100 | 76 | 114 | 99 | 54 | 59 | 125 | 75 | 82 | 65 | 109 | 88 | 76 | 84,92 | 85 | 84 | 1,002 | 0,003 | 0,012 | 0,99053273 | 0,99268067 | 0,061395 | 0,0190735 | 0,0388027 | 0,0388027 | TRUE | 0,8265122 | No |
| orf19.6305 | 1081 | 958 | 1029 | 1409 | 988 | 1124 | 1084 | 626 | 970 | 1220 | 1053 | 1162 | 845 | 1072 | 967 | 883 | 1021,5 | 1101 | 942 | 0,857 | -0,223 | -2,471 | 0,01346294 | 0,02621892 | 0,009562 | 0,0034007 | 0,0068267 | 0,0068267 | TRUE | 0,1046317 | No |
| orf19.6306 | 1464 | 1093 | 1326 | 1697 | 986 | 914 | 976 | 524 | 1313 | 1392 | 1357 | 1399 | 843 | 872 | 871 | 739 | 1098,3 | 1365 | 831 | 0,612 | -0,709 | -11,925 | 8,77E-33 | 3,74E-31 | 0,001544 | 0,0033013 | 0,0024275 | 0,0024275 | TRUE | 0,0574625 | No |
| orf19.6307 | 557 | 528 | 543 | 628 | 636 | 808 | 797 | 505 | 500 | 672 | 556 | 518 | 544 | 771 | 711 | 712 | 622,95 | 562 | 684 | 1,222 | 0,289 | 2,476 | 0,01330446 | 0,02594391 | 0,017582 | 0,0043099 | 0,011427 | 0,011427 | TRUE | 0,2329628 | No |
| orf19.6308 | 214 | 227 | 215 | 332 | 300 | 449 | 453 | 236 | 192 | 289 | 220 | 274 | 257 | 428 | 404 | 333 | 299,6 | 244 | 356 | 1,462 | 0,548 | 3,305 | 0,00094808 | 0,00239331 | 0,040291 | 0,006825 | 0,0229003 | 0,0229003 | TRUE | 0,3597475 | No |
| orf19.6309 | 1798 | 1270 | 1778 | 2057 | 1949 | 1668 | 1987 | 1227 | 1613 | 1617 | 1819 | 1696 | 1667 | 1592 | 1773 | 1731 | 1688,4 | 1686 | 1691 | 1,002 | 0,003 | 0,047 | 0,96254491 | 0,97258153 | 0,002118 | 0,0028393 | 0,0024144 | 0,0024144 | TRUE | 0,0334217 | No |
| orf19.6310 | 717 | 597 | 697 | 863 | 720 | 863 | 857 | 627 | 643 | 760 | 713 | 712 | 616 | 823 | 765 | 884 | 739,54 | 707 | 772 | 1,09 | 0,124 | 1,226 | 0,22018117 | 0,29894881 | 0,012814 | 0,0039425 | 0,0085233 | 0,0085233 | TRUE | 0,2248746 | No |
| orf19.6311 | 24109 | 17439 | 19825 | 25871 | 25777 | 23140 | 23162 | 14463 | 21628 | 22202 | 20283 | 21334 | 22045 | 22079 | 20668 | 20399 | 21330 | 21362 | 21298 | 0,997 | -0,004 | -0,094 | 0,92480324 | 0,94525653 | 0,00131 | 0,0020476 | 0,0016994 | 0,0016994 | TRUE | 0,014107 | No |
| orf19.6312 | 13194 | 8566 | 11199 | 13661 | 13314 | 11277 | 11986 | 8021 | 11836 | 10905 | 11458 | 11265 | 11387 | 10760 | 10696 | 11313 | 11202 | 11366 | 11039 | 0,971 | -0,043 | -1,099 | 0,27168068 | 0,35615481 | 0,001 | 0,0021091 | 0,001358 | 0,001358 | TRUE | 0,0094957 | No |
| orf19.6313 | 246 | 200 | 242 | 266 | 216 | 277 | 265 | 162 | 221 | 255 | 248 | 219 | 185 | 264 | 236 | 228 | 232,03 | 236 | 228 | 0,972 | -0,041 | -0,349 | 0,72716782 | 0,78604251 | 0,01011 | 0,0082361 | 0,0091415 | 0,0091415 | TRUE | 0,2010937 | No |
| orf19.6313.2 | 1965 | 1427 | 1653 | 2105 | 1899 | 1778 | 1924 | 973 | 1763 | 1817 | 1691 | 1736 | 1624 | 1697 | 1717 | 1372 | 1677 | 1752 | 1602 | 0,917 | -0,125 | -1,834 | 0,06664859 | 0,10825216 | 0,004544 | 0,0028452 | 0,0038248 | 0,0038248 | TRUE | 0,1057194 | No |
| orf19.6314 | 3194 | 2345 | 2694 | 3486 | 2833 | 2492 | 2519 | 1388 | 2865 | 2985 | 2756 | 2875 | 2423 | 2378 | 2248 | 1958 | 2561 | 2870 | 2252 | 0,786 | -0,347 | -5,397 | 6,76E-08 | 3,39E-07 | 0,004333 | 0,0025464 | 0,0035808 | 0,0035808 | TRUE | 0,0890774 | No |
| orf19.6315 | 187 | 118 | 133 | 187 | 91 | 74 | 75 | 54 | 168 | 150 | 136 | 154 | 78 | 71 | 67 | 76 | 112,47 | 152 | 73 | 0,477 | -1,067 | -7,691 | 1,46E-14 | 1,59E-13 | 1,00E-08 | 0,0148867 | 0,0083666 | 0,0083666 | TRUE | 0,0532984 | No |
| orf19.6316 | 383 | 298 | 333 | 401 | 381 | 415 | 422 | 228 | 344 | 379 | 341 | 331 | 326 | 396 | 377 | 322 | 351,79 | 349 | 355 | 1,024 | 0,035 | 0,38 | 0,70412029 | 0,76567505 | 0,004165 | 0,0061062 | 0,0051796 | 0,0051796 | TRUE | 0,0678138 | No |
| orf19.6316.4 | 3622 | 3256 | 3339 | 4743 | 3116 | 3229 | 3384 | 1781 | 3249 | 4145 | 3416 | 3911 | 2665 | 3081 | 3020 | 2512 | 3249,9 | 3680 | 2820 | 0,767 | -0,383 | -4,401 | 1,08E-05 | 3,90E-05 | 0,0108 | 0,0024262 | 0,0069688 | 0,0069688 | TRUE | 0,0868654 | No |
| orf19.6317 | 4266 | 3396 | 4077 | 4797 | 3590 | 3502 | 3677 | 2263 | 3827 | 4323 | 4171 | 3956 | 3070 | 3341 | 3281 | 3192 | 3645,3 | 4069 | 3221 | 0,792 | -0,336 | -6,787 | 1,15E-11 | 9,33E-11 | 0,001774 | 0,0023778 | 0,0020751 | 0,0020751 | TRUE | 0,0210404 | No |
| orf19.6318 | 1141 | 689 | 812 | 1120 | 1259 | 1075 | 983 | 780 | 1024 | 877 | 831 | 924 | 1077 | 1026 | 877 | 1100 | 966,86 | 914 | 1020 | 1,114 | 0,155 | 1,817 | 0,06914973 | 0,11153062 | 0,007505 | 0,003481 | 0,0059547 | 0,0059547 | TRUE | 0,1080727 | No |
| orf19.6319 | 6174 | 4005 | 5593 | 6985 | 7939 | 6679 | 7659 | 4805 | 5539 | 5099 | 5722 | 5760 | 6790 | 6373 | 6834 | 6777 | 6111,7 | 5530 | 6694 | 1,21 | 0,275 | 5,806 | 6,39E-09 | 3,64E-08 | 0,0018 | 0,0022171 | 0,0019822 | 0,0019822 | TRUE | 0,0327922 | No |
| orf19.6321 | 2693 | 1556 | 2223 | 2847 | 4179 | 3036 | 3266 | 2859 | 2416 | 1981 | 2274 | 2348 | 3574 | 2897 | 2914 | 4032 | 2804,6 | 2255 | 3354 | 1,486 | 0,571 | 5,581 | 2,39E-08 | 1,26E-07 | 0,016602 | 0,0024971 | 0,0096955 | 0,0096955 | TRUE | 0,2221933 | No |
| orf19.6322 | 31233 | 19043 | 26884 | 32855 | 37147 | 27718 | 31701 | 20491 | 28019 | 24244 | 27505 | 27093 | 31769 | 26448 | 28288 | 28901 | 27783 | 26715 | 28852 | 1,08 | 0,111 | 1,762 | 0,07804976 | 0,12382439 | 0,00488 | 0,0020318 | 0,0037691 | 0,0037691 | TRUE | 0,0569087 | No |
| orf19.6323 | 438 | 259 | 397 | 487 | 425 | 342 | 376 | 256 | 393 | 330 | 406 | 402 | 363 | 326 | 336 | 361 | 364,6 | 383 | 346 | 0,901 | -0,151 | -1,739 | 0,08206205 | 0,12930206 | 0,002632 | 0,0059611 | 0,0044473 | 0,0044473 | TRUE | 0,0878751 | No |
| orf19.6324 | 9826 | 6258 | 8680 | 10439 | 10438 | 8100 | 9353 | 5383 | 8815 | 7967 | 8880 | 8608 | 8927 | 7729 | 8346 | 7592 | 8358,1 | 8568 | 8148 | 0,951 | -0,072 | -1,226 | 0,22024259 | 0,29896464 | 0,003862 | 0,0021532 | 0,0032019 | 0,0032019 | TRUE | 0,0506537 | No |
| orf19.6325.1 | 196 | 100 | 118 | 175 | 153 | 104 | 130 | 113 | 176 | 127 | 121 | 144 | 131 | 99 | 116 | 159 | 134,2 | 142 | 126 | 0,879 | -0,186 | -1,127 | 0,25992072 | 0,34328505 | 0,025439 | 0,0127966 | 0,0187291 | 0,0187291 | TRUE | 0,2729893 | No |
| orf19.6326 | 915 | 550 | 678 | 825 | 880 | 652 | 759 | 473 | 821 | 700 | 694 | 680 | 753 | 622 | 677 | 667 | 701,77 | 724 | 680 | 0,94 | -0,09 | -1,095 | 0,27346721 | 0,3577163 | 0,005877 | 0,0040482 | 0,0050354 | 0,0050354 | TRUE | 0,0994893 | No |
| orf19.6327 | 844 | 552 | 782 | 922 | 932 | 720 | 820 | 577 | 757 | 703 | 800 | 760 | 797 | 687 | 732 | 814 | 756,23 | 755 | 758 | 1 | 0 | -0,002 | 0,99824974 | 0,99865955 | 0,002922 | 0,0038992 | 0,0033898 | 0,0033898 | TRUE | 0,0460793 | No |
| orf19.6328 | 185 | 114 | 159 | 240 | 207 | 143 | 172 | 121 | 166 | 145 | 163 | 198 | 177 | 136 | 153 | 171 | 163,66 | 168 | 159 | 0,942 | -0,087 | -0,665 | 0,50631017 | 0,59054776 | 0,009356 | 0,0108496 | 0,0101611 | 0,0101611 | TRUE | 0,1623355 | No |
| orf19.6329 | 450 | 306 | 430 | 495 | 445 | 368 | 439 | 270 | 404 | 390 | 440 | 408 | 381 | 351 | 392 | 381 | 393,21 | 410 | 376 | 0,916 | -0,127 | -1,673 | 0,09425599 | 0,14572568 | 1,00E-08 | 0,0056715 | 0,0030123 | 0,0030123 | TRUE | 0,0258787 | No |
| orf19.633 | 1395 | 1258 | 1399 | 1784 | 1405 | 1574 | 1661 | 922 | 1251 | 1602 | 1431 | 1471 | 1202 | 1502 | 1482 | 1300 | 1405,2 | 1439 | 1372 | 0,954 | -0,067 | -0,759 | 0,44795431 | 0,53452026 | 0,010162 | 0,0030126 | 0,0068399 | 0,0068399 | TRUE | 0,0931642 | No |
| orf19.6336 | 59 | 38 | 55 | 71 | 73 | 46 | 52 | 52 | 53 | 48 | 56 | 59 | 62 | 44 | 46 | 73 | 55,27 | 54 | 56 | 1,028 | 0,039 | 0,193 | 0,8472539 | 0,88395321 | 0,012795 | 0,0282433 | 0,0220394 | 0,0220394 | TRUE | 0,2594046 | No |
| orf19.6337 | 41 | 16 | 20 | 42 | 25 | 16 | 28 | 19 | 37 | 20 | 20 | 35 | 21 | 15 | 25 | 27 | 25,08 | 28 | 22 | 0,769 | -0,379 | -1,225 | 0,22071203 | 0,29933135 | 0,040269 | 0,0598516 | 0,0516887 | 0,0516887 | TRUE | 0,1417609 | No |
| orf19.6338 | 1 | 5 | 2 | 5 | 1 | 0 | 1 | 4 | 1 | 6 | 2 | 4 | 1 | 0 | 1 | 6 | 2,6 | 3 | 2 | 0,498 | -1,006 | -0,968 | 0,33285908 | NA | 0,608079 | 0,5597601 | 0,5777083 | 0,5777083 | TRUE | 0,7687912 | No |
| orf19.6339 | 0 | 0 | 0 | 0 | 0 | 0 | 0 | 0 | 0 | 0 | 0 | 0 | 0 | 0 | 0 | 0 | 0 | 0 | 0 | NA | NA | NA | NA | NA | NA | NA | NA | NA | NA | NA | NA |

| | | | | | | | | | | | | | | | | | | | | | | | | | | | | | | | |
|---|---|---|---|---|---|---|---|---|---|---|---|---|---|---|---|---|---|---|---|---|---|---|---|---|---|---|---|---|---|---|---|
| orf19.634 | 3 | 4 | 2 | 4 | 2 | 3 | 4 | 2 | 3 | 5 | 2 | 3 | 2 | 3 | 4 | 3 | 3,01 | 3 | 3 | 0,848 | -0,238 | -0,308 | 0,75782776 | NA | 1,00E-08 | 0,4840418 | 0,2363754 | 0,2363754 | TRUE | 0,136113 | No |
| orf19.6340 | 4990 | 4627 | 4580 | 5800 | 5127 | 6115 | 5563 | 3746 | 4477 | 5891 | 4686 | 4783 | 4385 | 5835 | 4964 | 5283 | 5037,8 | 4959 | 5117 | 1,032 | 0,045 | 0,474 | 0,63519893 | 0,70606387 | 0,013075 | 0,0022677 | 0,0086333 | 0,0086333 | TRUE | 0,1938133 | No |
| orf19.6341 | 2314 | 1784 | 2060 | 2665 | 2169 | 2020 | 2251 | 1372 | 2076 | 2271 | 2108 | 2198 | 1855 | 1927 | 2009 | 1935 | 2047,3 | 2163 | 1932 | 0,894 | -0,162 | -3,488 | 0,00048634 | 0,00131796 | 0,000899 | 0,0026886 | 0,0015897 | 0,0015897 | TRUE | 0,0128017 | No |
| orf19.6342 | 1169 | 879 | 969 | 1173 | 1003 | 879 | 932 | 428 | 1049 | 1119 | 991 | 967 | 858 | 839 | 832 | 604 | 907,28 | 1032 | 783 | 0,762 | -0,391 | -3,955 | 7,65E-05 | 0,00023829 | 0,012401 | 0,0035796 | 0,0082674 | 0,0082674 | TRUE | 0,2632116 | No |
| orf19.6343 | 8313 | 8001 | 8466 | 10930 | 10020 | 11251 | 11240 | 6409 | 7458 | 10186 | 8662 | 9013 | 8569 | 10735 | 10030 | 9039 | 9211,5 | 8830 | 9593 | 1,087 | 0,12 | 1,31 | 0,19009362 | 0,26319156 | 0,013024 | 0,0021371 | 0,007958 | 0,007958 | TRUE | 0,1338504 | No |
| orf19.6344 | 2787 | 2054 | 2483 | 3034 | 2584 | 2469 | 2593 | 1532 | 2500 | 2615 | 2540 | 2502 | 2210 | 2356 | 2314 | 2161 | 2399,7 | 2539 | 2260 | 0,892 | -0,165 | -3,909 | 9,27E-05 | 0,00028571 | 0,000464 | 0,0025845 | 0,0013006 | 0,0013006 | TRUE | 0,0095831 | No |
| orf19.6345 | 5986 | 5002 | 5750 | 7437 | 6169 | 6490 | 6728 | 4221 | 5370 | 6368 | 5883 | 6133 | 5276 | 6193 | 6004 | 5953 | 5897,4 | 5938 | 5856 | 0,986 | -0,02 | -0,309 | 0,75700886 | 0,81004889 | 0,004926 | 0,0022257 | 0,0038256 | 0,0038256 | TRUE | 0,0548746 | No |
| orf19.6346 | 2432 | 1984 | 2157 | 2657 | 2357 | 2545 | 2478 | 1708 | 2182 | 2526 | 2207 | 2191 | 2016 | 2428 | 2211 | 2409 | 2271,2 | 2276 | 2266 | 0,995 | -0,007 | -0,092 | 0,92660783 | 0,94693993 | 0,005826 | 0,0026187 | 0,0044369 | 0,0044369 | TRUE | 0,0678741 | No |
| orf19.6347 | 2403 | 2194 | 2336 | 3096 | 3110 | 3203 | 3347 | 2064 | 2156 | 2793 | 2390 | 2553 | 2660 | 3056 | 2987 | 2911 | 2688,2 | 2473 | 2904 | 1,175 | 0,232 | 3,05 | 0,00229065 | 0,00534096 | 0,007174 | 0,0025196 | 0,0052013 | 0,0052013 | TRUE | 0,0908281 | No |
| orf19.6348 | 826 | 651 | 747 | 1028 | 657 | 742 | 735 | 491 | 741 | 829 | 764 | 848 | 562 | 708 | 656 | 693 | 725 | 796 | 655 | 0,821 | -0,284 | -3,466 | 0,00052851 | 0,00141753 | 0,005996 | 0,0039819 | 0,005051 | 0,005051 | TRUE | 0,1118543 | No |
| orf19.6349 | 1401 | 1349 | 1384 | 1755 | 1797 | 2101 | 2138 | 1150 | 1257 | 1717 | 1416 | 1447 | 1537 | 2005 | 1908 | 1622 | 1613,6 | 1459 | 1768 | 1,213 | 0,279 | 2,72 | 0,00653557 | 0,01377261 | 0,015763 | 0,0028792 | 0,009462 | 0,009462 | TRUE | 0,1673113 | No |
| orf19.635 | 18 | 3 | 11 | 6 | 10 | 7 | 11 | 13 | 16 | 4 | 11 | 5 | 9 | 7 | 10 | 18 | 9,94 | 9 | 11 | 1,15 | 0,202 | 0,383 | 0,70189278 | NA | 0,188067 | 0,1479682 | 0,1650884 | 0,1650884 | TRUE | 0,5237209 | No |
| orf19.6350 | 189 | 106 | 142 | 209 | 216 | 165 | 184 | 138 | 170 | 135 | 145 | 172 | 185 | 157 | 164 | 195 | 165,39 | 156 | 175 | 1,114 | 0,156 | 1,274 | 0,20271095 | 0,27780278 | 0,005154 | 0,010757 | 0,008345 | 0,008345 | TRUE | 0,0732882 | No |
| orf19.6351 | 2684 | 1930 | 2335 | 2963 | 2970 | 2792 | 3080 | 1684 | 2408 | 2457 | 2389 | 2443 | 2540 | 2664 | 2748 | 2375 | 2503,1 | 2424 | 2582 | 1,067 | 0,093 | 1,878 | 0,06031301 | 0,09892454 | 0,001525 | 0,0025595 | 0,0019629 | 0,0019629 | TRUE | 0,032869 | No |
| orf19.6353 | 824 | 596 | 722 | 938 | 988 | 897 | 945 | 606 | 739 | 759 | 739 | 773 | 845 | 856 | 843 | 855 | 801,12 | 752 | 850 | 1,128 | 0,174 | 3,297 | 0,00097609 | 0,0024609 | 1,00E-08 | 0,0037916 | 0,0014341 | 0,0014341 | TRUE | 0,0047568 | No |
| orf19.6354 | 171 | 152 | 171 | 179 | 168 | 193 | 184 | 123 | 153 | 194 | 175 | 148 | 144 | 184 | 164 | 173 | 166,87 | 168 | 166 | 0,998 | -0,003 | -0,025 | 0,97995771 | 0,98508366 | 0,007536 | 0,010679 | 0,0092499 | 0,0092499 | TRUE | 0,1061849 | No |
| orf19.6355 | 15230 | 10471 | 12929 | 16298 | 16757 | 14504 | 15782 | 9142 | 13663 | 13331 | 13228 | 13440 | 14331 | 13839 | 14083 | 12894 | 13601 | 13416 | 13787 | 1,028 | 0,04 | 1,03 | 0,30318327 | 0,38910461 | 0,001037 | 0,0020863 | 0,0013728 | 0,0013728 | TRUE | 0,0227954 | No |
| orf19.6356 | 1032 | 974 | 1059 | 1341 | 1039 | 1224 | 1247 | 710 | 926 | 1240 | 1083 | 1106 | 889 | 1168 | 1113 | 1001 | 1065,7 | 1089 | 1043 | 0,959 | -0,06 | -0,606 | 0,54429613 | 0,6261563 | 0,013462 | 0,0033417 | 0,0086051 | 0,0086051 | TRUE | 0,1225266 | No |
| orf19.6357 | 390 | 354 | 402 | 488 | 492 | 547 | 590 | 367 | 350 | 451 | 411 | 402 | 421 | 522 | 526 | 518 | 450,13 | 404 | 497 | 1,232 | 0,301 | 3,059 | 0,00222344 | 0,00520391 | 0,008866 | 0,0052046 | 0,0070657 | 0,0070657 | TRUE | 0,129925 | No |
| orf19.6358 | 1017 | 1002 | 951 | 1361 | 1179 | 1550 | 1379 | 912 | 912 | 1276 | 973 | 1122 | 1008 | 1479 | 1231 | 1286 | 1160,9 | 1071 | 1251 | 1,169 | 0,225 | 1,881 | 0,05994354 | 0,09842601 | 0,022353 | 0,00323 | 0,0128756 | 0,0128756 | TRUE | 0,2082037 | No |
| orf19.6359 | 365 | 323 | 330 | 493 | 500 | 464 | 457 | 272 | 327 | 411 | 338 | 407 | 428 | 443 | 408 | 384 | 393,07 | 371 | 416 | 1,125 | 0,17 | 1,807 | 0,07073206 | 0,11371641 | 0,00625 | 0,0056727 | 0,0059518 | 0,0059518 | TRUE | 0,0734286 | No |
| orf19.6360 | 2358 | 1836 | 2031 | 2724 | 3006 | 2830 | 2918 | 1796 | 2115 | 2337 | 2078 | 2246 | 2571 | 2700 | 2604 | 2533 | 2398,1 | 2194 | 2602 | 1,187 | 0,248 | 5,13 | 2,89E-07 | 1,33E-06 | 0,001306 | 0,0025849 | 0,0018154 | 0,0018154 | TRUE | 0,0218843 | No |
| orf19.6362 | 5974 | 4284 | 5352 | 6477 | 6606 | 5978 | 6438 | 4056 | 5359 | 5454 | 5476 | 5341 | 5650 | 5704 | 5745 | 5721 | 5556,1 | 5408 | 5705 | 1,055 | 0,078 | 2,75 | 0,00596452 | 0,01267593 | 1,00E-08 | 0,0022408 | 0,0005832 | 0,0005832 | TRUE | 0,0009176 | No |
| orf19.6365 | 6831 | 4355 | 5893 | 7563 | 6681 | 5043 | 5703 | 3976 | 6128 | 5544 | 6029 | 6237 | 5714 | 4812 | 5089 | 5608 | 5645,1 | 5984 | 5306 | 0,886 | -0,175 | -2,805 | 0,00502665 | 0,01084775 | 0,004314 | 0,0022367 | 0,0035407 | 0,0035407 | TRUE | 0,0479439 | No |
| orf19.6366 | 90 | 58 | 76 | 103 | 84 | 70 | 64 | 38 | 81 | 74 | 78 | 85 | 72 | 67 | 57 | 54 | 70,83 | 80 | 62 | 0,79 | -0,34 | -2,05 | 0,04035185 | 0,06958493 | 1,00E-08 | 0,0224763 | 0,0123008 | 0,0123008 | TRUE | 0,0971281 | No |
| orf19.6367 | 44283 | 27537 | 38328 | 46881 | 43685 | 34842 | 38478 | 27585 | 39726 | 35058 | 39213 | 38659 | 37361 | 33245 | 34335 | 38906 | 37063 | 38164 | 35962 | 0,942 | -0,086 | -1,447 | 0,14784003 | 0,21305418 | 0,004163 | 0,0020187 | 0,0033604 | 0,0033604 | TRUE | 0,0370834 | No |
| orf19.6369 | 10747 | 6822 | 8961 | 11165 | 9898 | 8076 | 8840 | 5080 | 9641 | 8685 | 9168 | 9207 | 8465 | 7706 | 7888 | 7165 | 8490,6 | 9175 | 7806 | 0,851 | -0,233 | -4,259 | 2,06E-05 | 7,13E-05 | 0,003108 | 0,0021505 | 0,002749 | 0,002749 | TRUE | 0,039476 | No |
| orf19.637 | 4380 | 3014 | 3676 | 4612 | 4718 | 3955 | 4203 | 2885 | 3929 | 3837 | 3761 | 3803 | 4035 | 3774 | 3750 | 4069 | 3869,8 | 3832 | 3907 | 1,019 | 0,027 | 0,65 | 0,51544995 | 0,59969468 | 0,000757 | 0,0023547 | 0,0013571 | 0,0013571 | TRUE | 0,0088847 | No |
| orf19.6371 | 219 | 124 | 167 | 210 | 223 | 179 | 197 | 100 | 196 | 158 | 171 | 173 | 191 | 171 | 176 | 141 | 172,09 | 174 | 170 | 0,976 | -0,035 | -0,29 | 0,77176413 | 0,82174247 | 0,00472 | 0,0104153 | 0,0080152 | 0,0080152 | TRUE | 0,110546 | No |
| orf19.6373 | 6178 | 4126 | 5407 | 6778 | 8204 | 6875 | 7796 | 4820 | 5542 | 5253 | 5532 | 5589 | 7016 | 6560 | 6957 | 6798 | 6155,9 | 5479 | 6833 | 1,247 | 0,318 | 8,577 | 9,74E-18 | 1,47E-16 | 0,000615 | 0,0022153 | 0,001155 | 0,001155 | TRUE | 0,0090845 | No |
| orf19.6374 | 645 | 423 | 551 | 606 | 630 | 563 | 623 | 340 | 579 | 539 | 564 | 500 | 539 | 537 | 556 | 480 | 536,51 | 546 | 528 | 0,973 | -0,04 | -0,542 | 0,58787703 | 0,66615715 | 0,002113 | 0,0046854 | 0,0033993 | 0,0033993 | TRUE | 0,0407374 | No |
| orf19.6375 | 10056 | 6404 | 7689 | 10101 | 8818 | 7232 | 7436 | 4467 | 9021 | 8153 | 7867 | 8329 | 7541 | 6901 | 6635 | 6300 | 7593,5 | 8342 | 6844 | 0,821 | -0,285 | -4,61 | 4,02E-06 | 1,55E-05 | 0,004503 | 0,0021707 | 0,0035398 | 0,0035398 | TRUE | 0,057543 | No |
| orf19.6376 | 2196 | 1409 | 1967 | 2278 | 2530 | 2144 | 2409 | 1445 | 1970 | 1794 | 2012 | 1878 | 2164 | 2046 | 2150 | 2038 | 2006,5 | 1914 | 2100 | 1,097 | 0,134 | 2,726 | 0,00641848 | 0,01354487 | 0,001136 | 0,002703 | 0,0018075 | 0,0018075 | TRUE | 0,0199203 | No |
| orf19.6377 | 1264 | 860 | 1104 | 1317 | 1415 | 1177 | 1401 | 800 | 1134 | 1095 | 1130 | 1086 | 1210 | 1123 | 1250 | 1128 | 1144,5 | 1111 | 1178 | 1,062 | 0,087 | 1,621 | 0,10508202 | 0,15898584 | 0,000837 | 0,0032479 | 0,0018817 | 0,0018817 | TRUE | 0,0204705 | No |
| orf19.6378 | 3109 | 1831 | 2444 | 3096 | 2536 | 1996 | 2176 | 1321 | 2789 | 2331 | 2500 | 2553 | 2169 | 1905 | 1942 | 1863 | 2256,5 | 2543 | 1970 | 0,775 | -0,369 | -5,552 | 2,83E-08 | 1,48E-07 | 0,004575 | 0,0026229 | 0,00378 | 0,00378 | TRUE | 0,0569548 | No |
| orf19.6379 | 635 | 397 | 533 | 650 | 668 | 653 | 636 | 451 | 570 | 505 | 545 | 536 | 571 | 623 | 568 | 636 | 569,3 | 539 | 600 | 1,106 | 0,146 | 2,113 | 0,03456146 | 0,06078608 | 0,001056 | 0,0045295 | 0,0028111 | 0,0028111 | TRUE | 0,0182937 | No |
| orf19.638 | 9173 | 6361 | 7759 | 9989 | 14229 | 12308 | 12655 | 8815 | 8229 | 8098 | 7938 | 8237 | 12169 | 11744 | 11292 | 12433 | 10018 | 8126 | 11910 | 1,465 | 0,551 | 14,366 | 8,50E-47 | 7,30E-45 | 0,000891 | 0,0021244 | 0,0013087 | 0,0013087 | TRUE | 0,0149158 | No |
| orf19.6382 | 3079 | 2089 | 2820 | 3259 | 3383 | 2871 | 3196 | 2202 | 2762 | 2660 | 2885 | 2687 | 2893 | 2739 | 2852 | 3106 | 2823,1 | 2748 | 2898 | 1,053 | 0,074 | 1,517 | 0,12922771 | 0,18996822 | 0,001599 | 0,0024938 | 0,0019391 | 0,0019391 | TRUE | 0,0267403 | No |
| orf19.6385 | 20597 | 12435 | 17860 | 21656 | 16005 | 12914 | 14236 | 10114 | 18478 | 15831 | 18273 | 17858 | 13688 | 12322 | 12703 | 14265 | 15427 | 17610 | 13244 | 0,752 | -0,411 | -6,641 | 3,11E-11 | 2,42E-10 | 0,004549 | 0,0020736 | 0,0036205 | 0,0036205 | TRUE | 0,0563195 | No |
| orf19.6387 | 21747 | 13988 | 19188 | 23146 | 32317 | 26284 | 29280 | 19494 | 19509 | 17808 | 19631 | 19086 | 27639 | 25079 | 26128 | 27495 | 22797 | 19008 | 26585 | 1,398 | 0,484 | 10,476 | 1,11E-25 | 3,00E-24 | 0,001942 | 0,0020432 | 0,0020026 | 0,0020026 | TRUE | 0,021956 | No |
| orf19.639 | 133 | 135 | 156 | 203 | 164 | 140 | 144 | 108 | 119 | 172 | 160 | 167 | 140 | 134 | 128 | 152 | 146,61 | 154 | 138 | 0,896 | -0,158 | -1,16 | 0,2461914 | 0,32818174 | 0,00965 | 0,0118816 | 0,0108689 | 0,0108689 | TRUE | 0,2618999 | No |
| orf19.6390 | 4 | 2 | 3 | 2 | 1 | 1 | 4 | 3 | 4 | 3 | 3 | 2 | 1 | 1 | 4 | 4 | 2,56 | 3 | 2 | 0,847 | -0,239 | -0,273 | 0,7845959 | NA | 1,00E-08 | 0,5695181 | 0,3318327 | 0,3318327 | TRUE | 0,2000481 | No |
| orf19.6391 | 173 | 119 | 151 | 184 | 161 | 143 | 158 | 135 | 155 | 151 | 154 | 152 | 138 | 136 | 141 | 190 | 152,31 | 153 | 151 | 0,975 | -0,036 | -0,286 | 0,77482046 | 0,82367931 | 0,004649 | 0,011511 | 0,0087372 | 0,0087372 | TRUE | 0,2609213 | No |
| orf19.639.1 | 664 | 398 | 466 | 658 | 690 | 532 | 641 | 412 | 596 | 507 | 477 | 543 | 590 | 508 | 572 | 581 | 546,57 | 531 | 563 | 1,058 | 0,081 | 0,971 | 0,33133294 | 0,41811761 | 0,005157 | 0,0046356 | 0,0049048 | 0,0049048 | TRUE | 0,0812664 | No |
| orf19.6392 | 481 | 318 | 427 | 512 | 452 | 375 | 459 | 270 | 432 | 405 | 437 | 422 | 387 | 358 | 410 | 381 | 403,77 | 424 | 384 | 0,905 | -0,144 | -1,932 | 0,05334959 | 0,0889851 | 1,00E-08 | 0,0055749 | 0,002862 | 0,002862 | TRUE | 0,0255095 | No |
| orf19.6393 | 1201 | 786 | 1065 | 1307 | 1392 | 1146 | 1254 | 878 | 1077 | 1001 | 1090 | 1078 | 1190 | 1093 | 1119 | 1238 | 1110,8 | 1062 | 1160 | 1,09 | 0,124 | 2,181 | 0,02920728 | 0,05233443 | 0,001169 | 0,0032864 | 0,002191 | 0,002191 | TRUE | 0,0218298 | No |
| orf19.6396 | 2371 | 1549 | 2076 | 2472 | 2477 | 2253 | 2331 | 1504 | 2127 | 1972 | 2124 | 2038 | 2118 | 2150 | 2080 | 2121 | 2091,4 | 2065 | 2117 | 1,024 | 0,034 | 0,831 | 0,40586666 | 0,49411359 | 0,000191 | 0,0026737 | 0,001171 | 0,001171 | TRUE | 0,010189 | No |
| orf19.6398 | 589 | 372 | 496 | 629 | 410 | 344 | 392 | 256 | 528 | 474 | 507 | 519 | 351 | 328 | 350 | 361 | 427,23 | 507 | 348 | 0,682 | -0,552 | -7,562 | 3,97E-14 | 4,18E-13 | 1,00E-08 | 0,0053774 | 0,0026887 | 0,0026887 | TRUE | 0,0203924 | No |
| orf19.6399 | 587 | 345 | 456 | 597 | 577 | 448 | 500 | 314 | 527 | 439 | 467 | 492 | 493 | 427 | 446 | 443 | 466,83 | 481 | 452 | 0,939 | -0,091 | -1,138 | 0,2552379 | 0,33791743 | 0,002898 | 0,0050892 | 0,0039907 | 0,0039907 | TRUE | 0,0465589 | No |
| orf19.640 | 1149 | 850 | 988 | 1289 | 1010 | 1007 | 1027 | 621 | 1031 | 1082 | 1011 | 1063 | 864 | 961 | 916 | 876 | 975,45 | 1047 | 904 | 0,865 | -0,209 | -3,782 | 0,0001554 | 0,00046015 | 0,000557 | 0,0034678 | 0,0019024 | 0,0019024 | TRUE | 0,021386 | No |
| orf19.6400 | 1411 | 917 | 1275 | 1518 | 1539 | 1238 | 1309 | 989 | 1266 | 1167 | 1304 | 1252 | 1316 | 1181 | 1168 | 1395 | 1256,2 | 1247 | 1265 | 1,011 | 0,016 | 0,242 | 0,80887742 | 0,85183746 | 0,00381 | 0,0031351 | 0,0035072 | 0,0035072 | TRUE | 0,0530613 | No |
| orf19.6402 | 76418 | 48578 | 65096 | 80405 | 69240 | 54656 | 59709 | 43340 | 68555 | 61845 | 66600 | 66303 | 59216 | 52151 | 53280 | 61127 | 61135 | 65826 | 56444 | 0,857 | -0,222 | -3,83 | 0,0001282 | 0,00038453 | 0,003943 | 0,0020033 | 0,0032103 | 0,0032103 | TRUE | 0,0381428 | No |
| orf19.6403 | 3292 | 2083 | 2763 | 3330 | 3182 | 2713 | 3014 | 1637 | 2953 | 2652 | 2827 | 2746 | 2721 | 2589 | 2689 | 2309 | 2685,8 | 2794 | 2577 | 0,923 | -0,115 | -1,958 | 0,05028863 | 0,08460769 | 0,003196 | 0,00252 | 0,0029511 | 0,0029511 | TRUE | 0,056734 | No |
| orf19.6403.1 | 3399 | 2114 | 2701 | 3401 | 3499 | 2828 | 2957 | 2316 | 3049 | 2691 | 2763 | 2805 | 2992 | 2698 | 2639 | 3267 | 2863,1 | 2827 | 2899 | 1,024 | 0,034 | 0,483 | 0,62937386 | 0,70062594 | 0,005475 | 0,0024866 | 0,0044597 | 0,0044597 | TRUE | 0,0859082 | No |
| orf19.6404 | 1791 | 1214 | 1628 | 2031 | 2083 | 1659 | 1907 | 1122 | 1607 | 1546 | 1666 | 1675 | 1781 | 1583 | 1702 | 1582 | 1642,7 | 1624 | 1662 | 1,025 | 0,035 | 0,649 | 0,51657017 | 0,60041663 | 0,001583 | 0,0028633 | 0,0022169 | 0,0022169 | TRUE | 0,0286001 | No |
| orf19.6405 | 11 | 4 | 13 | 9 | 14 | 11 | 9 | 8 | 10 | 5 | 13 | 7 | 12 | 10 | 8 | 11 | 9,68 | 9 | 10 | 1,155 | 0,208 | 0,467 | 0,64026907 | NA | 1,00E-08 | 0,1518988 | 0,0873493 | 0,0873493 | TRUE | 0,318615 | No |
| orf19.6406 | 2035 | 1329 | 1743 | 2047 | 2173 | 1686 | 1861 | 1257 | 1826 | 1692 | 1783 | 1688 | 1858 | 1609 | 1661 | 1773 | 1736,2 | 1747 | 1725 | 0,987 | -0,019 | -0,335 | 0,73726697 | 0,79401475 | 0,002369 | 0,0028157 | 0,0025498 | 0,0025498 | TRUE | 0,0347561 | No |
| orf19.6407 | 3261 | 2089 | 2891 | 3443 | 2943 | 2403 | 2666 | 1588 | 2925 | 2660 | 2958 | 2839 | 2517 | 2293 | 2379 | 2240 | 2601,3 | 2846 | 2357 | 0,829 | -0,271 | -5,223 | 1,76E-07 | 8,32E-07 | 0,001905 | 0,0025376 | 0,0021925 | 0,0021925 | TRUE | 0,025513 | No |
| orf19.6408 | 4259 | 2802 | 3819 | 4584 | 6033 | 5022 | 5788 | 3760 | 3821 | 3567 | 3907 | 3780 | 5160 | 4792 | 5165 | 5303 | 4436,8 | 3769 | 5105 | 1,353 | 0,436 | 9,657 | 4,58E-22 | 9,59E-21 | 0,001347 | 0,0023067 | 0,0017299 | 0,0017299 | TRUE | 0,0207863 | No |

| | | | | | | | | | | | | | | | | | | | | | | | | | | | | | | | |
|---|---|---|---|---|---|---|---|---|---|---|---|---|---|---|---|---|---|---|---|---|---|---|---|---|---|---|---|---|---|---|---|
| orf19.641 | 964 | 768 | 864 | 1092 | 874 | 904 | 979 | 497 | 865 | 978 | 884 | 900 | 747 | 863 | 874 | 701 | 851,45 | 907 | 796 | 0,881 | -0,182 | -2,318 | 0,02043953 | 0,03799527 | 0,005661 | 0,0036845 | 0,0047377 | 0,0047377 | TRUE | 0,06952 | No |
| orf19.6411 | 1002 | 700 | 926 | 1068 | 1586 | 1304 | 1451 | 985 | 899 | 891 | 947 | 881 | 1356 | 1244 | 1295 | 1389 | 1112,9 | 904 | 1321 | 1,459 | 0,545 | 9,977 | 1,92E-23 | 4,40E-22 | 0,000874 | 0,003284 | 0,0019322 | 0,0019322 | TRUE | 0,0181862 | No |
| orf19.6413 | 675 | 466 | 615 | 741 | 798 | 764 | 881 | 504 | 606 | 593 | 629 | 611 | 682 | 729 | 786 | 711 | 668,44 | 610 | 727 | 1,193 | 0,254 | 3,955 | 7,66E-05 | 0,00023849 | 0,000993 | 0,0041513 | 0,0024588 | 0,0024588 | TRUE | 0,0375226 | No |
| orf19.6414 | 1153 | 717 | 1013 | 1236 | 1097 | 906 | 1089 | 699 | 1034 | 913 | 1036 | 1019 | 938 | 864 | 972 | 986 | 970,39 | 1000 | 940 | 0,936 | -0,095 | -1,495 | 0,13490227 | 0,19734425 | 0,002031 | 0,0034755 | 0,0028149 | 0,0028149 | TRUE | 0,0384922 | No |
| orf19.6414.3 | 2902 | 2198 | 2440 | 2996 | 3607 | 3151 | 3536 | 2123 | 2603 | 2798 | 2496 | 2471 | 3085 | 3007 | 3155 | 2994 | 2826,2 | 2592 | 3060 | 1,183 | 0,242 | 5,043 | 4,57E-07 | 2,04E-06 | 0,001477 | 0,0024932 | 0,0018508 | 0,0018508 | TRUE | 0,0332307 | No |
| orf19.6415.1 | 4121 | 2540 | 2950 | 3862 | 4106 | 2937 | 3079 | 2163 | 3697 | 3234 | 3018 | 3185 | 3512 | 2802 | 2747 | 3051 | 3155,7 | 3284 | 3028 | 0,922 | -0,117 | -1,388 | 0,16525911 | 0,23462569 | 0,010113 | 0,0024395 | 0,0064965 | 0,0064965 | TRUE | 0,1424547 | No |
| orf19.6416 | 4124 | 2760 | 3565 | 4580 | 4480 | 3712 | 4180 | 2844 | 3700 | 3514 | 3647 | 3777 | 3831 | 3542 | 3730 | 4011 | 3719 | 3660 | 3778 | 1,031 | 0,044 | 0,951 | 0,34161804 | 0,42839515 | 0,001445 | 0,0023699 | 0,0018056 | 0,0018056 | TRUE | 0,0214371 | No |
| orf19.6417 | 18106 | 12591 | 16384 | 19656 | 18389 | 16197 | 17757 | 10164 | 16243 | 16030 | 16762 | 16209 | 15727 | 15455 | 15845 | 14335 | 15826 | 16311 | 15340 | 0,941 | -0,088 | -2,24 | 0,0251089 | 0,04579668 | 0,001113 | 0,0020713 | 0,0014188 | 0,0014188 | TRUE | 0,0233905 | No |
| orf19.6418 | 6115 | 4081 | 5211 | 6365 | 5621 | 4717 | 5268 | 3052 | 5486 | 5196 | 5331 | 5249 | 4807 | 4501 | 4701 | 4305 | 4946,9 | 5316 | 4578 | 0,862 | -0,214 | -4,974 | 6,55E-07 | 2,85E-06 | 0,001192 | 0,002273 | 0,0015755 | 0,0015755 | TRUE | 0,0188569 | No |
| orf19.6419.1 | 809 | 580 | 650 | 843 | 828 | 714 | 696 | 399 | 726 | 738 | 665 | 695 | 708 | 681 | 621 | 563 | 674,69 | 706 | 643 | 0,915 | -0,128 | -1,656 | 0,09764744 | 0,14973604 | 0,004187 | 0,0041312 | 0,0042027 | 0,0042027 | TRUE | 0,0741799 | No |
| orf19.642 | 857 | 581 | 713 | 908 | 911 | 852 | 946 | 547 | 769 | 740 | 729 | 749 | 779 | 813 | 844 | 771 | 774,3 | 747 | 802 | 1,075 | 0,104 | 1,834 | 0,06661995 | 0,10823489 | 1,00E-08 | 0,0038544 | 0,0018129 | 0,0018129 | TRUE | 0,015254 | No |
| orf19.6420 | 7519 | 4899 | 6917 | 8396 | 8235 | 6767 | 7567 | 5932 | 6745 | 6237 | 7077 | 6923 | 7043 | 6457 | 6752 | 8367 | 6950,1 | 6746 | 7155 | 1,06 | 0,084 | 1,108 | 0,26807605 | 0,35219787 | 0,008208 | 0,0021884 | 0,0053746 | 0,0053746 | TRUE | 0,1572105 | No |
| orf19.6422 | 2657 | 1890 | 2447 | 3021 | 3310 | 3068 | 3249 | 2117 | 2384 | 2406 | 2504 | 2491 | 2831 | 2927 | 2899 | 2986 | 2678,5 | 2446 | 2911 | 1,189 | 0,249 | 6,545 | 5,94E-11 | 4,49E-10 | 0,000167 | 0,0025215 | 0,0010147 | 0,0010147 | TRUE | 0,0041914 | No |
| orf19.6423 | 1704 | 1181 | 1501 | 1833 | 2164 | 1874 | 2010 | 1324 | 1529 | 1504 | 1536 | 1512 | 1851 | 1788 | 1794 | 1867 | 1672,4 | 1520 | 1825 | 1,2 | 0,262 | 6,423 | 1,33E-10 | 9,53E-10 | 1,00E-08 | 0,0028476 | 0,0010005 | 0,0010005 | TRUE | 0,0025553 | No |
| orf19.6424 | 2013 | 1304 | 1650 | 1999 | 2645 | 1965 | 2109 | 1545 | 1806 | 1660 | 1688 | 1648 | 2262 | 1875 | 1882 | 2179 | 1875,1 | 1700 | 2050 | 1,204 | 0,268 | 3,822 | 0,00013212 | 0,00039569 | 0,004964 | 0,0027537 | 0,0041807 | 0,0041807 | TRUE | 0,0616891 | No |
| orf19.6425 | 1657 | 1193 | 1604 | 1766 | 1822 | 1657 | 1741 | 1030 | 1487 | 1519 | 1641 | 1456 | 1558 | 1581 | 1554 | 1453 | 1531 | 1526 | 1536 | 1,009 | 0,013 | 0,253 | 0,80061771 | 0,8452137 | 0,001374 | 0,0029277 | 0,0020243 | 0,0020243 | TRUE | 0,0315627 | No |
| orf19.643 | 466 | 338 | 409 | 531 | 485 | 436 | 437 | 279 | 418 | 430 | 418 | 438 | 415 | 416 | 390 | 394 | 414,87 | 426 | 404 | 0,948 | -0,077 | -1,089 | 0,27597736 | 0,36025364 | 1,00E-08 | 0,0054787 | 0,0023864 | 0,0023864 | TRUE | 0,0071378 | No |
| orf19.6431 | 1154 | 821 | 1046 | 1272 | 1197 | 1049 | 1090 | 674 | 1035 | 1045 | 1070 | 1049 | 1024 | 1001 | 973 | 951 | 1018,4 | 1050 | 987 | 0,942 | -0,086 | -1,738 | 0,08221858 | 0,12948079 | 1,00E-08 | 0,003405 | 0,0013865 | 0,0013865 | TRUE | 0,0075854 | No |
| orf19.6432 | 1675 | 1109 | 1541 | 1822 | 2036 | 1677 | 1915 | 1200 | 1503 | 1412 | 1577 | 1502 | 1741 | 1600 | 1709 | 1692 | 1592 | 1498 | 1686 | 1,124 | 0,169 | 3,377 | 0,00073177 | 0,0018975 | 0,000816 | 0,0028914 | 0,001765 | 0,001765 | TRUE | 0,0166094 | No |
| orf19.6434 | 6073 | 3840 | 5086 | 6423 | 5702 | 4666 | 5110 | 3392 | 5448 | 4889 | 5203 | 5296 | 4877 | 4452 | 4560 | 4784 | 4938,7 | 5209 | 4668 | 0,895 | -0,159 | -3,395 | 0,00068577 | 0,00178513 | 0,001639 | 0,0022735 | 0,0019095 | 0,0019095 | TRUE | 0,0204084 | No |
| orf19.6435 | 899 | 579 | 753 | 918 | 906 | 739 | 762 | 594 | 806 | 737 | 770 | 757 | 775 | 705 | 680 | 838 | 758,59 | 768 | 750 | 0,972 | -0,041 | -0,565 | 0,57202317 | 0,65101322 | 0,003576 | 0,0038932 | 0,0037284 | 0,0037284 | TRUE | 0,0669543 | No |
| orf19.6436 | 1234 | 901 | 1092 | 1256 | 1507 | 1257 | 1405 | 889 | 1107 | 1147 | 1117 | 1036 | 1289 | 1199 | 1254 | 1254 | 1175,4 | 1102 | 1249 | 1,136 | 0,184 | 3,541 | 0,00039801 | 0,00109691 | 0,000601 | 0,0032146 | 0,0017298 | 0,0017298 | TRUE | 0,0204833 | No |
| orf19.6437 | 515 | 341 | 450 | 587 | 725 | 644 | 678 | 481 | 462 | 434 | 460 | 484 | 620 | 614 | 605 | 678 | 544,81 | 460 | 629 | 1,36 | 0,443 | 6,536 | 6,33E-11 | 4,76E-10 | 1,00E-08 | 0,0046441 | 0,002538 | 0,002538 | TRUE | 0,0277091 | No |
| orf19.6438 | 3138 | 2056 | 2757 | 3261 | 2973 | 2538 | 2884 | 1750 | 2815 | 2618 | 2821 | 2689 | 2543 | 2422 | 2573 | 2468 | 2618,5 | 2736 | 2502 | 0,914 | -0,129 | -3,043 | 0,00234598 | 0,00545458 | 0,000651 | 0,0025339 | 0,0013445 | 0,0013445 | TRUE | 0,0095478 | No |
| orf19.644 | 129 | 90 | 132 | 164 | 217 | 195 | 198 | 131 | 116 | 115 | 135 | 135 | 186 | 186 | 177 | 185 | 154,21 | 125 | 184 | 1,458 | 0,544 | 4,763 | 1,90E-06 | 7,71E-06 | 1,00E-08 | 0,0113933 | 0,0058662 | 0,0058662 | TRUE | 0,0332903 | No |
| orf19.6440 | 4328 | 2899 | 3791 | 4554 | 6923 | 5801 | 6557 | 3893 | 3883 | 3691 | 3879 | 3755 | 5921 | 5535 | 5851 | 5491 | 4750,6 | 3802 | 5700 | 1,5 | 0,585 | 14,61 | 2,43E-48 | 2,24E-46 | 0,000749 | 0,0022851 | 0,0013175 | 0,0013175 | TRUE | 0,0083385 | No |
| orf19.6442 | 2630 | 1690 | 2399 | 2683 | 2875 | 2430 | 2725 | 1680 | 2359 | 2152 | 2454 | 2212 | 2459 | 2319 | 2432 | 2369 | 2344,5 | 2294 | 2395 | 1,044 | 0,061 | 1,214 | 0,22470546 | 0,30365056 | 0,00158 | 0,0025987 | 0,0020315 | 0,0020315 | TRUE | 0,0261243 | No |
| orf19.6443 | 765 | 525 | 652 | 849 | 852 | 703 | 712 | 591 | 686 | 668 | 667 | 700 | 729 | 671 | 635 | 834 | 698,77 | 680 | 717 | 1,049 | 0,069 | 0,849 | 0,39611152 | 0,48498838 | 0,00566 | 0,004057 | 0,0048467 | 0,0048467 | TRUE | 0,1291766 | No |
| orf19.6444 | 1953 | 1247 | 1713 | 2043 | 1966 | 1613 | 1873 | 1276 | 1752 | 1588 | 1753 | 1685 | 1681 | 1539 | 1671 | 1800 | 1683,5 | 1694 | 1673 | 0,985 | -0,022 | -0,379 | 0,70491877 | 0,76612735 | 0,002077 | 0,0028418 | 0,0025577 | 0,0025577 | TRUE | 0,0345275 | No |
| orf19.6445 | 1559 | 942 | 1200 | 1608 | 1076 | 850 | 960 | 673 | 1399 | 1199 | 1228 | 1326 | 920 | 811 | 857 | 949 | 1086,1 | 1288 | 884 | 0,684 | -0,547 | -7,87 | 3,54E-15 | 4,11E-14 | 0,003738 | 0,0033162 | 0,0036778 | 0,0036778 | TRUE | 0,0400678 | No |
| orf19.6447 | 7217 | 4573 | 6052 | 7375 | 9477 | 7534 | 8049 | 5073 | 6474 | 5822 | 6192 | 6082 | 8105 | 7189 | 7182 | 7155 | 6775,1 | 6142 | 7408 | 1,206 | 0,27 | 5,113 | 3,17E-07 | 1,44E-06 | 0,00274 | 0,0021938 | 0,0025341 | 0,0025341 | TRUE | 0,0494558 | No |
| orf19.6448 | 2919 | 1915 | 2533 | 2969 | 4123 | 3306 | 3813 | 2397 | 2619 | 2438 | 2592 | 2448 | 3526 | 3154 | 3402 | 3381 | 2945 | 2524 | 3366 | 1,333 | 0,415 | 8,686 | 3,76E-18 | 5,93E-17 | 0,001293 | 0,0024725 | 0,0018439 | 0,0018439 | TRUE | 0,0222005 | No |
| orf19.6449 | 319 | 237 | 287 | 357 | 321 | 237 | 270 | 172 | 286 | 302 | 294 | 294 | 275 | 226 | 241 | 243 | 270,01 | 294 | 246 | 0,84 | -0,252 | -2,786 | 0,00534041 | 0,01147539 | 1,00E-08 | 0,0073559 | 0,0041557 | 0,0041557 | TRUE | 0,0750591 | No |
| orf19.6450 | 2 | 2 | 5 | 2 | 3 | 1 | 1 | 1 | 2 | 3 | 5 | 2 | 3 | 1 | 1 | 1 | 2,12 | 3 | 2 | 0,538 | -0,895 | -0,926 | 0,35455092 | NA | 1,00E-08 | 0,6880489 | 0,3763451 | 0,3763451 | TRUE | 0,3950577 | No |
| orf19.645.1 | 1570 | 1390 | 1446 | 1933 | 1580 | 1656 | 1615 | 1087 | 1408 | 1770 | 1479 | 1594 | 1351 | 1580 | 1441 | 1533 | 1519,6 | 1563 | 1476 | 0,945 | -0,082 | -1,063 | 0,28774753 | 0,37229078 | 0,006568 | 0,0029348 | 0,0049951 | 0,0049951 | TRUE | 0,0924095 | No |
| orf19.6452 | 2369 | 1800 | 1956 | 2493 | 2712 | 2313 | 2276 | 1507 | 2125 | 2292 | 2001 | 2056 | 2319 | 2207 | 2031 | 2125 | 2144,6 | 2118 | 2170 | 1,026 | 0,037 | 0,639 | 0,5227719 | 0,60586681 | 0,002575 | 0,0026564 | 0,002767 | 0,002767 | TRUE | 0,0351237 | No |
| orf19.6453 | 845 | 767 | 792 | 1002 | 1337 | 1413 | 1419 | 897 | 758 | 976 | 810 | 826 | 1143 | 1348 | 1266 | 1265 | 1049,3 | 842 | 1256 | 1,493 | 0,578 | 7,17 | 7,51E-13 | 6,99E-12 | 0,007126 | 0,0033631 | 0,0052527 | 0,0052527 | TRUE | 0,1288331 | No |
| orf19.6455 | 1213 | 868 | 1033 | 1276 | 1716 | 1466 | 1561 | 1013 | 1088 | 1105 | 1057 | 1052 | 1468 | 1399 | 1393 | 1429 | 1248,8 | 1076 | 1422 | 1,323 | 0,404 | 8,653 | 5,01E-18 | 7,82E-17 | 1,00E-08 | 0,003142 | 0,0012793 | 0,0012793 | TRUE | 0,0061814 | No |
| orf19.6456 | 1077 | 761 | 1022 | 1225 | 964 | 960 | 1077 | 757 | 966 | 969 | 1046 | 1010 | 824 | 916 | 961 | 1068 | 969,99 | 998 | 942 | 0,941 | -0,088 | -1,164 | 0,24429142 | 0,32601129 | 0,005094 | 0,0034761 | 0,0043973 | 0,0043973 | TRUE | 0,0889691 | No |
| orf19.6457 | 1379 | 1001 | 1326 | 1485 | 1553 | 1517 | 1677 | 914 | 1237 | 1274 | 1357 | 1225 | 1328 | 1447 | 1496 | 1289 | 1331,7 | 1273 | 1390 | 1,094 | 0,13 | 2,108 | 0,0350011 | 0,06146935 | 0,002559 | 0,0030696 | 0,0029019 | 0,0029019 | TRUE | 0,0318101 | No |
| orf19.6458 | 22 | 25 | 25 | 37 | 19 | 40 | 39 | 30 | 20 | 32 | 26 | 31 | 16 | 38 | 35 | 42 | 29,9 | 27 | 33 | 1,21 | 0,275 | 0,883 | 0,37732658 | 0,46664844 | 0,069629 | 0,0505351 | 0,0589405 | 0,0589405 | TRUE | 0,9138521 | No |
| orf19.6458.1 | 2185 | 2055 | 1921 | 2532 | 2135 | 2488 | 2326 | 1107 | 1960 | 2616 | 1965 | 2088 | 1826 | 2374 | 2076 | 1561 | 2058,3 | 2157 | 1959 | 0,909 | -0,138 | -0,861 | 0,38934692 | 0,4784071 | 0,024167 | 0,0026848 | 0,0134139 | 0,0241675 | TRUE | 0,2464652 | No |
| orf19.6459 | 681 | 599 | 604 | 839 | 620 | 595 | 647 | 353 | 611 | 763 | 618 | 692 | 530 | 568 | 577 | 498 | 607,06 | 671 | 543 | 0,814 | -0,298 | -3,525 | 0,00042311 | 0,00116021 | 0,005678 | 0,0043709 | 0,0051775 | 0,0051775 | TRUE | 0,0930942 | No |
| orf19.646 | 1E+05 | 95625 | 1E+05 | 2E+05 | 1E+05 | 1E+05 | 1E+05 | 78054 | 1E+05 | 1E+05 | 1E+05 | 1E+05 | 1E+05 | 1E+05 | 1E+05 | 1E+05 | 122900 | 1E+05 | 1E+05 | 0,951 | -0,073 | -1,553 | 0,12049852 | 0,17888686 | 0,002171 | 0,0019913 | 0,0021295 | 0,0021295 | TRUE | 0,0363034 | No |
| orf19.6460 | 2120 | 1654 | 1905 | 2260 | 1909 | 1876 | 2080 | 1249 | 1902 | 2106 | 1949 | 1864 | 1633 | 1790 | 1856 | 1762 | 1857,6 | 1955 | 1760 | 0,901 | -0,15 | -2,65 | 0,00804519 | 0,01658164 | 0,002177 | 0,002761 | 0,0025307 | 0,0025307 | TRUE | 0,0312695 | No |
| orf19.6461 | 1150 | 721 | 928 | 1183 | 1135 | 957 | 1035 | 623 | 1032 | 918 | 949 | 976 | 971 | 913 | 924 | 879 | 945,08 | 969 | 922 | 0,952 | -0,071 | -1,239 | 0,21516185 | 0,29306108 | 0,000963 | 0,0035156 | 0,002135 | 0,002135 | TRUE | 0,0229029 | No |
| orf19.6462 | 1547 | 1174 | 1441 | 1772 | 1666 | 1394 | 1552 | 1020 | 1388 | 1495 | 1474 | 1461 | 1425 | 1330 | 1385 | 1439 | 1424,6 | 1454 | 1395 | 0,959 | -0,061 | -1,264 | 0,20636791 | 0,28249223 | 0,000418 | 0,0029986 | 0,0015088 | 0,0015088 | TRUE | 0,0116758 | No |
| orf19.6463 | 3111 | 2227 | 2787 | 3447 | 3383 | 3411 | 3532 | 2002 | 2791 | 2835 | 2851 | 2842 | 2893 | 3255 | 3152 | 2824 | 2930,4 | 2830 | 3031 | 1,072 | 0,1 | 1,963 | 0,04968011 | 0,0837011 | 0,001886 | 0,0024749 | 0,0021645 | 0,0021645 | TRUE | 0,029713 | No |
| orf19.6464 | 2114 | 1530 | 1915 | 2383 | 2484 | 2523 | 2667 | 1635 | 1896 | 1948 | 1959 | 1965 | 2124 | 2407 | 2380 | 2306 | 2123,3 | 1942 | 2304 | 1,186 | 0,246 | 4,988 | 6,09E-07 | 2,66E-06 | 0,001391 | 0,0026632 | 0,0018667 | 0,0018667 | TRUE | 0,0351574 | No |
| orf19.6465 | 164 | 137 | 179 | 213 | 213 | 173 | 202 | 156 | 147 | 174 | 183 | 176 | 182 | 165 | 180 | 220 | 178,48 | 170 | 187 | 1,092 | 0,127 | 1,063 | 0,28783134 | 0,37229078 | 0,005583 | 0,0101133 | 0,0081518 | 0,0081518 | TRUE | 0,1279615 | No |
| orf19.6469 | 575 | 381 | 557 | 645 | 670 | 623 | 681 | 467 | 516 | 485 | 570 | 532 | 573 | 594 | 608 | 659 | 567,05 | 526 | 608 | 1,152 | 0,204 | 2,834 | 0,00459387 | 0,00999636 | 0,00168 | 0,0045396 | 0,0031871 | 0,0031871 | TRUE | 0,0352835 | No |
| orf19.6470 | 790 | 525 | 715 | 859 | 820 | 726 | 808 | 496 | 709 | 668 | 732 | 708 | 701 | 693 | 721 | 700 | 703,94 | 704 | 704 | 0,998 | -0,003 | -0,049 | 0,96123526 | 0,97188311 | 1,00E-08 | 0,0040418 | 0,0017294 | 0,0017294 | TRUE | 0,0119957 | No |
| orf19.6472 | 3925 | 2560 | 3123 | 3950 | 5306 | 4112 | 4318 | 3078 | 3521 | 3259 | 3195 | 3257 | 4538 | 3924 | 3853 | 4341 | 3736,1 | 3308 | 4164 | 1,258 | 0,331 | 5,459 | 4,78E-08 | 2,44E-07 | 0,00373 | 0,0023681 | 0,0032641 | 0,0032641 | TRUE | 0,0457267 | No |
| orf19.647.3 | 164 | 133 | 154 | 213 | 173 | 184 | 187 | 100 | 147 | 169 | 158 | 176 | 148 | 176 | 167 | 141 | 160,13 | 162 | 158 | 0,977 | -0,034 | -0,286 | 0,77487275 | 0,82367931 | 1,00E-08 | 0,011045 | 0,007197 | 0,007197 | TRUE | 0,057844 | No |
| orf19.6474 | 3853 | 2454 | 3259 | 3981 | 3666 | 3045 | 3377 | 2001 | 3457 | 3124 | 3334 | 3283 | 3135 | 2905 | 3013 | 2822 | 3134,3 | 3300 | 2969 | 0,9 | -0,152 | -3,165 | 0,00155104 | 0,00375147 | 0,001488 | 0,0024427 | 0,0018802 | 0,0018802 | TRUE | 0,017314 | No |
| orf19.6475 | 32 | 20 | 29 | 38 | 32 | 26 | 29 | 20 | 29 | 25 | 30 | 31 | 27 | 25 | 26 | 28 | 27,68 | 29 | 26 | 0,914 | -0,13 | -0,515 | 0,60630141 | 0,68160006 | 1,00E-08 | 0,0544263 | 0,0250752 | 0,0250752 | TRUE | 0,0304031 | No |
| orf19.6476 | 1999 | 1624 | 1935 | 2310 | 2276 | 2351 | 2546 | 1415 | 1793 | 2068 | 1980 | 1905 | 1947 | 2243 | 2272 | 1996 | 2025,3 | 1936 | 2114 | 1,094 | 0,129 | 1,948 | 0,05145173 | 0,08619328 | 0,004262 | 0,0026963 | 0,003716 | 0,003716 | TRUE | 0,0362779 | No |
| orf19.6477 | 2466 | 2103 | 2278 | 2988 | 2895 | 3174 | 3291 | 1871 | 2212 | 2677 | 2331 | 2464 | 2476 | 3029 | 2937 | 2639 | 2595,5 | 2421 | 2770 | 1,145 | 0,195 | 2,56 | 0,01045549 | 0,02093886 | 0,007184 | 0,0025388 | 0,0052085 | 0,0052085 | TRUE | 0,0635498 | No |
| orf19.6478 | 10716 | 8246 | 9871 | 11674 | 13153 | 13445 | 13848 | 7578 | 9613 | 10498 | 10099 | 9627 | 11249 | 12829 | 12357 | 10688 | 10870 | 9959 | 11781 | 1,183 | 0,243 | 4,007 | 6,16E-05 | 0,00019578 | 0,004276 | 0,0021131 | 0,003438 | 0,003438 | TRUE | 0,0471464 | No |

| | | | | | | | | | | | | | | | | | | | | | | | | | | | | | | | |
|---|---|---|---|---|---|---|---|---|---|---|---|---|---|---|---|---|---|---|---|---|---|---|---|---|---|---|---|---|---|---|---|
| orf19.6479 | 679 | 536 | 638 | 833 | 906 | 860 | 841 | 497 | 609 | 682 | 653 | 687 | 775 | 821 | 750 | 701 | 709,75 | 658 | 762 | 1,162 | 0,216 | 3,176 | 0,00149367 | 0,00362586 | 0,002122 | 0,0040249 | 0,0030383 | 0,0030383 | TRUE | 0,0308662 | No |
| orf19.6480 | 407 | 282 | 359 | 450 | 508 | 581 | 564 | 321 | 365 | 359 | 367 | 371 | 434 | 554 | 503 | 453 | 425,92 | 366 | 486 | 1,331 | 0,412 | 4,715 | 2,42E-06 | 9,71E-06 | 0,004548 | 0,0053879 | 0,0049514 | 0,0049514 | TRUE | 0,1047297 | No |
| orf19.6481 | 2543 | 1942 | 2315 | 2871 | 2935 | 2803 | 3029 | 1765 | 2281 | 2472 | 2368 | 2367 | 2510 | 2675 | 2703 | 2489 | 2483,3 | 2372 | 2594 | 1,095 | 0,131 | 2,875 | 0,00403661 | 0,00889 | 0,000904 | 0,0025641 | 0,0015959 | 0,0015959 | TRUE | 0,0096728 | No |
| orf19.6482 | 375 | 366 | 345 | 465 | 304 | 349 | 348 | 192 | 336 | 466 | 353 | 383 | 260 | 333 | 311 | 271 | 339,14 | 384 | 294 | 0,767 | -0,383 | -3,266 | 0,00109077 | 0,0027283 | 0,01394 | 0,0062601 | 0,0101734 | 0,0101734 | TRUE | 0,217529 | No |
| orf19.6484 | 303 | 208 | 236 | 269 | 229 | 171 | 164 | 134 | 272 | 265 | 241 | 222 | 196 | 163 | 146 | 189 | 211,78 | 250 | 174 | 0,694 | -0,528 | -4,439 | 9,04E-06 | 3,31E-05 | 0,008447 | 0,0088342 | 0,0086728 | 0,0086728 | TRUE | 0,1269349 | No |
| orf19.6486 | 35 | 18 | 26 | 45 | 31 | 22 | 27 | 26 | 31 | 23 | 27 | 37 | 27 | 21 | 24 | 37 | 28,29 | 30 | 27 | 0,888 | -0,171 | -0,622 | 0,53384543 | 0,61608593 | 0,010145 | 0,0533013 | 0,0368093 | 0,0368093 | TRUE | 0,257181 | No |
| orf19.6487 | 33 | 23 | 23 | 29 | 16 | 19 | 19 | 19 | 30 | 29 | 24 | 24 | 14 | 18 | 17 | 27 | 22,74 | 27 | 19 | 0,693 | -0,529 | -1,774 | 0,07612018 | 0,12117912 | 1,00E-08 | 0,0658271 | 0,0395599 | 0,0395599 | TRUE | 0,2879642 | No |
| orf19.6488 | 24 | 11 | 11 | 22 | 8 | 7 | 11 | 5 | 22 | 14 | 11 | 18 | 7 | 7 | 10 | 7 | 11,91 | 16 | 8 | 0,466 | -1,103 | -2,631 | 0,00850174 | NA | 1,00E-08 | 0,1238189 | 0,0731503 | 0,0731503 | TRUE | 0,1542934 | No |
| orf19.6489 | 8917 | 5977 | 7718 | 9721 | 10048 | 8935 | 9554 | 6141 | 7999 | 7609 | 7896 | 8016 | 8593 | 8525 | 8525 | 8661 | 8228,3 | 7880 | 8576 | 1,088 | 0,121 | 4,116 | 3,86E-05 | 0,00012712 | 0,000152 | 0,0021559 | 0,0007095 | 0,0007095 | TRUE | 0,0062966 | No |
| orf19.649 | 7311 | 5967 | 7156 | 8555 | 8118 | 8269 | 8736 | 4802 | 6559 | 7597 | 7321 | 7055 | 6943 | 7890 | 7795 | 6773 | 7241,5 | 7133 | 7350 | 1,031 | 0,044 | 0,692 | 0,48913036 | 0,57452496 | 0,004409 | 0,00218 | 0,0037686 | 0,0037686 | TRUE | 0,0358929 | No |
| orf19.6491 | 25 | 30 | 48 | 55 | 29 | 41 | 42 | 18 | 22 | 38 | 49 | 45 | 25 | 39 | 37 | 25 | 35,23 | 38 | 32 | 0,824 | -0,279 | -0,976 | 0,32886146 | 0,41639614 | 0,05823 | 0,0431813 | 0,0498532 | 0,0498532 | TRUE | 0,6594607 | No |
| orf19.6492 | 1948 | 1770 | 1915 | 2408 | 1970 | 2270 | 2457 | 1481 | 1748 | 2253 | 1959 | 1986 | 1685 | 2166 | 2192 | 2089 | 2009,7 | 1986 | 2033 | 1,024 | 0,034 | 0,367 | 0,71394533 | 0,77439674 | 0,012225 | 0,0027019 | 0,0077388 | 0,0077388 | TRUE | 0,1634669 | No |
| orf19.6493 | 6 | 4 | 7 | 8 | 9 | 9 | 8 | 4 | 5 | 5 | 7 | 7 | 8 | 9 | 7 | 6 | 6,66 | 6 | 8 | 1,208 | 0,272 | 0,533 | 0,59404389 | NA | 1,00E-08 | 0,2198768 | 0,1017114 | 0,1017114 | TRUE | 0,0382086 | No |
| orf19.6494 | 2195 | 1412 | 1864 | 2449 | 2671 | 2199 | 2470 | 1730 | 1969 | 1798 | 1907 | 2019 | 2284 | 2098 | 2204 | 2440 | 2090 | 1923 | 2256 | 1,17 | 0,227 | 3,955 | 7,64E-05 | 0,00023823 | 0,00263 | 0,0026741 | 0,0026807 | 0,0026807 | TRUE | 0,0341872 | No |
| orf19.6496 | 1394 | 903 | 1096 | 1501 | 1691 | 1347 | 1410 | 907 | 1251 | 1150 | 1121 | 1238 | 1446 | 1285 | 1258 | 1279 | 1253,5 | 1190 | 1317 | 1,107 | 0,146 | 2,329 | 0,01987685 | 0,03700654 | 0,002886 | 0,0031376 | 0,0029927 | 0,0029927 | TRUE | 0,0542212 | No |
| orf19.6498 | 2985 | 1984 | 2630 | 3303 | 3065 | 2469 | 2731 | 1748 | 2678 | 2526 | 2691 | 2724 | 2621 | 2356 | 2437 | 2465 | 2562,2 | 2655 | 2470 | 0,93 | -0,105 | -2,249 | 0,02452672 | 0,0448301 | 0,001039 | 0,0025461 | 0,0016913 | 0,0016913 | TRUE | 0,0216159 | No |
| orf19.6499 | 1057 | 737 | 880 | 1079 | 1011 | 976 | 1076 | 698 | 948 | 938 | 900 | 890 | 865 | 931 | 960 | 984 | 927,14 | 919 | 935 | 1,016 | 0,022 | 0,385 | 0,69994144 | 0,76182032 | 0,001033 | 0,0035453 | 0,0021682 | 0,0021682 | TRUE | 0,0318882 | No |
| orf19.6500 | 2640 | 1766 | 2349 | 2861 | 3040 | 2487 | 2667 | 1888 | 2368 | 2248 | 2403 | 2359 | 2600 | 2373 | 2380 | 2663 | 2424,4 | 2344 | 2504 | 1,066 | 0,092 | 1,821 | 0,0686088 | 0,1108367 | 0,00164 | 0,0025783 | 0,0020569 | 0,0020569 | TRUE | 0,0206951 | No |
| orf19.6501 | 39 | 37 | 31 | 45 | 46 | 31 | 38 | 32 | 35 | 47 | 32 | 37 | 39 | 30 | 34 | 45 | 37,36 | 38 | 37 | 0,978 | -0,032 | -0,136 | 0,89156631 | 0,91815742 | 1,00E-08 | 0,040837 | 0,0255337 | 0,0255337 | TRUE | 0,1621247 | No |
| orf19.6502 | 34 | 22 | 42 | 40 | 23 | 38 | 18 | 16 | 31 | 28 | 43 | 33 | 20 | 36 | 16 | 23 | 28,63 | 34 | 24 | 0,7 | -0,515 | -1,727 | 0,08410146 | 0,1321343 | 0,044194 | 0,0526891 | 0,0491393 | 0,0491393 | TRUE | 0,8325121 | No |
| orf19.6503 | 1172 | 941 | 1044 | 1325 | 1357 | 1340 | 1368 | 1020 | 1051 | 1198 | 1068 | 1093 | 1161 | 1279 | 1221 | 1439 | 1188,6 | 1102 | 1275 | 1,154 | 0,207 | 2,867 | 0,00413802 | 0,00909004 | 0,00505 | 0,0032009 | 0,004162 | 0,004162 | TRUE | 0,0849812 | No |
| orf19.6506 | 2768 | 1858 | 2454 | 2874 | 3343 | 2800 | 2998 | 1995 | 2483 | 2365 | 2511 | 2370 | 2859 | 2672 | 2675 | 2814 | 2593,6 | 2432 | 2755 | 1,132 | 0,179 | 4,159 | 3,20E-05 | 0,00010735 | 0,000729 | 0,0025392 | 0,0013872 | 0,0013872 | TRUE | 0,0085092 | No |
| orf19.6507 | 9015 | 5882 | 7683 | 9252 | 8871 | 7361 | 8156 | 5350 | 8087 | 7488 | 7860 | 7629 | 7587 | 7024 | 7278 | 7546 | 7562,4 | 7766 | 7359 | 0,947 | -0,078 | -1,946 | 0,05168622 | 0,08649883 | 0,001032 | 0,0021715 | 0,001427 | 0,001427 | TRUE | 0,0114907 | No |
| orf19.6508 | 1204 | 791 | 1019 | 1298 | 1083 | 904 | 904 | 618 | 1080 | 1007 | 1043 | 1070 | 926 | 863 | 807 | 872 | 958,39 | 1050 | 867 | 0,824 | -0,278 | -4,793 | 1,64E-06 | 6,71E-06 | 0,001108 | 0,0034943 | 0,0021863 | 0,0021863 | TRUE | 0,0279867 | No |
| orf19.6509 | 627 | 510 | 509 | 669 | 634 | 606 | 612 | 445 | 562 | 649 | 521 | 552 | 542 | 578 | 546 | 628 | 572,3 | 571 | 574 | 1,004 | 0,006 | 0,069 | 0,94531473 | 0,96050244 | 0,00424 | 0,0045161 | 0,0045196 | 0,0045196 | TRUE | 0,0915967 | No |
| orf19.651 | 1083 | 926 | 997 | 1340 | 1197 | 1230 | 1214 | 691 | 972 | 1179 | 1020 | 1105 | 1024 | 1174 | 1083 | 975 | 1066,3 | 1069 | 1064 | 0,998 | -0,003 | -0,04 | 0,96808639 | 0,9775232 | 0,005783 | 0,0033409 | 0,0046602 | 0,0046602 | TRUE | 0,0572363 | No |
| orf19.6510 | 214 | 132 | 180 | 241 | 203 | 209 | 210 | 158 | 192 | 168 | 184 | 199 | 174 | 199 | 187 | 223 | 190,77 | 186 | 196 | 1,041 | 0,059 | 0,53 | 0,59588928 | 0,67281569 | 0,002228 | 0,0095891 | 0,0064642 | 0,0064642 | TRUE | 0,0781006 | No |
| orf19.6511 | 6815 | 4919 | 6216 | 7398 | 6903 | 6393 | 6768 | 4266 | 6114 | 6262 | 6360 | 6100 | 5904 | 6100 | 6039 | 6017 | 6112 | 6209 | 6015 | 0,969 | -0,045 | -1,487 | 0,13700914 | 0,19977588 | 0,000154 | 0,002217 | 0,0007324 | 0,0007324 | TRUE | 0,0033222 | No |
| orf19.6512 | 1240 | 938 | 1149 | 1340 | 1283 | 1249 | 1285 | 731 | 1112 | 1194 | 1176 | 1105 | 1097 | 1192 | 1147 | 1031 | 1131,7 | 1147 | 1117 | 0,977 | -0,033 | -0,572 | 0,56740083 | 0,64690024 | 0,001487 | 0,0032623 | 0,0023405 | 0,0023405 | TRUE | 0,0282396 | No |
| orf19.6514 | 30967 | 20320 | 27375 | 34472 | 39369 | 32758 | 36518 | 22629 | 27781 | 25870 | 28007 | 28426 | 33670 | 31257 | 32586 | 31916 | 29939 | 27521 | 32357 | 1,176 | 0,233 | 5,757 | 8,56E-09 | 4,78E-08 | 0,001309 | 0,002028 | 0,0015459 | 0,0015459 | TRUE | 0,0198561 | No |
| orf19.6515 | 10752 | 7210 | 9140 | 11132 | 14726 | 12804 | 13549 | 8381 | 9646 | 9179 | 9351 | 9180 | 12594 | 12217 | 12090 | 11821 | 10760 | 9339 | 12180 | 1,305 | 0,384 | 11,501 | 1,30E-30 | 4,77E-29 | 0,0005 | 0,0021144 | 0,000973 | 0,000973 | TRUE | 0,0064129 | No |
| orf19.6517 | 1045 | 629 | 912 | 1140 | 1026 | 847 | 1002 | 600 | 937 | 801 | 933 | 940 | 877 | 808 | 894 | 846 | 879,67 | 903 | 856 | 0,947 | -0,079 | -1,188 | 0,2348489 | 0,31579504 | 0,002589 | 0,0036298 | 0,0030805 | 0,0030805 | TRUE | 0,0636473 | No |
| orf19.6518 | 601 | 377 | 508 | 613 | 453 | 330 | 400 | 304 | 539 | 480 | 520 | 505 | 387 | 315 | 357 | 429 | 441,54 | 511 | 372 | 0,723 | -0,467 | -5,114 | 3,15E-07 | 1,44E-06 | 0,006463 | 0,0052673 | 0,005674 | 0,005674 | TRUE | 0,1202538 | No |
| orf19.652 | 580 | 418 | 440 | 552 | 553 | 535 | 550 | 310 | 520 | 532 | 450 | 455 | 473 | 510 | 491 | 437 | 483,66 | 489 | 478 | 0,981 | -0,027 | -0,336 | 0,73708438 | 0,79401475 | 0,003603 | 0,004981 | 0,0043033 | 0,0043033 | TRUE | 0,0363653 | No |
| orf19.6520 | 90 | 62 | 83 | 117 | 61 | 51 | 49 | 41 | 81 | 79 | 85 | 96 | 52 | 49 | 44 | 58 | 67,93 | 85 | 51 | 0,585 | -0,772 | -4,467 | 7,93E-06 | 2,94E-05 | 1,00E-08 | 0,0233496 | 0,0129521 | 0,0129521 | TRUE | 0,0825254 | No |
| orf19.6522 | 1418 | 908 | 1256 | 1505 | 1377 | 1125 | 1219 | 784 | 1272 | 1156 | 1285 | 1241 | 1178 | 1073 | 1088 | 1106 | 1174,8 | 1238 | 1111 | 0,896 | -0,158 | -2,883 | 0,00393392 | 0,00869253 | 0,001078 | 0,0032152 | 0,0020197 | 0,0020197 | TRUE | 0,0220749 | No |
| orf19.6524 | 5135 | 3357 | 4432 | 5545 | 5095 | 4206 | 4504 | 3204 | 4607 | 4274 | 4534 | 4572 | 4357 | 4013 | 4019 | 4519 | 4362 | 4497 | 4227 | 0,939 | -0,091 | -1,825 | 0,06802506 | 0,11004161 | 0,002045 | 0,0023123 | 0,0021534 | 0,0021534 | TRUE | 0,0252545 | No |
| orf19.6525 | 883 | 627 | 849 | 987 | 1414 | 1314 | 1296 | 842 | 792 | 798 | 869 | 814 | 1209 | 1254 | 1156 | 1188 | 1010 | 818 | 1202 | 1,469 | 0,555 | 10,213 | 1,74E-24 | 4,31E-23 | 0,000457 | 0,0034169 | 0,0018087 | 0,0018087 | TRUE | 0,0200154 | No |
| orf19.6526 | 1776 | 1199 | 1495 | 1975 | 1632 | 1357 | 1475 | 1000 | 1593 | 1526 | 1530 | 1629 | 1396 | 1295 | 1316 | 1410 | 1461,9 | 1570 | 1354 | 0,861 | -0,216 | -4,382 | 1,18E-05 | 4,24E-05 | 0,000684 | 0,0029726 | 0,0016452 | 0,0016452 | TRUE | 0,0102949 | No |
| orf19.6527 | 15420 | 10224 | 13942 | 16984 | 19284 | 16571 | 18573 | 11379 | 13833 | 13016 | 14264 | 14005 | 16492 | 15812 | 16573 | 16049 | 15006 | 13780 | 16232 | 1,178 | 0,236 | 6,303 | 2,91E-10 | 2,00E-09 | 0,000925 | 0,0020763 | 0,0012796 | 0,0012796 | TRUE | 0,0167457 | No |
| orf19.6528 | 829 | 544 | 744 | 888 | 960 | 821 | 913 | 563 | 744 | 693 | 761 | 732 | 821 | 783 | 815 | 794 | 767,86 | 732 | 803 | 1,096 | 0,132 | 2,354 | 0,01856609 | 0,03477092 | 1,00E-08 | 0,0038701 | 0,0017268 | 0,0017268 | TRUE | 0,0137494 | No |
| orf19.6529 | 17841 | 11534 | 15700 | 19060 | 17513 | 15009 | 16743 | 10314 | 16005 | 14684 | 16063 | 15717 | 14978 | 14321 | 14940 | 14547 | 15157 | 15617 | 14696 | 0,941 | -0,088 | -2,301 | 0,02141463 | 0,03967286 | 0,000985 | 0,0020753 | 0,0013342 | 0,0013342 | TRUE | 0,0195323 | No |
| orf19.653 | 2 | 1 | 1 | 0 | 1 | 2 | 1 | 1 | 2 | 1 | 1 | 0 | 1 | 2 | 1 | 1 | 1,14 | 1 | 1 | 1,254 | 0,326 | 0,258 | 0,79676455 | NA | 1,00E-08 | 1,270294 | 0,6462604 | 0,6462604 | TRUE | 0,3068692 | No |
| orf19.6530 | 3424 | 2459 | 3014 | 3835 | 3802 | 3413 | 3695 | 2511 | 3072 | 3131 | 3084 | 3162 | 3252 | 3257 | 3297 | 3542 | 3224,4 | 3112 | 3337 | 1,071 | 0,098 | 2,472 | 0,01343392 | 0,02617937 | 0,000533 | 0,0024297 | 0,0012096 | 0,0012096 | TRUE | 0,019278 | No |
| orf19.6531 | 6094 | 3971 | 5342 | 6442 | 6390 | 5629 | 5928 | 3877 | 5467 | 5056 | 5465 | 5312 | 5465 | 5371 | 5290 | 5468 | 5361,7 | 5325 | 5398 | 1,013 | 0,019 | 0,509 | 0,61061415 | 0,68517754 | 0,000532 | 0,0022503 | 0,0011059 | 0,0011059 | TRUE | 0,0135405 | No |
| orf19.6531.1 | 1614 | 1180 | 1245 | 1741 | 1859 | 1557 | 1436 | 1074 | 1448 | 1502 | 1274 | 1436 | 1590 | 1486 | 1281 | 1515 | 1441,4 | 1415 | 1468 | 1,037 | 0,053 | 0,705 | 0,48081601 | 0,56630907 | 0,006067 | 0,0029867 | 0,0047036 | 0,0047036 | TRUE | 0,0903012 | No |
| orf19.6532 | 578 | 362 | 481 | 579 | 601 | 501 | 544 | 356 | 519 | 461 | 492 | 477 | 514 | 478 | 485 | 502 | 491,07 | 487 | 495 | 1,014 | 0,019 | 0,283 | 0,77732088 | 0,82525956 | 1,00E-08 | 0,0049358 | 0,0025037 | 0,0025037 | TRUE | 0,0221999 | No |
| orf19.6533 | 658 | 433 | 508 | 633 | 739 | 746 | 775 | 497 | 590 | 551 | 520 | 522 | 632 | 712 | 692 | 701 | 614,95 | 546 | 684 | 1,252 | 0,324 | 4,645 | 3,41E-06 | 1,33E-05 | 0,001819 | 0,0043402 | 0,0030191 | 0,0030191 | TRUE | 0,0378159 | No |
| orf19.6534 | 4 | 9 | 2 | 9 | 7 | 6 | 9 | 9 | 4 | 11 | 2 | 7 | 6 | 6 | 8 | 13 | 7,12 | 6 | 8 | 1,32 | 0,4 | 0,692 | 0,48904519 | NA | 0,123055 | 0,2059044 | 0,1749111 | 0,1749111 | TRUE | 0,4208399 | No |
| orf19.6534.2 | 1517 | 993 | 1297 | 1684 | 1693 | 1433 | 1625 | 927 | 1361 | 1264 | 1327 | 1389 | 1448 | 1367 | 1450 | 1307 | 1364,2 | 1335 | 1393 | 1,044 | 0,062 | 1,169 | 0,24260212 | 0,32418895 | 0,001052 | 0,0030437 | 0,0019847 | 0,0019847 | TRUE | 0,0186286 | No |
| orf19.6536 | 364 | 267 | 344 | 370 | 432 | 436 | 466 | 263 | 327 | 340 | 352 | 305 | 369 | 416 | 416 | 371 | 361,97 | 331 | 393 | 1,193 | 0,255 | 3,056 | 0,0022443 | 0,00524456 | 0,001645 | 0,0059901 | 0,0039106 | 0,0039106 | TRUE | 0,0338907 | No |
| orf19.6537 | 238 | 148 | 201 | 237 | 188 | 213 | 201 | 116 | 214 | 188 | 206 | 195 | 161 | 203 | 179 | 164 | 188,75 | 201 | 177 | 0,881 | -0,183 | -1,659 | 0,09701712 | 0,14899766 | 0,001929 | 0,0096707 | 0,0063561 | 0,0063561 | TRUE | 0,1105746 | No |
| orf19.6538 | 769 | 551 | 705 | 900 | 843 | 687 | 732 | 476 | 690 | 701 | 721 | 742 | 721 | 656 | 653 | 671 | 694,48 | 714 | 675 | 0,946 | -0,08 | -1,305 | 0,19201861 | 0,26545049 | 1,00E-08 | 0,0040699 | 0,0021626 | 0,0021626 | TRUE | 0,0268765 | No |
| orf19.6539 | 6514 | 4255 | 5587 | 6991 | 6173 | 5162 | 5652 | 3594 | 5844 | 5417 | 5716 | 5765 | 5279 | 4925 | 5043 | 5069 | 5382,4 | 5686 | 5079 | 0,893 | -0,163 | -4,221 | 2,43E-05 | 8,34E-05 | 0,00071 | 0,0022492 | 0,0012482 | 0,0012482 | TRUE | 0,0119141 | No |
| orf19.654 | 12 | 12 | 15 | 13 | 10 | 8 | 8 | 3 | 11 | 15 | 15 | 11 | 9 | 8 | 7 | 4 | 9,96 | 13 | 7 | 0,55 | -0,862 | -1,956 | 0,05045773 | NA | 1,00E-08 | 0,1477609 | 0,077294 | 0,077294 | TRUE | 0,1039792 | No |
| orf19.6540 | 21817 | 14498 | 19590 | 23689 | 26616 | 22236 | 24327 | 15514 | 19572 | 18457 | 20042 | 19534 | 22763 | 21217 | 21708 | 21881 | 20647 | 19401 | 21892 | 1,128 | 0,174 | 4,653 | 3,27E-06 | 1,28E-05 | 0,000966 | 0,0020498 | 0,0012955 | 0,0012955 | TRUE | 0,0129812 | No |
| orf19.6541 | 19678 | 13002 | 16555 | 20823 | 19339 | 15612 | 16947 | 10505 | 17653 | 16553 | 16937 | 17171 | 16539 | 14896 | 15122 | 14816 | 16211 | 17078 | 15343 | 0,898 | -0,154 | -3,522 | 0,00042753 | 0,00117075 | 0,001665 | 0,0020691 | 0,0017837 | 0,0017837 | TRUE | 0,0338462 | No |
| orf19.6544 | 10233 | 6948 | 9100 | 10852 | 15501 | 13228 | 14532 | 8970 | 9180 | 8846 | 9310 | 8949 | 13257 | 12622 | 12967 | 12651 | 10973 | 9071 | 12874 | 1,419 | 0,505 | 15,683 | 1,99E-55 | 2,71E-53 | 0,000418 | 0,0021118 | 0,0009025 | 0,0009025 | TRUE | 0,0049329 | No |
| orf19.6547 | 11 | 8 | 8 | 11 | 15 | 5 | 6 | 8 | 10 | 10 | 8 | 9 | 13 | 5 | 5 | 11 | 8,94 | 9 | 8 | 0,908 | -0,138 | -0,29 | 0,77153393 | NA | 1,00E-08 | 0,1643052 | 0,1058378 | 0,1058378 | TRUE | 0,287619 | No |

| | | | | | | | | | | | | | | | | | | | | | | | | | | | | | | | |
|---|---|---|---|---|---|---|---|---|---|---|---|---|---|---|---|---|---|---|---|---|---|---|---|---|---|---|---|---|---|---|---|
| orf19.6548 | 12275 | 8098 | 10372 | 13072 | 13083 | 11008 | 11938 | 7706 | 11012 | 10310 | 10612 | 10779 | 11189 | 10503 | 10653 | 10869 | 10741 | 10678 | 10804 | 1,011 | 0,016 | 0,469 | 0,63940777 | 0,70969183 | 0,000598 | 0,0021147 | 0,0010867 | 0,0010867 | TRUE | 0,0072204 | No |
| orf19.655 | 46814 | 31658 | 42767 | 51441 | 55048 | 47273 | 51091 | 31929 | 41997 | 40304 | 43755 | 42419 | 47079 | 45106 | 45590 | 45033 | 43910 | 42119 | 45702 | 1,085 | 0,118 | 3,446 | 0,00056911 | 0,00151494 | 0,000757 | 0,0020126 | 0,0010993 | 0,0010993 | TRUE | 0,0102365 | No |
| orf19.6550 | 1196 | 867 | 1011 | 1373 | 1150 | 971 | 1021 | 592 | 1073 | 1104 | 1034 | 1132 | 984 | 926 | 911 | 835 | 999,91 | 1086 | 914 | 0,844 | -0,244 | -4,023 | 5,75E-05 | 0,00018406 | 0,001647 | 0,0034313 | 0,0025261 | 0,0025261 | TRUE | 0,0356998 | No |
| orf19.6551 | 697 | 626 | 598 | 848 | 641 | 693 | 675 | 379 | 625 | 797 | 612 | 699 | 548 | 661 | 602 | 535 | 634,96 | 683 | 586 | 0,861 | -0,215 | -2,223 | 0,02624063 | 0,04775937 | 0,010461 | 0,0042658 | 0,0074177 | 0,0074177 | TRUE | 0,1401274 | No |
| orf19.6551.1 | 140 | 113 | 117 | 135 | 163 | 130 | 163 | 109 | 126 | 144 | 120 | 111 | 139 | 124 | 145 | 154 | 132,89 | 125 | 140 | 1,127 | 0,173 | 1,345 | 0,17868227 | 0,25041359 | 1,00E-08 | 0,0129037 | 0,0082974 | 0,0082974 | TRUE | 0,0890949 | No |
| orf19.6552 | 2047 | 1534 | 1757 | 2236 | 2351 | 2197 | 2429 | 1601 | 1836 | 1953 | 1798 | 1844 | 2011 | 2096 | 2167 | 2258 | 1995,4 | 1858 | 2133 | 1,148 | 0,198 | 4,015 | 5,94E-05 | 0,00018948 | 0,001187 | 0,002707 | 0,0018403 | 0,0018403 | TRUE | 0,0182407 | No |
| orf19.6553 | 980 | 792 | 907 | 1122 | 1035 | 989 | 997 | 635 | 879 | 1008 | 928 | 925 | 885 | 944 | 890 | 896 | 919,34 | 935 | 904 | 0,968 | -0,046 | -0,798 | 0,42473781 | 0,51210612 | 0,000884 | 0,0035586 | 0,0021315 | 0,0021315 | TRUE | 0,0302758 | No |
| orf19.6554 | 6162 | 5707 | 5725 | 7528 | 6990 | 7925 | 7525 | 5061 | 5528 | 7266 | 5857 | 6208 | 5978 | 7562 | 6715 | 7138 | 6531,4 | 6215 | 6848 | 1,102 | 0,14 | 1,587 | 0,11250102 | 0,16847112 | 0,01201 | 0,0022018 | 0,0073492 | 0,0073492 | TRUE | 0,1572271 | No |
| orf19.6555 | 543 | 450 | 467 | 593 | 523 | 572 | 524 | 338 | 487 | 573 | 478 | 489 | 447 | 546 | 468 | 477 | 495,52 | 507 | 484 | 0,958 | -0,061 | -0,71 | 0,47767827 | 0,5629445 | 0,005046 | 0,0049092 | 0,0051112 | 0,0051112 | TRUE | 0,088157 | No |
| orf19.6556 | 8548 | 6346 | 7339 | 9527 | 11712 | 11539 | 11786 | 7553 | 7668 | 8079 | 7509 | 7856 | 10016 | 11010 | 10517 | 10653 | 9163,6 | 7778 | 10549 | 1,356 | 0,44 | 10,831 | 2,46E-27 | 7,73E-26 | 0,001112 | 0,0021379 | 0,0014707 | 0,0014707 | TRUE | 0,0143024 | No |
| orf19.6557 | 698 | 512 | 578 | 718 | 776 | 674 | 795 | 497 | 626 | 652 | 591 | 592 | 664 | 643 | 709 | 701 | 647,32 | 615 | 679 | 1,105 | 0,144 | 2,242 | 0,02499253 | 0,04562598 | 0,000578 | 0,0042221 | 0,002393 | 0,002393 | TRUE | 0,0166682 | No |
| orf19.6558 | 2215 | 1658 | 2066 | 2595 | 3002 | 2967 | 3064 | 1960 | 1987 | 2111 | 2114 | 2140 | 2567 | 2831 | 2734 | 2764 | 2406,1 | 2088 | 2724 | 1,304 | 0,383 | 8,27 | 1,34E-16 | 1,79E-15 | 0,00109 | 0,0025829 | 0,0016383 | 0,0016383 | TRUE | 0,0188806 | No |
| orf19.6559 | 1522 | 1049 | 1438 | 1688 | 2137 | 1772 | 1911 | 1297 | 1365 | 1335 | 1471 | 1392 | 1828 | 1691 | 1705 | 1829 | 1577,1 | 1391 | 1763 | 1,266 | 0,34 | 6,677 | 2,45E-11 | 1,92E-10 | 0,001102 | 0,0029 | 0,0018481 | 0,0018481 | TRUE | 0,0175956 | No |
| orf19.656 | 78 | 46 | 63 | 76 | 82 | 60 | 68 | 46 | 70 | 59 | 64 | 63 | 70 | 57 | 61 | 65 | 63,57 | 64 | 63 | 0,986 | -0,021 | -0,122 | 0,90328285 | 0,92752065 | 1,00E-08 | 0,024814 | 0,0124945 | 0,0124945 | TRUE | 0,0572987 | No |
| orf19.6561 | 6085 | 3878 | 5224 | 6303 | 6769 | 5740 | 6275 | 4286 | 5459 | 4937 | 5345 | 5198 | 5789 | 5477 | 5599 | 6045 | 5481,1 | 5235 | 5728 | 1,093 | 0,128 | 2,795 | 0,0051919 | 0,01118428 | 0,001616 | 0,0022444 | 0,0018457 | 0,0018457 | TRUE | 0,0173413 | No |
| orf19.6562 | 337 | 246 | 297 | 427 | 334 | 234 | 267 | 204 | 302 | 313 | 304 | 352 | 286 | 223 | 238 | 288 | 288,3 | 318 | 259 | 0,81 | -0,304 | -2,899 | 0,0037477 | 0,0083147 | 0,0071 | 0,007015 | 0,0070608 | 0,0070608 | TRUE | 0,0944675 | No |
| orf19.6563 | 2019 | 1462 | 1845 | 2239 | 2348 | 2046 | 2222 | 1481 | 1811 | 1861 | 1888 | 1846 | 2008 | 1952 | 1983 | 2089 | 1929,8 | 1852 | 2008 | 1,083 | 0,116 | 2,826 | 0,00471007 | 0,01023069 | 1,00E-08 | 0,0027318 | 0,0010842 | 0,0010842 | TRUE | 0,0078475 | No |
| orf19.6563.1 | 2839 | 1815 | 2175 | 2740 | 3385 | 2616 | 2717 | 1424 | 2547 | 2311 | 2225 | 2259 | 2895 | 2496 | 2424 | 2008 | 2395,8 | 2336 | 2456 | 1,053 | 0,074 | 0,803 | 0,42214348 | 0,50977463 | 0,011718 | 0,0025855 | 0,0077792 | 0,0077792 | TRUE | 0,1782165 | No |
| orf19.6565 | 1070 | 719 | 916 | 1141 | 1117 | 946 | 1056 | 664 | 960 | 915 | 937 | 941 | 955 | 903 | 942 | 937 | 936,26 | 938 | 934 | 0,995 | -0,007 | -0,147 | 0,88294078 | 0,91146189 | 1,00E-08 | 0,0035301 | 0,0013968 | 0,0013968 | TRUE | 0,0065099 | No |
| orf19.6566 | 1044 | 852 | 942 | 1215 | 1115 | 979 | 1113 | 685 | 937 | 1085 | 964 | 1002 | 954 | 934 | 993 | 966 | 979,24 | 997 | 962 | 0,967 | -0,049 | -0,831 | 0,40586727 | 0,49411359 | 0,001234 | 0,003462 | 0,0022608 | 0,0022608 | TRUE | 0,0385678 | No |
| orf19.6568 | 5524 | 3925 | 4835 | 5770 | 5365 | 5018 | 5270 | 2883 | 4956 | 4997 | 4947 | 4758 | 4588 | 4788 | 4703 | 4066 | 4725,3 | 4914 | 4536 | 0,924 | -0,114 | -2,16 | 0,0307848 | 0,05485026 | 0,002501 | 0,0022867 | 0,0024503 | 0,0024503 | TRUE | 0,0572061 | No |
| orf19.6569 | 674 | 520 | 628 | 757 | 449 | 463 | 493 | 244 | 605 | 662 | 643 | 624 | 384 | 442 | 440 | 344 | 517,91 | 634 | 402 | 0,64 | -0,645 | -7,786 | 6,91E-15 | 7,88E-14 | 0,004269 | 0,0047825 | 0,00455 | 0,00455 | TRUE | 0,0923971 | No |
| orf19.657 | 1338 | 841 | 1088 | 1347 | 1246 | 1058 | 1103 | 803 | 1200 | 1071 | 1113 | 1111 | 1066 | 1010 | 984 | 1133 | 1085,9 | 1124 | 1048 | 0,929 | -0,106 | -1,743 | 0,08132606 | 0,12837808 | 0,001927 | 0,0033164 | 0,0026073 | 0,0026073 | TRUE | 0,0316929 | No |
| orf19.6570 | 208 | 147 | 187 | 215 | 142 | 130 | 154 | 99 | 187 | 187 | 191 | 177 | 121 | 124 | 137 | 140 | 158,11 | 186 | 130 | 0,702 | -0,511 | -4,583 | 4,58E-06 | 1,75E-05 | 1,00E-08 | 0,011161 | 0,0054231 | 0,0054231 | TRUE | 0,0243959 | No |
| orf19.6573 | 2950 | 2451 | 2952 | 3469 | 3758 | 3829 | 3931 | 2303 | 2646 | 3120 | 3020 | 2861 | 3214 | 3654 | 3508 | 3248 | 3158,9 | 2912 | 3406 | 1,171 | 0,228 | 3,636 | 0,00027726 | 0,00078797 | 0,004014 | 0,0024391 | 0,0034432 | 0,0034432 | TRUE | 0,0458662 | No |
| orf19.6574 | 22 | 14 | 10 | 33 | 29 | 33 | 28 | 18 | 20 | 18 | 10 | 27 | 25 | 31 | 25 | 25 | 22,71 | 19 | 26 | 1,403 | 0,488 | 1,547 | 0,12181797 | 0,18075633 | 0,025884 | 0,0659082 | 0,0507917 | 0,0507917 | TRUE | 0,3707782 | No |
| orf19.6575 | 1 | 1 | 2 | 2 | 2 | 1 | 5 | 5 | 1 | 1 | 2 | 2 | 2 | 1 | 4 | 7 | 2,51 | 2 | 4 | 2,303 | 1,204 | 1,262 | 0,2069272 | NA | 1,00E-08 | 0,5813857 | 0,3883747 | 0,3883747 | TRUE | 0,209151 | No |
| orf19.6577 | 11967 | 8400 | 10789 | 13583 | 12358 | 11112 | 11530 | 7942 | 10736 | 10694 | 11038 | 11201 | 10569 | 10603 | 10289 | 11201 | 10791 | 10917 | 10666 | 0,976 | -0,035 | -0,952 | 0,34101579 | 0,42799725 | 0,000773 | 0,002114 | 0,0011712 | 0,0011712 | TRUE | 0,0136154 | No |
| orf19.6578 | 13146 | 9369 | 12078 | 14438 | 18536 | 17412 | 18835 | 11809 | 11793 | 11928 | 12357 | 11906 | 15853 | 16614 | 16807 | 16656 | 14239 | 11996 | 16482 | 1,374 | 0,458 | 14,276 | 3,10E-46 | 2,58E-44 | 0,000454 | 0,0020815 | 0,0009174 | 0,0009174 | TRUE | 0,0081966 | No |
| orf19.6579 | 812 | 732 | 764 | 909 | 715 | 853 | 932 | 487 | 728 | 932 | 782 | 750 | 611 | 814 | 832 | 687 | 766,94 | 798 | 736 | 0,925 | -0,113 | -1,052 | 0,29293894 | 0,37759647 | 0,01475 | 0,0038724 | 0,0097735 | 0,0097735 | TRUE | 0,1608406 | No |
| orf19.658 | 251 | 208 | 227 | 258 | 245 | 234 | 250 | 151 | 225 | 265 | 232 | 213 | 210 | 223 | 223 | 213 | 225,48 | 234 | 217 | 0,936 | -0,096 | -0,969 | 0,33264332 | 0,41941681 | 1,00E-08 | 0,0084178 | 0,0049729 | 0,0049729 | TRUE | 0,0783681 | No |
| orf19.6580 | 4643 | 3365 | 4082 | 5233 | 5318 | 4788 | 5040 | 3317 | 4165 | 4284 | 4176 | 4315 | 4548 | 4569 | 4497 | 4678 | 4404,1 | 4235 | 4573 | 1,079 | 0,11 | 3,423 | 0,00061836 | 0,00162657 | 1,00E-08 | 0,0023091 | 0,0007613 | 0,0007613 | TRUE | 0,0026927 | No |
| orf19.6581 | 939 | 692 | 839 | 1036 | 875 | 824 | 874 | 489 | 842 | 881 | 858 | 854 | 748 | 786 | 780 | 690 | 805,03 | 859 | 751 | 0,878 | -0,188 | -3,229 | 0,00124317 | 0,00307365 | 3,47E-05 | 0,0037828 | 0,0019918 | 0,0019918 | TRUE | 0,0301466 | No |
| orf19.6582 | 5904 | 4284 | 5135 | 6479 | 5855 | 5244 | 5503 | 3248 | 5297 | 5454 | 5254 | 5343 | 5007 | 5004 | 4911 | 4581 | 5106,2 | 5337 | 4876 | 0,915 | -0,129 | -3,338 | 0,00084441 | 0,00215973 | 0,000723 | 0,0022638 | 0,0012268 | 0,0012268 | TRUE | 0,0190385 | No |
| orf19.6583 | 246 | 225 | 258 | 349 | 266 | 291 | 317 | 205 | 221 | 286 | 264 | 288 | 227 | 278 | 283 | 289 | 267,01 | 265 | 269 | 1,016 | 0,022 | 0,197 | 0,84387649 | 0,8811279 | 0,009907 | 0,0074165 | 0,0086304 | 0,0086304 | TRUE | 0,1466576 | No |
| orf19.6584 | 9868 | 7518 | 8847 | 10850 | 9405 | 8861 | 9341 | 5707 | 8853 | 9571 | 9051 | 8947 | 8043 | 8455 | 8335 | 8049 | 8663,1 | 9106 | 8220 | 0,903 | -0,147 | -3,949 | 7,85E-05 | 0,00024348 | 0,000784 | 0,0021471 | 0,0012077 | 0,0012077 | TRUE | 0,0141866 | No |
| orf19.6585 | 653 | 522 | 593 | 728 | 660 | 713 | 660 | 421 | 586 | 665 | 607 | 600 | 564 | 680 | 589 | 594 | 610,61 | 614 | 607 | 0,99 | -0,015 | -0,192 | 0,84746513 | 0,88395321 | 0,003326 | 0,004357 | 0,0038195 | 0,0038195 | TRUE | 0,0807538 | No |
| orf19.6586 | 2522 | 1868 | 2352 | 2571 | 3178 | 3100 | 3298 | 2015 | 2262 | 2378 | 2406 | 2120 | 2718 | 2958 | 2943 | 2842 | 2578,5 | 2292 | 2865 | 1,252 | 0,324 | 6,184 | 6,23E-10 | 4,08E-09 | 0,001841 | 0,0025425 | 0,0022357 | 0,0022357 | TRUE | 0,031604 | No |
| orf19.6587 | 1541 | 1260 | 1392 | 1870 | 1829 | 1883 | 1905 | 1131 | 1382 | 1604 | 1424 | 1542 | 1564 | 1797 | 1700 | 1595 | 1576,1 | 1488 | 1664 | 1,12 | 0,163 | 2,523 | 0,01162255 | 0,02302284 | 0,003553 | 0,0029006 | 0,0033685 | 0,0033685 | TRUE | 0,0349497 | No |
| orf19.6588 | 1763 | 1372 | 1718 | 2122 | 2563 | 2467 | 2531 | 1472 | 1582 | 1747 | 1758 | 1750 | 2192 | 2354 | 2258 | 2076 | 1964,5 | 1709 | 2220 | 1,301 | 0,38 | 6,938 | 3,99E-12 | 3,42E-11 | 0,001931 | 0,0027185 | 0,0023586 | 0,0023586 | TRUE | 0,0311328 | No |
| orf19.6589 | 7250 | 5670 | 6326 | 8048 | 6977 | 7157 | 7339 | 4236 | 6504 | 7219 | 6472 | 6636 | 5967 | 6829 | 6549 | 5974 | 6518,8 | 6708 | 6330 | 0,944 | -0,083 | -1,457 | 0,1450731 | 0,20963741 | 0,003449 | 0,0022022 | 0,0029503 | 0,0029503 | TRUE | 0,0343688 | No |
| orf19.6590 | 94 | 78 | 63 | 88 | 123 | 128 | 101 | 68 | 84 | 99 | 64 | 73 | 105 | 122 | 90 | 96 | 91,75 | 80 | 103 | 1,301 | 0,38 | 2,26 | 0,02380185 | 0,04366458 | 0,013605 | 0,0178016 | 0,0160342 | 0,0160342 | TRUE | 0,201733 | No |
| orf19.6591 | 2001 | 1598 | 1903 | 2374 | 2300 | 2136 | 2236 | 1532 | 1795 | 2034 | 1947 | 1958 | 1967 | 2038 | 1995 | 2161 | 1986,9 | 1934 | 2040 | 1,055 | 0,077 | 1,481 | 0,13858718 | 0,20155695 | 0,001548 | 0,0027102 | 0,0020751 | 0,0020751 | TRUE | 0,0284215 | No |
| orf19.6592 | 415 | 272 | 362 | 435 | 581 | 552 | 598 | 358 | 372 | 346 | 370 | 359 | 497 | 527 | 534 | 505 | 438,72 | 362 | 516 | 1,423 | 0,509 | 7,25 | 4,16E-13 | 4,04E-12 | 1,00E-08 | 0,0052884 | 0,0024 | 0,0024 | TRUE | 0,0081732 | No |
| orf19.6594 | 20720 | 15777 | 18863 | 23296 | 24334 | 23595 | 24880 | 14880 | 18588 | 20086 | 19299 | 19210 | 20811 | 22514 | 22201 | 20987 | 20462 | 19296 | 21628 | 1,121 | 0,165 | 4,128 | 3,66E-05 | 0,00012122 | 0,001222 | 0,0020505 | 0,0014846 | 0,0014846 | TRUE | 0,0092928 | No |
| orf19.6595 | 88 | 81 | 75 | 116 | 65 | 55 | 57 | 35 | 79 | 103 | 77 | 96 | 56 | 52 | 51 | 49 | 70,34 | 89 | 52 | 0,592 | -0,756 | -4,364 | 1,28E-05 | 4,58E-05 | 1,00E-08 | 0,0226167 | 0,0136417 | 0,0136417 | TRUE | 0,0949935 | No |
| orf19.6596 | 1287 | 1162 | 1360 | 1713 | 1805 | 2004 | 1915 | 1355 | 1155 | 1479 | 1391 | 1413 | 1544 | 1912 | 1709 | 1911 | 1564,2 | 1360 | 1769 | 1,301 | 0,38 | 4,289 | 1,80E-05 | 6,29E-05 | 0,009625 | 0,0029076 | 0,006869 | 0,006869 | TRUE | 0,1251587 | No |
| orf19.6597 | 923 | 878 | 865 | 1102 | 918 | 1188 | 1174 | 606 | 828 | 1118 | 885 | 909 | 785 | 1134 | 1048 | 855 | 945,06 | 935 | 956 | 1,024 | 0,034 | 0,283 | 0,77699152 | 0,82505568 | 0,02208 | 0,0035156 | 0,0127282 | 0,0127282 | TRUE | 0,2018958 | No |
| orf19.6598 | 2443 | 1552 | 2255 | 2694 | 3418 | 2989 | 3315 | 2202 | 2192 | 1976 | 2307 | 2222 | 2923 | 2852 | 2958 | 3106 | 2566,9 | 2174 | 2960 | 1,359 | 0,442 | 8,306 | 9,93E-17 | 1,36E-15 | 0,002219 | 0,0025451 | 0,0023243 | 0,0023243 | TRUE | 0,0432518 | No |
| orf19.6599.1 | 2965 | 1798 | 2171 | 2687 | 2579 | 1903 | 1927 | 969 | 2660 | 2289 | 2221 | 2216 | 2206 | 1816 | 1720 | 1367 | 2061,7 | 2346 | 1777 | 0,758 | -0,4 | -2,6 | 0,00931866 | 0,01891437 | 0,022246 | 0,0026837 | 0,0122436 | 0,0222465 | TRUE | 0,319272 | No |
| orf19.660 | 1350 | 881 | 1213 | 1415 | 1450 | 1230 | 1385 | 899 | 1211 | 1122 | 1241 | 1167 | 1240 | 1174 | 1236 | 1268 | 1207,3 | 1185 | 1230 | 1,035 | 0,05 | 0,974 | 0,33027333 | 0,41730435 | 0,000506 | 0,003182 | 0,0017167 | 0,0017167 | TRUE | 0,0139418 | No |
| orf19.6600 | 1717 | 1473 | 1571 | 1937 | 2029 | 2151 | 2092 | 1165 | 1540 | 1875 | 1607 | 1597 | 1735 | 2052 | 1867 | 1643 | 1739,7 | 1655 | 1824 | 1,105 | 0,144 | 1,795 | 0,07265261 | 0,11636981 | 0,008062 | 0,002814 | 0,0055631 | 0,0055631 | TRUE | 0,0942853 | No |
| orf19.6601 | 9968 | 7185 | 8671 | 11541 | 11864 | 11490 | 11880 | 6829 | 8942 | 9147 | 8871 | 9517 | 10146 | 10963 | 10601 | 9632 | 9727,5 | 9119 | 10336 | 1,134 | 0,181 | 3,858 | 0,00011422 | 0,0003457 | 0,00192 | 0,0021288 | 0,0020104 | 0,0020104 | TRUE | 0,0251864 | No |
| orf19.6601.1 | 17727 | 13622 | 14865 | 18798 | 23400 | 22578 | 22798 | 13295 | 15903 | 17342 | 15208 | 15501 | 20012 | 21543 | 20343 | 18751 | 18076 | 15988 | 20162 | 1,261 | 0,335 | 6,135 | 8,50E-10 | 5,46E-09 | 0,003297 | 0,0020598 | 0,0028087 | 0,0028087 | TRUE | 0,0395068 | No |
| orf19.6602 | 735 | 855 | 788 | 1105 | 933 | 1189 | 1122 | 727 | 659 | 1089 | 806 | 911 | 798 | 1135 | 1001 | 1025 | 928,04 | 866 | 990 | 1,143 | 0,193 | 1,072 | 0,28363949 | 0,36789855 | 0,029996 | 0,0035438 | 0,0168505 | 0,0299963 | TRUE | 0,3130741 | No |
| orf19.6604 | 3565 | 3222 | 3439 | 4291 | 5319 | 6054 | 6020 | 3740 | 3198 | 4102 | 3518 | 3538 | 4549 | 5777 | 5372 | 5275 | 4416,2 | 3589 | 5243 | 1,461 | 0,547 | 6,542 | 6,06E-11 | 4,57E-10 | 0,009918 | 0,0023082 | 0,0064884 | 0,0064884 | TRUE | 0,1112636 | No |
| orf19.6605 | 931 | 878 | 854 | 1140 | 638 | 679 | 703 | 461 | 835 | 1118 | 874 | 940 | 546 | 648 | 627 | 650 | 779,73 | 942 | 618 | 0,656 | -0,608 | -6,391 | 1,65E-10 | 1,15E-09 | 0,010208 | 0,0038413 | 0,0073283 | 0,0073283 | TRUE | 0,1819693 | No |
| orf19.6606 | 338 | 346 | 360 | 436 | 239 | 271 | 230 | 194 | 303 | 440 | 368 | 360 | 204 | 259 | 205 | 274 | 301,67 | 368 | 236 | 0,638 | -0,647 | -4,951 | 7,38E-07 | 3,18E-06 | 0,018715 | 0,0067917 | 0,0128893 | 0,0128893 | TRUE | 0,1916913 | No |
| orf19.6607 | 3618 | 2880 | 3052 | 3780 | 3111 | 3198 | 2979 | 1500 | 3246 | 3667 | 3122 | 3117 | 2661 | 3051 | 2658 | 2116 | 2954,7 | 3288 | 2622 | 0,798 | -0,325 | -3,435 | 0,0005917 | 0,00156743 | 0,013412 | 0,0024708 | 0,0082447 | 0,0082447 | TRUE | 0,2001468 | No |

| | | | | | | | | | | | | | | | | | | | | | | | | | | | | | | | |
|---|---|---|---|---|---|---|---|---|---|---|---|---|---|---|---|---|---|---|---|---|---|---|---|---|---|---|---|---|---|---|---|
| orf19.6608 | 1961 | 1467 | 1816 | 2303 | 2937 | 2812 | 3061 | 1752 | 1759 | 1868 | 1858 | 1899 | 2512 | 2683 | 2731 | 2471 | 2222,7 | 1846 | 2599 | 1,41 | 0,495 | 10,117 | 4,66E-24 | 1,09E-22 | 0,001345 | 0,0026327 | 0,001837 | 0,001837 | TRUE | 0,014212 | No |
| orf19.661 | 9801 | 6972 | 8207 | 10308 | 10545 | 9457 | 9686 | 6395 | 8793 | 8876 | 8397 | 8500 | 9018 | 9024 | 8643 | 9020 | 8783,8 | 8642 | 8926 | 1,033 | 0,047 | 1,414 | 0,1573306 | 0,22436321 | 0,000424 | 0,0021448 | 0,0009408 | 0,0009408 | TRUE | 0,0056518 | No |
| orf19.6610 | 281 | 194 | 279 | 307 | 334 | 315 | 332 | 227 | 252 | 247 | 285 | 253 | 286 | 301 | 296 | 320 | 280,04 | 259 | 301 | 1,154 | 0,207 | 2,354 | 0,01859113 | 0,03480695 | 1,00E-08 | 0,0071635 | 0,0038582 | 0,0038582 | TRUE | 0,0493513 | No |
| orf19.6612 | 949 | 862 | 916 | 1224 | 1074 | 1177 | 1201 | 674 | 851 | 1097 | 937 | 1009 | 919 | 1123 | 1072 | 951 | 994,89 | 974 | 1016 | 1,045 | 0,064 | 0,721 | 0,47060588 | 0,55690371 | 0,009329 | 0,0034387 | 0,006563 | 0,006563 | TRUE | 0,0863135 | No |
| orf19.6621 | 11927 | 8241 | 10880 | 12573 | 15333 | 14187 | 15491 | 8977 | 10700 | 10492 | 11131 | 10368 | 13113 | 13537 | 13823 | 12661 | 11978 | 10673 | 13284 | 1,245 | 0,316 | 7,96 | 1,72E-15 | 2,08E-14 | 0,001109 | 0,0021007 | 0,0014303 | 0,0014303 | TRUE | 0,0119185 | No |
| orf19.6623 | 1776 | 1297 | 1487 | 1995 | 1682 | 1489 | 1422 | 929 | 1593 | 1651 | 1521 | 1645 | 1438 | 1421 | 1269 | 1310 | 1481,2 | 1602 | 1360 | 0,849 | -0,236 | -4,197 | 2,71E-05 | 9,19E-05 | 0,001902 | 0,0029596 | 0,0023459 | 0,0023459 | TRUE | 0,02632 | No |
| orf19.6624 | 1001 | 734 | 983 | 1125 | 887 | 749 | 782 | 521 | 898 | 934 | 1006 | 928 | 759 | 715 | 698 | 735 | 833,97 | 942 | 727 | 0,772 | -0,373 | -6,307 | 2,85E-10 | 1,96E-09 | 0,000823 | 0,0037202 | 0,0021454 | 0,0021454 | TRUE | 0,0252735 | No |
| orf19.6625 | 1183 | 832 | 986 | 1252 | 961 | 874 | 921 | 513 | 1061 | 1059 | 1009 | 1032 | 822 | 834 | 822 | 724 | 920,36 | 1040 | 800 | 0,773 | -0,372 | -6,456 | 1,07E-10 | 7,78E-10 | 0,000512 | 0,0035568 | 0,0020886 | 0,0020886 | TRUE | 0,0422173 | No |
| orf19.6626 | 241 | 149 | 192 | 296 | 237 | 186 | 220 | 147 | 216 | 190 | 196 | 244 | 203 | 177 | 196 | 207 | 203,78 | 212 | 196 | 0,919 | -0,122 | -1,102 | 0,27057131 | 0,35493285 | 0,004374 | 0,0091035 | 0,0069249 | 0,0069249 | TRUE | 0,12302 | No |
| orf19.6627 | 1494 | 930 | 1236 | 1478 | 1362 | 1090 | 1110 | 771 | 1340 | 1184 | 1265 | 1219 | 1165 | 1040 | 990 | 1087 | 1161,3 | 1252 | 1070 | 0,854 | -0,227 | -3,515 | 0,00043984 | 0,00120117 | 0,003059 | 0,0032296 | 0,0031326 | 0,0031326 | TRUE | 0,0450076 | No |
| orf19.6628 | 664 | 429 | 516 | 691 | 617 | 433 | 463 | 321 | 596 | 546 | 528 | 570 | 528 | 413 | 413 | 453 | 505,79 | 560 | 452 | 0,807 | -0,31 | -3,468 | 0,00052385 | 0,00140692 | 0,006449 | 0,0048497 | 0,0056756 | 0,0056756 | TRUE | 0,1604395 | No |
| orf19.6629 | 418 | 275 | 359 | 483 | 462 | 369 | 402 | 265 | 375 | 350 | 367 | 398 | 395 | 352 | 359 | 374 | 371,29 | 372 | 370 | 0,99 | -0,015 | -0,191 | 0,84828953 | 0,88415642 | 1,00E-08 | 0,0058894 | 0,0032988 | 0,0032988 | TRUE | 0,0273665 | No |
| orf19.663 | 1362 | 974 | 1150 | 1418 | 1315 | 1220 | 1321 | 755 | 1222 | 1240 | 1177 | 1169 | 1125 | 1164 | 1179 | 1065 | 1167,5 | 1202 | 1133 | 0,946 | -0,081 | -1,569 | 0,11654458 | 0,17370375 | 0,000276 | 0,0032229 | 0,0016696 | 0,0016696 | TRUE | 0,01703 | No |
| orf19.6630 | 311 | 196 | 242 | 288 | 364 | 246 | 252 | 142 | 279 | 250 | 248 | 237 | 311 | 235 | 225 | 200 | 248,1 | 254 | 243 | 0,964 | -0,053 | -0,409 | 0,68222606 | 0,74605281 | 0,016603 | 0,0078308 | 0,0121095 | 0,0121095 | TRUE | 0,4120149 | No |
| orf19.6632 | 1560 | 1046 | 1434 | 1685 | 2119 | 1702 | 1848 | 1252 | 1399 | 1332 | 1467 | 1389 | 1812 | 1624 | 1649 | 1766 | 1554,9 | 1397 | 1713 | 1,225 | 0,292 | 5,442 | 5,27E-08 | 2,68E-07 | 0,001568 | 0,0029132 | 0,0021206 | 0,0021206 | TRUE | 0,0200164 | No |
| orf19.6634 | 1586 | 1059 | 1320 | 1690 | 1720 | 1502 | 1685 | 1017 | 1423 | 1348 | 1350 | 1394 | 1471 | 1433 | 1504 | 1434 | 1419,7 | 1379 | 1460 | 1,059 | 0,083 | 1,845 | 0,06497882 | 0,10579749 | 1,00E-08 | 0,0030021 | 0,0012315 | 0,0012315 | TRUE | 0,0055334 | No |
| orf19.6635 | 1152 | 855 | 1009 | 1261 | 866 | 829 | 886 | 510 | 1033 | 1089 | 1032 | 1040 | 741 | 791 | 791 | 719 | 904,46 | 1048 | 760 | 0,728 | -0,459 | -8,346 | 7,07E-17 | 9,90E-16 | 1,00E-08 | 0,0035846 | 0,0017665 | 0,0017665 | TRUE | 0,0130166 | No |
| orf19.6636 | 108 | 63 | 93 | 119 | 76 | 86 | 84 | 48 | 97 | 80 | 95 | 98 | 65 | 82 | 75 | 68 | 82,51 | 92 | 72 | 0,779 | -0,36 | -2,316 | 0,02053855 | 0,03816752 | 1,00E-08 | 0,0195737 | 0,0109352 | 0,0109352 | TRUE | 0,0745857 | No |
| orf19.6637 | 3628 | 2505 | 3168 | 3911 | 3699 | 3198 | 3517 | 2209 | 3255 | 3189 | 3241 | 3225 | 3164 | 3051 | 3138 | 3116 | 3172,4 | 3228 | 3117 | 0,966 | -0,05 | -1,525 | 0,12735584 | 0,18776782 | 1,00E-08 | 0,0024371 | 0,0007296 | 0,0007296 | TRUE | 0,0025721 | No |
| orf19.6638 | 978 | 732 | 783 | 1067 | 985 | 880 | 849 | 560 | 877 | 932 | 801 | 880 | 842 | 840 | 758 | 790 | 839,97 | 872 | 808 | 0,927 | -0,11 | -1,7 | 0,08909914 | 0,13875263 | 0,001992 | 0,0037078 | 0,0027925 | 0,0027925 | TRUE | 0,0349794 | No |
| orf19.6639 | 3148 | 2149 | 2752 | 3407 | 3986 | 3582 | 3768 | 2397 | 2824 | 2736 | 2816 | 2809 | 3409 | 3418 | 3362 | 3381 | 3094,4 | 2796 | 3392 | 1,213 | 0,278 | 8,461 | 2,64E-17 | 3,86E-16 | 1,00E-08 | 0,0024487 | 0,0007108 | 0,0007108 | TRUE | 0,0023348 | No |
| orf19.664 | 87 | 57 | 72 | 93 | 85 | 93 | 87 | 63 | 78 | 73 | 74 | 77 | 73 | 89 | 78 | 89 | 78,61 | 76 | 82 | 1,08 | 0,111 | 0,716 | 0,47394575 | 0,5597535 | 1,00E-08 | 0,0204464 | 0,0103233 | 0,0103233 | TRUE | 0,0586211 | No |
| orf19.6640 | 5560 | 3987 | 4893 | 5998 | 9078 | 8090 | 8555 | 5240 | 4988 | 5076 | 5006 | 4946 | 7764 | 7719 | 7634 | 7391 | 6315,4 | 5004 | 7627 | 1,526 | 0,609 | 19,996 | 5,95E-89 | 2,55E-86 | 0,00013 | 0,0022094 | 0,0007257 | 0,0007257 | TRUE | 0,0050078 | No |
| orf19.6641 | 695 | 458 | 612 | 675 | 1179 | 1032 | 1030 | 733 | 623 | 583 | 626 | 557 | 1008 | 985 | 919 | 1034 | 791,91 | 597 | 986 | 1,65 | 0,722 | 11,197 | 4,21E-29 | 1,44E-27 | 0,001621 | 0,0038127 | 0,0026482 | 0,0026482 | TRUE | 0,0277538 | No |
| orf19.6642 | 1916 | 1370 | 1701 | 2164 | 1921 | 1789 | 1939 | 1360 | 1719 | 1744 | 1740 | 1784 | 1643 | 1707 | 1730 | 1918 | 1748,3 | 1747 | 1750 | 0,999 | -0,002 | -0,033 | 0,9735786 | 0,98139755 | 0,001487 | 0,0028099 | 0,0021014 | 0,0021014 | TRUE | 0,046975 | No |
| orf19.6643 | 3374 | 2293 | 2974 | 3604 | 3384 | 2846 | 3357 | 2300 | 3027 | 2919 | 3043 | 2972 | 2894 | 2716 | 2996 | 3244 | 2976,2 | 2990 | 2962 | 0,989 | -0,016 | -0,289 | 0,77247454 | 0,82214828 | 0,00219 | 0,0024673 | 0,0024746 | 0,0024746 | TRUE | 0,047441 | No |
| orf19.6644 | 198 | 128 | 170 | 211 | 182 | 192 | 245 | 167 | 178 | 163 | 174 | 174 | 156 | 183 | 219 | 236 | 185,19 | 172 | 198 | 1,142 | 0,191 | 1,455 | 0,14554373 | 0,21019873 | 0,012493 | 0,0098185 | 0,0110807 | 0,0110807 | TRUE | 0,2386923 | No |
| orf19.6645 | 18201 | 11406 | 14864 | 19535 | 18657 | 15509 | 16704 | 11931 | 16328 | 14521 | 15207 | 16109 | 15956 | 14798 | 14906 | 16828 | 15582 | 15541 | 15622 | 1,005 | 0,007 | 0,127 | 0,89916263 | 0,92360559 | 0,002797 | 0,0020727 | 0,0028036 | 0,0028036 | TRUE | 0,0327386 | No |
| orf19.6648 | 22616 | 15333 | 19744 | 24136 | 19444 | 16582 | 18279 | 10578 | 20289 | 19521 | 20200 | 19903 | 16629 | 15822 | 16311 | 14919 | 17949 | 19978 | 15920 | 0,797 | -0,327 | -8,259 | 1,47E-16 | 1,95E-15 | 0,001171 | 0,0020604 | 0,0014501 | 0,0014501 | TRUE | 0,0215751 | No |
| orf19.6649 | 4783 | 3056 | 4064 | 4750 | 4787 | 4057 | 4538 | 2736 | 4291 | 3891 | 4158 | 3917 | 4094 | 3871 | 4049 | 3859 | 4016,2 | 4064 | 3968 | 0,977 | -0,034 | -0,757 | 0,44909931 | 0,53561462 | 0,001261 | 0,002341 | 0,0016956 | 0,0016956 | TRUE | 0,0173789 | No |
| orf19.665 | 5406 | 3866 | 4495 | 5717 | 4785 | 4350 | 4703 | 2777 | 4850 | 4922 | 4599 | 4714 | 4092 | 4151 | 4197 | 3917 | 4430,1 | 4771 | 4089 | 0,858 | -0,221 | -5,754 | 8,71E-09 | 4,85E-08 | 0,000599 | 0,0023072 | 0,001187 | 0,001187 | TRUE | 0,0091504 | No |
| orf19.6650 | 652 | 408 | 505 | 659 | 588 | 528 | 577 | 291 | 585 | 519 | 517 | 543 | 503 | 504 | 515 | 410 | 512,05 | 541 | 483 | 0,896 | -0,158 | -1,928 | 0,05382258 | 0,08962475 | 0,00417 | 0,0048146 | 0,0044959 | 0,0044959 | TRUE | 0,1028727 | No |
| orf19.6652 | 6950 | 4564 | 6122 | 7328 | 6616 | 5646 | 6209 | 4006 | 6235 | 5810 | 6263 | 6043 | 5658 | 5387 | 5541 | 5650 | 5823,4 | 6088 | 5559 | 0,913 | -0,132 | -3,543 | 0,00039631 | 0,00109273 | 0,00062 | 0,0022288 | 0,0011589 | 0,0011589 | TRUE | 0,0110414 | No |
| orf19.6653 | 2861 | 1979 | 2714 | 3249 | 3067 | 2478 | 2892 | 1589 | 2567 | 2519 | 2777 | 2679 | 2623 | 2364 | 2581 | 2241 | 2543,9 | 2636 | 2452 | 0,932 | -0,102 | -1,736 | 0,08248454 | 0,1298656 | 0,00315 | 0,0025502 | 0,0029347 | 0,0029347 | TRUE | 0,0388163 | No |
| orf19.6654 | 6317 | 4343 | 5783 | 7187 | 6560 | 5384 | 5778 | 3756 | 5667 | 5529 | 5917 | 5926 | 5610 | 5137 | 5156 | 5298 | 5530 | 5760 | 5300 | 0,92 | -0,12 | -2,807 | 0,00500255 | 0,0108035 | 0,001184 | 0,002242 | 0,0015808 | 0,0015808 | TRUE | 0,0194334 | No |
| orf19.6656 | 2084 | 1343 | 1802 | 2243 | 2558 | 2051 | 2311 | 1471 | 1870 | 1710 | 1844 | 1850 | 2188 | 1957 | 2062 | 2075 | 1944,3 | 1818 | 2070 | 1,138 | 0,186 | 3,715 | 0,00020345 | 0,0005928 | 0,001343 | 0,0027262 | 0,0018909 | 0,0018909 | TRUE | 0,0185071 | No |
| orf19.6658 | 5189 | 3386 | 4512 | 5353 | 5295 | 4428 | 4720 | 3067 | 4655 | 4311 | 4616 | 4414 | 4528 | 4225 | 4212 | 4326 | 4410,9 | 4499 | 4323 | 0,961 | -0,058 | -1,386 | 0,16563057 | 0,23502798 | 0,000935 | 0,0023086 | 0,0014543 | 0,0014543 | TRUE | 0,0129512 | No |
| orf19.6659 | 39369 | 25911 | 35663 | 43286 | 41019 | 34069 | 38302 | 24353 | 35318 | 32987 | 36487 | 35694 | 35081 | 32508 | 34178 | 34348 | 34575 | 35122 | 34029 | 0,969 | -0,046 | -1,109 | 0,26755881 | 0,3518427 | 0,001388 | 0,0020215 | 0,0016061 | 0,0016061 | TRUE | 0,0203879 | No |
| orf19.666 | 1045 | 779 | 993 | 1147 | 976 | 891 | 1022 | 627 | 937 | 992 | 1016 | 946 | 835 | 850 | 912 | 884 | 921,52 | 973 | 870 | 0,895 | -0,16 | -2,848 | 0,00439862 | 0,00960978 | 0,000581 | 0,0035549 | 0,0019296 | 0,0019296 | TRUE | 0,013831 | No |
| orf19.6660 | 13181 | 8817 | 11917 | 14776 | 17193 | 13334 | 15370 | 10221 | 11825 | 11225 | 12192 | 12184 | 14704 | 12723 | 13715 | 14416 | 12873 | 11856 | 13890 | 1,171 | 0,228 | 4,402 | 1,07E-05 | 3,89E-05 | 0,002645 | 0,0020923 | 0,0024981 | 0,0024981 | TRUE | 0,0391842 | No |
| orf19.6661 | 379 | 224 | 330 | 391 | 353 | 307 | 353 | 214 | 340 | 285 | 338 | 322 | 302 | 293 | 315 | 302 | 312,11 | 321 | 303 | 0,94 | -0,09 | -1,057 | 0,29031032 | 0,37501277 | 1,00E-08 | 0,0066308 | 0,0037321 | 0,0037321 | TRUE | 0,0557611 | No |
| orf19.6662 | 429 | 264 | 335 | 416 | 391 | 306 | 334 | 196 | 385 | 336 | 343 | 343 | 334 | 292 | 298 | 276 | 325,95 | 352 | 300 | 0,856 | -0,224 | -2,529 | 0,01145106 | 0,02271433 | 0,002256 | 0,0064333 | 0,0044597 | 0,0044597 | TRUE | 0,0687976 | No |
| orf19.6663 | 12589 | 8107 | 10146 | 13119 | 12813 | 10175 | 10635 | 8442 | 11294 | 10321 | 10380 | 10818 | 10958 | 9709 | 9490 | 11907 | 10610 | 10703 | 10516 | 0,982 | -0,026 | -0,372 | 0,70976993 | 0,77070265 | 0,005636 | 0,0021164 | 0,0046186 | 0,0046186 | TRUE | 0,0961365 | No |
| orf19.6665 | 3736 | 2477 | 3462 | 4137 | 4202 | 3566 | 4119 | 2740 | 3352 | 3153 | 3542 | 3411 | 3594 | 3403 | 3676 | 3865 | 3499,3 | 3364 | 3634 | 1,079 | 0,109 | 2,12 | 0,03402348 | 0,05998036 | 0,002161 | 0,0023944 | 0,0022655 | 0,0022655 | TRUE | 0,0221705 | No |
| orf19.6667 | 1253 | 844 | 956 | 1314 | 1496 | 1280 | 1378 | 829 | 1124 | 1075 | 978 | 1084 | 1279 | 1221 | 1230 | 1169 | 1145 | 1065 | 1225 | 1,151 | 0,203 | 3,56 | 0,00037152 | 0,00103099 | 0,001444 | 0,0032474 | 0,0022477 | 0,0022477 | TRUE | 0,0362181 | No |
| orf19.6668 | 1407 | 888 | 1105 | 1326 | 1493 | 1224 | 1394 | 949 | 1262 | 1131 | 1131 | 1093 | 1277 | 1168 | 1244 | 1338 | 1205,5 | 1154 | 1257 | 1,087 | 0,12 | 1,913 | 0,05576317 | 0,09229343 | 0,002552 | 0,0031838 | 0,002966 | 0,002966 | TRUE | 0,0492554 | No |
| orf19.6670 | 249 | 206 | 218 | 258 | 269 | 262 | 241 | 168 | 223 | 262 | 223 | 213 | 230 | 250 | 215 | 237 | 231,68 | 230 | 233 | 1,016 | 0,023 | 0,225 | 0,82187854 | 0,86279209 | 0,001651 | 0,0082454 | 0,0053781 | 0,0053781 | TRUE | 0,0845869 | No |
| orf19.6671 | 773 | 523 | 684 | 759 | 757 | 721 | 780 | 455 | 693 | 666 | 700 | 626 | 647 | 688 | 696 | 642 | 669,76 | 671 | 668 | 0,998 | -0,003 | -0,049 | 0,96068177 | 0,97151592 | 0,000835 | 0,004147 | 0,0024189 | 0,0024189 | TRUE | 0,0269888 | No |
| orf19.667.1 | 13190 | 8441 | 10402 | 13634 | 13216 | 10336 | 10359 | 8168 | 11833 | 10746 | 10642 | 11243 | 11303 | 9862 | 9244 | 11520 | 10799 | 11116 | 10482 | 0,943 | -0,085 | -1,198 | 0,23085336 | 0,31104889 | 0,006476 | 0,002114 | 0,0047573 | 0,0047573 | TRUE | 0,0774865 | No |
| orf19.6672 | 6301 | 4177 | 5419 | 6709 | 8119 | 7010 | 7630 | 4685 | 5653 | 5318 | 5544 | 5532 | 6944 | 6689 | 6809 | 6608 | 6136,9 | 5512 | 6762 | 1,227 | 0,295 | 8,75 | 2,14E-18 | 3,45E-17 | 0,000338 | 0,0022161 | 0,0009253 | 0,0009253 | TRUE | 0,0065428 | No |
| orf19.6673 | 457 | 333 | 462 | 495 | 531 | 431 | 510 | 344 | 410 | 424 | 473 | 408 | 454 | 411 | 455 | 485 | 440,05 | 429 | 451 | 1,051 | 0,072 | 0,908 | 0,36411518 | 0,45282633 | 0,002304 | 0,0052784 | 0,0038394 | 0,0038394 | TRUE | 0,055638 | No |
| orf19.6674 | 434 | 278 | 359 | 426 | 344 | 275 | 321 | 141 | 389 | 354 | 367 | 351 | 294 | 262 | 286 | 199 | 312,97 | 365 | 260 | 0,719 | -0,476 | -4,385 | 1,16E-05 | 4,19E-05 | 0,009902 | 0,006618 | 0,0080201 | 0,0080201 | TRUE | 0,2396363 | No |
| orf19.6675 | 861 | 615 | 775 | 989 | 839 | 823 | 821 | 574 | 772 | 783 | 793 | 816 | 718 | 785 | 733 | 810 | 776,1 | 791 | 762 | 0,959 | -0,061 | -1,016 | 0,30939374 | 0,39541323 | 0,000543 | 0,00385 | 0,0021338 | 0,0021338 | TRUE | 0,0189616 | No |
| orf19.6676 | 199 | 115 | 162 | 163 | 162 | 114 | 138 | 102 | 179 | 146 | 166 | 134 | 139 | 109 | 123 | 144 | 142,43 | 156 | 129 | 0,819 | -0,288 | -2,108 | 0,03499582 | 0,06146935 | 0,009205 | 0,0121722 | 0,0108246 | 0,0108246 | TRUE | 0,1100167 | No |
| orf19.6678 | 223 | 148 | 183 | 220 | 367 | 332 | 352 | 246 | 200 | 188 | 187 | 181 | 314 | 317 | 314 | 347 | 256,1 | 189 | 323 | 1,7 | 0,765 | 8,445 | 3,03E-17 | 4,37E-16 | 1,00E-08 | 0,0076479 | 0,0037111 | 0,0037111 | TRUE | 0,0236596 | No |
| orf19.6679 | 1555 | 975 | 1291 | 1580 | 1407 | 1131 | 1230 | 865 | 1395 | 1241 | 1321 | 1303 | 1203 | 1079 | 1098 | 1220 | 1232,5 | 1315 | 1150 | 0,872 | -0,197 | -3,294 | 0,00098738 | 0,00248625 | 0,002026 | 0,0031574 | 0,0026181 | 0,0026181 | TRUE | 0,0202561 | No |
| orf19.668 | 1102 | 742 | 1052 | 1181 | 978 | 964 | 976 | 762 | 989 | 945 | 1076 | 974 | 836 | 920 | 871 | 1075 | 960,66 | 996 | 926 | 0,926 | -0,112 | -1,405 | 0,16006802 | 0,22788831 | 0,006692 | 0,0034907 | 0,0049992 | 0,0049992 | TRUE | 0,131451 | No |
| orf19.6680 | 2951 | 2058 | 2717 | 3299 | 2747 | 2466 | 2747 | 1778 | 2647 | 2620 | 2780 | 2720 | 2349 | 2353 | 2451 | 2508 | 2553,6 | 2692 | 2415 | 0,896 | -0,159 | -3,876 | 0,00010598 | 0,0003232 | 0,000353 | 0,002548 | 0,0012124 | 0,0012124 | TRUE | 0,0072947 | No |
| orf19.6681 | 1817 | 1180 | 1559 | 1961 | 2003 | 1889 | 2047 | 1253 | 1630 | 1502 | 1595 | 1617 | 1713 | 1802 | 1827 | 1767 | 1681,7 | 1586 | 1777 | 1,119 | 0,162 | 3,544 | 0,00039378 | 0,00108724 | 0,000418 | 0,0028428 | 0,0014105 | 0,0014105 | TRUE | 0,0140383 | No |

| | | | | | | | | | | | | | | | | | | | | | | | | | | | | | | | |
|---|---|---|---|---|---|---|---|---|---|---|---|---|---|---|---|---|---|---|---|---|---|---|---|---|---|---|---|---|---|---|---|
| orf19.6684 | 487 | 333 | 405 | 552 | 612 | 556 | 583 | 353 | 437 | 424 | 414 | 455 | 523 | 531 | 520 | 498 | 475,3 | 432 | 518 | 1,197 | 0,26 | 3,833 | 0,00012666 | 0,0003803 | 1,00E-08 | 0,0050338 | 0,0022971 | 0,0022971 | TRUE | 0,0153979 | No |
| orf19.6685 | 2227 | 1575 | 1891 | 2360 | 1903 | 1765 | 1831 | 965 | 1998 | 2005 | 1935 | 1946 | 1628 | 1684 | 1634 | 1361 | 1773,8 | 1971 | 1577 | 0,802 | -0,318 | -5,038 | 4,71E-07 | 2,09E-06 | 0,003687 | 0,0027979 | 0,0032523 | 0,0032523 | TRUE | 0,0949454 | No |
| orf19.6686 | 11358 | 8568 | 10365 | 12381 | 11994 | 11727 | 12382 | 7567 | 10189 | 10908 | 10604 | 10210 | 10258 | 11190 | 11049 | 10673 | 10635 | 10478 | 10792 | 1,03 | 0,043 | 1,06 | 0,28929296 | 0,37393978 | 0,001165 | 0,002116 | 0,001493 | 0,001493 | TRUE | 0,0138407 | No |
| orf19.6687 | 22 | 24 | 24 | 34 | 9 | 16 | 7 | 13 | 20 | 31 | 25 | 28 | 8 | 15 | 6 | 18 | 18,8 | 26 | 12 | 0,446 | -1,164 | -3,156 | 0,0016015 | 0,00385489 | 0,043797 | 0,0791835 | 0,0674462 | 0,0674462 | TRUE | 0,3777737 | No |
| orf19.6688 | 10299 | 7799 | 9449 | 12005 | 8126 | 7806 | 8399 | 4907 | 9239 | 9929 | 9667 | 9899 | 6950 | 7448 | 7495 | 6921 | 8443,6 | 9684 | 7204 | 0,744 | -0,426 | -10,033 | 1,09E-23 | 2,52E-22 | 0,001308 | 0,0021515 | 0,0016115 | 0,0016115 | TRUE | 0,0117456 | No |
| orf19.6689 | 3704 | 2811 | 3325 | 4134 | 4526 | 4341 | 4373 | 2606 | 3323 | 3579 | 3402 | 3409 | 3871 | 4142 | 3902 | 3676 | 3662,9 | 3428 | 3898 | 1,139 | 0,187 | 4,099 | 4,15E-05 | 0,00013623 | 0,001333 | 0,0023759 | 0,0017262 | 0,0017262 | TRUE | 0,0214516 | No |
| orf19.669 | 206 | 136 | 203 | 251 | 158 | 157 | 134 | 83 | 185 | 173 | 208 | 207 | 135 | 150 | 120 | 117 | 161,77 | 193 | 130 | 0,676 | -0,565 | -4,648 | 3,35E-06 | 1,31E-05 | 0,003316 | 0,0109532 | 0,0077818 | 0,0077818 | TRUE | 0,1010567 | No |
| orf19.6690 | 2643 | 1936 | 2374 | 2912 | 2460 | 2267 | 2436 | 1374 | 2371 | 2465 | 2429 | 2401 | 2104 | 2163 | 2174 | 1938 | 2255,6 | 2416 | 2095 | 0,869 | -0,203 | -4,443 | 8,87E-06 | 3,26E-05 | 0,000894 | 0,0026231 | 0,0015498 | 0,0015498 | TRUE | 0,0278452 | No |
| orf19.6691 | 10236 | 8059 | 9497 | 11737 | 11639 | 11692 | 12095 | 7275 | 9183 | 10260 | 9716 | 9678 | 9954 | 11156 | 10793 | 10261 | 10125 | 9709 | 10541 | 1,086 | 0,119 | 2,438 | 0,01474919 | 0,0284381 | 0,002193 | 0,0021229 | 0,0021887 | 0,0021887 | TRUE | 0,0189196 | No |
| orf19.6692 | 1650 | 1280 | 1498 | 1764 | 2060 | 2162 | 2237 | 1276 | 1480 | 1630 | 1533 | 1455 | 1762 | 2063 | 1996 | 1800 | 1714,7 | 1524 | 1905 | 1,252 | 0,324 | 5,078 | 3,81E-07 | 1,72E-06 | 0,003478 | 0,0028261 | 0,0033271 | 0,0033271 | TRUE | 0,0375564 | No |
| orf19.6693 | 1680 | 1480 | 1670 | 2052 | 1943 | 2074 | 2237 | 1191 | 1507 | 1884 | 1709 | 1692 | 1662 | 1979 | 1996 | 1680 | 1763,6 | 1698 | 1829 | 1,079 | 0,11 | 1,331 | 0,18304168 | 0,25521338 | 0,008376 | 0,0028027 | 0,0059462 | 0,0059462 | TRUE | 0,0693467 | No |
| orf19.6694 | 1681 | 1115 | 1403 | 1881 | 2147 | 1828 | 2078 | 1237 | 1508 | 1420 | 1435 | 1551 | 1836 | 1744 | 1854 | 1745 | 1636,7 | 1478 | 1795 | 1,213 | 0,279 | 5,76 | 8,40E-09 | 4,71E-08 | 0,000785 | 0,0028665 | 0,0016334 | 0,0016334 | TRUE | 0,0134389 | No |
| orf19.6696 | 900 | 856 | 852 | 1182 | 969 | 979 | 917 | 504 | 807 | 1090 | 872 | 975 | 829 | 934 | 818 | 711 | 879,44 | 936 | 823 | 0,882 | -0,181 | -1,792 | 0,07306816 | 0,11697135 | 0,013344 | 0,0036302 | 0,0086261 | 0,0086261 | TRUE | 0,141001 | No |
| orf19.6698 | 709 | 474 | 621 | 756 | 857 | 688 | 738 | 467 | 636 | 603 | 635 | 623 | 733 | 656 | 659 | 659 | 650,61 | 624 | 677 | 1,084 | 0,117 | 1,846 | 0,06494972 | 0,10577879 | 1,00E-08 | 0,0042108 | 0,0023006 | 0,0023006 | TRUE | 0,0395537 | No |
| orf19.6699 | 299 | 264 | 270 | 414 | 339 | 260 | 314 | 200 | 268 | 336 | 276 | 341 | 290 | 248 | 280 | 282 | 290,28 | 305 | 275 | 0,901 | -0,15 | -1,427 | 0,15354127 | 0,2200038 | 0,007355 | 0,0069806 | 0,0071632 | 0,0071632 | TRUE | 0,0805847 | No |
| orf19.670 | 5539 | 3521 | 4323 | 5336 | 5624 | 4298 | 4510 | 2821 | 4969 | 4483 | 4423 | 4400 | 4810 | 4101 | 4024 | 3979 | 4398,6 | 4569 | 4228 | 0,926 | -0,111 | -1,638 | 0,10144774 | 0,15449886 | 0,005703 | 0,0023096 | 0,0041869 | 0,0041869 | TRUE | 0,1052025 | No |
| orf19.6701 | 3897 | 2656 | 3443 | 4109 | 3832 | 3334 | 3639 | 2305 | 3496 | 3381 | 3523 | 3388 | 3277 | 3181 | 3247 | 3251 | 3343,1 | 3447 | 3239 | 0,94 | -0,09 | -2,639 | 0,00831185 | 0,01707268 | 1,00E-08 | 0,0024138 | 0,0008166 | 0,0008166 | TRUE | 0,0025701 | No |
| orf19.6702 | 3038 | 2067 | 2671 | 3276 | 3634 | 3114 | 3228 | 2304 | 2725 | 2632 | 2733 | 2701 | 3108 | 2971 | 2880 | 3250 | 2875 | 2698 | 3052 | 1,13 | 0,176 | 3,86 | 0,00011321 | 0,00034282 | 0,001078 | 0,0024845 | 0,0016418 | 0,0016418 | TRUE | 0,0215487 | No |
| orf19.6702.2 | 19391 | 12505 | 15251 | 19637 | 22344 | 17276 | 18283 | 8704 | 17396 | 15920 | 15603 | 16193 | 19109 | 16484 | 16315 | 12276 | 16162 | 16278 | 16046 | 0,986 | -0,02 | -0,201 | 0,84061353 | 0,8783315 | 0,014051 | 0,0020693 | 0,0098724 | 0,0098724 | TRUE | 0,3051508 | No |
| orf19.6703 | 791 | 598 | 742 | 855 | 978 | 903 | 1051 | 526 | 710 | 761 | 759 | 705 | 836 | 862 | 938 | 742 | 789,11 | 734 | 844 | 1,156 | 0,209 | 2,843 | 0,00446419 | 0,00973887 | 0,003754 | 0,0038192 | 0,0039302 | 0,0039302 | TRUE | 0,071629 | No |
| orf19.6704 | 153 | 104 | 118 | 132 | 150 | 132 | 152 | 90 | 137 | 132 | 121 | 109 | 128 | 126 | 136 | 127 | 127,01 | 125 | 129 | 1,04 | 0,057 | 0,449 | 0,6535231 | 0,72242259 | 1,00E-08 | 0,0134098 | 0,0074972 | 0,0074972 | TRUE | 0,0865167 | No |
| orf19.6705 | 4896 | 3580 | 4609 | 5581 | 5736 | 5259 | 5666 | 3651 | 4392 | 4558 | 4715 | 4602 | 4906 | 5018 | 5056 | 5149 | 4799,6 | 4567 | 5032 | 1,101 | 0,139 | 3,909 | 9,28E-05 | 0,00028589 | 0,000353 | 0,002282 | 0,0010121 | 0,0010121 | TRUE | 0,0082868 | No |
| orf19.6706 | 10659 | 7992 | 9701 | 11769 | 12393 | 11890 | 12588 | 6971 | 9562 | 10175 | 9925 | 9705 | 10599 | 11345 | 11233 | 9832 | 10297 | 9842 | 10752 | 1,093 | 0,128 | 2,578 | 0,00993791 | 0,02001574 | 0,002334 | 0,0021205 | 0,0022862 | 0,0022862 | TRUE | 0,0398966 | No |
| orf19.6707 | 3676 | 2642 | 3053 | 3854 | 4011 | 3772 | 3939 | 2328 | 3298 | 3364 | 3124 | 3178 | 3430 | 3599 | 3515 | 3283 | 3348,8 | 3241 | 3457 | 1,068 | 0,095 | 2,206 | 0,02741831 | 0,04958736 | 0,000931 | 0,002413 | 0,0014863 | 0,0014863 | TRUE | 0,012955 | No |
| orf19.6709 | 151 | 113 | 162 | 198 | 164 | 136 | 165 | 101 | 135 | 144 | 166 | 163 | 140 | 130 | 147 | 142 | 146,01 | 152 | 140 | 0,918 | -0,123 | -1,026 | 0,30483192 | 0,39058012 | 1,00E-08 | 0,0119223 | 0,0069782 | 0,0069782 | TRUE | 0,0642424 | No |
| orf19.671 | 2798 | 1988 | 2549 | 3212 | 2355 | 2282 | 2418 | 1571 | 2510 | 2531 | 2608 | 2649 | 2014 | 2177 | 2158 | 2216 | 2357,8 | 2574 | 2141 | 0,83 | -0,268 | -6,06 | 1,36E-09 | 8,55E-09 | 0,000762 | 0,0025952 | 0,0014494 | 0,0014494 | TRUE | 0,019941 | No |
| orf19.6710 | 19824 | 12968 | 16921 | 20426 | 16102 | 13506 | 14858 | 9156 | 17784 | 16510 | 17312 | 16844 | 13771 | 12887 | 13258 | 12914 | 15160 | 17112 | 13208 | 0,772 | -0,374 | -9,983 | 1,80E-23 | 4,13E-22 | 0,000924 | 0,0020753 | 0,0012772 | 0,0012772 | TRUE | 0,0101558 | No |
| orf19.6712 | 2202 | 1538 | 2007 | 2348 | 2851 | 2428 | 2708 | 1846 | 1975 | 1958 | 2053 | 1936 | 2438 | 2317 | 2416 | 2604 | 2212,3 | 1980 | 2444 | 1,232 | 0,301 | 6,461 | 1,04E-10 | 7,57E-10 | 0,000907 | 0,0026357 | 0,0016273 | 0,0016273 | TRUE | 0,0217021 | No |
| orf19.6713 | 8629 | 5750 | 8128 | 9582 | 9617 | 8243 | 9408 | 7386 | 7741 | 7320 | 8316 | 7901 | 8225 | 7865 | 8395 | 10417 | 8272,6 | 7820 | 8726 | 1,115 | 0,157 | 1,957 | 0,05035859 | 0,08467798 | 0,009897 | 0,002155 | 0,0060921 | 0,0060921 | TRUE | 0,2066853 | No |
| orf19.6715 | 16 | 20 | 18 | 17 | 9 | 8 | 10 | 10 | 14 | 25 | 18 | 14 | 8 | 8 | 9 | 14 | 13,83 | 18 | 10 | 0,524 | -0,932 | -2,352 | 0,01868321 | NA | 1,00E-08 | 0,1069776 | 0,069045 | 0,069045 | TRUE | 0,3117405 | No |
| orf19.6716 | 2760 | 1747 | 2252 | 2906 | 3313 | 2660 | 2906 | 1814 | 2476 | 2224 | 2304 | 2396 | 2833 | 2538 | 2593 | 2558 | 2490,4 | 2350 | 2630 | 1,119 | 0,162 | 3,098 | 0,00194774 | 0,00461071 | 0,001904 | 0,0025624 | 0,0022333 | 0,0022333 | TRUE | 0,0333985 | No |
| orf19.6717 | 1459 | 973 | 1219 | 1561 | 1531 | 1254 | 1355 | 927 | 1309 | 1239 | 1247 | 1287 | 1309 | 1197 | 1209 | 1307 | 1263,1 | 1270 | 1256 | 0,986 | -0,02 | -0,388 | 0,69795548 | 0,76034751 | 0,000739 | 0,0031289 | 0,001768 | 0,001768 | TRUE | 0,0118758 | No |
| orf19.6718 | 2551 | 1727 | 2150 | 2640 | 2965 | 2439 | 2542 | 1694 | 2289 | 2199 | 2200 | 2177 | 2536 | 2327 | 2268 | 2389 | 2298 | 2216 | 2380 | 1,074 | 0,103 | 2,213 | 0,02692782 | 0,04881784 | 0,001088 | 0,0026112 | 0,0016466 | 0,0016466 | TRUE | 0,0248228 | No |
| orf19.6719 | 1105 | 665 | 905 | 1143 | 1131 | 1060 | 1148 | 658 | 991 | 847 | 926 | 943 | 967 | 1011 | 1024 | 928 | 954,69 | 927 | 982 | 1,06 | 0,084 | 1,361 | 0,17346388 | 0,24418251 | 0,001705 | 0,0035001 | 0,0026065 | 0,0026065 | TRUE | 0,0370154 | No |
| orf19.672 | 1241 | 821 | 1092 | 1355 | 1291 | 1138 | 1306 | 799 | 1113 | 1045 | 1117 | 1117 | 1104 | 1086 | 1165 | 1127 | 1109,4 | 1098 | 1120 | 1,019 | 0,027 | 0,53 | 0,59599286 | 0,67281569 | 1,00E-08 | 0,003288 | 0,0015207 | 0,0015207 | TRUE | 0,0113694 | No |
| orf19.6720 | 27451 | 17369 | 22431 | 29140 | 22975 | 18692 | 20027 | 13189 | 24626 | 22113 | 22949 | 24029 | 19649 | 17835 | 17871 | 18602 | 20959 | 23429 | 18489 | 0,789 | -0,342 | -7,212 | 5,53E-13 | 5,25E-12 | 0,002133 | 0,0020488 | 0,0021093 | 0,0021093 | TRUE | 0,0219821 | No |
| orf19.6722 | 3078 | 2055 | 2815 | 3419 | 3421 | 2930 | 3259 | 2042 | 2761 | 2616 | 2880 | 2819 | 2926 | 2796 | 2908 | 2880 | 2823,3 | 2769 | 2878 | 1,038 | 0,054 | 1,302 | 0,19289519 | 0,26611121 | 0,00056 | 0,0024937 | 0,0013061 | 0,0013061 | TRUE | 0,01571 | No |
| orf19.6723 | 1113 | 805 | 1044 | 1236 | 1445 | 1348 | 1424 | 940 | 998 | 1025 | 1068 | 1019 | 1236 | 1286 | 1271 | 1326 | 1153,7 | 1028 | 1280 | 1,244 | 0,315 | 6,424 | 1,33E-10 | 9,51E-10 | 1,00E-08 | 0,0032379 | 0,001426 | 0,001426 | TRUE | 0,0084441 | No |
| orf19.6724 | 2314 | 1466 | 2033 | 2428 | 2556 | 1877 | 2189 | 1615 | 2076 | 1866 | 2080 | 2002 | 2186 | 1791 | 1953 | 2278 | 2029,1 | 2006 | 2052 | 1,021 | 0,03 | 0,408 | 0,68339071 | 0,74678272 | 0,006587 | 0,002695 | 0,0048621 | 0,0048621 | TRUE | 0,0885535 | No |
| orf19.6726 | 1938 | 1272 | 1841 | 2060 | 1833 | 1512 | 1749 | 1095 | 1739 | 1619 | 1884 | 1699 | 1568 | 1443 | 1561 | 1544 | 1632 | 1735 | 1529 | 0,881 | -0,183 | -3,257 | 0,00112464 | 0,00280367 | 0,001941 | 0,0028691 | 0,0024243 | 0,0024243 | TRUE | 0,0392175 | No |
| orf19.6727 | 2699 | 1693 | 2285 | 2828 | 2860 | 2336 | 2557 | 1612 | 2421 | 2155 | 2338 | 2332 | 2446 | 2229 | 2282 | 2274 | 2309,6 | 2312 | 2308 | 0,998 | -0,003 | -0,066 | 0,94706897 | 0,96195949 | 0,001566 | 0,0026081 | 0,0019715 | 0,0019715 | TRUE | 0,0238217 | No |
| orf19.6729 | 1301 | 889 | 1181 | 1322 | 1776 | 1583 | 1826 | 1006 | 1167 | 1132 | 1208 | 1090 | 1519 | 1510 | 1629 | 1419 | 1334,4 | 1149 | 1519 | 1,325 | 0,406 | 7,127 | 1,03E-12 | 9,43E-12 | 0,001811 | 0,0030675 | 0,002347 | 0,002347 | TRUE | 0,028494 | No |
| orf19.673 | 621 | 425 | 549 | 687 | 750 | 625 | 634 | 449 | 557 | 541 | 562 | 567 | 641 | 596 | 566 | 633 | 582,89 | 557 | 609 | 1,092 | 0,127 | 1,931 | 0,0534435 | 0,08907667 | 1,00E-08 | 0,00447 | 0,0024065 | 0,0024065 | TRUE | 0,0284548 | No |
| orf19.6730 | 2706 | 1786 | 2313 | 2841 | 2576 | 2208 | 2402 | 1501 | 2428 | 2274 | 2366 | 2343 | 2203 | 2107 | 2143 | 2117 | 2247,6 | 2353 | 2142 | 0,911 | -0,135 | -3,405 | 0,00066151 | 0,00172949 | 1,00E-08 | 0,0026254 | 0,0010664 | 0,0010664 | TRUE | 0,0056863 | No |
| orf19.6731 | 1657 | 1105 | 1300 | 1628 | 1604 | 1348 | 1472 | 975 | 1487 | 1407 | 1330 | 1342 | 1372 | 1286 | 1314 | 1375 | 1364,1 | 1392 | 1337 | 0,96 | -0,059 | -1,108 | 0,26790332 | 0,35204794 | 0,001218 | 0,0030438 | 0,0019749 | 0,0019749 | TRUE | 0,0267582 | No |
| orf19.6731.1 | 1116 | 794 | 851 | 1137 | 1048 | 867 | 861 | 462 | 1001 | 1011 | 871 | 938 | 896 | 827 | 768 | 652 | 870,46 | 955 | 786 | 0,826 | -0,276 | -3,045 | 0,00233038 | 0,00542041 | 0,009469 | 0,0036473 | 0,0067134 | 0,0067134 | TRUE | 0,1454006 | No |
| orf19.6732 | 1442 | 1047 | 1299 | 1596 | 1341 | 1262 | 1344 | 926 | 1294 | 1333 | 1329 | 1316 | 1147 | 1204 | 1199 | 1306 | 1266 | 1318 | 1214 | 0,919 | -0,122 | -2,399 | 0,01643214 | 0,03128147 | 0,000538 | 0,0031262 | 0,0016985 | 0,0016985 | TRUE | 0,0277177 | No |
| orf19.6734 | 7502 | 5030 | 7156 | 8359 | 8968 | 7828 | 9103 | 6171 | 6730 | 6404 | 7321 | 6893 | 7670 | 7469 | 8123 | 8704 | 7414,2 | 6837 | 7992 | 1,168 | 0,224 | 3,862 | 0,00011238 | 0,00034066 | 0,003727 | 0,0021753 | 0,0031013 | 0,0031013 | TRUE | 0,043314 | No |
| orf19.6736 | 7584 | 5558 | 7046 | 8551 | 8969 | 8462 | 9094 | 5878 | 6804 | 7076 | 7209 | 7051 | 7671 | 8074 | 8115 | 8290 | 7536,2 | 7035 | 8038 | 1,142 | 0,192 | 5,249 | 1,53E-07 | 7,29E-07 | 0,000642 | 0,0021722 | 0,0011478 | 0,0011478 | TRUE | 0,0116558 | No |
| orf19.6737 | 923 | 701 | 845 | 1054 | 1027 | 880 | 939 | 672 | 828 | 892 | 865 | 869 | 878 | 840 | 838 | 948 | 869,73 | 864 | 876 | 1,012 | 0,017 | 0,299 | 0,7652406 | 0,81676418 | 0,000666 | 0,0036487 | 0,0021355 | 0,0021355 | TRUE | 0,0316406 | No |
| orf19.6738 | 3355 | 2225 | 2870 | 3663 | 3387 | 2802 | 3081 | 1994 | 3010 | 2833 | 2936 | 3021 | 2897 | 2674 | 2749 | 2812 | 2866,4 | 2950 | 2783 | 0,943 | -0,085 | -2,044 | 0,04091389 | 0,07037236 | 0,000672 | 0,002486 | 0,0013217 | 0,0013217 | TRUE | 0,0097921 | No |
| orf19.6739 | 1585 | 1032 | 1349 | 1608 | 2218 | 1824 | 2071 | 1219 | 1422 | 1314 | 1380 | 1326 | 1897 | 1740 | 1848 | 1719 | 1580,8 | 1360 | 1801 | 1,325 | 0,406 | 7,909 | 2,60E-15 | 3,06E-14 | 0,00119 | 0,0028978 | 0,0018857 | 0,0018857 | TRUE | 0,0155564 | No |
| orf19.674 | 2920 | 1958 | 2507 | 3151 | 2673 | 2369 | 2611 | 1773 | 2620 | 2493 | 2565 | 2598 | 2286 | 2260 | 2330 | 2501 | 2456,6 | 2569 | 2344 | 0,91 | -0,136 | -3,1 | 0,00193228 | 0,0045774 | 0,000754 | 0,0025705 | 0,0014246 | 0,0014246 | TRUE | 0,0223957 | No |
| orf19.6740 | 2318 | 1539 | 1992 | 2487 | 2241 | 1877 | 2159 | 1388 | 2079 | 1959 | 2038 | 2051 | 1917 | 1791 | 1927 | 1958 | 1964,9 | 2032 | 1898 | 0,933 | -0,101 | -2,266 | 0,02347927 | 0,04316507 | 0,000502 | 0,0027183 | 0,0013876 | 0,0013876 | TRUE | 0,017469 | No |
| orf19.6741 | 240 | 150 | 196 | 246 | 227 | 187 | 194 | 147 | 215 | 191 | 201 | 203 | 194 | 178 | 173 | 207 | 195,33 | 202 | 188 | 0,923 | -0,116 | -1,122 | 0,26167152 | 0,34529376 | 1,00E-08 | 0,0094115 | 0,0051289 | 0,0051289 | TRUE | 0,0398558 | No |
| orf19.6742 | 1732 | 1109 | 1390 | 1739 | 1762 | 1537 | 1784 | 1015 | 1554 | 1412 | 1422 | 1434 | 1507 | 1467 | 1592 | 1432 | 1477,3 | 1456 | 1500 | 1,031 | 0,044 | 0,821 | 0,41187667 | 0,50011218 | 0,001397 | 0,0029622 | 0,0020461 | 0,0020461 | TRUE | 0,0254256 | No |
| orf19.6744 | 132 | 115 | 129 | 146 | 146 | 103 | 150 | 75 | 118 | 146 | 132 | 120 | 125 | 98 | 134 | 106 | 122,5 | 129 | 116 | 0,905 | -0,144 | -1,023 | 0,30650451 | 0,39238847 | 0,006698 | 0,0138305 | 0,0107695 | 0,0107695 | TRUE | 0,1160621 | No |
| orf19.6745 | 6704 | 4433 | 5554 | 7025 | 8296 | 6371 | 6859 | 4490 | 6014 | 5644 | 5682 | 5793 | 7095 | 6079 | 6121 | 6333 | 6095 | 5783 | 6407 | 1,108 | 0,148 | 2,728 | 0,00637401 | 0,01345575 | 0,002909 | 0,0022177 | 0,0026517 | 0,0026517 | TRUE | 0,0646621 | No |
| orf19.6747 | 4188 | 2614 | 3410 | 4414 | 4698 | 3815 | 4213 | 2714 | 3757 | 3328 | 3489 | 3640 | 4018 | 3640 | 3759 | 3828 | 3682,4 | 3554 | 3811 | 1,072 | 0,1 | 2,008 | 0,04463292 | 0,07620192 | 0,00187 | 0,0023738 | 0,0021096 | 0,0021096 | TRUE | 0,0215551 | No |

| | | | | | | | | | | | | | | | | | | | | | | | | | | | | | | | |
|---|---|---|---|---|---|---|---|---|---|---|---|---|---|---|---|---|---|---|---|---|---|---|---|---|---|---|---|---|---|---|---|
| orf19.6748 | 6308 | 4279 | 5427 | 6629 | 5720 | 5076 | 5336 | 3192 | 5659 | 5448 | 5552 | 5466 | 4892 | 4843 | 4761 | 4502 | 5140,5 | 5531 | 4750 | 0,859 | -0,219 | -5,916 | 3,30E-09 | 1,95E-08 | 0,00056 | 0,0022619 | 0,0011133 | 0,0011133 | TRUE | 0,0141006 | No |
| orf19.6749 | 10497 | 6805 | 8938 | 10649 | 10253 | 8869 | 9814 | 6448 | 9417 | 8663 | 9144 | 8781 | 8769 | 8463 | 8757 | 9094 | 8886,1 | 9001 | 8771 | 0,974 | -0,038 | -0,968 | 0,33317894 | 0,41999932 | 0,001015 | 0,0021429 | 0,0013833 | 0,0013833 | TRUE | 0,0118539 | No |
| orf19.675 | 11955 | 7805 | 10325 | 12469 | 10778 | 8632 | 9305 | 6927 | 10725 | 9937 | 10563 | 10282 | 9218 | 8236 | 8303 | 9770 | 9629,3 | 10377 | 8882 | 0,856 | -0,225 | -3,842 | 0,00012194 | 0,00036687 | 0,003908 | 0,0021303 | 0,0031929 | 0,0031929 | TRUE | 0,0546101 | No |
| orf19.6751 | 3078 | 1997 | 2612 | 3263 | 3364 | 2729 | 2949 | 2222 | 2761 | 2542 | 2672 | 2691 | 2877 | 2604 | 2631 | 3134 | 2739,1 | 2666 | 2812 | 1,053 | 0,074 | 1,189 | 0,23429872 | 0,31512574 | 0,003973 | 0,0025095 | 0,0033374 | 0,0033374 | TRUE | 0,0695725 | No |
| orf19.675.1 | 244 | 168 | 215 | 301 | 196 | 152 | 169 | 125 | 219 | 214 | 220 | 248 | 168 | 145 | 151 | 176 | 192,59 | 225 | 160 | 0,704 | -0,506 | -4,675 | 2,94E-06 | 1,16E-05 | 1,00E-08 | 0,0095173 | 0,005882 | 0,005882 | TRUE | 0,0561895 | No |
| orf19.6752 | 2298 | 1574 | 1840 | 2392 | 2759 | 2394 | 2519 | 1659 | 2062 | 2004 | 1883 | 1972 | 2360 | 2284 | 2248 | 2340 | 2144 | 1980 | 2308 | 1,165 | 0,22 | 5,099 | 3,42E-07 | 1,55E-06 | 0,000496 | 0,0026566 | 0,001323 | 0,001323 | TRUE | 0,0132247 | No |
| orf19.6753 | 1608 | 955 | 1376 | 1771 | 1225 | 1021 | 1089 | 632 | 1443 | 1216 | 1408 | 1460 | 1048 | 974 | 972 | 891 | 1176,4 | 1382 | 971 | 0,703 | -0,508 | -7,143 | 9,16E-13 | 8,44E-12 | 0,004317 | 0,0032135 | 0,0039727 | 0,0039727 | TRUE | 0,074991 | No |
| orf19.6754 | 3487 | 2204 | 2849 | 3662 | 3860 | 3055 | 3480 | 2610 | 3128 | 2806 | 2915 | 3020 | 3301 | 2915 | 3105 | 3681 | 3108,9 | 2967 | 3250 | 1,094 | 0,13 | 1,87 | 0,06149328 | 0,10066802 | 0,005731 | 0,0024465 | 0,0042868 | 0,0042868 | TRUE | 0,0940178 | No |
| orf19.6755 | 647 | 416 | 588 | 684 | 544 | 483 | 520 | 358 | 580 | 530 | 602 | 564 | 465 | 461 | 464 | 505 | 521,34 | 569 | 474 | 0,829 | -0,271 | -3,963 | 7,40E-05 | 0,00023106 | 1,00E-08 | 0,0047641 | 0,0025611 | 0,0025611 | TRUE | 0,0220161 | No |
| orf19.6756 | 1872 | 1182 | 1637 | 1959 | 1931 | 1554 | 1721 | 1020 | 1679 | 1505 | 1675 | 1615 | 1651 | 1483 | 1536 | 1439 | 1572,9 | 1618 | 1527 | 0,944 | -0,083 | -1,434 | 0,15162819 | 0,21762594 | 0,002222 | 0,0029025 | 0,0025742 | 0,0025742 | TRUE | 0,0367004 | No |
| orf19.6757 | 1140 | 733 | 946 | 1209 | 1175 | 1026 | 1075 | 664 | 1023 | 933 | 968 | 997 | 1005 | 979 | 959 | 937 | 975,04 | 980 | 970 | 0,989 | -0,016 | -0,293 | 0,76941797 | 0,8201283 | 1,00E-08 | 0,0034684 | 0,0016841 | 0,0016841 | TRUE | 0,0113058 | No |
| orf19.6758 | 855 | 552 | 706 | 793 | 946 | 818 | 902 | 531 | 767 | 703 | 722 | 654 | 809 | 781 | 805 | 749 | 748,67 | 712 | 786 | 1,107 | 0,147 | 2,265 | 0,02350438 | 0,04318867 | 0,001612 | 0,0039186 | 0,0026852 | 0,0026852 | TRUE | 0,0388649 | No |
| orf19.676 | 1052 | 685 | 853 | 1088 | 773 | 551 | 682 | 415 | 944 | 872 | 873 | 897 | 661 | 526 | 609 | 585 | 745,81 | 896 | 595 | 0,664 | -0,59 | -8,033 | 9,54E-16 | 1,18E-14 | 0,00363 | 0,003926 | 0,0037727 | 0,0037727 | TRUE | 0,0767396 | No |
| orf19.6760 | 1779 | 1273 | 1585 | 1843 | 1688 | 1539 | 1719 | 1034 | 1596 | 1621 | 1622 | 1520 | 1444 | 1468 | 1534 | 1458 | 1532,8 | 1590 | 1476 | 0,93 | -0,105 | -2,29 | 0,0220381 | 0,04075252 | 1,00E-08 | 0,0029266 | 0,0013513 | 0,0013513 | TRUE | 0,0110839 | No |
| orf19.6763 | 35835 | 24039 | 30743 | 36391 | 39002 | 34465 | 37600 | 22679 | 32148 | 30604 | 31453 | 30008 | 33356 | 32885 | 33552 | 31987 | 31999 | 31053 | 32945 | 1,061 | 0,085 | 2,587 | 0,00966985 | 0,01954793 | 0,00064 | 0,0020249 | 0,0010163 | 0,0010163 | TRUE | 0,0069283 | No |
| orf19.6766 | 10200 | 6777 | 8227 | 10436 | 10484 | 8734 | 9241 | 5479 | 9150 | 8628 | 8417 | 8606 | 8966 | 8334 | 8246 | 7728 | 8509,3 | 8700 | 8318 | 0,957 | -0,064 | -1,282 | 0,19989183 | 0,27450289 | 0,002362 | 0,0021501 | 0,0022877 | 0,0022877 | TRUE | 0,0336328 | No |
| orf19.6769 | 9046 | 6254 | 8139 | 9751 | 8081 | 7497 | 8132 | 4722 | 8115 | 7962 | 8327 | 8041 | 6911 | 7153 | 7256 | 6660 | 7553,2 | 8111 | 6995 | 0,863 | -0,213 | -5,746 | 9,12E-09 | 5,06E-08 | 0,000734 | 0,0021717 | 0,0011827 | 0,0011827 | TRUE | 0,0121557 | No |
| orf19.677 | 3765 | 2770 | 3355 | 4269 | 2922 | 2524 | 2857 | 1722 | 3378 | 3527 | 3432 | 3520 | 2499 | 2408 | 2549 | 2429 | 2967,8 | 3464 | 2471 | 0,714 | -0,486 | -12,915 | 3,68E-38 | 2,30E-36 | 0,000225 | 0,0024687 | 0,0010104 | 0,0010104 | TRUE | 0,0055487 | No |
| orf19.6770 | 9866 | 6908 | 8644 | 10984 | 8877 | 7978 | 8670 | 6032 | 8851 | 8795 | 8844 | 9058 | 7592 | 7612 | 7737 | 8508 | 8374,4 | 8887 | 7862 | 0,884 | -0,178 | -4,156 | 3,24E-05 | 0,00010846 | 0,001457 | 0,0021529 | 0,0016396 | 0,0016396 | TRUE | 0,036329 | No |
| orf19.6771 | 1E+05 | 69051 | 86638 | 1E+05 | 1E+05 | 80824 | 85226 | 54254 | 94132 | 87909 | 88639 | 87673 | 86167 | 77120 | 76050 | 76520 | 84276 | 89588 | 78964 | 0,881 | -0,182 | -3,768 | 0,00016433 | 0,00048516 | 0,002431 | 0,0019967 | 0,0022313 | 0,0022313 | TRUE | 0,0462542 | No |
| orf19.6773 | 2580 | 1876 | 2331 | 2631 | 2549 | 2362 | 2587 | 1526 | 2315 | 2388 | 2385 | 2170 | 2180 | 2254 | 2308 | 2152 | 2269 | 2314 | 2224 | 0,963 | -0,055 | -1,173 | 0,2408618 | 0,32207828 | 0,001104 | 0,0026193 | 0,001676 | 0,001676 | TRUE | 0,0221933 | No |
| orf19.6776 | 7482 | 5002 | 6427 | 7998 | 7797 | 6954 | 7453 | 4957 | 6712 | 6368 | 6575 | 6595 | 6668 | 6635 | 6651 | 6991 | 6649,6 | 6562 | 6736 | 1,025 | 0,036 | 1,089 | 0,27624643 | 0,36025364 | 0,000358 | 0,0021978 | 0,0009088 | 0,0009088 | TRUE | 0,0075181 | No |
| orf19.6777 | 94 | 86 | 97 | 103 | 85 | 73 | 83 | 64 | 84 | 109 | 99 | 85 | 73 | 70 | 74 | 90 | 85,58 | 94 | 77 | 0,81 | -0,304 | -1,92 | 0,0548574 | 0,09104509 | 1,00E-08 | 0,0189419 | 0,0122015 | 0,0122015 | TRUE | 0,100254 | No |
| orf19.6778 | 1470 | 1181 | 1437 | 1754 | 1181 | 1149 | 1223 | 804 | 1319 | 1504 | 1470 | 1446 | 1010 | 1096 | 1091 | 1134 | 1258,8 | 1435 | 1083 | 0,754 | -0,407 | -6,906 | 4,99E-12 | 4,20E-11 | 0,002056 | 0,0031328 | 0,0025168 | 0,0025168 | TRUE | 0,0361856 | No |
| orf19.6779 | 1215 | 961 | 1101 | 1328 | 1438 | 1514 | 1517 | 915 | 1090 | 1223 | 1126 | 1095 | 1230 | 1445 | 1354 | 1291 | 1231,7 | 1134 | 1330 | 1,175 | 0,233 | 3,648 | 0,00026402 | 0,00075284 | 0,002787 | 0,0031581 | 0,00309 | 0,00309 | TRUE | 0,0413586 | No |
| orf19.6780 | 417 | 345 | 345 | 499 | 417 | 391 | 401 | 251 | 374 | 439 | 353 | 411 | 357 | 373 | 358 | 354 | 377,41 | 394 | 360 | 0,916 | -0,126 | -1,504 | 0,13255 | 0,1944714 | 0,002161 | 0,005826 | 0,0041112 | 0,0041112 | TRUE | 0,0586249 | No |
| orf19.6781 | 2604 | 2657 | 2646 | 3447 | 3003 | 3587 | 3714 | 2160 | 2336 | 3383 | 2707 | 2842 | 2568 | 3423 | 3314 | 3046 | 2952,5 | 2817 | 3088 | 1,097 | 0,133 | 1,23 | 0,21880442 | 0,29755024 | 0,01843 | 0,0024712 | 0,0109202 | 0,0109202 | TRUE | 0,2197723 | No |
| orf19.6782 | 230 | 175 | 199 | 243 | 170 | 162 | 158 | 111 | 206 | 223 | 204 | 200 | 145 | 155 | 141 | 157 | 178,83 | 208 | 150 | 0,717 | -0,48 | -4,582 | 4,61E-06 | 1,76E-05 | 1,00E-08 | 0,0100974 | 0,004809 | 0,004809 | TRUE | 0,0207137 | No |
| orf19.6783 | 3185 | 2444 | 2848 | 3563 | 3530 | 3429 | 3751 | 2228 | 2857 | 3111 | 2914 | 2938 | 3019 | 3272 | 3347 | 3142 | 3075,1 | 2955 | 3195 | 1,082 | 0,114 | 2,43 | 0,0150886 | 0,02899018 | 0,001279 | 0,0024516 | 0,0017818 | 0,0017818 | TRUE | 0,0175878 | No |
| orf19.6784 | 183 | 141 | 166 | 179 | 213 | 184 | 181 | 129 | 164 | 180 | 170 | 148 | 182 | 176 | 162 | 182 | 170,29 | 166 | 176 | 1,064 | 0,09 | 0,813 | 0,41639776 | 0,50437858 | 1,00E-08 | 0,0105045 | 0,0058672 | 0,0058672 | TRUE | 0,0598586 | No |
| orf19.6785 | 10180 | 6703 | 7894 | 10376 | 9560 | 8163 | 8326 | 4930 | 9133 | 8534 | 8076 | 8556 | 8176 | 7789 | 7430 | 6953 | 8080,8 | 8575 | 7587 | 0,885 | -0,176 | -3,096 | 0,00195905 | 0,00463203 | 0,003464 | 0,0021592 | 0,0029789 | 0,0029789 | TRUE | 0,0379353 | No |
| orf19.6786 | 328 | 230 | 323 | 403 | 419 | 361 | 392 | 254 | 294 | 293 | 330 | 332 | 358 | 344 | 350 | 358 | 332,59 | 312 | 352 | 1,125 | 0,171 | 2,098 | 0,03587349 | 0,06283612 | 1,00E-08 | 0,0063444 | 0,0033343 | 0,0033343 | TRUE | 0,0224378 | No |
| orf19.6787 | 1677 | 1619 | 1474 | 2141 | 1470 | 1724 | 1634 | 936 | 1504 | 2061 | 1508 | 1765 | 1257 | 1645 | 1458 | 1320 | 1564,9 | 1710 | 1420 | 0,832 | -0,266 | -2,469 | 0,01353662 | 0,02633679 | 0,018399 | 0,0029072 | 0,0105135 | 0,0105135 | TRUE | 0,2270597 | No |
| orf19.6788 | 1430 | 1063 | 1136 | 1505 | 906 | 942 | 939 | 563 | 1283 | 1353 | 1162 | 1241 | 775 | 899 | 838 | 794 | 1043,1 | 1260 | 826 | 0,657 | -0,606 | -9,098 | 9,24E-20 | 1,63E-18 | 0,002891 | 0,0033712 | 0,003253 | 0,003253 | TRUE | 0,041876 | No |
| orf19.6789 | 419 | 359 | 376 | 466 | 518 | 512 | 519 | 265 | 376 | 457 | 385 | 384 | 443 | 489 | 463 | 374 | 421,29 | 400 | 442 | 1,113 | 0,154 | 1,613 | 0,10674898 | 0,1612237 | 0,007512 | 0,0054254 | 0,0063741 | 0,0063741 | TRUE | 0,1100881 | No |
| orf19.679 | 811 | 556 | 662 | 792 | 759 | 622 | 657 | 450 | 728 | 708 | 677 | 653 | 649 | 593 | 586 | 635 | 653,67 | 692 | 616 | 0,891 | -0,167 | -2,552 | 0,01072439 | 0,02143452 | 0,001108 | 0,0042004 | 0,0025738 | 0,0025738 | TRUE | 0,0179944 | No |
| orf19.6790 | 465 | 301 | 417 | 485 | 554 | 454 | 506 | 315 | 417 | 383 | 427 | 400 | 474 | 433 | 452 | 444 | 428,71 | 407 | 451 | 1,108 | 0,148 | 2,05 | 0,04039924 | 0,06964666 | 1,00E-08 | 0,0053657 | 0,0026628 | 0,0026628 | TRUE | 0,0149202 | No |
| orf19.6791 | 15567 | 10902 | 12819 | 15565 | 17656 | 16147 | 17569 | 8313 | 13965 | 13879 | 13115 | 12835 | 15100 | 15407 | 15677 | 11725 | 13963 | 13448 | 14477 | 1,077 | 0,107 | 1,316 | 0,18825528 | 0,26102879 | 0,008936 | 0,0020835 | 0,0062524 | 0,0062524 | TRUE | 0,1993938 | No |
| orf19.6792 | 1325 | 1024 | 1193 | 1479 | 872 | 812 | 876 | 599 | 1189 | 1304 | 1221 | 1220 | 746 | 775 | 782 | 845 | 1009,9 | 1234 | 787 | 0,637 | -0,651 | -11,488 | 1,52E-30 | 5,53E-29 | 0,000818 | 0,0034169 | 0,0020387 | 0,0020387 | TRUE | 0,0253972 | No |
| orf19.6793 | 5237 | 3517 | 4461 | 5759 | 3638 | 3431 | 3608 | 2057 | 4698 | 4478 | 4564 | 4749 | 3111 | 3274 | 3220 | 2901 | 3874,3 | 4622 | 3126 | 0,677 | -0,563 | -12,379 | 3,38E-35 | 1,69E-33 | 0,001389 | 0,0023542 | 0,0017148 | 0,0017148 | TRUE | 0,0267318 | No |
| orf19.6794 | 4813 | 4019 | 4400 | 5791 | 5076 | 4893 | 4914 | 3020 | 4318 | 5117 | 4502 | 4775 | 4341 | 4669 | 4385 | 4259 | 4545,7 | 4678 | 4414 | 0,944 | -0,083 | -1,452 | 0,14637157 | 0,21119126 | 0,003281 | 0,0022989 | 0,0029022 | 0,0029022 | TRUE | 0,0476301 | No |
| orf19.6795 | 590 | 458 | 477 | 623 | 457 | 540 | 503 | 293 | 529 | 583 | 488 | 514 | 391 | 515 | 449 | 413 | 485,29 | 528 | 442 | 0,839 | -0,254 | -2,689 | 0,0071678 | 0,01491658 | 0,00767 | 0,0049709 | 0,0064701 | 0,0064701 | TRUE | 0,1463214 | No |
| orf19.6796 | 458 | 336 | 361 | 510 | 437 | 378 | 423 | 280 | 411 | 428 | 369 | 421 | 374 | 361 | 377 | 395 | 391,91 | 407 | 377 | 0,923 | -0,115 | -1,493 | 0,1355053 | 0,19793724 | 1,00E-08 | 0,0056837 | 0,0031338 | 0,0031338 | TRUE | 0,0437628 | No |
| orf19.6797 | 1134 | 884 | 965 | 1336 | 1328 | 1251 | 1341 | 873 | 1017 | 1125 | 987 | 1102 | 1136 | 1194 | 1197 | 1231 | 1123,6 | 1058 | 1190 | 1,124 | 0,168 | 2,92 | 0,00349839 | 0,0078021 | 0,001316 | 0,0032715 | 0,0022901 | 0,0022901 | TRUE | 0,0233357 | No |
| orf19.6798 | 22320 | 14039 | 19699 | 23422 | 27587 | 23611 | 26920 | 17883 | 20023 | 17873 | 20154 | 19314 | 23593 | 22529 | 24022 | 25222 | 21591 | 19341 | 23842 | 1,232 | 0,301 | 6,027 | 1,67E-09 | 1,02E-08 | 0,002492 | 0,0020468 | 0,0023562 | 0,0023562 | TRUE | 0,0317429 | No |
| orf19.680 | 4270 | 2769 | 3642 | 4439 | 3636 | 3160 | 3491 | 2119 | 3831 | 3525 | 3726 | 3660 | 3110 | 3015 | 3115 | 2989 | 3371,4 | 3686 | 3057 | 0,83 | -0,27 | -6,889 | 5,64E-12 | 4,72E-11 | 0,000427 | 0,0024101 | 0,0011709 | 0,0011709 | TRUE | 0,0098809 | No |
| orf19.6800 | 1377 | 942 | 1248 | 1522 | 1513 | 1318 | 1488 | 947 | 1235 | 1199 | 1277 | 1255 | 1294 | 1258 | 1328 | 1336 | 1272,7 | 1242 | 1304 | 1,048 | 0,068 | 1,451 | 0,1468465 | 0,21174475 | 1,00E-08 | 0,0031202 | 0,0013183 | 0,0013183 | TRUE | 0,0067517 | No |
| orf19.6801 | 2607 | 1796 | 2222 | 2772 | 2730 | 2534 | 2683 | 1673 | 2339 | 2286 | 2273 | 2286 | 2335 | 2418 | 2394 | 2360 | 2336,4 | 2296 | 2377 | 1,035 | 0,05 | 1,364 | 0,17258196 | 0,24299799 | 1,00E-08 | 0,0026009 | 0,0008411 | 0,0008411 | TRUE | 0,001965 | No |
| orf19.6802 | 191 | 170 | 163 | 221 | 298 | 350 | 344 | 163 | 171 | 216 | 167 | 182 | 255 | 334 | 307 | 230 | 232,81 | 184 | 282 | 1,54 | 0,623 | 4,727 | 2,28E-06 | 9,20E-06 | 0,01659 | 0,0082152 | 0,0121893 | 0,0121893 | TRUE | 0,135828 | No |
| orf19.6803 | 632 | 548 | 553 | 744 | 598 | 559 | 570 | 326 | 567 | 698 | 566 | 614 | 511 | 533 | 509 | 460 | 557,14 | 611 | 503 | 0,828 | -0,272 | -3,312 | 0,00092769 | 0,00234478 | 0,004622 | 0,0045851 | 0,0046444 | 0,0046444 | TRUE | 0,0989954 | No |
| orf19.6804 | 969 | 1149 | 1058 | 1383 | 1115 | 1430 | 1334 | 845 | 869 | 1463 | 1082 | 1140 | 954 | 1364 | 1190 | 1192 | 1156,9 | 1138 | 1175 | 1,032 | 0,046 | 0,247 | 0,80515581 | 0,84881139 | 0,032551 | 0,0032343 | 0,0171893 | 0,0325514 | TRUE | 0,4302973 | No |
| orf19.6805 | 881 | 814 | 885 | 1121 | 912 | 1027 | 1128 | 666 | 790 | 1036 | 905 | 924 | 780 | 980 | 1007 | 939 | 920,28 | 914 | 926 | 1,014 | 0,021 | 0,216 | 0,82891421 | 0,86837255 | 0,011481 | 0,003557 | 0,0077164 | 0,0077164 | TRUE | 0,1397514 | No |
| orf19.6807 | 64 | 33 | 35 | 37 | 52 | 43 | 45 | 25 | 57 | 42 | 36 | 31 | 44 | 41 | 40 | 35 | 40,83 | 42 | 40 | 0,978 | -0,032 | -0,132 | 0,895112 | 0,92070414 | 0,023274 | 0,0375318 | 0,0312137 | 0,0312137 | TRUE | 0,5796434 | No |
| orf19.6809 | 621 | 452 | 507 | 696 | 774 | 718 | 750 | 411 | 557 | 575 | 519 | 574 | 662 | 685 | 669 | 580 | 602,64 | 556 | 649 | 1,171 | 0,228 | 3,242 | 0,00118809 | 0,00295205 | 0,001546 | 0,0043884 | 0,0030903 | 0,0030903 | TRUE | 0,0525634 | No |
| orf19.681 | 3178 | 2064 | 2802 | 3513 | 3496 | 2988 | 3377 | 2115 | 2851 | 2628 | 2867 | 2897 | 2990 | 2851 | 3013 | 2983 | 2885 | 2811 | 2959 | 1,052 | 0,073 | 1,671 | 0,09476726 | 0,14629026 | 0,000809 | 0,0024827 | 0,0014688 | 0,0014688 | TRUE | 0,0220575 | No |
| orf19.6810 | 1387 | 1183 | 1304 | 1641 | 1508 | 1522 | 1573 | 1061 | 1244 | 1506 | 1334 | 1353 | 1290 | 1452 | 1404 | 1496 | 1385 | 1359 | 1410 | 1,038 | 0,053 | 0,78 | 0,43516937 | 0,52227279 | 0,004065 | 0,0030277 | 0,0037695 | 0,0037695 | TRUE | 0,06066 | No |
| orf19.6811 | 3991 | 3069 | 3492 | 4488 | 5432 | 4757 | 5145 | 2627 | 3580 | 3907 | 3573 | 3701 | 4646 | 4539 | 4591 | 3705 | 4030,2 | 3690 | 4370 | 1,186 | 0,246 | 3,5 | 0,00046454 | 0,00126403 | 0,005757 | 0,0023397 | 0,0044812 | 0,0044812 | TRUE | 0,12508 | No |
| orf19.6812 | 1073 | 765 | 1004 | 1217 | 1505 | 1239 | 1336 | 773 | 963 | 974 | 1027 | 1004 | 1287 | 1182 | 1192 | 1090 | 1089,9 | 992 | 1188 | 1,201 | 0,264 | 4,448 | 8,66E-06 | 3,19E-05 | 0,001776 | 0,0033115 | 0,0024561 | 0,0024561 | TRUE | 0,0379454 | No |
| orf19.6813 | 7 | 3 | 8 | 8 | 6 | 4 | 9 | 5 | 6 | 4 | 8 | 7 | 5 | 4 | 8 | 7 | 6,11 | 6 | 6 | 0,945 | -0,082 | -0,149 | 0,88176842 | NA | 1,00E-08 | 0,239419 | 0,1267702 | 0,1267702 | TRUE | 0,1585434 | No |

| | | | | | | | | | | | | | | | | | | | | | | | | | | | | | | | |
|---|---|---|---|---|---|---|---|---|---|---|---|---|---|---|---|---|---|---|---|---|---|---|---|---|---|---|---|---|---|---|---|
| orf19.6814 | 84554 | 53348 | 72042 | 90233 | 1E+05 | 81646 | 89182 | 67659 | 75854 | 67918 | 73706 | 74407 | 90110 | 77904 | 79580 | 95427 | 79363 | 72971 | 85755 | 1,175 | 0,233 | 3,472 | 0,00051749 | 0,00139168 | 0,005935 | 0,0019978 | 0,004309 | 0,004309 | TRUE | 0,0705643 | No |
| orf19.6816 | 693 | 501 | 609 | 697 | 762 | 659 | 640 | 467 | 622 | 638 | 623 | 575 | 652 | 629 | 571 | 659 | 620,95 | 614 | 628 | 1,021 | 0,031 | 0,445 | 0,65621738 | 0,72486744 | 0,001621 | 0,0043174 | 0,0029174 | 0,0029174 | TRUE | 0,0448774 | No |
| orf19.6817 | 3149 | 2136 | 2753 | 3375 | 2736 | 2468 | 2695 | 1732 | 2825 | 2719 | 2817 | 2783 | 2340 | 2355 | 2405 | 2443 | 2585,8 | 2786 | 2386 | 0,855 | -0,226 | -6,211 | 5,26E-10 | 3,49E-09 | 1,00E-08 | 0,0025409 | 0,0008797 | 0,0008797 | TRUE | 0,0028877 | No |
| orf19.6818 | 2473 | 1741 | 2264 | 2623 | 2157 | 1976 | 2162 | 1292 | 2219 | 2216 | 2316 | 2163 | 1845 | 1885 | 1929 | 1822 | 2049,5 | 2228 | 1870 | 0,84 | -0,251 | -5,992 | 2,08E-09 | 1,26E-08 | 0,000274 | 0,0026878 | 0,0011914 | 0,0011914 | TRUE | 0,0085754 | No |
| orf19.68.2 | 361 | 393 | 396 | 518 | 389 | 457 | 429 | 309 | 324 | 500 | 405 | 427 | 333 | 436 | 383 | 436 | 405,48 | 414 | 397 | 0,959 | -0,061 | -0,476 | 0,6343225 | 0,70522013 | 0,021339 | 0,0055597 | 0,0133356 | 0,0133356 | TRUE | 0,2520931 | No |
| orf19.6820 | 2923 | 1896 | 2590 | 3174 | 4509 | 3916 | 4405 | 2884 | 2622 | 2414 | 2650 | 2617 | 3856 | 3737 | 3931 | 4068 | 3236,8 | 2576 | 3898 | 1,511 | 0,595 | 13,316 | 1,86E-40 | 1,27E-38 | 0,001097 | 0,002428 | 0,001595 | 0,001595 | TRUE | 0,0205149 | No |
| orf19.6821 | 201 | 153 | 197 | 205 | 208 | 177 | 188 | 121 | 180 | 195 | 202 | 169 | 178 | 169 | 168 | 171 | 178,86 | 186 | 172 | 0,923 | -0,115 | -1,081 | 0,27990737 | 0,3641584 | 1,00E-08 | 0,0100959 | 0,0053011 | 0,0053011 | TRUE | 0,0463518 | No |
| orf19.6822 | 1477 | 1182 | 1336 | 1660 | 1869 | 1680 | 1850 | 966 | 1325 | 1505 | 1367 | 1369 | 1598 | 1603 | 1651 | 1362 | 1472,5 | 1392 | 1554 | 1,12 | 0,164 | 2,439 | 0,0147331 | 0,0284162 | 0,004156 | 0,0029654 | 0,0036634 | 0,0036634 | TRUE | 0,077441 | No |
| orf19.6824 | 1104 | 758 | 900 | 1236 | 1104 | 1085 | 1100 | 675 | 990 | 965 | 921 | 1019 | 944 | 1035 | 982 | 952 | 976,06 | 974 | 978 | 1,004 | 0,006 | 0,101 | 0,91925711 | 0,94038766 | 0,000875 | 0,0034668 | 0,0020352 | 0,0020352 | TRUE | 0,0186069 | No |
| orf19.6826 | 53 | 32 | 37 | 38 | 53 | 35 | 37 | 24 | 48 | 41 | 38 | 31 | 45 | 33 | 33 | 34 | 37,88 | 40 | 36 | 0,933 | -0,1 | -0,428 | 0,6689854 | 0,7355901 | 1,00E-08 | 0,0403 | 0,0264649 | 0,0264649 | TRUE | 0,2270747 | No |
| orf19.6827 | 1149 | 844 | 1091 | 1228 | 1158 | 1051 | 1137 | 688 | 1031 | 1075 | 1116 | 1013 | 990 | 1003 | 1015 | 970 | 1026,5 | 1059 | 994 | 0,942 | -0,086 | -1,648 | 0,09937834 | 0,15207974 | 1,00E-08 | 0,0033937 | 0,0016656 | 0,0016656 | TRUE | 0,0167231 | No |
| orf19.6828 | 9611 | 6807 | 8090 | 9746 | 8307 | 7338 | 7699 | 4094 | 8622 | 8666 | 8277 | 8037 | 7104 | 7002 | 6870 | 5774 | 7544 | 8400 | 6688 | 0,797 | -0,328 | -5,132 | 2,86E-07 | 1,31E-06 | 0,0047 | 0,002172 | 0,0037852 | 0,0037852 | TRUE | 0,1014799 | No |
| orf19.6828.1 | 679 | 449 | 549 | 635 | 679 | 588 | 564 | 276 | 609 | 572 | 562 | 524 | 581 | 561 | 503 | 389 | 537,55 | 567 | 508 | 0,902 | -0,148 | -1,364 | 0,17252204 | 0,24297056 | 0,013752 | 0,0046801 | 0,0094461 | 0,0094461 | TRUE | 0,2655358 | No |
| orf19.6829 | 3125 | 2353 | 2885 | 3302 | 3725 | 3527 | 3694 | 2282 | 2803 | 2996 | 2952 | 2723 | 3186 | 3365 | 3296 | 3219 | 3067,4 | 2868 | 3266 | 1,14 | 0,189 | 4,346 | 1,39E-05 | 4,94E-05 | 0,00085 | 0,0024528 | 0,0014941 | 0,0014941 | TRUE | 0,0143964 | No |
| orf19.6830 | 443 | 328 | 406 | 519 | 456 | 436 | 438 | 268 | 397 | 418 | 415 | 428 | 390 | 416 | 391 | 378 | 404,15 | 414 | 394 | 0,951 | -0,072 | -0,995 | 0,3199736 | 0,40659928 | 1,00E-08 | 0,0055715 | 0,0025692 | 0,0025692 | TRUE | 0,0164519 | No |
| orf19.6831 | 10660 | 7221 | 9481 | 11430 | 11484 | 9887 | 10793 | 6193 | 9563 | 9193 | 9700 | 9425 | 9821 | 9434 | 9631 | 8735 | 9437,8 | 9470 | 9405 | 0,994 | -0,009 | -0,217 | 0,82796141 | 0,86782835 | 0,001213 | 0,0021333 | 0,0016509 | 0,0016509 | TRUE | 0,0274881 | No |
| orf19.6832 | 1041 | 756 | 953 | 1148 | 1084 | 932 | 1014 | 576 | 934 | 962 | 975 | 947 | 927 | 889 | 905 | 812 | 918,95 | 954 | 883 | 0,929 | -0,106 | -1,892 | 0,05847224 | 0,09629967 | 0,000314 | 0,0035593 | 0,0019014 | 0,0019014 | TRUE | 0,0299919 | No |
| orf19.6833.2 | 56 | 39 | 43 | 55 | 54 | 36 | 42 | 60 | 50 | 50 | 44 | 45 | 46 | 34 | 37 | 85 | 48,98 | 47 | 50 | 1,048 | 0,067 | 0,251 | 0,80162995 | 0,84555346 | 0,067133 | 0,0316158 | 0,0474759 | 0,0474759 | TRUE | 1,36274 | No |
| orf19.6834.1( | 21 | 12 | 17 | 14 | 27 | 21 | 24 | 6 | 19 | 15 | 17 | 12 | 23 | 20 | 21 | 8 | 17,01 | 16 | 18 | 1,195 | 0,257 | 0,719 | 0,47220415 | 0,55868502 | 0,021499 | 0,0873354 | 0,064074 | 0,064074 | TRUE | 0,4455964 | No |
| orf19.6837 | 264 | 167 | 238 | 259 | 279 | 213 | 250 | 183 | 237 | 213 | 243 | 214 | 239 | 203 | 223 | 258 | 228,69 | 227 | 231 | 1,012 | 0,018 | 0,173 | 0,86244741 | 0,89566363 | 0,002588 | 0,0083273 | 0,005837 | 0,005837 | TRUE | 0,0691792 | No |
| orf19.6838 | 2149 | 1379 | 1767 | 2215 | 1739 | 1501 | 1552 | 1057 | 1928 | 1756 | 1808 | 1827 | 1487 | 1432 | 1385 | 1491 | 1639,1 | 1830 | 1449 | 0,79 | -0,339 | -7,011 | 2,36E-12 | 2,08E-11 | 0,000787 | 0,0028652 | 0,001628 | 0,001628 | TRUE | 0,0159357 | No |
| orf19.684 | 4184 | 3104 | 3765 | 4733 | 5090 | 4576 | 5043 | 3183 | 3753 | 3952 | 3852 | 3903 | 4353 | 4366 | 4500 | 4489 | 4146,1 | 3865 | 4427 | 1,146 | 0,196 | 5,815 | 6,05E-09 | 3,46E-08 | 0,000157 | 0,0023297 | 0,0008473 | 0,0008473 | TRUE | 0,0046575 | No |
| orf19.6840 | 1249 | 860 | 1032 | 1302 | 996 | 987 | 1005 | 788 | 1120 | 1095 | 1056 | 1074 | 852 | 942 | 897 | 1111 | 1018,3 | 1086 | 950 | 0,871 | -0,198 | -2,598 | 0,00936547 | 0,01899654 | 0,006134 | 0,0034051 | 0,0046078 | 0,0046078 | TRUE | 0,1447474 | No |
| orf19.6842 | 9054 | 6363 | 8652 | 10069 | 11970 | 10749 | 12105 | 7345 | 8122 | 8101 | 8852 | 8303 | 10237 | 10256 | 10802 | 10359 | 9379,1 | 8344 | 10414 | 1,248 | 0,32 | 8,067 | 7,20E-16 | 9,03E-15 | 0,001058 | 0,0021343 | 0,0013987 | 0,0013987 | TRUE | 0,0203787 | No |
| orf19.6843 | 8906 | 5820 | 7480 | 9256 | 7817 | 6687 | 7302 | 3982 | 7990 | 7409 | 7653 | 7633 | 6685 | 6381 | 6516 | 5616 | 6985,3 | 7671 | 6300 | 0,822 | -0,283 | -5,121 | 3,04E-07 | 1,39E-06 | 0,003104 | 0,0021873 | 0,0027941 | 0,0027941 | TRUE | 0,0636128 | No |
| orf19.6844 | 36176 | 22671 | 31161 | 37615 | 30395 | 23819 | 25861 | 18126 | 32454 | 28863 | 31881 | 31018 | 25995 | 22727 | 23077 | 25565 | 27697 | 31054 | 24341 | 0,784 | -0,352 | -6,216 | 5,11E-10 | 3,40E-09 | 0,003611 | 0,0020319 | 0,0030367 | 0,0030367 | TRUE | 0,0275446 | No |
| orf19.6845 | 4413 | 3176 | 3909 | 4863 | 4082 | 3767 | 4111 | 2555 | 3959 | 4043 | 3999 | 4010 | 3491 | 3594 | 3668 | 3604 | 3796,1 | 4003 | 3589 | 0,897 | -0,157 | -4,816 | 1,46E-06 | 6,03E-06 | 1,00E-08 | 0,0023619 | 0,0007594 | 0,0007594 | TRUE | 0,0043934 | No |
| orf19.6846 | 3537 | 2421 | 3080 | 3632 | 3512 | 3252 | 3335 | 2270 | 3173 | 3082 | 3151 | 2995 | 3004 | 3103 | 2976 | 3202 | 3085,7 | 3100 | 3071 | 0,99 | -0,015 | -0,379 | 0,70486312 | 0,76612735 | 0,000461 | 0,00245 | 0,0012233 | 0,0012233 | TRUE | 0,0092429 | No |
| orf19.6847 | 1832 | 1294 | 1726 | 2056 | 1906 | 1678 | 1891 | 1158 | 1643 | 1647 | 1766 | 1695 | 1630 | 1601 | 1687 | 1633 | 1663 | 1688 | 1638 | 0,97 | -0,043 | -0,98 | 0,32729426 | 0,41449914 | 0,000218 | 0,0028525 | 0,0012898 | 0,0012898 | TRUE | 0,0113449 | No |
| orf19.6848 | 634 | 451 | 560 | 655 | 848 | 769 | 864 | 538 | 569 | 574 | 573 | 540 | 725 | 734 | 771 | 759 | 655,6 | 564 | 747 | 1,326 | 0,407 | 6,835 | 8,19E-12 | 6,75E-11 | 1,00E-08 | 0,0041938 | 0,0018479 | 0,0018479 | TRUE | 0,0103926 | No |
| orf19.6849 | 878 | 658 | 709 | 883 | 850 | 742 | 818 | 542 | 788 | 838 | 725 | 728 | 727 | 708 | 730 | 764 | 751,02 | 770 | 732 | 0,952 | -0,071 | -1,104 | 0,26944176 | 0,3538374 | 0,001376 | 0,0039125 | 0,0026068 | 0,0026068 | TRUE | 0,0381464 | No |
| orf19.685 | 2219 | 1478 | 1871 | 2321 | 2513 | 2060 | 2181 | 1597 | 1991 | 1882 | 1914 | 1914 | 2149 | 1966 | 1946 | 2252 | 2001,7 | 1925 | 2078 | 1,078 | 0,108 | 1,951 | 0,05108305 | 0,08568033 | 0,00215 | 0,0027047 | 0,0024264 | 0,0024264 | TRUE | 0,0359696 | No |
| orf19.6850 | 812 | 604 | 792 | 904 | 940 | 871 | 912 | 540 | 728 | 769 | 810 | 745 | 804 | 831 | 814 | 762 | 782,95 | 763 | 803 | 1,055 | 0,077 | 1,31 | 0,19012173 | 0,26319156 | 0,000365 | 0,0038337 | 0,0020538 | 0,0020538 | TRUE | 0,0208517 | No |
| orf19.6851 | 183 | 133 | 188 | 208 | 154 | 150 | 183 | 95 | 164 | 169 | 192 | 172 | 132 | 143 | 163 | 134 | 158,68 | 174 | 143 | 0,823 | -0,28 | -2,379 | 0,01735529 | 0,03278949 | 1,00E-08 | 0,0111279 | 0,0069812 | 0,0069812 | TRUE | 0,1011966 | No |
| orf19.685.1 | 1319 | 1149 | 1250 | 1558 | 1292 | 1309 | 1284 | 916 | 1183 | 1463 | 1279 | 1285 | 1105 | 1249 | 1146 | 1292 | 1250,2 | 1302 | 1198 | 0,92 | -0,121 | -1,624 | 0,10427889 | 0,15789002 | 0,005588 | 0,0031407 | 0,0045103 | 0,0045103 | TRUE | 0,0793733 | No |
| orf19.6852 | 646 | 440 | 600 | 704 | 515 | 443 | 433 | 306 | 580 | 560 | 614 | 581 | 440 | 423 | 386 | 432 | 501,9 | 584 | 420 | 0,719 | -0,476 | -6,785 | 1,16E-11 | 9,39E-11 | 1,00E-08 | 0,0048719 | 0,0026834 | 0,0026834 | TRUE | 0,0366044 | No |
| orf19.6852.1 | 1927 | 1535 | 1698 | 2339 | 2360 | 2037 | 1879 | 1197 | 1729 | 1954 | 1737 | 1929 | 2018 | 1944 | 1677 | 1688 | 1834,5 | 1837 | 1832 | 0,998 | -0,002 | -0,032 | 0,97457531 | 0,98199121 | 0,006175 | 0,0027709 | 0,0046788 | 0,0046788 | TRUE | 0,057402 | No |
| orf19.6853 | 2391 | 1589 | 1980 | 2319 | 2720 | 2305 | 2627 | 1625 | 2145 | 2023 | 2026 | 1912 | 2326 | 2199 | 2344 | 2292 | 2158,5 | 2026 | 2290 | 1,131 | 0,177 | 3,727 | 0,00019364 | 0,00056558 | 0,001127 | 0,0026521 | 0,0017018 | 0,0017018 | TRUE | 0,0196691 | No |
| orf19.6854 | 12000 | 7541 | 10389 | 12343 | 13796 | 11253 | 11982 | 8155 | 10765 | 9600 | 10629 | 10178 | 11799 | 10737 | 10692 | 11502 | 10738 | 10293 | 11182 | 1,086 | 0,119 | 2,373 | 0,01765263 | 0,03322588 | 0,002406 | 0,0021147 | 0,0023269 | 0,0023269 | TRUE | 0,0247791 | No |
| orf19.6855 | 2506 | 1762 | 2136 | 2715 | 2219 | 2111 | 2232 | 1462 | 2248 | 2243 | 2185 | 2239 | 1898 | 2014 | 1992 | 2062 | 2110,2 | 2229 | 1992 | 0,892 | -0,165 | -4,009 | 6,11E-05 | 0,0001947 | 0,000173 | 0,0026675 | 0,0011442 | 0,0011442 | TRUE | 0,0124302 | No |
| orf19.6858 | 1518 | 994 | 1262 | 1562 | 1679 | 1551 | 1529 | 944 | 1362 | 1265 | 1291 | 1288 | 1436 | 1480 | 1364 | 1331 | 1352,3 | 1302 | 1403 | 1,079 | 0,109 | 2,121 | 0,03391601 | 0,05980845 | 0,000875 | 0,0030531 | 0,0018061 | 0,0018061 | TRUE | 0,0159488 | No |
| orf19.6859 | 73 | 49 | 59 | 74 | 57 | 54 | 47 | 47 | 65 | 62 | 60 | 61 | 49 | 52 | 42 | 66 | 57,22 | 62 | 52 | 0,822 | -0,283 | -1,505 | 0,13234706 | 0,19422109 | 1,00E-08 | 0,0273502 | 0,0161578 | 0,0161578 | TRUE | 0,2066493 | No |
| orf19.686 | 1679 | 1134 | 1455 | 1813 | 1664 | 1476 | 1645 | 977 | 1506 | 1444 | 1489 | 1495 | 1423 | 1408 | 1468 | 1378 | 1451,4 | 1484 | 1419 | 0,957 | -0,063 | -1,437 | 0,15070055 | 0,21660258 | 1,00E-08 | 0,0029798 | 0,0011577 | 0,0011577 | TRUE | 0,0063625 | No |
| orf19.6860 | 1252 | 957 | 1082 | 1406 | 1165 | 1212 | 1192 | 736 | 1123 | 1218 | 1107 | 1159 | 996 | 1156 | 1064 | 1038 | 1107,8 | 1152 | 1064 | 0,924 | -0,113 | -1,885 | 0,0594395 | 0,09767847 | 0,001998 | 0,00329 | 0,0025673 | 0,0025673 | TRUE | 0,0422995 | No |
| orf19.6861 | 885 | 892 | 922 | 1102 | 1176 | 1418 | 1325 | 793 | 794 | 1136 | 943 | 909 | 1006 | 1353 | 1182 | 1118 | 1055,1 | 946 | 1165 | 1,234 | 0,304 | 2,774 | 0,00552933 | 0,01183899 | 0,017758 | 0,0033554 | 0,0105571 | 0,0105571 | TRUE | 0,2133636 | No |
| orf19.6862 | 3416 | 3394 | 3316 | 4460 | 3476 | 4253 | 4349 | 2460 | 3065 | 4321 | 3393 | 3678 | 2973 | 4058 | 3881 | 3470 | 3604,6 | 3614 | 3596 | 0,995 | -0,007 | -0,049 | 0,96061446 | 0,97151592 | 0,0197 | 0,0023823 | 0,011047 | 0,0197003 | TRUE | 0,210017 | No |
| orf19.6863 | 3566 | 2651 | 3196 | 3849 | 4219 | 3950 | 4351 | 2480 | 3199 | 3375 | 3270 | 3174 | 3608 | 3769 | 3883 | 3498 | 3471,9 | 3254 | 3690 | 1,135 | 0,183 | 4,15 | 3,32E-05 | 0,0001111 | 0,001062 | 0,0023976 | 0,0015797 | 0,0015797 | TRUE | 0,0150865 | No |
| orf19.6864 | 165 | 139 | 172 | 180 | 232 | 206 | 267 | 155 | 148 | 177 | 176 | 148 | 198 | 197 | 238 | 219 | 187,65 | 162 | 213 | 1,318 | 0,398 | 3,459 | 0,00054317 | 0,00145297 | 0,004402 | 0,0097156 | 0,0072971 | 0,0072971 | TRUE | 0,0782516 | No |
| orf19.6866 | 1783 | 1267 | 1677 | 1970 | 1891 | 1637 | 1827 | 1146 | 1600 | 1613 | 1716 | 1624 | 1617 | 1562 | 1630 | 1616 | 1622,3 | 1638 | 1606 | 0,981 | -0,028 | -0,642 | 0,52098298 | 0,60413048 | 1,00E-08 | 0,0028743 | 0,0012318 | 0,0012318 | TRUE | 0,0121263 | No |
| orf19.6867 | 507 | 354 | 434 | 521 | 658 | 565 | 634 | 298 | 455 | 451 | 444 | 430 | 563 | 539 | 566 | 420 | 483,38 | 445 | 522 | 1,181 | 0,24 | 2,634 | 0,0084499 | 0,0173148 | 0,006863 | 0,0049827 | 0,0058891 | 0,0058891 | TRUE | 0,1784279 | No |
| orf19.6868 | 639 | 397 | 481 | 646 | 695 | 548 | 673 | 469 | 573 | 505 | 492 | 533 | 594 | 523 | 601 | 661 | 560,35 | 526 | 595 | 1,126 | 0,171 | 2,07 | 0,03841603 | 0,06676411 | 0,004925 | 0,0045702 | 0,0047505 | 0,0047505 | TRUE | 0,0767514 | No |
| orf19.6869 | 845 | 655 | 749 | 903 | 997 | 942 | 1013 | 562 | 758 | 834 | 766 | 745 | 853 | 899 | 904 | 793 | 818,87 | 776 | 862 | 1,117 | 0,159 | 2,523 | 0,01164808 | 0,02306581 | 0,00148 | 0,0037523 | 0,0026109 | 0,0026109 | TRUE | 0,0305183 | No |
| orf19.687 | 205 | 167 | 198 | 241 | 219 | 183 | 185 | 140 | 184 | 213 | 203 | 199 | 187 | 175 | 165 | 197 | 190,28 | 200 | 181 | 0,907 | -0,142 | -1,346 | 0,17821397 | 0,24981557 | 1,00E-08 | 0,0096087 | 0,0053484 | 0,0053484 | TRUE | 0,0387262 | No |
| orf19.6871 | 2603 | 1852 | 2417 | 2925 | 2568 | 2379 | 2612 | 1736 | 2335 | 2358 | 2473 | 2412 | 2196 | 2270 | 2331 | 2448 | 2352,9 | 2394 | 2311 | 0,964 | -0,053 | -1,181 | 0,23771593 | 0,31858071 | 0,000827 | 0,0025965 | 0,0015407 | 0,0015407 | TRUE | 0,0178322 | No |
| orf19.687.1 | 16159 | 11112 | 13195 | 17294 | 16456 | 13868 | 14372 | 11335 | 14496 | 14147 | 13500 | 14261 | 14074 | 13232 | 12825 | 15987 | 14065 | 14101 | 14030 | 0,995 | -0,008 | -0,121 | 0,90373829 | 0,92768408 | 0,004605 | 0,0020827 | 0,0039294 | 0,0039294 | TRUE | 0,1072095 | No |
| orf19.6872 | 2493 | 1791 | 2216 | 2835 | 2337 | 2180 | 2260 | 1418 | 2236 | 2280 | 2267 | 2338 | 1999 | 2080 | 2017 | 2000 | 2152,1 | 2280 | 2024 | 0,888 | -0,172 | -4,464 | 8,05E-06 | 2,98E-05 | 1,00E-08 | 0,0026541 | 0,0009599 | 0,0009599 | TRUE | 0,004294 | No |
| orf19.6873 | 22807 | 13869 | 18607 | 22549 | 23255 | 18224 | 19882 | 12889 | 20460 | 17657 | 19037 | 18594 | 19888 | 17389 | 17741 | 18179 | 18618 | 18937 | 18299 | 0,966 | -0,049 | -0,875 | 0,38180223 | 0,4709948 | 0,003482 | 0,0020575 | 0,0030234 | 0,0030234 | TRUE | 0,0420128 | No |
| orf19.6873.1 | 1696 | 1193 | 1374 | 1824 | 1836 | 1448 | 1394 | 938 | 1521 | 1519 | 1406 | 1504 | 1570 | 1382 | 1244 | 1323 | 1433,6 | 1488 | 1380 | 0,928 | -0,107 | -1,508 | 0,13152975 | 0,19316322 | 0,004828 | 0,0029921 | 0,004151 | 0,004151 | TRUE | 0,1070647 | No |
| orf19.6874 | 2807 | 1900 | 2596 | 3101 | 3680 | 3316 | 3663 | 2503 | 2518 | 2419 | 2656 | 2557 | 3147 | 3164 | 3269 | 3530 | 2907,5 | 2538 | 3278 | 1,289 | 0,367 | 7,384 | 1,53E-13 | 1,52E-12 | 0,00172 | 0,0024788 | 0,0020165 | 0,0020165 | TRUE | 0,0311696 | No |

| | | | | | | | | | | | | | | | | | | | | | | | | | | | | | | | |
|---|---|---|---|---|---|---|---|---|---|---|---|---|---|---|---|---|---|---|---|---|---|---|---|---|---|---|---|---|---|---|---|
| orf19.6875 | 1156 | 854 | 1073 | 1273 | 1793 | 1700 | 1745 | 1047 | 1037 | 1087 | 1098 | 1050 | 1533 | 1622 | 1557 | 1477 | 1307,6 | 1068 | 1547 | 1,452 | 0,538 | 10,937 | 7,65E-28 | 2,48E-26 | 0,000341 | 0,0030897 | 0,0015359 | 0,0015359 | TRUE | 0,0123939 | No |
| orf19.6877 | 21685 | 15482 | 19767 | 24200 | 23697 | 21380 | 22949 | 15369 | 19454 | 19710 | 20224 | 19956 | 20266 | 20400 | 20478 | 21677 | 20271 | 19836 | 20705 | 1,044 | 0,061 | 1,895 | 0,05804948 | 0,09565585 | 0,00057 | 0,0020511 | 0,0009609 | 0,0009609 | TRUE | 0,0119921 | No |
| orf19.6879 | 1708 | 1172 | 1499 | 1905 | 2211 | 1930 | 2051 | 1402 | 1532 | 1492 | 1534 | 1571 | 1891 | 1842 | 1830 | 1977 | 1708,6 | 1532 | 1885 | 1,227 | 0,295 | 6,696 | 2,14E-11 | 1,69E-10 | 0,000173 | 0,0028292 | 0,0012746 | 0,0012746 | TRUE | 0,0117341 | No |
| orf19.688 | 2194 | 1472 | 1903 | 2357 | 2193 | 1911 | 2091 | 1440 | 1968 | 1874 | 1947 | 1944 | 1876 | 1823 | 1866 | 2031 | 1916,1 | 1933 | 1899 | 0,98 | -0,03 | -0,65 | 0,51596278 | 0,60005891 | 0,000674 | 0,0027372 | 0,00149 | 0,00149 | TRUE | 0,0239455 | No |
| orf19.6880 | 556 | 451 | 524 | 631 | 604 | 575 | 624 | 433 | 499 | 574 | 536 | 520 | 517 | 549 | 557 | 611 | 545,27 | 532 | 558 | 1,047 | 0,066 | 0,905 | 0,36568073 | 0,45458511 | 0,00179 | 0,0046419 | 0,0033432 | 0,0033432 | TRUE | 0,0411513 | No |
| orf19.6881 | 6782 | 4860 | 5630 | 7313 | 8603 | 7504 | 8042 | 5410 | 6084 | 6187 | 5760 | 6030 | 7358 | 7160 | 7176 | 7630 | 6673,3 | 6015 | 7331 | 1,218 | 0,285 | 7,643 | 2,13E-14 | 2,28E-13 | 0,000702 | 0,0021971 | 0,0011818 | 0,0011818 | TRUE | 0,0098612 | No |
| orf19.6882 | 30687 | 20025 | 26558 | 32325 | 38975 | 32439 | 36365 | 21310 | 27529 | 25494 | 27171 | 26656 | 33333 | 30952 | 32450 | 30056 | 29205 | 26712 | 31698 | 1,187 | 0,247 | 5,752 | 8,79E-09 | 4,89E-08 | 0,001583 | 0,0020292 | 0,0017357 | 0,0017357 | TRUE | 0,014792 | No |
| orf19.6882.1 | 13675 | 9244 | 10733 | 13675 | 13889 | 11557 | 11907 | 7298 | 12268 | 11769 | 10981 | 11277 | 11878 | 11027 | 10625 | 10293 | 11265 | 11574 | 10956 | 0,947 | -0,079 | -1,466 | 0,14261244 | 0,20656112 | 0,003026 | 0,0021084 | 0,0026778 | 0,0026778 | TRUE | 0,0393868 | No |
| orf19.6883 | 2408 | 1610 | 2019 | 2585 | 2161 | 1884 | 2014 | 1365 | 2160 | 2050 | 2066 | 2132 | 1848 | 1798 | 1797 | 1925 | 1971,9 | 2102 | 1842 | 0,874 | -0,194 | -4,555 | 5,25E-06 | 1,99E-05 | 0,000172 | 0,0027157 | 0,0012263 | 0,0012263 | TRUE | 0,0098963 | No |
| orf19.6884 | 277 | 195 | 269 | 296 | 355 | 297 | 358 | 225 | 248 | 248 | 275 | 244 | 304 | 283 | 319 | 317 | 279,98 | 254 | 306 | 1,204 | 0,267 | 3,037 | 0,00238805 | 0,00553313 | 1,00E-08 | 0,0071645 | 0,0038513 | 0,0038513 | TRUE | 0,034924 | No |
| orf19.6885 | 1927 | 1410 | 1770 | 2174 | 2705 | 2348 | 2631 | 1655 | 1729 | 1795 | 1811 | 1793 | 2313 | 2240 | 2348 | 2334 | 2045,4 | 1782 | 2309 | 1,296 | 0,374 | 9,419 | 4,56E-21 | 8,90E-20 | 1,00E-08 | 0,0026893 | 0,0010166 | 0,0010166 | TRUE | 0,0050495 | No |
| orf19.6886 | 12331 | 8471 | 10408 | 12434 | 12232 | 11181 | 12189 | 6313 | 11062 | 10784 | 10648 | 10253 | 10461 | 10669 | 10877 | 8904 | 10457 | 10687 | 10228 | 0,957 | -0,063 | -1,024 | 0,30588156 | 0,39175282 | 0,004063 | 0,0021183 | 0,0035078 | 0,0035078 | TRUE | 0,0916927 | No |
| orf19.6887 | 4500 | 2917 | 3526 | 4485 | 4539 | 3465 | 3852 | 2234 | 4037 | 3714 | 3607 | 3698 | 3882 | 3306 | 3437 | 3151 | 3604,1 | 3764 | 3444 | 0,916 | -0,127 | -1,929 | 0,0537086 | 0,08945974 | 0,005044 | 0,0023823 | 0,003892 | 0,003892 | TRUE | 0,089604 | No |
| orf19.6888 | 13484 | 9419 | 12254 | 15283 | 15625 | 13967 | 15316 | 10006 | 12097 | 11991 | 12537 | 12603 | 13363 | 13327 | 13667 | 14113 | 12962 | 12307 | 13618 | 1,106 | 0,145 | 4,342 | 1,41E-05 | 5,03E-05 | 0,000566 | 0,0020915 | 0,0009978 | 0,0009978 | TRUE | 0,007142 | No |
| orf19.6889 | 2481 | 1667 | 2100 | 2547 | 2618 | 2321 | 2461 | 1586 | 2226 | 2122 | 2149 | 2100 | 2239 | 2215 | 2196 | 2237 | 2185,4 | 2149 | 2222 | 1,033 | 0,047 | 1,227 | 0,21995924 | 0,29871499 | 1,00E-08 | 0,0026438 | 0,0009629 | 0,0009629 | TRUE | 0,0072683 | No |
| orf19.689 | 13 | 10 | 8 | 23 | 25 | 19 | 18 | 12 | 12 | 13 | 8 | 19 | 21 | 18 | 16 | 17 | 15,51 | 13 | 18 | 1,395 | 0,48 | 1,33 | 0,18353548 | 0,25572383 | 1,00E-08 | 0,0956065 | 0,0596859 | 0,0596859 | TRUE | 0,5224736 | No |
| orf19.6891 | 1583 | 1110 | 1500 | 1780 | 2047 | 1788 | 1886 | 1259 | 1420 | 1413 | 1535 | 1468 | 1751 | 1706 | 1683 | 1776 | 1593,9 | 1459 | 1729 | 1,184 | 0,243 | 5,294 | 1,20E-07 | 5,80E-07 | 0,000293 | 0,0028903 | 0,0013912 | 0,0013912 | TRUE | 0,0142442 | No |
| orf19.6893 | 1586 | 1082 | 1345 | 1594 | 1274 | 1116 | 1291 | 809 | 1423 | 1378 | 1376 | 1314 | 1090 | 1065 | 1152 | 1141 | 1242,3 | 1373 | 1112 | 0,81 | -0,305 | -6,027 | 1,67E-09 | 1,02E-08 | 0,000533 | 0,0031481 | 0,0016396 | 0,0016396 | TRUE | 0,0106252 | No |
| orf19.6896 | 10 | 8 | 18 | 18 | 18 | 16 | 27 | 9 | 9 | 10 | 18 | 15 | 15 | 15 | 24 | 13 | 14,98 | 13 | 17 | 1,298 | 0,376 | 0,994 | 0,32040916 | NA | 0,028284 | 0,0988708 | 0,0706155 | 0,0706155 | TRUE | 0,4438931 | No |
| orf19.6897 | 0 | 0 | 0 | 0 | 0 | 0 | 0 | 0 | 0 | 0 | 0 | 0 | 0 | 0 | 0 | 0 | 0 | 0 | 0 | NA | NA | NA | NA | NA | NA | NA | NA | NA | NA | NA | NA |
| orf19.6898 | 1865 | 1326 | 1623 | 1992 | 2104 | 1913 | 2111 | 1262 | 1673 | 1688 | 1660 | 1643 | 1799 | 1825 | 1884 | 1780 | 1744,1 | 1666 | 1822 | 1,095 | 0,131 | 3,207 | 0,0013415 | 0,00328478 | 1,00E-08 | 0,0028119 | 0,0010308 | 0,0010308 | TRUE | 0,0063772 | No |
| orf19.6898.1 | 1436 | 1013 | 1077 | 1440 | 1379 | 1182 | 1201 | 891 | 1288 | 1290 | 1102 | 1187 | 1179 | 1128 | 1072 | 1257 | 1187,9 | 1217 | 1159 | 0,951 | -0,072 | -1,049 | 0,29432781 | 0,37906141 | 0,003935 | 0,0032017 | 0,0037116 | 0,0037116 | TRUE | 0,0477067 | No |
| orf19.6899 | 5589 | 3799 | 5027 | 6297 | 6608 | 5600 | 6041 | 3743 | 5014 | 4837 | 5143 | 5193 | 5651 | 5343 | 5391 | 5279 | 5231,3 | 5047 | 5416 | 1,073 | 0,102 | 2,645 | 0,0081635 | 0,01679094 | 0,000696 | 0,002257 | 0,0012333 | 0,0012333 | TRUE | 0,0105746 | No |
| orf19.69 | 563 | 388 | 471 | 618 | 420 | 411 | 452 | 275 | 505 | 494 | 482 | 510 | 359 | 392 | 403 | 388 | 441,64 | 498 | 386 | 0,774 | -0,37 | -5,213 | 1,86E-07 | 8,78E-07 | 1,00E-08 | 0,0052666 | 0,002551 | 0,002551 | TRUE | 0,0276539 | No |
| orf19.690 | 66 | 58 | 64 | 73 | 76 | 84 | 76 | 56 | 59 | 74 | 65 | 60 | 65 | 80 | 68 | 79 | 68,83 | 64 | 73 | 1,13 | 0,176 | 1,043 | 0,29707795 | 0,38227551 | 1,00E-08 | 0,0230694 | 0,012854 | 0,012854 | TRUE | 0,0665808 | No |
| orf19.6900 | 1208 | 813 | 1026 | 1317 | 1347 | 1247 | 1395 | 790 | 1084 | 1035 | 1050 | 1086 | 1152 | 1190 | 1245 | 1114 | 1119,4 | 1064 | 1175 | 1,106 | 0,145 | 2,775 | 0,00551228 | 0,01181512 | 0,000486 | 0,0032764 | 0,0017274 | 0,0017274 | TRUE | 0,0190975 | No |
| orf19.6901 | 605 | 433 | 557 | 667 | 720 | 646 | 626 | 424 | 543 | 551 | 570 | 550 | 616 | 616 | 559 | 598 | 575,33 | 554 | 597 | 1,079 | 0,11 | 1,723 | 0,08495389 | 0,13329929 | 1,00E-08 | 0,0045028 | 0,0021734 | 0,0021734 | TRUE | 0,0242944 | No |
| orf19.6902 | 15137 | 9868 | 13114 | 16032 | 12966 | 11138 | 12346 | 7281 | 13579 | 12563 | 13417 | 13220 | 11089 | 10628 | 11017 | 10269 | 11973 | 13195 | 10751 | 0,815 | -0,295 | -7,481 | 7,37E-14 | 7,54E-13 | 0,001082 | 0,0021008 | 0,0014115 | 0,0014115 | TRUE | 0,0125247 | No |
| orf19.6903 | 3948 | 2532 | 3243 | 4252 | 4714 | 3893 | 4116 | 2693 | 3542 | 3224 | 3318 | 3506 | 4032 | 3715 | 3673 | 3798 | 3600,8 | 3398 | 3804 | 1,119 | 0,162 | 3,411 | 0,00064735 | 0,00169763 | 0,001515 | 0,0023827 | 0,0018926 | 0,0018926 | TRUE | 0,0203022 | No |
| orf19.6904 | 2699 | 1783 | 2208 | 2869 | 3356 | 2617 | 2904 | 1876 | 2421 | 2270 | 2259 | 2366 | 2870 | 2497 | 2591 | 2646 | 2490,1 | 2329 | 2651 | 1,138 | 0,186 | 3,567 | 0,00036097 | 0,00100451 | 0,001809 | 0,0025625 | 0,0022185 | 0,0022185 | TRUE | 0,0388931 | No |
| orf19.6905 | 3399 | 2395 | 3046 | 3815 | 4254 | 3622 | 3913 | 2791 | 3049 | 3049 | 3116 | 3146 | 3638 | 3456 | 3492 | 3936 | 3360,4 | 3090 | 3630 | 1,173 | 0,23 | 4,857 | 1,19E-06 | 4,99E-06 | 0,001538 | 0,0024115 | 0,0018605 | 0,0018605 | TRUE | 0,0372194 | No |
| orf19.6906 | 16554 | 10879 | 14136 | 17512 | 16576 | 13650 | 14234 | 9265 | 14851 | 13850 | 14463 | 14441 | 14176 | 13024 | 12701 | 13067 | 13822 | 14401 | 13242 | 0,92 | -0,121 | -2,826 | 0,00471867 | 0,01024566 | 0,001495 | 0,0020846 | 0,0016896 | 0,0016896 | TRUE | 0,0278313 | No |
| orf19.6907 | 1797 | 1296 | 1677 | 2055 | 2524 | 2217 | 2488 | 1386 | 1612 | 1650 | 1716 | 1695 | 2159 | 2115 | 2220 | 1955 | 1890,2 | 1668 | 2112 | 1,269 | 0,343 | 6,873 | 6,28E-12 | 5,22E-11 | 0,001111 | 0,0027476 | 0,0018559 | 0,0018559 | TRUE | 0,0280641 | No |
| orf19.6908 | 854 | 646 | 775 | 994 | 1112 | 1018 | 1106 | 775 | 766 | 822 | 793 | 820 | 951 | 971 | 987 | 1093 | 900,43 | 800 | 1000 | 1,247 | 0,318 | 5,299 | 1,16E-07 | 5,65E-07 | 0,001228 | 0,0035917 | 0,0023363 | 0,0023363 | TRUE | 0,0413115 | No |
| orf19.6909 | 187 | 179 | 152 | 216 | 253 | 268 | 252 | 167 | 168 | 228 | 156 | 178 | 216 | 256 | 225 | 236 | 207,72 | 182 | 233 | 1,286 | 0,363 | 2,963 | 0,00304398 | 0,00690643 | 0,010098 | 0,0089682 | 0,0094639 | 0,0094639 | TRUE | 0,2686337 | No |
| orf19.691 | 2E+05 | 1E+05 | 1E+05 | 2E+05 | 2E+05 | 1E+05 | 2E+05 | 99615 | 2E+05 | 1E+05 | 2E+05 | 1E+05 | 1E+05 | 1E+05 | 1E+05 | 1E+05 | 146450 | 1E+05 | 1E+05 | 0,964 | -0,053 | -1,732 | 0,08325105 | 0,13093515 | 0,000537 | 0,0019894 | 0,0008949 | 0,0008949 | TRUE | 0,0056009 | No |
| orf19.6910 | 68 | 59 | 48 | 83 | 77 | 61 | 62 | 44 | 61 | 75 | 49 | 68 | 66 | 58 | 55 | 62 | 61,89 | 63 | 60 | 0,954 | -0,067 | -0,367 | 0,7132972 | 0,77397319 | 1,00E-08 | 0,0254362 | 0,0162264 | 0,0162264 | TRUE | 0,1914448 | No |
| orf19.6911 | 4 | 0 | 2 | 1 | 1 | 1 | 3 | 0 | 4 | 0 | 2 | 1 | 1 | 1 | 3 | 0 | 1,37 | 2 | 1 | 0,714 | -0,485 | -0,393 | 0,69406229 | NA | 1,00E-08 | 1,0630066 | 0,7425155 | 0,7425155 | TRUE | 0,314407 | No |
| orf19.6912 | 676 | 536 | 652 | 785 | 1075 | 1070 | 1116 | 705 | 606 | 682 | 667 | 647 | 919 | 1021 | 996 | 994 | 816,72 | 650 | 982 | 1,512 | 0,596 | 9,681 | 3,62E-22 | 7,68E-21 | 0,001162 | 0,003757 | 0,002363 | 0,002363 | TRUE | 0,0251607 | No |
| orf19.6913 | 742 | 602 | 706 | 847 | 1222 | 1221 | 1248 | 692 | 666 | 766 | 722 | 698 | 1045 | 1165 | 1114 | 976 | 894,07 | 713 | 1075 | 1,514 | 0,598 | 8,568 | 1,06E-17 | 1,59E-16 | 0,003139 | 0,0036032 | 0,003507 | 0,003507 | TRUE | 0,041691 | No |
| orf19.6915 | 1303 | 1000 | 1262 | 1501 | 1507 | 1411 | 1513 | 917 | 1169 | 1273 | 1291 | 1238 | 1289 | 1346 | 1350 | 1293 | 1281,2 | 1243 | 1320 | 1,064 | 0,089 | 1,777 | 0,07549807 | 0,12028439 | 0,00052 | 0,0031126 | 0,0016326 | 0,0016326 | TRUE | 0,0196025 | No |
| orf19.6916 | 537 | 447 | 457 | 587 | 816 | 785 | 786 | 429 | 482 | 569 | 468 | 484 | 698 | 749 | 701 | 605 | 594,47 | 501 | 688 | 1,383 | 0,468 | 5,588 | 2,29E-08 | 1,21E-07 | 0,00579 | 0,0044215 | 0,0050107 | 0,0050107 | TRUE | 0,0907652 | No |
| orf19.6917 | 487 | 467 | 485 | 715 | 704 | 802 | 821 | 445 | 437 | 595 | 496 | 590 | 602 | 765 | 733 | 628 | 605,6 | 530 | 682 | 1,291 | 0,368 | 3,315 | 0,00091613 | 0,00231653 | 0,01494 | 0,0043767 | 0,0101662 | 0,0101662 | TRUE | 0,1651942 | No |
| orf19.6918 | 702 | 759 | 761 | 897 | 788 | 1040 | 981 | 554 | 630 | 966 | 779 | 740 | 674 | 992 | 875 | 781 | 804,66 | 779 | 830 | 1,069 | 0,096 | 0,73 | 0,46508853 | 0,55122252 | 0,027844 | 0,0037836 | 0,0154864 | 0,0154864 | TRUE | 0,3053782 | No |
| orf19.6919 | 182 | 153 | 190 | 209 | 126 | 152 | 142 | 80 | 163 | 195 | 194 | 172 | 108 | 145 | 127 | 113 | 152,14 | 181 | 123 | 0,684 | -0,549 | -4,273 | 1,93E-05 | 6,71E-05 | 0,005703 | 0,0115213 | 0,0090204 | 0,0090204 | TRUE | 0,1494287 | No |
| orf19.692 | 3478 | 2186 | 3054 | 3975 | 3940 | 3187 | 3665 | 2297 | 3120 | 2783 | 3125 | 3278 | 3370 | 3041 | 3270 | 3240 | 3153,3 | 3076 | 3230 | 1,049 | 0,069 | 1,225 | 0,22066411 | 0,29933135 | 0,002792 | 0,0024399 | 0,0027338 | 0,0027338 | TRUE | 0,0486268 | No |
| orf19.69.2 | 15 | 12 | 15 | 24 | 17 | 32 | 15 | 13 | 13 | 15 | 15 | 20 | 15 | 31 | 13 | 18 | 17,58 | 16 | 19 | 1,193 | 0,255 | 0,703 | 0,48235075 | 0,56778278 | 0,049762 | 0,0845425 | 0,0688957 | 0,0688957 | TRUE | 0,9120894 | No |
| orf19.6920 | 1590 | 1629 | 1563 | 1978 | 1485 | 2036 | 1871 | 1111 | 1426 | 2074 | 1599 | 1631 | 1270 | 1943 | 1670 | 1567 | 1647,5 | 1682 | 1612 | 0,959 | -0,061 | -0,358 | 0,72021122 | 0,78034789 | 0,027245 | 0,0028607 | 0,0146581 | 0,0272452 | TRUE | 0,293407 | No |
| orf19.6921 | 1624 | 1220 | 1424 | 1819 | 1978 | 1912 | 2069 | 1124 | 1457 | 1553 | 1457 | 1500 | 1692 | 1824 | 1846 | 1585 | 1614,3 | 1492 | 1737 | 1,167 | 0,223 | 3,907 | 9,35E-05 | 0,00028769 | 0,002105 | 0,0028788 | 0,0025029 | 0,0025029 | TRUE | 0,0383806 | No |
| orf19.6922 | 0 | 0 | 0 | 0 | 0 | 0 | 0 | 0 | 0 | 0 | 0 | 0 | 0 | 0 | 0 | 0 | 0 | 0 | 0 | NA | NA | NA | NA | NA | NA | NA | NA | NA | NA | NA | NA |
| orf19.6923 | 2360 | 1470 | 1961 | 2384 | 2143 | 1832 | 1942 | 1149 | 2117 | 1871 | 2006 | 1966 | 1833 | 1748 | 1733 | 1621 | 1861,9 | 1990 | 1734 | 0,872 | -0,198 | -3,657 | 0,00025539 | 0,00072995 | 0,001853 | 0,0027592 | 0,0022733 | 0,0022733 | TRUE | 0,0223849 | No |
| orf19.6923.1 | 2278 | 1354 | 1683 | 2145 | 1820 | 1375 | 1383 | 860 | 2044 | 1724 | 1722 | 1769 | 1557 | 1312 | 1234 | 1213 | 1571,7 | 1815 | 1329 | 0,733 | -0,448 | -5,213 | 1,86E-07 | 8,78E-07 | 0,009778 | 0,0029032 | 0,0064404 | 0,0064404 | TRUE | 0,1622346 | No |
| orf19.6924 | 5909 | 3770 | 4633 | 5904 | 4832 | 3699 | 3959 | 2726 | 5301 | 4800 | 4740 | 4869 | 4132 | 3529 | 3533 | 3845 | 4343,6 | 4928 | 3760 | 0,763 | -0,39 | -6,416 | 1,40E-10 | 9,95E-10 | 0,003978 | 0,0023137 | 0,0033217 | 0,0033217 | TRUE | 0,0554968 | No |
| orf19.6925 | 7476 | 5093 | 6468 | 8174 | 6157 | 5565 | 6051 | 4462 | 6707 | 6484 | 6617 | 6740 | 5266 | 5310 | 5400 | 6293 | 6102,1 | 6637 | 5567 | 0,838 | -0,255 | -4,442 | 8,92E-06 | 3,27E-05 | 0,00376 | 0,0022174 | 0,0030029 | 0,0030029 | TRUE | 0,0921048 | No |
| orf19.6926 | 2949 | 2020 | 2713 | 2993 | 2978 | 2603 | 2920 | 1802 | 2646 | 2572 | 2776 | 2468 | 2547 | 2484 | 2606 | 2542 | 2579,8 | 2616 | 2545 | 0,973 | -0,039 | -0,853 | 0,39346979 | 0,48224531 | 0,001086 | 0,0025422 | 0,0016188 | 0,0016188 | TRUE | 0,0207517 | No |
| orf19.6927 | 1388 | 940 | 1272 | 1608 | 1413 | 1200 | 1336 | 838 | 1245 | 1197 | 1301 | 1326 | 1208 | 1145 | 1192 | 1182 | 1224,6 | 1267 | 1182 | 0,931 | -0,103 | -2,021 | 0,04332099 | 0,0741306 | 0,000503 | 0,003165 | 0,0016601 | 0,0016601 | TRUE | 0,0154345 | No |
| orf19.6928 | 23351 | 15903 | 21051 | 25920 | 19667 | 17258 | 18386 | 12115 | 20948 | 20246 | 21537 | 21374 | 16820 | 16467 | 16406 | 17087 | 18861 | 21026 | 16695 | 0,794 | -0,333 | -10,528 | 6,45E-26 | 1,80E-24 | 0,000486 | 0,0020565 | 0,0009082 | 0,0009082 | TRUE | 0,0075282 | No |
| orf19.6929 | 1326 | 827 | 1139 | 1363 | 1182 | 978 | 1052 | 714 | 1190 | 1053 | 1165 | 1124 | 1011 | 933 | 939 | 1007 | 1052,7 | 1133 | 972 | 0,856 | -0,224 | -3,881 | 0,00010402 | 0,00031757 | 0,001311 | 0,0033586 | 0,0022383 | 0,0022383 | TRUE | 0,0248025 | No |
| orf19.693 | 198 | 143 | 164 | 200 | 169 | 176 | 216 | 119 | 178 | 182 | 168 | 165 | 145 | 168 | 193 | 168 | 170,68 | 173 | 168 | 0,974 | -0,039 | -0,332 | 0,73986194 | 0,79646847 | 0,003234 | 0,0104849 | 0,0072069 | 0,0072069 | TRUE | 0,1145884 | No |

| | | | | | | | | | | | | | | | | | | | | | | | | | | | | | | | |
|---|---|---|---|---|---|---|---|---|---|---|---|---|---|---|---|---|---|---|---|---|---|---|---|---|---|---|---|---|---|---|---|
| orf19.6930 | 2485 | 1666 | 2002 | 2533 | 2568 | 2235 | 2450 | 1664 | 2229 | 2121 | 2048 | 2089 | 2196 | 2133 | 2186 | 2347 | 2168,7 | 2122 | 2216 | 1,042 | 0,06 | 1,283 | 0,19939339 | 0,273881 | 0,000915 | 0,0026489 | 0,0016308 | 0,0016308 | TRUE | 0,0177433 | No |
| orf19.6931 | 2432 | 1686 | 2095 | 2528 | 2477 | 2190 | 2373 | 1422 | 2182 | 2146 | 2143 | 2085 | 2118 | 2090 | 2118 | 2006 | 2110,9 | 2139 | 2083 | 0,975 | -0,036 | -0,909 | 0,36341221 | 0,45204566 | 1,00E-08 | 0,0026672 | 0,0010309 | 0,0010309 | TRUE | 0,0066684 | No |
| orf19.6933 | 805 | 511 | 733 | 856 | 988 | 861 | 887 | 646 | 722 | 651 | 750 | 706 | 845 | 822 | 792 | 911 | 774,71 | 707 | 842 | 1,186 | 0,246 | 3,744 | 0,00018132 | 0,00053193 | 0,001841 | 0,0038534 | 0,0028385 | 0,0028385 | TRUE | 0,0320176 | No |
| orf19.6934 | 1479 | 985 | 1207 | 1537 | 1421 | 1196 | 1428 | 834 | 1327 | 1254 | 1235 | 1267 | 1215 | 1141 | 1274 | 1176 | 1236,3 | 1271 | 1202 | 0,946 | -0,08 | -1,519 | 0,12863795 | 0,18937927 | 0,000873 | 0,0031538 | 0,0018445 | 0,0018445 | TRUE | 0,0203812 | No |
| orf19.6936 | 1901 | 1157 | 1627 | 1990 | 1736 | 1451 | 1728 | 1025 | 1705 | 1473 | 1665 | 1641 | 1485 | 1384 | 1542 | 1446 | 1542,6 | 1621 | 1464 | 0,902 | -0,148 | -2,549 | 0,01079596 | 0,02156321 | 0,002144 | 0,0029206 | 0,0025888 | 0,0025888 | TRUE | 0,0429793 | No |
| orf19.6937 | 65474 | 43193 | 59746 | 72393 | 65122 | 55033 | 60206 | 40419 | 58737 | 54989 | 61126 | 59696 | 55694 | 52511 | 53724 | 57007 | 56686 | 58637 | 54734 | 0,933 | -0,1 | -2,304 | 0,02122534 | 0,03935859 | 0,001647 | 0,0020051 | 0,0017745 | 0,0017745 | TRUE | 0,0214275 | No |
| orf19.6938 | 3558 | 2373 | 3040 | 3807 | 2529 | 1977 | 2112 | 1392 | 3192 | 3021 | 3110 | 3139 | 2163 | 1886 | 1885 | 1963 | 2545 | 3116 | 1974 | 0,634 | -0,658 | -12,635 | 1,36E-36 | 7,47E-35 | 0,001912 | 0,0025499 | 0,0021884 | 0,0021884 | TRUE | 0,052315 | No |
| orf19.6939 | 4271 | 2816 | 3777 | 4456 | 4261 | 3633 | 3906 | 2331 | 3832 | 3585 | 3864 | 3674 | 3644 | 3466 | 3485 | 3288 | 3604,9 | 3739 | 3471 | 0,929 | -0,106 | -2,397 | 0,01652426 | 0,03142696 | 0,001134 | 0,0023822 | 0,0016055 | 0,0016055 | TRUE | 0,0144976 | No |
| orf19.6940 | 55 | 33 | 40 | 47 | 28 | 19 | 22 | 9 | 49 | 42 | 41 | 39 | 24 | 18 | 20 | 13 | 30,68 | 43 | 19 | 0,446 | -1,166 | -4,44 | 9,00E-06 | 3,30E-05 | 1,00E-08 | 0,049298 | 0,0284346 | 0,0284346 | TRUE | 0,2033563 | No |
| orf19.6941 | 2475 | 1640 | 2290 | 2620 | 2489 | 2183 | 2492 | 1526 | 2220 | 2088 | 2343 | 2160 | 2129 | 2083 | 2224 | 2152 | 2174,9 | 2203 | 2147 | 0,974 | -0,038 | -0,801 | 0,42316095 | 0,51061478 | 0,000961 | 0,002647 | 0,001681 | 0,001681 | TRUE | 0,0216233 | No |
| orf19.6942 | 886 | 626 | 776 | 971 | 772 | 675 | 770 | 464 | 795 | 797 | 794 | 801 | 660 | 644 | 687 | 654 | 729,03 | 797 | 661 | 0,831 | -0,267 | -4,835 | 1,33E-06 | 5,52E-06 | 1,00E-08 | 0,0039708 | 0,0015517 | 0,0015517 | TRUE | 0,0085229 | No |
| orf19.6943 | 777 | 547 | 683 | 899 | 799 | 717 | 747 | 412 | 697 | 696 | 699 | 741 | 683 | 684 | 667 | 581 | 681,08 | 708 | 654 | 0,926 | -0,111 | -1,671 | 0,09474655 | 0,14629026 | 0,001345 | 0,004111 | 0,0027335 | 0,0027335 | TRUE | 0,0565325 | No |
| orf19.6944 | 1842 | 1162 | 1561 | 1895 | 2050 | 1709 | 1869 | 1156 | 1652 | 1479 | 1597 | 1563 | 1753 | 1631 | 1668 | 1630 | 1621,7 | 1573 | 1670 | 1,062 | 0,086 | 1,734 | 0,08289048 | 0,13043635 | 0,000991 | 0,0028747 | 0,0017688 | 0,0017688 | TRUE | 0,0178424 | No |
| orf19.6947 | 707 | 519 | 644 | 841 | 647 | 605 | 673 | 453 | 634 | 661 | 659 | 693 | 553 | 577 | 601 | 639 | 627,18 | 662 | 592 | 0,891 | -0,166 | -2,517 | 0,01182413 | 0,02336821 | 0,000845 | 0,0042942 | 0,0025581 | 0,0025581 | TRUE | 0,0280648 | No |
| orf19.6948 | 1849 | 1262 | 1636 | 1991 | 1612 | 1383 | 1507 | 982 | 1659 | 1607 | 1674 | 1642 | 1379 | 1320 | 1345 | 1385 | 1501,1 | 1646 | 1357 | 0,824 | -0,28 | -6,523 | 6,88E-11 | 5,12E-10 | 1,00E-08 | 0,0029466 | 0,0010919 | 0,0010919 | TRUE | 0,0040884 | No |
| orf19.6949 | 4 | 3 | 5 | 4 | 1 | 2 | 2 | 2 | 4 | 4 | 5 | 3 | 1 | 2 | 2 | 3 | 2,9 | 4 | 2 | 0,447 | -1,162 | -1,412 | 0,15793984 | NA | 1,00E-08 | 0,502766 | 0,240611 | 0,240611 | TRUE | 0,1337754 | No |
| orf19.695 | 1281 | 1173 | 1162 | 1540 | 1440 | 1590 | 1509 | 982 | 1149 | 1493 | 1189 | 1270 | 1232 | 1517 | 1347 | 1385 | 1322,7 | 1275 | 1370 | 1,075 | 0,105 | 1,195 | 0,2322746 | 0,31286911 | 0,009888 | 0,0030771 | 0,0066421 | 0,0066421 | TRUE | 0,1541928 | No |
| orf19.6950 | 1592 | 1043 | 1413 | 1753 | 1693 | 1497 | 1688 | 1152 | 1428 | 1328 | 1446 | 1446 | 1448 | 1428 | 1506 | 1625 | 1456,8 | 1412 | 1502 | 1,06 | 0,084 | 1,536 | 0,12462047 | 0,1841869 | 0,001619 | 0,002976 | 0,0021964 | 0,0021964 | TRUE | 0,0334375 | No |
| orf19.6951 | 2725 | 1823 | 2461 | 2838 | 3202 | 2864 | 3096 | 1850 | 2445 | 2321 | 2518 | 2340 | 2738 | 2733 | 2763 | 2609 | 2558,3 | 2406 | 2711 | 1,128 | 0,173 | 4,095 | 4,22E-05 | 0,00013816 | 0,000536 | 0,002547 | 0,0013231 | 0,0013231 | TRUE | 0,0116351 | No |
| orf19.6952 | 2350 | 1752 | 2052 | 2555 | 2076 | 1931 | 2008 | 1193 | 2108 | 2230 | 2099 | 2107 | 1775 | 1842 | 1792 | 1683 | 1954,7 | 2136 | 1773 | 0,832 | -0,265 | -5,977 | 2,27E-09 | 1,37E-08 | 0,000454 | 0,0027222 | 0,0013716 | 0,0013716 | TRUE | 0,0126113 | No |
| orf19.6953 | 3384 | 2168 | 2959 | 3600 | 3352 | 2806 | 3238 | 2090 | 3036 | 2760 | 3027 | 2969 | 2867 | 2677 | 2889 | 2948 | 2896,6 | 2948 | 2845 | 0,964 | -0,053 | -1,118 | 0,26361142 | 0,34731961 | 0,001339 | 0,0024807 | 0,0018187 | 0,0018187 | TRUE | 0,0213222 | No |
| orf19.6955 | 14675 | 9434 | 12539 | 15448 | 13289 | 11010 | 12433 | 7685 | 13165 | 12010 | 12829 | 12739 | 11365 | 10505 | 11094 | 10839 | 11818 | 12686 | 10951 | 0,863 | -0,212 | -5,231 | 1,69E-07 | 8,01E-07 | 0,001181 | 0,0021024 | 0,0014963 | 0,0014963 | TRUE | 0,0154813 | No |
| orf19.6956 | 691 | 481 | 544 | 720 | 765 | 686 | 728 | 394 | 620 | 612 | 557 | 594 | 654 | 655 | 650 | 556 | 612,09 | 596 | 629 | 1,06 | 0,084 | 1,184 | 0,23651274 | 0,31739302 | 0,001937 | 0,0043513 | 0,0032217 | 0,0032217 | TRUE | 0,061935 | No |
| orf19.6957 | 0 | 0 | 0 | 0 | 0 | 0 | 0 | 0 | 0 | 0 | 0 | 0 | 0 | 0 | 0 | 0 | 0 | 0 | 0 | NA | NA | NA | NA | NA | NA | NA | NA | NA | NA | NA | NA |
| orf19.6958 | 76 | 52 | 64 | 80 | 61 | 40 | 57 | 33 | 68 | 66 | 65 | 66 | 52 | 38 | 51 | 47 | 56,7 | 66 | 47 | 0,708 | -0,498 | -2,716 | 0,00661166 | 0,0139183 | 1,00E-08 | 0,0275843 | 0,014174 | 0,014174 | TRUE | 0,1389387 | No |
| orf19.696 | 29 | 17 | 22 | 21 | 25 | 23 | 24 | 12 | 26 | 22 | 23 | 17 | 21 | 22 | 21 | 17 | 21,14 | 22 | 20 | 0,946 | -0,081 | -0,274 | 0,7843819 | 0,83114047 | 1,00E-08 | 0,0706376 | 0,036319 | 0,036319 | TRUE | 0,1434305 | No |
| orf19.6966 | 825 | 906 | 887 | 1177 | 1052 | 1299 | 1290 | 783 | 740 | 1153 | 907 | 971 | 900 | 1239 | 1151 | 1104 | 1020,8 | 943 | 1098 | 1,166 | 0,222 | 1,79 | 0,07343354 | 0,11743112 | 0,023671 | 0,0034017 | 0,0137972 | 0,0137972 | TRUE | 0,2634239 | No |
| orf19.6967 | 3117 | 2390 | 2873 | 3414 | 3407 | 3291 | 3669 | 2126 | 2796 | 3043 | 2939 | 2815 | 2914 | 3140 | 3274 | 2999 | 2990 | 2898 | 3082 | 1,064 | 0,09 | 1,809 | 0,07042554 | 0,11331459 | 0,001669 | 0,002465 | 0,0020445 | 0,0020445 | TRUE | 0,0218803 | No |
| orf19.6968 | 3402 | 2689 | 3294 | 4005 | 4307 | 4604 | 4610 | 3037 | 3052 | 3423 | 3370 | 3303 | 3683 | 4393 | 4114 | 4283 | 3702,7 | 3287 | 4118 | 1,253 | 0,325 | 5,284 | 1,26E-07 | 6,11E-07 | 0,003752 | 0,0023716 | 0,0033618 | 0,0033618 | TRUE | 0,062315 | No |
| orf19.697 | 2996 | 2804 | 2884 | 3628 | 3172 | 3564 | 3619 | 2107 | 2688 | 3570 | 2951 | 2992 | 2713 | 3401 | 3229 | 2972 | 3064,3 | 3050 | 3079 | 1,01 | 0,015 | 0,163 | 0,8707176 | 0,90111898 | 0,011637 | 0,0024533 | 0,0073449 | 0,0073449 | TRUE | 0,1581097 | No |
| orf19.6970 | 32 | 43 | 28 | 42 | 31 | 36 | 49 | 30 | 29 | 55 | 29 | 35 | 27 | 34 | 44 | 42 | 36,7 | 37 | 37 | 1,008 | 0,012 | 0,044 | 0,96519271 | 0,97492902 | 0,045809 | 0,0415313 | 0,0433733 | 0,0433733 | TRUE | 0,6609185 | No |
| orf19.6971 | 60 | 62 | 60 | 96 | 102 | 137 | 107 | 82 | 54 | 79 | 61 | 79 | 87 | 131 | 95 | 116 | 87,8 | 68 | 107 | 1,567 | 0,648 | 3,494 | 0,00047546 | 0,00129211 | 0,024092 | 0,0185138 | 0,0210532 | 0,0210532 | TRUE | 0,2286527 | No |
| orf19.6972 | 612 | 548 | 497 | 664 | 838 | 812 | 936 | 434 | 549 | 698 | 508 | 548 | 717 | 775 | 835 | 612 | 655,19 | 576 | 735 | 1,282 | 0,359 | 3,241 | 0,00119056 | 0,00295695 | 0,0168 | 0,0041952 | 0,0102081 | 0,0102081 | TRUE | 0,2278346 | No |
| orf19.6973 | 9660 | 7325 | 8817 | 10764 | 12171 | 12133 | 12924 | 7151 | 8666 | 9326 | 9021 | 8876 | 10409 | 11577 | 11533 | 10086 | 9936,6 | 8972 | 10901 | 1,216 | 0,282 | 5,324 | 1,01E-07 | 4,98E-07 | 0,002813 | 0,0021256 | 0,0025853 | 0,0025853 | TRUE | 0,0305173 | No |
| orf19.6975 | 18923 | 11961 | 15697 | 19710 | 18719 | 14823 | 16028 | 11352 | 16976 | 15228 | 16060 | 16253 | 16009 | 14144 | 14302 | 16011 | 15623 | 16129 | 15116 | 0,937 | -0,094 | -1,713 | 0,08667522 | 0,13546652 | 0,003246 | 0,0020724 | 0,0028221 | 0,0028221 | TRUE | 0,0229454 | No |
| orf19.6976 | 1707 | 1472 | 1596 | 1970 | 1776 | 1790 | 1768 | 1032 | 1531 | 1874 | 1633 | 1624 | 1519 | 1708 | 1578 | 1456 | 1615,4 | 1666 | 1565 | 0,942 | -0,087 | -1,217 | 0,22348621 | 0,30241107 | 0,005393 | 0,0028782 | 0,0042333 | 0,0042333 | TRUE | 0,0827796 | No |
| orf19.6977 | 271 | 291 | 256 | 307 | 257 | 375 | 332 | 174 | 243 | 370 | 262 | 253 | 220 | 358 | 296 | 245 | 280,99 | 282 | 280 | 0,997 | -0,005 | -0,028 | 0,97736288 | 0,98378316 | 0,039138 | 0,0071459 | 0,0224475 | 0,0224475 | TRUE | 0,4753476 | No |
| orf19.6978 | 97 | 83 | 89 | 114 | 98 | 112 | 106 | 67 | 87 | 106 | 91 | 94 | 84 | 107 | 95 | 94 | 94,69 | 94 | 95 | 1,009 | 0,012 | 0,086 | 0,93162235 | 0,95060921 | 1,00E-08 | 0,0173108 | 0,0098044 | 0,0098044 | TRUE | 0,0751945 | No |
| orf19.6979 | 2167 | 1613 | 1929 | 2447 | 2282 | 2084 | 2277 | 1363 | 1944 | 2054 | 1974 | 2018 | 1952 | 1988 | 2032 | 1922 | 1985,4 | 1998 | 1974 | 0,99 | -0,015 | -0,368 | 0,71304984 | 0,77384455 | 1,00E-08 | 0,0027107 | 0,0010927 | 0,0010927 | TRUE | 0,0047936 | No |
| orf19.698 | 4778 | 3975 | 4689 | 5992 | 5070 | 5069 | 5631 | 3211 | 4286 | 5061 | 4797 | 4941 | 4336 | 4837 | 5025 | 4529 | 4726,5 | 4771 | 4682 | 0,982 | -0,027 | -0,421 | 0,67395849 | 0,73943437 | 0,003935 | 0,0022867 | 0,0036748 | 0,0036748 | TRUE | 0,0573921 | No |
| orf19.6980 | 1483 | 1151 | 1333 | 1712 | 1564 | 1609 | 1778 | 1139 | 1330 | 1465 | 1364 | 1412 | 1338 | 1535 | 1587 | 1606 | 1454,6 | 1393 | 1516 | 1,088 | 0,122 | 1,864 | 0,06232239 | 0,10174808 | 0,003704 | 0,0029775 | 0,0033953 | 0,0033953 | TRUE | 0,0780996 | No |
| orf19.6981 | 540 | 420 | 441 | 554 | 931 | 974 | 974 | 565 | 484 | 535 | 451 | 457 | 796 | 929 | 869 | 797 | 664,84 | 482 | 848 | 1,767 | 0,821 | 10,614 | 2,58E-26 | 7,59E-25 | 0,00383 | 0,004163 | 0,0041202 | 0,0041202 | TRUE | 0,0576193 | No |
| orf19.6982 | 891 | 732 | 770 | 1030 | 1135 | 1267 | 1175 | 728 | 799 | 932 | 788 | 849 | 971 | 1209 | 1048 | 1027 | 952,91 | 842 | 1064 | 1,265 | 0,339 | 4,251 | 2,13E-05 | 7,37E-05 | 0,006229 | 0,003503 | 0,0050531 | 0,0050531 | TRUE | 0,1026889 | No |
| orf19.6983 | 4130 | 3631 | 3809 | 4920 | 5226 | 5546 | 5779 | 3226 | 3705 | 4623 | 3897 | 4057 | 4469 | 5292 | 5157 | 4550 | 4468,7 | 4070 | 4867 | 1,197 | 0,259 | 3,364 | 0,0007696 | 0,00198274 | 0,008094 | 0,0023044 | 0,0054638 | 0,0054638 | TRUE | 0,1003429 | No |
| orf19.6984 | 2286 | 1963 | 2261 | 2872 | 2520 | 2794 | 2896 | 1709 | 2051 | 2499 | 2313 | 2368 | 2155 | 2666 | 2584 | 2410 | 2380,9 | 2308 | 2454 | 1,064 | 0,089 | 1,158 | 0,24676519 | 0,32872843 | 0,007004 | 0,0025893 | 0,0052878 | 0,0052878 | TRUE | 0,0832845 | No |
| orf19.6985 | 1394 | 1133 | 1236 | 1629 | 1824 | 1919 | 2034 | 1231 | 1251 | 1442 | 1265 | 1343 | 1560 | 1831 | 1815 | 1736 | 1530,4 | 1325 | 1736 | 1,311 | 0,39 | 5,822 | 5,81E-09 | 3,34E-08 | 0,003886 | 0,0029281 | 0,0036483 | 0,0036483 | TRUE | 0,0581163 | No |
| orf19.6986 | 379 | 298 | 366 | 450 | 389 | 445 | 438 | 290 | 340 | 379 | 374 | 371 | 333 | 425 | 391 | 409 | 377,76 | 366 | 390 | 1,061 | 0,086 | 0,946 | 0,34425574 | 0,43143273 | 0,004599 | 0,0058225 | 0,005209 | 0,005209 | TRUE | 0,1175967 | No |
| orf19.6987 | 6801 | 5137 | 6330 | 7523 | 7097 | 6899 | 7230 | 4493 | 6101 | 6540 | 6476 | 6204 | 6070 | 6583 | 6452 | 6337 | 6345,2 | 6330 | 6360 | 1,005 | 0,007 | 0,183 | 0,85485834 | 0,88950275 | 0,000828 | 0,0022083 | 0,0013945 | 0,0013945 | TRUE | 0,011873 | No |
| orf19.6988 | 439 | 352 | 420 | 528 | 483 | 449 | 523 | 288 | 394 | 448 | 430 | 435 | 413 | 428 | 467 | 406 | 427,68 | 427 | 428 | 1,008 | 0,012 | 0,156 | 0,87604884 | 0,90543634 | 0,001255 | 0,0053739 | 0,0033817 | 0,0033817 | TRUE | 0,0427598 | No |
| orf19.6989 | 964 | 690 | 811 | 1023 | 947 | 858 | 974 | 585 | 865 | 878 | 830 | 844 | 810 | 819 | 869 | 825 | 842,42 | 854 | 831 | 0,974 | -0,039 | -0,713 | 0,47584361 | 0,56124065 | 1,00E-08 | 0,0037028 | 0,0016237 | 0,0016237 | TRUE | 0,0124316 | No |
| orf19.699 | 789 | 1225 | 901 | 1283 | 805 | 1362 | 1277 | 641 | 708 | 1560 | 922 | 1058 | 688 | 1300 | 1140 | 904 | 1034,9 | 1062 | 1008 | 0,95 | -0,074 | -0,242 | 0,80905547 | 0,85183746 | 0,090232 | 0,0033823 | 0,0403537 | 0,0902319 | TRUE | 0,6147755 | No |
| orf19.6990 | 1367 | 1000 | 1125 | 1474 | 1165 | 1159 | 1166 | 720 | 1226 | 1273 | 1151 | 1215 | 996 | 1106 | 1040 | 1015 | 1128 | 1216 | 1039 | 0,856 | -0,225 | -4,106 | 4,02E-05 | 0,00013204 | 0,000965 | 0,0032665 | 0,0019909 | 0,0019909 | TRUE | 0,0225228 | No |
| orf19.6991 | 4331 | 2966 | 3656 | 4659 | 4592 | 4115 | 4259 | 2290 | 3885 | 3776 | 3740 | 3842 | 3927 | 3926 | 3800 | 3230 | 3766 | 3811 | 3721 | 0,977 | -0,033 | -0,539 | 0,58982669 | 0,66748517 | 0,003954 | 0,002365 | 0,0032988 | 0,0032988 | TRUE | 0,092942 | No |
| orf19.6992 | 9941 | 7076 | 8834 | 11087 | 11882 | 10421 | 11239 | 7027 | 8918 | 9009 | 9038 | 9142 | 10162 | 9943 | 10029 | 9911 | 9519 | 9027 | 10011 | 1,109 | 0,15 | 5,805 | 6,44E-09 | 3,66E-08 | 1,00E-08 | 0,002132 | 0,0005318 | 0,0005318 | TRUE | 0,0012693 | No |
| orf19.6993 | 19760 | 12639 | 17401 | 21186 | 12993 | 10874 | 11723 | 7632 | 17727 | 16091 | 17803 | 17470 | 11112 | 10376 | 10461 | 10764 | 13975 | 17273 | 10678 | 0,618 | -0,694 | -16,128 | 1,62E-58 | 2,43E-56 | 0,001462 | 0,0020834 | 0,0017027 | 0,0017027 | TRUE | 0,0257148 | No |
| orf19.6994 | 5133 | 3552 | 4645 | 5654 | 5397 | 4502 | 4859 | 3383 | 4605 | 4522 | 4752 | 4662 | 4616 | 4296 | 4336 | 4771 | 4570 | 4635 | 4505 | 0,971 | -0,043 | -0,982 | 0,32626191 | 0,413366 | 0,001205 | 0,0022972 | 0,0015949 | 0,0015949 | TRUE | 0,0184233 | No |
| orf19.6995 | 457 | 298 | 424 | 499 | 406 | 350 | 396 | 270 | 410 | 379 | 434 | 411 | 347 | 334 | 353 | 381 | 381,25 | 408 | 354 | 0,861 | -0,216 | -2,784 | 0,00537378 | 0,01153883 | 1,00E-08 | 0,0057873 | 0,0031487 | 0,0031487 | TRUE | 0,0246634 | No |
| orf19.6996 | 5154 | 3966 | 4898 | 5875 | 4568 | 4594 | 4642 | 2832 | 4624 | 5049 | 5011 | 4845 | 3907 | 4383 | 4142 | 3994 | 4494,4 | 4882 | 4106 | 0,842 | -0,249 | -5,149 | 2,62E-07 | 1,21E-06 | 0,001779 | 0,0023025 | 0,0020149 | 0,0020149 | TRUE | 0,0252384 | No |
| orf19.6997 | 319 | 230 | 293 | 396 | 270 | 247 | 240 | 146 | 286 | 293 | 300 | 327 | 231 | 236 | 214 | 206 | 261,5 | 302 | 222 | 0,737 | -0,44 | -4,78 | 1,76E-06 | 7,14E-06 | 1,00E-08 | 0,007531 | 0,0042465 | 0,0042465 | TRUE | 0,0394799 | No |

| | | | | | | | | | | | | | | | | | | | | | | | | | | | | | | | |
|---|---|---|---|---|---|---|---|---|---|---|---|---|---|---|---|---|---|---|---|---|---|---|---|---|---|---|---|---|---|---|---|
| orf19.6998 | 36 | 20 | 29 | 40 | 17 | 11 | 25 | 16 | 32 | 25 | 30 | 33 | 15 | 10 | 22 | 23 | 23,79 | 30 | 18 | 0,567 | -0,818 | -2,7 | 0,00692647 | 0,01449467 | 1,00E-08 | 0,0630004 | 0,042563 | 0,042563 | TRUE | 0,3708842 | No |
| orf19.6999 | 13 | 11 | 16 | 12 | 3 | 14 | 13 | 5 | 12 | 14 | 16 | 10 | 3 | 13 | 12 | 7 | 10,81 | 13 | 9 | 0,673 | -0,572 | -1,198 | 0,23099611 | NA | 0,099458 | 0,1362286 | 0,122717 | 0,122717 | TRUE | 0,7089962 | No |
| orf19.700 | 2245 | 1696 | 1970 | 2343 | 2001 | 1942 | 2045 | 1130 | 2014 | 2159 | 2016 | 1932 | 1711 | 1853 | 1825 | 1594 | 1888 | 2030 | 1746 | 0,862 | -0,214 | -3,683 | 0,00023081 | 0,00066381 | 0,002722 | 0,0027484 | 0,0027045 | 0,0027045 | TRUE | 0,0383592 | No |
| orf19.7001 | 13813 | 9740 | 12708 | 15248 | 13878 | 12731 | 13760 | 8770 | 12392 | 12400 | 13002 | 12574 | 11869 | 12148 | 12279 | 12369 | 12379 | 12592 | 12166 | 0,966 | -0,05 | -1,657 | 0,09759756 | 0,14971904 | 0,00031 | 0,0020968 | 0,0007927 | 0,0007927 | TRUE | 0,0058414 | No |
| orf19.7002 | 3 | 5 | 3 | 9 | 11 | 10 | 9 | 4 | 3 | 6 | 3 | 7 | 9 | 10 | 8 | 6 | 6,52 | 5 | 8 | 1,699 | 0,764 | 1,385 | 0,16599777 | NA | 1,00E-08 | 0,2245924 | 0,1306745 | 0,1306745 | TRUE | 0,3459707 | No |
| orf19.7006 | 4638 | 3095 | 4267 | 5018 | 7566 | 6403 | 7318 | 4580 | 4161 | 3940 | 4366 | 4138 | 6471 | 6110 | 6530 | 6460 | 5271,8 | 4151 | 6393 | 1,539 | 0,622 | 14,883 | 4,28E-50 | 4,59E-48 | 0,00106 | 0,0022549 | 0,0014791 | 0,0014791 | TRUE | 0,0144104 | No |
| orf19.7007 | 15 | 12 | 11 | 20 | 23 | 17 | 21 | 12 | 13 | 15 | 11 | 16 | 20 | 16 | 19 | 17 | 16 | 14 | 18 | 1,27 | 0,345 | 1,032 | 0,30211221 | 0,38800559 | 1,00E-08 | 0,0926858 | 0,0447903 | 0,0447903 | TRUE | 0,0798989 | No |
| orf19.7009 | 93 | 70 | 85 | 99 | 110 | 106 | 99 | 66 | 83 | 89 | 87 | 82 | 94 | 101 | 88 | 93 | 89,72 | 85 | 94 | 1,108 | 0,147 | 1,033 | 0,30178363 | 0,38774937 | 1,00E-08 | 0,0181591 | 0,0084294 | 0,0084294 | TRUE | 0,0255244 | No |
| orf19.701 | 32 | 34 | 40 | 50 | 31 | 38 | 27 | 19 | 29 | 43 | 41 | 41 | 27 | 36 | 24 | 27 | 33,48 | 38 | 28 | 0,742 | -0,431 | -1,721 | 0,08519666 | 0,13357556 | 1,00E-08 | 0,0453448 | 0,0296756 | 0,0296756 | TRUE | 0,2397704 | No |
| orf19.7010 | 1387 | 910 | 1255 | 1534 | 1810 | 1549 | 1705 | 1130 | 1244 | 1159 | 1284 | 1265 | 1548 | 1478 | 1521 | 1594 | 1386,6 | 1238 | 1535 | 1,237 | 0,307 | 6,169 | 6,87E-10 | 4,47E-09 | 0,000526 | 0,0030265 | 0,0016418 | 0,0016418 | TRUE | 0,0202488 | No |
| orf19.7011 | 15509 | 11152 | 14289 | 16672 | 13436 | 12240 | 13300 | 8077 | 13913 | 14198 | 14619 | 13748 | 11491 | 11679 | 11868 | 11392 | 12863 | 14120 | 11608 | 0,822 | -0,282 | -8,867 | 7,50E-19 | 1,24E-17 | 0,000425 | 0,0020924 | 0,0008932 | 0,0008932 | TRUE | 0,0069884 | No |
| orf19.7012 | 1695 | 1143 | 1374 | 1818 | 2124 | 1866 | 1799 | 1226 | 1521 | 1455 | 1406 | 1499 | 1817 | 1780 | 1605 | 1729 | 1601,5 | 1470 | 1733 | 1,178 | 0,236 | 4,435 | 9,20E-06 | 3,37E-05 | 0,00157 | 0,002886 | 0,0020954 | 0,0020954 | TRUE | 0,0311675 | No |
| orf19.7013 | 218 | 137 | 184 | 213 | 235 | 278 | 296 | 203 | 196 | 174 | 188 | 176 | 201 | 265 | 264 | 286 | 218,82 | 184 | 254 | 1,376 | 0,461 | 3,851 | 0,00011762 | 0,00035509 | 0,00952 | 0,0086138 | 0,0090536 | 0,0090536 | TRUE | 0,2337992 | No |
| orf19.7014 | 24 | 9 | 11 | 10 | 20 | 6 | 9 | 18 | 22 | 11 | 11 | 8 | 17 | 6 | 8 | 25 | 13,59 | 13 | 14 | 1,041 | 0,057 | 0,116 | 0,90775373 | NA | 0,22251 | 0,1087844 | 0,1607763 | 0,1607763 | TRUE | 0,3642116 | No |
| orf19.7015 | 12676 | 7826 | 10529 | 12967 | 13605 | 10792 | 11269 | 8168 | 11372 | 9963 | 10772 | 10693 | 11635 | 10297 | 10056 | 11520 | 10789 | 10700 | 10877 | 1,016 | 0,023 | 0,386 | 0,69922385 | 0,76145313 | 0,003663 | 0,0021141 | 0,0033764 | 0,0033764 | TRUE | 0,031685 | No |
| orf19.7016 | 2113 | 1558 | 2001 | 2238 | 2767 | 2553 | 2753 | 1710 | 1896 | 1984 | 2047 | 1845 | 2366 | 2436 | 2457 | 2412 | 2180,3 | 1943 | 2418 | 1,246 | 0,317 | 7,073 | 1,52E-12 | 1,36E-11 | 0,000728 | 0,0026453 | 0,0014641 | 0,0014641 | TRUE | 0,0163582 | No |
| orf19.7017 | 90 | 61 | 69 | 129 | 86 | 70 | 92 | 52 | 81 | 78 | 71 | 106 | 74 | 67 | 82 | 73 | 78,89 | 84 | 74 | 0,877 | -0,19 | -1,099 | 0,27187521 | 0,35633208 | 0,010888 | 0,0203804 | 0,0160562 | 0,0160562 | TRUE | 0,3414864 | No |
| orf19.7018 | 17432 | 11833 | 14045 | 17744 | 16107 | 13885 | 14334 | 10241 | 15638 | 15065 | 14369 | 14632 | 13775 | 13249 | 12791 | 14444 | 14245 | 14926 | 13565 | 0,909 | -0,138 | -2,975 | 0,00292971 | 0,00667237 | 0,00197 | 0,0020814 | 0,0020076 | 0,0020076 | TRUE | 0,0229787 | No |
| orf19.7019 | 2252 | 1586 | 2064 | 2425 | 2283 | 2036 | 2273 | 1533 | 2020 | 2019 | 2112 | 2000 | 1952 | 1943 | 2028 | 2162 | 2029,6 | 2038 | 2021 | 0,99 | -0,014 | -0,3 | 0,76389152 | 0,81569566 | 0,000848 | 0,0026948 | 0,0016238 | 0,0016238 | TRUE | 0,0243223 | No |
| orf19.702 | 6120 | 4315 | 5479 | 6655 | 6542 | 5993 | 6461 | 3833 | 5490 | 5493 | 5606 | 5488 | 5595 | 5718 | 5765 | 5406 | 5570,2 | 5519 | 5621 | 1,019 | 0,028 | 0,84 | 0,40079205 | 0,48947984 | 0,000222 | 0,0022401 | 0,0008512 | 0,0008512 | TRUE | 0,0075456 | No |
| orf19.7020 | 7518 | 5519 | 6846 | 8360 | 7749 | 7253 | 7700 | 4772 | 6744 | 7026 | 7004 | 6894 | 6627 | 6921 | 6871 | 6730 | 6852,2 | 6917 | 6787 | 0,982 | -0,027 | -0,861 | 0,38938261 | 0,4784071 | 0,000237 | 0,0021914 | 0,0007838 | 0,0007838 | TRUE | 0,0034653 | No |
| orf19.7021 | 21644 | 15208 | 19167 | 23356 | 33380 | 30412 | 31617 | 19542 | 19417 | 19361 | 19610 | 19260 | 28548 | 29018 | 28213 | 27562 | 23874 | 19412 | 28335 | 1,46 | 0,546 | 20,561 | 6,12E-94 | 3,68E-91 | 0,000204 | 0,0020403 | 0,0006335 | 0,0006335 | TRUE | 0,0040421 | No |
| orf19.7022 | 892 | 702 | 818 | 1037 | 1326 | 1250 | 1287 | 738 | 800 | 894 | 837 | 855 | 1134 | 1193 | 1148 | 1041 | 987,75 | 846 | 1129 | 1,339 | 0,421 | 7,071 | 1,53E-12 | 1,37E-11 | 0,001276 | 0,0034492 | 0,002368 | 0,002368 | TRUE | 0,0293818 | No |
| orf19.7023 | 217 | 153 | 171 | 245 | 272 | 264 | 266 | 176 | 195 | 195 | 175 | 202 | 233 | 252 | 237 | 248 | 217,07 | 192 | 242 | 1,262 | 0,335 | 3,475 | 0,00051101 | 0,00137675 | 1,00E-08 | 0,0086673 | 0,0042896 | 0,0042896 | TRUE | 0,0370115 | No |
| orf19.7024 | 8 | 2 | 14 | 11 | 6 | 3 | 4 | 4 | 7 | 3 | 14 | 9 | 5 | 3 | 4 | 6 | 6,29 | 8 | 4 | 0,503 | -0,99 | -1,633 | 0,10237879 | NA | 1,00E-08 | 0,2327653 | 0,1755311 | 0,1755311 | TRUE | 0,1505893 | No |
| orf19.7025 | 2078 | 1374 | 1742 | 2148 | 2166 | 1981 | 2106 | 1430 | 1864 | 1749 | 1782 | 1771 | 1852 | 1890 | 1879 | 2017 | 1850,7 | 1792 | 1910 | 1,063 | 0,088 | 1,966 | 0,04930719 | 0,08320357 | 0,000406 | 0,0027639 | 0,0013994 | 0,0013994 | TRUE | 0,0156157 | No |
| orf19.7027 | 7722 | 5252 | 6866 | 8216 | 14477 | 12915 | 14901 | 8440 | 6927 | 6686 | 7025 | 6775 | 12381 | 12323 | 13297 | 11904 | 9664,8 | 6853 | 12476 | 1,821 | 0,865 | 20,901 | 5,24E-97 | 3,94E-94 | 0,0012 | 0,0021297 | 0,00153 | 0,00153 | TRUE | 0,0240175 | No |
| orf19.7029 | 158 | 130 | 170 | 201 | 168 | 182 | 178 | 123 | 142 | 166 | 174 | 166 | 144 | 174 | 159 | 173 | 162,07 | 162 | 162 | 1,001 | 0,002 | 0,016 | 0,98705444 | 0,9896901 | 1,00E-08 | 0,0109367 | 0,0071506 | 0,0071506 | TRUE | 0,0782855 | No |
| orf19.703 | 1358 | 899 | 1102 | 1496 | 1682 | 1396 | 1479 | 1009 | 1218 | 1145 | 1127 | 1234 | 1438 | 1332 | 1320 | 1423 | 1279,7 | 1181 | 1378 | 1,164 | 0,22 | 4,045 | 5,22E-05 | 0,00016852 | 0,001258 | 0,003114 | 0,0020442 | 0,0020442 | TRUE | 0,0117237 | No |
| orf19.7030 | 1921 | 1248 | 1803 | 2425 | 2105 | 1646 | 1944 | 1404 | 1723 | 1589 | 1845 | 2000 | 1800 | 1571 | 1735 | 1980 | 1780,3 | 1789 | 1772 | 0,988 | -0,017 | -0,21 | 0,83335942 | 0,87226889 | 0,008689 | 0,002795 | 0,0059864 | 0,0059864 | TRUE | 0,0765821 | No |
| orf19.7032 | 998 | 948 | 1044 | 1230 | 1045 | 1307 | 1229 | 780 | 895 | 1207 | 1068 | 1014 | 894 | 1247 | 1097 | 1100 | 1065,3 | 1046 | 1084 | 1,037 | 0,053 | 0,501 | 0,61654034 | 0,68979039 | 0,016008 | 0,0033423 | 0,0098388 | 0,0098388 | TRUE | 0,171796 | No |
| orf19.7033 | 1309 | 1133 | 1247 | 1544 | 1349 | 1591 | 1550 | 943 | 1174 | 1442 | 1276 | 1273 | 1154 | 1518 | 1383 | 1330 | 1318,8 | 1291 | 1346 | 1,043 | 0,061 | 0,708 | 0,47921055 | 0,56452883 | 0,009331 | 0,0030803 | 0,0064501 | 0,0064501 | TRUE | 0,1148935 | No |
| orf19.7034 | 2258 | 1550 | 2001 | 2494 | 2684 | 2573 | 2920 | 1789 | 2026 | 1973 | 2047 | 2057 | 2295 | 2455 | 2606 | 2523 | 2247,8 | 2026 | 2470 | 1,218 | 0,284 | 5,888 | 3,92E-09 | 2,31E-08 | 0,001296 | 0,0026254 | 0,0017897 | 0,0017897 | TRUE | 0,0284561 | No |
| orf19.7035 | 1015 | 889 | 938 | 1193 | 1186 | 1168 | 1241 | 759 | 911 | 1132 | 960 | 984 | 1014 | 1114 | 1107 | 1071 | 1036,6 | 997 | 1076 | 1,083 | 0,115 | 1,632 | 0,10276375 | 0,15618634 | 0,004185 | 0,00338 | 0,003801 | 0,003801 | TRUE | 0,0942667 | No |
| orf19.7036 | 3729 | 2979 | 3392 | 4252 | 3835 | 3811 | 4125 | 2352 | 3345 | 3793 | 3470 | 3506 | 3280 | 3636 | 3681 | 3317 | 3503,6 | 3528 | 3478 | 0,987 | -0,019 | -0,342 | 0,73257209 | 0,79100437 | 0,002544 | 0,0023939 | 0,0027068 | 0,0027068 | TRUE | 0,0300688 | No |
| orf19.7037 | 1153 | 799 | 950 | 1192 | 954 | 904 | 902 | 616 | 1034 | 1017 | 972 | 983 | 816 | 863 | 805 | 869 | 919,83 | 1002 | 838 | 0,835 | -0,26 | -4,806 | 1,54E-06 | 6,32E-06 | 1,00E-08 | 0,0035578 | 0,0017084 | 0,0017084 | TRUE | 0,0082993 | No |
| orf19.7038 | 830 | 731 | 705 | 945 | 785 | 913 | 915 | 559 | 745 | 931 | 721 | 779 | 671 | 871 | 816 | 788 | 790,4 | 794 | 786 | 0,992 | -0,011 | -0,113 | 0,90995326 | 0,93341287 | 0,01131 | 0,0038162 | 0,0077081 | 0,0077081 | TRUE | 0,1525991 | No |
| orf19.704 | 2214 | 1446 | 1788 | 2208 | 2418 | 1993 | 2135 | 1417 | 1986 | 1841 | 1829 | 1821 | 2068 | 1902 | 1905 | 1999 | 1918,8 | 1869 | 1968 | 1,052 | 0,074 | 1,489 | 0,13660922 | 0,19935589 | 0,001262 | 0,0027361 | 0,0018307 | 0,0018307 | TRUE | 0,0220523 | No |
| orf19.7041 | 3330 | 2565 | 3159 | 3724 | 4271 | 4055 | 4240 | 2704 | 2987 | 3266 | 3232 | 3071 | 3653 | 3869 | 3783 | 3814 | 3459,4 | 3139 | 3780 | 1,205 | 0,269 | 6,378 | 1,79E-10 | 1,25E-09 | 0,000791 | 0,0023992 | 0,0014142 | 0,0014142 | TRUE | 0,0127952 | No |
| orf19.7042 | 980 | 847 | 879 | 1175 | 1119 | 1105 | 1079 | 712 | 879 | 1078 | 899 | 969 | 957 | 1054 | 963 | 1004 | 975,51 | 956 | 994 | 1,041 | 0,059 | 0,825 | 0,40927418 | 0,49735423 | 0,003949 | 0,0034677 | 0,0038157 | 0,0038157 | TRUE | 0,0823352 | No |
| orf19.7043 | 29 | 60 | 37 | 47 | 38 | 65 | 40 | 20 | 26 | 76 | 38 | 39 | 32 | 62 | 36 | 28 | 42,18 | 45 | 40 | 0,901 | -0,15 | -0,442 | 0,65829637 | 0,72676306 | 0,142353 | 0,0363968 | 0,0860464 | 0,0860464 | TRUE | 1,280955 | No |
| orf19.7043.1 | 917 | 788 | 778 | 1117 | 739 | 749 | 662 | 450 | 823 | 1003 | 796 | 921 | 632 | 715 | 591 | 635 | 764,38 | 886 | 643 | 0,727 | -0,46 | -5,321 | 1,03E-07 | 5,07E-07 | 0,007214 | 0,0038787 | 0,0058316 | 0,0058316 | TRUE | 0,0898248 | No |
| orf19.7044 | 3022 | 2111 | 2703 | 3196 | 4141 | 3632 | 4062 | 2444 | 2711 | 2688 | 2765 | 2635 | 3542 | 3466 | 3625 | 3447 | 3109,8 | 2700 | 3520 | 1,305 | 0,384 | 10,542 | 5,50E-26 | 1,55E-24 | 0,000173 | 0,0024463 | 0,0009448 | 0,0009448 | TRUE | 0,0049273 | No |
| orf19.7046 | 2678 | 1962 | 2211 | 2894 | 3637 | 3502 | 3506 | 2123 | 2402 | 2498 | 2262 | 2386 | 3110 | 3341 | 3129 | 2994 | 2765,5 | 2387 | 3144 | 1,318 | 0,399 | 8,3 | 1,04E-16 | 1,41E-15 | 0,00139 | 0,0025045 | 0,001847 | 0,001847 | TRUE | 0,0216625 | No |
| orf19.7047 | 3683 | 2784 | 3233 | 3829 | 3249 | 3131 | 3359 | 1943 | 3304 | 3544 | 3308 | 3157 | 2779 | 2988 | 2997 | 2740 | 3102,2 | 3328 | 2876 | 0,865 | -0,209 | -4,147 | 3,36E-05 | 0,00011217 | 0,001833 | 0,0024475 | 0,0021015 | 0,0021015 | TRUE | 0,0225312 | No |
| orf19.7048.1 | 6078 | 3852 | 4652 | 6233 | 5817 | 4405 | 4654 | 3228 | 5453 | 4904 | 4759 | 5140 | 4975 | 4203 | 4153 | 4553 | 4767,5 | 5064 | 4471 | 0,883 | -0,18 | -2,734 | 0,00625769 | 0,01322879 | 0,005122 | 0,002284 | 0,0039548 | 0,0039548 | TRUE | 0,071355 | No |
| orf19.7049 | 1237 | 931 | 1124 | 1316 | 911 | 902 | 898 | 592 | 1110 | 1185 | 1150 | 1085 | 779 | 861 | 801 | 835 | 975,77 | 1132 | 819 | 0,723 | -0,467 | -8,316 | 9,12E-17 | 1,25E-15 | 0,000733 | 0,0034673 | 0,0019741 | 0,0019741 | TRUE | 0,0153181 | No |
| orf19.705 | 1396 | 868 | 1129 | 1438 | 1553 | 1231 | 1459 | 816 | 1252 | 1105 | 1155 | 1186 | 1328 | 1175 | 1302 | 1151 | 1206,7 | 1174 | 1239 | 1,056 | 0,078 | 1,217 | 0,2236171 | 0,30252005 | 0,002857 | 0,0031825 | 0,0031196 | 0,0031196 | TRUE | 0,0281842 | No |
| orf19.7050 | 8853 | 6504 | 7507 | 9419 | 9265 | 8639 | 8712 | 5089 | 7942 | 8280 | 7680 | 7767 | 7924 | 8243 | 7774 | 7178 | 7848,5 | 7917 | 7780 | 0,983 | -0,024 | -0,505 | 0,61343786 | 0,68695893 | 0,001835 | 0,0021645 | 0,0020901 | 0,0020901 | TRUE | 0,0323335 | No |
| orf19.7051 | 130 | 113 | 115 | 134 | 98 | 123 | 142 | 70 | 117 | 144 | 118 | 110 | 84 | 117 | 127 | 99 | 114,41 | 122 | 107 | 0,88 | -0,185 | -1,176 | 0,2394339 | 0,3205256 | 0,015027 | 0,0146685 | 0,0148442 | 0,0148442 | TRUE | 0,2389588 | No |
| orf19.7052 | 701 | 558 | 652 | 777 | 786 | 782 | 817 | 454 | 629 | 710 | 667 | 641 | 672 | 746 | 729 | 640 | 679,35 | 662 | 697 | 1,058 | 0,082 | 1,179 | 0,23838372 | 0,31926202 | 0,002021 | 0,0041164 | 0,0031546 | 0,0031546 | TRUE | 0,030596 | No |
| orf19.7053 | 31197 | 23142 | 29299 | 36053 | 41900 | 40319 | 43666 | 27996 | 27987 | 29462 | 29976 | 29730 | 35834 | 38471 | 38965 | 39486 | 33739 | 29289 | 38189 | 1,304 | 0,383 | 9,368 | 7,38E-21 | 1,43E-19 | 0,001328 | 0,0020226 | 0,0015732 | 0,0015732 | TRUE | 0,0211566 | No |
| orf19.7054 | 63 | 28 | 43 | 54 | 54 | 43 | 43 | 38 | 57 | 36 | 44 | 45 | 46 | 41 | 38 | 54 | 44,98 | 46 | 45 | 0,972 | -0,04 | -0,187 | 0,85163594 | 0,88661042 | 1,00E-08 | 0,0342517 | 0,0226182 | 0,0226182 | TRUE | 0,2317108 | No |
| orf19.7055 | 3 | 3 | 5 | 5 | 0 | 5 | 2 | 0 | 3 | 4 | 5 | 4 | 0 | 5 | 2 | 0 | 2,79 | 4 | 2 | 0,434 | -1,205 | -1,291 | 0,19669701 | NA | 0,10972 | 0,5226594 | 0,4158766 | 0,4158766 | TRUE | 1,2584916 | No |
| orf19.7056 | 267 | 199 | 275 | 316 | 316 | 257 | 295 | 199 | 240 | 253 | 281 | 261 | 270 | 245 | 263 | 281 | 261,77 | 259 | 265 | 1,022 | 0,031 | 0,336 | 0,73680818 | 0,79401475 | 1,00E-08 | 0,0075252 | 0,0042549 | 0,0042549 | TRUE | 0,038202 | No |
| orf19.7057 | 6033 | 3997 | 5083 | 6318 | 6279 | 5103 | 5628 | 3570 | 5412 | 5089 | 5200 | 5210 | 5370 | 4869 | 5022 | 5035 | 5150,9 | 5228 | 5074 | 0,971 | -0,043 | -1,049 | 0,29404875 | 0,37878321 | 0,000939 | 0,0022614 | 0,0014257 | 0,0014257 | TRUE | 0,0192858 | No |
| orf19.7058 | 1297 | 950 | 1122 | 1415 | 1344 | 1142 | 1214 | 863 | 1164 | 1209 | 1148 | 1167 | 1149 | 1090 | 1083 | 1217 | 1153,4 | 1172 | 1135 | 0,967 | -0,049 | -0,926 | 0,35458737 | 0,44216707 | 0,000609 | 0,0032381 | 0,0018183 | 0,0018183 | TRUE | 0,0250732 | No |
| orf19.7059 | 948 | 564 | 831 | 877 | 872 | 740 | 826 | 510 | 850 | 718 | 850 | 723 | 746 | 706 | 737 | 719 | 756,26 | 785 | 727 | 0,925 | -0,112 | -1,539 | 0,12382251 | 0,18327806 | 0,003636 | 0,0038991 | 0,0037577 | 0,0037577 | TRUE | 0,0377032 | No |
| orf19.706 | 24580 | 15879 | 21446 | 26112 | 21533 | 17923 | 19676 | 12211 | 22051 | 20216 | 21941 | 21532 | 18416 | 17102 | 17558 | 17223 | 19505 | 21435 | 17575 | 0,82 | -0,287 | -7,059 | 1,68E-12 | 1,50E-11 | 0,001274 | 0,002054 | 0,0015305 | 0,0015305 | TRUE | 0,0177975 | No |

| | | | | | | | | | | | | | | | | | | | | | | | | | | | | | | | |
|---|---|---|---|---|---|---|---|---|---|---|---|---|---|---|---|---|---|---|---|---|---|---|---|---|---|---|---|---|---|---|---|
| orf19.7060 | 1167 | 755 | 1021 | 1220 | 1566 | 1322 | 1503 | 908 | 1047 | 961 | 1045 | 1006 | 1339 | 1261 | 1341 | 1281 | 1160,2 | 1015 | 1306 | 1,286 | 0,363 | 7,116 | 1,11E-12 | 1,01E-11 | 0,000307 | 0,0032308 | 0,0016266 | 0,0016266 | TRUE | 0,0133335 | No |
| orf19.7061 | 135 | 86 | 123 | 138 | 150 | 107 | 141 | 84 | 121 | 109 | 126 | 114 | 128 | 102 | 126 | 118 | 118,11 | 118 | 118 | 1,01 | 0,014 | 0,105 | 0,91598211 | 0,93815556 | 1,00E-08 | 0,0142702 | 0,0081345 | 0,0081345 | TRUE | 0,0963486 | No |
| orf19.7062 | 14913 | 9809 | 13397 | 16047 | 14493 | 12405 | 13546 | 8571 | 13379 | 12488 | 13706 | 13233 | 12395 | 11836 | 12088 | 12089 | 12652 | 13202 | 12102 | 0,917 | -0,126 | -3,41 | 0,00065031 | 0,0017039 | 0,000834 | 0,0020943 | 0,0012259 | 0,0012259 | TRUE | 0,015898 | No |
| orf19.7063 | 685 | 439 | 580 | 680 | 709 | 580 | 624 | 429 | 615 | 559 | 593 | 561 | 606 | 553 | 557 | 605 | 581,15 | 582 | 580 | 0,995 | -0,007 | -0,105 | 0,91651796 | 0,93854438 | 0,000727 | 0,0044775 | 0,0025837 | 0,0025837 | TRUE | 0,0174095 | No |
| orf19.7064 | 8522 | 5665 | 7406 | 9151 | 7770 | 6871 | 7364 | 4869 | 7645 | 7212 | 7577 | 7546 | 6645 | 6556 | 6571 | 6867 | 7077,5 | 7495 | 6660 | 0,888 | -0,172 | -5,169 | 2,35E-07 | 1,09E-06 | 0,000375 | 0,0021846 | 0,0009178 | 0,0009178 | TRUE | 0,0076114 | No |
| orf19.7065 | 535 | 341 | 506 | 592 | 746 | 654 | 647 | 378 | 480 | 434 | 518 | 488 | 638 | 624 | 577 | 533 | 536,56 | 480 | 593 | 1,237 | 0,307 | 3,889 | 0,00010058 | 0,00030828 | 0,003315 | 0,0046851 | 0,0041137 | 0,0041137 | TRUE | 0,0478487 | No |
| orf19.7067 | 3071 | 2051 | 2794 | 3275 | 2841 | 2562 | 2777 | 1720 | 2755 | 2611 | 2859 | 2701 | 2430 | 2445 | 2478 | 2426 | 2587,9 | 2732 | 2445 | 0,895 | -0,16 | -4,005 | 6,19E-05 | 0,00019659 | 0,000336 | 0,0025405 | 0,0011525 | 0,0011525 | TRUE | 0,0115312 | No |
| orf19.7068 | 500 | 295 | 409 | 555 | 486 | 389 | 438 | 293 | 449 | 376 | 418 | 458 | 416 | 371 | 391 | 413 | 411,39 | 425 | 398 | 0,93 | -0,104 | -1,258 | 0,20856426 | 0,285109 | 0,002625 | 0,0055083 | 0,0041315 | 0,0041315 | TRUE | 0,0640516 | No |
| orf19.7069 | 3835 | 2443 | 3143 | 4099 | 3156 | 2370 | 2554 | 1822 | 3440 | 3110 | 3216 | 3380 | 2699 | 2261 | 2279 | 2570 | 2869,4 | 3286 | 2452 | 0,746 | -0,424 | -6,517 | 7,19E-11 | 5,34E-10 | 0,004496 | 0,0024854 | 0,0037005 | 0,0037005 | TRUE | 0,0578482 | No |
| orf19.707 | 859 | 555 | 722 | 898 | 1045 | 889 | 956 | 619 | 771 | 707 | 739 | 741 | 894 | 848 | 853 | 873 | 803,06 | 740 | 867 | 1,171 | 0,228 | 4,126 | 3,69E-05 | 0,00012216 | 1,00E-08 | 0,0037872 | 0,001675 | 0,001675 | TRUE | 0,0096746 | No |
| orf19.7070 | 9 | 3 | 10 | 6 | 6 | 8 | 10 | 4 | 8 | 4 | 10 | 5 | 5 | 8 | 9 | 6 | 6,8 | 7 | 7 | 1,009 | 0,014 | 0,025 | 0,97969819 | NA | 1,00E-08 | 0,2154586 | 0,1261995 | 0,1261995 | TRUE | 0,272163 | No |
| orf19.7071 | 16 | 6 | 16 | 12 | 11 | 10 | 5 | 8 | 14 | 8 | 16 | 10 | 9 | 10 | 4 | 11 | 10,37 | 12 | 8 | 0,7 | -0,515 | -1,144 | 0,25253877 | NA | 1,00E-08 | 0,1419845 | 0,0949391 | 0,0949391 | TRUE | 0,2700174 | No |
| orf19.7072 | 5662 | 3923 | 4989 | 6085 | 7753 | 6593 | 7525 | 4759 | 5079 | 4994 | 5104 | 5018 | 6631 | 6291 | 6715 | 6712 | 5818 | 5049 | 6587 | 1,304 | 0,383 | 11,285 | 1,55E-29 | 5,44E-28 | 0,000388 | 0,002229 | 0,0009314 | 0,0009314 | TRUE | 0,011268 | No |
| orf19.7073 | 303 | 219 | 287 | 333 | 229 | 251 | 231 | 153 | 272 | 279 | 294 | 275 | 196 | 239 | 206 | 216 | 247,02 | 280 | 214 | 0,765 | -0,386 | -4,057 | 4,96E-05 | 0,00016053 | 1,00E-08 | 0,0078565 | 0,0045624 | 0,0045624 | TRUE | 0,0746898 | No |
| orf19.7074 | 826 | 541 | 670 | 825 | 867 | 796 | 804 | 538 | 741 | 689 | 685 | 680 | 741 | 760 | 717 | 759 | 721,6 | 699 | 744 | 1,063 | 0,088 | 1,511 | 0,1307838 | 0,19211469 | 1,00E-08 | 0,0039913 | 0,0019028 | 0,0019028 | TRUE | 0,020561 | No |
| orf19.7076 | 4962 | 3390 | 4399 | 5463 | 5436 | 4698 | 5131 | 2995 | 4451 | 4316 | 4501 | 4505 | 4649 | 4483 | 4579 | 4224 | 4463,4 | 4443 | 4484 | 1,01 | 0,014 | 0,356 | 0,72215946 | 0,78217668 | 0,000707 | 0,0023048 | 0,0012936 | 0,0012936 | TRUE | 0,017463 | No |
| orf19.7077 | 3414 | 2144 | 2964 | 3574 | 2406 | 2040 | 2245 | 1513 | 3063 | 2730 | 3032 | 2947 | 2058 | 1947 | 2003 | 2134 | 2489,2 | 2943 | 2036 | 0,69 | -0,535 | -10,685 | 1,20E-26 | 3,62E-25 | 0,001523 | 0,0025627 | 0,0019858 | 0,0019858 | TRUE | 0,027684 | No |
| orf19.7078 | 462 | 313 | 385 | 409 | 565 | 443 | 464 | 275 | 414 | 398 | 394 | 337 | 483 | 423 | 414 | 388 | 406,49 | 386 | 427 | 1,112 | 0,153 | 1,644 | 0,10011141 | 0,15292889 | 0,006163 | 0,0055508 | 0,0058615 | 0,0058615 | TRUE | 0,0933753 | No |
| orf19.7079 | 196 | 107 | 160 | 201 | 165 | 140 | 187 | 128 | 176 | 136 | 164 | 166 | 141 | 134 | 167 | 181 | 157,95 | 160 | 156 | 0,958 | -0,062 | -0,476 | 0,63403919 | 0,70516613 | 0,007996 | 0,0111705 | 0,009812 | 0,009812 | TRUE | 0,1018575 | No |
| orf19.708 | 261 | 184 | 252 | 287 | 399 | 360 | 426 | 226 | 234 | 234 | 258 | 237 | 341 | 344 | 380 | 319 | 293,31 | 241 | 346 | 1,443 | 0,529 | 5,953 | 2,64E-09 | 1,58E-08 | 1,00E-08 | 0,0069288 | 0,0040694 | 0,0040694 | TRUE | 0,0510289 | No |
| orf19.7080 | 1239 | 862 | 1089 | 1380 | 788 | 682 | 644 | 579 | 1112 | 1097 | 1114 | 1138 | 674 | 651 | 575 | 817 | 897,13 | 1115 | 679 | 0,606 | -0,723 | -8,306 | 9,93E-17 | 1,36E-15 | 0,00862 | 0,0035977 | 0,0060821 | 0,0060821 | TRUE | 0,2034398 | No |
| orf19.7081 | 16446 | 10735 | 14060 | 17587 | 18219 | 14872 | 16153 | 11126 | 14754 | 13667 | 14385 | 14502 | 15581 | 14190 | 14414 | 15692 | 14648 | 14327 | 14969 | 1,045 | 0,063 | 1,387 | 0,16530801 | 0,23462569 | 0,001787 | 0,0020786 | 0,0019008 | 0,0019008 | TRUE | 0,0149929 | No |
| orf19.7082 | 134 | 106 | 103 | 130 | 114 | 97 | 107 | 78 | 120 | 135 | 105 | 107 | 97 | 93 | 95 | 110 | 107,91 | 117 | 99 | 0,845 | -0,243 | -1,747 | 0,08059026 | 0,12735042 | 1,00E-08 | 0,0154324 | 0,009138 | 0,009138 | TRUE | 0,0915751 | No |
| orf19.7083 | 77 | 53 | 83 | 88 | 78 | 82 | 61 | 65 | 69 | 67 | 85 | 73 | 67 | 78 | 54 | 92 | 73,14 | 74 | 73 | 0,975 | -0,036 | -0,195 | 0,8454924 | 0,8825084 | 0,014308 | 0,0218287 | 0,0187107 | 0,0187107 | TRUE | 0,2309778 | No |
| orf19.7084 | 15692 | 10382 | 13689 | 16935 | 19530 | 16783 | 18836 | 11159 | 14077 | 13217 | 14005 | 13965 | 16703 | 16014 | 16808 | 15739 | 15066 | 13816 | 16316 | 1,181 | 0,24 | 6,561 | 5,33E-11 | 4,06E-10 | 0,000845 | 0,0020759 | 0,0012174 | 0,0012174 | TRUE | 0,0102505 | No |
| orf19.7085 | 10223 | 6780 | 8942 | 11105 | 13245 | 11224 | 12349 | 8677 | 9171 | 8632 | 9149 | 9157 | 11328 | 10710 | 11019 | 12238 | 10175 | 9027 | 11324 | 1,254 | 0,326 | 6,934 | 4,10E-12 | 3,51E-11 | 0,001992 | 0,0021222 | 0,0020251 | 0,0020251 | TRUE | 0,0355512 | No |
| orf19.7086 | 6605 | 4243 | 5588 | 6834 | 7339 | 6452 | 6870 | 4375 | 5925 | 5402 | 5717 | 5635 | 6277 | 6156 | 6130 | 6171 | 5926,7 | 5670 | 6184 | 1,09 | 0,124 | 3,422 | 0,00062078 | 0,00163153 | 0,000557 | 0,0022245 | 0,0010984 | 0,0010984 | TRUE | 0,0118801 | No |
| orf19.7088 | 2993 | 1973 | 2656 | 3229 | 3252 | 2623 | 2987 | 1920 | 2685 | 2512 | 2717 | 2663 | 2781 | 2503 | 2665 | 2708 | 2654,3 | 2644 | 2664 | 1,007 | 0,01 | 0,21 | 0,83356939 | 0,87233668 | 0,0012 | 0,0025265 | 0,0016977 | 0,0016977 | TRUE | 0,0205791 | No |
| orf19.7089 | 1604 | 1096 | 1408 | 1725 | 1800 | 1515 | 1735 | 1065 | 1439 | 1395 | 1441 | 1422 | 1539 | 1446 | 1548 | 1502 | 1466,6 | 1424 | 1509 | 1,059 | 0,083 | 1,866 | 0,06205096 | 0,10149801 | 1,00E-08 | 0,0029694 | 0,0012231 | 0,0012231 | TRUE | 0,0101745 | No |
| orf19.709 | 4746 | 3196 | 4041 | 4972 | 4441 | 3740 | 3988 | 2446 | 4258 | 4069 | 4134 | 4100 | 3798 | 3569 | 3559 | 3450 | 3867 | 4140 | 3594 | 0,869 | -0,203 | -5,054 | 4,33E-07 | 1,93E-06 | 0,000683 | 0,0023549 | 0,0012892 | 0,0012892 | TRUE | 0,0180655 | No |
| orf19.7091 | 1505 | 941 | 1276 | 1490 | 999 | 744 | 874 | 660 | 1350 | 1198 | 1305 | 1229 | 854 | 710 | 780 | 931 | 1044,7 | 1270 | 819 | 0,642 | -0,639 | -7,986 | 1,40E-15 | 1,70E-14 | 0,006927 | 0,0033692 | 0,0051287 | 0,0051287 | TRUE | 0,093959 | No |
| orf19.7092 | 1689 | 1095 | 1456 | 1707 | 1665 | 1359 | 1464 | 972 | 1515 | 1394 | 1490 | 1408 | 1424 | 1297 | 1306 | 1371 | 1400,6 | 1452 | 1350 | 0,929 | -0,106 | -2,022 | 0,04318124 | 0,0739756 | 0,001177 | 0,003016 | 0,001939 | 0,001939 | TRUE | 0,0181061 | No |
| orf19.7093 | 298 | 217 | 268 | 359 | 333 | 282 | 316 | 190 | 267 | 276 | 274 | 296 | 285 | 269 | 282 | 268 | 277,21 | 278 | 276 | 0,992 | -0,012 | -0,139 | 0,88921785 | 0,91621006 | 1,00E-08 | 0,0072164 | 0,0033799 | 0,0033799 | TRUE | 0,0229969 | No |
| orf19.7094 | 47 | 25 | 28 | 41 | 12 | 16 | 15 | 10 | 42 | 32 | 29 | 34 | 10 | 15 | 13 | 14 | 23,68 | 34 | 13 | 0,384 | -1,383 | -4,552 | 5,32E-06 | 2,02E-05 | 1,00E-08 | 0,063276 | 0,0361918 | 0,0361918 | TRUE | 0,1912765 | No |
| orf19.7095 | 2471 | 1594 | 2167 | 2607 | 2618 | 2072 | 2335 | 1401 | 2217 | 2029 | 2217 | 2150 | 2239 | 1977 | 2084 | 1976 | 2111,1 | 2153 | 2069 | 0,961 | -0,057 | -1,051 | 0,29326915 | 0,37794102 | 0,002012 | 0,0026672 | 0,0023397 | 0,0023397 | TRUE | 0,0377579 | No |
| orf19.7096 | 2208 | 1573 | 1989 | 2412 | 3187 | 2767 | 2979 | 1850 | 1981 | 2003 | 2035 | 1989 | 2726 | 2640 | 2658 | 2609 | 2330,1 | 2002 | 2658 | 1,329 | 0,411 | 11,134 | 8,54E-29 | 2,88E-27 | 1,00E-08 | 0,0026025 | 0,0008684 | 0,0008684 | TRUE | 0,0035981 | No |
| orf19.7097 | 2606 | 1643 | 2094 | 2758 | 2445 | 1923 | 2161 | 1366 | 2338 | 2092 | 2142 | 2274 | 2091 | 1835 | 1928 | 1927 | 2078,4 | 2212 | 1945 | 0,879 | -0,186 | -3,338 | 0,000844 | 0,0021596 | 0,002187 | 0,002678 | 0,0024918 | 0,0024918 | TRUE | 0,0319489 | No |
| orf19.7098 | 1415 | 964 | 1150 | 1543 | 1507 | 1291 | 1447 | 869 | 1269 | 1227 | 1177 | 1272 | 1289 | 1232 | 1291 | 1226 | 1247,9 | 1236 | 1260 | 1,019 | 0,027 | 0,549 | 0,58331209 | 0,66160828 | 1,00E-08 | 0,0031428 | 0,0015282 | 0,0015282 | TRUE | 0,0130273 | No |
| orf19.71 | 963 | 663 | 924 | 1099 | 1203 | 1035 | 1156 | 762 | 864 | 844 | 945 | 906 | 1029 | 988 | 1032 | 1075 | 960,28 | 890 | 1031 | 1,155 | 0,208 | 3,714 | 0,00020384 | 0,00059363 | 0,000488 | 0,0034913 | 0,0019705 | 0,0019705 | TRUE | 0,0198718 | No |
| orf19.7100 | 270 | 127 | 198 | 277 | 190 | 195 | 194 | 133 | 242 | 162 | 203 | 228 | 162 | 186 | 173 | 188 | 193,02 | 209 | 177 | 0,841 | -0,25 | -1,951 | 0,05111041 | 0,0857023 | 0,011766 | 0,0095006 | 0,0105357 | 0,0105357 | TRUE | 0,2246332 | No |
| orf19.7101 | 427 | 332 | 413 | 505 | 445 | 431 | 438 | 252 | 383 | 423 | 423 | 416 | 381 | 411 | 391 | 355 | 397,85 | 411 | 384 | 0,94 | -0,09 | -1,173 | 0,24081246 | 0,32207828 | 1,00E-08 | 0,0056284 | 0,0031266 | 0,0031266 | TRUE | 0,0254967 | No |
| orf19.7102 | 457 | 281 | 347 | 439 | 377 | 304 | 310 | 235 | 410 | 358 | 355 | 362 | 322 | 290 | 277 | 331 | 338,16 | 371 | 305 | 0,818 | -0,29 | -3,23 | 0,00123724 | 0,00306151 | 0,003062 | 0,0062724 | 0,0047439 | 0,0047439 | TRUE | 0,058888 | No |
| orf19.7103 | 1224 | 817 | 1061 | 1319 | 1256 | 1056 | 1156 | 736 | 1098 | 1040 | 1086 | 1088 | 1074 | 1008 | 1032 | 1038 | 1057,8 | 1078 | 1038 | 0,962 | -0,056 | -1,133 | 0,25735425 | 0,34026943 | 1,00E-08 | 0,0033518 | 0,0013875 | 0,0013875 | TRUE | 0,0073005 | No |
| orf19.7104 | 581 | 322 | 506 | 590 | 712 | 534 | 581 | 416 | 521 | 410 | 518 | 487 | 609 | 510 | 518 | 587 | 519,87 | 484 | 556 | 1,144 | 0,195 | 2,116 | 0,0343221 | 0,06041817 | 0,007601 | 0,0047719 | 0,0061861 | 0,0061861 | TRUE | 0,1125843 | No |
| orf19.7105 | 683 | 519 | 654 | 775 | 848 | 766 | 907 | 628 | 613 | 661 | 669 | 639 | 725 | 731 | 809 | 886 | 716,61 | 646 | 788 | 1,217 | 0,284 | 3,813 | 0,00013714 | 0,0004089 | 0,003504 | 0,0040053 | 0,0039032 | 0,0039032 | TRUE | 0,0754401 | No |
| orf19.7106 | 563 | 410 | 517 | 615 | 590 | 467 | 561 | 342 | 505 | 522 | 529 | 507 | 505 | 446 | 501 | 482 | 499,53 | 516 | 484 | 0,939 | -0,091 | -1,328 | 0,18417892 | 0,25632313 | 1,00E-08 | 0,0048856 | 0,0025271 | 0,0025271 | TRUE | 0,0342546 | No |
| orf19.7107 | 8064 | 4836 | 6390 | 8430 | 6998 | 5489 | 6247 | 4540 | 7234 | 6157 | 6538 | 6951 | 5985 | 5237 | 5574 | 6403 | 6260 | 6720 | 5800 | 0,862 | -0,213 | -3,086 | 0,00202735 | 0,00478035 | 0,006046 | 0,0022114 | 0,0044343 | 0,0044343 | TRUE | 0,0588747 | No |
| orf19.7108 | 2109 | 1381 | 1738 | 2206 | 2356 | 1904 | 1896 | 1429 | 1892 | 1758 | 1778 | 1819 | 2015 | 1817 | 1692 | 2015 | 1848,3 | 1812 | 1885 | 1,039 | 0,055 | 0,868 | 0,38534728 | 0,47432216 | 0,003371 | 0,0027649 | 0,0032696 | 0,0032696 | TRUE | 0,0582045 | No |
| orf19.7109 | 845 | 493 | 618 | 722 | 770 | 590 | 648 | 479 | 758 | 628 | 632 | 595 | 659 | 563 | 578 | 676 | 636,08 | 653 | 619 | 0,945 | -0,082 | -0,892 | 0,37229086 | 0,46080006 | 0,008069 | 0,0042618 | 0,0064778 | 0,0064778 | TRUE | 0,1414663 | No |
| orf19.711 | 29291 | 20849 | 25409 | 34080 | 23492 | 19694 | 21152 | 14136 | 26277 | 26543 | 25996 | 28103 | 20091 | 18791 | 18875 | 19938 | 23077 | 26730 | 19424 | 0,727 | -0,461 | -11,679 | 1,62E-31 | 6,51E-30 | 0,001185 | 0,0020424 | 0,0014509 | 0,0014509 | TRUE | 0,0147225 | No |
| orf19.7110 | 1694 | 1115 | 1440 | 1840 | 1562 | 1283 | 1452 | 1093 | 1520 | 1420 | 1473 | 1517 | 1336 | 1224 | 1296 | 1542 | 1415,9 | 1482 | 1350 | 0,907 | -0,141 | -2,062 | 0,03918195 | 0,0678401 | 0,004657 | 0,0030048 | 0,0037512 | 0,0037512 | TRUE | 0,1035585 | No |
| orf19.7111 | 1466 | 973 | 1209 | 1484 | 1238 | 933 | 1075 | 696 | 1315 | 1239 | 1237 | 1224 | 1059 | 890 | 959 | 982 | 1113,1 | 1254 | 972 | 0,776 | -0,366 | -5,977 | 2,27E-09 | 1,37E-08 | 0,002222 | 0,0032838 | 0,0026882 | 0,0026882 | TRUE | 0,045836 | No |
| orf19.7111.1 | 3473 | 2249 | 3047 | 3689 | 4858 | 3765 | 4106 | 3008 | 3116 | 2863 | 3117 | 3042 | 4155 | 3592 | 3664 | 4243 | 3474 | 3034 | 3914 | 1,288 | 0,365 | 5,877 | 4,17E-09 | 2,45E-08 | 0,003996 | 0,0023974 | 0,00342 | 0,00342 | TRUE | 0,0378672 | No |
| orf19.7112 | 415 | 297 | 374 | 487 | 382 | 357 | 340 | 235 | 372 | 378 | 383 | 402 | 327 | 341 | 303 | 331 | 354,6 | 384 | 326 | 0,846 | -0,241 | -3,092 | 0,00198798 | 0,00469674 | 1,00E-08 | 0,0060734 | 0,0029797 | 0,0029797 | TRUE | 0,0255746 | No |
| orf19.7114 | 1033 | 673 | 964 | 1092 | 1733 | 1464 | 1472 | 1278 | 927 | 857 | 986 | 900 | 1482 | 1397 | 1314 | 1803 | 1208,2 | 918 | 1499 | 1,629 | 0,704 | 7,734 | 1,04E-14 | 1,16E-13 | 0,01075 | 0,0031811 | 0,007078 | 0,007078 | TRUE | 0,2165652 | No |
| orf19.7115 | 2596 | 1643 | 2368 | 2698 | 2911 | 2417 | 2780 | 2002 | 2329 | 2092 | 2423 | 2225 | 2490 | 2306 | 2481 | 2824 | 2396 | 2267 | 2525 | 1,112 | 0,153 | 2,265 | 0,02351367 | 0,04318867 | 0,005073 | 0,0025854 | 0,0039628 | 0,0039628 | TRUE | 0,0740995 | No |
| orf19.7116 | 2582 | 1543 | 2058 | 2722 | 2166 | 1637 | 1901 | 1097 | 2316 | 1964 | 2106 | 2245 | 1852 | 1562 | 1696 | 1547 | 1911,1 | 2158 | 1664 | 0,772 | -0,374 | -5,233 | 1,67E-07 | 7,93E-07 | 0,005542 | 0,0027391 | 0,0043666 | 0,0043666 | TRUE | 0,0704943 | No |
| orf19.7118 | 1322 | 817 | 1114 | 1360 | 1157 | 897 | 936 | 574 | 1186 | 1040 | 1140 | 1121 | 990 | 856 | 835 | 810 | 997,19 | 1122 | 873 | 0,779 | -0,361 | -5,001 | 5,70E-07 | 2,50E-06 | 0,004341 | 0,0034353 | 0,0039744 | 0,0039744 | TRUE | 0,0996137 | No |
| orf19.7119 | 5285 | 3398 | 4837 | 5638 | 7380 | 5735 | 6526 | 3678 | 4741 | 4326 | 4949 | 4649 | 6312 | 5472 | 5823 | 5187 | 5182,5 | 4666 | 5698 | 1,222 | 0,289 | 4,464 | 8,05E-06 | 2,98E-05 | 0,004895 | 0,0022596 | 0,0038248 | 0,0038248 | TRUE | 0,0640763 | No |

| | | | | | | | | | | | | | | | | | | | | | | | | | | | | | | | |
|---|---|---|---|---|---|---|---|---|---|---|---|---|---|---|---|---|---|---|---|---|---|---|---|---|---|---|---|---|---|---|---|
| orf19.712 | 552 | 484 | 482 | 681 | 476 | 534 | 521 | 341 | 495 | 616 | 493 | 562 | 407 | 510 | 465 | 481 | 503,57 | 542 | 466 | 0,86 | -0,217 | -2,295 | 0,02170791 | 0,04017904 | 0,008279 | 0,0048623 | 0,0066059 | 0,0066059 | TRUE | 0,0935704 | No |
| orf19.7121 | 850 | 549 | 688 | 968 | 881 | 646 | 733 | 483 | 763 | 699 | 704 | 798 | 753 | 616 | 654 | 681 | 708,59 | 741 | 676 | 0,911 | -0,134 | -1,74 | 0,08192107 | 0,12914767 | 0,004471 | 0,0040282 | 0,0042659 | 0,0042659 | TRUE | 0,0746373 | No |
| orf19.7123 | 3078 | 1984 | 2677 | 3362 | 3697 | 2947 | 3330 | 2656 | 2761 | 2526 | 2739 | 2772 | 3162 | 2812 | 2971 | 3746 | 2936,2 | 2700 | 3173 | 1,174 | 0,231 | 2,9 | 0,00372831 | 0,00827473 | 0,008746 | 0,0024739 | 0,0057429 | 0,0057429 | TRUE | 0,176084 | No |
| orf19.7124 | 13126 | 8180 | 10817 | 13571 | 16015 | 12375 | 13844 | 8237 | 11775 | 10414 | 11067 | 11191 | 13697 | 11808 | 12353 | 11618 | 11740 | 11112 | 12369 | 1,113 | 0,155 | 2,627 | 0,0086075 | 0,01760772 | 0,004027 | 0,0021032 | 0,0032457 | 0,0032457 | TRUE | 0,0640413 | No |
| orf19.7125 | 1332 | 979 | 1330 | 1606 | 1421 | 1136 | 1285 | 738 | 1195 | 1246 | 1361 | 1324 | 1215 | 1084 | 1147 | 1041 | 1201,6 | 1282 | 1122 | 0,877 | -0,189 | -2,936 | 0,00332083 | 0,00746131 | 0,002891 | 0,0031876 | 0,0031546 | 0,0031546 | TRUE | 0,0380435 | No |
| orf19.7127 | 24 | 20 | 21 | 21 | 26 | 19 | 28 | 16 | 22 | 25 | 21 | 17 | 22 | 18 | 25 | 23 | 21,71 | 21 | 22 | 1,037 | 0,052 | 0,179 | 0,8582967 | 0,89261668 | 1,00E-08 | 0,0688352 | 0,0355435 | 0,0355435 | TRUE | 0,1120284 | No |
| orf19.7127.1 | 98 | 55 | 77 | 71 | 58 | 47 | 40 | 35 | 88 | 70 | 79 | 59 | 50 | 45 | 36 | 49 | 59,35 | 74 | 45 | 0,605 | -0,725 | -3,617 | 0,00029788 | 0,00084259 | 0,013168 | 0,0264411 | 0,0205684 | 0,0205684 | TRUE | 0,2079124 | No |
| orf19.7128 | 608 | 323 | 489 | 615 | 436 | 333 | 356 | 259 | 545 | 411 | 500 | 507 | 373 | 318 | 318 | 365 | 417,21 | 491 | 344 | 0,696 | -0,523 | -5,312 | 1,08E-07 | 5,29E-07 | 0,00823 | 0,0054591 | 0,0068284 | 0,0068284 | TRUE | 0,1290627 | No |
| orf19.713 | 3177 | 2205 | 2550 | 3184 | 3530 | 3025 | 3351 | 2260 | 2850 | 2807 | 2609 | 2626 | 3019 | 2886 | 2990 | 3188 | 2871,9 | 2723 | 3021 | 1,109 | 0,149 | 3,104 | 0,00190859 | 0,00452339 | 0,001418 | 0,002485 | 0,0018589 | 0,0018589 | TRUE | 0,0157863 | No |
| orf19.7130 | 46 | 24 | 44 | 58 | 41 | 42 | 33 | 30 | 41 | 31 | 45 | 48 | 35 | 40 | 29 | 42 | 38,95 | 41 | 36 | 0,873 | -0,195 | -0,85 | 0,39553656 | 0,48438314 | 1,00E-08 | 0,0392547 | 0,0250396 | 0,0250396 | TRUE | 0,1805564 | No |
| orf19.7131 | 549 | 309 | 481 | 613 | 582 | 443 | 543 | 407 | 493 | 393 | 492 | 505 | 498 | 423 | 485 | 574 | 482,81 | 471 | 495 | 1,044 | 0,062 | 0,599 | 0,5494874 | 0,63050426 | 0,01161 | 0,0049863 | 0,008229 | 0,008229 | TRUE | 0,1328466 | No |
| orf19.7136 | 1776 | 1130 | 1580 | 1940 | 1671 | 1298 | 1486 | 935 | 1593 | 1439 | 1616 | 1600 | 1429 | 1239 | 1326 | 1319 | 1445,1 | 1562 | 1328 | 0,85 | -0,235 | -3,966 | 7,32E-05 | 0,0002294 | 0,002209 | 0,0029841 | 0,0026734 | 0,0026734 | TRUE | 0,0330516 | No |
| orf19.7139 | 659 | 413 | 567 | 758 | 657 | 473 | 497 | 369 | 591 | 526 | 580 | 625 | 562 | 451 | 443 | 520 | 537,41 | 580 | 494 | 0,849 | -0,237 | -2,61 | 0,0090596 | 0,01846338 | 0,007265 | 0,0046808 | 0,0060101 | 0,0060101 | TRUE | 0,1084169 | No |
| orf19.714 | 409 | 359 | 403 | 517 | 497 | 499 | 510 | 341 | 367 | 457 | 412 | 426 | 425 | 476 | 455 | 481 | 437,48 | 416 | 459 | 1,106 | 0,145 | 1,747 | 0,08064447 | 0,12740257 | 0,003267 | 0,0052979 | 0,0043058 | 0,0043058 | TRUE | 0,0736012 | No |
| orf19.7140 | 1611 | 995 | 1326 | 1669 | 1777 | 1422 | 1599 | 1073 | 1445 | 1267 | 1357 | 1376 | 1520 | 1357 | 1427 | 1513 | 1407,7 | 1361 | 1454 | 1,066 | 0,092 | 1,59 | 0,1118195 | 0,16761766 | 0,001933 | 0,0030108 | 0,0025067 | 0,0025067 | TRUE | 0,024643 | No |
| orf19.7141 | 2607 | 1587 | 2328 | 2776 | 3192 | 2519 | 2783 | 1726 | 2339 | 2020 | 2382 | 2289 | 2730 | 2404 | 2483 | 2434 | 2385,2 | 2258 | 2513 | 1,113 | 0,154 | 2,46 | 0,0139109 | 0,02700378 | 0,003776 | 0,0025882 | 0,0033426 | 0,0033426 | TRUE | 0,0583219 | No |
| orf19.7144 | 3704 | 2363 | 3335 | 4123 | 4068 | 3368 | 3790 | 2468 | 3323 | 3008 | 3412 | 3400 | 3479 | 3214 | 3382 | 3481 | 3337,3 | 3286 | 3389 | 1,03 | 0,043 | 0,845 | 0,39831976 | 0,48709651 | 0,001944 | 0,0024145 | 0,0021757 | 0,0021757 | TRUE | 0,0378711 | No |
| orf19.7148 | 192 | 125 | 185 | 206 | 201 | 162 | 167 | 111 | 172 | 159 | 189 | 170 | 172 | 155 | 149 | 157 | 165,32 | 172 | 158 | 0,915 | -0,128 | -1,156 | 0,24780691 | 0,32975059 | 1,00E-08 | 0,0107606 | 0,0057945 | 0,0057945 | TRUE | 0,0421621 | No |
| orf19.7149 | 1353 | 943 | 1229 | 1512 | 921 | 776 | 818 | 597 | 1214 | 1201 | 1257 | 1247 | 788 | 740 | 730 | 842 | 1002,3 | 1230 | 775 | 0,628 | -0,672 | -11,81 | 3,48E-32 | 1,45E-30 | 0,000624 | 0,0034279 | 0,0020501 | 0,0020501 | TRUE | 0,0344378 | No |
| orf19.715 | 1027 | 826 | 881 | 1129 | 1894 | 1897 | 1908 | 1257 | 921 | 1052 | 901 | 931 | 1620 | 1810 | 1703 | 1773 | 1338,8 | 951 | 1726 | 1,817 | 0,862 | 14,082 | 4,91E-45 | 3,93E-43 | 0,002646 | 0,0030638 | 0,0027783 | 0,0027783 | TRUE | 0,0555158 | No |
| orf19.7150 | 10338 | 6420 | 8852 | 11211 | 12441 | 10435 | 12253 | 8258 | 9274 | 8173 | 9056 | 9245 | 10640 | 9957 | 10934 | 11647 | 9865,8 | 8937 | 10794 | 1,207 | 0,272 | 4,703 | 2,57E-06 | 1,02E-05 | 0,003651 | 0,0021267 | 0,0031012 | 0,0031012 | TRUE | 0,0400598 | No |
| orf19.7151 | 3 | 7 | 16 | 11 | 3 | 5 | 8 | 3 | 3 | 9 | 16 | 9 | 3 | 5 | 7 | 4 | 6,97 | 9 | 5 | 0,51 | -0,97 | -1,573 | 0,11581633 | NA | 0,196725 | 0,2102961 | 0,2046531 | 0,2046531 | TRUE | 0,1832446 | No |
| orf19.7152 | 9195 | 5771 | 8015 | 9827 | 8504 | 6615 | 7470 | 5170 | 8249 | 7347 | 8200 | 8103 | 7273 | 6312 | 6666 | 7292 | 7430,2 | 7975 | 6886 | 0,863 | -0,213 | -3,642 | 0,00027068 | 0,00077035 | 0,00366 | 0,0021749 | 0,0031381 | 0,0031381 | TRUE | 0,0384286 | No |
| orf19.7153 | 4514 | 3415 | 4268 | 4967 | 4767 | 4695 | 4888 | 3131 | 4050 | 4348 | 4367 | 4096 | 4077 | 4480 | 4362 | 4416 | 4274,3 | 4215 | 4334 | 1,028 | 0,04 | 0,889 | 0,37399796 | 0,46272235 | 0,001301 | 0,0023192 | 0,0017351 | 0,0017351 | TRUE | 0,0197936 | No |
| orf19.7154 | 10880 | 7480 | 9587 | 11624 | 10386 | 9263 | 10313 | 5981 | 9760 | 9523 | 9808 | 9585 | 8882 | 8838 | 9203 | 8436 | 9254,5 | 9669 | 8840 | 0,915 | -0,129 | -3,709 | 0,00020824 | 0,00060498 | 0,000589 | 0,0021364 | 0,001047 | 0,001047 | TRUE | 0,0111401 | No |
| orf19.7156 | 3097 | 1880 | 2631 | 3327 | 1801 | 1614 | 1753 | 1117 | 2778 | 2393 | 2692 | 2743 | 1540 | 1540 | 1564 | 1575 | 2103,4 | 2652 | 1555 | 0,585 | -0,772 | -14,433 | 3,23E-47 | 2,81E-45 | 0,001957 | 0,0026697 | 0,0022368 | 0,0022368 | TRUE | 0,0500798 | No |
| orf19.7157 | 163 | 118 | 156 | 159 | 143 | 138 | 143 | 111 | 146 | 150 | 160 | 131 | 122 | 132 | 128 | 157 | 140,66 | 147 | 135 | 0,911 | -0,135 | -1,076 | 0,28170749 | 0,36570843 | 0,00172 | 0,0122999 | 0,0078827 | 0,0078827 | TRUE | 0,1008097 | No |
| orf19.7158 | 1233 | 815 | 1074 | 1349 | 1533 | 1302 | 1457 | 896 | 1106 | 1038 | 1099 | 1112 | 1311 | 1242 | 1300 | 1264 | 1184 | 1089 | 1279 | 1,174 | 0,231 | 4,786 | 1,71E-06 | 6,95E-06 | 1,00E-08 | 0,0032056 | 0,0013961 | 0,0013961 | TRUE | 0,0107436 | No |
| orf19.7159 | 1127 | 690 | 972 | 1148 | 935 | 822 | 977 | 601 | 1011 | 878 | 994 | 947 | 800 | 784 | 872 | 848 | 891,75 | 958 | 826 | 0,86 | -0,217 | -3,424 | 0,0006179 | 0,00162657 | 0,001975 | 0,0036074 | 0,002732 | 0,002732 | TRUE | 0,0338104 | No |
| orf19.716 | 599 | 569 | 572 | 726 | 1044 | 1115 | 1084 | 575 | 537 | 724 | 585 | 599 | 893 | 1064 | 967 | 811 | 772,58 | 611 | 934 | 1,534 | 0,617 | 6,092 | 1,11E-09 | 7,03E-09 | 0,013363 | 0,0038585 | 0,0084856 | 0,0084856 | TRUE | 0,1735115 | No |
| orf19.7160 | 3205 | 2060 | 2623 | 3203 | 3004 | 2454 | 2738 | 1813 | 2875 | 2623 | 2684 | 2641 | 2569 | 2342 | 2443 | 2557 | 2591,7 | 2706 | 2478 | 0,915 | -0,128 | -2,641 | 0,00826919 | 0,01699669 | 0,001335 | 0,0025397 | 0,0018766 | 0,0018766 | TRUE | 0,0219415 | No |
| orf19.7161 | 9164 | 5746 | 7519 | 9407 | 9260 | 7413 | 8109 | 4743 | 8221 | 7315 | 7693 | 7757 | 7919 | 7073 | 7236 | 6690 | 7488 | 7746 | 7230 | 0,933 | -0,099 | -1,745 | 0,08097964 | 0,12786483 | 0,003498 | 0,0021734 | 0,0029747 | 0,0029747 | TRUE | 0,050525 | No |
| orf19.7163 | 141 | 114 | 114 | 151 | 170 | 127 | 126 | 88 | 126 | 145 | 117 | 125 | 145 | 121 | 112 | 124 | 126,99 | 128 | 126 | 0,987 | -0,019 | -0,141 | 0,88775192 | 0,91519397 | 0,003053 | 0,0134114 | 0,0089564 | 0,0089564 | TRUE | 0,1274752 | No |
| orf19.7164 | 1066 | 720 | 934 | 1208 | 1511 | 1300 | 1494 | 943 | 956 | 917 | 956 | 996 | 1292 | 1240 | 1333 | 1330 | 1127,6 | 956 | 1299 | 1,355 | 0,438 | 8,586 | 9,03E-18 | 1,37E-16 | 1,00E-08 | 0,003267 | 0,001597 | 0,001597 | TRUE | 0,011105 | No |
| orf19.7165 | 520 | 307 | 404 | 545 | 548 | 493 | 469 | 327 | 466 | 391 | 413 | 449 | 469 | 470 | 419 | 461 | 442,36 | 430 | 455 | 1,053 | 0,074 | 0,927 | 0,35403675 | 0,4415721 | 0,002374 | 0,0052613 | 0,0038528 | 0,0038528 | TRUE | 0,0393524 | No |
| orf19.7166 | 8852 | 5857 | 7555 | 9311 | 10195 | 8453 | 9545 | 5909 | 7941 | 7457 | 7730 | 7678 | 8719 | 8066 | 8517 | 8334 | 8055,2 | 7702 | 8409 | 1,092 | 0,127 | 3,428 | 0,00060732 | 0,00160244 | 0,000713 | 0,0021597 | 0,001187 | 0,001187 | TRUE | 0,0094238 | No |
| orf19.7167 | 68 | 38 | 39 | 46 | 51 | 30 | 31 | 29 | 61 | 48 | 40 | 38 | 44 | 29 | 28 | 41 | 41 | 47 | 36 | 0,748 | -0,418 | -1,682 | 0,09253722 | 0,14340046 | 0,030714 | 0,0373847 | 0,0344291 | 0,0344291 | TRUE | 0,3707913 | No |
| orf19.717 | 5934 | 4116 | 5261 | 6345 | 9116 | 8064 | 8737 | 5169 | 5323 | 5240 | 5383 | 5232 | 7796 | 7694 | 7796 | 7290 | 6469,5 | 5294 | 7644 | 1,445 | 0,531 | 15,545 | 1,72E-54 | 2,19E-52 | 0,000375 | 0,0022039 | 0,0009594 | 0,0009594 | TRUE | 0,0113242 | No |
| orf19.7170 | 11 | 2 | 11 | 18 | 18 | 6 | 11 | 15 | 10 | 3 | 11 | 15 | 15 | 6 | 10 | 21 | 11,33 | 10 | 13 | 1,29 | 0,367 | 0,718 | 0,47291316 | NA | 0,208637 | 0,1301615 | 0,1611612 | 0,1611612 | TRUE | 0,2357284 | No |
| orf19.7173 | 432 | 290 | 372 | 415 | 404 | 313 | 371 | 235 | 388 | 369 | 381 | 342 | 346 | 299 | 331 | 331 | 348,28 | 370 | 327 | 0,884 | -0,178 | -2,147 | 0,03183015 | 0,056495 | 1,00E-08 | 0,0061477 | 0,0036952 | 0,0036952 | TRUE | 0,0398056 | No |
| orf19.7175 | 1321 | 840 | 1122 | 1378 | 1062 | 858 | 1049 | 613 | 1185 | 1069 | 1148 | 1136 | 908 | 819 | 936 | 865 | 1008,3 | 1134 | 882 | 0,777 | -0,364 | -6,098 | 1,07E-09 | 6,81E-09 | 0,001559 | 0,0034193 | 0,002406 | 0,002406 | TRUE | 0,029608 | No |
| orf19.7176 | 882 | 595 | 775 | 956 | 797 | 752 | 815 | 511 | 791 | 757 | 793 | 788 | 682 | 718 | 727 | 721 | 747,14 | 782 | 712 | 0,908 | -0,14 | -2,508 | 0,01214412 | 0,02394547 | 1,00E-08 | 0,0039225 | 0,0016351 | 0,0016351 | TRUE | 0,0105005 | No |
| orf19.7177 | 4770 | 3525 | 4396 | 5133 | 5185 | 4989 | 5360 | 3372 | 4279 | 4488 | 4498 | 4233 | 4434 | 4760 | 4783 | 4756 | 4528,8 | 4374 | 4683 | 1,071 | 0,099 | 2,402 | 0,01630546 | 0,03106984 | 0,000834 | 0,0023001 | 0,0014013 | 0,0014013 | TRUE | 0,0160199 | No |
| orf19.7178 | 7432 | 4947 | 6332 | 7822 | 7268 | 6044 | 6510 | 4373 | 6667 | 6298 | 6478 | 6450 | 6216 | 5767 | 5809 | 6168 | 6231,7 | 6473 | 5990 | 0,925 | -0,113 | -2,89 | 0,00385073 | 0,00853067 | 0,000826 | 0,0022125 | 0,0013016 | 0,0013016 | TRUE | 0,0082735 | No |
| orf19.7179 | 1152 | 706 | 1014 | 1197 | 1606 | 1307 | 1479 | 933 | 1033 | 899 | 1037 | 987 | 1374 | 1247 | 1320 | 1316 | 1151,6 | 989 | 1314 | 1,326 | 0,407 | 6,908 | 4,93E-12 | 4,15E-11 | 0,001718 | 0,0032401 | 0,0024519 | 0,0024519 | TRUE | 0,0410969 | No |
| orf19.718 | 5755 | 4543 | 5205 | 6344 | 6031 | 6082 | 6381 | 3709 | 5163 | 5784 | 5325 | 5231 | 5158 | 5803 | 5694 | 5231 | 5423,7 | 5376 | 5472 | 1,019 | 0,027 | 0,49 | 0,62422569 | 0,6970907 | 0,002577 | 0,0022472 | 0,0026373 | 0,0026373 | TRUE | 0,031305 | No |
| orf19.7181 | 1098 | 678 | 941 | 1055 | 878 | 807 | 881 | 605 | 985 | 863 | 963 | 870 | 751 | 770 | 786 | 853 | 855,16 | 920 | 790 | 0,855 | -0,225 | -3,374 | 0,00073984 | 0,00191674 | 0,002591 | 0,0036771 | 0,0030938 | 0,0030938 | TRUE | 0,0307614 | No |
| orf19.7182 | 2430 | 1618 | 2043 | 2533 | 1881 | 1629 | 1705 | 963 | 2180 | 2060 | 2090 | 2089 | 1609 | 1554 | 1521 | 1358 | 1807,7 | 2105 | 1510 | 0,72 | -0,475 | -8,667 | 4,42E-18 | 6,92E-17 | 0,002095 | 0,0027826 | 0,0023118 | 0,0023118 | TRUE | 0,0506682 | No |
| orf19.7183 | 973 | 603 | 757 | 978 | 1297 | 1039 | 1114 | 739 | 873 | 768 | 774 | 806 | 1109 | 991 | 994 | 1042 | 919,81 | 805 | 1034 | 1,282 | 0,359 | 5,628 | 1,83E-08 | 9,79E-08 | 0,002177 | 0,0035578 | 0,0027931 | 0,0027931 | TRUE | 0,0382804 | No |
| orf19.7184 | 2483 | 1664 | 2083 | 2561 | 2812 | 2509 | 2696 | 1952 | 2228 | 2118 | 2131 | 2112 | 2405 | 2394 | 2406 | 2753 | 2318,3 | 2147 | 2490 | 1,157 | 0,21 | 3,912 | 9,16E-05 | 0,00028238 | 0,002235 | 0,0026057 | 0,0023398 | 0,0023398 | TRUE | 0,0583525 | No |
| orf19.7185 | 2814 | 1813 | 2437 | 2887 | 3018 | 2526 | 2602 | 1645 | 2524 | 2308 | 2493 | 2381 | 2581 | 2410 | 2322 | 2320 | 2417,5 | 2426 | 2408 | 0,993 | -0,01 | -0,206 | 0,83679538 | 0,87510299 | 0,00172 | 0,00258 | 0,0020511 | 0,0020511 | TRUE | 0,0289298 | No |
| orf19.7186 | 185 | 129 | 162 | 174 | 246 | 223 | 272 | 147 | 166 | 164 | 166 | 143 | 210 | 213 | 243 | 207 | 189,08 | 160 | 218 | 1,373 | 0,458 | 4,243 | 2,21E-05 | 7,60E-05 | 1,00E-08 | 0,0096573 | 0,0057672 | 0,0057672 | TRUE | 0,0653997 | No |
| orf19.7187 | 6236 | 4532 | 5359 | 6734 | 6229 | 5563 | 5660 | 3662 | 5594 | 5770 | 5483 | 5553 | 5327 | 5308 | 5051 | 5165 | 5406,3 | 5600 | 5213 | 0,931 | -0,103 | -3 | 0,00270398 | 0,00621006 | 0,000302 | 0,002248 | 0,0009371 | 0,0009371 | TRUE | 0,0056107 | No |
| orf19.7188 | 2107 | 1216 | 1815 | 2160 | 2064 | 1674 | 1782 | 1568 | 1890 | 1548 | 1857 | 1781 | 1765 | 1597 | 1590 | 2212 | 1780,1 | 1769 | 1791 | 1,01 | 0,015 | 0,144 | 0,88533624 | 0,91330703 | 0,014204 | 0,0027951 | 0,0092984 | 0,0092984 | TRUE | 0,2946805 | No |
| orf19.719 | 847 | 677 | 845 | 980 | 1294 | 1295 | 1304 | 755 | 760 | 862 | 865 | 808 | 1107 | 1236 | 1164 | 1065 | 983,14 | 824 | 1143 | 1,392 | 0,477 | 7,239 | 4,51E-13 | 4,32E-12 | 0,002843 | 0,0034561 | 0,0031218 | 0,0031218 | TRUE | 0,0354391 | No |
| orf19.7190 | 630 | 433 | 552 | 667 | 584 | 523 | 533 | 346 | 565 | 551 | 565 | 550 | 499 | 499 | 476 | 488 | 524,16 | 558 | 490 | 0,879 | -0,185 | -2,94 | 0,00328614 | 0,00739997 | 1,00E-08 | 0,0047491 | 0,0019048 | 0,0019048 | TRUE | 0,0055234 | No |
| orf19.7192 | 25 | 21 | 28 | 37 | 18 | 15 | 22 | 18 | 22 | 27 | 29 | 31 | 15 | 14 | 20 | 25 | 22,88 | 27 | 18 | 0,674 | -0,57 | -1,926 | 0,05411414 | 0,09003538 | 1,00E-08 | 0,065426 | 0,0383087 | 0,0383087 | TRUE | 0,2053323 | No |
| orf19.7193 | 8061 | 5743 | 7052 | 8727 | 8825 | 7760 | 8425 | 5315 | 7232 | 7311 | 7215 | 7196 | 7547 | 7404 | 7518 | 7496 | 7365 | 7238 | 7491 | 1,035 | 0,05 | 1,931 | 0,05349359 | 0,08912632 | 1,00E-08 | 0,0021766 | 0,0005045 | 0,0005045 | TRUE | 0,0007715 | No |
| orf19.7194 | 1344 | 1063 | 1207 | 1498 | 1541 | 1590 | 1584 | 973 | 1206 | 1353 | 1235 | 1235 | 1318 | 1517 | 1413 | 1372 | 1331,3 | 1257 | 1405 | 1,12 | 0,163 | 2,736 | 0,00621365 | 0,013144 | 0,00215 | 0,00307 | 0,0026498 | 0,0026498 | TRUE | 0,0352514 | No |

| | | | | | | | | | | | | | | | | | | | | | | | | | | | | | | | |
|---|---|---|---|---|---|---|---|---|---|---|---|---|---|---|---|---|---|---|---|---|---|---|---|---|---|---|---|---|---|---|---|
| orf19.7195 | 2143 | 1505 | 1873 | 2485 | 1952 | 1715 | 1796 | 1040 | 1922 | 1916 | 1916 | 2049 | 1669 | 1636 | 1603 | 1467 | 1772,4 | 1951 | 1594 | 0,818 | -0,289 | -5,62 | 1,90E-08 | 1,02E-07 | 0,001361 | 0,0027986 | 0,0019716 | 0,0019716 | TRUE | 0,0314186 | No |
| orf19.7196 | 16303 | 11754 | 14594 | 17888 | 14158 | 12463 | 13205 | 9155 | 14625 | 14964 | 14931 | 14751 | 12108 | 11892 | 11783 | 12912 | 13496 | 14818 | 12174 | 0,821 | -0,284 | -7,856 | 3,96E-15 | 4,59E-14 | 0,000837 | 0,0020871 | 0,0011809 | 0,0011809 | TRUE | 0,0198913 | No |
| orf19.7197 | 9166 | 7381 | 8060 | 10451 | 8747 | 9018 | 9558 | 5369 | 8223 | 9397 | 8246 | 8618 | 7481 | 8605 | 8529 | 7572 | 8333,8 | 8621 | 8047 | 0,934 | -0,099 | -1,58 | 0,11402832 | 0,1703337 | 0,004663 | 0,0021537 | 0,0036382 | 0,0036382 | TRUE | 0,0444481 | No |
| orf19.7198 | 1317 | 1046 | 1287 | 1571 | 1169 | 1116 | 1104 | 674 | 1181 | 1332 | 1317 | 1295 | 1000 | 1065 | 985 | 951 | 1140,7 | 1281 | 1000 | 0,782 | -0,354 | -6,06 | 1,36E-09 | 8,52E-09 | 0,001703 | 0,0032521 | 0,0023814 | 0,0023814 | TRUE | 0,0337348 | No |
| orf19.7199 | 436 | 420 | 435 | 557 | 474 | 533 | 496 | 249 | 391 | 535 | 445 | 459 | 405 | 509 | 443 | 351 | 442,24 | 458 | 427 | 0,938 | -0,092 | -0,776 | 0,43758851 | 0,5244428 | 0,017361 | 0,0052621 | 0,0111835 | 0,0111835 | TRUE | 0,1884365 | No |
| orf19.720 | 293 | 180 | 222 | 287 | 235 | 208 | 244 | 132 | 263 | 229 | 227 | 237 | 201 | 198 | 218 | 186 | 219,89 | 239 | 201 | 0,842 | -0,248 | -2,5 | 0,01243667 | 0,02441017 | 1,00E-08 | 0,0085813 | 0,0048826 | 0,0048826 | TRUE | 0,0520761 | No |
| orf19.7200 | 437 | 374 | 367 | 504 | 377 | 426 | 423 | 270 | 392 | 476 | 375 | 416 | 322 | 406 | 377 | 381 | 393,3 | 415 | 372 | 0,897 | -0,156 | -1,603 | 0,10904282 | 0,16406943 | 0,007516 | 0,0056706 | 0,0065839 | 0,0065839 | TRUE | 0,1046185 | No |
| orf19.7201 | 2676 | 2144 | 2582 | 3225 | 3327 | 3383 | 3621 | 2103 | 2401 | 2730 | 2642 | 2659 | 2845 | 3228 | 3231 | 2966 | 2837,7 | 2608 | 3068 | 1,177 | 0,235 | 4,016 | 5,93E-05 | 0,00018925 | 0,003022 | 0,0024911 | 0,002943 | 0,002943 | TRUE | 0,0350534 | No |
| orf19.7202 | 364 | 266 | 302 | 354 | 298 | 325 | 356 | 173 | 327 | 339 | 309 | 292 | 255 | 310 | 318 | 244 | 299,09 | 317 | 282 | 0,896 | -0,159 | -1,529 | 0,12624029 | 0,18639165 | 0,007182 | 0,0068333 | 0,006997 | 0,006997 | TRUE | 0,0833051 | No |
| orf19.7203 | 2625 | 2067 | 2312 | 2769 | 2864 | 2780 | 2863 | 1564 | 2355 | 2632 | 2365 | 2283 | 2449 | 2653 | 2555 | 2206 | 2437,2 | 2409 | 2466 | 1,026 | 0,037 | 0,562 | 0,57402414 | 0,65242849 | 0,004086 | 0,0025752 | 0,0036782 | 0,0036782 | TRUE | 0,0583004 | No |
| orf19.7204 | 109 | 104 | 108 | 166 | 113 | 104 | 104 | 54 | 98 | 132 | 110 | 137 | 97 | 99 | 93 | 76 | 105,3 | 119 | 91 | 0,772 | -0,374 | -2,432 | 0,01503221 | 0,02892139 | 0,00973 | 0,0157657 | 0,0131116 | 0,0131116 | TRUE | 0,1609567 | No |
| orf19.7205 | 1319 | 851 | 1174 | 1442 | 1336 | 1225 | 1301 | 832 | 1183 | 1083 | 1201 | 1189 | 1143 | 1169 | 1161 | 1173 | 1162,8 | 1164 | 1162 | 0,995 | -0,007 | -0,141 | 0,88779312 | 0,91519397 | 1,30E-05 | 0,0032279 | 0,0015457 | 0,0015457 | TRUE | 0,0235291 | No |
| orf19.7206 | 138 | 93 | 114 | 170 | 161 | 121 | 147 | 118 | 124 | 118 | 117 | 140 | 138 | 115 | 131 | 166 | 131,22 | 125 | 138 | 1,089 | 0,124 | 0,903 | 0,36655771 | 0,45548681 | 0,006688 | 0,0130426 | 0,0103474 | 0,0103474 | TRUE | 0,1666045 | No |
| orf19.7207 | 5010 | 3527 | 4416 | 5491 | 4751 | 4363 | 4696 | 2915 | 4494 | 4490 | 4518 | 4528 | 4063 | 4163 | 4190 | 4111 | 4319,8 | 4508 | 4132 | 0,917 | -0,126 | -4,197 | 2,70E-05 | 9,18E-05 | 1,00E-08 | 0,0023156 | 0,0006265 | 0,0006265 | TRUE | 0,0016709 | No |
| orf19.7208 | 14 | 13 | 14 | 13 | 16 | 8 | 10 | 3 | 13 | 17 | 14 | 11 | 14 | 8 | 9 | 4 | 11,08 | 14 | 9 | 0,668 | -0,581 | -1,355 | 0,1755028 | NA | 1,00E-08 | 0,1330219 | 0,0836991 | 0,0836991 | TRUE | 0,5363139 | No |
| orf19.7209 | 130 | 65 | 94 | 122 | 95 | 81 | 89 | 47 | 117 | 83 | 96 | 101 | 81 | 77 | 79 | 66 | 87,55 | 99 | 76 | 0,769 | -0,38 | -2,431 | 0,01503835 | 0,02892139 | 1,00E-08 | 0,0185611 | 0,0119209 | 0,0119209 | TRUE | 0,1399811 | No |
| orf19.721 | 2163 | 1872 | 2108 | 2611 | 2236 | 2360 | 2505 | 1459 | 1940 | 2383 | 2157 | 2153 | 1912 | 2252 | 2235 | 2058 | 2136,3 | 2158 | 2114 | 0,981 | -0,028 | -0,391 | 0,69583707 | 0,75867659 | 0,005997 | 0,0026591 | 0,004549 | 0,004549 | TRUE | 0,0581365 | No |
| orf19.7210 | 2726 | 1736 | 2398 | 3047 | 2910 | 2389 | 2588 | 1939 | 2446 | 2210 | 2453 | 2513 | 2489 | 2280 | 2309 | 2735 | 2429,3 | 2406 | 2453 | 1,018 | 0,025 | 0,386 | 0,69982919 | 0,76182032 | 0,004169 | 0,0025771 | 0,00373 | 0,00373 | TRUE | 0,0696283 | No |
| orf19.7212 | 510 | 382 | 462 | 532 | 701 | 603 | 665 | 433 | 458 | 486 | 473 | 439 | 600 | 575 | 593 | 611 | 529,28 | 464 | 595 | 1,285 | 0,361 | 5,531 | 3,19E-08 | 1,66E-07 | 1,00E-08 | 0,0047223 | 0,0021853 | 0,0021853 | TRUE | 0,0162423 | No |
| orf19.7213 | 2405 | 1573 | 2074 | 2539 | 2203 | 1974 | 2179 | 1360 | 2158 | 2003 | 2122 | 2094 | 1884 | 1884 | 1944 | 1918 | 2000,7 | 2094 | 1908 | 0,91 | -0,136 | -3,318 | 0,00090794 | 0,00230067 | 1,00E-08 | 0,0027051 | 0,0011179 | 0,0011179 | TRUE | 0,0095624 | No |
| orf19.7214 | 3698 | 2321 | 3190 | 3891 | 2781 | 2356 | 2598 | 1519 | 3317 | 2955 | 3264 | 3209 | 2378 | 2248 | 2318 | 2142 | 2729 | 3186 | 2272 | 0,713 | -0,487 | -9,59 | 8,77E-22 | 1,80E-20 | 0,001761 | 0,0025115 | 0,0020989 | 0,0020989 | TRUE | 0,0279156 | No |
| orf19.7215 | 8154 | 5806 | 7718 | 8895 | 7056 | 6549 | 6851 | 4570 | 7315 | 7392 | 7896 | 7335 | 6035 | 6249 | 6113 | 6446 | 6847,5 | 7484 | 6211 | 0,829 | -0,27 | -6,874 | 6,26E-12 | 5,21E-11 | 0,000905 | 0,0021915 | 0,0013315 | 0,0013315 | TRUE | 0,0167646 | No |
| orf19.7215.3 | 1814 | 1099 | 1382 | 1697 | 2742 | 2200 | 2368 | 1428 | 1627 | 1399 | 1414 | 1399 | 2345 | 2099 | 2113 | 2014 | 1801,4 | 1460 | 2143 | 1,468 | 0,554 | 8,293 | 1,10E-16 | 1,49E-15 | 0,004353 | 0,0027854 | 0,0037088 | 0,0037088 | TRUE | 0,0730111 | No |
| orf19.7216 | 2423 | 1667 | 2007 | 2428 | 2814 | 2425 | 2668 | 1563 | 2174 | 2122 | 2053 | 2002 | 2407 | 2314 | 2381 | 2204 | 2207,2 | 2088 | 2326 | 1,116 | 0,159 | 3,438 | 0,00058526 | 0,00155174 | 0,000799 | 0,0026373 | 0,0015912 | 0,0015912 | TRUE | 0,0139037 | No |
| orf19.7217 | 21156 | 13144 | 18802 | 22607 | 22091 | 16748 | 18556 | 13687 | 18979 | 16734 | 19236 | 18642 | 18893 | 15980 | 16558 | 19304 | 18041 | 18398 | 17684 | 0,961 | -0,057 | -0,83 | 0,4063731 | 0,49462917 | 0,005714 | 0,00206 | 0,0045345 | 0,0045345 | TRUE | 0,0514472 | No |
| orf19.7218 | 52 | 44 | 47 | 58 | 99 | 54 | 60 | 76 | 47 | 56 | 48 | 48 | 85 | 52 | 54 | 107 | 61,94 | 50 | 74 | 1,482 | 0,568 | 2,362 | 0,01815872 | 0,03411433 | 0,053454 | 0,0254175 | 0,0385568 | 0,0385568 | TRUE | 0,296071 | No |
| orf19.7219 | 14821 | 9898 | 13469 | 15943 | 12335 | 10440 | 11780 | 7694 | 13296 | 12601 | 13780 | 13147 | 10549 | 9962 | 10512 | 10852 | 11837 | 13206 | 10469 | 0,792 | -0,336 | -8,285 | 1,18E-16 | 1,60E-15 | 0,001188 | 0,0021022 | 0,0014903 | 0,0014903 | TRUE | 0,0129771 | No |
| orf19.7221 | 2202 | 1446 | 1965 | 2403 | 2214 | 1928 | 2126 | 1260 | 1975 | 1841 | 2010 | 1982 | 1893 | 1840 | 1897 | 1777 | 1902 | 1952 | 1852 | 0,949 | -0,076 | -1,67 | 0,09498529 | 0,14655153 | 0,000512 | 0,0027428 | 0,0014461 | 0,0014461 | TRUE | 0,0164298 | No |
| orf19.7222 | 2805 | 1815 | 2370 | 2788 | 3209 | 2487 | 2751 | 1715 | 2516 | 2311 | 2425 | 2299 | 2744 | 2373 | 2455 | 2419 | 2442,7 | 2388 | 2498 | 1,047 | 0,066 | 1,156 | 0,24751858 | 0,32951288 | 0,002638 | 0,0025738 | 0,0026914 | 0,0026914 | TRUE | 0,0548008 | No |
| orf19.7223 | 2565 | 1753 | 2205 | 2709 | 2492 | 2380 | 2410 | 1660 | 2301 | 2232 | 2256 | 2234 | 2131 | 2271 | 2151 | 2341 | 2239,6 | 2256 | 2224 | 0,984 | -0,024 | -0,54 | 0,58952121 | 0,66748517 | 0,000516 | 0,0026277 | 0,0013798 | 0,0013798 | TRUE | 0,0141553 | No |
| orf19.7224 | 2984 | 2002 | 2498 | 3104 | 2945 | 2524 | 2728 | 1688 | 2677 | 2549 | 2556 | 2560 | 2519 | 2408 | 2434 | 2381 | 2510,4 | 2586 | 2436 | 0,943 | -0,085 | -2,192 | 0,02839373 | 0,0510747 | 0,000214 | 0,0025578 | 0,0010558 | 0,0010558 | TRUE | 0,0070906 | No |
| orf19.7225 | 127 | 85 | 110 | 131 | 85 | 82 | 104 | 74 | 114 | 108 | 113 | 108 | 73 | 78 | 93 | 104 | 98,85 | 111 | 87 | 0,777 | -0,365 | -2,476 | 0,01330023 | 0,02594391 | 0,000258 | 0,016665 | 0,0105274 | 0,0105274 | TRUE | 0,1325057 | No |
| orf19.7227 | 1611 | 1140 | 1303 | 1645 | 2577 | 2082 | 2284 | 1559 | 1445 | 1451 | 1333 | 1356 | 2204 | 1987 | 2038 | 2199 | 1751,7 | 1396 | 2107 | 1,509 | 0,593 | 10,985 | 4,49E-28 | 1,47E-26 | 0,001788 | 0,0028083 | 0,0022018 | 0,0022018 | TRUE | 0,017826 | No |
| orf19.7228 | 1113 | 815 | 924 | 1170 | 1167 | 1147 | 1117 | 756 | 998 | 1038 | 945 | 965 | 998 | 1094 | 997 | 1066 | 1012,7 | 986 | 1039 | 1,053 | 0,074 | 1,317 | 0,18797643 | 0,2607023 | 0,000963 | 0,003413 | 0,0020659 | 0,0020659 | TRUE | 0,0169322 | No |
| orf19.7229 | 6578 | 4491 | 5943 | 7080 | 6771 | 5778 | 6212 | 4225 | 5901 | 5718 | 6080 | 5838 | 5791 | 5513 | 5543 | 5959 | 5792,9 | 5884 | 5702 | 0,968 | -0,047 | -1,205 | 0,22827288 | 0,30812479 | 0,000791 | 0,0022301 | 0,0012652 | 0,0012652 | TRUE | 0,010731 | No |
| orf19.723 | 5893 | 4116 | 5499 | 6780 | 7405 | 6761 | 7347 | 5297 | 5287 | 5240 | 5626 | 5591 | 6333 | 6451 | 6556 | 7471 | 6069,3 | 5436 | 6703 | 1,232 | 0,301 | 5,309 | 1,10E-07 | 5,37E-07 | 0,00347 | 0,0022187 | 0,0029193 | 0,0029193 | TRUE | 0,071368 | No |
| orf19.7231 | 1845 | 1254 | 1726 | 2045 | 1200 | 1038 | 1119 | 716 | 1655 | 1596 | 1766 | 1686 | 1026 | 990 | 999 | 1010 | 1341,1 | 1676 | 1006 | 0,6 | -0,737 | -15,038 | 4,12E-51 | 4,59E-49 | 1,00E-08 | 0,003062 | 0,0015091 | 0,0015091 | TRUE | 0,0150977 | No |
| orf19.7232 | 404 | 326 | 408 | 442 | 515 | 430 | 447 | 284 | 362 | 415 | 417 | 364 | 440 | 410 | 399 | 401 | 401,19 | 390 | 412 | 1,064 | 0,089 | 1,109 | 0,26757145 | 0,3518427 | 0,001798 | 0,005598 | 0,0037589 | 0,0037589 | TRUE | 0,0285968 | No |
| orf19.7234 | 2720 | 1839 | 2278 | 2791 | 2746 | 2135 | 2472 | 1617 | 2440 | 2341 | 2331 | 2301 | 2348 | 2037 | 2206 | 2281 | 2285,7 | 2353 | 2218 | 0,942 | -0,086 | -1,67 | 0,09483548 | 0,14635797 | 0,001566 | 0,0026146 | 0,0021018 | 0,0021018 | TRUE | 0,0370273 | No |
| orf19.7235 | 4230 | 2747 | 3655 | 4321 | 5037 | 4383 | 4809 | 2862 | 3795 | 3497 | 3739 | 3563 | 4308 | 4182 | 4291 | 4037 | 3926,5 | 3648 | 4204 | 1,153 | 0,205 | 4,939 | 7,86E-07 | 3,37E-06 | 0,00085 | 0,0023492 | 0,0013983 | 0,0013983 | TRUE | 0,008996 | No |
| orf19.7236 | 7543 | 4557 | 6133 | 7632 | 6701 | 5229 | 5720 | 3568 | 6767 | 5802 | 6275 | 6293 | 5731 | 4989 | 5104 | 5032 | 5749,2 | 6284 | 5214 | 0,83 | -0,269 | -4,498 | 6,86E-06 | 2,56E-05 | 0,003957 | 0,002232 | 0,0032655 | 0,0032655 | TRUE | 0,0547916 | No |
| orf19.7237 | 598 | 406 | 546 | 666 | 451 | 374 | 378 | 295 | 536 | 517 | 559 | 549 | 386 | 357 | 337 | 416 | 457,14 | 540 | 374 | 0,688 | -0,54 | -6,971 | 3,15E-12 | 2,72E-11 | 0,001349 | 0,0051552 | 0,0034873 | 0,0034873 | TRUE | 0,0549504 | No |
| orf19.7238 | 22941 | 15297 | 20113 | 24563 | 20168 | 16087 | 17492 | 10768 | 20580 | 19475 | 20578 | 20255 | 17248 | 15350 | 15609 | 15187 | 18035 | 20222 | 15848 | 0,784 | -0,351 | -7,547 | 4,44E-14 | 4,65E-13 | 0,00205 | 0,00206 | 0,0020256 | 0,0020256 | TRUE | 0,0434407 | No |
| orf19.7239 | 25400 | 15878 | 21810 | 26111 | 23539 | 19510 | 22024 | 14145 | 22786 | 20214 | 22314 | 21531 | 20131 | 18616 | 19653 | 19950 | 20650 | 21711 | 19588 | 0,902 | -0,149 | -3,252 | 0,00114785 | 0,00285325 | 0,001874 | 0,0020498 | 0,0019618 | 0,0019618 | TRUE | 0,0261964 | No |
| orf19.7242 | 3017 | 2061 | 2862 | 3270 | 4252 | 3574 | 3883 | 2448 | 2707 | 2624 | 2928 | 2696 | 3636 | 3410 | 3465 | 3453 | 3114,9 | 2739 | 3491 | 1,275 | 0,35 | 7,756 | 8,78E-15 | 9,90E-14 | 0,001126 | 0,0024456 | 0,0016312 | 0,0016312 | TRUE | 0,0259251 | No |
| orf19.7243 | 769 | 569 | 728 | 915 | 747 | 705 | 690 | 473 | 690 | 724 | 745 | 755 | 639 | 673 | 616 | 667 | 688,5 | 728 | 649 | 0,889 | -0,17 | -2,775 | 0,00552568 | 0,01183899 | 1,00E-08 | 0,004088 | 0,0021523 | 0,0021523 | TRUE | 0,0161942 | No |
| orf19.7244 | 1000 | 662 | 862 | 1076 | 1028 | 793 | 883 | 623 | 897 | 843 | 882 | 887 | 879 | 757 | 788 | 879 | 851,44 | 877 | 826 | 0,939 | -0,092 | -1,423 | 0,15473925 | 0,22156182 | 0,002014 | 0,0036845 | 0,0027939 | 0,0027939 | TRUE | 0,0373787 | No |
| orf19.7245 | 2265 | 1469 | 1990 | 2528 | 2640 | 2138 | 2387 | 1630 | 2032 | 1870 | 2036 | 2085 | 2258 | 2040 | 2130 | 2299 | 2093,7 | 2006 | 2182 | 1,086 | 0,118 | 2,203 | 0,0275976 | 0,04974689 | 0,001928 | 0,0026729 | 0,0022944 | 0,0022944 | TRUE | 0,0238084 | No |
| orf19.7247 | 12600 | 8080 | 10989 | 13433 | 14525 | 12081 | 13291 | 8649 | 11304 | 10287 | 11243 | 11077 | 12422 | 11527 | 11860 | 12199 | 11490 | 10978 | 12002 | 1,093 | 0,128 | 3,052 | 0,00227227 | 0,00530372 | 0,001335 | 0,0021059 | 0,001608 | 0,001608 | TRUE | 0,0216005 | No |
| orf19.725 | 184 | 194 | 175 | 253 | 208 | 248 | 234 | 187 | 165 | 247 | 179 | 209 | 178 | 237 | 209 | 264 | 210,85 | 200 | 222 | 1,107 | 0,147 | 0,992 | 0,32100679 | 0,40768399 | 0,023908 | 0,0088645 | 0,0163127 | 0,0163127 | TRUE | 0,2495981 | No |
| orf19.7250 | 2544 | 1483 | 2072 | 2617 | 2314 | 1742 | 1857 | 1330 | 2282 | 1888 | 2120 | 2158 | 1979 | 1662 | 1657 | 1876 | 1952,8 | 2112 | 1794 | 0,848 | -0,238 | -3,176 | 0,00149269 | 0,00362493 | 0,006344 | 0,0027229 | 0,0048586 | 0,0048586 | TRUE | 0,0609405 | No |
| orf19.7251 | 816 | 546 | 714 | 806 | 768 | 651 | 698 | 573 | 732 | 695 | 730 | 665 | 657 | 621 | 623 | 808 | 691,41 | 706 | 677 | 0,955 | -0,066 | -0,766 | 0,4439389 | 0,53067746 | 0,007009 | 0,0040791 | 0,0056122 | 0,0056122 | TRUE | 0,1838691 | No |
| orf19.7252 | 6 | 2 | 5 | 13 | 9 | 3 | 7 | 5 | 5 | 3 | 5 | 11 | 8 | 3 | 6 | 7 | 5,95 | 6 | 6 | 0,967 | -0,049 | -0,082 | 0,93440627 | NA | 0,071772 | 0,2458488 | 0,1761312 | 0,1761312 | TRUE | 0,8164502 | No |
| orf19.7254 | 1421 | 852 | 1260 | 1585 | 1609 | 1424 | 1581 | 1057 | 1275 | 1085 | 1289 | 1307 | 1376 | 1359 | 1411 | 1491 | 1324 | 1239 | 1409 | 1,134 | 0,181 | 2,795 | 0,00519536 | 0,01118772 | 0,003354 | 0,003076 | 0,0032567 | 0,0032567 | TRUE | 0,0793183 | No |
| orf19.7255 | 2021 | 1288 | 1501 | 1923 | 2065 | 1308 | 1466 | 915 | 1813 | 1640 | 1536 | 1586 | 1766 | 1248 | 1308 | 1291 | 1523,4 | 1644 | 1403 | 0,855 | -0,227 | -2,203 | 0,02757453 | 0,04972397 | 0,016839 | 0,0029325 | 0,0095118 | 0,0095118 | TRUE | 0,3696204 | No |
| orf19.7256 | 1327 | 781 | 1065 | 1360 | 1394 | 961 | 1019 | 549 | 1190 | 994 | 1090 | 1121 | 1192 | 917 | 909 | 774 | 1023,6 | 1099 | 948 | 0,864 | -0,211 | -1,913 | 0,05577938 | 0,09229482 | 0,018542 | 0,0033978 | 0,0106522 | 0,0106522 | TRUE | 0,3606859 | No |
| orf19.7257 | 2018 | 1293 | 1870 | 2231 | 2537 | 1847 | 2275 | 1628 | 1810 | 1646 | 1913 | 1840 | 2170 | 1762 | 2030 | 2296 | 1933,5 | 1802 | 2064 | 1,144 | 0,194 | 2,447 | 0,01438905 | 0,02785995 | 0,007412 | 0,0027304 | 0,0054976 | 0,0054976 | TRUE | 0,1175231 | No |
| orf19.7258 | 1845 | 1182 | 1540 | 1990 | 1715 | 1278 | 1428 | 927 | 1655 | 1505 | 1576 | 1641 | 1467 | 1219 | 1274 | 1307 | 1455,6 | 1594 | 1317 | 0,826 | -0,276 | -4,3 | 1,71E-05 | 5,99E-05 | 0,003263 | 0,0029769 | 0,003261 | 0,003261 | TRUE | 0,0736375 | No |
| orf19.726 | 1990 | 1390 | 1667 | 2118 | 2216 | 2143 | 2196 | 1294 | 1785 | 1770 | 1706 | 1747 | 1895 | 2045 | 1960 | 1825 | 1841,4 | 1752 | 1931 | 1,104 | 0,143 | 3,041 | 0,00235749 | 0,00547704 | 0,000644 | 0,0027679 | 0,0015688 | 0,0015688 | TRUE | 0,0185504 | No |

| | | | | | | | | | | | | | | | | | | | | | | | | | | | | | | | |
|---|---|---|---|---|---|---|---|---|---|---|---|---|---|---|---|---|---|---|---|---|---|---|---|---|---|---|---|---|---|---|---|
| orf19.7260 | 300 | 204 | 296 | 345 | 422 | 323 | 396 | 280 | 269 | 260 | 303 | 284 | 361 | 308 | 353 | 395 | 316,69 | 279 | 354 | 1,263 | 0,337 | 3,571 | 0,00035593 | 0,00099094 | 0,003932 | 0,0065634 | 0,0053424 | 0,0053424 | TRUE | 0,0866118 | No |
| orf19.7261 | 2453 | 1539 | 2190 | 2678 | 2947 | 2187 | 2579 | 1669 | 2201 | 1959 | 2241 | 2208 | 2520 | 2087 | 2301 | 2354 | 2233,9 | 2152 | 2316 | 1,075 | 0,104 | 1,592 | 0,11137777 | 0,16712229 | 0,004203 | 0,0026294 | 0,0036807 | 0,0036807 | TRUE | 0,0543909 | No |
| orf19.7263 | 2584 | 1531 | 2258 | 2678 | 2535 | 1966 | 2188 | 1497 | 2318 | 1949 | 2310 | 2208 | 2168 | 1876 | 1952 | 2111 | 2111,7 | 2196 | 2027 | 0,922 | -0,118 | -1,735 | 0,08282339 | 0,13036493 | 0,004691 | 0,002667 | 0,0039466 | 0,0039466 | TRUE | 0,0672759 | No |
| orf19.7264 | 5823 | 3463 | 4714 | 5979 | 5749 | 4015 | 4693 | 2899 | 5224 | 4409 | 4823 | 4930 | 4917 | 3831 | 4188 | 4089 | 4551,3 | 4846 | 4256 | 0,878 | -0,187 | -2,434 | 0,01493935 | 0,02878083 | 0,008109 | 0,0022985 | 0,0054732 | 0,0054732 | TRUE | 0,1341146 | No |
| orf19.7265 | 906 | 577 | 817 | 1003 | 978 | 768 | 834 | 563 | 813 | 735 | 836 | 827 | 836 | 733 | 744 | 794 | 789,73 | 803 | 777 | 0,966 | -0,05 | -0,755 | 0,45028738 | 0,53623917 | 0,002257 | 0,0038178 | 0,0029909 | 0,0029909 | TRUE | 0,0352381 | No |
| orf19.7266 | 524 | 357 | 429 | 591 | 630 | 509 | 576 | 316 | 470 | 454 | 439 | 487 | 539 | 486 | 514 | 446 | 479,37 | 462 | 496 | 1,075 | 0,105 | 1,37 | 0,17081827 | 0,24102322 | 0,002002 | 0,0050079 | 0,0035443 | 0,0035443 | TRUE | 0,0467511 | No |
| orf19.7267 | 934 | 531 | 862 | 991 | 1024 | 743 | 948 | 714 | 838 | 676 | 882 | 817 | 876 | 709 | 846 | 1007 | 831,34 | 803 | 860 | 1,066 | 0,093 | 0,877 | 0,38058044 | 0,4698022 | 0,014272 | 0,0037257 | 0,0094944 | 0,0094944 | TRUE | 0,1645672 | No |
| orf19.7269 | 6000 | 3452 | 4972 | 6305 | 6009 | 4459 | 5104 | 3643 | 5383 | 4395 | 5087 | 5199 | 5139 | 4255 | 4554 | 5138 | 4893,7 | 5016 | 4772 | 0,951 | -0,073 | -0,951 | 0,34141213 | 0,42838202 | 0,007321 | 0,0022762 | 0,0054539 | 0,0054539 | TRUE | 0,0838151 | No |
| orf19.727 | 58 | 26 | 50 | 52 | 46 | 41 | 46 | 39 | 52 | 33 | 51 | 43 | 39 | 39 | 41 | 55 | 44,21 | 45 | 44 | 0,952 | -0,071 | -0,321 | 0,7479722 | 0,80214164 | 0,004879 | 0,0348158 | 0,0238512 | 0,0238512 | TRUE | 0,1891394 | No |
| orf19.7270 | 3829 | 2430 | 3554 | 4358 | 3741 | 2872 | 3242 | 1766 | 3435 | 3094 | 3636 | 3594 | 3199 | 2740 | 2893 | 2491 | 3135,2 | 3440 | 2831 | 0,824 | -0,28 | -3,646 | 0,00026635 | 0,00075875 | 0,007616 | 0,0024425 | 0,0053444 | 0,0053444 | TRUE | 0,0933832 | No |
| orf19.7271 | 2472 | 1615 | 2164 | 2468 | 2637 | 2290 | 2499 | 1450 | 2218 | 2056 | 2214 | 2035 | 2255 | 2185 | 2230 | 2045 | 2154,8 | 2131 | 2179 | 1,024 | 0,034 | 0,68 | 0,49656454 | 0,58155161 | 0,001379 | 0,0026532 | 0,0019458 | 0,0019458 | TRUE | 0,0191643 | No |
| orf19.7272 | 1393 | 997 | 1153 | 1435 | 1411 | 1155 | 1205 | 829 | 1250 | 1269 | 1180 | 1183 | 1207 | 1102 | 1075 | 1169 | 1179,4 | 1220 | 1138 | 0,933 | -0,1 | -1,797 | 0,07239426 | 0,11604714 | 0,001319 | 0,0032104 | 0,0021335 | 0,0021335 | TRUE | 0,0218064 | No |
| orf19.7274 | 244 | 178 | 187 | 268 | 162 | 116 | 135 | 98 | 219 | 227 | 191 | 221 | 139 | 111 | 120 | 138 | 170,72 | 214 | 127 | 0,59 | -0,761 | -6,496 | 8,23E-11 | 6,06E-10 | 0,001533 | 0,0104831 | 0,0068946 | 0,0068946 | TRUE | 0,0766069 | No |
| orf19.7275 | 18 | 13 | 15 | 9 | 7 | 6 | 11 | 7 | 16 | 17 | 15 | 7 | 6 | 6 | 10 | 10 | 10,86 | 14 | 8 | 0,565 | -0,824 | -1,838 | 0,06601457 | NA | 1,00E-08 | 0,1356745 | 0,0920667 | 0,0920667 | TRUE | 0,5293904 | No |
| orf19.7276.1 | 147 | 98 | 112 | 151 | 136 | 132 | 112 | 78 | 132 | 125 | 115 | 125 | 116 | 126 | 100 | 110 | 118,49 | 124 | 113 | 0,912 | -0,133 | -1,02 | 0,30791657 | 0,39386049 | 1,00E-08 | 0,0142307 | 0,0079014 | 0,0079014 | TRUE | 0,0700353 | No |
| orf19.7277 | 15 | 4 | 8 | 11 | 10 | 0 | 8 | 7 | 13 | 5 | 8 | 9 | 9 | 0 | 7 | 10 | 7,67 | 9 | 6 | 0,692 | -0,531 | -0,857 | 0,3913185 | NA | 0,335562 | 0,1912237 | 0,233152 | 0,233152 | TRUE | 1,0386795 | No |
| orf19.7278 | 0 | 0 | 0 | 0 | 0 | 0 | 0 | 0 | 0 | 0 | 0 | 0 | 0 | 0 | 0 | 0 | 0 | 0 | 0 | NA | NA | NA | NA | NA | NA | NA | NA | NA | NA | NA | NA |
| orf19.7279 | 136 | 97 | 97 | 124 | 136 | 120 | 120 | 112 | 122 | 123 | 99 | 102 | 116 | 115 | 107 | 158 | 117,86 | 112 | 124 | 1,099 | 0,137 | 0,903 | 0,36645002 | 0,45544719 | 0,01243 | 0,0142971 | 0,013398 | 0,013398 | TRUE | 0,2888547 | No |
| orf19.7279.1 | 3646 | 2537 | 2938 | 3696 | 2912 | 2468 | 2654 | 1710 | 3271 | 3230 | 3006 | 3048 | 2490 | 2355 | 2368 | 2412 | 2772,5 | 3139 | 2406 | 0,767 | -0,383 | -8,726 | 2,63E-18 | 4,19E-17 | 0,000909 | 0,0025031 | 0,0014771 | 0,0014771 | TRUE | 0,0102655 | No |
| orf19.728 | 590 | 428 | 511 | 608 | 658 | 752 | 786 | 418 | 529 | 545 | 523 | 501 | 563 | 718 | 701 | 590 | 583,69 | 524 | 643 | 1,229 | 0,297 | 3,389 | 0,00070248 | 0,00182547 | 0,006671 | 0,0044666 | 0,005636 | 0,005636 | TRUE | 0,0904594 | No |
| orf19.7280 | 4 | 5 | 5 | 3 | 4 | 2 | 2 | 3 | 4 | 6 | 5 | 2 | 3 | 2 | 2 | 4 | 3,61 | 4 | 3 | 0,646 | -0,631 | -0,862 | 0,38886004 | NA | 1,00E-08 | 0,4039953 | 0,2169268 | 0,2169268 | TRUE | 0,1957965 | No |
| orf19.7281 | 2442 | 1625 | 2171 | 2647 | 2115 | 1813 | 1971 | 1252 | 2191 | 2069 | 2221 | 2183 | 1809 | 1730 | 1759 | 1766 | 1965,9 | 2166 | 1766 | 0,815 | -0,296 | -7,119 | 1,08E-12 | 9,91E-12 | 1,00E-08 | 0,002718 | 0,0011412 | 0,0011412 | TRUE | 0,0101022 | No |
| orf19.7282 | 3395 | 2159 | 2799 | 3549 | 2788 | 2166 | 2466 | 1625 | 3046 | 2749 | 2864 | 2927 | 2384 | 2067 | 2200 | 2292 | 2566 | 2896 | 2236 | 0,771 | -0,375 | -6,895 | 5,39E-12 | 4,52E-11 | 0,002223 | 0,0025453 | 0,0024339 | 0,0024339 | TRUE | 0,0318555 | No |
| orf19.7283 | 17373 | 11290 | 14325 | 17699 | 20935 | 16851 | 18239 | 12063 | 15585 | 14373 | 14656 | 14595 | 17904 | 16079 | 16275 | 17014 | 15810 | 14802 | 16818 | 1,136 | 0,184 | 4,088 | 4,36E-05 | 0,00014243 | 0,001784 | 0,0020713 | 0,0018834 | 0,0018834 | TRUE | 0,0233559 | No |
| orf19.7284 | 9309 | 5862 | 7585 | 9668 | 13003 | 10369 | 11428 | 6671 | 8351 | 7463 | 7760 | 7972 | 11121 | 9894 | 10198 | 9409 | 9020,9 | 7886 | 10156 | 1,288 | 0,365 | 6,441 | 1,18E-10 | 8,55E-10 | 0,003491 | 0,0021404 | 0,0029695 | 0,0029695 | TRUE | 0,0501618 | No |
| orf19.7285 | 2346 | 1459 | 1900 | 2454 | 2522 | 2080 | 2310 | 1700 | 2105 | 1857 | 1944 | 2024 | 2157 | 1985 | 2061 | 2398 | 2066,3 | 1982 | 2150 | 1,082 | 0,114 | 1,754 | 0,07945005 | 0,12574707 | 0,004267 | 0,0026821 | 0,003552 | 0,003552 | TRUE | 0,0694196 | No |
| orf19.7286 | 3461 | 2344 | 3034 | 3854 | 2845 | 2646 | 2858 | 1885 | 3105 | 2984 | 3104 | 3178 | 2433 | 2525 | 2550 | 2659 | 2817,2 | 3093 | 2542 | 0,82 | -0,286 | -6,932 | 4,14E-12 | 3,54E-11 | 0,000525 | 0,0024948 | 0,001276 | 0,001276 | TRUE | 0,0106886 | No |
| orf19.7288 | 881 | 518 | 709 | 901 | 616 | 507 | 560 | 389 | 790 | 659 | 725 | 743 | 527 | 484 | 500 | 549 | 622,14 | 729 | 515 | 0,702 | -0,51 | -7,01 | 2,39E-12 | 2,09E-11 | 0,002575 | 0,0043129 | 0,0034273 | 0,0034273 | TRUE | 0,0455344 | No |
| orf19.729 | 1007 | 735 | 903 | 1126 | 1109 | 1030 | 1042 | 595 | 903 | 936 | 924 | 929 | 948 | 983 | 930 | 839 | 923,97 | 923 | 925 | 1,006 | 0,009 | 0,153 | 0,87820572 | 0,90735339 | 0,000866 | 0,0035507 | 0,0021551 | 0,0021551 | TRUE | 0,0403033 | No |
| orf19.7290 | 1779 | 1145 | 1307 | 1865 | 1321 | 1161 | 1169 | 720 | 1596 | 1458 | 1337 | 1538 | 1130 | 1108 | 1043 | 1015 | 1278,1 | 1482 | 1074 | 0,725 | -0,464 | -7,082 | 1,42E-12 | 1,28E-11 | 0,003463 | 0,0031153 | 0,0033125 | 0,0033125 | TRUE | 0,0527914 | No |
| orf19.7291 | 3853 | 2615 | 3282 | 4294 | 4096 | 3213 | 3769 | 2474 | 3457 | 3329 | 3358 | 3541 | 3503 | 3066 | 3363 | 3489 | 3388,2 | 3421 | 3355 | 0,98 | -0,029 | -0,579 | 0,56246353 | 0,64232672 | 0,00199 | 0,002408 | 0,0021464 | 0,0021464 | TRUE | 0,0413187 | No |
| orf19.729.1 | 1070 | 730 | 938 | 1135 | 1528 | 1315 | 1444 | 1011 | 960 | 929 | 960 | 936 | 1307 | 1255 | 1289 | 1426 | 1132,6 | 946 | 1319 | 1,39 | 0,475 | 8,83 | 1,05E-18 | 1,71E-17 | 0,000735 | 0,0032613 | 0,0018693 | 0,0018693 | TRUE | 0,0319722 | No |
| orf19.7292 | 2301 | 1486 | 1994 | 2409 | 2406 | 1944 | 2107 | 1329 | 2064 | 1892 | 2040 | 1986 | 2058 | 1855 | 1880 | 1874 | 1956,2 | 1996 | 1917 | 0,961 | -0,058 | -1,15 | 0,25027172 | 0,33236792 | 0,001269 | 0,0027216 | 0,0019408 | 0,0019408 | TRUE | 0,0307201 | No |
| orf19.7293 | 408 | 226 | 352 | 432 | 446 | 413 | 437 | 278 | 366 | 288 | 360 | 356 | 381 | 394 | 390 | 392 | 365,96 | 342 | 389 | 1,13 | 0,177 | 2,071 | 0,03840251 | 0,06675991 | 0,001922 | 0,0059464 | 0,0042494 | 0,0042494 | TRUE | 0,115506 | No |
| orf19.7295 | 1068 | 743 | 925 | 1205 | 846 | 719 | 858 | 610 | 958 | 946 | 946 | 994 | 724 | 686 | 766 | 860 | 859,95 | 961 | 759 | 0,786 | -0,348 | -5,016 | 5,28E-07 | 2,34E-06 | 0,003184 | 0,0036676 | 0,0034241 | 0,0034241 | TRUE | 0,0859379 | No |
| orf19.7296 | 15002 | 10380 | 12850 | 16222 | 7941 | 6914 | 7200 | 4954 | 13458 | 13215 | 13147 | 13377 | 6791 | 6597 | 6425 | 6987 | 9999,7 | 13299 | 6700 | 0,503 | -0,99 | -29,039 | 2,10E-185 | 6,32E-182 | 0,000566 | 0,0021247 | 0,0010031 | 0,0010031 | TRUE | 0,0096547 | No |
| orf19.7297 | 2364 | 1650 | 2131 | 2661 | 2203 | 1940 | 2009 | 1207 | 2121 | 2101 | 2180 | 2194 | 1884 | 1851 | 1793 | 1702 | 1978,3 | 2149 | 1808 | 0,842 | -0,247 | -5,528 | 3,24E-08 | 1,69E-07 | 0,000514 | 0,0027133 | 0,0014136 | 0,0014136 | TRUE | 0,0165759 | No |
| orf19.7298 | 2387 | 1784 | 2380 | 2688 | 1607 | 1624 | 1714 | 1002 | 2141 | 2271 | 2435 | 2217 | 1374 | 1550 | 1529 | 1413 | 1866,4 | 2266 | 1466 | 0,648 | -0,626 | -10,702 | 9,91E-27 | 3,01E-25 | 0,002482 | 0,0027574 | 0,0027215 | 0,0027215 | TRUE | 0,030741 | No |
| orf19.7299.1 | 12 | 8 | 11 | 6 | 6 | 7 | 6 | 5 | 11 | 10 | 11 | 5 | 5 | 7 | 5 | 7 | 7,67 | 9 | 6 | 0,651 | -0,618 | -1,224 | 0,22085566 | NA | 1,00E-08 | 0,1912234 | 0,1072666 | 0,1072666 | TRUE | 0,4213721 | No |
| orf19.73 | 4104 | 3545 | 3752 | 4850 | 5017 | 5057 | 5243 | 3071 | 3682 | 4513 | 3839 | 3999 | 4291 | 4825 | 4679 | 4331 | 4269,8 | 4008 | 4532 | 1,131 | 0,178 | 2,668 | 0,00762926 | 0,01578395 | 0,005408 | 0,0023195 | 0,0040478 | 0,0040478 | TRUE | 0,0861627 | No |
| orf19.730 | 1566 | 1025 | 1340 | 1710 | 2085 | 1697 | 1934 | 1258 | 1405 | 1305 | 1371 | 1410 | 1783 | 1619 | 1726 | 1774 | 1549,2 | 1373 | 1726 | 1,255 | 0,327 | 6,527 | 6,71E-11 | 5,01E-10 | 0,000944 | 0,0029166 | 0,0017581 | 0,0017581 | TRUE | 0,0210391 | No |
| orf19.7300 | 2325 | 1910 | 2199 | 2700 | 2112 | 2121 | 2280 | 1400 | 2086 | 2432 | 2250 | 2226 | 1806 | 2024 | 2035 | 1975 | 2104,1 | 2248 | 1960 | 0,872 | -0,197 | -3,334 | 0,00085677 | 0,00218575 | 0,002811 | 0,0026695 | 0,0028791 | 0,0028791 | TRUE | 0,0352317 | No |
| orf19.7301 | 247 | 147 | 204 | 258 | 198 | 165 | 157 | 109 | 222 | 187 | 209 | 213 | 169 | 157 | 140 | 154 | 181,35 | 208 | 155 | 0,745 | -0,424 | -3,884 | 0,00010282 | 0,00031423 | 1,00E-08 | 0,0099845 | 0,0058491 | 0,0058491 | TRUE | 0,0515665 | No |
| orf19.7303 | 18 | 15 | 8 | 16 | 10 | 11 | 9 | 3 | 16 | 19 | 8 | 13 | 9 | 10 | 8 | 4 | 10,99 | 14 | 8 | 0,573 | -0,804 | -1,832 | 0,06694843 | NA | 1,00E-08 | 0,1340522 | 0,0874315 | 0,0874315 | TRUE | 0,2526932 | No |
| orf19.7304 | 803 | 791 | 796 | 1063 | 1088 | 1166 | 1251 | 719 | 720 | 1007 | 814 | 877 | 930 | 1113 | 1116 | 1014 | 948,97 | 854 | 1043 | 1,223 | 0,291 | 2,983 | 0,00285316 | 0,00650789 | 0,012103 | 0,0035093 | 0,0080551 | 0,0080551 | TRUE | 0,1650401 | No |
| orf19.7305 | 525 | 432 | 486 | 612 | 504 | 480 | 435 | 230 | 471 | 550 | 497 | 505 | 431 | 458 | 388 | 324 | 453,06 | 506 | 400 | 0,797 | -0,327 | -3,29 | 0,00100149 | 0,00251965 | 0,009714 | 0,0051838 | 0,0072229 | 0,0072229 | TRUE | 0,1649285 | No |
| orf19.7306 | 1075 | 805 | 910 | 1091 | 681 | 622 | 663 | 407 | 964 | 1025 | 931 | 900 | 582 | 593 | 592 | 574 | 770,18 | 955 | 585 | 0,615 | -0,702 | -11,557 | 6,77E-31 | 2,53E-29 | 1,00E-08 | 0,0038644 | 0,0021582 | 0,0021582 | TRUE | 0,0266462 | No |
| orf19.7307 | 3250 | 2433 | 2920 | 3823 | 2994 | 2678 | 2864 | 2183 | 2916 | 3097 | 2987 | 3152 | 2561 | 2555 | 2556 | 3079 | 2862,9 | 3038 | 2688 | 0,883 | -0,179 | -2,771 | 0,00559253 | 0,011953 | 0,004861 | 0,0024866 | 0,0036665 | 0,0036665 | TRUE | 0,1126275 | No |
| orf19.7308 | 4872 | 3206 | 4169 | 5280 | 4789 | 3920 | 4311 | 2869 | 4371 | 4082 | 4265 | 4354 | 4096 | 3740 | 3847 | 4046 | 4100,1 | 4268 | 3932 | 0,921 | -0,119 | -2,76 | 0,00577464 | 0,01231158 | 0,001061 | 0,0023336 | 0,0015528 | 0,0015528 | TRUE | 0,0131954 | No |
| orf19.731 | 2192 | 1528 | 1890 | 2467 | 2220 | 1870 | 2056 | 1372 | 1966 | 1945 | 1934 | 2034 | 1899 | 1784 | 1835 | 1935 | 1916,6 | 1970 | 1863 | 0,944 | -0,083 | -1,905 | 0,05681776 | 0,09383208 | 0,000358 | 0,002737 | 0,0012967 | 0,0012967 | TRUE | 0,0096263 | No |
| orf19.7310 | 5181 | 3799 | 4610 | 5562 | 8335 | 7769 | 8074 | 5074 | 4648 | 4837 | 4716 | 4586 | 7128 | 7413 | 7205 | 7156 | 5961,2 | 4697 | 7226 | 1,54 | 0,623 | 19,331 | 2,97E-83 | 9,92E-81 | 0,000185 | 0,002223 | 0,0008196 | 0,0008196 | TRUE | 0,0046699 | No |
| orf19.7311 | 570 | 393 | 493 | 625 | 754 | 712 | 757 | 538 | 511 | 500 | 504 | 515 | 645 | 679 | 675 | 759 | 598,75 | 508 | 690 | 1,351 | 0,434 | 6,465 | 1,01E-10 | 7,37E-10 | 0,000844 | 0,0044041 | 0,0026151 | 0,0026151 | TRUE | 0,0471657 | No |
| orf19.7312 | 2891 | 1909 | 2550 | 3051 | 2362 | 2122 | 2121 | 1380 | 2594 | 2430 | 2609 | 2516 | 2020 | 2025 | 1893 | 1946 | 2254,1 | 2537 | 1971 | 0,777 | -0,365 | -8,388 | 4,96E-17 | 7,07E-16 | 0,000584 | 0,0026236 | 0,0013627 | 0,0013627 | TRUE | 0,0093383 | No |
| orf19.7313 | 3621 | 2505 | 3041 | 3816 | 3293 | 2965 | 3225 | 2156 | 3248 | 3189 | 3111 | 3147 | 2816 | 2829 | 2878 | 3041 | 3032,4 | 3174 | 2891 | 0,91 | -0,137 | -3,51 | 0,0004488 | 0,00122396 | 0,000371 | 0,0024582 | 0,0011265 | 0,0011265 | TRUE | 0,0135421 | No |
| orf19.7314 | 4259 | 2852 | 3770 | 4554 | 4952 | 4213 | 4436 | 3023 | 3821 | 3631 | 3857 | 3755 | 4235 | 4020 | 3958 | 4264 | 3942,6 | 3766 | 4119 | 1,093 | 0,128 | 3,174 | 0,00150262 | 0,00364168 | 0,000697 | 0,0023477 | 0,0013039 | 0,0013039 | TRUE | 0,0084718 | No |
| orf19.7316 | 1210 | 880 | 1113 | 1328 | 1517 | 1263 | 1532 | 884 | 1085 | 1120 | 1139 | 1095 | 1297 | 1205 | 1367 | 1247 | 1194,5 | 1110 | 1279 | 1,154 | 0,207 | 3,843 | 0,00012164 | 0,00036633 | 0,001047 | 0,0031948 | 0,0019503 | 0,0019503 | TRUE | 0,0264224 | No |
| orf19.7317 | 1558 | 1177 | 1409 | 1780 | 1787 | 1563 | 1757 | 1004 | 1398 | 1498 | 1442 | 1468 | 1528 | 1491 | 1568 | 1416 | 1476,1 | 1452 | 1501 | 1,037 | 0,052 | 1,079 | 0,28040504 | 0,36449015 | 0,000455 | 0,002963 | 0,0015901 | 0,0015901 | TRUE | 0,0155345 | No |
| orf19.7319 | 1233 | 825 | 1021 | 1354 | 1590 | 1427 | 1570 | 957 | 1106 | 1050 | 1045 | 1117 | 1360 | 1362 | 1401 | 1350 | 1223,7 | 1080 | 1368 | 1,266 | 0,34 | 7,152 | 8,56E-13 | 7,90E-12 | 1,00E-08 | 0,0031658 | 0,0013464 | 0,0013464 | TRUE | 0,00657 | No |
| orf19.732 | 4032 | 2593 | 3389 | 4425 | 4191 | 3278 | 3569 | 2526 | 3617 | 3301 | 3467 | 3649 | 3584 | 3128 | 3185 | 3563 | 3436,7 | 3508 | 3365 | 0,958 | -0,062 | -1,064 | 0,28725438 | 0,37178465 | 0,003233 | 0,0024019 | 0,0029605 | 0,0029605 | TRUE | 0,0273032 | No |

| | | | | | | | | | | | | | | | | | | | | | | | | | | | | | | | |
|---|---|---|---|---|---|---|---|---|---|---|---|---|---|---|---|---|---|---|---|---|---|---|---|---|---|---|---|---|---|---|---|
| orf19.7320 | 98 | 59 | 70 | 90 | 105 | 81 | 94 | 77 | 88 | 75 | 72 | 74 | 90 | 77 | 84 | 109 | 83,55 | 77 | 90 | 1,15 | 0,202 | 1,261 | 0,20738435 | 0,2836543 | 1,00E-08 | 0,019354 | 0,0125252 | 0,0125252 | TRUE | 0,1466086 | No |
| orf19.7321 | 616 | 393 | 546 | 600 | 621 | 553 | 670 | 369 | 553 | 500 | 559 | 495 | 531 | 528 | 598 | 520 | 535,42 | 527 | 544 | 1,035 | 0,05 | 0,662 | 0,5082095 | 0,59253303 | 0,002461 | 0,0046908 | 0,0035478 | 0,0035478 | TRUE | 0,0532535 | No |
| orf19.7322 | 5808 | 3841 | 4771 | 6023 | 5513 | 4619 | 4705 | 3432 | 5210 | 4890 | 4881 | 4967 | 4715 | 4407 | 4198 | 4841 | 4763,7 | 4987 | 4540 | 0,91 | -0,137 | -2,673 | 0,0075275 | 0,01560028 | 0,002258 | 0,0022843 | 0,0022983 | 0,0022983 | TRUE | 0,0314364 | No |
| orf19.7323 | 6020 | 4198 | 5318 | 6372 | 5907 | 5677 | 5986 | 3727 | 5401 | 5345 | 5441 | 5254 | 5052 | 5417 | 5342 | 5257 | 5313,4 | 5360 | 5267 | 0,983 | -0,025 | -0,73 | 0,46516281 | 0,55122252 | 0,000417 | 0,0022527 | 0,0009689 | 0,0009689 | TRUE | 0,0095939 | No |
| orf19.7324 | 286 | 196 | 224 | 320 | 386 | 317 | 330 | 226 | 257 | 250 | 229 | 264 | 330 | 302 | 294 | 319 | 280,62 | 250 | 311 | 1,243 | 0,314 | 3,552 | 0,0003828 | 0,00105984 | 1,00E-08 | 0,0071527 | 0,0039316 | 0,0039316 | TRUE | 0,0349056 | No |
| orf19.7325 | 3742 | 2406 | 3130 | 3832 | 4315 | 3563 | 3964 | 2487 | 3357 | 3063 | 3202 | 3160 | 3690 | 3400 | 3537 | 3508 | 3364,7 | 3196 | 3534 | 1,105 | 0,145 | 3,334 | 0,00085619 | 0,0021852 | 0,000912 | 0,002411 | 0,0015093 | 0,0015093 | TRUE | 0,0141367 | No |
| orf19.7326 | 2688 | 1905 | 2272 | 2756 | 2863 | 2642 | 2776 | 1943 | 2411 | 2425 | 2324 | 2273 | 2449 | 2521 | 2477 | 2740 | 2452,6 | 2358 | 2547 | 1,079 | 0,109 | 2,28 | 0,02258199 | 0,0416813 | 0,001282 | 0,0025714 | 0,0017868 | 0,0017868 | TRUE | 0,0297645 | No |
| orf19.7327 | 6467 | 4196 | 5399 | 6683 | 5006 | 4125 | 4486 | 3409 | 5802 | 5342 | 5524 | 5511 | 4281 | 3936 | 4003 | 4808 | 4900,8 | 5545 | 4257 | 0,767 | -0,383 | -6,117 | 9,54E-10 | 6,10E-09 | 0,004652 | 0,0022757 | 0,0035523 | 0,0035523 | TRUE | 0,0902403 | No |
| orf19.7328 | 669 | 442 | 624 | 712 | 730 | 708 | 680 | 485 | 600 | 563 | 638 | 587 | 624 | 676 | 607 | 684 | 622,39 | 597 | 648 | 1,081 | 0,112 | 1,647 | 0,09945581 | 0,15215953 | 0,001355 | 0,004312 | 0,002826 | 0,002826 | TRUE | 0,024328 | No |
| orf19.7329 | 4394 | 2751 | 3463 | 4578 | 5413 | 4499 | 4563 | 3312 | 3942 | 3502 | 3543 | 3775 | 4629 | 4293 | 4072 | 4671 | 4053,4 | 3690 | 4416 | 1,195 | 0,258 | 4,434 | 9,26E-06 | 3,39E-05 | 0,003317 | 0,0023377 | 0,0029911 | 0,0029911 | TRUE | 0,0338137 | No |
| orf19.733 | 2196 | 1322 | 1935 | 2235 | 2122 | 1678 | 1940 | 1338 | 1970 | 1683 | 1980 | 1843 | 1815 | 1601 | 1731 | 1887 | 1813,7 | 1869 | 1758 | 0,939 | -0,09 | -1,341 | 0,17984278 | 0,25180504 | 0,004456 | 0,0027799 | 0,003815 | 0,003815 | TRUE | 0,0519892 | No |
| orf19.7330 | 37 | 27 | 36 | 36 | 32 | 43 | 41 | 28 | 33 | 34 | 37 | 30 | 27 | 41 | 37 | 39 | 34,82 | 34 | 36 | 1,073 | 0,102 | 0,432 | 0,66579489 | 0,73328977 | 1,00E-08 | 0,0436714 | 0,0247662 | 0,0247662 | TRUE | 0,2230512 | No |
| orf19.7331 | 168 | 107 | 141 | 203 | 198 | 180 | 237 | 141 | 151 | 136 | 144 | 167 | 169 | 172 | 211 | 199 | 168,75 | 150 | 188 | 1,246 | 0,317 | 2,626 | 0,00863013 | 0,017648 | 0,004949 | 0,010582 | 0,0079961 | 0,0079961 | TRUE | 0,0891253 | No |
| orf19.7332 | 34968 | 25365 | 31249 | 37841 | 35068 | 32443 | 34566 | 20422 | 31370 | 32292 | 31971 | 31204 | 29991 | 30956 | 30844 | 28803 | 30929 | 31709 | 30148 | 0,951 | -0,072 | -2,204 | 0,02754587 | 0,04969837 | 0,000631 | 0,0020265 | 0,0010062 | 0,0010062 | TRUE | 0,0109072 | No |
| orf19.7335 | 5725 | 4117 | 5002 | 6334 | 5784 | 4940 | 5316 | 3115 | 5136 | 5241 | 5118 | 5223 | 4947 | 4714 | 4744 | 4393 | 4939,4 | 5180 | 4700 | 0,908 | -0,139 | -3,366 | 0,00076264 | 0,0019665 | 0,000985 | 0,0022734 | 0,0014251 | 0,0014251 | TRUE | 0,0222487 | No |
| orf19.7336 | 18 | 17 | 12 | 16 | 5 | 16 | 13 | 3 | 16 | 22 | 12 | 13 | 4 | 15 | 12 | 4 | 12,33 | 16 | 9 | 0,577 | -0,794 | -1,736 | 0,08247663 | NA | 0,10235 | 0,1197215 | 0,1134265 | 0,1134265 | TRUE | 0,5816674 | No |
| orf19.7337 | 1756 | 1330 | 1620 | 1995 | 2352 | 2429 | 2416 | 1497 | 1575 | 1693 | 1657 | 1645 | 2012 | 2318 | 2156 | 2111 | 1895,9 | 1642 | 2149 | 1,309 | 0,389 | 7,313 | 2,61E-13 | 2,56E-12 | 0,001807 | 0,0027452 | 0,0021725 | 0,0021725 | TRUE | 0,0344135 | No |
| orf19.7339 | 460 | 281 | 399 | 454 | 510 | 456 | 472 | 285 | 413 | 358 | 408 | 374 | 436 | 435 | 421 | 402 | 405,93 | 388 | 424 | 1,092 | 0,126 | 1,657 | 0,09756073 | 0,14971771 | 1,00E-08 | 0,0055558 | 0,0031307 | 0,0031307 | TRUE | 0,0273705 | No |
| orf19.734 | 8769 | 5605 | 7625 | 9117 | 14428 | 12173 | 13318 | 8421 | 7867 | 7136 | 7801 | 7518 | 12339 | 11615 | 11884 | 11877 | 9754,6 | 7580 | 11929 | 1,573 | 0,654 | 16,181 | 6,85E-59 | 1,08E-56 | 0,001138 | 0,0021283 | 0,0014592 | 0,0014592 | TRUE | 0,0185598 | No |
| orf19.7340 | 19 | 14 | 15 | 24 | 13 | 26 | 19 | 26 | 17 | 18 | 15 | 20 | 11 | 25 | 17 | 37 | 19,94 | 18 | 22 | 1,238 | 0,308 | 0,836 | 0,4031438 | 0,49149512 | 0,083725 | 0,0747661 | 0,0787043 | 0,0787043 | TRUE | 0,3092021 | No |
| orf19.7341 | 3693 | 2491 | 3232 | 4110 | 4699 | 3927 | 4271 | 2649 | 3313 | 3171 | 3307 | 3389 | 4019 | 3747 | 3811 | 3736 | 3561,7 | 3295 | 3828 | 1,162 | 0,216 | 5,379 | 7,47E-08 | 3,73E-07 | 0,000589 | 0,0023871 | 0,0012684 | 0,0012684 | TRUE | 0,0139534 | No |
| orf19.7341.1 | 24624 | 20091 | 21946 | 28819 | 30955 | 32246 | 32201 | 17266 | 22090 | 25578 | 22453 | 23765 | 26474 | 30768 | 28734 | 24352 | 25527 | 23472 | 27582 | 1,175 | 0,233 | 3,212 | 0,00131722 | 0,00323129 | 0,007276 | 0,0020364 | 0,0050174 | 0,0050174 | TRUE | 0,0758425 | No |
| orf19.7342 | 554 | 449 | 527 | 691 | 795 | 813 | 882 | 568 | 497 | 572 | 539 | 570 | 680 | 776 | 787 | 801 | 652,68 | 544 | 761 | 1,396 | 0,482 | 6,418 | 1,38E-10 | 9,82E-10 | 0,003499 | 0,0042038 | 0,0038303 | 0,0038303 | TRUE | 0,0635882 | No |
| orf19.7343 | 1614 | 1194 | 1311 | 1722 | 1816 | 1808 | 1946 | 994 | 1448 | 1520 | 1341 | 1420 | 1553 | 1725 | 1736 | 1402 | 1518,2 | 1432 | 1604 | 1,123 | 0,167 | 2,306 | 0,02111402 | 0,03916426 | 0,005349 | 0,0029357 | 0,00436 | 0,00436 | TRUE | 0,0819084 | No |
| orf19.7344 | 826 | 781 | 799 | 1021 | 1231 | 1462 | 1522 | 872 | 741 | 994 | 817 | 842 | 1053 | 1395 | 1358 | 1230 | 1053,8 | 848 | 1259 | 1,486 | 0,571 | 5,581 | 2,39E-08 | 1,26E-07 | 0,014049 | 0,0033571 | 0,0090726 | 0,0090726 | TRUE | 0,1533328 | No |
| orf19.7345 | 1038 | 868 | 947 | 1205 | 1240 | 1343 | 1306 | 747 | 931 | 1105 | 969 | 994 | 1060 | 1281 | 1165 | 1054 | 1070 | 1000 | 1140 | 1,144 | 0,193 | 2,504 | 0,01228816 | 0,02418187 | 0,005962 | 0,0033363 | 0,0047892 | 0,0047892 | TRUE | 0,0833169 | No |
| orf19.7347 | 845 | 618 | 721 | 1022 | 794 | 737 | 841 | 587 | 758 | 787 | 738 | 843 | 679 | 703 | 750 | 828 | 760,74 | 782 | 740 | 0,943 | -0,084 | -1,137 | 0,25557779 | 0,33821836 | 0,003879 | 0,0038878 | 0,0039569 | 0,0039569 | TRUE | 0,0683086 | No |
| orf19.7348 | 3 | 1 | 5 | 7 | 1 | 2 | 2 | 4 | 3 | 1 | 5 | 6 | 1 | 2 | 2 | 6 | 3,13 | 4 | 3 | 0,616 | -0,698 | -0,835 | 0,40352474 | NA | 0,08749 | 0,4657433 | 0,326522 | 0,326522 | TRUE | 0,4193562 | No |
| orf19.7349 | 5868 | 4183 | 5448 | 6606 | 6439 | 5910 | 6496 | 4123 | 5264 | 5325 | 5574 | 5447 | 5507 | 5639 | 5797 | 5815 | 5546,1 | 5402 | 5690 | 1,053 | 0,074 | 2,092 | 0,03642473 | 0,06372737 | 0,000407 | 0,0022413 | 0,0010158 | 0,0010158 | TRUE | 0,0057879 | No |
| orf19.7350 | 32074 | 19462 | 27228 | 33945 | 24412 | 18996 | 21408 | 11558 | 28774 | 24777 | 27857 | 27991 | 20878 | 18125 | 19103 | 16302 | 22976 | 27350 | 18602 | 0,68 | -0,556 | -7,608 | 2,79E-14 | 2,96E-13 | 0,007289 | 0,0020427 | 0,0050844 | 0,0050844 | TRUE | 0,0844597 | No |
| orf19.7353 | 878 | 591 | 792 | 1014 | 746 | 709 | 739 | 472 | 788 | 752 | 810 | 836 | 638 | 677 | 659 | 666 | 728,27 | 796 | 660 | 0,826 | -0,276 | -4,68 | 2,87E-06 | 1,13E-05 | 1,00E-08 | 0,0039729 | 0,0019497 | 0,0019497 | TRUE | 0,0149112 | No |
| orf19.7354 | 1268 | 1125 | 1227 | 1559 | 1131 | 1243 | 1305 | 672 | 1138 | 1432 | 1255 | 1286 | 967 | 1186 | 1164 | 948 | 1172 | 1278 | 1066 | 0,837 | -0,257 | -2,834 | 0,00459593 | 0,00999724 | 0,010365 | 0,0032181 | 0,0070519 | 0,0070519 | TRUE | 0,0773825 | No |
| orf19.7355 | 542 | 483 | 498 | 585 | 651 | 699 | 660 | 453 | 486 | 615 | 510 | 482 | 557 | 667 | 589 | 639 | 568,08 | 523 | 613 | 1,174 | 0,231 | 2,533 | 0,01129706 | 0,0224447 | 0,007968 | 0,004535 | 0,0062328 | 0,0062328 | TRUE | 0,152591 | No |
| orf19.7356 | 452 | 368 | 444 | 532 | 414 | 418 | 445 | 273 | 405 | 469 | 454 | 439 | 354 | 399 | 397 | 385 | 412,75 | 442 | 384 | 0,87 | -0,2 | -2,562 | 0,0103992 | 0,02084002 | 0,001219 | 0,0054967 | 0,0034385 | 0,0034385 | TRUE | 0,0358755 | No |
| orf19.7357 | 1591 | 1289 | 1369 | 1847 | 1907 | 1816 | 1927 | 1257 | 1427 | 1641 | 1401 | 1523 | 1631 | 1733 | 1720 | 1773 | 1606 | 1498 | 1714 | 1,145 | 0,195 | 3,355 | 0,00079308 | 0,00204061 | 0,002418 | 0,0028834 | 0,0026114 | 0,0026114 | TRUE | 0,0466938 | No |
| orf19.7358 | 732 | 823 | 726 | 1044 | 881 | 1070 | 1104 | 549 | 657 | 1048 | 743 | 861 | 753 | 1021 | 985 | 774 | 855,25 | 827 | 883 | 1,07 | 0,097 | 0,526 | 0,59878685 | 0,6752086 | 0,031449 | 0,0036769 | 0,0169753 | 0,0314492 | TRUE | 0,3776191 | No |
| orf19.7359 | 2573 | 1899 | 2343 | 2851 | 2808 | 2569 | 2752 | 1798 | 2308 | 2418 | 2397 | 2351 | 2401 | 2451 | 2456 | 2536 | 2414,8 | 2368 | 2461 | 1,039 | 0,055 | 1,439 | 0,15018123 | 0,21591023 | 1,00E-08 | 0,0025807 | 0,0009745 | 0,0009745 | TRUE | 0,0045668 | No |
| orf19.736 | 1318 | 1037 | 1139 | 1408 | 1859 | 1875 | 1984 | 1221 | 1182 | 1320 | 1165 | 1161 | 1590 | 1789 | 1770 | 1722 | 1462,6 | 1207 | 1718 | 1,425 | 0,511 | 8,559 | 1,14E-17 | 1,71E-16 | 0,002403 | 0,0029721 | 0,0027173 | 0,0027173 | TRUE | 0,0447028 | No |
| orf19.7360 | 502 | 357 | 411 | 516 | 498 | 588 | 575 | 313 | 450 | 454 | 420 | 426 | 426 | 561 | 513 | 441 | 461,54 | 438 | 485 | 1,112 | 0,153 | 1,635 | 0,10209528 | 0,15536689 | 0,007231 | 0,0051248 | 0,0062087 | 0,0062087 | TRUE | 0,1286184 | No |
| orf19.7361 | 3010 | 2159 | 2518 | 3175 | 3184 | 3080 | 3278 | 1946 | 2700 | 2749 | 2576 | 2618 | 2723 | 2939 | 2925 | 2745 | 2746,9 | 2661 | 2833 | 1,066 | 0,092 | 2,096 | 0,036041 | 0,06311113 | 0,000766 | 0,002508 | 0,0014783 | 0,0014783 | TRUE | 0,0089879 | No |
| orf19.7362 | 1187 | 839 | 1075 | 1338 | 1051 | 1066 | 1091 | 726 | 1065 | 1068 | 1100 | 1103 | 899 | 1017 | 974 | 1024 | 1031,2 | 1084 | 978 | 0,9 | -0,152 | -2,702 | 0,00688737 | 0,01442794 | 0,000974 | 0,0033873 | 0,0020569 | 0,0020569 | TRUE | 0,037275 | No |
| orf19.7363 | 9280 | 6585 | 8697 | 10323 | 10888 | 10306 | 11175 | 6791 | 8325 | 8383 | 8898 | 8512 | 9312 | 9834 | 9972 | 9578 | 9101,8 | 8530 | 9674 | 1,134 | 0,182 | 4,906 | 9,29E-07 | 3,94E-06 | 0,000762 | 0,002139 | 0,0012055 | 0,0012055 | TRUE | 0,0102788 | No |
| orf19.7364 | 54 | 29 | 50 | 45 | 36 | 31 | 28 | 20 | 48 | 37 | 51 | 37 | 31 | 30 | 25 | 28 | 35,9 | 43 | 28 | 0,653 | -0,615 | -2,607 | 0,00912112 | 0,01854473 | 1,00E-08 | 0,0424187 | 0,0243545 | 0,0243545 | TRUE | 0,1057667 | No |
| orf19.7365 | 829 | 735 | 796 | 1028 | 1185 | 1239 | 1310 | 827 | 744 | 936 | 814 | 848 | 1013 | 1182 | 1169 | 1166 | 984,07 | 836 | 1132 | 1,358 | 0,441 | 5,645 | 1,65E-08 | 8,92E-08 | 0,005651 | 0,0034547 | 0,0048136 | 0,0048136 | TRUE | 0,0733674 | No |
| orf19.7366 | 2991 | 2359 | 2641 | 3222 | 2973 | 2787 | 3005 | 1482 | 2683 | 3003 | 2702 | 2657 | 2543 | 2659 | 2681 | 2090 | 2627,4 | 2761 | 2493 | 0,905 | -0,145 | -1,879 | 0,06026105 | 0,0988663 | 0,007327 | 0,0025321 | 0,005298 | 0,005298 | TRUE | 0,1390016 | No |
| orf19.7367 | 2686 | 2236 | 2518 | 3112 | 2879 | 3039 | 3096 | 1890 | 2410 | 2847 | 2576 | 2566 | 2462 | 2900 | 2763 | 2666 | 2648,6 | 2600 | 2698 | 1,038 | 0,054 | 0,846 | 0,39781774 | 0,48658163 | 0,004086 | 0,0025276 | 0,0036073 | 0,0036073 | TRUE | 0,0484008 | No |
| orf19.7368 | 1773 | 1308 | 1656 | 1949 | 1624 | 1425 | 1577 | 1046 | 1591 | 1665 | 1694 | 1607 | 1389 | 1360 | 1407 | 1475 | 1523,5 | 1639 | 1408 | 0,858 | -0,221 | -4,759 | 1,94E-06 | 7,86E-06 | 0,000281 | 0,0029324 | 0,0014069 | 0,0014069 | TRUE | 0,0109934 | No |
| orf19.7369 | 664 | 530 | 582 | 787 | 613 | 705 | 663 | 415 | 596 | 675 | 595 | 649 | 524 | 673 | 592 | 585 | 611,09 | 629 | 594 | 0,945 | -0,082 | -0,985 | 0,32471899 | 0,41175848 | 0,005703 | 0,0043551 | 0,0050642 | 0,0050642 | TRUE | 0,098358 | No |
| orf19.7370 | 3234 | 2308 | 2717 | 3445 | 3240 | 3039 | 3196 | 1952 | 2901 | 2938 | 2780 | 2841 | 2771 | 2900 | 2852 | 2753 | 2842 | 2865 | 2819 | 0,985 | -0,022 | -0,581 | 0,56095206 | 0,64096614 | 0,000203 | 0,0024903 | 0,0010264 | 0,0010264 | TRUE | 0,0047045 | No |
| orf19.7371 | 2213 | 1750 | 2068 | 2505 | 2356 | 2379 | 2527 | 1546 | 1985 | 2228 | 2116 | 2066 | 2015 | 2270 | 2255 | 2180 | 2139,4 | 2099 | 2180 | 1,04 | 0,056 | 1,028 | 0,30375824 | 0,38961987 | 0,001984 | 0,0026581 | 0,002384 | 0,002384 | TRUE | 0,0328487 | No |
| orf19.7372 | 1628 | 1127 | 1456 | 1693 | 1580 | 1420 | 1548 | 1001 | 1460 | 1435 | 1490 | 1396 | 1351 | 1355 | 1381 | 1412 | 1410 | 1445 | 1375 | 0,951 | -0,073 | -1,629 | 0,103318 | 0,15690964 | 1,00E-08 | 0,0030091 | 0,0012271 | 0,0012271 | TRUE | 0,0069112 | No |
| orf19.7374 | 8344 | 6178 | 7923 | 9351 | 8162 | 7571 | 8377 | 5243 | 7485 | 7865 | 8106 | 7711 | 6980 | 7224 | 7475 | 7395 | 7530,2 | 7792 | 7268 | 0,933 | -0,1 | -2,601 | 0,00930116 | 0,01888522 | 0,000837 | 0,0021723 | 0,0012957 | 0,0012957 | TRUE | 0,0090674 | No |
| orf19.7375 | 253 | 188 | 194 | 258 | 361 | 328 | 307 | 201 | 227 | 239 | 198 | 213 | 309 | 313 | 274 | 283 | 257,09 | 219 | 295 | 1,349 | 0,432 | 4,53 | 5,91E-06 | 2,23E-05 | 2,88E-05 | 0,0076263 | 0,0047813 | 0,0047813 | TRUE | 0,0418101 | No |
| orf19.7376 | 438 | 288 | 397 | 443 | 494 | 540 | 518 | 346 | 393 | 367 | 406 | 365 | 422 | 515 | 462 | 488 | 427,38 | 383 | 472 | 1,23 | 0,299 | 3,612 | 0,00030386 | 0,00085871 | 0,003055 | 0,0053763 | 0,0041966 | 0,0041966 | TRUE | 0,0616664 | No |
| orf19.7377 | 180 | 105 | 147 | 171 | 256 | 241 | 216 | 153 | 161 | 134 | 150 | 141 | 219 | 230 | 193 | 216 | 180,5 | 146 | 214 | 1,456 | 0,542 | 4,892 | 1,00E-06 | 4,22E-06 | 1,00E-08 | 0,0100222 | 0,0060816 | 0,0060816 | TRUE | 0,0550561 | No |
| orf19.7378 | 3077 | 2075 | 2845 | 3437 | 3691 | 3379 | 3597 | 2264 | 2760 | 2642 | 2911 | 2834 | 3157 | 3224 | 3210 | 3193 | 2991,3 | 2787 | 3196 | 1,146 | 0,196 | 4,934 | 8,05E-07 | 3,44E-06 | 0,000438 | 0,0024648 | 0,001184 | 0,001184 | TRUE | 0,0138562 | No |
| orf19.7379 | 7356 | 5273 | 6865 | 8198 | 6918 | 6502 | 6870 | 4523 | 6599 | 6713 | 7024 | 6760 | 5916 | 6204 | 6130 | 6379 | 6465,8 | 6774 | 6157 | 0,908 | -0,138 | -3,765 | 0,00016663 | 0,00049146 | 0,000629 | 0,002204 | 0,0011433 | 0,0011433 | TRUE | 0,0085194 | No |
| orf19.738 | 4060 | 2897 | 3736 | 4508 | 4165 | 3847 | 4302 | 2598 | 3642 | 3688 | 3822 | 3717 | 3562 | 3671 | 3839 | 3664 | 3700,7 | 3717 | 3684 | 0,991 | -0,013 | -0,343 | 0,73161545 | 0,7904236 | 0,000468 | 0,0023718 | 0,0010917 | 0,0010917 | TRUE | 0,0101446 | No |
| orf19.7380 | 287 | 197 | 257 | 303 | 233 | 193 | 227 | 145 | 257 | 251 | 263 | 250 | 199 | 184 | 203 | 205 | 226,44 | 255 | 198 | 0,773 | -0,372 | -4,004 | 6,23E-05 | 0,00019768 | 1,00E-08 | 0,0083904 | 0,0037912 | 0,0037912 | TRUE | 0,0231514 | No |

| | | | | | | | | | | | | | | | | | | | | | | | | | | | | | | | |
|---|---|---|---|---|---|---|---|---|---|---|---|---|---|---|---|---|---|---|---|---|---|---|---|---|---|---|---|---|---|---|---|
| orf19.7381 | 7871 | 6061 | 7347 | 8958 | 10097 | 10064 | 10954 | 6822 | 7061 | 7716 | 7517 | 7387 | 8635 | 9603 | 9775 | 9622 | 8414,4 | 7420 | 9409 | 1,268 | 0,343 | 7,008 | 2,42E-12 | 2,12E-11 | 0,002061 | 0,002152 | 0,0021746 | 0,0021746 | TRUE | 0,0377929 | No |
| orf19.7382 | 10425 | 7419 | 8980 | 11712 | 11169 | 9580 | 10161 | 6894 | 9352 | 9445 | 9187 | 9658 | 9552 | 9141 | 9067 | 9723 | 9390,8 | 9410 | 9371 | 0,995 | -0,007 | -0,191 | 0,84890398 | 0,88437925 | 0,000574 | 0,0021341 | 0,0011049 | 0,0011049 | TRUE | 0,0075635 | No |
| orf19.7383 | 2715 | 2001 | 2479 | 3029 | 2855 | 2672 | 2733 | 1778 | 2436 | 2547 | 2536 | 2498 | 2442 | 2550 | 2439 | 2508 | 2494,4 | 2504 | 2485 | 0,992 | -0,011 | -0,301 | 0,76350982 | 0,81549635 | 1,00E-08 | 0,0025615 | 0,0009695 | 0,0009695 | TRUE | 0,0041676 | No |
| orf19.7384 | 27029 | 18684 | 23642 | 28847 | 24913 | 22133 | 23988 | 13916 | 24248 | 23787 | 24188 | 23788 | 21306 | 21119 | 21405 | 19627 | 22433 | 24003 | 20864 | 0,87 | -0,202 | -5,709 | 1,14E-08 | 6,24E-08 | 0,000792 | 0,0020442 | 0,0011543 | 0,0011543 | TRUE | 0,0191966 | No |
| orf19.7385 | 376 | 264 | 363 | 396 | 414 | 316 | 370 | 223 | 337 | 336 | 371 | 327 | 354 | 302 | 330 | 315 | 333,95 | 343 | 325 | 0,951 | -0,072 | -0,844 | 0,39882056 | 0,48760967 | 0,001367 | 0,0063266 | 0,0039811 | 0,0039811 | TRUE | 0,0437671 | No |
| orf19.7386 | 2238 | 1663 | 2052 | 2524 | 2722 | 2540 | 2682 | 1853 | 2008 | 2117 | 2099 | 2081 | 2328 | 2424 | 2393 | 2613 | 2258 | 2076 | 2440 | 1,173 | 0,231 | 4,961 | 7,01E-07 | 3,03E-06 | 0,000937 | 0,0026224 | 0,001626 | 0,001626 | TRUE | 0,0259685 | No |
| orf19.7387 | 8349 | 5798 | 7375 | 8989 | 10073 | 9012 | 9484 | 5945 | 7490 | 7381 | 7545 | 7412 | 8615 | 8599 | 8463 | 8385 | 7986,3 | 7457 | 8516 | 1,142 | 0,192 | 7,172 | 7,41E-13 | 6,90E-12 | 1,00E-08 | 0,0021613 | 0,0005602 | 0,0005602 | TRUE | 0,0012159 | No |
| orf19.7388 | 9901 | 6743 | 8731 | 10895 | 10961 | 9606 | 10507 | 6582 | 8882 | 8585 | 8933 | 8984 | 9374 | 9166 | 9376 | 9283 | 9072,8 | 8846 | 9300 | 1,051 | 0,072 | 2,527 | 0,01151938 | 0,02283354 | 0,000115 | 0,0021395 | 0,0006629 | 0,0006629 | TRUE | 0,0046486 | No |
| orf19.7389 | 827 | 604 | 733 | 883 | 707 | 647 | 700 | 451 | 742 | 769 | 750 | 728 | 605 | 617 | 625 | 636 | 683,96 | 747 | 621 | 0,831 | -0,268 | -4,701 | 2,59E-06 | 1,03E-05 | 1,00E-08 | 0,004102 | 0,0016423 | 0,0016423 | TRUE | 0,0040999 | No |
| orf19.7391 | 3581 | 2507 | 3196 | 3912 | 4073 | 3494 | 3880 | 2489 | 3213 | 3192 | 3270 | 3226 | 3483 | 3334 | 3462 | 3511 | 3336,2 | 3225 | 3448 | 1,068 | 0,095 | 2,794 | 0,00519989 | 0,01118963 | 1,00E-08 | 0,0024147 | 0,0008198 | 0,0008198 | TRUE | 0,006011 | No |
| orf19.7392 | 43101 | 28855 | 39236 | 46989 | 39678 | 32854 | 36561 | 24303 | 38666 | 36735 | 40142 | 38748 | 33934 | 31348 | 32625 | 34277 | 35809 | 38573 | 33046 | 0,857 | -0,223 | -5,371 | 7,83E-08 | 3,90E-07 | 0,00144 | 0,0020201 | 0,0016322 | 0,0016322 | TRUE | 0,0146481 | No |
| orf19.7393 | 3099 | 2197 | 2858 | 3467 | 3289 | 3002 | 3243 | 2161 | 2780 | 2797 | 2924 | 2859 | 2813 | 2864 | 2894 | 3048 | 2872,4 | 2840 | 2905 | 1,021 | 0,03 | 0,759 | 0,44785269 | 0,53450516 | 0,000407 | 0,0024849 | 0,0011849 | 0,0011849 | TRUE | 0,0123036 | No |
| orf19.7394 | 11866 | 8367 | 10702 | 13407 | 10600 | 9323 | 10321 | 6443 | 10645 | 10652 | 10949 | 11056 | 9065 | 8896 | 9210 | 9087 | 9945 | 10826 | 9064 | 0,837 | -0,256 | -8,916 | 4,84E-19 | 8,19E-18 | 0,000153 | 0,0021255 | 0,000692 | 0,000692 | TRUE | 0,0025509 | No |
| orf19.7395 | 4 | 2 | 2 | 4 | 0 | 2 | 2 | 3 | 4 | 3 | 2 | 3 | 0 | 2 | 2 | 4 | 2,43 | 3 | 2 | 0,627 | -0,674 | -0,733 | 0,46372081 | NA | 1,00E-08 | 0,6005107 | 0,3658753 | 0,3658753 | TRUE | 0,4353649 | No |
| orf19.7396 | 658 | 435 | 575 | 772 | 674 | 624 | 689 | 464 | 590 | 554 | 588 | 637 | 576 | 595 | 615 | 654 | 601,26 | 592 | 610 | 1,024 | 0,035 | 0,506 | 0,61306736 | 0,68680006 | 0,001306 | 0,004394 | 0,0028437 | 0,0028437 | TRUE | 0,0310207 | No |
| orf19.7397 | 1439 | 996 | 1247 | 1461 | 1396 | 1312 | 1473 | 859 | 1291 | 1268 | 1276 | 1205 | 1194 | 1252 | 1314 | 1212 | 1251,4 | 1260 | 1243 | 0,988 | -0,018 | -0,342 | 0,73267965 | 0,79100437 | 0,000732 | 0,0031396 | 0,0017377 | 0,0017377 | TRUE | 0,0187161 | No |
| orf19.7397.1 | 40 | 22 | 28 | 56 | 37 | 37 | 52 | 70 | 36 | 28 | 29 | 46 | 32 | 35 | 46 | 99 | 43,85 | 35 | 53 | 1,49 | 0,575 | 1,674 | 0,09406195 | 0,14553804 | 0,148985 | 0,035086 | 0,0892745 | 0,0892745 | TRUE | 2,3909672 | No |
| orf19.7397.3 | 5 | 1 | 1 | 1 | 4 | 4 | 5 | 0 | 4 | 1 | 1 | 1 | 3 | 4 | 4 | 0 | 2,41 | 2 | 3 | 1,584 | 0,663 | 0,695 | 0,48700597 | NA | 0,151375 | 0,6035461 | 0,4548792 | 0,4548792 | TRUE | 0,8368688 | No |
| orf19.7398 | 0 | 0 | 0 | 0 | 0 | 0 | 0 | 0 | 0 | 0 | 0 | 0 | 0 | 0 | 0 | 0 | 0 | 0 | 0 | NA | NA | NA | NA | NA | NA | NA | NA | NA | NA | NA | NA |
| orf19.7398.1 | 1 | 1 | 2 | 2 | 2 | 1 | 0 | 0 | 1 | 1 | 2 | 2 | 2 | 1 | 0 | 0 | 1,07 | 2 | 1 | 0,49 | -1,03 | -0,762 | 0,44614211 | NA | 1,00E-08 | 1,3634389 | 0,7220686 | 0,7220686 | TRUE | 0,4085716 | No |
| orf19.74 | 1091 | 1046 | 980 | 1326 | 1188 | 1462 | 1366 | 843 | 979 | 1332 | 1003 | 1093 | 1016 | 1395 | 1219 | 1189 | 1153,2 | 1102 | 1205 | 1,095 | 0,131 | 1,203 | 0,22884525 | 0,30882801 | 0,017533 | 0,0032384 | 0,0104654 | 0,0104654 | TRUE | 0,2307707 | No |
| orf19.740 | 35257 | 25461 | 31646 | 40234 | 45082 | 42743 | 46629 | 28225 | 31629 | 32415 | 32377 | 33177 | 38556 | 40784 | 41609 | 39809 | 36294 | 32400 | 40190 | 1,24 | 0,311 | 9,243 | 2,39E-20 | 4,47E-19 | 0,000688 | 0,0020195 | 0,0010584 | 0,0010584 | TRUE | 0,0092422 | No |
| orf19.7400 | 1556 | 1057 | 1379 | 1717 | 1611 | 1342 | 1483 | 1002 | 1396 | 1346 | 1411 | 1416 | 1378 | 1280 | 1323 | 1413 | 1370,4 | 1392 | 1348 | 0,966 | -0,049 | -1,014 | 0,31044816 | 0,39667642 | 0,000334 | 0,0030389 | 0,0015254 | 0,0015254 | TRUE | 0,0136122 | No |
| orf19.7401 | 1547 | 1058 | 1432 | 1663 | 1744 | 1560 | 1758 | 1077 | 1388 | 1347 | 1465 | 1371 | 1492 | 1489 | 1569 | 1519 | 1454,9 | 1393 | 1517 | 1,089 | 0,123 | 2,628 | 0,00859471 | 0,01758755 | 0,000244 | 0,0029773 | 0,0014043 | 0,0014043 | TRUE | 0,014277 | No |
| orf19.7402 | 1655 | 1153 | 1461 | 1780 | 1897 | 1649 | 1688 | 1083 | 1485 | 1468 | 1495 | 1468 | 1622 | 1573 | 1506 | 1527 | 1518,1 | 1479 | 1557 | 1,054 | 0,076 | 1,716 | 0,08610572 | 0,13478978 | 1,00E-08 | 0,0029358 | 0,0012175 | 0,0012175 | TRUE | 0,0100317 | No |
| orf19.7403 | 2359 | 1560 | 2023 | 2311 | 2844 | 2535 | 2761 | 1654 | 2116 | 1986 | 2070 | 1906 | 2432 | 2419 | 2464 | 2333 | 2215,7 | 2020 | 2412 | 1,195 | 0,258 | 5,622 | 1,89E-08 | 1,01E-07 | 0,000895 | 0,0026347 | 0,0015596 | 0,0015596 | TRUE | 0,0186889 | No |
| orf19.7404 | 2271 | 1346 | 1876 | 2333 | 2683 | 2087 | 2234 | 1589 | 2037 | 1714 | 1919 | 1924 | 2295 | 1991 | 1993 | 2241 | 2014,3 | 1898 | 2130 | 1,12 | 0,164 | 2,408 | 0,0160187 | 0,03060106 | 0,004697 | 0,0027002 | 0,0039423 | 0,0039423 | TRUE | 0,0501631 | No |
| orf19.7405 | 1283 | 890 | 1139 | 1347 | 1399 | 1229 | 1264 | 841 | 1151 | 1133 | 1165 | 1111 | 1196 | 1173 | 1128 | 1186 | 1155,4 | 1140 | 1171 | 1,027 | 0,038 | 0,798 | 0,42460303 | 0,51210612 | 1,00E-08 | 0,003236 | 0,0013176 | 0,0013176 | TRUE | 0,0077075 | No |
| orf19.7406 | 1333 | 922 | 1136 | 1414 | 1254 | 1145 | 1307 | 739 | 1196 | 1174 | 1162 | 1166 | 1072 | 1093 | 1166 | 1042 | 1133,9 | 1174 | 1093 | 0,933 | -0,101 | -1,967 | 0,04922889 | 0,08312737 | 0,000307 | 0,0032598 | 0,0016276 | 0,0016276 | TRUE | 0,0244771 | No |
| orf19.7409 | 771 | 542 | 652 | 814 | 832 | 698 | 725 | 492 | 692 | 690 | 667 | 671 | 712 | 666 | 647 | 694 | 679,8 | 680 | 680 | 0,999 | -0,001 | -0,021 | 0,98315498 | 0,98759323 | 1,00E-08 | 0,004115 | 0,0019148 | 0,0019148 | TRUE | 0,0137961 | No |
| orf19.7409.1 | 339 | 238 | 258 | 356 | 374 | 340 | 331 | 221 | 304 | 303 | 264 | 294 | 320 | 324 | 295 | 312 | 302 | 291 | 313 | 1,075 | 0,104 | 1,219 | 0,2226993 | 0,301686 | 1,00E-08 | 0,0067865 | 0,0036552 | 0,0036552 | TRUE | 0,0433107 | No |
| orf19.741 | 12 | 11 | 7 | 15 | 8 | 11 | 11 | 5 | 11 | 14 | 7 | 12 | 7 | 10 | 10 | 7 | 9,81 | 11 | 8 | 0,782 | -0,355 | -0,811 | 0,41727049 | NA | 1,00E-08 | 0,1499128 | 0,0808735 | 0,0808735 | TRUE | 0,199466 | No |
| orf19.7410 | 5193 | 3754 | 4801 | 5844 | 4638 | 4449 | 4637 | 2964 | 4659 | 4779 | 4912 | 4819 | 3967 | 4245 | 4138 | 4180 | 4462,3 | 4792 | 4132 | 0,862 | -0,214 | -5,881 | 4,08E-09 | 2,40E-08 | 0,000463 | 0,0023048 | 0,0010471 | 0,0010471 | TRUE | 0,0091972 | No |
| orf19.7411 | 1031 | 694 | 910 | 1117 | 665 | 497 | 579 | 379 | 925 | 884 | 931 | 921 | 569 | 474 | 517 | 535 | 719,34 | 915 | 524 | 0,572 | -0,807 | -12,299 | 9,13E-35 | 4,43E-33 | 0,001096 | 0,0039976 | 0,0026273 | 0,0026273 | TRUE | 0,0490363 | No |
| orf19.7412 | 4741 | 3043 | 4149 | 5086 | 4149 | 3729 | 4087 | 2410 | 4253 | 3874 | 4245 | 4194 | 3548 | 3558 | 3647 | 3399 | 3839,8 | 4142 | 3538 | 0,854 | -0,227 | -5,266 | 1,40E-07 | 6,71E-07 | 0,001019 | 0,0023576 | 0,0015232 | 0,0015232 | TRUE | 0,0221075 | No |
| orf19.7413 | 928 | 685 | 805 | 959 | 777 | 758 | 745 | 542 | 833 | 872 | 824 | 791 | 665 | 723 | 665 | 764 | 767 | 830 | 704 | 0,847 | -0,24 | -3,712 | 0,00020542 | 0,00059795 | 0,001455 | 0,0038722 | 0,0026874 | 0,0026874 | TRUE | 0,0345539 | No |
| orf19.7414 | 1970 | 1312 | 1782 | 2150 | 2323 | 2073 | 2245 | 1470 | 1767 | 1670 | 1823 | 1773 | 1987 | 1978 | 2003 | 2073 | 1884,4 | 1758 | 2010 | 1,141 | 0,19 | 4,42 | 9,85E-06 | 3,59E-05 | 0,000181 | 0,0027499 | 0,0012414 | 0,0012414 | TRUE | 0,0124485 | No |
| orf19.7417 | 4 | 0 | 0 | 0 | 3 | 1 | 1 | 1 | 4 | 0 | 0 | 0 | 3 | 1 | 1 | 1 | 1,18 | 1 | 2 | 1,558 | 0,64 | 0,445 | 0,65635527 | NA | 0,88406 | 1,2360224 | 1,1143841 | 1,1143841 | TRUE | 1,9195246 | No |
| orf19.7419 | 78 | 47 | 62 | 57 | 48 | 37 | 39 | 18 | 70 | 60 | 63 | 47 | 41 | 35 | 35 | 25 | 47,1 | 60 | 34 | 0,579 | -0,788 | -3,588 | 0,00033287 | 0,00093235 | 0,010795 | 0,0328021 | 0,0235469 | 0,0235469 | TRUE | 0,2171507 | No |
| orf19.742 | 2938 | 2068 | 2562 | 3247 | 3265 | 2797 | 2989 | 1939 | 2636 | 2633 | 2621 | 2678 | 2792 | 2669 | 2667 | 2735 | 2678,8 | 2642 | 2716 | 1,028 | 0,039 | 1,096 | 0,27293818 | 0,35725765 | 1,00E-08 | 0,0025214 | 0,0008639 | 0,0008639 | TRUE | 0,0047817 | No |
| orf19.7420 | 0 | 0 | 1 | 1 | 1 | 1 | 1 | 1 | 0 | 0 | 1 | 1 | 1 | 1 | 1 | 1 | 0,74 | 0 | 1 | 2,047 | 1,033 | 0,66 | 0,50920861 | NA | 1,00E-08 | 1,9506189 | 0,9201543 | 0,9201543 | TRUE | 0,1221952 | No |
| orf19.7421 | 2160 | 1381 | 1866 | 2289 | 2425 | 1898 | 2094 | 1500 | 1938 | 1758 | 1909 | 1888 | 2074 | 1811 | 1869 | 2116 | 1920,2 | 1873 | 1968 | 1,048 | 0,068 | 1,118 | 0,26339097 | 0,34718143 | 0,003112 | 0,0027355 | 0,0030308 | 0,0030308 | TRUE | 0,0343069 | No |
| orf19.7422 | 5769 | 3548 | 4676 | 6232 | 4606 | 3435 | 3945 | 2529 | 5175 | 4517 | 4784 | 5139 | 3939 | 3278 | 3520 | 3567 | 4239,9 | 4904 | 3576 | 0,729 | -0,456 | -7,089 | 1,35E-12 | 1,22E-11 | 0,00463 | 0,0023219 | 0,0037303 | 0,0037303 | TRUE | 0,0578855 | No |
| orf19.7424 | 0 | 0 | 0 | 1 | 1 | 0 | 2 | 1 | 0 | 0 | 0 | 1 | 1 | 0 | 2 | 1 | 0,61 | 0 | 1 | 3,33 | 1,736 | 0,961 | 0,33668038 | NA | 1,00E-08 | 2,3842963 | 1,6306473 | 1,6306473 | TRUE | 0,2428649 | No |
| orf19.7425 | 833 | 569 | 805 | 936 | 914 | 870 | 929 | 594 | 747 | 724 | 824 | 772 | 782 | 830 | 829 | 838 | 793,21 | 767 | 820 | 1,067 | 0,093 | 1,557 | 0,11946399 | 0,17752651 | 0,000621 | 0,0038097 | 0,0021664 | 0,0021664 | TRUE | 0,0281663 | No |
| orf19.7426 | 798 | 585 | 781 | 888 | 919 | 788 | 911 | 521 | 716 | 745 | 799 | 732 | 786 | 752 | 813 | 735 | 759,69 | 748 | 772 | 1,034 | 0,049 | 0,796 | 0,42576623 | 0,51283149 | 0,000841 | 0,0038904 | 0,0022981 | 0,0022981 | TRUE | 0,0252284 | No |
| orf19.7427 | 7063 | 4888 | 6374 | 7722 | 6879 | 6008 | 6528 | 4271 | 6336 | 6223 | 6521 | 6368 | 5883 | 5733 | 5825 | 6024 | 6114,1 | 6362 | 5866 | 0,921 | -0,118 | -3,73 | 0,00019157 | 0,00056008 | 0,000185 | 0,002217 | 0,0007977 | 0,0007977 | TRUE | 0,0037208 | No |
| orf19.7428 | 4067 | 2627 | 3490 | 4211 | 3791 | 3264 | 3658 | 1990 | 3649 | 3344 | 3571 | 3472 | 3242 | 3114 | 3264 | 2807 | 3307,9 | 3509 | 3107 | 0,886 | -0,174 | -3,197 | 0,00138799 | 0,00338849 | 0,002595 | 0,0024184 | 0,0025424 | 0,0025424 | TRUE | 0,0490636 | No |
| orf19.7433 | 5952 | 3818 | 5408 | 6328 | 6234 | 5462 | 6141 | 3799 | 5340 | 4861 | 5533 | 5218 | 5332 | 5212 | 5480 | 5358 | 5291,6 | 5238 | 5346 | 1,02 | 0,029 | 0,64 | 0,52230412 | 0,60544147 | 0,001336 | 0,0022539 | 0,0017313 | 0,0017313 | TRUE | 0,0277299 | No |
| orf19.7434 | 839 | 680 | 778 | 972 | 1040 | 1065 | 1070 | 627 | 753 | 866 | 796 | 802 | 889 | 1016 | 955 | 884 | 870,08 | 804 | 936 | 1,168 | 0,224 | 3,371 | 0,00074997 | 0,00193798 | 0,002609 | 0,003648 | 0,0030885 | 0,0030885 | TRUE | 0,0393643 | No |
| orf19.7436 | 6612 | 4578 | 6250 | 7497 | 9072 | 8106 | 9347 | 6214 | 5932 | 5828 | 6394 | 6182 | 7759 | 7734 | 8341 | 8764 | 7116,8 | 6084 | 8150 | 1,339 | 0,421 | 8,153 | 3,55E-16 | 4,58E-15 | 0,00254 | 0,0021835 | 0,0024142 | 0,0024142 | TRUE | 0,0309242 | No |
| orf19.7436.1 | 608 | 394 | 469 | 610 | 607 | 514 | 532 | 393 | 545 | 502 | 480 | 503 | 519 | 490 | 475 | 554 | 508,56 | 508 | 510 | 1 | 0 | -0,005 | 0,99572934 | 0,99717303 | 0,001645 | 0,0048341 | 0,003273 | 0,003273 | TRUE | 0,0347508 | No |
| orf19.7437 | 942 | 632 | 803 | 1011 | 777 | 643 | 708 | 468 | 845 | 805 | 822 | 834 | 665 | 614 | 632 | 660 | 734,35 | 826 | 643 | 0,776 | -0,366 | -6,414 | 1,42E-10 | 1,01E-09 | 1,00E-08 | 0,0039564 | 0,001742 | 0,001742 | TRUE | 0,0108834 | No |
| orf19.7438 | 5162 | 3522 | 4682 | 5555 | 5535 | 4887 | 5339 | 3549 | 4631 | 4484 | 4790 | 4581 | 4734 | 4663 | 4764 | 5006 | 4706,5 | 4622 | 4792 | 1,036 | 0,051 | 1,348 | 0,17780918 | 0,24953925 | 0,000582 | 0,002288 | 0,0011452 | 0,0011452 | TRUE | 0,0103736 | No |
| orf19.744 | 4468 | 3294 | 4222 | 5074 | 4888 | 4289 | 4834 | 3059 | 4008 | 4194 | 4320 | 4184 | 4180 | 4092 | 4314 | 4314 | 4200,8 | 4176 | 4225 | 1,012 | 0,017 | 0,436 | 0,66290895 | 0,73104937 | 0,000481 | 0,0023251 | 0,0011498 | 0,0011498 | TRUE | 0,0091158 | No |
| orf19.7440 | 59 | 46 | 50 | 64 | 44 | 66 | 58 | 38 | 53 | 59 | 51 | 53 | 38 | 63 | 52 | 54 | 52,67 | 54 | 52 | 0,954 | -0,068 | -0,339 | 0,73481924 | 0,79288685 | 1,00E-08 | 0,0295405 | 0,0201384 | 0,0201384 | TRUE | 0,3139897 | No |
| orf19.7441 | 1501 | 1059 | 1286 | 1719 | 1527 | 1532 | 1600 | 1046 | 1347 | 1348 | 1316 | 1418 | 1306 | 1462 | 1428 | 1475 | 1387,3 | 1357 | 1418 | 1,042 | 0,06 | 1,11 | 0,26704603 | 0,35130558 | 0,001408 | 0,0030259 | 0,0020753 | 0,0020753 | TRUE | 0,0351385 | No |
| orf19.7442 | 1602 | 1091 | 1328 | 1631 | 1214 | 1101 | 1176 | 694 | 1437 | 1389 | 1359 | 1345 | 1038 | 1051 | 1049 | 979 | 1205,8 | 1382 | 1029 | 0,746 | -0,422 | -8,6 | 7,98E-18 | 1,22E-16 | 1,00E-08 | 0,0031834 | 0,0014673 | 0,0014673 | TRUE | 0,0110542 | No |
| orf19.7443 | 406 | 451 | 421 | 595 | 477 | 580 | 575 | 360 | 364 | 574 | 431 | 491 | 408 | 553 | 513 | 508 | 480,25 | 465 | 496 | 1,067 | 0,094 | 0,714 | 0,47512911 | 0,56082038 | 0,024587 | 0,0050024 | 0,0146113 | 0,0146113 | TRUE | 0,2778293 | No |

| | | | | | | | | | | | | | | | | | | | | | | | | | | | | | | | |
|---|---|---|---|---|---|---|---|---|---|---|---|---|---|---|---|---|---|---|---|---|---|---|---|---|---|---|---|---|---|---|---|
| orf19.7444 | 669 | 558 | 679 | 881 | 784 | 788 | 786 | 483 | 600 | 710 | 695 | 726 | 671 | 752 | 701 | 681 | 692,09 | 683 | 701 | 1,028 | 0,04 | 0,539 | 0,58961914 | 0,66748517 | 0,003646 | 0,0040771 | 0,0038467 | 0,0038467 | TRUE | 0,081827 | No |
| orf19.7445 | 7691 | 5822 | 7278 | 9123 | 8828 | 8407 | 9260 | 5898 | 6900 | 7412 | 7446 | 7523 | 7550 | 8022 | 8263 | 8319 | 7679,3 | 7320 | 8038 | 1,098 | 0,135 | 2,997 | 0,00272544 | 0,00625695 | 0,001528 | 0,0021686 | 0,0018079 | 0,0018079 | TRUE | 0,0206219 | No |
| orf19.7446 | 181 | 185 | 172 | 263 | 214 | 208 | 211 | 126 | 162 | 236 | 176 | 217 | 183 | 198 | 188 | 178 | 192,28 | 198 | 187 | 0,95 | -0,073 | -0,579 | 0,5623544 | 0,64232419 | 0,010909 | 0,0095295 | 0,010167 | 0,010167 | TRUE | 0,1623444 | No |
| orf19.7447 | 97 | 75 | 101 | 142 | 39 | 28 | 39 | 18 | 87 | 95 | 103 | 117 | 33 | 27 | 35 | 25 | 65,4 | 100 | 30 | 0,302 | -1,729 | -8,731 | 2,52E-18 | 4,04E-17 | 1,00E-08 | 0,0241772 | 0,0163541 | 0,0163541 | TRUE | 0,1310804 | No |
| orf19.7448 | 1480 | 1026 | 1258 | 1528 | 1618 | 1380 | 1466 | 961 | 1328 | 1306 | 1287 | 1260 | 1384 | 1317 | 1308 | 1355 | 1318,1 | 1295 | 1341 | 1,036 | 0,05 | 1,093 | 0,27453975 | 0,35880679 | 1,00E-08 | 0,0030809 | 0,00128 | 0,00128 | TRUE | 0,0062323 | No |
| orf19.7449 | 2067 | 1545 | 1726 | 2179 | 2298 | 2246 | 2315 | 1452 | 1854 | 1967 | 1766 | 1797 | 1965 | 2143 | 2066 | 2048 | 1950,8 | 1846 | 2056 | 1,115 | 0,157 | 3,185 | 0,00144524 | 0,00351966 | 0,001095 | 0,0027237 | 0,0018037 | 0,0018037 | TRUE | 0,0220052 | No |
| orf19.745 | 6466 | 4644 | 5926 | 7534 | 6994 | 6542 | 6985 | 4995 | 5801 | 5912 | 6063 | 6213 | 5981 | 6242 | 6233 | 7045 | 6186,3 | 5997 | 6375 | 1,062 | 0,087 | 1,631 | 0,10298209 | 0,15643903 | 0,002862 | 0,0022142 | 0,0025695 | 0,0025695 | TRUE | 0,0597498 | No |
| orf19.7450 | 1292 | 921 | 1125 | 1399 | 1042 | 1020 | 1028 | 606 | 1159 | 1173 | 1151 | 1154 | 891 | 973 | 917 | 855 | 1034,1 | 1159 | 909 | 0,786 | -0,347 | -6,538 | 6,25E-11 | 4,71E-10 | 0,000103 | 0,0033834 | 0,0017205 | 0,0017205 | TRUE | 0,0261389 | No |
| orf19.7451 | 4529 | 3209 | 4141 | 4936 | 6407 | 5970 | 6432 | 4274 | 4063 | 4085 | 4237 | 4070 | 5479 | 5696 | 5739 | 6028 | 4924,9 | 4114 | 5736 | 1,393 | 0,478 | 12,242 | 1,86E-34 | 8,80E-33 | 0,000696 | 0,0022743 | 0,0012572 | 0,0012572 | TRUE | 0,0137368 | No |
| orf19.7452 | 45 | 34 | 45 | 69 | 113 | 125 | 120 | 65 | 40 | 43 | 46 | 57 | 97 | 119 | 107 | 92 | 75,16 | 46 | 104 | 2,217 | 1,148 | 6,51 | 7,52E-11 | 5,56E-10 | 1,00E-08 | 0,0212949 | 0,0145267 | 0,0145267 | TRUE | 0,2062719 | No |
| orf19.7453 | 21 | 14 | 18 | 27 | 65 | 56 | 70 | 42 | 19 | 18 | 18 | 22 | 56 | 53 | 62 | 59 | 38,51 | 19 | 58 | 2,955 | 1,563 | 6,62 | 3,60E-11 | 2,78E-10 | 1,00E-08 | 0,0396778 | 0,0195964 | 0,0195964 | TRUE | 0,0674984 | No |
| orf19.7454 | 857 | 665 | 791 | 901 | 1238 | 1094 | 1165 | 759 | 769 | 847 | 809 | 743 | 1059 | 1044 | 1040 | 1071 | 922,55 | 792 | 1054 | 1,333 | 0,415 | 7,415 | 1,22E-13 | 1,22E-12 | 0,000269 | 0,0035531 | 0,0019019 | 0,0019019 | TRUE | 0,0236367 | No |
| orf19.7455 | 221 | 168 | 211 | 212 | 343 | 300 | 351 | 206 | 198 | 214 | 216 | 175 | 293 | 286 | 313 | 291 | 248,27 | 201 | 296 | 1,484 | 0,569 | 5,859 | 4,64E-09 | 2,71E-08 | 1,00E-08 | 0,0078267 | 0,0048804 | 0,0048804 | TRUE | 0,0902753 | No |
| orf19.7456 | 43 | 52 | 48 | 58 | 56 | 52 | 38 | 31 | 39 | 66 | 49 | 48 | 48 | 50 | 34 | 44 | 47,11 | 50 | 44 | 0,878 | -0,187 | -0,837 | 0,40262005 | 0,49095622 | 0,017975 | 0,0327968 | 0,026634 | 0,026634 | TRUE | 0,3042172 | No |
| orf19.7457 | 1273 | 915 | 1088 | 1428 | 1015 | 852 | 940 | 544 | 1142 | 1165 | 1113 | 1178 | 868 | 813 | 839 | 767 | 985,58 | 1150 | 822 | 0,717 | -0,479 | -8,785 | 1,57E-18 | 2,54E-17 | 0,000265 | 0,0034525 | 0,0018154 | 0,0018154 | TRUE | 0,0202797 | No |
| orf19.7458 | 454 | 296 | 386 | 532 | 393 | 322 | 328 | 209 | 407 | 377 | 395 | 439 | 336 | 307 | 293 | 295 | 356,07 | 404 | 308 | 0,76 | -0,395 | -4,7 | 2,60E-06 | 1,04E-05 | 0,001644 | 0,0060566 | 0,003938 | 0,003938 | TRUE | 0,0469027 | No |
| orf19.7459 | 16511 | 11815 | 14586 | 18279 | 16260 | 15324 | 16376 | 11299 | 14812 | 15042 | 14923 | 15073 | 13906 | 14622 | 14613 | 15936 | 14866 | 14962 | 14769 | 0,987 | -0,019 | -0,447 | 0,6550276 | 0,72382012 | 0,001399 | 0,0020772 | 0,0017273 | 0,0017273 | TRUE | 0,0341557 | No |
| orf19.746 | 2558 | 2403 | 2591 | 3128 | 2492 | 2880 | 2943 | 1722 | 2295 | 3059 | 2651 | 2579 | 2131 | 2748 | 2626 | 2429 | 2564,8 | 2646 | 2484 | 0,939 | -0,09 | -0,974 | 0,33004358 | 0,41718953 | 0,011957 | 0,0025455 | 0,0078958 | 0,0078958 | TRUE | 0,1325029 | No |
| orf19.7460 | 1968 | 1409 | 1663 | 2212 | 2177 | 1798 | 1961 | 1235 | 1766 | 1794 | 1701 | 1824 | 1862 | 1716 | 1750 | 1742 | 1769,2 | 1771 | 1768 | 0,998 | -0,002 | -0,051 | 0,95903612 | 0,97028049 | 0,000624 | 0,0028 | 0,0014713 | 0,0014713 | TRUE | 0,0164526 | No |
| orf19.7463 | 419 | 280 | 366 | 494 | 510 | 431 | 518 | 337 | 376 | 356 | 374 | 407 | 436 | 411 | 462 | 475 | 412,39 | 378 | 446 | 1,172 | 0,229 | 2,931 | 0,00338047 | 0,00757548 | 0,001151 | 0,0054997 | 0,0034364 | 0,0034364 | TRUE | 0,0318239 | No |
| orf19.7464 | 13 | 6 | 9 | 11 | 16 | 8 | 11 | 3 | 12 | 8 | 9 | 9 | 14 | 8 | 10 | 4 | 9,12 | 10 | 9 | 0,966 | -0,05 | -0,107 | 0,91470496 | NA | 1,00E-08 | 0,1611931 | 0,0983324 | 0,0983324 | TRUE | 0,2807213 | No |
| orf19.7466 | 9500 | 6442 | 8884 | 9793 | 6592 | 5967 | 6412 | 3819 | 8522 | 8201 | 9089 | 8075 | 5638 | 5694 | 5722 | 5386 | 7041 | 8472 | 5610 | 0,662 | -0,594 | -13,078 | 4,38E-39 | 2,86E-37 | 0,0016 | 0,0021857 | 0,0018327 | 0,0018327 | TRUE | 0,0293349 | No |
| orf19.7468 | 3348 | 2294 | 3008 | 3760 | 3510 | 3095 | 3249 | 2171 | 3004 | 2921 | 3077 | 3101 | 3002 | 2953 | 2899 | 3062 | 3002,3 | 3026 | 2979 | 0,983 | -0,025 | -0,648 | 0,51702709 | 0,60071529 | 0,000188 | 0,0024631 | 0,0010391 | 0,0010391 | TRUE | 0,0062065 | No |
| orf19.7469 | 29934 | 19623 | 25170 | 30804 | 30259 | 25123 | 27164 | 15885 | 26854 | 24982 | 25751 | 25401 | 25878 | 23972 | 24239 | 22404 | 24935 | 25747 | 24123 | 0,937 | -0,094 | -1,976 | 0,04820463 | 0,08153531 | 0,002171 | 0,0020377 | 0,0021241 | 0,0021241 | TRUE | 0,0293742 | No |
| orf19.747 | 3751 | 2586 | 3395 | 4014 | 3951 | 3509 | 3896 | 2307 | 3365 | 3292 | 3473 | 3310 | 3379 | 3348 | 3477 | 3254 | 3362,3 | 3360 | 3364 | 1,002 | 0,003 | 0,079 | 0,93668825 | 0,95415785 | 0,00027 | 0,0024113 | 0,0010477 | 0,0010477 | TRUE | 0,0061662 | No |
| orf19.7472 | 1022 | 746 | 976 | 1168 | 1645 | 1446 | 1651 | 994 | 917 | 950 | 999 | 963 | 1407 | 1380 | 1473 | 1402 | 1186,3 | 957 | 1416 | 1,48 | 0,566 | 11,296 | 1,37E-29 | 4,83E-28 | 1,00E-08 | 0,0032033 | 0,001532 | 0,001532 | TRUE | 0,0101415 | No |
| orf19.7473 | 1265 | 1148 | 1234 | 1560 | 1862 | 1861 | 2018 | 1295 | 1135 | 1462 | 1263 | 1286 | 1592 | 1776 | 1801 | 1826 | 1517,6 | 1286 | 1749 | 1,361 | 0,445 | 5,849 | 4,95E-09 | 2,88E-08 | 0,006409 | 0,0029361 | 0,0048678 | 0,0048678 | TRUE | 0,0972582 | No |
| orf19.7475 | 709 | 664 | 704 | 855 | 940 | 1016 | 1033 | 670 | 636 | 845 | 720 | 705 | 804 | 969 | 922 | 945 | 818,35 | 726 | 910 | 1,254 | 0,327 | 3,647 | 0,00026529 | 0,00075611 | 0,008797 | 0,0037535 | 0,0064741 | 0,0064741 | TRUE | 0,1364252 | No |
| orf19.7477 | 5430 | 3889 | 4505 | 5520 | 5167 | 4813 | 5093 | 2718 | 4871 | 4951 | 4609 | 4552 | 4419 | 4592 | 4545 | 3833 | 4546,6 | 4746 | 4347 | 0,917 | -0,125 | -2,074 | 0,03808689 | 0,06626877 | 0,003798 | 0,0022988 | 0,0032568 | 0,0032568 | TRUE | 0,074916 | No |
| orf19.7478 | 2680 | 2086 | 2458 | 2938 | 3094 | 2933 | 3107 | 1952 | 2404 | 2656 | 2515 | 2423 | 2646 | 2799 | 2772 | 2753 | 2621 | 2500 | 2742 | 1,099 | 0,136 | 3,045 | 0,00232529 | 0,00541068 | 0,000845 | 0,0025334 | 0,0015195 | 0,0015195 | TRUE | 0,0204971 | No |
| orf19.7479 | 5226 | 4048 | 4959 | 5910 | 5064 | 4775 | 5134 | 3062 | 4688 | 5154 | 5074 | 4873 | 4331 | 4556 | 4581 | 4319 | 4697 | 4947 | 4447 | 0,9 | -0,153 | -3,543 | 0,00039622 | 0,00109273 | 0,001112 | 0,0022886 | 0,0015648 | 0,0015648 | TRUE | 0,015142 | No |
| orf19.748 | 3575 | 2766 | 3326 | 3924 | 3843 | 3840 | 4160 | 2340 | 3207 | 3521 | 3403 | 3236 | 3287 | 3664 | 3712 | 3300 | 3416,3 | 3342 | 3491 | 1,046 | 0,065 | 1,165 | 0,24386141 | 0,32565201 | 0,002731 | 0,0024045 | 0,0026565 | 0,0026565 | TRUE | 0,0222631 | No |
| orf19.7480 | 2 | 7 | 4 | 6 | 6 | 5 | 2 | 4 | 2 | 9 | 4 | 5 | 5 | 5 | 2 | 6 | 4,63 | 5 | 4 | 0,89 | -0,168 | -0,248 | 0,80400494 | NA | 1,00E-08 | 0,3152296 | 0,218117 | 0,218117 | TRUE | 0,3538667 | No |
| orf19.7481 | 2467 | 1582 | 2138 | 2554 | 2810 | 2452 | 2714 | 1891 | 2213 | 2014 | 2187 | 2106 | 2403 | 2340 | 2422 | 2667 | 2294 | 2130 | 2458 | 1,151 | 0,203 | 3,85 | 0,00011817 | 0,0003566 | 0,002017 | 0,0026123 | 0,0022363 | 0,0022363 | TRUE | 0,0374856 | No |
| orf19.7482 | 36 | 45 | 28 | 45 | 38 | 62 | 44 | 20 | 32 | 57 | 29 | 37 | 32 | 59 | 39 | 28 | 39,31 | 39 | 40 | 1,044 | 0,062 | 0,216 | 0,82862483 | 0,86822078 | 0,073309 | 0,0389105 | 0,0542564 | 0,0542564 | TRUE | 0,8127129 | No |
| orf19.7483 | 4157 | 3275 | 3935 | 4846 | 6166 | 6127 | 6502 | 3869 | 3729 | 4169 | 4026 | 3996 | 5273 | 5846 | 5802 | 5457 | 4787,4 | 3980 | 5594 | 1,407 | 0,492 | 9,873 | 5,43E-23 | 1,21E-21 | 0,001994 | 0,0022828 | 0,0021697 | 0,0021697 | TRUE | 0,0220338 | No |
| orf19.7484 | 2176 | 1424 | 1768 | 2248 | 1973 | 1606 | 1766 | 1127 | 1952 | 1813 | 1809 | 1854 | 1687 | 1532 | 1576 | 1590 | 1726,6 | 1857 | 1596 | 0,859 | -0,219 | -4,48 | 7,48E-06 | 2,78E-05 | 0,000971 | 0,0028203 | 0,0017002 | 0,0017002 | TRUE | 0,0188902 | No |
| orf19.7485 | 3503 | 2360 | 2905 | 3581 | 3583 | 3114 | 3342 | 1922 | 3143 | 3005 | 2972 | 2953 | 3064 | 2971 | 2982 | 2711 | 2975,1 | 3018 | 2932 | 0,973 | -0,04 | -0,843 | 0,39937739 | 0,48809181 | 0,001321 | 0,0024675 | 0,001776 | 0,001776 | TRUE | 0,0293943 | No |
| orf19.7486 | 1020 | 814 | 934 | 1193 | 947 | 941 | 976 | 602 | 915 | 1036 | 956 | 984 | 810 | 898 | 871 | 849 | 914,81 | 973 | 857 | 0,882 | -0,181 | -3,031 | 0,00243958 | 0,00564817 | 0,001257 | 0,0035664 | 0,0023096 | 0,0023096 | TRUE | 0,0211734 | No |
| orf19.7487 | 16 | 16 | 24 | 25 | 13 | 16 | 14 | 7 | 14 | 20 | 25 | 21 | 11 | 15 | 12 | 10 | 16,08 | 20 | 12 | 0,619 | -0,693 | -1,982 | 0,04752417 | 0,0804523 | 1,00E-08 | 0,0922571 | 0,0518098 | 0,0518098 | TRUE | 0,2191453 | No |
| orf19.7488 | 8384 | 5911 | 6916 | 8641 | 7342 | 6906 | 6739 | 4565 | 7521 | 7525 | 7076 | 7125 | 6279 | 6589 | 6013 | 6439 | 6821,1 | 7312 | 6330 | 0,866 | -0,208 | -5,005 | 5,59E-07 | 2,46E-06 | 0,001121 | 0,0021923 | 0,0015127 | 0,0015127 | TRUE | 0,0140025 | No |
| orf19.7489 | 2416 | 1583 | 2132 | 2505 | 2466 | 2133 | 2424 | 1515 | 2167 | 2015 | 2181 | 2066 | 2109 | 2035 | 2163 | 2137 | 2109,2 | 2107 | 2111 | 1,001 | 0,001 | 0,028 | 0,97765205 | 0,98378316 | 0,000561 | 0,0026678 | 0,0013786 | 0,0013786 | TRUE | 0,0095992 | No |
| orf19.7489.3 | 989 | 693 | 764 | 944 | 1259 | 1048 | 1087 | 613 | 887 | 882 | 782 | 778 | 1077 | 1000 | 970 | 865 | 905,1 | 832 | 978 | 1,179 | 0,238 | 3,079 | 0,00207373 | 0,00488204 | 0,005196 | 0,0035834 | 0,0046004 | 0,0046004 | TRUE | 0,0669789 | No |
| orf19.7490 | 291 | 204 | 285 | 360 | 298 | 279 | 303 | 162 | 261 | 260 | 292 | 297 | 255 | 266 | 270 | 228 | 266,14 | 278 | 255 | 0,922 | -0,117 | -1,252 | 0,21039438 | 0,28734936 | 0,001026 | 0,0074341 | 0,0046394 | 0,0046394 | TRUE | 0,0439204 | No |
| orf19.7491 | 90 | 49 | 65 | 91 | 72 | 68 | 82 | 47 | 81 | 62 | 67 | 75 | 62 | 65 | 73 | 66 | 68,82 | 71 | 66 | 0,927 | -0,109 | -0,649 | 0,51611877 | 0,60012416 | 1,00E-08 | 0,0230728 | 0,0125518 | 0,0125518 | TRUE | 0,0734465 | No |
| orf19.7492 | 2051 | 1308 | 1653 | 2217 | 2033 | 1863 | 1958 | 1220 | 1840 | 1665 | 1691 | 1828 | 1739 | 1778 | 1747 | 1721 | 1751,1 | 1756 | 1746 | 0,993 | -0,01 | -0,214 | 0,83065128 | 0,86991559 | 0,000842 | 0,0028085 | 0,0016342 | 0,0016342 | TRUE | 0,0138981 | No |
| orf19.7494 | 275 | 193 | 242 | 316 | 215 | 214 | 240 | 141 | 247 | 246 | 248 | 261 | 184 | 204 | 214 | 199 | 225,21 | 250 | 200 | 0,799 | -0,323 | -3,4 | 0,00067379 | 0,00175852 | 1,00E-08 | 0,0084255 | 0,0041887 | 0,0041887 | TRUE | 0,0386646 | No |
| orf19.7495 | 227 | 140 | 200 | 244 | 173 | 177 | 180 | 85 | 204 | 178 | 205 | 201 | 148 | 169 | 161 | 120 | 173,13 | 197 | 150 | 0,763 | -0,39 | -3,296 | 0,00098013 | 0,00247006 | 0,002843 | 0,0103645 | 0,0075819 | 0,0075819 | TRUE | 0,1434947 | No |
| orf19.7497 | 1909 | 1381 | 1781 | 2009 | 1791 | 1629 | 1824 | 1011 | 1713 | 1758 | 1822 | 1657 | 1532 | 1554 | 1628 | 1426 | 1636,1 | 1738 | 1535 | 0,886 | -0,175 | -3,279 | 0,00104251 | 0,00261738 | 0,001472 | 0,0028668 | 0,0021135 | 0,0021135 | TRUE | 0,0249542 | No |
| orf19.7498 | 5540 | 3663 | 4793 | 6055 | 3053 | 2794 | 3039 | 1982 | 4970 | 4663 | 4904 | 4993 | 2611 | 2666 | 2712 | 2795 | 3789,3 | 4882 | 2696 | 0,551 | -0,859 | -21,834 | 1,09E-105 | 1,09E-102 | 0,000513 | 0,0023626 | 0,0011971 | 0,0011971 | TRUE | 0,0107117 | No |
| orf19.7499 | 3105 | 2096 | 2821 | 3343 | 2746 | 2220 | 2476 | 1642 | 2786 | 2668 | 2886 | 2757 | 2348 | 2118 | 2209 | 2316 | 2511,1 | 2774 | 2248 | 0,81 | -0,305 | -6,502 | 7,94E-11 | 5,87E-10 | 0,001075 | 0,0025576 | 0,0017029 | 0,0017029 | TRUE | 0,0184314 | No |
| orf19.7499.1 | 662 | 407 | 499 | 680 | 564 | 486 | 514 | 338 | 594 | 518 | 511 | 561 | 482 | 464 | 459 | 477 | 508,09 | 546 | 470 | 0,859 | -0,219 | -3,052 | 0,00227076 | 0,00530226 | 0,001147 | 0,0048367 | 0,0029751 | 0,0029751 | TRUE | 0,0411778 | No |
| orf19.750 | 145 | 101 | 163 | 183 | 234 | 160 | 177 | 143 | 130 | 129 | 167 | 151 | 200 | 153 | 158 | 202 | 161,09 | 144 | 178 | 1,229 | 0,297 | 2,167 | 0,0302159 | 0,05392857 | 0,01271 | 0,010991 | 0,0117801 | 0,0117801 | TRUE | 0,1115603 | No |
| orf19.7500 | 4601 | 3596 | 4268 | 5351 | 5575 | 5261 | 5527 | 3518 | 4128 | 4578 | 4367 | 4413 | 4768 | 5020 | 4932 | 4962 | 4645,8 | 4372 | 4920 | 1,126 | 0,171 | 4,194 | 2,75E-05 | 9,31E-05 | 0,000859 | 0,002292 | 0,0013847 | 0,0013847 | TRUE | 0,0172483 | No |
| orf19.7501 | 31097 | 21003 | 26709 | 32358 | 28628 | 24688 | 26588 | 15327 | 27897 | 26739 | 27326 | 26683 | 24484 | 23557 | 23725 | 21617 | 25253 | 27161 | 23346 | 0,86 | -0,218 | -5,105 | 3,31E-07 | 1,50E-06 | 0,001534 | 0,002037 | 0,001713 | 0,001713 | TRUE | 0,0301162 | No |
| orf19.7502 | 9163 | 8842 | 8593 | 11588 | 9371 | 11307 | 11023 | 6778 | 8220 | 11257 | 8791 | 9556 | 8014 | 10789 | 9836 | 9560 | 9502,9 | 9456 | 9550 | 1,01 | 0,014 | 0,143 | 0,88589683 | 0,91372844 | 0,014469 | 0,0021323 | 0,0095976 | 0,0095976 | TRUE | 0,2002513 | No |
| orf19.7503 | 1306 | 895 | 1215 | 1509 | 1135 | 1070 | 1123 | 789 | 1172 | 1139 | 1243 | 1244 | 971 | 1021 | 1002 | 1113 | 1113,1 | 1200 | 1027 | 0,852 | -0,231 | -3,973 | 7,10E-05 | 0,00022294 | 0,001533 | 0,0032837 | 0,0023289 | 0,0023289 | TRUE | 0,0342479 | No |
| orf19.7504 | 21606 | 15421 | 19002 | 23505 | 19456 | 18081 | 19554 | 12525 | 19383 | 19633 | 19441 | 19383 | 16639 | 17252 | 17449 | 17665 | 18356 | 19460 | 17251 | 0,886 | -0,174 | -6,127 | 8,94E-10 | 5,74E-09 | 0,000273 | 0,0020586 | 0,0007199 | 0,0007199 | TRUE | 0,0070365 | No |
| orf19.7506 | 2581 | 1797 | 2262 | 2657 | 2010 | 1888 | 2010 | 1168 | 2315 | 2288 | 2314 | 2191 | 1719 | 1801 | 1794 | 1647 | 2008,7 | 2277 | 1740 | 0,766 | -0,385 | -8,559 | 1,14E-17 | 1,71E-16 | 0,00052 | 0,0027022 | 0,0014331 | 0,0014331 | TRUE | 0,0138459 | No |

| | | | | | | | | | | | | | | | | | | | | | | | | | | | | | | | |
|---|---|---|---|---|---|---|---|---|---|---|---|---|---|---|---|---|---|---|---|---|---|---|---|---|---|---|---|---|---|---|---|
| orf19.7507 | 1153 | 750 | 1032 | 1232 | 1100 | 893 | 961 | 621 | 1034 | 955 | 1056 | 1016 | 941 | 852 | 858 | 876 | 948,4 | 1015 | 882 | 0,868 | -0,205 | -3,562 | 0,0003674 | 0,00102051 | 0,000912 | 0,0035102 | 0,0021044 | 0,0021044 | TRUE | 0,026307 | No |
| orf19.7509.1 | 3378 | 2210 | 2596 | 3281 | 2937 | 2151 | 2217 | 1565 | 3030 | 2814 | 2656 | 2706 | 2512 | 2052 | 1978 | 2207 | 2494,4 | 2802 | 2187 | 0,781 | -0,357 | -4,725 | 2,30E-06 | 9,26E-06 | 0,007056 | 0,0025615 | 0,0050659 | 0,0050659 | TRUE | 0,1236485 | No |
| orf19.7509.2 | 2743 | 1894 | 2122 | 2833 | 2736 | 2060 | 2188 | 1100 | 2461 | 2411 | 2171 | 2336 | 2340 | 1966 | 1952 | 1551 | 2148,6 | 2345 | 1952 | 0,834 | -0,262 | -2,663 | 0,00773479 | 0,01599127 | 0,014437 | 0,0026552 | 0,0088234 | 0,0088234 | TRUE | 0,2249204 | No |
| orf19.751 | 58 | 40 | 62 | 80 | 38 | 32 | 40 | 15 | 52 | 51 | 63 | 66 | 32 | 31 | 36 | 21 | 44,03 | 58 | 30 | 0,524 | -0,933 | -4,22 | 2,44E-05 | 8,36E-05 | 1,00E-08 | 0,0349503 | 0,0220674 | 0,0220674 | TRUE | 0,1733025 | No |
| orf19.7510 | 3398 | 2240 | 3087 | 3901 | 4325 | 3399 | 4002 | 2645 | 3048 | 2852 | 3158 | 3217 | 3699 | 3243 | 3571 | 3731 | 3314,9 | 3069 | 3561 | 1,159 | 0,213 | 3,735 | 0,00018742 | 0,000549 | 0,002922 | 0,0024175 | 0,002819 | 0,002819 | TRUE | 0,0440484 | No |
| orf19.7511 | 4477 | 3097 | 4179 | 4687 | 5344 | 4792 | 5180 | 3184 | 4016 | 3943 | 4276 | 3865 | 4570 | 4572 | 4622 | 4491 | 4294,4 | 4025 | 4564 | 1,134 | 0,182 | 4,549 | 5,38E-06 | 2,04E-05 | 0,000739 | 0,0023176 | 0,0013007 | 0,0013007 | TRUE | 0,0213737 | No |
| orf19.7512 | 1140 | 691 | 1041 | 1174 | 1485 | 1222 | 1324 | 879 | 1023 | 880 | 1065 | 968 | 1270 | 1166 | 1181 | 1240 | 1099,1 | 984 | 1214 | 1,232 | 0,301 | 4,649 | 3,33E-06 | 1,31E-05 | 0,002888 | 0,0033003 | 0,0030959 | 0,0030959 | TRUE | 0,0556571 | No |
| orf19.7513 | 2051 | 1302 | 1766 | 2192 | 1876 | 1443 | 1585 | 970 | 1840 | 1658 | 1807 | 1808 | 1604 | 1377 | 1414 | 1368 | 1609,5 | 1778 | 1441 | 0,811 | -0,302 | -4,853 | 1,21E-06 | 5,06E-06 | 0,003176 | 0,0028815 | 0,0030975 | 0,0030975 | TRUE | 0,0720322 | No |
| orf19.7514 | 28421 | 18538 | 24856 | 30550 | 40286 | 32257 | 35024 | 23935 | 25497 | 23601 | 25430 | 25192 | 34454 | 30779 | 31253 | 33758 | 28745 | 24930 | 32561 | 1,306 | 0,385 | 8,111 | 5,02E-16 | 6,34E-15 | 0,002155 | 0,00203 | 0,00213 | 0,00213 | TRUE | 0,0188729 | No |
| orf19.7516 | 2165 | 1462 | 1886 | 2252 | 2078 | 1697 | 1914 | 1288 | 1942 | 1861 | 1930 | 1857 | 1777 | 1619 | 1708 | 1817 | 1813,9 | 1898 | 1730 | 0,911 | -0,135 | -2,809 | 0,00496295 | 0,01072182 | 0,000792 | 0,0027799 | 0,0016679 | 0,0016679 | TRUE | 0,0223509 | No |
| orf19.7517 | 337 | 216 | 287 | 362 | 273 | 224 | 209 | 143 | 302 | 275 | 294 | 299 | 233 | 214 | 186 | 202 | 250,61 | 292 | 209 | 0,714 | -0,485 | -5,019 | 5,18E-07 | 2,30E-06 | 1,00E-08 | 0,0077723 | 0,0048803 | 0,0048803 | TRUE | 0,0764378 | No |
| orf19.7518 | 1326 | 883 | 1192 | 1456 | 1308 | 1104 | 1149 | 836 | 1190 | 1124 | 1220 | 1201 | 1119 | 1053 | 1025 | 1179 | 1138,8 | 1184 | 1094 | 0,921 | -0,118 | -2,064 | 0,03898319 | 0,06755438 | 0,00142 | 0,0032543 | 0,0022693 | 0,0022693 | TRUE | 0,0291731 | No |
| orf19.7519 | 2168 | 1501 | 1761 | 2202 | 2354 | 1799 | 1963 | 1330 | 1945 | 1911 | 1802 | 1816 | 2013 | 1717 | 1752 | 1876 | 1853,8 | 1868 | 1840 | 0,985 | -0,022 | -0,376 | 0,70665513 | 0,76745915 | 0,002904 | 0,0027626 | 0,0028456 | 0,0028456 | TRUE | 0,051438 | No |
| orf19.752 | 2069 | 1617 | 1889 | 2257 | 1849 | 1675 | 1835 | 1136 | 1856 | 2059 | 1933 | 1861 | 1581 | 1598 | 1637 | 1602 | 1766 | 1927 | 1604 | 0,834 | -0,262 | -5,554 | 2,79E-08 | 1,46E-07 | 0,000728 | 0,0028016 | 0,0015549 | 0,0015549 | TRUE | 0,0240592 | No |
| orf19.7520 | 11570 | 7058 | 9655 | 11692 | 7323 | 5773 | 6383 | 4397 | 10379 | 8986 | 9878 | 9641 | 6263 | 5508 | 5696 | 6202 | 7819,1 | 9721 | 5917 | 0,608 | -0,717 | -12,418 | 2,10E-35 | 1,08E-33 | 0,003575 | 0,0021652 | 0,0030641 | 0,0030641 | TRUE | 0,0314045 | No |
| orf19.7521 | 1391 | 907 | 1247 | 1539 | 1009 | 819 | 980 | 628 | 1248 | 1155 | 1276 | 1269 | 863 | 781 | 874 | 886 | 1044 | 1237 | 851 | 0,686 | -0,543 | -9,209 | 3,31E-20 | 6,07E-19 | 0,001444 | 0,00337 | 0,0023411 | 0,0023411 | TRUE | 0,0365086 | No |
| orf19.7522 | 931 | 674 | 773 | 985 | 793 | 669 | 703 | 471 | 835 | 858 | 791 | 812 | 678 | 638 | 627 | 664 | 738,07 | 824 | 652 | 0,792 | -0,337 | -5,776 | 7,64E-09 | 4,31E-08 | 1,00E-08 | 0,0039464 | 0,0018994 | 0,0018994 | TRUE | 0,0100803 | No |
| orf19.7523 | 2907 | 1929 | 2476 | 3164 | 3170 | 2637 | 3015 | 1830 | 2608 | 2456 | 2533 | 2609 | 2711 | 2516 | 2690 | 2581 | 2588,1 | 2552 | 2624 | 1,028 | 0,04 | 0,952 | 0,3410029 | 0,42799725 | 0,000653 | 0,0025404 | 0,0013433 | 0,0013433 | TRUE | 0,0099363 | No |
| orf19.7527 | 2068 | 1365 | 1947 | 2220 | 2967 | 2497 | 2714 | 1793 | 1855 | 1738 | 1992 | 1831 | 2537 | 2383 | 2422 | 2529 | 2160,8 | 1854 | 2468 | 1,33 | 0,411 | 8,199 | 2,43E-16 | 3,17E-15 | 0,001466 | 0,0026514 | 0,0019398 | 0,0019398 | TRUE | 0,0295496 | No |
| orf19.7529 | 15290 | 9600 | 13410 | 16447 | 18782 | 15228 | 17404 | 10995 | 13717 | 12222 | 13720 | 13562 | 16063 | 14530 | 15530 | 15507 | 14356 | 13305 | 15408 | 1,158 | 0,211 | 4,309 | 1,64E-05 | 5,79E-05 | 0,002255 | 0,0020806 | 0,0022411 | 0,0022411 | TRUE | 0,0364014 | No |
| orf19.753 | 1760 | 1362 | 1618 | 1896 | 1363 | 1332 | 1514 | 854 | 1579 | 1734 | 1655 | 1563 | 1166 | 1271 | 1351 | 1204 | 1440,5 | 1633 | 1248 | 0,766 | -0,385 | -6,431 | 1,27E-10 | 9,10E-10 | 0,00233 | 0,0029873 | 0,0027288 | 0,0027288 | TRUE | 0,0382188 | No |
| orf19.7531 | 2909 | 1827 | 2500 | 3055 | 3731 | 3036 | 3158 | 2298 | 2610 | 2326 | 2558 | 2519 | 3191 | 2897 | 2818 | 3241 | 2769,9 | 2503 | 3037 | 1,211 | 0,277 | 4,705 | 2,54E-06 | 1,02E-05 | 0,00314 | 0,0025036 | 0,0029554 | 0,0029554 | TRUE | 0,0286995 | No |
| orf19.7534 | 19695 | 12082 | 17470 | 21087 | 18264 | 14646 | 16874 | 11800 | 17668 | 15382 | 17874 | 17389 | 15620 | 13975 | 15057 | 16643 | 16201 | 17078 | 15324 | 0,897 | -0,157 | -2,481 | 0,01310334 | 0,02560992 | 0,004817 | 0,0020691 | 0,0037751 | 0,0037751 | TRUE | 0,0544646 | No |
| orf19.7537 | 153 | 111 | 150 | 181 | 151 | 98 | 146 | 103 | 137 | 141 | 153 | 149 | 129 | 94 | 130 | 145 | 134,94 | 145 | 124 | 0,851 | -0,232 | -1,673 | 0,0942522 | 0,14572568 | 0,008534 | 0,0127379 | 0,0110145 | 0,0110145 | TRUE | 0,2895928 | No |
| orf19.7538 | 1038 | 648 | 823 | 1074 | 902 | 792 | 925 | 552 | 931 | 825 | 842 | 886 | 771 | 756 | 825 | 779 | 826,86 | 871 | 783 | 0,897 | -0,156 | -2,579 | 0,00991538 | 0,01997706 | 0,001104 | 0,0037352 | 0,0023175 | 0,0023175 | TRUE | 0,0257104 | No |
| orf19.7539 | 784 | 484 | 745 | 888 | 629 | 442 | 602 | 410 | 703 | 616 | 762 | 732 | 538 | 422 | 537 | 578 | 611,14 | 703 | 519 | 0,735 | -0,445 | -4,5 | 6,81E-06 | 2,55E-05 | 0,01104 | 0,0043549 | 0,0076899 | 0,0076899 | TRUE | 0,1863519 | No |
| orf19.7539.1 | 1745 | 1138 | 1493 | 1873 | 1549 | 1253 | 1334 | 913 | 1565 | 1449 | 1527 | 1544 | 1325 | 1196 | 1190 | 1288 | 1385,6 | 1521 | 1250 | 0,82 | -0,287 | -5,369 | 7,92E-08 | 3,93E-07 | 0,001246 | 0,0030272 | 0,0020052 | 0,0020052 | TRUE | 0,0218756 | No |
| orf19.754 | 5440 | 3770 | 4653 | 5850 | 5302 | 4624 | 4932 | 3056 | 4880 | 4800 | 4760 | 4824 | 4534 | 4412 | 4401 | 4310 | 4615,3 | 4816 | 4414 | 0,917 | -0,125 | -3,964 | 7,39E-05 | 0,00023088 | 1,00E-08 | 0,0022941 | 0,0007338 | 0,0007338 | TRUE | 0,0041202 | No |
| orf19.7544 | 253 | 150 | 196 | 252 | 279 | 213 | 260 | 142 | 227 | 191 | 201 | 208 | 239 | 203 | 232 | 200 | 212,55 | 207 | 218 | 1,06 | 0,084 | 0,8 | 0,42364597 | 0,51109739 | 0,001929 | 0,0088095 | 0,0058223 | 0,0058223 | TRUE | 0,0486903 | No |
| orf19.7545 | 72 | 52 | 62 | 87 | 36 | 41 | 41 | 30 | 65 | 66 | 63 | 72 | 31 | 39 | 37 | 42 | 51,85 | 66 | 37 | 0,552 | -0,857 | -4,408 | 1,04E-05 | 3,78E-05 | 1,00E-08 | 0,0299794 | 0,015275 | 0,015275 | TRUE | 0,1100574 | No |
| orf19.7546 | 6853 | 5158 | 6121 | 7640 | 7478 | 7132 | 7512 | 4854 | 6148 | 6567 | 6262 | 6300 | 6395 | 6805 | 6703 | 6846 | 6503,4 | 6319 | 6687 | 1,058 | 0,082 | 2,204 | 0,02754192 | 0,04969837 | 0,000652 | 0,0022027 | 0,0011646 | 0,0011646 | TRUE | 0,0107076 | No |
| orf19.7547 | 3618 | 2633 | 3228 | 3915 | 3309 | 2993 | 3012 | 1910 | 3246 | 3352 | 3303 | 3228 | 2830 | 2856 | 2688 | 2694 | 3024,5 | 3282 | 2767 | 0,844 | -0,245 | -6,416 | 1,40E-10 | 9,95E-10 | 0,000292 | 0,0024595 | 0,0010606 | 0,0010606 | TRUE | 0,0055763 | No |
| orf19.7548 | 292 | 197 | 229 | 271 | 278 | 245 | 246 | 197 | 262 | 251 | 234 | 223 | 238 | 234 | 220 | 278 | 242,43 | 242 | 242 | 0,993 | -0,01 | -0,103 | 0,91780839 | 0,93954555 | 0,002991 | 0,0079678 | 0,0057566 | 0,0057566 | TRUE | 0,0910086 | No |
| orf19.7549 | 178 | 159 | 172 | 238 | 216 | 208 | 211 | 135 | 160 | 202 | 176 | 196 | 185 | 198 | 188 | 190 | 187,03 | 184 | 190 | 1,04 | 0,057 | 0,522 | 0,60178798 | 0,67757537 | 1,00E-08 | 0,0097415 | 0,0059946 | 0,0059946 | TRUE | 0,0871406 | No |
| orf19.755 | 855 | 743 | 778 | 1117 | 1047 | 939 | 969 | 537 | 767 | 946 | 796 | 921 | 895 | 896 | 865 | 757 | 855,43 | 858 | 853 | 0,999 | -0,002 | -0,025 | 0,97998079 | 0,98508366 | 0,007102 | 0,0036766 | 0,005478 | 0,005478 | TRUE | 0,0628435 | No |
| orf19.7550 | 439 | 319 | 416 | 527 | 652 | 556 | 640 | 397 | 394 | 406 | 426 | 435 | 558 | 531 | 571 | 560 | 484,91 | 415 | 555 | 1,335 | 0,417 | 6,084 | 1,18E-09 | 7,40E-09 | 1,00E-08 | 0,0049733 | 0,002415 | 0,002415 | TRUE | 0,0154821 | No |
| orf19.7551 | 5582 | 3923 | 5009 | 6055 | 5826 | 5170 | 5529 | 3329 | 5008 | 4994 | 5125 | 4993 | 4983 | 4933 | 4934 | 4695 | 4958,1 | 5030 | 4886 | 0,972 | -0,04 | -1,232 | 0,21778265 | 0,29629488 | 0,000161 | 0,0022723 | 0,000831 | 0,000831 | TRUE | 0,0078109 | No |
| orf19.7552 | 17309 | 11804 | 15254 | 18678 | 16602 | 14540 | 16299 | 8957 | 15528 | 15028 | 15606 | 15402 | 14199 | 13874 | 14544 | 12633 | 14602 | 15391 | 13812 | 0,898 | -0,156 | -3,386 | 0,00071033 | 0,00184507 | 0,001864 | 0,0020789 | 0,0019611 | 0,0019611 | TRUE | 0,0398315 | No |
| orf19.7553 | 224 | 187 | 178 | 200 | 232 | 245 | 285 | 140 | 201 | 238 | 182 | 165 | 198 | 234 | 254 | 197 | 208,75 | 196 | 221 | 1,135 | 0,182 | 1,384 | 0,16644077 | 0,23584343 | 0,015086 | 0,0089337 | 0,0118363 | 0,0118363 | TRUE | 0,2004307 | No |
| orf19.7554 | 1077 | 784 | 934 | 1163 | 860 | 834 | 969 | 563 | 966 | 998 | 956 | 959 | 735 | 796 | 865 | 794 | 883,61 | 970 | 798 | 0,823 | -0,281 | -4,643 | 3,43E-06 | 1,34E-05 | 0,00132 | 0,0036224 | 0,0023748 | 0,0023748 | TRUE | 0,0412495 | No |
| orf19.7555 | 40 | 32 | 41 | 53 | 38 | 19 | 34 | 25 | 36 | 41 | 42 | 44 | 32 | 18 | 30 | 35 | 34,81 | 41 | 29 | 0,71 | -0,495 | -2,001 | 0,04540767 | 0,07730498 | 1,00E-08 | 0,0436797 | 0,029044 | 0,029044 | TRUE | 0,4367323 | No |
| orf19.7556 | 413 | 358 | 448 | 557 | 439 | 428 | 470 | 267 | 371 | 456 | 458 | 459 | 375 | 408 | 419 | 377 | 415,47 | 436 | 395 | 0,909 | -0,138 | -1,555 | 0,11985647 | 0,17806569 | 0,004904 | 0,0054737 | 0,0051862 | 0,0051862 | TRUE | 0,1234821 | No |
| orf19.7557 | 25 | 15 | 12 | 19 | 20 | 17 | 12 | 6 | 22 | 19 | 12 | 16 | 17 | 16 | 11 | 8 | 15,25 | 17 | 13 | 0,772 | -0,373 | -1,011 | 0,31192732 | NA | 1,00E-08 | 0,0972004 | 0,0649219 | 0,0649219 | TRUE | 0,2053064 | No |
| orf19.7558 | 3963 | 2819 | 3746 | 4324 | 4315 | 3892 | 4308 | 2736 | 3555 | 3589 | 3833 | 3566 | 3690 | 3714 | 3844 | 3859 | 3706,2 | 3636 | 3777 | 1,039 | 0,055 | 1,38 | 0,16772289 | 0,23743618 | 0,000566 | 0,0023712 | 0,0012291 | 0,0012291 | TRUE | 0,0161813 | No |
| orf19.756 | 9 | 6 | 10 | 18 | 10 | 8 | 8 | 8 | 8 | 8 | 10 | 15 | 9 | 8 | 7 | 11 | 9,42 | 10 | 9 | 0,819 | -0,287 | -0,638 | 0,52333701 | NA | 1,00E-08 | 0,156018 | 0,0877551 | 0,0877551 | TRUE | 0,4058745 | No |
| orf19.7561 | 19015 | 12217 | 17109 | 20835 | 27732 | 22638 | 25991 | 17695 | 17058 | 15554 | 17504 | 17181 | 23717 | 21600 | 23193 | 24957 | 20096 | 16824 | 23367 | 1,389 | 0,474 | 8,835 | 1,00E-18 | 1,64E-17 | 0,003033 | 0,0020518 | 0,0027092 | 0,0027092 | TRUE | 0,031686 | No |
| orf19.7563 | 714 | 456 | 730 | 809 | 734 | 652 | 723 | 488 | 641 | 581 | 747 | 667 | 628 | 622 | 645 | 688 | 652,29 | 659 | 646 | 0,977 | -0,034 | -0,432 | 0,66562537 | 0,73323749 | 0,004254 | 0,0042051 | 0,0043882 | 0,0043882 | TRUE | 0,0918932 | No |
| orf19.7564 | 2602 | 1936 | 2349 | 2830 | 2143 | 2059 | 2305 | 1340 | 2334 | 2465 | 2403 | 2334 | 1833 | 1965 | 2057 | 1890 | 2160 | 2384 | 1936 | 0,813 | -0,298 | -6,248 | 4,15E-10 | 2,80E-09 | 0,000979 | 0,0026516 | 0,0017206 | 0,0017206 | TRUE | 0,0220741 | No |
| orf19.7565 | 2413 | 1783 | 2223 | 2749 | 2321 | 2056 | 2172 | 1353 | 2165 | 2270 | 2274 | 2267 | 1985 | 1962 | 1938 | 1908 | 2096,1 | 2244 | 1948 | 0,869 | -0,202 | -5,136 | 2,80E-07 | 1,29E-06 | 1,00E-08 | 0,0026721 | 0,0010054 | 0,0010054 | TRUE | 0,0070731 | No |
| orf19.7566 | 32765 | 22277 | 29994 | 35992 | 33293 | 28720 | 31850 | 20347 | 29394 | 28361 | 30687 | 29679 | 28473 | 27404 | 28421 | 28698 | 28890 | 29530 | 28249 | 0,956 | -0,064 | -1,904 | 0,05687007 | 0,09389265 | 0,000687 | 0,0020298 | 0,0010565 | 0,0010565 | TRUE | 0,0086135 | No |
| orf19.7567 | 3830 | 2632 | 3390 | 4124 | 4165 | 3736 | 4026 | 2721 | 3436 | 3351 | 3468 | 3401 | 3562 | 3565 | 3593 | 3838 | 3526,6 | 3414 | 3640 | 1,064 | 0,09 | 2,38 | 0,01732623 | 0,03277582 | 0,000398 | 0,0023912 | 0,0010869 | 0,0010869 | TRUE | 0,0152056 | No |
| orf19.7568 | 123 | 58 | 100 | 127 | 102 | 89 | 96 | 47 | 110 | 74 | 102 | 105 | 87 | 85 | 86 | 66 | 89,42 | 98 | 81 | 0,829 | -0,27 | -1,664 | 0,09601435 | 0,14771653 | 0,007637 | 0,0182149 | 0,0141841 | 0,0141841 | TRUE | 0,2224498 | No |
| orf19.7569 | 27608 | 18769 | 23748 | 29564 | 24272 | 21260 | 22155 | 15441 | 24767 | 23895 | 24297 | 24379 | 20758 | 20286 | 19770 | 21778 | 22491 | 24334 | 20648 | 0,848 | -0,237 | -6,529 | 6,64E-11 | 4,96E-10 | 0,000905 | 0,0020441 | 0,0012256 | 0,0012256 | TRUE | 0,0163826 | No |
| orf19.757 | 2246 | 1634 | 2010 | 2485 | 2388 | 2221 | 2423 | 1454 | 2015 | 2080 | 2056 | 2049 | 2042 | 2119 | 2162 | 2051 | 2071,9 | 2050 | 2094 | 1,022 | 0,032 | 0,801 | 0,42296802 | 0,51048451 | 1,00E-08 | 0,0026802 | 0,0010281 | 0,0010281 | TRUE | 0,0060423 | No |
| orf19.7570 | 15649 | 11031 | 14001 | 16998 | 15452 | 14110 | 15778 | 8905 | 14039 | 14044 | 14324 | 14017 | 13215 | 13463 | 14079 | 12560 | 13718 | 14106 | 13329 | 0,945 | -0,081 | -2,084 | 0,03717709 | 0,06477957 | 0,001069 | 0,0020854 | 0,0013833 | 0,0013833 | TRUE | 0,018097 | No |
| orf19.7571 | 9250 | 6404 | 7625 | 9700 | 8293 | 7668 | 8056 | 5354 | 8298 | 8153 | 7801 | 7999 | 7092 | 7317 | 7189 | 7551 | 7675 | 8063 | 7287 | 0,903 | -0,147 | -4,185 | 2,85E-05 | 9,64E-05 | 0,000546 | 0,0021687 | 0,0010483 | 0,0010483 | TRUE | 0,0069751 | No |
| orf19.757.1 | 628 | 442 | 507 | 619 | 677 | 556 | 584 | 481 | 563 | 563 | 519 | 510 | 579 | 531 | 521 | 678 | 558,04 | 539 | 577 | 1,067 | 0,094 | 1,07 | 0,28441899 | 0,36867082 | 0,006764 | 0,004581 | 0,0055931 | 0,0055931 | TRUE | 0,148051 | No |
| orf19.7572 | 1718 | 1247 | 1586 | 1814 | 1742 | 1606 | 1762 | 1049 | 1541 | 1588 | 1623 | 1496 | 1490 | 1532 | 1572 | 1480 | 1540,2 | 1562 | 1518 | 0,974 | -0,038 | -0,808 | 0,41885281 | 0,50647075 | 0,000435 | 0,0029221 | 0,0014442 | 0,0014442 | TRUE | 0,0101942 | No |
| orf19.7574 | 2570 | 1861 | 2383 | 2800 | 3034 | 2838 | 3122 | 1975 | 2306 | 2369 | 2438 | 2309 | 2595 | 2708 | 2786 | 2786 | 2537 | 2356 | 2719 | 1,154 | 0,206 | 4,938 | 7,89E-07 | 3,38E-06 | 0,000566 | 0,0025517 | 0,0012807 | 0,0012807 | TRUE | 0,0117695 | No |

| | | | | | | | | | | | | | | | | | | | | | | | | | | | | | | | |
|---|---|---|---|---|---|---|---|---|---|---|---|---|---|---|---|---|---|---|---|---|---|---|---|---|---|---|---|---|---|---|---|
| orf19.7576 | 1129 | 722 | 948 | 1191 | 1261 | 1012 | 1127 | 662 | 1013 | 919 | 970 | 982 | 1078 | 966 | 1006 | 934 | 983,43 | 971 | 996 | 1,027 | 0,038 | 0,629 | 0,52909122 | 0,61201002 | 0,001669 | 0,0034557 | 0,0024783 | 0,0024783 | TRUE | 0,0378536 | No |
| orf19.7577 | 3692 | 2531 | 3352 | 3963 | 3839 | 3440 | 3641 | 2435 | 3312 | 3222 | 3429 | 3268 | 3283 | 3282 | 3249 | 3434 | 3310,1 | 3308 | 3312 | 1 | 0 | 0,003 | 0,9972932 | 0,99812386 | 0,000272 | 0,0024181 | 0,0010396 | 0,0010396 | TRUE | 0,0072806 | No |
| orf19.7578 | 116 | 81 | 76 | 98 | 93 | 75 | 71 | 63 | 104 | 103 | 78 | 81 | 80 | 72 | 63 | 89 | 83,63 | 92 | 76 | 0,825 | -0,277 | -1,637 | 0,10168145 | 0,15477633 | 0,010011 | 0,0193375 | 0,0153936 | 0,0153936 | TRUE | 0,1163178 | No |
| orf19.7579 | 930 | 614 | 757 | 1063 | 791 | 651 | 717 | 517 | 834 | 782 | 774 | 877 | 676 | 621 | 640 | 729 | 741,71 | 817 | 666 | 0,812 | -0,3 | -4,326 | 1,52E-05 | 5,40E-05 | 0,002457 | 0,0039368 | 0,0032571 | 0,0032571 | TRUE | 0,0412228 | No |
| orf19.758 | 1425 | 941 | 1233 | 1502 | 1645 | 1355 | 1517 | 957 | 1278 | 1198 | 1261 | 1239 | 1407 | 1293 | 1354 | 1350 | 1297,5 | 1244 | 1351 | 1,085 | 0,118 | 2,435 | 0,01488581 | 0,02869231 | 1,00E-08 | 0,0030984 | 0,0014781 | 0,0014781 | TRUE | 0,0105533 | No |
| orf19.7580 | 543 | 335 | 468 | 549 | 660 | 522 | 501 | 290 | 487 | 426 | 479 | 453 | 564 | 498 | 447 | 409 | 470,47 | 461 | 480 | 1,043 | 0,061 | 0,619 | 0,53576892 | 0,61783103 | 0,009147 | 0,0050652 | 0,0071178 | 0,0071178 | TRUE | 0,1670359 | No |
| orf19.7581 | 6757 | 4372 | 5522 | 6813 | 7330 | 6267 | 6980 | 4019 | 6062 | 5566 | 5650 | 5618 | 6269 | 5980 | 6228 | 5668 | 5880,1 | 5724 | 6036 | 1,055 | 0,077 | 1,687 | 0,09159049 | 0,14215336 | 0,001615 | 0,0022264 | 0,0018476 | 0,0018476 | TRUE | 0,0198981 | No |
| orf19.7583 | 2425 | 1758 | 2254 | 2797 | 1847 | 1794 | 1901 | 1146 | 2175 | 2238 | 2306 | 2306 | 1580 | 1712 | 1696 | 1616 | 1953,8 | 2256 | 1651 | 0,732 | -0,45 | -9,919 | 3,46E-23 | 7,78E-22 | 0,000646 | 0,0027225 | 0,0014522 | 0,0014522 | TRUE | 0,0113345 | No |
| orf19.7585 | 1340 | 903 | 1214 | 1408 | 1523 | 1271 | 1408 | 864 | 1202 | 1150 | 1242 | 1161 | 1303 | 1213 | 1256 | 1219 | 1218,1 | 1189 | 1248 | 1,05 | 0,071 | 1,417 | 0,15645617 | 0,22359389 | 0,000359 | 0,0031713 | 0,0015877 | 0,0015877 | TRUE | 0,0109455 | No |
| orf19.7586 | 594 | 404 | 538 | 653 | 630 | 563 | 608 | 429 | 533 | 514 | 550 | 538 | 539 | 537 | 543 | 605 | 544,96 | 534 | 556 | 1,035 | 0,05 | 0,757 | 0,44928362 | 0,53563278 | 1,00E-08 | 0,0046434 | 0,002381 | 0,002381 | TRUE | 0,0351247 | No |
| orf19.7588 | 642 | 549 | 553 | 804 | 721 | 721 | 760 | 415 | 576 | 699 | 566 | 663 | 617 | 688 | 678 | 585 | 633,96 | 626 | 642 | 1,029 | 0,042 | 0,488 | 0,6258609 | 0,69774954 | 0,006551 | 0,0042694 | 0,0054399 | 0,0054399 | TRUE | 0,0669545 | No |
| orf19.7589 | 743 | 543 | 603 | 804 | 789 | 712 | 710 | 435 | 667 | 691 | 617 | 663 | 675 | 679 | 634 | 614 | 654,87 | 660 | 650 | 0,989 | -0,015 | -0,242 | 0,80916052 | 0,85183746 | 0,000519 | 0,0041963 | 0,0023995 | 0,0023995 | TRUE | 0,0213057 | No |
| orf19.759 | 1145 | 748 | 961 | 1182 | 1321 | 1182 | 1236 | 797 | 1027 | 952 | 983 | 975 | 1130 | 1128 | 1103 | 1124 | 1052,8 | 984 | 1121 | 1,138 | 0,186 | 3,797 | 0,00014617 | 0,00043369 | 1,00E-08 | 0,0033585 | 0,0013487 | 0,0013487 | TRUE | 0,0104421 | No |
| orf19.7590 | 3875 | 2821 | 3507 | 4439 | 4896 | 4438 | 4748 | 2802 | 3476 | 3591 | 3588 | 3660 | 4187 | 4235 | 4237 | 3952 | 3865,8 | 3579 | 4153 | 1,161 | 0,216 | 5,719 | 1,07E-08 | 5,90E-08 | 0,000414 | 0,002355 | 0,0011076 | 0,0011076 | TRUE | 0,0119924 | No |
| orf19.7592 | 37868 | 27651 | 34913 | 42135 | 35265 | 33549 | 36495 | 22061 | 33972 | 35203 | 35719 | 34745 | 30160 | 32011 | 32566 | 31115 | 33186 | 34910 | 31463 | 0,901 | -0,15 | -4,354 | 1,34E-05 | 4,77E-05 | 0,000747 | 0,0020233 | 0,001108 | 0,001108 | TRUE | 0,0096113 | No |
| orf19.7593 | 2406 | 1838 | 2107 | 2756 | 2854 | 2709 | 2849 | 1744 | 2158 | 2340 | 2156 | 2273 | 2441 | 2585 | 2542 | 2460 | 2369,3 | 2232 | 2507 | 1,125 | 0,169 | 3,855 | 0,00011596 | 0,00035064 | 0,000626 | 0,0025922 | 0,0014269 | 0,0014269 | TRUE | 0,0119726 | No |
| orf19.7594 | 498 | 420 | 489 | 604 | 707 | 701 | 683 | 430 | 447 | 535 | 500 | 498 | 605 | 669 | 609 | 606 | 558,66 | 495 | 622 | 1,262 | 0,335 | 4,598 | 4,26E-06 | 1,64E-05 | 0,001982 | 0,0045781 | 0,0032863 | 0,0032863 | TRUE | 0,0509395 | No |
| orf19.7595 | 273 | 202 | 216 | 283 | 258 | 245 | 292 | 166 | 245 | 257 | 221 | 233 | 221 | 234 | 261 | 234 | 238,19 | 239 | 238 | 0,995 | -0,007 | -0,075 | 0,94001394 | 0,9568966 | 1,00E-08 | 0,0080742 | 0,0047853 | 0,0047853 | TRUE | 0,0530985 | No |
| orf19.7596 | 219 | 157 | 163 | 204 | 164 | 142 | 151 | 89 | 196 | 200 | 167 | 168 | 140 | 135 | 135 | 126 | 158,42 | 183 | 134 | 0,738 | -0,438 | -3,768 | 0,00016465 | 0,00048585 | 1,00E-08 | 0,0111432 | 0,0065267 | 0,0065267 | TRUE | 0,0363522 | No |
| orf19.7597 | 2 | 3 | 4 | 4 | 4 | 1 | 6 | 4 | 2 | 4 | 4 | 3 | 3 | 1 | 5 | 6 | 3,55 | 3 | 4 | 1,172 | 0,229 | 0,306 | 0,75951279 | NA | 1,00E-08 | 0,4112681 | 0,2477286 | 0,2477286 | TRUE | 0,4405539 | No |
| orf19.7598 | 7761 | 5208 | 6891 | 8268 | 5774 | 4958 | 5529 | 3549 | 6962 | 6630 | 7050 | 6818 | 4938 | 4731 | 4934 | 5006 | 5883,6 | 6865 | 4902 | 0,714 | -0,487 | -13,869 | 9,81E-44 | 7,46E-42 | 0,000418 | 0,0022263 | 0,001007 | 0,001007 | TRUE | 0,0067485 | No |
| orf19.7599 | 15226 | 10310 | 13478 | 16279 | 13512 | 11795 | 12869 | 8855 | 13659 | 13126 | 13789 | 13424 | 11556 | 11254 | 11483 | 12489 | 12598 | 13500 | 11696 | 0,866 | -0,208 | -5,185 | 2,16E-07 | 1,01E-06 | 0,00121 | 0,0020948 | 0,0014612 | 0,0014612 | TRUE | 0,0249702 | No |
| orf19.76 | 17622 | 13149 | 16115 | 19339 | 17714 | 16596 | 17809 | 10622 | 15809 | 16740 | 16487 | 15947 | 15150 | 15835 | 15892 | 14981 | 15855 | 16246 | 15464 | 0,952 | -0,071 | -2,004 | 0,04510144 | 0,07687077 | 0,000739 | 0,0020711 | 0,0011279 | 0,0011279 | TRUE | 0,0052876 | No |
| orf19.760 | 688 | 473 | 518 | 643 | 781 | 619 | 689 | 420 | 617 | 602 | 530 | 530 | 668 | 591 | 615 | 592 | 593,17 | 570 | 616 | 1,085 | 0,118 | 1,542 | 0,12298223 | 0,18225882 | 0,003427 | 0,0044269 | 0,0039049 | 0,0039049 | TRUE | 0,0401454 | No |
| orf19.7600 | 9658 | 5973 | 8262 | 10163 | 11623 | 9376 | 10329 | 6532 | 8664 | 7604 | 8453 | 8381 | 9940 | 8946 | 9217 | 9213 | 8802,3 | 8276 | 9329 | 1,127 | 0,173 | 3,389 | 0,0007015 | 0,00182372 | 0,002462 | 0,0021444 | 0,0023762 | 0,0023762 | TRUE | 0,0359092 | No |
| orf19.7601 | 2397 | 1610 | 2223 | 2739 | 2451 | 2138 | 2331 | 1580 | 2150 | 2050 | 2274 | 2259 | 2096 | 2040 | 2080 | 2228 | 2147,2 | 2183 | 2111 | 0,965 | -0,052 | -1,055 | 0,29140198 | 0,37601871 | 0,001277 | 0,0026556 | 0,0018395 | 0,0018395 | TRUE | 0,0193397 | No |
| orf19.7602 | 1110 | 708 | 914 | 1231 | 1674 | 1510 | 1530 | 953 | 996 | 901 | 935 | 1015 | 1432 | 1441 | 1365 | 1344 | 1178,7 | 962 | 1396 | 1,449 | 0,535 | 9,73 | 2,25E-22 | 4,84E-21 | 0,000927 | 0,0032112 | 0,0020229 | 0,0020229 | TRUE | 0,0197901 | No |
| orf19.7603 | 1367 | 931 | 1059 | 1366 | 1382 | 1144 | 1277 | 719 | 1226 | 1185 | 1083 | 1126 | 1182 | 1092 | 1140 | 1014 | 1131,1 | 1155 | 1107 | 0,961 | -0,058 | -0,922 | 0,35653241 | 0,44422371 | 0,002426 | 0,003263 | 0,0029049 | 0,0029049 | TRUE | 0,0345782 | No |
| orf19.7604 | 674 | 516 | 625 | 758 | 849 | 730 | 811 | 554 | 605 | 657 | 639 | 625 | 726 | 697 | 724 | 781 | 681,72 | 632 | 732 | 1,158 | 0,211 | 3,453 | 0,00055463 | 0,00148032 | 1,00E-08 | 0,004109 | 0,0021211 | 0,0021211 | TRUE | 0,0208993 | No |
| orf19.7605 | 10862 | 6574 | 8798 | 11058 | 10849 | 8330 | 9551 | 6309 | 9744 | 8369 | 9001 | 9119 | 9278 | 7948 | 8523 | 8898 | 8860,1 | 9058 | 8662 | 0,956 | -0,065 | -1,095 | 0,27339142 | 0,35769505 | 0,003926 | 0,0021434 | 0,0032693 | 0,0032693 | TRUE | 0,0377205 | No |
| orf19.7606 | 48 | 28 | 43 | 41 | 25 | 47 | 32 | 29 | 43 | 36 | 44 | 34 | 21 | 45 | 29 | 41 | 36,52 | 39 | 34 | 0,854 | -0,227 | -0,862 | 0,38856983 | 0,47760384 | 0,035463 | 0,0417258 | 0,0390811 | 0,0390811 | TRUE | 0,3375538 | No |
| orf19.7608 | 3 | 0 | 5 | 5 | 4 | 3 | 3 | 3 | 3 | 0 | 5 | 4 | 3 | 3 | 3 | 4 | 3,14 | 3 | 3 | 1,049 | 0,069 | 0,087 | 0,93041668 | NA | 1,00E-08 | 0,4642765 | 0,2773737 | 0,2773737 | TRUE | 0,3915703 | No |
| orf19.7609 | 28 | 17 | 20 | 26 | 20 | 12 | 11 | 11 | 25 | 22 | 20 | 21 | 17 | 11 | 10 | 16 | 17,82 | 22 | 14 | 0,602 | -0,731 | -2,217 | 0,02660973 | 0,04832867 | 1,00E-08 | 0,0834508 | 0,0448526 | 0,0448526 | TRUE | 0,195619 | No |
| orf19.761 | 3265 | 2051 | 2830 | 3592 | 4390 | 3520 | 4209 | 2864 | 2929 | 2611 | 2895 | 2962 | 3754 | 3359 | 3756 | 4039 | 3288,2 | 2849 | 3727 | 1,306 | 0,385 | 6,166 | 6,99E-10 | 4,54E-09 | 0,004004 | 0,002421 | 0,0034404 | 0,0034404 | TRUE | 0,0537539 | No |
| orf19.7610 | 3771 | 2657 | 3514 | 4232 | 3386 | 3101 | 3416 | 2127 | 3383 | 3383 | 3595 | 3490 | 2896 | 2959 | 3048 | 3000 | 3219,2 | 3463 | 2976 | 0,859 | -0,219 | -5,759 | 8,45E-09 | 4,74E-08 | 0,00038 | 0,0024305 | 0,0010788 | 0,0010788 | TRUE | 0,0079157 | No |
| orf19.7611 | 6486 | 4999 | 5544 | 7358 | 6296 | 6450 | 6126 | 4224 | 5819 | 6364 | 5672 | 6067 | 5385 | 6154 | 5466 | 5958 | 5860,7 | 5980 | 5741 | 0,96 | -0,059 | -1,059 | 0,28979693 | 0,37443009 | 0,003185 | 0,0022272 | 0,0028397 | 0,0028397 | TRUE | 0,0292486 | No |
| orf19.7612 | 17706 | 12708 | 15766 | 19150 | 22724 | 20731 | 21695 | 13760 | 15884 | 16179 | 16130 | 15791 | 19434 | 19781 | 19359 | 19407 | 17746 | 15996 | 19495 | 1,219 | 0,286 | 11,972 | 5,01E-33 | 2,18E-31 | 4,19E-05 | 0,0020613 | 0,0004898 | 0,0004898 | TRUE | 0,001198 | No |
| orf19.7613 | 5120 | 3237 | 4127 | 5354 | 5032 | 4132 | 4557 | 2812 | 4593 | 4121 | 4222 | 4415 | 4304 | 3943 | 4066 | 3966 | 4203,8 | 4338 | 4070 | 0,938 | -0,092 | -1,934 | 0,05310256 | 0,08859766 | 0,001668 | 0,0023249 | 0,0019468 | 0,0019468 | TRUE | 0,019074 | No |
| orf19.7614 | 10 | 4 | 9 | 17 | 4 | 3 | 5 | 2 | 9 | 5 | 9 | 14 | 3 | 3 | 4 | 3 | 6,36 | 9 | 3 | 0,361 | -1,47 | -2,494 | 0,01262071 | NA | 1,00E-08 | 0,2303435 | 0,1362351 | 0,1362351 | TRUE | 0,1890963 | No |
| orf19.7615 | 487 | 304 | 486 | 512 | 527 | 441 | 436 | 276 | 437 | 387 | 497 | 422 | 451 | 421 | 389 | 389 | 424,15 | 436 | 412 | 0,947 | -0,078 | -0,871 | 0,38371175 | 0,47279331 | 0,005429 | 0,0054022 | 0,0054116 | 0,0054116 | TRUE | 0,1003501 | No |
| orf19.7617 | 1941 | 1308 | 1651 | 2075 | 2015 | 1565 | 1748 | 1112 | 1741 | 1665 | 1689 | 1711 | 1723 | 1493 | 1560 | 1568 | 1643,9 | 1702 | 1586 | 0,932 | -0,101 | -1,918 | 0,05506094 | 0,09130723 | 0,001512 | 0,0028626 | 0,0020531 | 0,0020531 | TRUE | 0,0428931 | No |
| orf19.7618 | 9196 | 5876 | 7814 | 9827 | 8933 | 7456 | 7980 | 5337 | 8250 | 7481 | 7994 | 8103 | 7640 | 7114 | 7121 | 7527 | 7653,8 | 7957 | 7350 | 0,923 | -0,115 | -2,647 | 0,00811193 | 0,016702 | 0,001398 | 0,0021692 | 0,0016869 | 0,0016869 | TRUE | 0,0194705 | No |
| orf19.7619 | 8367 | 5792 | 7339 | 9148 | 9827 | 8924 | 9659 | 5677 | 7506 | 7374 | 7509 | 7544 | 8404 | 8515 | 8619 | 8007 | 7934,7 | 7483 | 8386 | 1,121 | 0,165 | 5,007 | 5,53E-07 | 2,44E-06 | 0,000391 | 0,0021625 | 0,0009167 | 0,0009167 | TRUE | 0,0108214 | No |
| orf19.762 | 32 | 18 | 20 | 23 | 32 | 30 | 21 | 23 | 29 | 23 | 20 | 19 | 27 | 29 | 19 | 32 | 24,78 | 23 | 27 | 1,163 | 0,218 | 0,757 | 0,44933243 | 0,53563278 | 1,00E-08 | 0,0605684 | 0,0389575 | 0,0389575 | TRUE | 0,283104 | No |
| orf19.7620 | 787 | 527 | 637 | 806 | 1062 | 866 | 925 | 550 | 706 | 671 | 652 | 665 | 908 | 826 | 825 | 776 | 753,62 | 674 | 834 | 1,242 | 0,312 | 4,852 | 1,22E-06 | 5,10E-06 | 0,001544 | 0,0039058 | 0,0026363 | 0,0026363 | TRUE | 0,0435874 | No |
| orf19.7621 | 385 | 298 | 389 | 501 | 417 | 376 | 392 | 233 | 345 | 379 | 398 | 413 | 357 | 359 | 350 | 329 | 366,21 | 384 | 349 | 0,909 | -0,137 | -1,664 | 0,09607957 | 0,14771653 | 0,001533 | 0,0059436 | 0,0037682 | 0,0037682 | TRUE | 0,057303 | No |
| orf19.7622 | 1001 | 699 | 921 | 1049 | 1214 | 1059 | 1123 | 669 | 898 | 890 | 942 | 865 | 1038 | 1010 | 1002 | 944 | 948,7 | 899 | 998 | 1,114 | 0,156 | 2,862 | 0,00420965 | 0,00921752 | 0,000307 | 0,0035097 | 0,0018018 | 0,0018018 | TRUE | 0,014264 | No |
| orf19.7624 | 8941 | 6194 | 7852 | 9805 | 9807 | 8206 | 9061 | 5570 | 8021 | 7886 | 8033 | 8085 | 8387 | 7830 | 8085 | 7856 | 8023 | 8006 | 8040 | 1,004 | 0,006 | 0,191 | 0,84844562 | 0,88415642 | 0,000372 | 0,0021605 | 0,0009397 | 0,0009397 | TRUE | 0,0104836 | No |
| orf19.7625 | 2369 | 1595 | 2069 | 2547 | 1893 | 1733 | 1803 | 1165 | 2125 | 2031 | 2117 | 2100 | 1619 | 1654 | 1609 | 1643 | 1862,2 | 2093 | 1631 | 0,778 | -0,362 | -9,131 | 6,77E-20 | 1,21E-18 | 1,00E-08 | 0,0027591 | 0,0009599 | 0,0009599 | TRUE | 0,0044601 | No |
| orf19.7626 | 4296 | 2828 | 3564 | 4445 | 3779 | 3203 | 3297 | 2139 | 3854 | 3600 | 3646 | 3665 | 3232 | 3056 | 2942 | 3017 | 3376,6 | 3691 | 3062 | 0,829 | -0,27 | -6,258 | 3,91E-10 | 2,64E-09 | 0,000865 | 0,0024094 | 0,0014844 | 0,0014844 | TRUE | 0,0176303 | No |
| orf19.7627 | 707 | 496 | 636 | 823 | 953 | 885 | 879 | 654 | 634 | 631 | 651 | 679 | 815 | 844 | 784 | 922 | 745,16 | 649 | 841 | 1,291 | 0,368 | 5,66 | 1,52E-08 | 8,22E-08 | 0,001592 | 0,0039277 | 0,0027023 | 0,0027023 | TRUE | 0,0443728 | No |
| orf19.7629 | 529 | 382 | 493 | 550 | 530 | 505 | 538 | 353 | 475 | 486 | 504 | 454 | 453 | 482 | 480 | 498 | 478,99 | 480 | 478 | 0,996 | -0,005 | -0,075 | 0,93982941 | 0,95687088 | 1,00E-08 | 0,0050103 | 0,0025358 | 0,0025358 | TRUE | 0,0170075 | No |
| orf19.763 | 3612 | 2360 | 2947 | 3617 | 3687 | 3331 | 3395 | 1769 | 3240 | 3005 | 3015 | 2983 | 3153 | 3178 | 3029 | 2495 | 3012,3 | 3061 | 2964 | 0,97 | -0,045 | -0,616 | 0,53809579 | 0,62039523 | 0,006458 | 0,0024614 | 0,0046943 | 0,0046943 | TRUE | 0,1336329 | No |
| orf19.7631 | 144 | 116 | 146 | 141 | 126 | 110 | 144 | 88 | 129 | 148 | 149 | 116 | 108 | 105 | 128 | 124 | 125,98 | 136 | 116 | 0,86 | -0,217 | -1,595 | 0,11064619 | 0,16623213 | 0,005082 | 0,0135028 | 0,009799 | 0,009799 | TRUE | 0,103577 | No |
| orf19.7632 | 541 | 371 | 500 | 522 | 666 | 550 | 625 | 388 | 485 | 472 | 512 | 430 | 570 | 525 | 558 | 547 | 512,37 | 475 | 550 | 1,16 | 0,214 | 2,948 | 0,00320151 | 0,00722293 | 0,001439 | 0,0048128 | 0,0031217 | 0,0031217 | TRUE | 0,05102 | No |
| orf19.7634 | 77 | 45 | 56 | 74 | 97 | 88 | 106 | 50 | 69 | 57 | 57 | 61 | 83 | 84 | 95 | 71 | 72,09 | 61 | 83 | 1,363 | 0,446 | 2,685 | 0,00725413 | 0,01508055 | 1,00E-08 | 0,0221171 | 0,0124835 | 0,0124835 | TRUE | 0,0818082 | No |
| orf19.7635 | 13950 | 8918 | 12201 | 14656 | 13785 | 11738 | 13103 | 8562 | 12515 | 11354 | 12483 | 12086 | 11789 | 11200 | 11692 | 12076 | 11899 | 12110 | 11689 | 0,965 | -0,051 | -1,22 | 0,22259708 | 0,30162093 | 0,00137 | 0,0021015 | 0,0016268 | 0,0016268 | TRUE | 0,021261 | No |
| orf19.7637 | 1710 | 1071 | 1483 | 1783 | 1879 | 1650 | 1748 | 1033 | 1534 | 1363 | 1517 | 1470 | 1607 | 1574 | 1560 | 1457 | 1510,4 | 1471 | 1550 | 1,054 | 0,075 | 1,415 | 0,15700843 | 0,22416983 | 0,001351 | 0,0029407 | 0,0020542 | 0,0020542 | TRUE | 0,0271059 | No |
| orf19.7638 | 2191 | 1391 | 1848 | 2384 | 2796 | 2197 | 2387 | 1727 | 1966 | 1771 | 1891 | 1966 | 2391 | 2096 | 2130 | 2436 | 2080,8 | 1898 | 2263 | 1,19 | 0,251 | 4,021 | 5,80E-05 | 0,00018571 | 0,003514 | 0,0026772 | 0,0032563 | 0,0032563 | TRUE | 0,0302635 | No |

| | | | | | | | | | | | | | | | | | | | | | | | | | | | | | | | |
|---|---|---|---|---|---|---|---|---|---|---|---|---|---|---|---|---|---|---|---|---|---|---|---|---|---|---|---|---|---|---|---|
| orf19.764 | 296 | 219 | 247 | 311 | 376 | 415 | 439 | 272 | 266 | 279 | 253 | 256 | 322 | 396 | 392 | 384 | 318,3 | 264 | 374 | 1,417 | 0,503 | 5,432 | 5,56E-08 | 2,82E-07 | 0,003438 | 0,0065403 | 0,0049846 | 0,0049846 | TRUE | 0,1079681 | No |
| orf19.7642 | 579 | 360 | 505 | 576 | 645 | 577 | 598 | 400 | 519 | 458 | 517 | 475 | 552 | 551 | 534 | 564 | 521,17 | 492 | 550 | 1,114 | 0,156 | 2,287 | 0,02222046 | 0,04107699 | 1,00E-08 | 0,004765 | 0,0025219 | 0,0025219 | TRUE | 0,0215108 | No |
| orf19.7643 | 1507 | 936 | 1261 | 1672 | 1641 | 1380 | 1514 | 996 | 1352 | 1192 | 1290 | 1379 | 1403 | 1317 | 1351 | 1405 | 1336,1 | 1303 | 1369 | 1,048 | 0,067 | 1,193 | 0,23298934 | 0,31350503 | 0,001524 | 0,0030661 | 0,0022808 | 0,0022808 | TRUE | 0,0373353 | No |
| orf19.7644 | 383 | 267 | 356 | 418 | 397 | 338 | 350 | 254 | 344 | 340 | 364 | 345 | 340 | 323 | 312 | 358 | 340,63 | 348 | 333 | 0,953 | -0,069 | -0,871 | 0,38398091 | 0,47293098 | 1,00E-08 | 0,0062414 | 0,0031253 | 0,0031253 | TRUE | 0,0238033 | No |
| orf19.7645 | 1852 | 1226 | 1599 | 1890 | 1703 | 1423 | 1510 | 971 | 1661 | 1561 | 1636 | 1559 | 1456 | 1358 | 1347 | 1370 | 1493,5 | 1604 | 1383 | 0,862 | -0,214 | -4,433 | 9,31E-06 | 3,40E-05 | 0,000613 | 0,0029515 | 0,0015599 | 0,0015599 | TRUE | 0,0165129 | No |
| orf19.7646 | 424 | 295 | 404 | 442 | 443 | 348 | 424 | 241 | 380 | 376 | 413 | 364 | 379 | 332 | 378 | 340 | 370,37 | 383 | 357 | 0,935 | -0,096 | -1,175 | 0,23985127 | 0,3210128 | 0,001337 | 0,0058992 | 0,0037439 | 0,0037439 | TRUE | 0,0316285 | No |
| orf19.7648 | 439 | 257 | 396 | 470 | 575 | 496 | 542 | 392 | 394 | 327 | 405 | 388 | 492 | 473 | 484 | 553 | 439,41 | 378 | 500 | 1,312 | 0,392 | 4,674 | 2,95E-06 | 1,16E-05 | 0,003726 | 0,0052833 | 0,004442 | 0,004442 | TRUE | 0,0839326 | No |
| orf19.7650 | 6197 | 3926 | 5319 | 6800 | 5963 | 4866 | 5582 | 3385 | 5559 | 4998 | 5442 | 5607 | 5100 | 4643 | 4981 | 4774 | 5138,1 | 5402 | 4874 | 0,902 | -0,149 | -3,029 | 0,00245476 | 0,00568111 | 0,001944 | 0,0022621 | 0,0021133 | 0,0021133 | TRUE | 0,0301733 | No |
| orf19.7652 | 8994 | 5855 | 8022 | 9738 | 9060 | 7461 | 8625 | 5245 | 8069 | 7454 | 8207 | 8030 | 7748 | 7119 | 7696 | 7398 | 7715,2 | 7940 | 7490 | 0,943 | -0,084 | -1,911 | 0,05598596 | 0,09258565 | 0,001464 | 0,0021677 | 0,0017381 | 0,0017381 | TRUE | 0,0203157 | No |
| orf19.7654 | 3667 | 2252 | 3042 | 3814 | 5214 | 4071 | 4576 | 3111 | 3290 | 2867 | 3112 | 3145 | 4459 | 3884 | 4083 | 4388 | 3653,6 | 3104 | 4204 | 1,353 | 0,436 | 7,486 | 7,09E-14 | 7,29E-13 | 0,003259 | 0,0023769 | 0,0029812 | 0,0029812 | TRUE | 0,0319216 | No |
| orf19.7655 | 4551 | 2903 | 3961 | 4825 | 4447 | 3755 | 4085 | 2527 | 4083 | 3696 | 4052 | 3979 | 3803 | 3583 | 3645 | 3564 | 3800,7 | 3952 | 3649 | 0,923 | -0,116 | -2,636 | 0,00839279 | 0,01720951 | 0,001089 | 0,0023615 | 0,0015817 | 0,0015817 | TRUE | 0,0223216 | No |
| orf19.7657 | 3569 | 2261 | 2791 | 3607 | 3291 | 2509 | 2725 | 1852 | 3202 | 2878 | 2855 | 2974 | 2815 | 2394 | 2432 | 2612 | 2770,3 | 2977 | 2563 | 0,86 | -0,217 | -3,51 | 0,00044797 | 0,00122225 | 0,003755 | 0,0025035 | 0,0033011 | 0,0033011 | TRUE | 0,0547134 | No |
| orf19.7658 | 1896 | 1220 | 1599 | 1908 | 1920 | 1623 | 1793 | 1198 | 1701 | 1553 | 1636 | 1573 | 1642 | 1549 | 1600 | 1690 | 1618 | 1616 | 1620 | 1,001 | 0,001 | 0,022 | 0,98210363 | 0,9867021 | 0,000872 | 0,0028768 | 0,0016973 | 0,0016973 | TRUE | 0,015456 | No |
| orf19.7660 | 567 | 374 | 511 | 583 | 630 | 556 | 539 | 417 | 509 | 476 | 523 | 481 | 539 | 531 | 481 | 588 | 515,85 | 497 | 535 | 1,071 | 0,098 | 1,303 | 0,19259986 | 0,26581301 | 0,002224 | 0,0047937 | 0,0035303 | 0,0035303 | TRUE | 0,0535259 | No |
| orf19.7661 | 1090 | 677 | 911 | 1143 | 1002 | 890 | 924 | 597 | 978 | 862 | 932 | 943 | 857 | 849 | 825 | 842 | 885,87 | 929 | 843 | 0,906 | -0,143 | -2,545 | 0,01093834 | 0,02181134 | 1,00E-08 | 0,0036182 | 0,0018805 | 0,0018805 | TRUE | 0,025147 | No |
| orf19.7662 | 3709 | 2420 | 3136 | 3696 | 3772 | 3143 | 3445 | 2066 | 3327 | 3081 | 3208 | 3048 | 3226 | 2999 | 3074 | 2914 | 3109,7 | 3166 | 3053 | 0,965 | -0,051 | -1,089 | 0,27605685 | 0,36025364 | 0,001327 | 0,0024464 | 0,0018009 | 0,0018009 | TRUE | 0,01775 | No |
| orf19.7663 | 675 | 445 | 547 | 716 | 756 | 588 | 718 | 420 | 606 | 567 | 560 | 590 | 647 | 561 | 641 | 592 | 595,35 | 581 | 610 | 1,052 | 0,073 | 1,053 | 0,29224831 | 0,37686797 | 0,001462 | 0,0044179 | 0,0028917 | 0,0028917 | TRUE | 0,0374854 | No |
| orf19.7664 | 2046 | 1391 | 1831 | 2360 | 1677 | 1495 | 1650 | 1137 | 1835 | 1771 | 1873 | 1946 | 1434 | 1426 | 1472 | 1604 | 1670,3 | 1856 | 1484 | 0,797 | -0,328 | -6,238 | 4,42E-10 | 2,97E-09 | 0,0015 | 0,0028486 | 0,00204 | 0,00204 | TRUE | 0,0321573 | No |
| orf19.7665 | 1198 | 718 | 944 | 1156 | 1224 | 910 | 986 | 677 | 1075 | 914 | 966 | 953 | 1047 | 868 | 880 | 955 | 957,21 | 977 | 938 | 0,959 | -0,061 | -0,805 | 0,42102786 | 0,50885846 | 0,005294 | 0,0034961 | 0,0044885 | 0,0044885 | TRUE | 0,0779658 | No |
| orf19.7666 | 23 | 9 | 18 | 25 | 18 | 13 | 16 | 11 | 21 | 11 | 18 | 21 | 15 | 12 | 14 | 16 | 16,09 | 18 | 14 | 0,793 | -0,334 | -0,98 | 0,32712726 | 0,41437499 | 1,00E-08 | 0,0922085 | 0,0493683 | 0,0493683 | TRUE | 0,215971 | No |
| orf19.7667 | 1239 | 807 | 1076 | 1337 | 1528 | 1313 | 1358 | 990 | 1112 | 1027 | 1101 | 1103 | 1307 | 1253 | 1212 | 1396 | 1188,8 | 1086 | 1292 | 1,186 | 0,246 | 4,309 | 1,64E-05 | 5,79E-05 | 0,001421 | 0,0032007 | 0,0022738 | 0,0022738 | TRUE | 0,0321565 | No |
| orf19.7668 | 325 | 180 | 279 | 291 | 416 | 306 | 351 | 218 | 292 | 229 | 285 | 240 | 356 | 292 | 313 | 307 | 289,32 | 262 | 317 | 1,212 | 0,277 | 2,635 | 0,00840262 | 0,0172238 | 0,00734 | 0,0069972 | 0,0071599 | 0,0071599 | TRUE | 0,0812671 | No |
| orf19.767 | 998 | 785 | 779 | 1187 | 1189 | 1295 | 935 | 708 | 895 | 999 | 797 | 979 | 1017 | 1236 | 834 | 999 | 969,49 | 918 | 1022 | 1,113 | 0,155 | 1,415 | 0,15719533 | 0,22427679 | 0,016675 | 0,0034769 | 0,0104642 | 0,0104642 | TRUE | 0,2414952 | No |
| orf19.7670 | 1251 | 809 | 1138 | 1441 | 1873 | 1434 | 1706 | 1074 | 1122 | 1030 | 1164 | 1188 | 1602 | 1368 | 1522 | 1515 | 1314 | 1126 | 1502 | 1,332 | 0,413 | 6,407 | 1,49E-10 | 1,05E-09 | 0,003312 | 0,0030843 | 0,003213 | 0,003213 | TRUE | 0,0442111 | No |
| orf19.7672 | 2066 | 1175 | 1782 | 2141 | 2284 | 1796 | 1983 | 1400 | 1853 | 1496 | 1823 | 1765 | 1953 | 1714 | 1769 | 1975 | 1793,6 | 1734 | 1853 | 1,066 | 0,092 | 1,237 | 0,21605117 | 0,294139 | 0,006169 | 0,0027889 | 0,004802 | 0,004802 | TRUE | 0,0999622 | No |
| orf19.7673 | 478 | 306 | 341 | 499 | 425 | 307 | 338 | 196 | 429 | 390 | 349 | 411 | 363 | 293 | 302 | 276 | 351,65 | 395 | 308 | 0,784 | -0,35 | -3,423 | 0,00061825 | 0,00162657 | 0,008257 | 0,0061078 | 0,0071665 | 0,0071665 | TRUE | 0,1679184 | No |
| orf19.7675 | 723 | 448 | 657 | 785 | 751 | 545 | 602 | 453 | 649 | 570 | 672 | 647 | 642 | 520 | 537 | 639 | 609,61 | 634 | 584 | 0,918 | -0,124 | -1,419 | 0,15583461 | 0,2228116 | 0,006441 | 0,0043605 | 0,0056452 | 0,0056452 | TRUE | 0,0646223 | No |
| orf19.7676 | 12674 | 7544 | 10820 | 12842 | 19956 | 15522 | 16642 | 12942 | 11370 | 9604 | 11070 | 10590 | 17067 | 14811 | 14850 | 18254 | 13452 | 10658 | 16246 | 1,524 | 0,608 | 8,017 | 1,09E-15 | 1,33E-14 | 0,008027 | 0,0020874 | 0,0054412 | 0,0054412 | TRUE | 0,0843522 | No |
| orf19.7678 | 6307 | 3634 | 5035 | 6074 | 5979 | 4593 | 4954 | 3449 | 5658 | 4626 | 5151 | 5009 | 5113 | 4382 | 4421 | 4865 | 4903,2 | 5111 | 4695 | 0,918 | -0,123 | -1,764 | 0,07764958 | 0,12335237 | 0,00606 | 0,0022756 | 0,0044705 | 0,0044705 | TRUE | 0,0633509 | No |
| orf19.768 | 588 | 450 | 535 | 633 | 559 | 546 | 578 | 316 | 527 | 573 | 547 | 522 | 478 | 521 | 516 | 446 | 516,28 | 542 | 490 | 0,909 | -0,138 | -1,915 | 0,05551586 | 0,09201084 | 0,000901 | 0,0047914 | 0,0030181 | 0,0030181 | TRUE | 0,0361033 | No |
| orf19.7680 | 449 | 252 | 360 | 438 | 523 | 396 | 447 | 293 | 403 | 321 | 368 | 361 | 447 | 378 | 399 | 413 | 386,3 | 363 | 409 | 1,123 | 0,168 | 1,891 | 0,05868351 | 0,09656821 | 0,004118 | 0,0057375 | 0,0049508 | 0,0049508 | TRUE | 0,0629986 | No |
| orf19.769 | 89 | 75 | 89 | 96 | 91 | 76 | 69 | 42 | 80 | 95 | 91 | 79 | 78 | 73 | 62 | 59 | 77,09 | 86 | 68 | 0,796 | -0,33 | -2,021 | 0,04331821 | 0,0741306 | 1,00E-08 | 0,0208115 | 0,0124354 | 0,0124354 | TRUE | 0,094422 | No |
| orf19.771 | 710 | 561 | 555 | 757 | 662 | 641 | 599 | 386 | 637 | 714 | 568 | 624 | 566 | 612 | 535 | 544 | 599,99 | 636 | 564 | 0,89 | -0,168 | -2,114 | 0,03450445 | 0,06070359 | 0,004112 | 0,0043991 | 0,0043966 | 0,0043966 | TRUE | 0,0745041 | No |
| orf19.77.1 | 6 | 5 | 5 | 10 | 3 | 7 | 10 | 3 | 5 | 6 | 5 | 8 | 3 | 7 | 9 | 4 | 5,94 | 6 | 6 | 0,896 | -0,159 | -0,276 | 0,7825494 | NA | 1,00E-08 | 0,2464291 | 0,1510104 | 0,1510104 | TRUE | 0,3309297 | No |
| orf19.772 | 2669 | 1848 | 2349 | 2844 | 2789 | 2453 | 2566 | 1584 | 2394 | 2353 | 2403 | 2345 | 2385 | 2341 | 2290 | 2234 | 2343,2 | 2374 | 2312 | 0,975 | -0,036 | -0,941 | 0,34675617 | 0,43400127 | 1,00E-08 | 0,0025991 | 0,000985 | 0,000985 | TRUE | 0,0056071 | No |
| orf19.772.1 | 1824 | 1431 | 1648 | 2213 | 2162 | 1971 | 1981 | 1374 | 1636 | 1822 | 1686 | 1825 | 1849 | 1881 | 1768 | 1938 | 1800,6 | 1742 | 1859 | 1,066 | 0,093 | 1,748 | 0,08046233 | 0,12718172 | 0,0015 | 0,0027858 | 0,0021303 | 0,0021303 | TRUE | 0,0207338 | No |
| orf19.773 | 3413 | 2705 | 3054 | 3886 | 3965 | 3905 | 3989 | 2551 | 3062 | 3444 | 3125 | 3204 | 3391 | 3726 | 3560 | 3598 | 3388,6 | 3209 | 3569 | 1,113 | 0,154 | 3,154 | 0,0016117 | 0,00387781 | 0,001714 | 0,0024079 | 0,001997 | 0,001997 | TRUE | 0,0284819 | No |
| orf19.775 | 2607 | 1962 | 2342 | 2944 | 3400 | 3349 | 3393 | 2121 | 2339 | 2498 | 2396 | 2428 | 2908 | 3196 | 3028 | 2991 | 2722,9 | 2415 | 3031 | 1,256 | 0,329 | 7,553 | 4,25E-14 | 4,47E-13 | 0,000844 | 0,0025127 | 0,0014429 | 0,0014429 | TRUE | 0,0166131 | No |
| orf19.776 | 140 | 96 | 115 | 130 | 117 | 110 | 110 | 65 | 126 | 122 | 118 | 107 | 100 | 105 | 98 | 92 | 108,44 | 118 | 99 | 0,841 | -0,25 | -1,886 | 0,05926321 | 0,09744209 | 1,00E-08 | 0,0153666 | 0,0076316 | 0,0076316 | TRUE | 0,0453208 | No |
| orf19.778 | 8963 | 5876 | 7706 | 8961 | 10178 | 8599 | 8989 | 6013 | 8041 | 7481 | 7884 | 7389 | 8705 | 8205 | 8021 | 8481 | 8025,8 | 7699 | 8353 | 1,085 | 0,117 | 2,748 | 0,00599818 | 0,01273845 | 0,001311 | 0,0021604 | 0,0016277 | 0,0016277 | TRUE | 0,0110337 | No |
| orf19.779 | 769 | 572 | 729 | 837 | 849 | 735 | 796 | 451 | 690 | 728 | 746 | 690 | 726 | 701 | 710 | 636 | 703,49 | 714 | 693 | 0,977 | -0,034 | -0,535 | 0,59253305 | 0,67004302 | 0,000679 | 0,0040431 | 0,0023719 | 0,0023719 | TRUE | 0,031343 | No |
| orf19.780 | 3117 | 2146 | 2889 | 3331 | 2115 | 1923 | 2251 | 1285 | 2796 | 2732 | 2956 | 2747 | 1809 | 1835 | 2009 | 1812 | 2337 | 2808 | 1866 | 0,665 | -0,588 | -11,86 | 1,90E-32 | 7,99E-31 | 0,001385 | 0,0026007 | 0,0019136 | 0,0019136 | TRUE | 0,0328023 | No |
| orf19.781 | 239 | 137 | 197 | 229 | 170 | 158 | 157 | 113 | 214 | 174 | 202 | 189 | 145 | 151 | 140 | 159 | 171,85 | 195 | 149 | 0,758 | -0,399 | -3,59 | 0,00033062 | 0,0009282 | 1,00E-08 | 0,0104268 | 0,0059437 | 0,0059437 | TRUE | 0,0504779 | No |
| orf19.782 | 5616 | 3979 | 5094 | 6149 | 4160 | 3853 | 4334 | 2745 | 5038 | 5066 | 5212 | 5071 | 3558 | 3676 | 3867 | 3872 | 4419,9 | 5097 | 3743 | 0,734 | -0,446 | -11,407 | 3,87E-30 | 1,39E-28 | 0,00065 | 0,002308 | 0,0012365 | 0,0012365 | TRUE | 0,0138308 | No |
| orf19.783 | 3412 | 2393 | 3101 | 3820 | 3769 | 3518 | 3611 | 2497 | 3061 | 3047 | 3173 | 3150 | 3223 | 3357 | 3222 | 3522 | 3219,3 | 3108 | 3331 | 1,07 | 0,098 | 2,358 | 0,01837345 | 0,03446385 | 0,0007 | 0,0024305 | 0,0013402 | 0,0013402 | TRUE | 0,0169322 | No |
| orf19.787 | 21 | 17 | 24 | 45 | 17 | 24 | 30 | 21 | 19 | 22 | 25 | 37 | 15 | 23 | 27 | 30 | 24,5 | 26 | 24 | 0,897 | -0,157 | -0,488 | 0,62527238 | 0,69768216 | 0,055028 | 0,0612412 | 0,0584833 | 0,0584833 | TRUE | 0,6712384 | No |
| orf19.787.1 | 4683 | 3750 | 3985 | 5625 | 6930 | 6727 | 6584 | 4589 | 4201 | 4774 | 4077 | 4638 | 5927 | 6419 | 5875 | 6472 | 5298 | 4422 | 6173 | 1,396 | 0,481 | 8 | 1,25E-15 | 1,53E-14 | 0,003936 | 0,0022535 | 0,0032774 | 0,0032774 | TRUE | 0,0340846 | No |
| orf19.789 | 4882 | 3404 | 4351 | 5222 | 5776 | 5511 | 5687 | 3568 | 4380 | 4334 | 4452 | 4306 | 4940 | 5258 | 5075 | 5032 | 4722 | 4368 | 5076 | 1,162 | 0,217 | 6,407 | 1,49E-10 | 1,05E-09 | 0,000281 | 0,002287 | 0,0008878 | 0,0008878 | TRUE | 0,0072405 | No |
| orf19.789.1 | 592 | 439 | 526 | 690 | 837 | 734 | 771 | 427 | 531 | 559 | 538 | 569 | 716 | 700 | 688 | 602 | 612,94 | 549 | 676 | 1,237 | 0,307 | 4,447 | 8,70E-06 | 3,20E-05 | 0,00147 | 0,0043479 | 0,0029338 | 0,0029338 | TRUE | 0,0559305 | No |
| orf19.791 | 0 | 0 | 0 | 0 | 0 | 0 | 0 | 0 | 0 | 0 | 0 | 0 | 0 | 0 | 0 | 0 | 0 | 0 | 0 | NA | NA | NA | NA | NA | NA | NA | NA | NA | NA | NA | NA |
| orf19.792 | 2815 | 2189 | 2552 | 3319 | 2084 | 2296 | 2190 | 1822 | 2525 | 2787 | 2611 | 2737 | 1782 | 2191 | 1954 | 2570 | 2394,6 | 2665 | 2124 | 0,796 | -0,329 | -3,521 | 0,00043059 | 0,00117698 | 0,013313 | 0,0025858 | 0,0079764 | 0,0079764 | TRUE | 0,2363666 | No |
| orf19.793 | 1424 | 1401 | 1379 | 1994 | 1452 | 1588 | 1708 | 1013 | 1277 | 1784 | 1411 | 1644 | 1242 | 1515 | 1524 | 1429 | 1478,3 | 1529 | 1428 | 0,934 | -0,098 | -0,972 | 0,33122324 | 0,41811761 | 0,014185 | 0,0029616 | 0,0091479 | 0,0091479 | TRUE | 0,1487961 | No |
| orf19.794 | 1013 | 755 | 912 | 1189 | 1193 | 1182 | 1234 | 828 | 909 | 961 | 933 | 980 | 1020 | 1128 | 1101 | 1168 | 1025,1 | 946 | 1104 | 1,165 | 0,22 | 3,809 | 0,00013932 | 0,0004148 | 0,001305 | 0,0033957 | 0,0022226 | 0,0022226 | TRUE | 0,0322228 | No |
| orf19.795 | 2080 | 1514 | 1927 | 2340 | 2424 | 2347 | 2415 | 1365 | 1866 | 1927 | 1972 | 1930 | 2073 | 2239 | 2155 | 1925 | 2010,9 | 1924 | 2098 | 1,093 | 0,128 | 2,456 | 0,01402999 | 0,02721736 | 0,001625 | 0,0027014 | 0,0021045 | 0,0021045 | TRUE | 0,0343735 | No |
| orf19.796 | 1158 | 824 | 1010 | 1276 | 1238 | 1188 | 1213 | 784 | 1039 | 1049 | 1033 | 1052 | 1059 | 1134 | 1082 | 1106 | 1069,2 | 1043 | 1095 | 1,049 | 0,069 | 1,417 | 0,15659307 | 0,22373631 | 1,00E-08 | 0,0033372 | 0,0013191 | 0,0013191 | TRUE | 0,0073065 | No |
| orf19.797 | 1710 | 1174 | 1557 | 1918 | 2262 | 1902 | 2027 | 1331 | 1534 | 1495 | 1593 | 1582 | 1935 | 1815 | 1809 | 1877 | 1704,8 | 1551 | 1859 | 1,197 | 0,26 | 5,764 | 8,23E-09 | 4,62E-08 | 0,000393 | 0,002831 | 0,0013609 | 0,0013609 | TRUE | 0,0098996 | No |
| orf19.798 | 2831 | 1949 | 2485 | 2944 | 2777 | 2479 | 2548 | 1631 | 2540 | 2481 | 2542 | 2428 | 2375 | 2365 | 2274 | 2300 | 2413,2 | 2498 | 2328 | 0,933 | -0,1 | -2,61 | 0,00906425 | 0,0184666 | 1,00E-08 | 0,0025811 | 0,0010037 | 0,0010037 | TRUE | 0,0046754 | No |
| orf19.799 | 202 | 142 | 178 | 210 | 184 | 131 | 181 | 131 | 181 | 181 | 182 | 173 | 157 | 125 | 162 | 185 | 168,24 | 179 | 157 | 0,871 | -0,2 | -1,64 | 0,10108489 | 0,15418076 | 0,004557 | 0,0106083 | 0,0082759 | 0,0082759 | TRUE | 0,1981088 | No |
| orf19.8 | 101 | 87 | 122 | 128 | 116 | 124 | 117 | 88 | 91 | 111 | 125 | 106 | 99 | 118 | 104 | 124 | 109,72 | 108 | 111 | 1,03 | 0,042 | 0,292 | 0,77024837 | 0,82065122 | 0,005274 | 0,0152102 | 0,0109971 | 0,0109971 | TRUE | 0,122927 | No |

| | | | | | | | | | | | | | | | | | | | | | | | | | | | | | | | |
|---|---|---|---|---|---|---|---|---|---|---|---|---|---|---|---|---|---|---|---|---|---|---|---|---|---|---|---|---|---|---|---|
| orf19.801 | 5045 | 3230 | 4356 | 5357 | 6166 | 5090 | 5890 | 3765 | 4526 | 4112 | 4457 | 4417 | 5273 | 4857 | 5256 | 5310 | 4776 | 4378 | 5174 | 1,181 | 0,24 | 5,26 | 1,44E-07 | 6,91E-07 | 0,001436 | 0,0022835 | 0,0017847 | 0,0017847 | TRUE | 0,0208987 | No |
| orf19.802 | 2277 | 1390 | 1960 | 2208 | 1813 | 1467 | 1656 | 1126 | 2043 | 1770 | 2005 | 1821 | 1551 | 1400 | 1478 | 1588 | 1706,8 | 1910 | 1504 | 0,786 | -0,347 | -5,534 | 3,13E-08 | 1,63E-07 | 0,003225 | 0,0028301 | 0,0031701 | 0,0031701 | TRUE | 0,028 | No |
| orf19.804 | 612 | 426 | 559 | 694 | 462 | 410 | 464 | 263 | 549 | 542 | 572 | 572 | 395 | 391 | 414 | 371 | 475,86 | 559 | 393 | 0,705 | -0,505 | -7,404 | 1,32E-13 | 1,32E-12 | 1,00E-08 | 0,0050302 | 0,0023138 | 0,0023138 | TRUE | 0,0148408 | No |
| orf19.804.1 | 4767 | 2798 | 3511 | 4638 | 6019 | 4147 | 4227 | 2955 | 4276 | 3562 | 3592 | 3825 | 5148 | 3957 | 3772 | 4168 | 4037,4 | 3814 | 4261 | 1,117 | 0,16 | 1,722 | 0,08503548 | 0,13335765 | 0,013753 | 0,0023391 | 0,008026 | 0,008026 | TRUE | 0,2407957 | No |
| orf19.805 | 765 | 496 | 597 | 771 | 904 | 743 | 840 | 510 | 686 | 631 | 611 | 636 | 773 | 709 | 750 | 719 | 689,41 | 641 | 738 | 1,151 | 0,203 | 3,238 | 0,00120307 | 0,00298557 | 0,000727 | 0,0040852 | 0,002326 | 0,002326 | TRUE | 0,027602 | No |
| orf19.806 | 266 | 189 | 203 | 311 | 279 | 200 | 252 | 146 | 239 | 241 | 208 | 256 | 239 | 191 | 225 | 206 | 225,45 | 236 | 215 | 0,913 | -0,131 | -1,214 | 0,22456121 | 0,30358213 | 0,004636 | 0,0084186 | 0,0066679 | 0,0066679 | TRUE | 0,071667 | No |
| orf19.807 | 2608 | 1657 | 2175 | 2620 | 2730 | 2289 | 2475 | 1478 | 2340 | 2110 | 2225 | 2160 | 2335 | 2184 | 2209 | 2085 | 2205,9 | 2209 | 2203 | 0,998 | -0,003 | -0,054 | 0,9573245 | 0,96942619 | 0,001595 | 0,0026376 | 0,0020051 | 0,0020051 | TRUE | 0,0197472 | No |
| orf19.808 | 3858 | 2672 | 3502 | 4157 | 3821 | 3470 | 3855 | 2349 | 3461 | 3402 | 3583 | 3428 | 3268 | 3311 | 3440 | 3313 | 3400,7 | 3468 | 3333 | 0,961 | -0,057 | -1,587 | 0,11253621 | 0,16848182 | 0,000237 | 0,0024064 | 0,0009608 | 0,0009608 | TRUE | 0,0059225 | No |
| orf19.809 | 11698 | 8292 | 10444 | 12564 | 10894 | 9526 | 10956 | 6318 | 10494 | 10557 | 10685 | 10360 | 9317 | 9089 | 9776 | 8911 | 9898,8 | 10524 | 9273 | 0,882 | -0,182 | -4,984 | 6,24E-07 | 2,72E-06 | 0,000761 | 0,0021262 | 0,0011755 | 0,0011755 | TRUE | 0,0163239 | No |
| orf19.81 | 2307 | 2286 | 2216 | 2978 | 2468 | 3131 | 3019 | 1723 | 2070 | 2910 | 2267 | 2456 | 2111 | 2988 | 2694 | 2430 | 2490,6 | 2426 | 2556 | 1,054 | 0,076 | 0,677 | 0,49837237 | 0,58310052 | 0,018954 | 0,0025624 | 0,011745 | 0,011745 | TRUE | 0,217022 | No |
| orf19.810 | 703 | 546 | 687 | 727 | 713 | 615 | 699 | 426 | 631 | 695 | 703 | 599 | 610 | 587 | 624 | 601 | 631,16 | 657 | 606 | 0,925 | -0,113 | -1,629 | 0,10337744 | 0,15693621 | 0,001941 | 0,0042796 | 0,0030458 | 0,0030458 | TRUE | 0,0427596 | No |
| orf19.811 | 719 | 535 | 592 | 755 | 739 | 649 | 725 | 412 | 645 | 681 | 606 | 623 | 632 | 619 | 647 | 581 | 629,21 | 639 | 620 | 0,975 | -0,036 | -0,559 | 0,57602137 | 0,65420347 | 0,000218 | 0,0042867 | 0,0023908 | 0,0023908 | TRUE | 0,0209692 | No |
| orf19.812 | 61 | 35 | 50 | 56 | 42 | 20 | 42 | 18 | 55 | 45 | 51 | 46 | 36 | 19 | 37 | 25 | 39,31 | 49 | 29 | 0,606 | -0,723 | -2,969 | 0,00299077 | 0,00679597 | 0,014128 | 0,0389086 | 0,030141 | 0,030141 | TRUE | 0,4197304 | No |
| orf19.813 | 40844 | 28601 | 35737 | 42869 | 37436 | 34497 | 36870 | 21246 | 36641 | 36412 | 36562 | 35350 | 32016 | 32916 | 32900 | 29966 | 34096 | 36241 | 31950 | 0,882 | -0,182 | -4,804 | 1,56E-06 | 6,37E-06 | 0,001042 | 0,0020221 | 0,0013427 | 0,0013427 | TRUE | 0,0212019 | No |
| orf19.814 | 4807 | 3064 | 4335 | 5085 | 4982 | 4400 | 4903 | 2967 | 4312 | 3901 | 4435 | 4193 | 4261 | 4198 | 4375 | 4185 | 4232,6 | 4210 | 4255 | 1,01 | 0,015 | 0,323 | 0,74650623 | 0,80106007 | 0,001224 | 0,0023225 | 0,0017336 | 0,0017336 | TRUE | 0,0286917 | No |
| orf19.815 | 8624 | 6536 | 8265 | 9487 | 11540 | 11105 | 11905 | 7181 | 7737 | 8321 | 8456 | 7823 | 9869 | 10596 | 10623 | 10128 | 9194,2 | 8084 | 10304 | 1,275 | 0,351 | 8,032 | 9,62E-16 | 1,19E-14 | 0,001469 | 0,0021374 | 0,0017195 | 0,0017195 | TRUE | 0,0118546 | No |
| orf19.816 | 496 | 326 | 430 | 494 | 709 | 634 | 696 | 416 | 445 | 415 | 440 | 407 | 606 | 605 | 621 | 587 | 515,8 | 427 | 605 | 1,419 | 0,505 | 7,647 | 2,06E-14 | 2,22E-13 | 1,00E-08 | 0,004794 | 0,0021915 | 0,0021915 | TRUE | 0,0129702 | No |
| orf19.819 | 1025 | 665 | 847 | 1022 | 904 | 773 | 838 | 559 | 920 | 847 | 867 | 843 | 773 | 738 | 748 | 788 | 815,3 | 869 | 762 | 0,875 | -0,193 | -3,398 | 0,00067877 | 0,00177075 | 1,00E-08 | 0,0037601 | 0,0018603 | 0,0018603 | TRUE | 0,0191522 | No |
| orf19.820 | 1322 | 962 | 1220 | 1384 | 1352 | 1142 | 1205 | 742 | 1186 | 1225 | 1248 | 1141 | 1156 | 1090 | 1075 | 1047 | 1146 | 1200 | 1092 | 0,913 | -0,132 | -2,455 | 0,01410658 | 0,02734828 | 0,000858 | 0,0032463 | 0,0018907 | 0,0018907 | TRUE | 0,0193533 | No |
| orf19.822 | 2E+05 | 1E+05 | 2E+05 | 2E+05 | 3E+05 | 2E+05 | 2E+05 | 1E+05 | 2E+05 | 2E+05 | 2E+05 | 2E+05 | 2E+05 | 2E+05 | 2E+05 | 2E+05 | 194296 | 2E+05 | 2E+05 | 1,206 | 0,271 | 5,224 | 1,75E-07 | 8,31E-07 | 0,002969 | 0,001987 | 0,0025759 | 0,0025759 | TRUE | 0,0460337 | No |
| orf19.823 | 967 | 773 | 785 | 954 | 988 | 1262 | 1242 | 1826 | 868 | 984 | 803 | 787 | 845 | 1204 | 1108 | 2575 | 1146,8 | 860 | 1433 | 1,665 | 0,736 | 2,027 | 0,04262996 | 0,07313558 | 0,125559 | 0,0032454 | 0,0533243 | 0,1255587 | TRUE | 2,0074161 | No |
| orf19.825 | 2310 | 1517 | 1971 | 2342 | 2018 | 1730 | 1857 | 1217 | 2072 | 1931 | 2017 | 1931 | 1726 | 1651 | 1657 | 1716 | 1837,7 | 1988 | 1688 | 0,848 | -0,238 | -5,384 | 7,29E-08 | 3,64E-07 | 0,000408 | 0,0027695 | 0,0013197 | 0,0013197 | TRUE | 0,0101793 | No |
| orf19.826 | 600 | 445 | 554 | 657 | 820 | 781 | 852 | 480 | 538 | 567 | 567 | 542 | 701 | 745 | 760 | 677 | 637,14 | 554 | 721 | 1,308 | 0,387 | 6,104 | 1,03E-09 | 6,57E-09 | 1,00E-08 | 0,004258 | 0,0022709 | 0,0022709 | TRUE | 0,0168781 | No |
| orf19.827 | 1057 | 781 | 979 | 1210 | 1449 | 1207 | 1374 | 942 | 948 | 994 | 1002 | 998 | 1239 | 1152 | 1226 | 1329 | 1110,9 | 986 | 1236 | 1,252 | 0,325 | 5,827 | 5,65E-09 | 3,25E-08 | 0,001079 | 0,0032863 | 0,002065 | 0,002065 | TRUE | 0,0269659 | No |
| orf19.827.1 | 12121 | 7946 | 9221 | 11960 | 11957 | 9751 | 9817 | 6475 | 10874 | 10116 | 9434 | 9862 | 10226 | 9304 | 8760 | 9132 | 9713,6 | 10072 | 9356 | 0,929 | -0,106 | -1,814 | 0,06963354 | 0,11228081 | 0,003881 | 0,002129 | 0,0031906 | 0,0031906 | TRUE | 0,0482717 | No |
| orf19.8278 | 1935 | 1447 | 1585 | 2310 | 2106 | 2043 | 2095 | 1289 | 1736 | 1842 | 1622 | 1905 | 1801 | 1949 | 1869 | 1818 | 1817,8 | 1776 | 1859 | 1,047 | 0,066 | 1,147 | 0,25154672 | 0,33398734 | 0,00236 | 0,0027781 | 0,0026648 | 0,0026648 | TRUE | 0,0412865 | No |
| orf19.828 | 1813 | 1230 | 1553 | 1983 | 2450 | 2219 | 2289 | 1472 | 1626 | 1566 | 1589 | 1635 | 2095 | 2117 | 2043 | 2076 | 1843,5 | 1604 | 2083 | 1,297 | 0,375 | 9,398 | 5,57E-21 | 1,08E-19 | 1,00E-08 | 0,002767 | 0,0009797 | 0,0009797 | TRUE | 0,0028491 | No |
| orf19.829 | 5213 | 3745 | 4857 | 5633 | 5573 | 5088 | 5406 | 3553 | 4677 | 4768 | 4969 | 4645 | 4766 | 4855 | 4824 | 5011 | 4814,4 | 4765 | 4864 | 1,021 | 0,029 | 0,809 | 0,41828294 | 0,50625381 | 0,000424 | 0,0022811 | 0,0010507 | 0,0010507 | TRUE | 0,010274 | No |
| orf19.831 | 2384 | 1767 | 2024 | 2648 | 2854 | 2710 | 2919 | 1813 | 2139 | 2250 | 2071 | 2184 | 2441 | 2586 | 2605 | 2557 | 2353,9 | 2161 | 2547 | 1,179 | 0,238 | 5,537 | 3,08E-08 | 1,60E-07 | 0,000588 | 0,0025963 | 0,001342 | 0,001342 | TRUE | 0,0100917 | No |
| orf19.832 | 797 | 616 | 817 | 937 | 1183 | 1155 | 1277 | 711 | 715 | 784 | 836 | 773 | 1012 | 1102 | 1140 | 1003 | 920,48 | 777 | 1064 | 1,373 | 0,457 | 6,781 | 1,19E-11 | 9,64E-11 | 0,002986 | 0,0035566 | 0,0032448 | 0,0032448 | TRUE | 0,0352219 | No |
| orf19.834 | 3441 | 2743 | 3186 | 3900 | 3889 | 3854 | 4106 | 2434 | 3087 | 3492 | 3260 | 3216 | 3326 | 3677 | 3664 | 3433 | 3394,4 | 3264 | 3525 | 1,081 | 0,113 | 2,165 | 0,03041677 | 0,05425889 | 0,002163 | 0,0024072 | 0,0023023 | 0,0023023 | TRUE | 0,026215 | No |
| orf19.835 | 1299 | 871 | 1151 | 1436 | 1152 | 1124 | 1186 | 719 | 1165 | 1109 | 1178 | 1184 | 985 | 1072 | 1058 | 1014 | 1095,8 | 1159 | 1032 | 0,89 | -0,168 | -3,255 | 0,0011328 | 0,00282049 | 1,00E-08 | 0,0033044 | 0,0016418 | 0,0016418 | TRUE | 0,0128241 | No |
| orf19.836 | 111 | 124 | 110 | 168 | 209 | 214 | 197 | 133 | 100 | 158 | 113 | 139 | 179 | 204 | 176 | 188 | 156,85 | 128 | 187 | 1,475 | 0,561 | 3,971 | 7,16E-05 | 0,00022475 | 0,01463 | 0,0112347 | 0,0125421 | 0,0125421 | TRUE | 0,2335817 | No |
| orf19.836.1 | 378 | 259 | 269 | 356 | 339 | 249 | 258 | 118 | 339 | 330 | 275 | 294 | 290 | 238 | 230 | 166 | 270,22 | 310 | 231 | 0,753 | -0,409 | -2,928 | 0,00341266 | 0,00763622 | 0,023684 | 0,0073518 | 0,0149366 | 0,0149366 | TRUE | 0,3428949 | No |
| orf19.837 | 1215 | 929 | 1058 | 1378 | 1214 | 1133 | 1126 | 611 | 1090 | 1183 | 1082 | 1136 | 1038 | 1081 | 1005 | 862 | 1059,7 | 1123 | 996 | 0,891 | -0,167 | -2,385 | 0,01708477 | 0,03237001 | 0,004146 | 0,0033495 | 0,0037338 | 0,0037338 | TRUE | 0,0898034 | No |
| orf19.837.1 | 1074 | 703 | 915 | 1057 | 919 | 856 | 945 | 499 | 963 | 895 | 936 | 872 | 786 | 817 | 843 | 704 | 852 | 916 | 788 | 0,862 | -0,214 | -3,228 | 0,00124566 | 0,00307853 | 0,002382 | 0,0036834 | 0,0030412 | 0,0030412 | TRUE | 0,0532864 | No |
| orf19.838.1 | 21939 | 13973 | 17751 | 22128 | 19928 | 16350 | 17684 | 10478 | 19682 | 17789 | 18161 | 18247 | 17043 | 15601 | 15780 | 14778 | 17135 | 18470 | 15800 | 0,856 | -0,225 | -4,406 | 1,05E-05 | 3,81E-05 | 0,002692 | 0,0020642 | 0,0024455 | 0,0024455 | TRUE | 0,0343384 | No |
| orf19.839 | 2710 | 1927 | 2378 | 2864 | 3240 | 2873 | 3104 | 1856 | 2431 | 2453 | 2433 | 2362 | 2771 | 2741 | 2770 | 2618 | 2572,4 | 2420 | 2725 | 1,128 | 0,174 | 4,614 | 3,96E-06 | 1,53E-05 | 1,00E-08 | 0,0025439 | 0,0009758 | 0,0009758 | TRUE | 0,0076646 | No |
| orf19.84 | 4154 | 2738 | 3871 | 4756 | 5341 | 4742 | 5175 | 3394 | 3727 | 3486 | 3960 | 3922 | 4568 | 4525 | 4618 | 4787 | 4199 | 3774 | 4624 | 1,224 | 0,292 | 6,178 | 6,47E-10 | 4,22E-09 | 0,001649 | 0,0023253 | 0,0018966 | 0,0018966 | TRUE | 0,0309317 | No |
| orf19.840 | 17875 | 11871 | 14314 | 18740 | 18678 | 14805 | 15522 | 11308 | 16036 | 15113 | 14645 | 15453 | 15974 | 14126 | 13851 | 15949 | 15143 | 15312 | 14975 | 0,978 | -0,032 | -0,572 | 0,56758055 | 0,64690024 | 0,003163 | 0,0020754 | 0,0030143 | 0,0030143 | TRUE | 0,0313031 | No |
| orf19.841 | 1709 | 1191 | 1505 | 1724 | 1848 | 1658 | 1763 | 1135 | 1533 | 1516 | 1540 | 1422 | 1580 | 1582 | 1573 | 1601 | 1543,4 | 1503 | 1584 | 1,055 | 0,077 | 1,713 | 0,08669587 | 0,13546652 | 1,00E-08 | 0,0029201 | 0,0012823 | 0,0012823 | TRUE | 0,0170979 | No |
| orf19.842 | 15653 | 10785 | 12914 | 15760 | 16568 | 15340 | 15962 | 9439 | 14042 | 13730 | 13212 | 12996 | 14169 | 14637 | 14243 | 13313 | 13793 | 13495 | 14090 | 1,044 | 0,063 | 1,517 | 0,12932608 | 0,19002542 | 0,001312 | 0,0020848 | 0,0015737 | 0,0015737 | TRUE | 0,0165903 | No |
| orf19.843 | 1101 | 792 | 1064 | 1240 | 1277 | 1212 | 1236 | 723 | 988 | 1008 | 1089 | 1023 | 1092 | 1156 | 1103 | 1020 | 1059,8 | 1027 | 1093 | 1,067 | 0,093 | 1,646 | 0,09982001 | 0,15267786 | 0,000968 | 0,0033493 | 0,0021308 | 0,0021308 | TRUE | 0,0212813 | No |
| orf19.844 | 6620 | 4883 | 6033 | 7247 | 10225 | 9579 | 10630 | 6246 | 5939 | 6217 | 6172 | 5976 | 8745 | 9140 | 9486 | 8809 | 7560,4 | 6076 | 9045 | 1,489 | 0,575 | 15,055 | 3,21E-51 | 3,64E-49 | 0,000788 | 0,0021715 | 0,0012618 | 0,0012618 | TRUE | 0,0132557 | No |
| orf19.846 | 6613 | 4570 | 5961 | 7167 | 6823 | 6156 | 6779 | 3942 | 5933 | 5818 | 6099 | 5910 | 5835 | 5874 | 6049 | 5560 | 5884,7 | 5940 | 5830 | 0,982 | -0,026 | -0,718 | 0,4728309 | 0,55898623 | 0,00057 | 0,0022262 | 0,0011008 | 0,0011008 | TRUE | 0,0112131 | No |
| orf19.847 | 6170 | 4441 | 5417 | 6646 | 4618 | 4469 | 4563 | 2951 | 5535 | 5654 | 5542 | 5480 | 3949 | 4264 | 4072 | 4162 | 4832,4 | 5553 | 4112 | 0,74 | -0,434 | -12,245 | 1,78E-34 | 8,51E-33 | 0,000333 | 0,0022799 | 0,0009916 | 0,0009916 | TRUE | 0,008893 | No |
| orf19.848 | 0 | 1 | 1 | 3 | 0 | 1 | 0 | 0 | 0 | 1 | 1 | 2 | 0 | 1 | 0 | 0 | 0,72 | 1 | 0 | 0,254 | -1,976 | -1,131 | 0,25796246 | NA | 1,00E-08 | 2,0308369 | 1,517841 | 1,517841 | TRUE | 0,2418566 | No |
| orf19.849 | 81 | 87 | 102 | 98 | 84 | 103 | 88 | 64 | 73 | 111 | 104 | 81 | 72 | 98 | 79 | 90 | 88,44 | 92 | 85 | 0,924 | -0,115 | -0,647 | 0,51774336 | 0,60131492 | 0,019203 | 0,0183944 | 0,0187228 | 0,0187228 | TRUE | 0,1815265 | No |
| orf19.85 | 2808 | 2014 | 2305 | 2931 | 2040 | 2072 | 2058 | 1183 | 2519 | 2564 | 2358 | 2417 | 1745 | 1977 | 1836 | 1669 | 2135,6 | 2464 | 1807 | 0,734 | -0,446 | -7,738 | 1,01E-14 | 1,13E-13 | 0,002784 | 0,0026593 | 0,0027016 | 0,0027016 | TRUE | 0,0482038 | No |
| orf19.850 | 2023 | 1473 | 1809 | 2294 | 1828 | 1711 | 1925 | 1125 | 1815 | 1875 | 1851 | 1892 | 1563 | 1633 | 1718 | 1587 | 1741,6 | 1858 | 1625 | 0,875 | -0,192 | -4,213 | 2,52E-05 | 8,60E-05 | 0,000506 | 0,0028131 | 0,0014192 | 0,0014192 | TRUE | 0,0184658 | No |
| orf19.851 | 2620 | 1976 | 2344 | 2936 | 3812 | 3786 | 4011 | 2491 | 2350 | 2516 | 2398 | 2421 | 3260 | 3612 | 3579 | 3513 | 2956,3 | 2421 | 3491 | 1,442 | 0,528 | 11,53 | 9,36E-31 | 3,45E-29 | 0,001219 | 0,0024706 | 0,0016662 | 0,0016662 | TRUE | 0,0250125 | No |
| orf19.852 | 633 | 464 | 677 | 814 | 875 | 783 | 854 | 640 | 568 | 591 | 693 | 671 | 748 | 747 | 762 | 903 | 710,33 | 631 | 790 | 1,248 | 0,319 | 3,706 | 0,00021052 | 0,00061102 | 0,007273 | 0,0040232 | 0,0057004 | 0,0057004 | TRUE | 0,1015197 | No |
| orf19.853 | 129 | 66 | 85 | 105 | 105 | 86 | 102 | 58 | 116 | 84 | 87 | 87 | 90 | 82 | 91 | 82 | 89,75 | 94 | 86 | 0,922 | -0,116 | -0,746 | 0,45575837 | 0,5415273 | 1,00E-08 | 0,0181549 | 0,0123205 | 0,0123205 | TRUE | 0,2662855 | No |
| orf19.854 | 3082 | 2121 | 2729 | 3340 | 3364 | 2845 | 3273 | 1984 | 2765 | 2700 | 2792 | 2754 | 2877 | 2715 | 2921 | 2798 | 2790,2 | 2753 | 2828 | 1,027 | 0,039 | 1,011 | 0,31215424 | 0,3984327 | 0,000288 | 0,0024998 | 0,0010658 | 0,0010658 | TRUE | 0,0092573 | No |
| orf19.855 | 16 | 21 | 22 | 26 | 42 | 28 | 29 | 16 | 14 | 27 | 23 | 21 | 36 | 27 | 26 | 23 | 24,51 | 21 | 28 | 1,339 | 0,421 | 1,437 | 0,15080301 | 0,21660258 | 1,00E-08 | 0,0611971 | 0,0409972 | 0,0409972 | TRUE | 0,2924885 | No |
| orf19.856 | 917 | 710 | 884 | 1075 | 1225 | 1129 | 1222 | 738 | 823 | 904 | 904 | 886 | 1048 | 1077 | 1090 | 1041 | 971,71 | 879 | 1064 | 1,213 | 0,278 | 5,198 | 2,01E-07 | 9,45E-07 | 1,00E-08 | 0,0034735 | 0,0017086 | 0,0017086 | TRUE | 0,023183 | No |
| orf19.857 | 1204 | 706 | 989 | 1181 | 1140 | 981 | 1060 | 671 | 1080 | 899 | 1012 | 974 | 975 | 936 | 946 | 946 | 970,99 | 991 | 951 | 0,958 | -0,062 | -1,008 | 0,31334347 | 0,39986567 | 0,001957 | 0,0034746 | 0,0026498 | 0,0026498 | TRUE | 0,0436282 | No |
| orf19.86 | 4146 | 2845 | 3384 | 4122 | 3072 | 2647 | 2765 | 1596 | 3719 | 3622 | 3462 | 3399 | 2627 | 2526 | 2467 | 2251 | 3009,2 | 3550 | 2468 | 0,696 | -0,522 | -9,642 | 5,30E-22 | 1,10E-20 | 0,002467 | 0,0024619 | 0,0024717 | 0,0024717 | TRUE | 0,0400154 | No |
| orf19.860 | 488 | 348 | 458 | 582 | 462 | 383 | 387 | 281 | 438 | 443 | 469 | 480 | 395 | 365 | 345 | 396 | 416,44 | 458 | 375 | 0,818 | -0,289 | -3,763 | 0,00016808 | 0,00049525 | 1,00E-08 | 0,0054655 | 0,0032558 | 0,0032558 | TRUE | 0,0351218 | No |

| | | | | | | | | | | | | | | | | | | | | | | | | | | | | | | | |
|---|---|---|---|---|---|---|---|---|---|---|---|---|---|---|---|---|---|---|---|---|---|---|---|---|---|---|---|---|---|---|---|
| orf19.861 | 592 | 385 | 498 | 577 | 751 | 596 | 589 | 434 | 531 | 490 | 510 | 476 | 642 | 569 | 526 | 612 | 544,4 | 502 | 587 | 1,169 | 0,225 | 2,867 | 0,00414447 | 0,00910086 | 0,003574 | 0,0046461 | 0,00408 | 0,00408 | TRUE | 0,0614566 | No |
| orf19.862 | 313 | 195 | 241 | 340 | 380 | 337 | 340 | 229 | 281 | 248 | 247 | 280 | 325 | 322 | 303 | 323 | 291,11 | 264 | 318 | 1,199 | 0,262 | 3,027 | 0,00247251 | 0,00572 | 1,00E-08 | 0,0069662 | 0,0037607 | 0,0037607 | TRUE | 0,0241941 | No |
| orf19.863 | 3000 | 2134 | 2639 | 3100 | 2798 | 2557 | 2854 | 1599 | 2691 | 2717 | 2700 | 2556 | 2393 | 2440 | 2547 | 2255 | 2537,4 | 2666 | 2409 | 0,905 | -0,144 | -3,072 | 0,00212468 | 0,00499611 | 0,001117 | 0,0025516 | 0,0017003 | 0,0017003 | TRUE | 0,0205607 | No |
| orf19.864 | 2280 | 1661 | 2068 | 2329 | 2896 | 2773 | 3084 | 1874 | 2045 | 2115 | 2116 | 1921 | 2477 | 2646 | 2752 | 2643 | 2339,3 | 2049 | 2630 | 1,284 | 0,361 | 7,179 | 7,02E-13 | 6,56E-12 | 0,001619 | 0,0026001 | 0,0019909 | 0,0019909 | TRUE | 0,0222683 | No |
| orf19.866 | 2356 | 1642 | 2127 | 2517 | 1467 | 1376 | 1473 | 895 | 2114 | 2090 | 2176 | 2076 | 1255 | 1313 | 1314 | 1262 | 1700 | 2114 | 1286 | 0,609 | -0,716 | -16,753 | 5,34E-63 | 9,72E-61 | 1,00E-08 | 0,0028335 | 0,0011267 | 0,0011267 | TRUE | 0,0047209 | No |
| orf19.867 | 846 | 564 | 712 | 904 | 971 | 828 | 917 | 526 | 759 | 718 | 728 | 745 | 830 | 790 | 818 | 742 | 766,44 | 738 | 795 | 1,08 | 0,111 | 1,912 | 0,05587042 | 0,09242001 | 1,00E-08 | 0,0038736 | 0,001942 | 0,001942 | TRUE | 0,0206847 | No |
| orf19.868 | 21172 | 14882 | 19046 | 23664 | 29071 | 25846 | 28612 | 19574 | 18993 | 18946 | 19486 | 19514 | 24862 | 24661 | 25531 | 27607 | 22450 | 19235 | 25665 | 1,334 | 0,416 | 9,999 | 1,54E-23 | 3,53E-22 | 0,001443 | 0,0020442 | 0,001615 | 0,001615 | TRUE | 0,03148 | No |
| orf19.87 | 368 | 247 | 312 | 367 | 212 | 196 | 186 | 112 | 330 | 314 | 319 | 303 | 181 | 187 | 166 | 158 | 244,84 | 316 | 173 | 0,55 | -0,864 | -9,146 | 5,90E-20 | 1,05E-18 | 1,00E-08 | 0,0079088 | 0,0041163 | 0,0041163 | TRUE | 0,0288665 | No |
| orf19.871 | 5561 | 3927 | 5173 | 6284 | 5993 | 5496 | 5894 | 3703 | 4989 | 4999 | 5292 | 5182 | 5125 | 5244 | 5259 | 5223 | 5164,3 | 5116 | 5213 | 1,019 | 0,027 | 0,796 | 0,42630152 | 0,51337333 | 0,000224 | 0,0022606 | 0,0008893 | 0,0008893 | TRUE | 0,0064724 | No |
| orf19.872 | 1434 | 919 | 1248 | 1543 | 1356 | 1165 | 1305 | 913 | 1286 | 1170 | 1277 | 1272 | 1160 | 1112 | 1164 | 1288 | 1216,1 | 1251 | 1181 | 0,94 | -0,09 | -1,541 | 0,12337727 | 0,1827541 | 0,001878 | 0,0031732 | 0,0024136 | 0,0024136 | TRUE | 0,0401328 | No |
| orf19.873 | 170 | 102 | 119 | 147 | 140 | 131 | 147 | 77 | 153 | 130 | 122 | 121 | 120 | 125 | 131 | 109 | 126,23 | 132 | 121 | 0,927 | -0,11 | -0,84 | 0,40081904 | 0,48947984 | 1,00E-08 | 0,01348 | 0,0085915 | 0,0085915 | TRUE | 0,1316283 | No |
| orf19.873.1 | 4532 | 3538 | 3883 | 4839 | 4854 | 4656 | 4764 | 2960 | 4066 | 4504 | 3973 | 3990 | 4151 | 4443 | 4251 | 4175 | 4194,1 | 4133 | 4255 | 1,03 | 0,043 | 0,877 | 0,38030027 | 0,4697862 | 0,001954 | 0,0023257 | 0,0020922 | 0,0020922 | TRUE | 0,043256 | No |
| orf19.874 | 1758 | 1140 | 1499 | 1875 | 2007 | 1636 | 1886 | 1177 | 1577 | 1451 | 1534 | 1546 | 1716 | 1561 | 1683 | 1660 | 1591,1 | 1527 | 1655 | 1,083 | 0,115 | 2,342 | 0,01920032 | 0,03584697 | 0,000827 | 0,0028919 | 0,0016724 | 0,0016724 | TRUE | 0,0183423 | No |
| orf19.875 | 1526 | 1100 | 1363 | 1604 | 1649 | 1588 | 1635 | 988 | 1369 | 1400 | 1394 | 1323 | 1410 | 1515 | 1459 | 1393 | 1408,1 | 1372 | 1444 | 1,055 | 0,078 | 1,627 | 0,10366328 | 0,15723529 | 0,000355 | 0,0030105 | 0,0014718 | 0,0014718 | TRUE | 0,0129491 | No |
| orf19.876 | 1291 | 758 | 1094 | 1305 | 1326 | 1025 | 1168 | 975 | 1158 | 965 | 1119 | 1076 | 1134 | 978 | 1042 | 1375 | 1106 | 1080 | 1132 | 1,045 | 0,064 | 0,653 | 0,51349223 | 0,59764846 | 0,012784 | 0,0032921 | 0,0083286 | 0,0083286 | TRUE | 0,2395818 | No |
| orf19.878 | 2382 | 1505 | 2094 | 2521 | 3098 | 2417 | 2554 | 1727 | 2137 | 1916 | 2142 | 2079 | 2650 | 2306 | 2279 | 2436 | 2243,1 | 2068 | 2418 | 1,168 | 0,224 | 3,718 | 0,00020099 | 0,00058589 | 0,003151 | 0,0026267 | 0,0030343 | 0,0030343 | TRUE | 0,0522966 | No |
| orf19.879 | 1195 | 877 | 1041 | 1275 | 971 | 879 | 992 | 658 | 1072 | 1117 | 1065 | 1051 | 830 | 839 | 885 | 928 | 973,42 | 1076 | 870 | 0,807 | -0,308 | -5,634 | 1,76E-08 | 9,47E-08 | 0,000309 | 0,0034709 | 0,0018364 | 0,0018364 | TRUE | 0,020276 | No |
| orf19.88 | 12235 | 7850 | 10478 | 12761 | 6833 | 5704 | 5885 | 3957 | 10976 | 9994 | 10720 | 10523 | 5844 | 5443 | 5251 | 5581 | 8041,5 | 10553 | 5530 | 0,524 | -0,933 | -20,578 | 4,32E-94 | 2,88E-91 | 0,001611 | 0,0021601 | 0,0018343 | 0,0018343 | TRUE | 0,0183406 | No |
| orf19.880 | 3291 | 2296 | 3047 | 3583 | 3606 | 3163 | 3394 | 2377 | 2952 | 2923 | 3117 | 2955 | 3084 | 3018 | 3029 | 3353 | 3053,8 | 2987 | 3121 | 1,043 | 0,061 | 1,335 | 0,18203191 | 0,25398228 | 0,001268 | 0,0024549 | 0,0016877 | 0,0016877 | TRUE | 0,0285754 | No |
| orf19.881 | 1043 | 701 | 952 | 1133 | 1011 | 892 | 925 | 640 | 936 | 892 | 974 | 934 | 865 | 851 | 825 | 903 | 897,53 | 934 | 861 | 0,919 | -0,122 | -2,238 | 0,0252179 | 0,04598153 | 1,00E-08 | 0,003597 | 0,0017443 | 0,0017443 | TRUE | 0,0108226 | No |
| orf19.882 | 7017 | 4610 | 6238 | 7490 | 11494 | 10177 | 11057 | 6781 | 6295 | 5869 | 6382 | 6176 | 9830 | 9711 | 9867 | 9564 | 7961,7 | 6180 | 9743 | 1,576 | 0,656 | 18,73 | 2,81E-78 | 7,03E-76 | 0,00058 | 0,0021619 | 0,001046 | 0,001046 | TRUE | 0,013481 | No |
| orf19.886 | 5238 | 3519 | 4719 | 5408 | 6730 | 5711 | 6645 | 4130 | 4699 | 4480 | 4828 | 4460 | 5756 | 5449 | 5930 | 5825 | 5178,3 | 4617 | 5740 | 1,243 | 0,314 | 7,337 | 2,19E-13 | 2,16E-12 | 0,001128 | 0,0022599 | 0,0015616 | 0,0015616 | TRUE | 0,0142921 | No |
| orf19.889 | 562 | 417 | 607 | 600 | 428 | 399 | 399 | 295 | 504 | 531 | 621 | 495 | 366 | 381 | 356 | 416 | 458,71 | 538 | 380 | 0,704 | -0,505 | -5,618 | 1,94E-08 | 1,03E-07 | 0,005859 | 0,0051442 | 0,005512 | 0,005512 | TRUE | 0,1284959 | No |
| orf19.89 | 2686 | 1787 | 2322 | 2691 | 1715 | 1387 | 1494 | 1020 | 2410 | 2275 | 2376 | 2219 | 1467 | 1323 | 1333 | 1439 | 1855,2 | 2320 | 1390 | 0,599 | -0,74 | -14,201 | 9,05E-46 | 7,45E-44 | 0,001525 | 0,002762 | 0,0020252 | 0,0020252 | TRUE | 0,0181952 | No |
| orf19.891 | 6098 | 4380 | 5640 | 6727 | 5248 | 4880 | 5190 | 3338 | 5471 | 5576 | 5770 | 5547 | 4488 | 4656 | 4631 | 4708 | 5106 | 5591 | 4621 | 0,826 | -0,275 | -8,205 | 2,31E-16 | 3,01E-15 | 0,000284 | 0,0022638 | 0,0008835 | 0,0008835 | TRUE | 0,0057028 | No |
| orf19.893 | 45 | 33 | 36 | 48 | 40 | 44 | 40 | 36 | 40 | 42 | 37 | 40 | 34 | 42 | 36 | 51 | 40,18 | 40 | 41 | 1,009 | 0,013 | 0,059 | 0,95308913 | 0,96659106 | 1,00E-08 | 0,0381077 | 0,0209041 | 0,0209041 | TRUE | 0,1563068 | No |
| orf19.894 | 613 | 450 | 584 | 681 | 598 | 494 | 522 | 324 | 550 | 573 | 597 | 562 | 511 | 471 | 466 | 457 | 523,43 | 570 | 476 | 0,838 | -0,255 | -3,764 | 0,00016721 | 0,00049293 | 1,00E-08 | 0,004753 | 0,0024827 | 0,0024827 | TRUE | 0,0301163 | No |
| orf19.895 | 4998 | 3444 | 4531 | 5344 | 5031 | 4450 | 4769 | 3210 | 4484 | 4385 | 4636 | 4407 | 4303 | 4246 | 4256 | 4527 | 4405,3 | 4478 | 4333 | 0,967 | -0,049 | -1,318 | 0,1874708 | 0,26012114 | 0,000483 | 0,0023091 | 0,0010911 | 0,0010911 | TRUE | 0,0104032 | No |
| orf19.896 | 3724 | 2662 | 3599 | 4125 | 4679 | 4308 | 4739 | 2811 | 3341 | 3389 | 3682 | 3402 | 4002 | 4111 | 4229 | 3965 | 3764,9 | 3454 | 4077 | 1,181 | 0,24 | 5,503 | 3,74E-08 | 1,93E-07 | 0,001073 | 0,0023651 | 0,0015587 | 0,0015587 | TRUE | 0,0241376 | No |
| orf19.897 | 1917 | 1227 | 1648 | 2047 | 2370 | 1987 | 2142 | 1511 | 1720 | 1562 | 1686 | 1688 | 2027 | 1896 | 1911 | 2131 | 1827,7 | 1664 | 1991 | 1,194 | 0,255 | 4,807 | 1,53E-06 | 6,28E-06 | 0,001666 | 0,0027738 | 0,0021541 | 0,0021541 | TRUE | 0,0250211 | No |
| orf19.898 | 1099 | 797 | 966 | 1209 | 1106 | 860 | 934 | 682 | 986 | 1015 | 988 | 997 | 946 | 821 | 833 | 962 | 943,46 | 996 | 890 | 0,892 | -0,165 | -2,584 | 0,00977111 | 0,01972609 | 0,002096 | 0,0035182 | 0,0028437 | 0,0028437 | TRUE | 0,0325366 | No |
| orf19.899 | 740 | 488 | 621 | 766 | 680 | 620 | 578 | 406 | 664 | 621 | 635 | 632 | 582 | 592 | 516 | 573 | 601,71 | 638 | 566 | 0,885 | -0,177 | -2,671 | 0,0075612 | 0,01565933 | 1,00E-08 | 0,0043922 | 0,0025385 | 0,0025385 | TRUE | 0,0439733 | No |
| orf19.9 | 110 | 108 | 103 | 150 | 128 | 119 | 108 | 73 | 99 | 137 | 105 | 124 | 109 | 114 | 96 | 103 | 110,95 | 116 | 106 | 0,913 | -0,131 | -0,909 | 0,36339543 | 0,45204566 | 0,005059 | 0,0150639 | 0,0108491 | 0,0108491 | TRUE | 0,1275931 | No |
| orf19.90 | 1453 | 1043 | 1385 | 1695 | 1234 | 1148 | 1274 | 751 | 1303 | 1328 | 1417 | 1398 | 1055 | 1095 | 1137 | 1059 | 1224,1 | 1362 | 1086 | 0,799 | -0,325 | -6,274 | 3,53E-10 | 2,39E-09 | 0,000682 | 0,0031655 | 0,0017413 | 0,0017413 | TRUE | 0,011945 | No |
| orf19.900 | 4506 | 2874 | 3957 | 4819 | 5177 | 4131 | 4581 | 3143 | 4042 | 3659 | 4048 | 3974 | 4428 | 3942 | 4088 | 4433 | 4076,7 | 3931 | 4223 | 1,073 | 0,102 | 1,913 | 0,05571755 | 0,09224334 | 0,002487 | 0,0023356 | 0,0024759 | 0,0024759 | TRUE | 0,0256715 | No |
| orf19.903 | 10195 | 6535 | 8528 | 10447 | 13840 | 11224 | 11723 | 8984 | 9146 | 8320 | 8725 | 8615 | 11836 | 10710 | 10461 | 12671 | 10060 | 8702 | 11420 | 1,312 | 0,392 | 6,269 | 3,63E-10 | 2,45E-09 | 0,004682 | 0,0021238 | 0,0036461 | 0,0036461 | TRUE | 0,0659519 | No |
| orf19.904 | 912 | 706 | 746 | 983 | 1222 | 1124 | 1165 | 747 | 818 | 899 | 763 | 811 | 1045 | 1072 | 1040 | 1054 | 937,69 | 823 | 1053 | 1,282 | 0,359 | 6,199 | 5,68E-10 | 3,75E-09 | 0,000703 | 0,0035277 | 0,0021293 | 0,0021293 | TRUE | 0,04121 | No |
| orf19.905 | 2019 | 1468 | 1829 | 2322 | 2095 | 1813 | 1982 | 1259 | 1811 | 1869 | 1871 | 1915 | 1792 | 1730 | 1769 | 1776 | 1816,5 | 1866 | 1767 | 0,946 | -0,08 | -1,963 | 0,04967232 | 0,0837011 | 1,00E-08 | 0,0027787 | 0,001032 | 0,001032 | TRUE | 0,0051371 | No |
| orf19.906 | 4434 | 3168 | 4017 | 4889 | 5158 | 4870 | 5354 | 3189 | 3978 | 4033 | 4110 | 4032 | 4411 | 4647 | 4778 | 4498 | 4310,7 | 4038 | 4584 | 1,135 | 0,183 | 4,918 | 8,75E-07 | 3,72E-06 | 0,000536 | 0,0023163 | 0,0010997 | 0,0010997 | TRUE | 0,0101416 | No |
| orf19.908 | 4891 | 3916 | 4346 | 5453 | 4697 | 4526 | 4748 | 2804 | 4388 | 4985 | 4446 | 4497 | 4017 | 4319 | 4237 | 3955 | 4355,4 | 4579 | 4132 | 0,903 | -0,147 | -2,841 | 0,0045009 | 0,00981184 | 0,002374 | 0,0023128 | 0,0023305 | 0,0023305 | TRUE | 0,0425174 | No |
| orf19.909 | 11044 | 7925 | 9993 | 12576 | 12697 | 11384 | 12262 | 7703 | 9908 | 10089 | 10224 | 10370 | 10859 | 10862 | 10942 | 10864 | 10515 | 10148 | 10882 | 1,072 | 0,101 | 3,733 | 0,00018944 | 0,00055438 | 0,000117 | 0,0021176 | 0,0006025 | 0,0006025 | TRUE | 0,0032078 | No |
| orf19.909.1 | 894 | 637 | 773 | 922 | 887 | 755 | 739 | 567 | 802 | 811 | 791 | 760 | 759 | 720 | 659 | 800 | 762,78 | 791 | 734 | 0,927 | -0,11 | -1,638 | 0,10142462 | 0,15449886 | 0,002228 | 0,0038827 | 0,0030192 | 0,0030192 | TRUE | 0,0568775 | No |
| orf19.91 | 1022 | 804 | 884 | 1178 | 1132 | 1030 | 1110 | 664 | 917 | 1024 | 904 | 971 | 968 | 983 | 990 | 937 | 961,77 | 954 | 970 | 1,019 | 0,027 | 0,483 | 0,62879475 | 0,70045065 | 0,000561 | 0,0034889 | 0,0019979 | 0,0019979 | TRUE | 0,0257063 | No |
| orf19.910 | 4814 | 3357 | 4235 | 5211 | 4263 | 4033 | 4210 | 2780 | 4319 | 4274 | 4333 | 4297 | 3646 | 3848 | 3757 | 3921 | 4049,3 | 4306 | 3793 | 0,88 | -0,185 | -5,294 | 1,20E-07 | 5,80E-07 | 0,000178 | 0,002338 | 0,0009176 | 0,0009176 | TRUE | 0,0083782 | No |
| orf19.913.2 | 3047 | 2064 | 2555 | 3189 | 2776 | 2468 | 2610 | 1578 | 2733 | 2628 | 2614 | 2630 | 2374 | 2355 | 2329 | 2226 | 2486,1 | 2651 | 2321 | 0,876 | -0,19 | -4,88 | 1,06E-06 | 4,45E-06 | 0,000145 | 0,0025635 | 0,0010551 | 0,0010551 | TRUE | 0,0082631 | No |
| orf19.915 | 789 | 775 | 784 | 937 | 1236 | 1418 | 1373 | 946 | 708 | 987 | 802 | 773 | 1057 | 1353 | 1225 | 1334 | 1029,8 | 818 | 1242 | 1,522 | 0,606 | 5,826 | 5,67E-09 | 3,26E-08 | 0,015322 | 0,0033892 | 0,00937 | 0,00937 | TRUE | 0,2238868 | No |
| orf19.916 | 3627 | 2952 | 3237 | 4169 | 3922 | 4065 | 4182 | 2162 | 3254 | 3758 | 3312 | 3438 | 3354 | 3879 | 3732 | 3049 | 3471,9 | 3440 | 3504 | 1,019 | 0,028 | 0,367 | 0,71394479 | 0,77439674 | 0,006602 | 0,0023976 | 0,0052568 | 0,0052568 | TRUE | 0,0903789 | No |
| orf19.918 | 3557 | 2465 | 3291 | 4096 | 3911 | 3299 | 3748 | 2269 | 3191 | 3138 | 3367 | 3378 | 3345 | 3148 | 3344 | 3200 | 3263,9 | 3268 | 3259 | 0,997 | -0,004 | -0,097 | 0,92288994 | 0,94362199 | 0,00087 | 0,0024243 | 0,0014194 | 0,0014194 | TRUE | 0,0083124 | No |
| orf19.92 | 1962 | 1314 | 1674 | 1965 | 2071 | 1863 | 2096 | 1236 | 1760 | 1673 | 1713 | 1620 | 1771 | 1778 | 1870 | 1743 | 1741,1 | 1692 | 1790 | 1,06 | 0,084 | 1,811 | 0,07009136 | 0,11292811 | 0,000598 | 0,0028133 | 0,0014646 | 0,0014646 | TRUE | 0,0110558 | No |
| orf19.920 | 3024 | 2318 | 2661 | 3394 | 2694 | 2665 | 2678 | 1653 | 2713 | 2951 | 2722 | 2799 | 2304 | 2543 | 2390 | 2331 | 2594,1 | 2796 | 2392 | 0,856 | -0,224 | -4,704 | 2,55E-06 | 1,02E-05 | 0,001254 | 0,0025391 | 0,001783 | 0,001783 | TRUE | 0,0220918 | No |
| orf19.921 | 4224 | 3246 | 3722 | 4874 | 5304 | 5323 | 5801 | 3505 | 3789 | 4133 | 3808 | 4019 | 4536 | 5079 | 5176 | 4943 | 4435,5 | 3937 | 4934 | 1,253 | 0,326 | 6,284 | 3,30E-10 | 2,25E-09 | 0,002223 | 0,0023068 | 0,0023515 | 0,0023515 | TRUE | 0,0366496 | No |
| orf19.922 | 3075 | 2575 | 3040 | 3686 | 2653 | 2655 | 2782 | 1663 | 2759 | 3278 | 3110 | 3040 | 2269 | 2533 | 2482 | 2346 | 2727,1 | 3047 | 2408 | 0,791 | -0,338 | -5,61 | 2,02E-08 | 1,08E-07 | 0,00331 | 0,0025118 | 0,0031164 | 0,0031164 | TRUE | 0,0494246 | No |
| orf19.923 | 607 | 476 | 539 | 681 | 775 | 674 | 758 | 515 | 545 | 606 | 551 | 562 | 663 | 643 | 676 | 726 | 621,53 | 566 | 677 | 1,196 | 0,258 | 3,953 | 7,71E-05 | 0,00023948 | 0,000407 | 0,0043152 | 0,0024661 | 0,0024661 | TRUE | 0,0242807 | No |
| orf19.925 | 425 | 411 | 439 | 515 | 527 | 652 | 670 | 354 | 381 | 523 | 449 | 425 | 451 | 622 | 598 | 499 | 493,54 | 444 | 542 | 1,225 | 0,292 | 2,459 | 0,01394945 | 0,02706986 | 0,017376 | 0,0049209 | 0,011528 | 0,011528 | TRUE | 0,1603407 | No |
| orf19.926 | 1270 | 1063 | 1204 | 1437 | 1310 | 1151 | 1244 | 787 | 1139 | 1353 | 1232 | 1185 | 1120 | 1098 | 1110 | 1110 | 1168,5 | 1227 | 1110 | 0,906 | -0,142 | -2,407 | 0,01608457 | 0,03069761 | 0,001703 | 0,0032219 | 0,0024666 | 0,0024666 | TRUE | 0,0539466 | No |
| orf19.927 | 142 | 159 | 132 | 202 | 215 | 242 | 219 | 144 | 127 | 202 | 135 | 167 | 184 | 231 | 195 | 203 | 180,59 | 158 | 203 | 1,295 | 0,373 | 2,561 | 0,01044748 | 0,02092979 | 0,020823 | 0,0100181 | 0,0146738 | 0,0146738 | TRUE | 0,3403611 | No |
| orf19.928 | 767 | 523 | 642 | 840 | 1209 | 1051 | 1168 | 814 | 688 | 666 | 657 | 693 | 1034 | 1003 | 1042 | 1148 | 866,32 | 676 | 1057 | 1,557 | 0,639 | 10,739 | 6,70E-27 | 2,05E-25 | 0,000889 | 0,0036552 | 0,0021879 | 0,0021879 | TRUE | 0,0359437 | No |
| orf19.929 | 96 | 65 | 72 | 113 | 102 | 102 | 111 | 73 | 86 | 83 | 74 | 93 | 87 | 97 | 99 | 103 | 90,29 | 84 | 96 | 1,142 | 0,191 | 1,298 | 0,19438799 | 0,26787477 | 1,00E-08 | 0,0180585 | 0,0097451 | 0,0097451 | TRUE | 0,0655656 | No |
| orf19.93 | 5431 | 3229 | 4337 | 5530 | 5490 | 4324 | 4780 | 3957 | 4872 | 4111 | 4437 | 4560 | 4695 | 4126 | 4265 | 5581 | 4581 | 4495 | 4667 | 1,037 | 0,053 | 0,599 | 0,54921033 | 0,63030672 | 0,010989 | 0,0022964 | 0,0072347 | 0,0072347 | TRUE | 0,2094448 | No |

| | | | | | | | | | | | | | | | | | | | | | | | | | | | | | | | |
|---|---|---|---|---|---|---|---|---|---|---|---|---|---|---|---|---|---|---|---|---|---|---|---|---|---|---|---|---|---|---|---|
| orf19.930 | 73279 | 46176 | 62542 | 76887 | 64742 | 53137 | 56479 | 37722 | 65739 | 58787 | 63987 | 63402 | 55369 | 50702 | 50398 | 53204 | 57698 | 62979 | 52418 | 0,832 | -0,265 | -5,687 | 1,29E-08 | 7,06E-08 | 0,002077 | 0,0020047 | 0,0020668 | 0,0020668 | TRUE | 0,0245446 | No |
| orf19.931 | 1038 | 776 | 922 | 1065 | 1163 | 1141 | 1110 | 719 | 931 | 988 | 943 | 878 | 995 | 1089 | 990 | 1014 | 978,57 | 935 | 1022 | 1,095 | 0,131 | 2,256 | 0,02404959 | 0,04403839 | 0,001199 | 0,003463 | 0,0022086 | 0,0022086 | TRUE | 0,0239666 | No |
| orf19.932 | 4941 | 3379 | 4330 | 5112 | 6300 | 5670 | 6128 | 3893 | 4433 | 4302 | 4430 | 4215 | 5388 | 5410 | 5468 | 5491 | 4892,1 | 4345 | 5439 | 1,252 | 0,324 | 10,077 | 7,01E-24 | 1,64E-22 | 0,000133 | 0,0022763 | 0,0007839 | 0,0007839 | TRUE | 0,0052358 | No |
| orf19.933 | 4726 | 3376 | 3994 | 4965 | 4326 | 3750 | 4215 | 2403 | 4240 | 4298 | 4086 | 4094 | 3700 | 3578 | 3761 | 3389 | 3893,3 | 4180 | 3607 | 0,864 | -0,21 | -4,926 | 8,40E-07 | 3,59E-06 | 0,00101 | 0,0023524 | 0,0014902 | 0,0014902 | TRUE | 0,0189181 | No |
| orf19.934 | 955 | 623 | 735 | 953 | 832 | 764 | 793 | 509 | 857 | 793 | 752 | 786 | 712 | 729 | 708 | 718 | 756,72 | 797 | 717 | 0,898 | -0,155 | -2,596 | 0,00942649 | 0,01911385 | 1,00E-08 | 0,0038979 | 0,0020907 | 0,0020907 | TRUE | 0,0317123 | No |
| orf19.935 | 42 | 49 | 65 | 63 | 24 | 12 | 15 | 24 | 38 | 62 | 67 | 52 | 21 | 11 | 13 | 34 | 37,22 | 55 | 20 | 0,352 | -1,508 | -4,785 | 1,71E-06 | 6,96E-06 | 0,090995 | 0,0409876 | 0,0598603 | 0,0598603 | TRUE | 0,8108122 | No |
| orf19.936 | 6 | 6 | 10 | 14 | 8 | 9 | 7 | 4 | 5 | 8 | 10 | 12 | 7 | 9 | 6 | 6 | 7,76 | 9 | 7 | 0,787 | -0,345 | -0,703 | 0,48207894 | NA | 1,00E-08 | 0,1889515 | 0,1025179 | 0,1025179 | TRUE | 0,2517532 | No |
| orf19.937 | 6 | 1 | 3 | 3 | 7 | 11 | 4 | 2 | 5 | 1 | 3 | 2 | 6 | 10 | 4 | 3 | 4,38 | 3 | 6 | 1,862 | 0,897 | 1,265 | 0,20569708 | NA | 0,107268 | 0,3331205 | 0,2379884 | 0,2379884 | TRUE | 0,7279268 | No |
| orf19.938 | 293 | 213 | 290 | 333 | 358 | 316 | 362 | 228 | 263 | 271 | 297 | 275 | 306 | 302 | 323 | 322 | 294,7 | 276 | 313 | 1,132 | 0,179 | 2,134 | 0,03281102 | 0,05799606 | 1,00E-08 | 0,0069055 | 0,0033478 | 0,0033478 | TRUE | 0,0272354 | No |
| orf19.939 | 7162 | 5398 | 6697 | 8215 | 7891 | 7226 | 7977 | 4994 | 6425 | 6872 | 6852 | 6774 | 6749 | 6895 | 7118 | 7044 | 6841,1 | 6731 | 6952 | 1,033 | 0,047 | 1,292 | 0,19651259 | 0,27048056 | 0,000561 | 0,0021917 | 0,0011036 | 0,0011036 | TRUE | 0,0115254 | No |
| orf19.94 | 4565 | 3095 | 3764 | 4767 | 4167 | 3740 | 3905 | 2130 | 4095 | 3940 | 3851 | 3931 | 3564 | 3569 | 3485 | 3004 | 3679,8 | 3954 | 3406 | 0,862 | -0,214 | -3,747 | 0,00017883 | 0,0005254 | 0,003116 | 0,002374 | 0,0028525 | 0,0028525 | TRUE | 0,0734295 | No |
| orf19.940 | 3043 | 2624 | 2882 | 3684 | 3815 | 4108 | 4219 | 2433 | 2730 | 3341 | 2949 | 3038 | 3263 | 3920 | 3765 | 3432 | 3304,5 | 3014 | 3595 | 1,194 | 0,255 | 3,487 | 0,00048861 | 0,00132292 | 0,006621 | 0,0024188 | 0,004838 | 0,004838 | TRUE | 0,0634072 | No |
| orf19.941 | 23142 | 16823 | 20091 | 25231 | 24587 | 23159 | 23541 | 14967 | 20761 | 21417 | 20555 | 20806 | 21028 | 22098 | 21006 | 21110 | 21098 | 20885 | 21310 | 1,021 | 0,029 | 0,975 | 0,3297637 | 0,41692346 | 0,0004 | 0,0020483 | 0,00082 | 0,00082 | TRUE | 0,0076113 | No |
| orf19.942 | 101 | 82 | 90 | 104 | 96 | 86 | 107 | 66 | 91 | 104 | 92 | 86 | 82 | 82 | 95 | 93 | 90,7 | 93 | 88 | 0,949 | -0,076 | -0,517 | 0,60507176 | 0,68076238 | 1,00E-08 | 0,0179857 | 0,0095027 | 0,0095027 | TRUE | 0,0531302 | No |
| orf19.943 | 3940 | 3172 | 3933 | 4706 | 3300 | 3148 | 3234 | 2042 | 3535 | 4038 | 4024 | 3881 | 2822 | 3004 | 2886 | 2880 | 3383,7 | 3870 | 2898 | 0,749 | -0,416 | -8,224 | 1,97E-16 | 2,60E-15 | 0,001775 | 0,0024086 | 0,0021562 | 0,0021562 | TRUE | 0,0416255 | No |
| orf19.944 | 2976 | 2223 | 2520 | 3141 | 3359 | 3230 | 3471 | 2090 | 2670 | 2830 | 2578 | 2590 | 2873 | 3082 | 3097 | 2948 | 2833,5 | 2667 | 3000 | 1,126 | 0,171 | 3,723 | 0,00019649 | 0,00057362 | 0,001125 | 0,0024919 | 0,001677 | 0,001677 | TRUE | 0,0195502 | No |
| orf19.945 | 1306 | 785 | 1074 | 1347 | 1239 | 1035 | 1109 | 706 | 1172 | 999 | 1099 | 1111 | 1060 | 988 | 990 | 996 | 1051,6 | 1095 | 1008 | 0,919 | -0,122 | -2,041 | 0,04124528 | 0,07090175 | 0,001714 | 0,0033599 | 0,0024537 | 0,0024537 | TRUE | 0,0383573 | No |
| orf19.946 | 2373 | 1640 | 1982 | 2499 | 1787 | 1564 | 1554 | 970 | 2129 | 2088 | 2028 | 2061 | 1528 | 1492 | 1387 | 1368 | 1760,1 | 2076 | 1444 | 0,697 | -0,522 | -10,634 | 2,08E-26 | 6,17E-25 | 0,000907 | 0,0028043 | 0,00172 | 0,00172 | TRUE | 0,0189515 | No |
| orf19.947 | 1516 | 955 | 1160 | 1586 | 1690 | 1376 | 1491 | 875 | 1360 | 1216 | 1187 | 1308 | 1445 | 1313 | 1330 | 1234 | 1299,2 | 1268 | 1330 | 1,05 | 0,071 | 1,1 | 0,27114404 | 0,35552889 | 0,003127 | 0,0030969 | 0,0032245 | 0,0032245 | TRUE | 0,0407566 | No |
| orf19.948 | 3 | 1 | 1 | 2 | 1 | 1 | 1 | 4 | 3 | 1 | 1 | 2 | 1 | 1 | 1 | 6 | 1,87 | 2 | 2 | 1,115 | 0,157 | 0,149 | 0,88165594 | NA | 0,151375 | 0,7772476 | 0,5070468 | 0,5070468 | TRUE | 0,833042 | No |
| orf19.949 | 840 | 656 | 822 | 954 | 918 | 901 | 955 | 556 | 754 | 835 | 841 | 787 | 785 | 860 | 852 | 784 | 812,2 | 804 | 820 | 1,023 | 0,033 | 0,533 | 0,59426907 | 0,67156469 | 0,001401 | 0,0037669 | 0,0025033 | 0,0025033 | TRUE | 0,0213833 | No |
| orf19.95 | 125 | 77 | 114 | 156 | 81 | 75 | 100 | 54 | 112 | 98 | 117 | 129 | 69 | 72 | 89 | 76 | 95,21 | 114 | 76 | 0,669 | -0,581 | -3,818 | 0,00013431 | 0,00040126 | 1,00E-08 | 0,017227 | 0,0113497 | 0,0113497 | TRUE | 0,1303014 | No |
| orf19.951 | 8182 | 6272 | 7370 | 9149 | 9358 | 9665 | 10208 | 6020 | 7340 | 7985 | 7540 | 7544 | 8003 | 9222 | 9109 | 8491 | 8154,3 | 7602 | 8706 | 1,146 | 0,196 | 3,755 | 0,00017365 | 0,00051143 | 0,002607 | 0,0021576 | 0,0024942 | 0,0024942 | TRUE | 0,0366426 | No |
| orf19.952 | 457 | 341 | 439 | 510 | 635 | 567 | 679 | 442 | 410 | 434 | 449 | 421 | 543 | 541 | 606 | 623 | 503,4 | 428 | 578 | 1,347 | 0,43 | 5,789 | 7,09E-09 | 4,01E-08 | 0,001705 | 0,0048633 | 0,0032712 | 0,0032712 | TRUE | 0,0280407 | No |
| orf19.953.1 | 1617 | 1156 | 1441 | 1795 | 1617 | 1375 | 1494 | 1010 | 1451 | 1472 | 1474 | 1480 | 1383 | 1312 | 1333 | 1425 | 1416,2 | 1469 | 1363 | 0,926 | -0,11 | -2,442 | 0,01461193 | 0,02820059 | 1,00E-08 | 0,0030046 | 0,001256 | 0,001256 | TRUE | 0,0095045 | No |
| orf19.954 | 2884 | 1789 | 2395 | 2886 | 2724 | 2284 | 2516 | 1543 | 2587 | 2278 | 2450 | 2380 | 2330 | 2179 | 2245 | 2176 | 2328,2 | 2424 | 2232 | 0,921 | -0,119 | -2,39 | 0,01685797 | 0,03197054 | 0,001387 | 0,0026031 | 0,0019358 | 0,0019358 | TRUE | 0,0251369 | No |
| orf19.955 | 1817 | 1272 | 1471 | 1869 | 2170 | 1918 | 1944 | 1298 | 1630 | 1619 | 1505 | 1541 | 1856 | 1830 | 1735 | 1831 | 1693,4 | 1574 | 1813 | 1,152 | 0,204 | 4,36 | 1,30E-05 | 4,66E-05 | 0,000621 | 0,0028368 | 0,00151 | 0,00151 | TRUE | 0,0106445 | No |
| orf19.956 | 674 | 529 | 658 | 738 | 991 | 923 | 1028 | 611 | 605 | 673 | 673 | 609 | 848 | 881 | 917 | 862 | 758,4 | 640 | 877 | 1,375 | 0,459 | 7,357 | 1,88E-13 | 1,86E-12 | 0,000986 | 0,0038937 | 0,0023883 | 0,0023883 | TRUE | 0,0158289 | No |
| orf19.96 | 13200 | 9906 | 12033 | 14626 | 11272 | 11367 | 11968 | 6858 | 11842 | 12611 | 12311 | 12061 | 9640 | 10846 | 10679 | 9673 | 11208 | 12206 | 10210 | 0,837 | -0,257 | -5,273 | 1,34E-07 | 6,48E-07 | 0,002228 | 0,002109 | 0,0021948 | 0,0021948 | TRUE | 0,0214863 | No |
| orf19.961.2 | 1350 | 854 | 1073 | 1345 | 1284 | 1148 | 1106 | 664 | 1211 | 1087 | 1098 | 1109 | 1098 | 1095 | 987 | 937 | 1077,8 | 1126 | 1029 | 0,915 | -0,128 | -1,924 | 0,05434335 | 0,09036669 | 0,00312 | 0,0033265 | 0,0032948 | 0,0032948 | TRUE | 0,0398058 | No |
| orf19.962 | 5598 | 3409 | 4224 | 5549 | 4907 | 3809 | 4174 | 2530 | 5022 | 4340 | 4322 | 4576 | 4197 | 3634 | 3725 | 3568 | 4172,9 | 4565 | 3781 | 0,828 | -0,271 | -4,142 | 3,44E-05 | 0,00011457 | 0,005056 | 0,0023274 | 0,0038824 | 0,0038824 | TRUE | 0,0672073 | No |
| orf19.964 | 3820 | 2692 | 3462 | 3982 | 4202 | 3877 | 4125 | 2474 | 3427 | 3427 | 3542 | 3284 | 3594 | 3699 | 3681 | 3489 | 3517,9 | 3420 | 3616 | 1,058 | 0,082 | 2,102 | 0,03552756 | 0,06232104 | 0,000452 | 0,0023922 | 0,0011757 | 0,0011757 | TRUE | 0,0091249 | No |
| orf19.966 | 3914 | 2511 | 3408 | 4250 | 3866 | 2964 | 3206 | 2233 | 3511 | 3197 | 3487 | 3505 | 3306 | 2828 | 2861 | 3149 | 3230,5 | 3425 | 3036 | 0,886 | -0,175 | -2,929 | 0,00340012 | 0,00761101 | 0,003466 | 0,0024289 | 0,0031116 | 0,0031116 | TRUE | 0,0451639 | No |
| orf19.967 | 2510 | 1531 | 2153 | 2580 | 2165 | 1721 | 1887 | 1308 | 2252 | 1949 | 2203 | 2128 | 1852 | 1642 | 1684 | 1845 | 1944,2 | 2133 | 1756 | 0,822 | -0,283 | -4,637 | 3,54E-06 | 1,38E-05 | 0,003146 | 0,0027262 | 0,0030583 | 0,0030583 | TRUE | 0,0390123 | No |
| orf19.968 | 18341 | 11160 | 14898 | 18524 | 16506 | 12580 | 13337 | 9678 | 16454 | 14208 | 15242 | 15275 | 14116 | 12003 | 11901 | 13650 | 14106 | 15295 | 12918 | 0,844 | -0,244 | -3,691 | 0,00022351 | 0,0006456 | 0,005544 | 0,0020824 | 0,0041262 | 0,0041262 | TRUE | 0,0481259 | No |
| orf19.969 | 12727 | 8356 | 11258 | 13976 | 12714 | 10275 | 11122 | 7444 | 11417 | 10638 | 11518 | 11525 | 10873 | 9804 | 9925 | 10499 | 10775 | 11274 | 10275 | 0,911 | -0,134 | -2,934 | 0,00334648 | 0,00751095 | 0,001782 | 0,0021143 | 0,0019196 | 0,0019196 | TRUE | 0,0190973 | No |
| orf19.97 | 64944 | 45430 | 58339 | 71131 | 48269 | 43659 | 46590 | 27529 | 58262 | 57837 | 59687 | 58656 | 41281 | 41658 | 41574 | 38827 | 49723 | 58610 | 40835 | 0,697 | -0,521 | -16,186 | 6,34E-59 | 1,03E-56 | 0,000609 | 0,0020087 | 0,000975 | 0,000975 | TRUE | 0,0132898 | No |
| orf19.970 | 708 | 476 | 609 | 730 | 764 | 664 | 694 | 463 | 635 | 606 | 623 | 602 | 653 | 634 | 619 | 653 | 628,18 | 616 | 640 | 1,036 | 0,052 | 0,869 | 0,38464236 | 0,47355149 | 1,00E-08 | 0,0042905 | 0,0018025 | 0,0018025 | TRUE | 0,005734 | No |
| orf19.971 | 403 | 260 | 372 | 478 | 547 | 485 | 533 | 336 | 362 | 331 | 381 | 394 | 468 | 463 | 476 | 474 | 418,42 | 367 | 470 | 1,276 | 0,351 | 4,698 | 2,63E-06 | 1,05E-05 | 1,00E-08 | 0,005449 | 0,0029484 | 0,0029484 | TRUE | 0,0428508 | No |
| orf19.972 | 265 | 162 | 232 | 283 | 338 | 247 | 270 | 224 | 238 | 206 | 237 | 233 | 289 | 236 | 241 | 316 | 249,54 | 228 | 270 | 1,173 | 0,23 | 2,062 | 0,03921092 | 0,06787069 | 0,007973 | 0,0077971 | 0,007878 | 0,007878 | TRUE | 0,1238336 | No |
| orf19.974 | 630 | 385 | 546 | 650 | 638 | 545 | 659 | 353 | 565 | 490 | 559 | 536 | 546 | 520 | 588 | 498 | 537,69 | 538 | 538 | 1,002 | 0,003 | 0,034 | 0,97261797 | 0,9809449 | 0,002394 | 0,0046794 | 0,0035589 | 0,0035589 | TRUE | 0,0460515 | No |
| orf19.976 | 10869 | 7436 | 9650 | 11438 | 12047 | 10710 | 11657 | 7107 | 9751 | 9467 | 9873 | 9432 | 10303 | 10219 | 10402 | 10024 | 9933,8 | 9631 | 10237 | 1,063 | 0,088 | 2,957 | 0,00310822 | 0,00703094 | 0,000239 | 0,0021257 | 0,0007584 | 0,0007584 | TRUE | 0,0034798 | No |
| orf19.978 | 4788 | 3210 | 4225 | 5012 | 4798 | 4030 | 4590 | 2919 | 4295 | 4087 | 4323 | 4133 | 4103 | 3845 | 4096 | 4117 | 4124,9 | 4210 | 4040 | 0,959 | -0,06 | -1,53 | 0,1261177 | 0,18626233 | 0,000584 | 0,0023315 | 0,0012243 | 0,0012243 | TRUE | 0,0129918 | No |
| orf19.979 | 8848 | 5899 | 8030 | 9374 | 6339 | 5623 | 5955 | 4092 | 7938 | 7510 | 8215 | 7730 | 5421 | 5365 | 5314 | 5771 | 6658,1 | 7848 | 5468 | 0,696 | -0,523 | -12,383 | 3,24E-35 | 1,64E-33 | 0,001244 | 0,0021976 | 0,0015544 | 0,0015544 | TRUE | 0,016351 | No |
| orf19.980 | 6555 | 4335 | 6112 | 7115 | 7867 | 6686 | 7536 | 4950 | 5881 | 5519 | 6253 | 5867 | 6728 | 6380 | 6725 | 6982 | 6291,7 | 5880 | 6704 | 1,139 | 0,188 | 4,036 | 5,43E-05 | 0,0001746 | 0,001768 | 0,0022103 | 0,001928 | 0,001928 | TRUE | 0,0220066 | No |
| orf19.984 | 103 | 73 | 125 | 135 | 87 | 75 | 87 | 65 | 92 | 93 | 128 | 111 | 74 | 72 | 78 | 92 | 92,48 | 106 | 79 | 0,735 | -0,445 | -2,784 | 0,00536258 | 0,01151891 | 0,007686 | 0,0176773 | 0,0134297 | 0,0134297 | TRUE | 0,179586 | No |
| orf19.985 | 2725 | 1995 | 2527 | 3015 | 2473 | 2321 | 2626 | 1637 | 2445 | 2540 | 2585 | 2486 | 2115 | 2215 | 2343 | 2309 | 2379,7 | 2514 | 2246 | 0,893 | -0,164 | -3,589 | 0,00033142 | 0,00093001 | 0,000964 | 0,0025896 | 0,0015709 | 0,0015709 | TRUE | 0,0192106 | No |
| orf19.986 | 33856 | 22958 | 29113 | 36615 | 38637 | 32894 | 35259 | 22100 | 30372 | 29228 | 29785 | 30193 | 33044 | 31386 | 31463 | 31170 | 30830 | 29894 | 31766 | 1,063 | 0,088 | 2,869 | 0,00411194 | 0,00904266 | 0,000468 | 0,0020266 | 0,0008626 | 0,0008626 | TRUE | 0,0090263 | No |
| orf19.988 | 2635 | 1808 | 2281 | 3006 | 2580 | 2203 | 2371 | 1621 | 2364 | 2302 | 2334 | 2479 | 2206 | 2102 | 2116 | 2286 | 2273,6 | 2370 | 2178 | 0,917 | -0,125 | -2,776 | 0,00550051 | 0,0117941 | 0,000722 | 0,002618 | 0,0015012 | 0,0015012 | TRUE | 0,0124271 | No |
| orf19.989 | 3804 | 2649 | 3250 | 4151 | 4212 | 3871 | 4153 | 2976 | 3413 | 3372 | 3325 | 3423 | 3602 | 3694 | 3706 | 4197 | 3591,5 | 3383 | 3800 | 1,121 | 0,165 | 3,232 | 0,00123094 | 0,00304717 | 0,002221 | 0,0023837 | 0,0022313 | 0,0022313 | TRUE | 0,058102 | No |
| orf19.99 | 1210 | 735 | 1012 | 1271 | 1124 | 908 | 926 | 690 | 1085 | 936 | 1035 | 1048 | 961 | 866 | 826 | 973 | 966,48 | 1026 | 906 | 0,881 | -0,183 | -2,591 | 0,00956396 | 0,01937299 | 0,003695 | 0,0034816 | 0,0037629 | 0,0037629 | TRUE | 0,0432428 | No |
| orf19.991 | 3274 | 2255 | 2718 | 3325 | 3328 | 2850 | 3137 | 1928 | 2937 | 2871 | 2781 | 2742 | 2846 | 2719 | 2799 | 2719 | 2801,8 | 2833 | 2771 | 0,979 | -0,03 | -0,764 | 0,44489235 | 0,53146187 | 0,000437 | 0,0024976 | 0,0011553 | 0,0011553 | TRUE | 0,0079331 | No |
| orf19.992 | 3403 | 2406 | 3139 | 3671 | 3853 | 3421 | 3667 | 2197 | 3053 | 3063 | 3212 | 3027 | 3295 | 3264 | 3272 | 3099 | 3160,6 | 3089 | 3232 | 1,048 | 0,068 | 1,754 | 0,07944822 | 0,12574707 | 0,000331 | 0,0024388 | 0,0011088 | 0,0011088 | TRUE | 0,0087293 | No |
| orf19.993 | 229 | 186 | 159 | 233 | 241 | 207 | 216 | 131 | 205 | 237 | 163 | 192 | 206 | 198 | 193 | 185 | 197,27 | 199 | 196 | 0,987 | -0,019 | -0,162 | 0,87097169 | 0,90111898 | 0,006852 | 0,0093384 | 0,0082577 | 0,0082577 | TRUE | 0,1566615 | No |
| orf19.994 | 2 | 4 | 2 | 5 | 3 | 1 | 2 | 3 | 2 | 5 | 2 | 4 | 3 | 1 | 2 | 4 | 2,82 | 3 | 2 | 0,709 | -0,497 | -0,596 | 0,55098219 | NA | 1,00E-08 | 0,5160458 | 0,2927444 | 0,2927444 | TRUE | 0,1758112 | No |
| orf19.996 | 114 | 91 | 106 | 101 | 118 | 135 | 112 | 73 | 102 | 116 | 108 | 83 | 101 | 129 | 100 | 103 | 105,31 | 102 | 108 | 1,064 | 0,09 | 0,59 | 0,55520313 | 0,63540861 | 0,008977 | 0,0157644 | 0,0127656 | 0,0127656 | TRUE | 0,1723322 | No |
| orf19.997 | 578 | 451 | 488 | 674 | 824 | 796 | 695 | 453 | 519 | 574 | 499 | 556 | 705 | 760 | 620 | 639 | 608,88 | 537 | 681 | 1,271 | 0,346 | 4,281 | 1,86E-05 | 6,49E-05 | 0,004818 | 0,0043637 | 0,0046019 | 0,0046019 | TRUE | 0,0715178 | No |
| orf19.998 | 139 | 104 | 128 | 185 | 175 | 183 | 157 | 122 | 125 | 132 | 131 | 153 | 150 | 175 | 140 | 172 | 147,13 | 135 | 159 | 1,168 | 0,224 | 1,786 | 0,07405515 | 0,11823634 | 0,00397 | 0,0118462 | 0,0083312 | 0,0083312 | TRUE | 0,0878468 | No |
| orf19.999 | 3956 | 2607 | 3507 | 4276 | 6022 | 4984 | 5414 | 3877 | 3549 | 3319 | 3588 | 3526 | 5150 | 4756 | 4831 | 5468 | 4273,4 | 3496 | 5051 | 1,443 | 0,529 | 10,152 | 3,24E-24 | 7,69E-23 | 0,002358 | 0,0023192 | 0,0023691 | 0,0023691 | TRUE | 0,0364856 | No |

| | | | | | | | | | | | | | | | | | | | | | | | | | | | | | | | |
|---|---|---|---|---|---|---|---|---|---|---|---|---|---|---|---|---|---|---|---|---|---|---|---|---|---|---|---|---|---|---|---|
| RDN18 | 65162 | 45858 | 59529 | 75212 | 80311 | 73414 | 80379 | 46846 | 58457 | 58382 | 60904 | 62021 | 68684 | 70049 | 71725 | 66072 | 64537 | 59941 | 69132 | 1,153 | 0,206 | 5,527 | 3,25E-08 | 1,69E-07 | 0,001035 | 0,002002 | 0,0013171 | 0,0013171 | TRUE | 0,0108359 | No |
| RDN25 | 63461 | 49009 | 58692 | 79446 | 73724 | 71932 | 77944 | 51278 | 56931 | 62394 | 60048 | 65512 | 63051 | 68635 | 69552 | 72323 | 64806 | 61221 | 68390 | 1,117 | 0,16 | 2,908 | 0,00363431 | 0,00807645 | 0,003356 | 0,0020019 | 0,0028818 | 0,0028818 | TRUE | 0,0338666 | No |
| RDN5 | 312 | 160 | 467 | 536 | 276 | 705 | 672 | 479 | 280 | 204 | 478 | 442 | 236 | 673 | 600 | 676 | 448,42 | 351 | 546 | 1,555 | 0,637 | 1,483 | 0,13814015 | 0,20100412 | 0,174812 | 0,0052169 | 0,0751978 | 0,1748119 | TRUE | 0,773014 | No |
| RDN58 | 2181 | 1251 | 1966 | 2654 | 2598 | 2571 | 2991 | 2394 | 1957 | 1593 | 2011 | 2189 | 2222 | 2453 | 2669 | 3377 | 2308,7 | 1938 | 2680 | 1,382 | 0,467 | 2,866 | 0,00415833 | 0,0091213 | 0,025068 | 0,0026083 | 0,0132902 | 0,0250675 | TRUE | 0,3685274 | No |
| RPR1 | 144 | 87 | 139 | 167 | 169 | 131 | 143 | 95 | 129 | 111 | 142 | 138 | 145 | 125 | 128 | 134 | 131,37 | 130 | 133 | 1,017 | 0,024 | 0,188 | 0,85112705 | 0,88638781 | 1,00E-08 | 0,0130297 | 0,0076771 | 0,0076771 | TRUE | 0,0838383 | No |
| SCR1 | 407 | 236 | 343 | 465 | 501 | 410 | 394 | 243 | 365 | 300 | 351 | 383 | 428 | 391 | 352 | 343 | 364,24 | 350 | 378 | 1,081 | 0,112 | 1,121 | 0,2621118 | 0,34572287 | 0,007647 | 0,0059651 | 0,0067836 | 0,0067836 | TRUE | 0,0944204 | No |
| snR10a | 513 | 421 | 455 | 559 | 588 | 563 | 540 | 408 | 460 | 536 | 466 | 461 | 503 | 537 | 482 | 575 | 502,51 | 481 | 524 | 1,09 | 0,125 | 1,557 | 0,11939248 | 0,17746413 | 0,003353 | 0,0048685 | 0,0041492 | 0,0041492 | TRUE | 0,0636278 | No |
| snR10b | 0 | 0 | 0 | 0 | 0 | 0 | 1 | 1 | 0 | 0 | 0 | 0 | 0 | 0 | 1 | 1 | 0,29 | 0 | 0 | 2,833 | 1,502 | 0,536 | 0,59180467 | NA | 1,00E-08 | 5,0453571 | 5,6379511 | 5,6379511 | TRUE | 0,1714431 | No |
| snR10c | 1051 | 625 | 857 | 1083 | 1360 | 1192 | 1128 | 847 | 943 | 796 | 877 | 893 | 1163 | 1137 | 1007 | 1195 | 1001,3 | 877 | 1126 | 1,279 | 0,355 | 4,993 | 5,96E-07 | 2,61E-06 | 0,004166 | 0,0034294 | 0,0038299 | 0,0038299 | TRUE | 0,0617789 | No |
| snR128 | 2602 | 1843 | 2388 | 2851 | 2875 | 2539 | 2784 | 1733 | 2334 | 2346 | 2443 | 2351 | 2459 | 2423 | 2484 | 2444 | 2410,6 | 2368 | 2452 | 1,036 | 0,051 | 1,381 | 0,16720839 | 0,23676361 | 1,00E-08 | 0,0025817 | 0,0008727 | 0,0008727 | TRUE | 0,0053896 | No |
| snR13 | 792 | 551 | 659 | 820 | 890 | 688 | 701 | 526 | 711 | 701 | 674 | 676 | 761 | 656 | 626 | 742 | 693,43 | 690 | 696 | 1,007 | 0,01 | 0,133 | 0,89431292 | 0,92035492 | 0,003503 | 0,004073 | 0,0037682 | 0,0037682 | TRUE | 0,0571329 | No |
| snR161a | 115 | 77 | 89 | 112 | 130 | 76 | 88 | 65 | 103 | 98 | 91 | 92 | 111 | 73 | 79 | 92 | 92,31 | 96 | 89 | 0,921 | -0,119 | -0,729 | 0,46588997 | 0,55186651 | 0,010674 | 0,0177053 | 0,0145764 | 0,0145764 | TRUE | 0,329247 | No |
| snR17A | 5000 | 3605 | 3844 | 5315 | 3673 | 3282 | 3188 | 2409 | 4486 | 4590 | 3933 | 4383 | 3141 | 3132 | 2845 | 3398 | 3738,3 | 4348 | 3129 | 0,719 | -0,476 | -7,423 | 1,14E-13 | 1,15E-12 | 0,004453 | 0,0023679 | 0,0036669 | 0,0036669 | TRUE | 0,05008 | No |
| snR18 | 0 | 0 | 0 | 0 | 0 | 0 | 0 | 0 | 0 | 0 | 0 | 0 | 0 | 0 | 0 | 0 | 0 | 0 | 0 | NA | NA | NA | NA | NA | NA | NA | NA | NA | NA | NA | NA |
| snR189a | 0 | 2 | 1 | 2 | 3 | 3 | 1 | 1 | 0 | 3 | 1 | 2 | 3 | 3 | 1 | 1 | 1,62 | 2 | 2 | 1,566 | 0,647 | 0,588 | 0,55670623 | NA | 1,00E-08 | 0,8988192 | 0,5131308 | 0,5131308 | TRUE | 0,3232073 | No |
| snR189b | 124 | 104 | 110 | 118 | 118 | 99 | 92 | 88 | 111 | 132 | 113 | 97 | 101 | 94 | 82 | 124 | 106,88 | 113 | 100 | 0,882 | -0,182 | -1,165 | 0,24418683 | 0,32594412 | 0,011551 | 0,0155614 | 0,0138705 | 0,0138705 | TRUE | 0,1999179 | No |
| snR189c | 185 | 139 | 165 | 194 | 183 | 205 | 203 | 144 | 166 | 177 | 169 | 160 | 157 | 196 | 181 | 203 | 176,01 | 168 | 184 | 1,092 | 0,127 | 1,119 | 0,26320145 | 0,34700775 | 1,00E-08 | 0,0102274 | 0,0066811 | 0,0066811 | TRUE | 0,116738 | No |
| snR189d | 0 | 0 | 1 | 0 | 0 | 1 | 1 | 0 | 0 | 0 | 1 | 0 | 0 | 1 | 1 | 0 | 0,36 | 0 | 0 | 1,674 | 0,744 | 0,324 | 0,74622179 | NA | 1,00E-08 | 4,0490961 | 3,1539764 | 3,1539764 | TRUE | 0,3677882 | No |
| snR190 | 971 | 1180 | 987 | 1525 | 1277 | 1862 | 1795 | 872 | 871 | 1502 | 1010 | 1258 | 1092 | 1777 | 1602 | 1230 | 1292,6 | 1160 | 1425 | 1,229 | 0,297 | 1,261 | 0,20740568 | 0,2836543 | 0,052699 | 0,0031026 | 0,0254034 | 0,0526994 | TRUE | 0,2470884 | No |
| snR24 | 0 | 0 | 0 | 0 | 0 | 0 | 0 | 0 | 0 | 0 | 0 | 0 | 0 | 0 | 0 | 0 | 0 | 0 | 0 | NA | NA | NA | NA | NA | NA | NA | NA | NA | NA | NA | NA |
| snR31a | 35 | 20 | 47 | 50 | 44 | 29 | 34 | 25 | 31 | 25 | 48 | 41 | 38 | 28 | 30 | 35 | 34,63 | 36 | 33 | 0,886 | -0,175 | -0,686 | 0,4926406 | 0,57763168 | 0,017612 | 0,0438941 | 0,0333909 | 0,0333909 | TRUE | 0,1851676 | No |
| snR32a | 47 | 32 | 35 | 43 | 37 | 39 | 30 | 10 | 42 | 41 | 36 | 35 | 32 | 37 | 27 | 14 | 32,99 | 38 | 28 | 0,731 | -0,452 | -1,712 | 0,08686371 | 0,13569349 | 0,01499 | 0,0459871 | 0,0361407 | 0,0361407 | TRUE | 0,4583566 | No |
| snR33a | 41513 | 24214 | 29874 | 39017 | 53991 | 35181 | 38180 | 32840 | 37241 | 30827 | 30564 | 32174 | 46175 | 33569 | 34069 | 46318 | 36367 | 32702 | 40033 | 1,224 | 0,292 | 1,994 | 0,04619482 | 0,07842285 | 0,020554 | 0,0020194 | 0,0111583 | 0,0205537 | TRUE | 0,1098266 | No |
| snR35a | 14 | 25 | 18 | 24 | 15 | 19 | 25 | 15 | 13 | 32 | 18 | 20 | 13 | 18 | 22 | 21 | 19,63 | 21 | 18 | 0,911 | -0,134 | -0,385 | 0,70031046 | 0,7620839 | 0,050546 | 0,0759441 | 0,0653825 | 0,0653825 | TRUE | 0,5184163 | No |
| snR35c | 42 | 25 | 40 | 47 | 36 | 17 | 38 | 19 | 38 | 32 | 41 | 39 | 31 | 16 | 34 | 27 | 32,11 | 38 | 27 | 0,722 | -0,47 | -1,792 | 0,07310225 | 0,11699476 | 0,010795 | 0,0471858 | 0,0342506 | 0,0342506 | TRUE | 0,4949954 | No |
| snR39b | 0 | 0 | 0 | 0 | 0 | 0 | 0 | 0 | 0 | 0 | 0 | 0 | 0 | 0 | 0 | 0 | 0 | 0 | 0 | NA | NA | NA | NA | NA | NA | NA | NA | NA | NA | NA | NA |
| snR3a | 104 | 77 | 87 | 116 | 87 | 83 | 88 | 44 | 93 | 98 | 89 | 96 | 74 | 79 | 79 | 62 | 83,77 | 94 | 74 | 0,791 | -0,339 | -2,263 | 0,02366171 | 0,04343402 | 1,00E-08 | 0,0193087 | 0,0095626 | 0,0095626 | TRUE | 0,0724023 | No |
| snR40 | 32 | 25 | 32 | 40 | 30 | 36 | 25 | 23 | 29 | 32 | 33 | 33 | 26 | 34 | 22 | 32 | 30,13 | 32 | 28 | 0,9 | -0,153 | -0,606 | 0,54462265 | 0,62623787 | 1,00E-08 | 0,0501661 | 0,0276276 | 0,0276276 | TRUE | 0,1588002 | No |
| snR41 | 1804 | 1361 | 1611 | 2160 | 1747 | 1609 | 1700 | 1006 | 1618 | 1733 | 1648 | 1781 | 1494 | 1535 | 1517 | 1419 | 1593,2 | 1695 | 1491 | 0,881 | -0,183 | -3,672 | 0,0002405 | 0,0006897 | 0,000953 | 0,0028907 | 0,0017423 | 0,0017423 | TRUE | 0,0152723 | No |
| snR41b | 674 | 504 | 529 | 775 | 602 | 568 | 520 | 495 | 605 | 642 | 541 | 639 | 515 | 542 | 464 | 698 | 580,7 | 607 | 555 | 0,91 | -0,136 | -1,214 | 0,22460429 | 0,30358213 | 0,016256 | 0,0044795 | 0,010284 | 0,010284 | TRUE | 0,3333525 | No |
| snR42a | 181 | 101 | 151 | 163 | 157 | 131 | 117 | 125 | 162 | 129 | 154 | 134 | 134 | 125 | 104 | 176 | 139,98 | 145 | 135 | 0,919 | -0,122 | -0,779 | 0,43584448 | 0,52276974 | 0,021085 | 0,0123504 | 0,0162541 | 0,0162541 | TRUE | 0,3737968 | No |
| snR43a | 8 | 0 | 7 | 5 | 3 | 4 | 1 | 3 | 7 | 0 | 7 | 4 | 3 | 4 | 1 | 4 | 3,75 | 4 | 3 | 0,589 | -0,764 | -0,944 | 0,34503889 | NA | 0,24198 | 0,3895194 | 0,3440836 | 0,3440836 | TRUE | 0,6064088 | No |
| snR43b | 30 | 15 | 26 | 36 | 29 | 25 | 17 | 19 | 27 | 19 | 27 | 30 | 25 | 24 | 15 | 27 | 24,11 | 26 | 23 | 0,866 | -0,208 | -0,718 | 0,4727975 | 0,55898623 | 1,00E-08 | 0,0621787 | 0,0389847 | 0,0389847 | TRUE | 0,3224643 | No |
| snR44a | 33 | 37 | 25 | 35 | 20 | 22 | 24 | 18 | 30 | 47 | 26 | 29 | 17 | 21 | 21 | 25 | 27,01 | 33 | 21 | 0,648 | -0,625 | -2,176 | 0,02958249 | 0,05291207 | 0,016974 | 0,0557343 | 0,039982 | 0,039982 | TRUE | 0,4867166 | No |
| snR44b | 82 | 53 | 47 | 67 | 62 | 65 | 64 | 42 | 74 | 67 | 48 | 55 | 53 | 62 | 57 | 59 | 59,47 | 61 | 58 | 0,946 | -0,079 | -0,419 | 0,67546873 | 0,74068556 | 1,00E-08 | 0,02639 | 0,0176538 | 0,0176538 | TRUE | 0,1873609 | No |
| snR44c | 9 | 20 | 11 | 9 | 12 | 18 | 17 | 17 | 8 | 25 | 11 | 7 | 10 | 17 | 15 | 24 | 14,85 | 13 | 16 | 1,296 | 0,374 | 0,835 | 0,40347669 | NA | 0,151711 | 0,0997404 | 0,1223544 | 0,1223544 | TRUE | 1,0548739 | No |
| snR44d | 239 | 275 | 275 | 415 | 382 | 414 | 339 | 254 | 214 | 350 | 281 | 342 | 327 | 395 | 303 | 358 | 321,32 | 297 | 346 | 1,165 | 0,22 | 1,524 | 0,12757407 | 0,18804343 | 0,027873 | 0,0064975 | 0,0168478 | 0,0168478 | TRUE | 0,4070746 | No |
| snR46a | 1 | 0 | 1 | 0 | 0 | 0 | 0 | 0 | 1 | 0 | 1 | 0 | 0 | 0 | 0 | 0 | 0,24 | 0 | 0 | 0,367 | -1,446 | -0,49 | 0,62399902 | NA | 1,00E-08 | 6,0500945 | 6,4379451 | 6,4379451 | TRUE | 0,0586 | No |
| snR47 | 523 | 390 | 401 | 641 | 706 | 490 | 535 | 317 | 469 | 497 | 410 | 529 | 604 | 468 | 477 | 447 | 487,55 | 476 | 499 | 1,05 | 0,071 | 0,654 | 0,51288159 | 0,59716911 | 0,013307 | 0,0049571 | 0,0092219 | 0,0092219 | TRUE | 0,2392225 | No |
| snR48 | 727 | 763 | 769 | 1068 | 1252 | 1381 | 1302 | 581 | 652 | 971 | 787 | 881 | 1071 | 1318 | 1162 | 819 | 957,59 | 823 | 1092 | 1,33 | 0,411 | 2,265 | 0,02351183 | 0,04318867 | 0,030558 | 0,0034955 | 0,017121 | 0,0305584 | TRUE | 0,3147737 | No |
| snR49a | 0 | 0 | 0 | 0 | 0 | 0 | 0 | 0 | 0 | 0 | 0 | 0 | 0 | 0 | 0 | 0 | 0 | 0 | 0 | NA | NA | NA | NA | NA | NA | NA | NA | NA | NA | NA | NA |
| snR50 | 5 | 3 | 5 | 3 | 2 | 5 | 5 | 2 | 4 | 4 | 5 | 2 | 2 | 5 | 4 | 3 | 3,71 | 4 | 4 | 0,877 | -0,19 | -0,269 | 0,78793855 | NA | 1,00E-08 | 0,3935643 | 0,2068713 | 0,2068713 | TRUE | 0,2301405 | No |
| snR52 | 18 | 13 | 13 | 13 | 11 | 8 | 11 | 4 | 16 | 17 | 13 | 11 | 9 | 8 | 10 | 6 | 11,15 | 14 | 8 | 0,592 | -0,757 | -1,831 | 0,06717358 | NA | 1,00E-08 | 0,1321525 | 0,069113 | 0,069113 | TRUE | 0,1473283 | No |
| snR54 | 0 | 0 | 0 | 0 | 0 | 0 | 0 | 0 | 0 | 0 | 0 | 0 | 0 | 0 | 0 | 0 | 0 | 0 | 0 | NA | NA | NA | NA | NA | NA | NA | NA | NA | NA | NA | NA |
| snR55 | 12 | 2 | 8 | 8 | 13 | 7 | 13 | 11 | 11 | 3 | 8 | 7 | 11 | 7 | 12 | 16 | 9,13 | 7 | 12 | 1,532 | 0,615 | 1,262 | 0,20688948 | NA | 0,021074 | 0,1610587 | 0,1139083 | 0,1139083 | TRUE | 0,283525 | No |
| snR56 | 554 | 373 | 449 | 601 | 570 | 388 | 434 | 344 | 497 | 475 | 459 | 496 | 487 | 370 | 387 | 485 | 457,12 | 482 | 432 | 0,895 | -0,16 | -1,639 | 0,10119107 | 0,15426438 | 0,008828 | 0,0051553 | 0,0069937 | 0,0069937 | TRUE | 0,1084816 | No |
| snR57 | 67 | 64 | 56 | 75 | 99 | 126 | 129 | 35 | 60 | 81 | 57 | 62 | 85 | 120 | 115 | 49 | 78,76 | 65 | 92 | 1,439 | 0,525 | 2,182 | 0,02912456 | 0,05221735 | 0,070115 | 0,0204111 | 0,0424872 | 0,0424872 | TRUE | 0,4391381 | No |
| snR57b | 422 | 616 | 481 | 716 | 833 | 853 | 790 | 348 | 379 | 784 | 492 | 590 | 712 | 814 | 705 | 491 | 620,93 | 561 | 680 | 1,214 | 0,28 | 1,086 | 0,27740281 | 0,36144785 | 0,062287 | 0,0043175 | 0,0317697 | 0,0622872 | TRUE | 0,4951713 | No |
| snR58 | 227 | 153 | 201 | 266 | 287 | 234 | 196 | 152 | 204 | 195 | 206 | 219 | 245 | 223 | 175 | 214 | 210,18 | 206 | 214 | 1,04 | 0,057 | 0,488 | 0,62573964 | 0,69774954 | 0,007335 | 0,0088865 | 0,0081451 | 0,0081451 | TRUE | 0,1853248 | No |
| snR5a | 245 | 210 | 185 | 288 | 362 | 285 | 254 | 229 | 220 | 267 | 189 | 237 | 310 | 272 | 227 | 323 | 255,63 | 228 | 283 | 1,236 | 0,306 | 2,322 | 0,02024645 | 0,03765965 | 0,018203 | 0,0076583 | 0,0127055 | 0,0127055 | TRUE | 0,2066168 | No |
| snR5b | 28 | 21 | 16 | 27 | 29 | 9 | 20 | 11 | 25 | 27 | 16 | 22 | 25 | 9 | 18 | 16 | 19,65 | 22 | 17 | 0,747 | -0,42 | -1,201 | 0,22966888 | 0,3095918 | 0,049956 | 0,0758393 | 0,0656774 | 0,0656774 | TRUE | 0,3485156 | No |
| snR5c | 5 | 3 | 5 | 10 | 6 | 6 | 7 | 0 | 4 | 4 | 5 | 8 | 5 | 6 | 6 | 0 | 4,85 | 5 | 4 | 0,819 | -0,287 | -0,44 | 0,66001731 | NA | 0,01202 | 0,3015371 | 0,2086843 | 0,2086843 | TRUE | 0,4672094 | No |
| snR5d | 226 | 176 | 179 | 256 | 215 | 195 | 175 | 171 | 203 | 224 | 183 | 211 | 184 | 186 | 156 | 241 | 198,54 | 205 | 192 | 0,926 | -0,11 | -0,848 | 0,39628121 | 0,48509729 | 0,013182 | 0,0092914 | 0,0111026 | 0,0111026 | TRUE | 0,2810688 | No |
| snR60 | 1 | 0 | 0 | 4 | 0 | 0 | 3 | 2 | 1 | 0 | 0 | 3 | 0 | 0 | 3 | 3 | 1,21 | 1 | 2 | 1,185 | 0,245 | 0,158 | 0,87481106 | NA | 2,273913 | 1,2000768 | 1,4979009 | 1,4979009 | TRUE | 0,4379107 | No |
| snR61 | 298 | 279 | 264 | 369 | 427 | 505 | 463 | 348 | 267 | 355 | 270 | 304 | 365 | 482 | 413 | 491 | 368,49 | 299 | 438 | 1,463 | 0,549 | 4,646 | 3,38E-06 | 1,32E-05 | 0,014848 | 0,0059191 | 0,0105701 | 0,0105701 | TRUE | 0,1646501 | No |
| snR62 | 208 | 224 | 164 | 272 | 200 | 223 | 176 | 126 | 187 | 285 | 168 | 224 | 171 | 213 | 157 | 178 | 197,81 | 216 | 180 | 0,835 | -0,26 | -1,625 | 0,10412582 | 0,15773775 | 0,03065 | 0,0093186 | 0,0195088 | 0,0195088 | TRUE | 0,4723761 | No |
| snR63 | 184 | 138 | 142 | 193 | 226 | 182 | 156 | 80 | 165 | 176 | 145 | 159 | 193 | 174 | 139 | 113 | 158,02 | 161 | 155 | 0,97 | -0,044 | -0,29 | 0,77217917 | 0,82197952 | 0,022359 | 0,0111663 | 0,0162006 | 0,0162006 | TRUE | 0,2242381 | No |
| snR64 | 8 | 15 | 9 | 16 | 18 | 12 | 7 | 13 | 7 | 19 | 9 | 13 | 15 | 11 | 6 | 18 | 12,51 | 12 | 12 | 1,055 | 0,077 | 0,172 | 0,86328497 | NA | 0,097305 | 0,1179984 | 0,1095142 | 0,1095142 | TRUE | 0,3914812 | No |
| snR65 | 11 | 4 | 8 | 14 | 20 | 7 | 16 | 6 | 10 | 5 | 8 | 12 | 17 | 7 | 14 | 8 | 10,15 | 9 | 12 | 1,336 | 0,418 | 0,904 | 0,36581063 | NA | 0,050859 | 0,1449804 | 0,107182 | 0,107182 | TRUE | 0,1975071 | No |

| | | | | | | | | | | | | | | | | | | | | | | | | | | | | | | | |
|---|---|---|---|---|---|---|---|---|---|---|---|---|---|---|---|---|---|---|---|---|---|---|---|---|---|---|---|---|---|---|---|
| snR66 | 49 | 79 | 64 | 111 | 82 | 93 | 132 | 113 | 44 | 101 | 65 | 92 | 70 | 89 | 118 | 159 | 92,2 | 76 | 109 | 1,441 | 0,527 | 1,908 | 0,05637764 | 0,09318208 | 0,113578 | 0,0177252 | 0,0618622 | 0,0618622 | TRUE | 0,3156943 | No |
| snR67 | 6 | 8 | 7 | 2 | 2 | 2 | 8 | 5 | 5 | 10 | 7 | 2 | 2 | 2 | 7 | 7 | 5,27 | 6 | 4 | 0,731 | -0,452 | -0,637 | 0,52440382 | NA | 0,291391 | 0,2772623 | 0,2829188 | 0,2829188 | TRUE | 0,2645199 | No |
| snR68 | 38 | 42 | 29 | 54 | 36 | 57 | 40 | 46 | 34 | 53 | 30 | 45 | 31 | 54 | 36 | 65 | 43,44 | 40 | 46 | 1,136 | 0,185 | 0,665 | 0,50609403 | 0,59052494 | 0,069137 | 0,0353993 | 0,0505368 | 0,0505368 | TRUE | 0,2164376 | No |
| snR69 | 409 | 250 | 298 | 433 | 488 | 389 | 422 | 372 | 367 | 318 | 305 | 357 | 417 | 371 | 377 | 525 | 379,61 | 337 | 422 | 1,245 | 0,316 | 2,758 | 0,00581705 | 0,0123932 | 0,014585 | 0,0058037 | 0,00993 | 0,00993 | TRUE | 0,2815788 | No |
| snR70 | 1552 | 1185 | 1319 | 1811 | 1546 | 1457 | 1509 | 921 | 1392 | 1509 | 1349 | 1493 | 1322 | 1390 | 1347 | 1299 | 1387,7 | 1436 | 1340 | 0,934 | -0,098 | -1,871 | 0,06129278 | 0,10040563 | 0,001129 | 0,0030256 | 0,001934 | 0,001934 | TRUE | 0,0192739 | No |
| snR71 | 8 | 3 | 7 | 6 | 11 | 13 | 14 | 6 | 7 | 4 | 7 | 5 | 9 | 12 | 12 | 8 | 8,23 | 6 | 10 | 1,848 | 0,886 | 1,844 | 0,06517093 | NA | 1,00E-08 | 0,1782842 | 0,0908151 | 0,0908151 | TRUE | 0,0784347 | No |
| snR72 | 17 | 8 | 11 | 8 | 5 | 11 | 7 | 8 | 15 | 10 | 11 | 7 | 4 | 10 | 6 | 11 | 9,45 | 11 | 8 | 0,726 | -0,462 | -0,96 | 0,33726099 | NA | 1,00E-08 | 0,1556226 | 0,1130192 | 0,1130192 | TRUE | 0,2883313 | No |
| snR73 | 470 | 329 | 392 | 478 | 419 | 407 | 397 | 299 | 422 | 419 | 401 | 394 | 358 | 388 | 354 | 422 | 394,8 | 409 | 380 | 0,926 | -0,11 | -1,396 | 0,16274059 | 0,23120016 | 0,000819 | 0,0056566 | 0,0034623 | 0,0034623 | TRUE | 0,0514147 | No |
| snR74 | 119 | 176 | 118 | 159 | 82 | 172 | 112 | 106 | 107 | 224 | 121 | 131 | 70 | 164 | 100 | 150 | 133,29 | 146 | 121 | 0,83 | -0,269 | -0,748 | 0,45472693 | 0,54056181 | 0,116374 | 0,0128705 | 0,0615459 | 0,1163736 | TRUE | 0,5559622 | No |
| snR75 | 174 | 146 | 150 | 163 | 163 | 226 | 207 | 171 | 156 | 186 | 153 | 134 | 139 | 216 | 185 | 241 | 176,35 | 157 | 195 | 1,237 | 0,307 | 1,879 | 0,06024971 | 0,0988663 | 0,030218 | 0,0102115 | 0,0197933 | 0,0197933 | TRUE | 0,330913 | No |
| snR76 | 492 | 410 | 415 | 552 | 409 | 399 | 420 | 228 | 441 | 522 | 425 | 455 | 350 | 381 | 375 | 322 | 408,75 | 461 | 357 | 0,779 | -0,36 | -4,134 | 3,56E-05 | 0,00011831 | 0,003834 | 0,0055311 | 0,0047713 | 0,0047713 | TRUE | 0,0841272 | No |
| snR77 | 191 | 275 | 256 | 336 | 259 | 321 | 356 | 231 | 171 | 350 | 262 | 277 | 222 | 306 | 318 | 326 | 278,96 | 265 | 293 | 1,106 | 0,146 | 0,797 | 0,42570852 | 0,51283149 | 0,052666 | 0,0071834 | 0,0284406 | 0,0284406 | TRUE | 0,3586166 | No |
| snR77b | 80 | 168 | 116 | 243 | 103 | 237 | 220 | 127 | 72 | 214 | 119 | 200 | 88 | 226 | 196 | 179 | 161,8 | 151 | 172 | 1,141 | 0,19 | 0,429 | 0,66779472 | 0,73454974 | 0,182661 | 0,0109519 | 0,0866427 | 0,1826615 | TRUE | 0,3755944 | No |
| snR78 | 28 | 51 | 39 | 59 | 51 | 96 | 100 | 32 | 25 | 65 | 40 | 49 | 44 | 92 | 89 | 45 | 56,02 | 45 | 68 | 1,53 | 0,613 | 1,942 | 0,05218578 | 0,08726194 | 0,134461 | 0,0278921 | 0,0770411 | 0,0770411 | TRUE | 0,1391246 | No |
| snR79 | 70 | 115 | 79 | 151 | 118 | 170 | 154 | 126 | 63 | 146 | 81 | 125 | 101 | 162 | 137 | 178 | 124,1 | 104 | 144 | 1,395 | 0,481 | 1,967 | 0,04919329 | 0,08311397 | 0,090859 | 0,0136773 | 0,0489705 | 0,0489705 | TRUE | 0,280982 | No |
| snR87 | 254 | 222 | 197 | 327 | 280 | 271 | 268 | 168 | 228 | 283 | 202 | 270 | 239 | 259 | 239 | 237 | 244,48 | 246 | 244 | 0,995 | -0,007 | -0,065 | 0,94781715 | 0,96239402 | 0,008248 | 0,0079175 | 0,00807 | 0,00807 | TRUE | 0,1554552 | No |
| snR8a | 277 | 218 | 224 | 261 | 235 | 234 | 215 | 133 | 248 | 278 | 229 | 215 | 201 | 223 | 192 | 188 | 221,77 | 242 | 201 | 0,834 | -0,261 | -2,408 | 0,01606138 | 0,0306631 | 0,004611 | 0,0085256 | 0,0067418 | 0,0067418 | TRUE | 0,093386 | No |
| snR8b | 22 | 21 | 18 | 40 | 18 | 28 | 23 | 22 | 20 | 27 | 18 | 33 | 15 | 27 | 21 | 31 | 23,94 | 24 | 24 | 0,936 | -0,096 | -0,303 | 0,76212953 | 0,81445645 | 0,043167 | 0,0626138 | 0,0543175 | 0,0543175 | TRUE | 0,3097842 | No |
| snR8c | 78 | 73 | 71 | 111 | 101 | 122 | 113 | 87 | 70 | 93 | 73 | 92 | 86 | 116 | 101 | 123 | 94,18 | 82 | 106 | 1,296 | 0,374 | 2,266 | 0,02342659 | 0,04308141 | 0,012727 | 0,0173943 | 0,0153537 | 0,0153537 | TRUE | 0,1714703 | No |
| SNRNAU1 | 136 | 95 | 102 | 135 | 163 | 137 | 121 | 68 | 122 | 121 | 104 | 111 | 139 | 131 | 108 | 96 | 116,58 | 114 | 118 | 1,044 | 0,063 | 0,432 | 0,66556094 | 0,73323749 | 0,007662 | 0,014432 | 0,0115594 | 0,0115594 | TRUE | 0,144506 | No |
| SNRNAU2 | 41 | 40 | 33 | 46 | 39 | 37 | 34 | 22 | 37 | 51 | 34 | 38 | 33 | 35 | 30 | 31 | 36,18 | 40 | 32 | 0,826 | -0,275 | -1,193 | 0,23281015 | 0,31334032 | 1,00E-08 | 0,0421058 | 0,0232678 | 0,0232678 | TRUE | 0,2089374 | No |
| SNRNAU4 | 170 | 94 | 120 | 151 | 151 | 162 | 180 | 78 | 153 | 120 | 123 | 125 | 129 | 155 | 161 | 110 | 134,23 | 130 | 139 | 1,072 | 0,101 | 0,694 | 0,48752115 | 0,57297087 | 0,012764 | 0,0127948 | 0,012779 | 0,012779 | TRUE | 0,161323 | No |
| SNRNAU5 | 24 | 22 | 17 | 31 | 20 | 20 | 19 | 16 | 22 | 28 | 17 | 26 | 17 | 19 | 17 | 23 | 21,03 | 23 | 19 | 0,811 | -0,302 | -1,005 | 0,31502449 | 0,4016696 | 1,00E-08 | 0,0710245 | 0,0384689 | 0,0384689 | TRUE | 0,1530317 | No |
| SNRNAU6 | 39 | 16 | 27 | 39 | 31 | 21 | 35 | 17 | 35 | 20 | 28 | 32 | 27 | 20 | 31 | 24 | 27,11 | 29 | 26 | 0,876 | -0,191 | -0,694 | 0,48774927 | 0,57312686 | 1,00E-08 | 0,0555235 | 0,0359809 | 0,0359809 | TRUE | 0,197094 | No |
| tA(AGC)1 | 0 | 0 | 0 | 0 | 0 | 0 | 0 | 1 | 0 | 0 | 0 | 0 | 0 | 0 | 0 | 1 | 0,18 | 0 | 0 | 1,675 | 0,744 | 0,211 | 0,83317474 | NA | 1,00E-08 | 8,2361844 | 10 | 10 | TRUE | 0,6248756 | No |
| tA(AGC)2 | 0 | 0 | 0 | 0 | 0 | 0 | 0 | 0 | 0 | 0 | 0 | 0 | 0 | 0 | 0 | 0 | 0 | 0 | 0 | NA | NA | NA | NA | NA | NA | NA | NA | NA | NA | NA | NA |
| tA(AGC)3 | 0 | 0 | 0 | 0 | 0 | 0 | 0 | 0 | 0 | 0 | 0 | 0 | 0 | 0 | 0 | 0 | 0 | 0 | 0 | NA | NA | NA | NA | NA | NA | NA | NA | NA | NA | NA | NA |
| tA(AGC)4 | 0 | 0 | 0 | 0 | 0 | 0 | 1 | 0 | 0 | 0 | 0 | 0 | 0 | 0 | 1 | 0 | 0,11 | 0 | 0 | 1,675 | 0,744 | 0,211 | 0,83317474 | NA | 1,00E-08 | 13,01685 | 10 | 10 | TRUE | 0,2773144 | No |
| tA(AGC)5 | 0 | 1 | 0 | 1 | 0 | 0 | 0 | 0 | 0 | 1 | 0 | 1 | 0 | 0 | 0 | 0 | 0,26 | 0 | 0 | 0,378 | -1,402 | -0,489 | 0,62500522 | NA | 1,00E-08 | 5,5382821 | 5,9647107 | 5,9647107 | TRUE | 0,1641193 | No |
| tA(AGC)7 | 0 | 0 | 0 | 0 | 0 | 0 | 0 | 1 | 0 | 0 | 0 | 0 | 0 | 0 | 0 | 1 | 0,18 | 0 | 0 | 1,675 | 0,744 | 0,211 | 0,83317474 | NA | 1,00E-08 | 8,2361844 | 10 | 10 | TRUE | 0,6248756 | No |
| tA(UGC)1 | 0 | 0 | 0 | 0 | 1 | 0 | 0 | 0 | 0 | 0 | 0 | 0 | 1 | 0 | 0 | 0 | 0,11 | 0 | 0 | 1,675 | 0,744 | 0,211 | 0,83317474 | NA | 1,00E-08 | 13,581472 | 10 | 10 | TRUE | 0,2537777 | No |
| tA(UGC)2 | 0 | 0 | 0 | 0 | 0 | 0 | 0 | 0 | 0 | 0 | 0 | 0 | 0 | 0 | 0 | 0 | 0 | 0 | 0 | NA | NA | NA | NA | NA | NA | NA | NA | NA | NA | NA | NA |
| tC(GCA)1 | 0 | 0 | 0 | 0 | 0 | 0 | 0 | 0 | 0 | 0 | 0 | 0 | 0 | 0 | 0 | 0 | 0 | 0 | 0 | NA | NA | NA | NA | NA | NA | NA | NA | NA | NA | NA | NA |
| tC(GCA)2 | 0 | 0 | 0 | 1 | 0 | 0 | 0 | 0 | 0 | 0 | 0 | 1 | 0 | 0 | 0 | 0 | 0,1 | 0 | 0 | 0,616 | -0,698 | -0,198 | 0,84334499 | NA | 1,00E-08 | 14,085693 | 10 | 10 | TRUE | 0,2428614 | No |
| tD(GUC)2 | 0 | 0 | 0 | 0 | 0 | 0 | 1 | 0 | 0 | 0 | 0 | 0 | 0 | 0 | 1 | 0 | 0,11 | 0 | 0 | 1,675 | 0,744 | 0,211 | 0,83317474 | NA | 1,00E-08 | 13,01685 | 10 | 10 | TRUE | 0,2773144 | No |
| tD(GUC)3 | 1 | 0 | 1 | 0 | 0 | 0 | 0 | 0 | 1 | 0 | 1 | 0 | 0 | 0 | 0 | 0 | 0,24 | 0 | 0 | 0,367 | -1,446 | -0,49 | 0,62399902 | NA | 1,00E-08 | 6,0500945 | 6,4379451 | 6,4379451 | TRUE | 0,0586 | No |
| tD(GUC)4 | 0 | 0 | 2 | 1 | 0 | 0 | 0 | 1 | 0 | 0 | 2 | 1 | 0 | 0 | 0 | 1 | 0,54 | 1 | 0 | 0,421 | -1,25 | -0,603 | 0,54664991 | NA | 1,00E-08 | 2,7146662 | 2,4445877 | 2,4445877 | TRUE | 0,6248746 | No |
| tD(GUC)5 | 1 | 0 | 0 | 0 | 1 | 1 | 0 | 0 | 1 | 0 | 0 | 0 | 1 | 1 | 0 | 0 | 0,34 | 0 | 0 | 1,656 | 0,728 | 0,312 | 0,75473354 | NA | 1,00E-08 | 4,2929789 | 3,2837448 | 3,2837448 | TRUE | 0,2895953 | No |
| tD(GUC)6 | 0 | 0 | 1 | 1 | 0 | 0 | 0 | 0 | 0 | 0 | 1 | 1 | 0 | 0 | 0 | 0 | 0,23 | 0 | 0 | 0,379 | -1,402 | -0,471 | 0,63792379 | NA | 1,00E-08 | 6,2873821 | 6,584019 | 6,584019 | TRUE | 0,0700646 | No |
| tD(GUC)7 | 0 | 0 | 0 | 0 | 0 | 0 | 0 | 0 | 0 | 0 | 0 | 0 | 0 | 0 | 0 | 0 | 0 | 0 | 0 | NA | NA | NA | NA | NA | NA | NA | NA | NA | NA | NA | NA |
| tE(CUC)1 | 0 | 0 | 0 | 0 | 1 | 0 | 0 | 0 | 0 | 0 | 0 | 0 | 1 | 0 | 0 | 0 | 0,11 | 0 | 0 | 1,675 | 0,744 | 0,211 | 0,83317474 | NA | 1,00E-08 | 13,581472 | 10 | 10 | TRUE | 0,2537777 | No |
| TER1 | 2764 | 2056 | 2510 | 3170 | 3391 | 2952 | 3071 | 2110 | 2480 | 2618 | 2568 | 2614 | 2900 | 2817 | 2740 | 2976 | 2714 | 2570 | 2858 | 1,111 | 0,153 | 3,68 | 0,00023362 | 0,00067061 | 0,000574 | 0,0025144 | 0,0012782 | 0,0012782 | TRUE | 0,0093698 | No |
| tE(UUC)1 | 3 | 2 | 4 | 1 | 8 | 4 | 6 | 3 | 3 | 3 | 4 | 1 | 7 | 4 | 5 | 4 | 3,8 | 3 | 5 | 2,067 | 1,047 | 1,442 | 0,14928018 | NA | 1,00E-08 | 0,3840276 | 0,2098585 | 0,2098585 | TRUE | 0,3232021 | No |
| tE(UUC)2 | 0 | 0 | 0 | 0 | 0 | 0 | 0 | 0 | 0 | 0 | 0 | 0 | 0 | 0 | 0 | 0 | 0 | 0 | 0 | NA | NA | NA | NA | NA | NA | NA | NA | NA | NA | NA | NA |
| tE(UUC)3 | 0 | 0 | 0 | 0 | 1 | 0 | 0 | 1 | 0 | 0 | 0 | 0 | 1 | 0 | 0 | 1 | 0,28 | 0 | 0 | 2,78 | 1,475 | 0,525 | 0,59977115 | NA | 1,00E-08 | 5,1279479 | 5,6799432 | 5,6799432 | TRUE | 0,1950519 | No |
| tE(UUC)4 | 0 | 0 | 0 | 0 | 0 | 0 | 0 | 0 | 0 | 0 | 0 | 0 | 0 | 0 | 0 | 0 | 0 | 0 | 0 | NA | NA | NA | NA | NA | NA | NA | NA | NA | NA | NA | NA |
| tE(UUC)5 | 0 | 0 | 0 | 0 | 0 | 0 | 0 | 0 | 0 | 0 | 0 | 0 | 0 | 0 | 0 | 0 | 0 | 0 | 0 | NA | NA | NA | NA | NA | NA | NA | NA | NA | NA | NA | NA |
| tE(UUC)6 | 0 | 0 | 3 | 0 | 2 | 1 | 0 | 2 | 0 | 0 | 3 | 0 | 2 | 1 | 0 | 3 | 1,07 | 1 | 2 | 1,742 | 0,8 | 0,495 | 0,62085719 | NA | 1,793258 | 1,3595425 | 1,4756124 | 1,4756124 | TRUE | 1,0262325 | No |
| tE(UUC)7 | 6 | 4 | 4 | 5 | 7 | 7 | 4 | 1 | 5 | 5 | 4 | 4 | 6 | 7 | 4 | 1 | 4,54 | 4 | 4 | 0,985 | -0,022 | -0,034 | 0,97268181 | NA | 1,00E-08 | 0,3215962 | 0,1745837 | 0,1745837 | TRUE | 0,2222215 | No |
| tF(GAA)1 | 2 | 3 | 6 | 6 | 3 | 3 | 2 | 3 | 2 | 4 | 6 | 5 | 3 | 3 | 2 | 4 | 3,52 | 4 | 3 | 0,664 | -0,591 | -0,798 | 0,42481437 | NA | 1,00E-08 | 0,4146306 | 0,227068 | 0,227068 | TRUE | 0,306608 | No |
| tF(GAA)2 | 0 | 0 | 0 | 0 | 0 | 0 | 1 | 0 | 0 | 0 | 0 | 0 | 0 | 0 | 1 | 0 | 0,11 | 0 | 0 | 1,675 | 0,744 | 0,211 | 0,83317474 | NA | 1,00E-08 | 13,01685 | 10 | 10 | TRUE | 0,2773144 | No |
| tF(GAA)3 | 0 | 1 | 0 | 1 | 1 | 2 | 0 | 0 | 0 | 1 | 0 | 1 | 1 | 2 | 0 | 0 | 0,61 | 0 | 1 | 1,465 | 0,551 | 0,297 | 0,76678388 | NA | 1,00E-08 | 2,390978 | 1,6785847 | 1,6785847 | TRUE | 0,4640956 | No |
| tF(GAA)4 | 1 | 0 | 0 | 0 | 0 | 0 | 0 | 0 | 1 | 0 | 0 | 0 | 0 | 0 | 0 | 0 | 0,11 | 0 | 0 | 0,616 | -0,698 | -0,198 | 0,84334499 | NA | 1,00E-08 | 12,947638 | 10 | 10 | TRUE | 0,289592 | No |
| tF(GAA)5 | 0 | 0 | 0 | 0 | 0 | 0 | 0 | 0 | 0 | 0 | 0 | 0 | 0 | 0 | 0 | 0 | 0 | 0 | 0 | NA | NA | NA | NA | NA | NA | NA | NA | NA | NA | NA | NA |
| tG(CCC)1 | 1 | 0 | 0 | 0 | 2 | 1 | 0 | 0 | 1 | 0 | 0 | 0 | 2 | 1 | 0 | 0 | 0,45 | 0 | 1 | 2,318 | 1,213 | 0,562 | 0,57406983 | NA | 1,00E-08 | 3,2626393 | 2,7485048 | 2,7485048 | TRUE | 0,4071564 | No |
| tG(GCC)1 | 0 | 0 | 1 | 1 | 0 | 0 | 0 | 0 | 0 | 0 | 1 | 1 | 0 | 0 | 0 | 0 | 0,23 | 0 | 0 | 0,379 | -1,402 | -0,471 | 0,63792379 | NA | 1,00E-08 | 6,2873821 | 6,584019 | 6,584019 | TRUE | 0,0700646 | No |
| tG(GCC)2 | 0 | 0 | 0 | 0 | 0 | 0 | 0 | 1 | 0 | 0 | 0 | 0 | 0 | 0 | 0 | 1 | 0,18 | 0 | 0 | 1,675 | 0,744 | 0,211 | 0,83317474 | NA | 1,00E-08 | 8,2361844 | 10 | 10 | TRUE | 0,6248756 | No |
| tG(GCC)3 | 0 | 0 | 0 | 0 | 0 | 0 | 0 | 0 | 0 | 0 | 0 | 0 | 0 | 0 | 0 | 0 | 0 | 0 | 0 | NA | NA | NA | NA | NA | NA | NA | NA | NA | NA | NA | NA |
| tG(GCC)4 | 0 | 0 | 0 | 0 | 0 | 1 | 2 | 0 | 0 | 0 | 0 | 0 | 0 | 1 | 2 | 0 | 0,34 | 0 | 1 | 3,85 | 1,945 | 0,739 | 0,45970473 | NA | 1,00E-08 | 4,2423243 | 4,9244377 | 4,9244377 | TRUE | 0,2828214 | No |

| | | | | | | | | | | | | | | | | | | | | | | | | | | | | | | | | |
|---|---|---|---|---|---|---|---|---|---|---|---|---|---|---|---|---|---|---|---|---|---|---|---|---|---|---|---|---|---|---|---|---|
| tG(GCC)5 | 0 | 0 | 0 | 0 | 0 | 0 | 1 | 0 | 0 | 0 | 0 | 0 | 0 | 0 | 1 | 0 | 0,11 | 0 | 0 | 1,675 | 0,744 | 0,211 | 0,83317474 | | NA | 1,00E-08 | 13,01685 | 10 | 10 | TRUE | 0,2773144 | No |
| tG(GCC)6 | 2 | 1 | 2 | 3 | 3 | 0 | 0 | 0 | 2 | 1 | 2 | 2 | 3 | 0 | 0 | 0 | 1,27 | 2 | 1 | 0,368 | -1,442 | -1,066 | 0,28651754 | | NA | 1,00E-08 | 1,1458343 | 0,7924557 | 0,7924557 | TRUE | 1,4519435 | No |
| tG(UCC)1 | 0 | 0 | 0 | 1 | 0 | 0 | 0 | 0 | 0 | 0 | 0 | 1 | 0 | 0 | 0 | 0 | 0,1 | 0 | 0 | 0,616 | -0,698 | -0,198 | 0,84334499 | | NA | 1,00E-08 | 14,085693 | 10 | 10 | TRUE | 0,2428614 | No |
| tG(UCC)2 | 0 | 0 | 1 | 1 | 0 | 2 | 0 | 0 | 0 | 0 | 1 | 1 | 0 | 2 | 0 | 0 | 0,47 | 0 | 0 | 1,044 | 0,063 | 0,028 | 0,97728768 | | NA | 1,00E-08 | 3,0939551 | 2,7861685 | 2,7861685 | TRUE | 0,8544801 | No |
| tH(GUG)1 | 0 | 0 | 1 | 0 | 2 | 0 | 0 | 0 | 0 | 0 | 1 | 0 | 2 | 0 | 0 | 0 | 0,34 | 0 | 0 | 1,6 | 0,678 | 0,251 | 0,80201412 | | NA | 1,00E-08 | 4,2505063 | 5,0796615 | 5,0796615 | TRUE | 0,8236705 | No |
| tH(GUG)3 | 5 | 3 | 4 | 7 | 10 | 4 | 9 | 2 | 4 | 4 | 4 | 6 | 9 | 4 | 8 | 3 | 5,17 | 4 | 6 | 1,306 | 0,385 | 0,634 | 0,52595703 | | NA | 1,00E-08 | 0,282567 | 0,1629128 | 0,1629128 | TRUE | 0,1327023 | No |
| tI(AAU)1 | 0 | 0 | 0 | 0 | 0 | 0 | 0 | 0 | 0 | 0 | 0 | 0 | 0 | 0 | 0 | 0 | 0 | 0 | 0 | NA | NA | NA | NA | | NA | NA | NA | NA | NA | NA | NA | NA |
| tI(AAU)2 | 0 | 0 | 0 | 0 | 0 | 0 | 0 | 0 | 0 | 0 | 0 | 0 | 0 | 0 | 0 | 0 | 0 | 0 | 0 | NA | NA | NA | NA | | NA | NA | NA | NA | NA | NA | NA | NA |
| tI(AAU)3 | 0 | 1 | 0 | 0 | 0 | 0 | 0 | 0 | 0 | 1 | 0 | 0 | 0 | 0 | 0 | 0 | 0,16 | 0 | 0 | 0,616 | -0,698 | -0,198 | 0,84334499 | | NA | 1,00E-08 | 9,1242388 | 10 | 10 | TRUE | 0,5444342 | No |
| tI(AAU)4 | 1 | 1 | 0 | 0 | 0 | 0 | 0 | 0 | 1 | 1 | 0 | 0 | 0 | 0 | 0 | 0 | 0,27 | 0 | 0 | 0,367 | -1,445 | -0,508 | 0,61177232 | | NA | 1,00E-08 | 5,3533522 | 5,868165 | 5,868165 | TRUE | 0,1275109 | No |
| tI(AAU)5 | 0 | 0 | 1 | 0 | 0 | 0 | 0 | 0 | 0 | 0 | 1 | 0 | 0 | 0 | 0 | 0 | 0,13 | 0 | 0 | 0,616 | -0,698 | -0,198 | 0,84334499 | | NA | 1,00E-08 | 11,353388 | 10 | 10 | TRUE | 0,3731057 | No |
| tI(UAU)1 | 0 | 0 | 1 | 0 | 0 | 0 | 1 | 0 | 0 | 0 | 1 | 0 | 0 | 0 | 1 | 0 | 0,24 | 0 | 0 | 1,016 | 0,023 | 0,008 | 0,99379373 | | NA | 1,00E-08 | 6,0651585 | 6,4027826 | 6,4027826 | TRUE | 0,3731075 | No |
| tK(CUU)1 | 8 | 7 | 5 | 6 | 3 | 6 | 2 | 3 | 7 | 9 | 5 | 5 | 3 | 6 | 2 | 4 | 5,06 | 6 | 4 | 0,543 | -0,88 | -1,393 | 0,16361697 | | NA | 1,00E-08 | 0,2890305 | 0,1640262 | 0,1640262 | TRUE | 0,3020239 | No |
| tK(CUU)2 | 7 | 4 | 3 | 5 | 1 | 1 | 5 | 3 | 6 | 5 | 3 | 4 | 1 | 1 | 4 | 4 | 3,63 | 4 | 2 | 0,544 | -0,88 | -1,15 | 0,24999111 | | NA | 1,00E-08 | 0,4015282 | 0,2563599 | 0,2563599 | TRUE | 0,3071051 | No |
| tK(UUU)1 | 1 | 0 | 0 | 0 | 0 | 0 | 0 | 0 | 1 | 0 | 0 | 0 | 0 | 0 | 0 | 0 | 0,11 | 0 | 0 | 0,616 | -0,698 | -0,198 | 0,84334499 | | NA | 1,00E-08 | 12,947638 | 10 | 10 | TRUE | 0,289592 | No |
| tK(UUU)2 | 0 | 0 | 0 | 0 | 0 | 0 | 0 | 0 | 0 | 0 | 0 | 0 | 0 | 0 | 0 | 0 | 0 | 0 | 0 | NA | NA | NA | NA | | NA | NA | NA | NA | NA | NA | NA | NA |
| tK(UUU)3 | 0 | 0 | 0 | 0 | 1 | 0 | 0 | 0 | 0 | 0 | 0 | 0 | 1 | 0 | 0 | 0 | 0,11 | 0 | 0 | 1,675 | 0,744 | 0,211 | 0,83317474 | | NA | 1,00E-08 | 13,581472 | 10 | 10 | TRUE | 0,2537777 | No |
| tK(UUU)4 | 0 | 0 | 0 | 0 | 0 | 0 | 0 | 0 | 0 | 0 | 0 | 0 | 0 | 0 | 0 | 0 | 0 | 0 | 0 | NA | NA | NA | NA | | NA | NA | NA | NA | NA | NA | NA | NA |
| tK(UUU)5 | 0 | 0 | 1 | 0 | 0 | 0 | 0 | 0 | 0 | 0 | 1 | 0 | 0 | 0 | 0 | 0 | 0,13 | 0 | 0 | 0,616 | -0,698 | -0,198 | 0,84334499 | | NA | 1,00E-08 | 11,353388 | 10 | 10 | TRUE | 0,3731057 | No |
| tL(AAG)1 | 3 | 1 | 5 | 5 | 3 | 3 | 2 | 4 | 3 | 1 | 5 | 4 | 3 | 3 | 2 | 6 | 3,26 | 3 | 4 | 0,907 | -0,14 | -0,183 | 0,8545702 | | NA | 1,00E-08 | 0,4476714 | 0,2533495 | 0,2533495 | TRUE | 0,22794 | No |
| tL(AAG)2 | 1 | 0 | 0 | 0 | 0 | 0 | 1 | 1 | 1 | 0 | 0 | 0 | 0 | 0 | 1 | 1 | 0,4 | 0 | 0 | 1,706 | 0,771 | 0,342 | 0,73235422 | | NA | 1,00E-08 | 3,6314038 | 2,9744678 | 2,9744678 | TRUE | 0,2895956 | No |
| tL(CAA)1 | 0 | 0 | 2 | 0 | 0 | 0 | 2 | 0 | 0 | 0 | 2 | 0 | 0 | 0 | 2 | 0 | 0,48 | 0 | 0 | 0,942 | -0,086 | -0,034 | 0,97262439 | | NA | 1,00E-08 | 3,033569 | 4,2608228 | 4,2608228 | TRUE | 0,9041947 | No |
| tL(CAA)2 | 0 | 0 | 0 | 0 | 0 | 0 | 1 | 0 | 0 | 0 | 0 | 0 | 0 | 0 | 1 | 0 | 0,11 | 0 | 0 | 1,675 | 0,744 | 0,211 | 0,83317474 | | NA | 1,00E-08 | 13,01685 | 10 | 10 | TRUE | 0,2773144 | No |
| tL(CAA)3 | 0 | 0 | 0 | 0 | 0 | 1 | 0 | 0 | 0 | 0 | 0 | 0 | 0 | 1 | 0 | 0 | 0,12 | 0 | 0 | 1,675 | 0,744 | 0,211 | 0,83317474 | | NA | 1,00E-08 | 12,173425 | 10 | 10 | TRUE | 0,3171327 | No |
| tL(CAA)4 | 73 | 57 | 70 | 81 | 71 | 48 | 74 | 40 | 65 | 73 | 72 | 67 | 61 | 46 | 66 | 56 | 63,18 | 69 | 57 | 0,833 | -0,263 | -1,484 | 0,13784011 | 0,20066475 | 1,00E-08 | 0,0249569 | 0,0142377 | 0,0142377 | TRUE | 0,1684224 | No | |
| tL(CAA)5 | 0 | 0 | 0 | 0 | 0 | 0 | 0 | 0 | 0 | 0 | 0 | 0 | 0 | 0 | 0 | 0 | 0 | 0 | 0 | NA | NA | NA | NA | | NA | NA | NA | NA | NA | NA | NA | NA |
| tL(CAA)6 | 0 | 0 | 1 | 0 | 2 | 0 | 1 | 0 | 0 | 0 | 1 | 0 | 2 | 0 | 1 | 0 | 0,45 | 0 | 1 | 2,288 | 1,194 | 0,557 | 0,57776593 | | NA | 1,00E-08 | 3,2049427 | 2,6904033 | 2,6904033 | TRUE | 0,4198533 | No |
| tL(UAA)1 | 0 | 0 | 0 | 0 | 0 | 0 | 0 | 0 | 0 | 0 | 0 | 0 | 0 | 0 | 0 | 0 | 0 | 0 | 0 | NA | NA | NA | NA | | NA | NA | NA | NA | NA | NA | NA | NA |
| tL(UAA)2 | 0 | 0 | 0 | 2 | 0 | 0 | 1 | 0 | 0 | 0 | 0 | 2 | 0 | 0 | 1 | 0 | 0,32 | 0 | 0 | 0,653 | -0,614 | -0,222 | 0,8242023 | | NA | 1,00E-08 | 4,5714615 | 5,3916762 | 5,3916762 | TRUE | 0,8163204 | No |
| tL(UAA)3 | 0 | 1 | 0 | 0 | 1 | 0 | 1 | 0 | 0 | 1 | 0 | 0 | 1 | 0 | 1 | 0 | 0,38 | 0 | 0 | 1,627 | 0,702 | 0,31 | 0,75693144 | | NA | 1,00E-08 | 3,84669 | 3,0099701 | 3,0099701 | TRUE | 0,5444409 | No |
| tL(UAA)4 | 0 | 2 | 2 | 2 | 1 | 0 | 0 | 1 | 0 | 3 | 2 | 2 | 1 | 0 | 0 | 1 | 1,06 | 2 | 0 | 0,331 | -1,595 | -1,062 | 0,28833388 | | NA | 1,00E-08 | 1,3671176 | 0,9214629 | 0,9214629 | TRUE | 0,3773939 | No |
| tL(UAA)5 | 1 | 0 | 0 | 0 | 0 | 0 | 0 | 0 | 1 | 0 | 0 | 0 | 0 | 0 | 0 | 0 | 0,11 | 0 | 0 | 0,616 | -0,698 | -0,198 | 0,84334499 | | NA | 1,00E-08 | 12,947638 | 10 | 10 | TRUE | 0,289592 | No |
| tM(CAU)1 | 1 | 0 | 0 | 0 | 0 | 0 | 0 | 0 | 1 | 0 | 0 | 0 | 0 | 0 | 0 | 0 | 0,11 | 0 | 0 | 0,616 | -0,698 | -0,198 | 0,84334499 | | NA | 1,00E-08 | 12,947638 | 10 | 10 | TRUE | 0,289592 | No |
| tM(CAU)2 | 0 | 0 | 0 | 0 | 1 | 1 | 0 | 0 | 0 | 0 | 0 | 0 | 1 | 1 | 0 | 0 | 0,23 | 0 | 0 | 2,726 | 1,447 | 0,483 | 0,62927289 | | NA | 1,00E-08 | 6,4204661 | 6,7048567 | 6,7048567 | TRUE | 0,0559113 | No |
| tM(CAU)3 | 0 | 1 | 0 | 0 | 0 | 0 | 0 | 1 | 0 | 1 | 0 | 0 | 0 | 0 | 0 | 1 | 0,34 | 0 | 0 | 1,016 | 0,023 | 0,009 | 0,99310922 | | NA | 1,00E-08 | 4,3297361 | 4,817097 | 4,817097 | TRUE | 0,6248753 | No |
| tM(CAU)4 | 0 | 0 | 1 | 1 | 0 | 0 | 0 | 0 | 0 | 0 | 1 | 1 | 0 | 0 | 0 | 0 | 0,23 | 0 | 0 | 0,379 | -1,402 | -0,471 | 0,63792379 | | NA | 1,00E-08 | 6,2873821 | 6,584019 | 6,584019 | TRUE | 0,0700646 | No |
| tN(GUU)1 | 0 | 0 | 0 | 0 | 2 | 0 | 0 | 0 | 0 | 0 | 0 | 0 | 2 | 0 | 0 | 0 | 0,21 | 0 | 0 | 2,624 | 1,392 | 0,395 | 0,69317318 | | NA | 1,00E-08 | 6,7917257 | 10 | 10 | TRUE | 0,82123 | No |
| tN(GUU)2 | 0 | 0 | 0 | 0 | 0 | 0 | 0 | 0 | 0 | 0 | 0 | 0 | 0 | 0 | 0 | 0 | 0 | 0 | 0 | NA | NA | NA | NA | | NA | NA | NA | NA | NA | NA | NA | NA |
| tN(GUU)3 | 0 | 0 | 1 | 0 | 0 | 1 | 0 | 0 | 0 | 0 | 1 | 0 | 0 | 1 | 0 | 0 | 0,25 | 0 | 0 | 1,016 | 0,023 | 0,008 | 0,99373105 | | NA | 1,00E-08 | 5,8755459 | 6,2357025 | 6,2357025 | TRUE | 0,3731077 | No |
| tN(GUU)4 | 0 | 0 | 0 | 0 | 0 | 1 | 0 | 0 | 0 | 0 | 0 | 0 | 0 | 1 | 0 | 0 | 0,12 | 0 | 0 | 1,675 | 0,744 | 0,211 | 0,83317474 | | NA | 1,00E-08 | 12,173425 | 10 | 10 | TRUE | 0,3171327 | No |
| tP(AGG)1 | 0 | 2 | 0 | 0 | 1 | 0 | 2 | 1 | 0 | 3 | 0 | 0 | 1 | 0 | 2 | 1 | 0,82 | 1 | 1 | 1,94 | 0,956 | 0,543 | 0,58695886 | | NA | 1,00E-08 | 1,7625448 | 1,5583579 | 1,5583579 | TRUE | 1,2621409 | No |
| tP(UGG)1 | 0 | 0 | 0 | 0 | 0 | 0 | 1 | 0 | 0 | 0 | 0 | 0 | 0 | 0 | 1 | 0 | 0,11 | 0 | 0 | 1,675 | 0,744 | 0,211 | 0,83317474 | | NA | 1,00E-08 | 13,01685 | 10 | 10 | TRUE | 0,2773144 | No |
| tP(UGG)2 | 1 | 1 | 2 | 2 | 2 | 0 | 1 | 0 | 1 | 1 | 2 | 2 | 2 | 0 | 1 | 0 | 1,06 | 2 | 1 | 0,486 | -1,041 | -0,769 | 0,44171548 | | NA | 1,00E-08 | 1,3733799 | 0,7213286 | 0,7213286 | TRUE | 0,4168735 | No |
| tP(UGG)3 | 1 | 0 | 1 | 2 | 4 | 4 | 2 | 3 | 1 | 0 | 1 | 2 | 3 | 4 | 2 | 4 | 2,1 | 1 | 3 | 3,406 | 1,768 | 1,713 | 0,08666001 | | NA | 1,00E-08 | 0,6923237 | 0,3549712 | 0,3549712 | TRUE | 0,1607486 | No |
| tP(UGG)4 | 0 | 0 | 1 | 0 | 0 | 0 | 0 | 0 | 0 | 0 | 1 | 0 | 0 | 0 | 0 | 0 | 0,13 | 0 | 0 | 0,616 | -0,698 | -0,198 | 0,84334499 | | NA | 1,00E-08 | 11,353388 | 10 | 10 | TRUE | 0,3731057 | No |
| tP(UGG)5 | 0 | 0 | 1 | 0 | 0 | 0 | 0 | 1 | 0 | 0 | 1 | 0 | 0 | 0 | 0 | 1 | 0,3 | 0 | 0 | 1,016 | 0,023 | 0,008 | 0,99331204 | | NA | 1,00E-08 | 4,774351 | 5,236601 | 5,236601 | TRUE | 0,6248751 | No |
| tQ(CUG)1 | 1 | 1 | 1 | 4 | 0 | 0 | 0 | 0 | 1 | 1 | 1 | 3 | 0 | 0 | 0 | 0 | 0,81 | 2 | 0 | 0,112 | -3,155 | -1,924 | 0,05433643 | | NA | 1,00E-08 | 1,7909563 | 1,2824545 | 1,2824545 | TRUE | 0,4522828 | No |
| tQ(UUG)1 | 1 | 0 | 0 | 0 | 0 | 0 | 2 | 1 | 1 | 0 | 0 | 0 | 0 | 0 | 2 | 1 | 0,51 | 0 | 1 | 2,584 | 1,37 | 0,649 | 0,51625401 | | NA | 1,00E-08 | 2,8399782 | 2,6265336 | 2,6265336 | TRUE | 0,1919053 | No |
| tQ(UUG)3 | 0 | 0 | 0 | 0 | 0 | 2 | 0 | 1 | 0 | 0 | 0 | 0 | 0 | 2 | 0 | 1 | 0,41 | 0 | 1 | 4,438 | 2,15 | 0,854 | 0,3932329 | | NA | 1,00E-08 | 3,5013748 | 4,4636645 | 4,4636645 | TRUE | 0,1014416 | No |
| tQ(UUG)4 | 0 | 0 | 0 | 0 | 0 | 0 | 0 | 0 | 0 | 0 | 0 | 0 | 0 | 0 | 0 | 0 | 0 | 0 | 0 | NA | NA | NA | NA | | NA | NA | NA | NA | NA | NA | NA | NA |
| tQ(UUG)5 | 0 | 0 | 1 | 0 | 0 | 1 | 1 | 0 | 0 | 0 | 1 | 0 | 0 | 1 | 1 | 0 | 0,36 | 0 | 0 | 1,674 | 0,744 | 0,324 | 0,74622179 | | NA | 1,00E-08 | 4,0490961 | 3,1539764 | 3,1539764 | TRUE | 0,3677882 | No |
| tR(ACG)1 | 0 | 2 | 0 | 0 | 0 | 0 | 1 | 0 | 0 | 3 | 0 | 0 | 0 | 0 | 1 | 0 | 0,43 | 1 | 0 | 0,561 | -0,835 | -0,323 | 0,74683659 | | NA | 10 | 3,3794551 | 4,5547842 | 4,5547842 | TRUE | 1,1850919 | No |
| tR(ACG)2 | 0 | 0 | 1 | 0 | 0 | 0 | 0 | 0 | 0 | 0 | 1 | 0 | 0 | 0 | 0 | 0 | 0,13 | 0 | 0 | 0,616 | -0,698 | -0,198 | 0,84334499 | | NA | 1,00E-08 | 11,353388 | 10 | 10 | TRUE | 0,3731057 | No |
| tR(CCG)1 | 10 | 8 | 6 | 7 | 6 | 6 | 8 | 9 | 9 | 10 | 6 | 6 | 5 | 6 | 7 | 13 | 7,72 | 8 | 8 | 0,962 | -0,056 | -0,111 | 0,91161828 | | NA | 1,00E-08 | 0,1900373 | 0,1146452 | 0,1146452 | TRUE | 0,366037 | No |
| tR(CCU)1 | 98 | 68 | 95 | 112 | 53 | 78 | 59 | 50 | 88 | 87 | 97 | 92 | 45 | 74 | 53 | 71 | 75,87 | 91 | 61 | 0,659 | -0,603 | -3,354 | 0,00079512 | 0,00204499 | 0,01086 | 0,0211136 | 0,017181 | 0,017181 | TRUE | 0,2788771 | No | |
| tR(UCU)1 | 0 | 0 | 0 | 1 | 3 | 1 | 1 | 0 | 0 | 0 | 0 | 1 | 3 | 1 | 1 | 0 | 0,65 | 0 | 1 | 3,847 | 1,944 | 1,103 | 0,26999694 | | NA | 1,00E-08 | 2,2196675 | 1,5463516 | 1,5463516 | TRUE | 0,3423076 | No |
| tR(UCU)2 | 0 | 0 | 0 | 0 | 0 | 0 | 0 | 0 | 0 | 0 | 0 | 0 | 0 | 0 | 0 | 0 | 0 | 0 | 0 | NA | NA | NA | NA | | NA | NA | NA | NA | NA | NA | NA | NA |
| tR(UCU)3 | 0 | 0 | 0 | 0 | 0 | 0 | 0 | 0 | 0 | 0 | 0 | 0 | 0 | 0 | 0 | 0 | 0 | 0 | 0 | NA | NA | NA | NA | | NA | NA | NA | NA | NA | NA | NA | NA |
| tR(UCU)4 | 6 | 2 | 2 | 2 | 4 | 2 | 6 | 3 | 5 | 3 | 2 | 2 | 3 | 2 | 5 | 4 | 3,32 | 3 | 4 | 1,27 | 0,344 | 0,452 | 0,65161203 | | NA | 1,00E-08 | 0,4395889 | 0,2548642 | 0,2548642 | TRUE | 0,49624 | No |

| | | | | | | | | | | | | | | | | | | | | | | | | | | | | | | | |
|---|---|---|---|---|---|---|---|---|---|---|---|---|---|---|---|---|---|---|---|---|---|---|---|---|---|---|---|---|---|---|---|
| tR(UCU)5 | 0 | 0 | 0 | 0 | 0 | 0 | 0 | 0 | 0 | 0 | 0 | 0 | 0 | 0 | 0 | 0 | 0 | 0 | 0 | NA | NA | NA | NA | NA | NA | NA | NA | NA | NA | NA | NA |
| tS(AGA)1 | 2 | 0 | 3 | 4 | 2 | 4 | 2 | 1 | 2 | 0 | 3 | 3 | 2 | 4 | 2 | 1 | 2,11 | 2 | 2 | 1,037 | 0,052 | 0,055 | 0,95635425 | NA | 1,00E-08 | 0,6898196 | 0,3998502 | 0,3998502 | TRUE | 0,2770245 | No |
| tS(AGA)2 | 0 | 0 | 1 | 0 | 0 | 0 | 0 | 0 | 0 | 0 | 1 | 0 | 0 | 0 | 0 | 0 | 0,13 | 0 | 0 | 0,616 | -0,698 | -0,198 | 0,84334499 | NA | 1,00E-08 | 11,353388 | 10 | 10 | TRUE | 0,3731057 | No |
| tS(AGA)3 | 1 | 0 | 0 | 0 | 1 | 0 | 1 | 0 | 1 | 0 | 0 | 0 | 1 | 0 | 1 | 0 | 0,33 | 0 | 0 | 1,626 | 0,701 | 0,299 | 0,76496839 | NA | 1,00E-08 | 4,3933062 | 3,3459086 | 3,3459086 | TRUE | 0,2851494 | No |
| tS(AGA)4 | 1 | 0 | 1 | 0 | 2 | 0 | 0 | 1 | 1 | 0 | 1 | 0 | 2 | 0 | 0 | 1 | 0,63 | 0 | 1 | 1,516 | 0,601 | 0,324 | 0,74592404 | NA | 1,00E-08 | 2,305775 | 1,7097623 | 1,7097623 | TRUE | 0,2105179 | No |
| tS(CAG)1 | 1 | 0 | 1 | 0 | 0 | 0 | 1 | 0 | 1 | 0 | 1 | 0 | 0 | 0 | 1 | 0 | 0,35 | 0 | 0 | 0,608 | -0,719 | -0,311 | 0,75614239 | NA | 1,00E-08 | 4,1312086 | 3,2208255 | 3,2208255 | TRUE | 0,2645147 | No |
| tS(CGA)1 | 0 | 1 | 0 | 1 | 0 | 0 | 1 | 0 | 0 | 1 | 0 | 1 | 0 | 0 | 1 | 0 | 0,37 | 0 | 0 | 0,623 | -0,682 | -0,299 | 0,76478835 | NA | 1,00E-08 | 3,8860606 | 3,0564468 | 3,0564468 | TRUE | 0,2773135 | No |
| tS(GCU)1 | 24 | 25 | 28 | 29 | 10 | 13 | 8 | 5 | 22 | 32 | 29 | 24 | 9 | 12 | 7 | 7 | 17,63 | 27 | 9 | 0,338 | -1,563 | -4,351 | 1,36E-05 | 4,84E-05 | 1,00E-08 | 0,084307 | 0,0488555 | 0,0488555 | TRUE | 0,2515129 | No |
| tS(GCU)2 | 15 | 13 | 14 | 29 | 8 | 6 | 5 | 7 | 13 | 17 | 14 | 24 | 7 | 6 | 4 | 10 | 11,89 | 17 | 7 | 0,379 | -1,4 | -3,189 | 0,00142803 | NA | 1,00E-08 | 0,1240411 | 0,0794393 | 0,0794393 | TRUE | 0,4405206 | No |
| tS(UGA)1 | 0 | 0 | 0 | 1 | 0 | 0 | 0 | 0 | 0 | 0 | 0 | 1 | 0 | 0 | 0 | 0 | 0,1 | 0 | 0 | 0,616 | -0,698 | -0,198 | 0,84334499 | NA | 1,00E-08 | 14,085693 | 10 | 10 | TRUE | 0,2428614 | No |
| tS(UGA)2 | 0 | 1 | 2 | 4 | 0 | 0 | 1 | 0 | 0 | 1 | 2 | 3 | 0 | 0 | 1 | 0 | 0,94 | 2 | 0 | 0,183 | -2,453 | -1,499 | 0,13377921 | NA | 1,00E-08 | 1,5483806 | 1,2742618 | 1,2742618 | TRUE | 0,169429 | No |
| tS(UGA)3 | 0 | 0 | 0 | 0 | 0 | 0 | 0 | 0 | 0 | 0 | 0 | 0 | 0 | 0 | 0 | 0 | 0 | 0 | 0 | NA | NA | NA | NA | NA | NA | NA | NA | NA | NA | NA | NA |
| tT(AGU)1 | 1 | 0 | 1 | 0 | 0 | 0 | 0 | 0 | 1 | 0 | 1 | 0 | 0 | 0 | 0 | 0 | 0,24 | 0 | 0 | 0,367 | -1,446 | -0,49 | 0,62399902 | NA | 1,00E-08 | 6,0500945 | 6,4379451 | 6,4379451 | TRUE | 0,0586 | No |
| tT(AGU)3 | 0 | 0 | 0 | 0 | 0 | 0 | 0 | 0 | 0 | 0 | 0 | 0 | 0 | 0 | 0 | 0 | 0 | 0 | 0 | NA | NA | NA | NA | NA | NA | NA | NA | NA | NA | NA | NA |
| tT(AGU)4 | 0 | 0 | 1 | 0 | 0 | 0 | 0 | 0 | 0 | 0 | 1 | 0 | 0 | 0 | 0 | 0 | 0,13 | 0 | 0 | 0,616 | -0,698 | -0,198 | 0,84334499 | NA | 1,00E-08 | 11,353388 | 10 | 10 | TRUE | 0,3731057 | No |
| tT(AGU)6 | 0 | 0 | 0 | 0 | 1 | 0 | 0 | 0 | 0 | 0 | 0 | 0 | 1 | 0 | 0 | 0 | 0,11 | 0 | 0 | 1,675 | 0,744 | 0,211 | 0,83317474 | NA | 1,00E-08 | 13,581472 | 10 | 10 | TRUE | 0,2537777 | No |
| tT(CGU)1 | 1 | 0 | 2 | 0 | 0 | 0 | 0 | 0 | 1 | 0 | 2 | 0 | 0 | 0 | 0 | 0 | 0,37 | 1 | 0 | 0,251 | -1,995 | -0,771 | 0,44077307 | NA | 1,00E-08 | 3,9477556 | 4,7598899 | 4,7598899 | TRUE | 0,4826136 | No |
| tT(UGU)1 | 0 | 0 | 0 | 0 | 0 | 0 | 0 | 0 | 0 | 0 | 0 | 0 | 0 | 0 | 0 | 0 | 0 | 0 | 0 | NA | NA | NA | NA | NA | NA | NA | NA | NA | NA | NA | NA |
| tT(UGU)2 | 0 | 0 | 0 | 0 | 0 | 0 | 0 | 0 | 0 | 0 | 0 | 0 | 0 | 0 | 0 | 0 | 0 | 0 | 0 | NA | NA | NA | NA | NA | NA | NA | NA | NA | NA | NA | NA |
| tV(AAC)1 | 0 | 1 | 0 | 1 | 1 | 0 | 0 | 0 | 0 | 1 | 0 | 1 | 1 | 0 | 0 | 0 | 0,37 | 0 | 0 | 0,623 | -0,682 | -0,298 | 0,76549438 | NA | 1,00E-08 | 3,9348622 | 3,0870344 | 3,0870344 | TRUE | 0,2537768 | No |
| tV(AAC)2 | 0 | 0 | 0 | 1 | 0 | 0 | 0 | 0 | 0 | 0 | 0 | 1 | 0 | 0 | 0 | 0 | 0,1 | 0 | 0 | 0,616 | -0,698 | -0,198 | 0,84334499 | NA | 1,00E-08 | 14,085693 | 10 | 10 | TRUE | 0,2428614 | No |
| tV(AAC)3 | 0 | 1 | 0 | 0 | 0 | 0 | 0 | 0 | 0 | 1 | 0 | 0 | 0 | 0 | 0 | 0 | 0,16 | 0 | 0 | 0,616 | -0,698 | -0,198 | 0,84334499 | NA | 1,00E-08 | 9,1242388 | 10 | 10 | TRUE | 0,5444342 | No |
| tV(AAC)4 | 0 | 0 | 0 | 0 | 0 | 0 | 0 | 0 | 0 | 0 | 0 | 0 | 0 | 0 | 0 | 0 | 0 | 0 | 0 | NA | NA | NA | NA | NA | NA | NA | NA | NA | NA | NA | NA |
| tV(AAC)5 | 1 | 0 | 1 | 6 | 2 | 1 | 0 | 1 | 1 | 0 | 1 | 5 | 2 | 1 | 0 | 1 | 1,37 | 2 | 1 | 0,552 | -0,857 | -0,655 | 0,51256315 | NA | 0,590091 | 1,0632692 | 0,8673932 | 0,8673932 | TRUE | 1,5127092 | No |
| tV(AAC)6 | 1 | 2 | 2 | 0 | 1 | 2 | 0 | 0 | 1 | 3 | 2 | 0 | 1 | 2 | 0 | 0 | 1,03 | 2 | 1 | 0,554 | -0,852 | -0,575 | 0,56523071 | NA | 1,00E-08 | 1,4091638 | 1,0372503 | 1,0372503 | TRUE | 0,378485 | No |
| tV(CAC)1 | 0 | 0 | 5 | 3 | 1 | 0 | 2 | 1 | 0 | 0 | 5 | 2 | 1 | 0 | 2 | 1 | 1,45 | 2 | 1 | 0,519 | -0,947 | -0,693 | 0,48853978 | NA | 1,095837 | 0,9997431 | 1,0316714 | 1,0316714 | TRUE | 0,5071874 | No |
| tV(UAC)1 | 3 | 3 | 8 | 5 | 8 | 9 | 9 | 10 | 3 | 4 | 8 | 4 | 7 | 9 | 8 | 14 | 7,05 | 5 | 10 | 1,956 | 0,968 | 1,755 | 0,07932085 | NA | 1,00E-08 | 0,2079568 | 0,1314723 | 0,1314723 | TRUE | 0,4674952 | No |
| tW(CCA)1 | 0 | 0 | 0 | 0 | 0 | 0 | 1 | 1 | 0 | 0 | 0 | 0 | 0 | 0 | 1 | 1 | 0,29 | 0 | 0 | 2,833 | 1,502 | 0,536 | 0,59180467 | NA | 1,00E-08 | 5,0453571 | 5,6379511 | 5,6379511 | TRUE | 0,1714431 | No |
| tW(CCA)2 | 4 | 0 | 2 | 0 | 1 | 0 | 0 | 0 | 4 | 0 | 2 | 0 | 1 | 0 | 0 | 0 | 0,81 | 2 | 0 | 0,214 | -2,226 | -1,182 | 0,23704856 | NA | 3,064302 | 1,7914839 | 2,0581385 | 2,0581385 | TRUE | 0,2252711 | No |
| tY(GUA)1 | 0 | 0 | 0 | 1 | 0 | 1 | 0 | 0 | 0 | 0 | 0 | 1 | 0 | 1 | 0 | 0 | 0,22 | 0 | 0 | 1,016 | 0,023 | 0,008 | 0,99393821 | NA | 1,00E-08 | 6,5309496 | 6,8078458 | 6,8078458 | TRUE | 0,3171316 | No |
| tY(GUA)2 | 1 | 0 | 2 | 3 | 1 | 0 | 0 | 0 | 1 | 0 | 2 | 2 | 1 | 0 | 0 | 0 | 0,78 | 1 | 0 | 0,219 | -2,193 | -1,312 | 0,18938248 | NA | 1,00E-08 | 1,8535296 | 1,3280253 | 1,3280253 | TRUE | 0,2034151 | No |
| tY(GUA)3 | 5 | 1 | 3 | 2 | 8 | 4 | 5 | 5 | 4 | 1 | 3 | 2 | 7 | 4 | 4 | 7 | 4,08 | 2 | 6 | 2,052 | 1,037 | 1,477 | 0,13971184 | NA | 1,00E-08 | 0,3576862 | 0,1973999 | 0,1973999 | TRUE | 0,3027917 | No |
| tY(GUA)4 | 0 | 0 | 0 | 0 | 0 | 0 | 0 | 0 | 0 | 0 | 0 | 0 | 0 | 0 | 0 | 0 | 0 | 0 | 0 | NA | NA | NA | NA | NA | NA | NA | NA | NA | NA | NA | NA |

Wild type doxycycline-treated up-regulated genes

| Gene ID | WT 1 | WT 2 | WT 3 | WT 4 | WT dox 1 | WT dox 2 | WT dox 3 | WT dox 4 | norm. WT 1 | norm. WT 2 | norm. WT 3 | norm. WT 4 | norm. WT dox 1 | norm. WT dox 2 | norm. WT dox 3 | norm. WT dox 4 | base Mean | WT | WT dox | FC | log2FC | stat | pvalue | padj | disp. GeneEst | dispFit | dispMAP | dispersion | betaConv | maxCooks | outlier |
|---|---|---|---|---|---|---|---|---|---|---|---|---|---|---|---|---|---|---|---|---|---|---|---|---|---|---|---|---|---|---|---|
| orf19.7453 | 21 | 14 | 18 | 27 | 65 | 56 | 70 | 42 | 19 | 18 | 18 | 22 | 56 | 53 | 62 | 59 | 38,51 | 19 | 58 | 2,955 | 1,563 | 6,62 | 3,60E-11 | 2,78E-10 | 1,00E-08 | 0,0396778 | 0,0195964 | 0,0195964 | TRUE | 0,0674984 | No |

| | WT | | | | WT dox | | | | norm. WT | | | | norm. WT dox | | | | | | | | | | | | | | | | | | |
|---|---|---|---|---|---|---|---|---|---|---|---|---|---|---|---|---|---|---|---|---|---|---|---|---|---|---|---|---|---|---|---|
| Gene ID | 1 | 2 | 3 | 4 | 1 | 2 | 3 | 4 | 1 | 2 | 3 | 4 | 1 | 2 | 3 | 4 | base Mean | WT | WT dox | FC | log2FC | stat | pvalue | padj | disp GeneEst | dispFit | dispMAP | dispersion | betaConv | maxCooks | outlier |
| orf19.4335 | 1590 | 1145 | 1479 | 1815 | 278 | 296 | 240 | 175 | 1426 | 1458 | 1513 | 1497 | 238 | 282 | 214 | 247 | 859,39 | 1474 | 245 | 0,166 | -2,59 | -34,682 | 1,46E-263 | 8,75E-260 | 0,0014368 | 0,0036687 | 0,0029702 | 0,0029702 | TRUE | 0,1256677 | No |
| orf19.4583 | 616 | 346 | 492 | 625 | 167 | 159 | 161 | 98 | 553 | 440 | 503 | 515 | 143 | 152 | 144 | 138 | 323,54 | 503 | 144 | 0,286 | -1,805 | -18,172 | 8,53E-74 | 2,05E-71 | 1,00E-08 | 0,0064665 | 0,0050209 | 0,0050209 | TRUE | 0,0745945 | No |
| orf19.403 | 81 | 63 | 62 | 90 | 26 | 23 | 19 | 17 | 73 | 80 | 63 | 74 | 22 | 22 | 17 | 24 | 46,95 | 72 | 21 | 0,291 | -1,783 | -7,948 | 1,90E-15 | 2,28E-14 | 1,00E-08 | 0,0328971 | 0,0178259 | 0,0178259 | TRUE | 0,117728 | No |
| orf19.7447 | 97 | 75 | 101 | 142 | 39 | 28 | 39 | 18 | 87 | 95 | 103 | 117 | 33 | 27 | 35 | 25 | 65,4 | 100 | 30 | 0,302 | -1,729 | -8,731 | 2,52E-18 | 4,04E-17 | 1,00E-08 | 0,0241772 | 0,0163541 | 0,0163541 | TRUE | 0,1310804 | No |
| orf19.1409.3 | 43 | 48 | 30 | 50 | 19 | 14 | 18 | 9 | 39 | 61 | 31 | 41 | 16 | 13 | 16 | 13 | 28,75 | 43 | 14 | 0,347 | -1,526 | -5,045 | 4,54E-07 | 2,02E-06 | 1,00E-08 | 0,0524797 | 0,0422115 | 0,0422115 | TRUE | 0,3966769 | No |
| orf19.935 | 42 | 49 | 65 | 63 | 24 | 12 | 15 | 24 | 38 | 62 | 67 | 52 | 21 | 11 | 13 | 34 | 37,22 | 55 | 20 | 0,352 | -1,508 | -4,785 | 1,71E-06 | 6,96E-06 | 0,0909948 | 0,0409876 | 0,0598603 | 0,0598603 | TRUE | 0,8108122 | No |
| orf19.3735 | 36 | 42 | 27 | 39 | 12 | 11 | 9 | 18 | 32 | 53 | 28 | 32 | 10 | 10 | 8 | 25 | 24,97 | 36 | 13 | 0,357 | -1,485 | -4,037 | 5,42E-05 | 0,000174 | 0,1046151 | 0,060127 | 0,0770807 | 0,0770807 | TRUE | 1,046267 | No |
| orf19.6169 | 1316 | 844 | 1168 | 1426 | 450 | 443 | 490 | 339 | 1181 | 1075 | 1195 | 1176 | 385 | 423 | 437 | 478 | 793,61 | 1157 | 431 | 0,37 | -1,435 | -20,221 | 6,46E-91 | 3,23E-88 | 0,002565 | 0,0038088 | 0,0032293 | 0,0032293 | TRUE | 0,0608939 | No |
| orf19.218 | 66 | 53 | 55 | 58 | 28 | 20 | 24 | 15 | 59 | 67 | 56 | 48 | 24 | 19 | 21 | 21 | 39,55 | 58 | 21 | 0,375 | -1,415 | -5,974 | 2,32E-09 | 1,40E-08 | 1,00E-08 | 0,0386869 | 0,0219898 | 0,0219898 | TRUE | 0,1261572 | No |
| orf19.7094 | 47 | 25 | 28 | 41 | 12 | 16 | 15 | 10 | 42 | 32 | 29 | 34 | 10 | 15 | 13 | 14 | 23,68 | 34 | 13 | 0,384 | -1,383 | -4,552 | 5,32E-06 | 2,02E-05 | 1,00E-08 | 0,063276 | 0,0361918 | 0,0361918 | TRUE | 0,1912765 | No |
| orf19.1655 | 5852 | 3731 | 5021 | 6147 | 2404 | 2002 | 2213 | 1558 | 5250 | 4750 | 5137 | 5069 | 2056 | 1910 | 1975 | 2197 | 3543 | 5052 | 2034 | 0,402 | -1,315 | -25,179 | 6,74E-140 | 1,35E-136 | 0,0022936 | 0,0023893 | 0,00227 | 0,00227 | TRUE | 0,0326763 | No |
| orf19.1345 | 88 | 54 | 84 | 88 | 47 | 28 | 38 | 15 | 79 | 69 | 86 | 73 | 40 | 27 | 34 | 21 | 53,52 | 77 | 30 | 0,407 | -1,296 | -6,115 | 9,67E-10 | 6,17E-09 | 0,0021636 | 0,0291029 | 0,0206272 | 0,0206272 | TRUE | 0,318968 | No |
| orf19.5302 | 299 | 207 | 250 | 290 | 127 | 104 | 111 | 80 | 268 | 264 | 256 | 239 | 109 | 99 | 99 | 113 | 180,8 | 257 | 105 | 0,408 | -1,295 | -11,647 | 2,38E-31 | 9,11E-30 | 1,00E-08 | 0,0100088 | 0,0051505 | 0,0051505 | TRUE | 0,0261205 | No |
| orf19.1748 | 405 | 270 | 359 | 423 | 168 | 151 | 173 | 126 | 363 | 344 | 367 | 349 | 144 | 144 | 154 | 178 | 255,38 | 356 | 155 | 0,431 | -1,214 | -12,665 | 9,26E-37 | 5,15E-35 | 1,00E-08 | 0,0076641 | 0,0041713 | 0,0041713 | TRUE | 0,0794407 | No |

$P_{TET}$-*ZCF15* doxycycline-treated complete data

| | WT dox | | | | ZCF15 | | | | norm. WT dox | | | | norm. ZCF15 | | | | | | | | | | | | | | | | | | |
|---|---|---|---|---|---|---|---|---|---|---|---|---|---|---|---|---|---|---|---|---|---|---|---|---|---|---|---|---|---|---|---|
| Gene ID | 1 | 2 | 3 | 4 | 1 | 2 | 3 | 4 | 1 | 2 | 3 | 4 | 1 | 2 | n3 | 4 | base Mean | WT | ZCF15 | FC | log2FC | stat | pvalue | padj | dispGeneEst | dispFit | dispMAP | dispersion | betaConv | maxCooks | outlier |
| CaalfMp01 | 20 | 33 | 22 | 23 | 15 | 18 | 19 | 23 | 18 | 33 | 21 | 35 | 13 | 20 | 19 | 18 | 22,11 | 27 | 18 | 0,662 | -0,594 | -1,875 | 0,060741 | 0,074624 | 0,028646 | 0,061796 | 0,0492682 | 0,0492682 | TRUE | 0,1379523 | No |
| CaalfMp02 | 1 | 3 | 4 | 3 | 3 | 2 | 3 | 1 | 1 | 3 | 4 | 5 | 3 | 2 | 3 | 1 | 2,61 | 3 | 2 | 0,716 | -0,483 | -0,558 | 0,576766 | 0,617237 | 1,00E-08 | 0,507397 | 0,3195841 | 0,3195841 | TRUE | 0,3522586 | No |
| CaalfMp03 | 8 | 6 | 7 | 5 | 4 | 3 | 7 | 2 | 7 | 6 | 7 | 8 | 4 | 3 | 7 | 2 | 5,37 | 7 | 4 | 0,551 | -0,861 | -1,408 | 0,159141 | 0,186665 | 1,00E-08 | 0,24782 | 0,1575036 | 0,1575036 | TRUE | 0,5257083 | No |
| CaalfMp04 | 0 | 0 | 0 | 0 | 0 | 0 | 0 | 0 | 0 | 0 | 0 | 0 | 0 | 0 | 0 | 0 | 0 | 0 | 0 | NA | NA | NA | NA | NA | NA | NA | NA | NA | NA | NA | NA |
| CaalfMp05 | 23 | 40 | 27 | 32 | 11 | 14 | 8 | 12 | 21 | 40 | 26 | 48 | 10 | 15 | 8 | 9 | 22,17 | 34 | 10 | 0,316 | -1,662 | -4,571 | 4,86E-06 | 7,85E-06 | 0,073509 | 0,061626 | 0,0662982 | 0,0662982 | TRUE | 0,0975802 | No |
| CaalfMp06 | 5 | 19 | 13 | 13 | 7 | 7 | 8 | 11 | 5 | 19 | 12 | 20 | 6 | 8 | 8 | 8 | 10,75 | 14 | 8 | 0,561 | -0,835 | -1,739 | 0,081957 | 0,099254 | 0,1130683 | 0,124906 | 0,1205865 | 0,1205865 | TRUE | 0,5910115 | No |
| CaalfMp07 | 0 | 0 | 1 | 0 | 0 | 0 | 1 | 1 | 0 | 0 | 1 | 0 | 0 | 0 | 1 | 1 | 0,34 | 0 | 0 | 1,406 | 0,491 | 0,214 | 0,83033 | 0,853767 | 1,00E-08 | 3,868357 | 3,0967151 | 3,0967151 | TRUE | 0,2762674 | No |
| CaalfMp08 | 21 | 21 | 20 | 17 | 14 | 16 | 24 | 15 | 19 | 21 | 19 | 26 | 12 | 17 | 25 | 11 | 18,83 | 21 | 16 | 0,773 | -0,372 | -1,111 | 0,266776 | 0,302669 | 0,0170878 | 0,072177 | 0,0535567 | 0,0535567 | TRUE | 0,5321179 | No |
| CaalfMp08.1 | 0 | 0 | 0 | 0 | 0 | 0 | 0 | 0 | 0 | 0 | 0 | 0 | 0 | 0 | 0 | 0 | 0 | 0 | 0 | NA | NA | NA | NA | NA | NA | NA | NA | NA | NA | NA | NA |
| CaalfMp08.2 | 0 | 0 | 0 | 0 | 0 | 0 | 0 | 0 | 0 | 0 | 0 | 0 | 0 | 0 | 0 | 0 | 0 | 0 | 0 | NA | NA | NA | NA | NA | NA | NA | NA | NA | NA | NA | NA |
| CaalfMp08.3 | 0 | 0 | 0 | 0 | 0 | 0 | 0 | 0 | 0 | 0 | 0 | 0 | 0 | 0 | 0 | 0 | 0 | 0 | 0 | NA | NA | NA | NA | NA | NA | NA | NA | NA | NA | NA | NA |
| CaalfMp08.4 | 0 | 0 | 0 | 0 | 0 | 0 | 0 | 0 | 0 | 0 | 0 | 0 | 0 | 0 | 0 | 0 | 0 | 0 | 0 | NA | NA | NA | NA | NA | NA | NA | NA | NA | NA | NA | NA |
| CaalfMp09 | 5 | 5 | 3 | 2 | 4 | 3 | 5 | 2 | 5 | 5 | 3 | 3 | 4 | 3 | 5 | 2 | 3,61 | 4 | 4 | 0,842 | -0,248 | -0,348 | 0,728203 | 0,760409 | 1,00E-08 | 0,367533 | 0,2124399 | 0,2124399 | TRUE | 0,2839132 | No |
| CaalfMp10 | 2 | 1 | 1 | 0 | 3 | 1 | 1 | 0 | 2 | 1 | 1 | 0 | 3 | 1 | 1 | 0 | 1,07 | 1 | 1 | 1,167 | 0,223 | 0,166 | 0,868472 | 0,88783 | 1,00E-08 | 1,237051 | 0,8195685 | 0,8195685 | TRUE | 0,4650489 | No |
| CaalfMp11 | 80 | 76 | 97 | 82 | 74 | 54 | 77 | 82 | 73 | 77 | 92 | 123 | 66 | 59 | 79 | 63 | 78,82 | 91 | 67 | 0,739 | -0,436 | -2,334 | 0,019584 | 0,025136 | 0,0227538 | 0,018797 | 0,0204845 | 0,0204845 | TRUE | 0,4340947 | No |
| CaalfMp11.1 | 0 | 0 | 0 | 0 | 0 | 0 | 0 | 0 | 0 | 0 | 0 | 0 | 0 | 0 | 0 | 0 | 0 | 0 | 0 | NA | NA | NA | NA | NA | NA | NA | NA | NA | NA | NA | NA |
| CaalfMp12 | 0 | 0 | 0 | 0 | 0 | 1 | 0 | 0 | 0 | 0 | 0 | 0 | 0 | 1 | 0 | 0 | 0,14 | 0 | 0 | 1,436 | 0,523 | 0,148 | 0,882445 | 0,8988 | 1,00E-08 | 9,73564 | 10 | 10 | TRUE | 0,4622362 | No |
| CaalfMp13 | 5 | 6 | 10 | 6 | 6 | 2 | 7 | 1 | 5 | 6 | 9 | 9 | 5 | 2 | 7 | 1 | 5,57 | 7 | 4 | 0,526 | -0,928 | -1,426 | 0,15379 | 0,180655 | 0,1389773 | 0,239319 | 0,2089269 | 0,2089269 | TRUE | 0,4615478 | No |
| CaalfMp14 | 3 | 3 | 5 | 3 | 3 | 1 | 1 | 2 | 3 | 3 | 5 | 5 | 3 | 1 | 1 | 2 | 2,66 | 4 | 2 | 0,433 | -1,209 | -1,414 | 0,157423 | 0,184786 | 1,00E-08 | 0,498001 | 0,2687321 | 0,2687321 | TRUE | 0,1728937 | No |
| CaalfMp15 | 0 | 0 | 0 | 0 | 0 | 0 | 0 | 0 | 0 | 0 | 0 | 0 | 0 | 0 | 0 | 0 | 0 | 0 | 0 | NA | NA | NA | NA | NA | NA | NA | NA | NA | NA | NA | NA |
| CaalfMr16 | 1586 | 2242 | 2176 | 1673 | 2002 | 2267 | 2700 | 2419 | 1444 | 2268 | 2059 | 2513 | 1781 | 2461 | 2765 | 1845 | 2142,06 | 2071 | 2213 | 1,069 | 0,096 | 0,42 | 0,674575 | 0,71093 | 0,0497134 | 0,002654 | 0,0221889 | 0,0497134 | TRUE | 0,4308951 | No |
| CaalfMr17 | 399 | 590 | 654 | 470 | 239 | 295 | 399 | 299 | 363 | 597 | 619 | 706 | 213 | 320 | 409 | 228 | 431,82 | 571 | 292 | 0,512 | -0,967 | -3,259 | 0,001119 | 0,001576 | 0,0819684 | 0,005096 | 0,0364142 | 0,0819684 | TRUE | 0,7050163 | No |
| CaalfMt18 | 0 | 0 | 0 | 0 | 0 | 0 | 0 | 0 | 0 | 0 | 0 | 0 | 0 | 0 | 0 | 0 | 0 | 0 | 0 | NA | NA | NA | NA | NA | NA | NA | NA | NA | NA | NA | NA |
| CaalfMt19 | 0 | 0 | 0 | 0 | 0 | 0 | 0 | 0 | 0 | 0 | 0 | 0 | 0 | 0 | 0 | 0 | 0 | 0 | 0 | NA | NA | NA | NA | NA | NA | NA | NA | NA | NA | NA | NA |
| CaalfMt20 | 0 | 0 | 0 | 0 | 0 | 0 | 0 | 0 | 0 | 0 | 0 | 0 | 0 | 0 | 0 | 0 | 0 | 0 | 0 | NA | NA | NA | NA | NA | NA | NA | NA | NA | NA | NA | NA |
| CaalfMt21 | 0 | 0 | 0 | 0 | 0 | 0 | 0 | 0 | 0 | 0 | 0 | 0 | 0 | 0 | 0 | 0 | 0 | 0 | 0 | NA | NA | NA | NA | NA | NA | NA | NA | NA | NA | NA | NA |
| CaalfMt22 | 0 | 0 | 0 | 0 | 0 | 0 | 0 | 0 | 0 | 0 | 0 | 0 | 0 | 0 | 0 | 0 | 0 | 0 | 0 | NA | NA | NA | NA | NA | NA | NA | NA | NA | NA | NA | NA |
| CaalfMt23 | 0 | 0 | 0 | 0 | 0 | 0 | 0 | 0 | 0 | 0 | 0 | 0 | 0 | 0 | 0 | 0 | 0 | 0 | 0 | NA | NA | NA | NA | NA | NA | NA | NA | NA | NA | NA | NA |
| CaalfMt24 | 0 | 0 | 0 | 0 | 0 | 0 | 0 | 0 | 0 | 0 | 0 | 0 | 0 | 0 | 0 | 0 | 0 | 0 | 0 | NA | NA | NA | NA | NA | NA | NA | NA | NA | NA | NA | NA |
| CaalfMt25 | 0 | 0 | 0 | 0 | 0 | 0 | 0 | 0 | 0 | 0 | 0 | 0 | 0 | 0 | 0 | 0 | 0 | 0 | 0 | NA | NA | NA | NA | NA | NA | NA | NA | NA | NA | NA | NA |
| CaalfMt26 | 0 | 0 | 1 | 0 | 0 | 0 | 0 | 0 | 0 | 0 | 1 | 0 | 0 | 0 | 0 | 0 | 0,12 | 0 | 0 | 0,528 | -0,92 | -0,26 | 0,794572 | 0,821771 | 1,00E-08 | 11,16843 | 10 | 10 | TRUE | 0,2762646 | No |
| CaalfMt27 | 2 | 0 | 1 | 0 | 0 | 0 | 0 | 1 | 2 | 0 | 1 | 0 | 0 | 0 | 0 | 1 | 0,44 | 1 | 0 | 0,386 | -1,372 | -0,647 | 0,51789 | 0,559869 | 1,00E-08 | 2,995687 | 2,5970863 | 2,5970863 | TRUE | 0,4205281 | No |
| CaalfMt28 | 0 | 0 | 1 | 0 | 0 | 0 | 0 | 1 | 0 | 0 | 1 | 0 | 0 | 0 | 0 | 1 | 0,21 | 0 | 0 | 0,871 | -0,199 | -0,067 | 0,946611 | 0,953341 | 1,00E-08 | 6,184973 | 6,4683949 | 6,4683949 | TRUE | 0,2762655 | No |
| CaalfMt29 | 0 | 0 | 0 | 0 | 0 | 1 | 0 | 0 | 0 | 0 | 0 | 0 | 0 | 1 | 0 | 0 | 0,14 | 0 | 0 | 1,436 | 0,523 | 0,148 | 0,882445 | 0,8988 | 1,00E-08 | 9,73564 | 10 | 10 | TRUE | 0,4622362 | No |
| CaalfMt30 | 0 | 0 | 0 | 0 | 0 | 0 | 0 | 0 | 0 | 0 | 0 | 0 | 0 | 0 | 0 | 0 | 0 | 0 | 0 | NA | NA | NA | NA | NA | NA | NA | NA | NA | NA | NA | NA |
| CaalfMt31 | 0 | 0 | 0 | 0 | 0 | 0 | 0 | 0 | 0 | 0 | 0 | 0 | 0 | 0 | 0 | 0 | 0 | 0 | 0 | NA | NA | NA | NA | NA | NA | NA | NA | NA | NA | NA | NA |
| CaalfMt32 | 0 | 0 | 0 | 0 | 0 | 0 | 1 | 0 | 0 | 0 | 0 | 0 | 0 | 0 | 1 | 0 | 0,13 | 0 | 0 | 1,436 | 0,523 | 0,148 | 0,882445 | 0,8988 | 1,00E-08 | 10,32106 | 10 | 10 | TRUE | 0,417491 | No |
| CaalfMt33 | 0 | 0 | 0 | 0 | 0 | 0 | 0 | 0 | 0 | 0 | 0 | 0 | 0 | 0 | 0 | 0 | 0 | 0 | 0 | NA | NA | NA | NA | NA | NA | NA | NA | NA | NA | NA | NA |
| CaalfMt34 | 0 | 0 | 0 | 0 | 0 | 0 | 0 | 0 | 0 | 0 | 0 | 0 | 0 | 0 | 0 | 0 | 0 | 0 | 0 | NA | NA | NA | NA | NA | NA | NA | NA | NA | NA | NA | NA |
| CaalfMt35 | 0 | 0 | 0 | 0 | 0 | 0 | 0 | 0 | 0 | 0 | 0 | 0 | 0 | 0 | 0 | 0 | 0 | 0 | 0 | NA | NA | NA | NA | NA | NA | NA | NA | NA | NA | NA | NA |
| CaalfMt36 | 0 | 0 | 0 | 0 | 0 | 0 | 0 | 0 | 0 | 0 | 0 | 0 | 0 | 0 | 0 | 0 | 0 | 0 | 0 | NA | NA | NA | NA | NA | NA | NA | NA | NA | NA | NA | NA |
| CaalfMt37 | 0 | 0 | 0 | 0 | 0 | 0 | 0 | 0 | 0 | 0 | 0 | 0 | 0 | 0 | 0 | 0 | 0 | 0 | 0 | NA | NA | NA | NA | NA | NA | NA | NA | NA | NA | NA | NA |
| CaalfMt38 | 0 | 1 | 1 | 1 | 1 | 0 | 1 | 0 | 0 | 1 | 1 | 2 | 1 | 0 | 1 | 0 | 0,67 | 1 | 0 | 0,575 | -0,799 | -0,466 | 0,640971 | 0,678656 | 1,00E-08 | 1,968701 | 1,2394469 | 1,2394469 | TRUE | 0,2214554 | No |
| CaalfMt39 | 0 | 0 | 0 | 0 | 0 | 0 | 0 | 0 | 0 | 0 | 0 | 0 | 0 | 0 | 0 | 0 | 0 | 0 | 0 | NA | NA | NA | NA | NA | NA | NA | NA | NA | NA | NA | NA |
| CaalfMt40 | 0 | 0 | 0 | 0 | 0 | 0 | 0 | 0 | 0 | 0 | 0 | 0 | 0 | 0 | 0 | 0 | 0 | 0 | 0 | NA | NA | NA | NA | NA | NA | NA | NA | NA | NA | NA | NA |
| CaalfMt41 | 0 | 0 | 0 | 0 | 0 | 0 | 0 | 0 | 0 | 0 | 0 | 0 | 0 | 0 | 0 | 0 | 0 | 0 | 0 | NA | NA | NA | NA | NA | NA | NA | NA | NA | NA | NA | NA |
| CaalfMt42 | 0 | 0 | 0 | 0 | 0 | 0 | 0 | 0 | 0 | 0 | 0 | 0 | 0 | 0 | 0 | 0 | 0 | 0 | 0 | NA | NA | NA | NA | NA | NA | NA | NA | NA | NA | NA | NA |
| CaalfMt43 | 0 | 0 | 0 | 0 | 0 | 0 | 0 | 0 | 0 | 0 | 0 | 0 | 0 | 0 | 0 | 0 | 0 | 0 | 0 | NA | NA | NA | NA | NA | NA | NA | NA | NA | NA | NA | NA |
| CaalfMt44 | 0 | 0 | 0 | 0 | 0 | 0 | 0 | 0 | 0 | 0 | 0 | 0 | 0 | 0 | 0 | 0 | 0 | 0 | 0 | NA | NA | NA | NA | NA | NA | NA | NA | NA | NA | NA | NA |
| CaalfMt45 | 0 | 0 | 0 | 0 | 0 | 0 | 0 | 0 | 0 | 0 | 0 | 0 | 0 | 0 | 0 | 0 | 0 | 0 | 0 | NA | NA | NA | NA | NA | NA | NA | NA | NA | NA | NA | NA |
| CaalfMt46 | 0 | 0 | 0 | 0 | 0 | 0 | 0 | 0 | 0 | 0 | 0 | 0 | 0 | 0 | 0 | 0 | 0 | 0 | 0 | NA | NA | NA | NA | NA | NA | NA | NA | NA | NA | NA | NA |
| CaalfMt47 | 0 | 0 | 0 | 0 | 0 | 0 | 0 | 0 | 0 | 0 | 0 | 0 | 0 | 0 | 0 | 0 | 0 | 0 | 0 | NA | NA | NA | NA | NA | NA | NA | NA | NA | NA | NA | NA |
| CD39 | 0 | 0 | 0 | 0 | 0 | 0 | 0 | 0 | 0 | 0 | 0 | 0 | 0 | 0 | 0 | 0 | 0 | 0 | 0 | NA | NA | NA | NA | NA | NA | NA | NA | NA | NA | NA | NA |
| ITS1 | 192 | 224 | 222 | 233 | 170 | 203 | 160 | 205 | 175 | 227 | 210 | 350 | 151 | 220 | 164 | 156 | 206,66 | 240 | 173 | 0,724 | -0,467 | -2,505 | 0,012248 | 0,015961 | 0,0550922 | 0,008429 | 0,028377 | 0,028377 | TRUE | 0,9650425 | No |
| ITS2 | 444 | 314 | 315 | 222 | 163 | 103 | 154 | 191 | 404 | 318 | 298 | 333 | 145 | 112 | 158 | 146 | 239,21 | 338 | 140 | 0,415 | -1,27 | -10,044 | 9,76E-24 | 2,72E-23 | 0,0143422 | 0,007559 | 0,0104423 | 0,0104423 | TRUE | 0,2076339 | No |
| LSU-C2809 | 108 | 116 | 123 | 93 | 99 | 80 | 83 | 124 | 98 | 117 | 116 | 140 | 88 | 87 | 85 | 95 | 103,28 | 118 | 89 | 0,762 | -0,393 | -2,727 | 0,006395 | 0,00851 | 1,00E-08 | 0,014827 | 0,0101159 | 0,0101159 | TRUE | 0,1300951 | No |
| LSU-G1431 | 11 | 18 | 17 | 9 | 4 | 8 | 14 | 12 | 10 | 18 | 16 | 14 | 4 | 9 | 14 | 9 | 11,7 | 14 | 9 | 0,612 | -0,709 | -1,612 | 0,107014 | 0,128211 | 0,0560337 | 0,114987 | 0,0962414 | 0,0962414 | TRUE | 0,3919435 | No |
| orf19.10 | 54 | 62 | 48 | 40 | 58 | 61 | 58 | 83 | 49 | 63 | 45 | 60 | 52 | 66 | 59 | 63 | 57,24 | 54 | 60 | 1,114 | 0,156 | 0,829 | 0,407307 | 0,448382 | 1,00E-08 | 0,025115 | 0,0165541 | 0,0165541 | TRUE | 0,106289 | No |
| orf19.100 | 2501 | 2105 | 2433 | 1417 | 942 | 742 | 854 | 1006 | 2277 | 2129 | 2303 | 2129 | 838 | 806 | 875 | 767 | 1515,34 | 2210 | 822 | 0,371 | -1,431 | -25,443 | 8,48E-143 | 1,05E-141 | 0,0016037 | 0,002909 | 0,0022237 | 0,0022237 | TRUE | 0,0252233 | No |
| orf19.1002 | 38 | 55 | 61 | 34 | 55 | 49 | 49 | 65 | 35 | 56 | 58 | 51 | 49 | 53 | 50 | 50 | 50,11 | 50 | 50 | 1,017 | 0,024 | 0,118 | 0,906141 | 0,919409 | 1,00E-08 | 0,028397 | 0,0206463 | 0,0206463 | TRUE | 0,3965099 | No |
| orf19.1005 | 97 | 115 | 104 | 76 | 125 | 121 | 133 | 157 | 88 | 116 | 98 | 114 | 111 | 131 | 136 | 120 | 114,47 | 104 | 124 | 1,2 | 0,263 | 1,879 | 0,060206 | 0,074024 | 0,0036271 | 0,013577 | 0,0098653 | 0,0098653 | TRUE | 0,1126732 | No |
| orf19.1007 | 5607 | 5297 | 5561 | 3619 | 4561 | 3838 | 4147 | 5120 | 5105 | 5357 | 5263 | 5436 | 4057 | 4167 | 4247 | 3906 | 4692,26 | 5290 | 4094 | 0,774 | -0,37 | -9,12 | 7,53E-20 | 1,90E-19 | 0,0007266 | 0,002319 | 0,0013634 | 0,0013634 | TRUE | 0,0119963 | No |
| orf19.1008 | 57 | 54 | 43 | 41 | 47 | 34 | 39 | 66 | 52 | 55 | 41 | 62 | 42 | 37 | 40 | 50 | 47,23 | 52 | 42 | 0,825 | -0,278 | -1,33 | 0,183385 | 0,213103 | 1,00E-08 | 0,030009 | 0,0206533 | 0,0206533 | TRUE | 0,1890992 | No |

| | | | | | | | | | | | | | | | | | | | | | | | | | | | | | | | |
|---|---|---|---|---|---|---|---|---|---|---|---|---|---|---|---|---|---|---|---|---|---|---|---|---|---|---|---|---|---|---|---|
| orf19.1009 | 100 | 63 | 92 | 75 | 103 | 82 | 81 | 124 | 91 | 64 | 87 | 113 | 92 | 89 | 83 | 95 | 89,09 | 89 | 90 | 1,025 | 0,035 | 0,21 | 0,833781 | 0,8569 | 0,0125299 | 0,016865 | 0,015368 | 0,015368 | TRUE | 0,2603827 | No |
| orf19.101 | 1027 | 926 | 1006 | 703 | 529 | 453 | 425 | 575 | 935 | 937 | 952 | 1056 | 471 | 492 | 435 | 439 | 714,49 | 970 | 459 | 0,474 | -1,077 | -15,864 | 1,12E-56 | 5,80E-56 | 0,0016167 | 0,003886 | 0,0028528 | 0,0028528 | TRUE | 0,0381677 | No |
| orf19.1010 | 1853 | 1986 | 1805 | 1295 | 1941 | 1697 | 1677 | 2171 | 1687 | 2009 | 1708 | 1945 | 1726 | 1842 | 1717 | 1656 | 1786,46 | 1837 | 1735 | 0,945 | -0,081 | -1,224 | 0,220774 | 0,253958 | 0,004219 | 0,002776 | 0,0036471 | 0,0036471 | TRUE | 0,0499131 | No |
| orf19.1011 | 2041 | 1971 | 2098 | 1314 | 2212 | 1903 | 2022 | 2552 | 1858 | 1993 | 1986 | 1974 | 1968 | 2066 | 2071 | 1947 | 1982,78 | 1953 | 2013 | 1,03 | 0,043 | 0,947 | 0,343753 | 0,38372 | 0,0006259 | 0,002703 | 0,0014954 | 0,0014954 | TRUE | 0,013538 | No |
| orf19.1012 | 523 | 442 | 470 | 241 | 627 | 431 | 537 | 697 | 476 | 447 | 445 | 362 | 558 | 468 | 550 | 532 | 479,66 | 432 | 527 | 1,212 | 0,277 | 3,1 | 0,001935 | 0,002673 | 0,0066309 | 0,004791 | 0,0055653 | 0,0055653 | TRUE | 0,1199993 | No |
| orf19.102 | 766 | 742 | 784 | 457 | 1148 | 961 | 1011 | 1292 | 697 | 750 | 742 | 686 | 1021 | 1043 | 1035 | 986 | 870,22 | 719 | 1021 | 1,416 | 0,502 | 8,919 | 4,69E-19 | 1,16E-18 | 1,00E-08 | 0,003555 | 0,0018441 | 0,0018441 | TRUE | 0,0097114 | No |
| orf19.1026 | 4451 | 4136 | 4317 | 2865 | 4296 | 3635 | 3912 | 4946 | 4052 | 4183 | 4086 | 4304 | 3821 | 3947 | 4006 | 3773 | 4021,49 | 4156 | 3887 | 0,935 | -0,096 | -2,495 | 0,012596 | 0,016398 | 0,0004248 | 0,002366 | 0,0011776 | 0,0011776 | TRUE | 0,0065135 | No |
| orf19.1026.1 | 353 | 299 | 296 | 186 | 464 | 375 | 406 | 470 | 321 | 302 | 280 | 279 | 413 | 407 | 416 | 359 | 347,19 | 296 | 399 | 1,34 | 0,422 | 4,859 | 1,18E-06 | 1,96E-06 | 0,0023463 | 0,005842 | 0,0042757 | 0,0042757 | TRUE | 0,0575554 | No |
| orf19.1027 | 4527 | 4448 | 4524 | 2647 | 2976 | 2219 | 2380 | 3144 | 4122 | 4499 | 4282 | 3976 | 2647 | 2409 | 2437 | 2398 | 3346,26 | 4220 | 2473 | 0,586 | -0,772 | -14,819 | 1,10E-49 | 5,03E-49 | 0,002105 | 0,002432 | 0,0022861 | 0,0022861 | TRUE | 0,0275249 | No |
| orf19.1028 | 2052 | 2048 | 2180 | 1512 | 1286 | 1149 | 1139 | 1533 | 1868 | 2071 | 2063 | 2271 | 1144 | 1247 | 1166 | 1169 | 1625,16 | 2068 | 1182 | 0,572 | -0,806 | -12,882 | 5,71E-38 | 2,15E-37 | 0,0030116 | 0,00285 | 0,0030996 | 0,0030996 | TRUE | 0,0517127 | No |
| orf19.1029 | 4948 | 5425 | 5372 | 3297 | 3785 | 3336 | 3144 | 3912 | 4505 | 5487 | 5084 | 4953 | 3367 | 3622 | 3220 | 2984 | 4152,63 | 5007 | 3298 | 0,658 | -0,603 | -8,561 | 1,12E-17 | 2,67E-17 | 0,0062135 | 0,002355 | 0,0045146 | 0,0045146 | TRUE | 0,0562979 | No |
| orf19.103 | 153 | 151 | 165 | 119 | 197 | 179 | 154 | 234 | 139 | 153 | 156 | 179 | 175 | 194 | 158 | 178 | 166,59 | 157 | 176 | 1,135 | 0,182 | 1,567 | 0,117076 | 0,139506 | 0,0013955 | 0,009967 | 0,0069376 | 0,0069376 | TRUE | 0,0752455 | No |
| orf19.1030 | 26080 | 23650 | 25185 | 14868 | 29845 | 22779 | 25421 | 34430 | 23745 | 23919 | 23835 | 22334 | 26546 | 24732 | 26034 | 26263 | 24676,2 | 23458 | 25894 | 1,104 | 0,142 | 3,773 | 0,000162 | 0,000239 | 0,0009413 | 0,002091 | 0,0013261 | 0,0013261 | TRUE | 0,0125736 | No |
| orf19.1031 | 1617 | 1427 | 1672 | 1007 | 1824 | 1629 | 1617 | 2247 | 1472 | 1443 | 1582 | 1513 | 1622 | 1769 | 1656 | 1714 | 1596,45 | 1502 | 1690 | 1,125 | 0,17 | 3,372 | 0,000746 | 0,001061 | 0,0009308 | 0,002864 | 0,0017921 | 0,0017921 | TRUE | 0,0164297 | No |
| orf19.1032 | 4068 | 3561 | 3902 | 2678 | 7142 | 6029 | 6302 | 9040 | 3704 | 3602 | 3693 | 4023 | 6353 | 6546 | 6454 | 6896 | 5158,65 | 3756 | 6562 | 1,75 | 0,807 | 17,559 | 5,11E-69 | 3,16E-68 | 0,0015703 | 0,002293 | 0,0018143 | 0,0018143 | TRUE | 0,0265438 | No |
| orf19.1033 | 4691 | 4344 | 4561 | 2484 | 6937 | 5476 | 5892 | 7710 | 4271 | 4393 | 4317 | 3731 | 6170 | 5945 | 6034 | 5881 | 5092,93 | 4178 | 6008 | 1,436 | 0,522 | 9,976 | 1,95E-23 | 5,38E-23 | 0,0025602 | 0,002296 | 0,0024246 | 0,0024246 | TRUE | 0,0605193 | No |
| orf19.1034 | 2002 | 1872 | 1942 | 1091 | 764 | 558 | 616 | 825 | 1823 | 1893 | 1838 | 1639 | 680 | 606 | 631 | 629 | 1217,3 | 1798 | 636 | 0,354 | -1,5 | -24,302 | 1,88E-130 | 2,10E-129 | 0,0021471 | 0,003122 | 0,0026272 | 0,0026272 | TRUE | 0,0396347 | No |
| orf19.1035 | 3933 | 3545 | 3802 | 2767 | 2412 | 2198 | 2275 | 3047 | 3581 | 3585 | 3598 | 4156 | 2145 | 2386 | 2330 | 2324 | 3013,36 | 3730 | 2296 | 0,616 | -0,698 | -11,8 | 3,91E-32 | 1,32E-31 | 0,0034711 | 0,002475 | 0,0030158 | 0,0030158 | TRUE | 0,0696521 | No |
| orf19.1036 | 1184 | 1100 | 1068 | 666 | 1496 | 1183 | 1277 | 1739 | 1078 | 1113 | 1011 | 1000 | 1331 | 1284 | 1308 | 1327 | 1181,39 | 1050 | 1312 | 1,248 | 0,319 | 6,135 | 8,52E-10 | 1,60E-09 | 0,0002991 | 0,003155 | 0,0017348 | 0,0017348 | TRUE | 0,0184201 | No |
| orf19.1037 | 93 | 97 | 93 | 52 | 284 | 253 | 240 | 305 | 85 | 98 | 88 | 78 | 253 | 275 | 246 | 233 | 169,33 | 87 | 252 | 2,858 | 1,515 | 12,476 | 1,01E-35 | 3,65E-35 | 1,00E-08 | 0,009838 | 0,0062556 | 0,0062556 | TRUE | 0,0633599 | No |
| orf19.1039 | 259 | 390 | 313 | 181 | 331 | 264 | 321 | 356 | 236 | 394 | 296 | 272 | 294 | 287 | 329 | 272 | 297,46 | 300 | 296 | 0,984 | -0,023 | -0,171 | 0,864165 | 0,884278 | 0,0242825 | 0,006478 | 0,0144653 | 0,0144653 | TRUE | 0,5211367 | No |
| orf19.104 | 252 | 258 | 310 | 182 | 357 | 295 | 272 | 394 | 229 | 261 | 293 | 273 | 318 | 320 | 279 | 301 | 284,26 | 264 | 304 | 1,154 | 0,206 | 2,124 | 0,033629 | 0,042318 | 0,0037931 | 0,006684 | 0,0054758 | 0,0054758 | TRUE | 0,0967465 | No |
| orf19.1040 | 343 | 305 | 333 | 189 | 235 | 215 | 205 | 295 | 312 | 308 | 315 | 284 | 209 | 233 | 210 | 225 | 262,16 | 305 | 219 | 0,716 | -0,481 | -5,381 | 7,42E-08 | 1,29E-07 | 1,00E-08 | 0,007076 | 0,0038177 | 0,0038177 | TRUE | 0,0196589 | No |
| orf19.1041 | 428 | 512 | 467 | 354 | 776 | 675 | 736 | 889 | 390 | 518 | 442 | 532 | 690 | 733 | 754 | 678 | 592,03 | 470 | 714 | 1,524 | 0,608 | 6,573 | 4,93E-11 | 9,69E-11 | 0,0089391 | 0,004268 | 0,0064045 | 0,0064045 | TRUE | 0,1592287 | No |
| orf19.1042 | 16601 | 14194 | 14985 | 10665 | 8972 | 7838 | 7710 | 11024 | 15115 | 14356 | 14182 | 16021 | 7980 | 8510 | 7896 | 8409 | 11558,5 | 14918 | 8199 | 0,55 | -0,863 | -17,841 | 3,37E-71 | 2,14E-70 | 0,0021511 | 0,002151 | 0,0021552 | 0,0021552 | TRUE | 0,0298922 | No |
| orf19.1043 | 690 | 794 | 858 | 434 | 881 | 780 | 823 | 980 | 628 | 803 | 812 | 652 | 784 | 847 | 843 | 748 | 764,51 | 724 | 806 | 1,109 | 0,15 | 1,66 | 0,096815 | 0,116473 | 0,0090588 | 0,003765 | 0,0064877 | 0,0064877 | TRUE | 0,1011232 | No |
| orf19.1045 | 2228 | 2289 | 2250 | 1022 | 2789 | 1946 | 2368 | 2921 | 2028 | 2315 | 2129 | 1535 | 2481 | 2113 | 2425 | 2228 | 2156,88 | 2002 | 2312 | 1,153 | 0,205 | 2,032 | 0,04213 | 0,052581 | 0,0159627 | 0,002649 | 0,0093432 | 0,0093432 | TRUE | 0,2893853 | No |
| orf19.1047 | 8238 | 8502 | 8740 | 5130 | 12276 | 10205 | 10428 | 13912 | 7500 | 8599 | 8272 | 7706 | 10919 | 11080 | 10680 | 10612 | 9420,95 | 8019 | 10823 | 1,349 | 0,432 | 9,019 | 1,90E-19 | 4,76E-19 | 0,0020425 | 0,002177 | 0,0020949 | 0,0020949 | TRUE | 0,0289738 | No |
| orf19.1048 | 564 | 522 | 546 | 326 | 600 | 507 | 482 | 690 | 513 | 528 | 517 | 490 | 534 | 550 | 494 | 526 | 519 | 512 | 526 | 1,025 | 0,036 | 0,537 | 0,591528 | 0,632081 | 1,00E-08 | 0,004582 | 0,002379 | 0,002379 | TRUE | 0,0187589 | No |
| orf19.1049 | 71 | 101 | 91 | 59 | 138 | 118 | 133 | 130 | 65 | 102 | 86 | 89 | 123 | 128 | 136 | 99 | 103,47 | 86 | 122 | 1,423 | 0,509 | 3,101 | 0,001929 | 0,002665 | 0,0172273 | 0,014804 | 0,0157679 | 0,0157679 | TRUE | 0,2828325 | No |
| orf19.105 | 1075 | 747 | 801 | 537 | 1280 | 938 | 1183 | 1590 | 979 | 756 | 758 | 807 | 1139 | 1018 | 1212 | 1213 | 985,04 | 825 | 1146 | 1,388 | 0,473 | 5,304 | 1,13E-07 | 1,95E-07 | 0,0099532 | 0,003378 | 0,0065848 | 0,0065848 | TRUE | 0,1948981 | No |
| orf19.1050 | 176 | 207 | 215 | 110 | 184 | 160 | 134 | 183 | 160 | 209 | 203 | 165 | 164 | 174 | 137 | 140 | 169,06 | 184 | 154 | 0,827 | -0,274 | -2,117 | 0,034284 | 0,043117 | 0,0104762 | 0,009851 | 0,0101052 | 0,0101052 | TRUE | 0,0989183 | No |
| orf19.1051 | 3102 | 2566 | 2657 | 2028 | 3035 | 2360 | 2255 | 3407 | 2824 | 2595 | 2515 | 3046 | 2700 | 2562 | 2309 | 2599 | 2643,82 | 2745 | 2542 | 0,928 | -0,109 | -1,586 | 0,112691 | 0,134508 | 0,0054414 | 0,002537 | 0,0041183 | 0,0041183 | TRUE | 0,0639292 | No |
| orf19.1052 | 1519 | 1302 | 1421 | 1077 | 1255 | 1043 | 960 | 1363 | 1383 | 1317 | 1345 | 1618 | 1116 | 1132 | 983 | 1040 | 1241,76 | 1416 | 1068 | 0,756 | -0,403 | -5,496 | 3,88E-08 | 6,81E-08 | 0,0057379 | 0,003101 | 0,0043517 | 0,0043517 | TRUE | 0,104793 | No |
| orf19.1053 | 1174 | 1122 | 1222 | 701 | 1445 | 1284 | 1099 | 1587 | 1069 | 1135 | 1157 | 1053 | 1285 | 1394 | 1126 | 1211 | 1178,58 | 1104 | 1254 | 1,135 | 0,182 | 2,633 | 0,008465 | 0,011159 | 0,0040294 | 0,003158 | 0,003743 | 0,003743 | TRUE | 0,0658509 | No |
| orf19.1054 | 413 | 407 | 451 | 317 | 243 | 275 | 213 | 281 | 376 | 412 | 427 | 476 | 216 | 299 | 218 | 214 | 329,73 | 423 | 237 | 0,56 | -0,837 | -7,261 | 3,85E-13 | 8,11E-13 | 0,0141848 | 0,006043 | 0,009497 | 0,009497 | TRUE | 0,3268142 | No |
| orf19.1055 | 2136 | 2050 | 2055 | 1365 | 1730 | 1390 | 1367 | 1819 | 1945 | 2073 | 1945 | 2050 | 1539 | 1509 | 1400 | 1388 | 1731,11 | 2003 | 1459 | 0,728 | -0,457 | -8,8 | 1,37E-18 | 3,36E-18 | 0,001367 | 0,0028 | 0,0020023 | 0,0020023 | TRUE | 0,0170876 | No |
| orf19.1057 | 1437 | 1302 | 1294 | 859 | 1089 | 778 | 808 | 1136 | 1308 | 1317 | 1225 | 1290 | 969 | 845 | 827 | 867 | 1080,94 | 1285 | 877 | 0,683 | -0,55 | -9,002 | 2,21E-19 | 5,52E-19 | 0,0018498 | 0,003259 | 0,0026299 | 0,0026299 | TRUE | 0,059553 | No |
| orf19.1058 | 3888 | 3456 | 3695 | 2232 | 4111 | 2923 | 3511 | 4456 | 3540 | 3495 | 3497 | 3353 | 3657 | 3174 | 3596 | 3399 | 3463,75 | 3471 | 3456 | 0,995 | -0,007 | -0,132 | 0,894835 | 0,909965 | 0,0019873 | 0,002418 | 0,0021643 | 0,0021643 | TRUE | 0,0360263 | No |
| orf19.1059 | 3262 | 2761 | 2890 | 1699 | 7203 | 5062 | 5786 | 7834 | 2970 | 2792 | 2735 | 2552 | 6407 | 5496 | 5926 | 5976 | 4356,73 | 2762 | 5951 | 2,152 | 1,106 | 18,809 | 6,38E-79 | 4,47E-78 | 0,0035469 | 0,00234 | 0,0030463 | 0,0030463 | TRUE | 0,0324186 | No |
| orf19.1060 | 108 | 88 | 98 | 56 | 217 | 150 | 197 | 219 | 98 | 89 | 93 | 84 | 193 | 163 | 202 | 167 | 136,11 | 91 | 181 | 1,977 | 0,983 | 7,52 | 5,48E-14 | 1,18E-13 | 1,00E-08 | 0,011742 | 0,0080343 | 0,0080343 | TRUE | 0,0599275 | No |
| orf19.1061 | 2149 | 1731 | 1842 | 953 | 1873 | 1242 | 1426 | 2013 | 1957 | 1751 | 1743 | 1432 | 1666 | 1348 | 1460 | 1536 | 1611,56 | 1721 | 1502 | 0,872 | -0,197 | -2,212 | 0,026938 | 0,034169 | 0,0110232 | 0,002857 | 0,0070045 | 0,0070045 | TRUE | 0,1483498 | No |
| orf19.1062 | 92 | 93 | 78 | 47 | 123 | 79 | 104 | 149 | 84 | 94 | 74 | 71 | 109 | 86 | 107 | 114 | 92,2 | 81 | 104 | 1,287 | 0,364 | 2,354 | 0,018582 | 0,023903 | 0,0040683 | 0,016365 | 0,0119625 | 0,0119625 | TRUE | 0,1250079 | No |
| orf19.1063 | 409 | 484 | 453 | 284 | 466 | 390 | 390 | 497 | 372 | 490 | 429 | 427 | 414 | 423 | 399 | 379 | 416,71 | 430 | 404 | 0,941 | -0,088 | -0,983 | 0,325739 | 0,365274 | 0,0053411 | 0,005207 | 0,0052688 | 0,0052688 | TRUE | 0,1081858 | No |
| orf19.1064 | 4444 | 4108 | 4375 | 2750 | 14815 | 12917 | 13200 | 18570 | 4046 | 4155 | 4141 | 4131 | 13177 | 14024 | 13518 | 14165 | 8919,74 | 4118 | 13721 | 3,333 | 1,737 | 47,629 | 0 | 0 | 0,0006167 | 0,002185 | 0,0011117 | 0,0011117 | TRUE | 0,0087726 | No |
| orf19.1065 | 1289 | 1045 | 1228 | 786 | 1482 | 1230 | 1253 | 1782 | 1174 | 1057 | 1162 | 1181 | 1318 | 1335 | 1283 | 1359 | 1233,69 | 1144 | 1324 | 1,16 | 0,214 | 4,04 | 5,35E-05 | 8,13E-05 | 0,0006865 | 0,003108 | 0,0018599 | 0,0018599 | TRUE | 0,0315094 | No |
| orf19.1066 | 177 | 164 | 210 | 89 | 238 | 210 | 220 | 258 | 161 | 166 | 199 | 134 | 212 | 228 | 225 | 197 | 190,16 | 165 | 216 | 1,291 | 0,369 | 3,077 | 0,002094 | 0,002885 | 0,007457 | 0,008984 | 0,0084106 | 0,0084106 | TRUE | 0,200488 | No |
| orf19.1067 | 1172 | 1077 | 1095 | 633 | 582 | 444 | 512 | 616 | 1067 | 1089 | 1036 | 951 | 518 | 482 | 524 | 470 | 767,19 | 1036 | 498 | 0,48 | -1,06 | -15,984 | 1,65E-57 | 8,66E-57 | 0,0016479 | 0,003759 | 0,0027708 | 0,0027708 | TRUE | 0,0323817 | No |
| orf19.1069 | 44639 | 39590 | 43620 | 27657 | 27924 | 23666 | 25019 | 33555 | 40642 | 40041 | 41282 | 41545 | 24838 | 25695 | 25623 | 25596 | 33157,7 | 40878 | 25438 | 0,622 | -0,684 | -24,734 | 4,63E-135 | 5,38E-134 | 0,0002187 | 0,002077 | 0,0007037 | 0,0007037 | TRUE | 0,0030966 | No |
| orf19.107 | 7797 | 7215 | 7799 | 4674 | 6056 | 5126 | 5466 | 7088 | 7099 | 7297 | 7381 | 7021 | 5387 | 5565 | 5598 | 5407 | 6344,36 | 7200 | 5489 | 0,762 | -0,392 | -11,594 | 4,43E-31 | 1,46E-30 | 0,0002508 | 0,002245 | 0,0009393 | 0,0009393 | TRUE | 0,0035155 | No |
| orf19.1070 | 338 | 308 | 366 | 198 | 509 | 444 | 479 | 615 | 308 | 312 | 346 | 297 | 453 | 482 | 491 | 469 | 394,69 | 316 | 474 | 1,493 | 0,579 | 7,442 | 9,93E-14 | 2,13E-13 | 1,00E-08 | 0,005384 | 0,0031352 | 0,0031352 | TRUE | 0,0498178 | No |
| orf19.1075 | 116 | 114 | 110 | 86 | 207 | 186 | 170 | 259 | 106 | 115 | 104 | 129 | 184 | 202 | 174 | 198 | 151,49 | 114 | 190 | 1,685 | 0,753 | 6,272 | 3,56E-10 | 6,78E-10 | 1,00E-08 | 0,010757 | 0,0066538 | 0,0066538 | TRUE | 0,0655392 | No |
| orf19.1077 | 2593 | 2494 | 2552 | 1720 | 1994 | 1727 | 1661 | 2177 | 2361 | 2522 | 2415 | 2584 | 1774 | 1875 | 1701 | 1661 | 2111,57 | 2470 | 1753 | 0,71 | -0,495 | -9,484 | 2,46E-21 | 6,44E-21 | 0,0016257 | 0,002663 | 0,0021312 | 0,0021312 | TRUE | 0,0260505 | No |
| orf19.1078 | 2823 | 2775 | 2704 | 1813 | 2648 | 2157 | 2303 | 3075 | 2570 | 2807 | 2559 | 2723 | 2355 | 2342 | 2359 | 2346 | 2507,59 | 2665 | 2350 | 0,883 | -0,18 | -4,12 | 3,80E-05 | 5,84E-05 | 0,0006762 | 0,002564 | 0,0014312 | 0,0014312 | TRUE | 0,0164977 | No |
| orf19.1080 | 18 | 18 | 13 | 16 | 35 | 28 | 35 | 42 | 16 | 18 | 12 | 24 | 31 | 30 | 36 | 32 | 25,04 | 18 | 32 | 1,873 | 0,905 | 3,179 | 0,001477 | 0,002062 | 1,00E-08 | 0,054786 | 0,032745 | 0,032745 | TRUE | 0,2194015 | No |
| orf19.1082 | 1029 | 1053 | 1009 | 606 | 794 | 661 | 714 | 892 | 937 | 1065 | 955 | 910 | 706 | 718 | 731 | 680 | 837,83 | 967 | 709 | 0,731 | -0,451 | -7,071 | 1,54E-12 | 3,18E-12 | 0,0017969 | 0,003614 | 0,0026973 | 0,0026973 | TRUE | 0,0557774 | No |
| orf19.1082.1 | 6626 | 5220 | 5190 | 4152 | 8150 | 5834 | 6037 | 8568 | 6033 | 5279 | 4912 | 6237 | 7249 | 6334 | 6183 | 6536 | 6095,33 | 5615 | 6576 | 1,172 | 0,228 | 2,966 | 0,003019 | 0,004107 | 0,0085303 | 0,002254 | 0,0055343 | 0,0055343 | TRUE | 0,0869439 | No |
| orf19.1083 | 682 | 600 | 640 | 415 | 751 | 569 | 616 | 832 | 621 | 607 | 606 | 623 | 668 | 618 | 631 | 635 | 626,02 | 614 | 638 | 1,04 | 0,056 | 0,933 | 0,350994 | 0,390988 | 1,00E-08 | 0,004147 | 0,0019218 | 0,0019218 | TRUE | 0,0120126 | No |
| orf19.1084 | 8396 | 8020 | 8317 | 5363 | 5181 | 4592 | 4661 | 6244 | 7644 | 8111 | 7871 | 8056 | 4608 | 4986 | 4773 | 4763 | 6351,67 | 7920 | 4782 | 0,604 | -0,728 | -19,014 | 1,31E-80 | 9,34E-80 | 0,0006564 | 0,002245 | 0,0012414 | 0,0012414 | TRUE | 0,0097582 | No |
| orf19.1085 | 966 | 883 | 971 | 627 | 1559 | 1244 | 1332 | 1819 | 880 | 893 | 919 | 942 | 1387 | 1351 | 1364 | 1388 | 1140,3 | 908 | 1372 | 1,514 | 0,599 | 12,244 | 1,80E-34 | 6,37E-34 | 1,00E-08 | 0,003195 | 0,001367 | 0,001367 | TRUE | 0,0061573 | No |
| orf19.1086 | 2821 | 2880 | 3071 | 2000 | 2559 | 2197 | 2296 | 3087 | 2568 | 2913 | 2906 | 3004 | 2276 | 2385 | 2351 | 2355 | 2594,95 | 2848 | 2342 | 0,823 | -0,281 | -5,258 | 1,45E-07 | 2,49E-07 | 0,0019426 | 0,002546 | 0,0023497 | 0,0023497 | TRUE | 0,0540581 | No |
| orf19.1087 | 1773 | 1951 | 2091 | 1119 | 1373 | 1143 | 1292 | 1552 | 1614 | 1973 | 1979 | 1681 | 1221 | 1241 | 1323 | 1184 | 1527,07 | 1812 | 1242 | 0,685 | -0,546 | -7,342 | 2,10E-13 | 4,44E-13 | 0,005811 | 0,002902 | 0,0046421 | 0,0046421 | TRUE | 0,0681762 | No |
| orf19.1089 | 1271 | 1531 | 1466 | 870 | 3897 | 3679 | 3403 | 4278 | 1157 | 1548 | 1387 | 1307 | 3466 | 3994 | 3485 | 3263 | 2451,12 | 1350 | 3552 | 2,63 | 1,395 | 16,35 | 4,36E-60 | 2,39E-59 | 0,0104596 | 0,002576 | 0,0064625 | 0,0064625 | TRUE | 0,1192783 | No |
| orf19.109 | 1541 | 1287 | 1426 | 870 | 937 | 693 | 750 | 1044 | 1403 | 1302 | 1350 | 1307 | 833 | 752 | 768 | 796 | 1063,93 | 1340 | 787 | 0,588 | -0,767 | -13,902 | 6,14E-44 | 2,58E-43 | 0,0006157 | 0,003279 | 0,001929 | 0,001929 | TRUE | 0,0179767 | No |
| orf19.1090 | 257 | 213 | 220 | 170 | 117 | 98 | 110 | 142 | 234 | 215 | 208 | 255 | 104 | 106 | 113 | 108 | 168,05 | 228 | 108 | 0,475 | -1,074 | -9,314 | 1,24E-20 | 3,18E-20 | 1,00E-08 | 0,009898 | 0,0061031 | 0,0061031 | TRUE | 0,0575414 | No |
| orf19.1091 | 6769 | 6749 | 7154 | 3907 | 8631 | 7149 | 7622 | 9483 | 6163 | 6826 | 6771 | 5869 | 7677 | 7762 | 7806 | 7234 | 7013,34 | 6407 | 7620 | 1,188 | 0,249 | 4,549 | 5,40E-06 | 8,68E-06 | 0,0030697 | 0,002225 | 0,0027337 | 0,0027337 | TRUE | 0,0380194 | No |
| orf19.1092 | 910 | 1083 | 1107 | 656 | 627 | 524 | 511 | 622 | 829 | 1095 | 1048 | 985 | 558 | 569 | 523 | 474 | 760,17 | 989 | 531 | 0,536 | -0,899 | -9,786 | 1,30E-22 | 3,49E-22 | 0,0096428 | 0,003775 | 0,0066755 | 0,0066755 | TRUE | 0,1484306 | No |

| | | | | | | | | | | | | | | | | | | | | | | | | | | | | | | | |
|---|---|---|---|---|---|---|---|---|---|---|---|---|---|---|---|---|---|---|---|---|---|---|---|---|---|---|---|---|---|---|---|
| orf19.1093 | 8049 | 6936 | 7919 | 5419 | 8343 | 7299 | 7848 | 10758 | 7328 | 7015 | 7495 | 8140 | 7421 | 7925 | 8037 | 8206 | 7695,91 | 7494 | 7897 | 1,054 | 0,076 | 1,454 | 0,145972 | 0,17185 | 0,0027442 | 0,002209 | 0,0025174 | 0,0025174 | TRUE | 0,0402126 | No |
| orf19.1095 | 2590 | 2383 | 2527 | 1772 | 1704 | 1481 | 1464 | 1992 | 2358 | 2410 | 2392 | 2662 | 1516 | 1608 | 1499 | 1520 | 1995,51 | 2456 | 1536 | 0,626 | -0,675 | -13,152 | 1,65E-39 | 6,37E-39 | 0,0014128 | 0,002699 | 0,0020055 | 0,0020055 | TRUE | 0,0363204 | No |
| orf19.1096 | 1151 | 1313 | 1370 | 772 | 1386 | 1190 | 1256 | 1573 | 1048 | 1328 | 1297 | 1160 | 1233 | 1292 | 1286 | 1200 | 1230,39 | 1208 | 1253 | 1,036 | 0,051 | 0,689 | 0,490999 | 0,532605 | 0,0057223 | 0,003111 | 0,004458 | 0,004458 | TRUE | 0,1007395 | No |
| orf19.1097 | 2298 | 1849 | 2067 | 1588 | 1760 | 1583 | 1833 | 2533 | 2092 | 1870 | 1956 | 2385 | 1565 | 1719 | 1877 | 1932 | 1924,69 | 2076 | 1773 | 0,856 | -0,225 | -2,697 | 0,007007 | 0,009298 | 0,0095682 | 0,002723 | 0,0061661 | 0,0061661 | TRUE | 0,1178913 | No |
| orf19.11 | 480 | 348 | 412 | 230 | 227 | 156 | 180 | 277 | 437 | 352 | 390 | 345 | 202 | 169 | 184 | 211 | 286,42 | 381 | 192 | 0,503 | -0,99 | -9,343 | 9,33E-21 | 2,41E-20 | 0,0073656 | 0,006649 | 0,0069612 | 0,0069612 | TRUE | 0,1145785 | No |
| orf19.1105 | 44 | 47 | 64 | 34 | 52 | 55 | 50 | 67 | 40 | 48 | 61 | 51 | 46 | 60 | 51 | 51 | 50,94 | 50 | 52 | 1,044 | 0,062 | 0,309 | 0,757609 | 0,789176 | 1,00E-08 | 0,027969 | 0,0189217 | 0,0189217 | TRUE | 0,1999511 | No |
| orf19.1105.2 | 69796 | 53337 | 58691 | 41323 | 60535 | 45034 | 50153 | 69769 | 63546 | 53945 | 55546 | 62074 | 53844 | 48894 | 51363 | 53220 | 55304 | 58778 | 51830 | 0,882 | -0,181 | -3,091 | 0,001994 | 0,002752 | 0,0041474 | 0,002061 | 0,0032905 | 0,0032905 | TRUE | 0,0375452 | No |
| orf19.1105.3 | 13 | 8 | 7 | 13 | 23 | 18 | 14 | 24 | 12 | 8 | 7 | 20 | 20 | 20 | 14 | 18 | 14,84 | 12 | 18 | 1,65 | 0,722 | 1,881 | 0,059957 | 0,073746 | 0,0006481 | 0,091049 | 0,0677446 | 0,0677446 | TRUE | 0,5327943 | No |
| orf19.1106 | 277 | 302 | 300 | 195 | 167 | 176 | 165 | 222 | 252 | 305 | 284 | 293 | 149 | 191 | 169 | 169 | 226,55 | 284 | 170 | 0,597 | -0,744 | -6,975 | 3,06E-12 | 6,25E-12 | 0,0038815 | 0,007868 | 0,0062663 | 0,0062663 | TRUE | 0,0745677 | No |
| orf19.1107 | 4921 | 4095 | 4409 | 3251 | 5496 | 4645 | 4377 | 6423 | 4480 | 4142 | 4173 | 4884 | 4889 | 5043 | 4483 | 4900 | 4624,01 | 4420 | 4829 | 1,094 | 0,129 | 2,151 | 0,031509 | 0,039753 | 0,0040049 | 0,002323 | 0,0032344 | 0,0032344 | TRUE | 0,0591541 | No |
| orf19.1108 | 3164 | 2859 | 2984 | 2078 | 2622 | 2257 | 2041 | 2962 | 2881 | 2892 | 2824 | 3121 | 2332 | 2450 | 2090 | 2259 | 2606,27 | 2930 | 2283 | 0,78 | -0,358 | -6,41 | 1,46E-10 | 2,81E-10 | 0,0026662 | 0,002544 | 0,0026133 | 0,0026133 | TRUE | 0,0382726 | No |
| orf19.1109 | 2217 | 2192 | 2289 | 1250 | 1999 | 1543 | 1697 | 2156 | 2018 | 2217 | 2166 | 1878 | 1778 | 1675 | 1738 | 1645 | 1889,42 | 2070 | 1709 | 0,824 | -0,279 | -4,77 | 1,84E-06 | 3,02E-06 | 0,0026523 | 0,002736 | 0,0027507 | 0,0027507 | TRUE | 0,0440015 | No |
| orf19.111 | 25665 | 23125 | 24607 | 15595 | 25354 | 21829 | 21873 | 31043 | 23367 | 23388 | 23288 | 23426 | 22552 | 23700 | 22401 | 23680 | 23225,3 | 23367 | 23083 | 0,988 | -0,018 | -0,561 | 0,574479 | 0,615204 | 0,0003951 | 0,002094 | 0,0009013 | 0,0009013 | TRUE | 0,004826 | No |
| orf19.1110 | 426 | 377 | 409 | 270 | 517 | 388 | 431 | 540 | 388 | 381 | 387 | 406 | 460 | 421 | 441 | 412 | 412,03 | 390 | 434 | 1,112 | 0,154 | 2,08 | 0,037559 | 0,047041 | 1,00E-08 | 0,005243 | 0,002787 | 0,002787 | TRUE | 0,0203798 | No |
| orf19.1111 | 756 | 720 | 837 | 557 | 783 | 660 | 688 | 1014 | 688 | 728 | 792 | 837 | 696 | 717 | 705 | 773 | 742,06 | 761 | 723 | 0,954 | -0,069 | -0,951 | 0,341441 | 0,381492 | 0,0034608 | 0,003817 | 0,0036434 | 0,0036434 | TRUE | 0,0501525 | No |
| orf19.1112 | 1208 | 1156 | 1244 | 727 | 477 | 451 | 394 | 529 | 1100 | 1169 | 1177 | 1092 | 424 | 490 | 404 | 404 | 782,42 | 1134 | 430 | 0,378 | -1,404 | -19,943 | 1,71E-88 | 1,33E-87 | 0,0024824 | 0,003725 | 0,0032005 | 0,0032005 | TRUE | 0,0940468 | No |
| orf19.1113 | 535 | 444 | 511 | 279 | 763 | 576 | 617 | 890 | 487 | 449 | 484 | 419 | 679 | 625 | 632 | 679 | 556,71 | 460 | 654 | 1,417 | 0,503 | 7,101 | 1,24E-12 | 2,56E-12 | 0,0009512 | 0,00441 | 0,0029483 | 0,0029483 | TRUE | 0,0341083 | No |
| orf19.1114 | 5176 | 4457 | 4583 | 2909 | 5671 | 4277 | 5054 | 6678 | 4713 | 4508 | 4337 | 4370 | 5044 | 4644 | 5176 | 5094 | 4735,66 | 4482 | 4990 | 1,113 | 0,155 | 3,285 | 0,001022 | 0,001441 | 0,0015688 | 0,002316 | 0,0019136 | 0,0019136 | TRUE | 0,0261236 | No |
| orf19.1115 | 2402 | 2249 | 2357 | 1372 | 2709 | 2148 | 2195 | 2971 | 2187 | 2275 | 2231 | 2061 | 2410 | 2332 | 2248 | 2266 | 2251,14 | 2188 | 2314 | 1,056 | 0,078 | 1,698 | 0,089554 | 0,108044 | 0,0007946 | 0,002624 | 0,0015975 | 0,0015975 | TRUE | 0,0168197 | No |
| orf19.1116 | 192 | 165 | 175 | 110 | 197 | 160 | 170 | 235 | 175 | 167 | 166 | 165 | 175 | 174 | 174 | 179 | 171,86 | 168 | 176 | 1,044 | 0,062 | 0,592 | 0,554103 | 0,595483 | 1,00E-08 | 0,009724 | 0,0046236 | 0,0046236 | TRUE | 0,008299 | No |
| orf19.1117 | 299 | 264 | 271 | 152 | 179 | 167 | 199 | 258 | 272 | 267 | 256 | 228 | 159 | 181 | 204 | 197 | 220,65 | 256 | 185 | 0,72 | -0,474 | -4,451 | 8,56E-06 | 1,36E-05 | 0,0034593 | 0,008024 | 0,0062841 | 0,0062841 | TRUE | 0,1016446 | No |
| orf19.1119 | 2020 | 1955 | 2126 | 1329 | 1450 | 1257 | 1255 | 1567 | 1839 | 1977 | 2012 | 1996 | 1290 | 1365 | 1285 | 1195 | 1619,99 | 1956 | 1284 | 0,656 | -0,609 | -11,145 | 7,54E-29 | 2,35E-28 | 0,0017152 | 0,002852 | 0,002225 | 0,002225 | TRUE | 0,0270579 | No |
| orf19.1120 | 465 | 393 | 460 | 280 | 719 | 565 | 597 | 806 | 423 | 397 | 435 | 421 | 640 | 613 | 611 | 615 | 519,5 | 419 | 620 | 1,478 | 0,564 | 8,501 | 1,88E-17 | 4,45E-17 | 1,00E-08 | 0,00458 | 0,0022055 | 0,0022055 | TRUE | 0,0149256 | No |
| orf19.1121 | 4245 | 4242 | 4295 | 2583 | 5192 | 4382 | 4276 | 5702 | 3865 | 4290 | 4065 | 3880 | 4618 | 4758 | 4379 | 4350 | 4275,57 | 4025 | 4526 | 1,124 | 0,168 | 3,435 | 0,000594 | 0,00085 | 0,0018417 | 0,002346 | 0,0020708 | 0,0020708 | TRUE | 0,0240258 | No |
| orf19.1122 | 60 | 77 | 79 | 53 | 56 | 37 | 32 | 59 | 55 | 78 | 75 | 80 | 50 | 40 | 33 | 45 | 56,83 | 72 | 42 | 0,593 | -0,755 | -3,74 | 0,000184 | 0,000272 | 0,0123032 | 0,025281 | 0,0205561 | 0,0205561 | TRUE | 0,2556035 | No |
| orf19.1123 | 4 | 0 | 3 | 1 | 3 | 5 | 5 | 5 | 4 | 0 | 3 | 2 | 3 | 5 | 5 | 4 | 3,13 | 2 | 4 | 2,04 | 1,029 | 1,253 | 0,210166 | 0,242631 | 1,00E-08 | 0,424504 | 0,2793762 | 0,2793762 | TRUE | 0,442365 | No |
| orf19.1124 | 317 | 256 | 260 | 161 | 295 | 242 | 252 | 338 | 289 | 259 | 246 | 242 | 262 | 263 | 258 | 258 | 259,56 | 259 | 260 | 1,001 | 0,001 | 0,014 | 0,989186 | 0,990235 | 1,00E-08 | 0,007126 | 0,0041556 | 0,0041556 | TRUE | 0,0706398 | No |
| orf19.1124.2 | 374 | 215 | 270 | 183 | 501 | 311 | 425 | 603 | 341 | 217 | 256 | 275 | 446 | 338 | 435 | 460 | 345,86 | 272 | 420 | 1,543 | 0,626 | 4,724 | 2,31E-06 | 3,77E-06 | 0,0238109 | 0,005856 | 0,0137739 | 0,0137739 | TRUE | 0,3412254 | No |
| orf19.113 | 2155 | 2036 | 2074 | 1309 | 2092 | 1688 | 1596 | 2156 | 1962 | 2059 | 1963 | 1966 | 1861 | 1833 | 1635 | 1645 | 1865,37 | 1988 | 1744 | 0,877 | -0,19 | -3,431 | 0,000602 | 0,000861 | 0,0019495 | 0,002745 | 0,0024099 | 0,0024099 | TRUE | 0,0257259 | No |
| orf19.1133 | 957 | 908 | 946 | 699 | 791 | 758 | 703 | 1068 | 871 | 918 | 895 | 1050 | 704 | 823 | 720 | 815 | 849,52 | 934 | 766 | 0,822 | -0,282 | -3,671 | 0,000241 | 0,000355 | 0,0053818 | 0,003592 | 0,004485 | 0,004485 | TRUE | 0,0773182 | No |
| orf19.1135 | 411 | 507 | 497 | 265 | 194 | 175 | 194 | 217 | 374 | 513 | 470 | 398 | 173 | 190 | 199 | 166 | 310,27 | 439 | 182 | 0,412 | -1,279 | -11,102 | 1,23E-28 | 3,82E-28 | 0,0120323 | 0,006295 | 0,0089488 | 0,0089488 | TRUE | 0,1357955 | No |
| orf19.1136 | 197 | 202 | 173 | 113 | 91 | 49 | 61 | 82 | 179 | 204 | 164 | 170 | 81 | 53 | 62 | 63 | 122,04 | 179 | 65 | 0,363 | -1,463 | -9,863 | 6,04E-23 | 1,64E-22 | 0,0070667 | 0,012862 | 0,0110093 | 0,0110093 | TRUE | 0,2554803 | No |
| orf19.1137 | 1646 | 1506 | 1529 | 1031 | 1556 | 1370 | 1409 | 1785 | 1499 | 1523 | 1447 | 1549 | 1384 | 1487 | 1443 | 1362 | 1461,7 | 1504 | 1419 | 0,943 | -0,085 | -1,717 | 0,08589 | 0,103801 | 0,0005557 | 0,002941 | 0,0016379 | 0,0016379 | TRUE | 0,012443 | No |
| orf19.1139 | 7250 | 6628 | 7452 | 4614 | 6652 | 5469 | 5952 | 8063 | 6601 | 6704 | 7053 | 6931 | 5917 | 5938 | 6096 | 6151 | 6423,58 | 6822 | 6026 | 0,884 | -0,179 | -4,969 | 6,74E-07 | 1,13E-06 | 0,000434 | 0,002243 | 0,0010845 | 0,0010845 | TRUE | 0,0066094 | No |
| orf19.114 | 153 | 187 | 188 | 107 | 283 | 236 | 239 | 361 | 139 | 189 | 178 | 161 | 252 | 256 | 245 | 275 | 211,9 | 167 | 257 | 1,543 | 0,626 | 5,665 | 1,47E-08 | 2,63E-08 | 0,0041479 | 0,008271 | 0,0067139 | 0,0067139 | TRUE | 0,1510727 | No |
| orf19.1140 | 743 | 743 | 768 | 459 | 1385 | 1053 | 1125 | 1440 | 676 | 751 | 727 | 689 | 1232 | 1143 | 1152 | 1098 | 933,75 | 711 | 1156 | 1,625 | 0,7 | 11,516 | 1,09E-30 | 3,57E-30 | 0,0013996 | 0,003452 | 0,0023909 | 0,0023909 | TRUE | 0,0239295 | No |
| orf19.1142 | 1322 | 1274 | 1378 | 853 | 1953 | 1811 | 1831 | 2404 | 1204 | 1289 | 1304 | 1281 | 1737 | 1966 | 1875 | 1834 | 1561,25 | 1270 | 1853 | 1,459 | 0,545 | 10,185 | 2,32E-24 | 6,56E-24 | 0,0014387 | 0,002883 | 0,0020779 | 0,0020779 | TRUE | 0,0213565 | No |
| orf19.1143 | 429 | 394 | 422 | 221 | 634 | 511 | 519 | 725 | 391 | 398 | 399 | 332 | 564 | 555 | 532 | 553 | 465,47 | 380 | 551 | 1,44 | 0,526 | 7,11 | 1,16E-12 | 2,40E-12 | 1,00E-08 | 0,004875 | 0,0030128 | 0,0030128 | TRUE | 0,0668819 | No |
| orf19.1144 | 730 | 677 | 738 | 427 | 501 | 397 | 477 | 598 | 665 | 685 | 698 | 641 | 446 | 431 | 489 | 456 | 563,82 | 672 | 456 | 0,676 | -0,565 | -8,457 | 2,74E-17 | 6,47E-17 | 1,00E-08 | 0,00438 | 0,0024663 | 0,0024663 | TRUE | 0,0259392 | No |
| orf19.1146 | 54 | 42 | 49 | 38 | 50 | 38 | 55 | 70 | 49 | 42 | 46 | 57 | 44 | 41 | 56 | 53 | 48,82 | 48 | 48 | 1,014 | 0,021 | 0,102 | 0,918716 | 0,930096 | 1,00E-08 | 0,029096 | 0,018293 | 0,018293 | TRUE | 0,0784281 | No |
| orf19.1148 | 4 | 3 | 7 | 6 | 14 | 10 | 15 | 23 | 4 | 3 | 7 | 9 | 12 | 11 | 15 | 18 | 9,82 | 6 | 14 | 2,654 | 1,408 | 2,925 | 0,003442 | 0,004659 | 1,00E-08 | 0,136609 | 0,090197 | 0,090197 | TRUE | 0,221424 | No |
| orf19.1149 | 2892 | 2328 | 2534 | 1631 | 4076 | 3343 | 3447 | 4896 | 2633 | 2355 | 2398 | 2450 | 3625 | 3630 | 3530 | 3735 | 3044,46 | 2459 | 3630 | 1,476 | 0,562 | 12,265 | 1,40E-34 | 4,96E-34 | 0,0010502 | 0,002471 | 0,0016669 | 0,0016669 | TRUE | 0,0288835 | No |
| orf19.115 | 129 | 149 | 110 | 59 | 217 | 153 | 184 | 200 | 117 | 151 | 104 | 89 | 193 | 166 | 188 | 153 | 145,13 | 115 | 175 | 1,504 | 0,589 | 3,883 | 0,000103 | 0,000154 | 0,0206795 | 0,011139 | 0,0148136 | 0,0148136 | TRUE | 0,4174736 | No |
| orf19.1150 | 5702 | 5076 | 5481 | 3883 | 5395 | 4636 | 4995 | 6874 | 5191 | 5134 | 5187 | 5833 | 4799 | 5033 | 5116 | 5244 | 5192,07 | 5336 | 5048 | 0,947 | -0,079 | -1,547 | 0,121868 | 0,145 | 0,0023669 | 0,002291 | 0,0022959 | 0,0022959 | TRUE | 0,0463985 | No |
| orf19.1150.1 | 200 | 184 | 194 | 149 | 383 | 315 | 315 | 429 | 182 | 186 | 184 | 224 | 341 | 342 | 323 | 327 | 263,52 | 194 | 333 | 1,735 | 0,795 | 8,515 | 1,67E-17 | 3,96E-17 | 1,00E-08 | 0,00705 | 0,0041889 | 0,0041889 | TRUE | 0,0916825 | No |
| orf19.1152 | 55367 | 49905 | 53037 | 32479 | 9128 | 7416 | 8052 | 10179 | 50409 | 50474 | 50195 | 48789 | 8119 | 8052 | 8246 | 7765 | 29006 | 49967 | 8046 | 0,161 | -2,635 | -84,238 | 0 | 0 | 0,0003511 | 0,002083 | 0,0008711 | 0,0008711 | TRUE | 0,0069113 | No |
| orf19.1153 | 7748 | 6907 | 7682 | 4561 | 1356 | 1129 | 1154 | 1602 | 7054 | 6986 | 7270 | 6851 | 1206 | 1226 | 1182 | 1222 | 4124,67 | 7040 | 1209 | 0,172 | -2,542 | -64,175 | 0 | 0 | 1,00E-08 | 0,002357 | 0,0010449 | 0,0010449 | TRUE | 0,0059025 | No |
| orf19.1154 | 2071 | 1698 | 1779 | 1223 | 3514 | 2770 | 2635 | 3753 | 1886 | 1717 | 1684 | 1837 | 3126 | 3007 | 2699 | 2863 | 2352,27 | 1781 | 2924 | 1,643 | 0,716 | 12,164 | 4,85E-34 | 1,69E-33 | 0,0028406 | 0,002599 | 0,0028636 | 0,0028636 | TRUE | 0,0321808 | No |
| orf19.1155 | 274 | 256 | 256 | 177 | 460 | 419 | 441 | 644 | 249 | 259 | 242 | 266 | 409 | 455 | 452 | 491 | 352,94 | 254 | 452 | 1,785 | 0,836 | 9,665 | 4,26E-22 | 1,13E-21 | 1,00E-08 | 0,00578 | 0,0040399 | 0,0040399 | TRUE | 0,0492829 | No |
| orf19.1156 | 208 | 189 | 220 | 136 | 312 | 264 | 250 | 383 | 189 | 191 | 208 | 204 | 278 | 287 | 256 | 292 | 238,17 | 198 | 278 | 1,407 | 0,493 | 5,196 | 2,04E-07 | 3,48E-07 | 1,00E-08 | 0,007583 | 0,0042732 | 0,0042732 | TRUE | 0,030825 | No |
| orf19.1158 | 458 | 423 | 417 | 335 | 301 | 260 | 243 | 360 | 417 | 428 | 395 | 503 | 268 | 282 | 249 | 275 | 352,02 | 436 | 268 | 0,62 | -0,69 | -7,572 | 3,68E-14 | 7,96E-14 | 0,0039102 | 0,00579 | 0,0049885 | 0,0049885 | TRUE | 0,1101715 | No |
| orf19.1159 | 3963 | 3282 | 3733 | 2139 | 4615 | 3379 | 4089 | 5410 | 3608 | 3319 | 3533 | 3213 | 4105 | 3669 | 4188 | 4127 | 3720,2 | 3418 | 4022 | 1,176 | 0,234 | 4,194 | 2,74E-05 | 4,24E-05 | 0,0028337 | 0,002392 | 0,0027093 | 0,0027093 | TRUE | 0,0420037 | No |
| orf19.1160 | 1247 | 1013 | 1183 | 776 | 933 | 760 | 802 | 1069 | 1135 | 1025 | 1120 | 1166 | 830 | 825 | 821 | 815 | 967,12 | 1112 | 823 | 0,741 | -0,431 | -7,674 | 1,67E-14 | 3,64E-14 | 0,0006085 | 0,003403 | 0,0019851 | 0,0019851 | TRUE | 0,0330778 | No |
| orf19.1161 | 3654 | 3059 | 3446 | 2090 | 4373 | 3546 | 4161 | 5503 | 3327 | 3094 | 3261 | 3140 | 3890 | 3850 | 4261 | 4198 | 3627,53 | 3206 | 4050 | 1,263 | 0,337 | 6,973 | 3,09E-12 | 6,32E-12 | 0,0015334 | 0,002401 | 0,0019576 | 0,0019576 | TRUE | 0,014726 | No |
| orf19.1162 | 2002 | 1668 | 1764 | 971 | 1591 | 1228 | 1374 | 1812 | 1823 | 1687 | 1669 | 1459 | 1415 | 1333 | 1407 | 1382 | 1521,95 | 1660 | 1384 | 0,832 | -0,265 | -4,112 | 3,92E-05 | 6,02E-05 | 0,003579 | 0,002905 | 0,0033187 | 0,0033187 | TRUE | 0,074649 | No |
| orf19.1162.1 | 502 | 447 | 444 | 278 | 608 | 466 | 504 | 719 | 457 | 452 | 420 | 418 | 541 | 506 | 516 | 548 | 482,29 | 437 | 528 | 1,208 | 0,272 | 3,887 | 0,000102 | 0,000152 | 1,00E-08 | 0,004776 | 0,0026168 | 0,0026168 | TRUE | 0,011626 | No |
| orf19.1163 | 1732 | 1526 | 1683 | 1209 | 1518 | 1352 | 1379 | 1757 | 1577 | 1543 | 1593 | 1816 | 1350 | 1468 | 1412 | 1340 | 1512,48 | 1632 | 1392 | 0,855 | -0,227 | -3,745 | 0,00018 | 0,000266 | 0,0029206 | 0,00291 | 0,0028516 | 0,0028516 | TRUE | 0,0644205 | No |
| orf19.1164 | 9610 | 7379 | 8223 | 5485 | 16240 | 12027 | 13843 | 19763 | 8749 | 7463 | 7782 | 8239 | 14445 | 13058 | 14177 | 15075 | 11123,7 | 8058 | 14189 | 1,761 | 0,816 | 13,776 | 3,54E-43 | 1,46E-42 | 0,0040491 | 0,002156 | 0,0032738 | 0,0032738 | TRUE | 0,0412168 | No |
| orf19.1166 | 6185 | 5158 | 5779 | 3889 | 7668 | 6447 | 6841 | 9822 | 5631 | 5217 | 5469 | 5842 | 6820 | 7000 | 7006 | 7492 | 6309,7 | 5540 | 7080 | 1,279 | 0,355 | 7,556 | 4,17E-14 | 8,98E-14 | 0,0017491 | 0,002246 | 0,0019551 | 0,0019551 | TRUE | 0,0190791 | No |
| orf19.1167 | 898 | 786 | 808 | 550 | 812 | 599 | 656 | 923 | 818 | 795 | 765 | 826 | 722 | 650 | 672 | 704 | 743,99 | 801 | 687 | 0,861 | -0,217 | -3,585 | 0,000337 | 0,00049 | 1,00E-08 | 0,003813 | 0,002159 | 0,002159 | TRUE | 0,0147006 | No |
| orf19.1168 | 9210 | 7383 | 8350 | 5658 | 8096 | 6465 | 7064 | 9836 | 8385 | 7467 | 7903 | 8499 | 7201 | 7019 | 7234 | 7503 | 7651,49 | 8064 | 7239 | 0,898 | -0,155 | -3,231 | 0,001236 | 0,001735 | 0,0019353 | 0,00221 | 0,0020746 | 0,0020746 | TRUE | 0,0302728 | No |
| orf19.1169 | 6683 | 5818 | 6486 | 3468 | 7101 | 5763 | 6566 | 8568 | 6085 | 5884 | 6138 | 5209 | 6316 | 6257 | 6724 | 6536 | 6143,76 | 5829 | 6458 | 1,107 | 0,147 | 2,673 | 0,007519 | 0,009953 | 0,0030195 | 0,002252 | 0,0027263 | 0,0027263 | TRUE | 0,060835 | No |
| orf19.117 | 4006 | 3732 | 4081 | 2527 | 2873 | 2320 | 2506 | 3286 | 3647 | 3775 | 3862 | 3796 | 2555 | 2519 | 2566 | 2507 | 3153,43 | 3770 | 2537 | 0,673 | -0,571 | -15,53 | 2,17E-54 | 1,09E-53 | 1,00E-08 | 0,002456 | 0,0009727 | 0,0009727 | TRUE | 0,0061278 | No |
| orf19.1170 | 1777 | 1564 | 1681 | 1061 | 2040 | 1772 | 1764 | 2270 | 1618 | 1582 | 1591 | 1594 | 1815 | 1924 | 1807 | 1732 | 1707,62 | 1596 | 1820 | 1,138 | 0,187 | 3,996 | 6,44E-05 | 9,74E-05 | 0,0005447 | 0,002811 | 0,0015064 | 0,0015064 | TRUE | 0,0177097 | No |
| orf19.1171 | 30 | 31 | 29 | 22 | 74 | 64 | 56 | 69 | 27 | 31 | 27 | 33 | 66 | 69 | 57 | 53 | 45,56 | 30 | 61 | 2,068 | 1,048 | 4,879 | 1,07E-06 | 1,77E-06 | 1,00E-08 | 0,031034 | 0,0187439 | 0,0187439 | TRUE | 0,0931804 | No |
| orf19.1177 | 479 | 363 | 445 | 251 | 493 | 397 | 465 | 600 | 436 | 367 | 421 | 377 | 439 | 431 | 476 | 458 | 425,61 | 400 | 451 | 1,122 | 0,166 | 2,085 | 0,037048 | 0,046446 | 0,0019034 | 0,005141 | 0,0037485 | 0,0037485 | TRUE | 0,0426444 | No |
| orf19.1178 | 130 | 104 | 123 | 84 | 212 | 160 | 154 | 256 | 118 | 105 | 116 | 126 | 189 | 174 | 158 | 195 | 147,68 | 116 | 179 | 1,547 | 0,629 | 5,13 | 2,89E-07 | 4,91E-07 | 1,00E-08 | 0,010982 | 0,0072638 | 0,0072638 | TRUE | 0,0670427 | No |

| | | | | | | | | | | | | | | | | | | | | | | | | | | | | | | | |
|---|---|---|---|---|---|---|---|---|---|---|---|---|---|---|---|---|---|---|---|---|---|---|---|---|---|---|---|---|---|---|---|
| orf19.1179 | 1708 | 1222 | 1268 | 1048 | 2146 | 1616 | 1558 | 2463 | 1555 | 1236 | 1200 | 1574 | 1909 | 1755 | 1596 | 1879 | 1587,88 | 1391 | 1785 | 1,285 | 0,361 | 3,865 | 0,000111 | 0,000166 | 0,0127553 | 0,002869 | 0,007749 | 0,007749 | TRUE | 0,1036766 | No |
| orf19.118 | 18331 | 17865 | 18334 | 11007 | 62301 | 52894 | 53277 | 71001 | 16690 | 18069 | 17351 | 16534 | 55415 | 57428 | 54562 | 54160 | 36276,2 | 17161 | 55391 | 3,227 | 1,69 | 43,217 | 0 | 0 | 0,0011289 | 0,002073 | 0,0014296 | 0,0014296 | TRUE | 0,0155011 | No |
| orf19.1180 | 296 | 209 | 281 | 190 | 281 | 193 | 264 | 361 | 269 | 211 | 266 | 285 | 250 | 210 | 270 | 275 | 254,68 | 258 | 251 | 0,979 | -0,03 | -0,259 | 0,795441 | 0,822537 | 0,0113378 | 0,007224 | 0,0089597 | 0,0089597 | TRUE | 0,1664185 | No |
| orf19.1181 | 93 | 69 | 91 | 50 | 125 | 117 | 118 | 172 | 85 | 70 | 86 | 75 | 111 | 127 | 121 | 131 | 100,74 | 79 | 122 | 1,546 | 0,629 | 4,402 | 1,07E-05 | 1,70E-05 | 1,00E-08 | 0,015149 | 0,0090987 | 0,0090987 | TRUE | 0,0638536 | No |
| orf19.1182 | 707 | 608 | 642 | 405 | 703 | 564 | 631 | 853 | 644 | 615 | 608 | 608 | 625 | 612 | 646 | 651 | 626,14 | 619 | 634 | 1,024 | 0,034 | 0,561 | 0,574915 | 0,61543 | 1,00E-08 | 0,004147 | 0,001962 | 0,001962 | TRUE | 0,0093696 | No |
| orf19.1183 | 2597 | 2139 | 2513 | 1473 | 2882 | 2122 | 2598 | 3395 | 2364 | 2163 | 2378 | 2213 | 2563 | 2304 | 2661 | 2590 | 2404,57 | 2280 | 2530 | 1,109 | 0,15 | 2,658 | 0,007867 | 0,010396 | 0,0024629 | 0,002586 | 0,0026197 | 0,0026197 | TRUE | 0,0427109 | No |
| orf19.1185 | 4333 | 3438 | 3895 | 2614 | 5472 | 4657 | 4807 | 7035 | 3945 | 3477 | 3686 | 3927 | 4867 | 5056 | 4923 | 5366 | 4405,97 | 3759 | 5053 | 1,345 | 0,428 | 8,134 | 4,14E-16 | 9,44E-16 | 0,0024065 | 0,002337 | 0,0024208 | 0,0024208 | TRUE | 0,031033 | No |
| orf19.1186 | 949 | 727 | 807 | 481 | 1146 | 810 | 888 | 1295 | 864 | 735 | 764 | 723 | 1019 | 879 | 909 | 988 | 860,2 | 772 | 949 | 1,229 | 0,297 | 4,026 | 5,67E-05 | 8,61E-05 | 0,0042665 | 0,003573 | 0,0040495 | 0,0040495 | TRUE | 0,0796583 | No |
| orf19.1187 | 36280 | 33438 | 37018 | 23905 | 18494 | 16539 | 16589 | 22443 | 33031 | 33819 | 35034 | 35909 | 16450 | 17957 | 16989 | 17120 | 25788,7 | 34448 | 17129 | 0,497 | -1,008 | -24,636 | 5,25E-134 | 6,06E-133 | 0,0012896 | 0,002088 | 0,0015653 | 0,0015653 | TRUE | 0,012887 | No |
| orf19.1189 | 27688 | 22718 | 25701 | 17458 | 4329 | 3548 | 3812 | 5361 | 25209 | 22977 | 24324 | 26225 | 3851 | 3852 | 3904 | 4089 | 14303,8 | 24684 | 3924 | 0,159 | -2,653 | -55,912 | 0 | 0 | 0,0017754 | 0,002129 | 0,002021 | 0,002021 | TRUE | 0,0264985 | No |
| orf19.119 | 255 | 248 | 254 | 171 | 449 | 424 | 422 | 522 | 232 | 251 | 240 | 257 | 399 | 460 | 432 | 398 | 333,79 | 245 | 422 | 1,725 | 0,787 | 9,1 | 9,05E-20 | 2,28E-19 | 1,00E-08 | 0,005995 | 0,00389 | 0,00389 | TRUE | 0,0406413 | No |
| orf19.1190 | 1669 | 1433 | 1656 | 971 | 1547 | 1262 | 1395 | 1854 | 1520 | 1449 | 1567 | 1459 | 1376 | 1370 | 1429 | 1414 | 1447,98 | 1499 | 1397 | 0,931 | -0,103 | -2,167 | 0,030233 | 0,038226 | 1,00E-08 | 0,002949 | 0,0014615 | 0,0014615 | TRUE | 0,0115634 | No |
| orf19.1191 | 4888 | 4380 | 4864 | 2929 | 2449 | 1884 | 2096 | 2848 | 4450 | 4430 | 4603 | 4400 | 2178 | 2045 | 2147 | 2172 | 3303,28 | 4471 | 2136 | 0,478 | -1,065 | -27,536 | 6,54E-167 | 9,59E-166 | 0,0002318 | 0,002437 | 0,0010982 | 0,0010982 | TRUE | 0,0094008 | No |
| orf19.1192 | 745 | 680 | 680 | 471 | 1798 | 1534 | 1750 | 2268 | 678 | 688 | 644 | 708 | 1599 | 1665 | 1792 | 1730 | 1188,02 | 680 | 1696 | 2,503 | 1,324 | 22,874 | 8,50E-116 | 8,40E-115 | 0,0012817 | 0,003149 | 0,0021509 | 0,0021509 | TRUE | 0,0183782 | No |
| orf19.1193 | 1E+05 | 1E+05 | 1E+05 | 73853 | 54076 | 45277 | 46842 | 67146 | 1E+05 | 1E+05 | 1E+05 | 1E+05 | 48099 | 49158 | 47972 | 51220 | 79633,7 | 110155 | 49112 | 0,446 | -1,165 | -32,423 | 1,31E-230 | 2,53E-229 | 0,0008552 | 0,002054 | 0,0012268 | 0,0012268 | TRUE | 0,0102571 | No |
| orf19.1195 | 1529 | 1366 | 1531 | 922 | 2825 | 2246 | 2530 | 3344 | 1392 | 1382 | 1449 | 1385 | 2513 | 2439 | 2591 | 2551 | 1962,59 | 1402 | 2524 | 1,799 | 0,847 | 19,652 | 5,61E-86 | 4,27E-85 | 1,00E-08 | 0,00271 | 0,0012185 | 0,0012185 | TRUE | 0,006413 | No |
| orf19.1196 | 1441 | 1320 | 1443 | 915 | 1297 | 1063 | 1159 | 1589 | 1312 | 1335 | 1366 | 1374 | 1154 | 1154 | 1187 | 1212 | 1261,75 | 1347 | 1177 | 0,875 | -0,192 | -4,076 | 4,58E-05 | 6,99E-05 | 1,00E-08 | 0,003084 | 0,0013461 | 0,0013461 | TRUE | 0,0053279 | No |
| orf19.1198 | 1540 | 1364 | 1439 | 940 | 908 | 766 | 844 | 1067 | 1402 | 1380 | 1362 | 1412 | 808 | 832 | 864 | 814 | 1109,14 | 1389 | 830 | 0,597 | -0,744 | -14,967 | 1,20E-50 | 5,60E-50 | 1,00E-08 | 0,003228 | 0,0014263 | 0,0014263 | TRUE | 0,0093963 | No |
| orf19.1199 | 22316 | 19379 | 21051 | 14824 | 36182 | 31800 | 31908 | 43508 | 20318 | 19600 | 19923 | 22268 | 32183 | 34526 | 32678 | 33188 | 26835,4 | 20527 | 33144 | 1,615 | 0,692 | 14,81 | 1,27E-49 | 5,78E-49 | 0,0021113 | 0,002086 | 0,0020544 | 0,0020544 | TRUE | 0,039522 | No |
| orf19.120 | 197 | 233 | 206 | 138 | 1204 | 1083 | 996 | 1300 | 179 | 236 | 195 | 207 | 1071 | 1176 | 1020 | 992 | 634,46 | 204 | 1065 | 5,22 | 2,384 | 26,181 | 4,41E-151 | 5,78E-150 | 0,0058034 | 0,004119 | 0,0049058 | 0,0049058 | TRUE | 0,1269995 | No |
| orf19.1200 | 2386 | 2034 | 2188 | 1489 | 2612 | 2114 | 2173 | 2917 | 2172 | 2057 | 2071 | 2237 | 2323 | 2295 | 2225 | 2225 | 2200,75 | 2134 | 2267 | 1,064 | 0,089 | 2,02 | 0,043361 | 0,054021 | 0,0004373 | 0,002637 | 0,001403 | 0,001403 | TRUE | 0,011355 | No |
| orf19.1201 | 2793 | 2281 | 2559 | 1640 | 1613 | 1375 | 1447 | 1965 | 2543 | 2307 | 2422 | 2464 | 1435 | 1493 | 1482 | 1499 | 1955,46 | 2434 | 1477 | 0,607 | -0,72 | -15,602 | 7,03E-55 | 3,55E-54 | 0,0006721 | 0,002713 | 0,0015099 | 0,0015099 | TRUE | 0,0152734 | No |
| orf19.1202 | 127 | 142 | 146 | 113 | 72 | 84 | 73 | 106 | 116 | 144 | 138 | 170 | 64 | 91 | 75 | 81 | 109,75 | 142 | 78 | 0,552 | -0,858 | -5,488 | 4,07E-08 | 7,13E-08 | 0,013006 | 0,014073 | 0,0136642 | 0,0136642 | TRUE | 0,1623212 | No |
| orf19.1203 | 976 | 831 | 874 | 523 | 1090 | 880 | 971 | 1268 | 889 | 840 | 827 | 786 | 970 | 955 | 994 | 967 | 903,56 | 836 | 972 | 1,159 | 0,213 | 3,839 | 0,000124 | 0,000184 | 1,00E-08 | 0,003499 | 0,0018482 | 0,0018482 | TRUE | 0,0219647 | No |
| orf19.1203.1 | 380 | 314 | 313 | 242 | 308 | 265 | 263 | 329 | 346 | 318 | 296 | 364 | 274 | 288 | 269 | 251 | 300,66 | 331 | 270 | 0,819 | -0,288 | -3,137 | 0,001708 | 0,00237 | 0,0023694 | 0,006431 | 0,0047303 | 0,0047303 | TRUE | 0,0571699 | No |
| orf19.1204 | 1244 | 1075 | 1232 | 763 | 655 | 608 | 652 | 806 | 1133 | 1087 | 1166 | 1146 | 583 | 660 | 668 | 615 | 882,16 | 1133 | 632 | 0,556 | -0,846 | -13,947 | 3,27E-44 | 1,38E-43 | 0,0007985 | 0,003534 | 0,0023214 | 0,0023214 | TRUE | 0,0313575 | No |
| orf19.1207 | 9 | 3 | 9 | 4 | 16 | 14 | 11 | 25 | 8 | 3 | 9 | 6 | 14 | 15 | 11 | 19 | 10,69 | 6 | 15 | 2,309 | 1,208 | 2,679 | 0,00738 | 0,009775 | 1,00E-08 | 0,125608 | 0,0827516 | 0,0827516 | TRUE | 0,3437343 | No |
| orf19.1208 | 20 | 20 | 27 | 19 | 55 | 53 | 55 | 96 | 18 | 20 | 26 | 29 | 49 | 58 | 56 | 73 | 41,07 | 23 | 59 | 2,605 | 1,382 | 5,652 | 1,59E-08 | 2,83E-08 | 1,00E-08 | 0,034202 | 0,026165 | 0,026165 | TRUE | 0,2634204 | No |
| orf19.121 | 339 | 379 | 347 | 174 | 183 | 123 | 139 | 161 | 309 | 383 | 328 | 261 | 163 | 134 | 142 | 123 | 230,4 | 320 | 140 | 0,436 | -1,199 | -9,291 | 1,53E-20 | 3,94E-20 | 0,0152764 | 0,00777 | 0,0109843 | 0,0109843 | TRUE | 0,1439747 | No |
| orf19.1210 | 4505 | 4400 | 4617 | 2773 | 1838 | 1504 | 1575 | 2188 | 4102 | 4450 | 4370 | 4165 | 1635 | 1633 | 1613 | 1669 | 2954,57 | 4272 | 1638 | 0,383 | -1,383 | -32,164 | 5,56E-227 | 1,06E-225 | 0,0006946 | 0,002484 | 0,0013647 | 0,0013647 | TRUE | 0,0096506 | No |
| orf19.1212 | 804 | 711 | 753 | 557 | 2358 | 2027 | 1997 | 2825 | 732 | 719 | 713 | 837 | 2097 | 2201 | 2045 | 2155 | 1437,34 | 750 | 2124 | 2,847 | 1,509 | 26,223 | 1,44E-151 | 1,90E-150 | 0,0013611 | 0,002956 | 0,0022435 | 0,0022435 | TRUE | 0,0615302 | No |
| orf19.1214 | 6763 | 6081 | 6547 | 3806 | 6231 | 4907 | 5447 | 7120 | 6157 | 6150 | 6196 | 5717 | 5542 | 5328 | 5578 | 5431 | 5762,57 | 6055 | 5470 | 0,903 | -0,148 | -3,834 | 0,000126 | 0,000188 | 0,000682 | 0,002266 | 0,0012534 | 0,0012534 | TRUE | 0,0163956 | No |
| orf19.1215 | 11 | 14 | 3 | 12 | 4 | 7 | 8 | 4 | 10 | 14 | 3 | 18 | 4 | 8 | 8 | 3 | 8,43 | 11 | 6 | 0,498 | -1,005 | -1,736 | 0,082572 | 0,09998 | 0,2359452 | 0,158735 | 0,1876584 | 0,1876584 | TRUE | 0,2699401 | No |
| orf19.1217 | 763 | 821 | 858 | 466 | 1007 | 962 | 920 | 1107 | 695 | 830 | 812 | 700 | 896 | 1044 | 942 | 844 | 845,48 | 759 | 932 | 1,223 | 0,291 | 3,49 | 0,000482 | 0,000695 | 0,0070671 | 0,003599 | 0,0054561 | 0,0054561 | TRUE | 0,0781579 | No |
| orf19.1219 | 309 | 313 | 320 | 157 | 426 | 342 | 370 | 519 | 281 | 317 | 303 | 236 | 379 | 371 | 379 | 396 | 332,71 | 284 | 381 | 1,333 | 0,414 | 4,589 | 4,45E-06 | 7,19E-06 | 0,0023566 | 0,006007 | 0,0047372 | 0,0047372 | TRUE | 0,1177807 | No |
| orf19.122 | 707 | 678 | 716 | 445 | 419 | 373 | 411 | 538 | 644 | 686 | 678 | 668 | 373 | 405 | 421 | 410 | 535,56 | 669 | 402 | 0,601 | -0,734 | -10,842 | 2,17E-27 | 6,57E-27 | 1,00E-08 | 0,004504 | 0,0024431 | 0,0024431 | TRUE | 0,0290838 | No |
| orf19.1220 | 12106 | 10859 | 12070 | 7778 | 5994 | 4820 | 5210 | 6831 | 11022 | 10983 | 11423 | 11684 | 5331 | 5233 | 5336 | 5211 | 8277,85 | 11278 | 5278 | 0,468 | -1,095 | -32,081 | 8,05E-226 | 1,53E-224 | 0,0003342 | 0,002197 | 0,0009824 | 0,0009824 | TRUE | 0,0069365 | No |
| orf19.1221 | 900 | 862 | 925 | 540 | 503 | 409 | 421 | 600 | 819 | 872 | 875 | 811 | 447 | 444 | 431 | 458 | 644,77 | 844 | 445 | 0,527 | -0,924 | -14,734 | 3,92E-49 | 1,78E-48 | 1,00E-08 | 0,004086 | 0,0021015 | 0,0021015 | TRUE | 0,0072265 | No |
| orf19.1223 | 958 | 1086 | 1084 | 642 | 823 | 753 | 762 | 900 | 872 | 1098 | 1026 | 964 | 732 | 818 | 780 | 687 | 872,17 | 990 | 754 | 0,76 | -0,396 | -4,888 | 1,02E-06 | 1,69E-06 | 0,0066729 | 0,003552 | 0,0051211 | 0,0051211 | TRUE | 0,081009 | No |
| orf19.1224 | 5623 | 5785 | 5913 | 4173 | 15671 | 14463 | 13465 | 18321 | 5120 | 5851 | 5596 | 6269 | 13939 | 15703 | 13790 | 13975 | 10030,2 | 5709 | 14352 | 2,515 | 1,331 | 20,559 | 6,42E-94 | 5,19E-93 | 0,0053681 | 0,002169 | 0,003898 | 0,003898 | TRUE | 0,0590642 | No |
| orf19.1225 | 1597 | 1639 | 1637 | 937 | 3050 | 2421 | 2351 | 3078 | 1454 | 1658 | 1549 | 1408 | 2713 | 2629 | 2408 | 2348 | 2020,69 | 1517 | 2524 | 1,661 | 0,732 | 11,111 | 1,11E-28 | 3,44E-28 | 0,0043436 | 0,002691 | 0,0036327 | 0,0036327 | TRUE | 0,0464804 | No |
| orf19.1227 | 133 | 120 | 129 | 76 | 117 | 92 | 85 | 117 | 121 | 121 | 122 | 114 | 104 | 100 | 87 | 89 | 107,37 | 120 | 95 | 0,791 | -0,338 | -2,53 | 0,011414 | 0,014904 | 1,00E-08 | 0,01434 | 0,0077705 | 0,0077705 | TRUE | 0,0431848 | No |
| orf19.1228 | 3775 | 3579 | 3831 | 2404 | 1654 | 1410 | 1600 | 1922 | 3437 | 3620 | 3626 | 3611 | 1471 | 1531 | 1639 | 1466 | 2550,05 | 3574 | 1527 | 0,427 | -1,228 | -25,767 | 2,07E-146 | 2,66E-145 | 0,0010816 | 0,002555 | 0,001724 | 0,001724 | TRUE | 0,0291101 | No |
| orf19.1229 | 1634 | 2169 | 2077 | 1214 | 1248 | 1271 | 1156 | 1401 | 1488 | 2194 | 1966 | 1824 | 1110 | 1380 | 1184 | 1069 | 1526,66 | 1868 | 1186 | 0,634 | -0,657 | -6,095 | 1,10E-09 | 2,05E-09 | 0,0183861 | 0,002902 | 0,010467 | 0,010467 | TRUE | 0,2304078 | No |
| orf19.123 | 414 | 456 | 492 | 228 | 340 | 298 | 300 | 382 | 377 | 461 | 466 | 342 | 302 | 324 | 307 | 291 | 358,86 | 412 | 306 | 0,739 | -0,436 | -4,235 | 2,29E-05 | 3,57E-05 | 0,0093309 | 0,005718 | 0,00733 | 0,00733 | TRUE | 0,1279866 | No |
| orf19.1232 | 3432 | 3901 | 3708 | 2047 | 5559 | 4687 | 4622 | 6280 | 3125 | 3945 | 3509 | 3075 | 4945 | 5089 | 4734 | 4790 | 4151,45 | 3414 | 4890 | 1,431 | 0,517 | 7,143 | 9,13E-13 | 1,90E-12 | 0,0072156 | 0,002355 | 0,0047862 | 0,0047862 | TRUE | 0,1337074 | No |
| orf19.1233 | 9158 | 9322 | 9884 | 5906 | 15162 | 13320 | 13194 | 17119 | 8338 | 9428 | 9354 | 8872 | 13486 | 14462 | 13512 | 13059 | 11313,9 | 8998 | 13630 | 1,515 | 0,599 | 11,929 | 8,38E-33 | 2,86E-32 | 0,0023937 | 0,002154 | 0,0023267 | 0,0023267 | TRUE | 0,0302491 | No |
| orf19.1234 | 39 | 42 | 53 | 33 | 81 | 73 | 91 | 122 | 36 | 42 | 50 | 50 | 72 | 79 | 93 | 93 | 64,41 | 44 | 84 | 1,918 | 0,94 | 5,008 | 5,49E-07 | 9,23E-07 | 1,00E-08 | 0,022546 | 0,0162238 | 0,0162238 | TRUE | 0,1608817 | No |
| orf19.1235 | 2594 | 2448 | 2579 | 1626 | 1872 | 1772 | 1791 | 2378 | 2362 | 2476 | 2441 | 2443 | 1665 | 1924 | 1834 | 1814 | 2119,76 | 2430 | 1809 | 0,744 | -0,426 | -8,44 | 3,16E-17 | 7,44E-17 | 0,0013161 | 0,00266 | 0,0019709 | 0,0019709 | TRUE | 0,0348672 | No |
| orf19.1236 | 2785 | 2663 | 2654 | 1989 | 1298 | 1111 | 981 | 1378 | 2536 | 2693 | 2512 | 2988 | 1155 | 1206 | 1005 | 1051 | 1893,14 | 2682 | 1104 | 0,412 | -1,279 | -17,681 | 5,90E-70 | 3,71E-69 | 0,0059936 | 0,002735 | 0,0044034 | 0,0044034 | TRUE | 0,069192 | No |
| orf19.1237 | 823 | 778 | 772 | 528 | 1145 | 969 | 939 | 1246 | 749 | 787 | 731 | 793 | 1018 | 1052 | 962 | 950 | 880,32 | 765 | 996 | 1,303 | 0,382 | 6,44 | 1,19E-10 | 2,31E-10 | 0,0008787 | 0,003538 | 0,0022 | 0,0022 | TRUE | 0,0166683 | No |
| orf19.1238 | 235 | 230 | 226 | 144 | 251 | 210 | 253 | 305 | 214 | 233 | 214 | 216 | 223 | 228 | 259 | 233 | 227,47 | 219 | 236 | 1,074 | 0,103 | 1,075 | 0,282212 | 0,318764 | 1,00E-08 | 0,007844 | 0,0044492 | 0,0044492 | TRUE | 0,0467533 | No |
| orf19.1239 | 10 | 3 | 6 | 4 | 19 | 25 | 17 | 25 | 9 | 3 | 6 | 6 | 17 | 27 | 17 | 19 | 13,04 | 6 | 20 | 3,33 | 1,735 | 3,976 | 7,01E-05 | 0,000106 | 1,00E-08 | 0,103312 | 0,071527 | 0,071527 | TRUE | 0,3688113 | No |
| orf19.124 | 8265 | 8415 | 8613 | 5433 | 8795 | 7455 | 7623 | 9922 | 7525 | 8511 | 8151 | 8161 | 7823 | 8094 | 7807 | 7569 | 7955,11 | 8087 | 7823 | 0,967 | -0,048 | -1,067 | 0,28593 | 0,322563 | 0,001497 | 0,002203 | 0,0018077 | 0,0018077 | TRUE | 0,0271604 | No |
| orf19.1240 | 9978 | 9008 | 9729 | 6093 | 4028 | 3348 | 3596 | 4582 | 9085 | 9111 | 9208 | 9153 | 3583 | 3635 | 3683 | 3495 | 6368,9 | 9139 | 3599 | 0,394 | -1,345 | -42,439 | 0 | 0 | 8,11E-06 | 0,002244 | 0,000776 | 0,000776 | TRUE | 0,0047787 | No |
| orf19.1244 | 2597 | 2492 | 2626 | 1542 | 2063 | 1668 | 1755 | 2542 | 2364 | 2520 | 2485 | 2316 | 1835 | 1811 | 1797 | 1939 | 2133,6 | 2421 | 1846 | 0,762 | -0,392 | -8,307 | 9,85E-17 | 2,28E-16 | 0,00094 | 0,002656 | 0,0016632 | 0,0016632 | TRUE | 0,0146588 | No |
| orf19.1246 | 4766 | 4092 | 4683 | 3028 | 4180 | 3590 | 3691 | 5177 | 4339 | 4139 | 4432 | 4549 | 3718 | 3898 | 3780 | 3949 | 4100,41 | 4365 | 3836 | 0,88 | -0,185 | -4,39 | 1,13E-05 | 1,79E-05 | 0,0008428 | 0,002359 | 0,0014611 | 0,0014611 | TRUE | 0,0147003 | No |
| orf19.1247 | 266 | 232 | 249 | 144 | 331 | 242 | 315 | 394 | 242 | 235 | 236 | 216 | 294 | 263 | 323 | 301 | 263,64 | 232 | 295 | 1,267 | 0,342 | 3,645 | 0,000267 | 0,000391 | 1,00E-08 | 0,007048 | 0,0045655 | 0,0045655 | TRUE | 0,0560193 | No |
| orf19.1248 | 4050 | 3417 | 3707 | 2565 | 4159 | 3235 | 3533 | 4937 | 3687 | 3456 | 3508 | 3853 | 3699 | 3512 | 3618 | 3766 | 3637,56 | 3626 | 3649 | 1,008 | 0,011 | 0,241 | 0,809595 | 0,834465 | 0,0011777 | 0,0024 | 0,0017595 | 0,0017595 | TRUE | 0,0203572 | No |
| orf19.1249 | 738 | 587 | 640 | 414 | 1264 | 1006 | 1111 | 1446 | 672 | 594 | 606 | 622 | 1124 | 1092 | 1138 | 1103 | 868,82 | 624 | 1114 | 1,786 | 0,837 | 14,372 | 7,72E-47 | 3,38E-46 | 2,60E-05 | 0,003557 | 0,0019808 | 0,0019808 | TRUE | 0,0360108 | No |
| orf19.125 | 9690 | 9205 | 9602 | 5680 | 1741 | 1378 | 1427 | 1919 | 8822 | 9310 | 9087 | 8532 | 1549 | 1496 | 1461 | 1464 | 5215,23 | 8938 | 1492 | 0,167 | -2,583 | -59,625 | 0 | 0 | 0,0009055 | 0,00229 | 0,0014288 | 0,0014288 | TRUE | 0,0110358 | No |
| orf19.1250 | 3919 | 3250 | 3574 | 1957 | 7319 | 4983 | 6429 | 8330 | 3568 | 3287 | 3382 | 2940 | 6510 | 5410 | 6584 | 6354 | 4754,47 | 3294 | 6214 | 1,885 | 0,915 | 12,602 | 2,07E-36 | 7,61E-36 | 0,006735 | 0,002315 | 0,0048208 | 0,0048208 | TRUE | 0,0921671 | No |
| orf19.1251 | 1912 | 1739 | 1863 | 1230 | 1660 | 1337 | 1503 | 2019 | 1741 | 1759 | 1763 | 1848 | 1477 | 1452 | 1539 | 1540 | 1639,74 | 1778 | 1502 | 0,847 | -0,24 | -5,316 | 1,06E-07 | 1,83E-07 | 1,00E-08 | 0,002843 | 0,0013481 | 0,0013481 | TRUE | 0,0075618 | No |
| orf19.1252 | 20845 | 17709 | 19667 | 12803 | 11616 | 9827 | 10523 | 14227 | 18978 | 17911 | 18613 | 19232 | 10332 | 10669 | 10777 | 10852 | 14670,7 | 18684 | 10658 | 0,571 | -0,81 | -23,064 | 1,07E-117 | 1,07E-116 | 0,0006068 | 0,002127 | 0,0011109 | 0,0011109 | TRUE | 0,0094854 | No |
| orf19.1253 | 2165 | 1835 | 2106 | 1387 | 1766 | 1559 | 1710 | 2418 | 1971 | 1856 | 1993 | 2083 | 1571 | 1693 | 1751 | 1844 | 1845,36 | 1976 | 1715 | 0,869 | -0,203 | -3,423 | 0,000619 | 0,000886 | 0,0026278 | 0,002753 | 0,0028156 | 0,0028156 | TRUE | 0,039575 | No |
| orf19.1254 | 1002 | 837 | 839 | 532 | 879 | 747 | 824 | 1080 | 912 | 847 | 794 | 799 | 782 | 811 | 844 | 824 | 826,57 | 838 | 815 | 0,971 | -0,043 | -0,688 | 0,49145 | 0,533003 | 0,001481 | 0,003635 | 0,0025601 | 0,0025601 | TRUE | 0,0441507 | No |
| orf19.1255 | 61 | 42 | 58 | 43 | 730 | 599 | 627 | 864 | 56 | 42 | 55 | 65 | 649 | 650 | 642 | 659 | 352,29 | 54 | 650 | 12,126 | 3,6 | 30,058 | 1,74E-198 | 2,96E-197 | 1,00E-08 | 0,005787 | 0,0032315 | 0,0032315 | TRUE | 0,1735978 | No |

| | | | | | | | | | | | | | | | | | | | | | | | | | | | | | | | |
|---|---|---|---|---|---|---|---|---|---|---|---|---|---|---|---|---|---|---|---|---|---|---|---|---|---|---|---|---|---|---|---|
| orf19.1257 | 12 | 9 | 10 | 9 | 53 | 37 | 38 | 45 | 11 | 9 | 9 | 14 | 47 | 40 | 39 | 34 | 25,45 | 11 | 40 | 3,798 | 1,925 | 6,149 | 7,80E-10 | 1,46E-09 | 1,00E-08 | 0,053951 | 0,0323717 | 0,0323717 | TRUE | 0,115233 | No |
| orf19.1258 | 92 | 91 | 70 | 52 | 343 | 261 | 329 | 414 | 84 | 92 | 66 | 78 | 305 | 283 | 337 | 316 | 195,17 | 80 | 310 | 3,878 | 1,955 | 15,784 | 3,99E-56 | 2,06E-55 | 0,0021613 | 0,008805 | 0,0066061 | 0,0066061 | TRUE | 0,1453049 | No |
| orf19.1259 | 3926 | 3647 | 3837 | 2424 | 5144 | 4204 | 4785 | 6354 | 3574 | 3689 | 3631 | 3641 | 4575 | 4564 | 4900 | 4847 | 4177,84 | 3634 | 4722 | 1,3 | 0,378 | 9,64 | 5,42E-22 | 1,44E-21 | 0,0006088 | 0,002353 | 0,001231 | 0,001231 | TRUE | 0,0076705 | No |
| orf19.1260 | 681 | 614 | 598 | 379 | 761 | 522 | 580 | 769 | 620 | 621 | 566 | 569 | 677 | 567 | 594 | 587 | 600,06 | 594 | 606 | 1,019 | 0,028 | 0,375 | 0,707688 | 0,741658 | 0,002935 | 0,004238 | 0,0036108 | 0,0036108 | TRUE | 0,0738262 | No |
| orf19.1261 | 3116 | 2822 | 3221 | 1903 | 2253 | 1816 | 2087 | 2629 | 2837 | 2854 | 3048 | 2859 | 2004 | 1972 | 2137 | 2005 | 2464,57 | 2900 | 2030 | 0,7 | -0,515 | -11,432 | 2,90E-30 | 9,40E-30 | 0,0006882 | 0,002573 | 0,0015358 | 0,0015358 | TRUE | 0,0150421 | No |
| orf19.1263 | 2360 | 2188 | 2438 | 1560 | 2575 | 2138 | 2210 | 3031 | 2149 | 2213 | 2307 | 2343 | 2290 | 2321 | 2263 | 2312 | 2274,92 | 2253 | 2296 | 1,021 | 0,03 | 0,698 | 0,484893 | 0,526698 | 0,0003348 | 0,002618 | 0,001295 | 0,001295 | TRUE | 0,0120087 | No |
| orf19.1264 | 619 | 561 | 575 | 404 | 1097 | 851 | 975 | 1293 | 564 | 567 | 544 | 607 | 976 | 924 | 999 | 986 | 770,82 | 570 | 971 | 1,709 | 0,774 | 12,938 | 2,73E-38 | 1,03E-37 | 1,00E-08 | 0,003751 | 0,0020125 | 0,0020125 | TRUE | 0,0178857 | No |
| orf19.1265 | 351 | 354 | 319 | 208 | 317 | 236 | 241 | 350 | 320 | 358 | 302 | 312 | 282 | 256 | 247 | 267 | 292,99 | 323 | 263 | 0,815 | -0,295 | -3,275 | 0,001057 | 0,001491 | 1,00E-08 | 0,006546 | 0,0043705 | 0,0043705 | TRUE | 0,0621497 | No |
| orf19.1266 | 1542 | 1499 | 1509 | 802 | 1278 | 880 | 996 | 1281 | 1404 | 1516 | 1428 | 1205 | 1137 | 955 | 1020 | 977 | 1205,28 | 1388 | 1022 | 0,735 | -0,444 | -5,741 | 9,42E-09 | 1,70E-08 | 0,0064795 | 0,003133 | 0,0048965 | 0,0048965 | TRUE | 0,089211 | No |
| orf19.1267 | 7180 | 6286 | 7137 | 4305 | 4790 | 3892 | 4467 | 5621 | 6537 | 6358 | 6755 | 6467 | 4261 | 4226 | 4575 | 4288 | 5433,09 | 6529 | 4338 | 0,664 | -0,591 | -14,871 | 5,08E-50 | 2,34E-49 | 0,0007573 | 0,00228 | 0,0013241 | 0,0013241 | TRUE | 0,0163166 | No |
| orf19.1267.1 | 1501 | 1396 | 1383 | 871 | 915 | 678 | 652 | 940 | 1367 | 1412 | 1309 | 1308 | 814 | 736 | 668 | 717 | 1041,31 | 1349 | 734 | 0,544 | -0,879 | -13,522 | 1,17E-41 | 4,67E-41 | 0,0027113 | 0,003306 | 0,0030188 | 0,0030188 | TRUE | 0,0651263 | No |
| orf19.1268 | 190 | 156 | 158 | 138 | 162 | 170 | 146 | 217 | 173 | 158 | 150 | 207 | 144 | 185 | 150 | 166 | 166,41 | 172 | 161 | 0,943 | -0,084 | -0,658 | 0,510841 | 0,552999 | 0,009395 | 0,009975 | 0,0097062 | 0,0097062 | TRUE | 0,1692827 | No |
| orf19.1270 | 1179 | 1091 | 1183 | 738 | 1616 | 1449 | 1452 | 1992 | 1073 | 1103 | 1120 | 1109 | 1437 | 1573 | 1487 | 1520 | 1302,77 | 1101 | 1504 | 1,366 | 0,45 | 9,115 | 7,86E-20 | 1,99E-19 | 1,00E-08 | 0,003051 | 0,0015416 | 0,0015416 | TRUE | 0,0108791 | No |
| orf19.1272 | 1009 | 930 | 922 | 621 | 2475 | 1887 | 2019 | 2701 | 919 | 941 | 873 | 933 | 2201 | 2049 | 2068 | 2060 | 1505,37 | 916 | 2094 | 2,289 | 1,195 | 23,794 | 3,82E-125 | 4,09E-124 | 0,0005655 | 0,002915 | 0,0016134 | 0,0016134 | TRUE | 0,0144492 | No |
| orf19.1273 | 178 | 125 | 122 | 82 | 236 | 150 | 229 | 253 | 162 | 126 | 115 | 123 | 210 | 163 | 235 | 193 | 165,93 | 132 | 200 | 1,513 | 0,597 | 4,204 | 2,62E-05 | 4,08E-05 | 0,017186 | 0,009998 | 0,0130072 | 0,0130072 | TRUE | 0,2654139 | No |
| orf19.1274 | 12 | 8 | 15 | 13 | 132 | 124 | 113 | 154 | 11 | 8 | 14 | 20 | 117 | 135 | 116 | 117 | 67,25 | 13 | 121 | 9,56 | 3,257 | 13,054 | 6,02E-39 | 2,30E-38 | 1,00E-08 | 0,021681 | 0,0143089 | 0,0143089 | TRUE | 0,2570811 | No |
| orf19.1275 | 282 | 239 | 258 | 185 | 1529 | 1307 | 1407 | 1905 | 257 | 242 | 244 | 278 | 1360 | 1419 | 1441 | 1453 | 836,71 | 255 | 1418 | 5,593 | 2,484 | 36,468 | 3,58E-291 | 8,95E-290 | 1,00E-08 | 0,003616 | 0,002033 | 0,002033 | TRUE | 0,0301555 | No |
| orf19.1276 | 201 | 223 | 225 | 136 | 205 | 160 | 167 | 215 | 183 | 226 | 213 | 204 | 182 | 174 | 171 | 164 | 189,61 | 206 | 173 | 0,837 | -0,257 | -2,408 | 0,016058 | 0,020773 | 1,00E-08 | 0,009004 | 0,0056739 | 0,0056739 | TRUE | 0,0723798 | No |
| orf19.1277 | 15782 | 13278 | 14984 | 10449 | 4495 | 4260 | 4142 | 5730 | 14369 | 13429 | 14181 | 15696 | 3998 | 4625 | 4242 | 4371 | 9363,92 | 14419 | 4309 | 0,299 | -1,742 | -29,965 | 2,78E-197 | 4,68E-196 | 0,0037785 | 0,002178 | 0,0031021 | 0,0031021 | TRUE | 0,0430975 | No |
| orf19.1279 | 4956 | 4818 | 5094 | 3458 | 7789 | 6521 | 6769 | 8928 | 4512 | 4873 | 4821 | 5194 | 6928 | 7080 | 6932 | 6810 | 5893,91 | 4850 | 6938 | 1,432 | 0,518 | 11,424 | 3,17E-30 | 1,02E-29 | 0,0015518 | 0,002261 | 0,001794 | 0,001794 | TRUE | 0,0268975 | No |
| orf19.1280 | 17903 | 17965 | 17399 | 9922 | 25797 | 20046 | 19871 | 25800 | 16300 | 18170 | 16467 | 14904 | 22946 | 21764 | 20350 | 19680 | 18822,7 | 16460 | 21185 | 1,287 | 0,364 | 5,573 | 2,50E-08 | 4,42E-08 | 0,0056055 | 0,002107 | 0,0040389 | 0,0040389 | TRUE | 0,0597396 | No |
| orf19.1281 | 4474 | 3926 | 4186 | 2621 | 2297 | 1811 | 2090 | 2700 | 4073 | 3971 | 3962 | 3937 | 2043 | 1966 | 2140 | 2060 | 3019,04 | 3986 | 2052 | 0,515 | -0,958 | -24,224 | 1,24E-129 | 1,36E-128 | 0,0002275 | 0,002475 | 0,0011384 | 0,0011384 | TRUE | 0,0095727 | No |
| orf19.1282 | 1174 | 1140 | 1195 | 669 | 2338 | 1768 | 2010 | 2626 | 1069 | 1153 | 1131 | 1005 | 2080 | 1920 | 2058 | 2003 | 1552,32 | 1090 | 2015 | 1,845 | 0,884 | 16,263 | 1,81E-59 | 9,83E-59 | 0,0015567 | 0,002888 | 0,0021134 | 0,0021134 | TRUE | 0,0283468 | No |
| orf19.1283 | 669 | 661 | 667 | 467 | 845 | 721 | 737 | 991 | 609 | 669 | 631 | 702 | 752 | 783 | 755 | 756 | 706,94 | 653 | 762 | 1,171 | 0,227 | 3,664 | 0,000248 | 0,000365 | 0,0002376 | 0,003906 | 0,0022652 | 0,0022652 | TRUE | 0,0257758 | No |
| orf19.1285 | 5777 | 5533 | 5973 | 3978 | 1945 | 1665 | 1722 | 2314 | 5260 | 5596 | 5653 | 5976 | 1730 | 1808 | 1764 | 1765 | 3693,83 | 5621 | 1767 | 0,314 | -1,669 | -35,367 | 5,58E-274 | 1,29E-272 | 0,0013907 | 0,002395 | 0,0017781 | 0,0017781 | TRUE | 0,0229335 | No |
| orf19.1286 | 144 | 88 | 116 | 67 | 48 | 50 | 35 | 57 | 131 | 89 | 110 | 101 | 43 | 54 | 36 | 43 | 75,86 | 108 | 44 | 0,406 | -1,299 | -7,001 | 2,53E-12 | 5,20E-12 | 0,0144143 | 0,019452 | 0,0175037 | 0,0175037 | TRUE | 0,2284314 | No |
| orf19.1287 | 10769 | 10359 | 10676 | 6717 | 9506 | 8090 | 8242 | 11041 | 9805 | 10477 | 10104 | 10090 | 8455 | 8783 | 8441 | 8422 | 9322,17 | 10119 | 8525 | 0,842 | -0,247 | -7,251 | 4,15E-13 | 8,73E-13 | 0,0004319 | 0,002179 | 0,0010097 | 0,0010097 | TRUE | 0,0070377 | No |
| orf19.1288 | 6296 | 5354 | 5631 | 3891 | 61786 | 51815 | 54425 | 78199 | 5732 | 5415 | 5329 | 5845 | 54957 | 56257 | 55738 | 59651 | 31115,5 | 5580 | 56651 | 10,157 | 3,344 | 76,899 | 0 | 0 | 0,0015184 | 0,002079 | 0,0017118 | 0,0017118 | TRUE | 0,0156082 | No |
| orf19.1289 | 2030 | 1898 | 2044 | 1165 | 1310 | 1091 | 1117 | 1653 | 1848 | 1920 | 1934 | 1750 | 1165 | 1185 | 1144 | 1261 | 1525,87 | 1863 | 1189 | 0,638 | -0,649 | -12,278 | 1,18E-34 | 4,21E-34 | 0,0012399 | 0,002903 | 0,0020023 | 0,0020023 | TRUE | 0,0213296 | No |
| orf19.1290 | 4791 | 4396 | 4762 | 2926 | 842 | 696 | 656 | 832 | 4362 | 4446 | 4507 | 4395 | 749 | 756 | 672 | 635 | 2565,16 | 4428 | 703 | 0,158 | -2,659 | -47,278 | 0 | 0 | 0,0018437 | 0,002552 | 0,0022496 | 0,0022496 | TRUE | 0,0549964 | No |
| orf19.1291 | 2356 | 2071 | 2223 | 1355 | 1096 | 879 | 890 | 1232 | 2145 | 2095 | 2104 | 2035 | 975 | 954 | 911 | 940 | 1519,92 | 2095 | 945 | 0,451 | -1,149 | -24,903 | 6,89E-137 | 8,15E-136 | 1,00E-08 | 0,002906 | 0,001296 | 0,001296 | TRUE | 0,0064866 | No |
| orf19.1295 | 5607 | 5834 | 5999 | 3443 | 8437 | 7038 | 7126 | 9184 | 5105 | 5900 | 5678 | 5172 | 7504 | 7641 | 7298 | 7006 | 6413,02 | 5464 | 7362 | 1,347 | 0,43 | 7,872 | 3,49E-15 | 7,75E-15 | 0,003007 | 0,002243 | 0,002698 | 0,002698 | TRUE | 0,0353017 | No |
| orf19.1296 | 1436 | 1533 | 1586 | 901 | 927 | 783 | 783 | 1047 | 1307 | 1550 | 1501 | 1353 | 825 | 850 | 802 | 799 | 1123,44 | 1428 | 819 | 0,573 | -0,804 | -12,282 | 1,13E-34 | 4,03E-34 | 0,0031371 | 0,003213 | 0,0031721 | 0,0031721 | TRUE | 0,0418875 | No |
| orf19.1297 | 924 | 949 | 985 | 612 | 472 | 386 | 410 | 506 | 841 | 960 | 932 | 919 | 420 | 419 | 420 | 386 | 662,17 | 913 | 411 | 0,45 | -1,153 | -16,871 | 7,38E-64 | 4,27E-63 | 0,0014726 | 0,004032 | 0,0027584 | 0,0027584 | TRUE | 0,0358154 | No |
| orf19.1298 | 3162 | 3341 | 3549 | 1888 | 2356 | 2043 | 2050 | 2621 | 2879 | 3379 | 3359 | 2836 | 2096 | 2218 | 2099 | 1999 | 2608,16 | 3113 | 2103 | 0,675 | -0,568 | -8,463 | 2,60E-17 | 6,16E-17 | 0,0048936 | 0,002544 | 0,0039245 | 0,0039245 | TRUE | 0,0419906 | No |
| orf19.1299 | 1890 | 2400 | 2371 | 1527 | 1683 | 1458 | 1433 | 1760 | 1721 | 2427 | 2244 | 2294 | 1497 | 1583 | 1468 | 1343 | 1821,99 | 2172 | 1473 | 0,678 | -0,56 | -5,986 | 2,16E-09 | 3,99E-09 | 0,0130768 | 0,002762 | 0,0078466 | 0,0078466 | TRUE | 0,2396338 | No |
| orf19.130 | 406 | 444 | 465 | 277 | 450 | 396 | 379 | 513 | 370 | 449 | 440 | 416 | 400 | 430 | 388 | 391 | 410,57 | 419 | 402 | 0,961 | -0,058 | -0,706 | 0,480384 | 0,522245 | 0,0026142 | 0,005254 | 0,004074 | 0,004074 | TRUE | 0,0794902 | No |
| orf19.1300 | 2655 | 2469 | 2605 | 1538 | 1863 | 1472 | 1574 | 2059 | 2417 | 2497 | 2465 | 2310 | 1657 | 1598 | 1612 | 1571 | 2016 | 2422 | 1610 | 0,663 | -0,592 | -13,542 | 8,80E-42 | 3,54E-41 | 0,0002796 | 0,002692 | 0,0013232 | 0,0013232 | TRUE | 0,0106132 | No |
| orf19.1301 | 433 | 482 | 448 | 319 | 658 | 605 | 507 | 728 | 394 | 487 | 424 | 479 | 585 | 657 | 519 | 555 | 512,7 | 446 | 579 | 1,301 | 0,379 | 4,108 | 3,99E-05 | 6,13E-05 | 0,0079791 | 0,004614 | 0,0061778 | 0,0061778 | TRUE | 0,0919874 | No |
| orf19.1302 | 44 | 34 | 27 | 23 | 23 | 20 | 12 | 37 | 40 | 34 | 26 | 35 | 20 | 22 | 12 | 28 | 27,15 | 34 | 20 | 0,623 | -0,683 | -2,349 | 0,018825 | 0,024201 | 0,0282179 | 0,050685 | 0,0429429 | 0,0429429 | TRUE | 0,4149261 | No |
| orf19.1303 | 1904 | 1746 | 1887 | 1149 | 2152 | 1629 | 1758 | 2412 | 1734 | 1766 | 1786 | 1726 | 1914 | 1769 | 1800 | 1840 | 1791,79 | 1753 | 1831 | 1,045 | 0,063 | 1,429 | 0,153077 | 0,179949 | 1,00E-08 | 0,002774 | 0,0013088 | 0,0013088 | TRUE | 0,0112793 | No |
| orf19.1304 | 5115 | 4919 | 5232 | 2790 | 4272 | 3223 | 3783 | 4840 | 4657 | 4975 | 4952 | 4191 | 3800 | 3499 | 3874 | 3692 | 4205 | 4694 | 3716 | 0,791 | -0,338 | -5,687 | 1,30E-08 | 2,32E-08 | 0,0036632 | 0,002351 | 0,0031511 | 0,0031511 | TRUE | 0,0614432 | No |
| orf19.1305 | 1202 | 1419 | 1458 | 887 | 1113 | 907 | 925 | 1133 | 1094 | 1435 | 1380 | 1332 | 990 | 985 | 947 | 864 | 1128,52 | 1310 | 946 | 0,722 | -0,469 | -5,616 | 1,96E-08 | 3,48E-08 | 0,008309 | 0,003208 | 0,0058049 | 0,0058049 | TRUE | 0,1526828 | No |
| orf19.1306 | 2372 | 2332 | 2436 | 1514 | 2794 | 2342 | 2411 | 3146 | 2160 | 2359 | 2305 | 2274 | 2485 | 2543 | 2469 | 2400 | 2374,35 | 2274 | 2474 | 1,088 | 0,121 | 2,772 | 0,00558 | 0,007454 | 0,0005992 | 0,002593 | 0,0014069 | 0,0014069 | TRUE | 0,014873 | No |
| orf19.1307 | 764 | 711 | 698 | 501 | 1246 | 1093 | 1108 | 1489 | 696 | 719 | 661 | 753 | 1108 | 1187 | 1135 | 1136 | 924,17 | 707 | 1142 | 1,619 | 0,695 | 12,098 | 1,08E-33 | 3,77E-33 | 0,0003886 | 0,003466 | 0,0020022 | 0,0020022 | TRUE | 0,0228315 | No |
| orf19.1308 | 4137 | 3649 | 3881 | 2514 | 2284 | 1916 | 1860 | 2581 | 3767 | 3691 | 3673 | 3776 | 2032 | 2080 | 1905 | 1969 | 2861,51 | 3727 | 1996 | 0,536 | -0,901 | -22,148 | 1,09E-108 | 1,01E-107 | 0,0003139 | 0,002499 | 0,0012088 | 0,0012088 | TRUE | 0,0109059 | No |
| orf19.1309 | 747 | 542 | 604 | 380 | 246 | 166 | 157 | 258 | 680 | 548 | 572 | 571 | 219 | 180 | 161 | 197 | 390,92 | 593 | 189 | 0,32 | -1,644 | -15,703 | 1,45E-55 | 7,40E-55 | 0,0095124 | 0,005416 | 0,0071662 | 0,0071662 | TRUE | 0,1203226 | No |
| orf19.131 | 368 | 331 | 334 | 206 | 291 | 209 | 247 | 282 | 335 | 335 | 316 | 309 | 259 | 227 | 253 | 215 | 281,15 | 324 | 238 | 0,734 | -0,447 | -4,876 | 1,08E-06 | 1,80E-06 | 1,00E-08 | 0,006736 | 0,0044627 | 0,0044627 | TRUE | 0,0556906 | No |
| orf19.1310 | 745 | 673 | 819 | 493 | 621 | 490 | 472 | 644 | 678 | 681 | 775 | 741 | 552 | 532 | 483 | 491 | 616,7 | 719 | 514 | 0,717 | -0,481 | -6,568 | 5,11E-11 | 1,00E-10 | 0,0027419 | 0,004179 | 0,0034864 | 0,0034864 | TRUE | 0,036316 | No |
| orf19.1311 | 14 | 13 | 12 | 1 | 7 | 10 | 5 | 9 | 13 | 13 | 11 | 2 | 6 | 11 | 5 | 7 | 8,48 | 10 | 7 | 0,715 | -0,485 | -0,937 | 0,348993 | 0,389024 | 0,0861722 | 0,157854 | 0,1394411 | 0,1394411 | TRUE | 0,6594365 | No |
| orf19.131.2 | 3045 | 2581 | 2569 | 1887 | 2482 | 1956 | 1820 | 2631 | 2772 | 2610 | 2431 | 2835 | 2208 | 2124 | 1864 | 2007 | 2356,35 | 2662 | 2051 | 0,771 | -0,375 | -5,751 | 8,86E-09 | 1,60E-08 | 0,0043716 | 0,002598 | 0,0036609 | 0,0036609 | TRUE | 0,0447691 | No |
| orf19.1313 | 602 | 517 | 550 | 394 | 2889 | 2609 | 2624 | 3635 | 548 | 523 | 521 | 592 | 2570 | 2833 | 2687 | 2773 | 1630,72 | 546 | 2716 | 4,996 | 2,321 | 39,95 | 0 | 0 | 0,0012563 | 0,002847 | 0,0020861 | 0,0020861 | TRUE | 0,0309494 | No |
| orf19.1314 | 4584 | 4462 | 4678 | 2902 | 6888 | 5605 | 5787 | 7435 | 4174 | 4513 | 4427 | 4359 | 6127 | 6085 | 5927 | 5671 | 5160,4 | 4368 | 5952 | 1,362 | 0,446 | 10,684 | 1,21E-26 | 3,59E-26 | 0,0008808 | 0,002293 | 0,001473 | 0,001473 | TRUE | 0,0126019 | No |
| orf19.1317 | 1246 | 1141 | 1211 | 802 | 2992 | 2611 | 2717 | 3757 | 1134 | 1154 | 1146 | 1205 | 2661 | 2835 | 2783 | 2866 | 1972,97 | 1160 | 2786 | 2,407 | 1,267 | 27,536 | 6,58E-167 | 9,62E-166 | 0,0004915 | 0,002707 | 0,0014029 | 0,0014029 | TRUE | 0,0112662 | No |
| orf19.1318 | 1137 | 1183 | 1195 | 668 | 1668 | 1424 | 1417 | 1870 | 1035 | 1196 | 1131 | 1003 | 1484 | 1546 | 1451 | 1426 | 1284,18 | 1091 | 1477 | 1,35 | 0,433 | 6,967 | 3,24E-12 | 6,62E-12 | 0,0028396 | 0,003066 | 0,0028929 | 0,0028929 | TRUE | 0,0493027 | No |
| orf19.132 | 3268 | 3043 | 3222 | 2075 | 2232 | 1858 | 1842 | 2331 | 2975 | 3078 | 3049 | 3117 | 1985 | 2017 | 1886 | 1778 | 2485,81 | 3055 | 1916 | 0,627 | -0,673 | -13,824 | 1,84E-43 | 7,66E-43 | 0,0012107 | 0,002568 | 0,0018564 | 0,0018564 | TRUE | 0,0300068 | No |
| orf19.1321 | 13694 | 11416 | 13298 | 8965 | 834 | 762 | 765 | 1065 | 12468 | 11546 | 12585 | 13467 | 742 | 827 | 783 | 812 | 6653,88 | 12516 | 791 | 0,063 | -3,984 | -68,629 | 0 | 0 | 0,0028887 | 0,002236 | 0,0026028 | 0,0026028 | TRUE | 0,0332436 | No |
| orf19.1323 | 1728 | 1630 | 1735 | 1078 | 1599 | 1317 | 1445 | 1817 | 1573 | 1649 | 1642 | 1619 | 1422 | 1430 | 1480 | 1386 | 1525,15 | 1621 | 1430 | 0,881 | -0,182 | -4,029 | 5,60E-05 | 8,51E-05 | 1,00E-08 | 0,002903 | 0,001304 | 0,001304 | TRUE | 0,0067331 | No |
| orf19.1324 | 1809 | 1568 | 1755 | 1084 | 1403 | 1187 | 1331 | 1671 | 1647 | 1586 | 1661 | 1628 | 1248 | 1289 | 1363 | 1275 | 1462,08 | 1630 | 1294 | 0,793 | -0,335 | -7,074 | 1,50E-12 | 3,11E-12 | 3,32E-06 | 0,002941 | 0,0014673 | 0,0014673 | TRUE | 0,0153791 | No |
| orf19.1325 | 369 | 307 | 353 | 204 | 394 | 318 | 324 | 461 | 336 | 310 | 334 | 306 | 350 | 345 | 332 | 352 | 333,27 | 322 | 345 | 1,069 | 0,096 | 1,215 | 0,224441 | 0,257803 | 1,00E-08 | 0,006001 | 0,0030405 | 0,0030405 | TRUE | 0,0098455 | No |
| orf19.1326 | 267 | 212 | 204 | 136 | 739 | 618 | 600 | 824 | 243 | 214 | 193 | 204 | 657 | 671 | 614 | 629 | 428,27 | 214 | 643 | 2,996 | 1,583 | 18,669 | 8,92E-78 | 6,14E-77 | 0,0011318 | 0,005121 | 0,0036959 | 0,0036959 | TRUE | 0,1037056 | No |
| orf19.1327 | 159 | 138 | 150 | 109 | 140 | 112 | 130 | 186 | 145 | 140 | 142 | 164 | 125 | 122 | 133 | 142 | 138,9 | 148 | 130 | 0,891 | -0,166 | -1,379 | 0,167753 | 0,196153 | 1,00E-08 | 0,011548 | 0,0066608 | 0,0066608 | TRUE | 0,0445146 | No |
| orf19.133 | 0 | 0 | 0 | 0 | 5 | 1 | 1 | 3 | 0 | 0 | 0 | 0 | 4 | 1 | 1 | 2 | 1,11 | 0 | 2 | 11,397 | 3,511 | 2,338 | 0,019382 | 0,024891 | 1,00E-08 | 1,196751 | 0,9576772 | 0,9576772 | TRUE | 0,5045898 | No |
| orf19.1330 | 204 | 164 | 209 | 120 | 177 | 147 | 152 | 180 | 186 | 166 | 198 | 180 | 157 | 160 | 156 | 137 | 167,46 | 182 | 152 | 0,832 | -0,265 | -2,365 | 0,018039 | 0,023223 | 1,00E-08 | 0,009926 | 0,0061182 | 0,0061182 | TRUE | 0,0541637 | No |
| orf19.1331 | 5106 | 5566 | 5505 | 3189 | 1118 | 879 | 913 | 1161 | 4649 | 5629 | 5210 | 4790 | 994 | 954 | 935 | 886 | 3006 | 5070 | 942 | 0,186 | -2,429 | -36,076 | 5,49E-285 | 1,33E-283 | 0,0047978 | 0,002476 | 0,0037501 | 0,0037501 | TRUE | 0,0674986 | No |
| orf19.1332 | 966 | 826 | 936 | 623 | 987 | 839 | 902 | 1209 | 880 | 835 | 886 | 936 | 878 | 911 | 924 | 922 | 896,43 | 884 | 909 | 1,03 | 0,043 | 0,772 | 0,439864 | 0,481809 | 1,00E-08 | 0,003511 | 0,0018403 | 0,0018403 | TRUE | 0,0156909 | No |
| orf19.1333 | 987 | 898 | 1009 | 651 | 1520 | 1309 | 1318 | 1697 | 899 | 908 | 955 | 978 | 1352 | 1421 | 1350 | 1294 | 1144,65 | 935 | 1354 | 1,45 | 0,536 | 9,951 | 2,50E-23 | 6,88E-23 | 0,0006555 | 0,003191 | 0,0018672 | 0,0018672 | TRUE | 0,0129445 | No |

| | | | | | | | | | | | | | | | | | | | | | | | | | | | | | | | |
|---|---|---|---|---|---|---|---|---|---|---|---|---|---|---|---|---|---|---|---|---|---|---|---|---|---|---|---|---|---|---|---|
| orf19.1334 | 783 | 731 | 882 | 484 | 402 | 354 | 377 | 476 | 713 | 739 | 835 | 727 | 358 | 384 | 386 | 363 | 563,14 | 754 | 373 | 0,493 | -1,019 | -13,721 | 7,58E-43 | 3,10E-42 | 0,0022653 | 0,004383 | 0,0033355 | 0,0033355 | TRUE | 0,0650233 | No |
| orf19.1335 | 7704 | 7521 | 8033 | 4862 | 11415 | 9646 | 10084 | 13143 | 7014 | 7607 | 7603 | 7303 | 10153 | 10473 | 10327 | 10026 | 8813,24 | 7382 | 10245 | 1,388 | 0,473 | 12,416 | 2,15E-35 | 7,73E-35 | 0,0007635 | 0,002187 | 0,0012727 | 0,0012727 | TRUE | 0,014152 | No |
| orf19.1336 | 4453 | 3706 | 3812 | 2386 | 3434 | 2485 | 2539 | 3650 | 4054 | 3748 | 3608 | 3584 | 3054 | 2698 | 2600 | 2784 | 3266,41 | 3748 | 2784 | 0,743 | -0,429 | -7,142 | 9,18E-13 | 1,91E-12 | 0,0037043 | 0,002441 | 0,0031574 | 0,0031574 | TRUE | 0,0518976 | No |
| orf19.1336.2 | 1645 | 1464 | 1435 | 728 | 624 | 448 | 474 | 661 | 1498 | 1481 | 1358 | 1094 | 555 | 486 | 485 | 504 | 932,64 | 1358 | 508 | 0,373 | -1,421 | -15,099 | 1,64E-51 | 7,83E-51 | 0,0112028 | 0,003453 | 0,0071905 | 0,0071905 | TRUE | 0,1959215 | No |
| orf19.1338 | 1371 | 1212 | 1338 | 748 | 1010 | 762 | 894 | 1137 | 1248 | 1226 | 1266 | 1124 | 898 | 827 | 916 | 867 | 1046,57 | 1216 | 877 | 0,72 | -0,475 | -8,213 | 2,15E-16 | 4,93E-16 | 0,0010429 | 0,003299 | 0,0022332 | 0,0022332 | TRUE | 0,027895 | No |
| orf19.1339 | 10661 | 9007 | 9824 | 6833 | 19525 | 16988 | 16803 | 24027 | 9706 | 9110 | 9298 | 10264 | 17367 | 18444 | 17208 | 18328 | 13715,7 | 9594 | 17837 | 1,86 | 0,895 | 19,239 | 1,75E-82 | 1,28E-81 | 0,0019556 | 0,002133 | 0,0019974 | 0,0019974 | TRUE | 0,0264264 | No |
| orf19.134 | 0 | 1 | 0 | 0 | 1 | 1 | 0 | 1 | 0 | 1 | 0 | 0 | 1 | 1 | 0 | 1 | 0,47 | 0 | 1 | 2,119 | 1,084 | 0,565 | 0,572274 | 0,612944 | 1,00E-08 | 2,820623 | 1,8588363 | 1,8588363 | TRUE | 0,3156291 | No |
| orf19.1340 | 142 | 118 | 153 | 76 | 325 | 262 | 278 | 333 | 129 | 119 | 145 | 114 | 289 | 284 | 285 | 254 | 202,48 | 127 | 278 | 2,171 | 1,118 | 10,184 | 2,33E-24 | 6,59E-24 | 1,00E-08 | 0,008561 | 0,0057384 | 0,0057384 | TRUE | 0,0927175 | No |
| orf19.1341 | 445 | 391 | 401 | 252 | 753 | 565 | 687 | 836 | 405 | 395 | 380 | 379 | 670 | 613 | 704 | 638 | 522,89 | 390 | 656 | 1,681 | 0,749 | 10,482 | 1,04E-25 | 3,04E-25 | 1,00E-08 | 0,004563 | 0,0028261 | 0,0028261 | TRUE | 0,0266296 | No |
| orf19.1342 | 6082 | 5453 | 5850 | 3576 | 8089 | 6421 | 7089 | 9270 | 5537 | 5515 | 5536 | 5372 | 7195 | 6971 | 7260 | 7071 | 6307,29 | 5490 | 7124 | 1,297 | 0,375 | 11,952 | 6,31E-33 | 2,15E-32 | 5,31E-05 | 0,002246 | 0,0007824 | 0,0007824 | TRUE | 0,0024641 | No |
| orf19.1343 | 809 | 813 | 822 | 473 | 501 | 433 | 448 | 575 | 737 | 822 | 778 | 711 | 446 | 470 | 459 | 439 | 607,56 | 762 | 454 | 0,593 | -0,754 | -11,068 | 1,80E-28 | 5,57E-28 | 0,0010629 | 0,004211 | 0,0027283 | 0,0027283 | TRUE | 0,0329917 | No |
| orf19.1344 | 1589 | 1830 | 1816 | 1063 | 4867 | 4188 | 4248 | 5250 | 1447 | 1851 | 1719 | 1597 | 4329 | 4547 | 4350 | 4005 | 2980,54 | 1654 | 4308 | 2,605 | 1,381 | 19,357 | 1,79E-83 | 1,33E-82 | 0,0062385 | 0,00248 | 0,0044537 | 0,0044537 | TRUE | 0,0888075 | No |
| orf19.1345 | 47 | 28 | 38 | 15 | 181 | 150 | 142 | 217 | 43 | 28 | 36 | 23 | 161 | 163 | 145 | 166 | 95,55 | 32 | 159 | 4,768 | 2,253 | 12,725 | 4,29E-37 | 1,60E-36 | 1,00E-08 | 0,015862 | 0,0112917 | 0,0112917 | TRUE | 0,3215573 | No |
| orf19.1348 | 1 | 0 | 3 | 3 | 1 | 3 | 3 | 4 | 1 | 0 | 3 | 5 | 1 | 3 | 3 | 3 | 2,32 | 2 | 2 | 1,328 | 0,409 | 0,425 | 0,671092 | 0,707493 | 1,00E-08 | 0,572465 | 0,4319219 | 0,4319219 | TRUE | 0,2698167 | No |
| orf19.1349 | 95 | 60 | 88 | 62 | 105 | 74 | 88 | 125 | 86 | 61 | 83 | 93 | 93 | 80 | 90 | 95 | 85,35 | 81 | 90 | 1,121 | 0,165 | 1,022 | 0,306683 | 0,345181 | 0,0038815 | 0,017514 | 0,0130709 | 0,0130709 | TRUE | 0,2658968 | No |
| orf19.135 | 1168 | 1183 | 1288 | 664 | 1276 | 1102 | 1221 | 1489 | 1063 | 1196 | 1219 | 997 | 1135 | 1196 | 1250 | 1136 | 1149,25 | 1119 | 1179 | 1,051 | 0,071 | 1,017 | 0,309063 | 0,347553 | 0,0044501 | 0,003186 | 0,0038464 | 0,0038464 | TRUE | 0,0583866 | No |
| orf19.1350 | 7818 | 7918 | 8155 | 5370 | 6335 | 5490 | 5661 | 7219 | 7118 | 8008 | 7718 | 8067 | 5635 | 5961 | 5798 | 5507 | 6726,3 | 7728 | 5725 | 0,741 | -0,433 | -8,858 | 8,12E-19 | 2,00E-18 | 0,0019681 | 0,002233 | 0,0021388 | 0,0021388 | TRUE | 0,034738 | No |
| orf19.1351 | 430 | 395 | 453 | 262 | 453 | 280 | 396 | 510 | 391 | 400 | 429 | 394 | 403 | 304 | 406 | 389 | 389,35 | 404 | 376 | 0,932 | -0,101 | -1,074 | 0,282915 | 0,319444 | 0,0067176 | 0,00543 | 0,0059872 | 0,0059872 | TRUE | 0,1772195 | No |
| orf19.1352 | 4902 | 4264 | 4794 | 2897 | 1368 | 992 | 1163 | 1496 | 4463 | 4313 | 4537 | 4352 | 1217 | 1077 | 1191 | 1141 | 2786,32 | 4416 | 1156 | 0,262 | -1,932 | -41,168 | 0 | 0 | 0,0007529 | 0,002511 | 0,0015906 | 0,0015906 | TRUE | 0,0239147 | No |
| orf19.1353 | 244 | 173 | 205 | 176 | 48 | 59 | 41 | 63 | 222 | 175 | 194 | 264 | 43 | 64 | 42 | 48 | 131,54 | 214 | 49 | 0,23 | -2,12 | -12,327 | 6,44E-35 | 2,30E-34 | 0,0234528 | 0,01208 | 0,0163483 | 0,0163483 | TRUE | 0,220111 | No |
| orf19.1354 | 1E+05 | 1E+05 | 1E+05 | 82571 | 12531 | 10954 | 10498 | 14308 | 1E+05 | 1E+05 | 1E+05 | 1E+05 | 11146 | 11893 | 10751 | 10914 | 73609 | 136042 | 11176 | 0,082 | -3,606 | -65,323 | 0 | 0 | 0,0033919 | 0,002055 | 0,0028817 | 0,0028817 | TRUE | 0,0432144 | No |
| orf19.1355 | 3259 | 3300 | 3399 | 2180 | 4216 | 3536 | 3507 | 4684 | 2967 | 3338 | 3217 | 3275 | 3750 | 3839 | 3592 | 3573 | 3443,76 | 3199 | 3688 | 1,153 | 0,206 | 4,258 | 2,06E-05 | 3,22E-05 | 0,0015001 | 0,002421 | 0,0019491 | 0,0019491 | TRUE | 0,0297045 | No |
| orf19.1356 | 552 | 572 | 613 | 388 | 873 | 733 | 666 | 928 | 503 | 579 | 580 | 583 | 777 | 796 | 682 | 708 | 650,8 | 561 | 741 | 1,322 | 0,403 | 5,318 | 1,05E-07 | 1,81E-07 | 0,0037455 | 0,004067 | 0,0039122 | 0,0039122 | TRUE | 0,0611142 | No |
| orf19.1357 | 25327 | 23935 | 26421 | 15406 | 28088 | 23733 | 24790 | 33715 | 23059 | 24208 | 25005 | 23142 | 24983 | 25767 | 25388 | 25718 | 24658,9 | 23854 | 25464 | 1,067 | 0,094 | 2,605 | 0,009189 | 0,012088 | 0,0008052 | 0,002091 | 0,0012123 | 0,0012123 | TRUE | 0,0129716 | No |
| orf19.1358 | 19307 | 16929 | 18143 | 12938 | 15121 | 14062 | 12875 | 19179 | 17578 | 17122 | 17171 | 19435 | 13450 | 15267 | 13186 | 14630 | 15979,8 | 17826 | 14133 | 0,793 | -0,335 | -5,67 | 1,43E-08 | 2,55E-08 | 0,0042089 | 0,00212 | 0,0032833 | 0,0032833 | TRUE | 0,0448411 | No |
| orf19.1359 | 1236 | 1324 | 1410 | 945 | 1272 | 1088 | 1194 | 1611 | 1125 | 1339 | 1334 | 1420 | 1131 | 1181 | 1223 | 1229 | 1247,84 | 1304 | 1191 | 0,915 | -0,128 | -1,815 | 0,069559 | 0,084967 | 0,0047905 | 0,003096 | 0,0039758 | 0,0039758 | TRUE | 0,1051748 | No |
| orf19.136 | 3774 | 3525 | 3872 | 2522 | 6374 | 5456 | 5817 | 7486 | 3436 | 3565 | 3664 | 3788 | 5669 | 5924 | 5957 | 5710 | 4714,38 | 3613 | 5815 | 1,611 | 0,688 | 16,605 | 6,46E-62 | 3,62E-61 | 0,0008728 | 0,002317 | 0,001417 | 0,001417 | TRUE | 0,0133891 | No |
| orf19.1360 | 533 | 489 | 543 | 334 | 454 | 423 | 392 | 549 | 485 | 495 | 514 | 502 | 404 | 459 | 401 | 419 | 459,85 | 499 | 421 | 0,843 | -0,247 | -3,434 | 0,000594 | 0,000851 | 1,00E-08 | 0,00491 | 0,0027832 | 0,0027832 | TRUE | 0,0401028 | No |
| orf19.1360.1 | 274 | 204 | 196 | 176 | 211 | 210 | 189 | 293 | 249 | 206 | 185 | 264 | 188 | 228 | 194 | 224 | 217,3 | 226 | 208 | 0,925 | -0,113 | -0,905 | 0,365446 | 0,40531 | 0,0132828 | 0,008116 | 0,0103068 | 0,0103068 | TRUE | 0,1665837 | No |
| orf19.1361 | 4349 | 4166 | 4281 | 2433 | 4338 | 3605 | 3746 | 5009 | 3960 | 4213 | 4052 | 3655 | 3859 | 3914 | 3836 | 3821 | 3913,65 | 3970 | 3858 | 0,97 | -0,043 | -0,93 | 0,352308 | 0,39224 | 0,0014644 | 0,002375 | 0,0018198 | 0,0018198 | TRUE | 0,0328588 | No |
| orf19.1362 | 7737 | 7457 | 7946 | 4596 | 6651 | 5317 | 5717 | 7161 | 7044 | 7542 | 7520 | 6904 | 5916 | 5773 | 5855 | 5462 | 6502,04 | 7252 | 5752 | 0,793 | -0,335 | -7,49 | 6,88E-14 | 1,47E-13 | 0,0014203 | 0,00224 | 0,0017706 | 0,0017706 | TRUE | 0,0142462 | No |
| orf19.1363 | 34139 | 32582 | 33618 | 22204 | 17823 | 15136 | 15569 | 20040 | 31082 | 32953 | 31816 | 33354 | 15853 | 16433 | 15945 | 15287 | 24090,4 | 32301 | 15880 | 0,492 | -1,024 | -27,415 | 1,83E-165 | 2,65E-164 | 0,0008957 | 0,002092 | 0,0012954 | 0,0012954 | TRUE | 0,0079175 | No |
| orf19.1364 | 2693 | 2579 | 2780 | 1850 | 2924 | 2374 | 2421 | 3339 | 2452 | 2608 | 2631 | 2779 | 2601 | 2578 | 2479 | 2547 | 2584,37 | 2618 | 2551 | 0,976 | -0,035 | -0,747 | 0,455108 | 0,497055 | 0,0010054 | 0,002548 | 0,0016772 | 0,0016772 | TRUE | 0,0222047 | No |
| orf19.1365 | 1250 | 1059 | 1103 | 704 | 1344 | 1172 | 1190 | 1558 | 1138 | 1071 | 1044 | 1058 | 1195 | 1272 | 1219 | 1188 | 1148,2 | 1078 | 1218 | 1,129 | 0,175 | 3,348 | 0,000815 | 0,001157 | 0,0004094 | 0,003188 | 0,0017341 | 0,0017341 | TRUE | 0,0181355 | No |
| orf19.1366 | 94 | 64 | 81 | 50 | 146 | 125 | 139 | 181 | 86 | 65 | 77 | 75 | 130 | 136 | 142 | 138 | 106,01 | 76 | 136 | 1,802 | 0,85 | 6,075 | 1,24E-09 | 2,31E-09 | 1,00E-08 | 0,014498 | 0,0083482 | 0,0083482 | TRUE | 0,0925821 | No |
| orf19.1367 | 162 | 186 | 177 | 96 | 79 | 50 | 67 | 61 | 147 | 188 | 168 | 144 | 70 | 54 | 69 | 47 | 110,88 | 162 | 60 | 0,367 | -1,445 | -8,935 | 4,07E-19 | 1,01E-18 | 0,0140837 | 0,013951 | 0,0139905 | 0,0139905 | TRUE | 0,2415641 | No |
| orf19.1367.1 | 1521 | 1200 | 1250 | 639 | 1662 | 1079 | 1229 | 1638 | 1385 | 1214 | 1183 | 960 | 1478 | 1171 | 1259 | 1249 | 1237,41 | 1186 | 1289 | 1,086 | 0,119 | 1,188 | 0,234791 | 0,268773 | 0,014595 | 0,003105 | 0,0087537 | 0,0087537 | TRUE | 0,1874489 | No |
| orf19.1368 | 2545 | 2250 | 2518 | 1694 | 1461 | 1274 | 1282 | 1768 | 2317 | 2276 | 2383 | 2545 | 1300 | 1383 | 1313 | 1349 | 1858,1 | 2380 | 1336 | 0,562 | -0,831 | -16,767 | 4,23E-63 | 2,42E-62 | 0,0010161 | 0,002748 | 0,0017824 | 0,0017824 | TRUE | 0,0242633 | No |
| orf19.1369 | 457 | 422 | 505 | 245 | 489 | 353 | 417 | 532 | 416 | 427 | 478 | 368 | 435 | 383 | 427 | 406 | 417,49 | 422 | 413 | 0,973 | -0,039 | -0,454 | 0,649921 | 0,687334 | 0,0040346 | 0,005201 | 0,0047807 | 0,0047807 | TRUE | 0,0899894 | No |
| orf19.137 | 3162 | 3713 | 3615 | 2285 | 4375 | 3954 | 3930 | 4713 | 2879 | 3755 | 3421 | 3432 | 3891 | 4293 | 4025 | 3595 | 3661,52 | 3372 | 3951 | 1,172 | 0,229 | 2,93 | 0,003386 | 0,004586 | 0,0081375 | 0,002398 | 0,005566 | 0,005566 | TRUE | 0,1194113 | No |
| orf19.1370 | 5 | 14 | 5 | 4 | 7 | 13 | 12 | 12 | 5 | 14 | 5 | 6 | 6 | 14 | 12 | 9 | 8,9 | 8 | 10 | 1,401 | 0,486 | 0,951 | 0,341457 | 0,381492 | 0,1038354 | 0,150388 | 0,1335345 | 0,1335345 | TRUE | 1,0468612 | No |
| orf19.1371 | 571 | 538 | 541 | 344 | 443 | 395 | 411 | 541 | 520 | 544 | 512 | 517 | 394 | 429 | 421 | 413 | 468,66 | 523 | 414 | 0,79 | -0,339 | -4,947 | 7,54E-07 | 1,26E-06 | 1,00E-08 | 0,004856 | 0,0023756 | 0,0023756 | TRUE | 0,0122794 | No |
| orf19.1372 | 431 | 442 | 420 | 281 | 414 | 308 | 343 | 446 | 392 | 447 | 397 | 422 | 368 | 334 | 351 | 340 | 381,65 | 414 | 348 | 0,842 | -0,248 | -3,151 | 0,001627 | 0,002261 | 1,00E-08 | 0,005498 | 0,0033204 | 0,0033204 | TRUE | 0,0346209 | No |
| orf19.1373 | 170 | 141 | 163 | 106 | 202 | 154 | 167 | 244 | 155 | 143 | 154 | 159 | 180 | 167 | 171 | 186 | 164,36 | 153 | 176 | 1,158 | 0,212 | 1,945 | 0,051813 | 0,064076 | 1,00E-08 | 0,010074 | 0,0052601 | 0,0052601 | TRUE | 0,0207164 | No |
| orf19.1374 | 381 | 377 | 337 | 232 | 567 | 443 | 522 | 634 | 347 | 381 | 319 | 349 | 504 | 481 | 535 | 484 | 424,89 | 349 | 501 | 1,436 | 0,522 | 6,613 | 3,77E-11 | 7,42E-11 | 0,0012964 | 0,005146 | 0,0035137 | 0,0035137 | TRUE | 0,04738 | No |
| orf19.1375 | 27705 | 23957 | 25968 | 16330 | 9534 | 8040 | 8220 | 11705 | 25224 | 24230 | 24576 | 24530 | 8480 | 8729 | 8418 | 8929 | 16639,7 | 24640 | 8639 | 0,351 | -1,512 | -46,563 | 0 | 0 | 0,0004107 | 0,002116 | 0,0009366 | 0,0009366 | TRUE | 0,0063737 | No |
| orf19.1376 | 2669 | 2541 | 2555 | 1746 | 2861 | 2441 | 2453 | 3209 | 2430 | 2570 | 2418 | 2623 | 2545 | 2650 | 2512 | 2448 | 2524,48 | 2510 | 2539 | 1,012 | 0,017 | 0,379 | 0,704391 | 0,738811 | 0,0008129 | 0,00256 | 0,0016074 | 0,0016074 | TRUE | 0,0103854 | No |
| orf19.1377 | 2005 | 1824 | 1801 | 1227 | 1158 | 923 | 1035 | 1308 | 1825 | 1845 | 1704 | 1843 | 1030 | 1002 | 1060 | 998 | 1413,47 | 1804 | 1022 | 0,567 | -0,819 | -16,467 | 6,30E-61 | 3,49E-60 | 0,0005232 | 0,002972 | 0,0016207 | 0,0016207 | TRUE | 0,0173588 | No |
| orf19.1378 | 10508 | 8709 | 9980 | 6277 | 11428 | 9407 | 10419 | 13953 | 9567 | 8808 | 9445 | 9429 | 10165 | 10213 | 10670 | 10643 | 9867,7 | 9312 | 10423 | 1,119 | 0,163 | 4,187 | 2,83E-05 | 4,38E-05 | 0,0008758 | 0,002171 | 0,0013475 | 0,0013475 | TRUE | 0,01637 | No |
| orf19.138 | 100 | 83 | 90 | 74 | 213 | 213 | 204 | 259 | 91 | 84 | 85 | 111 | 189 | 231 | 209 | 198 | 149,82 | 93 | 207 | 2,249 | 1,169 | 9,078 | 1,11E-19 | 2,78E-19 | 0,0026669 | 0,010854 | 0,0078691 | 0,0078691 | TRUE | 0,1258352 | No |
| orf19.1381 | 2848 | 2370 | 2455 | 1564 | 566 | 416 | 482 | 697 | 2593 | 2397 | 2323 | 2349 | 503 | 452 | 494 | 532 | 1455,4 | 2416 | 495 | 0,205 | -2,283 | -36,412 | 2,74E-290 | 6,83E-289 | 0,0022804 | 0,002945 | 0,0026136 | 0,0026136 | TRUE | 0,0387301 | No |
| orf19.1382 | 574 | 493 | 494 | 316 | 376 | 347 | 375 | 477 | 523 | 499 | 468 | 475 | 334 | 377 | 384 | 364 | 427,82 | 491 | 365 | 0,741 | -0,433 | -5,648 | 1,63E-08 | 2,90E-08 | 1,00E-08 | 0,005125 | 0,0032763 | 0,0032763 | TRUE | 0,0361875 | No |
| orf19.1383 | 748 | 722 | 752 | 474 | 926 | 757 | 843 | 1054 | 681 | 730 | 712 | 712 | 824 | 822 | 863 | 804 | 768,48 | 709 | 828 | 1,168 | 0,224 | 3,924 | 8,72E-05 | 0,000131 | 1,00E-08 | 0,003756 | 0,001824 | 0,001824 | TRUE | 0,0095094 | No |
| orf19.1386 | 2390 | 2177 | 2332 | 1439 | 2651 | 1984 | 2176 | 2856 | 2176 | 2202 | 2207 | 2162 | 2358 | 2154 | 2228 | 2179 | 2208,2 | 2187 | 2230 | 1,02 | 0,028 | 0,645 | 0,518896 | 0,560767 | 0,0004918 | 0,002635 | 0,0013611 | 0,0013611 | TRUE | 0,0183991 | No |
| orf19.1387 | 4628 | 3768 | 4093 | 2744 | 3488 | 2485 | 2920 | 3770 | 4214 | 3811 | 3874 | 4122 | 3102 | 2698 | 2990 | 2876 | 3460,85 | 4005 | 2916 | 0,729 | -0,457 | -8,418 | 3,83E-17 | 9,00E-17 | 0,0025221 | 0,002419 | 0,0025322 | 0,0025322 | TRUE | 0,030259 | No |
| orf19.1388 | 5760 | 5207 | 5575 | 3128 | 6902 | 4987 | 5998 | 7652 | 5244 | 5266 | 5276 | 4699 | 6139 | 5414 | 6143 | 5837 | 5502,36 | 5121 | 5883 | 1,148 | 0,199 | 3,614 | 0,000301 | 0,00044 | 0,002981 | 0,002277 | 0,0027334 | 0,0027334 | TRUE | 0,0365017 | No |
| orf19.1389 | 3758 | 3363 | 3496 | 2040 | 3517 | 2570 | 2931 | 3888 | 3421 | 3401 | 3309 | 3064 | 3128 | 2790 | 3002 | 2966 | 3135,24 | 3299 | 2972 | 0,9 | -0,152 | -2,986 | 0,002826 | 0,003854 | 0,0019093 | 0,002458 | 0,0021662 | 0,0021662 | TRUE | 0,0263746 | No |
| orf19.139 | 1031 | 970 | 1069 | 682 | 1955 | 1637 | 1678 | 2236 | 939 | 981 | 1012 | 1024 | 1739 | 1777 | 1718 | 1706 | 1362,03 | 989 | 1735 | 1,757 | 0,813 | 16,786 | 3,09E-63 | 1,77E-62 | 1,00E-08 | 0,003007 | 0,0014424 | 0,0014424 | TRUE | 0,0146117 | No |
| orf19.1390 | 586 | 507 | 539 | 382 | 455 | 377 | 388 | 548 | 534 | 513 | 510 | 574 | 405 | 409 | 397 | 418 | 469,96 | 533 | 407 | 0,768 | -0,38 | -5,423 | 5,85E-08 | 1,02E-07 | 1,00E-08 | 0,004848 | 0,00257 | 0,00257 | TRUE | 0,0265345 | No |
| orf19.1391 | 115 | 100 | 86 | 70 | 104 | 87 | 87 | 132 | 105 | 101 | 81 | 105 | 93 | 94 | 89 | 101 | 96,14 | 98 | 94 | 0,966 | -0,049 | -0,341 | 0,733332 | 0,765263 | 1,00E-08 | 0,015777 | 0,0097055 | 0,0097055 | TRUE | 0,1381224 | No |
| orf19.1392 | 430 | 390 | 441 | 259 | 842 | 664 | 758 | 936 | 391 | 394 | 417 | 389 | 749 | 721 | 776 | 714 | 569,06 | 398 | 740 | 1,855 | 0,892 | 13,395 | 6,48E-41 | 2,56E-40 | 1,00E-08 | 0,004358 | 0,002287 | 0,002287 | TRUE | 0,0128732 | No |
| orf19.1393 | 3387 | 3010 | 3417 | 2147 | 2786 | 2282 | 2543 | 3296 | 3084 | 3044 | 3234 | 3225 | 2478 | 2478 | 2604 | 2514 | 2832,66 | 3147 | 2518 | 0,801 | -0,321 | -7,901 | 2,76E-15 | 6,16E-15 | 0,0003943 | 0,002503 | 0,0012237 | 0,0012237 | TRUE | 0,0061964 | No |
| orf19.1394 | 1485 | 1325 | 1348 | 656 | 1526 | 1017 | 1209 | 1636 | 1352 | 1340 | 1276 | 985 | 1357 | 1104 | 1238 | 1248 | 1237,62 | 1238 | 1237 | 0,997 | -0,005 | -0,049 | 0,961206 | 0,965751 | 0,0128208 | 0,003104 | 0,0076937 | 0,0076937 | TRUE | 0,2153834 | No |
| orf19.1395 | 356 | 269 | 296 | 193 | 281 | 219 | 208 | 321 | 324 | 272 | 280 | 290 | 250 | 238 | 213 | 245 | 263,98 | 292 | 236 | 0,811 | -0,301 | -3,149 | 0,001637 | 0,002275 | 0,0021437 | 0,007041 | 0,0049955 | 0,0049955 | TRUE | 0,070222 | No |
| orf19.1396 | 2146 | 1796 | 1962 | 1300 | 1465 | 1227 | 1278 | 1839 | 1954 | 1816 | 1857 | 1953 | 1303 | 1332 | 1309 | 1403 | 1615,86 | 1895 | 1337 | 0,706 | -0,501 | -10,274 | 9,24E-25 | 2,63E-24 | 0,0007158 | 0,002855 | 0,0016531 | 0,0016531 | TRUE | 0,0142293 | No |
| orf19.1397 | 1110 | 925 | 1019 | 688 | 1194 | 1214 | 1126 | 1495 | 1011 | 936 | 964 | 1033 | 1062 | 1318 | 1153 | 1140 | 1077,21 | 986 | 1168 | 1,185 | 0,245 | 3,489 | 0,000484 | 0,000698 | 0,0040004 | 0,003263 | 0,003775 | 0,003775 | TRUE | 0,0869699 | No |
| orf19.1400 | 1463 | 1455 | 1542 | 825 | 987 | 829 | 909 | 1027 | 1332 | 1472 | 1459 | 1239 | 878 | 900 | 931 | 783 | 1124,32 | 1376 | 873 | 0,633 | -0,661 | -9,075 | 1,14E-19 | 2,85E-19 | 0,0046398 | 0,003212 | 0,0041601 | 0,0041601 | TRUE | 0,0595791 | No |

| | | | | | | | | | | | | | | | | | | | | | | | | | | | | | | | |
|---|---|---|---|---|---|---|---|---|---|---|---|---|---|---|---|---|---|---|---|---|---|---|---|---|---|---|---|---|---|---|---|
| orf19.1401 | 1009 | 880 | 998 | 713 | 1864 | 1709 | 1748 | 2427 | 919 | 890 | 945 | 1071 | 1658 | 1855 | 1790 | 1851 | 1372,4 | 956 | 1788 | 1,878 | 0,909 | 14,035 | 9,47E-45 | 4,04E-44 | 0,0037423 | 0,003 | 0,0032001 | 0,0032001 | TRUE | 0,0704973 | No |
| orf19.1402 | 5678 | 4683 | 5075 | 3636 | 7056 | 5861 | 5966 | 8763 | 5170 | 4736 | 4803 | 5462 | 6276 | 6363 | 6110 | 6684 | 5700,59 | 5043 | 6358 | 1,262 | 0,336 | 6,338 | 2,33E-10 | 4,47E-10 | 0,0027324 | 0,002269 | 0,0025156 | 0,0025156 | TRUE | 0,0369756 | No |
| orf19.1403 | 5164 | 4503 | 4983 | 3275 | 4336 | 3471 | 3795 | 4965 | 4702 | 4554 | 4716 | 4920 | 3857 | 3769 | 3887 | 3787 | 4273,83 | 4723 | 3825 | 0,81 | -0,303 | -8,244 | 1,66E-16 | 3,83E-16 | 0,0002968 | 0,002346 | 0,0010618 | 0,0010618 | TRUE | 0,0089305 | No |
| orf19.1404 | 7218 | 5914 | 6527 | 4326 | 6030 | 5013 | 5497 | 7323 | 6572 | 5981 | 6177 | 6498 | 5363 | 5443 | 5630 | 5586 | 5906,31 | 6307 | 5506 | 0,873 | -0,196 | -4,701 | 2,58E-06 | 4,22E-06 | 0,0009879 | 0,002261 | 0,0014932 | 0,0014932 | TRUE | 0,0147932 | No |
| orf19.1405 | 722 | 655 | 633 | 435 | 498 | 427 | 434 | 610 | 657 | 662 | 599 | 653 | 443 | 464 | 444 | 465 | 548,58 | 643 | 454 | 0,707 | -0,5 | -7,515 | 5,69E-14 | 1,22E-13 | 1,00E-08 | 0,004445 | 0,0023977 | 0,0023977 | TRUE | 0,0266244 | No |
| orf19.1406 | 787 | 819 | 804 | 465 | 939 | 802 | 801 | 997 | 717 | 828 | 761 | 699 | 835 | 871 | 820 | 761 | 786,39 | 751 | 822 | 1,09 | 0,125 | 1,783 | 0,074541 | 0,09067 | 0,0031321 | 0,003717 | 0,0034298 | 0,0034298 | TRUE | 0,0563379 | No |
| orf19.1409 | 929 | 791 | 901 | 580 | 768 | 584 | 666 | 872 | 846 | 800 | 853 | 871 | 683 | 634 | 682 | 665 | 754,28 | 842 | 666 | 0,792 | -0,336 | -5,753 | 8,78E-09 | 1,59E-08 | 1,00E-08 | 0,003788 | 0,0019424 | 0,0019424 | TRUE | 0,0135646 | No |
| orf19.1409.1 | 9708 | 8118 | 8666 | 5476 | 12695 | 9680 | 9840 | 13685 | 8839 | 8210 | 8202 | 8226 | 11292 | 10510 | 10077 | 10439 | 9474,33 | 8369 | 10580 | 1,264 | 0,338 | 7,381 | 1,57E-13 | 3,34E-13 | 0,0017251 | 0,002176 | 0,0019049 | 0,0019049 | TRUE | 0,0251747 | No |
| orf19.1409.2 | 327 | 302 | 301 | 209 | 309 | 271 | 269 | 372 | 298 | 305 | 285 | 314 | 275 | 294 | 275 | 284 | 291,29 | 300 | 282 | 0,941 | -0,088 | -1,052 | 0,292914 | 0,330033 | 1,00E-08 | 0,006572 | 0,0032757 | 0,0032757 | TRUE | 0,0138756 | No |
| orf19.1409.3 | 19 | 14 | 18 | 9 | 290 | 225 | 245 | 323 | 17 | 14 | 17 | 14 | 258 | 244 | 251 | 246 | 132,69 | 16 | 250 | 15,887 | 3,99 | 19,092 | 2,92E-81 | 2,11E-80 | 1,00E-08 | 0,011992 | 0,0066214 | 0,0066214 | TRUE | 0,0282969 | No |
| orf19.1411 | 1185 | 1100 | 1151 | 770 | 1307 | 1175 | 1207 | 1623 | 1079 | 1113 | 1089 | 1157 | 1163 | 1276 | 1236 | 1238 | 1168,73 | 1110 | 1228 | 1,109 | 0,149 | 2,874 | 0,004053 | 0,005462 | 0,000365 | 0,003167 | 0,0017046 | 0,0017046 | TRUE | 0,0156255 | No |
| orf19.1412 | 2297 | 2042 | 2225 | 1329 | 1376 | 1012 | 1195 | 1671 | 2091 | 2065 | 2106 | 1996 | 1224 | 1099 | 1224 | 1275 | 1634,98 | 2064 | 1206 | 0,584 | -0,775 | -14,637 | 1,63E-48 | 7,29E-48 | 0,0013336 | 0,002845 | 0,0020469 | 0,0020469 | TRUE | 0,0409051 | No |
| orf19.1413 | 1225 | 1027 | 1099 | 702 | 686 | 502 | 513 | 731 | 1115 | 1039 | 1040 | 1055 | 610 | 545 | 525 | 558 | 810,85 | 1062 | 560 | 0,527 | -0,923 | -14,689 | 7,59E-49 | 3,43E-48 | 0,0010926 | 0,003666 | 0,002457 | 0,002457 | TRUE | 0,0437277 | No |
| orf19.1414 | 2293 | 1824 | 2049 | 1220 | 1979 | 1507 | 1641 | 2426 | 2088 | 1845 | 1939 | 1833 | 1760 | 1636 | 1681 | 1851 | 1828,99 | 1926 | 1732 | 0,899 | -0,154 | -2,609 | 0,009075 | 0,01194 | 0,0026217 | 0,002759 | 0,0027787 | 0,0027787 | TRUE | 0,0391883 | No |
| orf19.1415 | 16264 | 14770 | 15800 | 10398 | 36373 | 32176 | 32441 | 45401 | 14808 | 14938 | 14953 | 15619 | 32353 | 34934 | 33224 | 34632 | 24432,7 | 15080 | 33786 | 2,241 | 1,164 | 31,312 | 3,18E-215 | 5,76E-214 | 0,0008734 | 0,002091 | 0,0012782 | 0,0012782 | TRUE | 0,0100045 | No |
| orf19.1416 | 2652 | 2190 | 2213 | 1487 | 1582 | 1131 | 1248 | 1891 | 2415 | 2215 | 2094 | 2234 | 1407 | 1228 | 1278 | 1442 | 1789,16 | 2240 | 1339 | 0,599 | -0,74 | -11,389 | 4,72E-30 | 1,52E-29 | 0,0038616 | 0,002775 | 0,003466 | 0,003466 | TRUE | 0,0369699 | No |
| orf19.1417 | 1139 | 988 | 1012 | 600 | 434 | 309 | 338 | 446 | 1037 | 999 | 958 | 901 | 386 | 335 | 346 | 340 | 662,9 | 974 | 352 | 0,361 | -1,471 | -20,359 | 3,89E-92 | 3,10E-91 | 0,0020712 | 0,00403 | 0,0031365 | 0,0031365 | TRUE | 0,0498545 | No |
| orf19.1418 | 684 | 644 | 685 | 425 | 529 | 439 | 471 | 688 | 623 | 651 | 648 | 638 | 471 | 477 | 482 | 525 | 564,39 | 640 | 489 | 0,765 | -0,387 | -5,891 | 3,85E-09 | 7,05E-09 | 1,00E-08 | 0,004378 | 0,0023548 | 0,0023548 | TRUE | 0,0320562 | No |
| orf19.1420 | 124 | 109 | 121 | 78 | 147 | 118 | 130 | 164 | 113 | 110 | 115 | 117 | 131 | 128 | 133 | 125 | 121,49 | 114 | 129 | 1,138 | 0,186 | 1,514 | 0,1301 | 0,154219 | 1,00E-08 | 0,01291 | 0,0062579 | 0,0062579 | TRUE | 0,0056625 | No |
| orf19.1421 | 162 | 120 | 143 | 109 | 327 | 246 | 231 | 344 | 147 | 121 | 135 | 164 | 291 | 267 | 237 | 262 | 203,11 | 142 | 264 | 1,877 | 0,909 | 7,947 | 1,91E-15 | 4,29E-15 | 0,0043693 | 0,008541 | 0,0070023 | 0,0070023 | TRUE | 0,0948641 | No |
| orf19.1422 | 8541 | 7897 | 8494 | 5312 | 5836 | 4922 | 5188 | 6741 | 7776 | 7987 | 8039 | 7979 | 5191 | 5344 | 5313 | 5142 | 6596,45 | 7945 | 5248 | 0,66 | -0,599 | -19,037 | 8,49E-81 | 6,07E-80 | 0,0001361 | 0,002237 | 0,0007933 | 0,0007933 | TRUE | 0,0025855 | No |
| orf19.1424 | 901 | 819 | 868 | 529 | 1361 | 1169 | 1236 | 1695 | 820 | 828 | 821 | 795 | 1211 | 1269 | 1266 | 1293 | 1037,92 | 816 | 1260 | 1,541 | 0,624 | 12,19 | 3,51E-34 | 1,23E-33 | 1,00E-08 | 0,00331 | 0,0014944 | 0,0014944 | TRUE | 0,0085714 | No |
| orf19.1426 | 6851 | 6139 | 6695 | 4436 | 6144 | 5357 | 5758 | 8076 | 6238 | 6209 | 6336 | 6664 | 5465 | 5816 | 5897 | 6160 | 6098,09 | 6362 | 5834 | 0,918 | -0,124 | -2,711 | 0,006717 | 0,008925 | 0,0015281 | 0,002254 | 0,0018399 | 0,0018399 | TRUE | 0,0224723 | No |
| orf19.1427 | 296 | 269 | 295 | 210 | 387 | 378 | 382 | 478 | 269 | 272 | 279 | 315 | 344 | 410 | 391 | 365 | 330,83 | 284 | 378 | 1,334 | 0,416 | 4,673 | 2,97E-06 | 4,84E-06 | 0,0022915 | 0,00603 | 0,0044642 | 0,0044642 | TRUE | 0,0511791 | No |
| orf19.1428 | 168 | 156 | 168 | 87 | 181 | 132 | 148 | 204 | 153 | 158 | 159 | 131 | 161 | 143 | 152 | 156 | 151,49 | 150 | 153 | 1,013 | 0,018 | 0,161 | 0,87207 | 0,891365 | 1,00E-08 | 0,010757 | 0,0060772 | 0,0060772 | TRUE | 0,0600104 | No |
| orf19.1429 | 85 | 76 | 85 | 46 | 151 | 104 | 115 | 131 | 77 | 77 | 80 | 69 | 134 | 113 | 118 | 100 | 96,09 | 76 | 116 | 1,519 | 0,603 | 4,014 | 5,98E-05 | 9,07E-05 | 1,00E-08 | 0,015784 | 0,010683 | 0,010683 | TRUE | 0,1199298 | No |
| orf19.1430 | 76 | 69 | 93 | 53 | 39 | 31 | 38 | 47 | 69 | 70 | 88 | 80 | 35 | 34 | 39 | 36 | 56,22 | 77 | 36 | 0,468 | -1,097 | -5,734 | 9,84E-09 | 1,77E-08 | 1,00E-08 | 0,025536 | 0,015166 | 0,015166 | TRUE | 0,1056864 | No |
| orf19.1433 | 2255 | 2007 | 2046 | 1300 | 1363 | 1137 | 1140 | 1538 | 2053 | 2030 | 1936 | 1953 | 1212 | 1234 | 1168 | 1173 | 1594,95 | 1993 | 1197 | 0,6 | -0,737 | -16,016 | 9,90E-58 | 5,23E-57 | 1,00E-08 | 0,002865 | 0,0013761 | 0,0013761 | TRUE | 0,0051343 | No |
| orf19.1433.1 | 4 | 4 | 5 | 9 | 1 | 1 | 1 | 5 | 4 | 4 | 5 | 14 | 1 | 1 | 1 | 4 | 4,09 | 7 | 2 | 0,29 | -1,786 | -2,197 | 0,027992 | 0,035463 | 0,2254498 | 0,324701 | 0,2882503 | 0,2882503 | TRUE | 0,8852627 | No |
| orf19.1434 | 1169 | 1044 | 1184 | 733 | 1412 | 1103 | 1172 | 1522 | 1064 | 1056 | 1121 | 1101 | 1256 | 1198 | 1200 | 1161 | 1144,58 | 1086 | 1204 | 1,109 | 0,149 | 2,933 | 0,003356 | 0,004548 | 1,00E-08 | 0,003191 | 0,0016127 | 0,0016127 | TRUE | 0,0107093 | No |
| orf19.1435 | 11795 | 8193 | 9107 | 5622 | 15784 | 12046 | 13176 | 20726 | 10739 | 8286 | 8619 | 8445 | 14039 | 13079 | 13494 | 15810 | 11563,9 | 9022 | 14106 | 1,563 | 0,645 | 7,702 | 1,34E-14 | 2,94E-14 | 0,0116698 | 0,002151 | 0,0066356 | 0,0066356 | TRUE | 0,2010915 | No |
| orf19.1438 | 378 | 356 | 390 | 239 | 195 | 194 | 178 | 218 | 344 | 360 | 369 | 359 | 173 | 211 | 182 | 166 | 270,62 | 358 | 183 | 0,509 | -0,975 | -10,301 | 6,97E-25 | 1,99E-24 | 9,20E-05 | 0,006918 | 0,0045514 | 0,0045514 | TRUE | 0,1033488 | No |
| orf19.1439 | 1628 | 1784 | 1863 | 1137 | 1719 | 1512 | 1403 | 1865 | 1482 | 1804 | 1763 | 1708 | 1529 | 1642 | 1437 | 1423 | 1598,47 | 1689 | 1508 | 0,892 | -0,165 | -2,316 | 0,020574 | 0,026337 | 0,0050258 | 0,002863 | 0,0042212 | 0,0042212 | TRUE | 0,0849279 | No |
| orf19.144 | 2494 | 2821 | 2959 | 1674 | 2302 | 1988 | 1967 | 2488 | 2271 | 2853 | 2800 | 2515 | 2048 | 2158 | 2014 | 1898 | 2319,64 | 2610 | 2030 | 0,777 | -0,364 | -4,964 | 6,92E-07 | 1,16E-06 | 0,0061862 | 0,002606 | 0,0047171 | 0,0047171 | TRUE | 0,0952653 | No |
| orf19.1440 | 960 | 1021 | 1009 | 632 | 1182 | 974 | 1070 | 1310 | 874 | 1033 | 955 | 949 | 1051 | 1057 | 1096 | 999 | 1001,86 | 953 | 1051 | 1,103 | 0,141 | 2,264 | 0,023596 | 0,030079 | 0,002119 | 0,003356 | 0,0027166 | 0,0027166 | TRUE | 0,039739 | No |
| orf19.1440.1 | 9 | 23 | 9 | 9 | 20 | 8 | 17 | 20 | 8 | 23 | 9 | 14 | 18 | 9 | 17 | 15 | 14,08 | 14 | 15 | 1,121 | 0,165 | 0,392 | 0,694837 | 0,729631 | 0,1011698 | 0,095862 | 0,0978445 | 0,0978445 | TRUE | 1,1314824 | No |
| orf19.1440.2 | 240 | 212 | 213 | 101 | 544 | 351 | 407 | 542 | 219 | 214 | 202 | 152 | 484 | 381 | 417 | 413 | 310,18 | 197 | 424 | 2,136 | 1,095 | 9,763 | 1,62E-22 | 4,35E-22 | 0,0114368 | 0,006296 | 0,0082805 | 0,0082805 | TRUE | 0,2057754 | No |
| orf19.1441 | 4737 | 4366 | 4800 | 2814 | 5054 | 4191 | 4599 | 5879 | 4313 | 4416 | 4543 | 4227 | 4495 | 4550 | 4710 | 4485 | 4467,32 | 4375 | 4560 | 1,041 | 0,059 | 1,544 | 0,122626 | 0,145848 | 0,0004453 | 0,002333 | 0,0011553 | 0,0011553 | TRUE | 0,0081704 | No |
| orf19.1442 | 441 | 374 | 372 | 265 | 834 | 689 | 727 | 1067 | 402 | 378 | 352 | 398 | 742 | 748 | 745 | 814 | 572,28 | 382 | 762 | 1,999 | 0,999 | 14,061 | 6,56E-45 | 2,80E-44 | 0,0011271 | 0,004345 | 0,0028397 | 0,0028397 | TRUE | 0,0348605 | No |
| orf19.1444 | 2408 | 2138 | 2475 | 1734 | 2782 | 2396 | 2544 | 3486 | 2192 | 2162 | 2342 | 2605 | 2475 | 2601 | 2605 | 2659 | 2455,28 | 2325 | 2585 | 1,114 | 0,155 | 2,579 | 0,009914 | 0,013002 | 0,0036925 | 0,002575 | 0,0030778 | 0,0030778 | TRUE | 0,0752903 | No |
| orf19.1445 | 85 | 74 | 78 | 44 | 42 | 36 | 37 | 38 | 77 | 75 | 74 | 66 | 37 | 39 | 38 | 29 | 54,43 | 73 | 36 | 0,484 | -1,048 | -5,454 | 4,91E-08 | 8,58E-08 | 1,00E-08 | 0,026305 | 0,0151111 | 0,0151111 | TRUE | 0,1503225 | No |
| orf19.1446 | 3728 | 3390 | 3747 | 2246 | 1490 | 1360 | 1507 | 1907 | 3394 | 3429 | 3546 | 3374 | 1325 | 1477 | 1543 | 1455 | 2442,85 | 3436 | 1450 | 0,422 | -1,246 | -24,486 | 2,07E-132 | 2,37E-131 | 0,0015788 | 0,002578 | 0,0020074 | 0,0020074 | TRUE | 0,0406281 | No |
| orf19.1447 | 288 | 237 | 293 | 193 | 179 | 135 | 139 | 233 | 262 | 240 | 277 | 290 | 159 | 147 | 142 | 178 | 211,88 | 267 | 156 | 0,591 | -0,76 | -6,998 | 2,60E-12 | 5,33E-12 | 0,0033658 | 0,008272 | 0,0063483 | 0,0063483 | TRUE | 0,0974308 | No |
| orf19.1448 | 3386 | 2972 | 3084 | 1870 | 5392 | 4197 | 4410 | 6020 | 3083 | 3006 | 2919 | 2809 | 4796 | 4557 | 4516 | 4592 | 3784,72 | 2954 | 4615 | 1,561 | 0,642 | 15,287 | 9,28E-53 | 4,50E-52 | 0,0007863 | 0,002386 | 0,0014107 | 0,0014107 | TRUE | 0,0122423 | No |
| orf19.1448.1 | 542 | 536 | 495 | 380 | 620 | 476 | 462 | 700 | 493 | 542 | 468 | 571 | 551 | 517 | 473 | 534 | 518,78 | 518 | 519 | 1,006 | 0,008 | 0,105 | 0,916384 | 0,928178 | 0,0039133 | 0,004583 | 0,0042781 | 0,0042781 | TRUE | 0,0491444 | No |
| orf19.1449 | 221 | 238 | 260 | 138 | 3832 | 3034 | 3026 | 4227 | 201 | 241 | 246 | 207 | 3408 | 3294 | 3099 | 3224 | 1740,15 | 224 | 3256 | 14,494 | 3,857 | 54,028 | 0 | 0 | 0,0017569 | 0,002796 | 0,0023903 | 0,0023903 | TRUE | 0,0653596 | No |
| orf19.145 | 1372 | 1362 | 1420 | 861 | 1540 | 1322 | 1364 | 1676 | 1249 | 1378 | 1344 | 1293 | 1370 | 1435 | 1397 | 1278 | 1343,05 | 1316 | 1370 | 1,04 | 0,056 | 1,009 | 0,312805 | 0,351574 | 0,0015183 | 0,003021 | 0,0021981 | 0,0021981 | TRUE | 0,025495 | No |
| orf19.1450 | 13 | 12 | 19 | 9 | 4 | 8 | 10 | 6 | 12 | 12 | 18 | 14 | 4 | 9 | 10 | 5 | 10,32 | 14 | 7 | 0,474 | -1,077 | -2,356 | 0,018481 | 0,023777 | 1,00E-08 | 0,130078 | 0,0918112 | 0,0918112 | TRUE | 0,3184694 | No |
| orf19.1451 | 822 | 760 | 750 | 537 | 863 | 766 | 726 | 976 | 748 | 769 | 710 | 807 | 768 | 832 | 744 | 745 | 765,1 | 758 | 772 | 1,019 | 0,028 | 0,432 | 0,666051 | 0,70276 | 0,0013218 | 0,003764 | 0,0026026 | 0,0026026 | TRUE | 0,0307562 | No |
| orf19.1453 | 14603 | 11721 | 13429 | 7928 | 13334 | 10584 | 12336 | 16666 | 13295 | 11855 | 12709 | 11909 | 11860 | 11491 | 12634 | 12713 | 12308,3 | 12442 | 12174 | 0,978 | -0,032 | -0,615 | 0,538534 | 0,580219 | 0,0024054 | 0,002144 | 0,0024486 | 0,0024486 | TRUE | 0,0261446 | No |
| orf19.1456 | 20 | 6 | 2 | 4 | 9 | 4 | 5 | 11 | 18 | 6 | 2 | 6 | 8 | 4 | 5 | 8 | 7,25 | 8 | 6 | 0,801 | -0,32 | -0,515 | 0,606803 | 0,646344 | 0,3053639 | 0,184122 | 0,2342162 | 0,2342162 | TRUE | 1,3283147 | No |
| orf19.1457 | 1954 | 1750 | 1922 | 1222 | 2045 | 1690 | 1771 | 2424 | 1779 | 1770 | 1819 | 1836 | 1819 | 1835 | 1814 | 1849 | 1815,03 | 1801 | 1829 | 1,017 | 0,024 | 0,591 | 0,55463 | 0,595949 | 1,00E-08 | 0,002765 | 0,0010111 | 0,0010111 | TRUE | 0,0017589 | No |
| orf19.1458 | 446 | 437 | 417 | 277 | 608 | 492 | 520 | 677 | 406 | 442 | 395 | 416 | 541 | 534 | 533 | 516 | 472,84 | 415 | 531 | 1,281 | 0,357 | 5,14 | 2,75E-07 | 4,68E-07 | 1,00E-08 | 0,004831 | 0,0024667 | 0,0024667 | TRUE | 0,024402 | No |
| orf19.1459 | 554 | 513 | 595 | 387 | 721 | 587 | 656 | 857 | 504 | 519 | 563 | 581 | 641 | 637 | 672 | 654 | 596,48 | 542 | 651 | 1,206 | 0,27 | 4,053 | 5,06E-05 | 7,71E-05 | 0,0002678 | 0,004252 | 0,0025558 | 0,0025558 | TRUE | 0,0259184 | No |
| orf19.1460 | 2833 | 2316 | 2719 | 1620 | 5634 | 4498 | 5165 | 6854 | 2579 | 2342 | 2573 | 2433 | 5011 | 4884 | 5290 | 5228 | 3792,65 | 2482 | 5103 | 2,055 | 1,039 | 22,049 | 9,71E-108 | 8,99E-107 | 0,0013587 | 0,002385 | 0,0018239 | 0,0018239 | TRUE | 0,018227 | No |
| orf19.1461 | 1663 | 1352 | 1600 | 1061 | 2638 | 2270 | 2384 | 3290 | 1514 | 1367 | 1514 | 1594 | 2346 | 2465 | 2442 | 2510 | 1968,96 | 1497 | 2441 | 1,633 | 0,707 | 13,426 | 4,28E-41 | 1,70E-40 | 0,001668 | 0,002708 | 0,0021127 | 0,0021127 | TRUE | 0,0408841 | No |
| orf19.1462 | 2175 | 1851 | 2029 | 1372 | 5828 | 4875 | 5269 | 7105 | 1980 | 1872 | 1920 | 2061 | 5184 | 5293 | 5396 | 5420 | 3640,77 | 1958 | 5323 | 2,723 | 1,445 | 34,971 | 6,24E-268 | 1,42E-266 | 0,0004509 | 0,0024 | 0,001277 | 0,001277 | TRUE | 0,0133082 | No |
| orf19.1464 | 5196 | 4681 | 5211 | 2968 | 1834 | 1376 | 1582 | 2004 | 4731 | 4734 | 4932 | 4458 | 1631 | 1494 | 1620 | 1529 | 3141,16 | 4714 | 1568 | 0,333 | -1,588 | -33,023 | 3,74E-239 | 7,50E-238 | 0,0012162 | 0,002458 | 0,0018096 | 0,0018096 | TRUE | 0,0154921 | No |
| orf19.1465 | 85 | 76 | 73 | 48 | 69 | 54 | 51 | 61 | 77 | 77 | 69 | 72 | 61 | 59 | 52 | 47 | 64,28 | 74 | 55 | 0,736 | -0,442 | -2,546 | 0,010888 | 0,01425 | 1,00E-08 | 0,022589 | 0,0131553 | 0,0131553 | TRUE | 0,1047102 | No |
| orf19.1466 | 1958 | 1709 | 1790 | 951 | 1568 | 1140 | 1312 | 1675 | 1783 | 1728 | 1694 | 1429 | 1395 | 1238 | 1344 | 1278 | 1485,94 | 1658 | 1314 | 0,79 | -0,34 | -4,808 | 1,52E-06 | 2,51E-06 | 0,0050054 | 0,002926 | 0,0041101 | 0,0041101 | TRUE | 0,0987681 | No |
| orf19.1467 | 8783 | 7302 | 7533 | 5809 | 9833 | 7745 | 7927 | 10564 | 7997 | 7385 | 7129 | 8726 | 8746 | 8409 | 8118 | 8058 | 8071,09 | 7809 | 8333 | 1,068 | 0,094 | 1,524 | 0,127486 | 0,151345 | 0,0046979 | 0,002201 | 0,0035566 | 0,0035566 | TRUE | 0,0751687 | No |
| orf19.1468 | 4421 | 3907 | 4124 | 2494 | 2318 | 1827 | 1978 | 2735 | 4025 | 3952 | 3903 | 3746 | 2062 | 1984 | 2026 | 2086 | 2972,93 | 3906 | 2040 | 0,522 | -0,938 | -23,34 | 1,74E-120 | 1,79E-119 | 0,0003428 | 0,002481 | 0,0011848 | 0,0011848 | TRUE | 0,0086105 | No |
| orf19.147 | 13321 | 13905 | 15000 | 9272 | 1875 | 1705 | 1653 | 2193 | 12128 | 14063 | 14196 | 13928 | 1668 | 1851 | 1693 | 1673 | 7650,05 | 13579 | 1721 | 0,127 | -2,981 | -49,854 | 0 | 0 | 0,0034201 | 0,00221 | 0,003122 | 0,003122 | TRUE | 0,0636232 | No |
| orf19.1470 | 10651 | 8522 | 9081 | 8039 | 17432 | 14688 | 13276 | 19685 | 9697 | 8619 | 8594 | 12076 | 15505 | 15947 | 13596 | 15016 | 12381,4 | 9746 | 15016 | 1,541 | 0,624 | 4,78 | 1,75E-06 | 2,88E-06 | 0,0162773 | 0,002144 | 0,008522 | 0,0162773 | TRUE | 0,3153513 | No |
| orf19.1471 | 5897 | 5129 | 5138 | 3238 | 8819 | 6421 | 6798 | 9889 | 5369 | 5187 | 4863 | 4864 | 7844 | 6971 | 6962 | 7543 | 6200,51 | 5071 | 7330 | 1,445 | 0,531 | 9,906 | 3,91E-23 | 1,07E-22 | 0,0027804 | 0,00225 | 0,0025941 | 0,0025941 | TRUE | 0,0271345 | No |
| orf19.1473 | 356 | 357 | 380 | 202 | 1101 | 888 | 883 | 1268 | 324 | 361 | 360 | 303 | 979 | 964 | 904 | 967 | 645,4 | 337 | 954 | 2,816 | 1,494 | 21,152 | 2,63E-99 | 2,28E-98 | 0,000622 | 0,004084 | 0,0027262 | 0,0027262 | TRUE | 0,0398925 | No |
| orf19.1474 | 5234 | 4465 | 4970 | 3282 | 5356 | 4556 | 5039 | 6694 | 4765 | 4516 | 4704 | 4930 | 4764 | 4947 | 5161 | 5106 | 4861,54 | 4729 | 4994 | 1,057 | 0,08 | 1,871 | 0,061335 | 0,075339 | 0,0010182 | 0,002309 | 0,0015346 | 0,0015346 | TRUE | 0,0118828 | No |

| | | | | | | | | | | | | | | | | | | | | | | | | | | | | | | | |
|---|---|---|---|---|---|---|---|---|---|---|---|---|---|---|---|---|---|---|---|---|---|---|---|---|---|---|---|---|---|---|---|
| orf19.1476 | 33 | 29 | 32 | 20 | 43 | 39 | 40 | 58 | 30 | 29 | 30 | 30 | 38 | 42 | 41 | 44 | 35,69 | 30 | 41 | 1,387 | 0,472 | 2,095 | 0,036164 | 0,045401 | 1,00E-08 | 0,039052 | 0,0196821 | 0,0196821 | TRUE | 0,0230484 | No |
| orf19.1477 | 1899 | 1910 | 1981 | 1264 | 1233 | 1042 | 1010 | 1366 | 1729 | 1932 | 1875 | 1899 | 1097 | 1131 | 1034 | 1042 | 1467,34 | 1859 | 1076 | 0,579 | -0,788 | -14,434 | 3,16E-47 | 1,39E-46 | 0,0014815 | 0,002937 | 0,0021389 | 0,0021389 | TRUE | 0,0276651 | No |
| orf19.1478 | 1181 | 1104 | 1173 | 738 | 1169 | 964 | 987 | 1403 | 1075 | 1117 | 1110 | 1109 | 1040 | 1047 | 1011 | 1070 | 1072,25 | 1103 | 1042 | 0,946 | -0,08 | -1,627 | 0,10376 | 0,124476 | 1,00E-08 | 0,003269 | 0,0013962 | 0,0013962 | TRUE | 0,004745 | No |
| orf19.1479 | 3460 | 2979 | 3222 | 2238 | 2199 | 1917 | 2140 | 2928 | 3150 | 3013 | 3049 | 3362 | 1956 | 2081 | 2192 | 2234 | 2629,59 | 3144 | 2116 | 0,674 | -0,57 | -10,313 | 6,16E-25 | 1,77E-24 | 0,0024404 | 0,002539 | 0,0025371 | 0,0025371 | TRUE | 0,0318927 | No |
| orf19.1480 | 1974 | 1675 | 1770 | 1287 | 3849 | 3042 | 3030 | 4502 | 1797 | 1694 | 1675 | 1933 | 3424 | 3303 | 3103 | 3434 | 2545,42 | 1775 | 3316 | 1,872 | 0,905 | 16,087 | 3,17E-58 | 1,69E-57 | 0,0027255 | 0,002556 | 0,0025937 | 0,0025937 | TRUE | 0,0403701 | No |
| orf19.1481 | 7286 | 6788 | 7489 | 4549 | 2140 | 1962 | 2050 | 2882 | 6634 | 6865 | 7088 | 6833 | 1903 | 2130 | 2099 | 2198 | 4468,93 | 6855 | 2082 | 0,304 | -1,718 | -34,775 | 5,81E-265 | 1,30E-263 | 0,0017965 | 0,002333 | 0,0020421 | 0,0020421 | TRUE | 0,0415056 | No |
| orf19.1482 | 1737 | 1310 | 1469 | 792 | 661 | 442 | 525 | 667 | 1581 | 1325 | 1390 | 1190 | 588 | 480 | 538 | 509 | 950,08 | 1372 | 529 | 0,385 | -1,378 | -15,348 | 3,67E-53 | 1,80E-52 | 0,0091085 | 0,003427 | 0,0064617 | 0,0064617 | TRUE | 0,1279055 | No |
| orf19.1483 | 1290 | 1339 | 1284 | 737 | 876 | 613 | 656 | 835 | 1174 | 1354 | 1215 | 1107 | 779 | 666 | 672 | 637 | 950,56 | 1212 | 688 | 0,567 | -0,819 | -10,308 | 6,49E-25 | 1,86E-24 | 0,0062922 | 0,003427 | 0,0049385 | 0,0049385 | TRUE | 0,0954139 | No |
| orf19.1484 | 1039 | 1252 | 1224 | 773 | 410 | 369 | 387 | 421 | 946 | 1266 | 1158 | 1161 | 365 | 401 | 396 | 321 | 751,82 | 1133 | 371 | 0,326 | -1,615 | -16,525 | 2,42E-61 | 1,35E-60 | 0,0113766 | 0,003794 | 0,0074303 | 0,0074303 | TRUE | 0,1519585 | No |
| orf19.1485 | 923 | 822 | 823 | 465 | 1160 | 775 | 925 | 1189 | 840 | 831 | 779 | 699 | 1032 | 841 | 947 | 907 | 859,58 | 787 | 932 | 1,181 | 0,24 | 3,088 | 0,002018 | 0,002783 | 0,0055063 | 0,003574 | 0,0046054 | 0,0046054 | TRUE | 0,0625293 | No |
| orf19.1486 | 2013 | 2099 | 2271 | 1619 | 1098 | 956 | 961 | 1247 | 1833 | 2123 | 2149 | 2432 | 977 | 1038 | 984 | 951 | 1560,87 | 2134 | 988 | 0,463 | -1,11 | -14,42 | 3,85E-47 | 1,69E-46 | 0,0062774 | 0,002883 | 0,0049656 | 0,0049656 | TRUE | 0,1101815 | No |
| orf19.1488 | 460 | 504 | 478 | 317 | 210 | 165 | 160 | 217 | 419 | 510 | 452 | 476 | 187 | 179 | 164 | 166 | 319,06 | 464 | 174 | 0,375 | -1,416 | -14,872 | 5,03E-50 | 2,32E-49 | 0,0032606 | 0,006177 | 0,0048651 | 0,0048651 | TRUE | 0,0543288 | No |
| orf19.149 | 17 | 12 | 11 | 15 | 8 | 7 | 5 | 12 | 15 | 12 | 10 | 23 | 7 | 8 | 5 | 9 | 11,19 | 15 | 7 | 0,496 | -1,01 | -2,331 | 0,019762 | 0,02535 | 1,00E-08 | 0,120052 | 0,0808145 | 0,0808145 | TRUE | 0,328744 | No |
| orf19.1490 | 3578 | 3311 | 3619 | 2607 | 1776 | 1748 | 1710 | 2583 | 3258 | 3349 | 3425 | 3916 | 1580 | 1898 | 1751 | 1970 | 2643,33 | 3487 | 1800 | 0,517 | -0,953 | -12,476 | 1,01E-35 | 3,64E-35 | 0,0076035 | 0,002537 | 0,0051854 | 0,0051854 | TRUE | 0,0828793 | No |
| orf19.1490.1 | 43 | 36 | 30 | 15 | 42 | 42 | 41 | 49 | 39 | 36 | 28 | 23 | 37 | 46 | 42 | 37 | 36,1 | 32 | 40 | 1,254 | 0,327 | 1,384 | 0,166401 | 0,194644 | 1,00E-08 | 0,038629 | 0,0254415 | 0,0254415 | TRUE | 0,1757459 | No |
| orf19.1491 | 2251 | 2271 | 2313 | 1388 | 1550 | 1274 | 1380 | 1723 | 2049 | 2297 | 2189 | 2085 | 1379 | 1383 | 1413 | 1314 | 1763,73 | 2155 | 1372 | 0,636 | -0,653 | -12,643 | 1,22E-36 | 4,52E-36 | 0,0013559 | 0,002786 | 0,0019703 | 0,0019703 | TRUE | 0,0239071 | No |
| orf19.1492 | 856 | 992 | 930 | 686 | 486 | 439 | 410 | 492 | 779 | 1003 | 880 | 1030 | 432 | 477 | 420 | 375 | 674,68 | 923 | 426 | 0,461 | -1,116 | -11,328 | 9,59E-30 | 3,06E-29 | 0,0116098 | 0,003995 | 0,007632 | 0,007632 | TRUE | 0,1333714 | No |
| orf19.1493 | 3457 | 3088 | 3362 | 1999 | 5017 | 4003 | 4351 | 5894 | 3147 | 3123 | 3182 | 3003 | 4462 | 4346 | 4456 | 4496 | 3776,98 | 3114 | 4440 | 1,425 | 0,511 | 14,389 | 6,09E-47 | 2,68E-46 | 5,54E-05 | 0,002387 | 0,0009314 | 0,0009314 | TRUE | 0,0063036 | No |
| orf19.1494 | 9007 | 7641 | 8335 | 5984 | 11093 | 10092 | 10119 | 13814 | 8200 | 7728 | 7888 | 8989 | 9867 | 10957 | 10363 | 10537 | 9316,29 | 8201 | 10431 | 1,272 | 0,348 | 6,439 | 1,20E-10 | 2,33E-10 | 0,0031254 | 0,002179 | 0,0026889 | 0,0026889 | TRUE | 0,0501129 | No |
| orf19.1495 | 622 | 747 | 718 | 457 | 968 | 931 | 846 | 1044 | 566 | 756 | 680 | 686 | 861 | 1011 | 866 | 796 | 777,8 | 672 | 884 | 1,315 | 0,395 | 4,191 | 2,78E-05 | 4,31E-05 | 0,0102997 | 0,003735 | 0,0071899 | 0,0071899 | TRUE | 0,1379974 | No |
| orf19.1496 | 462 | 510 | 489 | 284 | 805 | 696 | 721 | 950 | 421 | 516 | 463 | 427 | 716 | 756 | 738 | 725 | 595,07 | 457 | 734 | 1,603 | 0,681 | 9,304 | 1,35E-20 | 3,49E-20 | 0,0020663 | 0,004257 | 0,0033365 | 0,0033365 | TRUE | 0,089485 | No |
| orf19.1497 | 489 | 462 | 509 | 294 | 769 | 606 | 654 | 824 | 445 | 467 | 482 | 442 | 684 | 658 | 670 | 629 | 559,51 | 459 | 660 | 1,434 | 0,52 | 7,846 | 4,28E-15 | 9,49E-15 | 1,00E-08 | 0,004398 | 0,0023566 | 0,0023566 | TRUE | 0,0134571 | No |
| orf19.1499 | 2920 | 2793 | 3238 | 2021 | 3984 | 3609 | 3623 | 4826 | 2659 | 2825 | 3064 | 3036 | 3544 | 3918 | 3710 | 3681 | 3304,68 | 2896 | 3713 | 1,283 | 0,359 | 6,527 | 6,72E-11 | 1,31E-10 | 0,0026331 | 0,002437 | 0,0026008 | 0,0026008 | TRUE | 0,0376594 | No |
| orf19.150 | 5012 | 4113 | 4425 | 2809 | 4213 | 3392 | 3646 | 4955 | 4563 | 4160 | 4188 | 4220 | 3747 | 3683 | 3734 | 3780 | 4009,28 | 4283 | 3736 | 0,872 | -0,197 | -4,787 | 1,70E-06 | 2,79E-06 | 0,0006942 | 0,002367 | 0,001382 | 0,001382 | TRUE | 0,0243951 | No |
| orf19.1500 | 945 | 925 | 987 | 613 | 656 | 580 | 594 | 766 | 860 | 936 | 934 | 921 | 583 | 630 | 608 | 584 | 757,09 | 913 | 601 | 0,658 | -0,603 | -10,055 | 8,73E-24 | 2,44E-23 | 1,00E-08 | 0,003782 | 0,0020895 | 0,0020895 | TRUE | 0,0191569 | No |
| orf19.1502 | 483 | 561 | 566 | 346 | 161 | 150 | 130 | 180 | 440 | 567 | 536 | 520 | 143 | 163 | 133 | 137 | 329,88 | 516 | 144 | 0,279 | -1,842 | -17,313 | 3,74E-67 | 2,26E-66 | 0,0072388 | 0,006041 | 0,0065818 | 0,0065818 | TRUE | 0,1216932 | No |
| orf19.1504 | 2559 | 2611 | 2706 | 1516 | 4592 | 3811 | 4087 | 5168 | 2330 | 2641 | 2561 | 2277 | 4084 | 4138 | 4186 | 3942 | 3269,85 | 2452 | 4088 | 1,665 | 0,735 | 13,771 | 3,83E-43 | 1,58E-42 | 0,0024731 | 0,002441 | 0,0024029 | 0,0024029 | TRUE | 0,0320577 | No |
| orf19.1505 | 2297 | 2270 | 2269 | 1430 | 3698 | 3156 | 3128 | 4331 | 2091 | 2296 | 2147 | 2148 | 3289 | 3427 | 3203 | 3304 | 2738,21 | 2170 | 3306 | 1,523 | 0,607 | 13,74 | 5,83E-43 | 2,39E-42 | 0,0006918 | 0,002519 | 0,0014825 | 0,0014825 | TRUE | 0,0187631 | No |
| orf19.1506 | 0 | 0 | 0 | 0 | 0 | 0 | 0 | 0 | 0 | 0 | 0 | 0 | 0 | 0 | 0 | 0 | 0 | 0 | 0 | NA | NA | NA | NA | NA | NA | NA | NA | NA | NA | NA | NA |
| orf19.1507 | 255 | 281 | 303 | 147 | 239 | 216 | 231 | 258 | 232 | 284 | 287 | 221 | 213 | 235 | 237 | 197 | 238,05 | 256 | 220 | 0,853 | -0,23 | -2,038 | 0,041557 | 0,051885 | 0,0084855 | 0,007586 | 0,0079662 | 0,0079662 | TRUE | 0,0788761 | No |
| orf19.1508 | 26 | 20 | 19 | 15 | 31 | 32 | 20 | 28 | 24 | 20 | 18 | 23 | 28 | 35 | 20 | 21 | 23,57 | 21 | 26 | 1,228 | 0,297 | 1,016 | 0,309692 | 0,348199 | 1,00E-08 | 0,058079 | 0,0386147 | 0,0386147 | TRUE | 0,2839979 | No |
| orf19.1509 | 9211 | 9071 | 9649 | 5932 | 5181 | 4564 | 4626 | 6490 | 8386 | 9174 | 9132 | 8911 | 4608 | 4955 | 4738 | 4951 | 6856,89 | 8901 | 4813 | 0,541 | -0,887 | -20,326 | 7,56E-92 | 5,99E-91 | 0,0012377 | 0,00223 | 0,0016708 | 0,0016708 | TRUE | 0,0188281 | No |
| orf19.151 | 1172 | 1129 | 1268 | 797 | 1119 | 1018 | 1011 | 1319 | 1067 | 1142 | 1200 | 1197 | 995 | 1105 | 1035 | 1006 | 1093,54 | 1152 | 1035 | 0,899 | -0,153 | -2,579 | 0,009904 | 0,012995 | 0,0017226 | 0,003245 | 0,002452 | 0,002452 | TRUE | 0,0304343 | No |
| orf19.1510 | 1222 | 1151 | 1251 | 742 | 1541 | 1300 | 1374 | 1908 | 1113 | 1164 | 1184 | 1115 | 1371 | 1411 | 1407 | 1455 | 1277,49 | 1144 | 1411 | 1,233 | 0,302 | 6,212 | 5,22E-10 | 9,87E-10 | 1,00E-08 | 0,003071 | 0,001474 | 0,001474 | TRUE | 0,0068583 | No |
| orf19.1513 | 6521 | 6281 | 6545 | 3767 | 2053 | 1700 | 1762 | 2364 | 5937 | 6353 | 6194 | 5659 | 1826 | 1846 | 1805 | 1803 | 3927,77 | 6036 | 1820 | 0,301 | -1,731 | -37,866 | 0 | 0 | 0,0012251 | 0,002373 | 0,0016603 | 0,0016603 | TRUE | 0,0206877 | No |
| orf19.1514 | 2396 | 2069 | 2312 | 1342 | 2578 | 2097 | 2203 | 2971 | 2181 | 2093 | 2188 | 2016 | 2293 | 2277 | 2256 | 2266 | 2196,29 | 2120 | 2273 | 1,07 | 0,098 | 2,344 | 0,019093 | 0,024535 | 0,0001583 | 0,002638 | 0,0012296 | 0,0012296 | TRUE | 0,0117349 | No |
| orf19.1515 | 259 | 257 | 237 | 151 | 185 | 125 | 157 | 184 | 236 | 260 | 224 | 227 | 165 | 136 | 161 | 140 | 193,53 | 237 | 150 | 0,634 | -0,657 | -6,038 | 1,56E-09 | 2,90E-09 | 1,00E-08 | 0,008863 | 0,0060133 | 0,0060133 | TRUE | 0,0487588 | No |
| orf19.1516 | 1185 | 1164 | 1198 | 653 | 1716 | 1393 | 1526 | 2003 | 1079 | 1177 | 1134 | 981 | 1526 | 1512 | 1563 | 1528 | 1312,54 | 1093 | 1532 | 1,398 | 0,483 | 8,433 | 3,37E-17 | 7,92E-17 | 0,0015905 | 0,003043 | 0,0023568 | 0,0023568 | TRUE | 0,0497665 | No |
| orf19.1517 | 2073 | 1709 | 1802 | 1143 | 2938 | 2220 | 2421 | 3391 | 1887 | 1728 | 1705 | 1717 | 2613 | 2410 | 2479 | 2587 | 2140,99 | 1759 | 2522 | 1,433 | 0,519 | 10,332 | 5,03E-25 | 1,45E-24 | 0,0012458 | 0,002654 | 0,0019331 | 0,0019331 | TRUE | 0,0300473 | No |
| orf19.1519 | 1148 | 1003 | 1108 | 713 | 1221 | 1030 | 1092 | 1396 | 1045 | 1014 | 1049 | 1071 | 1086 | 1118 | 1118 | 1065 | 1070,86 | 1045 | 1097 | 1,05 | 0,07 | 1,411 | 0,15814 | 0,185593 | 1,00E-08 | 0,003271 | 0,001434 | 0,001434 | TRUE | 0,0048924 | No |
| orf19.1522 | 72 | 73 | 73 | 42 | 53 | 69 | 70 | 84 | 66 | 74 | 69 | 63 | 47 | 75 | 72 | 64 | 66,17 | 68 | 64 | 0,941 | -0,088 | -0,481 | 0,630267 | 0,668767 | 0,0072231 | 0,022 | 0,0167153 | 0,0167153 | TRUE | 0,2990665 | No |
| orf19.1523 | 1491 | 1198 | 1390 | 898 | 797 | 634 | 670 | 926 | 1357 | 1212 | 1316 | 1349 | 709 | 688 | 686 | 706 | 1002,92 | 1308 | 697 | 0,534 | -0,906 | -15,946 | 3,03E-57 | 1,58E-56 | 1,00E-08 | 0,003354 | 0,0020189 | 0,0020189 | TRUE | 0,0302305 | No |
| orf19.1524 | 5 | 6 | 5 | 4 | 7 | 12 | 4 | 7 | 5 | 6 | 5 | 6 | 6 | 13 | 4 | 5 | 6,26 | 6 | 7 | 1,334 | 0,416 | 0,731 | 0,465037 | 0,50685 | 1,00E-08 | 0,213175 | 0,144826 | 0,144826 | TRUE | 0,727425 | No |
| orf19.1525 | 1543 | 1524 | 1580 | 1082 | 1191 | 1031 | 1078 | 1414 | 1405 | 1541 | 1495 | 1625 | 1059 | 1119 | 1104 | 1079 | 1303,53 | 1516 | 1090 | 0,72 | -0,474 | -8,552 | 1,21E-17 | 2,88E-17 | 0,0013894 | 0,00305 | 0,0021609 | 0,0021609 | TRUE | 0,0299682 | No |
| orf19.1526 | 3719 | 3285 | 3732 | 2354 | 3131 | 2697 | 2915 | 3920 | 3386 | 3322 | 3532 | 3536 | 2785 | 2928 | 2985 | 2990 | 3183,14 | 3444 | 2922 | 0,849 | -0,236 | -5,599 | 2,16E-08 | 3,83E-08 | 0,0007269 | 0,002452 | 0,0013947 | 0,0013947 | TRUE | 0,0123529 | No |
| orf19.1527 | 1130 | 1019 | 1074 | 723 | 1251 | 935 | 954 | 1423 | 1029 | 1031 | 1016 | 1086 | 1113 | 1015 | 977 | 1085 | 1044,04 | 1040 | 1048 | 1,01 | 0,014 | 0,248 | 0,804384 | 0,830436 | 0,0012666 | 0,003302 | 0,0022544 | 0,0022544 | TRUE | 0,0245457 | No |
| orf19.1528 | 3731 | 3727 | 3950 | 2253 | 3456 | 2818 | 3014 | 3844 | 3397 | 3769 | 3738 | 3384 | 3074 | 3060 | 3087 | 2932 | 3305,2 | 3572 | 3038 | 0,85 | -0,235 | -4,791 | 1,66E-06 | 2,73E-06 | 0,0016013 | 0,002437 | 0,0020068 | 0,0020068 | TRUE | 0,01662 | No |
| orf19.1529 | 17 | 9 | 12 | 14 | 15 | 6 | 2 | 10 | 15 | 9 | 11 | 21 | 13 | 7 | 2 | 8 | 10,81 | 14 | 8 | 0,537 | -0,898 | -1,837 | 0,066182 | 0,080965 | 0,139329 | 0,12421 | 0,1290169 | 0,1290169 | TRUE | 0,6105385 | No |
| orf19.1530 | 1 | 5 | 7 | 0 | 2 | 3 | 1 | 1 | 1 | 5 | 7 | 0 | 2 | 3 | 1 | 1 | 2,43 | 3 | 2 | 0,507 | -0,979 | -0,977 | 0,328817 | 0,368467 | 0,4721954 | 0,546348 | 0,5210266 | 0,5210266 | TRUE | 0,1289178 | No |
| orf19.1531 | 3032 | 3448 | 3332 | 2018 | 3469 | 3028 | 2850 | 3850 | 2761 | 3487 | 3153 | 3031 | 3086 | 3288 | 2919 | 2937 | 3082,66 | 3108 | 3058 | 0,983 | -0,024 | -0,348 | 0,727605 | 0,760282 | 0,0059097 | 0,002466 | 0,0043237 | 0,0043237 | TRUE | 0,0826965 | No |
| orf19.1532 | 466 | 494 | 499 | 355 | 603 | 512 | 514 | 598 | 424 | 500 | 472 | 533 | 536 | 556 | 526 | 456 | 500,53 | 482 | 518 | 1,078 | 0,108 | 1,222 | 0,221692 | 0,25483 | 0,0063188 | 0,004676 | 0,0054466 | 0,0054466 | TRUE | 0,0817728 | No |
| orf19.1533 | 2622 | 2660 | 2720 | 1558 | 1857 | 1466 | 1491 | 1950 | 2387 | 2690 | 2574 | 2340 | 1652 | 1592 | 1527 | 1487 | 2031,25 | 2498 | 1564 | 0,626 | -0,677 | -11,678 | 1,65E-31 | 5,51E-31 | 0,0025098 | 0,002687 | 0,002711 | 0,002711 | TRUE | 0,0323113 | No |
| orf19.1534 | 776 | 818 | 882 | 470 | 440 | 403 | 440 | 591 | 707 | 827 | 835 | 706 | 391 | 438 | 451 | 451 | 600,62 | 769 | 433 | 0,562 | -0,832 | -10,15 | 3,32E-24 | 9,34E-24 | 0,0051497 | 0,004236 | 0,004677 | 0,004677 | TRUE | 0,0490913 | No |
| orf19.1535 | 144 | 113 | 136 | 66 | 55 | 36 | 55 | 69 | 131 | 114 | 129 | 99 | 49 | 39 | 56 | 53 | 83,78 | 118 | 49 | 0,415 | -1,27 | -7,588 | 3,25E-14 | 7,03E-14 | 1,00E-08 | 0,017805 | 0,013018 | 0,013018 | TRUE | 0,1356154 | No |
| orf19.1536 | 1257 | 1403 | 1423 | 1030 | 2742 | 2475 | 2290 | 3009 | 1144 | 1419 | 1347 | 1547 | 2439 | 2687 | 2345 | 2295 | 1903 | 1364 | 2442 | 1,793 | 0,842 | 10,118 | 4,59E-24 | 1,29E-23 | 0,0094015 | 0,002731 | 0,0060672 | 0,0060672 | TRUE | 0,1432538 | No |
| orf19.1538 | 247 | 223 | 226 | 143 | 228 | 174 | 202 | 248 | 225 | 226 | 214 | 215 | 203 | 189 | 207 | 189 | 208,36 | 220 | 197 | 0,894 | -0,161 | -1,663 | 0,096341 | 0,115948 | 1,00E-08 | 0,008377 | 0,0042256 | 0,0042256 | TRUE | 0,0118187 | No |
| orf19.1539 | 277 | 318 | 302 | 212 | 135 | 164 | 137 | 168 | 252 | 322 | 286 | 318 | 120 | 178 | 140 | 128 | 218,09 | 294 | 142 | 0,48 | -1,06 | -8,171 | 3,06E-16 | 6,99E-16 | 0,0155345 | 0,008094 | 0,0110239 | 0,0110239 | TRUE | 0,2961901 | No |
| orf19.154 | 2489 | 2664 | 2685 | 1616 | 2087 | 1799 | 1822 | 2184 | 2266 | 2694 | 2541 | 2427 | 1856 | 1953 | 1866 | 1666 | 2158,82 | 2482 | 1835 | 0,739 | -0,437 | -6,654 | 2,85E-11 | 5,65E-11 | 0,0041981 | 0,002649 | 0,0036688 | 0,0036688 | TRUE | 0,0478614 | No |
| orf19.1541 | 30 | 40 | 40 | 23 | 109 | 95 | 91 | 127 | 27 | 40 | 38 | 35 | 97 | 103 | 93 | 97 | 66,29 | 35 | 98 | 2,788 | 1,479 | 8,006 | 1,18E-15 | 2,66E-15 | 1,00E-08 | 0,021964 | 0,012846 | 0,012846 | TRUE | 0,1927509 | No |
| orf19.1542 | 4469 | 3948 | 4337 | 2562 | 3407 | 2731 | 3201 | 4241 | 4069 | 3993 | 4105 | 3849 | 3030 | 2965 | 3278 | 3235 | 3565,47 | 4004 | 3127 | 0,781 | -0,357 | -7,818 | 5,35E-15 | 1,18E-14 | 0,0012162 | 0,002408 | 0,0017179 | 0,0017179 | TRUE | 0,0144098 | No |
| orf19.1543 | 1017 | 916 | 966 | 637 | 2096 | 1716 | 1820 | 2461 | 926 | 926 | 914 | 957 | 1864 | 1863 | 1864 | 1877 | 1399,01 | 931 | 1867 | 2,009 | 1,007 | 22,352 | 1,14E-110 | 1,08E-109 | 1,00E-08 | 0,002981 | 0,0011236 | 0,0011236 | TRUE | 0,0034737 | No |
| orf19.1544 | 312 | 271 | 294 | 219 | 846 | 626 | 685 | 924 | 284 | 274 | 278 | 329 | 752 | 680 | 702 | 705 | 500,49 | 291 | 710 | 2,456 | 1,296 | 16,973 | 1,30E-64 | 7,57E-64 | 0,0007529 | 0,004676 | 0,0031008 | 0,0031008 | TRUE | 0,0679399 | No |
| orf19.1545 | 1507 | 1230 | 1313 | 855 | 858 | 634 | 689 | 908 | 1372 | 1244 | 1243 | 1284 | 763 | 688 | 706 | 693 | 999,1 | 1286 | 712 | 0,554 | -0,852 | -14,294 | 2,39E-46 | 1,04E-45 | 0,0013941 | 0,003359 | 0,0023366 | 0,0023366 | TRUE | 0,0278094 | No |
| orf19.1546 | 1567 | 1420 | 1459 | 895 | 1256 | 1015 | 939 | 1243 | 1427 | 1436 | 1381 | 1344 | 1117 | 1102 | 962 | 948 | 1214,64 | 1397 | 1032 | 0,738 | -0,439 | -6,746 | 1,52E-11 | 3,04E-11 | 0,0031161 | 0,003125 | 0,003228 | 0,003228 | TRUE | 0,0383563 | No |
| orf19.1547 | 352 | 333 | 340 | 227 | 351 | 278 | 333 | 426 | 320 | 337 | 322 | 341 | 312 | 302 | 341 | 325 | 325,01 | 330 | 320 | 0,972 | -0,04 | -0,5 | 0,617219 | 0,655904 | 1,00E-08 | 0,006102 | 0,0032121 | 0,0032121 | TRUE | 0,0202532 | No |
| orf19.1548 | 1059 | 814 | 898 | 519 | 1131 | 910 | 945 | 1318 | 964 | 823 | 850 | 780 | 1006 | 988 | 968 | 1005 | 923,02 | 854 | 992 | 1,157 | 0,211 | 3,153 | 0,001614 | 0,002246 | 0,0029682 | 0,003468 | 0,0031987 | 0,0031987 | TRUE | 0,0909615 | No |
| orf19.1549 | 2575 | 2496 | 2742 | 1400 | 4362 | 3341 | 3631 | 4665 | 2344 | 2524 | 2595 | 2103 | 3880 | 3627 | 3719 | 3558 | 3043,91 | 2392 | 3696 | 1,543 | 0,626 | 9,885 | 4,82E-23 | 1,31E-22 | 0,0044589 | 0,002471 | 0,0034943 | 0,0034943 | TRUE | 0,0759981 | No |

| | | | | | | | | | | | | | | | | | | | | | | | | | | | | | | | |
|---|---|---|---|---|---|---|---|---|---|---|---|---|---|---|---|---|---|---|---|---|---|---|---|---|---|---|---|---|---|---|---|
| orf19.155 | 2285 | 2371 | 2402 | 1551 | 1588 | 1524 | 1386 | 1927 | 2080 | 2398 | 2273 | 2330 | 1412 | 1655 | 1419 | 1470 | 1879,75 | 2270 | 1489 | 0,656 | -0,609 | -9,451 | 3,35E-21 | 8,74E-21 | 0,0039603 | 0,00274 | 0,0034303 | 0,0034303 | TRUE | 0,0656726 | No |
| orf19.1551 | 10 | 13 | 12 | 4 | 24 | 19 | 15 | 16 | 9 | 13 | 11 | 6 | 21 | 21 | 15 | 12 | 13,65 | 10 | 17 | 1,703 | 0,768 | 1,967 | 0,04916 | 0,060925 | 1,00E-08 | 0,098848 | 0,0670056 | 0,0670056 | TRUE | 0,2615375 | No |
| orf19.1552 | 3135 | 2407 | 2418 | 1645 | 4795 | 3473 | 3543 | 5158 | 2854 | 2434 | 2288 | 2471 | 4265 | 3771 | 3628 | 3935 | 3205,87 | 2512 | 3900 | 1,552 | 0,634 | 8,815 | 1,20E-18 | 2,94E-18 | 0,0066459 | 0,002449 | 0,0046397 | 0,0046397 | TRUE | 0,1040766 | No |
| orf19.1553 | 6783 | 5957 | 6498 | 4109 | 7818 | 6692 | 7300 | 9532 | 6176 | 6025 | 6150 | 6172 | 6954 | 7266 | 7476 | 7271 | 6686,17 | 6131 | 7242 | 1,181 | 0,24 | 6,968 | 3,21E-12 | 6,55E-12 | 0,0003097 | 0,002235 | 0,0009884 | 0,0009884 | TRUE | 0,0087795 | No |
| orf19.1555 | 2182 | 2111 | 2191 | 1361 | 2615 | 2256 | 2306 | 3019 | 1987 | 2135 | 2074 | 2044 | 2326 | 2449 | 2362 | 2303 | 2209,95 | 2060 | 2360 | 1,145 | 0,195 | 4,539 | 5,64E-06 | 9,06E-06 | 0,0003922 | 0,002635 | 0,0013171 | 0,0013171 | TRUE | 0,0076954 | No |
| orf19.1557 | 149 | 106 | 141 | 95 | 157 | 120 | 119 | 183 | 136 | 107 | 133 | 143 | 140 | 130 | 122 | 140 | 131,3 | 130 | 133 | 1,031 | 0,044 | 0,341 | 0,733069 | 0,765114 | 1,00E-08 | 0,012098 | 0,0081894 | 0,0081894 | TRUE | 0,1412177 | No |
| orf19.1559 | 5067 | 4228 | 4786 | 3069 | 6050 | 4908 | 5349 | 7255 | 4613 | 4276 | 4530 | 4610 | 5381 | 5329 | 5478 | 5534 | 4968,92 | 4507 | 5430 | 1,205 | 0,27 | 7,108 | 1,18E-12 | 2,44E-12 | 0,0004963 | 0,002303 | 0,001175 | 0,001175 | TRUE | 0,014637 | No |
| orf19.156 | 3098 | 2892 | 3073 | 1769 | 2845 | 2488 | 2630 | 3353 | 2821 | 2925 | 2908 | 2657 | 2531 | 2701 | 2693 | 2558 | 2724,27 | 2828 | 2621 | 0,925 | -0,112 | -2,439 | 0,014739 | 0,019118 | 0,0010281 | 0,002522 | 0,0016712 | 0,0016712 | TRUE | 0,0184932 | No |
| orf19.1560 | 5786 | 5000 | 5449 | 3380 | 5855 | 4656 | 5231 | 7087 | 5268 | 5057 | 5157 | 5077 | 5208 | 5055 | 5357 | 5406 | 5198,17 | 5140 | 5256 | 1,023 | 0,032 | 0,889 | 0,374212 | 0,414383 | 0,0003717 | 0,002291 | 0,0010836 | 0,0010836 | TRUE | 0,0079525 | No |
| orf19.1562 | 32160 | 29870 | 31589 | 22185 | 2205 | 1994 | 1853 | 2562 | 29280 | 30210 | 29896 | 33325 | 1961 | 2165 | 1898 | 1954 | 16336,3 | 30678 | 1994 | 0,065 | -3,944 | -71,562 | 0 | 0 | 0,0031014 | 0,002118 | 0,002665 | 0,002665 | TRUE | 0,0411335 | No |
| orf19.1563 | 2834 | 2517 | 2683 | 1762 | 4705 | 3927 | 4195 | 5629 | 2580 | 2546 | 2539 | 2647 | 4185 | 4264 | 4296 | 4294 | 3418,82 | 2578 | 4260 | 1,654 | 0,726 | 20,765 | 9,01E-96 | 7,39E-95 | 1,00E-08 | 0,002423 | 0,0008552 | 0,0008552 | TRUE | 0,0034486 | No |
| orf19.1564 | 1483 | 1381 | 1438 | 1016 | 1833 | 1526 | 1486 | 2043 | 1350 | 1397 | 1361 | 1526 | 1630 | 1657 | 1522 | 1558 | 1500,19 | 1408 | 1592 | 1,133 | 0,18 | 3,324 | 0,000887 | 0,001257 | 0,0015298 | 0,002918 | 0,0021416 | 0,0021416 | TRUE | 0,034506 | No |
| orf19.1565 | 984 | 875 | 921 | 589 | 1720 | 1490 | 1626 | 2156 | 896 | 885 | 872 | 885 | 1530 | 1618 | 1665 | 1645 | 1249,34 | 884 | 1614 | 1,825 | 0,868 | 17,188 | 3,26E-66 | 1,95E-65 | 1,00E-08 | 0,003094 | 0,0015552 | 0,0015552 | TRUE | 0,0152682 | No |
| orf19.1566 | 18399 | 16883 | 17981 | 11461 | 21842 | 18123 | 19281 | 26074 | 16752 | 17075 | 17017 | 17216 | 19428 | 19677 | 19746 | 19889 | 18350,1 | 17015 | 19685 | 1,157 | 0,211 | 8,258 | 1,48E-16 | 3,41E-16 | 4,04E-05 | 0,002109 | 0,0005688 | 0,0005688 | TRUE | 0,0013402 | No |
| orf19.1567 | 3284 | 3156 | 3346 | 1915 | 2041 | 1744 | 1878 | 2428 | 2990 | 3192 | 3167 | 2877 | 1815 | 1893 | 1923 | 1852 | 2463,69 | 3056 | 1871 | 0,611 | -0,71 | -15,016 | 5,78E-51 | 2,72E-50 | 0,0011146 | 0,002573 | 0,0017215 | 0,0017215 | TRUE | 0,0176811 | No |
| orf19.1569 | 10173 | 9891 | 10562 | 6839 | 9749 | 8628 | 8882 | 11529 | 9262 | 10004 | 9996 | 10273 | 8671 | 9368 | 9096 | 8794 | 9433,09 | 9884 | 8982 | 0,909 | -0,138 | -3,148 | 0,001647 | 0,002287 | 0,0014286 | 0,002177 | 0,0017328 | 0,0017328 | TRUE | 0,0220687 | No |
| orf19.1570 | 1001 | 889 | 925 | 526 | 1139 | 843 | 984 | 1283 | 911 | 899 | 875 | 790 | 1013 | 915 | 1008 | 979 | 923,86 | 869 | 979 | 1,123 | 0,167 | 2,719 | 0,00655 | 0,00871 | 0,0015828 | 0,003467 | 0,0025446 | 0,0025446 | TRUE | 0,0386198 | No |
| orf19.1571 | 1 | 3 | 2 | 1 | 5 | 2 | 0 | 1 | 1 | 3 | 2 | 2 | 4 | 2 | 0 | 1 | 1,84 | 2 | 2 | 1,005 | 0,007 | 0,007 | 0,994482 | 0,99495 | 1,00E-08 | 0,719912 | 0,5506776 | 0,5506776 | TRUE | 0,6849389 | No |
| orf19.1573 | 2338 | 2099 | 2171 | 1376 | 2810 | 2430 | 2434 | 3398 | 2129 | 2123 | 2055 | 2067 | 2499 | 2638 | 2493 | 2592 | 2324,46 | 2094 | 2556 | 1,22 | 0,287 | 7,059 | 1,68E-12 | 3,47E-12 | 1,00E-08 | 0,002605 | 0,0011499 | 0,0011499 | TRUE | 0,0054294 | No |
| orf19.1574 | 2188 | 1819 | 2106 | 1194 | 1562 | 1106 | 1293 | 1787 | 1992 | 1840 | 1993 | 1794 | 1389 | 1201 | 1324 | 1363 | 1612 | 1905 | 1319 | 0,693 | -0,53 | -8,841 | 9,50E-19 | 2,33E-18 | 0,0026165 | 0,002856 | 0,0028142 | 0,0028142 | TRUE | 0,042703 | No |
| orf19.1575 | 11725 | 10114 | 10546 | 6747 | 11666 | 9241 | 9516 | 13867 | 10675 | 10229 | 9981 | 10135 | 10377 | 10033 | 9746 | 10578 | 10219,2 | 10255 | 10184 | 0,993 | -0,01 | -0,254 | 0,799507 | 0,826071 | 0,0008364 | 0,002166 | 0,0013896 | 0,0013896 | TRUE | 0,0102454 | No |
| orf19.1576 | 1574 | 1379 | 1547 | 967 | 1211 | 986 | 1002 | 1441 | 1433 | 1395 | 1464 | 1453 | 1077 | 1071 | 1026 | 1099 | 1252,19 | 1436 | 1068 | 0,745 | -0,425 | -8,86 | 8,02E-19 | 1,97E-18 | 1,00E-08 | 0,003092 | 0,0014012 | 0,0014012 | TRUE | 0,0082859 | No |
| orf19.1577 | 69 | 69 | 65 | 53 | 118 | 125 | 80 | 169 | 63 | 70 | 62 | 80 | 105 | 136 | 82 | 129 | 90,66 | 69 | 113 | 1,667 | 0,737 | 4,092 | 4,28E-05 | 6,55E-05 | 0,0227863 | 0,016608 | 0,0191497 | 0,0191497 | TRUE | 0,2965006 | No |
| orf19.1578 | 12235 | 12656 | 12802 | 7283 | 21147 | 17334 | 18154 | 23205 | 11139 | 12800 | 12116 | 10940 | 18810 | 18820 | 18592 | 17701 | 15114,8 | 11749 | 18481 | 1,573 | 0,653 | 12,25 | 1,68E-34 | 5,93E-34 | 0,003054 | 0,002124 | 0,0026591 | 0,0026591 | TRUE | 0,0443109 | No |
| orf19.158 | 5391 | 6152 | 6296 | 3163 | 7113 | 5616 | 6122 | 7321 | 4908 | 6222 | 5959 | 4751 | 6327 | 6097 | 6270 | 5585 | 5764,84 | 5460 | 6070 | 1,111 | 0,152 | 1,828 | 0,067517 | 0,082519 | 0,0104625 | 0,002266 | 0,0064614 | 0,0064614 | TRUE | 0,1073921 | No |
| orf19.1580 | 293 | 308 | 303 | 201 | 149 | 120 | 121 | 165 | 267 | 312 | 287 | 302 | 133 | 130 | 124 | 126 | 209,95 | 292 | 128 | 0,44 | -1,183 | -11,415 | 3,53E-30 | 1,14E-29 | 1,00E-08 | 0,008329 | 0,0048565 | 0,0048565 | TRUE | 0,0400246 | No |
| orf19.1581 | 21 | 12 | 18 | 7 | 16 | 14 | 11 | 10 | 19 | 12 | 17 | 11 | 14 | 15 | 11 | 8 | 13,39 | 15 | 12 | 0,796 | -0,33 | -0,85 | 0,395102 | 0,436 | 1,00E-08 | 0,100682 | 0,0698245 | 0,0698245 | TRUE | 0,3222363 | No |
| orf19.1582 | 89 | 55 | 84 | 51 | 172 | 179 | 154 | 205 | 81 | 56 | 79 | 77 | 153 | 194 | 158 | 156 | 119,27 | 73 | 165 | 2,253 | 1,172 | 7,779 | 7,33E-15 | 1,62E-14 | 0,0090732 | 0,013112 | 0,0116776 | 0,0116776 | TRUE | 0,2495383 | No |
| orf19.1584 | 17607 | 14400 | 15754 | 9476 | 16991 | 13555 | 15251 | 20968 | 16030 | 14564 | 14910 | 14234 | 15113 | 14717 | 15619 | 15995 | 15147,8 | 14934 | 15361 | 1,028 | 0,04 | 0,869 | 0,384732 | 0,425293 | 0,0018856 | 0,002124 | 0,0020057 | 0,0020057 | TRUE | 0,0299391 | No |
| orf19.1585 | 1219 | 1034 | 1158 | 797 | 934 | 818 | 846 | 1233 | 1110 | 1046 | 1096 | 1197 | 831 | 888 | 866 | 941 | 996,83 | 1112 | 882 | 0,795 | -0,331 | -5,393 | 6,95E-08 | 1,21E-07 | 0,0018733 | 0,003362 | 0,0026058 | 0,0026058 | TRUE | 0,0283608 | No |
| orf19.1586 | 149 | 117 | 111 | 73 | 142 | 127 | 125 | 162 | 136 | 118 | 105 | 110 | 126 | 138 | 128 | 124 | 123,06 | 117 | 129 | 1,093 | 0,128 | 0,987 | 0,323658 | 0,363068 | 1,00E-08 | 0,012772 | 0,0080546 | 0,0080546 | TRUE | 0,1242462 | No |
| orf19.1587 | 102 | 110 | 99 | 79 | 262 | 227 | 231 | 344 | 93 | 111 | 94 | 119 | 233 | 246 | 237 | 262 | 174,37 | 104 | 244 | 2,379 | 1,25 | 10,588 | 3,38E-26 | 9,90E-26 | 1,00E-08 | 0,009613 | 0,0063234 | 0,0063234 | TRUE | 0,0661202 | No |
| orf19.1588 | 1124 | 940 | 1089 | 598 | 769 | 547 | 640 | 815 | 1023 | 951 | 1031 | 898 | 684 | 594 | 655 | 622 | 807,25 | 976 | 639 | 0,653 | -0,614 | -9,062 | 1,28E-19 | 3,20E-19 | 0,0025547 | 0,003673 | 0,0031332 | 0,0031332 | TRUE | 0,0306293 | No |
| orf19.1589 | 1835 | 1628 | 1789 | 1134 | 2713 | 2108 | 2258 | 2935 | 1671 | 1647 | 1693 | 1703 | 2413 | 2289 | 2312 | 2239 | 1995,87 | 1678 | 2313 | 1,379 | 0,463 | 10,761 | 5,27E-27 | 1,58E-26 | 1,00E-08 | 0,002699 | 0,0012583 | 0,0012583 | TRUE | 0,0105368 | No |
| orf19.1589.1 | 172 | 139 | 155 | 75 | 343 | 219 | 342 | 443 | 157 | 141 | 147 | 113 | 305 | 238 | 350 | 338 | 223,45 | 140 | 308 | 2,197 | 1,135 | 8,342 | 7,31E-17 | 1,70E-16 | 0,0186461 | 0,007949 | 0,0124567 | 0,0124567 | TRUE | 0,2544495 | No |
| orf19.1590 | 695 | 744 | 813 | 452 | 652 | 545 | 611 | 731 | 633 | 752 | 769 | 679 | 580 | 592 | 626 | 558 | 648,58 | 708 | 589 | 0,829 | -0,27 | -3,535 | 0,000408 | 0,00059 | 0,0040176 | 0,004074 | 0,0040461 | 0,0040461 | TRUE | 0,0669873 | No |
| orf19.1591 | 1521 | 1327 | 1370 | 1009 | 4693 | 3962 | 3950 | 5868 | 1385 | 1342 | 1297 | 1516 | 4174 | 4302 | 4045 | 4476 | 2817,07 | 1385 | 4249 | 3,078 | 1,622 | 29,35 | 2,42E-189 | 3,91E-188 | 0,0026215 | 0,002506 | 0,0024323 | 0,0024323 | TRUE | 0,044299 | No |
| orf19.1592 | 1555 | 1505 | 1551 | 1165 | 2118 | 1756 | 1673 | 2190 | 1416 | 1522 | 1468 | 1750 | 1884 | 1907 | 1713 | 1671 | 1666,27 | 1539 | 1794 | 1,168 | 0,224 | 3,115 | 0,001843 | 0,002548 | 0,0059744 | 0,00283 | 0,0043465 | 0,0043465 | TRUE | 0,0974414 | No |
| orf19.1595 | 659 | 778 | 778 | 447 | 745 | 679 | 612 | 843 | 600 | 787 | 736 | 671 | 663 | 737 | 627 | 643 | 683,04 | 698 | 668 | 0,954 | -0,068 | -0,77 | 0,441072 | 0,482931 | 0,0080297 | 0,003971 | 0,0059413 | 0,0059413 | TRUE | 0,113932 | No |
| orf19.1596 | 447 | 419 | 491 | 318 | 580 | 522 | 473 | 770 | 407 | 424 | 465 | 478 | 516 | 567 | 484 | 587 | 490,94 | 444 | 538 | 1,22 | 0,287 | 3,411 | 0,000648 | 0,000926 | 0,004723 | 0,004728 | 0,004727 | 0,004727 | TRUE | 0,0525555 | No |
| orf19.1597 | 1744 | 1683 | 1667 | 1141 | 2231 | 1883 | 1927 | 2700 | 1588 | 1702 | 1578 | 1714 | 1984 | 2044 | 1973 | 2060 | 1830,44 | 1646 | 2015 | 1,227 | 0,295 | 6,334 | 2,39E-10 | 4,57E-10 | 0,0004644 | 0,002759 | 0,0015305 | 0,0015305 | TRUE | 0,0090916 | No |
| orf19.1598 | 649 | 489 | 570 | 408 | 529 | 421 | 478 | 616 | 591 | 495 | 539 | 613 | 471 | 457 | 490 | 470 | 515,6 | 560 | 472 | 0,846 | -0,242 | -3,119 | 0,001816 | 0,002514 | 0,0028539 | 0,004599 | 0,0038085 | 0,0038085 | TRUE | 0,0705916 | No |
| orf19.1599 | 13 | 15 | 8 | 5 | 14 | 11 | 17 | 19 | 12 | 15 | 8 | 8 | 12 | 12 | 17 | 14 | 12,3 | 11 | 14 | 1,318 | 0,398 | 1,002 | 0,3165 | 0,35554 | 1,00E-08 | 0,109448 | 0,0686685 | 0,0686685 | TRUE | 0,306264 | No |
| orf19.1600 | 1483 | 1348 | 1386 | 983 | 1843 | 1615 | 1839 | 2448 | 1350 | 1363 | 1312 | 1477 | 1639 | 1753 | 1883 | 1867 | 1580,67 | 1376 | 1786 | 1,301 | 0,38 | 6,351 | 2,15E-10 | 4,12E-10 | 0,002492 | 0,002873 | 0,0027784 | 0,0027784 | TRUE | 0,0375533 | No |
| orf19.1601 | 27325 | 20869 | 22874 | 15894 | 25104 | 19798 | 21062 | 30214 | 24878 | 21107 | 21648 | 23875 | 22329 | 21495 | 21570 | 23047 | 22493,8 | 22877 | 22110 | 0,967 | -0,049 | -0,876 | 0,380921 | 0,421299 | 0,0034608 | 0,002096 | 0,0029618 | 0,0029618 | TRUE | 0,0427147 | No |
| orf19.1604 | 2610 | 2321 | 2521 | 1608 | 2724 | 2410 | 2546 | 3216 | 2376 | 2347 | 2386 | 2415 | 2423 | 2617 | 2607 | 2453 | 2453,15 | 2381 | 2525 | 1,06 | 0,084 | 1,963 | 0,049616 | 0,061478 | 0,0005201 | 0,002576 | 0,0013407 | 0,0013407 | TRUE | 0,0087978 | No |
| orf19.1605 | 90 | 81 | 90 | 58 | 113 | 82 | 102 | 127 | 82 | 82 | 85 | 87 | 101 | 89 | 104 | 97 | 90,88 | 84 | 98 | 1,167 | 0,222 | 1,554 | 0,120251 | 0,143183 | 1,00E-08 | 0,016573 | 0,0085399 | 0,0085399 | TRUE | 0,0305422 | No |
| orf19.1606 | 1299 | 1369 | 1365 | 817 | 1151 | 873 | 1136 | 1265 | 1183 | 1385 | 1292 | 1227 | 1024 | 948 | 1163 | 965 | 1148,3 | 1272 | 1025 | 0,805 | -0,312 | -4,163 | 3,14E-05 | 4,85E-05 | 0,005757 | 0,003187 | 0,0045231 | 0,0045231 | TRUE | 0,0971671 | No |
| orf19.1607 | 11431 | 11329 | 12154 | 6959 | 7856 | 6525 | 6482 | 8845 | 10407 | 11458 | 11503 | 10454 | 6988 | 7084 | 6638 | 6747 | 8909,89 | 10956 | 6864 | 0,626 | -0,675 | -14,373 | 7,66E-47 | 3,36E-46 | 0,0018298 | 0,002185 | 0,0020003 | 0,0020003 | TRUE | 0,0141719 | No |
| orf19.1608 | 750 | 712 | 749 | 434 | 1296 | 1106 | 1072 | 1416 | 683 | 720 | 709 | 652 | 1153 | 1201 | 1098 | 1080 | 911,91 | 691 | 1133 | 1,634 | 0,709 | 11,867 | 1,76E-32 | 5,98E-32 | 0,0010021 | 0,003486 | 0,0022377 | 0,0022377 | TRUE | 0,0188249 | No |
| orf19.1609 | 10109 | 9374 | 10243 | 6100 | 13252 | 10836 | 11699 | 15089 | 9204 | 9481 | 9694 | 9163 | 11787 | 11765 | 11981 | 11510 | 10573,2 | 9386 | 11761 | 1,253 | 0,325 | 9,94 | 2,78E-23 | 7,65E-23 | 0,0003608 | 0,002162 | 0,0009288 | 0,0009288 | TRUE | 0,0060262 | No |
| orf19.1610 | 116 | 97 | 109 | 55 | 69 | 62 | 62 | 75 | 106 | 98 | 103 | 83 | 61 | 67 | 63 | 57 | 79,86 | 98 | 62 | 0,632 | -0,663 | -4,218 | 2,46E-05 | 3,83E-05 | 1,00E-08 | 0,018578 | 0,0107822 | 0,0107822 | TRUE | 0,0757992 | No |
| orf19.1611 | 328 | 358 | 337 | 215 | 146 | 131 | 120 | 141 | 299 | 362 | 319 | 323 | 130 | 142 | 123 | 108 | 225,64 | 326 | 126 | 0,384 | -1,382 | -12,352 | 4,73E-35 | 1,69E-34 | 0,0047434 | 0,007891 | 0,006645 | 0,006645 | TRUE | 0,1102604 | No |
| orf19.1613 | 5363 | 4999 | 5488 | 3636 | 12492 | 11531 | 11180 | 15029 | 4883 | 5056 | 5194 | 5462 | 11111 | 12519 | 11450 | 11464 | 8392,39 | 5149 | 11636 | 2,261 | 1,177 | 23,519 | 2,61E-122 | 2,70E-121 | 0,00232 | 0,002194 | 0,0022609 | 0,0022609 | TRUE | 0,0317753 | No |
| orf19.1614 | 13144 | 12209 | 13313 | 7459 | 12423 | 9579 | 10889 | 14383 | 11967 | 12348 | 12600 | 11205 | 11050 | 10400 | 11152 | 10971 | 11461,6 | 12030 | 10893 | 0,905 | -0,144 | -3,201 | 0,001372 | 0,001919 | 0,0016411 | 0,002152 | 0,0018464 | 0,0018464 | TRUE | 0,025579 | No |
| orf19.1616 | 2455 | 2233 | 2377 | 1584 | 1319 | 1326 | 1273 | 1777 | 2235 | 2258 | 2250 | 2379 | 1173 | 1440 | 1304 | 1356 | 1799,34 | 2280 | 1318 | 0,578 | -0,791 | -12,909 | 4,01E-38 | 1,51E-37 | 0,0032135 | 0,002771 | 0,0030133 | 0,0030133 | TRUE | 0,0663512 | No |
| orf19.1617 | 497 | 527 | 521 | 296 | 634 | 497 | 525 | 677 | 452 | 533 | 493 | 445 | 564 | 540 | 538 | 516 | 510,1 | 481 | 540 | 1,118 | 0,162 | 2,153 | 0,031286 | 0,0395 | 0,0019183 | 0,004627 | 0,0034244 | 0,0034244 | TRUE | 0,061337 | No |
| orf19.1618 | 8029 | 8034 | 8233 | 4951 | 6646 | 5581 | 5564 | 7574 | 7310 | 8126 | 7792 | 7437 | 5911 | 6059 | 5698 | 5778 | 6763,89 | 7666 | 5862 | 0,764 | -0,388 | -8,866 | 7,61E-19 | 1,87E-18 | 0,0013164 | 0,002232 | 0,0016858 | 0,0016858 | TRUE | 0,0199339 | No |
| orf19.1618.1 | 528 | 543 | 515 | 343 | 556 | 385 | 402 | 552 | 481 | 549 | 487 | 515 | 495 | 418 | 412 | 421 | 472,23 | 508 | 436 | 0,86 | -0,217 | -2,642 | 0,008251 | 0,010883 | 0,0037783 | 0,004834 | 0,0043421 | 0,0043421 | TRUE | 0,0951539 | No |
| orf19.1619 | 4620 | 4301 | 4516 | 2801 | 3308 | 2782 | 2908 | 3802 | 4206 | 4350 | 4274 | 4208 | 2942 | 3020 | 2978 | 2900 | 3609,88 | 4260 | 2960 | 0,694 | -0,526 | -15,133 | 9,75E-52 | 4,65E-51 | 1,00E-08 | 0,002403 | 0,0008759 | 0,0008759 | TRUE | 0,0025003 | No |
| orf19.1620 | 7 | 13 | 13 | 6 | 17 | 6 | 13 | 11 | 6 | 13 | 12 | 9 | 15 | 7 | 13 | 8 | 10,52 | 10 | 11 | 1,06 | 0,084 | 0,19 | 0,849532 | 0,871403 | 1,00E-08 | 0,127581 | 0,0946166 | 0,0946166 | TRUE | 0,1870203 | No |
| orf19.1621 | 745 | 689 | 725 | 483 | 955 | 851 | 868 | 1105 | 678 | 697 | 686 | 726 | 849 | 924 | 889 | 843 | 786,51 | 697 | 876 | 1,259 | 0,332 | 5,659 | 1,52E-08 | 2,72E-08 | 1,00E-08 | 0,003717 | 0,0019966 | 0,0019966 | TRUE | 0,0154875 | No |
| orf19.1622 | 2920 | 2744 | 3050 | 1790 | 2184 | 1825 | 1989 | 2474 | 2659 | 2775 | 2887 | 2689 | 1943 | 1981 | 2037 | 1887 | 2357,18 | 2752 | 1962 | 0,712 | -0,49 | -10,77 | 4,76E-27 | 1,43E-26 | 0,0008404 | 0,002597 | 0,0015541 | 0,0015541 | TRUE | 0,0134002 | No |
| orf19.1623 | 10715 | 9628 | 10333 | 6261 | 7124 | 5568 | 6036 | 8156 | 9756 | 9738 | 9779 | 9405 | 6337 | 6045 | 6182 | 6221 | 7932,81 | 9670 | 6196 | 0,641 | -0,642 | -20,293 | 1,49E-91 | 1,18E-90 | 0,0001792 | 0,002204 | 0,0008308 | 0,0008308 | TRUE | 0,0039541 | No |
| orf19.1624 | 1069 | 1218 | 1133 | 703 | 2260 | 1881 | 1936 | 2413 | 973 | 1232 | 1072 | 1056 | 2010 | 2042 | 1983 | 1841 | 1526,16 | 1083 | 1969 | 1,817 | 0,861 | 12,294 | 9,79E-35 | 3,48E-34 | 0,0050424 | 0,002903 | 0,0039775 | 0,0039775 | TRUE | 0,1042935 | No |
| orf19.1624.1 | 28 | 37 | 25 | 25 | 31 | 25 | 23 | 23 | 25 | 37 | 24 | 38 | 28 | 27 | 24 | 18 | 27,49 | 31 | 24 | 0,776 | -0,365 | -1,302 | 0,192762 | 0,22355 | 1,00E-08 | 0,050085 | 0,038413 | 0,038413 | TRUE | 0,2101793 | No |
| orf19.1625 | 2435 | 2137 | 2200 | 1443 | 3363 | 2509 | 2802 | 3684 | 2217 | 2161 | 2082 | 2168 | 2991 | 2724 | 2870 | 2810 | 2502,9 | 2157 | 2849 | 1,321 | 0,402 | 9,009 | 2,07E-19 | 5,18E-19 | 0,000786 | 0,002565 | 0,0014968 | 0,0014968 | TRUE | 0,0138484 | No |

| | | | | | | | | | | | | | | | | | | | | | | | | | | | | | | | |
|---|---|---|---|---|---|---|---|---|---|---|---|---|---|---|---|---|---|---|---|---|---|---|---|---|---|---|---|---|---|---|---|
| orf19.1626 | 2469 | 2107 | 2412 | 1416 | 4022 | 3175 | 3503 | 4717 | 2248 | 2131 | 2283 | 2127 | 3577 | 3447 | 3588 | 3598 | 2874,88 | 2197 | 3552 | 1,615 | 0,692 | 16,808 | 2,14E-63 | 1,23E-62 | 0,000365 | 0,002497 | 0,0012507 | 0,0012507 | TRUE | 0,0082524 | No |
| orf19.1628 | 1150 | 1005 | 1053 | 631 | 3894 | 3259 | 3208 | 4473 | 1047 | 1016 | 997 | 948 | 3464 | 3538 | 3285 | 3412 | 2213,41 | 1002 | 3425 | 3,409 | 1,769 | 37,111 | 1,90E-301 | 4,94E-300 | 0,0004857 | 0,002634 | 0,0015132 | 0,0015132 | TRUE | 0,01339 | No |
| orf19.163 | 2298 | 2164 | 2302 | 1443 | 1809 | 1549 | 1542 | 2081 | 2092 | 2189 | 2179 | 2168 | 1609 | 1682 | 1579 | 1587 | 1885,57 | 2157 | 1614 | 0,748 | -0,418 | -9,755 | 1,75E-22 | 4,70E-22 | 1,00E-08 | 0,002738 | 0,0012274 | 0,0012274 | TRUE | 0,0090492 | No |
| orf19.1630 | 3281 | 2978 | 3368 | 2246 | 2103 | 1727 | 1870 | 2502 | 2987 | 3012 | 3188 | 3374 | 1871 | 1875 | 1915 | 1909 | 2516,22 | 3140 | 1892 | 0,604 | -0,729 | -15,056 | 3,13E-51 | 1,48E-50 | 0,0011421 | 0,002562 | 0,0018291 | 0,0018291 | TRUE | 0,0287363 | No |
| orf19.1631 | 2288 | 2192 | 2248 | 1314 | 8037 | 6630 | 6773 | 8900 | 2083 | 2217 | 2128 | 1974 | 7149 | 7198 | 6936 | 6789 | 4559,23 | 2100 | 7018 | 3,336 | 1,738 | 39,509 | 0 | 0 | 0,001113 | 0,002327 | 0,0015356 | 0,0015356 | TRUE | 0,0178664 | No |
| orf19.1632 | 404 | 419 | 435 | 241 | 1084 | 947 | 1008 | 1183 | 368 | 424 | 412 | 362 | 964 | 1028 | 1032 | 902 | 686,55 | 392 | 982 | 2,498 | 1,321 | 17,522 | 9,72E-69 | 5,98E-68 | 0,0032241 | 0,003961 | 0,0036151 | 0,0036151 | TRUE | 0,0367948 | No |
| orf19.1633 | 10624 | 10652 | 11253 | 6640 | 13770 | 11501 | 12177 | 15944 | 9673 | 10773 | 10650 | 9974 | 12248 | 12487 | 12471 | 12162 | 11304,8 | 10268 | 12342 | 1,202 | 0,265 | 6,289 | 3,20E-10 | 6,10E-10 | 0,0012982 | 0,002154 | 0,0016174 | 0,0016174 | TRUE | 0,0190159 | No |
| orf19.1634 | 3381 | 3035 | 3201 | 1969 | 4180 | 3258 | 3442 | 4699 | 3078 | 3070 | 3029 | 2958 | 3718 | 3537 | 3525 | 3584 | 3312,47 | 3034 | 3591 | 1,183 | 0,243 | 6,544 | 5,99E-11 | 1,17E-10 | 0,0001689 | 0,002436 | 0,0010128 | 0,0010128 | TRUE | 0,0068905 | No |
| orf19.1635 | 17038 | 14160 | 14512 | 10374 | 28519 | 22798 | 22238 | 31701 | 15512 | 14321 | 13734 | 15583 | 25367 | 24752 | 22775 | 24182 | 19528,3 | 14788 | 24269 | 1,641 | 0,715 | 13,639 | 2,35E-42 | 9,58E-42 | 0,002863 | 0,002105 | 0,0025842 | 0,0025842 | TRUE | 0,0282064 | No |
| orf19.1636 | 1930 | 1569 | 1694 | 1150 | 2370 | 1966 | 2197 | 2981 | 1757 | 1587 | 1603 | 1727 | 2108 | 2135 | 2250 | 2274 | 1930,16 | 1668 | 2192 | 1,315 | 0,395 | 7,628 | 2,39E-14 | 5,19E-14 | 0,001338 | 0,002721 | 0,0020359 | 0,0020359 | TRUE | 0,016934 | No |
| orf19.1637 | 627 | 475 | 532 | 317 | 770 | 584 | 663 | 894 | 571 | 480 | 503 | 476 | 685 | 634 | 679 | 682 | 588,86 | 508 | 670 | 1,316 | 0,396 | 5,44 | 5,33E-08 | 9,31E-08 | 0,0020507 | 0,00428 | 0,0033424 | 0,0033424 | TRUE | 0,0852581 | No |
| orf19.1638 | 729 | 664 | 761 | 493 | 872 | 731 | 790 | 1084 | 664 | 672 | 720 | 741 | 776 | 794 | 809 | 827 | 750,16 | 699 | 802 | 1,15 | 0,202 | 3,34 | 0,000837 | 0,001188 | 1,02E-05 | 0,003798 | 0,0021541 | 0,0021541 | TRUE | 0,0160431 | No |
| orf19.1639 | 104 | 66 | 82 | 46 | 162 | 141 | 134 | 185 | 95 | 67 | 78 | 69 | 144 | 153 | 137 | 141 | 110,46 | 77 | 144 | 1,848 | 0,886 | 6,195 | 5,84E-10 | 1,10E-09 | 1,00E-08 | 0,013996 | 0,0095757 | 0,0095757 | TRUE | 0,2370731 | No |
| orf19.164 | 1170 | 1198 | 1195 | 734 | 2013 | 1783 | 1570 | 2230 | 1065 | 1212 | 1131 | 1103 | 1790 | 1936 | 1608 | 1701 | 1443,21 | 1128 | 1759 | 1,559 | 0,64 | 9,621 | 6,54E-22 | 1,73E-21 | 0,0036987 | 0,002952 | 0,0035112 | 0,0035112 | TRUE | 0,0540762 | No |
| orf19.1641 | 38 | 37 | 50 | 32 | 203 | 147 | 157 | 221 | 35 | 37 | 47 | 48 | 181 | 160 | 161 | 169 | 104,62 | 42 | 168 | 4,054 | 2,019 | 12,577 | 2,81E-36 | 1,03E-35 | 1,00E-08 | 0,014664 | 0,0091711 | 0,0091711 | TRUE | 0,1154556 | No |
| orf19.1642 | 7872 | 6899 | 7351 | 4468 | 8225 | 6157 | 7126 | 9390 | 7167 | 6978 | 6957 | 6712 | 7316 | 6685 | 7298 | 7163 | 7034,35 | 6954 | 7116 | 1,023 | 0,033 | 0,798 | 0,425051 | 0,466306 | 0,0010312 | 0,002225 | 0,0014891 | 0,0014891 | TRUE | 0,0199668 | No |
| orf19.1643 | 2767 | 2427 | 2667 | 1841 | 1268 | 1072 | 1090 | 1525 | 2519 | 2455 | 2524 | 2765 | 1128 | 1164 | 1116 | 1163 | 1854,34 | 2566 | 1143 | 0,446 | -1,164 | -23,054 | 1,35E-117 | 1,36E-116 | 0,001115 | 0,002749 | 0,0018297 | 0,0018297 | TRUE | 0,0309501 | No |
| orf19.1646 | 12545 | 10706 | 12504 | 7437 | 13233 | 10746 | 12342 | 16128 | 11422 | 10828 | 11834 | 11172 | 11770 | 11667 | 12640 | 12303 | 11704,4 | 11314 | 12095 | 1,069 | 0,096 | 2,283 | 0,022401 | 0,02859 | 0,0013028 | 0,00215 | 0,0016181 | 0,0016181 | TRUE | 0,011793 | No |
| orf19.1647 | 31 | 33 | 27 | 21 | 34 | 32 | 31 | 45 | 28 | 33 | 26 | 32 | 30 | 35 | 32 | 34 | 31,22 | 30 | 33 | 1,111 | 0,151 | 0,627 | 0,53036 | 0,571993 | 1,00E-08 | 0,04435 | 0,023627 | 0,023627 | TRUE | 0,0621416 | No |
| orf19.1648 | 443 | 420 | 453 | 268 | 323 | 275 | 256 | 369 | 403 | 425 | 429 | 403 | 287 | 299 | 262 | 281 | 348,62 | 415 | 282 | 0,68 | -0,557 | -7,035 | 2,00E-12 | 4,11E-12 | 1,00E-08 | 0,005826 | 0,0030878 | 0,0030878 | TRUE | 0,0238744 | No |
| orf19.1649 | 3494 | 3137 | 3161 | 2089 | 3572 | 3032 | 3127 | 4375 | 3181 | 3173 | 2992 | 3138 | 3177 | 3292 | 3202 | 3337 | 3186,54 | 3121 | 3252 | 1,042 | 0,06 | 1,523 | 0,127814 | 0,151678 | 0,000406 | 0,002452 | 0,0011724 | 0,0011724 | TRUE | 0,0098509 | No |
| orf19.1650 | 995 | 738 | 710 | 482 | 1605 | 1145 | 1157 | 1837 | 906 | 746 | 672 | 724 | 1428 | 1243 | 1185 | 1401 | 1038,16 | 762 | 1314 | 1,724 | 0,785 | 8,464 | 2,58E-17 | 6,11E-17 | 0,0114438 | 0,003309 | 0,0072068 | 0,0072068 | TRUE | 0,1976256 | No |
| orf19.1651 | 14 | 22 | 16 | 9 | 49 | 35 | 46 | 63 | 13 | 22 | 15 | 14 | 44 | 38 | 47 | 48 | 30,05 | 16 | 44 | 2,771 | 1,47 | 5,348 | 8,89E-08 | 1,54E-07 | 1,00E-08 | 0,045995 | 0,028956 | 0,028956 | TRUE | 0,3473984 | No |
| orf19.1652 | 11107 | 9726 | 10252 | 7484 | 1E+05 | 99219 | 95980 | 1E+05 | 10112 | 9837 | 9703 | 11242 | 1E+05 | 1E+05 | 98296 | 1E+05 | 56985,6 | 10224 | 103748 | 10,153 | 3,344 | 62,941 | 0 | 0 | 0,0031932 | 0,00206 | 0,0026541 | 0,0026541 | TRUE | 0,0542029 | No |
| orf19.1653 | 2277 | 2200 | 2223 | 1227 | 1529 | 1266 | 1406 | 1736 | 2073 | 2225 | 2104 | 1843 | 1360 | 1375 | 1440 | 1324 | 1717,98 | 2061 | 1375 | 0,665 | -0,588 | -9,726 | 2,33E-22 | 6,24E-22 | 0,0028057 | 0,002806 | 0,002907 | 0,002907 | TRUE | 0,0573513 | No |
| orf19.1654 | 1520 | 1294 | 1379 | 1024 | 2769 | 2211 | 2316 | 3078 | 1384 | 1309 | 1305 | 1538 | 2463 | 2401 | 2372 | 2348 | 1889,9 | 1384 | 2396 | 1,736 | 0,796 | 14,34 | 1,23E-46 | 5,36E-46 | 0,0020168 | 0,002736 | 0,0023744 | 0,0023744 | TRUE | 0,0614533 | No |
| orf19.1655 | 2404 | 2002 | 2213 | 1558 | 24385 | 20873 | 20954 | 30277 | 2189 | 2025 | 2094 | 2340 | 21690 | 22662 | 21460 | 23096 | 12194,4 | 2162 | 22227 | 10,299 | 3,364 | 68,288 | 0 | 0 | 0,0022699 | 0,002145 | 0,0020605 | 0,0020605 | TRUE | 0,0344876 | No |
| orf19.1655.3 | 3569 | 3178 | 3256 | 2046 | 2515 | 1819 | 2055 | 2770 | 3249 | 3214 | 3082 | 3073 | 2237 | 1975 | 2105 | 2113 | 2631,01 | 3154 | 2108 | 0,668 | -0,582 | -12,233 | 2,08E-34 | 7,32E-34 | 0,0011728 | 0,002539 | 0,0017795 | 0,0017795 | TRUE | 0,0209916 | No |
| orf19.1656 | 1069 | 1042 | 1055 | 714 | 2153 | 1821 | 1869 | 2495 | 973 | 1054 | 998 | 1073 | 1915 | 1977 | 1914 | 1903 | 1475,95 | 1024 | 1927 | 1,885 | 0,915 | 18,844 | 3,27E-79 | 2,30E-78 | 1,74E-05 | 0,002932 | 0,0014972 | 0,0014972 | TRUE | 0,0136849 | No |
| orf19.1658 | 11833 | 10175 | 11442 | 7319 | 12439 | 9747 | 10618 | 14701 | 10773 | 10291 | 10829 | 10994 | 11064 | 10583 | 10874 | 11214 | 10827,8 | 10722 | 10934 | 1,02 | 0,029 | 0,814 | 0,415476 | 0,456823 | 0,0005843 | 0,002159 | 0,0011096 | 0,0011096 | TRUE | 0,0089518 | No |
| orf19.1659 | 661 | 751 | 795 | 477 | 939 | 847 | 835 | 1176 | 602 | 760 | 752 | 717 | 835 | 920 | 855 | 897 | 792,16 | 708 | 877 | 1,24 | 0,311 | 4,007 | 6,14E-05 | 9,31E-05 | 0,0052745 | 0,003705 | 0,0044722 | 0,0044722 | TRUE | 0,1272996 | No |
| orf19.166 | 2737 | 3004 | 3160 | 1966 | 2461 | 2212 | 2070 | 2907 | 2492 | 3038 | 2991 | 2953 | 2189 | 2402 | 2120 | 2217 | 2550,26 | 2868 | 2232 | 0,778 | -0,362 | -5,342 | 9,22E-08 | 1,59E-07 | 0,004824 | 0,002555 | 0,0040028 | 0,0040028 | TRUE | 0,0966538 | No |
| orf19.1661 | 5015 | 4533 | 4883 | 2904 | 6419 | 5435 | 5623 | 7538 | 4566 | 4585 | 4621 | 4362 | 5709 | 5901 | 5759 | 5750 | 5156,66 | 4534 | 5780 | 1,273 | 0,349 | 10,286 | 8,17E-25 | 2,33E-24 | 0,0001721 | 0,002293 | 0,0009044 | 0,0009044 | TRUE | 0,0072592 | No |
| orf19.1662 | 3147 | 2860 | 3085 | 1904 | 2532 | 1944 | 2050 | 2752 | 2865 | 2893 | 2920 | 2860 | 2252 | 2111 | 2099 | 2099 | 2512,38 | 2884 | 2140 | 0,742 | -0,43 | -10,617 | 2,50E-26 | 7,34E-26 | 0,0002233 | 0,002563 | 0,0011738 | 0,0011738 | TRUE | 0,0152204 | No |
| orf19.1663 | 2194 | 2211 | 2411 | 1324 | 3433 | 2908 | 2961 | 3921 | 1998 | 2236 | 2282 | 1989 | 3054 | 3157 | 3032 | 2991 | 2592,33 | 2126 | 3058 | 1,437 | 0,523 | 9,625 | 6,30E-22 | 1,66E-21 | 0,0023805 | 0,002547 | 0,0024255 | 0,0024255 | TRUE | 0,0295687 | No |
| orf19.1664 | 363 | 326 | 298 | 238 | 393 | 297 | 335 | 429 | 330 | 330 | 282 | 358 | 350 | 322 | 343 | 327 | 330,26 | 325 | 336 | 1,038 | 0,054 | 0,614 | 0,539462 | 0,580826 | 0,0019639 | 0,006037 | 0,0043856 | 0,0043856 | TRUE | 0,0920437 | No |
| orf19.1665 | 9438 | 8913 | 9468 | 6098 | 14040 | 12170 | 12624 | 16221 | 8593 | 9015 | 8961 | 9160 | 12488 | 13213 | 12929 | 12374 | 10841,5 | 8932 | 12751 | 1,428 | 0,514 | 13,937 | 3,76E-44 | 1,58E-43 | 0,0007063 | 0,002159 | 0,0012078 | 0,0012078 | TRUE | 0,0080549 | No |
| orf19.1666 | 1296 | 1056 | 1256 | 859 | 1094 | 921 | 1042 | 1427 | 1180 | 1068 | 1189 | 1290 | 973 | 1000 | 1067 | 1089 | 1106,97 | 1182 | 1032 | 0,876 | -0,191 | -2,895 | 0,00379 | 0,005115 | 0,0031439 | 0,00323 | 0,0032906 | 0,0032906 | TRUE | 0,0495559 | No |
| orf19.1667 | 1016 | 989 | 1066 | 598 | 1109 | 1026 | 1025 | 1302 | 925 | 1000 | 1009 | 898 | 986 | 1114 | 1050 | 993 | 996,97 | 958 | 1036 | 1,078 | 0,108 | 1,721 | 0,085272 | 0,103073 | 0,0019394 | 0,003362 | 0,0027508 | 0,0027508 | TRUE | 0,0300922 | No |
| orf19.1667.1 | 599 | 476 | 474 | 244 | 524 | 344 | 435 | 525 | 545 | 481 | 449 | 367 | 466 | 373 | 445 | 400 | 440,93 | 460 | 421 | 0,911 | -0,135 | -1,206 | 0,227882 | 0,261474 | 0,0150121 | 0,005033 | 0,0097048 | 0,0097048 | TRUE | 0,1974611 | No |
| orf19.1668 | 849 | 814 | 861 | 534 | 565 | 493 | 455 | 599 | 773 | 823 | 815 | 802 | 503 | 535 | 466 | 457 | 646,75 | 803 | 490 | 0,609 | -0,716 | -10,635 | 2,06E-26 | 6,07E-26 | 0,0008748 | 0,00408 | 0,0027265 | 0,0027265 | TRUE | 0,042179 | No |
| orf19.1669 | 6266 | 5727 | 6336 | 3908 | 4211 | 3618 | 3914 | 5070 | 5705 | 5792 | 5996 | 5870 | 3746 | 3928 | 4008 | 3867 | 4864,2 | 5841 | 3887 | 0,665 | -0,588 | -15,988 | 1,55E-57 | 8,14E-57 | 0,0003449 | 0,002309 | 0,0010851 | 0,0010851 | TRUE | 0,0073391 | No |
| orf19.1670 | 1433 | 1482 | 1608 | 860 | 616 | 545 | 518 | 617 | 1305 | 1499 | 1522 | 1292 | 548 | 592 | 530 | 471 | 969,75 | 1404 | 535 | 0,38 | -1,397 | -17,031 | 4,84E-65 | 2,84E-64 | 0,0070884 | 0,003399 | 0,0051995 | 0,0051995 | TRUE | 0,0816683 | No |
| orf19.1671 | 8775 | 8378 | 8615 | 5593 | 5405 | 4591 | 4453 | 6295 | 7989 | 8473 | 8153 | 8402 | 4808 | 4985 | 4560 | 4802 | 6521,51 | 8254 | 4789 | 0,58 | -0,785 | -19,865 | 8,13E-88 | 6,28E-87 | 0,0008087 | 0,00224 | 0,0013375 | 0,0013375 | TRUE | 0,0122873 | No |
| orf19.1671.1 | 10 | 7 | 4 | 2 | 9 | 7 | 4 | 11 | 9 | 7 | 4 | 3 | 8 | 8 | 4 | 8 | 6,38 | 6 | 7 | 1,196 | 0,258 | 0,467 | 0,640814 | 0,678603 | 1,00E-08 | 0,208981 | 0,1380673 | 0,1380673 | TRUE | 0,3829502 | No |
| orf19.1672 | 3731 | 3582 | 3875 | 2278 | 5346 | 4545 | 4493 | 6030 | 3397 | 3623 | 3667 | 3422 | 4755 | 4935 | 4601 | 4600 | 4124,98 | 3527 | 4723 | 1,338 | 0,42 | 9,68 | 3,67E-22 | 9,78E-22 | 0,0009976 | 0,002357 | 0,001556 | 0,001556 | TRUE | 0,0109317 | No |
| orf19.1673 | 2705 | 2787 | 2733 | 1535 | 1759 | 1371 | 1465 | 1826 | 2463 | 2819 | 2587 | 2306 | 1565 | 1489 | 1500 | 1393 | 2015,03 | 2544 | 1487 | 0,584 | -0,777 | -11,916 | 9,76E-33 | 3,32E-32 | 0,0040477 | 0,002693 | 0,0035551 | 0,0035551 | TRUE | 0,0638655 | No |
| orf19.1674 | 2799 | 2830 | 2917 | 1787 | 2160 | 1849 | 1928 | 2414 | 2548 | 2862 | 2761 | 2684 | 1921 | 2007 | 1975 | 1841 | 2325,04 | 2714 | 1936 | 0,713 | -0,488 | -9,7 | 3,01E-22 | 8,04E-22 | 0,0015605 | 0,002605 | 0,0019939 | 0,0019939 | TRUE | 0,0214486 | No |
| orf19.1675 | 1765 | 1510 | 1707 | 989 | 2819 | 2347 | 2462 | 3280 | 1607 | 1527 | 1616 | 1486 | 2507 | 2548 | 2521 | 2502 | 2039,29 | 1559 | 2520 | 1,613 | 0,689 | 16,021 | 9,17E-58 | 4,84E-57 | 1,00E-08 | 0,002685 | 0,0012492 | 0,0012492 | TRUE | 0,0105462 | No |
| orf19.1676 | 1555 | 1546 | 1587 | 953 | 1496 | 1245 | 1211 | 1581 | 1416 | 1564 | 1502 | 1432 | 1331 | 1352 | 1240 | 1206 | 1380,18 | 1478 | 1282 | 0,866 | -0,207 | -3,641 | 0,000272 | 0,000398 | 0,0018725 | 0,002994 | 0,0023876 | 0,0023876 | TRUE | 0,0202388 | No |
| orf19.1677 | 2522 | 2048 | 2404 | 1529 | 3936 | 3423 | 3798 | 5194 | 2296 | 2071 | 2275 | 2297 | 3501 | 3716 | 3890 | 3962 | 3001,06 | 2235 | 3767 | 1,686 | 0,754 | 13,909 | 5,56E-44 | 2,33E-43 | 0,0022093 | 0,002477 | 0,0024563 | 0,0024563 | TRUE | 0,0297128 | No |
| orf19.1678 | 1147 | 972 | 1070 | 713 | 2102 | 1757 | 1862 | 2465 | 1044 | 983 | 1013 | 1071 | 1870 | 1908 | 1907 | 1880 | 1459,45 | 1028 | 1891 | 1,843 | 0,882 | 19,029 | 9,78E-81 | 6,98E-80 | 1,00E-08 | 0,002942 | 0,0012945 | 0,0012945 | TRUE | 0,0098675 | No |
| orf19.1679 | 64 | 39 | 51 | 37 | 480 | 394 | 435 | 542 | 58 | 39 | 48 | 56 | 427 | 428 | 445 | 413 | 239,4 | 50 | 428 | 8,534 | 3,093 | 23,768 | 7,21E-125 | 7,67E-124 | 1,00E-08 | 0,007555 | 0,0046528 | 0,0046528 | TRUE | 0,1789353 | No |
| orf19.168 | 6503 | 5851 | 6045 | 3561 | 4321 | 3233 | 3672 | 4753 | 5921 | 5918 | 5721 | 5349 | 3843 | 3510 | 3761 | 3626 | 4706,04 | 5727 | 3685 | 0,643 | -0,637 | -13,503 | 1,50E-41 | 6,01E-41 | 0,0016002 | 0,002318 | 0,0019164 | 0,0019164 | TRUE | 0,023203 | No |
| orf19.1680 | 2017 | 1821 | 1929 | 1198 | 2854 | 2348 | 2343 | 3276 | 1836 | 1842 | 1826 | 1800 | 2539 | 2549 | 2400 | 2499 | 2161,21 | 1826 | 2497 | 1,367 | 0,451 | 11,1 | 1,25E-28 | 3,87E-28 | 1,00E-08 | 0,002648 | 0,0011022 | 0,0011022 | TRUE | 0,0080261 | No |
| orf19.1681 | 241 | 264 | 267 | 191 | 413 | 311 | 337 | 436 | 219 | 267 | 253 | 287 | 367 | 338 | 345 | 333 | 301,09 | 256 | 346 | 1,357 | 0,441 | 4,699 | 2,62E-06 | 4,28E-06 | 0,0025988 | 0,006424 | 0,0050023 | 0,0050023 | TRUE | 0,1101508 | No |
| orf19.1682 | 5701 | 4614 | 5189 | 3426 | 8174 | 6721 | 7086 | 9630 | 5191 | 4667 | 4911 | 5146 | 7271 | 7297 | 7257 | 7346 | 6135,61 | 4979 | 7293 | 1,465 | 0,551 | 13,463 | 2,60E-41 | 1,03E-40 | 0,0009659 | 0,002252 | 0,0014378 | 0,0014378 | TRUE | 0,0217977 | No |
| orf19.1683 | 1890 | 1694 | 1851 | 1167 | 2692 | 2075 | 2281 | 3194 | 1721 | 1713 | 1752 | 1753 | 2394 | 2253 | 2336 | 2436 | 2044,83 | 1735 | 2355 | 1,359 | 0,443 | 10,315 | 6,03E-25 | 1,73E-24 | 1,00E-08 | 0,002683 | 0,0012619 | 0,0012619 | TRUE | 0,0100861 | No |
| orf19.1684 | 701 | 669 | 754 | 486 | 755 | 662 | 707 | 990 | 638 | 677 | 714 | 730 | 672 | 719 | 724 | 755 | 703,5 | 690 | 718 | 1,043 | 0,061 | 0,926 | 0,354586 | 0,394432 | 0,0016139 | 0,003915 | 0,0027955 | 0,0027955 | TRUE | 0,0307509 | No |
| orf19.1685 | 433 | 383 | 435 | 301 | 570 | 451 | 489 | 678 | 394 | 387 | 412 | 452 | 507 | 490 | 501 | 517 | 457,51 | 411 | 504 | 1,232 | 0,301 | 4,169 | 3,06E-05 | 4,73E-05 | 1,00E-08 | 0,004924 | 0,002782 | 0,002782 | TRUE | 0,0421543 | No |
| orf19.1686 | 680 | 586 | 735 | 387 | 494 | 438 | 451 | 618 | 619 | 593 | 696 | 581 | 439 | 476 | 462 | 471 | 542,12 | 622 | 462 | 0,74 | -0,434 | -5,815 | 6,07E-09 | 1,10E-08 | 0,002366 | 0,004474 | 0,0034721 | 0,0034721 | TRUE | 0,0756573 | No |
| orf19.1687 | 13647 | 12223 | 13389 | 8458 | 17469 | 14506 | 14950 | 20142 | 12425 | 12362 | 12671 | 12705 | 15538 | 15749 | 15311 | 15364 | 14015,9 | 12541 | 15490 | 1,235 | 0,305 | 11,158 | 6,56E-29 | 2,05E-28 | 8,02E-05 | 0,002131 | 0,000644 | 0,000644 | TRUE | 0,0015254 | No |
| orf19.169 | 1686 | 1865 | 1977 | 1049 | 4160 | 3457 | 3783 | 4538 | 1535 | 1886 | 1871 | 1576 | 3700 | 3753 | 3874 | 3462 | 2707,19 | 1717 | 3697 | 2,151 | 1,105 | 15,343 | 3,97E-53 | 1,94E-52 | 0,0064894 | 0,002525 | 0,0045399 | 0,0045399 | TRUE | 0,0647226 | No |
| orf19.1690 | 3664 | 3064 | 3341 | 2157 | 6531 | 5381 | 5748 | 8026 | 3336 | 3099 | 3162 | 3240 | 5809 | 5842 | 5887 | 6122 | 4562,16 | 3209 | 5915 | 1,844 | 0,883 | 22,697 | 4,82E-114 | 4,71E-113 | 0,000475 | 0,002327 | 0,0012052 | 0,0012052 | TRUE | 0,0091503 | No |
| orf19.1691 | 406 | 373 | 376 | 268 | 164 | 140 | 126 | 177 | 370 | 377 | 356 | 403 | 146 | 152 | 129 | 135 | 258,41 | 376 | 140 | 0,374 | -1,419 | -14,698 | 6,62E-49 | 2,99E-48 | 1,00E-08 | 0,007149 | 0,004211 | 0,004211 | TRUE | 0,0278902 | No |
| orf19.1693 | 815 | 737 | 821 | 488 | 1780 | 1656 | 1595 | 2146 | 742 | 745 | 777 | 733 | 1583 | 1798 | 1633 | 1637 | 1206,14 | 749 | 1663 | 2,215 | 1,147 | 20,001 | 5,38E-89 | 4,20E-88 | 0,0013612 | 0,003132 | 0,0021642 | 0,0021642 | TRUE | 0,0349466 | No |
| orf19.1697 | 3724 | 3316 | 3435 | 2043 | 3600 | 2612 | 3092 | 3845 | 3391 | 3354 | 3251 | 3069 | 3202 | 2836 | 3167 | 2933 | 3150,22 | 3266 | 3034 | 0,928 | -0,107 | -2,02 | 0,043408 | 0,054069 | 0,002274 | 0,002456 | 0,0023924 | 0,0023924 | TRUE | 0,0229263 | No |

| | | | | | | | | | | | | | | | | | | | | | | | | | | | | | | | |
|---|---|---|---|---|---|---|---|---|---|---|---|---|---|---|---|---|---|---|---|---|---|---|---|---|---|---|---|---|---|---|---|
| orf19.1698 | 764 | 671 | 776 | 508 | 1402 | 1140 | 1258 | 1677 | 696 | 679 | 734 | 763 | 1247 | 1238 | 1288 | 1279 | 990,51 | 718 | 1263 | 1,765 | 0,82 | 14,908 | 2,91E-50 | 1,35E-49 | 1,00E-08 | 0,003371 | 0,0017869 | 0,0017869 | TRUE | 0,0176432 | No |
| orf19.1699 | 870 | 882 | 891 | 563 | 1645 | 1411 | 1512 | 1805 | 792 | 892 | 843 | 846 | 1463 | 1532 | 1548 | 1377 | 1161,7 | 843 | 1480 | 1,754 | 0,811 | 13,523 | 1,14E-41 | 4,56E-41 | 0,001911 | 0,003174 | 0,0024987 | 0,0024987 | TRUE | 0,0275692 | No |
| orf19.17 | 738 | 850 | 827 | 469 | 871 | 739 | 697 | 933 | 672 | 860 | 783 | 705 | 775 | 802 | 714 | 712 | 752,67 | 755 | 751 | 0,993 | -0,01 | -0,126 | 0,899984 | 0,914327 | 0,0065786 | 0,003792 | 0,0051567 | 0,0051567 | TRUE | 0,1046657 | No |
| orf19.1700 | 10363 | 9784 | 9502 | 6754 | 19383 | 15961 | 14711 | 20532 | 9435 | 9895 | 8993 | 10146 | 17241 | 17329 | 15066 | 15662 | 12970,8 | 9617 | 16324 | 1,698 | 0,764 | 13,277 | 3,13E-40 | 1,22E-39 | 0,0037162 | 0,002139 | 0,0030933 | 0,0030933 | TRUE | 0,0328511 | No |
| orf19.1701 | 1628 | 1564 | 1584 | 1105 | 2788 | 2352 | 2391 | 2983 | 1482 | 1582 | 1499 | 1660 | 2480 | 2554 | 2449 | 2275 | 1997,58 | 1556 | 2440 | 1,57 | 0,651 | 11,965 | 5,43E-33 | 1,86E-32 | 0,0018512 | 0,002698 | 0,0023028 | 0,0023028 | TRUE | 0,025553 | No |
| orf19.1702 | 1072 | 969 | 942 | 687 | 1925 | 1548 | 1572 | 2254 | 976 | 980 | 892 | 1032 | 1712 | 1681 | 1610 | 1719 | 1325,22 | 970 | 1680 | 1,738 | 0,798 | 14,538 | 6,93E-48 | 3,08E-47 | 0,0011547 | 0,003034 | 0,002058 | 0,002058 | TRUE | 0,0357635 | No |
| orf19.1704 | 6422 | 5352 | 5903 | 3504 | 60387 | 47565 | 50334 | 70701 | 5847 | 5413 | 5587 | 5264 | 53712 | 51642 | 51548 | 53931 | 29118,1 | 5528 | 52708 | 9,528 | 3,252 | 80,384 | 0 | 0 | 0,0011931 | 0,002082 | 0,0014658 | 0,0014658 | TRUE | 0,0185318 | No |
| orf19.1706 | 324 | 414 | 398 | 245 | 411 | 364 | 349 | 452 | 295 | 419 | 377 | 368 | 366 | 395 | 357 | 345 | 365,17 | 365 | 366 | 1,003 | 0,005 | 0,045 | 0,963795 | 0,968199 | 0,0096192 | 0,005654 | 0,0074614 | 0,0074614 | TRUE | 0,2031663 | No |
| orf19.1707 | 20 | 48 | 34 | 12 | 27 | 41 | 43 | 39 | 18 | 49 | 32 | 18 | 24 | 45 | 44 | 30 | 32,41 | 29 | 36 | 1,196 | 0,258 | 0,776 | 0,437761 | 0,479588 | 0,1226638 | 0,042796 | 0,0747023 | 0,0747023 | TRUE | 0,9510321 | No |
| orf19.1708 | 3571 | 3682 | 3816 | 2282 | 2767 | 2148 | 2535 | 2824 | 3251 | 3724 | 3611 | 3428 | 2461 | 2332 | 2596 | 2154 | 2944,78 | 3504 | 2386 | 0,681 | -0,555 | -8,6 | 7,96E-18 | 1,91E-17 | 0,004407 | 0,002486 | 0,0036525 | 0,0036525 | TRUE | 0,0531054 | No |
| orf19.1709 | 1862 | 1714 | 1679 | 921 | 9256 | 6170 | 6941 | 9533 | 1695 | 1734 | 1589 | 1383 | 8233 | 6699 | 7108 | 7272 | 4464,18 | 1600 | 7328 | 4,571 | 2,193 | 28,5 | 1,17E-178 | 1,80E-177 | 0,0082311 | 0,002333 | 0,0052843 | 0,0052843 | TRUE | 0,0950902 | No |
| orf19.171 | 49630 | 46105 | 49665 | 30828 | 62626 | 51698 | 54516 | 74471 | 45186 | 46630 | 47003 | 46309 | 55704 | 56130 | 55831 | 56807 | 51200 | 46282 | 56118 | 1,213 | 0,278 | 10,819 | 2,81E-27 | 8,48E-27 | 0,0001557 | 0,002063 | 0,0006146 | 0,0006146 | TRUE | 0,0031426 | No |
| orf19.1710 | 1819 | 1910 | 1883 | 1190 | 3560 | 3084 | 2753 | 3660 | 1656 | 1932 | 1782 | 1788 | 3167 | 3348 | 2819 | 2792 | 2410,46 | 1790 | 3032 | 1,694 | 0,76 | 10,889 | 1,30E-27 | 3,97E-27 | 0,0053984 | 0,002585 | 0,0042268 | 0,0042268 | TRUE | 0,0595722 | No |
| orf19.1711 | 1458 | 1620 | 1653 | 803 | 957 | 707 | 829 | 985 | 1327 | 1638 | 1564 | 1206 | 851 | 768 | 849 | 751 | 1119,47 | 1434 | 805 | 0,56 | -0,836 | -9,075 | 1,14E-19 | 2,86E-19 | 0,0107442 | 0,003217 | 0,0071971 | 0,0071971 | TRUE | 0,1311301 | No |
| orf19.1712 | 929 | 1013 | 967 | 658 | 1702 | 1535 | 1517 | 1913 | 846 | 1025 | 915 | 988 | 1514 | 1667 | 1554 | 1459 | 1245,91 | 944 | 1548 | 1,643 | 0,716 | 10,41 | 2,23E-25 | 6,45E-25 | 0,0040154 | 0,003097 | 0,0036708 | 0,0036708 | TRUE | 0,0599571 | No |
| orf19.1714 | 96 | 125 | 114 | 111 | 54 | 52 | 55 | 51 | 87 | 126 | 108 | 167 | 48 | 56 | 56 | 39 | 86,02 | 122 | 50 | 0,41 | -1,285 | -6,173 | 6,70E-10 | 1,26E-09 | 0,0420204 | 0,017393 | 0,0277289 | 0,0277289 | TRUE | 0,2268522 | No |
| orf19.1715 | 740 | 715 | 814 | 515 | 556 | 453 | 451 | 623 | 674 | 723 | 770 | 774 | 495 | 492 | 462 | 475 | 608,05 | 735 | 481 | 0,656 | -0,609 | -8,898 | 5,67E-19 | 1,40E-18 | 0,0012898 | 0,00421 | 0,0028012 | 0,0028012 | TRUE | 0,0392383 | No |
| orf19.1716 | 2608 | 2200 | 2285 | 1392 | 3651 | 2601 | 2871 | 4091 | 2374 | 2225 | 2163 | 2091 | 3247 | 2824 | 2940 | 3121 | 2623,18 | 2213 | 3033 | 1,37 | 0,454 | 7,869 | 3,56E-15 | 7,91E-15 | 0,0028838 | 0,002541 | 0,0027963 | 0,0027963 | TRUE | 0,0294095 | No |
| orf19.1717 | 166 | 177 | 184 | 134 | 187 | 155 | 146 | 178 | 151 | 179 | 174 | 201 | 166 | 168 | 150 | 136 | 165,69 | 176 | 155 | 0,882 | -0,18 | -1,471 | 0,141189 | 0,166404 | 0,0058513 | 0,01001 | 0,0083529 | 0,0083529 | TRUE | 0,1026122 | No |
| orf19.1718 | 898 | 982 | 1004 | 584 | 770 | 653 | 662 | 797 | 818 | 993 | 950 | 877 | 685 | 709 | 678 | 608 | 789,75 | 910 | 670 | 0,735 | -0,444 | -5,83 | 5,56E-09 | 1,01E-08 | 0,0048442 | 0,00371 | 0,0042828 | 0,0042828 | TRUE | 0,0592258 | No |
| orf19.1719 | 2259 | 2591 | 2404 | 1616 | 2523 | 2277 | 2158 | 2867 | 2057 | 2621 | 2275 | 2427 | 2244 | 2472 | 2210 | 2187 | 2311,66 | 2345 | 2278 | 0,972 | -0,041 | -0,567 | 0,570527 | 0,611176 | 0,0057446 | 0,002608 | 0,0046296 | 0,0046296 | TRUE | 0,0844476 | No |
| orf19.172 | 4746 | 4901 | 4703 | 3027 | 5958 | 4800 | 4679 | 6240 | 4321 | 4957 | 4451 | 4547 | 5299 | 5211 | 4792 | 4760 | 4792,32 | 4569 | 5016 | 1,098 | 0,135 | 2,407 | 0,016099 | 0,020818 | 0,0030807 | 0,002313 | 0,0027889 | 0,0027889 | TRUE | 0,0402831 | No |
| orf19.1720 | 4388 | 4671 | 4879 | 2981 | 10492 | 9400 | 9308 | 11775 | 3995 | 4724 | 4618 | 4478 | 9332 | 10206 | 9533 | 8982 | 6983,44 | 4454 | 9513 | 2,136 | 1,095 | 18,415 | 1,00E-75 | 6,69E-75 | 0,0038567 | 0,002226 | 0,0032255 | 0,0032255 | TRUE | 0,0595757 | No |
| orf19.1721 | 21670 | 22389 | 22725 | 13009 | 37060 | 29095 | 30285 | 39118 | 19730 | 22644 | 21507 | 19542 | 32964 | 31589 | 31016 | 29840 | 26103,8 | 20856 | 31352 | 1,503 | 0,588 | 10,777 | 4,41E-27 | 1,32E-26 | 0,0033395 | 0,002088 | 0,0028186 | 0,0028186 | TRUE | 0,0407789 | No |
| orf19.1722 | 72 | 48 | 54 | 55 | 73 | 43 | 53 | 70 | 66 | 49 | 51 | 83 | 65 | 47 | 54 | 53 | 58,39 | 62 | 55 | 0,901 | -0,15 | -0,736 | 0,46174 | 0,503436 | 0,0195817 | 0,024661 | 0,0228328 | 0,0228328 | TRUE | 0,3329313 | No |
| orf19.1723 | 1457 | 1343 | 1434 | 965 | 2355 | 1797 | 1989 | 2609 | 1327 | 1358 | 1357 | 1450 | 2095 | 1951 | 2037 | 1990 | 1695,56 | 1373 | 2018 | 1,474 | 0,559 | 11,822 | 3,00E-32 | 1,01E-31 | 0,0004055 | 0,002816 | 0,0015273 | 0,0015273 | TRUE | 0,0149877 | No |
| orf19.1724 | 0 | 1 | 1 | 1 | 9 | 8 | 7 | 12 | 0 | 1 | 1 | 2 | 8 | 9 | 7 | 9 | 4,56 | 1 | 8 | 10,409 | 3,38 | 3,525 | 0,000424 | 0,000613 | 1,00E-08 | 0,291781 | 0,1641802 | 0,1641802 | TRUE | 0,2342945 | No |
| orf19.1725 | 1347 | 1075 | 1355 | 965 | 1606 | 1505 | 1688 | 2220 | 1226 | 1087 | 1282 | 1450 | 1428 | 1634 | 1729 | 1693 | 1441,28 | 1261 | 1621 | 1,288 | 0,365 | 4,279 | 1,88E-05 | 2,94E-05 | 0,0095508 | 0,002954 | 0,0062608 | 0,0062608 | TRUE | 0,1149257 | No |
| orf19.1727 | 14135 | 13832 | 14788 | 9072 | 7879 | 6802 | 6826 | 9352 | 12869 | 13990 | 13996 | 13628 | 7008 | 7385 | 6991 | 7134 | 10375 | 13621 | 7130 | 0,523 | -0,934 | -23,539 | 1,63E-122 | 1,69E-121 | 0,000939 | 0,002164 | 0,001407 | 0,001407 | TRUE | 0,0170927 | No |
| orf19.1728 | 509 | 500 | 543 | 328 | 729 | 587 | 594 | 825 | 463 | 506 | 514 | 493 | 648 | 637 | 608 | 629 | 562,39 | 494 | 630 | 1,278 | 0,354 | 5,41 | 6,31E-08 | 1,10E-07 | 1,00E-08 | 0,004386 | 0,002288 | 0,002288 | TRUE | 0,0227843 | No |
| orf19.1729 | 1054 | 1274 | 1251 | 700 | 986 | 797 | 829 | 1090 | 960 | 1289 | 1184 | 1052 | 877 | 865 | 849 | 831 | 988,3 | 1121 | 856 | 0,763 | -0,391 | -4,681 | 2,85E-06 | 4,65E-06 | 0,0078864 | 0,003374 | 0,0056601 | 0,0056601 | TRUE | 0,1223948 | No |
| orf19.173 | 24307 | 22932 | 25099 | 15373 | 7519 | 6429 | 7034 | 9267 | 22130 | 23193 | 23754 | 23093 | 6688 | 6980 | 7204 | 7069 | 15013,9 | 23042 | 6985 | 0,303 | -1,722 | -45,823 | 0 | 0 | 0,0007925 | 0,002125 | 0,0012661 | 0,0012661 | TRUE | 0,0100848 | No |
| orf19.1730 | 1097 | 920 | 1021 | 584 | 697 | 570 | 599 | 834 | 999 | 930 | 966 | 877 | 620 | 619 | 613 | 636 | 782,66 | 943 | 622 | 0,658 | -0,604 | -10,07 | 7,48E-24 | 2,09E-23 | 1,00E-08 | 0,003725 | 0,002138 | 0,002138 | TRUE | 0,0230128 | No |
| orf19.1734 | 2399 | 2498 | 2449 | 1590 | 3600 | 3277 | 3262 | 4335 | 2184 | 2526 | 2318 | 2388 | 3202 | 3558 | 3341 | 3307 | 2853,04 | 2354 | 3352 | 1,424 | 0,51 | 9,382 | 6,50E-21 | 1,68E-20 | 0,0023492 | 0,0025 | 0,0024692 | 0,0024692 | TRUE | 0,0304477 | No |
| orf19.1735 | 485 | 479 | 473 | 269 | 202 | 135 | 149 | 190 | 442 | 484 | 448 | 404 | 180 | 147 | 153 | 145 | 300,19 | 444 | 156 | 0,35 | -1,514 | -15,313 | 6,28E-53 | 3,05E-52 | 0,0032081 | 0,006438 | 0,005206 | 0,005206 | TRUE | 0,1163313 | No |
| orf19.1736 | 156 | 107 | 126 | 93 | 227 | 223 | 194 | 268 | 142 | 108 | 119 | 140 | 202 | 242 | 199 | 204 | 169,54 | 127 | 212 | 1,666 | 0,737 | 5,976 | 2,29E-09 | 4,23E-09 | 0,0054212 | 0,009829 | 0,008179 | 0,008179 | TRUE | 0,1042889 | No |
| orf19.1737 | 21 | 29 | 29 | 15 | 36 | 29 | 24 | 43 | 19 | 29 | 27 | 23 | 32 | 31 | 25 | 33 | 27,41 | 24 | 30 | 1,23 | 0,299 | 1,126 | 0,260134 | 0,295769 | 1,00E-08 | 0,050223 | 0,0306441 | 0,0306441 | TRUE | 0,1700264 | No |
| orf19.1738 | 8865 | 7398 | 8135 | 4875 | 10823 | 8430 | 8994 | 12726 | 8071 | 7482 | 7699 | 7323 | 9627 | 9153 | 9211 | 9707 | 8534,17 | 7644 | 9424 | 1,233 | 0,302 | 7,218 | 5,28E-13 | 1,11E-12 | 0,001191 | 0,002192 | 0,0015574 | 0,0015574 | TRUE | 0,0173771 | No |
| orf19.1738.1 | 374 | 318 | 299 | 180 | 283 | 178 | 208 | 260 | 341 | 322 | 283 | 270 | 252 | 193 | 213 | 198 | 258,98 | 304 | 214 | 0,702 | -0,51 | -4,575 | 4,76E-06 | 7,68E-06 | 0,0091294 | 0,007138 | 0,007993 | 0,007993 | TRUE | 0,1583253 | No |
| orf19.1741 | 3 | 4 | 8 | 1 | 4 | 3 | 6 | 7 | 3 | 4 | 8 | 2 | 4 | 3 | 6 | 5 | 4,27 | 4 | 4 | 1,118 | 0,16 | 0,238 | 0,811536 | 0,836331 | 1,00E-08 | 0,311496 | 0,2040747 | 0,2040747 | TRUE | 0,6507003 | No |
| orf19.1742 | 6501 | 7783 | 6691 | 4831 | 7761 | 7876 | 7458 | 11496 | 5919 | 7872 | 6332 | 7257 | 6903 | 8551 | 7638 | 8769 | 7405,17 | 6845 | 7965 | 1,164 | 0,219 | 2,37 | 0,017789 | 0,022915 | 0,0140018 | 0,002215 | 0,0080681 | 0,0080681 | TRUE | 0,1252315 | No |
| orf19.1743 | 3455 | 2959 | 3085 | 1981 | 23815 | 19054 | 19655 | 28437 | 3146 | 2993 | 2920 | 2976 | 21183 | 20687 | 20129 | 21692 | 11965,6 | 3009 | 20923 | 6,953 | 2,798 | 69,844 | 0 | 0 | 0,0007743 | 0,002147 | 0,0013394 | 0,0013394 | TRUE | 0,01193 | No |
| orf19.1744 | 2570 | 2766 | 2846 | 1631 | 1816 | 1497 | 1565 | 1885 | 2340 | 2798 | 2693 | 2450 | 1615 | 1625 | 1603 | 1438 | 2070,27 | 2570 | 1570 | 0,61 | -0,712 | -10,674 | 1,35E-26 | 4,02E-26 | 0,0043034 | 0,002675 | 0,003771 | 0,003771 | TRUE | 0,04582 | No |
| orf19.1745 | 726 | 749 | 774 | 524 | 452 | 452 | 396 | 612 | 661 | 758 | 733 | 787 | 402 | 491 | 406 | 467 | 587,92 | 735 | 442 | 0,602 | -0,732 | -8,733 | 2,47E-18 | 5,98E-18 | 0,0053349 | 0,004284 | 0,004954 | 0,004954 | TRUE | 0,0610834 | No |
| orf19.1746 | 96 | 96 | 109 | 69 | 166 | 130 | 153 | 177 | 87 | 97 | 103 | 104 | 148 | 141 | 157 | 135 | 121,48 | 98 | 145 | 1,487 | 0,573 | 4,426 | 9,61E-06 | 1,53E-05 | 1,00E-08 | 0,012912 | 0,007411 | 0,007411 | TRUE | 0,0551834 | No |
| orf19.1747 | 102 | 102 | 93 | 57 | 138 | 124 | 128 | 185 | 93 | 103 | 88 | 86 | 123 | 135 | 131 | 141 | 112,41 | 92 | 132 | 1,429 | 0,515 | 3,867 | 0,00011 | 0,000165 | 1,00E-08 | 0,013789 | 0,0077543 | 0,0077543 | TRUE | 0,0576989 | No |
| orf19.1748 | 168 | 151 | 173 | 126 | 237 | 190 | 201 | 272 | 153 | 153 | 164 | 189 | 211 | 206 | 206 | 207 | 186,14 | 165 | 208 | 1,273 | 0,348 | 3,294 | 0,000986 | 0,001393 | 1,00E-08 | 0,009134 | 0,0052025 | 0,0052025 | TRUE | 0,0835503 | No |
| orf19.175 | 167 | 166 | 203 | 105 | 146 | 123 | 121 | 172 | 152 | 168 | 192 | 158 | 130 | 134 | 124 | 131 | 148,54 | 168 | 130 | 0,772 | -0,373 | -3,12 | 0,001811 | 0,002509 | 1,00E-08 | 0,01093 | 0,0069484 | 0,0069484 | TRUE | 0,111474 | No |
| orf19.1750 | 23658 | 22443 | 24658 | 14735 | 18895 | 16062 | 16880 | 22387 | 21540 | 22699 | 23337 | 22134 | 16807 | 17439 | 17287 | 17077 | 19789,9 | 22428 | 17152 | 0,765 | -0,387 | -11,114 | 1,07E-28 | 3,32E-28 | 0,0006471 | 0,002104 | 0,0011133 | 0,0011133 | TRUE | 0,0091845 | No |
| orf19.1751 | 12327 | 12083 | 12800 | 8129 | 12375 | 10624 | 10700 | 14400 | 11223 | 12221 | 12114 | 12211 | 11007 | 11535 | 10958 | 10984 | 11531,7 | 11942 | 11121 | 0,931 | -0,103 | -2,582 | 0,009825 | 0,012896 | 0,0010069 | 0,002152 | 0,0014295 | 0,0014295 | TRUE | 0,020296 | No |
| orf19.1753 | 8119 | 7681 | 8093 | 4812 | 8642 | 7099 | 7512 | 9677 | 7392 | 7769 | 7659 | 7228 | 7687 | 7708 | 7693 | 7382 | 7564,68 | 7512 | 7618 | 1,013 | 0,019 | 0,525 | 0,599754 | 0,639587 | 0,0005615 | 0,002212 | 0,0011438 | 0,0011438 | TRUE | 0,0075256 | No |
| orf19.1754 | 1930 | 1775 | 1772 | 1146 | 1981 | 1603 | 1654 | 2228 | 1757 | 1795 | 1677 | 1721 | 1762 | 1740 | 1694 | 1700 | 1730,85 | 1738 | 1724 | 0,992 | -0,012 | -0,273 | 0,784681 | 0,814316 | 1,00E-08 | 0,0028 | 0,0012595 | 0,0012595 | TRUE | 0,0072491 | No |
| orf19.1755 | 3383 | 3287 | 3501 | 2094 | 2055 | 1860 | 1868 | 2451 | 3080 | 3324 | 3313 | 3146 | 1828 | 2019 | 1913 | 1870 | 2561,68 | 3216 | 1908 | 0,592 | -0,755 | -15,823 | 2,17E-56 | 1,12E-55 | 0,001095 | 0,002553 | 0,0017742 | 0,0017742 | TRUE | 0,0183738 | No |
| orf19.1756 | 7614 | 7039 | 7851 | 4780 | 14521 | 12740 | 12491 | 16877 | 6932 | 7119 | 7430 | 7180 | 12916 | 13832 | 12792 | 12874 | 10134,5 | 7165 | 13104 | 1,829 | 0,871 | 21,939 | 1,09E-106 | 1,00E-105 | 0,0009625 | 0,002167 | 0,0014017 | 0,0014017 | TRUE | 0,0170055 | No |
| orf19.1757 | 3194 | 3068 | 3278 | 1939 | 6128 | 5023 | 5480 | 7113 | 2908 | 3103 | 3102 | 2913 | 5451 | 5454 | 5612 | 5426 | 4246,03 | 3006 | 5486 | 1,823 | 0,866 | 22,44 | 1,59E-111 | 1,52E-110 | 0,0004598 | 0,002348 | 0,0011672 | 0,0011672 | TRUE | 0,0065708 | No |
| orf19.1759 | 2705 | 2442 | 2502 | 1677 | 2158 | 1813 | 1938 | 2601 | 2463 | 2470 | 2368 | 2519 | 1919 | 1968 | 1985 | 1984 | 2209,54 | 2455 | 1964 | 0,801 | -0,32 | -7,962 | 1,70E-15 | 3,81E-15 | 1,00E-08 | 0,002635 | 0,0010989 | 0,0010989 | TRUE | 0,0069751 | No |
| orf19.176 | 2920 | 2632 | 2665 | 1796 | 222 | 161 | 188 | 295 | 2659 | 2662 | 2522 | 2698 | 197 | 175 | 193 | 225 | 1416,3 | 2635 | 198 | 0,076 | -3,726 | -52,926 | 0 | 0 | 0,0008275 | 0,00297 | 0,002223 | 0,002223 | TRUE | 0,1077848 | No |
| orf19.1760 | 11030 | 10671 | 11070 | 7002 | 5522 | 4442 | 4719 | 6374 | 10042 | 10793 | 10477 | 10518 | 4912 | 4823 | 4833 | 4862 | 7657,4 | 10458 | 4858 | 0,465 | -1,106 | -32,347 | 1,52E-229 | 2,92E-228 | 0,0004233 | 0,00221 | 0,0009747 | 0,0009747 | TRUE | 0,0089044 | No |
| orf19.1761 | 273 | 300 | 253 | 169 | 168 | 138 | 131 | 195 | 249 | 303 | 239 | 254 | 149 | 150 | 134 | 149 | 203,43 | 261 | 146 | 0,557 | -0,843 | -7,618 | 2,57E-14 | 5,58E-14 | 0,0038438 | 0,008531 | 0,0065127 | 0,0065127 | TRUE | 0,1363601 | No |
| orf19.1762 | 1517 | 1333 | 1568 | 882 | 1057 | 799 | 830 | 1176 | 1381 | 1348 | 1484 | 1325 | 940 | 867 | 850 | 897 | 1136,62 | 1384 | 888 | 0,642 | -0,64 | -11,16 | 6,38E-29 | 2,00E-28 | 0,0013824 | 0,003199 | 0,0022483 | 0,0022483 | TRUE | 0,0284702 | No |
| orf19.1763 | 1395 | 1483 | 1516 | 938 | 973 | 849 | 930 | 1156 | 1270 | 1500 | 1435 | 1409 | 865 | 922 | 952 | 882 | 1154,41 | 1404 | 905 | 0,645 | -0,633 | -10,007 | 1,42E-23 | 3,94E-23 | 0,0027342 | 0,003181 | 0,0029423 | 0,0029423 | TRUE | 0,0517253 | No |
| orf19.1764 | 5674 | 5573 | 5979 | 3800 | 3094 | 2672 | 2723 | 3496 | 5166 | 5636 | 5659 | 5708 | 2752 | 2901 | 2789 | 2667 | 4159,72 | 5542 | 2777 | 0,501 | -0,997 | -21,407 | 1,15E-101 | 1,01E-100 | 0,0012906 | 0,002355 | 0,0018176 | 0,0018176 | TRUE | 0,0258878 | No |
| orf19.1765 | 880 | 822 | 921 | 461 | 1087 | 896 | 1010 | 1227 | 801 | 831 | 872 | 692 | 967 | 973 | 1034 | 936 | 888,34 | 799 | 978 | 1,217 | 0,284 | 4,02 | 5,82E-05 | 8,83E-05 | 0,0039841 | 0,003524 | 0,003629 | 0,003629 | TRUE | 0,0846952 | No |
| orf19.1766 | 138 | 136 | 127 | 69 | 167 | 173 | 136 | 180 | 126 | 138 | 120 | 104 | 149 | 188 | 139 | 137 | 137,5 | 122 | 153 | 1,245 | 0,316 | 2,281 | 0,022528 | 0,028741 | 0,0099351 | 0,011644 | 0,0109692 | 0,0109692 | TRUE | 0,2294743 | No |
| orf19.1767 | 485 | 622 | 652 | 350 | 369 | 338 | 304 | 400 | 442 | 629 | 617 | 526 | 328 | 367 | 311 | 305 | 440,64 | 554 | 328 | 0,591 | -0,758 | -6,735 | 1,64E-11 | 3,29E-11 | 0,0150367 | 0,005035 | 0,0097536 | 0,0097536 | TRUE | 0,2286375 | No |
| orf19.1768 | 293 | 334 | 368 | 219 | 255 | 247 | 213 | 255 | 267 | 338 | 348 | 329 | 227 | 268 | 218 | 195 | 273,68 | 320 | 227 | 0,706 | -0,502 | -4,292 | 1,77E-05 | 2,78E-05 | 0,0125121 | 0,006864 | 0,0093714 | 0,0093714 | TRUE | 0,1595507 | No |
| orf19.1769 | 886 | 852 | 915 | 507 | 999 | 788 | 885 | 1071 | 807 | 862 | 866 | 762 | 889 | 856 | 906 | 817 | 845,42 | 824 | 867 | 1,048 | 0,068 | 1,085 | 0,277803 | 0,314174 | 0,0015107 | 0,0036 | 0,0025802 | 0,0025802 | TRUE | 0,0265762 | No |

| | | | | | | | | | | | | | | | | | | | | | | | | | | | | | | | |
|---|---|---|---|---|---|---|---|---|---|---|---|---|---|---|---|---|---|---|---|---|---|---|---|---|---|---|---|---|---|---|---|
| orf19.177 | 496 | 532 | 509 | 372 | 993 | 894 | 865 | 1207 | 452 | 538 | 482 | 559 | 883 | 971 | 886 | 921 | 711,33 | 508 | 915 | 1,812 | 0,857 | 11,586 | 4,84E-31 | 1,59E-30 | 0,0037099 | 0,003894 | 0,0036839 | 0,0036839 | TRUE | 0,0654703 | No |
| orf19.1770 | 8255 | 6835 | 6983 | 4786 | 2683 | 2057 | 2084 | 2743 | 7516 | 6913 | 6609 | 7189 | 2386 | 2233 | 2134 | 2092 | 4634,16 | 7057 | 2211 | 0,313 | -1,674 | -30,009 | 7,56E-198 | 1,28E-196 | 0,002855 | 0,002322 | 0,0027008 | 0,0027008 | TRUE | 0,0349296 | No |
| orf19.1771 | 1740 | 1803 | 1813 | 1138 | 1225 | 986 | 1085 | 1394 | 1584 | 1824 | 1716 | 1709 | 1090 | 1071 | 1111 | 1063 | 1395,96 | 1708 | 1084 | 0,634 | -0,656 | -12,064 | 1,64E-33 | 5,69E-33 | 0,0014051 | 0,002983 | 0,0020966 | 0,0020966 | TRUE | 0,0304638 | No |
| orf19.1772 | 3054 | 3485 | 3620 | 2194 | 3100 | 2706 | 2686 | 3582 | 2781 | 3525 | 3426 | 3296 | 2757 | 2938 | 2751 | 2732 | 3025,68 | 3257 | 2794 | 0,858 | -0,221 | -3,188 | 0,001434 | 0,002004 | 0,0053143 | 0,002474 | 0,0042691 | 0,0042691 | TRUE | 0,1199373 | No |
| orf19.1773 | 2507 | 2391 | 2437 | 1586 | 3038 | 2714 | 2748 | 3537 | 2283 | 2418 | 2306 | 2382 | 2702 | 2947 | 2814 | 2698 | 2568,85 | 2347 | 2790 | 1,188 | 0,249 | 5,514 | 3,50E-08 | 6,16E-08 | 0,0008974 | 0,002551 | 0,001562 | 0,001562 | TRUE | 0,0169862 | No |
| orf19.1774 | 501 | 499 | 472 | 343 | 466 | 403 | 434 | 521 | 456 | 505 | 447 | 515 | 414 | 438 | 444 | 397 | 452,09 | 481 | 423 | 0,882 | -0,181 | -2,339 | 0,019331 | 0,024836 | 0,001681 | 0,004959 | 0,0034968 | 0,0034968 | TRUE | 0,0258694 | No |
| orf19.1776 | 311 | 305 | 334 | 188 | 466 | 396 | 446 | 512 | 283 | 308 | 316 | 282 | 414 | 430 | 457 | 391 | 360,24 | 297 | 423 | 1,415 | 0,501 | 5,971 | 2,36E-09 | 4,35E-09 | 0,0015332 | 0,005704 | 0,0038546 | 0,0038546 | TRUE | 0,033706 | No |
| orf19.1777 | 11727 | 12362 | 12818 | 7314 | 5478 | 4851 | 4958 | 6267 | 10677 | 12503 | 12131 | 10987 | 4873 | 5267 | 5078 | 4781 | 8286,89 | 11574 | 5000 | 0,432 | -1,212 | -20,878 | 8,51E-97 | 7,15E-96 | 0,0036432 | 0,002196 | 0,0030956 | 0,0030956 | TRUE | 0,0356668 | No |
| orf19.1778 | 93 | 97 | 84 | 58 | 101 | 73 | 85 | 114 | 85 | 98 | 79 | 87 | 90 | 79 | 87 | 87 | 86,56 | 87 | 86 | 0,985 | -0,022 | -0,146 | 0,883754 | 0,89999 | 1,00E-08 | 0,017298 | 0,009573 | 0,009573 | TRUE | 0,070057 | No |
| orf19.1779 | 20322 | 17443 | 19167 | 13593 | 44897 | 42662 | 41468 | 58421 | 18502 | 17642 | 18140 | 20419 | 39934 | 46319 | 42468 | 44564 | 30998,6 | 18676 | 43321 | 2,32 | 1,214 | 20,857 | 1,31E-96 | 1,09E-95 | 0,0040818 | 0,00208 | 0,0032163 | 0,0032163 | TRUE | 0,0479908 | No |
| orf19.178 | 959 | 1077 | 1094 | 670 | 1293 | 1114 | 1167 | 1421 | 873 | 1089 | 1035 | 1006 | 1150 | 1209 | 1195 | 1084 | 1080,36 | 1001 | 1160 | 1,158 | 0,212 | 2,951 | 0,003166 | 0,004302 | 0,0047109 | 0,00326 | 0,0040043 | 0,0040043 | TRUE | 0,0931111 | No |
| orf19.1780 | 24 | 34 | 47 | 27 | 68 | 57 | 58 | 72 | 22 | 34 | 44 | 41 | 60 | 62 | 59 | 55 | 47,25 | 35 | 59 | 1,691 | 0,758 | 3,406 | 0,00066 | 0,000942 | 0,0103862 | 0,029997 | 0,0243959 | 0,0243959 | TRUE | 0,5447932 | No |
| orf19.1782 | 535 | 472 | 474 | 352 | 669 | 558 | 518 | 753 | 487 | 477 | 449 | 529 | 595 | 606 | 530 | 574 | 530,95 | 486 | 576 | 1,192 | 0,254 | 3,455 | 0,00055 | 0,000789 | 0,0014731 | 0,004525 | 0,0032648 | 0,0032648 | TRUE | 0,0355648 | No |
| orf19.1782.1 | 26 | 19 | 29 | 10 | 23 | 14 | 26 | 21 | 24 | 19 | 27 | 15 | 20 | 15 | 27 | 16 | 20,46 | 21 | 20 | 0,897 | -0,157 | -0,494 | 0,621417 | 0,659721 | 0,0121088 | 0,06661 | 0,048463 | 0,048463 | TRUE | 0,2903905 | No |
| orf19.1783 | 12123 | 12235 | 12792 | 7193 | 12600 | 10178 | 11074 | 14154 | 11037 | 12374 | 12106 | 10805 | 11207 | 11050 | 11341 | 10797 | 11339,9 | 11580 | 11099 | 0,958 | -0,062 | -1,245 | 0,213307 | 0,245946 | 0,0023451 | 0,002154 | 0,0022799 | 0,0022799 | TRUE | 0,0259271 | No |
| orf19.1785 | 29607 | 27220 | 29944 | 20314 | 3587 | 3030 | 3115 | 4101 | 26956 | 27530 | 28339 | 30515 | 3191 | 3290 | 3190 | 3128 | 15767,4 | 28335 | 3200 | 0,113 | -3,147 | -68,534 | 0 | 0 | 0,0015017 | 0,002121 | 0,0018598 | 0,0018598 | TRUE | 0,0326023 | No |
| orf19.1789 | 273 | 394 | 338 | 182 | 261 | 264 | 219 | 289 | 249 | 398 | 320 | 273 | 232 | 287 | 224 | 220 | 275,48 | 310 | 241 | 0,774 | -0,37 | -2,62 | 0,008781 | 0,011564 | 0,0259132 | 0,006832 | 0,0154172 | 0,0154172 | TRUE | 0,4204207 | No |
| orf19.1789.1 | 1763 | 1697 | 1743 | 1186 | 2320 | 2082 | 1953 | 2668 | 1605 | 1716 | 1650 | 1782 | 2064 | 2260 | 2000 | 2035 | 1888,99 | 1688 | 2090 | 1,239 | 0,309 | 5,68 | 1,35E-08 | 2,42E-08 | 0,0018915 | 0,002736 | 0,0023066 | 0,0023066 | TRUE | 0,0356056 | No |
| orf19.1791 | 6886 | 6867 | 7214 | 4031 | 6022 | 4581 | 5255 | 6521 | 6269 | 6945 | 6827 | 6055 | 5356 | 4974 | 5382 | 4974 | 5847,92 | 6524 | 5172 | 0,792 | -0,336 | -6,172 | 6,76E-10 | 1,27E-09 | 0,0029132 | 0,002263 | 0,0026732 | 0,0026732 | TRUE | 0,0278581 | No |
| orf19.1792 | 488 | 513 | 583 | 320 | 544 | 489 | 435 | 597 | 444 | 519 | 552 | 481 | 484 | 531 | 445 | 455 | 488,91 | 499 | 479 | 0,958 | -0,062 | -0,718 | 0,472988 | 0,514643 | 0,0055696 | 0,004739 | 0,0051278 | 0,0051278 | TRUE | 0,0698125 | No |
| orf19.1793 | 1556 | 1416 | 1537 | 927 | 1250 | 1068 | 1065 | 1456 | 1417 | 1432 | 1455 | 1393 | 1112 | 1160 | 1091 | 1111 | 1271,08 | 1424 | 1118 | 0,784 | -0,351 | -7,497 | 6,53E-14 | 1,40E-13 | 1,00E-08 | 0,003076 | 0,0013088 | 0,0013088 | TRUE | 0,006896 | No |
| orf19.1794 | 183 | 200 | 175 | 83 | 112 | 103 | 139 | 159 | 167 | 202 | 166 | 125 | 100 | 112 | 142 | 121 | 141,79 | 165 | 119 | 0,714 | -0,485 | -3,157 | 0,001593 | 0,002218 | 0,0217193 | 0,011354 | 0,0154886 | 0,0154886 | TRUE | 0,1846706 | No |
| orf19.1795 | 2354 | 2365 | 2381 | 1364 | 1476 | 1158 | 1230 | 1620 | 2143 | 2392 | 2253 | 2049 | 1313 | 1257 | 1260 | 1236 | 1737,88 | 2209 | 1266 | 0,572 | -0,805 | -14,412 | 4,36E-47 | 1,92E-46 | 0,002052 | 0,002797 | 0,002382 | 0,002382 | TRUE | 0,036983 | No |
| orf19.1795.1 | 268 | 266 | 270 | 184 | 202 | 156 | 155 | 216 | 244 | 269 | 256 | 276 | 180 | 169 | 159 | 165 | 214,69 | 261 | 168 | 0,646 | -0,63 | -6,378 | 1,79E-10 | 3,44E-10 | 1,00E-08 | 0,00819 | 0,0045424 | 0,0045424 | TRUE | 0,0237416 | No |
| orf19.1796 | 269 | 268 | 293 | 153 | 272 | 240 | 246 | 308 | 245 | 271 | 277 | 230 | 242 | 261 | 252 | 235 | 251,56 | 256 | 248 | 0,96 | -0,059 | -0,622 | 0,534248 | 0,575893 | 1,00E-08 | 0,007288 | 0,0046163 | 0,0046163 | TRUE | 0,0410242 | No |
| orf19.1797 | 6377 | 6646 | 6891 | 4306 | 5988 | 5314 | 5301 | 6577 | 5806 | 6722 | 6522 | 6468 | 5326 | 5770 | 5429 | 5017 | 5882,41 | 6380 | 5386 | 0,844 | -0,245 | -4,263 | 2,02E-05 | 3,15E-05 | 0,0033724 | 0,002262 | 0,0029916 | 0,0029916 | TRUE | 0,0452372 | No |
| orf19.1798 | 724 | 759 | 839 | 463 | 560 | 469 | 492 | 561 | 659 | 768 | 794 | 695 | 498 | 509 | 504 | 428 | 606,93 | 729 | 485 | 0,663 | -0,593 | -7,16 | 8,08E-13 | 1,68E-12 | 0,0056079 | 0,004214 | 0,0048934 | 0,0048934 | TRUE | 0,077683 | No |
| orf19.1799 | 15248 | 13446 | 14503 | 8524 | 14655 | 11349 | 12850 | 17702 | 13883 | 13599 | 13726 | 12804 | 13035 | 12322 | 13160 | 13503 | 13254 | 13503 | 13005 | 0,963 | -0,054 | -1,309 | 0,19052 | 0,221031 | 0,0012601 | 0,002137 | 0,0015857 | 0,0015857 | TRUE | 0,0152158 | No |
| orf19.18 | 32285 | 27614 | 28971 | 18080 | 42172 | 33622 | 34403 | 49663 | 29394 | 27929 | 27418 | 27159 | 37511 | 36504 | 35233 | 37883 | 32378,9 | 27975 | 36783 | 1,315 | 0,395 | 10,103 | 5,34E-24 | 1,50E-23 | 0,0011151 | 0,002078 | 0,001435 | 0,001435 | TRUE | 0,0143237 | No |
| orf19.1800 | 10424 | 9833 | 10849 | 6225 | 2973 | 2491 | 2755 | 3447 | 9491 | 9945 | 10268 | 9351 | 2644 | 2705 | 2821 | 2629 | 6231,75 | 9764 | 2700 | 0,276 | -1,856 | -41,987 | 0 | 0 | 0,0011369 | 0,002249 | 0,0016473 | 0,0016473 | TRUE | 0,0147938 | No |
| orf19.1801 | 1830 | 1796 | 1931 | 1080 | 1714 | 1343 | 1397 | 1745 | 1666 | 1816 | 1828 | 1622 | 1525 | 1458 | 1431 | 1331 | 1584,62 | 1733 | 1436 | 0,827 | -0,274 | -4,558 | 5,18E-06 | 8,34E-06 | 0,0025257 | 0,002871 | 0,00283 | 0,00283 | TRUE | 0,0306493 | No |
| orf19.1802 | 6960 | 6655 | 6950 | 3922 | 6504 | 5046 | 5739 | 7100 | 6337 | 6731 | 6578 | 5891 | 5785 | 5479 | 5877 | 5416 | 6011,71 | 6384 | 5639 | 0,883 | -0,18 | -3,578 | 0,000346 | 0,000503 | 0,0022291 | 0,002257 | 0,0022634 | 0,0022634 | TRUE | 0,0319619 | No |
| orf19.1803 | 163 | 192 | 163 | 130 | 96 | 85 | 75 | 121 | 148 | 194 | 154 | 195 | 85 | 92 | 77 | 92 | 129,87 | 173 | 86 | 0,505 | -0,985 | -6,98 | 2,95E-12 | 6,03E-12 | 0,0083705 | 0,012209 | 0,0106315 | 0,0106315 | TRUE | 0,1007871 | No |
| orf19.1804 | 4051 | 3892 | 4245 | 2461 | 3985 | 3206 | 3543 | 4584 | 3688 | 3936 | 4018 | 3697 | 3545 | 3481 | 3628 | 3497 | 3686,18 | 3835 | 3538 | 0,922 | -0,117 | -2,779 | 0,005458 | 0,007302 | 0,0007679 | 0,002395 | 0,0014392 | 0,0014392 | TRUE | 0,012652 | No |
| orf19.1805 | 3067 | 2917 | 3158 | 2082 | 5250 | 4382 | 4466 | 5944 | 2792 | 2950 | 2989 | 3127 | 4670 | 4758 | 4574 | 4534 | 3799,26 | 2964 | 4634 | 1,565 | 0,646 | 15,01 | 6,31E-51 | 2,96E-50 | 0,000946 | 0,002385 | 0,0014959 | 0,0014959 | TRUE | 0,0187664 | No |
| orf19.1806 | 45 | 38 | 43 | 22 | 49 | 53 | 39 | 58 | 41 | 38 | 41 | 33 | 44 | 58 | 40 | 44 | 42,31 | 38 | 46 | 1,194 | 0,255 | 1,191 | 0,233645 | 0,267557 | 1,00E-08 | 0,033261 | 0,0202592 | 0,0202592 | TRUE | 0,1884882 | No |
| orf19.1808 | 349 | 389 | 434 | 248 | 231 | 214 | 206 | 222 | 318 | 393 | 411 | 373 | 205 | 232 | 211 | 169 | 289,07 | 374 | 204 | 0,545 | -0,876 | -7,691 | 1,46E-14 | 3,20E-14 | 0,0113665 | 0,006607 | 0,0087162 | 0,0087162 | TRUE | 0,1601052 | No |
| orf19.1813 | 3086 | 2757 | 2898 | 1770 | 4608 | 3970 | 4022 | 5548 | 2810 | 2788 | 2743 | 2659 | 4099 | 4310 | 4119 | 4232 | 3469,96 | 2750 | 4190 | 1,522 | 0,606 | 16,031 | 7,79E-58 | 4,12E-57 | 0,0001751 | 0,002418 | 0,0010648 | 0,0010648 | TRUE | 0,0054313 | No |
| orf19.1814 | 2185 | 2220 | 2239 | 1202 | 2079 | 1842 | 1752 | 2421 | 1989 | 2245 | 2119 | 1806 | 1849 | 2000 | 1794 | 1847 | 1956,17 | 2040 | 1872 | 0,916 | -0,126 | -1,891 | 0,058663 | 0,072197 | 0,0046787 | 0,002712 | 0,0037579 | 0,0037579 | TRUE | 0,0682348 | No |
| orf19.1815 | 12663 | 10467 | 10748 | 6838 | 13065 | 10033 | 10444 | 15295 | 11529 | 10586 | 10172 | 10272 | 11621 | 10893 | 10696 | 11667 | 10929,5 | 10640 | 11219 | 1,054 | 0,076 | 1,503 | 0,132865 | 0,157204 | 0,0025508 | 0,002158 | 0,0023918 | 0,0023918 | TRUE | 0,0389603 | No |
| orf19.1816 | 7074 | 6242 | 6875 | 4597 | 3980 | 3532 | 3879 | 5203 | 6441 | 6313 | 6507 | 6905 | 3540 | 3835 | 3973 | 3969 | 5185,25 | 6542 | 3829 | 0,586 | -0,772 | -15,799 | 3,15E-56 | 1,62E-55 | 0,0018935 | 0,002292 | 0,002088 | 0,002088 | TRUE | 0,0317435 | No |
| orf19.182 | 2718 | 2414 | 2571 | 1495 | 2299 | 1863 | 1832 | 2486 | 2475 | 2442 | 2433 | 2246 | 2045 | 2023 | 1876 | 1896 | 2179,4 | 2399 | 1960 | 0,816 | -0,294 | -5,909 | 3,45E-09 | 6,34E-09 | 0,0012779 | 0,002643 | 0,0019134 | 0,0019134 | TRUE | 0,0206907 | No |
| orf19.1821 | 0 | 0 | 2 | 1 | 0 | 0 | 1 | 0 | 0 | 0 | 2 | 2 | 0 | 0 | 1 | 0 | 0,55 | 1 | 0 | 0,343 | -1,542 | -0,76 | 0,447356 | 0,489511 | 1,00E-08 | 2,393468 | 2,3034767 | 2,3034767 | TRUE | 0,2912229 | No |
| orf19.1822 | 77 | 65 | 63 | 47 | 70 | 55 | 71 | 79 | 70 | 66 | 60 | 71 | 62 | 60 | 73 | 60 | 65,13 | 67 | 64 | 0,96 | -0,06 | -0,35 | 0,726294 | 0,759037 | 1,00E-08 | 0,02232 | 0,0124774 | 0,0124774 | TRUE | 0,0745584 | No |
| orf19.1823 | 167 | 180 | 180 | 122 | 142 | 133 | 125 | 168 | 152 | 182 | 170 | 183 | 126 | 144 | 128 | 128 | 151,82 | 172 | 132 | 0,768 | -0,381 | -3,246 | 0,001169 | 0,001643 | 1,00E-08 | 0,010738 | 0,0065765 | 0,0065765 | TRUE | 0,0692296 | No |
| orf19.1824 | 0 | 1 | 0 | 0 | 0 | 1 | 0 | 0 | 0 | 1 | 0 | 0 | 0 | 1 | 0 | 0 | 0,26 | 0 | 0 | 0,871 | -0,199 | -0,071 | 0,943001 | 0,950906 | 1,00E-08 | 5,041333 | 5,4278855 | 5,4278855 | TRUE | 0,4622358 | No |
| orf19.1825 | 62 | 49 | 66 | 47 | 163 | 107 | 111 | 184 | 56 | 50 | 62 | 71 | 145 | 116 | 114 | 140 | 94,28 | 60 | 129 | 2,188 | 1,13 | 6,974 | 3,08E-12 | 6,29E-12 | 0,0067509 | 0,016048 | 0,0125908 | 0,0125908 | TRUE | 0,1073307 | No |
| orf19.1826 | 20168 | 20208 | 21038 | 13363 | 8410 | 7798 | 7413 | 9735 | 18362 | 20438 | 19911 | 20073 | 7480 | 8466 | 7592 | 7426 | 13718,6 | 19696 | 7741 | 0,393 | -1,348 | -25,467 | 4,58E-143 | 5,71E-142 | 0,0029526 | 0,002133 | 0,0026019 | 0,0026019 | TRUE | 0,048322 | No |
| orf19.1827 | 17 | 18 | 17 | 8 | 19 | 17 | 13 | 20 | 15 | 18 | 16 | 12 | 17 | 18 | 13 | 15 | 15,71 | 15 | 16 | 1,019 | 0,028 | 0,081 | 0,935099 | 0,944582 | 1,00E-08 | 0,086099 | 0,0464049 | 0,0464049 | TRUE | 0,0623218 | No |
| orf19.1828 | 453 | 400 | 462 | 226 | 590 | 483 | 484 | 636 | 412 | 405 | 437 | 339 | 525 | 524 | 496 | 485 | 452,97 | 398 | 508 | 1,264 | 0,338 | 4,178 | 2,94E-05 | 4,55E-05 | 0,0026218 | 0,004953 | 0,0040279 | 0,0040279 | TRUE | 0,0947737 | No |
| orf19.183 | 1367 | 1214 | 1175 | 803 | 1685 | 1390 | 1370 | 1945 | 1245 | 1228 | 1112 | 1206 | 1499 | 1509 | 1403 | 1484 | 1335,66 | 1198 | 1474 | 1,231 | 0,3 | 5,625 | 1,86E-08 | 3,30E-08 | 0,0010238 | 0,003026 | 0,0019627 | 0,0019627 | TRUE | 0,0295246 | No |
| orf19.1830 | 3333 | 3492 | 3543 | 2149 | 3891 | 2812 | 3021 | 3788 | 3035 | 3532 | 3353 | 3228 | 3461 | 3053 | 3094 | 2890 | 3205,62 | 3287 | 3124 | 0,95 | -0,073 | -1,135 | 0,256203 | 0,291556 | 0,0045905 | 0,002449 | 0,0036937 | 0,0036937 | TRUE | 0,0645514 | No |
| orf19.1831 | 1 | 3 | 0 | 2 | 4 | 0 | 7 | 6 | 1 | 3 | 0 | 3 | 4 | 0 | 7 | 5 | 2,78 | 2 | 4 | 2,353 | 1,234 | 1,237 | 0,216265 | 0,249041 | 0,6134543 | 0,476948 | 0,5111648 | 0,5111648 | TRUE | 0,18436 | No |
| orf19.1832 | 2551 | 2236 | 2561 | 1485 | 3323 | 2680 | 2885 | 3785 | 2323 | 2261 | 2424 | 2231 | 2956 | 2910 | 2955 | 2887 | 2618,22 | 2310 | 2927 | 1,265 | 0,34 | 8,353 | 6,64E-17 | 1,55E-16 | 0,0002302 | 0,002542 | 0,0011958 | 0,0011958 | TRUE | 0,0134917 | No |
| orf19.1833 | 18265 | 17982 | 18976 | 10723 | 28409 | 22943 | 23843 | 31366 | 16630 | 18187 | 17959 | 16108 | 25269 | 24910 | 24418 | 23926 | 20925,8 | 17221 | 24631 | 1,43 | 0,516 | 11,178 | 5,25E-29 | 1,65E-28 | 0,0019399 | 0,0021 | 0,0019968 | 0,0019968 | TRUE | 0,0228646 | No |
| orf19.1834 | 299 | 341 | 385 | 227 | 229 | 217 | 222 | 263 | 272 | 345 | 364 | 341 | 204 | 236 | 227 | 201 | 273,72 | 330 | 217 | 0,655 | -0,611 | -5,623 | 1,88E-08 | 3,34E-08 | 0,008367 | 0,006863 | 0,007541 | 0,007541 | TRUE | 0,1728119 | No |
| orf19.1835 | 4228 | 4317 | 4578 | 2829 | 3961 | 3310 | 3431 | 4393 | 3849 | 4366 | 4333 | 4250 | 3523 | 3594 | 3514 | 3351 | 3847,45 | 4200 | 3496 | 0,832 | -0,265 | -5,345 | 9,05E-08 | 1,57E-07 | 0,0016941 | 0,00238 | 0,0020947 | 0,0020947 | TRUE | 0,0393021 | No |
| orf19.1836 | 152 | 152 | 139 | 85 | 237 | 204 | 199 | 283 | 138 | 154 | 132 | 128 | 211 | 221 | 204 | 216 | 175,42 | 138 | 213 | 1,539 | 0,622 | 5,74 | 9,47E-09 | 1,71E-08 | 1,00E-08 | 0,009568 | 0,0052591 | 0,0052591 | TRUE | 0,0610067 | No |
| orf19.1837 | 5851 | 5002 | 5690 | 3193 | 3936 | 2786 | 3224 | 4232 | 5327 | 5059 | 5385 | 4796 | 3501 | 3025 | 3302 | 3228 | 4202,91 | 5142 | 3264 | 0,635 | -0,656 | -11,888 | 1,37E-32 | 4,64E-32 | 0,0028666 | 0,002351 | 0,0026783 | 0,0026783 | TRUE | 0,0290938 | No |
| orf19.1839 | 13989 | 13377 | 14033 | 7808 | 20668 | 16896 | 18119 | 23789 | 12736 | 13529 | 13281 | 11729 | 18384 | 18344 | 18556 | 18146 | 15588,3 | 12819 | 18358 | 1,431 | 0,517 | 11,211 | 3,61E-29 | 1,14E-28 | 0,0018942 | 0,002122 | 0,0019793 | 0,0019793 | TRUE | 0,039382 | No |
| orf19.184 | 567 | 579 | 661 | 340 | 517 | 468 | 462 | 612 | 516 | 586 | 626 | 511 | 460 | 508 | 473 | 467 | 518,26 | 560 | 477 | 0,849 | -0,236 | -2,928 | 0,003409 | 0,004616 | 0,0039715 | 0,004586 | 0,0042988 | 0,0042988 | TRUE | 0,0751171 | No |
| orf19.1840 | 2296 | 2068 | 2174 | 1340 | 1716 | 1443 | 1647 | 2160 | 2090 | 2092 | 2057 | 2013 | 1526 | 1567 | 1687 | 1648 | 1834,97 | 2063 | 1607 | 0,778 | -0,361 | -7,732 | 1,06E-14 | 2,32E-14 | 0,0005771 | 0,002757 | 0,0015468 | 0,0015468 | TRUE | 0,014143 | No |
| orf19.1841 | 51 | 87 | 80 | 40 | 146 | 111 | 114 | 106 | 46 | 88 | 76 | 60 | 130 | 121 | 117 | 81 | 89,78 | 68 | 112 | 1,649 | 0,722 | 3,489 | 0,000485 | 0,000699 | 0,0464304 | 0,016751 | 0,0289721 | 0,0289721 | TRUE | 0,4304016 | No |
| orf19.1842 | 3217 | 3298 | 3462 | 1969 | 4362 | 3746 | 4156 | 5328 | 2929 | 3336 | 3276 | 2958 | 3880 | 4067 | 4256 | 4064 | 3595,78 | 3125 | 4067 | 1,301 | 0,379 | 6,977 | 3,01E-12 | 6,16E-12 | 0,0026189 | 0,002404 | 0,0025481 | 0,0025481 | TRUE | 0,0249681 | No |
| orf19.1843 | 209 | 265 | 252 | 174 | 460 | 413 | 348 | 447 | 190 | 268 | 238 | 261 | 409 | 448 | 356 | 341 | 314,14 | 239 | 388 | 1,627 | 0,702 | 5,81 | 6,25E-09 | 1,14E-08 | 0,0160656 | 0,006242 | 0,0105748 | 0,0105748 | TRUE | 0,2245469 | No |
| orf19.1844 | 11 | 17 | 16 | 13 | 11 | 12 | 13 | 7 | 10 | 17 | 15 | 20 | 10 | 13 | 13 | 5 | 12,92 | 16 | 10 | 0,666 | -0,586 | -1,43 | 0,152836 | 0,1797 | 1,00E-08 | 0,104296 | 0,080156 | 0,080156 | TRUE | 0,514136 | No |
| orf19.1847 | 1025 | 872 | 1033 | 685 | 3308 | 2999 | 3081 | 4176 | 933 | 882 | 978 | 1029 | 2942 | 3256 | 3155 | 3185 | 2045,13 | 956 | 3134 | 3,289 | 1,718 | 30,384 | 9,02E-203 | 1,56E-201 | 0,0021718 | 0,002683 | 0,0023563 | 0,0023563 | TRUE | 0,0310484 | No |

| | | | | | | | | | | | | | | | | | | | | | | | | | | | | | | | |
|---|---|---|---|---|---|---|---|---|---|---|---|---|---|---|---|---|---|---|---|---|---|---|---|---|---|---|---|---|---|---|---|
| orf19.1848.1 | 503 | 388 | 342 | 267 | 188 | 155 | 144 | 245 | 458 | 392 | 324 | 401 | 167 | 168 | 147 | 187 | 280,62 | 394 | 167 | 0,427 | -1,229 | -10,379 | 3,08E-25 | 8,88E-25 | 0,0123327 | 0,006744 | 0,0093148 | 0,0093148 | TRUE | 0,1704398 | No |
| orf19.1849 | 5328 | 5114 | 5436 | 3209 | 5389 | 4693 | 4680 | 6130 | 4851 | 5172 | 5145 | 4820 | 4793 | 5095 | 4793 | 4676 | 4918,23 | 4997 | 4839 | 0,968 | -0,047 | -1,098 | 0,272334 | 0,308372 | 0,0011164 | 0,002306 | 0,0015892 | 0,0015892 | TRUE | 0,0152532 | No |
| orf19.185 | 3020 | 2755 | 2755 | 1614 | 2452 | 1905 | 2056 | 2812 | 2750 | 2786 | 2607 | 2424 | 2181 | 2068 | 2106 | 2145 | 2383,46 | 2642 | 2125 | 0,803 | -0,316 | -6,183 | 6,30E-10 | 1,19E-09 | 0,0016295 | 0,002591 | 0,0020836 | 0,0020836 | TRUE | 0,0347521 | No |
| orf19.1850 | 1002 | 908 | 981 | 601 | 1088 | 886 | 1027 | 1270 | 912 | 918 | 928 | 903 | 968 | 962 | 1052 | 969 | 951,51 | 915 | 988 | 1,077 | 0,108 | 2,001 | 0,045359 | 0,056393 | 1,00E-08 | 0,003425 | 0,0017184 | 0,0017184 | TRUE | 0,0220961 | No |
| orf19.1852 | 1066 | 868 | 960 | 628 | 744 | 591 | 608 | 880 | 971 | 878 | 909 | 943 | 662 | 642 | 623 | 671 | 787,21 | 925 | 650 | 0,703 | -0,508 | -8,54 | 1,35E-17 | 3,21E-17 | 1,00E-08 | 0,003715 | 0,0021029 | 0,0021029 | TRUE | 0,0146316 | No |
| orf19.1853 | 1338 | 1030 | 1032 | 648 | 980 | 649 | 680 | 993 | 1218 | 1042 | 977 | 973 | 872 | 705 | 696 | 757 | 905,03 | 1052 | 758 | 0,719 | -0,475 | -5,254 | 1,49E-07 | 2,55E-07 | 0,0101223 | 0,003497 | 0,0067222 | 0,0067222 | TRUE | 0,1368213 | No |
| orf19.1854 | 3489 | 3062 | 3215 | 2048 | 2410 | 1821 | 1881 | 2611 | 3177 | 3097 | 3043 | 3076 | 2144 | 1977 | 1926 | 1992 | 2553,92 | 3098 | 2010 | 0,649 | -0,624 | -13,963 | 2,63E-44 | 1,12E-43 | 0,0007053 | 0,002554 | 0,0015118 | 0,0015118 | TRUE | 0,0245203 | No |
| orf19.1855 | 335 | 345 | 362 | 234 | 459 | 398 | 361 | 507 | 305 | 349 | 343 | 352 | 408 | 432 | 370 | 387 | 368,11 | 337 | 399 | 1,187 | 0,247 | 2,955 | 0,003123 | 0,004246 | 0,0016779 | 0,005626 | 0,0039512 | 0,0039512 | TRUE | 0,049924 | No |
| orf19.1856 | 2920 | 2761 | 2893 | 1716 | 2575 | 1915 | 2144 | 2841 | 2659 | 2792 | 2738 | 2578 | 2290 | 2079 | 2196 | 2167 | 2437,38 | 2692 | 2183 | 0,811 | -0,303 | -6,584 | 4,57E-11 | 8,98E-11 | 0,0008487 | 0,002579 | 0,0016169 | 0,0016169 | TRUE | 0,0132827 | No |
| orf19.1857 | 763 | 661 | 773 | 439 | 1852 | 1257 | 1563 | 2053 | 695 | 669 | 732 | 659 | 1647 | 1365 | 1601 | 1566 | 1116,63 | 689 | 1545 | 2,241 | 1,164 | 16,809 | 2,09E-63 | 1,21E-62 | 0,0038052 | 0,00322 | 0,0035257 | 0,0035257 | TRUE | 0,0721672 | No |
| orf19.1860 | 5006 | 3788 | 4333 | 3096 | 8542 | 7325 | 7787 | 10905 | 4558 | 3831 | 4101 | 4651 | 7598 | 7953 | 7975 | 8318 | 6123,05 | 4285 | 7961 | 1,859 | 0,895 | 14,429 | 3,39E-47 | 1,49E-46 | 0,0045107 | 0,002253 | 0,0035068 | 0,0035068 | TRUE | 0,0618714 | No |
| orf19.1860.1 | 233 | 242 | 235 | 137 | 258 | 208 | 205 | 272 | 212 | 245 | 222 | 206 | 229 | 226 | 210 | 207 | 219,73 | 221 | 218 | 0,982 | -0,026 | -0,264 | 0,792119 | 0,820965 | 1,00E-08 | 0,008049 | 0,0048323 | 0,0048323 | TRUE | 0,055278 | No |
| orf19.1861 | 15241 | 13677 | 14933 | 9177 | 1857 | 1495 | 1560 | 1986 | 13876 | 13833 | 14133 | 13785 | 1652 | 1623 | 1598 | 1515 | 7751,83 | 13907 | 1597 | 0,115 | -3,124 | -82,955 | 0 | 0 | 0,0001377 | 0,002207 | 0,0010308 | 0,0010308 | TRUE | 0,0156527 | No |
| orf19.1862 | 7477 | 6116 | 6479 | 3788 | 2077 | 1496 | 1578 | 2266 | 6807 | 6186 | 6132 | 5690 | 1847 | 1624 | 1616 | 1729 | 3953,92 | 6204 | 1704 | 0,275 | -1,864 | -29,461 | 9,08E-191 | 1,47E-189 | 0,0042699 | 0,002371 | 0,0034837 | 0,0034837 | TRUE | 0,0524708 | No |
| orf19.1863 | 22 | 21 | 17 | 12 | 25 | 23 | 18 | 23 | 20 | 21 | 16 | 18 | 22 | 25 | 18 | 18 | 19,82 | 19 | 21 | 1,094 | 0,13 | 0,426 | 0,670174 | 0,706759 | 1,00E-08 | 0,068682 | 0,0385968 | 0,0385968 | TRUE | 0,0908461 | No |
| orf19.1864 | 3094 | 2645 | 3026 | 2047 | 4315 | 3551 | 3806 | 5033 | 2817 | 2675 | 2864 | 3075 | 3838 | 3855 | 3898 | 3839 | 3357,67 | 2858 | 3858 | 1,352 | 0,435 | 9,563 | 1,14E-21 | 3,01E-21 | 0,001298 | 0,00243 | 0,0016769 | 0,0016769 | TRUE | 0,0295623 | No |
| orf19.1865 | 948 | 845 | 882 | 541 | 639 | 605 | 629 | 863 | 863 | 855 | 835 | 813 | 568 | 657 | 644 | 658 | 736,61 | 842 | 632 | 0,75 | -0,415 | -6,442 | 1,18E-10 | 2,29E-10 | 0,0012488 | 0,00383 | 0,0026161 | 0,0026161 | TRUE | 0,0554952 | No |
| orf19.1866 | 436 | 317 | 340 | 236 | 755 | 527 | 600 | 778 | 397 | 321 | 322 | 355 | 672 | 572 | 614 | 593 | 480,69 | 349 | 613 | 1,759 | 0,815 | 9,343 | 9,39E-21 | 2,43E-20 | 0,0052821 | 0,004785 | 0,0050096 | 0,0050096 | TRUE | 0,1113825 | No |
| orf19.1867 | 619 | 529 | 551 | 394 | 869 | 732 | 718 | 1046 | 564 | 535 | 521 | 592 | 773 | 795 | 735 | 798 | 664,1 | 553 | 775 | 1,407 | 0,493 | 7,677 | 1,63E-14 | 3,57E-14 | 0,0003236 | 0,004026 | 0,0023888 | 0,0023888 | TRUE | 0,021768 | No |
| orf19.1868 | 7428 | 6458 | 7075 | 4361 | 2256 | 1790 | 1891 | 2442 | 6763 | 6532 | 6696 | 6551 | 2007 | 1943 | 1937 | 1863 | 4286,34 | 6636 | 1938 | 0,292 | -1,777 | -46,828 | 0 | 0 | 0,0002498 | 0,002345 | 0,0010607 | 0,0010607 | TRUE | 0,0088019 | No |
| orf19.1871 | 3420 | 2924 | 3239 | 2041 | 3309 | 2808 | 3173 | 4176 | 3114 | 2957 | 3065 | 3066 | 2943 | 3049 | 3250 | 3185 | 3078,68 | 3050 | 3107 | 1,018 | 0,026 | 0,598 | 0,549508 | 0,591044 | 0,0009391 | 0,002466 | 0,0015342 | 0,0015342 | TRUE | 0,0155152 | No |
| orf19.1872 | 3379 | 2693 | 2876 | 2111 | 6377 | 5092 | 5700 | 7637 | 3076 | 2724 | 2722 | 3171 | 5672 | 5528 | 5838 | 5826 | 4319,6 | 2923 | 5716 | 1,958 | 0,969 | 17,295 | 5,15E-67 | 3,10E-66 | 0,0032041 | 0,002343 | 0,0027488 | 0,0027488 | TRUE | 0,037638 | No |
| orf19.1873 | 1242 | 1056 | 1152 | 807 | 2815 | 2320 | 2414 | 3338 | 1131 | 1068 | 1090 | 1212 | 2504 | 2519 | 2472 | 2546 | 1817,82 | 1125 | 2510 | 2,238 | 1,162 | 24,32 | 1,20E-130 | 1,34E-129 | 4,12E-05 | 0,002764 | 0,0015295 | 0,0015295 | TRUE | 0,0277887 | No |
| orf19.1874 | 4 | 2 | 3 | 0 | 10 | 4 | 9 | 3 | 4 | 2 | 3 | 0 | 9 | 4 | 9 | 2 | 4,16 | 2 | 6 | 2,679 | 1,422 | 1,838 | 0,066049 | 0,080849 | 0,1697919 | 0,3199 | 0,2663148 | 0,2663148 | TRUE | 0,186438 | No |
| orf19.1876 | 1881 | 1508 | 1691 | 1213 | 1764 | 1498 | 1631 | 2149 | 1713 | 1525 | 1600 | 1822 | 1569 | 1626 | 1670 | 1639 | 1645,66 | 1665 | 1626 | 0,978 | -0,031 | -0,535 | 0,592794 | 0,632903 | 0,0024985 | 0,00284 | 0,0026953 | 0,0026953 | TRUE | 0,0448781 | No |
| orf19.1877 | 632 | 583 | 644 | 402 | 200 | 166 | 183 | 253 | 575 | 590 | 609 | 604 | 178 | 180 | 187 | 193 | 389,62 | 594 | 184 | 0,311 | -1,684 | -21,072 | 1,44E-98 | 1,24E-97 | 1,00E-08 | 0,005428 | 0,0027196 | 0,0027196 | TRUE | 0,0118621 | No |
| orf19.1878 | 1323 | 1072 | 1233 | 786 | 759 | 571 | 687 | 820 | 1205 | 1084 | 1167 | 1181 | 675 | 620 | 704 | 626 | 907,56 | 1159 | 656 | 0,566 | -0,822 | -13,134 | 2,11E-39 | 8,14E-39 | 0,0016372 | 0,003493 | 0,002582 | 0,002582 | TRUE | 0,0269901 | No |
| orf19.188 | 2284 | 2117 | 2236 | 1384 | 1707 | 1316 | 1482 | 1949 | 2079 | 2141 | 2116 | 2079 | 1518 | 1429 | 1518 | 1487 | 1795,92 | 2104 | 1488 | 0,707 | -0,5 | -11,762 | 6,15E-32 | 2,07E-31 | 1,00E-08 | 0,002773 | 0,001164 | 0,001164 | TRUE | 0,00803 | No |
| orf19.1880 | 3800 | 3321 | 3640 | 2206 | 3712 | 3120 | 3349 | 4691 | 3460 | 3359 | 3445 | 3314 | 3302 | 3387 | 3430 | 3578 | 3409,32 | 3394 | 3424 | 1,009 | 0,012 | 0,307 | 0,758561 | 0,789781 | 0,0005467 | 0,002424 | 0,0012395 | 0,0012395 | TRUE | 0,0116327 | No |
| orf19.1881 | 1402 | 1276 | 1230 | 835 | 499 | 439 | 515 | 664 | 1276 | 1291 | 1164 | 1254 | 444 | 477 | 527 | 507 | 867,47 | 1246 | 489 | 0,392 | -1,351 | -20,143 | 3,09E-90 | 2,42E-89 | 0,0022202 | 0,00356 | 0,0029312 | 0,0029312 | TRUE | 0,0459471 | No |
| orf19.1883 | 1956 | 1663 | 1853 | 1157 | 1600 | 1405 | 1470 | 1966 | 1781 | 1682 | 1754 | 1738 | 1423 | 1525 | 1505 | 1500 | 1613,53 | 1739 | 1488 | 0,856 | -0,225 | -4,95 | 7,40E-07 | 1,24E-06 | 1,00E-08 | 0,002856 | 0,0013634 | 0,0013634 | TRUE | 0,0106022 | No |
| orf19.1885 | 3025 | 2545 | 2838 | 1638 | 2408 | 1857 | 2188 | 2916 | 2754 | 2574 | 2686 | 2461 | 2142 | 2016 | 2241 | 2224 | 2387,22 | 2619 | 2156 | 0,823 | -0,282 | -5,414 | 6,18E-08 | 1,08E-07 | 0,0017819 | 0,00259 | 0,0021786 | 0,0021786 | TRUE | 0,0223656 | No |
| orf19.1886 | 6333 | 5347 | 5799 | 3325 | 6964 | 5560 | 6198 | 8006 | 5766 | 5408 | 5488 | 4995 | 6194 | 6037 | 6348 | 6107 | 5792,77 | 5414 | 6172 | 1,139 | 0,187 | 3,998 | 6,38E-05 | 9,66E-05 | 0,00174 | 0,002265 | 0,0019353 | 0,0019353 | TRUE | 0,0319747 | No |
| orf19.1887 | 1359 | 1146 | 1254 | 768 | 1320 | 1079 | 1275 | 1814 | 1237 | 1159 | 1187 | 1154 | 1174 | 1171 | 1306 | 1384 | 1221,49 | 1184 | 1259 | 1,063 | 0,088 | 1,363 | 0,172798 | 0,201497 | 0,0032969 | 0,003118 | 0,0032138 | 0,0032138 | TRUE | 0,0559098 | No |
| orf19.1888 | 1198 | 1028 | 1091 | 703 | 1084 | 878 | 925 | 1252 | 1091 | 1040 | 1033 | 1056 | 964 | 953 | 947 | 955 | 1004,85 | 1055 | 955 | 0,905 | -0,143 | -2,872 | 0,004084 | 0,005503 | 1,00E-08 | 0,003352 | 0,0014003 | 0,0014003 | TRUE | 0,0069992 | No |
| orf19.1889 | 528 | 476 | 442 | 316 | 582 | 431 | 443 | 633 | 481 | 481 | 418 | 475 | 518 | 468 | 454 | 483 | 472,16 | 464 | 481 | 1,039 | 0,055 | 0,715 | 0,474618 | 0,516242 | 0,0018837 | 0,004835 | 0,0035017 | 0,0035017 | TRUE | 0,0539035 | No |
| orf19.1890 | 4080 | 3460 | 3845 | 2680 | 2954 | 2418 | 2476 | 3417 | 3715 | 3499 | 3639 | 4026 | 2627 | 2625 | 2536 | 2607 | 3159,23 | 3720 | 2599 | 0,7 | -0,515 | -10,689 | 1,14E-26 | 3,40E-26 | 0,0014934 | 0,002455 | 0,0019081 | 0,0019081 | TRUE | 0,0354434 | No |
| orf19.1891 | 12297 | 9732 | 10461 | 6787 | 16429 | 13045 | 14051 | 20473 | 11196 | 9843 | 9900 | 10195 | 14613 | 14163 | 14390 | 15617 | 12489,7 | 10284 | 14696 | 1,429 | 0,515 | 10,013 | 1,33E-23 | 3,70E-23 | 0,0027026 | 0,002143 | 0,0024576 | 0,0024576 | TRUE | 0,0440199 | No |
| orf19.1893 | 1426 | 1209 | 1309 | 803 | 1377 | 1131 | 1191 | 1683 | 1298 | 1223 | 1239 | 1206 | 1225 | 1228 | 1220 | 1284 | 1240,31 | 1242 | 1239 | 0,997 | -0,004 | -0,079 | 0,937055 | 0,946408 | 1,00E-08 | 0,003102 | 0,0014983 | 0,0014983 | TRUE | 0,0117431 | No |
| orf19.1896 | 22154 | 18958 | 20649 | 13991 | 19856 | 16965 | 17789 | 25146 | 20170 | 19174 | 19542 | 21017 | 17661 | 18419 | 18218 | 19182 | 19173 | 19976 | 18370 | 0,92 | -0,121 | -2,906 | 0,003661 | 0,004949 | 0,0013316 | 0,002106 | 0,0016005 | 0,0016005 | TRUE | 0,0148655 | No |
| orf19.1897 | 3151 | 2661 | 2890 | 1827 | 3167 | 2658 | 2797 | 3937 | 2869 | 2691 | 2735 | 2744 | 2817 | 2886 | 2864 | 3003 | 2826,27 | 2760 | 2892 | 1,048 | 0,068 | 1,656 | 0,097655 | 0,117418 | 0,000484 | 0,002504 | 0,001252 | 0,001252 | TRUE | 0,0089897 | No |
| orf19.190 | 502 | 570 | 499 | 294 | 483 | 346 | 360 | 500 | 457 | 576 | 472 | 442 | 430 | 376 | 369 | 381 | 437,85 | 487 | 389 | 0,798 | -0,326 | -3,448 | 0,000564 | 0,000808 | 0,0073111 | 0,005054 | 0,0062836 | 0,0062836 | TRUE | 0,1793387 | No |
| orf19.1900 | 3124 | 2732 | 2941 | 1864 | 2863 | 2286 | 2633 | 3449 | 2844 | 2763 | 2783 | 2800 | 2547 | 2482 | 2697 | 2631 | 2693,35 | 2798 | 2589 | 0,926 | -0,112 | -2,743 | 0,006097 | 0,008125 | 0,0003348 | 0,002527 | 0,0012154 | 0,0012154 | TRUE | 0,0090358 | No |
| orf19.1901 | 9793 | 8687 | 9435 | 6000 | 12549 | 11001 | 11557 | 15740 | 8916 | 8786 | 8929 | 9013 | 11162 | 11944 | 11836 | 12007 | 10324,1 | 8911 | 11737 | 1,317 | 0,398 | 11,456 | 2,19E-30 | 7,13E-30 | 0,0004755 | 0,002165 | 0,0010561 | 0,0010561 | TRUE | 0,0133834 | No |
| orf19.1902 | 10423 | 9557 | 10461 | 6067 | 9979 | 8201 | 8412 | 11268 | 9490 | 9666 | 9900 | 9114 | 8876 | 8904 | 8615 | 8595 | 9144,98 | 9542 | 8748 | 0,916 | -0,126 | -3,476 | 0,000508 | 0,000731 | 0,0006361 | 0,002181 | 0,0011573 | 0,0011573 | TRUE | 0,0107886 | No |
| orf19.1906 | 78 | 81 | 68 | 55 | 205 | 172 | 192 | 247 | 71 | 82 | 64 | 83 | 182 | 187 | 197 | 188 | 131,76 | 75 | 188 | 2,536 | 1,343 | 10,243 | 1,27E-24 | 3,62E-24 | 1,00E-08 | 0,012063 | 0,0068895 | 0,0068895 | TRUE | 0,085964 | No |
| orf19.1907 | 4181 | 3427 | 4085 | 2558 | 7953 | 6305 | 7159 | 9898 | 3807 | 3466 | 3866 | 3843 | 7074 | 6845 | 7332 | 7550 | 5472,84 | 3746 | 7200 | 1,923 | 0,944 | 19,243 | 1,63E-82 | 1,19E-81 | 0,0018916 | 0,002278 | 0,0021004 | 0,0021004 | TRUE | 0,030935 | No |
| orf19.191 | 4539 | 4218 | 4559 | 2940 | 2166 | 1881 | 2047 | 2726 | 4133 | 4266 | 4315 | 4416 | 1927 | 2042 | 2096 | 2079 | 3159,29 | 4282 | 2036 | 0,476 | -1,072 | -25,007 | 5,17E-138 | 6,18E-137 | 0,000601 | 0,002455 | 0,0014086 | 0,0014086 | TRUE | 0,0161851 | No |
| orf19.1910 | 648 | 477 | 585 | 381 | 640 | 516 | 554 | 855 | 590 | 482 | 554 | 572 | 569 | 560 | 567 | 652 | 568,43 | 550 | 587 | 1,071 | 0,099 | 1,218 | 0,223121 | 0,256396 | 0,0049054 | 0,004361 | 0,004623 | 0,004623 | TRUE | 0,0809971 | No |
| orf19.1911 | 4663 | 3646 | 3960 | 2731 | 6462 | 5166 | 5656 | 8245 | 4245 | 3688 | 3748 | 4102 | 5748 | 5609 | 5792 | 6289 | 4902,7 | 3946 | 5860 | 1,486 | 0,571 | 9,981 | 1,85E-23 | 5,10E-23 | 0,0033592 | 0,002306 | 0,0029299 | 0,0029299 | TRUE | 0,0328881 | No |
| orf19.1913 | 875 | 582 | 681 | 570 | 592 | 529 | 561 | 818 | 797 | 589 | 645 | 856 | 527 | 574 | 575 | 624 | 648,18 | 722 | 575 | 0,799 | -0,323 | -2,986 | 0,002825 | 0,003854 | 0,0154797 | 0,004075 | 0,0097007 | 0,0097007 | TRUE | 0,1460204 | No |
| orf19.1914 | 543 | 417 | 458 | 335 | 768 | 658 | 753 | 1043 | 494 | 422 | 433 | 503 | 683 | 714 | 771 | 796 | 602,14 | 463 | 741 | 1,606 | 0,683 | 8,531 | 1,46E-17 | 3,47E-17 | 0,0045188 | 0,004231 | 0,004365 | 0,004365 | TRUE | 0,0411917 | No |
| orf19.1915 | 6484 | 4823 | 5370 | 3078 | 10506 | 7591 | 9501 | 12711 | 5903 | 4878 | 5082 | 4624 | 9345 | 8242 | 9730 | 9696 | 7187,5 | 5122 | 9253 | 1,806 | 0,853 | 11,18 | 5,10E-29 | 1,60E-28 | 0,008476 | 0,002221 | 0,0054319 | 0,0054319 | TRUE | 0,1290996 | No |
| orf19.1917 | 3139 | 2434 | 2834 | 1814 | 3996 | 3183 | 3711 | 5131 | 2858 | 2462 | 2682 | 2725 | 3554 | 3456 | 3801 | 3914 | 3181,42 | 2682 | 3681 | 1,373 | 0,458 | 7,862 | 3,79E-15 | 8,40E-15 | 0,0031188 | 0,002452 | 0,0029246 | 0,0029246 | TRUE | 0,0375289 | No |
| orf19.192 | 1765 | 1785 | 1882 | 1271 | 1660 | 1493 | 1491 | 1814 | 1607 | 1805 | 1781 | 1909 | 1477 | 1621 | 1527 | 1384 | 1638,86 | 1776 | 1502 | 0,846 | -0,241 | -3,67 | 0,000242 | 0,000356 | 0,0038496 | 0,002843 | 0,0035181 | 0,0035181 | TRUE | 0,0500629 | No |
| orf19.1920 | 32 | 12 | 18 | 10 | 20 | 11 | 15 | 35 | 29 | 12 | 17 | 15 | 18 | 12 | 15 | 27 | 18,14 | 18 | 18 | 0,981 | -0,027 | -0,073 | 0,941491 | 0,950026 | 0,0868928 | 0,074859 | 0,0796689 | 0,0796689 | TRUE | 0,9006023 | No |
| orf19.1923 | 2683 | 2023 | 2429 | 1775 | 2672 | 2304 | 2535 | 3566 | 2443 | 2046 | 2299 | 2666 | 2377 | 2502 | 2596 | 2720 | 2456,06 | 2364 | 2549 | 1,08 | 0,111 | 1,477 | 0,139775 | 0,164829 | 0,0071403 | 0,002575 | 0,0049886 | 0,0049886 | TRUE | 0,0986617 | No |
| orf19.1925 | 37 | 37 | 32 | 28 | 30 | 40 | 28 | 60 | 34 | 37 | 30 | 42 | 27 | 43 | 29 | 46 | 36 | 36 | 36 | 1,021 | 0,03 | 0,121 | 0,903847 | 0,917374 | 0,0227923 | 0,03873 | 0,0327575 | 0,0327575 | TRUE | 0,1852751 | No |
| orf19.1926 | 247 | 234 | 217 | 189 | 665 | 586 | 596 | 747 | 225 | 237 | 205 | 284 | 591 | 636 | 610 | 570 | 419,84 | 238 | 602 | 2,557 | 1,355 | 14,951 | 1,54E-50 | 7,16E-50 | 0,0041709 | 0,005183 | 0,0048356 | 0,0048356 | TRUE | 0,1553523 | No |
| orf19.1927 | 1726 | 1644 | 1713 | 1062 | 1928 | 1504 | 1663 | 2124 | 1571 | 1663 | 1621 | 1595 | 1715 | 1633 | 1703 | 1620 | 1640,23 | 1612 | 1668 | 1,034 | 0,048 | 1,062 | 0,288429 | 0,32521 | 1,00E-08 | 0,002842 | 0,0013433 | 0,0013433 | TRUE | 0,0053871 | No |
| orf19.193 | 433 | 509 | 497 | 287 | 196 | 154 | 158 | 195 | 394 | 515 | 470 | 431 | 174 | 167 | 162 | 149 | 307,83 | 452 | 163 | 0,359 | -1,477 | -13,979 | 2,09E-44 | 8,86E-44 | 0,0070641 | 0,006328 | 0,0066696 | 0,0066696 | TRUE | 0,1010311 | No |
| orf19.1930 | 147 | 141 | 144 | 86 | 32 | 26 | 24 | 44 | 134 | 143 | 136 | 129 | 28 | 28 | 25 | 34 | 82,09 | 136 | 29 | 0,214 | -2,227 | -12,572 | 3,00E-36 | 1,10E-35 | 1,00E-08 | 0,018128 | 0,0102151 | 0,0102151 | TRUE | 0,1074917 | No |
| orf19.1932 | 1051 | 867 | 910 | 655 | 505 | 397 | 413 | 585 | 957 | 877 | 861 | 984 | 449 | 431 | 423 | 446 | 678,54 | 920 | 437 | 0,477 | -1,068 | -15,805 | 2,88E-56 | 1,49E-55 | 0,0014426 | 0,003984 | 0,0027316 | 0,0027316 | TRUE | 0,0231591 | No |
| orf19.1933 | 1685 | 1569 | 1660 | 997 | 1153 | 988 | 1072 | 1391 | 1534 | 1587 | 1571 | 1498 | 1026 | 1073 | 1098 | 1061 | 1305,86 | 1548 | 1064 | 0,687 | -0,542 | -11,35 | 7,44E-30 | 2,38E-29 | 1,00E-08 | 0,003049 | 0,0014086 | 0,0014086 | TRUE | 0,007192 | No |
| orf19.1934 | 2698 | 2438 | 2704 | 1821 | 3202 | 2609 | 3001 | 3954 | 2456 | 2466 | 2559 | 2735 | 2848 | 2833 | 3073 | 3016 | 2748,37 | 2554 | 2942 | 1,154 | 0,207 | 4,163 | 3,14E-05 | 4,85E-05 | 0,0016343 | 0,002518 | 0,0019953 | 0,0019953 | TRUE | 0,0258925 | No |
| orf19.1935 | 11 | 13 | 10 | 7 | 20 | 20 | 15 | 25 | 10 | 13 | 9 | 11 | 18 | 22 | 15 | 19 | 14,63 | 11 | 18 | 1,714 | 0,778 | 2,16 | 0,030774 | 0,038879 | 1,00E-08 | 0,092301 | 0,0500261 | 0,0500261 | TRUE | 0,0851292 | No |
| orf19.1936 | 8020 | 6740 | 7596 | 4880 | 4531 | 3892 | 4309 | 5750 | 7302 | 6817 | 7189 | 7331 | 4030 | 4226 | 4413 | 4386 | 5711,63 | 7160 | 4264 | 0,596 | -0,747 | -17,25 | 1,12E-66 | 6,75E-66 | 0,0011512 | 0,002268 | 0,0016181 | 0,0016181 | TRUE | 0,0167713 | No |

| | | | | | | | | | | | | | | | | | | | | | | | | | | | | | | | |
|---|---|---|---|---|---|---|---|---|---|---|---|---|---|---|---|---|---|---|---|---|---|---|---|---|---|---|---|---|---|---|---|
| orf19.1938 | 1076 | 963 | 1037 | 687 | 1850 | 1441 | 1532 | 2078 | 980 | 974 | 981 | 1032 | 1646 | 1565 | 1569 | 1585 | 1291,39 | 992 | 1591 | 1,608 | 0,685 | 14,335 | 1,32E-46 | 5,76E-46 | 1,00E-08 | 0,00306 | 0,0013611 | 0,0013611 | TRUE | 0,0074907 | No |
| orf19.1939 | 20 | 18 | 19 | 8 | 46 | 43 | 45 | 50 | 18 | 18 | 18 | 12 | 41 | 47 | 46 | 38 | 29,78 | 16 | 43 | 2,522 | 1,334 | 5,002 | 5,68E-07 | 9,54E-07 | 1,00E-08 | 0,046395 | 0,026179 | 0,026179 | TRUE | 0,0977659 | No |
| orf19.194 | 1356 | 1648 | 1508 | 982 | 1743 | 1615 | 1517 | 1823 | 1235 | 1667 | 1427 | 1475 | 1550 | 1753 | 1554 | 1391 | 1506,46 | 1451 | 1562 | 1,076 | 0,106 | 1,176 | 0,239495 | 0,273763 | 0,0109995 | 0,002914 | 0,0071398 | 0,0071398 | TRUE | 0,1240598 | No |
| orf19.1940 | 2851 | 2350 | 2646 | 1669 | 2376 | 1924 | 2179 | 2911 | 2596 | 2377 | 2504 | 2507 | 2113 | 2089 | 2232 | 2221 | 2329,78 | 2496 | 2164 | 0,867 | -0,206 | -4,522 | 6,12E-06 | 9,81E-06 | 0,0008279 | 0,002604 | 0,0015575 | 0,0015575 | TRUE | 0,012903 | No |
| orf19.1941 | 935 | 794 | 849 | 493 | 1009 | 864 | 930 | 1121 | 851 | 803 | 804 | 741 | 897 | 938 | 952 | 855 | 855,18 | 800 | 910 | 1,134 | 0,181 | 2,902 | 0,003712 | 0,005015 | 0,0014962 | 0,003582 | 0,0025534 | 0,0025534 | TRUE | 0,0254488 | No |
| orf19.1942 | 3049 | 2818 | 3009 | 1970 | 4083 | 3617 | 3708 | 5099 | 2776 | 2850 | 2848 | 2959 | 3632 | 3927 | 3797 | 3890 | 3334,86 | 2858 | 3812 | 1,335 | 0,416 | 10,049 | 9,26E-24 | 2,58E-23 | 0,000663 | 0,002433 | 0,001338 | 0,001338 | TRUE | 0,0123946 | No |
| orf19.1943 | 431 | 446 | 430 | 296 | 555 | 508 | 450 | 648 | 392 | 451 | 407 | 445 | 494 | 552 | 461 | 494 | 461,93 | 424 | 500 | 1,182 | 0,242 | 3,019 | 0,002535 | 0,003469 | 0,0027899 | 0,004897 | 0,0039585 | 0,0039585 | TRUE | 0,0523644 | No |
| orf19.1944 | 20649 | 17719 | 20142 | 12906 | 4990 | 4386 | 4708 | 6215 | 18800 | 17921 | 19063 | 19387 | 4438 | 4762 | 4822 | 4741 | 11741,7 | 18793 | 4691 | 0,25 | -2,002 | -48,873 | 0 | 0 | 0,0010514 | 0,00215 | 0,0014839 | 0,0014839 | TRUE | 0,0160648 | No |
| orf19.1945 | 19365 | 16882 | 18378 | 11219 | 13004 | 10492 | 10776 | 14618 | 17631 | 17074 | 17393 | 16853 | 11567 | 11391 | 11036 | 11151 | 14262 | 17238 | 11286 | 0,655 | -0,611 | -19,547 | 4,35E-85 | 3,29E-84 | 0,000324 | 0,00213 | 0,0008667 | 0,0008667 | TRUE | 0,0034531 | No |
| orf19.1946 | 2556 | 2191 | 2292 | 1374 | 2421 | 1836 | 1905 | 2629 | 2327 | 2216 | 2169 | 2064 | 2153 | 1993 | 1951 | 2005 | 2109,92 | 2194 | 2026 | 0,922 | -0,117 | -2,26 | 0,02383 | 0,030365 | 0,0015524 | 0,002663 | 0,002087 | 0,002087 | TRUE | 0,0219075 | No |
| orf19.1948 | 1151 | 939 | 1037 | 733 | 958 | 859 | 894 | 1197 | 1048 | 950 | 981 | 1101 | 852 | 933 | 916 | 913 | 961,69 | 1020 | 904 | 0,887 | -0,173 | -2,805 | 0,005035 | 0,006754 | 0,0015778 | 0,003411 | 0,0025887 | 0,0025887 | TRUE | 0,0302525 | No |
| orf19.1949 | 4098 | 3521 | 3831 | 2389 | 3435 | 2909 | 2988 | 4080 | 3731 | 3561 | 3626 | 3589 | 3055 | 3158 | 3060 | 3112 | 3361,57 | 3627 | 3096 | 0,853 | -0,229 | -6,4 | 1,56E-10 | 3,01E-10 | 1,00E-08 | 0,00243 | 0,0009278 | 0,0009278 | TRUE | 0,0047357 | No |
| orf19.1950 | 583 | 442 | 497 | 389 | 352 | 275 | 287 | 442 | 531 | 447 | 470 | 584 | 313 | 299 | 294 | 337 | 409,41 | 508 | 311 | 0,615 | -0,7 | -7,412 | 1,25E-13 | 2,66E-13 | 0,0069061 | 0,005264 | 0,0060135 | 0,0060135 | TRUE | 0,1056779 | No |
| orf19.1953 | 3206 | 2721 | 2961 | 1953 | 2655 | 2004 | 2243 | 3103 | 2919 | 2752 | 2802 | 2934 | 2362 | 2176 | 2297 | 2367 | 2576,05 | 2852 | 2300 | 0,808 | -0,308 | -6,89 | 5,59E-12 | 1,14E-11 | 0,0007074 | 0,00255 | 0,0015273 | 0,0015273 | TRUE | 0,0156863 | No |
| orf19.1954 | 4939 | 4418 | 4921 | 2826 | 5145 | 3866 | 4316 | 5668 | 4497 | 4468 | 4657 | 4245 | 4576 | 4197 | 4420 | 4324 | 4423,11 | 4467 | 4379 | 0,98 | -0,029 | -0,679 | 0,497103 | 0,538677 | 0,0010723 | 0,002336 | 0,0015845 | 0,0015845 | TRUE | 0,0129106 | No |
| orf19.1955 | 887 | 766 | 802 | 541 | 943 | 729 | 812 | 1012 | 808 | 775 | 759 | 813 | 839 | 791 | 832 | 772 | 798,48 | 789 | 808 | 1,026 | 0,037 | 0,636 | 0,524761 | 0,566528 | 1,00E-08 | 0,003691 | 0,0019922 | 0,0019922 | TRUE | 0,0121374 | No |
| orf19.1956 | 3764 | 3379 | 3537 | 2316 | 3473 | 2951 | 2963 | 4029 | 3427 | 3417 | 3347 | 3479 | 3089 | 3204 | 3034 | 3073 | 3258,98 | 3418 | 3100 | 0,907 | -0,14 | -3,858 | 0,000114 | 0,000171 | 1,00E-08 | 0,002442 | 0,0009621 | 0,0009621 | TRUE | 0,0059057 | No |
| orf19.1957 | 10981 | 8991 | 9929 | 5810 | 8808 | 6866 | 7976 | 10816 | 9998 | 9093 | 9397 | 8728 | 7834 | 7455 | 8168 | 8251 | 8615,45 | 9304 | 7927 | 0,852 | -0,231 | -4,496 | 6,93E-06 | 1,11E-05 | 0,0025596 | 0,00219 | 0,002428 | 0,002428 | TRUE | 0,0308535 | No |
| orf19.1958 | 13 | 13 | 17 | 13 | 37 | 23 | 35 | 40 | 12 | 13 | 16 | 20 | 33 | 25 | 36 | 31 | 23,1 | 15 | 31 | 2,098 | 1,069 | 3,561 | 0,00037 | 0,000537 | 1,00E-08 | 0,059211 | 0,0358185 | 0,0358185 | TRUE | 0,1077013 | No |
| orf19.1959 | 1320 | 1259 | 1266 | 894 | 2907 | 2266 | 2492 | 3269 | 1202 | 1273 | 1198 | 1343 | 2586 | 2460 | 2552 | 2494 | 1888,49 | 1254 | 2523 | 2,018 | 1,013 | 20,777 | 7,01E-96 | 5,77E-95 | 0,0006487 | 0,002737 | 0,0016684 | 0,0016684 | TRUE | 0,0241936 | No |
| orf19.1960 | 6339 | 5973 | 6402 | 4019 | 7865 | 6590 | 6786 | 9311 | 5771 | 6041 | 6059 | 6037 | 6996 | 7155 | 6950 | 7103 | 6513,92 | 5977 | 7051 | 1,18 | 0,239 | 7,35 | 1,98E-13 | 4,21E-13 | 0,0001482 | 0,00224 | 0,0008566 | 0,0008566 | TRUE | 0,006763 | No |
| orf19.1961 | 9144 | 8859 | 9615 | 5557 | 7221 | 5830 | 6366 | 8166 | 8325 | 8960 | 9100 | 8348 | 6423 | 6330 | 6520 | 6229 | 7529,21 | 8683 | 6376 | 0,734 | -0,446 | -10,706 | 9,51E-27 | 2,84E-26 | 0,0010946 | 0,002212 | 0,0015335 | 0,0015335 | TRUE | 0,0127978 | No |
| orf19.1963 | 5059 | 4545 | 5046 | 2765 | 10227 | 8200 | 9118 | 11913 | 4606 | 4597 | 4776 | 4153 | 9097 | 8903 | 9338 | 9087 | 6819,58 | 4533 | 9106 | 2,007 | 1,005 | 21,955 | 7,82E-107 | 7,18E-106 | 0,0016935 | 0,002231 | 0,001841 | 0,001841 | TRUE | 0,0369131 | No |
| orf19.1964 | 2680 | 2592 | 2703 | 1789 | 3762 | 3214 | 3248 | 4444 | 2440 | 2622 | 2558 | 2687 | 3346 | 3490 | 3326 | 3390 | 2982,38 | 2577 | 3388 | 1,316 | 0,397 | 9,356 | 8,31E-21 | 2,15E-20 | 0,0006101 | 0,00248 | 0,0013789 | 0,0013789 | TRUE | 0,0157084 | No |
| orf19.1966 | 4567 | 3888 | 4219 | 2598 | 5832 | 4283 | 4798 | 6522 | 4158 | 3932 | 3993 | 3903 | 5187 | 4650 | 4914 | 4975 | 4464,02 | 3996 | 4932 | 1,234 | 0,303 | 6,881 | 5,95E-12 | 1,21E-11 | 0,0011118 | 0,002333 | 0,0016338 | 0,0016338 | TRUE | 0,0177216 | No |
| orf19.1967 | 1565 | 1301 | 1449 | 789 | 1807 | 1233 | 1427 | 1931 | 1425 | 1316 | 1371 | 1185 | 1607 | 1339 | 1461 | 1473 | 1397,2 | 1324 | 1470 | 1,108 | 0,148 | 2,114 | 0,034493 | 0,043372 | 0,0048491 | 0,002982 | 0,0040001 | 0,0040001 | TRUE | 0,0558689 | No |
| orf19.1968 | 106 | 95 | 117 | 78 | 144 | 140 | 123 | 163 | 97 | 96 | 111 | 117 | 128 | 152 | 126 | 124 | 118,86 | 105 | 132 | 1,265 | 0,34 | 2,532 | 0,011354 | 0,014829 | 1,00E-08 | 0,013151 | 0,0086459 | 0,0086459 | TRUE | 0,0928401 | No |
| orf19.1968.1 | 245 | 233 | 210 | 137 | 250 | 183 | 206 | 266 | 223 | 236 | 199 | 206 | 222 | 199 | 211 | 203 | 212,27 | 216 | 209 | 0,966 | -0,05 | -0,502 | 0,615434 | 0,654555 | 1,00E-08 | 0,00826 | 0,0049837 | 0,0049837 | TRUE | 0,0418843 | No |
| orf19.1969 | 50 | 31 | 40 | 34 | 67 | 79 | 52 | 80 | 46 | 31 | 38 | 51 | 60 | 86 | 53 | 61 | 53,18 | 42 | 65 | 1,576 | 0,657 | 3,01 | 0,002613 | 0,003572 | 0,0230297 | 0,026876 | 0,0254585 | 0,0254585 | TRUE | 0,381375 | No |
| orf19.1970 | 2499 | 2112 | 2370 | 1632 | 2185 | 1664 | 1798 | 2391 | 2275 | 2136 | 2243 | 2452 | 1943 | 1807 | 1841 | 1824 | 2065,15 | 2276 | 1854 | 0,816 | -0,294 | -5,693 | 1,25E-08 | 2,25E-08 | 0,0015521 | 0,002677 | 0,0020667 | 0,0020667 | TRUE | 0,0301227 | No |
| orf19.1971 | 12433 | 10958 | 11911 | 7586 | 12525 | 10123 | 10877 | 14588 | 11320 | 11083 | 11273 | 11395 | 11141 | 10991 | 11139 | 11128 | 11183,7 | 11268 | 11100 | 0,985 | -0,021 | -0,81 | 0,417711 | 0,459043 | 1,00E-08 | 0,002155 | 0,000577 | 0,000577 | TRUE | 0,0014871 | No |
| orf19.1972 | 4432 | 3654 | 3903 | 2581 | 5046 | 4218 | 4517 | 6173 | 4035 | 3696 | 3694 | 3877 | 4488 | 4580 | 4626 | 4709 | 4213,04 | 3826 | 4601 | 1,203 | 0,267 | 6,386 | 1,71E-10 | 3,29E-10 | 0,0008014 | 0,002351 | 0,0014303 | 0,0014303 | TRUE | 0,0174383 | No |
| orf19.1973 | 1248 | 950 | 1152 | 692 | 1220 | 873 | 1086 | 1464 | 1136 | 961 | 1090 | 1039 | 1085 | 948 | 1112 | 1117 | 1061,1 | 1056 | 1066 | 1,009 | 0,012 | 0,172 | 0,863341 | 0,884144 | 0,0045283 | 0,003282 | 0,0039171 | 0,0039171 | TRUE | 0,0640888 | No |
| orf19.1974 | 4953 | 4094 | 4594 | 3278 | 3968 | 3399 | 3294 | 4586 | 4509 | 4141 | 4348 | 4924 | 3529 | 3690 | 3373 | 3498 | 4001,69 | 4480 | 3522 | 0,787 | -0,346 | -6,167 | 6,97E-10 | 1,31E-09 | 0,0030929 | 0,002367 | 0,0027658 | 0,0027658 | TRUE | 0,0523999 | No |
| orf19.1975 | 588 | 558 | 534 | 298 | 849 | 653 | 716 | 956 | 535 | 564 | 505 | 448 | 755 | 709 | 733 | 729 | 622,42 | 513 | 732 | 1,419 | 0,505 | 7,027 | 2,11E-12 | 4,33E-12 | 0,001953 | 0,004159 | 0,0032741 | 0,0032741 | TRUE | 0,0719617 | No |
| orf19.1976 | 451 | 432 | 469 | 270 | 559 | 459 | 473 | 643 | 411 | 437 | 444 | 406 | 497 | 498 | 484 | 490 | 458,43 | 424 | 492 | 1,159 | 0,212 | 3,075 | 0,002108 | 0,002902 | 1,00E-08 | 0,004919 | 0,0023792 | 0,0023792 | TRUE | 0,011277 | No |
| orf19.1977 | 1387 | 1203 | 1265 | 925 | 2213 | 1912 | 1912 | 2642 | 1263 | 1217 | 1197 | 1389 | 1968 | 2076 | 1958 | 2015 | 1635,5 | 1266 | 2004 | 1,587 | 0,667 | 12,39 | 2,97E-35 | 1,07E-34 | 0,0014009 | 0,002845 | 0,0021211 | 0,0021211 | TRUE | 0,0458735 | No |
| orf19.1978 | 955 | 873 | 937 | 575 | 2525 | 1973 | 2060 | 2997 | 869 | 883 | 887 | 864 | 2246 | 2142 | 2110 | 2286 | 1535,85 | 876 | 2196 | 2,508 | 1,327 | 26,779 | 5,65E-158 | 7,74E-157 | 1,00E-08 | 0,002897 | 0,001534 | 0,001534 | TRUE | 0,0095081 | No |
| orf19.1979 | 22144 | 20492 | 21615 | 12990 | 7416 | 5952 | 6352 | 8674 | 20161 | 20725 | 20457 | 19513 | 6596 | 6462 | 6505 | 6617 | 13379,6 | 20214 | 6545 | 0,324 | -1,627 | -51,304 | 0 | 0 | 0,0002604 | 0,002136 | 0,0008677 | 0,0008677 | TRUE | 0,0065148 | No |
| orf19.198 | 5538 | 4894 | 5401 | 3618 | 11345 | 9715 | 9926 | 13494 | 5042 | 4950 | 5112 | 5435 | 10091 | 10548 | 10165 | 10293 | 7704,48 | 5135 | 10274 | 2,003 | 1,002 | 26,054 | 1,21E-149 | 1,56E-148 | 0,0008389 | 0,002208 | 0,0012699 | 0,0012699 | TRUE | 0,0178906 | No |
| orf19.1980 | 111 | 107 | 110 | 54 | 68 | 51 | 57 | 76 | 101 | 108 | 104 | 81 | 60 | 55 | 58 | 58 | 78,34 | 98 | 58 | 0,584 | -0,776 | -4,87 | 1,11E-06 | 1,85E-06 | 1,00E-08 | 0,0189 | 0,011042 | 0,011042 | TRUE | 0,1034879 | No |
| orf19.1981 | 2014 | 1736 | 1842 | 1073 | 1112 | 781 | 915 | 1248 | 1834 | 1756 | 1743 | 1612 | 989 | 848 | 937 | 952 | 1333,83 | 1736 | 932 | 0,536 | -0,899 | -14,764 | 2,52E-49 | 1,14E-48 | 0,0023694 | 0,003027 | 0,0027471 | 0,0027471 | TRUE | 0,0416961 | No |
| orf19.1985 | 643 | 584 | 680 | 414 | 13105 | 11448 | 10233 | 14367 | 585 | 591 | 644 | 622 | 11656 | 12429 | 10480 | 10959 | 5995,81 | 610 | 11381 | 18,659 | 4,222 | 65,29 | 0 | 0 | 0,0037748 | 0,002257 | 0,0030952 | 0,0030952 | TRUE | 0,0467813 | No |
| orf19.1986 | 2656 | 2283 | 2489 | 1519 | 4480 | 3458 | 3784 | 5222 | 2418 | 2309 | 2356 | 2282 | 3985 | 3754 | 3875 | 3983 | 3120,31 | 2341 | 3899 | 1,665 | 0,735 | 18,328 | 4,95E-75 | 3,28E-74 | 0,0004089 | 0,00246 | 0,0011964 | 0,0011964 | TRUE | 0,0074961 | No |
| orf19.1988 | 1378 | 1139 | 1245 | 858 | 1269 | 1002 | 1071 | 1511 | 1255 | 1152 | 1178 | 1289 | 1129 | 1088 | 1097 | 1153 | 1167,47 | 1218 | 1117 | 0,919 | -0,122 | -2,262 | 0,023721 | 0,030232 | 0,0007808 | 0,003169 | 0,0019376 | 0,0019376 | TRUE | 0,0159086 | No |
| orf19.1989 | 4210 | 3733 | 4028 | 2580 | 7119 | 5935 | 6493 | 8652 | 3833 | 3776 | 3812 | 3876 | 6332 | 6444 | 6650 | 6600 | 5165,2 | 3824 | 6506 | 1,702 | 0,767 | 22,91 | 3,69E-116 | 3,67E-115 | 0,0001842 | 0,002293 | 0,0008644 | 0,0008644 | TRUE | 0,0040279 | No |
| orf19.199 | 2665 | 2790 | 2955 | 1650 | 2279 | 2090 | 2066 | 2508 | 2426 | 2822 | 2797 | 2479 | 2027 | 2269 | 2116 | 1913 | 2356,07 | 2631 | 2081 | 0,79 | -0,34 | -4,976 | 6,48E-07 | 1,09E-06 | 0,0051363 | 0,002598 | 0,0040513 | 0,0040513 | TRUE | 0,0443891 | No |
| orf19.1990 | 3063 | 3345 | 3462 | 2059 | 2636 | 2192 | 2375 | 3023 | 2789 | 3383 | 3276 | 3093 | 2345 | 2380 | 2432 | 2306 | 2750,51 | 3135 | 2366 | 0,754 | -0,407 | -6,761 | 1,37E-11 | 2,75E-11 | 0,003209 | 0,002517 | 0,0031042 | 0,0031042 | TRUE | 0,0692316 | No |
| orf19.1991 | 3881 | 3731 | 3837 | 2448 | 2937 | 2400 | 2427 | 3173 | 3533 | 3774 | 3631 | 3677 | 2612 | 2606 | 2486 | 2420 | 3092,46 | 3654 | 2531 | 0,693 | -0,53 | -12,382 | 3,26E-35 | 1,17E-34 | 0,0007678 | 0,002464 | 0,0014273 | 0,0014273 | TRUE | 0,0110049 | No |
| orf19.1992 | 3360 | 3456 | 3692 | 2316 | 4636 | 3987 | 3995 | 5207 | 3059 | 3495 | 3494 | 3479 | 4124 | 4329 | 4091 | 3972 | 3755,42 | 3382 | 4129 | 1,221 | 0,288 | 5,422 | 5,90E-08 | 1,03E-07 | 0,0023135 | 0,002389 | 0,0024429 | 0,0024429 | TRUE | 0,0513378 | No |
| orf19.1993 | 1265 | 1651 | 1531 | 935 | 1365 | 1278 | 1144 | 1365 | 1152 | 1670 | 1449 | 1405 | 1214 | 1388 | 1172 | 1041 | 1311,19 | 1419 | 1204 | 0,848 | -0,238 | -2,244 | 0,024804 | 0,031556 | 0,0172477 | 0,003044 | 0,0100427 | 0,0100427 | TRUE | 0,1971518 | No |
| orf19.1994 | 1492 | 1642 | 1744 | 1157 | 1806 | 1716 | 1686 | 2057 | 1358 | 1661 | 1651 | 1738 | 1606 | 1863 | 1727 | 1569 | 1646,61 | 1602 | 1691 | 1,056 | 0,079 | 0,988 | 0,323386 | 0,362917 | 0,0074195 | 0,002839 | 0,0055614 | 0,0055614 | TRUE | 0,1284486 | No |
| orf19.1995 | 544 | 543 | 525 | 302 | 722 | 644 | 630 | 803 | 495 | 549 | 497 | 454 | 642 | 699 | 645 | 613 | 574,27 | 499 | 650 | 1,296 | 0,374 | 5,077 | 3,84E-07 | 6,49E-07 | 0,0022577 | 0,004337 | 0,0034287 | 0,0034287 | TRUE | 0,0520718 | No |
| orf19.1996 | 7192 | 6588 | 7097 | 4291 | 7946 | 6344 | 6375 | 8460 | 6548 | 6663 | 6717 | 6446 | 7068 | 6888 | 6529 | 6453 | 6663,9 | 6594 | 6734 | 1,021 | 0,03 | 0,74 | 0,45908 | 0,50105 | 0,0009086 | 0,002235 | 0,0014208 | 0,0014208 | TRUE | 0,0137494 | No |
| orf19.1998 | 367 | 319 | 337 | 221 | 323 | 266 | 249 | 340 | 334 | 323 | 319 | 332 | 287 | 289 | 255 | 259 | 299,77 | 327 | 272 | 0,833 | -0,263 | -3,103 | 0,001915 | 0,002646 | 1,00E-08 | 0,006444 | 0,0035499 | 0,0035499 | TRUE | 0,0188969 | No |
| orf19.1999 | 1911 | 1601 | 1723 | 1055 | 660 | 496 | 606 | 715 | 1740 | 1619 | 1631 | 1585 | 587 | 539 | 621 | 545 | 1108,27 | 1644 | 573 | 0,348 | -1,523 | -25,007 | 5,17E-138 | 6,18E-137 | 0,0015052 | 0,003229 | 0,0024206 | 0,0024206 | TRUE | 0,0352696 | No |
| orf19.20 | 2552 | 2444 | 2486 | 1629 | 1773 | 1494 | 1626 | 2211 | 2323 | 2472 | 2353 | 2447 | 1577 | 1622 | 1665 | 1687 | 2018,25 | 2399 | 1638 | 0,683 | -0,549 | -12,349 | 4,94E-35 | 1,76E-34 | 0,0004526 | 0,002692 | 0,0013887 | 0,0013887 | TRUE | 0,007827 | No |
| orf19.200 | 1072 | 998 | 1118 | 599 | 2063 | 1565 | 1701 | 2321 | 976 | 1009 | 1058 | 900 | 1835 | 1699 | 1742 | 1770 | 1373,74 | 986 | 1762 | 1,782 | 0,834 | 14,808 | 1,29E-49 | 5,91E-49 | 0,0014348 | 0,002999 | 0,0022344 | 0,0022344 | TRUE | 0,0357764 | No |
| orf19.2000 | 3654 | 3792 | 4004 | 2245 | 3105 | 2516 | 2846 | 3453 | 3327 | 3835 | 3789 | 3372 | 2762 | 2732 | 2915 | 2634 | 3170,74 | 3581 | 2761 | 0,77 | -0,377 | -6,409 | 1,47E-10 | 2,84E-10 | 0,0032705 | 0,002454 | 0,002996 | 0,002996 | TRUE | 0,0289019 | No |
| orf19.2001 | 607 | 678 | 702 | 388 | 537 | 493 | 475 | 581 | 553 | 686 | 664 | 583 | 478 | 535 | 486 | 443 | 553,68 | 622 | 486 | 0,779 | -0,36 | -4,122 | 3,76E-05 | 5,79E-05 | 0,0067024 | 0,004424 | 0,0055155 | 0,0055155 | TRUE | 0,071318 | No |
| orf19.2002 | 3030 | 3061 | 3031 | 2009 | 3236 | 2790 | 2864 | 3608 | 2759 | 3096 | 2869 | 3018 | 2878 | 3029 | 2933 | 2752 | 2916,72 | 2936 | 2898 | 0,987 | -0,018 | -0,36 | 0,718563 | 0,75182 | 0,0018107 | 0,00249 | 0,0020899 | 0,0020899 | TRUE | 0,0203116 | No |
| orf19.2003 | 8360 | 7676 | 8230 | 4854 | 9982 | 8262 | 8726 | 11677 | 7611 | 7763 | 7789 | 7291 | 8879 | 8970 | 8937 | 8907 | 8268,51 | 7614 | 8923 | 1,171 | 0,228 | 7,002 | 2,52E-12 | 5,17E-12 | 0,0002987 | 0,002197 | 0,000894 | 0,000894 | TRUE | 0,0093872 | No |
| orf19.2005 | 18724 | 16446 | 17945 | 11023 | 9635 | 8272 | 8845 | 11890 | 17047 | 16633 | 16983 | 16558 | 8570 | 8981 | 9058 | 9070 | 12862,7 | 16805 | 8920 | 0,531 | -0,914 | -28,524 | 5,96E-179 | 9,17E-178 | 0,0003438 | 0,00214 | 0,0009019 | 0,0009019 | TRUE | 0,0085867 | No |
| orf19.2006 | 908 | 867 | 935 | 566 | 1335 | 1081 | 1136 | 1541 | 827 | 877 | 885 | 850 | 1187 | 1174 | 1163 | 1175 | 1017,34 | 860 | 1175 | 1,367 | 0,451 | 8,894 | 5,90E-19 | 1,46E-18 | 1,00E-08 | 0,003336 | 0,0014468 | 0,0014468 | TRUE | 0,009116 | No |
| orf19.2006.1 | 2255 | 1972 | 1955 | 1188 | 880 | 650 | 619 | 946 | 2053 | 1994 | 1850 | 1785 | 783 | 706 | 634 | 722 | 1315,79 | 1920 | 711 | 0,37 | -1,433 | -20,324 | 7,96E-92 | 6,31E-91 | 0,0045156 | 0,003041 | 0,0038405 | 0,0038405 | TRUE | 0,0618009 | No |
| orf19.2007 | 536 | 526 | 510 | 317 | 451 | 371 | 427 | 557 | 488 | 532 | 483 | 476 | 401 | 403 | 437 | 425 | 455,62 | 495 | 416 | 0,841 | -0,25 | -3,456 | 0,000548 | 0,000786 | 1,00E-08 | 0,004936 | 0,0028105 | 0,0028105 | TRUE | 0,0301404 | No |
| orf19.2008 | 1670 | 1670 | 1786 | 1042 | 1311 | 1099 | 1139 | 1460 | 1520 | 1689 | 1690 | 1565 | 1166 | 1193 | 1166 | 1114 | 1388,06 | 1616 | 1160 | 0,717 | -0,481 | -8,916 | 4,84E-19 | 1,20E-18 | 0,0013049 | 0,002989 | 0,0020574 | 0,0020574 | TRUE | 0,0209767 | No |

| | | | | | | | | | | | | | | | | | | | | | | | | | | | | | | | |
|---|---|---|---|---|---|---|---|---|---|---|---|---|---|---|---|---|---|---|---|---|---|---|---|---|---|---|---|---|---|---|---|
| orf19.2009 | 1934 | 1752 | 1864 | 1136 | 1591 | 1319 | 1343 | 1740 | 1761 | 1772 | 1764 | 1706 | 1415 | 1432 | 1375 | 1327 | 1569,16 | 1751 | 1387 | 0,791 | -0,338 | -7,44 | 1,01E-13 | 2,16E-13 | 1,00E-08 | 0,002879 | 0,0013422 | 0,0013422 | TRUE | 0,0109979 | No |
| orf19.201 | 262 | 213 | 254 | 130 | 233 | 228 | 222 | 288 | 239 | 215 | 240 | 195 | 207 | 248 | 227 | 220 | 223,93 | 222 | 226 | 1,004 | 0,006 | 0,057 | 0,954682 | 0,960208 | 0,0020566 | 0,007936 | 0,005702 | 0,005702 | TRUE | 0,0595964 | No |
| orf19.2010 | 14 | 20 | 23 | 9 | 11 | 20 | 11 | 19 | 13 | 20 | 22 | 14 | 10 | 22 | 11 | 14 | 15,69 | 17 | 14 | 0,824 | -0,279 | -0,752 | 0,451883 | 0,493787 | 0,0339259 | 0,086232 | 0,0681445 | 0,0681445 | TRUE | 0,4600982 | No |
| orf19.2012 | 3759 | 3403 | 3540 | 2238 | 3671 | 3209 | 3386 | 4457 | 3422 | 3442 | 3350 | 3362 | 3265 | 3484 | 3468 | 3400 | 3399,14 | 3394 | 3404 | 1,002 | 0,003 | 0,091 | 0,92789 | 0,938043 | 0,000249 | 0,002426 | 0,0010541 | 0,0010541 | TRUE | 0,0091893 | No |
| orf19.2013 | 10555 | 8738 | 9585 | 6144 | 6602 | 5447 | 5674 | 7938 | 9610 | 8838 | 9071 | 9229 | 5872 | 5914 | 5811 | 6055 | 7550,04 | 9187 | 5913 | 0,644 | -0,635 | -17,09 | 1,75E-65 | 1,04E-64 | 0,0005929 | 0,002212 | 0,0011901 | 0,0011901 | TRUE | 0,0120178 | No |
| orf19.2014 | 6808 | 5818 | 6304 | 4158 | 5233 | 4628 | 4768 | 6597 | 6198 | 5884 | 5966 | 6246 | 4655 | 5025 | 4883 | 5032 | 5486,18 | 6074 | 4899 | 0,807 | -0,31 | -7,642 | 2,14E-14 | 4,66E-14 | 0,0008226 | 0,002278 | 0,0013936 | 0,0013936 | TRUE | 0,0138565 | No |
| orf19.2016 | 219 | 247 | 207 | 138 | 236 | 224 | 170 | 279 | 199 | 250 | 196 | 207 | 210 | 243 | 174 | 213 | 211,56 | 213 | 210 | 0,985 | -0,022 | -0,179 | 0,857845 | 0,879151 | 0,0108088 | 0,008281 | 0,0093393 | 0,0093393 | TRUE | 0,1528059 | No |
| orf19.2017 | 6121 | 5487 | 5798 | 3879 | 8543 | 7446 | 7047 | 9653 | 5573 | 5550 | 5487 | 5827 | 7599 | 8084 | 7217 | 7363 | 6587,5 | 5609 | 7566 | 1,349 | 0,432 | 9,737 | 2,09E-22 | 5,60E-22 | 0,001408 | 0,002238 | 0,0017334 | 0,0017334 | TRUE | 0,0256987 | No |
| orf19.2018 | 1651 | 1361 | 1460 | 1108 | 2238 | 1723 | 2000 | 2599 | 1503 | 1377 | 1382 | 1664 | 1991 | 1871 | 2048 | 1983 | 1727,24 | 1482 | 1973 | 1,335 | 0,417 | 6,547 | 5,88E-11 | 1,15E-10 | 0,0040883 | 0,002802 | 0,0033002 | 0,0033002 | TRUE | 0,077371 | No |
| orf19.2018.1 | 116 | 106 | 122 | 83 | 133 | 140 | 124 | 165 | 106 | 107 | 115 | 125 | 118 | 152 | 127 | 126 | 122,01 | 113 | 131 | 1,157 | 0,211 | 1,597 | 0,110354 | 0,131891 | 1,00E-08 | 0,012864 | 0,0084341 | 0,0084341 | TRUE | 0,1135188 | No |
| orf19.2018.2 | 18 | 12 | 9 | 7 | 23 | 18 | 8 | 31 | 16 | 12 | 9 | 11 | 20 | 20 | 8 | 24 | 14,92 | 12 | 18 | 1,513 | 0,597 | 1,499 | 0,133808 | 0,15829 | 0,0677921 | 0,090547 | 0,0823287 | 0,0823287 | TRUE | 0,6526024 | No |
| orf19.2019 | 2861 | 2363 | 2529 | 1850 | 2255 | 1823 | 1795 | 2540 | 2605 | 2390 | 2393 | 2779 | 2006 | 1979 | 1838 | 1938 | 2241,01 | 2542 | 1940 | 0,764 | -0,387 | -6,688 | 2,27E-11 | 4,51E-11 | 0,0028284 | 0,002626 | 0,0027695 | 0,0027695 | TRUE | 0,045355 | No |
| orf19.202 | 5247 | 4506 | 4771 | 3194 | 6250 | 5143 | 5584 | 7673 | 4777 | 4557 | 4515 | 4798 | 5559 | 5584 | 5719 | 5853 | 5170,31 | 4662 | 5679 | 1,219 | 0,286 | 7,537 | 4,81E-14 | 1,04E-13 | 0,0005176 | 0,002293 | 0,0011813 | 0,0011813 | TRUE | 0,0054915 | No |
| orf19.2020 | 11044 | 9528 | 10493 | 6820 | 4820 | 4141 | 4265 | 5933 | 10055 | 9637 | 9931 | 10245 | 4287 | 4496 | 4368 | 4526 | 7192,99 | 9967 | 4419 | 0,444 | -1,173 | -32,71 | 1,12E-234 | 2,21E-233 | 0,0004284 | 0,002221 | 0,0010747 | 0,0010747 | TRUE | 0,0060533 | No |
| orf19.2021 | 30152 | 25220 | 27159 | 16389 | 48467 | 37656 | 40920 | 57259 | 27452 | 25507 | 25704 | 24619 | 43110 | 40884 | 41907 | 43678 | 34107,6 | 25820 | 42395 | 1,642 | 0,715 | 17,069 | 2,54E-65 | 1,50E-64 | 0,0014466 | 0,002076 | 0,0016549 | 0,0016549 | TRUE | 0,0221954 | No |
| orf19.2023 | 27155 | 21870 | 24432 | 15567 | 10013 | 8183 | 8589 | 12220 | 24723 | 22119 | 23123 | 23384 | 8906 | 8884 | 8796 | 9322 | 16157,2 | 23337 | 8977 | 0,385 | -1,378 | -32,814 | 3,77E-236 | 7,46E-235 | 0,0012951 | 0,002119 | 0,0016193 | 0,0016193 | TRUE | 0,0197121 | No |
| orf19.2024 | 39 | 32 | 26 | 17 | 31 | 28 | 29 | 39 | 36 | 32 | 25 | 26 | 28 | 30 | 30 | 30 | 29,43 | 30 | 30 | 0,985 | -0,021 | -0,085 | 0,932144 | 0,941746 | 1,00E-08 | 0,046923 | 0,0266839 | 0,0266839 | TRUE | 0,1359221 | No |
| orf19.2025 | 1 | 4 | 3 | 0 | 0 | 0 | 3 | 0 | 1 | 4 | 3 | 0 | 0 | 0 | 3 | 0 | 1,36 | 2 | 1 | 0,364 | -1,46 | -0,993 | 0,320681 | 0,360046 | 1,9500637 | 0,974464 | 1,1695656 | 1,1695656 | TRUE | 0,7237682 | No |
| orf19.2026 | 569 | 540 | 540 | 329 | 935 | 734 | 803 | 1097 | 518 | 546 | 511 | 494 | 832 | 797 | 822 | 837 | 669,65 | 517 | 822 | 1,587 | 0,666 | 10,979 | 4,80E-28 | 1,47E-27 | 1,00E-08 | 0,00401 | 0,0019363 | 0,0019363 | TRUE | 0,0158 | No |
| orf19.2028 | 4196 | 3596 | 3849 | 2568 | 3165 | 2461 | 2715 | 3684 | 3820 | 3637 | 3643 | 3858 | 2815 | 2672 | 2781 | 2810 | 3254,42 | 3740 | 2770 | 0,741 | -0,432 | -10,824 | 2,64E-27 | 7,97E-27 | 0,0003524 | 0,002443 | 0,0012138 | 0,0012138 | TRUE | 0,0066144 | No |
| orf19.2029 | 517 | 522 | 549 | 330 | 731 | 601 | 568 | 818 | 471 | 528 | 520 | 496 | 650 | 653 | 582 | 624 | 565,29 | 504 | 627 | 1,245 | 0,317 | 4,553 | 5,28E-06 | 8,49E-06 | 0,0009941 | 0,004374 | 0,0028377 | 0,0028377 | TRUE | 0,0265917 | No |
| orf19.203 | 3379 | 2955 | 3321 | 2300 | 2209 | 1840 | 1954 | 2608 | 3076 | 2989 | 3143 | 3455 | 1965 | 1998 | 2001 | 1989 | 2577,03 | 3166 | 1988 | 0,629 | -0,669 | -13,337 | 1,40E-40 | 5,49E-40 | 0,0015418 | 0,00255 | 0,0020097 | 0,0020097 | TRUE | 0,0434624 | No |
| orf19.2030 | 11158 | 8570 | 9276 | 6447 | 3318 | 2590 | 2577 | 3789 | 10159 | 8668 | 8779 | 9684 | 2951 | 2812 | 2639 | 2890 | 6072,82 | 9322 | 2823 | 0,303 | -1,723 | -28,614 | 4,53E-180 | 7,00E-179 | 0,003822 | 0,002255 | 0,0032586 | 0,0032586 | TRUE | 0,045161 | No |
| orf19.2031 | 2101 | 1922 | 2084 | 1208 | 1157 | 906 | 1059 | 1324 | 1913 | 1944 | 1972 | 1815 | 1029 | 984 | 1085 | 1010 | 1468,87 | 1911 | 1027 | 0,536 | -0,898 | -17,792 | 8,09E-71 | 5,13E-70 | 0,0005141 | 0,002936 | 0,0017125 | 0,0017125 | TRUE | 0,016332 | No |
| orf19.2032 | 8 | 4 | 9 | 3 | 9 | 7 | 7 | 4 | 7 | 4 | 9 | 5 | 8 | 8 | 7 | 3 | 6,27 | 6 | 6 | 1,022 | 0,031 | 0,056 | 0,955516 | 0,96052 | 1,00E-08 | 0,212644 | 0,1406272 | 0,1406272 | TRUE | 0,4419755 | No |
| orf19.2033 | 6 | 2 | 11 | 8 | 6 | 15 | 13 | 14 | 5 | 2 | 10 | 12 | 5 | 16 | 13 | 11 | 9,44 | 7 | 11 | 1,555 | 0,637 | 1,203 | 0,229102 | 0,262684 | 0,1789111 | 0,141957 | 0,1542356 | 0,1542356 | TRUE | 0,4462818 | No |
| orf19.2034 | 16 | 14 | 17 | 5 | 35 | 28 | 32 | 35 | 15 | 14 | 16 | 8 | 31 | 30 | 33 | 27 | 21,67 | 13 | 30 | 2,233 | 1,159 | 3,761 | 0,00017 | 0,000251 | 1,00E-08 | 0,063008 | 0,0365026 | 0,0365026 | TRUE | 0,1953494 | No |
| orf19.2035 | 141 | 128 | 111 | 89 | 225 | 180 | 186 | 260 | 128 | 129 | 105 | 134 | 200 | 195 | 190 | 198 | 160,12 | 124 | 196 | 1,589 | 0,668 | 5,812 | 6,17E-09 | 1,12E-08 | 1,00E-08 | 0,010287 | 0,0060122 | 0,0060122 | TRUE | 0,1165892 | No |
| orf19.2036 | 487 | 421 | 459 | 309 | 751 | 587 | 631 | 838 | 443 | 426 | 434 | 464 | 668 | 637 | 646 | 639 | 544,82 | 442 | 648 | 1,47 | 0,556 | 8,578 | 9,64E-18 | 2,30E-17 | 1,00E-08 | 0,004462 | 0,0020997 | 0,0020997 | TRUE | 0,0105767 | No |
| orf19.2037 | 545 | 464 | 474 | 306 | 1074 | 990 | 931 | 1231 | 496 | 469 | 449 | 460 | 955 | 1075 | 953 | 939 | 724,55 | 468 | 980 | 2,088 | 1,062 | 15,5 | 3,46E-54 | 1,73E-53 | 0,0019196 | 0,00386 | 0,0028924 | 0,0028924 | TRUE | 0,0482444 | No |
| orf19.2038 | 576 | 615 | 609 | 427 | 940 | 864 | 841 | 1082 | 524 | 622 | 576 | 641 | 836 | 938 | 861 | 825 | 728,13 | 591 | 865 | 1,468 | 0,553 | 7,428 | 1,10E-13 | 2,35E-13 | 0,0037174 | 0,003851 | 0,0038782 | 0,0038782 | TRUE | 0,0696251 | No |
| orf19.2039 | 2359 | 1901 | 2235 | 1324 | 2373 | 1925 | 2093 | 2880 | 2148 | 1923 | 2115 | 1989 | 2111 | 2090 | 2143 | 2197 | 2089,45 | 2044 | 2135 | 1,044 | 0,062 | 1,286 | 0,198546 | 0,229882 | 0,0011103 | 0,002669 | 0,001774 | 0,001774 | TRUE | 0,0203859 | No |
| orf19.204 | 880 | 746 | 862 | 532 | 594 | 412 | 484 | 682 | 801 | 754 | 816 | 799 | 528 | 447 | 496 | 520 | 645,28 | 792 | 498 | 0,63 | -0,667 | -9,812 | 1,00E-22 | 2,70E-22 | 0,0010609 | 0,004084 | 0,0028241 | 0,0028241 | TRUE | 0,0514705 | No |
| orf19.2040 | 2568 | 2253 | 2474 | 1488 | 3361 | 2682 | 2872 | 3893 | 2338 | 2279 | 2341 | 2235 | 2990 | 2912 | 2941 | 2970 | 2625,71 | 2298 | 2953 | 1,284 | 0,36 | 9,654 | 4,72E-22 | 1,25E-21 | 1,00E-08 | 0,00254 | 0,0009461 | 0,0009461 | TRUE | 0,0036681 | No |
| orf19.2041 | 1001 | 985 | 1039 | 582 | 824 | 668 | 696 | 1016 | 911 | 996 | 983 | 874 | 733 | 725 | 713 | 775 | 838,89 | 941 | 736 | 0,782 | -0,355 | -5,723 | 1,04E-08 | 1,88E-08 | 0,0013551 | 0,003612 | 0,0025004 | 0,0025004 | TRUE | 0,0237897 | No |
| orf19.2042 | 1669 | 1369 | 1496 | 876 | 1792 | 1439 | 1542 | 2114 | 1520 | 1385 | 1416 | 1316 | 1594 | 1562 | 1579 | 1613 | 1497,99 | 1409 | 1587 | 1,124 | 0,169 | 3,252 | 0,001144 | 0,00161 | 0,0010053 | 0,002919 | 0,0019032 | 0,0019032 | TRUE | 0,0338762 | No |
| orf19.2043 | 899 | 777 | 857 | 567 | 736 | 574 | 612 | 901 | 819 | 786 | 811 | 852 | 655 | 623 | 627 | 687 | 732,38 | 817 | 648 | 0,796 | -0,328 | -5,436 | 5,44E-08 | 9,49E-08 | 1,00E-08 | 0,003841 | 0,0021309 | 0,0021309 | TRUE | 0,0209648 | No |
| orf19.2044 | 488 | 489 | 471 | 321 | 711 | 625 | 654 | 863 | 444 | 495 | 446 | 482 | 632 | 679 | 670 | 658 | 563,24 | 467 | 660 | 1,416 | 0,502 | 7,49 | 6,87E-14 | 1,47E-13 | 1,00E-08 | 0,004382 | 0,002459 | 0,002459 | TRUE | 0,0214612 | No |
| orf19.2045 | 5803 | 5492 | 5705 | 3795 | 7127 | 5780 | 6321 | 8301 | 5283 | 5555 | 5399 | 5701 | 6339 | 6275 | 6474 | 6332 | 5919,77 | 5484 | 6355 | 1,16 | 0,214 | 5,992 | 2,08E-09 | 3,85E-09 | 0,0004113 | 0,00226 | 0,0010502 | 0,0010502 | TRUE | 0,0081346 | No |
| orf19.2046 | 798 | 645 | 749 | 438 | 1568 | 1197 | 1375 | 1865 | 727 | 652 | 709 | 658 | 1395 | 1300 | 1408 | 1423 | 1033,85 | 686 | 1382 | 2,009 | 1,006 | 17,12 | 1,06E-65 | 6,27E-65 | 0,0010522 | 0,003315 | 0,0022018 | 0,0022018 | TRUE | 0,0186423 | No |
| orf19.2047 | 1643 | 1426 | 1521 | 931 | 267 | 237 | 216 | 328 | 1496 | 1442 | 1439 | 1399 | 237 | 257 | 221 | 250 | 842,79 | 1444 | 241 | 0,167 | -2,58 | -38,448 | 0 | 0 | 1,00E-08 | 0,003604 | 0,0020321 | 0,0020321 | TRUE | 0,0325862 | No |
| orf19.2048 | 15558 | 12582 | 13136 | 9718 | 1149 | 842 | 799 | 1117 | 14165 | 12725 | 12432 | 14598 | 1022 | 914 | 818 | 852 | 7190,85 | 13480 | 902 | 0,067 | -3,902 | -52,684 | 0 | 0 | 0,0072484 | 0,002221 | 0,00471 | 0,00471 | TRUE | 0,0979584 | No |
| orf19.2049 | 681 | 658 | 623 | 377 | 174 | 148 | 153 | 195 | 620 | 665 | 590 | 566 | 155 | 161 | 157 | 149 | 382,79 | 610 | 156 | 0,253 | -1,982 | -22,28 | 5,78E-110 | 5,45E-109 | 1,00E-08 | 0,005488 | 0,0037216 | 0,0037216 | TRUE | 0,042492 | No |
| orf19.2050 | 7142 | 6226 | 6871 | 4275 | 1115 | 920 | 971 | 1315 | 6502 | 6297 | 6503 | 6422 | 992 | 999 | 994 | 1003 | 3714,01 | 6431 | 997 | 0,155 | -2,689 | -69,538 | 0 | 0 | 1,00E-08 | 0,002393 | 0,0008856 | 0,0008856 | TRUE | 0,0024703 | No |
| orf19.2051 | 406 | 387 | 437 | 292 | 974 | 891 | 903 | 1167 | 370 | 391 | 414 | 439 | 866 | 967 | 925 | 890 | 657,75 | 404 | 912 | 2,27 | 1,183 | 16,771 | 4,00E-63 | 2,29E-62 | 0,0014686 | 0,004045 | 0,0029337 | 0,0029337 | TRUE | 0,037115 | No |
| orf19.2054 | 2150 | 1905 | 2224 | 1356 | 2163 | 1845 | 2060 | 2567 | 1957 | 1927 | 2105 | 2037 | 1924 | 2003 | 2110 | 1958 | 2002,6 | 2006 | 1999 | 0,996 | -0,006 | -0,121 | 0,903356 | 0,917191 | 0,0011398 | 0,002697 | 0,001803 | 0,001803 | TRUE | 0,0167491 | No |
| orf19.2055 | 2806 | 2307 | 2498 | 1493 | 3369 | 2516 | 2973 | 3950 | 2555 | 2333 | 2364 | 2243 | 2997 | 2732 | 3045 | 3013 | 2660,12 | 2374 | 2947 | 1,24 | 0,311 | 5,766 | 8,12E-09 | 1,47E-08 | 0,0022254 | 0,002534 | 0,002402 | 0,002402 | TRUE | 0,0322765 | No |
| orf19.2057 | 6606 | 5900 | 6590 | 4203 | 6531 | 5477 | 5623 | 7857 | 6014 | 5967 | 6237 | 6314 | 5809 | 5946 | 5759 | 5993 | 6004,97 | 6133 | 5877 | 0,959 | -0,061 | -1,734 | 0,082832 | 0,100257 | 0,0003326 | 0,002257 | 0,0010067 | 0,0010067 | TRUE | 0,0045214 | No |
| orf19.2059 | 1442 | 1292 | 1403 | 957 | 3965 | 3179 | 3345 | 4493 | 1313 | 1307 | 1328 | 1438 | 3527 | 3452 | 3426 | 3427 | 2402,03 | 1346 | 3458 | 2,576 | 1,365 | 31,579 | 7,21E-219 | 1,33E-217 | 1,95E-05 | 0,002587 | 0,0012605 | 0,0012605 | TRUE | 0,0216415 | No |
| orf19.206 | 1439 | 1228 | 1302 | 945 | 5160 | 4436 | 4637 | 6244 | 1310 | 1242 | 1232 | 1420 | 4590 | 4816 | 4749 | 4763 | 3015,21 | 1301 | 4730 | 3,647 | 1,867 | 38,151 | 0 | 0 | 0,0010695 | 0,002475 | 0,0017856 | 0,0017856 | TRUE | 0,0399493 | No |
| orf19.2060 | 54 | 36 | 41 | 33 | 121 | 104 | 111 | 155 | 49 | 36 | 39 | 50 | 108 | 113 | 114 | 118 | 78,3 | 44 | 113 | 2,623 | 1,391 | 8,232 | 1,84E-16 | 4,24E-16 | 1,00E-08 | 0,018908 | 0,0110317 | 0,0110317 | TRUE | 0,0902988 | No |
| orf19.2061 | 7 | 4 | 1 | 1 | 13 | 18 | 14 | 7 | 6 | 4 | 1 | 2 | 12 | 20 | 14 | 5 | 7,96 | 3 | 13 | 3,733 | 1,9 | 3,019 | 0,00254 | 0,003474 | 0,2168446 | 0,168068 | 0,184675 | 0,184675 | TRUE | 0,2625376 | No |
| orf19.2062 | 340 | 282 | 279 | 250 | 575 | 485 | 538 | 722 | 310 | 285 | 264 | 376 | 511 | 527 | 551 | 551 | 421,76 | 309 | 535 | 1,75 | 0,807 | 8,55 | 1,23E-17 | 2,92E-17 | 0,0071684 | 0,005169 | 0,0059284 | 0,0059284 | TRUE | 0,2060642 | No |
| orf19.2063 | 3845 | 3511 | 3841 | 2453 | 3117 | 2612 | 2829 | 3782 | 3501 | 3551 | 3635 | 3685 | 2772 | 2836 | 2897 | 2885 | 3220,28 | 3593 | 2848 | 0,793 | -0,334 | -8,977 | 2,77E-19 | 6,91E-19 | 0,0001534 | 0,002447 | 0,0010177 | 0,0010177 | TRUE | 0,0039254 | No |
| orf19.2064 | 2696 | 2179 | 2351 | 1439 | 1230 | 1042 | 1078 | 1629 | 2455 | 2204 | 2225 | 2162 | 1094 | 1131 | 1104 | 1243 | 1702,13 | 2262 | 1143 | 0,506 | -0,984 | -16,329 | 6,15E-60 | 3,36E-59 | 0,0025973 | 0,002813 | 0,0028412 | 0,0028412 | TRUE | 0,0433891 | No |
| orf19.2065 | 111 | 88 | 134 | 54 | 150 | 122 | 123 | 165 | 101 | 89 | 127 | 81 | 133 | 132 | 126 | 126 | 114,46 | 100 | 129 | 1,286 | 0,363 | 2,495 | 0,012592 | 0,016396 | 0,0072376 | 0,013578 | 0,0113707 | 0,0113707 | TRUE | 0,336211 | No |
| orf19.2066 | 1060 | 944 | 987 | 654 | 1696 | 1364 | 1552 | 2017 | 965 | 955 | 934 | 982 | 1509 | 1481 | 1589 | 1539 | 1244,23 | 959 | 1530 | 1,597 | 0,675 | 13,796 | 2,70E-43 | 1,12E-42 | 1,00E-08 | 0,003099 | 0,0014359 | 0,0014359 | TRUE | 0,0079946 | No |
| orf19.2066.1 | 2408 | 1878 | 1897 | 1279 | 3017 | 2176 | 2214 | 3332 | 2192 | 1899 | 1795 | 1921 | 2684 | 2363 | 2267 | 2542 | 2207,94 | 1952 | 2464 | 1,262 | 0,336 | 4,693 | 2,69E-06 | 4,39E-06 | 0,0059835 | 0,002635 | 0,004457 | 0,004457 | TRUE | 0,0851848 | No |
| orf19.2067 | 4618 | 4025 | 4257 | 2472 | 1761 | 1202 | 1462 | 1922 | 4204 | 4071 | 4029 | 3713 | 1566 | 1305 | 1497 | 1466 | 2731,54 | 4004 | 1458 | 0,364 | -1,457 | -23,897 | 3,27E-126 | 3,52E-125 | 0,0036034 | 0,002521 | 0,0031141 | 0,0031141 | TRUE | 0,0586952 | No |
| orf19.2068 | 544 | 494 | 531 | 320 | 1033 | 899 | 830 | 1100 | 495 | 500 | 503 | 481 | 919 | 976 | 850 | 839 | 695,27 | 495 | 896 | 1,807 | 0,854 | 12,459 | 1,24E-35 | 4,49E-35 | 0,0019208 | 0,003937 | 0,0029045 | 0,0029045 | TRUE | 0,0415244 | No |
| orf19.2069 | 2614 | 2210 | 2411 | 1584 | 3540 | 2851 | 2887 | 4111 | 2380 | 2235 | 2282 | 2379 | 3149 | 3095 | 2957 | 3136 | 2701,62 | 2319 | 3084 | 1,331 | 0,413 | 9,797 | 1,16E-22 | 3,12E-22 | 0,0005101 | 0,002526 | 0,0013203 | 0,0013203 | TRUE | 0,009304 | No |
| orf19.207 | 2516 | 2078 | 2292 | 1647 | 4127 | 3735 | 3850 | 5341 | 2291 | 2102 | 2169 | 2474 | 3671 | 4055 | 3943 | 4074 | 3097,33 | 2259 | 3936 | 1,745 | 0,803 | 13,881 | 8,29E-44 | 3,47E-43 | 0,0032662 | 0,002464 | 0,0028573 | 0,0028573 | TRUE | 0,04698 | No |
| orf19.2070 | 3312 | 3040 | 3241 | 2039 | 3178 | 2544 | 2836 | 3689 | 3015 | 3075 | 3067 | 3063 | 2827 | 2762 | 2904 | 2814 | 2940,94 | 3055 | 2827 | 0,925 | -0,112 | -3,113 | 0,001852 | 0,002562 | 1,00E-08 | 0,002486 | 0,0008975 | 0,0008975 | TRUE | 0,0039798 | No |
| orf19.2071 | 1 | 1 | 3 | 1 | 5 | 3 | 0 | 5 | 1 | 1 | 3 | 2 | 4 | 3 | 0 | 4 | 2,22 | 2 | 3 | 1,863 | 0,898 | 0,908 | 0,363702 | 0,403656 | 1,00E-08 | 0,596352 | 0,4411462 | 0,4411462 | TRUE | 0,533967 | No |
| orf19.2072 | 6 | 9 | 10 | 17 | 41 | 31 | 48 | 44 | 5 | 9 | 9 | 26 | 36 | 34 | 49 | 34 | 25,3 | 12 | 38 | 3,305 | 1,725 | 4,738 | 2,16E-06 | 3,54E-06 | 0,1129647 | 0,054247 | 0,0688838 | 0,0688838 | TRUE | 1,3782883 | No |
| orf19.2073 | 10479 | 8744 | 9716 | 6082 | 5526 | 4762 | 5183 | 7110 | 9541 | 8844 | 9195 | 9136 | 4915 | 5170 | 5308 | 5424 | 7191,6 | 9179 | 5204 | 0,567 | -0,819 | -19,207 | 3,23E-82 | 2,36E-81 | 0,0011682 | 0,002221 | 0,001596 | 0,001596 | TRUE | 0,0172828 | No |
| orf19.2074 | 254 | 194 | 204 | 122 | 120 | 89 | 83 | 134 | 231 | 196 | 193 | 183 | 107 | 97 | 85 | 102 | 149,3 | 201 | 98 | 0,485 | -1,043 | -8,21 | 2,21E-16 | 5,08E-16 | 1,00E-08 | 0,010885 | 0,0080922 | 0,0080922 | TRUE | 0,1171346 | No |

| | | | | | | | | | | | | | | | | | | | | | | | | | | | | | | | |
|---|---|---|---|---|---|---|---|---|---|---|---|---|---|---|---|---|---|---|---|---|---|---|---|---|---|---|---|---|---|---|---|
| orf19.2075 | 3332 | 2879 | 3019 | 1780 | 3044 | 2321 | 2633 | 3643 | 3034 | 2912 | 2857 | 2674 | 2708 | 2520 | 2697 | 2779 | 2772,43 | 2869 | 2676 | 0,932 | -0,102 | -2,012 | 0,044226 | 0,055023 | 0,0017246 | 0,002513 | 0,0020933 | 0,0020933 | TRUE | 0,0239748 | No |
| orf19.2076 | 184 | 199 | 193 | 119 | 190 | 191 | 171 | 278 | 168 | 201 | 183 | 179 | 169 | 207 | 175 | 212 | 186,72 | 183 | 191 | 1,047 | 0,066 | 0,571 | 0,567794 | 0,60876 | 0,0051154 | 0,009112 | 0,0074511 | 0,0074511 | TRUE | 0,0691211 | No |
| orf19.2077 | 3291 | 2852 | 3063 | 2069 | 3552 | 2913 | 3273 | 4306 | 2996 | 2884 | 2899 | 3108 | 3159 | 3163 | 3352 | 3285 | 3105,79 | 2972 | 3240 | 1,091 | 0,126 | 3,012 | 0,002591 | 0,003543 | 0,0006042 | 0,002462 | 0,0013574 | 0,0013574 | TRUE | 0,0106073 | No |
| orf19.2078 | 1352 | 1238 | 1309 | 740 | 806 | 660 | 737 | 989 | 1231 | 1252 | 1239 | 1112 | 717 | 717 | 755 | 754 | 972,02 | 1208 | 736 | 0,607 | -0,719 | -12,525 | 5,42E-36 | 1,97E-35 | 0,0007023 | 0,003396 | 0,0020884 | 0,0020884 | TRUE | 0,0307065 | No |
| orf19.2079 | 4523 | 3582 | 3630 | 2955 | 868 | 688 | 650 | 953 | 4118 | 3623 | 3435 | 4439 | 772 | 747 | 666 | 727 | 2315,85 | 3904 | 728 | 0,187 | -2,422 | -29,586 | 2,25E-192 | 3,68E-191 | 0,0077544 | 0,002607 | 0,0056534 | 0,0056534 | TRUE | 0,0946079 | No |
| orf19.2081 | 2518 | 2298 | 2486 | 1523 | 1914 | 1622 | 1653 | 2295 | 2293 | 2324 | 2353 | 2288 | 1702 | 1761 | 1693 | 1751 | 2020,54 | 2314 | 1727 | 0,746 | -0,423 | -10,618 | 2,45E-26 | 7,21E-26 | 1,00E-08 | 0,002691 | 0,0010215 | 0,0010215 | TRUE | 0,0020098 | No |
| orf19.2082 | 4 | 4 | 1 | 3 | 3 | 4 | 6 | 8 | 4 | 4 | 1 | 5 | 3 | 4 | 6 | 6 | 4,05 | 4 | 5 | 1,511 | 0,595 | 0,851 | 0,394536 | 0,435527 | 1,00E-08 | 0,328221 | 0,2066141 | 0,2066141 | TRUE | 0,3691851 | No |
| orf19.2084 | 2196 | 1800 | 2087 | 1323 | 2758 | 2281 | 2415 | 3380 | 1999 | 1821 | 1975 | 1987 | 2453 | 2477 | 2473 | 2578 | 2220,45 | 1946 | 2495 | 1,283 | 0,36 | 7,914 | 2,49E-15 | 5,57E-15 | 0,0007085 | 0,002632 | 0,0015237 | 0,0015237 | TRUE | 0,0230836 | No |
| orf19.2087 | 902 | 761 | 833 | 570 | 621 | 488 | 538 | 764 | 821 | 770 | 788 | 856 | 552 | 530 | 551 | 583 | 681,43 | 809 | 554 | 0,688 | -0,54 | -8,588 | 8,89E-18 | 2,13E-17 | 1,00E-08 | 0,003976 | 0,0022963 | 0,0022963 | TRUE | 0,0157977 | No |
| orf19.2088 | 695 | 516 | 620 | 382 | 510 | 391 | 476 | 615 | 633 | 522 | 587 | 574 | 454 | 425 | 487 | 469 | 518,75 | 579 | 459 | 0,792 | -0,336 | -4,318 | 1,57E-05 | 2,47E-05 | 0,0030574 | 0,004584 | 0,003871 | 0,003871 | TRUE | 0,0543427 | No |
| orf19.2089 | 3928 | 3486 | 3597 | 2257 | 2540 | 2072 | 2158 | 2929 | 3576 | 3526 | 3404 | 3390 | 2259 | 2250 | 2210 | 2234 | 2856,22 | 3474 | 2238 | 0,644 | -0,635 | -16,784 | 3,20E-63 | 1,84E-62 | 1,00E-08 | 0,0025 | 0,0010119 | 0,0010119 | TRUE | 0,0048387 | No |
| orf19.209 | 350 | 304 | 269 | 218 | 142 | 136 | 128 | 201 | 319 | 307 | 255 | 327 | 126 | 148 | 131 | 153 | 220,82 | 302 | 140 | 0,464 | -1,107 | -9,682 | 3,58E-22 | 9,55E-22 | 0,0066397 | 0,008019 | 0,0074347 | 0,0074347 | TRUE | 0,1314349 | No |
| orf19.2090 | 8730 | 7576 | 8205 | 4715 | 11117 | 8681 | 10164 | 13442 | 7948 | 7662 | 7765 | 7083 | 9888 | 9425 | 10409 | 10254 | 8804,35 | 7614 | 9994 | 1,312 | 0,392 | 8,157 | 3,44E-16 | 7,86E-16 | 0,0020097 | 0,002187 | 0,0020964 | 0,0020964 | TRUE | 0,0263335 | No |
| orf19.2091 | 3481 | 3031 | 3331 | 2161 | 5738 | 4720 | 4904 | 6501 | 3169 | 3066 | 3152 | 3246 | 5104 | 5125 | 5022 | 4959 | 4105,4 | 3158 | 5052 | 1,601 | 0,679 | 19,409 | 6,42E-84 | 4,79E-83 | 4,65E-05 | 0,002359 | 0,0009109 | 0,0009109 | TRUE | 0,0046533 | No |
| orf19.2092 | 2851 | 2409 | 2570 | 1741 | 2839 | 2321 | 2422 | 3560 | 2596 | 2436 | 2432 | 2615 | 2525 | 2520 | 2480 | 2716 | 2540,11 | 2520 | 2560 | 1,017 | 0,025 | 0,525 | 0,599743 | 0,639587 | 0,0012516 | 0,002557 | 0,0017839 | 0,0017839 | TRUE | 0,0209063 | No |
| orf19.2093 | 3738 | 3094 | 3453 | 2388 | 4164 | 3665 | 3841 | 5101 | 3403 | 3129 | 3268 | 3587 | 3704 | 3979 | 3934 | 3891 | 3611,91 | 3347 | 3877 | 1,159 | 0,213 | 4,33 | 1,49E-05 | 2,34E-05 | 0,001819 | 0,002403 | 0,0020487 | 0,0020487 | TRUE | 0,026853 | No |
| orf19.2094 | 6578 | 6162 | 6594 | 4214 | 1513 | 1281 | 1392 | 1768 | 5989 | 6232 | 6241 | 6330 | 1346 | 1391 | 1426 | 1349 | 3787,84 | 6198 | 1378 | 0,222 | -2,17 | -53,957 | 0 | 0 | 0,0003364 | 0,002386 | 0,0011276 | 0,0011276 | TRUE | 0,0065645 | No |
| orf19.2095 | 13608 | 11724 | 12599 | 8669 | 6266 | 5061 | 5421 | 7318 | 12390 | 11858 | 11924 | 13022 | 5573 | 5495 | 5552 | 5582 | 8924,42 | 12298 | 5550 | 0,452 | -1,147 | -29,508 | 2,29E-191 | 3,74E-190 | 0,000806 | 0,002185 | 0,0013231 | 0,0013231 | TRUE | 0,0187434 | No |
| orf19.2097 | 701 | 615 | 703 | 402 | 1328 | 1159 | 1276 | 1618 | 638 | 622 | 665 | 604 | 1181 | 1258 | 1307 | 1234 | 938,75 | 632 | 1245 | 1,963 | 0,973 | 16,663 | 2,42E-62 | 1,37E-61 | 0,0006936 | 0,003444 | 0,0020545 | 0,0020545 | TRUE | 0,0143723 | No |
| orf19.2098 | 1986 | 1579 | 1793 | 1247 | 2619 | 2179 | 2356 | 3321 | 1808 | 1597 | 1697 | 1873 | 2330 | 2366 | 2413 | 2533 | 2077,09 | 1744 | 2410 | 1,385 | 0,47 | 8,317 | 9,03E-17 | 2,10E-16 | 0,0024268 | 0,002673 | 0,0025582 | 0,0025582 | TRUE | 0,0384258 | No |
| orf19.2099 | 1887 | 1552 | 1750 | 1128 | 1625 | 1327 | 1492 | 1960 | 1718 | 1570 | 1656 | 1694 | 1445 | 1441 | 1528 | 1495 | 1568,45 | 1660 | 1477 | 0,891 | -0,167 | -3,462 | 0,000536 | 0,00077 | 0,000553 | 0,002879 | 0,0015968 | 0,0015968 | TRUE | 0,0163324 | No |
| orf19.21 | 630 | 532 | 573 | 428 | 272 | 226 | 221 | 322 | 574 | 538 | 542 | 643 | 242 | 245 | 226 | 246 | 407,02 | 574 | 240 | 0,42 | -1,252 | -14,96 | 1,35E-50 | 6,29E-50 | 0,0020235 | 0,005283 | 0,0038489 | 0,0038489 | TRUE | 0,0663795 | No |
| orf19.210 | 146 | 152 | 128 | 97 | 61 | 55 | 70 | 83 | 133 | 154 | 121 | 146 | 54 | 60 | 72 | 63 | 100,31 | 138 | 62 | 0,451 | -1,15 | -7,611 | 2,73E-14 | 5,91E-14 | 1,00E-08 | 0,015206 | 0,0105551 | 0,0105551 | TRUE | 0,0844235 | No |
| orf19.2101 | 1135 | 920 | 1009 | 631 | 752 | 550 | 619 | 907 | 1033 | 930 | 955 | 948 | 669 | 597 | 634 | 692 | 807,31 | 966 | 648 | 0,671 | -0,575 | -8,831 | 1,04E-18 | 2,55E-18 | 0,0018961 | 0,003673 | 0,0027975 | 0,0027975 | TRUE | 0,0317896 | No |
| orf19.2102 | 4914 | 4087 | 4480 | 2726 | 3886 | 2991 | 3348 | 4631 | 4474 | 4134 | 4240 | 4095 | 3456 | 3247 | 3429 | 3533 | 3825,94 | 4236 | 3416 | 0,806 | -0,31 | -6,973 | 3,10E-12 | 6,32E-12 | 0,001082 | 0,002382 | 0,001638 | 0,001638 | TRUE | 0,0177111 | No |
| orf19.2103 | 25 | 23 | 32 | 16 | 36 | 46 | 41 | 50 | 23 | 23 | 30 | 24 | 32 | 50 | 42 | 38 | 32,8 | 25 | 40 | 1,599 | 0,678 | 2,696 | 0,007012 | 0,009303 | 1,00E-08 | 0,042306 | 0,0279429 | 0,0279429 | TRUE | 0,1713318 | No |
| orf19.2104 | 593 | 507 | 492 | 348 | 491 | 438 | 412 | 625 | 540 | 513 | 466 | 523 | 437 | 476 | 422 | 477 | 481,5 | 510 | 453 | 0,889 | -0,17 | -2,233 | 0,025561 | 0,032493 | 0,001981 | 0,00478 | 0,0035028 | 0,0035028 | TRUE | 0,0430506 | No |
| orf19.2105 | 1134 | 966 | 1052 | 779 | 1845 | 1579 | 1722 | 2293 | 1032 | 977 | 996 | 1170 | 1641 | 1714 | 1764 | 1749 | 1380,42 | 1044 | 1717 | 1,651 | 0,723 | 11,997 | 3,70E-33 | 1,27E-32 | 0,0023922 | 0,002994 | 0,0027027 | 0,0027027 | TRUE | 0,0712049 | No |
| orf19.2106 | 1498 | 1128 | 1307 | 805 | 2173 | 1707 | 2057 | 2691 | 1364 | 1141 | 1237 | 1209 | 1933 | 1853 | 2107 | 2053 | 1612,05 | 1238 | 1986 | 1,604 | 0,681 | 10,53 | 6,30E-26 | 1,84E-25 | 0,0036317 | 0,002856 | 0,0033511 | 0,0033511 | TRUE | 0,0586153 | No |
| orf19.2107 | 869 | 683 | 739 | 520 | 1535 | 1237 | 1397 | 1899 | 791 | 691 | 699 | 781 | 1365 | 1343 | 1431 | 1449 | 1068,77 | 740 | 1397 | 1,89 | 0,919 | 14,951 | 1,52E-50 | 7,09E-50 | 0,0018276 | 0,003273 | 0,0025652 | 0,0025652 | TRUE | 0,0291734 | No |
| orf19.2107.1 | 1E+05 | 78158 | 83372 | 44334 | 18323 | 10213 | 13622 | 18705 | 1E+05 | 79048 | 78904 | 66597 | 16298 | 11088 | 13951 | 14268 | 47612 | 81322 | 13901 | 0,171 | -2,548 | -15,168 | 5,77E-52 | 2,77E-51 | 0,0270846 | 0,002065 | 0,0131677 | 0,0270846 | TRUE | 0,2015577 | No |
| orf19.2108 | 4061 | 3383 | 3667 | 2304 | 1440 | 1160 | 1173 | 1642 | 3697 | 3422 | 3470 | 3461 | 1281 | 1259 | 1201 | 1253 | 2380,56 | 3512 | 1248 | 0,355 | -1,492 | -32,908 | 1,71E-237 | 3,41E-236 | 0,000682 | 0,002592 | 0,0014481 | 0,0014481 | TRUE | 0,0157238 | No |
| orf19.211 | 1916 | 1627 | 1802 | 1232 | 330 | 223 | 256 | 377 | 1744 | 1646 | 1705 | 1851 | 294 | 242 | 262 | 288 | 1003,93 | 1736 | 272 | 0,157 | -2,669 | -37,549 | 0 | 0 | 0,0022279 | 0,003353 | 0,0028353 | 0,0028353 | TRUE | 0,0550405 | No |
| orf19.2110 | 250 | 200 | 213 | 118 | 189 | 123 | 144 | 216 | 228 | 202 | 202 | 177 | 168 | 134 | 147 | 165 | 177,83 | 202 | 154 | 0,758 | -0,4 | -3,416 | 0,000636 | 0,000909 | 0,0043815 | 0,009466 | 0,0075209 | 0,0075209 | TRUE | 0,0783913 | No |
| orf19.2111 | 3763 | 2907 | 3351 | 2192 | 3220 | 2460 | 2877 | 3746 | 3426 | 2940 | 3171 | 3293 | 2864 | 2671 | 2946 | 2857 | 3021,15 | 3208 | 2834 | 0,884 | -0,178 | -3,235 | 0,001217 | 0,00171 | 0,0024718 | 0,002474 | 0,0025659 | 0,0025659 | TRUE | 0,0387043 | No |
| orf19.2111.2 | 6678 | 5169 | 5233 | 3337 | 9792 | 7474 | 7351 | 11410 | 6080 | 5228 | 4953 | 5013 | 8710 | 8115 | 7528 | 8704 | 6791,19 | 5318 | 8264 | 1,554 | 0,636 | 8,954 | 3,42E-19 | 8,49E-19 | 0,0069313 | 0,002232 | 0,0046817 | 0,0046817 | TRUE | 0,1141417 | No |
| orf19.2112 | 1015 | 782 | 957 | 671 | 1095 | 910 | 1044 | 1404 | 924 | 791 | 906 | 1008 | 974 | 988 | 1069 | 1071 | 966,35 | 907 | 1026 | 1,134 | 0,181 | 2,472 | 0,013428 | 0,017456 | 0,0047418 | 0,003404 | 0,0041035 | 0,0041035 | TRUE | 0,0878345 | No |
| orf19.2113 | 2533 | 2180 | 2340 | 1552 | 2641 | 2064 | 2180 | 3181 | 2306 | 2205 | 2215 | 2331 | 2349 | 2241 | 2233 | 2426 | 2288,26 | 2264 | 2312 | 1,023 | 0,033 | 0,719 | 0,472198 | 0,514047 | 0,0008014 | 0,002614 | 0,0015443 | 0,0015443 | TRUE | 0,0137947 | No |
| orf19.2114 | 201 | 157 | 183 | 154 | 616 | 576 | 563 | 788 | 183 | 159 | 173 | 231 | 548 | 625 | 577 | 601 | 387,16 | 186 | 588 | 3,189 | 1,673 | 16,903 | 4,27E-64 | 2,47E-63 | 0,0063021 | 0,005449 | 0,0057178 | 0,0057178 | TRUE | 0,2246305 | No |
| orf19.2115 | 2784 | 2207 | 2510 | 1548 | 4922 | 3630 | 4523 | 5895 | 2535 | 2232 | 2375 | 2325 | 4378 | 3941 | 4632 | 4497 | 3364,46 | 2367 | 4362 | 1,842 | 0,882 | 14,794 | 1,61E-49 | 7,31E-49 | 0,0033488 | 0,00243 | 0,0030759 | 0,0030759 | TRUE | 0,0507705 | No |
| orf19.2116 | 2491 | 1970 | 2162 | 1317 | 2330 | 1725 | 1873 | 2780 | 2268 | 1992 | 2046 | 1978 | 2072 | 1873 | 1918 | 2121 | 2033,63 | 2071 | 1996 | 0,964 | -0,054 | -0,88 | 0,379082 | 0,419339 | 0,00323 | 0,002687 | 0,0030602 | 0,0030602 | TRUE | 0,0503971 | No |
| orf19.2117 | 3896 | 3145 | 3511 | 2333 | 3105 | 2496 | 2858 | 3969 | 3547 | 3181 | 3323 | 3505 | 2762 | 2710 | 2927 | 3028 | 3122,71 | 3389 | 2857 | 0,844 | -0,245 | -4,649 | 3,34E-06 | 5,42E-06 | 0,00214 | 0,00246 | 0,0023511 | 0,0023511 | TRUE | 0,0208429 | No |
| orf19.2119 | 8972 | 7230 | 8601 | 6332 | 9066 | 7957 | 8702 | 12618 | 8169 | 7312 | 8140 | 9512 | 8064 | 8639 | 8912 | 9625 | 8546,6 | 8283 | 8810 | 1,064 | 0,09 | 1,176 | 0,239672 | 0,273885 | 0,0082928 | 0,002192 | 0,0054661 | 0,0054661 | TRUE | 0,1206281 | No |
| orf19.212 | 1767 | 1562 | 1724 | 1180 | 764 | 643 | 622 | 863 | 1609 | 1580 | 1632 | 1773 | 680 | 698 | 637 | 658 | 1158,21 | 1648 | 668 | 0,406 | -1,3 | -22,72 | 2,83E-114 | 2,77E-113 | 0,0010871 | 0,003178 | 0,0021189 | 0,0021189 | TRUE | 0,0283826 | No |
| orf19.2123 | 1 | 0 | 2 | 0 | 4 | 4 | 1 | 5 | 1 | 0 | 2 | 0 | 4 | 4 | 1 | 4 | 1,94 | 1 | 3 | 4,164 | 2,058 | 1,777 | 0,075495 | 0,091795 | 1,00E-08 | 0,681991 | 0,4536093 | 0,4536093 | TRUE | 0,4308411 | No |
| orf19.2124 | 2947 | 2637 | 2731 | 1792 | 13954 | 10504 | 11143 | 16099 | 2683 | 2667 | 2585 | 2692 | 12412 | 11404 | 11412 | 12280 | 7266,88 | 2657 | 11877 | 4,473 | 2,161 | 49,759 | 0 | 0 | 0,0012124 | 0,002219 | 0,0015695 | 0,0015695 | TRUE | 0,0111649 | No |
| orf19.2125 | 5095 | 3726 | 3907 | 3136 | 2459 | 1759 | 2007 | 2685 | 4639 | 3768 | 3698 | 4711 | 2187 | 1910 | 2055 | 2048 | 3127,02 | 4204 | 2050 | 0,488 | -1,035 | -12,535 | 4,82E-36 | 1,76E-35 | 0,0094555 | 0,002459 | 0,0061916 | 0,0061916 | TRUE | 0,0763593 | No |
| orf19.2127 | 449 | 376 | 422 | 266 | 373 | 275 | 316 | 428 | 409 | 380 | 399 | 400 | 332 | 299 | 324 | 326 | 358,56 | 397 | 320 | 0,808 | -0,308 | -3,989 | 6,64E-05 | 0,0001 | 1,00E-08 | 0,005721 | 0,0029317 | 0,0029317 | TRUE | 0,0214468 | No |
| orf19.2128 | 755 | 563 | 595 | 444 | 1213 | 893 | 1093 | 1440 | 687 | 569 | 563 | 667 | 1079 | 970 | 1119 | 1098 | 844,15 | 622 | 1066 | 1,72 | 0,782 | 9,928 | 3,14E-23 | 8,59E-23 | 0,0056643 | 0,003602 | 0,0046541 | 0,0046541 | TRUE | 0,0667419 | No |
| orf19.2129 | 706 | 667 | 642 | 477 | 1692 | 1384 | 1558 | 2043 | 643 | 675 | 608 | 717 | 1505 | 1503 | 1596 | 1558 | 1100,39 | 661 | 1540 | 2,343 | 1,229 | 20,897 | 5,67E-97 | 4,79E-96 | 0,000911 | 0,003237 | 0,0022016 | 0,0022016 | TRUE | 0,0335899 | No |
| orf19.213 | 2749 | 2736 | 2776 | 1906 | 2438 | 2168 | 1988 | 2711 | 2503 | 2767 | 2627 | 2863 | 2169 | 2354 | 2036 | 2068 | 2423,33 | 2690 | 2157 | 0,802 | -0,318 | -5,304 | 1,13E-07 | 1,95E-07 | 0,0033702 | 0,002582 | 0,0030369 | 0,0030369 | TRUE | 0,0450017 | No |
| orf19.2131 | 3005 | 2622 | 2809 | 1697 | 1793 | 1446 | 1624 | 2204 | 2736 | 2652 | 2658 | 2549 | 1595 | 1570 | 1663 | 1681 | 2138,08 | 2649 | 1627 | 0,614 | -0,704 | -15,95 | 2,87E-57 | 1,49E-56 | 0,0004165 | 0,002655 | 0,0013786 | 0,0013786 | TRUE | 0,0071405 | No |
| orf19.2132 | 1510 | 1166 | 1277 | 787 | 940 | 682 | 798 | 1045 | 1375 | 1179 | 1209 | 1182 | 836 | 740 | 817 | 797 | 1016,97 | 1236 | 798 | 0,645 | -0,633 | -9,446 | 3,52E-21 | 9,17E-21 | 0,0032521 | 0,003336 | 0,003292 | 0,003292 | TRUE | 0,0704629 | No |
| orf19.2133 | 57 | 49 | 56 | 41 | 65 | 49 | 51 | 74 | 52 | 50 | 53 | 62 | 58 | 53 | 52 | 56 | 54,47 | 54 | 55 | 1,028 | 0,04 | 0,22 | 0,825733 | 0,849588 | 1,00E-08 | 0,02629 | 0,0137682 | 0,0137682 | TRUE | 0,0523694 | No |
| orf19.2135 | 1967 | 1721 | 1805 | 1156 | 1922 | 1531 | 1910 | 2277 | 1791 | 1741 | 1708 | 1736 | 1710 | 1662 | 1956 | 1737 | 1755,13 | 1744 | 1766 | 1,012 | 0,018 | 0,312 | 0,75533 | 0,786931 | 0,0022556 | 0,00279 | 0,0024946 | 0,0024946 | TRUE | 0,0619403 | No |
| orf19.2137 | 814 | 671 | 708 | 498 | 511 | 456 | 463 | 635 | 741 | 679 | 670 | 748 | 455 | 495 | 474 | 484 | 593,26 | 710 | 477 | 0,673 | -0,571 | -8,449 | 2,94E-17 | 6,92E-17 | 1,00E-08 | 0,004264 | 0,0026437 | 0,0026437 | TRUE | 0,0167271 | No |
| orf19.2138 | 13730 | 11723 | 12719 | 8135 | 12095 | 10033 | 10631 | 14734 | 12501 | 11857 | 12037 | 12220 | 10758 | 10893 | 10887 | 11239 | 11549,1 | 12154 | 10944 | 0,901 | -0,151 | -4,731 | 2,23E-06 | 3,65E-06 | 0,0003173 | 0,002151 | 0,0008912 | 0,0008912 | TRUE | 0,0046355 | No |
| orf19.2143 | 4455 | 3651 | 4245 | 2821 | 5653 | 4906 | 5493 | 7373 | 4056 | 3693 | 4018 | 4238 | 5028 | 5327 | 5626 | 5624 | 4701,03 | 4001 | 5401 | 1,351 | 0,434 | 7,992 | 1,33E-15 | 2,98E-15 | 0,0027159 | 0,002318 | 0,0026079 | 0,0026079 | TRUE | 0,0327013 | No |
| orf19.2146 | 943 | 758 | 834 | 570 | 703 | 532 | 611 | 843 | 859 | 767 | 789 | 856 | 625 | 578 | 626 | 643 | 717,8 | 818 | 618 | 0,758 | -0,4 | -6,153 | 7,60E-10 | 1,43E-09 | 0,0013074 | 0,003877 | 0,0026439 | 0,0026439 | TRUE | 0,0215699 | No |
| orf19.2149 | 854 | 646 | 750 | 490 | 1598 | 1395 | 1569 | 2125 | 778 | 653 | 710 | 736 | 1421 | 1515 | 1607 | 1621 | 1130,07 | 719 | 1541 | 2,143 | 1,1 | 16,235 | 2,84E-59 | 1,54E-58 | 0,0035027 | 0,003206 | 0,0033568 | 0,0033568 | TRUE | 0,0462541 | No |
| orf19.215 | 10232 | 8792 | 9481 | 6411 | 4048 | 3352 | 3621 | 4984 | 9316 | 8892 | 8973 | 9630 | 3601 | 3639 | 3708 | 3802 | 6445,17 | 9203 | 3688 | 0,401 | -1,318 | -33,293 | 4,88E-243 | 9,94E-242 | 0,0007121 | 0,002242 | 0,0013207 | 0,0013207 | TRUE | 0,0115689 | No |
| orf19.2150 | 1079 | 842 | 907 | 639 | 1790 | 1404 | 1562 | 2123 | 982 | 852 | 858 | 960 | 1592 | 1524 | 1600 | 1619 | 1248,48 | 913 | 1584 | 1,738 | 0,797 | 13,479 | 2,08E-41 | 8,28E-41 | 0,0019413 | 0,003095 | 0,0024765 | 0,0024765 | TRUE | 0,0354302 | No |
| orf19.2151 | 12561 | 10014 | 11528 | 7117 | 5847 | 4616 | 5657 | 7552 | 11436 | 10128 | 10910 | 10691 | 5201 | 5012 | 5793 | 5761 | 8116,51 | 10791 | 5442 | 0,504 | -0,988 | -17,033 | 4,70E-65 | 2,76E-64 | 0,0037122 | 0,0022 | 0,0030936 | 0,0030936 | TRUE | 0,0342048 | No |
| orf19.2154 | 2297 | 1716 | 2033 | 1220 | 3191 | 2619 | 2961 | 4095 | 2091 | 1736 | 1924 | 1833 | 2838 | 2844 | 3032 | 3124 | 2427,69 | 1896 | 2960 | 1,56 | 0,642 | 10,329 | 5,18E-25 | 1,49E-24 | 0,0037583 | 0,002581 | 0,0032617 | 0,0032617 | TRUE | 0,0595415 | No |
| orf19.2156 | 2096 | 1576 | 1804 | 1201 | 1947 | 1425 | 1740 | 2396 | 1908 | 1594 | 1707 | 1804 | 1732 | 1547 | 1782 | 1828 | 1737,79 | 1753 | 1722 | 0,983 | -0,025 | -0,357 | 0,720776 | 0,753888 | 0,0050209 | 0,002797 | 0,0039766 | 0,0039766 | TRUE | 0,0554689 | No |
| orf19.2157 | 5435 | 4302 | 4715 | 3257 | 3363 | 2757 | 3188 | 4284 | 4948 | 4351 | 4462 | 4893 | 2991 | 2993 | 3265 | 3268 | 3896,45 | 4664 | 3129 | 0,671 | -0,575 | -10,159 | 3,03E-24 | 8,55E-24 | 0,0030048 | 0,002376 | 0,0028091 | 0,0028091 | TRUE | 0,0247976 | No |
| orf19.2158 | 272 | 212 | 237 | 166 | 451 | 403 | 433 | 565 | 248 | 214 | 224 | 249 | 401 | 438 | 443 | 431 | 331,11 | 234 | 428 | 1,835 | 0,876 | 10,228 | 1,49E-24 | 4,24E-24 | 1,00E-08 | 0,006027 | 0,0036672 | 0,0036672 | TRUE | 0,0342042 | No |

| | | | | | | | | | | | | | | | | | | | | | | | | | | | | | | | |
|---|---|---|---|---|---|---|---|---|---|---|---|---|---|---|---|---|---|---|---|---|---|---|---|---|---|---|---|---|---|---|---|
| orf19.216 | 5434 | 4831 | 5353 | 3051 | 2464 | 1972 | 2148 | 2820 | 4947 | 4886 | 5066 | 4583 | 2192 | 2141 | 2200 | 2151 | 3520,79 | 4870 | 2171 | 0,445 | -1,167 | -27,649 | 2,88E-168 | 4,24E-167 | 0,0007531 | 0,002412 | 0,0013858 | 0,0013858 | TRUE | 0,0182312 | No |
| orf19.2160 | 44 | 14 | 27 | 21 | 68 | 50 | 79 | 119 | 40 | 14 | 26 | 32 | 60 | 54 | 81 | 91 | 49,72 | 28 | 72 | 2,587 | 1,371 | 4,989 | 6,08E-07 | 1,02E-06 | 0,0797469 | 0,028605 | 0,0468034 | 0,0468034 | TRUE | 0,3558108 | No |
| orf19.216.1 | 2472 | 1853 | 1767 | 1275 | 2253 | 1774 | 1786 | 2712 | 2251 | 1874 | 1672 | 1915 | 2004 | 1926 | 1829 | 2069 | 1942,52 | 1928 | 1957 | 1,015 | 0,022 | 0,269 | 0,787731 | 0,816683 | 0,0085235 | 0,002717 | 0,005707 | 0,005707 | TRUE | 0,1570065 | No |
| orf19.2163 | 1074 | 945 | 1083 | 605 | 2676 | 2064 | 2581 | 3358 | 978 | 956 | 1025 | 909 | 2380 | 2241 | 2643 | 2562 | 1711,66 | 967 | 2456 | 2,537 | 1,343 | 21,067 | 1,60E-98 | 1,37E-97 | 0,0034543 | 0,002809 | 0,0031564 | 0,0031564 | TRUE | 0,0414539 | No |
| orf19.2164.1 | 0 | 0 | 1 | 2 | 0 | 2 | 2 | 1 | 0 | 0 | 1 | 3 | 0 | 2 | 2 | 1 | 1,12 | 1 | 1 | 1,391 | 0,476 | 0,328 | 0,743047 | 0,774641 | 0,5668428 | 1,185034 | 1,0268789 | 1,0268789 | TRUE | 0,554517 | No |
| orf19.2165 | 1430 | 1650 | 1664 | 938 | 2056 | 1645 | 1709 | 2130 | 1302 | 1669 | 1575 | 1409 | 1829 | 1786 | 1750 | 1625 | 1618,05 | 1489 | 1748 | 1,173 | 0,23 | 3,033 | 0,002418 | 0,003314 | 0,0065318 | 0,002853 | 0,0048923 | 0,0048923 | TRUE | 0,0897732 | No |
| orf19.2166 | 1339 | 1573 | 1587 | 980 | 1774 | 1531 | 1461 | 1790 | 1219 | 1591 | 1502 | 1472 | 1578 | 1662 | 1496 | 1365 | 1485,74 | 1446 | 1525 | 1,055 | 0,077 | 0,912 | 0,361902 | 0,402149 | 0,0093314 | 0,002926 | 0,0061626 | 0,0061626 | TRUE | 0,1379431 | No |
| orf19.2167 | 9837 | 9265 | 9602 | 5517 | 12713 | 10091 | 11114 | 14509 | 8956 | 9371 | 9087 | 8287 | 11308 | 10956 | 11382 | 11068 | 10051,9 | 8925 | 11178 | 1,252 | 0,324 | 7,614 | 2,65E-14 | 5,75E-14 | 0,0013488 | 0,002168 | 0,0016358 | 0,0016358 | TRUE | 0,0274796 | No |
| orf19.2168 | 459 | 435 | 466 | 270 | 732 | 593 | 605 | 797 | 418 | 440 | 441 | 406 | 651 | 644 | 620 | 608 | 528,37 | 426 | 631 | 1,475 | 0,561 | 8,378 | 5,40E-17 | 1,26E-16 | 1,00E-08 | 0,004537 | 0,002317 | 0,002317 | TRUE | 0,0096223 | No |
| orf19.2168.3 | 2647 | 2487 | 2522 | 1765 | 4683 | 4179 | 3647 | 5144 | 2410 | 2515 | 2387 | 2651 | 4165 | 4537 | 3735 | 3924 | 3290,62 | 2491 | 4090 | 1,643 | 0,717 | 11,307 | 1,21E-29 | 3,85E-29 | 0,0043228 | 0,002438 | 0,003526 | 0,003526 | TRUE | 0,0651572 | No |
| orf19.2169 | 12 | 7 | 13 | 14 | 21 | 10 | 23 | 22 | 11 | 7 | 12 | 21 | 19 | 11 | 24 | 17 | 15,15 | 13 | 18 | 1,414 | 0,499 | 1,276 | 0,202064 | 0,233743 | 0,0523725 | 0,089223 | 0,0774766 | 0,0774766 | TRUE | 0,4052835 | No |
| orf19.217 | 138 | 131 | 121 | 67 | 257 | 246 | 276 | 343 | 126 | 132 | 115 | 101 | 229 | 267 | 283 | 262 | 189,16 | 118 | 260 | 2,175 | 1,121 | 9,534 | 1,51E-21 | 3,96E-21 | 0,0037887 | 0,009021 | 0,0070199 | 0,0070199 | TRUE | 0,0751519 | No |
| orf19.2170 | 6274 | 5988 | 6388 | 4084 | 3178 | 2784 | 2735 | 3636 | 5712 | 6056 | 6046 | 6135 | 2827 | 3023 | 2801 | 2774 | 4421,61 | 5987 | 2856 | 0,477 | -1,068 | -24,996 | 6,72E-138 | 8,02E-137 | 0,0009318 | 0,002336 | 0,0014998 | 0,0014998 | TRUE | 0,0182831 | No |
| orf19.2172 | 11777 | 11357 | 11125 | 6229 | 5913 | 4522 | 4804 | 6186 | 10722 | 11486 | 10529 | 9357 | 5259 | 4910 | 4920 | 4719 | 7737,79 | 10524 | 4952 | 0,47 | -1,088 | -17,679 | 6,05E-70 | 3,79E-69 | 0,0043363 | 0,002208 | 0,0034937 | 0,0034937 | TRUE | 0,0672265 | No |
| orf19.2173 | 13142 | 11478 | 12638 | 7630 | 1756 | 1366 | 1525 | 2077 | 11965 | 11609 | 11961 | 11461 | 1562 | 1483 | 1562 | 1584 | 6648,42 | 11749 | 1548 | 0,132 | -2,923 | -76,905 | 0 | 0 | 0,0003295 | 0,002236 | 0,0010405 | 0,0010405 | TRUE | 0,0090676 | No |
| orf19.2174 | 575 | 484 | 516 | 290 | 688 | 490 | 559 | 741 | 524 | 490 | 488 | 436 | 612 | 532 | 572 | 565 | 527,34 | 484 | 570 | 1,173 | 0,231 | 3,085 | 0,002033 | 0,002803 | 0,0019177 | 0,004542 | 0,0034503 | 0,0034503 | TRUE | 0,044834 | No |
| orf19.2175 | 21385 | 18393 | 20029 | 12354 | 2267 | 1888 | 1923 | 2686 | 19470 | 18603 | 18956 | 18558 | 2016 | 2050 | 1969 | 2049 | 10458,8 | 18897 | 2021 | 0,107 | -3,225 | -91,872 | 0 | 0 | 0,0003321 | 0,002163 | 0,0009238 | 0,0009238 | TRUE | 0,0051403 | No |
| orf19.2176 | 1796 | 1396 | 1636 | 980 | 819 | 683 | 777 | 1041 | 1635 | 1412 | 1548 | 1472 | 728 | 742 | 796 | 794 | 1140,92 | 1517 | 765 | 0,504 | -0,989 | -15,856 | 1,29E-56 | 6,66E-56 | 0,0023936 | 0,003195 | 0,0027679 | 0,0027679 | TRUE | 0,0341334 | No |
| orf19.2177 | 520 | 436 | 495 | 388 | 659 | 667 | 643 | 852 | 473 | 441 | 468 | 583 | 586 | 724 | 659 | 650 | 573,06 | 491 | 655 | 1,338 | 0,42 | 4,566 | 4,97E-06 | 8,01E-06 | 0,0087187 | 0,004342 | 0,0063152 | 0,0063152 | TRUE | 0,1634704 | No |
| orf19.2178 | 3490 | 3007 | 3359 | 2511 | 4231 | 3892 | 3923 | 5215 | 3177 | 3041 | 3179 | 3772 | 3763 | 4226 | 4018 | 3978 | 3644,29 | 3292 | 3996 | 1,215 | 0,281 | 4,224 | 2,40E-05 | 3,73E-05 | 0,0056601 | 0,0024 | 0,0039726 | 0,0039726 | TRUE | 0,1134389 | No |
| orf19.2178.1 | 836 | 666 | 660 | 399 | 674 | 448 | 483 | 765 | 761 | 674 | 625 | 599 | 600 | 486 | 495 | 584 | 602,85 | 665 | 541 | 0,814 | -0,297 | -3,153 | 0,001618 | 0,00225 | 0,0094194 | 0,004228 | 0,0068726 | 0,0068726 | TRUE | 0,1145296 | No |
| orf19.2179 | 11681 | 9583 | 10435 | 6393 | 11048 | 8736 | 9697 | 13646 | 10635 | 9692 | 9876 | 9603 | 9827 | 9485 | 9931 | 10409 | 9932,28 | 9952 | 9913 | 0,996 | -0,006 | -0,126 | 0,89936 | 0,913838 | 0,0014932 | 0,00217 | 0,0018929 | 0,0018929 | TRUE | 0,0263012 | No |
| orf19.2179.2 | 15850 | 12553 | 12815 | 8071 | 22203 | 15737 | 18218 | 24412 | 14431 | 12696 | 12128 | 12124 | 19749 | 17086 | 18658 | 18622 | 15686,6 | 12845 | 18529 | 1,442 | 0,528 | 8,303 | 1,02E-16 | 2,36E-16 | 0,0052645 | 0,002121 | 0,003824 | 0,003824 | TRUE | 0,0847842 | No |
| orf19.218 | 28 | 20 | 24 | 15 | 84 | 58 | 75 | 82 | 25 | 20 | 23 | 23 | 75 | 63 | 77 | 63 | 46 | 23 | 70 | 3,033 | 1,601 | 7,147 | 8,85E-13 | 1,84E-12 | 1,00E-08 | 0,030753 | 0,0182078 | 0,0182078 | TRUE | 0,0462458 | No |
| orf19.2180 | 3070 | 2811 | 3147 | 2115 | 2012 | 1839 | 1779 | 2511 | 2795 | 2843 | 2978 | 3177 | 1790 | 1997 | 1822 | 1915 | 2414,64 | 2948 | 1881 | 0,639 | -0,647 | -11,76 | 6,30E-32 | 2,12E-31 | 0,0023463 | 0,002584 | 0,0024756 | 0,0024756 | TRUE | 0,031523 | No |
| orf19.2181 | 23 | 27 | 19 | 12 | 43 | 37 | 44 | 57 | 21 | 27 | 18 | 18 | 38 | 40 | 45 | 43 | 31,4 | 21 | 42 | 1,968 | 0,977 | 3,855 | 0,000116 | 0,000173 | 1,00E-08 | 0,044104 | 0,025491 | 0,025491 | TRUE | 0,2206613 | No |
| orf19.2182 | 7587 | 7146 | 7412 | 4787 | 6682 | 5852 | 5987 | 8226 | 6908 | 7227 | 7015 | 7191 | 5943 | 6354 | 6131 | 6275 | 6630,51 | 7085 | 6176 | 0,872 | -0,198 | -5,514 | 3,51E-08 | 6,17E-08 | 0,0004489 | 0,002236 | 0,0010847 | 0,0010847 | TRUE | 0,007891 | No |
| orf19.2183 | 24152 | 20980 | 22497 | 15119 | 36287 | 29584 | 31003 | 43980 | 21989 | 21219 | 21291 | 22711 | 32276 | 32120 | 31751 | 33548 | 27113,3 | 21802 | 32424 | 1,488 | 0,573 | 16,212 | 4,12E-59 | 2,22E-58 | 0,0007519 | 0,002086 | 0,0011604 | 0,0011604 | TRUE | 0,0094766 | No |
| orf19.2184 | 234 | 182 | 201 | 135 | 329 | 274 | 288 | 362 | 213 | 184 | 190 | 203 | 293 | 297 | 295 | 276 | 243,92 | 198 | 290 | 1,468 | 0,554 | 5,923 | 3,16E-09 | 5,81E-09 | 1,00E-08 | 0,007453 | 0,0040973 | 0,0040973 | TRUE | 0,0362237 | No |
| orf19.2185 | 4931 | 4107 | 4640 | 2764 | 5213 | 4218 | 4799 | 6279 | 4489 | 4154 | 4391 | 4152 | 4637 | 4580 | 4915 | 4790 | 4513,42 | 4296 | 4730 | 1,1 | 0,138 | 3,216 | 0,001298 | 0,00182 | 0,0010078 | 0,00233 | 0,0015378 | 0,0015378 | TRUE | 0,0111866 | No |
| orf19.2186 | 1090 | 917 | 1026 | 668 | 556 | 423 | 512 | 736 | 992 | 927 | 971 | 1003 | 495 | 459 | 524 | 561 | 741,74 | 973 | 510 | 0,526 | -0,928 | -13,413 | 5,05E-41 | 2,00E-40 | 0,0022331 | 0,003818 | 0,0031302 | 0,0031302 | TRUE | 0,0578529 | No |
| orf19.2187 | 835 | 707 | 859 | 562 | 1475 | 1294 | 1259 | 1855 | 760 | 715 | 813 | 844 | 1312 | 1405 | 1289 | 1415 | 1069,22 | 783 | 1355 | 1,736 | 0,796 | 12,51 | 6,55E-36 | 2,38E-35 | 0,0024263 | 0,003272 | 0,0028511 | 0,0028511 | TRUE | 0,0397073 | No |
| orf19.2190 | 1007 | 797 | 951 | 612 | 708 | 593 | 670 | 998 | 917 | 806 | 900 | 919 | 630 | 644 | 686 | 761 | 782,91 | 886 | 680 | 0,77 | -0,377 | -5,037 | 4,72E-07 | 7,95E-07 | 0,0044516 | 0,003724 | 0,004088 | 0,004088 | TRUE | 0,0805029 | No |
| orf19.2191 | 1661 | 1359 | 1447 | 944 | 2005 | 1595 | 1728 | 2285 | 1512 | 1374 | 1369 | 1418 | 1783 | 1732 | 1770 | 1743 | 1587,76 | 1418 | 1757 | 1,238 | 0,308 | 6,407 | 1,49E-10 | 2,87E-10 | 0,0005092 | 0,002869 | 0,0015776 | 0,0015776 | TRUE | 0,0257158 | No |
| orf19.2191.1 | 251 | 199 | 194 | 172 | 283 | 177 | 212 | 298 | 229 | 201 | 184 | 258 | 252 | 192 | 217 | 227 | 220,01 | 218 | 222 | 1,029 | 0,041 | 0,337 | 0,736397 | 0,76821 | 0,0118274 | 0,008041 | 0,0096025 | 0,0096025 | TRUE | 0,1448875 | No |
| orf19.2192 | 81189 | 72951 | 78116 | 50388 | 1E+05 | 95199 | 97245 | 1E+05 | 73919 | 73782 | 73930 | 75691 | 98139 | 1E+05 | 99591 | 1E+05 | 87597,9 | 74330 | 100866 | 1,357 | 0,44 | 15,223 | 2,47E-52 | 1,19E-51 | 0,0003468 | 0,002052 | 0,0007924 | 0,0007924 | TRUE | 0,0040622 | No |
| orf19.2193 | 9165 | 7676 | 8212 | 5366 | 8158 | 6909 | 7246 | 10097 | 8344 | 7763 | 7772 | 8061 | 7256 | 7501 | 7421 | 7702 | 7727,59 | 7985 | 7470 | 0,936 | -0,096 | -2,51 | 0,012056 | 0,015718 | 0,0007341 | 0,002208 | 0,0012737 | 0,0012737 | TRUE | 0,0115355 | No |
| orf19.2194 | 984 | 903 | 928 | 583 | 892 | 712 | 766 | 1009 | 896 | 913 | 878 | 876 | 793 | 773 | 784 | 770 | 835,47 | 891 | 780 | 0,875 | -0,193 | -3,626 | 0,000288 | 0,00042 | 1,00E-08 | 0,003618 | 0,0015099 | 0,0015099 | TRUE | 0,003387 | No |
| orf19.2195 | 7 | 9 | 3 | 5 | 10 | 6 | 3 | 10 | 6 | 9 | 3 | 8 | 9 | 7 | 3 | 8 | 6,49 | 6 | 7 | 1,038 | 0,053 | 0,096 | 0,923897 | 0,934303 | 1,00E-08 | 0,205521 | 0,1456709 | 0,1456709 | TRUE | 0,4023123 | No |
| orf19.2196 | 983 | 939 | 1045 | 589 | 293 | 275 | 246 | 351 | 895 | 950 | 989 | 885 | 261 | 299 | 252 | 268 | 599,66 | 930 | 270 | 0,289 | -1,79 | -23,944 | 1,07E-126 | 1,16E-125 | 0,0015257 | 0,00424 | 0,0030593 | 0,0030593 | TRUE | 0,052182 | No |
| orf19.2197 | 65 | 38 | 45 | 41 | 97 | 81 | 80 | 122 | 59 | 38 | 43 | 62 | 86 | 88 | 82 | 93 | 68,88 | 50 | 87 | 1,753 | 0,81 | 4,456 | 8,36E-06 | 1,33E-05 | 0,0012186 | 0,021217 | 0,0157188 | 0,0157188 | TRUE | 0,2044155 | No |
| orf19.2198 | 3973 | 3634 | 3934 | 2495 | 5330 | 4501 | 4726 | 6458 | 3617 | 3675 | 3723 | 3748 | 4741 | 4887 | 4840 | 4926 | 4269,71 | 3691 | 4848 | 1,314 | 0,394 | 11,762 | 6,16E-32 | 2,07E-31 | 1,00E-08 | 0,002346 | 0,0008372 | 0,0008372 | TRUE | 0,002779 | No |
| orf19.2199 | 3640 | 3281 | 3538 | 2438 | 4756 | 4068 | 3836 | 5511 | 3314 | 3318 | 3348 | 3662 | 4230 | 4417 | 3929 | 4204 | 3802,81 | 3410 | 4195 | 1,231 | 0,3 | 5,983 | 2,20E-09 | 4,06E-09 | 0,0020144 | 0,002384 | 0,0021472 | 0,0021472 | TRUE | 0,0285295 | No |
| orf19.22 | 3427 | 3108 | 3220 | 2059 | 4793 | 3925 | 4030 | 5413 | 3120 | 3143 | 3047 | 3093 | 4263 | 4261 | 4127 | 4129 | 3648,11 | 3101 | 4195 | 1,353 | 0,436 | 12,56 | 3,52E-36 | 1,29E-35 | 1,00E-08 | 0,002399 | 0,000871 | 0,000871 | TRUE | 0,0017537 | No |
| orf19.220 | 14471 | 12613 | 13578 | 9559 | 11356 | 10358 | 10631 | 14220 | 13175 | 12757 | 12850 | 14359 | 10101 | 11246 | 10887 | 10847 | 12027,8 | 13285 | 10770 | 0,811 | -0,302 | -6,062 | 1,34E-09 | 2,51E-09 | 0,002428 | 0,002147 | 0,0023067 | 0,0023067 | TRUE | 0,0357402 | No |
| orf19.2200 | 1503 | 1372 | 1509 | 924 | 1394 | 1214 | 1293 | 1695 | 1368 | 1388 | 1428 | 1388 | 1240 | 1318 | 1324 | 1293 | 1343,41 | 1393 | 1294 | 0,928 | -0,108 | -2,3 | 0,021451 | 0,027427 | 1,00E-08 | 0,00302 | 0,0013605 | 0,0013605 | TRUE | 0,0094604 | No |
| orf19.2201 | 1142 | 1015 | 999 | 620 | 977 | 672 | 720 | 917 | 1040 | 1027 | 945 | 931 | 869 | 730 | 737 | 699 | 872,32 | 986 | 759 | 0,769 | -0,379 | -5,002 | 5,68E-07 | 9,54E-07 | 0,0049298 | 0,003551 | 0,004359 | 0,004359 | TRUE | 0,1157702 | No |
| orf19.2202 | 128 | 122 | 125 | 86 | 267 | 209 | 231 | 325 | 117 | 123 | 118 | 129 | 237 | 227 | 237 | 248 | 179,54 | 122 | 237 | 1,96 | 0,971 | 9,017 | 1,94E-19 | 4,85E-19 | 1,00E-08 | 0,009395 | 0,0047878 | 0,0047878 | TRUE | 0,0124296 | No |
| orf19.2203 | 3 | 5 | 5 | 3 | 11 | 3 | 7 | 6 | 3 | 5 | 5 | 5 | 10 | 3 | 7 | 5 | 5,23 | 4 | 6 | 1,473 | 0,559 | 0,908 | 0,363878 | 0,403781 | 1,00E-08 | 0,254775 | 0,1633363 | 0,1633363 | TRUE | 0,313031 | No |
| orf19.2203.1 | 59 | 56 | 55 | 43 | 20 | 24 | 16 | 29 | 54 | 57 | 52 | 65 | 18 | 26 | 16 | 22 | 38,67 | 57 | 20 | 0,365 | -1,455 | -6,128 | 8,87E-10 | 1,66E-09 | 1,00E-08 | 0,036198 | 0,0220197 | 0,0220197 | TRUE | 0,144518 | No |
| orf19.2204 | 6978 | 5840 | 6374 | 4202 | 6450 | 5230 | 5529 | 7849 | 6353 | 5907 | 6032 | 6312 | 5737 | 5678 | 5662 | 5987 | 5958,66 | 6151 | 5766 | 0,938 | -0,092 | -2,35 | 0,018773 | 0,024138 | 0,0007375 | 0,002259 | 0,0013143 | 0,0013143 | TRUE | 0,0087443 | No |
| orf19.2204.2 | 1005 | 815 | 890 | 645 | 1120 | 910 | 883 | 1416 | 915 | 824 | 842 | 969 | 996 | 988 | 904 | 1080 | 939,89 | 888 | 992 | 1,122 | 0,166 | 2,312 | 0,020778 | 0,026593 | 0,0040389 | 0,003442 | 0,003862 | 0,003862 | TRUE | 0,0442082 | No |
| orf19.2208 | 7226 | 6154 | 6634 | 4384 | 4987 | 4419 | 4458 | 6103 | 6579 | 6224 | 6278 | 6585 | 4436 | 4798 | 4566 | 4655 | 5515,2 | 6416 | 4614 | 0,719 | -0,476 | -11,984 | 4,29E-33 | 1,47E-32 | 0,000746 | 0,002277 | 0,0013281 | 0,0013281 | TRUE | 0,0085852 | No |
| orf19.2209 | 1356 | 1086 | 1289 | 743 | 2295 | 1805 | 2044 | 2626 | 1235 | 1098 | 1220 | 1116 | 2041 | 1960 | 2093 | 2003 | 1595,81 | 1167 | 2024 | 1,731 | 0,791 | 14,89 | 3,83E-50 | 1,76E-49 | 0,0012871 | 0,002865 | 0,0020222 | 0,0020222 | TRUE | 0,0208804 | No |
| orf19.2210 | 2050 | 1824 | 1985 | 1351 | 1528 | 1281 | 1427 | 1914 | 1866 | 1845 | 1879 | 2029 | 1359 | 1391 | 1461 | 1460 | 1661,33 | 1905 | 1418 | 0,746 | -0,423 | -8,484 | 2,17E-17 | 5,15E-17 | 0,0008093 | 0,002832 | 0,0017736 | 0,0017736 | TRUE | 0,0213484 | No |
| orf19.2211 | 11 | 13 | 9 | 9 | 12 | 12 | 14 | 23 | 10 | 13 | 9 | 14 | 11 | 13 | 14 | 18 | 12,6 | 12 | 14 | 1,257 | 0,33 | 0,851 | 0,394925 | 0,43588 | 1,00E-08 | 0,106894 | 0,0633341 | 0,0633341 | TRUE | 0,1803324 | No |
| orf19.2212 | 35 | 23 | 42 | 15 | 59 | 57 | 47 | 73 | 32 | 23 | 40 | 23 | 52 | 62 | 48 | 56 | 41,95 | 30 | 54 | 1,824 | 0,867 | 3,765 | 0,000167 | 0,000247 | 0,0021636 | 0,033527 | 0,0245971 | 0,0245971 | TRUE | 0,379671 | No |
| orf19.2213 | 125 | 129 | 152 | 88 | 269 | 200 | 197 | 265 | 114 | 130 | 144 | 132 | 239 | 217 | 202 | 202 | 172,58 | 130 | 215 | 1,656 | 0,728 | 6,201 | 5,62E-10 | 1,06E-09 | 1,00E-08 | 0,009691 | 0,0069502 | 0,0069502 | TRUE | 0,0832113 | No |
| orf19.2214 | 2492 | 2131 | 2307 | 1546 | 1811 | 1422 | 1435 | 1947 | 2269 | 2155 | 2183 | 2322 | 1611 | 1544 | 1470 | 1485 | 1879,92 | 2232 | 1528 | 0,685 | -0,546 | -11,296 | 1,38E-29 | 4,37E-29 | 0,000881 | 0,00274 | 0,0016987 | 0,0016987 | TRUE | 0,0167456 | No |
| orf19.2215 | 5108 | 4387 | 4811 | 3150 | 3309 | 2839 | 2919 | 4005 | 4651 | 4437 | 4553 | 4732 | 2943 | 3082 | 2989 | 3055 | 3805,33 | 4593 | 3017 | 0,657 | -0,605 | -16,097 | 2,70E-58 | 1,44E-57 | 0,0003276 | 0,002384 | 0,0010861 | 0,0010861 | TRUE | 0,0063144 | No |
| orf19.2216 | 238 | 229 | 225 | 145 | 427 | 309 | 369 | 492 | 217 | 232 | 213 | 218 | 380 | 335 | 378 | 375 | 293,44 | 220 | 367 | 1,673 | 0,743 | 8,475 | 2,36E-17 | 5,57E-17 | 1,00E-08 | 0,006539 | 0,0036864 | 0,0036864 | TRUE | 0,0358161 | No |
| orf19.2219 | 3019 | 2689 | 3026 | 1971 | 1677 | 1446 | 1591 | 2045 | 2749 | 2720 | 2864 | 2961 | 1492 | 1570 | 1629 | 1560 | 2192,98 | 2824 | 1563 | 0,554 | -0,852 | -18,071 | 5,42E-73 | 3,53E-72 | 0,0007601 | 0,002639 | 0,0016466 | 0,0016466 | TRUE | 0,0121281 | No |
| orf19.2220 | 285 | 266 | 226 | 161 | 193 | 132 | 160 | 177 | 259 | 269 | 214 | 242 | 172 | 143 | 164 | 135 | 199,76 | 246 | 154 | 0,622 | -0,684 | -5,855 | 4,77E-09 | 8,71E-09 | 0,0068045 | 0,00865 | 0,0078821 | 0,0078821 | TRUE | 0,0937235 | No |
| orf19.2222 | 2727 | 2514 | 2842 | 1696 | 2772 | 2345 | 2503 | 3173 | 2483 | 2543 | 2690 | 2548 | 2466 | 2546 | 2563 | 2420 | 2532,28 | 2566 | 2499 | 0,973 | -0,039 | -0,911 | 0,362078 | 0,402273 | 0,000614 | 0,002559 | 0,0013842 | 0,0013842 | TRUE | 0,0134119 | No |
| orf19.2227 | 146 | 158 | 131 | 88 | 107 | 115 | 114 | 160 | 133 | 160 | 124 | 132 | 95 | 125 | 117 | 122 | 125,97 | 137 | 115 | 0,834 | -0,261 | -1,907 | 0,05656 | 0,069716 | 0,0061254 | 0,012524 | 0,0100426 | 0,0100426 | TRUE | 0,1381794 | No |
| orf19.2228 | 195 | 226 | 203 | 146 | 126 | 125 | 84 | 115 | 178 | 229 | 192 | 219 | 112 | 136 | 86 | 88 | 154,89 | 204 | 106 | 0,514 | -0,959 | -6,353 | 2,11E-10 | 4,04E-10 | 0,0218934 | 0,010566 | 0,0147814 | 0,0147814 | TRUE | 0,3478297 | No |
| orf19.2229 | 1259 | 1083 | 1175 | 727 | 1546 | 1356 | 1482 | 1892 | 1146 | 1095 | 1112 | 1092 | 1375 | 1472 | 1518 | 1443 | 1281,76 | 1111 | 1452 | 1,304 | 0,383 | 7,566 | 3,84E-14 | 8,28E-14 | 0,0004773 | 0,003068 | 0,0016607 | 0,0016607 | TRUE | 0,0153398 | No |

| | | | | | | | | | | | | | | | | | | | | | | | | | | | | | | | |
|---|---|---|---|---|---|---|---|---|---|---|---|---|---|---|---|---|---|---|---|---|---|---|---|---|---|---|---|---|---|---|---|
| orf19.223 | 10082 | 9243 | 9935 | 6231 | 11365 | 9536 | 10029 | 13293 | 9179 | 9348 | 9403 | 9360 | 10109 | 10353 | 10271 | 10140 | 9770,41 | 9322 | 10218 | 1,096 | 0,132 | 4,824 | 1,41E-06 | 2,33E-06 | 1,00E-08 | 0,002172 | 0,0006188 | 0,0006188 | TRUE | 0,0013484 | No |
| orf19.2230 | 550 | 490 | 534 | 290 | 631 | 444 | 525 | 678 | 501 | 496 | 505 | 436 | 561 | 482 | 538 | 517 | 504,44 | 484 | 524 | 1,079 | 0,11 | 1,461 | 0,144114 | 0,169776 | 0,0016811 | 0,004656 | 0,0034326 | 0,0034326 | TRUE | 0,0447631 | No |
| orf19.2231 | 2191 | 2044 | 2094 | 1567 | 2210 | 2071 | 1944 | 2568 | 1995 | 2067 | 1982 | 2354 | 1966 | 2249 | 1991 | 1959 | 2070,23 | 2100 | 2041 | 0,973 | -0,039 | -0,581 | 0,560947 | 0,602128 | 0,0046882 | 0,002675 | 0,0038318 | 0,0038318 | TRUE | 0,0770183 | No |
| orf19.2232 | 12829 | 10735 | 10913 | 6754 | 18372 | 12849 | 13178 | 19783 | 11680 | 10857 | 10328 | 10146 | 16341 | 13950 | 13496 | 15091 | 12736,2 | 10753 | 14720 | 1,369 | 0,453 | 6,889 | 5,63E-12 | 1,14E-11 | 0,0056756 | 0,002141 | 0,0040707 | 0,0040707 | TRUE | 0,0672914 | No |
| orf19.2233 | 4739 | 4026 | 4291 | 2560 | 3507 | 2964 | 3001 | 4312 | 4315 | 4072 | 4061 | 3846 | 3119 | 3218 | 3073 | 3289 | 3624,15 | 4074 | 3175 | 0,779 | -0,361 | -7,896 | 2,89E-15 | 6,44E-15 | 0,0012025 | 0,002402 | 0,0017223 | 0,0017223 | TRUE | 0,0194162 | No |
| orf19.2236 | 4447 | 4008 | 4436 | 2735 | 4006 | 3322 | 3566 | 4831 | 4049 | 4054 | 4198 | 4108 | 3563 | 3607 | 3652 | 3685 | 3864,54 | 4102 | 3627 | 0,884 | -0,177 | -5,218 | 1,81E-07 | 3,10E-07 | 1,00E-08 | 0,002379 | 0,0008511 | 0,0008511 | TRUE | 0,0031867 | No |
| orf19.2237 | 76 | 56 | 67 | 41 | 33 | 29 | 24 | 35 | 69 | 57 | 63 | 62 | 29 | 31 | 25 | 27 | 45,37 | 63 | 28 | 0,445 | -1,168 | -5,559 | 2,71E-08 | 4,78E-08 | 1,00E-08 | 0,031154 | 0,0173265 | 0,0173265 | TRUE | 0,0473011 | No |
| orf19.2237.1 | 2362 | 2450 | 2452 | 1556 | 2328 | 1900 | 1911 | 2494 | 2151 | 2478 | 2321 | 2337 | 2071 | 2063 | 1957 | 1902 | 2159,93 | 2322 | 1998 | 0,861 | -0,216 | -3,984 | 6,79E-05 | 0,000103 | 0,0021568 | 0,002649 | 0,0023668 | 0,0023668 | TRUE | 0,0309279 | No |
| orf19.2238 | 1040 | 1095 | 1113 | 658 | 643 | 560 | 548 | 644 | 947 | 1107 | 1053 | 988 | 572 | 608 | 561 | 491 | 791,07 | 1024 | 558 | 0,543 | -0,88 | -11,391 | 4,62E-30 | 1,49E-29 | 0,0050479 | 0,003707 | 0,0043651 | 0,0043651 | TRUE | 0,0811623 | No |
| orf19.2241 | 5701 | 4703 | 5235 | 3228 | 2257 | 1831 | 1971 | 2471 | 5191 | 4757 | 4954 | 4849 | 2008 | 1988 | 2019 | 1885 | 3456,18 | 4938 | 1975 | 0,4 | -1,324 | -29,925 | 9,30E-197 | 1,56E-195 | 0,0009813 | 0,002419 | 0,0015329 | 0,0015329 | TRUE | 0,0147647 | No |
| orf19.2242 | 10748 | 9808 | 10517 | 6796 | 15837 | 13061 | 13330 | 18770 | 9786 | 9920 | 9953 | 10209 | 14087 | 14181 | 13652 | 14318 | 12013 | 9967 | 14060 | 1,411 | 0,497 | 16,118 | 1,91E-58 | 1,02E-57 | 0,0002525 | 0,002147 | 0,0008256 | 0,0008256 | TRUE | 0,0046364 | No |
| orf19.2244 | 805 | 711 | 788 | 549 | 1695 | 1379 | 1335 | 1960 | 733 | 719 | 746 | 825 | 1508 | 1497 | 1367 | 1495 | 1111,21 | 756 | 1467 | 1,949 | 0,963 | 16,132 | 1,53E-58 | 8,22E-58 | 0,001655 | 0,003226 | 0,0023921 | 0,0023921 | TRUE | 0,0386921 | No |
| orf19.2245 | 642 | 597 | 603 | 365 | 1395 | 1128 | 1097 | 1460 | 585 | 604 | 571 | 548 | 1241 | 1225 | 1123 | 1114 | 876,24 | 577 | 1176 | 2,033 | 1,023 | 16,209 | 4,34E-59 | 2,34E-58 | 0,001536 | 0,003545 | 0,0025038 | 0,0025038 | TRUE | 0,0174694 | No |
| orf19.2246 | 1003 | 876 | 896 | 505 | 1186 | 838 | 938 | 1265 | 913 | 886 | 848 | 759 | 1055 | 910 | 961 | 965 | 912,01 | 852 | 973 | 1,139 | 0,188 | 2,697 | 0,006986 | 0,009273 | 0,0035906 | 0,003485 | 0,003555 | 0,003555 | TRUE | 0,0570626 | No |
| orf19.2247 | 102 | 96 | 90 | 81 | 28 | 25 | 35 | 41 | 93 | 97 | 85 | 122 | 25 | 27 | 36 | 31 | 64,5 | 99 | 30 | 0,303 | -1,721 | -8,636 | 5,80E-18 | 1,39E-17 | 1,00E-08 | 0,022518 | 0,0170804 | 0,0170804 | TRUE | 0,1826615 | No |
| orf19.2248 | 1205 | 1128 | 1244 | 804 | 1410 | 1211 | 1226 | 1593 | 1097 | 1141 | 1177 | 1208 | 1254 | 1315 | 1256 | 1215 | 1207,84 | 1156 | 1260 | 1,091 | 0,126 | 2,419 | 0,015557 | 0,020141 | 0,0005113 | 0,003131 | 0,0017624 | 0,0017624 | TRUE | 0,0142943 | No |
| orf19.2249 | 361 | 273 | 328 | 202 | 532 | 369 | 379 | 545 | 329 | 276 | 310 | 303 | 473 | 401 | 388 | 416 | 362,04 | 304 | 420 | 1,377 | 0,462 | 5,099 | 3,42E-07 | 5,78E-07 | 0,004178 | 0,005686 | 0,0050043 | 0,0050043 | TRUE | 0,0867558 | No |
| orf19.225 | 3602 | 3276 | 3568 | 2043 | 4861 | 3914 | 4243 | 5668 | 3279 | 3313 | 3377 | 3069 | 4324 | 4250 | 4345 | 4324 | 3785,08 | 3260 | 4311 | 1,321 | 0,401 | 10,238 | 1,34E-24 | 3,80E-24 | 0,0003707 | 0,002386 | 0,0012005 | 0,0012005 | TRUE | 0,0172822 | No |
| orf19.2250 | 4273 | 3530 | 3599 | 2541 | 5204 | 4174 | 3888 | 5986 | 3890 | 3570 | 3406 | 3817 | 4629 | 4532 | 3982 | 4566 | 4049,04 | 3671 | 4427 | 1,207 | 0,271 | 4,458 | 8,27E-06 | 1,32E-05 | 0,0038098 | 0,002363 | 0,0032967 | 0,0032967 | TRUE | 0,0555967 | No |
| orf19.2251 | 9027 | 7569 | 7943 | 5390 | 9009 | 7310 | 6958 | 10295 | 8219 | 7655 | 7517 | 8097 | 8013 | 7937 | 7126 | 7853 | 7802,09 | 7872 | 7732 | 0,983 | -0,025 | -0,514 | 0,607363 | 0,646833 | 0,0019479 | 0,002206 | 0,0022297 | 0,0022297 | TRUE | 0,0338686 | No |
| orf19.2252 | 5 | 8 | 3 | 3 | 14 | 4 | 10 | 17 | 5 | 8 | 3 | 5 | 12 | 4 | 10 | 13 | 7,5 | 5 | 10 | 2,033 | 1,024 | 1,869 | 0,061652 | 0,075714 | 1,00E-08 | 0,178189 | 0,1358023 | 0,1358023 | TRUE | 0,446606 | No |
| orf19.2253 | 273 | 201 | 231 | 152 | 506 | 404 | 466 | 662 | 249 | 203 | 219 | 228 | 450 | 439 | 477 | 505 | 346,21 | 225 | 468 | 2,084 | 1,06 | 11,75 | 7,07E-32 | 2,37E-31 | 0,0025415 | 0,005853 | 0,0044388 | 0,0044388 | TRUE | 0,0645693 | No |
| orf19.2256 | 4175 | 3294 | 3645 | 2261 | 5094 | 3815 | 4643 | 5890 | 3801 | 3332 | 3450 | 3396 | 4531 | 4142 | 4755 | 4493 | 3987,46 | 3495 | 4480 | 1,282 | 0,358 | 6,34 | 2,30E-10 | 4,41E-10 | 0,0030645 | 0,002368 | 0,0028015 | 0,0028015 | TRUE | 0,0430976 | No |
| orf19.2257 | 1604 | 1403 | 1507 | 898 | 2558 | 2218 | 2162 | 2915 | 1460 | 1419 | 1426 | 1349 | 2275 | 2408 | 2214 | 2224 | 1846,96 | 1414 | 2280 | 1,609 | 0,686 | 14,377 | 7,27E-47 | 3,18E-46 | 0,0007077 | 0,002752 | 0,0016044 | 0,0016044 | TRUE | 0,0167177 | No |
| orf19.2258 | 1115 | 1015 | 1138 | 602 | 1439 | 1072 | 1263 | 1540 | 1015 | 1027 | 1077 | 904 | 1280 | 1164 | 1293 | 1175 | 1116,88 | 1006 | 1228 | 1,217 | 0,283 | 4,387 | 1,15E-05 | 1,82E-05 | 0,0029526 | 0,00322 | 0,0030926 | 0,0030926 | TRUE | 0,0492204 | No |
| orf19.2259 | 1951 | 1698 | 1774 | 1044 | 1693 | 1271 | 1470 | 1767 | 1776 | 1717 | 1679 | 1568 | 1506 | 1380 | 1505 | 1348 | 1560 | 1685 | 1435 | 0,85 | -0,235 | -4,051 | 5,09E-05 | 7,76E-05 | 0,0021608 | 0,002884 | 0,0025894 | 0,0025894 | TRUE | 0,0242676 | No |
| orf19.2260 | 2216 | 1976 | 2275 | 1293 | 2990 | 2462 | 2686 | 3559 | 2018 | 1999 | 2153 | 1942 | 2660 | 2673 | 2751 | 2715 | 2363,71 | 2028 | 2700 | 1,329 | 0,411 | 9,396 | 5,65E-21 | 1,46E-20 | 0,0005139 | 0,002596 | 0,0013963 | 0,0013963 | TRUE | 0,0210964 | No |
| orf19.2261 | 1930 | 1745 | 1848 | 1167 | 1548 | 1179 | 1336 | 1848 | 1757 | 1765 | 1749 | 1753 | 1377 | 1280 | 1368 | 1410 | 1557,36 | 1756 | 1359 | 0,775 | -0,368 | -7,991 | 1,34E-15 | 3,02E-15 | 1,00E-08 | 0,002885 | 0,0013896 | 0,0013896 | TRUE | 0,0174954 | No |
| orf19.2262 | 2957 | 2610 | 2879 | 1698 | 1362 | 1073 | 1191 | 1494 | 2692 | 2640 | 2725 | 2551 | 1211 | 1165 | 1220 | 1140 | 1917,89 | 2652 | 1184 | 0,446 | -1,166 | -25,601 | 1,50E-144 | 1,89E-143 | 0,0003222 | 0,002726 | 0,0013942 | 0,0013942 | TRUE | 0,0083471 | No |
| orf19.2263 | 875 | 789 | 838 | 539 | 1171 | 993 | 973 | 1321 | 797 | 798 | 793 | 810 | 1042 | 1078 | 996 | 1008 | 915,15 | 800 | 1031 | 1,29 | 0,367 | 6,87 | 6,40E-12 | 1,30E-11 | 1,00E-08 | 0,00348 | 0,0016189 | 0,0016189 | TRUE | 0,0106593 | No |
| orf19.2264 | 5565 | 4724 | 5115 | 3406 | 4730 | 4090 | 4216 | 5824 | 5067 | 4778 | 4841 | 5116 | 4207 | 4441 | 4318 | 4443 | 4651,23 | 4950 | 4352 | 0,88 | -0,185 | -4,698 | 2,63E-06 | 4,29E-06 | 0,0006117 | 0,002321 | 0,0012752 | 0,0012752 | TRUE | 0,0066464 | No |
| orf19.2265 | 2284 | 2001 | 2199 | 1487 | 3002 | 2498 | 2623 | 3564 | 2079 | 2024 | 2081 | 2234 | 2670 | 2712 | 2686 | 2719 | 2400,67 | 2104 | 2697 | 1,284 | 0,361 | 8,597 | 8,17E-18 | 1,96E-17 | 0,0001421 | 0,002587 | 0,0012607 | 0,0012607 | TRUE | 0,0184744 | No |
| orf19.2266 | 1425 | 1305 | 1294 | 866 | 2197 | 1719 | 2018 | 2621 | 1297 | 1320 | 1225 | 1301 | 1954 | 1866 | 2067 | 1999 | 1628,66 | 1286 | 1972 | 1,535 | 0,618 | 12,341 | 5,47E-35 | 1,95E-34 | 0,0008763 | 0,002848 | 0,0017542 | 0,0017542 | TRUE | 0,0149583 | No |
| orf19.2267 | 2681 | 2058 | 2194 | 1545 | 3041 | 2431 | 2579 | 3782 | 2441 | 2081 | 2076 | 2321 | 2705 | 2639 | 2641 | 2885 | 2473,76 | 2230 | 2718 | 1,22 | 0,287 | 4,657 | 3,21E-06 | 5,23E-06 | 0,0036591 | 0,002571 | 0,0032237 | 0,0032237 | TRUE | 0,051553 | No |
| orf19.2268 | 7054 | 6247 | 6832 | 4505 | 9856 | 8343 | 9022 | 12124 | 6422 | 6318 | 6466 | 6767 | 8767 | 9058 | 9240 | 9248 | 7785,8 | 6493 | 9078 | 1,399 | 0,484 | 13,363 | 9,90E-41 | 3,89E-40 | 0,0005778 | 0,002207 | 0,001127 | 0,001127 | TRUE | 0,0093575 | No |
| orf19.2269 | 689 | 699 | 654 | 409 | 924 | 751 | 750 | 1059 | 627 | 707 | 619 | 614 | 822 | 815 | 768 | 808 | 722,59 | 642 | 803 | 1,25 | 0,322 | 5,013 | 5,35E-07 | 9,00E-07 | 0,0009014 | 0,003865 | 0,0025522 | 0,0025522 | TRUE | 0,0555467 | No |
| orf19.227 | 2901 | 2558 | 2673 | 1582 | 3980 | 3133 | 3371 | 4164 | 2641 | 2587 | 2530 | 2376 | 3540 | 3402 | 3452 | 3176 | 2963,11 | 2534 | 3392 | 1,337 | 0,419 | 8,391 | 4,83E-17 | 1,13E-16 | 0,0016025 | 0,002483 | 0,0020412 | 0,0020412 | TRUE | 0,0230695 | No |
| orf19.2270 | 8753 | 7895 | 8530 | 5228 | 6124 | 5130 | 5232 | 7198 | 7969 | 7985 | 8073 | 7853 | 5447 | 5570 | 5358 | 5491 | 6718,27 | 7970 | 5466 | 0,686 | -0,544 | -18,121 | 2,17E-73 | 1,42E-72 | 1,00E-08 | 0,002234 | 0,0007136 | 0,0007136 | TRUE | 0,0020997 | No |
| orf19.2272 | 2942 | 2668 | 3017 | 1954 | 2625 | 2540 | 2492 | 3446 | 2679 | 2698 | 2855 | 2935 | 2335 | 2758 | 2552 | 2629 | 2680,11 | 2792 | 2568 | 0,92 | -0,12 | -2,082 | 0,037302 | 0,046738 | 0,0028706 | 0,00253 | 0,002801 | 0,002801 | TRUE | 0,0456876 | No |
| orf19.2274 | 49 | 53 | 57 | 36 | 42 | 34 | 32 | 40 | 45 | 54 | 54 | 54 | 37 | 37 | 33 | 31 | 42,97 | 52 | 34 | 0,667 | -0,584 | -2,787 | 0,005316 | 0,007115 | 1,00E-08 | 0,032776 | 0,0182151 | 0,0182151 | TRUE | 0,0762266 | No |
| orf19.2275 | 1519 | 1271 | 1361 | 931 | 1571 | 1117 | 1222 | 1635 | 1383 | 1285 | 1288 | 1399 | 1397 | 1213 | 1251 | 1247 | 1307,98 | 1339 | 1277 | 0,956 | -0,066 | -1,103 | 0,270169 | 0,306137 | 0,002056 | 0,003047 | 0,0026324 | 0,0026324 | TRUE | 0,0486247 | No |
| orf19.2276 | 1020 | 824 | 846 | 475 | 1538 | 1092 | 1172 | 1631 | 929 | 833 | 801 | 714 | 1368 | 1186 | 1200 | 1244 | 1034,28 | 819 | 1250 | 1,521 | 0,605 | 7,714 | 1,22E-14 | 2,68E-14 | 0,0067358 | 0,003314 | 0,004877 | 0,004877 | TRUE | 0,0965272 | No |
| orf19.2277 | 2187 | 1956 | 2155 | 1416 | 2316 | 1880 | 2073 | 2752 | 1991 | 1978 | 2040 | 2127 | 2060 | 2041 | 2123 | 2099 | 2057,43 | 2034 | 2081 | 1,025 | 0,035 | 0,84 | 0,40076 | 0,441557 | 5,97E-05 | 0,002679 | 0,0012126 | 0,0012126 | TRUE | 0,0102074 | No |
| orf19.2278 | 2482 | 2255 | 2357 | 1588 | 1903 | 1634 | 1590 | 2118 | 2260 | 2281 | 2231 | 2385 | 1693 | 1774 | 1628 | 1616 | 1983,41 | 2289 | 1678 | 0,733 | -0,448 | -9,552 | 1,27E-21 | 3,34E-21 | 0,0006802 | 0,002703 | 0,0015964 | 0,0015964 | TRUE | 0,0174012 | No |
| orf19.2280 | 727 | 653 | 732 | 458 | 1475 | 1230 | 1313 | 1739 | 662 | 660 | 693 | 688 | 1312 | 1335 | 1345 | 1327 | 1002,71 | 676 | 1330 | 1,969 | 0,978 | 19,011 | 1,39E-80 | 9,90E-80 | 1,00E-08 | 0,003354 | 0,0013983 | 0,0013983 | TRUE | 0,0040857 | No |
| orf19.2281 | 125 | 99 | 107 | 60 | 397 | 312 | 337 | 473 | 114 | 100 | 101 | 90 | 353 | 339 | 345 | 361 | 225,39 | 101 | 350 | 3,42 | 1,774 | 16,673 | 2,06E-62 | 1,16E-61 | 1,00E-08 | 0,007898 | 0,0043594 | 0,0043594 | TRUE | 0,0688425 | No |
| orf19.2282 | 1179 | 1030 | 1047 | 618 | 575 | 464 | 472 | 683 | 1073 | 1042 | 991 | 928 | 511 | 504 | 483 | 521 | 756,75 | 1008 | 505 | 0,5 | -1,001 | -15,441 | 8,63E-54 | 4,28E-53 | 0,001322 | 0,003783 | 0,0025788 | 0,0025788 | TRUE | 0,030305 | No |
| orf19.2283 | 1933 | 1515 | 1650 | 1115 | 1764 | 1347 | 1445 | 2051 | 1760 | 1532 | 1562 | 1675 | 1569 | 1462 | 1480 | 1565 | 1575,57 | 1632 | 1519 | 0,932 | -0,102 | -1,791 | 0,073259 | 0,089229 | 0,0021631 | 0,002875 | 0,0024828 | 0,0024828 | TRUE | 0,0358612 | No |
| orf19.2284 | 1792 | 1466 | 1613 | 1136 | 794 | 619 | 649 | 958 | 1632 | 1483 | 1527 | 1706 | 706 | 672 | 665 | 731 | 1140,12 | 1587 | 694 | 0,438 | -1,19 | -19,198 | 3,84E-82 | 2,80E-81 | 0,0021957 | 0,003196 | 0,0026746 | 0,0026746 | TRUE | 0,0285196 | No |
| orf19.2285 | 111 | 108 | 112 | 70 | 130 | 130 | 113 | 172 | 101 | 109 | 106 | 105 | 116 | 141 | 116 | 131 | 115,64 | 105 | 126 | 1,195 | 0,257 | 1,953 | 0,050853 | 0,062913 | 1,00E-08 | 0,01346 | 0,0078501 | 0,0078501 | TRUE | 0,0596972 | No |
| orf19.2286 | 5580 | 4521 | 4802 | 3309 | 4915 | 3935 | 4094 | 5909 | 5080 | 4573 | 4545 | 4971 | 4372 | 4272 | 4193 | 4507 | 4564,05 | 4792 | 4336 | 0,905 | -0,143 | -2,93 | 0,003386 | 0,004586 | 0,0018256 | 0,002326 | 0,0020767 | 0,0020767 | TRUE | 0,0208107 | No |
| orf19.2287 | 1802 | 1401 | 1536 | 1042 | 2427 | 1834 | 1898 | 2593 | 1641 | 1417 | 1454 | 1565 | 2159 | 1991 | 1944 | 1978 | 1768,53 | 1519 | 2018 | 1,329 | 0,41 | 6,918 | 4,57E-12 | 9,30E-12 | 0,0026451 | 0,002784 | 0,0027935 | 0,0027935 | TRUE | 0,0374028 | No |
| orf19.2288 | 5162 | 4275 | 4932 | 2885 | 6696 | 5180 | 6118 | 8184 | 4700 | 4324 | 4668 | 4334 | 5956 | 5624 | 6266 | 6243 | 5264,16 | 4506 | 6022 | 1,336 | 0,418 | 8,404 | 4,32E-17 | 1,01E-16 | 0,0020252 | 0,002288 | 0,0021767 | 0,0021767 | TRUE | 0,0238961 | No |
| orf19.2289 | 1312 | 1199 | 1189 | 842 | 840 | 652 | 719 | 1032 | 1195 | 1213 | 1125 | 1265 | 747 | 708 | 736 | 787 | 971,99 | 1200 | 744 | 0,623 | -0,682 | -11,62 | 3,26E-31 | 1,08E-30 | 0,0010972 | 0,003396 | 0,00235 | 0,00235 | TRUE | 0,0207655 | No |
| orf19.229 | 1953 | 1910 | 1896 | 1086 | 2198 | 1682 | 1867 | 2463 | 1778 | 1932 | 1794 | 1631 | 1955 | 1826 | 1912 | 1879 | 1838,46 | 1784 | 1893 | 1,059 | 0,083 | 1,517 | 0,129306 | 0,153392 | 0,0020585 | 0,002756 | 0,0023442 | 0,0023442 | TRUE | 0,0366381 | No |
| orf19.2290 | 6307 | 5542 | 6064 | 3802 | 2984 | 2534 | 2767 | 3712 | 5742 | 5605 | 5739 | 5711 | 2654 | 2751 | 2834 | 2832 | 4233,54 | 5699 | 2768 | 0,486 | -1,042 | -28,612 | 4,78E-180 | 7,38E-179 | 0,0001759 | 0,002349 | 0,0010103 | 0,0010103 | TRUE | 0,0094978 | No |
| orf19.2296 | 1E+05 | 82301 | 92018 | 62502 | 16210 | 14304 | 14292 | 20205 | 92267 | 83239 | 87087 | 93888 | 14418 | 15530 | 14637 | 15413 | 52059,7 | 89120 | 15000 | 0,168 | -2,571 | -54,094 | 0 | 0 | 0,0021282 | 0,002062 | 0,0021328 | 0,0021328 | TRUE | 0,0241838 | No |
| orf19.2297 | 228 | 198 | 233 | 145 | 199 | 154 | 171 | 213 | 208 | 200 | 221 | 218 | 177 | 167 | 175 | 162 | 191 | 212 | 170 | 0,806 | -0,311 | -3,068 | 0,002157 | 0,002968 | 1,00E-08 | 0,008953 | 0,0046123 | 0,0046123 | TRUE | 0,0140773 | No |
| orf19.2298 | 1555 | 1439 | 1412 | 1060 | 1119 | 996 | 1066 | 1399 | 1416 | 1455 | 1336 | 1592 | 995 | 1081 | 1092 | 1067 | 1254,42 | 1450 | 1059 | 0,732 | -0,45 | -7,279 | 3,36E-13 | 7,08E-13 | 0,0024816 | 0,00309 | 0,0028626 | 0,0028626 | TRUE | 0,0486481 | No |
| orf19.2299 | 905 | 682 | 779 | 492 | 1908 | 1309 | 1491 | 2165 | 824 | 690 | 737 | 739 | 1697 | 1421 | 1527 | 1651 | 1160,85 | 748 | 1574 | 2,106 | 1,075 | 14,618 | 2,17E-48 | 9,71E-48 | 0,0051217 | 0,003175 | 0,0041731 | 0,0041731 | TRUE | 0,0592794 | No |
| orf19.23 | 3901 | 3437 | 3770 | 2275 | 4786 | 3955 | 4296 | 5902 | 3552 | 3476 | 3568 | 3417 | 4257 | 4294 | 4400 | 4502 | 3933,25 | 3503 | 4363 | 1,245 | 0,316 | 8,601 | 7,91E-18 | 1,90E-17 | 0,0003029 | 0,002373 | 0,0010355 | 0,0010355 | TRUE | 0,0058139 | No |
| orf19.230 | 670 | 811 | 691 | 405 | 660 | 645 | 617 | 704 | 610 | 820 | 654 | 608 | 587 | 700 | 632 | 537 | 643,6 | 673 | 614 | 0,91 | -0,137 | -1,286 | 0,198454 | 0,229818 | 0,0154 | 0,00409 | 0,0092981 | 0,0092981 | TRUE | 0,2567351 | No |
| orf19.2301 | 2001 | 1714 | 1891 | 1101 | 1754 | 1334 | 1441 | 2003 | 1822 | 1734 | 1790 | 1654 | 1560 | 1448 | 1476 | 1528 | 1626,38 | 1750 | 1503 | 0,858 | -0,221 | -4,498 | 6,87E-06 | 1,10E-05 | 0,0006212 | 0,002849 | 0,0016965 | 0,0016965 | TRUE | 0,014812 | No |
| orf19.2302 | 945 | 861 | 1002 | 616 | 2193 | 1975 | 2059 | 2667 | 860 | 871 | 948 | 925 | 1951 | 2144 | 2109 | 2034 | 1480,35 | 901 | 2060 | 2,286 | 1,193 | 21,895 | 2,88E-106 | 2,62E-105 | 0,0012496 | 0,002929 | 0,0020285 | 0,0020285 | TRUE | 0,0166854 | No |
| orf19.2303 | 233 | 225 | 211 | 153 | 362 | 303 | 314 | 397 | 212 | 228 | 200 | 230 | 322 | 329 | 322 | 303 | 268,07 | 218 | 319 | 1,471 | 0,557 | 6,192 | 5,95E-10 | 1,12E-09 | 1,00E-08 | 0,006965 | 0,003843 | 0,003843 | TRUE | 0,0332925 | No |
| orf19.2304 | 778 | 709 | 710 | 544 | 732 | 663 | 649 | 902 | 708 | 717 | 672 | 817 | 651 | 720 | 665 | 688 | 704,77 | 728 | 681 | 0,938 | -0,092 | -1,311 | 0,18993 | 0,220426 | 0,0027244 | 0,003911 | 0,0032964 | 0,0032964 | TRUE | 0,0702101 | No |
| orf19.2305 | 499 | 398 | 399 | 239 | 409 | 274 | 304 | 366 | 454 | 403 | 378 | 359 | 364 | 297 | 311 | 279 | 355,66 | 398 | 313 | 0,783 | -0,354 | -3,439 | 0,000584 | 0,000836 | 0,0091868 | 0,005751 | 0,0073205 | 0,0073205 | TRUE | 0,1409528 | No |

| | | | | | | | | | | | | | | | | | | | | | | | | | | | | | | |
|---|---|---|---|---|---|---|---|---|---|---|---|---|---|---|---|---|---|---|---|---|---|---|---|---|---|---|---|---|---|---|
| orf19.2306 | 702 | 625 | 666 | 341 | 1020 | 705 | 802 | 963 | 639 | 632 | 630 | 512 | 907 | 765 | 821 | 735 | 705,3 | 603 | 807 | 1,332 | 0,413 | 4,764 | 1,90E-06 | 3,11E-06 | 0,0075742 | 0,00391 | 0,0057543 | 0,0057543 | TRUE | 0,1085417 | No |
| orf19.2307 | 2298 | 1945 | 2264 | 1328 | 1915 | 1618 | 1728 | 2405 | 2092 | 1967 | 2143 | 1995 | 1703 | 1757 | 1770 | 1835 | 1907,65 | 2049 | 1766 | 0,861 | -0,215 | -4,548 | 5,42E-06 | 8,71E-06 | 0,0008189 | 0,002729 | 0,0016272 | 0,0016272 | TRUE | 0,0114884 | No |
| orf19.2308 | 30929 | 28640 | 30843 | 19118 | 24428 | 20291 | 21379 | 27764 | 28160 | 28966 | 29190 | 28718 | 21728 | 22030 | 21895 | 21179 | 25233,2 | 28758 | 21708 | 0,755 | -0,406 | -14,448 | 2,58E-47 | 1,14E-46 | 0,0002133 | 0,002089 | 0,0007179 | 0,0007179 | TRUE | 0,003331 | No |
| orf19.2309 | 5332 | 4940 | 5377 | 3548 | 7723 | 6893 | 7095 | 9291 | 4855 | 4996 | 5089 | 5330 | 6869 | 7484 | 7266 | 7087 | 6122 | 5068 | 7176 | 1,417 | 0,503 | 11,68 | 1,61E-31 | 5,37E-31 | 0,0012146 | 0,002253 | 0,0016084 | 0,0016084 | TRUE | 0,0141806 | No |
| orf19.2309.2 | 30325 | 24404 | 25995 | 17536 | 44909 | 33755 | 36763 | 54095 | 27610 | 24682 | 24602 | 26342 | 39945 | 36649 | 37650 | 41264 | 32342,9 | 25809 | 38877 | 1,506 | 0,591 | 11,235 | 2,75E-29 | 8,67E-29 | 0,0029895 | 0,002078 | 0,0026272 | 0,0026272 | TRUE | 0,0271596 | No |
| orf19.231 | 725 | 739 | 773 | 418 | 817 | 668 | 682 | 864 | 660 | 747 | 732 | 628 | 727 | 725 | 698 | 659 | 697,06 | 692 | 702 | 1,011 | 0,016 | 0,23 | 0,818436 | 0,843033 | 0,0027657 | 0,003932 | 0,0033792 | 0,0033792 | TRUE | 0,039308 | No |
| orf19.2310 | 1102 | 1055 | 1135 | 720 | 1192 | 979 | 1018 | 1410 | 1003 | 1067 | 1074 | 1082 | 1060 | 1063 | 1043 | 1076 | 1058,42 | 1056 | 1060 | 1,005 | 0,008 | 0,152 | 0,878951 | 0,897679 | 1,00E-08 | 0,003285 | 0,001507 | 0,001507 | TRUE | 0,0145553 | No |
| orf19.2310.1 | 9479 | 7163 | 7316 | 5824 | 15042 | 10896 | 11024 | 16504 | 8630 | 7245 | 6924 | 8749 | 13379 | 11830 | 11290 | 12589 | 10079,5 | 7887 | 12272 | 1,557 | 0,638 | 7,976 | 1,52E-15 | 3,41E-15 | 0,0096731 | 0,002168 | 0,0060483 | 0,0060483 | TRUE | 0,0826587 | No |
| orf19.2311 | 14631 | 12221 | 13426 | 9909 | 16862 | 15009 | 14544 | 20390 | 13321 | 12360 | 12706 | 14885 | 14998 | 16296 | 14895 | 15554 | 14376,9 | 13318 | 15436 | 1,159 | 0,213 | 3,606 | 0,000311 | 0,000453 | 0,0042925 | 0,002129 | 0,0032908 | 0,0032908 | TRUE | 0,0760004 | No |
| orf19.2312 | 69 | 46 | 65 | 47 | 136 | 120 | 116 | 154 | 63 | 47 | 62 | 71 | 121 | 130 | 119 | 117 | 91,12 | 61 | 122 | 2,034 | 1,025 | 6,604 | 4,00E-11 | 7,88E-11 | 1,00E-08 | 0,016534 | 0,0104037 | 0,0104037 | TRUE | 0,2031273 | No |
| orf19.2313 | 1326 | 1173 | 1215 | 884 | 1479 | 1171 | 1277 | 1712 | 1207 | 1186 | 1150 | 1328 | 1316 | 1271 | 1308 | 1306 | 1259,01 | 1218 | 1300 | 1,071 | 0,099 | 1,854 | 0,063788 | 0,078247 | 0,0008052 | 0,003086 | 0,0019631 | 0,0019631 | TRUE | 0,0394154 | No |
| orf19.2314 | 7229 | 6271 | 6724 | 4081 | 7797 | 5672 | 6483 | 8648 | 6582 | 6342 | 6364 | 6130 | 6935 | 6158 | 6639 | 6597 | 6468,46 | 6354 | 6582 | 1,036 | 0,051 | 1,136 | 0,255999 | 0,291375 | 0,0013994 | 0,002241 | 0,0017465 | 0,0017465 | TRUE | 0,0226871 | No |
| orf19.2315 | 3416 | 2831 | 3249 | 2193 | 1895 | 1674 | 1730 | 2379 | 3110 | 2863 | 3075 | 3294 | 1686 | 1817 | 1772 | 1815 | 2429 | 3086 | 1772 | 0,575 | -0,799 | -15,345 | 3,85E-53 | 1,88E-52 | 0,0016509 | 0,002581 | 0,0021619 | 0,0021619 | TRUE | 0,0283319 | No |
| orf19.2317 | 101 | 77 | 67 | 54 | 42 | 36 | 42 | 63 | 92 | 78 | 63 | 81 | 37 | 39 | 43 | 48 | 60,23 | 78 | 42 | 0,536 | -0,9 | -4,708 | 2,50E-06 | 4,09E-06 | 1,00E-08 | 0,023968 | 0,017258 | 0,017258 | TRUE | 0,1729237 | No |
| orf19.2318.1 | 1527 | 1269 | 1357 | 922 | 1827 | 1329 | 1531 | 2136 | 1390 | 1283 | 1284 | 1385 | 1625 | 1443 | 1568 | 1629 | 1451,03 | 1336 | 1566 | 1,175 | 0,233 | 4,108 | 4,00E-05 | 6,13E-05 | 0,0018924 | 0,002947 | 0,0023771 | 0,0023771 | TRUE | 0,0330067 | No |
| orf19.2319 | 7428 | 6436 | 7092 | 3844 | 6943 | 5366 | 6602 | 8208 | 6763 | 6509 | 6712 | 5774 | 6176 | 5826 | 6761 | 6261 | 6347,8 | 6440 | 6256 | 0,971 | -0,043 | -0,701 | 0,48356 | 0,525429 | 0,0038826 | 0,002245 | 0,0033932 | 0,0033932 | TRUE | 0,0578698 | No |
| orf19.232 | 384 | 395 | 418 | 277 | 251 | 173 | 226 | 270 | 350 | 400 | 396 | 416 | 223 | 188 | 231 | 206 | 301,16 | 390 | 212 | 0,546 | -0,874 | -9,234 | 2,61E-20 | 6,67E-20 | 0,0029675 | 0,006423 | 0,0050226 | 0,0050226 | TRUE | 0,0589526 | No |
| orf19.2320 | 10101 | 8433 | 9526 | 4994 | 11628 | 8877 | 10707 | 13673 | 9197 | 8529 | 9015 | 7502 | 10343 | 9638 | 10965 | 10430 | 9452,35 | 8561 | 10344 | 1,208 | 0,272 | 4,202 | 2,64E-05 | 4,10E-05 | 0,0051891 | 0,002177 | 0,0039257 | 0,0039257 | TRUE | 0,0835983 | No |
| orf19.2322.3 | 1122 | 930 | 1025 | 613 | 1404 | 1055 | 1196 | 1598 | 1022 | 941 | 970 | 921 | 1249 | 1145 | 1225 | 1219 | 1086,39 | 964 | 1210 | 1,254 | 0,326 | 5,946 | 2,75E-09 | 5,06E-09 | 0,0006734 | 0,003253 | 0,0019498 | 0,0019498 | TRUE | 0,0202118 | No |
| orf19.2324 | 1697 | 1393 | 1507 | 1115 | 1771 | 1622 | 1617 | 2243 | 1545 | 1409 | 1426 | 1675 | 1575 | 1761 | 1656 | 1711 | 1594,79 | 1514 | 1676 | 1,109 | 0,15 | 2,363 | 0,018151 | 0,023363 | 0,003607 | 0,002865 | 0,0032104 | 0,0032104 | TRUE | 0,0574285 | No |
| orf19.2325 | 952 | 795 | 927 | 607 | 706 | 580 | 650 | 926 | 867 | 804 | 877 | 912 | 628 | 630 | 666 | 706 | 761,21 | 865 | 658 | 0,762 | -0,391 | -6,013 | 1,82E-09 | 3,37E-09 | 0,0016464 | 0,003772 | 0,0027342 | 0,0027342 | TRUE | 0,0319501 | No |
| orf19.2326 | 2812 | 2444 | 2731 | 1548 | 2148 | 1723 | 1939 | 2511 | 2560 | 2472 | 2585 | 2325 | 1911 | 1871 | 1986 | 1915 | 2203,06 | 2486 | 1921 | 0,771 | -0,374 | -8,021 | 1,04E-15 | 2,36E-15 | 0,0008077 | 0,002637 | 0,0016328 | 0,0016328 | TRUE | 0,0210059 | No |
| orf19.2328 | 2428 | 2099 | 2363 | 1393 | 1236 | 949 | 1094 | 1430 | 2211 | 2123 | 2236 | 2093 | 1099 | 1030 | 1120 | 1091 | 1625,42 | 2166 | 1085 | 0,501 | -0,998 | -20,833 | 2,19E-96 | 1,82E-95 | 0,0003852 | 0,00285 | 0,0015238 | 0,0015238 | TRUE | 0,0129036 | No |
| orf19.2329.1 | 15691 | 13976 | 13706 | 8625 | 25900 | 18577 | 18513 | 25864 | 14286 | 14135 | 12971 | 12956 | 23037 | 20169 | 18960 | 19729 | 17030,6 | 13587 | 20474 | 1,507 | 0,591 | 9,392 | 5,89E-21 | 1,53E-20 | 0,0051137 | 0,002115 | 0,003747 | 0,003747 | TRUE | 0,0870426 | No |
| orf19.2330 | 5895 | 5040 | 5533 | 3286 | 6266 | 4984 | 5518 | 7523 | 5367 | 5097 | 5236 | 4936 | 5573 | 5411 | 5651 | 5739 | 5376,44 | 5159 | 5594 | 1,084 | 0,116 | 2,952 | 0,00316 | 0,004294 | 0,0007043 | 0,002283 | 0,0012895 | 0,0012895 | TRUE | 0,0098179 | No |
| orf19.2331 | 1727 | 1573 | 1774 | 1026 | 1776 | 1507 | 1583 | 2087 | 1572 | 1591 | 1679 | 1541 | 1580 | 1636 | 1621 | 1592 | 1601,56 | 1596 | 1607 | 1,005 | 0,008 | 0,169 | 0,865837 | 0,885719 | 1,00E-08 | 0,002862 | 0,0013716 | 0,0013716 | TRUE | 0,0150543 | No |
| orf19.233.1 | 2106 | 1818 | 1766 | 1177 | 3552 | 2614 | 2828 | 3818 | 1917 | 1839 | 1671 | 1768 | 3159 | 2838 | 2896 | 2912 | 2375,2 | 1799 | 2951 | 1,64 | 0,714 | 12,885 | 5,43E-38 | 2,05E-37 | 0,0022562 | 0,002593 | 0,0024911 | 0,0024911 | TRUE | 0,0290712 | No |
| orf19.2332 | 1 | 2 | 0 | 1 | 1 | 0 | 0 | 0 | 1 | 2 | 0 | 2 | 1 | 0 | 0 | 0 | 0,67 | 1 | 0 | 0,261 | -1,939 | -1,099 | 0,271891 | 0,307979 | 1,00E-08 | 1,986682 | 1,4980379 | 1,4980379 | TRUE | 0,3173692 | No |
| orf19.2333 | 9513 | 8095 | 9249 | 5565 | 5265 | 4455 | 4951 | 6659 | 8661 | 8187 | 8753 | 8360 | 4683 | 4837 | 5070 | 5080 | 6703,9 | 8490 | 4918 | 0,579 | -0,788 | -18,923 | 7,42E-80 | 5,24E-79 | 0,0010311 | 0,002234 | 0,0015071 | 0,0015071 | TRUE | 0,012744 | No |
| orf19.2334 | 943 | 959 | 883 | 526 | 575 | 498 | 447 | 603 | 859 | 970 | 836 | 790 | 511 | 541 | 458 | 460 | 678,02 | 864 | 492 | 0,568 | -0,815 | -10,072 | 7,38E-24 | 2,06E-23 | 0,0051846 | 0,003985 | 0,0047129 | 0,0047129 | TRUE | 0,081111 | No |
| orf19.2335 | 1251 | 1210 | 1298 | 769 | 1250 | 1007 | 1061 | 1442 | 1139 | 1224 | 1228 | 1155 | 1112 | 1093 | 1087 | 1100 | 1142,26 | 1186 | 1098 | 0,925 | -0,113 | -2,255 | 0,024104 | 0,030689 | 1,00E-08 | 0,003193 | 0,00153 | 0,00153 | TRUE | 0,0100778 | No |
| orf19.2336 | 229 | 195 | 190 | 136 | 263 | 214 | 198 | 292 | 208 | 197 | 180 | 204 | 234 | 232 | 203 | 223 | 210,2 | 197 | 223 | 1,132 | 0,178 | 1,767 | 0,077269 | 0,093899 | 1,00E-08 | 0,008321 | 0,0049978 | 0,0049978 | TRUE | 0,042333 | No |
| orf19.2337 | 14599 | 12810 | 13661 | 7770 | 15273 | 11877 | 13094 | 18408 | 13292 | 12956 | 12929 | 11672 | 13585 | 12895 | 13410 | 14042 | 13097,5 | 12712 | 13483 | 1,06 | 0,085 | 1,762 | 0,078005 | 0,094702 | 0,0020882 | 0,002138 | 0,0021326 | 0,0021326 | TRUE | 0,0365442 | No |
| orf19.234 | 727 | 653 | 719 | 469 | 1166 | 991 | 918 | 1301 | 662 | 660 | 680 | 705 | 1037 | 1076 | 940 | 992 | 844,12 | 677 | 1011 | 1,497 | 0,582 | 9,506 | 1,99E-21 | 5,21E-21 | 0,0010817 | 0,003602 | 0,0023401 | 0,0023401 | TRUE | 0,0254735 | No |
| orf19.2340 | 21992 | 17998 | 19671 | 12859 | 17702 | 15102 | 15499 | 22044 | 20023 | 18203 | 18617 | 19316 | 15745 | 16397 | 15873 | 16815 | 17623,6 | 19040 | 16208 | 0,851 | -0,232 | -5,635 | 1,75E-08 | 3,12E-08 | 0,001271 | 0,002112 | 0,0015746 | 0,0015746 | TRUE | 0,0149528 | No |
| orf19.2341 | 5624 | 4982 | 5407 | 3320 | 2102 | 1701 | 1844 | 2322 | 5120 | 5039 | 5117 | 4987 | 1870 | 1847 | 1888 | 1771 | 3454,97 | 5066 | 1844 | 0,364 | -1,46 | -38,808 | 0 | 0 | 1,00E-08 | 0,002419 | 0,0009987 | 0,0009987 | TRUE | 0,009186 | No |
| orf19.2342 | 1949 | 1623 | 1750 | 992 | 1485 | 1018 | 1207 | 1733 | 1774 | 1641 | 1656 | 1490 | 1321 | 1105 | 1236 | 1322 | 1443,32 | 1640 | 1246 | 0,759 | -0,397 | -5,625 | 1,86E-08 | 3,31E-08 | 0,0049428 | 0,002952 | 0,004091 | 0,004091 | TRUE | 0,0680294 | No |
| orf19.2343 | 270 | 267 | 253 | 195 | 223 | 205 | 226 | 306 | 246 | 270 | 239 | 293 | 198 | 223 | 231 | 233 | 241,75 | 262 | 221 | 0,85 | -0,234 | -2,323 | 0,020202 | 0,025882 | 0,0026504 | 0,007501 | 0,0055615 | 0,0055615 | TRUE | 0,0588095 | No |
| orf19.2343.1 | 369 | 312 | 345 | 241 | 797 | 618 | 661 | 874 | 336 | 316 | 327 | 362 | 709 | 671 | 677 | 667 | 507,95 | 335 | 681 | 2,042 | 1,03 | 14,648 | 1,38E-48 | 6,21E-48 | 1,00E-08 | 0,004638 | 0,0024653 | 0,0024653 | TRUE | 0,0263264 | No |
| orf19.2344 | 2352 | 1820 | 1963 | 1502 | 3289 | 2880 | 3054 | 4436 | 2141 | 1841 | 1858 | 2256 | 2925 | 3127 | 3128 | 3384 | 2582,5 | 2024 | 3141 | 1,554 | 0,636 | 8,828 | 1,06E-18 | 2,61E-18 | 0,0064907 | 0,002549 | 0,0045705 | 0,0045705 | TRUE | 0,0689866 | No |
| orf19.2346 | 989 | 891 | 962 | 692 | 2380 | 1862 | 2009 | 2765 | 900 | 901 | 910 | 1039 | 2117 | 2022 | 2057 | 2109 | 1507,09 | 938 | 2076 | 2,225 | 1,154 | 21,335 | 5,38E-101 | 4,72E-100 | 0,0009006 | 0,002914 | 0,0020086 | 0,0020086 | TRUE | 0,0552548 | No |
| orf19.2347 | 2887 | 2579 | 2664 | 1745 | 4506 | 3823 | 3728 | 5262 | 2628 | 2608 | 2521 | 2621 | 4008 | 4151 | 3818 | 4014 | 3296,23 | 2594 | 3998 | 1,541 | 0,624 | 15,451 | 7,48E-54 | 3,71E-53 | 0,0005161 | 0,002438 | 0,0012412 | 0,0012412 | TRUE | 0,0108702 | No |
| orf19.2350 | 528 | 413 | 437 | 277 | 238 | 210 | 221 | 295 | 481 | 418 | 414 | 416 | 212 | 228 | 226 | 225 | 327,4 | 432 | 223 | 0,514 | -0,96 | -10,975 | 5,04E-28 | 1,54E-27 | 1,00E-08 | 0,006072 | 0,0040165 | 0,0040165 | TRUE | 0,0709499 | No |
| orf19.2351 | 1290 | 1159 | 1164 | 827 | 529 | 387 | 432 | 661 | 1174 | 1172 | 1102 | 1242 | 471 | 420 | 442 | 504 | 815,99 | 1172 | 459 | 0,394 | -1,345 | -19,462 | 2,30E-84 | 1,72E-83 | 0,0023073 | 0,003656 | 0,0031115 | 0,0031115 | TRUE | 0,0532412 | No |
| orf19.2352 | 437 | 357 | 383 | 208 | 629 | 452 | 548 | 703 | 398 | 361 | 362 | 312 | 559 | 491 | 561 | 536 | 447,69 | 358 | 537 | 1,489 | 0,575 | 6,913 | 4,76E-12 | 9,69E-12 | 0,003557 | 0,004988 | 0,004283 | 0,004283 | TRUE | 0,0708358 | No |
| orf19.2356 | 28074 | 25541 | 28171 | 17897 | 5462 | 5080 | 5245 | 7000 | 25560 | 25832 | 26661 | 26884 | 4858 | 5515 | 5372 | 5340 | 15752,8 | 26234 | 5271 | 0,201 | -2,315 | -51,918 | 0 | 0 | 0,0016129 | 0,002121 | 0,0018014 | 0,0018014 | TRUE | 0,0340341 | No |
| orf19.236 | 10652 | 9144 | 9387 | 5525 | 16716 | 11510 | 12627 | 17130 | 9698 | 9248 | 8884 | 8299 | 14868 | 12497 | 12932 | 13067 | 11186,7 | 9032 | 13341 | 1,477 | 0,562 | 8,871 | 7,26E-19 | 1,79E-18 | 0,0050989 | 0,002155 | 0,0037653 | 0,0037653 | TRUE | 0,0727314 | No |
| orf19.2360 | 8826 | 8153 | 8552 | 5580 | 23573 | 20792 | 20818 | 28786 | 8036 | 8246 | 8094 | 8382 | 20967 | 22574 | 21320 | 21958 | 14947,2 | 8190 | 21705 | 2,651 | 1,407 | 39,653 | 0 | 0 | 0,0006044 | 0,002125 | 0,0011212 | 0,0011212 | TRUE | 0,0088328 | No |
| orf19.2361 | 3823 | 3544 | 3918 | 2484 | 1329 | 1161 | 1181 | 1550 | 3481 | 3584 | 3708 | 3731 | 1182 | 1261 | 1209 | 1182 | 2417,37 | 3626 | 1208 | 0,333 | -1,586 | -35,329 | 2,11E-273 | 4,86E-272 | 0,0005664 | 0,002583 | 0,0013991 | 0,0013991 | TRUE | 0,0096204 | No |
| orf19.2362 | 3635 | 3188 | 3316 | 2188 | 3456 | 2796 | 2993 | 3849 | 3310 | 3224 | 3138 | 3287 | 3074 | 3036 | 3065 | 2936 | 3133,72 | 3240 | 3028 | 0,935 | -0,098 | -2,572 | 0,010101 | 0,013241 | 0,000244 | 0,002459 | 0,0010632 | 0,0010632 | TRUE | 0,0055609 | No |
| orf19.2363 | 3989 | 3693 | 4079 | 2430 | 2675 | 2101 | 2257 | 2911 | 3632 | 3735 | 3860 | 3650 | 2379 | 2281 | 2311 | 2221 | 3008,74 | 3719 | 2298 | 0,617 | -0,696 | -16,949 | 1,96E-64 | 1,14E-63 | 0,0005195 | 0,002476 | 0,0012688 | 0,0012688 | TRUE | 0,0080912 | No |
| orf19.2364 | 1588 | 1419 | 1502 | 1014 | 2597 | 2151 | 2339 | 3280 | 1446 | 1435 | 1422 | 1523 | 2310 | 2335 | 2395 | 2502 | 1921,06 | 1456 | 2386 | 1,642 | 0,715 | 15,349 | 3,61E-53 | 1,77E-52 | 0,0006548 | 0,002725 | 0,0015203 | 0,0015203 | TRUE | 0,0136619 | No |
| orf19.2365 | 2473 | 2329 | 2400 | 1404 | 2658 | 2227 | 2394 | 3210 | 2252 | 2356 | 2271 | 2109 | 2364 | 2418 | 2452 | 2449 | 2333,75 | 2247 | 2421 | 1,076 | 0,105 | 2,378 | 0,01742 | 0,022476 | 0,0006061 | 0,002603 | 0,0014424 | 0,0014424 | TRUE | 0,0185876 | No |
| orf19.2366 | 1 | 3 | 2 | 2 | 2 | 1 | 0 | 2 | 1 | 3 | 2 | 3 | 2 | 1 | 0 | 2 | 1,65 | 2 | 1 | 0,527 | -0,924 | -0,853 | 0,393511 | 0,43447 | 1,00E-08 | 0,800703 | 0,4685944 | 0,4685944 | TRUE | 0,212464 | No |
| orf19.2367 | 1423 | 1439 | 1467 | 672 | 718 | 487 | 566 | 683 | 1296 | 1455 | 1388 | 1009 | 639 | 529 | 580 | 521 | 927,11 | 1287 | 567 | 0,44 | -1,186 | -11,316 | 1,09E-29 | 3,46E-29 | 0,0159482 | 0,003462 | 0,0093023 | 0,0093023 | TRUE | 0,2423672 | No |
| orf19.2368 | 2116 | 2133 | 2214 | 1125 | 1864 | 1358 | 1619 | 2034 | 1927 | 2157 | 2095 | 1690 | 1658 | 1474 | 1658 | 1552 | 1776,39 | 1967 | 1586 | 0,805 | -0,313 | -4,196 | 2,72E-05 | 4,22E-05 | 0,006478 | 0,002781 | 0,0047905 | 0,0047905 | TRUE | 0,103704 | No |
| orf19.2369 | 621 | 631 | 642 | 410 | 565 | 517 | 497 | 635 | 565 | 638 | 608 | 616 | 503 | 561 | 509 | 484 | 560,54 | 607 | 514 | 0,847 | -0,24 | -3,366 | 0,000762 | 0,001083 | 0,0015983 | 0,004394 | 0,0030956 | 0,0030956 | TRUE | 0,04236 | No |
| orf19.2369.1 | 1469 | 1332 | 1377 | 702 | 905 | 527 | 728 | 842 | 1337 | 1347 | 1303 | 1055 | 805 | 572 | 746 | 642 | 975,92 | 1260 | 691 | 0,548 | -0,869 | -8,304 | 1,01E-16 | 2,34E-16 | 0,0158789 | 0,003391 | 0,0094042 | 0,0094042 | TRUE | 0,1542253 | No |
| orf19.2370 | 342 | 346 | 367 | 204 | 312 | 242 | 246 | 365 | 311 | 350 | 347 | 306 | 278 | 263 | 252 | 278 | 298,21 | 328 | 268 | 0,814 | -0,298 | -3,408 | 0,000655 | 0,000935 | 1,00E-08 | 0,006467 | 0,0039739 | 0,0039739 | TRUE | 0,0202822 | No |
| orf19.2371 | 2275 | 1898 | 1992 | 1259 | 1496 | 1188 | 1220 | 1700 | 2071 | 1920 | 1885 | 1891 | 1331 | 1290 | 1249 | 1297 | 1616,76 | 1942 | 1292 | 0,665 | -0,589 | -11,883 | 1,45E-32 | 4,93E-32 | 0,0008647 | 0,002854 | 0,0017208 | 0,0017208 | TRUE | 0,0252718 | No |
| orf19.2372 | 2856 | 2500 | 2645 | 1648 | 3199 | 2627 | 2662 | 3653 | 2600 | 2528 | 2503 | 2476 | 2845 | 2852 | 2726 | 2787 | 2664,74 | 2527 | 2802 | 1,108 | 0,148 | 3,857 | 0,000115 | 0,000172 | 1,00E-08 | 0,002533 | 0,0010391 | 0,0010391 | TRUE | 0,0047461 | No |
| orf19.2376 | 65 | 67 | 69 | 45 | 62 | 62 | 66 | 89 | 59 | 68 | 65 | 68 | 55 | 67 | 68 | 68 | 64,72 | 65 | 64 | 0,995 | -0,007 | -0,041 | 0,967648 | 0,971611 | 1,00E-08 | 0,022447 | 0,0125492 | 0,0125492 | TRUE | 0,0891979 | No |
| orf19.2378 | 468 | 417 | 430 | 305 | 513 | 402 | 452 | 538 | 426 | 422 | 407 | 458 | 456 | 436 | 463 | 410 | 434,88 | 428 | 441 | 1,033 | 0,047 | 0,627 | 0,530602 | 0,572156 | 1,00E-08 | 0,005075 | 0,0030715 | 0,0030715 | TRUE | 0,0289829 | No |
| orf19.2379 | 4079 | 3420 | 3826 | 2559 | 2947 | 2635 | 2839 | 3861 | 3714 | 3459 | 3621 | 3844 | 2621 | 2861 | 2907 | 2945 | 3246,57 | 3660 | 2834 | 0,775 | -0,368 | -7,201 | 5,98E-13 | 1,25E-12 | 0,0018851 | 0,002444 | 0,0021958 | 0,0021958 | TRUE | 0,0313335 | No |
| orf19.238 | 7210 | 6274 | 6632 | 3910 | 1586 | 1318 | 1311 | 1823 | 6564 | 6345 | 6277 | 5873 | 1411 | 1431 | 1343 | 1391 | 3829,35 | 6265 | 1394 | 0,222 | -2,169 | -46,328 | 0 | 0 | 0,0012276 | 0,002382 | 0,0016848 | 0,0016848 | TRUE | 0,0208189 | No |
| orf19.2381 | 802 | 635 | 690 | 500 | 264 | 197 | 184 | 255 | 730 | 642 | 653 | 751 | 235 | 214 | 188 | 195 | 451,02 | 694 | 208 | 0,3 | -1,737 | -19,089 | 3,09E-81 | 2,23E-80 | 0,0049025 | 0,004966 | 0,0049306 | 0,0049306 | TRUE | 0,0876426 | No |
| orf19.2382 | 453 | 438 | 455 | 278 | 927 | 794 | 797 | 1147 | 412 | 443 | 431 | 418 | 825 | 862 | 816 | 875 | 635,18 | 426 | 844 | 1,982 | 0,987 | 15,481 | 4,63E-54 | 2,31E-53 | 1,00E-08 | 0,004117 | 0,0021022 | 0,0021022 | TRUE | 0,0085449 | No |

| | | | | | | | | | | | | | | | | | | | | | | | | | | | | | | | |
|---|---|---|---|---|---|---|---|---|---|---|---|---|---|---|---|---|---|---|---|---|---|---|---|---|---|---|---|---|---|---|---|
| orf19.2383 | 133 | 130 | 127 | 112 | 191 | 161 | 211 | 242 | 121 | 131 | 120 | 168 | 170 | 175 | 216 | 185 | 160,8 | 135 | 186 | 1,394 | 0,479 | 3,64 | 0,000273 | 0,0004 | 0,009935 | 0,010252 | 0,0101517 | 0,0101517 | TRUE | 0,2256301 | No |
| orf19.2384 | 2831 | 2343 | 2477 | 1600 | 2112 | 1635 | 1756 | 2576 | 2578 | 2370 | 2344 | 2403 | 1879 | 1775 | 1798 | 1965 | 2139 | 2424 | 1854 | 0,766 | -0,385 | -7,521 | 5,44E-14 | 1,17E-13 | 0,0016037 | 0,002655 | 0,0020466 | 0,0020466 | TRUE | 0,0229953 | No |
| orf19.2385 | 2872 | 2557 | 2624 | 1650 | 2490 | 1882 | 2057 | 2817 | 2615 | 2586 | 2483 | 2479 | 2215 | 2043 | 2107 | 2149 | 2334,56 | 2541 | 2128 | 0,838 | -0,256 | -5,895 | 3,74E-09 | 6,85E-09 | 0,0005036 | 0,002603 | 0,0013727 | 0,0013727 | TRUE | 0,0089315 | No |
| orf19.2386 | 5500 | 4865 | 5232 | 3364 | 3896 | 3307 | 3362 | 4637 | 5008 | 4920 | 4952 | 5053 | 3465 | 3590 | 3443 | 3537 | 4246,12 | 4983 | 3509 | 0,704 | -0,506 | -15,213 | 2,89E-52 | 1,39E-51 | 1,00E-08 | 0,002348 | 0,0008195 | 0,0008195 | TRUE | 0,002802 | No |
| orf19.2387 | 4580 | 4030 | 4499 | 2747 | 3576 | 2756 | 2981 | 4137 | 4170 | 4076 | 4258 | 4126 | 3181 | 2992 | 3053 | 3156 | 3626,47 | 4158 | 3096 | 0,745 | -0,425 | -11,317 | 1,08E-29 | 3,45E-29 | 0,0002871 | 0,002401 | 0,001074 | 0,001074 | TRUE | 0,0059946 | No |
| orf19.2389 | 1166 | 1001 | 1010 | 640 | 875 | 648 | 728 | 1038 | 1062 | 1012 | 956 | 961 | 778 | 704 | 746 | 792 | 876,31 | 998 | 755 | 0,757 | -0,402 | -6,507 | 7,67E-11 | 1,50E-10 | 0,0014804 | 0,003544 | 0,0025105 | 0,0025105 | TRUE | 0,023884 | No |
| orf19.239 | 4165 | 3995 | 4302 | 2513 | 3890 | 3230 | 3451 | 4390 | 3792 | 4041 | 4071 | 3775 | 3460 | 3507 | 3534 | 3349 | 3691,11 | 3920 | 3462 | 0,882 | -0,18 | -4,294 | 1,75E-05 | 2,75E-05 | 0,0007286 | 0,002395 | 0,0014227 | 0,0014227 | TRUE | 0,0082871 | No |
| orf19.2391 | 899 | 727 | 828 | 601 | 1236 | 972 | 1111 | 1422 | 819 | 735 | 784 | 903 | 1099 | 1055 | 1138 | 1085 | 952,18 | 810 | 1094 | 1,356 | 0,439 | 6,779 | 1,21E-11 | 2,44E-11 | 0,0023797 | 0,003424 | 0,0029424 | 0,0029424 | TRUE | 0,0620021 | No |
| orf19.2392 | 0 | 0 | 0 | 0 | 0 | 0 | 0 | 0 | 0 | 0 | 0 | 0 | 0 | 0 | 0 | 0 | 0 | 0 | 0 | NA | NA | NA | NA | NA | NA | NA | NA | NA | NA | NA | NA |
| orf19.2393 | 6773 | 5616 | 6314 | 3769 | 6206 | 5111 | 5352 | 7460 | 6167 | 5680 | 5976 | 5662 | 5520 | 5549 | 5481 | 5691 | 5715,58 | 5871 | 5560 | 0,947 | -0,079 | -1,998 | 0,045711 | 0,056783 | 0,0007648 | 0,002268 | 0,0013316 | 0,0013316 | TRUE | 0,0141123 | No |
| orf19.2394 | 28 | 31 | 35 | 32 | 93 | 64 | 94 | 105 | 25 | 31 | 33 | 48 | 83 | 69 | 96 | 80 | 58,33 | 34 | 82 | 2,448 | 1,292 | 6,17 | 6,81E-10 | 1,28E-09 | 0,0108437 | 0,024685 | 0,0205522 | 0,0205522 | TRUE | 0,3443784 | No |
| orf19.2395 | 14 | 13 | 17 | 12 | 14 | 19 | 24 | 14 | 13 | 13 | 16 | 18 | 12 | 21 | 25 | 11 | 16,04 | 15 | 17 | 1,131 | 0,178 | 0,48 | 0,630882 | 0,669197 | 0,0416358 | 0,084375 | 0,0681836 | 0,0681836 | TRUE | 0,2442958 | No |
| orf19.2396 | 643 | 568 | 628 | 396 | 1092 | 853 | 892 | 1273 | 585 | 574 | 594 | 595 | 971 | 926 | 914 | 971 | 766,39 | 587 | 946 | 1,613 | 0,69 | 12,004 | 3,38E-33 | 1,16E-32 | 1,00E-08 | 0,003761 | 0,0017667 | 0,0017667 | TRUE | 0,0062447 | No |
| orf19.2397 | 229 | 208 | 176 | 138 | 272 | 217 | 233 | 339 | 208 | 210 | 167 | 207 | 242 | 236 | 239 | 259 | 220,94 | 198 | 244 | 1,235 | 0,305 | 2,946 | 0,003224 | 0,004377 | 0,0019289 | 0,008016 | 0,0056853 | 0,0056853 | TRUE | 0,1350923 | No |
| orf19.2397.3 | 6566 | 6653 | 6954 | 3814 | 4189 | 3279 | 3711 | 4551 | 5978 | 6729 | 6581 | 5729 | 3726 | 3560 | 3801 | 3472 | 4946,95 | 6254 | 3640 | 0,582 | -0,782 | -13,423 | 4,43E-41 | 1,76E-40 | 0,0034797 | 0,002304 | 0,0030456 | 0,0030456 | TRUE | 0,038047 | No |
| orf19.2398 | 19972 | 17786 | 19466 | 12883 | 3121 | 2639 | 2804 | 3842 | 18184 | 17989 | 18423 | 19352 | 2776 | 2865 | 2872 | 2931 | 10673,9 | 18487 | 2861 | 0,155 | -2,691 | -70,988 | 0 | 0 | 0,0006931 | 0,002161 | 0,001188 | 0,001188 | TRUE | 0,0119402 | No |
| orf19.2399 | 463 | 489 | 520 | 253 | 548 | 466 | 476 | 628 | 422 | 495 | 492 | 380 | 487 | 506 | 487 | 479 | 468,52 | 447 | 490 | 1,09 | 0,124 | 1,444 | 0,148668 | 0,17496 | 0,0052811 | 0,004857 | 0,0049208 | 0,0049208 | TRUE | 0,1001989 | No |
| orf19.24 | 4698 | 4031 | 4267 | 2773 | 3921 | 3383 | 3591 | 4940 | 4277 | 4077 | 4038 | 4165 | 3488 | 3673 | 3678 | 3768 | 3895,57 | 4139 | 3652 | 0,882 | -0,181 | -4,518 | 6,24E-06 | 1,00E-05 | 0,000649 | 0,002376 | 0,0012782 | 0,0012782 | TRUE | 0,0113488 | No |
| orf19.240 | 1857 | 1820 | 1766 | 908 | 1230 | 945 | 1060 | 1414 | 1691 | 1841 | 1671 | 1364 | 1094 | 1026 | 1086 | 1079 | 1356,38 | 1642 | 1071 | 0,651 | -0,619 | -7,868 | 3,59E-15 | 7,98E-15 | 0,006805 | 0,003011 | 0,0051773 | 0,0051773 | TRUE | 0,1481167 | No |
| orf19.2400 | 4499 | 4240 | 4511 | 2486 | 3592 | 2955 | 3399 | 4289 | 4096 | 4288 | 4269 | 3734 | 3195 | 3208 | 3481 | 3272 | 3693,01 | 4097 | 3289 | 0,802 | -0,319 | -6,008 | 1,88E-09 | 3,48E-09 | 0,0024075 | 0,002395 | 0,0024259 | 0,0024259 | TRUE | 0,0414399 | No |
| orf19.2401 | 306 | 330 | 336 | 205 | 452 | 317 | 396 | 527 | 279 | 334 | 318 | 308 | 402 | 344 | 406 | 402 | 349,01 | 310 | 388 | 1,258 | 0,331 | 3,733 | 0,000189 | 0,00028 | 0,0030151 | 0,005822 | 0,0046267 | 0,0046267 | TRUE | 0,0635595 | No |
| orf19.2402 | 3380 | 2821 | 2988 | 2034 | 2477 | 1912 | 2088 | 2924 | 3077 | 2853 | 2828 | 3055 | 2203 | 2076 | 2138 | 2230 | 2557,71 | 2953 | 2162 | 0,733 | -0,448 | -9,551 | 1,28E-21 | 3,37E-21 | 0,0011378 | 0,002553 | 0,0017154 | 0,0017154 | TRUE | 0,0104817 | No |
| orf19.2404 | 3935 | 3729 | 4122 | 2436 | 3147 | 2694 | 2910 | 3730 | 3583 | 3771 | 3901 | 3659 | 2799 | 2925 | 2980 | 2845 | 3308,01 | 3728 | 2887 | 0,774 | -0,37 | -8,681 | 3,92E-18 | 9,46E-18 | 0,0008196 | 0,002436 | 0,0014371 | 0,0014371 | TRUE | 0,0120836 | No |
| orf19.2406 | 2487 | 2323 | 2451 | 1420 | 1515 | 1149 | 1343 | 1678 | 2264 | 2349 | 2320 | 2133 | 1348 | 1247 | 1375 | 1280 | 1789,62 | 2266 | 1312 | 0,578 | -0,79 | -15,481 | 4,68E-54 | 2,33E-53 | 0,0011032 | 0,002775 | 0,0019076 | 0,0019076 | TRUE | 0,0175108 | No |
| orf19.2407 | 4061 | 3657 | 3741 | 2374 | 3342 | 2747 | 2835 | 4017 | 3697 | 3699 | 3541 | 3566 | 2973 | 2982 | 2903 | 3064 | 3303,17 | 3626 | 2980 | 0,822 | -0,283 | -7,505 | 6,15E-14 | 1,32E-13 | 0,000245 | 0,002437 | 0,0010561 | 0,0010561 | TRUE | 0,0045179 | No |
| orf19.2408 | 1691 | 1617 | 1604 | 1030 | 689 | 594 | 564 | 744 | 1540 | 1635 | 1518 | 1547 | 613 | 645 | 578 | 568 | 1080,4 | 1560 | 601 | 0,384 | -1,38 | -23,786 | 4,62E-125 | 4,94E-124 | 0,0007383 | 0,00326 | 0,0021078 | 0,0021078 | TRUE | 0,027203 | No |
| orf19.2410 | 2360 | 2205 | 2397 | 1452 | 2788 | 2187 | 2406 | 3057 | 2149 | 2230 | 2269 | 2181 | 2480 | 2374 | 2464 | 2332 | 2309,84 | 2207 | 2412 | 1,092 | 0,128 | 3,029 | 0,002453 | 0,003361 | 0,0003777 | 0,002609 | 0,0012668 | 0,0012668 | TRUE | 0,0065388 | No |
| orf19.2411 | 1476 | 1352 | 1429 | 866 | 1331 | 1068 | 1052 | 1439 | 1344 | 1367 | 1352 | 1301 | 1184 | 1160 | 1077 | 1098 | 1235,38 | 1341 | 1130 | 0,841 | -0,249 | -4,906 | 9,32E-07 | 1,55E-06 | 0,0003281 | 0,003106 | 0,0016707 | 0,0016707 | TRUE | 0,0130547 | No |
| orf19.2412 | 383 | 345 | 369 | 189 | 267 | 236 | 235 | 331 | 349 | 349 | 349 | 284 | 237 | 256 | 241 | 252 | 289,71 | 333 | 246 | 0,736 | -0,441 | -4,833 | 1,34E-06 | 2,22E-06 | 0,0010472 | 0,006597 | 0,0045248 | 0,0045248 | TRUE | 0,0908543 | No |
| orf19.2414 | 9629 | 8265 | 8602 | 5697 | 1280 | 935 | 1111 | 1359 | 8767 | 8359 | 8141 | 8558 | 1139 | 1015 | 1138 | 1037 | 4769,11 | 8456 | 1082 | 0,128 | -2,966 | -59,778 | 0 | 0 | 0,0016343 | 0,002314 | 0,0018695 | 0,0018695 | TRUE | 0,0190417 | No |
| orf19.2415 | 602 | 492 | 570 | 331 | 939 | 746 | 789 | 1074 | 548 | 498 | 539 | 497 | 835 | 810 | 808 | 819 | 669,35 | 520 | 818 | 1,567 | 0,648 | 10,524 | 6,71E-26 | 1,96E-25 | 1,00E-08 | 0,004011 | 0,0020496 | 0,0020496 | TRUE | 0,0148584 | No |
| orf19.2416.1 | 401 | 308 | 322 | 243 | 305 | 183 | 240 | 344 | 365 | 312 | 305 | 365 | 271 | 199 | 246 | 262 | 290,67 | 337 | 244 | 0,731 | -0,452 | -4,126 | 3,69E-05 | 5,68E-05 | 0,0099747 | 0,006583 | 0,0080199 | 0,0080199 | TRUE | 0,1673017 | No |
| orf19.2417 | 811 | 720 | 810 | 504 | 1875 | 1607 | 1763 | 2209 | 738 | 728 | 767 | 757 | 1668 | 1745 | 1806 | 1685 | 1236,67 | 748 | 1726 | 2,308 | 1,207 | 23,118 | 3,03E-118 | 3,06E-117 | 1,00E-08 | 0,003105 | 0,00163 | 0,00163 | TRUE | 0,0116463 | No |
| orf19.2418 | 903 | 656 | 723 | 495 | 804 | 629 | 656 | 956 | 822 | 663 | 684 | 744 | 715 | 683 | 672 | 729 | 714,07 | 728 | 700 | 0,962 | -0,056 | -0,747 | 0,455332 | 0,497215 | 0,0042184 | 0,003887 | 0,0040517 | 0,0040517 | TRUE | 0,0959739 | No |
| orf19.2419 | 1732 | 1461 | 1542 | 985 | 1597 | 1201 | 1301 | 1707 | 1577 | 1478 | 1459 | 1480 | 1420 | 1304 | 1332 | 1302 | 1419,06 | 1498 | 1340 | 0,894 | -0,162 | -3,166 | 0,001548 | 0,002158 | 0,0008786 | 0,002968 | 0,0018123 | 0,0018123 | TRUE | 0,0202115 | No |
| orf19.242 | 156 | 154 | 165 | 99 | 334 | 277 | 305 | 407 | 142 | 156 | 156 | 149 | 297 | 301 | 312 | 310 | 227,91 | 151 | 305 | 2,026 | 1,018 | 10,548 | 5,20E-26 | 1,52E-25 | 1,00E-08 | 0,007833 | 0,0039005 | 0,0039005 | TRUE | 0,0188237 | No |
| orf19.2422 | 3373 | 2930 | 3210 | 1947 | 3769 | 3009 | 3382 | 4322 | 3071 | 2963 | 3038 | 2925 | 3352 | 3267 | 3464 | 3297 | 3172,1 | 2999 | 3345 | 1,114 | 0,156 | 4,07 | 4,69E-05 | 7,16E-05 | 0,0002441 | 0,002453 | 0,0010911 | 0,0010911 | TRUE | 0,0067702 | No |
| orf19.242.2 | 153 | 147 | 153 | 68 | 240 | 211 | 206 | 202 | 139 | 149 | 145 | 102 | 213 | 229 | 211 | 154 | 167,82 | 134 | 202 | 1,49 | 0,575 | 3,891 | 9,98E-05 | 0,00015 | 0,0213484 | 0,009909 | 0,0146737 | 0,0146737 | TRUE | 0,2993592 | No |
| orf19.2423 | 2322 | 1941 | 2221 | 1539 | 2126 | 1867 | 2014 | 2845 | 2114 | 1963 | 2102 | 2312 | 1891 | 2027 | 2063 | 2170 | 2080,23 | 2123 | 2038 | 0,962 | -0,057 | -0,935 | 0,349999 | 0,390078 | 0,00324 | 0,002672 | 0,0030286 | 0,0030286 | TRUE | 0,0410255 | No |
| orf19.2425 | 263 | 238 | 252 | 155 | 199 | 165 | 181 | 298 | 239 | 241 | 238 | 233 | 177 | 179 | 185 | 227 | 215,04 | 238 | 192 | 0,811 | -0,302 | -2,832 | 0,00463 | 0,00622 | 0,0035942 | 0,00818 | 0,0062399 | 0,0062399 | TRUE | 0,1819564 | No |
| orf19.2426 | 494 | 434 | 440 | 316 | 348 | 288 | 289 | 381 | 450 | 439 | 416 | 475 | 310 | 313 | 296 | 291 | 373,58 | 445 | 302 | 0,681 | -0,555 | -7,13 | 1,00E-12 | 2,08E-12 | 1,00E-08 | 0,005573 | 0,0030734 | 0,0030734 | TRUE | 0,0214933 | No |
| orf19.2429 | 35 | 29 | 32 | 17 | 34 | 38 | 43 | 38 | 32 | 29 | 30 | 26 | 30 | 41 | 44 | 29 | 32,69 | 29 | 36 | 1,211 | 0,277 | 1,114 | 0,26517 | 0,301061 | 1,00E-08 | 0,042444 | 0,0281527 | 0,0281527 | TRUE | 0,1615266 | No |
| orf19.243 | 1195 | 974 | 1087 | 678 | 818 | 627 | 655 | 952 | 1088 | 985 | 1029 | 1018 | 728 | 681 | 671 | 726 | 865,7 | 1030 | 702 | 0,681 | -0,553 | -9,397 | 5,60E-21 | 1,45E-20 | 0,0007382 | 0,003563 | 0,0021455 | 0,0021455 | TRUE | 0,0181747 | No |
| orf19.2430 | 9 | 8 | 4 | 1 | 15 | 5 | 13 | 15 | 8 | 8 | 4 | 2 | 13 | 5 | 13 | 11 | 8,14 | 6 | 10 | 1,944 | 0,959 | 1,809 | 0,070498 | 0,085997 | 0,050781 | 0,164373 | 0,1330926 | 0,1330926 | TRUE | 0,3682707 | No |
| orf19.2431 | 32 | 30 | 36 | 24 | 40 | 27 | 32 | 39 | 29 | 30 | 34 | 36 | 36 | 29 | 33 | 30 | 32,13 | 32 | 32 | 0,991 | -0,013 | -0,057 | 0,954786 | 0,960208 | 1,00E-08 | 0,043155 | 0,0230348 | 0,0230348 | TRUE | 0,0465012 | No |
| orf19.2432 | 6562 | 6629 | 7026 | 4325 | 4784 | 4024 | 4272 | 5469 | 5974 | 6705 | 6649 | 6497 | 4255 | 4369 | 4375 | 4172 | 5374,54 | 6456 | 4293 | 0,665 | -0,589 | -12,936 | 2,82E-38 | 1,07E-37 | 0,0013309 | 0,002283 | 0,0017982 | 0,0017982 | TRUE | 0,0314429 | No |
| orf19.2433 | 150 | 181 | 174 | 94 | 167 | 139 | 157 | 209 | 137 | 183 | 165 | 141 | 149 | 151 | 161 | 159 | 155,65 | 156 | 155 | 0,988 | -0,018 | -0,145 | 0,884754 | 0,900864 | 0,0035813 | 0,010524 | 0,0078768 | 0,0078768 | TRUE | 0,1389535 | No |
| orf19.2434 | 1819 | 1857 | 1904 | 1177 | 2031 | 1641 | 1700 | 2314 | 1656 | 1878 | 1802 | 1768 | 1807 | 1782 | 1741 | 1765 | 1774,83 | 1776 | 1774 | 0,999 | -0,002 | -0,032 | 0,974372 | 0,97713 | 0,0009758 | 0,002781 | 0,0017586 | 0,0017586 | TRUE | 0,026631 | No |
| orf19.2435 | 10574 | 10403 | 10638 | 6488 | 13382 | 11815 | 11689 | 15559 | 9627 | 10522 | 10068 | 9746 | 11903 | 12828 | 11971 | 11869 | 11066,6 | 9991 | 12143 | 1,215 | 0,281 | 6,568 | 5,10E-11 | 1,00E-10 | 0,0013933 | 0,002156 | 0,0016662 | 0,0016662 | TRUE | 0,0175459 | No |
| orf19.2436 | 8296 | 7833 | 8404 | 5187 | 3783 | 3168 | 3281 | 4461 | 7553 | 7922 | 7954 | 7792 | 3365 | 3440 | 3360 | 3403 | 5598,52 | 7805 | 3392 | 0,435 | -1,202 | -35,635 | 3,99E-278 | 9,32E-277 | 0,0002448 | 0,002273 | 0,0008864 | 0,0008864 | TRUE | 0,0059794 | No |
| orf19.2437 | 742 | 870 | 841 | 596 | 1006 | 902 | 851 | 1008 | 676 | 880 | 796 | 895 | 895 | 979 | 872 | 769 | 845,16 | 812 | 879 | 1,084 | 0,116 | 1,214 | 0,224837 | 0,258166 | 0,0118233 | 0,0036 | 0,0075654 | 0,0075654 | TRUE | 0,1546893 | No |
| orf19.2438 | 2572 | 2316 | 2340 | 1327 | 1791 | 1352 | 1508 | 1838 | 2342 | 2342 | 2215 | 1993 | 1593 | 1468 | 1544 | 1402 | 1862,43 | 2223 | 1502 | 0,674 | -0,569 | -8,968 | 3,03E-19 | 7,54E-19 | 0,0035623 | 0,002746 | 0,0033068 | 0,0033068 | TRUE | 0,0553719 | No |
| orf19.2439 | 1668 | 1518 | 1481 | 900 | 2113 | 1505 | 1492 | 2124 | 1519 | 1535 | 1402 | 1352 | 1879 | 1634 | 1528 | 1620 | 1558,65 | 1452 | 1665 | 1,146 | 0,197 | 2,793 | 0,005225 | 0,007002 | 0,0051019 | 0,002885 | 0,004112 | 0,004112 | TRUE | 0,0909336 | No |
| orf19.2439.1 | 2202 | 1831 | 1829 | 1080 | 2989 | 1940 | 2006 | 3069 | 2005 | 1852 | 1731 | 1622 | 2659 | 2106 | 2054 | 2341 | 2046,3 | 1802 | 2290 | 1,27 | 0,345 | 3,987 | 6,68E-05 | 0,000101 | 0,010729 | 0,002683 | 0,0066691 | 0,0066691 | TRUE | 0,142193 | No |
| orf19.244 | 79 | 66 | 58 | 42 | 63 | 42 | 47 | 80 | 72 | 67 | 55 | 63 | 56 | 46 | 48 | 61 | 58,43 | 64 | 53 | 0,827 | -0,274 | -1,475 | 0,140191 | 0,165289 | 1,00E-08 | 0,024644 | 0,0160553 | 0,0160553 | TRUE | 0,1069065 | No |
| orf19.2440 | 965 | 1037 | 1029 | 615 | 1039 | 905 | 886 | 1176 | 879 | 1049 | 974 | 924 | 924 | 983 | 907 | 897 | 942,03 | 956 | 928 | 0,969 | -0,046 | -0,695 | 0,487134 | 0,528862 | 0,0026952 | 0,003439 | 0,0030602 | 0,0030602 | TRUE | 0,0515806 | No |
| orf19.2442 | 408 | 474 | 492 | 296 | 308 | 241 | 260 | 319 | 371 | 479 | 466 | 445 | 274 | 262 | 266 | 243 | 350,8 | 440 | 261 | 0,593 | -0,753 | -7,842 | 4,43E-15 | 9,80E-15 | 0,0058802 | 0,005803 | 0,0058389 | 0,0058389 | TRUE | 0,1379265 | No |
| orf19.2443 | 1099 | 1112 | 1115 | 677 | 1702 | 1526 | 1451 | 1868 | 1001 | 1125 | 1055 | 1017 | 1514 | 1657 | 1486 | 1425 | 1284,89 | 1050 | 1520 | 1,447 | 0,533 | 8,59 | 8,67E-18 | 2,08E-17 | 0,0027001 | 0,003065 | 0,0028706 | 0,0028706 | TRUE | 0,0430664 | No |
| orf19.2444 | 3217 | 3034 | 3106 | 1945 | 4548 | 3792 | 3915 | 5234 | 2929 | 3069 | 2940 | 2922 | 4045 | 4117 | 4009 | 3993 | 3502,89 | 2965 | 4041 | 1,362 | 0,446 | 12,484 | 9,18E-36 | 3,33E-35 | 1,00E-08 | 0,002414 | 0,0009292 | 0,0009292 | TRUE | 0,0068422 | No |
| orf19.2445 | 92 | 79 | 84 | 68 | 199 | 160 | 183 | 288 | 84 | 80 | 79 | 102 | 177 | 174 | 187 | 220 | 137,89 | 86 | 190 | 2,228 | 1,156 | 8,451 | 2,89E-17 | 6,82E-17 | 0,0059162 | 0,011617 | 0,0092667 | 0,0092667 | TRUE | 0,1363353 | No |
| orf19.2446 | 678 | 715 | 682 | 473 | 762 | 616 | 648 | 771 | 617 | 723 | 645 | 711 | 678 | 669 | 664 | 588 | 661,84 | 674 | 650 | 0,965 | -0,052 | -0,696 | 0,486173 | 0,527909 | 0,003682 | 0,004033 | 0,0038607 | 0,0038607 | TRUE | 0,0511651 | No |
| orf19.2447 | 1484 | 1404 | 1483 | 907 | 2608 | 2143 | 2187 | 2860 | 1351 | 1420 | 1404 | 1362 | 2320 | 2327 | 2240 | 2182 | 1825,62 | 1384 | 2267 | 1,636 | 0,71 | 15,767 | 5,26E-56 | 2,70E-55 | 1,00E-08 | 0,002761 | 0,001355 | 0,001355 | TRUE | 0,0081848 | No |
| orf19.2449 | 10 | 7 | 12 | 3 | 8 | 10 | 13 | 11 | 9 | 7 | 11 | 5 | 7 | 11 | 13 | 8 | 8,97 | 8 | 10 | 1,188 | 0,249 | 0,535 | 0,592398 | 0,632691 | 1,00E-08 | 0,149377 | 0,0951294 | 0,0951294 | TRUE | 0,2272492 | No |
| orf19.245 | 541 | 562 | 600 | 377 | 829 | 744 | 712 | 952 | 493 | 568 | 568 | 566 | 737 | 808 | 729 | 726 | 649,45 | 549 | 750 | 1,368 | 0,452 | 6,431 | 1,27E-10 | 2,46E-10 | 0,0020959 | 0,004071 | 0,0031461 | 0,0031461 | TRUE | 0,059809 | No |
| orf19.2451 | 568 | 590 | 571 | 409 | 5283 | 5080 | 4358 | 6097 | 517 | 597 | 540 | 614 | 4699 | 5515 | 4463 | 4651 | 2699,64 | 567 | 4832 | 8,546 | 3,095 | 39,742 | 0 | 0 | 0,0066281 | 0,002526 | 0,0047777 | 0,0047777 | TRUE | 0,109782 | No |
| orf19.2452 | 974 | 1041 | 1057 | 629 | 3879 | 3398 | 3517 | 4463 | 887 | 1053 | 1000 | 945 | 3450 | 3689 | 3602 | 3404 | 2253,83 | 971 | 3536 | 3,639 | 1,863 | 32,229 | 6,98E-228 | 1,34E-226 | 0,0025348 | 0,002623 | 0,0025248 | 0,0025248 | TRUE | 0,0448409 | No |
| orf19.2454 | 12521 | 12228 | 12522 | 8083 | 16249 | 13660 | 13701 | 17937 | 11400 | 12367 | 11851 | 12142 | 14453 | 14831 | 14032 | 13682 | 13094,8 | 11940 | 14250 | 1,193 | 0,255 | 6,304 | 2,89E-10 | 5,52E-10 | 0,0011233 | 0,002138 | 0,0014957 | 0,0014957 | TRUE | 0,0114454 | No |

| | | | | | | | | | | | | | | | | | | | | | | | | | | | | | | | |
|---|---|---|---|---|---|---|---|---|---|---|---|---|---|---|---|---|---|---|---|---|---|---|---|---|---|---|---|---|---|---|---|
| orf19.2455 | 3630 | 3562 | 3736 | 2227 | 3626 | 3284 | 3242 | 4299 | 3305 | 3603 | 3536 | 3345 | 3225 | 3566 | 3320 | 3279 | 3397,36 | 3447 | 3348 | 0,97 | -0,044 | -0,908 | 0,363626 | 0,403642 | 0,001495 | 0,002426 | 0,0019082 | 0,0019082 | TRUE | 0,0229602 | No |
| orf19.2457 | 1179 | 1160 | 1173 | 742 | 864 | 701 | 717 | 908 | 1073 | 1173 | 1110 | 1115 | 769 | 761 | 734 | 693 | 928,49 | 1118 | 739 | 0,66 | -0,598 | -10,304 | 6,73E-25 | 1,93E-24 | 0,0008415 | 0,00346 | 0,0021258 | 0,0021258 | TRUE | 0,0232224 | No |
| orf19.2458 | 21292 | 19620 | 21289 | 14171 | 10434 | 9069 | 9652 | 12612 | 19385 | 19844 | 20148 | 21287 | 9281 | 9846 | 9885 | 9621 | 14912,1 | 20166 | 9658 | 0,479 | -1,062 | -26,426 | 6,84E-154 | 9,12E-153 | 0,0011177 | 0,002126 | 0,001476 | 0,001476 | TRUE | 0,0169241 | No |
| orf19.2459 | 5436 | 4999 | 5332 | 3531 | 4952 | 4373 | 4692 | 6189 | 4949 | 5056 | 5046 | 5304 | 4405 | 4748 | 4805 | 4721 | 4879,29 | 5089 | 4670 | 0,918 | -0,123 | -2,954 | 0,00314 | 0,004268 | 0,0009127 | 0,002308 | 0,0014673 | 0,0014673 | TRUE | 0,0178907 | No |
| orf19.246 | 481 | 428 | 457 | 287 | 1022 | 911 | 885 | 1204 | 438 | 433 | 433 | 431 | 909 | 989 | 906 | 918 | 682,17 | 434 | 930 | 2,144 | 1,1 | 17,561 | 4,87E-69 | 3,01E-68 | 1,00E-08 | 0,003973 | 0,0020379 | 0,0020379 | TRUE | 0,0200934 | No |
| orf19.2460 | 13 | 9 | 7 | 6 | 64 | 54 | 56 | 81 | 12 | 9 | 7 | 9 | 57 | 59 | 57 | 62 | 33,91 | 9 | 59 | 6,399 | 2,678 | 8,825 | 1,09E-18 | 2,68E-18 | 1,00E-08 | 0,040994 | 0,023016 | 0,023016 | TRUE | 0,164924 | No |
| orf19.2461 | 1161 | 1029 | 1132 | 678 | 1367 | 1051 | 1175 | 1548 | 1057 | 1041 | 1071 | 1018 | 1216 | 1141 | 1203 | 1181 | 1116,09 | 1047 | 1185 | 1,131 | 0,178 | 3,598 | 0,00032 | 0,000467 | 1,00E-08 | 0,003221 | 0,0014386 | 0,0014386 | TRUE | 0,007045 | No |
| orf19.2462 | 2895 | 2573 | 2811 | 1655 | 1657 | 1347 | 1352 | 1813 | 2636 | 2602 | 2660 | 2486 | 1474 | 1462 | 1385 | 1383 | 2011,05 | 2596 | 1426 | 0,548 | -0,867 | -19,181 | 5,32E-82 | 3,87E-81 | 0,0004155 | 0,002694 | 0,001426 | 0,001426 | TRUE | 0,0090433 | No |
| orf19.2463 | 939 | 942 | 979 | 523 | 714 | 489 | 570 | 793 | 855 | 953 | 927 | 786 | 635 | 531 | 584 | 605 | 734,31 | 880 | 589 | 0,668 | -0,581 | -7,525 | 5,28E-14 | 1,14E-13 | 0,0045455 | 0,003836 | 0,0043259 | 0,0043259 | TRUE | 0,0558073 | No |
| orf19.2465 | 3805 | 3248 | 3540 | 2109 | 3253 | 2481 | 2885 | 3846 | 3464 | 3285 | 3350 | 3168 | 2893 | 2694 | 2955 | 2934 | 3092,89 | 3317 | 2869 | 0,864 | -0,21 | -4,563 | 5,05E-06 | 8,13E-06 | 0,0011315 | 0,002464 | 0,0017106 | 0,0017106 | TRUE | 0,0199722 | No |
| orf19.2467 | 3007 | 2443 | 2694 | 1692 | 1290 | 1034 | 1116 | 1552 | 2738 | 2471 | 2550 | 2542 | 1147 | 1123 | 1143 | 1184 | 1862,09 | 2575 | 1149 | 0,446 | -1,164 | -23,812 | 2,50E-125 | 2,67E-124 | 0,0009346 | 0,002746 | 0,0016777 | 0,0016777 | TRUE | 0,0228068 | No |
| orf19.2468 | 8 | 8 | 3 | 3 | 8 | 4 | 8 | 14 | 7 | 8 | 3 | 5 | 7 | 4 | 8 | 11 | 6,63 | 6 | 8 | 1,342 | 0,424 | 0,771 | 0,440781 | 0,482731 | 1,00E-08 | 0,201239 | 0,1373023 | 0,1373023 | TRUE | 0,2975875 | No |
| orf19.2469 | 161 | 146 | 149 | 94 | 249 | 164 | 222 | 264 | 147 | 148 | 141 | 141 | 221 | 178 | 227 | 201 | 175,59 | 144 | 207 | 1,437 | 0,523 | 4,604 | 4,15E-06 | 6,72E-06 | 1,00E-08 | 0,00956 | 0,0064406 | 0,0064406 | TRUE | 0,0878038 | No |
| orf19.247 | 202 | 203 | 188 | 98 | 127 | 90 | 93 | 110 | 184 | 205 | 178 | 147 | 113 | 98 | 95 | 84 | 138,02 | 178 | 98 | 0,541 | -0,887 | -6,416 | 1,40E-10 | 2,71E-10 | 0,0088782 | 0,011608 | 0,0105674 | 0,0105674 | TRUE | 0,1226223 | No |
| orf19.2471 | 898 | 804 | 853 | 549 | 1138 | 882 | 833 | 1130 | 818 | 813 | 807 | 825 | 1012 | 958 | 853 | 862 | 868,45 | 816 | 921 | 1,129 | 0,175 | 2,633 | 0,00847 | 0,011164 | 0,0026766 | 0,003558 | 0,0030997 | 0,0030997 | TRUE | 0,0545408 | No |
| orf19.2472 | 814 | 695 | 734 | 502 | 943 | 719 | 781 | 1041 | 741 | 703 | 695 | 754 | 839 | 781 | 800 | 794 | 763,26 | 723 | 804 | 1,113 | 0,155 | 2,653 | 0,007973 | 0,010528 | 1,00E-08 | 0,003768 | 0,0019516 | 0,0019516 | TRUE | 0,0106227 | No |
| orf19.2472.1 | 1720 | 1359 | 1361 | 911 | 1019 | 599 | 666 | 1011 | 1566 | 1374 | 1288 | 1368 | 906 | 650 | 682 | 771 | 1075,87 | 1399 | 752 | 0,538 | -0,893 | -9,198 | 3,66E-20 | 9,34E-20 | 0,0137924 | 0,003265 | 0,008052 | 0,008052 | TRUE | 0,2242765 | No |
| orf19.2473 | 3305 | 3002 | 3155 | 2074 | 2704 | 2248 | 2351 | 3289 | 3009 | 3036 | 2986 | 3115 | 2405 | 2441 | 2408 | 2509 | 2738,64 | 3036 | 2441 | 0,805 | -0,313 | -8,34 | 7,41E-17 | 1,73E-16 | 1,00E-08 | 0,002519 | 0,0009874 | 0,0009874 | TRUE | 0,0045753 | No |
| orf19.2474 | 1256 | 1113 | 1202 | 813 | 3273 | 2765 | 2834 | 4076 | 1144 | 1126 | 1138 | 1221 | 2911 | 3002 | 2902 | 3109 | 2069,11 | 1157 | 2981 | 2,584 | 1,369 | 29,125 | 1,71E-186 | 2,71E-185 | 0,0006286 | 0,002675 | 0,0015019 | 0,0015019 | TRUE | 0,0144226 | No |
| orf19.2475 | 5 | 6 | 8 | 2 | 5 | 4 | 6 | 9 | 5 | 6 | 8 | 3 | 4 | 4 | 6 | 7 | 5,37 | 6 | 5 | 1,011 | 0,015 | 0,027 | 0,978665 | 0,980356 | 1,00E-08 | 0,247824 | 0,1393174 | 0,1393174 | TRUE | 0,1791526 | No |
| orf19.2476 | 1644 | 1575 | 1583 | 1012 | 1948 | 1697 | 1730 | 2265 | 1497 | 1593 | 1498 | 1520 | 1733 | 1842 | 1772 | 1728 | 1647,84 | 1527 | 1769 | 1,157 | 0,211 | 4,577 | 4,71E-06 | 7,61E-06 | 0,0002998 | 0,002839 | 0,0014231 | 0,0014231 | TRUE | 0,0105935 | No |
| orf19.2478 | 94 | 75 | 77 | 34 | 31 | 28 | 23 | 37 | 86 | 76 | 73 | 51 | 28 | 30 | 24 | 28 | 49,39 | 72 | 28 | 0,38 | -1,398 | -6,372 | 1,87E-10 | 3,59E-10 | 1,00E-08 | 0,028782 | 0,0218795 | 0,0218795 | TRUE | 0,2483254 | No |
| orf19.2478.1 | 22386 | 18853 | 19644 | 14457 | 29946 | 24487 | 24730 | 35142 | 20382 | 19068 | 18591 | 21717 | 26636 | 26586 | 25327 | 26807 | 23139,1 | 19940 | 26339 | 1,321 | 0,402 | 7,846 | 4,31E-15 | 9,55E-15 | 0,0027624 | 0,002094 | 0,0024768 | 0,0024768 | TRUE | 0,043712 | No |
| orf19.2479 | 161 | 155 | 163 | 98 | 418 | 404 | 428 | 548 | 147 | 157 | 154 | 147 | 372 | 439 | 438 | 418 | 283,95 | 151 | 417 | 2,748 | 1,459 | 14,93 | 2,09E-50 | 9,69E-50 | 1,00E-08 | 0,006689 | 0,0045316 | 0,0045316 | TRUE | 0,0613994 | No |
| orf19.248 | 1031 | 1087 | 1111 | 678 | 1156 | 936 | 952 | 1268 | 939 | 1099 | 1051 | 1018 | 1028 | 1016 | 975 | 967 | 1011,83 | 1027 | 996 | 0,97 | -0,043 | -0,715 | 0,474502 | 0,516204 | 0,0017991 | 0,003343 | 0,0025391 | 0,0025391 | TRUE | 0,0432349 | No |
| orf19.2480.1 | 17088 | 14277 | 15811 | 9393 | 7331 | 5533 | 6383 | 8384 | 15558 | 14440 | 14964 | 14110 | 6521 | 6007 | 6537 | 6395 | 10566,4 | 14768 | 6365 | 0,431 | -1,214 | -27,363 | 7,64E-165 | 1,10E-163 | 0,0015213 | 0,002162 | 0,0017818 | 0,0017818 | TRUE | 0,0172321 | No |
| orf19.2481 | 541 | 462 | 499 | 379 | 348 | 280 | 279 | 391 | 493 | 467 | 472 | 569 | 310 | 304 | 286 | 298 | 399,87 | 500 | 300 | 0,602 | -0,733 | -8,822 | 1,12E-18 | 2,75E-18 | 0,0021915 | 0,005341 | 0,003989 | 0,003989 | TRUE | 0,0868217 | No |
| orf19.2483 | 2182 | 1678 | 1694 | 1469 | 949 | 743 | 738 | 1101 | 1987 | 1697 | 1603 | 2207 | 844 | 807 | 756 | 840 | 1342,51 | 1874 | 812 | 0,434 | -1,204 | -13,128 | 2,29E-39 | 8,79E-39 | 0,0109113 | 0,003021 | 0,0072115 | 0,0072115 | TRUE | 0,1678934 | No |
| orf19.2484 | 2262 | 1830 | 1975 | 1572 | 2988 | 2555 | 2431 | 3474 | 2059 | 1851 | 1869 | 2361 | 2658 | 2774 | 2490 | 2650 | 2339,03 | 2035 | 2643 | 1,301 | 0,38 | 5,169 | 2,35E-07 | 4,00E-07 | 0,0071791 | 0,002602 | 0,004736 | 0,004736 | TRUE | 0,1352025 | No |
| orf19.2485 | 3296 | 3290 | 3556 | 2191 | 3354 | 3013 | 2845 | 3736 | 3001 | 3327 | 3365 | 3291 | 2983 | 3271 | 2914 | 2850 | 3125,38 | 3246 | 3004 | 0,925 | -0,112 | -1,99 | 0,046632 | 0,05787 | 0,0028024 | 0,00246 | 0,002721 | 0,002721 | TRUE | 0,042835 | No |
| orf19.2487 | 2634 | 2332 | 2555 | 1545 | 3000 | 2519 | 2454 | 3511 | 2398 | 2359 | 2418 | 2321 | 2668 | 2735 | 2513 | 2678 | 2511,3 | 2374 | 2648 | 1,115 | 0,157 | 3,752 | 0,000176 | 0,00026 | 0,0004004 | 0,002563 | 0,0012809 | 0,0012809 | TRUE | 0,0139905 | No |
| orf19.2488 | 4821 | 4260 | 4675 | 2668 | 5063 | 3928 | 4495 | 5952 | 4389 | 4309 | 4424 | 4008 | 4503 | 4265 | 4603 | 4540 | 4380,23 | 4282 | 4478 | 1,045 | 0,063 | 1,422 | 0,154922 | 0,181918 | 0,0012186 | 0,002339 | 0,0016628 | 0,0016628 | TRUE | 0,0215532 | No |
| orf19.2489 | 17737 | 16166 | 17228 | 11717 | 26091 | 23831 | 22453 | 31077 | 16149 | 16350 | 16305 | 17601 | 23207 | 25874 | 22995 | 23706 | 20273,2 | 16601 | 23946 | 1,443 | 0,529 | 10,988 | 4,35E-28 | 1,34E-27 | 0,0022701 | 0,002102 | 0,0021721 | 0,0021721 | TRUE | 0,0358821 | No |
| orf19.2492 | 38 | 22 | 24 | 23 | 85 | 76 | 73 | 102 | 35 | 22 | 23 | 35 | 76 | 83 | 75 | 78 | 53,1 | 29 | 78 | 2,753 | 1,461 | 6,985 | 2,85E-12 | 5,83E-12 | 1,00E-08 | 0,026915 | 0,0172123 | 0,0172123 | TRUE | 0,1894433 | No |
| orf19.2494 | 534 | 428 | 521 | 317 | 571 | 518 | 522 | 776 | 486 | 433 | 493 | 476 | 508 | 562 | 535 | 592 | 510,64 | 472 | 549 | 1,164 | 0,219 | 2,895 | 0,003787 | 0,005112 | 0,0022897 | 0,004624 | 0,0035191 | 0,0035191 | TRUE | 0,0379976 | No |
| orf19.2495 | 663 | 564 | 657 | 368 | 588 | 478 | 517 | 694 | 604 | 570 | 622 | 553 | 523 | 519 | 529 | 529 | 556,19 | 587 | 525 | 0,892 | -0,165 | -2,533 | 0,011324 | 0,014797 | 1,00E-08 | 0,004412 | 0,0022948 | 0,0022948 | TRUE | 0,0179706 | No |
| orf19.2496 | 2143 | 1778 | 1878 | 1371 | 7318 | 5940 | 6094 | 8663 | 1951 | 1798 | 1777 | 2059 | 6509 | 6449 | 6241 | 6608 | 4174,22 | 1896 | 6452 | 3,409 | 1,769 | 34,781 | 4,76E-265 | 1,07E-263 | 0,0023078 | 0,002353 | 0,0021283 | 0,0021283 | TRUE | 0,037135 | No |
| orf19.2498 | 11072 | 9143 | 10296 | 6861 | 4408 | 3516 | 4079 | 5490 | 10081 | 9247 | 9744 | 10306 | 3921 | 3817 | 4177 | 4188 | 6935,21 | 9844 | 4026 | 0,409 | -1,289 | -26,59 | 8,80E-156 | 1,20E-154 | 0,0019265 | 0,002227 | 0,0020872 | 0,0020872 | TRUE | 0,0203772 | No |
| orf19.2499 | 688 | 569 | 625 | 357 | 611 | 497 | 559 | 708 | 626 | 575 | 592 | 536 | 543 | 540 | 572 | 540 | 565,66 | 582 | 549 | 0,938 | -0,092 | -1,364 | 0,17255 | 0,201246 | 0,0004094 | 0,004372 | 0,0026242 | 0,0026242 | TRUE | 0,0300932 | No |
| orf19.25 | 3180 | 2748 | 2978 | 1896 | 2067 | 1658 | 1715 | 2307 | 2895 | 2779 | 2818 | 2848 | 1839 | 1800 | 1756 | 1760 | 2311,99 | 2835 | 1789 | 0,631 | -0,665 | -16,819 | 1,76E-63 | 1,02E-62 | 1,00E-08 | 0,002608 | 0,001049 | 0,001049 | TRUE | 0,0044048 | No |
| orf19.250 | 3790 | 4159 | 4322 | 2654 | 6486 | 5441 | 5425 | 7023 | 3451 | 4206 | 4090 | 3987 | 5769 | 5907 | 5556 | 5357 | 4790,46 | 3934 | 5647 | 1,436 | 0,522 | 8,363 | 6,13E-17 | 1,43E-16 | 0,0042363 | 0,002313 | 0,0035213 | 0,0035213 | TRUE | 0,0846016 | No |
| orf19.2500 | 524 | 495 | 494 | 324 | 437 | 340 | 387 | 472 | 477 | 501 | 468 | 487 | 389 | 369 | 396 | 360 | 430,77 | 483 | 378 | 0,783 | -0,352 | -4,887 | 1,02E-06 | 1,70E-06 | 1,00E-08 | 0,005104 | 0,0026432 | 0,0026432 | TRUE | 0,0141697 | No |
| orf19.2501 | 9940 | 9948 | 10472 | 6720 | 8253 | 7844 | 7340 | 9600 | 9050 | 10061 | 9911 | 10095 | 7341 | 8516 | 7517 | 7323 | 8726,73 | 9779 | 7674 | 0,785 | -0,35 | -6,02 | 1,74E-09 | 3,24E-09 | 0,0038548 | 0,002188 | 0,0031268 | 0,0031268 | TRUE | 0,066341 | No |
| orf19.2503 | 16 | 24 | 14 | 9 | 14 | 14 | 8 | 21 | 15 | 24 | 13 | 14 | 12 | 15 | 8 | 16 | 14,68 | 16 | 13 | 0,792 | -0,337 | -0,898 | 0,369124 | 0,409032 | 1,00E-08 | 0,091997 | 0,0672826 | 0,0672826 | TRUE | 0,510755 | No |
| orf19.2504 | 14899 | 14026 | 14711 | 8875 | 18522 | 15297 | 16111 | 21352 | 13565 | 14186 | 13923 | 13332 | 16475 | 16608 | 16500 | 16287 | 15109,4 | 13752 | 16468 | 1,197 | 0,26 | 8,419 | 3,81E-17 | 8,94E-17 | 0,0003219 | 0,002124 | 0,0008455 | 0,0008455 | TRUE | 0,0054938 | No |
| orf19.2505 | 21 | 13 | 17 | 11 | 17 | 17 | 21 | 25 | 19 | 13 | 16 | 17 | 15 | 18 | 22 | 19 | 17,38 | 16 | 18 | 1,138 | 0,187 | 0,576 | 0,5646 | 0,605539 | 1,00E-08 | 0,078046 | 0,0432519 | 0,0432519 | TRUE | 0,087516 | No |
| orf19.2506 | 782 | 710 | 778 | 478 | 1341 | 1051 | 1167 | 1509 | 712 | 718 | 736 | 718 | 1193 | 1141 | 1195 | 1151 | 945,56 | 721 | 1170 | 1,622 | 0,698 | 13,345 | 1,26E-40 | 4,94E-40 | 1,00E-08 | 0,003434 | 0,0014875 | 0,0014875 | TRUE | 0,0030245 | No |
| orf19.2507 | 1462 | 1582 | 1610 | 943 | 1604 | 1334 | 1409 | 1655 | 1331 | 1600 | 1524 | 1417 | 1427 | 1448 | 1443 | 1262 | 1431,48 | 1468 | 1395 | 0,949 | -0,075 | -1,082 | 0,279462 | 0,315993 | 0,0043429 | 0,00296 | 0,0039344 | 0,0039344 | TRUE | 0,0509882 | No |
| orf19.2508 | 154 | 173 | 159 | 106 | 79 | 66 | 65 | 88 | 140 | 175 | 150 | 159 | 70 | 72 | 67 | 67 | 112,56 | 156 | 69 | 0,442 | -1,179 | -8,48 | 2,25E-17 | 5,32E-17 | 1,00E-08 | 0,013773 | 0,0083862 | 0,0083862 | TRUE | 0,0750129 | No |
| orf19.2509 | 0 | 0 | 0 | 0 | 0 | 0 | 0 | 0 | 0 | 0 | 0 | 0 | 0 | 0 | 0 | 0 | 0 | 0 | 0 | NA | NA | NA | NA | NA | NA | NA | NA | NA | NA | NA | NA |
| orf19.2509.1 | 88 | 89 | 71 | 55 | 446 | 334 | 352 | 491 | 80 | 90 | 67 | 83 | 397 | 363 | 360 | 375 | 226,79 | 80 | 374 | 4,695 | 2,231 | 19,279 | 8,08E-83 | 5,96E-82 | 1,00E-08 | 0,007862 | 0,004966 | 0,004966 | TRUE | 0,1203113 | No |
| orf19.251 | 11165 | 9856 | 9932 | 5974 | 6788 | 5260 | 5293 | 7668 | 10165 | 9968 | 9400 | 8974 | 6038 | 5711 | 5421 | 5849 | 7690,71 | 9627 | 5755 | 0,598 | -0,743 | -14,522 | 8,82E-48 | 3,92E-47 | 0,0024245 | 0,002209 | 0,0023746 | 0,0023746 | TRUE | 0,0250077 | No |
| orf19.2510 | 113 | 172 | 192 | 83 | 255 | 236 | 233 | 250 | 103 | 174 | 182 | 125 | 227 | 256 | 239 | 191 | 186,95 | 146 | 228 | 1,556 | 0,638 | 3,857 | 0,000115 | 0,000171 | 0,0365307 | 0,009103 | 0,0205172 | 0,0205172 | TRUE | 0,4541529 | No |
| orf19.2511.1 | 1647 | 1315 | 1294 | 748 | 1751 | 1152 | 1273 | 1883 | 1500 | 1330 | 1225 | 1124 | 1557 | 1251 | 1304 | 1436 | 1340,76 | 1295 | 1387 | 1,07 | 0,098 | 1,078 | 0,280994 | 0,317557 | 0,0112837 | 0,003022 | 0,0072067 | 0,0072067 | TRUE | 0,1373174 | No |
| orf19.2511.2 | 911 | 1060 | 1015 | 559 | 978 | 889 | 885 | 1058 | 829 | 1072 | 961 | 840 | 870 | 965 | 906 | 807 | 906,29 | 926 | 887 | 0,956 | -0,065 | -0,737 | 0,461115 | 0,502927 | 0,0085174 | 0,003495 | 0,0062713 | 0,0062713 | TRUE | 0,1356397 | No |
| orf19.2512 | 170 | 174 | 196 | 128 | 185 | 189 | 175 | 220 | 155 | 176 | 185 | 192 | 165 | 205 | 179 | 168 | 178,17 | 177 | 179 | 1,013 | 0,018 | 0,156 | 0,875874 | 0,894824 | 0,0038934 | 0,009451 | 0,0072643 | 0,0072643 | TRUE | 0,0970609 | No |
| orf19.2513 | 431 | 611 | 587 | 365 | 444 | 402 | 418 | 514 | 392 | 618 | 556 | 548 | 395 | 436 | 428 | 392 | 470,72 | 528 | 413 | 0,782 | -0,356 | -3,034 | 0,002414 | 0,00331 | 0,0181125 | 0,004843 | 0,0110389 | 0,0110389 | TRUE | 0,3656647 | No |
| orf19.2514 | 1130 | 1128 | 1199 | 666 | 741 | 594 | 556 | 750 | 1029 | 1141 | 1135 | 1000 | 659 | 645 | 569 | 572 | 843,8 | 1076 | 611 | 0,567 | -0,819 | -11,346 | 7,78E-30 | 2,49E-29 | 0,0038853 | 0,003603 | 0,0037411 | 0,0037411 | TRUE | 0,0341686 | No |
| orf19.2515 | 11265 | 11190 | 11752 | 6926 | 7447 | 6508 | 6665 | 8992 | 10256 | 11317 | 11122 | 10404 | 6624 | 7066 | 6826 | 6859 | 8809,33 | 10775 | 6844 | 0,635 | -0,655 | -15,039 | 4,05E-51 | 1,91E-50 | 0,0013697 | 0,002187 | 0,0017049 | 0,0017049 | TRUE | 0,0140404 | No |
| orf19.2516 | 497 | 538 | 540 | 316 | 666 | 515 | 589 | 762 | 452 | 544 | 511 | 475 | 592 | 559 | 603 | 581 | 539,8 | 496 | 584 | 1,178 | 0,236 | 3,222 | 0,001271 | 0,001784 | 0,0017305 | 0,004484 | 0,0032606 | 0,0032606 | TRUE | 0,0515328 | No |
| orf19.2517 | 281 | 264 | 256 | 174 | 251 | 218 | 230 | 312 | 256 | 267 | 242 | 261 | 223 | 237 | 236 | 238 | 245 | 256 | 234 | 0,911 | -0,135 | -1,499 | 0,133837 | 0,158296 | 1,00E-08 | 0,007429 | 0,0037229 | 0,0037229 | TRUE | 0,0167737 | No |
| orf19.2518 | 3974 | 3679 | 3902 | 2350 | 2682 | 2348 | 2421 | 3151 | 3618 | 3721 | 3693 | 3530 | 2386 | 2549 | 2479 | 2404 | 3047,49 | 3640 | 2454 | 0,673 | -0,571 | -14,322 | 1,59E-46 | 6,94E-46 | 0,0002895 | 0,00247 | 0,0011866 | 0,0011866 | TRUE | 0,0080134 | No |
| orf19.2519 | 522 | 446 | 466 | 328 | 179 | 131 | 137 | 158 | 475 | 451 | 441 | 493 | 159 | 142 | 140 | 121 | 302,79 | 465 | 140 | 0,302 | -1,728 | -17,566 | 4,52E-69 | 2,80E-68 | 0,0017364 | 0,0064 | 0,0048181 | 0,0048181 | TRUE | 0,1151558 | No |
| orf19.252 | 445 | 515 | 518 | 236 | 620 | 427 | 453 | 617 | 405 | 521 | 490 | 355 | 551 | 464 | 464 | 471 | 465,05 | 443 | 488 | 1,097 | 0,134 | 1,197 | 0,231159 | 0,264805 | 0,0156883 | 0,004878 | 0,0097792 | 0,0097792 | TRUE | 0,1860157 | No |
| orf19.2520 | 2829 | 2561 | 2757 | 1755 | 2418 | 2004 | 2133 | 2799 | 2576 | 2590 | 2609 | 2636 | 2151 | 2176 | 2184 | 2135 | 2382,19 | 2603 | 2162 | 0,831 | -0,268 | -7,296 | 2,96E-13 | 6,26E-13 | 1,00E-08 | 0,002592 | 0,0008721 | 0,0008721 | TRUE | 0,0008595 | No |
| orf19.2521 | 430 | 392 | 445 | 257 | 896 | 780 | 807 | 1127 | 391 | 396 | 421 | 386 | 797 | 847 | 826 | 860 | 615,64 | 398 | 832 | 2,084 | 1,059 | 16,252 | 2,17E-59 | 1,18E-58 | 1,00E-08 | 0,004183 | 0,002186 | 0,002186 | TRUE | 0,0170606 | No |
| orf19.2524 | 7296 | 5970 | 6383 | 4267 | 3305 | 2522 | 2487 | 3621 | 6643 | 6038 | 6041 | 6410 | 2940 | 2738 | 2547 | 2762 | 4514,8 | 6283 | 2747 | 0,437 | -1,193 | -22,461 | 1,00E-111 | 9,57E-111 | 0,0024767 | 0,00233 | 0,0024535 | 0,0024535 | TRUE | 0,0288418 | No |

| | | | | | | | | | | | | | | | | | | | | | | | | | | | | | | | |
|---|---|---|---|---|---|---|---|---|---|---|---|---|---|---|---|---|---|---|---|---|---|---|---|---|---|---|---|---|---|---|---|
| orf19.2525 | 2070 | 1662 | 1791 | 1222 | 2220 | 1801 | 1958 | 2678 | 1885 | 1681 | 1695 | 1836 | 1975 | 1955 | 2005 | 2043 | 1884,29 | 1774 | 1994 | 1,125 | 0,17 | 3,381 | 0,000723 | 0,001031 | 0,0011117 | 0,002738 | 0,0019031 | 0,0019031 | TRUE | 0,0232921 | No |
| orf19.2527 | 4143 | 3418 | 3829 | 2335 | 2571 | 2044 | 2233 | 3126 | 3772 | 3457 | 3624 | 3508 | 2287 | 2219 | 2287 | 2385 | 2942,22 | 3590 | 2294 | 0,639 | -0,646 | -14,591 | 3,21E-48 | 1,43E-47 | 0,0009141 | 0,002486 | 0,0015278 | 0,0015278 | TRUE | 0,0144451 | No |
| orf19.2528 | 3618 | 2957 | 3330 | 2146 | 2796 | 2314 | 2667 | 3617 | 3294 | 2991 | 3152 | 3224 | 2487 | 2512 | 2731 | 2759 | 2893,7 | 3165 | 2622 | 0,829 | -0,271 | -5,243 | 1,58E-07 | 2,71E-07 | 0,0018563 | 0,002494 | 0,0022089 | 0,0022089 | TRUE | 0,0168875 | No |
| orf19.2529 | 1302 | 1190 | 1246 | 786 | 812 | 613 | 611 | 889 | 1185 | 1204 | 1179 | 1181 | 722 | 666 | 626 | 678 | 930,07 | 1187 | 673 | 0,568 | -0,817 | -14,38 | 6,91E-47 | 3,03E-46 | 3,89E-05 | 0,003457 | 0,0019578 | 0,0019578 | TRUE | 0,0286849 | No |
| orf19.2529.1 | 17392 | 12949 | 13002 | 7510 | 5280 | 3176 | 4278 | 5559 | 15835 | 13097 | 12305 | 11281 | 4696 | 3448 | 4381 | 4240 | 8660,49 | 13130 | 4191 | 0,319 | -1,647 | -11,713 | 1,09E-31 | 3,64E-31 | 0,0188491 | 0,00219 | 0,010008 | 0,0188491 | TRUE | 0,2355037 | No |
| orf19.2530 | 8 | 9 | 8 | 4 | 9 | 5 | 6 | 10 | 7 | 9 | 8 | 6 | 8 | 5 | 6 | 8 | 7,15 | 8 | 7 | 0,908 | -0,14 | -0,283 | 0,777375 | 0,807391 | 1,00E-08 | 0,18688 | 0,097001 | 0,097001 | TRUE | 0,0549712 | No |
| orf19.2531 | 11271 | 10210 | 10723 | 7177 | 11452 | 9593 | 9476 | 12899 | 10262 | 10326 | 10148 | 10781 | 10186 | 10415 | 9705 | 9839 | 10207,9 | 10379 | 10036 | 0,967 | -0,048 | -1,296 | 0,195007 | 0,225908 | 0,0007482 | 0,002166 | 0,0012229 | 0,0012229 | TRUE | 0,0080567 | No |
| orf19.2532 | 1132 | 994 | 1138 | 679 | 1009 | 856 | 906 | 1183 | 1031 | 1005 | 1077 | 1020 | 897 | 929 | 928 | 902 | 973,76 | 1033 | 914 | 0,884 | -0,179 | -3,455 | 0,00055 | 0,000789 | 1,00E-08 | 0,003394 | 0,0015388 | 0,0015388 | TRUE | 0,0101881 | No |
| orf19.2533.1 | 1152 | 997 | 1061 | 706 | 1550 | 1213 | 1285 | 1839 | 1049 | 1008 | 1004 | 1061 | 1379 | 1317 | 1316 | 1403 | 1192,04 | 1030 | 1354 | 1,317 | 0,397 | 7,889 | 3,05E-15 | 6,80E-15 | 1,00E-08 | 0,003145 | 0,0015649 | 0,0015649 | TRUE | 0,007478 | No |
| orf19.2534 | 3945 | 3523 | 3920 | 2587 | 2843 | 2556 | 2701 | 3650 | 3592 | 3563 | 3710 | 3886 | 2529 | 2775 | 2766 | 2784 | 3200,65 | 3688 | 2714 | 0,736 | -0,442 | -9,223 | 2,88E-20 | 7,36E-20 | 0,0013994 | 0,00245 | 0,0018817 | 0,0018817 | TRUE | 0,0258018 | No |
| orf19.2537 | 22 | 26 | 30 | 19 | 28 | 22 | 25 | 45 | 20 | 26 | 28 | 29 | 25 | 24 | 26 | 34 | 26,5 | 26 | 27 | 1,071 | 0,099 | 0,367 | 0,713971 | 0,747396 | 1,00E-08 | 0,051891 | 0,0327666 | 0,0327666 | TRUE | 0,2444375 | No |
| orf19.2538 | 22341 | 21422 | 23143 | 13111 | 33021 | 26274 | 28625 | 37646 | 20341 | 21666 | 21903 | 19695 | 29371 | 28526 | 29316 | 28717 | 24941,7 | 20901 | 28982 | 1,386 | 0,471 | 11,434 | 2,83E-30 | 9,16E-30 | 0,0013335 | 0,00209 | 0,0015903 | 0,0015903 | TRUE | 0,0182312 | No |
| orf19.2539 | 1 | 1 | 2 | 3 | 1 | 0 | 2 | 2 | 1 | 1 | 2 | 5 | 1 | 0 | 2 | 2 | 1,6 | 2 | 1 | 0,586 | -0,771 | -0,684 | 0,494275 | 0,535794 | 1,00E-08 | 0,828663 | 0,5382613 | 0,5382613 | TRUE | 0,3534974 | No |
| orf19.254 | 7588 | 6946 | 7546 | 5258 | 11655 | 11047 | 10989 | 15155 | 6909 | 7025 | 7142 | 7898 | 10367 | 11994 | 11254 | 11560 | 9268,61 | 7244 | 11294 | 1,56 | 0,641 | 11,266 | 1,94E-29 | 6,14E-29 | 0,0036105 | 0,00218 | 0,0029987 | 0,0029987 | TRUE | 0,0444379 | No |
| orf19.2540 | 1289 | 1168 | 1259 | 675 | 1653 | 1301 | 1415 | 1791 | 1174 | 1181 | 1192 | 1014 | 1470 | 1413 | 1449 | 1366 | 1282,31 | 1140 | 1424 | 1,245 | 0,316 | 5,356 | 8,52E-08 | 1,48E-07 | 0,0022013 | 0,003067 | 0,0025399 | 0,0025399 | TRUE | 0,0590901 | No |
| orf19.2541 | 726 | 769 | 790 | 431 | 582 | 475 | 433 | 621 | 661 | 778 | 748 | 647 | 518 | 516 | 443 | 474 | 598,05 | 708 | 488 | 0,687 | -0,542 | -6,678 | 2,42E-11 | 4,81E-11 | 0,0050231 | 0,004246 | 0,0046126 | 0,0046126 | TRUE | 0,0503177 | No |
| orf19.2542 | 526 | 535 | 538 | 339 | 389 | 311 | 325 | 358 | 479 | 541 | 509 | 509 | 346 | 338 | 333 | 273 | 416 | 510 | 322 | 0,631 | -0,665 | -7,558 | 4,09E-14 | 8,82E-14 | 0,0045676 | 0,005213 | 0,0049233 | 0,0049233 | TRUE | 0,1331222 | No |
| orf19.2544 | 396 | 358 | 351 | 234 | 311 | 231 | 245 | 333 | 361 | 362 | 332 | 352 | 277 | 251 | 251 | 254 | 304,83 | 352 | 258 | 0,735 | -0,445 | -5,31 | 1,10E-07 | 1,89E-07 | 1,00E-08 | 0,006371 | 0,0034088 | 0,0034088 | TRUE | 0,0267115 | No |
| orf19.2545 | 5566 | 4620 | 5235 | 3245 | 6443 | 5303 | 5955 | 8217 | 5068 | 4673 | 4954 | 4875 | 5731 | 5758 | 6099 | 6268 | 5428,04 | 4892 | 5964 | 1,219 | 0,286 | 6,39 | 1,66E-10 | 3,20E-10 | 0,0013035 | 0,00228 | 0,0017326 | 0,0017326 | TRUE | 0,014657 | No |
| orf19.2546 | 2366 | 2240 | 2432 | 1491 | 1851 | 1584 | 1695 | 2161 | 2154 | 2266 | 2302 | 2240 | 1646 | 1720 | 1736 | 1648 | 1963,95 | 2240 | 1688 | 0,753 | -0,41 | -9,329 | 1,07E-20 | 2,75E-20 | 0,0003493 | 0,00271 | 0,0013347 | 0,0013347 | TRUE | 0,0086725 | No |
| orf19.2547 | 3365 | 3428 | 3570 | 2131 | 3073 | 2561 | 2759 | 3521 | 3064 | 3467 | 3379 | 3201 | 2733 | 2781 | 2826 | 2686 | 3016,98 | 3278 | 2756 | 0,84 | -0,251 | -5,214 | 1,84E-07 | 3,15E-07 | 0,0013094 | 0,002475 | 0,0018856 | 0,0018856 | TRUE | 0,0246351 | No |
| orf19.2549 | 3335 | 2969 | 3134 | 1878 | 3039 | 2418 | 2619 | 3510 | 3036 | 3003 | 2966 | 2821 | 2703 | 2625 | 2682 | 2677 | 2814,29 | 2956 | 2672 | 0,903 | -0,148 | -3,803 | 0,000143 | 0,000212 | 0,0001158 | 0,002506 | 0,0010923 | 0,0010923 | TRUE | 0,010481 | No |
| orf19.255 | 11443 | 11114 | 11718 | 7165 | 20078 | 17209 | 18173 | 23757 | 10418 | 11241 | 11090 | 10763 | 17859 | 18684 | 18611 | 18122 | 14598,6 | 10878 | 18319 | 1,684 | 0,752 | 20,853 | 1,42E-96 | 1,18E-95 | 0,0007002 | 0,002128 | 0,0011731 | 0,0011731 | TRUE | 0,010131 | No |
| orf19.2551 | 7293 | 6391 | 6606 | 4724 | 24236 | 19944 | 20097 | 29306 | 6640 | 6464 | 6252 | 7096 | 21557 | 21654 | 20582 | 22355 | 14074,9 | 6613 | 21537 | 3,259 | 1,705 | 36,839 | 4,38E-297 | 1,12E-295 | 0,0019156 | 0,002131 | 0,0019529 | 0,0019529 | TRUE | 0,0286743 | No |
| orf19.2552 | 943 | 906 | 930 | 601 | 186 | 183 | 162 | 222 | 859 | 916 | 880 | 903 | 165 | 199 | 166 | 169 | 532,15 | 890 | 175 | 0,196 | -2,352 | -29,773 | 8,87E-195 | 1,47E-193 | 2,80E-05 | 0,004519 | 0,0027205 | 0,0027205 | TRUE | 0,0804854 | No |
| orf19.2553 | 329 | 348 | 357 | 200 | 64 | 73 | 64 | 97 | 300 | 352 | 338 | 300 | 57 | 79 | 66 | 74 | 195,69 | 322 | 69 | 0,213 | -2,231 | -17,562 | 4,83E-69 | 2,99E-68 | 0,0039513 | 0,008787 | 0,0070907 | 0,0070907 | TRUE | 0,1313509 | No |
| orf19.2555 | 1849 | 1680 | 1703 | 1148 | 2087 | 1645 | 1656 | 2291 | 1683 | 1699 | 1612 | 1724 | 1856 | 1786 | 1696 | 1748 | 1725,58 | 1680 | 1772 | 1,056 | 0,078 | 1,66 | 0,096909 | 0,116565 | 0,0005917 | 0,002803 | 0,001557 | 0,001557 | TRUE | 0,012772 | No |
| orf19.2557 | 2302 | 1944 | 2108 | 1303 | 1695 | 1379 | 1508 | 1956 | 2096 | 1966 | 1995 | 1957 | 1508 | 1497 | 1544 | 1492 | 1756,96 | 2004 | 1510 | 0,753 | -0,41 | -9,327 | 1,09E-20 | 2,80E-20 | 1,00E-08 | 0,002789 | 0,0012748 | 0,0012748 | TRUE | 0,0120349 | No |
| orf19.2558 | 881 | 821 | 880 | 560 | 954 | 864 | 780 | 975 | 802 | 830 | 833 | 841 | 849 | 938 | 799 | 744 | 829,46 | 826 | 832 | 1,005 | 0,008 | 0,108 | 0,913812 | 0,926308 | 0,0040311 | 0,00363 | 0,0038325 | 0,0038325 | TRUE | 0,0848952 | No |
| orf19.2559 | 4400 | 3795 | 4130 | 2542 | 3633 | 3062 | 3277 | 4496 | 4006 | 3838 | 3909 | 3818 | 3231 | 3324 | 3356 | 3430 | 3614,12 | 3893 | 3335 | 0,856 | -0,224 | -5,981 | 2,22E-09 | 4,10E-09 | 0,0003021 | 0,002403 | 0,0010628 | 0,0010628 | TRUE | 0,0055247 | No |
| orf19.2560 | 7977 | 6872 | 7414 | 4871 | 10991 | 9378 | 9795 | 13613 | 7263 | 6950 | 7017 | 7317 | 9776 | 10182 | 10031 | 10384 | 8615,02 | 7137 | 10093 | 1,415 | 0,501 | 14,241 | 5,08E-46 | 2,20E-45 | 0,0004818 | 0,00219 | 0,0010648 | 0,0010648 | TRUE | 0,0055352 | No |
| orf19.2563 | 1618 | 1406 | 1619 | 1016 | 1147 | 917 | 999 | 1334 | 1473 | 1422 | 1532 | 1526 | 1020 | 996 | 1023 | 1018 | 1251,26 | 1488 | 1014 | 0,682 | -0,552 | -11,363 | 6,40E-30 | 2,05E-29 | 1,00E-08 | 0,003093 | 0,0014451 | 0,0014451 | TRUE | 0,0108574 | No |
| orf19.2564 | 2040 | 1739 | 1959 | 1200 | 1721 | 1274 | 1519 | 1905 | 1857 | 1759 | 1854 | 1803 | 1531 | 1383 | 1556 | 1453 | 1649,44 | 1818 | 1481 | 0,814 | -0,297 | -5,849 | 4,95E-09 | 9,03E-09 | 0,0009129 | 0,002838 | 0,0018565 | 0,0018565 | TRUE | 0,0222895 | No |
| orf19.2565 | 2550 | 2343 | 2507 | 1446 | 1706 | 1478 | 1668 | 1995 | 2322 | 2370 | 2373 | 2172 | 1517 | 1605 | 1708 | 1522 | 1948,54 | 2309 | 1588 | 0,686 | -0,543 | -10,2 | 1,98E-24 | 5,62E-24 | 0,0017166 | 0,002715 | 0,0021959 | 0,0021959 | TRUE | 0,0311931 | No |
| orf19.2568 | 6834 | 5240 | 6051 | 4055 | 3829 | 2941 | 3535 | 4866 | 6222 | 5300 | 5727 | 6091 | 3406 | 3193 | 3620 | 3712 | 4658,84 | 5835 | 3483 | 0,597 | -0,744 | -11,805 | 3,70E-32 | 1,25E-31 | 0,0044244 | 0,002321 | 0,0035859 | 0,0035859 | TRUE | 0,0466582 | No |
| orf19.257 | 4 | 3 | 4 | 1 | 3 | 1 | 1 | 1 | 4 | 3 | 4 | 2 | 3 | 1 | 1 | 1 | 2,19 | 3 | 2 | 0,45 | -1,151 | -1,227 | 0,219723 | 0,252841 | 1,00E-08 | 0,605754 | 0,3369967 | 0,3369967 | TRUE | 0,2922841 | No |
| orf19.2570 | 1982 | 1728 | 1912 | 1149 | 2680 | 1959 | 2124 | 2772 | 1805 | 1748 | 1810 | 1726 | 2384 | 2127 | 2175 | 2115 | 1986,02 | 1772 | 2200 | 1,241 | 0,311 | 6,081 | 1,20E-09 | 2,23E-09 | 0,0014946 | 0,002702 | 0,0019997 | 0,0019997 | TRUE | 0,0387806 | No |
| orf19.2571 | 4179 | 3852 | 4319 | 2484 | 4271 | 3586 | 3639 | 4939 | 3805 | 3896 | 4088 | 3731 | 3799 | 3893 | 3727 | 3768 | 3838,28 | 3880 | 3797 | 0,978 | -0,033 | -0,807 | 0,419648 | 0,461013 | 0,000614 | 0,002381 | 0,0013082 | 0,0013082 | TRUE | 0,0159207 | No |
| orf19.2573 | 6129 | 5396 | 5837 | 3713 | 8370 | 7069 | 7444 | 9858 | 5580 | 5457 | 5524 | 5578 | 7445 | 7675 | 7624 | 7520 | 6550,32 | 5535 | 7566 | 1,367 | 0,451 | 15,162 | 6,32E-52 | 3,03E-51 | 1,00E-08 | 0,002239 | 0,0006901 | 0,0006901 | TRUE | 0,0014014 | No |
| orf19.2574 | 3906 | 3302 | 3693 | 2232 | 3985 | 3269 | 3758 | 5003 | 3556 | 3340 | 3495 | 3353 | 3545 | 3549 | 3849 | 3816 | 3562,82 | 3436 | 3690 | 1,073 | 0,102 | 2,258 | 0,023972 | 0,030534 | 0,0011128 | 0,002408 | 0,0016864 | 0,0016864 | TRUE | 0,0099393 | No |
| orf19.2575 | 4112 | 3627 | 3907 | 2502 | 4527 | 3516 | 3721 | 5215 | 3744 | 3668 | 3698 | 3758 | 4027 | 3817 | 3811 | 3978 | 3812,62 | 3717 | 3908 | 1,052 | 0,073 | 2,026 | 0,042771 | 0,053317 | 0,0002281 | 0,002383 | 0,0009982 | 0,0009982 | TRUE | 0,0049736 | No |
| orf19.2579 | 1081 | 948 | 1129 | 679 | 1079 | 932 | 1049 | 1360 | 984 | 959 | 1068 | 1020 | 960 | 1012 | 1074 | 1037 | 1014,35 | 1008 | 1021 | 1,013 | 0,019 | 0,316 | 0,751902 | 0,783615 | 0,0013304 | 0,003339 | 0,0023002 | 0,0023002 | TRUE | 0,0211887 | No |
| orf19.258 | 428 | 397 | 410 | 283 | 1113 | 905 | 912 | 1161 | 390 | 402 | 388 | 425 | 990 | 983 | 934 | 886 | 674,56 | 401 | 948 | 2,37 | 1,245 | 18,399 | 1,33E-75 | 8,89E-75 | 1,00E-08 | 0,003995 | 0,0025675 | 0,0025675 | TRUE | 0,0250644 | No |
| orf19.2580 | 827 | 744 | 843 | 449 | 655 | 508 | 620 | 794 | 753 | 752 | 798 | 674 | 583 | 552 | 635 | 606 | 669,06 | 744 | 594 | 0,795 | -0,331 | -4,719 | 2,37E-06 | 3,87E-06 | 0,0021233 | 0,004011 | 0,0032053 | 0,0032053 | TRUE | 0,0413755 | No |
| orf19.2581 | 95 | 80 | 100 | 62 | 71 | 57 | 49 | 78 | 86 | 81 | 95 | 93 | 63 | 62 | 50 | 59 | 73,74 | 89 | 58 | 0,663 | -0,592 | -3,652 | 0,00026 | 0,000382 | 1,00E-08 | 0,019952 | 0,011296 | 0,011296 | TRUE | 0,0754027 | No |
| orf19.2582 | 1939 | 1850 | 1916 | 1106 | 2446 | 2095 | 2053 | 2790 | 1765 | 1871 | 1813 | 1661 | 2176 | 2275 | 2103 | 2128 | 1974,02 | 1778 | 2170 | 1,218 | 0,285 | 5,744 | 9,24E-09 | 1,67E-08 | 0,0011769 | 0,002706 | 0,0018413 | 0,0018413 | TRUE | 0,0210779 | No |
| orf19.2583 | 9 | 7 | 13 | 4 | 38 | 37 | 34 | 36 | 8 | 7 | 12 | 6 | 34 | 40 | 35 | 27 | 21,23 | 8 | 34 | 3,932 | 1,975 | 5,656 | 1,55E-08 | 2,76E-08 | 1,00E-08 | 0,064261 | 0,0418286 | 0,0418286 | TRUE | 0,3150605 | No |
| orf19.2585 | 9 | 12 | 17 | 19 | 20 | 31 | 7 | 31 | 8 | 12 | 16 | 29 | 18 | 34 | 7 | 24 | 18,4 | 16 | 21 | 1,304 | 0,383 | 0,844 | 0,39885 | 0,439604 | 0,2426676 | 0,073819 | 0,1407129 | 0,1407129 | TRUE | 0,229111 | No |
| orf19.2587 | 6 | 7 | 5 | 6 | 42 | 23 | 32 | 44 | 5 | 7 | 5 | 9 | 37 | 25 | 33 | 34 | 19,37 | 6 | 32 | 5,091 | 2,348 | 6,112 | 9,81E-10 | 1,84E-09 | 1,00E-08 | 0,070239 | 0,0440727 | 0,0440727 | TRUE | 0,1409643 | No |
| orf19.259 | 1562 | 1439 | 1487 | 905 | 1886 | 1461 | 1696 | 2116 | 1422 | 1455 | 1407 | 1359 | 1678 | 1586 | 1737 | 1614 | 1532,39 | 1411 | 1654 | 1,17 | 0,227 | 4,665 | 3,09E-06 | 5,03E-06 | 0,0005293 | 0,002899 | 0,0016087 | 0,0016087 | TRUE | 0,0133937 | No |
| orf19.2590 | 13 | 4 | 4 | 10 | 51 | 51 | 48 | 66 | 12 | 4 | 4 | 15 | 45 | 55 | 49 | 50 | 29,37 | 9 | 50 | 6,073 | 2,602 | 7,615 | 2,64E-14 | 5,72E-14 | 1,00E-08 | 0,047021 | 0,0385889 | 0,0385889 | TRUE | 0,483341 | No |
| orf19.2591 | 51 | 33 | 47 | 29 | 138 | 116 | 118 | 167 | 46 | 33 | 44 | 44 | 123 | 126 | 121 | 127 | 83,1 | 42 | 124 | 2,961 | 1,566 | 9,446 | 3,52E-21 | 9,19E-21 | 1,00E-08 | 0,017934 | 0,010032 | 0,010032 | TRUE | 0,153983 | No |
| orf19.2593 | 94 | 84 | 93 | 69 | 252 | 175 | 195 | 296 | 86 | 85 | 88 | 104 | 224 | 190 | 200 | 226 | 150,23 | 91 | 210 | 2,349 | 1,232 | 9,755 | 1,75E-22 | 4,71E-22 | 1,00E-08 | 0,01083 | 0,0071824 | 0,0071824 | TRUE | 0,0663946 | No |
| orf19.2594 | 4970 | 4331 | 4753 | 2970 | 6472 | 5283 | 5579 | 7718 | 4525 | 4380 | 4498 | 4461 | 5757 | 5736 | 5714 | 5887 | 5119,81 | 4466 | 5774 | 1,293 | 0,371 | 11,706 | 1,18E-31 | 3,95E-31 | 1,00E-08 | 0,002295 | 0,0007604 | 0,0007604 | TRUE | 0,0022254 | No |
| orf19.2597 | 708 | 627 | 721 | 431 | 1108 | 934 | 995 | 1299 | 645 | 634 | 682 | 647 | 986 | 1014 | 1019 | 991 | 827,25 | 652 | 1002 | 1,535 | 0,618 | 11,194 | 4,38E-29 | 1,38E-28 | 1,00E-08 | 0,003634 | 0,0016482 | 0,0016482 | TRUE | 0,0123912 | No |
| orf19.2598 | 2272 | 2080 | 2118 | 1285 | 3180 | 2403 | 2535 | 3372 | 2069 | 2104 | 2004 | 1930 | 2829 | 2609 | 2596 | 2572 | 2339,11 | 2027 | 2652 | 1,307 | 0,386 | 7,944 | 1,95E-15 | 4,38E-15 | 0,0011359 | 0,002602 | 0,0018251 | 0,0018251 | TRUE | 0,0248451 | No |
| orf19.2599 | 2339 | 1889 | 2144 | 1456 | 9574 | 7940 | 8089 | 11740 | 2130 | 1911 | 2029 | 2187 | 8516 | 8621 | 8284 | 8955 | 5329,03 | 2064 | 8594 | 4,169 | 2,06 | 42,27 | 0 | 0 | 0,0018567 | 0,002285 | 0,0019647 | 0,0019647 | TRUE | 0,0301448 | No |
| orf19.26 | 2122 | 1983 | 2055 | 1365 | 1104 | 977 | 1016 | 1362 | 1932 | 2006 | 1945 | 2050 | 982 | 1061 | 1041 | 1039 | 1506,89 | 1983 | 1031 | 0,52 | -0,944 | -19,817 | 2,12E-87 | 1,63E-86 | 1,00E-08 | 0,002914 | 0,0014524 | 0,0014524 | TRUE | 0,012169 | No |
| orf19.260 | 6578 | 6064 | 6574 | 3882 | 3957 | 3374 | 3466 | 4577 | 5989 | 6133 | 6222 | 5831 | 3520 | 3663 | 3550 | 3491 | 4799,87 | 6044 | 3556 | 0,588 | -0,767 | -20,86 | 1,24E-96 | 1,04E-95 | 0,0003271 | 0,002312 | 0,0010764 | 0,0010764 | TRUE | 0,0064456 | No |
| orf19.2600 | 1203 | 1164 | 1234 | 807 | 764 | 763 | 694 | 975 | 1095 | 1177 | 1168 | 1212 | 680 | 828 | 711 | 744 | 951,89 | 1163 | 741 | 0,637 | -0,651 | -9,648 | 5,01E-22 | 1,33E-21 | 0,0031672 | 0,003425 | 0,0032794 | 0,0032794 | TRUE | 0,0717345 | No |
| orf19.2601 | 10697 | 9158 | 9865 | 6861 | 9967 | 9018 | 8809 | 12523 | 9739 | 9262 | 9336 | 10306 | 8865 | 9791 | 9022 | 9553 | 9484,34 | 9661 | 9308 | 0,964 | -0,053 | -1,093 | 0,274396 | 0,31054 | 0,0021848 | 0,002176 | 0,0021746 | 0,0021746 | TRUE | 0,0242353 | No |
| orf19.2602 | 994 | 801 | 915 | 609 | 11406 | 10209 | 10707 | 14885 | 905 | 810 | 866 | 915 | 10145 | 11084 | 10965 | 11354 | 5880,63 | 874 | 10887 | 12,475 | 3,641 | 67,028 | 0 | 0 | 0,0017974 | 0,002262 | 0,0021764 | 0,0021764 | TRUE | 0,0287884 | No |
| orf19.2604 | 6324 | 5295 | 5911 | 3611 | 4581 | 3715 | 4159 | 5537 | 5758 | 5355 | 5594 | 5424 | 4075 | 4033 | 4259 | 4224 | 4840,34 | 5533 | 4148 | 0,749 | -0,416 | -10,6 | 2,99E-26 | 8,79E-26 | 0,0006051 | 0,00231 | 0,0012696 | 0,0012696 | TRUE | 0,0092659 | No |
| orf19.2605 | 2641 | 2409 | 2564 | 1618 | 2561 | 2101 | 2329 | 3098 | 2405 | 2436 | 2427 | 2430 | 2278 | 2281 | 2385 | 2363 | 2375,68 | 2424 | 2327 | 0,96 | -0,059 | -1,535 | 0,124703 | 0,14818 | 1,00E-08 | 0,002593 | 0,0009878 | 0,0009878 | TRUE | 0,0032511 | No |
| orf19.2606 | 2151 | 1989 | 2107 | 1415 | 2534 | 2190 | 2357 | 3043 | 1958 | 2012 | 1994 | 2126 | 2254 | 2378 | 2414 | 2321 | 2182,05 | 2022 | 2342 | 1,159 | 0,213 | 4,807 | 1,53E-06 | 2,52E-06 | 0,0004683 | 0,002642 | 0,001424 | 0,001424 | TRUE | 0,0128987 | No |
| orf19.2607 | 939 | 773 | 895 | 633 | 1028 | 818 | 941 | 1234 | 855 | 782 | 847 | 951 | 914 | 888 | 964 | 941 | 892,77 | 859 | 927 | 1,084 | 0,116 | 1,787 | 0,073883 | 0,08992 | 0,0024203 | 0,003517 | 0,0028979 | 0,0028979 | TRUE | 0,0548703 | No |
| orf19.2608 | 2E+05 | 1E+05 | 1E+05 | 82223 | 45928 | 36647 | 41368 | 58390 | 1E+05 | 1E+05 | 1E+05 | 1E+05 | 40852 | 39788 | 42366 | 44540 | 84664,8 | 127443 | 41886 | 0,329 | -1,605 | -32,469 | 2,89E-231 | 5,63E-230 | 0,002537 | 0,002053 | 0,0023332 | 0,0023332 | TRUE | 0,0293409 | No |

| | | | | | | | | | | | | | | | | | | | | | | | | | | | | | | | |
|---|---|---|---|---|---|---|---|---|---|---|---|---|---|---|---|---|---|---|---|---|---|---|---|---|---|---|---|---|---|---|---|
| orf19.2609 | 9664 | 8023 | 9319 | 5670 | 6116 | 4658 | 5367 | 7127 | 8799 | 8114 | 8820 | 8517 | 5440 | 5057 | 5496 | 5437 | 6960,02 | 8562 | 5358 | 0,626 | -0,676 | -15,604 | 6,86E-55 | 3,46E-54 | 0,0012471 | 0,002227 | 0,0016538 | 0,0016538 | TRUE | 0,0170771 | No |
| orf19.261 | 1738 | 1831 | 1810 | 1080 | 2259 | 1932 | 1837 | 2546 | 1582 | 1852 | 1713 | 1622 | 2009 | 2098 | 1881 | 1942 | 1837,49 | 1692 | 1982 | 1,171 | 0,227 | 3,814 | 0,000137 | 0,000203 | 0,0028599 | 0,002756 | 0,0028505 | 0,0028505 | TRUE | 0,0489659 | No |
| orf19.2610 | 1320 | 1044 | 1093 | 766 | 1067 | 866 | 849 | 1311 | 1202 | 1056 | 1034 | 1151 | 949 | 940 | 869 | 1000 | 1025,2 | 1111 | 940 | 0,848 | -0,239 | -3,563 | 0,000367 | 0,000533 | 0,0033193 | 0,003326 | 0,0033246 | 0,0033246 | TRUE | 0,0396205 | No |
| orf19.2611 | 2317 | 1938 | 2134 | 1281 | 2907 | 2430 | 2664 | 3480 | 2110 | 1960 | 2020 | 1924 | 2586 | 2638 | 2728 | 2655 | 2327,54 | 2004 | 2652 | 1,321 | 0,402 | 9,158 | 5,27E-20 | 1,34E-19 | 0,0004792 | 0,002605 | 0,0014063 | 0,0014063 | TRUE | 0,0156145 | No |
| orf19.2612 | 2433 | 1736 | 2067 | 1535 | 2375 | 1866 | 2033 | 2778 | 2215 | 1756 | 1956 | 2306 | 2112 | 2026 | 2082 | 2119 | 2071,57 | 2058 | 2085 | 1,014 | 0,02 | 0,272 | 0,785986 | 0,815538 | 0,0071509 | 0,002675 | 0,0049821 | 0,0049821 | TRUE | 0,1182422 | No |
| orf19.2613 | 16506 | 13696 | 15090 | 9402 | 11056 | 8538 | 9423 | 12844 | 15028 | 13852 | 14281 | 14123 | 9834 | 9270 | 9650 | 9798 | 11979,6 | 14321 | 9638 | 0,673 | -0,571 | -14,979 | 1,01E-50 | 4,72E-50 | 0,0008602 | 0,002147 | 0,0013111 | 0,0013111 | TRUE | 0,013634 | No |
| orf19.2614 | 2263 | 1910 | 2120 | 1425 | 3182 | 2580 | 2863 | 3839 | 2060 | 1932 | 2006 | 2141 | 2830 | 2801 | 2932 | 2928 | 2453,88 | 2035 | 2873 | 1,414 | 0,5 | 11,304 | 1,26E-29 | 4,00E-29 | 0,0006504 | 0,002575 | 0,001452 | 0,001452 | TRUE | 0,0137013 | No |
| orf19.2615 | 743 | 626 | 743 | 405 | 483 | 407 | 464 | 531 | 676 | 633 | 703 | 608 | 430 | 442 | 475 | 405 | 546,61 | 655 | 438 | 0,665 | -0,589 | -7,852 | 4,11E-15 | 9,11E-15 | 0,0024228 | 0,004454 | 0,0035211 | 0,0035211 | TRUE | 0,0380031 | No |
| orf19.2616 | 4883 | 4392 | 4796 | 2839 | 4637 | 3774 | 4246 | 5721 | 4446 | 4442 | 4539 | 4265 | 4124 | 4098 | 4348 | 4364 | 4328,23 | 4423 | 4234 | 0,957 | -0,064 | -1,61 | 0,107324 | 0,128548 | 0,0005914 | 0,002342 | 0,0012718 | 0,0012718 | TRUE | 0,0066311 | No |
| orf19.2618 | 9361 | 7726 | 8456 | 5278 | 4142 | 3260 | 3645 | 4968 | 8523 | 7814 | 8003 | 7928 | 3684 | 3539 | 3733 | 3790 | 5876,78 | 8067 | 3686 | 0,457 | -1,13 | -27,141 | 3,23E-162 | 4,54E-161 | 0,0009143 | 0,002262 | 0,0014698 | 0,0014698 | TRUE | 0,0179222 | No |
| orf19.2619 | 423 | 416 | 429 | 310 | 340 | 311 | 306 | 399 | 385 | 421 | 406 | 466 | 302 | 338 | 313 | 304 | 366,92 | 420 | 314 | 0,752 | -0,411 | -4,922 | 8,56E-07 | 1,43E-06 | 0,0014939 | 0,005637 | 0,0039103 | 0,0039103 | TRUE | 0,0545698 | No |
| orf19.262 | 95 | 93 | 121 | 66 | 166 | 100 | 139 | 168 | 86 | 94 | 115 | 99 | 148 | 109 | 142 | 128 | 115,12 | 98 | 132 | 1,339 | 0,422 | 2,896 | 0,003782 | 0,005106 | 0,0080241 | 0,013512 | 0,0114008 | 0,0114008 | TRUE | 0,1308455 | No |
| orf19.2620 | 262 | 203 | 255 | 184 | 99 | 75 | 81 | 139 | 239 | 205 | 241 | 276 | 88 | 81 | 83 | 106 | 165,01 | 240 | 90 | 0,377 | -1,406 | -10,64 | 1,95E-26 | 5,76E-26 | 0,0082819 | 0,010043 | 0,0093596 | 0,0093596 | TRUE | 0,1655333 | No |
| orf19.2621 | 795 | 833 | 787 | 502 | 991 | 807 | 786 | 1083 | 724 | 842 | 745 | 754 | 881 | 876 | 805 | 826 | 806,74 | 766 | 847 | 1,105 | 0,145 | 2,207 | 0,02729 | 0,034601 | 0,002033 | 0,003674 | 0,0028632 | 0,0028632 | TRUE | 0,0550638 | No |
| orf19.2622 | 2251 | 2154 | 2175 | 1297 | 4399 | 3318 | 3678 | 4658 | 2049 | 2179 | 2058 | 1948 | 3913 | 3602 | 3767 | 3553 | 2883,73 | 2058 | 3709 | 1,799 | 0,847 | 17,097 | 1,55E-65 | 9,22E-65 | 0,0015193 | 0,002495 | 0,0019717 | 0,0019717 | TRUE | 0,0182357 | No |
| orf19.2623 | 15021 | 13342 | 14429 | 9492 | 4723 | 4107 | 4199 | 5653 | 13676 | 13494 | 13656 | 14258 | 4201 | 4459 | 4300 | 4312 | 9044,59 | 13771 | 4318 | 0,314 | -1,673 | -48,165 | 0 | 0 | 0,0003965 | 0,002183 | 0,0010114 | 0,0010114 | TRUE | 0,006755 | No |
| orf19.2624 | 5 | 14 | 6 | 3 | 10 | 15 | 14 | 15 | 5 | 14 | 6 | 5 | 9 | 16 | 14 | 11 | 9,98 | 8 | 12 | 1,726 | 0,787 | 1,636 | 0,101748 | 0,122179 | 0,0681703 | 0,134373 | 0,112547 | 0,112547 | TRUE | 1,1456063 | No |
| orf19.2625 | 1 | 2 | 2 | 0 | 0 | 0 | 0 | 0 | 1 | 2 | 2 | 0 | 0 | 0 | 0 | 0 | 0,6 | 1 | 0 | 0,137 | -2,863 | -1,54 | 0,123597 | 0,146919 | 1,00E-08 | 2,19181 | 1,8886372 | 1,8886372 | TRUE | 0,1477843 | No |
| orf19.2629 | 1181 | 1149 | 1264 | 736 | 1276 | 1015 | 1147 | 1349 | 1075 | 1162 | 1196 | 1106 | 1135 | 1102 | 1175 | 1029 | 1122,48 | 1135 | 1110 | 0,977 | -0,034 | -0,568 | 0,570199 | 0,610927 | 0,0019943 | 0,003214 | 0,0025643 | 0,0025643 | TRUE | 0,0307328 | No |
| orf19.2630 | 649 | 655 | 590 | 372 | 545 | 431 | 477 | 638 | 591 | 662 | 558 | 559 | 485 | 468 | 489 | 487 | 537,3 | 592 | 482 | 0,812 | -0,3 | -4,111 | 3,94E-05 | 6,05E-05 | 0,0018009 | 0,004496 | 0,0032411 | 0,0032411 | TRUE | 0,0742026 | No |
| orf19.2631 | 1311 | 1287 | 1251 | 815 | 1491 | 1298 | 1380 | 1749 | 1194 | 1302 | 1184 | 1224 | 1326 | 1409 | 1413 | 1334 | 1298,3 | 1226 | 1370 | 1,117 | 0,16 | 3,067 | 0,002162 | 0,002974 | 0,0007944 | 0,003054 | 0,0018426 | 0,0018426 | TRUE | 0,0217648 | No |
| orf19.263.1 | 7503 | 5863 | 6582 | 3739 | 4517 | 2898 | 3500 | 4829 | 6831 | 5930 | 6229 | 5617 | 4018 | 3146 | 3584 | 3684 | 4879,87 | 6152 | 3608 | 0,586 | -0,77 | -10,056 | 8,67E-24 | 2,42E-23 | 0,0082019 | 0,002308 | 0,005413 | 0,005413 | TRUE | 0,0896274 | No |
| orf19.2633 | 3499 | 2988 | 3282 | 2188 | 4867 | 4088 | 4422 | 6028 | 3186 | 3022 | 3106 | 3287 | 4329 | 4438 | 4529 | 4598 | 3811,87 | 3150 | 4474 | 1,422 | 0,508 | 12,478 | 9,90E-36 | 3,59E-35 | 0,0006209 | 0,002384 | 0,0013133 | 0,0013133 | TRUE | 0,0095125 | No |
| orf19.2633.1 | 236 | 217 | 230 | 140 | 149 | 127 | 122 | 181 | 215 | 219 | 218 | 210 | 133 | 138 | 125 | 138 | 174,47 | 216 | 134 | 0,619 | -0,693 | -6,572 | 4,98E-11 | 9,78E-11 | 1,00E-08 | 0,009609 | 0,0047294 | 0,0047294 | TRUE | 0,0174406 | No |
| orf19.2637 | 1217 | 1136 | 1164 | 694 | 739 | 554 | 684 | 821 | 1108 | 1149 | 1102 | 1042 | 657 | 601 | 701 | 626 | 873,33 | 1100 | 646 | 0,586 | -0,77 | -12,394 | 2,83E-35 | 1,01E-34 | 0,0013383 | 0,00355 | 0,0024967 | 0,0024967 | TRUE | 0,0355693 | No |
| orf19.2638 | 467 | 443 | 460 | 300 | 238 | 229 | 213 | 297 | 425 | 448 | 435 | 451 | 212 | 249 | 218 | 227 | 333,03 | 440 | 226 | 0,514 | -0,96 | -11,581 | 5,13E-31 | 1,69E-30 | 1,00E-08 | 0,006004 | 0,0033154 | 0,0033154 | TRUE | 0,0439022 | No |
| orf19.2638.1 | 2999 | 2620 | 2603 | 1476 | 1205 | 759 | 856 | 1208 | 2730 | 2650 | 2464 | 2217 | 1072 | 824 | 877 | 921 | 1719,37 | 2515 | 924 | 0,367 | -1,445 | -17,047 | 3,69E-65 | 2,18E-64 | 0,009782 | 0,002805 | 0,0061849 | 0,0061849 | TRUE | 0,1395536 | No |
| orf19.2639 | 1774 | 1532 | 1725 | 1026 | 1276 | 1082 | 1018 | 1446 | 1615 | 1549 | 1633 | 1541 | 1135 | 1175 | 1043 | 1103 | 1349,21 | 1584 | 1114 | 0,702 | -0,51 | -9,823 | 8,94E-23 | 2,42E-22 | 0,0006589 | 0,003016 | 0,0018347 | 0,0018347 | TRUE | 0,0210088 | No |
| orf19.2639.1 | 1255 | 1147 | 1199 | 625 | 1762 | 1167 | 1416 | 1787 | 1143 | 1160 | 1135 | 939 | 1567 | 1267 | 1450 | 1363 | 1252,98 | 1094 | 1412 | 1,288 | 0,365 | 4,538 | 5,67E-06 | 9,10E-06 | 0,0075316 | 0,003091 | 0,0053758 | 0,0053758 | TRUE | 0,1014395 | No |
| orf19.264 | 881 | 940 | 1004 | 603 | 828 | 763 | 824 | 995 | 802 | 951 | 950 | 906 | 736 | 828 | 844 | 759 | 847,07 | 902 | 792 | 0,877 | -0,189 | -2,589 | 0,009628 | 0,012642 | 0,004292 | 0,003597 | 0,0039482 | 0,0039482 | TRUE | 0,0704192 | No |
| orf19.2640 | 1693 | 1574 | 1560 | 913 | 1556 | 1183 | 1300 | 1683 | 1541 | 1592 | 1476 | 1371 | 1384 | 1284 | 1331 | 1284 | 1408,1 | 1495 | 1321 | 0,881 | -0,182 | -3,219 | 0,001287 | 0,001805 | 0,0017231 | 0,002975 | 0,0023574 | 0,0023574 | TRUE | 0,0338074 | No |
| orf19.2641 | 135 | 114 | 125 | 70 | 132 | 100 | 110 | 143 | 123 | 115 | 118 | 105 | 117 | 109 | 113 | 109 | 113,67 | 115 | 112 | 0,964 | -0,053 | -0,411 | 0,68073 | 0,716351 | 1,00E-08 | 0,013658 | 0,007042 | 0,007042 | TRUE | 0,0269144 | No |
| orf19.2642 | 35 | 49 | 51 | 28 | 61 | 44 | 54 | 82 | 32 | 50 | 48 | 42 | 54 | 48 | 55 | 63 | 48,95 | 43 | 55 | 1,289 | 0,366 | 1,764 | 0,077786 | 0,094491 | 1,00E-08 | 0,029021 | 0,0205208 | 0,0205208 | TRUE | 0,2759221 | No |
| orf19.2643 | 5342 | 4441 | 5065 | 3075 | 6256 | 4955 | 5133 | 6891 | 4864 | 4492 | 4794 | 4619 | 5565 | 5380 | 5257 | 5257 | 5028,2 | 4692 | 5365 | 1,143 | 0,193 | 4,78 | 1,76E-06 | 2,89E-06 | 0,0007468 | 0,0023 | 0,0013571 | 0,0013571 | TRUE | 0,010443 | No |
| orf19.2644 | 2837 | 2685 | 2665 | 1956 | 5536 | 5109 | 4848 | 6792 | 2583 | 2716 | 2522 | 2938 | 4924 | 5547 | 4965 | 5181 | 3922 | 2690 | 5154 | 1,919 | 0,94 | 16,212 | 4,14E-59 | 2,24E-58 | 0,0034991 | 0,002374 | 0,0029382 | 0,0029382 | TRUE | 0,0449001 | No |
| orf19.2645 | 40 | 50 | 35 | 27 | 32 | 34 | 36 | 43 | 36 | 51 | 33 | 41 | 28 | 37 | 37 | 33 | 36,96 | 40 | 34 | 0,839 | -0,253 | -1,086 | 0,277658 | 0,314065 | 1,00E-08 | 0,037774 | 0,0248871 | 0,0248871 | TRUE | 0,2469981 | No |
| orf19.2646 | 1871 | 2001 | 2141 | 1261 | 2046 | 1743 | 1914 | 2426 | 1703 | 2024 | 2026 | 1894 | 1820 | 1892 | 1960 | 1851 | 1896,35 | 1912 | 1881 | 0,983 | -0,024 | -0,393 | 0,694124 | 0,729122 | 0,0033246 | 0,002734 | 0,0030599 | 0,0030599 | TRUE | 0,0679968 | No |
| orf19.2647 | 131 | 141 | 135 | 82 | 208 | 163 | 157 | 181 | 119 | 143 | 128 | 123 | 185 | 177 | 161 | 138 | 146,71 | 128 | 165 | 1,283 | 0,36 | 2,771 | 0,00559 | 0,007465 | 0,0066624 | 0,011041 | 0,0091834 | 0,0091834 | TRUE | 0,1468946 | No |
| orf19.2649 | 267 | 273 | 334 | 169 | 550 | 543 | 455 | 599 | 243 | 276 | 316 | 254 | 489 | 590 | 466 | 457 | 386,35 | 272 | 500 | 1,83 | 0,872 | 8,155 | 3,48E-16 | 7,94E-16 | 0,0106658 | 0,005456 | 0,0080768 | 0,0080768 | TRUE | 0,1631473 | No |
| orf19.265 | 5820 | 5882 | 5786 | 3606 | 468 | 359 | 324 | 478 | 5299 | 5949 | 5476 | 5417 | 416 | 390 | 332 | 365 | 2955,38 | 5535 | 376 | 0,068 | -3,881 | -58,04 | 0 | 0 | 0,0040336 | 0,002484 | 0,0029563 | 0,0029563 | TRUE | 0,0663322 | No |
| orf19.2650 | 957 | 1128 | 1099 | 679 | 1447 | 1341 | 1275 | 1665 | 871 | 1141 | 1040 | 1020 | 1287 | 1456 | 1306 | 1270 | 1173,88 | 1018 | 1330 | 1,306 | 0,385 | 4,837 | 1,32E-06 | 2,18E-06 | 0,0068733 | 0,003162 | 0,0052058 | 0,0052058 | TRUE | 0,1174597 | No |
| orf19.2650.1 | 494 | 369 | 465 | 206 | 341 | 259 | 320 | 359 | 450 | 373 | 440 | 309 | 303 | 281 | 328 | 274 | 344,82 | 393 | 296 | 0,749 | -0,417 | -3,656 | 0,000257 | 0,000376 | 0,0142962 | 0,005868 | 0,0095662 | 0,0095662 | TRUE | 0,2100725 | No |
| orf19.2651 | 3548 | 3566 | 3520 | 2213 | 7291 | 6242 | 5967 | 8102 | 3230 | 3607 | 3331 | 3324 | 6485 | 6777 | 6111 | 6180 | 4880,75 | 3373 | 6388 | 1,893 | 0,921 | 18,465 | 3,97E-76 | 2,68E-75 | 0,0020023 | 0,002308 | 0,0021563 | 0,0021563 | TRUE | 0,0267789 | No |
| orf19.2652 | 95 | 106 | 118 | 62 | 908 | 818 | 734 | 983 | 86 | 107 | 112 | 93 | 808 | 888 | 752 | 750 | 449,48 | 100 | 800 | 7,99 | 2,998 | 28,102 | 9,20E-174 | 1,40E-172 | 0,005133 | 0,004976 | 0,0050384 | 0,0050384 | TRUE | 0,097445 | No |
| orf19.2653 | 2485 | 2546 | 2559 | 1619 | 5302 | 4514 | 4819 | 6102 | 2262 | 2575 | 2422 | 2432 | 4716 | 4901 | 4935 | 4655 | 3612,27 | 2423 | 4802 | 1,982 | 0,987 | 20,851 | 1,50E-96 | 1,25E-95 | 0,0013855 | 0,002403 | 0,0018318 | 0,0018318 | TRUE | 0,0251437 | No |
| orf19.2654 | 3420 | 3528 | 3554 | 2110 | 2523 | 2120 | 2155 | 2758 | 3114 | 3568 | 3364 | 3170 | 2244 | 2302 | 2207 | 2104 | 2758,97 | 3304 | 2214 | 0,67 | -0,579 | -10,741 | 6,54E-27 | 1,96E-26 | 0,0021525 | 0,002516 | 0,0024132 | 0,0024132 | TRUE | 0,0354303 | No |
| orf19.2655 | 24787 | 23324 | 24963 | 14754 | 2514 | 2046 | 2213 | 2855 | 22568 | 23590 | 23625 | 22163 | 2236 | 2221 | 2266 | 2178 | 12605,9 | 22986 | 2225 | 0,097 | -3,369 | -89,335 | 0 | 0 | 0,0006434 | 0,002142 | 0,0011328 | 0,0011328 | TRUE | 0,0070024 | No |
| orf19.2657 | 22 | 32 | 28 | 20 | 68 | 50 | 51 | 77 | 20 | 32 | 26 | 30 | 60 | 54 | 52 | 59 | 41,83 | 27 | 56 | 2,102 | 1,072 | 4,774 | 1,80E-06 | 2,96E-06 | 1,00E-08 | 0,033614 | 0,0204321 | 0,0204321 | TRUE | 0,2338603 | No |
| orf19.2660 | 3036 | 3255 | 3287 | 2209 | 3017 | 2598 | 2694 | 3496 | 2764 | 3292 | 3111 | 3318 | 2684 | 2821 | 2759 | 2667 | 2926,92 | 3121 | 2733 | 0,876 | -0,191 | -3,209 | 0,001331 | 0,001865 | 0,0032283 | 0,002488 | 0,0030551 | 0,0030551 | TRUE | 0,0731492 | No |
| orf19.2661 | 1402 | 1053 | 1151 | 699 | 746 | 549 | 686 | 1009 | 1276 | 1065 | 1089 | 1050 | 664 | 596 | 703 | 770 | 901,58 | 1120 | 683 | 0,61 | -0,713 | -8,214 | 2,14E-16 | 4,91E-16 | 0,0085146 | 0,003502 | 0,0060743 | 0,0060743 | TRUE | 0,1076356 | No |
| orf19.2662 | 212 | 180 | 194 | 114 | 145 | 111 | 110 | 216 | 193 | 182 | 184 | 171 | 129 | 121 | 113 | 165 | 157,1 | 182 | 132 | 0,725 | -0,463 | -3,539 | 0,000402 | 0,000582 | 0,0093834 | 0,010445 | 0,0100138 | 0,0100138 | TRUE | 0,3230136 | No |
| orf19.2663 | 794 | 668 | 815 | 465 | 548 | 488 | 445 | 652 | 723 | 676 | 771 | 699 | 487 | 530 | 456 | 497 | 604,84 | 717 | 492 | 0,686 | -0,544 | -7,713 | 1,23E-14 | 2,70E-14 | 0,0017627 | 0,004221 | 0,0030845 | 0,0030845 | TRUE | 0,0316956 | No |
| orf19.2664 | 2568 | 2423 | 2558 | 1618 | 1350 | 1206 | 1272 | 1601 | 2338 | 2451 | 2421 | 2430 | 1201 | 1309 | 1303 | 1221 | 1834,27 | 2410 | 1258 | 0,522 | -0,939 | -20,064 | 1,54E-89 | 1,20E-88 | 0,0003794 | 0,002757 | 0,0015078 | 0,0015078 | TRUE | 0,0111079 | No |
| orf19.2665 | 39545 | 35601 | 38783 | 23638 | 7568 | 6480 | 6862 | 9046 | 36004 | 36007 | 36705 | 35508 | 6731 | 7035 | 7028 | 6900 | 21489,8 | 36056 | 6924 | 0,192 | -2,381 | -80,993 | 0 | 0 | 0,0001896 | 0,002098 | 0,0007476 | 0,0007476 | TRUE | 0,0042569 | No |
| orf19.2667 | 11398 | 9230 | 10163 | 5968 | 12905 | 9420 | 11376 | 15049 | 10377 | 9335 | 9618 | 8965 | 11479 | 10227 | 11650 | 11479 | 10391,5 | 9574 | 11209 | 1,171 | 0,227 | 3,996 | 6,46E-05 | 9,77E-05 | 0,0035563 | 0,002164 | 0,0030068 | 0,0030068 | TRUE | 0,0422938 | No |
| orf19.2668 | 1099 | 946 | 996 | 707 | 1939 | 1643 | 1591 | 2236 | 1001 | 957 | 943 | 1062 | 1725 | 1784 | 1629 | 1706 | 1350,7 | 991 | 1711 | 1,732 | 0,792 | 14,651 | 1,32E-48 | 5,93E-48 | 0,0010999 | 0,003015 | 0,0019922 | 0,0019922 | TRUE | 0,0244995 | No |
| orf19.2669 | 1653 | 1511 | 1691 | 1034 | 3207 | 2766 | 2964 | 3831 | 1505 | 1528 | 1600 | 1553 | 2853 | 3003 | 3036 | 2922 | 2250,03 | 1546 | 2954 | 1,909 | 0,933 | 21,604 | 1,64E-103 | 1,48E-102 | 0,0003598 | 0,002624 | 0,0012849 | 0,0012849 | TRUE | 0,0071686 | No |
| orf19.267 | 3106 | 2788 | 2964 | 1951 | 3728 | 3152 | 3416 | 4757 | 2828 | 2820 | 2805 | 2931 | 3316 | 3422 | 3498 | 3629 | 3156,09 | 2846 | 3466 | 1,219 | 0,286 | 6,855 | 7,15E-12 | 1,45E-11 | 0,0006841 | 0,002456 | 0,0013489 | 0,0013489 | TRUE | 0,0125987 | No |
| orf19.2670 | 1383 | 1240 | 1299 | 815 | 866 | 761 | 831 | 1146 | 1259 | 1254 | 1229 | 1224 | 770 | 826 | 851 | 874 | 1036,08 | 1242 | 830 | 0,669 | -0,58 | -10,549 | 5,13E-26 | 1,50E-25 | 0,0004965 | 0,003312 | 0,0019135 | 0,0019135 | TRUE | 0,0296255 | No |
| orf19.2671 | 545 | 441 | 457 | 363 | 502 | 458 | 511 | 649 | 496 | 446 | 433 | 545 | 447 | 497 | 523 | 495 | 485,27 | 480 | 490 | 1,026 | 0,037 | 0,431 | 0,666782 | 0,703415 | 0,0052448 | 0,004759 | 0,0049762 | 0,0049762 | TRUE | 0,0851932 | No |
| orf19.2672 | 2730 | 2450 | 2582 | 1696 | 4611 | 3918 | 4030 | 5439 | 2486 | 2478 | 2444 | 2548 | 4101 | 4254 | 4127 | 4149 | 3323,26 | 2489 | 4158 | 1,672 | 0,741 | 20,867 | 1,08E-96 | 9,03E-96 | 1,00E-08 | 0,002435 | 0,0008841 | 0,0008841 | TRUE | 0,0028439 | No |
| orf19.2673 | 474 | 463 | 457 | 248 | 1313 | 955 | 1165 | 1426 | 432 | 468 | 433 | 373 | 1168 | 1037 | 1193 | 1088 | 773,81 | 426 | 1122 | 2,617 | 1,388 | 18,416 | 9,69E-76 | 6,49E-75 | 0,0042353 | 0,003744 | 0,003789 | 0,003789 | TRUE | 0,0686826 | No |
| orf19.2674 | 413 | 292 | 342 | 240 | 378 | 295 | 346 | 480 | 376 | 295 | 324 | 361 | 336 | 320 | 354 | 366 | 341,57 | 339 | 344 | 1,019 | 0,027 | 0,295 | 0,768239 | 0,798689 | 0,0046386 | 0,005904 | 0,0053533 | 0,0053533 | TRUE | 0,086649 | No |
| orf19.2675 | 1887 | 1698 | 1844 | 1167 | 2606 | 2136 | 2282 | 3088 | 1718 | 1717 | 1745 | 1753 | 2318 | 2319 | 2337 | 2356 | 2032,91 | 1733 | 2332 | 1,346 | 0,429 | 11,126 | 9,41E-29 | 2,93E-28 | 1,00E-08 | 0,002687 | 0,0009199 | 0,0009199 | TRUE | 0,0005994 | No |
| orf19.2676 | 1979 | 1749 | 1845 | 1267 | 1789 | 1603 | 1462 | 2055 | 1802 | 1769 | 1746 | 1903 | 1591 | 1740 | 1497 | 1568 | 1702,07 | 1805 | 1599 | 0,887 | -0,174 | -3,13 | 0,00175 | 0,002425 | 0,0019979 | 0,002813 | 0,0023672 | 0,0023672 | TRUE | 0,0411015 | No |
| orf19.2677 | 363 | 327 | 376 | 262 | 384 | 324 | 343 | 450 | 330 | 331 | 356 | 394 | 342 | 352 | 351 | 343 | 349,81 | 353 | 347 | 0,989 | -0,016 | -0,193 | 0,847175 | 0,869125 | 1,00E-08 | 0,005813 | 0,0034888 | 0,0034888 | TRUE | 0,0574416 | No |
| orf19.2678 | 726 | 573 | 659 | 438 | 612 | 483 | 521 | 659 | 661 | 580 | 624 | 658 | 544 | 524 | 534 | 503 | 578,4 | 631 | 526 | 0,835 | -0,26 | -3,82 | 0,000133 | 0,000198 | 0,0009146 | 0,004321 | 0,0027127 | 0,0027127 | TRUE | 0,035253 | No |

| | | | | | | | | | | | | | | | | | | | | | | | | | | | | | | | |
|---|---|---|---|---|---|---|---|---|---|---|---|---|---|---|---|---|---|---|---|---|---|---|---|---|---|---|---|---|---|---|---|
| orf19.268 | 566 | 461 | 530 | 333 | 671 | 532 | 583 | 723 | 515 | 466 | 502 | 500 | 597 | 578 | 597 | 552 | 538,3 | 496 | 581 | 1,17 | 0,226 | 3,368 | 0,000757 | 0,001077 | 1,00E-08 | 0,004491 | 0,0024551 | 0,0024551 | TRUE | 0,0197718 | No |
| orf19.2680 | 1992 | 1714 | 1821 | 1211 | 2141 | 1803 | 1884 | 2526 | 1814 | 1734 | 1723 | 1819 | 1904 | 1958 | 1929 | 1927 | 1850,99 | 1772 | 1930 | 1,089 | 0,123 | 2,942 | 0,00326 | 0,004422 | 1,00E-08 | 0,002751 | 0,0011465 | 0,0011465 | TRUE | 0,0042271 | No |
| orf19.2681 | 106 | 86 | 106 | 63 | 166 | 154 | 152 | 204 | 97 | 87 | 100 | 95 | 148 | 167 | 156 | 156 | 125,57 | 95 | 157 | 1,65 | 0,723 | 5,73 | 1,01E-08 | 1,81E-08 | 1,00E-08 | 0,012557 | 0,006687 | 0,006687 | TRUE | 0,0307308 | No |
| orf19.2682 | 2891 | 2683 | 2910 | 1583 | 2553 | 1972 | 2258 | 2980 | 2632 | 2714 | 2754 | 2378 | 2271 | 2141 | 2312 | 2273 | 2434,4 | 2620 | 2249 | 0,858 | -0,222 | -4,156 | 3,24E-05 | 5,00E-05 | 0,0020769 | 0,00258 | 0,0023188 | 0,0023188 | TRUE | 0,0438081 | No |
| orf19.2684 | 1866 | 1854 | 1941 | 1128 | 1223 | 981 | 1057 | 1314 | 1699 | 1875 | 1837 | 1694 | 1088 | 1065 | 1083 | 1002 | 1417,9 | 1776 | 1060 | 0,595 | -0,748 | -13,61 | 3,50E-42 | 1,42E-41 | 0,001492 | 0,002969 | 0,0021585 | 0,0021585 | TRUE | 0,0168779 | No |
| orf19.2685 | 321 | 307 | 299 | 238 | 114 | 113 | 93 | 157 | 292 | 310 | 283 | 358 | 101 | 123 | 95 | 120 | 210,29 | 311 | 110 | 0,356 | -1,491 | -12,557 | 3,66E-36 | 1,34E-35 | 0,0063387 | 0,008319 | 0,0075738 | 0,0075738 | TRUE | 0,1009928 | No |
| orf19.2686 | 2340 | 2121 | 2202 | 1516 | 5534 | 5191 | 4886 | 6850 | 2130 | 2145 | 2084 | 2277 | 4922 | 5636 | 5004 | 5225 | 3678,04 | 2159 | 5197 | 2,409 | 1,269 | 23,955 | 8,14E-127 | 8,81E-126 | 0,00219 | 0,002396 | 0,0023529 | 0,0023529 | TRUE | 0,0389459 | No |
| orf19.2687.1 | 4727 | 4066 | 4203 | 2461 | 5585 | 3761 | 4548 | 5751 | 4304 | 4112 | 3978 | 3697 | 4968 | 4083 | 4658 | 4387 | 4273,29 | 4023 | 4524 | 1,124 | 0,169 | 2,556 | 0,010589 | 0,013872 | 0,0051804 | 0,002346 | 0,0039456 | 0,0039456 | TRUE | 0,0532529 | No |
| orf19.2688 | 15612 | 15136 | 16085 | 10202 | 14829 | 12994 | 13285 | 17227 | 14214 | 15308 | 15223 | 15325 | 13190 | 14108 | 13606 | 13141 | 14264,3 | 15018 | 13511 | 0,9 | -0,153 | -3,803 | 0,000143 | 0,000212 | 0,0010995 | 0,00213 | 0,0014744 | 0,0014744 | TRUE | 0,0160011 | No |
| orf19.269 | 4359 | 3994 | 4151 | 2791 | 6323 | 4892 | 5159 | 7127 | 3969 | 4040 | 3929 | 4193 | 5624 | 5311 | 5283 | 5437 | 4723,09 | 4033 | 5414 | 1,344 | 0,427 | 11,087 | 1,46E-28 | 4,51E-28 | 0,0005378 | 0,002317 | 0,0012025 | 0,0012025 | TRUE | 0,0083242 | No |
| orf19.2690 | 9652 | 8819 | 9507 | 6244 | 6407 | 5614 | 5601 | 7687 | 8788 | 8919 | 8998 | 9379 | 5699 | 6095 | 5736 | 5864 | 7434,76 | 9021 | 5848 | 0,649 | -0,625 | -16,674 | 2,04E-62 | 1,15E-61 | 0,0006481 | 0,002215 | 0,0012088 | 0,0012088 | TRUE | 0,0096295 | No |
| orf19.2691 | 1018 | 972 | 979 | 731 | 2554 | 2144 | 2228 | 2996 | 927 | 983 | 927 | 1098 | 2272 | 2328 | 2282 | 2285 | 1637,64 | 984 | 2292 | 2,341 | 1,227 | 22,072 | 5,85E-108 | 5,43E-107 | 0,0014177 | 0,002844 | 0,0022147 | 0,0022147 | TRUE | 0,0649184 | No |
| orf19.2693 | 2146 | 1846 | 1969 | 1151 | 7687 | 6036 | 6426 | 8763 | 1954 | 1867 | 1863 | 1729 | 6837 | 6553 | 6581 | 6684 | 4258,7 | 1853 | 6664 | 3,587 | 1,843 | 42,886 | 0 | 0 | 0,0006418 | 0,002347 | 0,0014139 | 0,0014139 | TRUE | 0,0219865 | No |
| orf19.2694 | 6547 | 5720 | 6138 | 3697 | 5569 | 4439 | 4745 | 6559 | 5961 | 5785 | 5809 | 5553 | 4953 | 4820 | 4859 | 5003 | 5343,02 | 5777 | 4909 | 0,849 | -0,236 | -6,633 | 3,28E-11 | 6,48E-11 | 0,0003198 | 0,002284 | 0,001023 | 0,001023 | TRUE | 0,0078146 | No |
| orf19.2695 | 3680 | 3381 | 3602 | 2171 | 3446 | 2890 | 3127 | 4080 | 3350 | 3420 | 3409 | 3261 | 3065 | 3138 | 3202 | 3112 | 3244,71 | 3360 | 3129 | 0,93 | -0,104 | -2,864 | 0,004182 | 0,00563 | 1,00E-08 | 0,002444 | 0,0009609 | 0,0009609 | TRUE | 0,0043073 | No |
| orf19.2697 | 1148 | 1087 | 1141 | 687 | 1222 | 1050 | 1151 | 1471 | 1045 | 1099 | 1080 | 1032 | 1087 | 1140 | 1179 | 1122 | 1098,03 | 1064 | 1132 | 1,062 | 0,087 | 1,683 | 0,092418 | 0,111332 | 1,00E-08 | 0,00324 | 0,0016246 | 0,0016246 | TRUE | 0,0091445 | No |
| orf19.2698 | 539 | 487 | 534 | 304 | 393 | 334 | 315 | 421 | 491 | 493 | 505 | 457 | 350 | 363 | 323 | 321 | 412,66 | 486 | 339 | 0,694 | -0,528 | -6,948 | 3,70E-12 | 7,54E-12 | 1,00E-08 | 0,005238 | 0,0030615 | 0,0030615 | TRUE | 0,0233758 | No |
| orf19.2699 | 3394 | 2739 | 3107 | 1907 | 4565 | 3789 | 4135 | 5651 | 3090 | 2770 | 2940 | 2865 | 4060 | 4114 | 4235 | 4311 | 3548,13 | 2916 | 4180 | 1,433 | 0,519 | 11,558 | 6,72E-31 | 2,21E-30 | 0,0010554 | 0,002409 | 0,0016369 | 0,0016369 | TRUE | 0,0201068 | No |
| orf19.27 | 1896 | 1604 | 1809 | 1189 | 1080 | 913 | 992 | 1428 | 1726 | 1622 | 1712 | 1786 | 961 | 991 | 1016 | 1089 | 1362,97 | 1712 | 1014 | 0,594 | -0,752 | -13,531 | 1,03E-41 | 4,14E-41 | 0,0014752 | 0,003006 | 0,0021905 | 0,0021905 | TRUE | 0,0317461 | No |
| orf19.270 | 413 | 316 | 317 | 230 | 261 | 178 | 201 | 321 | 376 | 320 | 300 | 345 | 232 | 193 | 206 | 245 | 277,16 | 335 | 219 | 0,657 | -0,607 | -5,736 | 9,71E-09 | 1,75E-08 | 0,0073025 | 0,006803 | 0,0070214 | 0,0070214 | TRUE | 0,0844053 | No |
| orf19.2701 | 2932 | 2679 | 2535 | 1733 | 4016 | 3003 | 3107 | 4194 | 2669 | 2710 | 2399 | 2603 | 3572 | 3260 | 3182 | 3199 | 2949,38 | 2595 | 3303 | 1,273 | 0,348 | 6,314 | 2,71E-10 | 5,18E-10 | 0,0024758 | 0,002485 | 0,0025729 | 0,0025729 | TRUE | 0,0367389 | No |
| orf19.2703 | 5572 | 4691 | 5270 | 3273 | 4775 | 3922 | 4277 | 5651 | 5073 | 4744 | 4988 | 4917 | 4247 | 4258 | 4380 | 4311 | 4614,74 | 4930 | 4299 | 0,872 | -0,198 | -5,546 | 2,92E-08 | 5,15E-08 | 0,0003126 | 0,002323 | 0,001006 | 0,001006 | TRUE | 0,0081102 | No |
| orf19.2704 | 2 | 4 | 3 | 2 | 3 | 0 | 3 | 3 | 2 | 4 | 3 | 3 | 3 | 0 | 3 | 2 | 2,47 | 3 | 2 | 0,705 | -0,504 | -0,576 | 0,564424 | 0,605452 | 1,00E-08 | 0,537416 | 0,3252821 | 0,3252821 | TRUE | 0,3354438 | No |
| orf19.2706 | 608 | 553 | 570 | 400 | 345 | 267 | 262 | 428 | 554 | 559 | 539 | 601 | 307 | 290 | 268 | 326 | 430,59 | 563 | 298 | 0,532 | -0,909 | -11,387 | 4,83E-30 | 1,55E-29 | 0,0014045 | 0,005105 | 0,0036125 | 0,0036125 | TRUE | 0,0510451 | No |
| orf19.2707.1 | 2067 | 1418 | 1569 | 1108 | 3340 | 2288 | 2970 | 3644 | 1882 | 1434 | 1485 | 1664 | 2971 | 2484 | 3042 | 2780 | 2217,71 | 1616 | 2819 | 1,744 | 0,803 | 9,171 | 4,70E-20 | 1,20E-19 | 0,0111243 | 0,002633 | 0,0068594 | 0,0068594 | TRUE | 0,1516897 | No |
| orf19.2708 | 2343 | 2066 | 2185 | 1443 | 2754 | 2239 | 2282 | 3056 | 2133 | 2090 | 2068 | 2168 | 2450 | 2431 | 2337 | 2331 | 2250,87 | 2115 | 2387 | 1,13 | 0,176 | 4,303 | 1,69E-05 | 2,65E-05 | 1,00E-08 | 0,002624 | 0,0011524 | 0,0011524 | TRUE | 0,0039235 | No |
| orf19.2709 | 12457 | 10702 | 11615 | 7308 | 13204 | 10786 | 11574 | 15325 | 11342 | 10824 | 10993 | 10978 | 11745 | 11711 | 11853 | 11690 | 11391,8 | 11034 | 11750 | 1,065 | 0,09 | 3,164 | 0,001556 | 0,002168 | 9,86E-05 | 0,002153 | 0,0006953 | 0,0006953 | TRUE | 0,0044107 | No |
| orf19.271 | 71 | 63 | 84 | 49 | 66 | 56 | 55 | 66 | 65 | 64 | 79 | 74 | 59 | 61 | 56 | 50 | 63,46 | 70 | 56 | 0,802 | -0,318 | -1,823 | 0,068307 | 0,083452 | 1,00E-08 | 0,022855 | 0,0134266 | 0,0134266 | TRUE | 0,0810156 | No |
| orf19.2710 | 4553 | 4036 | 4442 | 2258 | 2250 | 1638 | 2016 | 2515 | 4145 | 4082 | 4204 | 3392 | 2001 | 1778 | 2065 | 1918 | 2948,24 | 3956 | 1940 | 0,49 | -1,029 | -14,272 | 3,27E-46 | 1,42E-45 | 0,0061126 | 0,002485 | 0,0046118 | 0,0046118 | TRUE | 0,1093009 | No |
| orf19.2711 | 7602 | 6535 | 7165 | 4133 | 7470 | 6078 | 6699 | 8901 | 6921 | 6609 | 6781 | 6208 | 6644 | 6599 | 6861 | 6790 | 6676,74 | 6630 | 6724 | 1,013 | 0,019 | 0,462 | 0,644123 | 0,681768 | 0,0009636 | 0,002235 | 0,0014822 | 0,0014822 | TRUE | 0,0215253 | No |
| orf19.2711.1 | 197 | 179 | 177 | 116 | 223 | 210 | 187 | 268 | 179 | 181 | 168 | 174 | 198 | 228 | 192 | 204 | 190,56 | 176 | 206 | 1,169 | 0,225 | 2,164 | 0,030439 | 0,038471 | 1,00E-08 | 0,008969 | 0,0050961 | 0,0050961 | TRUE | 0,0533186 | No |
| orf19.2712 | 10876 | 9470 | 10241 | 6073 | 9534 | 7481 | 8080 | 10910 | 9902 | 9578 | 9692 | 9123 | 8480 | 8122 | 8275 | 8322 | 8936,81 | 9574 | 8300 | 0,867 | -0,207 | -5,735 | 9,76E-09 | 1,76E-08 | 0,0005994 | 0,002185 | 0,001134 | 0,001134 | TRUE | 0,0118784 | No |
| orf19.2713 | 59 | 55 | 58 | 49 | 112 | 88 | 88 | 133 | 54 | 56 | 55 | 74 | 100 | 96 | 90 | 101 | 78,07 | 60 | 97 | 1,657 | 0,728 | 4,504 | 6,68E-06 | 1,07E-05 | 1,00E-08 | 0,018957 | 0,0112252 | 0,0112252 | TRUE | 0,1580578 | No |
| orf19.2715 | 5595 | 4991 | 5479 | 3214 | 3793 | 3151 | 3492 | 4498 | 5094 | 5048 | 5185 | 4828 | 3374 | 3421 | 3576 | 3431 | 4244,68 | 5039 | 3450 | 0,684 | -0,548 | -14,238 | 5,34E-46 | 2,31E-45 | 0,0004491 | 0,002348 | 0,0011785 | 0,0011785 | TRUE | 0,0091113 | No |
| orf19.2717 | 16771 | 15058 | 15637 | 8737 | 16961 | 12796 | 14701 | 19276 | 15269 | 15230 | 14799 | 13124 | 15086 | 13893 | 15056 | 14704 | 14645,1 | 14606 | 14685 | 1,005 | 0,007 | 0,137 | 0,891035 | 0,906535 | 0,0025948 | 0,002127 | 0,002715 | 0,002715 | TRUE | 0,0563817 | No |
| orf19.272 | 2394 | 2254 | 2424 | 1579 | 37475 | 32988 | 32808 | 44075 | 2180 | 2280 | 2294 | 2372 | 33333 | 35816 | 33600 | 33621 | 18186,8 | 2282 | 34092 | 14,963 | 3,903 | 94,557 | 0 | 0 | 0,0008628 | 0,00211 | 0,0013866 | 0,0013866 | TRUE | 0,0141466 | No |
| orf19.2720 | 6055 | 5245 | 5753 | 4136 | 6376 | 5576 | 5571 | 7619 | 5513 | 5305 | 5445 | 6213 | 5671 | 6054 | 5705 | 5812 | 5714,71 | 5619 | 5810 | 1,035 | 0,05 | 0,939 | 0,347642 | 0,387654 | 0,0026937 | 0,002268 | 0,0025025 | 0,0025025 | TRUE | 0,0601357 | No |
| orf19.2721 | 3544 | 3235 | 3305 | 2169 | 2789 | 2287 | 2298 | 3044 | 3227 | 3272 | 3128 | 3258 | 2481 | 2483 | 2353 | 2322 | 2815,47 | 3221 | 2410 | 0,748 | -0,419 | -10,166 | 2,82E-24 | 7,95E-24 | 0,0004643 | 0,002506 | 0,0012688 | 0,0012688 | TRUE | 0,0076927 | No |
| orf19.2722 | 723 | 668 | 696 | 398 | 249 | 224 | 193 | 291 | 658 | 676 | 659 | 598 | 221 | 243 | 198 | 222 | 434,34 | 648 | 221 | 0,34 | -1,555 | -19,071 | 4,41E-81 | 3,17E-80 | 0,0006518 | 0,005078 | 0,0034559 | 0,0034559 | TRUE | 0,0506773 | No |
| orf19.2723 | 2769 | 2725 | 2840 | 1980 | 3344 | 2447 | 2600 | 3450 | 2521 | 2756 | 2688 | 2974 | 2974 | 2657 | 2663 | 2632 | 2733,09 | 2735 | 2732 | 1 | 0 | 0,004 | 0,996765 | 0,997078 | 0,00364 | 0,00252 | 0,0031446 | 0,0031446 | TRUE | 0,0438827 | No |
| orf19.2724 | 58125 | 58917 | 61853 | 41939 | 27165 | 26463 | 24786 | 32830 | 52920 | 59588 | 58538 | 62999 | 24162 | 28731 | 25384 | 25043 | 42170,8 | 58511 | 25830 | 0,441 | -1,18 | -18,28 | 1,20E-74 | 7,89E-74 | 0,0055235 | 0,002068 | 0,0039743 | 0,0039743 | TRUE | 0,069929 | No |
| orf19.2725 | 6 | 12 | 5 | 10 | 5 | 9 | 6 | 9 | 5 | 12 | 5 | 15 | 4 | 10 | 6 | 7 | 8,07 | 9 | 7 | 0,75 | -0,415 | -0,79 | 0,42975 | 0,471136 | 0,0849823 | 0,165674 | 0,1368132 | 0,1368132 | TRUE | 0,1730635 | No |
| orf19.2726 | 4309 | 3895 | 4249 | 2632 | 4326 | 3707 | 3610 | 5160 | 3923 | 3939 | 4021 | 3954 | 3848 | 4025 | 3697 | 3936 | 3917,91 | 3959 | 3876 | 0,979 | -0,031 | -0,796 | 0,426168 | 0,467325 | 0,0003682 | 0,002374 | 0,0011567 | 0,0011567 | TRUE | 0,0115268 | No |
| orf19.2727 | 4548 | 4225 | 4265 | 2855 | 4127 | 3502 | 3359 | 4601 | 4141 | 4273 | 4036 | 4289 | 3671 | 3802 | 3440 | 3510 | 3895,22 | 4185 | 3606 | 0,862 | -0,215 | -4,855 | 1,20E-06 | 1,99E-06 | 0,0010801 | 0,002376 | 0,001617 | 0,001617 | TRUE | 0,0159815 | No |
| orf19.2728 | 881 | 889 | 918 | 506 | 1510 | 1222 | 1282 | 1575 | 802 | 899 | 869 | 760 | 1343 | 1327 | 1313 | 1201 | 1064,29 | 832 | 1296 | 1,551 | 0,634 | 9,726 | 2,35E-22 | 6,28E-22 | 0,0027819 | 0,003278 | 0,0030708 | 0,0030708 | TRUE | 0,0355339 | No |
| orf19.273 | 1362 | 1316 | 1394 | 939 | 900 | 905 | 764 | 1020 | 1240 | 1331 | 1319 | 1411 | 801 | 983 | 782 | 778 | 1080,56 | 1325 | 836 | 0,63 | -0,666 | -8,541 | 1,33E-17 | 3,18E-17 | 0,0069952 | 0,00326 | 0,004863 | 0,004863 | TRUE | 0,1615142 | No |
| orf19.2730 | 2418 | 1715 | 2096 | 1483 | 504 | 374 | 495 | 667 | 2201 | 1735 | 1984 | 2228 | 448 | 406 | 507 | 509 | 1252,19 | 2037 | 468 | 0,23 | -2,12 | -22,981 | 7,24E-117 | 7,23E-116 | 0,0102301 | 0,003092 | 0,0069105 | 0,0069105 | TRUE | 0,1210337 | No |
| orf19.2731 | 147 | 122 | 117 | 57 | 137 | 107 | 147 | 171 | 134 | 123 | 111 | 86 | 122 | 116 | 151 | 130 | 121,57 | 114 | 130 | 1,13 | 0,177 | 1,199 | 0,230461 | 0,264053 | 0,0125113 | 0,012903 | 0,0126026 | 0,0126026 | TRUE | 0,2098804 | No |
| orf19.2733 | 829 | 765 | 795 | 491 | 574 | 437 | 448 | 594 | 755 | 774 | 752 | 738 | 511 | 474 | 459 | 453 | 614,42 | 755 | 474 | 0,628 | -0,672 | -10,458 | 1,34E-25 | 3,91E-25 | 1,00E-08 | 0,004187 | 0,00227 | 0,00227 | TRUE | 0,0325237 | No |
| orf19.2734 | 114 | 103 | 100 | 68 | 274 | 245 | 264 | 380 | 104 | 104 | 95 | 102 | 244 | 266 | 270 | 290 | 184,34 | 101 | 268 | 2,649 | 1,405 | 12,235 | 2,02E-34 | 7,12E-34 | 1,00E-08 | 0,009203 | 0,0056763 | 0,0056763 | TRUE | 0,0427696 | No |
| orf19.2735 | 11850 | 11019 | 11883 | 7323 | 12525 | 10576 | 11234 | 14450 | 10789 | 11145 | 11246 | 11000 | 11141 | 11483 | 11505 | 11023 | 11166,3 | 11045 | 11288 | 1,022 | 0,031 | 0,987 | 0,323645 | 0,363068 | 0,0002735 | 0,002155 | 0,0008589 | 0,0008589 | TRUE | 0,0030915 | No |
| orf19.2736 | 903 | 801 | 882 | 557 | 1180 | 998 | 1027 | 1414 | 822 | 810 | 835 | 837 | 1050 | 1084 | 1052 | 1079 | 945,9 | 826 | 1066 | 1,291 | 0,369 | 7,248 | 4,23E-13 | 8,89E-13 | 1,00E-08 | 0,003434 | 0,0014024 | 0,0014024 | TRUE | 0,0019395 | No |
| orf19.2737 | 8395 | 6930 | 7605 | 5080 | 6464 | 5588 | 5714 | 7993 | 7643 | 7009 | 7197 | 7631 | 5750 | 6067 | 5852 | 6097 | 6655,77 | 7370 | 5942 | 0,806 | -0,31 | -7,333 | 2,26E-13 | 4,77E-13 | 0,0011275 | 0,002235 | 0,0015694 | 0,0015694 | TRUE | 0,0132838 | No |
| orf19.2738 | 3229 | 2865 | 3144 | 2100 | 7291 | 6545 | 6819 | 9359 | 2940 | 2898 | 2976 | 3155 | 6485 | 7106 | 6984 | 7139 | 4960,17 | 2992 | 6928 | 2,318 | 1,213 | 26,411 | 1,03E-153 | 1,37E-152 | 0,0013508 | 0,002303 | 0,0017777 | 0,0017777 | TRUE | 0,0227901 | No |
| orf19.2739 | 307 | 289 | 311 | 181 | 665 | 537 | 582 | 760 | 280 | 292 | 294 | 272 | 591 | 583 | 596 | 580 | 436,04 | 284 | 588 | 2,059 | 1,042 | 14,438 | 2,99E-47 | 1,32E-46 | 1,00E-08 | 0,005067 | 0,0023463 | 0,0023463 | TRUE | 0,0075597 | No |
| orf19.274 | 499 | 452 | 526 | 269 | 260 | 203 | 220 | 293 | 454 | 457 | 498 | 404 | 231 | 220 | 225 | 224 | 339,23 | 453 | 225 | 0,494 | -1,016 | -11,706 | 1,19E-31 | 3,97E-31 | 1,00E-08 | 0,005931 | 0,003992 | 0,003992 | TRUE | 0,0523439 | No |
| orf19.2740 | 659 | 605 | 668 | 416 | 1601 | 1349 | 1387 | 1945 | 600 | 612 | 632 | 625 | 1424 | 1465 | 1420 | 1484 | 1032,72 | 617 | 1448 | 2,349 | 1,232 | 23,373 | 7,99E-121 | 8,22E-120 | 1,00E-08 | 0,003316 | 0,0014814 | 0,0014814 | TRUE | 0,0045452 | No |
| orf19.2742 | 135 | 94 | 108 | 49 | 111 | 63 | 77 | 107 | 123 | 95 | 102 | 74 | 99 | 68 | 79 | 82 | 90,18 | 98 | 82 | 0,825 | -0,278 | -1,595 | 0,110646 | 0,13219 | 0,0202444 | 0,016686 | 0,0180428 | 0,0180428 | TRUE | 0,2000283 | No |
| orf19.2743 | 1197 | 922 | 1006 | 773 | 256 | 179 | 220 | 269 | 1090 | 933 | 952 | 1161 | 228 | 194 | 225 | 205 | 623,52 | 1034 | 213 | 0,207 | -2,275 | -24,615 | 8,72E-134 | 1,00E-132 | 0,0069958 | 0,004156 | 0,0054914 | 0,0054914 | TRUE | 0,0764878 | No |
| orf19.2745 | 20 | 17 | 21 | 11 | 23 | 15 | 17 | 20 | 18 | 17 | 20 | 17 | 20 | 16 | 17 | 15 | 17,65 | 18 | 17 | 0,96 | -0,059 | -0,185 | 0,85321 | 0,874894 | 1,00E-08 | 0,076875 | 0,0403372 | 0,0403372 | TRUE | 0,0824672 | No |
| orf19.2746 | 2523 | 2607 | 2658 | 1418 | 2160 | 1736 | 1849 | 2386 | 2297 | 2637 | 2516 | 2130 | 1921 | 1885 | 1894 | 1820 | 2137,39 | 2395 | 1880 | 0,784 | -0,351 | -5,472 | 4,46E-08 | 7,80E-08 | 0,0040649 | 0,002655 | 0,003489 | 0,003489 | TRUE | 0,0636643 | No |
| orf19.2747 | 923 | 1046 | 1125 | 739 | 759 | 796 | 714 | 983 | 840 | 1058 | 1065 | 1110 | 675 | 864 | 731 | 750 | 886,69 | 1018 | 755 | 0,742 | -0,43 | -4,524 | 6,07E-06 | 9,73E-06 | 0,0110319 | 0,003527 | 0,0075384 | 0,0075384 | TRUE | 0,1689607 | No |
| orf19.2748 | 1392 | 1582 | 1567 | 949 | 548 | 512 | 488 | 637 | 1267 | 1600 | 1483 | 1426 | 487 | 556 | 500 | 486 | 975,62 | 1444 | 507 | 0,351 | -1,511 | -19,102 | 2,41E-81 | 1,74E-80 | 0,0059869 | 0,003391 | 0,0047159 | 0,0047159 | TRUE | 0,0847824 | No |
| orf19.2749 | 5668 | 6031 | 6434 | 3841 | 1172 | 1036 | 978 | 1315 | 5160 | 6100 | 6089 | 5770 | 1042 | 1125 | 1002 | 1003 | 3411,39 | 5780 | 1043 | 0,18 | -2,471 | -38,445 | 0 | 0 | 0,0042469 | 0,002424 | 0,0034269 | 0,0034269 | TRUE | 0,0643743 | No |
| orf19.275 | 463 | 362 | 381 | 227 | 551 | 433 | 449 | 587 | 422 | 366 | 361 | 341 | 490 | 470 | 460 | 448 | 419,63 | 372 | 467 | 1,248 | 0,319 | 3,938 | 8,22E-05 | 0,000124 | 0,0021619 | 0,005185 | 0,0038797 | 0,0038797 | TRUE | 0,0942394 | No |
| orf19.2751 | 1022 | 1118 | 1151 | 637 | 1145 | 939 | 962 | 1258 | 930 | 1131 | 1089 | 957 | 1018 | 1019 | 985 | 960 | 1011,27 | 1027 | 996 | 0,968 | -0,047 | -0,674 | 0,50012 | 0,541761 | 0,0036486 | 0,003343 | 0,0036679 | 0,0036679 | TRUE | 0,0552552 | No |
| orf19.2752 | 2150 | 2164 | 2305 | 1391 | 6203 | 5419 | 5752 | 7383 | 1957 | 2189 | 2181 | 2090 | 5517 | 5884 | 5891 | 5632 | 3917,57 | 2104 | 5731 | 2,723 | 1,445 | 30,27 | 2,88E-201 | 4,95E-200 | 0,0014075 | 0,002374 | 0,0018514 | 0,0018514 | TRUE | 0,0283055 | No |

| | | | | | | | | | | | | | | | | | | | | | | | | | | | | | | | |
|---|---|---|---|---|---|---|---|---|---|---|---|---|---|---|---|---|---|---|---|---|---|---|---|---|---|---|---|---|---|---|---|
| orf19.2753 | 2106 | 2040 | 2206 | 1375 | 21217 | 18341 | 19199 | 25774 | 1917 | 2063 | 2088 | 2065 | 18872 | 19913 | 19662 | 19661 | 10780,2 | 2033 | 19527 | 9,61 | 3,265 | 81,574 | 0 | 0 | 0,0006453 | 0,00216 | 0,0012496 | 0,0012496 | TRUE | 0,0188859 | No |
| orf19.2754 | 527 | 494 | 473 | 375 | 646 | 549 | 572 | 710 | 480 | 500 | 448 | 563 | 575 | 596 | 586 | 542 | 536,06 | 498 | 575 | 1,16 | 0,214 | 2,773 | 0,005562 | 0,007432 | 0,0031473 | 0,004501 | 0,0038084 | 0,0038084 | TRUE | 0,0798978 | No |
| orf19.2755 | 5445 | 5081 | 5411 | 3402 | 4664 | 4081 | 3893 | 5257 | 4957 | 5139 | 5121 | 5110 | 4148 | 4431 | 3987 | 4010 | 4613 | 5082 | 4144 | 0,815 | -0,295 | -6,876 | 6,15E-12 | 1,25E-11 | 0,0010561 | 0,002323 | 0,0015455 | 0,0015455 | TRUE | 0,0259092 | No |
| orf19.2756 | 193 | 310 | 271 | 129 | 323 | 295 | 264 | 356 | 176 | 314 | 256 | 194 | 287 | 320 | 270 | 272 | 261,13 | 235 | 287 | 1,219 | 0,286 | 1,876 | 0,060619 | 0,074502 | 0,0338925 | 0,007096 | 0,0183618 | 0,0183618 | TRUE | 0,5282976 | No |
| orf19.2757 | 670 | 696 | 678 | 518 | 1089 | 1003 | 948 | 1228 | 610 | 704 | 642 | 778 | 969 | 1089 | 971 | 937 | 837,37 | 684 | 992 | 1,456 | 0,542 | 6,785 | 1,16E-11 | 2,33E-11 | 0,0065286 | 0,003615 | 0,0048534 | 0,0048534 | TRUE | 0,093586 | No |
| orf19.2758 | 414 | 335 | 360 | 269 | 167 | 161 | 168 | 223 | 377 | 339 | 341 | 404 | 149 | 175 | 172 | 170 | 265,75 | 365 | 166 | 0,457 | -1,13 | -11,398 | 4,29E-30 | 1,38E-29 | 0,0024282 | 0,007008 | 0,0051676 | 0,0051676 | TRUE | 0,0571558 | No |
| orf19.276 | 897 | 858 | 862 | 538 | 630 | 552 | 523 | 653 | 817 | 868 | 816 | 808 | 560 | 599 | 536 | 498 | 687,73 | 827 | 548 | 0,661 | -0,598 | -8,739 | 2,35E-18 | 5,70E-18 | 0,0018769 | 0,003958 | 0,0029878 | 0,0029878 | TRUE | 0,0480549 | No |
| orf19.2760 | 1950 | 2103 | 2125 | 1189 | 2366 | 1989 | 2127 | 2555 | 1775 | 2127 | 2011 | 1786 | 2104 | 2159 | 2178 | 1949 | 2011,35 | 1925 | 2098 | 1,088 | 0,122 | 1,827 | 0,067749 | 0,082787 | 0,0046095 | 0,002694 | 0,0037845 | 0,0037845 | TRUE | 0,0602348 | No |
| orf19.2761 | 253 | 291 | 290 | 185 | 510 | 433 | 415 | 520 | 230 | 294 | 274 | 278 | 454 | 470 | 425 | 397 | 352,8 | 269 | 436 | 1,622 | 0,698 | 7,371 | 1,69E-13 | 3,59E-13 | 0,0052387 | 0,005781 | 0,0055449 | 0,0055449 | TRUE | 0,1155799 | No |
| orf19.2762 | 15534 | 13772 | 14189 | 9355 | 7932 | 6245 | 6097 | 8799 | 14143 | 13929 | 13429 | 14053 | 7055 | 6780 | 6244 | 6712 | 10293,1 | 13888 | 6698 | 0,482 | -1,052 | -24,393 | 2,05E-131 | 2,31E-130 | 0,0013814 | 0,002165 | 0,0016772 | 0,0016772 | TRUE | 0,0252136 | No |
| orf19.2763 | 139 | 148 | 131 | 59 | 242 | 183 | 196 | 235 | 127 | 150 | 124 | 89 | 215 | 199 | 201 | 179 | 160,35 | 122 | 198 | 1,602 | 0,68 | 5,032 | 4,85E-07 | 8,17E-07 | 0,0119868 | 0,010275 | 0,0108458 | 0,0108458 | TRUE | 0,2641094 | No |
| orf19.2765 | 92176 | 76457 | 85080 | 71014 | 79444 | 78665 | 74729 | 1E+05 | 83922 | 77328 | 80521 | 1E+05 | 70663 | 85408 | 76532 | 82945 | 82999,1 | 87111 | 78887 | 0,906 | -0,143 | -1,137 | 0,255733 | 0,291142 | 0,015209 | 0,002053 | 0,0080866 | 0,015209 | TRUE | 0,2799924 | No |
| orf19.2766 | 9 | 12 | 12 | 4 | 7 | 5 | 5 | 8 | 8 | 12 | 11 | 6 | 6 | 5 | 5 | 6 | 7,57 | 9 | 6 | 0,599 | -0,739 | -1,496 | 0,134572 | 0,159134 | 1,00E-08 | 0,1765 | 0,0981301 | 0,0981301 | TRUE | 0,1160108 | No |
| orf19.2767 | 27267 | 22772 | 24868 | 18897 | 24979 | 22468 | 21603 | 31383 | 24825 | 23031 | 23535 | 28386 | 22218 | 24394 | 22124 | 23939 | 24056,7 | 24944 | 23169 | 0,929 | -0,106 | -1,619 | 0,105443 | 0,126449 | 0,0059033 | 0,002092 | 0,0040996 | 0,0040996 | TRUE | 0,1052055 | No |
| orf19.2768 | 40960 | 40990 | 42868 | 23944 | 6848 | 5857 | 5853 | 7667 | 37292 | 41457 | 40571 | 35968 | 6091 | 6359 | 5994 | 5848 | 22447,6 | 38822 | 6073 | 0,156 | -2,677 | -50,905 | 0 | 0 | 0,0027659 | 0,002096 | 0,0025655 | 0,0025655 | TRUE | 0,0298634 | No |
| orf19.2769 | 16305 | 13916 | 14030 | 8185 | 5242 | 3440 | 3936 | 5366 | 14845 | 14075 | 13278 | 12295 | 4663 | 3735 | 4031 | 4093 | 8876,82 | 13623 | 4130 | 0,303 | -1,722 | -23,689 | 4,69E-124 | 4,95E-123 | 0,0074092 | 0,002186 | 0,0049217 | 0,0049217 | TRUE | 0,0917014 | No |
| orf19.277 | 955 | 846 | 867 | 624 | 843 | 694 | 712 | 984 | 869 | 856 | 821 | 937 | 750 | 753 | 729 | 751 | 808,26 | 871 | 746 | 0,859 | -0,218 | -3,739 | 0,000185 | 0,000273 | 3,24E-05 | 0,003671 | 0,0020364 | 0,0020364 | TRUE | 0,0272268 | No |
| orf19.2770 | 5078 | 4615 | 4754 | 3049 | 613 | 424 | 485 | 652 | 4623 | 4668 | 4499 | 4580 | 545 | 460 | 497 | 497 | 2546,23 | 4592 | 500 | 0,109 | -3,196 | -61,663 | 0 | 0 | 0,0001341 | 0,002556 | 0,0015345 | 0,0015345 | TRUE | 0,0432724 | No |
| orf19.2770.1 | 7699 | 6124 | 6291 | 4266 | 1722 | 1294 | 1286 | 1873 | 7010 | 6194 | 5954 | 6408 | 1532 | 1405 | 1317 | 1429 | 3905,98 | 6392 | 1421 | 0,222 | -2,169 | -34,835 | 7,22E-266 | 1,63E-264 | 0,0038578 | 0,002375 | 0,0033099 | 0,0033099 | TRUE | 0,0524503 | No |
| orf19.2771 | 1508 | 1440 | 1549 | 919 | 2071 | 1742 | 1779 | 2436 | 1373 | 1456 | 1466 | 1380 | 1842 | 1891 | 1822 | 1858 | 1636,17 | 1419 | 1853 | 1,305 | 0,384 | 8,55 | 1,23E-17 | 2,93E-17 | 1,00E-08 | 0,002844 | 0,0013061 | 0,0013061 | TRUE | 0,0066814 | No |
| orf19.2772 | 6301 | 5902 | 6326 | 3990 | 1961 | 1749 | 1730 | 2346 | 5737 | 5969 | 5987 | 5994 | 1744 | 1899 | 1772 | 1790 | 3861,39 | 5922 | 1801 | 0,304 | -1,718 | -42,455 | 0 | 0 | 0,0004443 | 0,002379 | 0,0012216 | 0,0012216 | TRUE | 0,0153647 | No |
| orf19.2774 | 6012 | 5104 | 5308 | 3173 | 4102 | 3102 | 3352 | 4695 | 5474 | 5162 | 5024 | 4766 | 3649 | 3368 | 3433 | 3581 | 4307,06 | 5106 | 3508 | 0,687 | -0,542 | -10,77 | 4,78E-27 | 1,43E-26 | 0,0020274 | 0,002344 | 0,0021966 | 0,0021966 | TRUE | 0,0286382 | No |
| orf19.2775 | 4236 | 3852 | 4125 | 2520 | 4754 | 4114 | 3891 | 5444 | 3857 | 3896 | 3904 | 3785 | 4229 | 4467 | 3985 | 4153 | 4034,34 | 3860 | 4208 | 1,089 | 0,124 | 2,897 | 0,003768 | 0,00509 | 0,0008981 | 0,002364 | 0,0014968 | 0,0014968 | TRUE | 0,0202947 | No |
| orf19.2778 | 822 | 1053 | 924 | 618 | 1358 | 1223 | 1092 | 1381 | 748 | 1065 | 874 | 928 | 1208 | 1328 | 1118 | 1053 | 1040,47 | 904 | 1177 | 1,302 | 0,381 | 3,785 | 0,000154 | 0,000228 | 0,0143215 | 0,003307 | 0,0087235 | 0,0087235 | TRUE | 0,175807 | No |
| orf19.2779 | 2 | 1 | 2 | 1 | 3 | 1 | 1 | 2 | 2 | 1 | 2 | 2 | 3 | 1 | 1 | 2 | 1,57 | 2 | 2 | 1,02 | 0,029 | 0,028 | 0,977745 | 0,979743 | 1,00E-08 | 0,845374 | 0,426671 | 0,426671 | TRUE | 0,1736789 | No |
| orf19.278 | 1349 | 1157 | 1295 | 908 | 5013 | 4334 | 4504 | 6314 | 1228 | 1170 | 1226 | 1364 | 4459 | 4706 | 4613 | 4816 | 2947,68 | 1247 | 4648 | 3,74 | 1,903 | 36,96 | 5,08E-299 | 1,32E-297 | 0,0015725 | 0,002485 | 0,0020127 | 0,0020127 | TRUE | 0,0425215 | No |
| orf19.2781 | 1796 | 1995 | 2022 | 1200 | 2217 | 1998 | 1938 | 2428 | 1635 | 2018 | 1914 | 1803 | 1972 | 2169 | 1985 | 1852 | 1918,4 | 1842 | 1994 | 1,082 | 0,114 | 1,604 | 0,108807 | 0,130165 | 0,0054036 | 0,002726 | 0,0042856 | 0,0042856 | TRUE | 0,0717238 | No |
| orf19.2782 | 2141 | 2019 | 2099 | 1433 | 2631 | 2143 | 2147 | 2902 | 1949 | 2042 | 1987 | 2153 | 2340 | 2327 | 2199 | 2214 | 2151,22 | 2033 | 2270 | 1,119 | 0,162 | 3,438 | 0,000585 | 0,000838 | 0,0008584 | 0,002651 | 0,0016507 | 0,0016507 | TRUE | 0,017502 | No |
| orf19.2783 | 1 | 3 | 6 | 1 | 2 | 2 | 8 | 7 | 1 | 3 | 6 | 2 | 2 | 2 | 8 | 5 | 3,58 | 3 | 4 | 1,537 | 0,62 | 0,77 | 0,441115 | 0,482931 | 0,252621 | 0,37144 | 0,3281522 | 0,3281522 | TRUE | 0,4704054 | No |
| orf19.2784 | 22 | 31 | 26 | 33 | 26 | 22 | 18 | 22 | 20 | 31 | 25 | 50 | 23 | 24 | 18 | 17 | 25,97 | 32 | 20 | 0,67 | -0,577 | -1,838 | 0,06611 | 0,080893 | 0,0568224 | 0,052896 | 0,0544025 | 0,0544025 | TRUE | 0,6044201 | No |
| orf19.2785 | 2304 | 2291 | 2085 | 1316 | 3854 | 2888 | 2695 | 3918 | 2098 | 2317 | 1973 | 1977 | 3428 | 3136 | 2760 | 2989 | 2584,65 | 2091 | 3078 | 1,471 | 0,557 | 7,632 | 2,31E-14 | 5,01E-14 | 0,0065719 | 0,002548 | 0,0047067 | 0,0047067 | TRUE | 0,0715309 | No |
| orf19.2786 | 625 | 683 | 759 | 416 | 763 | 744 | 656 | 905 | 569 | 691 | 718 | 625 | 679 | 808 | 672 | 690 | 681,46 | 651 | 712 | 1,092 | 0,128 | 1,461 | 0,144107 | 0,169776 | 0,0074223 | 0,003976 | 0,0058433 | 0,0058433 | TRUE | 0,0937266 | No |
| orf19.2787 | 9 | 11 | 11 | 7 | 14 | 19 | 19 | 13 | 8 | 11 | 10 | 11 | 12 | 21 | 19 | 10 | 12,84 | 10 | 16 | 1,533 | 0,616 | 1,526 | 0,12694 | 0,150753 | 1,00E-08 | 0,104937 | 0,0728987 | 0,0728987 | TRUE | 0,1959804 | No |
| orf19.2788 | 601 | 557 | 627 | 398 | 526 | 407 | 479 | 648 | 547 | 563 | 593 | 598 | 468 | 442 | 491 | 494 | 524,55 | 575 | 474 | 0,826 | -0,276 | -3,997 | 6,41E-05 | 9,70E-05 | 1,00E-08 | 0,004555 | 0,0026565 | 0,0026565 | TRUE | 0,022115 | No |
| orf19.2789 | 1925 | 1914 | 1891 | 1257 | 1382 | 1035 | 1124 | 1467 | 1753 | 1936 | 1790 | 1888 | 1229 | 1124 | 1151 | 1119 | 1498,68 | 1842 | 1156 | 0,628 | -0,671 | -12,43 | 1,79E-35 | 6,44E-35 | 0,0014298 | 0,002918 | 0,0020974 | 0,0020974 | TRUE | 0,0224549 | No |
| orf19.279 | 45 | 26 | 28 | 19 | 80 | 40 | 60 | 85 | 41 | 26 | 26 | 29 | 71 | 43 | 61 | 65 | 45,4 | 30 | 60 | 1,966 | 0,976 | 4,143 | 3,42E-05 | 5,27E-05 | 0,0231248 | 0,031135 | 0,0282293 | 0,0282293 | TRUE | 0,402211 | No |
| orf19.2790 | 1718 | 1429 | 1545 | 965 | 2413 | 2088 | 2198 | 3094 | 1564 | 1445 | 1462 | 1450 | 2146 | 2267 | 2251 | 2360 | 1868,21 | 1480 | 2256 | 1,523 | 0,607 | 12,307 | 8,32E-35 | 2,97E-34 | 0,0010482 | 0,002744 | 0,0017655 | 0,0017655 | TRUE | 0,0182998 | No |
| orf19.2791 | 5422 | 4782 | 5346 | 3286 | 4033 | 3525 | 3720 | 4999 | 4937 | 4836 | 5060 | 4936 | 3587 | 3827 | 3810 | 3813 | 4350,75 | 4942 | 3759 | 0,76 | -0,395 | -10,569 | 4,16E-26 | 1,22E-25 | 0,0004203 | 0,002341 | 0,0011079 | 0,0011079 | TRUE | 0,011682 | No |
| orf19.2792 | 978 | 818 | 955 | 599 | 423 | 367 | 350 | 493 | 890 | 827 | 904 | 900 | 376 | 398 | 358 | 376 | 628,82 | 880 | 377 | 0,429 | -1,222 | -18,548 | 8,49E-77 | 5,76E-76 | 1,00E-08 | 0,004138 | 0,002327 | 0,002327 | TRUE | 0,0199491 | No |
| orf19.2794 | 1123 | 1018 | 1037 | 624 | 1271 | 987 | 985 | 1319 | 1022 | 1030 | 981 | 937 | 1131 | 1072 | 1009 | 1006 | 1023,48 | 992 | 1054 | 1,06 | 0,084 | 1,408 | 0,158992 | 0,186559 | 0,0015275 | 0,003328 | 0,0023981 | 0,0023981 | TRUE | 0,0293397 | No |
| orf19.2795 | 4382 | 4566 | 4507 | 2704 | 7231 | 5576 | 6045 | 7677 | 3990 | 4618 | 4265 | 4062 | 6432 | 6054 | 6191 | 5856 | 5183,45 | 4234 | 6133 | 1,448 | 0,534 | 9,994 | 1,61E-23 | 4,46E-23 | 0,0027229 | 0,002292 | 0,0025399 | 0,0025399 | TRUE | 0,045669 | No |
| orf19.2796 | 522 | 422 | 482 | 310 | 286 | 264 | 257 | 350 | 475 | 427 | 456 | 466 | 254 | 287 | 263 | 267 | 361,89 | 456 | 268 | 0,587 | -0,769 | -9,639 | 5,46E-22 | 1,45E-21 | 1,00E-08 | 0,005687 | 0,0032041 | 0,0032041 | TRUE | 0,0228433 | No |
| orf19.2797 | 89 | 82 | 73 | 54 | 62 | 44 | 53 | 78 | 81 | 83 | 69 | 81 | 55 | 48 | 54 | 59 | 66,36 | 78 | 54 | 0,695 | -0,525 | -3,086 | 0,002031 | 0,002801 | 1,00E-08 | 0,021944 | 0,0124161 | 0,0124161 | TRUE | 0,0663372 | No |
| orf19.2798 | 187 | 155 | 168 | 116 | 299 | 221 | 259 | 355 | 170 | 157 | 159 | 174 | 266 | 240 | 265 | 271 | 212,78 | 165 | 260 | 1,586 | 0,666 | 6,682 | 2,35E-11 | 4,68E-11 | 1,00E-08 | 0,008245 | 0,0045088 | 0,0045088 | TRUE | 0,0293077 | No |
| orf19.2799 | 1657 | 1367 | 1576 | 971 | 1200 | 1017 | 1051 | 1358 | 1509 | 1383 | 1492 | 1459 | 1067 | 1104 | 1076 | 1036 | 1265,64 | 1461 | 1071 | 0,732 | -0,449 | -8,887 | 6,30E-19 | 1,55E-18 | 1,00E-08 | 0,003081 | 0,0016525 | 0,0016525 | TRUE | 0,016215 | No |
| orf19.28 | 8061 | 7261 | 7685 | 4231 | 5843 | 4220 | 4921 | 6305 | 7339 | 7344 | 7273 | 6356 | 5197 | 4582 | 5040 | 4810 | 5992,48 | 7078 | 4907 | 0,693 | -0,529 | -9,076 | 1,13E-19 | 2,84E-19 | 0,0035973 | 0,002257 | 0,0030943 | 0,0030943 | TRUE | 0,0564683 | No |
| orf19.280 | 35 | 30 | 43 | 25 | 43 | 45 | 34 | 48 | 32 | 30 | 41 | 38 | 38 | 49 | 35 | 37 | 37,37 | 35 | 40 | 1,127 | 0,173 | 0,753 | 0,451189 | 0,493198 | 1,00E-08 | 0,037382 | 0,0233634 | 0,0233634 | TRUE | 0,165098 | No |
| orf19.2800 | 0 | 0 | 0 | 0 | 0 | 0 | 0 | 0 | 0 | 0 | 0 | 0 | 0 | 0 | 0 | 0 | 0 | 0 | 0 | NA | NA | NA | NA | NA | NA | NA | NA | NA | NA | NA | NA |
| orf19.2803 | 2875 | 2482 | 2689 | 1703 | 1399 | 1052 | 1236 | 1635 | 2618 | 2510 | 2545 | 2558 | 1244 | 1142 | 1266 | 1247 | 1891,31 | 2558 | 1225 | 0,479 | -1,061 | -22,982 | 7,09E-117 | 7,08E-116 | 0,0002697 | 0,002735 | 0,0014553 | 0,0014553 | TRUE | 0,0233042 | No |
| orf19.2804 | 5 | 1 | 6 | 1 | 3 | 4 | 3 | 7 | 5 | 1 | 6 | 2 | 3 | 4 | 3 | 5 | 3,52 | 4 | 4 | 1,17 | 0,227 | 0,304 | 0,761294 | 0,792238 | 1,00E-08 | 0,377219 | 0,2538868 | 0,2538868 | TRUE | 0,2605337 | No |
| orf19.2805 | 288 | 291 | 321 | 194 | 675 | 592 | 601 | 774 | 262 | 294 | 304 | 291 | 600 | 643 | 615 | 590 | 450,1 | 288 | 612 | 2,127 | 1,089 | 14,303 | 2,10E-46 | 9,12E-46 | 1,00E-08 | 0,004972 | 0,0029464 | 0,0029464 | TRUE | 0,0461055 | No |
| orf19.2808 | 754 | 832 | 853 | 493 | 1864 | 1553 | 1617 | 2089 | 686 | 841 | 807 | 741 | 1658 | 1686 | 1656 | 1594 | 1208,68 | 769 | 1648 | 2,142 | 1,099 | 17,042 | 4,04E-65 | 2,38E-64 | 0,0030509 | 0,00313 | 0,0030093 | 0,0030093 | TRUE | 0,0682595 | No |
| orf19.2809 | 1565 | 1381 | 1406 | 988 | 2619 | 2214 | 2376 | 3097 | 1425 | 1397 | 1331 | 1484 | 2330 | 2404 | 2433 | 2362 | 1895,68 | 1409 | 2382 | 1,694 | 0,76 | 16,504 | 3,41E-61 | 1,89E-60 | 0,0003242 | 0,002734 | 0,0014589 | 0,0014589 | TRUE | 0,016915 | No |
| orf19.281 | 205 | 183 | 176 | 134 | 243 | 227 | 258 | 339 | 187 | 185 | 167 | 201 | 216 | 246 | 264 | 259 | 215,63 | 185 | 246 | 1,339 | 0,421 | 3,986 | 6,73E-05 | 0,000102 | 0,0026166 | 0,008163 | 0,0059114 | 0,0059114 | TRUE | 0,0787632 | No |
| orf19.2810 | 387 | 320 | 361 | 241 | 1900 | 1722 | 1627 | 2341 | 352 | 324 | 342 | 362 | 1690 | 1870 | 1666 | 1786 | 1048,91 | 345 | 1753 | 5,094 | 2,349 | 35,263 | 2,14E-272 | 4,90E-271 | 0,0016232 | 0,003296 | 0,0024455 | 0,0024455 | TRUE | 0,0232906 | No |
| orf19.2812 | 357 | 297 | 351 | 205 | 857 | 761 | 728 | 1130 | 325 | 300 | 332 | 308 | 762 | 826 | 746 | 862 | 557,7 | 316 | 799 | 2,522 | 1,335 | 17,391 | 9,69E-68 | 5,90E-67 | 0,0023474 | 0,004406 | 0,0033897 | 0,0033897 | TRUE | 0,0356949 | No |
| orf19.2813 | 3 | 2 | 0 | 0 | 0 | 1 | 1 | 2 | 3 | 2 | 0 | 0 | 0 | 1 | 1 | 2 | 1,05 | 1 | 1 | 0,73 | -0,455 | -0,322 | 0,747298 | 0,778945 | 1,00E-08 | 1,261686 | 0,9760656 | 0,9760656 | TRUE | 0,2326836 | No |
| orf19.2818 | 1591 | 1498 | 1656 | 1037 | 1558 | 1334 | 1425 | 1875 | 1449 | 1515 | 1567 | 1558 | 1386 | 1448 | 1459 | 1430 | 1476,55 | 1522 | 1431 | 0,941 | -0,088 | -1,864 | 0,062318 | 0,076488 | 1,00E-08 | 0,002932 | 0,0014732 | 0,0014732 | TRUE | 0,0134062 | No |
| orf19.2819 | 138 | 181 | 180 | 107 | 178 | 138 | 126 | 159 | 126 | 183 | 170 | 161 | 158 | 150 | 129 | 121 | 149,78 | 160 | 140 | 0,873 | -0,197 | -1,386 | 0,165607 | 0,193857 | 0,0149396 | 0,010856 | 0,0125924 | 0,0125924 | TRUE | 0,2424135 | No |
| orf19.2820 | 4432 | 3959 | 4294 | 2769 | 3814 | 3143 | 3428 | 4601 | 4035 | 4004 | 4064 | 4159 | 3392 | 3412 | 3511 | 3510 | 3760,98 | 4066 | 3456 | 0,851 | -0,233 | -6,76 | 1,38E-11 | 2,76E-11 | 1,00E-08 | 0,002388 | 0,0008745 | 0,0008745 | TRUE | 0,0026844 | No |
| orf19.2821 | 2194 | 2140 | 2187 | 1407 | 2617 | 2130 | 2120 | 2969 | 1998 | 2164 | 2070 | 2114 | 2328 | 2313 | 2171 | 2265 | 2177,69 | 2086 | 2269 | 1,088 | 0,122 | 2,728 | 0,006369 | 0,008478 | 0,0006176 | 0,002644 | 0,0014643 | 0,0014643 | TRUE | 0,010358 | No |
| orf19.2822 | 255 | 258 | 239 | 154 | 159 | 106 | 94 | 134 | 232 | 261 | 226 | 231 | 141 | 115 | 96 | 102 | 175,7 | 238 | 114 | 0,479 | -1,063 | -8,348 | 6,92E-17 | 1,62E-16 | 0,0090678 | 0,009555 | 0,0092189 | 0,0092189 | TRUE | 0,2841113 | No |
| orf19.2823 | 3512 | 3236 | 3505 | 2598 | 2829 | 2620 | 2563 | 3446 | 3198 | 3273 | 3317 | 3903 | 2516 | 2845 | 2625 | 2629 | 3038,07 | 3423 | 2654 | 0,776 | -0,366 | -5,499 | 3,81E-08 | 6,68E-08 | 0,0053741 | 0,002472 | 0,003912 | 0,003912 | TRUE | 0,1054106 | No |
| orf19.2825 | 7112 | 6561 | 6923 | 4081 | 3669 | 3010 | 3047 | 3993 | 6475 | 6636 | 6552 | 6130 | 3263 | 3268 | 3121 | 3046 | 4811,39 | 6448 | 3174 | 0,492 | -1,024 | -24,372 | 3,38E-131 | 3,80E-130 | 0,000876 | 0,002312 | 0,0014637 | 0,0014637 | TRUE | 0,0128922 | No |
| orf19.2826 | 2118 | 1840 | 2048 | 1246 | 1478 | 1209 | 1357 | 1908 | 1928 | 1861 | 1938 | 1872 | 1315 | 1313 | 1390 | 1455 | 1633,96 | 1900 | 1368 | 0,721 | -0,473 | -9,537 | 1,46E-21 | 3,85E-21 | 0,0008469 | 0,002845 | 0,0017368 | 0,0017368 | TRUE | 0,0235736 | No |
| orf19.2827 | 296 | 348 | 374 | 194 | 269 | 250 | 233 | 328 | 269 | 352 | 354 | 291 | 239 | 271 | 239 | 250 | 283,29 | 316 | 250 | 0,787 | -0,346 | -3,254 | 0,001138 | 0,001602 | 0,0080624 | 0,0067 | 0,0073059 | 0,0073059 | TRUE | 0,128327 | No |
| orf19.2828 | 451 | 427 | 460 | 241 | 240 | 194 | 181 | 227 | 411 | 432 | 435 | 362 | 213 | 211 | 185 | 173 | 302,81 | 410 | 196 | 0,474 | -1,077 | -10,953 | 6,44E-28 | 1,97E-27 | 0,0043826 | 0,006399 | 0,0055989 | 0,0055989 | TRUE | 0,0742706 | No |

| | | | | | | | | | | | | | | | | | | | | | | | | | | | | | | | |
|---|---|---|---|---|---|---|---|---|---|---|---|---|---|---|---|---|---|---|---|---|---|---|---|---|---|---|---|---|---|---|---|
| orf19.2829 | 1627 | 1535 | 1548 | 907 | 2542 | 1759 | 1930 | 2430 | 1481 | 1552 | 1465 | 1362 | 2261 | 1910 | 1977 | 1854 | 1732,79 | 1465 | 2000 | 1,363 | 0,447 | 6,517 | 7,18E-11 | 1,40E-10 | 0,0047354 | 0,002799 | 0,0039207 | 0,0039207 | TRUE | 0,0942328 | No |
| orf19.2830 | 10887 | 10294 | 10555 | 6740 | 9762 | 8087 | 8377 | 11004 | 9912 | 10411 | 9989 | 10125 | 8683 | 8780 | 8579 | 8394 | 9359,2 | 10109 | 8609 | 0,852 | -0,232 | -7,201 | 5,99E-13 | 1,25E-12 | 0,0002768 | 0,002178 | 0,0008885 | 0,0008885 | TRUE | 0,0050367 | No |
| orf19.2831 | 2771 | 2448 | 2656 | 1440 | 3600 | 2723 | 3124 | 4101 | 2523 | 2476 | 2514 | 2163 | 3202 | 2956 | 3199 | 3128 | 2770,21 | 2419 | 3121 | 1,288 | 0,365 | 6,571 | 5,01E-11 | 9,83E-11 | 0,0027126 | 0,002514 | 0,0025979 | 0,0025979 | TRUE | 0,0577157 | No |
| orf19.2832 | 4022 | 3903 | 4244 | 2561 | 1386 | 1125 | 1206 | 1547 | 3662 | 3947 | 4017 | 3847 | 1233 | 1221 | 1235 | 1180 | 2542,79 | 3868 | 1217 | 0,315 | -1,669 | -36,108 | 1,73E-285 | 4,20E-284 | 0,0008754 | 0,002557 | 0,0015281 | 0,0015281 | TRUE | 0,0163902 | No |
| orf19.2833 | 88 | 84 | 100 | 75 | 83 | 76 | 61 | 77 | 80 | 85 | 95 | 113 | 74 | 83 | 62 | 59 | 81,24 | 93 | 70 | 0,749 | -0,417 | -2,45 | 0,014293 | 0,018558 | 0,0098687 | 0,018297 | 0,0152324 | 0,0152324 | TRUE | 0,1519218 | No |
| orf19.2834 | 3465 | 3230 | 3535 | 2148 | 1845 | 1464 | 1607 | 2063 | 3155 | 3267 | 3346 | 3227 | 1641 | 1589 | 1646 | 1574 | 2430,47 | 3249 | 1612 | 0,496 | -1,011 | -24,664 | 2,60E-134 | 3,02E-133 | 1,00E-08 | 0,002581 | 0,0011582 | 0,0011582 | TRUE | 0,0051327 | No |
| orf19.2835 | 591 | 618 | 639 | 359 | 687 | 665 | 607 | 797 | 538 | 625 | 605 | 539 | 611 | 722 | 622 | 608 | 608,73 | 577 | 641 | 1,107 | 0,146 | 1,835 | 0,066559 | 0,081395 | 0,004441 | 0,004207 | 0,0044585 | 0,0044585 | TRUE | 0,0831806 | No |
| orf19.2836 | 262 | 178 | 205 | 143 | 323 | 240 | 279 | 385 | 239 | 180 | 194 | 215 | 287 | 261 | 286 | 294 | 244,33 | 207 | 282 | 1,365 | 0,449 | 4,321 | 1,55E-05 | 2,44E-05 | 0,0041421 | 0,007444 | 0,0061424 | 0,0061424 | TRUE | 0,1333249 | No |
| orf19.2837 | 204 | 174 | 176 | 112 | 110 | 95 | 118 | 153 | 186 | 176 | 167 | 168 | 98 | 103 | 121 | 117 | 141,88 | 174 | 110 | 0,629 | -0,669 | -5,511 | 3,56E-08 | 6,25E-08 | 1,00E-08 | 0,011347 | 0,0068484 | 0,0068484 | TRUE | 0,0566977 | No |
| orf19.2838 | 2453 | 2390 | 2564 | 1499 | 596 | 493 | 475 | 678 | 2233 | 2417 | 2427 | 2252 | 530 | 535 | 486 | 517 | 1424,74 | 2332 | 517 | 0,222 | -2,173 | -38,049 | 0 | 0 | 0,0011589 | 0,002964 | 0,002002 | 0,002002 | TRUE | 0,0175036 | No |
| orf19.2839 | 4738 | 4177 | 4483 | 2617 | 492 | 411 | 490 | 641 | 4314 | 4225 | 4243 | 3931 | 438 | 446 | 502 | 489 | 2323,36 | 4178 | 469 | 0,112 | -3,156 | -54,464 | 0 | 0 | 0,0014965 | 0,002606 | 0,0021033 | 0,0021033 | TRUE | 0,0246764 | No |
| orf19.284 | 117 | 84 | 82 | 72 | 130 | 110 | 148 | 190 | 107 | 85 | 78 | 108 | 116 | 119 | 152 | 145 | 113,6 | 94 | 133 | 1,42 | 0,506 | 3,293 | 0,000991 | 0,001399 | 0,0131061 | 0,013665 | 0,0134481 | 0,0134481 | TRUE | 0,143555 | No |
| orf19.2841 | 8829 | 9009 | 9147 | 5089 | 3355 | 2649 | 2798 | 3672 | 8038 | 9112 | 8657 | 7644 | 2984 | 2876 | 2866 | 2801 | 5622,27 | 8363 | 2882 | 0,344 | -1,538 | -27,14 | 3,32E-162 | 4,67E-161 | 0,0030589 | 0,002272 | 0,0028568 | 0,0028568 | TRUE | 0,0442079 | No |
| orf19.2842 | 9774 | 8734 | 9780 | 6401 | 8312 | 7288 | 7901 | 10437 | 8899 | 8834 | 9256 | 9615 | 7393 | 7913 | 8092 | 7961 | 8495,32 | 9151 | 7840 | 0,857 | -0,223 | -5,112 | 3,19E-07 | 5,40E-07 | 0,0013875 | 0,002193 | 0,0017031 | 0,0017031 | TRUE | 0,0180315 | No |
| orf19.2843 | 4508 | 3923 | 3958 | 2346 | 8359 | 5667 | 6273 | 9336 | 4104 | 3968 | 3746 | 3524 | 7435 | 6153 | 6424 | 7122 | 5309,47 | 3836 | 6784 | 1,768 | 0,822 | 12,066 | 1,60E-33 | 5,54E-33 | 0,0057943 | 0,002286 | 0,0042491 | 0,0042491 | TRUE | 0,0509321 | No |
| orf19.2844 | 724 | 754 | 789 | 436 | 686 | 626 | 647 | 846 | 659 | 763 | 747 | 655 | 610 | 680 | 663 | 645 | 677,65 | 706 | 650 | 0,917 | -0,125 | -1,751 | 0,079942 | 0,096906 | 0,0027244 | 0,003986 | 0,0033832 | 0,0033832 | TRUE | 0,0338088 | No |
| orf19.2846 | 342 | 291 | 361 | 215 | 425 | 360 | 341 | 520 | 311 | 294 | 342 | 323 | 378 | 391 | 349 | 397 | 348,14 | 318 | 379 | 1,194 | 0,256 | 3,083 | 0,002051 | 0,002826 | 1,00E-08 | 0,005832 | 0,0037212 | 0,0037212 | TRUE | 0,0331084 | No |
| orf19.2847 | 3999 | 4431 | 4452 | 2800 | 3305 | 3021 | 2895 | 3635 | 3641 | 4481 | 4213 | 4206 | 2940 | 3280 | 2965 | 2773 | 3562,4 | 4135 | 2990 | 0,723 | -0,469 | -6,737 | 1,61E-11 | 3,22E-11 | 0,0057808 | 0,002408 | 0,0043587 | 0,0043587 | TRUE | 0,0799712 | No |
| orf19.2848 | 2954 | 2498 | 2742 | 1786 | 2598 | 2114 | 2332 | 3152 | 2689 | 2526 | 2595 | 2683 | 2311 | 2295 | 2388 | 2404 | 2486,57 | 2623 | 2350 | 0,896 | -0,158 | -3,833 | 0,000127 | 0,000189 | 0,0003198 | 0,002568 | 0,0012237 | 0,0012237 | TRUE | 0,0075349 | No |
| orf19.2849 | 24746 | 19196 | 20827 | 12641 | 23523 | 18168 | 19835 | 28887 | 22530 | 19415 | 19711 | 18989 | 20923 | 19725 | 20314 | 22035 | 20455,2 | 20161 | 20749 | 1,029 | 0,041 | 0,704 | 0,48127 | 0,52312 | 0,0038291 | 0,002102 | 0,003268 | 0,003268 | TRUE | 0,0767572 | No |
| orf19.285 | 6130 | 5272 | 5735 | 3081 | 1834 | 1319 | 1528 | 1929 | 5581 | 5332 | 5428 | 4628 | 1631 | 1432 | 1565 | 1471 | 3383,58 | 5242 | 1525 | 0,291 | -1,782 | -27,215 | 4,30E-163 | 6,06E-162 | 0,0044246 | 0,002427 | 0,0037102 | 0,0037102 | TRUE | 0,0741238 | No |
| orf19.2850 | 6961 | 6192 | 6785 | 4059 | 2944 | 2274 | 2693 | 3336 | 6338 | 6263 | 6421 | 6097 | 2619 | 2469 | 2758 | 2545 | 4438,64 | 6280 | 2598 | 0,413 | -1,274 | -29,813 | 2,66E-195 | 4,43E-194 | 0,0010023 | 0,002335 | 0,0014884 | 0,0014884 | TRUE | 0,0204187 | No |
| orf19.2851 | 857 | 884 | 908 | 602 | 1169 | 1044 | 1028 | 1388 | 780 | 894 | 859 | 904 | 1040 | 1133 | 1053 | 1059 | 965,35 | 859 | 1071 | 1,249 | 0,32 | 5,194 | 2,05E-07 | 3,50E-07 | 0,001761 | 0,003405 | 0,0025887 | 0,0025887 | TRUE | 0,0477309 | No |
| orf19.2852 | 3546 | 3193 | 3264 | 2106 | 3671 | 2936 | 3055 | 4053 | 3228 | 3229 | 3089 | 3164 | 3265 | 3188 | 3129 | 3092 | 3172,97 | 3178 | 3168 | 0,997 | -0,005 | -0,121 | 0,903379 | 0,917191 | 0,0002528 | 0,002453 | 0,0010652 | 0,0010652 | TRUE | 0,0053149 | No |
| orf19.2853 | 1457 | 1446 | 1509 | 1029 | 662 | 583 | 568 | 780 | 1327 | 1462 | 1428 | 1546 | 589 | 633 | 582 | 595 | 1020,17 | 1441 | 600 | 0,417 | -1,263 | -20,256 | 3,16E-91 | 2,50E-90 | 0,00188 | 0,003332 | 0,0025776 | 0,0025776 | TRUE | 0,0346586 | No |
| orf19.2857 | 8865 | 7774 | 8404 | 5119 | 10214 | 8129 | 9044 | 12108 | 8071 | 7863 | 7954 | 7690 | 9085 | 8826 | 9262 | 9236 | 8498,26 | 7894 | 9102 | 1,153 | 0,205 | 6,271 | 3,59E-10 | 6,83E-10 | 0,000292 | 0,002192 | 0,0009065 | 0,0009065 | TRUE | 0,005024 | No |
| orf19.2859 | 7255 | 6151 | 6816 | 4134 | 10742 | 8397 | 9363 | 12398 | 6605 | 6221 | 6451 | 6210 | 9555 | 9117 | 9589 | 9457 | 7900,59 | 6372 | 9430 | 1,479 | 0,565 | 15,695 | 1,64E-55 | 8,37E-55 | 0,0005367 | 0,002204 | 0,0011098 | 0,0011098 | TRUE | 0,0074957 | No |
| orf19.286 | 80 | 67 | 74 | 55 | 60 | 75 | 66 | 90 | 73 | 68 | 70 | 83 | 53 | 81 | 68 | 69 | 70,54 | 74 | 68 | 0,926 | -0,111 | -0,638 | 0,523726 | 0,565603 | 1,00E-08 | 0,020765 | 0,0146595 | 0,0146595 | TRUE | 0,1891148 | No |
| orf19.2862 | 4231 | 3440 | 3887 | 2436 | 4464 | 3714 | 4191 | 5451 | 3852 | 3479 | 3679 | 3659 | 3971 | 4032 | 4292 | 4158 | 3890,3 | 3667 | 4113 | 1,121 | 0,165 | 3,688 | 0,000226 | 0,000333 | 0,0011203 | 0,002377 | 0,001666 | 0,001666 | TRUE | 0,014895 | No |
| orf19.2863 | 447 | 365 | 409 | 250 | 433 | 347 | 342 | 555 | 407 | 369 | 387 | 376 | 385 | 377 | 350 | 423 | 384,28 | 385 | 384 | 0,999 | -0,001 | -0,013 | 0,989303 | 0,990235 | 0,0019525 | 0,005475 | 0,0038815 | 0,0038815 | TRUE | 0,059387 | No |
| orf19.2863.1 | 593 | 458 | 545 | 315 | 594 | 435 | 472 | 709 | 540 | 463 | 516 | 473 | 528 | 472 | 483 | 541 | 502,12 | 498 | 506 | 1,016 | 0,023 | 0,295 | 0,768245 | 0,798689 | 0,0029126 | 0,004668 | 0,0038555 | 0,0038555 | TRUE | 0,0384195 | No |
| orf19.2864.1 | 20529 | 16209 | 17035 | 10274 | 28321 | 18970 | 22670 | 30500 | 18691 | 16394 | 16122 | 15433 | 25191 | 20596 | 23217 | 23266 | 19863,6 | 16660 | 23068 | 1,384 | 0,469 | 6,701 | 2,06E-11 | 4,12E-11 | 0,0068719 | 0,002104 | 0,0046598 | 0,0046598 | TRUE | 0,0825358 | No |
| orf19.2866 | 109 | 74 | 80 | 59 | 110 | 72 | 79 | 137 | 99 | 75 | 76 | 89 | 98 | 78 | 81 | 105 | 87,48 | 85 | 90 | 1,076 | 0,106 | 0,649 | 0,516054 | 0,558168 | 0,0094884 | 0,017137 | 0,0141602 | 0,0141602 | TRUE | 0,1501013 | No |
| orf19.2867 | 4169 | 3762 | 4061 | 2608 | 1820 | 1639 | 1667 | 2305 | 3796 | 3805 | 3843 | 3918 | 1619 | 1779 | 1707 | 1758 | 2778,17 | 3840 | 1716 | 0,447 | -1,162 | -27,507 | 1,44E-166 | 2,10E-165 | 0,0004104 | 0,002513 | 0,0013008 | 0,0013008 | TRUE | 0,0178636 | No |
| orf19.2868 | 3 | 4 | 4 | 2 | 1 | 4 | 3 | 2 | 3 | 4 | 4 | 3 | 1 | 4 | 3 | 2 | 2,92 | 4 | 2 | 0,695 | -0,525 | -0,654 | 0,512921 | 0,554874 | 1,00E-08 | 0,453713 | 0,265049 | 0,265049 | TRUE | 0,2658433 | No |
| orf19.2869 | 5 | 2 | 5 | 0 | 1 | 2 | 2 | 2 | 5 | 2 | 5 | 0 | 1 | 2 | 2 | 2 | 2,24 | 3 | 2 | 0,545 | -0,876 | -0,926 | 0,354219 | 0,394092 | 1,00E-08 | 0,591048 | 0,3949487 | 0,3949487 | TRUE | 0,2278513 | No |
| orf19.287 | 4030 | 3267 | 3560 | 2283 | 6664 | 4855 | 5316 | 7198 | 3669 | 3304 | 3369 | 3429 | 5927 | 5271 | 5444 | 5491 | 4488,19 | 3443 | 5533 | 1,607 | 0,684 | 13,588 | 4,70E-42 | 1,90E-41 | 0,0020274 | 0,002331 | 0,0021956 | 0,0021956 | TRUE | 0,0280694 | No |
| orf19.2870 | 2828 | 2341 | 2453 | 1299 | 861 | 582 | 659 | 838 | 2575 | 2368 | 2322 | 1951 | 766 | 632 | 675 | 639 | 1490,89 | 2304 | 678 | 0,294 | -1,766 | -20,365 | 3,39E-92 | 2,70E-91 | 0,0091894 | 0,002923 | 0,0063009 | 0,0063009 | TRUE | 0,1244536 | No |
| orf19.2871 | 7784 | 6696 | 7059 | 4763 | 13208 | 11191 | 11810 | 16928 | 7087 | 6772 | 6681 | 7155 | 11748 | 12150 | 12095 | 12913 | 9575,12 | 6924 | 12226 | 1,767 | 0,821 | 19,49 | 1,33E-84 | 9,99E-84 | 0,0012187 | 0,002175 | 0,0015882 | 0,0015882 | TRUE | 0,0176521 | No |
| orf19.2873 | 2513 | 2349 | 2581 | 1551 | 3874 | 3219 | 3361 | 4428 | 2288 | 2376 | 2443 | 2330 | 3446 | 3495 | 3442 | 3378 | 2899,6 | 2359 | 3440 | 1,457 | 0,543 | 14,157 | 1,69E-45 | 7,29E-45 | 0,0001158 | 0,002493 | 0,0010508 | 0,0010508 | TRUE | 0,0072195 | No |
| orf19.2875 | 1788 | 1654 | 1710 | 961 | 2743 | 2022 | 2338 | 2880 | 1628 | 1673 | 1618 | 1444 | 2440 | 2195 | 2394 | 2197 | 1948,64 | 1591 | 2306 | 1,447 | 0,533 | 9,018 | 1,91E-19 | 4,78E-19 | 0,002866 | 0,002715 | 0,002814 | 0,002814 | TRUE | 0,0429148 | No |
| orf19.2876 | 3371 | 2850 | 3096 | 1994 | 1355 | 1044 | 1197 | 1772 | 3069 | 2882 | 2930 | 2995 | 1205 | 1133 | 1226 | 1352 | 2099,16 | 2969 | 1229 | 0,415 | -1,27 | -22,624 | 2,53E-113 | 2,44E-112 | 0,0023366 | 0,002666 | 0,0024642 | 0,0024642 | TRUE | 0,0564196 | No |
| orf19.2877 | 28896 | 24558 | 26727 | 18340 | 15352 | 12829 | 12543 | 18458 | 26309 | 24838 | 25295 | 27550 | 13655 | 13929 | 12846 | 14080 | 19812,5 | 25998 | 13628 | 0,524 | -0,932 | -20,497 | 2,30E-93 | 1,85E-92 | 0,0018113 | 0,002104 | 0,0019294 | 0,0019294 | TRUE | 0,0196085 | No |
| orf19.2878 | 81 | 72 | 95 | 48 | 40 | 36 | 30 | 56 | 74 | 73 | 90 | 72 | 36 | 39 | 31 | 43 | 57,09 | 77 | 37 | 0,481 | -1,056 | -5,501 | 3,79E-08 | 6,64E-08 | 1,00E-08 | 0,025178 | 0,0160309 | 0,0160309 | TRUE | 0,121279 | No |
| orf19.2879 | 230 | 193 | 211 | 154 | 276 | 231 | 252 | 301 | 209 | 195 | 200 | 231 | 245 | 251 | 258 | 230 | 227,45 | 209 | 246 | 1,182 | 0,241 | 2,481 | 0,013117 | 0,017058 | 1,00E-08 | 0,007845 | 0,0045869 | 0,0045869 | TRUE | 0,0447847 | No |
| orf19.288 | 2035 | 1854 | 2040 | 1341 | 5757 | 4796 | 5113 | 6719 | 1853 | 1875 | 1931 | 2014 | 5121 | 5207 | 5236 | 5125 | 3545,3 | 1918 | 5172 | 2,701 | 1,434 | 36,94 | 1,05E-298 | 2,70E-297 | 2,19E-05 | 0,00241 | 0,0010758 | 0,0010758 | TRUE | 0,0121108 | No |
| orf19.2881 | 8320 | 8986 | 8936 | 5199 | 10201 | 8927 | 8853 | 11360 | 7575 | 9088 | 8457 | 7810 | 9073 | 9692 | 9067 | 8665 | 8678,5 | 8232 | 9124 | 1,108 | 0,148 | 2,443 | 0,014586 | 0,018922 | 0,0043529 | 0,002189 | 0,0034087 | 0,0034087 | TRUE | 0,0599233 | No |
| orf19.2882 | 2708 | 2496 | 2670 | 1610 | 5478 | 4588 | 4894 | 6766 | 2466 | 2524 | 2527 | 2418 | 4873 | 4981 | 5012 | 5161 | 3745,3 | 2484 | 5007 | 2,014 | 1,01 | 26,991 | 1,89E-160 | 2,62E-159 | 0,0002895 | 0,00239 | 0,0010361 | 0,0010361 | TRUE | 0,0054527 | No |
| orf19.2883 | 1281 | 1229 | 1243 | 886 | 2974 | 2697 | 2659 | 3502 | 1166 | 1243 | 1176 | 1331 | 2645 | 2928 | 2723 | 2671 | 1985,57 | 1229 | 2742 | 2,236 | 1,161 | 20,816 | 3,12E-96 | 2,60E-95 | 0,0022154 | 0,002702 | 0,0023776 | 0,0023776 | TRUE | 0,033778 | No |
| orf19.2884 | 4127 | 4181 | 4262 | 2427 | 3851 | 3134 | 3327 | 4269 | 3757 | 4229 | 4034 | 3646 | 3425 | 3403 | 3407 | 3256 | 3644,64 | 3916 | 3373 | 0,86 | -0,217 | -4,184 | 2,86E-05 | 4,42E-05 | 0,0021806 | 0,002399 | 0,0023044 | 0,0023044 | TRUE | 0,0348147 | No |
| orf19.2885 | 54 | 63 | 52 | 33 | 32 | 43 | 35 | 56 | 49 | 64 | 49 | 50 | 28 | 47 | 36 | 43 | 45,67 | 53 | 38 | 0,724 | -0,467 | -2,149 | 0,031607 | 0,039869 | 0,0081019 | 0,03096 | 0,0230693 | 0,0230693 | TRUE | 0,2430944 | No |
| orf19.2886 | 964 | 1134 | 1136 | 773 | 1082 | 1024 | 916 | 1186 | 878 | 1147 | 1075 | 1161 | 962 | 1112 | 938 | 905 | 1022,23 | 1065 | 979 | 0,92 | -0,12 | -1,304 | 0,192337 | 0,223098 | 0,0105689 | 0,003329 | 0,0071926 | 0,0071926 | TRUE | 0,1716296 | No |
| orf19.2887 | 4514 | 3934 | 4215 | 2808 | 7423 | 5476 | 5695 | 8170 | 4110 | 3979 | 3989 | 4218 | 6603 | 5945 | 5832 | 6232 | 5113,53 | 4074 | 6153 | 1,512 | 0,596 | 12,514 | 6,23E-36 | 2,27E-35 | 0,0016719 | 0,002295 | 0,0019697 | 0,0019697 | TRUE | 0,029464 | No |
| orf19.2888 | 170 | 223 | 197 | 98 | 274 | 184 | 235 | 288 | 155 | 226 | 186 | 147 | 244 | 200 | 241 | 220 | 202,23 | 178 | 226 | 1,26 | 0,334 | 2,513 | 0,011972 | 0,015614 | 0,0173877 | 0,008569 | 0,0118806 | 0,0118806 | TRUE | 0,3330616 | No |
| orf19.2889 | 861 | 785 | 831 | 502 | 613 | 453 | 481 | 612 | 784 | 794 | 786 | 754 | 545 | 492 | 493 | 467 | 639,36 | 780 | 499 | 0,639 | -0,646 | -9,806 | 1,06E-22 | 2,85E-22 | 0,0006018 | 0,004103 | 0,0025444 | 0,0025444 | TRUE | 0,0475369 | No |
| orf19.2890 | 1012 | 986 | 919 | 623 | 1705 | 1353 | 1304 | 1870 | 921 | 997 | 870 | 936 | 1517 | 1469 | 1335 | 1426 | 1183,96 | 931 | 1437 | 1,544 | 0,627 | 10,3 | 7,01E-25 | 2,00E-24 | 0,0022038 | 0,003153 | 0,0026565 | 0,0026565 | TRUE | 0,0286813 | No |
| orf19.2891 | 1006 | 1074 | 1112 | 722 | 1809 | 1663 | 1619 | 2053 | 916 | 1086 | 1052 | 1085 | 1609 | 1806 | 1658 | 1566 | 1347,23 | 1035 | 1660 | 1,605 | 0,683 | 9,997 | 1,57E-23 | 4,36E-23 | 0,0039725 | 0,003018 | 0,0036762 | 0,0036762 | TRUE | 0,0741783 | No |
| orf19.2892 | 3694 | 4010 | 4122 | 2655 | 3755 | 3481 | 3386 | 4446 | 3363 | 4056 | 3901 | 3988 | 3340 | 3779 | 3468 | 3391 | 3660,84 | 3827 | 3494 | 0,913 | -0,131 | -2,018 | 0,04358 | 0,054273 | 0,0046244 | 0,002398 | 0,0037626 | 0,0037626 | TRUE | 0,0821035 | No |
| orf19.2893 | 903 | 1121 | 1176 | 642 | 1000 | 843 | 866 | 1083 | 822 | 1134 | 1113 | 964 | 889 | 915 | 887 | 826 | 943,88 | 1008 | 879 | 0,871 | -0,199 | -2,149 | 0,031637 | 0,039899 | 0,0109291 | 0,003437 | 0,0071274 | 0,0071274 | TRUE | 0,1924182 | No |
| orf19.2895 | 696 | 590 | 645 | 387 | 734 | 525 | 619 | 815 | 634 | 597 | 610 | 581 | 653 | 570 | 634 | 622 | 612,58 | 606 | 620 | 1,022 | 0,032 | 0,481 | 0,630388 | 0,668785 | 0,0005778 | 0,004193 | 0,0025605 | 0,0025605 | TRUE | 0,0319071 | No |
| orf19.2896 | 1669 | 1318 | 1522 | 956 | 1425 | 1104 | 1109 | 1589 | 1520 | 1333 | 1440 | 1436 | 1267 | 1199 | 1136 | 1212 | 1317,88 | 1432 | 1204 | 0,841 | -0,251 | -4,399 | 1,09E-05 | 1,73E-05 | 0,0017681 | 0,003039 | 0,002354 | 0,002354 | TRUE | 0,0271411 | No |
| orf19.2897 | 112 | 80 | 105 | 69 | 57 | 27 | 43 | 45 | 102 | 81 | 99 | 104 | 51 | 29 | 44 | 34 | 68,04 | 96 | 40 | 0,412 | -1,279 | -6,656 | 2,81E-11 | 5,58E-11 | 0,0108318 | 0,021453 | 0,0181564 | 0,0181564 | TRUE | 0,2867167 | No |
| orf19.2898 | 1419 | 1360 | 1449 | 910 | 1285 | 971 | 1082 | 1436 | 1292 | 1375 | 1371 | 1367 | 1143 | 1054 | 1108 | 1095 | 1225,81 | 1351 | 1100 | 0,815 | -0,295 | -5,886 | 3,95E-09 | 7,22E-09 | 1,00E-08 | 0,003115 | 0,0015892 | 0,0015892 | TRUE | 0,0112358 | No |
| orf19.2899 | 112 | 90 | 121 | 75 | 129 | 102 | 103 | 123 | 102 | 91 | 115 | 113 | 115 | 111 | 105 | 94 | 105,62 | 105 | 106 | 1,011 | 0,016 | 0,115 | 0,90861 | 0,921621 | 1,00E-08 | 0,014544 | 0,0094632 | 0,0094632 | TRUE | 0,0799778 | No |
| orf19.29 | 5432 | 5613 | 5711 | 3754 | 2476 | 2185 | 2161 | 2611 | 4946 | 5677 | 5405 | 5639 | 2202 | 2372 | 2213 | 1992 | 3805,76 | 5417 | 2195 | 0,405 | -1,304 | -20,924 | 3,27E-97 | 2,77E-96 | 0,0040312 | 0,002384 | 0,0034174 | 0,0034174 | TRUE | 0,048073 | No |
| orf19.290 | 1005 | 913 | 1046 | 578 | 856 | 764 | 809 | 991 | 915 | 923 | 990 | 868 | 761 | 829 | 829 | 756 | 858,99 | 924 | 794 | 0,856 | -0,225 | -3,605 | 0,000312 | 0,000456 | 0,0015486 | 0,003575 | 0,0025727 | 0,0025727 | TRUE | 0,0273745 | No |
| orf19.2901 | 2995 | 2899 | 3018 | 2073 | 1728 | 1487 | 1451 | 1935 | 2727 | 2932 | 2856 | 3114 | 1537 | 1614 | 1486 | 1476 | 2217,82 | 2907 | 1528 | 0,526 | -0,927 | -17,449 | 3,49E-68 | 2,13E-67 | 0,0017402 | 0,002633 | 0,0022164 | 0,0022164 | TRUE | 0,0264277 | No |

| | | | | | | | | | | | | | | | | | | | | | | | | | | | | | | | |
|---|---|---|---|---|---|---|---|---|---|---|---|---|---|---|---|---|---|---|---|---|---|---|---|---|---|---|---|---|---|---|---|
| orf19.2903 | 195 | 270 | 237 | 138 | 778 | 662 | 655 | 845 | 178 | 273 | 224 | 207 | 692 | 719 | 671 | 645 | 451,04 | 220 | 682 | 3,086 | 1,626 | 15,491 | 4,00E-54 | 1,99E-53 | 0,011694 | 0,004966 | 0,0074654 | 0,0074654 | TRUE | 0,2922038 | No |
| orf19.2905 | 16 | 18 | 15 | 4 | 33 | 36 | 35 | 44 | 15 | 18 | 14 | 6 | 29 | 39 | 36 | 34 | 23,85 | 13 | 34 | 2,5 | 1,322 | 4,309 | 1,64E-05 | 2,57E-05 | 1,00E-08 | 0,057418 | 0,0381736 | 0,0381736 | TRUE | 0,3202702 | No |
| orf19.2906 | 161 | 123 | 158 | 111 | 181 | 134 | 115 | 199 | 147 | 124 | 150 | 167 | 161 | 145 | 118 | 152 | 145,41 | 147 | 144 | 0,99 | -0,015 | -0,114 | 0,909268 | 0,922142 | 0,0081011 | 0,011121 | 0,0099707 | 0,0099707 | TRUE | 0,1514538 | No |
| orf19.2907 | 4 | 6 | 6 | 7 | 8 | 5 | 5 | 3 | 4 | 6 | 6 | 11 | 7 | 5 | 5 | 2 | 5,73 | 7 | 5 | 0,79 | -0,341 | -0,572 | 0,567608 | 0,608663 | 1,00E-08 | 0,232491 | 0,1607089 | 0,1607089 | TRUE | 0,3884772 | No |
| orf19.2907.1 | 353 | 258 | 294 | 150 | 143 | 89 | 122 | 171 | 321 | 261 | 278 | 225 | 127 | 97 | 125 | 130 | 195,64 | 271 | 120 | 0,441 | -1,182 | -8,989 | 2,49E-19 | 6,22E-19 | 0,0136484 | 0,008789 | 0,010762 | 0,010762 | TRUE | 0,1564881 | No |
| orf19.2908 | 353 | 318 | 333 | 244 | 282 | 276 | 297 | 387 | 321 | 322 | 315 | 367 | 251 | 300 | 304 | 295 | 309,32 | 331 | 288 | 0,871 | -0,199 | -2,17 | 0,030015 | 0,037966 | 0,0026493 | 0,006308 | 0,0048173 | 0,0048173 | TRUE | 0,0849803 | No |
| orf19.2909 | 942 | 877 | 919 | 582 | 944 | 864 | 794 | 1089 | 858 | 887 | 870 | 874 | 840 | 938 | 813 | 831 | 863,78 | 872 | 856 | 0,98 | -0,03 | -0,494 | 0,621432 | 0,659721 | 0,0009568 | 0,003566 | 0,0022907 | 0,0022907 | TRUE | 0,0477019 | No |
| orf19.291 | 1536 | 1329 | 1452 | 908 | 2481 | 1939 | 2268 | 2928 | 1398 | 1344 | 1374 | 1364 | 2207 | 2105 | 2323 | 2234 | 1793,62 | 1370 | 2217 | 1,618 | 0,694 | 14,901 | 3,25E-50 | 1,50E-49 | 0,0005222 | 0,002774 | 0,0014815 | 0,0014815 | TRUE | 0,0133156 | No |
| orf19.2910 | 130 | 98 | 107 | 81 | 173 | 125 | 149 | 183 | 118 | 99 | 101 | 122 | 154 | 136 | 153 | 140 | 127,77 | 110 | 146 | 1,328 | 0,409 | 3,208 | 0,001338 | 0,001874 | 1,00E-08 | 0,012375 | 0,0075771 | 0,0075771 | TRUE | 0,0427684 | No |
| orf19.2911 | 2162 | 1930 | 2137 | 1251 | 1190 | 919 | 1038 | 1387 | 1968 | 1952 | 2022 | 1879 | 1058 | 998 | 1063 | 1058 | 1499,92 | 1955 | 1044 | 0,534 | -0,906 | -19,127 | 1,51E-81 | 1,09E-80 | 1,00E-08 | 0,002918 | 0,0014324 | 0,0014324 | TRUE | 0,01008 | No |
| orf19.2914 | 688 | 542 | 637 | 376 | 244 | 191 | 204 | 277 | 626 | 548 | 603 | 565 | 217 | 207 | 209 | 211 | 398,36 | 586 | 211 | 0,36 | -1,473 | -18,067 | 5,81E-73 | 3,78E-72 | 1,00E-08 | 0,005353 | 0,0032731 | 0,0032731 | TRUE | 0,0268047 | No |
| orf19.2915 | 3275 | 3220 | 3381 | 2047 | 1801 | 1476 | 1624 | 2033 | 2982 | 3257 | 3200 | 3075 | 1602 | 1603 | 1663 | 1551 | 2366,45 | 3128 | 1605 | 0,512 | -0,964 | -20,946 | 2,04E-97 | 1,74E-96 | 0,0008811 | 0,002595 | 0,001572 | 0,001572 | TRUE | 0,0128104 | No |
| orf19.2916 | 63 | 79 | 85 | 32 | 94 | 52 | 65 | 87 | 57 | 80 | 80 | 48 | 84 | 56 | 67 | 66 | 67,35 | 66 | 68 | 1,019 | 0,027 | 0,136 | 0,891857 | 0,907227 | 0,0258603 | 0,021652 | 0,023213 | 0,023213 | TRUE | 0,2265803 | No |
| orf19.2917 | 15512 | 13669 | 14351 | 8884 | 24594 | 20172 | 21670 | 29934 | 14123 | 13825 | 13582 | 13345 | 21876 | 21901 | 22193 | 22834 | 17959,8 | 13719 | 22201 | 1,618 | 0,694 | 21,702 | 1,96E-104 | 1,78E-103 | 0,0004179 | 0,002111 | 0,0009223 | 0,0009223 | TRUE | 0,0048275 | No |
| orf19.2917.1 | 715 | 550 | 585 | 455 | 218 | 161 | 169 | 268 | 651 | 556 | 554 | 683 | 194 | 175 | 173 | 204 | 398,82 | 611 | 186 | 0,307 | -1,702 | -17,164 | 4,93E-66 | 2,95E-65 | 0,0069354 | 0,005349 | 0,006068 | 0,006068 | TRUE | 0,0674353 | No |
| orf19.2919 | 599 | 558 | 585 | 355 | 613 | 531 | 568 | 747 | 545 | 564 | 554 | 533 | 545 | 577 | 582 | 570 | 558,74 | 549 | 568 | 1,033 | 0,047 | 0,738 | 0,460756 | 0,502622 | 1,00E-08 | 0,004401 | 0,0020368 | 0,0020368 | TRUE | 0,008853 | No |
| orf19.292 | 106 | 77 | 70 | 60 | 167 | 124 | 150 | 209 | 97 | 78 | 66 | 90 | 149 | 135 | 154 | 159 | 115,87 | 83 | 149 | 1,815 | 0,86 | 5,982 | 2,21E-09 | 4,08E-09 | 0,0037405 | 0,013437 | 0,0102418 | 0,0102418 | TRUE | 0,1840349 | No |
| orf19.2920 | 2723 | 2173 | 2371 | 1479 | 2698 | 2134 | 2141 | 3156 | 2479 | 2198 | 2244 | 2222 | 2400 | 2317 | 2193 | 2407 | 2307,42 | 2286 | 2329 | 1,019 | 0,027 | 0,5 | 0,617037 | 0,655821 | 0,0021155 | 0,00261 | 0,0023264 | 0,0023264 | TRUE | 0,0404516 | No |
| orf19.2921 | 105 | 121 | 114 | 80 | 213 | 167 | 194 | 246 | 96 | 122 | 108 | 120 | 189 | 181 | 199 | 188 | 150,39 | 112 | 189 | 1,709 | 0,773 | 6,448 | 1,13E-10 | 2,19E-10 | 1,00E-08 | 0,010821 | 0,0065507 | 0,0065507 | TRUE | 0,1004004 | No |
| orf19.2922 | 31 | 14 | 17 | 12 | 21 | 21 | 18 | 24 | 28 | 14 | 16 | 18 | 19 | 23 | 18 | 18 | 19,34 | 19 | 20 | 1,01 | 0,014 | 0,044 | 0,965232 | 0,96949 | 1,00E-08 | 0,070341 | 0,0489854 | 0,0489854 | TRUE | 0,6294989 | No |
| orf19.2923 | 288 | 260 | 245 | 170 | 294 | 244 | 215 | 315 | 262 | 263 | 232 | 255 | 262 | 265 | 220 | 240 | 249,91 | 253 | 247 | 0,975 | -0,036 | -0,378 | 0,705316 | 0,739538 | 1,00E-08 | 0,007323 | 0,0049413 | 0,0049413 | TRUE | 0,0540374 | No |
| orf19.2925 | 289 | 316 | 315 | 174 | 298 | 254 | 268 | 332 | 263 | 320 | 298 | 261 | 265 | 276 | 274 | 253 | 276,35 | 286 | 267 | 0,931 | -0,103 | -1,103 | 0,269971 | 0,306022 | 0,001943 | 0,006817 | 0,0048263 | 0,0048263 | TRUE | 0,0713045 | No |
| orf19.2926 | 2136 | 2073 | 2309 | 1310 | 3373 | 2776 | 2934 | 3737 | 1945 | 2097 | 2185 | 1968 | 3000 | 3014 | 3005 | 2851 | 2508 | 2049 | 2968 | 1,447 | 0,533 | 10,877 | 1,48E-27 | 4,49E-27 | 0,0013166 | 0,002564 | 0,0018831 | 0,0018831 | TRUE | 0,0248931 | No |
| orf19.2927 | 1507 | 1968 | 1961 | 1166 | 1336 | 1215 | 1151 | 1439 | 1372 | 1990 | 1856 | 1752 | 1188 | 1319 | 1179 | 1098 | 1469,23 | 1742 | 1196 | 0,686 | -0,543 | -5,48 | 4,26E-08 | 7,45E-08 | 0,0149097 | 0,002936 | 0,0087438 | 0,0087438 | TRUE | 0,251851 | No |
| orf19.2928 | 778 | 857 | 909 | 521 | 1054 | 998 | 1040 | 1210 | 708 | 867 | 860 | 783 | 937 | 1084 | 1065 | 923 | 903,39 | 804 | 1002 | 1,244 | 0,315 | 3,875 | 0,000107 | 0,00016 | 0,0069035 | 0,003499 | 0,0052013 | 0,0052013 | TRUE | 0,0818868 | No |
| orf19.2929 | 9400 | 9250 | 9451 | 6113 | 27997 | 24986 | 24819 | 35803 | 8558 | 9355 | 8945 | 9183 | 24902 | 27128 | 25418 | 27311 | 17600 | 9010 | 26190 | 2,907 | 1,54 | 34,044 | 4,90E-254 | 1,05E-252 | 0,001701 | 0,002112 | 0,001887 | 0,001887 | TRUE | 0,0140647 | No |
| orf19.2930 | 4404 | 4052 | 4384 | 2995 | 5153 | 4559 | 4353 | 5971 | 4010 | 4098 | 4149 | 4499 | 4583 | 4950 | 4458 | 4555 | 4412,73 | 4189 | 4636 | 1,108 | 0,148 | 2,962 | 0,003052 | 0,004151 | 0,0020903 | 0,002336 | 0,0021523 | 0,0021523 | TRUE | 0,0290395 | No |
| orf19.2933 | 1767 | 1719 | 1840 | 1082 | 1347 | 1034 | 1240 | 1533 | 1609 | 1739 | 1741 | 1625 | 1198 | 1123 | 1270 | 1169 | 1434,27 | 1678 | 1190 | 0,708 | -0,497 | -9,11 | 8,22E-20 | 2,08E-19 | 0,0012884 | 0,002958 | 0,0021474 | 0,0021474 | TRUE | 0,0235542 | No |
| orf19.2934 | 6255 | 6512 | 6561 | 4012 | 7638 | 6067 | 6149 | 7934 | 5695 | 6586 | 6209 | 6027 | 6794 | 6587 | 6297 | 6052 | 6280,93 | 6129 | 6432 | 1,049 | 0,069 | 1,277 | 0,201701 | 0,233365 | 0,0029043 | 0,002247 | 0,0026818 | 0,0026818 | TRUE | 0,0309529 | No |
| orf19.2935 | 26374 | 21971 | 23633 | 14688 | 41790 | 31071 | 34172 | 48007 | 24012 | 22221 | 22366 | 22064 | 37171 | 33734 | 34996 | 36620 | 29148,2 | 22666 | 35630 | 1,572 | 0,653 | 14,731 | 4,05E-49 | 1,84E-48 | 0,0017157 | 0,002082 | 0,001848 | 0,001848 | TRUE | 0,0196485 | No |
| orf19.2936 | 311 | 324 | 340 | 177 | 347 | 294 | 314 | 449 | 283 | 328 | 322 | 266 | 309 | 319 | 322 | 343 | 311,3 | 300 | 323 | 1,074 | 0,103 | 1,143 | 0,252992 | 0,288468 | 0,0020535 | 0,00628 | 0,0045787 | 0,0045787 | TRUE | 0,0520763 | No |
| orf19.2937 | 1402 | 1385 | 1392 | 783 | 2354 | 1817 | 1833 | 2317 | 1276 | 1401 | 1317 | 1176 | 2094 | 1973 | 1877 | 1767 | 1610,26 | 1292 | 1928 | 1,488 | 0,573 | 8,489 | 2,09E-17 | 4,95E-17 | 0,0043392 | 0,002857 | 0,0037238 | 0,0037238 | TRUE | 0,0415197 | No |
| orf19.2938 | 1498 | 1581 | 1616 | 993 | 1910 | 1448 | 1642 | 1953 | 1364 | 1599 | 1529 | 1492 | 1699 | 1572 | 1682 | 1490 | 1553,29 | 1496 | 1611 | 1,076 | 0,106 | 1,665 | 0,095937 | 0,115483 | 0,0035756 | 0,002887 | 0,0032597 | 0,0032597 | TRUE | 0,0445944 | No |
| orf19.2939 | 457 | 375 | 407 | 230 | 698 | 505 | 545 | 727 | 416 | 379 | 385 | 345 | 621 | 548 | 558 | 555 | 475,99 | 381 | 570 | 1,488 | 0,574 | 7,369 | 1,72E-13 | 3,66E-13 | 0,0020884 | 0,004812 | 0,003608 | 0,003608 | TRUE | 0,0424873 | No |
| orf19.2940 | 523 | 485 | 556 | 315 | 520 | 430 | 423 | 647 | 476 | 491 | 526 | 473 | 463 | 467 | 433 | 494 | 477,78 | 492 | 464 | 0,944 | -0,083 | -1,141 | 0,253807 | 0,289203 | 1,00E-08 | 0,004802 | 0,0029843 | 0,0029843 | TRUE | 0,0272772 | No |
| orf19.2941 | 3 | 7 | 5 | 3 | 18 | 20 | 12 | 29 | 3 | 7 | 5 | 5 | 16 | 22 | 12 | 22 | 11,4 | 5 | 18 | 3,82 | 1,934 | 4,048 | 5,17E-05 | 7,87E-05 | 1,00E-08 | 0,117932 | 0,0816421 | 0,0816421 | TRUE | 0,2209575 | No |
| orf19.2942 | 43201 | 38206 | 41933 | 25897 | 46887 | 39465 | 41386 | 56729 | 39333 | 38641 | 39686 | 38901 | 41705 | 42848 | 42384 | 43273 | 40846,4 | 39140 | 42552 | 1,087 | 0,121 | 4,612 | 4,00E-06 | 6,47E-06 | 0,0001616 | 0,002069 | 0,0006316 | 0,0006316 | TRUE | 0,0022149 | No |
| orf19.2943.5 | 2 | 2 | 1 | 3 | 2 | 2 | 0 | 5 | 2 | 2 | 1 | 5 | 2 | 2 | 0 | 4 | 2,13 | 2 | 2 | 0,914 | -0,129 | -0,132 | 0,895247 | 0,910195 | 1,00E-08 | 0,62146 | 0,446911 | 0,446911 | TRUE | 0,3486184 | No |
| orf19.2945 | 56 | 54 | 52 | 17 | 90 | 76 | 72 | 97 | 51 | 55 | 49 | 26 | 80 | 83 | 74 | 74 | 61,33 | 45 | 78 | 1,675 | 0,744 | 3,821 | 0,000133 | 0,000198 | 1,00E-08 | 0,023576 | 0,0189079 | 0,0189079 | TRUE | 0,446192 | No |
| orf19.2946 | 245 | 257 | 250 | 166 | 543 | 457 | 462 | 671 | 223 | 260 | 237 | 249 | 483 | 496 | 473 | 512 | 366,64 | 242 | 491 | 2,035 | 1,025 | 12,575 | 2,89E-36 | 1,06E-35 | 1,00E-08 | 0,00564 | 0,0032283 | 0,0032283 | TRUE | 0,0343632 | No |
| orf19.2947 | 4150 | 3605 | 3792 | 2438 | 1416 | 1136 | 1252 | 1603 | 3778 | 3646 | 3589 | 3662 | 1259 | 1233 | 1282 | 1223 | 2459,17 | 3669 | 1249 | 0,34 | -1,555 | -37,761 | 0 | 0 | 1,00E-08 | 0,002574 | 0,0011079 | 0,0011079 | TRUE | 0,0051648 | No |
| orf19.2948 | 1707 | 1436 | 1615 | 1078 | 472 | 353 | 404 | 528 | 1554 | 1452 | 1528 | 1619 | 420 | 383 | 414 | 403 | 971,74 | 1538 | 405 | 0,264 | -1,923 | -31,471 | 2,18E-217 | 3,98E-216 | 1,00E-08 | 0,003396 | 0,0020859 | 0,0020859 | TRUE | 0,0168352 | No |
| orf19.2949 | 1273 | 1268 | 1302 | 805 | 848 | 735 | 771 | 1075 | 1159 | 1282 | 1232 | 1209 | 754 | 798 | 790 | 820 | 1005,6 | 1220 | 790 | 0,648 | -0,626 | -11,201 | 4,01E-29 | 1,26E-28 | 0,0007184 | 0,003351 | 0,0019728 | 0,0019728 | TRUE | 0,0151204 | No |
| orf19.295 | 158 | 119 | 141 | 88 | 189 | 132 | 137 | 214 | 144 | 120 | 133 | 132 | 168 | 143 | 140 | 163 | 143,1 | 132 | 154 | 1,164 | 0,219 | 1,793 | 0,072962 | 0,088889 | 1,00E-08 | 0,011268 | 0,0072559 | 0,0072559 | TRUE | 0,0419671 | No |
| orf19.2951 | 699 | 637 | 691 | 468 | 769 | 595 | 581 | 848 | 636 | 644 | 654 | 703 | 684 | 646 | 595 | 647 | 651,19 | 659 | 643 | 0,979 | -0,031 | -0,468 | 0,63978 | 0,677733 | 0,0008928 | 0,004066 | 0,0026137 | 0,0026137 | TRUE | 0,0287697 | No |
| orf19.2952 | 568 | 493 | 566 | 375 | 616 | 477 | 563 | 680 | 517 | 499 | 536 | 563 | 548 | 518 | 577 | 519 | 534,48 | 529 | 540 | 1,024 | 0,035 | 0,496 | 0,620162 | 0,658795 | 0,0005778 | 0,004509 | 0,0027801 | 0,0027801 | TRUE | 0,0230529 | No |
| orf19.2953 | 2016 | 1753 | 1780 | 1150 | 3561 | 2822 | 2879 | 4013 | 1835 | 1773 | 1685 | 1727 | 3167 | 3064 | 2948 | 3061 | 2407,68 | 1755 | 3060 | 1,743 | 0,801 | 17,903 | 1,11E-71 | 7,12E-71 | 0,0006691 | 0,002586 | 0,0014656 | 0,0014656 | TRUE | 0,0120852 | No |
| orf19.2954 | 4402 | 3651 | 3861 | 2482 | 7182 | 5453 | 5932 | 8175 | 4008 | 3693 | 3654 | 3728 | 6388 | 5920 | 6075 | 6236 | 4962,82 | 3771 | 6155 | 1,632 | 0,707 | 16,103 | 2,44E-58 | 1,31E-57 | 0,0011641 | 0,002303 | 0,001631 | 0,001631 | TRUE | 0,0224908 | No |
| orf19.2956 | 1360 | 1200 | 1286 | 876 | 1543 | 1315 | 1409 | 1912 | 1238 | 1214 | 1217 | 1316 | 1372 | 1428 | 1443 | 1458 | 1335,82 | 1246 | 1425 | 1,146 | 0,197 | 4,039 | 5,38E-05 | 8,17E-05 | 0,0001731 | 0,003026 | 0,0015312 | 0,0015312 | TRUE | 0,0147663 | No |
| orf19.2957 | 19 | 29 | 25 | 15 | 23 | 25 | 16 | 37 | 17 | 29 | 24 | 23 | 20 | 27 | 16 | 28 | 23,13 | 23 | 23 | 1 | 0 | 0 | 0,999897 | 0,999897 | 1,00E-08 | 0,059151 | 0,0432903 | 0,0432903 | TRUE | 0,209269 | No |
| orf19.2958 | 399 | 328 | 386 | 240 | 520 | 354 | 427 | 582 | 363 | 332 | 365 | 361 | 463 | 384 | 437 | 444 | 393,62 | 355 | 432 | 1,219 | 0,286 | 3,509 | 0,00045 | 0,00065 | 0,0017058 | 0,005393 | 0,0037948 | 0,0037948 | TRUE | 0,0600965 | No |
| orf19.2959 | 7 | 7 | 8 | 3 | 8 | 6 | 9 | 11 | 6 | 7 | 8 | 5 | 7 | 7 | 9 | 8 | 7,1 | 6 | 8 | 1,203 | 0,267 | 0,536 | 0,592062 | 0,632545 | 1,00E-08 | 0,188195 | 0,0976813 | 0,0976813 | TRUE | 0,053973 | No |
| orf19.2959.1 | 2842 | 2460 | 2334 | 1436 | 1155 | 811 | 930 | 1267 | 2588 | 2488 | 2209 | 2157 | 1027 | 881 | 952 | 966 | 1658,54 | 2360 | 956 | 0,405 | -1,303 | -18,291 | 9,85E-75 | 6,51E-74 | 0,0049599 | 0,002833 | 0,0041583 | 0,0041583 | TRUE | 0,0510965 | No |
| orf19.296 | 53 | 48 | 53 | 39 | 44 | 27 | 49 | 48 | 48 | 49 | 50 | 59 | 39 | 29 | 50 | 37 | 45,1 | 52 | 39 | 0,761 | -0,394 | -1,845 | 0,065037 | 0,079702 | 1,00E-08 | 0,031328 | 0,0212958 | 0,0212958 | TRUE | 0,2644315 | No |
| orf19.2960 | 3469 | 3054 | 3227 | 2007 | 5831 | 4516 | 5008 | 6858 | 3158 | 3089 | 3054 | 3015 | 5186 | 4903 | 5129 | 5231 | 4095,73 | 3079 | 5112 | 1,66 | 0,731 | 19,406 | 6,81E-84 | 5,07E-83 | 0,0003939 | 0,00236 | 0,001097 | 0,001097 | TRUE | 0,0090974 | No |
| orf19.2961 | 720 | 663 | 719 | 420 | 1288 | 1009 | 1099 | 1569 | 656 | 671 | 680 | 631 | 1146 | 1095 | 1126 | 1197 | 900,12 | 660 | 1141 | 1,729 | 0,79 | 13,93 | 4,14E-44 | 1,74E-43 | 1,00E-08 | 0,003505 | 0,0018691 | 0,0018691 | TRUE | 0,0137096 | No |
| orf19.2962 | 916 | 899 | 972 | 572 | 148 | 117 | 144 | 161 | 834 | 909 | 920 | 859 | 132 | 127 | 147 | 123 | 506,42 | 880 | 132 | 0,15 | -2,741 | -31,548 | 1,90E-218 | 3,49E-217 | 1,00E-08 | 0,004646 | 0,0031184 | 0,0031184 | TRUE | 0,058904 | No |
| orf19.2963 | 156 | 140 | 156 | 73 | 160 | 131 | 155 | 224 | 142 | 142 | 148 | 110 | 142 | 142 | 159 | 171 | 144,38 | 136 | 154 | 1,127 | 0,172 | 1,362 | 0,173119 | 0,201798 | 0,00363 | 0,011186 | 0,0084429 | 0,0084429 | TRUE | 0,1278991 | No |
| orf19.2964 | 5263 | 4679 | 4934 | 3174 | 3623 | 2993 | 3247 | 4394 | 4792 | 4732 | 4670 | 4768 | 3223 | 3250 | 3325 | 3352 | 4013,84 | 4740 | 3288 | 0,694 | -0,528 | -15,696 | 1,61E-55 | 8,21E-55 | 1,00E-08 | 0,002366 | 0,0008292 | 0,0008292 | TRUE | 0,0022525 | No |
| orf19.2965 | 2193 | 2016 | 2040 | 1416 | 3583 | 3002 | 3140 | 4287 | 1997 | 2039 | 1931 | 2127 | 3187 | 3259 | 3216 | 3270 | 2628,19 | 2024 | 3233 | 1,601 | 0,679 | 16,408 | 1,69E-60 | 9,30E-60 | 0,0001939 | 0,00254 | 0,0012338 | 0,0012338 | TRUE | 0,0129123 | No |
| orf19.2965.1 | 443 | 410 | 434 | 249 | 472 | 352 | 342 | 462 | 403 | 415 | 411 | 374 | 420 | 382 | 350 | 352 | 388,43 | 401 | 376 | 0,936 | -0,096 | -1,153 | 0,248993 | 0,284009 | 0,0022688 | 0,005438 | 0,0040444 | 0,0040444 | TRUE | 0,0732494 | No |
| orf19.2966 | 268 | 265 | 241 | 135 | 602 | 452 | 511 | 621 | 244 | 268 | 228 | 203 | 535 | 491 | 523 | 474 | 370,77 | 236 | 506 | 2,13 | 1,091 | 12,055 | 1,81E-33 | 6,27E-33 | 0,0032266 | 0,0056 | 0,0046875 | 0,0046875 | TRUE | 0,0876643 | No |
| orf19.2967 | 8542 | 7044 | 7680 | 4656 | 13329 | 10011 | 11903 | 15804 | 7777 | 7124 | 7268 | 6994 | 11856 | 10869 | 12190 | 12055 | 9516,79 | 7291 | 11742 | 1,61 | 0,687 | 13,836 | 1,55E-43 | 6,46E-43 | 0,002256 | 0,002176 | 0,0022558 | 0,0022558 | TRUE | 0,0304869 | No |
| orf19.2968 | 352 | 318 | 331 | 191 | 416 | 293 | 342 | 439 | 320 | 322 | 313 | 287 | 370 | 318 | 350 | 335 | 326,94 | 310 | 343 | 1,102 | 0,14 | 1,669 | 0,095176 | 0,114632 | 1,00E-08 | 0,006077 | 0,0036883 | 0,0036883 | TRUE | 0,0318737 | No |
| orf19.2969 | 22894 | 19524 | 21950 | 13394 | 9201 | 7482 | 8628 | 11204 | 20844 | 19746 | 20774 | 20120 | 8184 | 8123 | 8836 | 8547 | 14396,8 | 20371 | 8422 | 0,413 | -1,274 | -32,566 | 1,26E-232 | 2,46E-231 | 0,0010071 | 0,002129 | 0,0013887 | 0,0013887 | TRUE | 0,0131871 | No |
| orf19.297 | 900 | 618 | 696 | 432 | 783 | 640 | 647 | 963 | 819 | 625 | 659 | 649 | 696 | 695 | 663 | 735 | 692,57 | 688 | 697 | 1,012 | 0,017 | 0,196 | 0,844253 | 0,866406 | 0,0073944 | 0,003944 | 0,0056405 | 0,0056405 | TRUE | 0,2024273 | No |
| orf19.2970 | 2833 | 2542 | 2639 | 1669 | 3569 | 3076 | 3278 | 4273 | 2579 | 2571 | 2498 | 2507 | 3175 | 3340 | 3357 | 3259 | 2910,72 | 2539 | 3283 | 1,292 | 0,37 | 9,654 | 4,74E-22 | 1,26E-21 | 1,00E-08 | 0,002491 | 0,0010532 | 0,0010532 | TRUE | 0,0059543 | No |

| | | | | | | | | | | | | | | | | | | | | | | | | | | | | | | | |
|---|---|---|---|---|---|---|---|---|---|---|---|---|---|---|---|---|---|---|---|---|---|---|---|---|---|---|---|---|---|---|---|
| orf19.2971 | 739 | 781 | 815 | 511 | 584 | 432 | 503 | 650 | 673 | 790 | 771 | 768 | 519 | 469 | 515 | 496 | 625,14 | 750 | 500 | 0,667 | -0,583 | -8,152 | 3,57E-16 | 8,14E-16 | 0,0023383 | 0,00415 | 0,0032655 | 0,0032655 | TRUE | 0,0610696 | No |
| orf19.2972 | 11936 | 11120 | 11897 | 7501 | 9998 | 8541 | 8966 | 11855 | 10867 | 11247 | 11259 | 11268 | 8893 | 9273 | 9182 | 9043 | 10129,1 | 11160 | 9098 | 0,815 | -0,295 | -9,639 | 5,50E-22 | 1,46E-21 | 0,0001874 | 0,002167 | 0,0007985 | 0,0007985 | TRUE | 0,0038895 | No |
| orf19.2973 | 7081 | 6572 | 7268 | 4692 | 4989 | 4525 | 4359 | 6081 | 6447 | 6647 | 6879 | 7048 | 4438 | 4913 | 4464 | 4639 | 5684,21 | 6755 | 4614 | 0,683 | -0,55 | -11,843 | 2,33E-32 | 7,89E-32 | 0,0016396 | 0,002269 | 0,0018887 | 0,0018887 | TRUE | 0,02286 | No |
| orf19.2974 | 1466 | 1459 | 1523 | 850 | 1721 | 1309 | 1462 | 1797 | 1335 | 1476 | 1441 | 1277 | 1531 | 1421 | 1497 | 1371 | 1418,57 | 1382 | 1455 | 1,051 | 0,071 | 1,173 | 0,240621 | 0,274755 | 0,0025329 | 0,002968 | 0,0028184 | 0,0028184 | TRUE | 0,0286665 | No |
| orf19.2975 | 1002 | 909 | 932 | 565 | 1763 | 1444 | 1590 | 2069 | 912 | 919 | 882 | 849 | 1568 | 1568 | 1628 | 1578 | 1238,12 | 890 | 1586 | 1,776 | 0,829 | 16,908 | 3,93E-64 | 2,28E-63 | 1,00E-08 | 0,003104 | 0,0014128 | 0,0014128 | TRUE | 0,0098674 | No |
| orf19.2977 | 5616 | 4918 | 5560 | 3655 | 5199 | 4540 | 4763 | 6042 | 5113 | 4974 | 5262 | 5490 | 4624 | 4929 | 4878 | 4609 | 4984,99 | 5210 | 4760 | 0,914 | -0,13 | -2,947 | 0,003209 | 0,004359 | 0,001247 | 0,002302 | 0,0016629 | 0,0016629 | TRUE | 0,0153299 | No |
| orf19.2978 | 1714 | 1373 | 1509 | 937 | 2180 | 1561 | 1820 | 2358 | 1561 | 1389 | 1428 | 1408 | 1939 | 1695 | 1864 | 1799 | 1635,16 | 1446 | 1824 | 1,261 | 0,334 | 5,731 | 9,98E-09 | 1,80E-08 | 0,002235 | 0,002845 | 0,0026337 | 0,0026337 | TRUE | 0,0352909 | No |
| orf19.298 | 41 | 31 | 41 | 25 | 39 | 34 | 45 | 50 | 37 | 31 | 39 | 38 | 35 | 37 | 46 | 38 | 37,61 | 36 | 39 | 1,073 | 0,101 | 0,455 | 0,649463 | 0,686963 | 1,00E-08 | 0,037162 | 0,0209291 | 0,0209291 | TRUE | 0,1063138 | No |
| orf19.2980 | 1748 | 1491 | 1590 | 1055 | 1736 | 1378 | 1556 | 1990 | 1591 | 1508 | 1505 | 1585 | 1544 | 1496 | 1594 | 1518 | 1542,6 | 1547 | 1538 | 0,995 | -0,008 | -0,169 | 0,865851 | 0,885719 | 1,00E-08 | 0,002893 | 0,0014064 | 0,0014064 | TRUE | 0,0068602 | No |
| orf19.2982 | 300 | 305 | 303 | 180 | 355 | 293 | 352 | 453 | 273 | 308 | 287 | 270 | 316 | 318 | 360 | 346 | 309,84 | 284 | 335 | 1,175 | 0,232 | 2,67 | 0,007581 | 0,010028 | 1,00E-08 | 0,006301 | 0,0040166 | 0,0040166 | TRUE | 0,0354032 | No |
| orf19.2983 | 1205 | 1038 | 1141 | 802 | 1732 | 1477 | 1607 | 2191 | 1097 | 1050 | 1080 | 1205 | 1541 | 1604 | 1646 | 1671 | 1361,6 | 1108 | 1616 | 1,463 | 0,549 | 9,966 | 2,15E-23 | 5,93E-23 | 0,0014646 | 0,003007 | 0,0021352 | 0,0021352 | TRUE | 0,0366264 | No |
| orf19.2984 | 1743 | 1468 | 1625 | 991 | 1274 | 974 | 1176 | 1544 | 1587 | 1485 | 1538 | 1489 | 1133 | 1057 | 1204 | 1178 | 1333,88 | 1525 | 1143 | 0,75 | -0,415 | -7,661 | 1,85E-14 | 4,03E-14 | 0,0010568 | 0,003027 | 0,0020626 | 0,0020626 | TRUE | 0,0289502 | No |
| orf19.2985 | 1329 | 1076 | 1191 | 775 | 1008 | 779 | 916 | 1242 | 1210 | 1088 | 1127 | 1164 | 897 | 846 | 938 | 947 | 1027,18 | 1147 | 907 | 0,791 | -0,337 | -5,744 | 9,25E-09 | 1,67E-08 | 0,0013898 | 0,003323 | 0,00233 | 0,00233 | TRUE | 0,023426 | No |
| orf19.2986 | 547 | 411 | 447 | 335 | 308 | 266 | 280 | 397 | 498 | 416 | 423 | 503 | 274 | 289 | 287 | 303 | 374,04 | 460 | 288 | 0,629 | -0,67 | -7,508 | 6,01E-14 | 1,29E-13 | 0,0040087 | 0,005569 | 0,0048548 | 0,0048548 | TRUE | 0,0478018 | No |
| orf19.2987 | 4280 | 3590 | 3913 | 2598 | 2992 | 2537 | 2685 | 3643 | 3897 | 3631 | 3703 | 3903 | 2661 | 2754 | 2750 | 2779 | 3259,75 | 3784 | 2736 | 0,724 | -0,467 | -11,464 | 2,00E-30 | 6,49E-30 | 0,0005692 | 0,002442 | 0,0012784 | 0,0012784 | TRUE | 0,0089146 | No |
| orf19.2988 | 2356 | 2005 | 2120 | 1461 | 787 | 683 | 717 | 956 | 2145 | 2028 | 2006 | 2195 | 700 | 742 | 734 | 729 | 1409,88 | 2094 | 726 | 0,347 | -1,526 | -29,075 | 7,43E-186 | 1,17E-184 | 0,0007425 | 0,002974 | 0,0017469 | 0,0017469 | TRUE | 0,0116685 | No |
| orf19.2989 | 4404 | 3676 | 4081 | 2654 | 3020 | 2492 | 2767 | 3633 | 4010 | 3718 | 3862 | 3987 | 2686 | 2706 | 2834 | 2771 | 3321,68 | 3894 | 2749 | 0,706 | -0,502 | -12,237 | 1,98E-34 | 6,99E-34 | 0,0006122 | 0,002435 | 0,0013062 | 0,0013062 | TRUE | 0,0113667 | No |
| orf19.299 | 2983 | 2681 | 2891 | 2089 | 2518 | 2113 | 2132 | 2942 | 2716 | 2712 | 2736 | 3138 | 2240 | 2294 | 2183 | 2244 | 2532,87 | 2826 | 2240 | 0,794 | -0,332 | -6,146 | 7,97E-10 | 1,50E-09 | 0,0023985 | 0,002559 | 0,0024043 | 0,0024043 | TRUE | 0,0639809 | No |
| orf19.2990 | 2176 | 1913 | 2027 | 1418 | 4850 | 4174 | 4228 | 6036 | 1981 | 1935 | 1918 | 2130 | 4314 | 4532 | 4330 | 4604 | 3218,05 | 1991 | 4445 | 2,237 | 1,162 | 25,113 | 3,59E-139 | 4,37E-138 | 0,0012209 | 0,002448 | 0,0016787 | 0,0016787 | TRUE | 0,0243002 | No |
| orf19.2991 | 123 | 93 | 104 | 90 | 239 | 220 | 204 | 264 | 112 | 94 | 98 | 135 | 213 | 239 | 209 | 201 | 162,68 | 110 | 216 | 1,981 | 0,986 | 7,8 | 6,19E-15 | 1,37E-14 | 0,0043783 | 0,010157 | 0,0082635 | 0,0082635 | TRUE | 0,1816659 | No |
| orf19.2992 | 6796 | 5684 | 5906 | 4283 | 12009 | 9862 | 9484 | 13428 | 6187 | 5749 | 5589 | 6434 | 10682 | 10707 | 9713 | 10243 | 8163,03 | 5990 | 10336 | 1,727 | 0,788 | 14,652 | 1,31E-48 | 5,87E-48 | 0,0029564 | 0,002199 | 0,0026421 | 0,0026421 | TRUE | 0,0295912 | No |
| orf19.2994 | 21751 | 17635 | 19237 | 12741 | 37175 | 28119 | 30050 | 41593 | 19803 | 17836 | 18206 | 19139 | 33066 | 30529 | 30775 | 31727 | 25135,3 | 18746 | 31524 | 1,682 | 0,75 | 16,816 | 1,86E-63 | 1,07E-62 | 0,0017436 | 0,00209 | 0,0018674 | 0,0018674 | TRUE | 0,0178279 | No |
| orf19.2994.1 | 12694 | 10396 | 10736 | 7298 | 20284 | 14724 | 15286 | 22190 | 11557 | 10514 | 10161 | 10963 | 18042 | 15986 | 15655 | 16927 | 13725,6 | 10799 | 16652 | 1,542 | 0,625 | 11,126 | 9,33E-29 | 2,91E-28 | 0,0035273 | 0,002133 | 0,0029536 | 0,0029536 | TRUE | 0,0386076 | No |
| orf19.2995 | 1175 | 947 | 1102 | 722 | 793 | 577 | 673 | 892 | 1070 | 958 | 1043 | 1085 | 705 | 626 | 689 | 680 | 857,07 | 1039 | 675 | 0,652 | -0,618 | -9,831 | 8,31E-23 | 2,25E-22 | 0,0015468 | 0,003578 | 0,0025861 | 0,0025861 | TRUE | 0,0331017 | No |
| orf19.2996 | 1800 | 1702 | 1918 | 1109 | 2311 | 2047 | 2051 | 2735 | 1639 | 1721 | 1815 | 1666 | 2056 | 2222 | 2100 | 2086 | 1913,26 | 1710 | 2116 | 1,236 | 0,305 | 6,2 | 5,64E-10 | 1,06E-09 | 0,0009324 | 0,002727 | 0,0017944 | 0,0017944 | TRUE | 0,0212514 | No |
| orf19.2998 | 2590 | 2077 | 2170 | 1361 | 2539 | 1933 | 2203 | 3011 | 2358 | 2101 | 2054 | 2044 | 2258 | 2099 | 2256 | 2297 | 2183,37 | 2139 | 2228 | 1,041 | 0,058 | 1,011 | 0,311834 | 0,350545 | 0,0025595 | 0,002642 | 0,0026445 | 0,0026445 | TRUE | 0,0587479 | No |
| orf19.30 | 2565 | 2389 | 2494 | 1575 | 4425 | 3777 | 3900 | 5345 | 2335 | 2416 | 2360 | 2366 | 3936 | 4101 | 3994 | 4077 | 3198,22 | 2369 | 4027 | 1,7 | 0,765 | 21,102 | 7,61E-99 | 6,56E-98 | 1,00E-08 | 0,00245 | 0,0009201 | 0,0009201 | TRUE | 0,0028661 | No |
| orf19.300 | 1720 | 1360 | 1463 | 998 | 2844 | 2475 | 2589 | 3522 | 1566 | 1375 | 1385 | 1499 | 2530 | 2687 | 2651 | 2687 | 2047,51 | 1456 | 2639 | 1,813 | 0,858 | 16,17 | 8,18E-59 | 4,40E-58 | 0,0016963 | 0,002682 | 0,0021576 | 0,0021576 | TRUE | 0,0337322 | No |
| orf19.3000 | 715 | 605 | 672 | 469 | 740 | 578 | 656 | 797 | 651 | 612 | 636 | 705 | 658 | 628 | 672 | 608 | 646,11 | 651 | 642 | 0,988 | -0,018 | -0,265 | 0,791002 | 0,819941 | 0,0009917 | 0,004082 | 0,0026685 | 0,0026685 | TRUE | 0,0309734 | No |
| orf19.3001 | 216 | 199 | 247 | 140 | 172 | 126 | 133 | 206 | 197 | 201 | 234 | 210 | 153 | 137 | 136 | 157 | 178,14 | 210 | 146 | 0,695 | -0,524 | -4,712 | 2,46E-06 | 4,01E-06 | 1,00E-08 | 0,009452 | 0,0061573 | 0,0061573 | TRUE | 0,0667256 | No |
| orf19.3002 | 14698 | 12055 | 12543 | 8012 | 28102 | 21288 | 21847 | 31723 | 13382 | 12192 | 11871 | 12035 | 24996 | 23113 | 22374 | 24199 | 18020,2 | 12370 | 23670 | 1,913 | 0,936 | 18,372 | 2,22E-75 | 1,48E-74 | 0,0026589 | 0,00211 | 0,0024308 | 0,0024308 | TRUE | 0,0373127 | No |
| orf19.3003 | 2850 | 2656 | 2765 | 1658 | 1444 | 1186 | 1359 | 1794 | 2595 | 2686 | 2617 | 2491 | 1284 | 1288 | 1392 | 1368 | 1965,1 | 2597 | 1333 | 0,513 | -0,963 | -20,408 | 1,42E-92 | 1,14E-91 | 0,0005862 | 0,002709 | 0,0015807 | 0,0015807 | TRUE | 0,0099147 | No |
| orf19.3003.1 | 14572 | 12058 | 12889 | 9273 | 20206 | 16455 | 15973 | 22250 | 13267 | 12195 | 12198 | 13930 | 17973 | 17866 | 16358 | 16972 | 15094,9 | 12898 | 17292 | 1,341 | 0,423 | 7,908 | 2,61E-15 | 5,84E-15 | 0,0030819 | 0,002125 | 0,0026852 | 0,0026852 | TRUE | 0,0350535 | No |
| orf19.3004 | 6796 | 6407 | 6991 | 4045 | 7033 | 5567 | 6385 | 7739 | 6187 | 6480 | 6616 | 6076 | 6256 | 6044 | 6539 | 5903 | 6262,79 | 6340 | 6186 | 0,975 | -0,036 | -0,796 | 0,4262 | 0,467325 | 0,0015349 | 0,002248 | 0,0018509 | 0,0018509 | TRUE | 0,0179855 | No |
| orf19.3006 | 3130 | 2872 | 3002 | 1994 | 2031 | 1726 | 1754 | 2472 | 2850 | 2905 | 2841 | 2995 | 1807 | 1874 | 1796 | 1886 | 2369,17 | 2898 | 1841 | 0,636 | -0,653 | -16,015 | 1,01E-57 | 5,32E-57 | 0,0001376 | 0,002595 | 0,0011563 | 0,0011563 | TRUE | 0,005712 | No |
| orf19.3007 | 759 | 605 | 664 | 423 | 442 | 364 | 345 | 502 | 691 | 612 | 628 | 635 | 393 | 395 | 353 | 383 | 511,42 | 642 | 381 | 0,594 | -0,752 | -10,424 | 1,92E-25 | 5,56E-25 | 1,00E-08 | 0,00462 | 0,0029445 | 0,0029445 | TRUE | 0,0340273 | No |
| orf19.3007.2 | 912 | 802 | 859 | 441 | 1167 | 810 | 964 | 1254 | 830 | 811 | 813 | 662 | 1038 | 879 | 987 | 957 | 872,27 | 779 | 965 | 1,234 | 0,304 | 3,903 | 9,48E-05 | 0,000142 | 0,00587 | 0,003551 | 0,0046455 | 0,0046455 | TRUE | 0,1084759 | No |
| orf19.3008 | 1449 | 1221 | 1343 | 805 | 1114 | 829 | 980 | 1235 | 1319 | 1235 | 1271 | 1209 | 991 | 900 | 1004 | 942 | 1108,88 | 1258 | 959 | 0,761 | -0,394 | -7,093 | 1,31E-12 | 2,71E-12 | 0,0008933 | 0,003228 | 0,0020483 | 0,0020483 | TRUE | 0,0190244 | No |
| orf19.3009 | 655 | 672 | 730 | 470 | 737 | 600 | 636 | 770 | 596 | 680 | 691 | 706 | 656 | 651 | 651 | 587 | 652,32 | 668 | 636 | 0,953 | -0,069 | -0,952 | 0,340864 | 0,381013 | 0,002839 | 0,004062 | 0,0034644 | 0,0034644 | TRUE | 0,0648461 | No |
| orf19.301 | 2355 | 1843 | 2078 | 1646 | 1047 | 946 | 884 | 1298 | 2144 | 1864 | 1967 | 2473 | 931 | 1027 | 905 | 990 | 1537,64 | 2112 | 963 | 0,457 | -1,13 | -13,632 | 2,59E-42 | 1,05E-41 | 0,0086247 | 0,002896 | 0,0058577 | 0,0058577 | TRUE | 0,1547656 | No |
| orf19.3010 | 602 | 468 | 500 | 277 | 623 | 494 | 509 | 707 | 548 | 473 | 473 | 416 | 554 | 536 | 521 | 539 | 507,73 | 478 | 538 | 1,12 | 0,163 | 2,014 | 0,044013 | 0,054801 | 0,0040984 | 0,004639 | 0,0043231 | 0,0043231 | TRUE | 0,1153006 | No |
| orf19.3010.1 | 9256 | 7967 | 8283 | 6021 | 10195 | 9429 | 8885 | 12651 | 8427 | 8058 | 7839 | 9045 | 9068 | 10237 | 9099 | 9650 | 8927,95 | 8342 | 9514 | 1,141 | 0,19 | 3,374 | 0,00074 | 0,001053 | 0,0035286 | 0,002185 | 0,0029358 | 0,0029358 | TRUE | 0,0386004 | No |
| orf19.3012 | 1251 | 1127 | 1248 | 866 | 1207 | 1062 | 1062 | 1461 | 1139 | 1140 | 1181 | 1301 | 1074 | 1153 | 1088 | 1114 | 1148,69 | 1190 | 1107 | 0,933 | -0,1 | -1,761 | 0,078155 | 0,094866 | 0,001394 | 0,003187 | 0,0022367 | 0,0022367 | TRUE | 0,0420551 | No |
| orf19.3013 | 1270 | 1113 | 1134 | 715 | 1550 | 1198 | 1419 | 1761 | 1156 | 1126 | 1073 | 1074 | 1379 | 1301 | 1453 | 1343 | 1238,14 | 1107 | 1369 | 1,235 | 0,304 | 5,627 | 1,84E-08 | 3,27E-08 | 0,0009772 | 0,003104 | 0,0019804 | 0,0019804 | TRUE | 0,019916 | No |
| orf19.3014 | 9551 | 8316 | 8868 | 6199 | 7421 | 6499 | 6362 | 8978 | 8696 | 8411 | 8393 | 9312 | 6601 | 7056 | 6515 | 6848 | 7729 | 8703 | 6755 | 0,777 | -0,365 | -7,976 | 1,51E-15 | 3,39E-15 | 0,0016999 | 0,002208 | 0,0018778 | 0,0018778 | TRUE | 0,0264804 | No |
| orf19.3015 | 11540 | 9842 | 10909 | 7214 | 14507 | 12193 | 12859 | 17568 | 10507 | 9954 | 10324 | 10837 | 12904 | 13238 | 13169 | 13401 | 11791,7 | 10406 | 13178 | 1,267 | 0,341 | 9,624 | 6,30E-22 | 1,67E-21 | 0,0006345 | 0,002149 | 0,0011209 | 0,0011209 | TRUE | 0,010356 | No |
| orf19.3016 | 1434 | 1138 | 1319 | 717 | 1691 | 1061 | 1396 | 1785 | 1306 | 1151 | 1248 | 1077 | 1504 | 1152 | 1430 | 1362 | 1278,66 | 1196 | 1362 | 1,138 | 0,186 | 2,196 | 0,028118 | 0,035615 | 0,0087502 | 0,00307 | 0,006122 | 0,006122 | TRUE | 0,126919 | No |
| orf19.3018 | 1072 | 961 | 1032 | 645 | 1324 | 1046 | 1160 | 1541 | 976 | 972 | 977 | 969 | 1178 | 1136 | 1188 | 1175 | 1071,29 | 974 | 1169 | 1,201 | 0,265 | 5,473 | 4,43E-08 | 7,74E-08 | 1,00E-08 | 0,00327 | 0,0012975 | 0,0012975 | TRUE | 0,0041607 | No |
| orf19.3019 | 414 | 359 | 385 | 223 | 1060 | 932 | 984 | 1414 | 377 | 363 | 364 | 335 | 943 | 1012 | 1008 | 1079 | 685,06 | 360 | 1010 | 2,797 | 1,484 | 21,196 | 1,05E-99 | 9,09E-99 | 0,0015906 | 0,003965 | 0,0027697 | 0,0027697 | TRUE | 0,0265191 | No |
| orf19.302 | 344 | 306 | 314 | 185 | 376 | 274 | 282 | 392 | 313 | 309 | 297 | 278 | 334 | 297 | 289 | 299 | 302,19 | 299 | 305 | 1,015 | 0,022 | 0,255 | 0,798385 | 0,825045 | 1,00E-08 | 0,006408 | 0,0039104 | 0,0039104 | TRUE | 0,0490128 | No |
| orf19.3021 | 2105 | 1788 | 1903 | 1197 | 760 | 658 | 634 | 907 | 1917 | 1808 | 1801 | 1798 | 676 | 714 | 649 | 692 | 1256,94 | 1831 | 683 | 0,373 | -1,424 | -27,094 | 1,15E-161 | 1,60E-160 | 1,00E-08 | 0,003088 | 0,0016765 | 0,0016765 | TRUE | 0,0124268 | No |
| orf19.3022 | 3917 | 3129 | 3530 | 2247 | 3760 | 2980 | 3201 | 4395 | 3566 | 3165 | 3341 | 3375 | 3344 | 3235 | 3278 | 3353 | 3332,21 | 3362 | 3302 | 0,983 | -0,025 | -0,571 | 0,567961 | 0,608837 | 0,0009415 | 0,002433 | 0,0016014 | 0,0016014 | TRUE | 0,0212548 | No |
| orf19.3023 | 1908 | 1522 | 1653 | 988 | 1748 | 1446 | 1507 | 2065 | 1737 | 1539 | 1564 | 1484 | 1555 | 1570 | 1543 | 1575 | 1571,04 | 1581 | 1561 | 0,985 | -0,022 | -0,395 | 0,692597 | 0,727757 | 0,0015086 | 0,002878 | 0,0022253 | 0,0022253 | TRUE | 0,0541617 | No |
| orf19.3026 | 3352 | 2780 | 3176 | 2058 | 2439 | 1885 | 2241 | 2951 | 3052 | 2812 | 3006 | 3091 | 2169 | 2047 | 2295 | 2251 | 2590,36 | 2990 | 2190 | 0,733 | -0,448 | -8,878 | 6,82E-19 | 1,68E-18 | 0,0015752 | 0,002547 | 0,0020475 | 0,0020475 | TRUE | 0,0229583 | No |
| orf19.3027 | 5752 | 5138 | 5548 | 3635 | 2306 | 2008 | 2017 | 2692 | 5237 | 5197 | 5251 | 5460 | 2051 | 2180 | 2066 | 2053 | 3686,86 | 5286 | 2088 | 0,395 | -1,341 | -34,72 | 3,98E-264 | 8,88E-263 | 0,0002442 | 0,002395 | 0,0011055 | 0,0011055 | TRUE | 0,0103406 | No |
| orf19.3029 | 3221 | 3017 | 3105 | 2089 | 9013 | 7679 | 7963 | 10847 | 2933 | 3051 | 2939 | 3138 | 8017 | 8337 | 8155 | 8274 | 5605,49 | 3015 | 8196 | 2,722 | 1,445 | 39,533 | 0 | 0 | 0,0003793 | 0,002273 | 0,0010465 | 0,0010465 | TRUE | 0,0083612 | No |
| orf19.3030 | 2078 | 1852 | 2059 | 1314 | 2088 | 1833 | 1903 | 2459 | 1892 | 1873 | 1949 | 1974 | 1857 | 1990 | 1949 | 1876 | 1919,94 | 1922 | 1918 | 0,998 | -0,003 | -0,066 | 0,947232 | 0,953816 | 0,0002947 | 0,002725 | 0,0013402 | 0,0013402 | TRUE | 0,0075855 | No |
| orf19.3031 | 1859 | 1452 | 1532 | 934 | 2782 | 1984 | 2455 | 3314 | 1693 | 1469 | 1450 | 1403 | 2475 | 2154 | 2514 | 2528 | 1960,59 | 1504 | 2418 | 1,607 | 0,684 | 9,613 | 7,02E-22 | 1,85E-21 | 0,005648 | 0,002711 | 0,0043098 | 0,0043098 | TRUE | 0,0877527 | No |
| orf19.3034 | 10893 | 9671 | 10661 | 6499 | 15026 | 12752 | 13453 | 18226 | 9918 | 9781 | 10090 | 9763 | 13365 | 13845 | 13778 | 13903 | 11805,2 | 9888 | 13723 | 1,388 | 0,473 | 15,969 | 2,12E-57 | 1,11E-56 | 0,0001589 | 0,002149 | 0,0007522 | 0,0007522 | TRUE | 0,0037896 | No |
| orf19.3035 | 5029 | 3987 | 4634 | 2806 | 4318 | 3487 | 3903 | 5219 | 4579 | 4032 | 4386 | 4215 | 3841 | 3786 | 3997 | 3981 | 4102,09 | 4303 | 3901 | 0,906 | -0,142 | -2,999 | 0,002711 | 0,003702 | 0,0015048 | 0,002359 | 0,0019042 | 0,0019042 | TRUE | 0,0230651 | No |
| orf19.3037 | 25733 | 21208 | 23228 | 15940 | 39843 | 34420 | 35853 | 50340 | 23429 | 21450 | 21983 | 23944 | 35439 | 37371 | 36718 | 38400 | 29841,7 | 22702 | 36982 | 1,629 | 0,704 | 15,525 | 2,36E-54 | 1,18E-53 | 0,0018589 | 0,002081 | 0,0019406 | 0,0019406 | TRUE | 0,016844 | No |
| orf19.3038 | 70134 | 61043 | 66769 | 41592 | 12025 | 10147 | 11018 | 14894 | 63854 | 61738 | 63191 | 62478 | 10696 | 11017 | 11284 | 11361 | 36952,4 | 62815 | 11090 | 0,177 | -2,502 | -79,967 | 0 | 0 | 0,000403 | 0,002073 | 0,0008896 | 0,0008896 | TRUE | 0,0070565 | No |
| orf19.304 | 1662 | 1502 | 1561 | 959 | 2912 | 2540 | 2731 | 3674 | 1513 | 1519 | 1477 | 1441 | 2590 | 2758 | 2797 | 2803 | 2112,19 | 1488 | 2737 | 1,838 | 0,878 | 19,301 | 5,29E-83 | 3,92E-82 | 0,0006294 | 0,002662 | 0,0014548 | 0,0014548 | TRUE | 0,0160583 | No |
| orf19.3040 | 6462 | 5643 | 6012 | 3867 | 6759 | 5448 | 6000 | 7953 | 5883 | 5707 | 5690 | 5809 | 6012 | 5915 | 6145 | 6067 | 5903,45 | 5772 | 6035 | 1,046 | 0,064 | 2,043 | 0,041089 | 0,051341 | 8,19E-05 | 0,002261 | 0,0007822 | 0,0007822 | TRUE | 0,0021775 | No |
| orf19.3041 | 7375 | 6559 | 7263 | 4379 | 8482 | 6859 | 7530 | 9875 | 6715 | 6634 | 6874 | 6578 | 7544 | 7447 | 7712 | 7533 | 7129,49 | 6700 | 7559 | 1,128 | 0,173 | 5,571 | 2,53E-08 | 4,48E-08 | 0,000104 | 0,002222 | 0,0007882 | 0,0007882 | TRUE | 0,0037375 | No |
| orf19.3042 | 207 | 206 | 203 | 164 | 152 | 146 | 150 | 160 | 188 | 208 | 192 | 246 | 135 | 159 | 154 | 122 | 175,58 | 208 | 142 | 0,683 | -0,55 | -4,394 | 1,11E-05 | 1,76E-05 | 0,0085093 | 0,00956 | 0,0091421 | 0,0091421 | TRUE | 0,1356328 | No |
| orf19.3043 | 415 | 374 | 385 | 289 | 474 | 403 | 458 | 580 | 378 | 378 | 364 | 434 | 422 | 438 | 469 | 442 | 415,65 | 388 | 443 | 1,145 | 0,196 | 2,509 | 0,012122 | 0,0158 | 0,0006802 | 0,005215 | 0,0033986 | 0,0033986 | TRUE | 0,0594631 | No |

| | | | | | | | | | | | | | | | | | | | | | | | | | | | | | | | |
|---|---|---|---|---|---|---|---|---|---|---|---|---|---|---|---|---|---|---|---|---|---|---|---|---|---|---|---|---|---|---|---|
| orf19.3045 | 2070 | 1896 | 2045 | 1304 | 2870 | 2445 | 2645 | 3473 | 1885 | 1918 | 1935 | 1959 | 2553 | 2655 | 2709 | 2649 | 2282,74 | 1924 | 2642 | 1,373 | 0,458 | 11,408 | 3,83E-30 | 1,24E-29 | 1,00E-08 | 0,002616 | 0,0010905 | 0,0010905 | TRUE | 0,0062309 | No |
| orf19.3047 | 5448 | 5084 | 5647 | 3470 | 1158 | 949 | 1019 | 1354 | 4960 | 5142 | 5344 | 5212 | 1030 | 1030 | 1044 | 1033 | 3099,47 | 5164 | 1034 | 0,2 | -2,32 | -54,152 | 0 | 0 | 1,00E-08 | 0,002463 | 0,0012074 | 0,0012074 | TRUE | 0,0089444 | No |
| orf19.3048 | 161 | 133 | 169 | 117 | 457 | 436 | 452 | 625 | 147 | 135 | 160 | 176 | 406 | 473 | 463 | 477 | 304,54 | 154 | 455 | 2,973 | 1,572 | 15,522 | 2,45E-54 | 1,23E-53 | 0,0037631 | 0,006375 | 0,0053403 | 0,0053403 | TRUE | 0,072876 | No |
| orf19.3049 | 66 | 50 | 44 | 40 | 97 | 82 | 84 | 123 | 60 | 51 | 42 | 60 | 86 | 89 | 86 | 94 | 70,94 | 53 | 89 | 1,688 | 0,755 | 4,406 | 1,05E-05 | 1,67E-05 | 1,00E-08 | 0,020657 | 0,0128679 | 0,0128679 | TRUE | 0,1898151 | No |
| orf19.305 | 419 | 403 | 433 | 256 | 564 | 471 | 487 | 652 | 381 | 408 | 410 | 385 | 502 | 511 | 499 | 497 | 449,07 | 396 | 502 | 1,267 | 0,341 | 4,925 | 8,44E-07 | 1,41E-06 | 1,00E-08 | 0,004979 | 0,0023314 | 0,0023314 | TRUE | 0,00852 | No |
| orf19.3050 | 3777 | 3618 | 3968 | 2156 | 1963 | 1569 | 1797 | 2298 | 3439 | 3659 | 3755 | 3239 | 1746 | 1703 | 1840 | 1753 | 2641,85 | 3523 | 1760 | 0,499 | -1,002 | -18,426 | 8,15E-76 | 5,47E-75 | 0,002119 | 0,002537 | 0,0024223 | 0,0024223 | TRUE | 0,0342002 | No |
| orf19.3051 | 1054 | 947 | 975 | 623 | 4822 | 4056 | 4471 | 5830 | 960 | 958 | 923 | 936 | 4289 | 4404 | 4579 | 4447 | 2686,84 | 944 | 4430 | 4,69 | 2,23 | 50,456 | 0 | 0 | 1,00E-08 | 0,002529 | 0,0012041 | 0,0012041 | TRUE | 0,0061675 | No |
| orf19.3052 | 8718 | 7438 | 7966 | 4471 | 8811 | 6661 | 7405 | 10129 | 7937 | 7523 | 7539 | 6716 | 7837 | 7232 | 7584 | 7726 | 7511,82 | 7429 | 7595 | 1,022 | 0,031 | 0,584 | 0,558976 | 0,600315 | 0,0025239 | 0,002213 | 0,0025899 | 0,0025899 | TRUE | 0,0498295 | No |
| orf19.3053 | 1732 | 1474 | 1508 | 1156 | 1347 | 1029 | 1043 | 1398 | 1577 | 1491 | 1427 | 1736 | 1198 | 1117 | 1068 | 1066 | 1335,16 | 1558 | 1112 | 0,716 | -0,483 | -7,083 | 1,41E-12 | 2,92E-12 | 0,0042117 | 0,003026 | 0,0036942 | 0,0036942 | TRUE | 0,067417 | No |
| orf19.3054 | 7752 | 6709 | 7329 | 4716 | 6688 | 5952 | 6056 | 8286 | 7058 | 6785 | 6936 | 7084 | 5949 | 6462 | 6202 | 6321 | 6599,68 | 6966 | 6234 | 0,895 | -0,16 | -4,278 | 1,89E-05 | 2,95E-05 | 0,0005983 | 0,002237 | 0,0011945 | 0,0011945 | TRUE | 0,0115918 | No |
| orf19.3055 | 1848 | 1666 | 1752 | 1145 | 1681 | 1399 | 1439 | 1852 | 1683 | 1685 | 1658 | 1720 | 1495 | 1519 | 1474 | 1413 | 1580,77 | 1686 | 1475 | 0,875 | -0,193 | -4,321 | 1,55E-05 | 2,44E-05 | 1,00E-08 | 0,002873 | 0,0012879 | 0,0012879 | TRUE | 0,0104339 | No |
| orf19.3057 | 742 | 578 | 690 | 366 | 483 | 417 | 422 | 557 | 676 | 585 | 653 | 550 | 430 | 453 | 432 | 425 | 525,3 | 616 | 435 | 0,702 | -0,51 | -6,549 | 5,78E-11 | 1,13E-10 | 0,0030908 | 0,004552 | 0,0038721 | 0,0038721 | TRUE | 0,0536315 | No |
| orf19.3058 | 1142 | 970 | 1024 | 702 | 1115 | 907 | 929 | 1359 | 1040 | 981 | 969 | 1055 | 992 | 985 | 951 | 1037 | 1001,13 | 1011 | 991 | 0,983 | -0,025 | -0,452 | 0,651441 | 0,688713 | 0,0005339 | 0,003357 | 0,0019167 | 0,0019167 | TRUE | 0,0121312 | No |
| orf19.3059 | 32410 | 26084 | 28912 | 18704 | 15586 | 12422 | 13936 | 19264 | 29508 | 26381 | 27363 | 28096 | 13863 | 13487 | 14272 | 14695 | 20958,1 | 27837 | 14079 | 0,506 | -0,983 | -21,923 | 1,58E-106 | 1,44E-105 | 0,0017321 | 0,0021 | 0,0018801 | 0,0018801 | TRUE | 0,0201123 | No |
| orf19.3059.1 | 0 | 4 | 1 | 2 | 13 | 10 | 5 | 7 | 0 | 4 | 1 | 3 | 12 | 11 | 5 | 5 | 5,11 | 2 | 8 | 4,347 | 2,12 | 2,765 | 0,005691 | 0,007595 | 0,1331346 | 0,260569 | 0,2258807 | 0,2258807 | TRUE | 0,3351669 | No |
| orf19.306 | 211 | 191 | 198 | 133 | 233 | 194 | 191 | 226 | 192 | 193 | 187 | 200 | 207 | 211 | 196 | 172 | 194,79 | 193 | 196 | 1,016 | 0,022 | 0,212 | 0,831876 | 0,855218 | 1,00E-08 | 0,008819 | 0,0054673 | 0,0054673 | TRUE | 0,0837107 | No |
| orf19.3060 | 963 | 829 | 850 | 519 | 1227 | 969 | 926 | 1271 | 877 | 838 | 804 | 780 | 1091 | 1052 | 948 | 970 | 920,07 | 825 | 1015 | 1,228 | 0,296 | 4,543 | 5,55E-06 | 8,92E-06 | 0,0025038 | 0,003473 | 0,0029777 | 0,0029777 | TRUE | 0,0313525 | No |
| orf19.3061 | 402 | 300 | 331 | 203 | 344 | 241 | 319 | 398 | 366 | 303 | 313 | 305 | 306 | 262 | 327 | 304 | 310,69 | 322 | 300 | 0,929 | -0,107 | -1,105 | 0,269337 | 0,305412 | 0,0050369 | 0,006289 | 0,0057522 | 0,0057522 | TRUE | 0,1019716 | No |
| orf19.3061.1 | 7120 | 6400 | 6402 | 4724 | 6120 | 4970 | 4703 | 6636 | 6482 | 6473 | 6059 | 7096 | 5444 | 5396 | 4816 | 5062 | 5853,57 | 6528 | 5180 | 0,794 | -0,333 | -5,787 | 7,17E-09 | 1,30E-08 | 0,0035416 | 0,002263 | 0,0030067 | 0,0030067 | TRUE | 0,0411441 | No |
| orf19.3062 | 996 | 1070 | 1073 | 689 | 1864 | 1515 | 1608 | 2122 | 907 | 1082 | 1015 | 1035 | 1658 | 1645 | 1647 | 1619 | 1325,98 | 1010 | 1642 | 1,628 | 0,703 | 12,201 | 3,08E-34 | 1,08E-33 | 0,0016838 | 0,003033 | 0,0023743 | 0,0023743 | TRUE | 0,0598039 | No |
| orf19.3063 | 786 | 626 | 662 | 451 | 851 | 640 | 768 | 981 | 716 | 633 | 627 | 677 | 757 | 695 | 787 | 748 | 704,92 | 663 | 747 | 1,127 | 0,172 | 2,558 | 0,01052 | 0,013785 | 0,0018476 | 0,003911 | 0,0029234 | 0,0029234 | TRUE | 0,0373167 | No |
| orf19.3064 | 3040 | 2479 | 2638 | 1653 | 2952 | 2246 | 2428 | 3415 | 2768 | 2507 | 2497 | 2483 | 2626 | 2439 | 2487 | 2605 | 2551,32 | 2564 | 2539 | 0,99 | -0,014 | -0,286 | 0,774505 | 0,804541 | 0,0016519 | 0,002555 | 0,0020267 | 0,0020267 | TRUE | 0,0357512 | No |
| orf19.3064.1 | 465 | 369 | 398 | 292 | 677 | 522 | 507 | 695 | 423 | 373 | 377 | 439 | 602 | 567 | 519 | 530 | 478,77 | 403 | 554 | 1,381 | 0,465 | 5,752 | 8,84E-09 | 1,60E-08 | 0,0032806 | 0,004796 | 0,0041136 | 0,0041136 | TRUE | 0,0404387 | No |
| orf19.3065 | 1035 | 935 | 1024 | 648 | 2778 | 2370 | 2360 | 3309 | 942 | 946 | 969 | 973 | 2471 | 2573 | 2417 | 2524 | 1726,96 | 958 | 2496 | 2,609 | 1,383 | 30,278 | 2,21E-201 | 3,80E-200 | 1,00E-08 | 0,002802 | 0,0012593 | 0,0012593 | TRUE | 0,005362 | No |
| orf19.3066 | 2177 | 1998 | 2210 | 1464 | 1723 | 1438 | 1514 | 2056 | 1982 | 2021 | 2092 | 2199 | 1533 | 1561 | 1551 | 1568 | 1813,28 | 2074 | 1553 | 0,751 | -0,414 | -8,947 | 3,67E-19 | 9,09E-19 | 0,0004999 | 0,002766 | 0,0014953 | 0,0014953 | TRUE | 0,0183223 | No |
| orf19.307 | 2587 | 2168 | 2440 | 1674 | 1711 | 1487 | 1649 | 2226 | 2355 | 2193 | 2309 | 2515 | 1522 | 1614 | 1689 | 1698 | 1986,88 | 2343 | 1631 | 0,697 | -0,521 | -9,304 | 1,36E-20 | 3,50E-20 | 0,0021799 | 0,002702 | 0,0024917 | 0,0024917 | TRUE | 0,0276402 | No |
| orf19.3071 | 7979 | 6977 | 7846 | 5231 | 2888 | 2697 | 2888 | 3741 | 7265 | 7056 | 7426 | 7858 | 2569 | 2928 | 2958 | 2854 | 5114,08 | 7401 | 2827 | 0,382 | -1,388 | -25,681 | 1,91E-145 | 2,43E-144 | 0,0027035 | 0,002295 | 0,0025688 | 0,0025688 | TRUE | 0,0461418 | No |
| orf19.3073 | 0 | 0 | 0 | 0 | 0 | 0 | 0 | 0 | 0 | 0 | 0 | 0 | 0 | 0 | 0 | 0 | 0 | 0 | 0 | NA | NA | NA | NA | NA | NA | NA | NA | NA | NA | NA | NA |
| orf19.3074 | 131 | 105 | 135 | 84 | 178 | 102 | 140 | 201 | 119 | 106 | 128 | 126 | 158 | 111 | 143 | 153 | 130,65 | 120 | 141 | 1,189 | 0,249 | 1,802 | 0,071507 | 0,087212 | 0,0081971 | 0,012148 | 0,0106173 | 0,0106173 | TRUE | 0,2065652 | No |
| orf19.3076 | 859 | 680 | 728 | 410 | 1519 | 935 | 1184 | 1698 | 782 | 688 | 689 | 616 | 1351 | 1015 | 1213 | 1295 | 956,1 | 694 | 1218 | 1,754 | 0,81 | 8,674 | 4,19E-18 | 1,01E-17 | 0,0105953 | 0,003419 | 0,0072264 | 0,0072264 | TRUE | 0,1492329 | No |
| orf19.3077 | 1978 | 1515 | 1807 | 1255 | 2571 | 2058 | 2484 | 3440 | 1801 | 1532 | 1710 | 1885 | 2287 | 2234 | 2544 | 2624 | 2077,22 | 1732 | 2422 | 1,4 | 0,486 | 6,642 | 3,10E-11 | 6,14E-11 | 0,0062507 | 0,002673 | 0,004636 | 0,004636 | TRUE | 0,0727245 | No |
| orf19.308 | 1505 | 1358 | 1386 | 999 | 1470 | 1252 | 1325 | 1845 | 1370 | 1373 | 1312 | 1501 | 1308 | 1359 | 1357 | 1407 | 1373,41 | 1389 | 1358 | 0,98 | -0,028 | -0,531 | 0,595077 | 0,635127 | 0,0011242 | 0,002999 | 0,0020249 | 0,0020249 | TRUE | 0,0316995 | No |
| orf19.3080 | 1504 | 1116 | 1390 | 976 | 1392 | 1156 | 1291 | 1822 | 1369 | 1129 | 1316 | 1466 | 1238 | 1255 | 1322 | 1390 | 1310,61 | 1320 | 1301 | 0,988 | -0,017 | -0,229 | 0,818599 | 0,843064 | 0,0064858 | 0,003045 | 0,0047867 | 0,0047867 | TRUE | 0,1140791 | No |
| orf19.3083 | 2265 | 1788 | 2188 | 1480 | 620 | 503 | 602 | 803 | 2062 | 1808 | 2071 | 2223 | 551 | 546 | 617 | 613 | 1311,39 | 2041 | 582 | 0,286 | -1,808 | -24,803 | 8,32E-136 | 9,71E-135 | 0,0049928 | 0,003044 | 0,0040377 | 0,0040377 | TRUE | 0,0711196 | No |
| orf19.3084 | 2281 | 1752 | 1942 | 1114 | 539 | 413 | 518 | 677 | 2077 | 1772 | 1838 | 1673 | 479 | 448 | 530 | 516 | 1166,85 | 1840 | 493 | 0,268 | -1,899 | -24,294 | 2,29E-130 | 2,55E-129 | 0,0060949 | 0,003169 | 0,0046276 | 0,0046276 | TRUE | 0,0913238 | No |
| orf19.3086 | 2221 | 1752 | 1873 | 1160 | 2233 | 1811 | 2107 | 2730 | 2022 | 1772 | 1773 | 1743 | 1986 | 1966 | 2158 | 2082 | 1937,74 | 1828 | 2048 | 1,12 | 0,163 | 2,771 | 0,005592 | 0,007465 | 0,0028091 | 0,002719 | 0,0027997 | 0,0027997 | TRUE | 0,0637433 | No |
| orf19.3087 | 17946 | 13740 | 14644 | 10516 | 32172 | 25473 | 25989 | 38668 | 16339 | 13897 | 13859 | 15797 | 28616 | 27657 | 26616 | 29496 | 21534,6 | 14973 | 28096 | 1,877 | 0,908 | 14,908 | 2,92E-50 | 1,35E-49 | 0,0045815 | 0,002098 | 0,0035132 | 0,0035132 | TRUE | 0,0465596 | No |
| orf19.3087.1 | 195 | 125 | 132 | 90 | 178 | 144 | 146 | 206 | 178 | 126 | 125 | 135 | 158 | 156 | 150 | 157 | 148,18 | 141 | 155 | 1,098 | 0,135 | 1,02 | 0,307528 | 0,345949 | 0,0084662 | 0,010952 | 0,0099421 | 0,0099421 | TRUE | 0,3464426 | No |
| orf19.3087.2 | 417 | 348 | 341 | 281 | 460 | 423 | 373 | 556 | 380 | 352 | 323 | 422 | 409 | 459 | 382 | 424 | 393,87 | 369 | 418 | 1,14 | 0,189 | 2,037 | 0,04162 | 0,051953 | 0,0061249 | 0,005391 | 0,0056826 | 0,0056826 | TRUE | 0,0922607 | No |
| orf19.3088 | 2957 | 2450 | 2577 | 1644 | 3057 | 2241 | 2353 | 3383 | 2692 | 2478 | 2439 | 2470 | 2719 | 2433 | 2410 | 2581 | 2527,64 | 2520 | 2536 | 1,006 | 0,009 | 0,172 | 0,863625 | 0,884152 | 0,0022784 | 0,00256 | 0,0023964 | 0,0023964 | TRUE | 0,0286554 | No |
| orf19.3089 | 7642 | 6313 | 6747 | 3775 | 5608 | 4393 | 4945 | 6536 | 6958 | 6385 | 6385 | 5671 | 4988 | 4770 | 5064 | 4986 | 5650,81 | 6350 | 4952 | 0,779 | -0,36 | -6,214 | 5,18E-10 | 9,79E-10 | 0,0035336 | 0,002271 | 0,0030391 | 0,0030391 | TRUE | 0,0618735 | No |
| orf19.309 | 375 | 315 | 362 | 242 | 190 | 151 | 164 | 219 | 341 | 319 | 343 | 364 | 169 | 164 | 168 | 167 | 254,26 | 342 | 167 | 0,491 | -1,027 | -11,166 | 6,01E-29 | 1,88E-28 | 1,00E-08 | 0,007232 | 0,0037803 | 0,0037803 | TRUE | 0,0224446 | No |
| orf19.3091 | 4879 | 4260 | 4653 | 2808 | 6187 | 5174 | 5426 | 7714 | 4442 | 4309 | 4404 | 4218 | 5503 | 5618 | 5557 | 5884 | 4991,78 | 4343 | 5640 | 1,298 | 0,377 | 10,003 | 1,47E-23 | 4,09E-23 | 0,0004456 | 0,002302 | 0,0011534 | 0,0011534 | TRUE | 0,0106233 | No |
| orf19.3093 | 2796 | 2320 | 2559 | 1712 | 3397 | 2737 | 2994 | 4296 | 2546 | 2346 | 2422 | 2572 | 3022 | 2972 | 3066 | 3277 | 2777,75 | 2472 | 3084 | 1,25 | 0,321 | 6,582 | 4,63E-11 | 9,10E-11 | 0,0013801 | 0,002513 | 0,0019197 | 0,0019197 | TRUE | 0,0221203 | No |
| orf19.3097 | 14516 | 11428 | 12973 | 7962 | 16173 | 11857 | 13768 | 19187 | 13216 | 11558 | 12278 | 11960 | 14385 | 12873 | 14100 | 14636 | 13125,9 | 12253 | 13998 | 1,142 | 0,192 | 3,542 | 0,000397 | 0,000576 | 0,0031179 | 0,002138 | 0,0027497 | 0,0027497 | TRUE | 0,0357192 | No |
| orf19.3098 | 1577 | 1434 | 1503 | 972 | 2329 | 2009 | 2033 | 2855 | 1436 | 1450 | 1422 | 1460 | 2072 | 2181 | 2082 | 2178 | 1785,17 | 1442 | 2128 | 1,477 | 0,562 | 13,032 | 8,01E-39 | 3,06E-38 | 1,00E-08 | 0,002777 | 0,0011969 | 0,0011969 | TRUE | 0,0040031 | No |
| orf19.3099 | 1047 | 869 | 972 | 652 | 1098 | 933 | 980 | 1308 | 953 | 879 | 920 | 979 | 977 | 1013 | 1004 | 998 | 965,31 | 933 | 998 | 1,071 | 0,099 | 1,844 | 0,065189 | 0,079873 | 1,00E-08 | 0,003405 | 0,0017374 | 0,0017374 | TRUE | 0,0176873 | No |
| orf19.31 | 210 | 157 | 153 | 112 | 437 | 351 | 429 | 606 | 191 | 159 | 145 | 168 | 389 | 381 | 439 | 462 | 291,8 | 166 | 418 | 2,524 | 1,336 | 12,316 | 7,39E-35 | 2,64E-34 | 0,0074948 | 0,006564 | 0,0069595 | 0,0069595 | TRUE | 0,1301528 | No |
| orf19.310 | 41 | 25 | 34 | 21 | 259 | 189 | 217 | 312 | 37 | 25 | 32 | 32 | 230 | 205 | 222 | 238 | 127,77 | 32 | 224 | 7,076 | 2,823 | 16,938 | 2,35E-64 | 1,37E-63 | 1,00E-08 | 0,012376 | 0,0079516 | 0,0079516 | TRUE | 0,1351781 | No |
| orf19.3100 | 265 | 190 | 241 | 150 | 405 | 337 | 427 | 511 | 241 | 192 | 228 | 225 | 360 | 366 | 437 | 390 | 305,01 | 222 | 388 | 1,75 | 0,807 | 8,128 | 4,37E-16 | 9,96E-16 | 0,0051663 | 0,006368 | 0,0058578 | 0,0058578 | TRUE | 0,0930554 | No |
| orf19.3102 | 1733 | 1469 | 1581 | 989 | 897 | 658 | 819 | 1052 | 1578 | 1486 | 1496 | 1486 | 798 | 714 | 839 | 802 | 1149,87 | 1512 | 788 | 0,522 | -0,938 | -16,246 | 2,37E-59 | 1,29E-58 | 0,0013737 | 0,003186 | 0,0022557 | 0,0022557 | TRUE | 0,0441614 | No |
| orf19.3103 | 6955 | 5874 | 6537 | 3873 | 9890 | 8284 | 9318 | 12341 | 6332 | 5941 | 6187 | 5818 | 8797 | 8994 | 9543 | 9414 | 7628,16 | 6070 | 9187 | 1,513 | 0,597 | 13,865 | 1,03E-43 | 4,31E-43 | 0,001279 | 0,00221 | 0,0016422 | 0,0016422 | TRUE | 0,0103677 | No |
| orf19.3104 | 1316 | 1161 | 1297 | 706 | 1694 | 1384 | 1486 | 2029 | 1198 | 1174 | 1227 | 1061 | 1507 | 1503 | 1522 | 1548 | 1342,42 | 1165 | 1520 | 1,3 | 0,379 | 7,196 | 6,22E-13 | 1,30E-12 | 0,0006567 | 0,003021 | 0,0018932 | 0,0018932 | TRUE | 0,0382189 | No |
| orf19.3105 | 950 | 787 | 890 | 595 | 2518 | 2320 | 2193 | 3177 | 865 | 796 | 842 | 894 | 2240 | 2519 | 2246 | 2423 | 1603,11 | 849 | 2357 | 2,78 | 1,475 | 24,974 | 1,17E-137 | 1,39E-136 | 0,0022257 | 0,002861 | 0,0025179 | 0,0025179 | TRUE | 0,0251107 | No |
| orf19.3106 | 1358 | 1124 | 1185 | 878 | 1574 | 1296 | 1516 | 1941 | 1236 | 1137 | 1121 | 1319 | 1400 | 1407 | 1553 | 1481 | 1331,74 | 1203 | 1460 | 1,216 | 0,282 | 4,482 | 7,41E-06 | 1,18E-05 | 0,0031357 | 0,003029 | 0,0030447 | 0,0030447 | TRUE | 0,0456015 | No |
| orf19.3107 | 97 | 102 | 94 | 45 | 301 | 227 | 249 | 289 | 88 | 103 | 89 | 68 | 268 | 246 | 255 | 220 | 167,21 | 87 | 247 | 2,802 | 1,487 | 11,471 | 1,84E-30 | 5,99E-30 | 0,0050261 | 0,009937 | 0,008205 | 0,008205 | TRUE | 0,1499361 | No |
| orf19.3108 | 407 | 371 | 367 | 198 | 522 | 385 | 472 | 607 | 371 | 375 | 347 | 297 | 464 | 418 | 483 | 463 | 402,41 | 348 | 457 | 1,308 | 0,387 | 4,524 | 6,08E-06 | 9,75E-06 | 0,0031696 | 0,00532 | 0,0044761 | 0,0044761 | TRUE | 0,0886768 | No |
| orf19.3109 | 27 | 26 | 42 | 20 | 78 | 67 | 60 | 75 | 25 | 26 | 40 | 30 | 69 | 73 | 61 | 57 | 47,68 | 30 | 65 | 2,153 | 1,106 | 5,058 | 4,24E-07 | 7,15E-07 | 1,00E-08 | 0,029742 | 0,0211522 | 0,0211522 | TRUE | 0,3532649 | No |
| orf19.3110 | 3812 | 3332 | 3537 | 2156 | 5399 | 4500 | 4840 | 6382 | 3471 | 3370 | 3347 | 3239 | 4802 | 4886 | 4957 | 4868 | 4117,47 | 3357 | 4878 | 1,451 | 0,538 | 15,074 | 2,38E-51 | 1,13E-50 | 0,0001516 | 0,002358 | 0,000965 | 0,000965 | TRUE | 0,0063499 | No |
| orf19.3111 | 51 | 38 | 50 | 37 | 80 | 53 | 72 | 88 | 46 | 38 | 47 | 56 | 71 | 58 | 74 | 67 | 57,17 | 47 | 68 | 1,454 | 0,54 | 2,871 | 0,004087 | 0,005506 | 1,00E-08 | 0,025145 | 0,015563 | 0,015563 | TRUE | 0,1126936 | No |
| orf19.3112 | 513 | 486 | 535 | 348 | 578 | 505 | 558 | 786 | 467 | 492 | 506 | 523 | 514 | 548 | 571 | 600 | 527,64 | 497 | 558 | 1,128 | 0,173 | 2,374 | 0,017604 | 0,022691 | 0,0017244 | 0,004541 | 0,003204 | 0,003204 | TRUE | 0,035276 | No |
| orf19.3113 | 501 | 413 | 471 | 304 | 452 | 429 | 397 | 621 | 456 | 418 | 446 | 457 | 402 | 466 | 407 | 474 | 440,54 | 444 | 437 | 0,986 | -0,021 | -0,258 | 0,796457 | 0,82332 | 0,002654 | 0,005036 | 0,0039462 | 0,0039462 | TRUE | 0,041152 | No |
| orf19.3114 | 884 | 757 | 796 | 497 | 1022 | 696 | 842 | 1120 | 805 | 766 | 753 | 747 | 909 | 756 | 862 | 854 | 806,47 | 768 | 845 | 1,101 | 0,139 | 2,091 | 0,036491 | 0,045803 | 0,0023452 | 0,003675 | 0,0030173 | 0,0030173 | TRUE | 0,0578295 | No |
| orf19.3115 | 838 | 778 | 800 | 496 | 947 | 688 | 824 | 1048 | 763 | 787 | 757 | 745 | 842 | 747 | 844 | 799 | 785,58 | 763 | 808 | 1,059 | 0,083 | 1,371 | 0,170482 | 0,198906 | 0,0005309 | 0,003719 | 0,0022079 | 0,0022079 | TRUE | 0,0286259 | No |
| orf19.3116 | 1793 | 1444 | 1593 | 787 | 1092 | 781 | 1015 | 1277 | 1632 | 1460 | 1508 | 1182 | 971 | 848 | 1039 | 974 | 1201,95 | 1446 | 958 | 0,662 | -0,596 | -6,492 | 8,46E-11 | 1,65E-10 | 0,0109673 | 0,003136 | 0,007232 | 0,007232 | TRUE | 0,1731221 | No |
| orf19.3117 | 318 | 240 | 215 | 165 | 34 | 34 | 35 | 47 | 290 | 243 | 203 | 248 | 30 | 37 | 36 | 36 | 140,31 | 246 | 35 | 0,141 | -2,828 | -16,754 | 5,25E-63 | 3,00E-62 | 1,00E-08 | 0,011452 | 0,011775 | 0,011775 | TRUE | 0,0943788 | No |
| orf19.3120 | 40 | 29 | 41 | 26 | 37 | 27 | 36 | 43 | 36 | 29 | 39 | 39 | 33 | 29 | 37 | 33 | 34,44 | 36 | 33 | 0,923 | -0,116 | -0,503 | 0,615146 | 0,654545 | 1,00E-08 | 0,040396 | 0,0224156 | 0,0224156 | TRUE | 0,1107154 | No |

| | | | | | | | | | | | | | | | | | | | | | | | | | | | | | | | |
|---|---|---|---|---|---|---|---|---|---|---|---|---|---|---|---|---|---|---|---|---|---|---|---|---|---|---|---|---|---|---|---|
| orf19.3121 | 40 | 35 | 40 | 25 | 53 | 49 | 39 | 72 | 36 | 35 | 38 | 38 | 47 | 53 | 40 | 55 | 42,8 | 37 | 49 | 1,332 | 0,413 | 1,957 | 0,050372 | 0,062377 | 1,00E-08 | 0,032899 | 0,0188754 | 0,0188754 | TRUE | 0,1144285 | No |
| orf19.3122 | 1464 | 1229 | 1344 | 875 | 1692 | 1393 | 1385 | 1938 | 1333 | 1243 | 1272 | 1314 | 1505 | 1512 | 1418 | 1478 | 1384,55 | 1290 | 1478 | 1,146 | 0,197 | 4,094 | 4,24E-05 | 6,50E-05 | 1,00E-08 | 0,002991 | 0,0014907 | 0,0014907 | TRUE | 0,0086555 | No |
| orf19.3122.2 | 2489 | 1882 | 1914 | 1233 | 1590 | 1029 | 1136 | 1681 | 2266 | 1903 | 1811 | 1852 | 1414 | 1117 | 1163 | 1282 | 1601,29 | 1958 | 1244 | 0,636 | -0,654 | -7,524 | 5,30E-14 | 1,14E-13 | 0,0105078 | 0,002862 | 0,0066036 | 0,0066036 | TRUE | 0,1372909 | No |
| orf19.3123 | 9063 | 7261 | 7977 | 5172 | 6125 | 4978 | 5386 | 7352 | 8251 | 7344 | 7550 | 7769 | 5448 | 5405 | 5516 | 5608 | 6611,34 | 7728 | 5494 | 0,711 | -0,492 | -11,422 | 3,24E-30 | 1,05E-29 | 0,001193 | 0,002237 | 0,0016281 | 0,0016281 | TRUE | 0,0258864 | No |
| orf19.3123.2 | 308 | 242 | 253 | 182 | 309 | 241 | 266 | 356 | 280 | 245 | 239 | 273 | 275 | 262 | 272 | 272 | 264,81 | 259 | 270 | 1,044 | 0,062 | 0,684 | 0,493847 | 0,535421 | 1,00E-08 | 0,007025 | 0,0041047 | 0,0041047 | TRUE | 0,0401271 | No |
| orf19.3124 | 6407 | 5283 | 5760 | 3975 | 7311 | 6194 | 6505 | 8929 | 5833 | 5343 | 5451 | 5971 | 6503 | 6725 | 6662 | 6811 | 6162,48 | 5650 | 6675 | 1,182 | 0,242 | 5,468 | 4,56E-08 | 7,97E-08 | 0,0013836 | 0,002251 | 0,0017088 | 0,0017088 | TRUE | 0,0171534 | No |
| orf19.3125 | 540 | 491 | 508 | 315 | 337 | 294 | 311 | 381 | 492 | 497 | 481 | 473 | 300 | 319 | 319 | 291 | 396,29 | 486 | 307 | 0,63 | -0,667 | -8,971 | 2,94E-19 | 7,32E-19 | 1,00E-08 | 0,00537 | 0,0026909 | 0,0026909 | TRUE | 0,015985 | No |
| orf19.3126 | 5261 | 4631 | 5049 | 3184 | 5646 | 4670 | 4986 | 6787 | 4790 | 4684 | 4778 | 4783 | 5022 | 5070 | 5106 | 5177 | 4926,34 | 4759 | 5094 | 1,071 | 0,099 | 3,166 | 0,001544 | 0,002153 | 1,00E-08 | 0,002305 | 0,0007248 | 0,0007248 | TRUE | 0,001548 | No |
| orf19.3127 | 2096 | 1679 | 2029 | 1234 | 972 | 791 | 920 | 1272 | 1908 | 1698 | 1920 | 1854 | 865 | 859 | 942 | 970 | 1377,03 | 1845 | 909 | 0,493 | -1,02 | -16,526 | 2,38E-61 | 1,32E-60 | 0,0027264 | 0,002996 | 0,0028534 | 0,0028534 | TRUE | 0,0355796 | No |
| orf19.3128 | 1328 | 1207 | 1334 | 866 | 1206 | 1074 | 986 | 1459 | 1209 | 1221 | 1263 | 1301 | 1073 | 1166 | 1010 | 1113 | 1169,34 | 1248 | 1090 | 0,875 | -0,193 | -3,399 | 0,000678 | 0,000967 | 0,001246 | 0,003167 | 0,0022437 | 0,0022437 | TRUE | 0,0283793 | No |
| orf19.3129 | 5692 | 4711 | 5196 | 3219 | 4746 | 3841 | 4238 | 5717 | 5182 | 4765 | 4918 | 4835 | 4221 | 4170 | 4340 | 4361 | 4599,11 | 4925 | 4273 | 0,867 | -0,205 | -5,163 | 2,42E-07 | 4,13E-07 | 0,0006283 | 0,002324 | 0,0012974 | 0,0012974 | TRUE | 0,0154153 | No |
| orf19.313 | 7 | 8 | 4 | 5 | 31 | 36 | 31 | 41 | 6 | 8 | 4 | 8 | 28 | 39 | 32 | 31 | 19,43 | 6 | 32 | 5,103 | 2,351 | 6,127 | 8,97E-10 | 1,68E-09 | 1,00E-08 | 0,070023 | 0,0434942 | 0,0434942 | TRUE | 0,2191745 | No |
| orf19.3130 | 6433 | 5721 | 5947 | 3831 | 5625 | 4588 | 4882 | 6494 | 5857 | 5786 | 5628 | 5755 | 5003 | 4981 | 5000 | 4954 | 5370,53 | 5756 | 4984 | 0,866 | -0,208 | -6,76 | 1,38E-11 | 2,76E-11 | 1,00E-08 | 0,002283 | 0,0007206 | 0,0007206 | TRUE | 0,0028764 | No |
| orf19.3131 | 2124 | 1851 | 1877 | 1326 | 1256 | 1038 | 1064 | 1455 | 1934 | 1872 | 1776 | 1992 | 1117 | 1127 | 1090 | 1110 | 1502,24 | 1894 | 1111 | 0,588 | -0,767 | -15,284 | 9,78E-53 | 4,74E-52 | 0,0007667 | 0,002916 | 0,0017149 | 0,0017149 | TRUE | 0,021153 | No |
| orf19.3132 | 2856 | 2593 | 2639 | 1758 | 2000 | 1668 | 1774 | 2411 | 2600 | 2623 | 2498 | 2641 | 1779 | 1811 | 1817 | 1839 | 2200,88 | 2590 | 1812 | 0,7 | -0,515 | -12,817 | 1,31E-37 | 4,91E-37 | 1,00E-08 | 0,002637 | 0,0010833 | 0,0010833 | TRUE | 0,0071998 | No |
| orf19.3133 | 20477 | 17515 | 18699 | 12784 | 26695 | 23331 | 22650 | 33540 | 18643 | 17715 | 17697 | 19204 | 23744 | 25331 | 23196 | 25585 | 21389,4 | 18315 | 24464 | 1,336 | 0,418 | 9,093 | 9,63E-20 | 2,43E-19 | 0,0019019 | 0,002099 | 0,001981 | 0,001981 | TRUE | 0,0148747 | No |
| orf19.3134 | 4 | 4 | 5 | 5 | 1 | 3 | 2 | 5 | 4 | 4 | 5 | 8 | 1 | 3 | 2 | 4 | 3,74 | 5 | 2 | 0,521 | -0,94 | -1,291 | 0,19654 | 0,227643 | 1,00E-08 | 0,355019 | 0,2139356 | 0,2139356 | TRUE | 0,2483817 | No |
| orf19.3135 | 2420 | 2263 | 2402 | 1513 | 2095 | 1808 | 1850 | 2354 | 2203 | 2289 | 2273 | 2273 | 1863 | 1963 | 1895 | 1796 | 2069,35 | 2260 | 1879 | 0,831 | -0,267 | -6,129 | 8,83E-10 | 1,65E-09 | 0,0003781 | 0,002675 | 0,0013369 | 0,0013369 | TRUE | 0,0112841 | No |
| orf19.3136 | 6056 | 5488 | 5859 | 3512 | 3720 | 3044 | 3236 | 4483 | 5514 | 5551 | 5545 | 5276 | 3309 | 3305 | 3314 | 3420 | 4404,04 | 5472 | 3337 | 0,61 | -0,714 | -20,322 | 8,18E-92 | 6,47E-91 | 0,0001753 | 0,002337 | 0,0009465 | 0,0009465 | TRUE | 0,0065985 | No |
| orf19.3138 | 24277 | 17839 | 20810 | 13803 | 29957 | 23549 | 26970 | 38038 | 22103 | 18042 | 19695 | 20734 | 26646 | 25568 | 27621 | 29016 | 23678 | 20144 | 27213 | 1,351 | 0,434 | 6,893 | 5,46E-12 | 1,11E-11 | 0,00501 | 0,002093 | 0,0037655 | 0,0037655 | TRUE | 0,0604689 | No |
| orf19.3139 | 3333 | 2802 | 2991 | 1875 | 6860 | 5502 | 5822 | 8322 | 3035 | 2834 | 2831 | 2817 | 6102 | 5974 | 5962 | 6348 | 4487,71 | 2879 | 6096 | 2,117 | 1,082 | 26,061 | 1,02E-149 | 1,32E-148 | 0,0007441 | 0,002331 | 0,0013913 | 0,0013913 | TRUE | 0,0165894 | No |
| orf19.314 | 922 | 850 | 954 | 592 | 765 | 700 | 769 | 971 | 839 | 860 | 903 | 889 | 680 | 760 | 788 | 741 | 807,5 | 873 | 742 | 0,85 | -0,234 | -3,768 | 0,000164 | 0,000243 | 0,0012178 | 0,003673 | 0,0024728 | 0,0024728 | TRUE | 0,0373771 | No |
| orf19.3140 | 23 | 19 | 18 | 18 | 12 | 10 | 11 | 19 | 21 | 19 | 17 | 27 | 11 | 11 | 11 | 14 | 16,44 | 21 | 12 | 0,576 | -0,796 | -2,327 | 0,019948 | 0,025572 | 1,00E-08 | 0,082389 | 0,0477791 | 0,0477791 | TRUE | 0,1355917 | No |
| orf19.3140.1 | 4355 | 3321 | 3793 | 2510 | 4105 | 3065 | 3382 | 4761 | 3965 | 3359 | 3590 | 3770 | 3651 | 3328 | 3464 | 3632 | 3594,79 | 3671 | 3519 | 0,959 | -0,06 | -1,059 | 0,289783 | 0,326562 | 0,0030477 | 0,002404 | 0,0028506 | 0,0028506 | TRUE | 0,0402283 | No |
| orf19.3141 | 5012 | 4268 | 4808 | 3249 | 3403 | 3075 | 3117 | 4287 | 4563 | 4317 | 4550 | 4881 | 3027 | 3339 | 3192 | 3270 | 3892,31 | 4578 | 3207 | 0,701 | -0,513 | -10,407 | 2,30E-25 | 6,64E-25 | 0,0018002 | 0,002376 | 0,0020654 | 0,0020654 | TRUE | 0,023189 | No |
| orf19.3142 | 646 | 610 | 614 | 442 | 488 | 457 | 408 | 524 | 588 | 617 | 581 | 664 | 434 | 496 | 418 | 400 | 524,74 | 612 | 437 | 0,714 | -0,487 | -6,047 | 1,48E-09 | 2,76E-09 | 0,0038705 | 0,004554 | 0,0042769 | 0,0042769 | TRUE | 0,0923668 | No |
| orf19.3147 | 571 | 795 | 858 | 425 | 517 | 442 | 407 | 535 | 520 | 804 | 812 | 638 | 460 | 480 | 417 | 408 | 567,38 | 694 | 441 | 0,635 | -0,655 | -5,218 | 1,81E-07 | 3,09E-07 | 0,0236238 | 0,004365 | 0,0132758 | 0,0132758 | TRUE | 0,3335123 | No |
| orf19.3148 | 731 | 682 | 788 | 429 | 402 | 310 | 360 | 471 | 666 | 690 | 746 | 644 | 358 | 337 | 369 | 359 | 520,95 | 686 | 356 | 0,517 | -0,951 | -13,103 | 3,16E-39 | 1,21E-38 | 1,00E-08 | 0,004573 | 0,0029724 | 0,0029724 | TRUE | 0,0406587 | No |
| orf19.3149 | 7794 | 7369 | 7374 | 4847 | 5432 | 4373 | 4264 | 5667 | 7096 | 7453 | 6979 | 7281 | 4832 | 4748 | 4367 | 4323 | 5884,75 | 7202 | 4568 | 0,634 | -0,657 | -13,789 | 2,97E-43 | 1,23E-42 | 0,0017969 | 0,002261 | 0,0020044 | 0,0020044 | TRUE | 0,0188146 | No |
| orf19.315 | 93 | 66 | 66 | 59 | 58 | 45 | 36 | 67 | 85 | 67 | 62 | 89 | 52 | 49 | 37 | 51 | 61,37 | 76 | 47 | 0,63 | -0,666 | -3,462 | 0,000537 | 0,000771 | 0,0095962 | 0,023563 | 0,018547 | 0,018547 | TRUE | 0,1506373 | No |
| orf19.3150 | 2578 | 2455 | 2396 | 1612 | 1392 | 1115 | 1127 | 1462 | 2347 | 2483 | 2268 | 2421 | 1238 | 1211 | 1154 | 1115 | 1779,67 | 2380 | 1180 | 0,496 | -1,013 | -19,514 | 8,41E-85 | 6,33E-84 | 0,001317 | 0,002779 | 0,0019644 | 0,0019644 | TRUE | 0,0173732 | No |
| orf19.3151 | 421 | 450 | 402 | 259 | 475 | 423 | 371 | 512 | 383 | 455 | 380 | 389 | 422 | 459 | 380 | 391 | 407,53 | 402 | 413 | 1,026 | 0,036 | 0,41 | 0,682025 | 0,717595 | 0,0049868 | 0,005279 | 0,0051464 | 0,0051464 | TRUE | 0,0938719 | No |
| orf19.3152 | 1455 | 1354 | 1331 | 860 | 4647 | 4139 | 3917 | 5272 | 1325 | 1369 | 1260 | 1292 | 4133 | 4494 | 4012 | 4022 | 2738,23 | 1312 | 4165 | 3,174 | 1,666 | 31,848 | 1,38E-222 | 2,60E-221 | 0,0018107 | 0,002519 | 0,0021061 | 0,0021061 | TRUE | 0,0338939 | No |
| orf19.3153 | 6349 | 6114 | 6861 | 3927 | 6743 | 5523 | 5878 | 7737 | 5780 | 6184 | 6493 | 5899 | 5998 | 5996 | 6020 | 5902 | 6034,03 | 6089 | 5979 | 0,982 | -0,027 | -0,626 | 0,531296 | 0,572808 | 0,0011509 | 0,002256 | 0,0016082 | 0,0016082 | TRUE | 0,0246663 | No |
| orf19.3154 | 1949 | 1996 | 2060 | 1221 | 1543 | 1302 | 1430 | 1810 | 1774 | 2019 | 1950 | 1834 | 1372 | 1414 | 1465 | 1381 | 1651,03 | 1894 | 1408 | 0,742 | -0,43 | -7,988 | 1,37E-15 | 3,08E-15 | 0,0016337 | 0,002837 | 0,0021614 | 0,0021614 | TRUE | 0,0238606 | No |
| orf19.3156 | 1810 | 1628 | 1779 | 1091 | 879 | 751 | 771 | 1040 | 1648 | 1647 | 1684 | 1639 | 782 | 815 | 790 | 793 | 1224,64 | 1654 | 795 | 0,48 | -1,058 | -22,39 | 4,88E-111 | 4,65E-110 | 1,00E-08 | 0,003116 | 0,0012356 | 0,0012356 | TRUE | 0,0032371 | No |
| orf19.3158 | 2542 | 2333 | 2520 | 1614 | 1030 | 902 | 877 | 1146 | 2314 | 2360 | 2385 | 2424 | 916 | 979 | 898 | 874 | 1643,9 | 2371 | 917 | 0,386 | -1,372 | -28,118 | 5,87E-174 | 8,92E-173 | 0,0002365 | 0,002841 | 0,0015498 | 0,0015498 | TRUE | 0,0237923 | No |
| orf19.3159 | 3380 | 3547 | 3504 | 2001 | 7566 | 6697 | 6544 | 8331 | 3077 | 3587 | 3316 | 3006 | 6730 | 7271 | 6702 | 6355 | 5005,55 | 3246 | 6764 | 2,082 | 1,058 | 16,984 | 1,07E-64 | 6,29E-64 | 0,0045432 | 0,002301 | 0,0034901 | 0,0034901 | TRUE | 0,0605243 | No |
| orf19.3159.4 | 0 | 2 | 1 | 0 | 2 | 2 | 3 | 3 | 0 | 2 | 1 | 0 | 2 | 2 | 3 | 2 | 1,54 | 1 | 2 | 2,974 | 1,572 | 1,311 | 0,189985 | 0,220451 | 1,00E-08 | 0,862595 | 0,5024363 | 0,5024363 | TRUE | 0,4471253 | No |
| orf19.316 | 1212 | 939 | 1062 | 724 | 1101 | 840 | 964 | 1320 | 1103 | 950 | 1005 | 1088 | 979 | 912 | 987 | 1007 | 1003,91 | 1036 | 971 | 0,939 | -0,091 | -1,433 | 0,151769 | 0,178544 | 0,0023402 | 0,003353 | 0,0028363 | 0,0028363 | TRUE | 0,0381747 | No |
| orf19.3160 | 2415 | 1873 | 1853 | 1233 | 3004 | 2097 | 2322 | 3272 | 2199 | 1894 | 1754 | 1852 | 2672 | 2277 | 2378 | 2496 | 2190,2 | 1925 | 2456 | 1,276 | 0,351 | 4,751 | 2,02E-06 | 3,31E-06 | 0,0067439 | 0,00264 | 0,0047737 | 0,0047737 | TRUE | 0,1133709 | No |
| orf19.3161 | 6650 | 5573 | 5989 | 3475 | 7101 | 5282 | 6455 | 8388 | 6055 | 5636 | 5668 | 5220 | 6316 | 5735 | 6611 | 6398 | 5954,89 | 5645 | 6265 | 1,109 | 0,15 | 2,627 | 0,008609 | 0,011339 | 0,0033887 | 0,002259 | 0,0029497 | 0,0029497 | TRUE | 0,0392612 | No |
| orf19.3163 | 831 | 704 | 790 | 484 | 856 | 648 | 728 | 945 | 757 | 712 | 748 | 727 | 761 | 704 | 746 | 721 | 734,33 | 736 | 733 | 0,995 | -0,007 | -0,117 | 0,907152 | 0,920288 | 1,00E-08 | 0,003836 | 0,0018948 | 0,0018948 | TRUE | 0,0083087 | No |
| orf19.3166 | 619 | 578 | 599 | 385 | 654 | 545 | 590 | 750 | 564 | 585 | 567 | 578 | 582 | 592 | 604 | 572 | 580,39 | 574 | 588 | 1,024 | 0,035 | 0,561 | 0,574801 | 0,61543 | 1,00E-08 | 0,004313 | 0,0019173 | 0,0019173 | TRUE | 0,0043674 | No |
| orf19.3167 | 3675 | 2737 | 3090 | 2226 | 3424 | 2950 | 3194 | 4403 | 3346 | 2768 | 2924 | 3344 | 3046 | 3203 | 3271 | 3359 | 3157,56 | 3096 | 3220 | 1,041 | 0,058 | 0,872 | 0,383346 | 0,423908 | 0,004608 | 0,002455 | 0,0038858 | 0,0038858 | TRUE | 0,0615732 | No |
| orf19.3168 | 6312 | 5632 | 6024 | 3595 | 5286 | 3930 | 4277 | 5864 | 5747 | 5696 | 5701 | 5400 | 4702 | 4267 | 4380 | 4473 | 5045,79 | 5636 | 4456 | 0,79 | -0,339 | -8,084 | 6,27E-16 | 1,42E-15 | 0,0009873 | 0,002299 | 0,0014927 | 0,0014927 | TRUE | 0,0168352 | No |
| orf19.3169 | 7882 | 6592 | 7370 | 4604 | 5038 | 4060 | 4524 | 6035 | 7176 | 6667 | 6975 | 6916 | 4481 | 4408 | 4633 | 4604 | 5732,52 | 6934 | 4532 | 0,654 | -0,614 | -16,444 | 9,21E-61 | 5,09E-60 | 0,0004844 | 0,002267 | 0,001156 | 0,001156 | TRUE | 0,0083497 | No |
| orf19.317 | 1105 | 914 | 1027 | 634 | 482 | 348 | 420 | 565 | 1006 | 924 | 972 | 952 | 429 | 378 | 430 | 431 | 690,31 | 964 | 417 | 0,433 | -1,207 | -18,42 | 9,08E-76 | 6,09E-75 | 0,0005411 | 0,003951 | 0,002449 | 0,002449 | TRUE | 0,0423586 | No |
| orf19.3170 | 2908 | 2698 | 2999 | 1721 | 4220 | 3254 | 3883 | 4754 | 2648 | 2729 | 2838 | 2585 | 3754 | 3533 | 3977 | 3626 | 3211,18 | 2700 | 3722 | 1,377 | 0,462 | 9,254 | 2,16E-20 | 5,53E-20 | 0,0017261 | 0,002448 | 0,0020685 | 0,0020685 | TRUE | 0,0253921 | No |
| orf19.3171 | 9767 | 8394 | 8929 | 6676 | 8979 | 7823 | 7669 | 10826 | 8892 | 8490 | 8450 | 10028 | 7987 | 8494 | 7854 | 8258 | 8556,66 | 8965 | 8148 | 0,909 | -0,137 | -2,386 | 0,017021 | 0,02197 | 0,0038299 | 0,002191 | 0,0030543 | 0,0030543 | TRUE | 0,0767542 | No |
| orf19.3172 | 13 | 7 | 16 | 14 | 17 | 20 | 10 | 23 | 12 | 7 | 15 | 21 | 15 | 22 | 10 | 18 | 14,96 | 14 | 16 | 1,205 | 0,269 | 0,69 | 0,490367 | 0,53201 | 0,050969 | 0,090317 | 0,0777252 | 0,0777252 | TRUE | 0,2714854 | No |
| orf19.3173 | 613 | 494 | 531 | 295 | 1201 | 822 | 1056 | 1247 | 558 | 500 | 503 | 443 | 1068 | 892 | 1081 | 951 | 749,6 | 501 | 998 | 1,985 | 0,989 | 11,738 | 8,10E-32 | 2,71E-31 | 0,0066142 | 0,003799 | 0,0052815 | 0,0052815 | TRUE | 0,0689271 | No |
| orf19.3174 | 1202 | 1102 | 1231 | 811 | 1542 | 1379 | 1412 | 1942 | 1094 | 1115 | 1165 | 1218 | 1372 | 1497 | 1446 | 1481 | 1298,55 | 1148 | 1449 | 1,265 | 0,339 | 6,31 | 2,78E-10 | 5,32E-10 | 0,0008695 | 0,003054 | 0,0019799 | 0,0019799 | TRUE | 0,0179448 | No |
| orf19.3175 | 6107 | 5247 | 5637 | 3617 | 4406 | 3477 | 3840 | 5369 | 5560 | 5307 | 5335 | 5433 | 3919 | 3775 | 3933 | 4096 | 4669,67 | 5409 | 3931 | 0,727 | -0,46 | -11,983 | 4,36E-33 | 1,49E-32 | 0,0004874 | 0,00232 | 0,0011948 | 0,0011948 | TRUE | 0,0099748 | No |
| orf19.3176 | 2033 | 1734 | 1920 | 1278 | 1207 | 1026 | 1162 | 1528 | 1851 | 1754 | 1817 | 1920 | 1074 | 1114 | 1190 | 1166 | 1485,59 | 1836 | 1136 | 0,62 | -0,691 | -13,299 | 2,36E-40 | 9,20E-40 | 0,0009657 | 0,002926 | 0,0018852 | 0,0018852 | TRUE | 0,0167984 | No |
| orf19.3177 | 509 | 444 | 477 | 258 | 1091 | 875 | 929 | 1288 | 463 | 449 | 451 | 388 | 970 | 950 | 951 | 982 | 700,73 | 438 | 963 | 2,187 | 1,129 | 17,537 | 7,44E-69 | 4,58E-68 | 1,00E-08 | 0,003922 | 0,0022849 | 0,0022849 | TRUE | 0,0553375 | No |
| orf19.3178 | 2878 | 2583 | 2843 | 1554 | 2328 | 1848 | 1910 | 2545 | 2620 | 2612 | 2691 | 2334 | 2071 | 2006 | 1956 | 1941 | 2279,03 | 2564 | 1994 | 0,776 | -0,366 | -7,029 | 2,08E-12 | 4,29E-12 | 0,0017179 | 0,002617 | 0,0021589 | 0,0021589 | TRUE | 0,0413282 | No |
| orf19.318 | 1006 | 908 | 944 | 576 | 991 | 854 | 827 | 1187 | 916 | 918 | 893 | 865 | 881 | 927 | 847 | 905 | 894,25 | 898 | 890 | 0,99 | -0,015 | -0,275 | 0,783274 | 0,81312 | 1,00E-08 | 0,003514 | 0,001772 | 0,001772 | TRUE | 0,0121095 | No |
| orf19.3180 | 1346 | 1152 | 1249 | 778 | 954 | 877 | 905 | 1163 | 1225 | 1165 | 1182 | 1169 | 849 | 952 | 927 | 887 | 1044,51 | 1185 | 904 | 0,761 | -0,395 | -7,228 | 4,89E-13 | 1,03E-12 | 0,0004855 | 0,003302 | 0,0018929 | 0,0018929 | TRUE | 0,0198044 | No |
| orf19.3181.1 | 424 | 318 | 357 | 209 | 495 | 334 | 367 | 544 | 386 | 322 | 338 | 314 | 440 | 363 | 376 | 415 | 369,15 | 340 | 398 | 1,17 | 0,227 | 2,429 | 0,015149 | 0,019641 | 0,0056928 | 0,005615 | 0,0056498 | 0,0056498 | TRUE | 0,0988121 | No |
| orf19.3182 | 13072 | 10295 | 11098 | 7595 | 23482 | 19390 | 20936 | 29419 | 11902 | 10412 | 10503 | 11409 | 20886 | 21052 | 21441 | 22441 | 16255,8 | 11056 | 21455 | 1,941 | 0,957 | 18,954 | 4,06E-80 | 2,88E-79 | 0,0025624 | 0,002118 | 0,0023765 | 0,0023765 | TRUE | 0,0328352 | No |
| orf19.3183 | 1268 | 1043 | 1181 | 903 | 948 | 873 | 786 | 1068 | 1154 | 1055 | 1118 | 1356 | 843 | 948 | 805 | 815 | 1011,77 | 1171 | 853 | 0,73 | -0,455 | -5,573 | 2,51E-08 | 4,44E-08 | 0,0077965 | 0,003343 | 0,0053897 | 0,0053897 | TRUE | 0,1284138 | No |
| orf19.3184 | 2536 | 2111 | 2241 | 1405 | 1839 | 1372 | 1497 | 1993 | 2309 | 2135 | 2121 | 2111 | 1636 | 1490 | 1533 | 1520 | 1856,77 | 2169 | 1545 | 0,712 | -0,49 | -9,702 | 2,95E-22 | 7,89E-22 | 0,0012614 | 0,002748 | 0,0019019 | 0,0019019 | TRUE | 0,0235855 | No |
| orf19.3185 | 4943 | 4496 | 4768 | 3135 | 5415 | 4590 | 4851 | 6519 | 4500 | 4547 | 4512 | 4709 | 4816 | 4983 | 4968 | 4973 | 4751,26 | 4567 | 4935 | 1,081 | 0,113 | 3,361 | 0,000776 | 0,001103 | 7,24E-05 | 0,002315 | 0,000869 | 0,000869 | TRUE | 0,0049286 | No |
| orf19.3186 | 0 | 1 | 3 | 0 | 4 | 1 | 2 | 0 | 0 | 1 | 3 | 0 | 4 | 1 | 2 | 0 | 1,32 | 1 | 2 | 1,633 | 0,707 | 0,519 | 0,603664 | 0,643431 | 0,8963154 | 1,004457 | 0,974799 | 0,974799 | TRUE | 0,4942693 | No |
| orf19.3187 | 3880 | 3734 | 3933 | 2465 | 3895 | 3536 | 3628 | 4839 | 3533 | 3777 | 3722 | 3703 | 3464 | 3839 | 3716 | 3691 | 3680,56 | 3684 | 3678 | 0,998 | -0,003 | -0,057 | 0,954329 | 0,960052 | 0,0010826 | 0,002396 | 0,0015866 | 0,0015866 | TRUE | 0,018557 | No |
| orf19.3188 | 1955 | 1880 | 1967 | 1265 | 2184 | 1914 | 1928 | 2627 | 1780 | 1901 | 1862 | 1900 | 1943 | 2078 | 1975 | 2004 | 1930,28 | 1861 | 2000 | 1,075 | 0,105 | 2,354 | 0,018589 | 0,023907 | 0,0003699 | 0,002721 | 0,0013801 | 0,0013801 | TRUE | 0,0107436 | No |

| | | | | | | | | | | | | | | | | | | | | | | | | | | | | | | | |
|---|---|---|---|---|---|---|---|---|---|---|---|---|---|---|---|---|---|---|---|---|---|---|---|---|---|---|---|---|---|---|---|
| orf19.319 | 192 | 158 | 181 | 99 | 182 | 130 | 139 | 213 | 175 | 160 | 171 | 149 | 162 | 141 | 142 | 162 | 157,81 | 164 | 152 | 0,927 | -0,109 | -0,955 | 0,339735 | 0,3799 | 1,00E-08 | 0,010408 | 0,0063073 | 0,0063073 | TRUE | 0,0313207 | No |
| orf19.3190 | 7022 | 6486 | 6792 | 4490 | 5021 | 4338 | 4374 | 5895 | 6393 | 6560 | 6428 | 6745 | 4466 | 4710 | 4480 | 4497 | 5534,75 | 6532 | 4538 | 0,695 | -0,525 | -14,661 | 1,15E-48 | 5,17E-48 | 0,0003707 | 0,002276 | 0,0010453 | 0,0010453 | TRUE | 0,0077114 | No |
| orf19.3192 | 6510 | 6062 | 6419 | 4334 | 2554 | 2235 | 2200 | 2973 | 5927 | 6131 | 6075 | 6510 | 2272 | 2427 | 2253 | 2268 | 4232,84 | 6161 | 2305 | 0,374 | -1,418 | -32,02 | 5,67E-225 | 1,07E-223 | 0,0010417 | 0,002349 | 0,0015931 | 0,0015931 | TRUE | 0,0171646 | No |
| orf19.3193 | 1620 | 1598 | 1642 | 1104 | 2440 | 2052 | 2047 | 2609 | 1475 | 1616 | 1554 | 1658 | 2170 | 2228 | 2096 | 1990 | 1848,54 | 1576 | 2121 | 1,347 | 0,43 | 7,888 | 3,07E-15 | 6,82E-15 | 0,0017144 | 0,002752 | 0,0022924 | 0,0022924 | TRUE | 0,022748 | No |
| orf19.3195 | 10526 | 9494 | 10692 | 6442 | 9211 | 7631 | 8249 | 11302 | 9583 | 9602 | 10119 | 9677 | 8193 | 8285 | 8448 | 8621 | 9066,11 | 9745 | 8387 | 0,861 | -0,217 | -6,297 | 3,04E-10 | 5,80E-10 | 0,00044 | 0,002183 | 0,0010247 | 0,0010247 | TRUE | 0,00828 | No |
| orf19.3196 | 259 | 278 | 265 | 205 | 236 | 235 | 178 | 261 | 236 | 281 | 251 | 308 | 210 | 255 | 182 | 199 | 240,27 | 269 | 212 | 0,789 | -0,341 | -2,846 | 0,004433 | 0,005966 | 0,0123553 | 0,007535 | 0,0095743 | 0,0095743 | TRUE | 0,1975026 | No |
| orf19.3197 | 1435 | 1273 | 1349 | 825 | 1394 | 1157 | 1240 | 1586 | 1307 | 1288 | 1277 | 1239 | 1240 | 1256 | 1270 | 1210 | 1260,73 | 1278 | 1244 | 0,972 | -0,042 | -0,889 | 0,37383 | 0,414104 | 1,00E-08 | 0,003085 | 0,001307 | 0,001307 | TRUE | 0,0043716 | No |
| orf19.3198 | 2695 | 2190 | 2390 | 1464 | 1949 | 1585 | 1806 | 2354 | 2454 | 2215 | 2262 | 2199 | 1734 | 1721 | 1850 | 1796 | 2028,67 | 2282 | 1775 | 0,777 | -0,364 | -7,217 | 5,30E-13 | 1,11E-12 | 0,0013966 | 0,002688 | 0,0019466 | 0,0019466 | TRUE | 0,0316465 | No |
| orf19.3199 | 1411 | 1256 | 1344 | 784 | 1067 | 899 | 984 | 1225 | 1285 | 1270 | 1272 | 1178 | 949 | 976 | 1008 | 934 | 1108,99 | 1251 | 967 | 0,77 | -0,377 | -7,159 | 8,15E-13 | 1,70E-12 | 0,000274 | 0,003228 | 0,0017507 | 0,0017507 | TRUE | 0,0164046 | No |
| orf19.320 | 2226 | 1882 | 2062 | 1541 | 1579 | 1295 | 1403 | 1953 | 2027 | 1903 | 1951 | 2315 | 1404 | 1406 | 1437 | 1490 | 1741,69 | 2049 | 1434 | 0,702 | -0,511 | -8,111 | 5,04E-16 | 1,15E-15 | 0,0036396 | 0,002795 | 0,0032229 | 0,0032229 | TRUE | 0,0872481 | No |
| orf19.3200 | 103 | 124 | 106 | 92 | 116 | 111 | 104 | 110 | 94 | 125 | 100 | 138 | 103 | 121 | 107 | 84 | 108,98 | 114 | 104 | 0,909 | -0,138 | -0,853 | 0,393443 | 0,43447 | 0,018306 | 0,014159 | 0,015827 | 0,015827 | TRUE | 0,1797817 | No |
| orf19.3201 | 1140 | 967 | 1064 | 704 | 902 | 757 | 840 | 1008 | 1038 | 978 | 1007 | 1058 | 802 | 822 | 860 | 769 | 916,73 | 1020 | 813 | 0,797 | -0,327 | -5,721 | 1,06E-08 | 1,90E-08 | 0,0005932 | 0,003478 | 0,0020445 | 0,0020445 | TRUE | 0,017993 | No |
| orf19.3202 | 870 | 1042 | 978 | 628 | 599 | 578 | 532 | 623 | 792 | 1054 | 926 | 943 | 533 | 628 | 545 | 475 | 736,91 | 929 | 545 | 0,586 | -0,771 | -7,862 | 3,77E-15 | 8,36E-15 | 0,0113718 | 0,00383 | 0,0077918 | 0,0077918 | TRUE | 0,1206593 | No |
| orf19.3203 | 464 | 541 | 587 | 320 | 352 | 279 | 296 | 346 | 422 | 547 | 556 | 481 | 313 | 303 | 303 | 264 | 398,62 | 502 | 296 | 0,588 | -0,765 | -7,56 | 4,02E-14 | 8,68E-14 | 0,0091937 | 0,005351 | 0,0071786 | 0,0071786 | TRUE | 0,1411986 | No |
| orf19.3204 | 1935 | 2394 | 2341 | 1262 | 1563 | 1405 | 1484 | 1798 | 1762 | 2421 | 2216 | 1896 | 1390 | 1525 | 1520 | 1372 | 1762,66 | 2074 | 1452 | 0,699 | -0,516 | -5,736 | 9,67E-09 | 1,75E-08 | 0,011286 | 0,002786 | 0,0071812 | 0,0071812 | TRUE | 0,154565 | No |
| orf19.3205 | 4323 | 3781 | 4093 | 2300 | 2899 | 2204 | 2461 | 3246 | 3936 | 3824 | 3874 | 3455 | 2579 | 2393 | 2520 | 2476 | 3132,07 | 3772 | 2492 | 0,66 | -0,6 | -11,99 | 4,00E-33 | 1,37E-32 | 0,0016935 | 0,002459 | 0,0020713 | 0,0020713 | TRUE | 0,0371007 | No |
| orf19.3206 | 5872 | 5819 | 6254 | 3661 | 7254 | 6262 | 6474 | 8346 | 5346 | 5885 | 5919 | 5499 | 6452 | 6799 | 6630 | 6366 | 6112,17 | 5662 | 6562 | 1,158 | 0,212 | 4,687 | 2,77E-06 | 4,52E-06 | 0,0014683 | 0,002253 | 0,0017981 | 0,0017981 | TRUE | 0,0178676 | No |
| orf19.3207 | 135 | 137 | 160 | 90 | 188 | 182 | 183 | 228 | 123 | 139 | 151 | 135 | 167 | 198 | 187 | 174 | 159,28 | 137 | 182 | 1,321 | 0,401 | 3,428 | 0,000608 | 0,00087 | 1,00E-08 | 0,010331 | 0,006682 | 0,006682 | TRUE | 0,0583307 | No |
| orf19.3208 | 843 | 844 | 973 | 515 | 1399 | 1294 | 1272 | 1637 | 768 | 854 | 921 | 774 | 1244 | 1405 | 1303 | 1249 | 1064,54 | 829 | 1300 | 1,564 | 0,646 | 9,164 | 5,02E-20 | 1,28E-19 | 0,0041884 | 0,003278 | 0,0037579 | 0,0037579 | TRUE | 0,067712 | No |
| orf19.3209 | 23 | 18 | 22 | 15 | 33 | 24 | 23 | 36 | 21 | 18 | 21 | 23 | 29 | 26 | 24 | 27 | 23,62 | 21 | 26 | 1,301 | 0,38 | 1,373 | 0,169642 | 0,197999 | 1,00E-08 | 0,057975 | 0,0301035 | 0,0301035 | TRUE | 0,0360467 | No |
| orf19.321 | 1413 | 1290 | 1354 | 985 | 2045 | 1796 | 1844 | 2585 | 1286 | 1305 | 1281 | 1480 | 1819 | 1950 | 1888 | 1972 | 1622,69 | 1338 | 1907 | 1,43 | 0,516 | 9,128 | 6,96E-20 | 1,76E-19 | 0,0020201 | 0,002851 | 0,0024211 | 0,0024211 | TRUE | 0,0554917 | No |
| orf19.3210 | 23 | 35 | 36 | 18 | 39 | 30 | 41 | 47 | 21 | 35 | 34 | 27 | 35 | 33 | 42 | 36 | 32,82 | 29 | 36 | 1,233 | 0,302 | 1,214 | 0,22462 | 0,257963 | 1,00E-08 | 0,042288 | 0,0285341 | 0,0285341 | TRUE | 0,300286 | No |
| orf19.3211 | 1266 | 1187 | 1365 | 808 | 1250 | 1048 | 1089 | 1399 | 1153 | 1201 | 1292 | 1214 | 1112 | 1138 | 1115 | 1067 | 1161,36 | 1215 | 1108 | 0,911 | -0,134 | -2,495 | 0,012608 | 0,01641 | 0,0008009 | 0,003174 | 0,001899 | 0,001899 | TRUE | 0,0235423 | No |
| orf19.3212 | 1980 | 2007 | 2057 | 1261 | 940 | 771 | 810 | 1046 | 1803 | 2030 | 1947 | 1894 | 836 | 837 | 830 | 798 | 1371,77 | 1918 | 825 | 0,43 | -1,218 | -22,59 | 5,45E-113 | 5,25E-112 | 0,0011385 | 0,003 | 0,0019465 | 0,0019465 | TRUE | 0,0212298 | No |
| orf19.3213 | 2120 | 2136 | 2317 | 1444 | 3406 | 2896 | 3124 | 4025 | 1930 | 2160 | 2193 | 2169 | 3030 | 3144 | 3199 | 3070 | 2611,99 | 2113 | 3111 | 1,473 | 0,559 | 11,185 | 4,82E-29 | 1,52E-28 | 0,0014533 | 0,002543 | 0,0019916 | 0,0019916 | TRUE | 0,0427357 | No |
| orf19.3214 | 424 | 477 | 504 | 297 | 726 | 621 | 659 | 790 | 386 | 482 | 477 | 446 | 646 | 674 | 675 | 603 | 548,64 | 448 | 650 | 1,449 | 0,535 | 6,548 | 5,83E-11 | 1,14E-10 | 0,0045363 | 0,004445 | 0,0044868 | 0,0044868 | TRUE | 0,1097168 | No |
| orf19.3215 | 2122 | 2040 | 2101 | 1237 | 1442 | 1106 | 1261 | 1694 | 1932 | 2063 | 1988 | 1858 | 1283 | 1201 | 1291 | 1292 | 1613,61 | 1960 | 1267 | 0,646 | -0,631 | -12,556 | 3,70E-36 | 1,35E-35 | 0,0008688 | 0,002856 | 0,0017807 | 0,0017807 | TRUE | 0,0146918 | No |
| orf19.3216 | 1142 | 1277 | 1381 | 894 | 2087 | 1855 | 1844 | 2438 | 1040 | 1292 | 1307 | 1343 | 1856 | 2014 | 1888 | 1860 | 1574,97 | 1246 | 1904 | 1,532 | 0,615 | 8,208 | 2,24E-16 | 5,13E-16 | 0,0060806 | 0,002876 | 0,0047141 | 0,0047141 | TRUE | 0,1522766 | No |
| orf19.3218 | 766 | 781 | 804 | 466 | 587 | 511 | 520 | 704 | 697 | 790 | 761 | 700 | 522 | 555 | 533 | 537 | 636,84 | 737 | 537 | 0,727 | -0,461 | -6,999 | 2,58E-12 | 5,29E-12 | 0,0008676 | 0,004111 | 0,002569 | 0,002569 | TRUE | 0,0273192 | No |
| orf19.3219 | 250 | 372 | 366 | 200 | 583 | 482 | 467 | 577 | 228 | 376 | 346 | 300 | 519 | 523 | 478 | 440 | 401,37 | 312 | 490 | 1,566 | 0,647 | 5,052 | 4,38E-07 | 7,39E-07 | 0,0231104 | 0,005328 | 0,0130964 | 0,0130964 | TRUE | 0,4051912 | No |
| orf19.322 | 9 | 6 | 6 | 9 | 3 | 9 | 1 | 8 | 8 | 6 | 6 | 14 | 3 | 10 | 1 | 6 | 6,63 | 8 | 5 | 0,597 | -0,743 | -1,216 | 0,224061 | 0,257413 | 0,1872033 | 0,201332 | 0,1971443 | 0,1971443 | TRUE | 0,5333688 | No |
| orf19.3220 | 7007 | 6462 | 6648 | 3854 | 9002 | 6650 | 7230 | 9518 | 6380 | 6536 | 6292 | 5789 | 8007 | 7220 | 7404 | 7260 | 6861,02 | 6249 | 7473 | 1,195 | 0,257 | 5,103 | 3,35E-07 | 5,67E-07 | 0,002328 | 0,00223 | 0,0022923 | 0,0022923 | TRUE | 0,0290545 | No |
| orf19.3221 | 11925 | 11127 | 12024 | 7275 | 8849 | 7846 | 8069 | 10832 | 10857 | 11254 | 11380 | 10928 | 7871 | 8519 | 8264 | 8263 | 9666,84 | 11105 | 8229 | 0,741 | -0,433 | -11,879 | 1,53E-32 | 5,18E-32 | 0,0006438 | 0,002174 | 0,0011695 | 0,0011695 | TRUE | 0,0104975 | No |
| orf19.3222 | 811 | 824 | 930 | 521 | 871 | 704 | 737 | 947 | 738 | 833 | 880 | 783 | 775 | 764 | 755 | 722 | 781,35 | 808 | 754 | 0,931 | -0,103 | -1,54 | 0,123534 | 0,146872 | 0,0022909 | 0,003728 | 0,0030035 | 0,0030035 | TRUE | 0,0450289 | No |
| orf19.3223 | 5584 | 5052 | 5083 | 3465 | 8755 | 7576 | 6871 | 9720 | 5084 | 5110 | 4811 | 5205 | 7787 | 8225 | 7037 | 7414 | 6334,14 | 5052 | 7616 | 1,508 | 0,592 | 11,245 | 2,45E-29 | 7,73E-29 | 0,0025743 | 0,002246 | 0,0024972 | 0,0024972 | TRUE | 0,0351238 | No |
| orf19.3223.1 | 1620 | 1469 | 1384 | 934 | 3110 | 2339 | 2605 | 3241 | 1475 | 1486 | 1310 | 1403 | 2766 | 2540 | 2668 | 2472 | 2014,92 | 1418 | 2612 | 1,84 | 0,88 | 15,457 | 6,75E-54 | 3,35E-53 | 0,0021746 | 0,002693 | 0,0025544 | 0,0025544 | TRUE | 0,033778 | No |
| orf19.3225 | 815 | 814 | 888 | 500 | 1276 | 1066 | 1017 | 1467 | 742 | 823 | 840 | 751 | 1135 | 1157 | 1042 | 1119 | 951,21 | 789 | 1113 | 1,409 | 0,494 | 7,94 | 2,02E-15 | 4,53E-15 | 0,0017967 | 0,003426 | 0,0026222 | 0,0026222 | TRUE | 0,0231731 | No |
| orf19.3226 | 965 | 988 | 996 | 647 | 1410 | 1146 | 1143 | 1542 | 879 | 999 | 943 | 972 | 1254 | 1244 | 1171 | 1176 | 1079,7 | 948 | 1211 | 1,279 | 0,355 | 6,182 | 6,34E-10 | 1,19E-09 | 0,0012346 | 0,003261 | 0,0022179 | 0,0022179 | TRUE | 0,0306902 | No |
| orf19.3227 | 2375 | 2526 | 2660 | 1402 | 3281 | 2745 | 2859 | 3589 | 2162 | 2555 | 2517 | 2106 | 2918 | 2980 | 2928 | 2738 | 2613,12 | 2335 | 2891 | 1,236 | 0,306 | 4,555 | 5,24E-06 | 8,44E-06 | 0,0051611 | 0,002543 | 0,0039453 | 0,0039453 | TRUE | 0,0502962 | No |
| orf19.3228 | 1208 | 1410 | 1465 | 853 | 2435 | 2257 | 2123 | 2680 | 1100 | 1426 | 1386 | 1281 | 2166 | 2450 | 2174 | 2044 | 1753,58 | 1298 | 2208 | 1,7 | 0,766 | 9,322 | 1,14E-20 | 2,94E-20 | 0,0085131 | 0,00279 | 0,0058553 | 0,0058553 | TRUE | 0,1319644 | No |
| orf19.3229 | 4 | 8 | 2 | 0 | 7 | 3 | 3 | 13 | 4 | 8 | 2 | 0 | 6 | 3 | 3 | 10 | 4,51 | 4 | 6 | 1,613 | 0,69 | 0,92 | 0,357383 | 0,397404 | 0,335166 | 0,294794 | 0,3079458 | 0,3079458 | TRUE | 0,8407707 | No |
| orf19.323 | 2120 | 1786 | 1879 | 1158 | 2008 | 1583 | 1739 | 2190 | 1930 | 1806 | 1778 | 1740 | 1786 | 1719 | 1781 | 1671 | 1776,32 | 1814 | 1739 | 0,957 | -0,063 | -1,28 | 0,200467 | 0,232022 | 0,0009961 | 0,002781 | 0,0017687 | 0,0017687 | TRUE | 0,023157 | No |
| orf19.3230 | 6073 | 5393 | 6068 | 3870 | 4029 | 3518 | 3798 | 5025 | 5529 | 5454 | 5743 | 5813 | 3584 | 3820 | 3890 | 3833 | 4708,22 | 5635 | 3782 | 0,671 | -0,575 | -13,95 | 3,13E-44 | 1,32E-43 | 0,0008036 | 0,002318 | 0,0014123 | 0,0014123 | TRUE | 0,015225 | No |
| orf19.3231 | 476 | 584 | 547 | 331 | 570 | 521 | 490 | 663 | 433 | 591 | 518 | 497 | 507 | 566 | 502 | 506 | 514,89 | 510 | 520 | 1,02 | 0,028 | 0,303 | 0,761713 | 0,792508 | 0,0078969 | 0,004603 | 0,0061561 | 0,0061561 | TRUE | 0,1375498 | No |
| orf19.3232 | 438 | 422 | 411 | 290 | 112 | 86 | 82 | 154 | 399 | 427 | 389 | 436 | 100 | 93 | 84 | 117 | 255,58 | 413 | 98 | 0,242 | -2,048 | -18,596 | 3,48E-77 | 2,37E-76 | 0,0022467 | 0,007206 | 0,0056715 | 0,0056715 | TRUE | 0,1853269 | No |
| orf19.3233 | 2864 | 2531 | 2750 | 1820 | 1851 | 1448 | 1685 | 2264 | 2608 | 2560 | 2603 | 2734 | 1646 | 1572 | 1726 | 1727 | 2146,89 | 2626 | 1668 | 0,636 | -0,652 | -14,022 | 1,14E-44 | 4,87E-44 | 0,0007416 | 0,002652 | 0,0015935 | 0,0015935 | TRUE | 0,0172489 | No |
| orf19.3234 | 255 | 243 | 248 | 151 | 322 | 307 | 306 | 403 | 232 | 246 | 235 | 227 | 286 | 333 | 313 | 307 | 272,5 | 235 | 310 | 1,316 | 0,396 | 4,439 | 9,02E-06 | 1,43E-05 | 1,00E-08 | 0,006885 | 0,003865 | 0,003865 | TRUE | 0,0299322 | No |
| orf19.3234.1 | 749 | 644 | 629 | 363 | 253 | 167 | 178 | 231 | 682 | 651 | 595 | 545 | 225 | 181 | 182 | 176 | 404,84 | 618 | 191 | 0,308 | -1,697 | -16,785 | 3,13E-63 | 1,80E-62 | 0,0081521 | 0,0053 | 0,0065119 | 0,0065119 | TRUE | 0,1599216 | No |
| orf19.3235 | 673 | 728 | 743 | 462 | 628 | 495 | 482 | 646 | 613 | 736 | 703 | 694 | 559 | 537 | 494 | 493 | 603,58 | 686 | 521 | 0,759 | -0,399 | -5,209 | 1,90E-07 | 3,24E-07 | 0,0036394 | 0,004226 | 0,0039403 | 0,0039403 | TRUE | 0,0660273 | No |
| orf19.3237 | 2265 | 2207 | 2163 | 1261 | 3059 | 2635 | 2784 | 3643 | 2062 | 2232 | 2047 | 1894 | 2721 | 2861 | 2851 | 2779 | 2430,93 | 2059 | 2803 | 1,359 | 0,442 | 8,525 | 1,53E-17 | 3,65E-17 | 0,0019661 | 0,00258 | 0,0021569 | 0,0021569 | TRUE | 0,0380281 | No |
| orf19.3238 | 5 | 6 | 6 | 5 | 6 | 5 | 3 | 5 | 5 | 6 | 6 | 8 | 5 | 5 | 3 | 4 | 5,18 | 6 | 4 | 0,753 | -0,41 | -0,697 | 0,486112 | 0,527909 | 1,00E-08 | 0,256922 | 0,1363037 | 0,1363037 | TRUE | 0,066996 | No |
| orf19.3239 | 952 | 761 | 878 | 512 | 1067 | 823 | 943 | 1238 | 867 | 770 | 831 | 769 | 949 | 894 | 966 | 944 | 873,65 | 809 | 938 | 1,157 | 0,21 | 3,501 | 0,000464 | 0,000669 | 0,0009796 | 0,003549 | 0,0023095 | 0,0023095 | TRUE | 0,0278579 | No |
| orf19.3240 | 562 | 476 | 609 | 408 | 505 | 429 | 392 | 579 | 512 | 481 | 576 | 613 | 449 | 466 | 401 | 442 | 492,55 | 546 | 440 | 0,809 | -0,306 | -3,538 | 0,000404 | 0,000584 | 0,0056667 | 0,004719 | 0,0051487 | 0,0051487 | TRUE | 0,0714948 | No |
| orf19.3241 | 1411 | 1265 | 1350 | 889 | 1465 | 1287 | 1329 | 1815 | 1285 | 1279 | 1278 | 1335 | 1303 | 1397 | 1361 | 1384 | 1327,89 | 1294 | 1361 | 1,053 | 0,075 | 1,576 | 0,115042 | 0,137211 | 1,00E-08 | 0,003032 | 0,001397 | 0,001397 | TRUE | 0,0102331 | No |
| orf19.3242 | 1619 | 1311 | 1354 | 904 | 1434 | 1052 | 1247 | 1741 | 1474 | 1326 | 1281 | 1358 | 1275 | 1142 | 1277 | 1328 | 1307,77 | 1360 | 1256 | 0,924 | -0,114 | -1,789 | 0,073561 | 0,089563 | 0,003155 | 0,003047 | 0,0031044 | 0,0031044 | TRUE | 0,0432872 | No |
| orf19.3243 | 3528 | 3020 | 3361 | 2058 | 5107 | 4184 | 4715 | 6165 | 3212 | 3054 | 3181 | 3091 | 4543 | 4543 | 4829 | 4703 | 3894,43 | 3134 | 4654 | 1,484 | 0,569 | 14,654 | 1,27E-48 | 5,71E-48 | 0,0004873 | 0,002376 | 0,0011781 | 0,0011781 | TRUE | 0,0075895 | No |
| orf19.3244 | 389 | 336 | 339 | 214 | 554 | 403 | 459 | 598 | 354 | 340 | 321 | 321 | 493 | 438 | 470 | 456 | 399,1 | 334 | 464 | 1,387 | 0,472 | 6,128 | 8,92E-10 | 1,67E-09 | 1,00E-08 | 0,005347 | 0,003101 | 0,003101 | TRUE | 0,0203193 | No |
| orf19.3245 | 1839 | 1731 | 1774 | 1123 | 805 | 688 | 638 | 936 | 1674 | 1751 | 1679 | 1687 | 716 | 747 | 653 | 714 | 1202,66 | 1698 | 708 | 0,417 | -1,263 | -23,589 | 4,95E-123 | 5,19E-122 | 0,0001878 | 0,003135 | 0,0017751 | 0,0017751 | TRUE | 0,0294983 | No |
| orf19.3247 | 547 | 667 | 626 | 396 | 496 | 398 | 427 | 481 | 498 | 675 | 592 | 595 | 441 | 432 | 437 | 367 | 504,68 | 590 | 419 | 0,71 | -0,494 | -5,039 | 4,67E-07 | 7,87E-07 | 0,0099163 | 0,004655 | 0,0071846 | 0,0071846 | TRUE | 0,1359461 | No |
| orf19.3248 | 394 | 473 | 401 | 277 | 298 | 272 | 238 | 295 | 359 | 478 | 380 | 416 | 265 | 295 | 244 | 225 | 332,73 | 408 | 257 | 0,629 | -0,668 | -5,989 | 2,11E-09 | 3,90E-09 | 0,0121411 | 0,006007 | 0,0088059 | 0,0088059 | TRUE | 0,1617968 | No |
| orf19.3249 | 1883 | 1982 | 1977 | 1148 | 3569 | 2999 | 3159 | 3996 | 1714 | 2005 | 1871 | 1724 | 3175 | 3256 | 3235 | 3048 | 2503,56 | 1828 | 3178 | 1,736 | 0,796 | 14,198 | 9,39E-46 | 4,05E-45 | 0,0026824 | 0,002565 | 0,0025748 | 0,0025748 | TRUE | 0,0507237 | No |
| orf19.325 | 3062 | 2807 | 3029 | 1921 | 3159 | 2644 | 2902 | 3666 | 2788 | 2839 | 2867 | 2886 | 2810 | 2871 | 2972 | 2796 | 2853,51 | 2845 | 2862 | 1,006 | 0,009 | 0,227 | 0,820262 | 0,844641 | 0,0001816 | 0,0025 | 0,0010701 | 0,0010701 | TRUE | 0,0080478 | No |
| orf19.3250 | 1316 | 1340 | 1391 | 783 | 4638 | 3612 | 4075 | 5013 | 1198 | 1355 | 1316 | 1176 | 4125 | 3922 | 4173 | 3824 | 2636,29 | 1261 | 4011 | 3,174 | 1,666 | 29,671 | 1,84E-193 | 3,02E-192 | 0,0025783 | 0,002538 | 0,0024858 | 0,0024858 | TRUE | 0,0294373 | No |
| orf19.3251 | 964 | 965 | 958 | 641 | 839 | 762 | 709 | 871 | 878 | 976 | 907 | 963 | 746 | 827 | 726 | 664 | 835,91 | 931 | 741 | 0,795 | -0,33 | -4,539 | 5,65E-06 | 9,07E-06 | 0,0041336 | 0,003617 | 0,0038741 | 0,0038741 | TRUE | 0,071115 | No |
| orf19.3252 | 2365 | 2300 | 2501 | 1574 | 1951 | 1703 | 1827 | 2371 | 2153 | 2326 | 2367 | 2364 | 1735 | 1849 | 1871 | 1809 | 2059,35 | 2302 | 1816 | 0,789 | -0,342 | -7,044 | 1,86E-12 | 3,84E-12 | 0,0011139 | 0,002678 | 0,0017724 | 0,0017724 | TRUE | 0,0239452 | No |
| orf19.3254 | 573 | 536 | 593 | 355 | 745 | 657 | 750 | 889 | 522 | 542 | 561 | 533 | 663 | 713 | 768 | 678 | 622,56 | 540 | 706 | 1,305 | 0,384 | 5,617 | 1,95E-08 | 3,46E-08 | 0,0014462 | 0,004159 | 0,0028337 | 0,0028337 | TRUE | 0,0417209 | No |
| orf19.3255 | 495 | 570 | 525 | 262 | 553 | 446 | 503 | 542 | 451 | 576 | 497 | 394 | 492 | 484 | 515 | 413 | 477,79 | 480 | 476 | 0,988 | -0,017 | -0,156 | 0,876278 | 0,895092 | 0,0147793 | 0,004802 | 0,0093184 | 0,0093184 | TRUE | 0,2120498 | No |

| | | | | | | | | | | | | | | | | | | | | | | | | | | | | | | | |
|---|---|---|---|---|---|---|---|---|---|---|---|---|---|---|---|---|---|---|---|---|---|---|---|---|---|---|---|---|---|---|---|
| orf19.3256 | 7104 | 6884 | 7177 | 4526 | 5576 | 4704 | 4809 | 6483 | 6468 | 6962 | 6792 | 6799 | 4960 | 5107 | 4925 | 4945 | 5869,84 | 6755 | 4984 | 0,738 | -0,439 | -12,173 | 4,33E-34 | 1,51E-33 | 0,0004777 | 0,002262 | 0,0010728 | 0,0010728 | TRUE | 0,0102921 | No |
| orf19.3259 | 492 | 438 | 478 | 265 | 364 | 283 | 286 | 336 | 448 | 443 | 452 | 398 | 324 | 307 | 293 | 256 | 365,2 | 435 | 295 | 0,674 | -0,57 | -6,395 | 1,60E-10 | 3,09E-10 | 0,0036875 | 0,005654 | 0,0048143 | 0,0048143 | TRUE | 0,0987624 | No |
| orf19.326 | 372 | 302 | 304 | 205 | 230 | 203 | 198 | 306 | 339 | 305 | 288 | 308 | 205 | 220 | 203 | 233 | 262,62 | 310 | 215 | 0,695 | -0,524 | -5,53 | 3,20E-08 | 5,64E-08 | 1,00E-08 | 0,007067 | 0,0047396 | 0,0047396 | TRUE | 0,0488557 | No |
| orf19.3260 | 3857 | 3425 | 3825 | 2324 | 4826 | 3933 | 4426 | 5798 | 3512 | 3464 | 3620 | 3491 | 4293 | 4270 | 4533 | 4423 | 3950,62 | 3522 | 4380 | 1,243 | 0,314 | 8,433 | 3,38E-17 | 7,94E-17 | 0,0003436 | 0,002371 | 0,0010735 | 0,0010735 | TRUE | 0,0065667 | No |
| orf19.3261 | 102 | 70 | 86 | 50 | 127 | 128 | 135 | 186 | 93 | 71 | 81 | 75 | 113 | 139 | 138 | 142 | 106,53 | 80 | 133 | 1,651 | 0,724 | 4,946 | 7,59E-07 | 1,27E-06 | 1,00E-08 | 0,014437 | 0,0104418 | 0,0104418 | TRUE | 0,1180207 | No |
| orf19.3263 | 27 | 26 | 31 | 19 | 333 | 304 | 280 | 364 | 25 | 26 | 29 | 29 | 296 | 330 | 287 | 278 | 162,43 | 27 | 298 | 10,978 | 3,457 | 20,24 | 4,31E-91 | 3,40E-90 | 1,00E-08 | 0,01017 | 0,0069494 | 0,0069494 | TRUE | 0,0580025 | No |
| orf19.3264 | 340 | 381 | 377 | 237 | 145 | 132 | 122 | 141 | 310 | 385 | 357 | 356 | 129 | 143 | 125 | 108 | 239,06 | 352 | 126 | 0,357 | -1,487 | -13,16 | 1,48E-39 | 5,74E-39 | 0,006223 | 0,007563 | 0,007017 | 0,007017 | TRUE | 0,1156288 | No |
| orf19.3264.1 | 6478 | 4812 | 5025 | 2697 | 3544 | 2440 | 2471 | 4112 | 5898 | 4867 | 4756 | 4051 | 3152 | 2649 | 2531 | 3137 | 3880,06 | 4893 | 2867 | 0,586 | -0,771 | -5,575 | 2,47E-08 | 4,38E-08 | 0,01811 | 0,002377 | 0,0097699 | 0,01811 | TRUE | 0,2332924 | No |
| orf19.3265 | 1512 | 1421 | 1519 | 878 | 1610 | 1343 | 1449 | 1817 | 1377 | 1437 | 1438 | 1319 | 1432 | 1458 | 1484 | 1386 | 1416,31 | 1393 | 1440 | 1,031 | 0,044 | 0,901 | 0,36752 | 0,407397 | 0,000444 | 0,00297 | 0,0016264 | 0,0016264 | TRUE | 0,0133085 | No |
| orf19.3265.1 | 293 | 249 | 271 | 175 | 296 | 205 | 238 | 308 | 267 | 252 | 256 | 263 | 263 | 223 | 244 | 235 | 250,31 | 260 | 241 | 0,931 | -0,104 | -1,128 | 0,259409 | 0,295099 | 1,00E-08 | 0,007314 | 0,0041118 | 0,0041118 | TRUE | 0,0433007 | No |
| orf19.3266 | 188 | 157 | 202 | 164 | 521 | 428 | 393 | 567 | 171 | 159 | 191 | 246 | 463 | 465 | 402 | 433 | 316,32 | 192 | 441 | 2,326 | 1,218 | 10,659 | 1,58E-26 | 4,67E-26 | 0,0130923 | 0,006213 | 0,0086483 | 0,0086483 | TRUE | 0,3300265 | No |
| orf19.3267 | 83 | 74 | 89 | 65 | 146 | 124 | 128 | 172 | 76 | 75 | 84 | 98 | 130 | 135 | 131 | 131 | 107,38 | 83 | 132 | 1,603 | 0,681 | 4,968 | 6,75E-07 | 1,13E-06 | 1,00E-08 | 0,014339 | 0,0080264 | 0,0080264 | TRUE | 0,0954741 | No |
| orf19.3268 | 11881 | 9287 | 9628 | 7778 | 20591 | 17155 | 16826 | 24706 | 10817 | 9393 | 9112 | 11684 | 18315 | 18626 | 17232 | 18846 | 14253 | 10252 | 18255 | 1,781 | 0,833 | 11,436 | 2,78E-30 | 9,00E-30 | 0,0076564 | 0,00213 | 0,0050189 | 0,0050189 | TRUE | 0,1072492 | No |
| orf19.3269 | 445 | 371 | 431 | 262 | 441 | 309 | 339 | 440 | 405 | 375 | 408 | 394 | 392 | 335 | 347 | 336 | 374,05 | 396 | 352 | 0,892 | -0,166 | -2,038 | 0,041553 | 0,051885 | 0,0012186 | 0,005569 | 0,0036692 | 0,0036692 | TRUE | 0,0675319 | No |
| orf19.327 | 544 | 496 | 526 | 342 | 550 | 494 | 465 | 633 | 495 | 502 | 498 | 514 | 489 | 536 | 476 | 483 | 499,14 | 502 | 496 | 0,988 | -0,018 | -0,258 | 0,796126 | 0,823111 | 1,00E-08 | 0,004684 | 0,0024768 | 0,0024768 | TRUE | 0,0322204 | No |
| orf19.3271 | 2 | 5 | 2 | 1 | 1 | 3 | 2 | 4 | 2 | 5 | 2 | 2 | 1 | 3 | 2 | 3 | 2,44 | 3 | 2 | 0,884 | -0,177 | -0,202 | 0,840071 | 0,862531 | 1,00E-08 | 0,543458 | 0,3334996 | 0,3334996 | TRUE | 0,480997 | No |
| orf19.3272 | 1023 | 891 | 950 | 714 | 971 | 794 | 834 | 1074 | 931 | 901 | 899 | 1073 | 864 | 862 | 854 | 819 | 900,41 | 951 | 850 | 0,897 | -0,157 | -2,419 | 0,015549 | 0,020135 | 0,0025606 | 0,003504 | 0,002945 | 0,002945 | TRUE | 0,0791367 | No |
| orf19.3273 | 317 | 272 | 260 | 190 | 141 | 108 | 105 | 134 | 289 | 275 | 246 | 285 | 125 | 117 | 108 | 102 | 193,45 | 274 | 113 | 0,413 | -1,276 | -11,34 | 8,28E-30 | 2,65E-29 | 1,00E-08 | 0,008866 | 0,0060789 | 0,0060789 | TRUE | 0,0600247 | No |
| orf19.3275 | 2624 | 2125 | 2246 | 1380 | 1339 | 872 | 1103 | 1308 | 2389 | 2149 | 2126 | 2073 | 1191 | 947 | 1130 | 998 | 1625,25 | 2184 | 1066 | 0,488 | -1,035 | -13,666 | 1,62E-42 | 6,61E-42 | 0,0067811 | 0,00285 | 0,0048254 | 0,0048254 | TRUE | 0,0751471 | No |
| orf19.3276 | 8846 | 7907 | 8591 | 5343 | 9634 | 8051 | 8766 | 11439 | 8054 | 7997 | 8131 | 8026 | 8569 | 8741 | 8977 | 8726 | 8402,65 | 8052 | 8753 | 1,087 | 0,12 | 4,074 | 4,63E-05 | 7,07E-05 | 0,0001059 | 0,002194 | 0,0007158 | 0,0007158 | TRUE | 0,0035957 | No |
| orf19.3277 | 26 | 17 | 24 | 20 | 9 | 17 | 19 | 18 | 24 | 17 | 23 | 30 | 8 | 18 | 19 | 14 | 19,16 | 24 | 15 | 0,637 | -0,651 | -1,91 | 0,056117 | 0,069237 | 0,0254548 | 0,070985 | 0,056894 | 0,056894 | TRUE | 0,4786788 | No |
| orf19.3278 | 26585 | 22972 | 24811 | 16191 | 13233 | 10842 | 11233 | 15703 | 24205 | 23234 | 23481 | 24321 | 11770 | 11771 | 11504 | 11978 | 17783,2 | 23810 | 11756 | 0,494 | -1,018 | -33,222 | 5,27E-242 | 1,07E-240 | 0,0003041 | 0,002111 | 0,0008392 | 0,0008392 | TRUE | 0,0032442 | No |
| orf19.3279 | 2485 | 2175 | 2412 | 1536 | 2155 | 1825 | 1956 | 2623 | 2262 | 2200 | 2283 | 2307 | 1917 | 1981 | 2003 | 2001 | 2119,33 | 2263 | 1976 | 0,873 | -0,195 | -4,845 | 1,27E-06 | 2,09E-06 | 1,00E-08 | 0,00266 | 0,0010859 | 0,0010859 | TRUE | 0,004967 | No |
| orf19.328 | 533 | 518 | 559 | 313 | 477 | 422 | 403 | 591 | 485 | 524 | 529 | 470 | 424 | 458 | 413 | 451 | 469,3 | 502 | 436 | 0,867 | -0,205 | -2,81 | 0,004958 | 0,006652 | 1,56E-05 | 0,004852 | 0,003002 | 0,003002 | TRUE | 0,0176298 | No |
| orf19.3280 | 2 | 1 | 2 | 1 | 1 | 2 | 0 | 1 | 2 | 1 | 2 | 2 | 1 | 2 | 0 | 1 | 1,26 | 2 | 1 | 0,597 | -0,745 | -0,614 | 0,539267 | 0,580714 | 1,00E-08 | 1,053472 | 0,5777939 | 0,5777939 | TRUE | 0,2749703 | No |
| orf19.3281 | 829 | 745 | 752 | 543 | 164 | 131 | 160 | 186 | 755 | 753 | 712 | 816 | 146 | 142 | 164 | 142 | 453,68 | 759 | 148 | 0,196 | -2,353 | -27,24 | 2,21E-163 | 3,14E-162 | 1,00E-08 | 0,004949 | 0,0033221 | 0,0033221 | TRUE | 0,0480005 | No |
| orf19.3282 | 1894 | 1646 | 1888 | 1054 | 1917 | 1621 | 1688 | 2204 | 1724 | 1665 | 1787 | 1583 | 1705 | 1760 | 1729 | 1681 | 1704,28 | 1690 | 1719 | 1,014 | 0,021 | 0,422 | 0,67311 | 0,709504 | 0,0007272 | 0,002812 | 0,001692 | 0,001692 | TRUE | 0,019518 | No |
| orf19.3283 | 95 | 93 | 104 | 57 | 95 | 73 | 60 | 113 | 86 | 94 | 98 | 86 | 84 | 79 | 61 | 86 | 84,5 | 91 | 78 | 0,856 | -0,224 | -1,428 | 0,153166 | 0,179987 | 1,00E-08 | 0,01767 | 0,0118566 | 0,0118566 | TRUE | 0,1785052 | No |
| orf19.3284 | 0 | 1 | 0 | 1 | 0 | 1 | 1 | 0 | 0 | 1 | 0 | 2 | 0 | 1 | 1 | 0 | 0,58 | 1 | 0 | 0,878 | -0,188 | -0,098 | 0,922092 | 0,933069 | 1,00E-08 | 2,287797 | 1,7275167 | 1,7275167 | TRUE | 0,2184578 | No |
| orf19.3285 | 151 | 124 | 110 | 77 | 200 | 161 | 180 | 203 | 137 | 125 | 104 | 116 | 178 | 175 | 184 | 155 | 146,82 | 120 | 173 | 1,426 | 0,511 | 4,073 | 4,63E-05 | 7,08E-05 | 0,0032337 | 0,011034 | 0,008032 | 0,008032 | TRUE | 0,1006222 | No |
| orf19.3286 | 845 | 771 | 787 | 490 | 869 | 736 | 765 | 1100 | 769 | 780 | 745 | 736 | 773 | 799 | 783 | 839 | 778,07 | 758 | 798 | 1,054 | 0,076 | 1,319 | 0,187097 | 0,217218 | 1,00E-08 | 0,003735 | 0,0018875 | 0,0018875 | TRUE | 0,0151207 | No |
| orf19.3287 | 8810 | 7443 | 8325 | 4770 | 7778 | 6101 | 6924 | 8931 | 8021 | 7528 | 7879 | 7165 | 6918 | 6624 | 7091 | 6813 | 7254,88 | 7648 | 6862 | 0,897 | -0,157 | -3,536 | 0,000406 | 0,000587 | 0,0014753 | 0,002219 | 0,0017666 | 0,0017666 | TRUE | 0,0214511 | No |
| orf19.3288 | 9764 | 8266 | 9233 | 5676 | 12462 | 10273 | 11329 | 15590 | 8890 | 8360 | 8738 | 8526 | 11085 | 11154 | 11602 | 11892 | 10030,9 | 8628 | 11433 | 1,325 | 0,406 | 10,686 | 1,18E-26 | 3,50E-26 | 0,0007884 | 0,002169 | 0,0012824 | 0,0012824 | TRUE | 0,0090514 | No |
| orf19.3289 | 620 | 531 | 566 | 322 | 514 | 387 | 422 | 597 | 564 | 537 | 536 | 484 | 457 | 420 | 432 | 455 | 485,73 | 530 | 441 | 0,83 | -0,269 | -3,728 | 0,000193 | 0,000285 | 0,0004593 | 0,004757 | 0,0029295 | 0,0029295 | TRUE | 0,0342457 | No |
| orf19.329 | 3430 | 2897 | 3175 | 2006 | 2682 | 2048 | 2293 | 3129 | 3123 | 2930 | 3005 | 3013 | 2386 | 2224 | 2348 | 2387 | 2676,91 | 3018 | 2336 | 0,774 | -0,369 | -8,76 | 1,95E-18 | 4,75E-18 | 0,0005144 | 0,00253 | 0,0013223 | 0,0013223 | TRUE | 0,0123349 | No |
| orf19.3290 | 2577 | 2078 | 2241 | 1441 | 3450 | 2621 | 2952 | 3919 | 2346 | 2102 | 2121 | 2165 | 3069 | 2846 | 3023 | 2989 | 2582,56 | 2184 | 2982 | 1,365 | 0,449 | 9,209 | 3,29E-20 | 8,39E-20 | 0,0013027 | 0,002549 | 0,0018826 | 0,0018826 | TRUE | 0,0319391 | No |
| orf19.3291 | 10237 | 8696 | 9281 | 5296 | 14203 | 11468 | 12655 | 17154 | 9320 | 8795 | 8784 | 7955 | 12633 | 12451 | 12960 | 13085 | 10748 | 8714 | 12782 | 1,466 | 0,552 | 11,3 | 1,32E-29 | 4,18E-29 | 0,0022271 | 0,00216 | 0,002194 | 0,002194 | TRUE | 0,0410256 | No |
| orf19.3292 | 2618 | 2151 | 2270 | 1634 | 1862 | 1540 | 1654 | 2312 | 2384 | 2176 | 2148 | 2455 | 1656 | 1672 | 1694 | 1764 | 1993,46 | 2291 | 1696 | 0,742 | -0,431 | -7,88 | 3,28E-15 | 7,29E-15 | 0,001898 | 0,0027 | 0,0023576 | 0,0023576 | TRUE | 0,0262852 | No |
| orf19.3293 | 875 | 783 | 847 | 485 | 967 | 789 | 801 | 997 | 797 | 792 | 802 | 729 | 860 | 857 | 820 | 761 | 802,04 | 780 | 824 | 1,053 | 0,075 | 1,184 | 0,236545 | 0,270634 | 0,001408 | 0,003684 | 0,0025509 | 0,0025509 | TRUE | 0,0347823 | No |
| orf19.3294 | 15847 | 12436 | 13316 | 8048 | 12344 | 8622 | 10298 | 13973 | 14428 | 12578 | 12602 | 12089 | 10980 | 9361 | 10546 | 10659 | 11655,4 | 12924 | 10386 | 0,804 | -0,316 | -4,87 | 1,12E-06 | 1,85E-06 | 0,0053811 | 0,00215 | 0,0039457 | 0,0039457 | TRUE | 0,0752133 | No |
| orf19.3295 | 1248 | 1127 | 1172 | 731 | 798 | 632 | 680 | 848 | 1136 | 1140 | 1109 | 1098 | 710 | 686 | 696 | 647 | 902,83 | 1121 | 685 | 0,61 | -0,714 | -13,017 | 9,78E-39 | 3,73E-38 | 1,00E-08 | 0,0035 | 0,0017276 | 0,0017276 | TRUE | 0,0183569 | No |
| orf19.3296 | 197 | 164 | 171 | 100 | 74 | 37 | 62 | 79 | 179 | 166 | 162 | 150 | 66 | 40 | 63 | 60 | 110,88 | 164 | 57 | 0,351 | -1,511 | -9,934 | 2,97E-23 | 8,15E-23 | 0,0021636 | 0,013951 | 0,0109428 | 0,0109428 | TRUE | 0,3035444 | No |
| orf19.3297 | 2134 | 1984 | 2038 | 1426 | 1597 | 1332 | 1245 | 1746 | 1943 | 2007 | 1929 | 2142 | 1420 | 1446 | 1275 | 1332 | 1686,74 | 2005 | 1368 | 0,683 | -0,549 | -9,783 | 1,33E-22 | 3,59E-22 | 0,0020346 | 0,00282 | 0,0024172 | 0,0024172 | TRUE | 0,0243275 | No |
| orf19.3298 | 2271 | 2093 | 2338 | 1349 | 2399 | 1930 | 1996 | 2616 | 2068 | 2117 | 2213 | 2026 | 2134 | 2095 | 2044 | 1996 | 2086,57 | 2106 | 2067 | 0,98 | -0,029 | -0,632 | 0,527372 | 0,568962 | 0,0006807 | 0,00267 | 0,0015155 | 0,0015155 | TRUE | 0,01424 | No |
| orf19.33 | 4209 | 3677 | 3783 | 2480 | 1604 | 1120 | 1205 | 1719 | 3832 | 3719 | 3580 | 3725 | 1427 | 1216 | 1234 | 1311 | 2505,59 | 3714 | 1297 | 0,35 | -1,516 | -27,622 | 6,13E-168 | 9,03E-167 | 0,0024222 | 0,002564 | 0,0023878 | 0,0023878 | TRUE | 0,0541995 | No |
| orf19.3300 | 17731 | 15674 | 16858 | 10315 | 15254 | 12593 | 13190 | 17927 | 16143 | 15853 | 15955 | 15495 | 13568 | 13672 | 13508 | 13675 | 14733,6 | 15862 | 13606 | 0,858 | -0,222 | -8,227 | 1,93E-16 | 4,43E-16 | 7,46E-05 | 0,002127 | 0,000629 | 0,000629 | TRUE | 0,0028563 | No |
| orf19.3301 | 29617 | 26467 | 29114 | 17610 | 15460 | 13323 | 13932 | 18903 | 26965 | 26769 | 27554 | 26453 | 13751 | 14465 | 14268 | 14419 | 20580,5 | 26935 | 14226 | 0,528 | -0,921 | -30,001 | 9,50E-198 | 1,61E-196 | 0,0003413 | 0,002101 | 0,0008527 | 0,0008527 | TRUE | 0,0062039 | No |
| orf19.330.1 | 444 | 340 | 381 | 269 | 619 | 421 | 526 | 672 | 404 | 344 | 361 | 404 | 551 | 457 | 539 | 513 | 446,47 | 378 | 515 | 1,366 | 0,45 | 5,274 | 1,33E-07 | 2,29E-07 | 0,004217 | 0,004996 | 0,0046504 | 0,0046504 | TRUE | 0,0626476 | No |
| orf19.3302 | 23393 | 22188 | 23519 | 14726 | 2609 | 2296 | 2272 | 3024 | 21298 | 22441 | 22259 | 22121 | 2321 | 2493 | 2327 | 2307 | 12195,7 | 22030 | 2362 | 0,107 | -3,223 | -84,195 | 0 | 0 | 0,0006408 | 0,002145 | 0,0011846 | 0,0011846 | TRUE | 0,016335 | No |
| orf19.3303 | 3710 | 3263 | 3506 | 2095 | 2607 | 2091 | 2265 | 2931 | 3378 | 3300 | 3318 | 3147 | 2319 | 2270 | 2320 | 2236 | 2785,95 | 3286 | 2286 | 0,695 | -0,525 | -13,304 | 2,20E-40 | 8,60E-40 | 0,0002176 | 0,002511 | 0,0011297 | 0,0011297 | TRUE | 0,0090094 | No |
| orf19.3304 | 6525 | 5849 | 6098 | 3329 | 4588 | 3334 | 4008 | 4981 | 5941 | 5916 | 5771 | 5001 | 4081 | 3620 | 4105 | 3800 | 4779,15 | 5657 | 3902 | 0,689 | -0,537 | -8,449 | 2,94E-17 | 6,92E-17 | 0,0045775 | 0,002313 | 0,003666 | 0,003666 | TRUE | 0,0728179 | No |
| orf19.3305 | 2598 | 2352 | 2493 | 1549 | 2174 | 1801 | 1808 | 2376 | 2365 | 2379 | 2359 | 2327 | 1934 | 1955 | 1852 | 1812 | 2122,94 | 2358 | 1888 | 0,8 | -0,322 | -7,622 | 2,49E-14 | 5,42E-14 | 0,0002177 | 0,002659 | 0,0012368 | 0,0012368 | TRUE | 0,0093899 | No |
| orf19.3306 | 20 | 20 | 21 | 14 | 59 | 59 | 57 | 89 | 18 | 20 | 20 | 21 | 52 | 64 | 58 | 68 | 40,27 | 20 | 60 | 3,082 | 1,624 | 6,8 | 1,05E-11 | 2,11E-11 | 1,00E-08 | 0,034842 | 0,0202444 | 0,0202444 | TRUE | 0,0785146 | No |
| orf19.3307 | 261 | 242 | 276 | 177 | 285 | 265 | 241 | 328 | 238 | 245 | 261 | 266 | 253 | 288 | 247 | 250 | 255,96 | 252 | 260 | 1,029 | 0,042 | 0,448 | 0,65423 | 0,691317 | 1,00E-08 | 0,007198 | 0,0043796 | 0,0043796 | TRUE | 0,0551533 | No |
| orf19.3308 | 506 | 470 | 528 | 321 | 897 | 788 | 785 | 1116 | 461 | 475 | 500 | 482 | 798 | 856 | 804 | 851 | 653,32 | 480 | 827 | 1,726 | 0,787 | 12,486 | 8,88E-36 | 3,22E-35 | 1,00E-08 | 0,004059 | 0,0021425 | 0,0021425 | TRUE | 0,010654 | No |
| orf19.3309 | 3325 | 2736 | 2885 | 1842 | 3324 | 2725 | 2824 | 3911 | 3027 | 2767 | 2730 | 2767 | 2957 | 2959 | 2892 | 2983 | 2885,31 | 2823 | 2948 | 1,044 | 0,062 | 1,387 | 0,16534 | 0,19358 | 0,0008079 | 0,002495 | 0,0015453 | 0,0015453 | TRUE | 0,0298705 | No |
| orf19.331 | 1157 | 922 | 1006 | 710 | 1700 | 1444 | 1425 | 2079 | 1053 | 933 | 952 | 1067 | 1512 | 1568 | 1459 | 1586 | 1266,21 | 1001 | 1531 | 1,533 | 0,616 | 10,433 | 1,75E-25 | 5,06E-25 | 0,0019476 | 0,00308 | 0,0025077 | 0,0025077 | TRUE | 0,0249439 | No |
| orf19.3310 | 942 | 785 | 920 | 599 | 417 | 401 | 361 | 523 | 858 | 794 | 871 | 900 | 371 | 435 | 370 | 399 | 624,63 | 856 | 394 | 0,46 | -1,119 | -15,459 | 6,51E-54 | 3,24E-53 | 0,0019648 | 0,004152 | 0,0032179 | 0,0032179 | TRUE | 0,0533153 | No |
| orf19.3311 | 189 | 161 | 164 | 92 | 470 | 401 | 414 | 535 | 172 | 163 | 155 | 138 | 418 | 435 | 424 | 408 | 289,23 | 157 | 421 | 2,656 | 1,409 | 15,286 | 9,51E-53 | 4,61E-52 | 1,00E-08 | 0,006604 | 0,003704 | 0,003704 | TRUE | 0,0506692 | No |
| orf19.3312 | 545 | 517 | 524 | 403 | 715 | 586 | 595 | 818 | 496 | 523 | 496 | 605 | 636 | 636 | 609 | 624 | 578,24 | 530 | 626 | 1,189 | 0,25 | 3,453 | 0,000554 | 0,000795 | 0,0018074 | 0,004322 | 0,0032817 | 0,0032817 | TRUE | 0,0922441 | No |
| orf19.3314 | 244 | 217 | 229 | 173 | 155 | 134 | 122 | 190 | 222 | 219 | 217 | 260 | 138 | 145 | 125 | 145 | 183,93 | 230 | 138 | 0,607 | -0,719 | -6,559 | 5,43E-11 | 1,06E-10 | 1,00E-08 | 0,009219 | 0,005846 | 0,005846 | TRUE | 0,0714506 | No |
| orf19.3315 | 2353 | 2155 | 2460 | 1559 | 725 | 584 | 681 | 884 | 2142 | 2180 | 2328 | 2342 | 645 | 634 | 697 | 674 | 1455,32 | 2248 | 662 | 0,295 | -1,76 | -31,696 | 1,77E-220 | 3,28E-219 | 0,0012108 | 0,002945 | 0,0020185 | 0,0020185 | TRUE | 0,0135789 | No |
| orf19.3317 | 289 | 318 | 286 | 217 | 376 | 285 | 334 | 365 | 263 | 322 | 271 | 326 | 334 | 309 | 342 | 278 | 305,72 | 296 | 316 | 1,074 | 0,103 | 1 | 0,317089 | 0,356138 | 0,0073674 | 0,006358 | 0,0068037 | 0,0068037 | TRUE | 0,0797415 | No |
| orf19.3318 | 313 | 316 | 327 | 187 | 1306 | 1061 | 1233 | 1682 | 285 | 320 | 309 | 281 | 1162 | 1152 | 1263 | 1283 | 756,79 | 299 | 1215 | 4,058 | 2,021 | 27,821 | 2,44E-170 | 3,64E-169 | 0,0019535 | 0,003783 | 0,0029036 | 0,0029036 | TRUE | 0,0240076 | No |
| orf19.3319 | 1453 | 1316 | 1316 | 967 | 1800 | 1401 | 1310 | 1895 | 1323 | 1331 | 1245 | 1453 | 1601 | 1521 | 1342 | 1446 | 1407,65 | 1338 | 1478 | 1,106 | 0,146 | 2,194 | 0,028221 | 0,035731 | 0,0038948 | 0,002975 | 0,0035296 | 0,0035296 | TRUE | 0,0451679 | No |
| orf19.332 | 4369 | 3789 | 4280 | 2607 | 4780 | 3742 | 4309 | 5618 | 3978 | 3832 | 4051 | 3916 | 4252 | 4063 | 4413 | 4285 | 4098,7 | 3944 | 4253 | 1,078 | 0,109 | 2,75 | 0,005966 | 0,007957 | 0,0005295 | 0,002359 | 0,0012551 | 0,0012551 | TRUE | 0,0107331 | No |
| orf19.3320 | 25 | 11 | 20 | 20 | 36 | 50 | 31 | 46 | 23 | 11 | 19 | 30 | 32 | 54 | 32 | 35 | 29,5 | 21 | 38 | 1,878 | 0,909 | 2,971 | 0,002971 | 0,004044 | 0,0601394 | 0,046816 | 0,0513912 | 0,0513912 | TRUE | 0,428924 | No |

| | | | | | | | | | | | | | | | | | | | | | | | | | | | | | | | |
|---|---|---|---|---|---|---|---|---|---|---|---|---|---|---|---|---|---|---|---|---|---|---|---|---|---|---|---|---|---|---|---|
| orf19.3321 | 5751 | 4933 | 5504 | 3324 | 7460 | 6718 | 6701 | 9182 | 5236 | 4989 | 5209 | 4993 | 6635 | 7294 | 6863 | 7004 | 6027,94 | 5107 | 6949 | 1,36 | 0,444 | 10,842 | 2,18E-27 | 6,61E-27 | 0,000904 | 0,002256 | 0,0014342 | 0,0014342 | TRUE | 0,0134201 | No |
| orf19.332.1 | 1695 | 1366 | 1391 | 963 | 1936 | 1472 | 1528 | 2201 | 1543 | 1382 | 1316 | 1447 | 1722 | 1598 | 1565 | 1679 | 1531,48 | 1422 | 1641 | 1,155 | 0,208 | 3,478 | 0,000505 | 0,000727 | 0,0024065 | 0,0029 | 0,0027683 | 0,0027683 | TRUE | 0,0423754 | No |
| orf19.3322 | 579 | 389 | 469 | 355 | 410 | 378 | 339 | 502 | 527 | 393 | 444 | 533 | 365 | 410 | 347 | 383 | 425,36 | 474 | 376 | 0,795 | -0,331 | -3,24 | 0,001197 | 0,001682 | 0,0105988 | 0,005143 | 0,0076423 | 0,0076423 | TRUE | 0,1536119 | No |
| orf19.3323 | 704 | 725 | 703 | 482 | 916 | 770 | 819 | 1064 | 641 | 733 | 665 | 724 | 815 | 836 | 839 | 812 | 758,09 | 691 | 826 | 1,197 | 0,26 | 4,19 | 2,79E-05 | 4,32E-05 | 0,0006937 | 0,00378 | 0,0023471 | 0,0023471 | TRUE | 0,0289338 | No |
| orf19.3324 | 21151 | 18138 | 19788 | 11826 | 21694 | 17771 | 19100 | 25229 | 19257 | 18345 | 18728 | 17765 | 19296 | 19294 | 19561 | 19245 | 18936,3 | 18524 | 19349 | 1,044 | 0,062 | 1,878 | 0,060363 | 0,074203 | 0,0005339 | 0,002107 | 0,0010081 | 0,0010081 | TRUE | 0,0091247 | No |
| orf19.3325 | 1907 | 1657 | 1718 | 1132 | 4175 | 3545 | 4040 | 5187 | 1736 | 1676 | 1626 | 1700 | 3714 | 3849 | 4137 | 3957 | 2799,38 | 1684 | 3914 | 2,324 | 1,216 | 25,647 | 4,60E-145 | 5,84E-144 | 0,0012049 | 0,002509 | 0,0017214 | 0,0017214 | TRUE | 0,0177039 | No |
| orf19.3325.3 | 8301 | 6301 | 6284 | 3776 | 12662 | 8419 | 9410 | 13782 | 7558 | 6373 | 5947 | 5672 | 11262 | 9141 | 9637 | 10513 | 8262,88 | 6388 | 10138 | 1,587 | 0,666 | 7,637 | 2,22E-14 | 4,84E-14 | 0,0126049 | 0,002197 | 0,0071785 | 0,0071785 | TRUE | 0,1861305 | No |
| orf19.3327 | 5071 | 4450 | 4798 | 3124 | 5815 | 4666 | 5004 | 6829 | 4617 | 4501 | 4541 | 4693 | 5172 | 5066 | 5125 | 5209 | 4865,43 | 4588 | 5143 | 1,122 | 0,166 | 5,148 | 2,63E-07 | 4,46E-07 | 1,00E-08 | 0,002309 | 0,0007898 | 0,0007898 | TRUE | 0,0026331 | No |
| orf19.3328 | 2513 | 2197 | 2410 | 1584 | 1986 | 1697 | 1829 | 2503 | 2288 | 2222 | 2281 | 2379 | 1766 | 1842 | 1873 | 1909 | 2070,21 | 2292 | 1848 | 0,807 | -0,309 | -6,995 | 2,66E-12 | 5,45E-12 | 0,0004401 | 0,002675 | 0,0013898 | 0,0013898 | TRUE | 0,0109926 | No |
| orf19.3329 | 861 | 640 | 724 | 491 | 3698 | 3142 | 3358 | 4555 | 784 | 647 | 685 | 738 | 3289 | 3411 | 3439 | 3475 | 2058,52 | 714 | 3404 | 4,773 | 2,255 | 38,612 | 0 | 0 | 0,0020178 | 0,002679 | 0,0023865 | 0,0023865 | TRUE | 0,0588878 | No |
| orf19.333 | 560 | 489 | 514 | 302 | 1016 | 803 | 825 | 1120 | 510 | 495 | 486 | 454 | 904 | 872 | 845 | 854 | 677,41 | 486 | 869 | 1,78 | 0,832 | 13,345 | 1,27E-40 | 4,98E-40 | 1,00E-08 | 0,003987 | 0,0021012 | 0,0021012 | TRUE | 0,0190197 | No |
| orf19.3330 | 2407 | 1909 | 2264 | 1529 | 2021 | 1543 | 1795 | 2350 | 2191 | 1931 | 2143 | 2297 | 1798 | 1675 | 1838 | 1793 | 1958,19 | 2140 | 1776 | 0,831 | -0,267 | -4,557 | 5,19E-06 | 8,36E-06 | 0,0026777 | 0,002712 | 0,0027868 | 0,0027868 | TRUE | 0,0525137 | No |
| orf19.3330.3 | 155 | 160 | 131 | 108 | 208 | 139 | 182 | 245 | 141 | 162 | 124 | 162 | 185 | 151 | 186 | 187 | 162,3 | 147 | 177 | 1,215 | 0,281 | 2,252 | 0,024326 | 0,030966 | 0,0061801 | 0,010176 | 0,0086611 | 0,0086611 | TRUE | 0,1226629 | No |
| orf19.3331 | 23800 | 21332 | 23257 | 15506 | 16538 | 13947 | 14390 | 19492 | 21669 | 21575 | 22011 | 23292 | 14710 | 15143 | 14737 | 14869 | 18500,7 | 22137 | 14865 | 0,672 | -0,574 | -16,53 | 2,25E-61 | 1,25E-60 | 0,0006551 | 0,002108 | 0,0011034 | 0,0011034 | TRUE | 0,0148829 | No |
| orf19.3332 | 2421 | 2041 | 2272 | 1462 | 1533 | 1330 | 1348 | 1826 | 2204 | 2064 | 2150 | 2196 | 1364 | 1444 | 1381 | 1393 | 1774,48 | 2154 | 1396 | 0,648 | -0,626 | -14,025 | 1,10E-44 | 4,68E-44 | 1,00E-08 | 0,002781 | 0,0013312 | 0,0013312 | TRUE | 0,0095693 | No |
| orf19.3333 | 7684 | 6479 | 7065 | 5050 | 8336 | 6957 | 7141 | 9817 | 6996 | 6553 | 6686 | 7586 | 7415 | 7553 | 7313 | 7488 | 7198,85 | 6955 | 7442 | 1,071 | 0,099 | 2,048 | 0,040533 | 0,050656 | 0,0020935 | 0,002221 | 0,0020909 | 0,0020909 | TRUE | 0,0442732 | No |
| orf19.3334 | 20239 | 15571 | 16619 | 10620 | 36360 | 27200 | 29993 | 44156 | 18427 | 15748 | 15728 | 15953 | 32341 | 29532 | 30717 | 33683 | 24016 | 16464 | 31568 | 1,917 | 0,939 | 15,351 | 3,47E-53 | 1,70E-52 | 0,0047778 | 0,002092 | 0,0035476 | 0,0035476 | TRUE | 0,0790871 | No |
| orf19.3335 | 13244 | 11188 | 11997 | 8482 | 16018 | 12806 | 13321 | 18039 | 12058 | 11315 | 11354 | 12741 | 14248 | 13904 | 13642 | 13760 | 12877,9 | 11867 | 13888 | 1,171 | 0,228 | 5,101 | 3,38E-07 | 5,72E-07 | 0,0016932 | 0,00214 | 0,0018317 | 0,0018317 | TRUE | 0,0295092 | No |
| orf19.3336 | 6 | 7 | 6 | 2 | 24 | 10 | 11 | 27 | 5 | 7 | 6 | 3 | 21 | 11 | 11 | 21 | 10,66 | 5 | 16 | 2,976 | 1,573 | 3,289 | 0,001005 | 0,001419 | 1,00E-08 | 0,125942 | 0,0938647 | 0,0938647 | TRUE | 0,064698 | No |
| orf19.3337 | 28155 | 24424 | 27176 | 18157 | 12112 | 10967 | 11508 | 15179 | 25634 | 24702 | 25720 | 27275 | 10773 | 11907 | 11786 | 11579 | 18671,9 | 25833 | 11511 | 0,446 | -1,166 | -25,826 | 4,49E-147 | 5,78E-146 | 0,0017504 | 0,002108 | 0,001897 | 0,001897 | TRUE | 0,0228033 | No |
| orf19.334 | 6384 | 6225 | 6353 | 4352 | 4618 | 3948 | 3854 | 5046 | 5812 | 6296 | 6013 | 6537 | 4108 | 4286 | 3947 | 3849 | 5106,04 | 6164 | 4048 | 0,657 | -0,607 | -11,991 | 3,94E-33 | 1,35E-32 | 0,0021785 | 0,002296 | 0,0022545 | 0,0022545 | TRUE | 0,019725 | No |
| orf19.3340 | 2726 | 2290 | 2416 | 1834 | 3023 | 2493 | 2438 | 3414 | 2482 | 2316 | 2287 | 2755 | 2689 | 2707 | 2497 | 2604 | 2542,01 | 2460 | 2624 | 1,069 | 0,096 | 1,558 | 0,119135 | 0,14188 | 0,0039521 | 0,002557 | 0,0032215 | 0,0032215 | TRUE | 0,0754153 | No |
| orf19.3341 | 9456 | 8392 | 8889 | 5851 | 11590 | 9444 | 9703 | 13614 | 8609 | 8488 | 8413 | 8789 | 10309 | 10254 | 9937 | 10385 | 9397,89 | 8575 | 10221 | 1,193 | 0,254 | 8,106 | 5,22E-16 | 1,19E-15 | 0,0002262 | 0,002178 | 0,0008353 | 0,0008353 | TRUE | 0,0042562 | No |
| orf19.3342 | 2859 | 2643 | 2751 | 1700 | 2227 | 1868 | 1958 | 2518 | 2603 | 2673 | 2604 | 2554 | 1981 | 2028 | 2005 | 1921 | 2296,04 | 2608 | 1984 | 0,759 | -0,397 | -9,964 | 2,18E-23 | 6,02E-23 | 1,00E-08 | 0,002612 | 0,0010821 | 0,0010821 | TRUE | 0,0058175 | No |
| orf19.3344 | 2486 | 2210 | 2375 | 1503 | 2279 | 1818 | 1968 | 2570 | 2263 | 2235 | 2248 | 2258 | 2027 | 1974 | 2015 | 1960 | 2122,61 | 2251 | 1994 | 0,886 | -0,175 | -4,564 | 5,01E-06 | 8,08E-06 | 1,00E-08 | 0,002659 | 0,0009374 | 0,0009374 | TRUE | 0,0017266 | No |
| orf19.3345 | 3220 | 2805 | 3003 | 1943 | 2674 | 2152 | 2442 | 3328 | 2932 | 2837 | 2842 | 2919 | 2378 | 2336 | 2501 | 2539 | 2660,48 | 2882 | 2438 | 0,847 | -0,24 | -5,658 | 1,53E-08 | 2,74E-08 | 0,0005603 | 0,002534 | 0,0013488 | 0,0013488 | TRUE | 0,0096533 | No |
| orf19.3347 | 1078 | 983 | 1075 | 640 | 1069 | 836 | 935 | 1285 | 981 | 994 | 1017 | 961 | 951 | 908 | 958 | 980 | 968,84 | 988 | 949 | 0,96 | -0,059 | -1,126 | 0,260137 | 0,295769 | 1,00E-08 | 0,003401 | 0,0016022 | 0,0016022 | TRUE | 0,009805 | No |
| orf19.3348 | 2046 | 1669 | 1664 | 1056 | 1624 | 1252 | 1394 | 1810 | 1863 | 1688 | 1575 | 1586 | 1444 | 1359 | 1428 | 1381 | 1540,51 | 1678 | 1403 | 0,835 | -0,26 | -4,271 | 1,95E-05 | 3,04E-05 | 0,0026839 | 0,002895 | 0,0029059 | 0,0029059 | TRUE | 0,0679443 | No |
| orf19.3349 | 6569 | 5799 | 6459 | 3756 | 7629 | 6060 | 6977 | 9313 | 5981 | 5865 | 6113 | 5642 | 6786 | 6579 | 7145 | 7104 | 6401,93 | 5900 | 6904 | 1,17 | 0,226 | 5,347 | 8,93E-08 | 1,55E-07 | 0,0011236 | 0,002243 | 0,0015561 | 0,0015561 | TRUE | 0,0120369 | No |
| orf19.335 | 1305 | 1536 | 1503 | 999 | 316 | 288 | 276 | 368 | 1188 | 1553 | 1422 | 1501 | 281 | 313 | 283 | 281 | 852,74 | 1416 | 290 | 0,204 | -2,292 | -25,428 | 1,24E-142 | 1,53E-141 | 0,0079859 | 0,003586 | 0,0058051 | 0,0058051 | TRUE | 0,1445579 | No |
| orf19.3350 | 3404 | 3188 | 3288 | 2186 | 3093 | 2530 | 2478 | 3431 | 3099 | 3224 | 3112 | 3284 | 2751 | 2747 | 2538 | 2617 | 2921,5 | 3180 | 2663 | 0,838 | -0,255 | -5,869 | 4,39E-09 | 8,03E-09 | 0,0006995 | 0,002489 | 0,0014631 | 0,0014631 | TRUE | 0,0118054 | No |
| orf19.3351 | 1945 | 2047 | 2105 | 1246 | 1713 | 1326 | 1412 | 1832 | 1771 | 2070 | 1992 | 1872 | 1524 | 1440 | 1446 | 1397 | 1688,99 | 1926 | 1452 | 0,753 | -0,409 | -6,995 | 2,66E-12 | 5,44E-12 | 0,0025621 | 0,002819 | 0,0026758 | 0,0026758 | TRUE | 0,0377173 | No |
| orf19.3352 | 749 | 723 | 754 | 409 | 950 | 860 | 772 | 994 | 682 | 731 | 714 | 614 | 845 | 934 | 791 | 758 | 758,59 | 685 | 832 | 1,209 | 0,273 | 3,465 | 0,000531 | 0,000763 | 0,0053031 | 0,003778 | 0,0046323 | 0,0046323 | TRUE | 0,0787559 | No |
| orf19.3353 | 3046 | 2741 | 2832 | 1647 | 3976 | 3007 | 3246 | 4319 | 2773 | 2772 | 2680 | 2474 | 3537 | 3265 | 3324 | 3295 | 3014,99 | 2675 | 3355 | 1,252 | 0,325 | 6,628 | 3,41E-11 | 6,75E-11 | 0,0016148 | 0,002475 | 0,0019658 | 0,0019658 | TRUE | 0,0288922 | No |
| orf19.3354 | 9353 | 7345 | 7884 | 4287 | 18421 | 12694 | 14738 | 20387 | 8516 | 7429 | 7461 | 6440 | 16385 | 13782 | 15094 | 15551 | 11332,2 | 7462 | 15203 | 2,037 | 1,026 | 13,253 | 4,33E-40 | 1,68E-39 | 0,0088804 | 0,002154 | 0,0056578 | 0,0056578 | TRUE | 0,1104891 | No |
| orf19.3355 | 2775 | 2693 | 2808 | 1810 | 1751 | 1444 | 1455 | 2013 | 2527 | 2724 | 2658 | 2719 | 1557 | 1568 | 1490 | 1536 | 2097,19 | 2657 | 1538 | 0,579 | -0,788 | -17,8 | 7,09E-71 | 4,50E-70 | 0,0004759 | 0,002667 | 0,0013734 | 0,0013734 | TRUE | 0,0136059 | No |
| orf19.3356 | 468 | 590 | 518 | 293 | 328 | 294 | 312 | 332 | 426 | 597 | 490 | 440 | 292 | 319 | 320 | 253 | 392,11 | 488 | 296 | 0,603 | -0,729 | -6,304 | 2,89E-10 | 5,52E-10 | 0,0156188 | 0,005406 | 0,0101385 | 0,0101385 | TRUE | 0,2617947 | No |
| orf19.3357 | 1014 | 885 | 990 | 556 | 880 | 684 | 731 | 955 | 923 | 895 | 937 | 835 | 783 | 743 | 749 | 728 | 824,11 | 898 | 751 | 0,834 | -0,263 | -4,483 | 7,36E-06 | 1,18E-05 | 0,000365 | 0,00364 | 0,002081 | 0,002081 | TRUE | 0,0224834 | No |
| orf19.3358 | 2489 | 1927 | 2081 | 1468 | 6258 | 4844 | 5259 | 7451 | 2266 | 1949 | 1969 | 2205 | 5566 | 5259 | 5386 | 5684 | 3785,6 | 2097 | 5474 | 2,612 | 1,385 | 24,429 | 8,36E-132 | 9,46E-131 | 0,0031255 | 0,002386 | 0,0027453 | 0,0027453 | TRUE | 0,0378467 | No |
| orf19.3359 | 1665 | 1692 | 1727 | 938 | 1374 | 1136 | 1213 | 1512 | 1516 | 1711 | 1634 | 1409 | 1222 | 1233 | 1242 | 1153 | 1390,23 | 1568 | 1212 | 0,772 | -0,374 | -5,855 | 4,76E-09 | 8,68E-09 | 0,0031711 | 0,002987 | 0,0031889 | 0,0031889 | TRUE | 0,0515298 | No |
| orf19.336 | 2640 | 2257 | 2385 | 1426 | 1236 | 925 | 1035 | 1433 | 2404 | 2283 | 2257 | 2142 | 1099 | 1004 | 1060 | 1093 | 1667,79 | 2272 | 1064 | 0,468 | -1,094 | -20,771 | 7,97E-96 | 6,54E-95 | 0,0012888 | 0,002829 | 0,0019894 | 0,0019894 | TRUE | 0,0186346 | No |
| orf19.3360 | 9339 | 9626 | 9941 | 6252 | 3391 | 2805 | 2954 | 3636 | 8503 | 9736 | 9408 | 9392 | 3016 | 3045 | 3025 | 2774 | 6112,33 | 9260 | 2965 | 0,32 | -1,643 | -31,398 | 2,14E-216 | 3,90E-215 | 0,0022699 | 0,002253 | 0,0024151 | 0,0024151 | TRUE | 0,0373705 | No |
| orf19.3362 | 253 | 229 | 242 | 161 | 243 | 193 | 222 | 318 | 230 | 232 | 229 | 242 | 216 | 210 | 227 | 243 | 228,56 | 233 | 224 | 0,965 | -0,051 | -0,543 | 0,587266 | 0,627632 | 1,00E-08 | 0,007817 | 0,0042507 | 0,0042507 | TRUE | 0,038751 | No |
| orf19.3363 | 30621 | 27690 | 29468 | 17871 | 5348 | 4386 | 4611 | 6257 | 27879 | 28005 | 27889 | 26845 | 4757 | 4762 | 4722 | 4773 | 16204,1 | 27654 | 4754 | 0,172 | -2,541 | -86,302 | 0 | 0 | 0,0001901 | 0,002119 | 0,0007145 | 0,0007145 | TRUE | 0,0046592 | No |
| orf19.3364 | 834 | 790 | 864 | 488 | 493 | 409 | 371 | 515 | 759 | 799 | 818 | 733 | 439 | 444 | 380 | 393 | 595,55 | 777 | 414 | 0,531 | -0,914 | -12,586 | 2,51E-36 | 9,23E-36 | 0,0017682 | 0,004255 | 0,0032413 | 0,0032413 | TRUE | 0,0320914 | No |
| orf19.3365 | 3069 | 2566 | 2773 | 1751 | 4185 | 3373 | 3700 | 4714 | 2794 | 2595 | 2624 | 2630 | 3722 | 3662 | 3789 | 3596 | 3176,73 | 2661 | 3692 | 1,387 | 0,471 | 11,606 | 3,84E-31 | 1,27E-30 | 0,0005089 | 0,002453 | 0,0012556 | 0,0012556 | TRUE | 0,0144011 | No |
| orf19.3366 | 3792 | 3066 | 3231 | 2094 | 5207 | 3937 | 4456 | 5828 | 3452 | 3101 | 3058 | 3146 | 4631 | 4274 | 4564 | 4446 | 3833,98 | 3189 | 4479 | 1,404 | 0,49 | 9,888 | 4,68E-23 | 1,28E-22 | 0,0018286 | 0,002382 | 0,0020801 | 0,0020801 | TRUE | 0,0386562 | No |
| orf19.3366.1 | 1480 | 1317 | 1366 | 896 | 2710 | 2094 | 2125 | 2877 | 1347 | 1332 | 1293 | 1346 | 2410 | 2274 | 2176 | 2195 | 1796,63 | 1330 | 2264 | 1,703 | 0,768 | 15,6 | 7,31E-55 | 3,68E-54 | 0,0009802 | 0,002772 | 0,0017193 | 0,0017193 | TRUE | 0,023498 | No |
| orf19.3367 | 4334 | 4123 | 4139 | 2605 | 3848 | 3110 | 3472 | 4273 | 3946 | 4170 | 3917 | 3913 | 3423 | 3377 | 3556 | 3259 | 3695,09 | 3986 | 3404 | 0,853 | -0,229 | -5,417 | 6,05E-08 | 1,05E-07 | 0,0007617 | 0,002395 | 0,0014436 | 0,0014436 | TRUE | 0,0117728 | No |
| orf19.3368 | 319 | 333 | 307 | 241 | 209 | 171 | 197 | 207 | 290 | 337 | 291 | 362 | 186 | 186 | 202 | 158 | 251,38 | 320 | 183 | 0,572 | -0,805 | -7,343 | 2,09E-13 | 4,43E-13 | 0,0073289 | 0,007292 | 0,0073016 | 0,0073016 | TRUE | 0,1013747 | No |
| orf19.3369 | 22127 | 23586 | 24885 | 15033 | 5990 | 5110 | 5181 | 6396 | 20146 | 23855 | 23551 | 22582 | 5328 | 5548 | 5306 | 4879 | 13899,3 | 22534 | 5265 | 0,234 | -2,098 | -34,859 | 3,13E-266 | 7,10E-265 | 0,0040408 | 0,002132 | 0,0033668 | 0,0033668 | TRUE | 0,0625331 | No |
| orf19.337 | 3055 | 2932 | 2842 | 1826 | 1618 | 1320 | 1383 | 1800 | 2781 | 2965 | 2690 | 2743 | 1439 | 1433 | 1416 | 1373 | 2105,15 | 2795 | 1415 | 0,506 | -0,983 | -21,026 | 3,80E-98 | 3,26E-97 | 0,0008321 | 0,002665 | 0,0015745 | 0,0015745 | TRUE | 0,0207676 | No |
| orf19.3370 | 12294 | 11725 | 12505 | 7878 | 12056 | 10479 | 11165 | 14372 | 11193 | 11859 | 11835 | 11834 | 10723 | 11377 | 11434 | 10963 | 11402,4 | 11680 | 11124 | 0,952 | -0,07 | -1,899 | 0,057617 | 0,070964 | 0,0007369 | 0,002153 | 0,0012299 | 0,0012299 | TRUE | 0,0097568 | No |
| orf19.3371 | 2231 | 2097 | 2287 | 1256 | 1123 | 900 | 992 | 1327 | 2031 | 2121 | 2164 | 1887 | 999 | 977 | 1016 | 1012 | 1525,93 | 2051 | 1001 | 0,487 | -1,037 | -19,192 | 4,32E-82 | 3,14E-81 | 0,0013923 | 0,002903 | 0,0020743 | 0,0020743 | TRUE | 0,0322013 | No |
| orf19.3373 | 14416 | 12283 | 13125 | 9620 | 1051 | 892 | 866 | 1226 | 13125 | 12423 | 12422 | 14451 | 935 | 968 | 887 | 935 | 7018,24 | 13105 | 931 | 0,071 | -3,814 | -64,734 | 0 | 0 | 0,0032135 | 0,002225 | 0,0027918 | 0,0027918 | TRUE | 0,0577991 | No |
| orf19.3374 | 16501 | 13556 | 14242 | 10402 | 336 | 281 | 235 | 421 | 15023 | 13710 | 13479 | 15625 | 299 | 305 | 241 | 321 | 7375,49 | 14459 | 292 | 0,02 | -5,628 | -72,049 | 0 | 0 | 0,0074802 | 0,002216 | 0,004229 | 0,004229 | TRUE | 0,1494169 | No |
| orf19.3375 | 2 | 3 | 1 | 0 | 10 | 8 | 9 | 12 | 2 | 3 | 1 | 0 | 9 | 9 | 9 | 9 | 5,22 | 2 | 9 | 5,771 | 2,529 | 3,39 | 0,000699 | 0,000997 | 1,00E-08 | 0,255145 | 0,1439851 | 0,1439851 | TRUE | 0,3070413 | No |
| orf19.3376 | 135 | 149 | 134 | 89 | 51 | 48 | 60 | 73 | 123 | 151 | 127 | 134 | 45 | 52 | 61 | 56 | 93,59 | 134 | 54 | 0,402 | -1,316 | -8,428 | 3,52E-17 | 8,27E-17 | 1,00E-08 | 0,016152 | 0,0107153 | 0,0107153 | TRUE | 0,0966735 | No |
| orf19.3378 | 94 | 70 | 68 | 44 | 2 | 2 | 4 | 13 | 86 | 71 | 64 | 66 | 2 | 2 | 4 | 10 | 38,1 | 72 | 4 | 0,067 | -3,909 | -10,322 | 5,60E-25 | 1,61E-24 | 0,017338 | 0,036709 | 0,0336797 | 0,0336797 | TRUE | 1,6974797 | No |
| orf19.338 | 3947 | 3291 | 3542 | 2355 | 7576 | 6353 | 6869 | 9451 | 3594 | 3328 | 3352 | 3538 | 6739 | 6898 | 7035 | 7209 | 5211,51 | 3453 | 6970 | 2,02 | 1,014 | 24,45 | 5,04E-132 | 5,73E-131 | 0,0008588 | 0,00229 | 0,0014286 | 0,0014286 | TRUE | 0,0097991 | No |
| orf19.3380 | 23 | 18 | 13 | 11 | 19 | 12 | 12 | 23 | 21 | 18 | 12 | 17 | 17 | 13 | 12 | 18 | 15,97 | 17 | 15 | 0,886 | -0,174 | -0,506 | 0,612944 | 0,65234 | 1,00E-08 | 0,08477 | 0,0514275 | 0,0514275 | TRUE | 0,1998232 | No |
| orf19.339 | 8216 | 7279 | 7632 | 5111 | 8634 | 7274 | 7083 | 10069 | 7480 | 7362 | 7223 | 7678 | 7680 | 7898 | 7254 | 7681 | 7531,82 | 7436 | 7628 | 1,026 | 0,037 | 0,969 | 0,332409 | 0,372165 | 0,0007826 | 0,002212 | 0,0012968 | 0,0012968 | TRUE | 0,0131504 | No |
| orf19.3391 | 1260 | 1086 | 1165 | 810 | 2419 | 1930 | 2013 | 2790 | 1147 | 1098 | 1103 | 1217 | 2152 | 2095 | 2062 | 2128 | 1625,22 | 1141 | 2109 | 1,854 | 0,891 | 18,69 | 5,93E-78 | 4,10E-77 | 0,0001032 | 0,00285 | 0,0014881 | 0,0014881 | TRUE | 0,0203982 | No |
| orf19.3392 | 670 | 584 | 628 | 409 | 986 | 793 | 782 | 1126 | 610 | 591 | 594 | 614 | 877 | 861 | 801 | 859 | 725,9 | 602 | 850 | 1,412 | 0,498 | 8,418 | 3,83E-17 | 9,00E-17 | 1,00E-08 | 0,003857 | 0,0019264 | 0,0019264 | TRUE | 0,0170189 | No |
| orf19.3393 | 10852 | 10150 | 10835 | 6735 | 12194 | 10433 | 10611 | 14126 | 9880 | 10266 | 10254 | 10117 | 10846 | 11327 | 10867 | 10775 | 10541,7 | 10129 | 10954 | 1,081 | 0,113 | 3,536 | 0,000406 | 0,000587 | 0,0002944 | 0,002162 | 0,0008795 | 0,0008795 | TRUE | 0,0063616 | No |
| orf19.3394 | 483 | 445 | 462 | 350 | 556 | 444 | 471 | 590 | 440 | 450 | 437 | 526 | 495 | 482 | 482 | 450 | 470,23 | 463 | 477 | 1,036 | 0,051 | 0,656 | 0,511683 | 0,553722 | 0,0018558 | 0,004846 | 0,0035764 | 0,0035764 | TRUE | 0,0820854 | No |
| orf19.3395 | 2124 | 1850 | 1972 | 1211 | 2891 | 2253 | 2452 | 3514 | 1934 | 1871 | 1866 | 1819 | 2571 | 2446 | 2511 | 2681 | 2212,45 | 1872 | 2552 | 1,363 | 0,447 | 9,876 | 5,29E-23 | 1,44E-22 | 0,000718 | 0,002634 | 0,0014934 | 0,0014934 | TRUE | 0,0142728 | No |

| | | | | | | | | | | | | | | | | | | | | | | | | | | | | | | | |
|---|---|---|---|---|---|---|---|---|---|---|---|---|---|---|---|---|---|---|---|---|---|---|---|---|---|---|---|---|---|---|---|
| orf19.3396 | 1314 | 1025 | 1258 | 711 | 667 | 461 | 551 | 734 | 1196 | 1037 | 1191 | 1068 | 593 | 501 | 564 | 560 | 838,7 | 1123 | 554 | 0,494 | -1,018 | -13,995 | 1,68E-44 | 7,12E-44 | 0,003919 | 0,003612 | 0,0037629 | 0,0037629 | TRUE | 0,0475893 | No |
| orf19.3399 | 1922 | 1540 | 1770 | 1095 | 1755 | 1304 | 1533 | 2010 | 1750 | 1558 | 1675 | 1645 | 1561 | 1416 | 1570 | 1533 | 1588,44 | 1657 | 1520 | 0,917 | -0,124 | -2,29 | 0,022043 | 0,028161 | 0,0016435 | 0,002869 | 0,0021982 | 0,0021982 | TRUE | 0,0246372 | No |
| orf19.34 | 383 | 334 | 354 | 206 | 92 | 67 | 94 | 120 | 349 | 338 | 335 | 309 | 82 | 73 | 96 | 92 | 209,17 | 333 | 86 | 0,257 | -1,958 | -17,288 | 5,79E-67 | 3,48E-66 | 1,00E-08 | 0,008352 | 0,0053087 | 0,0053087 | TRUE | 0,0835098 | No |
| orf19.340 | 581 | 551 | 588 | 324 | 148 | 100 | 138 | 147 | 529 | 557 | 556 | 487 | 132 | 109 | 141 | 112 | 327,89 | 532 | 124 | 0,231 | -2,115 | -21,112 | 6,16E-99 | 5,32E-98 | 0,0023578 | 0,006066 | 0,0048617 | 0,0048617 | TRUE | 0,0901831 | No |
| orf19.3400 | 1551 | 1301 | 1400 | 750 | 1556 | 1154 | 1339 | 1795 | 1412 | 1316 | 1325 | 1127 | 1384 | 1253 | 1371 | 1369 | 1319,63 | 1295 | 1344 | 1,036 | 0,051 | 0,74 | 0,45904 | 0,50105 | 0,0041171 | 0,003038 | 0,0037107 | 0,0037107 | TRUE | 0,0848162 | No |
| orf19.3401 | 2327 | 1948 | 2170 | 1365 | 2345 | 1876 | 1919 | 2787 | 2119 | 1970 | 2054 | 2050 | 2086 | 2037 | 1965 | 2126 | 2050,86 | 2048 | 2054 | 1,003 | 0,004 | 0,099 | 0,920929 | 0,93204 | 0,0005882 | 0,002681 | 0,0014592 | 0,0014592 | TRUE | 0,0100876 | No |
| orf19.3402 | 2872 | 2334 | 2584 | 1850 | 2129 | 1550 | 1742 | 2379 | 2615 | 2361 | 2446 | 2779 | 1894 | 1683 | 1784 | 1815 | 2171,9 | 2550 | 1794 | 0,705 | -0,505 | -8,399 | 4,51E-17 | 1,06E-16 | 0,0031998 | 0,002645 | 0,0029978 | 0,0029978 | TRUE | 0,0421169 | No |
| orf19.3404 | 7 | 15 | 12 | 4 | 10 | 9 | 7 | 16 | 6 | 15 | 11 | 6 | 9 | 10 | 7 | 12 | 9,62 | 10 | 10 | 0,971 | -0,043 | -0,094 | 0,925206 | 0,935478 | 1,00E-08 | 0,139373 | 0,0977925 | 0,0977925 | TRUE | 0,4665002 | No |
| orf19.3405 | 470 | 424 | 477 | 343 | 1156 | 1016 | 1063 | 1400 | 428 | 429 | 451 | 515 | 1028 | 1103 | 1089 | 1068 | 763,91 | 456 | 1072 | 2,367 | 1,243 | 18,582 | 4,53E-77 | 3,09E-76 | 0,000952 | 0,003766 | 0,0026794 | 0,0026794 | TRUE | 0,0741802 | No |
| orf19.3406 | 1231 | 1091 | 1167 | 754 | 472 | 365 | 415 | 559 | 1121 | 1103 | 1104 | 1133 | 420 | 396 | 425 | 426 | 766,1 | 1115 | 417 | 0,375 | -1,417 | -24,257 | 5,54E-130 | 6,13E-129 | 1,00E-08 | 0,003761 | 0,0016808 | 0,0016808 | TRUE | 0,0116315 | No |
| orf19.3407 | 1384 | 1128 | 1301 | 874 | 743 | 574 | 665 | 941 | 1260 | 1141 | 1231 | 1313 | 661 | 623 | 681 | 718 | 953,5 | 1236 | 671 | 0,544 | -0,877 | -13,609 | 3,53E-42 | 1,43E-41 | 0,002297 | 0,003422 | 0,0028611 | 0,0028611 | TRUE | 0,0321319 | No |
| orf19.3409 | 1570 | 1330 | 1476 | 948 | 1308 | 1076 | 1187 | 1614 | 1429 | 1345 | 1397 | 1424 | 1163 | 1168 | 1216 | 1231 | 1296,75 | 1399 | 1194 | 0,855 | -0,226 | -4,685 | 2,79E-06 | 4,55E-06 | 1,00E-08 | 0,003056 | 0,001465 | 0,001465 | TRUE | 0,0081473 | No |
| orf19.341 | 376 | 291 | 346 | 213 | 2246 | 1835 | 1863 | 2701 | 342 | 294 | 327 | 320 | 1998 | 1992 | 1908 | 2060 | 1155,3 | 321 | 1990 | 6,191 | 2,63 | 41,438 | 0 | 0 | 0,0005998 | 0,00318 | 0,0019819 | 0,0019819 | TRUE | 0,0388258 | No |
| orf19.3411 | 204 | 166 | 179 | 102 | 440 | 371 | 355 | 520 | 186 | 168 | 169 | 153 | 391 | 403 | 364 | 397 | 278,83 | 169 | 389 | 2,283 | 1,191 | 12,82 | 1,26E-37 | 4,73E-37 | 1,00E-08 | 0,006775 | 0,0039686 | 0,0039686 | TRUE | 0,0474125 | No |
| orf19.3412 | 5005 | 4056 | 4652 | 2881 | 2552 | 2129 | 2436 | 3159 | 4557 | 4102 | 4403 | 4328 | 2270 | 2312 | 2495 | 2410 | 3359,42 | 4348 | 2372 | 0,545 | -0,875 | -18,134 | 1,72E-73 | 1,13E-72 | 0,0014189 | 0,00243 | 0,0019129 | 0,0019129 | TRUE | 0,0179289 | No |
| orf19.3413 | 3 | 7 | 0 | 1 | 3 | 2 | 1 | 6 | 3 | 7 | 0 | 2 | 3 | 2 | 1 | 5 | 2,72 | 3 | 3 | 0,933 | -0,099 | -0,106 | 0,915388 | 0,927463 | 0,4619683 | 0,487829 | 0,4797676 | 0,4797676 | TRUE | 0,8409953 | No |
| orf19.3414 | 9053 | 8538 | 9260 | 5360 | 3772 | 2981 | 3111 | 4284 | 8242 | 8635 | 8764 | 8052 | 3355 | 3237 | 3186 | 3268 | 5842,31 | 8423 | 3262 | 0,387 | -1,369 | -33,762 | 7,08E-250 | 1,47E-248 | 0,0007325 | 0,002263 | 0,001372 | 0,001372 | TRUE | 0,0103995 | No |
| orf19.3415 | 11953 | 10233 | 11804 | 7584 | 8748 | 7705 | 8160 | 11395 | 10883 | 10350 | 11171 | 11392 | 7781 | 8365 | 8357 | 8692 | 9623,96 | 10949 | 8299 | 0,758 | -0,399 | -8,638 | 5,74E-18 | 1,38E-17 | 0,0017486 | 0,002174 | 0,0019484 | 0,0019484 | TRUE | 0,0217128 | No |
| orf19.3415.1 | 15953 | 13458 | 13657 | 8936 | 24315 | 17970 | 17992 | 26052 | 14525 | 13611 | 12925 | 13423 | 21627 | 19510 | 18426 | 19873 | 16740,1 | 13621 | 19859 | 1,458 | 0,544 | 9,877 | 5,24E-23 | 1,42E-22 | 0,0033618 | 0,002116 | 0,0028507 | 0,0028507 | TRUE | 0,0439996 | No |
| orf19.3417 | 114 | 114 | 128 | 86 | 216 | 170 | 180 | 244 | 104 | 115 | 121 | 129 | 192 | 185 | 184 | 186 | 152,07 | 117 | 187 | 1,604 | 0,682 | 5,888 | 3,92E-09 | 7,16E-09 | 1,00E-08 | 0,010724 | 0,005825 | 0,005825 | TRUE | 0,0646279 | No |
| orf19.3418 | 132 | 120 | 133 | 57 | 278 | 238 | 247 | 366 | 120 | 121 | 126 | 86 | 247 | 258 | 253 | 279 | 186,36 | 113 | 259 | 2,259 | 1,176 | 9,994 | 1,63E-23 | 4,51E-23 | 0,0017842 | 0,009126 | 0,0068653 | 0,0068653 | TRUE | 0,1957639 | No |
| orf19.3419 | 913 | 845 | 859 | 585 | 823 | 663 | 707 | 987 | 831 | 855 | 813 | 879 | 732 | 720 | 724 | 753 | 788,3 | 844 | 732 | 0,87 | -0,201 | -3,596 | 0,000323 | 0,000471 | 1,00E-08 | 0,003713 | 0,001734 | 0,001734 | TRUE | 0,0076282 | No |
| orf19.342 | 35 | 33 | 24 | 23 | 79 | 69 | 63 | 80 | 32 | 33 | 23 | 35 | 70 | 75 | 65 | 61 | 49,15 | 31 | 68 | 2,225 | 1,154 | 5,488 | 4,06E-08 | 7,12E-08 | 1,00E-08 | 0,028912 | 0,0180497 | 0,0180497 | TRUE | 0,2228858 | No |
| orf19.3420 | 0 | 0 | 0 | 0 | 1 | 3 | 4 | 1 | 0 | 0 | 0 | 0 | 1 | 3 | 4 | 1 | 1,13 | 0 | 2 | 11,187 | 3,484 | 2,29 | 0,022033 | 0,028154 | 1,00E-08 | 1,175482 | 1,0116419 | 1,0116419 | TRUE | 0,0843906 | No |
| orf19.3421.1 | 1902 | 1763 | 1771 | 1216 | 1193 | 909 | 979 | 1267 | 1732 | 1783 | 1676 | 1827 | 1061 | 987 | 1003 | 966 | 1379,33 | 1754 | 1004 | 0,573 | -0,803 | -15,514 | 2,81E-54 | 1,40E-53 | 0,0007631 | 0,002995 | 0,001801 | 0,001801 | TRUE | 0,01801 | No |
| orf19.3422 | 824 | 771 | 868 | 502 | 2336 | 2093 | 2083 | 2736 | 750 | 780 | 821 | 754 | 2078 | 2272 | 2133 | 2087 | 1459,51 | 776 | 2142 | 2,754 | 1,462 | 26,937 | 8,05E-160 | 1,11E-158 | 0,0010407 | 0,002942 | 0,0019206 | 0,0019206 | TRUE | 0,0196745 | No |
| orf19.3423 | 10311 | 8900 | 10229 | 6156 | 13806 | 11559 | 12855 | 16436 | 9388 | 9001 | 9681 | 9247 | 12280 | 12550 | 13165 | 12538 | 10981,2 | 9329 | 12633 | 1,354 | 0,437 | 11,658 | 2,09E-31 | 6,96E-31 | 0,0007803 | 0,002157 | 0,001255 | 0,001255 | TRUE | 0,0097893 | No |
| orf19.3425 | 2533 | 2377 | 2564 | 1634 | 2993 | 2407 | 2659 | 3417 | 2306 | 2404 | 2427 | 2455 | 2662 | 2613 | 2723 | 2607 | 2524,57 | 2398 | 2651 | 1,106 | 0,146 | 3,657 | 0,000255 | 0,000375 | 0,0001731 | 0,00256 | 0,0011298 | 0,0011298 | TRUE | 0,0082846 | No |
| orf19.3426 | 13165 | 11876 | 12008 | 9136 | 21754 | 18754 | 17517 | 24499 | 11986 | 12011 | 11364 | 13724 | 19349 | 20362 | 17940 | 18688 | 15678,1 | 12271 | 19085 | 1,556 | 0,638 | 10,404 | 2,39E-25 | 6,89E-25 | 0,0047706 | 0,002121 | 0,0035401 | 0,0035401 | TRUE | 0,0769268 | No |
| orf19.3427 | 224 | 213 | 243 | 188 | 252 | 215 | 196 | 278 | 204 | 215 | 230 | 282 | 224 | 233 | 201 | 212 | 225,26 | 233 | 218 | 0,943 | -0,085 | -0,767 | 0,442887 | 0,484704 | 0,0066954 | 0,007901 | 0,0074291 | 0,0074291 | TRUE | 0,1907007 | No |
| orf19.3428 | 9425 | 8804 | 9674 | 5936 | 5874 | 4909 | 5074 | 6753 | 8581 | 8904 | 9156 | 8917 | 5225 | 5330 | 5196 | 5151 | 7057,49 | 8890 | 5226 | 0,588 | -0,767 | -22,659 | 1,13E-113 | 1,10E-112 | 0,0003663 | 0,002224 | 0,0009492 | 0,0009492 | TRUE | 0,0068497 | No |
| orf19.3429 | 298 | 318 | 317 | 216 | 1059 | 910 | 927 | 1244 | 271 | 322 | 300 | 324 | 942 | 988 | 949 | 949 | 630,71 | 304 | 957 | 3,161 | 1,66 | 23,552 | 1,19E-122 | 1,24E-121 | 1,00E-08 | 0,004131 | 0,0025251 | 0,0025251 | TRUE | 0,0645249 | No |
| orf19.3430 | 142 | 137 | 145 | 84 | 105 | 91 | 106 | 106 | 129 | 139 | 137 | 126 | 93 | 99 | 109 | 81 | 114,11 | 133 | 96 | 0,712 | -0,49 | -3,625 | 0,000289 | 0,000422 | 1,00E-08 | 0,013614 | 0,0086391 | 0,0086391 | TRUE | 0,1185546 | No |
| orf19.3431 | 1615 | 1689 | 1692 | 977 | 1744 | 1418 | 1492 | 1983 | 1470 | 1708 | 1601 | 1468 | 1551 | 1540 | 1528 | 1513 | 1547,37 | 1562 | 1533 | 0,98 | -0,029 | -0,511 | 0,60953 | 0,649032 | 0,0018928 | 0,002891 | 0,002435 | 0,002435 | TRUE | 0,0478266 | No |
| orf19.3432 | 354 | 335 | 350 | 192 | 221 | 185 | 205 | 264 | 322 | 339 | 331 | 288 | 197 | 201 | 210 | 201 | 261,19 | 320 | 202 | 0,628 | -0,671 | -7,39 | 1,47E-13 | 3,13E-13 | 1,00E-08 | 0,007095 | 0,0039567 | 0,0039567 | TRUE | 0,0407528 | No |
| orf19.3432.1 | 28 | 32 | 28 | 31 | 43 | 31 | 38 | 41 | 25 | 32 | 26 | 47 | 38 | 34 | 39 | 31 | 34,13 | 32 | 36 | 1,112 | 0,153 | 0,614 | 0,539259 | 0,580714 | 0,0077289 | 0,040745 | 0,0299732 | 0,0299732 | TRUE | 0,3985443 | No |
| orf19.3433 | 568 | 570 | 554 | 365 | 182 | 174 | 145 | 226 | 517 | 576 | 524 | 548 | 162 | 189 | 148 | 172 | 354,74 | 541 | 168 | 0,31 | -1,689 | -18,738 | 2,43E-78 | 1,69E-77 | 0,0011948 | 0,005761 | 0,004038 | 0,004038 | TRUE | 0,0650597 | No |
| orf19.3434 | 602 | 651 | 679 | 425 | 1493 | 1413 | 1426 | 1759 | 548 | 658 | 643 | 638 | 1328 | 1534 | 1460 | 1342 | 1018,98 | 622 | 1416 | 2,278 | 1,188 | 16,156 | 1,03E-58 | 5,55E-58 | 0,0046657 | 0,003333 | 0,003996 | 0,003996 | TRUE | 0,0799345 | No |
| orf19.3435 | 210 | 168 | 214 | 154 | 452 | 436 | 433 | 640 | 191 | 170 | 203 | 231 | 402 | 473 | 443 | 488 | 325,25 | 199 | 452 | 2,292 | 1,196 | 11,665 | 1,93E-31 | 6,42E-31 | 0,0070001 | 0,006098 | 0,0063571 | 0,0063571 | TRUE | 0,1065575 | No |
| orf19.3436 | 69 | 67 | 76 | 30 | 136 | 109 | 109 | 151 | 63 | 68 | 72 | 45 | 121 | 118 | 112 | 115 | 89,21 | 62 | 116 | 1,85 | 0,888 | 5,718 | 1,08E-08 | 1,94E-08 | 1,00E-08 | 0,016844 | 0,0107058 | 0,0107058 | TRUE | 0,1977707 | No |
| orf19.3437 | 88 | 89 | 89 | 69 | 310 | 247 | 244 | 335 | 80 | 90 | 84 | 104 | 276 | 268 | 250 | 256 | 175,92 | 90 | 262 | 2,968 | 1,57 | 13,275 | 3,22E-40 | 1,25E-39 | 1,00E-08 | 0,009546 | 0,0056446 | 0,0056446 | TRUE | 0,0788545 | No |
| orf19.3438 | 1087 | 975 | 1120 | 571 | 1761 | 1409 | 1613 | 2053 | 990 | 986 | 1060 | 858 | 1566 | 1530 | 1652 | 1566 | 1275,95 | 974 | 1578 | 1,615 | 0,692 | 11,063 | 1,90E-28 | 5,88E-28 | 0,0027255 | 0,003072 | 0,0029085 | 0,0029085 | TRUE | 0,0676876 | No |
| orf19.3439 | 3312 | 3561 | 3734 | 2208 | 1468 | 1209 | 1233 | 1627 | 3015 | 3602 | 3534 | 3317 | 1306 | 1313 | 1263 | 1241 | 2323,74 | 3367 | 1281 | 0,38 | -1,395 | -22,864 | 1,07E-115 | 1,05E-114 | 0,0033835 | 0,002605 | 0,0030504 | 0,0030504 | TRUE | 0,0617316 | No |
| orf19.344 | 939 | 864 | 852 | 630 | 1297 | 1150 | 1181 | 1650 | 855 | 874 | 806 | 946 | 1154 | 1249 | 1209 | 1259 | 1043,98 | 870 | 1218 | 1,404 | 0,489 | 8,129 | 4,32E-16 | 9,85E-16 | 0,0016786 | 0,003302 | 0,0024789 | 0,0024789 | TRUE | 0,0362296 | No |
| orf19.3440 | 118 | 133 | 119 | 81 | 158 | 143 | 157 | 172 | 107 | 135 | 113 | 122 | 141 | 155 | 161 | 131 | 133 | 119 | 147 | 1,232 | 0,301 | 2,331 | 0,019763 | 0,02535 | 1,00E-08 | 0,011969 | 0,00838 | 0,00838 | TRUE | 0,0850133 | No |
| orf19.3441 | 3189 | 2984 | 3111 | 2135 | 1920 | 1646 | 1588 | 2233 | 2903 | 3018 | 2944 | 3207 | 1708 | 1787 | 1626 | 1703 | 2362,17 | 3018 | 1706 | 0,566 | -0,821 | -17,001 | 8,08E-65 | 4,73E-64 | 0,0011229 | 0,002596 | 0,0017881 | 0,0017881 | TRUE | 0,0202949 | No |
| orf19.3442 | 3612 | 3387 | 3625 | 2371 | 23488 | 20333 | 19424 | 26813 | 3289 | 3426 | 3431 | 3562 | 20892 | 22076 | 19893 | 20453 | 12127,5 | 3427 | 20828 | 6,083 | 2,605 | 59,23 | 0 | 0 | 0,0012929 | 0,002146 | 0,0016775 | 0,0016775 | TRUE | 0,0198111 | No |
| orf19.3443 | 2896 | 2805 | 2867 | 1845 | 2532 | 2063 | 2173 | 2830 | 2637 | 2837 | 2713 | 2771 | 2252 | 2240 | 2225 | 2159 | 2479,33 | 2740 | 2219 | 0,81 | -0,304 | -7,386 | 1,52E-13 | 3,23E-13 | 0,0003489 | 0,00257 | 0,0012175 | 0,0012175 | TRUE | 0,0080716 | No |
| orf19.3444 | 509 | 349 | 373 | 289 | 114 | 115 | 92 | 134 | 463 | 353 | 353 | 434 | 101 | 125 | 94 | 102 | 253,28 | 401 | 106 | 0,263 | -1,926 | -15,085 | 2,04E-51 | 9,69E-51 | 0,0133059 | 0,007253 | 0,0098856 | 0,0098856 | TRUE | 0,1271922 | No |
| orf19.3445 | 807 | 855 | 853 | 581 | 1532 | 1304 | 1157 | 1717 | 735 | 865 | 807 | 873 | 1363 | 1416 | 1185 | 1310 | 1069,08 | 820 | 1318 | 1,612 | 0,688 | 9,426 | 4,27E-21 | 1,11E-20 | 0,0045234 | 0,003273 | 0,0041098 | 0,0041098 | TRUE | 0,0597604 | No |
| orf19.3446 | 667 | 692 | 703 | 348 | 321 | 254 | 270 | 321 | 607 | 700 | 665 | 523 | 286 | 276 | 277 | 245 | 447,24 | 624 | 271 | 0,431 | -1,213 | -12,491 | 8,35E-36 | 3,03E-35 | 0,008196 | 0,004991 | 0,0064613 | 0,0064613 | TRUE | 0,1256265 | No |
| orf19.3447 | 304 | 383 | 331 | 244 | 1165 | 1130 | 1024 | 1285 | 277 | 387 | 313 | 367 | 1036 | 1227 | 1049 | 980 | 704,49 | 336 | 1073 | 3,204 | 1,68 | 16,327 | 6,37E-60 | 3,48E-59 | 0,0135616 | 0,003912 | 0,0081242 | 0,0081242 | TRUE | 0,1674587 | No |
| orf19.3448 | 1776 | 2920 | 2753 | 1405 | 538 | 506 | 462 | 546 | 1617 | 2953 | 2605 | 2111 | 479 | 549 | 473 | 416 | 1400,47 | 2322 | 479 | 0,206 | -2,277 | -11,048 | 2,25E-28 | 6,94E-28 | 0,0395929 | 0,00298 | 0,0187939 | 0,0395929 | TRUE | 0,1044109 | No |
| orf19.3449 | 2699 | 2425 | 2747 | 1853 | 2001 | 1588 | 1625 | 2101 | 2457 | 2453 | 2600 | 2784 | 1780 | 1724 | 1664 | 1603 | 2133,01 | 2574 | 1693 | 0,658 | -0,603 | -10,865 | 1,69E-27 | 5,12E-27 | 0,0021997 | 0,002656 | 0,0024697 | 0,0024697 | TRUE | 0,0346715 | No |
| orf19.3449.2 | 909 | 962 | 936 | 629 | 920 | 826 | 781 | 1036 | 828 | 973 | 886 | 945 | 818 | 897 | 800 | 790 | 867,06 | 908 | 826 | 0,911 | -0,135 | -1,971 | 0,048687 | 0,06035 | 0,0031126 | 0,003561 | 0,0033349 | 0,0033349 | TRUE | 0,0437416 | No |
| orf19.345 | 203 | 194 | 208 | 132 | 271 | 239 | 244 | 259 | 185 | 196 | 197 | 198 | 241 | 259 | 250 | 198 | 215,52 | 194 | 237 | 1,217 | 0,284 | 2,603 | 0,009238 | 0,012148 | 0,0047753 | 0,008166 | 0,0066632 | 0,0066632 | TRUE | 0,1528181 | No |
| orf19.3450.1 | 356 | 337 | 368 | 221 | 443 | 380 | 382 | 528 | 324 | 341 | 348 | 332 | 394 | 413 | 391 | 403 | 368,23 | 336 | 400 | 1,189 | 0,25 | 3,325 | 0,000885 | 0,001254 | 1,00E-08 | 0,005624 | 0,0026697 | 0,0026697 | TRUE | 0,0080682 | No |
| orf19.3453 | 164 | 166 | 169 | 112 | 303 | 230 | 240 | 312 | 149 | 168 | 160 | 168 | 270 | 250 | 246 | 238 | 206,05 | 161 | 251 | 1,559 | 0,641 | 6,304 | 2,90E-10 | 5,53E-10 | 1,00E-08 | 0,008448 | 0,0047499 | 0,0047499 | TRUE | 0,0299392 | No |
| orf19.3455 | 2098 | 2156 | 2258 | 1292 | 1300 | 1137 | 1118 | 1462 | 1910 | 2181 | 2137 | 1941 | 1156 | 1234 | 1145 | 1115 | 1602,43 | 2042 | 1162 | 0,568 | -0,815 | -13,688 | 1,20E-42 | 4,92E-42 | 0,002636 | 0,002861 | 0,0027376 | 0,0027376 | TRUE | 0,025101 | No |
| orf19.3456 | 866 | 870 | 900 | 591 | 569 | 466 | 457 | 591 | 788 | 880 | 852 | 888 | 506 | 506 | 468 | 451 | 667,35 | 852 | 483 | 0,567 | -0,819 | -11,963 | 5,55E-33 | 1,90E-32 | 0,0016901 | 0,004016 | 0,0029023 | 0,0029023 | TRUE | 0,031116 | No |
| orf19.3457 | 542 | 539 | 525 | 314 | 689 | 594 | 628 | 764 | 493 | 545 | 497 | 472 | 613 | 645 | 643 | 583 | 561,36 | 502 | 621 | 1,232 | 0,301 | 4,3 | 1,71E-05 | 2,68E-05 | 0,0011247 | 0,00439 | 0,0028986 | 0,0028986 | TRUE | 0,0387903 | No |
| orf19.3458 | 1614 | 1531 | 1625 | 843 | 1201 | 972 | 965 | 1327 | 1469 | 1548 | 1538 | 1266 | 1068 | 1055 | 988 | 1012 | 1243,28 | 1455 | 1031 | 0,706 | -0,501 | -7,492 | 6,80E-14 | 1,46E-13 | 0,0035917 | 0,0031 | 0,0034784 | 0,0034784 | TRUE | 0,0851193 | No |
| orf19.3459 | 303 | 307 | 315 | 205 | 295 | 259 | 229 | 319 | 276 | 310 | 298 | 308 | 262 | 281 | 235 | 243 | 276,74 | 298 | 255 | 0,857 | -0,222 | -2,423 | 0,015376 | 0,019919 | 1,00E-08 | 0,006811 | 0,0044684 | 0,0044684 | TRUE | 0,0475485 | No |
| orf19.346 | 3617 | 3121 | 3398 | 2043 | 4878 | 3798 | 4031 | 5496 | 3293 | 3157 | 3216 | 3069 | 4339 | 4124 | 4128 | 4192 | 3689,69 | 3184 | 4196 | 1,317 | 0,397 | 10,262 | 1,04E-24 | 2,96E-24 | 0,0003608 | 0,002395 | 0,0011568 | 0,0011568 | TRUE | 0,0065779 | No |
| orf19.3460 | 416 | 461 | 474 | 271 | 530 | 417 | 454 | 562 | 379 | 466 | 449 | 407 | 471 | 453 | 465 | 429 | 439,81 | 425 | 454 | 1,067 | 0,093 | 1,14 | 0,254294 | 0,289694 | 0,0031149 | 0,005041 | 0,0041691 | 0,0041691 | TRUE | 0,0705401 | No |
| orf19.3461 | 11 | 10 | 18 | 11 | 50 | 39 | 45 | 54 | 10 | 10 | 17 | 17 | 44 | 42 | 46 | 41 | 28,47 | 14 | 43 | 3,296 | 1,721 | 5,975 | 2,30E-09 | 4,25E-09 | 1,00E-08 | 0,048432 | 0,0288019 | 0,0288019 | TRUE | 0,1943367 | No |
| orf19.3462 | 1679 | 1582 | 1494 | 993 | 1937 | 1607 | 1432 | 2010 | 1529 | 1600 | 1414 | 1492 | 1723 | 1745 | 1467 | 1533 | 1562,71 | 1509 | 1617 | 1,071 | 0,099 | 1,478 | 0,139345 | 0,164352 | 0,0041105 | 0,002882 | 0,0036939 | 0,0036939 | TRUE | 0,0459921 | No |

| | | | | | | | | | | | | | | | | | | | | | | | | | | | | | | | |
|---|---|---|---|---|---|---|---|---|---|---|---|---|---|---|---|---|---|---|---|---|---|---|---|---|---|---|---|---|---|---|---|
| orf19.3463 | 15181 | 14783 | 15533 | 9084 | 15193 | 12397 | 13197 | 16880 | 13822 | 14951 | 14701 | 13646 | 13514 | 13460 | 13515 | 12876 | 13810,5 | 14280 | 13341 | 0,934 | -0,099 | -2,413 | 0,01583 | 0,020486 | 0,0011889 | 0,002133 | 0,0015292 | 0,0015292 | TRUE | 0,0122174 | No |
| orf19.3465 | 20221 | 17666 | 17905 | 11984 | 37387 | 29470 | 28132 | 39763 | 18410 | 17867 | 16945 | 18002 | 33255 | 31996 | 28811 | 30332 | 24452,3 | 17806 | 31098 | 1,747 | 0,804 | 16,063 | 4,64E-58 | 2,47E-57 | 0,0024879 | 0,002091 | 0,0023646 | 0,0023646 | TRUE | 0,0299821 | No |
| orf19.3466 | 1273 | 1332 | 1372 | 821 | 982 | 882 | 799 | 1094 | 1159 | 1347 | 1298 | 1233 | 873 | 958 | 818 | 835 | 1065,22 | 1259 | 871 | 0,691 | -0,534 | -7,866 | 3,66E-15 | 8,12E-15 | 0,0033582 | 0,003277 | 0,0034585 | 0,0034585 | TRUE | 0,0513211 | No |
| orf19.3467 | 3079 | 3107 | 3430 | 1839 | 3455 | 2728 | 2983 | 3929 | 2803 | 3142 | 3246 | 2762 | 3073 | 2962 | 3055 | 2997 | 3005,17 | 2988 | 3022 | 1,01 | 0,015 | 0,257 | 0,797358 | 0,824117 | 0,0030138 | 0,002477 | 0,0027797 | 0,0027797 | TRUE | 0,0410105 | No |
| orf19.3468 | 3958 | 4238 | 4200 | 2689 | 3935 | 3501 | 3299 | 4362 | 3604 | 4286 | 3975 | 4039 | 3500 | 3801 | 3379 | 3327 | 3738,9 | 3976 | 3502 | 0,881 | -0,183 | -2,983 | 0,002851 | 0,003886 | 0,0039989 | 0,00239 | 0,0033573 | 0,0033573 | TRUE | 0,0491987 | No |
| orf19.3469 | 3772 | 3481 | 3908 | 2989 | 3197 | 3207 | 2931 | 4201 | 3434 | 3521 | 3699 | 4490 | 2844 | 3482 | 3002 | 3205 | 3459,4 | 3786 | 3133 | 0,828 | -0,272 | -3,178 | 0,001485 | 0,002073 | 0,0115885 | 0,002419 | 0,0067439 | 0,0067439 | TRUE | 0,1877394 | No |
| orf19.347 | 4986 | 4248 | 4594 | 3217 | 1107 | 959 | 1087 | 1451 | 4540 | 4296 | 4348 | 4832 | 985 | 1041 | 1113 | 1107 | 2782,76 | 4504 | 1062 | 0,236 | -2,084 | -36,849 | 3,04E-297 | 7,78E-296 | 0,0023096 | 0,002512 | 0,0025111 | 0,0025111 | TRUE | 0,0292924 | No |
| orf19.3470 | 5746 | 5429 | 5881 | 3561 | 6581 | 5384 | 5713 | 7480 | 5231 | 5491 | 5566 | 5349 | 5854 | 5846 | 5851 | 5706 | 5611,64 | 5409 | 5814 | 1,074 | 0,104 | 3,019 | 0,002534 | 0,003468 | 0,0002956 | 0,002272 | 0,0009507 | 0,0009507 | TRUE | 0,0063252 | No |
| orf19.3471 | 211 | 166 | 190 | 105 | 189 | 144 | 161 | 244 | 192 | 168 | 180 | 158 | 168 | 156 | 165 | 186 | 171,63 | 174 | 169 | 0,966 | -0,05 | -0,443 | 0,658123 | 0,69497 | 1,00E-08 | 0,009734 | 0,0062968 | 0,0062968 | TRUE | 0,0556978 | No |
| orf19.3473 | 498 | 461 | 508 | 366 | 586 | 502 | 496 | 734 | 453 | 466 | 481 | 550 | 521 | 545 | 508 | 560 | 510,54 | 488 | 534 | 1,102 | 0,14 | 1,85 | 0,064252 | 0,078755 | 0,0019673 | 0,004624 | 0,0034838 | 0,0034838 | TRUE | 0,0736832 | No |
| orf19.3474 | 1928 | 1942 | 2126 | 1201 | 377 | 270 | 333 | 401 | 1755 | 1964 | 2012 | 1804 | 335 | 293 | 341 | 306 | 1101,38 | 1884 | 319 | 0,169 | -2,564 | -35,453 | 2,57E-275 | 5,97E-274 | 0,0033141 | 0,003236 | 0,0032742 | 0,0032742 | TRUE | 0,0294575 | No |
| orf19.3475 | 1546 | 1442 | 1516 | 1079 | 460 | 407 | 430 | 599 | 1408 | 1458 | 1435 | 1621 | 409 | 442 | 440 | 457 | 958,74 | 1480 | 437 | 0,296 | -1,756 | -26,566 | 1,66E-155 | 2,24E-154 | 0,0021091 | 0,003415 | 0,0027642 | 0,0027642 | TRUE | 0,0453912 | No |
| orf19.3476 | 12550 | 11587 | 11980 | 7398 | 10595 | 8318 | 8926 | 12419 | 11426 | 11719 | 11338 | 11113 | 9424 | 9031 | 9141 | 9473 | 10333,2 | 11399 | 9267 | 0,813 | -0,299 | -9,048 | 1,46E-19 | 3,66E-19 | 0,0003807 | 0,002165 | 0,0009488 | 0,0009488 | TRUE | 0,0043344 | No |
| orf19.3477 | 8942 | 8989 | 9557 | 5114 | 11561 | 8915 | 8946 | 12112 | 8141 | 9091 | 9045 | 7682 | 10283 | 9679 | 9162 | 9239 | 9040,36 | 8490 | 9591 | 1,129 | 0,175 | 2,831 | 0,004644 | 0,006238 | 0,0046309 | 0,002183 | 0,0035748 | 0,0035748 | TRUE | 0,0493556 | No |
| orf19.3478 | 8103 | 7344 | 7716 | 4105 | 7434 | 5086 | 6158 | 7589 | 7377 | 7428 | 7302 | 6166 | 6612 | 5522 | 6307 | 5789 | 6562,97 | 7068 | 6058 | 0,857 | -0,223 | -3,121 | 0,001805 | 0,002501 | 0,0067519 | 0,002238 | 0,0047669 | 0,0047669 | TRUE | 0,0888702 | No |
| orf19.348 | 1110 | 934 | 1107 | 631 | 2574 | 2180 | 2379 | 3163 | 1011 | 945 | 1048 | 948 | 2289 | 2367 | 2436 | 2413 | 1682,04 | 988 | 2376 | 2,401 | 1,263 | 24,984 | 9,14E-138 | 1,09E-136 | 0,0007074 | 0,002822 | 0,0017176 | 0,0017176 | TRUE | 0,0200383 | No |
| orf19.3480 | 3799 | 3650 | 3723 | 2551 | 2982 | 2695 | 2552 | 3394 | 3459 | 3692 | 3523 | 3832 | 2652 | 2926 | 2614 | 2589 | 3160,86 | 3626 | 2695 | 0,743 | -0,428 | -8,078 | 6,59E-16 | 1,49E-15 | 0,0023247 | 0,002455 | 0,0023705 | 0,0023705 | TRUE | 0,0396151 | No |
| orf19.3481 | 6129 | 5967 | 6303 | 4220 | 7431 | 6666 | 6503 | 8463 | 5580 | 6035 | 5965 | 6339 | 6610 | 7237 | 6660 | 6456 | 6360,26 | 5980 | 6741 | 1,128 | 0,173 | 3,378 | 0,000731 | 0,001041 | 0,0024409 | 0,002245 | 0,0023666 | 0,0023666 | TRUE | 0,0298588 | No |
| orf19.3482 | 3365 | 3394 | 3530 | 2048 | 3852 | 3124 | 3279 | 4362 | 3064 | 3433 | 3341 | 3076 | 3426 | 3392 | 3358 | 3327 | 3302,14 | 3228 | 3376 | 1,045 | 0,063 | 1,347 | 0,177934 | 0,207108 | 0,0013484 | 0,002437 | 0,0018181 | 0,0018181 | TRUE | 0,0219494 | No |
| orf19.3483 | 10181 | 11277 | 11144 | 6422 | 8928 | 7168 | 6979 | 8962 | 9269 | 11405 | 10547 | 9647 | 7941 | 7782 | 7147 | 6836 | 8821,97 | 10217 | 7426 | 0,727 | -0,46 | -6,544 | 6,00E-11 | 1,17E-10 | 0,0066977 | 0,002187 | 0,0046415 | 0,0046415 | TRUE | 0,0750291 | No |
| orf19.3487 | 614 | 489 | 585 | 350 | 825 | 555 | 652 | 865 | 559 | 495 | 554 | 526 | 734 | 603 | 668 | 660 | 599,62 | 534 | 666 | 1,249 | 0,321 | 4,234 | 2,29E-05 | 3,57E-05 | 0,0033333 | 0,00424 | 0,0038045 | 0,0038045 | TRUE | 0,0555735 | No |
| orf19.349 | 1427 | 1304 | 1405 | 1001 | 1639 | 1377 | 1415 | 1838 | 1299 | 1319 | 1330 | 1504 | 1458 | 1495 | 1449 | 1402 | 1406,94 | 1363 | 1451 | 1,067 | 0,094 | 1,691 | 0,090776 | 0,109395 | 0,0015786 | 0,002976 | 0,0022607 | 0,0022607 | TRUE | 0,0527465 | No |
| orf19.3490 | 39 | 28 | 29 | 18 | 68 | 45 | 50 | 64 | 36 | 28 | 27 | 27 | 60 | 49 | 51 | 49 | 40,96 | 30 | 52 | 1,757 | 0,813 | 3,683 | 0,00023 | 0,000339 | 1,00E-08 | 0,034288 | 0,0201047 | 0,0201047 | TRUE | 0,1357293 | No |
| orf19.3494 | 75 | 69 | 63 | 26 | 45 | 40 | 49 | 56 | 68 | 70 | 60 | 39 | 40 | 43 | 50 | 43 | 51,64 | 59 | 44 | 0,731 | -0,453 | -2,181 | 0,029162 | 0,036894 | 0,0100351 | 0,027619 | 0,021942 | 0,021942 | TRUE | 0,3236768 | No |
| orf19.3496 | 2746 | 2782 | 2770 | 1811 | 4106 | 3688 | 3516 | 4788 | 2500 | 2814 | 2622 | 2720 | 3652 | 4004 | 3601 | 3652 | 3195,65 | 2664 | 3727 | 1,4 | 0,485 | 9,292 | 1,51E-20 | 3,88E-20 | 0,0020914 | 0,00245 | 0,0022878 | 0,0022878 | TRUE | 0,0299276 | No |
| orf19.3498 | 339 | 323 | 312 | 221 | 399 | 362 | 348 | 483 | 309 | 327 | 295 | 332 | 355 | 393 | 356 | 368 | 341,92 | 316 | 368 | 1,169 | 0,226 | 2,789 | 0,005292 | 0,007089 | 1,00E-08 | 0,005901 | 0,0033249 | 0,0033249 | TRUE | 0,0217041 | No |
| orf19.3499 | 2073 | 1863 | 1968 | 1327 | 438 | 389 | 363 | 469 | 1887 | 1884 | 1863 | 1993 | 390 | 422 | 372 | 358 | 1146,12 | 1907 | 386 | 0,202 | -2,31 | -37,766 | 0 | 0 | 0,0004987 | 0,00319 | 0,0020988 | 0,0020988 | TRUE | 0,0448728 | No |
| orf19.35 | 18923 | 20636 | 21636 | 11880 | 9919 | 8209 | 8919 | 11049 | 17229 | 20871 | 20477 | 17846 | 8823 | 8913 | 9134 | 8428 | 13965 | 19106 | 8824 | 0,462 | -1,115 | -17,45 | 3,46E-68 | 2,12E-67 | 0,0050624 | 0,002132 | 0,0038391 | 0,0038391 | TRUE | 0,0538585 | No |
| orf19.350 | 4484 | 3876 | 4153 | 2598 | 3284 | 2620 | 2624 | 3688 | 4082 | 3920 | 3930 | 3903 | 2921 | 2845 | 2687 | 2813 | 3387,73 | 3959 | 2816 | 0,711 | -0,491 | -12,283 | 1,12E-34 | 3,99E-34 | 0,0004126 | 0,002427 | 0,0012351 | 0,0012351 | TRUE | 0,0113282 | No |
| orf19.3501 | 1665 | 1511 | 1653 | 1052 | 2020 | 1759 | 1918 | 2436 | 1516 | 1528 | 1564 | 1580 | 1797 | 1910 | 1964 | 1858 | 1714,72 | 1547 | 1882 | 1,217 | 0,283 | 6,133 | 8,64E-10 | 1,62E-09 | 0,0004225 | 0,002807 | 0,0014486 | 0,0014486 | TRUE | 0,0112403 | No |
| orf19.3503 | 244 | 146 | 149 | 96 | 150 | 81 | 66 | 135 | 222 | 148 | 141 | 144 | 133 | 88 | 68 | 103 | 130,87 | 164 | 98 | 0,599 | -0,74 | -3,674 | 0,000239 | 0,000351 | 0,0575335 | 0,012131 | 0,0308903 | 0,0308903 | TRUE | 0,6342214 | No |
| orf19.3504 | 18030 | 14785 | 15592 | 9821 | 23115 | 16817 | 18402 | 25750 | 16416 | 14953 | 14756 | 14753 | 20560 | 18259 | 18846 | 19642 | 17273,1 | 15220 | 19327 | 1,27 | 0,345 | 6,778 | 1,22E-11 | 2,44E-11 | 0,0026484 | 0,002113 | 0,0024235 | 0,0024235 | TRUE | 0,0344006 | No |
| orf19.3505 | 8757 | 7587 | 8261 | 5476 | 3435 | 2915 | 3117 | 4273 | 7973 | 7673 | 7818 | 8226 | 3055 | 3165 | 3192 | 3259 | 5545,29 | 7922 | 3168 | 0,4 | -1,322 | -34,377 | 5,65E-259 | 1,23E-257 | 0,0005097 | 0,002275 | 0,0012036 | 0,0012036 | TRUE | 0,0077831 | No |
| orf19.3506 | 242 | 216 | 199 | 119 | 308 | 261 | 288 | 342 | 220 | 218 | 188 | 179 | 274 | 283 | 295 | 261 | 239,88 | 201 | 278 | 1,37 | 0,454 | 4,553 | 5,28E-06 | 8,49E-06 | 0,0014233 | 0,007544 | 0,0052357 | 0,0052357 | TRUE | 0,0487993 | No |
| orf19.3507 | 2570 | 2105 | 2141 | 1403 | 4327 | 3136 | 3596 | 4836 | 2340 | 2129 | 2026 | 2108 | 3849 | 3405 | 3683 | 3689 | 2903,49 | 2151 | 3656 | 1,7 | 0,765 | 13,565 | 6,43E-42 | 2,59E-41 | 0,0027114 | 0,002492 | 0,0026779 | 0,0026779 | TRUE | 0,0438435 | No |
| orf19.3508 | 482 | 405 | 414 | 306 | 850 | 699 | 719 | 911 | 439 | 410 | 392 | 460 | 756 | 759 | 736 | 695 | 580,77 | 425 | 736 | 1,738 | 0,798 | 11,323 | 1,01E-29 | 3,23E-29 | 0,0010511 | 0,004312 | 0,0028679 | 0,0028679 | TRUE | 0,0319672 | No |
| orf19.351 | 268 | 206 | 218 | 123 | 109 | 94 | 84 | 149 | 244 | 208 | 206 | 185 | 97 | 102 | 86 | 114 | 155,27 | 211 | 100 | 0,472 | -1,083 | -8,326 | 8,35E-17 | 1,94E-16 | 0,0067981 | 0,010545 | 0,0090634 | 0,0090634 | TRUE | 0,1250006 | No |
| orf19.35.1 | 15 | 5 | 8 | 4 | 12 | 9 | 3 | 6 | 14 | 5 | 8 | 6 | 11 | 10 | 3 | 5 | 7,55 | 8 | 7 | 0,849 | -0,237 | -0,429 | 0,667956 | 0,704537 | 0,1320255 | 0,177038 | 0,160355 | 0,160355 | TRUE | 0,3781471 | No |
| orf19.3512 | 350 | 306 | 334 | 197 | 340 | 298 | 318 | 385 | 319 | 309 | 316 | 296 | 302 | 324 | 326 | 294 | 310,69 | 310 | 312 | 0,999 | -0,002 | -0,024 | 0,980669 | 0,98221 | 1,00E-08 | 0,006289 | 0,0033213 | 0,0033213 | TRUE | 0,0178841 | No |
| orf19.3515 | 1651 | 1467 | 1567 | 904 | 976 | 690 | 724 | 970 | 1503 | 1484 | 1483 | 1358 | 868 | 749 | 741 | 740 | 1115,81 | 1457 | 774 | 0,531 | -0,913 | -14,021 | 1,15E-44 | 4,91E-44 | 0,0028829 | 0,003221 | 0,0030994 | 0,0030994 | TRUE | 0,079803 | No |
| orf19.3516 | 1382 | 1080 | 1140 | 681 | 1856 | 1387 | 1506 | 1992 | 1258 | 1092 | 1079 | 1023 | 1651 | 1506 | 1542 | 1520 | 1333,88 | 1113 | 1555 | 1,393 | 0,479 | 7,187 | 6,64E-13 | 1,39E-12 | 0,0040123 | 0,003027 | 0,0034767 | 0,0034767 | TRUE | 0,0940656 | No |
| orf19.3517 | 1195 | 1111 | 1155 | 613 | 839 | 642 | 715 | 979 | 1088 | 1124 | 1093 | 921 | 746 | 697 | 732 | 747 | 893,49 | 1056 | 730 | 0,69 | -0,536 | -7,901 | 2,78E-15 | 6,20E-15 | 0,0027385 | 0,003515 | 0,0032709 | 0,0032709 | TRUE | 0,0801791 | No |
| orf19.3518 | 2873 | 2570 | 2657 | 1717 | 3314 | 2731 | 2914 | 3886 | 2616 | 2599 | 2515 | 2579 | 2948 | 2965 | 2984 | 2964 | 2771,28 | 2577 | 2965 | 1,151 | 0,202 | 5,661 | 1,50E-08 | 2,69E-08 | 1,00E-08 | 0,002514 | 0,0008608 | 0,0008608 | TRUE | 0,0034678 | No |
| orf19.3519 | 534 | 482 | 539 | 362 | 1088 | 861 | 934 | 1279 | 486 | 487 | 510 | 544 | 968 | 935 | 957 | 976 | 732,79 | 507 | 959 | 1,901 | 0,926 | 15,403 | 1,57E-53 | 7,74E-53 | 1,00E-08 | 0,00384 | 0,001931 | 0,001931 | TRUE | 0,0227537 | No |
| orf19.352 | 1831 | 1739 | 1804 | 1169 | 1543 | 1409 | 1332 | 1766 | 1667 | 1759 | 1707 | 1756 | 1372 | 1530 | 1364 | 1347 | 1562,84 | 1722 | 1403 | 0,814 | -0,296 | -5,583 | 2,36E-08 | 4,18E-08 | 0,0012921 | 0,002882 | 0,0020566 | 0,0020566 | TRUE | 0,0427611 | No |
| orf19.3521 | 451 | 390 | 401 | 286 | 407 | 305 | 349 | 402 | 411 | 394 | 380 | 430 | 362 | 331 | 357 | 307 | 371,43 | 404 | 339 | 0,842 | -0,249 | -2,984 | 0,002843 | 0,003876 | 0,0018355 | 0,005594 | 0,0039564 | 0,0039564 | TRUE | 0,0538282 | No |
| orf19.3522 | 2 | 1 | 2 | 1 | 1 | 1 | 1 | 0 | 2 | 1 | 2 | 2 | 1 | 1 | 1 | 0 | 1,15 | 2 | 1 | 0,454 | -1,138 | -0,882 | 0,377586 | 0,417902 | 1,00E-08 | 1,147419 | 0,6054087 | 0,6054087 | TRUE | 0,2503288 | No |
| orf19.3523 | 2441 | 2073 | 2270 | 1392 | 2916 | 2338 | 2646 | 3521 | 2222 | 2097 | 2148 | 2091 | 2594 | 2538 | 2710 | 2686 | 2385,77 | 2140 | 2632 | 1,229 | 0,298 | 6,972 | 3,12E-12 | 6,37E-12 | 0,0004495 | 0,002591 | 0,0013244 | 0,0013244 | TRUE | 0,0084323 | No |
| orf19.3526 | 8574 | 7359 | 8023 | 5125 | 8914 | 7579 | 8026 | 11037 | 7806 | 7443 | 7593 | 7699 | 7929 | 8229 | 8220 | 8419 | 7917,11 | 7635 | 8199 | 1,074 | 0,103 | 3,037 | 0,002392 | 0,003281 | 0,0003419 | 0,002204 | 0,0009766 | 0,0009766 | TRUE | 0,0061027 | No |
| orf19.3527 | 4427 | 3683 | 3989 | 2647 | 6998 | 5606 | 6345 | 8639 | 4031 | 3725 | 3775 | 3976 | 6224 | 6087 | 6498 | 6590 | 5113,25 | 3877 | 6350 | 1,639 | 0,713 | 16,284 | 1,29E-59 | 7,01E-59 | 0,0011456 | 0,002295 | 0,0016265 | 0,0016265 | TRUE | 0,0093969 | No |
| orf19.3528 | 746 | 695 | 738 | 402 | 638 | 453 | 526 | 671 | 679 | 703 | 698 | 604 | 567 | 492 | 539 | 512 | 599,29 | 671 | 528 | 0,784 | -0,352 | -4,869 | 1,12E-06 | 1,86E-06 | 0,0020327 | 0,004241 | 0,0033357 | 0,0033357 | TRUE | 0,0463692 | No |
| orf19.3529 | 231 | 221 | 249 | 152 | 95 | 89 | 85 | 104 | 210 | 224 | 236 | 228 | 84 | 97 | 87 | 79 | 155,67 | 224 | 87 | 0,385 | -1,376 | -11,489 | 1,50E-30 | 4,91E-30 | 1,00E-08 | 0,010523 | 0,006016 | 0,006016 | TRUE | 0,0504482 | No |
| orf19.353 | 1340 | 1123 | 1271 | 736 | 1783 | 1451 | 1573 | 2069 | 1220 | 1136 | 1203 | 1106 | 1586 | 1575 | 1611 | 1578 | 1376,85 | 1166 | 1588 | 1,358 | 0,441 | 9,083 | 1,05E-19 | 2,65E-19 | 1,00E-08 | 0,002996 | 0,0015137 | 0,0015137 | TRUE | 0,0126911 | No |
| orf19.3530 | 3933 | 4027 | 4170 | 2384 | 2501 | 2134 | 2358 | 2910 | 3581 | 4073 | 3947 | 3581 | 2225 | 2317 | 2415 | 2220 | 3044,69 | 3796 | 2294 | 0,604 | -0,728 | -13,108 | 2,95E-39 | 1,13E-38 | 0,0025594 | 0,002471 | 0,0026144 | 0,0026144 | TRUE | 0,0293853 | No |
| orf19.3531 | 538 | 499 | 568 | 339 | 582 | 468 | 505 | 606 | 490 | 505 | 538 | 509 | 518 | 508 | 517 | 462 | 505,82 | 510 | 501 | 0,98 | -0,028 | -0,403 | 0,687207 | 0,722451 | 1,00E-08 | 0,004649 | 0,0028176 | 0,0028176 | TRUE | 0,0353097 | No |
| orf19.3532 | 2157 | 1954 | 2084 | 1352 | 2119 | 1585 | 1694 | 2403 | 1964 | 1976 | 1972 | 2031 | 1885 | 1721 | 1735 | 1833 | 1889,61 | 1986 | 1794 | 0,905 | -0,144 | -3,153 | 0,001615 | 0,002247 | 0,0005258 | 0,002736 | 0,0014849 | 0,0014849 | TRUE | 0,013947 | No |
| orf19.3533 | 940 | 753 | 846 | 540 | 1634 | 1302 | 1306 | 1722 | 856 | 762 | 801 | 811 | 1453 | 1414 | 1338 | 1314 | 1093,41 | 808 | 1380 | 1,707 | 0,772 | 13,125 | 2,36E-39 | 9,09E-39 | 0,0014875 | 0,003245 | 0,0023182 | 0,0023182 | TRUE | 0,0212593 | No |
| orf19.3534 | 703 | 623 | 686 | 421 | 1233 | 970 | 1061 | 1426 | 640 | 630 | 649 | 632 | 1097 | 1053 | 1087 | 1088 | 859,5 | 638 | 1081 | 1,694 | 0,761 | 14,198 | 9,49E-46 | 4,09E-45 | 1,00E-08 | 0,003574 | 0,0014904 | 0,0014904 | TRUE | 0,0034087 | No |
| orf19.3535 | 4274 | 3367 | 3892 | 2412 | 2969 | 2293 | 2588 | 3489 | 3891 | 3405 | 3683 | 3623 | 2641 | 2490 | 2650 | 2661 | 3130,69 | 3650 | 2610 | 0,715 | -0,484 | -9,774 | 1,46E-22 | 3,93E-22 | 0,001544 | 0,002459 | 0,002026 | 0,002026 | TRUE | 0,0255406 | No |
| orf19.3536 | 1091 | 1031 | 1148 | 714 | 1957 | 1722 | 1641 | 2256 | 993 | 1043 | 1086 | 1073 | 1741 | 1870 | 1681 | 1721 | 1400,86 | 1049 | 1753 | 1,672 | 0,742 | 13,785 | 3,12E-43 | 1,29E-42 | 0,0011685 | 0,00298 | 0,0020015 | 0,0020015 | TRUE | 0,023353 | No |
| orf19.3537 | 2762 | 2208 | 2140 | 1683 | 410 | 343 | 343 | 458 | 2515 | 2233 | 2025 | 2528 | 365 | 372 | 351 | 349 | 1342,38 | 2325 | 359 | 0,155 | -2,693 | -33,497 | 5,26E-246 | 1,08E-244 | 0,0061627 | 0,003021 | 0,0046775 | 0,0046775 | TRUE | 0,0922839 | No |
| orf19.3538 | 5906 | 5193 | 5902 | 3730 | 1305 | 1190 | 1258 | 1692 | 5377 | 5252 | 5586 | 5603 | 1161 | 1292 | 1288 | 1291 | 3356,23 | 5454 | 1258 | 0,231 | -2,117 | -44,005 | 0 | 0 | 0,001158 | 0,002431 | 0,0017512 | 0,0017512 | TRUE | 0,0330969 | No |
| orf19.3539 | 2769 | 2420 | 2535 | 1680 | 2648 | 2075 | 2109 | 2892 | 2521 | 2448 | 2399 | 2524 | 2355 | 2253 | 2160 | 2206 | 2358,19 | 2473 | 2244 | 0,908 | -0,139 | -3,193 | 0,00141 | 0,001971 | 0,0005875 | 0,002597 | 0,0014073 | 0,0014073 | TRUE | 0,0138551 | No |
| orf19.354 | 1130 | 966 | 1085 | 695 | 815 | 717 | 737 | 1056 | 1029 | 977 | 1027 | 1044 | 725 | 778 | 755 | 806 | 892,55 | 1019 | 766 | 0,753 | -0,41 | -7,231 | 4,79E-13 | 1,01E-12 | 1,00E-08 | 0,003517 | 0,0019542 | 0,0019542 | TRUE | 0,016312 | No |
| orf19.3540 | 7946 | 6403 | 6933 | 4422 | 10317 | 8121 | 9147 | 12340 | 7234 | 6476 | 6561 | 6643 | 9177 | 8817 | 9368 | 9413 | 7961,12 | 6728 | 9194 | 1,366 | 0,45 | 10,009 | 1,39E-23 | 3,86E-23 | 0,0015704 | 0,002203 | 0,0018118 | 0,0018118 | TRUE | 0,0318015 | No |
| orf19.3541 | 11370 | 9837 | 10769 | 6355 | 14930 | 12260 | 13122 | 17513 | 10352 | 9949 | 10192 | 9546 | 13280 | 13311 | 13439 | 13359 | 11678,4 | 10010 | 13347 | 1,333 | 0,414 | 12,208 | 2,82E-34 | 9,90E-34 | 0,0005103 | 0,00215 | 0,0010169 | 0,0010169 | TRUE | 0,0114581 | No |
| orf19.3542 | 6455 | 5412 | 5729 | 3709 | 4956 | 3969 | 4108 | 5723 | 5877 | 5474 | 5422 | 5572 | 4408 | 4309 | 4207 | 4366 | 4954,28 | 5586 | 4322 | 0,774 | -0,37 | -9,449 | 3,41E-21 | 8,89E-21 | 0,0005934 | 0,002304 | 0,0012664 | 0,0012664 | TRUE | 0,0154785 | No |

| | | | | | | | | | | | | | | | | | | | | | | | | | | | | | | | |
|---|---|---|---|---|---|---|---|---|---|---|---|---|---|---|---|---|---|---|---|---|---|---|---|---|---|---|---|---|---|---|---|
| orf19.3543 | 2 | 0 | 1 | 2 | 2 | 2 | 2 | 0 | 2 | 0 | 1 | 3 | 2 | 2 | 2 | 0 | 1,47 | 2 | 2 | 1,061 | 0,086 | 0,072 | 0,942977 | 0,950906 | 1,00E-08 | 0,89989 | 0,6697927 | 0,6697927 | TRUE | 0,3463843 | No |
| orf19.3544 | 292 | 317 | 319 | 196 | 767 | 612 | 726 | 896 | 266 | 321 | 302 | 294 | 682 | 664 | 744 | 683 | 494,56 | 296 | 693 | 2,347 | 1,231 | 15,771 | 4,95E-56 | 2,54E-55 | 0,0014837 | 0,004708 | 0,003368 | 0,003368 | TRUE | 0,0591254 | No |
| orf19.3545 | 1592 | 1480 | 1473 | 975 | 2279 | 1934 | 2035 | 2735 | 1449 | 1497 | 1394 | 1465 | 2027 | 2100 | 2084 | 2086 | 1762,78 | 1451 | 2074 | 1,43 | 0,516 | 11,887 | 1,38E-32 | 4,68E-32 | 1,00E-08 | 0,002786 | 0,0012145 | 0,0012145 | TRUE | 0,008904 | No |
| orf19.3546 | 2610 | 2213 | 2320 | 1375 | 2129 | 1713 | 1775 | 2403 | 2376 | 2238 | 2196 | 2065 | 1894 | 1860 | 1818 | 1833 | 2035 | 2219 | 1851 | 0,833 | -0,264 | -5,303 | 1,14E-07 | 1,96E-07 | 0,0011674 | 0,002686 | 0,0018866 | 0,0018866 | TRUE | 0,0276569 | No |
| orf19.3547 | 14660 | 13512 | 14167 | 8585 | 21498 | 17380 | 18827 | 24737 | 13347 | 13666 | 13408 | 12896 | 19122 | 18870 | 19281 | 18870 | 16182,4 | 13329 | 19036 | 1,428 | 0,514 | 17,29 | 5,62E-67 | 3,38E-66 | 0,0002599 | 0,002119 | 0,0007824 | 0,0007824 | TRUE | 0,0056355 | No |
| orf19.3548 | 2531 | 2099 | 2335 | 1413 | 3398 | 2521 | 2695 | 3781 | 2304 | 2123 | 2210 | 2123 | 3022 | 2737 | 2760 | 2884 | 2520,43 | 2190 | 2851 | 1,301 | 0,38 | 7,725 | 1,12E-14 | 2,45E-14 | 0,0013054 | 0,002561 | 0,0019127 | 0,0019127 | TRUE | 0,0198265 | No |
| orf19.3548.1 | 82538 | 59849 | 60697 | 46945 | 97506 | 73826 | 65845 | 1E+05 | 75147 | 60531 | 57444 | 70519 | 86728 | 80154 | 67434 | 81005 | 72370,3 | 65910 | 78830 | 1,196 | 0,258 | 2,88 | 0,003978 | 0,005364 | 0,0132603 | 0,002055 | 0,0077144 | 0,0077144 | TRUE | 0,1160661 | No |
| orf19.3549 | 1433 | 1191 | 1257 | 790 | 2686 | 2273 | 2498 | 3269 | 1305 | 1205 | 1190 | 1187 | 2389 | 2468 | 2558 | 2494 | 1849,31 | 1222 | 2477 | 2,025 | 1,018 | 20,884 | 7,52E-97 | 6,32E-96 | 0,0006378 | 0,002751 | 0,0016538 | 0,0016538 | TRUE | 0,0265638 | No |
| orf19.355 | 99 | 88 | 81 | 62 | 577 | 472 | 495 | 703 | 90 | 89 | 77 | 93 | 513 | 512 | 507 | 536 | 302,23 | 87 | 517 | 5,968 | 2,577 | 24,663 | 2,67E-134 | 3,09E-133 | 1,00E-08 | 0,006408 | 0,0034923 | 0,0034923 | TRUE | 0,0681858 | No |
| orf19.3550 | 1176 | 1035 | 1117 | 694 | 551 | 456 | 436 | 584 | 1071 | 1047 | 1057 | 1042 | 490 | 495 | 447 | 445 | 761,79 | 1054 | 469 | 0,444 | -1,171 | -19,364 | 1,56E-83 | 1,16E-82 | 1,00E-08 | 0,003771 | 0,002007 | 0,002007 | TRUE | 0,0152867 | No |
| orf19.3551 | 311 | 243 | 257 | 140 | 43 | 30 | 39 | 45 | 283 | 246 | 243 | 210 | 38 | 33 | 40 | 34 | 140,94 | 246 | 36 | 0,147 | -2,769 | -17,345 | 2,14E-67 | 1,30E-66 | 1,00E-08 | 0,01141 | 0,0095026 | 0,0095026 | TRUE | 0,1004608 | No |
| orf19.3552 | 2964 | 2904 | 3115 | 1866 | 2714 | 2504 | 2425 | 3191 | 2699 | 2937 | 2948 | 2803 | 2414 | 2719 | 2484 | 2434 | 2679,63 | 2847 | 2513 | 0,882 | -0,181 | -3,478 | 0,000505 | 0,000726 | 0,0019325 | 0,00253 | 0,0022342 | 0,0022342 | TRUE | 0,0362953 | No |
| orf19.3553 | 9901 | 9452 | 9631 | 5028 | 14840 | 10209 | 12062 | 15219 | 9014 | 9560 | 9115 | 7553 | 13200 | 11084 | 12353 | 11609 | 10436 | 8810 | 12062 | 1,368 | 0,453 | 6,099 | 1,07E-09 | 2,00E-09 | 0,0076687 | 0,002164 | 0,0051915 | 0,0051915 | TRUE | 0,1116169 | No |
| orf19.3554 | 3454 | 3385 | 3386 | 2089 | 8655 | 7387 | 6922 | 9860 | 3145 | 3424 | 3205 | 3138 | 7698 | 8020 | 7089 | 7521 | 5404,96 | 3228 | 7582 | 2,348 | 1,232 | 25,022 | 3,51E-138 | 4,21E-137 | 0,001878 | 0,002281 | 0,0020978 | 0,0020978 | TRUE | 0,0231987 | No |
| orf19.3555 | 10230 | 8880 | 9872 | 5584 | 2805 | 2324 | 2621 | 3430 | 9314 | 8981 | 9343 | 8388 | 2495 | 2523 | 2684 | 2616 | 5793,13 | 9006 | 2580 | 0,286 | -1,805 | -38,471 | 0 | 0 | 0,0014487 | 0,002265 | 0,001872 | 0,001872 | TRUE | 0,025497 | No |
| orf19.3556 | 3688 | 3937 | 4117 | 2720 | 4384 | 4077 | 3855 | 5194 | 3358 | 3982 | 3896 | 4086 | 3899 | 4426 | 3948 | 3962 | 3944,73 | 3830 | 4059 | 1,06 | 0,084 | 1,271 | 0,203866 | 0,235785 | 0,005094 | 0,002372 | 0,0039622 | 0,0039622 | TRUE | 0,084858 | No |
| orf19.3557 | 281 | 246 | 254 | 137 | 129 | 96 | 110 | 164 | 256 | 249 | 240 | 206 | 115 | 104 | 113 | 125 | 175,94 | 238 | 114 | 0,48 | -1,06 | -9,201 | 3,53E-20 | 9,01E-20 | 1,00E-08 | 0,009545 | 0,0064586 | 0,0064586 | TRUE | 0,0735108 | No |
| orf19.3558 | 749 | 848 | 788 | 481 | 847 | 742 | 723 | 934 | 682 | 858 | 746 | 723 | 753 | 806 | 740 | 712 | 752,47 | 752 | 753 | 1 | 0 | -0,004 | 0,997171 | 0,997328 | 0,004896 | 0,003793 | 0,0043376 | 0,0043376 | TRUE | 0,1083032 | No |
| orf19.3559 | 6055 | 5602 | 5562 | 3670 | 4682 | 3762 | 3928 | 5169 | 5513 | 5666 | 5264 | 5513 | 4164 | 4084 | 4023 | 3943 | 4771,27 | 5489 | 4054 | 0,739 | -0,437 | -11,491 | 1,47E-30 | 4,78E-30 | 0,0005686 | 0,002314 | 0,0011773 | 0,0011773 | TRUE | 0,0095199 | No |
| orf19.356 | 208 | 195 | 193 | 105 | 677 | 540 | 558 | 766 | 189 | 197 | 183 | 158 | 602 | 586 | 571 | 584 | 383,9 | 182 | 586 | 3,197 | 1,677 | 20,018 | 3,85E-89 | 3,01E-88 | 1,00E-08 | 0,005478 | 0,0030467 | 0,0030467 | TRUE | 0,0618247 | No |
| orf19.3561 | 2966 | 2920 | 3228 | 1978 | 2588 | 2198 | 2307 | 3065 | 2700 | 2953 | 3055 | 2971 | 2302 | 2386 | 2363 | 2338 | 2633,62 | 2920 | 2347 | 0,804 | -0,315 | -6,652 | 2,89E-11 | 5,73E-11 | 0,0012475 | 0,002539 | 0,0017634 | 0,0017634 | TRUE | 0,0322668 | No |
| orf19.3563 | 3398 | 3019 | 3348 | 2390 | 2348 | 2129 | 2140 | 3127 | 3094 | 3053 | 3169 | 3590 | 2088 | 2312 | 2192 | 2385 | 2735,35 | 3226 | 2244 | 0,697 | -0,522 | -8,333 | 7,86E-17 | 1,83E-16 | 0,004132 | 0,00252 | 0,0033885 | 0,0033885 | TRUE | 0,0675938 | No |
| orf19.3564 | 8036 | 7118 | 7665 | 4263 | 8420 | 6569 | 6969 | 9084 | 7316 | 7199 | 7254 | 6404 | 7489 | 7132 | 7137 | 6929 | 7107,67 | 7043 | 7172 | 1,017 | 0,025 | 0,503 | 0,615221 | 0,654545 | 0,0019719 | 0,002223 | 0,0022317 | 0,0022317 | TRUE | 0,044433 | No |
| orf19.3565 | 480 | 427 | 457 | 302 | 420 | 323 | 367 | 500 | 437 | 432 | 433 | 454 | 374 | 351 | 376 | 381 | 404,57 | 439 | 370 | 0,847 | -0,239 | -3,316 | 0,000912 | 0,00129 | 1,00E-08 | 0,005302 | 0,0025335 | 0,0025335 | TRUE | 0,0136198 | No |
| orf19.3567 | 2209 | 2099 | 2057 | 1385 | 996 | 886 | 880 | 1117 | 2011 | 2123 | 1947 | 2080 | 886 | 962 | 901 | 852 | 1470,31 | 2040 | 900 | 0,441 | -1,182 | -21,915 | 1,88E-106 | 1,72E-105 | 0,0011907 | 0,002935 | 0,0020104 | 0,0020104 | TRUE | 0,0245352 | No |
| orf19.3568 | 1758 | 1674 | 1869 | 1115 | 1134 | 878 | 980 | 1350 | 1601 | 1693 | 1769 | 1675 | 1009 | 953 | 1004 | 1030 | 1341,59 | 1684 | 999 | 0,594 | -0,752 | -14,52 | 9,03E-48 | 4,01E-47 | 0,0007889 | 0,003022 | 0,0017916 | 0,0017916 | TRUE | 0,0145769 | No |
| orf19.3569 | 2642 | 2606 | 2672 | 1556 | 2733 | 2146 | 2385 | 3046 | 2405 | 2636 | 2529 | 2337 | 2431 | 2330 | 2443 | 2324 | 2429,28 | 2477 | 2382 | 0,96 | -0,058 | -1,203 | 0,229153 | 0,262695 | 0,001235 | 0,002581 | 0,0018361 | 0,0018361 | TRUE | 0,0222919 | No |
| orf19.357 | 1917 | 1532 | 1748 | 1066 | 1876 | 1456 | 1789 | 2268 | 1745 | 1549 | 1654 | 1601 | 1669 | 1581 | 1832 | 1730 | 1670,26 | 1637 | 1703 | 1,039 | 0,056 | 0,947 | 0,343617 | 0,383635 | 0,0024318 | 0,002828 | 0,0027096 | 0,0027096 | TRUE | 0,0305039 | No |
| orf19.3571 | 255 | 224 | 222 | 180 | 59 | 35 | 34 | 56 | 232 | 227 | 210 | 270 | 52 | 38 | 35 | 43 | 138,4 | 235 | 42 | 0,181 | -2,464 | -15,89 | 7,44E-57 | 3,86E-56 | 0,0061167 | 0,011582 | 0,0098206 | 0,0098206 | TRUE | 0,219955 | No |
| orf19.3572 | 322 | 376 | 347 | 205 | 171 | 152 | 164 | 176 | 293 | 380 | 328 | 308 | 152 | 165 | 168 | 134 | 241,14 | 327 | 155 | 0,47 | -1,089 | -9,569 | 1,08E-21 | 2,85E-21 | 0,0081229 | 0,007515 | 0,0077794 | 0,0077794 | TRUE | 0,1363755 | No |
| orf19.3572.3 | 11419 | 9967 | 10117 | 6395 | 18858 | 13629 | 14738 | 20012 | 10397 | 10081 | 9575 | 9606 | 16774 | 14797 | 15094 | 15265 | 12698,5 | 9915 | 15482 | 1,561 | 0,643 | 13,065 | 5,24E-39 | 2,00E-38 | 0,0023105 | 0,002141 | 0,0022411 | 0,0022411 | TRUE | 0,0386008 | No |
| orf19.3573 | 1537 | 1469 | 1540 | 916 | 8706 | 7457 | 7475 | 10195 | 1399 | 1486 | 1457 | 1376 | 7744 | 8096 | 7655 | 7777 | 4623,83 | 1430 | 7818 | 5,46 | 2,449 | 59,276 | 0 | 0 | 0,0003806 | 0,002323 | 0,001205 | 0,001205 | TRUE | 0,0081074 | No |
| orf19.3574 | 628 | 577 | 589 | 411 | 954 | 713 | 795 | 1028 | 572 | 584 | 557 | 617 | 849 | 774 | 814 | 784 | 693,9 | 582 | 805 | 1,387 | 0,472 | 7,569 | 3,77E-14 | 8,14E-14 | 1,00E-08 | 0,003941 | 0,0022351 | 0,0022351 | TRUE | 0,015977 | No |
| orf19.3575 | 4633 | 3803 | 4074 | 2877 | 2942 | 2568 | 2501 | 3729 | 4218 | 3846 | 3856 | 4322 | 2617 | 2788 | 2561 | 2845 | 3381,58 | 4060 | 2703 | 0,666 | -0,586 | -10,607 | 2,78E-26 | 8,16E-26 | 0,0026675 | 0,002428 | 0,0026252 | 0,0026252 | TRUE | 0,021976 | No |
| orf19.3577 | 1807 | 1458 | 1520 | 1020 | 1996 | 1511 | 1660 | 2308 | 1645 | 1475 | 1439 | 1532 | 1775 | 1641 | 1700 | 1761 | 1620,88 | 1523 | 1719 | 1,13 | 0,176 | 3,201 | 0,001369 | 0,001915 | 0,0018097 | 0,002852 | 0,0022757 | 0,0022757 | TRUE | 0,0377089 | No |
| orf19.3577.1 | 97 | 86 | 84 | 69 | 305 | 224 | 223 | 337 | 88 | 87 | 79 | 104 | 271 | 243 | 228 | 257 | 169,8 | 90 | 250 | 2,826 | 1,499 | 12,232 | 2,09E-34 | 7,38E-34 | 1,00E-08 | 0,009817 | 0,006567 | 0,006567 | TRUE | 0,0772845 | No |
| orf19.3578 | 721 | 579 | 633 | 502 | 928 | 778 | 844 | 1103 | 656 | 586 | 599 | 754 | 825 | 845 | 864 | 841 | 746,38 | 649 | 844 | 1,307 | 0,387 | 5,123 | 3,00E-07 | 5,08E-07 | 0,0044245 | 0,003807 | 0,0040801 | 0,0040801 | TRUE | 0,1248176 | No |
| orf19.3579 | 5120 | 4222 | 4582 | 3379 | 5816 | 5097 | 5073 | 6896 | 4662 | 4270 | 4336 | 5076 | 5173 | 5534 | 5195 | 5260 | 4938,33 | 4586 | 5290 | 1,155 | 0,207 | 3,656 | 0,000256 | 0,000376 | 0,0034534 | 0,002305 | 0,002885 | 0,002885 | TRUE | 0,06113 | No |
| orf19.3581 | 466 | 455 | 487 | 309 | 717 | 533 | 610 | 809 | 424 | 460 | 461 | 464 | 638 | 579 | 625 | 617 | 533,47 | 452 | 615 | 1,362 | 0,446 | 6,572 | 4,95E-11 | 9,73E-11 | 1,00E-08 | 0,004513 | 0,0024841 | 0,0024841 | TRUE | 0,0219014 | No |
| orf19.3582 | 1188 | 1011 | 1047 | 599 | 1426 | 1004 | 1099 | 1465 | 1082 | 1023 | 991 | 900 | 1268 | 1090 | 1126 | 1118 | 1074,54 | 999 | 1150 | 1,149 | 0,201 | 2,879 | 0,003989 | 0,005377 | 0,0040674 | 0,003266 | 0,0037168 | 0,0037168 | TRUE | 0,0577023 | No |
| orf19.3583 | 3846 | 3414 | 3707 | 2183 | 2177 | 1796 | 2062 | 2560 | 3502 | 3453 | 3508 | 3279 | 1936 | 1950 | 2112 | 1953 | 2711,62 | 3436 | 1988 | 0,578 | -0,791 | -17,613 | 1,95E-69 | 1,21E-68 | 0,0008154 | 0,002524 | 0,0015469 | 0,0015469 | TRUE | 0,0209148 | No |
| orf19.3585 | 988 | 913 | 889 | 531 | 1445 | 1039 | 1105 | 1346 | 900 | 923 | 841 | 798 | 1285 | 1128 | 1132 | 1027 | 1004,21 | 866 | 1143 | 1,318 | 0,398 | 5,235 | 1,65E-07 | 2,82E-07 | 0,0051564 | 0,003352 | 0,0045145 | 0,0045145 | TRUE | 0,0862689 | No |
| orf19.3586 | 2153 | 1827 | 2092 | 1295 | 3736 | 2938 | 3333 | 4507 | 1960 | 1848 | 1980 | 1945 | 3323 | 3190 | 3413 | 3438 | 2637,19 | 1933 | 3341 | 1,729 | 0,79 | 18,012 | 1,57E-72 | 1,02E-71 | 0,0006796 | 0,002538 | 0,0014284 | 0,0014284 | TRUE | 0,0110558 | No |
| orf19.3589 | 99 | 78 | 90 | 59 | 117 | 102 | 102 | 110 | 90 | 79 | 85 | 89 | 104 | 111 | 104 | 84 | 93,25 | 86 | 101 | 1,17 | 0,227 | 1,523 | 0,127721 | 0,151596 | 1,00E-08 | 0,016203 | 0,0104522 | 0,0104522 | TRUE | 0,1437717 | No |
| orf19.359 | 28 | 18 | 31 | 22 | 570 | 435 | 431 | 644 | 25 | 18 | 29 | 33 | 507 | 472 | 441 | 491 | 252,25 | 26 | 478 | 18,382 | 4,2 | 25,28 | 5,31E-141 | 6,51E-140 | 0,0015631 | 0,007274 | 0,0053043 | 0,0053043 | TRUE | 0,2709133 | No |
| orf19.3590 | 2096 | 1812 | 1956 | 1446 | 5327 | 4359 | 4250 | 6081 | 1908 | 1833 | 1851 | 2172 | 4738 | 4733 | 4353 | 4639 | 3278,29 | 1941 | 4616 | 2,383 | 1,253 | 21,695 | 2,29E-104 | 2,07E-103 | 0,0035689 | 0,00244 | 0,0028239 | 0,0028239 | TRUE | 0,0728391 | No |
| orf19.3591 | 1787 | 1663 | 1750 | 1123 | 3762 | 3140 | 3331 | 4362 | 1627 | 1682 | 1656 | 1687 | 3346 | 3409 | 3411 | 3327 | 2518,27 | 1663 | 3373 | 2,03 | 1,021 | 26,816 | 2,12E-158 | 2,92E-157 | 1,00E-08 | 0,002562 | 0,0009319 | 0,0009319 | TRUE | 0,0026521 | No |
| orf19.3592 | 1408 | 1407 | 1444 | 928 | 1043 | 917 | 1017 | 1285 | 1282 | 1423 | 1367 | 1394 | 928 | 996 | 1042 | 980 | 1176,33 | 1366 | 986 | 0,722 | -0,47 | -8,318 | 8,97E-17 | 2,09E-16 | 0,0013404 | 0,00316 | 0,0021994 | 0,0021994 | TRUE | 0,0217735 | No |
| orf19.3593 | 6056 | 5226 | 5400 | 3536 | 5489 | 4460 | 4732 | 6627 | 5514 | 5286 | 5111 | 5312 | 4882 | 4842 | 4846 | 5055 | 5105,93 | 5306 | 4906 | 0,925 | -0,112 | -3,008 | 0,002629 | 0,003593 | 0,0005422 | 0,002296 | 0,0011432 | 0,0011432 | TRUE | 0,0088745 | No |
| orf19.3594 | 632 | 565 | 551 | 363 | 416 | 365 | 342 | 489 | 575 | 571 | 521 | 545 | 370 | 396 | 350 | 373 | 462,9 | 553 | 372 | 0,672 | -0,573 | -7,895 | 2,91E-15 | 6,48E-15 | 1,00E-08 | 0,004891 | 0,0028421 | 0,0028421 | TRUE | 0,0196658 | No |
| orf19.3599 | 15362 | 13806 | 15036 | 9075 | 17582 | 14535 | 16070 | 21081 | 13986 | 13963 | 14230 | 13632 | 15639 | 15781 | 16458 | 16081 | 14971,3 | 13953 | 15990 | 1,146 | 0,196 | 6,327 | 2,50E-10 | 4,79E-10 | 0,0003225 | 0,002125 | 0,0008556 | 0,0008556 | TRUE | 0,0047139 | No |
| orf19.36 | 276 | 326 | 253 | 158 | 276 | 224 | 243 | 327 | 251 | 330 | 239 | 237 | 245 | 243 | 249 | 249 | 255,6 | 264 | 246 | 0,931 | -0,103 | -0,929 | 0,352956 | 0,392892 | 0,0086544 | 0,007205 | 0,0079637 | 0,0079637 | TRUE | 0,3113858 | No |
| orf19.360 | 3 | 3 | 0 | 5 | 28 | 27 | 25 | 34 | 3 | 3 | 0 | 8 | 25 | 29 | 26 | 26 | 14,88 | 4 | 26 | 8,979 | 3,167 | 6,095 | 1,09E-09 | 2,04E-09 | 1,00E-08 | 0,090818 | 0,0621341 | 0,0621341 | TRUE | 0,7183618 | No |
| orf19.3600 | 121 | 136 | 141 | 94 | 127 | 113 | 122 | 164 | 110 | 138 | 133 | 141 | 113 | 123 | 125 | 125 | 126,01 | 130 | 122 | 0,935 | -0,097 | -0,753 | 0,451458 | 0,493407 | 1,00E-08 | 0,012521 | 0,0078698 | 0,0078698 | TRUE | 0,1237579 | No |
| orf19.3601 | 1260 | 1346 | 1335 | 788 | 1240 | 971 | 1069 | 1366 | 1147 | 1361 | 1263 | 1184 | 1103 | 1054 | 1095 | 1042 | 1156,2 | 1239 | 1074 | 0,866 | -0,208 | -3,355 | 0,000795 | 0,001129 | 0,0025129 | 0,00318 | 0,0028269 | 0,0028269 | TRUE | 0,0536237 | No |
| orf19.3603 | 1338 | 1250 | 1356 | 858 | 1064 | 962 | 986 | 1287 | 1218 | 1264 | 1283 | 1289 | 946 | 1044 | 1010 | 982 | 1129,62 | 1264 | 996 | 0,788 | -0,344 | -6,622 | 3,55E-11 | 7,01E-11 | 2,19E-08 | 0,003206 | 0,0017042 | 0,0017042 | TRUE | 0,0130144 | No |
| orf19.3604 | 881 | 875 | 898 | 478 | 1430 | 1148 | 1339 | 1752 | 802 | 885 | 850 | 718 | 1272 | 1246 | 1371 | 1336 | 1060,14 | 814 | 1306 | 1,6 | 0,678 | 10,185 | 2,31E-24 | 6,53E-24 | 0,0031937 | 0,003283 | 0,0032417 | 0,0032417 | TRUE | 0,065109 | No |
| orf19.3605 | 1349 | 1561 | 1492 | 820 | 2129 | 1721 | 1794 | 2296 | 1228 | 1579 | 1412 | 1232 | 1894 | 1869 | 1837 | 1751 | 1600,21 | 1363 | 1838 | 1,347 | 0,429 | 5,603 | 2,11E-08 | 3,75E-08 | 0,0072884 | 0,002863 | 0,0049916 | 0,0049916 | TRUE | 0,1369358 | No |
| orf19.3606 | 5398 | 4769 | 5342 | 3079 | 4380 | 3335 | 3896 | 5188 | 4915 | 4823 | 5056 | 4625 | 3896 | 3621 | 3990 | 3957 | 4360,38 | 4855 | 3866 | 0,796 | -0,329 | -7,243 | 4,39E-13 | 9,24E-13 | 0,0013256 | 0,00234 | 0,0017495 | 0,0017495 | TRUE | 0,0217294 | No |
| orf19.3607 | 538 | 587 | 578 | 318 | 530 | 489 | 476 | 557 | 490 | 594 | 547 | 478 | 471 | 531 | 487 | 425 | 502,87 | 527 | 476 | 0,904 | -0,146 | -1,607 | 0,107998 | 0,129221 | 0,0073196 | 0,004664 | 0,0059065 | 0,0059065 | TRUE | 0,0836755 | No |
| orf19.3608 | 611 | 727 | 782 | 413 | 1217 | 987 | 1057 | 1390 | 556 | 735 | 740 | 620 | 1082 | 1072 | 1083 | 1060 | 868,62 | 663 | 1074 | 1,618 | 0,695 | 8,26 | 1,46E-16 | 3,36E-16 | 0,0076625 | 0,003558 | 0,0055423 | 0,0055423 | TRUE | 0,1496447 | No |
| orf19.3609 | 8219 | 7901 | 8259 | 5478 | 7394 | 6270 | 6090 | 8360 | 7483 | 7991 | 7816 | 8229 | 6577 | 6807 | 6237 | 6377 | 7189,69 | 7880 | 6500 | 0,825 | -0,277 | -6,372 | 1,87E-10 | 3,59E-10 | 0,0013352 | 0,002221 | 0,0016791 | 0,0016791 | TRUE | 0,0140888 | No |
| orf19.36.1 | 196 | 118 | 120 | 91 | 663 | 418 | 414 | 684 | 178 | 119 | 114 | 137 | 590 | 454 | 424 | 522 | 317,17 | 137 | 498 | 3,628 | 1,859 | 12,997 | 1,27E-38 | 4,85E-38 | 0,0277428 | 0,006202 | 0,0148013 | 0,0148013 | TRUE | 0,4238926 | No |
| orf19.3610 | 1407 | 1484 | 1533 | 878 | 1873 | 1494 | 1587 | 2050 | 1281 | 1501 | 1451 | 1319 | 1666 | 1622 | 1625 | 1564 | 1503,59 | 1388 | 1619 | 1,165 | 0,221 | 3,744 | 0,000181 | 0,000268 | 0,0022935 | 0,002916 | 0,0026605 | 0,0026605 | TRUE | 0,0361619 | No |
| orf19.3611 | 3032 | 2681 | 2717 | 1710 | 3642 | 2782 | 2967 | 3877 | 2761 | 2712 | 2571 | 2569 | 3239 | 3020 | 3039 | 2957 | 2858,51 | 2653 | 3064 | 1,154 | 0,207 | 4,48 | 7,48E-06 | 1,20E-05 | 0,0010052 | 0,002499 | 0,0016875 | 0,0016875 | TRUE | 0,018351 | No |
| orf19.3612 | 896 | 862 | 847 | 598 | 563 | 457 | 382 | 598 | 816 | 872 | 802 | 898 | 501 | 496 | 391 | 456 | 653,98 | 847 | 461 | 0,546 | -0,874 | -10,817 | 2,85E-27 | 8,61E-27 | 0,0052729 | 0,004057 | 0,0046201 | 0,0046201 | TRUE | 0,1157178 | No |
| orf19.3613 | 2823 | 2887 | 3141 | 1666 | 2778 | 2215 | 2466 | 3043 | 2570 | 2920 | 2973 | 2503 | 2471 | 2405 | 2525 | 2321 | 2585,99 | 2742 | 2430 | 0,886 | -0,175 | -2,8 | 0,005107 | 0,00685 | 0,003946 | 0,002548 | 0,0033754 | 0,0033754 | TRUE | 0,0397628 | No |

| | | | | | | | | | | | | | | | | | | | | | | | | | | | | | | |
|---|---|---|---|---|---|---|---|---|---|---|---|---|---|---|---|---|---|---|---|---|---|---|---|---|---|---|---|---|---|---|
| orf19.3615 | 1254 | 1325 | 1316 | 844 | 927 | 827 | 760 | 1025 | 1142 | 1340 | 1245 | 1268 | 825 | 898 | 778 | 782 | 1034,72 | 1249 | 821 | 0,657 | -0,606 | -8,965 | 3,11E-19 | 7,72E-19 | 0,0034618 | 0,003314 | 0,0033896 | 0,0033896 | TRUE | 0,0458554 | No |
| orf19.3616 | 1593 | 1687 | 1678 | 937 | 1255 | 1084 | 1080 | 1340 | 1450 | 1706 | 1588 | 1408 | 1116 | 1177 | 1106 | 1022 | 1321,7 | 1538 | 1105 | 0,717 | -0,48 | -6,839 | 7,99E-12 | 1,62E-11 | 0,0045335 | 0,003036 | 0,0039556 | 0,0039556 | TRUE | 0,064806 | No |
| orf19.3617 | 823 | 1036 | 1074 | 595 | 739 | 701 | 656 | 868 | 749 | 1048 | 1016 | 894 | 657 | 761 | 672 | 662 | 807,46 | 927 | 688 | 0,742 | -0,431 | -4,371 | 1,23E-05 | 1,95E-05 | 0,012746 | 0,003673 | 0,0080771 | 0,0080771 | TRUE | 0,2061346 | No |
| orf19.3618 | 2010 | 1675 | 1842 | 1420 | 5079 | 4863 | 4568 | 6424 | 1830 | 1694 | 1743 | 2133 | 4518 | 5280 | 4678 | 4900 | 3347,05 | 1850 | 4844 | 2,623 | 1,391 | 19,02 | 1,16E-80 | 8,26E-80 | 0,0074849 | 0,002432 | 0,0047497 | 0,0047497 | TRUE | 0,12254 | No |
| orf19.362 | 70 | 55 | 57 | 44 | 149 | 110 | 130 | 209 | 64 | 56 | 54 | 66 | 133 | 119 | 133 | 159 | 97,99 | 60 | 136 | 2,3 | 1,201 | 7,642 | 2,14E-14 | 4,66E-14 | 1,00E-08 | 0,015518 | 0,0113905 | 0,0113905 | TRUE | 0,1504116 | No |
| orf19.3621 | 747 | 688 | 755 | 504 | 33 | 38 | 31 | 37 | 680 | 696 | 715 | 757 | 29 | 41 | 32 | 28 | 372,27 | 712 | 32 | 0,045 | -4,466 | -31,837 | 2,00E-222 | 3,75E-221 | 1,00E-08 | 0,005586 | 0,0037558 | 0,0037558 | TRUE | 0,1923731 | No |
| orf19.3622 | 1240 | 1275 | 1280 | 757 | 1167 | 1016 | 1015 | 1465 | 1129 | 1290 | 1211 | 1137 | 1038 | 1103 | 1039 | 1118 | 1133,14 | 1192 | 1074 | 0,901 | -0,15 | -2,534 | 0,011283 | 0,014746 | 0,001855 | 0,003203 | 0,0024961 | 0,0024961 | TRUE | 0,0366796 | No |
| orf19.3623 | 1211 | 1152 | 1326 | 809 | 1174 | 989 | 966 | 1306 | 1103 | 1165 | 1255 | 1215 | 1044 | 1074 | 989 | 996 | 1105,18 | 1184 | 1026 | 0,866 | -0,207 | -3,562 | 0,000368 | 0,000534 | 0,0015164 | 0,003232 | 0,0023275 | 0,0023275 | TRUE | 0,0273742 | No |
| orf19.3624 | 4243 | 4595 | 4628 | 2674 | 5346 | 4766 | 4581 | 6021 | 3863 | 4647 | 4380 | 4017 | 4755 | 5175 | 4692 | 4593 | 4515,15 | 4227 | 4804 | 1,136 | 0,184 | 2,906 | 0,003665 | 0,004953 | 0,0046404 | 0,00233 | 0,003618 | 0,003618 | TRUE | 0,0547838 | No |
| orf19.3625 | 492 | 493 | 521 | 265 | 257 | 210 | 260 | 280 | 448 | 499 | 493 | 398 | 229 | 228 | 266 | 214 | 346,77 | 460 | 234 | 0,506 | -0,982 | -10,015 | 1,31E-23 | 3,65E-23 | 0,006387 | 0,005846 | 0,0060757 | 0,0060757 | TRUE | 0,0928103 | No |
| orf19.3626 | 1586 | 1742 | 1679 | 1029 | 1407 | 1318 | 1266 | 1588 | 1444 | 1762 | 1589 | 1546 | 1251 | 1431 | 1297 | 1211 | 1441,37 | 1585 | 1298 | 0,818 | -0,29 | -4,043 | 5,28E-05 | 8,02E-05 | 0,0052707 | 0,002954 | 0,0042564 | 0,0042564 | TRUE | 0,0689276 | No |
| orf19.3626.1 | 46 | 61 | 51 | 24 | 40 | 38 | 41 | 51 | 42 | 62 | 48 | 36 | 36 | 41 | 42 | 39 | 43,2 | 47 | 40 | 0,829 | -0,271 | -1,235 | 0,216829 | 0,249645 | 1,00E-08 | 0,032614 | 0,0230615 | 0,0230615 | TRUE | 0,3419095 | No |
| orf19.3627 | 1186 | 1125 | 1240 | 647 | 2396 | 1854 | 1986 | 2436 | 1080 | 1138 | 1174 | 972 | 2131 | 2013 | 2034 | 1858 | 1549,91 | 1091 | 2009 | 1,836 | 0,877 | 13,374 | 8,55E-41 | 3,37E-40 | 0,0039048 | 0,002889 | 0,0033999 | 0,0033999 | TRUE | 0,0581967 | No |
| orf19.3628 | 6245 | 5797 | 6014 | 3660 | 4018 | 3189 | 3415 | 4502 | 5686 | 5863 | 5692 | 5498 | 3574 | 3462 | 3497 | 3434 | 4588,28 | 5685 | 3492 | 0,614 | -0,704 | -19,742 | 9,36E-87 | 7,18E-86 | 0,0002674 | 0,002325 | 0,0009921 | 0,0009921 | TRUE | 0,0055775 | No |
| orf19.3629 | 1706 | 1740 | 1741 | 1081 | 1309 | 1148 | 1152 | 1492 | 1553 | 1760 | 1648 | 1624 | 1164 | 1246 | 1180 | 1138 | 1414,15 | 1646 | 1182 | 0,717 | -0,479 | -8,749 | 2,14E-18 | 5,21E-18 | 0,0014919 | 0,002971 | 0,0021585 | 0,0021585 | TRUE | 0,0267487 | No |
| orf19.3630 | 7191 | 7372 | 7583 | 4734 | 9473 | 7955 | 8311 | 10577 | 6547 | 7456 | 7177 | 7111 | 8426 | 8637 | 8512 | 8068 | 7741,69 | 7073 | 8411 | 1,189 | 0,25 | 5,376 | 7,61E-08 | 1,32E-07 | 0,0016987 | 0,002208 | 0,0019438 | 0,0019438 | TRUE | 0,0310895 | No |
| orf19.3631 | 229 | 247 | 267 | 165 | 495 | 444 | 447 | 601 | 208 | 250 | 253 | 248 | 440 | 482 | 458 | 458 | 349,68 | 240 | 460 | 1,922 | 0,943 | 10,966 | 5,59E-28 | 1,71E-27 | 0,0002679 | 0,005815 | 0,0038482 | 0,0038482 | TRUE | 0,0948759 | No |
| orf19.3633 | 360 | 366 | 352 | 228 | 313 | 289 | 276 | 348 | 328 | 370 | 333 | 342 | 278 | 314 | 283 | 265 | 314,23 | 343 | 285 | 0,828 | -0,272 | -3,133 | 0,001728 | 0,002396 | 1,00E-08 | 0,006241 | 0,0040579 | 0,0040579 | TRUE | 0,0491411 | No |
| orf19.3635 | 1546 | 1394 | 1479 | 931 | 2217 | 1886 | 2056 | 2716 | 1408 | 1410 | 1400 | 1399 | 1972 | 2048 | 2106 | 2072 | 1726,59 | 1404 | 2050 | 1,459 | 0,545 | 12,619 | 1,66E-36 | 6,13E-36 | 1,00E-08 | 0,002802 | 0,0011829 | 0,0011829 | TRUE | 0,007943 | No |
| orf19.3637 | 0 | 1 | 0 | 1 | 2 | 1 | 1 | 1 | 0 | 1 | 0 | 2 | 2 | 1 | 1 | 1 | 0,9 | 1 | 1 | 2,134 | 1,094 | 0,732 | 0,464303 | 0,506144 | 1,00E-08 | 1,476952 | 0,8375768 | 0,8375768 | TRUE | 0,2207644 | No |
| orf19.3638 | 41 | 42 | 42 | 23 | 22 | 19 | 12 | 21 | 37 | 42 | 40 | 35 | 20 | 21 | 12 | 16 | 27,83 | 38 | 17 | 0,442 | -1,179 | -4,371 | 1,23E-05 | 1,95E-05 | 1,00E-08 | 0,04951 | 0,0289182 | 0,0289182 | TRUE | 0,1582075 | No |
| orf19.3639 | 879 | 825 | 943 | 561 | 772 | 630 | 662 | 823 | 800 | 834 | 892 | 843 | 687 | 684 | 678 | 628 | 755,79 | 842 | 669 | 0,793 | -0,334 | -5,404 | 6,53E-08 | 1,14E-07 | 0,0009013 | 0,003785 | 0,0023403 | 0,0023403 | TRUE | 0,0222267 | No |
| orf19.364 | 1595 | 1317 | 1392 | 913 | 3141 | 2549 | 2693 | 3819 | 1452 | 1332 | 1317 | 1371 | 2794 | 2768 | 2758 | 2913 | 2088,19 | 1368 | 2808 | 2,053 | 1,038 | 22,037 | 1,27E-107 | 1,17E-106 | 0,0007162 | 0,00267 | 0,0015702 | 0,0015702 | TRUE | 0,0222892 | No |
| orf19.3642 | 9544 | 8061 | 8767 | 5478 | 18041 | 14803 | 15984 | 22751 | 8689 | 8153 | 8297 | 8229 | 16047 | 16072 | 16370 | 17355 | 12401,4 | 8342 | 16461 | 1,973 | 0,98 | 24,83 | 4,30E-136 | 5,03E-135 | 0,0009848 | 0,002144 | 0,0014045 | 0,0014045 | TRUE | 0,0164842 | No |
| orf19.3643 | 557 | 460 | 535 | 362 | 1014 | 944 | 980 | 1358 | 507 | 465 | 506 | 544 | 902 | 1025 | 1004 | 1036 | 748,61 | 506 | 992 | 1,967 | 0,976 | 13,921 | 4,69E-44 | 1,97E-43 | 0,0025367 | 0,003802 | 0,0031847 | 0,0031847 | TRUE | 0,045193 | No |
| orf19.3644 | 4364 | 3774 | 4106 | 2778 | 3200 | 2727 | 2793 | 4074 | 3973 | 3817 | 3886 | 4173 | 2846 | 2961 | 2860 | 3108 | 3453,04 | 3962 | 2944 | 0,744 | -0,427 | -9,305 | 1,34E-20 | 3,45E-20 | 0,0012025 | 0,00242 | 0,001726 | 0,001726 | TRUE | 0,0176985 | No |
| orf19.3646 | 41323 | 33937 | 35671 | 23957 | 35409 | 27526 | 28786 | 41351 | 37623 | 34324 | 33759 | 35987 | 31495 | 29886 | 29480 | 31543 | 33012,2 | 35423 | 30601 | 0,864 | -0,211 | -4,715 | 2,42E-06 | 3,95E-06 | 0,0017873 | 0,002077 | 0,0018938 | 0,0018938 | TRUE | 0,0215141 | No |
| orf19.3647 | 537 | 490 | 546 | 335 | 734 | 543 | 609 | 836 | 489 | 496 | 517 | 503 | 653 | 590 | 624 | 638 | 563,53 | 501 | 626 | 1,251 | 0,323 | 4,969 | 6,73E-07 | 1,13E-06 | 1,00E-08 | 0,004381 | 0,0022588 | 0,0022588 | TRUE | 0,0170492 | No |
| orf19.3648 | 425 | 335 | 360 | 291 | 555 | 531 | 573 | 695 | 387 | 339 | 341 | 437 | 494 | 577 | 587 | 530 | 461,34 | 376 | 547 | 1,461 | 0,547 | 5,89 | 3,87E-09 | 7,08E-09 | 0,0077703 | 0,0049 | 0,0059839 | 0,0059839 | TRUE | 0,1193828 | No |
| orf19.3649 | 2262 | 2236 | 2263 | 1504 | 2534 | 2043 | 1986 | 2676 | 2059 | 2261 | 2142 | 2259 | 2254 | 2218 | 2034 | 2041 | 2158,64 | 2180 | 2137 | 0,981 | -0,028 | -0,525 | 0,599757 | 0,639587 | 0,0020469 | 0,002649 | 0,002306 | 0,002306 | TRUE | 0,0171895 | No |
| orf19.3651 | 20804 | 16377 | 17502 | 11579 | 4940 | 3836 | 3991 | 5785 | 18941 | 16564 | 16564 | 17394 | 4394 | 4165 | 4087 | 4413 | 10815,2 | 17366 | 4265 | 0,246 | -2,025 | -38,686 | 0 | 0 | 0,0026223 | 0,002159 | 0,0024921 | 0,0024921 | TRUE | 0,045952 | No |
| orf19.3653 | 5454 | 4726 | 5023 | 3349 | 19669 | 16366 | 16976 | 24091 | 4966 | 4780 | 4754 | 5031 | 17495 | 17769 | 17386 | 18377 | 11319,5 | 4883 | 17757 | 3,639 | 1,864 | 52,186 | 0 | 0 | 0,0005297 | 0,002154 | 0,0010878 | 0,0010878 | TRUE | 0,0068354 | No |
| orf19.3655 | 41 | 44 | 39 | 29 | 59 | 46 | 43 | 56 | 37 | 45 | 37 | 44 | 52 | 50 | 44 | 43 | 43,93 | 41 | 47 | 1,171 | 0,227 | 1,099 | 0,271782 | 0,30791 | 1,00E-08 | 0,032104 | 0,0179798 | 0,0179798 | TRUE | 0,0493107 | No |
| orf19.3656 | 4086 | 3935 | 4167 | 2923 | 3227 | 2853 | 2804 | 3813 | 3720 | 3980 | 3944 | 4391 | 2870 | 3098 | 2872 | 2909 | 3472,82 | 4009 | 2937 | 0,733 | -0,447 | -8,173 | 3,02E-16 | 6,89E-16 | 0,0027194 | 0,002417 | 0,0025843 | 0,0025843 | TRUE | 0,0484657 | No |
| orf19.3658 | 237 | 218 | 204 | 143 | 449 | 410 | 396 | 540 | 216 | 220 | 193 | 215 | 399 | 445 | 406 | 412 | 313,27 | 211 | 416 | 1,969 | 0,978 | 11,233 | 2,82E-29 | 8,90E-29 | 1,00E-08 | 0,006254 | 0,003628 | 0,003628 | TRUE | 0,0393021 | No |
| orf19.3659 | 1756 | 1592 | 1751 | 1030 | 1214 | 976 | 1047 | 1405 | 1599 | 1610 | 1657 | 1547 | 1080 | 1060 | 1072 | 1072 | 1337,1 | 1603 | 1071 | 0,667 | -0,584 | -12,663 | 9,42E-37 | 3,50E-36 | 1,00E-08 | 0,003025 | 0,0012727 | 0,0012727 | TRUE | 0,0060732 | No |
| orf19.366 | 1549 | 1091 | 1257 | 650 | 2863 | 1933 | 2605 | 3461 | 1410 | 1103 | 1190 | 976 | 2547 | 2099 | 2668 | 2640 | 1829,12 | 1170 | 2488 | 2,123 | 1,086 | 10,627 | 2,24E-26 | 6,60E-26 | 0,0170087 | 0,002759 | 0,0093854 | 0,0093854 | TRUE | 0,2300335 | No |
| orf19.3660 | 2320 | 2008 | 2273 | 1338 | 1646 | 1385 | 1495 | 2054 | 2112 | 2031 | 2151 | 2010 | 1464 | 1504 | 1531 | 1567 | 1796,24 | 2076 | 1516 | 0,73 | -0,454 | -9,959 | 2,31E-23 | 6,37E-23 | 0,0004144 | 0,002772 | 0,0014316 | 0,0014316 | TRUE | 0,0071149 | No |
| orf19.3661 | 4994 | 4921 | 5164 | 3355 | 3448 | 3120 | 3032 | 4182 | 4547 | 4977 | 4887 | 5040 | 3067 | 3387 | 3105 | 3190 | 4025,06 | 4863 | 3187 | 0,656 | -0,609 | -12,548 | 4,09E-36 | 1,49E-35 | 0,0016803 | 0,002365 | 0,0020045 | 0,0020045 | TRUE | 0,0236124 | No |
| orf19.3663 | 7744 | 7396 | 7846 | 4857 | 10104 | 8288 | 8777 | 11705 | 7051 | 7480 | 7426 | 7296 | 8987 | 8998 | 8989 | 8929 | 8144,43 | 7313 | 8976 | 1,227 | 0,296 | 9,354 | 8,45E-21 | 2,19E-20 | 0,000183 | 0,002199 | 0,000833 | 0,000833 | TRUE | 0,0074142 | No |
| orf19.3663.1 | 873 | 754 | 784 | 504 | 478 | 375 | 348 | 501 | 795 | 763 | 742 | 757 | 425 | 407 | 356 | 382 | 578,42 | 764 | 392 | 0,514 | -0,961 | -13,684 | 1,27E-42 | 5,18E-42 | 0,0007625 | 0,004321 | 0,0028455 | 0,0028455 | TRUE | 0,041679 | No |
| orf19.3664 | 10653 | 9513 | 9835 | 6309 | 430 | 324 | 339 | 445 | 9699 | 9621 | 9308 | 9477 | 382 | 352 | 347 | 339 | 4940,8 | 9526 | 355 | 0,037 | -4,746 | -91,825 | 0 | 0 | 1,00E-08 | 0,002304 | 0,0012017 | 0,0012017 | TRUE | 0,0326558 | No |
| orf19.3665 | 1607 | 1467 | 1502 | 959 | 1645 | 1301 | 1333 | 1792 | 1463 | 1484 | 1422 | 1441 | 1463 | 1413 | 1365 | 1367 | 1427,09 | 1452 | 1402 | 0,965 | -0,051 | -1,097 | 0,272548 | 0,308504 | 1,00E-08 | 0,002963 | 0,0013993 | 0,0013993 | TRUE | 0,0107703 | No |
| orf19.3666 | 635 | 626 | 661 | 385 | 1372 | 1135 | 1200 | 1537 | 578 | 633 | 626 | 578 | 1220 | 1232 | 1229 | 1172 | 908,65 | 604 | 1213 | 2,005 | 1,003 | 17,605 | 2,24E-69 | 1,39E-68 | 1,00E-08 | 0,003491 | 0,0018519 | 0,0018519 | TRUE | 0,0121813 | No |
| orf19.3667 | 56 | 49 | 46 | 33 | 57 | 43 | 47 | 75 | 51 | 50 | 44 | 50 | 51 | 47 | 48 | 57 | 49,55 | 49 | 51 | 1,054 | 0,077 | 0,397 | 0,691544 | 0,726891 | 1,00E-08 | 0,028698 | 0,0156066 | 0,0156066 | TRUE | 0,0721865 | No |
| orf19.3668 | 4258 | 3577 | 3770 | 2602 | 16387 | 14344 | 14074 | 20508 | 3877 | 3618 | 3568 | 3909 | 14576 | 15574 | 14414 | 15644 | 9397,2 | 3743 | 15052 | 4,024 | 2,009 | 42,541 | 0 | 0 | 0,0017923 | 0,002178 | 0,0019659 | 0,0019659 | TRUE | 0,0120808 | No |
| orf19.3669 | 17247 | 15782 | 17267 | 11387 | 3294 | 2887 | 2911 | 3939 | 15703 | 15962 | 16342 | 17105 | 2930 | 3134 | 2981 | 3005 | 9645,19 | 16278 | 3012 | 0,185 | -2,434 | -59,811 | 0 | 0 | 0,0008593 | 0,002174 | 0,001402 | 0,001402 | TRUE | 0,0140855 | No |
| orf19.367 | 4440 | 3473 | 4148 | 2873 | 6510 | 5877 | 6342 | 9143 | 4042 | 3513 | 3926 | 4316 | 5790 | 6381 | 6495 | 6974 | 5179,62 | 3949 | 6410 | 1,624 | 0,7 | 10,034 | 1,09E-23 | 3,02E-23 | 0,0061805 | 0,002292 | 0,0044628 | 0,0044628 | TRUE | 0,0673017 | No |
| orf19.3670 | 6097 | 5715 | 6019 | 3658 | 2338 | 1936 | 2039 | 2672 | 5551 | 5780 | 5696 | 5495 | 2080 | 2102 | 2088 | 2038 | 3853,81 | 5630 | 2077 | 0,369 | -1,44 | -39,515 | 0 | 0 | 1,00E-08 | 0,00238 | 0,0009548 | 0,0009548 | TRUE | 0,0038484 | No |
| orf19.3671 | 1 | 1 | 0 | 2 | 4 | 9 | 3 | 8 | 1 | 1 | 0 | 3 | 4 | 10 | 3 | 6 | 3,43 | 1 | 6 | 5,152 | 2,365 | 2,491 | 0,012749 | 0,01659 | 0,0956334 | 0,387303 | 0,2951937 | 0,2951937 | TRUE | 0,411922 | No |
| orf19.3672 | 6920 | 6218 | 6519 | 4217 | 2670 | 2371 | 2307 | 3160 | 6300 | 6289 | 6170 | 6335 | 2375 | 2574 | 2363 | 2410 | 4351,96 | 6274 | 2430 | 0,387 | -1,369 | -35,396 | 1,98E-274 | 4,59E-273 | 0,0003103 | 0,002341 | 0,0011572 | 0,0011572 | TRUE | 0,0183764 | No |
| orf19.3673 | 182 | 139 | 163 | 104 | 177 | 104 | 127 | 155 | 166 | 141 | 154 | 156 | 157 | 113 | 130 | 118 | 141,93 | 154 | 130 | 0,842 | -0,248 | -1,888 | 0,059025 | 0,072613 | 0,0068825 | 0,011345 | 0,0095714 | 0,0095714 | TRUE | 0,2218358 | No |
| orf19.3674 | 2682 | 2505 | 2592 | 1716 | 1512 | 1223 | 1228 | 1593 | 2442 | 2534 | 2453 | 2578 | 1345 | 1328 | 1258 | 1215 | 1893,96 | 2502 | 1286 | 0,514 | -0,96 | -19,834 | 1,51E-87 | 1,17E-86 | 0,0006664 | 0,002734 | 0,0016686 | 0,0016686 | TRUE | 0,0179574 | No |
| orf19.3675 | 7745 | 7553 | 7704 | 5487 | 3580 | 3107 | 2929 | 3917 | 7051 | 7639 | 7291 | 8242 | 3184 | 3373 | 3000 | 2988 | 5346,16 | 7556 | 3136 | 0,415 | -1,268 | -21,548 | 5,59E-103 | 5,00E-102 | 0,0037227 | 0,002284 | 0,0031071 | 0,0031071 | TRUE | 0,0449473 | No |
| orf19.3676 | 3237 | 3287 | 3270 | 2101 | 4101 | 3178 | 3277 | 4279 | 2947 | 3324 | 3095 | 3156 | 3648 | 3450 | 3356 | 3264 | 3280,08 | 3130 | 3430 | 1,096 | 0,132 | 2,553 | 0,010687 | 0,013998 | 0,001982 | 0,00244 | 0,0022538 | 0,0022538 | TRUE | 0,022646 | No |
| orf19.3677 | 302 | 270 | 355 | 202 | 191 | 186 | 161 | 239 | 275 | 273 | 336 | 303 | 170 | 202 | 165 | 182 | 238,31 | 297 | 180 | 0,605 | -0,724 | -6,794 | 1,09E-11 | 2,20E-11 | 0,0050643 | 0,00758 | 0,0065151 | 0,0065151 | TRUE | 0,0973514 | No |
| orf19.3678 | 2926 | 2855 | 3033 | 1925 | 1573 | 1418 | 1496 | 1957 | 2664 | 2888 | 2870 | 2892 | 1399 | 1540 | 1532 | 1493 | 2159,65 | 2828 | 1491 | 0,527 | -0,924 | -18,661 | 1,02E-77 | 7,00E-77 | 0,0012497 | 0,002649 | 0,0018492 | 0,0018492 | TRUE | 0,0207602 | No |
| orf19.3679 | 1802 | 1721 | 1785 | 1095 | 638 | 574 | 515 | 708 | 1641 | 1741 | 1689 | 1645 | 567 | 623 | 527 | 540 | 1121,7 | 1679 | 564 | 0,336 | -1,576 | -26,377 | 2,52E-153 | 3,34E-152 | 0,0010521 | 0,003215 | 0,0022784 | 0,0022784 | TRUE | 0,0534209 | No |
| orf19.3680 | 388 | 428 | 425 | 261 | 775 | 670 | 656 | 871 | 353 | 433 | 402 | 392 | 689 | 727 | 672 | 664 | 541,68 | 395 | 688 | 1,742 | 0,8 | 10,67 | 1,41E-26 | 4,19E-26 | 0,0018989 | 0,004476 | 0,0033706 | 0,0033706 | TRUE | 0,0654489 | No |
| orf19.3681 | 2377 | 2202 | 2398 | 1691 | 2396 | 2132 | 2047 | 2821 | 2164 | 2227 | 2269 | 2540 | 2131 | 2315 | 2096 | 2152 | 2236,88 | 2300 | 2174 | 0,946 | -0,079 | -1,376 | 0,168725 | 0,197072 | 0,0029916 | 0,002628 | 0,002751 | 0,002751 | TRUE | 0,0565533 | No |
| orf19.3682 | 1634 | 1943 | 2008 | 1208 | 1495 | 1435 | 1326 | 1630 | 1488 | 1965 | 1900 | 1815 | 1330 | 1558 | 1358 | 1243 | 1582,12 | 1792 | 1372 | 0,765 | -0,386 | -4,283 | 1,85E-05 | 2,89E-05 | 0,0108426 | 0,002872 | 0,0071478 | 0,0071478 | TRUE | 0,1610572 | No |
| orf19.3683 | 1814 | 1819 | 1977 | 1047 | 2923 | 2381 | 2521 | 3214 | 1652 | 1840 | 1871 | 1573 | 2600 | 2585 | 2582 | 2452 | 2144,2 | 1734 | 2555 | 1,47 | 0,556 | 9,385 | 6,27E-21 | 1,62E-20 | 0,0031996 | 0,002653 | 0,0028793 | 0,0028793 | TRUE | 0,0435374 | No |
| orf19.3684 | 2405 | 2012 | 2137 | 1482 | 14144 | 12590 | 12154 | 17424 | 2190 | 2035 | 2022 | 2226 | 12581 | 13669 | 12447 | 13291 | 7557,69 | 2118 | 12997 | 6,142 | 2,619 | 53,282 | 0 | 0 | 0,0018552 | 0,002212 | 0,0020292 | 0,0020292 | TRUE | 0,0147057 | No |
| orf19.3685 | 620 | 650 | 696 | 392 | 764 | 607 | 719 | 803 | 564 | 657 | 659 | 589 | 680 | 659 | 736 | 613 | 644,61 | 617 | 672 | 1,086 | 0,119 | 1,514 | 0,130144 | 0,154242 | 0,0045734 | 0,004086 | 0,0043242 | 0,0043242 | TRUE | 0,0489743 | No |
| orf19.3686 | 780 | 778 | 730 | 487 | 686 | 525 | 531 | 780 | 710 | 787 | 691 | 732 | 610 | 570 | 544 | 595 | 654,8 | 730 | 580 | 0,796 | -0,329 | -4,887 | 1,02E-06 | 1,70E-06 | 0,0014744 | 0,004054 | 0,0028181 | 0,0028181 | TRUE | 0,0344857 | No |
| orf19.3687 | 238 | 225 | 227 | 144 | 298 | 210 | 256 | 315 | 217 | 228 | 215 | 216 | 265 | 228 | 262 | 240 | 233,87 | 219 | 249 | 1,137 | 0,186 | 1,956 | 0,050429 | 0,062412 | 1,00E-08 | 0,007686 | 0,0043516 | 0,0043516 | TRUE | 0,0315977 | No |
| orf19.3688 | 296 | 278 | 303 | 135 | 323 | 201 | 230 | 334 | 269 | 281 | 287 | 203 | 287 | 218 | 236 | 255 | 254,51 | 260 | 249 | 0,953 | -0,07 | -0,586 | 0,557636 | 0,598977 | 0,0135197 | 0,007227 | 0,0097913 | 0,0097913 | TRUE | 0,2076424 | No |

| | | | | | | | | | | | | | | | | | | | | | | | | | | | | | | | |
|---|---|---|---|---|---|---|---|---|---|---|---|---|---|---|---|---|---|---|---|---|---|---|---|---|---|---|---|---|---|---|---|
| orf19.3689 | 3880 | 3766 | 4007 | 2278 | 4725 | 3953 | 3942 | 5295 | 3533 | 3809 | 3792 | 3422 | 4203 | 4292 | 4037 | 4039 | 3890,8 | 3639 | 4143 | 1,137 | 0,186 | 3,942 | 8,07E-05 | 0,000122 | 0,0015142 | 0,002377 | 0,0018651 | 0,0018651 | TRUE | 0,0184688 | No |
| orf19.3690.2 | 6008 | 5918 | 5344 | 3462 | 12331 | 9617 | 9158 | 12384 | 5470 | 5985 | 5058 | 5200 | 10968 | 10441 | 9379 | 9447 | 7743,56 | 5428 | 10059 | 1,853 | 0,89 | 13,414 | 5,00E-41 | 1,98E-40 | 0,0056287 | 0,002208 | 0,0040785 | 0,0040785 | TRUE | 0,0583352 | No |
| orf19.3691 | 941 | 966 | 1014 | 496 | 749 | 584 | 686 | 780 | 857 | 977 | 960 | 745 | 666 | 634 | 703 | 595 | 767,04 | 885 | 650 | 0,731 | -0,451 | -5,149 | 2,62E-07 | 4,45E-07 | 0,0080407 | 0,003759 | 0,0060502 | 0,0060502 | TRUE | 0,1234373 | No |
| orf19.3693 | 33 | 32 | 27 | 13 | 41 | 43 | 35 | 53 | 30 | 32 | 26 | 20 | 36 | 47 | 36 | 40 | 33,36 | 27 | 40 | 1,455 | 0,541 | 2,224 | 0,026181 | 0,033234 | 1,00E-08 | 0,04163 | 0,0257231 | 0,0257231 | TRUE | 0,1345439 | No |
| orf19.3694 | 411 | 408 | 451 | 227 | 349 | 300 | 271 | 409 | 374 | 413 | 427 | 341 | 310 | 326 | 278 | 312 | 347,54 | 389 | 306 | 0,785 | -0,349 | -3,851 | 0,000118 | 0,000175 | 0,0038367 | 0,005838 | 0,0050045 | 0,0050045 | TRUE | 0,0658336 | No |
| orf19.3695 | 2220 | 2456 | 2459 | 1161 | 2308 | 1897 | 2053 | 2496 | 2021 | 2484 | 2327 | 1744 | 2053 | 2060 | 2103 | 1904 | 2086,93 | 2144 | 2030 | 0,945 | -0,081 | -0,898 | 0,369017 | 0,408985 | 0,0117453 | 0,00267 | 0,0073914 | 0,0073914 | TRUE | 0,1848059 | No |
| orf19.3696 | 1753 | 1579 | 1540 | 890 | 1670 | 1310 | 1159 | 1788 | 1596 | 1597 | 1457 | 1337 | 1485 | 1422 | 1187 | 1364 | 1430,75 | 1497 | 1364 | 0,911 | -0,135 | -1,731 | 0,083401 | 0,100926 | 0,00699 | 0,00296 | 0,0051598 | 0,0051598 | TRUE | 0,0911741 | No |
| orf19.3697 | 243 | 314 | 333 | 191 | 432 | 392 | 382 | 489 | 221 | 318 | 315 | 287 | 384 | 426 | 391 | 373 | 339,37 | 285 | 394 | 1,38 | 0,465 | 4,25 | 2,14E-05 | 3,33E-05 | 0,0116223 | 0,00593 | 0,0083971 | 0,0083971 | TRUE | 0,2785816 | No |
| orf19.3698 | 338 | 436 | 437 | 232 | 102 | 78 | 96 | 113 | 308 | 441 | 414 | 349 | 91 | 85 | 98 | 86 | 233,84 | 378 | 90 | 0,238 | -2,072 | -15,407 | 1,48E-53 | 7,30E-53 | 0,0143218 | 0,007686 | 0,0107719 | 0,0107719 | TRUE | 0,1055258 | No |
| orf19.3699 | 522 | 615 | 657 | 359 | 408 | 345 | 372 | 447 | 475 | 622 | 622 | 539 | 363 | 375 | 381 | 341 | 464,72 | 564 | 365 | 0,645 | -0,633 | -6,675 | 2,48E-11 | 4,93E-11 | 0,0080156 | 0,00488 | 0,0063975 | 0,0063975 | TRUE | 0,1431228 | No |
| orf19.3700 | 3232 | 3644 | 3679 | 2364 | 4248 | 3998 | 3698 | 4795 | 2943 | 3686 | 3482 | 3551 | 3778 | 4341 | 3787 | 3658 | 3653,14 | 3416 | 3891 | 1,139 | 0,188 | 2,519 | 0,011772 | 0,01536 | 0,0072801 | 0,002399 | 0,0050901 | 0,0050901 | TRUE | 0,1068619 | No |
| orf19.3701 | 1029 | 1056 | 1074 | 624 | 1003 | 829 | 886 | 1157 | 937 | 1068 | 1016 | 937 | 892 | 900 | 907 | 883 | 942,6 | 990 | 896 | 0,903 | -0,147 | -2,5 | 0,012431 | 0,016189 | 0,0010842 | 0,003438 | 0,002253 | 0,002253 | TRUE | 0,0337182 | No |
| orf19.3704 | 7712 | 7132 | 7592 | 4801 | 7472 | 5915 | 6569 | 8452 | 7021 | 7213 | 7185 | 7212 | 6646 | 6422 | 6727 | 6447 | 6859,32 | 7158 | 6560 | 0,917 | -0,126 | -3,915 | 9,04E-05 | 0,000136 | 0,0001479 | 0,00223 | 0,0008402 | 0,0008402 | TRUE | 0,003515 | No |
| orf19.3704.1 | 1256 | 928 | 949 | 588 | 926 | 572 | 724 | 993 | 1144 | 939 | 898 | 883 | 824 | 621 | 741 | 757 | 850,89 | 966 | 736 | 0,762 | -0,393 | -3,998 | 6,38E-05 | 9,66E-05 | 0,0127974 | 0,00359 | 0,0080771 | 0,0080771 | TRUE | 0,1861294 | No |
| orf19.3705 | 2870 | 2492 | 2680 | 1966 | 4319 | 3724 | 3991 | 5456 | 2613 | 2520 | 2536 | 2953 | 3842 | 4043 | 4087 | 4162 | 3344,63 | 2656 | 4034 | 1,522 | 0,606 | 10,909 | 1,04E-27 | 3,18E-27 | 0,0030646 | 0,002432 | 0,0026393 | 0,0026393 | TRUE | 0,0655402 | No |
| orf19.3706 | 3454 | 3379 | 3557 | 1954 | 2533 | 2086 | 2265 | 2841 | 3145 | 3417 | 3366 | 2935 | 2253 | 2265 | 2320 | 2167 | 2733,55 | 3216 | 2251 | 0,699 | -0,517 | -9,529 | 1,59E-21 | 4,18E-21 | 0,0022603 | 0,00252 | 0,0024498 | 0,0024498 | TRUE | 0,0397964 | No |
| orf19.3707 | 29758 | 26245 | 28293 | 17328 | 11098 | 8824 | 9101 | 12336 | 27093 | 26544 | 26777 | 26029 | 9871 | 9580 | 9321 | 9410 | 18078,2 | 26611 | 9546 | 0,359 | -1,479 | -46,809 | 0 | 0 | 0,0003695 | 0,00211 | 0,00089 | 0,00089 | TRUE | 0,006505 | No |
| orf19.3708 | 15 | 13 | 17 | 5 | 42 | 39 | 41 | 42 | 14 | 13 | 16 | 8 | 37 | 42 | 42 | 32 | 25,52 | 13 | 38 | 2,939 | 1,555 | 5,182 | 2,19E-07 | 3,74E-07 | 1,00E-08 | 0,053807 | 0,0335287 | 0,0335287 | TRUE | 0,1697446 | No |
| orf19.371 | 1115 | 911 | 947 | 695 | 1335 | 1181 | 1209 | 1803 | 1015 | 921 | 896 | 1044 | 1187 | 1282 | 1238 | 1375 | 1120 | 969 | 1270 | 1,315 | 0,395 | 5,824 | 5,75E-09 | 1,05E-08 | 0,0034735 | 0,003216 | 0,0034836 | 0,0034836 | TRUE | 0,0385651 | No |
| orf19.3710 | 3704 | 3854 | 3906 | 2143 | 2437 | 2120 | 2177 | 2646 | 3372 | 3898 | 3697 | 3219 | 2168 | 2302 | 2230 | 2018 | 2862,92 | 3546 | 2180 | 0,614 | -0,704 | -10,642 | 1,91E-26 | 5,64E-26 | 0,0047913 | 0,002498 | 0,0038421 | 0,0038421 | TRUE | 0,0537723 | No |
| orf19.3711 | 59 | 50 | 61 | 29 | 1540 | 1306 | 1353 | 1862 | 54 | 51 | 58 | 44 | 1370 | 1418 | 1386 | 1420 | 724,91 | 52 | 1398 | 26,804 | 4,744 | 41,618 | 0 | 0 | 1,00E-08 | 0,003859 | 0,00206 | 0,00206 | TRUE | 0,0531348 | No |
| orf19.3712 | 15372 | 16687 | 16874 | 9718 | 9610 | 8317 | 8409 | 10374 | 13996 | 16877 | 15970 | 14598 | 8548 | 9030 | 8612 | 7913 | 11942,9 | 15360 | 8526 | 0,555 | -0,85 | -13,41 | 5,27E-41 | 2,08E-40 | 0,0049531 | 0,002148 | 0,0037666 | 0,0037666 | TRUE | 0,0541338 | No |
| orf19.3713 | 4025 | 5711 | 4514 | 3380 | 3506 | 3022 | 2243 | 3008 | 3665 | 5776 | 4272 | 5077 | 3118 | 3281 | 2297 | 2295 | 3722,65 | 4698 | 2748 | 0,585 | -0,774 | -3,914 | 9,06E-05 | 0,000136 | 0,037245 | 0,002392 | 0,0174408 | 0,037245 | TRUE | 0,1480015 | No |
| orf19.3714 | 868 | 818 | 897 | 547 | 1422 | 1239 | 1315 | 1786 | 790 | 827 | 849 | 822 | 1265 | 1345 | 1347 | 1362 | 1075,92 | 822 | 1330 | 1,618 | 0,694 | 13,098 | 3,39E-39 | 1,30E-38 | 1,00E-08 | 0,003265 | 0,0016947 | 0,0016947 | TRUE | 0,0133377 | No |
| orf19.3715 | 1176 | 921 | 997 | 582 | 1085 | 782 | 968 | 1209 | 1071 | 931 | 944 | 874 | 965 | 849 | 991 | 922 | 943,46 | 955 | 932 | 0,974 | -0,038 | -0,525 | 0,599727 | 0,639587 | 0,0047294 | 0,003437 | 0,0040822 | 0,0040822 | TRUE | 0,0805342 | No |
| orf19.3719 | 14 | 9 | 12 | 8 | 6 | 16 | 10 | 11 | 13 | 9 | 11 | 12 | 5 | 17 | 10 | 8 | 10,82 | 11 | 10 | 0,897 | -0,157 | -0,356 | 0,721545 | 0,754569 | 0,0360402 | 0,12412 | 0,0931851 | 0,0931851 | TRUE | 0,6789303 | No |
| orf19.372 | 717 | 524 | 617 | 394 | 543 | 423 | 408 | 616 | 653 | 530 | 584 | 592 | 483 | 459 | 418 | 470 | 523,57 | 590 | 458 | 0,776 | -0,365 | -4,549 | 5,40E-06 | 8,68E-06 | 0,0039283 | 0,00456 | 0,0042599 | 0,0042599 | TRUE | 0,0656469 | No |
| orf19.3720 | 4107 | 3550 | 3899 | 2297 | 2972 | 2325 | 2755 | 3703 | 3739 | 3590 | 3690 | 3450 | 2643 | 2524 | 2821 | 2825 | 3160,52 | 3617 | 2703 | 0,747 | -0,421 | -8,515 | 1,67E-17 | 3,97E-17 | 0,0016561 | 0,002455 | 0,0020214 | 0,0020214 | TRUE | 0,0235973 | No |
| orf19.3721 | 251 | 222 | 239 | 182 | 276 | 226 | 258 | 352 | 229 | 225 | 226 | 273 | 245 | 245 | 264 | 269 | 247,03 | 238 | 256 | 1,084 | 0,117 | 1,218 | 0,223135 | 0,256396 | 0,000274 | 0,007385 | 0,0047745 | 0,0047745 | TRUE | 0,0889551 | No |
| orf19.3722 | 3306 | 2793 | 2978 | 2003 | 2835 | 2358 | 2575 | 3555 | 3010 | 2825 | 2818 | 3009 | 2522 | 2560 | 2637 | 2712 | 2761,59 | 2916 | 2608 | 0,895 | -0,16 | -3,591 | 0,00033 | 0,00048 | 0,0008835 | 0,002515 | 0,0015319 | 0,0015319 | TRUE | 0,0091621 | No |
| orf19.3724 | 3361 | 3008 | 3138 | 2012 | 3193 | 2549 | 2719 | 3788 | 3060 | 3042 | 2970 | 3022 | 2840 | 2768 | 2785 | 2890 | 2922,02 | 3024 | 2821 | 0,933 | -0,1 | -2,737 | 0,006191 | 0,008249 | 1,00E-08 | 0,002489 | 0,0009258 | 0,0009258 | TRUE | 0,0034243 | No |
| orf19.3726 | 895 | 816 | 917 | 523 | 1802 | 1442 | 1640 | 2245 | 815 | 825 | 868 | 786 | 1603 | 1566 | 1680 | 1713 | 1231,77 | 824 | 1640 | 1,989 | 0,992 | 18,161 | 1,05E-73 | 6,86E-73 | 0,0008909 | 0,003109 | 0,0019318 | 0,0019318 | TRUE | 0,0157102 | No |
| orf19.3727 | 115 | 112 | 108 | 71 | 124 | 85 | 95 | 122 | 105 | 113 | 102 | 107 | 110 | 92 | 97 | 93 | 102,47 | 107 | 98 | 0,922 | -0,118 | -0,858 | 0,390639 | 0,431598 | 1,00E-08 | 0,014928 | 0,0082623 | 0,0082623 | TRUE | 0,0702345 | No |
| orf19.3728 | 2122 | 2011 | 2183 | 1454 | 2296 | 1995 | 2175 | 2748 | 1932 | 2034 | 2066 | 2184 | 2042 | 2166 | 2227 | 2096 | 2093,49 | 2054 | 2133 | 1,04 | 0,056 | 1,111 | 0,266732 | 0,302669 | 0,0013601 | 0,002668 | 0,0019686 | 0,0019686 | TRUE | 0,0203727 | No |
| orf19.3730 | 413 | 432 | 411 | 277 | 912 | 716 | 745 | 1028 | 376 | 437 | 389 | 416 | 811 | 777 | 763 | 784 | 594,22 | 404 | 784 | 1,944 | 0,959 | 14,063 | 6,45E-45 | 2,76E-44 | 0,0001134 | 0,00426 | 0,0025492 | 0,0025492 | TRUE | 0,0377823 | No |
| orf19.3732 | 4912 | 4720 | 4657 | 3050 | 5492 | 4871 | 4594 | 6450 | 4472 | 4774 | 4407 | 4582 | 4885 | 5289 | 4705 | 4920 | 4754,18 | 4559 | 4950 | 1,086 | 0,119 | 2,542 | 0,011015 | 0,014407 | 0,0015613 | 0,002315 | 0,0018795 | 0,0018795 | TRUE | 0,0254476 | No |
| orf19.3733 | 1623 | 1452 | 1525 | 996 | 6759 | 5931 | 5719 | 8082 | 1478 | 1469 | 1443 | 1496 | 6012 | 6439 | 5857 | 6165 | 3794,87 | 1472 | 6118 | 4,16 | 2,057 | 46,624 | 0 | 0 | 0,0008652 | 0,002385 | 0,0014273 | 0,0014273 | TRUE | 0,01491 | No |
| orf19.3734 | 3928 | 3761 | 3996 | 2289 | 2991 | 2497 | 2635 | 3434 | 3576 | 3804 | 3782 | 3438 | 2660 | 2711 | 2699 | 2619 | 3161,24 | 3650 | 2672 | 0,731 | -0,452 | -10,279 | 8,80E-25 | 2,51E-24 | 0,0008289 | 0,002455 | 0,0015325 | 0,0015325 | TRUE | 0,0173014 | No |
| orf19.3735 | 12 | 11 | 9 | 18 | 44 | 41 | 37 | 58 | 11 | 11 | 9 | 27 | 39 | 45 | 38 | 44 | 27,92 | 14 | 42 | 3,069 | 1,618 | 5,162 | 2,45E-07 | 4,17E-07 | 0,0107955 | 0,049343 | 0,0437884 | 0,0437884 | TRUE | 1,0168811 | No |
| orf19.3736 | 6 | 9 | 11 | 3 | 5 | 8 | 2 | 7 | 5 | 9 | 10 | 5 | 4 | 9 | 2 | 5 | 6,25 | 7 | 5 | 0,679 | -0,558 | -0,974 | 0,32995 | 0,369651 | 1,00E-08 | 0,213383 | 0,1530451 | 0,1530451 | TRUE | 0,3032829 | No |
| orf19.3737 | 4730 | 4397 | 4693 | 3076 | 3565 | 3024 | 3150 | 4270 | 4306 | 4447 | 4441 | 4621 | 3171 | 3283 | 3226 | 3257 | 3844,13 | 4454 | 3234 | 0,727 | -0,46 | -12,53 | 5,12E-36 | 1,87E-35 | 0,0002529 | 0,002381 | 0,0010308 | 0,0010308 | TRUE | 0,007226 | No |
| orf19.3738 | 34 | 37 | 29 | 18 | 47 | 42 | 59 | 82 | 31 | 37 | 27 | 27 | 42 | 46 | 60 | 63 | 41,66 | 30 | 53 | 1,709 | 0,773 | 3,321 | 0,000897 | 0,00127 | 1,00E-08 | 0,03375 | 0,0259293 | 0,0259293 | TRUE | 0,1396124 | No |
| orf19.374 | 6878 | 5462 | 6148 | 3701 | 4956 | 4049 | 4799 | 6455 | 6262 | 5524 | 5819 | 5559 | 4408 | 4396 | 4915 | 4924 | 5225,92 | 5791 | 4661 | 0,805 | -0,314 | -5,42 | 5,97E-08 | 1,04E-07 | 0,003522 | 0,00229 | 0,0030215 | 0,0030215 | TRUE | 0,0368374 | No |
| orf19.3740 | 370 | 281 | 289 | 262 | 310 | 279 | 267 | 403 | 337 | 284 | 274 | 394 | 276 | 303 | 273 | 307 | 305,96 | 322 | 290 | 0,907 | -0,141 | -1,258 | 0,208292 | 0,240511 | 0,0121702 | 0,006355 | 0,008811 | 0,008811 | TRUE | 0,2233147 | No |
| orf19.3742 | 808 | 655 | 702 | 485 | 325 | 252 | 285 | 358 | 736 | 662 | 664 | 729 | 289 | 274 | 292 | 273 | 489,83 | 698 | 282 | 0,404 | -1,306 | -17,493 | 1,63E-68 | 1,00E-67 | 1,00E-08 | 0,004734 | 0,0029338 | 0,0029338 | TRUE | 0,018319 | No |
| orf19.3743 | 510 | 414 | 514 | 370 | 594 | 562 | 509 | 766 | 464 | 419 | 486 | 556 | 528 | 610 | 521 | 584 | 521,18 | 481 | 561 | 1,171 | 0,228 | 2,543 | 0,011002 | 0,014394 | 0,0074315 | 0,004572 | 0,0057493 | 0,0057493 | TRUE | 0,1115702 | No |
| orf19.3744 | 485 | 473 | 524 | 315 | 722 | 581 | 639 | 784 | 442 | 478 | 496 | 473 | 642 | 631 | 654 | 598 | 551,81 | 472 | 631 | 1,336 | 0,418 | 6,148 | 7,84E-10 | 1,47E-09 | 1,00E-08 | 0,004431 | 0,002565 | 0,002565 | TRUE | 0,0249678 | No |
| orf19.3746 | 3145 | 3152 | 3323 | 2078 | 1461 | 1302 | 1340 | 1804 | 2863 | 3188 | 3145 | 3121 | 1300 | 1414 | 1372 | 1376 | 2222,41 | 3079 | 1366 | 0,443 | -1,173 | -23,195 | 5,16E-119 | 5,23E-118 | 0,0014462 | 0,002631 | 0,0019399 | 0,0019399 | TRUE | 0,0279795 | No |
| orf19.3748 | 59 | 51 | 76 | 34 | 23 | 20 | 16 | 25 | 54 | 52 | 72 | 51 | 20 | 22 | 16 | 19 | 38,24 | 57 | 19 | 0,338 | -1,566 | -6,38 | 1,77E-10 | 3,40E-10 | 1,00E-08 | 0,036581 | 0,0245548 | 0,0245548 | TRUE | 0,2750113 | No |
| orf19.3749 | 15278 | 14480 | 15053 | 9386 | 6720 | 5463 | 5767 | 7584 | 13910 | 14645 | 14246 | 14099 | 5977 | 5931 | 5906 | 5785 | 10062,5 | 14225 | 5900 | 0,415 | -1,27 | -40,454 | 0 | 0 | 0,0002751 | 0,002168 | 0,0008292 | 0,0008292 | TRUE | 0,0048658 | No |
| orf19.3751 | 2854 | 2557 | 2802 | 1745 | 1427 | 1259 | 1344 | 1800 | 2598 | 2586 | 2652 | 2621 | 1269 | 1367 | 1376 | 1373 | 1980,42 | 2614 | 1346 | 0,515 | -0,958 | -21,892 | 3,10E-106 | 2,82E-105 | 4,47E-05 | 0,002704 | 0,0012852 | 0,0012852 | TRUE | 0,0182871 | No |
| orf19.3752 | 6264 | 5149 | 5566 | 3660 | 3921 | 3268 | 3301 | 4878 | 5703 | 5208 | 5268 | 5498 | 3488 | 3548 | 3381 | 3721 | 4476,72 | 5419 | 3534 | 0,653 | -0,616 | -13,35 | 1,18E-40 | 4,63E-40 | 0,0013768 | 0,002332 | 0,0018122 | 0,0018122 | TRUE | 0,0157877 | No |
| orf19.3753 | 9456 | 8391 | 9476 | 6216 | 5156 | 4839 | 4674 | 6677 | 8609 | 8487 | 8968 | 9337 | 4586 | 5254 | 4787 | 5093 | 6890,18 | 8850 | 4930 | 0,557 | -0,844 | -16,209 | 4,35E-59 | 2,35E-58 | 0,0025825 | 0,002229 | 0,0024473 | 0,0024473 | TRUE | 0,0270186 | No |
| orf19.3755 | 2450 | 2307 | 2254 | 1392 | 1458 | 1056 | 1147 | 1485 | 2231 | 2333 | 2133 | 2091 | 1297 | 1147 | 1175 | 1133 | 1692,37 | 2197 | 1188 | 0,54 | -0,888 | -15,239 | 1,94E-52 | 9,34E-52 | 0,0023284 | 0,002818 | 0,0026246 | 0,0026246 | TRUE | 0,0468307 | No |
| orf19.3756 | 13268 | 12851 | 13770 | 8438 | 10635 | 9572 | 9702 | 12486 | 12080 | 12997 | 13032 | 12675 | 9459 | 10393 | 9936 | 9524 | 11262,1 | 12696 | 9828 | 0,774 | -0,37 | -8,515 | 1,66E-17 | 3,95E-17 | 0,0014466 | 0,002154 | 0,0017205 | 0,0017205 | TRUE | 0,01818 | No |
| orf19.3757 | 1652 | 1673 | 1621 | 1262 | 3589 | 3009 | 2542 | 3571 | 1504 | 1692 | 1534 | 1896 | 3192 | 3267 | 2603 | 2724 | 2301,57 | 1656 | 2946 | 1,781 | 0,833 | 9,431 | 4,08E-21 | 1,06E-20 | 0,0115865 | 0,002611 | 0,0070095 | 0,0070095 | TRUE | 0,1102613 | No |
| orf19.3758 | 4932 | 4944 | 5002 | 3324 | 1888 | 1752 | 1709 | 2205 | 4490 | 5000 | 4734 | 4993 | 1679 | 1902 | 1750 | 1682 | 3278,94 | 4804 | 1753 | 0,365 | -1,455 | -26,424 | 7,19E-154 | 9,57E-153 | 0,0025379 | 0,00244 | 0,0025321 | 0,0025321 | TRUE | 0,0386108 | No |
| orf19.3759 | 6728 | 6236 | 6900 | 4342 | 3194 | 2899 | 2795 | 3881 | 6126 | 6307 | 6530 | 6522 | 2841 | 3148 | 2862 | 2960 | 4662,07 | 6371 | 2953 | 0,463 | -1,11 | -24,891 | 9,34E-137 | 1,10E-135 | 0,0012186 | 0,00232 | 0,0016652 | 0,0016652 | TRUE | 0,0232207 | No |
| orf19.376 | 1601 | 1269 | 1512 | 882 | 2885 | 2289 | 2730 | 3741 | 1458 | 1283 | 1431 | 1325 | 2566 | 2485 | 2796 | 2854 | 2024,73 | 1374 | 2675 | 1,945 | 0,96 | 15,38 | 2,22E-53 | 1,09E-52 | 0,0033041 | 0,002689 | 0,0031743 | 0,0031743 | TRUE | 0,0273475 | No |
| orf19.3760 | 111 | 95 | 88 | 52 | 64 | 51 | 61 | 100 | 101 | 96 | 83 | 78 | 57 | 55 | 62 | 76 | 76,2 | 90 | 62 | 0,701 | -0,513 | -3,024 | 0,002495 | 0,003417 | 0,0057412 | 0,019173 | 0,0143998 | 0,0143998 | TRUE | 0,2149853 | No |
| orf19.3761 | 3014 | 2889 | 3052 | 1760 | 2109 | 1688 | 1904 | 2485 | 2744 | 2922 | 2888 | 2644 | 1876 | 1833 | 1950 | 1896 | 2344,05 | 2800 | 1889 | 0,674 | -0,569 | -12,257 | 1,54E-34 | 5,46E-34 | 0,0008081 | 0,002601 | 0,0016328 | 0,0016328 | TRUE | 0,0156881 | No |
| orf19.3762 | 713 | 732 | 654 | 473 | 468 | 383 | 358 | 568 | 649 | 740 | 619 | 711 | 416 | 416 | 367 | 433 | 543,87 | 680 | 408 | 0,602 | -0,731 | -9,05 | 1,43E-19 | 3,58E-19 | 0,0041834 | 0,004466 | 0,0043326 | 0,0043326 | TRUE | 0,0524683 | No |
| orf19.3763 | 961 | 985 | 1041 | 598 | 835 | 633 | 687 | 890 | 875 | 996 | 985 | 898 | 743 | 687 | 704 | 679 | 820,89 | 938 | 703 | 0,748 | -0,418 | -6,483 | 9,00E-11 | 1,75E-10 | 0,0019065 | 0,003646 | 0,0027647 | 0,0027647 | TRUE | 0,0278248 | No |
| orf19.3764 | 656 | 780 | 790 | 422 | 363 | 320 | 356 | 469 | 597 | 789 | 748 | 634 | 323 | 347 | 365 | 358 | 520,05 | 692 | 348 | 0,502 | -0,993 | -10,364 | 3,62E-25 | 1,04E-24 | 0,0089253 | 0,004577 | 0,0066954 | 0,0066954 | TRUE | 0,107758 | No |
| orf19.3765 | 1131 | 1151 | 1179 | 682 | 2280 | 1946 | 2008 | 2508 | 1030 | 1164 | 1116 | 1024 | 2028 | 2113 | 2056 | 1913 | 1555,56 | 1084 | 2028 | 1,867 | 0,901 | 15,792 | 3,55E-56 | 1,83E-55 | 0,0018912 | 0,002886 | 0,0023989 | 0,0023989 | TRUE | 0,0296924 | No |
| orf19.3767 | 740 | 657 | 659 | 384 | 807 | 650 | 755 | 941 | 674 | 664 | 624 | 577 | 718 | 706 | 773 | 718 | 681,66 | 635 | 729 | 1,143 | 0,193 | 2,903 | 0,003701 | 0,005001 | 0,0013306 | 0,003975 | 0,0027586 | 0,0027586 | TRUE | 0,0375762 | No |
| orf19.3769 | 1209 | 1129 | 1135 | 715 | 2030 | 1783 | 1837 | 2403 | 1101 | 1142 | 1074 | 1074 | 1806 | 1936 | 1881 | 1833 | 1480,83 | 1098 | 1864 | 1,696 | 0,762 | 15,902 | 6,15E-57 | 3,19E-56 | 1,00E-08 | 0,002929 | 0,0014649 | 0,0014649 | TRUE | 0,0087833 | No |

| | | | | | | | | | | | | | | | | | | | | | | | | | | | | | | | |
|---|---|---|---|---|---|---|---|---|---|---|---|---|---|---|---|---|---|---|---|---|---|---|---|---|---|---|---|---|---|---|---|
| orf19.377 | 143 | 125 | 91 | 80 | 207 | 143 | 157 | 248 | 130 | 126 | 86 | 120 | 184 | 155 | 161 | 189 | 144,03 | 116 | 172 | 1,497 | 0,582 | 4,102 | 4,09E-05 | 6,27E-05 | 0,0133305 | 0,011209 | 0,0120245 | 0,0120245 | TRUE | 0,325626 | No |
| orf19.3770 | 7402 | 7204 | 7481 | 4199 | 3519 | 2762 | 2942 | 3889 | 6739 | 7286 | 7080 | 6308 | 3130 | 2999 | 3013 | 2967 | 4940,16 | 6853 | 3027 | 0,441 | -1,18 | -23,569 | 8,09E-123 | 8,47E-122 | 0,0019103 | 0,002304 | 0,0021732 | 0,0021732 | TRUE | 0,034052 | No |
| orf19.3773 | 2334 | 2196 | 2490 | 1560 | 1916 | 1646 | 1805 | 2384 | 2125 | 2221 | 2357 | 2343 | 1704 | 1787 | 1849 | 1819 | 2025,54 | 2262 | 1790 | 0,792 | -0,336 | -6,7 | 2,09E-11 | 4,16E-11 | 0,0013542 | 0,002689 | 0,0019219 | 0,0019219 | TRUE | 0,0205373 | No |
| orf19.3774 | 1724 | 1535 | 1621 | 981 | 1646 | 1305 | 1473 | 1989 | 1570 | 1552 | 1534 | 1474 | 1464 | 1417 | 1509 | 1517 | 1504,57 | 1532 | 1477 | 0,963 | -0,055 | -1,175 | 0,239832 | 0,274002 | 1,00E-08 | 0,002915 | 0,0014118 | 0,0014118 | TRUE | 0,0085408 | No |
| orf19.3774.1 | 93 | 87 | 108 | 68 | 65 | 59 | 47 | 81 | 85 | 88 | 102 | 102 | 58 | 64 | 48 | 62 | 76,1 | 94 | 58 | 0,619 | -0,692 | -4,212 | 2,53E-05 | 3,94E-05 | 1,00E-08 | 0,019395 | 0,0122551 | 0,0122551 | TRUE | 0,1028675 | No |
| orf19.3775 | 4353 | 3920 | 4215 | 2584 | 4386 | 3670 | 4185 | 5561 | 3963 | 3965 | 3989 | 3882 | 3901 | 3985 | 4286 | 4242 | 4026,54 | 3950 | 4104 | 1,039 | 0,055 | 1,299 | 0,193902 | 0,224668 | 0,0008036 | 0,002365 | 0,0014609 | 0,0014609 | TRUE | 0,0136567 | No |
| orf19.3777 | 293 | 503 | 488 | 193 | 258 | 283 | 276 | 283 | 267 | 509 | 462 | 290 | 229 | 307 | 283 | 216 | 320,32 | 382 | 259 | 0,676 | -0,564 | -2,195 | 0,028162 | 0,035664 | 0,0601859 | 0,006161 | 0,0302124 | 0,0601859 | TRUE | 0,1403666 | No |
| orf19.3778 | 8931 | 7877 | 8274 | 5299 | 8874 | 7351 | 7611 | 10677 | 8131 | 7967 | 7831 | 7960 | 7893 | 7981 | 7795 | 8145 | 7962,75 | 7972 | 7954 | 0,998 | -0,003 | -0,104 | 0,916994 | 0,928593 | 0,0001952 | 0,002203 | 0,0007949 | 0,0007949 | TRUE | 0,0032855 | No |
| orf19.3780 | 354 | 361 | 349 | 235 | 675 | 652 | 563 | 750 | 322 | 365 | 330 | 353 | 600 | 708 | 577 | 572 | 478,46 | 342 | 614 | 1,793 | 0,842 | 9,637 | 5,58E-22 | 1,48E-21 | 0,0052266 | 0,004798 | 0,0050031 | 0,0050031 | TRUE | 0,1197662 | No |
| orf19.3781 | 1433 | 1333 | 1449 | 978 | 1420 | 1307 | 1338 | 1812 | 1305 | 1348 | 1371 | 1469 | 1263 | 1419 | 1370 | 1382 | 1365,99 | 1373 | 1358 | 0,991 | -0,013 | -0,231 | 0,817032 | 0,841722 | 0,0016635 | 0,003004 | 0,0022945 | 0,0022945 | TRUE | 0,0272681 | No |
| orf19.3782 | 549 | 490 | 559 | 369 | 1134 | 940 | 994 | 1240 | 500 | 496 | 529 | 554 | 1009 | 1021 | 1018 | 946 | 758,98 | 520 | 998 | 1,926 | 0,945 | 15,22 | 2,59E-52 | 1,25E-51 | 0,0005151 | 0,003778 | 0,0022043 | 0,0022043 | TRUE | 0,0192807 | No |
| orf19.3782.2 | 1261 | 1009 | 1120 | 559 | 2215 | 1450 | 1814 | 2457 | 1148 | 1020 | 1060 | 840 | 1970 | 1574 | 1858 | 1874 | 1418,09 | 1017 | 1819 | 1,784 | 0,835 | 9,165 | 4,93E-20 | 1,26E-19 | 0,0115068 | 0,002969 | 0,0071908 | 0,0071908 | TRUE | 0,1546922 | No |
| orf19.3783 | 149 | 138 | 175 | 89 | 347 | 253 | 318 | 381 | 136 | 140 | 166 | 134 | 309 | 275 | 326 | 291 | 221,77 | 144 | 300 | 2,079 | 1,056 | 9,821 | 9,14E-23 | 2,47E-22 | 0,0023316 | 0,007994 | 0,0058528 | 0,0058528 | TRUE | 0,1168509 | No |
| orf19.3784 | 280 | 296 | 325 | 164 | 450 | 354 | 362 | 477 | 255 | 299 | 308 | 246 | 400 | 384 | 371 | 364 | 328,43 | 277 | 380 | 1,364 | 0,447 | 4,921 | 8,59E-07 | 1,43E-06 | 0,002949 | 0,006059 | 0,0047894 | 0,0047894 | TRUE | 0,0616197 | No |
| orf19.3785 | 1005 | 921 | 918 | 636 | 1132 | 910 | 977 | 1254 | 915 | 931 | 869 | 955 | 1007 | 988 | 1001 | 957 | 952,84 | 918 | 988 | 1,078 | 0,109 | 2,02 | 0,04335 | 0,054018 | 1,00E-08 | 0,003423 | 0,0017216 | 0,0017216 | TRUE | 0,0154803 | No |
| orf19.3787 | 1233 | 977 | 1036 | 727 | 2268 | 1907 | 2052 | 2768 | 1123 | 988 | 980 | 1092 | 2017 | 2070 | 2102 | 2111 | 1560,5 | 1046 | 2075 | 1,987 | 0,99 | 17,976 | 3,00E-72 | 1,93E-71 | 0,0016313 | 0,002884 | 0,0021755 | 0,0021755 | TRUE | 0,0330627 | No |
| orf19.3788 | 893 | 733 | 808 | 480 | 1213 | 924 | 1059 | 1335 | 813 | 741 | 765 | 721 | 1079 | 1003 | 1085 | 1018 | 903,14 | 760 | 1046 | 1,373 | 0,457 | 7,72 | 1,17E-14 | 2,56E-14 | 0,0008858 | 0,0035 | 0,0022171 | 0,0022171 | TRUE | 0,0266979 | No |
| orf19.3788.1 | 19120 | 15439 | 15834 | 11064 | 19269 | 14903 | 14823 | 21269 | 17408 | 15615 | 14985 | 16620 | 17139 | 16181 | 15181 | 16224 | 16169,1 | 16157 | 16181 | 1,002 | 0,002 | 0,042 | 0,966781 | 0,970893 | 0,00287 | 0,002119 | 0,0028586 | 0,0028586 | TRUE | 0,0335205 | No |
| orf19.3789 | 7622 | 6689 | 6565 | 4638 | 14636 | 11676 | 10254 | 15107 | 6940 | 6765 | 6213 | 6967 | 13018 | 12677 | 10501 | 11524 | 9325,65 | 6721 | 11930 | 1,775 | 0,828 | 12,203 | 3,01E-34 | 1,06E-33 | 0,0058263 | 0,002179 | 0,0043041 | 0,0043041 | TRUE | 0,0792876 | No |
| orf19.3791 | 565 | 463 | 522 | 327 | 837 | 659 | 765 | 991 | 514 | 468 | 494 | 491 | 744 | 715 | 783 | 756 | 620,91 | 492 | 750 | 1,524 | 0,608 | 9,504 | 2,01E-21 | 5,27E-21 | 1,00E-08 | 0,004165 | 0,0022172 | 0,0022172 | TRUE | 0,013195 | No |
| orf19.3792 | 4616 | 4119 | 4475 | 2872 | 4095 | 3337 | 3576 | 4694 | 4203 | 4166 | 4235 | 4314 | 3642 | 3623 | 3662 | 3581 | 3928,29 | 4230 | 3627 | 0,858 | -0,221 | -6,765 | 1,33E-11 | 2,67E-11 | 1,00E-08 | 0,002373 | 0,0007699 | 0,0007699 | TRUE | 0,0020084 | No |
| orf19.3793 | 1609 | 1591 | 1677 | 1057 | 1278 | 1092 | 1160 | 1442 | 1465 | 1609 | 1587 | 1588 | 1137 | 1186 | 1188 | 1100 | 1357,41 | 1562 | 1153 | 0,737 | -0,439 | -8,371 | 5,71E-17 | 1,34E-16 | 0,0009818 | 0,00301 | 0,0018957 | 0,0018957 | TRUE | 0,0222846 | No |
| orf19.3794 | 446 | 447 | 484 | 317 | 686 | 539 | 567 | 804 | 406 | 452 | 458 | 476 | 610 | 585 | 581 | 613 | 522,72 | 448 | 597 | 1,34 | 0,422 | 6,007 | 1,89E-09 | 3,51E-09 | 7,51E-06 | 0,004564 | 0,0027593 | 0,0027593 | TRUE | 0,048725 | No |
| orf19.3795 | 3281 | 2867 | 3182 | 2048 | 5214 | 4496 | 4732 | 6318 | 2987 | 2900 | 3011 | 3076 | 4638 | 4881 | 4846 | 4819 | 3894,93 | 2994 | 4796 | 1,603 | 0,681 | 18,339 | 4,06E-75 | 2,69E-74 | 0,0002729 | 0,002376 | 0,0010459 | 0,0010459 | TRUE | 0,006058 | No |
| orf19.3796 | 1726 | 1643 | 1716 | 1167 | 3314 | 2863 | 3021 | 4025 | 1571 | 1662 | 1624 | 1753 | 2948 | 3108 | 3094 | 3070 | 2353,82 | 1652 | 3055 | 1,853 | 0,89 | 19,565 | 3,08E-85 | 2,33E-84 | 0,0007383 | 0,002598 | 0,0015064 | 0,0015064 | TRUE | 0,0181385 | No |
| orf19.3797 | 1969 | 1877 | 1854 | 1155 | 1686 | 1329 | 1507 | 1832 | 1793 | 1898 | 1755 | 1735 | 1500 | 1443 | 1543 | 1397 | 1633,01 | 1795 | 1471 | 0,818 | -0,289 | -5,643 | 1,67E-08 | 2,97E-08 | 0,0011593 | 0,002846 | 0,0019095 | 0,0019095 | TRUE | 0,0180712 | No |
| orf19.3798 | 5657 | 4686 | 5077 | 2970 | 4876 | 3605 | 4167 | 5425 | 5150 | 4739 | 4805 | 4461 | 4337 | 3914 | 4268 | 4138 | 4476,63 | 4789 | 4164 | 0,869 | -0,203 | -3,858 | 0,000114 | 0,000171 | 0,0024475 | 0,002332 | 0,0024235 | 0,0024235 | TRUE | 0,0315104 | No |
| orf19.3799 | 3643 | 3440 | 3721 | 2526 | 5534 | 5199 | 4892 | 6699 | 3317 | 3479 | 3522 | 3794 | 4922 | 5645 | 5010 | 5110 | 4349,89 | 3528 | 5172 | 1,467 | 0,553 | 9,787 | 1,29E-22 | 3,46E-22 | 0,0032305 | 0,002341 | 0,0028211 | 0,0028211 | TRUE | 0,0457592 | No |
| orf19.3801 | 92 | 66 | 98 | 42 | 46 | 55 | 62 | 47 | 84 | 67 | 93 | 63 | 41 | 60 | 63 | 36 | 63,29 | 77 | 50 | 0,64 | -0,644 | -2,976 | 0,002918 | 0,003975 | 0,0374816 | 0,022909 | 0,028551 | 0,028551 | TRUE | 0,3143193 | No |
| orf19.3802 | 601 | 519 | 646 | 382 | 697 | 568 | 585 | 840 | 547 | 525 | 611 | 574 | 620 | 617 | 599 | 641 | 591,73 | 564 | 619 | 1,099 | 0,136 | 2,002 | 0,045313 | 0,056354 | 0,0008848 | 0,004269 | 0,0027133 | 0,0027133 | TRUE | 0,0408025 | No |
| orf19.3803 | 1660 | 1484 | 1654 | 1036 | 2657 | 2332 | 2375 | 3209 | 1511 | 1501 | 1565 | 1556 | 2363 | 2532 | 2432 | 2448 | 1988,65 | 1533 | 2444 | 1,594 | 0,673 | 15,622 | 5,12E-55 | 2,59E-54 | 1,00E-08 | 0,002701 | 0,0012388 | 0,0012388 | TRUE | 0,0068602 | No |
| orf19.3804 | 1827 | 1733 | 1759 | 988 | 1388 | 1103 | 1152 | 1552 | 1663 | 1753 | 1665 | 1484 | 1235 | 1198 | 1180 | 1184 | 1420,1 | 1641 | 1199 | 0,729 | -0,456 | -8,106 | 5,22E-16 | 1,19E-15 | 0,0016151 | 0,002967 | 0,0023263 | 0,0023263 | TRUE | 0,0454854 | No |
| orf19.3806 | 2227 | 1906 | 2095 | 1285 | 1465 | 1058 | 1256 | 1657 | 2028 | 1928 | 1983 | 1930 | 1303 | 1149 | 1286 | 1264 | 1608,79 | 1967 | 1250 | 0,636 | -0,653 | -12,75 | 3,13E-37 | 1,17E-36 | 0,0009765 | 0,002858 | 0,0018724 | 0,0018724 | TRUE | 0,0342486 | No |
| orf19.3809 | 3798 | 3160 | 3604 | 2272 | 3680 | 3059 | 3504 | 4865 | 3458 | 3196 | 3411 | 3413 | 3273 | 3321 | 3589 | 3711 | 3421,47 | 3370 | 3474 | 1,031 | 0,045 | 0,856 | 0,391769 | 0,432697 | 0,0020563 | 0,002423 | 0,0023008 | 0,0023008 | TRUE | 0,0264149 | No |
| orf19.3810 | 1757 | 1552 | 1686 | 1048 | 1558 | 1267 | 1339 | 1846 | 1600 | 1570 | 1596 | 1574 | 1386 | 1376 | 1371 | 1408 | 1485,01 | 1585 | 1385 | 0,874 | -0,194 | -4,548 | 5,41E-06 | 8,70E-06 | 1,00E-08 | 0,002927 | 0,0010735 | 0,0010735 | TRUE | 0,0015966 | No |
| orf19.3811 | 978 | 858 | 931 | 543 | 1383 | 1190 | 1364 | 1686 | 890 | 868 | 881 | 816 | 1230 | 1292 | 1397 | 1286 | 1082,52 | 864 | 1301 | 1,502 | 0,587 | 10,172 | 2,65E-24 | 7,48E-24 | 0,0012619 | 0,003257 | 0,0022171 | 0,0022171 | TRUE | 0,0288777 | No |
| orf19.3812 | 13382 | 11210 | 12271 | 8501 | 19166 | 16055 | 16957 | 24039 | 12184 | 11338 | 11613 | 12770 | 17048 | 17431 | 17366 | 18337 | 14760,9 | 11976 | 17546 | 1,466 | 0,552 | 12,139 | 6,52E-34 | 2,27E-33 | 0,001813 | 0,002127 | 0,001911 | 0,001911 | TRUE | 0,0239024 | No |
| orf19.3813 | 1342 | 1058 | 1230 | 782 | 1245 | 1088 | 1254 | 1626 | 1222 | 1070 | 1164 | 1175 | 1107 | 1181 | 1284 | 1240 | 1180,49 | 1158 | 1203 | 1,039 | 0,056 | 0,882 | 0,377836 | 0,418105 | 0,0028198 | 0,003156 | 0,0029788 | 0,0029788 | TRUE | 0,0352179 | No |
| orf19.3815 | 7932 | 6947 | 7849 | 5236 | 5338 | 4581 | 5005 | 6672 | 7222 | 7026 | 7428 | 7865 | 4748 | 4974 | 5126 | 5089 | 6184,81 | 7385 | 4984 | 0,675 | -0,567 | -12,409 | 2,34E-35 | 8,41E-35 | 0,0015454 | 0,002251 | 0,0018349 | 0,0018349 | TRUE | 0,0227036 | No |
| orf19.3817 | 717 | 568 | 652 | 432 | 886 | 721 | 726 | 1005 | 653 | 574 | 617 | 649 | 788 | 783 | 744 | 767 | 696,78 | 623 | 770 | 1,237 | 0,307 | 4,905 | 9,32E-07 | 1,55E-06 | 0,0003073 | 0,003933 | 0,002305 | 0,002305 | TRUE | 0,0331132 | No |
| orf19.3818 | 295 | 245 | 306 | 191 | 611 | 500 | 502 | 739 | 269 | 248 | 290 | 287 | 543 | 543 | 514 | 564 | 407,13 | 274 | 541 | 1,987 | 0,991 | 12,535 | 4,80E-36 | 1,75E-35 | 1,00E-08 | 0,005282 | 0,0031822 | 0,0031822 | TRUE | 0,044135 | No |
| orf19.3819 | 98 | 77 | 93 | 61 | 174 | 130 | 156 | 219 | 89 | 78 | 88 | 92 | 155 | 141 | 160 | 167 | 121,18 | 87 | 156 | 1,806 | 0,853 | 6,498 | 8,16E-11 | 1,59E-10 | 1,00E-08 | 0,012938 | 0,0074129 | 0,0074129 | TRUE | 0,0445656 | No |
| orf19.382 | 10247 | 7690 | 8246 | 6829 | 14622 | 11544 | 11585 | 18690 | 9329 | 7778 | 7804 | 10258 | 13006 | 12534 | 11865 | 14257 | 10853,8 | 8792 | 12916 | 1,469 | 0,555 | 6,369 | 1,91E-10 | 3,66E-10 | 0,0125084 | 0,002159 | 0,0072075 | 0,0072075 | TRUE | 0,1528369 | No |
| orf19.3820 | 43 | 34 | 38 | 23 | 71 | 45 | 48 | 93 | 39 | 34 | 36 | 35 | 63 | 49 | 49 | 71 | 47,02 | 36 | 58 | 1,62 | 0,696 | 3,261 | 0,00111 | 0,001564 | 1,00E-08 | 0,030131 | 0,0211178 | 0,0211178 | TRUE | 0,2150208 | No |
| orf19.3821 | 550 | 476 | 496 | 300 | 1282 | 1036 | 1232 | 1551 | 501 | 481 | 469 | 451 | 1140 | 1125 | 1262 | 1183 | 826,52 | 476 | 1178 | 2,468 | 1,304 | 20,373 | 2,89E-92 | 2,31E-91 | 0,0011754 | 0,003635 | 0,0024152 | 0,0024152 | TRUE | 0,0268079 | No |
| orf19.3822 | 17957 | 14290 | 15634 | 10440 | 17018 | 14026 | 14246 | 20664 | 16349 | 14453 | 14796 | 15683 | 15137 | 15228 | 14590 | 15763 | 15249,8 | 15320 | 15180 | 0,991 | -0,013 | -0,28 | 0,779676 | 0,809517 | 0,0017447 | 0,002124 | 0,0020506 | 0,0020506 | TRUE | 0,0252988 | No |
| orf19.3823 | 3874 | 3202 | 3586 | 2208 | 4647 | 3713 | 4372 | 5861 | 3527 | 3238 | 3394 | 3317 | 4133 | 4031 | 4477 | 4471 | 3823,64 | 3369 | 4278 | 1,27 | 0,345 | 6,955 | 3,53E-12 | 7,20E-12 | 0,0017583 | 0,002382 | 0,002087 | 0,002087 | TRUE | 0,0181093 | No |
| orf19.3825 | 1059 | 948 | 1043 | 692 | 2202 | 1816 | 1798 | 2522 | 964 | 959 | 987 | 1039 | 1959 | 1972 | 1841 | 1924 | 1455,63 | 987 | 1924 | 1,953 | 0,966 | 19,655 | 5,22E-86 | 3,98E-85 | 0,0002567 | 0,002945 | 0,0015329 | 0,0015329 | TRUE | 0,0128128 | No |
| orf19.3826 | 5473 | 4596 | 4910 | 3452 | 4883 | 4277 | 4365 | 6264 | 4983 | 4648 | 4647 | 5185 | 4343 | 4644 | 4470 | 4778 | 4712,38 | 4866 | 4559 | 0,938 | -0,093 | -1,853 | 0,063825 | 0,078277 | 0,0020819 | 0,002317 | 0,0021988 | 0,0021988 | TRUE | 0,0229518 | No |
| orf19.3827 | 2671 | 2191 | 2503 | 1599 | 3093 | 2556 | 2883 | 3673 | 2432 | 2216 | 2369 | 2402 | 2751 | 2775 | 2953 | 2802 | 2587,4 | 2355 | 2820 | 1,198 | 0,26 | 5,708 | 1,15E-08 | 2,06E-08 | 0,0008279 | 0,002548 | 0,0016038 | 0,0016038 | TRUE | 0,0193675 | No |
| orf19.3828 | 10 | 12 | 12 | 7 | 36 | 22 | 26 | 30 | 9 | 12 | 11 | 11 | 32 | 24 | 27 | 23 | 18,57 | 11 | 26 | 2,446 | 1,291 | 3,835 | 0,000125 | 0,000187 | 1,00E-08 | 0,073187 | 0,0418513 | 0,0418513 | TRUE | 0,1448789 | No |
| orf19.3829 | 3002 | 2551 | 2704 | 1736 | 6031 | 5537 | 5234 | 7614 | 2733 | 2580 | 2559 | 2608 | 5364 | 6012 | 5360 | 5808 | 4128,05 | 2620 | 5636 | 2,151 | 1,105 | 22,039 | 1,22E-107 | 1,12E-106 | 0,0017524 | 0,002357 | 0,0021243 | 0,0021243 | TRUE | 0,024131 | No |
| orf19.3831 | 7764 | 6118 | 6643 | 4026 | 7272 | 5509 | 6576 | 8681 | 7069 | 6188 | 6287 | 6048 | 6468 | 5981 | 6735 | 6622 | 6424,65 | 6398 | 6452 | 1,008 | 0,012 | 0,201 | 0,840571 | 0,862905 | 0,0031839 | 0,002243 | 0,0030581 | 0,0030581 | TRUE | 0,0612206 | No |
| orf19.3833 | 4956 | 4323 | 4882 | 3053 | 4320 | 3604 | 4091 | 5277 | 4512 | 4372 | 4620 | 4586 | 3843 | 3913 | 4190 | 4025 | 4257,68 | 4522 | 3993 | 0,883 | -0,18 | -4,414 | 1,02E-05 | 1,61E-05 | 0,0007007 | 0,002347 | 0,0013552 | 0,0013552 | TRUE | 0,0131894 | No |
| orf19.3836 | 2599 | 2209 | 2409 | 1592 | 2867 | 2355 | 2602 | 3389 | 2366 | 2234 | 2280 | 2391 | 2550 | 2557 | 2665 | 2585 | 2453,59 | 2318 | 2589 | 1,118 | 0,161 | 3,924 | 8,73E-05 | 0,000131 | 0,0001639 | 0,002575 | 0,0011993 | 0,0011993 | TRUE | 0,0070613 | No |
| orf19.3838 | 14581 | 12035 | 12630 | 8657 | 14925 | 11217 | 11190 | 16575 | 13275 | 12172 | 11953 | 13004 | 13275 | 12179 | 11460 | 12644 | 12495,3 | 12601 | 12390 | 0,983 | -0,024 | -0,447 | 0,654977 | 0,691992 | 0,0027263 | 0,002143 | 0,0027275 | 0,0027275 | TRUE | 0,0311602 | No |
| orf19.3839 | 2254 | 2082 | 2253 | 1386 | 6107 | 4985 | 5481 | 7493 | 2052 | 2106 | 2132 | 2082 | 5432 | 5412 | 5613 | 5716 | 3818,18 | 2093 | 5543 | 2,649 | 1,405 | 36,771 | 5,38E-296 | 1,37E-294 | 0,0003503 | 0,002383 | 0,0010618 | 0,0010618 | TRUE | 0,0055006 | No |
| orf19.384 | 1991 | 1665 | 1842 | 1240 | 2183 | 1902 | 2111 | 2909 | 1813 | 1684 | 1743 | 1863 | 1942 | 2065 | 2162 | 2219 | 1936,29 | 1776 | 2097 | 1,182 | 0,242 | 4,357 | 1,32E-05 | 2,08E-05 | 0,0019561 | 0,002719 | 0,0024298 | 0,0024298 | TRUE | 0,0307316 | No |
| orf19.3840 | 1750 | 1486 | 1655 | 1033 | 2513 | 2078 | 2223 | 2774 | 1593 | 1503 | 1566 | 1552 | 2235 | 2256 | 2277 | 2116 | 1887,29 | 1554 | 2221 | 1,428 | 0,514 | 11,383 | 5,09E-30 | 1,64E-29 | 0,0004728 | 0,002737 | 0,0014025 | 0,0014025 | TRUE | 0,012835 | No |
| orf19.3841 | 7339 | 6369 | 7080 | 4522 | 2377 | 1973 | 2066 | 2732 | 6682 | 6442 | 6701 | 6793 | 2114 | 2142 | 2116 | 2084 | 4384,12 | 6654 | 2114 | 0,318 | -1,654 | -46,416 | 0 | 0 | 1,00E-08 | 0,002338 | 0,000918 | 0,000918 | TRUE | 0,005642 | No |
| orf19.3843 | 1542 | 1382 | 1494 | 843 | 2177 | 1658 | 1817 | 2394 | 1404 | 1398 | 1414 | 1266 | 1936 | 1800 | 1861 | 1826 | 1613,18 | 1370 | 1856 | 1,351 | 0,434 | 8,586 | 9,04E-18 | 2,16E-17 | 0,0009577 | 0,002856 | 0,0018083 | 0,0018083 | TRUE | 0,0277779 | No |
| orf19.3844 | 1038 | 968 | 977 | 651 | 614 | 493 | 542 | 721 | 945 | 979 | 925 | 978 | 546 | 535 | 555 | 550 | 751,64 | 957 | 546 | 0,572 | -0,805 | -14,119 | 2,89E-45 | 1,24E-44 | 1,00E-08 | 0,003795 | 0,001711 | 0,001711 | TRUE | 0,0061256 | No |
| orf19.3845 | 2399 | 1990 | 2398 | 1534 | 1232 | 929 | 1035 | 1582 | 2184 | 2013 | 2269 | 2304 | 1096 | 1009 | 1060 | 1207 | 1642,73 | 2192 | 1093 | 0,5 | -1,001 | -15,275 | 1,12E-52 | 5,42E-52 | 0,0038011 | 0,002841 | 0,0034525 | 0,0034525 | TRUE | 0,0609798 | No |
| orf19.3846 | 1735 | 1571 | 1630 | 1012 | 3409 | 2928 | 3304 | 4408 | 1580 | 1589 | 1543 | 1520 | 3032 | 3179 | 3384 | 3362 | 2398,59 | 1558 | 3239 | 2,078 | 1,055 | 21,431 | 6,80E-102 | 6,03E-101 | 0,0013405 | 0,002588 | 0,0018386 | 0,0018386 | TRUE | 0,0228947 | No |
| orf19.3848 | 573 | 517 | 528 | 309 | 791 | 616 | 711 | 935 | 522 | 523 | 500 | 464 | 704 | 669 | 728 | 713 | 602,78 | 502 | 704 | 1,396 | 0,482 | 7,381 | 1,57E-13 | 3,35E-13 | 1,00E-08 | 0,004229 | 0,002366 | 0,002366 | TRUE | 0,0245595 | No |
| orf19.385 | 7046 | 6168 | 6720 | 4617 | 11077 | 9571 | 9827 | 13360 | 6415 | 6238 | 6360 | 6935 | 9853 | 10391 | 10064 | 10191 | 8306 | 6487 | 10125 | 1,562 | 0,643 | 15,557 | 1,42E-54 | 7,12E-54 | 0,0012066 | 0,002196 | 0,0015129 | 0,0015129 | TRUE | 0,0254273 | No |
| orf19.3851 | 735 | 627 | 702 | 460 | 1419 | 1128 | 1314 | 1661 | 669 | 634 | 664 | 691 | 1262 | 1225 | 1346 | 1267 | 969,78 | 664 | 1275 | 1,921 | 0,942 | 16,71 | 1,11E-62 | 6,29E-62 | 1,00E-08 | 0,003399 | 0,0018805 | 0,0018805 | TRUE | 0,0161216 | No |

| | | | | | | | | | | | | | | | | | | | | | | | | | | | | | | | |
|---|---|---|---|---|---|---|---|---|---|---|---|---|---|---|---|---|---|---|---|---|---|---|---|---|---|---|---|---|---|---|---|
| orf19.3852 | 181 | 186 | 200 | 122 | 153 | 146 | 167 | 223 | 165 | 188 | 189 | 183 | 136 | 159 | 171 | 170 | 170,15 | 181 | 159 | 0,877 | -0,189 | -1,638 | 0,101443 | 0,121881 | 1,00E-08 | 0,009801 | 0,0068491 | 0,0068491 | TRUE | 0,1044849 | No |
| orf19.3854 | 8583 | 7519 | 8436 | 5254 | 7997 | 6891 | 7617 | 10040 | 7814 | 7605 | 7984 | 7892 | 7113 | 7482 | 7801 | 7659 | 7668,69 | 7824 | 7514 | 0,96 | -0,058 | -1,489 | 0,136484 | 0,161156 | 0,0008218 | 0,002209 | 0,0013437 | 0,0013437 | TRUE | 0,0158308 | No |
| orf19.3856 | 654 | 646 | 647 | 338 | 1154 | 876 | 978 | 1197 | 595 | 653 | 612 | 508 | 1026 | 951 | 1002 | 913 | 782,63 | 592 | 973 | 1,635 | 0,709 | 9,42 | 4,52E-21 | 1,18E-20 | 0,0044671 | 0,003725 | 0,004056 | 0,004056 | TRUE | 0,0934182 | No |
| orf19.3858 | 2774 | 2401 | 2560 | 1454 | 2286 | 1568 | 2029 | 2506 | 2526 | 2428 | 2423 | 2184 | 2033 | 1702 | 2078 | 1912 | 2160,77 | 2390 | 1931 | 0,807 | -0,309 | -4,533 | 5,82E-06 | 9,33E-06 | 0,0050073 | 0,002648 | 0,0039863 | 0,0039863 | TRUE | 0,0752102 | No |
| orf19.3859 | 1562 | 1419 | 1511 | 1010 | 2242 | 1888 | 1861 | 2727 | 1422 | 1435 | 1430 | 1517 | 1994 | 2050 | 1906 | 2080 | 1729,33 | 1451 | 2008 | 1,386 | 0,471 | 9,915 | 3,59E-23 | 9,79E-23 | 0,000607 | 0,002801 | 0,0015676 | 0,0015676 | TRUE | 0,0138097 | No |
| orf19.386 | 1943 | 1654 | 1725 | 1236 | 3303 | 2748 | 2718 | 3826 | 1769 | 1673 | 1633 | 1857 | 2938 | 2984 | 2784 | 2919 | 2319,33 | 1733 | 2906 | 1,68 | 0,749 | 14,983 | 9,50E-51 | 4,45E-50 | 0,0015055 | 0,002607 | 0,001926 | 0,001926 | TRUE | 0,0253758 | No |
| orf19.3861 | 7648 | 6665 | 7311 | 4313 | 2043 | 1680 | 1918 | 2503 | 6963 | 6741 | 6919 | 6479 | 1817 | 1824 | 1964 | 1909 | 4327,11 | 6776 | 1878 | 0,277 | -1,851 | -42,989 | 0 | 0 | 0,0007835 | 0,002342 | 0,0014526 | 0,0014526 | TRUE | 0,0108777 | No |
| orf19.3862 | 3247 | 2623 | 2902 | 1923 | 2322 | 1831 | 2130 | 2860 | 2956 | 2653 | 2746 | 2889 | 2065 | 1988 | 2181 | 2182 | 2457,57 | 2811 | 2104 | 0,749 | -0,417 | -8,09 | 5,96E-16 | 1,35E-15 | 0,0016488 | 0,002575 | 0,0021343 | 0,0021343 | TRUE | 0,0175513 | No |
| orf19.3863 | 97 | 107 | 119 | 68 | 232 | 200 | 213 | 253 | 88 | 108 | 113 | 102 | 206 | 217 | 218 | 193 | 155,74 | 103 | 208 | 2,026 | 1,019 | 8,405 | 4,27E-17 | 1,00E-16 | 1,00E-08 | 0,010519 | 0,0066867 | 0,0066867 | TRUE | 0,1048048 | No |
| orf19.3865 | 2295 | 2101 | 2282 | 1473 | 2164 | 1833 | 2045 | 2754 | 2090 | 2125 | 2160 | 2213 | 1925 | 1990 | 2094 | 2101 | 2087,11 | 2147 | 2028 | 0,946 | -0,081 | -1,756 | 0,079043 | 0,095908 | 0,000737 | 0,00267 | 0,0015468 | 0,0015468 | TRUE | 0,0145094 | No |
| orf19.3867 | 8785 | 8491 | 9122 | 5490 | 11709 | 9285 | 9565 | 12412 | 7998 | 8588 | 8633 | 8247 | 10415 | 10081 | 9796 | 9468 | 9153,19 | 8366 | 9940 | 1,188 | 0,248 | 5,765 | 8,18E-09 | 1,48E-08 | 0,0013205 | 0,002181 | 0,001671 | 0,001671 | TRUE | 0,01277 | No |
| orf19.3868 | 1 | 1 | 1 | 1 | 4 | 0 | 3 | 2 | 1 | 1 | 1 | 2 | 4 | 0 | 3 | 2 | 1,57 | 1 | 2 | 1,938 | 0,955 | 0,833 | 0,404639 | 0,445599 | 1,00E-08 | 0,845701 | 0,5352536 | 0,5352536 | TRUE | 0,2755193 | No |
| orf19.3869 | 16240 | 15531 | 16956 | 11323 | 17783 | 16485 | 16383 | 21789 | 14786 | 15708 | 16047 | 17009 | 15817 | 17898 | 16778 | 16621 | 16333,1 | 15888 | 16778 | 1,056 | 0,079 | 1,505 | 0,132365 | 0,156699 | 0,0028919 | 0,002118 | 0,0025836 | 0,0025836 | TRUE | 0,0273726 | No |
| orf19.387 | 2184 | 1978 | 2263 | 1419 | 3643 | 3274 | 3230 | 4393 | 1988 | 2001 | 2142 | 2132 | 3240 | 3555 | 3308 | 3351 | 2714,52 | 2066 | 3364 | 1,629 | 0,704 | 14,907 | 2,99E-50 | 1,38E-49 | 0,0010829 | 0,002524 | 0,0017428 | 0,0017428 | TRUE | 0,0173996 | No |
| orf19.3870 | 6592 | 6038 | 6279 | 4048 | 7482 | 6109 | 6380 | 8726 | 6002 | 6107 | 5943 | 6081 | 6655 | 6633 | 6534 | 6656 | 6326,2 | 6033 | 6620 | 1,098 | 0,134 | 4,587 | 4,49E-06 | 7,25E-06 | 1,00E-08 | 0,002246 | 0,0006639 | 0,0006639 | TRUE | 0,0012579 | No |
| orf19.3871 | 31 | 29 | 43 | 21 | 39 | 37 | 49 | 55 | 28 | 29 | 41 | 32 | 35 | 40 | 50 | 42 | 37,1 | 32 | 42 | 1,281 | 0,357 | 1,526 | 0,127099 | 0,150913 | 1,00E-08 | 0,037644 | 0,025115 | 0,025115 | TRUE | 0,2277085 | No |
| orf19.3872 | 211 | 184 | 217 | 125 | 181 | 125 | 158 | 187 | 192 | 186 | 205 | 188 | 161 | 136 | 162 | 143 | 171,56 | 193 | 150 | 0,778 | -0,362 | -3,292 | 0,000993 | 0,001402 | 1,00E-08 | 0,009737 | 0,0057491 | 0,0057491 | TRUE | 0,0387654 | No |
| orf19.3873 | 1406 | 1184 | 1287 | 822 | 846 | 654 | 697 | 977 | 1280 | 1197 | 1218 | 1235 | 752 | 710 | 714 | 745 | 981,5 | 1232 | 730 | 0,593 | -0,753 | -14,163 | 1,55E-45 | 6,65E-45 | 1,00E-08 | 0,003383 | 0,0016445 | 0,0016445 | TRUE | 0,0087878 | No |
| orf19.3874 | 4173 | 3962 | 4226 | 2632 | 3933 | 3443 | 3431 | 4666 | 3799 | 4007 | 4000 | 3954 | 3498 | 3738 | 3514 | 3559 | 3758,64 | 3940 | 3577 | 0,908 | -0,139 | -3,562 | 0,000368 | 0,000535 | 0,0004203 | 0,002388 | 0,001205 | 0,001205 | TRUE | 0,0107996 | No |
| orf19.3876 | 79 | 63 | 91 | 48 | 255 | 226 | 240 | 312 | 72 | 64 | 86 | 72 | 227 | 245 | 246 | 238 | 156,23 | 74 | 239 | 3,241 | 1,696 | 13,366 | 9,57E-41 | 3,77E-40 | 1,00E-08 | 0,010492 | 0,0063206 | 0,0063206 | TRUE | 0,1370624 | No |
| orf19.3877 | 486 | 513 | 573 | 324 | 784 | 654 | 656 | 903 | 442 | 519 | 542 | 487 | 697 | 710 | 672 | 689 | 594,8 | 498 | 692 | 1,39 | 0,475 | 6,561 | 5,34E-11 | 1,05E-10 | 0,0020769 | 0,004258 | 0,0032861 | 0,0032861 | TRUE | 0,0727485 | No |
| orf19.3878 | 1015 | 962 | 1011 | 575 | 1149 | 1033 | 1038 | 1379 | 924 | 973 | 957 | 864 | 1022 | 1122 | 1063 | 1052 | 997,02 | 930 | 1065 | 1,141 | 0,191 | 3,347 | 0,000818 | 0,001162 | 0,0007727 | 0,003362 | 0,0021066 | 0,0021066 | TRUE | 0,0231073 | No |
| orf19.3879 | 69 | 69 | 72 | 64 | 333 | 298 | 324 | 388 | 63 | 70 | 68 | 96 | 296 | 324 | 332 | 296 | 193,05 | 74 | 312 | 4,291 | 2,101 | 16,441 | 9,70E-61 | 5,36E-60 | 0,0019503 | 0,00888 | 0,006877 | 0,006877 | TRUE | 0,2618649 | No |
| orf19.388 | 2042 | 1646 | 1855 | 1246 | 877 | 754 | 789 | 1043 | 1859 | 1665 | 1756 | 1872 | 780 | 819 | 808 | 796 | 1294,19 | 1788 | 801 | 0,448 | -1,159 | -20,985 | 8,98E-98 | 7,67E-97 | 0,0012152 | 0,003058 | 0,0020421 | 0,0020421 | TRUE | 0,0259035 | No |
| orf19.3881 | 1512 | 1140 | 1272 | 845 | 1283 | 1018 | 1169 | 1652 | 1377 | 1153 | 1204 | 1269 | 1141 | 1105 | 1197 | 1260 | 1213,32 | 1251 | 1176 | 0,941 | -0,088 | -1,302 | 0,192918 | 0,22361 | 0,0035344 | 0,003126 | 0,0035239 | 0,0035239 | TRUE | 0,0582833 | No |
| orf19.3884 | 338 | 270 | 299 | 202 | 1089 | 853 | 1041 | 1407 | 308 | 273 | 283 | 303 | 969 | 926 | 1066 | 1073 | 650,17 | 292 | 1008 | 3,465 | 1,793 | 22,927 | 2,49E-116 | 2,48E-115 | 0,0030603 | 0,004069 | 0,0035766 | 0,0035766 | TRUE | 0,0349624 | No |
| orf19.3885 | 173 | 182 | 156 | 118 | 178 | 140 | 134 | 195 | 158 | 184 | 148 | 177 | 158 | 152 | 137 | 149 | 157,85 | 167 | 149 | 0,9 | -0,153 | -1,302 | 0,19283 | 0,223588 | 1,00E-08 | 0,010406 | 0,00684 | 0,00684 | TRUE | 0,0645648 | No |
| orf19.3886 | 2852 | 2453 | 2654 | 1779 | 1915 | 1570 | 1666 | 2069 | 2597 | 2481 | 2512 | 2672 | 1703 | 1705 | 1706 | 1578 | 2119,26 | 2566 | 1673 | 0,652 | -0,617 | -13,22 | 6,71E-40 | 2,60E-39 | 0,0008172 | 0,00266 | 0,0016 | 0,0016 | TRUE | 0,0185711 | No |
| orf19.3887 | 2821 | 2414 | 2691 | 1398 | 2345 | 1607 | 1948 | 2478 | 2568 | 2442 | 2547 | 2100 | 2086 | 1745 | 1995 | 1890 | 2171,56 | 2414 | 1929 | 0,798 | -0,325 | -4,453 | 8,48E-06 | 1,35E-05 | 0,0062893 | 0,002645 | 0,0046639 | 0,0046639 | TRUE | 0,089454 | No |
| orf19.3888 | 4151 | 3525 | 3802 | 2528 | 1025 | 804 | 859 | 1229 | 3779 | 3565 | 3598 | 3797 | 912 | 873 | 880 | 937 | 2292,75 | 3685 | 900 | 0,245 | -2,03 | -43,043 | 0 | 0 | 0,0006197 | 0,002613 | 0,0014724 | 0,0014724 | TRUE | 0,0096811 | No |
| orf19.3888.2 | 1 | 1 | 3 | 0 | 4 | 2 | 0 | 5 | 1 | 1 | 3 | 0 | 4 | 2 | 0 | 4 | 1,79 | 1 | 2 | 1,926 | 0,945 | 0,848 | 0,396228 | 0,437017 | 1,00E-08 | 0,740828 | 0,5828991 | 0,5828991 | TRUE | 0,4856627 | No |
| orf19.389 | 168 | 147 | 136 | 104 | 283 | 178 | 204 | 267 | 153 | 149 | 129 | 156 | 252 | 193 | 209 | 204 | 180,52 | 147 | 214 | 1,469 | 0,555 | 4,626 | 3,73E-06 | 6,05E-06 | 0,0061432 | 0,009355 | 0,0079982 | 0,0079982 | TRUE | 0,1540519 | No |
| orf19.3893 | 156 | 129 | 162 | 112 | 74 | 62 | 64 | 108 | 142 | 130 | 153 | 168 | 66 | 67 | 66 | 82 | 109,39 | 148 | 70 | 0,479 | -1,062 | -7,303 | 2,82E-13 | 5,95E-13 | 1,00E-08 | 0,014113 | 0,0101154 | 0,0101154 | TRUE | 0,1435336 | No |
| orf19.3894 | 3210 | 2833 | 2999 | 1850 | 2414 | 1900 | 2152 | 2837 | 2923 | 2865 | 2838 | 2779 | 2147 | 2063 | 2204 | 2164 | 2497,9 | 2851 | 2144 | 0,752 | -0,412 | -10,283 | 8,39E-25 | 2,39E-24 | 0,0001032 | 0,002566 | 0,0011322 | 0,0011322 | TRUE | 0,0075345 | No |
| orf19.3895 | 825 | 659 | 728 | 566 | 1313 | 1304 | 1213 | 1747 | 751 | 667 | 689 | 850 | 1168 | 1416 | 1242 | 1333 | 1014,42 | 739 | 1290 | 1,75 | 0,808 | 9,717 | 2,55E-22 | 6,81E-22 | 0,0082553 | 0,003339 | 0,0055374 | 0,0055374 | TRUE | 0,11061 | No |
| orf19.3897 | 24 | 15 | 31 | 10 | 30 | 24 | 23 | 33 | 22 | 15 | 29 | 15 | 27 | 26 | 24 | 25 | 22,86 | 20 | 26 | 1,224 | 0,291 | 0,98 | 0,326976 | 0,366597 | 1,00E-08 | 0,059833 | 0,0407538 | 0,0407538 | TRUE | 0,4982228 | No |
| orf19.3898 | 298 | 267 | 258 | 158 | 488 | 370 | 385 | 566 | 271 | 270 | 244 | 237 | 434 | 402 | 394 | 432 | 335,59 | 256 | 416 | 1,621 | 0,697 | 8,257 | 1,50E-16 | 3,46E-16 | 1,00E-08 | 0,005973 | 0,0036434 | 0,0036434 | TRUE | 0,0203478 | No |
| orf19.3899 | 324 | 289 | 287 | 164 | 448 | 303 | 352 | 475 | 295 | 292 | 272 | 246 | 398 | 329 | 360 | 362 | 319,44 | 276 | 362 | 1,307 | 0,387 | 4,246 | 2,18E-05 | 3,39E-05 | 0,0027102 | 0,006172 | 0,0047485 | 0,0047485 | TRUE | 0,0508689 | No |
| orf19.390 | 1875 | 1591 | 1858 | 1088 | 2890 | 2360 | 2606 | 3547 | 1707 | 1609 | 1758 | 1634 | 2571 | 2562 | 2669 | 2706 | 2152,05 | 1677 | 2627 | 1,565 | 0,646 | 14,106 | 3,48E-45 | 1,49E-44 | 0,0006663 | 0,002651 | 0,0015163 | 0,0015163 | TRUE | 0,0129841 | No |
| orf19.3900 | 9397 | 7973 | 9040 | 5511 | 4251 | 3493 | 3964 | 5298 | 8556 | 8064 | 8556 | 8278 | 3781 | 3792 | 4060 | 4041 | 6140,99 | 8364 | 3918 | 0,469 | -1,094 | -26,639 | 2,40E-156 | 3,27E-155 | 0,0009145 | 0,002252 | 0,0014349 | 0,0014349 | TRUE | 0,007238 | No |
| orf19.3901 | 897 | 741 | 822 | 569 | 733 | 603 | 634 | 849 | 817 | 749 | 778 | 855 | 652 | 655 | 649 | 648 | 725,3 | 800 | 651 | 0,816 | -0,293 | -4,846 | 1,26E-06 | 2,08E-06 | 1,00E-08 | 0,003858 | 0,0021319 | 0,0021319 | TRUE | 0,0216301 | No |
| orf19.3902 | 36 | 48 | 44 | 38 | 17 | 18 | 7 | 20 | 33 | 49 | 42 | 57 | 15 | 20 | 7 | 15 | 29,64 | 45 | 14 | 0,322 | -1,637 | -5,414 | 6,18E-08 | 1,08E-07 | 0,0355018 | 0,046602 | 0,0432029 | 0,0432029 | TRUE | 0,2472006 | No |
| orf19.3903 | 1 | 1 | 1 | 0 | 0 | 0 | 0 | 2 | 1 | 1 | 1 | 0 | 0 | 0 | 0 | 2 | 0,55 | 1 | 0 | 0,587 | -0,768 | -0,408 | 0,683503 | 0,719031 | 1,00E-08 | 2,407198 | 1,7322424 | 1,7322424 | TRUE | 0,8316477 | No |
| orf19.3904 | 10 | 19 | 20 | 6 | 0 | 3 | 1 | 2 | 9 | 19 | 19 | 9 | 0 | 3 | 1 | 2 | 7,76 | 14 | 2 | 0,098 | -3,349 | -4,545 | 5,48E-06 | 8,81E-06 | 0,103643 | 0,172299 | 0,1519162 | 0,1519162 | TRUE | 0,2272579 | No |
| orf19.3905 | 2 | 7 | 8 | 2 | 3 | 4 | 2 | 4 | 2 | 7 | 8 | 3 | 3 | 4 | 2 | 3 | 3,95 | 5 | 3 | 0,609 | -0,715 | -0,997 | 0,31881 | 0,358009 | 1,00E-08 | 0,336604 | 0,2310673 | 0,2310673 | TRUE | 0,1795777 | No |
| orf19.3906 | 5 | 9 | 15 | 9 | 5 | 3 | 11 | 7 | 5 | 9 | 14 | 14 | 4 | 3 | 11 | 5 | 8,21 | 10 | 6 | 0,594 | -0,752 | -1,391 | 0,164292 | 0,192423 | 0,130396 | 0,162938 | 0,1512025 | 0,1512025 | TRUE | 0,6670317 | No |
| orf19.3907 | 1 | 0 | 0 | 0 | 1 | 1 | 1 | 0 | 1 | 0 | 0 | 0 | 1 | 1 | 1 | 0 | 0,49 | 0 | 1 | 2,243 | 1,165 | 0,61 | 0,542032 | 0,583494 | 1,00E-08 | 2,705002 | 1,8687726 | 1,8687726 | TRUE | 0,2548036 | No |
| orf19.3908 | 23 | 20 | 26 | 19 | 68 | 72 | 82 | 101 | 21 | 20 | 25 | 29 | 60 | 78 | 84 | 77 | 49,25 | 24 | 75 | 3,22 | 1,687 | 7,497 | 6,52E-14 | 1,40E-13 | 1,00E-08 | 0,02886 | 0,0195611 | 0,0195611 | TRUE | 0,1611493 | No |
| orf19.391 | 1017 | 937 | 1019 | 718 | 1964 | 1760 | 1808 | 2485 | 926 | 948 | 964 | 1079 | 1747 | 1911 | 1852 | 1896 | 1415,19 | 979 | 1852 | 1,898 | 0,924 | 15,962 | 2,36E-57 | 1,24E-56 | 0,0018624 | 0,00297 | 0,002416 | 0,002416 | TRUE | 0,04951 | No |
| orf19.3910 | 328 | 274 | 310 | 206 | 553 | 479 | 450 | 622 | 299 | 277 | 293 | 309 | 492 | 520 | 461 | 474 | 390,73 | 294 | 487 | 1,655 | 0,727 | 9,291 | 1,52E-20 | 3,92E-20 | 1,00E-08 | 0,005418 | 0,0031004 | 0,0031004 | TRUE | 0,0228998 | No |
| orf19.3911 | 7965 | 6358 | 6457 | 4885 | 19486 | 15951 | 15855 | 23941 | 7252 | 6430 | 6111 | 7338 | 17332 | 17318 | 16238 | 18262 | 12035,2 | 6783 | 17288 | 2,55 | 1,35 | 21,317 | 7,88E-101 | 6,91E-100 | 0,0050082 | 0,002147 | 0,0037485 | 0,0037485 | TRUE | 0,0544458 | No |
| orf19.3912 | 3820 | 3473 | 3842 | 2564 | 3215 | 2882 | 2905 | 3852 | 3478 | 3513 | 3636 | 3852 | 2860 | 3129 | 2975 | 2938 | 3297,53 | 3620 | 2976 | 0,823 | -0,282 | -5,976 | 2,29E-09 | 4,23E-09 | 0,0013669 | 0,002438 | 0,0018279 | 0,0018279 | TRUE | 0,0214783 | No |
| orf19.3914 | 8283 | 7991 | 8517 | 5103 | 7313 | 6238 | 6597 | 8291 | 7541 | 8082 | 8061 | 7666 | 6505 | 6773 | 6756 | 6324 | 7213,43 | 7838 | 6590 | 0,84 | -0,251 | -6,172 | 6,74E-10 | 1,27E-09 | 0,0009621 | 0,00222 | 0,0014475 | 0,0014475 | TRUE | 0,0090749 | No |
| orf19.3915 | 4783 | 4081 | 4368 | 2734 | 5632 | 4553 | 4588 | 6519 | 4355 | 4127 | 4134 | 4107 | 5009 | 4943 | 4699 | 4973 | 4543,41 | 4181 | 4906 | 1,173 | 0,23 | 5,952 | 2,66E-09 | 4,89E-09 | 0,0005054 | 0,002328 | 0,0012151 | 0,0012151 | TRUE | 0,0097962 | No |
| orf19.3916 | 1058 | 1069 | 1104 | 700 | 1765 | 1462 | 1579 | 1969 | 963 | 1081 | 1045 | 1052 | 1570 | 1587 | 1617 | 1502 | 1302,14 | 1035 | 1569 | 1,516 | 0,601 | 11,249 | 2,34E-29 | 7,38E-29 | 0,0009183 | 0,003052 | 0,0019198 | 0,0019198 | TRUE | 0,0279592 | No |
| orf19.3919 | 1483 | 1374 | 1464 | 960 | 886 | 821 | 783 | 1022 | 1350 | 1390 | 1386 | 1442 | 788 | 891 | 802 | 780 | 1103,55 | 1392 | 815 | 0,585 | -0,773 | -13,611 | 3,43E-42 | 1,39E-41 | 0,0010756 | 0,003234 | 0,0021355 | 0,0021355 | TRUE | 0,0448967 | No |
| orf19.3919.1 | 792 | 635 | 662 | 457 | 693 | 517 | 549 | 728 | 721 | 642 | 627 | 686 | 616 | 561 | 562 | 555 | 621,45 | 669 | 574 | 0,858 | -0,22 | -3,151 | 0,001625 | 0,002259 | 0,001936 | 0,004163 | 0,0030819 | 0,0030819 | TRUE | 0,0361523 | No |
| orf19.392 | 731 | 588 | 714 | 433 | 1081 | 796 | 879 | 1174 | 666 | 595 | 676 | 650 | 962 | 864 | 900 | 896 | 775,99 | 647 | 906 | 1,401 | 0,486 | 7,633 | 2,30E-14 | 5,00E-14 | 0,0012717 | 0,003739 | 0,0025498 | 0,0025498 | TRUE | 0,0360972 | No |
| orf19.3920 | 639 | 529 | 570 | 368 | 1176 | 864 | 865 | 1230 | 582 | 535 | 539 | 553 | 1046 | 938 | 886 | 938 | 752,16 | 552 | 952 | 1,724 | 0,786 | 11,474 | 1,79E-30 | 5,82E-30 | 0,00234 | 0,003793 | 0,0030476 | 0,0030476 | TRUE | 0,0531935 | No |
| orf19.3921 | 1590 | 1209 | 1418 | 895 | 1382 | 1314 | 1429 | 1933 | 1448 | 1223 | 1342 | 1344 | 1229 | 1427 | 1463 | 1475 | 1368,84 | 1339 | 1398 | 1,044 | 0,062 | 0,863 | 0,388127 | 0,428972 | 0,0052969 | 0,003002 | 0,0042094 | 0,0042094 | TRUE | 0,0809076 | No |
| orf19.3922 | 4759 | 3962 | 4227 | 2873 | 2313 | 1934 | 1954 | 2652 | 4333 | 4007 | 4000 | 4316 | 2057 | 2100 | 2001 | 2023 | 3104,68 | 4164 | 2045 | 0,491 | -1,025 | -22,992 | 5,59E-117 | 5,60E-116 | 0,0009784 | 0,002463 | 0,0015512 | 0,0015512 | TRUE | 0,0096974 | No |
| orf19.3923 | 420 | 374 | 371 | 297 | 124 | 81 | 110 | 132 | 382 | 378 | 351 | 446 | 110 | 88 | 113 | 101 | 246,19 | 389 | 103 | 0,266 | -1,911 | -16,749 | 5,79E-63 | 3,30E-62 | 0,0053197 | 0,007403 | 0,0065975 | 0,0065975 | TRUE | 0,0969927 | No |
| orf19.3924 | 26 | 27 | 36 | 12 | 504 | 408 | 429 | 581 | 24 | 27 | 34 | 18 | 448 | 443 | 439 | 443 | 234,61 | 26 | 443 | 16,764 | 4,067 | 25,044 | 2,02E-138 | 2,44E-137 | 1,00E-08 | 0,007668 | 0,0043809 | 0,0043809 | TRUE | 0,2863188 | No |
| orf19.3925 | 8 | 12 | 10 | 5 | 88 | 65 | 61 | 68 | 7 | 12 | 9 | 8 | 78 | 71 | 62 | 52 | 37,45 | 9 | 66 | 7,147 | 2,837 | 9,081 | 1,08E-19 | 2,71E-19 | 1,00E-08 | 0,037313 | 0,0290833 | 0,0290833 | TRUE | 0,1317936 | No |
| orf19.3926 | 51 | 33 | 29 | 29 | 96 | 80 | 107 | 118 | 46 | 33 | 27 | 44 | 85 | 87 | 110 | 90 | 65,33 | 38 | 93 | 2,482 | 1,311 | 6,627 | 3,44E-11 | 6,79E-11 | 0,0099572 | 0,022257 | 0,0183652 | 0,0183652 | TRUE | 0,2715124 | No |
| orf19.3928 | 652 | 516 | 602 | 356 | 568 | 430 | 527 | 701 | 594 | 522 | 570 | 535 | 505 | 467 | 540 | 535 | 533,32 | 555 | 512 | 0,921 | -0,118 | -1,603 | 0,108979 | 0,130321 | 0,00199 | 0,004514 | 0,0033652 | 0,0033652 | TRUE | 0,0380184 | No |
| orf19.3929 | 778 | 667 | 694 | 480 | 1342 | 1105 | 1259 | 1726 | 708 | 675 | 657 | 721 | 1194 | 1200 | 1289 | 1317 | 970,02 | 690 | 1250 | 1,815 | 0,86 | 14,346 | 1,13E-46 | 4,92E-46 | 0,0012926 | 0,003399 | 0,0023028 | 0,0023028 | TRUE | 0,0164308 | No |

| | | | | | | | | | | | | | | | | | | | | | | | | | | | | | | | |
|---|---|---|---|---|---|---|---|---|---|---|---|---|---|---|---|---|---|---|---|---|---|---|---|---|---|---|---|---|---|---|---|
| orf19.393 | 1059 | 863 | 908 | 651 | 966 | 720 | 818 | 1088 | 964 | 873 | 859 | 978 | 859 | 782 | 838 | 830 | 872,86 | 918 | 827 | 0,903 | -0,147 | -2,336 | 0,019474 | 0,025005 | 0,0017627 | 0,00355 | 0,0026672 | 0,0026672 | TRUE | 0,0226354 | No |
| orf19.3930 | 3271 | 2821 | 3061 | 2032 | 3456 | 2953 | 3075 | 4052 | 2978 | 2853 | 2897 | 3052 | 3074 | 3206 | 3149 | 3091 | 3037,6 | 2945 | 3130 | 1,063 | 0,089 | 2,286 | 0,022248 | 0,028409 | 0,000194 | 0,002472 | 0,001111 | 0,001111 | TRUE | 0,0065976 | No |
| orf19.3931 | 1858 | 1574 | 1686 | 1158 | 6181 | 5290 | 5405 | 7473 | 1692 | 1592 | 1596 | 1740 | 5498 | 5743 | 5535 | 5700 | 3636,98 | 1655 | 5619 | 3,402 | 1,766 | 41,467 | 0 | 0 | 0,0005058 | 0,0024 | 0,0013341 | 0,0013341 | TRUE | 0,0125706 | No |
| orf19.3931.2 | 12 | 9 | 15 | 16 | 27 | 28 | 24 | 35 | 11 | 9 | 14 | 24 | 24 | 30 | 25 | 27 | 20,49 | 14 | 26 | 1,89 | 0,919 | 2,788 | 0,005308 | 0,007107 | 1,00E-08 | 0,066495 | 0,0484862 | 0,0484862 | TRUE | 0,551725 | No |
| orf19.3932 | 11765 | 9706 | 10734 | 6896 | 3723 | 2892 | 2998 | 4267 | 10712 | 9817 | 10159 | 10359 | 3311 | 3140 | 3070 | 3255 | 6727,8 | 10262 | 3194 | 0,311 | -1,683 | -39,935 | 0 | 0 | 0,0009757 | 0,002233 | 0,0015069 | 0,0015069 | TRUE | 0,0108658 | No |
| orf19.3932.1 | 1946 | 1361 | 1406 | 1212 | 808 | 753 | 640 | 976 | 1772 | 1377 | 1331 | 1821 | 719 | 818 | 655 | 745 | 1154,46 | 1575 | 734 | 0,467 | -1,1 | -10,458 | 1,34E-25 | 3,90E-25 | 0,016158 | 0,003181 | 0,0096438 | 0,0096438 | TRUE | 0,078386 | No |
| orf19.3934 | 13578 | 11100 | 12103 | 7436 | 18822 | 14469 | 16436 | 22784 | 12362 | 11226 | 11454 | 11170 | 16742 | 15709 | 16833 | 17380 | 14109,5 | 11553 | 16666 | 1,442 | 0,528 | 11,357 | 6,82E-30 | 2,18E-29 | 0,0019233 | 0,002131 | 0,0020047 | 0,0020047 | TRUE | 0,0273253 | No |
| orf19.3935 | 4 | 3 | 2 | 2 | 12 | 5 | 7 | 11 | 4 | 3 | 2 | 3 | 11 | 5 | 7 | 8 | 5,4 | 3 | 8 | 2,767 | 1,468 | 2,329 | 0,019841 | 0,025444 | 1,00E-08 | 0,246467 | 0,1399983 | 0,1399983 | TRUE | 0,1713653 | No |
| orf19.3936 | 2512 | 2332 | 2419 | 1497 | 4745 | 4049 | 4354 | 5658 | 2287 | 2359 | 2289 | 2249 | 4221 | 4396 | 4459 | 4316 | 3321,92 | 2296 | 4348 | 1,892 | 0,92 | 24,293 | 2,34E-130 | 2,59E-129 | 0,0002145 | 0,002435 | 0,0010356 | 0,0010356 | TRUE | 0,0047198 | No |
| orf19.3937 | 621 | 604 | 634 | 406 | 880 | 715 | 748 | 1095 | 565 | 611 | 600 | 610 | 783 | 776 | 766 | 835 | 693,31 | 596 | 790 | 1,328 | 0,41 | 6,646 | 3,01E-11 | 5,96E-11 | 1,00E-08 | 0,003942 | 0,0021624 | 0,0021624 | TRUE | 0,0188467 | No |
| orf19.3938 | 2352 | 2134 | 2281 | 1782 | 1484 | 1305 | 1249 | 1720 | 2141 | 2158 | 2159 | 2677 | 1320 | 1417 | 1279 | 1312 | 1807,91 | 2284 | 1332 | 0,584 | -0,775 | -10,442 | 1,59E-25 | 4,63E-25 | 0,0067849 | 0,002768 | 0,0047089 | 0,0047089 | TRUE | 0,1570007 | No |
| orf19.3939 | 652 | 613 | 676 | 435 | 717 | 557 | 539 | 737 | 594 | 620 | 640 | 653 | 638 | 605 | 552 | 562 | 607,94 | 627 | 589 | 0,942 | -0,087 | -1,25 | 0,211322 | 0,243744 | 0,0015803 | 0,00421 | 0,0029659 | 0,0029659 | TRUE | 0,037712 | No |
| orf19.394 | 1906 | 1538 | 1738 | 1173 | 1397 | 1246 | 1228 | 1696 | 1735 | 1556 | 1645 | 1762 | 1243 | 1353 | 1258 | 1294 | 1480,56 | 1674 | 1287 | 0,769 | -0,379 | -6,909 | 4,89E-12 | 9,96E-12 | 0,0015875 | 0,002929 | 0,0022044 | 0,0022044 | TRUE | 0,0274715 | No |
| orf19.3940 | 80 | 74 | 70 | 53 | 105 | 96 | 85 | 120 | 73 | 75 | 66 | 80 | 93 | 104 | 87 | 92 | 83,72 | 74 | 94 | 1,287 | 0,364 | 2,4 | 0,01639 | 0,021176 | 1,00E-08 | 0,017816 | 0,0098165 | 0,0098165 | TRUE | 0,0460448 | No |
| orf19.3940.1 | 74206 | 42907 | 49512 | 40372 | 17793 | 12679 | 12480 | 20786 | 67561 | 43396 | 46859 | 60645 | 15826 | 13766 | 12781 | 15856 | 34586,3 | 54615 | 14557 | 0,267 | -1,908 | -11,308 | 1,20E-29 | 3,80E-29 | 0,0273 | 0,002075 | 0,0133377 | 0,0273 | TRUE | 0,1038138 | No |
| orf19.3941 | 18056 | 15293 | 16580 | 10269 | 22482 | 18401 | 19518 | 27548 | 16439 | 15467 | 15691 | 15426 | 19997 | 19978 | 19989 | 21014 | 18000,2 | 15756 | 20244 | 1,285 | 0,362 | 10,234 | 1,39E-24 | 3,95E-24 | 0,0006912 | 0,00211 | 0,0011412 | 0,0011412 | TRUE | 0,010505 | No |
| orf19.3942 | 386 | 357 | 361 | 206 | 356 | 281 | 324 | 497 | 351 | 361 | 342 | 309 | 317 | 305 | 332 | 379 | 337,03 | 341 | 333 | 0,977 | -0,034 | -0,369 | 0,712054 | 0,745744 | 0,0040581 | 0,005956 | 0,0051152 | 0,0051152 | TRUE | 0,1059428 | No |
| orf19.3942.1 | 15321 | 12228 | 12211 | 7621 | 22127 | 15872 | 16631 | 24280 | 13949 | 12367 | 11557 | 11448 | 19681 | 17233 | 17032 | 18521 | 15223,5 | 12330 | 18117 | 1,469 | 0,555 | 8,067 | 7,23E-16 | 1,64E-15 | 0,0066132 | 0,002124 | 0,0044774 | 0,0044774 | TRUE | 0,0957799 | No |
| orf19.3944 | 10661 | 9403 | 10300 | 6634 | 8701 | 7275 | 7933 | 10557 | 9706 | 9510 | 9748 | 9965 | 7739 | 7899 | 8124 | 8053 | 8843,14 | 9732 | 7954 | 0,817 | -0,291 | -9,043 | 1,52E-19 | 3,81E-19 | 0,0002635 | 0,002186 | 0,0008782 | 0,0008782 | TRUE | 0,0040739 | No |
| orf19.3945 | 4851 | 4673 | 4942 | 3022 | 3525 | 3031 | 3035 | 4140 | 4417 | 4726 | 4677 | 4540 | 3135 | 3291 | 3108 | 3158 | 3881,5 | 4590 | 3173 | 0,691 | -0,533 | -13,558 | 7,15E-42 | 2,88E-41 | 0,0005473 | 0,002377 | 0,0012197 | 0,0012197 | TRUE | 0,0082719 | No |
| orf19.3946 | 461 | 417 | 420 | 312 | 520 | 454 | 480 | 575 | 420 | 422 | 397 | 469 | 463 | 493 | 492 | 439 | 449,16 | 427 | 472 | 1,107 | 0,147 | 1,907 | 0,05655 | 0,069716 | 0,0014152 | 0,004978 | 0,0034273 | 0,0034273 | TRUE | 0,042222 | No |
| orf19.3947 | 1341 | 1088 | 1106 | 688 | 1149 | 823 | 901 | 1223 | 1221 | 1100 | 1047 | 1033 | 1022 | 894 | 923 | 933 | 1021,59 | 1100 | 943 | 0,856 | -0,225 | -3,281 | 0,001036 | 0,001461 | 0,003424 | 0,00333 | 0,0035184 | 0,0035184 | TRUE | 0,0667601 | No |
| orf19.3949 | 4641 | 3976 | 4353 | 2756 | 3340 | 2753 | 3020 | 4046 | 4225 | 4021 | 4120 | 4140 | 2971 | 2989 | 3093 | 3086 | 3580,68 | 4126 | 3035 | 0,736 | -0,443 | -12,189 | 3,58E-34 | 1,25E-33 | 0,0001778 | 0,002406 | 0,0009851 | 0,0009851 | TRUE | 0,0036924 | No |
| orf19.395 | 14978 | 11641 | 12454 | 8741 | 4495 | 3535 | 3555 | 5348 | 13637 | 11774 | 11787 | 13130 | 3998 | 3838 | 3641 | 4079 | 8235,48 | 12582 | 3889 | 0,309 | -1,693 | -28,493 | 1,43E-178 | 2,19E-177 | 0,0038566 | 0,002197 | 0,0032293 | 0,0032293 | TRUE | 0,039375 | No |
| orf19.3950 | 337 | 301 | 336 | 207 | 608 | 524 | 533 | 801 | 307 | 304 | 318 | 311 | 541 | 569 | 546 | 611 | 438,35 | 310 | 567 | 1,83 | 0,872 | 11,502 | 1,29E-30 | 4,22E-30 | 1,00E-08 | 0,005051 | 0,0029794 | 0,0029794 | TRUE | 0,0355277 | No |
| orf19.3951 | 1364 | 1142 | 1326 | 807 | 1753 | 1378 | 1503 | 2027 | 1242 | 1155 | 1255 | 1212 | 1559 | 1496 | 1539 | 1546 | 1375,61 | 1216 | 1535 | 1,262 | 0,336 | 7,048 | 1,82E-12 | 3,75E-12 | 1,00E-08 | 0,002997 | 0,00144 | 0,00144 | TRUE | 0,0145052 | No |
| orf19.3952 | 2684 | 2275 | 2537 | 1576 | 3750 | 3049 | 3354 | 4452 | 2444 | 2301 | 2401 | 2367 | 3336 | 3310 | 3435 | 3396 | 2873,73 | 2378 | 3369 | 1,416 | 0,502 | 13,144 | 1,85E-39 | 7,12E-39 | 1,00E-08 | 0,002497 | 0,0010366 | 0,0010366 | TRUE | 0,0061432 | No |
| orf19.3954 | 1285 | 1301 | 1324 | 804 | 1387 | 1104 | 1199 | 1569 | 1170 | 1316 | 1253 | 1208 | 1234 | 1199 | 1228 | 1197 | 1225,46 | 1237 | 1214 | 0,981 | -0,027 | -0,518 | 0,60467 | 0,644287 | 0,0006315 | 0,003115 | 0,0017998 | 0,0017998 | TRUE | 0,0227677 | No |
| orf19.3954.1 | 39 | 38 | 52 | 32 | 58 | 59 | 43 | 75 | 36 | 38 | 49 | 48 | 52 | 64 | 44 | 57 | 48,51 | 43 | 54 | 1,276 | 0,352 | 1,688 | 0,091376 | 0,110097 | 1,00E-08 | 0,029266 | 0,0206027 | 0,0206027 | TRUE | 0,1237673 | No |
| orf19.3955 | 4608 | 4169 | 4278 | 3020 | 6730 | 5787 | 5652 | 8050 | 4195 | 4216 | 4049 | 4537 | 5986 | 6283 | 5788 | 6141 | 5149,41 | 4249 | 6050 | 1,425 | 0,511 | 11,209 | 3,67E-29 | 1,16E-28 | 0,0015175 | 0,002294 | 0,0017923 | 0,0017923 | TRUE | 0,024076 | No |
| orf19.3956 | 1275 | 1107 | 1163 | 810 | 1043 | 868 | 975 | 1348 | 1161 | 1120 | 1101 | 1217 | 928 | 942 | 999 | 1028 | 1061,85 | 1150 | 974 | 0,85 | -0,235 | -4,095 | 4,22E-05 | 6,47E-05 | 0,001186 | 0,003281 | 0,0022045 | 0,0022045 | TRUE | 0,0178911 | No |
| orf19.3957 | 2248 | 1887 | 2067 | 1230 | 2097 | 1632 | 1903 | 2409 | 2047 | 1908 | 1956 | 1848 | 1865 | 1772 | 1949 | 1838 | 1897,84 | 1940 | 1856 | 0,955 | -0,066 | -1,332 | 0,182811 | 0,212474 | 0,0010152 | 0,002733 | 0,001821 | 0,001821 | TRUE | 0,0167595 | No |
| orf19.3959 | 4464 | 3652 | 3919 | 2647 | 3597 | 2849 | 2942 | 4226 | 4064 | 3694 | 3709 | 3976 | 3199 | 3093 | 3013 | 3224 | 3496,54 | 3861 | 3132 | 0,812 | -0,301 | -6,431 | 1,26E-10 | 2,45E-10 | 0,0012827 | 0,002415 | 0,001808 | 0,001808 | TRUE | 0,0161443 | No |
| orf19.396 | 314 | 286 | 310 | 185 | 669 | 457 | 582 | 748 | 286 | 289 | 293 | 278 | 595 | 496 | 596 | 571 | 425,53 | 286 | 564 | 1,969 | 0,977 | 11,842 | 2,38E-32 | 8,04E-32 | 0,0023684 | 0,005141 | 0,0038569 | 0,0038569 | TRUE | 0,0732275 | No |
| orf19.3960 | 2927 | 2767 | 2918 | 1773 | 2598 | 2037 | 2360 | 3155 | 2665 | 2799 | 2762 | 2663 | 2311 | 2212 | 2417 | 2407 | 2529,3 | 2722 | 2337 | 0,858 | -0,22 | -4,974 | 6,56E-07 | 1,10E-06 | 0,0006316 | 0,002559 | 0,0014839 | 0,0014839 | TRUE | 0,0151445 | No |
| orf19.3962 | 16540 | 14753 | 16086 | 10102 | 19366 | 15972 | 16973 | 22787 | 15059 | 14921 | 15224 | 15175 | 17225 | 17341 | 17382 | 17382 | 16213,8 | 15095 | 17332 | 1,148 | 0,2 | 8,198 | 2,45E-16 | 5,62E-16 | 1,00E-08 | 0,002118 | 0,0005063 | 0,0005063 | TRUE | 0,0007364 | No |
| orf19.3963 | 1465 | 1343 | 1417 | 1012 | 1143 | 936 | 944 | 1330 | 1334 | 1358 | 1341 | 1520 | 1017 | 1016 | 967 | 1015 | 1195,95 | 1388 | 1004 | 0,725 | -0,463 | -8,264 | 1,41E-16 | 3,26E-16 | 0,0011858 | 0,003142 | 0,0021656 | 0,0021656 | TRUE | 0,0445195 | No |
| orf19.3964 | 716 | 580 | 638 | 397 | 494 | 395 | 456 | 603 | 652 | 587 | 604 | 596 | 439 | 429 | 467 | 460 | 529,24 | 610 | 449 | 0,736 | -0,443 | -6,474 | 9,52E-11 | 1,85E-10 | 1,00E-08 | 0,004533 | 0,0025821 | 0,0025821 | TRUE | 0,0272524 | No |
| orf19.3965 | 406 | 381 | 370 | 252 | 295 | 227 | 270 | 345 | 370 | 385 | 350 | 379 | 262 | 246 | 277 | 263 | 316,53 | 371 | 262 | 0,708 | -0,498 | -6,046 | 1,49E-09 | 2,76E-09 | 1,00E-08 | 0,00621 | 0,0032955 | 0,0032955 | TRUE | 0,0171082 | No |
| orf19.3966 | 25 | 32 | 26 | 13 | 17 | 19 | 21 | 25 | 23 | 32 | 25 | 20 | 15 | 21 | 22 | 19 | 21,95 | 25 | 19 | 0,758 | -0,4 | -1,351 | 0,176823 | 0,20589 | 1,00E-08 | 0,062224 | 0,0383965 | 0,0383965 | TRUE | 0,2467033 | No |
| orf19.3967 | 33248 | 29380 | 31722 | 19913 | 13681 | 11449 | 12394 | 16618 | 30271 | 29715 | 30022 | 29913 | 12169 | 12430 | 12693 | 12676 | 21236,1 | 29980 | 12492 | 0,417 | -1,263 | -45,845 | 0 | 0 | 0,0001494 | 0,002099 | 0,0006735 | 0,0006735 | TRUE | 0,0037536 | No |
| orf19.3968 | 6 | 14 | 9 | 8 | 6 | 3 | 5 | 7 | 5 | 14 | 9 | 12 | 5 | 3 | 5 | 5 | 7,4 | 10 | 4 | 0,488 | -1,036 | -1,957 | 0,050395 | 0,062382 | 1,00E-08 | 0,180516 | 0,1188834 | 0,1188834 | TRUE | 0,1890232 | No |
| orf19.3969 | 6451 | 5738 | 6325 | 3964 | 2119 | 1814 | 1988 | 2681 | 5873 | 5803 | 5986 | 5955 | 1885 | 1969 | 2036 | 2045 | 3944,08 | 5904 | 1984 | 0,336 | -1,573 | -39,969 | 0 | 0 | 0,0003465 | 0,002372 | 0,0011607 | 0,0011607 | TRUE | 0,0141145 | No |
| orf19.397 | 1138 | 984 | 1031 | 796 | 1212 | 1031 | 924 | 1236 | 1036 | 995 | 976 | 1196 | 1078 | 1119 | 946 | 943 | 1036,16 | 1051 | 1022 | 0,974 | -0,038 | -0,471 | 0,63759 | 0,67575 | 0,0065036 | 0,003312 | 0,0051524 | 0,0051524 | TRUE | 0,0964265 | No |
| orf19.3970 | 1983 | 1706 | 1635 | 1011 | 978 | 802 | 897 | 1130 | 1805 | 1725 | 1547 | 1519 | 870 | 871 | 919 | 862 | 1264,77 | 1649 | 880 | 0,533 | -0,909 | -13,91 | 5,50E-44 | 2,31E-43 | 0,0033768 | 0,003081 | 0,0032406 | 0,0032406 | TRUE | 0,0495177 | No |
| orf19.3971 | 367 | 329 | 355 | 232 | 222 | 193 | 160 | 264 | 334 | 333 | 336 | 349 | 197 | 210 | 164 | 201 | 265,45 | 338 | 193 | 0,573 | -0,803 | -8,55 | 1,23E-17 | 2,94E-17 | 1,00E-08 | 0,007013 | 0,004479 | 0,004479 | TRUE | 0,1050059 | No |
| orf19.3972 | 687 | 781 | 836 | 567 | 747 | 660 | 623 | 737 | 625 | 790 | 791 | 852 | 664 | 717 | 638 | 562 | 704,94 | 764 | 645 | 0,845 | -0,243 | -2,461 | 0,013852 | 0,017999 | 0,0123622 | 0,003911 | 0,0079537 | 0,0079537 | TRUE | 0,1819971 | No |
| orf19.3973 | 1606 | 1351 | 1375 | 966 | 690 | 576 | 610 | 841 | 1462 | 1366 | 1301 | 1451 | 614 | 625 | 625 | 642 | 1010,79 | 1395 | 626 | 0,45 | -1,153 | -19,706 | 1,92E-86 | 1,46E-85 | 0,0011511 | 0,003344 | 0,0021592 | 0,0021592 | TRUE | 0,0254031 | No |
| orf19.3974 | 34189 | 31460 | 33836 | 22477 | 45635 | 40313 | 37911 | 54329 | 31128 | 31818 | 32023 | 33764 | 40591 | 43769 | 38826 | 41443 | 36670,1 | 32183 | 41157 | 1,279 | 0,355 | 7,922 | 2,35E-15 | 5,25E-15 | 0,0018155 | 0,002073 | 0,0019011 | 0,0019011 | TRUE | 0,022298 | No |
| orf19.3976 | 423 | 386 | 443 | 240 | 432 | 372 | 375 | 521 | 385 | 390 | 419 | 361 | 384 | 404 | 384 | 397 | 390,61 | 389 | 392 | 1,005 | 0,007 | 0,097 | 0,922503 | 0,933337 | 1,00E-08 | 0,005419 | 0,0029586 | 0,0029586 | TRUE | 0,0316138 | No |
| orf19.3977 | 3682 | 3088 | 3297 | 1965 | 3117 | 2328 | 2521 | 3272 | 3352 | 3123 | 3120 | 2952 | 2772 | 2528 | 2582 | 2496 | 2865,66 | 3137 | 2594 | 0,826 | -0,275 | -5,259 | 1,45E-07 | 2,49E-07 | 0,0020448 | 0,002498 | 0,0022815 | 0,0022815 | TRUE | 0,0262289 | No |
| orf19.3978 | 5622 | 5602 | 5736 | 2606 | 4643 | 3289 | 3655 | 4448 | 5119 | 5666 | 5429 | 3915 | 4130 | 3571 | 3743 | 3393 | 4370,57 | 5032 | 3709 | 0,737 | -0,441 | -4,494 | 6,98E-06 | 1,12E-05 | 0,0152351 | 0,002339 | 0,0090142 | 0,0090142 | TRUE | 0,2690843 | No |
| orf19.398 | 124 | 113 | 127 | 96 | 323 | 303 | 331 | 430 | 113 | 114 | 120 | 144 | 287 | 329 | 339 | 328 | 221,86 | 123 | 321 | 2,639 | 1,4 | 12,701 | 5,84E-37 | 2,17E-36 | 0,0023022 | 0,007991 | 0,0058304 | 0,0058304 | TRUE | 0,1047614 | No |
| orf19.3980 | 985 | 924 | 1079 | 635 | 2284 | 1826 | 1979 | 2601 | 897 | 935 | 1021 | 954 | 2032 | 1983 | 2027 | 1984 | 1478,91 | 952 | 2006 | 2,109 | 1,076 | 21,112 | 6,13E-99 | 5,30E-98 | 0,0003902 | 0,00293 | 0,0016998 | 0,0016998 | TRUE | 0,0316418 | No |
| orf19.3981 | 412 | 344 | 390 | 260 | 583 | 460 | 522 | 734 | 375 | 348 | 369 | 391 | 519 | 499 | 535 | 560 | 449,4 | 371 | 528 | 1,431 | 0,517 | 7,014 | 2,32E-12 | 4,76E-12 | 1,00E-08 | 0,004977 | 0,0028893 | 0,0028893 | TRUE | 0,0206403 | No |
| orf19.3982 | 234 | 195 | 219 | 124 | 761 | 678 | 661 | 881 | 213 | 197 | 207 | 186 | 677 | 736 | 677 | 672 | 445,72 | 201 | 690 | 3,412 | 1,771 | 22,01 | 2,33E-107 | 2,14E-106 | 1,00E-08 | 0,005001 | 0,0029122 | 0,0029122 | TRUE | 0,0223051 | No |
| orf19.3983 | 992 | 951 | 974 | 642 | 2281 | 2036 | 1913 | 2608 | 903 | 962 | 922 | 964 | 2029 | 2211 | 1959 | 1989 | 1492,4 | 938 | 2047 | 2,185 | 1,127 | 20,393 | 1,95E-92 | 1,55E-91 | 0,0014991 | 0,002922 | 0,0021331 | 0,0021331 | TRUE | 0,0340081 | No |
| orf19.3984 | 11600 | 12418 | 12609 | 7504 | 5112 | 4335 | 4367 | 5604 | 10561 | 12559 | 11933 | 11272 | 4547 | 4707 | 4472 | 4275 | 8040,87 | 11581 | 4500 | 0,388 | -1,364 | -23,97 | 5,78E-127 | 6,28E-126 | 0,0033418 | 0,002201 | 0,0029611 | 0,0029611 | TRUE | 0,0434361 | No |
| orf19.3986 | 1116 | 1270 | 1269 | 737 | 550 | 484 | 489 | 651 | 1016 | 1284 | 1201 | 1107 | 489 | 525 | 501 | 497 | 827,59 | 1152 | 503 | 0,436 | -1,197 | -15,428 | 1,05E-53 | 5,22E-53 | 0,0050793 | 0,003633 | 0,0043858 | 0,0043858 | TRUE | 0,0801655 | No |
| orf19.3988 | 9452 | 8183 | 8926 | 5710 | 301 | 243 | 268 | 376 | 8606 | 8276 | 8448 | 8577 | 268 | 264 | 274 | 287 | 4374,96 | 8477 | 273 | 0,032 | -4,951 | -90,418 | 0 | 0 | 1,00E-08 | 0,002339 | 0,0011243 | 0,0011243 | TRUE | 0,0147275 | No |
| orf19.399 | 7580 | 6709 | 7297 | 4881 | 3516 | 2834 | 3035 | 4189 | 6901 | 6785 | 6906 | 7332 | 3127 | 3077 | 3108 | 3195 | 5054,08 | 6981 | 3127 | 0,448 | -1,157 | -30,543 | 6,90E-205 | 1,20E-203 | 0,0005246 | 0,002298 | 0,0011515 | 0,0011515 | TRUE | 0,0133547 | No |
| orf19.3990 | 668 | 743 | 719 | 424 | 817 | 590 | 657 | 780 | 608 | 751 | 680 | 637 | 727 | 641 | 673 | 595 | 664,02 | 669 | 659 | 0,983 | -0,025 | -0,294 | 0,768582 | 0,798909 | 0,0065094 | 0,004026 | 0,0052471 | 0,0052471 | TRUE | 0,0817495 | No |
| orf19.3991 | 2627 | 2320 | 2628 | 1517 | 2215 | 1959 | 2028 | 2669 | 2392 | 2346 | 2487 | 2279 | 1970 | 2127 | 2077 | 2036 | 2214,26 | 2376 | 2052 | 0,862 | -0,214 | -4,718 | 2,38E-06 | 3,88E-06 | 0,0005934 | 0,002634 | 0,0015168 | 0,0015168 | TRUE | 0,0119651 | No |
| orf19.3992 | 67 | 40 | 46 | 41 | 82 | 38 | 66 | 82 | 61 | 40 | 44 | 62 | 73 | 41 | 68 | 63 | 56,36 | 52 | 61 | 1,197 | 0,26 | 1,182 | 0,237147 | 0,271225 | 0,0329444 | 0,025474 | 0,0282553 | 0,0282553 | TRUE | 0,3780107 | No |
| orf19.3994 | 464 | 421 | 426 | 307 | 431 | 330 | 356 | 495 | 422 | 426 | 403 | 461 | 383 | 358 | 365 | 378 | 399,55 | 428 | 371 | 0,871 | -0,199 | -2,654 | 0,007959 | 0,010511 | 1,00E-08 | 0,005343 | 0,002883 | 0,002883 | TRUE | 0,0258425 | No |
| orf19.3995 | 382 | 379 | 363 | 197 | 612 | 528 | 530 | 755 | 348 | 383 | 344 | 296 | 544 | 573 | 543 | 576 | 450,86 | 343 | 559 | 1,622 | 0,698 | 8,78 | 1,63E-18 | 3,98E-18 | 0,0014094 | 0,004967 | 0,0036756 | 0,0036756 | TRUE | 0,0769008 | No |
| orf19.3996 | 103 | 83 | 102 | 58 | 119 | 111 | 122 | 176 | 94 | 84 | 97 | 87 | 106 | 121 | 125 | 134 | 105,87 | 90 | 122 | 1,341 | 0,424 | 3,034 | 0,002412 | 0,003308 | 1,00E-08 | 0,014515 | 0,0090275 | 0,0090275 | TRUE | 0,0815062 | No |

| | | | | | | | | | | | | | | | | | | | | | | | | | | | | | | | |
|---|---|---|---|---|---|---|---|---|---|---|---|---|---|---|---|---|---|---|---|---|---|---|---|---|---|---|---|---|---|---|---|
| orf19.3997 | 75794 | 57465 | 64327 | 42099 | 22165 | 17219 | 18973 | 27419 | 69007 | 58120 | 60880 | 63239 | 19715 | 18695 | 19431 | 20915 | 41250,3 | 62812 | 19689 | 0,313 | -1,674 | -29,521 | 1,54E-191 | 2,51E-190 | 0,0037729 | 0,002069 | 0,0030554 | 0,0030554 | TRUE | 0,0541128 | No |
| orf19.3998 | 158 | 173 | 160 | 122 | 122 | 77 | 77 | 121 | 144 | 175 | 151 | 183 | 109 | 84 | 79 | 92 | 127,1 | 163 | 91 | 0,562 | -0,831 | -5,904 | 3,55E-09 | 6,52E-09 | 0,0076673 | 0,012431 | 0,0106427 | 0,0106427 | TRUE | 0,166609 | No |
| orf19.3999 | 966 | 909 | 976 | 612 | 865 | 669 | 732 | 974 | 880 | 919 | 924 | 919 | 769 | 726 | 750 | 743 | 828,78 | 910 | 747 | 0,821 | -0,284 | -5,221 | 1,78E-07 | 3,04E-07 | 1,00E-08 | 0,003631 | 0,001623 | 0,001623 | TRUE | 0,0067406 | No |
| orf19.400 | 2649 | 2312 | 2435 | 1652 | 1821 | 1520 | 1520 | 2048 | 2412 | 2338 | 2305 | 2482 | 1620 | 1650 | 1557 | 1562 | 1990,64 | 2384 | 1597 | 0,67 | -0,577 | -12,94 | 2,68E-38 | 1,02E-37 | 0,0003866 | 0,002701 | 0,0013902 | 0,0013902 | TRUE | 0,0083362 | No |
| orf19.4000 | 15040 | 13746 | 15383 | 9817 | 9892 | 8607 | 8974 | 12110 | 13693 | 13903 | 14559 | 14747 | 8799 | 9345 | 9191 | 9238 | 11684,1 | 14226 | 9143 | 0,643 | -0,637 | -16,686 | 1,67E-62 | 9,44E-62 | 0,0008545 | 0,00215 | 0,0013129 | 0,0013129 | TRUE | 0,0078797 | No |
| orf19.4001 | 377 | 366 | 393 | 233 | 181 | 147 | 177 | 230 | 343 | 370 | 372 | 350 | 161 | 160 | 181 | 175 | 264,08 | 359 | 169 | 0,472 | -1,083 | -11,773 | 5,40E-32 | 1,82E-31 | 1,00E-08 | 0,007039 | 0,0038983 | 0,0038983 | TRUE | 0,0211785 | No |
| orf19.4002 | 462 | 395 | 481 | 247 | 386 | 322 | 321 | 426 | 421 | 400 | 455 | 371 | 343 | 350 | 329 | 325 | 374,13 | 412 | 337 | 0,813 | -0,299 | -3,626 | 0,000288 | 0,000421 | 0,0014604 | 0,005568 | 0,0038273 | 0,0038273 | TRUE | 0,0577102 | No |
| orf19.4003 | 778 | 748 | 744 | 483 | 296 | 225 | 243 | 292 | 708 | 757 | 704 | 726 | 263 | 244 | 249 | 223 | 484,21 | 724 | 245 | 0,338 | -1,566 | -20,614 | 2,04E-94 | 1,66E-93 | 1,00E-08 | 0,004765 | 0,0028983 | 0,0028983 | TRUE | 0,0480849 | No |
| orf19.4004 | 5834 | 4850 | 5512 | 3351 | 6395 | 4957 | 5677 | 7840 | 5312 | 4905 | 5217 | 5034 | 5688 | 5382 | 5814 | 5980 | 5416,46 | 5117 | 5716 | 1,117 | 0,16 | 3,544 | 0,000394 | 0,000572 | 0,0013657 | 0,002281 | 0,0017662 | 0,0017662 | TRUE | 0,0186938 | No |
| orf19.4005 | 407 | 378 | 409 | 282 | 509 | 403 | 472 | 611 | 371 | 382 | 387 | 424 | 453 | 438 | 483 | 466 | 425,41 | 391 | 460 | 1,183 | 0,242 | 3,26 | 0,001114 | 0,00157 | 1,00E-08 | 0,005142 | 0,0029153 | 0,0029153 | TRUE | 0,0300676 | No |
| orf19.4006 | 1278 | 1143 | 1126 | 802 | 737 | 610 | 652 | 849 | 1164 | 1156 | 1066 | 1205 | 656 | 662 | 668 | 648 | 902,89 | 1148 | 658 | 0,574 | -0,8 | -13,876 | 8,89E-44 | 3,72E-43 | 1,00E-08 | 0,0035 | 0,0020129 | 0,0020129 | TRUE | 0,0281417 | No |
| orf19.4007 | 259 | 200 | 231 | 164 | 371 | 274 | 298 | 446 | 236 | 202 | 219 | 246 | 330 | 297 | 305 | 340 | 271,99 | 226 | 318 | 1,419 | 0,505 | 5,341 | 9,22E-08 | 1,59E-07 | 0,0015106 | 0,006894 | 0,0047556 | 0,0047556 | TRUE | 0,0525907 | No |
| orf19.4009 | 1113 | 983 | 984 | 618 | 1315 | 1032 | 1166 | 1456 | 1013 | 994 | 931 | 928 | 1170 | 1120 | 1194 | 1111 | 1057,75 | 966 | 1149 | 1,186 | 0,246 | 4,479 | 7,51E-06 | 1,20E-05 | 0,0006328 | 0,003286 | 0,0019301 | 0,0019301 | TRUE | 0,0128606 | No |
| orf19.401 | 3427 | 2824 | 3035 | 2541 | 5072 | 4668 | 4480 | 6339 | 3120 | 2856 | 2872 | 3817 | 4511 | 5068 | 4588 | 4835 | 3958,59 | 3166 | 4750 | 1,502 | 0,587 | 7,061 | 1,66E-12 | 3,42E-12 | 0,0111732 | 0,002371 | 0,0063712 | 0,0063712 | TRUE | 0,22783 | No |
| orf19.4010 | 1985 | 1807 | 1893 | 1426 | 1292 | 1236 | 1208 | 1587 | 1807 | 1828 | 1792 | 2142 | 1149 | 1342 | 1237 | 1211 | 1563,42 | 1892 | 1235 | 0,653 | -0,614 | -8,828 | 1,07E-18 | 2,61E-18 | 0,0051375 | 0,002882 | 0,0039816 | 0,0039816 | TRUE | 0,0910807 | No |
| orf19.4011 | 46 | 34 | 53 | 29 | 189 | 158 | 154 | 206 | 42 | 34 | 50 | 44 | 168 | 172 | 158 | 157 | 103,06 | 42 | 164 | 3,845 | 1,943 | 12,247 | 1,75E-34 | 6,20E-34 | 1,00E-08 | 0,014854 | 0,0088715 | 0,0088715 | TRUE | 0,1346819 | No |
| orf19.4012 | 9852 | 10704 | 11112 | 6808 | 5840 | 4938 | 4955 | 6368 | 8970 | 10826 | 10517 | 10227 | 5194 | 5361 | 5075 | 4858 | 7628,36 | 10135 | 5122 | 0,505 | -0,985 | -16,379 | 2,70E-60 | 1,49E-59 | 0,0038872 | 0,00221 | 0,0033274 | 0,0033274 | TRUE | 0,0737617 | No |
| orf19.4013 | 522 | 460 | 504 | 302 | 904 | 729 | 771 | 1049 | 475 | 465 | 477 | 454 | 804 | 791 | 790 | 800 | 632,06 | 468 | 796 | 1,699 | 0,765 | 12,707 | 5,42E-37 | 2,02E-36 | 1,00E-08 | 0,004127 | 0,0017594 | 0,0017594 | TRUE | 0,0038425 | No |
| orf19.4014 | 1087 | 980 | 1063 | 699 | 1186 | 944 | 1014 | 1343 | 990 | 991 | 1006 | 1050 | 1055 | 1025 | 1038 | 1024 | 1022,45 | 1009 | 1036 | 1,028 | 0,04 | 0,806 | 0,420353 | 0,461618 | 1,00E-08 | 0,003329 | 0,0014104 | 0,0014104 | TRUE | 0,0074886 | No |
| orf19.4015 | 918 | 826 | 917 | 590 | 1143 | 877 | 1027 | 1218 | 836 | 835 | 868 | 886 | 1017 | 952 | 1052 | 929 | 921,88 | 856 | 988 | 1,154 | 0,206 | 3,477 | 0,000506 | 0,000729 | 0,0011704 | 0,00347 | 0,002283 | 0,002283 | TRUE | 0,0225393 | No |
| orf19.4016 | 6671 | 5375 | 5913 | 4318 | 11101 | 9307 | 9756 | 13925 | 6074 | 5436 | 5596 | 6486 | 9874 | 10105 | 9991 | 10622 | 8023,08 | 5898 | 10148 | 1,722 | 0,784 | 13,822 | 1,88E-43 | 7,82E-43 | 0,0036136 | 0,002202 | 0,0029522 | 0,0029522 | TRUE | 0,0537059 | No |
| orf19.4017 | 417 | 338 | 376 | 267 | 559 | 449 | 523 | 683 | 380 | 342 | 356 | 401 | 497 | 487 | 536 | 521 | 439,97 | 370 | 510 | 1,387 | 0,472 | 6,235 | 4,53E-10 | 8,58E-10 | 0,0003183 | 0,005039 | 0,0031366 | 0,0031366 | TRUE | 0,030409 | No |
| orf19.4018 | 1836 | 1551 | 1788 | 1125 | 1721 | 1377 | 1357 | 1976 | 1672 | 1569 | 1692 | 1690 | 1531 | 1495 | 1390 | 1507 | 1568,16 | 1656 | 1481 | 0,895 | -0,16 | -3,173 | 0,001509 | 0,002106 | 0,0008847 | 0,002879 | 0,0017882 | 0,0017882 | TRUE | 0,0201756 | No |
| orf19.4019 | 53 | 48 | 60 | 30 | 64 | 53 | 76 | 95 | 48 | 49 | 57 | 45 | 57 | 58 | 78 | 72 | 57,93 | 50 | 66 | 1,327 | 0,409 | 2,152 | 0,031361 | 0,039581 | 1,00E-08 | 0,024841 | 0,0169133 | 0,0169133 | TRUE | 0,1126785 | No |
| orf19.4021 | 1228 | 907 | 1086 | 651 | 1971 | 1462 | 1748 | 2368 | 1118 | 917 | 1028 | 978 | 1753 | 1587 | 1790 | 1806 | 1372,26 | 1010 | 1734 | 1,715 | 0,778 | 11,372 | 5,77E-30 | 1,85E-29 | 0,0041612 | 0,003 | 0,0036972 | 0,0036972 | TRUE | 0,0637973 | No |
| orf19.4022 | 1077 | 845 | 930 | 606 | 1454 | 1047 | 1165 | 1592 | 981 | 855 | 880 | 910 | 1293 | 1137 | 1193 | 1214 | 1057,9 | 906 | 1209 | 1,335 | 0,417 | 6,657 | 2,79E-11 | 5,54E-11 | 0,0023167 | 0,003286 | 0,0027834 | 0,0027834 | TRUE | 0,0389471 | No |
| orf19.4023 | 1273 | 1005 | 1033 | 756 | 1139 | 981 | 951 | 1449 | 1159 | 1016 | 978 | 1136 | 1013 | 1065 | 974 | 1105 | 1055,77 | 1072 | 1039 | 0,971 | -0,042 | -0,606 | 0,544295 | 0,585832 | 0,0040768 | 0,003288 | 0,0036952 | 0,0036952 | TRUE | 0,0431307 | No |
| orf19.4024 | 1087 | 967 | 1055 | 647 | 589 | 474 | 498 | 690 | 990 | 978 | 998 | 972 | 524 | 515 | 510 | 526 | 751,62 | 984 | 519 | 0,527 | -0,924 | -16,482 | 4,97E-61 | 2,76E-60 | 1,00E-08 | 0,003795 | 0,00158 | 0,00158 | TRUE | 0,0015052 | No |
| orf19.4025 | 4929 | 3940 | 4086 | 2573 | 4006 | 3016 | 3346 | 4544 | 4488 | 3985 | 3867 | 3865 | 3563 | 3275 | 3427 | 3466 | 3741,91 | 4051 | 3433 | 0,847 | -0,24 | -4,295 | 1,75E-05 | 2,74E-05 | 0,0029278 | 0,00239 | 0,0027179 | 0,0027179 | TRUE | 0,0649038 | No |
| orf19.4026 | 1179 | 863 | 981 | 521 | 1889 | 1468 | 1570 | 2101 | 1073 | 873 | 928 | 783 | 1680 | 1594 | 1608 | 1603 | 1267,74 | 914 | 1621 | 1,767 | 0,821 | 10,378 | 3,13E-25 | 9,03E-25 | 0,0080485 | 0,003079 | 0,0051343 | 0,0051343 | TRUE | 0,1641876 | No |
| orf19.4028 | 2811 | 2373 | 2570 | 1619 | 3102 | 2395 | 2769 | 3706 | 2559 | 2400 | 2432 | 2432 | 2759 | 2600 | 2836 | 2827 | 2605,73 | 2456 | 2756 | 1,122 | 0,166 | 3,721 | 0,000199 | 0,000293 | 0,0008365 | 0,002544 | 0,0015271 | 0,0015271 | TRUE | 0,0169441 | No |
| orf19.4029 | 17983 | 14420 | 16288 | 10090 | 18623 | 15016 | 17335 | 23104 | 16373 | 14584 | 15415 | 15157 | 16565 | 16303 | 17753 | 17624 | 16221,7 | 15382 | 17061 | 1,109 | 0,149 | 3,174 | 0,001505 | 0,0021 | 0,0020194 | 0,002118 | 0,0020657 | 0,0020657 | TRUE | 0,0231342 | No |
| orf19.403 | 26 | 23 | 19 | 17 | 89 | 51 | 60 | 78 | 24 | 23 | 18 | 26 | 79 | 55 | 61 | 59 | 43,24 | 23 | 64 | 2,858 | 1,515 | 6,407 | 1,48E-10 | 2,87E-10 | 1,00E-08 | 0,032586 | 0,0227328 | 0,0227328 | TRUE | 0,2385409 | No |
| orf19.4030 | 813 | 672 | 774 | 466 | 695 | 502 | 553 | 801 | 740 | 680 | 733 | 700 | 618 | 545 | 566 | 611 | 649,12 | 713 | 585 | 0,822 | -0,284 | -4,284 | 1,83E-05 | 2,87E-05 | 0,0010555 | 0,004072 | 0,0026633 | 0,0026633 | TRUE | 0,0241033 | No |
| orf19.4031 | 4283 | 3484 | 3914 | 2647 | 4408 | 3662 | 4152 | 5696 | 3899 | 3524 | 3704 | 3976 | 3921 | 3976 | 4252 | 4345 | 3949,68 | 3776 | 4124 | 1,093 | 0,128 | 2,427 | 0,015232 | 0,01974 | 0,0023723 | 0,002371 | 0,0024221 | 0,0024221 | TRUE | 0,0245386 | No |
| orf19.4032 | 5117 | 4407 | 4531 | 2936 | 3847 | 2935 | 3242 | 4282 | 4659 | 4457 | 4288 | 4410 | 3422 | 3187 | 3320 | 3266 | 3876,18 | 4454 | 3299 | 0,741 | -0,433 | -10,412 | 2,17E-25 | 6,29E-25 | 0,0006955 | 0,002378 | 0,0013988 | 0,0013988 | TRUE | 0,012086 | No |
| orf19.4033 | 587 | 459 | 619 | 335 | 1522 | 1214 | 1417 | 1957 | 534 | 464 | 586 | 503 | 1354 | 1318 | 1451 | 1493 | 962,94 | 522 | 1404 | 2,686 | 1,425 | 18,978 | 2,59E-80 | 1,84E-79 | 0,0047856 | 0,003409 | 0,0040579 | 0,0040579 | TRUE | 0,0821648 | No |
| orf19.4035 | 9465 | 7588 | 8482 | 5375 | 2652 | 1938 | 2161 | 3046 | 8617 | 7674 | 8027 | 8074 | 2359 | 2104 | 2213 | 2324 | 5174,14 | 8098 | 2250 | 0,278 | -1,847 | -36,138 | 5,82E-286 | 1,42E-284 | 0,0020907 | 0,002292 | 0,0022339 | 0,0022339 | TRUE | 0,0230487 | No |
| orf19.4036 | 762 | 643 | 732 | 470 | 1179 | 893 | 1102 | 1377 | 694 | 650 | 693 | 706 | 1049 | 970 | 1129 | 1050 | 867,51 | 686 | 1050 | 1,532 | 0,616 | 9,844 | 7,29E-23 | 1,97E-22 | 0,0015002 | 0,00356 | 0,0025281 | 0,0025281 | TRUE | 0,0294767 | No |
| orf19.4040 | 3025 | 2327 | 2603 | 1809 | 5698 | 4858 | 5309 | 7586 | 2754 | 2354 | 2464 | 2717 | 5068 | 5274 | 5437 | 5787 | 3981,87 | 2572 | 5392 | 2,098 | 1,069 | 17,366 | 1,49E-67 | 9,03E-67 | 0,0040697 | 0,002369 | 0,0033413 | 0,0033413 | TRUE | 0,0397719 | No |
| orf19.4041 | 793 | 618 | 691 | 460 | 1842 | 1325 | 1477 | 2071 | 722 | 625 | 654 | 691 | 1638 | 1439 | 1513 | 1580 | 1107,67 | 673 | 1542 | 2,295 | 1,198 | 18,619 | 2,24E-77 | 1,53E-76 | 0,002536 | 0,00323 | 0,0028778 | 0,0028778 | TRUE | 0,0317678 | No |
| orf19.4043 | 3422 | 2750 | 3215 | 2187 | 3599 | 2943 | 3385 | 4719 | 3116 | 2781 | 3043 | 3285 | 3201 | 3195 | 3467 | 3600 | 3210,96 | 3056 | 3366 | 1,102 | 0,141 | 2,324 | 0,020102 | 0,025764 | 0,003715 | 0,002448 | 0,0032056 | 0,0032056 | TRUE | 0,0443878 | No |
| orf19.4044 | 535 | 430 | 456 | 364 | 991 | 637 | 808 | 1160 | 487 | 435 | 432 | 547 | 881 | 692 | 827 | 885 | 648,22 | 475 | 821 | 1,738 | 0,798 | 8,311 | 9,49E-17 | 2,20E-16 | 0,0103363 | 0,004075 | 0,0071417 | 0,0071417 | TRUE | 0,1315447 | No |
| orf19.4045 | 6662 | 5359 | 5995 | 3656 | 6446 | 5320 | 6118 | 8012 | 6065 | 5420 | 5674 | 5492 | 5734 | 5776 | 6266 | 6112 | 5817,24 | 5663 | 5972 | 1,054 | 0,076 | 1,553 | 0,120355 | 0,14328 | 0,0020113 | 0,002264 | 0,0021319 | 0,0021319 | TRUE | 0,0282671 | No |
| orf19.4046 | 1070 | 804 | 824 | 575 | 749 | 622 | 689 | 944 | 974 | 813 | 780 | 864 | 666 | 675 | 706 | 720 | 774,77 | 858 | 692 | 0,807 | -0,31 | -4,112 | 3,91E-05 | 6,02E-05 | 0,0045361 | 0,003742 | 0,0041482 | 0,0041482 | TRUE | 0,1057704 | No |
| orf19.4048 | 11704 | 9201 | 10285 | 6809 | 8004 | 6067 | 7164 | 9757 | 10656 | 9306 | 9734 | 10228 | 7119 | 6587 | 7337 | 7443 | 8551,22 | 9981 | 7122 | 0,714 | -0,487 | -8,953 | 3,45E-19 | 8,56E-19 | 0,0029991 | 0,002191 | 0,0027179 | 0,0027179 | TRUE | 0,0309503 | No |
| orf19.405 | 974 | 818 | 911 | 595 | 2199 | 1752 | 1885 | 2597 | 887 | 827 | 862 | 894 | 1956 | 1902 | 1930 | 1981 | 1404,96 | 868 | 1942 | 2,243 | 1,165 | 24,293 | 2,33E-130 | 2,59E-129 | 1,00E-08 | 0,002977 | 0,001353 | 0,001353 | TRUE | 0,0115249 | No |
| orf19.4051 | 4386 | 3332 | 3890 | 2249 | 6350 | 5075 | 5229 | 7346 | 3993 | 3370 | 3682 | 3378 | 5648 | 5510 | 5355 | 5604 | 4567,5 | 3606 | 5529 | 1,532 | 0,616 | 10,854 | 1,90E-27 | 5,78E-27 | 0,0033431 | 0,002326 | 0,0028549 | 0,0028549 | TRUE | 0,0641979 | No |
| orf19.4054 | 560 | 399 | 521 | 310 | 786 | 612 | 703 | 933 | 510 | 404 | 493 | 466 | 699 | 664 | 720 | 712 | 583,42 | 468 | 699 | 1,492 | 0,577 | 7,46 | 8,68E-14 | 1,86E-13 | 0,00321 | 0,004301 | 0,0039351 | 0,0039351 | TRUE | 0,1055803 | No |
| orf19.4055 | 243 | 172 | 178 | 117 | 586 | 432 | 527 | 720 | 221 | 174 | 168 | 176 | 521 | 469 | 540 | 549 | 352,33 | 185 | 520 | 2,804 | 1,487 | 14,833 | 9,02E-50 | 4,14E-49 | 0,0062323 | 0,005786 | 0,0058835 | 0,0058835 | TRUE | 0,2073318 | No |
| orf19.4056 | 26344 | 22994 | 25417 | 17877 | 12151 | 10964 | 11834 | 16120 | 23985 | 23256 | 24055 | 26854 | 10808 | 11904 | 12120 | 12296 | 18159,7 | 24538 | 11782 | 0,48 | -1,058 | -18,756 | 1,73E-78 | 1,20E-77 | 0,0036169 | 0,00211 | 0,0029964 | 0,0029964 | TRUE | 0,0491826 | No |
| orf19.4057 | 14 | 11 | 10 | 7 | 15 | 18 | 11 | 24 | 13 | 11 | 9 | 11 | 13 | 20 | 11 | 18 | 13,29 | 11 | 16 | 1,422 | 0,508 | 1,338 | 0,181047 | 0,210463 | 1,00E-08 | 0,101445 | 0,0603476 | 0,0603476 | TRUE | 0,1525858 | No |
| orf19.4059 | 2198 | 1932 | 2172 | 1262 | 1763 | 1443 | 1687 | 2160 | 2001 | 1954 | 2056 | 1896 | 1568 | 1567 | 1728 | 1648 | 1802,09 | 1977 | 1628 | 0,822 | -0,282 | -5,673 | 1,41E-08 | 2,52E-08 | 0,0009576 | 0,00277 | 0,0018145 | 0,0018145 | TRUE | 0,0199335 | No |
| orf19.406 | 2738 | 2393 | 2712 | 1778 | 3016 | 2688 | 2658 | 3714 | 2493 | 2420 | 2567 | 2671 | 2683 | 2918 | 2722 | 2833 | 2663,36 | 2538 | 2789 | 1,1 | 0,138 | 2,935 | 0,00333 | 0,004515 | 0,0011265 | 0,002533 | 0,0017336 | 0,0017336 | TRUE | 0,013994 | No |
| orf19.4060 | 6784 | 5642 | 5990 | 4353 | 9384 | 7352 | 7985 | 10942 | 6177 | 5706 | 5669 | 6539 | 8347 | 7982 | 8178 | 8347 | 7118 | 6023 | 8214 | 1,365 | 0,449 | 8,926 | 4,42E-19 | 1,09E-18 | 0,0023991 | 0,002223 | 0,0022809 | 0,0022809 | TRUE | 0,0394624 | No |
| orf19.4061 | 612 | 591 | 574 | 425 | 893 | 776 | 769 | 1000 | 557 | 598 | 543 | 638 | 794 | 843 | 788 | 763 | 690,47 | 584 | 797 | 1,369 | 0,453 | 6,724 | 1,77E-11 | 3,53E-11 | 0,001599 | 0,00395 | 0,002847 | 0,002847 | TRUE | 0,0396159 | No |
| orf19.4062 | 3227 | 2851 | 2980 | 1732 | 2812 | 2208 | 2486 | 3268 | 2938 | 2883 | 2820 | 2602 | 2501 | 2397 | 2546 | 2493 | 2647,61 | 2811 | 2484 | 0,883 | -0,18 | -3,814 | 0,000137 | 0,000203 | 0,0012186 | 0,002536 | 0,0017658 | 0,0017658 | TRUE | 0,0283199 | No |
| orf19.4063 | 243 | 196 | 228 | 150 | 498 | 450 | 445 | 675 | 221 | 198 | 216 | 225 | 443 | 489 | 456 | 515 | 345,34 | 215 | 476 | 2,217 | 1,149 | 12,98 | 1,59E-38 | 6,06E-38 | 1,00E-08 | 0,005862 | 0,0040597 | 0,0040597 | TRUE | 0,0395616 | No |
| orf19.4064 | 715 | 670 | 714 | 415 | 1007 | 891 | 894 | 1131 | 651 | 678 | 676 | 623 | 896 | 967 | 916 | 863 | 783,64 | 657 | 910 | 1,38 | 0,465 | 7,635 | 2,26E-14 | 4,91E-14 | 0,0007065 | 0,003723 | 0,0022353 | 0,0022353 | TRUE | 0,0206399 | No |
| orf19.4066 | 1124 | 1439 | 1401 | 873 | 3950 | 3736 | 3737 | 4524 | 1023 | 1455 | 1326 | 1311 | 3513 | 4056 | 3827 | 3451 | 2495,48 | 1279 | 3712 | 2,903 | 1,538 | 16,745 | 6,21E-63 | 3,54E-62 | 0,0123543 | 0,002566 | 0,0075523 | 0,0075523 | TRUE | 0,2229753 | No |
| orf19.4067 | 3 | 6 | 3 | 0 | 2 | 5 | 3 | 8 | 3 | 6 | 3 | 0 | 2 | 5 | 3 | 6 | 3,5 | 3 | 4 | 1,345 | 0,428 | 0,549 | 0,583065 | 0,62356 | 0,105134 | 0,379177 | 0,2957347 | 0,2957347 | TRUE | 0,356225 | No |
| orf19.4068 | 561 | 524 | 563 | 362 | 975 | 824 | 811 | 1115 | 511 | 530 | 533 | 544 | 867 | 895 | 831 | 851 | 695,04 | 530 | 861 | 1,628 | 0,703 | 11,757 | 6,50E-32 | 2,18E-31 | 1,00E-08 | 0,003938 | 0,0018853 | 0,0018853 | TRUE | 0,0077729 | No |
| orf19.4069 | 55 | 67 | 54 | 46 | 125 | 100 | 104 | 146 | 50 | 68 | 51 | 69 | 111 | 109 | 107 | 111 | 84,46 | 60 | 110 | 1,865 | 0,899 | 5,642 | 1,68E-08 | 2,99E-08 | 1,00E-08 | 0,017678 | 0,0110971 | 0,0110971 | TRUE | 0,1014998 | No |
| orf19.407 | 4134 | 3390 | 3744 | 2269 | 3291 | 2697 | 3146 | 4158 | 3764 | 3429 | 3543 | 3408 | 2927 | 2928 | 3222 | 3172 | 3299,16 | 3536 | 3062 | 0,866 | -0,208 | -4,068 | 4,74E-05 | 7,23E-05 | 0,0019666 | 0,002437 | 0,0022105 | 0,0022105 | TRUE | 0,0231619 | No |
| orf19.4070 | 258 | 234 | 222 | 146 | 383 | 284 | 307 | 483 | 235 | 237 | 210 | 219 | 341 | 308 | 314 | 368 | 279,1 | 225 | 333 | 1,482 | 0,567 | 5,983 | 2,19E-09 | 4,04E-09 | 0,0024268 | 0,00677 | 0,004863 | 0,004863 | TRUE | 0,062311 | No |
| orf19.4072 | 3314 | 2573 | 2971 | 2006 | 3137 | 2584 | 3156 | 4607 | 3017 | 2602 | 2812 | 3013 | 2790 | 2806 | 3232 | 3514 | 2973,36 | 2861 | 3086 | 1,079 | 0,11 | 1,394 | 0,163266 | 0,191292 | 0,0084251 | 0,002481 | 0,0056321 | 0,0056321 | TRUE | 0,1076295 | No |
| orf19.4076 | 8342 | 7612 | 8011 | 5007 | 20752 | 16659 | 17834 | 24208 | 7595 | 7699 | 7582 | 7521 | 18458 | 18087 | 18264 | 18466 | 12959 | 7599 | 18319 | 2,41 | 1,269 | 47,769 | 0 | 0 | 1,00E-08 | 0,002139 | 0,0005815 | 0,0005815 | TRUE | 0,0009403 | No |

| | | | | | | | | | | | | | | | | | | | | | | | | | | | | | | | |
|---|---|---|---|---|---|---|---|---|---|---|---|---|---|---|---|---|---|---|---|---|---|---|---|---|---|---|---|---|---|---|---|
| orf19.4077 | 12385 | 11968 | 12832 | 7787 | 9261 | 7847 | 8282 | 10619 | 11276 | 12104 | 12144 | 11697 | 8237 | 8520 | 8482 | 8100 | 10070,1 | 11805 | 8335 | 0,706 | -0,503 | -13,356 | 1,09E-40 | 4,28E-40 | 0,0007455 | 0,002168 | 0,0012583 | 0,0012583 | TRUE | 0,0113789 | No |
| orf19.4078 | 3819 | 3672 | 3739 | 2374 | 2881 | 2441 | 2603 | 3377 | 3477 | 3714 | 3539 | 3566 | 2563 | 2650 | 2666 | 2576 | 3093,78 | 3574 | 2614 | 0,731 | -0,452 | -11,617 | 3,37E-31 | 1,11E-30 | 0,0003214 | 0,002464 | 0,0011236 | 0,0011236 | TRUE | 0,0087134 | No |
| orf19.4079 | 1517 | 1427 | 1537 | 1010 | 1260 | 1209 | 1136 | 1539 | 1381 | 1443 | 1455 | 1517 | 1121 | 1313 | 1163 | 1174 | 1320,87 | 1449 | 1193 | 0,823 | -0,281 | -4,743 | 2,11E-06 | 3,45E-06 | 0,0021543 | 0,003037 | 0,0025956 | 0,0025956 | TRUE | 0,0530251 | No |
| orf19.4080 | 781 | 859 | 993 | 523 | 883 | 807 | 805 | 1020 | 711 | 869 | 940 | 786 | 785 | 876 | 824 | 778 | 821,17 | 826 | 816 | 0,986 | -0,021 | -0,244 | 0,80751 | 0,832721 | 0,0076745 | 0,003646 | 0,0056418 | 0,0056418 | TRUE | 0,1117399 | No |
| orf19.4081 | 3 | 0 | 0 | 0 | 1 | 0 | 4 | 0 | 3 | 0 | 0 | 0 | 1 | 0 | 4 | 0 | 0,96 | 1 | 1 | 1,676 | 0,745 | 0,407 | 0,683771 | 0,719194 | 5,2940903 | 1,371417 | 2,1068165 | 2,1068165 | TRUE | 1,4381607 | No |
| orf19.4082 | 7723 | 6095 | 6396 | 4586 | 2508 | 1987 | 2055 | 2939 | 7031 | 6164 | 6053 | 6889 | 2231 | 2157 | 2105 | 2242 | 4359,08 | 6534 | 2184 | 0,334 | -1,58 | -27,527 | 8,44E-167 | 1,23E-165 | 0,0029245 | 0,00234 | 0,0028692 | 0,0028692 | TRUE | 0,0327798 | No |
| orf19.4084 | 4276 | 3680 | 4179 | 3006 | 2554 | 2312 | 2415 | 3312 | 3893 | 3722 | 3955 | 4515 | 2272 | 2510 | 2473 | 2526 | 3233,39 | 4021 | 2445 | 0,609 | -0,716 | -11,336 | 8,70E-30 | 2,78E-29 | 0,0043793 | 0,002446 | 0,0035082 | 0,0035082 | TRUE | 0,081004 | No |
| orf19.4085 | 345 | 379 | 342 | 186 | 282 | 239 | 239 | 339 | 314 | 383 | 324 | 279 | 251 | 259 | 245 | 259 | 289,27 | 325 | 254 | 0,776 | -0,366 | -3,675 | 0,000238 | 0,00035 | 0,0052658 | 0,006604 | 0,0060378 | 0,0060378 | TRUE | 0,1606917 | No |
| orf19.4086 | 716 | 613 | 557 | 411 | 311 | 228 | 239 | 346 | 652 | 620 | 527 | 617 | 277 | 248 | 245 | 264 | 431,16 | 604 | 258 | 0,429 | -1,222 | -13,924 | 4,53E-44 | 1,90E-43 | 0,0042759 | 0,005101 | 0,004708 | 0,004708 | TRUE | 0,0910999 | No |
| orf19.4088 | 852 | 768 | 777 | 548 | 813 | 668 | 730 | 969 | 776 | 777 | 735 | 823 | 723 | 725 | 748 | 739 | 755,77 | 778 | 734 | 0,946 | -0,08 | -1,376 | 0,168881 | 0,197218 | 1,00E-08 | 0,003785 | 0,0018947 | 0,0018947 | TRUE | 0,0157488 | No |
| orf19.4089 | 4572 | 3903 | 4305 | 2609 | 4126 | 3152 | 3649 | 4898 | 4163 | 3947 | 4074 | 3919 | 3670 | 3422 | 3737 | 3736 | 3833,61 | 4026 | 3641 | 0,904 | -0,145 | -3,376 | 0,000736 | 0,001048 | 0,0008903 | 0,002382 | 0,0015101 | 0,0015101 | TRUE | 0,0195091 | No |
| orf19.409 | 53612 | 43511 | 47000 | 30771 | 19595 | 15689 | 16259 | 22959 | 48811 | 44007 | 44481 | 46223 | 17429 | 17034 | 16651 | 17513 | 31518,8 | 45880 | 17157 | 0,374 | -1,419 | -34,321 | 3,76E-258 | 8,12E-257 | 0,0013363 | 0,002079 | 0,0016032 | 0,0016032 | TRUE | 0,0227492 | No |
| orf19.4090 | 132 | 120 | 135 | 102 | 245 | 223 | 196 | 262 | 120 | 121 | 128 | 153 | 218 | 242 | 201 | 200 | 172,89 | 130 | 215 | 1,66 | 0,731 | 6,156 | 7,44E-10 | 1,40E-09 | 0,0030034 | 0,009678 | 0,0072492 | 0,0072492 | TRUE | 0,1079773 | No |
| orf19.4090.1 | 523 | 385 | 422 | 270 | 747 | 488 | 508 | 776 | 476 | 389 | 399 | 406 | 664 | 530 | 520 | 592 | 497,12 | 418 | 576 | 1,381 | 0,465 | 4,834 | 1,33E-06 | 2,20E-06 | 0,009129 | 0,004694 | 0,0068063 | 0,0068063 | TRUE | 0,123018 | No |
| orf19.4091 | 767 | 631 | 754 | 427 | 453 | 320 | 394 | 494 | 698 | 638 | 714 | 641 | 403 | 347 | 404 | 377 | 527,78 | 673 | 383 | 0,568 | -0,816 | -10,926 | 8,70E-28 | 2,66E-27 | 0,0018311 | 0,00454 | 0,0033444 | 0,0033444 | TRUE | 0,0401572 | No |
| orf19.4092 | 846 | 754 | 743 | 482 | 759 | 534 | 579 | 874 | 770 | 763 | 703 | 724 | 675 | 580 | 593 | 667 | 684,33 | 740 | 629 | 0,851 | -0,233 | -3,31 | 0,000934 | 0,00132 | 0,0025463 | 0,003967 | 0,003283 | 0,003283 | TRUE | 0,0315045 | No |
| orf19.4093 | 18677 | 16417 | 18002 | 10855 | 28994 | 22670 | 25517 | 33760 | 17005 | 16604 | 17037 | 16306 | 25789 | 24613 | 26133 | 25752 | 21154,9 | 16738 | 25572 | 1,528 | 0,611 | 18,711 | 4,02E-78 | 2,79E-77 | 0,0004861 | 0,002099 | 0,0009744 | 0,0009744 | TRUE | 0,007746 | No |
| orf19.4094 | 60 | 65 | 59 | 55 | 22 | 26 | 19 | 28 | 55 | 66 | 56 | 83 | 20 | 28 | 19 | 21 | 43,43 | 65 | 22 | 0,345 | -1,536 | -6,459 | 1,05E-10 | 2,04E-10 | 1,00E-08 | 0,032454 | 0,024804 | 0,024804 | TRUE | 0,2387998 | No |
| orf19.4095 | 6 | 7 | 1 | 7 | 5 | 3 | 5 | 1 | 5 | 7 | 1 | 11 | 4 | 3 | 5 | 1 | 4,7 | 6 | 3 | 0,574 | -0,801 | -1,08 | 0,280128 | 0,316634 | 0,319283 | 0,28316 | 0,2954264 | 0,2954264 | TRUE | 0,2284054 | No |
| orf19.4096 | 1983 | 1653 | 1886 | 1175 | 1640 | 1327 | 1435 | 1991 | 1805 | 1672 | 1785 | 1765 | 1459 | 1441 | 1470 | 1519 | 1614,39 | 1757 | 1472 | 0,838 | -0,254 | -5,515 | 3,48E-08 | 6,12E-08 | 1,00E-08 | 0,002855 | 0,0014202 | 0,0014202 | TRUE | 0,0132946 | No |
| orf19.4097 | 939 | 961 | 942 | 579 | 1425 | 1227 | 1380 | 1685 | 855 | 972 | 892 | 870 | 1267 | 1332 | 1413 | 1285 | 1110,8 | 897 | 1324 | 1,475 | 0,56 | 9,286 | 1,61E-20 | 4,13E-20 | 0,0019078 | 0,003226 | 0,0025443 | 0,0025443 | TRUE | 0,0383495 | No |
| orf19.4099 | 14727 | 12743 | 13838 | 8735 | 27010 | 22930 | 24295 | 32859 | 13408 | 12888 | 13096 | 13121 | 24025 | 24896 | 24881 | 25065 | 18922,6 | 13128 | 24717 | 1,883 | 0,913 | 30,992 | 6,89E-211 | 1,23E-209 | 0,0002348 | 0,002107 | 0,0007729 | 0,0007729 | TRUE | 0,0043679 | No |
| orf19.4101 | 265 | 169 | 207 | 134 | 191 | 158 | 163 | 236 | 241 | 171 | 196 | 201 | 170 | 172 | 167 | 180 | 187,22 | 202 | 172 | 0,85 | -0,235 | -1,988 | 0,046848 | 0,058127 | 0,0066394 | 0,009093 | 0,0080446 | 0,0080446 | TRUE | 0,2003982 | No |
| orf19.4102 | 5541 | 4865 | 5162 | 3595 | 3860 | 3027 | 3134 | 4540 | 5045 | 4920 | 4885 | 5400 | 3433 | 3286 | 3210 | 3463 | 4205,44 | 5062 | 3348 | 0,662 | -0,595 | -12,92 | 3,45E-38 | 1,30E-37 | 0,0013948 | 0,002351 | 0,0017877 | 0,0017877 | TRUE | 0,0235692 | No |
| orf19.410.3 | 1153 | 931 | 1045 | 717 | 328 | 266 | 267 | 411 | 1050 | 942 | 989 | 1077 | 292 | 289 | 273 | 314 | 653,11 | 1014 | 292 | 0,289 | -1,791 | -24,407 | 1,43E-131 | 1,61E-130 | 0,0018925 | 0,00406 | 0,0030479 | 0,0030479 | TRUE | 0,0311243 | No |
| orf19.4104 | 148 | 131 | 137 | 74 | 129 | 109 | 133 | 161 | 135 | 132 | 130 | 111 | 115 | 118 | 136 | 123 | 125,02 | 127 | 123 | 0,96 | -0,058 | -0,461 | 0,645129 | 0,682605 | 1,00E-08 | 0,012603 | 0,00738 | 0,00738 | TRUE | 0,0542943 | No |
| orf19.4105 | 769 | 666 | 698 | 442 | 629 | 520 | 509 | 728 | 700 | 674 | 661 | 664 | 559 | 565 | 521 | 555 | 612,37 | 675 | 550 | 0,815 | -0,295 | -4,761 | 1,92E-06 | 3,16E-06 | 1,00E-08 | 0,004194 | 0,0020456 | 0,0020456 | TRUE | 0,0138232 | No |
| orf19.4106 | 1269 | 1164 | 1303 | 794 | 1245 | 1114 | 1051 | 1350 | 1155 | 1177 | 1233 | 1193 | 1107 | 1209 | 1076 | 1030 | 1147,69 | 1190 | 1106 | 0,928 | -0,108 | -1,815 | 0,06958 | 0,084975 | 0,0018875 | 0,003188 | 0,0025025 | 0,0025025 | TRUE | 0,0463875 | No |
| orf19.4107 | 1744 | 1442 | 1621 | 952 | 1633 | 1267 | 1407 | 1818 | 1588 | 1458 | 1534 | 1430 | 1453 | 1376 | 1441 | 1387 | 1458,29 | 1502 | 1414 | 0,939 | -0,091 | -1,789 | 0,073551 | 0,089563 | 0,000776 | 0,002943 | 0,001771 | 0,001771 | TRUE | 0,0176677 | No |
| orf19.4109 | 1223 | 1071 | 1108 | 754 | 856 | 715 | 781 | 1082 | 1113 | 1083 | 1049 | 1133 | 761 | 776 | 800 | 825 | 942,6 | 1094 | 790 | 0,724 | -0,466 | -8,48 | 2,26E-17 | 5,35E-17 | 1,00E-08 | 0,003438 | 0,0018184 | 0,0018184 | TRUE | 0,0113543 | No |
| orf19.411 | 19940 | 17605 | 18706 | 10985 | 8349 | 6677 | 7423 | 9610 | 18155 | 17806 | 17704 | 16501 | 7426 | 7249 | 7602 | 7331 | 12471,6 | 17542 | 7402 | 0,422 | -1,245 | -31,806 | 5,32E-222 | 9,92E-221 | 0,0009236 | 0,002143 | 0,0013784 | 0,0013784 | TRUE | 0,0191674 | No |
| orf19.4110 | 1805 | 1613 | 1806 | 1209 | 2499 | 2139 | 2358 | 3077 | 1643 | 1631 | 1709 | 1816 | 2223 | 2322 | 2415 | 2347 | 2013,41 | 1700 | 2327 | 1,371 | 0,456 | 9,242 | 2,43E-20 | 6,22E-20 | 0,0011455 | 0,002693 | 0,0018148 | 0,0018148 | TRUE | 0,0230718 | No |
| orf19.4112 | 1806 | 1475 | 1580 | 1034 | 2075 | 1626 | 1875 | 2450 | 1644 | 1492 | 1495 | 1553 | 1846 | 1765 | 1920 | 1869 | 1698,1 | 1546 | 1850 | 1,197 | 0,259 | 5,133 | 2,86E-07 | 4,85E-07 | 0,0010724 | 0,002815 | 0,0018457 | 0,0018457 | TRUE | 0,0236148 | No |
| orf19.4114 | 2387 | 2152 | 2370 | 1423 | 14572 | 11951 | 12870 | 17627 | 2173 | 2177 | 2243 | 2138 | 12961 | 12975 | 13181 | 13446 | 7661,71 | 2183 | 13141 | 6,016 | 2,589 | 75,133 | 0 | 0 | 1,00E-08 | 0,002209 | 0,0008591 | 0,0008591 | TRUE | 0,0042435 | No |
| orf19.4115 | 16 | 8 | 8 | 4 | 19 | 18 | 6 | 6 | 15 | 8 | 8 | 6 | 17 | 20 | 6 | 5 | 10,43 | 9 | 12 | 1,261 | 0,335 | 0,634 | 0,526242 | 0,567936 | 0,2291351 | 0,128747 | 0,1702574 | 0,1702574 | TRUE | 0,1441194 | No |
| orf19.4116 | 716 | 589 | 641 | 421 | 638 | 529 | 539 | 775 | 652 | 596 | 607 | 632 | 567 | 574 | 552 | 591 | 596,46 | 622 | 571 | 0,92 | -0,12 | -1,89 | 0,058701 | 0,072229 | 1,00E-08 | 0,004252 | 0,0021995 | 0,0021995 | TRUE | 0,0144602 | No |
| orf19.4117 | 1557 | 1450 | 1581 | 898 | 1018 | 770 | 811 | 1155 | 1418 | 1467 | 1496 | 1349 | 905 | 836 | 831 | 881 | 1147,8 | 1432 | 863 | 0,602 | -0,731 | -13,235 | 5,49E-40 | 2,13E-39 | 0,0008782 | 0,003188 | 0,0020159 | 0,0020159 | TRUE | 0,0164747 | No |
| orf19.4118 | 28153 | 26406 | 27868 | 17511 | 40476 | 35268 | 35078 | 48111 | 25632 | 26707 | 26375 | 26304 | 36002 | 38291 | 35924 | 36699 | 31491,9 | 26254 | 36729 | 1,399 | 0,484 | 14,749 | 3,12E-49 | 1,42E-48 | 0,0005506 | 0,002079 | 0,0010027 | 0,0010027 | TRUE | 0,0099933 | No |
| orf19.4119 | 5369 | 5083 | 5394 | 3191 | 3550 | 3013 | 3149 | 4115 | 4888 | 5141 | 5105 | 4793 | 3158 | 3271 | 3225 | 3139 | 4090,04 | 4982 | 3198 | 0,641 | -0,641 | -16,548 | 1,65E-61 | 9,21E-61 | 0,0004094 | 0,00236 | 0,0011861 | 0,0011861 | TRUE | 0,0074048 | No |
| orf19.412 | 2540 | 2879 | 2684 | 1470 | 3577 | 3267 | 3018 | 3966 | 2313 | 2912 | 2540 | 2208 | 3182 | 3547 | 3091 | 3025 | 2852,19 | 2493 | 3211 | 1,287 | 0,364 | 4,434 | 9,27E-06 | 1,47E-05 | 0,0099657 | 0,0025 | 0,0060998 | 0,0060998 | TRUE | 0,1546792 | No |
| orf19.4120 | 6174 | 5406 | 5653 | 3595 | 5028 | 4174 | 4314 | 6009 | 5621 | 5468 | 5350 | 5400 | 4472 | 4532 | 4418 | 4584 | 4980,61 | 5460 | 4502 | 0,824 | -0,279 | -8,249 | 1,60E-16 | 3,68E-16 | 0,0001948 | 0,002302 | 0,0008927 | 0,0008927 | TRUE | 0,0049792 | No |
| orf19.4121 | 1907 | 1599 | 1766 | 1102 | 4051 | 3345 | 3396 | 4856 | 1736 | 1617 | 1671 | 1655 | 3603 | 3632 | 3478 | 3704 | 2637,16 | 1670 | 3604 | 2,157 | 1,109 | 26,215 | 1,81E-151 | 2,38E-150 | 0,0004162 | 0,002538 | 0,0012687 | 0,0012687 | TRUE | 0,0091053 | No |
| orf19.4122 | 1979 | 1839 | 1890 | 1267 | 5772 | 4578 | 4900 | 6728 | 1802 | 1860 | 1789 | 1903 | 5134 | 4970 | 5018 | 5132 | 3451,07 | 1838 | 5064 | 2,76 | 1,465 | 38,337 | 0 | 0 | 5,63E-05 | 0,00242 | 0,0010169 | 0,0010169 | TRUE | 0,0059812 | No |
| orf19.4123 | 3394 | 2973 | 3285 | 2003 | 3251 | 2654 | 2859 | 3751 | 3090 | 3007 | 3109 | 3009 | 2892 | 2882 | 2928 | 2861 | 2972,15 | 3054 | 2891 | 0,946 | -0,08 | -2,271 | 0,023173 | 0,029551 | 1,00E-08 | 0,002481 | 0,0008642 | 0,0008642 | TRUE | 0,001789 | No |
| orf19.4125 | 3310 | 3100 | 3241 | 2025 | 3926 | 3400 | 3425 | 4453 | 3014 | 3135 | 3067 | 3042 | 3492 | 3691 | 3508 | 3397 | 3293,26 | 3064 | 3522 | 1,148 | 0,2 | 5,004 | 5,62E-07 | 9,45E-07 | 0,0004955 | 0,002438 | 0,0012205 | 0,0012205 | TRUE | 0,0125317 | No |
| orf19.4127 | 10413 | 8902 | 9841 | 5774 | 2911 | 2423 | 2551 | 3480 | 9481 | 9003 | 9314 | 8673 | 2589 | 2631 | 2613 | 2655 | 5869,77 | 9118 | 2622 | 0,287 | -1,799 | -45,341 | 0 | 0 | 0,0007545 | 0,002262 | 0,0012742 | 0,0012742 | TRUE | 0,0127375 | No |
| orf19.4128 | 1045 | 955 | 1083 | 594 | 7869 | 6273 | 6630 | 8535 | 951 | 966 | 1025 | 892 | 6999 | 6811 | 6790 | 6511 | 3868,13 | 958 | 6778 | 7,047 | 2,817 | 57,813 | 0 | 0 | 0,0012629 | 0,002379 | 0,0016541 | 0,0016541 | TRUE | 0,0254812 | No |
| orf19.413 | 4678 | 4447 | 4784 | 2895 | 734 | 640 | 621 | 877 | 4259 | 4498 | 4528 | 4349 | 653 | 695 | 636 | 669 | 2535,73 | 4408 | 663 | 0,15 | -2,733 | -56,769 | 0 | 0 | 1,00E-08 | 0,002558 | 0,0014018 | 0,0014018 | TRUE | 0,0108457 | No |
| orf19.4131 | 975 | 1049 | 1024 | 602 | 1141 | 915 | 960 | 1267 | 888 | 1061 | 969 | 904 | 1015 | 993 | 983 | 966 | 972,5 | 956 | 989 | 1,034 | 0,048 | 0,757 | 0,44894 | 0,490991 | 0,0022195 | 0,003395 | 0,002902 | 0,002902 | TRUE | 0,0663353 | No |
| orf19.413.1 | 4665 | 4937 | 4999 | 2167 | 3189 | 2267 | 2573 | 3098 | 4247 | 4993 | 4731 | 3255 | 2837 | 2461 | 2635 | 2363 | 3440,36 | 4306 | 2574 | 0,597 | -0,743 | -5,287 | 1,24E-07 | 2,13E-07 | 0,0186722 | 0,002421 | 0,0106404 | 0,0186722 | TRUE | 0,3263644 | No |
| orf19.4132 | 2179 | 1969 | 1975 | 1178 | 2086 | 1515 | 1687 | 2216 | 1984 | 1991 | 1869 | 1770 | 1855 | 1645 | 1728 | 1690 | 1816,55 | 1904 | 1730 | 0,908 | -0,14 | -2,489 | 0,012805 | 0,016659 | 0,0021649 | 0,002764 | 0,0024843 | 0,0024843 | TRUE | 0,0292117 | No |
| orf19.4133 | 2890 | 2478 | 2771 | 1673 | 2477 | 1861 | 2246 | 2958 | 2631 | 2506 | 2623 | 2513 | 2203 | 2021 | 2300 | 2256 | 2381,67 | 2568 | 2195 | 0,855 | -0,226 | -4,566 | 4,97E-06 | 8,02E-06 | 0,0013536 | 0,002592 | 0,00194 | 0,00194 | TRUE | 0,0335901 | No |
| orf19.4134 | 912 | 880 | 937 | 587 | 812 | 632 | 648 | 889 | 830 | 890 | 887 | 882 | 722 | 686 | 664 | 678 | 779,89 | 872 | 688 | 0,789 | -0,342 | -5,883 | 4,02E-09 | 7,34E-09 | 1,00E-08 | 0,003731 | 0,0019446 | 0,0019446 | TRUE | 0,0139678 | No |
| orf19.4135 | 946 | 863 | 814 | 562 | 708 | 608 | 568 | 884 | 861 | 873 | 770 | 844 | 630 | 660 | 582 | 674 | 736,83 | 837 | 636 | 0,761 | -0,393 | -5,754 | 8,71E-09 | 1,58E-08 | 0,0023597 | 0,00383 | 0,0031111 | 0,0031111 | TRUE | 0,0384917 | No |
| orf19.4136 | 385 | 397 | 415 | 231 | 315 | 293 | 297 | 394 | 351 | 402 | 393 | 347 | 280 | 318 | 304 | 301 | 336,85 | 373 | 301 | 0,803 | -0,316 | -3,711 | 0,000206 | 0,000304 | 0,0012186 | 0,005959 | 0,003993 | 0,003993 | TRUE | 0,0294928 | No |
| orf19.414 | 6693 | 5998 | 6511 | 3678 | 5976 | 4813 | 5249 | 6822 | 6094 | 6066 | 6162 | 5525 | 5315 | 5226 | 5376 | 5204 | 5620,95 | 5962 | 5280 | 0,885 | -0,177 | -4,171 | 3,03E-05 | 4,68E-05 | 0,0010923 | 0,002272 | 0,0015429 | 0,0015429 | TRUE | 0,0284426 | No |
| orf19.4142 | 681 | 597 | 578 | 403 | 1019 | 823 | 933 | 1164 | 620 | 604 | 547 | 605 | 906 | 894 | 956 | 888 | 752,44 | 594 | 911 | 1,534 | 0,617 | 9,93 | 3,08E-23 | 8,43E-23 | 0,0005611 | 0,003793 | 0,0022951 | 0,0022951 | TRUE | 0,0358376 | No |
| orf19.4143 | 1748 | 1455 | 1615 | 981 | 1473 | 1050 | 1199 | 1613 | 1591 | 1472 | 1528 | 1474 | 1310 | 1140 | 1228 | 1230 | 1371,71 | 1516 | 1227 | 0,809 | -0,305 | -5,518 | 3,43E-08 | 6,03E-08 | 0,0015087 | 0,003 | 0,0022064 | 0,0022064 | TRUE | 0,0262822 | No |
| orf19.4144 | 3381 | 2954 | 3185 | 2139 | 1571 | 1373 | 1337 | 1871 | 3078 | 2988 | 3014 | 3213 | 1397 | 1491 | 1369 | 1427 | 2247,23 | 3073 | 1421 | 0,463 | -1,112 | -24,519 | 9,26E-133 | 1,06E-131 | 0,0005213 | 0,002625 | 0,0014702 | 0,0014702 | TRUE | 0,0122253 | No |
| orf19.4145 | 2624 | 2462 | 2615 | 1556 | 2986 | 2424 | 2557 | 3378 | 2389 | 2490 | 2475 | 2337 | 2656 | 2632 | 2619 | 2577 | 2521,81 | 2423 | 2621 | 1,08 | 0,111 | 2,839 | 0,00452 | 0,006076 | 1,00E-08 | 0,002561 | 0,0010806 | 0,0010806 | TRUE | 0,0060475 | No |
| orf19.4146 | 1416 | 1204 | 1192 | 617 | 1311 | 887 | 977 | 1423 | 1289 | 1218 | 1128 | 927 | 1166 | 963 | 1001 | 1085 | 1097,13 | 1140 | 1054 | 0,922 | -0,117 | -1,239 | 0,215176 | 0,247921 | 0,011955 | 0,003241 | 0,007595 | 0,007595 | TRUE | 0,1802195 | No |
| orf19.4147 | 2664 | 2430 | 2554 | 1686 | 2870 | 2381 | 2361 | 3367 | 2425 | 2458 | 2417 | 2533 | 2553 | 2585 | 2418 | 2568 | 2494,64 | 2458 | 2531 | 1,031 | 0,044 | 1,078 | 0,281143 | 0,317612 | 0,000233 | 0,002567 | 0,0011919 | 0,0011919 | TRUE | 0,0106994 | No |
| orf19.4148 | 557 | 522 | 558 | 327 | 378 | 330 | 351 | 455 | 507 | 528 | 528 | 491 | 336 | 358 | 359 | 347 | 431,93 | 514 | 350 | 0,68 | -0,557 | -7,821 | 5,26E-15 | 1,16E-14 | 1,00E-08 | 0,005095 | 0,0025005 | 0,0025005 | TRUE | 0,0082249 | No |
| orf19.4149 | 17 | 16 | 11 | 14 | 22 | 18 | 19 | 19 | 15 | 16 | 10 | 21 | 20 | 20 | 19 | 14 | 17,02 | 16 | 18 | 1,176 | 0,234 | 0,692 | 0,488924 | 0,530625 | 1,00E-08 | 0,079649 | 0,0494547 | 0,0494547 | TRUE | 0,2597515 | No |
| orf19.4149.1 | 14018 | 11936 | 12490 | 8860 | 23322 | 19195 | 19102 | 27056 | 12763 | 12072 | 11821 | 13309 | 20744 | 20840 | 19563 | 20639 | 16468,8 | 12491 | 20446 | 1,637 | 0,711 | 15,712 | 1,25E-55 | 6,40E-55 | 0,0017986 | 0,002117 | 0,0019036 | 0,0019036 | TRUE | 0,0233645 | No |
| orf19.415 | 1405 | 1155 | 1247 | 853 | 1623 | 1245 | 1390 | 1947 | 1279 | 1168 | 1180 | 1281 | 1444 | 1352 | 1424 | 1485 | 1326,61 | 1227 | 1426 | 1,165 | 0,22 | 4,047 | 5,19E-05 | 7,90E-05 | 0,0012161 | 0,003033 | 0,0020678 | 0,0020678 | TRUE | 0,0144221 | No |
| orf19.4150 | 7482 | 6617 | 6985 | 4566 | 6485 | 5372 | 5693 | 7575 | 6812 | 6692 | 6611 | 6859 | 5768 | 5832 | 5830 | 5778 | 6272,91 | 6744 | 5802 | 0,861 | -0,217 | -7,243 | 4,40E-13 | 9,25E-13 | 1,00E-08 | 0,002248 | 0,0006976 | 0,0006976 | TRUE | 0,0021576 | No |

| | | | | | | | | | | | | | | | | | | | | | | | | | | | | | | | |
|---|---|---|---|---|---|---|---|---|---|---|---|---|---|---|---|---|---|---|---|---|---|---|---|---|---|---|---|---|---|---|---|
| orf19.4151 | 111 | 107 | 103 | 66 | 164 | 134 | 151 | 215 | 101 | 108 | 97 | 99 | 146 | 145 | 155 | 164 | 126,99 | 101 | 152 | 1,505 | 0,59 | 4,729 | 2,26E-06 | 3,69E-06 | 1,00E-08 | 0,01244 | 0,0066512 | 0,0066512 | TRUE | 0,0307763 | No |
| orf19.4152 | 59685 | 52446 | 56865 | 37633 | 91135 | 79350 | 81025 | 1E+05 | 54341 | 53044 | 53818 | 56531 | 81062 | 86152 | 82980 | 87069 | 69374,4 | 54434 | 84316 | 1,549 | 0,631 | 17,301 | 4,59E-67 | 2,77E-66 | 0,0009046 | 0,002056 | 0,0012643 | 0,0012643 | TRUE | 0,0082807 | No |
| orf19.4153 | 898 | 1074 | 1080 | 581 | 464 | 428 | 444 | 486 | 818 | 1086 | 1022 | 873 | 413 | 465 | 455 | 371 | 687,69 | 950 | 426 | 0,447 | -1,162 | -11,479 | 1,69E-30 | 5,51E-30 | 0,0127897 | 0,003958 | 0,0081783 | 0,0081783 | TRUE | 0,1051895 | No |
| orf19.4157 | 403 | 331 | 348 | 237 | 2060 | 1795 | 1729 | 2636 | 367 | 335 | 329 | 356 | 1832 | 1949 | 1771 | 2011 | 1118,71 | 347 | 1891 | 5,46 | 2,449 | 35,798 | 1,18E-280 | 2,78E-279 | 0,0022294 | 0,003218 | 0,0027083 | 0,0027083 | TRUE | 0,0229646 | No |
| orf19.4159 | 1746 | 1500 | 1655 | 1075 | 2168 | 1875 | 1885 | 2724 | 1590 | 1517 | 1566 | 1615 | 1928 | 2036 | 1930 | 2078 | 1782,54 | 1572 | 1993 | 1,269 | 0,344 | 7,352 | 1,95E-13 | 4,13E-13 | 0,0006061 | 0,002778 | 0,0015273 | 0,0015273 | TRUE | 0,0105429 | No |
| orf19.416 | 126 | 104 | 134 | 80 | 162 | 167 | 130 | 217 | 115 | 105 | 127 | 120 | 144 | 181 | 133 | 166 | 136,37 | 117 | 156 | 1,338 | 0,42 | 3,144 | 0,001666 | 0,002312 | 0,006189 | 0,011724 | 0,0095348 | 0,0095348 | TRUE | 0,1130387 | No |
| orf19.4160 | 2536 | 2238 | 2407 | 1712 | 2776 | 2406 | 2451 | 3252 | 2309 | 2263 | 2278 | 2572 | 2469 | 2612 | 2510 | 2481 | 2436,79 | 2356 | 2518 | 1,071 | 0,099 | 1,974 | 0,048412 | 0,060021 | 0,0016905 | 0,002579 | 0,0019844 | 0,0019844 | TRUE | 0,0430581 | No |
| orf19.4161 | 727 | 698 | 682 | 460 | 933 | 771 | 807 | 994 | 662 | 706 | 645 | 691 | 830 | 837 | 826 | 758 | 744,5 | 676 | 813 | 1,202 | 0,266 | 4,288 | 1,81E-05 | 2,83E-05 | 0,0008154 | 0,003811 | 0,0023219 | 0,0023219 | TRUE | 0,0261962 | No |
| orf19.4162 | 295 | 307 | 342 | 185 | 309 | 247 | 262 | 330 | 269 | 310 | 324 | 278 | 275 | 268 | 268 | 252 | 280,47 | 295 | 266 | 0,897 | -0,156 | -1,7 | 0,089198 | 0,107635 | 1,00E-08 | 0,006747 | 0,0045622 | 0,0045622 | TRUE | 0,0498461 | No |
| orf19.4163 | 1234 | 1208 | 1325 | 830 | 2251 | 2032 | 1847 | 2472 | 1124 | 1222 | 1254 | 1247 | 2002 | 2206 | 1892 | 1886 | 1603,96 | 1212 | 1996 | 1,648 | 0,721 | 11,338 | 8,48E-30 | 2,71E-29 | 0,0031677 | 0,002861 | 0,0032029 | 0,0032029 | TRUE | 0,0594254 | No |
| orf19.4164 | 1547 | 1559 | 1556 | 922 | 1576 | 1181 | 1321 | 1577 | 1408 | 1577 | 1473 | 1385 | 1402 | 1282 | 1353 | 1203 | 1385,34 | 1461 | 1310 | 0,895 | -0,159 | -2,514 | 0,011934 | 0,015567 | 0,0029437 | 0,002991 | 0,0031278 | 0,0031278 | TRUE | 0,0383016 | No |
| orf19.4166 | 5157 | 4319 | 4909 | 3594 | 5609 | 5132 | 5188 | 7429 | 4695 | 4368 | 4646 | 5399 | 4989 | 5572 | 5313 | 5667 | 5081,14 | 4777 | 5385 | 1,128 | 0,174 | 2,635 | 0,008409 | 0,011087 | 0,0055431 | 0,002297 | 0,0039936 | 0,0039936 | TRUE | 0,0915849 | No |
| orf19.4167 | 800 | 731 | 781 | 443 | 1554 | 1013 | 1370 | 1669 | 728 | 739 | 739 | 665 | 1382 | 1100 | 1403 | 1273 | 1003,82 | 718 | 1290 | 1,793 | 0,843 | 10,631 | 2,14E-26 | 6,31E-26 | 0,0064568 | 0,003353 | 0,0049207 | 0,0049207 | TRUE | 0,1146137 | No |
| orf19.4168 | 1753 | 1343 | 1471 | 895 | 1427 | 1050 | 1190 | 1595 | 1596 | 1358 | 1392 | 1344 | 1269 | 1140 | 1219 | 1217 | 1316,95 | 1422 | 1211 | 0,85 | -0,234 | -3,59 | 0,000331 | 0,000482 | 0,0033462 | 0,00304 | 0,0033074 | 0,0033074 | TRUE | 0,0828826 | No |
| orf19.4169 | 87 | 76 | 71 | 52 | 105 | 126 | 135 | 183 | 79 | 77 | 67 | 78 | 93 | 137 | 138 | 140 | 101,18 | 75 | 127 | 1,687 | 0,755 | 4,616 | 3,91E-06 | 6,33E-06 | 0,0145837 | 0,015093 | 0,0148803 | 0,0148803 | TRUE | 0,3381461 | No |
| orf19.417 | 3621 | 3201 | 3499 | 2264 | 2679 | 2302 | 2407 | 3154 | 3297 | 3237 | 3311 | 3401 | 2383 | 2499 | 2465 | 2406 | 2874,97 | 3312 | 2438 | 0,736 | -0,441 | -11,689 | 1,46E-31 | 4,86E-31 | 1,00E-08 | 0,002496 | 0,0010156 | 0,0010156 | TRUE | 0,0036319 | No |
| orf19.4170 | 1398 | 1276 | 1297 | 824 | 231 | 193 | 227 | 280 | 1273 | 1291 | 1227 | 1238 | 205 | 210 | 232 | 214 | 736,21 | 1257 | 215 | 0,171 | -2,548 | -36,868 | 1,50E-297 | 3,87E-296 | 1,00E-08 | 0,003831 | 0,0020025 | 0,0020025 | TRUE | 0,0293459 | No |
| orf19.4171 | 354 | 340 | 403 | 188 | 401 | 337 | 352 | 475 | 322 | 344 | 381 | 282 | 357 | 366 | 360 | 362 | 346,92 | 332 | 361 | 1,079 | 0,11 | 1,223 | 0,221498 | 0,254653 | 0,0035171 | 0,005845 | 0,0049214 | 0,0049214 | TRUE | 0,1095075 | No |
| orf19.4172 | 1343 | 1353 | 1282 | 731 | 1548 | 1246 | 1284 | 1751 | 1223 | 1368 | 1213 | 1098 | 1377 | 1353 | 1315 | 1336 | 1285,36 | 1226 | 1345 | 1,095 | 0,131 | 2,066 | 0,038808 | 0,048548 | 0,0032357 | 0,003065 | 0,0030601 | 0,0030601 | TRUE | 0,0722444 | No |
| orf19.4173 | 1257 | 929 | 1087 | 609 | 1157 | 883 | 1084 | 1514 | 1144 | 940 | 1029 | 915 | 1029 | 959 | 1110 | 1155 | 1035,06 | 1007 | 1063 | 1,055 | 0,077 | 0,938 | 0,348041 | 0,388031 | 0,0072112 | 0,003313 | 0,005427 | 0,005427 | TRUE | 0,1021728 | No |
| orf19.4174 | 2658 | 2222 | 2304 | 1676 | 812 | 677 | 697 | 1029 | 2420 | 2247 | 2181 | 2518 | 722 | 735 | 714 | 785 | 1540,19 | 2342 | 739 | 0,316 | -1,66 | -27,102 | 9,45E-162 | 1,32E-160 | 0,0026129 | 0,002895 | 0,002742 | 0,002742 | TRUE | 0,029342 | No |
| orf19.4175 | 4573 | 3997 | 4360 | 2816 | 6076 | 5415 | 5573 | 7516 | 4164 | 4043 | 4126 | 4230 | 5404 | 5879 | 5707 | 5733 | 4910,85 | 4141 | 5681 | 1,372 | 0,457 | 11,819 | 3,13E-32 | 1,06E-31 | 0,0005156 | 0,002306 | 0,0012207 | 0,0012207 | TRUE | 0,0131608 | No |
| orf19.4176 | 1335 | 1201 | 1245 | 743 | 1485 | 1060 | 1161 | 1629 | 1215 | 1215 | 1178 | 1116 | 1321 | 1151 | 1189 | 1243 | 1203,48 | 1181 | 1226 | 1,037 | 0,053 | 0,911 | 0,362454 | 0,402481 | 0,0016843 | 0,003135 | 0,0023764 | 0,0023764 | TRUE | 0,0324188 | No |
| orf19.4177 | 2146 | 1951 | 2048 | 1394 | 2105 | 1711 | 1832 | 2495 | 1954 | 1973 | 1938 | 2094 | 1872 | 1858 | 1876 | 1903 | 1933,59 | 1990 | 1877 | 0,946 | -0,081 | -1,901 | 0,057259 | 0,07055 | 1,00E-08 | 0,00272 | 0,0012105 | 0,0012105 | TRUE | 0,0133954 | No |
| orf19.4179 | 56 | 40 | 47 | 31 | 71 | 51 | 43 | 64 | 51 | 40 | 44 | 47 | 63 | 55 | 44 | 49 | 49,23 | 46 | 53 | 1,158 | 0,212 | 1,051 | 0,293423 | 0,330548 | 1,00E-08 | 0,028868 | 0,0185782 | 0,0185782 | TRUE | 0,1539666 | No |
| orf19.418 | 3209 | 3003 | 3226 | 1865 | 2078 | 1848 | 1955 | 2581 | 2922 | 3037 | 3053 | 2802 | 1848 | 2006 | 2002 | 1969 | 2454,9 | 2954 | 1956 | 0,661 | -0,596 | -12,714 | 4,93E-37 | 1,84E-36 | 0,0009332 | 0,002575 | 0,0016904 | 0,0016904 | TRUE | 0,0168451 | No |
| orf19.4180 | 1146 | 1014 | 1015 | 686 | 1293 | 942 | 988 | 1426 | 1043 | 1026 | 961 | 1030 | 1150 | 1023 | 1012 | 1088 | 1041,56 | 1015 | 1068 | 1,054 | 0,076 | 1,282 | 0,199979 | 0,2315 | 0,0016137 | 0,003305 | 0,002422 | 0,002422 | TRUE | 0,0316417 | No |
| orf19.4181 | 569 | 421 | 452 | 309 | 627 | 473 | 479 | 705 | 518 | 426 | 428 | 464 | 558 | 514 | 491 | 538 | 491,92 | 459 | 525 | 1,145 | 0,195 | 2,383 | 0,017154 | 0,022137 | 0,0039664 | 0,004722 | 0,0043703 | 0,0043703 | TRUE | 0,0957602 | No |
| orf19.4182 | 1777 | 1580 | 1576 | 1069 | 1525 | 1263 | 1278 | 1752 | 1618 | 1598 | 1492 | 1606 | 1356 | 1371 | 1309 | 1336 | 1460,78 | 1578 | 1343 | 0,852 | -0,232 | -4,875 | 1,09E-06 | 1,80E-06 | 1,00E-08 | 0,002941 | 0,0014843 | 0,0014843 | TRUE | 0,0172744 | No |
| orf19.4183 | 4016 | 3370 | 3730 | 2399 | 2378 | 2059 | 2222 | 3054 | 3656 | 3408 | 3530 | 3604 | 2115 | 2236 | 2276 | 2330 | 2894,31 | 3550 | 2239 | 0,631 | -0,664 | -14,785 | 1,82E-49 | 8,30E-49 | 0,0009675 | 0,002493 | 0,0015779 | 0,0015779 | TRUE | 0,0172404 | No |
| orf19.4184 | 2389 | 2104 | 2475 | 1717 | 1771 | 1648 | 1638 | 2273 | 2175 | 2128 | 2342 | 2579 | 1575 | 1789 | 1678 | 1734 | 2000,07 | 2306 | 1694 | 0,736 | -0,443 | -6,641 | 3,12E-11 | 6,17E-11 | 0,0046826 | 0,002697 | 0,0037657 | 0,0037657 | TRUE | 0,0736759 | No |
| orf19.4185 | 230 | 237 | 222 | 133 | 252 | 192 | 260 | 285 | 209 | 240 | 210 | 200 | 224 | 208 | 266 | 217 | 221,91 | 215 | 229 | 1,063 | 0,088 | 0,816 | 0,414383 | 0,455778 | 0,0044782 | 0,00799 | 0,0065637 | 0,0065637 | TRUE | 0,1262556 | No |
| orf19.4186 | 1179 | 1043 | 1211 | 728 | 1062 | 803 | 884 | 1146 | 1073 | 1055 | 1146 | 1094 | 945 | 872 | 905 | 874 | 995,49 | 1092 | 899 | 0,823 | -0,281 | -5,113 | 3,18E-07 | 5,38E-07 | 0,0004593 | 0,003364 | 0,0018816 | 0,0018816 | TRUE | 0,0143213 | No |
| orf19.4187 | 1380 | 1159 | 1379 | 803 | 730 | 630 | 630 | 945 | 1256 | 1172 | 1305 | 1206 | 649 | 684 | 645 | 721 | 954,92 | 1235 | 675 | 0,547 | -0,872 | -14,296 | 2,32E-46 | 1,01E-45 | 0,0015076 | 0,00342 | 0,0024432 | 0,0024432 | TRUE | 0,027153 | No |
| orf19.4188 | 6954 | 6801 | 6959 | 4124 | 7687 | 6449 | 6566 | 8378 | 6331 | 6878 | 6586 | 6195 | 6837 | 7002 | 6724 | 6391 | 6618,14 | 6498 | 6738 | 1,036 | 0,052 | 1,124 | 0,26095 | 0,296587 | 0,0016077 | 0,002237 | 0,0018701 | 0,0018701 | TRUE | 0,0189239 | No |
| orf19.4189 | 22 | 15 | 15 | 11 | 44 | 31 | 36 | 48 | 20 | 15 | 14 | 17 | 39 | 34 | 37 | 37 | 26,52 | 16 | 37 | 2,218 | 1,149 | 4,175 | 2,98E-05 | 4,61E-05 | 1,00E-08 | 0,051839 | 0,0279553 | 0,0279553 | TRUE | 0,1262923 | No |
| orf19.419 | 7407 | 6750 | 6900 | 4020 | 2489 | 1942 | 2057 | 2700 | 6744 | 6827 | 6530 | 6039 | 2214 | 2108 | 2107 | 2060 | 4328,52 | 6535 | 2122 | 0,324 | -1,624 | -33,176 | 2,37E-241 | 4,77E-240 | 0,0015201 | 0,002342 | 0,0019977 | 0,0019977 | TRUE | 0,0308869 | No |
| orf19.4190 | 3479 | 2861 | 3195 | 2078 | 3316 | 2532 | 2991 | 3720 | 3167 | 2894 | 3024 | 3121 | 2949 | 2749 | 3063 | 2838 | 2975,71 | 3052 | 2900 | 0,951 | -0,073 | -1,507 | 0,131806 | 0,156096 | 0,0014449 | 0,002481 | 0,0019291 | 0,0019291 | TRUE | 0,0171018 | No |
| orf19.4191 | 11630 | 10606 | 10870 | 6369 | 15752 | 11292 | 12740 | 18187 | 10589 | 10727 | 10287 | 9567 | 14011 | 12260 | 13047 | 13873 | 11795,2 | 10292 | 13298 | 1,292 | 0,369 | 6,852 | 7,30E-12 | 1,48E-11 | 0,003013 | 0,002149 | 0,0027042 | 0,0027042 | TRUE | 0,0336786 | No |
| orf19.4191.1 | 1438 | 1479 | 1591 | 929 | 704 | 637 | 694 | 803 | 1309 | 1496 | 1506 | 1396 | 626 | 692 | 711 | 613 | 1043,43 | 1427 | 660 | 0,462 | -1,115 | -16,015 | 1,00E-57 | 5,30E-57 | 0,0038229 | 0,003303 | 0,0035673 | 0,0035673 | TRUE | 0,0389939 | No |
| orf19.4192 | 3207 | 2828 | 3000 | 2033 | 2008 | 1719 | 1900 | 2503 | 2920 | 2860 | 2839 | 3054 | 1786 | 1866 | 1946 | 1909 | 2397,59 | 2918 | 1877 | 0,644 | -0,635 | -14,133 | 2,39E-45 | 1,03E-44 | 0,0006349 | 0,002588 | 0,0015072 | 0,0015072 | TRUE | 0,0130814 | No |
| orf19.4192.1 | 864 | 655 | 734 | 434 | 709 | 506 | 564 | 736 | 787 | 662 | 695 | 652 | 631 | 549 | 578 | 561 | 639,34 | 699 | 580 | 0,828 | -0,273 | -3,542 | 0,000397 | 0,000575 | 0,004155 | 0,004103 | 0,0041285 | 0,0041285 | TRUE | 0,0867749 | No |
| orf19.4193 | 552 | 501 | 512 | 371 | 527 | 407 | 435 | 578 | 503 | 507 | 485 | 557 | 469 | 442 | 445 | 441 | 481,02 | 513 | 449 | 0,88 | -0,184 | -2,612 | 0,00901 | 0,01186 | 1,00E-08 | 0,004783 | 0,0026899 | 0,0026899 | TRUE | 0,0335162 | No |
| orf19.4193.1 | 18375 | 14749 | 15131 | 10531 | 23422 | 17230 | 16585 | 25361 | 16730 | 14917 | 14320 | 15819 | 20833 | 18707 | 16985 | 19346 | 17207,1 | 15446 | 18968 | 1,228 | 0,296 | 4,465 | 8,01E-06 | 1,28E-05 | 0,0057815 | 0,002114 | 0,0041744 | 0,0041744 | TRUE | 0,0605706 | No |
| orf19.4194 | 634 | 567 | 610 | 393 | 582 | 509 | 503 | 695 | 577 | 573 | 577 | 590 | 518 | 553 | 515 | 530 | 554,24 | 579 | 529 | 0,913 | -0,131 | -2,088 | 0,036756 | 0,046099 | 1,00E-08 | 0,00442 | 0,0020028 | 0,0020028 | TRUE | 0,009683 | No |
| orf19.4195 | 264 | 221 | 241 | 137 | 274 | 242 | 237 | 318 | 240 | 224 | 228 | 206 | 244 | 263 | 243 | 243 | 236,19 | 224 | 248 | 1,097 | 0,134 | 1,433 | 0,15193 | 0,1787 | 1,00E-08 | 0,00763 | 0,0041172 | 0,0041172 | TRUE | 0,027007 | No |
| orf19.4195.1 | 2467 | 2025 | 2161 | 1220 | 8082 | 5566 | 6822 | 9028 | 2246 | 2048 | 2045 | 1833 | 7189 | 6043 | 6987 | 6887 | 4409,63 | 2043 | 6776 | 3,312 | 1,728 | 25,103 | 4,66E-139 | 5,64E-138 | 0,0057638 | 0,002337 | 0,0042175 | 0,0042175 | TRUE | 0,0644127 | No |
| orf19.4196 | 46 | 23 | 38 | 26 | 82 | 64 | 60 | 75 | 42 | 23 | 36 | 39 | 73 | 69 | 61 | 57 | 50,16 | 35 | 65 | 1,864 | 0,899 | 4,193 | 2,76E-05 | 4,28E-05 | 0,0072143 | 0,028375 | 0,0217175 | 0,0217175 | TRUE | 0,378316 | No |
| orf19.4197 | 2265 | 1711 | 1969 | 1269 | 4087 | 3307 | 3639 | 5007 | 2062 | 1730 | 1863 | 1906 | 3635 | 3590 | 3727 | 3819 | 2791,79 | 1890 | 3693 | 1,953 | 0,966 | 17,618 | 1,80E-69 | 1,12E-68 | 0,002459 | 0,00251 | 0,0024751 | 0,0024751 | TRUE | 0,0473516 | No |
| orf19.4199 | 10509 | 9793 | 10586 | 7065 | 5206 | 4876 | 4606 | 6437 | 9568 | 9905 | 10019 | 10613 | 4631 | 5294 | 4717 | 4910 | 7456,99 | 10026 | 4888 | 0,488 | -1,036 | -20,161 | 2,16E-90 | 1,69E-89 | 0,0025594 | 0,002214 | 0,0023882 | 0,0023882 | TRUE | 0,0376056 | No |
| orf19.420 | 32 | 23 | 12 | 13 | 4 | 8 | 9 | 14 | 29 | 23 | 11 | 20 | 4 | 9 | 9 | 11 | 14,43 | 21 | 8 | 0,385 | -1,375 | -3,213 | 0,001314 | 0,001841 | 0,0903168 | 0,093597 | 0,0925558 | 0,0925558 | TRUE | 0,2225741 | No |
| orf19.4200 | 22 | 26 | 35 | 36 | 68 | 57 | 54 | 69 | 20 | 26 | 33 | 54 | 60 | 62 | 55 | 53 | 45,48 | 33 | 58 | 1,783 | 0,834 | 3,318 | 0,000905 | 0,001281 | 0,0474316 | 0,031083 | 0,0360229 | 0,0360229 | TRUE | 0,8883416 | No |
| orf19.4201 | 2689 | 2275 | 2637 | 1770 | 2697 | 2318 | 2426 | 3204 | 2448 | 2301 | 2496 | 2659 | 2399 | 2517 | 2485 | 2444 | 2468,48 | 2476 | 2461 | 0,995 | -0,007 | -0,141 | 0,887798 | 0,90353 | 0,0013109 | 0,002572 | 0,0019211 | 0,0019211 | TRUE | 0,0278345 | No |
| orf19.4203 | 3341 | 3449 | 3659 | 2388 | 2185 | 1958 | 1909 | 2489 | 3042 | 3488 | 3463 | 3587 | 1943 | 2126 | 1955 | 1899 | 2687,9 | 3395 | 1981 | 0,584 | -0,777 | -12,888 | 5,28E-38 | 1,99E-37 | 0,0032648 | 0,002528 | 0,0030969 | 0,0030969 | TRUE | 0,0604655 | No |
| orf19.4204 | 3618 | 3217 | 3350 | 2150 | 3267 | 2686 | 2876 | 3804 | 3294 | 3254 | 3170 | 3230 | 2906 | 2916 | 2945 | 2902 | 3077,13 | 3237 | 2917 | 0,901 | -0,15 | -4,331 | 1,48E-05 | 2,33E-05 | 1,00E-08 | 0,002466 | 0,0008307 | 0,0008307 | TRUE | 0,0024937 | No |
| orf19.4205.1 | 886 | 684 | 759 | 359 | 567 | 378 | 496 | 655 | 807 | 692 | 718 | 539 | 504 | 410 | 508 | 500 | 584,8 | 689 | 480 | 0,696 | -0,524 | -4,756 | 1,98E-06 | 3,24E-06 | 0,0166613 | 0,004296 | 0,0098932 | 0,0098932 | TRUE | 0,2351827 | No |
| orf19.4206 | 603 | 572 | 623 | 364 | 511 | 445 | 437 | 632 | 549 | 579 | 590 | 547 | 455 | 483 | 448 | 482 | 516,4 | 566 | 467 | 0,824 | -0,279 | -4,145 | 3,39E-05 | 5,23E-05 | 1,00E-08 | 0,004595 | 0,0024191 | 0,0024191 | TRUE | 0,0094442 | No |
| orf19.4207 | 235 | 195 | 220 | 160 | 274 | 194 | 233 | 300 | 214 | 197 | 208 | 240 | 244 | 211 | 239 | 229 | 222,69 | 215 | 231 | 1,08 | 0,111 | 1,116 | 0,264412 | 0,300307 | 1,00E-08 | 0,007969 | 0,0050179 | 0,0050179 | TRUE | 0,0551443 | No |
| orf19.4208 | 1178 | 960 | 1082 | 686 | 1288 | 1054 | 1189 | 1595 | 1073 | 971 | 1024 | 1030 | 1146 | 1144 | 1218 | 1217 | 1102,79 | 1024 | 1181 | 1,153 | 0,206 | 3,814 | 0,000137 | 0,000203 | 0,000616 | 0,003235 | 0,001872 | 0,001872 | TRUE | 0,0154746 | No |
| orf19.4209 | 1391 | 1410 | 1495 | 852 | 916 | 731 | 831 | 1026 | 1266 | 1426 | 1415 | 1280 | 815 | 794 | 851 | 783 | 1078,67 | 1347 | 811 | 0,601 | -0,735 | -12,115 | 8,78E-34 | 3,05E-33 | 0,0019319 | 0,003262 | 0,0025593 | 0,0025593 | TRUE | 0,0214647 | No |
| orf19.4210 | 1055 | 974 | 1005 | 649 | 2531 | 2040 | 2145 | 2747 | 961 | 985 | 951 | 975 | 2251 | 2215 | 2197 | 2095 | 1578,75 | 968 | 2190 | 2,262 | 1,178 | 24,927 | 3,79E-137 | 4,49E-136 | 1,00E-08 | 0,002874 | 0,0013772 | 0,0013772 | TRUE | 0,010725 | No |
| orf19.4211 | 1934 | 1835 | 1865 | 1212 | 25508 | 21802 | 22277 | 30863 | 1761 | 1856 | 1765 | 1821 | 22689 | 23671 | 22814 | 23543 | 12489,9 | 1801 | 23179 | 12,883 | 3,687 | 100,011 | 0 | 0 | 0,0003748 | 0,002143 | 0,0009874 | 0,0009874 | TRUE | 0,0056423 | No |
| orf19.4212 | 370 | 318 | 344 | 206 | 1551 | 1320 | 1329 | 1861 | 337 | 322 | 326 | 309 | 1380 | 1433 | 1361 | 1420 | 860,86 | 324 | 1398 | 4,31 | 2,108 | 33,931 | 2,31E-252 | 4,85E-251 | 1,00E-08 | 0,003572 | 0,0017351 | 0,0017351 | TRUE | 0,0089366 | No |
| orf19.4213 | 975 | 818 | 1000 | 585 | 942 | 796 | 797 | 1081 | 888 | 827 | 946 | 879 | 838 | 864 | 816 | 825 | 860,39 | 885 | 836 | 0,943 | -0,084 | -1,433 | 0,151993 | 0,178741 | 0,0007427 | 0,003572 | 0,0021678 | 0,0021678 | TRUE | 0,0274187 | No |
| orf19.4214 | 42 | 38 | 44 | 26 | 16 | 21 | 19 | 42 | 38 | 38 | 42 | 39 | 14 | 23 | 19 | 32 | 30,74 | 39 | 22 | 0,568 | -0,815 | -3,015 | 0,002567 | 0,00351 | 0,0165632 | 0,045014 | 0,0357798 | 0,0357798 | TRUE | 0,6676358 | No |
| orf19.4215 | 16848 | 14225 | 15404 | 9726 | 7716 | 6636 | 7352 | 9884 | 15339 | 14387 | 14578 | 14610 | 6863 | 7205 | 7529 | 7540 | 11006,5 | 14728 | 7284 | 0,495 | -1,016 | -24,234 | 9,74E-130 | 1,08E-128 | 0,0012379 | 0,002157 | 0,0015869 | 0,0015869 | TRUE | 0,0186596 | No |
| orf19.4216 | 2318 | 1924 | 1917 | 1335 | 3104 | 2203 | 2266 | 3422 | 2110 | 1946 | 1814 | 2005 | 2761 | 2392 | 2321 | 2610 | 2244,97 | 1969 | 2521 | 1,281 | 0,358 | 5,365 | 8,10E-08 | 1,40E-07 | 0,0045747 | 0,002625 | 0,003808 | 0,003808 | TRUE | 0,0496063 | No |

| | | | | | | | | | | | | | | | | | | | | | | | | | | | | | | | |
|---|---|---|---|---|---|---|---|---|---|---|---|---|---|---|---|---|---|---|---|---|---|---|---|---|---|---|---|---|---|---|---|
| orf19.4217 | 7 | 3 | 3 | 0 | 7 | 4 | 1 | 1 | 6 | 3 | 3 | 0 | 6 | 4 | 1 | 1 | 3,08 | 3 | 3 | 0,948 | -0,077 | -0,085 | 0,932065 | 0,941746 | 0,509571 | 0,431582 | 0,4578077 | 0,4578077 | TRUE | 0,3022459 | No |
| orf19.4219 | 6798 | 5688 | 6356 | 3968 | 7156 | 5824 | 6333 | 8699 | 6189 | 5753 | 6015 | 5961 | 6365 | 6323 | 6486 | 6636 | 6215,97 | 5980 | 6452 | 1,079 | 0,11 | 3,006 | 0,002646 | 0,003616 | 0,0004738 | 0,00225 | 0,0011199 | 0,0011199 | TRUE | 0,0081536 | No |
| orf19.422 | 947 | 797 | 899 | 652 | 747 | 704 | 750 | 975 | 862 | 806 | 851 | 979 | 664 | 764 | 768 | 744 | 804,89 | 874 | 735 | 0,843 | -0,247 | -3,368 | 0,000756 | 0,001076 | 0,0040663 | 0,003678 | 0,0039139 | 0,0039139 | TRUE | 0,0701061 | No |
| orf19.4220 | 1520 | 1306 | 1424 | 959 | 909 | 703 | 830 | 1031 | 1384 | 1321 | 1348 | 1441 | 809 | 763 | 850 | 786 | 1087,66 | 1374 | 802 | 0,585 | -0,774 | -14,094 | 4,12E-45 | 1,76E-44 | 0,000454 | 0,003252 | 0,0019211 | 0,0019211 | TRUE | 0,0182429 | No |
| orf19.4221 | 2397 | 2052 | 2278 | 1491 | 2109 | 1906 | 1851 | 2590 | 2182 | 2075 | 2156 | 2240 | 1876 | 2069 | 1896 | 1976 | 2058,75 | 2163 | 1954 | 0,904 | -0,146 | -3,062 | 0,002202 | 0,003029 | 0,0008653 | 0,002679 | 0,0016968 | 0,0016968 | TRUE | 0,0182237 | No |
| orf19.4222 | 2136 | 2065 | 2138 | 1438 | 1404 | 1282 | 1296 | 1758 | 1945 | 2089 | 2023 | 2160 | 1249 | 1392 | 1327 | 1341 | 1690,72 | 2054 | 1327 | 0,647 | -0,629 | -12,02 | 2,79E-33 | 9,59E-33 | 0,0013511 | 0,002818 | 0,0020131 | 0,0020131 | TRUE | 0,0191634 | No |
| orf19.4223 | 13098 | 10504 | 11788 | 7254 | 15294 | 11870 | 13684 | 18948 | 11925 | 10624 | 11156 | 10897 | 13604 | 12888 | 14014 | 14454 | 12445,1 | 11150 | 13740 | 1,232 | 0,301 | 6,084 | 1,17E-09 | 2,19E-09 | 0,0023287 | 0,002143 | 0,0022716 | 0,0022716 | TRUE | 0,0269286 | No |
| orf19.4225 | 5405 | 4861 | 5354 | 3318 | 5159 | 4353 | 4629 | 6432 | 4921 | 4916 | 5067 | 4984 | 4589 | 4726 | 4741 | 4906 | 4856,33 | 4972 | 4740 | 0,954 | -0,068 | -1,938 | 0,052592 | 0,065001 | 0,0003273 | 0,002309 | 0,0009905 | 0,0009905 | TRUE | 0,0070478 | No |
| orf19.4225.1 | 4465 | 3724 | 4043 | 2785 | 3047 | 2321 | 2679 | 3478 | 4065 | 3766 | 3826 | 4184 | 2710 | 2520 | 2744 | 2653 | 3308,54 | 3960 | 2657 | 0,671 | -0,575 | -11,891 | 1,32E-32 | 4,48E-32 | 0,0015051 | 0,002436 | 0,00193 | 0,00193 | TRUE | 0,0166399 | No |
| orf19.4227 | 94 | 76 | 98 | 67 | 87 | 72 | 59 | 89 | 86 | 77 | 93 | 101 | 77 | 78 | 60 | 68 | 79,96 | 89 | 71 | 0,802 | -0,318 | -1,978 | 0,047949 | 0,059458 | 1,00E-08 | 0,018557 | 0,0121621 | 0,0121621 | TRUE | 0,0822272 | No |
| orf19.4228 | 1165 | 982 | 936 | 665 | 1944 | 1560 | 1656 | 2246 | 1061 | 993 | 886 | 999 | 1729 | 1694 | 1696 | 1713 | 1346,34 | 985 | 1708 | 1,736 | 0,795 | 13,818 | 1,98E-43 | 8,21E-43 | 0,0016994 | 0,003018 | 0,002362 | 0,002362 | TRUE | 0,0579853 | No |
| orf19.4229 | 850 | 731 | 812 | 495 | 2055 | 1634 | 1780 | 2302 | 774 | 739 | 768 | 744 | 1828 | 1774 | 1823 | 1756 | 1275,77 | 756 | 1795 | 2,37 | 1,245 | 25,442 | 8,77E-143 | 1,09E-141 | 1,00E-08 | 0,003072 | 0,0013347 | 0,0013347 | TRUE | 0,0031717 | No |
| orf19.423 | 1197 | 990 | 1053 | 873 | 1330 | 1164 | 1078 | 1496 | 1090 | 1001 | 997 | 1311 | 1183 | 1264 | 1104 | 1141 | 1136,37 | 1100 | 1173 | 1,07 | 0,097 | 1,154 | 0,248631 | 0,283647 | 0,0090694 | 0,003199 | 0,00592 | 0,00592 | TRUE | 0,1895004 | No |
| orf19.4230 | 6338 | 5515 | 5827 | 3761 | 5257 | 4234 | 4478 | 6248 | 5770 | 5578 | 5515 | 5650 | 4676 | 4597 | 4586 | 4766 | 5142,2 | 5628 | 4656 | 0,828 | -0,273 | -8,142 | 3,89E-16 | 8,86E-16 | 0,0001947 | 0,002294 | 0,0008829 | 0,0008829 | TRUE | 0,0037204 | No |
| orf19.4231 | 608 | 540 | 599 | 420 | 1188 | 1013 | 1103 | 1399 | 554 | 546 | 567 | 631 | 1057 | 1100 | 1130 | 1067 | 831,35 | 574 | 1088 | 1,903 | 0,928 | 15,197 | 3,69E-52 | 1,77E-51 | 0,000403 | 0,003626 | 0,0022178 | 0,0022178 | TRUE | 0,0429657 | No |
| orf19.4232 | 2001 | 1896 | 1994 | 1188 | 2004 | 1682 | 1879 | 2383 | 1822 | 1918 | 1887 | 1785 | 1782 | 1826 | 1924 | 1818 | 1845,24 | 1853 | 1838 | 0,99 | -0,014 | -0,304 | 0,760817 | 0,791871 | 0,0004704 | 0,002753 | 0,0014749 | 0,0014749 | TRUE | 0,0119415 | No |
| orf19.4233 | 4160 | 3747 | 3954 | 2433 | 5838 | 4851 | 5078 | 6798 | 3788 | 3790 | 3742 | 3655 | 5193 | 5267 | 5201 | 5186 | 4477,46 | 3744 | 5212 | 1,391 | 0,476 | 14,935 | 1,94E-50 | 9,01E-50 | 1,00E-08 | 0,002332 | 0,0007421 | 0,0007421 | TRUE | 0,0027798 | No |
| orf19.4234 | 5399 | 4839 | 5320 | 3373 | 5563 | 4851 | 4776 | 6259 | 4916 | 4894 | 5035 | 5067 | 4948 | 5267 | 4891 | 4774 | 4974 | 4978 | 4970 | 0,998 | -0,003 | -0,062 | 0,950602 | 0,956605 | 0,0006919 | 0,002303 | 0,0013661 | 0,0013661 | TRUE | 0,0194032 | No |
| orf19.4235 | 2049 | 1933 | 2172 | 1449 | 2306 | 2161 | 2092 | 2899 | 1866 | 1955 | 2056 | 2177 | 2051 | 2346 | 2142 | 2211 | 2100,5 | 2014 | 2188 | 1,088 | 0,122 | 2,03 | 0,042338 | 0,05282 | 0,0031879 | 0,002666 | 0,0029732 | 0,0029732 | TRUE | 0,0340898 | No |
| orf19.4236 | 1352 | 1245 | 1246 | 771 | 1530 | 1268 | 1388 | 1855 | 1231 | 1259 | 1179 | 1158 | 1361 | 1377 | 1421 | 1415 | 1300,2 | 1207 | 1394 | 1,153 | 0,205 | 4,215 | 2,50E-05 | 3,89E-05 | 1,00E-08 | 0,003053 | 0,0015017 | 0,0015017 | TRUE | 0,0098182 | No |
| orf19.4239 | 11 | 16 | 19 | 15 | 30 | 20 | 24 | 25 | 10 | 16 | 18 | 23 | 27 | 22 | 25 | 19 | 19,84 | 17 | 23 | 1,411 | 0,497 | 1,54 | 0,123441 | 0,146789 | 1,00E-08 | 0,068603 | 0,0469126 | 0,0469126 | TRUE | 0,3973012 | No |
| orf19.424 | 1760 | 1415 | 1473 | 1008 | 1628 | 1280 | 1204 | 1756 | 1602 | 1431 | 1394 | 1514 | 1448 | 1390 | 1233 | 1339 | 1419,01 | 1485 | 1352 | 0,911 | -0,135 | -2,083 | 0,037211 | 0,046633 | 0,003281 | 0,002968 | 0,0032955 | 0,0032955 | TRUE | 0,0416809 | No |
| orf19.4240 | 709 | 622 | 663 | 453 | 474 | 392 | 413 | 553 | 646 | 629 | 627 | 680 | 422 | 426 | 423 | 422 | 534,32 | 646 | 423 | 0,657 | -0,607 | -9,353 | 8,50E-21 | 2,20E-20 | 1,00E-08 | 0,004509 | 0,0021085 | 0,0021085 | TRUE | 0,0129923 | No |
| orf19.4241 | 780 | 707 | 744 | 545 | 600 | 497 | 478 | 678 | 710 | 715 | 704 | 819 | 534 | 540 | 490 | 517 | 628,5 | 737 | 520 | 0,709 | -0,497 | -7,166 | 7,70E-13 | 1,60E-12 | 0,00149 | 0,004139 | 0,0029892 | 0,0029892 | TRUE | 0,0579595 | No |
| orf19.4242 | 1901 | 1708 | 1824 | 1217 | 3829 | 3292 | 3398 | 4552 | 1731 | 1727 | 1726 | 1828 | 3406 | 3574 | 3480 | 3472 | 2618,11 | 1753 | 3483 | 1,99 | 0,993 | 24,838 | 3,51E-136 | 4,11E-135 | 6,52E-05 | 0,002542 | 0,0010928 | 0,0010928 | TRUE | 0,0087307 | No |
| orf19.4243 | 215 | 198 | 181 | 105 | 291 | 280 | 251 | 376 | 196 | 200 | 171 | 158 | 259 | 304 | 257 | 287 | 228,97 | 181 | 277 | 1,516 | 0,6 | 5,637 | 1,73E-08 | 3,09E-08 | 0,0035012 | 0,007806 | 0,0062622 | 0,0062622 | TRUE | 0,0637823 | No |
| orf19.4244 | 83 | 113 | 115 | 58 | 167 | 101 | 129 | 150 | 76 | 114 | 109 | 87 | 149 | 110 | 132 | 114 | 111,32 | 96 | 126 | 1,306 | 0,385 | 2,378 | 0,017426 | 0,022479 | 0,0188773 | 0,013904 | 0,0158943 | 0,0158943 | TRUE | 0,2395204 | No |
| orf19.4245 | 1241 | 1045 | 1139 | 769 | 803 | 699 | 745 | 987 | 1130 | 1057 | 1078 | 1155 | 714 | 759 | 763 | 753 | 926,12 | 1105 | 747 | 0,677 | -0,563 | -10,119 | 4,56E-24 | 1,28E-23 | 1,00E-08 | 0,003463 | 0,0018603 | 0,0018603 | TRUE | 0,0106405 | No |
| orf19.4246 | 18310 | 15829 | 17288 | 10706 | 11435 | 9247 | 10016 | 13479 | 16670 | 16009 | 16362 | 16082 | 10171 | 10040 | 10258 | 10282 | 13234,2 | 16281 | 10188 | 0,626 | -0,676 | -23,697 | 3,88E-124 | 4,10E-123 | 0,0001221 | 0,002137 | 0,0007036 | 0,0007036 | TRUE | 0,0032061 | No |
| orf19.4247 | 726 | 585 | 654 | 382 | 554 | 403 | 473 | 599 | 661 | 592 | 619 | 574 | 493 | 438 | 484 | 457 | 539,63 | 612 | 468 | 0,763 | -0,39 | -5,406 | 6,46E-08 | 1,12E-07 | 0,0014736 | 0,004485 | 0,0031163 | 0,0031163 | TRUE | 0,035761 | No |
| orf19.4248 | 1263 | 1090 | 1164 | 714 | 1465 | 1091 | 1168 | 1635 | 1150 | 1102 | 1102 | 1073 | 1303 | 1185 | 1196 | 1247 | 1169,68 | 1107 | 1233 | 1,114 | 0,155 | 2,944 | 0,00324 | 0,004396 | 0,000583 | 0,003166 | 0,0018038 | 0,0018038 | TRUE | 0,0175199 | No |
| orf19.425 | 326 | 274 | 313 | 201 | 388 | 303 | 365 | 430 | 297 | 277 | 296 | 302 | 345 | 329 | 374 | 328 | 318,5 | 293 | 344 | 1,174 | 0,231 | 2,765 | 0,005687 | 0,007591 | 1,00E-08 | 0,006185 | 0,0035475 | 0,0035475 | TRUE | 0,0373655 | No |
| orf19.4250 | 156 | 121 | 113 | 89 | 137 | 126 | 129 | 163 | 142 | 122 | 107 | 134 | 122 | 137 | 132 | 124 | 127,52 | 126 | 129 | 1,02 | 0,029 | 0,22 | 0,826025 | 0,849752 | 1,00E-08 | 0,012396 | 0,0083211 | 0,0083211 | TRUE | 0,1177014 | No |
| orf19.4251 | 193 | 189 | 198 | 149 | 313 | 276 | 291 | 369 | 176 | 191 | 187 | 224 | 278 | 300 | 298 | 281 | 241,96 | 194 | 289 | 1,5 | 0,585 | 6,005 | 1,91E-09 | 3,54E-09 | 1,00E-08 | 0,007497 | 0,0047251 | 0,0047251 | TRUE | 0,0887563 | No |
| orf19.4252 | 3116 | 2755 | 2905 | 1929 | 2007 | 1680 | 1708 | 2365 | 2837 | 2786 | 2749 | 2898 | 1785 | 1824 | 1749 | 1804 | 2304,1 | 2818 | 1790 | 0,636 | -0,653 | -16,466 | 6,44E-61 | 3,57E-60 | 1,00E-08 | 0,00261 | 0,0010572 | 0,0010572 | TRUE | 0,0040043 | No |
| orf19.4253 | 524 | 555 | 555 | 307 | 876 | 806 | 855 | 1040 | 477 | 561 | 525 | 461 | 779 | 875 | 876 | 793 | 668,5 | 506 | 831 | 1,634 | 0,709 | 9,269 | 1,88E-20 | 4,81E-20 | 0,0039056 | 0,004013 | 0,0039926 | 0,0039926 | TRUE | 0,0612545 | No |
| orf19.4255 | 434 | 339 | 381 | 292 | 126 | 123 | 102 | 160 | 395 | 343 | 361 | 439 | 112 | 134 | 104 | 122 | 251,17 | 384 | 118 | 0,308 | -1,698 | -15 | 7,33E-51 | 3,44E-50 | 0,006433 | 0,007296 | 0,0069516 | 0,0069516 | TRUE | 0,0910016 | No |
| orf19.4257 | 3042 | 2815 | 3159 | 1848 | 3041 | 2682 | 2768 | 3626 | 2770 | 2847 | 2990 | 2776 | 2705 | 2912 | 2835 | 2766 | 2824,99 | 2846 | 2804 | 0,984 | -0,023 | -0,518 | 0,604551 | 0,644268 | 0,0006798 | 0,002505 | 0,0014757 | 0,0014757 | TRUE | 0,0144212 | No |
| orf19.4258 | 3397 | 2832 | 3266 | 2035 | 3119 | 2484 | 2904 | 3977 | 3093 | 2864 | 3091 | 3057 | 2774 | 2697 | 2974 | 3034 | 2947,99 | 3026 | 2870 | 0,949 | -0,076 | -1,49 | 0,136094 | 0,160726 | 0,001752 | 0,002485 | 0,0021448 | 0,0021448 | TRUE | 0,0195899 | No |
| orf19.426 | 1107 | 1018 | 1119 | 666 | 990 | 817 | 885 | 1127 | 1008 | 1030 | 1059 | 1000 | 881 | 887 | 906 | 860 | 953,82 | 1024 | 884 | 0,861 | -0,216 | -4,171 | 3,04E-05 | 4,69E-05 | 1,00E-08 | 0,003422 | 0,0015275 | 0,0015275 | TRUE | 0,006398 | No |
| orf19.4261 | 4059 | 4250 | 4304 | 2625 | 5951 | 5068 | 5142 | 6729 | 3696 | 4298 | 4073 | 3943 | 5293 | 5502 | 5266 | 5133 | 4650,64 | 4002 | 5298 | 1,324 | 0,404 | 8,004 | 1,21E-15 | 2,72E-15 | 0,0021036 | 0,002321 | 0,0022283 | 0,0022283 | TRUE | 0,0335029 | No |
| orf19.4262 | 986 | 840 | 953 | 684 | 1717 | 1609 | 1494 | 2189 | 898 | 850 | 902 | 1027 | 1527 | 1747 | 1530 | 1670 | 1268,83 | 919 | 1618 | 1,766 | 0,821 | 11,901 | 1,17E-32 | 3,97E-32 | 0,0045266 | 0,003078 | 0,0036898 | 0,0036898 | TRUE | 0,0679396 | No |
| orf19.4263 | 592 | 610 | 724 | 418 | 1416 | 1192 | 1237 | 1720 | 539 | 617 | 685 | 628 | 1259 | 1294 | 1267 | 1312 | 950,2 | 617 | 1283 | 2,081 | 1,058 | 15,319 | 5,70E-53 | 2,78E-52 | 0,0033292 | 0,003427 | 0,0033415 | 0,0033415 | TRUE | 0,0929878 | No |
| orf19.4264 | 473 | 491 | 495 | 322 | 620 | 515 | 489 | 651 | 431 | 497 | 468 | 484 | 551 | 559 | 501 | 497 | 498,43 | 470 | 527 | 1,123 | 0,167 | 2,207 | 0,027307 | 0,034616 | 0,0020262 | 0,004687 | 0,0034609 | 0,0034609 | TRUE | 0,0391687 | No |
| orf19.4265 | 4080 | 4514 | 4450 | 2716 | 3916 | 3618 | 3371 | 4272 | 3715 | 4565 | 4212 | 4080 | 3483 | 3928 | 3452 | 3259 | 3836,73 | 4143 | 3530 | 0,852 | -0,231 | -3,252 | 0,001145 | 0,00161 | 0,0064624 | 0,002381 | 0,0045952 | 0,0045952 | TRUE | 0,0694995 | No |
| orf19.4266 | 860 | 943 | 896 | 592 | 1481 | 1399 | 1247 | 1714 | 783 | 954 | 848 | 889 | 1317 | 1519 | 1277 | 1307 | 1111,85 | 868 | 1355 | 1,561 | 0,643 | 8,539 | 1,35E-17 | 3,21E-17 | 0,0054736 | 0,003225 | 0,0044702 | 0,0044702 | TRUE | 0,0778369 | No |
| orf19.4268 | 12445 | 11899 | 12805 | 6819 | 17586 | 14133 | 15648 | 19660 | 11331 | 12035 | 12119 | 10243 | 15642 | 15345 | 16026 | 14997 | 13467 | 11432 | 15502 | 1,355 | 0,439 | 8,046 | 8,56E-16 | 1,93E-15 | 0,0031765 | 0,002135 | 0,0027797 | 0,0027797 | TRUE | 0,0590476 | No |
| orf19.4269 | 2088 | 2070 | 2149 | 1339 | 1216 | 1104 | 1099 | 1467 | 1901 | 2094 | 2034 | 2011 | 1082 | 1199 | 1126 | 1119 | 1570,58 | 2010 | 1132 | 0,562 | -0,83 | -16,023 | 8,88E-58 | 4,70E-57 | 0,0010963 | 0,002878 | 0,0018977 | 0,0018977 | TRUE | 0,0184586 | No |
| orf19.427 | 2706 | 2368 | 2671 | 1513 | 3021 | 2470 | 2645 | 3502 | 2464 | 2395 | 2528 | 2273 | 2687 | 2682 | 2709 | 2671 | 2551,04 | 2415 | 2687 | 1,111 | 0,151 | 3,553 | 0,000381 | 0,000552 | 0,0005146 | 0,002555 | 0,0013477 | 0,0013477 | TRUE | 0,0172541 | No |
| orf19.4270 | 1055 | 1112 | 1134 | 710 | 457 | 406 | 401 | 487 | 961 | 1125 | 1073 | 1067 | 406 | 441 | 411 | 371 | 731,8 | 1056 | 407 | 0,385 | -1,378 | -18,666 | 9,40E-78 | 6,46E-77 | 0,0032925 | 0,003842 | 0,0035746 | 0,0035746 | TRUE | 0,0467932 | No |
| orf19.4271 | 1913 | 1795 | 1869 | 864 | 1969 | 1222 | 1783 | 2011 | 1742 | 1815 | 1769 | 1298 | 1751 | 1327 | 1826 | 1534 | 1632,75 | 1656 | 1610 | 0,971 | -0,042 | -0,281 | 0,778869 | 0,808811 | 0,0209569 | 0,002846 | 0,0111357 | 0,0209569 | TRUE | 0,2507171 | No |
| orf19.4273 | 5223 | 6084 | 5950 | 3470 | 4971 | 4354 | 4274 | 5140 | 4755 | 6153 | 5631 | 5212 | 4422 | 4727 | 4377 | 3921 | 4899,87 | 5438 | 4362 | 0,802 | -0,319 | -4,067 | 4,76E-05 | 7,26E-05 | 0,0086385 | 0,002307 | 0,0056936 | 0,0056936 | TRUE | 0,0959338 | No |
| orf19.4274 | 36042 | 40494 | 41408 | 26188 | 34555 | 31159 | 30651 | 38408 | 32815 | 40955 | 39189 | 39339 | 30736 | 33830 | 31390 | 29298 | 34693,9 | 38074 | 31314 | 0,822 | -0,282 | -4,107 | 4,01E-05 | 6,15E-05 | 0,0062472 | 0,002075 | 0,0045025 | 0,0045025 | TRUE | 0,1060865 | No |
| orf19.4275 | 835 | 776 | 851 | 496 | 613 | 439 | 538 | 657 | 760 | 785 | 805 | 745 | 545 | 477 | 551 | 501 | 646,19 | 774 | 518 | 0,669 | -0,579 | -8,674 | 4,17E-18 | 1,01E-17 | 0,0008158 | 0,004081 | 0,0026927 | 0,0026927 | TRUE | 0,0312122 | No |
| orf19.4278 | 4571 | 4079 | 4295 | 2659 | 6776 | 5353 | 5719 | 7615 | 4162 | 4125 | 4065 | 3994 | 6027 | 5812 | 5857 | 5809 | 4981,36 | 4086 | 5876 | 1,437 | 0,523 | 15,752 | 6,67E-56 | 3,42E-55 | 0,0001153 | 0,002302 | 0,0008478 | 0,0008478 | TRUE | 0,0036859 | No |
| orf19.4279 | 21197 | 19906 | 21241 | 12835 | 1556 | 1299 | 1353 | 1802 | 19299 | 20133 | 20103 | 19280 | 1384 | 1410 | 1386 | 1375 | 10546,2 | 19704 | 1389 | 0,07 | -3,828 | -103,51 | 0 | 0 | 1,00E-08 | 0,002162 | 0,00095 | 0,00095 | TRUE | 0,0026192 | No |
| orf19.428 | 1758 | 1659 | 1708 | 1038 | 571 | 494 | 491 | 677 | 1601 | 1678 | 1616 | 1559 | 508 | 536 | 503 | 516 | 1064,71 | 1614 | 516 | 0,319 | -1,647 | -30,257 | 4,21E-201 | 7,22E-200 | 1,00E-08 | 0,003278 | 0,0016112 | 0,0016112 | TRUE | 0,0085218 | No |
| orf19.4280 | 8023 | 7385 | 7803 | 4774 | 755 | 653 | 659 | 854 | 7305 | 7469 | 7385 | 7171 | 672 | 709 | 675 | 651 | 4004,59 | 7332 | 677 | 0,092 | -3,441 | -78,556 | 0 | 0 | 1,00E-08 | 0,002367 | 0,0010805 | 0,0010805 | TRUE | 0,011483 | No |
| orf19.4282 | 4752 | 4237 | 4647 | 2752 | 8683 | 6982 | 7865 | 10623 | 4326 | 4285 | 4398 | 4134 | 7723 | 7581 | 8055 | 8103 | 6075,69 | 4286 | 7866 | 1,834 | 0,875 | 22,74 | 1,82E-114 | 1,78E-113 | 0,0006203 | 0,002254 | 0,001237 | 0,001237 | TRUE | 0,0071721 | No |
| orf19.4283 | 5800 | 5222 | 5522 | 3320 | 6727 | 5510 | 5933 | 7541 | 5281 | 5281 | 5226 | 4987 | 5983 | 5982 | 6076 | 5752 | 5571,21 | 5194 | 5948 | 1,144 | 0,194 | 5,386 | 7,22E-08 | 1,25E-07 | 0,000393 | 0,002274 | 0,0010662 | 0,0010662 | TRUE | 0,0082013 | No |
| orf19.4284 | 545 | 515 | 559 | 303 | 852 | 714 | 771 | 1016 | 496 | 521 | 529 | 455 | 758 | 775 | 790 | 775 | 637,36 | 500 | 774 | 1,54 | 0,623 | 9,733 | 2,18E-22 | 5,83E-22 | 1,00E-08 | 0,00411 | 0,0022717 | 0,0022717 | TRUE | 0,034709 | No |
| orf19.4286 | 4476 | 4169 | 4432 | 2616 | 2209 | 1807 | 1823 | 2586 | 4075 | 4216 | 4194 | 3930 | 1965 | 1962 | 1867 | 1973 | 3022,77 | 4104 | 1942 | 0,473 | -1,08 | -26,106 | 3,08E-150 | 4,01E-149 | 0,0004729 | 0,002474 | 0,0012728 | 0,0012728 | TRUE | 0,0092705 | No |
| orf19.4287 | 808 | 629 | 729 | 432 | 2804 | 2200 | 2451 | 3210 | 736 | 636 | 690 | 649 | 2494 | 2389 | 2510 | 2449 | 1569,01 | 678 | 2460 | 3,62 | 1,856 | 33,952 | 1,15E-252 | 2,43E-251 | 0,0006308 | 0,002879 | 0,0018931 | 0,0018931 | TRUE | 0,0409908 | No |
| orf19.4288 | 2319 | 2049 | 2223 | 1550 | 3940 | 3389 | 3464 | 4839 | 2111 | 2072 | 2104 | 2328 | 3505 | 3680 | 3548 | 3691 | 2879,84 | 2154 | 3606 | 1,678 | 0,746 | 15,988 | 1,56E-57 | 8,18E-57 | 0,001096 | 0,002496 | 0,0017129 | 0,0017129 | TRUE | 0,032975 | No |
| orf19.429 | 3300 | 3023 | 3207 | 1923 | 3364 | 2723 | 2880 | 3980 | 3005 | 3057 | 3035 | 2889 | 2992 | 2956 | 2949 | 3036 | 2989,97 | 2996 | 2983 | 0,995 | -0,007 | -0,198 | 0,843432 | 0,865702 | 1,00E-08 | 0,002479 | 0,0009689 | 0,0009689 | TRUE | 0,0063838 | No |
| orf19.4290 | 4759 | 3993 | 4653 | 2749 | 4628 | 3691 | 3891 | 5494 | 4333 | 4038 | 4404 | 4129 | 4116 | 4007 | 3985 | 4191 | 4150,5 | 4226 | 4075 | 0,964 | -0,053 | -1,27 | 0,204198 | 0,236126 | 0,0007742 | 0,002355 | 0,001422 | 0,001422 | TRUE | 0,0113482 | No |
| orf19.4292 | 1419 | 1210 | 1231 | 832 | 1265 | 1041 | 1092 | 1460 | 1292 | 1224 | 1165 | 1250 | 1125 | 1130 | 1118 | 1114 | 1177,25 | 1233 | 1122 | 0,91 | -0,136 | -2,676 | 0,007445 | 0,009856 | 1,00E-08 | 0,003159 | 0,0016123 | 0,0016123 | TRUE | 0,0174012 | No |

| | | | | | | | | | | | | | | | | | | | | | | | | | | | | | | | |
|---|---|---|---|---|---|---|---|---|---|---|---|---|---|---|---|---|---|---|---|---|---|---|---|---|---|---|---|---|---|---|---|
| orf19.4293 | 3879 | 3097 | 3442 | 2222 | 2423 | 2011 | 2050 | 2949 | 3532 | 3132 | 3258 | 3338 | 2155 | 2183 | 2099 | 2250 | 2743,35 | 3315 | 2172 | 0,655 | -0,61 | -12,62 | 1,64E-36 | 6,05E-36 | 0,0014256 | 0,002519 | 0,0018621 | 0,0018621 | TRUE | 0,0245288 | No |
| orf19.4294 | 2734 | 2541 | 2822 | 1642 | 2640 | 2119 | 2298 | 3012 | 2489 | 2570 | 2671 | 2467 | 2348 | 2301 | 2353 | 2298 | 2437,04 | 2549 | 2325 | 0,911 | -0,134 | -3,238 | 0,001203 | 0,001691 | 0,0003269 | 0,002579 | 0,0012378 | 0,0012378 | TRUE | 0,0127068 | No |
| orf19.4295 | 1366 | 1213 | 1382 | 778 | 1455 | 1347 | 1433 | 1950 | 1244 | 1227 | 1308 | 1169 | 1294 | 1462 | 1468 | 1487 | 1332,35 | 1237 | 1428 | 1,152 | 0,205 | 3,407 | 0,000658 | 0,000939 | 0,0024262 | 0,003028 | 0,0027055 | 0,0027055 | TRUE | 0,0486627 | No |
| orf19.4297 | 2319 | 2366 | 2443 | 1542 | 1817 | 1422 | 1408 | 1910 | 2111 | 2393 | 2312 | 2316 | 1616 | 1544 | 1442 | 1457 | 1898,96 | 2283 | 1515 | 0,664 | -0,591 | -10,439 | 1,64E-25 | 4,77E-25 | 0,0023841 | 0,002733 | 0,0025371 | 0,0025371 | TRUE | 0,0321248 | No |
| orf19.4299 | 1788 | 1584 | 1745 | 1145 | 925 | 869 | 860 | 1177 | 1628 | 1602 | 1651 | 1720 | 823 | 943 | 881 | 898 | 1268,28 | 1650 | 886 | 0,537 | -0,897 | -16,471 | 5,96E-61 | 3,31E-60 | 0,0008998 | 0,003079 | 0,0019922 | 0,0019922 | TRUE | 0,0279206 | No |
| orf19.430 | 2152 | 1795 | 1993 | 1257 | 772 | 634 | 751 | 934 | 1959 | 1815 | 1886 | 1888 | 687 | 688 | 769 | 712 | 1300,72 | 1887 | 714 | 0,378 | -1,403 | -25,837 | 3,40E-147 | 4,39E-146 | 0,0006502 | 0,003053 | 0,0018928 | 0,0018928 | TRUE | 0,0301427 | No |
| orf19.4301 | 435 | 367 | 414 | 248 | 238 | 190 | 203 | 293 | 396 | 371 | 392 | 373 | 212 | 206 | 208 | 224 | 297,62 | 383 | 212 | 0,555 | -0,85 | -10,066 | 7,82E-24 | 2,19E-23 | 1,00E-08 | 0,006476 | 0,0032653 | 0,0032653 | TRUE | 0,0153898 | No |
| orf19.4304 | 36040 | 32669 | 34838 | 21350 | 34658 | 29114 | 30192 | 40690 | 32813 | 33041 | 32971 | 32071 | 30827 | 31610 | 30920 | 31039 | 31911,5 | 32724 | 31099 | 0,95 | -0,074 | -2,896 | 0,003783 | 0,005107 | 0,0001147 | 0,002078 | 0,000591 | 0,000591 | TRUE | 0,0021641 | No |
| orf19.4305 | 154 | 144 | 149 | 94 | 167 | 175 | 172 | 224 | 140 | 146 | 141 | 141 | 149 | 190 | 176 | 171 | 156,7 | 142 | 172 | 1,203 | 0,267 | 2,294 | 0,021768 | 0,027821 | 1,00E-08 | 0,010467 | 0,0065099 | 0,0065099 | TRUE | 0,0870839 | No |
| orf19.4305.1 | 1232 | 1018 | 1050 | 615 | 1410 | 1032 | 1285 | 1631 | 1122 | 1030 | 994 | 924 | 1254 | 1120 | 1316 | 1244 | 1125,45 | 1018 | 1234 | 1,21 | 0,275 | 3,932 | 8,41E-05 | 0,000127 | 0,004284 | 0,003211 | 0,0037981 | 0,0037981 | TRUE | 0,057483 | No |
| orf19.4306 | 1329 | 1154 | 1247 | 846 | 1531 | 1314 | 1217 | 1725 | 1210 | 1167 | 1180 | 1271 | 1362 | 1427 | 1246 | 1316 | 1272,35 | 1207 | 1338 | 1,11 | 0,15 | 2,681 | 0,007336 | 0,009719 | 0,0012927 | 0,003075 | 0,0022141 | 0,0022141 | TRUE | 0,0242239 | No |
| orf19.4307 | 3301 | 2955 | 2888 | 1883 | 2047 | 1577 | 1752 | 2209 | 3005 | 2989 | 2733 | 2829 | 1821 | 1712 | 1794 | 1685 | 2321,02 | 2889 | 1753 | 0,606 | -0,722 | -14,642 | 1,52E-48 | 6,84E-48 | 0,001366 | 0,002606 | 0,001879 | 0,001879 | TRUE | 0,0168306 | No |
| orf19.4308 | 397 | 353 | 373 | 261 | 723 | 611 | 662 | 909 | 361 | 357 | 353 | 392 | 643 | 663 | 678 | 693 | 517,67 | 366 | 669 | 1,838 | 0,879 | 12,757 | 2,86E-37 | 1,07E-36 | 1,00E-08 | 0,004589 | 0,0023957 | 0,0023957 | TRUE | 0,0211983 | No |
| orf19.4309 | 3683 | 3306 | 3350 | 2226 | 1117 | 911 | 915 | 1260 | 3353 | 3344 | 3170 | 3344 | 994 | 989 | 937 | 961 | 2136,5 | 3303 | 970 | 0,294 | -1,767 | -39,355 | 0 | 0 | 1,00E-08 | 0,002655 | 0,001292 | 0,001292 | TRUE | 0,0091037 | No |
| orf19.431 | 7604 | 6593 | 7260 | 4616 | 3311 | 2763 | 2939 | 4070 | 6923 | 6668 | 6871 | 6934 | 2945 | 3000 | 3010 | 3105 | 4931,94 | 6849 | 3015 | 0,44 | -1,183 | -34,001 | 2,15E-253 | 4,59E-252 | 0,0002035 | 0,002305 | 0,000929 | 0,000929 | TRUE | 0,0051781 | No |
| orf19.4310 | 1 | 0 | 1 | 1 | 0 | 1 | 0 | 0 | 1 | 0 | 1 | 2 | 0 | 1 | 0 | 0 | 0,56 | 1 | 0 | 0,348 | -1,523 | -0,818 | 0,41331 | 0,454677 | 1,00E-08 | 2,37967 | 1,6701323 | 1,6701323 | TRUE | 0,4622362 | No |
| orf19.4311 | 2458 | 2068 | 2212 | 1327 | 4048 | 2911 | 3117 | 4124 | 2238 | 2092 | 2093 | 1993 | 3601 | 3161 | 3192 | 3146 | 2689,43 | 2104 | 3275 | 1,555 | 0,637 | 11,09 | 1,41E-28 | 4,36E-28 | 0,0028422 | 0,002528 | 0,0027675 | 0,0027675 | TRUE | 0,0549126 | No |
| orf19.4312 | 913 | 771 | 847 | 487 | 945 | 779 | 889 | 1096 | 831 | 780 | 802 | 732 | 841 | 846 | 910 | 836 | 822,13 | 786 | 858 | 1,087 | 0,121 | 2 | 0,045533 | 0,056576 | 0,0007804 | 0,003644 | 0,0022825 | 0,0022825 | TRUE | 0,0222299 | No |
| orf19.4315 | 2033 | 2201 | 2308 | 1282 | 2819 | 2344 | 2366 | 3100 | 1851 | 2226 | 2184 | 1926 | 2507 | 2545 | 2423 | 2365 | 2253,41 | 2047 | 2460 | 1,201 | 0,264 | 4,126 | 3,68E-05 | 5,67E-05 | 0,0040911 | 0,002623 | 0,0034762 | 0,0034762 | TRUE | 0,0528209 | No |
| orf19.4316 | 700 | 694 | 722 | 461 | 3293 | 2793 | 2904 | 3740 | 637 | 702 | 683 | 692 | 2929 | 3032 | 2974 | 2853 | 1812,93 | 678 | 2947 | 4,347 | 2,12 | 41,967 | 0 | 0 | 0,0004094 | 0,002766 | 0,0015049 | 0,0015049 | TRUE | 0,0216682 | No |
| orf19.4317 | 2151 | 2085 | 2080 | 1282 | 903 | 655 | 667 | 857 | 1958 | 2109 | 1969 | 1926 | 803 | 711 | 683 | 654 | 1351,58 | 1990 | 713 | 0,358 | -1,483 | -22,384 | 5,67E-111 | 5,38E-110 | 0,0036734 | 0,003014 | 0,0032868 | 0,0032868 | TRUE | 0,0893769 | No |
| orf19.4318 | 17510 | 16362 | 18446 | 11314 | 9441 | 8311 | 8411 | 11529 | 15942 | 16548 | 17457 | 16995 | 8397 | 9023 | 8614 | 8794 | 12721,6 | 16736 | 8707 | 0,52 | -0,943 | -23,294 | 5,12E-120 | 5,23E-119 | 0,0010959 | 0,002141 | 0,001487 | 0,001487 | TRUE | 0,0125789 | No |
| orf19.432 | 2978 | 2591 | 2827 | 1748 | 2644 | 2207 | 2432 | 3281 | 2711 | 2621 | 2675 | 2626 | 2352 | 2396 | 2491 | 2503 | 2546,82 | 2658 | 2436 | 0,916 | -0,127 | -3,159 | 0,001585 | 0,002208 | 0,0002406 | 0,002556 | 0,0011512 | 0,0011512 | TRUE | 0,0067188 | No |
| orf19.4320 | 11 | 5 | 6 | 3 | 6 | 4 | 5 | 13 | 10 | 5 | 6 | 5 | 5 | 4 | 5 | 10 | 6,25 | 6 | 6 | 0,979 | -0,031 | -0,056 | 0,955511 | 0,96052 | 1,00E-08 | 0,213508 | 0,1429935 | 0,1429935 | TRUE | 0,5295345 | No |
| orf19.4321 | 34 | 10 | 20 | 20 | 28 | 35 | 20 | 54 | 31 | 10 | 19 | 30 | 25 | 38 | 20 | 41 | 26,83 | 22 | 31 | 1,403 | 0,488 | 1,4 | 0,161658 | 0,189512 | 0,1214492 | 0,051277 | 0,0779432 | 0,0779432 | TRUE | 0,3697715 | No |
| orf19.4322 | 7371 | 7081 | 7496 | 4323 | 9260 | 7400 | 7885 | 10190 | 6711 | 7162 | 7094 | 6494 | 8236 | 8034 | 8075 | 7773 | 7447,48 | 6865 | 8030 | 1,169 | 0,225 | 5,36 | 8,33E-08 | 1,44E-07 | 0,0011743 | 0,002214 | 0,0015577 | 0,0015577 | TRUE | 0,0155633 | No |
| orf19.4323 | 1469 | 1444 | 1402 | 781 | 378 | 293 | 319 | 316 | 1337 | 1460 | 1327 | 1173 | 336 | 318 | 327 | 241 | 815,01 | 1324 | 306 | 0,23 | -2,123 | -21,374 | 2,36E-101 | 2,07E-100 | 0,0126351 | 0,003658 | 0,0075339 | 0,0075339 | TRUE | 0,2487844 | No |
| orf19.4324 | 498 | 438 | 438 | 291 | 236 | 176 | 184 | 277 | 453 | 443 | 415 | 437 | 210 | 191 | 188 | 211 | 318,6 | 437 | 200 | 0,46 | -1,121 | -13,129 | 2,24E-39 | 8,61E-39 | 1,00E-08 | 0,006183 | 0,0034626 | 0,0034626 | TRUE | 0,0173624 | No |
| orf19.4325 | 746 | 814 | 838 | 496 | 430 | 413 | 395 | 488 | 679 | 823 | 793 | 745 | 382 | 448 | 405 | 372 | 581,04 | 760 | 402 | 0,527 | -0,924 | -11,103 | 1,21E-28 | 3,75E-28 | 0,005277 | 0,004311 | 0,0047717 | 0,0047717 | TRUE | 0,0675771 | No |
| orf19.4326 | 1519 | 1563 | 1636 | 957 | 1852 | 1533 | 1599 | 2025 | 1383 | 1581 | 1548 | 1438 | 1647 | 1664 | 1638 | 1545 | 1555,46 | 1488 | 1624 | 1,09 | 0,125 | 2,23 | 0,025729 | 0,032693 | 0,0018947 | 0,002886 | 0,0023425 | 0,0023425 | TRUE | 0,0292017 | No |
| orf19.4328 | 9486 | 9638 | 10104 | 6236 | 7540 | 6763 | 6542 | 8720 | 8637 | 9748 | 9563 | 9367 | 6707 | 7343 | 6700 | 6652 | 8089,4 | 9329 | 6850 | 0,734 | -0,446 | -8,829 | 1,06E-18 | 2,59E-18 | 0,0023477 | 0,0022 | 0,0023218 | 0,0023218 | TRUE | 0,0308355 | No |
| orf19.433 | 212 | 169 | 199 | 120 | 378 | 311 | 266 | 436 | 193 | 171 | 188 | 180 | 336 | 338 | 272 | 333 | 251,43 | 183 | 320 | 1,745 | 0,803 | 7,841 | 4,47E-15 | 9,89E-15 | 0,0035952 | 0,007291 | 0,0057128 | 0,0057128 | TRUE | 0,1084578 | No |
| orf19.4330 | 485 | 362 | 418 | 238 | 339 | 292 | 352 | 484 | 442 | 366 | 396 | 358 | 302 | 317 | 360 | 369 | 363,63 | 390 | 337 | 0,862 | -0,215 | -2,225 | 0,026072 | 0,033103 | 0,0068292 | 0,00567 | 0,006194 | 0,006194 | TRUE | 0,0925768 | No |
| orf19.4332 | 822 | 713 | 782 | 494 | 740 | 603 | 701 | 870 | 748 | 721 | 740 | 742 | 658 | 655 | 718 | 664 | 705,77 | 738 | 674 | 0,912 | -0,132 | -2,236 | 0,02533 | 0,032205 | 1,00E-08 | 0,003909 | 0,001945 | 0,001945 | TRUE | 0,0221149 | No |
| orf19.4334 | 94 | 71 | 87 | 68 | 101 | 99 | 100 | 143 | 86 | 72 | 82 | 102 | 90 | 107 | 102 | 109 | 93,84 | 86 | 102 | 1,208 | 0,273 | 1,81 | 0,07024 | 0,085749 | 0,0001457 | 0,016115 | 0,0109843 | 0,0109843 | TRUE | 0,1214397 | No |
| orf19.4335 | 278 | 296 | 240 | 175 | 1847 | 1573 | 1633 | 2228 | 253 | 299 | 227 | 263 | 1643 | 1708 | 1672 | 1700 | 970,64 | 260 | 1681 | 6,463 | 2,692 | 37,491 | 1,28E-307 | 3,38E-306 | 1,00E-08 | 0,003398 | 0,0026293 | 0,0026293 | TRUE | 0,1220873 | No |
| orf19.4336 | 12170 | 10109 | 10141 | 7408 | 20827 | 16634 | 16534 | 23664 | 11080 | 10224 | 9598 | 11128 | 18525 | 18060 | 16933 | 18051 | 14199,9 | 10508 | 17892 | 1,703 | 0,768 | 14,258 | 3,97E-46 | 1,72E-45 | 0,0030622 | 0,00213 | 0,0027119 | 0,0027119 | TRUE | 0,0416971 | No |
| orf19.4337 | 3537 | 3275 | 3482 | 2133 | 3465 | 3001 | 3102 | 4231 | 3220 | 3312 | 3295 | 3204 | 3082 | 3258 | 3177 | 3227 | 3222,08 | 3258 | 3186 | 0,977 | -0,033 | -0,893 | 0,372116 | 0,412277 | 0,0001376 | 0,002447 | 0,0009987 | 0,0009987 | TRUE | 0,0059596 | No |
| orf19.4339 | 3345 | 2920 | 3231 | 2192 | 3789 | 3344 | 3454 | 4750 | 3045 | 2953 | 3058 | 3293 | 3370 | 3631 | 3537 | 3623 | 3313,86 | 3087 | 3540 | 1,148 | 0,199 | 4,315 | 1,59E-05 | 2,50E-05 | 0,0012787 | 0,002436 | 0,0017415 | 0,0017415 | TRUE | 0,0228293 | No |
| orf19.434 | 546 | 500 | 559 | 326 | 430 | 301 | 375 | 456 | 497 | 506 | 529 | 490 | 382 | 327 | 384 | 348 | 432,84 | 506 | 360 | 0,712 | -0,49 | -6,392 | 1,64E-10 | 3,16E-10 | 0,0005937 | 0,005089 | 0,003282 | 0,003282 | TRUE | 0,0401766 | No |
| orf19.4340 | 1014 | 929 | 925 | 551 | 879 | 666 | 734 | 997 | 923 | 940 | 875 | 828 | 782 | 723 | 752 | 761 | 822,88 | 892 | 754 | 0,845 | -0,244 | -4,064 | 4,82E-05 | 7,35E-05 | 0,0006596 | 0,003642 | 0,0022368 | 0,0022368 | TRUE | 0,0237535 | No |
| orf19.4340.1 | 1132 | 1056 | 1083 | 566 | 1330 | 953 | 1109 | 1342 | 1031 | 1068 | 1025 | 850 | 1183 | 1035 | 1136 | 1024 | 1043,87 | 994 | 1094 | 1,098 | 0,135 | 1,752 | 0,079711 | 0,096682 | 0,006047 | 0,003302 | 0,0047074 | 0,0047074 | TRUE | 0,1029833 | No |
| orf19.4341 | 369 | 413 | 403 | 261 | 632 | 512 | 575 | 678 | 336 | 418 | 381 | 392 | 562 | 556 | 589 | 517 | 468,9 | 382 | 556 | 1,458 | 0,544 | 6,664 | 2,66E-11 | 5,28E-11 | 0,0032797 | 0,004854 | 0,0041481 | 0,0041481 | TRUE | 0,0811945 | No |
| orf19.4342 | 2662 | 2589 | 2835 | 1674 | 2644 | 2269 | 2368 | 3024 | 2424 | 2618 | 2683 | 2515 | 2352 | 2464 | 2425 | 2307 | 2473,37 | 2560 | 2387 | 0,932 | -0,102 | -2,189 | 0,028611 | 0,03621 | 0,0010934 | 0,002571 | 0,0016938 | 0,0016938 | TRUE | 0,0166417 | No |
| orf19.4346 | 1880 | 1623 | 1763 | 1198 | 2232 | 1941 | 2058 | 2861 | 1712 | 1641 | 1669 | 1800 | 1985 | 2107 | 2108 | 2182 | 1900,5 | 1706 | 2096 | 1,231 | 0,3 | 6,132 | 8,66E-10 | 1,62E-09 | 0,000872 | 0,002732 | 0,0017588 | 0,0017588 | TRUE | 0,0156296 | No |
| orf19.4347 | 8411 | 7558 | 8546 | 5481 | 3075 | 2514 | 2863 | 3725 | 7658 | 7644 | 8088 | 8233 | 2735 | 2730 | 2932 | 2841 | 5357,68 | 7906 | 2810 | 0,356 | -1,492 | -34,77 | 6,93E-265 | 1,55E-263 | 0,0009903 | 0,002284 | 0,0015334 | 0,0015334 | TRUE | 0,0101647 | No |
| orf19.4348 | 843 | 992 | 1054 | 584 | 842 | 726 | 766 | 848 | 768 | 1003 | 998 | 877 | 749 | 788 | 784 | 647 | 826,76 | 912 | 742 | 0,813 | -0,299 | -3,179 | 0,001479 | 0,002065 | 0,0110396 | 0,003635 | 0,0072649 | 0,0072649 | TRUE | 0,140754 | No |
| orf19.4349 | 732 | 687 | 786 | 407 | 506 | 377 | 425 | 605 | 666 | 695 | 744 | 611 | 450 | 409 | 435 | 461 | 559,08 | 679 | 439 | 0,646 | -0,631 | -8,346 | 7,07E-17 | 1,65E-16 | 0,0027629 | 0,0044 | 0,0036464 | 0,0036464 | TRUE | 0,0479576 | No |
| orf19.4353 | 2075 | 2111 | 2270 | 1345 | 991 | 852 | 839 | 1091 | 1889 | 2135 | 2148 | 2020 | 881 | 925 | 859 | 832 | 1461,37 | 2048 | 874 | 0,426 | -1,23 | -20,79 | 5,29E-96 | 4,36E-95 | 0,0022504 | 0,002941 | 0,0025615 | 0,0025615 | TRUE | 0,0347612 | No |
| orf19.4354 | 3915 | 3629 | 3917 | 2299 | 4581 | 3751 | 4184 | 5321 | 3564 | 3670 | 3707 | 3453 | 4075 | 4073 | 4285 | 4059 | 3860,8 | 3598 | 4123 | 1,144 | 0,194 | 4,938 | 7,87E-07 | 1,31E-06 | 0,0005021 | 0,002379 | 0,0012263 | 0,0012263 | TRUE | 0,0084353 | No |
| orf19.4355 | 1715 | 1749 | 1803 | 1096 | 1043 | 900 | 910 | 1217 | 1561 | 1769 | 1706 | 1646 | 928 | 977 | 932 | 928 | 1306,03 | 1670 | 941 | 0,563 | -0,829 | -15,196 | 3,75E-52 | 1,80E-51 | 0,0012304 | 0,003048 | 0,0020385 | 0,0020385 | TRUE | 0,0250979 | No |
| orf19.4356 | 188 | 219 | 196 | 136 | 397 | 352 | 344 | 449 | 171 | 221 | 185 | 204 | 353 | 382 | 352 | 343 | 276,57 | 195 | 358 | 1,833 | 0,874 | 8,952 | 3,51E-19 | 8,70E-19 | 0,0021697 | 0,006813 | 0,0051122 | 0,0051122 | TRUE | 0,0971476 | No |
| orf19.4357 | 1706 | 1619 | 1652 | 1019 | 2398 | 2036 | 1984 | 2663 | 1553 | 1637 | 1563 | 1531 | 2133 | 2211 | 2032 | 2031 | 1836,44 | 1571 | 2102 | 1,336 | 0,418 | 8,753 | 2,08E-18 | 5,05E-18 | 0,0007675 | 0,002756 | 0,0016239 | 0,0016239 | TRUE | 0,0142438 | No |
| orf19.4358 | 1458 | 1217 | 1269 | 766 | 595 | 437 | 470 | 712 | 1327 | 1231 | 1201 | 1151 | 529 | 474 | 481 | 543 | 867,26 | 1228 | 507 | 0,413 | -1,274 | -18,58 | 4,65E-77 | 3,17E-76 | 0,002738 | 0,00356 | 0,0031598 | 0,0031598 | TRUE | 0,0365534 | No |
| orf19.4361 | 75 | 49 | 50 | 34 | 46 | 26 | 32 | 45 | 68 | 50 | 47 | 51 | 41 | 28 | 33 | 34 | 44,06 | 54 | 34 | 0,631 | -0,665 | -3,023 | 0,002505 | 0,00343 | 1,00E-08 | 0,032019 | 0,0231069 | 0,0231069 | TRUE | 0,2923777 | No |
| orf19.4362 | 2260 | 2206 | 2436 | 1335 | 2157 | 1759 | 1837 | 2382 | 2058 | 2231 | 2305 | 2005 | 1919 | 1910 | 1881 | 1817 | 2015,79 | 2150 | 1882 | 0,874 | -0,195 | -3,625 | 0,00029 | 0,000423 | 0,0018043 | 0,002692 | 0,0022711 | 0,0022711 | TRUE | 0,0287508 | No |
| orf19.4363 | 8027 | 7439 | 8204 | 4695 | 9289 | 6988 | 8127 | 10538 | 7308 | 7524 | 7764 | 7053 | 8262 | 7587 | 8323 | 8038 | 7732,48 | 7412 | 8052 | 1,086 | 0,119 | 2,657 | 0,007882 | 0,010414 | 0,0015248 | 0,002208 | 0,0017986 | 0,0017986 | TRUE | 0,0182919 | No |
| orf19.4365 | 1836 | 1874 | 1941 | 1259 | 1416 | 1144 | 1236 | 1572 | 1672 | 1895 | 1837 | 1891 | 1259 | 1242 | 1266 | 1199 | 1532,71 | 1824 | 1242 | 0,681 | -0,554 | -10,243 | 1,27E-24 | 3,60E-24 | 0,0015363 | 0,002899 | 0,0021325 | 0,0021325 | TRUE | 0,0394017 | No |
| orf19.4366 | 44 | 33 | 42 | 34 | 25 | 19 | 26 | 34 | 40 | 33 | 40 | 51 | 22 | 21 | 27 | 26 | 32,46 | 41 | 24 | 0,59 | -0,76 | -3,075 | 0,002106 | 0,0029 | 1,00E-08 | 0,042732 | 0,0260957 | 0,0260957 | TRUE | 0,1446819 | No |
| orf19.4367 | 617 | 594 | 603 | 395 | 1058 | 815 | 920 | 1187 | 562 | 601 | 571 | 593 | 941 | 885 | 942 | 905 | 750,02 | 582 | 918 | 1,582 | 0,661 | 11,288 | 1,51E-29 | 4,79E-29 | 1,00E-08 | 0,003798 | 0,0018707 | 0,0018707 | TRUE | 0,0066398 | No |
| orf19.4368 | 1220 | 1074 | 1216 | 826 | 1388 | 1283 | 1274 | 1620 | 1111 | 1086 | 1151 | 1241 | 1235 | 1393 | 1305 | 1236 | 1219,58 | 1147 | 1292 | 1,128 | 0,173 | 2,845 | 0,004442 | 0,005977 | 0,002332 | 0,00312 | 0,0027276 | 0,0027276 | TRUE | 0,0326435 | No |
| orf19.4369 | 2076 | 1987 | 2095 | 1332 | 1529 | 1341 | 1469 | 1962 | 1890 | 2010 | 1983 | 2001 | 1360 | 1456 | 1504 | 1497 | 1712,55 | 1971 | 1454 | 0,738 | -0,438 | -8,89 | 6,10E-19 | 1,51E-18 | 0,0008848 | 0,002808 | 0,0017336 | 0,0017336 | TRUE | 0,0234713 | No |
| orf19.437 | 4873 | 4353 | 4774 | 2780 | 8629 | 7073 | 7639 | 10346 | 4437 | 4403 | 4518 | 4176 | 7675 | 7679 | 7823 | 7892 | 6075,41 | 4384 | 7767 | 1,77 | 0,824 | 23,17 | 9,19E-119 | 9,28E-118 | 0,000417 | 0,002254 | 0,0010313 | 0,0010313 | TRUE | 0,0114562 | No |
| orf19.4370 | 2691 | 2490 | 2641 | 1615 | 1019 | 859 | 899 | 1179 | 2450 | 2518 | 2499 | 2426 | 906 | 933 | 921 | 899 | 1694,11 | 2473 | 915 | 0,369 | -1,437 | -32,749 | 3,11E-235 | 6,14E-234 | 1,00E-08 | 0,002817 | 0,0011223 | 0,0011223 | TRUE | 0,001992 | No |
| orf19.4371 | 2963 | 2517 | 2659 | 1830 | 3252 | 2638 | 2631 | 3892 | 2698 | 2546 | 2517 | 2749 | 2893 | 2864 | 2694 | 2969 | 2741,1 | 2628 | 2855 | 1,088 | 0,122 | 2,539 | 0,011113 | 0,014533 | 0,0012695 | 0,002519 | 0,001847 | 0,001847 | TRUE | 0,0172942 | No |
| orf19.4372 | 5333 | 5067 | 5410 | 3454 | 1394 | 1308 | 1293 | 1713 | 4855 | 5125 | 5120 | 5188 | 1240 | 1420 | 1324 | 1307 | 3197,46 | 5072 | 1323 | 0,261 | -1,94 | -40,017 | 0 | 0 | 0,0013707 | 0,00245 | 0,0017977 | 0,0017977 | TRUE | 0,0283998 | No |
| orf19.4373 | 1735 | 1617 | 1577 | 1130 | 1558 | 1369 | 1436 | 1824 | 1580 | 1635 | 1492 | 1697 | 1386 | 1486 | 1471 | 1391 | 1517,39 | 1601 | 1434 | 0,896 | -0,158 | -2,946 | 0,003222 | 0,004374 | 0,0012623 | 0,002908 | 0,0021072 | 0,0021072 | TRUE | 0,0252341 | No |

| | | | | | | | | | | | | | | | | | | | | | | | | | | | | | | | |
|---|---|---|---|---|---|---|---|---|---|---|---|---|---|---|---|---|---|---|---|---|---|---|---|---|---|---|---|---|---|---|---|
| orf19.4374 | 598 | 599 | 632 | 354 | 800 | 624 | 656 | 814 | 544 | 606 | 598 | 532 | 712 | 677 | 672 | 621 | 620,25 | 570 | 670 | 1,172 | 0,229 | 3,226 | 0,001254 | 0,00176 | 0,0021633 | 0,004167 | 0,0032094 | 0,0032094 | TRUE | 0,0316878 | No |
| orf19.4375 | 2839 | 2354 | 2562 | 1656 | 2646 | 2147 | 2276 | 2986 | 2585 | 2381 | 2425 | 2488 | 2354 | 2331 | 2331 | 2278 | 2396,39 | 2470 | 2324 | 0,941 | -0,088 | -2,111 | 0,034733 | 0,043665 | 0,0003915 | 0,002588 | 0,0012642 | 0,0012642 | TRUE | 0,012851 | No |
| orf19.4375.1 | 10056 | 7792 | 8021 | 6341 | 13012 | 9462 | 9048 | 14513 | 9156 | 7881 | 7591 | 9525 | 11574 | 10273 | 9266 | 11071 | 9542,06 | 8538 | 10546 | 1,236 | 0,305 | 3,679 | 0,000234 | 0,000345 | 0,010557 | 0,002175 | 0,0065044 | 0,0065044 | TRUE | 0,0815536 | No |
| orf19.4376 | 384 | 371 | 379 | 256 | 214 | 195 | 194 | 256 | 350 | 375 | 359 | 385 | 190 | 212 | 199 | 195 | 283,01 | 367 | 199 | 0,543 | -0,882 | -10,085 | 6,44E-24 | 1,80E-23 | 1,00E-08 | 0,006705 | 0,0035435 | 0,0035435 | TRUE | 0,0184791 | No |
| orf19.4377 | 2574 | 2458 | 2626 | 1636 | 695 | 582 | 631 | 779 | 2344 | 2486 | 2485 | 2458 | 618 | 632 | 646 | 594 | 1532,86 | 2443 | 622 | 0,254 | -1,975 | -38,706 | 0 | 0 | 1,00E-08 | 0,002899 | 0,0015293 | 0,0015293 | TRUE | 0,0121557 | No |
| orf19.4378 | 715 | 561 | 600 | 357 | 862 | 656 | 840 | 1058 | 651 | 567 | 568 | 536 | 767 | 712 | 860 | 807 | 683,59 | 580 | 786 | 1,351 | 0,434 | 5,495 | 3,91E-08 | 6,86E-08 | 0,0047619 | 0,003969 | 0,0044805 | 0,0044805 | TRUE | 0,0801273 | No |
| orf19.4379 | 1701 | 1283 | 1587 | 1019 | 1045 | 920 | 941 | 1431 | 1549 | 1298 | 1502 | 1531 | 929 | 999 | 964 | 1092 | 1232,82 | 1470 | 996 | 0,678 | -0,56 | -7,802 | 6,11E-15 | 1,35E-14 | 0,0050196 | 0,003109 | 0,0041054 | 0,0041054 | TRUE | 0,0757172 | No |
| orf19.438 | 720 | 765 | 741 | 541 | 958 | 786 | 849 | 1137 | 656 | 774 | 701 | 813 | 852 | 853 | 869 | 867 | 798,19 | 736 | 860 | 1,175 | 0,232 | 3,352 | 0,000803 | 0,00114 | 0,0029868 | 0,003692 | 0,0033409 | 0,0033409 | TRUE | 0,0647047 | No |
| orf19.4380.1 | 1803 | 1455 | 1626 | 1059 | 1217 | 942 | 1013 | 1468 | 1642 | 1472 | 1539 | 1591 | 1082 | 1023 | 1037 | 1120 | 1313,16 | 1561 | 1066 | 0,684 | -0,548 | -10,018 | 1,27E-23 | 3,53E-23 | 0,001291 | 0,003043 | 0,0020916 | 0,0020916 | TRUE | 0,0181629 | No |
| orf19.4381 | 2444 | 2317 | 2488 | 1717 | 2266 | 2004 | 1925 | 2592 | 2225 | 2343 | 2355 | 2579 | 2016 | 2176 | 1971 | 1977 | 2205,3 | 2376 | 2035 | 0,858 | -0,222 | -3,987 | 6,70E-05 | 0,000101 | 0,002438 | 0,002636 | 0,0025099 | 0,0025099 | TRUE | 0,0381212 | No |
| orf19.4382 | 1278 | 1228 | 1202 | 720 | 1102 | 916 | 1014 | 1307 | 1164 | 1242 | 1138 | 1082 | 980 | 995 | 1038 | 997 | 1079,36 | 1156 | 1002 | 0,865 | -0,209 | -3,751 | 0,000176 | 0,00026 | 0,0007333 | 0,003261 | 0,0020549 | 0,0020549 | TRUE | 0,029131 | No |
| orf19.4383 | 151 | 149 | 162 | 128 | 198 | 160 | 171 | 228 | 137 | 151 | 153 | 192 | 176 | 174 | 175 | 174 | 166,58 | 158 | 175 | 1,116 | 0,159 | 1,36 | 0,173896 | 0,202667 | 0,0013392 | 0,009967 | 0,0070435 | 0,0070435 | TRUE | 0,1753398 | No |
| orf19.4384 | 28 | 11 | 28 | 18 | 24 | 20 | 25 | 33 | 25 | 11 | 26 | 27 | 21 | 22 | 26 | 25 | 23 | 22 | 24 | 1,049 | 0,069 | 0,226 | 0,821247 | 0,845267 | 0,0191667 | 0,059474 | 0,047195 | 0,047195 | TRUE | 0,6914408 | No |
| orf19.4384.1 | 33 | 20 | 22 | 12 | 36 | 33 | 28 | 43 | 30 | 20 | 21 | 18 | 32 | 36 | 29 | 33 | 27,31 | 22 | 32 | 1,428 | 0,514 | 1,921 | 0,054686 | 0,067536 | 1,00E-08 | 0,050415 | 0,0307605 | 0,0307605 | TRUE | 0,3471292 | No |
| orf19.4388 | 1920 | 1551 | 1835 | 1084 | 1769 | 1393 | 1641 | 2100 | 1748 | 1569 | 1737 | 1628 | 1573 | 1512 | 1681 | 1602 | 1631,26 | 1670 | 1592 | 0,952 | -0,071 | -1,3 | 0,193544 | 0,224295 | 0,0014902 | 0,002847 | 0,0022083 | 0,0022083 | TRUE | 0,0214683 | No |
| orf19.439 | 2395 | 1919 | 2177 | 1469 | 2130 | 1620 | 1878 | 2396 | 2181 | 1941 | 2060 | 2207 | 1895 | 1759 | 1923 | 1828 | 1974,11 | 2097 | 1851 | 0,884 | -0,179 | -3,31 | 0,000932 | 0,001317 | 0,0018031 | 0,002706 | 0,0022826 | 0,0022826 | TRUE | 0,0304085 | No |
| orf19.4390 | 624 | 592 | 606 | 377 | 731 | 650 | 679 | 944 | 568 | 599 | 574 | 566 | 650 | 706 | 695 | 720 | 634,76 | 577 | 693 | 1,201 | 0,264 | 4,2 | 2,67E-05 | 4,14E-05 | 1,00E-08 | 0,004118 | 0,0021984 | 0,0021984 | TRUE | 0,0211263 | No |
| orf19.4391 | 65 | 52 | 50 | 39 | 49 | 39 | 39 | 49 | 59 | 53 | 47 | 59 | 44 | 42 | 40 | 37 | 47,62 | 54 | 41 | 0,751 | -0,413 | -2,086 | 0,037018 | 0,046419 | 1,00E-08 | 0,02978 | 0,0164266 | 0,0164266 | TRUE | 0,0694643 | No |
| orf19.4392 | 407 | 365 | 381 | 225 | 492 | 410 | 451 | 548 | 371 | 369 | 361 | 338 | 438 | 445 | 462 | 418 | 400,12 | 360 | 441 | 1,219 | 0,286 | 3,828 | 0,000129 | 0,000193 | 1,00E-08 | 0,005339 | 0,0028189 | 0,0028189 | TRUE | 0,0151193 | No |
| orf19.4393 | 12760 | 10656 | 11509 | 8198 | 14121 | 12165 | 12094 | 17702 | 11617 | 10777 | 10892 | 12315 | 12560 | 13208 | 12386 | 13503 | 12157,4 | 11400 | 12914 | 1,133 | 0,18 | 3,51 | 0,000447 | 0,000646 | 0,0026883 | 0,002146 | 0,0024546 | 0,0024546 | TRUE | 0,0351126 | No |
| orf19.4394 | 17 | 11 | 18 | 19 | 60 | 61 | 61 | 84 | 15 | 11 | 17 | 29 | 53 | 66 | 62 | 64 | 39,79 | 18 | 61 | 3,552 | 1,829 | 7,051 | 1,78E-12 | 3,67E-12 | 1,00E-08 | 0,035236 | 0,0263917 | 0,0263917 | TRUE | 0,4953001 | No |
| orf19.4395 | 1295 | 1164 | 1246 | 823 | 2525 | 2127 | 2206 | 2914 | 1179 | 1177 | 1179 | 1236 | 2246 | 2309 | 2259 | 2223 | 1726,14 | 1193 | 2259 | 1,896 | 0,923 | 21,42 | 8,66E-102 | 7,67E-101 | 1,00E-08 | 0,002802 | 0,0011283 | 0,0011283 | TRUE | 0,0060473 | No |
| orf19.4396 | 3583 | 3083 | 3534 | 2146 | 4829 | 3989 | 4342 | 5800 | 3262 | 3118 | 3345 | 3224 | 4295 | 4331 | 4447 | 4424 | 3805,72 | 3237 | 4374 | 1,351 | 0,434 | 11,666 | 1,91E-31 | 6,36E-31 | 0,0002744 | 0,002384 | 0,001056 | 0,001056 | TRUE | 0,0077577 | No |
| orf19.4398 | 2151 | 1853 | 1975 | 1378 | 2715 | 2249 | 2232 | 3093 | 1958 | 1874 | 1869 | 2070 | 2415 | 2442 | 2286 | 2359 | 2159,19 | 1943 | 2376 | 1,225 | 0,293 | 6,185 | 6,22E-10 | 1,17E-09 | 0,0009695 | 0,002649 | 0,001674 | 0,001674 | TRUE | 0,0212403 | No |
| orf19.4399 | 9811 | 8410 | 9100 | 5659 | 9673 | 7691 | 8646 | 11437 | 8932 | 8506 | 8612 | 8501 | 8604 | 8350 | 8855 | 8724 | 8635,54 | 8638 | 8633 | 0,999 | -0,001 | -0,029 | 0,976611 | 0,97876 | 0,0003841 | 0,00219 | 0,0010167 | 0,0010167 | TRUE | 0,0065352 | No |
| orf19.440 | 499 | 430 | 457 | 341 | 418 | 376 | 379 | 560 | 454 | 435 | 433 | 512 | 372 | 408 | 388 | 427 | 428,66 | 458 | 399 | 0,875 | -0,193 | -2,389 | 0,016904 | 0,021823 | 0,0023204 | 0,005119 | 0,0039055 | 0,0039055 | TRUE | 0,0617698 | No |
| orf19.4400 | 471 | 358 | 422 | 287 | 455 | 364 | 362 | 586 | 429 | 362 | 399 | 431 | 405 | 395 | 371 | 447 | 404,88 | 405 | 404 | 1,002 | 0,003 | 0,037 | 0,970864 | 0,974379 | 0,0039959 | 0,0053 | 0,0047093 | 0,0047093 | TRUE | 0,0618919 | No |
| orf19.4401 | 3461 | 2937 | 3128 | 2080 | 3432 | 2872 | 2816 | 4023 | 3151 | 2970 | 2960 | 3124 | 3053 | 3118 | 2884 | 3069 | 3041,25 | 3051 | 3031 | 0,994 | -0,009 | -0,21 | 0,833734 | 0,8569 | 0,0007677 | 0,002471 | 0,0014411 | 0,0014411 | TRUE | 0,0126677 | No |
| orf19.4403 | 1007 | 1012 | 1077 | 615 | 556 | 553 | 472 | 621 | 917 | 1024 | 1019 | 924 | 495 | 600 | 483 | 474 | 741,94 | 971 | 513 | 0,527 | -0,925 | -11,414 | 3,56E-30 | 1,15E-29 | 0,0064131 | 0,003817 | 0,0048423 | 0,0048423 | TRUE | 0,1477479 | No |
| orf19.4404 | 15 | 15 | 16 | 8 | 8 | 10 | 7 | 10 | 14 | 15 | 15 | 12 | 7 | 11 | 7 | 8 | 11,09 | 14 | 8 | 0,575 | -0,797 | -1,957 | 0,050384 | 0,062381 | 1,00E-08 | 0,121103 | 0,0644174 | 0,0644174 | TRUE | 0,1234335 | No |
| orf19.4405 | 36 | 33 | 30 | 31 | 55 | 56 | 28 | 58 | 33 | 33 | 28 | 47 | 49 | 61 | 29 | 44 | 40,47 | 35 | 46 | 1,314 | 0,394 | 1,545 | 0,122315 | 0,145504 | 0,0405468 | 0,034679 | 0,0368445 | 0,0368445 | TRUE | 0,2500044 | No |
| orf19.4406 | 426 | 397 | 366 | 332 | 514 | 488 | 518 | 588 | 388 | 402 | 346 | 499 | 457 | 530 | 530 | 449 | 450,07 | 409 | 492 | 1,208 | 0,273 | 2,581 | 0,00986 | 0,012939 | 0,0136105 | 0,004972 | 0,0084775 | 0,0084775 | TRUE | 0,2294089 | No |
| orf19.4407 | 3110 | 2827 | 3012 | 1886 | 3709 | 3121 | 3466 | 4587 | 2832 | 2859 | 2851 | 2833 | 3299 | 3389 | 3550 | 3499 | 3138,82 | 2844 | 3434 | 1,208 | 0,272 | 7,047 | 1,83E-12 | 3,76E-12 | 0,0003218 | 0,002458 | 0,0011064 | 0,0011064 | TRUE | 0,0086903 | No |
| orf19.4409 | 2378 | 2019 | 2249 | 1525 | 2102 | 1823 | 1920 | 2674 | 2165 | 2042 | 2128 | 2291 | 1870 | 1979 | 1966 | 2040 | 2060,17 | 2156 | 1964 | 0,912 | -0,133 | -2,685 | 0,007257 | 0,009617 | 0,0011412 | 0,002678 | 0,0018614 | 0,0018614 | TRUE | 0,0194957 | No |
| orf19.441 | 9876 | 7984 | 8973 | 5326 | 7572 | 5915 | 6739 | 9374 | 8992 | 8075 | 8492 | 8001 | 6735 | 6422 | 6902 | 7151 | 7596,06 | 8390 | 6802 | 0,811 | -0,303 | -6,031 | 1,63E-09 | 3,03E-09 | 0,0023081 | 0,002211 | 0,0022884 | 0,0022884 | TRUE | 0,0286144 | No |
| orf19.4410 | 1201 | 1088 | 1140 | 704 | 1403 | 1244 | 1260 | 1711 | 1093 | 1100 | 1079 | 1058 | 1248 | 1351 | 1290 | 1305 | 1190,55 | 1082 | 1298 | 1,198 | 0,26 | 5,309 | 1,10E-07 | 1,90E-07 | 1,00E-08 | 0,003147 | 0,0014521 | 0,0014521 | TRUE | 0,0083002 | No |
| orf19.4411 | 1246 | 1099 | 1242 | 755 | 1128 | 920 | 1005 | 1373 | 1134 | 1112 | 1175 | 1134 | 1003 | 999 | 1029 | 1047 | 1079,29 | 1139 | 1020 | 0,896 | -0,159 | -3,203 | 0,00136 | 0,001904 | 1,00E-08 | 0,003261 | 0,0014394 | 0,0014394 | TRUE | 0,005952 | No |
| orf19.4412 | 1220 | 1032 | 1192 | 679 | 1492 | 1183 | 1334 | 1824 | 1111 | 1044 | 1128 | 1020 | 1327 | 1284 | 1366 | 1391 | 1208,96 | 1076 | 1342 | 1,246 | 0,317 | 5,896 | 3,71E-09 | 6,81E-09 | 0,0008108 | 0,00313 | 0,0019327 | 0,0019327 | TRUE | 0,0127611 | No |
| orf19.4413 | 9297 | 7643 | 8344 | 5335 | 3939 | 2860 | 3153 | 4390 | 8465 | 7730 | 7897 | 8014 | 3504 | 3105 | 3229 | 3349 | 5661,5 | 8026 | 3297 | 0,411 | -1,283 | -26,687 | 6,67E-157 | 9,12E-156 | 0,0018132 | 0,00227 | 0,0020111 | 0,0020111 | TRUE | 0,0215956 | No |
| orf19.4414 | 929 | 792 | 849 | 594 | 1192 | 947 | 960 | 1308 | 846 | 801 | 804 | 892 | 1060 | 1028 | 983 | 998 | 926,5 | 836 | 1017 | 1,221 | 0,288 | 5,036 | 4,75E-07 | 8,00E-07 | 0,0004352 | 0,003463 | 0,0020344 | 0,0020344 | TRUE | 0,0211119 | No |
| orf19.4415 | 1534 | 1327 | 1496 | 1008 | 762 | 631 | 650 | 895 | 1397 | 1342 | 1416 | 1514 | 678 | 685 | 666 | 683 | 1047,5 | 1417 | 678 | 0,479 | -1,061 | -19,157 | 8,42E-82 | 6,10E-81 | 1,00E-08 | 0,003298 | 0,0018762 | 0,0018762 | TRUE | 0,0227556 | No |
| orf19.4416 | 1284 | 1103 | 1271 | 769 | 837 | 674 | 805 | 1027 | 1169 | 1116 | 1203 | 1155 | 744 | 732 | 824 | 783 | 965,84 | 1161 | 771 | 0,664 | -0,59 | -10,306 | 6,60E-25 | 1,89E-24 | 0,0006853 | 0,003405 | 0,0020824 | 0,0020824 | TRUE | 0,0241587 | No |
| orf19.4418 | 1114 | 963 | 1123 | 704 | 1014 | 873 | 908 | 1223 | 1014 | 974 | 1063 | 1058 | 902 | 948 | 930 | 933 | 977,64 | 1027 | 928 | 0,904 | -0,145 | -2,72 | 0,006531 | 0,008687 | 1,00E-08 | 0,003388 | 0,0017119 | 0,0017119 | TRUE | 0,014511 | No |
| orf19.4418.1 | 326 | 246 | 281 | 237 | 200 | 174 | 157 | 202 | 297 | 249 | 266 | 356 | 178 | 189 | 161 | 154 | 231,16 | 292 | 170 | 0,586 | -0,77 | -6,315 | 2,70E-10 | 5,15E-10 | 0,0123031 | 0,007752 | 0,0096902 | 0,0096902 | TRUE | 0,2158616 | No |
| orf19.442 | 85 | 52 | 58 | 45 | 68 | 52 | 38 | 86 | 77 | 53 | 55 | 68 | 60 | 56 | 39 | 66 | 59,24 | 63 | 55 | 0,884 | -0,177 | -0,857 | 0,391646 | 0,432637 | 0,0242972 | 0,024336 | 0,0243161 | 0,0243161 | TRUE | 0,325176 | No |
| orf19.4420 | 1769 | 1612 | 1751 | 1121 | 2714 | 2328 | 2588 | 3276 | 1611 | 1630 | 1657 | 1684 | 2414 | 2528 | 2650 | 2499 | 2084,13 | 1646 | 2523 | 1,534 | 0,617 | 13,718 | 7,96E-43 | 3,26E-42 | 0,0005773 | 0,002671 | 0,0014311 | 0,0014311 | TRUE | 0,0139172 | No |
| orf19.4423 | 1507 | 1412 | 1548 | 925 | 2439 | 2045 | 2213 | 2929 | 1372 | 1428 | 1465 | 1389 | 2169 | 2220 | 2266 | 2234 | 1818,13 | 1414 | 2222 | 1,571 | 0,652 | 14,998 | 7,57E-51 | 3,55E-50 | 1,00E-08 | 0,002764 | 0,0012244 | 0,0012244 | TRUE | 0,0075397 | No |
| orf19.4424 | 43 | 42 | 63 | 37 | 37 | 31 | 50 | 57 | 39 | 42 | 60 | 56 | 33 | 34 | 51 | 43 | 44,76 | 49 | 40 | 0,825 | -0,277 | -1,211 | 0,226044 | 0,259505 | 0,0215254 | 0,031549 | 0,0278349 | 0,0278349 | TRUE | 0,2196447 | No |
| orf19.4426 | 1976 | 1999 | 2118 | 1219 | 2050 | 1815 | 1734 | 2379 | 1799 | 2022 | 2004 | 1831 | 1823 | 1971 | 1776 | 1815 | 1880,13 | 1914 | 1846 | 0,963 | -0,054 | -0,959 | 0,337722 | 0,377848 | 0,0021774 | 0,00274 | 0,0024974 | 0,0024974 | TRUE | 0,024164 | No |
| orf19.4427 | 6930 | 5924 | 6384 | 3598 | 5973 | 4374 | 5107 | 6547 | 6309 | 5991 | 6042 | 5405 | 5313 | 4749 | 5230 | 4994 | 5504,21 | 5937 | 5072 | 0,854 | -0,228 | -4,11 | 3,95E-05 | 6,07E-05 | 0,0030713 | 0,002277 | 0,0027773 | 0,0027773 | TRUE | 0,0433074 | No |
| orf19.4428 | 1415 | 1355 | 1442 | 872 | 1111 | 1018 | 1127 | 1265 | 1288 | 1370 | 1365 | 1310 | 988 | 1105 | 1154 | 965 | 1193,24 | 1333 | 1053 | 0,788 | -0,344 | -5,26 | 1,44E-07 | 2,48E-07 | 0,0030752 | 0,003144 | 0,003247 | 0,003247 | TRUE | 0,0504537 | No |
| orf19.443 | 863 | 769 | 804 | 493 | 1287 | 956 | 993 | 1282 | 786 | 778 | 761 | 741 | 1145 | 1038 | 1017 | 978 | 905,32 | 766 | 1044 | 1,361 | 0,444 | 7,04 | 1,92E-12 | 3,96E-12 | 0,0019656 | 0,003496 | 0,0026807 | 0,0026807 | TRUE | 0,0515995 | No |
| orf19.4430 | 4595 | 4147 | 4472 | 2718 | 9081 | 7582 | 8338 | 10924 | 4184 | 4194 | 4232 | 4083 | 8077 | 8232 | 8539 | 8333 | 6234,29 | 4173 | 8295 | 1,986 | 0,99 | 29,488 | 4,11E-191 | 6,68E-190 | 0,0002572 | 0,002249 | 0,0008975 | 0,0008975 | TRUE | 0,0047241 | No |
| orf19.4432 | 18400 | 16196 | 17266 | 10464 | 3030 | 2430 | 2631 | 3501 | 16752 | 16381 | 16341 | 15719 | 2695 | 2638 | 2694 | 2671 | 9486,34 | 16298 | 2674 | 0,164 | -2,608 | -75,784 | 0 | 0 | 0,000367 | 0,002176 | 0,0009293 | 0,0009293 | TRUE | 0,0068347 | No |
| orf19.4433 | 314 | 248 | 282 | 208 | 591 | 535 | 536 | 771 | 286 | 251 | 267 | 312 | 526 | 581 | 549 | 588 | 419,95 | 279 | 561 | 2,023 | 1,016 | 12,238 | 1,95E-34 | 6,88E-34 | 0,0019214 | 0,005183 | 0,0038656 | 0,0038656 | TRUE | 0,0583872 | No |
| orf19.4435 | 4059 | 3940 | 4004 | 2733 | 1500 | 1275 | 1304 | 1810 | 3696 | 3985 | 3789 | 4105 | 1334 | 1384 | 1335 | 1381 | 2626,24 | 3894 | 1358 | 0,349 | -1,517 | -31,969 | 2,90E-224 | 5,47E-223 | 0,0011094 | 0,00254 | 0,0016808 | 0,0016808 | TRUE | 0,0155114 | No |
| orf19.4436 | 412 | 392 | 394 | 271 | 60 | 45 | 44 | 76 | 375 | 396 | 373 | 407 | 53 | 49 | 45 | 58 | 219,6 | 388 | 51 | 0,134 | -2,901 | -23,096 | 5,08E-118 | 5,12E-117 | 1,00E-08 | 0,008052 | 0,0048261 | 0,0048261 | TRUE | 0,0750598 | No |
| orf19.4437 | 2715 | 2615 | 2826 | 1727 | 4038 | 3275 | 3719 | 4607 | 2472 | 2645 | 2675 | 2594 | 3592 | 3556 | 3809 | 3514 | 3106,98 | 2596 | 3618 | 1,393 | 0,478 | 10,79 | 3,82E-27 | 1,15E-26 | 0,0009636 | 0,002462 | 0,00155 | 0,00155 | TRUE | 0,0152407 | No |
| orf19.4438 | 911 | 811 | 931 | 654 | 1044 | 886 | 1008 | 1289 | 829 | 820 | 881 | 982 | 929 | 962 | 1032 | 983 | 927,41 | 878 | 976 | 1,116 | 0,158 | 2,368 | 0,017865 | 0,023008 | 0,0031367 | 0,003461 | 0,003188 | 0,003188 | TRUE | 0,0679039 | No |
| orf19.4439 | 17 | 11 | 11 | 5 | 17 | 27 | 26 | 28 | 15 | 11 | 10 | 8 | 15 | 29 | 27 | 21 | 17,12 | 11 | 23 | 2,008 | 1,006 | 2,771 | 0,005582 | 0,007455 | 1,00E-08 | 0,079206 | 0,059512 | 0,059512 | TRUE | 0,2870128 | No |
| orf19.4439.1 | 425 | 345 | 358 | 287 | 442 | 351 | 312 | 455 | 387 | 349 | 339 | 431 | 393 | 381 | 320 | 347 | 368,33 | 376 | 360 | 0,962 | -0,057 | -0,583 | 0,559714 | 0,600906 | 0,0068733 | 0,005623 | 0,0062862 | 0,0062862 | TRUE | 0,0952651 | No |
| orf19.444 | 468 | 394 | 419 | 265 | 476 | 397 | 389 | 530 | 426 | 398 | 397 | 398 | 423 | 431 | 398 | 404 | 409,54 | 405 | 414 | 1,021 | 0,03 | 0,417 | 0,676887 | 0,713131 | 1,00E-08 | 0,005263 | 0,0026737 | 0,0026737 | TRUE | 0,0157134 | No |
| orf19.4440 | 893 | 739 | 832 | 509 | 677 | 587 | 598 | 790 | 813 | 747 | 787 | 765 | 602 | 637 | 612 | 603 | 695,88 | 778 | 614 | 0,787 | -0,346 | -5,805 | 6,45E-09 | 1,17E-08 | 1,00E-08 | 0,003935 | 0,0019656 | 0,0019656 | TRUE | 0,0114175 | No |
| orf19.4441 | 417 | 376 | 436 | 264 | 565 | 465 | 490 | 633 | 380 | 380 | 413 | 397 | 503 | 505 | 502 | 483 | 445,15 | 392 | 498 | 1,269 | 0,344 | 4,892 | 9,97E-07 | 1,66E-06 | 1,00E-08 | 0,005005 | 0,0024453 | 0,0024453 | TRUE | 0,0160037 | No |
| orf19.4442 | 1075 | 955 | 1039 | 701 | 1034 | 893 | 999 | 1353 | 979 | 966 | 983 | 1053 | 920 | 970 | 1023 | 1032 | 990,67 | 995 | 986 | 0,993 | -0,01 | -0,166 | 0,868076 | 0,887567 | 0,0011559 | 0,00337 | 0,0022415 | 0,0022415 | TRUE | 0,0255024 | No |
| orf19.4443 | 1474 | 1230 | 1377 | 787 | 2474 | 1802 | 2076 | 2757 | 1342 | 1244 | 1303 | 1182 | 2201 | 1956 | 2126 | 2103 | 1682,2 | 1268 | 2096 | 1,65 | 0,723 | 12,933 | 2,92E-38 | 1,11E-37 | 0,0018644 | 0,002822 | 0,0023538 | 0,0023538 | TRUE | 0,0238139 | No |
| orf19.4444 | 982 | 913 | 924 | 593 | 1563 | 1147 | 1259 | 1732 | 894 | 923 | 874 | 891 | 1390 | 1245 | 1289 | 1321 | 1103,61 | 896 | 1311 | 1,466 | 0,551 | 10,187 | 2,28E-24 | 6,44E-24 | 0,0006613 | 0,003234 | 0,0018607 | 0,0018607 | TRUE | 0,0194654 | No |

| | | | | | | | | | | | | | | | | | | | | | | | | | | | | | | | |
|---|---|---|---|---|---|---|---|---|---|---|---|---|---|---|---|---|---|---|---|---|---|---|---|---|---|---|---|---|---|---|---|
| orf19.4445 | 23556 | 21849 | 22926 | 15876 | 13638 | 12095 | 11350 | 15382 | 21447 | 22098 | 21697 | 23848 | 12131 | 13132 | 11624 | 11734 | 17213,8 | 22272 | 12155 | 0,546 | -0,874 | -17,167 | 4,71E-66 | 2,82E-65 | 0,0026957 | 0,002114 | 0,0024255 | 0,0024255 | TRUE | 0,0356323 | No |
| orf19.4446 | 4959 | 4347 | 4732 | 2975 | 5114 | 4197 | 4732 | 6394 | 4515 | 4397 | 4478 | 4469 | 4549 | 4557 | 4846 | 4877 | 4585,99 | 4465 | 4707 | 1,055 | 0,077 | 1,983 | 0,047421 | 0,058826 | 0,0006096 | 0,002325 | 0,0012153 | 0,0012153 | TRUE | 0,0074455 | No |
| orf19.4447 | 1600 | 1246 | 1444 | 880 | 3271 | 2492 | 2847 | 3948 | 1457 | 1260 | 1367 | 1322 | 2909 | 2706 | 2916 | 3012 | 2118,47 | 1352 | 2886 | 2,134 | 1,094 | 19,526 | 6,63E-85 | 4,99E-84 | 0,0021434 | 0,002661 | 0,0024526 | 0,0024526 | TRUE | 0,0344467 | No |
| orf19.4448 | 2370 | 2010 | 2175 | 1492 | 1771 | 1570 | 1658 | 2188 | 2158 | 2033 | 2058 | 2241 | 1575 | 1705 | 1698 | 1669 | 1892,15 | 2122 | 1662 | 0,784 | -0,352 | -7,133 | 9,82E-13 | 2,04E-12 | 0,0009459 | 0,002735 | 0,0018018 | 0,0018018 | TRUE | 0,015664 | No |
| orf19.4449 | 3557 | 2996 | 3294 | 2121 | 1640 | 1302 | 1402 | 1950 | 3238 | 3030 | 3117 | 3186 | 1459 | 1414 | 1436 | 1487 | 2295,98 | 3143 | 1449 | 0,461 | -1,116 | -26,273 | 3,93E-152 | 5,19E-151 | 1,00E-08 | 0,002612 | 0,0012384 | 0,0012384 | TRUE | 0,0072131 | No |
| orf19.445 | 3621 | 3191 | 3452 | 2040 | 3191 | 2569 | 2924 | 3765 | 3297 | 3227 | 3267 | 3064 | 2838 | 2789 | 2995 | 2872 | 3043,7 | 3214 | 2874 | 0,893 | -0,163 | -3,928 | 8,57E-05 | 0,000129 | 0,0005446 | 0,002471 | 0,0013278 | 0,0013278 | TRUE | 0,0110146 | No |
| orf19.4450 | 493 | 397 | 446 | 295 | 1084 | 918 | 1059 | 1459 | 449 | 402 | 422 | 443 | 964 | 997 | 1085 | 1113 | 734,25 | 429 | 1040 | 2,428 | 1,28 | 17,848 | 2,98E-71 | 1,89E-70 | 0,0026584 | 0,003836 | 0,0032392 | 0,0032392 | TRUE | 0,0296547 | No |
| orf19.4450.1 | 4231 | 3527 | 3514 | 2520 | 438 | 338 | 301 | 447 | 3852 | 3567 | 3326 | 3785 | 390 | 367 | 308 | 341 | 1992,03 | 3632 | 352 | 0,097 | -3,369 | -45,579 | 0 | 0 | 0,0050198 | 0,0027 | 0,0037709 | 0,0037709 | TRUE | 0,0737259 | No |
| orf19.4450.2 | 580 | 458 | 436 | 344 | 519 | 416 | 443 | 612 | 528 | 463 | 413 | 517 | 462 | 452 | 454 | 467 | 469,31 | 480 | 459 | 0,958 | -0,062 | -0,744 | 0,457087 | 0,499046 | 0,0043401 | 0,004852 | 0,0046165 | 0,0046165 | TRUE | 0,1087041 | No |
| orf19.4451 | 6857 | 6024 | 6743 | 3823 | 7064 | 5521 | 6463 | 8428 | 6243 | 6093 | 6382 | 5743 | 6283 | 5994 | 6619 | 6429 | 6223,17 | 6115 | 6331 | 1,035 | 0,049 | 1,065 | 0,28684 | 0,323475 | 0,0016609 | 0,002249 | 0,0019008 | 0,0019008 | TRUE | 0,019811 | No |
| orf19.4455 | 3509 | 2887 | 3357 | 2192 | 4023 | 3449 | 3765 | 4949 | 3195 | 2920 | 3177 | 3293 | 3578 | 3745 | 3856 | 3775 | 3442,31 | 3146 | 3738 | 1,189 | 0,25 | 5,296 | 1,18E-07 | 2,04E-07 | 0,0013761 | 0,002421 | 0,0018391 | 0,0018391 | TRUE | 0,0284837 | No |
| orf19.4456 | 37945 | 31914 | 34616 | 21680 | 23829 | 20319 | 21637 | 31033 | 34547 | 32278 | 32761 | 32567 | 21195 | 22061 | 22159 | 23672 | 27655 | 33038 | 22272 | 0,674 | -0,569 | -13,376 | 8,39E-41 | 3,31E-40 | 0,0014993 | 0,002085 | 0,0017006 | 0,0017006 | TRUE | 0,0220643 | No |
| orf19.4457 | 2268 | 2179 | 2362 | 1324 | 2710 | 2317 | 2370 | 3181 | 2065 | 2204 | 2235 | 1989 | 2410 | 2516 | 2427 | 2426 | 2284,1 | 2123 | 2445 | 1,149 | 0,201 | 4,209 | 2,56E-05 | 3,98E-05 | 0,0011027 | 0,002615 | 0,0017427 | 0,0017427 | TRUE | 0,019946 | No |
| orf19.4459 | 2428 | 2452 | 2565 | 1583 | 5092 | 4357 | 4481 | 5702 | 2211 | 2480 | 2428 | 2378 | 4529 | 4730 | 4589 | 4350 | 3461,79 | 2374 | 4550 | 1,916 | 0,938 | 19,562 | 3,25E-85 | 2,46E-84 | 0,0013811 | 0,002419 | 0,0018793 | 0,0018793 | TRUE | 0,0272742 | No |
| orf19.446 | 868 | 851 | 868 | 543 | 913 | 763 | 818 | 1026 | 790 | 861 | 821 | 816 | 812 | 828 | 838 | 783 | 818,62 | 822 | 815 | 0,991 | -0,014 | -0,241 | 0,809469 | 0,834465 | 1,00E-08 | 0,003651 | 0,0018664 | 0,0018664 | TRUE | 0,0124605 | No |
| orf19.4461 | 0 | 0 | 1 | 0 | 3 | 2 | 2 | 1 | 0 | 0 | 1 | 0 | 3 | 2 | 2 | 1 | 1,07 | 0 | 2 | 5,824 | 2,542 | 1,751 | 0,079967 | 0,096918 | 1,00E-08 | 1,231255 | 0,7782076 | 0,7782076 | TRUE | 0,2762684 | No |
| orf19.446.1 | 533 | 467 | 462 | 294 | 1251 | 923 | 935 | 1282 | 485 | 472 | 437 | 442 | 1113 | 1002 | 958 | 978 | 735,85 | 459 | 1013 | 2,202 | 1,139 | 16,059 | 4,96E-58 | 2,64E-57 | 0,0025895 | 0,003832 | 0,0032039 | 0,0032039 | TRUE | 0,0538591 | No |
| orf19.446.2 | 470 | 431 | 397 | 312 | 897 | 658 | 680 | 1060 | 428 | 436 | 376 | 469 | 798 | 714 | 696 | 809 | 590,68 | 427 | 754 | 1,777 | 0,829 | 10,108 | 5,07E-24 | 1,42E-23 | 0,0049441 | 0,004273 | 0,0045828 | 0,0045828 | TRUE | 0,0768411 | No |
| orf19.4463 | 9 | 4 | 8 | 3 | 39 | 26 | 25 | 37 | 8 | 4 | 8 | 5 | 35 | 28 | 26 | 28 | 17,63 | 6 | 29 | 4,674 | 2,225 | 5,681 | 1,34E-08 | 2,39E-08 | 1,00E-08 | 0,076954 | 0,0468341 | 0,0468341 | TRUE | 0,145965 | No |
| orf19.4464 | 804 | 740 | 726 | 439 | 494 | 388 | 472 | 600 | 732 | 748 | 687 | 659 | 439 | 421 | 483 | 458 | 578,59 | 706 | 450 | 0,636 | -0,653 | -9,256 | 2,12E-20 | 5,42E-20 | 0,0012974 | 0,00432 | 0,0029824 | 0,0029824 | TRUE | 0,0259866 | No |
| orf19.4465 | 1520 | 1582 | 1634 | 1039 | 1333 | 1041 | 1260 | 1530 | 1384 | 1600 | 1546 | 1561 | 1186 | 1130 | 1290 | 1167 | 1358,06 | 1523 | 1193 | 0,784 | -0,351 | -5,656 | 1,55E-08 | 2,76E-08 | 0,0029392 | 0,00301 | 0,0029452 | 0,0029452 | TRUE | 0,0471396 | No |
| orf19.4466 | 371 | 373 | 339 | 249 | 474 | 442 | 416 | 610 | 338 | 377 | 321 | 374 | 422 | 480 | 426 | 465 | 400,34 | 352 | 448 | 1,276 | 0,352 | 4,219 | 2,45E-05 | 3,82E-05 | 0,0025074 | 0,005337 | 0,0041112 | 0,0041112 | TRUE | 0,0423666 | No |
| orf19.4467 | 648 | 709 | 653 | 433 | 752 | 587 | 684 | 759 | 590 | 717 | 618 | 650 | 669 | 637 | 701 | 579 | 645,15 | 644 | 646 | 1,004 | 0,005 | 0,063 | 0,950078 | 0,956228 | 0,0055381 | 0,004085 | 0,0048001 | 0,0048001 | TRUE | 0,0724043 | No |
| orf19.4468 | 2860 | 2540 | 2581 | 1404 | 4607 | 3463 | 3850 | 5076 | 2604 | 2569 | 2443 | 2109 | 4098 | 3760 | 3943 | 3872 | 3174,64 | 2431 | 3918 | 1,609 | 0,686 | 10,817 | 2,86E-27 | 8,64E-27 | 0,0045144 | 0,002453 | 0,0035222 | 0,0035222 | TRUE | 0,0917572 | No |
| orf19.4470 | 298 | 330 | 299 | 171 | 250 | 178 | 193 | 261 | 271 | 334 | 283 | 257 | 222 | 193 | 198 | 199 | 244,66 | 286 | 203 | 0,708 | -0,498 | -4,772 | 1,83E-06 | 3,00E-06 | 0,0047653 | 0,007436 | 0,0063097 | 0,0063097 | TRUE | 0,1393943 | No |
| orf19.4471 | 907 | 899 | 974 | 571 | 1278 | 1138 | 1143 | 1524 | 826 | 909 | 922 | 858 | 1137 | 1236 | 1171 | 1163 | 1027,49 | 879 | 1177 | 1,337 | 0,419 | 7,359 | 1,85E-13 | 3,93E-13 | 0,0009057 | 0,003323 | 0,002105 | 0,002105 | TRUE | 0,0220316 | No |
| orf19.4472 | 41 | 34 | 34 | 29 | 93 | 66 | 62 | 86 | 37 | 34 | 32 | 44 | 83 | 72 | 63 | 66 | 53,87 | 37 | 71 | 1,946 | 0,961 | 4,837 | 1,32E-06 | 2,18E-06 | 1,00E-08 | 0,026561 | 0,016718 | 0,016718 | TRUE | 0,1222273 | No |
| orf19.4473 | 396 | 469 | 389 | 260 | 333 | 294 | 279 | 345 | 361 | 474 | 368 | 391 | 296 | 319 | 286 | 263 | 344,74 | 398 | 291 | 0,729 | -0,456 | -4,402 | 1,07E-05 | 1,70E-05 | 0,0090239 | 0,005869 | 0,007332 | 0,007332 | TRUE | 0,196324 | No |
| orf19.4474 | 2004 | 2266 | 2273 | 1275 | 2167 | 1777 | 1923 | 2222 | 1825 | 2292 | 2151 | 1915 | 1927 | 1929 | 1969 | 1695 | 1963 | 2046 | 1880 | 0,918 | -0,123 | -1,61 | 0,107487 | 0,128682 | 0,0070122 | 0,00271 | 0,0051119 | 0,0051119 | TRUE | 0,0795816 | No |
| orf19.4475 | 2123 | 2460 | 2509 | 1427 | 2034 | 1838 | 1965 | 2360 | 1933 | 2488 | 2375 | 2144 | 1809 | 1996 | 2012 | 1800 | 2069,55 | 2235 | 1904 | 0,852 | -0,232 | -2,995 | 0,002749 | 0,003752 | 0,007296 | 0,002675 | 0,0052721 | 0,0052721 | TRUE | 0,1029522 | No |
| orf19.4476 | 510 | 431 | 444 | 352 | 1060 | 975 | 923 | 1351 | 464 | 436 | 420 | 529 | 943 | 1059 | 945 | 1031 | 728,31 | 462 | 994 | 2,163 | 1,113 | 14,493 | 1,35E-47 | 5,96E-47 | 0,0042582 | 0,003851 | 0,0040295 | 0,0040295 | TRUE | 0,0934155 | No |
| orf19.4477 | 956 | 825 | 795 | 503 | 986 | 762 | 804 | 1145 | 870 | 834 | 752 | 756 | 877 | 827 | 823 | 873 | 826,74 | 803 | 850 | 1,057 | 0,08 | 1,24 | 0,21487 | 0,247613 | 0,0019494 | 0,003635 | 0,0028057 | 0,0028057 | TRUE | 0,0387967 | No |
| orf19.4478 | 282 | 233 | 277 | 176 | 448 | 328 | 416 | 508 | 257 | 236 | 262 | 264 | 398 | 356 | 426 | 388 | 323,39 | 255 | 392 | 1,542 | 0,625 | 7,149 | 8,75E-13 | 1,82E-12 | 1,00E-08 | 0,006122 | 0,0040482 | 0,0040482 | TRUE | 0,0399624 | No |
| orf19.4479 | 14988 | 13470 | 14449 | 8285 | 19194 | 15172 | 16229 | 21242 | 13646 | 13623 | 13675 | 12445 | 17072 | 16473 | 16621 | 16204 | 14969,8 | 13347 | 16592 | 1,243 | 0,313 | 7,747 | 9,41E-15 | 2,07E-14 | 0,0011661 | 0,002125 | 0,0015026 | 0,0015026 | TRUE | 0,0247951 | No |
| orf19.448 | 2735 | 2327 | 2446 | 1532 | 2684 | 2017 | 2311 | 3244 | 2490 | 2354 | 2315 | 2301 | 2387 | 2190 | 2367 | 2475 | 2359,8 | 2365 | 2355 | 0,996 | -0,006 | -0,121 | 0,903318 | 0,917191 | 0,0015546 | 0,002597 | 0,0019929 | 0,0019929 | TRUE | 0,0263496 | No |
| orf19.4481 | 10 | 8 | 10 | 6 | 57 | 43 | 50 | 69 | 9 | 8 | 9 | 9 | 51 | 47 | 51 | 53 | 29,61 | 9 | 50 | 5,649 | 2,498 | 8,014 | 1,11E-15 | 2,51E-15 | 1,00E-08 | 0,046647 | 0,0249301 | 0,0249301 | TRUE | 0,017629 | No |
| orf19.4482 | 358 | 357 | 352 | 203 | 334 | 315 | 304 | 388 | 326 | 361 | 333 | 305 | 297 | 342 | 311 | 296 | 321,43 | 331 | 312 | 0,935 | -0,097 | -1,117 | 0,263857 | 0,299731 | 0,0012177 | 0,006147 | 0,0041592 | 0,0041592 | TRUE | 0,0464801 | No |
| orf19.4488 | 11215 | 10307 | 11046 | 6873 | 12041 | 9767 | 10019 | 13448 | 10211 | 10424 | 10454 | 10324 | 10710 | 10604 | 10261 | 10258 | 10405,9 | 10353 | 10458 | 1,01 | 0,014 | 0,476 | 0,634413 | 0,672607 | 0,0001779 | 0,002164 | 0,0007864 | 0,0007864 | TRUE | 0,0032619 | No |
| orf19.449 | 38067 | 35063 | 37919 | 24321 | 3742 | 2934 | 3286 | 4249 | 34658 | 35462 | 35887 | 36534 | 3328 | 3186 | 3365 | 3241 | 19457,8 | 35635 | 3280 | 0,092 | -3,441 | -100,91 | 0 | 0 | 0,0003321 | 0,002105 | 0,0009595 | 0,0009595 | TRUE | 0,0043263 | No |
| orf19.4490 | 14942 | 12863 | 12830 | 9295 | 23516 | 18612 | 16911 | 25940 | 13604 | 13010 | 12142 | 13963 | 20917 | 20207 | 17319 | 19787 | 16368,6 | 13180 | 19558 | 1,484 | 0,57 | 8,97 | 2,95E-19 | 7,35E-19 | 0,0049318 | 0,002118 | 0,00381 | 0,00381 | TRUE | 0,072596 | No |
| orf19.4490.2 | 2873 | 2472 | 2541 | 1731 | 4766 | 3886 | 3683 | 5392 | 2616 | 2500 | 2405 | 2600 | 4239 | 4219 | 3772 | 4113 | 3308,03 | 2530 | 4086 | 1,616 | 0,692 | 13,665 | 1,65E-42 | 6,73E-42 | 0,0017281 | 0,002436 | 0,0021368 | 0,0021368 | TRUE | 0,0322222 | No |
| orf19.4491 | 3871 | 4162 | 4377 | 2259 | 4155 | 3423 | 3739 | 4269 | 3524 | 4209 | 4142 | 3393 | 3696 | 3716 | 3829 | 3256 | 3720,93 | 3817 | 3624 | 0,949 | -0,076 | -0,987 | 0,323846 | 0,363216 | 0,0078817 | 0,002392 | 0,0054481 | 0,0054481 | TRUE | 0,0664303 | No |
| orf19.4492 | 14470 | 12945 | 13404 | 8915 | 15637 | 12947 | 13358 | 18073 | 13174 | 13092 | 12686 | 13392 | 13909 | 14057 | 13680 | 13786 | 13472 | 13086 | 13858 | 1,059 | 0,083 | 2,783 | 0,005381 | 0,007201 | 0,000224 | 0,002135 | 0,0007797 | 0,0007797 | TRUE | 0,0052095 | No |
| orf19.4494 | 888 | 918 | 917 | 597 | 1386 | 1115 | 1163 | 1543 | 808 | 928 | 868 | 897 | 1233 | 1211 | 1191 | 1177 | 1039,13 | 875 | 1203 | 1,376 | 0,461 | 8,158 | 3,40E-16 | 7,78E-16 | 0,0007871 | 0,003308 | 0,0020596 | 0,0020596 | TRUE | 0,033513 | No |
| orf19.4495 | 5836 | 5236 | 5610 | 3384 | 10261 | 8703 | 8790 | 12265 | 5313 | 5296 | 5309 | 5083 | 9127 | 9449 | 9002 | 9356 | 7241,94 | 5250 | 9234 | 1,757 | 0,813 | 24,229 | 1,09E-129 | 1,21E-128 | 0,0002741 | 0,002219 | 0,0009291 | 0,0009291 | TRUE | 0,0052178 | No |
| orf19.4496 | 2247 | 2178 | 2312 | 1421 | 1678 | 1464 | 1447 | 1872 | 2046 | 2203 | 2188 | 2135 | 1493 | 1589 | 1482 | 1428 | 1820,4 | 2143 | 1498 | 0,698 | -0,518 | -10,503 | 8,33E-26 | 2,43E-25 | 0,0010068 | 0,002763 | 0,0017722 | 0,0017722 | TRUE | 0,0198111 | No |
| orf19.4497 | 1167 | 1199 | 1086 | 607 | 852 | 690 | 754 | 877 | 1063 | 1213 | 1028 | 912 | 758 | 749 | 772 | 669 | 895,37 | 1054 | 737 | 0,697 | -0,52 | -6,21 | 5,30E-10 | 1,00E-09 | 0,0073681 | 0,003512 | 0,005598 | 0,005598 | TRUE | 0,1214647 | No |
| orf19.4498 | 388 | 463 | 480 | 247 | 532 | 415 | 457 | 546 | 353 | 468 | 454 | 371 | 473 | 451 | 468 | 416 | 431,89 | 412 | 452 | 1,095 | 0,13 | 1,325 | 0,185033 | 0,214939 | 0,0091452 | 0,005096 | 0,0069432 | 0,0069432 | TRUE | 0,1174443 | No |
| orf19.4499 | 2846 | 2643 | 2772 | 1721 | 2074 | 1673 | 1791 | 2392 | 2591 | 2673 | 2623 | 2585 | 1845 | 1816 | 1834 | 1825 | 2224,12 | 2618 | 1830 | 0,699 | -0,517 | -13,633 | 2,53E-42 | 1,03E-41 | 1,00E-08 | 0,002631 | 0,0009215 | 0,0009215 | TRUE | 0,0024157 | No |
| orf19.450 | 51 | 51 | 59 | 58 | 17 | 14 | 9 | 17 | 46 | 52 | 56 | 87 | 15 | 15 | 9 | 13 | 36,69 | 60 | 13 | 0,222 | -2,172 | -7,384 | 1,54E-13 | 3,28E-13 | 0,0397012 | 0,038046 | 0,0386766 | 0,0386766 | TRUE | 0,6225518 | No |
| orf19.4502 | 1347 | 1212 | 1311 | 722 | 2513 | 2152 | 2198 | 3049 | 1226 | 1226 | 1241 | 1085 | 2235 | 2336 | 2251 | 2326 | 1740,75 | 1194 | 2287 | 1,908 | 0,932 | 18,348 | 3,42E-75 | 2,27E-74 | 0,0007838 | 0,002796 | 0,0018264 | 0,0018264 | TRUE | 0,0400109 | No |
| orf19.4503 | 967 | 720 | 864 | 581 | 1696 | 1295 | 1514 | 1962 | 880 | 728 | 818 | 873 | 1509 | 1406 | 1551 | 1497 | 1157,6 | 825 | 1491 | 1,81 | 0,856 | 13,018 | 9,61E-39 | 3,67E-38 | 0,0031973 | 0,003178 | 0,0031849 | 0,0031849 | TRUE | 0,0746737 | No |
| orf19.4504 | 510 | 443 | 489 | 273 | 11643 | 8656 | 10024 | 13371 | 464 | 448 | 463 | 410 | 10356 | 9398 | 10266 | 10200 | 5250,59 | 446 | 10055 | 22,414 | 4,486 | 78,199 | 0 | 0 | 0,0016198 | 0,002289 | 0,0019272 | 0,0019272 | TRUE | 0,0274543 | No |
| orf19.4505 | 162 | 138 | 129 | 99 | 612 | 496 | 537 | 786 | 147 | 140 | 122 | 149 | 544 | 539 | 550 | 600 | 348,78 | 140 | 558 | 4,026 | 2,009 | 21,282 | 1,65E-100 | 1,44E-99 | 1,00E-08 | 0,005824 | 0,0039022 | 0,0039022 | TRUE | 0,0785305 | No |
| orf19.4506 | 3545 | 3177 | 3357 | 1939 | 2790 | 2202 | 2221 | 3150 | 3228 | 3213 | 3177 | 2913 | 2482 | 2391 | 2275 | 2403 | 2760,04 | 3133 | 2388 | 0,761 | -0,393 | -8,237 | 1,77E-16 | 4,06E-16 | 0,0012655 | 0,002516 | 0,0018226 | 0,0018226 | TRUE | 0,0254318 | No |
| orf19.4507 | 79 | 91 | 70 | 48 | 442 | 352 | 332 | 488 | 72 | 92 | 66 | 72 | 393 | 382 | 340 | 372 | 223,74 | 76 | 372 | 4,92 | 2,299 | 18,915 | 8,54E-80 | 6,03E-79 | 0,0017346 | 0,007941 | 0,0059214 | 0,0059214 | TRUE | 0,2074357 | No |
| orf19.4508 | 31 | 43 | 29 | 26 | 57 | 42 | 32 | 51 | 28 | 43 | 27 | 39 | 51 | 46 | 33 | 39 | 38,27 | 34 | 42 | 1,229 | 0,297 | 1,224 | 0,220829 | 0,253976 | 0,0180496 | 0,036551 | 0,0299169 | 0,0299169 | TRUE | 0,2479275 | No |
| orf19.4509 | 8 | 9 | 10 | 4 | 7 | 11 | 3 | 5 | 7 | 9 | 9 | 6 | 6 | 12 | 3 | 4 | 7,11 | 8 | 6 | 0,76 | -0,396 | -0,73 | 0,465109 | 0,50685 | 0,041177 | 0,187719 | 0,1400786 | 0,1400786 | TRUE | 0,805884 | No |
| orf19.451 | 18265 | 17169 | 18307 | 12512 | 12866 | 11962 | 11421 | 15799 | 16630 | 17365 | 17326 | 18795 | 11444 | 12987 | 11697 | 12052 | 14786,8 | 17529 | 12045 | 0,687 | -0,541 | -10,428 | 1,86E-25 | 5,38E-25 | 0,0028268 | 0,002126 | 0,0025174 | 0,0025174 | TRUE | 0,0337651 | No |
| orf19.4510 | 375 | 364 | 385 | 215 | 225 | 171 | 156 | 243 | 341 | 368 | 364 | 323 | 200 | 186 | 160 | 185 | 265,98 | 349 | 183 | 0,523 | -0,936 | -9,743 | 1,97E-22 | 5,30E-22 | 0,0009146 | 0,007004 | 0,0047859 | 0,0047859 | TRUE | 0,0730666 | No |
| orf19.4511 | 684 | 604 | 754 | 444 | 333 | 263 | 326 | 385 | 623 | 611 | 714 | 667 | 296 | 286 | 334 | 294 | 477,93 | 654 | 302 | 0,462 | -1,113 | -13,722 | 7,45E-43 | 3,05E-42 | 0,002945 | 0,004801 | 0,0039479 | 0,0039479 | TRUE | 0,0529963 | No |
| orf19.4513 | 7353 | 7474 | 8078 | 4696 | 6063 | 5381 | 5638 | 7502 | 6695 | 7559 | 7645 | 7054 | 5393 | 5842 | 5774 | 5723 | 6460,59 | 7238 | 5683 | 0,785 | -0,349 | -6,833 | 8,31E-12 | 1,68E-11 | 0,002309 | 0,002241 | 0,0023536 | 0,0023536 | TRUE | 0,0318537 | No |
| orf19.4515 | 202 | 168 | 207 | 114 | 250 | 228 | 209 | 299 | 184 | 170 | 196 | 171 | 222 | 248 | 214 | 228 | 204,13 | 180 | 228 | 1,259 | 0,332 | 3,258 | 0,001123 | 0,001582 | 1,00E-08 | 0,008508 | 0,0050096 | 0,0050096 | TRUE | 0,0375762 | No |
| orf19.4516 | 1258 | 1102 | 1265 | 691 | 1036 | 900 | 1015 | 1268 | 1145 | 1115 | 1197 | 1038 | 921 | 977 | 1039 | 967 | 1050,06 | 1124 | 976 | 0,866 | -0,208 | -3,424 | 0,000617 | 0,000883 | 0,0017173 | 0,003295 | 0,0025758 | 0,0025758 | TRUE | 0,0282678 | No |
| orf19.4517 | 2900 | 3679 | 3368 | 2042 | 3379 | 3404 | 3062 | 3807 | 2640 | 3721 | 3188 | 3067 | 3006 | 3696 | 3136 | 2904 | 3169,67 | 3154 | 3186 | 1,01 | 0,014 | 0,142 | 0,887305 | 0,903317 | 0,015435 | 0,002454 | 0,0087268 | 0,0087268 | TRUE | 0,1789113 | No |
| orf19.4518 | 1121 | 982 | 1107 | 670 | 1253 | 1080 | 1133 | 1634 | 1021 | 993 | 1048 | 1006 | 1115 | 1173 | 1160 | 1246 | 1095,22 | 1017 | 1174 | 1,154 | 0,207 | 3,819 | 0,000134 | 0,000199 | 0,0007613 | 0,003243 | 0,0018904 | 0,0018904 | TRUE | 0,0226202 | No |
| orf19.4519 | 6786 | 6707 | 6934 | 4591 | 5843 | 5235 | 4914 | 6692 | 6178 | 6783 | 6562 | 6896 | 5197 | 5684 | 5033 | 5105 | 5929,85 | 6605 | 5255 | 0,796 | -0,33 | -6,375 | 1,83E-10 | 3,51E-10 | 0,0024854 | 0,00226 | 0,0023995 | 0,0023995 | TRUE | 0,0364708 | No |

| | | | | | | | | | | | | | | | | | | | | | | | | | | | | | | | |
|---|---|---|---|---|---|---|---|---|---|---|---|---|---|---|---|---|---|---|---|---|---|---|---|---|---|---|---|---|---|---|---|
| orf19.4520 | 165 | 163 | 169 | 84 | 218 | 168 | 192 | 260 | 150 | 165 | 160 | 126 | 194 | 182 | 197 | 198 | 171,56 | 150 | 193 | 1,273 | 0,348 | 3,135 | 0,001717 | 0,002382 | 1,00E-08 | 0,009737 | 0,0059331 | 0,0059331 | TRUE | 0,0915058 | No |
| orf19.4521 | 458 | 436 | 486 | 276 | 493 | 448 | 413 | 535 | 417 | 441 | 460 | 415 | 439 | 486 | 423 | 408 | 436,06 | 433 | 439 | 1,009 | 0,013 | 0,169 | 0,866151 | 0,885883 | 0,0019452 | 0,005066 | 0,0036733 | 0,0036733 | TRUE | 0,0583557 | No |
| orf19.4522 | 962 | 757 | 809 | 596 | 632 | 532 | 483 | 730 | 876 | 766 | 766 | 895 | 562 | 578 | 495 | 557 | 686,71 | 826 | 548 | 0,665 | -0,588 | -7,69 | 1,47E-14 | 3,23E-14 | 0,0042796 | 0,003961 | 0,0041137 | 0,0041137 | TRUE | 0,048102 | No |
| orf19.4523 | 322 | 292 | 304 | 203 | 342 | 257 | 275 | 401 | 293 | 295 | 288 | 305 | 304 | 279 | 282 | 306 | 293,99 | 295 | 293 | 0,996 | -0,006 | -0,073 | 0,94183 | 0,950026 | 1,00E-08 | 0,00653 | 0,0033318 | 0,0033318 | TRUE | 0,0109723 | No |
| orf19.4524 | 918 | 886 | 986 | 580 | 1515 | 1270 | 1246 | 1677 | 836 | 896 | 933 | 871 | 1348 | 1379 | 1276 | 1279 | 1102,25 | 884 | 1320 | 1,492 | 0,577 | 10,319 | 5,76E-25 | 1,65E-24 | 0,0009343 | 0,003235 | 0,0020464 | 0,0020464 | TRUE | 0,0180606 | No |
| orf19.4525 | 2829 | 2566 | 2690 | 1725 | 2670 | 2221 | 2188 | 3151 | 2576 | 2595 | 2546 | 2591 | 2375 | 2411 | 2241 | 2404 | 2467,33 | 2577 | 2358 | 0,915 | -0,128 | -3,16 | 0,001579 | 0,0022 | 0,0001731 | 0,002572 | 0,0011578 | 0,0011578 | TRUE | 0,0131529 | No |
| orf19.4526 | 39009 | 35560 | 36958 | 23720 | 2030 | 1703 | 1735 | 2343 | 35516 | 35965 | 34977 | 35631 | 1806 | 1849 | 1777 | 1787 | 18663,6 | 35522 | 1805 | 0,051 | -4,3 | -134,08 | 0 | 0 | 1,00E-08 | 0,002108 | 0,0007139 | 0,0007139 | TRUE | 0,0030688 | No |
| orf19.4527 | 9545 | 7787 | 8528 | 5721 | 8254 | 7031 | 7128 | 10511 | 8690 | 7876 | 8071 | 8594 | 7342 | 7634 | 7300 | 8018 | 7940,51 | 8308 | 7574 | 0,912 | -0,133 | -2,787 | 0,005312 | 0,007111 | 0,0019269 | 0,002203 | 0,0020606 | 0,0020606 | TRUE | 0,0193273 | No |
| orf19.4528 | 1355 | 1300 | 1306 | 892 | 1677 | 1360 | 1335 | 1918 | 1234 | 1315 | 1236 | 1340 | 1492 | 1477 | 1367 | 1463 | 1365,36 | 1281 | 1450 | 1,134 | 0,182 | 3,489 | 0,000484 | 0,000697 | 0,0008418 | 0,003005 | 0,0018591 | 0,0018591 | TRUE | 0,0171647 | No |
| orf19.4528.1 | 3 | 2 | 2 | 4 | 4 | 5 | 4 | 6 | 3 | 2 | 2 | 6 | 4 | 5 | 4 | 5 | 3,79 | 3 | 4 | 1,481 | 0,566 | 0,798 | 0,424636 | 0,46593 | 1,00E-08 | 0,350638 | 0,1984475 | 0,1984475 | TRUE | 0,2940421 | No |
| orf19.4529 | 910 | 819 | 828 | 580 | 977 | 800 | 854 | 1100 | 829 | 828 | 784 | 871 | 869 | 869 | 875 | 839 | 845,38 | 828 | 863 | 1,044 | 0,062 | 1,117 | 0,263856 | 0,299731 | 1,00E-08 | 0,0036 | 0,0017919 | 0,0017919 | TRUE | 0,015349 | No |
| orf19.4530.1 | 8342 | 6990 | 7902 | 5093 | 4339 | 3635 | 4015 | 5314 | 7595 | 7070 | 7479 | 7650 | 3859 | 3947 | 4112 | 4054 | 5720,64 | 7448 | 3993 | 0,536 | -0,899 | -22,352 | 1,15E-110 | 1,09E-109 | 0,0007597 | 0,002268 | 0,0013645 | 0,0013645 | TRUE | 0,0143552 | No |
| orf19.4531 | 26415 | 24471 | 26708 | 15803 | 8489 | 7168 | 7627 | 10017 | 24050 | 24750 | 25277 | 23739 | 7551 | 7782 | 7811 | 7641 | 16075 | 24454 | 7696 | 0,315 | -1,668 | -50,387 | 0 | 0 | 0,0004135 | 0,002119 | 0,0009701 | 0,0009701 | TRUE | 0,0062902 | No |
| orf19.4532 | 2466 | 2173 | 2272 | 1497 | 5737 | 4556 | 4620 | 6465 | 2245 | 2198 | 2150 | 2249 | 5103 | 4947 | 4731 | 4932 | 3569,3 | 2210 | 4928 | 2,231 | 1,158 | 28,936 | 4,24E-184 | 6,66E-183 | 0,0004972 | 0,002407 | 0,0011989 | 0,0011989 | TRUE | 0,0086915 | No |
| orf19.4533 | 961 | 853 | 956 | 644 | 1265 | 1042 | 1047 | 1394 | 875 | 863 | 905 | 967 | 1125 | 1131 | 1072 | 1063 | 1000,24 | 902 | 1098 | 1,22 | 0,287 | 5,136 | 2,81E-07 | 4,76E-07 | 0,0005092 | 0,003358 | 0,0019682 | 0,0019682 | TRUE | 0,0240823 | No |
| orf19.4534 | 11102 | 9086 | 10092 | 6261 | 5155 | 4179 | 4779 | 6439 | 10108 | 9190 | 9551 | 9405 | 4585 | 4537 | 4894 | 4912 | 7147,76 | 9564 | 4732 | 0,495 | -1,015 | -22,435 | 1,79E-111 | 1,71E-110 | 0,0015096 | 0,002222 | 0,0018106 | 0,0018106 | TRUE | 0,0181364 | No |
| orf19.4535 | 520 | 436 | 475 | 270 | 455 | 394 | 411 | 515 | 473 | 441 | 450 | 406 | 405 | 428 | 421 | 393 | 426,97 | 442 | 412 | 0,924 | -0,114 | -1,517 | 0,129362 | 0,15343 | 1,00E-08 | 0,005131 | 0,0030456 | 0,0030456 | TRUE | 0,0301025 | No |
| orf19.4536 | 7762 | 6685 | 7329 | 4656 | 12220 | 9687 | 9887 | 13962 | 7067 | 6761 | 6936 | 6994 | 10869 | 10517 | 10126 | 10650 | 8740,12 | 6940 | 10540 | 1,519 | 0,603 | 17,207 | 2,34E-66 | 1,40E-65 | 0,0004567 | 0,002188 | 0,0010585 | 0,0010585 | TRUE | 0,0085563 | No |
| orf19.4537 | 3559 | 2940 | 3216 | 2012 | 3870 | 3158 | 3385 | 4814 | 3240 | 2973 | 3044 | 3022 | 3442 | 3429 | 3467 | 3672 | 3286,2 | 3070 | 3502 | 1,141 | 0,19 | 4,321 | 1,55E-05 | 2,44E-05 | 0,0008544 | 0,002439 | 0,0015484 | 0,0015484 | TRUE | 0,017502 | No |
| orf19.4538 | 3732 | 2967 | 3206 | 1869 | 3078 | 2279 | 2640 | 3453 | 3398 | 3001 | 3034 | 2808 | 2738 | 2474 | 2704 | 2634 | 2848,77 | 3060 | 2638 | 0,861 | -0,216 | -3,541 | 0,000399 | 0,000577 | 0,0037532 | 0,002501 | 0,003212 | 0,003212 | TRUE | 0,0674545 | No |
| orf19.4539 | 1829 | 1469 | 1699 | 959 | 2009 | 1547 | 1790 | 2335 | 1665 | 1486 | 1608 | 1441 | 1787 | 1680 | 1833 | 1781 | 1660,05 | 1550 | 1770 | 1,14 | 0,189 | 3,335 | 0,000853 | 0,00121 | 0,0021336 | 0,002833 | 0,0024752 | 0,0024752 | TRUE | 0,0302091 | No |
| orf19.454 | 7286 | 6316 | 7187 | 4852 | 10451 | 9211 | 9691 | 13145 | 6634 | 6388 | 6802 | 7288 | 9296 | 10001 | 9925 | 10027 | 8295,03 | 6778 | 9812 | 1,449 | 0,535 | 11,157 | 6,58E-29 | 2,06E-28 | 0,0020336 | 0,002196 | 0,0020775 | 0,0020775 | TRUE | 0,030519 | No |
| orf19.4540 | 6306 | 5452 | 5987 | 3765 | 3704 | 3134 | 3217 | 4313 | 5741 | 5514 | 5666 | 5656 | 3295 | 3403 | 3295 | 3290 | 4482,39 | 5644 | 3321 | 0,588 | -0,766 | -22,867 | 9,87E-116 | 9,74E-115 | 1,00E-08 | 0,002332 | 0,0008411 | 0,0008411 | TRUE | 0,0032515 | No |
| orf19.4542 | 894 | 859 | 923 | 568 | 841 | 768 | 760 | 973 | 814 | 869 | 874 | 853 | 748 | 834 | 778 | 742 | 813,99 | 852 | 776 | 0,908 | -0,139 | -2,302 | 0,021332 | 0,02728 | 0,0007799 | 0,00366 | 0,0022439 | 0,0022439 | TRUE | 0,0291852 | No |
| orf19.4543 | 2325 | 1920 | 2186 | 1403 | 1765 | 1451 | 1538 | 2157 | 2117 | 1942 | 2069 | 2108 | 1570 | 1575 | 1575 | 1645 | 1825,11 | 2059 | 1591 | 0,774 | -0,37 | -7,991 | 1,34E-15 | 3,02E-15 | 0,0005733 | 0,002761 | 0,0015036 | 0,0015036 | TRUE | 0,0179216 | No |
| orf19.4544 | 1125 | 1002 | 1010 | 639 | 901 | 716 | 797 | 1065 | 1024 | 1013 | 956 | 960 | 801 | 777 | 816 | 812 | 895,11 | 988 | 802 | 0,811 | -0,303 | -5,58 | 2,40E-08 | 4,26E-08 | 1,00E-08 | 0,003513 | 0,0017041 | 0,0017041 | TRUE | 0,0073752 | No |
| orf19.4545 | 2386 | 2102 | 2341 | 1504 | 3064 | 2515 | 2668 | 3597 | 2172 | 2126 | 2216 | 2259 | 2725 | 2731 | 2732 | 2744 | 2463,15 | 2193 | 2733 | 1,247 | 0,319 | 8,397 | 4,59E-17 | 1,08E-16 | 1,00E-08 | 0,002573 | 0,0009703 | 0,0009703 | TRUE | 0,0050009 | No |
| orf19.4546 | 6584 | 5836 | 6460 | 3871 | 6650 | 5619 | 5960 | 7980 | 5994 | 5902 | 6114 | 5815 | 5915 | 6101 | 6104 | 6087 | 6004,03 | 5956 | 6052 | 1,015 | 0,022 | 0,676 | 0,499103 | 0,540752 | 0,000123 | 0,002257 | 0,0008478 | 0,0008478 | TRUE | 0,0038441 | No |
| orf19.4548 | 321 | 293 | 339 | 206 | 251 | 206 | 247 | 307 | 292 | 296 | 321 | 309 | 223 | 224 | 253 | 234 | 269,12 | 304 | 234 | 0,767 | -0,383 | -4,307 | 1,66E-05 | 2,60E-05 | 1,00E-08 | 0,006946 | 0,0038591 | 0,0038591 | TRUE | 0,0323285 | No |
| orf19.4549 | 525 | 461 | 513 | 310 | 371 | 328 | 376 | 431 | 478 | 466 | 486 | 466 | 330 | 356 | 385 | 329 | 411,92 | 474 | 350 | 0,736 | -0,443 | -5,761 | 8,37E-09 | 1,51E-08 | 1,00E-08 | 0,005244 | 0,0032132 | 0,0032132 | TRUE | 0,0517139 | No |
| orf19.4550 | 1022 | 926 | 974 | 634 | 2072 | 1796 | 1817 | 2380 | 930 | 937 | 922 | 952 | 1843 | 1950 | 1861 | 1815 | 1401,31 | 935 | 1867 | 1,997 | 0,998 | 20,838 | 1,97E-96 | 1,64E-95 | 1,00E-08 | 0,00298 | 0,0013789 | 0,0013789 | TRUE | 0,010411 | No |
| orf19.4551 | 2484 | 2417 | 2567 | 1702 | 7890 | 6853 | 6721 | 8842 | 2262 | 2445 | 2429 | 2557 | 7018 | 7440 | 6883 | 6745 | 4722,31 | 2423 | 7022 | 2,901 | 1,537 | 31,245 | 2,60E-214 | 4,70E-213 | 0,0018082 | 0,002317 | 0,0020332 | 0,0020332 | TRUE | 0,0247108 | No |
| orf19.4553 | 6165 | 5595 | 6243 | 4119 | 3252 | 2946 | 3042 | 4173 | 5613 | 5659 | 5908 | 6187 | 2893 | 3199 | 3115 | 3183 | 4469,65 | 5842 | 3098 | 0,53 | -0,915 | -18,831 | 4,24E-79 | 2,98E-78 | 0,0017351 | 0,002333 | 0,0020224 | 0,0020224 | TRUE | 0,0243235 | No |
| orf19.4555 | 4196 | 3375 | 3868 | 2814 | 2894 | 2784 | 2819 | 3835 | 3820 | 3413 | 3661 | 4227 | 2574 | 3023 | 2887 | 2925 | 3316,33 | 3780 | 2852 | 0,755 | -0,405 | -5,806 | 6,39E-09 | 1,16E-08 | 0,0059865 | 0,002435 | 0,0043776 | 0,0043776 | TRUE | 0,0750696 | No |
| orf19.4557 | 576 | 539 | 535 | 320 | 224 | 161 | 176 | 231 | 524 | 545 | 506 | 481 | 199 | 175 | 180 | 176 | 348,39 | 514 | 182 | 0,355 | -1,496 | -17,362 | 1,61E-67 | 9,76E-67 | 1,00E-08 | 0,005829 | 0,0035459 | 0,0035459 | TRUE | 0,0421372 | No |
| orf19.4560 | 4915 | 4294 | 4683 | 2990 | 5304 | 4640 | 4567 | 6170 | 4475 | 4343 | 4432 | 4491 | 4718 | 5038 | 4677 | 4707 | 4610,06 | 4435 | 4785 | 1,079 | 0,109 | 2,885 | 0,003913 | 0,005279 | 0,0004502 | 0,002324 | 0,001156 | 0,001156 | TRUE | 0,015087 | No |
| orf19.4563 | 5851 | 4982 | 5502 | 3469 | 5520 | 4560 | 4695 | 6198 | 5327 | 5039 | 5207 | 5211 | 4910 | 4951 | 4808 | 4728 | 5022,61 | 5196 | 4849 | 0,933 | -0,1 | -2,856 | 0,004287 | 0,005771 | 0,0003071 | 0,0023 | 0,0009798 | 0,0009798 | TRUE | 0,0051829 | No |
| orf19.4565 | 5431 | 4383 | 4582 | 3334 | 10222 | 8413 | 8040 | 12213 | 4945 | 4433 | 4336 | 5008 | 9092 | 9134 | 8234 | 9316 | 6812,34 | 4680 | 8944 | 1,912 | 0,935 | 15,69 | 1,78E-55 | 9,06E-55 | 0,0039578 | 0,002231 | 0,0032445 | 0,0032445 | TRUE | 0,0348393 | No |
| orf19.4566 | 27 | 23 | 26 | 13 | 67 | 32 | 49 | 68 | 25 | 23 | 25 | 20 | 60 | 35 | 50 | 52 | 36,05 | 23 | 49 | 2,126 | 1,088 | 4,343 | 1,40E-05 | 2,21E-05 | 1,00E-08 | 0,038684 | 0,0279797 | 0,0279797 | TRUE | 0,2848335 | No |
| orf19.4567 | 41 | 41 | 36 | 23 | 360 | 305 | 316 | 432 | 37 | 41 | 34 | 35 | 320 | 331 | 324 | 330 | 181,49 | 37 | 326 | 8,814 | 3,14 | 21,43 | 7,00E-102 | 6,21E-101 | 1,00E-08 | 0,009316 | 0,0049357 | 0,0049357 | TRUE | 0,051163 | No |
| orf19.4568 | 274 | 233 | 261 | 164 | 313 | 241 | 263 | 350 | 249 | 236 | 247 | 246 | 278 | 262 | 269 | 267 | 256,86 | 244 | 269 | 1,101 | 0,138 | 1,585 | 0,112948 | 0,134789 | 1,00E-08 | 0,00718 | 0,0034078 | 0,0034078 | TRUE | 0,0071094 | No |
| orf19.4569 | 66 | 48 | 54 | 36 | 3048 | 2634 | 2511 | 3506 | 60 | 49 | 51 | 54 | 2711 | 2860 | 2572 | 2674 | 1378,84 | 54 | 2704 | 50,5 | 5,658 | 50,004 | 0 | 0 | 1,00E-08 | 0,002995 | 0,0022902 | 0,0022902 | TRUE | 0,0730836 | No |
| orf19.457 | 3486 | 3493 | 3749 | 2089 | 2291 | 1862 | 1987 | 2583 | 3174 | 3533 | 3548 | 3138 | 2038 | 2022 | 2035 | 1970 | 2682,18 | 3348 | 2016 | 0,602 | -0,733 | -13,974 | 2,23E-44 | 9,46E-44 | 0,0018005 | 0,00253 | 0,002251 | 0,002251 | TRUE | 0,0205259 | No |
| orf19.4570 | 140 | 129 | 135 | 91 | 1287 | 926 | 1011 | 1457 | 127 | 130 | 128 | 137 | 1145 | 1005 | 1035 | 1111 | 602,42 | 130 | 1074 | 8,268 | 3,047 | 34,149 | 1,36E-255 | 2,93E-254 | 1,00E-08 | 0,00423 | 0,0031429 | 0,0031429 | TRUE | 0,0229248 | No |
| orf19.4571 | 86 | 65 | 82 | 42 | 5807 | 5012 | 5341 | 7123 | 78 | 66 | 78 | 63 | 5165 | 5442 | 5470 | 5433 | 2724,36 | 71 | 5378 | 74,571 | 6,221 | 64,481 | 0 | 0 | 1,00E-08 | 0,002522 | 0,0015517 | 0,0015517 | TRUE | 0,0390507 | No |
| orf19.4573 | 239 | 168 | 220 | 165 | 401 | 365 | 401 | 484 | 218 | 170 | 208 | 248 | 357 | 396 | 411 | 369 | 297,05 | 211 | 383 | 1,828 | 0,87 | 8,262 | 1,43E-16 | 3,31E-16 | 0,0075282 | 0,006484 | 0,0068706 | 0,0068706 | TRUE | 0,1834122 | No |
| orf19.4574 | 1416 | 1270 | 1372 | 857 | 1466 | 1161 | 1184 | 1787 | 1289 | 1284 | 1298 | 1287 | 1304 | 1261 | 1213 | 1363 | 1287,46 | 1290 | 1285 | 0,998 | -0,003 | -0,065 | 0,947874 | 0,954311 | 0,0006493 | 0,003063 | 0,0017393 | 0,0017393 | TRUE | 0,0210578 | No |
| orf19.4575 | 999 | 952 | 1030 | 683 | 1765 | 1508 | 1636 | 2066 | 910 | 963 | 975 | 1026 | 1570 | 1637 | 1675 | 1576 | 1291,47 | 968 | 1614 | 1,671 | 0,741 | 14,024 | 1,10E-44 | 4,71E-44 | 0,0006092 | 0,00306 | 0,0018344 | 0,0018344 | TRUE | 0,0201923 | No |
| orf19.4577 | 381 | 326 | 342 | 195 | 495 | 386 | 461 | 601 | 347 | 330 | 324 | 293 | 440 | 419 | 472 | 458 | 385,39 | 324 | 447 | 1,378 | 0,462 | 5,794 | 6,85E-09 | 1,24E-08 | 0,000192 | 0,005465 | 0,0034284 | 0,0034284 | TRUE | 0,0365083 | No |
| orf19.4577.3 | 1602 | 1355 | 1379 | 922 | 2037 | 1536 | 1476 | 2199 | 1459 | 1370 | 1305 | 1385 | 1812 | 1668 | 1512 | 1677 | 1523,45 | 1380 | 1667 | 1,209 | 0,274 | 4,376 | 1,21E-05 | 1,91E-05 | 0,0032366 | 0,002904 | 0,0030858 | 0,0030858 | TRUE | 0,0470118 | No |
| orf19.4578 | 1655 | 1451 | 1495 | 887 | 1329 | 1000 | 1103 | 1486 | 1507 | 1468 | 1415 | 1332 | 1182 | 1086 | 1130 | 1134 | 1281,58 | 1430 | 1133 | 0,791 | -0,339 | -6,251 | 4,07E-10 | 7,73E-10 | 0,00094 | 0,003068 | 0,0020351 | 0,0020351 | TRUE | 0,0229132 | No |
| orf19.4579 | 476 | 417 | 452 | 290 | 713 | 593 | 627 | 821 | 433 | 422 | 428 | 436 | 634 | 644 | 642 | 626 | 533,12 | 430 | 636 | 1,482 | 0,567 | 8,885 | 6,38E-19 | 1,57E-18 | 1,00E-08 | 0,004515 | 0,0019423 | 0,0019423 | TRUE | 0,0017253 | No |
| orf19.458 | 1700 | 1432 | 1565 | 970 | 2083 | 1627 | 1861 | 2501 | 1548 | 1448 | 1481 | 1457 | 1853 | 1766 | 1906 | 1908 | 1670,9 | 1484 | 1858 | 1,252 | 0,324 | 6,846 | 7,57E-12 | 1,53E-11 | 0,0005292 | 0,002828 | 0,0015431 | 0,0015431 | TRUE | 0,0128364 | No |
| orf19.4580 | 1319 | 1236 | 1238 | 705 | 1343 | 982 | 1119 | 1366 | 1201 | 1250 | 1172 | 1059 | 1195 | 1066 | 1146 | 1042 | 1141,3 | 1170 | 1112 | 0,948 | -0,077 | -1,172 | 0,241355 | 0,275544 | 0,0032433 | 0,003194 | 0,0033065 | 0,0033065 | TRUE | 0,044616 | No |
| orf19.4581 | 194 | 202 | 205 | 118 | 169 | 158 | 150 | 180 | 177 | 204 | 194 | 177 | 150 | 172 | 154 | 137 | 170,62 | 188 | 153 | 0,809 | -0,306 | -2,696 | 0,007024 | 0,009314 | 1,00E-08 | 0,009779 | 0,0064746 | 0,0064746 | TRUE | 0,0652311 | No |
| orf19.458.1 | 294 | 201 | 233 | 141 | 224 | 161 | 191 | 209 | 268 | 203 | 221 | 212 | 199 | 175 | 196 | 159 | 204,04 | 226 | 182 | 0,803 | -0,317 | -2,654 | 0,007945 | 0,010495 | 0,0091415 | 0,008511 | 0,0087854 | 0,0087854 | TRUE | 0,1814722 | No |
| orf19.4582 | 1948 | 1685 | 1812 | 1136 | 2468 | 1804 | 2008 | 2628 | 1774 | 1704 | 1715 | 1706 | 2195 | 1959 | 2056 | 2005 | 1889,26 | 1725 | 2054 | 1,191 | 0,252 | 5,173 | 2,30E-07 | 3,92E-07 | 0,0010076 | 0,002736 | 0,0017326 | 0,0017326 | TRUE | 0,0262938 | No |
| orf19.4583 | 167 | 159 | 161 | 98 | 392 | 303 | 299 | 443 | 152 | 161 | 152 | 147 | 349 | 329 | 306 | 338 | 241,78 | 153 | 330 | 2,157 | 1,109 | 11,445 | 2,50E-30 | 8,11E-30 | 1,00E-08 | 0,007501 | 0,0041445 | 0,0041445 | TRUE | 0,0270072 | No |
| orf19.4584 | 383 | 359 | 357 | 182 | 347 | 324 | 389 | 437 | 349 | 363 | 338 | 273 | 309 | 352 | 398 | 333 | 339,4 | 331 | 348 | 1,044 | 0,062 | 0,597 | 0,550706 | 0,592232 | 0,0089375 | 0,005929 | 0,0074353 | 0,0074353 | TRUE | 0,132266 | No |
| orf19.4585 | 10863 | 9315 | 10334 | 6040 | 5930 | 4691 | 5553 | 7337 | 9890 | 9421 | 9780 | 9073 | 5275 | 5093 | 5687 | 5597 | 7477 | 9541 | 5413 | 0,567 | -0,818 | -17,247 | 1,18E-66 | 7,11E-66 | 0,0018724 | 0,002214 | 0,0020179 | 0,0020179 | TRUE | 0,0190124 | No |
| orf19.4587 | 4524 | 4038 | 4209 | 2528 | 6772 | 5430 | 5796 | 7563 | 4119 | 4084 | 3983 | 3797 | 6023 | 5895 | 5936 | 5769 | 4950,96 | 3996 | 5906 | 1,476 | 0,562 | 14,826 | 9,88E-50 | 4,53E-49 | 0,0005322 | 0,002304 | 0,001165 | 0,001165 | TRUE | 0,0126323 | No |
| orf19.4589 | 1129 | 1007 | 1102 | 652 | 1283 | 1099 | 1094 | 1404 | 1028 | 1018 | 1043 | 979 | 1141 | 1193 | 1120 | 1071 | 1074,31 | 1017 | 1131 | 1,109 | 0,149 | 2,778 | 0,00547 | 0,007317 | 0,0005292 | 0,003267 | 0,0018332 | 0,0018332 | TRUE | 0,0164207 | No |
| orf19.459 | 2550 | 2306 | 2492 | 1436 | 2670 | 2199 | 2386 | 3208 | 2322 | 2332 | 2358 | 2157 | 2375 | 2388 | 2444 | 2447 | 2352,82 | 2292 | 2414 | 1,051 | 0,072 | 1,693 | 0,090418 | 0,108984 | 0,0003497 | 0,002598 | 0,0012926 | 0,0012926 | TRUE | 0,0171663 | No |
| orf19.4590 | 418 | 424 | 455 | 327 | 266 | 273 | 245 | 351 | 381 | 429 | 431 | 491 | 237 | 296 | 251 | 268 | 347,86 | 433 | 263 | 0,61 | -0,714 | -7,267 | 3,67E-13 | 7,73E-13 | 0,0067963 | 0,005835 | 0,0062488 | 0,0062488 | TRUE | 0,0848979 | No |
| orf19.4591 | 2955 | 2653 | 2862 | 1929 | 25281 | 22606 | 21890 | 30630 | 2690 | 2683 | 2709 | 2898 | 22487 | 24544 | 22418 | 23365 | 12974,2 | 2745 | 23204 | 8,463 | 3,081 | 69,413 | 0 | 0 | 0,0013377 | 0,002139 | 0,0016757 | 0,0016757 | TRUE | 0,0184323 | No |
| orf19.4592 | 2153 | 2011 | 2093 | 1331 | 5586 | 4739 | 4925 | 6685 | 1960 | 2034 | 1981 | 1999 | 4969 | 5145 | 5044 | 5099 | 3528,92 | 1994 | 5064 | 2,541 | 1,345 | 37,496 | 1,07E-307 | 2,85E-306 | 1,00E-08 | 0,002411 | 0,0008749 | 0,0008749 | TRUE | 0,0024058 | No |
| orf19.4593 | 10418 | 9199 | 10176 | 6298 | 3070 | 2538 | 2936 | 3801 | 9485 | 9304 | 9631 | 9461 | 2731 | 2756 | 3007 | 2899 | 6159,09 | 9470 | 2848 | 0,301 | -1,733 | -43,319 | 0 | 0 | 0,0008265 | 0,002251 | 0,0013165 | 0,0013165 | TRUE | 0,016544 | No |

| | | | | | | | | | | | | | | | | | | | | | | | | | | | | | | | |
|---|---|---|---|---|---|---|---|---|---|---|---|---|---|---|---|---|---|---|---|---|---|---|---|---|---|---|---|---|---|---|---|
| orf19.4593.1 | 334 | 306 | 310 | 193 | 277 | 234 | 223 | 278 | 304 | 309 | 293 | 290 | 246 | 254 | 228 | 212 | 267,22 | 299 | 235 | 0,782 | -0,354 | -3,875 | 0,000107 | 0,00016 | 1,00E-08 | 0,00698 | 0,0042461 | 0,0042461 | TRUE | 0,0548683 | No |
| orf19.4594 | 929 | 818 | 924 | 514 | 1305 | 920 | 1036 | 1424 | 846 | 827 | 874 | 772 | 1161 | 999 | 1061 | 1086 | 953,32 | 830 | 1077 | 1,295 | 0,373 | 5,892 | 3,81E-09 | 6,98E-09 | 0,0021311 | 0,003423 | 0,0027738 | 0,0027738 | TRUE | 0,0328377 | No |
| orf19.4595 | 2087 | 2132 | 2130 | 1564 | 1434 | 1238 | 1243 | 1603 | 1900 | 2156 | 2016 | 2349 | 1275 | 1344 | 1273 | 1223 | 1692,13 | 2105 | 1279 | 0,608 | -0,717 | -10,754 | 5,68E-27 | 1,70E-26 | 0,0041425 | 0,002818 | 0,0036513 | 0,0036513 | TRUE | 0,0704526 | No |
| orf19.4596 | 1822 | 1512 | 1664 | 1097 | 1330 | 1169 | 1326 | 1785 | 1659 | 1529 | 1575 | 1648 | 1183 | 1269 | 1358 | 1362 | 1447,82 | 1603 | 1293 | 0,807 | -0,309 | -5,285 | 1,26E-07 | 2,17E-07 | 0,0022662 | 0,002949 | 0,0025805 | 0,0025805 | TRUE | 0,0404426 | No |
| orf19.4597 | 2418 | 1982 | 2207 | 1448 | 1284 | 977 | 1063 | 1449 | 2201 | 2005 | 2089 | 2175 | 1142 | 1061 | 1089 | 1105 | 1608,34 | 2118 | 1099 | 0,52 | -0,944 | -18,848 | 3,03E-79 | 2,13E-78 | 0,0008468 | 0,002858 | 0,0017305 | 0,0017305 | TRUE | 0,0157253 | No |
| orf19.4599 | 238 | 242 | 234 | 179 | 565 | 459 | 505 | 727 | 217 | 245 | 221 | 269 | 503 | 498 | 517 | 555 | 378,05 | 238 | 518 | 2,199 | 1,137 | 13,13 | 2,21E-39 | 8,51E-39 | 0,0017796 | 0,005531 | 0,004043 | 0,004043 | TRUE | 0,0687032 | No |
| orf19.460 | 1593 | 1358 | 1528 | 1067 | 773 | 690 | 715 | 942 | 1450 | 1373 | 1446 | 1603 | 688 | 749 | 732 | 719 | 1095,03 | 1468 | 722 | 0,492 | -1,022 | -16,854 | 9,87E-64 | 5,71E-63 | 0,0018273 | 0,003243 | 0,0025171 | 0,0025171 | TRUE | 0,041898 | No |
| orf19.4600 | 2641 | 2135 | 2400 | 1407 | 2635 | 1894 | 2252 | 3000 | 2405 | 2159 | 2271 | 2114 | 2344 | 2056 | 2306 | 2288 | 2242,95 | 2237 | 2248 | 1,004 | 0,006 | 0,112 | 0,911077 | 0,923829 | 0,0028989 | 0,002626 | 0,0027763 | 0,0027763 | TRUE | 0,0392852 | No |
| orf19.4600.1 | 662 | 530 | 533 | 372 | 1001 | 612 | 759 | 1053 | 603 | 536 | 504 | 559 | 890 | 664 | 777 | 803 | 667,17 | 550 | 784 | 1,426 | 0,512 | 5,64 | 1,70E-08 | 3,02E-08 | 0,0087929 | 0,004017 | 0,0063411 | 0,0063411 | TRUE | 0,1214745 | No |
| orf19.4601 | 552 | 480 | 558 | 366 | 668 | 548 | 618 | 770 | 503 | 485 | 528 | 550 | 594 | 595 | 633 | 587 | 559,42 | 516 | 602 | 1,169 | 0,225 | 3,372 | 0,000746 | 0,001062 | 1,00E-08 | 0,004398 | 0,002469 | 0,002469 | TRUE | 0,0181546 | No |
| orf19.4602 | 2671 | 2047 | 2192 | 1783 | 3074 | 2810 | 2732 | 3958 | 2432 | 2070 | 2075 | 2678 | 2734 | 3051 | 2798 | 3019 | 2607,16 | 2314 | 2900 | 1,255 | 0,328 | 4,087 | 4,36E-05 | 6,68E-05 | 0,0091643 | 0,002544 | 0,0057907 | 0,0057907 | TRUE | 0,1321112 | No |
| orf19.4603 | 2547 | 2344 | 2541 | 1482 | 1736 | 1395 | 1473 | 2135 | 2319 | 2371 | 2405 | 2226 | 1544 | 1515 | 1509 | 1629 | 1939,56 | 2330 | 1549 | 0,665 | -0,589 | -12,674 | 8,23E-37 | 3,06E-36 | 0,0006427 | 0,002718 | 0,0015436 | 0,0015436 | TRUE | 0,0151706 | No |
| orf19.4605 | 1126 | 1013 | 1121 | 708 | 1822 | 1465 | 1602 | 2011 | 1025 | 1025 | 1061 | 1064 | 1621 | 1591 | 1641 | 1534 | 1320 | 1044 | 1597 | 1,53 | 0,614 | 12,754 | 2,96E-37 | 1,11E-36 | 1,00E-08 | 0,003038 | 0,0014176 | 0,0014176 | TRUE | 0,0089386 | No |
| orf19.4606 | 1023 | 942 | 1024 | 623 | 1195 | 1063 | 1116 | 1525 | 931 | 953 | 969 | 936 | 1063 | 1154 | 1143 | 1163 | 1039,04 | 947 | 1131 | 1,193 | 0,255 | 4,801 | 1,58E-06 | 2,59E-06 | 1,00E-08 | 0,003308 | 0,0017233 | 0,0017233 | TRUE | 0,0200964 | No |
| orf19.4607 | 263 | 251 | 303 | 133 | 243 | 228 | 220 | 281 | 239 | 254 | 287 | 200 | 216 | 248 | 225 | 214 | 235,4 | 245 | 226 | 0,913 | -0,132 | -1,182 | 0,237096 | 0,271216 | 0,0074309 | 0,007649 | 0,0076563 | 0,0076563 | TRUE | 0,1429958 | No |
| orf19.4608 | 244 | 240 | 262 | 158 | 353 | 300 | 272 | 429 | 222 | 243 | 248 | 237 | 314 | 326 | 279 | 327 | 274,46 | 238 | 312 | 1,313 | 0,393 | 4,285 | 1,83E-05 | 2,86E-05 | 1,00E-08 | 0,00685 | 0,0043368 | 0,0043368 | TRUE | 0,0547153 | No |
| orf19.4609 | 1641 | 1387 | 1465 | 1077 | 1616 | 1369 | 1379 | 1871 | 1494 | 1403 | 1386 | 1618 | 1437 | 1486 | 1412 | 1427 | 1458,05 | 1475 | 1440 | 0,979 | -0,031 | -0,547 | 0,584251 | 0,624618 | 0,001922 | 0,002943 | 0,0023515 | 0,0023515 | TRUE | 0,0463609 | No |
| orf19.4610 | 344 | 334 | 341 | 183 | 242 | 215 | 196 | 307 | 313 | 338 | 323 | 275 | 215 | 233 | 201 | 234 | 266,53 | 312 | 221 | 0,705 | -0,505 | -5,266 | 1,39E-07 | 2,39E-07 | 0,0017484 | 0,006993 | 0,005001 | 0,005001 | TRUE | 0,0597926 | No |
| orf19.4611 | 4824 | 3868 | 4228 | 2772 | 4331 | 3417 | 3527 | 5080 | 4392 | 3912 | 4001 | 4164 | 3852 | 3710 | 3612 | 3875 | 3939,86 | 4117 | 3762 | 0,914 | -0,13 | -2,724 | 0,006455 | 0,00859 | 0,0015136 | 0,002372 | 0,0019175 | 0,0019175 | TRUE | 0,0254226 | No |
| orf19.4612 | 66 | 47 | 65 | 39 | 88 | 77 | 70 | 107 | 60 | 48 | 62 | 59 | 78 | 84 | 72 | 82 | 67,86 | 57 | 79 | 1,385 | 0,47 | 2,777 | 0,005491 | 0,007343 | 1,00E-08 | 0,021502 | 0,0122563 | 0,0122563 | TRUE | 0,1103865 | No |
| orf19.4614 | 1481 | 1438 | 1441 | 862 | 1379 | 1203 | 1225 | 1634 | 1348 | 1454 | 1364 | 1295 | 1227 | 1306 | 1255 | 1246 | 1311,89 | 1365 | 1258 | 0,92 | -0,121 | -2,346 | 0,018969 | 0,02438 | 0,0005237 | 0,003044 | 0,0017813 | 0,0017813 | TRUE | 0,0227569 | No |
| orf19.4615 | 2359 | 2136 | 2192 | 1213 | 2704 | 1937 | 2311 | 2939 | 2148 | 2160 | 2075 | 1822 | 2405 | 2103 | 2367 | 2242 | 2165,2 | 2051 | 2279 | 1,109 | 0,15 | 2,339 | 0,019344 | 0,024847 | 0,004119 | 0,002647 | 0,0034733 | 0,0034733 | TRUE | 0,0645111 | No |
| orf19.4616 | 489 | 443 | 543 | 353 | 654 | 549 | 557 | 749 | 445 | 448 | 514 | 530 | 582 | 596 | 570 | 571 | 532,12 | 484 | 580 | 1,202 | 0,265 | 3,587 | 0,000334 | 0,000487 | 0,001828 | 0,00452 | 0,0033338 | 0,0033338 | TRUE | 0,0401772 | No |
| orf19.4617 | 1383 | 1431 | 1331 | 836 | 504 | 382 | 368 | 532 | 1259 | 1447 | 1260 | 1256 | 448 | 415 | 377 | 406 | 858,46 | 1306 | 412 | 0,315 | -1,665 | -22,63 | 2,19E-113 | 2,13E-112 | 0,0037382 | 0,003576 | 0,0036553 | 0,0036553 | TRUE | 0,0649991 | No |
| orf19.4618 | 10564 | 8608 | 9382 | 6692 | 3194 | 2864 | 2741 | 4104 | 9618 | 8706 | 8879 | 10052 | 2841 | 3110 | 2807 | 3131 | 6143 | 9314 | 2972 | 0,319 | -1,647 | -27,988 | 2,27E-172 | 3,39E-171 | 0,0036707 | 0,002252 | 0,0031122 | 0,0031122 | TRUE | 0,0343051 | No |
| orf19.4620 | 388 | 396 | 361 | 236 | 404 | 327 | 312 | 453 | 353 | 401 | 342 | 355 | 359 | 355 | 320 | 346 | 353,67 | 363 | 345 | 0,952 | -0,072 | -0,861 | 0,389362 | 0,430262 | 0,0010865 | 0,005772 | 0,0038086 | 0,0038086 | TRUE | 0,0592687 | No |
| orf19.4621 | 998 | 1071 | 1093 | 670 | 1493 | 1182 | 1224 | 1610 | 909 | 1083 | 1034 | 1006 | 1328 | 1283 | 1254 | 1228 | 1140,71 | 1008 | 1273 | 1,263 | 0,337 | 5,444 | 5,22E-08 | 9,11E-08 | 0,0024006 | 0,003195 | 0,0027828 | 0,0027828 | TRUE | 0,0565345 | No |
| orf19.4622 | 8890 | 7445 | 8149 | 5254 | 5554 | 4458 | 5069 | 6695 | 8094 | 7530 | 7712 | 7892 | 4940 | 4840 | 5191 | 5107 | 6413,37 | 7807 | 5020 | 0,643 | -0,637 | -16,213 | 4,07E-59 | 2,20E-58 | 0,0007618 | 0,002243 | 0,0013198 | 0,0013198 | TRUE | 0,0076987 | No |
| orf19.4623 | 3540 | 2850 | 3449 | 2585 | 2035 | 1828 | 2037 | 2828 | 3223 | 2882 | 3264 | 3883 | 1810 | 1985 | 2086 | 2157 | 2661,36 | 3313 | 2010 | 0,607 | -0,72 | -8,572 | 1,02E-17 | 2,44E-17 | 0,0101848 | 0,002533 | 0,0063771 | 0,0063771 | TRUE | 0,1597572 | No |
| orf19.4623.3 | 8847 | 8470 | 8249 | 4941 | 7639 | 5601 | 5714 | 7755 | 8055 | 8567 | 7807 | 7422 | 6795 | 6081 | 5852 | 5916 | 7061,7 | 7963 | 6161 | 0,774 | -0,37 | -6,223 | 4,87E-10 | 9,20E-10 | 0,0040958 | 0,002224 | 0,0032607 | 0,0032607 | TRUE | 0,0587666 | No |
| orf19.4624 | 422 | 378 | 351 | 207 | 702 | 562 | 537 | 765 | 384 | 382 | 332 | 311 | 624 | 610 | 550 | 584 | 472,22 | 352 | 592 | 1,671 | 0,741 | 8,924 | 4,48E-19 | 1,11E-18 | 0,0035791 | 0,004834 | 0,0043152 | 0,0043152 | TRUE | 0,0589444 | No |
| orf19.4625 | 3173 | 3345 | 3356 | 1820 | 2781 | 2148 | 2285 | 2859 | 2889 | 3383 | 3176 | 2734 | 2474 | 2332 | 2340 | 2181 | 2688,6 | 3046 | 2332 | 0,765 | -0,387 | -5,658 | 1,53E-08 | 2,74E-08 | 0,0053521 | 0,002528 | 0,0041122 | 0,0041122 | TRUE | 0,0671823 | No |
| orf19.4626 | 2090 | 2037 | 2088 | 1100 | 2071 | 1563 | 1843 | 2313 | 1903 | 2060 | 1976 | 1652 | 1842 | 1697 | 1887 | 1764 | 1847,81 | 1898 | 1798 | 0,945 | -0,081 | -1,204 | 0,228423 | 0,262 | 0,0046678 | 0,002752 | 0,0037907 | 0,0037907 | TRUE | 0,0863032 | No |
| orf19.4627 | 1438 | 1575 | 1697 | 980 | 1342 | 1104 | 1170 | 1510 | 1309 | 1593 | 1606 | 1472 | 1194 | 1199 | 1198 | 1152 | 1340,34 | 1495 | 1186 | 0,793 | -0,335 | -4,986 | 6,17E-07 | 1,04E-06 | 0,0040654 | 0,003023 | 0,0035822 | 0,0035822 | TRUE | 0,0884863 | No |
| orf19.4628 | 1657 | 1523 | 1697 | 970 | 1013 | 826 | 814 | 1090 | 1509 | 1540 | 1606 | 1457 | 901 | 897 | 834 | 831 | 1196,88 | 1528 | 866 | 0,565 | -0,823 | -15,052 | 3,36E-51 | 1,58E-50 | 0,0008879 | 0,003141 | 0,0019779 | 0,0019779 | TRUE | 0,0142168 | No |
| orf19.4629 | 2561 | 2841 | 2799 | 1800 | 3795 | 3542 | 3326 | 4249 | 2332 | 2873 | 2649 | 2704 | 3376 | 3846 | 3406 | 3241 | 3053,31 | 2640 | 3467 | 1,314 | 0,394 | 5,555 | 2,78E-08 | 4,91E-08 | 0,0060935 | 0,00247 | 0,0044819 | 0,0044819 | TRUE | 0,0761348 | No |
| orf19.4630 | 11614 | 10251 | 10895 | 7145 | 5556 | 4916 | 4876 | 6620 | 10574 | 10368 | 10311 | 10733 | 4942 | 5337 | 4994 | 5050 | 7788,58 | 10496 | 5081 | 0,484 | -1,047 | -28,813 | 1,47E-182 | 2,31E-181 | 0,000581 | 0,002207 | 0,0011245 | 0,0011245 | TRUE | 0,0137822 | No |
| orf19.4631 | 3360 | 3463 | 3468 | 2520 | 900 | 1031 | 770 | 1144 | 3059 | 3502 | 3282 | 3785 | 801 | 1119 | 789 | 873 | 2151,29 | 3407 | 896 | 0,263 | -1,929 | -19,215 | 2,78E-82 | 2,04E-81 | 0,0179911 | 0,002651 | 0,0089984 | 0,0089984 | TRUE | 0,3321476 | No |
| orf19.4632 | 11607 | 10271 | 10175 | 7688 | 18811 | 15250 | 13342 | 19428 | 10568 | 10388 | 9630 | 11549 | 16732 | 16557 | 13664 | 14820 | 12988,4 | 10534 | 15443 | 1,466 | 0,552 | 7,665 | 1,79E-14 | 3,91E-14 | 0,0072136 | 0,002139 | 0,0049073 | 0,0049073 | TRUE | 0,0734787 | No |
| orf19.4633 | 1580 | 1408 | 1566 | 960 | 2133 | 1808 | 1824 | 2439 | 1439 | 1424 | 1482 | 1442 | 1897 | 1963 | 1868 | 1860 | 1671,93 | 1447 | 1897 | 1,31 | 0,39 | 8,983 | 2,64E-19 | 6,59E-19 | 1,00E-08 | 0,002827 | 0,0011934 | 0,0011934 | TRUE | 0,0063167 | No |
| orf19.4634 | 4596 | 4248 | 4624 | 2807 | 3732 | 3041 | 3241 | 4251 | 4184 | 4296 | 4376 | 4217 | 3319 | 3302 | 3319 | 3243 | 3782,09 | 4268 | 3296 | 0,772 | -0,374 | -10,875 | 1,51E-27 | 4,60E-27 | 1,00E-08 | 0,002386 | 0,0008665 | 0,0008665 | TRUE | 0,003616 | No |
| orf19.4635 | 7645 | 7972 | 7982 | 4619 | 12418 | 10418 | 10518 | 13667 | 6960 | 8063 | 7554 | 6938 | 11045 | 11311 | 10772 | 10425 | 9133,69 | 7379 | 10888 | 1,475 | 0,561 | 10,347 | 4,33E-25 | 1,25E-24 | 0,0031104 | 0,002182 | 0,0027052 | 0,0027052 | TRUE | 0,0475175 | No |
| orf19.4636 | 1 | 5 | 2 | 0 | 1 | 1 | 1 | 1 | 1 | 5 | 2 | 0 | 1 | 1 | 1 | 1 | 1,45 | 2 | 1 | 0,456 | -1,134 | -0,946 | 0,344109 | 0,383983 | 1,00E-08 | 0,911317 | 0,6189526 | 0,6189526 | TRUE | 0,6498953 | No |
| orf19.4637 | 445 | 463 | 489 | 285 | 363 | 359 | 342 | 453 | 405 | 468 | 463 | 428 | 323 | 390 | 350 | 346 | 396,6 | 441 | 352 | 0,796 | -0,329 | -3,908 | 9,31E-05 | 0,00014 | 0,0028744 | 0,005368 | 0,0042715 | 0,0042715 | TRUE | 0,0559611 | No |
| orf19.4639 | 738 | 697 | 753 | 435 | 773 | 671 | 579 | 745 | 672 | 705 | 713 | 653 | 688 | 729 | 593 | 568 | 665,04 | 686 | 644 | 0,937 | -0,094 | -1,129 | 0,258843 | 0,294507 | 0,0063644 | 0,004023 | 0,0051611 | 0,0051611 | TRUE | 0,088634 | No |
| orf19.4640 | 11180 | 9625 | 10689 | 6320 | 12157 | 9868 | 10591 | 14427 | 10179 | 9735 | 10116 | 9494 | 10813 | 10714 | 10847 | 11005 | 10362,8 | 9881 | 10845 | 1,097 | 0,134 | 3,933 | 8,37E-05 | 0,000126 | 0,0004718 | 0,002164 | 0,0010111 | 0,0010111 | TRUE | 0,0082069 | No |
| orf19.4641 | 2845 | 3015 | 2946 | 1904 | 1531 | 1370 | 1401 | 1775 | 2590 | 3049 | 2788 | 2860 | 1362 | 1487 | 1435 | 1354 | 2115,73 | 2822 | 1410 | 0,499 | -1,002 | -17,014 | 6,48E-65 | 3,80E-64 | 0,0029254 | 0,002661 | 0,0028089 | 0,0028089 | TRUE | 0,0380295 | No |
| orf19.4642 | 891 | 1006 | 992 | 504 | 822 | 698 | 696 | 874 | 811 | 1017 | 939 | 757 | 731 | 758 | 713 | 667 | 799,13 | 881 | 717 | 0,811 | -0,302 | -3,399 | 0,000676 | 0,000964 | 0,0088014 | 0,00369 | 0,0063071 | 0,0063071 | TRUE | 0,1271375 | No |
| orf19.4643 | 1842 | 1777 | 1910 | 1030 | 1254 | 976 | 1106 | 1381 | 1677 | 1797 | 1808 | 1547 | 1115 | 1060 | 1133 | 1053 | 1398,79 | 1707 | 1090 | 0,637 | -0,65 | -10,836 | 2,32E-27 | 7,02E-27 | 0,002216 | 0,002981 | 0,0027141 | 0,0027141 | TRUE | 0,0442168 | No |
| orf19.4645 | 5027 | 5704 | 5826 | 3310 | 1783 | 1492 | 1535 | 2004 | 4577 | 5769 | 5514 | 4972 | 1586 | 1620 | 1572 | 1529 | 3392,29 | 5208 | 1577 | 0,303 | -1,725 | -24,935 | 3,08E-137 | 3,65E-136 | 0,005034 | 0,002426 | 0,0041936 | 0,0041936 | TRUE | 0,0824685 | No |
| orf19.4646 | 315 | 251 | 295 | 260 | 82 | 92 | 74 | 135 | 287 | 254 | 279 | 391 | 73 | 100 | 76 | 103 | 195,25 | 303 | 88 | 0,293 | -1,772 | -11,308 | 1,19E-29 | 3,80E-29 | 0,0276026 | 0,008803 | 0,0164988 | 0,0164988 | TRUE | 0,3943722 | No |
| orf19.4647 | 323 | 236 | 289 | 176 | 230 | 205 | 216 | 308 | 294 | 239 | 274 | 264 | 205 | 223 | 221 | 235 | 244,25 | 268 | 221 | 0,824 | -0,279 | -2,835 | 0,004583 | 0,006159 | 1,00E-08 | 0,007445 | 0,0051872 | 0,0051872 | TRUE | 0,0638713 | No |
| orf19.4649 | 740 | 887 | 801 | 563 | 934 | 842 | 833 | 1134 | 674 | 897 | 758 | 846 | 831 | 914 | 853 | 865 | 829,71 | 794 | 866 | 1,093 | 0,128 | 1,536 | 0,124648 | 0,148142 | 0,006908 | 0,003629 | 0,0054698 | 0,0054698 | TRUE | 0,126859 | No |
| orf19.465 | 36 | 18 | 28 | 32 | 23 | 20 | 21 | 22 | 33 | 18 | 26 | 48 | 20 | 22 | 22 | 17 | 25,75 | 31 | 20 | 0,651 | -0,618 | -1,991 | 0,04644 | 0,057654 | 0,049303 | 0,053335 | 0,0518348 | 0,0518348 | TRUE | 0,2704478 | No |
| orf19.4650 | 1129 | 969 | 1088 | 649 | 1952 | 1573 | 1663 | 2173 | 1028 | 980 | 1030 | 975 | 1736 | 1708 | 1703 | 1658 | 1352,17 | 1003 | 1701 | 1,692 | 0,759 | 16,034 | 7,41E-58 | 3,93E-57 | 1,00E-08 | 0,003014 | 0,0013454 | 0,0013454 | TRUE | 0,0038624 | No |
| orf19.4651 | 1459 | 1492 | 1505 | 992 | 3131 | 2503 | 2424 | 3427 | 1328 | 1509 | 1424 | 1490 | 2785 | 2718 | 2482 | 2614 | 2043,87 | 1438 | 2650 | 1,845 | 0,884 | 15,693 | 1,68E-55 | 8,56E-55 | 0,0021314 | 0,002683 | 0,0024948 | 0,0024948 | TRUE | 0,0327234 | No |
| orf19.4652 | 1 | 0 | 1 | 0 | 4 | 1 | 4 | 2 | 1 | 0 | 1 | 0 | 4 | 1 | 4 | 2 | 1,52 | 0 | 3 | 4,855 | 2,279 | 1,73 | 0,083584 | 0,10111 | 1,00E-08 | 0,873797 | 0,5574325 | 0,5574325 | TRUE | 0,0839782 | No |
| orf19.4653 | 12 | 8 | 14 | 14 | 13 | 19 | 16 | 20 | 11 | 8 | 13 | 21 | 12 | 21 | 16 | 15 | 14,64 | 13 | 16 | 1,229 | 0,298 | 0,78 | 0,43529 | 0,477046 | 0,0184885 | 0,092261 | 0,0688653 | 0,0688653 | TRUE | 0,4127555 | No |
| orf19.4654 | 28 | 15 | 23 | 14 | 18 | 21 | 11 | 25 | 25 | 15 | 22 | 21 | 16 | 23 | 11 | 19 | 19,08 | 21 | 17 | 0,826 | -0,275 | -0,843 | 0,399199 | 0,439912 | 1,00E-08 | 0,071286 | 0,0500374 | 0,0500374 | TRUE | 0,2503646 | No |
| orf19.4655 | 412 | 470 | 473 | 306 | 419 | 466 | 372 | 478 | 375 | 475 | 448 | 460 | 373 | 506 | 381 | 365 | 422,75 | 440 | 406 | 0,923 | -0,115 | -1,021 | 0,307089 | 0,345577 | 0,0148383 | 0,005162 | 0,0098928 | 0,0098928 | TRUE | 0,3081977 | No |
| orf19.4656 | 195 | 250 | 236 | 136 | 428 | 347 | 377 | 474 | 178 | 253 | 223 | 204 | 381 | 377 | 386 | 362 | 295,39 | 214 | 376 | 1,752 | 0,809 | 7,904 | 2,71E-15 | 6,04E-15 | 0,0064476 | 0,006509 | 0,0063304 | 0,0063304 | TRUE | 0,1682987 | No |
| orf19.4657 | 11544 | 11204 | 12027 | 6889 | 5830 | 4874 | 5171 | 6792 | 10510 | 11332 | 11382 | 10348 | 5186 | 5292 | 5296 | 5181 | 8065,87 | 10893 | 5239 | 0,481 | -1,057 | -25,053 | 1,61E-138 | 1,94E-137 | 0,0011148 | 0,002201 | 0,0015703 | 0,0015703 | TRUE | 0,0135085 | No |
| orf19.4658 | 3247 | 3301 | 3505 | 1850 | 4307 | 3308 | 3826 | 4741 | 2956 | 3339 | 3317 | 2779 | 3831 | 3592 | 3918 | 3616 | 3418,54 | 3098 | 3739 | 1,206 | 0,27 | 4,265 | 2,00E-05 | 3,13E-05 | 0,0044847 | 0,002423 | 0,0035512 | 0,0035512 | TRUE | 0,055954 | No |
| orf19.4659 | 3284 | 3059 | 3296 | 2038 | 4330 | 3550 | 3651 | 4992 | 2990 | 3094 | 3119 | 3061 | 3851 | 3854 | 3739 | 3808 | 3439,66 | 3066 | 3813 | 1,244 | 0,315 | 8,985 | 2,60E-19 | 6,48E-19 | 1,00E-08 | 0,002421 | 0,00088 | 0,00088 | TRUE | 0,0036625 | No |
| orf19.4660 | 20674 | 17816 | 18599 | 11675 | 31326 | 24640 | 25191 | 35094 | 18823 | 18019 | 17602 | 17538 | 27863 | 26752 | 25799 | 26770 | 22395,8 | 17996 | 26796 | 1,489 | 0,574 | 15,082 | 2,11E-51 | 1,00E-50 | 0,000974 | 0,002096 | 0,001345 | 0,001345 | TRUE | 0,0117862 | No |
| orf19.4662 | 1378 | 1111 | 1245 | 909 | 1791 | 1636 | 1690 | 2316 | 1255 | 1124 | 1178 | 1365 | 1593 | 1776 | 1731 | 1767 | 1473,59 | 1230 | 1717 | 1,398 | 0,484 | 7,417 | 1,20E-13 | 2,56E-13 | 0,0039133 | 0,002933 | 0,003373 | 0,003373 | TRUE | 0,0600312 | No |
| orf19.4664 | 3626 | 3316 | 3423 | 1968 | 273 | 199 | 248 | 288 | 3301 | 3354 | 3240 | 2956 | 243 | 216 | 254 | 220 | 1722,93 | 3213 | 233 | 0,072 | -3,788 | -53,422 | 0 | 0 | 0,0024993 | 0,002804 | 0,0026641 | 0,0026641 | TRUE | 0,0372901 | No |

| | | | | | | | | | | | | | | | | | | | | | | | | | | | | | | | |
|---|---|---|---|---|---|---|---|---|---|---|---|---|---|---|---|---|---|---|---|---|---|---|---|---|---|---|---|---|---|---|---|
| orf19.4665 | 2799 | 2139 | 2429 | 1335 | 790 | 561 | 704 | 890 | 2548 | 2163 | 2299 | 2005 | 703 | 609 | 721 | 679 | 1465,95 | 2254 | 678 | 0,301 | -1,734 | -22,019 | 1,89E-107 | 1,74E-106 | 0,00654 | 0,002938 | 0,0050298 | 0,0050298 | TRUE | 0,0938641 | No |
| orf19.4666 | 3481 | 3208 | 3108 | 1984 | 2209 | 1743 | 1723 | 2361 | 3169 | 3245 | 2941 | 2980 | 1965 | 1892 | 1765 | 1801 | 2469,8 | 3084 | 1856 | 0,601 | -0,734 | -14,073 | 5,53E-45 | 2,37E-44 | 0,001737 | 0,002572 | 0,0021824 | 0,0021824 | TRUE | 0,019336 | No |
| orf19.4668 | 1407 | 1270 | 1440 | 919 | 1207 | 971 | 1035 | 1383 | 1281 | 1284 | 1363 | 1380 | 1074 | 1054 | 1060 | 1055 | 1193,94 | 1327 | 1061 | 0,801 | -0,321 | -6,495 | 8,29E-11 | 1,62E-10 | 1,00E-08 | 0,003143 | 0,0015008 | 0,0015008 | TRUE | 0,0076547 | No |
| orf19.4669 | 485 | 582 | 534 | 334 | 2483 | 2112 | 2067 | 2609 | 442 | 589 | 505 | 502 | 2209 | 2293 | 2117 | 1990 | 1330,74 | 510 | 2152 | 4,226 | 2,079 | 26,124 | 1,92E-150 | 2,51E-149 | 0,0072362 | 0,00303 | 0,0048106 | 0,0048106 | TRUE | 0,1335996 | No |
| orf19.467 | 11616 | 10141 | 11279 | 7755 | 2592 | 2256 | 2504 | 3410 | 10576 | 10257 | 10675 | 11649 | 2306 | 2449 | 2564 | 2601 | 6634,59 | 10789 | 2480 | 0,23 | -2,12 | -39,46 | 0 | 0 | 0,0026601 | 0,002236 | 0,0025347 | 0,0025347 | TRUE | 0,0347018 | No |
| orf19.4670 | 5949 | 5515 | 6067 | 3678 | 3826 | 3307 | 3621 | 4709 | 5416 | 5578 | 5742 | 5525 | 3403 | 3590 | 3708 | 3592 | 4569,37 | 5565 | 3573 | 0,642 | -0,64 | -16,077 | 3,69E-58 | 1,97E-57 | 0,000615 | 0,002326 | 0,0012919 | 0,0012919 | TRUE | 0,0126172 | No |
| orf19.4672 | 3 | 4 | 6 | 4 | 11 | 5 | 7 | 9 | 3 | 4 | 6 | 6 | 10 | 5 | 7 | 7 | 5,96 | 5 | 7 | 1,63 | 0,705 | 1,244 | 0,213432 | 0,246044 | 1,00E-08 | 0,223537 | 0,1274652 | 0,1274652 | TRUE | 0,1688664 | No |
| orf19.4673 | 1537 | 1419 | 1564 | 941 | 292 | 238 | 225 | 298 | 1399 | 1435 | 1480 | 1414 | 260 | 258 | 230 | 227 | 838,02 | 1432 | 244 | 0,17 | -2,557 | -37,784 | 0 | 0 | 1,00E-08 | 0,003613 | 0,0021153 | 0,0021153 | TRUE | 0,0277728 | No |
| orf19.4674.1 | 310 | 276 | 255 | 149 | 566 | 501 | 465 | 603 | 282 | 279 | 241 | 224 | 503 | 544 | 476 | 460 | 376,26 | 256 | 496 | 1,917 | 0,939 | 10,093 | 5,91E-24 | 1,66E-23 | 0,0050569 | 0,005548 | 0,0052947 | 0,0052947 | TRUE | 0,0664668 | No |
| orf19.4675 | 709 | 600 | 712 | 390 | 738 | 590 | 682 | 836 | 646 | 607 | 674 | 586 | 656 | 641 | 698 | 638 | 643,15 | 628 | 658 | 1,044 | 0,062 | 0,933 | 0,351 | 0,390988 | 0,0010578 | 0,004091 | 0,002701 | 0,002701 | TRUE | 0,0279079 | No |
| orf19.4676 | 370 | 364 | 370 | 178 | 551 | 431 | 468 | 566 | 337 | 368 | 350 | 267 | 490 | 468 | 479 | 432 | 398,96 | 330 | 467 | 1,401 | 0,487 | 5,252 | 1,51E-07 | 2,58E-07 | 0,006084 | 0,005348 | 0,0056344 | 0,0056344 | TRUE | 0,1565404 | No |
| orf19.4677 | 1119 | 1133 | 1149 | 709 | 1241 | 836 | 965 | 1370 | 1019 | 1146 | 1087 | 1065 | 1104 | 908 | 988 | 1045 | 1045,25 | 1079 | 1011 | 0,938 | -0,092 | -1,348 | 0,177786 | 0,206974 | 0,0036958 | 0,003301 | 0,0035025 | 0,0035025 | TRUE | 0,0551846 | No |
| orf19.4678 | 1182 | 1196 | 1231 | 703 | 1333 | 1135 | 1166 | 1496 | 1076 | 1210 | 1165 | 1056 | 1186 | 1232 | 1194 | 1141 | 1157,51 | 1127 | 1188 | 1,052 | 0,073 | 1,262 | 0,206949 | 0,239046 | 0,0014298 | 0,003178 | 0,0023478 | 0,0023478 | TRUE | 0,0288339 | No |
| orf19.4679 | 23209 | 19566 | 21758 | 14503 | 55419 | 46508 | 49834 | 69361 | 21131 | 19789 | 20592 | 21786 | 49293 | 50495 | 51036 | 52909 | 35878,9 | 20824 | 50933 | 2,446 | 1,291 | 32,271 | 1,79E-228 | 3,44E-227 | 0,0012157 | 0,002074 | 0,0015015 | 0,0015015 | TRUE | 0,0136874 | No |
| orf19.4680 | 435 | 408 | 437 | 279 | 420 | 331 | 369 | 515 | 396 | 413 | 414 | 419 | 374 | 359 | 378 | 393 | 393,13 | 410 | 376 | 0,919 | -0,122 | -1,659 | 0,097114 | 0,11679 | 1,00E-08 | 0,005397 | 0,0026307 | 0,0026307 | TRUE | 0,011517 | No |
| orf19.4681 | 7015 | 6617 | 7222 | 4585 | 9149 | 7794 | 8517 | 10878 | 6387 | 6692 | 6835 | 6887 | 8138 | 8462 | 8722 | 8298 | 7552,72 | 6700 | 8405 | 1,255 | 0,327 | 8,358 | 6,36E-17 | 1,49E-16 | 0,0008245 | 0,002212 | 0,001337 | 0,001337 | TRUE | 0,0122782 | No |
| orf19.4682 | 273 | 288 | 317 | 200 | 325 | 297 | 260 | 376 | 249 | 291 | 300 | 300 | 289 | 322 | 266 | 287 | 288,11 | 285 | 291 | 1,024 | 0,034 | 0,35 | 0,726009 | 0,758863 | 0,003522 | 0,006622 | 0,0053311 | 0,0053311 | TRUE | 0,0900451 | No |
| orf19.4683 | 5045 | 4886 | 5143 | 3165 | 3058 | 2702 | 2647 | 3461 | 4593 | 4942 | 4867 | 4754 | 2720 | 2934 | 2711 | 2640 | 3770,15 | 4789 | 2751 | 0,574 | -0,801 | -17,68 | 6,03E-70 | 3,78E-69 | 0,0011816 | 0,002387 | 0,0016875 | 0,0016875 | TRUE | 0,0237195 | No |
| orf19.4684.2 | 4860 | 3794 | 3905 | 2553 | 9195 | 6198 | 7538 | 9584 | 4425 | 3837 | 3696 | 3835 | 8179 | 6729 | 7720 | 7311 | 5716,42 | 3948 | 7485 | 1,895 | 0,922 | 13,125 | 2,37E-39 | 9,10E-39 | 0,0065061 | 0,002268 | 0,0045457 | 0,0045457 | TRUE | 0,0813433 | No |
| orf19.4685 | 11 | 10 | 18 | 6 | 45 | 25 | 27 | 32 | 10 | 10 | 17 | 9 | 40 | 27 | 28 | 24 | 20,68 | 12 | 30 | 2,541 | 1,346 | 3,958 | 7,56E-05 | 0,000114 | 1,00E-08 | 0,065928 | 0,0500722 | 0,0500722 | TRUE | 0,4264209 | No |
| orf19.4686 | 3317 | 3273 | 3465 | 2046 | 4234 | 3498 | 3665 | 4650 | 3020 | 3310 | 3279 | 3073 | 3766 | 3798 | 3753 | 3547 | 3443,42 | 3170 | 3716 | 1,171 | 0,228 | 4,982 | 6,29E-07 | 1,06E-06 | 0,0011348 | 0,002421 | 0,0017108 | 0,0017108 | TRUE | 0,0132842 | No |
| orf19.4688 | 1842 | 1626 | 1834 | 1243 | 8312 | 7321 | 7364 | 9676 | 1677 | 1645 | 1736 | 1867 | 7393 | 7949 | 7542 | 7381 | 4648,61 | 1731 | 7566 | 4,38 | 2,131 | 44,156 | 0 | 0 | 0,0017516 | 0,002321 | 0,0018629 | 0,0018629 | TRUE | 0,0306615 | No |
| orf19.4689 | 936 | 781 | 897 | 590 | 897 | 825 | 824 | 1142 | 852 | 790 | 849 | 886 | 798 | 896 | 844 | 871 | 848,23 | 844 | 852 | 1,01 | 0,015 | 0,247 | 0,804697 | 0,830626 | 0,0010936 | 0,003594 | 0,0023739 | 0,0023739 | TRUE | 0,0222584 | No |
| orf19.469 | 3398 | 2984 | 3327 | 2128 | 1852 | 1515 | 1610 | 2215 | 3094 | 3018 | 3149 | 3197 | 1647 | 1645 | 1649 | 1690 | 2385,96 | 3114 | 1658 | 0,533 | -0,908 | -22,887 | 6,30E-116 | 6,25E-115 | 1,00E-08 | 0,002591 | 0,0010575 | 0,0010575 | TRUE | 0,0051771 | No |
| orf19.4690 | 181 | 158 | 171 | 96 | 62 | 66 | 64 | 83 | 165 | 160 | 162 | 144 | 55 | 72 | 66 | 63 | 110,79 | 158 | 64 | 0,402 | -1,316 | -9,414 | 4,77E-21 | 1,24E-20 | 1,00E-08 | 0,013961 | 0,0080882 | 0,0080882 | TRUE | 0,0765404 | No |
| orf19.4691 | 183 | 170 | 192 | 107 | 75 | 60 | 65 | 77 | 167 | 172 | 182 | 161 | 67 | 65 | 67 | 59 | 117,27 | 170 | 64 | 0,375 | -1,414 | -10,453 | 1,42E-25 | 4,13E-25 | 1,00E-08 | 0,013302 | 0,0072021 | 0,0072021 | TRUE | 0,0368536 | No |
| orf19.4697 | 982 | 930 | 1015 | 564 | 4687 | 3812 | 4264 | 5339 | 894 | 941 | 961 | 847 | 4169 | 4139 | 4367 | 4073 | 2548,71 | 911 | 4187 | 4,583 | 2,196 | 44,114 | 0 | 0 | 0,0008726 | 0,002555 | 0,0016822 | 0,0016822 | TRUE | 0,0219429 | No |
| orf19.4698 | 2517 | 2627 | 2843 | 1456 | 1255 | 1042 | 1061 | 1258 | 2292 | 2657 | 2691 | 2187 | 1116 | 1131 | 1087 | 960 | 1765,02 | 2457 | 1074 | 0,436 | -1,197 | -15,202 | 3,43E-52 | 1,65E-51 | 0,0071347 | 0,002785 | 0,0053001 | 0,0053001 | TRUE | 0,0636984 | No |
| orf19.4699 | 2994 | 3125 | 3198 | 1923 | 2474 | 2122 | 2043 | 2676 | 2726 | 3161 | 3027 | 2889 | 2201 | 2304 | 2092 | 2041 | 2554,97 | 2951 | 2160 | 0,731 | -0,451 | -7,732 | 1,06E-14 | 2,33E-14 | 0,0029268 | 0,002554 | 0,0028705 | 0,0028705 | TRUE | 0,0330193 | No |
| orf19.470 | 2629 | 2213 | 2484 | 1560 | 1376 | 1241 | 1374 | 1881 | 2394 | 2238 | 2351 | 2343 | 1224 | 1347 | 1407 | 1435 | 1842,42 | 2332 | 1353 | 0,581 | -0,785 | -13,951 | 3,11E-44 | 1,32E-43 | 0,002018 | 0,002754 | 0,00246 | 0,00246 | TRUE | 0,0511239 | No |
| orf19.4701 | 344 | 445 | 403 | 266 | 242 | 238 | 228 | 254 | 313 | 450 | 381 | 400 | 215 | 258 | 234 | 194 | 305,64 | 386 | 225 | 0,582 | -0,781 | -6,432 | 1,26E-10 | 2,44E-10 | 0,0157997 | 0,006359 | 0,010661 | 0,010661 | TRUE | 0,1947192 | No |
| orf19.4702 | 117 | 93 | 116 | 78 | 162 | 134 | 130 | 164 | 107 | 94 | 110 | 117 | 144 | 145 | 133 | 125 | 121,92 | 107 | 137 | 1,285 | 0,361 | 2,788 | 0,005304 | 0,007104 | 1,00E-08 | 0,012872 | 0,0077077 | 0,0077077 | TRUE | 0,0629537 | No |
| orf19.4703 | 410 | 367 | 379 | 224 | 458 | 453 | 465 | 628 | 373 | 371 | 359 | 336 | 407 | 492 | 476 | 479 | 411,76 | 360 | 464 | 1,283 | 0,359 | 4,363 | 1,28E-05 | 2,03E-05 | 0,0025592 | 0,005245 | 0,0040271 | 0,0040271 | TRUE | 0,0790837 | No |
| orf19.4704 | 7364 | 6644 | 7070 | 4433 | 6071 | 4986 | 5375 | 7173 | 6705 | 6720 | 6691 | 6659 | 5400 | 5413 | 5505 | 5472 | 6070,52 | 6694 | 5448 | 0,814 | -0,297 | -10,43 | 1,81E-25 | 5,24E-25 | 1,00E-08 | 0,002255 | 0,0006142 | 0,0006142 | TRUE | 0,0005883 | No |
| orf19.4705 | 1562 | 1414 | 1510 | 1013 | 1565 | 1261 | 1232 | 1850 | 1422 | 1430 | 1429 | 1522 | 1392 | 1369 | 1262 | 1411 | 1404,63 | 1451 | 1358 | 0,939 | -0,091 | -1,745 | 0,081063 | 0,098191 | 0,0009563 | 0,002977 | 0,0019012 | 0,0019012 | TRUE | 0,0272217 | No |
| orf19.4706 | 1293 | 1163 | 1224 | 815 | 879 | 753 | 798 | 1134 | 1177 | 1176 | 1158 | 1224 | 782 | 818 | 817 | 865 | 1002,23 | 1184 | 820 | 0,694 | -0,526 | -9,792 | 1,22E-22 | 3,30E-22 | 1,00E-08 | 0,003355 | 0,0017493 | 0,0017493 | TRUE | 0,0175879 | No |
| orf19.4707 | 181 | 132 | 167 | 105 | 132 | 112 | 107 | 149 | 165 | 134 | 158 | 158 | 117 | 122 | 110 | 114 | 134,54 | 154 | 116 | 0,752 | -0,411 | -3,338 | 0,000843 | 0,001195 | 1,00E-08 | 0,011856 | 0,0070226 | 0,0070226 | TRUE | 0,0847049 | No |
| orf19.4711 | 3544 | 3118 | 3431 | 2066 | 2162 | 1703 | 1858 | 2646 | 3227 | 3154 | 3247 | 3103 | 1923 | 1849 | 1903 | 2018 | 2553 | 3183 | 1923 | 0,605 | -0,726 | -17,145 | 6,86E-66 | 4,08E-65 | 0,0004355 | 0,002554 | 0,0013102 | 0,0013102 | TRUE | 0,0139771 | No |
| orf19.4712 | 25 | 17 | 15 | 15 | 27 | 24 | 23 | 28 | 23 | 17 | 14 | 23 | 24 | 26 | 24 | 21 | 21,46 | 19 | 24 | 1,245 | 0,317 | 1,067 | 0,28612 | 0,322721 | 1,00E-08 | 0,063597 | 0,0368901 | 0,0368901 | TRUE | 0,1770518 | No |
| orf19.4713 | 392 | 412 | 410 | 297 | 291 | 239 | 264 | 330 | 357 | 417 | 388 | 446 | 259 | 259 | 270 | 252 | 331,02 | 402 | 260 | 0,65 | -0,622 | -6,994 | 2,67E-12 | 5,47E-12 | 0,0024244 | 0,006028 | 0,0044499 | 0,0044499 | TRUE | 0,0667149 | No |
| orf19.4715 | 1818 | 1685 | 1831 | 1117 | 1798 | 1534 | 1647 | 2090 | 1655 | 1704 | 1733 | 1678 | 1599 | 1665 | 1687 | 1594 | 1664,5 | 1692 | 1636 | 0,966 | -0,05 | -1,136 | 0,255748 | 0,291142 | 1,00E-08 | 0,002831 | 0,0012774 | 0,0012774 | TRUE | 0,0052296 | No |
| orf19.4716 | 16048 | 13178 | 14286 | 9270 | 45594 | 36142 | 38772 | 54422 | 14611 | 13328 | 13520 | 13925 | 40554 | 39240 | 39707 | 41514 | 27050 | 13846 | 40254 | 2,907 | 1,54 | 39,435 | 0 | 0 | 0,0010837 | 0,002086 | 0,0014138 | 0,0014138 | TRUE | 0,0171631 | No |
| orf19.4718 | 10656 | 9119 | 9850 | 6451 | 11875 | 9787 | 10464 | 14696 | 9702 | 9223 | 9322 | 9690 | 10562 | 10626 | 10716 | 11210 | 10131,6 | 9484 | 10778 | 1,137 | 0,185 | 5,198 | 2,01E-07 | 3,44E-07 | 0,0005829 | 0,002167 | 0,0011176 | 0,0011176 | TRUE | 0,0090127 | No |
| orf19.4719 | 798 | 758 | 784 | 431 | 851 | 723 | 817 | 1074 | 727 | 767 | 742 | 647 | 757 | 785 | 837 | 819 | 760,06 | 721 | 800 | 1,105 | 0,144 | 2,194 | 0,028254 | 0,035766 | 0,0017876 | 0,003775 | 0,0028298 | 0,0028298 | TRUE | 0,0473811 | No |
| orf19.4720 | 3542 | 3073 | 3343 | 2143 | 2499 | 1951 | 2217 | 2987 | 3225 | 3108 | 3164 | 3219 | 2223 | 2118 | 2270 | 2279 | 2700,73 | 3179 | 2222 | 0,7 | -0,515 | -12,788 | 1,91E-37 | 7,16E-37 | 0,0001887 | 0,002526 | 0,0011782 | 0,0011782 | TRUE | 0,0115929 | No |
| orf19.4721 | 4739 | 4327 | 4485 | 2515 | 3238 | 2454 | 2876 | 3748 | 4315 | 4376 | 4245 | 3778 | 2880 | 2664 | 2945 | 2859 | 3507,8 | 4178 | 2837 | 0,678 | -0,56 | -10,19 | 2,20E-24 | 6,23E-24 | 0,0026383 | 0,002414 | 0,0026031 | 0,0026031 | TRUE | 0,0487605 | No |
| orf19.4722 | 2758 | 2324 | 2631 | 1528 | 656 | 514 | 610 | 781 | 2511 | 2350 | 2490 | 2295 | 583 | 558 | 625 | 596 | 1501,11 | 2412 | 590 | 0,245 | -2,031 | -36,35 | 2,65E-289 | 6,58E-288 | 0,0011331 | 0,002917 | 0,0019863 | 0,0019863 | TRUE | 0,0163892 | No |
| orf19.4723 | 1040 | 864 | 932 | 593 | 1790 | 1423 | 1384 | 1939 | 947 | 874 | 882 | 891 | 1592 | 1545 | 1417 | 1479 | 1203,39 | 898 | 1508 | 1,678 | 0,747 | 13,225 | 6,30E-40 | 2,44E-39 | 0,0013166 | 0,003135 | 0,0021546 | 0,0021546 | TRUE | 0,0192467 | No |
| orf19.4724 | 1179 | 1017 | 1145 | 657 | 2419 | 1867 | 2048 | 2778 | 1073 | 1029 | 1084 | 987 | 2152 | 2027 | 2097 | 2119 | 1570,97 | 1043 | 2099 | 2,007 | 1,005 | 20,793 | 5,01E-96 | 4,14E-95 | 0,0002676 | 0,002878 | 0,0015104 | 0,0015104 | TRUE | 0,0134571 | No |
| orf19.4725 | 1147 | 1030 | 1033 | 753 | 1961 | 1699 | 1724 | 2326 | 1044 | 1042 | 978 | 1131 | 1744 | 1845 | 1766 | 1774 | 1415,45 | 1049 | 1782 | 1,704 | 0,769 | 14,608 | 2,51E-48 | 1,12E-47 | 0,0007432 | 0,00297 | 0,0018875 | 0,0018875 | TRUE | 0,0292806 | No |
| orf19.4726 | 1102 | 1020 | 1109 | 609 | 622 | 456 | 516 | 716 | 1003 | 1032 | 1050 | 915 | 553 | 495 | 528 | 546 | 765,29 | 1000 | 530 | 0,53 | -0,915 | -14,018 | 1,20E-44 | 5,11E-44 | 0,0014247 | 0,003763 | 0,0026784 | 0,0026784 | TRUE | 0,034544 | No |
| orf19.4727 | 967 | 855 | 848 | 540 | 1417 | 1001 | 1077 | 1464 | 880 | 865 | 803 | 811 | 1260 | 1087 | 1103 | 1117 | 990,72 | 840 | 1142 | 1,359 | 0,442 | 6,829 | 8,55E-12 | 1,73E-11 | 0,0026287 | 0,00337 | 0,0029816 | 0,0029816 | TRUE | 0,0591487 | No |
| orf19.4728 | 4659 | 4068 | 4648 | 2625 | 3714 | 2969 | 3411 | 4497 | 4242 | 4114 | 4399 | 3943 | 3303 | 3224 | 3493 | 3430 | 3768,61 | 4174 | 3362 | 0,805 | -0,313 | -6,742 | 1,57E-11 | 3,13E-11 | 0,0013695 | 0,002388 | 0,0018026 | 0,0018026 | TRUE | 0,0161435 | No |
| orf19.473 | 1877 | 1494 | 1689 | 1054 | 1896 | 1592 | 1704 | 2430 | 1709 | 1511 | 1598 | 1583 | 1686 | 1728 | 1745 | 1854 | 1676,92 | 1600 | 1753 | 1,095 | 0,132 | 2,458 | 0,013957 | 0,018129 | 0,0016249 | 0,002825 | 0,0021485 | 0,0021485 | TRUE | 0,0262978 | No |
| orf19.4730 | 2151 | 1831 | 2053 | 1225 | 2335 | 1833 | 1976 | 2684 | 1958 | 1852 | 1943 | 1840 | 2077 | 1990 | 2024 | 2047 | 1966,43 | 1898 | 2034 | 1,071 | 0,099 | 2,308 | 0,020973 | 0,026832 | 1,00E-08 | 0,002709 | 0,0012391 | 0,0012391 | TRUE | 0,0054572 | No |
| orf19.4731 | 1040 | 934 | 1113 | 619 | 1920 | 1693 | 1678 | 2177 | 947 | 945 | 1053 | 930 | 1708 | 1838 | 1718 | 1661 | 1349,97 | 969 | 1731 | 1,783 | 0,834 | 14,521 | 8,88E-48 | 3,94E-47 | 0,0016074 | 0,003016 | 0,0023456 | 0,0023456 | TRUE | 0,0425671 | No |
| orf19.4732 | 2160 | 1889 | 2080 | 1402 | 1308 | 1323 | 1307 | 1726 | 1967 | 1911 | 1969 | 2106 | 1163 | 1436 | 1339 | 1317 | 1650,83 | 1988 | 1314 | 0,661 | -0,598 | -9,281 | 1,68E-20 | 4,31E-20 | 0,0037284 | 0,002837 | 0,003353 | 0,003353 | TRUE | 0,0715501 | No |
| orf19.4733 | 1265 | 1132 | 1201 | 621 | 1468 | 1099 | 1264 | 1769 | 1152 | 1145 | 1137 | 933 | 1306 | 1193 | 1294 | 1349 | 1188,62 | 1092 | 1286 | 1,175 | 0,232 | 3,268 | 0,001083 | 0,001526 | 0,0051185 | 0,003148 | 0,0040003 | 0,0040003 | TRUE | 0,1047174 | No |
| orf19.4734 | 768 | 639 | 717 | 489 | 1079 | 789 | 854 | 1115 | 699 | 646 | 679 | 735 | 960 | 857 | 875 | 851 | 787,52 | 690 | 886 | 1,287 | 0,364 | 5,606 | 2,07E-08 | 3,68E-08 | 0,0017511 | 0,003714 | 0,0027441 | 0,0027441 | TRUE | 0,0389405 | No |
| orf19.4735 | 1643 | 1432 | 1570 | 1052 | 2925 | 2526 | 2546 | 3479 | 1496 | 1448 | 1486 | 1580 | 2602 | 2743 | 2607 | 2654 | 2076,97 | 1502 | 2652 | 1,768 | 0,822 | 18,653 | 1,19E-77 | 8,15E-77 | 0,0002932 | 0,002673 | 0,0013297 | 0,0013297 | TRUE | 0,0127003 | No |
| orf19.4736 | 1177 | 1053 | 1164 | 729 | 1972 | 1807 | 1838 | 2429 | 1072 | 1065 | 1102 | 1095 | 1754 | 1962 | 1882 | 1853 | 1473,05 | 1084 | 1863 | 1,719 | 0,782 | 15,432 | 1,00E-53 | 4,96E-53 | 0,0007872 | 0,002934 | 0,0017166 | 0,0017166 | TRUE | 0,0186196 | No |
| orf19.4737 | 1674 | 1540 | 1589 | 1033 | 846 | 787 | 718 | 996 | 1524 | 1558 | 1504 | 1552 | 752 | 854 | 735 | 760 | 1154,91 | 1534 | 775 | 0,505 | -0,986 | -17,54 | 7,13E-69 | 4,40E-68 | 0,0007552 | 0,003181 | 0,0020807 | 0,0020807 | TRUE | 0,0522456 | No |
| orf19.4738 | 1356 | 1173 | 1314 | 811 | 707 | 598 | 611 | 869 | 1235 | 1186 | 1244 | 1218 | 629 | 649 | 626 | 663 | 931,19 | 1221 | 642 | 0,526 | -0,927 | -17,389 | 9,91E-68 | 6,03E-67 | 1,00E-08 | 0,003456 | 0,0015651 | 0,0015651 | TRUE | 0,0065686 | No |
| orf19.4739 | 21910 | 18915 | 20381 | 13354 | 26745 | 22182 | 23240 | 31910 | 19948 | 19131 | 19289 | 20060 | 23789 | 24083 | 23801 | 24341 | 21805,2 | 19607 | 24004 | 1,224 | 0,292 | 9,942 | 2,73E-23 | 7,52E-23 | 0,00028 | 0,002098 | 0,0007825 | 0,0007825 | TRUE | 0,0032586 | No |
| orf19.474 | 1350 | 1332 | 1308 | 829 | 1987 | 1722 | 1871 | 2408 | 1229 | 1347 | 1238 | 1245 | 1767 | 1870 | 1916 | 1837 | 1556,18 | 1265 | 1848 | 1,46 | 0,546 | 10,81 | 3,07E-27 | 9,27E-27 | 0,0008394 | 0,002886 | 0,0017688 | 0,0017688 | TRUE | 0,0237054 | No |
| orf19.4740 | 577 | 485 | 557 | 354 | 415 | 355 | 394 | 486 | 525 | 491 | 527 | 532 | 369 | 385 | 404 | 371 | 450,45 | 519 | 382 | 0,736 | -0,442 | -6,21 | 5,28E-10 | 9,98E-10 | 1,00E-08 | 0,00497 | 0,0026076 | 0,0026076 | TRUE | 0,0159753 | No |
| orf19.4741 | 18 | 13 | 16 | 7 | 193 | 172 | 142 | 263 | 16 | 13 | 15 | 11 | 172 | 187 | 145 | 201 | 94,96 | 14 | 176 | 12,496 | 3,643 | 15,354 | 3,35E-53 | 1,64E-52 | 1,00E-08 | 0,015949 | 0,0140938 | 0,0140938 | TRUE | 0,136426 | No |

| | | | | | | | | | | | | | | | | | | | | | | | | | | | | | | | |
|---|---|---|---|---|---|---|---|---|---|---|---|---|---|---|---|---|---|---|---|---|---|---|---|---|---|---|---|---|---|---|---|
| orf19.4742 | 1149 | 973 | 1053 | 690 | 697 | 683 | 615 | 852 | 1046 | 984 | 997 | 1036 | 620 | 742 | 630 | 650 | 838,06 | 1016 | 660 | 0,649 | -0,623 | -9,47 | 2,79E-21 | 7,29E-21 | 0,0023152 | 0,003613 | 0,0029202 | 0,0029202 | TRUE | 0,0765286 | No |
| orf19.4743 | 443 | 456 | 488 | 289 | 412 | 339 | 385 | 526 | 403 | 461 | 462 | 434 | 366 | 368 | 394 | 401 | 411,32 | 440 | 382 | 0,87 | -0,201 | -2,577 | 0,009978 | 0,013083 | 1,00E-08 | 0,005249 | 0,0034008 | 0,0034008 | TRUE | 0,0412368 | No |
| orf19.4744 | 2931 | 2607 | 2733 | 1804 | 1935 | 1793 | 1834 | 2409 | 2669 | 2637 | 2587 | 2710 | 1721 | 1947 | 1878 | 1838 | 2248,17 | 2651 | 1846 | 0,696 | -0,522 | -11,074 | 1,68E-28 | 5,18E-28 | 0,0008934 | 0,002625 | 0,0016796 | 0,0016796 | TRUE | 0,0250055 | No |
| orf19.4746 | 9692 | 8253 | 9329 | 5456 | 10703 | 8743 | 10104 | 13271 | 8824 | 8347 | 8829 | 8196 | 9520 | 9492 | 10348 | 10123 | 9209,93 | 8549 | 9871 | 1,154 | 0,207 | 4,652 | 3,29E-06 | 5,34E-06 | 0,001537 | 0,00218 | 0,0017903 | 0,0017903 | TRUE | 0,012806 | No |
| orf19.4747 | 339 | 290 | 343 | 171 | 439 | 412 | 357 | 492 | 309 | 293 | 325 | 257 | 390 | 447 | 366 | 375 | 345,27 | 296 | 394 | 1,323 | 0,404 | 4,275 | 1,91E-05 | 2,99E-05 | 0,0051398 | 0,005863 | 0,0056003 | 0,0056003 | TRUE | 0,088215 | No |
| orf19.4748 | 341 | 391 | 355 | 268 | 355 | 318 | 253 | 314 | 310 | 395 | 336 | 403 | 316 | 345 | 259 | 240 | 325,52 | 361 | 290 | 0,803 | -0,316 | -2,524 | 0,011606 | 0,015152 | 0,0190742 | 0,006095 | 0,0119146 | 0,0119146 | TRUE | 0,1809277 | No |
| orf19.4749 | 969 | 862 | 982 | 656 | 987 | 908 | 870 | 1251 | 882 | 872 | 929 | 985 | 878 | 986 | 891 | 954 | 922,23 | 917 | 927 | 1,013 | 0,019 | 0,306 | 0,759301 | 0,790422 | 0,0018242 | 0,003469 | 0,0026574 | 0,0026574 | TRUE | 0,0264022 | No |
| orf19.475 | 4586 | 4108 | 4425 | 2932 | 5649 | 4736 | 4667 | 6160 | 4175 | 4155 | 4188 | 4404 | 5025 | 5142 | 4780 | 4699 | 4570,93 | 4230 | 4912 | 1,161 | 0,216 | 5,068 | 4,03E-07 | 6,80E-07 | 0,0009746 | 0,002326 | 0,0015182 | 0,0015182 | TRUE | 0,0120215 | No |
| orf19.4750 | 839 | 883 | 954 | 545 | 526 | 465 | 462 | 569 | 764 | 893 | 903 | 819 | 468 | 505 | 473 | 434 | 657,3 | 845 | 470 | 0,555 | -0,85 | -11,18 | 5,12E-29 | 1,61E-28 | 0,0037805 | 0,004047 | 0,0039149 | 0,0039149 | TRUE | 0,0533276 | No |
| orf19.4751 | 2558 | 2528 | 2564 | 1548 | 1879 | 1619 | 1527 | 2228 | 2329 | 2557 | 2427 | 2325 | 1671 | 1758 | 1564 | 1700 | 2041,27 | 2410 | 1673 | 0,694 | -0,527 | -10,126 | 4,25E-24 | 1,20E-23 | 0,0015253 | 0,002684 | 0,0020965 | 0,0020965 | TRUE | 0,0226431 | No |
| orf19.4751.1 | 1268 | 1008 | 996 | 702 | 1707 | 1040 | 1251 | 1661 | 1154 | 1019 | 943 | 1055 | 1518 | 1129 | 1281 | 1267 | 1170,85 | 1043 | 1299 | 1,246 | 0,318 | 3,567 | 0,000361 | 0,000525 | 0,0101349 | 0,003165 | 0,0067463 | 0,0067463 | TRUE | 0,155831 | No |
| orf19.4752 | 1643 | 1398 | 1548 | 979 | 1787 | 1536 | 1601 | 2141 | 1496 | 1414 | 1465 | 1471 | 1589 | 1668 | 1640 | 1633 | 1546,93 | 1462 | 1632 | 1,117 | 0,159 | 3,596 | 0,000324 | 0,000472 | 1,00E-08 | 0,002891 | 0,0012293 | 0,0012293 | TRUE | 0,0059728 | No |
| orf19.4753 | 8483 | 7877 | 8677 | 4862 | 2422 | 1973 | 2150 | 2700 | 7723 | 7967 | 8212 | 7303 | 2154 | 2142 | 2202 | 2060 | 4970,44 | 7801 | 2140 | 0,274 | -1,868 | -40,076 | 0 | 0 | 0,001484 | 0,002303 | 0,0017987 | 0,0017987 | TRUE | 0,0218862 | No |
| orf19.4754 | 1540 | 1661 | 1687 | 978 | 3291 | 2927 | 2786 | 3642 | 1402 | 1680 | 1597 | 1469 | 2927 | 3178 | 2853 | 2778 | 2235,53 | 1537 | 2934 | 1,907 | 0,932 | 14,251 | 4,44E-46 | 1,92E-45 | 0,0043891 | 0,002628 | 0,003593 | 0,003593 | TRUE | 0,0475972 | No |
| orf19.4755 | 3667 | 3409 | 3733 | 2299 | 4858 | 4085 | 4320 | 5674 | 3339 | 3448 | 3533 | 3453 | 4321 | 4435 | 4424 | 4328 | 3910,19 | 3443 | 4377 | 1,271 | 0,346 | 9,836 | 7,84E-23 | 2,12E-22 | 0,0001226 | 0,002375 | 0,0009255 | 0,0009255 | TRUE | 0,0053937 | No |
| orf19.4756 | 1106 | 1097 | 1072 | 689 | 820 | 626 | 643 | 855 | 1007 | 1109 | 1015 | 1035 | 729 | 680 | 659 | 652 | 860,72 | 1042 | 680 | 0,653 | -0,615 | -10,034 | 1,08E-23 | 3,00E-23 | 0,0012551 | 0,003572 | 0,0024064 | 0,0024064 | TRUE | 0,0293991 | No |
| orf19.4757 | 7216 | 6665 | 7218 | 4237 | 6292 | 5208 | 5501 | 7245 | 6570 | 6741 | 6831 | 6365 | 5597 | 5654 | 5634 | 5527 | 6114,73 | 6627 | 5603 | 0,845 | -0,243 | -7,034 | 2,01E-12 | 4,13E-12 | 0,0003118 | 0,002253 | 0,0009841 | 0,0009841 | TRUE | 0,0081625 | No |
| orf19.4758 | 3194 | 3180 | 3266 | 2118 | 3412 | 3258 | 3003 | 3982 | 2908 | 3216 | 3091 | 3182 | 3035 | 3537 | 3075 | 3038 | 3135,23 | 3099 | 3171 | 1,023 | 0,033 | 0,556 | 0,578365 | 0,618845 | 0,0032675 | 0,002458 | 0,0030675 | 0,0030675 | TRUE | 0,0725044 | No |
| orf19.4759 | 4186 | 3699 | 3632 | 2978 | 5656 | 4212 | 3803 | 5920 | 3811 | 3741 | 3437 | 4473 | 5031 | 4573 | 3895 | 4516 | 4184,7 | 3866 | 4504 | 1,166 | 0,222 | 2,579 | 0,009908 | 0,012997 | 0,0114221 | 0,002353 | 0,0068497 | 0,0068497 | TRUE | 0,1341497 | No |
| orf19.4760 | 4783 | 5144 | 5289 | 2714 | 5841 | 4502 | 4974 | 6157 | 4355 | 5203 | 5006 | 4077 | 5195 | 4888 | 5094 | 4697 | 4814,21 | 4660 | 4968 | 1,065 | 0,092 | 1,25 | 0,211156 | 0,243597 | 0,0070938 | 0,002311 | 0,0049354 | 0,0049354 | TRUE | 0,0846085 | No |
| orf19.4761 | 1440 | 1461 | 1529 | 915 | 2373 | 1898 | 2044 | 2673 | 1311 | 1478 | 1447 | 1374 | 2111 | 2061 | 2093 | 2039 | 1739,24 | 1402 | 2076 | 1,479 | 0,565 | 11,459 | 2,11E-30 | 6,86E-30 | 0,0007454 | 0,002797 | 0,0017271 | 0,0017271 | TRUE | 0,0254804 | No |
| orf19.4763 | 203 | 183 | 217 | 143 | 324 | 287 | 246 | 345 | 185 | 185 | 205 | 215 | 288 | 312 | 252 | 263 | 238,12 | 198 | 279 | 1,415 | 0,501 | 4,867 | 1,13E-06 | 1,88E-06 | 0,0031817 | 0,007585 | 0,0057889 | 0,0057889 | TRUE | 0,0661357 | No |
| orf19.4764 | 1214 | 1403 | 1364 | 841 | 950 | 915 | 866 | 1082 | 1105 | 1419 | 1291 | 1263 | 845 | 993 | 887 | 825 | 1078,65 | 1270 | 888 | 0,698 | -0,518 | -6,273 | 3,55E-10 | 6,76E-10 | 0,0074461 | 0,003262 | 0,0055921 | 0,0055921 | TRUE | 0,094421 | No |
| orf19.4765 | 159 | 146 | 132 | 116 | 100 | 85 | 72 | 110 | 145 | 148 | 125 | 174 | 89 | 92 | 74 | 84 | 116,31 | 148 | 85 | 0,578 | -0,791 | -5,571 | 2,53E-08 | 4,48E-08 | 0,0048223 | 0,013395 | 0,0102329 | 0,0102329 | TRUE | 0,1227404 | No |
| orf19.4766 | 3187 | 3210 | 3423 | 2020 | 2143 | 1883 | 1834 | 2350 | 2902 | 3247 | 3240 | 3034 | 1906 | 2044 | 1878 | 1793 | 2505,44 | 3106 | 1905 | 0,613 | -0,706 | -12,65 | 1,12E-36 | 4,16E-36 | 0,0024009 | 0,002564 | 0,0025744 | 0,0025744 | TRUE | 0,0287046 | No |
| orf19.4767 | 511 | 450 | 526 | 319 | 776 | 601 | 700 | 1004 | 465 | 455 | 498 | 479 | 690 | 653 | 717 | 766 | 590,36 | 474 | 706 | 1,492 | 0,578 | 8,101 | 5,46E-16 | 1,24E-15 | 0,0018836 | 0,004275 | 0,0030952 | 0,0030952 | TRUE | 0,0401839 | No |
| orf19.4768 | 659 | 808 | 803 | 432 | 1028 | 917 | 978 | 1065 | 600 | 817 | 760 | 649 | 914 | 996 | 1002 | 812 | 818,76 | 706 | 931 | 1,315 | 0,395 | 3,962 | 7,44E-05 | 0,000112 | 0,0128001 | 0,00365 | 0,0082764 | 0,0082764 | TRUE | 0,1320441 | No |
| orf19.4769 | 3245 | 3134 | 3270 | 2001 | 2662 | 2413 | 2381 | 3102 | 2954 | 3170 | 3095 | 3006 | 2368 | 2620 | 2438 | 2366 | 2752,13 | 3056 | 2448 | 0,8 | -0,322 | -6,809 | 9,85E-12 | 1,99E-11 | 0,0011848 | 0,002517 | 0,0017742 | 0,0017742 | TRUE | 0,0264958 | No |
| orf19.477 | 1123 | 1069 | 1134 | 685 | 1506 | 1211 | 1192 | 1667 | 1022 | 1081 | 1073 | 1029 | 1340 | 1315 | 1221 | 1272 | 1169,07 | 1051 | 1287 | 1,223 | 0,29 | 5,569 | 2,56E-08 | 4,52E-08 | 0,0004475 | 0,003167 | 0,0017386 | 0,0017386 | TRUE | 0,0137028 | No |
| orf19.4771 | 5948 | 5715 | 6135 | 3878 | 1869 | 1666 | 1642 | 2046 | 5415 | 5780 | 5806 | 5825 | 1662 | 1809 | 1682 | 1561 | 3692,58 | 5706 | 1678 | 0,294 | -1,767 | -34,672 | 2,11E-263 | 4,64E-262 | 0,0019121 | 0,002395 | 0,0021214 | 0,0021214 | TRUE | 0,0324805 | No |
| orf19.4772 | 1200 | 1128 | 1179 | 813 | 403 | 349 | 335 | 514 | 1093 | 1141 | 1116 | 1221 | 358 | 379 | 343 | 392 | 755,38 | 1143 | 368 | 0,323 | -1,628 | -24,278 | 3,35E-130 | 3,71E-129 | 0,0011287 | 0,003786 | 0,0025848 | 0,0025848 | TRUE | 0,0248279 | No |
| orf19.4773 | 7716 | 6976 | 7287 | 4438 | 15624 | 12883 | 13049 | 18029 | 7025 | 7055 | 6896 | 6667 | 13897 | 13987 | 13364 | 13753 | 10330,6 | 6911 | 13750 | 1,988 | 0,992 | 29,851 | 8,54E-196 | 1,43E-194 | 0,0003827 | 0,002165 | 0,0009478 | 0,0009478 | TRUE | 0,006536 | No |
| orf19.4774 | 1072 | 938 | 992 | 700 | 734 | 640 | 663 | 866 | 976 | 949 | 939 | 1052 | 653 | 695 | 679 | 661 | 825,3 | 979 | 672 | 0,688 | -0,54 | -9,19 | 3,93E-20 | 1,00E-19 | 0,0003447 | 0,003638 | 0,0020742 | 0,0020742 | TRUE | 0,026062 | No |
| orf19.4775 | 10028 | 8998 | 9702 | 6360 | 8515 | 7397 | 8214 | 10848 | 9130 | 9101 | 9182 | 9554 | 7574 | 8031 | 8412 | 8275 | 8657,3 | 9242 | 8073 | 0,874 | -0,195 | -4,679 | 2,88E-06 | 4,70E-06 | 0,0011456 | 0,00219 | 0,001546 | 0,001546 | TRUE | 0,0212977 | No |
| orf19.4776 | 1721 | 1652 | 1711 | 1129 | 2336 | 1878 | 2127 | 2804 | 1567 | 1671 | 1619 | 1696 | 2078 | 2039 | 2178 | 2139 | 1873,37 | 1638 | 2108 | 1,289 | 0,367 | 8,022 | 1,04E-15 | 2,36E-15 | 0,0004451 | 0,002742 | 0,001456 | 0,001456 | TRUE | 0,0105478 | No |
| orf19.4777 | 3414 | 3062 | 3315 | 1916 | 1061 | 908 | 1041 | 1335 | 3108 | 3097 | 3137 | 2878 | 944 | 986 | 1066 | 1018 | 2029,34 | 3055 | 1004 | 0,328 | -1,608 | -31,018 | 3,04E-211 | 5,45E-210 | 0,0012075 | 0,002688 | 0,0019395 | 0,0019395 | TRUE | 0,0202253 | No |
| orf19.4778 | 422 | 367 | 423 | 237 | 366 | 340 | 303 | 414 | 384 | 371 | 400 | 356 | 326 | 369 | 310 | 316 | 354,07 | 378 | 330 | 0,869 | -0,202 | -2,422 | 0,015431 | 0,019986 | 0,0011175 | 0,005768 | 0,0038838 | 0,0038838 | TRUE | 0,0668338 | No |
| orf19.4779 | 7221 | 6279 | 6822 | 4356 | 2643 | 2165 | 2381 | 3243 | 6574 | 6351 | 6456 | 6543 | 2351 | 2351 | 2438 | 2474 | 4442,31 | 6481 | 2404 | 0,371 | -1,43 | -39,855 | 0 | 0 | 0,0001544 | 0,002334 | 0,0009593 | 0,0009593 | TRUE | 0,0050102 | No |
| orf19.478 | 1565 | 1347 | 1466 | 986 | 1070 | 905 | 938 | 1284 | 1425 | 1362 | 1387 | 1481 | 952 | 983 | 961 | 979 | 1191,27 | 1414 | 969 | 0,686 | -0,543 | -10,997 | 3,94E-28 | 1,21E-27 | 1,00E-08 | 0,003146 | 0,0014823 | 0,0014823 | TRUE | 0,0107245 | No |
| orf19.4780 | 1804 | 1482 | 1721 | 1090 | 2571 | 2196 | 2351 | 3229 | 1642 | 1499 | 1629 | 1637 | 2287 | 2384 | 2408 | 2463 | 1993,67 | 1602 | 2386 | 1,49 | 0,575 | 12,005 | 3,34E-33 | 1,15E-32 | 0,000891 | 0,0027 | 0,0016737 | 0,0016737 | TRUE | 0,0230235 | No |
| orf19.4781 | 433 | 336 | 407 | 314 | 355 | 324 | 326 | 430 | 394 | 340 | 385 | 472 | 316 | 352 | 334 | 328 | 365,04 | 398 | 332 | 0,84 | -0,252 | -2,618 | 0,008853 | 0,011655 | 0,0068066 | 0,005656 | 0,0061208 | 0,0061208 | TRUE | 0,1574066 | No |
| orf19.4783 | 115 | 103 | 134 | 77 | 52 | 51 | 64 | 62 | 105 | 104 | 127 | 116 | 46 | 55 | 66 | 47 | 83,23 | 113 | 54 | 0,472 | -1,084 | -6,533 | 6,45E-11 | 1,26E-10 | 5,88E-05 | 0,017909 | 0,0129408 | 0,0129408 | TRUE | 0,1847441 | No |
| orf19.4784 | 1763 | 1507 | 1579 | 1077 | 686 | 555 | 591 | 779 | 1605 | 1524 | 1494 | 1618 | 610 | 603 | 605 | 594 | 1081,72 | 1560 | 603 | 0,387 | -1,37 | -25,412 | 1,85E-142 | 2,28E-141 | 1,00E-08 | 0,003258 | 0,0016752 | 0,0016752 | TRUE | 0,0098927 | No |
| orf19.4785 | 1951 | 1572 | 1798 | 1090 | 1208 | 992 | 1138 | 1575 | 1776 | 1590 | 1702 | 1637 | 1074 | 1077 | 1165 | 1201 | 1402,95 | 1676 | 1129 | 0,674 | -0,569 | -9,815 | 9,72E-23 | 2,63E-22 | 0,00209 | 0,002979 | 0,0024979 | 0,0024979 | TRUE | 0,0232252 | No |
| orf19.4786 | 588 | 575 | 589 | 371 | 383 | 360 | 319 | 467 | 535 | 582 | 557 | 557 | 341 | 391 | 327 | 356 | 455,76 | 558 | 354 | 0,633 | -0,659 | -8,751 | 2,11E-18 | 5,14E-18 | 0,0005393 | 0,004935 | 0,0031635 | 0,0031635 | TRUE | 0,0520096 | No |
| orf19.4787 | 1885 | 1624 | 1875 | 1191 | 1176 | 1080 | 1139 | 1580 | 1716 | 1643 | 1775 | 1789 | 1046 | 1173 | 1166 | 1205 | 1439,08 | 1731 | 1148 | 0,664 | -0,592 | -10,382 | 3,00E-25 | 8,66E-25 | 0,0019153 | 0,002955 | 0,0023995 | 0,0023995 | TRUE | 0,0436332 | No |
| orf19.4788 | 13751 | 11957 | 13040 | 8093 | 12543 | 10394 | 10815 | 15036 | 12520 | 12093 | 12341 | 12157 | 11157 | 11285 | 11076 | 11470 | 11762,3 | 12278 | 11247 | 0,916 | -0,127 | -4,379 | 1,19E-05 | 1,89E-05 | 0,0001259 | 0,002149 | 0,0007177 | 0,0007177 | TRUE | 0,0022245 | No |
| orf19.4789 | 62 | 60 | 76 | 50 | 362 | 301 | 294 | 439 | 56 | 61 | 72 | 75 | 322 | 327 | 301 | 335 | 193,62 | 66 | 321 | 4,923 | 2,3 | 18,349 | 3,37E-75 | 2,24E-74 | 1,00E-08 | 0,00886 | 0,0055307 | 0,0055307 | TRUE | 0,0915868 | No |
| orf19.4791 | 1776 | 1962 | 2039 | 1151 | 1281 | 1061 | 1123 | 1467 | 1617 | 1984 | 1930 | 1729 | 1139 | 1152 | 1150 | 1119 | 1477,57 | 1815 | 1140 | 0,628 | -0,672 | -9,998 | 1,56E-23 | 4,31E-23 | 0,0042145 | 0,002931 | 0,0036297 | 0,0036297 | TRUE | 0,0684428 | No |
| orf19.4792 | 17765 | 15331 | 17092 | 11488 | 7290 | 6461 | 6735 | 9069 | 16174 | 15506 | 16176 | 17257 | 6484 | 7015 | 6897 | 6918 | 11553,4 | 16278 | 6828 | 0,42 | -1,253 | -28,72 | 2,17E-181 | 3,37E-180 | 0,0014458 | 0,002151 | 0,0017269 | 0,0017269 | TRUE | 0,0197514 | No |
| orf19.479.2 | 449 | 384 | 417 | 256 | 260 | 232 | 220 | 282 | 409 | 388 | 395 | 385 | 231 | 252 | 225 | 215 | 312,49 | 394 | 231 | 0,583 | -0,778 | -9,235 | 2,59E-20 | 6,62E-20 | 1,00E-08 | 0,006264 | 0,0034458 | 0,0034458 | TRUE | 0,0387133 | No |
| orf19.4793 | 5251 | 4956 | 5108 | 3095 | 4840 | 3605 | 3902 | 5053 | 4781 | 5012 | 4834 | 4649 | 4305 | 3914 | 3996 | 3854 | 4418,3 | 4819 | 4017 | 0,833 | -0,263 | -5,711 | 1,13E-08 | 2,02E-08 | 0,0014269 | 0,002336 | 0,00181 | 0,00181 | TRUE | 0,0285915 | No |
| orf19.4795 | 2047 | 2018 | 2077 | 1296 | 2577 | 2171 | 2184 | 2905 | 1864 | 2041 | 1966 | 1947 | 2292 | 2357 | 2237 | 2216 | 2114,89 | 1954 | 2276 | 1,164 | 0,219 | 4,848 | 1,25E-06 | 2,06E-06 | 0,0006273 | 0,002662 | 0,0014805 | 0,0014805 | TRUE | 0,0126026 | No |
| orf19.4796 | 2698 | 2322 | 2497 | 1495 | 3286 | 2533 | 2824 | 3629 | 2456 | 2348 | 2363 | 2246 | 2923 | 2750 | 2892 | 2768 | 2593,38 | 2353 | 2833 | 1,202 | 0,266 | 6,057 | 1,38E-09 | 2,58E-09 | 0,0006229 | 0,002546 | 0,0014534 | 0,0014534 | TRUE | 0,010502 | No |
| orf19.4798 | 4024 | 3799 | 3874 | 2680 | 6672 | 5918 | 5698 | 7942 | 3664 | 3842 | 3666 | 4026 | 5935 | 6425 | 5835 | 6058 | 4931,46 | 3800 | 6063 | 1,597 | 0,676 | 14,398 | 5,34E-47 | 2,35E-46 | 0,0016491 | 0,002305 | 0,0018952 | 0,0018952 | TRUE | 0,0194178 | No |
| orf19.4799 | 5708 | 5207 | 5770 | 3627 | 4588 | 3965 | 4175 | 5715 | 5197 | 5266 | 5461 | 5448 | 4081 | 4305 | 4276 | 4359 | 4799,16 | 5343 | 4255 | 0,797 | -0,328 | -8,694 | 3,48E-18 | 8,42E-18 | 0,0004296 | 0,002312 | 0,0011548 | 0,0011548 | TRUE | 0,0094143 | No |
| orf19.48 | 4019 | 4635 | 4748 | 3005 | 1737 | 1661 | 1636 | 2022 | 3659 | 4688 | 4494 | 4514 | 1545 | 1803 | 1675 | 1542 | 2990,09 | 4339 | 1641 | 0,378 | -1,403 | -17,658 | 8,87E-70 | 5,54E-69 | 0,0079682 | 0,002479 | 0,0056522 | 0,0056522 | TRUE | 0,1366825 | No |
| orf19.480 | 4638 | 3914 | 4286 | 2741 | 3900 | 3235 | 3559 | 4810 | 4223 | 3959 | 4056 | 4117 | 3469 | 3512 | 3645 | 3669 | 3831,28 | 4089 | 3574 | 0,874 | -0,194 | -4,972 | 6,63E-07 | 1,11E-06 | 0,0005202 | 0,002382 | 0,0011978 | 0,0011978 | TRUE | 0,0062345 | No |
| orf19.4800 | 2528 | 2253 | 2426 | 1470 | 911 | 700 | 811 | 1020 | 2302 | 2279 | 2296 | 2208 | 810 | 760 | 831 | 778 | 1532,93 | 2271 | 795 | 0,35 | -1,516 | -31,631 | 1,40E-219 | 2,60E-218 | 1,00E-08 | 0,002899 | 0,0013836 | 0,0013836 | TRUE | 0,0102191 | No |
| orf19.4801 | 634 | 598 | 605 | 363 | 726 | 599 | 646 | 864 | 577 | 605 | 573 | 545 | 646 | 650 | 662 | 659 | 614,58 | 575 | 654 | 1,135 | 0,183 | 2,982 | 0,002864 | 0,003903 | 1,00E-08 | 0,004186 | 0,0019671 | 0,0019671 | TRUE | 0,0135211 | No |
| orf19.4802 | 5052 | 4436 | 4921 | 2801 | 1581 | 1350 | 1432 | 1781 | 4600 | 4487 | 4657 | 4208 | 1406 | 1466 | 1467 | 1359 | 2956,01 | 4488 | 1424 | 0,317 | -1,659 | -34,29 | 1,12E-257 | 2,41E-256 | 0,0013071 | 0,002484 | 0,0017987 | 0,0017987 | TRUE | 0,0205352 | No |
| orf19.4803 | 14 | 11 | 14 | 9 | 19 | 18 | 18 | 32 | 13 | 11 | 13 | 14 | 17 | 20 | 18 | 24 | 16,24 | 13 | 20 | 1,581 | 0,661 | 1,94 | 0,052412 | 0,064791 | 1,00E-08 | 0,083374 | 0,0459428 | 0,0459428 | TRUE | 0,1680087 | No |
| orf19.4804 | 69 | 68 | 82 | 34 | 112 | 101 | 97 | 151 | 63 | 69 | 78 | 51 | 100 | 110 | 99 | 115 | 85,51 | 65 | 106 | 1,61 | 0,687 | 4,323 | 1,54E-05 | 2,42E-05 | 0,0006528 | 0,017486 | 0,0117654 | 0,0117654 | TRUE | 0,143569 | No |
| orf19.4805 | 5039 | 4495 | 4949 | 3108 | 9452 | 7957 | 8307 | 11175 | 4588 | 4546 | 4684 | 4669 | 8407 | 8639 | 8507 | 8524 | 6570,58 | 4622 | 8519 | 1,844 | 0,883 | 29,283 | 1,69E-188 | 2,73E-187 | 1,00E-08 | 0,002238 | 0,000701 | 0,000701 | TRUE | 0,0014642 | No |
| orf19.4807 | 1352 | 1273 | 1380 | 860 | 3647 | 2913 | 3085 | 4060 | 1231 | 1288 | 1306 | 1292 | 3244 | 3163 | 3159 | 3097 | 2222,42 | 1279 | 3166 | 2,476 | 1,308 | 30,978 | 1,06E-210 | 1,88E-209 | 1,00E-08 | 0,002631 | 0,0011463 | 0,0011463 | TRUE | 0,0083477 | No |
| orf19.4808 | 4085 | 4000 | 3968 | 2509 | 3708 | 3220 | 3246 | 4330 | 3719 | 4046 | 3755 | 3769 | 3298 | 3496 | 3324 | 3303 | 3588,81 | 3822 | 3355 | 0,877 | -0,189 | -4,429 | 9,45E-06 | 1,50E-05 | 0,0007967 | 0,002405 | 0,0014613 | 0,0014613 | TRUE | 0,0191217 | No |
| orf19.4809 | 1293 | 1139 | 1123 | 736 | 1073 | 896 | 982 | 1298 | 1177 | 1152 | 1063 | 1106 | 954 | 973 | 1006 | 990 | 1052,58 | 1124 | 981 | 0,872 | -0,198 | -3,688 | 0,000226 | 0,000332 | 1,00E-08 | 0,003292 | 0,0018218 | 0,0018218 | TRUE | 0,0179508 | No |

| | | | | | | | | | | | | | | | | | | | | | | | | | | | | | | | |
|---|---|---|---|---|---|---|---|---|---|---|---|---|---|---|---|---|---|---|---|---|---|---|---|---|---|---|---|---|---|---|---|
| orf19.481 | 3700 | 2997 | 3298 | 2215 | 4972 | 4075 | 4563 | 5881 | 3369 | 3031 | 3121 | 3327 | 4422 | 4424 | 4673 | 4486 | 3856,79 | 3212 | 4501 | 1,402 | 0,488 | 10,702 | 9,91E-27 | 2,95E-26 | 0,0012616 | 0,00238 | 0,0017208 | 0,0017208 | TRUE | 0,0175158 | No |
| orf19.4811 | 2435 | 2068 | 2112 | 1599 | 2560 | 2116 | 2150 | 3009 | 2217 | 2092 | 1999 | 2402 | 2277 | 2297 | 2202 | 2295 | 2222,61 | 2178 | 2268 | 1,044 | 0,062 | 1,08 | 0,280078 | 0,316634 | 0,0028043 | 0,002631 | 0,0026754 | 0,0026754 | TRUE | 0,0549279 | No |
| orf19.4812 | 476 | 386 | 382 | 341 | 568 | 488 | 458 | 684 | 433 | 390 | 362 | 512 | 505 | 530 | 469 | 522 | 465,43 | 424 | 506 | 1,201 | 0,265 | 2,714 | 0,006655 | 0,008846 | 0,0096239 | 0,004875 | 0,0069354 | 0,0069354 | TRUE | 0,199744 | No |
| orf19.4813 | 22379 | 18635 | 19951 | 12995 | 32141 | 26094 | 27290 | 38648 | 20375 | 18847 | 18882 | 19521 | 28588 | 28331 | 27948 | 29481 | 23996,7 | 19406 | 28587 | 1,473 | 0,559 | 15,097 | 1,69E-51 | 8,05E-51 | 0,0008831 | 0,002092 | 0,0012725 | 0,0012725 | TRUE | 0,0139931 | No |
| orf19.4814 | 82 | 70 | 65 | 39 | 103 | 65 | 68 | 78 | 75 | 71 | 62 | 59 | 92 | 71 | 70 | 59 | 69,61 | 67 | 73 | 1,088 | 0,122 | 0,669 | 0,503179 | 0,544889 | 0,0120959 | 0,021014 | 0,0174521 | 0,0174521 | TRUE | 0,2972748 | No |
| orf19.4815 | 10958 | 9181 | 10036 | 6101 | 10458 | 8315 | 9114 | 12603 | 9977 | 9286 | 9498 | 9165 | 9302 | 9028 | 9334 | 9614 | 9400,33 | 9482 | 9320 | 0,983 | -0,025 | -0,645 | 0,518719 | 0,56067 | 0,0008763 | 0,002178 | 0,0013517 | 0,0013517 | TRUE | 0,0152262 | No |
| orf19.4816 | 2145 | 1845 | 1983 | 1301 | 1410 | 1183 | 1200 | 1612 | 1953 | 1866 | 1877 | 1954 | 1254 | 1284 | 1229 | 1230 | 1580,89 | 1912 | 1249 | 0,653 | -0,614 | -13,796 | 2,68E-43 | 1,11E-42 | 1,00E-08 | 0,002873 | 0,0012486 | 0,0012486 | TRUE | 0,0040541 | No |
| orf19.4817 | 562 | 551 | 560 | 384 | 672 | 596 | 563 | 734 | 512 | 557 | 530 | 577 | 598 | 647 | 577 | 560 | 569,63 | 544 | 596 | 1,096 | 0,132 | 1,857 | 0,063298 | 0,077675 | 0,0016089 | 0,004356 | 0,0030853 | 0,0030853 | TRUE | 0,0385373 | No |
| orf19.4818 | 3642 | 3003 | 3281 | 2218 | 6537 | 5410 | 5969 | 8052 | 3316 | 3037 | 3105 | 3332 | 5814 | 5874 | 6113 | 6142 | 4591,68 | 3198 | 5986 | 1,874 | 0,906 | 20,625 | 1,64E-94 | 1,33E-93 | 0,0011443 | 0,002325 | 0,0016052 | 0,0016052 | TRUE | 0,0137507 | No |
| orf19.4819 | 211 | 257 | 196 | 126 | 454 | 329 | 346 | 451 | 192 | 260 | 185 | 189 | 404 | 357 | 354 | 344 | 285,77 | 206 | 365 | 1,762 | 0,817 | 7,193 | 6,35E-13 | 1,33E-12 | 0,0111792 | 0,00666 | 0,00852 | 0,00852 | TRUE | 0,3344867 | No |
| orf19.482 | 5045 | 4295 | 4796 | 2717 | 4807 | 3821 | 4153 | 5377 | 4593 | 4344 | 4539 | 4081 | 4276 | 4149 | 4253 | 4102 | 4292,07 | 4389 | 4195 | 0,954 | -0,067 | -1,496 | 0,134767 | 0,159306 | 0,0012777 | 0,002345 | 0,0017067 | 0,0017067 | TRUE | 0,0258769 | No |
| orf19.4820 | 2598 | 2283 | 2503 | 1544 | 2510 | 2017 | 2107 | 2816 | 2365 | 2309 | 2369 | 2319 | 2233 | 2190 | 2158 | 2148 | 2261,37 | 2340 | 2182 | 0,932 | -0,102 | -2,658 | 0,007858 | 0,010387 | 1,00E-08 | 0,002621 | 0,0009706 | 0,0009706 | TRUE | 0,0030471 | No |
| orf19.4821 | 15 | 19 | 14 | 8 | 80 | 48 | 38 | 75 | 14 | 19 | 13 | 12 | 71 | 52 | 39 | 57 | 34,69 | 14 | 55 | 3,758 | 1,91 | 6,587 | 4,49E-11 | 8,84E-11 | 1,00E-08 | 0,040114 | 0,0360966 | 0,0360966 | TRUE | 0,1053154 | No |
| orf19.4822 | 5 | 5 | 7 | 3 | 7 | 6 | 5 | 8 | 5 | 5 | 7 | 5 | 6 | 7 | 5 | 6 | 5,59 | 6 | 6 | 1,146 | 0,197 | 0,353 | 0,724034 | 0,756923 | 1,00E-08 | 0,238434 | 0,1203524 | 0,1203524 | TRUE | 0,0808099 | No |
| orf19.4823 | 282 | 248 | 260 | 178 | 369 | 292 | 306 | 398 | 257 | 251 | 246 | 267 | 328 | 317 | 313 | 304 | 285,41 | 255 | 316 | 1,239 | 0,309 | 3,657 | 0,000256 | 0,000375 | 1,00E-08 | 0,006665 | 0,0032904 | 0,0032904 | TRUE | 0,0089721 | No |
| orf19.4824 | 1610 | 1504 | 1591 | 1027 | 2430 | 2244 | 2246 | 3139 | 1466 | 1521 | 1506 | 1543 | 2161 | 2436 | 2300 | 2394 | 1915,98 | 1509 | 2323 | 1,541 | 0,624 | 12,344 | 5,24E-35 | 1,87E-34 | 0,0012952 | 0,002726 | 0,0018922 | 0,0018922 | TRUE | 0,0269296 | No |
| orf19.4825 | 1407 | 1184 | 1251 | 853 | 1008 | 790 | 740 | 988 | 1281 | 1197 | 1184 | 1281 | 897 | 858 | 758 | 754 | 1026,2 | 1236 | 817 | 0,661 | -0,598 | -8,736 | 2,42E-18 | 5,87E-18 | 0,0034226 | 0,003324 | 0,0034924 | 0,0034924 | TRUE | 0,0540162 | No |
| orf19.4826 | 14758 | 11921 | 12683 | 8864 | 17999 | 14381 | 15105 | 21807 | 13437 | 12057 | 12003 | 13315 | 16010 | 15614 | 15469 | 16635 | 14317,4 | 12703 | 15932 | 1,255 | 0,327 | 6,651 | 2,91E-11 | 5,77E-11 | 0,0023204 | 0,002129 | 0,0022523 | 0,0022523 | TRUE | 0,0188052 | No |
| orf19.4827 | 11444 | 9282 | 9855 | 6333 | 15475 | 12049 | 12905 | 18320 | 10419 | 9388 | 9327 | 9513 | 13765 | 13082 | 13216 | 13975 | 11585,6 | 9662 | 13510 | 1,398 | 0,484 | 10,593 | 3,22E-26 | 9,44E-26 | 0,0017823 | 0,002151 | 0,0019111 | 0,0019111 | TRUE | 0,0344381 | No |
| orf19.4828 | 12077 | 10530 | 11767 | 6704 | 3417 | 2643 | 2942 | 4012 | 10996 | 10650 | 11136 | 10070 | 3039 | 2870 | 3013 | 3060 | 6854,34 | 10713 | 2996 | 0,28 | -1,839 | -42,108 | 0 | 0 | 0,0011109 | 0,00223 | 0,0016246 | 0,0016246 | TRUE | 0,019485 | No |
| orf19.4829 | 1904 | 1548 | 1683 | 1150 | 1871 | 1497 | 1730 | 2227 | 1734 | 1566 | 1593 | 1727 | 1664 | 1625 | 1772 | 1699 | 1672,43 | 1655 | 1690 | 1,022 | 0,032 | 0,605 | 0,545184 | 0,586689 | 0,0012963 | 0,002827 | 0,0020602 | 0,0020602 | TRUE | 0,0157714 | No |
| orf19.4830 | 510 | 406 | 381 | 289 | 457 | 355 | 349 | 504 | 464 | 411 | 361 | 434 | 406 | 385 | 357 | 384 | 400,43 | 418 | 383 | 0,92 | -0,12 | -1,345 | 0,178731 | 0,207883 | 0,0048412 | 0,005336 | 0,0051096 | 0,0051096 | TRUE | 0,1029532 | No |
| orf19.4831 | 3253 | 2724 | 2989 | 1867 | 6322 | 5199 | 5520 | 7849 | 2962 | 2755 | 2829 | 2805 | 5623 | 5645 | 5653 | 5987 | 4282,31 | 2838 | 5727 | 2,018 | 1,013 | 24,846 | 2,89E-136 | 3,39E-135 | 0,000736 | 0,002345 | 0,0013246 | 0,0013246 | TRUE | 0,0116695 | No |
| orf19.4833 | 11923 | 9778 | 10533 | 7209 | 57142 | 46905 | 50194 | 69756 | 10855 | 9889 | 9969 | 10829 | 50826 | 50926 | 51405 | 53210 | 30988,7 | 10386 | 51592 | 4,969 | 2,313 | 54,858 | 0 | 0 | 0,0014478 | 0,00208 | 0,0016466 | 0,0016466 | TRUE | 0,0126123 | No |
| orf19.4834 | 26 | 31 | 24 | 16 | 95 | 74 | 87 | 102 | 24 | 31 | 23 | 24 | 84 | 80 | 89 | 78 | 54,19 | 26 | 83 | 3,252 | 1,701 | 8,188 | 2,66E-16 | 6,10E-16 | 1,00E-08 | 0,026414 | 0,0150499 | 0,0150499 | TRUE | 0,157608 | No |
| orf19.4835 | 13975 | 11804 | 12981 | 8113 | 13957 | 11420 | 13006 | 17004 | 12724 | 11938 | 12285 | 12187 | 12414 | 12399 | 13320 | 12971 | 12529,8 | 12284 | 12776 | 1,04 | 0,057 | 1,489 | 0,136594 | 0,161257 | 0,0008687 | 0,002142 | 0,001307 | 0,001307 | TRUE | 0,0099527 | No |
| orf19.4836 | 2272 | 1798 | 1930 | 1347 | 3118 | 2565 | 2761 | 3723 | 2069 | 1818 | 1827 | 2023 | 2773 | 2785 | 2828 | 2840 | 2370,35 | 1934 | 2806 | 1,452 | 0,538 | 10,429 | 1,83E-25 | 5,30E-25 | 0,0017552 | 0,002594 | 0,002113 | 0,002113 | TRUE | 0,028762 | No |
| orf19.4837 | 514 | 383 | 466 | 349 | 264 | 200 | 258 | 336 | 468 | 387 | 441 | 524 | 235 | 217 | 264 | 256 | 349,14 | 455 | 243 | 0,537 | -0,897 | -8,717 | 2,85E-18 | 6,91E-18 | 0,0086361 | 0,005821 | 0,0070583 | 0,0070583 | TRUE | 0,1145267 | No |
| orf19.4837.1 | 666 | 577 | 612 | 319 | 270 | 176 | 257 | 292 | 606 | 584 | 579 | 479 | 240 | 191 | 263 | 223 | 395,69 | 562 | 229 | 0,407 | -1,298 | -12,597 | 2,19E-36 | 8,06E-36 | 0,0097461 | 0,005375 | 0,0072063 | 0,0072063 | TRUE | 0,126622 | No |
| orf19.4839 | 1240 | 933 | 1082 | 738 | 2245 | 1799 | 1993 | 2630 | 1129 | 944 | 1024 | 1109 | 1997 | 1953 | 2041 | 2006 | 1525,32 | 1052 | 1999 | 1,904 | 0,929 | 15,781 | 4,18E-56 | 2,15E-55 | 0,0024611 | 0,002903 | 0,0025832 | 0,0025832 | TRUE | 0,057395 | No |
| orf19.484 | 6909 | 5819 | 6356 | 3942 | 5392 | 4489 | 4915 | 6618 | 6290 | 5885 | 6015 | 5922 | 4796 | 4874 | 5034 | 5048 | 5483,02 | 6028 | 4938 | 0,819 | -0,288 | -7,629 | 2,37E-14 | 5,15E-14 | 0,0005277 | 0,002278 | 0,0011865 | 0,0011865 | TRUE | 0,010657 | No |
| orf19.4841 | 3360 | 2828 | 3135 | 1863 | 1243 | 975 | 1121 | 1498 | 3059 | 2860 | 2967 | 2799 | 1106 | 1059 | 1148 | 1143 | 2017,47 | 2921 | 1114 | 0,381 | -1,392 | -28,484 | 1,85E-178 | 2,81E-177 | 0,0009077 | 0,002692 | 0,0016888 | 0,0016888 | TRUE | 0,0125775 | No |
| orf19.4842 | 1438 | 1104 | 1183 | 940 | 474 | 377 | 416 | 546 | 1309 | 1117 | 1120 | 1412 | 422 | 409 | 426 | 416 | 828,86 | 1240 | 418 | 0,338 | -1,563 | -18,76 | 1,61E-78 | 1,13E-77 | 0,0065396 | 0,003631 | 0,0051173 | 0,0051173 | TRUE | 0,0991751 | No |
| orf19.4843 | 6186 | 5757 | 6168 | 4012 | 1979 | 1755 | 1688 | 2374 | 5632 | 5823 | 5837 | 6027 | 1760 | 1905 | 1729 | 1811 | 3815,52 | 5830 | 1801 | 0,309 | -1,694 | -39,195 | 0 | 0 | 0,0008167 | 0,002383 | 0,001442 | 0,001442 | TRUE | 0,0172925 | No |
| orf19.4844 | 3863 | 3033 | 3467 | 2089 | 2272 | 1841 | 2149 | 2852 | 3517 | 3068 | 3281 | 3138 | 2021 | 1999 | 2201 | 2176 | 2674,99 | 3251 | 2099 | 0,645 | -0,632 | -11,256 | 2,15E-29 | 6,81E-29 | 0,0025649 | 0,002531 | 0,0026357 | 0,0026357 | TRUE | 0,0374665 | No |
| orf19.4845 | 499 | 497 | 512 | 270 | 465 | 396 | 352 | 456 | 454 | 503 | 485 | 406 | 414 | 430 | 360 | 348 | 424,88 | 462 | 388 | 0,835 | -0,259 | -2,792 | 0,005238 | 0,007018 | 0,0068865 | 0,005146 | 0,0059218 | 0,0059218 | TRUE | 0,0676325 | No |
| orf19.4846 | 1795 | 1459 | 1644 | 1030 | 1532 | 1227 | 1398 | 1892 | 1634 | 1476 | 1556 | 1547 | 1363 | 1332 | 1432 | 1443 | 1472,85 | 1553 | 1392 | 0,897 | -0,157 | -3,052 | 0,002274 | 0,003123 | 0,0010073 | 0,002934 | 0,0018721 | 0,0018721 | TRUE | 0,0157221 | No |
| orf19.4847 | 50 | 31 | 35 | 22 | 29 | 35 | 39 | 39 | 46 | 31 | 33 | 33 | 26 | 38 | 40 | 30 | 34,57 | 36 | 34 | 0,919 | -0,122 | -0,495 | 0,620675 | 0,659137 | 1,00E-08 | 0,040253 | 0,0296089 | 0,0296089 | TRUE | 0,2714722 | No |
| orf19.4848 | 2511 | 2226 | 2367 | 1497 | 2101 | 1804 | 1951 | 2531 | 2286 | 2251 | 2240 | 2249 | 1869 | 1959 | 1998 | 1931 | 2097,82 | 2256 | 1939 | 0,859 | -0,22 | -5,423 | 5,86E-08 | 1,02E-07 | 1,00E-08 | 0,002667 | 0,0010988 | 0,0010988 | TRUE | 0,007188 | No |
| orf19.4849 | 7368 | 6896 | 7254 | 4868 | 5713 | 4626 | 4794 | 6253 | 6708 | 6975 | 6865 | 7313 | 5082 | 5023 | 4910 | 4770 | 5955,52 | 6965 | 4946 | 0,71 | -0,493 | -12,301 | 8,96E-35 | 3,19E-34 | 0,0008309 | 0,002259 | 0,0013721 | 0,0013721 | TRUE | 0,0132696 | No |
| orf19.485 | 893 | 794 | 891 | 530 | 693 | 526 | 607 | 814 | 813 | 803 | 843 | 796 | 616 | 571 | 622 | 621 | 710,69 | 814 | 608 | 0,747 | -0,422 | -7,124 | 1,05E-12 | 2,18E-12 | 1,00E-08 | 0,003896 | 0,0019398 | 0,0019398 | TRUE | 0,0177744 | No |
| orf19.4850 | 10183 | 9749 | 10635 | 6082 | 4947 | 4147 | 4440 | 5405 | 9271 | 9860 | 10065 | 9136 | 4400 | 4502 | 4547 | 4123 | 6988,16 | 9583 | 4393 | 0,458 | -1,126 | -23,549 | 1,29E-122 | 1,35E-121 | 0,0018173 | 0,002226 | 0,0020342 | 0,0020342 | TRUE | 0,0212771 | No |
| orf19.4851 | 1782 | 1578 | 1693 | 1194 | 3408 | 2820 | 3052 | 4270 | 1622 | 1596 | 1602 | 1794 | 3031 | 3062 | 3126 | 3257 | 2386,27 | 1654 | 3119 | 1,892 | 0,92 | 18,572 | 5,39E-77 | 3,67E-76 | 0,0015155 | 0,002591 | 0,0018784 | 0,0018784 | TRUE | 0,0355089 | No |
| orf19.4853 | 169 | 145 | 167 | 118 | 335 | 318 | 354 | 443 | 154 | 147 | 158 | 177 | 298 | 345 | 363 | 338 | 247,44 | 159 | 336 | 2,125 | 1,087 | 10,571 | 4,04E-26 | 1,18E-25 | 0,0026042 | 0,007376 | 0,0053932 | 0,0053932 | TRUE | 0,0669275 | No |
| orf19.4855 | 930 | 730 | 829 | 518 | 743 | 579 | 589 | 796 | 847 | 738 | 785 | 778 | 661 | 629 | 603 | 607 | 705,96 | 787 | 625 | 0,793 | -0,334 | -5,13 | 2,90E-07 | 4,92E-07 | 0,0013453 | 0,003908 | 0,0026426 | 0,0026426 | TRUE | 0,0331561 | No |
| orf19.4856 | 21 | 9 | 21 | 8 | 25 | 14 | 14 | 20 | 19 | 9 | 20 | 12 | 22 | 15 | 14 | 15 | 15,89 | 15 | 16 | 1,1 | 0,137 | 0,379 | 0,704948 | 0,739273 | 0,0214986 | 0,085155 | 0,0638275 | 0,0638275 | TRUE | 0,3227642 | No |
| orf19.4857 | 2683 | 2392 | 2679 | 1914 | 4116 | 3595 | 3640 | 4969 | 2443 | 2419 | 2535 | 2875 | 3661 | 3903 | 3728 | 3790 | 3169,38 | 2568 | 3770 | 1,471 | 0,557 | 9,845 | 7,22E-23 | 1,96E-22 | 0,0033024 | 0,002454 | 0,0027363 | 0,0027363 | TRUE | 0,0746663 | No |
| orf19.4858 | 634 | 564 | 614 | 366 | 1279 | 1048 | 1103 | 1443 | 577 | 570 | 581 | 550 | 1138 | 1138 | 1130 | 1101 | 848,04 | 570 | 1127 | 1,972 | 0,98 | 17,858 | 2,48E-71 | 1,58E-70 | 1,00E-08 | 0,003595 | 0,0015426 | 0,0015426 | TRUE | 0,0051767 | No |
| orf19.486 | 592 | 508 | 553 | 320 | 86 | 86 | 105 | 126 | 539 | 514 | 523 | 481 | 76 | 93 | 108 | 96 | 303,79 | 514 | 93 | 0,181 | -2,469 | -23,474 | 7,52E-122 | 7,74E-121 | 0,000274 | 0,006385 | 0,0045978 | 0,0045978 | TRUE | 0,1501343 | No |
| orf19.4861 | 15 | 27 | 32 | 26 | 48 | 56 | 42 | 67 | 14 | 27 | 30 | 39 | 43 | 61 | 43 | 51 | 38,49 | 28 | 50 | 1,835 | 0,876 | 3,219 | 0,001286 | 0,001804 | 0,0534344 | 0,036357 | 0,0417807 | 0,0417807 | TRUE | 0,8103507 | No |
| orf19.4861.1 | 8 | 10 | 15 | 1 | 14 | 14 | 18 | 29 | 7 | 10 | 14 | 2 | 12 | 15 | 18 | 22 | 12,66 | 8 | 17 | 1,975 | 0,982 | 2,226 | 0,026012 | 0,03304 | 0,0753672 | 0,106356 | 0,0980882 | 0,0980882 | TRUE | 0,462171 | No |
| orf19.4862 | 1066 | 1006 | 1144 | 749 | 1002 | 799 | 977 | 1192 | 971 | 1017 | 1083 | 1125 | 891 | 867 | 1001 | 909 | 983,05 | 1049 | 917 | 0,876 | -0,191 | -2,891 | 0,003841 | 0,005182 | 0,0027828 | 0,003381 | 0,0031594 | 0,0031594 | TRUE | 0,0432729 | No |
| orf19.4862.2 | 2195 | 1999 | 1972 | 1292 | 1553 | 1188 | 1136 | 1637 | 1998 | 2022 | 1866 | 1941 | 1381 | 1290 | 1163 | 1249 | 1613,83 | 1957 | 1271 | 0,65 | -0,622 | -10,668 | 1,43E-26 | 4,24E-26 | 0,0023005 | 0,002856 | 0,0026254 | 0,0026254 | TRUE | 0,0416977 | No |
| orf19.4863 | 1858 | 1699 | 1870 | 1156 | 1789 | 1471 | 1651 | 2152 | 1692 | 1718 | 1770 | 1736 | 1591 | 1597 | 1691 | 1642 | 1679,63 | 1729 | 1630 | 0,943 | -0,085 | -1,93 | 0,053665 | 0,066301 | 1,00E-08 | 0,002823 | 0,0012526 | 0,0012526 | TRUE | 0,0072147 | No |
| orf19.4864 | 993 | 914 | 956 | 562 | 1812 | 1485 | 1466 | 1816 | 904 | 924 | 905 | 844 | 1612 | 1612 | 1501 | 1385 | 1211,01 | 894 | 1528 | 1,703 | 0,768 | 12,144 | 6,15E-34 | 2,14E-33 | 0,0027777 | 0,003128 | 0,0029377 | 0,0029377 | TRUE | 0,0494471 | No |
| orf19.4865 | 1019 | 1125 | 1138 | 646 | 1786 | 1591 | 1589 | 1971 | 928 | 1138 | 1077 | 970 | 1589 | 1727 | 1627 | 1503 | 1319,97 | 1028 | 1612 | 1,565 | 0,646 | 9,035 | 1,64E-19 | 4,10E-19 | 0,0049847 | 0,003038 | 0,0040945 | 0,0040945 | TRUE | 0,0618228 | No |
| orf19.4866 | 4864 | 4501 | 5017 | 3522 | 5206 | 4852 | 4601 | 6477 | 4428 | 4552 | 4748 | 5291 | 4631 | 5268 | 4712 | 4941 | 4821,34 | 4755 | 4888 | 1,029 | 0,041 | 0,655 | 0,512614 | 0,554636 | 0,0041792 | 0,002311 | 0,003528 | 0,003528 | TRUE | 0,0685228 | No |
| orf19.4867 | 11634 | 10837 | 11881 | 7471 | 1485 | 1230 | 1195 | 1681 | 10592 | 10960 | 11244 | 11223 | 1321 | 1335 | 1224 | 1282 | 6147,76 | 11005 | 1290 | 0,117 | -3,092 | -73,018 | 0 | 0 | 0,0005498 | 0,002252 | 0,0013113 | 0,0013113 | TRUE | 0,0137668 | No |
| orf19.4869 | 3752 | 3035 | 3591 | 2069 | 3848 | 3044 | 3464 | 4778 | 3416 | 3070 | 3399 | 3108 | 3423 | 3305 | 3548 | 3645 | 3364 | 3248 | 3480 | 1,071 | 0,099 | 1,898 | 0,057751 | 0,071115 | 0,0021232 | 0,00243 | 0,0023047 | 0,0023047 | TRUE | 0,0173775 | No |
| orf19.487 | 1847 | 1627 | 1827 | 1228 | 993 | 764 | 822 | 1094 | 1682 | 1646 | 1729 | 1845 | 883 | 829 | 842 | 835 | 1286,25 | 1726 | 847 | 0,492 | -1,023 | -19,018 | 1,22E-80 | 8,66E-80 | 0,0008866 | 0,003064 | 0,0019139 | 0,0019139 | TRUE | 0,0237991 | No |
| orf19.4870 | 13486 | 12435 | 13531 | 8673 | 15266 | 13701 | 13619 | 18312 | 12278 | 12577 | 12806 | 13028 | 13579 | 14875 | 13948 | 13969 | 13382,4 | 12672 | 14093 | 1,112 | 0,153 | 3,941 | 8,11E-05 | 0,000122 | 0,0009872 | 0,002136 | 0,001379 | 0,001379 | TRUE | 0,0169954 | No |
| orf19.4871 | 18055 | 17150 | 18083 | 11063 | 12690 | 11034 | 10837 | 14852 | 16438 | 17345 | 17114 | 16618 | 11287 | 11980 | 11098 | 11329 | 14151,4 | 16879 | 11424 | 0,677 | -0,563 | -15,357 | 3,19E-53 | 1,57E-52 | 0,0007803 | 0,00213 | 0,0012202 | 0,0012202 | TRUE | 0,0129863 | No |
| orf19.4872 | 139 | 154 | 163 | 112 | 263 | 296 | 283 | 356 | 127 | 156 | 154 | 168 | 234 | 321 | 290 | 272 | 215,19 | 151 | 279 | 1,854 | 0,89 | 7,194 | 6,31E-13 | 1,32E-12 | 0,0112109 | 0,008176 | 0,0094685 | 0,0094685 | TRUE | 0,1343772 | No |
| orf19.4873 | 12611 | 11711 | 11972 | 7825 | 2746 | 2320 | 2150 | 2982 | 11482 | 11844 | 11330 | 11754 | 2442 | 2519 | 2202 | 2275 | 6981,12 | 11602 | 2360 | 0,203 | -2,298 | -49,011 | 0 | 0 | 0,0017964 | 0,002226 | 0,0018675 | 0,0018675 | TRUE | 0,0242703 | No |
| orf19.4874 | 5444 | 5082 | 5469 | 3197 | 4114 | 3616 | 3614 | 4829 | 4957 | 5140 | 5176 | 4802 | 3659 | 3926 | 3701 | 3684 | 4380,6 | 5019 | 3742 | 0,745 | -0,425 | -10,256 | 1,12E-24 | 3,18E-24 | 0,0008295 | 0,002339 | 0,0014153 | 0,0014153 | TRUE | 0,0129709 | No |
| orf19.4875 | 923 | 757 | 857 | 453 | 577 | 414 | 478 | 644 | 840 | 766 | 811 | 680 | 513 | 449 | 490 | 491 | 630,13 | 774 | 486 | 0,626 | -0,676 | -8,819 | 1,16E-18 | 2,83E-18 | 0,0038411 | 0,004133 | 0,0039975 | 0,0039975 | TRUE | 0,0705767 | No |
| orf19.4878 | 1139 | 921 | 1094 | 658 | 939 | 752 | 883 | 1215 | 1037 | 931 | 1035 | 988 | 835 | 816 | 904 | 927 | 934,39 | 998 | 870 | 0,873 | -0,196 | -3,098 | 0,00195 | 0,002693 | 0,0021462 | 0,003451 | 0,002784 | 0,002784 | TRUE | 0,0253362 | No |

| | | | | | | | | | | | | | | | | | | | | | | | | | | | | | | | |
|---|---|---|---|---|---|---|---|---|---|---|---|---|---|---|---|---|---|---|---|---|---|---|---|---|---|---|---|---|---|---|---|
| orf19.4879.2 | 3664 | 3277 | 3239 | 2201 | 2287 | 1806 | 1824 | 2551 | 3336 | 3314 | 3065 | 3306 | 2034 | 1961 | 1868 | 1946 | 2603,86 | 3255 | 1952 | 0,6 | -0,737 | -15,915 | 4,98E-57 | 2,59E-56 | 0,0010461 | 0,002544 | 0,0016535 | 0,0016535 | TRUE | 0,0192814 | No |
| orf19.488 | 4631 | 4238 | 4526 | 2871 | 4395 | 3754 | 3987 | 5358 | 4216 | 4286 | 4283 | 4313 | 3909 | 4076 | 4083 | 4087 | 4156,76 | 4274 | 4039 | 0,945 | -0,082 | -2,414 | 0,015772 | 0,020416 | 1,00E-08 | 0,002355 | 0,0008555 | 0,0008555 | TRUE | 0,0057627 | No |
| orf19.4880 | 263 | 207 | 252 | 126 | 461 | 391 | 421 | 643 | 239 | 209 | 238 | 189 | 410 | 425 | 431 | 490 | 329,1 | 219 | 439 | 1,991 | 0,994 | 10,145 | 3,50E-24 | 9,84E-24 | 0,0052921 | 0,006051 | 0,0057304 | 0,0057304 | TRUE | 0,0770374 | No |
| orf19.4882 | 5917 | 4942 | 5415 | 2990 | 3530 | 2695 | 3265 | 4438 | 5387 | 4998 | 5125 | 4491 | 3140 | 2926 | 3344 | 3385 | 4099,59 | 5000 | 3199 | 0,639 | -0,645 | -10,045 | 9,68E-24 | 2,70E-23 | 0,004663 | 0,002359 | 0,0037078 | 0,0037078 | TRUE | 0,0559253 | No |
| orf19.4883 | 3483 | 3258 | 3473 | 2209 | 7049 | 5964 | 6173 | 8114 | 3171 | 3295 | 3287 | 3318 | 6270 | 6475 | 6322 | 6189 | 4790,98 | 3268 | 6314 | 1,933 | 0,951 | 27,42 | 1,60E-165 | 2,32E-164 | 0,0001851 | 0,002313 | 0,0009156 | 0,0009156 | TRUE | 0,0049897 | No |
| orf19.4884 | 111 | 120 | 118 | 89 | 763 | 660 | 678 | 971 | 101 | 121 | 112 | 134 | 679 | 717 | 694 | 741 | 412,26 | 117 | 708 | 6,132 | 2,616 | 27,484 | 2,69E-166 | 3,91E-165 | 0,0005364 | 0,005241 | 0,0034563 | 0,0034563 | TRUE | 0,0867725 | No |
| orf19.4885 | 15225 | 12508 | 13116 | 9831 | 16803 | 13584 | 13924 | 20082 | 13862 | 12651 | 12413 | 14768 | 14946 | 14748 | 14260 | 15319 | 14120,7 | 13424 | 14818 | 1,104 | 0,143 | 2,524 | 0,011611 | 0,015156 | 0,0036941 | 0,002131 | 0,0030193 | 0,0030193 | TRUE | 0,0550408 | No |
| orf19.4886 | 86 | 82 | 80 | 42 | 360 | 351 | 355 | 416 | 78 | 83 | 76 | 63 | 320 | 381 | 364 | 317 | 210,28 | 75 | 346 | 4,544 | 2,184 | 17,441 | 4,06E-68 | 2,48E-67 | 1,00E-08 | 0,008319 | 0,0067079 | 0,0067079 | TRUE | 0,0711232 | No |
| orf19.4887 | 13227 | 12112 | 12871 | 8651 | 3452 | 3137 | 3146 | 4298 | 12043 | 12250 | 12181 | 12995 | 3070 | 3406 | 3222 | 3279 | 7805,73 | 12367 | 3244 | 0,262 | -1,931 | -44,484 | 0 | 0 | 0,0012565 | 0,002206 | 0,0016211 | 0,0016211 | TRUE | 0,0158702 | No |
| orf19.4888 | 3555 | 3353 | 3399 | 2097 | 1432 | 1072 | 1200 | 1629 | 3237 | 3391 | 3217 | 3150 | 1274 | 1164 | 1229 | 1243 | 2237,99 | 3249 | 1228 | 0,378 | -1,404 | -30,518 | 1,51E-204 | 2,64E-203 | 0,0005914 | 0,002627 | 0,0014858 | 0,0014858 | TRUE | 0,0137943 | No |
| orf19.4889 | 5898 | 5561 | 5937 | 3608 | 3407 | 2894 | 3131 | 3909 | 5370 | 5624 | 5619 | 5420 | 3030 | 3142 | 3207 | 2982 | 4299,21 | 5508 | 3090 | 0,561 | -0,835 | -21,248 | 3,45E-100 | 3,01E-99 | 0,0005032 | 0,002344 | 0,0012342 | 0,0012342 | TRUE | 0,0078631 | No |
| orf19.489 | 553 | 498 | 535 | 345 | 1708 | 1376 | 1406 | 1844 | 503 | 504 | 506 | 518 | 1519 | 1494 | 1440 | 1407 | 986,43 | 508 | 1465 | 2,886 | 1,529 | 26,962 | 4,10E-160 | 5,67E-159 | 1,00E-08 | 0,003376 | 0,0017198 | 0,0017198 | TRUE | 0,0092508 | No |
| orf19.4890 | 1905 | 1875 | 1924 | 1301 | 2838 | 2462 | 2516 | 3302 | 1734 | 1896 | 1821 | 1954 | 2524 | 2673 | 2577 | 2519 | 2212,35 | 1851 | 2573 | 1,392 | 0,477 | 9,867 | 5,78E-23 | 1,57E-22 | 0,0010905 | 0,002634 | 0,0017705 | 0,0017705 | TRUE | 0,0221576 | No |
| orf19.4892 | 12078 | 11737 | 12118 | 7205 | 11856 | 9703 | 10467 | 13119 | 10997 | 11871 | 11469 | 10823 | 10546 | 10535 | 10720 | 10007 | 10870,8 | 11290 | 10452 | 0,925 | -0,112 | -2,705 | 0,006829 | 0,00907 | 0,0011923 | 0,002159 | 0,0015493 | 0,0015493 | TRUE | 0,0146359 | No |
| orf19.4893 | 2906 | 2713 | 2949 | 1773 | 1503 | 1243 | 1312 | 1697 | 2646 | 2744 | 2791 | 2663 | 1337 | 1350 | 1344 | 1294 | 2021,07 | 2711 | 1331 | 0,49 | -1,028 | -24,219 | 1,41E-129 | 1,55E-128 | 1,00E-08 | 0,002691 | 0,0011806 | 0,0011806 | TRUE | 0,0049022 | No |
| orf19.4894 | 530 | 460 | 456 | 365 | 714 | 619 | 643 | 963 | 483 | 465 | 432 | 548 | 635 | 672 | 659 | 735 | 578,48 | 482 | 675 | 1,41 | 0,495 | 6,132 | 8,67E-10 | 1,62E-09 | 0,0047938 | 0,004321 | 0,0044551 | 0,0044551 | TRUE | 0,0871204 | No |
| orf19.4895 | 1604 | 1318 | 1542 | 893 | 1714 | 1366 | 1578 | 2186 | 1460 | 1333 | 1459 | 1341 | 1525 | 1483 | 1616 | 1667 | 1485,67 | 1398 | 1573 | 1,124 | 0,168 | 2,945 | 0,003229 | 0,004382 | 0,0020715 | 0,002926 | 0,0024607 | 0,0024607 | TRUE | 0,0205866 | No |
| orf19.4896 | 4036 | 3590 | 3811 | 2489 | 4647 | 3638 | 3817 | 5230 | 3675 | 3631 | 3607 | 3739 | 4133 | 3950 | 3909 | 3989 | 3829,12 | 3663 | 3995 | 1,092 | 0,127 | 3,541 | 0,000398 | 0,000577 | 0,0001698 | 0,002382 | 0,0009628 | 0,0009628 | TRUE | 0,0065919 | No |
| orf19.4897 | 682 | 627 | 707 | 410 | 614 | 548 | 531 | 692 | 621 | 634 | 669 | 616 | 546 | 595 | 544 | 528 | 594,11 | 635 | 553 | 0,868 | -0,204 | -3,091 | 0,001994 | 0,002752 | 1,00E-08 | 0,004261 | 0,0024885 | 0,0024885 | TRUE | 0,028767 | No |
| orf19.4898 | 2257 | 1904 | 2154 | 1244 | 1102 | 825 | 982 | 1196 | 2055 | 1926 | 2039 | 1869 | 980 | 896 | 1006 | 912 | 1460,22 | 1972 | 948 | 0,48 | -1,058 | -18,781 | 1,09E-78 | 7,60E-78 | 0,0015436 | 0,002942 | 0,0022835 | 0,0022835 | TRUE | 0,0189251 | No |
| orf19.4899 | 1082 | 863 | 936 | 660 | 889 | 681 | 745 | 1037 | 985 | 873 | 886 | 991 | 791 | 739 | 763 | 791 | 852,42 | 934 | 771 | 0,828 | -0,272 | -4,325 | 1,53E-05 | 2,40E-05 | 0,0016529 | 0,003587 | 0,0026229 | 0,0026229 | TRUE | 0,0225998 | No |
| orf19.490 | 1169 | 1014 | 1066 | 675 | 1974 | 1578 | 1585 | 2257 | 1064 | 1026 | 1009 | 1014 | 1756 | 1713 | 1623 | 1722 | 1365,84 | 1028 | 1704 | 1,656 | 0,728 | 14,87 | 5,16E-50 | 2,37E-49 | 1,00E-08 | 0,003004 | 0,0015067 | 0,0015067 | TRUE | 0,0118824 | No |
| orf19.4900 | 1156 | 1224 | 1241 | 755 | 583 | 482 | 486 | 602 | 1052 | 1238 | 1174 | 1134 | 519 | 523 | 498 | 459 | 824,73 | 1150 | 500 | 0,434 | -1,205 | -17,089 | 1,78E-65 | 1,06E-64 | 0,0031072 | 0,003639 | 0,0033688 | 0,0033688 | TRUE | 0,0413136 | No |
| orf19.4901 | 700 | 636 | 649 | 455 | 545 | 474 | 483 | 619 | 637 | 643 | 614 | 683 | 485 | 515 | 495 | 472 | 568,06 | 644 | 492 | 0,763 | -0,389 | -5,939 | 2,86E-09 | 5,27E-09 | 1,00E-08 | 0,004362 | 0,0023437 | 0,0023437 | TRUE | 0,0165437 | No |
| orf19.4903 | 847 | 994 | 936 | 589 | 1702 | 1534 | 1468 | 1755 | 771 | 1005 | 886 | 885 | 1514 | 1665 | 1503 | 1339 | 1196,08 | 887 | 1505 | 1,697 | 0,763 | 8,966 | 3,06E-19 | 7,62E-19 | 0,0085519 | 0,003141 | 0,0060334 | 0,0060334 | TRUE | 0,0992155 | No |
| orf19.4904 | 979 | 967 | 1003 | 668 | 872 | 731 | 708 | 916 | 891 | 978 | 949 | 1003 | 776 | 794 | 725 | 699 | 851,89 | 955 | 748 | 0,784 | -0,352 | -5,507 | 3,65E-08 | 6,41E-08 | 0,0018688 | 0,003588 | 0,002729 | 0,002729 | TRUE | 0,0252591 | No |
| orf19.4905 | 1129 | 1021 | 1181 | 679 | 796 | 672 | 721 | 938 | 1028 | 1033 | 1118 | 1020 | 708 | 730 | 738 | 716 | 886,22 | 1050 | 723 | 0,687 | -0,541 | -9,654 | 4,72E-22 | 1,25E-21 | 1,00E-08 | 0,003528 | 0,0018557 | 0,0018557 | TRUE | 0,0236891 | No |
| orf19.4906 | 7828 | 6904 | 7642 | 4890 | 6239 | 5421 | 6004 | 8068 | 7127 | 6983 | 7232 | 7346 | 5549 | 5886 | 6149 | 6154 | 6553,25 | 7172 | 5934 | 0,828 | -0,273 | -6,352 | 2,12E-10 | 4,08E-10 | 0,0011861 | 0,002239 | 0,0016178 | 0,0016178 | TRUE | 0,0235245 | No |
| orf19.4907 | 12038 | 11315 | 11786 | 7531 | 557 | 517 | 535 | 714 | 10960 | 11444 | 11154 | 11313 | 495 | 561 | 548 | 545 | 5877,56 | 11218 | 537 | 0,048 | -4,386 | -92,709 | 0 | 0 | 0,0003137 | 0,002262 | 0,0012347 | 0,0012347 | TRUE | 0,0327823 | No |
| orf19.4908 | 117 | 89 | 109 | 74 | 7 | 6 | 12 | 9 | 107 | 90 | 103 | 111 | 6 | 7 | 12 | 7 | 55,34 | 103 | 8 | 0,077 | -3,703 | -12,731 | 3,95E-37 | 1,48E-36 | 1,00E-08 | 0,025906 | 0,017211 | 0,017211 | TRUE | 0,3575953 | No |
| orf19.4909 | 2528 | 2258 | 2474 | 1572 | 2841 | 2455 | 2604 | 3457 | 2302 | 2284 | 2341 | 2361 | 2527 | 2665 | 2667 | 2637 | 2473,06 | 2322 | 2624 | 1,13 | 0,177 | 4,512 | 6,43E-06 | 1,03E-05 | 1,00E-08 | 0,002571 | 0,0010646 | 0,0010646 | TRUE | 0,0075816 | No |
| orf19.4909.1 | 10814 | 8262 | 8737 | 5351 | 16547 | 11306 | 12766 | 17870 | 9846 | 8356 | 8269 | 8038 | 14718 | 12275 | 13074 | 13631 | 11025,9 | 8627 | 13424 | 1,556 | 0,638 | 8,91 | 5,08E-19 | 1,26E-18 | 0,0073494 | 0,002157 | 0,0048229 | 0,0048229 | TRUE | 0,1108526 | No |
| orf19.491 | 3241 | 2838 | 3188 | 1882 | 4099 | 3284 | 3718 | 4950 | 2951 | 2870 | 3017 | 2827 | 3646 | 3566 | 3808 | 3776 | 3307,55 | 2916 | 3699 | 1,267 | 0,342 | 8,389 | 4,88E-17 | 1,14E-16 | 0,0004786 | 0,002436 | 0,0012843 | 0,0012843 | TRUE | 0,0069532 | No |
| orf19.4910 | 178 | 164 | 153 | 101 | 104 | 101 | 85 | 98 | 162 | 166 | 145 | 152 | 93 | 110 | 87 | 75 | 123,55 | 156 | 91 | 0,578 | -0,792 | -5,73 | 1,00E-08 | 1,81E-08 | 0,0042078 | 0,012729 | 0,0097599 | 0,0097599 | TRUE | 0,1725679 | No |
| orf19.4911 | 478 | 426 | 472 | 301 | 96 | 69 | 75 | 109 | 435 | 431 | 447 | 452 | 85 | 75 | 77 | 83 | 260,65 | 441 | 80 | 0,182 | -2,454 | -23,266 | 9,75E-120 | 9,94E-119 | 1,00E-08 | 0,007105 | 0,0037076 | 0,0037076 | TRUE | 0,0174188 | No |
| orf19.4912 | 2198 | 1832 | 2055 | 1327 | 2005 | 1682 | 1864 | 2527 | 2001 | 1853 | 1945 | 1993 | 1783 | 1826 | 1909 | 1928 | 1904,81 | 1948 | 1862 | 0,956 | -0,064 | -1,359 | 0,174108 | 0,20284 | 0,0008134 | 0,002731 | 0,0016315 | 0,0016315 | TRUE | 0,0132213 | No |
| orf19.4913 | 1526 | 1596 | 1654 | 979 | 1124 | 1068 | 1033 | 1293 | 1389 | 1614 | 1565 | 1471 | 1000 | 1160 | 1058 | 986 | 1280,38 | 1510 | 1051 | 0,695 | -0,525 | -7,717 | 1,19E-14 | 2,61E-14 | 0,0039324 | 0,003069 | 0,0036475 | 0,0036475 | TRUE | 0,0568385 | No |
| orf19.4914 | 206 | 171 | 176 | 111 | 271 | 199 | 209 | 277 | 188 | 173 | 167 | 167 | 241 | 216 | 214 | 211 | 197,03 | 174 | 220 | 1,269 | 0,343 | 3,323 | 0,00089 | 0,00126 | 1,00E-08 | 0,008742 | 0,0050652 | 0,0050652 | TRUE | 0,0446481 | No |
| orf19.4914.1 | 42186 | 39346 | 39854 | 21332 | 12767 | 8487 | 10179 | 12537 | 38409 | 39794 | 37718 | 32044 | 11356 | 9215 | 10425 | 9563 | 23565,4 | 36991 | 10140 | 0,274 | -1,867 | -24,331 | 9,24E-131 | 1,03E-129 | 0,0086186 | 0,002093 | 0,0055984 | 0,0055984 | TRUE | 0,0990521 | No |
| orf19.4916 | 843 | 774 | 866 | 535 | 635 | 527 | 536 | 749 | 768 | 783 | 820 | 804 | 565 | 572 | 549 | 571 | 678,86 | 794 | 564 | 0,712 | -0,49 | -8,32 | 8,83E-17 | 2,05E-16 | 1,00E-08 | 0,003983 | 0,0018319 | 0,0018319 | TRUE | 0,0066863 | No |
| orf19.4917 | 93 | 83 | 103 | 51 | 74 | 61 | 66 | 128 | 85 | 84 | 97 | 77 | 66 | 66 | 68 | 98 | 80 | 86 | 74 | 0,87 | -0,2 | -1,141 | 0,253741 | 0,289203 | 0,0151897 | 0,01855 | 0,0171794 | 0,0171794 | TRUE | 0,464643 | No |
| orf19.4918 | 253 | 238 | 301 | 209 | 512 | 381 | 439 | 545 | 230 | 241 | 285 | 314 | 455 | 414 | 450 | 416 | 350,53 | 268 | 434 | 1,634 | 0,708 | 7,208 | 5,68E-13 | 1,19E-12 | 0,0067568 | 0,005806 | 0,0061765 | 0,0061765 | TRUE | 0,1289982 | No |
| orf19.4919 | 376 | 295 | 320 | 226 | 607 | 465 | 499 | 682 | 342 | 298 | 303 | 339 | 540 | 505 | 511 | 520 | 419,88 | 320 | 519 | 1,623 | 0,699 | 9,116 | 7,79E-20 | 1,97E-19 | 1,00E-08 | 0,005183 | 0,003081 | 0,003081 | TRUE | 0,0290846 | No |
| orf19.492 | 7265 | 6248 | 6879 | 4822 | 8873 | 7619 | 7811 | 10908 | 6614 | 6319 | 6510 | 7243 | 7892 | 8272 | 7999 | 8321 | 7396,49 | 6672 | 8121 | 1,218 | 0,285 | 6,041 | 1,54E-09 | 2,86E-09 | 0,0019449 | 0,002216 | 0,0019953 | 0,0019953 | TRUE | 0,0394333 | No |
| orf19.4921 | 1125 | 1002 | 1019 | 657 | 1214 | 1076 | 1055 | 1475 | 1024 | 1013 | 964 | 987 | 1080 | 1168 | 1080 | 1125 | 1055,33 | 997 | 1113 | 1,116 | 0,158 | 2,999 | 0,002707 | 0,003696 | 1,00E-08 | 0,003289 | 0,0017013 | 0,0017013 | TRUE | 0,0122539 | No |
| orf19.4921.1 | 977 | 919 | 865 | 471 | 1695 | 1248 | 1174 | 1686 | 890 | 929 | 819 | 708 | 1508 | 1355 | 1202 | 1286 | 1087,02 | 836 | 1338 | 1,595 | 0,674 | 7,591 | 3,16E-14 | 6,84E-14 | 0,0101471 | 0,003252 | 0,0065719 | 0,0065719 | TRUE | 0,1176692 | No |
| orf19.4922 | 764 | 760 | 799 | 461 | 1012 | 883 | 886 | 1177 | 696 | 769 | 756 | 692 | 900 | 959 | 907 | 898 | 822,12 | 728 | 916 | 1,255 | 0,327 | 5,49 | 4,03E-08 | 7,05E-08 | 0,0004895 | 0,003644 | 0,0021654 | 0,0021654 | TRUE | 0,0160656 | No |
| orf19.4923 | 239 | 195 | 229 | 139 | 166 | 157 | 146 | 191 | 218 | 197 | 217 | 209 | 148 | 170 | 150 | 146 | 181,71 | 210 | 154 | 0,727 | -0,461 | -4,34 | 1,42E-05 | 2,25E-05 | 1,00E-08 | 0,009307 | 0,0052471 | 0,0052471 | TRUE | 0,0548948 | No |
| orf19.4923.1 | 33 | 28 | 22 | 15 | 32 | 15 | 20 | 26 | 30 | 28 | 21 | 23 | 28 | 16 | 20 | 20 | 23,35 | 26 | 21 | 0,835 | -0,26 | -0,888 | 0,374311 | 0,41442 | 1,00E-08 | 0,058616 | 0,0394355 | 0,0394355 | TRUE | 0,311028 | No |
| orf19.4924 | 2550 | 2534 | 2645 | 1596 | 1848 | 1472 | 1677 | 2031 | 2322 | 2563 | 2503 | 2397 | 1644 | 1598 | 1717 | 1549 | 2036,73 | 2446 | 1627 | 0,665 | -0,589 | -11,538 | 8,53E-31 | 2,80E-30 | 0,001486 | 0,002686 | 0,0019989 | 0,0019989 | TRUE | 0,016664 | No |
| orf19.4927 | 3915 | 3605 | 3949 | 2350 | 2307 | 1889 | 2016 | 2614 | 3564 | 3646 | 3737 | 3530 | 2052 | 2051 | 2065 | 1994 | 2829,94 | 3619 | 2040 | 0,563 | -0,828 | -21,461 | 3,62E-102 | 3,23E-101 | 1,00E-08 | 0,002504 | 0,0010534 | 0,0010534 | TRUE | 0,0059018 | No |
| orf19.4928 | 4362 | 4309 | 4628 | 2668 | 3270 | 2690 | 2965 | 3753 | 3971 | 4358 | 4380 | 4008 | 2909 | 2921 | 3037 | 2863 | 3555,72 | 4179 | 2932 | 0,701 | -0,512 | -10,905 | 1,09E-27 | 3,33E-27 | 0,0013134 | 0,002409 | 0,001832 | 0,001832 | TRUE | 0,0144163 | No |
| orf19.4929 | 3113 | 2998 | 3193 | 1845 | 1949 | 1669 | 1714 | 2359 | 2834 | 3032 | 3022 | 2771 | 1734 | 1812 | 1755 | 1799 | 2345,03 | 2915 | 1775 | 0,608 | -0,717 | -15,634 | 4,27E-55 | 2,16E-54 | 0,0008539 | 0,0026 | 0,0015719 | 0,0015719 | TRUE | 0,0122757 | No |
| orf19.493 | 25663 | 20052 | 20963 | 12759 | 38402 | 27439 | 31442 | 43289 | 23365 | 20280 | 19840 | 19166 | 34157 | 29791 | 32201 | 33021 | 26477,7 | 20663 | 32292 | 1,563 | 0,644 | 9,942 | 2,73E-23 | 7,52E-23 | 0,0056358 | 0,002087 | 0,0039912 | 0,0039912 | TRUE | 0,0950326 | No |
| orf19.4930 | 502 | 464 | 481 | 342 | 309 | 236 | 226 | 389 | 457 | 469 | 455 | 514 | 275 | 256 | 231 | 297 | 369,32 | 474 | 265 | 0,563 | -0,828 | -9,278 | 1,72E-20 | 4,41E-20 | 0,0035156 | 0,005614 | 0,00476 | 0,00476 | TRUE | 0,0799963 | No |
| orf19.4931 | 4585 | 4225 | 4546 | 2897 | 5529 | 4841 | 5023 | 6806 | 4174 | 4273 | 4302 | 4352 | 4918 | 5256 | 5144 | 5192 | 4701,43 | 4275 | 5128 | 1,2 | 0,263 | 7,216 | 5,36E-13 | 1,12E-12 | 0,0003885 | 0,002318 | 0,0010547 | 0,0010547 | TRUE | 0,009296 | No |
| orf19.4931.1 | 11232 | 9633 | 9728 | 6462 | 18868 | 13677 | 14149 | 19550 | 10226 | 9743 | 9207 | 9707 | 16782 | 14849 | 14490 | 14913 | 12489,7 | 9721 | 15258 | 1,57 | 0,651 | 12,083 | 1,30E-33 | 4,51E-33 | 0,003096 | 0,002143 | 0,0026986 | 0,0026986 | TRUE | 0,0553613 | No |
| orf19.4932 | 4548 | 3951 | 4094 | 2596 | 5184 | 4442 | 4501 | 6289 | 4141 | 3996 | 3875 | 3900 | 4611 | 4823 | 4610 | 4797 | 4343,96 | 3978 | 4710 | 1,184 | 0,243 | 6,321 | 2,59E-10 | 4,96E-10 | 0,0004528 | 0,002341 | 0,0011858 | 0,0011858 | TRUE | 0,0095132 | No |
| orf19.4933 | 5888 | 5212 | 5547 | 3822 | 16476 | 15116 | 14966 | 20737 | 5361 | 5271 | 5250 | 5741 | 14655 | 16412 | 15327 | 15818 | 10479,4 | 5406 | 15553 | 2,879 | 1,526 | 32,468 | 3,05E-231 | 5,91E-230 | 0,0018767 | 0,002163 | 0,0019905 | 0,0019905 | TRUE | 0,020547 | No |
| orf19.4934 | 25 | 21 | 12 | 14 | 327 | 229 | 270 | 392 | 23 | 21 | 11 | 21 | 291 | 249 | 277 | 299 | 148,93 | 19 | 279 | 14,768 | 3,884 | 19,511 | 8,80E-85 | 6,60E-84 | 0,0031718 | 0,010907 | 0,0085406 | 0,0085406 | TRUE | 0,4626867 | No |
| orf19.4936 | 96 | 87 | 101 | 59 | 67 | 46 | 51 | 65 | 87 | 88 | 96 | 89 | 60 | 50 | 52 | 50 | 71,37 | 90 | 53 | 0,587 | -0,767 | -4,693 | 2,69E-06 | 4,39E-06 | 1,00E-08 | 0,020546 | 0,0109671 | 0,0109671 | TRUE | 0,066452 | No |
| orf19.4936.1 | 17231 | 15923 | 16570 | 9585 | 3314 | 2504 | 2731 | 3450 | 15688 | 16104 | 15682 | 14398 | 2948 | 2719 | 2797 | 2632 | 9120,95 | 15468 | 2774 | 0,179 | -2,48 | -49,942 | 0 | 0 | 0,0020316 | 0,002182 | 0,002165 | 0,002165 | TRUE | 0,026185 | No |
| orf19.4937 | 8210 | 7445 | 7933 | 4811 | 6997 | 6017 | 6383 | 8408 | 7475 | 7530 | 7508 | 7227 | 6224 | 6533 | 6537 | 6414 | 6930,81 | 7435 | 6427 | 0,864 | -0,211 | -6,371 | 1,87E-10 | 3,60E-10 | 0,0002467 | 0,002228 | 0,0009096 | 0,0009096 | TRUE | 0,005527 | No |
| orf19.4939 | 382 | 436 | 419 | 275 | 171 | 158 | 174 | 195 | 348 | 441 | 397 | 413 | 152 | 172 | 178 | 149 | 281,12 | 400 | 163 | 0,406 | -1,3 | -12,39 | 2,97E-35 | 1,07E-34 | 0,0058642 | 0,006736 | 0,0063529 | 0,0063529 | TRUE | 0,0930816 | No |
| orf19.494 | 7920 | 6339 | 7163 | 4856 | 7087 | 5778 | 6538 | 8780 | 7211 | 6411 | 6779 | 7294 | 6304 | 6273 | 6696 | 6697 | 6708,23 | 6924 | 6492 | 0,938 | -0,092 | -1,841 | 0,065568 | 0,080306 | 0,0022117 | 0,002234 | 0,0022562 | 0,0022562 | TRUE | 0,0303259 | No |
| orf19.4940 | 11721 | 9842 | 10929 | 7392 | 7820 | 6724 | 6921 | 9763 | 10671 | 9954 | 10343 | 11104 | 6956 | 7300 | 7088 | 7447 | 8858,02 | 10518 | 7198 | 0,685 | -0,547 | -12,575 | 2,90E-36 | 1,06E-35 | 0,0013832 | 0,002186 | 0,0016988 | 0,0016988 | TRUE | 0,0167858 | No |
| orf19.4941 | 28783 | 25073 | 28101 | 17479 | 11923 | 9743 | 10670 | 13972 | 26206 | 25359 | 26595 | 26256 | 10605 | 10578 | 10927 | 10658 | 18398 | 26104 | 10692 | 0,41 | -1,288 | -43,57 | 0 | 0 | 0,0002275 | 0,002109 | 0,0007746 | 0,0007746 | TRUE | 0,0045475 | No |

| | | | | | | | | | | | | | | | | | | | | | | | | | | | | | | | |
|---|---|---|---|---|---|---|---|---|---|---|---|---|---|---|---|---|---|---|---|---|---|---|---|---|---|---|---|---|---|---|---|
| orf19.4942 | 41 | 46 | 40 | 34 | 86 | 87 | 69 | 97 | 37 | 47 | 38 | 51 | 76 | 94 | 71 | 74 | 61,05 | 43 | 79 | 1,846 | 0,884 | 4,652 | 3,29E-06 | 5,35E-06 | 1,00E-08 | 0,023675 | 0,0162919 | 0,0162919 | TRUE | 0,1503221 | No |
| orf19.4943 | 8732 | 7984 | 8528 | 5467 | 1704 | 1321 | 1365 | 2050 | 7950 | 8075 | 8071 | 8212 | 1516 | 1434 | 1398 | 1564 | 4777,49 | 8077 | 1478 | 0,183 | -2,448 | -57,1 | 0 | 0 | 0,0005541 | 0,002314 | 0,0013822 | 0,0013822 | TRUE | 0,0192056 | No |
| orf19.4945 | 547 | 516 | 525 | 276 | 966 | 863 | 854 | 1161 | 498 | 522 | 497 | 415 | 859 | 937 | 875 | 886 | 685,97 | 483 | 889 | 1,829 | 0,871 | 12,239 | 1,92E-34 | 6,81E-34 | 0,0020932 | 0,003963 | 0,0032349 | 0,0032349 | TRUE | 0,0871969 | No |
| orf19.4946 | 6 | 5 | 5 | 5 | 11 | 11 | 16 | 11 | 5 | 5 | 5 | 8 | 10 | 12 | 16 | 8 | 8,66 | 6 | 12 | 2,065 | 1,046 | 2,137 | 0,032595 | 0,04105 | 1,00E-08 | 0,154607 | 0,0941468 | 0,0941468 | TRUE | 0,2913173 | No |
| orf19.4947 | 7612 | 6886 | 6472 | 4453 | 2793 | 2421 | 1956 | 3049 | 6930 | 6964 | 6125 | 6689 | 2484 | 2629 | 2003 | 2326 | 4518,87 | 6677 | 2360 | 0,353 | -1,5 | -19,825 | 1,82E-87 | 1,40E-86 | 0,0077678 | 0,002329 | 0,0052224 | 0,0052224 | TRUE | 0,1244751 | No |
| orf19.4948 | 609 | 614 | 638 | 323 | 939 | 759 | 822 | 1053 | 554 | 621 | 604 | 485 | 835 | 824 | 842 | 803 | 696,1 | 566 | 826 | 1,451 | 0,537 | 7,363 | 1,80E-13 | 3,82E-13 | 0,0031254 | 0,003935 | 0,0036018 | 0,0036018 | TRUE | 0,092117 | No |
| orf19.4949 | 1 | 4 | 1 | 5 | 8 | 3 | 4 | 2 | 1 | 4 | 1 | 8 | 7 | 3 | 4 | 2 | 3,68 | 4 | 4 | 1,282 | 0,359 | 0,439 | 0,660934 | 0,697476 | 0,3381403 | 0,361391 | 0,3529503 | 0,3529503 | TRUE | 0,6106831 | No |
| orf19.4950 | 6524 | 6414 | 6792 | 4498 | 4282 | 3765 | 3862 | 5011 | 5940 | 6487 | 6428 | 6757 | 3809 | 4088 | 3955 | 3822 | 5160,71 | 6403 | 3918 | 0,612 | -0,708 | -14,797 | 1,54E-49 | 7,03E-49 | 0,0016943 | 0,002293 | 0,0019956 | 0,0019956 | TRUE | 0,0291303 | No |
| orf19.4951 | 2724 | 2847 | 2993 | 1878 | 1877 | 1592 | 1542 | 2160 | 2480 | 2879 | 2833 | 2821 | 1670 | 1728 | 1579 | 1648 | 2204,76 | 2753 | 1656 | 0,602 | -0,733 | -12,812 | 1,41E-37 | 5,27E-37 | 0,0026814 | 0,002636 | 0,0026614 | 0,0026614 | TRUE | 0,0556012 | No |
| orf19.4952 | 6275 | 5791 | 6262 | 3664 | 6890 | 6037 | 5993 | 7717 | 5713 | 5857 | 5926 | 5504 | 6128 | 6555 | 6138 | 5887 | 5963,44 | 5750 | 6177 | 1,073 | 0,102 | 2,32 | 0,020324 | 0,026027 | 0,0013109 | 0,002259 | 0,0016861 | 0,0016861 | TRUE | 0,0205142 | No |
| orf19.4952.1 | 1304 | 1078 | 1146 | 632 | 767 | 601 | 599 | 883 | 1187 | 1090 | 1085 | 949 | 682 | 653 | 613 | 674 | 866,65 | 1078 | 656 | 0,607 | -0,721 | -9,968 | 2,10E-23 | 5,79E-23 | 0,004067 | 0,003561 | 0,003811 | 0,003811 | TRUE | 0,0704012 | No |
| orf19.4953 | 1588 | 1808 | 1646 | 1046 | 990 | 820 | 831 | 1057 | 1446 | 1829 | 1558 | 1571 | 881 | 890 | 851 | 806 | 1228,96 | 1601 | 857 | 0,535 | -0,902 | -12,219 | 2,47E-34 | 8,70E-34 | 0,005459 | 0,003112 | 0,0043545 | 0,0043545 | TRUE | 0,1125865 | No |
| orf19.4954 | 835 | 788 | 820 | 536 | 1171 | 1002 | 957 | 1266 | 760 | 797 | 776 | 805 | 1042 | 1088 | 980 | 966 | 901,71 | 784 | 1019 | 1,299 | 0,377 | 6,383 | 1,73E-10 | 3,33E-10 | 0,0009971 | 0,003502 | 0,0022095 | 0,0022095 | TRUE | 0,0239179 | No |
| orf19.4955 | 46 | 66 | 69 | 47 | 61 | 61 | 52 | 59 | 42 | 67 | 65 | 71 | 54 | 66 | 53 | 45 | 57,91 | 61 | 54 | 0,897 | -0,157 | -0,755 | 0,450331 | 0,492344 | 0,0222232 | 0,024848 | 0,024002 | 0,024002 | TRUE | 0,4294562 | No |
| orf19.4956 | 8033 | 7704 | 7989 | 4956 | 9594 | 8176 | 8647 | 11378 | 7314 | 7792 | 7561 | 7445 | 8534 | 8877 | 8856 | 8679 | 8132,04 | 7528 | 8736 | 1,16 | 0,214 | 6,319 | 2,63E-10 | 5,04E-10 | 0,0003695 | 0,002199 | 0,0009811 | 0,0009811 | TRUE | 0,0068815 | No |
| orf19.4958 | 995 | 1088 | 1181 | 666 | 765 | 778 | 744 | 861 | 906 | 1100 | 1118 | 1000 | 680 | 845 | 762 | 657 | 883,54 | 1031 | 736 | 0,712 | -0,49 | -5,393 | 6,93E-08 | 1,21E-07 | 0,009704 | 0,003532 | 0,0067583 | 0,0067583 | TRUE | 0,1157706 | No |
| orf19.4959 | 6207 | 5421 | 5885 | 3898 | 3365 | 2995 | 3129 | 4326 | 5651 | 5483 | 5570 | 5855 | 2993 | 3252 | 3204 | 3300 | 4413,53 | 5640 | 3187 | 0,565 | -0,823 | -19,117 | 1,81E-81 | 1,30E-80 | 0,0009719 | 0,002336 | 0,0015356 | 0,0015356 | TRUE | 0,0207773 | No |
| orf19.496 | 551 | 498 | 587 | 360 | 711 | 640 | 614 | 864 | 502 | 504 | 556 | 541 | 632 | 695 | 629 | 659 | 589,6 | 526 | 654 | 1,245 | 0,317 | 4,695 | 2,66E-06 | 4,34E-06 | 0,0006481 | 0,004278 | 0,0026321 | 0,0026321 | TRUE | 0,0200811 | No |
| orf19.4960 | 11514 | 10913 | 11334 | 7025 | 5771 | 4772 | 4666 | 6347 | 10483 | 11037 | 10727 | 10553 | 5133 | 5181 | 4779 | 4842 | 7841,74 | 10700 | 4984 | 0,466 | -1,103 | -27,791 | 5,52E-170 | 8,20E-169 | 0,0009 | 0,002205 | 0,0013678 | 0,0013678 | TRUE | 0,0091025 | No |
| orf19.4961 | 39259 | 35803 | 38734 | 25804 | 23063 | 20111 | 20516 | 28494 | 35744 | 36211 | 36658 | 38762 | 20514 | 21835 | 21011 | 21735 | 29058,7 | 36844 | 21274 | 0,577 | -0,792 | -20,645 | 1,09E-94 | 8,85E-94 | 0,0010368 | 0,002082 | 0,0013778 | 0,0013778 | TRUE | 0,014929 | No |
| orf19.4962 | 1159 | 1266 | 1347 | 814 | 1020 | 810 | 798 | 1023 | 1055 | 1280 | 1275 | 1223 | 907 | 879 | 817 | 780 | 1027,19 | 1208 | 846 | 0,7 | -0,514 | -6,799 | 1,05E-11 | 2,12E-11 | 0,0055977 | 0,003323 | 0,004495 | 0,004495 | TRUE | 0,0908528 | No |
| orf19.4963 | 1926 | 2286 | 2214 | 1145 | 2786 | 2202 | 2316 | 2761 | 1754 | 2312 | 2095 | 1720 | 2478 | 2391 | 2372 | 2106 | 2153,46 | 1970 | 2337 | 1,185 | 0,244 | 2,717 | 0,006579 | 0,008746 | 0,0121298 | 0,00265 | 0,0072997 | 0,0072997 | TRUE | 0,1645378 | No |
| orf19.4964 | 1876 | 1723 | 1846 | 1046 | 1111 | 945 | 967 | 1303 | 1708 | 1743 | 1747 | 1571 | 988 | 1026 | 990 | 994 | 1345,93 | 1692 | 1000 | 0,589 | -0,763 | -14,957 | 1,40E-50 | 6,53E-50 | 0,0005264 | 0,003018 | 0,0017168 | 0,0017168 | TRUE | 0,0249669 | No |
| orf19.4965 | 1215 | 1400 | 1350 | 769 | 864 | 738 | 772 | 936 | 1106 | 1416 | 1278 | 1155 | 769 | 801 | 791 | 714 | 1003,67 | 1239 | 769 | 0,619 | -0,691 | -8,661 | 4,69E-18 | 1,13E-17 | 0,0067147 | 0,003353 | 0,0050738 | 0,0050738 | TRUE | 0,1119255 | No |
| orf19.4966 | 5207 | 5879 | 5875 | 3746 | 7627 | 7112 | 6577 | 8632 | 4741 | 5946 | 5560 | 5627 | 6784 | 7722 | 6736 | 6585 | 6212,48 | 5468 | 6957 | 1,272 | 0,347 | 4,808 | 1,52E-06 | 2,51E-06 | 0,0069549 | 0,00225 | 0,0048463 | 0,0048463 | TRUE | 0,0987483 | No |
| orf19.4967 | 2609 | 2352 | 2310 | 1396 | 2833 | 2228 | 2183 | 2999 | 2375 | 2379 | 2186 | 2097 | 2520 | 2419 | 2236 | 2288 | 2312,45 | 2259 | 2366 | 1,046 | 0,064 | 1,112 | 0,266138 | 0,302053 | 0,0028411 | 0,002608 | 0,0027846 | 0,0027846 | TRUE | 0,0266152 | No |
| orf19.4969 | 2959 | 3033 | 3107 | 1813 | 2568 | 2327 | 2263 | 2889 | 2694 | 3068 | 2940 | 2723 | 2284 | 2526 | 2318 | 2204 | 2594,69 | 2856 | 2333 | 0,816 | -0,294 | -4,988 | 6,11E-07 | 1,03E-06 | 0,0031739 | 0,002546 | 0,0029412 | 0,0029412 | TRUE | 0,037307 | No |
| orf19.497 | 2315 | 1901 | 2228 | 1353 | 2253 | 1714 | 2139 | 2755 | 2108 | 1923 | 2109 | 2032 | 2004 | 1861 | 2191 | 2102 | 2041,05 | 2043 | 2040 | 0,998 | -0,002 | -0,04 | 0,967802 | 0,971612 | 0,0028559 | 0,002684 | 0,0027789 | 0,0027789 | TRUE | 0,0407727 | No |
| orf19.4970 | 2740 | 2574 | 2731 | 1583 | 2392 | 2018 | 1960 | 2597 | 2495 | 2603 | 2585 | 2378 | 2128 | 2191 | 2007 | 1981 | 2295,93 | 2515 | 2077 | 0,824 | -0,279 | -5,571 | 2,54E-08 | 4,49E-08 | 0,0013982 | 0,002612 | 0,0019656 | 0,0019656 | TRUE | 0,0161921 | No |
| orf19.4972 | 258 | 263 | 275 | 168 | 37 | 28 | 42 | 52 | 235 | 266 | 260 | 252 | 33 | 30 | 43 | 40 | 144,94 | 253 | 36 | 0,145 | -2,789 | -18,486 | 2,65E-76 | 1,79E-75 | 1,00E-08 | 0,011151 | 0,0070993 | 0,0070993 | TRUE | 0,0866673 | No |
| orf19.4975 | 5466 | 4689 | 5034 | 3614 | 401 | 398 | 341 | 505 | 4977 | 4742 | 4764 | 5429 | 357 | 432 | 349 | 385 | 2679,41 | 4978 | 381 | 0,076 | -3,71 | -52,67 | 0 | 0 | 0,0050264 | 0,00253 | 0,0034296 | 0,0034296 | TRUE | 0,0865393 | No |
| orf19.4976 | 2 | 5 | 7 | 0 | 4 | 11 | 5 | 13 | 2 | 5 | 7 | 0 | 4 | 12 | 5 | 10 | 5,51 | 4 | 8 | 2,139 | 1,097 | 1,59 | 0,111932 | 0,133701 | 0,2577519 | 0,241997 | 0,2465388 | 0,2465388 | TRUE | 0,2468423 | No |
| orf19.4979 | 526 | 511 | 534 | 249 | 469 | 370 | 385 | 501 | 479 | 517 | 505 | 374 | 417 | 402 | 394 | 382 | 433,81 | 469 | 399 | 0,845 | -0,243 | -2,585 | 0,009732 | 0,012776 | 0,0072313 | 0,005082 | 0,0061487 | 0,0061487 | TRUE | 0,1871996 | No |
| orf19.498 | 3763 | 3372 | 3682 | 2370 | 3111 | 2647 | 2779 | 3623 | 3426 | 3410 | 3485 | 3560 | 2767 | 2874 | 2846 | 2764 | 3141,5 | 3470 | 2813 | 0,811 | -0,303 | -8,281 | 1,22E-16 | 2,82E-16 | 1,00E-08 | 0,002458 | 0,0009629 | 0,0009629 | TRUE | 0,0033584 | No |
| orf19.4980 | 61165 | 56134 | 59582 | 36289 | 14496 | 12330 | 12194 | 17364 | 55688 | 56774 | 56389 | 54512 | 12894 | 13387 | 12488 | 13245 | 34422,1 | 55841 | 13004 | 0,233 | -2,102 | -63,084 | 0 | 0 | 0,0005712 | 0,002075 | 0,0010214 | 0,0010214 | TRUE | 0,0086384 | No |
| orf19.4981 | 688 | 804 | 798 | 480 | 996 | 962 | 939 | 1135 | 626 | 813 | 755 | 721 | 886 | 1044 | 962 | 866 | 834,21 | 729 | 940 | 1,287 | 0,365 | 4,203 | 2,63E-05 | 4,09E-05 | 0,0078768 | 0,003621 | 0,0059854 | 0,0059854 | TRUE | 0,1120951 | No |
| orf19.4982 | 1212 | 1257 | 1341 | 732 | 802 | 769 | 780 | 994 | 1103 | 1271 | 1269 | 1100 | 713 | 835 | 799 | 758 | 981,1 | 1186 | 776 | 0,653 | -0,615 | -8,462 | 2,63E-17 | 6,21E-17 | 0,0046347 | 0,003383 | 0,0040161 | 0,0040161 | TRUE | 0,0351635 | No |
| orf19.4983 | 1643 | 1579 | 1663 | 988 | 1411 | 1046 | 1118 | 1504 | 1496 | 1597 | 1574 | 1484 | 1255 | 1136 | 1145 | 1147 | 1354,23 | 1538 | 1171 | 0,761 | -0,394 | -7,422 | 1,15E-13 | 2,46E-13 | 0,0010425 | 0,003012 | 0,0019577 | 0,0019577 | TRUE | 0,0285686 | No |
| orf19.4984 | 2986 | 2802 | 3027 | 1676 | 4092 | 3348 | 3502 | 4605 | 2719 | 2834 | 2865 | 2518 | 3640 | 3635 | 3586 | 3513 | 3163,61 | 2734 | 3594 | 1,312 | 0,392 | 8,407 | 4,22E-17 | 9,89E-17 | 0,0013588 | 0,002455 | 0,0017594 | 0,0017594 | TRUE | 0,0318119 | No |
| orf19.4985 | 594 | 657 | 644 | 317 | 714 | 636 | 681 | 817 | 541 | 664 | 609 | 476 | 635 | 691 | 697 | 623 | 617,15 | 572 | 662 | 1,149 | 0,201 | 2,167 | 0,030204 | 0,038197 | 0,009682 | 0,004177 | 0,0065984 | 0,0065984 | TRUE | 0,1344841 | No |
| orf19.4987 | 3956 | 3551 | 3934 | 2449 | 3082 | 2531 | 2911 | 3726 | 3602 | 3591 | 3723 | 3679 | 2741 | 2748 | 2981 | 2842 | 3238,49 | 3649 | 2828 | 0,775 | -0,367 | -8,965 | 3,11E-19 | 7,72E-19 | 0,0005143 | 0,002445 | 0,0012997 | 0,0012997 | TRUE | 0,0156669 | No |
| orf19.4988 | 1659 | 1646 | 1677 | 1023 | 1059 | 945 | 942 | 1184 | 1510 | 1665 | 1587 | 1537 | 942 | 1026 | 965 | 903 | 1266,86 | 1575 | 959 | 0,608 | -0,719 | -12,679 | 7,73E-37 | 2,87E-36 | 0,0015192 | 0,00308 | 0,0022559 | 0,0022559 | TRUE | 0,0257738 | No |
| orf19.499 | 1866 | 1576 | 1729 | 1004 | 1608 | 1201 | 1368 | 1850 | 1699 | 1594 | 1636 | 1508 | 1430 | 1304 | 1401 | 1411 | 1497,97 | 1609 | 1386 | 0,861 | -0,217 | -4,077 | 4,57E-05 | 6,98E-05 | 0,0011699 | 0,002919 | 0,002042 | 0,002042 | TRUE | 0,0196085 | No |
| orf19.4991 | 974 | 937 | 1036 | 692 | 3207 | 3095 | 3052 | 4089 | 887 | 948 | 980 | 1039 | 2853 | 3360 | 3126 | 3119 | 2039 | 964 | 3114 | 3,239 | 1,696 | 26,437 | 5,09E-154 | 6,80E-153 | 0,0035037 | 0,002685 | 0,0032402 | 0,0032402 | TRUE | 0,0389917 | No |
| orf19.4993 | 2522 | 2523 | 2644 | 1485 | 2463 | 1974 | 2171 | 2790 | 2296 | 2552 | 2502 | 2231 | 2191 | 2143 | 2223 | 2128 | 2283,31 | 2395 | 2171 | 0,905 | -0,144 | -2,759 | 0,005795 | 0,007732 | 0,0017334 | 0,002616 | 0,0021579 | 0,0021579 | TRUE | 0,023987 | No |
| orf19.4995 | 53 | 49 | 49 | 30 | 33 | 42 | 41 | 49 | 48 | 50 | 46 | 45 | 29 | 46 | 42 | 37 | 42,95 | 47 | 38 | 0,807 | -0,309 | -1,452 | 0,146363 | 0,172279 | 1,00E-08 | 0,032796 | 0,0199767 | 0,0199767 | TRUE | 0,1999366 | No |
| orf19.4996 | 30 | 42 | 39 | 26 | 39 | 24 | 23 | 43 | 27 | 42 | 37 | 39 | 35 | 26 | 24 | 33 | 32,86 | 36 | 30 | 0,816 | -0,294 | -1,174 | 0,240325 | 0,274466 | 1,00E-08 | 0,042241 | 0,0295075 | 0,0295075 | TRUE | 0,2341538 | No |
| orf19.4997 | 2097 | 1907 | 2072 | 1305 | 2891 | 2457 | 2547 | 3362 | 1909 | 1929 | 1961 | 1960 | 2571 | 2668 | 2608 | 2565 | 2271,41 | 1940 | 2603 | 1,342 | 0,424 | 10,979 | 4,81E-28 | 1,48E-27 | 1,00E-08 | 0,002619 | 0,0009775 | 0,0009775 | TRUE | 0,0032917 | No |
| orf19.4998 | 3675 | 3554 | 3875 | 2293 | 2990 | 2376 | 2585 | 3320 | 3346 | 3594 | 3667 | 3444 | 2660 | 2580 | 2647 | 2533 | 3058,91 | 3513 | 2605 | 0,741 | -0,432 | -9,936 | 2,91E-23 | 7,99E-23 | 0,0008722 | 0,002469 | 0,0014847 | 0,0014847 | TRUE | 0,0132379 | No |
| orf19.50 | 1863 | 1929 | 1919 | 1129 | 1390 | 1113 | 1189 | 1506 | 1696 | 1951 | 1816 | 1696 | 1236 | 1208 | 1218 | 1149 | 1496,31 | 1790 | 1203 | 0,671 | -0,575 | -9,927 | 3,19E-23 | 8,72E-23 | 0,0022255 | 0,00292 | 0,0025372 | 0,0025372 | TRUE | 0,0444203 | No |
| orf19.500 | 6365 | 5734 | 6214 | 3698 | 7782 | 6103 | 6490 | 8642 | 5795 | 5799 | 5881 | 5555 | 6922 | 6626 | 6647 | 6592 | 6227,14 | 5758 | 6697 | 1,162 | 0,217 | 6,238 | 4,44E-10 | 8,41E-10 | 0,0003402 | 0,002249 | 0,0009993 | 0,0009993 | TRUE | 0,0064343 | No |
| orf19.5000 | 4355 | 3924 | 4197 | 2534 | 16458 | 13149 | 14825 | 19103 | 3965 | 3969 | 3972 | 3806 | 14639 | 14276 | 15183 | 14572 | 9297,74 | 3928 | 14668 | 3,731 | 1,9 | 54,735 | 0 | 0 | 0,00033 | 0,002179 | 0,0009877 | 0,0009877 | TRUE | 0,0067845 | No |
| orf19.5001 | 2765 | 2430 | 2654 | 1619 | 1747 | 1450 | 1648 | 2061 | 2517 | 2458 | 2512 | 2432 | 1554 | 1574 | 1688 | 1572 | 2038,37 | 2480 | 1597 | 0,643 | -0,637 | -14,618 | 2,17E-48 | 9,71E-48 | 0,0001887 | 0,002685 | 0,0013116 | 0,0013116 | TRUE | 0,017188 | No |
| orf19.5003 | 1762 | 1798 | 1819 | 988 | 1374 | 1072 | 1171 | 1519 | 1604 | 1818 | 1722 | 1484 | 1222 | 1164 | 1199 | 1159 | 1421,54 | 1657 | 1186 | 0,714 | -0,485 | -7,605 | 2,86E-14 | 6,19E-14 | 0,0031755 | 0,002966 | 0,0031922 | 0,0031922 | TRUE | 0,0550637 | No |
| orf19.5004 | 373 | 350 | 368 | 194 | 359 | 329 | 324 | 420 | 340 | 354 | 348 | 291 | 319 | 357 | 332 | 320 | 332,75 | 333 | 332 | 0,989 | -0,017 | -0,195 | 0,845666 | 0,867716 | 0,0008728 | 0,006007 | 0,0040521 | 0,0040521 | TRUE | 0,0660208 | No |
| orf19.5005 | 3667 | 3324 | 3587 | 2231 | 18992 | 15313 | 15967 | 21705 | 3339 | 3362 | 3395 | 3351 | 16893 | 16626 | 16352 | 16557 | 9984,25 | 3362 | 16607 | 4,94 | 2,304 | 77,032 | 0 | 0 | 1,00E-08 | 0,002169 | 0,0006715 | 0,0006715 | TRUE | 0,0016502 | No |
| orf19.5006 | 2633 | 2420 | 2428 | 1608 | 2710 | 2178 | 2234 | 2997 | 2397 | 2448 | 2298 | 2415 | 2410 | 2365 | 2288 | 2286 | 2363,42 | 2390 | 2337 | 0,978 | -0,031 | -0,755 | 0,450244 | 0,492333 | 0,0003352 | 0,002596 | 0,0012435 | 0,0012435 | TRUE | 0,008372 | No |
| orf19.5006.1 | 4834 | 4694 | 4879 | 2610 | 4822 | 3946 | 3942 | 5260 | 4401 | 4747 | 4618 | 3921 | 4289 | 4284 | 4037 | 4012 | 4288,69 | 4422 | 4156 | 0,939 | -0,091 | -1,529 | 0,126167 | 0,149863 | 0,0037293 | 0,002345 | 0,0031689 | 0,0031689 | TRUE | 0,0686498 | No |
| orf19.5007 | 5516 | 4594 | 4905 | 3441 | 7628 | 6082 | 6247 | 8871 | 5022 | 4646 | 4642 | 5169 | 6785 | 6603 | 6398 | 6767 | 5754,04 | 4870 | 6638 | 1,364 | 0,448 | 9,699 | 3,04E-22 | 8,11E-22 | 0,001637 | 0,002267 | 0,0018692 | 0,0018692 | TRUE | 0,0199478 | No |
| orf19.5008.1 | 245 | 218 | 238 | 159 | 367 | 312 | 278 | 381 | 223 | 220 | 225 | 239 | 326 | 339 | 285 | 291 | 268,52 | 227 | 310 | 1,369 | 0,453 | 4,828 | 1,38E-06 | 2,28E-06 | 1,00E-08 | 0,006957 | 0,004576 | 0,004576 | TRUE | 0,0407681 | No |
| orf19.5009 | 1782 | 1729 | 1799 | 1030 | 2438 | 1958 | 2165 | 2808 | 1622 | 1749 | 1703 | 1547 | 2169 | 2126 | 2217 | 2142 | 1909,31 | 1655 | 2164 | 1,304 | 0,383 | 7,985 | 1,40E-15 | 3,15E-15 | 0,0007419 | 0,002729 | 0,0016737 | 0,0016737 | TRUE | 0,0206052 | No |
| orf19.501 | 16976 | 14359 | 15808 | 9554 | 25461 | 20083 | 22410 | 30155 | 15456 | 14523 | 14961 | 14352 | 22647 | 21805 | 22951 | 23002 | 18711,9 | 14823 | 22601 | 1,524 | 0,608 | 16,758 | 4,98E-63 | 2,84E-62 | 0,000781 | 0,002108 | 0,0012086 | 0,0012086 | TRUE | 0,0101496 | No |
| orf19.5010 | 3305 | 3009 | 3145 | 1814 | 3228 | 2466 | 2474 | 3388 | 3009 | 3043 | 2976 | 2725 | 2871 | 2677 | 2534 | 2584 | 2802,55 | 2938 | 2666 | 0,907 | -0,142 | -2,632 | 0,008478 | 0,011172 | 0,0022958 | 0,002508 | 0,0024186 | 0,0024186 | TRUE | 0,0327164 | No |
| orf19.5011 | 922 | 891 | 928 | 574 | 926 | 840 | 858 | 1054 | 839 | 901 | 878 | 862 | 824 | 912 | 879 | 804 | 862,43 | 870 | 855 | 0,98 | -0,029 | -0,487 | 0,626237 | 0,664712 | 0,001118 | 0,003569 | 0,00233 | 0,00233 | TRUE | 0,0239 | No |
| orf19.5012 | 185 | 193 | 160 | 98 | 311 | 161 | 250 | 281 | 168 | 195 | 151 | 147 | 277 | 175 | 256 | 214 | 198,01 | 165 | 230 | 1,389 | 0,474 | 3,249 | 0,001158 | 0,001629 | 0,0236601 | 0,008708 | 0,0152029 | 0,0152029 | TRUE | 0,2440675 | No |
| orf19.5013 | 277 | 286 | 300 | 248 | 335 | 290 | 257 | 362 | 252 | 289 | 284 | 373 | 298 | 315 | 263 | 276 | 293,76 | 300 | 288 | 0,969 | -0,045 | -0,393 | 0,693964 | 0,729074 | 0,0126925 | 0,006534 | 0,0090494 | 0,0090494 | TRUE | 0,2692999 | No |
| orf19.5014 | 203 | 172 | 179 | 118 | 544 | 390 | 408 | 558 | 185 | 174 | 169 | 177 | 484 | 423 | 418 | 426 | 307,03 | 176 | 438 | 2,484 | 1,312 | 14,181 | 1,21E-45 | 5,20E-45 | 1,00E-08 | 0,00634 | 0,004144 | 0,004144 | TRUE | 0,0595329 | No |

| | | | | | | | | | | | | | | | | | | | | | | | | | | | | | | | |
|---|---|---|---|---|---|---|---|---|---|---|---|---|---|---|---|---|---|---|---|---|---|---|---|---|---|---|---|---|---|---|---|
| orf19.5015 | 1362 | 1312 | 1292 | 798 | 1773 | 1472 | 1502 | 1996 | 1240 | 1327 | 1223 | 1199 | 1577 | 1598 | 1538 | 1523 | 1403,06 | 1247 | 1559 | 1,248 | 0,32 | 6,483 | 8,98E-11 | 1,75E-10 | 0,0002888 | 0,002979 | 0,0016043 | 0,0016043 | TRUE | 0,0221807 | No |
| orf19.5016 | 1137 | 1023 | 1016 | 765 | 533 | 412 | 416 | 531 | 1035 | 1035 | 962 | 1149 | 474 | 447 | 426 | 405 | 741,63 | 1045 | 438 | 0,42 | -1,252 | -16,903 | 4,26E-64 | 2,47E-63 | 0,0035407 | 0,003818 | 0,0036854 | 0,0036854 | TRUE | 0,0492247 | No |
| orf19.5017 | 70 | 63 | 61 | 42 | 98 | 103 | 101 | 146 | 64 | 64 | 58 | 63 | 87 | 112 | 103 | 111 | 82,76 | 62 | 103 | 1,668 | 0,738 | 4,648 | 3,35E-06 | 5,44E-06 | 1,00E-08 | 0,017999 | 0,0111289 | 0,0111289 | TRUE | 0,1166996 | No |
| orf19.5019 | 2799 | 2295 | 2572 | 1660 | 1866 | 1592 | 1653 | 2318 | 2548 | 2321 | 2434 | 2494 | 1660 | 1728 | 1693 | 1768 | 2080,82 | 2449 | 1712 | 0,699 | -0,516 | -11,138 | 8,18E-29 | 2,55E-28 | 0,0007886 | 0,002672 | 0,0015657 | 0,0015657 | TRUE | 0,0152986 | No |
| orf19.5020 | 6710 | 5853 | 6368 | 4469 | 7360 | 6098 | 6496 | 9143 | 6109 | 5920 | 6027 | 6713 | 6546 | 6621 | 6653 | 6974 | 6445,38 | 6192 | 6698 | 1,083 | 0,115 | 2,452 | 0,014224 | 0,018472 | 0,001843 | 0,002242 | 0,0019496 | 0,0019496 | TRUE | 0,0379298 | No |
| orf19.5021 | 5121 | 4428 | 4828 | 3118 | 5766 | 4804 | 5011 | 6824 | 4662 | 4478 | 4569 | 4684 | 5129 | 5216 | 5132 | 5205 | 4884,46 | 4598 | 5170 | 1,125 | 0,17 | 5,247 | 1,55E-07 | 2,65E-07 | 1,00E-08 | 0,002307 | 0,0007972 | 0,0007972 | TRUE | 0,0037466 | No |
| orf19.5022 | 4509 | 3939 | 4295 | 2809 | 3492 | 3115 | 3365 | 4595 | 4105 | 3984 | 4065 | 4220 | 3106 | 3382 | 3446 | 3505 | 3726,61 | 4094 | 3360 | 0,821 | -0,284 | -6,101 | 1,06E-09 | 1,97E-09 | 0,0013059 | 0,002391 | 0,0018133 | 0,0018133 | TRUE | 0,0318898 | No |
| orf19.5023 | 205 | 168 | 167 | 117 | 354 | 333 | 328 | 392 | 187 | 170 | 158 | 176 | 315 | 362 | 336 | 299 | 250,21 | 173 | 328 | 1,894 | 0,922 | 9,088 | 1,01E-19 | 2,55E-19 | 1,00E-08 | 0,007316 | 0,0053614 | 0,0053614 | TRUE | 0,053079 | No |
| orf19.5024 | 4833 | 4186 | 4469 | 2966 | 9077 | 7430 | 7828 | 11119 | 4400 | 4234 | 4230 | 4455 | 8074 | 8067 | 8017 | 8482 | 6244,75 | 4330 | 8160 | 1,886 | 0,915 | 24,861 | 1,99E-136 | 2,34E-135 | 0,0004709 | 0,002249 | 0,0011201 | 0,0011201 | TRUE | 0,0087711 | No |
| orf19.5025 | 14259 | 13157 | 13528 | 8291 | 30613 | 24334 | 24949 | 34463 | 12982 | 13307 | 12803 | 12454 | 27229 | 26420 | 25551 | 26289 | 19629,4 | 12886 | 26372 | 2,046 | 1,033 | 29,696 | 8,71E-194 | 1,44E-192 | 0,0006438 | 0,002104 | 0,0011022 | 0,0011022 | TRUE | 0,0060556 | No |
| orf19.5026 | 19789 | 18176 | 19551 | 12735 | 20611 | 18530 | 18675 | 25983 | 18017 | 18383 | 18503 | 19130 | 18333 | 20118 | 19126 | 19820 | 18928,8 | 18508 | 19349 | 1,046 | 0,064 | 1,629 | 0,103282 | 0,123927 | 0,0010869 | 0,002107 | 0,0014434 | 0,0014434 | TRUE | 0,0153412 | No |
| orf19.5027 | 3227 | 3039 | 3103 | 1829 | 6951 | 5764 | 6135 | 8400 | 2938 | 3074 | 2937 | 2747 | 6183 | 6258 | 6283 | 6408 | 4603,4 | 2924 | 6283 | 2,146 | 1,102 | 27,136 | 3,67E-162 | 5,15E-161 | 0,0006105 | 0,002324 | 0,001324 | 0,001324 | TRUE | 0,0182925 | No |
| orf19.5029 | 2356 | 2207 | 2459 | 1489 | 2646 | 2181 | 2194 | 2921 | 2145 | 2232 | 2327 | 2237 | 2354 | 2368 | 2247 | 2228 | 2267,21 | 2235 | 2299 | 1,028 | 0,04 | 0,903 | 0,366694 | 0,406552 | 0,0006681 | 0,00262 | 0,001462 | 0,001462 | TRUE | 0,0098536 | No |
| orf19.5030 | 1773 | 1568 | 1688 | 949 | 701 | 539 | 610 | 809 | 1614 | 1586 | 1598 | 1426 | 624 | 585 | 625 | 617 | 1084,22 | 1556 | 613 | 0,393 | -1,347 | -22,977 | 8,01E-117 | 7,98E-116 | 0,0011617 | 0,003255 | 0,0021947 | 0,0021947 | TRUE | 0,0346412 | No |
| orf19.5031 | 10198 | 8895 | 9985 | 6676 | 4893 | 4542 | 4568 | 6281 | 9285 | 8996 | 9450 | 10028 | 4352 | 4931 | 4678 | 4791 | 7064,05 | 9440 | 4688 | 0,497 | -1,009 | -20,162 | 2,12E-90 | 1,66E-89 | 0,0022338 | 0,002224 | 0,0022507 | 0,0022507 | TRUE | 0,0285053 | No |
| orf19.5032 | 33778 | 27708 | 29922 | 20112 | 79699 | 62769 | 70002 | 99986 | 30753 | 28024 | 28318 | 30211 | 70890 | 68150 | 71691 | 76270 | 50538,4 | 29326 | 71750 | 2,447 | 1,291 | 27,365 | 7,22E-165 | 1,04E-163 | 0,0021453 | 0,002063 | 0,0021131 | 0,0021131 | TRUE | 0,0220628 | No |
| orf19.5033 | 189 | 169 | 159 | 100 | 194 | 148 | 166 | 216 | 172 | 171 | 150 | 150 | 173 | 161 | 170 | 165 | 163,96 | 161 | 167 | 1,034 | 0,048 | 0,44 | 0,659649 | 0,696235 | 1,00E-08 | 0,010094 | 0,0055079 | 0,0055079 | TRUE | 0,025373 | No |
| orf19.5034 | 1087 | 1093 | 1144 | 787 | 1213 | 997 | 1069 | 1292 | 990 | 1105 | 1083 | 1182 | 1079 | 1082 | 1095 | 986 | 1075,22 | 1090 | 1060 | 0,975 | -0,037 | -0,577 | 0,563679 | 0,604755 | 0,0028513 | 0,003266 | 0,0030521 | 0,0030521 | TRUE | 0,0466078 | No |
| orf19.5035 | 1539 | 1335 | 1496 | 913 | 2478 | 1959 | 2349 | 3123 | 1401 | 1350 | 1416 | 1371 | 2204 | 2127 | 2406 | 2382 | 1832,21 | 1384 | 2280 | 1,646 | 0,719 | 13,502 | 1,51E-41 | 6,04E-41 | 0,0016693 | 0,002758 | 0,0021325 | 0,0021325 | TRUE | 0,0239869 | No |
| orf19.5037 | 6654 | 7434 | 7287 | 4734 | 5047 | 4496 | 4275 | 5476 | 6058 | 7519 | 6896 | 7111 | 4489 | 4881 | 4378 | 4177 | 5688,8 | 6896 | 4481 | 0,65 | -0,622 | -9,037 | 1,61E-19 | 4,02E-19 | 0,0059209 | 0,002269 | 0,0043665 | 0,0043665 | TRUE | 0,0822904 | No |
| orf19.5038 | 4898 | 5528 | 5744 | 3301 | 4912 | 4644 | 4368 | 5500 | 4459 | 5591 | 5436 | 4959 | 4369 | 5042 | 4473 | 4195 | 4815,65 | 5111 | 4520 | 0,884 | -0,178 | -2,335 | 0,019542 | 0,025086 | 0,0080094 | 0,002311 | 0,0053654 | 0,0053654 | TRUE | 0,0909831 | No |
| orf19.5039 | 3020 | 2804 | 2676 | 1682 | 2466 | 1918 | 1946 | 2717 | 2750 | 2836 | 2533 | 2527 | 2193 | 2082 | 1993 | 2073 | 2373,26 | 2662 | 2085 | 0,783 | -0,353 | -6,627 | 3,43E-11 | 6,78E-11 | 0,0019292 | 0,002594 | 0,0022929 | 0,0022929 | TRUE | 0,023497 | No |
| orf19.504 | 3426 | 3010 | 3318 | 2183 | 3064 | 2399 | 2776 | 3697 | 3119 | 3044 | 3140 | 3279 | 2725 | 2605 | 2843 | 2820 | 2947 | 3146 | 2748 | 0,875 | -0,193 | -4,391 | 1,13E-05 | 1,78E-05 | 0,0008098 | 0,002485 | 0,0015126 | 0,0015126 | TRUE | 0,0145551 | No |
| orf19.5040 | 4089 | 4330 | 4573 | 2453 | 2443 | 2052 | 2287 | 2752 | 3723 | 4379 | 4328 | 3685 | 2173 | 2228 | 2342 | 2099 | 3119,65 | 4029 | 2210 | 0,548 | -0,867 | -12,976 | 1,68E-38 | 6,38E-38 | 0,0049617 | 0,00246 | 0,0039455 | 0,0039455 | TRUE | 0,0415451 | No |
| orf19.5041 | 225 | 202 | 176 | 125 | 70 | 71 | 55 | 85 | 205 | 204 | 167 | 188 | 62 | 77 | 56 | 65 | 128 | 191 | 65 | 0,34 | -1,556 | -11,022 | 2,98E-28 | 9,18E-28 | 1,00E-08 | 0,012357 | 0,0091764 | 0,0091764 | TRUE | 0,1168016 | No |
| orf19.5042 | 59 | 87 | 77 | 45 | 23 | 22 | 19 | 29 | 54 | 88 | 73 | 68 | 20 | 24 | 19 | 22 | 46,01 | 71 | 21 | 0,304 | -1,716 | -7,249 | 4,19E-13 | 8,81E-13 | 1,00E-08 | 0,030746 | 0,0245529 | 0,0245529 | TRUE | 0,122815 | No |
| orf19.5043 | 203 | 195 | 209 | 112 | 124 | 89 | 112 | 123 | 185 | 197 | 198 | 168 | 110 | 97 | 115 | 94 | 145,44 | 187 | 104 | 0,551 | -0,859 | -7,01 | 2,38E-12 | 4,88E-12 | 1,00E-08 | 0,01112 | 0,0070703 | 0,0070703 | TRUE | 0,0504531 | No |
| orf19.5045 | 8709 | 9723 | 10201 | 5931 | 2096 | 1863 | 1896 | 2351 | 7929 | 9834 | 9654 | 8909 | 1864 | 2023 | 1942 | 1793 | 5493,59 | 9082 | 1906 | 0,21 | -2,254 | -32,547 | 2,34E-232 | 4,56E-231 | 0,0054724 | 0,002277 | 0,0043006 | 0,0043006 | TRUE | 0,0899202 | No |
| orf19.5046 | 270 | 284 | 293 | 180 | 266 | 205 | 202 | 278 | 246 | 287 | 277 | 270 | 237 | 223 | 207 | 212 | 244,86 | 270 | 220 | 0,813 | -0,298 | -3,13 | 0,001751 | 0,002427 | 1,00E-08 | 0,007432 | 0,0046184 | 0,0046184 | TRUE | 0,0461818 | No |
| orf19.5049 | 5125 | 4417 | 4635 | 2984 | 5117 | 3731 | 4166 | 5576 | 4666 | 4467 | 4387 | 4482 | 4551 | 4051 | 4267 | 4253 | 4390,58 | 4500 | 4280 | 0,951 | -0,072 | -1,616 | 0,106029 | 0,127079 | 0,0011961 | 0,002338 | 0,0016794 | 0,0016794 | TRUE | 0,0221251 | No |
| orf19.505 | 3397 | 2905 | 3202 | 2081 | 1918 | 1626 | 1705 | 2429 | 3093 | 2938 | 3030 | 3126 | 1706 | 1765 | 1746 | 1853 | 2407,21 | 3047 | 1768 | 0,581 | -0,784 | -17,91 | 9,83E-72 | 6,29E-71 | 0,0005704 | 0,002586 | 0,0013985 | 0,0013985 | TRUE | 0,013597 | No |
| orf19.5050 | 2426 | 2217 | 2349 | 1467 | 1836 | 1481 | 1598 | 2174 | 2209 | 2242 | 2223 | 2204 | 1633 | 1608 | 1637 | 1658 | 1926,72 | 2220 | 1634 | 0,736 | -0,441 | -11,243 | 2,52E-29 | 7,95E-29 | 1,00E-08 | 0,002723 | 0,0009532 | 0,0009532 | TRUE | 0,0013352 | No |
| orf19.5051 | 2815 | 2485 | 2680 | 1768 | 2492 | 2142 | 2245 | 3051 | 2563 | 2513 | 2536 | 2656 | 2217 | 2326 | 2299 | 2327 | 2429,64 | 2567 | 2292 | 0,894 | -0,162 | -4,061 | 4,90E-05 | 7,46E-05 | 3,61E-06 | 0,002581 | 0,0011155 | 0,0011155 | TRUE | 0,0060941 | No |
| orf19.5052 | 646 | 559 | 551 | 410 | 573 | 454 | 478 | 635 | 588 | 565 | 521 | 616 | 510 | 493 | 490 | 484 | 533,42 | 572 | 494 | 0,866 | -0,208 | -2,971 | 0,002967 | 0,004039 | 0,0008154 | 0,004513 | 0,0028378 | 0,0028378 | TRUE | 0,0432386 | No |
| orf19.5053 | 1292 | 1213 | 1340 | 844 | 318 | 329 | 283 | 376 | 1176 | 1227 | 1268 | 1268 | 283 | 357 | 290 | 287 | 769,48 | 1235 | 304 | 0,245 | -2,027 | -27,034 | 5,85E-161 | 8,13E-160 | 0,0028273 | 0,003754 | 0,0034215 | 0,0034215 | TRUE | 0,1452991 | No |
| orf19.5054 | 2484 | 2270 | 2239 | 1322 | 1980 | 1380 | 1530 | 2056 | 2262 | 2296 | 2119 | 1986 | 1761 | 1498 | 1567 | 1568 | 1882,12 | 2166 | 1598 | 0,738 | -0,439 | -6,842 | 7,79E-12 | 1,58E-11 | 0,0039352 | 0,002739 | 0,0034115 | 0,0034115 | TRUE | 0,0567259 | No |
| orf19.5056 | 162 | 131 | 131 | 91 | 172 | 136 | 158 | 151 | 147 | 132 | 124 | 137 | 153 | 148 | 162 | 115 | 139,79 | 135 | 144 | 1,063 | 0,088 | 0,656 | 0,511528 | 0,553649 | 0,0082934 | 0,011487 | 0,0101424 | 0,0101424 | TRUE | 0,2189137 | No |
| orf19.5057 | 18 | 22 | 14 | 8 | 20 | 12 | 14 | 29 | 16 | 22 | 13 | 12 | 18 | 13 | 14 | 22 | 16,4 | 16 | 17 | 1,055 | 0,077 | 0,221 | 0,825017 | 0,848989 | 1,00E-08 | 0,082596 | 0,0570292 | 0,0570292 | TRUE | 0,3222772 | No |
| orf19.5058 | 21958 | 19774 | 21530 | 12778 | 7462 | 6266 | 6634 | 8608 | 19992 | 19999 | 20376 | 19195 | 6637 | 6803 | 6794 | 6566 | 13295,3 | 19890 | 6700 | 0,337 | -1,571 | -48,294 | 0 | 0 | 0,0003331 | 0,002136 | 0,0009188 | 0,0009188 | TRUE | 0,0066432 | No |
| orf19.5059 | 9309 | 9485 | 10118 | 5908 | 10828 | 9341 | 9783 | 12314 | 8475 | 9593 | 9576 | 8875 | 9631 | 10142 | 10019 | 9393 | 9463,02 | 9130 | 9796 | 1,073 | 0,101 | 2,029 | 0,042469 | 0,052972 | 0,0022949 | 0,002177 | 0,0022862 | 0,0022862 | TRUE | 0,0289369 | No |
| orf19.506 | 9252 | 7847 | 8248 | 5234 | 14032 | 11325 | 12134 | 17058 | 8424 | 7936 | 7806 | 7862 | 12481 | 12296 | 12427 | 13012 | 10280,5 | 8007 | 12554 | 1,568 | 0,649 | 17,13 | 8,81E-66 | 5,24E-65 | 0,0008022 | 0,002166 | 0,0012723 | 0,0012723 | TRUE | 0,0152218 | No |
| orf19.5061 | 8545 | 7739 | 8296 | 5017 | 12015 | 10203 | 10384 | 14186 | 7780 | 7827 | 7851 | 7536 | 10687 | 11078 | 10635 | 10821 | 9276,88 | 7748 | 10805 | 1,394 | 0,479 | 15,475 | 5,11E-54 | 2,54E-53 | 0,0001944 | 0,002179 | 0,0008068 | 0,0008068 | TRUE | 0,003923 | No |
| orf19.5063 | 4013 | 3818 | 3650 | 2698 | 6664 | 5578 | 5123 | 7332 | 3654 | 3861 | 3454 | 4053 | 5927 | 6056 | 5247 | 5593 | 4730,68 | 3756 | 5706 | 1,52 | 0,605 | 9,907 | 3,87E-23 | 1,06E-22 | 0,0041015 | 0,002316 | 0,0033503 | 0,0033503 | TRUE | 0,0355312 | No |
| orf19.5064 | 4917 | 4810 | 4772 | 2908 | 3981 | 3240 | 3123 | 4303 | 4477 | 4865 | 4516 | 4368 | 3541 | 3518 | 3198 | 3282 | 3970,69 | 4556 | 3385 | 0,742 | -0,43 | -8,491 | 2,05E-17 | 4,87E-17 | 0,0020385 | 0,00237 | 0,0022016 | 0,0022016 | TRUE | 0,0252701 | No |
| orf19.5064.1 | 725 | 577 | 568 | 340 | 1353 | 881 | 986 | 1404 | 660 | 584 | 538 | 511 | 1203 | 957 | 1010 | 1071 | 816,59 | 573 | 1060 | 1,846 | 0,884 | 9,727 | 2,30E-22 | 6,17E-22 | 0,0097506 | 0,003655 | 0,0065597 | 0,0065597 | TRUE | 0,1249234 | No |
| orf19.5065 | 457 | 596 | 544 | 318 | 427 | 397 | 380 | 443 | 416 | 603 | 515 | 478 | 380 | 431 | 389 | 338 | 443,67 | 503 | 384 | 0,763 | -0,391 | -3,494 | 0,000475 | 0,000685 | 0,0150693 | 0,005014 | 0,0097204 | 0,0097204 | TRUE | 0,2125057 | No |
| orf19.5066 | 4123 | 3803 | 3849 | 2223 | 4653 | 3820 | 3986 | 4943 | 3754 | 3846 | 3643 | 3339 | 4139 | 4147 | 4082 | 3771 | 3840,13 | 3646 | 4035 | 1,105 | 0,144 | 2,705 | 0,006828 | 0,00907 | 0,002459 | 0,002381 | 0,0024685 | 0,0024685 | TRUE | 0,0371561 | No |
| orf19.5067 | 3528 | 3413 | 3382 | 1884 | 2979 | 2214 | 2381 | 3064 | 3212 | 3452 | 3201 | 2830 | 2650 | 2404 | 2438 | 2337 | 2815,5 | 3174 | 2457 | 0,773 | -0,371 | -5,82 | 5,90E-09 | 1,07E-08 | 0,0042983 | 0,002506 | 0,0035364 | 0,0035364 | TRUE | 0,062043 | No |
| orf19.5068 | 3256 | 3356 | 3443 | 2040 | 2263 | 1918 | 1985 | 2788 | 2964 | 3394 | 3258 | 3064 | 2013 | 2082 | 2033 | 2127 | 2617,06 | 3170 | 2064 | 0,651 | -0,62 | -12,022 | 2,72E-33 | 9,36E-33 | 0,0018841 | 0,002542 | 0,0021542 | 0,0021542 | TRUE | 0,027622 | No |
| orf19.5069 | 10880 | 10053 | 10907 | 6945 | 1602 | 1290 | 1429 | 1824 | 9906 | 10168 | 10322 | 10432 | 1425 | 1401 | 1463 | 1391 | 5813,58 | 10207 | 1420 | 0,139 | -2,846 | -74,783 | 0 | 0 | 1,00E-08 | 0,002264 | 0,0010092 | 0,0010092 | TRUE | 0,0049363 | No |
| orf19.5070 | 2148 | 1979 | 2207 | 1497 | 986 | 945 | 915 | 1316 | 1956 | 2002 | 2089 | 2249 | 877 | 1026 | 937 | 1004 | 1517,33 | 2074 | 961 | 0,464 | -1,108 | -17,203 | 2,50E-66 | 1,50E-65 | 0,0033931 | 0,002908 | 0,0032351 | 0,0032351 | TRUE | 0,0420814 | No |
| orf19.5071 | 218 | 190 | 241 | 161 | 286 | 263 | 310 | 319 | 198 | 192 | 228 | 242 | 254 | 286 | 317 | 243 | 245,17 | 215 | 275 | 1,281 | 0,358 | 3,148 | 0,001646 | 0,002285 | 0,0091973 | 0,007425 | 0,0081909 | 0,0081909 | TRUE | 0,1206413 | No |
| orf19.5073 | 733 | 629 | 674 | 383 | 617 | 467 | 515 | 666 | 667 | 636 | 638 | 575 | 549 | 507 | 527 | 508 | 576 | 629 | 523 | 0,828 | -0,273 | -4,024 | 5,71E-05 | 8,68E-05 | 0,0005609 | 0,00433 | 0,0026765 | 0,0026765 | TRUE | 0,0331667 | No |
| orf19.5074 | 984 | 913 | 896 | 590 | 879 | 700 | 807 | 1070 | 896 | 923 | 848 | 886 | 782 | 760 | 826 | 816 | 842,26 | 888 | 796 | 0,897 | -0,157 | -2,745 | 0,006051 | 0,008067 | 1,00E-08 | 0,003605 | 0,0019531 | 0,0019531 | TRUE | 0,012053 | No |
| orf19.5076 | 1550 | 1368 | 1384 | 924 | 2183 | 1739 | 1776 | 2422 | 1411 | 1384 | 1310 | 1388 | 1942 | 1888 | 1819 | 1848 | 1623,6 | 1373 | 1874 | 1,365 | 0,449 | 9,609 | 7,34E-22 | 1,93E-21 | 0,0003553 | 0,002851 | 0,0014593 | 0,0014593 | TRUE | 0,0122789 | No |
| orf19.5076.1 | 902 | 832 | 884 | 713 | 1411 | 1280 | 1184 | 1627 | 821 | 841 | 837 | 1071 | 1255 | 1390 | 1213 | 1241 | 1083,6 | 892 | 1275 | 1,434 | 0,52 | 6,231 | 4,63E-10 | 8,77E-10 | 0,0084256 | 0,003256 | 0,0057022 | 0,0057022 | TRUE | 0,2012819 | No |
| orf19.5077 | 3045 | 2617 | 2909 | 1806 | 5903 | 4847 | 5253 | 6952 | 2772 | 2647 | 2753 | 2713 | 5251 | 5262 | 5380 | 5303 | 4010,12 | 2721 | 5299 | 1,947 | 0,961 | 27,99 | 2,17E-172 | 3,25E-171 | 1,00E-08 | 0,002366 | 0,0008463 | 0,0008463 | TRUE | 0,0043549 | No |
| orf19.5078 | 792 | 657 | 713 | 417 | 694 | 530 | 624 | 799 | 721 | 664 | 675 | 626 | 617 | 575 | 639 | 609 | 641 | 672 | 610 | 0,906 | -0,142 | -2,164 | 0,030499 | 0,038539 | 0,0008005 | 0,004098 | 0,0026022 | 0,0026022 | TRUE | 0,0291008 | No |
| orf19.5079 | 81762 | 70731 | 78184 | 51925 | 8264 | 7290 | 7675 | 10382 | 74441 | 71537 | 73994 | 78000 | 7351 | 7915 | 7860 | 7919 | 41127,1 | 74493 | 7761 | 0,104 | -3,263 | -80,555 | 0 | 0 | 0,0011728 | 0,002069 | 0,0015085 | 0,0015085 | TRUE | 0,015583 | No |
| orf19.5079.1 | 2261 | 1808 | 2037 | 1200 | 4678 | 3407 | 3925 | 4994 | 2059 | 1829 | 1928 | 1803 | 4161 | 3699 | 4020 | 3809 | 2913,34 | 1905 | 3922 | 2,057 | 1,04 | 18,42 | 9,07E-76 | 6,09E-75 | 0,0027351 | 0,00249 | 0,0026622 | 0,0026622 | TRUE | 0,0363533 | No |
| orf19.508 | 2538 | 2303 | 2476 | 1642 | 514 | 449 | 473 | 627 | 2311 | 2329 | 2343 | 2467 | 457 | 487 | 484 | 478 | 1419,65 | 2362 | 476 | 0,202 | -2,308 | -43,559 | 0 | 0 | 1,00E-08 | 0,002968 | 0,0014921 | 0,0014921 | TRUE | 0,0097568 | No |
| orf19.5081 | 10991 | 9044 | 9989 | 6339 | 21437 | 17554 | 20083 | 27225 | 10007 | 9147 | 9454 | 9522 | 19068 | 19059 | 20568 | 20767 | 14698,9 | 9532 | 19866 | 2,084 | 1,059 | 23,474 | 7,52E-122 | 7,74E-121 | 0,0016888 | 0,002127 | 0,0018761 | 0,0018761 | TRUE | 0,0139071 | No |
| orf19.5083 | 3301 | 2642 | 3015 | 1933 | 4637 | 3649 | 4013 | 5646 | 3005 | 2672 | 2853 | 2904 | 4124 | 3962 | 4110 | 4307 | 3492,19 | 2858 | 4126 | 1,444 | 0,53 | 11,171 | 5,66E-29 | 1,77E-28 | 0,0013829 | 0,002415 | 0,0018598 | 0,0018598 | TRUE | 0,0238488 | No |
| orf19.5085 | 4489 | 4023 | 4373 | 2870 | 7656 | 6581 | 7070 | 9364 | 4087 | 4069 | 4139 | 4311 | 6810 | 7145 | 7241 | 7143 | 5618,02 | 4152 | 7085 | 1,708 | 0,772 | 20,885 | 7,38E-97 | 6,22E-96 | 0,0004549 | 0,002272 | 0,0011168 | 0,0011168 | TRUE | 0,0083708 | No |
| orf19.5087 | 3086 | 2535 | 2895 | 1682 | 1323 | 1033 | 1171 | 1637 | 2810 | 2564 | 2740 | 2527 | 1177 | 1122 | 1199 | 1249 | 1923,29 | 2660 | 1187 | 0,446 | -1,165 | -21,594 | 2,04E-103 | 1,83E-102 | 0,0017805 | 0,002724 | 0,0021973 | 0,0021973 | TRUE | 0,0174348 | No |
| orf19.5089 | 427 | 438 | 476 | 309 | 511 | 358 | 442 | 538 | 389 | 443 | 450 | 464 | 455 | 389 | 453 | 410 | 431,58 | 436 | 427 | 0,98 | -0,029 | -0,342 | 0,732714 | 0,764868 | 0,0035411 | 0,005098 | 0,0044012 | 0,0044012 | TRUE | 0,0662405 | No |
| orf19.5090 | 566 | 471 | 515 | 362 | 510 | 339 | 442 | 620 | 515 | 476 | 487 | 544 | 454 | 368 | 453 | 473 | 471,27 | 506 | 437 | 0,868 | -0,204 | -2,36 | 0,018255 | 0,023491 | 0,0052468 | 0,00484 | 0,0050198 | 0,0050198 | TRUE | 0,1237646 | No |

| | | | | | | | | | | | | | | | | | | | | | | | | | | | | | | | |
|---|---|---|---|---|---|---|---|---|---|---|---|---|---|---|---|---|---|---|---|---|---|---|---|---|---|---|---|---|---|---|---|
| orf19.5092 | 19 | 19 | 16 | 9 | 7 | 4 | 4 | 8 | 17 | 19 | 15 | 14 | 6 | 4 | 4 | 6 | 10,74 | 16 | 5 | 0,321 | -1,639 | -3,689 | 0,000225 | 0,000332 | 1,00E-08 | 0,124999 | 0,0691756 | 0,0691756 | TRUE | 0,0658099 | No |
| orf19.5093 | 3272 | 2830 | 3066 | 2003 | 2989 | 2531 | 2709 | 3469 | 2979 | 2862 | 2902 | 3009 | 2659 | 2748 | 2774 | 2646 | 2822,36 | 2938 | 2707 | 0,921 | -0,118 | -3,046 | 0,00232 | 0,003185 | 0,000184 | 0,002505 | 0,0010905 | 0,0010905 | TRUE | 0,0035882 | No |
| orf19.5094 | 1565 | 1401 | 1487 | 890 | 466 | 415 | 463 | 539 | 1425 | 1417 | 1407 | 1337 | 414 | 451 | 474 | 411 | 917,06 | 1396 | 438 | 0,312 | -1,681 | -27,13 | 4,33E-162 | 6,05E-161 | 0,0004048 | 0,003477 | 0,0022333 | 0,0022333 | TRUE | 0,0368857 | No |
| orf19.5095 | 5115 | 4358 | 4944 | 2903 | 5226 | 4514 | 4962 | 6611 | 4657 | 4408 | 4679 | 4361 | 4648 | 4901 | 5082 | 5043 | 4722,3 | 4526 | 4918 | 1,086 | 0,119 | 2,691 | 0,007125 | 0,009445 | 0,0011785 | 0,002317 | 0,0016632 | 0,0016632 | TRUE | 0,0168257 | No |
| orf19.5097 | 740 | 683 | 731 | 424 | 1553 | 1257 | 1446 | 1936 | 674 | 691 | 692 | 637 | 1381 | 1365 | 1481 | 1477 | 1049,63 | 674 | 1426 | 2,113 | 1,08 | 19,034 | 8,88E-81 | 6,35E-80 | 0,0007446 | 0,003296 | 0,0019695 | 0,0019695 | TRUE | 0,0129121 | No |
| orf19.5098 | 1832 | 1465 | 1633 | 1049 | 3330 | 2507 | 2724 | 3821 | 1668 | 1482 | 1545 | 1576 | 2962 | 2722 | 2790 | 2915 | 2207,39 | 1568 | 2847 | 1,817 | 0,861 | 16,921 | 3,15E-64 | 1,83E-63 | 0,0013168 | 0,002635 | 0,0019806 | 0,0019806 | TRUE | 0,0238292 | No |
| orf19.51 | 2375 | 2738 | 2647 | 1457 | 2623 | 2314 | 2213 | 2826 | 2162 | 2769 | 2505 | 2189 | 2333 | 2512 | 2266 | 2156 | 2361,61 | 2406 | 2317 | 0,962 | -0,056 | -0,703 | 0,481906 | 0,523722 | 0,0080572 | 0,002596 | 0,0056856 | 0,0056856 | TRUE | 0,1249576 | No |
| orf19.510 | 44 | 41 | 42 | 19 | 59 | 47 | 49 | 66 | 40 | 41 | 40 | 29 | 52 | 51 | 50 | 50 | 44,23 | 38 | 51 | 1,341 | 0,423 | 2,064 | 0,039061 | 0,048845 | 1,00E-08 | 0,031903 | 0,0172809 | 0,0172809 | TRUE | 0,1219956 | No |
| orf19.5100 | 3541 | 3111 | 3497 | 2178 | 5669 | 4919 | 5282 | 6749 | 3224 | 3146 | 3310 | 3272 | 5042 | 5341 | 5409 | 5148 | 4236,54 | 3238 | 5235 | 1,617 | 0,693 | 17,725 | 2,67E-70 | 1,69E-69 | 0,0005786 | 0,002349 | 0,0012117 | 0,0012117 | TRUE | 0,0074431 | No |
| orf19.5101 | 4760 | 4154 | 4725 | 2954 | 4788 | 3873 | 4009 | 5558 | 4334 | 4201 | 4472 | 4437 | 4259 | 4205 | 4106 | 4240 | 4281,68 | 4361 | 4202 | 0,964 | -0,053 | -1,46 | 0,144235 | 0,169868 | 0,0002851 | 0,002346 | 0,0010188 | 0,0010188 | TRUE | 0,0074472 | No |
| orf19.5102 | 1683 | 1473 | 1590 | 971 | 2334 | 1989 | 2096 | 2916 | 1532 | 1490 | 1505 | 1459 | 2076 | 2159 | 2147 | 2224 | 1823,99 | 1496 | 2152 | 1,437 | 0,523 | 11,927 | 8,60E-33 | 2,93E-32 | 1,00E-08 | 0,002761 | 0,0012691 | 0,0012691 | TRUE | 0,0070425 | No |
| orf19.5103 | 569 | 470 | 517 | 356 | 599 | 445 | 534 | 744 | 518 | 475 | 489 | 535 | 533 | 483 | 547 | 568 | 518,48 | 504 | 533 | 1,061 | 0,085 | 1,151 | 0,249563 | 0,284609 | 0,0017114 | 0,004585 | 0,0032906 | 0,0032906 | TRUE | 0,043405 | No |
| orf19.5104 | 1753 | 1438 | 1462 | 1260 | 2810 | 2267 | 2130 | 3169 | 1596 | 1454 | 1384 | 1893 | 2499 | 2461 | 2181 | 2417 | 1985,78 | 1582 | 2390 | 1,514 | 0,599 | 6,963 | 3,32E-12 | 6,77E-12 | 0,0112401 | 0,002702 | 0,0065644 | 0,0065644 | TRUE | 0,2032235 | No |
| orf19.5105 | 3309 | 2854 | 3186 | 2153 | 2980 | 2674 | 2887 | 4114 | 3013 | 2887 | 3015 | 3234 | 2651 | 2903 | 2957 | 3138 | 2974,66 | 3037 | 2912 | 0,96 | -0,059 | -1,018 | 0,308506 | 0,346987 | 0,0030711 | 0,002481 | 0,0028987 | 0,0028987 | TRUE | 0,0450775 | No |
| orf19.5106 | 16350 | 14711 | 16180 | 9220 | 16027 | 13153 | 14868 | 19259 | 14886 | 14879 | 15313 | 13850 | 14256 | 14280 | 15227 | 14691 | 14672,6 | 14732 | 14614 | 0,992 | -0,012 | -0,292 | 0,769926 | 0,800176 | 0,0011317 | 0,002127 | 0,0015851 | 0,0015851 | TRUE | 0,0195203 | No |
| orf19.5107 | 5907 | 4916 | 5484 | 3731 | 5129 | 4288 | 4663 | 6364 | 5378 | 4972 | 5190 | 5605 | 4562 | 4656 | 4775 | 4855 | 4999,05 | 5286 | 4712 | 0,892 | -0,165 | -3,635 | 0,000278 | 0,000407 | 0,0014064 | 0,002301 | 0,001772 | 0,001772 | TRUE | 0,019352 | No |
| orf19.511 | 2728 | 2347 | 2506 | 1613 | 2444 | 1761 | 1959 | 2672 | 2484 | 2374 | 2372 | 2423 | 2174 | 1912 | 2006 | 2038 | 2222,81 | 2413 | 2032 | 0,843 | -0,246 | -5,095 | 3,48E-07 | 5,89E-07 | 0,0010562 | 0,002631 | 0,0017941 | 0,0017941 | TRUE | 0,0264121 | No |
| orf19.5110 | 6078 | 5183 | 5939 | 3857 | 8963 | 7703 | 8520 | 11453 | 5534 | 5242 | 5621 | 5794 | 7972 | 8363 | 8726 | 8736 | 6998,5 | 5548 | 8449 | 1,524 | 0,608 | 13,257 | 4,12E-40 | 1,60E-39 | 0,0015839 | 0,002226 | 0,0018654 | 0,0018654 | TRUE | 0,0177726 | No |
| orf19.5112 | 5727 | 4746 | 5123 | 3670 | 9088 | 7770 | 7726 | 11569 | 5214 | 4800 | 4848 | 5513 | 8083 | 8436 | 7912 | 8825 | 6704,06 | 5094 | 8314 | 1,633 | 0,708 | 12,896 | 4,75E-38 | 1,79E-37 | 0,0031334 | 0,002234 | 0,0027325 | 0,0027325 | TRUE | 0,0363072 | No |
| orf19.5113 | 4356 | 3074 | 3294 | 2418 | 1941 | 1448 | 1562 | 2311 | 3966 | 3109 | 3117 | 3632 | 1726 | 1572 | 1600 | 1763 | 2560,72 | 3456 | 1665 | 0,482 | -1,052 | -13,177 | 1,19E-39 | 4,59E-39 | 0,0083245 | 0,002553 | 0,0056884 | 0,0056884 | TRUE | 0,1221979 | No |
| orf19.5114 | 1064 | 894 | 926 | 509 | 580 | 364 | 500 | 657 | 969 | 904 | 876 | 765 | 516 | 395 | 512 | 501 | 679,78 | 878 | 481 | 0,547 | -0,87 | -9,274 | 1,80E-20 | 4,62E-20 | 0,0097243 | 0,00398 | 0,0068629 | 0,0068629 | TRUE | 0,1610033 | No |
| orf19.5114.1 | 22816 | 16304 | 17523 | 9896 | 3679 | 2444 | 2891 | 4106 | 20773 | 16490 | 16584 | 14865 | 3272 | 2654 | 2961 | 3132 | 10091,3 | 17178 | 3005 | 0,175 | -2,515 | -27,232 | 2,70E-163 | 3,83E-162 | 0,0144613 | 0,002168 | 0,0080081 | 0,0080081 | TRUE | 0,243099 | No |
| orf19.5117 | 2E+05 | 1E+05 | 1E+05 | 95503 | 2E+05 | 2E+05 | 2E+05 | 2E+05 | 1E+05 | 1E+05 | 1E+05 | 1E+05 | 2E+05 | 2E+05 | 2E+05 | 2E+05 | 156350 | 138302 | 174398 | 1,261 | 0,335 | 8,124 | 4,50E-16 | 1,02E-15 | 0,0014012 | 0,002045 | 0,0016234 | 0,0016234 | TRUE | 0,013431 | No |
| orf19.5118 | 48886 | 42179 | 46387 | 31412 | 7728 | 6805 | 6954 | 10014 | 44509 | 42660 | 43901 | 47186 | 6874 | 7388 | 7122 | 7639 | 25909,7 | 44564 | 7256 | 0,163 | -2,618 | -56,992 | 0 | 0 | 0,0018503 | 0,002088 | 0,0019513 | 0,0019513 | TRUE | 0,019127 | No |
| orf19.512 | 9834 | 8762 | 9776 | 6272 | 12824 | 11317 | 11605 | 15610 | 8953 | 8862 | 9252 | 9422 | 11407 | 12287 | 11885 | 11907 | 10496,9 | 9122 | 11872 | 1,302 | 0,38 | 10,245 | 1,24E-24 | 3,53E-24 | 0,0007385 | 0,002163 | 0,0012253 | 0,0012253 | TRUE | 0,0085082 | No |
| orf19.5121 | 54 | 68 | 69 | 24 | 41 | 42 | 39 | 61 | 49 | 69 | 65 | 36 | 36 | 46 | 40 | 47 | 48,48 | 55 | 42 | 0,758 | -0,4 | -1,797 | 0,072386 | 0,088217 | 0,022292 | 0,029286 | 0,0268172 | 0,0268172 | TRUE | 0,2529029 | No |
| orf19.5124 | 1514 | 1207 | 1404 | 1028 | 1961 | 1714 | 1720 | 2711 | 1378 | 1221 | 1329 | 1544 | 1744 | 1861 | 1761 | 2068 | 1613,35 | 1368 | 1858 | 1,362 | 0,446 | 5,809 | 6,28E-09 | 1,14E-08 | 0,007198 | 0,002856 | 0,0050071 | 0,0050071 | TRUE | 0,0852692 | No |
| orf19.5125 | 6690 | 6414 | 6442 | 4394 | 6995 | 6337 | 5895 | 8200 | 6091 | 6487 | 6097 | 6600 | 6222 | 6880 | 6037 | 6255 | 6333,7 | 6319 | 6348 | 1,005 | 0,007 | 0,141 | 0,887485 | 0,903355 | 0,0020297 | 0,002246 | 0,0022881 | 0,0022881 | TRUE | 0,0383778 | No |
| orf19.5126 | 2656 | 2442 | 2515 | 1882 | 2141 | 2043 | 1889 | 2828 | 2418 | 2470 | 2380 | 2827 | 1904 | 2218 | 1935 | 2157 | 2288,69 | 2524 | 2054 | 0,815 | -0,295 | -4,263 | 2,01E-05 | 3,15E-05 | 0,0056596 | 0,002614 | 0,0041715 | 0,0041715 | TRUE | 0,0765517 | No |
| orf19.5129 | 423 | 408 | 448 | 260 | 768 | 644 | 624 | 810 | 385 | 413 | 424 | 391 | 683 | 699 | 639 | 618 | 531,45 | 403 | 660 | 1,633 | 0,707 | 9,801 | 1,11E-22 | 2,99E-22 | 0,0012186 | 0,004523 | 0,0029661 | 0,0029661 | TRUE | 0,0232381 | No |
| orf19.513 | 0 | 2 | 4 | 1 | 9 | 3 | 3 | 5 | 0 | 2 | 4 | 2 | 8 | 3 | 3 | 4 | 3,18 | 2 | 4 | 2,489 | 1,316 | 1,533 | 0,125398 | 0,148977 | 1,00E-08 | 0,417129 | 0,316646 | 0,316646 | TRUE | 0,5512728 | No |
| orf19.5130 | 3746 | 3192 | 3354 | 2272 | 4712 | 3997 | 4011 | 5712 | 3411 | 3228 | 3174 | 3413 | 4191 | 4340 | 4108 | 4357 | 3777,73 | 3306 | 4249 | 1,286 | 0,363 | 8,665 | 4,52E-18 | 1,09E-17 | 0,0007377 | 0,002387 | 0,001411 | 0,001411 | TRUE | 0,0089183 | No |
| orf19.5131 | 5690 | 5102 | 5669 | 3449 | 1460 | 1271 | 1263 | 1790 | 5181 | 5160 | 5365 | 5181 | 1299 | 1380 | 1293 | 1365 | 3278,03 | 5222 | 1334 | 0,256 | -1,968 | -48,478 | 0 | 0 | 0,0002005 | 0,00244 | 0,0011305 | 0,0011305 | TRUE | 0,0057257 | No |
| orf19.5132 | 112 | 126 | 116 | 72 | 93 | 66 | 76 | 115 | 102 | 127 | 110 | 108 | 83 | 72 | 78 | 88 | 95,91 | 112 | 80 | 0,719 | -0,476 | -3,271 | 0,001073 | 0,001513 | 1,00E-08 | 0,01581 | 0,0098181 | 0,0098181 | TRUE | 0,0917897 | No |
| orf19.5133 | 4115 | 3688 | 4043 | 2614 | 2443 | 2127 | 2193 | 3062 | 3747 | 3730 | 3826 | 3927 | 2173 | 2309 | 2246 | 2336 | 3036,68 | 3808 | 2266 | 0,596 | -0,748 | -18,444 | 5,85E-76 | 3,93E-75 | 0,0003225 | 0,002472 | 0,0012295 | 0,0012295 | TRUE | 0,0095171 | No |
| orf19.5134 | 2455 | 2342 | 2499 | 1475 | 2334 | 1882 | 2125 | 2700 | 2235 | 2369 | 2365 | 2216 | 2076 | 2043 | 2176 | 2060 | 2192,48 | 2296 | 2089 | 0,909 | -0,138 | -3,116 | 0,001835 | 0,002539 | 0,0005425 | 0,00264 | 0,0014286 | 0,0014286 | TRUE | 0,009375 | No |
| orf19.5136 | 1439 | 1312 | 1395 | 824 | 1332 | 954 | 1094 | 1464 | 1310 | 1327 | 1320 | 1238 | 1185 | 1036 | 1120 | 1117 | 1206,6 | 1299 | 1114 | 0,858 | -0,222 | -4,049 | 5,15E-05 | 7,84E-05 | 0,0008925 | 0,003132 | 0,0020456 | 0,0020456 | TRUE | 0,0257271 | No |
| orf19.5137 | 12 | 19 | 14 | 9 | 59 | 42 | 33 | 71 | 11 | 19 | 13 | 14 | 52 | 46 | 34 | 54 | 30,37 | 14 | 46 | 3,295 | 1,72 | 5,861 | 4,59E-09 | 8,38E-09 | 1,00E-08 | 0,045536 | 0,0354014 | 0,0354014 | TRUE | 0,2633815 | No |
| orf19.5137.1 | 1841 | 1524 | 1549 | 1120 | 1908 | 1706 | 1625 | 2221 | 1676 | 1541 | 1466 | 1682 | 1697 | 1852 | 1664 | 1694 | 1659,21 | 1591 | 1727 | 1,086 | 0,119 | 2 | 0,045536 | 0,056576 | 0,0026196 | 0,002833 | 0,0027808 | 0,0027808 | TRUE | 0,0345211 | No |
| orf19.5138 | 336 | 350 | 342 | 206 | 540 | 466 | 495 | 646 | 306 | 354 | 324 | 309 | 480 | 506 | 507 | 493 | 409,87 | 323 | 496 | 1,533 | 0,616 | 8,1 | 5,52E-16 | 1,25E-15 | 1,00E-08 | 0,00526 | 0,0029656 | 0,0029656 | TRUE | 0,0472161 | No |
| orf19.5139 | 2678 | 2458 | 2651 | 1496 | 2208 | 1779 | 1977 | 2554 | 2438 | 2486 | 2509 | 2247 | 1964 | 1931 | 2025 | 1948 | 2193,59 | 2420 | 1967 | 0,811 | -0,302 | -6,487 | 8,76E-11 | 1,70E-10 | 0,0008068 | 0,002639 | 0,0016179 | 0,0016179 | TRUE | 0,0256262 | No |
| orf19.514 | 861 | 806 | 796 | 514 | 1147 | 922 | 971 | 1336 | 784 | 815 | 753 | 772 | 1020 | 1001 | 994 | 1019 | 894,92 | 781 | 1008 | 1,292 | 0,369 | 6,929 | 4,23E-12 | 8,61E-12 | 1,00E-08 | 0,003513 | 0,0015783 | 0,0015783 | TRUE | 0,0105256 | No |
| orf19.5140 | 1597 | 1337 | 1486 | 903 | 115 | 112 | 109 | 135 | 1454 | 1352 | 1406 | 1356 | 102 | 122 | 112 | 103 | 750,94 | 1392 | 110 | 0,078 | -3,678 | -43,119 | 0 | 0 | 1,00E-08 | 0,003796 | 0,0023463 | 0,0023463 | TRUE | 0,0493289 | No |
| orf19.5141 | 15 | 25 | 20 | 18 | 61 | 54 | 63 | 62 | 14 | 25 | 19 | 27 | 54 | 59 | 65 | 47 | 38,7 | 21 | 56 | 2,698 | 1,432 | 5,663 | 1,48E-08 | 2,65E-08 | 0,007787 | 0,03617 | 0,0274347 | 0,0274347 | TRUE | 0,3632959 | No |
| orf19.5142 | 81 | 86 | 82 | 52 | 120 | 92 | 112 | 145 | 74 | 87 | 78 | 78 | 107 | 100 | 115 | 111 | 93,55 | 79 | 108 | 1,367 | 0,451 | 3,151 | 0,001625 | 0,002259 | 1,00E-08 | 0,016158 | 0,0086774 | 0,0086774 | TRUE | 0,0444092 | No |
| orf19.5143 | 3716 | 2939 | 3394 | 2010 | 5987 | 4847 | 5416 | 7225 | 3383 | 2972 | 3212 | 3019 | 5325 | 5262 | 5547 | 5511 | 4279,11 | 3146 | 5411 | 1,718 | 0,781 | 16,1 | 2,56E-58 | 1,37E-57 | 0,0017921 | 0,002346 | 0,0020015 | 0,0020015 | TRUE | 0,0315401 | No |
| orf19.5144 | 182 | 142 | 166 | 112 | 147 | 136 | 136 | 182 | 166 | 144 | 157 | 168 | 131 | 148 | 139 | 139 | 148,9 | 159 | 139 | 0,878 | -0,188 | -1,635 | 0,102143 | 0,122607 | 1,00E-08 | 0,010909 | 0,006019 | 0,006019 | TRUE | 0,0429913 | No |
| orf19.5145 | 482 | 429 | 468 | 289 | 349 | 279 | 281 | 376 | 439 | 434 | 443 | 434 | 310 | 303 | 288 | 287 | 367,21 | 438 | 297 | 0,678 | -0,56 | -7,427 | 1,11E-13 | 2,37E-13 | 1,00E-08 | 0,005634 | 0,0026788 | 0,0026788 | TRUE | 0,0113697 | No |
| orf19.5147 | 1647 | 1644 | 1602 | 1001 | 2410 | 2042 | 1979 | 2657 | 1500 | 1663 | 1516 | 1504 | 2144 | 2217 | 2027 | 2027 | 1824,53 | 1546 | 2104 | 1,36 | 0,443 | 8,33 | 8,06E-17 | 1,87E-16 | 0,0015302 | 0,002761 | 0,0021501 | 0,0021501 | TRUE | 0,0318759 | No |
| orf19.5148 | 4093 | 3575 | 3891 | 2390 | 3066 | 2544 | 2648 | 3689 | 3727 | 3616 | 3682 | 3590 | 2727 | 2762 | 2712 | 2814 | 3203,74 | 3654 | 2754 | 0,753 | -0,408 | -11,456 | 2,20E-30 | 7,16E-30 | 1,00E-08 | 0,002449 | 0,0009034 | 0,0009034 | TRUE | 0,0027955 | No |
| orf19.515 | 489 | 393 | 450 | 282 | 317 | 207 | 250 | 324 | 445 | 397 | 426 | 424 | 282 | 225 | 256 | 247 | 337,76 | 423 | 252 | 0,598 | -0,743 | -8,478 | 2,29E-17 | 5,41E-17 | 0,0016874 | 0,005948 | 0,0042592 | 0,0042592 | TRUE | 0,0695992 | No |
| orf19.5155 | 340 | 282 | 292 | 203 | 309 | 239 | 273 | 388 | 310 | 285 | 276 | 305 | 275 | 259 | 280 | 296 | 285,74 | 294 | 278 | 0,948 | -0,078 | -0,886 | 0,375806 | 0,416004 | 1,00E-08 | 0,00666 | 0,0038753 | 0,0038753 | TRUE | 0,0247228 | No |
| orf19.5156 | 708 | 652 | 604 | 442 | 1095 | 830 | 955 | 1256 | 645 | 659 | 572 | 664 | 974 | 901 | 978 | 958 | 793,86 | 635 | 953 | 1,505 | 0,59 | 9,157 | 5,36E-20 | 1,36E-19 | 0,0014191 | 0,003701 | 0,0026476 | 0,0026476 | TRUE | 0,0552581 | No |
| orf19.5157 | 260 | 266 | 262 | 161 | 443 | 364 | 322 | 500 | 237 | 269 | 248 | 242 | 394 | 395 | 330 | 381 | 312 | 249 | 375 | 1,507 | 0,592 | 6,502 | 7,93E-11 | 1,55E-10 | 0,0022586 | 0,006271 | 0,0045709 | 0,0045709 | TRUE | 0,0727201 | No |
| orf19.5158 | 1254 | 1225 | 1186 | 810 | 703 | 608 | 595 | 824 | 1142 | 1239 | 1122 | 1217 | 625 | 660 | 609 | 629 | 905,4 | 1180 | 631 | 0,535 | -0,902 | -15,37 | 2,59E-53 | 1,28E-52 | 0,0007858 | 0,003496 | 0,0021121 | 0,0021121 | TRUE | 0,0149008 | No |
| orf19.5159 | 3966 | 3631 | 3816 | 2338 | 1410 | 1043 | 1133 | 1568 | 3611 | 3672 | 3611 | 3512 | 1254 | 1132 | 1160 | 1196 | 2393,72 | 3602 | 1186 | 0,329 | -1,602 | -35,988 | 1,30E-283 | 3,14E-282 | 0,0002127 | 0,002589 | 0,0013599 | 0,0013599 | TRUE | 0,0178665 | No |
| orf19.516 | 299 | 300 | 339 | 178 | 387 | 320 | 344 | 422 | 272 | 303 | 321 | 267 | 344 | 347 | 352 | 322 | 316,22 | 291 | 341 | 1,167 | 0,223 | 2,552 | 0,010711 | 0,014026 | 0,0010865 | 0,006215 | 0,0041406 | 0,0041406 | TRUE | 0,054811 | No |
| orf19.5160 | 2369 | 2014 | 2272 | 1312 | 3196 | 2553 | 2871 | 3609 | 2157 | 2037 | 2150 | 1971 | 2843 | 2772 | 2940 | 2753 | 2452,84 | 2079 | 2827 | 1,357 | 0,441 | 9,624 | 6,32E-22 | 1,67E-21 | 0,0008431 | 0,002576 | 0,0015901 | 0,0015901 | TRUE | 0,0134617 | No |
| orf19.5161 | 1797 | 1459 | 1456 | 1054 | 1216 | 1108 | 1113 | 1471 | 1636 | 1476 | 1378 | 1583 | 1082 | 1203 | 1140 | 1122 | 1327,43 | 1518 | 1137 | 0,749 | -0,418 | -6,546 | 5,91E-11 | 1,16E-10 | 0,0032356 | 0,003032 | 0,0031412 | 0,0031412 | TRUE | 0,0478553 | No |
| orf19.5162 | 1214 | 1168 | 1243 | 754 | 1164 | 987 | 984 | 1439 | 1105 | 1181 | 1176 | 1133 | 1035 | 1072 | 1008 | 1098 | 1101 | 1149 | 1053 | 0,917 | -0,125 | -2,37 | 0,017772 | 0,022897 | 1,00E-08 | 0,003237 | 0,001761 | 0,001761 | TRUE | 0,0105075 | No |
| orf19.5163 | 1342 | 1263 | 1325 | 825 | 1610 | 1363 | 1482 | 1892 | 1222 | 1277 | 1254 | 1239 | 1432 | 1480 | 1518 | 1443 | 1358,17 | 1248 | 1468 | 1,175 | 0,233 | 5,023 | 5,10E-07 | 8,58E-07 | 1,00E-08 | 0,00301 | 0,0013214 | 0,0013214 | TRUE | 0,0061747 | No |
| orf19.5164 | 358 | 341 | 351 | 271 | 812 | 671 | 672 | 914 | 326 | 345 | 332 | 407 | 722 | 729 | 688 | 697 | 530,79 | 352 | 709 | 2,029 | 1,02 | 13,542 | 8,82E-42 | 3,54E-41 | 0,0008615 | 0,004526 | 0,0032661 | 0,0032661 | TRUE | 0,1027245 | No |
| orf19.5165 | 1155 | 1057 | 1172 | 650 | 924 | 716 | 811 | 1165 | 1052 | 1069 | 1109 | 976 | 822 | 777 | 831 | 889 | 940,59 | 1052 | 830 | 0,788 | -0,343 | -5,551 | 2,84E-08 | 5,00E-08 | 0,001735 | 0,003441 | 0,0025916 | 0,0025916 | TRUE | 0,028763 | No |
| orf19.5166 | 1661 | 1379 | 1626 | 1060 | 2516 | 2118 | 2362 | 3232 | 1512 | 1395 | 1539 | 1592 | 2238 | 2300 | 2419 | 2465 | 1932,5 | 1510 | 2356 | 1,563 | 0,644 | 11,851 | 2,12E-32 | 7,17E-32 | 0,0017973 | 0,002721 | 0,0022816 | 0,0022816 | TRUE | 0,0314241 | No |
| orf19.5167 | 5764 | 4837 | 5481 | 3292 | 5569 | 4495 | 5258 | 6915 | 5248 | 4892 | 5187 | 4945 | 4953 | 4880 | 5385 | 5275 | 5095,73 | 5068 | 5123 | 1,011 | 0,015 | 0,332 | 0,73955 | 0,771247 | 0,0013299 | 0,002296 | 0,0018315 | 0,0018315 | TRUE | 0,0141176 | No |
| orf19.5168 | 4005 | 3439 | 3767 | 2363 | 3480 | 2709 | 3025 | 4191 | 3646 | 3478 | 3565 | 3550 | 3095 | 2941 | 3098 | 3197 | 3321,35 | 3560 | 3083 | 0,866 | -0,207 | -5,184 | 2,17E-07 | 3,70E-07 | 0,0004861 | 0,002435 | 0,0012251 | 0,0012251 | TRUE | 0,011366 | No |
| orf19.5169 | 201 | 179 | 175 | 131 | 1250 | 1079 | 1076 | 1468 | 183 | 181 | 166 | 197 | 1112 | 1171 | 1102 | 1120 | 653,94 | 182 | 1126 | 6,24 | 2,641 | 34,661 | 3,07E-263 | 6,74E-262 | 1,00E-08 | 0,004057 | 0,002228 | 0,002228 | TRUE | 0,0379613 | No |

| | | | | | | | | | | | | | | | | | | | | | | | | | | | | | | |
|---|---|---|---|---|---|---|---|---|---|---|---|---|---|---|---|---|---|---|---|---|---|---|---|---|---|---|---|---|---|---|
| orf19.517 | 1288 | 1097 | 1225 | 870 | 2094 | 1715 | 1694 | 2227 | 1173 | 1109 | 1159 | 1307 | 1863 | 1862 | 1735 | 1699 | 1488,32 | 1187 | 1790 | 1,511 | 0,596 | 9,937 | 2,87E-23 | 7,89E-23 | 0,0027256 | 0,002925 | 0,0027336 | 0,0027336 | TRUE | 0,0500542 | No |
| orf19.5170 | 6910 | 6510 | 6886 | 4192 | 7674 | 6369 | 6680 | 9130 | 6291 | 6584 | 6517 | 6297 | 6826 | 6915 | 6841 | 6964 | 6654,47 | 6422 | 6886 | 1,072 | 0,1 | 3,156 | 0,001599 | 0,002226 | 0,0001186 | 0,002236 | 0,0008184 | 0,0008184 | TRUE | 0,0034902 | No |
| orf19.5171 | 1216 | 1195 | 1325 | 876 | 1421 | 1176 | 1267 | 1686 | 1107 | 1209 | 1254 | 1316 | 1264 | 1277 | 1298 | 1286 | 1251,25 | 1222 | 1281 | 1,052 | 0,073 | 1,263 | 0,206645 | 0,238739 | 0,0015892 | 0,003093 | 0,0023771 | 0,0023771 | TRUE | 0,0487007 | No |
| orf19.5172 | 13 | 7 | 8 | 4 | 7 | 6 | 6 | 7 | 12 | 7 | 8 | 6 | 6 | 7 | 6 | 5 | 7,09 | 8 | 6 | 0,727 | -0,46 | -0,909 | 0,363187 | 0,403225 | 1,00E-08 | 0,188353 | 0,1042414 | 0,1042414 | TRUE | 0,3142806 | No |
| orf19.5174 | 925 | 672 | 694 | 452 | 1015 | 703 | 859 | 1077 | 842 | 680 | 657 | 679 | 903 | 763 | 880 | 822 | 778,12 | 714 | 842 | 1,177 | 0,235 | 2,696 | 0,007017 | 0,009308 | 0,0082542 | 0,003735 | 0,0059807 | 0,0059807 | TRUE | 0,1778973 | No |
| orf19.5175 | 3734 | 3341 | 3649 | 2526 | 5188 | 4680 | 4726 | 6431 | 3400 | 3379 | 3453 | 3794 | 4615 | 5081 | 4840 | 4906 | 4183,5 | 3506 | 4860 | 1,388 | 0,473 | 9,57 | 1,07E-21 | 2,81E-21 | 0,0019972 | 0,002353 | 0,0020913 | 0,0020913 | TRUE | 0,0353572 | No |
| orf19.5177 | 2857 | 2534 | 2720 | 1662 | 848 | 693 | 718 | 976 | 2601 | 2563 | 2574 | 2497 | 754 | 752 | 735 | 745 | 1652,67 | 2559 | 746 | 0,292 | -1,778 | -39,334 | 0 | 0 | 1,00E-08 | 0,002836 | 0,0011291 | 0,0011291 | TRUE | 0,0029237 | No |
| orf19.5178 | 3075 | 2641 | 2897 | 1801 | 5141 | 4248 | 4419 | 6173 | 2800 | 2671 | 2742 | 2705 | 4573 | 4612 | 4526 | 4709 | 3667,15 | 2730 | 4605 | 1,687 | 0,754 | 21,115 | 5,84E-99 | 5,06E-98 | 1,00E-08 | 0,002397 | 0,0009268 | 0,0009268 | TRUE | 0,0037955 | No |
| orf19.5179 | 23 | 19 | 29 | 25 | 59 | 60 | 59 | 64 | 21 | 19 | 27 | 38 | 52 | 65 | 60 | 49 | 41,5 | 26 | 56 | 2,206 | 1,142 | 4,74 | 2,14E-06 | 3,51E-06 | 0,0099283 | 0,033866 | 0,026639 | 0,026639 | TRUE | 0,3479341 | No |
| orf19.518 | 7857 | 7052 | 7651 | 4481 | 9920 | 7861 | 8541 | 11551 | 7153 | 7132 | 7241 | 6731 | 8824 | 8535 | 8747 | 8811 | 7896,83 | 7064 | 8729 | 1,235 | 0,304 | 8,752 | 2,10E-18 | 5,11E-18 | 0,0004636 | 0,002204 | 0,0010316 | 0,0010316 | TRUE | 0,0117065 | No |
| orf19.5180 | 1165 | 862 | 1007 | 680 | 803 | 659 | 706 | 930 | 1061 | 872 | 953 | 1021 | 714 | 715 | 723 | 709 | 846,15 | 977 | 715 | 0,733 | -0,448 | -6,625 | 3,48E-11 | 6,87E-11 | 0,0028025 | 0,003598 | 0,0031892 | 0,0031892 | TRUE | 0,0634019 | No |
| orf19.5181 | 9089 | 7678 | 8503 | 4947 | 12029 | 9283 | 11232 | 14914 | 8275 | 7765 | 8047 | 7431 | 10699 | 10079 | 11503 | 11377 | 9397,1 | 7880 | 10914 | 1,385 | 0,47 | 8,913 | 4,95E-19 | 1,22E-18 | 0,0027496 | 0,002178 | 0,0025549 | 0,0025549 | TRUE | 0,0323042 | No |
| orf19.5182 | 3613 | 3139 | 3584 | 2115 | 4265 | 3447 | 3783 | 5074 | 3289 | 3175 | 3392 | 3177 | 3794 | 3742 | 3874 | 3870 | 3539,26 | 3258 | 3820 | 1,172 | 0,229 | 5,988 | 2,13E-09 | 3,94E-09 | 0,000269 | 0,00241 | 0,0011124 | 0,0011124 | TRUE | 0,0093444 | No |
| orf19.5184 | 4994 | 3980 | 4524 | 2704 | 4426 | 3199 | 4005 | 5245 | 4547 | 4025 | 4282 | 4062 | 3937 | 3473 | 4102 | 4001 | 4053,52 | 4229 | 3878 | 0,917 | -0,125 | -2,064 | 0,039026 | 0,048811 | 0,0038601 | 0,002363 | 0,0032826 | 0,0032826 | TRUE | 0,059398 | No |
| orf19.5188 | 217 | 212 | 214 | 137 | 127 | 107 | 108 | 152 | 198 | 214 | 203 | 206 | 113 | 116 | 111 | 116 | 159,5 | 205 | 114 | 0,556 | -0,846 | -7,641 | 2,16E-14 | 4,71E-14 | 1,00E-08 | 0,010319 | 0,0051031 | 0,0051031 | TRUE | 0,0111156 | No |
| orf19.519 | 1112 | 1039 | 1080 | 610 | 1105 | 943 | 1034 | 1318 | 1012 | 1051 | 1022 | 916 | 983 | 1024 | 1059 | 1005 | 1009,09 | 1000 | 1018 | 1,013 | 0,019 | 0,339 | 0,734367 | 0,766218 | 0,0008817 | 0,003346 | 0,0021252 | 0,0021252 | TRUE | 0,0330797 | No |
| orf19.5190 | 1 | 2 | 1 | 1 | 2 | 4 | 2 | 9 | 1 | 2 | 1 | 2 | 2 | 4 | 2 | 7 | 2,55 | 2 | 4 | 2,889 | 1,531 | 1,61 | 0,107415 | 0,128619 | 1,00E-08 | 0,51964 | 0,3451802 | 0,3451802 | TRUE | 0,572082 | No |
| orf19.5191 | 41 | 32 | 25 | 23 | 67 | 62 | 59 | 67 | 37 | 32 | 24 | 35 | 60 | 67 | 60 | 51 | 45,79 | 32 | 60 | 1,863 | 0,898 | 4,138 | 3,51E-05 | 5,41E-05 | 1,00E-08 | 0,030884 | 0,0206375 | 0,0206375 | TRUE | 0,2366404 | No |
| orf19.5194.1 | 876 | 703 | 791 | 491 | 550 | 453 | 451 | 696 | 798 | 711 | 749 | 738 | 489 | 492 | 462 | 531 | 621,07 | 749 | 494 | 0,66 | -0,6 | -8,761 | 1,94E-18 | 4,72E-18 | 0,0014006 | 0,004164 | 0,0028441 | 0,0028441 | TRUE | 0,0330161 | No |
| orf19.5195 | 2321 | 1915 | 2105 | 1367 | 2562 | 2038 | 2143 | 3171 | 2113 | 1937 | 1992 | 2053 | 2279 | 2213 | 2195 | 2419 | 2150,09 | 2024 | 2276 | 1,126 | 0,171 | 3,46 | 0,00054 | 0,000776 | 0,0013235 | 0,002651 | 0,0018751 | 0,0018751 | TRUE | 0,0222147 | No |
| orf19.5196 | 2115 | 1632 | 1841 | 1094 | 1839 | 1343 | 1517 | 2073 | 1926 | 1651 | 1742 | 1643 | 1636 | 1458 | 1554 | 1581 | 1648,83 | 1740 | 1557 | 0,894 | -0,162 | -2,601 | 0,0093 | 0,012224 | 0,0031777 | 0,002838 | 0,0031021 | 0,0031021 | TRUE | 0,0630217 | No |
| orf19.5197 | 9023 | 7635 | 8235 | 5573 | 8320 | 7112 | 7363 | 10274 | 8215 | 7722 | 7794 | 8372 | 7400 | 7722 | 7541 | 7837 | 7825,25 | 8026 | 7625 | 0,951 | -0,073 | -1,838 | 0,06608 | 0,080872 | 0,0009305 | 0,002206 | 0,0013952 | 0,0013952 | TRUE | 0,0099214 | No |
| orf19.5198 | 24411 | 21059 | 22778 | 13004 | 26215 | 19861 | 23184 | 30578 | 22225 | 21299 | 21557 | 19534 | 23317 | 21564 | 23743 | 23325 | 22070,6 | 21154 | 22987 | 1,086 | 0,12 | 2,444 | 0,014533 | 0,018858 | 0,0023026 | 0,002097 | 0,0022576 | 0,0022576 | TRUE | 0,0322397 | No |
| orf19.52 | 2521 | 2346 | 2439 | 1556 | 2890 | 2512 | 2461 | 3485 | 2295 | 2373 | 2308 | 2337 | 2571 | 2727 | 2520 | 2658 | 2473,79 | 2328 | 2619 | 1,125 | 0,17 | 4,104 | 4,06E-05 | 6,22E-05 | 0,0003439 | 0,002571 | 0,0012387 | 0,0012387 | TRUE | 0,0088751 | No |
| orf19.520 | 1483 | 1169 | 1360 | 947 | 1930 | 1724 | 1717 | 2418 | 1350 | 1182 | 1287 | 1423 | 1717 | 1872 | 1758 | 1844 | 1554,19 | 1310 | 1798 | 1,374 | 0,459 | 7,521 | 5,43E-14 | 1,17E-13 | 0,002953 | 0,002887 | 0,0029006 | 0,0029006 | TRUE | 0,0515903 | No |
| orf19.5200 | 3239 | 2810 | 3164 | 1967 | 4334 | 3752 | 3871 | 5164 | 2949 | 2842 | 2994 | 2955 | 3855 | 4074 | 3964 | 3939 | 3446,54 | 2935 | 3958 | 1,348 | 0,431 | 11,529 | 9,45E-31 | 3,09E-30 | 0,0002234 | 0,00242 | 0,0010408 | 0,0010408 | TRUE | 0,0056657 | No |
| orf19.5201 | 2906 | 2511 | 2676 | 1745 | 1956 | 1705 | 1625 | 2225 | 2646 | 2540 | 2533 | 2621 | 1740 | 1851 | 1664 | 1697 | 2161,46 | 2585 | 1738 | 0,672 | -0,573 | -12,447 | 1,46E-35 | 5,26E-35 | 0,0007021 | 0,002648 | 0,0015587 | 0,0015587 | TRUE | 0,0221435 | No |
| orf19.5201.1 | 7393 | 6652 | 6500 | 3999 | 5245 | 3622 | 4001 | 5339 | 6731 | 6728 | 6152 | 6007 | 4665 | 3932 | 4098 | 4073 | 5298,19 | 6404 | 4192 | 0,654 | -0,612 | -9,882 | 4,99E-23 | 1,36E-22 | 0,004479 | 0,002286 | 0,0034846 | 0,0034846 | TRUE | 0,0705665 | No |
| orf19.5203 | 507 | 418 | 497 | 298 | 977 | 802 | 945 | 1215 | 462 | 423 | 470 | 448 | 869 | 871 | 968 | 927 | 679,59 | 451 | 909 | 2,015 | 1,011 | 15,07 | 2,55E-51 | 1,20E-50 | 0,001145 | 0,003981 | 0,0026206 | 0,0026206 | TRUE | 0,02186 | No |
| orf19.5204 | 2145 | 1739 | 1910 | 1462 | 1665 | 1488 | 1624 | 2338 | 1953 | 1759 | 1808 | 2196 | 1481 | 1616 | 1663 | 1783 | 1782,34 | 1929 | 1636 | 0,85 | -0,235 | -3,056 | 0,002244 | 0,003083 | 0,0073501 | 0,002778 | 0,0051156 | 0,0051156 | TRUE | 0,100958 | No |
| orf19.5205 | 727 | 696 | 758 | 551 | 653 | 621 | 499 | 789 | 662 | 704 | 717 | 828 | 581 | 674 | 511 | 602 | 659,86 | 728 | 592 | 0,816 | -0,294 | -3,236 | 0,00121 | 0,0017 | 0,009117 | 0,004039 | 0,0063698 | 0,0063698 | TRUE | 0,0979259 | No |
| orf19.5206 | 1856 | 1730 | 1828 | 1255 | 1769 | 1540 | 1526 | 1999 | 1690 | 1750 | 1730 | 1885 | 1573 | 1672 | 1563 | 1525 | 1673,49 | 1764 | 1583 | 0,899 | -0,154 | -3,003 | 0,002669 | 0,003647 | 0,0011404 | 0,002826 | 0,0019211 | 0,0019211 | TRUE | 0,0237571 | No |
| orf19.5207 | 4564 | 3931 | 4190 | 2407 | 4116 | 3263 | 3485 | 4631 | 4155 | 3976 | 3965 | 3616 | 3661 | 3543 | 3569 | 3533 | 3752,21 | 3928 | 3576 | 0,909 | -0,137 | -2,938 | 0,0033 | 0,004476 | 0,0014293 | 0,002389 | 0,0018267 | 0,0018267 | TRUE | 0,0330969 | No |
| orf19.5209 | 364 | 406 | 409 | 233 | 347 | 223 | 246 | 358 | 331 | 411 | 387 | 350 | 309 | 242 | 252 | 273 | 319,36 | 370 | 269 | 0,728 | -0,457 | -4,502 | 6,73E-06 | 1,08E-05 | 0,0074089 | 0,006173 | 0,0067174 | 0,0067174 | TRUE | 0,1102746 | No |
| orf19.5210 | 395 | 315 | 393 | 291 | 974 | 879 | 874 | 1206 | 360 | 319 | 372 | 437 | 866 | 954 | 895 | 920 | 640,38 | 372 | 909 | 2,462 | 1,3 | 15,913 | 5,17E-57 | 2,69E-56 | 0,0050087 | 0,0041 | 0,004441 | 0,004441 | TRUE | 0,1353766 | No |
| orf19.5210.1 | 59 | 59 | 71 | 65 | 48 | 62 | 35 | 60 | 54 | 60 | 67 | 98 | 43 | 67 | 36 | 46 | 58,73 | 70 | 48 | 0,696 | -0,523 | -2,237 | 0,025269 | 0,032135 | 0,0511635 | 0,024529 | 0,0348821 | 0,0348821 | TRUE | 0,5471876 | No |
| orf19.5211 | 628 | 489 | 505 | 377 | 714 | 668 | 619 | 943 | 572 | 495 | 478 | 566 | 635 | 725 | 634 | 719 | 603,02 | 528 | 678 | 1,289 | 0,366 | 4,474 | 7,68E-06 | 1,23E-05 | 0,0052566 | 0,004228 | 0,0047152 | 0,0047152 | TRUE | 0,048238 | No |
| orf19.5212 | 11 | 17 | 17 | 13 | 40 | 26 | 24 | 36 | 10 | 17 | 16 | 20 | 36 | 28 | 25 | 27 | 22,33 | 16 | 29 | 1,886 | 0,916 | 2,977 | 0,002915 | 0,003972 | 1,00E-08 | 0,061184 | 0,0402755 | 0,0402755 | TRUE | 0,3214028 | No |
| orf19.5213 | 2701 | 2517 | 2825 | 1812 | 3660 | 3172 | 3338 | 4547 | 2459 | 2546 | 2674 | 2722 | 3255 | 3444 | 3419 | 3468 | 2998,34 | 2600 | 3396 | 1,308 | 0,387 | 8,503 | 1,85E-17 | 4,40E-17 | 0,0010109 | 0,002478 | 0,0016462 | 0,0016462 | TRUE | 0,016393 | No |
| orf19.5213.1 | 7334 | 5872 | 6094 | 3668 | 8751 | 6133 | 6286 | 9142 | 6677 | 5939 | 5767 | 5510 | 7784 | 6659 | 6438 | 6974 | 6468,41 | 5973 | 6964 | 1,165 | 0,221 | 3,119 | 0,001813 | 0,002511 | 0,0068966 | 0,002241 | 0,0046587 | 0,0046587 | TRUE | 0,0770589 | No |
| orf19.5213.2 | 4602 | 3548 | 3554 | 2991 | 4594 | 3347 | 3352 | 4719 | 4190 | 3588 | 3364 | 4493 | 4086 | 3634 | 3433 | 3600 | 3798,44 | 3909 | 3688 | 0,944 | -0,083 | -0,957 | 0,338666 | 0,378771 | 0,0111376 | 0,002385 | 0,0069131 | 0,0069131 | TRUE | 0,1212329 | No |
| orf19.5214 | 979 | 892 | 1021 | 611 | 604 | 461 | 511 | 637 | 891 | 902 | 966 | 918 | 537 | 501 | 523 | 486 | 715,57 | 919 | 512 | 0,556 | -0,846 | -13,613 | 3,33E-42 | 1,35E-41 | 1,00E-08 | 0,003883 | 0,0022176 | 0,0022176 | TRUE | 0,0153812 | No |
| orf19.5215 | 2342 | 1940 | 2077 | 1465 | 12971 | 10281 | 11248 | 15704 | 2132 | 1962 | 1966 | 2201 | 11537 | 11162 | 11519 | 11979 | 6807,36 | 2065 | 11549 | 5,602 | 2,486 | 52,866 | 0 | 0 | 0,001734 | 0,002231 | 0,0018218 | 0,0018218 | TRUE | 0,0215357 | No |
| orf19.5216 | 677 | 661 | 633 | 427 | 571 | 492 | 476 | 750 | 616 | 669 | 599 | 641 | 508 | 534 | 487 | 572 | 578,38 | 631 | 525 | 0,835 | -0,261 | -3,626 | 0,000288 | 0,000421 | 0,0020445 | 0,004321 | 0,0032363 | 0,0032363 | TRUE | 0,0454378 | No |
| orf19.5217 | 494 | 443 | 447 | 301 | 4637 | 3673 | 3853 | 5328 | 450 | 448 | 423 | 452 | 4124 | 3988 | 3946 | 4064 | 2236,94 | 443 | 4030 | 9,109 | 3,187 | 61,889 | 0 | 0 | 1,00E-08 | 0,002628 | 0,0012322 | 0,0012322 | TRUE | 0,011682 | No |
| orf19.5219 | 4002 | 3935 | 4095 | 2418 | 6112 | 5466 | 5580 | 7170 | 3644 | 3980 | 3876 | 3632 | 5436 | 5935 | 5715 | 5469 | 4710,77 | 3783 | 5639 | 1,489 | 0,575 | 12,198 | 3,19E-34 | 1,12E-33 | 0,0016013 | 0,002317 | 0,0019063 | 0,0019063 | TRUE | 0,0151094 | No |
| orf19.522 | 6315 | 5637 | 5977 | 3673 | 7521 | 6268 | 6564 | 8809 | 5750 | 5701 | 5657 | 5517 | 6690 | 6805 | 6722 | 6720 | 6195,23 | 5656 | 6734 | 1,19 | 0,251 | 8,315 | 9,14E-17 | 2,12E-16 | 1,00E-08 | 0,00225 | 0,0007076 | 0,0007076 | TRUE | 0,0030694 | No |
| orf19.5220 | 3641 | 3213 | 3570 | 2191 | 2889 | 2343 | 2467 | 3312 | 3315 | 3250 | 3379 | 3291 | 2570 | 2544 | 2527 | 2526 | 2925,12 | 3309 | 2542 | 0,768 | -0,381 | -10,767 | 4,94E-27 | 1,48E-26 | 1,00E-08 | 0,002489 | 0,0008555 | 0,0008555 | TRUE | 0,0025562 | No |
| orf19.5221 | 1654 | 1556 | 1734 | 1039 | 885 | 751 | 813 | 1010 | 1506 | 1574 | 1641 | 1561 | 787 | 815 | 833 | 770 | 1185,88 | 1570 | 801 | 0,51 | -0,973 | -18,413 | 1,04E-75 | 6,93E-75 | 1,00E-08 | 0,003151 | 0,0017551 | 0,0017551 | TRUE | 0,011722 | No |
| orf19.5223 | 29 | 28 | 29 | 15 | 26 | 23 | 20 | 21 | 26 | 28 | 27 | 23 | 23 | 25 | 20 | 16 | 23,66 | 26 | 21 | 0,794 | -0,333 | -1,186 | 0,235603 | 0,269653 | 1,00E-08 | 0,057864 | 0,0334159 | 0,0334159 | TRUE | 0,1936601 | No |
| orf19.5224 | 2674 | 2587 | 2788 | 1771 | 3896 | 3290 | 3434 | 4577 | 2435 | 2616 | 2639 | 2660 | 3465 | 3572 | 3517 | 3491 | 3049,45 | 2588 | 3511 | 1,358 | 0,442 | 10,731 | 7,27E-27 | 2,17E-26 | 0,0004368 | 0,00247 | 0,0012852 | 0,0012852 | TRUE | 0,0198989 | No |
| orf19.5225 | 556 | 561 | 519 | 346 | 685 | 529 | 596 | 799 | 506 | 567 | 491 | 520 | 609 | 574 | 610 | 609 | 561 | 521 | 600 | 1,155 | 0,207 | 3,041 | 0,002361 | 0,003238 | 0,0004593 | 0,004392 | 0,0026675 | 0,0026675 | TRUE | 0,0441782 | No |
| orf19.5225.2 | 14223 | 11965 | 12432 | 9059 | 20634 | 16881 | 16328 | 22817 | 12949 | 12101 | 11766 | 13608 | 18353 | 18328 | 16722 | 17405 | 15154,1 | 12606 | 17702 | 1,405 | 0,49 | 9,145 | 5,98E-20 | 1,52E-19 | 0,0030857 | 0,002124 | 0,0026913 | 0,0026913 | TRUE | 0,0345853 | No |
| orf19.5226 | 2211 | 2020 | 2267 | 1319 | 3254 | 2464 | 2924 | 3644 | 2013 | 2043 | 2146 | 1981 | 2894 | 2675 | 2995 | 2780 | 2440,83 | 2046 | 2836 | 1,385 | 0,47 | 9,573 | 1,04E-21 | 2,73E-21 | 0,0012364 | 0,002578 | 0,0018876 | 0,0018876 | TRUE | 0,0170277 | No |
| orf19.5227 | 1789 | 1614 | 1781 | 993 | 934 | 778 | 815 | 1080 | 1629 | 1632 | 1686 | 1492 | 831 | 845 | 835 | 824 | 1221,54 | 1610 | 834 | 0,516 | -0,953 | -17,955 | 4,40E-72 | 2,82E-71 | 0,0006382 | 0,003118 | 0,0018148 | 0,0018148 | TRUE | 0,0262588 | No |
| orf19.5228 | 6173 | 5548 | 6109 | 3945 | 2965 | 2362 | 2434 | 3326 | 5620 | 5611 | 5782 | 5926 | 2637 | 2564 | 2493 | 2537 | 4146,33 | 5735 | 2558 | 0,446 | -1,164 | -30,872 | 2,80E-209 | 4,96E-208 | 0,000347 | 0,002356 | 0,0010878 | 0,0010878 | TRUE | 0,0058098 | No |
| orf19.5229 | 9449 | 9124 | 9410 | 5797 | 11672 | 10091 | 10153 | 13581 | 8603 | 9228 | 8906 | 8708 | 10382 | 10956 | 10398 | 10360 | 9692,52 | 8861 | 10524 | 1,187 | 0,248 | 6,685 | 2,31E-11 | 4,60E-11 | 0,0007183 | 0,002173 | 0,0012134 | 0,0012134 | TRUE | 0,0095389 | No |
| orf19.5230 | 5030 | 4569 | 4987 | 2798 | 4698 | 3620 | 4067 | 5424 | 4580 | 4621 | 4720 | 4203 | 4179 | 3930 | 4165 | 4137 | 4316,88 | 4531 | 4103 | 0,905 | -0,144 | -3,18 | 0,001471 | 0,002055 | 0,0013398 | 0,002343 | 0,0017453 | 0,0017453 | TRUE | 0,0275078 | No |
| orf19.5231 | 38 | 49 | 39 | 18 | 84 | 68 | 78 | 107 | 35 | 50 | 37 | 27 | 75 | 74 | 80 | 82 | 57,27 | 37 | 78 | 2,066 | 1,047 | 5,332 | 9,69E-08 | 1,67E-07 | 1,00E-08 | 0,025104 | 0,0168468 | 0,0168468 | TRUE | 0,3559319 | No |
| orf19.5231.2 | 1311 | 1117 | 1193 | 685 | 2903 | 1902 | 2359 | 2983 | 1194 | 1130 | 1129 | 1029 | 2582 | 2065 | 2416 | 2275 | 1727,49 | 1120 | 2334 | 2,081 | 1,057 | 14,586 | 3,47E-48 | 1,55E-47 | 0,0053669 | 0,002802 | 0,0043627 | 0,0043627 | TRUE | 0,0720133 | No |
| orf19.5232 | 5019 | 5436 | 5175 | 3463 | 5232 | 4800 | 4369 | 5881 | 4570 | 5498 | 4898 | 5202 | 4654 | 5211 | 4474 | 4486 | 4874,1 | 5042 | 4706 | 0,934 | -0,099 | -1,484 | 0,137903 | 0,162681 | 0,0055672 | 0,002308 | 0,0040902 | 0,0040902 | TRUE | 0,0631789 | No |
| orf19.5233 | 0 | 0 | 1 | 1 | 6 | 2 | 5 | 3 | 0 | 0 | 1 | 2 | 5 | 2 | 5 | 2 | 2,17 | 1 | 4 | 6,952 | 2,798 | 2,282 | 0,022465 | 0,028665 | 1,00E-08 | 0,610586 | 0,4008308 | 0,4008308 | TRUE | 0,2014173 | No |
| orf19.5234 | 1540 | 1373 | 1507 | 894 | 2845 | 2365 | 2422 | 3277 | 1402 | 1389 | 1426 | 1343 | 2531 | 2568 | 2480 | 2500 | 1954,79 | 1390 | 2520 | 1,809 | 0,855 | 20,598 | 2,85E-94 | 2,31E-93 | 1,00E-08 | 0,002713 | 0,0010863 | 0,0010863 | TRUE | 0,0053188 | No |
| orf19.5235 | 797 | 1044 | 1025 | 537 | 1080 | 1002 | 948 | 1043 | 726 | 1056 | 970 | 807 | 961 | 1088 | 971 | 796 | 921,66 | 890 | 954 | 1,07 | 0,098 | 0,844 | 0,39842 | 0,439282 | 0,0215966 | 0,00347 | 0,0118445 | 0,0118445 | TRUE | 0,1898813 | No |
| orf19.5236 | 484 | 528 | 537 | 330 | 280 | 278 | 238 | 370 | 441 | 534 | 508 | 496 | 249 | 302 | 244 | 282 | 381,93 | 495 | 269 | 0,544 | -0,877 | -9,426 | 4,27E-21 | 1,11E-20 | 0,005499 | 0,005496 | 0,0054954 | 0,0054954 | TRUE | 0,0675069 | No |
| orf19.5237 | 679 | 592 | 645 | 378 | 283 | 287 | 275 | 343 | 618 | 599 | 610 | 568 | 252 | 312 | 282 | 262 | 437,72 | 599 | 277 | 0,459 | -1,123 | -14,01 | 1,35E-44 | 5,74E-44 | 0,0010588 | 0,005055 | 0,0035892 | 0,0035892 | TRUE | 0,076877 | No |
| orf19.5238 | 3024 | 2814 | 2967 | 1979 | 2317 | 2036 | 2186 | 2829 | 2753 | 2846 | 2808 | 2973 | 2061 | 2211 | 2239 | 2158 | 2506,03 | 2845 | 2167 | 0,762 | -0,392 | -8,815 | 1,20E-18 | 2,93E-18 | 0,0006482 | 0,002564 | 0,0014914 | 0,0014914 | TRUE | 0,0131712 | No |

| | | | | | | | | | | | | | | | | | | | | | | | | | | | | | | | |
|---|---|---|---|---|---|---|---|---|---|---|---|---|---|---|---|---|---|---|---|---|---|---|---|---|---|---|---|---|---|---|---|
| orf19.5239 | 535 | 513 | 513 | 327 | 316 | 243 | 244 | 350 | 487 | 519 | 486 | 491 | 281 | 264 | 250 | 267 | 380,55 | 496 | 266 | 0,536 | -0,899 | -11,669 | 1,84E-31 | 6,12E-31 | 1,00E-08 | 0,005508 | 0,0028706 | 0,0028706 | TRUE | 0,0175987 | No |
| orf19.524 | 1362 | 1123 | 1324 | 757 | 545 | 356 | 504 | 636 | 1240 | 1136 | 1253 | 1137 | 485 | 387 | 516 | 485 | 829,83 | 1192 | 468 | 0,393 | -1,347 | -16,621 | 4,94E-62 | 2,78E-61 | 0,0066992 | 0,003629 | 0,0048549 | 0,0048549 | TRUE | 0,1507804 | No |
| orf19.5241 | 7932 | 6839 | 7505 | 4650 | 3406 | 2800 | 3112 | 3964 | 7222 | 6917 | 7103 | 6985 | 3030 | 3040 | 3187 | 3024 | 5063,36 | 7057 | 3070 | 0,435 | -1,202 | -33,899 | 6,97E-252 | 1,46E-250 | 0,0001943 | 0,002298 | 0,0009777 | 0,0009777 | TRUE | 0,0078529 | No |
| orf19.5242 | 75 | 97 | 81 | 42 | 103 | 95 | 102 | 97 | 68 | 98 | 77 | 63 | 92 | 103 | 104 | 74 | 84,92 | 76 | 93 | 1,204 | 0,268 | 1,51 | 0,131012 | 0,155242 | 0,0191488 | 0,017593 | 0,0182035 | 0,0182035 | TRUE | 0,326518 | No |
| orf19.5243 | 1473 | 1538 | 1596 | 854 | 1379 | 1128 | 1228 | 1632 | 1341 | 1556 | 1510 | 1283 | 1227 | 1225 | 1258 | 1245 | 1330,47 | 1422 | 1239 | 0,869 | -0,203 | -3,135 | 0,001719 | 0,002385 | 0,0033205 | 0,00303 | 0,0032538 | 0,0032538 | TRUE | 0,048101 | No |
| orf19.5244 | 450 | 517 | 552 | 323 | 614 | 549 | 644 | 752 | 410 | 523 | 522 | 485 | 546 | 596 | 660 | 574 | 539,45 | 485 | 594 | 1,224 | 0,291 | 3,2 | 0,001374 | 0,001922 | 0,0078266 | 0,004486 | 0,0060633 | 0,0060633 | TRUE | 0,1365799 | No |
| orf19.5245 | 1781 | 1677 | 1871 | 1222 | 630 | 568 | 546 | 766 | 1622 | 1696 | 1771 | 1836 | 560 | 617 | 559 | 584 | 1155,57 | 1731 | 580 | 0,335 | -1,576 | -26,214 | 1,82E-151 | 2,39E-150 | 0,0015799 | 0,00318 | 0,0023538 | 0,0023538 | TRUE | 0,0221297 | No |
| orf19.5246 | 327 | 285 | 328 | 220 | 145 | 130 | 123 | 162 | 298 | 288 | 310 | 330 | 129 | 141 | 126 | 124 | 218,32 | 306 | 130 | 0,424 | -1,239 | -12,154 | 5,43E-34 | 1,89E-33 | 1,00E-08 | 0,008088 | 0,0046347 | 0,0046347 | TRUE | 0,0316542 | No |
| orf19.5247 | 1863 | 1666 | 1678 | 1053 | 1189 | 979 | 1006 | 1372 | 1696 | 1685 | 1588 | 1582 | 1058 | 1063 | 1030 | 1047 | 1343,54 | 1638 | 1050 | 0,64 | -0,644 | -13,314 | 1,92E-40 | 7,49E-40 | 1,00E-08 | 0,00302 | 0,0014749 | 0,0014749 | TRUE | 0,0070504 | No |
| orf19.5248 | 4183 | 3305 | 3756 | 2379 | 2260 | 1755 | 1965 | 2650 | 3808 | 3343 | 3555 | 3574 | 2010 | 1905 | 2012 | 2021 | 2778,62 | 3570 | 1987 | 0,557 | -0,845 | -17,15 | 6,27E-66 | 3,74E-65 | 0,0015601 | 0,002512 | 0,001943 | 0,001943 | TRUE | 0,02543 | No |
| orf19.5249 | 778 | 612 | 677 | 432 | 602 | 448 | 501 | 655 | 708 | 619 | 641 | 649 | 535 | 486 | 513 | 500 | 581,44 | 654 | 508 | 0,777 | -0,364 | -5,298 | 1,17E-07 | 2,02E-07 | 0,0011299 | 0,004309 | 0,0027941 | 0,0027941 | TRUE | 0,0394956 | No |
| orf19.5250 | 1216 | 1095 | 1135 | 751 | 1674 | 1291 | 1315 | 1836 | 1107 | 1107 | 1074 | 1128 | 1489 | 1402 | 1347 | 1401 | 1256,85 | 1104 | 1410 | 1,278 | 0,354 | 6,989 | 2,76E-12 | 5,66E-12 | 1,00E-08 | 0,003088 | 0,0016487 | 0,0016487 | TRUE | 0,01751 | No |
| orf19.5251 | 5538 | 4921 | 5500 | 3773 | 4639 | 3942 | 4029 | 5498 | 5042 | 4977 | 5205 | 5668 | 4126 | 4280 | 4126 | 4194 | 4702,29 | 5223 | 4182 | 0,801 | -0,319 | -6,832 | 8,39E-12 | 1,70E-11 | 0,0016442 | 0,002318 | 0,0018848 | 0,0018848 | TRUE | 0,0385681 | No |
| orf19.5253 | 869 | 815 | 905 | 529 | 1602 | 1399 | 1436 | 1964 | 791 | 824 | 857 | 795 | 1425 | 1519 | 1471 | 1498 | 1147,41 | 817 | 1478 | 1,807 | 0,854 | 16,4 | 1,90E-60 | 1,05E-59 | 1,00E-08 | 0,003188 | 0,0016337 | 0,0016337 | TRUE | 0,0133552 | No |
| orf19.5254 | 429 | 423 | 439 | 312 | 573 | 465 | 434 | 621 | 391 | 428 | 415 | 469 | 510 | 505 | 444 | 474 | 454,41 | 426 | 483 | 1,141 | 0,19 | 2,407 | 0,016096 | 0,020818 | 0,0022243 | 0,004944 | 0,0037685 | 0,0037685 | TRUE | 0,0457818 | No |
| orf19.5255 | 3542 | 3163 | 3491 | 2224 | 5282 | 4328 | 4537 | 6114 | 3225 | 3199 | 3304 | 3341 | 4698 | 4699 | 4646 | 4664 | 3972 | 3267 | 4677 | 1,432 | 0,519 | 15,579 | 1,01E-54 | 5,07E-54 | 1,00E-08 | 0,00237 | 0,0007991 | 0,0007991 | TRUE | 0,0024969 | No |
| orf19.5256 | 72 | 57 | 78 | 43 | 100 | 84 | 88 | 92 | 66 | 58 | 74 | 65 | 89 | 91 | 90 | 70 | 75,26 | 66 | 85 | 1,292 | 0,369 | 2,21 | 0,027087 | 0,034351 | 1,00E-08 | 0,01959 | 0,0131565 | 0,0131565 | TRUE | 0,1534038 | No |
| orf19.5257 | 979 | 834 | 905 | 607 | 917 | 771 | 767 | 1052 | 891 | 844 | 857 | 912 | 816 | 837 | 786 | 802 | 842,98 | 876 | 810 | 0,926 | -0,11 | -1,996 | 0,045948 | 0,057066 | 1,00E-08 | 0,003604 | 0,0017546 | 0,0017546 | TRUE | 0,0076638 | No |
| orf19.5258 | 1249 | 1257 | 1330 | 791 | 545 | 459 | 466 | 623 | 1137 | 1271 | 1259 | 1188 | 485 | 498 | 477 | 475 | 848,87 | 1214 | 484 | 0,398 | -1,328 | -21,57 | 3,43E-103 | 3,09E-102 | 1,00E-08 | 0,003593 | 0,0022362 | 0,0022362 | TRUE | 0,0237392 | No |
| orf19.5259 | 664 | 619 | 667 | 433 | 240 | 227 | 210 | 290 | 605 | 626 | 631 | 650 | 213 | 246 | 215 | 221 | 426,06 | 628 | 224 | 0,356 | -1,488 | -19,196 | 4,02E-82 | 2,93E-81 | 1,00E-08 | 0,005137 | 0,0028369 | 0,0028369 | TRUE | 0,0447381 | No |
| orf19.526 | 10843 | 9646 | 9999 | 5503 | 16071 | 12020 | 13629 | 17509 | 9872 | 9756 | 9463 | 8266 | 14295 | 13050 | 13958 | 13356 | 11502 | 9339 | 13665 | 1,462 | 0,548 | 9,403 | 5,32E-21 | 1,38E-20 | 0,0038356 | 0,002152 | 0,0031753 | 0,0031753 | TRUE | 0,0720033 | No |
| orf19.5260 | 6356 | 5629 | 6119 | 4134 | 6470 | 5554 | 5879 | 8006 | 5787 | 5693 | 5791 | 6210 | 5755 | 6030 | 6021 | 6107 | 5924,23 | 5870 | 5978 | 1,019 | 0,027 | 0,685 | 0,493079 | 0,534679 | 0,0009146 | 0,00226 | 0,0013725 | 0,0013725 | TRUE | 0,0176925 | No |
| orf19.5262 | 43 | 21 | 24 | 14 | 59 | 72 | 51 | 84 | 39 | 21 | 23 | 21 | 52 | 78 | 52 | 64 | 43,89 | 26 | 62 | 2,329 | 1,22 | 4,817 | 1,46E-06 | 2,41E-06 | 0,0370099 | 0,032138 | 0,0338538 | 0,0338538 | TRUE | 0,7923798 | No |
| orf19.5263 | 7556 | 6613 | 7013 | 4695 | 10049 | 8767 | 8593 | 12320 | 6879 | 6688 | 6637 | 7053 | 8938 | 9519 | 8800 | 9398 | 7989,06 | 6814 | 9164 | 1,345 | 0,428 | 10,703 | 9,81E-27 | 2,92E-26 | 0,0009397 | 0,002202 | 0,0014051 | 0,0014051 | TRUE | 0,0086328 | No |
| orf19.5264 | 3 | 11 | 2 | 7 | 11 | 7 | 6 | 5 | 3 | 11 | 2 | 11 | 10 | 8 | 6 | 4 | 6,7 | 7 | 7 | 1,075 | 0,104 | 0,167 | 0,867403 | 0,887022 | 0,2470301 | 0,199173 | 0,2168897 | 0,2168897 | TRUE | 0,180475 | No |
| orf19.5265 | 3359 | 3061 | 3214 | 1910 | 2592 | 2008 | 2325 | 2981 | 3058 | 3096 | 3042 | 2869 | 2306 | 2180 | 2381 | 2274 | 2650,71 | 3016 | 2285 | 0,757 | -0,402 | -9,124 | 7,23E-20 | 1,83E-19 | 0,0007072 | 0,002535 | 0,0014796 | 0,0014796 | TRUE | 0,0120942 | No |
| orf19.5266 | 64 | 45 | 47 | 22 | 19 | 18 | 13 | 17 | 58 | 46 | 44 | 33 | 17 | 20 | 13 | 13 | 30,5 | 45 | 16 | 0,338 | -1,564 | -5,566 | 2,61E-08 | 4,61E-08 | 1,00E-08 | 0,045342 | 0,0342798 | 0,0342798 | TRUE | 0,1365796 | No |
| orf19.5267 | 138 | 120 | 122 | 95 | 83 | 90 | 83 | 112 | 126 | 121 | 115 | 143 | 74 | 98 | 85 | 85 | 105,89 | 126 | 86 | 0,679 | -0,558 | -3,957 | 7,60E-05 | 0,000115 | 1,00E-08 | 0,014512 | 0,0094026 | 0,0094026 | TRUE | 0,0815195 | No |
| orf19.5268 | 169 | 139 | 163 | 122 | 119 | 99 | 109 | 114 | 154 | 141 | 154 | 183 | 106 | 107 | 112 | 87 | 130,49 | 158 | 103 | 0,653 | -0,615 | -4,604 | 4,13E-06 | 6,69E-06 | 0,0040098 | 0,012161 | 0,0091702 | 0,0091702 | TRUE | 0,1254015 | No |
| orf19.5269 | 658 | 545 | 599 | 353 | 1264 | 1045 | 1200 | 1579 | 599 | 551 | 567 | 530 | 1124 | 1135 | 1229 | 1204 | 867,47 | 562 | 1173 | 2,081 | 1,057 | 17,164 | 4,98E-66 | 2,97E-65 | 0,0009825 | 0,00356 | 0,0022989 | 0,0022989 | TRUE | 0,0236198 | No |
| orf19.5269.1 | 374 | 382 | 320 | 215 | 523 | 380 | 343 | 468 | 341 | 386 | 303 | 323 | 465 | 413 | 351 | 357 | 367,34 | 338 | 396 | 1,171 | 0,228 | 2,095 | 0,036151 | 0,045393 | 0,0120097 | 0,005633 | 0,008589 | 0,008589 | TRUE | 0,1614998 | No |
| orf19.527 | 1662 | 1487 | 1582 | 1111 | 1362 | 1130 | 1088 | 1603 | 1513 | 1504 | 1497 | 1669 | 1211 | 1227 | 1114 | 1223 | 1369,82 | 1546 | 1194 | 0,774 | -0,369 | -6,672 | 2,52E-11 | 5,02E-11 | 0,0014308 | 0,003001 | 0,0021934 | 0,0021934 | TRUE | 0,0315526 | No |
| orf19.5270 | 182 | 170 | 196 | 113 | 458 | 371 | 348 | 462 | 166 | 172 | 185 | 170 | 407 | 403 | 356 | 352 | 276,48 | 173 | 380 | 2,187 | 1,129 | 11,659 | 2,06E-31 | 6,85E-31 | 1,00E-08 | 0,006815 | 0,0047052 | 0,0047052 | TRUE | 0,029657 | No |
| orf19.5271 | 213 | 232 | 253 | 141 | 304 | 231 | 213 | 310 | 194 | 235 | 239 | 212 | 270 | 251 | 218 | 236 | 231,95 | 220 | 244 | 1,109 | 0,149 | 1,398 | 0,162197 | 0,19011 | 0,0048555 | 0,007732 | 0,0065322 | 0,0065322 | TRUE | 0,0799889 | No |
| orf19.5274 | 3597 | 3374 | 3837 | 2271 | 2129 | 1760 | 1846 | 2599 | 3275 | 3412 | 3631 | 3411 | 1894 | 1911 | 1891 | 1983 | 2675,97 | 3432 | 1920 | 0,559 | -0,838 | -18,536 | 1,05E-76 | 7,13E-76 | 0,0009519 | 0,002531 | 0,0015624 | 0,0015624 | TRUE | 0,0191442 | No |
| orf19.5275 | 488 | 517 | 510 | 333 | 153 | 136 | 132 | 163 | 444 | 523 | 483 | 500 | 136 | 148 | 135 | 124 | 311,67 | 488 | 136 | 0,278 | -1,848 | -19,064 | 4,99E-81 | 3,57E-80 | 1,00E-08 | 0,006275 | 0,0044821 | 0,0044821 | TRUE | 0,0440009 | No |
| orf19.5276 | 805 | 866 | 867 | 538 | 532 | 427 | 402 | 495 | 733 | 876 | 821 | 808 | 473 | 464 | 412 | 378 | 620,45 | 810 | 432 | 0,532 | -0,91 | -10,624 | 2,30E-26 | 6,77E-26 | 0,006532 | 0,004166 | 0,0052885 | 0,0052885 | TRUE | 0,0887752 | No |
| orf19.5277 | 1814 | 1676 | 1873 | 1058 | 1247 | 954 | 1037 | 1318 | 1652 | 1695 | 1773 | 1589 | 1109 | 1036 | 1062 | 1005 | 1365,11 | 1677 | 1053 | 0,627 | -0,674 | -12,483 | 9,27E-36 | 3,36E-35 | 0,0012013 | 0,003005 | 0,0020352 | 0,0020352 | TRUE | 0,017691 | No |
| orf19.5278 | 1940 | 1784 | 1824 | 1071 | 3142 | 2618 | 2639 | 3703 | 1766 | 1804 | 1726 | 1609 | 2795 | 2842 | 2703 | 2825 | 2258,77 | 1726 | 2791 | 1,613 | 0,69 | 15,053 | 3,29E-51 | 1,55E-50 | 0,0005984 | 0,002622 | 0,0015396 | 0,0015396 | TRUE | 0,0225012 | No |
| orf19.5279 | 2962 | 2606 | 2608 | 1482 | 2314 | 1466 | 1860 | 2303 | 2697 | 2636 | 2468 | 2226 | 2058 | 1592 | 1905 | 1757 | 2167,3 | 2507 | 1828 | 0,729 | -0,457 | -5,611 | 2,01E-08 | 3,57E-08 | 0,0088733 | 0,002647 | 0,0058876 | 0,0058876 | TRUE | 0,0899224 | No |
| orf19.528 | 3213 | 3089 | 3182 | 1957 | 3665 | 3143 | 3214 | 4298 | 2925 | 3124 | 3011 | 2940 | 3260 | 3412 | 3292 | 3279 | 3155,39 | 3000 | 3311 | 1,103 | 0,141 | 3,605 | 0,000312 | 0,000455 | 0,0002529 | 0,002456 | 0,0011484 | 0,0011484 | TRUE | 0,0094787 | No |
| orf19.5280 | 68553 | 57925 | 63100 | 41383 | 68040 | 56943 | 57750 | 85111 | 62415 | 58585 | 59718 | 62164 | 60519 | 61824 | 59143 | 64923 | 61161,5 | 60720 | 61602 | 1,015 | 0,021 | 0,522 | 0,60184 | 0,641594 | 0,0011433 | 0,002059 | 0,0015237 | 0,0015237 | TRUE | 0,0161702 | No |
| orf19.5281 | 1603 | 1477 | 1600 | 1002 | 2084 | 1751 | 1848 | 2405 | 1459 | 1494 | 1514 | 1505 | 1854 | 1901 | 1893 | 1835 | 1681,82 | 1493 | 1871 | 1,252 | 0,325 | 7,701 | 1,35E-14 | 2,96E-14 | 1,00E-08 | 0,002822 | 0,0010996 | 0,0010996 | TRUE | 0,0029766 | No |
| orf19.5282 | 9071 | 7863 | 8643 | 5760 | 4159 | 3520 | 3514 | 4970 | 8259 | 7953 | 8180 | 8652 | 3699 | 3822 | 3599 | 3791 | 5994,32 | 8261 | 3728 | 0,452 | -1,147 | -28,973 | 1,44E-184 | 2,27E-183 | 0,0007265 | 0,002257 | 0,0013148 | 0,0013148 | TRUE | 0,011998 | No |
| orf19.5284 | 63 | 58 | 60 | 51 | 100 | 110 | 66 | 146 | 57 | 59 | 57 | 77 | 89 | 119 | 68 | 111 | 79,59 | 62 | 97 | 1,573 | 0,653 | 3,341 | 0,000834 | 0,001183 | 0,0299156 | 0,018634 | 0,0232136 | 0,0232136 | TRUE | 0,274495 | No |
| orf19.5285 | 1366 | 1089 | 1071 | 848 | 1630 | 1305 | 1250 | 1866 | 1244 | 1101 | 1014 | 1274 | 1450 | 1417 | 1280 | 1423 | 1275,35 | 1158 | 1392 | 1,205 | 0,269 | 3,566 | 0,000362 | 0,000526 | 0,0060142 | 0,003073 | 0,0046649 | 0,0046649 | TRUE | 0,0856578 | No |
| orf19.5286 | 3661 | 2881 | 3100 | 2443 | 3513 | 2892 | 3007 | 4181 | 3333 | 2914 | 2934 | 3670 | 3125 | 3140 | 3080 | 3189 | 3173,01 | 3213 | 3134 | 0,977 | -0,034 | -0,496 | 0,620042 | 0,658795 | 0,0054862 | 0,002453 | 0,0042627 | 0,0042627 | TRUE | 0,1082441 | No |
| orf19.5287 | 128 | 103 | 116 | 98 | 170 | 150 | 162 | 195 | 117 | 104 | 110 | 147 | 151 | 163 | 166 | 149 | 138,3 | 120 | 157 | 1,33 | 0,411 | 3,204 | 0,001355 | 0,001897 | 0,0016739 | 0,011588 | 0,0083402 | 0,0083402 | TRUE | 0,1920509 | No |
| orf19.5288 | 2E+05 | 1E+05 | 2E+05 | 1E+05 | 16814 | 13499 | 13675 | 20038 | 2E+05 | 1E+05 | 1E+05 | 2E+05 | 14956 | 14656 | 14005 | 15285 | 83467,5 | 152210 | 14726 | 0,097 | -3,369 | -77,4 | 0 | 0 | 0,0016082 | 0,002053 | 0,0017856 | 0,0017856 | TRUE | 0,0132679 | No |
| orf19.5288.1 | 796 | 765 | 759 | 441 | 1141 | 886 | 900 | 1099 | 725 | 774 | 718 | 662 | 1015 | 962 | 922 | 838 | 827,01 | 720 | 934 | 1,294 | 0,371 | 5,078 | 3,81E-07 | 6,43E-07 | 0,0041401 | 0,003634 | 0,0038892 | 0,0038892 | TRUE | 0,0602188 | No |
| orf19.5289 | 5 | 3 | 11 | 4 | 10 | 11 | 14 | 22 | 5 | 3 | 10 | 6 | 9 | 12 | 14 | 17 | 9,5 | 6 | 13 | 2,166 | 1,115 | 2,278 | 0,02274 | 0,029005 | 0,0360402 | 0,141158 | 0,1062668 | 0,1062668 | TRUE | 0,654509 | No |
| orf19.529 | 2659 | 2208 | 2530 | 1680 | 1541 | 1350 | 1425 | 1969 | 2421 | 2233 | 2394 | 2524 | 1371 | 1466 | 1459 | 1502 | 1921,23 | 2393 | 1450 | 0,606 | -0,722 | -13,89 | 7,24E-44 | 3,04E-43 | 0,0015161 | 0,002725 | 0,0020445 | 0,0020445 | TRUE | 0,0243381 | No |
| orf19.5290 | 0 | 0 | 0 | 0 | 0 | 0 | 0 | 0 | 0 | 0 | 0 | 0 | 0 | 0 | 0 | 0 | 0 | 0 | 0 | NA | NA | NA | NA | NA | NA | NA | NA | NA | NA | NA | NA |
| orf19.5291 | 919 | 1049 | 1032 | 703 | 1488 | 1352 | 1269 | 1609 | 837 | 1061 | 977 | 1056 | 1324 | 1468 | 1300 | 1227 | 1156,1 | 983 | 1330 | 1,355 | 0,438 | 5,371 | 7,83E-08 | 1,36E-07 | 0,0073673 | 0,00318 | 0,0054837 | 0,0054837 | TRUE | 0,1224666 | No |
| orf19.529.1 | 64 | 77 | 97 | 58 | 47 | 50 | 49 | 63 | 58 | 78 | 92 | 87 | 42 | 54 | 50 | 48 | 63,68 | 79 | 48 | 0,618 | -0,694 | -3,606 | 0,000311 | 0,000454 | 0,0133882 | 0,022783 | 0,0190886 | 0,0190886 | TRUE | 0,3143113 | No |
| orf19.5292 | 914 | 945 | 873 | 538 | 1626 | 1367 | 1381 | 1789 | 832 | 956 | 826 | 808 | 1446 | 1484 | 1414 | 1365 | 1141,47 | 856 | 1427 | 1,665 | 0,735 | 11,805 | 3,67E-32 | 1,24E-31 | 0,0025011 | 0,003194 | 0,0027723 | 0,0027723 | TRUE | 0,074399 | No |
| orf19.5293 | 2404 | 2684 | 2742 | 1651 | 4465 | 4173 | 3866 | 4830 | 2189 | 2715 | 2595 | 2480 | 3971 | 4531 | 3959 | 3684 | 3265,53 | 2495 | 4036 | 1,618 | 0,694 | 9,152 | 5,58E-20 | 1,42E-19 | 0,0075162 | 0,002442 | 0,0051894 | 0,0051894 | TRUE | 0,0843989 | No |
| orf19.5294 | 9527 | 8395 | 8787 | 6011 | 12684 | 10628 | 10268 | 14661 | 8674 | 8491 | 8316 | 9029 | 11282 | 11539 | 10516 | 11184 | 9878,81 | 8628 | 11130 | 1,291 | 0,368 | 8,748 | 2,17E-18 | 5,28E-18 | 0,0012563 | 0,002171 | 0,0015956 | 0,0015956 | TRUE | 0,0167681 | No |
| orf19.5295 | 1104 | 1055 | 1023 | 670 | 2330 | 1913 | 1952 | 2628 | 1005 | 1067 | 968 | 1006 | 2072 | 2077 | 1999 | 2005 | 1525 | 1012 | 2038 | 2,015 | 1,01 | 20,978 | 1,04E-97 | 8,88E-97 | 0,0001376 | 0,002903 | 0,0014697 | 0,0014697 | TRUE | 0,016972 | No |
| orf19.5296 | 1687 | 1473 | 1589 | 915 | 1161 | 896 | 924 | 1249 | 1536 | 1490 | 1504 | 1374 | 1033 | 973 | 946 | 953 | 1226,07 | 1476 | 976 | 0,66 | -0,6 | -10,962 | 5,79E-28 | 1,77E-27 | 0,0010099 | 0,003114 | 0,0020311 | 0,0020311 | TRUE | 0,0231943 | No |
| orf19.5297 | 1103 | 1141 | 1159 | 712 | 1011 | 895 | 865 | 1092 | 1004 | 1154 | 1097 | 1070 | 899 | 972 | 886 | 833 | 989,31 | 1081 | 898 | 0,829 | -0,271 | -4,156 | 3,24E-05 | 5,00E-05 | 0,0027674 | 0,003372 | 0,0030594 | 0,0030594 | TRUE | 0,0360537 | No |
| orf19.5299 | 3697 | 3624 | 3606 | 1957 | 4856 | 3463 | 3673 | 4915 | 3366 | 3665 | 3413 | 2940 | 4319 | 3760 | 3762 | 3749 | 3621,71 | 3346 | 3898 | 1,164 | 0,219 | 3,115 | 0,001837 | 0,002541 | 0,0062804 | 0,002402 | 0,0044611 | 0,0044611 | TRUE | 0,0786931 | No |
| orf19.5299.1 | 1261 | 1184 | 1348 | 588 | 649 | 495 | 556 | 690 | 1148 | 1197 | 1276 | 883 | 577 | 537 | 569 | 526 | 839,38 | 1126 | 552 | 0,489 | -1,032 | -10,636 | 2,03E-26 | 5,99E-26 | 0,0122119 | 0,003611 | 0,0077097 | 0,0077097 | TRUE | 0,238473 | No |
| orf19.53 | 942 | 815 | 957 | 532 | 1259 | 933 | 1106 | 1446 | 858 | 824 | 906 | 799 | 1120 | 1013 | 1133 | 1103 | 969,42 | 847 | 1092 | 1,288 | 0,365 | 6,024 | 1,70E-09 | 3,16E-09 | 0,001496 | 0,0034 | 0,0024597 | 0,0024597 | TRUE | 0,0271212 | No |
| orf19.530 | 284 | 314 | 326 | 190 | 169 | 119 | 162 | 159 | 259 | 318 | 309 | 285 | 150 | 129 | 166 | 121 | 217,1 | 293 | 142 | 0,483 | -1,051 | -8,858 | 8,16E-19 | 2,01E-18 | 0,0087488 | 0,008122 | 0,0083735 | 0,0083735 | TRUE | 0,1338598 | No |
| orf19.5300 | 2077 | 1980 | 2055 | 1318 | 3195 | 2819 | 2683 | 3670 | 1891 | 2003 | 1945 | 1980 | 2842 | 3061 | 2748 | 2800 | 2408,5 | 1955 | 2863 | 1,465 | 0,551 | 11,654 | 2,20E-31 | 7,29E-31 | 0,0010939 | 0,002585 | 0,0017038 | 0,0017038 | TRUE | 0,0256821 | No |
| orf19.5302 | 127 | 104 | 111 | 80 | 1146 | 1130 | 1064 | 1522 | 116 | 105 | 105 | 120 | 1019 | 1227 | 1090 | 1161 | 617,86 | 112 | 1124 | 10,128 | 3,34 | 33,776 | 4,48E-250 | 9,30E-249 | 1,00E-08 | 0,004175 | 0,0041996 | 0,0041996 | TRUE | 0,0477225 | No |

| | | | | | | | | | | | | | | | | | | | | | | | | | | | | | | | |
|---|---|---|---|---|---|---|---|---|---|---|---|---|---|---|---|---|---|---|---|---|---|---|---|---|---|---|---|---|---|---|---|
| orf19.5303 | 1 | 4 | 1 | 0 | 2 | 2 | 2 | 7 | 1 | 4 | 1 | 0 | 2 | 2 | 2 | 5 | 2,16 | 2 | 3 | 1,875 | 0,907 | 0,907 | 0,364461 | 0,404287 | 0,1671392 | 0,615003 | 0,4559355 | 0,4559355 | TRUE | 0,881301 | No |
| orf19.5304 | 0 | 0 | 0 | 0 | 0 | 0 | 1 | 1 | 0 | 0 | 0 | 0 | 0 | 0 | 1 | 1 | 0,22 | 0 | 0 | 2,309 | 1,207 | 0,414 | 0,678623 | 0,714487 | 1,00E-08 | 5,916067 | 6,2007652 | 6,2007652 | TRUE | 0,1072431 | No |
| orf19.5305 | 3624 | 2964 | 3130 | 2256 | 1938 | 1511 | 1501 | 2258 | 3299 | 2998 | 2962 | 3389 | 1724 | 1641 | 1537 | 1722 | 2409,04 | 3162 | 1656 | 0,524 | -0,931 | -15,565 | 1,25E-54 | 6,28E-54 | 0,0030938 | 0,002585 | 0,0029826 | 0,0029826 | TRUE | 0,0279787 | No |
| orf19.5306 | 0 | 0 | 0 | 0 | 2 | 1 | 5 | 8 | 0 | 0 | 0 | 0 | 2 | 1 | 5 | 6 | 1,76 | 0 | 4 | 18,126 | 4,18 | 3,041 | 0,002356 | 0,003233 | 0,4678609 | 0,752191 | 0,6840427 | 0,6840427 | TRUE | 0,0568797 | No |
| orf19.5307 | 1372 | 1154 | 1221 | 781 | 4781 | 3952 | 4379 | 5937 | 1249 | 1167 | 1156 | 1173 | 4253 | 4291 | 4485 | 4529 | 2787,73 | 1186 | 4390 | 3,698 | 1,887 | 41,057 | 0 | 0 | 0,0007364 | 0,002511 | 0,0014699 | 0,0014699 | TRUE | 0,0164304 | No |
| orf19.5308 | 92 | 81 | 82 | 48 | 129 | 110 | 114 | 167 | 84 | 82 | 78 | 72 | 115 | 119 | 117 | 127 | 99,21 | 79 | 120 | 1,51 | 0,595 | 4,272 | 1,94E-05 | 3,03E-05 | 1,00E-08 | 0,015352 | 0,0080231 | 0,0080231 | TRUE | 0,0227213 | No |
| orf19.5311 | 2132 | 1880 | 2105 | 1252 | 2597 | 2047 | 2298 | 3048 | 1941 | 1901 | 1992 | 1881 | 2310 | 2222 | 2353 | 2325 | 2115,79 | 1929 | 2302 | 1,193 | 0,255 | 6,099 | 1,07E-09 | 2,00E-09 | 1,00E-08 | 0,002661 | 0,0011925 | 0,0011925 | TRUE | 0,0063644 | No |
| orf19.5312 | 2645 | 2323 | 2580 | 1684 | 3579 | 3363 | 3411 | 4746 | 2408 | 2349 | 2442 | 2530 | 3183 | 3651 | 3493 | 3620 | 2959,66 | 2432 | 3487 | 1,435 | 0,521 | 9,923 | 3,31E-23 | 9,04E-23 | 0,0018912 | 0,002483 | 0,0022872 | 0,0022872 | TRUE | 0,0421418 | No |
| orf19.5316 | 165 | 147 | 128 | 88 | 197 | 166 | 200 | 277 | 150 | 149 | 121 | 132 | 175 | 180 | 205 | 211 | 165,48 | 138 | 193 | 1,397 | 0,482 | 4,004 | 6,22E-05 | 9,43E-05 | 0,004066 | 0,01002 | 0,00764 | 0,00764 | TRUE | 0,0813003 | No |
| orf19.5318 | 1453 | 1342 | 1474 | 937 | 1025 | 863 | 937 | 1163 | 1323 | 1357 | 1395 | 1408 | 912 | 937 | 960 | 887 | 1147,27 | 1371 | 924 | 0,674 | -0,57 | -11,159 | 6,47E-29 | 2,02E-28 | 1,00E-08 | 0,003188 | 0,0016038 | 0,0016038 | TRUE | 0,0091021 | No |
| orf19.532 | 723 | 605 | 558 | 386 | 3373 | 2640 | 2503 | 3830 | 658 | 612 | 528 | 580 | 3000 | 2866 | 2563 | 2922 | 1716,19 | 594 | 2838 | 4,77 | 2,254 | 30,699 | 5,87E-207 | 1,03E-205 | 0,00515 | 0,002807 | 0,004109 | 0,004109 | TRUE | 0,0722982 | No |
| orf19.5320 | 271 | 227 | 274 | 184 | 121 | 101 | 101 | 145 | 247 | 230 | 259 | 276 | 108 | 110 | 103 | 111 | 180,42 | 253 | 108 | 0,428 | -1,223 | -10,976 | 4,99E-28 | 1,53E-27 | 1,00E-08 | 0,009359 | 0,0054955 | 0,0054955 | TRUE | 0,0415189 | No |
| orf19.5321 | 2304 | 1929 | 2211 | 1375 | 2757 | 2352 | 2582 | 3497 | 2098 | 1951 | 2093 | 2065 | 2452 | 2554 | 2644 | 2668 | 2315,55 | 2052 | 2580 | 1,257 | 0,33 | 7,128 | 1,02E-12 | 2,11E-12 | 0,0009377 | 0,002608 | 0,0016196 | 0,0016196 | TRUE | 0,0136788 | No |
| orf19.5322 | 776 | 604 | 696 | 388 | 1212 | 981 | 1136 | 1441 | 707 | 611 | 659 | 583 | 1078 | 1065 | 1163 | 1099 | 870,58 | 640 | 1101 | 1,714 | 0,777 | 11,598 | 4,21E-31 | 1,39E-30 | 0,0027576 | 0,003554 | 0,0030561 | 0,0030561 | TRUE | 0,058539 | No |
| orf19.5323 | 1022 | 784 | 868 | 609 | 2883 | 2402 | 2424 | 3541 | 930 | 793 | 821 | 915 | 2564 | 2608 | 2482 | 2701 | 1726,94 | 865 | 2589 | 2,999 | 1,584 | 26,541 | 3,29E-155 | 4,45E-154 | 0,0026776 | 0,002802 | 0,0026199 | 0,0026199 | TRUE | 0,0373385 | No |
| orf19.5325 | 3061 | 2351 | 2700 | 1573 | 2680 | 1909 | 2362 | 3237 | 2787 | 2378 | 2555 | 2363 | 2384 | 2073 | 2419 | 2469 | 2428,44 | 2521 | 2336 | 0,926 | -0,11 | -1,586 | 0,112652 | 0,134486 | 0,0054693 | 0,002581 | 0,0042175 | 0,0042175 | TRUE | 0,0689368 | No |
| orf19.5326 | 2953 | 2411 | 2639 | 1916 | 2383 | 1979 | 2053 | 2881 | 2689 | 2438 | 2498 | 2878 | 2120 | 2149 | 2103 | 2198 | 2383,9 | 2626 | 2142 | 0,817 | -0,291 | -5,228 | 1,71E-07 | 2,93E-07 | 0,0025042 | 0,002591 | 0,0025561 | 0,0025561 | TRUE | 0,048051 | No |
| orf19.5328 | 1885 | 1605 | 1722 | 1045 | 3470 | 3017 | 3357 | 4407 | 1716 | 1623 | 1630 | 1570 | 3086 | 3276 | 3438 | 3362 | 2462,59 | 1635 | 3290 | 2,01 | 1,007 | 20,34 | 5,68E-92 | 4,51E-91 | 0,0013949 | 0,002573 | 0,0018833 | 0,0018833 | TRUE | 0,0214001 | No |
| orf19.533 | 0 | 0 | 0 | 1 | 0 | 0 | 0 | 0 | 0 | 0 | 0 | 2 | 0 | 0 | 0 | 0 | 0,19 | 0 | 0 | 0,528 | -0,92 | -0,26 | 0,794572 | 0,821771 | 1,00E-08 | 7,03723 | 10 | 10 | TRUE | 0,6271619 | No |
| orf19.5334 | 3949 | 3043 | 3312 | 2275 | 1914 | 1545 | 1672 | 2540 | 3595 | 3078 | 3135 | 3417 | 1702 | 1677 | 1712 | 1938 | 2531,84 | 3306 | 1757 | 0,532 | -0,91 | -13,873 | 9,17E-44 | 3,83E-43 | 0,004497 | 0,002559 | 0,0037046 | 0,0037046 | TRUE | 0,0589638 | No |
| orf19.5335 | 2468 | 2020 | 2303 | 1477 | 3006 | 2469 | 2699 | 3687 | 2247 | 2043 | 2180 | 2219 | 2674 | 2681 | 2764 | 2812 | 2452,41 | 2172 | 2733 | 1,259 | 0,332 | 7,428 | 1,10E-13 | 2,36E-13 | 0,0007425 | 0,002576 | 0,0015017 | 0,0015017 | TRUE | 0,0197018 | No |
| orf19.5337 | 416 | 316 | 335 | 300 | 429 | 351 | 317 | 492 | 379 | 320 | 317 | 451 | 382 | 381 | 325 | 375 | 366,08 | 367 | 366 | 1,005 | 0,007 | 0,06 | 0,952107 | 0,957968 | 0,0133571 | 0,005645 | 0,0089979 | 0,0089979 | TRUE | 0,2441314 | No |
| orf19.5338 | 1709 | 1466 | 1610 | 1124 | 2434 | 1971 | 2065 | 2943 | 1556 | 1483 | 1524 | 1688 | 2165 | 2140 | 2115 | 2245 | 1864,44 | 1563 | 2166 | 1,39 | 0,475 | 9,531 | 1,56E-21 | 4,11E-21 | 0,0010214 | 0,002746 | 0,0018285 | 0,0018285 | TRUE | 0,03173 | No |
| orf19.5339 | 2825 | 1961 | 2175 | 1511 | 1838 | 1338 | 1623 | 2164 | 2572 | 1983 | 2058 | 2270 | 1635 | 1453 | 1662 | 1651 | 1910,5 | 2221 | 1600 | 0,721 | -0,472 | -5,882 | 4,07E-09 | 7,43E-09 | 0,0082394 | 0,002728 | 0,0056605 | 0,0056605 | TRUE | 0,1404892 | No |
| orf19.5340 | 5155 | 4336 | 4802 | 2916 | 2995 | 2312 | 2612 | 3492 | 4693 | 4385 | 4545 | 4380 | 2664 | 2510 | 2675 | 2664 | 3564,58 | 4501 | 2628 | 0,584 | -0,776 | -18,7 | 4,92E-78 | 3,41E-77 | 0,0005985 | 0,002408 | 0,0013576 | 0,0013576 | TRUE | 0,0107119 | No |
| orf19.5341 | 7571 | 5535 | 5844 | 4185 | 12987 | 10083 | 10312 | 15751 | 6893 | 5598 | 5531 | 6287 | 11552 | 10947 | 10561 | 12015 | 8672,89 | 6077 | 11269 | 1,855 | 0,891 | 12,508 | 6,71E-36 | 2,44E-35 | 0,0070455 | 0,002189 | 0,0047454 | 0,0047454 | TRUE | 0,1009172 | No |
| orf19.5342 | 1211 | 1098 | 1182 | 841 | 2251 | 1984 | 2117 | 2860 | 1103 | 1111 | 1119 | 1263 | 2002 | 2154 | 2168 | 2182 | 1637,63 | 1149 | 2126 | 1,857 | 0,893 | 15,954 | 2,67E-57 | 1,39E-56 | 0,0018263 | 0,002844 | 0,0023205 | 0,0023205 | TRUE | 0,0483846 | No |
| orf19.5342.1 | 1495 | 1066 | 1269 | 800 | 1401 | 915 | 1122 | 1569 | 1361 | 1078 | 1201 | 1202 | 1246 | 993 | 1149 | 1197 | 1178,44 | 1210 | 1146 | 0,948 | -0,078 | -0,933 | 0,351077 | 0,391005 | 0,0078298 | 0,003158 | 0,0058225 | 0,0058225 | TRUE | 0,0948286 | No |
| orf19.5342.2 | 190 | 123 | 157 | 101 | 594 | 332 | 471 | 596 | 173 | 124 | 149 | 152 | 528 | 360 | 482 | 455 | 302,94 | 150 | 456 | 3,056 | 1,612 | 12,354 | 4,66E-35 | 1,67E-34 | 0,0185962 | 0,006398 | 0,0117365 | 0,0117365 | TRUE | 0,2253574 | No |
| orf19.5343 | 323 | 288 | 345 | 231 | 230 | 191 | 208 | 274 | 294 | 291 | 327 | 347 | 205 | 207 | 213 | 209 | 261,61 | 315 | 208 | 0,666 | -0,587 | -6,32 | 2,62E-10 | 5,00E-10 | 1,00E-08 | 0,007087 | 0,0043471 | 0,0043471 | TRUE | 0,0446548 | No |
| orf19.5345 | 3409 | 2671 | 2985 | 2179 | 3085 | 2837 | 2931 | 3998 | 3104 | 2701 | 2825 | 3273 | 2744 | 3080 | 3002 | 3050 | 2972,38 | 2976 | 2969 | 0,999 | -0,002 | -0,033 | 0,973664 | 0,976574 | 0,0047756 | 0,002481 | 0,0037309 | 0,0037309 | TRUE | 0,0529317 | No |
| orf19.5346 | 1446 | 1213 | 1423 | 821 | 1053 | 803 | 949 | 1223 | 1317 | 1227 | 1347 | 1233 | 937 | 872 | 972 | 933 | 1104,58 | 1281 | 928 | 0,724 | -0,466 | -8,239 | 1,73E-16 | 3,98E-16 | 0,0011108 | 0,003233 | 0,0021478 | 0,0021478 | TRUE | 0,0186905 | No |
| orf19.5348 | 11880 | 9871 | 11245 | 7228 | 1430 | 1129 | 1197 | 1773 | 10816 | 9983 | 10642 | 10858 | 1272 | 1226 | 1226 | 1352 | 5921,97 | 10575 | 1269 | 0,12 | -3,056 | -63,691 | 0 | 0 | 0,0012532 | 0,00226 | 0,0017933 | 0,0017933 | TRUE | 0,0246981 | No |
| orf19.535 | 58 | 38 | 39 | 18 | 1348 | 1096 | 1101 | 1710 | 53 | 38 | 37 | 27 | 1199 | 1190 | 1128 | 1304 | 622,01 | 39 | 1205 | 30,153 | 4,914 | 36,265 | 5,78E-288 | 1,42E-286 | 0,0038782 | 0,004161 | 0,0040755 | 0,0040755 | TRUE | 0,4504749 | No |
| orf19.5350 | 907 | 749 | 869 | 549 | 1261 | 1021 | 1067 | 1449 | 826 | 758 | 822 | 825 | 1122 | 1109 | 1093 | 1105 | 957,33 | 808 | 1107 | 1,372 | 0,456 | 8,578 | 9,69E-18 | 2,32E-17 | 1,00E-08 | 0,003417 | 0,0016294 | 0,0016294 | TRUE | 0,0212527 | No |
| orf19.5351 | 5765 | 4410 | 4902 | 2748 | 9890 | 6703 | 8380 | 11345 | 5249 | 4460 | 4639 | 4128 | 8797 | 7278 | 8582 | 8654 | 6473,37 | 4619 | 8328 | 1,802 | 0,85 | 10,987 | 4,41E-28 | 1,35E-27 | 0,0085485 | 0,002241 | 0,0055717 | 0,0055717 | TRUE | 0,1028544 | No |
| orf19.5352 | 2701 | 2219 | 2469 | 1659 | 2047 | 1787 | 1798 | 2567 | 2459 | 2244 | 2337 | 2492 | 1821 | 1940 | 1841 | 1958 | 2136,58 | 2383 | 1890 | 0,794 | -0,333 | -6,682 | 2,36E-11 | 4,69E-11 | 0,0011827 | 0,002655 | 0,0019094 | 0,0019094 | TRUE | 0,0185232 | No |
| orf19.5353 | 13 | 11 | 14 | 6 | 29 | 35 | 33 | 44 | 12 | 11 | 13 | 9 | 26 | 38 | 34 | 34 | 22,05 | 11 | 33 | 2,843 | 1,507 | 4,741 | 2,13E-06 | 3,49E-06 | 1,00E-08 | 0,061954 | 0,0367406 | 0,0367406 | TRUE | 0,1516263 | No |
| orf19.5356 | 6200 | 5202 | 5729 | 3921 | 7504 | 6634 | 6907 | 9704 | 5645 | 5261 | 5422 | 5890 | 6675 | 7203 | 7074 | 7402 | 6321,4 | 5554 | 7088 | 1,277 | 0,353 | 7,301 | 2,87E-13 | 6,05E-13 | 0,0019363 | 0,002246 | 0,002078 | 0,002078 | TRUE | 0,0194656 | No |
| orf19.5357 | 2319 | 1975 | 2249 | 1587 | 2259 | 1909 | 2157 | 2881 | 2111 | 1998 | 2128 | 2384 | 2009 | 2073 | 2209 | 2198 | 2138,74 | 2155 | 2122 | 0,986 | -0,02 | -0,328 | 0,74265 | 0,774352 | 0,0029743 | 0,002655 | 0,0029594 | 0,0029594 | TRUE | 0,0582712 | No |
| orf19.5358 | 1585 | 1302 | 1351 | 1011 | 1871 | 1541 | 1658 | 2355 | 1443 | 1317 | 1279 | 1519 | 1664 | 1673 | 1698 | 1796 | 1548,61 | 1390 | 1708 | 1,233 | 0,302 | 4,949 | 7,46E-07 | 1,25E-06 | 0,0029682 | 0,00289 | 0,0029067 | 0,0029067 | TRUE | 0,0433553 | No |
| orf19.536 | 2944 | 2502 | 2835 | 1698 | 2828 | 2286 | 2685 | 3473 | 2680 | 2531 | 2683 | 2551 | 2515 | 2482 | 2750 | 2649 | 2605,13 | 2611 | 2599 | 0,995 | -0,007 | -0,157 | 0,87549 | 0,894574 | 0,0012259 | 0,002544 | 0,0017696 | 0,0017696 | TRUE | 0,0179181 | No |
| orf19.5360 | 1095 | 799 | 843 | 612 | 1142 | 859 | 913 | 1412 | 997 | 808 | 798 | 919 | 1016 | 933 | 935 | 1077 | 935,34 | 880 | 990 | 1,126 | 0,172 | 2,101 | 0,035646 | 0,044777 | 0,0068798 | 0,003449 | 0,005334 | 0,005334 | TRUE | 0,1005325 | No |
| orf19.5362 | 261 | 180 | 225 | 159 | 231 | 198 | 172 | 305 | 238 | 182 | 213 | 239 | 205 | 215 | 176 | 233 | 212,59 | 218 | 207 | 0,957 | -0,064 | -0,536 | 0,592244 | 0,632634 | 0,0097393 | 0,008251 | 0,0088734 | 0,0088734 | TRUE | 0,1354248 | No |
| orf19.5363 | 1149 | 881 | 967 | 642 | 989 | 713 | 803 | 1080 | 1046 | 891 | 915 | 964 | 880 | 774 | 822 | 824 | 889,59 | 954 | 825 | 0,865 | -0,209 | -3,106 | 0,001894 | 0,002618 | 0,0029177 | 0,003522 | 0,0032126 | 0,0032126 | TRUE | 0,0538412 | No |
| orf19.5364 | 1712 | 1361 | 1529 | 997 | 2280 | 1750 | 2032 | 2800 | 1559 | 1377 | 1447 | 1498 | 2028 | 1900 | 2081 | 2136 | 1753,1 | 1470 | 2036 | 1,386 | 0,471 | 8,447 | 3,00E-17 | 7,06E-17 | 0,0020744 | 0,002791 | 0,0023946 | 0,0023946 | TRUE | 0,0239988 | No |
| orf19.5365 | 841 | 782 | 861 | 562 | 2318 | 2094 | 2193 | 3094 | 766 | 791 | 815 | 844 | 2062 | 2274 | 2246 | 2360 | 1519,63 | 804 | 2236 | 2,787 | 1,479 | 25,29 | 4,15E-141 | 5,10E-140 | 0,0020032 | 0,002906 | 0,0024053 | 0,0024053 | TRUE | 0,0336938 | No |
| orf19.5365.1 | 743 | 559 | 556 | 424 | 824 | 638 | 597 | 890 | 676 | 565 | 526 | 637 | 733 | 693 | 611 | 679 | 640,11 | 601 | 679 | 1,131 | 0,178 | 2,026 | 0,042721 | 0,053265 | 0,0076751 | 0,004101 | 0,0058139 | 0,0058139 | TRUE | 0,091303 | No |
| orf19.5367 | 559 | 493 | 517 | 341 | 1020 | 838 | 940 | 1243 | 509 | 499 | 489 | 512 | 907 | 910 | 963 | 948 | 717,13 | 502 | 932 | 1,858 | 0,894 | 15,05 | 3,47E-51 | 1,63E-50 | 1,00E-08 | 0,003879 | 0,0018239 | 0,0018239 | TRUE | 0,0054665 | No |
| orf19.5368 | 828 | 657 | 732 | 617 | 870 | 733 | 846 | 1193 | 754 | 664 | 693 | 927 | 774 | 796 | 866 | 910 | 798,01 | 760 | 836 | 1,106 | 0,146 | 1,51 | 0,131122 | 0,155344 | 0,0131497 | 0,003692 | 0,007699 | 0,007699 | TRUE | 0,2437871 | No |
| orf19.5369 | 1293 | 1064 | 1173 | 901 | 3293 | 2709 | 2893 | 3999 | 1177 | 1076 | 1110 | 1353 | 2929 | 2941 | 2963 | 3050 | 2075,05 | 1179 | 2971 | 2,529 | 1,339 | 21,089 | 9,96E-99 | 8,57E-98 | 0,0038899 | 0,002674 | 0,0032526 | 0,0032526 | TRUE | 0,108689 | No |
| orf19.537 | 260 | 190 | 239 | 147 | 300 | 194 | 239 | 319 | 237 | 192 | 226 | 221 | 267 | 211 | 245 | 243 | 230,18 | 219 | 242 | 1,104 | 0,143 | 1,352 | 0,17639 | 0,205424 | 0,0042845 | 0,007776 | 0,0063593 | 0,0063593 | TRUE | 0,0786965 | No |
| orf19.5370 | 746 | 603 | 637 | 429 | 1057 | 750 | 864 | 1163 | 679 | 610 | 603 | 644 | 940 | 814 | 885 | 887 | 757,85 | 634 | 882 | 1,392 | 0,477 | 7,138 | 9,46E-13 | 1,96E-12 | 0,0020262 | 0,00378 | 0,0029189 | 0,0029189 | TRUE | 0,0301491 | No |
| orf19.5376 | 648 | 605 | 674 | 404 | 465 | 422 | 432 | 570 | 590 | 612 | 638 | 607 | 414 | 458 | 442 | 435 | 524,45 | 612 | 437 | 0,714 | -0,487 | -7,252 | 4,12E-13 | 8,67E-13 | 1,00E-08 | 0,004556 | 0,0023837 | 0,0023837 | TRUE | 0,0152526 | No |
| orf19.5377 | 2526 | 2198 | 2396 | 1439 | 1482 | 1236 | 1268 | 1823 | 2300 | 2223 | 2268 | 2162 | 1318 | 1342 | 1299 | 1391 | 1787,67 | 2238 | 1338 | 0,597 | -0,743 | -16,556 | 1,44E-61 | 8,02E-61 | 1,95E-06 | 0,002776 | 0,0013473 | 0,0013473 | TRUE | 0,0093043 | No |
| orf19.5378 | 2644 | 2321 | 2545 | 1377 | 2499 | 1973 | 2237 | 2946 | 2407 | 2347 | 2409 | 2068 | 2223 | 2142 | 2291 | 2247 | 2266,86 | 2308 | 2226 | 0,963 | -0,055 | -1,007 | 0,314012 | 0,352869 | 0,0022746 | 0,00262 | 0,0024268 | 0,0024268 | TRUE | 0,0552272 | No |
| orf19.5379 | 1985 | 1766 | 1751 | 1284 | 1225 | 908 | 951 | 1336 | 1807 | 1786 | 1657 | 1929 | 1090 | 986 | 974 | 1019 | 1405,98 | 1795 | 1017 | 0,568 | -0,816 | -13,595 | 4,27E-42 | 1,73E-41 | 0,0021987 | 0,002977 | 0,0026979 | 0,0026979 | TRUE | 0,032228 | No |
| orf19.538 | 314 | 279 | 297 | 234 | 200 | 187 | 142 | 203 | 286 | 282 | 281 | 352 | 178 | 203 | 145 | 155 | 235,23 | 300 | 170 | 0,569 | -0,814 | -6,839 | 8,00E-12 | 1,62E-11 | 0,0111734 | 0,007653 | 0,0090472 | 0,0090472 | TRUE | 0,1664562 | No |
| orf19.5380 | 761 | 734 | 767 | 432 | 914 | 747 | 799 | 1058 | 693 | 742 | 726 | 649 | 813 | 811 | 818 | 807 | 757,42 | 702 | 812 | 1,152 | 0,204 | 3,46 | 0,00054 | 0,000775 | 1,00E-08 | 0,003781 | 0,0020204 | 0,0020204 | TRUE | 0,0257504 | No |
| orf19.5381 | 1352 | 1333 | 1412 | 971 | 1024 | 894 | 964 | 1257 | 1231 | 1348 | 1336 | 1459 | 911 | 971 | 987 | 959 | 1150,2 | 1344 | 957 | 0,714 | -0,486 | -8,041 | 8,94E-16 | 2,02E-15 | 0,002111 | 0,003186 | 0,0026249 | 0,0026249 | TRUE | 0,0385481 | No |
| orf19.5382 | 0 | 0 | 0 | 0 | 0 | 0 | 0 | 0 | 0 | 0 | 0 | 0 | 0 | 0 | 0 | 0 | 0 | 0 | 0 | NA | NA | NA | NA | NA | NA | NA | NA | NA | NA | NA | NA |
| orf19.5383 | 4131 | 3571 | 3894 | 2657 | 9305 | 7793 | 7831 | 11393 | 3761 | 3612 | 3685 | 3991 | 8277 | 8461 | 8020 | 8691 | 6062,18 | 3762 | 8362 | 2,225 | 1,154 | 26,477 | 1,80E-154 | 2,42E-153 | 0,0012743 | 0,002255 | 0,0016247 | 0,0016247 | TRUE | 0,0192396 | No |
| orf19.5384 | 969 | 910 | 962 | 626 | 1845 | 1517 | 1603 | 2204 | 882 | 920 | 910 | 940 | 1641 | 1647 | 1642 | 1681 | 1283,05 | 913 | 1653 | 1,814 | 0,859 | 18,226 | 3,19E-74 | 2,10E-73 | 1,00E-08 | 0,003067 | 0,0012637 | 0,0012637 | TRUE | 0,0064063 | No |
| orf19.5387 | 777 | 834 | 806 | 527 | 1114 | 918 | 895 | 1118 | 707 | 844 | 763 | 792 | 991 | 997 | 917 | 853 | 857,79 | 776 | 940 | 1,21 | 0,275 | 3,74 | 0,000184 | 0,000272 | 0,0043997 | 0,003577 | 0,0039956 | 0,0039956 | TRUE | 0,04807 | No |
| orf19.5388 | 1108 | 958 | 1065 | 669 | 1038 | 851 | 965 | 1290 | 1009 | 969 | 1008 | 1005 | 923 | 924 | 988 | 984 | 976,26 | 998 | 955 | 0,958 | -0,062 | -1,176 | 0,239687 | 0,273885 | 1,00E-08 | 0,00339 | 0,0016645 | 0,0016645 | TRUE | 0,0063352 | No |
| orf19.5389 | 1226 | 1116 | 1192 | 866 | 625 | 596 | 640 | 839 | 1116 | 1129 | 1128 | 1301 | 556 | 647 | 655 | 640 | 896,55 | 1168 | 624 | 0,536 | -0,901 | -12,418 | 2,09E-35 | 7,54E-35 | 0,0039658 | 0,00351 | 0,0038427 | 0,0038427 | TRUE | 0,06561 | No |
| orf19.539 | 831 | 739 | 780 | 491 | 2182 | 1780 | 1856 | 2519 | 757 | 747 | 738 | 738 | 1941 | 1933 | 1901 | 1922 | 1334,43 | 745 | 1924 | 2,581 | 1,368 | 29,132 | 1,40E-186 | 2,23E-185 | 1,00E-08 | 0,003027 | 0,0011601 | 0,0011601 | TRUE | 0,0013918 | No |

| | | | | | | | | | | | | | | | | | | | | | | | | | | | | | | | |
|---|---|---|---|---|---|---|---|---|---|---|---|---|---|---|---|---|---|---|---|---|---|---|---|---|---|---|---|---|---|---|---|
| orf19.5390 | 0 | 0 | 0 | 0 | 0 | 0 | 0 | 0 | 0 | 0 | 0 | 0 | 0 | 0 | 0 | 0 | 0 | 0 | 0 | NA | NA | NA | NA | NA | NA | NA | NA | NA | NA | NA | NA |
| orf19.5391 | 1707 | 1737 | 1818 | 1133 | 1723 | 1429 | 1555 | 2031 | 1554 | 1757 | 1721 | 1702 | 1533 | 1551 | 1593 | 1549 | 1619,91 | 1684 | 1556 | 0,925 | -0,112 | -2,219 | 0,026454 | 0,033568 | 0,0010538 | 0,002852 | 0,0018411 | 0,0018411 | TRUE | 0,0340473 | No |
| orf19.5392 | 54 | 34 | 35 | 22 | 51 | 69 | 50 | 72 | 49 | 34 | 33 | 33 | 45 | 75 | 51 | 55 | 47,02 | 37 | 56 | 1,491 | 0,576 | 2,513 | 0,011976 | 0,015616 | 0,0248152 | 0,030134 | 0,0280837 | 0,0280837 | TRUE | 0,3719498 | No |
| orf19.5393 | 5202 | 4989 | 5375 | 3109 | 8553 | 7163 | 7430 | 9794 | 4736 | 5046 | 5087 | 4670 | 7608 | 7777 | 7609 | 7471 | 6250,51 | 4885 | 7616 | 1,558 | 0,64 | 16,097 | 2,66E-58 | 1,42E-57 | 0,0008537 | 0,002248 | 0,0013444 | 0,0013444 | TRUE | 0,010034 | No |
| orf19.5394.1 | 1696 | 1395 | 1450 | 998 | 885 | 662 | 696 | 956 | 1544 | 1411 | 1372 | 1499 | 787 | 719 | 713 | 729 | 1096,81 | 1456 | 737 | 0,507 | -0,981 | -16,349 | 4,39E-60 | 2,41E-59 | 0,0017516 | 0,003241 | 0,0024549 | 0,0024549 | TRUE | 0,0251626 | No |
| orf19.5395 | 2957 | 2706 | 2877 | 1838 | 1494 | 1189 | 1346 | 1646 | 2692 | 2737 | 2723 | 2761 | 1329 | 1291 | 1378 | 1256 | 2020,84 | 2728 | 1314 | 0,481 | -1,055 | -24,07 | 5,10E-128 | 5,57E-127 | 5,18E-05 | 0,002691 | 0,0012928 | 0,0012928 | TRUE | 0,0131152 | No |
| orf19.5397 | 324 | 312 | 327 | 202 | 308 | 289 | 270 | 385 | 295 | 316 | 309 | 303 | 274 | 314 | 277 | 294 | 297,67 | 306 | 290 | 0,945 | -0,081 | -0,952 | 0,34091 | 0,381013 | 1,00E-08 | 0,006475 | 0,0035671 | 0,0035671 | TRUE | 0,032436 | No |
| orf19.5399 | 31 | 31 | 15 | 18 | 70 | 73 | 57 | 96 | 28 | 31 | 14 | 27 | 62 | 79 | 58 | 73 | 46,74 | 25 | 68 | 2,727 | 1,448 | 6,031 | 1,63E-09 | 3,03E-09 | 0,018762 | 0,030298 | 0,0272494 | 0,0272494 | TRUE | 0,6027793 | No |
| orf19.54 | 302 | 250 | 282 | 203 | 3334 | 2988 | 3013 | 3968 | 275 | 253 | 267 | 305 | 2965 | 3244 | 3086 | 3027 | 1677,72 | 275 | 3080 | 11,28 | 3,496 | 53,279 | 0 | 0 | 0,0013161 | 0,002824 | 0,002038 | 0,002038 | TRUE | 0,0459039 | No |
| orf19.540 | 393 | 303 | 385 | 215 | 1000 | 712 | 813 | 1041 | 358 | 306 | 364 | 323 | 889 | 773 | 833 | 794 | 580,1 | 338 | 822 | 2,426 | 1,279 | 16,293 | 1,11E-59 | 6,04E-59 | 0,0031076 | 0,004314 | 0,003771 | 0,003771 | TRUE | 0,0497744 | No |
| orf19.5402 | 0 | 0 | 0 | 0 | 0 | 0 | 0 | 0 | 0 | 0 | 0 | 0 | 0 | 0 | 0 | 0 | 0 | 0 | 0 | NA | NA | NA | NA | NA | NA | NA | NA | NA | NA | NA | NA |
| orf19.5404.1 | 1129 | 993 | 1111 | 744 | 1567 | 1383 | 1519 | 1970 | 1028 | 1004 | 1051 | 1118 | 1394 | 1502 | 1556 | 1503 | 1269,38 | 1050 | 1489 | 1,42 | 0,506 | 9,163 | 5,03E-20 | 1,28E-19 | 0,0012132 | 0,003078 | 0,0020989 | 0,0020989 | TRUE | 0,0223021 | No |
| orf19.5406 | 4036 | 3538 | 3864 | 2249 | 3936 | 3062 | 3589 | 4700 | 3675 | 3578 | 3657 | 3378 | 3501 | 3324 | 3676 | 3585 | 3546,8 | 3572 | 3522 | 0,985 | -0,022 | -0,472 | 0,636682 | 0,6749 | 0,0011604 | 0,002409 | 0,0017339 | 0,0017339 | TRUE | 0,01683 | No |
| orf19.5407 | 14021 | 11945 | 12862 | 7879 | 14695 | 11617 | 13102 | 17890 | 12766 | 12081 | 12173 | 11836 | 13071 | 12613 | 13418 | 13647 | 12700,4 | 12214 | 13187 | 1,08 | 0,11 | 2,815 | 0,004883 | 0,006556 | 0,000999 | 0,002141 | 0,0013991 | 0,0013991 | TRUE | 0,0113667 | No |
| orf19.5408 | 7561 | 6412 | 7223 | 4628 | 5744 | 4822 | 5266 | 7101 | 6884 | 6485 | 6836 | 6952 | 5109 | 5235 | 5393 | 5417 | 6038,89 | 6789 | 5288 | 0,779 | -0,36 | -9,416 | 4,68E-21 | 1,22E-20 | 0,0006181 | 0,002256 | 0,001235 | 0,001235 | TRUE | 0,0111401 | No |
| orf19.5409 | 501 | 418 | 490 | 308 | 247 | 180 | 192 | 261 | 456 | 423 | 464 | 463 | 220 | 195 | 197 | 199 | 327,02 | 452 | 203 | 0,45 | -1,152 | -13,588 | 4,69E-42 | 1,90E-41 | 1,00E-08 | 0,006077 | 0,0034227 | 0,0034227 | TRUE | 0,0355562 | No |
| orf19.541 | 2366 | 2025 | 2264 | 1499 | 3819 | 3172 | 3350 | 4456 | 2154 | 2048 | 2143 | 2252 | 3397 | 3444 | 3431 | 3399 | 2783,41 | 2149 | 3418 | 1,592 | 0,671 | 16,798 | 2,50E-63 | 1,44E-62 | 1,00E-08 | 0,002512 | 0,0011459 | 0,0011459 | TRUE | 0,0117843 | No |
| orf19.5410 | 1293 | 1044 | 1253 | 759 | 912 | 706 | 848 | 1079 | 1177 | 1056 | 1186 | 1140 | 811 | 767 | 868 | 823 | 978,54 | 1140 | 817 | 0,717 | -0,48 | -7,895 | 2,91E-15 | 6,48E-15 | 0,0016433 | 0,003387 | 0,0025043 | 0,0025043 | TRUE | 0,0306029 | No |
| orf19.5411 | 967 | 799 | 963 | 639 | 913 | 727 | 757 | 994 | 880 | 808 | 911 | 960 | 812 | 789 | 775 | 758 | 836,84 | 890 | 784 | 0,883 | -0,18 | -2,842 | 0,004481 | 0,006028 | 0,0016542 | 0,003616 | 0,0026622 | 0,0026622 | TRUE | 0,0448747 | No |
| orf19.5412 | 1549 | 1305 | 1470 | 941 | 1236 | 1022 | 1086 | 1386 | 1410 | 1320 | 1391 | 1414 | 1099 | 1110 | 1112 | 1057 | 1239,17 | 1384 | 1094 | 0,791 | -0,339 | -6,914 | 4,72E-12 | 9,60E-12 | 1,00E-08 | 0,003103 | 0,0014894 | 0,0014894 | TRUE | 0,0117549 | No |
| orf19.5413 | 1467 | 1196 | 1324 | 859 | 967 | 769 | 860 | 1132 | 1336 | 1210 | 1253 | 1290 | 860 | 835 | 881 | 863 | 1065,99 | 1272 | 860 | 0,676 | -0,565 | -10,599 | 3,02E-26 | 8,88E-26 | 1,00E-08 | 0,003276 | 0,0017601 | 0,0017601 | TRUE | 0,014914 | No |
| orf19.5416 | 7673 | 6941 | 7685 | 4787 | 5015 | 4094 | 4567 | 5942 | 6986 | 7020 | 7273 | 7191 | 4461 | 4445 | 4677 | 4533 | 5823,18 | 7118 | 4529 | 0,636 | -0,652 | -19,125 | 1,57E-81 | 1,13E-80 | 0,0002186 | 0,002264 | 0,0009371 | 0,0009371 | TRUE | 0,0057775 | No |
| orf19.5417 | 13564 | 10673 | 11939 | 7925 | 15799 | 12465 | 13388 | 18500 | 12349 | 10795 | 11299 | 11905 | 14053 | 13534 | 13711 | 14112 | 12719,6 | 11587 | 13852 | 1,196 | 0,258 | 5,602 | 2,12E-08 | 3,77E-08 | 0,0018226 | 0,002141 | 0,0019552 | 0,0019552 | TRUE | 0,0260062 | No |
| orf19.5418 | 1564 | 1341 | 1413 | 937 | 2193 | 1730 | 1865 | 2619 | 1424 | 1356 | 1337 | 1408 | 1951 | 1878 | 1910 | 1998 | 1657,72 | 1381 | 1934 | 1,402 | 0,488 | 10,652 | 1,71E-26 | 5,05E-26 | 1,00E-08 | 0,002834 | 0,0013836 | 0,0013836 | TRUE | 0,0061457 | No |
| orf19.5419 | 4880 | 4043 | 4373 | 2986 | 8583 | 7074 | 7442 | 9921 | 4443 | 4089 | 4139 | 4485 | 7634 | 7680 | 7622 | 7568 | 5957,53 | 4289 | 7626 | 1,779 | 0,831 | 20,652 | 9,30E-95 | 7,59E-94 | 0,0009072 | 0,002259 | 0,0013685 | 0,0013685 | TRUE | 0,0118645 | No |
| orf19.542 | 4983 | 4098 | 4554 | 2720 | 1263 | 939 | 1087 | 1513 | 4537 | 4145 | 4310 | 4086 | 1123 | 1019 | 1113 | 1154 | 2685,95 | 4270 | 1102 | 0,258 | -1,952 | -36,491 | 1,52E-291 | 3,81E-290 | 0,001744 | 0,002529 | 0,0021986 | 0,0021986 | TRUE | 0,0296667 | No |
| orf19.5420 | 5162 | 4070 | 4609 | 3058 | 4917 | 4048 | 4355 | 5920 | 4700 | 4116 | 4362 | 4594 | 4374 | 4395 | 4460 | 4516 | 4439,52 | 4443 | 4436 | 0,999 | -0,002 | -0,035 | 0,971778 | 0,974837 | 0,0015999 | 0,002335 | 0,0018867 | 0,0018867 | TRUE | 0,0300189 | No |
| orf19.5422 | 56 | 43 | 66 | 46 | 68 | 46 | 51 | 81 | 51 | 43 | 62 | 69 | 60 | 50 | 52 | 62 | 56,31 | 56 | 56 | 1,01 | 0,014 | 0,073 | 0,941707 | 0,950026 | 0,0057412 | 0,025496 | 0,0188116 | 0,0188116 | TRUE | 0,1964995 | No |
| orf19.542.2 | 1253 | 1019 | 1130 | 741 | 719 | 489 | 558 | 727 | 1141 | 1031 | 1069 | 1113 | 640 | 531 | 571 | 555 | 831,3 | 1088 | 574 | 0,528 | -0,921 | -13,493 | 1,73E-41 | 6,90E-41 | 0,0025228 | 0,003626 | 0,0031679 | 0,0031679 | TRUE | 0,0701025 | No |
| orf19.5423 | 42 | 49 | 35 | 32 | 59 | 50 | 51 | 75 | 38 | 50 | 33 | 48 | 52 | 54 | 52 | 57 | 48,15 | 42 | 54 | 1,296 | 0,374 | 1,843 | 0,065311 | 0,080007 | 1,00E-08 | 0,029472 | 0,0182184 | 0,0182184 | TRUE | 0,1703872 | No |
| orf19.5424 | 3 | 0 | 3 | 0 | 5 | 5 | 3 | 5 | 3 | 0 | 3 | 0 | 4 | 5 | 3 | 4 | 2,79 | 2 | 4 | 2,74 | 1,454 | 1,618 | 0,105644 | 0,126666 | 1,19E-08 | 0,475238 | 0,3161634 | 0,3161634 | TRUE | 0,3128519 | No |
| orf19.5425 | 6491 | 5568 | 6298 | 3975 | 7248 | 6061 | 6334 | 9053 | 5910 | 5631 | 5960 | 5971 | 6447 | 6581 | 6487 | 6906 | 6236,59 | 5868 | 6605 | 1,126 | 0,171 | 4,458 | 8,28E-06 | 1,32E-05 | 0,0006542 | 0,002249 | 0,0012547 | 0,0012547 | TRUE | 0,0117264 | No |
| orf19.5426 | 1874 | 1631 | 1751 | 1118 | 3100 | 2526 | 2984 | 3858 | 1706 | 1650 | 1657 | 1679 | 2757 | 2743 | 3056 | 2943 | 2273,89 | 1673 | 2875 | 1,718 | 0,781 | 16,026 | 8,44E-58 | 4,46E-57 | 0,0012397 | 0,002618 | 0,0017972 | 0,0017972 | TRUE | 0,0212923 | No |
| orf19.5428 | 134 | 150 | 122 | 89 | 136 | 100 | 85 | 148 | 122 | 152 | 115 | 134 | 121 | 109 | 87 | 113 | 119,04 | 131 | 108 | 0,826 | -0,275 | -1,918 | 0,055125 | 0,068052 | 0,008493 | 0,013134 | 0,0113441 | 0,0113441 | TRUE | 0,1547054 | No |
| orf19.5429 | 2080 | 1836 | 2032 | 1303 | 1037 | 811 | 811 | 1091 | 1894 | 1857 | 1923 | 1957 | 922 | 881 | 831 | 832 | 1387,1 | 1908 | 866 | 0,454 | -1,138 | -22,142 | 1,24E-108 | 1,15E-107 | 0,0004161 | 0,002989 | 0,0017166 | 0,0017166 | TRUE | 0,0235505 | No |
| orf19.543 | 774 | 689 | 713 | 465 | 332 | 283 | 275 | 380 | 705 | 697 | 675 | 699 | 295 | 307 | 282 | 290 | 493,61 | 694 | 294 | 0,423 | -1,242 | -17,967 | 3,56E-72 | 2,28E-71 | 1,00E-08 | 0,004713 | 0,0022292 | 0,0022292 | TRUE | 0,0099243 | No |
| orf19.5430 | 5163 | 4137 | 4540 | 2727 | 6356 | 4683 | 5339 | 7390 | 4701 | 4184 | 4297 | 4096 | 5653 | 5084 | 5468 | 5637 | 4890,1 | 4320 | 5460 | 1,264 | 0,338 | 6,225 | 4,80E-10 | 9,09E-10 | 0,0028078 | 0,002307 | 0,0026152 | 0,0026152 | TRUE | 0,0433676 | No |
| orf19.5431 | 851 | 781 | 854 | 629 | 1043 | 818 | 854 | 1255 | 775 | 790 | 808 | 945 | 928 | 888 | 875 | 957 | 870,69 | 830 | 912 | 1,106 | 0,145 | 2,089 | 0,036726 | 0,04607 | 0,0033466 | 0,003554 | 0,0034561 | 0,0034561 | TRUE | 0,0937904 | No |
| orf19.5432 | 1224 | 1102 | 1183 | 780 | 1585 | 1187 | 1388 | 1790 | 1114 | 1115 | 1120 | 1172 | 1410 | 1289 | 1421 | 1365 | 1250,71 | 1130 | 1371 | 1,216 | 0,282 | 5,514 | 3,51E-08 | 6,16E-08 | 0,0004171 | 0,003093 | 0,0016975 | 0,0016975 | TRUE | 0,0186264 | No |
| orf19.5433 | 556 | 474 | 475 | 300 | 499 | 352 | 422 | 517 | 506 | 479 | 450 | 451 | 444 | 382 | 432 | 394 | 442,3 | 472 | 413 | 0,875 | -0,193 | -2,46 | 0,013889 | 0,018045 | 0,0019749 | 0,005024 | 0,003666 | 0,003666 | TRUE | 0,0299465 | No |
| orf19.5436 | 12961 | 11348 | 12340 | 7960 | 9109 | 7468 | 7847 | 10951 | 11800 | 11477 | 11679 | 11957 | 8102 | 8108 | 8036 | 8354 | 9939,22 | 11728 | 8150 | 0,695 | -0,525 | -17,345 | 2,15E-67 | 1,30E-66 | 0,0001569 | 0,00217 | 0,0007748 | 0,0007748 | TRUE | 0,0035374 | No |
| orf19.5437 | 86361 | 67816 | 73595 | 49806 | 2E+05 | 1E+05 | 1E+05 | 2E+05 | 78628 | 68589 | 69651 | 74817 | 1E+05 | 1E+05 | 1E+05 | 1E+05 | 104167 | 72921 | 135414 | 1,857 | 0,893 | 18,142 | 1,49E-73 | 9,78E-73 | 0,0025057 | 0,00205 | 0,0023173 | 0,0023173 | TRUE | 0,0340886 | No |
| orf19.5438 | 1702 | 1341 | 1437 | 933 | 3769 | 2805 | 3259 | 4351 | 1550 | 1356 | 1360 | 1402 | 3352 | 3045 | 3338 | 3319 | 2340,23 | 1417 | 3264 | 2,303 | 1,203 | 21,335 | 5,38E-101 | 4,72E-100 | 0,0023826 | 0,002601 | 0,0025317 | 0,0025317 | TRUE | 0,0503297 | No |
| orf19.5439 | 462 | 454 | 461 | 314 | 729 | 656 | 749 | 957 | 421 | 459 | 436 | 472 | 648 | 712 | 767 | 730 | 580,69 | 447 | 714 | 1,603 | 0,681 | 9,24 | 2,47E-20 | 6,31E-20 | 0,0023182 | 0,004312 | 0,0033536 | 0,0033536 | TRUE | 0,0466528 | No |
| orf19.544 | 888 | 793 | 802 | 546 | 1003 | 802 | 859 | 1101 | 808 | 802 | 759 | 820 | 892 | 871 | 880 | 840 | 834,02 | 797 | 871 | 1,093 | 0,128 | 2,297 | 0,021607 | 0,027621 | 1,00E-08 | 0,003621 | 0,0017644 | 0,0017644 | TRUE | 0,012909 | No |
| orf19.5440 | 17491 | 13971 | 15724 | 9461 | 11837 | 9354 | 10560 | 14923 | 15925 | 14130 | 14881 | 14212 | 10529 | 10156 | 10815 | 11383 | 12753,9 | 14787 | 10721 | 0,725 | -0,464 | -9,065 | 1,25E-19 | 3,14E-19 | 0,0026331 | 0,002141 | 0,0024368 | 0,0024368 | TRUE | 0,032938 | No |
| orf19.5441 | 1554 | 1406 | 1593 | 986 | 2069 | 1764 | 1887 | 2620 | 1415 | 1422 | 1508 | 1481 | 1840 | 1915 | 1933 | 1999 | 1689,03 | 1456 | 1922 | 1,321 | 0,401 | 8,464 | 2,58E-17 | 6,09E-17 | 0,0006005 | 0,002819 | 0,0015465 | 0,0015465 | TRUE | 0,0101947 | No |
| orf19.544.1 | 7242 | 5679 | 6078 | 3804 | 4931 | 3757 | 4086 | 5795 | 6594 | 5744 | 5752 | 5714 | 4386 | 4079 | 4185 | 4420 | 5109,23 | 5951 | 4268 | 0,717 | -0,48 | -8,664 | 4,55E-18 | 1,10E-17 | 0,0030572 | 0,002296 | 0,0027458 | 0,0027458 | TRUE | 0,0651156 | No |
| orf19.5442 | 2757 | 2331 | 2642 | 1564 | 927 | 729 | 808 | 1101 | 2510 | 2358 | 2500 | 2349 | 825 | 791 | 827 | 840 | 1625,11 | 2429 | 821 | 0,338 | -1,565 | -31,846 | 1,50E-222 | 2,81E-221 | 1,00E-08 | 0,00285 | 0,0015318 | 0,0015318 | TRUE | 0,0065003 | No |
| orf19.5443 | 6086 | 5043 | 5542 | 3193 | 2858 | 2275 | 2625 | 3562 | 5541 | 5100 | 5245 | 4796 | 2542 | 2470 | 2688 | 2717 | 3887,56 | 5170 | 2604 | 0,503 | -0,99 | -18,462 | 4,19E-76 | 2,82E-75 | 0,002433 | 0,002377 | 0,0024778 | 0,0024778 | TRUE | 0,028323 | No |
| orf19.5444 | 3661 | 3043 | 3342 | 2071 | 6032 | 4816 | 5497 | 7337 | 3333 | 3078 | 3163 | 3111 | 5365 | 5229 | 5630 | 5597 | 4313,15 | 3171 | 5455 | 1,72 | 0,782 | 18,187 | 6,56E-74 | 4,31E-73 | 0,0009235 | 0,002343 | 0,0015199 | 0,0015199 | TRUE | 0,0147656 | No |
| orf19.5445 | 2890 | 2208 | 2669 | 1741 | 4412 | 3582 | 4220 | 5627 | 2631 | 2233 | 2526 | 2615 | 3924 | 3889 | 4322 | 4292 | 3304,14 | 2501 | 4107 | 1,643 | 0,716 | 11,593 | 4,49E-31 | 1,48E-30 | 0,0038926 | 0,002437 | 0,0033352 | 0,0033352 | TRUE | 0,0636631 | No |
| orf19.5446 | 0 | 2 | 2 | 4 | 4 | 2 | 2 | 2 | 0 | 2 | 2 | 6 | 4 | 2 | 2 | 2 | 2,4 | 2 | 2 | 1,026 | 0,037 | 0,039 | 0,968832 | 0,972493 | 0,1349565 | 0,551668 | 0,4249132 | 0,4249132 | TRUE | 0,518063 | No |
| orf19.5447 | 3376 | 2733 | 2974 | 2074 | 3465 | 2915 | 3093 | 4407 | 3074 | 2764 | 2815 | 3115 | 3082 | 3165 | 3168 | 3362 | 3068,02 | 2942 | 3194 | 1,087 | 0,12 | 2,302 | 0,02133 | 0,02728 | 0,0020893 | 0,002468 | 0,002289 | 0,002289 | TRUE | 0,0199272 | No |
| orf19.5449 | 60 | 51 | 54 | 44 | 2867 | 2275 | 2342 | 3313 | 55 | 52 | 51 | 66 | 2550 | 2470 | 2399 | 2527 | 1271,15 | 56 | 2486 | 45,282 | 5,501 | 50,058 | 0 | 0 | 1,00E-08 | 0,003076 | 0,0018305 | 0,0018305 | TRUE | 0,0820559 | No |
| orf19.5450 | 2658 | 1985 | 2180 | 1502 | 2951 | 2286 | 2573 | 3686 | 2420 | 2008 | 2063 | 2256 | 2625 | 2482 | 2635 | 2812 | 2412,57 | 2187 | 2638 | 1,207 | 0,272 | 4,166 | 3,10E-05 | 4,78E-05 | 0,0044674 | 0,002585 | 0,0036649 | 0,0036649 | TRUE | 0,0649172 | No |
| orf19.5451 | 681 | 529 | 652 | 574 | 136 | 147 | 134 | 207 | 620 | 535 | 617 | 862 | 121 | 160 | 137 | 158 | 401,26 | 658 | 144 | 0,219 | -2,189 | -15,527 | 2,28E-54 | 1,14E-53 | 0,0273955 | 0,005329 | 0,01502 | 0,01502 | TRUE | 0,487116 | No |
| orf19.5454 | 603 | 469 | 520 | 363 | 1252 | 1010 | 1115 | 1590 | 549 | 474 | 492 | 545 | 1114 | 1097 | 1142 | 1213 | 828,22 | 515 | 1142 | 2,222 | 1,152 | 17,216 | 2,00E-66 | 1,20E-65 | 0,0017995 | 0,003632 | 0,0028478 | 0,0028478 | TRUE | 0,033022 | No |
| orf19.5455 | 2806 | 2170 | 2244 | 1369 | 1160 | 854 | 941 | 1393 | 2555 | 2195 | 2124 | 2056 | 1032 | 927 | 964 | 1063 | 1614,37 | 2232 | 996 | 0,446 | -1,164 | -15,647 | 3,51E-55 | 1,78E-54 | 0,0059139 | 0,002855 | 0,0046024 | 0,0046024 | TRUE | 0,1156938 | No |
| orf19.5457 | 2483 | 1887 | 2130 | 1363 | 2330 | 1987 | 2252 | 3105 | 2261 | 1908 | 2016 | 2047 | 2072 | 2157 | 2306 | 2369 | 2142,14 | 2058 | 2226 | 1,082 | 0,113 | 1,784 | 0,074499 | 0,090636 | 0,003798 | 0,002654 | 0,0033917 | 0,0033917 | TRUE | 0,0547859 | No |
| orf19.5459 | 6804 | 4955 | 5986 | 3946 | 3985 | 3255 | 3600 | 4998 | 6195 | 5011 | 5665 | 5928 | 3545 | 3534 | 3687 | 3813 | 4672,11 | 5700 | 3645 | 0,64 | -0,645 | -10,221 | 1,59E-24 | 4,51E-24 | 0,0043152 | 0,00232 | 0,0035988 | 0,0035988 | TRUE | 0,0809979 | No |
| orf19.5461 | 51 | 23 | 28 | 24 | 28 | 26 | 21 | 30 | 46 | 23 | 26 | 36 | 25 | 28 | 22 | 23 | 28,72 | 33 | 24 | 0,736 | -0,442 | -1,569 | 0,116716 | 0,139103 | 0,030268 | 0,04803 | 0,0410507 | 0,0410507 | TRUE | 0,4520275 | No |
| orf19.5462 | 5 | 1 | 2 | 0 | 3 | 4 | 1 | 1 | 5 | 1 | 2 | 0 | 3 | 4 | 1 | 1 | 2,03 | 2 | 2 | 1,077 | 0,107 | 0,104 | 0,917086 | 0,928593 | 0,2866476 | 0,652185 | 0,5260799 | 0,5260799 | TRUE | 0,8428713 | No |
| orf19.5463 | 1533 | 1236 | 1466 | 816 | 1252 | 909 | 1202 | 1532 | 1396 | 1250 | 1387 | 1226 | 1114 | 987 | 1231 | 1169 | 1219,9 | 1315 | 1125 | 0,855 | -0,226 | -3,057 | 0,002238 | 0,003077 | 0,0054665 | 0,00312 | 0,0044251 | 0,0044251 | TRUE | 0,0792577 | No |
| orf19.5464 | 103 | 76 | 103 | 65 | 187 | 148 | 172 | 199 | 94 | 77 | 97 | 98 | 166 | 161 | 176 | 152 | 127,59 | 92 | 164 | 1,792 | 0,842 | 6,448 | 1,13E-10 | 2,20E-10 | 1,00E-08 | 0,01239 | 0,0076842 | 0,0076842 | TRUE | 0,1129095 | No |
| orf19.5465 | 1283 | 1053 | 1248 | 782 | 2423 | 1912 | 2191 | 3014 | 1168 | 1065 | 1181 | 1175 | 2155 | 2076 | 2244 | 2299 | 1670,37 | 1147 | 2194 | 1,914 | 0,937 | 17,312 | 3,79E-67 | 2,29E-66 | 0,0015504 | 0,002828 | 0,0021292 | 0,0021292 | TRUE | 0,0284766 | No |
| orf19.5466 | 12074 | 9100 | 9664 | 6308 | 23561 | 15906 | 18214 | 25720 | 10993 | 9204 | 9146 | 9476 | 20957 | 17269 | 18653 | 19619 | 14414,7 | 9705 | 19124 | 1,971 | 0,979 | 13,735 | 6,30E-43 | 2,59E-42 | 0,0072329 | 0,002129 | 0,0047979 | 0,0047979 | TRUE | 0,0981121 | No |

| | | | | | | | | | | | | | | | | | | | | | | | | | | | | | | | |
|---|---|---|---|---|---|---|---|---|---|---|---|---|---|---|---|---|---|---|---|---|---|---|---|---|---|---|---|---|---|---|---|
| orf19.5467 | 12 | 12 | 17 | 9 | 44 | 20 | 23 | 32 | 11 | 12 | 16 | 14 | 39 | 22 | 24 | 24 | 20,19 | 13 | 27 | 2,079 | 1,056 | 3,147 | 0,001649 | 0,00229 | 1,00E-08 | 0,067479 | 0,0506602 | 0,0506602 | TRUE | 0,5910928 | No |
| orf19.5468 | 94 | 80 | 77 | 44 | 303 | 276 | 250 | 376 | 86 | 81 | 73 | 66 | 270 | 300 | 256 | 287 | 177,18 | 76 | 278 | 3,603 | 1,849 | 14,905 | 3,08E-50 | 1,42E-49 | 1,00E-08 | 0,009492 | 0,0062346 | 0,0062346 | TRUE | 0,061086 | No |
| orf19.5469 | 363 | 321 | 300 | 197 | 1357 | 1137 | 1452 | 1897 | 330 | 325 | 284 | 296 | 1207 | 1234 | 1487 | 1447 | 826,32 | 309 | 1344 | 4,346 | 2,12 | 23,711 | 2,75E-124 | 2,92E-123 | 0,0076146 | 0,003636 | 0,0055946 | 0,0055946 | TRUE | 0,0603981 | No |
| orf19.547 | 1083 | 887 | 995 | 602 | 942 | 803 | 851 | 1192 | 986 | 897 | 942 | 904 | 838 | 872 | 872 | 909 | 902,45 | 932 | 873 | 0,935 | -0,097 | -1,702 | 0,088689 | 0,107062 | 0,0005778 | 0,003501 | 0,002011 | 0,002011 | TRUE | 0,0186409 | No |
| orf19.5472 | 2 | 3 | 4 | 2 | 8 | 3 | 7 | 9 | 2 | 3 | 4 | 3 | 7 | 3 | 7 | 7 | 4,51 | 3 | 6 | 2,129 | 1,09 | 1,637 | 0,101645 | 0,122078 | 1,00E-08 | 0,295168 | 0,1674888 | 0,1674888 | TRUE | 0,2023426 | No |
| orf19.5474 | 359 | 275 | 274 | 141 | 838 | 619 | 836 | 1111 | 327 | 278 | 259 | 212 | 745 | 672 | 856 | 847 | 524,65 | 269 | 780 | 2,88 | 1,526 | 13,272 | 3,38E-40 | 1,32E-39 | 0,0186689 | 0,004555 | 0,0101186 | 0,0101186 | TRUE | 0,2341424 | No |
| orf19.5475 | 15 | 14 | 22 | 20 | 40 | 51 | 41 | 67 | 14 | 14 | 21 | 30 | 36 | 55 | 42 | 51 | 32,84 | 20 | 46 | 2,419 | 1,274 | 4,482 | 7,38E-06 | 1,18E-05 | 0,0333456 | 0,042261 | 0,0394458 | 0,0394458 | TRUE | 0,3988709 | No |
| orf19.5479 | 4 | 3 | 1 | 0 | 7 | 4 | 3 | 11 | 4 | 3 | 1 | 0 | 6 | 4 | 3 | 8 | 3,71 | 2 | 5 | 2,742 | 1,455 | 1,841 | 0,065638 | 0,080376 | 0,0286949 | 0,358403 | 0,2573721 | 0,2573721 | TRUE | 0,2838254 | No |
| orf19.548 | 579 | 553 | 587 | 364 | 595 | 467 | 540 | 693 | 527 | 559 | 556 | 547 | 529 | 507 | 553 | 529 | 538,34 | 547 | 530 | 0,968 | -0,047 | -0,721 | 0,471023 | 0,512857 | 1,00E-08 | 0,004491 | 0,0021808 | 0,0021808 | TRUE | 0,0096687 | No |
| orf19.5480 | 1810 | 1497 | 1718 | 1057 | 2240 | 1827 | 1911 | 2675 | 1648 | 1514 | 1626 | 1588 | 1992 | 1984 | 1957 | 2041 | 1793,67 | 1594 | 1994 | 1,251 | 0,323 | 7,182 | 6,88E-13 | 1,44E-12 | 0,0002527 | 0,002773 | 0,001366 | 0,001366 | TRUE | 0,0144847 | No |
| orf19.5483 | 1524 | 1363 | 1598 | 922 | 1330 | 1073 | 1194 | 1625 | 1388 | 1379 | 1512 | 1385 | 1183 | 1165 | 1223 | 1240 | 1309,22 | 1416 | 1203 | 0,849 | -0,236 | -4,554 | 5,27E-06 | 8,48E-06 | 0,0007723 | 0,003046 | 0,0018081 | 0,0018081 | TRUE | 0,0263014 | No |
| orf19.5484 | 2515 | 2300 | 2363 | 1492 | 1313 | 1088 | 1196 | 1643 | 2290 | 2326 | 2236 | 2241 | 1168 | 1181 | 1225 | 1253 | 1740,11 | 2273 | 1207 | 0,531 | -0,913 | -20,408 | 1,41E-92 | 1,13E-91 | 1,00E-08 | 0,002796 | 0,0012995 | 0,0012995 | TRUE | 0,008791 | No |
| orf19.5485 | 605 | 527 | 551 | 316 | 793 | 654 | 621 | 929 | 551 | 533 | 521 | 475 | 705 | 710 | 636 | 709 | 605 | 520 | 690 | 1,322 | 0,403 | 5,825 | 5,72E-09 | 1,04E-08 | 0,0010755 | 0,00422 | 0,002896 | 0,002896 | TRUE | 0,0335685 | No |
| orf19.5486 | 4142 | 3734 | 4158 | 2319 | 4371 | 3640 | 4006 | 5242 | 3771 | 3777 | 3935 | 3484 | 3888 | 3952 | 4103 | 3999 | 3863,44 | 3742 | 3986 | 1,064 | 0,089 | 2,001 | 0,045362 | 0,056393 | 0,0011762 | 0,002379 | 0,0016431 | 0,0016431 | TRUE | 0,0246889 | No |
| orf19.5486.1 | 2202 | 1891 | 1791 | 1056 | 1614 | 1135 | 1239 | 1652 | 2005 | 1913 | 1695 | 1586 | 1436 | 1232 | 1269 | 1260 | 1549,45 | 1800 | 1299 | 0,721 | -0,472 | -6,014 | 1,81E-09 | 3,37E-09 | 0,0073649 | 0,00289 | 0,0052524 | 0,0052524 | TRUE | 0,0734291 | No |
| orf19.5487 | 4707 | 4159 | 4413 | 2724 | 4932 | 4115 | 4478 | 6074 | 4286 | 4206 | 4177 | 4092 | 4387 | 4468 | 4586 | 4633 | 4354,28 | 4190 | 4518 | 1,078 | 0,108 | 3,022 | 0,002513 | 0,003441 | 0,0002828 | 0,00234 | 0,0009994 | 0,0009994 | TRUE | 0,0048127 | No |
| orf19.5488 | 321 | 276 | 298 | 200 | 588 | 427 | 470 | 610 | 292 | 279 | 282 | 300 | 523 | 464 | 481 | 465 | 385,89 | 288 | 483 | 1,68 | 0,748 | 9,465 | 2,94E-21 | 7,69E-21 | 1,00E-08 | 0,00546 | 0,0031838 | 0,0031838 | TRUE | 0,0366314 | No |
| orf19.5489 | 137 | 150 | 151 | 98 | 312 | 240 | 262 | 362 | 125 | 152 | 143 | 147 | 278 | 261 | 268 | 276 | 206,14 | 142 | 271 | 1,921 | 0,942 | 9,121 | 7,47E-20 | 1,89E-19 | 1,00E-08 | 0,008445 | 0,0047535 | 0,0047535 | TRUE | 0,0750372 | No |
| orf19.549 | 2955 | 2689 | 2748 | 2061 | 3453 | 2787 | 2886 | 3868 | 2690 | 2720 | 2601 | 3096 | 3071 | 3026 | 2956 | 2951 | 2888,77 | 2777 | 3001 | 1,083 | 0,115 | 2,084 | 0,037196 | 0,046623 | 0,00283 | 0,002494 | 0,0025575 | 0,0025575 | TRUE | 0,0692498 | No |
| orf19.5491 | 66 | 107 | 84 | 64 | 76 | 62 | 53 | 53 | 60 | 108 | 79 | 96 | 68 | 67 | 54 | 40 | 71,7 | 86 | 57 | 0,668 | -0,583 | -2,665 | 0,007697 | 0,01018 | 0,0467918 | 0,020462 | 0,0314796 | 0,0314796 | TRUE | 0,2503509 | No |
| orf19.5491.1 | 3179 | 2682 | 2762 | 2170 | 4932 | 4070 | 4042 | 5553 | 2894 | 2713 | 2614 | 3260 | 4387 | 4419 | 4140 | 4236 | 3582,71 | 2870 | 4296 | 1,499 | 0,584 | 9,128 | 6,98E-20 | 1,77E-19 | 0,0049893 | 0,002406 | 0,003631 | 0,003631 | TRUE | 0,097713 | No |
| orf19.5492 | 2253 | 1846 | 1926 | 1120 | 3021 | 2137 | 2499 | 3371 | 2051 | 1867 | 1823 | 1682 | 2687 | 2320 | 2559 | 2571 | 2195,19 | 1856 | 2534 | 1,364 | 0,448 | 6,8 | 1,05E-11 | 2,11E-11 | 0,004587 | 0,002639 | 0,0036895 | 0,0036895 | TRUE | 0,06093 | No |
| orf19.5493 | 9408 | 8328 | 8640 | 5398 | 14022 | 10529 | 10355 | 14919 | 8566 | 8423 | 8177 | 8109 | 12472 | 11432 | 10605 | 11380 | 9895,37 | 8319 | 11472 | 1,379 | 0,464 | 9,165 | 4,96E-20 | 1,26E-19 | 0,002421 | 0,002171 | 0,0023526 | 0,0023526 | TRUE | 0,0421574 | No |
| orf19.5495 | 2099 | 1920 | 2177 | 1340 | 3285 | 2812 | 3032 | 4087 | 1911 | 1942 | 2060 | 2013 | 2922 | 3053 | 3105 | 3118 | 2515,48 | 1982 | 3050 | 1,54 | 0,623 | 14,197 | 9,52E-46 | 4,10E-45 | 0,0006515 | 0,002562 | 0,0014224 | 0,0014224 | TRUE | 0,0097997 | No |
| orf19.5496 | 1897 | 1673 | 1959 | 1312 | 2889 | 2545 | 2588 | 3631 | 1727 | 1692 | 1854 | 1971 | 2570 | 2763 | 2650 | 2770 | 2249,63 | 1811 | 2688 | 1,487 | 0,573 | 10,311 | 6,25E-25 | 1,79E-24 | 0,0024922 | 0,002624 | 0,0024905 | 0,0024905 | TRUE | 0,0395108 | No |
| orf19.5498 | 282 | 260 | 299 | 173 | 496 | 396 | 448 | 682 | 257 | 263 | 283 | 260 | 441 | 430 | 459 | 520 | 364,09 | 266 | 462 | 1,744 | 0,802 | 9,059 | 1,32E-19 | 3,30E-19 | 0,0032733 | 0,005665 | 0,0045154 | 0,0045154 | TRUE | 0,0874559 | No |
| orf19.5499 | 160 | 182 | 179 | 98 | 184 | 187 | 185 | 206 | 146 | 184 | 169 | 147 | 164 | 203 | 189 | 157 | 169,96 | 162 | 178 | 1,095 | 0,131 | 1,043 | 0,297101 | 0,334514 | 0,0081667 | 0,00981 | 0,0091369 | 0,0091369 | TRUE | 0,0921649 | No |
| orf19.55 | 3222 | 3467 | 3505 | 2029 | 8385 | 7022 | 6853 | 9075 | 2933 | 3507 | 3317 | 3048 | 7458 | 7624 | 7018 | 6922 | 5228,5 | 3201 | 7256 | 2,265 | 1,18 | 19,532 | 5,88E-85 | 4,43E-84 | 0,0040712 | 0,00229 | 0,0032713 | 0,0032713 | TRUE | 0,0502536 | No |
| orf19.550 | 23973 | 20045 | 22250 | 14692 | 5836 | 4531 | 5121 | 6602 | 21826 | 20273 | 21058 | 22070 | 5191 | 4919 | 5245 | 5036 | 13202,3 | 21307 | 5098 | 0,239 | -2,063 | -51,259 | 0 | 0 | 0,0009879 | 0,002137 | 0,001439 | 0,001439 | TRUE | 0,0130444 | No |
| orf19.5500 | 17716 | 15895 | 17628 | 9014 | 17342 | 12583 | 15478 | 19800 | 16130 | 16076 | 16683 | 13540 | 15425 | 13662 | 15851 | 15104 | 15308,9 | 15607 | 15010 | 0,962 | -0,057 | -0,831 | 0,405712 | 0,446703 | 0,0056281 | 0,002123 | 0,0043831 | 0,0043831 | TRUE | 0,0965828 | No |
| orf19.5501 | 1088 | 1050 | 1074 | 704 | 645 | 542 | 540 | 688 | 991 | 1062 | 1016 | 1058 | 574 | 588 | 553 | 525 | 795,81 | 1032 | 560 | 0,542 | -0,882 | -14,603 | 2,70E-48 | 1,20E-47 | 1,00E-08 | 0,003697 | 0,0021517 | 0,0021517 | TRUE | 0,0230378 | No |
| orf19.5502 | 336 | 397 | 420 | 334 | 455 | 399 | 337 | 488 | 306 | 402 | 397 | 502 | 405 | 433 | 345 | 372 | 395,24 | 402 | 389 | 0,973 | -0,039 | -0,312 | 0,754853 | 0,786562 | 0,0218189 | 0,005379 | 0,0126648 | 0,0126648 | TRUE | 0,245811 | No |
| orf19.5503 | 51 | 43 | 45 | 27 | 19 | 18 | 25 | 35 | 46 | 43 | 43 | 41 | 17 | 20 | 26 | 27 | 32,73 | 43 | 22 | 0,513 | -0,962 | -3,895 | 9,82E-05 | 0,000147 | 1,00E-08 | 0,042401 | 0,0254068 | 0,0254068 | TRUE | 0,1709518 | No |
| orf19.5504 | 2270 | 2688 | 2726 | 1554 | 1466 | 1242 | 1276 | 1660 | 2067 | 2719 | 2580 | 2334 | 1304 | 1348 | 1307 | 1266 | 1865,64 | 2425 | 1306 | 0,538 | -0,893 | -11,58 | 5,21E-31 | 1,71E-30 | 0,0072608 | 0,002745 | 0,0051312 | 0,0051312 | TRUE | 0,1229458 | No |
| orf19.5505 | 2317 | 2231 | 2531 | 1360 | 1948 | 1794 | 1816 | 2310 | 2110 | 2256 | 2395 | 2043 | 1733 | 1948 | 1860 | 1762 | 2013,33 | 2201 | 1826 | 0,828 | -0,273 | -4,456 | 8,36E-06 | 1,33E-05 | 0,0033231 | 0,002693 | 0,0030917 | 0,0030917 | TRUE | 0,0429185 | No |
| orf19.5506 | 1084 | 1194 | 1105 | 713 | 972 | 834 | 879 | 1118 | 987 | 1208 | 1046 | 1071 | 865 | 905 | 900 | 853 | 979,31 | 1078 | 881 | 0,817 | -0,292 | -4,325 | 1,53E-05 | 2,40E-05 | 0,0033035 | 0,003386 | 0,0033424 | 0,0033424 | TRUE | 0,0813263 | No |
| orf19.5507 | 9368 | 9190 | 9617 | 5654 | 7629 | 6438 | 6410 | 8485 | 8529 | 9295 | 9102 | 8493 | 6786 | 6990 | 6565 | 6472 | 7778,93 | 8855 | 6703 | 0,757 | -0,402 | -9,054 | 1,37E-19 | 3,45E-19 | 0,0014683 | 0,002207 | 0,001765 | 0,001765 | TRUE | 0,0136338 | No |
| orf19.5508 | 527 | 497 | 506 | 349 | 377 | 323 | 364 | 448 | 480 | 503 | 479 | 524 | 335 | 351 | 373 | 342 | 423,27 | 496 | 350 | 0,706 | -0,501 | -6,795 | 1,09E-11 | 2,19E-11 | 1,00E-08 | 0,005158 | 0,0028179 | 0,0028179 | TRUE | 0,0211174 | No |
| orf19.551 | 18926 | 17083 | 18210 | 11867 | 6014 | 4910 | 5101 | 6877 | 17231 | 17278 | 17234 | 17826 | 5349 | 5331 | 5224 | 5246 | 11339,9 | 17392 | 5288 | 0,304 | -1,718 | -58,743 | 0 | 0 | 0,0001163 | 0,002154 | 0,0007015 | 0,0007015 | TRUE | 0,0033474 | No |
| orf19.5510 | 2157 | 2075 | 2314 | 1261 | 5189 | 4268 | 4566 | 5928 | 1964 | 2099 | 2190 | 1894 | 4615 | 4634 | 4676 | 4522 | 3324,26 | 2037 | 4612 | 2,26 | 1,177 | 24,157 | 6,24E-129 | 6,83E-128 | 0,0017549 | 0,002434 | 0,0019122 | 0,0019122 | TRUE | 0,0310519 | No |
| orf19.5513 | 2189 | 1966 | 2118 | 1531 | 1593 | 1329 | 1261 | 1720 | 1993 | 1988 | 2004 | 2300 | 1417 | 1443 | 1291 | 1312 | 1718,62 | 2071 | 1366 | 0,66 | -0,599 | -9,628 | 6,09E-22 | 1,61E-21 | 0,0034142 | 0,002806 | 0,0031095 | 0,0031095 | TRUE | 0,0630777 | No |
| orf19.5514 | 793 | 748 | 765 | 488 | 973 | 772 | 941 | 1073 | 722 | 757 | 724 | 733 | 865 | 838 | 964 | 818 | 802,67 | 734 | 871 | 1,186 | 0,246 | 3,794 | 0,000149 | 0,00022 | 0,0019256 | 0,003683 | 0,0027866 | 0,0027866 | TRUE | 0,0592942 | No |
| orf19.5515 | 2828 | 2756 | 2798 | 1789 | 2925 | 2285 | 2428 | 3214 | 2575 | 2787 | 2648 | 2687 | 2602 | 2481 | 2487 | 2452 | 2589,8 | 2674 | 2506 | 0,937 | -0,094 | -2,205 | 0,02742 | 0,034753 | 0,0005679 | 0,002547 | 0,0013474 | 0,0013474 | TRUE | 0,0102576 | No |
| orf19.5516 | 1274 | 1387 | 1474 | 884 | 1646 | 1347 | 1359 | 1674 | 1160 | 1403 | 1395 | 1328 | 1464 | 1462 | 1392 | 1277 | 1360,11 | 1322 | 1399 | 1,058 | 0,082 | 1,141 | 0,253818 | 0,289203 | 0,0052331 | 0,003008 | 0,0041804 | 0,0041804 | TRUE | 0,0847734 | No |
| orf19.5517 | 2123 | 1805 | 1901 | 1279 | 5297 | 4339 | 4286 | 6310 | 1933 | 1826 | 1799 | 1921 | 4712 | 4711 | 4389 | 4813 | 3263 | 1870 | 4656 | 2,493 | 1,318 | 28,5 | 1,18E-178 | 1,81E-177 | 0,001147 | 0,002442 | 0,0016646 | 0,0016646 | TRUE | 0,0180256 | No |
| orf19.5518 | 2492 | 2431 | 2613 | 1700 | 1574 | 1406 | 1340 | 1782 | 2269 | 2459 | 2473 | 2554 | 1400 | 1527 | 1372 | 1359 | 1926,55 | 2439 | 1414 | 0,58 | -0,786 | -14,25 | 4,47E-46 | 1,93E-45 | 0,0019428 | 0,002723 | 0,002366 | 0,002366 | TRUE | 0,0331307 | No |
| orf19.5519 | 1282 | 1161 | 1245 | 725 | 1324 | 1134 | 1155 | 1524 | 1167 | 1174 | 1178 | 1089 | 1178 | 1231 | 1183 | 1163 | 1170,38 | 1152 | 1189 | 1,028 | 0,04 | 0,806 | 0,420292 | 0,461618 | 1,00E-08 | 0,003166 | 0,0015359 | 0,0015359 | TRUE | 0,0139652 | No |
| orf19.552 | 4 | 2 | 1 | 2 | 2 | 1 | 0 | 2 | 4 | 2 | 1 | 3 | 2 | 1 | 0 | 2 | 1,75 | 2 | 1 | 0,477 | -1,069 | -1,004 | 0,315228 | 0,354173 | 1,00E-08 | 0,75659 | 0,4545754 | 0,4545754 | TRUE | 0,2122797 | No |
| orf19.5520 | 3 | 2 | 2 | 2 | 2 | 0 | 2 | 5 | 3 | 2 | 2 | 3 | 2 | 0 | 2 | 4 | 2,16 | 2 | 2 | 0,84 | -0,252 | -0,27 | 0,787415 | 0,816489 | 1,00E-08 | 0,613167 | 0,3818147 | 0,3818147 | TRUE | 0,3411039 | No |
| orf19.5521 | 4509 | 3933 | 4260 | 2598 | 2971 | 2259 | 2731 | 3571 | 4105 | 3978 | 4032 | 3903 | 2643 | 2453 | 2797 | 2724 | 3329,19 | 4004 | 2654 | 0,663 | -0,593 | -12,621 | 1,63E-36 | 6,00E-36 | 0,0013722 | 0,002434 | 0,0018118 | 0,0018118 | TRUE | 0,0307116 | No |
| orf19.5522 | 505 | 527 | 508 | 257 | 1080 | 721 | 811 | 1070 | 460 | 533 | 481 | 386 | 961 | 783 | 831 | 816 | 656,23 | 465 | 848 | 1,816 | 0,86 | 9,145 | 5,95E-20 | 1,51E-19 | 0,0105984 | 0,00405 | 0,0067967 | 0,0067967 | TRUE | 0,1325487 | No |
| orf19.5523 | 443 | 499 | 416 | 320 | 560 | 506 | 451 | 637 | 403 | 505 | 394 | 481 | 498 | 549 | 462 | 486 | 472,21 | 446 | 499 | 1,122 | 0,167 | 1,775 | 0,075936 | 0,092314 | 0,0076291 | 0,004834 | 0,0063119 | 0,0063119 | TRUE | 0,1008702 | No |
| orf19.5524 | 49 | 58 | 56 | 37 | 22 | 19 | 16 | 36 | 45 | 59 | 53 | 56 | 20 | 21 | 16 | 27 | 36,99 | 53 | 21 | 0,404 | -1,309 | -5,366 | 8,05E-08 | 1,40E-07 | 1,00E-08 | 0,037752 | 0,0252066 | 0,0252066 | TRUE | 0,3036038 | No |
| orf19.5525 | 1285 | 1065 | 1175 | 796 | 808 | 645 | 657 | 978 | 1170 | 1077 | 1112 | 1196 | 719 | 700 | 673 | 746 | 924,08 | 1139 | 710 | 0,625 | -0,678 | -11,473 | 1,80E-30 | 5,86E-30 | 0,0009792 | 0,003467 | 0,0022257 | 0,0022257 | TRUE | 0,0155653 | No |
| orf19.5526 | 130 | 115 | 119 | 90 | 56 | 48 | 42 | 48 | 118 | 116 | 113 | 135 | 50 | 52 | 43 | 37 | 83 | 120 | 46 | 0,375 | -1,414 | -8,496 | 1,97E-17 | 4,67E-17 | 1,00E-08 | 0,017952 | 0,0118085 | 0,0118085 | TRUE | 0,1673476 | No |
| orf19.5527 | 335 | 327 | 326 | 238 | 273 | 242 | 234 | 338 | 305 | 331 | 309 | 358 | 243 | 263 | 240 | 258 | 288,1 | 326 | 251 | 0,775 | -0,368 | -4,156 | 3,24E-05 | 5,00E-05 | 1,00E-08 | 0,006622 | 0,0040157 | 0,0040157 | TRUE | 0,0417542 | No |
| orf19.5528 | 1002 | 886 | 972 | 624 | 314 | 232 | 254 | 395 | 912 | 896 | 920 | 937 | 279 | 252 | 260 | 301 | 594,78 | 916 | 273 | 0,3 | -1,737 | -24,406 | 1,47E-131 | 1,66E-130 | 1,00E-08 | 0,004258 | 0,0025865 | 0,0025865 | TRUE | 0,0590193 | No |
| orf19.553 | 11 | 16 | 18 | 9 | 3 | 7 | 7 | 12 | 10 | 16 | 17 | 14 | 3 | 8 | 7 | 9 | 10,42 | 14 | 7 | 0,47 | -1,09 | -2,394 | 0,016686 | 0,021546 | 1,00E-08 | 0,128838 | 0,0916429 | 0,0916429 | TRUE | 0,4822949 | No |
| orf19.5530 | 6582 | 5652 | 6443 | 4001 | 5254 | 4706 | 5051 | 6767 | 5993 | 5716 | 6098 | 6010 | 4673 | 5109 | 5173 | 5162 | 5491,79 | 5954 | 5029 | 0,845 | -0,244 | -5,474 | 4,40E-08 | 7,69E-08 | 0,001269 | 0,002278 | 0,0017181 | 0,0017181 | TRUE | 0,0279016 | No |
| orf19.5531 | 2360 | 2050 | 2154 | 1444 | 525 | 426 | 432 | 591 | 2149 | 2073 | 2039 | 2169 | 467 | 463 | 442 | 451 | 1281,56 | 2108 | 456 | 0,216 | -2,208 | -40,89 | 0 | 0 | 1,00E-08 | 0,003068 | 0,001525 | 0,001525 | TRUE | 0,0059032 | No |
| orf19.5532 | 182 | 155 | 143 | 94 | 205 | 224 | 161 | 231 | 166 | 157 | 135 | 141 | 182 | 243 | 165 | 176 | 170,71 | 150 | 192 | 1,272 | 0,347 | 2,544 | 0,010953 | 0,014333 | 0,0146785 | 0,009775 | 0,0118779 | 0,0118779 | TRUE | 0,3368322 | No |
| orf19.5533 | 1586 | 1421 | 1579 | 961 | 969 | 759 | 845 | 1147 | 1444 | 1437 | 1494 | 1444 | 862 | 824 | 865 | 875 | 1155,68 | 1455 | 856 | 0,589 | -0,763 | -15,633 | 4,32E-55 | 2,19E-54 | 1,00E-08 | 0,00318 | 0,0013765 | 0,0013765 | TRUE | 0,0072872 | No |
| orf19.5534 | 1593 | 1341 | 1370 | 935 | 1491 | 1168 | 1289 | 1847 | 1450 | 1356 | 1297 | 1405 | 1326 | 1268 | 1320 | 1409 | 1353,88 | 1377 | 1331 | 0,968 | -0,047 | -0,849 | 0,395759 | 0,436574 | 0,0014796 | 0,003013 | 0,0021738 | 0,0021738 | TRUE | 0,0195332 | No |
| orf19.5535 | 1008 | 930 | 1012 | 699 | 2051 | 1855 | 1866 | 2558 | 918 | 941 | 958 | 1050 | 1824 | 2014 | 1911 | 1951 | 1445,84 | 967 | 1925 | 1,998 | 0,999 | 17,885 | 1,54E-71 | 9,84E-71 | 0,0016178 | 0,002951 | 0,0021939 | 0,0021939 | TRUE | 0,0354958 | No |
| orf19.5536 | 521 | 451 | 504 | 334 | 260 | 267 | 249 | 347 | 474 | 456 | 477 | 502 | 231 | 290 | 255 | 265 | 368,76 | 477 | 260 | 0,545 | -0,875 | -10,42 | 2,01E-25 | 5,83E-25 | 0,0008333 | 0,005619 | 0,0038582 | 0,0038582 | TRUE | 0,0635953 | No |
| orf19.5537 | 8943 | 7389 | 8145 | 5499 | 4859 | 4007 | 4448 | 6369 | 8142 | 7473 | 7709 | 8260 | 4322 | 4350 | 4555 | 4858 | 6208,79 | 7896 | 4521 | 0,573 | -0,803 | -15,711 | 1,27E-55 | 6,52E-55 | 0,0023543 | 0,00225 | 0,0023402 | 0,0023402 | TRUE | 0,0311863 | No |
| orf19.5539 | 644 | 622 | 694 | 420 | 739 | 654 | 644 | 864 | 586 | 629 | 657 | 631 | 657 | 710 | 660 | 659 | 648,64 | 626 | 672 | 1,073 | 0,101 | 1,587 | 0,112428 | 0,134269 | 1,00E-08 | 0,004074 | 0,0023458 | 0,0023458 | TRUE | 0,0233346 | No |

| | | | | | | | | | | | | | | | | | | | | | | | | | | | | | | | |
|---|---|---|---|---|---|---|---|---|---|---|---|---|---|---|---|---|---|---|---|---|---|---|---|---|---|---|---|---|---|---|---|
| orf19.554 | 652 | 580 | 610 | 408 | 144 | 119 | 136 | 145 | 594 | 587 | 577 | 613 | 128 | 129 | 139 | 111 | 359,7 | 593 | 127 | 0,213 | -2,232 | -24,484 | 2,19E-132 | 2,50E-131 | 1,00E-08 | 0,00571 | 0,0033905 | 0,0033905 | TRUE | 0,0899571 | No |
| orf19.5541 | 4160 | 3858 | 4315 | 2692 | 5002 | 4272 | 4454 | 6001 | 3788 | 3902 | 4084 | 4044 | 4449 | 4638 | 4561 | 4578 | 4255,43 | 3954 | 4556 | 1,153 | 0,205 | 5,355 | 8,55E-08 | 1,48E-07 | 0,0004248 | 0,002347 | 0,0011707 | 0,0011707 | TRUE | 0,0100829 | No |
| orf19.5542 | 67 | 55 | 82 | 58 | 295 | 232 | 238 | 337 | 61 | 56 | 78 | 87 | 262 | 252 | 244 | 257 | 162,06 | 70 | 254 | 3,67 | 1,876 | 14,136 | 2,28E-45 | 9,82E-45 | 1,00E-08 | 0,010189 | 0,0074082 | 0,0074082 | TRUE | 0,1652704 | No |
| orf19.5543 | 1582 | 1472 | 1642 | 932 | 1460 | 1150 | 1238 | 1759 | 1440 | 1489 | 1554 | 1400 | 1299 | 1249 | 1268 | 1342 | 1380 | 1471 | 1290 | 0,876 | -0,191 | -3,723 | 0,000196 | 0,00029 | 0,0007695 | 0,002994 | 0,0017952 | 0,0017952 | TRUE | 0,0175343 | No |
| orf19.5544 | 7138 | 6360 | 6793 | 3988 | 4936 | 3965 | 4483 | 5799 | 6499 | 6432 | 6429 | 5991 | 4390 | 4305 | 4591 | 4424 | 5382,61 | 6338 | 4428 | 0,698 | -0,519 | -12,935 | 2,87E-38 | 1,09E-37 | 0,0007911 | 0,002282 | 0,0013528 | 0,0013528 | TRUE | 0,0158668 | No |
| orf19.5547 | 2470 | 2066 | 2220 | 1274 | 3236 | 2331 | 2596 | 3633 | 2249 | 2090 | 2101 | 1914 | 2878 | 2531 | 2659 | 2771 | 2399,02 | 2088 | 2710 | 1,296 | 0,374 | 6,359 | 2,03E-10 | 3,89E-10 | 0,0031072 | 0,002588 | 0,0028945 | 0,0028945 | TRUE | 0,0360067 | No |
| orf19.5548 | 720 | 690 | 710 | 457 | 783 | 657 | 702 | 898 | 656 | 698 | 672 | 686 | 696 | 713 | 719 | 685 | 690,69 | 678 | 703 | 1,038 | 0,053 | 0,914 | 0,360944 | 0,401224 | 1,00E-08 | 0,00395 | 0,0018061 | 0,0018061 | TRUE | 0,0062482 | No |
| orf19.5549 | 28 | 32 | 38 | 25 | 36 | 38 | 46 | 56 | 25 | 32 | 36 | 38 | 32 | 41 | 47 | 43 | 36,81 | 33 | 41 | 1,252 | 0,324 | 1,379 | 0,167989 | 0,196393 | 1,00E-08 | 0,037924 | 0,0252353 | 0,0252353 | TRUE | 0,1762227 | No |
| orf19.555 | 2 | 1 | 2 | 3 | 7 | 9 | 4 | 6 | 2 | 1 | 2 | 5 | 6 | 10 | 4 | 5 | 4,24 | 2 | 6 | 2,836 | 1,504 | 2,03 | 0,042326 | 0,052814 | 1,00E-08 | 0,313752 | 0,2038866 | 0,2038866 | TRUE | 0,3315092 | No |
| orf19.5550 | 9470 | 9208 | 9334 | 4899 | 12317 | 8692 | 10019 | 12258 | 8622 | 9313 | 8834 | 7359 | 10956 | 9437 | 10261 | 9350 | 9266,46 | 8532 | 10001 | 1,172 | 0,229 | 3,108 | 0,001881 | 0,0026 | 0,0074942 | 0,00218 | 0,0050869 | 0,0050869 | TRUE | 0,1034526 | No |
| orf19.5551 | 6529 | 5747 | 6293 | 3782 | 1917 | 1537 | 1704 | 2321 | 5944 | 5812 | 5956 | 5681 | 1705 | 1669 | 1745 | 1770 | 3785,4 | 5848 | 1722 | 0,295 | -1,763 | -45,708 | 0 | 0 | 0,0002308 | 0,002386 | 0,0010664 | 0,0010664 | TRUE | 0,0050764 | No |
| orf19.5552 | 4808 | 4416 | 4769 | 2861 | 1542 | 1170 | 1208 | 1627 | 4377 | 4466 | 4513 | 4298 | 1372 | 1270 | 1237 | 1241 | 2846,87 | 4414 | 1280 | 0,29 | -1,786 | -39,2 | 0 | 0 | 0,0007665 | 0,002501 | 0,0015071 | 0,0015071 | TRUE | 0,0285577 | No |
| orf19.5553 | 1033 | 969 | 1052 | 603 | 612 | 483 | 535 | 728 | 941 | 980 | 996 | 906 | 544 | 524 | 548 | 555 | 749,24 | 956 | 543 | 0,568 | -0,817 | -13,75 | 5,07E-43 | 2,08E-42 | 1,00E-08 | 0,0038 | 0,0019706 | 0,0019706 | TRUE | 0,0125098 | No |
| orf19.5555 | 2266 | 2032 | 2197 | 1396 | 3066 | 2639 | 3021 | 3883 | 2063 | 2055 | 2079 | 2097 | 2727 | 2865 | 3094 | 2962 | 2492,84 | 2074 | 2912 | 1,405 | 0,49 | 10,314 | 6,08E-25 | 1,74E-24 | 0,0012053 | 0,002567 | 0,0017479 | 0,0017479 | TRUE | 0,0224568 | No |
| orf19.5556 | 1 | 2 | 1 | 0 | 8 | 5 | 3 | 4 | 1 | 2 | 1 | 0 | 7 | 5 | 3 | 3 | 2,82 | 1 | 4 | 4,486 | 2,165 | 2,232 | 0,025582 | 0,032514 | 1,00E-08 | 0,470733 | 0,2933168 | 0,2933168 | TRUE | 0,2791562 | No |
| orf19.5557 | 630 | 685 | 640 | 444 | 7315 | 6352 | 6430 | 8304 | 574 | 693 | 606 | 667 | 6506 | 6897 | 6585 | 6334 | 3607,69 | 635 | 6580 | 10,4 | 3,379 | 56,745 | 0 | 0 | 0,0025903 | 0,002403 | 0,0024868 | 0,0024868 | TRUE | 0,0519943 | No |
| orf19.5558 | 1625 | 1262 | 1508 | 1082 | 1120 | 1010 | 1057 | 1509 | 1479 | 1276 | 1427 | 1625 | 996 | 1097 | 1083 | 1151 | 1266,84 | 1452 | 1082 | 0,746 | -0,422 | -5,666 | 1,46E-08 | 2,62E-08 | 0,0056044 | 0,00308 | 0,0045231 | 0,0045231 | TRUE | 0,0791459 | No |
| orf19.5559 | 621 | 555 | 559 | 350 | 854 | 766 | 702 | 983 | 565 | 561 | 529 | 526 | 760 | 832 | 719 | 750 | 655,2 | 545 | 765 | 1,399 | 0,484 | 7,248 | 4,24E-13 | 8,93E-13 | 0,0011947 | 0,004053 | 0,0026992 | 0,0026992 | TRUE | 0,0384572 | No |
| orf19.556 | 995 | 1094 | 991 | 651 | 2641 | 2395 | 2054 | 2982 | 906 | 1106 | 938 | 978 | 2349 | 2600 | 2104 | 2275 | 1656,98 | 982 | 2332 | 2,375 | 1,248 | 16,13 | 1,56E-58 | 8,36E-58 | 0,0069677 | 0,002834 | 0,0049973 | 0,0049973 | TRUE | 0,0893269 | No |
| orf19.5561 | 3914 | 3929 | 4145 | 2547 | 5958 | 5251 | 5104 | 6651 | 3564 | 3974 | 3923 | 3826 | 5299 | 5701 | 5227 | 5073 | 4573,41 | 3822 | 5325 | 1,393 | 0,478 | 9,405 | 5,23E-21 | 1,36E-20 | 0,0021054 | 0,002326 | 0,0022564 | 0,0022564 | TRUE | 0,0272976 | No |
| orf19.5563 | 1649 | 1344 | 1471 | 896 | 2340 | 1861 | 2135 | 2842 | 1501 | 1359 | 1392 | 1346 | 2081 | 2021 | 2187 | 2168 | 1756,88 | 1400 | 2114 | 1,508 | 0,593 | 11,499 | 1,33E-30 | 4,34E-30 | 0,0011468 | 0,002789 | 0,0019499 | 0,0019499 | TRUE | 0,0296464 | No |
| orf19.5564 | 437 | 422 | 427 | 267 | 971 | 778 | 928 | 1138 | 398 | 427 | 404 | 401 | 864 | 845 | 950 | 868 | 644,59 | 408 | 882 | 2,162 | 1,113 | 16,575 | 1,06E-61 | 5,94E-61 | 1,00E-08 | 0,004086 | 0,002488 | 0,002488 | TRUE | 0,0315896 | No |
| orf19.5565 | 298 | 302 | 264 | 199 | 908 | 711 | 773 | 1100 | 271 | 305 | 250 | 299 | 808 | 772 | 792 | 839 | 541,98 | 281 | 803 | 2,869 | 1,521 | 19,82 | 2,00E-87 | 1,54E-86 | 0,001041 | 0,004474 | 0,0031687 | 0,0031687 | TRUE | 0,0662581 | No |
| orf19.5566 | 1986 | 1748 | 1947 | 1198 | 1833 | 1518 | 1645 | 2059 | 1808 | 1768 | 1843 | 1800 | 1630 | 1648 | 1685 | 1571 | 1719,02 | 1805 | 1634 | 0,904 | -0,145 | -3,323 | 0,000892 | 0,001263 | 1,00E-08 | 0,002805 | 0,0012552 | 0,0012552 | TRUE | 0,0085742 | No |
| orf19.5567 | 2333 | 2020 | 2251 | 1339 | 1692 | 1317 | 1377 | 1838 | 2124 | 2043 | 2130 | 2011 | 1505 | 1430 | 1410 | 1402 | 1757 | 2077 | 1437 | 0,691 | -0,533 | -11,655 | 2,16E-31 | 7,18E-31 | 0,0003588 | 0,002789 | 0,001428 | 0,001428 | TRUE | 0,012645 | No |
| orf19.5568 | 363 | 348 | 435 | 243 | 255 | 189 | 198 | 253 | 330 | 352 | 412 | 365 | 227 | 205 | 203 | 193 | 285,87 | 365 | 207 | 0,567 | -0,819 | -8,308 | 9,70E-17 | 2,25E-16 | 0,0043053 | 0,006658 | 0,0055967 | 0,0055967 | TRUE | 0,0928098 | No |
| orf19.5569 | 10795 | 9605 | 10421 | 6263 | 5352 | 4465 | 4760 | 6335 | 9828 | 9714 | 9863 | 9408 | 4760 | 4848 | 4875 | 4832 | 7266,1 | 9703 | 4829 | 0,497 | -1,008 | -32,143 | 1,09E-226 | 2,07E-225 | 0,0001446 | 0,002219 | 0,0007913 | 0,0007913 | TRUE | 0,0048957 | No |
| orf19.557 | 51 | 58 | 49 | 28 | 58 | 32 | 47 | 61 | 46 | 59 | 46 | 42 | 52 | 35 | 48 | 47 | 46,82 | 48 | 46 | 0,935 | -0,097 | -0,468 | 0,64007 | 0,677928 | 1,00E-08 | 0,030254 | 0,020372 | 0,020372 | TRUE | 0,1704459 | No |
| orf19.5572 | 412 | 378 | 398 | 228 | 1460 | 1217 | 1214 | 1699 | 375 | 382 | 377 | 342 | 1299 | 1321 | 1243 | 1296 | 829,48 | 369 | 1290 | 3,478 | 1,798 | 29,263 | 3,11E-188 | 4,99E-187 | 1,00E-08 | 0,00363 | 0,0018312 | 0,0018312 | TRUE | 0,0207341 | No |
| orf19.5573 | 266 | 264 | 294 | 167 | 534 | 459 | 474 | 597 | 242 | 267 | 278 | 251 | 475 | 498 | 485 | 455 | 369,06 | 260 | 478 | 1,838 | 0,878 | 10,872 | 1,57E-27 | 4,76E-27 | 1,00E-08 | 0,005616 | 0,0032373 | 0,0032373 | TRUE | 0,028127 | No |
| orf19.5574 | 1238 | 1139 | 1203 | 731 | 965 | 830 | 805 | 1049 | 1127 | 1152 | 1139 | 1098 | 858 | 901 | 824 | 800 | 987,48 | 1129 | 846 | 0,747 | -0,42 | -7,604 | 2,87E-14 | 6,22E-14 | 0,0003549 | 0,003375 | 0,0019011 | 0,0019011 | TRUE | 0,0218932 | No |
| orf19.5575 | 1616 | 1734 | 1692 | 1163 | 2834 | 2298 | 2515 | 3299 | 1471 | 1754 | 1601 | 1747 | 2521 | 2495 | 2576 | 2517 | 2085,16 | 1643 | 2527 | 1,541 | 0,623 | 10,736 | 6,88E-27 | 2,06E-26 | 0,0028829 | 0,002671 | 0,0027248 | 0,0027248 | TRUE | 0,0612962 | No |
| orf19.5576 | 4774 | 4731 | 4807 | 3119 | 12960 | 11066 | 11282 | 15044 | 4347 | 4785 | 4549 | 4685 | 11528 | 12015 | 11554 | 11476 | 8117,25 | 4592 | 11643 | 2,537 | 1,343 | 33,851 | 3,51E-251 | 7,32E-250 | 0,0008815 | 0,0022 | 0,001354 | 0,001354 | TRUE | 0,016078 | No |
| orf19.5578 | 464 | 430 | 477 | 321 | 1390 | 1169 | 1288 | 1694 | 422 | 435 | 451 | 482 | 1236 | 1269 | 1319 | 1292 | 863,48 | 448 | 1279 | 2,871 | 1,521 | 25,095 | 5,69E-139 | 6,88E-138 | 3,30E-05 | 0,003567 | 0,0019686 | 0,0019686 | TRUE | 0,0251051 | No |
| orf19.5579 | 306 | 330 | 322 | 222 | 262 | 252 | 229 | 305 | 279 | 334 | 305 | 333 | 233 | 274 | 235 | 233 | 278,05 | 313 | 244 | 0,78 | -0,359 | -3,717 | 0,000202 | 0,000297 | 0,0032882 | 0,006788 | 0,0053289 | 0,0053289 | TRUE | 0,0722424 | No |
| orf19.558 | 6477 | 5626 | 6252 | 3885 | 10338 | 8902 | 9595 | 13051 | 5897 | 5690 | 5917 | 5836 | 9195 | 9665 | 9826 | 9955 | 7747,79 | 5835 | 9660 | 1,656 | 0,727 | 19,597 | 1,64E-85 | 1,24E-84 | 0,0005622 | 0,002208 | 0,001182 | 0,001182 | TRUE | 0,0129588 | No |
| orf19.5580 | 707 | 626 | 696 | 365 | 1008 | 815 | 947 | 1181 | 644 | 633 | 659 | 548 | 897 | 885 | 970 | 901 | 767 | 621 | 913 | 1,462 | 0,548 | 8,227 | 1,92E-16 | 4,41E-16 | 0,001774 | 0,003759 | 0,0028955 | 0,0028955 | TRUE | 0,0619588 | No |
| orf19.5584 | 611 | 615 | 629 | 371 | 688 | 515 | 555 | 711 | 556 | 622 | 595 | 557 | 612 | 559 | 568 | 542 | 576,59 | 582 | 570 | 0,977 | -0,033 | -0,477 | 0,633713 | 0,671976 | 0,0012584 | 0,004328 | 0,0028867 | 0,0028867 | TRUE | 0,029348 | No |
| orf19.5585 | 920 | 792 | 833 | 591 | 373 | 391 | 337 | 501 | 838 | 801 | 788 | 888 | 332 | 425 | 345 | 382 | 599,8 | 829 | 371 | 0,448 | -1,159 | -14,308 | 1,94E-46 | 8,45E-46 | 0,0045999 | 0,004239 | 0,0043912 | 0,0043912 | TRUE | 0,1005776 | No |
| orf19.5586 | 801 | 656 | 727 | 497 | 839 | 622 | 767 | 967 | 729 | 663 | 688 | 747 | 746 | 675 | 786 | 738 | 721,51 | 707 | 736 | 1,044 | 0,062 | 0,932 | 0,35119 | 0,391064 | 0,0017585 | 0,003868 | 0,0028798 | 0,0028798 | TRUE | 0,0342741 | No |
| orf19.5587 | 931 | 894 | 889 | 580 | 1226 | 1010 | 1011 | 1318 | 848 | 904 | 841 | 871 | 1090 | 1097 | 1035 | 1005 | 961,53 | 866 | 1057 | 1,22 | 0,287 | 5,127 | 2,94E-07 | 4,99E-07 | 0,0005778 | 0,003411 | 0,001945 | 0,001945 | TRUE | 0,0138572 | No |
| orf19.5588 | 86 | 72 | 63 | 43 | 36 | 37 | 39 | 42 | 78 | 73 | 60 | 65 | 32 | 40 | 40 | 32 | 52,44 | 69 | 36 | 0,517 | -0,95 | -4,774 | 1,80E-06 | 2,96E-06 | 1,00E-08 | 0,027228 | 0,0173232 | 0,0173232 | TRUE | 0,0852161 | No |
| orf19.5589 | 217 | 211 | 243 | 136 | 409 | 380 | 368 | 514 | 198 | 213 | 230 | 204 | 364 | 413 | 377 | 392 | 298,82 | 211 | 386 | 1,824 | 0,867 | 9,68 | 3,67E-22 | 9,78E-22 | 1,00E-08 | 0,006458 | 0,0039794 | 0,0039794 | TRUE | 0,0417852 | No |
| orf19.559 | 2942 | 2859 | 3148 | 1886 | 4375 | 3922 | 3978 | 5129 | 2679 | 2892 | 2979 | 2833 | 3891 | 4258 | 4074 | 3912 | 3439,82 | 2846 | 4034 | 1,417 | 0,503 | 10,407 | 2,31E-25 | 6,66E-25 | 0,00148 | 0,002421 | 0,0019366 | 0,0019366 | TRUE | 0,0198442 | No |
| orf19.5591 | 2676 | 2392 | 2503 | 1782 | 4930 | 4485 | 3994 | 5761 | 2436 | 2419 | 2369 | 2677 | 4385 | 4869 | 4090 | 4395 | 3455,1 | 2475 | 4435 | 1,793 | 0,843 | 14,002 | 1,52E-44 | 6,46E-44 | 0,0037611 | 0,002419 | 0,0031556 | 0,0031556 | TRUE | 0,0523681 | No |
| orf19.5592 | 20 | 19 | 23 | 8 | 23 | 21 | 14 | 7 | 18 | 19 | 22 | 12 | 20 | 23 | 14 | 5 | 16,77 | 18 | 16 | 0,857 | -0,222 | -0,545 | 0,585761 | 0,626128 | 0,1319566 | 0,080817 | 0,0998474 | 0,0998474 | TRUE | 0,9187898 | No |
| orf19.5595 | 750 | 667 | 729 | 486 | 773 | 711 | 724 | 929 | 683 | 675 | 690 | 730 | 688 | 772 | 741 | 709 | 710,88 | 694 | 728 | 1,048 | 0,068 | 1,097 | 0,272443 | 0,30844 | 0,0004094 | 0,003895 | 0,0022667 | 0,0022667 | TRUE | 0,0193906 | No |
| orf19.5596 | 1744 | 1591 | 1619 | 1175 | 1139 | 945 | 987 | 1267 | 1588 | 1609 | 1532 | 1765 | 1013 | 1026 | 1011 | 966 | 1313,83 | 1624 | 1004 | 0,62 | -0,691 | -12,431 | 1,78E-35 | 6,41E-35 | 0,0013967 | 0,003042 | 0,0021677 | 0,0021677 | TRUE | 0,0380048 | No |
| orf19.5597 | 3895 | 3773 | 4036 | 2318 | 3947 | 3138 | 3430 | 4363 | 3546 | 3816 | 3820 | 3482 | 3511 | 3407 | 3513 | 3328 | 3552,82 | 3666 | 3440 | 0,937 | -0,093 | -2,066 | 0,038803 | 0,048548 | 0,0011307 | 0,002409 | 0,0016775 | 0,0016775 | TRUE | 0,0130134 | No |
| orf19.5597.1 | 1238 | 1097 | 1089 | 900 | 1640 | 1493 | 1330 | 1870 | 1127 | 1109 | 1031 | 1352 | 1459 | 1621 | 1362 | 1426 | 1310,93 | 1155 | 1467 | 1,274 | 0,349 | 4,263 | 2,02E-05 | 3,15E-05 | 0,0089253 | 0,003045 | 0,0056487 | 0,0056487 | TRUE | 0,1495025 | No |
| orf19.5599 | 3638 | 3169 | 3542 | 2221 | 2885 | 2509 | 2504 | 3388 | 3312 | 3205 | 3352 | 3336 | 2566 | 2724 | 2564 | 2584 | 2955,6 | 3301 | 2610 | 0,79 | -0,34 | -8,685 | 3,80E-18 | 9,17E-18 | 0,0001767 | 0,002484 | 0,0011262 | 0,0011262 | TRUE | 0,0101397 | No |
| orf19.56 | 2318 | 2141 | 2291 | 1440 | 1557 | 1322 | 1331 | 1797 | 2110 | 2165 | 2168 | 2163 | 1385 | 1435 | 1363 | 1371 | 1770,16 | 2152 | 1388 | 0,645 | -0,632 | -15,09 | 1,88E-51 | 8,95E-51 | 1,00E-08 | 0,002783 | 0,0011008 | 0,0011008 | TRUE | 0,0058125 | No |
| orf19.5601 | 1288 | 1141 | 1272 | 879 | 1296 | 1082 | 1129 | 1521 | 1173 | 1154 | 1204 | 1320 | 1153 | 1175 | 1156 | 1160 | 1186,86 | 1213 | 1161 | 0,96 | -0,058 | -1,092 | 0,274989 | 0,311102 | 0,0006096 | 0,00315 | 0,001906 | 0,001906 | TRUE | 0,0379028 | No |
| orf19.5602 | 2277 | 2055 | 2197 | 1428 | 984 | 845 | 850 | 1209 | 2073 | 2078 | 2079 | 2145 | 875 | 917 | 871 | 922 | 1495,16 | 2094 | 896 | 0,429 | -1,222 | -26,468 | 2,24E-154 | 3,01E-153 | 1,00E-08 | 0,002921 | 0,0012713 | 0,0012713 | TRUE | 0,0050933 | No |
| orf19.5604 | 1404 | 1236 | 1370 | 847 | 1717 | 1391 | 1460 | 1977 | 1278 | 1250 | 1297 | 1272 | 1527 | 1510 | 1495 | 1508 | 1392,25 | 1274 | 1510 | 1,185 | 0,245 | 5,58 | 2,40E-08 | 4,26E-08 | 1,00E-08 | 0,002986 | 0,0011199 | 0,0011199 | TRUE | 0,0021011 | No |
| orf19.5605 | 14609 | 12284 | 14029 | 9124 | 10296 | 8801 | 9492 | 13230 | 13301 | 12424 | 13277 | 13706 | 9158 | 9555 | 9721 | 10092 | 11404,3 | 13177 | 9632 | 0,731 | -0,452 | -10,227 | 1,50E-24 | 4,25E-24 | 0,0015258 | 0,002153 | 0,0017849 | 0,0017849 | TRUE | 0,0180939 | No |
| orf19.5606 | 55 | 53 | 55 | 43 | 70 | 66 | 62 | 79 | 50 | 54 | 52 | 65 | 62 | 72 | 63 | 60 | 59,75 | 55 | 64 | 1,179 | 0,237 | 1,33 | 0,183425 | 0,21311 | 1,00E-08 | 0,024146 | 0,0135375 | 0,0135375 | TRUE | 0,0812268 | No |
| orf19.5607 | 10441 | 9627 | 10041 | 6245 | 4940 | 3703 | 4194 | 5427 | 9506 | 9737 | 9503 | 9381 | 4394 | 4020 | 4295 | 4140 | 6872 | 9532 | 4212 | 0,442 | -1,178 | -31,117 | 1,41E-212 | 2,55E-211 | 0,000663 | 0,002229 | 0,0012089 | 0,0012089 | TRUE | 0,0110818 | No |
| orf19.5608 | 3616 | 3465 | 3508 | 1902 | 2939 | 2223 | 2413 | 2937 | 3292 | 3504 | 3320 | 2857 | 2614 | 2414 | 2471 | 2240 | 2839,14 | 3243 | 2435 | 0,75 | -0,416 | -6,153 | 7,59E-10 | 1,43E-09 | 0,0051124 | 0,002502 | 0,004023 | 0,004023 | TRUE | 0,0754699 | No |
| orf19.5609 | 2229 | 1848 | 2033 | 1299 | 2527 | 2021 | 2245 | 2877 | 2029 | 1869 | 1924 | 1951 | 2248 | 2194 | 2299 | 2195 | 2088,69 | 1943 | 2234 | 1,149 | 0,2 | 4,599 | 4,25E-06 | 6,87E-06 | 0,0003753 | 0,002669 | 0,0013389 | 0,0013389 | TRUE | 0,0115366 | No |
| orf19.5610 | 3599 | 3539 | 3692 | 2163 | 1229 | 1044 | 976 | 1408 | 3277 | 3579 | 3494 | 3249 | 1093 | 1133 | 1000 | 1074 | 2237,45 | 3400 | 1075 | 0,316 | -1,662 | -30,825 | 1,23E-208 | 2,16E-207 | 0,0016844 | 0,002627 | 0,0021985 | 0,0021985 | TRUE | 0,0255664 | No |
| orf19.5611 | 236 | 236 | 240 | 118 | 133 | 103 | 111 | 130 | 215 | 239 | 227 | 177 | 118 | 112 | 114 | 99 | 162,62 | 214 | 111 | 0,511 | -0,968 | -7,82 | 5,28E-15 | 1,17E-14 | 0,0044698 | 0,01016 | 0,0080165 | 0,0080165 | TRUE | 0,1212618 | No |
| orf19.5612 | 1036 | 1070 | 1138 | 602 | 1599 | 1329 | 1360 | 1903 | 943 | 1082 | 1077 | 904 | 1422 | 1443 | 1393 | 1452 | 1214,54 | 1002 | 1428 | 1,422 | 0,507 | 7,97 | 1,59E-15 | 3,56E-15 | 0,0031971 | 0,003125 | 0,0030294 | 0,0030294 | TRUE | 0,0452984 | No |
| orf19.5614 | 7951 | 6518 | 7279 | 4557 | 4455 | 3647 | 3989 | 5419 | 7239 | 6592 | 6889 | 6845 | 3963 | 3960 | 4085 | 4134 | 5463,33 | 6891 | 4036 | 0,586 | -0,772 | -19,224 | 2,35E-82 | 1,72E-81 | 0,000721 | 0,002279 | 0,0013551 | 0,0013551 | TRUE | 0,0143808 | No |
| orf19.5615 | 1827 | 1525 | 1615 | 1061 | 2750 | 2163 | 2147 | 3161 | 1663 | 1542 | 1528 | 1594 | 2446 | 2348 | 2199 | 2411 | 1966,56 | 1582 | 2351 | 1,487 | 0,572 | 11,207 | 3,78E-29 | 1,19E-28 | 0,001412 | 0,002709 | 0,0019663 | 0,0019663 | TRUE | 0,0228442 | No |
| orf19.5616 | 42 | 40 | 39 | 35 | 143 | 143 | 114 | 176 | 38 | 40 | 37 | 53 | 127 | 155 | 117 | 134 | 87,7 | 42 | 133 | 3,228 | 1,691 | 9,722 | 2,43E-22 | 6,51E-22 | 0,0045633 | 0,017099 | 0,0126851 | 0,0126851 | TRUE | 0,1489173 | No |
| orf19.5617 | 4949 | 4286 | 4572 | 2695 | 2622 | 1976 | 2341 | 3013 | 4506 | 4335 | 4327 | 4048 | 2332 | 2145 | 2397 | 2298 | 3298,67 | 4304 | 2293 | 0,533 | -0,909 | -18,527 | 1,24E-76 | 8,41E-76 | 0,0016084 | 0,002437 | 0,0019827 | 0,0019827 | TRUE | 0,0220623 | No |

| | | | | | | | | | | | | | | | | | | | | | | | | | | | | | | | |
|---|---|---|---|---|---|---|---|---|---|---|---|---|---|---|---|---|---|---|---|---|---|---|---|---|---|---|---|---|---|---|---|
| orf19.5618 | 1455 | 1489 | 1539 | 980 | 516 | 430 | 404 | 549 | 1325 | 1506 | 1457 | 1472 | 459 | 467 | 414 | 419 | 939,71 | 1440 | 440 | 0,305 | -1,712 | -25,618 | 9,53E-145 | 1,20E-143 | 0,0022541 | 0,003443 | 0,0028531 | 0,0028531 | TRUE | 0,0362583 | No |
| orf19.5619 | 1628 | 1463 | 1536 | 966 | 1086 | 899 | 969 | 1271 | 1482 | 1480 | 1454 | 1451 | 966 | 976 | 992 | 970 | 1221,32 | 1467 | 976 | 0,665 | -0,589 | -12,797 | 1,70E-37 | 6,36E-37 | 1,00E-08 | 0,003119 | 0,0011901 | 0,0011901 | TRUE | 0,0014898 | No |
| orf19.562 | 698 | 617 | 730 | 452 | 1460 | 1337 | 1445 | 1741 | 635 | 624 | 691 | 679 | 1299 | 1452 | 1480 | 1328 | 1023,44 | 657 | 1390 | 2,114 | 1,08 | 16,606 | 6,30E-62 | 3,54E-61 | 0,0025246 | 0,003328 | 0,0029077 | 0,0029077 | TRUE | 0,0232616 | No |
| orf19.5620 | 795 | 777 | 751 | 485 | 757 | 648 | 617 | 798 | 724 | 786 | 711 | 729 | 673 | 704 | 632 | 609 | 695,81 | 738 | 654 | 0,886 | -0,175 | -2,603 | 0,009243 | 0,012152 | 0,0017573 | 0,003936 | 0,0028773 | 0,0028773 | TRUE | 0,0290185 | No |
| orf19.5621 | 681 | 730 | 763 | 434 | 699 | 584 | 621 | 713 | 620 | 738 | 722 | 652 | 622 | 634 | 636 | 544 | 646,01 | 683 | 609 | 0,889 | -0,17 | -2,148 | 0,031715 | 0,039981 | 0,0048389 | 0,004082 | 0,0044536 | 0,0044536 | TRUE | 0,0645731 | No |
| orf19.5622 | 3019 | 2963 | 3198 | 1763 | 5722 | 4539 | 4680 | 6318 | 2749 | 2997 | 3027 | 2648 | 5090 | 4928 | 4793 | 4819 | 3881,29 | 2855 | 4908 | 1,717 | 0,78 | 15,395 | 1,75E-53 | 8,65E-53 | 0,0021252 | 0,002377 | 0,0021796 | 0,0021796 | TRUE | 0,0270478 | No |
| orf19.5623 | 1560 | 1568 | 1648 | 999 | 1731 | 1344 | 1491 | 2019 | 1420 | 1586 | 1560 | 1501 | 1540 | 1459 | 1527 | 1540 | 1516,56 | 1517 | 1516 | 1 | 0,001 | 0,014 | 0,988901 | 0,990144 | 0,0009034 | 0,002908 | 0,0018034 | 0,0018034 | TRUE | 0,0238128 | No |
| orf19.5625 | 703 | 763 | 767 | 438 | 1049 | 883 | 898 | 1128 | 640 | 772 | 726 | 658 | 933 | 959 | 920 | 860 | 808,43 | 699 | 918 | 1,31 | 0,39 | 5,503 | 3,73E-08 | 6,54E-08 | 0,0031585 | 0,003671 | 0,0035422 | 0,0035422 | TRUE | 0,0582817 | No |
| orf19.5626 | 27896 | 26829 | 28764 | 19222 | 3555 | 3101 | 3131 | 4256 | 25398 | 27135 | 27223 | 28875 | 3162 | 3367 | 3207 | 3247 | 15201,5 | 27158 | 3246 | 0,12 | -3,065 | -66,757 | 0 | 0 | 0,0015039 | 0,002124 | 0,0018607 | 0,0018607 | TRUE | 0,0233109 | No |
| orf19.5627 | 10552 | 10280 | 11084 | 6798 | 8788 | 7253 | 7763 | 10425 | 9607 | 10397 | 10490 | 10212 | 7817 | 7875 | 7950 | 7952 | 9037,49 | 10176 | 7898 | 0,776 | -0,365 | -9,855 | 6,50E-23 | 1,76E-22 | 0,0006446 | 0,002183 | 0,0012082 | 0,0012082 | TRUE | 0,0176829 | No |
| orf19.5628 | 1875 | 1727 | 1822 | 1156 | 2400 | 2028 | 2092 | 2851 | 1707 | 1747 | 1724 | 1736 | 2135 | 2202 | 2142 | 2175 | 1946,06 | 1728 | 2164 | 1,252 | 0,324 | 8,198 | 2,45E-16 | 5,61E-16 | 1,00E-08 | 0,002716 | 0,0009739 | 0,0009739 | TRUE | 0,0016166 | No |
| orf19.5629 | 2815 | 2579 | 2543 | 1579 | 4271 | 3335 | 3361 | 4349 | 2563 | 2608 | 2407 | 2372 | 3799 | 3621 | 3442 | 3317 | 3016,16 | 2488 | 3545 | 1,424 | 0,51 | 9,331 | 1,05E-20 | 2,72E-20 | 0,0023852 | 0,002475 | 0,0025159 | 0,0025159 | TRUE | 0,0287905 | No |
| orf19.563 | 8292 | 6797 | 7622 | 4362 | 9235 | 6759 | 8163 | 10634 | 7550 | 6874 | 7214 | 6552 | 8214 | 7338 | 8360 | 8112 | 7526,77 | 7048 | 8006 | 1,136 | 0,183 | 3,269 | 0,001081 | 0,001524 | 0,0033098 | 0,002213 | 0,002891 | 0,002891 | TRUE | 0,0382281 | No |
| orf19.5630 | 1680 | 1761 | 1795 | 1096 | 1184 | 1018 | 982 | 1299 | 1530 | 1781 | 1699 | 1646 | 1053 | 1105 | 1006 | 991 | 1351,35 | 1664 | 1039 | 0,624 | -0,681 | -11,15 | 7,15E-29 | 2,23E-28 | 0,0026303 | 0,003015 | 0,0028073 | 0,0028073 | TRUE | 0,0375417 | No |
| orf19.5633 | 6258 | 5481 | 5717 | 3714 | 5214 | 4364 | 4769 | 6455 | 5698 | 5543 | 5411 | 5579 | 4638 | 4738 | 4884 | 4924 | 5176,81 | 5558 | 4796 | 0,863 | -0,212 | -5,84 | 5,21E-09 | 9,51E-09 | 0,0004477 | 0,002292 | 0,0010759 | 0,0010759 | TRUE | 0,0061515 | No |
| orf19.5634 | 4214 | 3498 | 3830 | 2331 | 2264 | 1835 | 2055 | 2648 | 3837 | 3538 | 3625 | 3502 | 2014 | 1992 | 2105 | 2020 | 2828,92 | 3626 | 2033 | 0,56 | -0,836 | -18,922 | 7,59E-80 | 5,35E-79 | 0,0008469 | 0,002504 | 0,0014963 | 0,0014963 | TRUE | 0,0190605 | No |
| orf19.5635 | 3366 | 2570 | 2830 | 2270 | 1645 | 1457 | 1470 | 2258 | 3065 | 2599 | 2678 | 3410 | 1463 | 1582 | 1505 | 1722 | 2253,13 | 2938 | 1568 | 0,535 | -0,904 | -10,661 | 1,55E-26 | 4,58E-26 | 0,0101129 | 0,002623 | 0,0064199 | 0,0064199 | TRUE | 0,1387385 | No |
| orf19.5636 | 23644 | 18043 | 19957 | 15914 | 5169 | 4345 | 4350 | 6587 | 21527 | 18249 | 18887 | 23905 | 4598 | 4717 | 4455 | 5025 | 12670,4 | 20642 | 4699 | 0,228 | -2,135 | -27,311 | 3,16E-164 | 4,53E-163 | 0,0090726 | 0,002141 | 0,0057449 | 0,0057449 | TRUE | 0,1381384 | No |
| orf19.5639 | 5998 | 5118 | 5728 | 3708 | 4565 | 3824 | 4186 | 5814 | 5461 | 5176 | 5421 | 5570 | 4060 | 4152 | 4287 | 4435 | 4820,31 | 5407 | 4234 | 0,783 | -0,352 | -8,348 | 6,97E-17 | 1,63E-16 | 0,0009365 | 0,002311 | 0,0014979 | 0,0014979 | TRUE | 0,0128801 | No |
| orf19.564 | 435 | 401 | 492 | 268 | 254 | 192 | 232 | 309 | 396 | 406 | 466 | 403 | 226 | 208 | 238 | 236 | 322,19 | 418 | 227 | 0,544 | -0,879 | -9,81 | 1,02E-22 | 2,76E-22 | 0,0021541 | 0,006137 | 0,004379 | 0,004379 | TRUE | 0,0733236 | No |
| orf19.5640 | 13546 | 10993 | 12318 | 8335 | 14214 | 11728 | 13150 | 17864 | 12333 | 11118 | 11658 | 12520 | 12643 | 12733 | 13467 | 13627 | 12512,5 | 11907 | 13118 | 1,102 | 0,14 | 2,922 | 0,003474 | 0,004701 | 0,0020841 | 0,002143 | 0,0021249 | 0,0021249 | TRUE | 0,0243251 | No |
| orf19.5641 | 17672 | 13823 | 15433 | 9742 | 52250 | 40579 | 46903 | 64973 | 16090 | 13980 | 14606 | 14634 | 46475 | 44057 | 48035 | 49562 | 30929,9 | 14828 | 47032 | 3,172 | 1,665 | 31,565 | 1,10E-218 | 2,03E-217 | 0,0030059 | 0,00208 | 0,0026277 | 0,0026277 | TRUE | 0,0404029 | No |
| orf19.5642 | 7754 | 6023 | 6412 | 4436 | 11223 | 8021 | 8838 | 12538 | 7060 | 6092 | 6068 | 6664 | 9983 | 8709 | 9051 | 9564 | 7898,71 | 6471 | 9327 | 1,442 | 0,528 | 8,644 | 5,43E-18 | 1,31E-17 | 0,0043802 | 0,002204 | 0,0034475 | 0,0034475 | TRUE | 0,0465526 | No |
| orf19.5643 | 980 | 835 | 943 | 600 | 2068 | 1692 | 1874 | 2547 | 892 | 845 | 892 | 901 | 1839 | 1837 | 1919 | 1943 | 1383,63 | 882 | 1884 | 2,138 | 1,096 | 22,229 | 1,81E-109 | 1,70E-108 | 1,00E-08 | 0,002992 | 0,001481 | 0,001481 | TRUE | 0,0101701 | No |
| orf19.5644 | 274 | 251 | 275 | 187 | 690 | 667 | 635 | 818 | 249 | 254 | 260 | 281 | 614 | 724 | 650 | 624 | 457,09 | 261 | 653 | 2,507 | 1,326 | 16,049 | 5,77E-58 | 3,07E-57 | 0,0024989 | 0,004927 | 0,0037886 | 0,0037886 | TRUE | 0,0612681 | No |
| orf19.5645 | 3024 | 2384 | 2672 | 1892 | 7610 | 6296 | 6571 | 9466 | 2753 | 2411 | 2529 | 2842 | 6769 | 6836 | 6730 | 7221 | 4761,26 | 2634 | 6889 | 2,619 | 1,389 | 25,256 | 9,72E-141 | 1,19E-139 | 0,0030273 | 0,002314 | 0,0026313 | 0,0026313 | TRUE | 0,0389977 | No |
| orf19.5646 | 407 | 311 | 342 | 233 | 685 | 481 | 595 | 831 | 371 | 315 | 324 | 350 | 609 | 522 | 609 | 634 | 466,69 | 340 | 594 | 1,752 | 0,809 | 9,483 | 2,46E-21 | 6,44E-21 | 0,0043449 | 0,004868 | 0,0046279 | 0,0046279 | TRUE | 0,0741468 | No |
| orf19.5647 | 5038 | 4306 | 4677 | 2995 | 7548 | 6520 | 6435 | 9021 | 4587 | 4355 | 4426 | 4499 | 6714 | 7079 | 6590 | 6881 | 5641,43 | 4467 | 6816 | 1,526 | 0,61 | 16,284 | 1,27E-59 | 6,93E-59 | 0,0004836 | 0,002271 | 0,0011567 | 0,0011567 | TRUE | 0,0080296 | No |
| orf19.5648 | 531 | 454 | 444 | 279 | 1412 | 948 | 1166 | 1473 | 483 | 459 | 420 | 419 | 1256 | 1029 | 1194 | 1124 | 798,11 | 445 | 1151 | 2,578 | 1,366 | 17,428 | 5,04E-68 | 3,08E-67 | 0,0049129 | 0,003692 | 0,004293 | 0,004293 | TRUE | 0,0585548 | No |
| orf19.5649 | 45 | 42 | 30 | 27 | 90 | 105 | 88 | 117 | 41 | 42 | 28 | 41 | 80 | 114 | 90 | 89 | 65,73 | 38 | 93 | 2,453 | 1,294 | 6,589 | 4,43E-11 | 8,73E-11 | 0,01072 | 0,022135 | 0,0180184 | 0,0180184 | TRUE | 0,2425876 | No |
| orf19.5650 | 6629 | 5221 | 5722 | 3631 | 13886 | 11199 | 11934 | 16803 | 6035 | 5280 | 5415 | 5454 | 12351 | 12159 | 12222 | 12817 | 8966,89 | 5546 | 12387 | 2,233 | 1,159 | 24,622 | 7,42E-134 | 8,56E-133 | 0,0019324 | 0,002184 | 0,0019929 | 0,0019929 | TRUE | 0,0437392 | No |
| orf19.5651 | 7528 | 6128 | 6905 | 4362 | 6985 | 5835 | 6377 | 8755 | 6854 | 6198 | 6535 | 6552 | 6213 | 6335 | 6531 | 6678 | 6487,06 | 6535 | 6439 | 0,985 | -0,021 | -0,495 | 0,620249 | 0,658795 | 0,0010707 | 0,002241 | 0,0015945 | 0,0015945 | TRUE | 0,0149238 | No |
| orf19.5653 | 13174 | 10753 | 11642 | 7883 | 18377 | 15223 | 15706 | 23712 | 11994 | 10876 | 11018 | 11842 | 16346 | 16528 | 16085 | 18088 | 14097 | 11432 | 16762 | 1,467 | 0,552 | 10,88 | 1,43E-27 | 4,36E-27 | 0,0025822 | 0,002131 | 0,002401 | 0,002401 | TRUE | 0,0348807 | No |
| orf19.5654 | 2799 | 2492 | 2610 | 1758 | 3017 | 2560 | 2678 | 3612 | 2548 | 2520 | 2470 | 2641 | 2684 | 2779 | 2743 | 2755 | 2642,57 | 2545 | 2740 | 1,078 | 0,108 | 2,8 | 0,005108 | 0,00685 | 1,74E-05 | 0,002537 | 0,0010536 | 0,0010536 | TRUE | 0,0069915 | No |
| orf19.5655 | 135 | 166 | 123 | 74 | 243 | 177 | 221 | 316 | 123 | 168 | 116 | 111 | 216 | 192 | 226 | 241 | 174,26 | 130 | 219 | 1,685 | 0,753 | 5,536 | 3,10E-08 | 5,46E-08 | 0,0145625 | 0,009618 | 0,0115304 | 0,0115304 | TRUE | 0,405824 | No |
| orf19.5657 | 3759 | 3125 | 3593 | 2251 | 2609 | 2264 | 2361 | 3226 | 3422 | 3161 | 3400 | 3381 | 2321 | 2458 | 2418 | 2461 | 2877,79 | 3341 | 2414 | 0,723 | -0,469 | -10,862 | 1,74E-27 | 5,29E-27 | 0,0007508 | 0,002496 | 0,0014329 | 0,0014329 | TRUE | 0,0164007 | No |
| orf19.5658 | 1039 | 805 | 850 | 584 | 1900 | 1440 | 1645 | 2184 | 946 | 814 | 804 | 877 | 1690 | 1563 | 1685 | 1666 | 1255,74 | 860 | 1651 | 1,92 | 0,941 | 15,265 | 1,31E-52 | 6,31E-52 | 0,0023641 | 0,003089 | 0,0027416 | 0,0027416 | TRUE | 0,0583563 | No |
| orf19.5660 | 828 | 695 | 715 | 495 | 1370 | 1005 | 1143 | 1570 | 754 | 703 | 677 | 744 | 1219 | 1091 | 1171 | 1198 | 944,37 | 720 | 1170 | 1,63 | 0,705 | 11,632 | 2,83E-31 | 9,38E-31 | 0,0013219 | 0,003436 | 0,0023783 | 0,0023783 | TRUE | 0,0235087 | No |
| orf19.5660.1 | 2592 | 2101 | 2156 | 1572 | 5239 | 4089 | 4102 | 5930 | 2360 | 2125 | 2040 | 2361 | 4660 | 4440 | 4201 | 4523 | 3338,82 | 2222 | 4456 | 2,008 | 1,006 | 17,369 | 1,41E-67 | 8,59E-67 | 0,0032151 | 0,002433 | 0,0028719 | 0,0028719 | TRUE | 0,0369334 | No |
| orf19.5661 | 2061 | 1649 | 1765 | 1210 | 2944 | 2700 | 2593 | 3833 | 1876 | 1668 | 1670 | 1818 | 2619 | 2931 | 2656 | 2924 | 2270,21 | 1758 | 2782 | 1,583 | 0,663 | 11,105 | 1,18E-28 | 3,68E-28 | 0,003015 | 0,002619 | 0,002949 | 0,002949 | TRUE | 0,0264687 | No |
| orf19.5662 | 1436 | 1225 | 1328 | 834 | 1181 | 950 | 1063 | 1418 | 1307 | 1239 | 1257 | 1253 | 1050 | 1031 | 1089 | 1082 | 1163,53 | 1264 | 1063 | 0,841 | -0,25 | -5,125 | 2,97E-07 | 5,03E-07 | 1,00E-08 | 0,003172 | 0,0014219 | 0,0014219 | TRUE | 0,0068452 | No |
| orf19.5663 | 1534 | 1349 | 1391 | 856 | 2388 | 1857 | 1999 | 2808 | 1397 | 1364 | 1316 | 1286 | 2124 | 2016 | 2047 | 2142 | 1711,59 | 1341 | 2082 | 1,552 | 0,634 | 13,493 | 1,72E-41 | 6,89E-41 | 0,0004296 | 0,002809 | 0,0014943 | 0,0014943 | TRUE | 0,0095112 | No |
| orf19.5664 | 144 | 159 | 170 | 86 | 190 | 131 | 153 | 234 | 131 | 161 | 161 | 129 | 169 | 142 | 157 | 178 | 153,55 | 146 | 162 | 1,11 | 0,151 | 1,203 | 0,228988 | 0,262601 | 0,0053699 | 0,01064 | 0,0086015 | 0,0086015 | TRUE | 0,065721 | No |
| orf19.5665 | 546 | 452 | 498 | 322 | 792 | 582 | 635 | 806 | 497 | 457 | 471 | 484 | 704 | 632 | 650 | 615 | 563,84 | 477 | 650 | 1,363 | 0,446 | 6,397 | 1,59E-10 | 3,06E-10 | 3,72E-06 | 0,00438 | 0,0028367 | 0,0028367 | TRUE | 0,0383786 | No |
| orf19.5666 | 899 | 729 | 768 | 436 | 1710 | 1053 | 1383 | 1805 | 819 | 737 | 727 | 655 | 1521 | 1143 | 1416 | 1377 | 1049,39 | 734 | 1364 | 1,854 | 0,891 | 9,902 | 4,07E-23 | 1,11E-22 | 0,00959 | 0,003296 | 0,0066985 | 0,0066985 | TRUE | 0,1406587 | No |
| orf19.5667 | 6904 | 5628 | 6325 | 3781 | 4148 | 3366 | 4019 | 4999 | 6286 | 5692 | 5986 | 5680 | 3690 | 3655 | 4116 | 3813 | 4864,61 | 5911 | 3818 | 0,646 | -0,631 | -12,181 | 3,91E-34 | 1,37E-33 | 0,0024027 | 0,002309 | 0,0023646 | 0,0023646 | TRUE | 0,0331136 | No |
| orf19.5669 | 730 | 592 | 649 | 391 | 783 | 568 | 654 | 842 | 665 | 599 | 614 | 587 | 696 | 617 | 670 | 642 | 636,27 | 616 | 656 | 1,063 | 0,088 | 1,302 | 0,192886 | 0,22361 | 0,0013249 | 0,004113 | 0,0028006 | 0,0028006 | TRUE | 0,0340647 | No |
| orf19.567 | 927 | 1030 | 1070 | 670 | 494 | 436 | 422 | 489 | 844 | 1042 | 1013 | 1006 | 439 | 473 | 432 | 373 | 702,85 | 976 | 429 | 0,439 | -1,187 | -13,433 | 3,89E-41 | 1,55E-40 | 0,0079369 | 0,003917 | 0,0058569 | 0,0058569 | TRUE | 0,1027383 | No |
| orf19.5671 | 5360 | 4573 | 4938 | 3464 | 1643 | 1342 | 1431 | 1917 | 4880 | 4625 | 4673 | 5203 | 1461 | 1457 | 1466 | 1462 | 3153,53 | 4845 | 1462 | 0,302 | -1,728 | -36,142 | 5,01E-286 | 1,23E-284 | 0,0013186 | 0,002456 | 0,0017634 | 0,0017634 | TRUE | 0,0288478 | No |
| orf19.5672 | 42 | 51 | 43 | 40 | 327 | 282 | 296 | 441 | 38 | 52 | 41 | 60 | 291 | 306 | 303 | 336 | 178,4 | 48 | 309 | 6,657 | 2,735 | 18,934 | 5,98E-80 | 4,23E-79 | 0,0023141 | 0,009442 | 0,0071543 | 0,0071543 | TRUE | 0,1703452 | No |
| orf19.5673 | 2127 | 1841 | 2087 | 1499 | 1709 | 1652 | 1667 | 2403 | 1937 | 1862 | 1975 | 2252 | 1520 | 1794 | 1707 | 1833 | 1859,92 | 2006 | 1714 | 0,855 | -0,225 | -3,088 | 0,002014 | 0,002778 | 0,0061776 | 0,002747 | 0,004574 | 0,004574 | TRUE | 0,0785672 | No |
| orf19.5674 | 460 | 332 | 410 | 308 | 302 | 306 | 277 | 417 | 419 | 336 | 388 | 463 | 269 | 332 | 284 | 318 | 350,99 | 402 | 301 | 0,752 | -0,411 | -3,871 | 0,000108 | 0,000162 | 0,010651 | 0,005801 | 0,0079467 | 0,0079467 | TRUE | 0,1374545 | No |
| orf19.5675 | 5733 | 4972 | 5471 | 3400 | 4475 | 3776 | 4131 | 5700 | 5220 | 5029 | 5178 | 5107 | 3980 | 4100 | 4231 | 4348 | 4649,02 | 5134 | 4165 | 0,811 | -0,302 | -7,672 | 1,70E-14 | 3,71E-14 | 0,0005751 | 0,002321 | 0,0012662 | 0,0012662 | TRUE | 0,0111003 | No |
| orf19.5676 | 2752 | 2123 | 2554 | 1503 | 2588 | 1964 | 2416 | 2986 | 2506 | 2147 | 2417 | 2258 | 2302 | 2132 | 2474 | 2278 | 2314,25 | 2332 | 2296 | 0,984 | -0,023 | -0,37 | 0,711033 | 0,744797 | 0,0037828 | 0,002608 | 0,0032554 | 0,0032554 | TRUE | 0,0355821 | No |
| orf19.5677 | 80 | 81 | 86 | 47 | 135 | 113 | 133 | 169 | 73 | 82 | 81 | 71 | 120 | 123 | 136 | 129 | 101,83 | 77 | 127 | 1,648 | 0,721 | 5,168 | 2,37E-07 | 4,03E-07 | 1,00E-08 | 0,01501 | 0,0081132 | 0,0081132 | TRUE | 0,0227899 | No |
| orf19.5678 | 1741 | 1309 | 1566 | 981 | 1455 | 1201 | 1501 | 2016 | 1585 | 1324 | 1482 | 1474 | 1294 | 1304 | 1537 | 1538 | 1442,23 | 1466 | 1418 | 0,968 | -0,048 | -0,619 | 0,536036 | 0,577723 | 0,0068393 | 0,002953 | 0,0049619 | 0,0049619 | TRUE | 0,0523914 | No |
| orf19.568 | 3187 | 3160 | 3311 | 1847 | 3259 | 2572 | 2647 | 3624 | 2902 | 3196 | 3134 | 2774 | 2899 | 2792 | 2711 | 2764 | 2896,53 | 3002 | 2792 | 0,929 | -0,106 | -2,023 | 0,043086 | 0,0537 | 0,0021005 | 0,002493 | 0,0022897 | 0,0022897 | TRUE | 0,0296254 | No |
| orf19.5680 | 1929 | 1591 | 1725 | 1075 | 2432 | 1840 | 2160 | 2952 | 1756 | 1609 | 1633 | 1615 | 2163 | 1998 | 2212 | 2252 | 1904,7 | 1653 | 2156 | 1,304 | 0,383 | 7,195 | 6,25E-13 | 1,31E-12 | 0,0017537 | 0,002731 | 0,0021794 | 0,0021794 | TRUE | 0,0289234 | No |
| orf19.5681 | 1139 | 965 | 969 | 642 | 1049 | 809 | 1010 | 1315 | 1037 | 976 | 917 | 964 | 933 | 878 | 1034 | 1003 | 967,92 | 974 | 962 | 0,989 | -0,017 | -0,249 | 0,803178 | 0,829326 | 0,0029553 | 0,003402 | 0,0031737 | 0,0031737 | TRUE | 0,0392663 | No |
| orf19.5682 | 2944 | 2365 | 2582 | 1812 | 4237 | 3545 | 3733 | 5302 | 2680 | 2392 | 2444 | 2722 | 3769 | 3849 | 3823 | 4044 | 3215,36 | 2560 | 3871 | 1,514 | 0,599 | 11,519 | 1,06E-30 | 3,46E-30 | 0,0021479 | 0,002448 | 0,0022627 | 0,0022627 | TRUE | 0,0233048 | No |
| orf19.5683 | 1761 | 1460 | 1616 | 923 | 3127 | 2527 | 2852 | 3945 | 1603 | 1477 | 1529 | 1386 | 2781 | 2744 | 2921 | 3009 | 2181,36 | 1499 | 2864 | 1,907 | 0,931 | 17,072 | 2,40E-65 | 1,42E-64 | 0,002102 | 0,002643 | 0,0023351 | 0,0023351 | TRUE | 0,0279903 | No |
| orf19.5684 | 2145 | 1556 | 1731 | 1068 | 2043 | 1544 | 1869 | 2460 | 1953 | 1574 | 1638 | 1604 | 1817 | 1676 | 1914 | 1877 | 1756,67 | 1692 | 1821 | 1,075 | 0,105 | 1,429 | 0,153144 | 0,179987 | 0,0062747 | 0,002789 | 0,0045829 | 0,0045829 | TRUE | 0,1322963 | No |
| orf19.5684.1 | 110 | 95 | 90 | 78 | 131 | 71 | 106 | 173 | 100 | 96 | 85 | 117 | 117 | 77 | 109 | 132 | 104,09 | 100 | 109 | 1,105 | 0,145 | 0,85 | 0,395243 | 0,436081 | 0,0231187 | 0,014728 | 0,0180886 | 0,0180886 | TRUE | 0,3561247 | No |
| orf19.5685 | 4357 | 3552 | 3982 | 2430 | 8032 | 5961 | 7290 | 10008 | 3967 | 3592 | 3769 | 3650 | 7144 | 6472 | 7466 | 7634 | 5461,8 | 3744 | 7179 | 1,917 | 0,939 | 16,334 | 5,68E-60 | 3,11E-59 | 0,0032606 | 0,002279 | 0,0029638 | 0,0029638 | TRUE | 0,0533487 | No |
| orf19.5686 | 5857 | 4399 | 4676 | 2874 | 584 | 451 | 486 | 725 | 5333 | 4449 | 4425 | 4317 | 519 | 490 | 498 | 553 | 2573,02 | 4631 | 515 | 0,111 | -3,167 | -41,669 | 0 | 0 | 0,0063091 | 0,00255 | 0,0045259 | 0,0045259 | TRUE | 0,1275848 | No |
| orf19.5688 | 2757 | 2157 | 2306 | 1372 | 1739 | 1101 | 1351 | 1844 | 2510 | 2182 | 2182 | 2061 | 1547 | 1195 | 1384 | 1407 | 1808,43 | 2234 | 1383 | 0,619 | -0,692 | -8,583 | 9,20E-18 | 2,20E-17 | 0,0083999 | 0,002767 | 0,0056571 | 0,0056571 | TRUE | 0,0986484 | No |
| orf19.5689 | 1096 | 845 | 957 | 619 | 1220 | 1045 | 1131 | 1549 | 998 | 855 | 906 | 930 | 1085 | 1135 | 1158 | 1182 | 1030,96 | 922 | 1140 | 1,236 | 0,306 | 5,049 | 4,44E-07 | 7,49E-07 | 0,0017982 | 0,003318 | 0,0025348 | 0,0025348 | TRUE | 0,0396308 | No |

| | | | | | | | | | | | | | | | | | | | | | | | | | | | | | | | |
|---|---|---|---|---|---|---|---|---|---|---|---|---|---|---|---|---|---|---|---|---|---|---|---|---|---|---|---|---|---|---|---|
| orf19.5691 | 742 | 597 | 670 | 367 | 529 | 353 | 467 | 641 | 676 | 604 | 634 | 551 | 471 | 383 | 478 | 489 | 535,72 | 616 | 455 | 0,738 | -0,438 | -4,907 | 9,26E-07 | 1,54E-06 | 0,0072809 | 0,004503 | 0,0057564 | 0,0057564 | TRUE | 0,1256103 | No |
| orf19.5692 | 484 | 398 | 413 | 320 | 567 | 430 | 552 | 721 | 441 | 403 | 391 | 481 | 504 | 467 | 565 | 550 | 475,16 | 429 | 522 | 1,223 | 0,291 | 3,339 | 0,000841 | 0,001192 | 0,0053494 | 0,004817 | 0,0051394 | 0,0051394 | TRUE | 0,0667311 | No |
| orf19.5693 | 1036 | 792 | 922 | 560 | 1566 | 1236 | 1408 | 2059 | 943 | 801 | 873 | 841 | 1393 | 1342 | 1442 | 1571 | 1150,69 | 864 | 1437 | 1,661 | 0,732 | 10,688 | 1,16E-26 | 3,45E-26 | 0,0039082 | 0,003185 | 0,0035634 | 0,0035634 | TRUE | 0,0487355 | No |
| orf19.5694 | 153 | 129 | 171 | 118 | 353 | 246 | 304 | 378 | 139 | 130 | 162 | 177 | 314 | 267 | 311 | 288 | 223,7 | 152 | 295 | 1,958 | 0,969 | 8,692 | 3,57E-18 | 8,63E-18 | 0,0048026 | 0,007942 | 0,0068239 | 0,0068239 | TRUE | 0,1008412 | No |
| orf19.5698 | 2260 | 1775 | 2001 | 1481 | 1594 | 1302 | 1440 | 2132 | 2058 | 1795 | 1894 | 2225 | 1418 | 1414 | 1475 | 1626 | 1737,97 | 1993 | 1483 | 0,746 | -0,423 | -5,823 | 5,79E-09 | 1,05E-08 | 0,0059471 | 0,002797 | 0,0044849 | 0,0044849 | TRUE | 0,0708727 | No |
| orf19.57 | 693 | 625 | 649 | 405 | 756 | 581 | 619 | 805 | 631 | 632 | 614 | 608 | 672 | 631 | 634 | 614 | 629,61 | 621 | 638 | 1,025 | 0,036 | 0,581 | 0,561532 | 0,602654 | 1,00E-08 | 0,004135 | 0,0020354 | 0,0020354 | TRUE | 0,0165293 | No |
| orf19.570 | 194 | 166 | 190 | 111 | 129 | 82 | 103 | 137 | 177 | 168 | 180 | 167 | 115 | 89 | 105 | 105 | 138,1 | 173 | 104 | 0,6 | -0,738 | -6,054 | 1,42E-09 | 2,64E-09 | 1,00E-08 | 0,011602 | 0,0067049 | 0,0067049 | TRUE | 0,0771735 | No |
| orf19.5700 | 57 | 49 | 59 | 33 | 148 | 110 | 148 | 164 | 52 | 50 | 56 | 50 | 132 | 119 | 152 | 125 | 91,83 | 52 | 132 | 2,54 | 1,345 | 8,505 | 1,82E-17 | 4,32E-17 | 1,00E-08 | 0,016423 | 0,0102436 | 0,0102436 | TRUE | 0,0981289 | No |
| orf19.5701 | 26 | 25 | 30 | 13 | 35 | 24 | 23 | 38 | 24 | 25 | 28 | 20 | 31 | 26 | 24 | 29 | 25,83 | 24 | 28 | 1,125 | 0,169 | 0,635 | 0,525367 | 0,567087 | 1,00E-08 | 0,053187 | 0,0297685 | 0,0297685 | TRUE | 0,0792842 | No |
| orf19.5702 | 1671 | 1657 | 1690 | 1033 | 1600 | 1379 | 1378 | 1919 | 1521 | 1676 | 1599 | 1552 | 1423 | 1497 | 1411 | 1464 | 1517,98 | 1587 | 1449 | 0,912 | -0,132 | -2,696 | 0,007023 | 0,009314 | 0,0006163 | 0,002907 | 0,0016496 | 0,0016496 | TRUE | 0,0173677 | No |
| orf19.5704 | 4242 | 3745 | 4335 | 2622 | 3074 | 2452 | 2661 | 3438 | 3862 | 3788 | 4103 | 3939 | 2734 | 2662 | 2725 | 2623 | 3304,42 | 3923 | 2686 | 0,685 | -0,547 | -13,527 | 1,08E-41 | 4,33E-41 | 0,0005644 | 0,002437 | 0,0012567 | 0,0012567 | TRUE | 0,0120804 | No |
| orf19.5705 | 938 | 889 | 960 | 583 | 1172 | 1110 | 1156 | 1581 | 854 | 899 | 909 | 876 | 1042 | 1205 | 1184 | 1206 | 1021,87 | 884 | 1159 | 1,31 | 0,39 | 6,282 | 3,34E-10 | 6,37E-10 | 0,0021281 | 0,00333 | 0,0026893 | 0,0026893 | TRUE | 0,0562207 | No |
| orf19.5710 | 5921 | 5328 | 5784 | 3667 | 3808 | 3289 | 3432 | 4762 | 5391 | 5389 | 5474 | 5508 | 3387 | 3571 | 3515 | 3632 | 4483,41 | 5440 | 3526 | 0,648 | -0,625 | -17,477 | 2,14E-68 | 1,32E-67 | 0,0001891 | 0,002332 | 0,0009966 | 0,0009966 | TRUE | 0,0088113 | No |
| orf19.5711 | 958 | 929 | 1002 | 663 | 4739 | 3964 | 3987 | 5657 | 872 | 940 | 948 | 996 | 4215 | 4304 | 4083 | 4315 | 2584,18 | 939 | 4229 | 4,519 | 2,176 | 44,682 | 0 | 0 | 0,0006813 | 0,002548 | 0,0015953 | 0,0015953 | TRUE | 0,0281036 | No |
| orf19.5713 | 6403 | 5970 | 6462 | 4157 | 5403 | 4475 | 4592 | 6164 | 5830 | 6038 | 6116 | 6244 | 4806 | 4859 | 4703 | 4702 | 5412,12 | 6057 | 4768 | 0,787 | -0,345 | -9,746 | 1,93E-22 | 5,17E-22 | 0,0002957 | 0,002281 | 0,0010146 | 0,0010146 | TRUE | 0,0078665 | No |
| orf19.5714 | 842 | 713 | 777 | 476 | 1013 | 924 | 885 | 1337 | 767 | 721 | 735 | 715 | 901 | 1003 | 906 | 1020 | 846,07 | 734 | 958 | 1,303 | 0,381 | 6,002 | 1,95E-09 | 3,61E-09 | 0,0017883 | 0,003598 | 0,0026602 | 0,0026602 | TRUE | 0,0246947 | No |
| orf19.5716 | 74 | 80 | 60 | 51 | 122 | 122 | 134 | 166 | 67 | 81 | 57 | 77 | 109 | 132 | 137 | 127 | 98,31 | 70 | 126 | 1,802 | 0,849 | 5,479 | 4,29E-08 | 7,51E-08 | 0,0048446 | 0,015474 | 0,0117539 | 0,0117539 | TRUE | 0,1635018 | No |
| orf19.5717 | 2 | 11 | 5 | 4 | 30 | 17 | 35 | 42 | 2 | 11 | 5 | 6 | 27 | 18 | 36 | 32 | 17,09 | 6 | 28 | 4,869 | 2,284 | 5,154 | 2,55E-07 | 4,34E-07 | 0,0864629 | 0,079339 | 0,081076 | 0,081076 | TRUE | 0,7156669 | No |
| orf19.5718 | 541 | 547 | 553 | 375 | 491 | 444 | 437 | 558 | 493 | 553 | 523 | 563 | 437 | 482 | 448 | 426 | 490,56 | 533 | 448 | 0,841 | -0,25 | -3,387 | 0,000707 | 0,001008 | 0,0012495 | 0,00473 | 0,0031696 | 0,0031696 | TRUE | 0,0316569 | No |
| orf19.5719 | 1905 | 1592 | 1728 | 1074 | 3235 | 2486 | 2603 | 3709 | 1734 | 1610 | 1635 | 1613 | 2877 | 2699 | 2666 | 2829 | 2208,11 | 1648 | 2768 | 1,679 | 0,747 | 15,956 | 2,59E-57 | 1,35E-56 | 0,0008806 | 0,002635 | 0,0016113 | 0,0016113 | TRUE | 0,0155296 | No |
| orf19.5720 | 4265 | 3693 | 3969 | 2277 | 3209 | 2439 | 2730 | 3570 | 3883 | 3735 | 3756 | 3420 | 2854 | 2648 | 2796 | 2723 | 3227,04 | 3698 | 2755 | 0,744 | -0,426 | -8,8 | 1,37E-18 | 3,35E-18 | 0,0015183 | 0,002446 | 0,0019389 | 0,0019389 | TRUE | 0,0296158 | No |
| orf19.5722 | 1100 | 872 | 938 | 611 | 2305 | 1823 | 2095 | 2806 | 1002 | 882 | 888 | 918 | 2050 | 1979 | 2146 | 2140 | 1500,56 | 922 | 2079 | 2,253 | 1,172 | 20,811 | 3,43E-96 | 2,84E-95 | 0,0014749 | 0,002917 | 0,0022398 | 0,0022398 | TRUE | 0,043097 | No |
| orf19.5723 | 3726 | 3173 | 3397 | 2376 | 34673 | 31097 | 30148 | 43630 | 3392 | 3209 | 3215 | 3569 | 30841 | 33763 | 30875 | 33281 | 17768,2 | 3346 | 32190 | 9,629 | 3,267 | 65,922 | 0 | 0 | 0,0022476 | 0,002111 | 0,0021835 | 0,0021835 | TRUE | 0,0231791 | No |
| orf19.5724 | 8 | 5 | 11 | 4 | 57 | 49 | 48 | 62 | 7 | 5 | 10 | 6 | 51 | 53 | 49 | 47 | 28,64 | 7 | 50 | 6,823 | 2,77 | 8,221 | 2,02E-16 | 4,63E-16 | 1,00E-08 | 0,048163 | 0,0277525 | 0,0277525 | TRUE | 0,2716088 | No |
| orf19.5725 | 379 | 391 | 477 | 294 | 497 | 392 | 443 | 581 | 345 | 395 | 451 | 442 | 442 | 426 | 454 | 443 | 424,77 | 408 | 441 | 1,084 | 0,117 | 1,318 | 0,187643 | 0,217812 | 0,0052025 | 0,005147 | 0,0051715 | 0,0051715 | TRUE | 0,1331389 | No |
| orf19.5727 | 1374 | 1123 | 1189 | 705 | 1097 | 764 | 899 | 1112 | 1251 | 1136 | 1125 | 1059 | 976 | 829 | 921 | 848 | 1018,15 | 1143 | 894 | 0,78 | -0,358 | -5,076 | 3,86E-07 | 6,51E-07 | 0,0039542 | 0,003334 | 0,0037707 | 0,0037707 | TRUE | 0,0492376 | No |
| orf19.5728 | 269 | 262 | 243 | 173 | 6816 | 5446 | 5756 | 7510 | 245 | 265 | 230 | 260 | 6063 | 5913 | 5895 | 5729 | 3074,85 | 250 | 5900 | 23,701 | 4,567 | 74,484 | 0 | 0 | 1,00E-08 | 0,002467 | 0,0014057 | 0,0014057 | TRUE | 0,0337691 | No |
| orf19.5729 | 418 | 383 | 404 | 277 | 718 | 646 | 625 | 810 | 381 | 387 | 382 | 416 | 639 | 701 | 640 | 618 | 520,54 | 392 | 650 | 1,662 | 0,733 | 10,297 | 7,27E-25 | 2,08E-24 | 1,00E-08 | 0,004575 | 0,0027724 | 0,0027724 | TRUE | 0,0327031 | No |
| orf19.5730 | 2136 | 1966 | 2092 | 1379 | 2010 | 1854 | 1912 | 2509 | 1945 | 1988 | 1980 | 2071 | 1788 | 2013 | 1958 | 1914 | 1957,16 | 1996 | 1918 | 0,961 | -0,057 | -1,161 | 0,245735 | 0,280394 | 0,0010897 | 0,002712 | 0,0017933 | 0,0017933 | TRUE | 0,0253257 | No |
| orf19.5731 | 2012 | 1847 | 1969 | 1175 | 2447 | 1831 | 2072 | 2796 | 1832 | 1868 | 1863 | 1765 | 2177 | 1988 | 2122 | 2133 | 1968,46 | 1832 | 2105 | 1,148 | 0,2 | 4,382 | 1,18E-05 | 1,86E-05 | 0,0005282 | 0,002708 | 0,0014762 | 0,0014762 | TRUE | 0,0163094 | No |
| orf19.5732 | 29322 | 26514 | 28239 | 16621 | 31489 | 24425 | 26508 | 35220 | 26696 | 26816 | 26726 | 24967 | 28008 | 26519 | 27148 | 26866 | 26718,3 | 26301 | 27135 | 1,032 | 0,045 | 1,243 | 0,213977 | 0,246628 | 0,0007987 | 0,002086 | 0,0012086 | 0,0012086 | TRUE | 0,0140881 | No |
| orf19.5734 | 3364 | 2896 | 3246 | 2088 | 1669 | 1363 | 1447 | 1920 | 3063 | 2929 | 3072 | 3137 | 1485 | 1480 | 1482 | 1465 | 2263,9 | 3050 | 1478 | 0,485 | -1,045 | -25,524 | 1,06E-143 | 1,33E-142 | 1,00E-08 | 0,002621 | 0,0011165 | 0,0011165 | TRUE | 0,008627 | No |
| orf19.5735 | 1926 | 1627 | 1746 | 982 | 2159 | 1667 | 1871 | 2494 | 1754 | 1646 | 1652 | 1475 | 1920 | 1810 | 1916 | 1902 | 1759,43 | 1632 | 1887 | 1,154 | 0,207 | 3,72 | 0,000199 | 0,000294 | 0,0021224 | 0,002788 | 0,0023836 | 0,0023836 | TRUE | 0,0460024 | No |
| orf19.5735.3 | 3054 | 2483 | 2673 | 1687 | 281 | 228 | 247 | 335 | 2781 | 2511 | 2530 | 2534 | 250 | 248 | 253 | 256 | 1420,21 | 2589 | 252 | 0,097 | -3,364 | -50,953 | 0 | 0 | 1,00E-08 | 0,002967 | 0,0021311 | 0,0021311 | TRUE | 0,0309915 | No |
| orf19.5736 | 4827 | 4138 | 4711 | 3143 | 4446 | 4025 | 4253 | 5795 | 4395 | 4185 | 4459 | 4721 | 3955 | 4370 | 4356 | 4420 | 4357,55 | 4440 | 4275 | 0,963 | -0,054 | -1,043 | 0,296889 | 0,334334 | 0,002203 | 0,00234 | 0,0023096 | 0,0023096 | TRUE | 0,0312758 | No |
| orf19.5741 | 51254 | 43385 | 48418 | 33794 | 88799 | 81230 | 86271 | 1E+05 | 46665 | 43879 | 45823 | 50764 | 78984 | 88193 | 88352 | 89776 | 66554,6 | 46783 | 86326 | 1,845 | 0,884 | 15,903 | 6,07E-57 | 3,15E-56 | 0,0035398 | 0,002057 | 0,0029517 | 0,0029517 | TRUE | 0,0401926 | No |
| orf19.5742 | 1742 | 1472 | 1669 | 1051 | 1482 | 1437 | 1430 | 1913 | 1586 | 1489 | 1580 | 1579 | 1318 | 1560 | 1465 | 1459 | 1504,41 | 1558 | 1450 | 0,93 | -0,104 | -1,809 | 0,070419 | 0,085934 | 0,0021935 | 0,002915 | 0,0025242 | 0,0025242 | TRUE | 0,0455938 | No |
| orf19.5746 | 8669 | 8205 | 8664 | 5168 | 10162 | 8326 | 8853 | 11834 | 7893 | 8298 | 8200 | 7763 | 9039 | 9040 | 9067 | 9027 | 8540,78 | 8038 | 9043 | 1,124 | 0,169 | 5,126 | 2,96E-07 | 5,02E-07 | 0,0003367 | 0,002192 | 0,0009275 | 0,0009275 | TRUE | 0,0061559 | No |
| orf19.5747 | 2620 | 2536 | 2663 | 1764 | 2574 | 2212 | 2224 | 2768 | 2385 | 2565 | 2520 | 2650 | 2289 | 2402 | 2278 | 2111 | 2400,08 | 2530 | 2270 | 0,897 | -0,156 | -2,974 | 0,002938 | 0,004001 | 0,0017826 | 0,002587 | 0,0022291 | 0,0022291 | TRUE | 0,0276156 | No |
| orf19.5748 | 17 | 18 | 20 | 12 | 50 | 21 | 22 | 29 | 15 | 18 | 19 | 18 | 44 | 23 | 23 | 22 | 22,82 | 18 | 28 | 1,593 | 0,672 | 2,057 | 0,039649 | 0,049571 | 0,0495704 | 0,059924 | 0,0554797 | 0,0554797 | TRUE | 1,0911438 | No |
| orf19.5749 | 4462 | 4053 | 4311 | 2718 | 1085 | 863 | 941 | 1255 | 4062 | 4099 | 4080 | 4083 | 965 | 937 | 964 | 957 | 2518,44 | 4081 | 956 | 0,234 | -2,094 | -52,754 | 0 | 0 | 1,00E-08 | 0,002562 | 0,0008937 | 0,0008937 | TRUE | 0,0018962 | No |
| orf19.575 | 9303 | 7467 | 8420 | 5451 | 14635 | 12406 | 13662 | 19476 | 8470 | 7552 | 7969 | 8188 | 13017 | 13469 | 13992 | 14856 | 10939,3 | 8045 | 13834 | 1,72 | 0,782 | 15,065 | 2,74E-51 | 1,29E-50 | 0,0026562 | 0,002158 | 0,0024891 | 0,0024891 | TRUE | 0,0304963 | No |
| orf19.5750 | 4209 | 3815 | 4109 | 2438 | 7815 | 6744 | 6732 | 9140 | 3832 | 3858 | 3889 | 3662 | 6951 | 7322 | 6894 | 6972 | 5422,68 | 3810 | 7035 | 1,844 | 0,883 | 23,687 | 4,96E-124 | 5,23E-123 | 0,0004658 | 0,002281 | 0,0011267 | 0,0011267 | TRUE | 0,0090558 | No |
| orf19.5751 | 4615 | 4492 | 4752 | 3051 | 1973 | 1812 | 1738 | 2327 | 4202 | 4543 | 4497 | 4583 | 1755 | 1967 | 1780 | 1775 | 3137,82 | 4456 | 1819 | 0,408 | -1,293 | -25,743 | 3,85E-146 | 4,93E-145 | 0,0017197 | 0,002458 | 0,0020445 | 0,0020445 | TRUE | 0,0351896 | No |
| orf19.5752 | 2129 | 2086 | 2203 | 1253 | 1755 | 1461 | 1538 | 1976 | 1938 | 2110 | 2085 | 1882 | 1561 | 1586 | 1575 | 1507 | 1780,62 | 2004 | 1557 | 0,776 | -0,366 | -7,225 | 5,01E-13 | 1,05E-12 | 0,0012087 | 0,002779 | 0,0019016 | 0,0019016 | TRUE | 0,0183193 | No |
| orf19.5753 | 5390 | 4940 | 5312 | 3376 | 30012 | 25770 | 25461 | 35691 | 4907 | 4996 | 5027 | 5071 | 26695 | 27979 | 26075 | 27225 | 15997,1 | 5000 | 26994 | 5,4 | 2,433 | 70,719 | 0 | 0 | 0,0003858 | 0,00212 | 0,0010115 | 0,0010115 | TRUE | 0,0073388 | No |
| orf19.5754 | 10340 | 10691 | 11027 | 6505 | 8522 | 7439 | 7596 | 9426 | 9414 | 10813 | 10436 | 9772 | 7580 | 8077 | 7779 | 7190 | 8882,59 | 10109 | 7656 | 0,757 | -0,401 | -7,417 | 1,20E-13 | 2,55E-13 | 0,0029821 | 0,002186 | 0,0026977 | 0,0026977 | TRUE | 0,0269345 | No |
| orf19.5755 | 3292 | 3064 | 3368 | 1981 | 1076 | 848 | 890 | 1142 | 2997 | 3099 | 3188 | 2976 | 957 | 921 | 911 | 871 | 1989,97 | 3065 | 915 | 0,298 | -1,746 | -35,955 | 4,19E-283 | 9,98E-282 | 0,00068 | 0,002701 | 0,0015781 | 0,0015781 | TRUE | 0,0138312 | No |
| orf19.5757 | 1916 | 1884 | 2030 | 1243 | 1600 | 1250 | 1406 | 1714 | 1744 | 1905 | 1921 | 1867 | 1423 | 1357 | 1440 | 1307 | 1620,75 | 1859 | 1382 | 0,743 | -0,429 | -8,113 | 4,93E-16 | 1,12E-15 | 0,00143 | 0,002852 | 0,0020535 | 0,0020535 | TRUE | 0,0221233 | No |
| orf19.5758 | 7575 | 7146 | 7461 | 4425 | 9442 | 7945 | 7982 | 10683 | 6897 | 7227 | 7061 | 6647 | 8398 | 8626 | 8175 | 8149 | 7647,55 | 6958 | 8337 | 1,197 | 0,26 | 6,723 | 1,78E-11 | 3,56E-11 | 0,0008138 | 0,00221 | 0,0013012 | 0,0013012 | TRUE | 0,0105679 | No |
| orf19.5759 | 4075 | 3883 | 4187 | 2456 | 6865 | 5975 | 6619 | 8470 | 3710 | 3927 | 3963 | 3689 | 6106 | 6487 | 6779 | 6461 | 5140,29 | 3822 | 6458 | 1,689 | 0,756 | 16,716 | 1,01E-62 | 5,71E-62 | 0,0013269 | 0,002294 | 0,0017495 | 0,0017495 | TRUE | 0,0164777 | No |
| orf19.576 | 270 | 223 | 240 | 165 | 213 | 171 | 188 | 234 | 246 | 226 | 227 | 248 | 189 | 186 | 193 | 178 | 211,56 | 237 | 186 | 0,789 | -0,342 | -3,523 | 0,000427 | 0,000617 | 1,00E-08 | 0,008281 | 0,0042793 | 0,0042793 | TRUE | 0,0104889 | No |
| orf19.5760 | 5180 | 3905 | 4443 | 3473 | 755 | 647 | 640 | 912 | 4716 | 3949 | 4205 | 5217 | 672 | 702 | 655 | 696 | 2601,59 | 4522 | 681 | 0,151 | -2,729 | -33,647 | 3,43E-248 | 7,05E-247 | 0,0082418 | 0,002545 | 0,0055146 | 0,0055146 | TRUE | 0,1153981 | No |
| orf19.5762 | 106 | 63 | 71 | 59 | 60 | 62 | 41 | 91 | 97 | 64 | 67 | 89 | 53 | 67 | 42 | 69 | 68,52 | 79 | 58 | 0,739 | -0,436 | -2,13 | 0,033172 | 0,041768 | 0,0316097 | 0,021317 | 0,0253709 | 0,0253709 | TRUE | 0,2477113 | No |
| orf19.5763 | 195 | 136 | 143 | 98 | 100 | 59 | 69 | 104 | 178 | 138 | 135 | 147 | 89 | 64 | 71 | 79 | 112,58 | 150 | 76 | 0,509 | -0,974 | -6,46 | 1,05E-10 | 2,03E-10 | 0,0096553 | 0,013771 | 0,012135 | 0,012135 | TRUE | 0,184527 | No |
| orf19.5764 | 1259 | 1007 | 1085 | 669 | 1189 | 933 | 1032 | 1447 | 1146 | 1018 | 1027 | 1005 | 1058 | 1013 | 1057 | 1104 | 1053,47 | 1049 | 1058 | 1,007 | 0,011 | 0,179 | 0,857913 | 0,879151 | 0,0016168 | 0,003291 | 0,0024171 | 0,0024171 | TRUE | 0,0482674 | No |
| orf19.5765 | 1644 | 1503 | 1666 | 1005 | 1087 | 862 | 971 | 1273 | 1497 | 1520 | 1577 | 1510 | 967 | 936 | 994 | 971 | 1246,44 | 1526 | 967 | 0,634 | -0,658 | -13,731 | 6,62E-43 | 2,71E-42 | 1,00E-08 | 0,003097 | 0,0013718 | 0,0013718 | TRUE | 0,0063738 | No |
| orf19.5767 | 730 | 677 | 703 | 410 | 726 | 502 | 620 | 802 | 665 | 685 | 665 | 616 | 646 | 545 | 635 | 612 | 633,51 | 658 | 610 | 0,926 | -0,112 | -1,597 | 0,110224 | 0,131761 | 0,001849 | 0,004122 | 0,0031016 | 0,0031016 | TRUE | 0,0556777 | No |
| orf19.5768 | 7483 | 6481 | 6947 | 4334 | 5748 | 4544 | 4992 | 6611 | 6813 | 6555 | 6575 | 6510 | 5113 | 4934 | 5112 | 5043 | 5831,8 | 6613 | 5050 | 0,763 | -0,389 | -11,86 | 1,91E-32 | 6,46E-32 | 0,0001832 | 0,002264 | 0,0008606 | 0,0008606 | TRUE | 0,0051435 | No |
| orf19.577 | 4089 | 3534 | 3854 | 2133 | 9296 | 7165 | 8289 | 11023 | 3723 | 3574 | 3647 | 3204 | 8268 | 7779 | 8489 | 8408 | 5886,72 | 3537 | 8236 | 2,326 | 1,218 | 23,263 | 1,05E-119 | 1,07E-118 | 0,0025936 | 0,00261 | 0,002422 | 0,002422 | TRUE | 0,0466623 | No |
| orf19.5770 | 7355 | 6628 | 7127 | 4140 | 9668 | 7858 | 8515 | 12026 | 6696 | 6704 | 6745 | 6219 | 8599 | 8532 | 8720 | 9174 | 7673,61 | 6591 | 8756 | 1,328 | 0,409 | 9,935 | 2,92E-23 | 8,01E-23 | 0,0010806 | 0,002209 | 0,0014933 | 0,0014933 | TRUE | 0,0169352 | No |
| orf19.5771 | 8586 | 7005 | 7871 | 5502 | 8579 | 7127 | 8026 | 10626 | 7817 | 7085 | 7449 | 8265 | 7631 | 7738 | 8220 | 8106 | 7788,75 | 7654 | 7924 | 1,036 | 0,051 | 0,974 | 0,32999 | 0,369651 | 0,0025822 | 0,002207 | 0,0024672 | 0,0024672 | TRUE | 0,0344814 | No |
| orf19.5772 | 1302 | 1125 | 1213 | 759 | 1353 | 1061 | 1279 | 1575 | 1185 | 1138 | 1148 | 1140 | 1203 | 1152 | 1310 | 1201 | 1184,76 | 1153 | 1216 | 1,055 | 0,077 | 1,427 | 0,153639 | 0,180511 | 0,0008084 | 0,003152 | 0,0019265 | 0,0019265 | TRUE | 0,0306597 | No |
| orf19.5773 | 1921 | 1831 | 1916 | 1247 | 2556 | 2183 | 2233 | 3052 | 1749 | 1852 | 1813 | 1873 | 2273 | 2370 | 2287 | 2328 | 2068,24 | 1822 | 2314 | 1,272 | 0,347 | 8,266 | 1,38E-16 | 3,20E-16 | 1,00E-08 | 0,002676 | 0,001196 | 0,001196 | TRUE | 0,0089881 | No |
| orf19.5775 | 47 | 57 | 50 | 22 | 87 | 92 | 86 | 97 | 43 | 58 | 47 | 33 | 77 | 100 | 88 | 74 | 65,02 | 45 | 85 | 1,84 | 0,88 | 4,566 | 4,98E-06 | 8,03E-06 | 0,0111007 | 0,022354 | 0,0185211 | 0,0185211 | TRUE | 0,2496219 | No |
| orf19.5776 | 4457 | 4037 | 4286 | 2571 | 3889 | 3376 | 3457 | 4575 | 4058 | 4083 | 4056 | 3862 | 3459 | 3665 | 3540 | 3490 | 3776,76 | 4015 | 3538 | 0,88 | -0,184 | -4,878 | 1,07E-06 | 1,78E-06 | 0,0003016 | 0,002387 | 0,0011007 | 0,0011007 | TRUE | 0,0073855 | No |
| orf19.5777 | 286 | 213 | 300 | 188 | 1225 | 1059 | 1015 | 1441 | 260 | 215 | 284 | 282 | 1090 | 1150 | 1039 | 1099 | 677,53 | 260 | 1094 | 4,215 | 2,076 | 25,647 | 4,61E-145 | 5,85E-144 | 0,0034408 | 0,003987 | 0,0038085 | 0,0038085 | TRUE | 0,1537936 | No |

| | | | | | | | | | | | | | | | | | | | | | | | | | | | | | | | |
|---|---|---|---|---|---|---|---|---|---|---|---|---|---|---|---|---|---|---|---|---|---|---|---|---|---|---|---|---|---|---|---|
| orf19.5779 | 4492 | 3859 | 4266 | 2823 | 4872 | 4125 | 4387 | 5980 | 4090 | 3903 | 4037 | 4241 | 4333 | 4479 | 4493 | 4562 | 4267,16 | 4068 | 4467 | 1,099 | 0,136 | 3,519 | 0,000434 | 0,000626 | 0,0005058 | 0,002347 | 0,0012025 | 0,0012025 | TRUE | 0,0092658 | No |
| orf19.578 | 3785 | 3431 | 3620 | 2075 | 3524 | 2790 | 3055 | 4079 | 3446 | 3470 | 3426 | 3117 | 3134 | 3029 | 3129 | 3111 | 3232,88 | 3365 | 3101 | 0,92 | -0,12 | -2,731 | 0,00632 | 0,008414 | 0,0009306 | 0,002446 | 0,0015361 | 0,0015361 | TRUE | 0,0278357 | No |
| orf19.5780 | 1378 | 1218 | 1336 | 896 | 1729 | 1450 | 1636 | 2140 | 1255 | 1232 | 1264 | 1346 | 1538 | 1574 | 1675 | 1632 | 1439,61 | 1274 | 1605 | 1,262 | 0,336 | 6,669 | 2,57E-11 | 5,10E-11 | 0,0005451 | 0,002955 | 0,0017238 | 0,0017238 | TRUE | 0,0151472 | No |
| orf19.5782 | 8804 | 7776 | 8659 | 4841 | 5718 | 4472 | 5178 | 6717 | 8016 | 7865 | 8195 | 7272 | 5086 | 4855 | 5303 | 5124 | 6464,4 | 7837 | 5092 | 0,649 | -0,623 | -13,24 | 5,13E-40 | 1,99E-39 | 0,0017396 | 0,002241 | 0,0019642 | 0,0019642 | TRUE | 0,0280023 | No |
| orf19.5783 | 1112 | 909 | 1003 | 652 | 1066 | 851 | 830 | 1235 | 1012 | 919 | 949 | 979 | 948 | 924 | 850 | 942 | 940,58 | 965 | 916 | 0,95 | -0,074 | -1,252 | 0,210533 | 0,243011 | 0,0010836 | 0,003441 | 0,0022505 | 0,0022505 | TRUE | 0,0274384 | No |
| orf19.5784 | 786 | 710 | 707 | 463 | 2310 | 2033 | 2027 | 2808 | 716 | 718 | 669 | 695 | 2055 | 2207 | 2076 | 2142 | 1409,77 | 700 | 2120 | 3,029 | 1,599 | 30,842 | 7,16E-209 | 1,26E-207 | 1,00E-08 | 0,002974 | 0,0015971 | 0,0015971 | TRUE | 0,0113525 | No |
| orf19.5785 | 1876 | 1758 | 1884 | 1175 | 645 | 623 | 584 | 791 | 1708 | 1778 | 1783 | 1765 | 574 | 676 | 598 | 603 | 1185,71 | 1758 | 613 | 0,348 | -1,523 | -26,585 | 1,01E-155 | 1,37E-154 | 0,0006485 | 0,003151 | 0,0020846 | 0,0020846 | TRUE | 0,0535062 | No |
| orf19.5788 | 41699 | 33997 | 37247 | 24690 | 54487 | 45462 | 48510 | 69464 | 37965 | 34384 | 35251 | 37088 | 48464 | 49359 | 49680 | 52988 | 43147,5 | 36172 | 50123 | 1,386 | 0,471 | 10,578 | 3,77E-26 | 1,11E-25 | 0,0017786 | 0,002068 | 0,0018781 | 0,0018781 | TRUE | 0,0181781 | No |
| orf19.5789 | 3918 | 3191 | 3447 | 2275 | 3417 | 2898 | 3101 | 4152 | 3567 | 3227 | 3262 | 3417 | 3039 | 3146 | 3176 | 3167 | 3250,37 | 3368 | 3132 | 0,93 | -0,105 | -2,375 | 0,017551 | 0,022626 | 0,0008656 | 0,002443 | 0,0015636 | 0,0015636 | TRUE | 0,0201748 | No |
| orf19.579 | 1371 | 1244 | 1277 | 834 | 1152 | 975 | 929 | 1358 | 1248 | 1258 | 1209 | 1253 | 1025 | 1059 | 951 | 1036 | 1129,79 | 1242 | 1018 | 0,82 | -0,286 | -5,522 | 3,36E-08 | 5,91E-08 | 1,11E-05 | 0,003206 | 0,0016913 | 0,0016913 | TRUE | 0,0219334 | No |
| orf19.5791 | 7603 | 6005 | 6648 | 4725 | 8720 | 7506 | 7474 | 11028 | 6922 | 6073 | 6292 | 7098 | 7756 | 8149 | 7654 | 8412 | 7294,65 | 6596 | 7993 | 1,212 | 0,278 | 4,858 | 1,18E-06 | 1,96E-06 | 0,0035559 | 0,002218 | 0,0030007 | 0,0030007 | TRUE | 0,0346534 | No |
| orf19.5792 | 477 | 457 | 497 | 335 | 409 | 390 | 374 | 521 | 434 | 462 | 470 | 503 | 364 | 423 | 383 | 397 | 429,72 | 467 | 392 | 0,841 | -0,251 | -3,221 | 0,001277 | 0,001792 | 0,0012606 | 0,005111 | 0,0034736 | 0,0034736 | TRUE | 0,0311413 | No |
| orf19.5793 | 9273 | 7706 | 8210 | 4965 | 8355 | 6545 | 7230 | 9805 | 8443 | 7794 | 7770 | 7458 | 7432 | 7106 | 7404 | 7479 | 7610,75 | 7866 | 7355 | 0,935 | -0,097 | -2,201 | 0,027736 | 0,035145 | 0,0014712 | 0,002211 | 0,00175 | 0,00175 | TRUE | 0,0298895 | No |
| orf19.5796 | 1222 | 1010 | 1139 | 654 | 1131 | 948 | 1041 | 1455 | 1113 | 1022 | 1078 | 982 | 1006 | 1029 | 1066 | 1110 | 1050,71 | 1049 | 1053 | 1,002 | 0,003 | 0,053 | 0,958019 | 0,962852 | 0,0014773 | 0,003294 | 0,0023571 | 0,0023571 | TRUE | 0,0200262 | No |
| orf19.5797 | 2083 | 1866 | 1854 | 1217 | 1946 | 1532 | 1554 | 2401 | 1896 | 1887 | 1755 | 1828 | 1731 | 1663 | 1591 | 1832 | 1772,97 | 1842 | 1704 | 0,926 | -0,11 | -2,013 | 0,044067 | 0,054857 | 0,0019524 | 0,002782 | 0,0023152 | 0,0023152 | TRUE | 0,0314244 | No |
| orf19.5798 | 2431 | 2190 | 2350 | 1408 | 1687 | 1366 | 1482 | 2022 | 2213 | 2215 | 2224 | 2115 | 1501 | 1483 | 1518 | 1542 | 1851,39 | 2192 | 1511 | 0,689 | -0,538 | -12,885 | 5,47E-38 | 2,06E-37 | 1,00E-08 | 0,002751 | 0,0011189 | 0,0011189 | TRUE | 0,0059722 | No |
| orf19.5799 | 2656 | 2254 | 2404 | 1746 | 3863 | 3173 | 3358 | 4710 | 2418 | 2280 | 2275 | 2623 | 3436 | 3445 | 3439 | 3593 | 2938,58 | 2399 | 3478 | 1,453 | 0,539 | 10,564 | 4,38E-26 | 1,28E-25 | 0,0020895 | 0,002487 | 0,0021398 | 0,0021398 | TRUE | 0,0446697 | No |
| orf19.58 | 14747 | 13186 | 14392 | 9359 | 15006 | 13100 | 13192 | 18016 | 13427 | 13336 | 13621 | 14059 | 13347 | 14223 | 13510 | 13743 | 13658,2 | 13611 | 13706 | 1,007 | 0,01 | 0,301 | 0,763117 | 0,793746 | 0,0005105 | 0,002134 | 0,001058 | 0,001058 | TRUE | 0,0078055 | No |
| orf19.580 | 1155 | 1018 | 1056 | 607 | 652 | 515 | 530 | 728 | 1052 | 1030 | 999 | 912 | 580 | 559 | 543 | 555 | 778,7 | 998 | 559 | 0,559 | -0,84 | -13,37 | 9,03E-41 | 3,56E-40 | 0,000985 | 0,003733 | 0,0024175 | 0,0024175 | TRUE | 0,0355067 | No |
| orf19.5801 | 10033 | 8863 | 9426 | 5789 | 6112 | 4926 | 5073 | 7027 | 9135 | 8964 | 8921 | 8696 | 5436 | 5348 | 5195 | 5360 | 7131,97 | 8929 | 5335 | 0,597 | -0,743 | -22,877 | 7,92E-116 | 7,84E-115 | 0,0001841 | 0,002222 | 0,0008658 | 0,0008658 | TRUE | 0,00374 | No |
| orf19.5802 | 1579 | 1217 | 1411 | 841 | 2026 | 1493 | 1921 | 2476 | 1438 | 1231 | 1335 | 1263 | 1802 | 1621 | 1967 | 1889 | 1568,29 | 1317 | 1820 | 1,381 | 0,466 | 6,622 | 3,55E-11 | 7,01E-11 | 0,0048882 | 0,002879 | 0,0040849 | 0,0040849 | TRUE | 0,0639361 | No |
| orf19.5804 | 1701 | 1502 | 1696 | 1081 | 1297 | 1051 | 1110 | 1597 | 1549 | 1519 | 1605 | 1624 | 1154 | 1141 | 1137 | 1218 | 1368,31 | 1574 | 1162 | 0,74 | -0,434 | -8,88 | 6,67E-19 | 1,64E-18 | 0,0003447 | 0,003002 | 0,0015538 | 0,0015538 | TRUE | 0,0133288 | No |
| orf19.5805 | 773 | 610 | 726 | 475 | 1900 | 1605 | 1898 | 2468 | 704 | 617 | 687 | 714 | 1690 | 1743 | 1944 | 1883 | 1247,54 | 680 | 1815 | 2,671 | 1,418 | 21,292 | 1,35E-100 | 1,18E-99 | 0,0033157 | 0,003096 | 0,0032084 | 0,0032084 | TRUE | 0,0468667 | No |
| orf19.5806 | 60096 | 48214 | 53462 | 36525 | 2E+05 | 1E+05 | 1E+05 | 2E+05 | 54715 | 48763 | 50597 | 54866 | 1E+05 | 1E+05 | 1E+05 | 1E+05 | 97388,6 | 52235 | 142542 | 2,729 | 1,448 | 30,163 | 7,27E-200 | 1,24E-198 | 0,0022959 | 0,002051 | 0,0022018 | 0,0022018 | TRUE | 0,0245296 | No |
| orf19.5808 | 3724 | 3232 | 3539 | 2332 | 3310 | 2693 | 2897 | 4025 | 3391 | 3269 | 3349 | 3503 | 2944 | 2924 | 2967 | 3070 | 3177,12 | 3378 | 2976 | 0,882 | -0,181 | -4,651 | 3,31E-06 | 5,37E-06 | 0,0002345 | 0,002453 | 0,0011329 | 0,0011329 | TRUE | 0,0069161 | No |
| orf19.5809 | 66 | 55 | 67 | 57 | 158 | 144 | 137 | 198 | 60 | 56 | 63 | 86 | 141 | 156 | 140 | 151 | 106,62 | 66 | 147 | 2,265 | 1,18 | 7,928 | 2,23E-15 | 4,99E-15 | 1,00E-08 | 0,014427 | 0,0099006 | 0,0099006 | TRUE | 0,2499609 | No |
| orf19.581 | 14233 | 12148 | 13839 | 9161 | 15676 | 13870 | 14762 | 20071 | 12959 | 12286 | 13097 | 13761 | 13943 | 15059 | 15118 | 15310 | 13941,8 | 13026 | 14858 | 1,141 | 0,19 | 4,118 | 3,83E-05 | 5,88E-05 | 0,0018495 | 0,002132 | 0,0019764 | 0,0019764 | TRUE | 0,0210626 | No |
| orf19.5811 | 6038 | 4433 | 5083 | 3639 | 4268 | 3636 | 4165 | 5972 | 5497 | 4484 | 4811 | 5466 | 3796 | 3948 | 4265 | 4555 | 4602,84 | 5064 | 4141 | 0,818 | -0,29 | -3,808 | 0,00014 | 0,000208 | 0,007925 | 0,002324 | 0,0053372 | 0,0053372 | TRUE | 0,072751 | No |
| orf19.5812 | 5939 | 5284 | 5749 | 3719 | 4670 | 3861 | 4160 | 5402 | 5407 | 5344 | 5441 | 5587 | 4154 | 4192 | 4260 | 4121 | 4813,21 | 5445 | 4182 | 0,768 | -0,38 | -11,602 | 4,02E-31 | 1,33E-30 | 1,00E-08 | 0,002311 | 0,0008211 | 0,0008211 | TRUE | 0,0034758 | No |
| orf19.5813 | 15116 | 13131 | 14448 | 9098 | 9197 | 7017 | 7528 | 10755 | 13762 | 13281 | 13674 | 13667 | 8180 | 7619 | 7710 | 8204 | 10762 | 13596 | 7928 | 0,583 | -0,778 | -20,958 | 1,58E-97 | 1,34E-96 | 0,0007429 | 0,00216 | 0,0012234 | 0,0012234 | TRUE | 0,0084066 | No |
| orf19.5814 | 963 | 863 | 872 | 573 | 1008 | 824 | 798 | 999 | 877 | 873 | 825 | 861 | 897 | 895 | 817 | 762 | 850,77 | 859 | 843 | 0,98 | -0,03 | -0,446 | 0,655366 | 0,692288 | 0,0025249 | 0,00359 | 0,003041 | 0,003041 | TRUE | 0,052635 | No |
| orf19.5814.1 | 574 | 493 | 535 | 310 | 1092 | 776 | 899 | 1162 | 523 | 499 | 506 | 466 | 971 | 843 | 921 | 886 | 701,76 | 498 | 905 | 1,811 | 0,857 | 12,692 | 6,54E-37 | 2,43E-36 | 0,0016048 | 0,003919 | 0,0027932 | 0,0027932 | TRUE | 0,0290462 | No |
| orf19.5815 | 1175 | 1092 | 1127 | 722 | 2091 | 1734 | 1747 | 2465 | 1070 | 1104 | 1067 | 1085 | 1860 | 1883 | 1789 | 1880 | 1467,17 | 1082 | 1853 | 1,715 | 0,778 | 17,088 | 1,83E-65 | 1,08E-64 | 1,00E-08 | 0,002937 | 0,0012436 | 0,0012436 | TRUE | 0,0063182 | No |
| orf19.5816 | 1102 | 961 | 1098 | 715 | 1803 | 1479 | 1586 | 2144 | 1003 | 972 | 1039 | 1074 | 1604 | 1606 | 1624 | 1635 | 1319,71 | 1022 | 1617 | 1,586 | 0,666 | 13,77 | 3,87E-43 | 1,59E-42 | 1,00E-08 | 0,003038 | 0,0014297 | 0,0014297 | TRUE | 0,0124459 | No |
| orf19.5817 | 989 | 950 | 988 | 593 | 1033 | 851 | 869 | 1235 | 900 | 961 | 935 | 891 | 919 | 924 | 890 | 942 | 920,24 | 922 | 919 | 0,996 | -0,006 | -0,105 | 0,916063 | 0,927999 | 1,00E-08 | 0,003473 | 0,0016685 | 0,0016685 | TRUE | 0,0094923 | No |
| orf19.5818 | 49848 | 43607 | 47803 | 28523 | 22164 | 17971 | 18980 | 25778 | 45384 | 44104 | 45241 | 42846 | 19714 | 19511 | 19438 | 19664 | 31987,9 | 44394 | 19582 | 0,441 | -1,181 | -39,56 | 0 | 0 | 0,000326 | 0,002078 | 0,0008201 | 0,0008201 | TRUE | 0,0066766 | No |
| orf19.5820 | 21 | 8 | 25 | 16 | 37 | 34 | 29 | 40 | 19 | 8 | 24 | 24 | 33 | 37 | 30 | 31 | 25,62 | 19 | 33 | 1,754 | 0,811 | 2,671 | 0,007574 | 0,010021 | 0,0187866 | 0,053603 | 0,0452919 | 0,0452919 | TRUE | 0,7798715 | No |
| orf19.5821 | 1290 | 1225 | 1298 | 773 | 763 | 626 | 629 | 904 | 1174 | 1239 | 1228 | 1161 | 679 | 680 | 644 | 690 | 936,89 | 1200 | 673 | 0,56 | -0,836 | -15,355 | 3,29E-53 | 1,62E-52 | 1,00E-08 | 0,003447 | 0,0017055 | 0,0017055 | TRUE | 0,0095209 | No |
| orf19.5823 | 5255 | 4252 | 4555 | 3078 | 5549 | 4708 | 4832 | 6844 | 4784 | 4300 | 4311 | 4624 | 4936 | 5112 | 4949 | 5221 | 4779,49 | 4505 | 5054 | 1,122 | 0,166 | 3,591 | 0,00033 | 0,00048 | 0,0014969 | 0,002313 | 0,0018522 | 0,0018522 | TRUE | 0,022155 | No |
| orf19.5824 | 7128 | 6190 | 6573 | 4555 | 7228 | 6066 | 6109 | 8379 | 6490 | 6261 | 6221 | 6842 | 6429 | 6586 | 6256 | 6392 | 6434,55 | 6454 | 6416 | 0,995 | -0,007 | -0,183 | 0,854666 | 0,876105 | 0,0008836 | 0,002242 | 0,0014289 | 0,0014289 | TRUE | 0,0192962 | No |
| orf19.5825 | 2571 | 2102 | 2238 | 1429 | 1630 | 1332 | 1499 | 1920 | 2341 | 2126 | 2118 | 2147 | 1450 | 1446 | 1535 | 1465 | 1828,39 | 2183 | 1474 | 0,675 | -0,568 | -11,397 | 4,35E-30 | 1,40E-29 | 0,0011327 | 0,00276 | 0,0018195 | 0,0018195 | TRUE | 0,0299134 | No |
| orf19.5825.1 | 899 | 732 | 737 | 509 | 761 | 502 | 595 | 802 | 819 | 740 | 698 | 765 | 677 | 545 | 609 | 612 | 683 | 756 | 611 | 0,81 | -0,304 | -3,918 | 8,92E-05 | 0,000134 | 0,0046597 | 0,003971 | 0,0043049 | 0,0043049 | TRUE | 0,0623097 | No |
| orf19.5826 | 16 | 16 | 17 | 10 | 29 | 26 | 37 | 51 | 15 | 16 | 16 | 15 | 26 | 28 | 38 | 39 | 24,08 | 16 | 33 | 2,12 | 1,084 | 3,656 | 0,000256 | 0,000376 | 1,00E-08 | 0,056885 | 0,0359569 | 0,0359569 | TRUE | 0,135831 | No |
| orf19.5827 | 153 | 154 | 159 | 126 | 66 | 62 | 58 | 82 | 139 | 156 | 150 | 189 | 59 | 67 | 59 | 63 | 110,35 | 158 | 62 | 0,394 | -1,345 | -9,107 | 8,49E-20 | 2,14E-19 | 1,00E-08 | 0,014008 | 0,0100178 | 0,0100178 | TRUE | 0,1466972 | No |
| orf19.5828 | 8705 | 7485 | 8333 | 5177 | 8535 | 6892 | 7898 | 10289 | 7926 | 7570 | 7886 | 7777 | 7592 | 7483 | 8089 | 7849 | 7771,3 | 7790 | 7753 | 0,995 | -0,007 | -0,185 | 0,853545 | 0,875097 | 0,0005779 | 0,002207 | 0,0012009 | 0,0012009 | TRUE | 0,0102078 | No |
| orf19.583 | 12120 | 10405 | 11307 | 7465 | 10404 | 8456 | 9496 | 12799 | 11035 | 10524 | 10701 | 11214 | 9254 | 9181 | 9725 | 9763 | 10174,5 | 10868 | 9481 | 0,873 | -0,197 | -5,192 | 2,08E-07 | 3,54E-07 | 0,0008035 | 0,002167 | 0,0012766 | 0,0012766 | TRUE | 0,0055398 | No |
| orf19.5830 | 1376 | 1189 | 1405 | 833 | 2548 | 2032 | 2290 | 2977 | 1253 | 1203 | 1330 | 1251 | 2266 | 2206 | 2345 | 2271 | 1765,63 | 1259 | 2272 | 1,804 | 0,851 | 17,963 | 3,79E-72 | 2,43E-71 | 0,0003621 | 0,002785 | 0,0015241 | 0,0015241 | TRUE | 0,0181241 | No |
| orf19.5831 | 37 | 41 | 42 | 27 | 51 | 33 | 52 | 60 | 34 | 41 | 40 | 41 | 45 | 36 | 53 | 46 | 41,96 | 39 | 45 | 1,167 | 0,222 | 1,034 | 0,300981 | 0,338823 | 1,00E-08 | 0,03352 | 0,0203543 | 0,0203543 | TRUE | 0,1298926 | No |
| orf19.5832 | 10298 | 8643 | 9182 | 5574 | 11647 | 8618 | 9423 | 12896 | 9376 | 8741 | 8690 | 8373 | 10360 | 9357 | 9650 | 9837 | 9298,03 | 8795 | 9801 | 1,114 | 0,156 | 3,311 | 0,000929 | 0,001314 | 0,0019224 | 0,002179 | 0,0020194 | 0,0020194 | TRUE | 0,024249 | No |
| orf19.5833 | 5304 | 4432 | 4877 | 3085 | 4291 | 3392 | 3740 | 5075 | 4829 | 4482 | 4616 | 4634 | 3817 | 3683 | 3830 | 3871 | 4220,29 | 4640 | 3800 | 0,819 | -0,288 | -7,542 | 4,62E-14 | 9,93E-14 | 0,0004995 | 0,00235 | 0,0011599 | 0,0011599 | TRUE | 0,0094867 | No |
| orf19.5834 | 6435 | 5757 | 6201 | 4021 | 7529 | 6476 | 7102 | 9177 | 5859 | 5823 | 5869 | 6040 | 6697 | 7031 | 7273 | 7000 | 6448,98 | 5898 | 7000 | 1,187 | 0,248 | 6,764 | 1,34E-11 | 2,68E-11 | 0,0004974 | 0,002242 | 0,0011288 | 0,0011288 | TRUE | 0,0104332 | No |
| orf19.5835 | 8778 | 7343 | 8215 | 5251 | 7627 | 6224 | 6596 | 8730 | 7992 | 7427 | 7775 | 7888 | 6784 | 6758 | 6755 | 6659 | 7254,65 | 7770 | 6739 | 0,867 | -0,205 | -5,971 | 2,36E-09 | 4,35E-09 | 0,0003481 | 0,002219 | 0,0009974 | 0,0009974 | TRUE | 0,0109307 | No |
| orf19.5837 | 1129 | 936 | 1090 | 643 | 1227 | 909 | 1047 | 1395 | 1028 | 947 | 1032 | 966 | 1091 | 987 | 1072 | 1064 | 1023,34 | 993 | 1054 | 1,06 | 0,085 | 1,496 | 0,134761 | 0,159306 | 0,000888 | 0,003328 | 0,0020966 | 0,0020966 | TRUE | 0,0206542 | No |
| orf19.5838 | 924 | 791 | 839 | 547 | 997 | 784 | 869 | 1141 | 841 | 800 | 794 | 822 | 887 | 851 | 890 | 870 | 844,42 | 814 | 874 | 1,075 | 0,104 | 1,922 | 0,05459 | 0,067431 | 1,00E-08 | 0,003601 | 0,0016072 | 0,0016072 | TRUE | 0,0067846 | No |
| orf19.5839 | 634 | 493 | 565 | 393 | 1775 | 1422 | 1669 | 2027 | 577 | 499 | 535 | 590 | 1579 | 1544 | 1709 | 1546 | 1072,39 | 550 | 1594 | 2,905 | 1,538 | 23,532 | 1,94E-122 | 2,02E-121 | 0,0024568 | 0,003269 | 0,0028353 | 0,0028353 | TRUE | 0,0461404 | No |
| orf19.584 | 7 | 10 | 11 | 7 | 5 | 4 | 5 | 9 | 6 | 10 | 10 | 11 | 4 | 4 | 5 | 7 | 7,27 | 9 | 5 | 0,568 | -0,815 | -1,603 | 0,108977 | 0,130321 | 1,00E-08 | 0,183652 | 0,1025301 | 0,1025301 | TRUE | 0,179408 | No |
| orf19.5840 | 47 | 42 | 36 | 44 | 43 | 30 | 23 | 46 | 43 | 42 | 34 | 66 | 38 | 33 | 24 | 35 | 39,36 | 46 | 32 | 0,716 | -0,481 | -1,918 | 0,05505 | 0,067973 | 0,0318931 | 0,035597 | 0,0342603 | 0,0342603 | TRUE | 0,4937611 | No |
| orf19.5841 | 534 | 435 | 473 | 332 | 734 | 578 | 580 | 850 | 486 | 440 | 448 | 499 | 653 | 628 | 594 | 648 | 549,41 | 468 | 631 | 1,353 | 0,436 | 6,255 | 3,97E-10 | 7,55E-10 | 0,0006167 | 0,004441 | 0,0027768 | 0,0027768 | TRUE | 0,0183824 | No |
| orf19.5842 | 655 | 612 | 682 | 409 | 632 | 503 | 597 | 746 | 596 | 619 | 645 | 614 | 562 | 546 | 611 | 569 | 595,48 | 618 | 572 | 0,924 | -0,113 | -1,753 | 0,079687 | 0,096671 | 1,00E-08 | 0,004255 | 0,0023467 | 0,0023467 | TRUE | 0,0235333 | No |
| orf19.5843 | 2802 | 2790 | 2926 | 1856 | 1610 | 1407 | 1470 | 1970 | 2551 | 2822 | 2769 | 2788 | 1432 | 1528 | 1505 | 1503 | 2112,24 | 2732 | 1492 | 0,546 | -0,873 | -18,022 | 1,31E-72 | 8,45E-72 | 0,0010948 | 0,002662 | 0,0017403 | 0,0017403 | TRUE | 0,0250694 | No |
| orf19.5844 | 6 | 3 | 4 | 4 | 5 | 8 | 6 | 6 | 5 | 3 | 4 | 6 | 4 | 9 | 6 | 5 | 5,27 | 4 | 6 | 1,304 | 0,383 | 0,641 | 0,521512 | 0,563307 | 1,00E-08 | 0,252784 | 0,1452665 | 0,1452665 | TRUE | 0,2000298 | No |
| orf19.5845 | 4843 | 4288 | 4752 | 2993 | 5250 | 4220 | 4487 | 6017 | 4409 | 4337 | 4497 | 4496 | 4670 | 4582 | 4595 | 4590 | 4522 | 4435 | 4609 | 1,04 | 0,056 | 1,746 | 0,080726 | 0,0978 | 1,00E-08 | 0,002329 | 0,0007699 | 0,0007699 | TRUE | 0,0026893 | No |
| orf19.5846 | 3361 | 2888 | 3216 | 2093 | 1342 | 1135 | 1134 | 1576 | 3060 | 2921 | 3044 | 3144 | 1194 | 1232 | 1161 | 1202 | 2119,77 | 3042 | 1197 | 0,394 | -1,344 | -30,57 | 3,03E-205 | 5,31E-204 | 1,00E-08 | 0,00266 | 0,00129 | 0,00129 | TRUE | 0,0086477 | No |
| orf19.5847 | 10411 | 8906 | 9811 | 6183 | 10594 | 9207 | 9485 | 13180 | 9479 | 9007 | 9285 | 9288 | 9423 | 9996 | 9714 | 10054 | 9530,78 | 9265 | 9797 | 1,057 | 0,081 | 2,286 | 0,022251 | 0,028409 | 0,0005176 | 0,002176 | 0,0010852 | 0,0010852 | TRUE | 0,0081285 | No |
| orf19.5848 | 6486 | 5569 | 5956 | 4021 | 825 | 735 | 773 | 1036 | 5905 | 5632 | 5637 | 6040 | 734 | 798 | 792 | 790 | 3291,05 | 5804 | 778 | 0,134 | -2,898 | -60,145 | 0 | 0 | 0,0008561 | 0,002438 | 0,0015382 | 0,0015382 | TRUE | 0,0180356 | No |
| orf19.5849 | 3485 | 2757 | 3144 | 2030 | 2346 | 1983 | 2144 | 2918 | 3173 | 2788 | 2976 | 3049 | 2087 | 2153 | 2196 | 2226 | 2580,94 | 2996 | 2166 | 0,723 | -0,468 | -9,457 | 3,18E-21 | 8,30E-21 | 0,0015552 | 0,002549 | 0,0019599 | 0,0019599 | TRUE | 0,0269903 | No |
| orf19.585 | 3872 | 3501 | 3660 | 2382 | 3140 | 2554 | 2621 | 3501 | 3525 | 3541 | 3464 | 3578 | 2793 | 2773 | 2684 | 2671 | 3128,61 | 3527 | 2730 | 0,774 | -0,369 | -10,149 | 3,34E-24 | 9,40E-24 | 1,00E-08 | 0,002459 | 0,0009469 | 0,0009469 | TRUE | 0,0030474 | No |

| | | | | | | | | | | | | | | | | | | | | | | | | | | | | | | | |
|---|---|---|---|---|---|---|---|---|---|---|---|---|---|---|---|---|---|---|---|---|---|---|---|---|---|---|---|---|---|---|---|
| orf19.5850 | 16022 | 14594 | 16116 | 10022 | 22219 | 18929 | 19608 | 26089 | 14587 | 14760 | 15252 | 15055 | 19763 | 20552 | 20081 | 19901 | 17493,9 | 14914 | 20074 | 1,346 | 0,429 | 14,371 | 7,84E-47 | 3,43E-46 | 0,0002671 | 0,002113 | 0,0007953 | 0,0007953 | TRUE | 0,0031168 | No |
| orf19.5851 | 771 | 703 | 747 | 513 | 1574 | 1281 | 1322 | 1874 | 702 | 711 | 707 | 771 | 1400 | 1391 | 1354 | 1430 | 1058,1 | 723 | 1394 | 1,937 | 0,954 | 17,981 | 2,73E-72 | 1,76E-71 | 1,00E-08 | 0,003285 | 0,0016239 | 0,0016239 | TRUE | 0,0196897 | No |
| orf19.5852 | 259 | 281 | 271 | 168 | 417 | 373 | 414 | 526 | 236 | 284 | 256 | 252 | 371 | 405 | 424 | 401 | 328,74 | 257 | 400 | 1,555 | 0,637 | 7,275 | 3,47E-13 | 7,33E-13 | 0,0013669 | 0,006055 | 0,0041203 | 0,0041203 | TRUE | 0,0585443 | No |
| orf19.5854 | 4547 | 4216 | 4487 | 2757 | 4756 | 3785 | 4007 | 5324 | 4140 | 4264 | 4247 | 4141 | 4230 | 4109 | 4104 | 4061 | 4162,06 | 4198 | 4126 | 0,983 | -0,025 | -0,75 | 0,452977 | 0,494854 | 1,00E-08 | 0,002354 | 0,0008552 | 0,0008552 | TRUE | 0,0035941 | No |
| orf19.5854.1 | 126 | 122 | 134 | 90 | 128 | 93 | 112 | 125 | 115 | 123 | 127 | 135 | 114 | 101 | 115 | 95 | 115,62 | 125 | 106 | 0,852 | -0,231 | -1,751 | 0,0799 | 0,096874 | 1,00E-08 | 0,013462 | 0,0080437 | 0,0080437 | TRUE | 0,0561672 | No |
| orf19.5855 | 4107 | 3523 | 4013 | 2631 | 2413 | 2093 | 2308 | 3223 | 3739 | 3563 | 3798 | 3952 | 2146 | 2272 | 2364 | 2459 | 3036,68 | 3763 | 2310 | 0,614 | -0,703 | -13,306 | 2,12E-40 | 8,29E-40 | 0,0021066 | 0,002472 | 0,0023311 | 0,0023311 | TRUE | 0,0283164 | No |
| orf19.5856 | 2142 | 2040 | 2143 | 1280 | 2056 | 1745 | 1800 | 2400 | 1950 | 2063 | 2028 | 1923 | 1829 | 1895 | 1843 | 1831 | 1920,23 | 1991 | 1850 | 0,927 | -0,109 | -2,532 | 0,011329 | 0,0148 | 1,00E-08 | 0,002725 | 0,0012439 | 0,0012439 | TRUE | 0,0070108 | No |
| orf19.5857 | 1231 | 1070 | 1207 | 761 | 916 | 737 | 821 | 993 | 1121 | 1082 | 1142 | 1143 | 815 | 800 | 841 | 757 | 962,7 | 1122 | 803 | 0,715 | -0,483 | -8,797 | 1,40E-18 | 3,43E-18 | 1,00E-08 | 0,003409 | 0,0018392 | 0,0018392 | TRUE | 0,0191156 | No |
| orf19.5858 | 2808 | 2122 | 2282 | 1372 | 3068 | 2026 | 2330 | 3306 | 2557 | 2146 | 2160 | 2061 | 2729 | 2200 | 2386 | 2522 | 2345 | 2231 | 2459 | 1,102 | 0,14 | 1,77 | 0,076757 | 0,093293 | 0,0084749 | 0,0026 | 0,0055727 | 0,0055727 | TRUE | 0,1181494 | No |
| orf19.5859 | 759 | 589 | 695 | 430 | 845 | 763 | 748 | 1153 | 691 | 596 | 658 | 646 | 752 | 828 | 766 | 880 | 727 | 648 | 806 | 1,245 | 0,317 | 4,356 | 1,33E-05 | 2,09E-05 | 0,0034831 | 0,003854 | 0,0036669 | 0,0036669 | TRUE | 0,0471933 | No |
| orf19.586 | 1429 | 1282 | 1308 | 772 | 1974 | 1571 | 1687 | 2296 | 1301 | 1297 | 1238 | 1160 | 1756 | 1706 | 1728 | 1751 | 1491,98 | 1249 | 1735 | 1,386 | 0,471 | 9,599 | 8,10E-22 | 2,14E-21 | 0,0002385 | 0,002922 | 0,0016134 | 0,0016134 | TRUE | 0,024001 | No |
| orf19.5860 | 134 | 128 | 128 | 77 | 113 | 95 | 107 | 168 | 122 | 129 | 121 | 116 | 101 | 103 | 110 | 128 | 116,21 | 122 | 110 | 0,906 | -0,142 | -1,069 | 0,285142 | 0,321731 | 1,00E-08 | 0,013405 | 0,008475 | 0,008475 | TRUE | 0,1353175 | No |
| orf19.5861 | 4917 | 4308 | 4513 | 3276 | 6644 | 6031 | 5689 | 8456 | 4477 | 4357 | 4271 | 4921 | 5910 | 6548 | 5826 | 6450 | 5345,02 | 4506 | 6184 | 1,373 | 0,458 | 7,95 | 1,87E-15 | 4,19E-15 | 0,0035724 | 0,002284 | 0,0029878 | 0,0029878 | TRUE | 0,0453954 | No |
| orf19.5861.1 | 451 | 397 | 460 | 261 | 796 | 603 | 615 | 789 | 411 | 402 | 435 | 392 | 708 | 655 | 630 | 602 | 529,24 | 410 | 649 | 1,578 | 0,658 | 8,776 | 1,69E-18 | 4,13E-18 | 0,0021649 | 0,004533 | 0,0033728 | 0,0033728 | TRUE | 0,0468965 | No |
| orf19.5862 | 744 | 635 | 710 | 470 | 1229 | 1097 | 1187 | 1571 | 677 | 642 | 672 | 706 | 1093 | 1191 | 1216 | 1198 | 924,47 | 674 | 1174 | 1,744 | 0,803 | 13,569 | 6,09E-42 | 2,45E-41 | 0,0009364 | 0,003466 | 0,0021683 | 0,0021683 | TRUE | 0,0265424 | No |
| orf19.5863 | 71 | 79 | 73 | 32 | 128 | 115 | 114 | 159 | 65 | 80 | 69 | 48 | 114 | 125 | 117 | 121 | 92,31 | 66 | 119 | 1,793 | 0,843 | 5,445 | 5,17E-08 | 9,03E-08 | 1,00E-08 | 0,016348 | 0,0110787 | 0,0110787 | TRUE | 0,1923325 | No |
| orf19.5864 | 5885 | 5804 | 6192 | 3443 | 3875 | 3110 | 3438 | 4331 | 5358 | 5870 | 5860 | 5172 | 3447 | 3377 | 3521 | 3304 | 4488,53 | 5565 | 3412 | 0,613 | -0,707 | -13,876 | 8,83E-44 | 3,69E-43 | 0,0020896 | 0,002331 | 0,0022587 | 0,0022587 | TRUE | 0,0266382 | No |
| orf19.5865 | 2181 | 2142 | 2415 | 1499 | 2486 | 2215 | 2253 | 2854 | 1986 | 2166 | 2286 | 2252 | 2211 | 2405 | 2307 | 2177 | 2223,74 | 2172 | 2275 | 1,047 | 0,067 | 1,191 | 0,233785 | 0,267668 | 0,0023237 | 0,002631 | 0,0025791 | 0,0025791 | TRUE | 0,0416643 | No |
| orf19.5866 | 27 | 24 | 20 | 14 | 17 | 13 | 12 | 9 | 25 | 24 | 19 | 21 | 15 | 14 | 12 | 7 | 17,15 | 22 | 12 | 0,535 | -0,902 | -2,603 | 0,009228 | 0,012137 | 1,00E-08 | 0,079061 | 0,0523395 | 0,0523395 | TRUE | 0,4664691 | No |
| orf19.5867 | 4279 | 3874 | 4088 | 2645 | 2639 | 2344 | 2444 | 3296 | 3896 | 3918 | 3869 | 3973 | 2347 | 2545 | 2503 | 2514 | 3195,69 | 3914 | 2477 | 0,633 | -0,66 | -16,774 | 3,76E-63 | 2,15E-62 | 0,0002626 | 0,00245 | 0,001159 | 0,001159 | TRUE | 0,0153228 | No |
| orf19.5869 | 4730 | 4341 | 4722 | 3001 | 3220 | 2684 | 2904 | 3943 | 4306 | 4390 | 4469 | 4508 | 2864 | 2914 | 2974 | 3008 | 3679,23 | 4418 | 2940 | 0,666 | -0,587 | -16,147 | 1,19E-58 | 6,41E-58 | 0,000185 | 0,002396 | 0,0009867 | 0,0009867 | TRUE | 0,0038119 | No |
| orf19.587 | 357 | 310 | 304 | 195 | 393 | 296 | 355 | 403 | 325 | 314 | 288 | 293 | 350 | 321 | 364 | 307 | 320,14 | 305 | 336 | 1,097 | 0,133 | 1,508 | 0,131589 | 0,155867 | 0,0019715 | 0,006163 | 0,0043405 | 0,0043405 | TRUE | 0,040343 | No |
| orf19.5870 | 484 | 449 | 450 | 266 | 918 | 734 | 798 | 1090 | 441 | 454 | 426 | 400 | 817 | 797 | 817 | 831 | 622,8 | 430 | 816 | 1,89 | 0,918 | 14,44 | 2,90E-47 | 1,28E-46 | 1,00E-08 | 0,004158 | 0,0020785 | 0,0020785 | TRUE | 0,0206987 | No |
| orf19.5871 | 640 | 539 | 597 | 367 | 614 | 530 | 538 | 732 | 583 | 545 | 565 | 551 | 546 | 575 | 551 | 558 | 559,38 | 561 | 558 | 0,992 | -0,011 | -0,178 | 0,859073 | 0,880198 | 1,00E-08 | 0,004399 | 0,0020451 | 0,0020451 | TRUE | 0,0083815 | No |
| orf19.5873 | 539 | 527 | 518 | 328 | 1328 | 1090 | 1261 | 1660 | 491 | 533 | 490 | 493 | 1181 | 1183 | 1291 | 1266 | 866,13 | 502 | 1230 | 2,453 | 1,294 | 20,9 | 5,41E-97 | 4,57E-96 | 0,0009337 | 0,003562 | 0,0022402 | 0,0022402 | TRUE | 0,0212912 | No |
| orf19.5874 | 2 | 1 | 1 | 1 | 6 | 5 | 3 | 7 | 2 | 1 | 1 | 2 | 5 | 5 | 3 | 5 | 3,06 | 2 | 4 | 3,668 | 1,875 | 2,149 | 0,031661 | 0,039921 | 1,00E-08 | 0,434118 | 0,2321376 | 0,2321376 | TRUE | 0,1074016 | No |
| orf19.5875 | 254 | 198 | 222 | 151 | 719 | 600 | 651 | 892 | 231 | 200 | 210 | 227 | 640 | 651 | 667 | 680 | 438,32 | 217 | 660 | 3,045 | 1,606 | 20,427 | 9,60E-93 | 7,71E-92 | 1,00E-08 | 0,005051 | 0,0027812 | 0,0027812 | TRUE | 0,0301899 | No |
| orf19.5876 | 21643 | 17430 | 18729 | 11093 | 16179 | 11409 | 13076 | 18422 | 19705 | 17629 | 17725 | 16663 | 14391 | 12387 | 13391 | 14052 | 15743 | 17930 | 13555 | 0,756 | -0,404 | -6,61 | 3,85E-11 | 7,58E-11 | 0,0045669 | 0,002121 | 0,003518 | 0,003518 | TRUE | 0,0543892 | No |
| orf19.5877 | 13681 | 12497 | 13790 | 8256 | 7625 | 6229 | 6810 | 8828 | 12456 | 12639 | 13051 | 12402 | 6782 | 6763 | 6974 | 6734 | 9725,21 | 12637 | 6813 | 0,539 | -0,892 | -27,92 | 1,52E-171 | 2,27E-170 | 0,0002484 | 0,002173 | 0,0008682 | 0,0008682 | TRUE | 0,0059928 | No |
| orf19.5879 | 150 | 110 | 112 | 66 | 201 | 147 | 146 | 211 | 137 | 111 | 106 | 99 | 179 | 160 | 150 | 161 | 137,73 | 113 | 162 | 1,422 | 0,508 | 3,888 | 0,000101 | 0,000151 | 0,0041019 | 0,011628 | 0,0088634 | 0,0088634 | TRUE | 0,2032767 | No |
| orf19.588 | 600 | 556 | 527 | 372 | 919 | 840 | 774 | 1049 | 546 | 562 | 499 | 559 | 817 | 912 | 793 | 800 | 686,06 | 542 | 830 | 1,534 | 0,618 | 8,836 | 9,90E-19 | 2,43E-18 | 0,0022505 | 0,003963 | 0,0031374 | 0,0031374 | TRUE | 0,0496546 | No |
| orf19.5880 | 552 | 437 | 524 | 341 | 533 | 460 | 507 | 639 | 503 | 442 | 496 | 512 | 474 | 499 | 519 | 487 | 491,61 | 488 | 495 | 1,015 | 0,021 | 0,291 | 0,771064 | 0,801097 | 0,0006481 | 0,004724 | 0,0029726 | 0,0029726 | TRUE | 0,0477134 | No |
| orf19.5884 | 1836 | 1650 | 1800 | 1050 | 3224 | 2647 | 2781 | 3630 | 1672 | 1669 | 1704 | 1577 | 2868 | 2874 | 2848 | 2769 | 2247,48 | 1656 | 2840 | 1,711 | 0,775 | 18,696 | 5,31E-78 | 3,67E-77 | 4,68E-05 | 0,002625 | 0,0011631 | 0,0011631 | TRUE | 0,0105391 | No |
| orf19.5885 | 15771 | 13780 | 14068 | 9321 | 18347 | 14058 | 13524 | 19757 | 14359 | 13937 | 13314 | 14002 | 16319 | 15263 | 13850 | 15071 | 14514,4 | 13903 | 15126 | 1,088 | 0,122 | 2,368 | 0,017884 | 0,023027 | 0,002634 | 0,002128 | 0,0024695 | 0,0024695 | TRUE | 0,0393468 | No |
| orf19.5886 | 5306 | 4320 | 4492 | 3243 | 4909 | 4127 | 3998 | 5800 | 4831 | 4369 | 4251 | 4872 | 4366 | 4481 | 4094 | 4424 | 4461,1 | 4581 | 4341 | 0,948 | -0,076 | -1,389 | 0,164811 | 0,192996 | 0,0028643 | 0,002333 | 0,002686 | 0,002686 | TRUE | 0,0286902 | No |
| orf19.5887 | 3704 | 3473 | 3611 | 2287 | 2499 | 2200 | 2323 | 3080 | 3372 | 3513 | 3417 | 3435 | 2223 | 2389 | 2379 | 2349 | 2884,71 | 3434 | 2335 | 0,68 | -0,557 | -13,945 | 3,38E-44 | 1,43E-43 | 0,0003052 | 0,002495 | 0,0011756 | 0,0011756 | TRUE | 0,0127668 | No |
| orf19.589 | 20209 | 16518 | 18226 | 11740 | 13813 | 11209 | 12664 | 17497 | 18399 | 16706 | 17249 | 17635 | 12286 | 12170 | 12970 | 13347 | 15095,3 | 17497 | 12693 | 0,726 | -0,463 | -10,29 | 7,80E-25 | 2,23E-24 | 0,0017126 | 0,002125 | 0,0018767 | 0,0018767 | TRUE | 0,0148698 | No |
| orf19.5890 | 4013 | 3530 | 3759 | 2387 | 3409 | 2849 | 3006 | 3956 | 3654 | 3570 | 3558 | 3586 | 3032 | 3093 | 3079 | 3018 | 3323,59 | 3592 | 3056 | 0,85 | -0,234 | -6,893 | 5,47E-12 | 1,11E-11 | 1,00E-08 | 0,002434 | 0,0008045 | 0,0008045 | TRUE | 0,0017382 | No |
| orf19.5892 | 249 | 230 | 257 | 175 | 513 | 389 | 444 | 583 | 227 | 233 | 243 | 263 | 456 | 422 | 455 | 445 | 342,94 | 242 | 444 | 1,854 | 0,89 | 10,842 | 2,18E-27 | 6,61E-27 | 1,00E-08 | 0,005889 | 0,003212 | 0,003212 | TRUE | 0,031293 | No |
| orf19.5893 | 6832 | 5490 | 5945 | 3696 | 13200 | 9856 | 10912 | 14928 | 6220 | 5553 | 5626 | 5552 | 11741 | 10701 | 11175 | 11387 | 8494,44 | 5738 | 11251 | 1,96 | 0,971 | 19,866 | 7,99E-88 | 6,18E-87 | 0,0021607 | 0,002193 | 0,0021593 | 0,0021593 | TRUE | 0,0395656 | No |
| orf19.5894 | 30 | 25 | 27 | 22 | 5 | 9 | 16 | 14 | 27 | 25 | 26 | 33 | 4 | 10 | 16 | 11 | 19,06 | 28 | 10 | 0,371 | -1,432 | -4,045 | 5,22E-05 | 7,95E-05 | 1,00E-08 | 0,071343 | 0,0554324 | 0,0554324 | TRUE | 0,5837237 | No |
| orf19.5896 | 121 | 108 | 102 | 75 | 104 | 102 | 95 | 153 | 110 | 109 | 97 | 113 | 93 | 111 | 97 | 117 | 105,73 | 107 | 104 | 0,978 | -0,032 | -0,226 | 0,821201 | 0,845267 | 1,00E-08 | 0,014531 | 0,0092401 | 0,0092401 | TRUE | 0,0751355 | No |
| orf19.5897 | 550 | 492 | 579 | 380 | 412 | 345 | 333 | 454 | 501 | 498 | 548 | 571 | 366 | 375 | 341 | 346 | 443,19 | 530 | 357 | 0,676 | -0,564 | -7,39 | 1,47E-13 | 3,12E-13 | 0,0010258 | 0,005018 | 0,0032744 | 0,0032744 | TRUE | 0,0278133 | No |
| orf19.59 | 11987 | 10379 | 11078 | 7019 | 11172 | 8938 | 10093 | 12921 | 10914 | 10497 | 10484 | 10544 | 9937 | 9704 | 10337 | 9856 | 10284,1 | 10610 | 9958 | 0,938 | -0,092 | -2,731 | 0,006314 | 0,008408 | 0,0004226 | 0,002165 | 0,0009824 | 0,0009824 | TRUE | 0,0078937 | No |
| orf19.590 | 1739 | 1356 | 1516 | 1012 | 1329 | 1138 | 1280 | 1721 | 1583 | 1371 | 1435 | 1520 | 1182 | 1236 | 1311 | 1313 | 1368,88 | 1477 | 1260 | 0,854 | -0,228 | -3,763 | 0,000168 | 0,000249 | 0,0026322 | 0,003002 | 0,0028028 | 0,0028028 | TRUE | 0,0300468 | No |
| orf19.5901 | 4463 | 3720 | 4137 | 2726 | 3000 | 2439 | 2694 | 3645 | 4063 | 3762 | 3915 | 4095 | 2668 | 2648 | 2759 | 2780 | 3336,48 | 3959 | 2714 | 0,686 | -0,544 | -12,878 | 5,98E-38 | 2,25E-37 | 0,0007587 | 0,002433 | 0,0014021 | 0,0014021 | TRUE | 0,0135885 | No |
| orf19.5902 | 5768 | 5316 | 5793 | 3986 | 2447 | 2133 | 2283 | 2971 | 5252 | 5377 | 5483 | 5988 | 2177 | 2316 | 2338 | 2266 | 3899,37 | 5525 | 2274 | 0,412 | -1,28 | -25,558 | 4,53E-144 | 5,67E-143 | 0,0017902 | 0,002376 | 0,0021043 | 0,0021043 | TRUE | 0,0376241 | No |
| orf19.5903 | 3698 | 3340 | 3896 | 2379 | 4228 | 3816 | 4017 | 5156 | 3367 | 3378 | 3687 | 3574 | 3761 | 4143 | 4114 | 3933 | 3744,56 | 3502 | 3988 | 1,139 | 0,188 | 3,856 | 0,000115 | 0,000172 | 0,0016272 | 0,00239 | 0,0019996 | 0,0019996 | TRUE | 0,0178437 | No |
| orf19.5904 | 23427 | 19785 | 20655 | 13114 | 36072 | 26594 | 28515 | 39008 | 21329 | 20010 | 19548 | 19699 | 32085 | 28874 | 29203 | 29756 | 25063 | 20146 | 29980 | 1,488 | 0,573 | 12,527 | 5,34E-36 | 1,95E-35 | 0,0019258 | 0,00209 | 0,0019702 | 0,0019702 | TRUE | 0,0273902 | No |
| orf19.5905 | 4868 | 4107 | 4387 | 2411 | 5925 | 4375 | 5101 | 6512 | 4432 | 4154 | 4152 | 3622 | 5270 | 4750 | 5224 | 4967 | 4571,39 | 4090 | 5053 | 1,234 | 0,304 | 4,914 | 8,93E-07 | 1,49E-06 | 0,0042807 | 0,002326 | 0,0034427 | 0,0034427 | TRUE | 0,0701381 | No |
| orf19.5906 | 9784 | 8326 | 8706 | 6151 | 10863 | 8955 | 9372 | 13169 | 8908 | 8421 | 8239 | 9240 | 9662 | 9723 | 9598 | 10045 | 9229,56 | 8702 | 9757 | 1,122 | 0,166 | 3,82 | 0,000133 | 0,000198 | 0,0014374 | 0,00218 | 0,0017025 | 0,0017025 | TRUE | 0,0205867 | No |
| orf19.5908 | 10566 | 9139 | 10326 | 6845 | 1842 | 1566 | 1678 | 2368 | 9620 | 9243 | 9773 | 10282 | 1638 | 1700 | 1718 | 1806 | 5722,67 | 9730 | 1716 | 0,177 | -2,502 | -52,63 | 0 | 0 | 0,001339 | 0,002268 | 0,001843 | 0,001843 | TRUE | 0,0174803 | No |
| orf19.5910 | 834 | 684 | 757 | 495 | 864 | 687 | 833 | 1026 | 759 | 692 | 716 | 744 | 769 | 746 | 853 | 783 | 757,66 | 728 | 788 | 1,082 | 0,114 | 1,801 | 0,071691 | 0,08742 | 0,0012081 | 0,003781 | 0,0025394 | 0,0025394 | TRUE | 0,0356445 | No |
| orf19.5911 | 1957 | 1728 | 1852 | 1318 | 3333 | 2787 | 2910 | 4049 | 1782 | 1748 | 1753 | 1980 | 2965 | 3026 | 2980 | 3089 | 2415,17 | 1816 | 3015 | 1,665 | 0,736 | 15,313 | 6,30E-53 | 3,06E-52 | 0,0010426 | 0,002584 | 0,0017648 | 0,0017648 | TRUE | 0,0407543 | No |
| orf19.5912 | 24050 | 21002 | 23064 | 13614 | 30169 | 24042 | 27269 | 35781 | 21897 | 21241 | 21828 | 20450 | 26834 | 26103 | 27927 | 27294 | 24196,8 | 21354 | 27040 | 1,266 | 0,34 | 9,301 | 1,39E-20 | 3,56E-20 | 0,0008407 | 0,002092 | 0,0012431 | 0,0012431 | TRUE | 0,0097881 | No |
| orf19.5915 | 280 | 220 | 259 | 174 | 529 | 411 | 450 | 626 | 255 | 223 | 245 | 261 | 471 | 446 | 461 | 478 | 354,88 | 246 | 464 | 1,893 | 0,921 | 11,311 | 1,16E-29 | 3,69E-29 | 1,00E-08 | 0,005759 | 0,0031919 | 0,0031919 | TRUE | 0,0455755 | No |
| orf19.5917 | 2916 | 2522 | 2873 | 1721 | 5555 | 5115 | 5372 | 7122 | 2655 | 2551 | 2719 | 2585 | 4941 | 5553 | 5502 | 5433 | 3992,33 | 2628 | 5357 | 2,038 | 1,027 | 21,244 | 3,72E-100 | 3,24E-99 | 0,0016861 | 0,002368 | 0,0019513 | 0,0019513 | TRUE | 0,0334813 | No |
| orf19.5917.3 | 7939 | 7433 | 7900 | 5492 | 10180 | 9089 | 8596 | 11956 | 7228 | 7518 | 7477 | 8250 | 9055 | 9868 | 8803 | 9120 | 8414,84 | 7618 | 9212 | 1,21 | 0,275 | 5,283 | 1,27E-07 | 2,19E-07 | 0,0027481 | 0,002194 | 0,002476 | 0,002476 | TRUE | 0,0373156 | No |
| orf19.5918 | 9 | 10 | 7 | 2 | 6 | 7 | 3 | 9 | 8 | 10 | 7 | 3 | 5 | 8 | 3 | 7 | 6,35 | 7 | 6 | 0,798 | -0,326 | -0,592 | 0,553718 | 0,59527 | 1,00E-08 | 0,21002 | 0,1364335 | 0,1364335 | TRUE | 0,2236553 | No |
| orf19.5919 | 464 | 553 | 567 | 376 | 764 | 757 | 629 | 851 | 422 | 559 | 537 | 565 | 680 | 822 | 644 | 649 | 609,74 | 521 | 699 | 1,343 | 0,426 | 4,089 | 4,34E-05 | 6,64E-05 | 0,0131237 | 0,004203 | 0,0087124 | 0,0087124 | TRUE | 0,1956555 | No |
| orf19.592 | 2635 | 2036 | 2442 | 1536 | 965 | 737 | 827 | 1150 | 2399 | 2059 | 2311 | 2307 | 858 | 800 | 847 | 877 | 1557,42 | 2269 | 846 | 0,373 | -1,422 | -23,715 | 2,52E-124 | 2,68E-123 | 0,0024863 | 0,002885 | 0,0026646 | 0,0026646 | TRUE | 0,0478467 | No |
| orf19.5920 | 2437 | 2071 | 2191 | 1059 | 1544 | 1148 | 1277 | 1713 | 2219 | 2095 | 2074 | 1591 | 1373 | 1246 | 1308 | 1307 | 1651,5 | 1995 | 1308 | 0,655 | -0,61 | -7,033 | 2,03E-12 | 4,17E-12 | 0,0095596 | 0,002837 | 0,0066093 | 0,0066093 | TRUE | 0,2163461 | No |
| orf19.5921 | 1046 | 1063 | 993 | 732 | 2111 | 2030 | 1838 | 2601 | 952 | 1075 | 940 | 1100 | 1878 | 2204 | 1882 | 1984 | 1501,86 | 1017 | 1987 | 1,958 | 0,97 | 13,577 | 5,51E-42 | 2,22E-41 | 0,0052435 | 0,002917 | 0,0041304 | 0,0041304 | TRUE | 0,0641622 | No |
| orf19.5924 | 802 | 810 | 906 | 500 | 913 | 840 | 857 | 1182 | 730 | 819 | 857 | 751 | 812 | 912 | 878 | 902 | 832,67 | 789 | 876 | 1,108 | 0,147 | 2,152 | 0,031409 | 0,039635 | 0,0029651 | 0,003623 | 0,0032955 | 0,0032955 | TRUE | 0,0410844 | No |
| orf19.5925 | 712 | 700 | 730 | 417 | 676 | 581 | 534 | 794 | 648 | 708 | 691 | 626 | 601 | 631 | 547 | 606 | 632,27 | 668 | 596 | 0,89 | -0,168 | -2,449 | 0,014326 | 0,018597 | 0,0014725 | 0,004126 | 0,0029265 | 0,0029265 | TRUE | 0,0346256 | No |
| orf19.5926 | 1050 | 963 | 1052 | 640 | 1000 | 871 | 906 | 1236 | 956 | 974 | 996 | 961 | 889 | 946 | 928 | 943 | 949,1 | 972 | 926 | 0,953 | -0,07 | -1,343 | 0,179169 | 0,208355 | 1,00E-08 | 0,003429 | 0,0015339 | 0,0015339 | TRUE | 0,0088527 | No |
| orf19.5927 | 13795 | 11278 | 11473 | 8004 | 19934 | 13939 | 14673 | 20976 | 12560 | 11406 | 10858 | 12023 | 17731 | 15134 | 15027 | 16001 | 13842,5 | 11712 | 15973 | 1,364 | 0,448 | 7,146 | 8,96E-13 | 1,86E-12 | 0,0049547 | 0,002132 | 0,0036987 | 0,0036987 | TRUE | 0,0671634 | No |

| | | | | | | | | | | | | | | | | | | | | | | | | | | | | | | | |
|---|---|---|---|---|---|---|---|---|---|---|---|---|---|---|---|---|---|---|---|---|---|---|---|---|---|---|---|---|---|---|---|
| orf19.5928 | 7115 | 5470 | 5372 | 4482 | 11093 | 8528 | 7549 | 12490 | 6478 | 5532 | 5084 | 6733 | 9867 | 9259 | 7731 | 9527 | 7526,44 | 5957 | 9096 | 1,528 | 0,611 | 6,617 | 3,65E-11 | 7,20E-11 | 0,0137195 | 0,002213 | 0,0080567 | 0,0080567 | TRUE | 0,1246287 | No |
| orf19.5929 | 860 | 769 | 803 | 567 | 821 | 643 | 668 | 972 | 783 | 778 | 760 | 852 | 730 | 698 | 684 | 741 | 753,3 | 793 | 713 | 0,904 | -0,146 | -2,4 | 0,016392 | 0,021176 | 0,0003769 | 0,003791 | 0,0022146 | 0,0022146 | TRUE | 0,0253119 | No |
| orf19.593 | 2883 | 2439 | 2722 | 1724 | 2272 | 1967 | 2179 | 3125 | 2625 | 2467 | 2576 | 2590 | 2021 | 2136 | 2232 | 2384 | 2378,66 | 2564 | 2193 | 0,856 | -0,225 | -4,063 | 4,83E-05 | 7,37E-05 | 0,0022437 | 0,002592 | 0,0025168 | 0,0025168 | TRUE | 0,0430785 | No |
| orf19.5930 | 1929 | 1481 | 1650 | 1180 | 1331 | 1135 | 1284 | 1727 | 1756 | 1498 | 1562 | 1773 | 1184 | 1232 | 1315 | 1317 | 1454,6 | 1647 | 1262 | 0,767 | -0,382 | -5,693 | 1,25E-08 | 2,25E-08 | 0,0039288 | 0,002945 | 0,0036277 | 0,0036277 | TRUE | 0,0446427 | No |
| orf19.5931 | 166 | 183 | 165 | 74 | 121 | 102 | 129 | 132 | 151 | 185 | 156 | 111 | 108 | 111 | 132 | 101 | 131,84 | 151 | 113 | 0,738 | -0,437 | -2,869 | 0,00412 | 0,00555 | 0,0187227 | 0,012057 | 0,0146554 | 0,0146554 | TRUE | 0,2397918 | No |
| orf19.5932 | 2542 | 2238 | 2471 | 1486 | 1452 | 1167 | 1234 | 1638 | 2314 | 2263 | 2339 | 2232 | 1292 | 1267 | 1264 | 1249 | 1777,56 | 2287 | 1268 | 0,554 | -0,853 | -20,202 | 9,35E-91 | 7,36E-90 | 1,00E-08 | 0,00278 | 0,001108 | 0,001108 | TRUE | 0,0028413 | No |
| orf19.5933 | 72 | 71 | 86 | 45 | 121 | 126 | 127 | 149 | 66 | 72 | 81 | 68 | 108 | 137 | 130 | 114 | 96,81 | 72 | 122 | 1,692 | 0,758 | 5,018 | 5,22E-07 | 8,78E-07 | 1,00E-08 | 0,015682 | 0,010687 | 0,010687 | TRUE | 0,0833396 | No |
| orf19.5934 | 281 | 275 | 281 | 156 | 998 | 847 | 933 | 1243 | 256 | 278 | 266 | 234 | 888 | 920 | 956 | 948 | 593,15 | 258 | 928 | 3,568 | 1,835 | 25,354 | 8,19E-142 | 1,01E-140 | 1,00E-08 | 0,004264 | 0,0024844 | 0,0024844 | TRUE | 0,0330548 | No |
| orf19.5935 | 1090 | 970 | 1076 | 708 | 1186 | 1012 | 1074 | 1377 | 992 | 981 | 1018 | 1064 | 1055 | 1099 | 1100 | 1050 | 1044,91 | 1014 | 1076 | 1,063 | 0,088 | 1,696 | 0,089948 | 0,108463 | 1,00E-08 | 0,003301 | 0,0015982 | 0,0015982 | TRUE | 0,0111138 | No |
| orf19.5938 | 2342 | 2185 | 2308 | 1478 | 3823 | 3357 | 3449 | 4631 | 2132 | 2210 | 2184 | 2220 | 3400 | 3645 | 3532 | 3533 | 2857,08 | 2186 | 3528 | 1,614 | 0,69 | 17,423 | 5,56E-68 | 3,40E-67 | 0,0002892 | 0,002499 | 0,0011295 | 0,0011295 | TRUE | 0,0071546 | No |
| orf19.5940 | 1169 | 996 | 1083 | 749 | 1673 | 1549 | 1601 | 2086 | 1064 | 1007 | 1025 | 1125 | 1488 | 1682 | 1640 | 1591 | 1327,81 | 1055 | 1600 | 1,518 | 0,602 | 10,531 | 6,19E-26 | 1,81E-25 | 0,0015328 | 0,003032 | 0,0023397 | 0,0023397 | TRUE | 0,0268188 | No |
| orf19.5941 | 642 | 546 | 572 | 333 | 918 | 706 | 788 | 1014 | 585 | 552 | 541 | 500 | 817 | 767 | 807 | 773 | 667,73 | 544 | 791 | 1,446 | 0,532 | 8,255 | 1,52E-16 | 3,50E-16 | 0,0004571 | 0,004015 | 0,0024201 | 0,0024201 | TRUE | 0,0291696 | No |
| orf19.5942 | 2792 | 2452 | 2670 | 1548 | 2821 | 2191 | 2448 | 3335 | 2542 | 2480 | 2527 | 2325 | 2509 | 2379 | 2507 | 2544 | 2476,65 | 2468 | 2485 | 1,005 | 0,008 | 0,177 | 0,85965 | 0,880648 | 0,0006559 | 0,00257 | 0,0014956 | 0,0014956 | TRUE | 0,0169122 | No |
| orf19.5943 | 1312 | 1289 | 1359 | 768 | 1422 | 1292 | 1344 | 1773 | 1195 | 1304 | 1286 | 1154 | 1265 | 1403 | 1376 | 1352 | 1291,81 | 1235 | 1349 | 1,09 | 0,125 | 2,166 | 0,030291 | 0,038291 | 0,0016541 | 0,00306 | 0,0023967 | 0,0023967 | TRUE | 0,0213563 | No |
| orf19.5943.1 | 7773 | 6414 | 6865 | 4467 | 8997 | 7341 | 7776 | 10699 | 7077 | 6487 | 6497 | 6710 | 8003 | 7970 | 7964 | 8161 | 7358,63 | 6693 | 8024 | 1,199 | 0,262 | 6,848 | 7,46E-12 | 1,51E-11 | 0,0007345 | 0,002217 | 0,0012648 | 0,0012648 | TRUE | 0,0187692 | No |
| orf19.5947 | 2604 | 2335 | 2558 | 1658 | 2863 | 2348 | 2468 | 3258 | 2371 | 2362 | 2421 | 2491 | 2547 | 2549 | 2528 | 2485 | 2469,07 | 2411 | 2527 | 1,049 | 0,069 | 1,81 | 0,070237 | 0,085749 | 1,00E-08 | 0,002572 | 0,0009936 | 0,0009936 | TRUE | 0,0053148 | No |
| orf19.5949 | 6066 | 5619 | 6144 | 3856 | 50021 | 41473 | 43371 | 58689 | 5523 | 5683 | 5815 | 5792 | 44492 | 45028 | 44417 | 44768 | 25189,9 | 5703 | 44676 | 7,837 | 2,97 | 101,501 | 0 | 0 | 5,37E-05 | 0,002089 | 0,0007175 | 0,0007175 | TRUE | 0,0057006 | No |
| orf19.5952 | 272 | 249 | 295 | 159 | 670 | 566 | 515 | 730 | 248 | 252 | 279 | 239 | 596 | 615 | 527 | 557 | 414,03 | 254 | 574 | 2,246 | 1,167 | 13,888 | 7,51E-44 | 3,14E-43 | 0,0021534 | 0,005228 | 0,0038814 | 0,0038814 | TRUE | 0,0494257 | No |
| orf19.5953 | 7255 | 5963 | 6990 | 4468 | 7840 | 6724 | 7728 | 10407 | 6605 | 6031 | 6615 | 6712 | 6973 | 7300 | 7914 | 7939 | 7011,27 | 6491 | 7532 | 1,161 | 0,215 | 3,937 | 8,24E-05 | 0,000124 | 0,0029412 | 0,002225 | 0,002717 | 0,002717 | TRUE | 0,0306074 | No |
| orf19.5954 | 100 | 71 | 63 | 41 | 70 | 41 | 57 | 85 | 91 | 72 | 60 | 62 | 62 | 45 | 58 | 65 | 64,26 | 71 | 58 | 0,809 | -0,306 | -1,57 | 0,116371 | 0,138744 | 0,0181117 | 0,022595 | 0,0208638 | 0,0208638 | TRUE | 0,349542 | No |
| orf19.5956 | 12060 | 9884 | 10840 | 7587 | 8180 | 6686 | 7461 | 10416 | 10980 | 9997 | 10259 | 11397 | 7276 | 7259 | 7641 | 7945 | 9094,26 | 10658 | 7530 | 0,707 | -0,501 | -9,665 | 4,24E-22 | 1,13E-21 | 0,0026263 | 0,002182 | 0,0024633 | 0,0024633 | TRUE | 0,0261844 | No |
| orf19.5958 | 3771 | 3399 | 3634 | 2015 | 2700 | 2293 | 2346 | 3229 | 3433 | 3438 | 3439 | 3027 | 2402 | 2490 | 2403 | 2463 | 2886,75 | 3334 | 2440 | 0,73 | -0,453 | -9,128 | 6,95E-20 | 1,76E-19 | 0,0015806 | 0,002495 | 0,0020142 | 0,0020142 | TRUE | 0,044203 | No |
| orf19.5959 | 13492 | 13161 | 13982 | 7954 | 14164 | 11350 | 12235 | 16078 | 12284 | 13311 | 13233 | 11948 | 12598 | 12323 | 12530 | 12264 | 12561,5 | 12694 | 12429 | 0,979 | -0,031 | -0,721 | 0,471024 | 0,512857 | 0,0013793 | 0,002142 | 0,0016845 | 0,0016845 | TRUE | 0,0187354 | No |
| orf19.5960 | 38765 | 35479 | 38516 | 28931 | 12406 | 11230 | 10545 | 14476 | 35294 | 35883 | 36452 | 43459 | 11035 | 12193 | 10799 | 11042 | 24519,7 | 37772 | 11267 | 0,298 | -1,745 | -25,652 | 3,98E-145 | 5,06E-144 | 0,0065879 | 0,002091 | 0,0043908 | 0,0043908 | TRUE | 0,1255355 | No |
| orf19.5961 | 4536 | 3928 | 4210 | 2650 | 1145 | 869 | 926 | 1300 | 4130 | 3973 | 3984 | 3981 | 1018 | 943 | 948 | 992 | 2496,2 | 4017 | 975 | 0,243 | -2,04 | -46,649 | 0 | 0 | 0,0001731 | 0,002566 | 0,0012256 | 0,0012256 | TRUE | 0,0101846 | No |
| orf19.596.1 | 2835 | 2267 | 2588 | 1696 | 4220 | 2951 | 3451 | 4655 | 2581 | 2293 | 2449 | 2548 | 3754 | 3204 | 3534 | 3551 | 2989,2 | 2468 | 3511 | 1,424 | 0,51 | 8,769 | 1,81E-18 | 4,40E-18 | 0,0030126 | 0,002479 | 0,0028917 | 0,0028917 | TRUE | 0,0415093 | No |
| orf19.5962 | 2440 | 2397 | 2366 | 1416 | 3008 | 2407 | 2654 | 3470 | 2222 | 2424 | 2239 | 2127 | 2676 | 2613 | 2718 | 2647 | 2458,24 | 2253 | 2664 | 1,181 | 0,24 | 5,098 | 3,43E-07 | 5,79E-07 | 0,0011158 | 0,002574 | 0,0017083 | 0,0017083 | TRUE | 0,0314281 | No |
| orf19.596.2 | 549 | 430 | 494 | 276 | 571 | 432 | 557 | 676 | 500 | 435 | 468 | 415 | 508 | 469 | 570 | 516 | 484,98 | 454 | 516 | 1,131 | 0,178 | 2,143 | 0,032107 | 0,040467 | 0,0039142 | 0,004761 | 0,0045241 | 0,0045241 | TRUE | 0,05665 | No |
| orf19.5963 | 7248 | 6400 | 6908 | 3731 | 4071 | 3013 | 3397 | 4553 | 6599 | 6473 | 6538 | 5605 | 3621 | 3271 | 3479 | 3473 | 4882,32 | 6304 | 3461 | 0,549 | -0,866 | -14,94 | 1,81E-50 | 8,40E-50 | 0,0033312 | 0,002308 | 0,0030044 | 0,0030044 | TRUE | 0,0664568 | No |
| orf19.5964 | 4698 | 4202 | 4580 | 2603 | 5883 | 4674 | 5149 | 6653 | 4277 | 4250 | 4335 | 3910 | 5233 | 5075 | 5273 | 5075 | 4678,43 | 4193 | 5164 | 1,23 | 0,299 | 7,148 | 8,78E-13 | 1,83E-12 | 0,0009621 | 0,002319 | 0,0014567 | 0,0014567 | TRUE | 0,0236375 | No |
| orf19.5964.2 | 10104 | 9677 | 9414 | 5548 | 15209 | 11459 | 11749 | 15236 | 9199 | 9787 | 8909 | 8334 | 13528 | 12441 | 12032 | 11622 | 10731,7 | 9057 | 12406 | 1,369 | 0,453 | 7,555 | 4,18E-14 | 9,01E-14 | 0,0042842 | 0,00216 | 0,0033619 | 0,0033619 | TRUE | 0,0454773 | No |
| orf19.5965 | 1689 | 1691 | 1772 | 1060 | 2682 | 2164 | 2267 | 2959 | 1538 | 1710 | 1677 | 1592 | 2386 | 2350 | 2322 | 2257 | 1978,91 | 1629 | 2329 | 1,428 | 0,514 | 10,728 | 7,52E-27 | 2,25E-26 | 0,0007427 | 0,002705 | 0,0016746 | 0,0016746 | TRUE | 0,0189335 | No |
| orf19.5966 | 12 | 7 | 9 | 8 | 12 | 18 | 11 | 20 | 11 | 7 | 9 | 12 | 11 | 20 | 11 | 15 | 11,91 | 10 | 14 | 1,486 | 0,571 | 1,397 | 0,162349 | 0,190252 | 1,00E-08 | 0,112954 | 0,0716535 | 0,0716535 | TRUE | 0,2471026 | No |
| orf19.5967 | 1233 | 1009 | 1199 | 777 | 1469 | 1081 | 1218 | 1658 | 1123 | 1020 | 1135 | 1167 | 1307 | 1174 | 1247 | 1265 | 1179,68 | 1111 | 1248 | 1,126 | 0,171 | 2,933 | 0,003358 | 0,00455 | 0,0014806 | 0,003157 | 0,0023946 | 0,0023946 | TRUE | 0,0362257 | No |
| orf19.5968 | 3161 | 2990 | 2907 | 1975 | 2215 | 1762 | 1841 | 2518 | 2878 | 3024 | 2751 | 2967 | 1970 | 1913 | 1885 | 1921 | 2413,67 | 2905 | 1922 | 0,662 | -0,594 | -13,437 | 3,65E-41 | 1,45E-40 | 0,0007011 | 0,002584 | 0,0014501 | 0,0014501 | TRUE | 0,0157435 | No |
| orf19.597 | 3467 | 2676 | 3094 | 2125 | 4458 | 3795 | 4014 | 5850 | 3157 | 2706 | 2928 | 3192 | 3965 | 4120 | 4111 | 4462 | 3580,27 | 2996 | 4164 | 1,391 | 0,477 | 7,909 | 2,59E-15 | 5,78E-15 | 0,0037328 | 0,002406 | 0,0031933 | 0,0031933 | TRUE | 0,0513886 | No |
| orf19.5970 | 696 | 628 | 718 | 373 | 934 | 795 | 831 | 1070 | 634 | 635 | 680 | 560 | 831 | 863 | 851 | 816 | 733,73 | 627 | 840 | 1,332 | 0,414 | 6,372 | 1,86E-10 | 3,58E-10 | 0,0010447 | 0,003837 | 0,0026433 | 0,0026433 | TRUE | 0,0510784 | No |
| orf19.5971 | 1174 | 1046 | 1066 | 681 | 656 | 574 | 576 | 773 | 1069 | 1058 | 1009 | 1023 | 583 | 623 | 590 | 590 | 818,11 | 1040 | 596 | 0,573 | -0,804 | -14,296 | 2,33E-46 | 1,01E-45 | 1,00E-08 | 0,003652 | 0,0017401 | 0,0017401 | TRUE | 0,0098561 | No |
| orf19.5973 | 3931 | 3704 | 3813 | 2424 | 2549 | 1981 | 2000 | 2775 | 3579 | 3746 | 3609 | 3641 | 2267 | 2151 | 2048 | 2117 | 2894,78 | 3644 | 2146 | 0,589 | -0,763 | -17,736 | 2,19E-70 | 1,39E-69 | 0,000616 | 0,002493 | 0,0014135 | 0,0014135 | TRUE | 0,0178175 | No |
| orf19.5974 | 1379 | 1557 | 1646 | 946 | 2218 | 1886 | 2046 | 2490 | 1256 | 1575 | 1558 | 1421 | 1973 | 2048 | 2095 | 1899 | 1728,04 | 1452 | 2004 | 1,379 | 0,464 | 6,49 | 8,58E-11 | 1,67E-10 | 0,0053295 | 0,002801 | 0,0042984 | 0,0042984 | TRUE | 0,1045773 | No |
| orf19.5975 | 1678 | 1391 | 1493 | 898 | 3190 | 2469 | 2835 | 3708 | 1528 | 1407 | 1413 | 1349 | 2837 | 2681 | 2903 | 2828 | 2118,31 | 1424 | 2812 | 1,971 | 0,979 | 19,551 | 4,08E-85 | 3,09E-84 | 0,0012353 | 0,002661 | 0,0018655 | 0,0018655 | TRUE | 0,0294028 | No |
| orf19.5976 | 1505 | 1421 | 1548 | 969 | 1495 | 1374 | 1433 | 1821 | 1370 | 1437 | 1465 | 1456 | 1330 | 1492 | 1468 | 1389 | 1425,78 | 1432 | 1420 | 0,991 | -0,013 | -0,251 | 0,801825 | 0,828064 | 0,0011841 | 0,002964 | 0,001991 | 0,001991 | TRUE | 0,0216002 | No |
| orf19.5977 | 1607 | 1323 | 1451 | 912 | 1690 | 1391 | 1390 | 2046 | 1463 | 1338 | 1373 | 1370 | 1503 | 1510 | 1424 | 1561 | 1442,76 | 1386 | 1500 | 1,082 | 0,113 | 2,223 | 0,026231 | 0,033292 | 0,0008642 | 0,002953 | 0,0018001 | 0,0018001 | TRUE | 0,017784 | No |
| orf19.5978 | 264 | 248 | 281 | 209 | 331 | 288 | 300 | 400 | 240 | 251 | 266 | 314 | 294 | 313 | 307 | 305 | 286,32 | 268 | 305 | 1,148 | 0,199 | 2,127 | 0,033444 | 0,042102 | 0,0018403 | 0,006651 | 0,0048939 | 0,0048939 | TRUE | 0,1249183 | No |
| orf19.598 | 5 | 6 | 1 | 1 | 1 | 0 | 0 | 6 | 5 | 6 | 1 | 2 | 1 | 0 | 0 | 5 | 2,32 | 4 | 2 | 0,445 | -1,167 | -1,059 | 0,289599 | 0,326413 | 0,9445427 | 0,572184 | 0,6898332 | 0,6898332 | TRUE | 0,9181911 | No |
| orf19.5980 | 1288 | 1224 | 1287 | 806 | 1294 | 1042 | 1120 | 1519 | 1173 | 1238 | 1218 | 1211 | 1151 | 1131 | 1147 | 1159 | 1178,43 | 1210 | 1147 | 0,949 | -0,076 | -1,609 | 0,107537 | 0,128717 | 1,00E-08 | 0,003158 | 0,0012874 | 0,0012874 | TRUE | 0,0056262 | No |
| orf19.5982 | 17187 | 13356 | 14325 | 8991 | 24282 | 18377 | 19873 | 28129 | 15648 | 13508 | 13557 | 13506 | 21598 | 19952 | 20352 | 21457 | 17447,4 | 14055 | 20840 | 1,483 | 0,568 | 10,257 | 1,10E-24 | 3,14E-24 | 0,0035376 | 0,002113 | 0,0028868 | 0,0028868 | TRUE | 0,0715493 | No |
| orf19.5984 | 979 | 839 | 898 | 606 | 1170 | 916 | 1016 | 1337 | 891 | 849 | 850 | 910 | 1041 | 995 | 1041 | 1020 | 949,46 | 875 | 1024 | 1,172 | 0,23 | 4,352 | 1,35E-05 | 2,12E-05 | 1,00E-08 | 0,003428 | 0,0016039 | 0,0016039 | TRUE | 0,0073172 | No |
| orf19.5985 | 1665 | 1386 | 1477 | 944 | 1822 | 1313 | 1554 | 1962 | 1516 | 1402 | 1398 | 1418 | 1621 | 1426 | 1591 | 1497 | 1483,48 | 1434 | 1534 | 1,07 | 0,097 | 1,735 | 0,082717 | 0,100137 | 0,001832 | 0,002927 | 0,0023282 | 0,0023282 | TRUE | 0,0258681 | No |
| orf19.5986 | 764 | 631 | 656 | 545 | 950 | 921 | 855 | 1225 | 696 | 638 | 621 | 819 | 845 | 1000 | 876 | 934 | 803,54 | 694 | 914 | 1,323 | 0,404 | 4,618 | 3,87E-06 | 6,28E-06 | 0,0094365 | 0,003681 | 0,0060538 | 0,0060538 | TRUE | 0,1605676 | No |
| orf19.5987 | 3805 | 3187 | 3415 | 2167 | 5765 | 4621 | 5192 | 6973 | 3464 | 3223 | 3232 | 3255 | 5128 | 5017 | 5317 | 5319 | 4244,5 | 3294 | 5195 | 1,577 | 0,657 | 16,044 | 6,27E-58 | 3,33E-57 | 0,0006844 | 0,002348 | 0,0013575 | 0,0013575 | TRUE | 0,0153345 | No |
| orf19.5989 | 5557 | 4555 | 5195 | 3959 | 8993 | 8199 | 8503 | 11812 | 5059 | 4607 | 4917 | 5947 | 7999 | 8902 | 8708 | 9010 | 6893,65 | 5132 | 8655 | 1,688 | 0,755 | 10,479 | 1,08E-25 | 3,14E-25 | 0,0074478 | 0,002229 | 0,004827 | 0,004827 | TRUE | 0,1368744 | No |
| orf19.599 | 1 | 0 | 0 | 0 | 1 | 0 | 0 | 0 | 1 | 0 | 0 | 0 | 1 | 0 | 0 | 0 | 0,22 | 0 | 0 | 0,871 | -0,199 | -0,068 | 0,945712 | 0,952989 | 1,00E-08 | 5,873379 | 6,1895949 | 6,1895949 | TRUE | 0,3214243 | No |
| orf19.5991 | 19515 | 17617 | 19336 | 11538 | 23565 | 19160 | 20829 | 28168 | 17768 | 17818 | 18300 | 17332 | 20960 | 20802 | 21332 | 21487 | 19474,8 | 17804 | 21145 | 1,187 | 0,248 | 8,316 | 9,07E-17 | 2,11E-16 | 0,0002912 | 0,002105 | 0,0008007 | 0,0008007 | TRUE | 0,0042901 | No |
| orf19.5992 | 3279 | 2860 | 3159 | 2280 | 1523 | 1454 | 1450 | 2065 | 2985 | 2893 | 2990 | 3425 | 1355 | 1579 | 1485 | 1575 | 2285,76 | 3073 | 1498 | 0,488 | -1,035 | -15,437 | 9,17E-54 | 4,54E-53 | 0,004819 | 0,002615 | 0,0038272 | 0,0038272 | TRUE | 0,0697869 | No |
| orf19.5994 | 2362 | 2254 | 2221 | 1393 | 4282 | 3661 | 3610 | 4843 | 2151 | 2280 | 2102 | 2093 | 3809 | 3975 | 3697 | 3694 | 2974,95 | 2156 | 3794 | 1,758 | 0,814 | 17,982 | 2,71E-72 | 1,75E-71 | 0,0009187 | 0,002481 | 0,0015938 | 0,0015938 | TRUE | 0,0180608 | No |
| orf19.5995 | 3682 | 3173 | 3311 | 2256 | 3039 | 2522 | 2651 | 3612 | 3352 | 3209 | 3134 | 3389 | 2703 | 2738 | 2715 | 2755 | 2999,42 | 3271 | 2728 | 0,835 | -0,261 | -6,533 | 6,45E-11 | 1,26E-10 | 0,0003865 | 0,002477 | 0,001192 | 0,001192 | TRUE | 0,0098069 | No |
| orf19.5996.1 | 17196 | 13051 | 13830 | 8931 | 28143 | 20649 | 22032 | 32446 | 15656 | 13200 | 13089 | 13416 | 25032 | 22419 | 22564 | 24750 | 18765,7 | 13840 | 23691 | 1,712 | 0,775 | 12,05 | 1,94E-33 | 6,69E-33 | 0,0055233 | 0,002107 | 0,0039198 | 0,0039198 | TRUE | 0,09584 | No |
| orf19.5999 | 1264 | 1134 | 1196 | 712 | 812 | 709 | 743 | 946 | 1151 | 1147 | 1132 | 1070 | 722 | 770 | 761 | 722 | 934,22 | 1125 | 744 | 0,659 | -0,602 | -11,041 | 2,43E-28 | 7,50E-28 | 1,00E-08 | 0,003451 | 0,0017522 | 0,0017522 | TRUE | 0,0113388 | No |
| orf19.600 | 3736 | 3053 | 3602 | 2016 | 4061 | 3299 | 3823 | 5186 | 3401 | 3088 | 3409 | 3028 | 3612 | 3582 | 3915 | 3956 | 3498,96 | 3232 | 3766 | 1,165 | 0,22 | 3,907 | 9,33E-05 | 0,00014 | 0,0029309 | 0,002415 | 0,002748 | 0,002748 | TRUE | 0,0209136 | No |
| orf19.6000 | 75534 | 71159 | 76660 | 44869 | 45433 | 37874 | 40077 | 54238 | 68771 | 71970 | 72552 | 67400 | 40411 | 41121 | 41044 | 41373 | 55580,2 | 70173 | 40987 | 0,584 | -0,776 | -22,995 | 5,18E-117 | 5,19E-116 | 0,0006516 | 0,002061 | 0,0010745 | 0,0010745 | TRUE | 0,0086186 | No |
| orf19.6001 | 612 | 631 | 672 | 378 | 446 | 380 | 426 | 539 | 557 | 638 | 636 | 568 | 397 | 413 | 436 | 411 | 506,99 | 600 | 414 | 0,689 | -0,538 | -7,234 | 4,68E-13 | 9,83E-13 | 0,0017308 | 0,004643 | 0,003286 | 0,003286 | TRUE | 0,030939 | No |
| orf19.6002 | 10673 | 10072 | 10401 | 7337 | 16488 | 14185 | 13333 | 18045 | 9717 | 10187 | 9844 | 11021 | 14666 | 15401 | 13655 | 13765 | 12281,9 | 10192 | 14372 | 1,41 | 0,496 | 9,167 | 4,85E-20 | 1,24E-19 | 0,0031846 | 0,002145 | 0,0027291 | 0,0027291 | TRUE | 0,0361598 | No |
| orf19.6003 | 3127 | 3122 | 3481 | 2140 | 4051 | 3433 | 3415 | 4776 | 2847 | 3158 | 3294 | 3215 | 3603 | 3727 | 3497 | 3643 | 3373,09 | 3128 | 3618 | 1,157 | 0,21 | 4,111 | 3,94E-05 | 6,04E-05 | 0,0019099 | 0,002429 | 0,0022156 | 0,0022156 | TRUE | 0,0457651 | No |
| orf19.6005 | 348 | 350 | 361 | 229 | 740 | 685 | 692 | 854 | 317 | 354 | 342 | 344 | 658 | 744 | 709 | 651 | 514,82 | 339 | 690 | 2,035 | 1,025 | 13,574 | 5,74E-42 | 2,31E-41 | 0,0016949 | 0,004603 | 0,0032227 | 0,0032227 | TRUE | 0,031104 | No |
| orf19.6007 | 2445 | 2961 | 2822 | 1801 | 8011 | 7153 | 6646 | 8212 | 2226 | 2995 | 2671 | 2705 | 7126 | 7766 | 6806 | 6264 | 4819,9 | 2649 | 6990 | 2,639 | 1,4 | 16,393 | 2,15E-60 | 1,18E-59 | 0,0108549 | 0,002311 | 0,0067363 | 0,0067363 | TRUE | 0,142275 | No |
| orf19.6008 | 312 | 392 | 401 | 249 | 677 | 574 | 646 | 811 | 284 | 396 | 380 | 374 | 602 | 623 | 662 | 619 | 492,46 | 358 | 626 | 1,751 | 0,808 | 8,527 | 1,51E-17 | 3,58E-17 | 0,00859 | 0,004719 | 0,0063844 | 0,0063844 | TRUE | 0,2402222 | No |

| | | | | | | | | | | | | | | | | | | | | | | | | | | | | | | | |
|---|---|---|---|---|---|---|---|---|---|---|---|---|---|---|---|---|---|---|---|---|---|---|---|---|---|---|---|---|---|---|---|
| orf19.6008.3 | 295 | 265 | 292 | 168 | 328 | 262 | 293 | 374 | 269 | 268 | 276 | 252 | 292 | 284 | 300 | 285 | 278,36 | 266 | 290 | 1,086 | 0,119 | 1,414 | 0,157393 | 0,184785 | 1,00E-08 | 0,006783 | 0,0032616 | 0,0032616 | TRUE | 0,0108891 | No |
| orf19.6008.4 | 2460 | 1903 | 1999 | 1306 | 1527 | 1073 | 1198 | 1655 | 2240 | 1925 | 1892 | 1962 | 1358 | 1165 | 1227 | 1262 | 1628,83 | 2005 | 1253 | 0,625 | -0,677 | -9,993 | 1,63E-23 | 4,52E-23 | 0,0043755 | 0,002848 | 0,0037705 | 0,0037705 | TRUE | 0,0774796 | No |
| orf19.6009 | 1345 | 1278 | 1342 | 846 | 1916 | 1573 | 1836 | 2314 | 1225 | 1293 | 1270 | 1271 | 1704 | 1708 | 1880 | 1765 | 1514,44 | 1265 | 1764 | 1,396 | 0,481 | 9,583 | 9,41E-22 | 2,48E-21 | 0,0007971 | 0,002909 | 0,0017289 | 0,0017289 | TRUE | 0,0229221 | No |
| orf19.6010 | 557 | 531 | 549 | 339 | 249 | 214 | 226 | 310 | 507 | 537 | 520 | 509 | 221 | 232 | 231 | 236 | 374,34 | 518 | 230 | 0,445 | -1,17 | -15,137 | 9,23E-52 | 4,40E-51 | 1,00E-08 | 0,005566 | 0,0026858 | 0,0026858 | TRUE | 0,0081198 | No |
| orf19.6010.1 | 1383 | 1257 | 1241 | 833 | 1261 | 1008 | 1027 | 1331 | 1259 | 1271 | 1174 | 1251 | 1122 | 1094 | 1052 | 1015 | 1154,92 | 1239 | 1071 | 0,864 | -0,211 | -3,89 | 0,0001 | 0,00015 | 0,0008888 | 0,003181 | 0,0019555 | 0,0019555 | TRUE | 0,0157502 | No |
| orf19.6011 | 5898 | 5257 | 5762 | 3514 | 8932 | 7641 | 7990 | 10655 | 5370 | 5317 | 5453 | 5279 | 7945 | 8296 | 8183 | 8128 | 6746,22 | 5355 | 8138 | 1,519 | 0,603 | 19,279 | 8,00E-83 | 5,91E-82 | 0,0001282 | 0,002233 | 0,0007815 | 0,0007815 | TRUE | 0,0031125 | No |
| orf19.6012 | 1435 | 1402 | 1498 | 902 | 1335 | 1258 | 1186 | 1584 | 1307 | 1418 | 1418 | 1355 | 1187 | 1366 | 1215 | 1208 | 1309,17 | 1374 | 1244 | 0,904 | -0,146 | -2,482 | 0,013047 | 0,016971 | 0,0019258 | 0,003046 | 0,0025365 | 0,0025365 | TRUE | 0,0504384 | No |
| orf19.6013 | 1262 | 1068 | 1205 | 743 | 1530 | 1208 | 1302 | 1655 | 1149 | 1080 | 1140 | 1116 | 1361 | 1312 | 1333 | 1262 | 1219,25 | 1121 | 1317 | 1,173 | 0,23 | 4,607 | 4,08E-06 | 6,61E-06 | 1,00E-08 | 0,00312 | 0,0015708 | 0,0015708 | TRUE | 0,010154 | No |
| orf19.6014 | 7947 | 6759 | 7524 | 3949 | 9258 | 6715 | 7864 | 9959 | 7235 | 6836 | 7121 | 5932 | 8235 | 7291 | 8054 | 7597 | 7287,51 | 6781 | 7794 | 1,149 | 0,2 | 3,08 | 0,002072 | 0,002856 | 0,0051263 | 0,002218 | 0,0039111 | 0,0039111 | TRUE | 0,0852701 | No |
| orf19.6016 | 1869 | 1828 | 1954 | 1200 | 1020 | 888 | 844 | 1064 | 1702 | 1849 | 1849 | 1803 | 907 | 964 | 864 | 812 | 1343,71 | 1801 | 887 | 0,492 | -1,024 | -16,84 | 1,23E-63 | 7,13E-63 | 0,0022828 | 0,00302 | 0,0027241 | 0,0027241 | TRUE | 0,0411009 | No |
| orf19.6017 | 506 | 451 | 494 | 313 | 247 | 219 | 228 | 274 | 461 | 456 | 468 | 470 | 220 | 238 | 234 | 209 | 344,31 | 464 | 225 | 0,484 | -1,048 | -12,983 | 1,53E-38 | 5,81E-38 | 1,00E-08 | 0,005874 | 0,0030275 | 0,0030275 | TRUE | 0,0280487 | No |
| orf19.6018 | 1476 | 1357 | 1524 | 906 | 2603 | 2055 | 2291 | 3069 | 1344 | 1372 | 1442 | 1361 | 2315 | 2231 | 2346 | 2341 | 1844,17 | 1380 | 2308 | 1,673 | 0,742 | 16,685 | 1,68E-62 | 9,52E-62 | 1,00E-08 | 0,002753 | 0,0013091 | 0,0013091 | TRUE | 0,0117704 | No |
| orf19.6020 | 1226 | 1367 | 1357 | 736 | 1155 | 946 | 1001 | 1290 | 1116 | 1383 | 1284 | 1106 | 1027 | 1027 | 1025 | 984 | 1119,03 | 1222 | 1016 | 0,83 | -0,269 | -3,66 | 0,000253 | 0,000371 | 0,0050879 | 0,003217 | 0,0043023 | 0,0043023 | TRUE | 0,093092 | No |
| orf19.6021 | 283 | 247 | 251 | 169 | 363 | 270 | 282 | 399 | 258 | 250 | 238 | 254 | 323 | 293 | 289 | 304 | 276,01 | 250 | 302 | 1,213 | 0,279 | 3,201 | 0,00137 | 0,001917 | 1,00E-08 | 0,006823 | 0,0036262 | 0,0036262 | TRUE | 0,023687 | No |
| orf19.6022 | 96 | 83 | 110 | 70 | 27 | 28 | 29 | 34 | 87 | 84 | 104 | 105 | 24 | 30 | 30 | 26 | 61,33 | 95 | 28 | 0,288 | -1,793 | -9,054 | 1,38E-19 | 3,45E-19 | 1,00E-08 | 0,023575 | 0,0150456 | 0,0150456 | TRUE | 0,0589165 | No |
| orf19.6023 | 1500 | 1423 | 1564 | 922 | 3755 | 3155 | 3219 | 4493 | 1366 | 1439 | 1480 | 1385 | 3340 | 3425 | 3297 | 3427 | 2394,93 | 1418 | 3372 | 2,377 | 1,249 | 28,972 | 1,48E-184 | 2,34E-183 | 0,0001923 | 0,002589 | 0,0012693 | 0,0012693 | TRUE | 0,0111061 | No |
| orf19.6024 | 282 | 308 | 323 | 183 | 176 | 141 | 139 | 194 | 257 | 312 | 306 | 275 | 157 | 153 | 142 | 148 | 218,6 | 288 | 150 | 0,522 | -0,939 | -8,98 | 2,71E-19 | 6,75E-19 | 1,00E-08 | 0,00808 | 0,0055259 | 0,0055259 | TRUE | 0,0656089 | No |
| orf19.6025 | 36 | 50 | 51 | 27 | 32 | 28 | 36 | 36 | 33 | 51 | 48 | 41 | 28 | 30 | 37 | 27 | 36,92 | 43 | 30 | 0,711 | -0,492 | -2,081 | 0,037475 | 0,046946 | 1,00E-08 | 0,037817 | 0,0261352 | 0,0261352 | TRUE | 0,2360697 | No |
| orf19.6026 | 166 | 188 | 183 | 112 | 273 | 250 | 245 | 337 | 151 | 190 | 173 | 168 | 243 | 271 | 251 | 257 | 213,12 | 170 | 256 | 1,497 | 0,582 | 5,648 | 1,62E-08 | 2,90E-08 | 1,00E-08 | 0,008235 | 0,0052513 | 0,0052513 | TRUE | 0,0731781 | No |
| orf19.6027 | 146 | 140 | 152 | 80 | 92 | 89 | 98 | 124 | 133 | 142 | 144 | 120 | 82 | 97 | 100 | 95 | 114 | 135 | 94 | 0,688 | -0,539 | -4,05 | 5,12E-05 | 7,79E-05 | 1,00E-08 | 0,013625 | 0,0081001 | 0,0081001 | TRUE | 0,0691055 | No |
| orf19.6028 | 332 | 342 | 333 | 214 | 31 | 22 | 33 | 30 | 302 | 346 | 315 | 321 | 28 | 24 | 34 | 23 | 174,12 | 321 | 27 | 0,084 | -3,581 | -22,04 | 1,20E-107 | 1,11E-106 | 1,00E-08 | 0,009624 | 0,0064305 | 0,0064305 | TRUE | 0,1672742 | No |
| orf19.6029 | 558 | 570 | 574 | 364 | 780 | 635 | 616 | 762 | 508 | 576 | 543 | 547 | 694 | 689 | 631 | 581 | 596,24 | 544 | 649 | 1,192 | 0,254 | 3,313 | 0,000924 | 0,001307 | 0,0036112 | 0,004253 | 0,0039355 | 0,0039355 | TRUE | 0,0620563 | No |
| orf19.603 | 12142 | 10075 | 10608 | 7014 | 13287 | 10172 | 10912 | 15488 | 11055 | 10190 | 10040 | 10536 | 11818 | 11044 | 11175 | 11814 | 10959 | 10455 | 11463 | 1,097 | 0,133 | 3,059 | 0,002221 | 0,003053 | 0,0014516 | 0,002158 | 0,0017245 | 0,0017245 | TRUE | 0,0185338 | No |
| orf19.6030 | 2492 | 2098 | 2276 | 1358 | 1048 | 779 | 924 | 1242 | 2269 | 2122 | 2154 | 2040 | 932 | 846 | 946 | 947 | 1532,05 | 2146 | 918 | 0,428 | -1,225 | -22,23 | 1,76E-109 | 1,65E-108 | 0,0013406 | 0,002899 | 0,0021587 | 0,0021587 | TRUE | 0,0316195 | No |
| orf19.6031 | 5005 | 4838 | 5076 | 3124 | 2496 | 2215 | 2264 | 2901 | 4557 | 4893 | 4804 | 4693 | 2220 | 2405 | 2319 | 2213 | 3512,9 | 4737 | 2289 | 0,483 | -1,05 | -24,077 | 4,32E-128 | 4,72E-127 | 0,0008281 | 0,002413 | 0,0015095 | 0,0015095 | TRUE | 0,0137886 | No |
| orf19.6032 | 4484 | 4487 | 4773 | 3010 | 6138 | 5433 | 5210 | 6846 | 4082 | 4538 | 4517 | 4521 | 5460 | 5899 | 5336 | 5222 | 4946,94 | 4414 | 5479 | 1,241 | 0,312 | 5,93 | 3,02E-09 | 5,56E-09 | 0,002449 | 0,002304 | 0,0024451 | 0,0024451 | TRUE | 0,0321659 | No |
| orf19.6033 | 1539 | 1396 | 1539 | 1004 | 2998 | 2428 | 2426 | 3401 | 1401 | 1412 | 1457 | 1508 | 2667 | 2636 | 2485 | 2594 | 2019,92 | 1444 | 2596 | 1,8 | 0,848 | 18,702 | 4,81E-78 | 3,33E-77 | 0,0004468 | 0,002691 | 0,001423 | 0,001423 | TRUE | 0,0097958 | No |
| orf19.6034 | 2073 | 1876 | 2014 | 1274 | 2202 | 1785 | 1833 | 2372 | 1887 | 1897 | 1906 | 1914 | 1959 | 1938 | 1877 | 1809 | 1898,47 | 1901 | 1896 | 0,997 | -0,005 | -0,107 | 0,914488 | 0,926797 | 1,00E-08 | 0,002733 | 0,0012838 | 0,0012838 | TRUE | 0,0121456 | No |
| orf19.6035 | 971 | 933 | 919 | 508 | 1676 | 1143 | 1279 | 1615 | 884 | 944 | 870 | 763 | 1491 | 1241 | 1310 | 1232 | 1091,76 | 865 | 1318 | 1,52 | 0,604 | 7,675 | 1,65E-14 | 3,61E-14 | 0,0066456 | 0,003247 | 0,0049738 | 0,0049738 | TRUE | 0,0942173 | No |
| orf19.6036 | 177 | 213 | 215 | 126 | 118 | 95 | 114 | 127 | 161 | 215 | 203 | 189 | 105 | 103 | 117 | 97 | 148,88 | 192 | 106 | 0,547 | -0,871 | -6,718 | 1,84E-11 | 3,68E-11 | 0,0061473 | 0,01091 | 0,0089107 | 0,0089107 | TRUE | 0,1437235 | No |
| orf19.6037 | 930 | 796 | 900 | 624 | 1606 | 1288 | 1350 | 1825 | 847 | 805 | 852 | 937 | 1428 | 1398 | 1383 | 1392 | 1130,31 | 860 | 1400 | 1,634 | 0,709 | 13,017 | 9,80E-39 | 3,73E-38 | 0,0001158 | 0,003206 | 0,0018879 | 0,0018879 | TRUE | 0,0369634 | No |
| orf19.6038 | 518 | 395 | 434 | 301 | 810 | 624 | 706 | 935 | 472 | 400 | 411 | 452 | 720 | 677 | 723 | 713 | 571,03 | 434 | 708 | 1,637 | 0,711 | 10,012 | 1,35E-23 | 3,76E-23 | 0,0010472 | 0,00435 | 0,0029564 | 0,0029564 | TRUE | 0,0475015 | No |
| orf19.6039 | 1325 | 1182 | 1298 | 851 | 1076 | 800 | 860 | 1245 | 1206 | 1195 | 1228 | 1278 | 957 | 869 | 881 | 950 | 1070,59 | 1227 | 914 | 0,747 | -0,42 | -7,668 | 1,75E-14 | 3,81E-14 | 0,0005886 | 0,003271 | 0,0019366 | 0,0019366 | TRUE | 0,0130571 | No |
| orf19.604 | 32 | 33 | 36 | 36 | 69 | 57 | 57 | 116 | 29 | 33 | 34 | 54 | 61 | 62 | 58 | 88 | 52,6 | 38 | 67 | 1,851 | 0,888 | 3,853 | 0,000117 | 0,000174 | 0,033624 | 0,027152 | 0,0296117 | 0,0296117 | TRUE | 0,4581508 | No |
| orf19.6040 | 5253 | 4686 | 5237 | 3037 | 4114 | 3100 | 3771 | 4710 | 4783 | 4739 | 4956 | 4562 | 3659 | 3366 | 3862 | 3593 | 4190,03 | 4760 | 3620 | 0,76 | -0,396 | -8,076 | 6,67E-16 | 1,51E-15 | 0,0018504 | 0,002352 | 0,002062 | 0,002062 | TRUE | 0,0265001 | No |
| orf19.6041 | 11197 | 10288 | 10978 | 7332 | 11931 | 10138 | 10708 | 14749 | 10194 | 10405 | 10390 | 11014 | 10612 | 11007 | 10966 | 11251 | 10729,9 | 10501 | 10959 | 1,044 | 0,062 | 1,696 | 0,089844 | 0,108374 | 0,0007361 | 0,00216 | 0,0012059 | 0,0012059 | TRUE | 0,0128503 | No |
| orf19.6043 | 525 | 426 | 481 | 326 | 883 | 695 | 769 | 1090 | 478 | 431 | 455 | 490 | 785 | 755 | 788 | 831 | 626,6 | 464 | 790 | 1,711 | 0,774 | 11,632 | 2,83E-31 | 9,37E-31 | 0,0005778 | 0,004145 | 0,0025115 | 0,0025115 | TRUE | 0,0255083 | No |
| orf19.6044 | 2775 | 2381 | 2564 | 1874 | 2550 | 2323 | 2312 | 3211 | 2527 | 2408 | 2427 | 2815 | 2268 | 2522 | 2368 | 2449 | 2472,96 | 2544 | 2402 | 0,945 | -0,081 | -1,384 | 0,166392 | 0,194644 | 0,0032237 | 0,002571 | 0,0028749 | 0,0028749 | TRUE | 0,05929 | No |
| orf19.6045 | 5504 | 4507 | 5259 | 3024 | 3137 | 2601 | 2944 | 3867 | 5011 | 4558 | 4977 | 4543 | 2790 | 2824 | 3015 | 2950 | 3833,53 | 4772 | 2895 | 0,606 | -0,722 | -14,609 | 2,46E-48 | 1,10E-47 | 0,0017408 | 0,002382 | 0,0020722 | 0,0020722 | TRUE | 0,0138159 | No |
| orf19.6046 | 984 | 800 | 816 | 565 | 664 | 572 | 633 | 810 | 896 | 809 | 772 | 849 | 591 | 621 | 648 | 618 | 725,47 | 832 | 620 | 0,745 | -0,425 | -6,449 | 1,13E-10 | 2,19E-10 | 0,0016903 | 0,003858 | 0,0027723 | 0,0027723 | TRUE | 0,0357349 | No |
| orf19.6047 | 8705 | 7055 | 8010 | 5150 | 12300 | 9884 | 10927 | 15257 | 7926 | 7135 | 7581 | 7736 | 10940 | 10731 | 11191 | 11638 | 9359,79 | 7594 | 11125 | 1,465 | 0,551 | 12,526 | 5,41E-36 | 1,97E-35 | 0,0014516 | 0,002178 | 0,0017464 | 0,0017464 | TRUE | 0,0203645 | No |
| orf19.6048 | 1440 | 1248 | 1314 | 871 | 1060 | 932 | 948 | 1251 | 1311 | 1262 | 1244 | 1308 | 943 | 1012 | 971 | 954 | 1125,64 | 1281 | 970 | 0,757 | -0,402 | -7,989 | 1,36E-15 | 3,07E-15 | 1,00E-08 | 0,003211 | 0,0015329 | 0,0015329 | TRUE | 0,0096226 | No |
| orf19.6049 | 2375 | 1921 | 2124 | 1461 | 1352 | 1170 | 1219 | 1662 | 2162 | 1943 | 2010 | 2195 | 1203 | 1270 | 1248 | 1268 | 1662,39 | 2078 | 1247 | 0,601 | -0,735 | -13,795 | 2,74E-43 | 1,13E-42 | 0,0015075 | 0,002832 | 0,0020891 | 0,0020891 | TRUE | 0,0228188 | No |
| orf19.605 | 85 | 79 | 85 | 76 | 174 | 176 | 138 | 251 | 77 | 80 | 80 | 114 | 155 | 191 | 141 | 191 | 128,82 | 88 | 170 | 1,961 | 0,972 | 6,221 | 4,94E-10 | 9,34E-10 | 0,0180524 | 0,012292 | 0,0145186 | 0,0145186 | TRUE | 0,2992949 | No |
| orf19.6052 | 3722 | 3277 | 3596 | 2358 | 3134 | 2568 | 2707 | 3583 | 3389 | 3314 | 3403 | 3542 | 2788 | 2788 | 2772 | 2733 | 3091,2 | 3412 | 2770 | 0,813 | -0,299 | -8,099 | 5,54E-16 | 1,26E-15 | 1,00E-08 | 0,002464 | 0,0009861 | 0,0009861 | TRUE | 0,0072704 | No |
| orf19.6053 | 2990 | 2660 | 3051 | 1801 | 7366 | 6135 | 6824 | 9104 | 2722 | 2690 | 2887 | 2705 | 6552 | 6661 | 6989 | 6945 | 4768,93 | 2751 | 6787 | 2,465 | 1,302 | 31,502 | 8,04E-218 | 1,47E-216 | 0,0007094 | 0,002314 | 0,0013755 | 0,0013755 | TRUE | 0,0138338 | No |
| orf19.6054 | 10249 | 8847 | 9859 | 6110 | 10254 | 8327 | 9039 | 12113 | 9331 | 8948 | 9331 | 9178 | 9121 | 9041 | 9257 | 9240 | 9180,79 | 9197 | 9165 | 0,996 | -0,005 | -0,172 | 0,863534 | 0,884152 | 0,000111 | 0,002181 | 0,0007482 | 0,0007482 | TRUE | 0,0041635 | No |
| orf19.6055 | 2592 | 2210 | 2410 | 1590 | 1244 | 1061 | 1128 | 1620 | 2360 | 2235 | 2281 | 2388 | 1106 | 1152 | 1155 | 1236 | 1739,22 | 2316 | 1162 | 0,503 | -0,992 | -19,916 | 2,96E-88 | 2,30E-87 | 0,0008922 | 0,002797 | 0,00175 | 0,00175 | TRUE | 0,0233626 | No |
| orf19.6056 | 2443 | 2053 | 2243 | 1248 | 1919 | 1459 | 1684 | 2209 | 2224 | 2076 | 2123 | 1875 | 1707 | 1584 | 1725 | 1685 | 1874,84 | 2074 | 1675 | 0,806 | -0,311 | -5,393 | 6,92E-08 | 1,20E-07 | 0,0024894 | 0,002742 | 0,0026536 | 0,0026536 | TRUE | 0,0475591 | No |
| orf19.6057 | 876 | 638 | 783 | 499 | 1540 | 1211 | 1334 | 1815 | 798 | 645 | 741 | 750 | 1370 | 1315 | 1366 | 1384 | 1046,09 | 734 | 1359 | 1,854 | 0,89 | 13,752 | 4,92E-43 | 2,02E-42 | 0,0026323 | 0,0033 | 0,0029485 | 0,0029485 | TRUE | 0,0802048 | No |
| orf19.6058 | 1234 | 956 | 1007 | 591 | 1257 | 915 | 1071 | 1446 | 1124 | 967 | 953 | 888 | 1118 | 993 | 1097 | 1103 | 1030,32 | 983 | 1078 | 1,095 | 0,13 | 1,757 | 0,078967 | 0,095833 | 0,0052708 | 0,003319 | 0,0043018 | 0,0043018 | TRUE | 0,1125961 | No |
| orf19.6059 | 7134 | 5375 | 5420 | 4169 | 3933 | 3270 | 2952 | 4556 | 6495 | 5436 | 5130 | 6263 | 3498 | 3550 | 3023 | 3475 | 4608,83 | 5831 | 3386 | 0,581 | -0,783 | -9,967 | 2,13E-23 | 5,88E-23 | 0,0085439 | 0,002324 | 0,0057029 | 0,0057029 | TRUE | 0,0803363 | No |
| orf19.606 | 5343 | 4866 | 5310 | 3504 | 4202 | 3849 | 3906 | 5228 | 4865 | 4921 | 5025 | 5264 | 3738 | 4179 | 4000 | 3988 | 4497,46 | 5019 | 3976 | 0,793 | -0,335 | -7,383 | 1,55E-13 | 3,30E-13 | 0,0013576 | 0,002331 | 0,0017575 | 0,0017575 | TRUE | 0,0199009 | No |
| orf19.6060 | 2821 | 2402 | 2661 | 1613 | 3894 | 3082 | 3679 | 4814 | 2568 | 2429 | 2518 | 2423 | 3464 | 3346 | 3768 | 3672 | 3023,61 | 2484 | 3562 | 1,433 | 0,519 | 10,629 | 2,19E-26 | 6,46E-26 | 0,001404 | 0,002474 | 0,0019428 | 0,0019428 | TRUE | 0,0198543 | No |
| orf19.6061 | 1810 | 1587 | 1735 | 1030 | 1871 | 1504 | 1484 | 2131 | 1648 | 1605 | 1642 | 1547 | 1664 | 1633 | 1520 | 1626 | 1610,59 | 1610 | 1611 | 0,999 | -0,001 | -0,031 | 0,975645 | 0,977946 | 0,0004849 | 0,002857 | 0,0015668 | 0,0015668 | TRUE | 0,0169244 | No |
| orf19.6062 | 5416 | 4374 | 4552 | 2898 | 2237 | 1555 | 1682 | 2387 | 4931 | 4424 | 4308 | 4353 | 1990 | 1688 | 1723 | 1821 | 3154,7 | 4504 | 1806 | 0,401 | -1,318 | -20,814 | 3,20E-96 | 2,66E-95 | 0,0043376 | 0,002456 | 0,0034748 | 0,0034748 | TRUE | 0,0570417 | No |
| orf19.6062.3 | 1198 | 929 | 1033 | 725 | 1643 | 1296 | 1390 | 1901 | 1091 | 940 | 978 | 1089 | 1461 | 1407 | 1424 | 1450 | 1229,89 | 1024 | 1436 | 1,404 | 0,49 | 8,344 | 7,16E-17 | 1,67E-16 | 0,0018864 | 0,003111 | 0,0024598 | 0,0024598 | TRUE | 0,0367936 | No |
| orf19.6063 | 7959 | 7195 | 7891 | 4519 | 6169 | 4790 | 5487 | 6995 | 7246 | 7277 | 7468 | 6788 | 5487 | 5201 | 5619 | 5336 | 6302,83 | 7195 | 5411 | 0,752 | -0,412 | -9,701 | 2,98E-22 | 7,97E-22 | 0,0011514 | 0,002247 | 0,0015709 | 0,0015709 | TRUE | 0,0170649 | No |
| orf19.6064 | 1716 | 1499 | 1623 | 1091 | 2635 | 2052 | 2185 | 3089 | 1562 | 1516 | 1536 | 1639 | 2344 | 2228 | 2238 | 2356 | 1927,37 | 1563 | 2292 | 1,469 | 0,555 | 12,275 | 1,23E-34 | 4,38E-34 | 0,000315 | 0,002722 | 0,0014164 | 0,0014164 | TRUE | 0,0111963 | No |
| orf19.6065 | 1117 | 1020 | 1061 | 615 | 4151 | 3318 | 3417 | 4817 | 1017 | 1032 | 1004 | 924 | 3692 | 3602 | 3499 | 3674 | 2305,63 | 994 | 3617 | 3,627 | 1,859 | 39,711 | 0 | 0 | 0,0003829 | 0,00261 | 0,0014386 | 0,0014386 | TRUE | 0,0226471 | No |
| orf19.6066 | 1586 | 1433 | 1488 | 980 | 10185 | 8946 | 8537 | 11955 | 1444 | 1449 | 1408 | 1472 | 9059 | 9713 | 8743 | 9119 | 5301,01 | 1443 | 9158 | 6,349 | 2,667 | 58,799 | 0 | 0 | 0,0011229 | 0,002286 | 0,0015518 | 0,0015518 | TRUE | 0,0200373 | No |
| orf19.6068 | 9829 | 8271 | 8910 | 5979 | 10712 | 8976 | 9040 | 12780 | 8949 | 8365 | 8433 | 8981 | 9528 | 9745 | 9258 | 9749 | 9126,03 | 8682 | 9570 | 1,103 | 0,141 | 3,641 | 0,000271 | 0,000397 | 0,0008631 | 0,002182 | 0,0013293 | 0,0013293 | TRUE | 0,0073318 | No |
| orf19.607 | 4544 | 3904 | 4436 | 2690 | 4892 | 4041 | 4721 | 6242 | 4137 | 3948 | 4198 | 4041 | 4351 | 4387 | 4835 | 4761 | 4332,46 | 4081 | 4584 | 1,123 | 0,167 | 3,535 | 0,000408 | 0,00059 | 0,0015376 | 0,002342 | 0,0019207 | 0,0019207 | TRUE | 0,0162496 | No |
| orf19.6070 | 9478 | 8431 | 9024 | 6144 | 21102 | 18243 | 19188 | 27211 | 8629 | 8527 | 8540 | 9229 | 18770 | 19807 | 19651 | 20757 | 14238,8 | 8731 | 19746 | 2,263 | 1,178 | 27,256 | 1,43E-163 | 2,03E-162 | 0,0014753 | 0,00213 | 0,0017088 | 0,0017088 | TRUE | 0,0175359 | No |
| orf19.6071 | 2626 | 2530 | 2726 | 1726 | 3027 | 2597 | 2725 | 3478 | 2391 | 2559 | 2580 | 2593 | 2692 | 2820 | 2791 | 2653 | 2634,77 | 2531 | 2739 | 1,083 | 0,114 | 2,599 | 0,00934 | 0,012274 | 0,0007691 | 0,002538 | 0,0014777 | 0,0014777 | TRUE | 0,0173395 | No |
| orf19.6072 | 370 | 333 | 346 | 232 | 793 | 610 | 670 | 989 | 337 | 337 | 327 | 349 | 705 | 662 | 686 | 754 | 519,73 | 338 | 702 | 2,089 | 1,063 | 14,703 | 6,17E-49 | 2,79E-48 | 1,00E-08 | 0,004579 | 0,0027727 | 0,0027727 | TRUE | 0,031435 | No |
| orf19.6072.1 | 852 | 823 | 833 | 627 | 378 | 286 | 286 | 336 | 776 | 832 | 788 | 942 | 336 | 311 | 293 | 256 | 566,78 | 834 | 299 | 0,358 | -1,48 | -15,998 | 1,32E-57 | 6,94E-57 | 0,0080269 | 0,004368 | 0,0060056 | 0,0060056 | TRUE | 0,1156934 | No |

| | | | | | | | | | | | | | | | | | | | | | | | | | | | | | | | |
|---|---|---|---|---|---|---|---|---|---|---|---|---|---|---|---|---|---|---|---|---|---|---|---|---|---|---|---|---|---|---|---|
| orf19.6073 | 25710 | 21227 | 23025 | 14898 | 16866 | 13717 | 14300 | 20124 | 23408 | 21469 | 21791 | 22379 | 15002 | 14893 | 14645 | 15351 | 18617,2 | 22262 | 14973 | 0,673 | -0,572 | -15,396 | 1,75E-53 | 8,64E-53 | 0,0008493 | 0,002108 | 0,0012711 | 0,0012711 | TRUE | 0,0148557 | No |
| orf19.6074 | 5195 | 4332 | 4447 | 2619 | 4817 | 3472 | 4308 | 5494 | 4730 | 4381 | 4209 | 3934 | 4285 | 3770 | 4412 | 4191 | 4238,88 | 4314 | 4164 | 0,965 | -0,052 | -0,796 | 0,425791 | 0,467037 | 0,0046072 | 0,002349 | 0,0037956 | 0,0037956 | TRUE | 0,0515156 | No |
| orf19.6075 | 1355 | 1126 | 1171 | 928 | 1285 | 1111 | 992 | 1542 | 1234 | 1139 | 1108 | 1394 | 1143 | 1206 | 1016 | 1176 | 1177,02 | 1219 | 1135 | 0,934 | -0,098 | -1,247 | 0,212356 | 0,244893 | 0,0071419 | 0,003159 | 0,0051055 | 0,0051055 | TRUE | 0,1058456 | No |
| orf19.6076 | 569 | 469 | 532 | 399 | 343 | 300 | 303 | 466 | 518 | 474 | 503 | 599 | 305 | 326 | 310 | 355 | 423,98 | 524 | 324 | 0,623 | -0,683 | -7,715 | 1,21E-14 | 2,65E-14 | 0,0049519 | 0,005153 | 0,0050625 | 0,0050625 | TRUE | 0,0972638 | No |
| orf19.6077 | 2331 | 1919 | 2112 | 1586 | 2062 | 1889 | 1872 | 2615 | 2122 | 1941 | 1999 | 2382 | 1834 | 2051 | 1917 | 1995 | 2030,16 | 2111 | 1949 | 0,925 | -0,113 | -1,696 | 0,089952 | 0,108463 | 0,0048264 | 0,002688 | 0,0037491 | 0,0037491 | TRUE | 0,0863255 | No |
| orf19.6078 | 3114 | 2668 | 2897 | 1646 | 1492 | 1164 | 1281 | 1631 | 2835 | 2698 | 2742 | 2473 | 1327 | 1264 | 1312 | 1244 | 1986,85 | 2687 | 1287 | 0,478 | -1,065 | -20,172 | 1,73E-90 | 1,36E-89 | 0,0016544 | 0,002702 | 0,0021131 | 0,0021131 | TRUE | 0,0328305 | No |
| orf19.6078.1 | 0 | 4 | 3 | 3 | 1 | 1 | 0 | 0 | 0 | 4 | 3 | 5 | 1 | 1 | 0 | 0 | 1,67 | 3 | 0 | 0,17 | -2,556 | -1,852 | 0,064068 | 0,078561 | 1,00E-08 | 0,792671 | 0,7009287 | 0,7009287 | TRUE | 0,6611441 | No |
| orf19.6079 | 468 | 366 | 354 | 245 | 301 | 289 | 284 | 368 | 426 | 370 | 335 | 368 | 268 | 314 | 291 | 281 | 331,55 | 375 | 288 | 0,766 | -0,384 | -4,071 | 4,68E-05 | 7,15E-05 | 0,0048279 | 0,006021 | 0,0054801 | 0,0054801 | TRUE | 0,1051856 | No |
| orf19.6080 | 3012 | 2491 | 2717 | 1655 | 3303 | 2793 | 3028 | 4115 | 2742 | 2519 | 2571 | 2486 | 2938 | 3032 | 3101 | 3139 | 2816,19 | 2580 | 3052 | 1,182 | 0,241 | 5,317 | 1,05E-07 | 1,82E-07 | 0,0009155 | 0,002506 | 0,0016168 | 0,0016168 | TRUE | 0,0222584 | No |
| orf19.6081 | 13887 | 11419 | 12710 | 8623 | 15340 | 13016 | 13806 | 19821 | 12644 | 11549 | 12029 | 12953 | 13644 | 14132 | 14139 | 15120 | 13276,2 | 12294 | 14259 | 1,16 | 0,214 | 4,453 | 8,45E-06 | 1,35E-05 | 0,0021441 | 0,002137 | 0,0021492 | 0,0021492 | TRUE | 0,0203557 | No |
| orf19.6082 | 4258 | 3623 | 3947 | 2541 | 6373 | 5170 | 5557 | 7380 | 3877 | 3664 | 3735 | 3817 | 5669 | 5613 | 5691 | 5630 | 4711,98 | 3773 | 5651 | 1,498 | 0,583 | 17,399 | 8,33E-68 | 5,08E-67 | 2,86E-05 | 0,002317 | 0,0008519 | 0,0008519 | TRUE | 0,0046831 | No |
| orf19.6083 | 613 | 539 | 564 | 357 | 803 | 623 | 682 | 899 | 558 | 545 | 534 | 536 | 714 | 676 | 698 | 686 | 618,52 | 543 | 694 | 1,276 | 0,352 | 5,805 | 6,46E-09 | 1,17E-08 | 1,00E-08 | 0,004173 | 0,0018714 | 0,0018714 | TRUE | 0,0047907 | No |
| orf19.6084 | 11794 | 10222 | 10864 | 7148 | 3937 | 3315 | 3471 | 4668 | 10738 | 10338 | 10282 | 10737 | 3502 | 3599 | 3555 | 3561 | 7039,02 | 10524 | 3554 | 0,338 | -1,566 | -47,278 | 0 | 0 | 0,0002628 | 0,002225 | 0,0008708 | 0,0008708 | TRUE | 0,0029464 | No |
| orf19.6085 | 18708 | 15845 | 16961 | 10695 | 32848 | 25488 | 27249 | 36783 | 17033 | 16026 | 16052 | 16066 | 29217 | 27673 | 27906 | 28058 | 22253,9 | 16294 | 28214 | 1,731 | 0,792 | 22,625 | 2,48E-113 | 2,41E-112 | 0,0006898 | 0,002096 | 0,0011272 | 0,0011272 | TRUE | 0,0115131 | No |
| orf19.6086 | 486 | 386 | 406 | 278 | 1059 | 876 | 925 | 1349 | 442 | 390 | 384 | 418 | 942 | 951 | 947 | 1029 | 688,01 | 408 | 967 | 2,371 | 1,245 | 18,112 | 2,57E-73 | 1,68E-72 | 0,0013793 | 0,003957 | 0,0027494 | 0,0027494 | TRUE | 0,0414295 | No |
| orf19.609 | 48 | 45 | 45 | 37 | 71 | 60 | 39 | 68 | 44 | 46 | 43 | 56 | 63 | 65 | 40 | 52 | 50,94 | 47 | 55 | 1,187 | 0,247 | 1,184 | 0,236482 | 0,270611 | 0,0106346 | 0,027971 | 0,0218689 | 0,0218689 | TRUE | 0,2649309 | No |
| orf19.6090 | 31343 | 25417 | 27699 | 18183 | 57383 | 45314 | 50422 | 69995 | 28536 | 25707 | 26215 | 27314 | 51040 | 49198 | 51638 | 53393 | 39130,2 | 26943 | 51317 | 1,905 | 0,93 | 21,445 | 5,07E-102 | 4,51E-101 | 0,0016178 | 0,002071 | 0,0017762 | 0,0017762 | TRUE | 0,019549 | No |
| orf19.6091 | 4651 | 3889 | 4425 | 2969 | 3736 | 3082 | 3346 | 4679 | 4235 | 3933 | 4188 | 4460 | 3323 | 3346 | 3427 | 3569 | 3810,1 | 4204 | 3416 | 0,814 | -0,298 | -6,284 | 3,30E-10 | 6,29E-10 | 0,0014958 | 0,002384 | 0,0018901 | 0,0018901 | TRUE | 0,0226754 | No |
| orf19.6092 | 1430 | 1173 | 1358 | 933 | 2913 | 2445 | 2718 | 3674 | 1302 | 1186 | 1285 | 1402 | 2591 | 2655 | 2784 | 2803 | 2000,85 | 1294 | 2708 | 2,098 | 1,069 | 19,258 | 1,21E-82 | 8,87E-82 | 0,0022659 | 0,002697 | 0,00237 | 0,00237 | TRUE | 0,0366977 | No |
| orf19.6094 | 1082 | 863 | 927 | 545 | 1850 | 1326 | 1588 | 2228 | 985 | 873 | 877 | 819 | 1646 | 1440 | 1626 | 1700 | 1245,62 | 888 | 1603 | 1,801 | 0,849 | 12,048 | 1,99E-33 | 6,87E-33 | 0,0044 | 0,003098 | 0,0038695 | 0,0038695 | TRUE | 0,0648188 | No |
| orf19.6096 | 768 | 604 | 663 | 410 | 1259 | 961 | 1056 | 1587 | 699 | 611 | 627 | 616 | 1120 | 1043 | 1081 | 1211 | 876,09 | 638 | 1114 | 1,744 | 0,802 | 11,645 | 2,44E-31 | 8,09E-31 | 0,0030679 | 0,003545 | 0,0033014 | 0,0033014 | TRUE | 0,0510233 | No |
| orf19.6099 | 5230 | 3867 | 4519 | 3373 | 6789 | 5635 | 5992 | 8822 | 4762 | 3911 | 4277 | 5067 | 6039 | 6118 | 6137 | 6729 | 5379,88 | 4504 | 6256 | 1,39 | 0,475 | 6,442 | 1,18E-10 | 2,29E-10 | 0,0075034 | 0,002283 | 0,0050283 | 0,0050283 | TRUE | 0,0955304 | No |
| orf19.610 | 7007 | 5678 | 6610 | 4667 | 3473 | 3209 | 3472 | 4858 | 6380 | 5743 | 6256 | 7011 | 3089 | 3484 | 3556 | 3706 | 4902,91 | 6348 | 3459 | 0,545 | -0,875 | -12,658 | 1,02E-36 | 3,76E-36 | 0,0059669 | 0,002306 | 0,0043719 | 0,0043719 | TRUE | 0,0634134 | No |
| orf19.6100 | 689 | 486 | 517 | 380 | 1222 | 936 | 1026 | 1582 | 627 | 492 | 489 | 571 | 1087 | 1016 | 1051 | 1207 | 817,45 | 545 | 1090 | 2,005 | 1,003 | 11,392 | 4,57E-30 | 1,47E-29 | 0,0085843 | 0,003653 | 0,0060333 | 0,0060333 | TRUE | 0,1324109 | No |
| orf19.6102 | 1897 | 1533 | 1858 | 1059 | 1324 | 882 | 1196 | 1633 | 1727 | 1550 | 1758 | 1591 | 1178 | 958 | 1225 | 1246 | 1404,08 | 1656 | 1152 | 0,695 | -0,524 | -6,52 | 7,03E-11 | 1,37E-10 | 0,0075005 | 0,002978 | 0,0054745 | 0,0054745 | TRUE | 0,1505398 | No |
| orf19.6103 | 353 | 300 | 332 | 223 | 285 | 254 | 291 | 411 | 321 | 303 | 314 | 335 | 253 | 276 | 298 | 314 | 301,85 | 318 | 285 | 0,899 | -0,154 | -1,693 | 0,090381 | 0,10896 | 0,0022023 | 0,006413 | 0,0046147 | 0,0046147 | TRUE | 0,0674946 | No |
| orf19.6105 | 1705 | 1139 | 1382 | 1107 | 1222 | 1033 | 1040 | 1556 | 1552 | 1152 | 1308 | 1663 | 1087 | 1122 | 1065 | 1187 | 1266,95 | 1419 | 1115 | 0,787 | -0,345 | -3,543 | 0,000396 | 0,000573 | 0,0133926 | 0,00308 | 0,0082952 | 0,0082952 | TRUE | 0,192421 | No |
| orf19.6109 | 6045 | 4454 | 5209 | 3291 | 7005 | 5464 | 6571 | 9332 | 5504 | 4505 | 4930 | 4944 | 6231 | 5932 | 6730 | 7119 | 5736,63 | 4971 | 6503 | 1,308 | 0,388 | 5,529 | 3,23E-08 | 5,68E-08 | 0,0064319 | 0,002267 | 0,0045451 | 0,0045451 | TRUE | 0,0642808 | No |
| orf19.6110 | 10 | 5 | 5 | 4 | 20 | 8 | 12 | 8 | 9 | 5 | 5 | 6 | 18 | 9 | 12 | 6 | 8,72 | 6 | 11 | 1,783 | 0,834 | 1,631 | 0,102867 | 0,123453 | 1,00E-08 | 0,153507 | 0,1247137 | 0,1247137 | TRUE | 0,3351673 | No |
| orf19.6112 | 322 | 290 | 304 | 180 | 440 | 330 | 429 | 556 | 293 | 293 | 288 | 270 | 391 | 358 | 439 | 424 | 344,71 | 286 | 403 | 1,407 | 0,493 | 5,653 | 1,58E-08 | 2,82E-08 | 0,0022511 | 0,005869 | 0,0042908 | 0,0042908 | TRUE | 0,0606406 | No |
| orf19.6113 | 90 | 64 | 77 | 37 | 242 | 154 | 237 | 284 | 82 | 65 | 73 | 56 | 215 | 167 | 243 | 217 | 139,62 | 69 | 210 | 3,024 | 1,597 | 10,358 | 3,83E-25 | 1,10E-24 | 0,0152822 | 0,011499 | 0,0129502 | 0,0129502 | TRUE | 0,1969574 | No |
| orf19.6114 | 2421 | 1762 | 2145 | 1200 | 4022 | 2848 | 3501 | 4909 | 2204 | 1782 | 2030 | 1803 | 3577 | 3092 | 3585 | 3745 | 2727,32 | 1955 | 3500 | 1,789 | 0,839 | 10,787 | 3,96E-27 | 1,19E-26 | 0,0079994 | 0,002521 | 0,0054039 | 0,0054039 | TRUE | 0,0899557 | No |
| orf19.6115 | 4 | 0 | 2 | 0 | 10 | 7 | 2 | 2 | 4 | 0 | 2 | 0 | 9 | 8 | 2 | 2 | 3,2 | 2 | 5 | 3,31 | 1,727 | 1,751 | 0,079864 | 0,096848 | 0,6387435 | 0,414796 | 0,4856286 | 0,4856286 | TRUE | 0,2649989 | No |
| orf19.6116 | 2015 | 1700 | 1939 | 1119 | 269 | 201 | 240 | 301 | 1835 | 1719 | 1835 | 1681 | 239 | 218 | 246 | 230 | 1000,35 | 1768 | 233 | 0,132 | -2,923 | -42,588 | 0 | 0 | 1,00E-08 | 0,003358 | 0,0022302 | 0,0022302 | TRUE | 0,0176329 | No |
| orf19.6117 | 8545 | 6833 | 7600 | 5667 | 2451 | 2154 | 2243 | 3236 | 7780 | 6911 | 7193 | 8513 | 2180 | 2339 | 2297 | 2468 | 4960,06 | 7599 | 2321 | 0,306 | -1,71 | -25,56 | 4,29E-144 | 5,38E-143 | 0,0052866 | 0,002303 | 0,0040272 | 0,0040272 | TRUE | 0,0789149 | No |
| orf19.6118 | 350 | 301 | 279 | 176 | 312 | 239 | 228 | 324 | 319 | 304 | 264 | 264 | 278 | 259 | 234 | 247 | 271,15 | 288 | 254 | 0,881 | -0,183 | -1,864 | 0,062294 | 0,076473 | 0,0038042 | 0,006909 | 0,0056025 | 0,0056025 | TRUE | 0,0603655 | No |
| orf19.6119 | 243 | 191 | 239 | 164 | 249 | 246 | 252 | 396 | 221 | 193 | 226 | 246 | 221 | 267 | 258 | 302 | 241,96 | 222 | 262 | 1,189 | 0,25 | 2,184 | 0,028936 | 0,036616 | 0,0094303 | 0,007497 | 0,0083488 | 0,0083488 | TRUE | 0,1289662 | No |
| orf19.612 | 532 | 501 | 447 | 325 | 618 | 452 | 488 | 732 | 484 | 507 | 423 | 488 | 550 | 491 | 500 | 558 | 500,11 | 476 | 525 | 1,107 | 0,147 | 1,851 | 0,064227 | 0,078739 | 0,0032951 | 0,004678 | 0,0040386 | 0,0040386 | TRUE | 0,0679308 | No |
| orf19.6121 | 8452 | 7340 | 8294 | 5299 | 1852 | 1480 | 1661 | 2207 | 7695 | 7424 | 7850 | 7960 | 1647 | 1607 | 1701 | 1684 | 4695,88 | 7732 | 1660 | 0,215 | -2,219 | -55,15 | 0 | 0 | 0,0005477 | 0,002318 | 0,0012035 | 0,0012035 | TRUE | 0,0087814 | No |
| orf19.6124 | 1746 | 1630 | 1700 | 1077 | 840 | 662 | 727 | 974 | 1590 | 1649 | 1609 | 1618 | 747 | 719 | 745 | 743 | 1177,3 | 1616 | 738 | 0,457 | -1,129 | -23,334 | 2,00E-120 | 2,05E-119 | 1,00E-08 | 0,003159 | 0,001286 | 0,001286 | TRUE | 0,0034786 | No |
| orf19.6126 | 5981 | 5076 | 5667 | 3410 | 11030 | 8665 | 9173 | 12843 | 5445 | 5134 | 5363 | 5122 | 9811 | 9408 | 9394 | 9797 | 7434,33 | 5266 | 9602 | 1,823 | 0,866 | 23,484 | 6,00E-122 | 6,20E-121 | 0,0005769 | 0,002215 | 0,001155 | 0,001155 | TRUE | 0,0064298 | No |
| orf19.6127 | 19612 | 15662 | 17550 | 12195 | 20377 | 16587 | 18405 | 25127 | 17856 | 15840 | 16609 | 18319 | 18125 | 18009 | 18849 | 19167 | 17846,8 | 17156 | 18538 | 1,081 | 0,112 | 2,214 | 0,02682 | 0,034026 | 0,0025749 | 0,002111 | 0,0024032 | 0,0024032 | TRUE | 0,032586 | No |
| orf19.6129 | 2108 | 1831 | 1970 | 1176 | 2000 | 1475 | 1609 | 2208 | 1919 | 1852 | 1864 | 1767 | 1779 | 1601 | 1648 | 1684 | 1764,32 | 1850 | 1678 | 0,906 | -0,142 | -2,881 | 0,00396 | 0,00534 | 0,0008312 | 0,002786 | 0,0017664 | 0,0017664 | TRUE | 0,0196678 | No |
| orf19.6131 | 781 | 596 | 715 | 513 | 546 | 558 | 549 | 769 | 711 | 603 | 677 | 771 | 486 | 606 | 562 | 587 | 625,18 | 690 | 560 | 0,813 | -0,299 | -3,355 | 0,000794 | 0,001128 | 0,0080674 | 0,00415 | 0,0059997 | 0,0059997 | TRUE | 0,0954162 | No |
| orf19.6132 | 1463 | 1224 | 1291 | 744 | 1341 | 983 | 1047 | 1453 | 1332 | 1238 | 1222 | 1118 | 1193 | 1067 | 1072 | 1108 | 1168,75 | 1228 | 1110 | 0,903 | -0,147 | -2,317 | 0,020493 | 0,026238 | 0,0027829 | 0,003167 | 0,0030323 | 0,0030323 | TRUE | 0,0395853 | No |
| orf19.6133 | 4407 | 3780 | 4345 | 2519 | 3607 | 3012 | 3385 | 4562 | 4012 | 3823 | 4112 | 3784 | 3208 | 3270 | 3467 | 3480 | 3644,58 | 3933 | 3356 | 0,853 | -0,229 | -4,981 | 6,34E-07 | 1,06E-06 | 0,0012553 | 0,002399 | 0,0017621 | 0,0017621 | TRUE | 0,01143 | No |
| orf19.6134 | 805 | 720 | 793 | 506 | 313 | 281 | 253 | 347 | 733 | 728 | 751 | 760 | 278 | 305 | 259 | 265 | 509,88 | 743 | 277 | 0,372 | -1,427 | -19,769 | 5,48E-87 | 4,21E-86 | 1,00E-08 | 0,004628 | 0,0025941 | 0,0025941 | TRUE | 0,0481022 | No |
| orf19.6135 | 1003 | 840 | 941 | 598 | 1159 | 962 | 1039 | 1391 | 913 | 850 | 891 | 898 | 1031 | 1044 | 1064 | 1061 | 969,01 | 888 | 1050 | 1,183 | 0,242 | 4,695 | 2,66E-06 | 4,34E-06 | 1,00E-08 | 0,0034 | 0,0015131 | 0,0015131 | TRUE | 0,0104247 | No |
| orf19.6135.1 | 981 | 836 | 792 | 465 | 776 | 478 | 614 | 834 | 893 | 846 | 750 | 699 | 690 | 519 | 629 | 636 | 707,62 | 797 | 618 | 0,776 | -0,366 | -3,8 | 0,000145 | 0,000215 | 0,0110087 | 0,003904 | 0,0074994 | 0,0074994 | TRUE | 0,1340958 | No |
| orf19.6136 | 4031 | 3178 | 3607 | 2111 | 2292 | 1758 | 1936 | 2714 | 3670 | 3214 | 3414 | 3171 | 2039 | 1909 | 1983 | 2070 | 2683,67 | 3367 | 2000 | 0,594 | -0,752 | -13,571 | 5,95E-42 | 2,40E-41 | 0,002386 | 0,002529 | 0,0025553 | 0,0025553 | TRUE | 0,0449353 | No |
| orf19.6137 | 2891 | 2544 | 2778 | 1702 | 3424 | 2713 | 2919 | 3703 | 2632 | 2573 | 2629 | 2557 | 3046 | 2946 | 2989 | 2825 | 2774,52 | 2598 | 2952 | 1,135 | 0,183 | 4,555 | 5,24E-06 | 8,43E-06 | 0,0003814 | 0,002513 | 0,0011794 | 0,0011794 | TRUE | 0,0106802 | No |
| orf19.6138 | 44 | 37 | 67 | 27 | 34 | 46 | 33 | 56 | 40 | 37 | 63 | 41 | 30 | 50 | 34 | 43 | 42,27 | 45 | 39 | 0,857 | -0,223 | -0,908 | 0,364027 | 0,403876 | 0,036032 | 0,03329 | 0,0343666 | 0,0343666 | TRUE | 0,6041142 | No |
| orf19.6139 | 2189 | 1950 | 2103 | 1322 | 4018 | 3264 | 3672 | 5148 | 1993 | 1972 | 1990 | 1986 | 3574 | 3544 | 3761 | 3927 | 2843,32 | 1985 | 3702 | 1,865 | 0,899 | 19,723 | 1,38E-86 | 1,05E-85 | 0,0010555 | 0,002502 | 0,0015994 | 0,0015994 | TRUE | 0,0210716 | No |
| orf19.6140 | 889 | 857 | 955 | 620 | 1892 | 1627 | 1586 | 2238 | 809 | 867 | 904 | 931 | 1683 | 1766 | 1624 | 1707 | 1286,51 | 878 | 1695 | 1,936 | 0,953 | 16,952 | 1,87E-64 | 1,09E-63 | 0,0012556 | 0,003064 | 0,0021489 | 0,0021489 | TRUE | 0,0338829 | No |
| orf19.6141 | 867 | 764 | 847 | 549 | 2209 | 1766 | 1861 | 2647 | 789 | 773 | 802 | 825 | 1965 | 1917 | 1906 | 2019 | 1374,45 | 797 | 1952 | 2,455 | 1,296 | 26,324 | 1,03E-152 | 1,36E-151 | 1,00E-08 | 0,002998 | 0,001416 | 0,001416 | TRUE | 0,0067841 | No |
| orf19.6142 | 13 | 6 | 5 | 4 | 18 | 18 | 22 | 33 | 12 | 6 | 5 | 6 | 16 | 20 | 23 | 25 | 13,99 | 7 | 21 | 2,863 | 1,518 | 3,683 | 0,00023 | 0,000339 | 1,00E-08 | 0,096477 | 0,0679955 | 0,0679955 | TRUE | 0,6232164 | No |
| orf19.6143 | 4771 | 3909 | 4233 | 2864 | 9524 | 8087 | 8747 | 12128 | 4344 | 3954 | 4006 | 4302 | 8471 | 8780 | 8958 | 9251 | 6508,32 | 4152 | 8865 | 2,137 | 1,095 | 23,677 | 6,22E-124 | 6,54E-123 | 0,0016035 | 0,00224 | 0,0018724 | 0,0018724 | TRUE | 0,0125152 | No |
| orf19.6146 | 12158 | 9752 | 10461 | 7078 | 10248 | 8122 | 8337 | 11941 | 11069 | 9863 | 9900 | 10632 | 9115 | 8818 | 8538 | 9109 | 9630,68 | 10366 | 8895 | 0,858 | -0,22 | -4,647 | 3,36E-06 | 5,46E-06 | 0,0019455 | 0,002174 | 0,0020577 | 0,0020577 | TRUE | 0,0259202 | No |
| orf19.6147 | 729 | 632 | 621 | 317 | 976 | 559 | 791 | 981 | 664 | 639 | 588 | 476 | 868 | 607 | 810 | 748 | 675,03 | 592 | 758 | 1,278 | 0,354 | 3,072 | 0,002129 | 0,002931 | 0,0188396 | 0,003994 | 0,0112471 | 0,0112471 | TRUE | 0,2097376 | No |
| orf19.6148 | 352 | 341 | 403 | 245 | 243 | 193 | 250 | 247 | 320 | 345 | 381 | 368 | 216 | 210 | 256 | 188 | 285,62 | 354 | 218 | 0,614 | -0,704 | -6,701 | 2,07E-11 | 4,12E-11 | 0,0072915 | 0,006662 | 0,0069295 | 0,0069295 | TRUE | 0,1537025 | No |
| orf19.6151 | 2333 | 1833 | 1963 | 1225 | 1485 | 1149 | 1202 | 1687 | 2124 | 1854 | 1858 | 1840 | 1321 | 1247 | 1231 | 1287 | 1595,27 | 1919 | 1272 | 0,662 | -0,595 | -10,123 | 4,36E-24 | 1,22E-23 | 0,0025064 | 0,002865 | 0,0026647 | 0,0026647 | TRUE | 0,0642487 | No |
| orf19.6152 | 2487 | 2149 | 2511 | 1445 | 2347 | 1956 | 2069 | 2758 | 2264 | 2173 | 2376 | 2171 | 2088 | 2124 | 2119 | 2104 | 2177,36 | 2246 | 2109 | 0,937 | -0,093 | -2,124 | 0,033647 | 0,042332 | 0,0004883 | 0,002644 | 0,0013882 | 0,0013882 | TRUE | 0,0186678 | No |
| orf19.6155 | 4935 | 4521 | 4708 | 2965 | 4233 | 3559 | 3620 | 4863 | 4493 | 4573 | 4456 | 4454 | 3765 | 3864 | 3707 | 3710 | 4127,66 | 4494 | 3762 | 0,837 | -0,257 | -7,672 | 1,69E-14 | 3,69E-14 | 1,00E-08 | 0,002357 | 0,0008362 | 0,0008362 | TRUE | 0,0039575 | No |
| orf19.6156 | 1496 | 1119 | 1200 | 708 | 1167 | 925 | 1093 | 1393 | 1362 | 1132 | 1136 | 1064 | 1038 | 1004 | 1119 | 1063 | 1114,66 | 1174 | 1056 | 0,898 | -0,155 | -2,039 | 0,041404 | 0,051715 | 0,0060076 | 0,003222 | 0,0046375 | 0,0046375 | TRUE | 0,1429404 | No |
| orf19.6160 | 8309 | 7308 | 7834 | 5127 | 5816 | 5046 | 5374 | 7194 | 7565 | 7391 | 7414 | 7702 | 5173 | 5479 | 5504 | 5488 | 6464,37 | 7518 | 5411 | 0,72 | -0,474 | -13,349 | 1,20E-40 | 4,69E-40 | 0,0004035 | 0,002241 | 0,0010536 | 0,0010536 | TRUE | 0,0107504 | No |
| orf19.6163 | 1205 | 975 | 1071 | 601 | 697 | 490 | 619 | 816 | 1097 | 986 | 1014 | 903 | 620 | 532 | 634 | 622 | 801 | 1000 | 602 | 0,601 | -0,734 | -9,66 | 4,45E-22 | 1,18E-21 | 0,0044978 | 0,003686 | 0,0042249 | 0,0042249 | TRUE | 0,0686542 | No |
| orf19.6164 | 17 | 10 | 6 | 2 | 8 | 2 | 4 | 13 | 15 | 10 | 6 | 3 | 7 | 2 | 4 | 10 | 7,2 | 8 | 6 | 0,679 | -0,558 | -0,933 | 0,350999 | 0,390988 | 0,238699 | 0,18559 | 0,2050494 | 0,2050494 | TRUE | 0,2888322 | No |

| | | | | | | | | | | | | | | | | | | | | | | | | | | | | | | | |
|---|---|---|---|---|---|---|---|---|---|---|---|---|---|---|---|---|---|---|---|---|---|---|---|---|---|---|---|---|---|---|---|
| orf19.6165 | 15448 | 12889 | 14048 | 9354 | 15392 | 12885 | 13217 | 19209 | 14065 | 13036 | 13295 | 14051 | 13691 | 13990 | 13536 | 14653 | 13789,5 | 13612 | 13968 | 1,026 | 0,038 | 0,903 | 0,366366 | 0,406259 | 0,0012763 | 0,002133 | 0,0015885 | 0,0015885 | TRUE | 0,0134756 | No |
| orf19.6166 | 1062 | 1014 | 1037 | 624 | 904 | 711 | 756 | 1176 | 967 | 1026 | 981 | 937 | 804 | 772 | 774 | 897 | 894,82 | 978 | 812 | 0,831 | -0,267 | -4,143 | 3,43E-05 | 5,28E-05 | 0,0022274 | 0,003513 | 0,0028567 | 0,0028567 | TRUE | 0,0625014 | No |
| orf19.6167 | 1000 | 810 | 977 | 656 | 1330 | 1029 | 1153 | 1565 | 910 | 819 | 925 | 985 | 1183 | 1117 | 1181 | 1194 | 1039,32 | 910 | 1169 | 1,289 | 0,366 | 5,969 | 2,39E-09 | 4,40E-09 | 0,0019868 | 0,003308 | 0,0026132 | 0,0026132 | TRUE | 0,0528985 | No |
| orf19.6168 | 771 | 733 | 777 | 440 | 689 | 517 | 629 | 827 | 702 | 741 | 735 | 661 | 613 | 561 | 644 | 631 | 661,1 | 710 | 612 | 0,861 | -0,215 | -3,223 | 0,001269 | 0,00178 | 0,001087 | 0,004035 | 0,0027724 | 0,0027724 | TRUE | 0,0345648 | No |
| orf19.6169 | 450 | 443 | 490 | 339 | 1184 | 959 | 995 | 1436 | 410 | 448 | 464 | 509 | 1053 | 1041 | 1019 | 1095 | 754,93 | 458 | 1052 | 2,315 | 1,211 | 17,727 | 2,57E-70 | 1,63E-69 | 0,0015074 | 0,003787 | 0,002862 | 0,002862 | TRUE | 0,058709 | No |
| orf19.6170 | 907 | 756 | 797 | 494 | 836 | 625 | 719 | 929 | 826 | 765 | 754 | 742 | 744 | 679 | 736 | 709 | 744,24 | 772 | 717 | 0,927 | -0,109 | -1,763 | 0,077878 | 0,094584 | 0,0006668 | 0,003812 | 0,002304 | 0,002304 | TRUE | 0,0275525 | No |
| orf19.6171 | 1320 | 1222 | 1212 | 863 | 1564 | 1293 | 1459 | 2043 | 1202 | 1236 | 1147 | 1296 | 1391 | 1404 | 1494 | 1558 | 1341,09 | 1220 | 1462 | 1,201 | 0,265 | 4,553 | 5,28E-06 | 8,49E-06 | 0,0020196 | 0,003022 | 0,0024831 | 0,0024831 | TRUE | 0,0249216 | No |
| orf19.6173 | 2791 | 2320 | 2548 | 2083 | 1727 | 1565 | 1574 | 2191 | 2541 | 2346 | 2411 | 3129 | 1536 | 1699 | 1612 | 1671 | 2118,32 | 2607 | 1630 | 0,626 | -0,676 | -8,214 | 2,14E-16 | 4,92E-16 | 0,0097073 | 0,002661 | 0,0060055 | 0,0060055 | TRUE | 0,2148301 | No |
| orf19.6175 | 2810 | 2401 | 2628 | 1719 | 3181 | 2333 | 2633 | 3593 | 2558 | 2428 | 2487 | 2582 | 2829 | 2533 | 2697 | 2741 | 2606,97 | 2514 | 2700 | 1,075 | 0,105 | 2,259 | 0,023884 | 0,030428 | 0,0009338 | 0,002544 | 0,0016722 | 0,0016722 | TRUE | 0,0205105 | No |
| orf19.6176 | 1976 | 1580 | 1684 | 1158 | 2422 | 1992 | 2085 | 2986 | 1799 | 1598 | 1594 | 1740 | 2154 | 2163 | 2135 | 2278 | 1932,55 | 1683 | 2182 | 1,299 | 0,377 | 7,098 | 1,27E-12 | 2,62E-12 | 0,0016103 | 0,002721 | 0,0021728 | 0,0021728 | TRUE | 0,028482 | No |
| orf19.6177 | 599 | 472 | 494 | 371 | 1158 | 893 | 946 | 1461 | 545 | 477 | 468 | 557 | 1030 | 970 | 969 | 1114 | 766,3 | 512 | 1021 | 2,003 | 1,002 | 13,072 | 4,77E-39 | 1,83E-38 | 0,0043476 | 0,003761 | 0,004133 | 0,004133 | TRUE | 0,0471643 | No |
| orf19.6178 | 3497 | 2647 | 2813 | 2131 | 5147 | 4192 | 4425 | 6592 | 3184 | 2677 | 2662 | 3201 | 4578 | 4551 | 4532 | 5028 | 3801,75 | 2931 | 4672 | 1,596 | 0,674 | 9,638 | 5,53E-22 | 1,47E-21 | 0,0062353 | 0,002384 | 0,0044145 | 0,0044145 | TRUE | 0,0464737 | No |
| orf19.6179 | 7 | 4 | 6 | 6 | 19 | 18 | 12 | 14 | 6 | 4 | 6 | 9 | 17 | 20 | 12 | 11 | 10,57 | 6 | 15 | 2,414 | 1,271 | 2,774 | 0,005544 | 0,00741 | 1,00E-08 | 0,127069 | 0,0831226 | 0,0831226 | TRUE | 0,1493082 | No |
| orf19.6180 | 1063 | 817 | 905 | 571 | 1170 | 871 | 1034 | 1449 | 968 | 826 | 857 | 858 | 1041 | 946 | 1059 | 1105 | 957,37 | 877 | 1038 | 1,183 | 0,243 | 3,514 | 0,000441 | 0,000637 | 0,0036097 | 0,003417 | 0,0035152 | 0,0035152 | TRUE | 0,0606011 | No |
| orf19.6181 | 23 | 17 | 22 | 18 | 31 | 27 | 28 | 41 | 21 | 17 | 21 | 27 | 28 | 29 | 29 | 31 | 25,35 | 22 | 29 | 1,383 | 0,468 | 1,728 | 0,084039 | 0,101622 | 1,00E-08 | 0,054139 | 0,0294583 | 0,0294583 | TRUE | 0,1077285 | No |
| orf19.6182 | 1685 | 1325 | 1459 | 1056 | 2102 | 1907 | 2140 | 2935 | 1534 | 1340 | 1381 | 1586 | 1870 | 2070 | 2192 | 2239 | 1776,49 | 1460 | 2093 | 1,435 | 0,521 | 7,227 | 4,92E-13 | 1,03E-12 | 0,0056116 | 0,002781 | 0,0043984 | 0,0043984 | TRUE | 0,0630537 | No |
| orf19.6183 | 450 | 319 | 354 | 270 | 500 | 319 | 452 | 559 | 410 | 323 | 335 | 406 | 445 | 346 | 463 | 426 | 394,17 | 368 | 420 | 1,145 | 0,195 | 1,797 | 0,072279 | 0,088103 | 0,0128889 | 0,005388 | 0,008749 | 0,008749 | TRUE | 0,1543682 | No |
| orf19.6184 | 52 | 38 | 60 | 24 | 63 | 37 | 56 | 74 | 47 | 38 | 57 | 36 | 56 | 40 | 57 | 56 | 48,58 | 44 | 52 | 1,167 | 0,223 | 1,045 | 0,296229 | 0,33365 | 0,0106832 | 0,029231 | 0,022963 | 0,022963 | TRUE | 0,2640534 | No |
| orf19.6185 | 0 | 0 | 0 | 0 | 0 | 0 | 0 | 0 | 0 | 0 | 0 | 0 | 0 | 0 | 0 | 0 | 0 | 0 | 0 | NA | NA | NA | NA | NA | NA | NA | NA | NA | NA | NA | NA |
| orf19.6186 | 571 | 458 | 565 | 385 | 387 | 287 | 344 | 498 | 520 | 463 | 535 | 578 | 344 | 312 | 352 | 380 | 435,52 | 524 | 347 | 0,666 | -0,586 | -6,683 | 2,34E-11 | 4,65E-11 | 0,0049519 | 0,00507 | 0,0050169 | 0,0050169 | TRUE | 0,0701372 | No |
| orf19.6187 | 1285 | 1022 | 1133 | 737 | 1172 | 991 | 1168 | 1654 | 1170 | 1034 | 1072 | 1107 | 1042 | 1076 | 1196 | 1262 | 1119,9 | 1096 | 1144 | 1,045 | 0,063 | 0,889 | 0,374118 | 0,41435 | 0,0046951 | 0,003217 | 0,0039934 | 0,0039934 | TRUE | 0,0598543 | No |
| orf19.6188 | 9 | 3 | 3 | 1 | 8 | 4 | 3 | 4 | 8 | 3 | 3 | 2 | 7 | 4 | 3 | 3 | 4,14 | 4 | 4 | 1,081 | 0,113 | 0,16 | 0,872864 | 0,892034 | 0,1048608 | 0,320811 | 0,24066 | 0,24066 | TRUE | 0,9964159 | No |
| orf19.6189 | 2163 | 1688 | 1866 | 1145 | 1503 | 1086 | 1359 | 1849 | 1969 | 1707 | 1766 | 1720 | 1337 | 1179 | 1392 | 1410 | 1560,09 | 1790 | 1330 | 0,743 | -0,429 | -6,25 | 4,10E-10 | 7,79E-10 | 0,00456 | 0,002884 | 0,0038803 | 0,0038803 | TRUE | 0,0680282 | No |
| orf19.6190 | 5399 | 4050 | 4371 | 2805 | 7184 | 5492 | 6150 | 8802 | 4916 | 4096 | 4137 | 4214 | 6390 | 5963 | 6298 | 6714 | 5340,92 | 4341 | 6341 | 1,461 | 0,547 | 8,622 | 6,58E-18 | 1,58E-17 | 0,0048925 | 0,002284 | 0,0036638 | 0,0036638 | TRUE | 0,0981273 | No |
| orf19.6191 | 22 | 14 | 18 | 9 | 73 | 53 | 76 | 71 | 20 | 14 | 17 | 14 | 65 | 58 | 78 | 54 | 39,9 | 16 | 64 | 3,855 | 1,947 | 7,508 | 5,98E-14 | 1,28E-13 | 1,00E-08 | 0,035144 | 0,0250082 | 0,0250082 | TRUE | 0,1931335 | No |
| orf19.6192 | 394 | 267 | 288 | 279 | 475 | 454 | 443 | 671 | 359 | 270 | 273 | 419 | 422 | 493 | 454 | 512 | 400,17 | 330 | 470 | 1,433 | 0,519 | 4,083 | 4,44E-05 | 6,79E-05 | 0,0241273 | 0,005338 | 0,0129025 | 0,0129025 | TRUE | 0,3383444 | No |
| orf19.6193 | 3691 | 3229 | 3538 | 2219 | 2370 | 2055 | 2338 | 3031 | 3360 | 3266 | 3348 | 3333 | 2108 | 2231 | 2394 | 2312 | 2794,21 | 3327 | 2261 | 0,68 | -0,557 | -12,036 | 2,30E-33 | 7,93E-33 | 0,0010422 | 0,00251 | 0,0016892 | 0,0016892 | TRUE | 0,025652 | No |
| orf19.6194 | 178 | 188 | 175 | 108 | 139 | 125 | 150 | 158 | 162 | 190 | 166 | 162 | 124 | 136 | 154 | 121 | 151,69 | 170 | 134 | 0,78 | -0,359 | -2,935 | 0,003331 | 0,004516 | 0,0027221 | 0,010745 | 0,0077065 | 0,0077065 | TRUE | 0,1067985 | No |
| orf19.6195 | 6493 | 5972 | 6584 | 3848 | 5362 | 4222 | 4505 | 5947 | 5912 | 6040 | 6231 | 5780 | 4769 | 4584 | 4614 | 4536 | 5308,31 | 5991 | 4626 | 0,772 | -0,374 | -9,956 | 2,37E-23 | 6,54E-23 | 0,0004965 | 0,002286 | 0,0011629 | 0,0011629 | TRUE | 0,0089122 | No |
| orf19.6196 | 1090 | 939 | 992 | 725 | 1993 | 1572 | 1712 | 2252 | 992 | 950 | 939 | 1089 | 1773 | 1707 | 1753 | 1718 | 1365,08 | 992 | 1738 | 1,758 | 0,814 | 15,086 | 2,01E-51 | 9,53E-51 | 0,0007705 | 0,003005 | 0,0019815 | 0,0019815 | TRUE | 0,0446131 | No |
| orf19.6197 | 591 | 517 | 549 | 359 | 634 | 547 | 489 | 732 | 538 | 523 | 520 | 539 | 564 | 594 | 501 | 558 | 542,1 | 530 | 554 | 1,047 | 0,066 | 0,942 | 0,346286 | 0,386278 | 0,0007269 | 0,004474 | 0,0028235 | 0,0028235 | TRUE | 0,0466124 | No |
| orf19.6198.1 | 1742 | 1516 | 1605 | 899 | 3195 | 2094 | 2480 | 3336 | 1586 | 1533 | 1519 | 1350 | 2842 | 2274 | 2540 | 2545 | 2023,58 | 1497 | 2550 | 1,701 | 0,766 | 10,675 | 1,34E-26 | 3,96E-26 | 0,0057595 | 0,00269 | 0,0044068 | 0,0044068 | TRUE | 0,0720318 | No |
| orf19.6199 | 731 | 604 | 716 | 398 | 1462 | 1219 | 1366 | 1737 | 666 | 611 | 678 | 598 | 1300 | 1323 | 1399 | 1325 | 987,47 | 638 | 1337 | 2,087 | 1,062 | 18 | 1,96E-72 | 1,27E-71 | 0,0008843 | 0,003375 | 0,0021552 | 0,0021552 | TRUE | 0,0198694 | No |
| orf19.6200 | 27 | 17 | 22 | 16 | 6 | 4 | 8 | 7 | 25 | 17 | 21 | 24 | 5 | 4 | 8 | 5 | 13,73 | 22 | 6 | 0,268 | -1,9 | -4,612 | 3,99E-06 | 6,47E-06 | 1,00E-08 | 0,098246 | 0,0580792 | 0,0580792 | TRUE | 0,1581242 | No |
| orf19.6202 | 1446 | 1319 | 1317 | 1027 | 3276 | 2842 | 2850 | 4302 | 1317 | 1334 | 1246 | 1543 | 2914 | 3086 | 2919 | 3282 | 2204,95 | 1360 | 3050 | 2,25 | 1,17 | 17,382 | 1,13E-67 | 6,89E-67 | 0,0053179 | 0,002636 | 0,0037986 | 0,0037986 | TRUE | 0,0921239 | No |
| orf19.6203 | 2545 | 2226 | 2445 | 1549 | 1830 | 1518 | 1672 | 2174 | 2317 | 2251 | 2314 | 2327 | 1628 | 1648 | 1712 | 1658 | 1981,98 | 2302 | 1662 | 0,722 | -0,47 | -11,612 | 3,58E-31 | 1,18E-30 | 1,00E-08 | 0,002704 | 0,001062 | 0,001062 | TRUE | 0,0049119 | No |
| orf19.6205 | 323 | 286 | 319 | 222 | 246 | 216 | 228 | 321 | 294 | 289 | 302 | 333 | 219 | 235 | 234 | 245 | 268,8 | 304 | 233 | 0,769 | -0,379 | -4,225 | 2,39E-05 | 3,72E-05 | 1,00E-08 | 0,006951 | 0,0039789 | 0,0039789 | TRUE | 0,0373881 | No |
| orf19.6208 | 2809 | 2225 | 2468 | 1794 | 2521 | 2203 | 2461 | 3273 | 2557 | 2250 | 2336 | 2695 | 2242 | 2392 | 2520 | 2497 | 2436,21 | 2460 | 2413 | 0,982 | -0,026 | -0,407 | 0,684139 | 0,719463 | 0,003832 | 0,002579 | 0,003526 | 0,003526 | TRUE | 0,0479705 | No |
| orf19.6209 | 1278 | 1045 | 1179 | 731 | 756 | 619 | 679 | 923 | 1164 | 1057 | 1116 | 1098 | 672 | 672 | 695 | 704 | 897,29 | 1109 | 686 | 0,619 | -0,693 | -12,37 | 3,79E-35 | 1,36E-34 | 1,00E-08 | 0,003509 | 0,0018484 | 0,0018484 | TRUE | 0,0141815 | No |
| orf19.6211 | 5670 | 4992 | 5503 | 3288 | 1460 | 1202 | 1299 | 1789 | 5162 | 5049 | 5208 | 4939 | 1299 | 1305 | 1330 | 1365 | 3207,13 | 5090 | 1325 | 0,26 | -1,942 | -48,248 | 0 | 0 | 1,00E-08 | 0,002449 | 0,0010978 | 0,0010978 | TRUE | 0,0054096 | No |
| orf19.6213 | 7472 | 5913 | 6336 | 4333 | 9290 | 7157 | 7605 | 10827 | 6803 | 5980 | 5996 | 6509 | 8263 | 7771 | 7788 | 8259 | 7171,21 | 6322 | 8020 | 1,269 | 0,344 | 6,712 | 1,92E-11 | 3,83E-11 | 0,0024514 | 0,002221 | 0,0023748 | 0,0023748 | TRUE | 0,0327573 | No |
| orf19.6214 | 1834 | 1572 | 1705 | 1163 | 4803 | 4272 | 4515 | 6134 | 1670 | 1590 | 1614 | 1747 | 4272 | 4638 | 4624 | 4679 | 3104,21 | 1655 | 4553 | 2,755 | 1,462 | 30,308 | 9,13E-202 | 1,58E-200 | 0,0012019 | 0,002463 | 0,001807 | 0,001807 | TRUE | 0,0212409 | No |
| orf19.6217 | 1285 | 1010 | 1223 | 832 | 2059 | 1865 | 1850 | 2571 | 1170 | 1022 | 1157 | 1250 | 1831 | 2025 | 1895 | 1961 | 1538,85 | 1150 | 1928 | 1,68 | 0,749 | 11,858 | 1,95E-32 | 6,60E-32 | 0,0034074 | 0,002895 | 0,0031144 | 0,0031144 | TRUE | 0,0670733 | No |
| orf19.6219 | 134 | 128 | 93 | 75 | 725 | 625 | 628 | 905 | 122 | 129 | 88 | 113 | 645 | 679 | 643 | 690 | 388,63 | 113 | 664 | 5,884 | 2,557 | 25,19 | 5,18E-140 | 6,33E-139 | 0,001218 | 0,005436 | 0,0044936 | 0,0044936 | TRUE | 0,2546358 | No |
| orf19.6220 | 3015 | 2781 | 2881 | 1967 | 2472 | 2092 | 2296 | 3046 | 2745 | 2813 | 2727 | 2955 | 2199 | 2271 | 2351 | 2324 | 2548,01 | 2810 | 2286 | 0,815 | -0,296 | -6,77 | 1,29E-11 | 2,59E-11 | 0,0005803 | 0,002555 | 0,0014331 | 0,0014331 | TRUE | 0,0135594 | No |
| orf19.6220.3 | 1528 | 1173 | 1331 | 969 | 684 | 530 | 596 | 793 | 1391 | 1186 | 1260 | 1456 | 608 | 575 | 610 | 605 | 961,49 | 1323 | 600 | 0,454 | -1,138 | -15,959 | 2,46E-57 | 1,29E-56 | 0,0039626 | 0,003411 | 0,0036971 | 0,0036971 | TRUE | 0,0575919 | No |
| orf19.6220.4 | 14906 | 11699 | 12207 | 8082 | 22050 | 15855 | 17060 | 24495 | 13571 | 11832 | 11553 | 12140 | 19613 | 17214 | 17472 | 18685 | 15260 | 12274 | 18246 | 1,487 | 0,572 | 9,501 | 2,07E-21 | 5,43E-21 | 0,0044469 | 0,002124 | 0,0034117 | 0,0034117 | TRUE | 0,0622693 | No |
| orf19.6222 | 106 | 103 | 97 | 67 | 250 | 218 | 232 | 312 | 97 | 104 | 92 | 101 | 222 | 237 | 238 | 238 | 165,97 | 98 | 234 | 2,383 | 1,253 | 10,954 | 6,37E-28 | 1,95E-27 | 1,00E-08 | 0,009996 | 0,0051578 | 0,0051578 | TRUE | 0,0205459 | No |
| orf19.6222.1 | 3104 | 2416 | 2458 | 1473 | 5852 | 3682 | 4138 | 6029 | 2826 | 2444 | 2326 | 2213 | 5205 | 3998 | 4238 | 4599 | 3481,02 | 2452 | 4510 | 1,838 | 0,878 | 9,947 | 2,60E-23 | 7,15E-23 | 0,0122693 | 0,002417 | 0,007165 | 0,007165 | TRUE | 0,1312379 | No |
| orf19.6224 | 155 | 132 | 173 | 84 | 152 | 140 | 136 | 193 | 141 | 134 | 164 | 126 | 135 | 152 | 139 | 147 | 142,28 | 141 | 143 | 1,009 | 0,013 | 0,107 | 0,914585 | 0,926797 | 1,00E-08 | 0,011322 | 0,0073346 | 0,0073346 | TRUE | 0,1225294 | No |
| orf19.6225 | 1549 | 1433 | 1644 | 1067 | 1772 | 1654 | 1591 | 2238 | 1410 | 1449 | 1556 | 1603 | 1576 | 1796 | 1629 | 1707 | 1590,85 | 1504 | 1677 | 1,116 | 0,159 | 2,659 | 0,007832 | 0,010355 | 0,0025699 | 0,002867 | 0,002774 | 0,002774 | TRUE | 0,0264773 | No |
| orf19.6225.1 | 1282 | 1076 | 1107 | 687 | 655 | 503 | 565 | 772 | 1167 | 1088 | 1048 | 1032 | 583 | 546 | 579 | 589 | 828,92 | 1084 | 574 | 0,529 | -0,918 | -14,828 | 9,62E-50 | 4,41E-49 | 0,0011631 | 0,003631 | 0,0023717 | 0,0023717 | TRUE | 0,0330977 | No |
| orf19.6227 | 4893 | 4025 | 4614 | 2789 | 4352 | 3623 | 4070 | 5418 | 4455 | 4071 | 4367 | 4190 | 3871 | 3934 | 4168 | 4133 | 4148,45 | 4271 | 4026 | 0,943 | -0,085 | -1,909 | 0,056234 | 0,069355 | 0,001158 | 0,002355 | 0,0016796 | 0,0016796 | TRUE | 0,0124805 | No |
| orf19.6229 | 7493 | 6494 | 7055 | 5035 | 21449 | 18944 | 18835 | 27107 | 6822 | 6568 | 6677 | 7563 | 19078 | 20568 | 19289 | 20677 | 13405,4 | 6908 | 19903 | 2,883 | 1,528 | 29,39 | 7,44E-190 | 1,20E-188 | 0,0028694 | 0,002136 | 0,00249 | 0,00249 | TRUE | 0,0487438 | No |
| orf19.6230 | 4745 | 4696 | 5005 | 2758 | 5471 | 4488 | 4869 | 6067 | 4320 | 4750 | 4737 | 4143 | 4866 | 4873 | 4986 | 4628 | 4662,85 | 4488 | 4838 | 1,077 | 0,107 | 2,04 | 0,04136 | 0,051669 | 0,0024959 | 0,00232 | 0,0024379 | 0,0024379 | TRUE | 0,0312479 | No |
| orf19.6231 | 2310 | 1747 | 1922 | 1478 | 2093 | 1751 | 1738 | 2498 | 2103 | 1767 | 1819 | 2220 | 1862 | 1901 | 1780 | 1905 | 1919,68 | 1977 | 1862 | 0,943 | -0,084 | -1,175 | 0,239884 | 0,274011 | 0,0059384 | 0,002725 | 0,0044242 | 0,0044242 | TRUE | 0,0788841 | No |
| orf19.6232 | 5510 | 4758 | 5183 | 3157 | 3634 | 3134 | 3530 | 4523 | 5017 | 4812 | 4905 | 4742 | 3232 | 3403 | 3615 | 3450 | 4147,09 | 4869 | 3425 | 0,703 | -0,508 | -11,698 | 1,31E-31 | 4,36E-31 | 0,0010236 | 0,002356 | 0,001567 | 0,001567 | TRUE | 0,0175865 | No |
| orf19.6233 | 3894 | 3135 | 3293 | 2039 | 2976 | 2318 | 2729 | 3691 | 3545 | 3171 | 3117 | 3063 | 2647 | 2517 | 2795 | 2816 | 2958,7 | 3224 | 2694 | 0,835 | -0,26 | -4,457 | 8,32E-06 | 1,33E-05 | 0,0031939 | 0,002483 | 0,0029233 | 0,0029233 | TRUE | 0,0554897 | No |
| orf19.6234 | 9983 | 8670 | 9419 | 5888 | 9463 | 8014 | 8556 | 11733 | 9089 | 8769 | 8914 | 8845 | 8417 | 8701 | 8762 | 8950 | 8805,91 | 8904 | 8708 | 0,978 | -0,032 | -0,987 | 0,323408 | 0,362917 | 0,0002881 | 0,002187 | 0,0009133 | 0,0009133 | TRUE | 0,006258 | No |
| orf19.6235 | 923 | 714 | 816 | 600 | 537 | 411 | 435 | 601 | 840 | 722 | 772 | 901 | 478 | 446 | 445 | 458 | 632,98 | 809 | 457 | 0,567 | -0,819 | -10,654 | 1,67E-26 | 4,93E-26 | 0,0038331 | 0,004124 | 0,0039833 | 0,0039833 | TRUE | 0,0627791 | No |
| orf19.6236 | 6354 | 5751 | 5984 | 3488 | 5641 | 4244 | 4860 | 6211 | 5785 | 5817 | 5663 | 5240 | 5017 | 4608 | 4977 | 4738 | 5230,6 | 5626 | 4835 | 0,859 | -0,22 | -4,661 | 3,15E-06 | 5,12E-06 | 0,0016788 | 0,00229 | 0,0019391 | 0,0019391 | TRUE | 0,0251399 | No |
| orf19.6237 | 2774 | 2491 | 2682 | 1722 | 3131 | 2499 | 2640 | 3531 | 2526 | 2519 | 2538 | 2587 | 2785 | 2713 | 2704 | 2693 | 2633,16 | 2542 | 2724 | 1,072 | 0,1 | 2,758 | 0,005824 | 0,007769 | 1,00E-08 | 0,002539 | 0,0008886 | 0,0008886 | TRUE | 0,0028337 | No |
| orf19.6238 | 337 | 330 | 337 | 202 | 434 | 353 | 403 | 473 | 307 | 334 | 319 | 303 | 386 | 383 | 413 | 361 | 350,72 | 316 | 386 | 1,217 | 0,284 | 3,516 | 0,000438 | 0,000633 | 1,00E-08 | 0,005804 | 0,0033518 | 0,0033518 | TRUE | 0,025357 | No |
| orf19.6239 | 1614 | 1377 | 1477 | 1020 | 1215 | 989 | 1030 | 1392 | 1469 | 1393 | 1398 | 1532 | 1081 | 1074 | 1055 | 1062 | 1257,92 | 1448 | 1068 | 0,739 | -0,437 | -8,712 | 2,99E-18 | 7,24E-18 | 0,0002442 | 0,003087 | 0,0016041 | 0,0016041 | TRUE | 0,0162094 | No |
| orf19.6240 | 665 | 581 | 600 | 394 | 360 | 277 | 308 | 429 | 605 | 588 | 568 | 592 | 320 | 301 | 315 | 327 | 452,05 | 588 | 316 | 0,538 | -0,894 | -12,629 | 1,47E-36 | 5,43E-36 | 1,00E-08 | 0,004959 | 0,0024281 | 0,0024281 | TRUE | 0,0110937 | No |
| orf19.6242 | 270 | 242 | 275 | 153 | 161 | 142 | 166 | 201 | 246 | 245 | 260 | 230 | 143 | 154 | 170 | 153 | 200,17 | 245 | 155 | 0,63 | -0,668 | -6,516 | 7,20E-11 | 1,41E-10 | 1,00E-08 | 0,008636 | 0,0049035 | 0,0049035 | TRUE | 0,0416517 | No |
| orf19.6243 | 814 | 775 | 871 | 500 | 665 | 587 | 550 | 766 | 741 | 784 | 824 | 751 | 591 | 637 | 563 | 584 | 684,59 | 775 | 594 | 0,765 | -0,387 | -5,955 | 2,61E-09 | 4,80E-09 | 0,0009986 | 0,003967 | 0,0025694 | 0,0025694 | TRUE | 0,0261892 | No |

| | | | | | | | | | | | | | | | | | | | | | | | | | | | | | | | |
|---|---|---|---|---|---|---|---|---|---|---|---|---|---|---|---|---|---|---|---|---|---|---|---|---|---|---|---|---|---|---|---|
| orf19.6244 | 1352 | 1173 | 1324 | 905 | 1543 | 1368 | 1447 | 2016 | 1231 | 1186 | 1253 | 1359 | 1372 | 1485 | 1482 | 1538 | 1363,41 | 1257 | 1469 | 1,171 | 0,228 | 3,967 | 7,27E-05 | 0,00011 | 0,0018401 | 0,003006 | 0,0024342 | 0,0024342 | TRUE | 0,0322511 | No |
| orf19.6245 | 14100 | 12871 | 13791 | 8222 | 3594 | 2873 | 3132 | 4033 | 12837 | 13018 | 13052 | 12351 | 3197 | 3119 | 3208 | 3076 | 7982,22 | 12814 | 3150 | 0,246 | -2,025 | -57,8 | 0 | 0 | 0,0003862 | 0,002203 | 0,0009885 | 0,0009885 | TRUE | 0,0070321 | No |
| orf19.6246 | 1461 | 1196 | 1269 | 759 | 685 | 543 | 559 | 777 | 1330 | 1210 | 1201 | 1140 | 609 | 590 | 572 | 593 | 905,62 | 1220 | 591 | 0,483 | -1,048 | -16,734 | 7,35E-63 | 4,18E-62 | 0,0016709 | 0,003496 | 0,0025407 | 0,0025407 | TRUE | 0,0448821 | No |
| orf19.6247 | 80 | 63 | 81 | 49 | 43 | 35 | 37 | 50 | 73 | 64 | 77 | 74 | 38 | 38 | 38 | 38 | 54,89 | 72 | 38 | 0,531 | -0,912 | -4,894 | 9,87E-07 | 1,64E-06 | 1,00E-08 | 0,026104 | 0,0137322 | 0,0137322 | TRUE | 0,052954 | No |
| orf19.6247.1 | 4590 | 3806 | 3990 | 1710 | 1038 | 604 | 738 | 1043 | 4179 | 3849 | 3776 | 2569 | 923 | 656 | 756 | 796 | 2187,96 | 3593 | 783 | 0,218 | -2,199 | -12,663 | 9,48E-37 | 3,52E-36 | 0,0282225 | 0,002641 | 0,0150998 | 0,0282225 | TRUE | 0,4449436 | No |
| orf19.6248 | 45 | 66 | 60 | 32 | 45 | 45 | 65 | 62 | 41 | 67 | 57 | 48 | 40 | 49 | 67 | 47 | 51,92 | 53 | 51 | 0,947 | -0,078 | -0,354 | 0,723389 | 0,756373 | 0,0273642 | 0,027482 | 0,0274472 | 0,0274472 | TRUE | 0,3533699 | No |
| orf19.6249 | 2885 | 2568 | 2819 | 1622 | 507 | 439 | 477 | 623 | 2627 | 2597 | 2668 | 2437 | 451 | 477 | 489 | 475 | 1527,46 | 2582 | 473 | 0,183 | -2,452 | -44,464 | 0 | 0 | 1,00E-08 | 0,002902 | 0,0017274 | 0,0017274 | TRUE | 0,0161622 | No |
| orf19.6250 | 6215 | 4632 | 5435 | 3645 | 4569 | 3486 | 4104 | 5672 | 5658 | 4685 | 5144 | 5475 | 4064 | 3785 | 4203 | 4327 | 4667,6 | 5240 | 4095 | 0,782 | -0,355 | -5,547 | 2,91E-08 | 5,14E-08 | 0,0046514 | 0,00232 | 0,0037249 | 0,0037249 | TRUE | 0,0624025 | No |
| orf19.6252 | 3544 | 2996 | 3344 | 2096 | 4197 | 3345 | 3659 | 4918 | 3227 | 3030 | 3165 | 3149 | 3733 | 3632 | 3747 | 3751 | 3429,21 | 3143 | 3716 | 1,183 | 0,242 | 6,571 | 5,01E-11 | 9,83E-11 | 0,0001501 | 0,002422 | 0,0010047 | 0,0010047 | TRUE | 0,0073021 | No |
| orf19.6253 | 16846 | 13493 | 14563 | 9070 | 27984 | 20771 | 22230 | 31427 | 15338 | 13647 | 13783 | 13625 | 24891 | 22552 | 22766 | 23973 | 18821,6 | 14098 | 23546 | 1,67 | 0,74 | 14,459 | 2,19E-47 | 9,70E-47 | 0,0027336 | 0,002107 | 0,002457 | 0,002457 | TRUE | 0,0430765 | No |
| orf19.6254 | 3570 | 3022 | 3291 | 2158 | 26110 | 22164 | 23298 | 32011 | 3250 | 3056 | 3115 | 3242 | 23224 | 24064 | 23860 | 24418 | 13528,7 | 3166 | 23892 | 7,551 | 2,917 | 81,476 | 0 | 0 | 0,0004386 | 0,002135 | 0,001041 | 0,001041 | TRUE | 0,0065548 | No |
| orf19.6255 | 1307 | 1219 | 1289 | 751 | 1671 | 1329 | 1515 | 2030 | 1190 | 1233 | 1220 | 1128 | 1486 | 1443 | 1552 | 1549 | 1350,02 | 1193 | 1508 | 1,262 | 0,336 | 6,66 | 2,74E-11 | 5,45E-11 | 0,0003688 | 0,003016 | 0,00168 | 0,00168 | TRUE | 0,0137501 | No |
| orf19.6257 | 4608 | 4220 | 4502 | 2739 | 15666 | 12979 | 13802 | 18703 | 4195 | 4268 | 4261 | 4114 | 13934 | 14092 | 14135 | 14267 | 9158,3 | 4210 | 14107 | 3,349 | 1,744 | 58,774 | 0 | 0 | 1,00E-08 | 0,002181 | 0,0006844 | 0,0006844 | TRUE | 0,0025314 | No |
| orf19.6259 | 4470 | 4158 | 4464 | 2610 | 4894 | 3863 | 4311 | 5337 | 4070 | 4205 | 4225 | 3921 | 4353 | 4194 | 4415 | 4071 | 4181,73 | 4105 | 4258 | 1,036 | 0,051 | 1,201 | 0,229932 | 0,263493 | 0,0009392 | 0,002353 | 0,0015185 | 0,0015185 | TRUE | 0,011004 | No |
| orf19.6260 | 3616 | 3299 | 3566 | 2277 | 3790 | 3187 | 3428 | 4451 | 3292 | 3337 | 3375 | 3420 | 3371 | 3460 | 3511 | 3395 | 3395,17 | 3356 | 3434 | 1,024 | 0,034 | 0,953 | 0,340356 | 0,380527 | 1,00E-08 | 0,002426 | 0,0008991 | 0,0008991 | TRUE | 0,002723 | No |
| orf19.6261 | 2125 | 2013 | 2199 | 1245 | 1164 | 1025 | 1059 | 1358 | 1935 | 2036 | 2081 | 1870 | 1035 | 1113 | 1085 | 1036 | 1523,83 | 1980 | 1067 | 0,537 | -0,896 | -17,038 | 4,32E-65 | 2,54E-64 | 0,0011429 | 0,002904 | 0,0019445 | 0,0019445 | TRUE | 0,0154842 | No |
| orf19.6263 | 4770 | 3944 | 4101 | 2890 | 4077 | 3152 | 3383 | 4920 | 4343 | 3989 | 3881 | 4341 | 3626 | 3422 | 3465 | 3753 | 3852,56 | 4138 | 3566 | 0,863 | -0,213 | -4,086 | 4,38E-05 | 6,70E-05 | 0,0022318 | 0,00238 | 0,0023513 | 0,0023513 | TRUE | 0,0214945 | No |
| orf19.6264 | 425 | 362 | 401 | 278 | 239 | 195 | 198 | 269 | 387 | 366 | 380 | 418 | 213 | 212 | 203 | 205 | 297,81 | 388 | 208 | 0,539 | -0,892 | -10,412 | 2,18E-25 | 6,31E-25 | 1,00E-08 | 0,006473 | 0,0034353 | 0,0034353 | TRUE | 0,025798 | No |
| orf19.6264.3 | 5554 | 5495 | 5505 | 2746 | 4594 | 3371 | 3785 | 4626 | 5057 | 5558 | 5210 | 4125 | 4086 | 3660 | 3876 | 3529 | 4387,56 | 4988 | 3788 | 0,759 | -0,398 | -4,89 | 1,01E-06 | 1,68E-06 | 0,0093123 | 0,002338 | 0,0061314 | 0,0061314 | TRUE | 0,1624476 | No |
| orf19.6264.4 | 7119 | 5447 | 5871 | 3676 | 9997 | 6801 | 8308 | 11240 | 6482 | 5509 | 5556 | 5522 | 8892 | 7384 | 8508 | 8574 | 7053,42 | 5767 | 8340 | 1,446 | 0,532 | 7,667 | 1,76E-14 | 3,84E-14 | 0,0063391 | 0,002224 | 0,0044741 | 0,0044741 | TRUE | 0,0854035 | No |
| orf19.6265 | 8804 | 6569 | 6532 | 4459 | 13100 | 9124 | 10030 | 14334 | 8016 | 6644 | 6182 | 6698 | 11652 | 9906 | 10272 | 10934 | 8787,98 | 6885 | 10691 | 1,553 | 0,635 | 8,235 | 1,79E-16 | 4,12E-16 | 0,0089882 | 0,002187 | 0,0055864 | 0,0055864 | TRUE | 0,1501981 | No |
| orf19.6265.1 | 16128 | 12924 | 13608 | 8399 | 24013 | 16959 | 19748 | 26789 | 14684 | 13071 | 12879 | 12617 | 21359 | 18413 | 20224 | 20435 | 16710,2 | 13313 | 20108 | 1,51 | 0,595 | 10,008 | 1,40E-23 | 3,90E-23 | 0,0042516 | 0,002116 | 0,0033294 | 0,0033294 | TRUE | 0,0589991 | No |
| orf19.6266 | 2490 | 2635 | 2700 | 1599 | 2840 | 2446 | 2514 | 3081 | 2267 | 2665 | 2555 | 2402 | 2526 | 2656 | 2575 | 2350 | 2499,49 | 2472 | 2527 | 1,021 | 0,03 | 0,501 | 0,616162 | 0,655109 | 0,0034745 | 0,002566 | 0,0030758 | 0,0030758 | TRUE | 0,0394533 | No |
| orf19.6267 | 537 | 500 | 534 | 309 | 91 | 81 | 71 | 104 | 489 | 506 | 505 | 464 | 81 | 88 | 73 | 79 | 285,64 | 491 | 80 | 0,163 | -2,619 | -24,876 | 1,37E-136 | 1,61E-135 | 1,00E-08 | 0,006662 | 0,0037686 | 0,0037686 | TRUE | 0,0325343 | No |
| orf19.6268 | 1758 | 1647 | 1757 | 1080 | 1721 | 1491 | 1564 | 2059 | 1601 | 1666 | 1663 | 1622 | 1531 | 1619 | 1602 | 1571 | 1609,18 | 1638 | 1581 | 0,964 | -0,053 | -1,202 | 0,229421 | 0,262956 | 1,00E-08 | 0,002858 | 0,001223 | 0,001223 | TRUE | 0,0053526 | No |
| orf19.6269 | 686 | 766 | 804 | 465 | 600 | 498 | 514 | 660 | 625 | 775 | 761 | 699 | 534 | 541 | 526 | 503 | 620,37 | 715 | 526 | 0,735 | -0,443 | -5,712 | 1,11E-08 | 2,00E-08 | 0,0041271 | 0,004166 | 0,0041468 | 0,0041468 | TRUE | 0,092058 | No |
| orf19.6271 | 3313 | 3188 | 3529 | 2311 | 5188 | 4436 | 4622 | 6067 | 3016 | 3224 | 3340 | 3471 | 4615 | 4816 | 4734 | 4628 | 3980,54 | 3263 | 4698 | 1,441 | 0,527 | 11,096 | 1,31E-28 | 4,06E-28 | 0,0016072 | 0,002369 | 0,0019042 | 0,0019042 | TRUE | 0,031842 | No |
| orf19.6272 | 879 | 1258 | 1200 | 705 | 892 | 917 | 840 | 997 | 800 | 1272 | 1136 | 1059 | 793 | 996 | 860 | 761 | 959,64 | 1067 | 852 | 0,798 | -0,325 | -2,663 | 0,007752 | 0,01025 | 0,0233363 | 0,003414 | 0,0132218 | 0,0132218 | TRUE | 0,3457062 | No |
| orf19.6273 | 696 | 532 | 570 | 361 | 738 | 546 | 617 | 834 | 634 | 538 | 539 | 542 | 656 | 593 | 632 | 636 | 596,35 | 563 | 629 | 1,116 | 0,158 | 2,153 | 0,03129 | 0,0395 | 0,0026006 | 0,004252 | 0,0034713 | 0,0034713 | TRUE | 0,0877883 | No |
| orf19.6274 | 201 | 164 | 156 | 131 | 143 | 101 | 110 | 144 | 183 | 166 | 148 | 197 | 127 | 110 | 113 | 110 | 144,08 | 174 | 115 | 0,667 | -0,584 | -4,594 | 4,34E-06 | 7,02E-06 | 0,0038398 | 0,011205 | 0,0083567 | 0,0083567 | TRUE | 0,1087308 | No |
| orf19.6275 | 1265 | 1314 | 1339 | 721 | 1070 | 835 | 888 | 1146 | 1152 | 1329 | 1267 | 1083 | 952 | 907 | 909 | 874 | 1059,11 | 1208 | 910 | 0,752 | -0,411 | -5,987 | 2,14E-09 | 3,95E-09 | 0,0035447 | 0,003284 | 0,0035681 | 0,0035681 | TRUE | 0,0534657 | No |
| orf19.6276 | 1667 | 1613 | 1804 | 1009 | 906 | 717 | 731 | 973 | 1518 | 1631 | 1707 | 1516 | 806 | 778 | 749 | 742 | 1180,91 | 1593 | 769 | 0,482 | -1,053 | -17,787 | 8,90E-71 | 5,64E-70 | 0,0017893 | 0,003156 | 0,0024215 | 0,0024215 | TRUE | 0,0285454 | No |
| orf19.6277 | 50 | 46 | 31 | 26 | 13 | 11 | 22 | 24 | 46 | 47 | 29 | 39 | 12 | 12 | 23 | 18 | 28,1 | 40 | 16 | 0,402 | -1,316 | -4,462 | 8,12E-06 | 1,30E-05 | 0,0258317 | 0,049051 | 0,0413289 | 0,0413289 | TRUE | 0,3087043 | No |
| orf19.6280 | 5 | 4 | 3 | 2 | 10 | 12 | 3 | 3 | 5 | 4 | 3 | 3 | 9 | 13 | 3 | 2 | 5,22 | 4 | 7 | 1,835 | 0,875 | 1,266 | 0,205555 | 0,237609 | 0,2270214 | 0,255311 | 0,2435771 | 0,2435771 | TRUE | 0,2491157 | No |
| orf19.6282 | 798 | 773 | 761 | 529 | 526 | 404 | 429 | 539 | 727 | 782 | 720 | 795 | 468 | 439 | 439 | 411 | 597,53 | 756 | 439 | 0,582 | -0,781 | -11,343 | 8,06E-30 | 2,58E-29 | 0,001084 | 0,004248 | 0,0027788 | 0,0027788 | TRUE | 0,0243299 | No |
| orf19.6283 | 3410 | 3145 | 3277 | 2005 | 2413 | 1769 | 1819 | 2458 | 3105 | 3181 | 3101 | 3012 | 2146 | 1921 | 1863 | 1875 | 2525,44 | 3100 | 1951 | 0,629 | -0,668 | -12,723 | 4,39E-37 | 1,64E-36 | 0,0019289 | 0,00256 | 0,0022342 | 0,0022342 | TRUE | 0,0555293 | No |
| orf19.6284 | 3713 | 3568 | 3822 | 2065 | 2072 | 1696 | 1819 | 2418 | 3381 | 3609 | 3617 | 3102 | 1843 | 1841 | 1863 | 1844 | 2637,5 | 3427 | 1848 | 0,539 | -0,893 | -16,612 | 5,74E-62 | 3,22E-61 | 0,0019709 | 0,002538 | 0,0023646 | 0,0023646 | TRUE | 0,0471169 | No |
| orf19.6285 | 6181 | 5510 | 6224 | 3629 | 8397 | 6726 | 7385 | 9902 | 5628 | 5573 | 5890 | 5451 | 7469 | 7303 | 7563 | 7553 | 6553,75 | 5636 | 7472 | 1,325 | 0,406 | 11,357 | 6,83E-30 | 2,19E-29 | 0,0004628 | 0,002239 | 0,0010701 | 0,0010701 | TRUE | 0,0113567 | No |
| orf19.6286 | 474 | 369 | 431 | 242 | 971 | 688 | 803 | 1109 | 432 | 373 | 408 | 364 | 864 | 747 | 822 | 846 | 606,9 | 394 | 820 | 2,074 | 1,052 | 13,788 | 3,03E-43 | 1,25E-42 | 0,0030115 | 0,004214 | 0,0036707 | 0,0036707 | TRUE | 0,0478226 | No |
| orf19.6286.2 | 9817 | 8388 | 8621 | 7149 | 13998 | 12748 | 11060 | 16362 | 8938 | 8484 | 8159 | 10739 | 12451 | 13841 | 11327 | 12481 | 10802,4 | 9080 | 12525 | 1,38 | 0,465 | 5,593 | 2,23E-08 | 3,95E-08 | 0,0112841 | 0,002159 | 0,0065324 | 0,0065324 | TRUE | 0,1836013 | No |
| orf19.6287 | 4746 | 4065 | 4354 | 3002 | 13325 | 11273 | 11402 | 16066 | 4321 | 4111 | 4121 | 4509 | 11852 | 12239 | 11677 | 12255 | 8135,79 | 4266 | 12006 | 2,817 | 1,494 | 36,71 | 5,14E-295 | 1,30E-293 | 0,0010588 | 0,002199 | 0,0014259 | 0,0014259 | TRUE | 0,0169923 | No |
| orf19.6288 | 158 | 127 | 130 | 92 | 42 | 33 | 45 | 52 | 144 | 128 | 123 | 138 | 37 | 36 | 46 | 40 | 86,56 | 133 | 40 | 0,298 | -1,746 | -10,628 | 2,21E-26 | 6,50E-26 | 1,00E-08 | 0,017298 | 0,0102243 | 0,0102243 | TRUE | 0,0777311 | No |
| orf19.6291 | 3531 | 3142 | 3291 | 2054 | 6600 | 5481 | 6277 | 8233 | 3215 | 3178 | 3115 | 3085 | 5870 | 5951 | 6428 | 6280 | 4640,33 | 3148 | 6132 | 1,947 | 0,961 | 22,755 | 1,29E-114 | 1,26E-113 | 0,0009827 | 0,002322 | 0,0014667 | 0,0014667 | TRUE | 0,0126697 | No |
| orf19.6293 | 832 | 690 | 785 | 445 | 697 | 549 | 624 | 792 | 758 | 698 | 743 | 668 | 620 | 596 | 639 | 604 | 665,75 | 717 | 615 | 0,855 | -0,226 | -3,56 | 0,000371 | 0,000539 | 0,0004593 | 0,004021 | 0,0023813 | 0,0023813 | TRUE | 0,0208341 | No |
| orf19.6294 | 310 | 387 | 370 | 229 | 914 | 832 | 846 | 1074 | 282 | 391 | 350 | 344 | 813 | 903 | 866 | 819 | 596,22 | 342 | 850 | 2,489 | 1,316 | 14,815 | 1,16E-49 | 5,33E-49 | 0,0071874 | 0,004253 | 0,0054523 | 0,0054523 | TRUE | 0,1722643 | No |
| orf19.6295 | 3489 | 3209 | 3505 | 2096 | 3381 | 2808 | 2869 | 3871 | 3177 | 3246 | 3317 | 3149 | 3007 | 3049 | 2938 | 2953 | 3104,36 | 3222 | 2987 | 0,926 | -0,111 | -2,991 | 0,00278 | 0,003795 | 1,00E-08 | 0,002463 | 0,0009891 | 0,0009891 | TRUE | 0,0048574 | No |
| orf19.6296 | 325 | 355 | 342 | 189 | 252 | 164 | 190 | 244 | 296 | 359 | 324 | 284 | 224 | 178 | 195 | 186 | 255,68 | 316 | 196 | 0,619 | -0,692 | -6,466 | 1,01E-10 | 1,96E-10 | 0,0065367 | 0,007204 | 0,0069251 | 0,0069251 | TRUE | 0,106643 | No |
| orf19.6297 | 3003 | 2718 | 2874 | 1740 | 3007 | 2551 | 2592 | 3599 | 2734 | 2749 | 2720 | 2614 | 2675 | 2770 | 2655 | 2745 | 2707,62 | 2704 | 2711 | 1,001 | 0,002 | 0,056 | 0,955548 | 0,96052 | 1,00E-08 | 0,002525 | 0,0010306 | 0,0010306 | TRUE | 0,0055054 | No |
| orf19.6298 | 3120 | 3013 | 3036 | 1780 | 3308 | 2563 | 2665 | 3376 | 2841 | 3047 | 2873 | 2674 | 2942 | 2783 | 2729 | 2575 | 2808,09 | 2859 | 2757 | 0,963 | -0,054 | -0,978 | 0,327936 | 0,367545 | 0,0024671 | 0,002507 | 0,0025459 | 0,0025459 | TRUE | 0,0252811 | No |
| orf19.6302 | 31 | 23 | 11 | 7 | 11 | 2 | 9 | 13 | 28 | 23 | 10 | 11 | 10 | 2 | 9 | 10 | 12,94 | 18 | 8 | 0,431 | -1,215 | -2,538 | 0,011157 | 0,014587 | 0,1837585 | 0,104142 | 0,1321474 | 0,1321474 | TRUE | 0,1997813 | No |
| orf19.6304 | 76 | 114 | 99 | 54 | 124 | 140 | 120 | 145 | 69 | 115 | 94 | 81 | 110 | 152 | 123 | 111 | 106,89 | 90 | 124 | 1,371 | 0,456 | 2,643 | 0,008221 | 0,010849 | 0,0253311 | 0,014396 | 0,0188076 | 0,0188076 | TRUE | 0,3564816 | No |
| orf19.6305 | 988 | 1124 | 1084 | 626 | 2887 | 2278 | 2462 | 3042 | 900 | 1137 | 1026 | 940 | 2568 | 2473 | 2521 | 2320 | 1735,7 | 1001 | 2470 | 2,466 | 1,302 | 18,508 | 1,76E-76 | 1,19E-75 | 0,0055075 | 0,002798 | 0,0040274 | 0,0040274 | TRUE | 0,1011239 | No |
| orf19.6306 | 986 | 914 | 976 | 524 | 1539 | 1320 | 1371 | 1851 | 898 | 924 | 924 | 787 | 1369 | 1433 | 1404 | 1412 | 1143,88 | 883 | 1404 | 1,583 | 0,663 | 11,582 | 5,06E-31 | 1,66E-30 | 0,0009153 | 0,003192 | 0,0022057 | 0,0022057 | TRUE | 0,0547189 | No |
| orf19.6307 | 636 | 808 | 797 | 505 | 1142 | 973 | 1005 | 1245 | 579 | 817 | 754 | 759 | 1016 | 1056 | 1029 | 950 | 870,03 | 727 | 1013 | 1,394 | 0,479 | 5,231 | 1,69E-07 | 2,89E-07 | 0,0104737 | 0,003555 | 0,0068539 | 0,0068539 | TRUE | 0,2319897 | No |
| orf19.6308 | 300 | 449 | 453 | 236 | 575 | 566 | 585 | 605 | 273 | 454 | 429 | 355 | 511 | 615 | 599 | 461 | 462,13 | 378 | 546 | 1,445 | 0,531 | 3,782 | 0,000156 | 0,000231 | 0,0316234 | 0,004895 | 0,0166586 | 0,0166586 | TRUE | 0,4182544 | No |
| orf19.6309 | 1949 | 1668 | 1987 | 1227 | 1631 | 1363 | 1481 | 2088 | 1774 | 1687 | 1881 | 1843 | 1451 | 1480 | 1517 | 1593 | 1653,15 | 1796 | 1510 | 0,842 | -0,249 | -4,726 | 2,29E-06 | 3,74E-06 | 0,0014372 | 0,002836 | 0,0020508 | 0,0020508 | TRUE | 0,0204477 | No |
| orf19.6310 | 720 | 863 | 857 | 627 | 447 | 427 | 377 | 546 | 656 | 873 | 811 | 942 | 398 | 464 | 386 | 416 | 618,13 | 820 | 416 | 0,508 | -0,977 | -9,361 | 7,90E-21 | 2,05E-20 | 0,0136458 | 0,004174 | 0,0086703 | 0,0086703 | TRUE | 0,2211835 | No |
| orf19.6311 | 25777 | 23140 | 23162 | 14463 | 14078 | 10529 | 11133 | 14808 | 23469 | 23404 | 21921 | 21726 | 12522 | 11432 | 11402 | 11296 | 17146,2 | 22630 | 11663 | 0,515 | -0,956 | -20,512 | 1,69E-93 | 1,36E-92 | 0,0019857 | 0,002114 | 0,0020248 | 0,0020248 | TRUE | 0,0301758 | No |
| orf19.6312 | 13314 | 11277 | 11986 | 8021 | 23098 | 19124 | 17766 | 26738 | 12122 | 11405 | 11344 | 12049 | 20545 | 20763 | 18195 | 20396 | 15852,3 | 11730 | 19975 | 1,703 | 0,768 | 15,428 | 1,06E-53 | 5,26E-53 | 0,0023206 | 0,00212 | 0,0023123 | 0,0023123 | TRUE | 0,0439678 | No |
| orf19.6313 | 216 | 277 | 265 | 162 | 639 | 564 | 564 | 768 | 197 | 280 | 251 | 243 | 568 | 612 | 578 | 586 | 414,39 | 243 | 586 | 2,418 | 1,274 | 13,485 | 1,92E-41 | 7,67E-41 | 0,0061546 | 0,005225 | 0,0055654 | 0,0055654 | TRUE | 0,2021344 | No |
| orf19.6313.2 | 1899 | 1778 | 1924 | 973 | 2577 | 1777 | 2032 | 2611 | 1729 | 1798 | 1821 | 1462 | 2292 | 1929 | 2081 | 1992 | 1887,99 | 1702 | 2074 | 1,216 | 0,282 | 3,739 | 0,000185 | 0,000273 | 0,0069184 | 0,002737 | 0,0049198 | 0,0049198 | TRUE | 0,1037404 | No |
| orf19.6314 | 2833 | 2492 | 2519 | 1388 | 4120 | 2926 | 3411 | 4441 | 2579 | 2520 | 2384 | 2085 | 3665 | 3177 | 3493 | 3388 | 2911,38 | 2392 | 3431 | 1,432 | 0,518 | 7,57 | 3,73E-14 | 8,05E-14 | 0,0055453 | 0,002491 | 0,0041373 | 0,0041373 | TRUE | 0,0866641 | No |
| orf19.6315 | 91 | 74 | 75 | 54 | 331 | 257 | 259 | 331 | 83 | 75 | 71 | 81 | 294 | 279 | 265 | 252 | 175,12 | 78 | 272 | 3,528 | 1,819 | 14,774 | 2,17E-49 | 9,87E-49 | 1,00E-08 | 0,00958 | 0,0059833 | 0,0059833 | TRUE | 0,0341987 | No |
| orf19.6316 | 381 | 415 | 422 | 228 | 580 | 493 | 539 | 624 | 347 | 420 | 399 | 342 | 516 | 535 | 552 | 476 | 448,45 | 377 | 520 | 1,372 | 0,456 | 5,304 | 1,13E-07 | 1,95E-07 | 0,0045128 | 0,004983 | 0,0047736 | 0,0047736 | TRUE | 0,0648739 | No |
| orf19.6316.4 | 3116 | 3229 | 3384 | 1781 | 5847 | 4323 | 4840 | 5973 | 2837 | 3266 | 3203 | 2675 | 5201 | 4694 | 4957 | 4556 | 3923,51 | 2995 | 4852 | 1,618 | 0,694 | 10,113 | 4,85E-24 | 1,36E-23 | 0,0059011 | 0,002374 | 0,0042535 | 0,0042535 | TRUE | 0,060503 | No |
| orf19.6317 | 3590 | 3502 | 3677 | 2263 | 8439 | 7558 | 7397 | 9690 | 3269 | 3542 | 3480 | 3399 | 7506 | 8206 | 7575 | 7392 | 5546,12 | 3422 | 7670 | 2,24 | 1,164 | 25,176 | 7,37E-140 | 8,98E-139 | 0,0014475 | 0,002275 | 0,0018335 | 0,0018335 | TRUE | 0,0268396 | No |
| orf19.6318 | 1259 | 1075 | 983 | 780 | 930 | 675 | 625 | 990 | 1146 | 1087 | 930 | 1172 | 827 | 733 | 640 | 755 | 911,36 | 1084 | 739 | 0,683 | -0,55 | -6,172 | 6,74E-10 | 1,27E-09 | 0,00893 | 0,003487 | 0,0064846 | 0,0064846 | TRUE | 0,110598 | No |

| | | | | | | | | | | | | | | | | | | | | | | | | | | | | | | | |
|---|---|---|---|---|---|---|---|---|---|---|---|---|---|---|---|---|---|---|---|---|---|---|---|---|---|---|---|---|---|---|---|
| orf19.6319 | 7939 | 6679 | 7659 | 4805 | 3519 | 2946 | 3421 | 4474 | 7228 | 6755 | 7249 | 7218 | 3130 | 3199 | 3504 | 3413 | 5211,82 | 7112 | 3312 | 0,466 | -1,103 | -23,247 | 1,54E-119 | 1,56E-118 | 0,0016785 | 0,00229 | 0,001944 | 0,001944 | TRUE | 0,018178 | No |
| orf19.6321 | 4179 | 3036 | 3266 | 2859 | 2057 | 1975 | 1839 | 2756 | 3805 | 3071 | 3091 | 4295 | 1830 | 2144 | 1883 | 2102 | 2777,58 | 3566 | 1990 | 0,559 | -0,84 | -8,435 | 3,31E-17 | 7,79E-17 | 0,0164584 | 0,002513 | 0,0091455 | 0,0091455 | TRUE | 0,2275212 | No |
| orf19.6322 | 37147 | 27718 | 31701 | 20491 | 35723 | 27976 | 33410 | 46513 | 33821 | 28034 | 30002 | 30781 | 31774 | 30374 | 34216 | 35480 | 31810,3 | 30660 | 32961 | 1,075 | 0,104 | 1,617 | 0,105901 | 0,12695 | 0,0054895 | 0,002079 | 0,00398 | 0,00398 | TRUE | 0,0591529 | No |
| orf19.6323 | 425 | 342 | 376 | 256 | 682 | 528 | 622 | 844 | 387 | 346 | 356 | 385 | 607 | 573 | 637 | 644 | 491,74 | 368 | 615 | 1,675 | 0,744 | 10,081 | 6,69E-24 | 1,87E-23 | 0,0009146 | 0,004723 | 0,0030271 | 0,0030271 | TRUE | 0,0234715 | No |
| orf19.6324 | 10438 | 8100 | 9353 | 5383 | 8868 | 6672 | 8328 | 11201 | 9503 | 8192 | 8852 | 8086 | 7888 | 7244 | 8529 | 8544 | 8354,8 | 8658 | 8051 | 0,93 | -0,105 | -1,578 | 0,114658 | 0,136779 | 0,0056776 | 0,002195 | 0,0041443 | 0,0041443 | TRUE | 0,0554163 | No |
| orf19.6325.1 | 153 | 104 | 130 | 113 | 343 | 187 | 226 | 318 | 139 | 105 | 123 | 170 | 305 | 203 | 231 | 243 | 189,93 | 134 | 246 | 1,848 | 0,886 | 5,771 | 7,88E-09 | 1,43E-08 | 0,0285339 | 0,008992 | 0,0167079 | 0,0167079 | TRUE | 0,2984892 | No |
| orf19.6326 | 880 | 652 | 759 | 473 | 1386 | 1066 | 1209 | 1690 | 801 | 659 | 718 | 711 | 1233 | 1157 | 1238 | 1289 | 975,87 | 722 | 1229 | 1,701 | 0,766 | 11,433 | 2,86E-30 | 9,27E-30 | 0,0028105 | 0,003391 | 0,0031923 | 0,0031923 | TRUE | 0,0679057 | No |
| orf19.6327 | 932 | 720 | 820 | 577 | 1553 | 1227 | 1271 | 1891 | 849 | 728 | 776 | 867 | 1381 | 1332 | 1302 | 1442 | 1084,65 | 805 | 1364 | 1,7 | 0,766 | 11,741 | 7,90E-32 | 2,65E-31 | 0,0029013 | 0,003255 | 0,0030747 | 0,0030747 | TRUE | 0,0483851 | No |
| orf19.6328 | 207 | 143 | 172 | 121 | 303 | 254 | 293 | 390 | 188 | 145 | 163 | 182 | 270 | 276 | 300 | 297 | 227,56 | 170 | 286 | 1,692 | 0,758 | 7,217 | 5,30E-13 | 1,11E-12 | 0,0022654 | 0,007842 | 0,0058162 | 0,0058162 | TRUE | 0,1052301 | No |
| orf19.6329 | 445 | 368 | 439 | 270 | 1191 | 1022 | 1101 | 1484 | 405 | 372 | 415 | 406 | 1059 | 1110 | 1128 | 1132 | 753,37 | 400 | 1107 | 2,771 | 1,471 | 23,318 | 2,93E-120 | 3,00E-119 | 1,00E-08 | 0,00379 | 0,0020648 | 0,0020648 | TRUE | 0,0256654 | No |
| orf19.633 | 1405 | 1574 | 1661 | 922 | 783 | 662 | 622 | 804 | 1279 | 1592 | 1572 | 1385 | 696 | 719 | 637 | 613 | 1061,7 | 1457 | 666 | 0,457 | -1,131 | -13,768 | 3,95E-43 | 1,62E-42 | 0,0074726 | 0,003281 | 0,0054055 | 0,0054055 | TRUE | 0,0847784 | No |
| orf19.6336 | 73 | 46 | 52 | 52 | 433 | 382 | 439 | 605 | 66 | 47 | 49 | 78 | 385 | 415 | 450 | 461 | 243,91 | 60 | 428 | 7,264 | 2,861 | 20,8 | 4,32E-96 | 3,58E-95 | 0,0099256 | 0,007453 | 0,0080768 | 0,0080768 | TRUE | 0,2425753 | No |
| orf19.6337 | 25 | 16 | 28 | 19 | 68 | 61 | 64 | 98 | 23 | 16 | 26 | 29 | 60 | 66 | 66 | 75 | 45,12 | 24 | 67 | 2,885 | 1,528 | 6,697 | 2,13E-11 | 4,24E-11 | 1,00E-08 | 0,031312 | 0,0201938 | 0,0201938 | TRUE | 0,2567313 | No |
| orf19.6338 | 1 | 0 | 1 | 4 | 5 | 7 | 1 | 4 | 1 | 0 | 1 | 6 | 4 | 8 | 1 | 3 | 3 | 2 | 4 | 2,32 | 1,214 | 1,25 | 0,211122 | 0,243597 | 0,5644014 | 0,442585 | 0,4794977 | 0,4794977 | TRUE | 0,7309125 | No |
| orf19.6339 | 0 | 0 | 0 | 0 | 0 | 0 | 0 | 0 | 0 | 0 | 0 | 0 | 0 | 0 | 0 | 0 | 0 | 0 | 0 | NA | NA | NA | NA | NA | NA | NA | NA | NA | NA | NA | NA |
| orf19.634 | 2 | 3 | 4 | 2 | 2 | 2 | 2 | 0 | 2 | 3 | 4 | 3 | 2 | 2 | 2 | 0 | 2,21 | 3 | 2 | 0,493 | -1,021 | -1,069 | 0,285099 | 0,321731 | 1,00E-08 | 0,601004 | 0,3676809 | 0,3676809 | TRUE | 0,477626 | No |
| orf19.6340 | 5127 | 6115 | 5563 | 3746 | 5239 | 4372 | 4156 | 5402 | 4668 | 6185 | 5265 | 5627 | 4660 | 4747 | 4256 | 4121 | 4941,03 | 5436 | 4446 | 0,818 | -0,29 | -3,656 | 0,000256 | 0,000376 | 0,0089315 | 0,002304 | 0,005839 | 0,005839 | TRUE | 0,1111607 | No |
| orf19.6341 | 2169 | 2020 | 2251 | 1372 | 3212 | 2899 | 3015 | 3930 | 1975 | 2043 | 2130 | 2061 | 2857 | 3148 | 3088 | 2998 | 2537,4 | 2052 | 3023 | 1,472 | 0,558 | 12 | 3,57E-33 | 1,23E-32 | 0,0010532 | 0,002558 | 0,0016598 | 0,0016598 | TRUE | 0,0164811 | No |
| orf19.6342 | 1003 | 879 | 932 | 428 | 2821 | 1763 | 2380 | 2822 | 913 | 889 | 882 | 643 | 2509 | 1914 | 2437 | 2153 | 1542,57 | 832 | 2253 | 2,701 | 1,433 | 13,458 | 2,78E-41 | 1,11E-40 | 0,0178594 | 0,002893 | 0,0100407 | 0,0100407 | TRUE | 0,2609707 | No |
| orf19.6343 | 10020 | 11251 | 11240 | 6409 | 8721 | 8071 | 7951 | 10181 | 9123 | 11379 | 10638 | 9627 | 7757 | 8763 | 8143 | 7766 | 9149,48 | 10192 | 8107 | 0,795 | -0,33 | -4,762 | 1,92E-06 | 3,14E-06 | 0,0064776 | 0,002181 | 0,0045162 | 0,0045162 | TRUE | 0,075295 | No |
| orf19.6344 | 2584 | 2469 | 2593 | 1532 | 2286 | 1836 | 1892 | 2625 | 2353 | 2497 | 2454 | 2301 | 2033 | 1993 | 1938 | 2002 | 2196,48 | 2401 | 1992 | 0,829 | -0,271 | -6,251 | 4,08E-10 | 7,74E-10 | 0,0003983 | 0,002638 | 0,0013453 | 0,0013453 | TRUE | 0,0085453 | No |
| orf19.6345 | 6169 | 6490 | 6728 | 4221 | 9967 | 9071 | 8576 | 11502 | 5617 | 6564 | 6367 | 6341 | 8865 | 9849 | 8783 | 8774 | 7644,91 | 6222 | 9068 | 1,457 | 0,543 | 9,331 | 1,05E-20 | 2,70E-20 | 0,0037106 | 0,00221 | 0,0031194 | 0,0031194 | TRUE | 0,0529897 | No |
| orf19.6346 | 2357 | 2545 | 2478 | 1708 | 3275 | 2911 | 2643 | 3623 | 2146 | 2574 | 2345 | 2566 | 2913 | 3161 | 2707 | 2764 | 2646,85 | 2408 | 2886 | 1,199 | 0,262 | 3,79 | 0,000151 | 0,000223 | 0,005552 | 0,002536 | 0,0042171 | 0,0042171 | TRUE | 0,0657281 | No |
| orf19.6347 | 3110 | 3203 | 3347 | 2064 | 2703 | 2391 | 2465 | 2945 | 2832 | 3239 | 3168 | 3100 | 2404 | 2596 | 2524 | 2246 | 2763,78 | 3085 | 2442 | 0,791 | -0,338 | -5,689 | 1,28E-08 | 2,29E-08 | 0,0031377 | 0,002515 | 0,0030133 | 0,0030133 | TRUE | 0,0380669 | No |
| orf19.6348 | 657 | 742 | 735 | 491 | 1741 | 1539 | 1485 | 1959 | 598 | 750 | 696 | 738 | 1549 | 1671 | 1521 | 1494 | 1127,06 | 696 | 1559 | 2,246 | 1,167 | 16,039 | 6,78E-58 | 3,60E-57 | 0,004782 | 0,003209 | 0,0040118 | 0,0040118 | TRUE | 0,1096665 | No |
| orf19.6349 | 1797 | 2101 | 2138 | 1150 | 579 | 479 | 550 | 565 | 1636 | 2125 | 2023 | 1727 | 515 | 520 | 563 | 431 | 1192,66 | 1878 | 507 | 0,269 | -1,892 | -19,439 | 3,58E-84 | 2,67E-83 | 0,0130778 | 0,003145 | 0,0078848 | 0,0078848 | TRUE | 0,0996576 | No |
| orf19.635 | 10 | 7 | 11 | 13 | 2 | 4 | 6 | 5 | 9 | 7 | 10 | 20 | 2 | 4 | 6 | 4 | 7,78 | 12 | 4 | 0,355 | -1,496 | -2,678 | 0,007416 | 0,009821 | 1,00E-08 | 0,171931 | 0,1344415 | 0,1344415 | TRUE | 0,558177 | No |
| orf19.6350 | 216 | 165 | 184 | 138 | 41 | 44 | 43 | 58 | 197 | 167 | 174 | 207 | 36 | 48 | 44 | 44 | 114,69 | 186 | 43 | 0,232 | -2,109 | -13,583 | 5,06E-42 | 2,04E-41 | 1,00E-08 | 0,013555 | 0,0094739 | 0,0094739 | TRUE | 0,0883984 | No |
| orf19.6351 | 2970 | 2792 | 3080 | 1684 | 2030 | 1681 | 1710 | 2334 | 2704 | 2824 | 2915 | 2530 | 1806 | 1825 | 1751 | 1780 | 2266,85 | 2743 | 1790 | 0,652 | -0,618 | -12,184 | 3,77E-34 | 1,32E-33 | 0,0013855 | 0,00262 | 0,0020101 | 0,0020101 | TRUE | 0,0310738 | No |
| orf19.6353 | 988 | 897 | 945 | 606 | 798 | 738 | 662 | 927 | 900 | 907 | 894 | 910 | 710 | 801 | 678 | 707 | 813,45 | 903 | 724 | 0,801 | -0,32 | -5,16 | 2,47E-07 | 4,19E-07 | 0,0010992 | 0,003661 | 0,0024502 | 0,0024502 | TRUE | 0,0573226 | No |
| orf19.6354 | 168 | 193 | 184 | 123 | 217 | 206 | 185 | 239 | 153 | 195 | 174 | 185 | 193 | 224 | 189 | 182 | 186,94 | 177 | 197 | 1,116 | 0,158 | 1,378 | 0,168071 | 0,196452 | 0,0043314 | 0,009103 | 0,0071951 | 0,0071951 | TRUE | 0,095859 | No |
| orf19.6355 | 16757 | 14504 | 15782 | 9142 | 16663 | 12471 | 14134 | 18548 | 15257 | 14669 | 14936 | 13733 | 14821 | 13540 | 14475 | 14149 | 14447,5 | 14649 | 14246 | 0,972 | -0,041 | -0,911 | 0,362314 | 0,402396 | 0,0016427 | 0,002128 | 0,0018354 | 0,0018354 | TRUE | 0,0213497 | No |
| orf19.6356 | 1039 | 1224 | 1247 | 710 | 1631 | 1324 | 1357 | 1746 | 946 | 1238 | 1180 | 1067 | 1451 | 1437 | 1390 | 1332 | 1255,05 | 1108 | 1402 | 1,265 | 0,34 | 4,367 | 1,26E-05 | 1,99E-05 | 0,0064617 | 0,00309 | 0,0049875 | 0,0049875 | TRUE | 0,1216487 | No |
| orf19.6357 | 492 | 547 | 590 | 367 | 558 | 521 | 489 | 596 | 448 | 553 | 558 | 551 | 496 | 566 | 501 | 455 | 516,03 | 528 | 504 | 0,955 | -0,066 | -0,718 | 0,472561 | 0,514355 | 0,0077682 | 0,004597 | 0,0061044 | 0,0061044 | TRUE | 0,1274701 | No |
| orf19.6358 | 1179 | 1550 | 1379 | 912 | 968 | 910 | 755 | 1013 | 1073 | 1568 | 1305 | 1370 | 861 | 988 | 773 | 773 | 1088,89 | 1329 | 849 | 0,638 | -0,647 | -5,987 | 2,13E-09 | 3,95E-09 | 0,0176415 | 0,00325 | 0,0102675 | 0,0102675 | TRUE | 0,1981086 | No |
| orf19.6359 | 500 | 464 | 457 | 272 | 278 | 227 | 220 | 320 | 455 | 469 | 433 | 409 | 247 | 246 | 225 | 244 | 341,09 | 442 | 240 | 0,544 | -0,878 | -10,606 | 2,80E-26 | 8,23E-26 | 1,00E-08 | 0,00591 | 0,0034392 | 0,0034392 | TRUE | 0,0239837 | No |
| orf19.6360 | 3006 | 2830 | 2918 | 1796 | 1669 | 1361 | 1484 | 1844 | 2737 | 2862 | 2762 | 2698 | 1485 | 1478 | 1520 | 1407 | 2118,4 | 2765 | 1472 | 0,532 | -0,911 | -20,79 | 5,28E-96 | 4,36E-95 | 0,0003442 | 0,002661 | 0,0013328 | 0,0013328 | TRUE | 0,0115681 | No |
| orf19.6362 | 6606 | 5978 | 6438 | 4056 | 8559 | 7230 | 7813 | 10281 | 6014 | 6046 | 6093 | 6093 | 7613 | 7850 | 8001 | 7842 | 6944,12 | 6062 | 7826 | 1,291 | 0,369 | 11,995 | 3,79E-33 | 1,30E-32 | 0,0001269 | 0,002227 | 0,0007587 | 0,0007587 | TRUE | 0,0041439 | No |
| orf19.6365 | 6681 | 5043 | 5703 | 3976 | 5587 | 4566 | 5153 | 7172 | 6083 | 5100 | 5397 | 5973 | 4969 | 4957 | 5277 | 5471 | 5403,53 | 5638 | 5168 | 0,917 | -0,125 | -2,012 | 0,044224 | 0,055023 | 0,0043646 | 0,002281 | 0,0035094 | 0,0035094 | TRUE | 0,0503747 | No |
| orf19.6366 | 84 | 70 | 64 | 38 | 132 | 99 | 112 | 171 | 76 | 71 | 61 | 57 | 117 | 107 | 115 | 130 | 91,87 | 66 | 117 | 1,764 | 0,819 | 5,345 | 9,06E-08 | 1,57E-07 | 1,00E-08 | 0,016416 | 0,0106569 | 0,0106569 | TRUE | 0,1001431 | No |
| orf19.6367 | 43685 | 34842 | 38478 | 27585 | 72801 | 62477 | 65109 | 93225 | 39773 | 35239 | 36416 | 41437 | 64754 | 67833 | 66680 | 71113 | 52905,6 | 38216 | 67595 | 1,769 | 0,823 | 14,776 | 2,08E-49 | 9,47E-49 | 0,003593 | 0,002062 | 0,0029587 | 0,0029587 | TRUE | 0,039274 | No |
| orf19.6369 | 9898 | 8076 | 8840 | 5080 | 12530 | 9144 | 10888 | 14649 | 9012 | 8168 | 8366 | 7631 | 11145 | 9928 | 11151 | 11174 | 9571,86 | 8294 | 10850 | 1,308 | 0,387 | 6,634 | 3,26E-11 | 6,44E-11 | 0,0038265 | 0,002175 | 0,0031602 | 0,0031602 | TRUE | 0,0414545 | No |
| orf19.637 | 4718 | 3955 | 4203 | 2885 | 5752 | 4862 | 4586 | 6392 | 4296 | 4000 | 3978 | 4334 | 5116 | 5279 | 4697 | 4876 | 4571,82 | 4152 | 4992 | 1,203 | 0,266 | 5,295 | 1,19E-07 | 2,05E-07 | 0,0020681 | 0,002326 | 0,0022069 | 0,0022069 | TRUE | 0,019028 | No |
| orf19.6371 | 223 | 179 | 197 | 100 | 243 | 245 | 225 | 313 | 203 | 181 | 186 | 150 | 216 | 266 | 230 | 239 | 209,01 | 180 | 238 | 1,306 | 0,385 | 3,453 | 0,000555 | 0,000796 | 0,0048807 | 0,008357 | 0,0070162 | 0,0070162 | TRUE | 0,1064498 | No |
| orf19.6373 | 8204 | 6875 | 7796 | 4820 | 2847 | 2379 | 2644 | 3637 | 7469 | 6953 | 7378 | 7240 | 2532 | 2583 | 2708 | 2774 | 4954,84 | 7260 | 2649 | 0,365 | -1,454 | -33,389 | 2,00E-244 | 4,08E-243 | 0,0010199 | 0,002304 | 0,0015704 | 0,0015704 | TRUE | 0,0127892 | No |
| orf19.6374 | 630 | 563 | 623 | 340 | 1119 | 855 | 940 | 1335 | 574 | 569 | 590 | 511 | 995 | 928 | 963 | 1018 | 768,5 | 561 | 976 | 1,735 | 0,795 | 12,593 | 2,31E-36 | 8,49E-36 | 0,0008305 | 0,003756 | 0,0023985 | 0,0023985 | TRUE | 0,0350289 | No |
| orf19.6375 | 8818 | 7232 | 7436 | 4467 | 14141 | 9846 | 11473 | 15698 | 8028 | 7314 | 7038 | 6710 | 12578 | 10690 | 11750 | 11975 | 9510,35 | 7272 | 11748 | 1,615 | 0,691 | 10,836 | 2,32E-27 | 7,02E-27 | 0,0050738 | 0,002176 | 0,0037976 | 0,0037976 | TRUE | 0,0599332 | No |
| orf19.6376 | 2530 | 2144 | 2409 | 1445 | 2433 | 1937 | 2108 | 2975 | 2303 | 2168 | 2280 | 2171 | 2164 | 2103 | 2159 | 2269 | 2202,22 | 2230 | 2174 | 0,974 | -0,037 | -0,844 | 0,398806 | 0,439604 | 0,0006038 | 0,002637 | 0,001439 | 0,001439 | TRUE | 0,0110154 | No |
| orf19.6377 | 1415 | 1177 | 1401 | 800 | 1455 | 1136 | 1296 | 1761 | 1288 | 1190 | 1326 | 1202 | 1294 | 1233 | 1327 | 1343 | 1275,56 | 1252 | 1299 | 1,037 | 0,053 | 0,962 | 0,336195 | 0,376206 | 0,0012693 | 0,003073 | 0,00212 | 0,00212 | TRUE | 0,019227 | No |
| orf19.6378 | 2536 | 1996 | 2176 | 1321 | 3655 | 2598 | 3199 | 4253 | 2309 | 2019 | 2059 | 1984 | 3251 | 2821 | 3276 | 3244 | 2620,44 | 2093 | 3148 | 1,503 | 0,588 | 9,091 | 9,84E-20 | 2,48E-19 | 0,0043577 | 0,002541 | 0,0036132 | 0,0036132 | TRUE | 0,0593584 | No |
| orf19.6379 | 668 | 653 | 636 | 451 | 532 | 437 | 404 | 542 | 608 | 660 | 602 | 677 | 473 | 474 | 414 | 413 | 540,36 | 637 | 444 | 0,698 | -0,52 | -6,802 | 1,03E-11 | 2,09E-11 | 0,0027815 | 0,004482 | 0,0037075 | 0,0037075 | TRUE | 0,0268728 | No |
| orf19.638 | 14229 | 12308 | 12655 | 8815 | 7575 | 6349 | 6011 | 8929 | 12955 | 12448 | 11977 | 13242 | 6738 | 6893 | 6156 | 6811 | 9652,45 | 12656 | 6650 | 0,526 | -0,928 | -19,075 | 4,06E-81 | 2,92E-80 | 0,0020914 | 0,002174 | 0,0021605 | 0,0021605 | TRUE | 0,0302779 | No |
| orf19.6382 | 3383 | 2871 | 3196 | 2202 | 3984 | 3497 | 3852 | 5133 | 3080 | 2904 | 3025 | 3308 | 3544 | 3797 | 3945 | 3915 | 3439,64 | 3079 | 3800 | 1,236 | 0,305 | 5,807 | 6,36E-09 | 1,16E-08 | 0,0022749 | 0,002421 | 0,0023527 | 0,0023527 | TRUE | 0,0288263 | No |
| orf19.6385 | 16005 | 12914 | 14236 | 10114 | 35735 | 30692 | 32422 | 46637 | 14572 | 13061 | 13473 | 15193 | 31785 | 33323 | 33204 | 35575 | 23773,3 | 14075 | 33472 | 2,379 | 1,25 | 22,629 | 2,24E-113 | 2,18E-112 | 0,0034566 | 0,002093 | 0,0028802 | 0,0028802 | TRUE | 0,0346136 | No |
| orf19.6387 | 32317 | 26284 | 29280 | 19494 | 3649 | 3040 | 3288 | 4385 | 29423 | 26583 | 27711 | 29283 | 3246 | 3301 | 3367 | 3345 | 15782,4 | 28250 | 3315 | 0,117 | -3,091 | -72,215 | 0 | 0 | 0,0010575 | 0,002121 | 0,0015999 | 0,0015999 | TRUE | 0,0193267 | No |
| orf19.639 | 164 | 140 | 144 | 108 | 483 | 374 | 418 | 498 | 149 | 142 | 136 | 162 | 430 | 406 | 428 | 380 | 279,13 | 147 | 411 | 2,803 | 1,487 | 15,302 | 7,41E-53 | 3,59E-52 | 1,00E-08 | 0,00677 | 0,0043045 | 0,0043045 | TRUE | 0,0360347 | No |
| orf19.6390 | 1 | 1 | 4 | 3 | 0 | 7 | 1 | 2 | 1 | 1 | 4 | 5 | 0 | 8 | 1 | 2 | 2,55 | 3 | 3 | 0,993 | -0,01 | -0,009 | 0,992448 | 0,993071 | 0,8303191 | 0,520997 | 0,6367187 | 0,6367187 | TRUE | 1,3861381 | No |
| orf19.6391 | 161 | 143 | 158 | 135 | 33 | 27 | 49 | 54 | 147 | 145 | 150 | 203 | 29 | 29 | 50 | 41 | 99,2 | 161 | 37 | 0,235 | -2,087 | -11,01 | 3,42E-28 | 1,05E-27 | 0,0259004 | 0,015354 | 0,018745 | 0,018745 | TRUE | 0,3399697 | No |
| orf19.639.1 | 690 | 532 | 641 | 412 | 623 | 465 | 485 | 726 | 628 | 538 | 607 | 619 | 554 | 505 | 497 | 554 | 562,66 | 598 | 528 | 0,884 | -0,177 | -2,396 | 0,016597 | 0,021436 | 0,0024366 | 0,004385 | 0,0034751 | 0,0034751 | TRUE | 0,0546552 | No |
| orf19.6392 | 452 | 375 | 459 | 270 | 898 | 784 | 827 | 1094 | 412 | 379 | 434 | 406 | 799 | 851 | 847 | 835 | 620,27 | 408 | 833 | 2,04 | 1,028 | 15,557 | 1,42E-54 | 7,11E-54 | 1,00E-08 | 0,004167 | 0,0023289 | 0,0023289 | TRUE | 0,0273628 | No |
| orf19.6393 | 1392 | 1146 | 1254 | 878 | 1414 | 1156 | 1289 | 1842 | 1267 | 1159 | 1187 | 1319 | 1258 | 1255 | 1320 | 1405 | 1271,26 | 1233 | 1310 | 1,065 | 0,091 | 1,504 | 0,132647 | 0,157004 | 0,0021693 | 0,003076 | 0,0027004 | 0,0027004 | TRUE | 0,030355 | No |
| orf19.6396 | 2477 | 2253 | 2331 | 1504 | 1998 | 1780 | 1828 | 2526 | 2255 | 2279 | 2206 | 2259 | 1777 | 1933 | 1872 | 1927 | 2063,49 | 2250 | 1877 | 0,834 | -0,261 | -5,979 | 2,24E-09 | 4,15E-09 | 0,0003756 | 0,002677 | 0,0013438 | 0,0013438 | TRUE | 0,0158744 | No |
| orf19.6398 | 410 | 344 | 392 | 256 | 1622 | 1344 | 1405 | 1847 | 373 | 348 | 371 | 385 | 1443 | 1459 | 1439 | 1409 | 903,31 | 369 | 1438 | 3,901 | 1,964 | 33,052 | 1,46E-239 | 2,94E-238 | 1,00E-08 | 0,003499 | 0,0016231 | 0,0016231 | TRUE | 0,0168709 | No |
| orf19.6399 | 577 | 448 | 500 | 314 | 716 | 574 | 696 | 932 | 525 | 453 | 473 | 472 | 637 | 623 | 713 | 711 | 575,89 | 481 | 671 | 1,395 | 0,48 | 6,325 | 2,53E-10 | 4,84E-10 | 0,0030522 | 0,004331 | 0,0037187 | 0,0037187 | TRUE | 0,0484382 | No |
| orf19.640 | 1010 | 1007 | 1027 | 621 | 1176 | 936 | 963 | 1250 | 920 | 1018 | 972 | 933 | 1046 | 1016 | 986 | 954 | 980,6 | 961 | 1000 | 1,04 | 0,057 | 0,988 | 0,32312 | 0,362721 | 0,0009546 | 0,003384 | 0,0021295 | 0,0021295 | TRUE | 0,0197564 | No |
| orf19.6400 | 1539 | 1238 | 1309 | 989 | 1265 | 942 | 1063 | 1444 | 1401 | 1252 | 1239 | 1486 | 1125 | 1023 | 1089 | 1101 | 1214,48 | 1344 | 1084 | 0,809 | -0,306 | -4,566 | 4,96E-06 | 8,01E-06 | 0,003657 | 0,003125 | 0,0034763 | 0,0034763 | TRUE | 0,0555555 | No |
| orf19.6402 | 69240 | 54656 | 59709 | 43340 | 69089 | 58028 | 62310 | 86851 | 63040 | 55279 | 56509 | 65104 | 61452 | 63002 | 63813 | 66251 | 61806,3 | 59983 | 63630 | 1,061 | 0,085 | 1,514 | 0,130003 | 0,154161 | 0,0037056 | 0,002058 | 0,0030291 | 0,0030291 | TRUE | 0,0403693 | No |

| | | | | | | | | | | | | | | | | | | | | | | | | | | | | | | | |
|---|---|---|---|---|---|---|---|---|---|---|---|---|---|---|---|---|---|---|---|---|---|---|---|---|---|---|---|---|---|---|---|
| orf19.6403 | 3182 | 2713 | 3014 | 1637 | 3419 | 2617 | 3126 | 4048 | 2897 | 2744 | 2852 | 2459 | 3041 | 2841 | 3201 | 3088 | 2890,52 | 2738 | 3043 | 1,11 | 0,15 | 2,551 | 0,01073 | 0,014048 | 0,0033421 | 0,002494 | 0,0029836 | 0,0029836 | TRUE | 0,0544128 | No |
| orf19.6403.1 | 3499 | 2828 | 2957 | 2316 | 5592 | 4169 | 4192 | 6224 | 3186 | 2860 | 2799 | 3479 | 4974 | 4526 | 4293 | 4748 | 3858,07 | 3081 | 4635 | 1,506 | 0,591 | 8,237 | 1,76E-16 | 4,05E-16 | 0,0068582 | 0,002379 | 0,0046699 | 0,0046699 | TRUE | 0,0893582 | No |
| orf19.6404 | 2083 | 1659 | 1907 | 1122 | 2294 | 1787 | 2087 | 2758 | 1896 | 1678 | 1805 | 1685 | 2040 | 1940 | 2137 | 2104 | 1910,8 | 1766 | 2055 | 1,163 | 0,217 | 3,964 | 7,37E-05 | 0,000111 | 0,0019399 | 0,002728 | 0,0023536 | 0,0023536 | TRUE | 0,030275 | No |
| orf19.6405 | 14 | 11 | 9 | 8 | 13 | 14 | 14 | 12 | 13 | 11 | 9 | 12 | 12 | 15 | 14 | 9 | 11,83 | 11 | 12 | 1,121 | 0,165 | 0,416 | 0,677332 | 0,713246 | 1,00E-08 | 0,113677 | 0,0656416 | 0,0656416 | TRUE | 0,1777337 | No |
| orf19.6406 | 2173 | 1686 | 1861 | 1257 | 1594 | 1106 | 1299 | 1777 | 1978 | 1705 | 1761 | 1888 | 1418 | 1201 | 1330 | 1356 | 1579,7 | 1833 | 1326 | 0,724 | -0,465 | -7,095 | 1,30E-12 | 2,69E-12 | 0,0037671 | 0,002873 | 0,0034871 | 0,0034871 | TRUE | 0,0473785 | No |
| orf19.6407 | 2943 | 2403 | 2666 | 1588 | 3844 | 3114 | 3766 | 4843 | 2679 | 2430 | 2523 | 2385 | 3419 | 3381 | 3857 | 3694 | 3046,2 | 2504 | 3588 | 1,431 | 0,517 | 9,086 | 1,03E-19 | 2,60E-19 | 0,0028999 | 0,002471 | 0,0027679 | 0,0027679 | TRUE | 0,0305231 | No |
| orf19.6408 | 6033 | 5022 | 5788 | 3760 | 2244 | 1937 | 2214 | 2887 | 5493 | 5079 | 5478 | 5648 | 1996 | 2103 | 2267 | 2202 | 3783,32 | 5424 | 2142 | 0,395 | -1,34 | -25,761 | 2,44E-146 | 3,13E-145 | 0,0020996 | 0,002386 | 0,0022806 | 0,0022806 | TRUE | 0,0259508 | No |
| orf19.641 | 874 | 904 | 979 | 497 | 1335 | 999 | 1113 | 1428 | 796 | 914 | 927 | 747 | 1187 | 1085 | 1140 | 1089 | 985,55 | 846 | 1125 | 1,327 | 0,408 | 5,565 | 2,62E-08 | 4,62E-08 | 0,0050334 | 0,003377 | 0,0041052 | 0,0041052 | TRUE | 0,0662218 | No |
| orf19.6411 | 1586 | 1304 | 1451 | 985 | 702 | 593 | 646 | 819 | 1444 | 1319 | 1373 | 1480 | 624 | 644 | 662 | 625 | 1021,28 | 1404 | 639 | 0,455 | -1,136 | -19,698 | 2,23E-86 | 1,70E-85 | 0,0009239 | 0,00333 | 0,0020783 | 0,0020783 | TRUE | 0,0197169 | No |
| orf19.6413 | 798 | 764 | 881 | 504 | 1134 | 1002 | 1002 | 1436 | 727 | 773 | 834 | 757 | 1009 | 1088 | 1026 | 1095 | 913,53 | 773 | 1054 | 1,364 | 0,448 | 7,306 | 2,75E-13 | 5,81E-13 | 0,0014923 | 0,003483 | 0,0024784 | 0,0024784 | TRUE | 0,0358582 | No |
| orf19.6414 | 1097 | 906 | 1089 | 699 | 1251 | 1097 | 1245 | 1713 | 999 | 916 | 1031 | 1050 | 1113 | 1191 | 1275 | 1307 | 1110,15 | 999 | 1222 | 1,225 | 0,292 | 4,356 | 1,32E-05 | 2,09E-05 | 0,0035626 | 0,003227 | 0,003402 | 0,003402 | TRUE | 0,0442239 | No |
| orf19.6414.3 | 3607 | 3151 | 3536 | 2123 | 2653 | 1963 | 2128 | 2765 | 3284 | 3187 | 3347 | 3189 | 2360 | 2131 | 2179 | 2109 | 2723,26 | 3252 | 2195 | 0,675 | -0,568 | -12,064 | 1,64E-33 | 5,69E-33 | 0,0011204 | 0,002522 | 0,0017466 | 0,0017466 | TRUE | 0,0314976 | No |
| orf19.6415.1 | 4106 | 2937 | 3079 | 2163 | 6389 | 4127 | 4482 | 6381 | 3738 | 2970 | 2914 | 3249 | 5683 | 4481 | 4590 | 4867 | 4061,65 | 3218 | 4905 | 1,525 | 0,608 | 6,875 | 6,19E-12 | 1,26E-11 | 0,0126374 | 0,002362 | 0,0072582 | 0,0072582 | TRUE | 0,1459084 | No |
| orf19.6416 | 4480 | 3712 | 4180 | 2844 | 2507 | 2115 | 2282 | 3097 | 4079 | 3754 | 3956 | 4272 | 2230 | 2296 | 2337 | 2362 | 3160,87 | 4015 | 2306 | 0,575 | -0,798 | -16,619 | 5,11E-62 | 2,87E-61 | 0,0013235 | 0,002455 | 0,0018796 | 0,0018796 | TRUE | 0,0230926 | No |
| orf19.6417 | 18389 | 16197 | 17757 | 10164 | 23704 | 19356 | 22323 | 28952 | 16742 | 16382 | 16805 | 15268 | 21084 | 21015 | 22862 | 22085 | 19030,4 | 16299 | 21762 | 1,335 | 0,417 | 9,336 | 9,98E-21 | 2,58E-20 | 0,0017041 | 0,002106 | 0,0018579 | 0,0018579 | TRUE | 0,0219006 | No |
| orf19.6418 | 5621 | 4717 | 5268 | 3052 | 5182 | 3864 | 4283 | 5811 | 5118 | 4771 | 4986 | 4585 | 4609 | 4195 | 4386 | 4433 | 4635,27 | 4865 | 4406 | 0,905 | -0,144 | -3,044 | 0,002335 | 0,003206 | 0,0016257 | 0,002322 | 0,0019233 | 0,0019233 | TRUE | 0,0176027 | No |
| orf19.6419.1 | 828 | 714 | 696 | 399 | 891 | 627 | 646 | 893 | 754 | 722 | 659 | 599 | 793 | 681 | 662 | 681 | 693,76 | 684 | 704 | 1,027 | 0,038 | 0,458 | 0,64721 | 0,684694 | 0,0068711 | 0,003941 | 0,0053398 | 0,0053398 | TRUE | 0,0857543 | No |
| orf19.642 | 911 | 852 | 946 | 547 | 1054 | 888 | 893 | 1201 | 829 | 862 | 895 | 822 | 937 | 964 | 915 | 916 | 892,55 | 852 | 933 | 1,093 | 0,128 | 2,332 | 0,01968 | 0,025254 | 1,00E-08 | 0,003517 | 0,0017643 | 0,0017643 | TRUE | 0,0142216 | No |
| orf19.6420 | 8235 | 6767 | 7567 | 5932 | 2416 | 2257 | 2358 | 3406 | 7498 | 6844 | 7161 | 8911 | 2149 | 2450 | 2415 | 2598 | 5003,31 | 7604 | 2403 | 0,316 | -1,661 | -20,398 | 1,73E-92 | 1,38E-91 | 0,0098505 | 0,002301 | 0,0061036 | 0,0061036 | TRUE | 0,1621312 | No |
| orf19.6422 | 3310 | 3068 | 3249 | 2117 | 2826 | 2491 | 2505 | 3397 | 3014 | 3103 | 3075 | 3180 | 2514 | 2705 | 2565 | 2591 | 2843,3 | 3093 | 2594 | 0,839 | -0,253 | -6,317 | 2,67E-10 | 5,10E-10 | 0,0002598 | 0,002502 | 0,0011898 | 0,0011898 | TRUE | 0,0096484 | No |
| orf19.6423 | 2164 | 1874 | 2010 | 1324 | 1992 | 1538 | 1706 | 2316 | 1970 | 1895 | 1902 | 1989 | 1772 | 1670 | 1747 | 1767 | 1839,03 | 1939 | 1739 | 0,898 | -0,155 | -3,582 | 0,000341 | 0,000496 | 1,00E-08 | 0,002755 | 0,001255 | 0,001255 | TRUE | 0,008262 | No |
| orf19.6424 | 2645 | 1965 | 2109 | 1545 | 1948 | 1512 | 1677 | 2345 | 2408 | 1987 | 1996 | 2321 | 1733 | 1642 | 1717 | 1789 | 1949,11 | 2178 | 1720 | 0,791 | -0,339 | -4,913 | 8,98E-07 | 1,50E-06 | 0,0049466 | 0,002715 | 0,0040517 | 0,0040517 | TRUE | 0,0640827 | No |
| orf19.6425 | 1822 | 1657 | 1741 | 1030 | 1964 | 1515 | 1801 | 2384 | 1659 | 1676 | 1648 | 1547 | 1747 | 1645 | 1844 | 1819 | 1698,05 | 1632 | 1764 | 1,079 | 0,11 | 2,142 | 0,032199 | 0,040574 | 0,0011262 | 0,002815 | 0,0019522 | 0,0019522 | TRUE | 0,0239107 | No |
| orf19.643 | 485 | 436 | 437 | 279 | 591 | 486 | 544 | 635 | 442 | 441 | 414 | 419 | 526 | 528 | 557 | 484 | 476,26 | 429 | 524 | 1,217 | 0,284 | 3,917 | 8,98E-05 | 0,000135 | 1,00E-08 | 0,004811 | 0,0029064 | 0,0029064 | TRUE | 0,0324633 | No |
| orf19.6431 | 1197 | 1049 | 1090 | 674 | 1105 | 882 | 1014 | 1434 | 1090 | 1061 | 1032 | 1012 | 983 | 958 | 1038 | 1094 | 1033,45 | 1049 | 1018 | 0,971 | -0,042 | -0,723 | 0,469388 | 0,51125 | 0,0014443 | 0,003315 | 0,0023347 | 0,0023347 | TRUE | 0,0316417 | No |
| orf19.6432 | 2036 | 1677 | 1915 | 1200 | 2244 | 1874 | 2059 | 2836 | 1854 | 1696 | 1812 | 1803 | 1996 | 2035 | 2109 | 2163 | 1933,42 | 1791 | 2076 | 1,159 | 0,213 | 4,443 | 8,85E-06 | 1,41E-05 | 0,0009243 | 0,00272 | 0,001683 | 0,001683 | TRUE | 0,0159347 | No |
| orf19.6434 | 5702 | 4666 | 5110 | 3392 | 9156 | 7187 | 7723 | 10704 | 5191 | 4719 | 4836 | 5095 | 8144 | 7803 | 7909 | 8165 | 6482,94 | 4960 | 8005 | 1,615 | 0,691 | 16,734 | 7,36E-63 | 4,19E-62 | 0,0010126 | 0,002241 | 0,001471 | 0,001471 | TRUE | 0,013133 | No |
| orf19.6435 | 906 | 739 | 762 | 594 | 1526 | 1170 | 1239 | 1812 | 825 | 747 | 721 | 892 | 1357 | 1270 | 1269 | 1382 | 1058,06 | 796 | 1320 | 1,664 | 0,735 | 10,539 | 5,70E-26 | 1,67E-25 | 0,0043942 | 0,003286 | 0,0036376 | 0,0036376 | TRUE | 0,0696085 | No |
| orf19.6436 | 1507 | 1257 | 1405 | 889 | 1202 | 931 | 1034 | 1448 | 1372 | 1271 | 1330 | 1335 | 1069 | 1011 | 1059 | 1105 | 1193,99 | 1327 | 1061 | 0,8 | -0,321 | -6,254 | 4,01E-10 | 7,63E-10 | 0,0004094 | 0,003143 | 0,0016899 | 0,0016899 | TRUE | 0,011651 | No |
| orf19.6437 | 725 | 644 | 678 | 481 | 525 | 438 | 505 | 673 | 660 | 651 | 642 | 723 | 467 | 476 | 517 | 513 | 581,09 | 669 | 493 | 0,74 | -0,434 | -6,334 | 2,39E-10 | 4,58E-10 | 0,0007701 | 0,00431 | 0,0027514 | 0,0027514 | TRUE | 0,0296478 | No |
| orf19.6438 | 2973 | 2538 | 2884 | 1750 | 4997 | 4168 | 4547 | 6061 | 2707 | 2567 | 2729 | 2629 | 4445 | 4525 | 4657 | 4623 | 3610,25 | 2658 | 4562 | 1,716 | 0,779 | 20,341 | 5,62E-92 | 4,47E-91 | 0,0003186 | 0,002403 | 0,0011028 | 0,0011028 | TRUE | 0,0068533 | No |
| orf19.644 | 217 | 195 | 198 | 131 | 137 | 106 | 119 | 155 | 198 | 197 | 187 | 197 | 122 | 115 | 122 | 118 | 157 | 195 | 119 | 0,613 | -0,706 | -6,392 | 1,64E-10 | 3,15E-10 | 1,00E-08 | 0,010451 | 0,0050766 | 0,0050766 | TRUE | 0,0075673 | No |
| orf19.6440 | 6923 | 5801 | 6557 | 3893 | 2422 | 1993 | 2284 | 3134 | 6303 | 5867 | 6206 | 5848 | 2154 | 2164 | 2339 | 2391 | 4158,95 | 6056 | 2262 | 0,374 | -1,421 | -28,811 | 1,54E-182 | 2,41E-181 | 0,0017578 | 0,002355 | 0,0020392 | 0,0020392 | TRUE | 0,0184579 | No |
| orf19.6442 | 2875 | 2430 | 2725 | 1680 | 4707 | 3992 | 4443 | 5820 | 2618 | 2458 | 2579 | 2524 | 4187 | 4334 | 4550 | 4440 | 3461,06 | 2545 | 4378 | 1,72 | 0,782 | 18,521 | 1,40E-76 | 9,49E-76 | 0,0007717 | 0,002419 | 0,0013941 | 0,0013941 | TRUE | 0,0106134 | No |
| orf19.6443 | 852 | 703 | 712 | 591 | 1560 | 1231 | 1334 | 1870 | 776 | 711 | 674 | 888 | 1388 | 1337 | 1366 | 1426 | 1070,63 | 762 | 1379 | 1,819 | 0,863 | 11,53 | 9,32E-31 | 3,05E-30 | 0,0056047 | 0,003271 | 0,0043325 | 0,0043325 | TRUE | 0,1322099 | No |
| orf19.6444 | 1966 | 1613 | 1873 | 1276 | 2066 | 1786 | 2052 | 2661 | 1790 | 1631 | 1773 | 1917 | 1838 | 1939 | 2102 | 2030 | 1877,35 | 1778 | 1977 | 1,114 | 0,155 | 2,558 | 0,010514 | 0,01378 | 0,0031155 | 0,002741 | 0,0029975 | 0,0029975 | TRUE | 0,0366398 | No |
| orf19.6445 | 1076 | 850 | 960 | 673 | 22443 | 16400 | 18367 | 26464 | 980 | 860 | 909 | 1011 | 19962 | 17806 | 18810 | 20187 | 10065,5 | 940 | 19191 | 20,468 | 4,355 | 72,188 | 0 | 0 | 0,0035588 | 0,002168 | 0,0028981 | 0,0028981 | TRUE | 0,0386224 | No |
| orf19.6447 | 9477 | 7534 | 8049 | 5073 | 4127 | 2943 | 3578 | 4887 | 8628 | 7620 | 7618 | 7620 | 3671 | 3195 | 3664 | 3728 | 5718,08 | 7872 | 3564 | 0,453 | -1,143 | -18,679 | 7,40E-78 | 5,11E-77 | 0,0041997 | 0,002268 | 0,0033948 | 0,0033948 | TRUE | 0,0585214 | No |
| orf19.6448 | 4123 | 3306 | 3813 | 2397 | 2299 | 1846 | 2024 | 2697 | 3754 | 3344 | 3609 | 3601 | 2045 | 2004 | 2073 | 2057 | 2810,76 | 3577 | 2045 | 0,572 | -0,807 | -17,706 | 3,78E-70 | 2,38E-69 | 0,0010484 | 0,002507 | 0,0016134 | 0,0016134 | TRUE | 0,0239084 | No |
| orf19.6449 | 321 | 237 | 270 | 172 | 443 | 378 | 397 | 558 | 292 | 240 | 256 | 258 | 394 | 410 | 407 | 426 | 335,32 | 262 | 409 | 1,563 | 0,644 | 7,545 | 4,53E-14 | 9,77E-14 | 1,00E-08 | 0,005977 | 0,0038222 | 0,0038222 | TRUE | 0,0781013 | No |
| orf19.6450 | 3 | 1 | 1 | 1 | 2 | 2 | 3 | 0 | 3 | 1 | 1 | 2 | 2 | 2 | 3 | 0 | 1,65 | 2 | 2 | 1,08 | 0,112 | 0,102 | 0,919004 | 0,93024 | 1,00E-08 | 0,801788 | 0,5417139 | 0,5417139 | TRUE | 0,5325085 | No |
| orf19.645.1 | 1580 | 1656 | 1615 | 1087 | 1330 | 1328 | 1157 | 1385 | 1439 | 1675 | 1528 | 1633 | 1183 | 1442 | 1185 | 1056 | 1392,62 | 1569 | 1216 | 0,775 | -0,368 | -4,24 | 2,24E-05 | 3,48E-05 | 0,0101668 | 0,002986 | 0,0064977 | 0,0064977 | TRUE | 0,1845826 | No |
| orf19.6452 | 2712 | 2313 | 2276 | 1507 | 2095 | 1555 | 1541 | 2190 | 2469 | 2339 | 2154 | 2264 | 1863 | 1688 | 1578 | 1671 | 2003,34 | 2306 | 1700 | 0,737 | -0,44 | -7,099 | 1,26E-12 | 2,61E-12 | 0,0034209 | 0,002696 | 0,0031828 | 0,0031828 | TRUE | 0,0508627 | No |
| orf19.6453 | 1337 | 1413 | 1419 | 897 | 852 | 705 | 810 | 946 | 1217 | 1429 | 1343 | 1347 | 758 | 765 | 830 | 722 | 1051,4 | 1334 | 769 | 0,576 | -0,797 | -12,062 | 1,67E-33 | 5,79E-33 | 0,0030688 | 0,003293 | 0,0031762 | 0,0031762 | TRUE | 0,0437453 | No |
| orf19.6455 | 1716 | 1466 | 1561 | 1013 | 928 | 746 | 755 | 1086 | 1562 | 1483 | 1477 | 1522 | 825 | 810 | 773 | 828 | 1160,13 | 1511 | 809 | 0,536 | -0,899 | -17,732 | 2,38E-70 | 1,50E-69 | 1,00E-08 | 0,003176 | 0,001539 | 0,001539 | TRUE | 0,0104311 | No |
| orf19.6456 | 964 | 960 | 1077 | 757 | 1039 | 889 | 829 | 1100 | 878 | 971 | 1019 | 1137 | 924 | 965 | 849 | 839 | 947,81 | 1001 | 894 | 0,895 | -0,16 | -1,991 | 0,046519 | 0,057741 | 0,0066998 | 0,003431 | 0,0051092 | 0,0051092 | TRUE | 0,0931284 | No |
| orf19.6457 | 1553 | 1517 | 1677 | 914 | 1802 | 1456 | 1523 | 2030 | 1414 | 1534 | 1587 | 1373 | 1603 | 1581 | 1560 | 1549 | 1525,03 | 1477 | 1573 | 1,063 | 0,088 | 1,61 | 0,107335 | 0,128548 | 0,0015487 | 0,002903 | 0,002199 | 0,002199 | TRUE | 0,0302884 | No |
| orf19.6458 | 19 | 40 | 39 | 30 | 65 | 51 | 38 | 56 | 17 | 40 | 37 | 45 | 58 | 55 | 39 | 43 | 41,82 | 35 | 49 | 1,417 | 0,502 | 1,867 | 0,06197 | 0,07609 | 0,0634709 | 0,033626 | 0,0443868 | 0,0443868 | TRUE | 0,9177825 | No |
| orf19.6458.1 | 2135 | 2488 | 2326 | 1107 | 2465 | 1718 | 1930 | 2303 | 1944 | 2516 | 2201 | 1663 | 2193 | 1865 | 1977 | 1757 | 2014,44 | 2081 | 1948 | 0,935 | -0,097 | -0,911 | 0,36229 | 0,402396 | 0,018408 | 0,002693 | 0,0104168 | 0,0104168 | TRUE | 0,2285986 | No |
| orf19.6459 | 620 | 595 | 647 | 353 | 818 | 709 | 636 | 900 | 564 | 602 | 612 | 530 | 728 | 770 | 651 | 687 | 643,01 | 577 | 709 | 1,223 | 0,291 | 3,985 | 6,74E-05 | 0,000102 | 0,0028846 | 0,004091 | 0,0035301 | 0,0035301 | TRUE | 0,0375705 | No |
| orf19.646 | 1E+05 | 1E+05 | 1E+05 | 78054 | 2E+05 | 2E+05 | 2E+05 | 3E+05 | 1E+05 | 1E+05 | 1E+05 | 1E+05 | 2E+05 | 2E+05 | 2E+05 | 2E+05 | 156041 | 127269 | 184812 | 1,452 | 0,538 | 10,243 | 1,28E-24 | 3,63E-24 | 0,0030143 | 0,002045 | 0,0026457 | 0,0026457 | TRUE | 0,0343801 | No |
| orf19.6460 | 1909 | 1876 | 2080 | 1249 | 5070 | 4404 | 4239 | 5607 | 1738 | 1897 | 1969 | 1876 | 4510 | 4782 | 4341 | 4277 | 3173,7 | 1870 | 4478 | 2,394 | 1,259 | 23,565 | 8,76E-123 | 9,15E-122 | 0,0020897 | 0,002453 | 0,0023503 | 0,0023503 | TRUE | 0,028659 | No |
| orf19.6461 | 1135 | 957 | 1035 | 623 | 1538 | 1042 | 1330 | 1673 | 1033 | 968 | 980 | 936 | 1368 | 1131 | 1362 | 1276 | 1131,78 | 979 | 1284 | 1,311 | 0,39 | 5,733 | 9,84E-09 | 1,77E-08 | 0,0038469 | 0,003204 | 0,0035377 | 0,0035377 | TRUE | 0,0746476 | No |
| orf19.6462 | 1666 | 1394 | 1552 | 1020 | 1512 | 1282 | 1227 | 1733 | 1517 | 1410 | 1469 | 1532 | 1345 | 1392 | 1257 | 1322 | 1405,38 | 1482 | 1329 | 0,897 | -0,156 | -3,047 | 0,002312 | 0,003175 | 0,000777 | 0,002977 | 0,0018128 | 0,0018128 | TRUE | 0,0153225 | No |
| orf19.6463 | 3383 | 3411 | 3532 | 2002 | 2976 | 2400 | 2449 | 3239 | 3080 | 3450 | 3343 | 3007 | 2647 | 2606 | 2508 | 2471 | 2888,95 | 3220 | 2558 | 0,794 | -0,334 | -6,26 | 3,86E-10 | 7,34E-10 | 0,002183 | 0,002494 | 0,002377 | 0,002377 | TRUE | 0,0277881 | No |
| orf19.6464 | 2484 | 2523 | 2667 | 1635 | 1222 | 1004 | 1052 | 1328 | 2262 | 2552 | 2524 | 2456 | 1087 | 1090 | 1077 | 1013 | 1757,6 | 2448 | 1067 | 0,435 | -1,199 | -22,051 | 9,35E-108 | 8,66E-107 | 0,0017302 | 0,002789 | 0,0021825 | 0,0021825 | TRUE | 0,0333088 | No |
| orf19.6465 | 213 | 173 | 202 | 156 | 173 | 183 | 149 | 253 | 194 | 175 | 191 | 234 | 154 | 199 | 153 | 193 | 186,57 | 198 | 175 | 0,885 | -0,176 | -1,385 | 0,166059 | 0,194315 | 0,0109276 | 0,009117 | 0,0100613 | 0,0100613 | TRUE | 0,1337955 | No |
| orf19.6469 | 670 | 623 | 681 | 467 | 490 | 538 | 481 | 709 | 610 | 630 | 645 | 702 | 436 | 584 | 493 | 541 | 579,94 | 647 | 514 | 0,795 | -0,331 | -3,717 | 0,000202 | 0,000298 | 0,0072334 | 0,004315 | 0,0058552 | 0,0058552 | TRUE | 0,1226328 | No |
| orf19.6470 | 820 | 726 | 808 | 496 | 770 | 639 | 610 | 837 | 747 | 734 | 765 | 745 | 685 | 694 | 625 | 638 | 704,06 | 748 | 660 | 0,882 | -0,18 | -2,976 | 0,002921 | 0,003978 | 1,00E-08 | 0,003913 | 0,0021051 | 0,0021051 | TRUE | 0,0143272 | No |
| orf19.6472 | 5306 | 4112 | 4318 | 3078 | 3382 | 2560 | 2543 | 3744 | 4831 | 4159 | 4087 | 4624 | 3008 | 2779 | 2604 | 2856 | 3618,49 | 4425 | 2812 | 0,636 | -0,653 | -10,065 | 7,91E-24 | 2,21E-23 | 0,0047112 | 0,002402 | 0,0037584 | 0,0037584 | TRUE | 0,0475753 | No |
| orf19.647.3 | 173 | 184 | 187 | 100 | 273 | 224 | 200 | 304 | 158 | 186 | 177 | 150 | 243 | 243 | 205 | 232 | 199,19 | 168 | 231 | 1,37 | 0,454 | 4,173 | 3,01E-05 | 4,65E-05 | 0,0019138 | 0,008669 | 0,0061771 | 0,0061771 | TRUE | 0,0596038 | No |
| orf19.6474 | 3666 | 3045 | 3377 | 2001 | 2801 | 2032 | 2201 | 3164 | 3338 | 3080 | 3196 | 3006 | 2491 | 2206 | 2254 | 2414 | 2748,06 | 3155 | 2341 | 0,742 | -0,43 | -8,048 | 8,38E-16 | 1,90E-15 | 0,0021888 | 0,002518 | 0,0023751 | 0,0023751 | TRUE | 0,0223178 | No |
| orf19.6475 | 32 | 26 | 29 | 20 | 129 | 90 | 91 | 140 | 29 | 26 | 27 | 30 | 115 | 98 | 93 | 107 | 65,67 | 28 | 103 | 3,679 | 1,879 | 9,658 | 4,56E-22 | 1,21E-21 | 1,00E-08 | 0,022153 | 0,0130286 | 0,0130286 | TRUE | 0,056596 | No |
| orf19.6476 | 2276 | 2351 | 2546 | 1415 | 2250 | 1965 | 2021 | 2544 | 2072 | 2378 | 2410 | 2126 | 2001 | 2133 | 2070 | 1941 | 2141,27 | 2246 | 2036 | 0,905 | -0,144 | -2,375 | 0,017527 | 0,0226 | 0,0031518 | 0,002654 | 0,0030347 | 0,0030347 | TRUE | 0,0348645 | No |
| orf19.6477 | 2895 | 3174 | 3291 | 1871 | 3576 | 2974 | 2988 | 3880 | 2636 | 3210 | 3115 | 2811 | 3181 | 3229 | 3060 | 2960 | 3025,07 | 2943 | 3108 | 1,055 | 0,078 | 1,209 | 0,226638 | 0,26014 | 0,0044168 | 0,002474 | 0,0036348 | 0,0036348 | TRUE | 0,0618685 | No |
| orf19.6478 | 13153 | 13445 | 13848 | 7578 | 5034 | 4035 | 4357 | 5406 | 11975 | 13598 | 13106 | 11383 | 4478 | 4381 | 4462 | 4124 | 8438,37 | 12516 | 4361 | 0,348 | -1,522 | -25,73 | 5,44E-146 | 6,94E-145 | 0,0037917 | 0,002194 | 0,0032093 | 0,0032093 | TRUE | 0,0448279 | No |
| orf19.6479 | 906 | 860 | 841 | 497 | 844 | 784 | 717 | 975 | 825 | 870 | 796 | 747 | 751 | 851 | 734 | 744 | 789,64 | 810 | 770 | 0,948 | -0,077 | -1,115 | 0,264688 | 0,300568 | 0,0028608 | 0,00371 | 0,0033247 | 0,0033247 | TRUE | 0,0572757 | No |

| | | | | | | | | | | | | | | | | | | | | | | | | | | | | | | | |
|---|---|---|---|---|---|---|---|---|---|---|---|---|---|---|---|---|---|---|---|---|---|---|---|---|---|---|---|---|---|---|---|
| orf19.6480 | 508 | 581 | 564 | 321 | 511 | 409 | 427 | 512 | 463 | 588 | 534 | 482 | 455 | 444 | 437 | 391 | 474,07 | 517 | 432 | 0,833 | -0,263 | -2,955 | 0,003123 | 0,004246 | 0,0062422 | 0,004824 | 0,0054952 | 0,0054952 | TRUE | 0,1010331 | No |
| orf19.6481 | 2935 | 2803 | 3029 | 1765 | 2891 | 2376 | 2458 | 3292 | 2672 | 2835 | 2867 | 2651 | 2571 | 2580 | 2517 | 2511 | 2650,59 | 2756 | 2545 | 0,922 | -0,117 | -2,776 | 0,005498 | 0,007352 | 0,0004958 | 0,002535 | 0,0013164 | 0,0013164 | TRUE | 0,0088512 | No |
| orf19.6482 | 304 | 349 | 348 | 192 | 306 | 252 | 236 | 310 | 277 | 353 | 329 | 288 | 272 | 274 | 242 | 236 | 283,93 | 312 | 256 | 0,818 | -0,29 | -2,84 | 0,004516 | 0,006072 | 0,0062024 | 0,00669 | 0,0064763 | 0,0064763 | TRUE | 0,0892906 | No |
| orf19.6484 | 229 | 171 | 164 | 134 | 128 | 98 | 110 | 155 | 208 | 173 | 155 | 201 | 114 | 106 | 113 | 118 | 148,64 | 184 | 113 | 0,615 | -0,702 | -5,479 | 4,28E-08 | 7,50E-08 | 0,0056793 | 0,010925 | 0,0087225 | 0,0087225 | TRUE | 0,1297312 | No |
| orf19.6486 | 31 | 22 | 27 | 26 | 68 | 55 | 65 | 64 | 28 | 22 | 26 | 39 | 60 | 60 | 67 | 49 | 43,83 | 29 | 59 | 2,084 | 1,06 | 4,653 | 3,28E-06 | 5,33E-06 | 0,0047697 | 0,032174 | 0,0229562 | 0,0229562 | TRUE | 0,2509452 | No |
| orf19.6487 | 16 | 19 | 19 | 19 | 216 | 210 | 167 | 237 | 15 | 19 | 18 | 29 | 192 | 228 | 171 | 181 | 106,53 | 20 | 193 | 9,973 | 3,318 | 15,677 | 2,17E-55 | 1,10E-54 | 0,0107981 | 0,014437 | 0,0132509 | 0,0132509 | TRUE | 0,2736019 | No |
| orf19.6488 | 8 | 7 | 11 | 5 | 26 | 17 | 19 | 26 | 7 | 7 | 10 | 8 | 23 | 18 | 19 | 20 | 14,14 | 8 | 20 | 2,495 | 1,319 | 3,459 | 0,000541 | 0,000777 | 1,00E-08 | 0,095427 | 0,0514609 | 0,0514609 | TRUE | 0,1292553 | No |
| orf19.6489 | 10048 | 8935 | 9554 | 6141 | 2171 | 1751 | 1833 | 2380 | 9148 | 9037 | 9042 | 9225 | 1931 | 1901 | 1877 | 1815 | 5497,08 | 9113 | 1881 | 0,206 | -2,277 | -64,859 | 0 | 0 | 1,00E-08 | 0,002277 | 0,0008761 | 0,0008761 | TRUE | 0,0072839 | No |
| orf19.649 | 8118 | 8269 | 8736 | 4802 | 1908 | 1600 | 1682 | 2217 | 7391 | 8363 | 8268 | 7213 | 1697 | 1737 | 1723 | 1691 | 4760,44 | 7809 | 1712 | 0,219 | -2,19 | -38,955 | 0 | 0 | 0,0029579 | 0,002315 | 0,0026943 | 0,0026943 | TRUE | 0,0314872 | No |
| orf19.6491 | 29 | 41 | 42 | 18 | 76 | 67 | 62 | 90 | 26 | 41 | 40 | 27 | 68 | 73 | 63 | 69 | 50,89 | 34 | 68 | 2,003 | 1,002 | 4,846 | 1,26E-06 | 2,08E-06 | 1,00E-08 | 0,027993 | 0,0185983 | 0,0185983 | TRUE | 0,1953357 | No |
| orf19.6492 | 1970 | 2270 | 2457 | 1481 | 1509 | 1343 | 1338 | 1748 | 1794 | 2296 | 2325 | 2225 | 1342 | 1458 | 1370 | 1333 | 1767,94 | 2160 | 1376 | 0,637 | -0,65 | -8,309 | 9,67E-17 | 2,24E-16 | 0,007587 | 0,002784 | 0,005294 | 0,005294 | TRUE | 0,1610765 | No |
| orf19.6493 | 9 | 9 | 8 | 4 | 4 | 3 | 7 | 12 | 8 | 9 | 8 | 6 | 4 | 3 | 7 | 9 | 6,75 | 8 | 6 | 0,755 | -0,405 | -0,757 | 0,448876 | 0,490991 | 1,00E-08 | 0,197689 | 0,1275463 | 0,1275463 | TRUE | 0,4787749 | No |
| orf19.6494 | 2671 | 2199 | 2470 | 1730 | 1729 | 1394 | 1599 | 2333 | 2432 | 2224 | 2338 | 2599 | 1538 | 1513 | 1638 | 1780 | 2007,61 | 2398 | 1617 | 0,676 | -0,566 | -8,647 | 5,29E-18 | 1,27E-17 | 0,0042875 | 0,002695 | 0,0035954 | 0,0035954 | TRUE | 0,0568186 | No |
| orf19.6496 | 1691 | 1347 | 1410 | 907 | 994 | 736 | 793 | 1135 | 1540 | 1362 | 1334 | 1362 | 884 | 799 | 812 | 866 | 1120 | 1400 | 840 | 0,601 | -0,736 | -11,587 | 4,80E-31 | 1,58E-30 | 0,0026795 | 0,003216 | 0,0029309 | 0,0029309 | TRUE | 0,056575 | No |
| orf19.6498 | 3065 | 2469 | 2731 | 1748 | 2933 | 2239 | 2487 | 3444 | 2791 | 2497 | 2585 | 2626 | 2609 | 2431 | 2547 | 2627 | 2588,99 | 2625 | 2554 | 0,973 | -0,039 | -0,811 | 0,417128 | 0,458481 | 0,0011849 | 0,002547 | 0,0018385 | 0,0018385 | TRUE | 0,0229906 | No |
| orf19.6499 | 1011 | 976 | 1076 | 698 | 1330 | 1109 | 1114 | 1420 | 920 | 987 | 1018 | 1049 | 1183 | 1204 | 1141 | 1083 | 1073,19 | 994 | 1153 | 1,162 | 0,216 | 3,629 | 0,000285 | 0,000416 | 0,0017089 | 0,003268 | 0,002457 | 0,002457 | TRUE | 0,0302001 | No |
| orf19.6500 | 3040 | 2487 | 2667 | 1888 | 2286 | 1969 | 2013 | 2885 | 2768 | 2515 | 2524 | 2836 | 2033 | 2138 | 2062 | 2201 | 2384,58 | 2661 | 2108 | 0,793 | -0,334 | -6,224 | 4,84E-10 | 9,16E-10 | 0,0020095 | 0,002591 | 0,0023389 | 0,0023389 | TRUE | 0,0224244 | No |
| orf19.6501 | 46 | 31 | 38 | 32 | 179 | 144 | 165 | 235 | 42 | 31 | 36 | 48 | 159 | 156 | 169 | 179 | 102,63 | 39 | 166 | 4,293 | 2,102 | 12,772 | 2,35E-37 | 8,80E-37 | 1,00E-08 | 0,014908 | 0,0095426 | 0,0095426 | TRUE | 0,1292059 | No |
| orf19.6502 | 23 | 38 | 18 | 16 | 29 | 28 | 28 | 33 | 21 | 38 | 17 | 24 | 26 | 30 | 29 | 25 | 26,31 | 25 | 28 | 1,094 | 0,129 | 0,441 | 0,658991 | 0,695656 | 0,0297788 | 0,052244 | 0,0440869 | 0,0440869 | TRUE | 0,8268413 | No |
| orf19.6503 | 1357 | 1340 | 1368 | 1020 | 868 | 708 | 743 | 956 | 1235 | 1355 | 1295 | 1532 | 772 | 769 | 761 | 729 | 1056,07 | 1354 | 758 | 0,561 | -0,834 | -12,07 | 1,52E-33 | 5,29E-33 | 0,0038362 | 0,003288 | 0,0035716 | 0,0035716 | TRUE | 0,0877689 | No |
| orf19.6506 | 3343 | 2800 | 2998 | 1995 | 1991 | 1670 | 1848 | 2475 | 3044 | 2832 | 2837 | 2997 | 1771 | 1813 | 1893 | 1888 | 2384,29 | 2928 | 1841 | 0,629 | -0,668 | -14,668 | 1,03E-48 | 4,63E-48 | 0,0008418 | 0,002591 | 0,0015548 | 0,0015548 | TRUE | 0,0093465 | No |
| orf19.6507 | 8871 | 7361 | 8156 | 5350 | 7941 | 6661 | 6990 | 9400 | 8077 | 7445 | 7719 | 8037 | 7063 | 7232 | 7159 | 7170 | 7487,66 | 7820 | 7156 | 0,915 | -0,128 | -3,467 | 0,000526 | 0,000756 | 0,0005879 | 0,002213 | 0,0011654 | 0,0011654 | TRUE | 0,0127209 | No |
| orf19.6508 | 1083 | 904 | 904 | 618 | 1174 | 921 | 1020 | 1447 | 986 | 914 | 856 | 928 | 1044 | 1000 | 1045 | 1104 | 984,6 | 921 | 1048 | 1,139 | 0,188 | 3,117 | 0,001824 | 0,002525 | 0,0016286 | 0,003379 | 0,0024727 | 0,0024727 | TRUE | 0,0294515 | No |
| orf19.6509 | 634 | 606 | 612 | 445 | 762 | 618 | 581 | 809 | 577 | 613 | 579 | 668 | 678 | 671 | 595 | 617 | 624,83 | 609 | 640 | 1,054 | 0,077 | 1,061 | 0,288816 | 0,325588 | 0,0023152 | 0,004151 | 0,0033826 | 0,0033826 | TRUE | 0,0437667 | No |
| orf19.651 | 1197 | 1230 | 1214 | 691 | 1578 | 1323 | 1377 | 1729 | 1090 | 1244 | 1149 | 1038 | 1404 | 1436 | 1410 | 1319 | 1261,23 | 1130 | 1392 | 1,229 | 0,297 | 4,81 | 1,51E-06 | 2,49E-06 | 0,0026586 | 0,003084 | 0,002848 | 0,002848 | TRUE | 0,0541835 | No |
| orf19.6510 | 203 | 209 | 210 | 158 | 250 | 216 | 245 | 332 | 185 | 211 | 199 | 237 | 222 | 235 | 251 | 253 | 224,17 | 208 | 240 | 1,165 | 0,221 | 2,138 | 0,032506 | 0,040946 | 0,0020856 | 0,00793 | 0,0056939 | 0,0056939 | TRUE | 0,0801497 | No |
| orf19.6511 | 6903 | 6393 | 6768 | 4266 | 6104 | 5158 | 5559 | 7041 | 6285 | 6466 | 6405 | 6408 | 5429 | 5600 | 5693 | 5371 | 5957,22 | 6391 | 5523 | 0,864 | -0,211 | -6,243 | 4,29E-10 | 8,14E-10 | 0,0002107 | 0,002259 | 0,0009276 | 0,0009276 | TRUE | 0,0052497 | No |
| orf19.6512 | 1283 | 1249 | 1285 | 731 | 514 | 474 | 447 | 624 | 1168 | 1263 | 1216 | 1098 | 457 | 515 | 458 | 476 | 831,39 | 1186 | 476 | 0,4 | -1,321 | -19,963 | 1,15E-88 | 8,94E-88 | 0,0018202 | 0,003626 | 0,0027705 | 0,0027705 | TRUE | 0,0313287 | No |
| orf19.6514 | 39369 | 32758 | 36518 | 22629 | 23421 | 19128 | 21820 | 29900 | 35844 | 33131 | 34561 | 33992 | 20832 | 20768 | 22346 | 22808 | 28035,3 | 34382 | 21688 | 0,631 | -0,665 | -15,151 | 7,49E-52 | 3,58E-51 | 0,0016552 | 0,002084 | 0,0018119 | 0,0018119 | TRUE | 0,0148543 | No |
| orf19.6515 | 14726 | 12804 | 13549 | 8381 | 6526 | 5287 | 5770 | 7831 | 13407 | 12950 | 12823 | 12590 | 5805 | 5740 | 5909 | 5974 | 9399,68 | 12942 | 5857 | 0,452 | -1,144 | -33,987 | 3,50E-253 | 7,44E-252 | 0,0003467 | 0,002178 | 0,0009667 | 0,0009667 | TRUE | 0,0071922 | No |
| orf19.6517 | 1026 | 847 | 1002 | 600 | 1324 | 1111 | 1205 | 1595 | 934 | 857 | 948 | 901 | 1178 | 1206 | 1234 | 1217 | 1059,38 | 910 | 1209 | 1,327 | 0,408 | 7,697 | 1,40E-14 | 3,06E-14 | 1,00E-08 | 0,003284 | 0,0017213 | 0,0017213 | TRUE | 0,0199872 | No |
| orf19.6518 | 453 | 330 | 400 | 304 | 1227 | 1075 | 1132 | 1649 | 412 | 334 | 379 | 457 | 1091 | 1167 | 1159 | 1258 | 782,14 | 396 | 1169 | 2,974 | 1,572 | 18,601 | 3,15E-77 | 2,16E-76 | 0,0070259 | 0,003726 | 0,0050975 | 0,0050975 | TRUE | 0,1236433 | No |
| orf19.652 | 553 | 535 | 550 | 310 | 786 | 647 | 686 | 899 | 503 | 541 | 521 | 466 | 699 | 702 | 703 | 686 | 602,58 | 508 | 698 | 1,368 | 0,452 | 7,025 | 2,13E-12 | 4,38E-12 | 1,00E-08 | 0,004229 | 0,0022531 | 0,0022531 | TRUE | 0,0291138 | No |
| orf19.6520 | 61 | 51 | 49 | 41 | 41 | 55 | 61 | 60 | 56 | 52 | 46 | 62 | 36 | 60 | 62 | 46 | 52,44 | 54 | 51 | 0,949 | -0,076 | -0,358 | 0,720686 | 0,753888 | 0,0177462 | 0,027229 | 0,0237521 | 0,0237521 | TRUE | 0,3134689 | No |
| orf19.6522 | 1377 | 1125 | 1219 | 784 | 1930 | 1610 | 1781 | 2472 | 1254 | 1138 | 1154 | 1178 | 1717 | 1748 | 1824 | 1886 | 1487,15 | 1181 | 1794 | 1,519 | 0,603 | 11,311 | 1,15E-29 | 3,67E-29 | 0,0012904 | 0,002925 | 0,0020162 | 0,0020162 | TRUE | 0,0222866 | No |
| orf19.6524 | 5095 | 4206 | 4504 | 3204 | 8895 | 7388 | 7820 | 10915 | 4639 | 4254 | 4263 | 4813 | 7912 | 8021 | 8009 | 8326 | 6279,51 | 4492 | 8067 | 1,797 | 0,846 | 17,709 | 3,58E-70 | 2,25E-69 | 0,0019479 | 0,002247 | 0,0020136 | 0,0020136 | TRUE | 0,0269726 | No |
| orf19.6525 | 1414 | 1314 | 1296 | 842 | 920 | 806 | 830 | 1160 | 1287 | 1329 | 1227 | 1265 | 818 | 875 | 850 | 885 | 1067 | 1277 | 857 | 0,671 | -0,575 | -10,804 | 3,31E-27 | 9,98E-27 | 1,00E-08 | 0,003275 | 0,0017576 | 0,0017576 | TRUE | 0,0115031 | No |
| orf19.6526 | 1632 | 1357 | 1475 | 1000 | 1277 | 1061 | 1168 | 1509 | 1486 | 1372 | 1396 | 1502 | 1136 | 1152 | 1196 | 1151 | 1298,94 | 1439 | 1159 | 0,806 | -0,311 | -6,147 | 7,91E-10 | 1,48E-09 | 0,0004785 | 0,003054 | 0,0016851 | 0,0016851 | TRUE | 0,0114539 | No |
| orf19.6527 | 19284 | 16571 | 18573 | 11379 | 13088 | 10808 | 12452 | 16465 | 17557 | 16760 | 17578 | 17093 | 11641 | 11734 | 12752 | 12560 | 14709,5 | 17247 | 12172 | 0,706 | -0,503 | -12,194 | 3,34E-34 | 1,17E-33 | 0,0012481 | 0,002127 | 0,0015634 | 0,0015634 | TRUE | 0,0124867 | No |
| orf19.6528 | 960 | 821 | 913 | 563 | 680 | 575 | 580 | 824 | 874 | 830 | 864 | 846 | 605 | 624 | 594 | 629 | 733,23 | 854 | 613 | 0,718 | -0,478 | -8,364 | 6,07E-17 | 1,42E-16 | 1,00E-08 | 0,003839 | 0,0017511 | 0,0017511 | TRUE | 0,0048816 | No |
| orf19.6529 | 17513 | 15009 | 16743 | 10314 | 16399 | 12997 | 14425 | 19887 | 15945 | 15180 | 15846 | 15493 | 14586 | 14111 | 14773 | 15170 | 15138 | 15616 | 14660 | 0,939 | -0,091 | -2,611 | 0,009036 | 0,011892 | 0,0006114 | 0,002124 | 0,0011014 | 0,0011014 | TRUE | 0,0077294 | No |
| orf19.653 | 1 | 2 | 1 | 1 | 1 | 6 | 2 | 2 | 1 | 2 | 1 | 2 | 1 | 7 | 2 | 2 | 2,04 | 2 | 3 | 2,003 | 1,002 | 0,966 | 0,334083 | 0,373973 | 1,00E-08 | 0,648022 | 0,457693 | 0,457693 | TRUE | 0,9364271 | No |
| orf19.6530 | 3802 | 3413 | 3695 | 2511 | 2249 | 1934 | 2005 | 2725 | 3462 | 3452 | 3497 | 3772 | 2000 | 2100 | 2053 | 2079 | 2801,82 | 3546 | 2058 | 0,581 | -0,783 | -17,968 | 3,48E-72 | 2,23E-71 | 0,0006254 | 0,002508 | 0,0014443 | 0,0014443 | TRUE | 0,0210341 | No |
| orf19.6531 | 6390 | 5629 | 5928 | 3877 | 9086 | 7353 | 7894 | 11226 | 5818 | 5693 | 5610 | 5824 | 8082 | 7983 | 8084 | 8563 | 6957,24 | 5736 | 8178 | 1,426 | 0,512 | 14,134 | 2,34E-45 | 1,01E-44 | 0,0004739 | 0,002227 | 0,0011111 | 0,0011111 | TRUE | 0,0124882 | No |
| orf19.6531.1 | 1859 | 1557 | 1436 | 1074 | 1758 | 1371 | 1306 | 2097 | 1693 | 1575 | 1359 | 1613 | 1564 | 1489 | 1338 | 1600 | 1528,62 | 1560 | 1498 | 0,961 | -0,057 | -0,75 | 0,453015 | 0,494854 | 0,006868 | 0,002901 | 0,0049546 | 0,0049546 | TRUE | 0,0926353 | No |
| orf19.6532 | 601 | 501 | 544 | 356 | 736 | 609 | 729 | 905 | 547 | 507 | 515 | 535 | 655 | 661 | 747 | 690 | 607,04 | 526 | 688 | 1,309 | 0,388 | 5,736 | 9,69E-09 | 1,75E-08 | 0,0010208 | 0,004213 | 0,0027037 | 0,0027037 | TRUE | 0,0368947 | No |
| orf19.6533 | 739 | 746 | 775 | 497 | 933 | 739 | 850 | 1067 | 673 | 754 | 733 | 747 | 830 | 802 | 871 | 814 | 778 | 727 | 829 | 1,142 | 0,192 | 3,155 | 0,001604 | 0,002232 | 0,0006587 | 0,003735 | 0,0022615 | 0,0022615 | TRUE | 0,0315898 | No |
| orf19.6534 | 7 | 6 | 9 | 9 | 5 | 4 | 2 | 3 | 6 | 6 | 9 | 14 | 4 | 4 | 2 | 2 | 5,95 | 9 | 3 | 0,389 | -1,364 | -2,257 | 0,024039 | 0,030613 | 1,00E-08 | 0,224027 | 0,144074 | 0,144074 | TRUE | 0,2913987 | No |
| orf19.6534.2 | 1693 | 1433 | 1625 | 927 | 2129 | 1721 | 2036 | 2644 | 1541 | 1449 | 1538 | 1393 | 1894 | 1869 | 2085 | 2017 | 1723,17 | 1480 | 1966 | 1,326 | 0,407 | 7,402 | 1,34E-13 | 2,86E-13 | 0,0017649 | 0,002804 | 0,0023027 | 0,0023027 | TRUE | 0,0194464 | No |
| orf19.6536 | 432 | 436 | 466 | 263 | 207 | 156 | 164 | 226 | 393 | 441 | 441 | 395 | 184 | 169 | 168 | 172 | 295,53 | 418 | 173 | 0,415 | -1,269 | -13,95 | 3,14E-44 | 1,32E-43 | 1,00E-08 | 0,006507 | 0,0039831 | 0,0039831 | TRUE | 0,0210013 | No |
| orf19.6537 | 188 | 213 | 201 | 116 | 291 | 242 | 256 | 303 | 171 | 215 | 190 | 174 | 259 | 263 | 262 | 231 | 220,74 | 188 | 254 | 1,345 | 0,427 | 4,072 | 4,65E-05 | 7,11E-05 | 0,0026428 | 0,008021 | 0,0058927 | 0,0058927 | TRUE | 0,1070021 | No |
| orf19.6538 | 843 | 687 | 732 | 476 | 1172 | 936 | 913 | 1328 | 768 | 695 | 693 | 715 | 1042 | 1016 | 935 | 1013 | 859,61 | 718 | 1002 | 1,396 | 0,481 | 7,872 | 3,49E-15 | 7,75E-15 | 0,0011689 | 0,003574 | 0,0023766 | 0,0023766 | TRUE | 0,0284134 | No |
| orf19.6539 | 6173 | 5162 | 5652 | 3594 | 6071 | 4902 | 5497 | 7398 | 5620 | 5221 | 5349 | 5399 | 5400 | 5322 | 5630 | 5643 | 5448 | 5397 | 5499 | 1,019 | 0,027 | 0,691 | 0,489812 | 0,531498 | 0,0006663 | 0,002279 | 0,0012847 | 0,0012847 | TRUE | 0,0097475 | No |
| orf19.654 | 10 | 8 | 8 | 3 | 8 | 13 | 11 | 18 | 9 | 8 | 8 | 5 | 7 | 14 | 11 | 14 | 9,44 | 8 | 12 | 1,536 | 0,619 | 1,359 | 0,174106 | 0,20284 | 1,00E-08 | 0,142011 | 0,0879717 | 0,0879717 | TRUE | 0,2896093 | No |
| orf19.6540 | 26616 | 22236 | 24327 | 15514 | 7214 | 6149 | 6632 | 9268 | 24233 | 22489 | 23023 | 23305 | 6417 | 6676 | 6792 | 7070 | 15000,5 | 23262 | 6739 | 0,29 | -1,787 | -43,516 | 0 | 0 | 0,0011723 | 0,002125 | 0,0015278 | 0,0015278 | TRUE | 0,0136406 | No |
| orf19.6541 | 19339 | 15612 | 16947 | 10505 | 29087 | 22655 | 25023 | 34571 | 17607 | 15790 | 16039 | 15780 | 25872 | 24597 | 25627 | 26371 | 20960,4 | 16304 | 25617 | 1,571 | 0,652 | 14,494 | 1,33E-47 | 5,89E-47 | 0,0018052 | 0,0021 | 0,0018909 | 0,0018909 | TRUE | 0,0356152 | No |
| orf19.6544 | 15501 | 13228 | 14532 | 8970 | 4243 | 3648 | 4067 | 5225 | 14113 | 13379 | 13753 | 13474 | 3774 | 3961 | 4165 | 3986 | 8825,6 | 13680 | 3972 | 0,29 | -1,785 | -45,005 | 0 | 0 | 0,0008853 | 0,002187 | 0,0013534 | 0,0013534 | TRUE | 0,0137061 | No |
| orf19.6547 | 15 | 5 | 6 | 8 | 26 | 16 | 20 | 33 | 14 | 5 | 6 | 12 | 23 | 17 | 20 | 25 | 15,32 | 9 | 21 | 2,419 | 1,274 | 3,258 | 0,001124 | 0,001582 | 0,0032337 | 0,088262 | 0,0661956 | 0,0661956 | TRUE | 0,5065467 | No |
| orf19.6548 | 13083 | 11008 | 11938 | 7706 | 11494 | 9874 | 10022 | 14057 | 11912 | 11133 | 11298 | 11576 | 10224 | 10720 | 10264 | 10723 | 10981,2 | 11480 | 10483 | 0,913 | -0,131 | -3,609 | 0,000307 | 0,000448 | 0,0006668 | 0,002157 | 0,001174 | 0,001174 | TRUE | 0,0080174 | No |
| orf19.655 | 55048 | 47273 | 51091 | 31929 | 47250 | 38778 | 40816 | 59262 | 50119 | 47812 | 48353 | 47962 | 42027 | 42102 | 41801 | 45206 | 45672,7 | 48562 | 42784 | 0,881 | -0,183 | -4,965 | 6,86E-07 | 1,15E-06 | 0,0009267 | 0,002066 | 0,0012788 | 0,0012788 | TRUE | 0,0178447 | No |
| orf19.6550 | 1150 | 971 | 1021 | 592 | 1531 | 1199 | 1237 | 1770 | 1047 | 982 | 966 | 889 | 1362 | 1302 | 1267 | 1350 | 1145,65 | 971 | 1320 | 1,356 | 0,439 | 7,543 | 4,59E-14 | 9,88E-14 | 0,0015915 | 0,00319 | 0,0023479 | 0,0023479 | TRUE | 0,0337135 | No |
| orf19.6551 | 641 | 693 | 675 | 379 | 885 | 648 | 736 | 929 | 584 | 701 | 639 | 569 | 787 | 704 | 754 | 709 | 680,72 | 623 | 738 | 1,182 | 0,241 | 3,171 | 0,001518 | 0,002117 | 0,0040835 | 0,003978 | 0,0040664 | 0,0040664 | TRUE | 0,0821698 | No |
| orf19.6551.1 | 163 | 130 | 163 | 109 | 253 | 224 | 196 | 271 | 148 | 131 | 154 | 164 | 225 | 243 | 201 | 207 | 184,2 | 149 | 219 | 1,469 | 0,555 | 4,924 | 8,46E-07 | 1,41E-06 | 0,0019289 | 0,009209 | 0,0064642 | 0,0064642 | TRUE | 0,0672261 | No |
| orf19.6552 | 2351 | 2197 | 2429 | 1601 | 2022 | 1601 | 1622 | 2274 | 2140 | 2222 | 2299 | 2405 | 1799 | 1738 | 1661 | 1735 | 1999,85 | 2266 | 1733 | 0,766 | -0,384 | -7,745 | 9,52E-15 | 2,10E-14 | 0,0010827 | 0,002698 | 0,0018579 | 0,0018579 | TRUE | 0,0189706 | No |
| orf19.6553 | 1035 | 989 | 997 | 635 | 1636 | 1376 | 1417 | 1843 | 942 | 1000 | 944 | 954 | 1455 | 1494 | 1451 | 1406 | 1205,77 | 960 | 1452 | 1,511 | 0,595 | 12,05 | 1,93E-33 | 6,67E-33 | 1,00E-08 | 0,003133 | 0,001465 | 0,001465 | TRUE | 0,0099473 | No |
| orf19.6554 | 6990 | 7925 | 7525 | 5061 | 1784 | 1579 | 1508 | 1892 | 6364 | 8015 | 7122 | 7602 | 1587 | 1714 | 1544 | 1443 | 4424,04 | 7276 | 1572 | 0,216 | -2,211 | -29,837 | 1,28E-195 | 2,13E-194 | 0,0067841 | 0,002336 | 0,0049029 | 0,0049029 | TRUE | 0,0873922 | No |
| orf19.6555 | 523 | 572 | 524 | 338 | 308 | 223 | 232 | 332 | 476 | 579 | 496 | 508 | 274 | 242 | 238 | 253 | 383,16 | 515 | 252 | 0,49 | -1,028 | -11,602 | 4,04E-31 | 1,33E-30 | 0,0036805 | 0,005485 | 0,0046535 | 0,0046535 | TRUE | 0,0849176 | No |

| | | | | | | | | | | | | | | | | | | | | | | | | | | | | | | | |
|---|---|---|---|---|---|---|---|---|---|---|---|---|---|---|---|---|---|---|---|---|---|---|---|---|---|---|---|---|---|---|---|
| orf19.6556 | 11712 | 11539 | 11786 | 7553 | 6101 | 5073 | 5064 | 6868 | 10663 | 11670 | 11154 | 11346 | 5427 | 5508 | 5186 | 5239 | 8274,2 | 11208 | 5340 | 0,476 | -1,07 | -26,54 | 3,31E-155 | 4,46E-154 | 0,0009286 | 0,002197 | 0,0014243 | 0,0014243 | TRUE | 0,0132657 | No |
| orf19.6557 | 776 | 674 | 795 | 497 | 850 | 768 | 837 | 1093 | 707 | 682 | 752 | 747 | 756 | 834 | 857 | 834 | 771 | 722 | 820 | 1,137 | 0,185 | 2,91 | 0,00361 | 0,004883 | 0,0013735 | 0,00375 | 0,002582 | 0,002582 | TRUE | 0,0334944 | No |
| orf19.6558 | 3002 | 2967 | 3064 | 1960 | 1825 | 1472 | 1562 | 1998 | 2733 | 3001 | 2900 | 2944 | 1623 | 1598 | 1600 | 1524 | 2240,41 | 2894 | 1586 | 0,548 | -0,868 | -18,714 | 3,83E-78 | 2,67E-77 | 0,0008773 | 0,002627 | 0,001583 | 0,001583 | TRUE | 0,0176661 | No |
| orf19.6559 | 2137 | 1772 | 1911 | 1297 | 1115 | 866 | 956 | 1325 | 1946 | 1792 | 1809 | 1948 | 992 | 940 | 979 | 1011 | 1427,06 | 1874 | 980 | 0,524 | -0,931 | -17,937 | 6,11E-72 | 3,91E-71 | 0,0008972 | 0,002963 | 0,0018265 | 0,0018265 | TRUE | 0,0102186 | No |
| orf19.656 | 82 | 60 | 68 | 46 | 64 | 55 | 60 | 84 | 75 | 61 | 64 | 69 | 57 | 60 | 61 | 64 | 63,87 | 67 | 60 | 0,902 | -0,148 | -0,869 | 0,384609 | 0,425231 | 1,00E-08 | 0,02272 | 0,012305 | 0,012305 | TRUE | 0,0595151 | No |
| orf19.6561 | 6769 | 5740 | 6275 | 4286 | 7946 | 6498 | 6847 | 9452 | 6163 | 5805 | 5939 | 6438 | 7068 | 7055 | 7012 | 7210 | 6586,28 | 6086 | 7086 | 1,165 | 0,221 | 5,551 | 2,85E-08 | 5,02E-08 | 0,0009066 | 0,002238 | 0,0013632 | 0,0013632 | TRUE | 0,0176572 | No |
| orf19.6562 | 334 | 234 | 267 | 204 | 611 | 544 | 547 | 769 | 304 | 237 | 253 | 306 | 543 | 591 | 560 | 587 | 422,6 | 275 | 570 | 2,083 | 1,059 | 11,84 | 2,43E-32 | 8,21E-32 | 0,0044698 | 0,005163 | 0,0049037 | 0,0049037 | TRUE | 0,0973675 | No |
| orf19.6563 | 2348 | 2046 | 2222 | 1481 | 2380 | 1989 | 2103 | 2829 | 2138 | 2069 | 2103 | 2225 | 2117 | 2159 | 2154 | 2158 | 2140,36 | 2134 | 2147 | 1,008 | 0,011 | 0,271 | 0,786219 | 0,815542 | 1,00E-08 | 0,002654 | 0,0011088 | 0,0011088 | TRUE | 0,0088187 | No |
| orf19.6563.1 | 3385 | 2616 | 2717 | 1424 | 1535 | 949 | 1268 | 1552 | 3082 | 2646 | 2571 | 2139 | 1365 | 1030 | 1299 | 1184 | 1914,54 | 2610 | 1220 | 0,467 | -1,098 | -10,544 | 5,41E-26 | 1,58E-25 | 0,0173796 | 0,002727 | 0,0098336 | 0,0098336 | TRUE | 0,1295632 | No |
| orf19.6565 | 1117 | 946 | 1056 | 664 | 1583 | 1278 | 1304 | 1827 | 1017 | 957 | 999 | 997 | 1408 | 1388 | 1335 | 1394 | 1186,91 | 992 | 1381 | 1,392 | 0,477 | 9,77 | 1,51E-22 | 4,07E-22 | 1,00E-08 | 0,00315 | 0,0014119 | 0,0014119 | TRUE | 0,0074196 | No |
| orf19.6566 | 1115 | 979 | 1113 | 685 | 1102 | 962 | 927 | 1249 | 1015 | 990 | 1053 | 1029 | 980 | 1044 | 949 | 953 | 1001,8 | 1022 | 982 | 0,96 | -0,059 | -1,093 | 0,274527 | 0,310634 | 0,0003073 | 0,003356 | 0,0018264 | 0,0018264 | TRUE | 0,0209634 | No |
| orf19.6568 | 5365 | 5018 | 5270 | 2883 | 6751 | 5380 | 6110 | 7759 | 4885 | 5075 | 4988 | 4331 | 6005 | 5841 | 6257 | 5919 | 5412,51 | 4820 | 6006 | 1,245 | 0,316 | 5,996 | 2,02E-09 | 3,74E-09 | 0,0026199 | 0,002281 | 0,0024759 | 0,0024759 | TRUE | 0,0548061 | No |
| orf19.6569 | 449 | 463 | 493 | 244 | 1094 | 926 | 903 | 1208 | 409 | 468 | 467 | 367 | 973 | 1005 | 925 | 921 | 691,86 | 428 | 956 | 2,223 | 1,152 | 14,93 | 2,11E-50 | 9,77E-50 | 0,0040417 | 0,003946 | 0,003984 | 0,003984 | TRUE | 0,0884356 | No |
| orf19.657 | 1246 | 1058 | 1103 | 803 | 2071 | 1805 | 1580 | 2461 | 1134 | 1070 | 1044 | 1206 | 1842 | 1960 | 1618 | 1877 | 1468,98 | 1114 | 1824 | 1,641 | 0,715 | 10,37 | 3,38E-25 | 9,74E-25 | 0,004359 | 0,002936 | 0,0038237 | 0,0038237 | TRUE | 0,0688972 | No |
| orf19.6570 | 142 | 130 | 154 | 99 | 446 | 412 | 415 | 554 | 129 | 131 | 146 | 149 | 397 | 447 | 425 | 423 | 280,86 | 139 | 423 | 3,059 | 1,613 | 16,719 | 9,52E-63 | 5,41E-62 | 1,00E-08 | 0,00674 | 0,0039911 | 0,0039911 | TRUE | 0,0230383 | No |
| orf19.6573 | 3758 | 3829 | 3931 | 2303 | 5012 | 4105 | 4583 | 5881 | 3421 | 3873 | 3720 | 3459 | 4458 | 4457 | 4694 | 4486 | 4071,06 | 3618 | 4524 | 1,249 | 0,321 | 6,627 | 3,43E-11 | 6,78E-11 | 0,0017564 | 0,002362 | 0,0020045 | 0,0020045 | TRUE | 0,0272004 | No |
| orf19.6574 | 29 | 33 | 28 | 18 | 37 | 40 | 35 | 44 | 26 | 33 | 26 | 27 | 33 | 43 | 36 | 34 | 32,38 | 28 | 36 | 1,277 | 0,353 | 1,461 | 0,14413 | 0,169776 | 1,00E-08 | 0,04283 | 0,0244763 | 0,0244763 | TRUE | 0,1104274 | No |
| orf19.6575 | 2 | 1 | 5 | 5 | 1 | 1 | 3 | 1 | 2 | 1 | 5 | 8 | 1 | 1 | 3 | 1 | 2,61 | 4 | 2 | 0,393 | -1,348 | -1,406 | 0,159593 | 0,187161 | 1,00E-08 | 0,508032 | 0,4048852 | 0,4048852 | TRUE | 0,2089597 | No |
| orf19.6577 | 12358 | 11112 | 11530 | 7942 | 5746 | 5146 | 4952 | 7005 | 11251 | 11239 | 10912 | 11930 | 5111 | 5587 | 5071 | 5343 | 8305,65 | 11333 | 5278 | 0,466 | -1,102 | -24,673 | 2,07E-134 | 2,40E-133 | 0,0015472 | 0,002196 | 0,0017801 | 0,0017801 | TRUE | 0,0185638 | No |
| orf19.6578 | 18536 | 17412 | 18835 | 11809 | 3671 | 3123 | 3278 | 4360 | 16876 | 17610 | 17826 | 17739 | 3265 | 3391 | 3357 | 3326 | 10423,8 | 17513 | 3335 | 0,19 | -2,393 | -70,522 | 0 | 0 | 0,0003884 | 0,002164 | 0,0009348 | 0,0009348 | TRUE | 0,0074083 | No |
| orf19.6579 | 715 | 853 | 932 | 487 | 917 | 754 | 802 | 982 | 651 | 863 | 882 | 732 | 816 | 819 | 821 | 749 | 791,5 | 782 | 801 | 1,023 | 0,033 | 0,363 | 0,716421 | 0,749825 | 0,0099214 | 0,003706 | 0,00673 | 0,00673 | TRUE | 0,1597178 | No |
| orf19.658 | 245 | 234 | 250 | 151 | 514 | 375 | 414 | 526 | 223 | 237 | 237 | 227 | 457 | 407 | 424 | 401 | 326,59 | 231 | 422 | 1,829 | 0,871 | 10,222 | 1,58E-24 | 4,48E-24 | 1,00E-08 | 0,006082 | 0,0035614 | 0,0035614 | TRUE | 0,0370091 | No |
| orf19.6580 | 5318 | 4788 | 5040 | 3317 | 6291 | 5345 | 5397 | 7135 | 4842 | 4843 | 4770 | 4983 | 5596 | 5803 | 5527 | 5443 | 5225,7 | 4860 | 5592 | 1,151 | 0,203 | 5,726 | 1,03E-08 | 1,85E-08 | 0,0002906 | 0,00229 | 0,001012 | 0,001012 | TRUE | 0,0077357 | No |
| orf19.6581 | 875 | 824 | 874 | 489 | 1403 | 1164 | 1279 | 1588 | 797 | 833 | 827 | 735 | 1248 | 1264 | 1310 | 1211 | 1028,08 | 798 | 1258 | 1,57 | 0,651 | 11,5 | 1,32E-30 | 4,30E-30 | 0,0007657 | 0,003322 | 0,0020363 | 0,0020363 | TRUE | 0,0285525 | No |
| orf19.6582 | 5855 | 5244 | 5503 | 3248 | 4198 | 3221 | 3587 | 4693 | 5331 | 5304 | 5208 | 4879 | 3734 | 3497 | 3674 | 3580 | 4400,76 | 5180 | 3621 | 0,698 | -0,518 | -12,266 | 1,39E-34 | 4,92E-34 | 0,0009049 | 0,002337 | 0,0014777 | 0,0014777 | TRUE | 0,0177905 | No |
| orf19.6583 | 266 | 291 | 317 | 205 | 427 | 392 | 359 | 509 | 242 | 294 | 300 | 308 | 380 | 426 | 368 | 388 | 338,22 | 286 | 390 | 1,369 | 0,453 | 4,877 | 1,08E-06 | 1,79E-06 | 0,0042106 | 0,005943 | 0,0052248 | 0,0052248 | TRUE | 0,1287941 | No |
| orf19.6584 | 9405 | 8861 | 9341 | 5707 | 11348 | 9830 | 9613 | 12966 | 8563 | 8962 | 8840 | 8573 | 10094 | 10673 | 9845 | 9891 | 9429,98 | 8734 | 10126 | 1,159 | 0,213 | 5,567 | 2,59E-08 | 4,58E-08 | 0,0008228 | 0,002177 | 0,0012943 | 0,0012943 | TRUE | 0,0160131 | No |
| orf19.6585 | 660 | 713 | 660 | 421 | 482 | 351 | 349 | 448 | 601 | 721 | 625 | 632 | 429 | 381 | 357 | 342 | 511 | 645 | 377 | 0,585 | -0,774 | -8,765 | 1,88E-18 | 4,57E-18 | 0,0063141 | 0,004622 | 0,0054189 | 0,0054189 | TRUE | 0,1013645 | No |
| orf19.6586 | 3178 | 3100 | 3298 | 2015 | 1402 | 1138 | 1155 | 1385 | 2893 | 3135 | 3121 | 3027 | 1247 | 1236 | 1183 | 1056 | 2112,35 | 3044 | 1180 | 0,387 | -1,369 | -23,243 | 1,68E-119 | 1,70E-118 | 0,0026184 | 0,002662 | 0,0027561 | 0,0027561 | TRUE | 0,063268 | No |
| orf19.6587 | 1829 | 1883 | 1905 | 1131 | 1095 | 908 | 1012 | 1160 | 1665 | 1904 | 1803 | 1699 | 974 | 986 | 1036 | 885 | 1369,08 | 1768 | 970 | 0,548 | -0,868 | -13,531 | 1,02E-41 | 4,11E-41 | 0,0033152 | 0,003002 | 0,0031683 | 0,0031683 | TRUE | 0,0440989 | No |
| orf19.6588 | 2563 | 2467 | 2531 | 1472 | 1080 | 910 | 952 | 1327 | 2334 | 2495 | 2395 | 2211 | 961 | 988 | 975 | 1012 | 1671,37 | 2359 | 984 | 0,417 | -1,263 | -24,583 | 1,93E-133 | 2,22E-132 | 0,0010501 | 0,002827 | 0,0018313 | 0,0018313 | TRUE | 0,0197148 | No |
| orf19.6589 | 6977 | 7157 | 7339 | 4236 | 8058 | 6082 | 6969 | 8676 | 6352 | 7239 | 6946 | 6363 | 7167 | 6603 | 7137 | 6618 | 6803,2 | 6725 | 6881 | 1,023 | 0,033 | 0,603 | 0,546836 | 0,588367 | 0,002706 | 0,002231 | 0,0026891 | 0,0026891 | TRUE | 0,0322403 | No |
| orf19.6590 | 123 | 128 | 101 | 68 | 37 | 25 | 27 | 37 | 112 | 129 | 96 | 102 | 33 | 27 | 28 | 28 | 69,39 | 110 | 29 | 0,264 | -1,921 | -9,985 | 1,77E-23 | 4,89E-23 | 1,00E-08 | 0,021075 | 0,0147064 | 0,0147064 | TRUE | 0,1464037 | No |
| orf19.6591 | 2300 | 2136 | 2236 | 1532 | 1672 | 1420 | 1338 | 1826 | 2094 | 2160 | 2116 | 2301 | 1487 | 1542 | 1370 | 1393 | 1807,99 | 2168 | 1448 | 0,668 | -0,581 | -10,825 | 2,62E-27 | 7,92E-27 | 0,0016954 | 0,002768 | 0,002195 | 0,002195 | TRUE | 0,0219804 | No |
| orf19.6592 | 581 | 552 | 598 | 358 | 303 | 221 | 235 | 332 | 529 | 558 | 566 | 538 | 270 | 240 | 241 | 253 | 399,3 | 548 | 251 | 0,459 | -1,124 | -14,586 | 3,45E-48 | 1,54E-47 | 1,00E-08 | 0,005345 | 0,0028797 | 0,0028797 | TRUE | 0,0276345 | No |
| orf19.6594 | 24334 | 23595 | 24880 | 14880 | 6168 | 5320 | 5536 | 7167 | 22155 | 23864 | 23547 | 22352 | 5486 | 5776 | 5670 | 5467 | 14289,6 | 22980 | 5600 | 0,244 | -2,037 | -52,069 | 0 | 0 | 0,0008922 | 0,002129 | 0,0013638 | 0,0013638 | TRUE | 0,0082151 | No |
| orf19.6595 | 65 | 55 | 57 | 35 | 63 | 42 | 52 | 81 | 59 | 56 | 54 | 53 | 56 | 46 | 53 | 62 | 54,75 | 56 | 54 | 0,983 | -0,025 | -0,135 | 0,892594 | 0,907831 | 1,00E-08 | 0,026164 | 0,0148885 | 0,0148885 | TRUE | 0,0873116 | No |
| orf19.6596 | 1805 | 2004 | 1915 | 1355 | 2686 | 2469 | 2270 | 2963 | 1643 | 2027 | 1812 | 2035 | 2389 | 2681 | 2325 | 2260 | 2146,59 | 1879 | 2414 | 1,285 | 0,362 | 4,734 | 2,21E-06 | 3,61E-06 | 0,0073397 | 0,002652 | 0,0051371 | 0,0051371 | TRUE | 0,0875017 | No |
| orf19.6597 | 918 | 1188 | 1174 | 606 | 1000 | 878 | 945 | 1034 | 836 | 1202 | 1111 | 910 | 889 | 953 | 968 | 789 | 957,25 | 1015 | 900 | 0,885 | -0,176 | -1,635 | 0,101992 | 0,122448 | 0,0168026 | 0,003417 | 0,0100564 | 0,0100564 | TRUE | 0,1829308 | No |
| orf19.6598 | 3418 | 2989 | 3315 | 2202 | 1078 | 825 | 890 | 1291 | 3112 | 3023 | 3137 | 3308 | 959 | 896 | 911 | 985 | 2041,37 | 3145 | 938 | 0,299 | -1,742 | -34,689 | 1,16E-263 | 2,57E-262 | 0,0009629 | 0,002684 | 0,0017528 | 0,0017528 | TRUE | 0,0142588 | No |
| orf19.6599.1 | 2579 | 1903 | 1927 | 969 | 3575 | 2020 | 2541 | 3767 | 2348 | 1925 | 1824 | 1456 | 3180 | 2193 | 2602 | 2873 | 2300,11 | 1888 | 2712 | 1,436 | 0,522 | 2,921 | 0,003487 | 0,004717 | 0,030218 | 0,002611 | 0,0149074 | 0,030218 | TRUE | 0,3257065 | No |
| orf19.660 | 1450 | 1230 | 1385 | 899 | 1372 | 1171 | 1238 | 1617 | 1320 | 1244 | 1311 | 1350 | 1220 | 1271 | 1268 | 1233 | 1277,31 | 1306 | 1248 | 0,956 | -0,065 | -1,346 | 0,178246 | 0,207433 | 1,00E-08 | 0,003071 | 0,0014492 | 0,0014492 | TRUE | 0,0122935 | No |
| orf19.6600 | 2029 | 2151 | 2092 | 1165 | 1603 | 1266 | 1128 | 1650 | 1847 | 2176 | 1980 | 1750 | 1426 | 1375 | 1155 | 1259 | 1620,86 | 1938 | 1304 | 0,672 | -0,574 | -7,167 | 7,67E-13 | 1,60E-12 | 0,0079743 | 0,002852 | 0,005514 | 0,005514 | TRUE | 0,0817422 | No |
| orf19.6601 | 11864 | 11490 | 11880 | 6829 | 6232 | 4724 | 5423 | 6817 | 10802 | 11621 | 11243 | 10258 | 5543 | 5129 | 5554 | 5200 | 8168,77 | 10981 | 5356 | 0,488 | -1,036 | -21,095 | 8,91E-99 | 7,68E-98 | 0,0021139 | 0,002199 | 0,0021809 | 0,0021809 | TRUE | 0,0235578 | No |
| orf19.6601.1 | 23400 | 22578 | 22798 | 13295 | 5815 | 4422 | 4550 | 5711 | 21305 | 22835 | 21576 | 19971 | 5172 | 4801 | 4660 | 4356 | 13084,6 | 21422 | 4747 | 0,222 | -2,174 | -37,242 | 1,43E-303 | 3,73E-302 | 0,0038754 | 0,002138 | 0,0031508 | 0,0031508 | TRUE | 0,0447029 | No |
| orf19.6602 | 933 | 1189 | 1122 | 727 | 752 | 685 | 613 | 792 | 849 | 1203 | 1062 | 1092 | 669 | 744 | 628 | 604 | 856,31 | 1052 | 661 | 0,629 | -0,67 | -6,67 | 2,55E-11 | 5,07E-11 | 0,0136755 | 0,00358 | 0,0084571 | 0,0084571 | TRUE | 0,2052028 | No |
| orf19.6604 | 5319 | 6054 | 6020 | 3740 | 909 | 807 | 742 | 938 | 4843 | 6123 | 5697 | 5618 | 809 | 876 | 760 | 716 | 3180,16 | 5570 | 790 | 0,142 | -2,82 | -35,814 | 6,61E-281 | 1,56E-279 | 0,0074179 | 0,002452 | 0,0052638 | 0,0052638 | TRUE | 0,0951692 | No |
| orf19.6605 | 638 | 679 | 703 | 461 | 946 | 836 | 779 | 971 | 581 | 687 | 665 | 692 | 841 | 908 | 798 | 741 | 739,13 | 656 | 822 | 1,253 | 0,325 | 4,072 | 4,67E-05 | 7,13E-05 | 0,0056822 | 0,003824 | 0,0047427 | 0,0047427 | TRUE | 0,0736044 | No |
| orf19.6606 | 239 | 271 | 230 | 194 | 304 | 221 | 224 | 293 | 218 | 274 | 218 | 291 | 270 | 240 | 229 | 224 | 245,5 | 250 | 241 | 0,969 | -0,045 | -0,385 | 0,699869 | 0,73443 | 0,0109392 | 0,007418 | 0,0089286 | 0,0089286 | TRUE | 0,119135 | No |
| orf19.6607 | 3111 | 3198 | 2979 | 1500 | 3877 | 2813 | 3017 | 3737 | 2832 | 3234 | 2819 | 2253 | 3448 | 3054 | 3090 | 2851 | 2947,81 | 2784 | 3111 | 1,116 | 0,158 | 1,712 | 0,0869 | 0,104962 | 0,0132303 | 0,002485 | 0,007846 | 0,007846 | TRUE | 0,1956592 | No |
| orf19.6608 | 2937 | 2812 | 3061 | 1752 | 352 | 328 | 314 | 438 | 2674 | 2844 | 2897 | 2632 | 313 | 356 | 322 | 334 | 1546,46 | 2762 | 331 | 0,12 | -3,063 | -48,828 | 0 | 0 | 0,001476 | 0,002891 | 0,0021772 | 0,0021772 | TRUE | 0,0260229 | No |
| orf19.661 | 10545 | 9457 | 9686 | 6395 | 9402 | 7392 | 7587 | 9992 | 9601 | 9565 | 9167 | 9606 | 8363 | 8026 | 7770 | 7622 | 8714,9 | 9485 | 7945 | 0,838 | -0,255 | -6,438 | 1,21E-10 | 2,35E-10 | 0,0009314 | 0,002189 | 0,0013969 | 0,0013969 | TRUE | 0,0154709 | No |
| orf19.6610 | 334 | 315 | 332 | 227 | 336 | 289 | 267 | 384 | 304 | 319 | 314 | 341 | 299 | 314 | 273 | 293 | 307,11 | 320 | 295 | 0,926 | -0,111 | -1,319 | 0,187002 | 0,217148 | 1,00E-08 | 0,006338 | 0,0035456 | 0,0035456 | TRUE | 0,0245189 | No |
| orf19.6612 | 1074 | 1177 | 1201 | 674 | 1528 | 1246 | 1236 | 1623 | 978 | 1190 | 1137 | 1012 | 1359 | 1353 | 1266 | 1238 | 1191,64 | 1079 | 1304 | 1,206 | 0,271 | 3,868 | 0,00011 | 0,000164 | 0,0043205 | 0,003146 | 0,0038437 | 0,0038437 | TRUE | 0,0574911 | No |
| orf19.6621 | 15333 | 14187 | 15491 | 8977 | 2391 | 1903 | 2172 | 2825 | 13960 | 14349 | 14661 | 13485 | 2127 | 2066 | 2224 | 2155 | 8128,32 | 14114 | 2143 | 0,152 | -2,72 | -65,542 | 0 | 0 | 0,0007586 | 0,0022 | 0,0013973 | 0,0013973 | TRUE | 0,0107696 | No |
| orf19.6623 | 1682 | 1489 | 1422 | 929 | 1925 | 1295 | 1541 | 2049 | 1531 | 1506 | 1346 | 1396 | 1712 | 1406 | 1578 | 1563 | 1504,76 | 1445 | 1565 | 1,083 | 0,115 | 1,7 | 0,089112 | 0,107552 | 0,0041947 | 0,002915 | 0,0037443 | 0,0037443 | TRUE | 0,054951 | No |
| orf19.6624 | 887 | 749 | 782 | 521 | 1688 | 1316 | 1486 | 2017 | 808 | 758 | 740 | 783 | 1501 | 1429 | 1522 | 1539 | 1134,81 | 772 | 1498 | 1,942 | 0,958 | 17,893 | 1,33E-71 | 8,46E-71 | 1,00E-08 | 0,003201 | 0,0017463 | 0,0017463 | TRUE | 0,0131384 | No |
| orf19.6625 | 961 | 874 | 921 | 513 | 1836 | 1299 | 1593 | 2055 | 875 | 884 | 872 | 771 | 1633 | 1410 | 1631 | 1568 | 1205,45 | 850 | 1560 | 1,831 | 0,872 | 13,355 | 1,11E-40 | 4,35E-40 | 0,0030098 | 0,003133 | 0,003168 | 0,003168 | TRUE | 0,0490886 | No |
| orf19.6626 | 237 | 186 | 220 | 147 | 531 | 422 | 443 | 608 | 216 | 188 | 208 | 221 | 472 | 458 | 454 | 464 | 335,11 | 208 | 462 | 2,226 | 1,155 | 13,821 | 1,90E-43 | 7,89E-43 | 1,00E-08 | 0,005979 | 0,0031333 | 0,0031333 | TRUE | 0,0460244 | No |
| orf19.6627 | 1362 | 1090 | 1110 | 771 | 2852 | 2022 | 2318 | 3102 | 1240 | 1102 | 1051 | 1158 | 2537 | 2195 | 2374 | 2366 | 1752,92 | 1138 | 2368 | 2,083 | 1,058 | 16,594 | 7,72E-62 | 4,33E-61 | 0,0034076 | 0,002791 | 0,0032354 | 0,0032354 | TRUE | 0,0470137 | No |
| orf19.6628 | 617 | 433 | 463 | 321 | 1509 | 1041 | 1369 | 1705 | 562 | 438 | 438 | 482 | 1342 | 1130 | 1402 | 1301 | 886,89 | 480 | 1294 | 2,695 | 1,43 | 15,722 | 1,07E-55 | 5,50E-55 | 0,0095464 | 0,003526 | 0,0064639 | 0,0064639 | TRUE | 0,1634995 | No |
| orf19.6629 | 462 | 369 | 402 | 265 | 1055 | 839 | 949 | 1299 | 421 | 373 | 380 | 398 | 938 | 911 | 972 | 991 | 673,06 | 393 | 953 | 2,426 | 1,279 | 19,262 | 1,12E-82 | 8,22E-82 | 1,00E-08 | 0,004 | 0,0023872 | 0,0023872 | TRUE | 0,0295827 | No |
| orf19.663 | 1315 | 1220 | 1321 | 755 | 1888 | 1589 | 1783 | 2270 | 1197 | 1234 | 1250 | 1134 | 1679 | 1725 | 1826 | 1732 | 1472,2 | 1204 | 1740 | 1,442 | 0,528 | 10,468 | 1,21E-25 | 3,53E-25 | 0,0006037 | 0,002934 | 0,0017314 | 0,0017314 | TRUE | 0,0158094 | No |
| orf19.6630 | 364 | 246 | 252 | 142 | 549 | 337 | 440 | 590 | 331 | 249 | 238 | 213 | 488 | 366 | 451 | 450 | 348,36 | 258 | 439 | 1,692 | 0,759 | 5,924 | 3,14E-09 | 5,77E-09 | 0,0225111 | 0,005829 | 0,0126288 | 0,0126288 | TRUE | 0,4187112 | No |
| orf19.6632 | 2119 | 1702 | 1848 | 1252 | 2624 | 2112 | 2297 | 3209 | 1929 | 1721 | 1749 | 1881 | 2334 | 2293 | 2352 | 2448 | 2088,45 | 1820 | 2357 | 1,296 | 0,374 | 7,438 | 1,03E-13 | 2,19E-13 | 0,0012669 | 0,00267 | 0,0019394 | 0,0019394 | TRUE | 0,0215028 | No |
| orf19.6634 | 1720 | 1502 | 1685 | 1017 | 1980 | 1727 | 1710 | 2427 | 1566 | 1519 | 1595 | 1528 | 1761 | 1875 | 1751 | 1851 | 1680,78 | 1552 | 1810 | 1,165 | 0,221 | 4,819 | 1,44E-06 | 2,38E-06 | 0,0003113 | 0,002823 | 0,0014101 | 0,0014101 | TRUE | 0,0066448 | No |
| orf19.6635 | 866 | 829 | 886 | 510 | 1310 | 1217 | 1203 | 1552 | 788 | 838 | 839 | 766 | 1165 | 1321 | 1232 | 1184 | 1016,74 | 808 | 1226 | 1,512 | 0,597 | 9,878 | 5,17E-23 | 1,41E-22 | 0,0016316 | 0,003336 | 0,0024599 | 0,0024599 | TRUE | 0,0322935 | No |

| | | | | | | | | | | | | | | | | | | | | | | | | | | | | | | | |
|---|---|---|---|---|---|---|---|---|---|---|---|---|---|---|---|---|---|---|---|---|---|---|---|---|---|---|---|---|---|---|---|
| orf19.6636 | 76 | 86 | 84 | 48 | 136 | 110 | 143 | 180 | 69 | 87 | 79 | 72 | 121 | 119 | 146 | 137 | 103,99 | 77 | 131 | 1,701 | 0,766 | 5,281 | 1,28E-07 | 2,21E-07 | 1,00E-08 | 0,01474 | 0,0097533 | 0,0097533 | TRUE | 0,0719455 | No |
| orf19.6637 | 3699 | 3198 | 3517 | 2209 | 4622 | 3808 | 3909 | 5368 | 3368 | 3234 | 3329 | 3318 | 4111 | 4134 | 4003 | 4095 | 3699,08 | 3312 | 4086 | 1,234 | 0,303 | 8,852 | 8,57E-19 | 2,11E-18 | 1,00E-08 | 0,002394 | 0,0008466 | 0,0008466 | TRUE | 0,0031559 | No |
| orf19.6638 | 985 | 880 | 849 | 560 | 986 | 807 | 889 | 1219 | 897 | 890 | 804 | 841 | 877 | 876 | 910 | 930 | 878,13 | 858 | 898 | 1,047 | 0,066 | 1,138 | 0,255299 | 0,290735 | 0,0007148 | 0,003541 | 0,0021266 | 0,0021266 | TRUE | 0,0239914 | No |
| orf19.6639 | 3986 | 3582 | 3768 | 2397 | 1527 | 1266 | 1324 | 1767 | 3629 | 3623 | 3566 | 3601 | 1358 | 1375 | 1356 | 1348 | 2481,9 | 3605 | 1359 | 0,377 | -1,408 | -37,411 | 2,63E-306 | 6,94E-305 | 1,00E-08 | 0,002569 | 0,0008674 | 0,0008674 | TRUE | 0,0006714 | No |
| orf19.664 | 85 | 93 | 87 | 63 | 109 | 86 | 97 | 94 | 77 | 94 | 82 | 95 | 97 | 93 | 99 | 72 | 88,72 | 87 | 90 | 1,037 | 0,053 | 0,333 | 0,739094 | 0,770897 | 1,00E-08 | 0,016926 | 0,0129496 | 0,0129496 | TRUE | 0,2153997 | No |
| orf19.6640 | 9078 | 8090 | 8555 | 5240 | 2457 | 1982 | 2049 | 2783 | 8265 | 8182 | 8097 | 7871 | 2185 | 2152 | 2098 | 2123 | 5121,73 | 8104 | 2140 | 0,264 | -1,922 | -54,36 | 0 | 0 | 1,00E-08 | 0,002295 | 0,0009155 | 0,0009155 | TRUE | 0,004338 | No |
| orf19.6641 | 1179 | 1032 | 1030 | 733 | 431 | 367 | 337 | 487 | 1073 | 1044 | 975 | 1101 | 383 | 398 | 345 | 371 | 711,44 | 1048 | 374 | 0,358 | -1,483 | -21,662 | 4,68E-104 | 4,23E-103 | 0,0014012 | 0,003894 | 0,0027417 | 0,0027417 | TRUE | 0,0296272 | No |
| orf19.6642 | 1921 | 1789 | 1939 | 1360 | 1922 | 1695 | 1688 | 2256 | 1749 | 1809 | 1835 | 2043 | 1710 | 1840 | 1729 | 1721 | 1804,48 | 1859 | 1750 | 0,943 | -0,085 | -1,522 | 0,128073 | 0,151957 | 0,0022254 | 0,002769 | 0,0024143 | 0,0024143 | TRUE | 0,0498105 | No |
| orf19.6643 | 3384 | 2846 | 3357 | 2300 | 2192 | 1950 | 1994 | 2710 | 3081 | 2878 | 3177 | 3455 | 1950 | 2117 | 2042 | 2067 | 2595,96 | 3148 | 2044 | 0,65 | -0,622 | -10,714 | 8,72E-27 | 2,60E-26 | 0,0029487 | 0,002546 | 0,0028308 | 0,0028308 | TRUE | 0,0501855 | No |
| orf19.6644 | 182 | 192 | 245 | 167 | 184 | 190 | 215 | 234 | 166 | 194 | 232 | 251 | 164 | 206 | 220 | 178 | 201,41 | 211 | 192 | 0,915 | -0,128 | -0,913 | 0,36125 | 0,401493 | 0,0205873 | 0,008596 | 0,0137511 | 0,0137511 | TRUE | 0,2339393 | No |
| orf19.6645 | 18657 | 15509 | 16704 | 11931 | 12853 | 10307 | 10502 | 15338 | 16986 | 15686 | 15809 | 17922 | 11432 | 11191 | 10755 | 11700 | 13935,2 | 16601 | 11270 | 0,679 | -0,558 | -11,009 | 3,44E-28 | 1,06E-27 | 0,0025655 | 0,002132 | 0,0023966 | 0,0023966 | TRUE | 0,0347713 | No |
| orf19.6648 | 19444 | 16582 | 18279 | 10578 | 28964 | 22775 | 26490 | 34432 | 17703 | 16771 | 17299 | 15890 | 25763 | 24727 | 27129 | 26265 | 21443,4 | 16916 | 25971 | 1,535 | 0,618 | 13,775 | 3,60E-43 | 1,49E-42 | 0,0017525 | 0,002099 | 0,0018848 | 0,0018848 | TRUE | 0,0201424 | No |
| orf19.6649 | 4787 | 4057 | 4538 | 2736 | 3753 | 2940 | 3538 | 4477 | 4358 | 4103 | 4295 | 4110 | 3338 | 3192 | 3623 | 3415 | 3804,37 | 4216 | 3392 | 0,804 | -0,314 | -6,644 | 3,06E-11 | 6,05E-11 | 0,0015225 | 0,002384 | 0,0018845 | 0,0018845 | TRUE | 0,0250385 | No |
| orf19.665 | 4785 | 4350 | 4703 | 2777 | 5372 | 4245 | 4616 | 6113 | 4357 | 4400 | 4451 | 4171 | 4778 | 4609 | 4727 | 4663 | 4519,51 | 4345 | 4694 | 1,08 | 0,11 | 3,118 | 0,00182 | 0,002519 | 0,000221 | 0,002329 | 0,000981 | 0,000981 | TRUE | 0,0080892 | No |
| orf19.6650 | 588 | 528 | 577 | 291 | 404 | 291 | 293 | 393 | 535 | 534 | 546 | 437 | 359 | 316 | 300 | 300 | 415,96 | 513 | 319 | 0,618 | -0,694 | -7,558 | 4,08E-14 | 8,80E-14 | 0,0060516 | 0,005213 | 0,0055773 | 0,0055773 | TRUE | 0,1017048 | No |
| orf19.6652 | 6616 | 5646 | 6209 | 4006 | 6507 | 5501 | 5837 | 8319 | 6024 | 5710 | 5876 | 6018 | 5788 | 5973 | 5978 | 6346 | 5963,97 | 5907 | 6021 | 1,02 | 0,028 | 0,699 | 0,48478 | 0,526665 | 0,0008435 | 0,002259 | 0,0013989 | 0,0013989 | TRUE | 0,0164729 | No |
| orf19.6653 | 3067 | 2478 | 2892 | 1589 | 2947 | 2327 | 2677 | 3543 | 2792 | 2506 | 2737 | 2387 | 2621 | 2526 | 2742 | 2703 | 2626,81 | 2606 | 2648 | 1,015 | 0,021 | 0,375 | 0,707495 | 0,741579 | 0,0025251 | 0,00254 | 0,0027249 | 0,0027249 | TRUE | 0,0367659 | No |
| orf19.6654 | 6560 | 5384 | 5778 | 3756 | 9776 | 8287 | 8569 | 11904 | 5973 | 5445 | 5468 | 5642 | 8695 | 8997 | 8776 | 9080 | 7259,68 | 5632 | 8887 | 1,578 | 0,658 | 16,314 | 7,86E-60 | 4,28E-59 | 0,0009487 | 0,002219 | 0,0014138 | 0,0014138 | TRUE | 0,0208299 | No |
| orf19.6656 | 2558 | 2051 | 2311 | 1471 | 3299 | 2788 | 2940 | 4207 | 2329 | 2074 | 2187 | 2210 | 2934 | 3027 | 3011 | 3209 | 2622,69 | 2200 | 3045 | 1,384 | 0,469 | 9,522 | 1,70E-21 | 4,45E-21 | 0,0013339 | 0,002541 | 0,0019346 | 0,0019346 | TRUE | 0,0198209 | No |
| orf19.6658 | 5295 | 4428 | 4720 | 3067 | 5634 | 4416 | 4809 | 6677 | 4821 | 4478 | 4467 | 4607 | 5011 | 4795 | 4925 | 5093 | 4774,7 | 4593 | 4956 | 1,079 | 0,11 | 2,744 | 0,006066 | 0,008085 | 0,000693 | 0,002314 | 0,0013325 | 0,0013325 | TRUE | 0,0140897 | No |
| orf19.6659 | 41019 | 34069 | 38302 | 24353 | 27814 | 23193 | 25076 | 34284 | 37346 | 34457 | 36249 | 36582 | 24740 | 25181 | 25681 | 26152 | 30798,6 | 36158 | 25438 | 0,704 | -0,507 | -13,994 | 1,71E-44 | 7,24E-44 | 0,0008172 | 0,00208 | 0,0012291 | 0,0012291 | TRUE | 0,0123234 | No |
| orf19.666 | 976 | 891 | 1022 | 627 | 1529 | 1266 | 1423 | 1784 | 889 | 901 | 967 | 942 | 1360 | 1375 | 1457 | 1361 | 1156,44 | 925 | 1388 | 1,502 | 0,587 | 11,084 | 1,50E-28 | 4,63E-28 | 0,0004664 | 0,003179 | 0,0017725 | 0,0017725 | TRUE | 0,0132758 | No |
| orf19.6660 | 17193 | 13334 | 15370 | 10221 | 23779 | 20073 | 22143 | 30412 | 15653 | 13486 | 14546 | 15354 | 21151 | 21794 | 22677 | 23199 | 18482,4 | 14760 | 22205 | 1,505 | 0,589 | 11,139 | 8,09E-29 | 2,52E-28 | 0,0029239 | 0,002108 | 0,002632 | 0,002632 | TRUE | 0,0413731 | No |
| orf19.6661 | 353 | 307 | 353 | 214 | 1531 | 1217 | 1271 | 1742 | 321 | 310 | 334 | 321 | 1362 | 1321 | 1302 | 1329 | 825,13 | 322 | 1328 | 4,126 | 2,045 | 32,99 | 1,15E-238 | 2,29E-237 | 1,00E-08 | 0,003638 | 0,0016906 | 0,0016906 | TRUE | 0,0082597 | No |
| orf19.6662 | 391 | 306 | 334 | 196 | 726 | 634 | 586 | 813 | 356 | 309 | 316 | 294 | 646 | 688 | 600 | 620 | 478,8 | 319 | 638 | 1,99 | 0,993 | 12,424 | 1,93E-35 | 6,96E-35 | 0,0023301 | 0,004796 | 0,0037368 | 0,0037368 | TRUE | 0,0716557 | No |
| orf19.6663 | 12813 | 10175 | 10635 | 8442 | 21404 | 16966 | 17132 | 22805 | 11666 | 10291 | 10065 | 12681 | 19038 | 18420 | 17545 | 17396 | 14637,8 | 11176 | 18100 | 1,62 | 0,696 | 9,967 | 2,13E-23 | 5,87E-23 | 0,006848 | 0,002127 | 0,0046118 | 0,0046118 | TRUE | 0,0997282 | No |
| orf19.6665 | 4202 | 3566 | 4119 | 2740 | 5772 | 4823 | 5266 | 7137 | 3826 | 3607 | 3898 | 4116 | 5134 | 5236 | 5393 | 5444 | 4581,78 | 3862 | 5302 | 1,374 | 0,459 | 9,907 | 3,89E-23 | 1,06E-22 | 0,0015422 | 0,002325 | 0,0018318 | 0,0018318 | TRUE | 0,0237796 | No |
| orf19.6667 | 1496 | 1280 | 1378 | 829 | 997 | 698 | 799 | 1012 | 1362 | 1295 | 1304 | 1245 | 887 | 758 | 818 | 772 | 1055,12 | 1302 | 809 | 0,621 | -0,688 | -11,147 | 7,40E-29 | 2,31E-28 | 0,0019214 | 0,003289 | 0,0026673 | 0,0026673 | TRUE | 0,0512853 | No |
| orf19.6668 | 1493 | 1224 | 1394 | 949 | 1360 | 1074 | 1196 | 1607 | 1359 | 1238 | 1319 | 1426 | 1210 | 1166 | 1225 | 1226 | 1271,07 | 1336 | 1207 | 0,906 | -0,143 | -2,642 | 0,008251 | 0,010883 | 0,0009425 | 0,003076 | 0,0020221 | 0,0020221 | TRUE | 0,0285368 | No |
| orf19.6670 | 269 | 262 | 241 | 168 | 332 | 285 | 300 | 414 | 245 | 265 | 228 | 252 | 295 | 309 | 307 | 316 | 277,26 | 248 | 307 | 1,242 | 0,313 | 3,575 | 0,00035 | 0,000508 | 1,00E-08 | 0,006801 | 0,0036935 | 0,0036935 | TRUE | 0,0334848 | No |
| orf19.6671 | 757 | 721 | 780 | 455 | 889 | 631 | 827 | 1025 | 689 | 729 | 738 | 683 | 791 | 685 | 847 | 782 | 743,1 | 710 | 776 | 1,093 | 0,128 | 1,799 | 0,071971 | 0,087744 | 0,003159 | 0,003815 | 0,0034985 | 0,0034985 | TRUE | 0,0699819 | No |
| orf19.667.1 | 13216 | 10336 | 10359 | 8168 | 20265 | 15758 | 15431 | 23016 | 12033 | 10454 | 9804 | 12270 | 18025 | 17109 | 15803 | 17557 | 14131,7 | 11140 | 17124 | 1,538 | 0,621 | 8,594 | 8,40E-18 | 2,01E-17 | 0,0072748 | 0,00213 | 0,0049346 | 0,0049346 | TRUE | 0,0798366 | No |
| orf19.6672 | 8119 | 7010 | 7630 | 4685 | 4274 | 3406 | 3712 | 5006 | 7392 | 7090 | 7221 | 7038 | 3802 | 3698 | 3802 | 3819 | 5482,54 | 7185 | 3780 | 0,526 | -0,927 | -27,72 | 3,98E-169 | 5,91E-168 | 0,000192 | 0,002278 | 0,000875 | 0,000875 | TRUE | 0,0046295 | No |
| orf19.6673 | 531 | 431 | 510 | 344 | 997 | 806 | 827 | 1211 | 483 | 436 | 483 | 517 | 887 | 875 | 847 | 924 | 681,42 | 480 | 883 | 1,849 | 0,887 | 13,359 | 1,05E-40 | 4,15E-40 | 0,0009038 | 0,003976 | 0,0025875 | 0,0025875 | TRUE | 0,0428395 | No |
| orf19.6674 | 344 | 275 | 321 | 141 | 728 | 536 | 533 | 729 | 313 | 278 | 304 | 212 | 648 | 582 | 546 | 556 | 429,79 | 277 | 583 | 2,086 | 1,061 | 10,136 | 3,81E-24 | 1,07E-23 | 0,012038 | 0,005111 | 0,0077983 | 0,0077983 | TRUE | 0,2349541 | No |
| orf19.6675 | 839 | 823 | 821 | 574 | 1071 | 953 | 883 | 1237 | 764 | 832 | 777 | 862 | 953 | 1035 | 904 | 944 | 883,84 | 809 | 959 | 1,188 | 0,249 | 3,905 | 9,41E-05 | 0,000141 | 0,0017528 | 0,003532 | 0,0027355 | 0,0027355 | TRUE | 0,0322129 | No |
| orf19.6676 | 162 | 114 | 138 | 102 | 193 | 130 | 170 | 263 | 147 | 115 | 131 | 153 | 172 | 141 | 174 | 201 | 154,27 | 136 | 172 | 1,27 | 0,344 | 2,534 | 0,011273 | 0,014735 | 0,0118147 | 0,0106 | 0,0110953 | 0,0110953 | TRUE | 0,1473739 | No |
| orf19.6678 | 367 | 332 | 352 | 246 | 266 | 259 | 265 | 334 | 334 | 336 | 333 | 370 | 237 | 281 | 271 | 255 | 302,07 | 343 | 261 | 0,761 | -0,393 | -4,494 | 7,00E-06 | 1,12E-05 | 1,00E-08 | 0,00641 | 0,0040058 | 0,0040058 | TRUE | 0,0438027 | No |
| orf19.6679 | 1407 | 1131 | 1230 | 865 | 2560 | 2192 | 2356 | 3231 | 1281 | 1144 | 1164 | 1299 | 2277 | 2380 | 2413 | 2465 | 1802,85 | 1222 | 2384 | 1,954 | 0,966 | 17,658 | 8,86E-70 | 5,54E-69 | 0,0018379 | 0,00277 | 0,0022399 | 0,0022399 | TRUE | 0,021806 | No |
| orf19.668 | 978 | 964 | 976 | 762 | 403 | 349 | 382 | 485 | 890 | 975 | 924 | 1145 | 358 | 379 | 391 | 370 | 679,04 | 984 | 374 | 0,382 | -1,388 | -16,56 | 1,36E-61 | 7,60E-61 | 0,005945 | 0,003982 | 0,0049501 | 0,0049501 | TRUE | 0,135631 | No |
| orf19.6680 | 2747 | 2466 | 2747 | 1778 | 6071 | 5306 | 5476 | 7432 | 2501 | 2494 | 2600 | 2671 | 5400 | 5761 | 5608 | 5669 | 4087,98 | 2566 | 5610 | 2,188 | 1,13 | 28,074 | 2,01E-173 | 3,04E-172 | 0,0005388 | 0,00236 | 0,0012614 | 0,0012614 | TRUE | 0,0082606 | No |
| orf19.6681 | 2003 | 1889 | 2047 | 1253 | 2531 | 2157 | 2326 | 3042 | 1824 | 1911 | 1937 | 1882 | 2251 | 2342 | 2382 | 2320 | 2106,17 | 1888 | 2324 | 1,23 | 0,299 | 7,106 | 1,19E-12 | 2,47E-12 | 1,00E-08 | 0,002664 | 0,001214 | 0,001214 | TRUE | 0,0070215 | No |
| orf19.6684 | 612 | 556 | 583 | 353 | 296 | 254 | 289 | 354 | 557 | 562 | 552 | 530 | 263 | 276 | 296 | 270 | 413,33 | 550 | 276 | 0,5 | -0,999 | -13,423 | 4,43E-41 | 1,76E-40 | 1,00E-08 | 0,005233 | 0,0026671 | 0,0026671 | TRUE | 0,0249843 | No |
| orf19.6685 | 1903 | 1765 | 1831 | 965 | 1500 | 1154 | 1252 | 1599 | 1733 | 1785 | 1733 | 1450 | 1334 | 1253 | 1282 | 1220 | 1473,65 | 1675 | 1272 | 0,758 | -0,401 | -6,012 | 1,84E-09 | 3,41E-09 | 0,0039405 | 0,002933 | 0,003574 | 0,003574 | TRUE | 0,0924354 | No |
| orf19.6686 | 11994 | 11727 | 12382 | 7567 | 10881 | 9079 | 9462 | 12099 | 10920 | 11861 | 11718 | 11367 | 9678 | 9857 | 9690 | 9229 | 10540,1 | 11466 | 9614 | 0,838 | -0,255 | -6,471 | 9,71E-11 | 1,89E-10 | 0,0009327 | 0,002162 | 0,0013903 | 0,0013903 | TRUE | 0,0128221 | No |
| orf19.6687 | 9 | 16 | 7 | 13 | 30 | 25 | 21 | 16 | 8 | 16 | 7 | 20 | 27 | 27 | 22 | 12 | 17,26 | 13 | 22 | 1,779 | 0,831 | 2,057 | 0,039686 | 0,049607 | 0,1133613 | 0,078579 | 0,0913419 | 0,0913419 | TRUE | 0,3809802 | No |
| orf19.6688 | 8126 | 7806 | 8399 | 4907 | 10141 | 8669 | 9607 | 12068 | 7398 | 7895 | 7949 | 7371 | 9020 | 9412 | 9839 | 9206 | 8511,23 | 7653 | 9369 | 1,224 | 0,291 | 6,714 | 1,89E-11 | 3,78E-11 | 0,0013724 | 0,002192 | 0,0016862 | 0,0016862 | TRUE | 0,0138902 | No |
| orf19.6689 | 4526 | 4341 | 4373 | 2606 | 1708 | 1383 | 1514 | 1861 | 4121 | 4390 | 4139 | 3915 | 1519 | 1502 | 1551 | 1420 | 2819,42 | 4141 | 1498 | 0,361 | -1,469 | -29,783 | 6,54E-195 | 1,09E-193 | 0,00146 | 0,002506 | 0,0018952 | 0,0018952 | TRUE | 0,0198 | No |
| orf19.669 | 158 | 157 | 134 | 83 | 380 | 318 | 379 | 458 | 144 | 159 | 127 | 125 | 338 | 345 | 388 | 349 | 246,86 | 139 | 355 | 2,548 | 1,349 | 12,898 | 4,61E-38 | 1,74E-37 | 0,0019782 | 0,007388 | 0,0053761 | 0,0053761 | TRUE | 0,0970365 | No |
| orf19.6690 | 2460 | 2267 | 2436 | 1374 | 871 | 663 | 737 | 965 | 2240 | 2293 | 2305 | 2064 | 775 | 720 | 755 | 736 | 1485,93 | 2226 | 746 | 0,335 | -1,578 | -29,098 | 3,78E-186 | 5,99E-185 | 0,0011147 | 0,002926 | 0,001958 | 0,001958 | TRUE | 0,0265975 | No |
| orf19.6691 | 11639 | 11692 | 12095 | 7275 | 12383 | 10349 | 11093 | 13871 | 10597 | 11825 | 11447 | 10928 | 11014 | 11236 | 11361 | 10581 | 11123,6 | 11199 | 11048 | 0,986 | -0,02 | -0,451 | 0,652044 | 0,689235 | 0,0013953 | 0,002156 | 0,0017932 | 0,0017932 | TRUE | 0,017364 | No |
| orf19.6692 | 2060 | 2162 | 2237 | 1276 | 2974 | 2388 | 2584 | 3257 | 1876 | 2187 | 2117 | 1917 | 2645 | 2593 | 2646 | 2484 | 2308,1 | 2024 | 2592 | 1,279 | 0,355 | 6,25 | 4,11E-10 | 7,80E-10 | 0,0026373 | 0,002609 | 0,0026596 | 0,0026596 | TRUE | 0,0353473 | No |
| orf19.6693 | 1943 | 2074 | 2237 | 1191 | 2311 | 1999 | 2021 | 2517 | 1769 | 2098 | 2117 | 1789 | 2056 | 2170 | 2070 | 1920 | 1998,56 | 1943 | 2054 | 1,055 | 0,078 | 1,113 | 0,265523 | 0,301408 | 0,0053619 | 0,002698 | 0,0041863 | 0,0041863 | TRUE | 0,0464335 | No |
| orf19.6694 | 2147 | 1828 | 2078 | 1237 | 2216 | 1734 | 2085 | 2647 | 1955 | 1849 | 1967 | 1858 | 1971 | 1883 | 2135 | 2019 | 1954,57 | 1907 | 2002 | 1,049 | 0,069 | 1,363 | 0,172878 | 0,201554 | 0,0012326 | 0,002713 | 0,0019545 | 0,0019545 | TRUE | 0,0234552 | No |
| orf19.6696 | 969 | 979 | 917 | 504 | 1580 | 1190 | 1187 | 1586 | 882 | 990 | 868 | 757 | 1405 | 1292 | 1216 | 1210 | 1077,52 | 874 | 1281 | 1,461 | 0,547 | 6,878 | 6,09E-12 | 1,24E-11 | 0,0071941 | 0,003263 | 0,0050838 | 0,0050838 | TRUE | 0,0922359 | No |
| orf19.6698 | 857 | 688 | 738 | 467 | 1007 | 883 | 868 | 1247 | 780 | 696 | 698 | 702 | 896 | 959 | 889 | 951 | 821,33 | 719 | 924 | 1,283 | 0,359 | 5,807 | 6,36E-09 | 1,16E-08 | 0,0011652 | 0,003645 | 0,002425 | 0,002425 | TRUE | 0,0415079 | No |
| orf19.6699 | 339 | 260 | 314 | 200 | 461 | 395 | 370 | 588 | 309 | 263 | 297 | 300 | 410 | 429 | 379 | 449 | 354,45 | 292 | 417 | 1,428 | 0,514 | 5,914 | 3,34E-09 | 6,13E-09 | 0,0024329 | 0,005764 | 0,0043084 | 0,0043084 | TRUE | 0,0533373 | No |
| orf19.670 | 5624 | 4298 | 4510 | 2821 | 6481 | 4193 | 4848 | 6585 | 5120 | 4347 | 4268 | 4238 | 5765 | 4552 | 4965 | 5023 | 4784,8 | 4493 | 5076 | 1,13 | 0,176 | 2,245 | 0,024738 | 0,031478 | 0,0090392 | 0,002313 | 0,0056714 | 0,0056714 | TRUE | 0,1082477 | No |
| orf19.6701 | 3832 | 3334 | 3639 | 2305 | 4914 | 4177 | 4377 | 6084 | 3489 | 3372 | 3444 | 3462 | 4371 | 4535 | 4483 | 4641 | 3974,59 | 3442 | 4508 | 1,31 | 0,389 | 10,838 | 2,27E-27 | 6,88E-27 | 0,0002552 | 0,002369 | 0,0009797 | 0,0009797 | TRUE | 0,0052013 | No |
| orf19.6702 | 3634 | 3114 | 3228 | 2304 | 4423 | 3681 | 3734 | 5197 | 3309 | 3149 | 3055 | 3461 | 3934 | 3997 | 3824 | 3964 | 3586,64 | 3244 | 3930 | 1,213 | 0,279 | 6,077 | 1,22E-09 | 2,28E-09 | 0,0013259 | 0,002405 | 0,0017345 | 0,0017345 | TRUE | 0,0232429 | No |
| orf19.670.2 | 22344 | 17276 | 18283 | 8704 | 30308 | 18386 | 24590 | 32155 | 20343 | 17473 | 17303 | 13075 | 26958 | 19962 | 25183 | 24528 | 20603,2 | 17048 | 24158 | 1,417 | 0,503 | 3,249 | 0,001159 | 0,00163 | 0,0229593 | 0,002101 | 0,0121547 | 0,0229593 | TRUE | 0,3008047 | No |
| orf19.6703 | 978 | 903 | 1051 | 526 | 674 | 537 | 586 | 772 | 890 | 913 | 995 | 790 | 600 | 583 | 600 | 589 | 745,01 | 897 | 593 | 0,659 | -0,603 | -8,452 | 2,85E-17 | 6,73E-17 | 0,0031652 | 0,00381 | 0,0034964 | 0,0034964 | TRUE | 0,0684545 | No |
| orf19.6704 | 150 | 132 | 152 | 90 | 102 | 86 | 103 | 133 | 137 | 134 | 144 | 135 | 91 | 93 | 105 | 101 | 117,52 | 138 | 98 | 0,712 | -0,491 | -3,85 | 0,000118 | 0,000176 | 1,00E-08 | 0,013278 | 0,0069702 | 0,0069702 | TRUE | 0,024805 | No |
| orf19.6705 | 5736 | 5259 | 5666 | 3651 | 5303 | 4525 | 4886 | 6306 | 5222 | 5319 | 5362 | 5484 | 4717 | 4913 | 5004 | 4810 | 5103,99 | 5347 | 4861 | 0,909 | -0,137 | -3,877 | 0,000106 | 0,000158 | 0,0002724 | 0,002296 | 0,0010045 | 0,0010045 | TRUE | 0,0048279 | No |
| orf19.6706 | 12393 | 11890 | 12588 | 6971 | 5002 | 4017 | 4441 | 5536 | 11283 | 12025 | 11913 | 10472 | 4449 | 4361 | 4548 | 4223 | 7909,4 | 11423 | 4395 | 0,385 | -1,379 | -27,22 | 3,76E-163 | 5,31E-162 | 0,0022347 | 0,002204 | 0,0023108 | 0,0023108 | TRUE | 0,0378264 | No |
| orf19.6707 | 4011 | 3772 | 3939 | 2328 | 2180 | 1597 | 1786 | 2300 | 3652 | 3815 | 3728 | 3497 | 1939 | 1734 | 1829 | 1754 | 2743,53 | 3673 | 1814 | 0,494 | -1,018 | -20,689 | 4,37E-95 | 3,57E-94 | 0,0014332 | 0,002519 | 0,0019226 | 0,0019226 | TRUE | 0,026358 | No |
| orf19.6709 | 164 | 136 | 165 | 101 | 236 | 167 | 211 | 301 | 149 | 138 | 156 | 152 | 210 | 181 | 216 | 230 | 178,96 | 149 | 209 | 1,413 | 0,499 | 4,429 | 9,45E-06 | 1,50E-05 | 1,00E-08 | 0,009419 | 0,0063829 | 0,0063829 | TRUE | 0,0826496 | No |

| | | | | | | | | | | | | | | | | | | | | | | | | | | | | | | | |
|---|---|---|---|---|---|---|---|---|---|---|---|---|---|---|---|---|---|---|---|---|---|---|---|---|---|---|---|---|---|---|---|
| orf19.671 | 2355 | 2282 | 2418 | 1571 | 3207 | 2939 | 2896 | 3862 | 2144 | 2308 | 2288 | 2360 | 2853 | 3191 | 2966 | 2946 | 2631,97 | 2275 | 2989 | 1,314 | 0,394 | 7,927 | 2,24E-15 | 5,02E-15 | 0,0014612 | 0,002539 | 0,0019831 | 0,0019831 | TRUE | 0,0246954 | No |
| orf19.6710 | 16102 | 13506 | 14858 | 9156 | 18923 | 14305 | 16623 | 21950 | 14660 | 13660 | 14062 | 13754 | 16831 | 15531 | 17024 | 16744 | 15283,2 | 14034 | 16532 | 1,178 | 0,236 | 5,69 | 1,27E-08 | 2,27E-08 | 0,0012698 | 0,002123 | 0,0015898 | 0,0015898 | TRUE | 0,0202212 | No |
| orf19.6712 | 2851 | 2428 | 2708 | 1846 | 1512 | 1280 | 1427 | 1827 | 2596 | 2456 | 2563 | 2773 | 1345 | 1390 | 1461 | 1394 | 1997,11 | 2597 | 1398 | 0,539 | -0,893 | -17,633 | 1,37E-69 | 8,53E-69 | 0,0011458 | 0,002698 | 0,0019181 | 0,0019181 | TRUE | 0,023556 | No |
| orf19.6713 | 9617 | 8243 | 9408 | 7386 | 6774 | 6607 | 6157 | 9041 | 8756 | 8337 | 8904 | 11095 | 6025 | 7173 | 6306 | 6897 | 7936,53 | 9273 | 6600 | 0,712 | -0,49 | -5,764 | 8,21E-09 | 1,49E-08 | 0,011932 | 0,002203 | 0,006813 | 0,006813 | TRUE | 0,2124262 | No |
| orf19.6715 | 9 | 8 | 10 | 10 | 20 | 17 | 24 | 22 | 8 | 8 | 9 | 15 | 18 | 18 | 25 | 17 | 14,8 | 10 | 20 | 1,959 | 0,97 | 2,597 | 0,009401 | 0,01235 | 1,00E-08 | 0,09131 | 0,0558777 | 0,0558777 | TRUE | 0,2177329 | No |
| orf19.6716 | 3313 | 2660 | 2906 | 1814 | 2429 | 1834 | 2209 | 2762 | 3016 | 2690 | 2750 | 2725 | 2161 | 1991 | 2262 | 2107 | 2462,84 | 2795 | 2130 | 0,762 | -0,393 | -7,18 | 6,95E-13 | 1,45E-12 | 0,0022788 | 0,002573 | 0,002461 | 0,002461 | TRUE | 0,0351337 | No |
| orf19.6717 | 1531 | 1254 | 1355 | 927 | 1245 | 940 | 1070 | 1452 | 1394 | 1268 | 1282 | 1393 | 1107 | 1021 | 1096 | 1108 | 1208,56 | 1334 | 1083 | 0,813 | -0,298 | -5,372 | 7,80E-08 | 1,35E-07 | 0,0012146 | 0,00313 | 0,0021252 | 0,0021252 | TRUE | 0,0171607 | No |
| orf19.6718 | 2965 | 2439 | 2542 | 1694 | 1449 | 1119 | 1130 | 1622 | 2700 | 2467 | 2406 | 2545 | 1289 | 1215 | 1157 | 1237 | 1876,88 | 2530 | 1224 | 0,485 | -1,045 | -19,402 | 7,47E-84 | 5,55E-83 | 0,0017715 | 0,002741 | 0,0021935 | 0,0021935 | TRUE | 0,0260807 | No |
| orf19.6719 | 1131 | 1060 | 1148 | 658 | 917 | 742 | 851 | 1060 | 1030 | 1072 | 1086 | 988 | 816 | 806 | 872 | 809 | 934,76 | 1044 | 826 | 0,788 | -0,343 | -6,108 | 1,01E-09 | 1,88E-09 | 0,0003326 | 0,00345 | 0,0019494 | 0,0019494 | TRUE | 0,0162777 | No |
| orf19.672 | 1291 | 1138 | 1306 | 799 | 991 | 789 | 870 | 1126 | 1175 | 1151 | 1236 | 1200 | 881 | 857 | 891 | 859 | 1031,33 | 1190 | 872 | 0,732 | -0,449 | -8,775 | 1,70E-18 | 4,16E-18 | 1,00E-08 | 0,003318 | 0,0015326 | 0,0015326 | TRUE | 0,0086241 | No |
| orf19.6720 | 22975 | 18692 | 20027 | 13189 | 18351 | 15035 | 15517 | 22442 | 20918 | 18905 | 18954 | 19812 | 16323 | 16324 | 15891 | 17119 | 18030,7 | 19647 | 16414 | 0,836 | -0,259 | -5,955 | 2,60E-09 | 4,80E-09 | 0,0015663 | 0,00211 | 0,0017649 | 0,0017649 | TRUE | 0,0234062 | No |
| orf19.6722 | 3421 | 2930 | 3259 | 2042 | 3441 | 2687 | 3224 | 4125 | 3115 | 2963 | 3084 | 3067 | 3061 | 2917 | 3302 | 3147 | 3082,02 | 3057 | 3107 | 1,016 | 0,023 | 0,502 | 0,615687 | 0,654713 | 0,001105 | 0,002466 | 0,0017298 | 0,0017298 | TRUE | 0,0211257 | No |
| orf19.6723 | 1445 | 1348 | 1424 | 940 | 1252 | 1071 | 1019 | 1428 | 1316 | 1363 | 1348 | 1412 | 1114 | 1163 | 1044 | 1089 | 1231 | 1360 | 1102 | 0,812 | -0,301 | -5,821 | 5,86E-09 | 1,07E-08 | 0,000357 | 0,00311 | 0,0017504 | 0,0017504 | TRUE | 0,0152535 | No |
| orf19.6724 | 2556 | 1877 | 2189 | 1615 | 5657 | 4995 | 5016 | 7184 | 2327 | 1898 | 2072 | 2426 | 5032 | 5423 | 5137 | 5480 | 3724,39 | 2181 | 5268 | 2,418 | 1,274 | 18,035 | 1,04E-72 | 6,74E-72 | 0,0064901 | 0,002392 | 0,0044571 | 0,0044571 | TRUE | 0,0917486 | No |
| orf19.6726 | 1833 | 1512 | 1749 | 1095 | 2388 | 1854 | 2193 | 2978 | 1669 | 1529 | 1655 | 1645 | 2124 | 2013 | 2246 | 2272 | 1894,09 | 1624 | 2164 | 1,333 | 0,414 | 7,683 | 1,55E-14 | 3,39E-14 | 0,0018692 | 0,002734 | 0,0022467 | 0,0022467 | TRUE | 0,0259735 | No |
| orf19.6727 | 2860 | 2336 | 2557 | 1612 | 3654 | 2648 | 2995 | 4105 | 2604 | 2363 | 2420 | 2421 | 3250 | 2875 | 3067 | 3131 | 2766,46 | 2452 | 3081 | 1,257 | 0,33 | 6,439 | 1,21E-10 | 2,34E-10 | 0,0017223 | 0,002515 | 0,0021435 | 0,0021435 | TRUE | 0,0241748 | No |
| orf19.6729 | 1776 | 1583 | 1826 | 1006 | 951 | 776 | 847 | 1116 | 1617 | 1601 | 1728 | 1511 | 846 | 843 | 867 | 851 | 1233,06 | 1614 | 852 | 0,527 | -0,925 | -17,039 | 4,19E-65 | 2,47E-64 | 0,0009715 | 0,003108 | 0,0019514 | 0,0019514 | TRUE | 0,0270021 | No |
| orf19.673 | 750 | 625 | 634 | 449 | 639 | 513 | 573 | 850 | 683 | 632 | 600 | 674 | 568 | 557 | 587 | 648 | 618,75 | 647 | 590 | 0,915 | -0,129 | -1,763 | 0,07792 | 0,094617 | 0,0027806 | 0,004172 | 0,0034915 | 0,0034915 | TRUE | 0,0553237 | No |
| orf19.6730 | 2576 | 2208 | 2402 | 1501 | 4331 | 3235 | 3694 | 4858 | 2345 | 2233 | 2273 | 2255 | 3852 | 3512 | 3783 | 3706 | 2994,99 | 2276 | 3713 | 1,631 | 0,706 | 16,36 | 3,68E-60 | 2,02E-59 | 0,0007602 | 0,002478 | 0,0014243 | 0,0014243 | TRUE | 0,0157197 | No |
| orf19.6731 | 1604 | 1348 | 1472 | 975 | 2432 | 1942 | 1942 | 2565 | 1460 | 1363 | 1393 | 1465 | 2163 | 2108 | 1989 | 1957 | 1737,32 | 1420 | 2054 | 1,447 | 0,533 | 10,351 | 4,13E-25 | 1,19E-24 | 0,0012903 | 0,002797 | 0,0019396 | 0,0019396 | TRUE | 0,0159286 | No |
| orf19.6731.1 | 1048 | 867 | 861 | 462 | 1866 | 1213 | 1525 | 1922 | 954 | 877 | 815 | 694 | 1660 | 1317 | 1562 | 1466 | 1168,07 | 835 | 1501 | 1,793 | 0,842 | 9,004 | 2,18E-19 | 5,45E-19 | 0,0121527 | 0,003168 | 0,0074469 | 0,0074469 | TRUE | 0,1427107 | No |
| orf19.6732 | 1341 | 1262 | 1344 | 926 | 1588 | 1228 | 1339 | 1795 | 1221 | 1276 | 1272 | 1391 | 1412 | 1333 | 1371 | 1369 | 1330,82 | 1290 | 1371 | 1,067 | 0,093 | 1,787 | 0,07401 | 0,090058 | 0,0008264 | 0,00303 | 0,0018579 | 0,0018579 | TRUE | 0,0297243 | No |
| orf19.6734 | 8968 | 7828 | 9103 | 6171 | 8269 | 7487 | 7409 | 10744 | 8165 | 7917 | 8615 | 9270 | 7355 | 8129 | 7588 | 8196 | 8154,29 | 8492 | 7817 | 0,921 | -0,119 | -2,071 | 0,038322 | 0,047958 | 0,0036621 | 0,002199 | 0,0030375 | 0,0030375 | TRUE | 0,0457513 | No |
| orf19.6736 | 8969 | 8462 | 9094 | 5878 | 6754 | 5985 | 5857 | 7977 | 8166 | 8558 | 8607 | 8830 | 6007 | 6498 | 5998 | 6085 | 7343,67 | 8540 | 6147 | 0,72 | -0,474 | -11,49 | 1,48E-30 | 4,84E-30 | 0,0010799 | 0,002217 | 0,0014961 | 0,0014961 | TRUE | 0,0177414 | No |
| orf19.6737 | 1027 | 880 | 939 | 672 | 2798 | 2466 | 2283 | 3176 | 935 | 890 | 889 | 1009 | 2489 | 2677 | 2338 | 2423 | 1706,26 | 931 | 2482 | 2,673 | 1,418 | 23,656 | 1,02E-123 | 1,07E-122 | 0,0025598 | 0,002811 | 0,0026848 | 0,0026848 | TRUE | 0,0341381 | No |
| orf19.6738 | 3387 | 2802 | 3081 | 1994 | 4635 | 3887 | 4193 | 5686 | 3084 | 2834 | 2916 | 2995 | 4123 | 4220 | 4294 | 4337 | 3600,4 | 2957 | 4244 | 1,435 | 0,521 | 12,836 | 1,04E-37 | 3,89E-37 | 0,0005177 | 0,002404 | 0,0012919 | 0,0012919 | TRUE | 0,0107733 | No |
| orf19.6739 | 2218 | 1824 | 2071 | 1219 | 1251 | 1034 | 1101 | 1514 | 2019 | 1845 | 1960 | 1831 | 1113 | 1123 | 1128 | 1155 | 1521,64 | 1914 | 1130 | 0,589 | -0,763 | -15,246 | 1,75E-52 | 8,45E-52 | 0,0007888 | 0,002905 | 0,0017082 | 0,0017082 | TRUE | 0,0168132 | No |
| orf19.674 | 2673 | 2369 | 2611 | 1773 | 2535 | 2017 | 2269 | 2933 | 2434 | 2396 | 2471 | 2663 | 2255 | 2190 | 2324 | 2237 | 2371,23 | 2491 | 2252 | 0,905 | -0,143 | -3,132 | 0,001735 | 0,002405 | 0,0008787 | 0,002594 | 0,0015879 | 0,0015879 | TRUE | 0,0242185 | No |
| orf19.6740 | 2241 | 1877 | 2159 | 1388 | 2898 | 2559 | 2625 | 3420 | 2040 | 1898 | 2043 | 2085 | 2578 | 2778 | 2688 | 2609 | 2340,02 | 2016 | 2663 | 1,321 | 0,401 | 8,658 | 4,80E-18 | 1,16E-17 | 0,0008036 | 0,002602 | 0,0016216 | 0,0016216 | TRUE | 0,018941 | No |
| orf19.6741 | 227 | 187 | 194 | 147 | 279 | 237 | 253 | 372 | 207 | 189 | 184 | 221 | 248 | 257 | 259 | 284 | 231,07 | 200 | 262 | 1,32 | 0,4 | 4,039 | 5,36E-05 | 8,15E-05 | 1,00E-08 | 0,007754 | 0,0049742 | 0,0049742 | TRUE | 0,0416338 | No |
| orf19.6742 | 1762 | 1537 | 1784 | 1015 | 1782 | 1503 | 1727 | 2220 | 1604 | 1555 | 1688 | 1525 | 1585 | 1632 | 1769 | 1693 | 1631,35 | 1593 | 1670 | 1,047 | 0,066 | 1,237 | 0,216134 | 0,24898 | 0,0013254 | 0,002847 | 0,0021006 | 0,0021006 | TRUE | 0,0196995 | No |
| orf19.6744 | 146 | 103 | 150 | 75 | 179 | 159 | 145 | 216 | 133 | 104 | 142 | 113 | 159 | 173 | 148 | 165 | 142,1 | 123 | 161 | 1,304 | 0,382 | 2,988 | 0,00281 | 0,003834 | 0,0033698 | 0,011333 | 0,0085279 | 0,0085279 | TRUE | 0,120643 | No |
| orf19.6745 | 8296 | 6371 | 6859 | 4490 | 7373 | 5379 | 6183 | 8670 | 7553 | 6444 | 6491 | 6745 | 6558 | 5840 | 6332 | 6614 | 6572,09 | 6808 | 6336 | 0,931 | -0,104 | -1,713 | 0,086621 | 0,104644 | 0,0041912 | 0,002238 | 0,0033558 | 0,0033558 | TRUE | 0,0669516 | No |
| orf19.6747 | 4698 | 3815 | 4213 | 2714 | 4398 | 3502 | 3820 | 5242 | 4277 | 3858 | 3987 | 4077 | 3912 | 3802 | 3912 | 3999 | 3978,09 | 4050 | 3906 | 0,965 | -0,052 | -1,223 | 0,221438 | 0,25463 | 0,0008222 | 0,002369 | 0,0014701 | 0,0014701 | TRUE | 0,0181172 | No |
| orf19.6748 | 5720 | 5076 | 5336 | 3192 | 8476 | 6390 | 6920 | 9277 | 5208 | 5134 | 5050 | 4795 | 7539 | 6938 | 7087 | 7077 | 6103,37 | 5047 | 7160 | 1,418 | 0,504 | 11,994 | 3,83E-33 | 1,31E-32 | 0,0010772 | 0,002253 | 0,0015219 | 0,0015219 | TRUE | 0,0155518 | No |
| orf19.6749 | 10253 | 8869 | 9814 | 6448 | 14105 | 12111 | 12308 | 17344 | 9335 | 8970 | 9288 | 9686 | 12546 | 13149 | 12605 | 13230 | 11101,2 | 9320 | 12882 | 1,383 | 0,468 | 12,617 | 1,70E-36 | 6,27E-36 | 0,0007549 | 0,002156 | 0,0012252 | 0,0012252 | TRUE | 0,0082568 | No |
| orf19.675 | 10778 | 8632 | 9305 | 6927 | 2173 | 1893 | 1950 | 2748 | 9813 | 8730 | 8806 | 10405 | 1933 | 2055 | 1997 | 2096 | 5729,55 | 9438 | 2020 | 0,214 | -2,223 | -35,958 | 3,76E-283 | 8,99E-282 | 0,0039163 | 0,002268 | 0,0033835 | 0,0033835 | TRUE | 0,0573273 | No |
| orf19.6751 | 3364 | 2729 | 2949 | 2222 | 2860 | 2333 | 2405 | 3507 | 3063 | 2760 | 2791 | 3338 | 2544 | 2533 | 2463 | 2675 | 2770,84 | 2988 | 2554 | 0,856 | -0,224 | -3,576 | 0,000349 | 0,000508 | 0,0041996 | 0,002514 | 0,0034115 | 0,0034115 | TRUE | 0,0726206 | No |
| orf19.675.1 | 196 | 152 | 169 | 125 | 264 | 227 | 213 | 296 | 178 | 154 | 160 | 188 | 235 | 246 | 218 | 226 | 200,64 | 170 | 231 | 1,367 | 0,451 | 4,317 | 1,58E-05 | 2,49E-05 | 1,00E-08 | 0,008621 | 0,0053046 | 0,0053046 | TRUE | 0,0417341 | No |
| orf19.6752 | 2759 | 2394 | 2519 | 1659 | 2447 | 1856 | 1957 | 2781 | 2512 | 2421 | 2384 | 2492 | 2177 | 2015 | 2004 | 2121 | 2265,82 | 2452 | 2079 | 0,849 | -0,236 | -5,284 | 1,27E-07 | 2,18E-07 | 0,0006651 | 0,00262 | 0,0014769 | 0,0014769 | TRUE | 0,0118032 | No |
| orf19.6753 | 1225 | 1021 | 1089 | 632 | 1755 | 1250 | 1545 | 2138 | 1115 | 1033 | 1031 | 949 | 1561 | 1357 | 1582 | 1631 | 1282,41 | 1032 | 1533 | 1,483 | 0,569 | 8,181 | 2,82E-16 | 6,45E-16 | 0,0041753 | 0,003067 | 0,0038185 | 0,0038185 | TRUE | 0,070236 | No |
| orf19.6754 | 3860 | 3055 | 3480 | 2610 | 6792 | 5540 | 6066 | 8191 | 3514 | 3090 | 3294 | 3921 | 6041 | 6015 | 6212 | 6248 | 4791,87 | 3455 | 6129 | 1,776 | 0,829 | 12,932 | 2,97E-38 | 1,12E-37 | 0,0051879 | 0,002313 | 0,003714 | 0,003714 | TRUE | 0,0972062 | No |
| orf19.6755 | 544 | 483 | 520 | 358 | 1064 | 960 | 950 | 1321 | 495 | 489 | 492 | 538 | 946 | 1042 | 973 | 1008 | 747,87 | 504 | 992 | 1,978 | 0,984 | 15,794 | 3,43E-56 | 1,77E-55 | 1,00E-08 | 0,003803 | 0,002191 | 0,002191 | TRUE | 0,0201565 | No |
| orf19.6756 | 1931 | 1554 | 1721 | 1020 | 1389 | 1126 | 1162 | 1645 | 1758 | 1572 | 1629 | 1532 | 1235 | 1223 | 1190 | 1255 | 1424,2 | 1623 | 1226 | 0,754 | -0,407 | -7,477 | 7,62E-14 | 1,63E-13 | 0,0014461 | 0,002965 | 0,0021297 | 0,0021297 | TRUE | 0,0386998 | No |
| orf19.6757 | 1175 | 1026 | 1075 | 664 | 2383 | 1901 | 1940 | 2626 | 1070 | 1038 | 1017 | 997 | 2120 | 2064 | 1987 | 2003 | 1536,97 | 1030 | 2044 | 1,979 | 0,985 | 20,575 | 4,56E-94 | 3,70E-93 | 1,00E-08 | 0,002896 | 0,0014536 | 0,0014536 | TRUE | 0,008008 | No |
| orf19.6758 | 946 | 818 | 902 | 531 | 727 | 634 | 648 | 892 | 861 | 827 | 854 | 798 | 647 | 688 | 664 | 680 | 752,37 | 835 | 670 | 0,8 | -0,322 | -5,59 | 2,27E-08 | 4,03E-08 | 1,00E-08 | 0,003793 | 0,0018455 | 0,0018455 | TRUE | 0,0091766 | No |
| orf19.676 | 773 | 551 | 682 | 415 | 883 | 678 | 706 | 1035 | 704 | 557 | 645 | 623 | 785 | 736 | 723 | 790 | 695,5 | 632 | 758 | 1,199 | 0,262 | 3,46 | 0,00054 | 0,000776 | 0,004133 | 0,003936 | 0,0040312 | 0,0040312 | TRUE | 0,0794921 | No |
| orf19.6760 | 1688 | 1539 | 1719 | 1034 | 2674 | 2310 | 2575 | 3206 | 1537 | 1557 | 1627 | 1553 | 2378 | 2508 | 2637 | 2446 | 2030,33 | 1568 | 2492 | 1,588 | 0,667 | 13,959 | 2,76E-44 | 1,17E-43 | 0,0009393 | 0,002688 | 0,0016616 | 0,0016616 | TRUE | 0,0184311 | No |
| orf19.6763 | 39002 | 34465 | 37600 | 22679 | 7781 | 6224 | 6902 | 9121 | 35510 | 34858 | 35585 | 34067 | 6921 | 6758 | 7069 | 6958 | 20965,6 | 35005 | 6926 | 0,198 | -2,337 | -76,15 | 0 | 0 | 0,0002603 | 0,0021 | 0,0008221 | 0,0008221 | TRUE | 0,0039279 | No |
| orf19.6766 | 10484 | 8734 | 9241 | 5479 | 8964 | 6927 | 7687 | 9976 | 9545 | 8834 | 8746 | 8230 | 7973 | 7521 | 7872 | 7610 | 8291,38 | 8839 | 7744 | 0,876 | -0,191 | -3,949 | 7,85E-05 | 0,000118 | 0,0020919 | 0,002196 | 0,0021365 | 0,0021365 | TRUE | 0,0354516 | No |
| orf19.6769 | 8081 | 7497 | 8132 | 4722 | 10403 | 8314 | 9198 | 12136 | 7357 | 7582 | 7696 | 7093 | 9253 | 9027 | 9420 | 9257 | 8335,8 | 7432 | 9239 | 1,242 | 0,313 | 8,59 | 8,70E-18 | 2,08E-17 | 0,0006331 | 0,002195 | 0,0011534 | 0,0011534 | TRUE | 0,0110043 | No |
| orf19.677 | 2922 | 2524 | 2857 | 1722 | 3582 | 2818 | 3013 | 3906 | 2660 | 2553 | 2704 | 2587 | 3186 | 3060 | 3086 | 2980 | 2851,82 | 2626 | 3078 | 1,171 | 0,228 | 5,612 | 1,99E-08 | 3,54E-08 | 0,0004394 | 0,0025 | 0,0012267 | 0,0012267 | TRUE | 0,0070298 | No |
| orf19.6770 | 8877 | 7978 | 8670 | 6032 | 6112 | 5299 | 5322 | 7587 | 8082 | 8069 | 8205 | 9061 | 5436 | 5753 | 5450 | 5787 | 6980,61 | 8354 | 5606 | 0,672 | -0,574 | -11,968 | 5,23E-33 | 1,79E-32 | 0,0019886 | 0,002226 | 0,0020642 | 0,0020642 | TRUE | 0,0387453 | No |
| orf19.6771 | 1E+05 | 80824 | 85226 | 54254 | 86382 | 61631 | 69827 | 99028 | 91731 | 81745 | 80659 | 81498 | 76834 | 66914 | 71512 | 75539 | 78304 | 83908 | 72700 | 0,866 | -0,207 | -3,648 | 0,000264 | 0,000387 | 0,0038233 | 0,002054 | 0,0030759 | 0,0030759 | TRUE | 0,0483124 | No |
| orf19.6773 | 2549 | 2362 | 2587 | 1526 | 3344 | 2902 | 3016 | 4004 | 2321 | 2389 | 2448 | 2292 | 2974 | 3151 | 3089 | 3054 | 2714,81 | 2362 | 3067 | 1,297 | 0,375 | 9,231 | 2,68E-20 | 6,85E-20 | 0,0003148 | 0,002524 | 0,0012021 | 0,0012021 | TRUE | 0,0072969 | No |
| orf19.6776 | 7797 | 6954 | 7453 | 4957 | 6248 | 5338 | 5361 | 7222 | 7099 | 7033 | 7054 | 7446 | 5557 | 5796 | 5490 | 5509 | 6373,02 | 7158 | 5588 | 0,781 | -0,357 | -9,928 | 3,13E-23 | 8,58E-23 | 0,0004744 | 0,002244 | 0,0010803 | 0,0010803 | TRUE | 0,0085625 | No |
| orf19.6777 | 85 | 73 | 83 | 64 | 115 | 106 | 110 | 125 | 77 | 74 | 79 | 96 | 102 | 115 | 113 | 95 | 93,91 | 82 | 106 | 1,313 | 0,393 | 2,652 | 0,008008 | 0,010572 | 1,00E-08 | 0,016103 | 0,0101166 | 0,0101166 | TRUE | 0,1014167 | No |
| orf19.6778 | 1181 | 1149 | 1223 | 804 | 2399 | 2190 | 2148 | 2879 | 1075 | 1162 | 1157 | 1208 | 2134 | 2378 | 2200 | 2196 | 1688,76 | 1150 | 2227 | 1,938 | 0,955 | 17,626 | 1,55E-69 | 9,66E-69 | 0,0013816 | 0,002819 | 0,0021391 | 0,0021391 | TRUE | 0,0245856 | No |
| orf19.6779 | 1438 | 1514 | 1517 | 915 | 1424 | 1139 | 1167 | 1433 | 1309 | 1531 | 1436 | 1374 | 1267 | 1237 | 1195 | 1093 | 1305,27 | 1412 | 1198 | 0,847 | -0,24 | -3,647 | 0,000265 | 0,000389 | 0,0036524 | 0,003049 | 0,003372 | 0,003372 | TRUE | 0,0436127 | No |
| orf19.6780 | 417 | 391 | 401 | 251 | 596 | 436 | 548 | 626 | 380 | 395 | 380 | 377 | 530 | 473 | 561 | 478 | 446,74 | 383 | 510 | 1,332 | 0,413 | 5,256 | 1,47E-07 | 2,52E-07 | 0,0020627 | 0,004994 | 0,0036242 | 0,0036242 | TRUE | 0,0502568 | No |
| orf19.6781 | 3003 | 3587 | 3714 | 2160 | 1936 | 1845 | 1653 | 2157 | 2734 | 3628 | 3515 | 3245 | 1722 | 2003 | 1693 | 1645 | 2523,13 | 3280 | 1766 | 0,538 | -0,894 | -10,175 | 2,57E-24 | 7,25E-24 | 0,0109563 | 0,002561 | 0,0069893 | 0,0069893 | TRUE | 0,1548311 | No |
| orf19.6782 | 170 | 162 | 158 | 111 | 1587 | 1320 | 1273 | 1626 | 155 | 164 | 150 | 167 | 1412 | 1433 | 1304 | 1240 | 752,96 | 159 | 1347 | 8,517 | 3,09 | 35,971 | 2,40E-283 | 5,76E-282 | 1,00E-08 | 0,003791 | 0,0033836 | 0,0033836 | TRUE | 0,0356916 | No |
| orf19.6783 | 3530 | 3429 | 3751 | 2228 | 2751 | 2384 | 2380 | 3137 | 3214 | 3468 | 3550 | 3347 | 2447 | 2588 | 2437 | 2393 | 2930,55 | 3395 | 2466 | 0,726 | -0,462 | -9,937 | 2,86E-23 | 7,86E-23 | 0,0010546 | 0,002488 | 0,0017262 | 0,0017262 | TRUE | 0,0164173 | No |
| orf19.6784 | 213 | 184 | 181 | 129 | 180 | 137 | 143 | 223 | 194 | 186 | 171 | 194 | 160 | 149 | 146 | 170 | 171,31 | 186 | 156 | 0,845 | -0,244 | -2,226 | 0,026042 | 0,033071 | 1,00E-08 | 0,009748 | 0,0056779 | 0,0056779 | TRUE | 0,040834 | No |
| orf19.6785 | 9560 | 8163 | 8326 | 4930 | 14426 | 10320 | 10644 | 15220 | 8704 | 8256 | 7880 | 7406 | 12831 | 11205 | 10901 | 11610 | 9849,03 | 8062 | 11637 | 1,443 | 0,529 | 8,452 | 2,86E-17 | 6,74E-17 | 0,0048576 | 0,002171 | 0,0036584 | 0,0036584 | TRUE | 0,0584536 | No |
| orf19.6786 | 419 | 361 | 392 | 254 | 619 | 513 | 545 | 633 | 381 | 365 | 371 | 382 | 551 | 557 | 558 | 483 | 455,96 | 375 | 537 | 1,431 | 0,517 | 6,791 | 1,11E-11 | 2,24E-11 | 1,00E-08 | 0,004934 | 0,003262 | 0,003262 | TRUE | 0,058677 | No |
| orf19.6787 | 1470 | 1724 | 1634 | 936 | 2405 | 2011 | 2007 | 2444 | 1338 | 1744 | 1546 | 1406 | 2139 | 2183 | 2055 | 1864 | 1784,59 | 1508 | 2060 | 1,364 | 0,448 | 5,518 | 3,43E-08 | 6,03E-08 | 0,0086252 | 0,002777 | 0,0057553 | 0,0057553 | TRUE | 0,1332097 | No |

| | | | | | | | | | | | | | | | | | | | | | | | | | | | | | | | |
|---|---|---|---|---|---|---|---|---|---|---|---|---|---|---|---|---|---|---|---|---|---|---|---|---|---|---|---|---|---|---|---|
| orf19.6788 | 906 | 942 | 939 | 563 | 2574 | 1921 | 2048 | 2719 | 825 | 953 | 889 | 846 | 2289 | 2086 | 2097 | 2074 | 1507,33 | 878 | 2136 | 2,432 | 1,282 | 21,514 | 1,14E-102 | 1,02E-101 | 0,0021432 | 0,002913 | 0,0025802 | 0,0025802 | TRUE | 0,0396258 | No |
| orf19.6789 | 518 | 512 | 519 | 265 | 688 | 582 | 616 | 750 | 472 | 518 | 491 | 398 | 612 | 632 | 631 | 572 | 540,69 | 470 | 612 | 1,293 | 0,371 | 4,623 | 3,77E-06 | 6,12E-06 | 0,0040049 | 0,00448 | 0,0042822 | 0,0042822 | TRUE | 0,1033781 | No |
| orf19.679 | 759 | 622 | 657 | 450 | 748 | 517 | 565 | 755 | 691 | 629 | 622 | 676 | 665 | 561 | 579 | 576 | 624,88 | 654 | 595 | 0,911 | -0,134 | -1,828 | 0,067508 | 0,082519 | 0,002957 | 0,004151 | 0,0035732 | 0,0035732 | TRUE | 0,0748138 | No |
| orf19.6790 | 554 | 454 | 506 | 315 | 625 | 484 | 546 | 672 | 504 | 459 | 479 | 473 | 556 | 525 | 559 | 513 | 508,6 | 479 | 538 | 1,122 | 0,166 | 2,411 | 0,015905 | 0,020579 | 1,00E-08 | 0,004634 | 0,002559 | 0,002559 | TRUE | 0,0161875 | No |
| orf19.6791 | 17656 | 16147 | 17569 | 8313 | 12032 | 7825 | 9291 | 11483 | 16075 | 16331 | 16627 | 12487 | 10702 | 8496 | 9515 | 8759 | 12374,2 | 15380 | 9368 | 0,609 | -0,716 | -7,822 | 5,21E-15 | 1,15E-14 | 0,0134239 | 0,002144 | 0,0079555 | 0,0079555 | TRUE | 0,195321 | No |
| orf19.6792 | 872 | 812 | 876 | 599 | 1726 | 1527 | 1497 | 1935 | 794 | 821 | 829 | 900 | 1535 | 1658 | 1533 | 1476 | 1193,29 | 836 | 1550 | 1,859 | 0,895 | 15,317 | 5,88E-53 | 2,86E-52 | 0,0014498 | 0,003144 | 0,0023295 | 0,0023295 | TRUE | 0,0273779 | No |
| orf19.6793 | 3638 | 3431 | 3608 | 2057 | 8685 | 6664 | 7123 | 9984 | 3312 | 3470 | 3415 | 3090 | 7725 | 7235 | 7295 | 7616 | 5394,74 | 3322 | 7468 | 2,246 | 1,167 | 25,626 | 7,74E-145 | 9,79E-144 | 0,0015001 | 0,002282 | 0,0017677 | 0,0017677 | TRUE | 0,0251127 | No |
| orf19.6794 | 5076 | 4893 | 4914 | 3020 | 7039 | 6291 | 6424 | 8508 | 4621 | 4949 | 4651 | 4537 | 6261 | 6830 | 6579 | 6490 | 5614,7 | 4690 | 6540 | 1,394 | 0,479 | 11,177 | 5,26E-29 | 1,65E-28 | 0,0011365 | 0,002272 | 0,0015772 | 0,0015772 | TRUE | 0,0169035 | No |
| orf19.6795 | 457 | 540 | 503 | 293 | 652 | 494 | 541 | 639 | 416 | 546 | 476 | 440 | 580 | 536 | 554 | 487 | 504,52 | 470 | 539 | 1,146 | 0,197 | 2,139 | 0,032442 | 0,040873 | 0,0078262 | 0,004655 | 0,0061488 | 0,0061488 | TRUE | 0,1419894 | No |
| orf19.6796 | 437 | 378 | 423 | 280 | 619 | 564 | 551 | 721 | 398 | 382 | 400 | 421 | 551 | 612 | 564 | 550 | 484,79 | 400 | 569 | 1,423 | 0,509 | 7,157 | 8,25E-13 | 1,72E-12 | 1,00E-08 | 0,004762 | 0,002705 | 0,002705 | TRUE | 0,0289196 | No |
| orf19.6797 | 1328 | 1251 | 1341 | 873 | 1384 | 1209 | 1193 | 1577 | 1209 | 1265 | 1269 | 1311 | 1231 | 1313 | 1222 | 1203 | 1252,91 | 1264 | 1242 | 0,983 | -0,024 | -0,47 | 0,638008 | 0,675969 | 0,0004592 | 0,003091 | 0,0017098 | 0,0017098 | TRUE | 0,0168453 | No |
| orf19.6798 | 27587 | 23611 | 26920 | 17883 | 12656 | 10617 | 11535 | 15874 | 25117 | 23880 | 25477 | 26863 | 11257 | 11527 | 11813 | 12109 | 18505,5 | 25334 | 11676 | 0,461 | -1,117 | -25,444 | 8,21E-143 | 1,02E-141 | 0,001584 | 0,002108 | 0,0017906 | 0,0017906 | TRUE | 0,02002 | No |
| orf19.680 | 3636 | 3160 | 3491 | 2119 | 5017 | 4060 | 4451 | 5877 | 3310 | 3196 | 3304 | 3183 | 4462 | 4408 | 4558 | 4483 | 3863,16 | 3248 | 4478 | 1,378 | 0,462 | 13,366 | 9,54E-41 | 3,76E-40 | 1,00E-08 | 0,002379 | 0,0008778 | 0,0008778 | TRUE | 0,0020245 | No |
| orf19.6800 | 1513 | 1318 | 1488 | 947 | 2465 | 2120 | 2221 | 2820 | 1378 | 1333 | 1408 | 1423 | 2193 | 2302 | 2275 | 2151 | 1807,66 | 1386 | 2230 | 1,61 | 0,687 | 14,996 | 7,76E-51 | 3,63E-50 | 0,0004106 | 0,002768 | 0,0014177 | 0,0014177 | TRUE | 0,007687 | No |
| orf19.6801 | 2730 | 2534 | 2683 | 1673 | 2665 | 2091 | 2515 | 3172 | 2486 | 2563 | 2539 | 2513 | 2370 | 2270 | 2576 | 2420 | 2467,09 | 2525 | 2409 | 0,954 | -0,068 | -1,459 | 0,144693 | 0,170376 | 0,0009432 | 0,002572 | 0,0016703 | 0,0016703 | TRUE | 0,025475 | No |
| orf19.6802 | 298 | 350 | 344 | 163 | 254 | 217 | 241 | 309 | 271 | 354 | 326 | 245 | 226 | 236 | 247 | 236 | 267,47 | 299 | 236 | 0,785 | -0,349 | -3,096 | 0,001959 | 0,002704 | 0,0104301 | 0,006976 | 0,0084554 | 0,0084554 | TRUE | 0,1559563 | No |
| orf19.6803 | 598 | 559 | 570 | 326 | 600 | 497 | 556 | 661 | 544 | 565 | 539 | 490 | 534 | 540 | 569 | 504 | 535,74 | 534 | 537 | 0,998 | -0,003 | -0,036 | 0,97151 | 0,974721 | 0,0008811 | 0,004503 | 0,0029081 | 0,0029081 | TRUE | 0,0312762 | No |
| orf19.6804 | 1115 | 1430 | 1334 | 845 | 1415 | 1401 | 1304 | 1511 | 1015 | 1446 | 1263 | 1269 | 1259 | 1521 | 1335 | 1153 | 1282,63 | 1248 | 1317 | 1,055 | 0,077 | 0,74 | 0,459169 | 0,501061 | 0,0165892 | 0,003067 | 0,0095092 | 0,0095092 | TRUE | 0,1939278 | No |
| orf19.6805 | 912 | 1027 | 1128 | 666 | 992 | 949 | 918 | 1146 | 830 | 1039 | 1068 | 1000 | 882 | 1030 | 940 | 874 | 958,01 | 984 | 932 | 0,946 | -0,08 | -0,945 | 0,344673 | 0,384545 | 0,0081808 | 0,003416 | 0,0057971 | 0,0057971 | TRUE | 0,1376941 | No |
| orf19.6807 | 52 | 43 | 45 | 25 | 78 | 53 | 36 | 65 | 47 | 43 | 43 | 38 | 69 | 58 | 37 | 50 | 48,04 | 43 | 54 | 1,241 | 0,312 | 1,404 | 0,160358 | 0,188023 | 0,0209722 | 0,029533 | 0,026241 | 0,026241 | TRUE | 0,1983183 | No |
| orf19.6809 | 774 | 718 | 750 | 411 | 821 | 599 | 634 | 855 | 705 | 726 | 710 | 617 | 730 | 650 | 649 | 652 | 680,02 | 690 | 670 | 0,969 | -0,045 | -0,642 | 0,520925 | 0,562864 | 0,0023239 | 0,00398 | 0,0032619 | 0,0032619 | TRUE | 0,0506294 | No |
| orf19.681 | 3496 | 2988 | 3377 | 2115 | 3076 | 2572 | 2723 | 3660 | 3183 | 3022 | 3196 | 3177 | 2736 | 2792 | 2789 | 2792 | 2960,89 | 3144 | 2777 | 0,883 | -0,179 | -4,802 | 1,57E-06 | 2,58E-06 | 1,00E-08 | 0,002483 | 0,0009957 | 0,0009957 | TRUE | 0,0085368 | No |
| orf19.6810 | 1508 | 1522 | 1573 | 1061 | 2081 | 1832 | 1705 | 2213 | 1373 | 1539 | 1489 | 1594 | 1851 | 1989 | 1746 | 1688 | 1658,63 | 1499 | 1818 | 1,214 | 0,28 | 4,298 | 1,72E-05 | 2,70E-05 | 0,0038023 | 0,002833 | 0,0034631 | 0,0034631 | TRUE | 0,0473579 | No |
| orf19.6811 | 5432 | 4757 | 5145 | 2627 | 2671 | 1869 | 2293 | 2809 | 4946 | 4811 | 4869 | 3946 | 2376 | 2029 | 2348 | 2143 | 3433,54 | 4643 | 2224 | 0,479 | -1,063 | -13,94 | 3,62E-44 | 1,52E-43 | 0,0074311 | 0,002422 | 0,0052584 | 0,0052584 | TRUE | 0,122006 | No |
| orf19.6812 | 1505 | 1239 | 1336 | 773 | 1404 | 1067 | 1201 | 1537 | 1370 | 1253 | 1264 | 1161 | 1249 | 1158 | 1230 | 1172 | 1232,33 | 1262 | 1202 | 0,95 | -0,074 | -1,251 | 0,210763 | 0,243232 | 0,0018773 | 0,003109 | 0,0025271 | 0,0025271 | TRUE | 0,0399405 | No |
| orf19.6813 | 6 | 4 | 9 | 5 | 0 | 3 | 5 | 3 | 5 | 4 | 9 | 8 | 0 | 3 | 5 | 2 | 4,53 | 6 | 2 | 0,408 | -1,293 | -1,809 | 0,070466 | 0,085975 | 0,0150903 | 0,293946 | 0,2269674 | 0,2269674 | TRUE | 0,605935 | No |
| orf19.6814 | 1E+05 | 81646 | 89182 | 67659 | 19142 | 16441 | 16840 | 24508 | 95929 | 82576 | 84403 | 1E+05 | 17026 | 17850 | 17246 | 18695 | 54420 | 91136 | 17704 | 0,194 | -2,364 | -35,651 | 2,28E-278 | 5,35E-277 | 0,0058969 | 0,002061 | 0,0041918 | 0,0041918 | TRUE | 0,0736226 | No |
| orf19.6816 | 762 | 659 | 640 | 467 | 740 | 607 | 643 | 827 | 694 | 667 | 606 | 702 | 658 | 659 | 659 | 631 | 659,26 | 667 | 652 | 0,979 | -0,031 | -0,479 | 0,632225 | 0,670509 | 0,0008989 | 0,004041 | 0,0025659 | 0,0025659 | TRUE | 0,0467406 | No |
| orf19.6817 | 2736 | 2468 | 2695 | 1732 | 1377 | 1225 | 1240 | 1643 | 2491 | 2496 | 2551 | 2602 | 1225 | 1330 | 1270 | 1253 | 1902,18 | 2535 | 1270 | 0,501 | -0,998 | -22,605 | 3,90E-113 | 3,77E-112 | 3,55E-05 | 0,002731 | 0,0012916 | 0,0012916 | TRUE | 0,0117736 | No |
| orf19.6818 | 2157 | 1976 | 2162 | 1292 | 4313 | 3649 | 4034 | 5273 | 1964 | 1999 | 2046 | 1941 | 3836 | 3962 | 4131 | 4022 | 2987,62 | 1988 | 3988 | 2,005 | 1,004 | 24,389 | 2,21E-131 | 2,49E-130 | 0,0004728 | 0,002479 | 0,0012383 | 0,0012383 | TRUE | 0,0080577 | No |
| orf19.68.2 | 389 | 457 | 429 | 309 | 550 | 472 | 482 | 634 | 354 | 462 | 406 | 464 | 489 | 512 | 494 | 484 | 458,18 | 422 | 495 | 1,178 | 0,237 | 2,686 | 0,007228 | 0,00958 | 0,0056382 | 0,00492 | 0,0052376 | 0,0052376 | TRUE | 0,1397723 | No |
| orf19.6820 | 4509 | 3916 | 4405 | 2884 | 1717 | 1512 | 1615 | 2147 | 4105 | 3961 | 4169 | 4332 | 1527 | 1642 | 1654 | 1638 | 2878,45 | 4142 | 1615 | 0,39 | -1,358 | -29,675 | 1,63E-193 | 2,68E-192 | 0,0009488 | 0,002496 | 0,0015914 | 0,0015914 | TRUE | 0,016423 | No |
| orf19.6821 | 208 | 177 | 188 | 121 | 208 | 190 | 162 | 264 | 189 | 179 | 178 | 182 | 185 | 206 | 166 | 201 | 185,83 | 182 | 190 | 1,043 | 0,06 | 0,561 | 0,57498 | 0,61543 | 1,00E-08 | 0,009146 | 0,0056864 | 0,0056864 | TRUE | 0,0723854 | No |
| orf19.6822 | 1869 | 1680 | 1850 | 966 | 1792 | 1417 | 1500 | 1848 | 1702 | 1699 | 1751 | 1451 | 1594 | 1538 | 1536 | 1410 | 1585,12 | 1651 | 1520 | 0,918 | -0,124 | -1,891 | 0,058662 | 0,072197 | 0,0038271 | 0,00287 | 0,0034668 | 0,0034668 | TRUE | 0,0745103 | No |
| orf19.6824 | 1104 | 1085 | 1100 | 675 | 1136 | 863 | 957 | 1283 | 1005 | 1097 | 1041 | 1014 | 1010 | 937 | 980 | 979 | 1007,96 | 1039 | 976 | 0,94 | -0,09 | -1,666 | 0,095707 | 0,115228 | 1,00E-08 | 0,003348 | 0,0017995 | 0,0017995 | TRUE | 0,0171026 | No |
| orf19.6826 | 53 | 35 | 37 | 24 | 63 | 28 | 31 | 63 | 48 | 35 | 35 | 36 | 56 | 30 | 32 | 48 | 40,12 | 38 | 42 | 1,08 | 0,112 | 0,445 | 0,656464 | 0,693333 | 0,0365115 | 0,034963 | 0,0355689 | 0,0355689 | TRUE | 0,2946591 | No |
| orf19.6827 | 1158 | 1051 | 1137 | 688 | 1433 | 1080 | 1237 | 1577 | 1054 | 1063 | 1076 | 1033 | 1275 | 1173 | 1267 | 1203 | 1142,98 | 1056 | 1230 | 1,162 | 0,217 | 4,257 | 2,07E-05 | 3,24E-05 | 1,00E-08 | 0,003193 | 0,00161 | 0,00161 | TRUE | 0,0106984 | No |
| orf19.6828 | 8307 | 7338 | 7699 | 4094 | 8891 | 6168 | 7322 | 9312 | 7563 | 7422 | 7286 | 6150 | 7908 | 6697 | 7499 | 7103 | 7203,49 | 7105 | 7302 | 1,027 | 0,039 | 0,548 | 0,58367 | 0,624102 | 0,0057633 | 0,00222 | 0,0046426 | 0,0046426 | TRUE | 0,0986332 | No |
| orf19.6828.1 | 679 | 588 | 564 | 276 | 1050 | 671 | 811 | 1035 | 618 | 595 | 534 | 415 | 934 | 729 | 831 | 790 | 680,48 | 540 | 821 | 1,512 | 0,597 | 5,395 | 6,85E-08 | 1,19E-07 | 0,0176944 | 0,003978 | 0,0101888 | 0,0101888 | TRUE | 0,2622426 | No |
| orf19.6829 | 3725 | 3527 | 3694 | 2282 | 3322 | 2837 | 3056 | 3634 | 3391 | 3567 | 3496 | 3428 | 2955 | 3080 | 3130 | 2772 | 3227,42 | 3470 | 2984 | 0,859 | -0,219 | -4,639 | 3,50E-06 | 5,69E-06 | 0,0012618 | 0,002446 | 0,0018339 | 0,0018339 | TRUE | 0,0285748 | No |
| orf19.6830 | 456 | 436 | 438 | 268 | 4107 | 3154 | 3274 | 4505 | 415 | 441 | 415 | 403 | 3653 | 3424 | 3353 | 3436 | 1942,51 | 418 | 3466 | 8,273 | 3,048 | 53,581 | 0 | 0 | 1,00E-08 | 0,002717 | 0,0017047 | 0,0017047 | TRUE | 0,0160098 | No |
| orf19.6831 | 11484 | 9887 | 10793 | 6193 | 11291 | 8780 | 9968 | 13159 | 10456 | 10000 | 10215 | 9303 | 10043 | 9533 | 10208 | 10038 | 9974,34 | 9994 | 9956 | 0,996 | -0,006 | -0,138 | 0,890431 | 0,906065 | 0,0013334 | 0,002169 | 0,0017802 | 0,0017802 | TRUE | 0,0258433 | No |
| orf19.6832 | 1084 | 932 | 1014 | 576 | 1905 | 1492 | 1667 | 2145 | 987 | 943 | 960 | 865 | 1694 | 1620 | 1707 | 1636 | 1301,53 | 939 | 1664 | 1,767 | 0,821 | 15,613 | 5,90E-55 | 2,98E-54 | 0,0004545 | 0,003052 | 0,0018058 | 0,0018058 | TRUE | 0,0282289 | No |
| orf19.6833.2 | 54 | 36 | 42 | 60 | 27 | 31 | 23 | 38 | 49 | 36 | 40 | 90 | 24 | 34 | 24 | 29 | 40,71 | 54 | 28 | 0,522 | -0,939 | -3,221 | 0,001279 | 0,001795 | 0,0825347 | 0,034487 | 0,0544434 | 0,0544434 | TRUE | 1,383059 | No |
| orf19.6834.10 | 27 | 21 | 24 | 6 | 10 | 12 | 8 | 18 | 25 | 21 | 23 | 9 | 9 | 13 | 8 | 14 | 15,17 | 20 | 11 | 0,553 | -0,854 | -2,241 | 0,025022 | 0,031826 | 0,028646 | 0,089091 | 0,0704585 | 0,0704585 | TRUE | 0,4464696 | No |
| orf19.6837 | 279 | 213 | 250 | 183 | 803 | 625 | 655 | 883 | 254 | 215 | 237 | 275 | 714 | 679 | 671 | 674 | 464,77 | 245 | 684 | 2,809 | 1,49 | 18,658 | 1,08E-77 | 7,43E-77 | 6,52E-05 | 0,004879 | 0,0032574 | 0,0032574 | TRUE | 0,0710103 | No |
| orf19.6838 | 1739 | 1501 | 1552 | 1057 | 25022 | 20622 | 20373 | 29763 | 1583 | 1518 | 1469 | 1588 | 22256 | 22390 | 20865 | 22703 | 11796,5 | 1540 | 22054 | 14,341 | 3,842 | 86,118 | 0 | 0 | 0,000901 | 0,002149 | 0,001541 | 0,001541 | TRUE | 0,0160973 | No |
| orf19.684 | 5090 | 4576 | 5043 | 3183 | 3212 | 2834 | 2987 | 3901 | 4634 | 4628 | 4773 | 4781 | 2857 | 3077 | 3059 | 2976 | 3848,15 | 4704 | 2992 | 0,636 | -0,653 | -16,949 | 1,94E-64 | 1,13E-63 | 0,0003585 | 0,00238 | 0,0011544 | 0,0011544 | TRUE | 0,0112138 | No |
| orf19.6840 | 996 | 987 | 1005 | 788 | 1305 | 1119 | 1127 | 1480 | 907 | 998 | 951 | 1184 | 1161 | 1215 | 1154 | 1129 | 1087,34 | 1010 | 1165 | 1,158 | 0,211 | 2,797 | 0,005164 | 0,006924 | 0,0058904 | 0,003252 | 0,0045308 | 0,0045308 | TRUE | 0,1490928 | No |
| orf19.6842 | 11970 | 10749 | 12105 | 7345 | 1392 | 1191 | 1237 | 1779 | 10898 | 10871 | 11456 | 11033 | 1238 | 1293 | 1267 | 1357 | 6176,8 | 11064 | 1289 | 0,117 | -3,101 | -73,673 | 0 | 0 | 0,0004961 | 0,002251 | 0,0012907 | 0,0012907 | TRUE | 0,0166622 | No |
| orf19.6843 | 7817 | 6687 | 7302 | 3982 | 3880 | 2682 | 3382 | 4235 | 7117 | 6763 | 6911 | 5982 | 3451 | 2912 | 3464 | 3230 | 4978,71 | 6693 | 3264 | 0,488 | -1,036 | -15,304 | 7,16E-53 | 3,48E-52 | 0,0055701 | 0,002302 | 0,0041812 | 0,0041812 | TRUE | 0,0633557 | No |
| orf19.6844 | 30395 | 23819 | 25861 | 18126 | 88736 | 73809 | 79342 | 1E+05 | 27673 | 24090 | 24475 | 27228 | 78928 | 80136 | 81256 | 85613 | 53675 | 25866 | 81483 | 3,15 | 1,656 | 31,052 | 1,07E-211 | 1,92E-210 | 0,0031481 | 0,002062 | 0,0027045 | 0,0027045 | TRUE | 0,0272572 | No |
| orf19.6845 | 4082 | 3767 | 4111 | 2555 | 4171 | 3504 | 3728 | 4969 | 3716 | 3810 | 3891 | 3838 | 3710 | 3804 | 3818 | 3790 | 3797,22 | 3814 | 3780 | 0,991 | -0,012 | -0,362 | 0,717105 | 0,750418 | 1,00E-08 | 0,002385 | 0,0008628 | 0,0008628 | TRUE | 0,0037712 | No |
| orf19.6846 | 3512 | 3252 | 3335 | 2270 | 3937 | 3178 | 3289 | 4494 | 3198 | 3289 | 3156 | 3410 | 3502 | 3450 | 3368 | 3428 | 3350,18 | 3263 | 3437 | 1,055 | 0,077 | 1,981 | 0,047614 | 0,059054 | 0,000325 | 0,002431 | 0,0011469 | 0,0011469 | TRUE | 0,0102773 | No |
| orf19.6847 | 1906 | 1678 | 1891 | 1158 | 2091 | 1750 | 1907 | 2593 | 1735 | 1697 | 1790 | 1740 | 1860 | 1900 | 1953 | 1978 | 1831,56 | 1740 | 1923 | 1,105 | 0,144 | 3,309 | 0,000937 | 0,001325 | 1,00E-08 | 0,002758 | 0,0012663 | 0,0012663 | TRUE | 0,0061105 | No |
| orf19.6848 | 848 | 769 | 864 | 538 | 615 | 543 | 549 | 640 | 772 | 778 | 818 | 808 | 547 | 590 | 562 | 488 | 670,34 | 794 | 547 | 0,687 | -0,542 | -7,842 | 4,43E-15 | 9,81E-15 | 0,0019803 | 0,004008 | 0,0030508 | 0,0030508 | TRUE | 0,0654899 | No |
| orf19.6849 | 850 | 742 | 818 | 542 | 1037 | 810 | 951 | 1172 | 774 | 750 | 774 | 814 | 922 | 879 | 974 | 894 | 847,81 | 778 | 917 | 1,181 | 0,24 | 4,147 | 3,36E-05 | 5,18E-05 | 0,0003447 | 0,003595 | 0,0020146 | 0,0020146 | TRUE | 0,0196888 | No |
| orf19.685 | 2513 | 2060 | 2181 | 1597 | 1581 | 1294 | 1254 | 1885 | 2288 | 2083 | 2064 | 2399 | 1406 | 1405 | 1284 | 1438 | 1795,98 | 2208 | 1383 | 0,628 | -0,672 | -10,897 | 1,19E-27 | 3,63E-27 | 0,0031182 | 0,002773 | 0,0030728 | 0,0030728 | TRUE | 0,0385018 | No |
| orf19.6850 | 940 | 871 | 912 | 540 | 877 | 662 | 746 | 910 | 856 | 881 | 863 | 811 | 780 | 719 | 764 | 694 | 796 | 853 | 739 | 0,865 | -0,21 | -3,449 | 0,000563 | 0,000807 | 0,00082 | 0,003697 | 0,0022947 | 0,0022947 | TRUE | 0,0220917 | No |
| orf19.6851 | 154 | 150 | 183 | 95 | 313 | 255 | 281 | 360 | 140 | 152 | 173 | 143 | 278 | 277 | 288 | 275 | 215,68 | 152 | 280 | 1,831 | 0,873 | 8,574 | 9,97E-18 | 2,38E-17 | 1,00E-08 | 0,008162 | 0,0047936 | 0,0047936 | TRUE | 0,0994046 | No |
| orf19.685.1 | 1292 | 1309 | 1284 | 916 | 826 | 728 | 665 | 938 | 1176 | 1324 | 1215 | 1376 | 735 | 790 | 681 | 716 | 1001,63 | 1273 | 730 | 0,575 | -0,799 | -11,704 | 1,22E-31 | 4,08E-31 | 0,0031874 | 0,003356 | 0,0034115 | 0,0034115 | TRUE | 0,0342837 | No |
| orf19.6852 | 515 | 443 | 433 | 306 | 916 | 672 | 724 | 977 | 469 | 448 | 410 | 460 | 815 | 730 | 741 | 745 | 602,18 | 447 | 758 | 1,7 | 0,766 | 10,922 | 9,01E-28 | 2,75E-27 | 0,0013895 | 0,004231 | 0,0029038 | 0,0029038 | TRUE | 0,0380894 | No |
| orf19.6852.1 | 2360 | 2037 | 1879 | 1197 | 2808 | 1994 | 2201 | 3064 | 2149 | 2060 | 1778 | 1798 | 2498 | 2165 | 2254 | 2337 | 2129,9 | 1946 | 2314 | 1,188 | 0,248 | 3,498 | 0,000469 | 0,000676 | 0,0058209 | 0,002657 | 0,0043611 | 0,0043611 | TRUE | 0,0598513 | No |
| orf19.6853 | 2720 | 2305 | 2627 | 1625 | 2525 | 2195 | 2237 | 2998 | 2476 | 2331 | 2486 | 2441 | 2246 | 2383 | 2291 | 2287 | 2367,73 | 2434 | 2302 | 0,945 | -0,081 | -1,944 | 0,051913 | 0,064186 | 0,0003295 | 0,002595 | 0,001255 | 0,001255 | TRUE | 0,0100782 | No |
| orf19.6854 | 13796 | 11253 | 11982 | 8155 | 18874 | 15603 | 15781 | 23314 | 12561 | 11381 | 11340 | 12250 | 16788 | 16941 | 16162 | 17784 | 14400,8 | 11883 | 16919 | 1,424 | 0,51 | 10,778 | 4,36E-27 | 1,31E-26 | 0,002031 | 0,002129 | 0,0020778 | 0,0020778 | TRUE | 0,0183276 | No |
| orf19.6855 | 2219 | 2111 | 2232 | 1462 | 4058 | 3224 | 3482 | 4822 | 2020 | 2135 | 2112 | 2196 | 3609 | 3500 | 3566 | 3678 | 2852,25 | 2116 | 3588 | 1,699 | 0,765 | 18,675 | 7,89E-78 | 5,44E-77 | 0,0003103 | 0,0025 | 0,001226 | 0,001226 | TRUE | 0,011401 | No |
| orf19.6858 | 1679 | 1551 | 1529 | 944 | 1290 | 995 | 1117 | 1419 | 1529 | 1569 | 1447 | 1418 | 1147 | 1080 | 1144 | 1082 | 1302,06 | 1491 | 1113 | 0,746 | -0,423 | -8,022 | 1,04E-15 | 2,35E-15 | 0,0008665 | 0,003052 | 0,0018913 | 0,0018913 | TRUE | 0,0145413 | No |

| | | | | | | | | | | | | | | | | | | | | | | | | | | | | | | | |
|---|---|---|---|---|---|---|---|---|---|---|---|---|---|---|---|---|---|---|---|---|---|---|---|---|---|---|---|---|---|---|---|
| orf19.6859 | 57 | 54 | 47 | 47 | 38 | 22 | 24 | 34 | 52 | 55 | 44 | 71 | 34 | 24 | 25 | 26 | 41,22 | 56 | 27 | 0,497 | -1,007 | -4,336 | 1,45E-05 | 2,29E-05 | 1,00E-08 | 0,034081 | 0,024931 | 0,024931 | TRUE | 0,2174059 | No |
| orf19.686 | 1664 | 1476 | 1645 | 977 | 1469 | 1218 | 1304 | 1774 | 1515 | 1493 | 1557 | 1468 | 1307 | 1322 | 1335 | 1353 | 1418,75 | 1508 | 1329 | 0,881 | -0,183 | -4,075 | 4,61E-05 | 7,04E-05 | 1,00E-08 | 0,002968 | 0,0012366 | 0,0012366 | TRUE | 0,0056942 | No |
| orf19.6860 | 1165 | 1212 | 1192 | 736 | 1239 | 1076 | 961 | 1460 | 1061 | 1226 | 1128 | 1106 | 1102 | 1168 | 984 | 1114 | 1111,05 | 1130 | 1092 | 0,966 | -0,049 | -0,74 | 0,459421 | 0,501251 | 0,0035243 | 0,003226 | 0,003379 | 0,003379 | TRUE | 0,0514651 | No |
| orf19.6861 | 1176 | 1418 | 1325 | 793 | 857 | 763 | 721 | 939 | 1071 | 1434 | 1254 | 1191 | 762 | 828 | 738 | 716 | 999,43 | 1238 | 761 | 0,615 | -0,702 | -8,178 | 2,88E-16 | 6,60E-16 | 0,008114 | 0,003359 | 0,0060329 | 0,0060329 | TRUE | 0,1392251 | No |
| orf19.6862 | 3476 | 4253 | 4349 | 2460 | 2766 | 2556 | 2597 | 3096 | 3165 | 4301 | 4116 | 3695 | 2460 | 2775 | 2660 | 2362 | 3191,77 | 3819 | 2564 | 0,671 | -0,575 | -6,583 | 4,63E-11 | 9,09E-11 | 0,0109697 | 0,002451 | 0,0070183 | 0,0070183 | TRUE | 0,1640428 | No |
| orf19.6863 | 4219 | 3950 | 4351 | 2480 | 4144 | 3505 | 3725 | 4844 | 3841 | 3995 | 4118 | 3725 | 3686 | 3805 | 3815 | 3695 | 3835,09 | 3920 | 3750 | 0,956 | -0,066 | -1,565 | 0,11753 | 0,139995 | 0,0007867 | 0,002381 | 0,0014201 | 0,0014201 | TRUE | 0,014038 | No |
| orf19.6864 | 232 | 206 | 267 | 155 | 244 | 248 | 266 | 327 | 211 | 208 | 253 | 233 | 217 | 269 | 272 | 249 | 239,16 | 226 | 252 | 1,112 | 0,154 | 1,438 | 0,150568 | 0,177163 | 0,0056686 | 0,007561 | 0,0067595 | 0,0067595 | TRUE | 0,0983812 | No |
| orf19.6866 | 1891 | 1637 | 1827 | 1146 | 1604 | 1244 | 1487 | 1903 | 1722 | 1656 | 1729 | 1721 | 1427 | 1351 | 1523 | 1452 | 1572,47 | 1707 | 1438 | 0,843 | -0,247 | -5,014 | 5,34E-07 | 8,98E-07 | 0,0005699 | 0,002877 | 0,0016849 | 0,0016849 | TRUE | 0,018929 | No |
| orf19.6867 | 658 | 565 | 634 | 298 | 373 | 234 | 241 | 386 | 599 | 571 | 600 | 448 | 332 | 254 | 247 | 294 | 418,16 | 554 | 282 | 0,508 | -0,978 | -8,627 | 6,32E-18 | 1,52E-17 | 0,0155951 | 0,005196 | 0,0097323 | 0,0097323 | TRUE | 0,18049 | No |
| orf19.6868 | 695 | 548 | 673 | 469 | 491 | 394 | 444 | 633 | 633 | 554 | 637 | 705 | 437 | 428 | 455 | 483 | 541,32 | 632 | 451 | 0,716 | -0,482 | -5,954 | 2,61E-09 | 4,81E-09 | 0,0042971 | 0,004477 | 0,0043936 | 0,0043936 | TRUE | 0,0795923 | No |
| orf19.6869 | 997 | 942 | 1013 | 562 | 991 | 805 | 833 | 1058 | 908 | 953 | 959 | 844 | 881 | 874 | 853 | 807 | 884,88 | 916 | 854 | 0,929 | -0,107 | -1,764 | 0,077655 | 0,094349 | 0,0011541 | 0,00353 | 0,0023706 | 0,0023706 | TRUE | 0,0285817 | No |
| orf19.687 | 219 | 183 | 185 | 140 | 429 | 363 | 362 | 475 | 199 | 185 | 175 | 210 | 382 | 394 | 371 | 362 | 284,83 | 192 | 377 | 1,968 | 0,977 | 10,791 | 3,81E-27 | 1,15E-26 | 1,00E-08 | 0,006675 | 0,0038547 | 0,0038547 | TRUE | 0,0403065 | No |
| orf19.6871 | 2568 | 2379 | 2612 | 1736 | 2769 | 2406 | 2591 | 3397 | 2338 | 2406 | 2472 | 2608 | 2463 | 2612 | 2654 | 2591 | 2517,99 | 2456 | 2580 | 1,052 | 0,073 | 1,552 | 0,120573 | 0,143512 | 0,0010743 | 0,002562 | 0,001719 | 0,001719 | TRUE | 0,0194515 | No |
| orf19.687.1 | 16456 | 13868 | 14372 | 11335 | 23460 | 19905 | 18729 | 26656 | 14982 | 14026 | 13602 | 17027 | 20867 | 21611 | 19181 | 20333 | 17703,7 | 14909 | 20498 | 1,375 | 0,46 | 6,784 | 1,17E-11 | 2,35E-11 | 0,0064005 | 0,002112 | 0,0043521 | 0,0043521 | TRUE | 0,1111533 | No |
| orf19.6872 | 2337 | 2180 | 2260 | 1418 | 1264 | 1081 | 1052 | 1444 | 2128 | 2205 | 2139 | 2130 | 1124 | 1174 | 1077 | 1101 | 1634,79 | 2150 | 1119 | 0,52 | -0,943 | -20,725 | 2,06E-95 | 1,69E-94 | 1,00E-08 | 0,002845 | 0,0013224 | 0,0013224 | TRUE | 0,0119542 | No |
| orf19.6873 | 23255 | 18224 | 19882 | 12889 | 31852 | 23421 | 24749 | 35773 | 21173 | 18432 | 18817 | 19361 | 28331 | 25429 | 25346 | 27288 | 23022 | 19446 | 26598 | 1,368 | 0,452 | 8,241 | 1,71E-16 | 3,93E-16 | 0,0033974 | 0,002094 | 0,0028441 | 0,0028441 | TRUE | 0,0439689 | No |
| orf19.6873.1 | 1836 | 1448 | 1394 | 938 | 1449 | 978 | 1170 | 1634 | 1672 | 1464 | 1319 | 1409 | 1289 | 1062 | 1198 | 1246 | 1332,47 | 1466 | 1199 | 0,818 | -0,29 | -3,598 | 0,000321 | 0,000468 | 0,0075662 | 0,003028 | 0,0054862 | 0,0054862 | TRUE | 0,1097436 | No |
| orf19.6874 | 3680 | 3316 | 3663 | 2503 | 1459 | 1219 | 1342 | 1783 | 3350 | 3354 | 3467 | 3760 | 1298 | 1323 | 1374 | 1360 | 2410,82 | 3483 | 1339 | 0,385 | -1,377 | -27,304 | 3,78E-164 | 5,41E-163 | 0,0014698 | 0,002585 | 0,0019388 | 0,0019388 | TRUE | 0,0331378 | No |
| orf19.6875 | 1793 | 1700 | 1745 | 1047 | 638 | 533 | 550 | 777 | 1632 | 1719 | 1651 | 1573 | 567 | 579 | 563 | 593 | 1109,78 | 1644 | 576 | 0,35 | -1,515 | -28,031 | 6,79E-173 | 1,02E-171 | 1,00E-08 | 0,003227 | 0,0016697 | 0,0016697 | TRUE | 0,0111788 | No |
| orf19.6877 | 23697 | 21380 | 22949 | 15369 | 13952 | 12372 | 12330 | 16911 | 21575 | 21624 | 21719 | 23087 | 12410 | 13433 | 12627 | 12900 | 17421,8 | 22001 | 12842 | 0,584 | -0,776 | -19,909 | 3,42E-88 | 2,65E-87 | 0,0010535 | 0,002113 | 0,0014003 | 0,0014003 | TRUE | 0,013336 | No |
| orf19.6879 | 2211 | 1930 | 2051 | 1402 | 1344 | 1104 | 1174 | 1525 | 2013 | 1952 | 1941 | 2106 | 1195 | 1199 | 1202 | 1163 | 1596,48 | 2003 | 1190 | 0,595 | -0,75 | -16,174 | 7,71E-59 | 4,15E-58 | 1,00E-08 | 0,002864 | 0,0014027 | 0,0014027 | TRUE | 0,0129976 | No |
| orf19.688 | 2193 | 1911 | 2091 | 1440 | 2556 | 2274 | 2301 | 3115 | 1997 | 1933 | 1979 | 2163 | 2273 | 2469 | 2357 | 2376 | 2193,32 | 2018 | 2369 | 1,176 | 0,234 | 4,816 | 1,46E-06 | 2,41E-06 | 0,0012023 | 0,002639 | 0,001793 | 0,001793 | TRUE | 0,0259621 | No |
| orf19.6880 | 604 | 575 | 624 | 433 | 487 | 498 | 459 | 544 | 550 | 582 | 591 | 650 | 433 | 541 | 470 | 415 | 528,92 | 593 | 465 | 0,783 | -0,352 | -3,942 | 8,09E-05 | 0,000122 | 0,0071029 | 0,004535 | 0,0057597 | 0,0057597 | TRUE | 0,1373296 | No |
| orf19.6881 | 8603 | 7504 | 8042 | 5410 | 1352 | 1049 | 1045 | 1425 | 7833 | 7589 | 7611 | 8127 | 1203 | 1139 | 1070 | 1087 | 4457,32 | 7790 | 1125 | 0,144 | -2,792 | -58,385 | 0 | 0 | 0,0012403 | 0,002333 | 0,0017112 | 0,0017112 | TRUE | 0,0268804 | No |
| orf19.6882 | 38975 | 32439 | 36365 | 21310 | 10111 | 7881 | 8753 | 11886 | 35485 | 32809 | 34416 | 32011 | 8993 | 8557 | 8964 | 9067 | 21287,7 | 33680 | 8895 | 0,264 | -1,921 | -45,693 | 0 | 0 | 0,0013183 | 0,002099 | 0,001629 | 0,001629 | TRUE | 0,0159424 | No |
| orf19.6882.1 | 13889 | 11557 | 11907 | 7298 | 20378 | 14314 | 15922 | 21722 | 12645 | 11689 | 11269 | 10963 | 18126 | 15541 | 16306 | 16570 | 14138,5 | 11642 | 16636 | 1,429 | 0,515 | 8,837 | 9,85E-19 | 2,42E-18 | 0,0040085 | 0,00213 | 0,0031858 | 0,0031858 | TRUE | 0,0445011 | No |
| orf19.6883 | 2161 | 1884 | 2014 | 1365 | 3071 | 2728 | 2660 | 3733 | 1967 | 1905 | 1906 | 2050 | 2732 | 2962 | 2724 | 2848 | 2386,83 | 1957 | 2816 | 1,44 | 0,527 | 11,386 | 4,90E-30 | 1,57E-29 | 0,0008306 | 0,00259 | 0,0016122 | 0,0016122 | TRUE | 0,0141729 | No |
| orf19.6884 | 355 | 297 | 358 | 225 | 126 | 75 | 112 | 141 | 323 | 300 | 339 | 338 | 112 | 81 | 115 | 108 | 214,52 | 325 | 104 | 0,322 | -1,636 | -14,466 | 1,99E-47 | 8,78E-47 | 0,0014808 | 0,008195 | 0,0061829 | 0,0061829 | TRUE | 0,1839196 | No |
| orf19.6885 | 2705 | 2348 | 2631 | 1655 | 1760 | 1506 | 1661 | 2142 | 2463 | 2375 | 2490 | 2486 | 1565 | 1635 | 1701 | 1634 | 2043,65 | 2454 | 1634 | 0,666 | -0,587 | -13,489 | 1,81E-41 | 7,20E-41 | 0,0002638 | 0,002683 | 0,0013114 | 0,0013114 | TRUE | 0,0096542 | No |
| orf19.6886 | 12232 | 11181 | 12189 | 6313 | 14195 | 10349 | 12500 | 15772 | 11137 | 11308 | 11536 | 9483 | 12626 | 11236 | 12802 | 12031 | 11519,8 | 10866 | 12174 | 1,12 | 0,163 | 2,504 | 0,012283 | 0,016003 | 0,0052812 | 0,002152 | 0,0040052 | 0,0040052 | TRUE | 0,0887812 | No |
| orf19.6887 | 4539 | 3465 | 3852 | 2234 | 4872 | 3106 | 3992 | 5275 | 4133 | 3504 | 3646 | 3356 | 4333 | 3372 | 4088 | 4024 | 3807,04 | 3660 | 3954 | 1,08 | 0,111 | 1,377 | 0,168524 | 0,19691 | 0,0092694 | 0,002384 | 0,0060162 | 0,0060162 | TRUE | 0,118808 | No |
| orf19.6888 | 15625 | 13967 | 15316 | 10006 | 3237 | 2714 | 2830 | 3783 | 14226 | 14126 | 14495 | 15031 | 2879 | 2947 | 2898 | 2886 | 8685,96 | 14470 | 2902 | 0,201 | -2,318 | -66,742 | 0 | 0 | 0,000394 | 0,002189 | 0,0009596 | 0,0009596 | TRUE | 0,0081115 | No |
| orf19.6889 | 2618 | 2321 | 2461 | 1586 | 2179 | 1874 | 1826 | 2524 | 2384 | 2347 | 2329 | 2382 | 1938 | 2035 | 1870 | 1925 | 2151,34 | 2360 | 1942 | 0,823 | -0,282 | -6,748 | 1,50E-11 | 3,00E-11 | 6,01E-05 | 0,002651 | 0,0012064 | 0,0012064 | TRUE | 0,0117782 | No |
| orf19.689 | 25 | 19 | 18 | 12 | 22 | 15 | 9 | 19 | 23 | 19 | 17 | 18 | 20 | 16 | 9 | 14 | 17,08 | 19 | 15 | 0,773 | -0,372 | -1,103 | 0,270046 | 0,306052 | 1,00E-08 | 0,079399 | 0,0504922 | 0,0504922 | TRUE | 0,2760621 | No |
| orf19.6891 | 2047 | 1788 | 1886 | 1259 | 1765 | 1473 | 1551 | 2100 | 1864 | 1808 | 1785 | 1891 | 1570 | 1599 | 1588 | 1602 | 1713,47 | 1837 | 1590 | 0,866 | -0,207 | -4,882 | 1,05E-06 | 1,74E-06 | 1,00E-08 | 0,002808 | 0,0011395 | 0,0011395 | TRUE | 0,004401 | No |
| orf19.6893 | 1274 | 1116 | 1291 | 809 | 2049 | 1673 | 1907 | 2448 | 1160 | 1129 | 1222 | 1215 | 1823 | 1816 | 1953 | 1867 | 1523,12 | 1182 | 1865 | 1,58 | 0,66 | 13,357 | 1,08E-40 | 4,23E-40 | 0,000556 | 0,002904 | 0,0016385 | 0,0016385 | TRUE | 0,0118603 | No |
| orf19.6896 | 18 | 16 | 27 | 9 | 23 | 28 | 21 | 35 | 16 | 16 | 26 | 14 | 20 | 30 | 22 | 27 | 21,34 | 18 | 25 | 1,358 | 0,442 | 1,424 | 0,154492 | 0,181445 | 1,00E-08 | 0,063945 | 0,044378 | 0,044378 | TRUE | 0,4408432 | No |
| orf19.6897 | 0 | 0 | 0 | 0 | 0 | 0 | 0 | 0 | 0 | 0 | 0 | 0 | 0 | 0 | 0 | 0 | 0 | 0 | 0 | NA | NA | NA | NA | NA | NA | NA | NA | NA | NA | NA | NA |
| orf19.6898 | 2104 | 1913 | 2111 | 1262 | 1425 | 1257 | 1239 | 1687 | 1916 | 1935 | 1998 | 1896 | 1267 | 1365 | 1269 | 1287 | 1616,5 | 1936 | 1297 | 0,669 | -0,58 | -12,631 | 1,42E-36 | 5,24E-36 | 0,0001226 | 0,002854 | 0,0013868 | 0,0013868 | TRUE | 0,0140016 | No |
| orf19.6898.1 | 1379 | 1182 | 1201 | 891 | 2068 | 1538 | 1473 | 2039 | 1256 | 1195 | 1137 | 1338 | 1839 | 1670 | 1509 | 1555 | 1437,4 | 1232 | 1643 | 1,337 | 0,419 | 5,749 | 8,97E-09 | 1,62E-08 | 0,0055199 | 0,002956 | 0,0043686 | 0,0043686 | TRUE | 0,0788363 | No |
| orf19.6899 | 6608 | 5600 | 6041 | 3743 | 5702 | 4500 | 4922 | 6695 | 6016 | 5664 | 5717 | 5623 | 5072 | 4886 | 5041 | 5107 | 5390,65 | 5755 | 5026 | 0,873 | -0,196 | -5,336 | 9,51E-08 | 1,64E-07 | 0,0004207 | 0,002282 | 0,0011044 | 0,0011044 | TRUE | 0,0115976 | No |
| orf19.69 | 420 | 411 | 452 | 275 | 744 | 746 | 730 | 926 | 382 | 416 | 428 | 413 | 662 | 810 | 748 | 706 | 570,58 | 410 | 732 | 1,783 | 0,834 | 10,598 | 3,06E-26 | 8,98E-26 | 0,0036432 | 0,004352 | 0,004005 | 0,004005 | TRUE | 0,0601956 | No |
| orf19.690 | 76 | 84 | 76 | 56 | 68 | 54 | 50 | 89 | 69 | 85 | 72 | 84 | 60 | 59 | 51 | 68 | 68,55 | 78 | 60 | 0,777 | -0,363 | -2,123 | 0,03372 | 0,042416 | 1,00E-08 | 0,021307 | 0,0134284 | 0,0134284 | TRUE | 0,0891615 | No |
| orf19.6900 | 1347 | 1247 | 1395 | 790 | 1334 | 1068 | 1185 | 1540 | 1226 | 1261 | 1320 | 1187 | 1187 | 1160 | 1214 | 1175 | 1216,12 | 1248 | 1184 | 0,946 | -0,08 | -1,575 | 0,11537 | 0,137576 | 0,000274 | 0,003123 | 0,001654 | 0,001654 | TRUE | 0,0179251 | No |
| orf19.6901 | 720 | 646 | 626 | 424 | 597 | 519 | 508 | 678 | 656 | 653 | 592 | 637 | 531 | 563 | 520 | 517 | 583,77 | 634 | 533 | 0,839 | -0,253 | -3,836 | 0,000125 | 0,000186 | 1,00E-08 | 0,0043 | 0,002464 | 0,002464 | TRUE | 0,0254539 | No |
| orf19.6902 | 12966 | 11138 | 12346 | 7281 | 15902 | 12750 | 15061 | 19428 | 11805 | 11265 | 11684 | 10937 | 14144 | 13843 | 15424 | 14820 | 12990,4 | 11423 | 14558 | 1,274 | 0,35 | 7,778 | 7,35E-15 | 1,62E-14 | 0,0016837 | 0,002139 | 0,0018602 | 0,0018602 | TRUE | 0,0195251 | No |
| orf19.6903 | 4714 | 3893 | 4116 | 2693 | 3419 | 2728 | 3106 | 4131 | 4292 | 3937 | 3895 | 4045 | 3041 | 2962 | 3181 | 3151 | 3563,13 | 4042 | 3084 | 0,763 | -0,39 | -8,585 | 9,06E-18 | 2,17E-17 | 0,0011242 | 0,002408 | 0,0017002 | 0,0017002 | TRUE | 0,0217403 | No |
| orf19.6904 | 3356 | 2617 | 2904 | 1876 | 1823 | 1419 | 1595 | 2212 | 3055 | 2647 | 2748 | 2818 | 1622 | 1541 | 1633 | 1687 | 2218,96 | 2817 | 1621 | 0,576 | -0,797 | -14,516 | 9,54E-48 | 4,23E-47 | 0,0022532 | 0,002632 | 0,0024142 | 0,0024142 | TRUE | 0,0407152 | No |
| orf19.6905 | 4254 | 3622 | 3913 | 2791 | 983 | 750 | 838 | 1121 | 3873 | 3663 | 3703 | 4193 | 874 | 814 | 858 | 855 | 2354,27 | 3858 | 850 | 0,221 | -2,179 | -39,029 | 0 | 0 | 0,0020813 | 0,002598 | 0,0023047 | 0,0023047 | TRUE | 0,039728 | No |
| orf19.6906 | 16576 | 13650 | 14234 | 9265 | 30999 | 24272 | 25402 | 36925 | 15092 | 13806 | 13471 | 13918 | 27573 | 26353 | 26015 | 28167 | 20549,1 | 14072 | 27027 | 1,921 | 0,942 | 20,562 | 6,00E-94 | 4,85E-93 | 0,0018888 | 0,002101 | 0,0019588 | 0,0019588 | TRUE | 0,0293918 | No |
| orf19.6907 | 2524 | 2217 | 2488 | 1386 | 1938 | 1518 | 1626 | 2155 | 2298 | 2242 | 2355 | 2082 | 1724 | 1648 | 1665 | 1644 | 1957,24 | 2244 | 1670 | 0,743 | -0,429 | -8,783 | 1,59E-18 | 3,89E-18 | 0,0009178 | 0,002712 | 0,0017701 | 0,0017701 | TRUE | 0,026324 | No |
| orf19.6908 | 1112 | 1018 | 1106 | 775 | 610 | 549 | 521 | 767 | 1012 | 1030 | 1047 | 1164 | 543 | 596 | 534 | 585 | 813,78 | 1063 | 564 | 0,532 | -0,909 | -13,731 | 6,63E-43 | 2,72E-42 | 0,0020164 | 0,00366 | 0,0028774 | 0,0028774 | TRUE | 0,0441678 | No |
| orf19.6909 | 253 | 268 | 252 | 167 | 310 | 224 | 254 | 312 | 230 | 271 | 238 | 251 | 276 | 243 | 260 | 238 | 250,98 | 248 | 254 | 1,028 | 0,04 | 0,413 | 0,679552 | 0,715347 | 1,00E-08 | 0,0073 | 0,0048656 | 0,0048656 | TRUE | 0,0487817 | No |
| orf19.691 | 2E+05 | 1E+05 | 2E+05 | 99615 | 2E+05 | 2E+05 | 2E+05 | 2E+05 | 2E+05 | 2E+05 | 2E+05 | 1E+05 | 2E+05 | 2E+05 | 2E+05 | 2E+05 | 161961 | 152749 | 171174 | 1,121 | 0,164 | 5,342 | 9,20E-08 | 1,59E-07 | 0,0004767 | 0,002045 | 0,0009025 | 0,0009025 | TRUE | 0,0049845 | No |
| orf19.6910 | 77 | 61 | 62 | 44 | 218 | 146 | 163 | 209 | 70 | 62 | 59 | 66 | 194 | 159 | 167 | 159 | 116,92 | 64 | 170 | 2,65 | 1,406 | 9,764 | 1,61E-22 | 4,32E-22 | 1,00E-08 | 0,013335 | 0,0088857 | 0,0088857 | TRUE | 0,1020767 | No |
| orf19.6911 | 1 | 1 | 3 | 0 | 2 | 1 | 6 | 1 | 1 | 1 | 3 | 0 | 2 | 1 | 6 | 1 | 1,82 | 1 | 2 | 1,887 | 0,916 | 0,814 | 0,41566 | 0,456946 | 0,4065655 | 0,729193 | 0,6024349 | 0,6024349 | TRUE | 1,0966999 | No |
| orf19.6912 | 1075 | 1070 | 1116 | 705 | 835 | 693 | 720 | 974 | 979 | 1082 | 1056 | 1059 | 743 | 752 | 737 | 743 | 893,95 | 1044 | 744 | 0,713 | -0,488 | -8,794 | 1,45E-18 | 3,53E-18 | 1,00E-08 | 0,003515 | 0,0018132 | 0,0018132 | TRUE | 0,0226982 | No |
| orf19.6913 | 1222 | 1221 | 1248 | 692 | 490 | 447 | 449 | 555 | 1113 | 1235 | 1181 | 1039 | 436 | 485 | 460 | 423 | 796,56 | 1142 | 451 | 0,393 | -1,347 | -18,695 | 5,42E-78 | 3,75E-77 | 0,0032397 | 0,003695 | 0,0034752 | 0,0034752 | TRUE | 0,0396308 | No |
| orf19.6915 | 1507 | 1411 | 1513 | 917 | 1311 | 1053 | 1244 | 1509 | 1372 | 1427 | 1432 | 1377 | 1166 | 1143 | 1274 | 1151 | 1292,87 | 1402 | 1184 | 0,843 | -0,246 | -4,73 | 2,25E-06 | 3,68E-06 | 0,0006955 | 0,003059 | 0,0018197 | 0,0018197 | TRUE | 0,0307847 | No |
| orf19.6916 | 816 | 785 | 786 | 429 | 532 | 501 | 470 | 592 | 743 | 794 | 744 | 644 | 473 | 544 | 481 | 452 | 609,41 | 731 | 488 | 0,663 | -0,593 | -7,371 | 1,70E-13 | 3,60E-13 | 0,0046964 | 0,004205 | 0,0045248 | 0,0045248 | TRUE | 0,0683601 | No |
| orf19.6917 | 704 | 802 | 821 | 445 | 423 | 324 | 347 | 447 | 641 | 811 | 777 | 668 | 376 | 352 | 355 | 341 | 540,24 | 724 | 356 | 0,491 | -1,027 | -11,675 | 1,71E-31 | 5,70E-31 | 0,0062915 | 0,004482 | 0,0053763 | 0,0053763 | TRUE | 0,0774313 | No |
| orf19.6918 | 788 | 1040 | 981 | 554 | 601 | 549 | 559 | 714 | 717 | 1052 | 928 | 832 | 535 | 596 | 572 | 545 | 722,21 | 882 | 562 | 0,636 | -0,653 | -6,412 | 1,44E-10 | 2,78E-10 | 0,0134185 | 0,003866 | 0,0085093 | 0,0085093 | TRUE | 0,1616263 | No |
| orf19.6919 | 126 | 152 | 142 | 80 | 30 | 23 | 18 | 33 | 115 | 154 | 134 | 120 | 27 | 25 | 18 | 25 | 77,28 | 131 | 24 | 0,183 | -2,452 | -12,298 | 9,29E-35 | 3,31E-34 | 1,00E-08 | 0,01913 | 0,0147086 | 0,0147086 | TRUE | 0,1451303 | No |
| orf19.692 | 3940 | 3187 | 3665 | 2297 | 6218 | 5052 | 5984 | 7929 | 3587 | 3223 | 3469 | 3450 | 5531 | 5485 | 6128 | 6048 | 4615,25 | 3432 | 5798 | 1,689 | 0,756 | 14,346 | 1,12E-46 | 4,90E-46 | 0,0023628 | 0,002323 | 0,0024323 | 0,0024323 | TRUE | 0,0207879 | No |
| orf19.69.2 | 17 | 32 | 15 | 13 | 9 | 10 | 15 | 13 | 15 | 32 | 14 | 20 | 8 | 11 | 15 | 10 | 15,71 | 20 | 11 | 0,538 | -0,895 | -2,287 | 0,022168 | 0,028316 | 0,0658696 | 0,086105 | 0,0780102 | 0,0780102 | TRUE | 0,9013804 | No |
| orf19.6920 | 1485 | 2036 | 1871 | 1111 | 913 | 707 | 744 | 877 | 1352 | 2059 | 1771 | 1669 | 812 | 768 | 762 | 669 | 1232,68 | 1713 | 753 | 0,439 | -1,187 | -11,06 | 1,96E-28 | 6,06E-28 | 0,0164278 | 0,003109 | 0,0101249 | 0,0101249 | TRUE | 0,1394177 | No |
| orf19.6921 | 1978 | 1912 | 2069 | 1124 | 1166 | 939 | 1020 | 1275 | 1801 | 1934 | 1958 | 1688 | 1037 | 1019 | 1045 | 973 | 1431,88 | 1845 | 1018 | 0,551 | -0,861 | -14,685 | 8,00E-49 | 3,61E-48 | 0,0022007 | 0,00296 | 0,0025504 | 0,0025504 | TRUE | 0,0364511 | No |

| | | | | | | | | | | | | | | | | | | | | | | | | | | | | | | | |
|---|---|---|---|---|---|---|---|---|---|---|---|---|---|---|---|---|---|---|---|---|---|---|---|---|---|---|---|---|---|---|---|
| orf19.6922 | 0 | 0 | 0 | 0 | 0 | 0 | 0 | 0 | 0 | 0 | 0 | 0 | 0 | 0 | 0 | 0 | 0 | 0 | 0 | NA | NA | NA | NA | NA | NA | NA | NA | NA | NA | NA | NA |
| orf19.6923 | 2143 | 1832 | 1942 | 1149 | 2072 | 1488 | 1815 | 2455 | 1951 | 1853 | 1838 | 1726 | 1843 | 1616 | 1859 | 1873 | 1819,74 | 1842 | 1798 | 0,975 | -0,036 | -0,602 | 0,547228 | 0,588691 | 0,0029993 | 0,002763 | 0,002891 | 0,002891 | TRUE | 0,0547127 | No |
| orf19.6923.1 | 1820 | 1375 | 1383 | 860 | 2223 | 1370 | 1723 | 2475 | 1657 | 1391 | 1309 | 1292 | 1977 | 1487 | 1765 | 1888 | 1595,71 | 1412 | 1779 | 1,26 | 0,333 | 3,487 | 0,000488 | 0,000702 | 0,0134856 | 0,002865 | 0,0081063 | 0,0081063 | TRUE | 0,1660225 | No |
| orf19.6924 | 4832 | 3699 | 3959 | 2726 | 5086 | 3799 | 3760 | 5710 | 4399 | 3741 | 3747 | 4095 | 4524 | 4125 | 3851 | 4356 | 4104,63 | 3996 | 4214 | 1,055 | 0,077 | 1,166 | 0,243718 | 0,278192 | 0,0052048 | 0,002359 | 0,0039693 | 0,0039693 | TRUE | 0,0575305 | No |
| orf19.6925 | 6157 | 5565 | 6051 | 4462 | 6385 | 5518 | 5118 | 7265 | 5606 | 5628 | 5727 | 6703 | 5679 | 5991 | 5241 | 5542 | 5764,62 | 5916 | 5613 | 0,95 | -0,075 | -1,161 | 0,2457 | 0,280394 | 0,0052744 | 0,002266 | 0,0038044 | 0,0038044 | TRUE | 0,0961127 | No |
| orf19.6926 | 2978 | 2603 | 2920 | 1802 | 2622 | 2179 | 2498 | 3423 | 2711 | 2633 | 2764 | 2707 | 2332 | 2366 | 2558 | 2611 | 2585,22 | 2704 | 2467 | 0,913 | -0,132 | -2,7 | 0,006929 | 0,0092 | 0,0012565 | 0,002548 | 0,0018923 | 0,0018923 | TRUE | 0,0195447 | No |
| orf19.6927 | 1413 | 1200 | 1336 | 838 | 1373 | 1106 | 1085 | 1532 | 1286 | 1214 | 1264 | 1259 | 1221 | 1201 | 1111 | 1169 | 1215,65 | 1256 | 1176 | 0,936 | -0,095 | -1,875 | 0,060728 | 0,074623 | 0,0003073 | 0,003124 | 0,0016586 | 0,0016586 | TRUE | 0,015575 | No |
| orf19.6928 | 19667 | 17258 | 18386 | 12115 | 11870 | 10364 | 10893 | 14840 | 17906 | 17455 | 17401 | 18199 | 10558 | 11252 | 11156 | 11320 | 14405,8 | 17740 | 11072 | 0,624 | -0,68 | -19,287 | 6,93E-83 | 5,13E-82 | 0,000629 | 0,002129 | 0,0011212 | 0,0011212 | TRUE | 0,011993 | No |
| orf19.6929 | 1182 | 978 | 1052 | 714 | 1803 | 1422 | 1568 | 2046 | 1076 | 989 | 996 | 1073 | 1604 | 1544 | 1606 | 1561 | 1305,95 | 1034 | 1579 | 1,53 | 0,613 | 12,193 | 3,37E-34 | 1,18E-33 | 0,0001731 | 0,003049 | 0,0016133 | 0,0016133 | TRUE | 0,011044 | No |
| orf19.693 | 169 | 176 | 216 | 119 | 70 | 61 | 56 | 64 | 154 | 178 | 204 | 179 | 62 | 66 | 57 | 49 | 118,71 | 179 | 58 | 0,326 | -1,617 | -10,704 | 9,78E-27 | 2,91E-26 | 1,00E-08 | 0,013165 | 0,0109409 | 0,0109409 | TRUE | 0,1330856 | No |
| orf19.6930 | 2568 | 2235 | 2450 | 1664 | 2469 | 1973 | 2106 | 2866 | 2338 | 2260 | 2319 | 2500 | 2196 | 2142 | 2157 | 2186 | 2262,26 | 2354 | 2170 | 0,924 | -0,114 | -2,647 | 0,008125 | 0,010724 | 0,0004223 | 0,002621 | 0,0013499 | 0,0013499 | TRUE | 0,0189736 | No |
| orf19.6931 | 2477 | 2190 | 2373 | 1422 | 2115 | 1721 | 1938 | 2551 | 2255 | 2215 | 2246 | 2136 | 1881 | 1869 | 1985 | 1946 | 2066,56 | 2213 | 1920 | 0,867 | -0,206 | -4,875 | 1,09E-06 | 1,80E-06 | 0,0001226 | 0,002676 | 0,0012339 | 0,0012339 | TRUE | 0,0059512 | No |
| orf19.6933 | 988 | 861 | 887 | 646 | 710 | 583 | 591 | 849 | 900 | 871 | 839 | 970 | 632 | 633 | 605 | 648 | 762,2 | 895 | 630 | 0,706 | -0,502 | -8,044 | 8,70E-16 | 1,97E-15 | 0,0008973 | 0,00377 | 0,0024029 | 0,0024029 | TRUE | 0,0335035 | No |
| orf19.6934 | 1421 | 1196 | 1428 | 834 | 1512 | 1195 | 1290 | 1669 | 1294 | 1210 | 1351 | 1253 | 1345 | 1297 | 1321 | 1273 | 1293,03 | 1277 | 1309 | 1,024 | 0,034 | 0,661 | 0,508534 | 0,550595 | 0,0006734 | 0,003059 | 0,0017791 | 0,0017791 | TRUE | 0,0191454 | No |
| orf19.6936 | 1736 | 1451 | 1728 | 1025 | 1321 | 1019 | 1209 | 1682 | 1581 | 1468 | 1635 | 1540 | 1175 | 1106 | 1238 | 1283 | 1378,22 | 1556 | 1200 | 0,772 | -0,373 | -6,267 | 3,69E-10 | 7,02E-10 | 0,0023735 | 0,002995 | 0,0026632 | 0,0026632 | TRUE | 0,0325084 | No |
| orf19.6937 | 65122 | 55033 | 60206 | 40419 | 86831 | 74492 | 76269 | 1E+05 | 59291 | 55660 | 56980 | 60716 | 77233 | 80878 | 78109 | 83721 | 69073,4 | 58162 | 79985 | 1,375 | 0,46 | 11,105 | 1,19E-28 | 3,68E-28 | 0,001415 | 0,002056 | 0,0016318 | 0,0016318 | TRUE | 0,0121372 | No |
| orf19.6938 | 2529 | 1977 | 2112 | 1392 | 5321 | 4259 | 4612 | 6454 | 2303 | 2000 | 1999 | 2091 | 4733 | 4624 | 4723 | 4923 | 3424,41 | 2098 | 4751 | 2,264 | 1,179 | 22,576 | 7,45E-113 | 7,17E-112 | 0,0022137 | 0,002423 | 0,0022637 | 0,0022637 | TRUE | 0,0545079 | No |
| orf19.6939 | 4261 | 3633 | 3906 | 2331 | 3502 | 2748 | 3199 | 4153 | 3879 | 3674 | 3697 | 3502 | 3115 | 2984 | 3276 | 3168 | 3411,83 | 3688 | 3136 | 0,85 | -0,235 | -5,112 | 3,19E-07 | 5,40E-07 | 0,0012369 | 0,002424 | 0,001738 | 0,001738 | TRUE | 0,0148724 | No |
| orf19.6940 | 28 | 19 | 22 | 9 | 31 | 19 | 22 | 35 | 25 | 19 | 21 | 14 | 28 | 21 | 23 | 27 | 22,06 | 20 | 25 | 1,214 | 0,279 | 0,948 | 0,34332 | 0,383371 | 1,00E-08 | 0,061919 | 0,0381803 | 0,0381803 | TRUE | 0,2077086 | No |
| orf19.6941 | 2489 | 2183 | 2492 | 1526 | 3778 | 3177 | 3323 | 4476 | 2266 | 2208 | 2358 | 2292 | 3360 | 3449 | 3403 | 3414 | 2844 | 2281 | 3406 | 1,493 | 0,579 | 15,25 | 1,64E-52 | 7,92E-52 | 1,00E-08 | 0,002502 | 0,0010093 | 0,0010093 | TRUE | 0,0067574 | No |
| orf19.6942 | 772 | 675 | 770 | 464 | 946 | 834 | 884 | 1147 | 703 | 683 | 729 | 697 | 841 | 905 | 905 | 875 | 792,31 | 703 | 882 | 1,253 | 0,325 | 5,686 | 1,30E-08 | 2,34E-08 | 1,00E-08 | 0,003704 | 0,0018477 | 0,0018477 | TRUE | 0,0112381 | No |
| orf19.6943 | 799 | 717 | 747 | 412 | 834 | 585 | 726 | 857 | 727 | 725 | 707 | 619 | 742 | 635 | 744 | 654 | 694,09 | 694 | 694 | 0,995 | -0,007 | -0,096 | 0,923592 | 0,934143 | 0,0044484 | 0,00394 | 0,0041798 | 0,0041798 | TRUE | 0,0562594 | No |
| orf19.6944 | 2050 | 1709 | 1869 | 1156 | 1718 | 1348 | 1646 | 2144 | 1866 | 1728 | 1769 | 1736 | 1528 | 1464 | 1686 | 1635 | 1676,63 | 1775 | 1578 | 0,889 | -0,17 | -3,042 | 0,002348 | 0,003222 | 0,0018795 | 0,002825 | 0,0023944 | 0,0023944 | TRUE | 0,0277461 | No |
| orf19.6947 | 647 | 605 | 673 | 453 | 511 | 458 | 435 | 593 | 589 | 612 | 637 | 680 | 455 | 497 | 445 | 452 | 546 | 630 | 462 | 0,736 | -0,442 | -6,242 | 4,32E-10 | 8,18E-10 | 0,0010479 | 0,004456 | 0,0029519 | 0,0029519 | TRUE | 0,0301771 | No |
| orf19.6948 | 1612 | 1383 | 1507 | 982 | 4814 | 3866 | 4152 | 5663 | 1468 | 1399 | 1426 | 1475 | 4282 | 4197 | 4252 | 4320 | 2852,38 | 1442 | 4263 | 2,96 | 1,565 | 40,285 | 0 | 0 | 1,00E-08 | 0,0025 | 0,0009691 | 0,0009691 | TRUE | 0,0048195 | No |
| orf19.6949 | 1 | 2 | 2 | 2 | 3 | 5 | 3 | 4 | 1 | 2 | 2 | 3 | 3 | 5 | 3 | 3 | 2,76 | 2 | 4 | 1,875 | 0,907 | 1,078 | 0,2811 | 0,317612 | 1,00E-08 | 0,481289 | 0,2632885 | 0,2632885 | TRUE | 0,1675479 | No |
| orf19.695 | 1440 | 1590 | 1509 | 982 | 2016 | 1651 | 1583 | 2113 | 1311 | 1608 | 1428 | 1475 | 1793 | 1793 | 1621 | 1612 | 1580,14 | 1456 | 1705 | 1,171 | 0,228 | 3,33 | 0,000868 | 0,00123 | 0,0044995 | 0,002873 | 0,003869 | 0,003869 | TRUE | 0,0619131 | No |
| orf19.6950 | 1693 | 1497 | 1688 | 1152 | 825 | 678 | 723 | 1012 | 1541 | 1514 | 1598 | 1730 | 734 | 736 | 740 | 772 | 1170,73 | 1596 | 746 | 0,469 | -1,093 | -19,066 | 4,85E-81 | 3,48E-80 | 0,0013257 | 0,003165 | 0,0021949 | 0,0021949 | TRUE | 0,0354654 | No |
| orf19.6951 | 3202 | 2864 | 3096 | 1850 | 1669 | 1364 | 1604 | 2056 | 2915 | 2897 | 2930 | 2779 | 1485 | 1481 | 1643 | 1568 | 2212,18 | 2880 | 1544 | 0,536 | -0,9 | -19,171 | 6,51E-82 | 4,73E-81 | 0,0008704 | 0,002634 | 0,0016289 | 0,0016289 | TRUE | 0,0212893 | No |
| orf19.6952 | 2076 | 1931 | 2008 | 1193 | 2353 | 1908 | 2165 | 2854 | 1890 | 1953 | 1900 | 1792 | 2093 | 2072 | 2217 | 2177 | 2011,79 | 1884 | 2140 | 1,134 | 0,182 | 4,004 | 6,23E-05 | 9,43E-05 | 0,0004877 | 0,002694 | 0,0014776 | 0,0014776 | TRUE | 0,011571 | No |
| orf19.6953 | 3352 | 2806 | 3238 | 2090 | 3069 | 2715 | 2982 | 4002 | 3052 | 2838 | 3064 | 3140 | 2730 | 2948 | 3054 | 3053 | 2984,75 | 3024 | 2946 | 0,975 | -0,037 | -0,715 | 0,474726 | 0,51627 | 0,0017785 | 0,00248 | 0,0021901 | 0,0021901 | TRUE | 0,0300404 | No |
| orf19.6955 | 13289 | 11010 | 12433 | 7685 | 14880 | 11325 | 13749 | 17882 | 12099 | 11135 | 11767 | 11544 | 13235 | 12296 | 14081 | 13641 | 12474,7 | 11636 | 13313 | 1,144 | 0,194 | 4,009 | 6,10E-05 | 9,24E-05 | 0,0021302 | 0,002143 | 0,0021729 | 0,0021729 | TRUE | 0,0321782 | No |
| orf19.6956 | 765 | 686 | 728 | 394 | 972 | 805 | 831 | 1148 | 697 | 694 | 689 | 592 | 865 | 874 | 851 | 876 | 767,06 | 668 | 866 | 1,291 | 0,368 | 5,914 | 3,34E-09 | 6,13E-09 | 0,000274 | 0,003759 | 0,002386 | 0,002386 | TRUE | 0,0580319 | No |
| orf19.6957 | 0 | 0 | 0 | 0 | 1 | 0 | 1 | 0 | 0 | 0 | 0 | 0 | 1 | 0 | 1 | 0 | 0,24 | 0 | 0 | 2,441 | 1,287 | 0,45 | 0,652795 | 0,689915 | 1,00E-08 | 5,524619 | 5,9419166 | 5,9419166 | TRUE | 0,0630912 | No |
| orf19.6958 | 61 | 40 | 57 | 33 | 517 | 437 | 417 | 583 | 56 | 40 | 54 | 50 | 460 | 474 | 427 | 445 | 250,7 | 50 | 452 | 9,011 | 3,172 | 24,437 | 6,89E-132 | 7,82E-131 | 1,00E-08 | 0,007306 | 0,0046066 | 0,0046066 | TRUE | 0,1449372 | No |
| orf19.696 | 25 | 23 | 24 | 12 | 27 | 22 | 29 | 33 | 23 | 23 | 23 | 18 | 24 | 24 | 30 | 25 | 23,69 | 22 | 26 | 1,17 | 0,226 | 0,821 | 0,411532 | 0,452798 | 1,00E-08 | 0,057794 | 0,0304498 | 0,0304498 | TRUE | 0,06338 | No |
| orf19.6966 | 1052 | 1299 | 1290 | 783 | 1518 | 1395 | 1443 | 1733 | 958 | 1314 | 1221 | 1176 | 1350 | 1515 | 1478 | 1322 | 1291,65 | 1167 | 1416 | 1,213 | 0,279 | 3,169 | 0,001528 | 0,002131 | 0,0095343 | 0,00306 | 0,0066363 | 0,0066363 | TRUE | 0,1804468 | No |
| orf19.6967 | 3407 | 3291 | 3669 | 2126 | 2719 | 2244 | 2588 | 3170 | 3102 | 3328 | 3472 | 3194 | 2418 | 2436 | 2650 | 2418 | 2877,47 | 3274 | 2480 | 0,757 | -0,401 | -7,86 | 3,85E-15 | 8,54E-15 | 0,001791 | 0,002496 | 0,0021493 | 0,0021493 | TRUE | 0,0254059 | No |
| orf19.6968 | 4307 | 4604 | 4610 | 3037 | 5756 | 5122 | 4942 | 6714 | 3921 | 4656 | 4363 | 4562 | 5120 | 5561 | 5061 | 5121 | 4795,8 | 4376 | 5216 | 1,192 | 0,254 | 4,352 | 1,35E-05 | 2,12E-05 | 0,0034943 | 0,002312 | 0,0030565 | 0,0030565 | TRUE | 0,0602725 | No |
| orf19.697 | 3172 | 3564 | 3619 | 2107 | 2509 | 2273 | 2161 | 2695 | 2888 | 3605 | 3425 | 3165 | 2232 | 2468 | 2213 | 2056 | 2756,39 | 3271 | 2242 | 0,685 | -0,546 | -7,318 | 2,51E-13 | 5,32E-13 | 0,0069353 | 0,002516 | 0,0049674 | 0,0049674 | TRUE | 0,0769689 | No |
| orf19.6970 | 31 | 36 | 49 | 30 | 32 | 37 | 27 | 30 | 28 | 36 | 46 | 45 | 28 | 40 | 28 | 23 | 34,41 | 39 | 30 | 0,762 | -0,393 | -1,514 | 0,130087 | 0,154219 | 0,0260175 | 0,040432 | 0,0350907 | 0,0350907 | TRUE | 0,3382807 | No |
| orf19.6971 | 102 | 137 | 107 | 82 | 110 | 91 | 90 | 86 | 93 | 139 | 101 | 123 | 98 | 99 | 92 | 66 | 101,29 | 114 | 89 | 0,777 | -0,365 | -2,079 | 0,037573 | 0,04705 | 0,0256421 | 0,015079 | 0,0195146 | 0,0195146 | TRUE | 0,3332062 | No |
| orf19.6972 | 838 | 812 | 936 | 434 | 1033 | 858 | 938 | 1076 | 763 | 821 | 886 | 652 | 919 | 932 | 961 | 821 | 844,22 | 780 | 908 | 1,158 | 0,212 | 2,426 | 0,015253 | 0,019764 | 0,0087418 | 0,003602 | 0,0061063 | 0,0061063 | TRUE | 0,1331158 | No |
| orf19.6973 | 12171 | 12133 | 12924 | 7151 | 8439 | 7231 | 7376 | 9388 | 11081 | 12271 | 12231 | 10742 | 7506 | 7851 | 7554 | 7161 | 9549,75 | 11581 | 7518 | 0,649 | -0,624 | -11,708 | 1,15E-31 | 3,87E-31 | 0,0028628 | 0,002175 | 0,0026201 | 0,0026201 | TRUE | 0,0286747 | No |
| orf19.6975 | 18719 | 14823 | 16028 | 11352 | 30899 | 24812 | 25986 | 37116 | 17043 | 14992 | 15169 | 17053 | 27484 | 26939 | 26613 | 28312 | 21700,5 | 16064 | 27337 | 1,702 | 0,767 | 14,817 | 1,13E-49 | 5,17E-49 | 0,002813 | 0,002098 | 0,0025256 | 0,0025256 | TRUE | 0,0246756 | No |
| orf19.6976 | 1776 | 1790 | 1768 | 1032 | 3636 | 3012 | 3078 | 3877 | 1617 | 1810 | 1673 | 1550 | 3234 | 3270 | 3152 | 2957 | 2408,1 | 1662 | 3153 | 1,893 | 0,921 | 16,367 | 3,27E-60 | 1,80E-59 | 0,0025797 | 0,002586 | 0,0025678 | 0,0025678 | TRUE | 0,04271 | No |
| orf19.6977 | 257 | 375 | 332 | 174 | 198 | 193 | 222 | 215 | 234 | 379 | 314 | 261 | 176 | 210 | 227 | 164 | 245,73 | 297 | 194 | 0,65 | -0,621 | -4,103 | 4,07E-05 | 6,24E-05 | 0,0305672 | 0,007413 | 0,017732 | 0,017732 | TRUE | 0,3116287 | No |
| orf19.6978 | 98 | 112 | 106 | 67 | 85 | 92 | 66 | 118 | 89 | 113 | 100 | 101 | 76 | 100 | 68 | 90 | 92,07 | 101 | 84 | 0,827 | -0,275 | -1,731 | 0,083509 | 0,101038 | 0,0079355 | 0,016385 | 0,0133167 | 0,0133167 | TRUE | 0,1483781 | No |
| orf19.6979 | 2282 | 2084 | 2277 | 1363 | 3418 | 2809 | 3007 | 4054 | 2078 | 2108 | 2155 | 2047 | 3040 | 3050 | 3080 | 3092 | 2581,22 | 2097 | 3066 | 1,461 | 0,547 | 14,657 | 1,22E-48 | 5,49E-48 | 1,00E-08 | 0,002549 | 0,0009277 | 0,0009277 | TRUE | 0,0042178 | No |
| orf19.698 | 5070 | 5069 | 5631 | 3211 | 2679 | 2534 | 2514 | 3312 | 4616 | 5127 | 5329 | 4823 | 2383 | 2751 | 2575 | 2526 | 3766,33 | 4974 | 2559 | 0,514 | -0,96 | -16,332 | 5,85E-60 | 3,20E-59 | 0,0033921 | 0,002388 | 0,0030262 | 0,0030262 | TRUE | 0,0306603 | No |
| orf19.6980 | 1564 | 1609 | 1778 | 1139 | 1544 | 1424 | 1479 | 1924 | 1424 | 1627 | 1683 | 1711 | 1373 | 1546 | 1515 | 1468 | 1543,34 | 1611 | 1476 | 0,917 | -0,126 | -1,907 | 0,056505 | 0,069675 | 0,0040797 | 0,002893 | 0,0035254 | 0,0035254 | TRUE | 0,0756748 | No |
| orf19.6981 | 931 | 974 | 974 | 565 | 284 | 259 | 243 | 341 | 848 | 985 | 922 | 849 | 253 | 281 | 249 | 260 | 580,76 | 901 | 261 | 0,289 | -1,792 | -22,792 | 5,56E-115 | 5,46E-114 | 0,0027263 | 0,004312 | 0,0035545 | 0,0035545 | TRUE | 0,0471883 | No |
| orf19.6982 | 1135 | 1267 | 1175 | 728 | 557 | 461 | 484 | 581 | 1033 | 1281 | 1112 | 1094 | 495 | 501 | 496 | 443 | 806,9 | 1130 | 484 | 0,427 | -1,227 | -15,698 | 1,55E-55 | 7,91E-55 | 0,0051387 | 0,003674 | 0,0044228 | 0,0044228 | TRUE | 0,0990667 | No |
| orf19.6983 | 5226 | 5546 | 5779 | 3226 | 3553 | 2976 | 3102 | 4067 | 4758 | 5609 | 5469 | 4846 | 3160 | 3231 | 3177 | 3102 | 4169,13 | 5170 | 3168 | 0,612 | -0,708 | -12,189 | 3,56E-34 | 1,25E-33 | 0,003274 | 0,002354 | 0,002986 | 0,002986 | TRUE | 0,0396875 | No |
| orf19.6984 | 2520 | 2794 | 2896 | 1709 | 3354 | 2832 | 2976 | 3764 | 2294 | 2826 | 2741 | 2567 | 2983 | 3075 | 3048 | 2871 | 2800,65 | 2607 | 2994 | 1,148 | 0,199 | 3,149 | 0,001639 | 0,002277 | 0,0040282 | 0,002509 | 0,0034894 | 0,0034894 | TRUE | 0,0815343 | No |
| orf19.6985 | 1824 | 1919 | 2034 | 1231 | 927 | 744 | 741 | 943 | 1661 | 1941 | 1925 | 1849 | 825 | 808 | 759 | 719 | 1310,78 | 1844 | 778 | 0,421 | -1,247 | -18,519 | 1,45E-76 | 9,82E-76 | 0,0038217 | 0,003045 | 0,0034574 | 0,0034574 | TRUE | 0,0560978 | No |
| orf19.6986 | 389 | 445 | 438 | 290 | 643 | 511 | 522 | 715 | 354 | 450 | 415 | 436 | 572 | 555 | 535 | 545 | 482,64 | 414 | 552 | 1,339 | 0,421 | 5,203 | 1,96E-07 | 3,35E-07 | 0,0033041 | 0,004774 | 0,0041355 | 0,0041355 | TRUE | 0,1154853 | No |
| orf19.6987 | 7097 | 6899 | 7230 | 4493 | 9122 | 7822 | 7781 | 10360 | 6462 | 6978 | 6843 | 6749 | 8114 | 8493 | 7969 | 7903 | 7438,56 | 6758 | 8120 | 1,201 | 0,265 | 6,675 | 2,48E-11 | 4,93E-11 | 0,0008706 | 0,002215 | 0,0013724 | 0,0013724 | TRUE | 0,011567 | No |
| orf19.6988 | 483 | 449 | 523 | 288 | 494 | 480 | 444 | 606 | 440 | 454 | 495 | 433 | 439 | 521 | 455 | 462 | 462,37 | 456 | 469 | 1,027 | 0,038 | 0,484 | 0,628372 | 0,666868 | 0,0024309 | 0,004894 | 0,0037919 | 0,0037919 | TRUE | 0,0604924 | No |
| orf19.6989 | 947 | 858 | 974 | 585 | 800 | 685 | 725 | 867 | 862 | 868 | 922 | 879 | 712 | 744 | 742 | 661 | 798,71 | 883 | 715 | 0,808 | -0,308 | -5,061 | 4,18E-07 | 7,05E-07 | 0,0008789 | 0,003691 | 0,0022914 | 0,0022914 | TRUE | 0,0321316 | No |
| orf19.699 | 805 | 1362 | 1277 | 641 | 485 | 476 | 447 | 478 | 733 | 1378 | 1209 | 963 | 431 | 517 | 458 | 365 | 756,56 | 1071 | 443 | 0,413 | -1,275 | -5,719 | 1,07E-08 | 1,93E-08 | 0,0462018 | 0,003783 | 0,022049 | 0,0462018 | TRUE | 0,1369651 | No |
| orf19.6990 | 1165 | 1159 | 1166 | 720 | 1164 | 921 | 986 | 1332 | 1061 | 1172 | 1104 | 1082 | 1035 | 1000 | 1010 | 1016 | 1059,89 | 1105 | 1015 | 0,919 | -0,122 | -2,321 | 0,020291 | 0,02599 | 1,00E-08 | 0,003283 | 0,0016899 | 0,0016899 | TRUE | 0,0208834 | No |
| orf19.6991 | 4592 | 4115 | 4259 | 2290 | 3047 | 2281 | 2409 | 3357 | 4181 | 4162 | 4031 | 3440 | 2710 | 2477 | 2467 | 2561 | 3253,5 | 3954 | 2554 | 0,645 | -0,632 | -9,808 | 1,04E-22 | 2,80E-22 | 0,0044317 | 0,002443 | 0,0036651 | 0,0036651 | TRUE | 0,0902157 | No |
| orf19.6992 | 11882 | 10421 | 11239 | 7027 | 4468 | 4065 | 4083 | 5564 | 10818 | 10540 | 10637 | 10556 | 3974 | 4413 | 4182 | 4244 | 7420,44 | 10638 | 4203 | 0,395 | -1,34 | -34,629 | 9,23E-263 | 2,02E-261 | 0,000783 | 0,002215 | 0,0012763 | 0,0012763 | TRUE | 0,0164694 | No |
| orf19.6993 | 12993 | 10874 | 11723 | 7632 | 32117 | 26707 | 28136 | 40643 | 11830 | 10998 | 11095 | 11464 | 28567 | 28996 | 28815 | 31003 | 20346 | 11347 | 29345 | 2,586 | 1,371 | 33,731 | 2,04E-249 | 4,20E-248 | 0,001205 | 0,002102 | 0,0015234 | 0,0015234 | TRUE | 0,0177852 | No |
| orf19.6994 | 5397 | 4502 | 4859 | 3383 | 6628 | 5391 | 5536 | 7873 | 4914 | 4553 | 4599 | 5082 | 5895 | 5853 | 5670 | 6006 | 5321,79 | 4787 | 5856 | 1,224 | 0,292 | 6,493 | 8,43E-11 | 1,64E-10 | 0,0014366 | 0,002285 | 0,001752 | 0,001752 | TRUE | 0,0200168 | No |
| orf19.6995 | 406 | 350 | 396 | 270 | 678 | 586 | 620 | 850 | 370 | 354 | 375 | 406 | 603 | 636 | 635 | 648 | 503,33 | 376 | 630 | 1,685 | 0,753 | 10,778 | 4,37E-27 | 1,31E-26 | 1,00E-08 | 0,004662 | 0,0025185 | 0,0025185 | TRUE | 0,0257194 | No |

| | | | | | | | | | | | | | | | | | | | | | | | | | | | | | | | |
|---|---|---|---|---|---|---|---|---|---|---|---|---|---|---|---|---|---|---|---|---|---|---|---|---|---|---|---|---|---|---|---|
| orf19.6996 | 4568 | 4594 | 4642 | 2832 | 2138 | 1824 | 1892 | 2363 | 4159 | 4646 | 4393 | 4254 | 1902 | 1980 | 1938 | 1803 | 3134,36 | 4363 | 1906 | 0,436 | -1,197 | -24,064 | 6,03E-128 | 6,57E-127 | 0,0014685 | 0,002458 | 0,0020059 | 0,0020059 | TRUE | 0,0234212 | No |
| orf19.6997 | 270 | 247 | 240 | 146 | 163 | 141 | 149 | 195 | 246 | 250 | 227 | 219 | 145 | 153 | 153 | 149 | 192,69 | 236 | 150 | 0,633 | -0,659 | -6,429 | 1,29E-10 | 2,49E-10 | 1,00E-08 | 0,008893 | 0,0047285 | 0,0047285 | TRUE | 0,0187091 | No |
| orf19.6998 | 17 | 11 | 25 | 16 | 26 | 24 | 14 | 32 | 15 | 11 | 24 | 24 | 23 | 26 | 14 | 24 | 20,28 | 18 | 22 | 1,205 | 0,269 | 0,806 | 0,420417 | 0,461618 | 0,0343744 | 0,067179 | 0,0564363 | 0,0564363 | TRUE | 0,3758754 | No |
| orf19.6999 | 3 | 14 | 13 | 5 | 4 | 3 | 6 | 9 | 3 | 14 | 12 | 8 | 4 | 3 | 6 | 7 | 7,07 | 9 | 5 | 0,545 | -0,875 | -1,494 | 0,1353 | 0,159906 | 0,1610633 | 0,188981 | 0,1784958 | 0,1784958 | TRUE | 0,3040416 | No |
| orf19.700 | 2001 | 1942 | 2045 | 1130 | 7471 | 5868 | 6690 | 8474 | 1822 | 1964 | 1935 | 1697 | 6645 | 6371 | 6851 | 6464 | 4218,81 | 1854 | 6583 | 3,542 | 1,825 | 35,928 | 1,13E-282 | 2,68E-281 | 0,0022048 | 0,00235 | 0,002116 | 0,002116 | TRUE | 0,035819 | No |
| orf19.7001 | 13878 | 12731 | 13760 | 8770 | 9996 | 8495 | 9199 | 12235 | 12635 | 12876 | 13023 | 13174 | 8891 | 9223 | 9421 | 9333 | 11072 | 12927 | 9217 | 0,713 | -0,488 | -15,008 | 6,50E-51 | 3,05E-50 | 0,0003498 | 0,002156 | 0,0009219 | 0,0009219 | TRUE | 0,0069764 | No |
| orf19.7002 | 11 | 10 | 9 | 4 | 24 | 12 | 9 | 13 | 10 | 10 | 9 | 6 | 21 | 13 | 9 | 10 | 11,02 | 9 | 13 | 1,517 | 0,601 | 1,366 | 0,17184 | 0,200453 | 1,00E-08 | 0,121903 | 0,0907981 | 0,0907981 | TRUE | 0,7626179 | No |
| orf19.7006 | 7566 | 6403 | 7318 | 4580 | 1357 | 1115 | 1368 | 1737 | 6889 | 6476 | 6926 | 6880 | 1207 | 1211 | 1401 | 1325 | 4039,22 | 6793 | 1286 | 0,189 | -2,401 | -44,756 | 0 | 0 | 0,0025591 | 0,002364 | 0,0023207 | 0,0023207 | TRUE | 0,0418598 | No |
| orf19.7007 | 23 | 17 | 21 | 12 | 7 | 7 | 11 | 5 | 21 | 17 | 20 | 18 | 6 | 8 | 11 | 4 | 13,12 | 19 | 7 | 0,368 | -1,441 | -3,524 | 0,000426 | 0,000615 | 1,00E-08 | 0,102742 | 0,0664841 | 0,0664841 | TRUE | 0,3900042 | No |
| orf19.7009 | 110 | 106 | 99 | 66 | 92 | 95 | 93 | 103 | 100 | 107 | 94 | 99 | 82 | 103 | 95 | 79 | 94,87 | 100 | 90 | 0,89 | -0,168 | -1,142 | 0,253313 | 0,288782 | 1,00E-08 | 0,015961 | 0,0102126 | 0,0102126 | TRUE | 0,0936515 | No |
| orf19.701 | 31 | 38 | 27 | 19 | 22 | 23 | 21 | 26 | 28 | 38 | 26 | 29 | 20 | 25 | 22 | 20 | 25,83 | 30 | 22 | 0,706 | -0,502 | -1,839 | 0,065847 | 0,080617 | 1,00E-08 | 0,053181 | 0,031987 | 0,031987 | TRUE | 0,2353093 | No |
| orf19.7010 | 1810 | 1549 | 1705 | 1130 | 1423 | 1188 | 1291 | 1698 | 1648 | 1567 | 1614 | 1697 | 1266 | 1290 | 1322 | 1295 | 1462,32 | 1632 | 1293 | 0,794 | -0,334 | -7,213 | 5,47E-13 | 1,15E-12 | 1,00E-08 | 0,00294 | 0,0013624 | 0,0013624 | TRUE | 0,0083474 | No |
| orf19.7011 | 13436 | 12240 | 13300 | 8077 | 21763 | 18929 | 19527 | 26168 | 12233 | 12379 | 12587 | 12133 | 19357 | 20552 | 19998 | 19961 | 16150,1 | 12333 | 19967 | 1,619 | 0,695 | 22,182 | 5,17E-109 | 4,84E-108 | 0,0003392 | 0,002119 | 0,0008749 | 0,0008749 | TRUE | 0,0051694 | No |
| orf19.7012 | 2124 | 1866 | 1799 | 1226 | 1889 | 1483 | 1593 | 2113 | 1934 | 1887 | 1703 | 1842 | 1680 | 1610 | 1631 | 1612 | 1737,36 | 1842 | 1633 | 0,887 | -0,173 | -3,393 | 0,000691 | 0,000985 | 0,0012404 | 0,002797 | 0,0019115 | 0,0019115 | TRUE | 0,0327362 | No |
| orf19.7013 | 235 | 278 | 296 | 203 | 538 | 479 | 450 | 652 | 214 | 281 | 280 | 305 | 479 | 520 | 461 | 497 | 379,63 | 270 | 489 | 1,824 | 0,867 | 8,712 | 3,00E-18 | 7,25E-18 | 0,0080042 | 0,005517 | 0,0065529 | 0,0065529 | TRUE | 0,2318837 | No |
| orf19.7014 | 20 | 6 | 9 | 18 | 20 | 16 | 11 | 18 | 18 | 6 | 9 | 27 | 18 | 17 | 11 | 14 | 15 | 15 | 15 | 1,039 | 0,055 | 0,128 | 0,898065 | 0,912668 | 0,1485966 | 0,09011 | 0,1111444 | 0,1111444 | TRUE | 0,3737938 | No |
| orf19.7015 | 13605 | 10792 | 11269 | 8168 | 22564 | 18237 | 18175 | 27812 | 12387 | 10915 | 10665 | 12270 | 20070 | 19800 | 18613 | 21215 | 15741,9 | 11559 | 19924 | 1,724 | 0,786 | 13,063 | 5,35E-39 | 2,05E-38 | 0,004344 | 0,002121 | 0,0034069 | 0,0034069 | TRUE | 0,0332287 | No |
| orf19.7016 | 2767 | 2553 | 2753 | 1710 | 3522 | 2958 | 3167 | 4357 | 2519 | 2582 | 2605 | 2569 | 3133 | 3212 | 3243 | 3324 | 2898,34 | 2569 | 3228 | 1,257 | 0,33 | 8,699 | 3,36E-18 | 8,13E-18 | 1,00E-08 | 0,002493 | 0,0010271 | 0,0010271 | TRUE | 0,0051235 | No |
| orf19.7017 | 86 | 70 | 92 | 52 | 57 | 45 | 37 | 70 | 78 | 71 | 87 | 78 | 51 | 49 | 38 | 53 | 63,14 | 78 | 48 | 0,61 | -0,712 | -3,977 | 6,96E-05 | 0,000105 | 1,00E-08 | 0,022959 | 0,0143326 | 0,0143326 | TRUE | 0,1466142 | No |
| orf19.7018 | 16107 | 13885 | 14334 | 10241 | 26115 | 19706 | 20568 | 27897 | 14665 | 14043 | 13566 | 15384 | 23228 | 21395 | 21064 | 21280 | 18078,2 | 14414 | 21742 | 1,509 | 0,593 | 11,946 | 6,85E-33 | 2,34E-32 | 0,0024737 | 0,00211 | 0,0023115 | 0,0023115 | TRUE | 0,0259663 | No |
| orf19.7019 | 2283 | 2036 | 2273 | 1533 | 2960 | 2303 | 2354 | 3408 | 2079 | 2059 | 2151 | 2303 | 2633 | 2500 | 2411 | 2600 | 2341,93 | 2148 | 2536 | 1,183 | 0,243 | 4,831 | 1,36E-06 | 2,25E-06 | 0,0015389 | 0,002601 | 0,0019899 | 0,0019899 | TRUE | 0,02638 | No |
| orf19.702 | 6542 | 5993 | 6461 | 3833 | 6296 | 5432 | 5746 | 7472 | 5956 | 6061 | 6115 | 5758 | 5600 | 5898 | 5885 | 5700 | 5871,51 | 5972 | 5771 | 0,965 | -0,051 | -1,408 | 0,159104 | 0,186655 | 0,0004374 | 0,002262 | 0,0010849 | 0,0010849 | TRUE | 0,0067466 | No |
| orf19.7020 | 7749 | 7253 | 7700 | 4772 | 10067 | 8465 | 9119 | 12080 | 7055 | 7336 | 7287 | 7168 | 8954 | 9191 | 9339 | 9215 | 8193,13 | 7212 | 9175 | 1,272 | 0,347 | 11,172 | 5,60E-29 | 1,75E-28 | 0,0001404 | 0,002198 | 0,0008016 | 0,0008016 | TRUE | 0,0032125 | No |
| orf19.7021 | 33380 | 30412 | 31617 | 19542 | 16862 | 13832 | 14098 | 19618 | 30391 | 30758 | 29923 | 29355 | 14998 | 15018 | 14438 | 14965 | 22480,8 | 30107 | 14855 | 0,493 | -1,019 | -33,875 | 1,53E-251 | 3,21E-250 | 0,0003119 | 0,002096 | 0,0008202 | 0,0008202 | TRUE | 0,0043297 | No |
| orf19.7022 | 1326 | 1250 | 1287 | 738 | 451 | 378 | 349 | 456 | 1207 | 1264 | 1218 | 1109 | 401 | 410 | 357 | 348 | 789,37 | 1200 | 379 | 0,315 | -1,667 | -23,245 | 1,58E-119 | 1,61E-118 | 0,0025522 | 0,003711 | 0,0032601 | 0,0032601 | TRUE | 0,0395554 | No |
| orf19.7023 | 272 | 264 | 266 | 176 | 329 | 264 | 298 | 368 | 248 | 267 | 252 | 264 | 293 | 287 | 305 | 281 | 274,49 | 258 | 292 | 1,132 | 0,178 | 2,073 | 0,038217 | 0,047836 | 1,00E-08 | 0,00685 | 0,0034375 | 0,0034375 | TRUE | 0,011292 | No |
| orf19.7024 | 6 | 3 | 4 | 4 | 27 | 21 | 18 | 17 | 5 | 3 | 4 | 6 | 24 | 23 | 18 | 13 | 12,06 | 4 | 20 | 4,331 | 2,115 | 4,426 | 9,58E-06 | 1,52E-05 | 1,00E-08 | 0,11154 | 0,0771525 | 0,0771525 | TRUE | 0,3497532 | No |
| orf19.7025 | 2166 | 1981 | 2106 | 1430 | 1722 | 1460 | 1551 | 2041 | 1972 | 2004 | 1993 | 2148 | 1532 | 1585 | 1588 | 1557 | 1797,37 | 2029 | 1566 | 0,773 | -0,372 | -8,281 | 1,23E-16 | 2,84E-16 | 0,000174 | 0,002772 | 0,001374 | 0,001374 | TRUE | 0,0169657 | No |
| orf19.7027 | 14477 | 12915 | 14901 | 8440 | 1159 | 902 | 1030 | 1361 | 13181 | 13062 | 14102 | 12678 | 1031 | 979 | 1055 | 1038 | 7140,84 | 13256 | 1026 | 0,077 | -3,691 | -76,935 | 0 | 0 | 0,0013894 | 0,002222 | 0,0017156 | 0,0017156 | TRUE | 0,0226513 | No |
| orf19.7029 | 168 | 182 | 178 | 123 | 337 | 324 | 299 | 383 | 153 | 184 | 168 | 185 | 300 | 352 | 306 | 292 | 242,52 | 172 | 312 | 1,815 | 0,86 | 8,278 | 1,26E-16 | 2,90E-16 | 0,0032976 | 0,007484 | 0,0057687 | 0,0057687 | TRUE | 0,0777011 | No |
| orf19.703 | 1682 | 1396 | 1479 | 1009 | 702 | 556 | 588 | 756 | 1531 | 1412 | 1400 | 1516 | 624 | 604 | 602 | 577 | 1033,21 | 1465 | 602 | 0,411 | -1,283 | -22,183 | 5,05E-109 | 4,73E-108 | 0,0009624 | 0,003316 | 0,0020708 | 0,0020708 | TRUE | 0,0126596 | No |
| orf19.7030 | 2105 | 1646 | 1944 | 1404 | 3863 | 4036 | 3318 | 5206 | 1917 | 1665 | 1840 | 2109 | 3436 | 4382 | 3398 | 3971 | 2839,67 | 1883 | 3797 | 2,019 | 1,013 | 11,431 | 2,92E-30 | 9,44E-30 | 0,0119072 | 0,002502 | 0,0071387 | 0,0071387 | TRUE | 0,1300767 | No |
| orf19.7032 | 1045 | 1307 | 1229 | 780 | 1143 | 1067 | 1052 | 1273 | 951 | 1322 | 1163 | 1172 | 1017 | 1158 | 1077 | 971 | 1103,96 | 1152 | 1056 | 0,916 | -0,126 | -1,375 | 0,168982 | 0,1973 | 0,0104785 | 0,003234 | 0,0071592 | 0,0071592 | TRUE | 0,1694477 | No |
| orf19.7033 | 1349 | 1591 | 1550 | 943 | 2376 | 2027 | 2119 | 2711 | 1228 | 1609 | 1467 | 1417 | 2113 | 2201 | 2170 | 2068 | 1784,13 | 1430 | 2138 | 1,495 | 0,58 | 8,066 | 7,27E-16 | 1,65E-15 | 0,0056624 | 0,002777 | 0,004371 | 0,004371 | TRUE | 0,1131784 | No |
| orf19.7034 | 2684 | 2573 | 2920 | 1789 | 1524 | 1276 | 1376 | 1853 | 2444 | 2602 | 2764 | 2687 | 1356 | 1385 | 1409 | 1413 | 2007,56 | 2624 | 1391 | 0,53 | -0,915 | -18,235 | 2,75E-74 | 1,81E-73 | 0,0013078 | 0,002695 | 0,0018757 | 0,0018757 | TRUE | 0,0269274 | No |
| orf19.7035 | 1186 | 1168 | 1241 | 759 | 1392 | 1192 | 1201 | 1486 | 1080 | 1181 | 1174 | 1140 | 1238 | 1294 | 1230 | 1134 | 1183,95 | 1144 | 1224 | 1,069 | 0,096 | 1,655 | 0,097921 | 0,117716 | 0,0016901 | 0,003153 | 0,0023632 | 0,0023632 | TRUE | 0,0311601 | No |
| orf19.7036 | 3835 | 3811 | 4125 | 2352 | 3474 | 2876 | 3165 | 4002 | 3492 | 3854 | 3904 | 3533 | 3090 | 3123 | 3241 | 3053 | 3411,21 | 3696 | 3127 | 0,845 | -0,242 | -4,948 | 7,49E-07 | 1,25E-06 | 0,0016145 | 0,002424 | 0,0020095 | 0,0020095 | TRUE | 0,0177785 | No |
| orf19.7037 | 954 | 904 | 902 | 616 | 937 | 663 | 682 | 977 | 869 | 914 | 854 | 925 | 833 | 720 | 698 | 745 | 819,86 | 890 | 749 | 0,844 | -0,245 | -3,65 | 0,000262 | 0,000384 | 0,0025517 | 0,003648 | 0,0031056 | 0,0031056 | TRUE | 0,0681098 | No |
| orf19.7038 | 785 | 913 | 915 | 559 | 1014 | 858 | 893 | 1115 | 715 | 923 | 866 | 840 | 902 | 932 | 915 | 851 | 867,79 | 836 | 900 | 1,076 | 0,106 | 1,377 | 0,168664 | 0,197037 | 0,0054728 | 0,003559 | 0,0045159 | 0,0045159 | TRUE | 0,1197289 | No |
| orf19.704 | 2418 | 1993 | 2135 | 1417 | 2094 | 1546 | 1644 | 2374 | 2201 | 2016 | 2021 | 2129 | 1863 | 1679 | 1684 | 1811 | 1925,25 | 2092 | 1759 | 0,842 | -0,248 | -4,66 | 3,16E-06 | 5,14E-06 | 0,0016033 | 0,002723 | 0,0022012 | 0,0022012 | TRUE | 0,0187068 | No |
| orf19.7041 | 4271 | 4055 | 4240 | 2704 | 3140 | 2610 | 2671 | 3402 | 3889 | 4101 | 4013 | 4062 | 2793 | 2834 | 2735 | 2595 | 3377,69 | 4016 | 2739 | 0,682 | -0,553 | -13,212 | 7,51E-40 | 2,91E-39 | 0,0007212 | 0,002428 | 0,0013745 | 0,0013745 | TRUE | 0,0158817 | No |
| orf19.7042 | 1119 | 1105 | 1079 | 712 | 404 | 347 | 350 | 398 | 1019 | 1118 | 1021 | 1070 | 359 | 377 | 358 | 304 | 703,15 | 1057 | 350 | 0,33 | -1,601 | -21,477 | 2,54E-102 | 2,27E-101 | 0,0029315 | 0,003916 | 0,0034899 | 0,0034899 | TRUE | 0,0989289 | No |
| orf19.7043 | 38 | 65 | 40 | 20 | 55 | 45 | 38 | 47 | 35 | 66 | 38 | 30 | 49 | 49 | 39 | 36 | 42,6 | 42 | 43 | 1,013 | 0,019 | 0,074 | 0,941034 | 0,949975 | 0,0511221 | 0,033048 | 0,0401651 | 0,0401651 | TRUE | 1,091573 | No |
| orf19.7043.1 | 739 | 749 | 662 | 450 | 2743 | 1967 | 2076 | 2638 | 673 | 758 | 627 | 676 | 2440 | 2136 | 2126 | 2012 | 1430,83 | 684 | 2178 | 3,189 | 1,673 | 22,255 | 1,01E-109 | 9,50E-109 | 0,0054246 | 0,00296 | 0,0044246 | 0,0044246 | TRUE | 0,0798826 | No |
| orf19.7044 | 4141 | 3632 | 4062 | 2444 | 3874 | 3076 | 3345 | 4352 | 3770 | 3673 | 3844 | 3671 | 3446 | 3340 | 3426 | 3320 | 3561,26 | 3740 | 3383 | 0,904 | -0,146 | -4,029 | 5,59E-05 | 8,49E-05 | 0,000155 | 0,002408 | 0,0009746 | 0,0009746 | TRUE | 0,0043328 | No |
| orf19.7046 | 3637 | 3502 | 3506 | 2123 | 4008 | 2977 | 3028 | 4142 | 3311 | 3542 | 3318 | 3189 | 3565 | 3232 | 3101 | 3160 | 3302,28 | 3340 | 3264 | 0,977 | -0,034 | -0,618 | 0,536778 | 0,578425 | 0,0023823 | 0,002437 | 0,0025433 | 0,0025433 | TRUE | 0,0470555 | No |
| orf19.7047 | 3249 | 3131 | 3359 | 1943 | 4864 | 3703 | 4112 | 5382 | 2958 | 3167 | 3179 | 2919 | 4326 | 4020 | 4211 | 4105 | 3610,73 | 3056 | 4166 | 1,362 | 0,446 | 9,925 | 3,23E-23 | 8,82E-23 | 0,0011235 | 0,002403 | 0,0016515 | 0,0016515 | TRUE | 0,0102389 | No |
| orf19.7048.1 | 5817 | 4405 | 4654 | 3228 | 10275 | 7559 | 8131 | 11303 | 5296 | 4455 | 4405 | 4849 | 9139 | 8207 | 8327 | 8622 | 6662,54 | 4751 | 8574 | 1,805 | 0,852 | 13,592 | 4,46E-42 | 1,80E-41 | 0,0047521 | 0,002235 | 0,0036048 | 0,0036048 | TRUE | 0,073996 | No |
| orf19.7049 | 911 | 902 | 898 | 592 | 3801 | 3051 | 2812 | 3984 | 829 | 912 | 850 | 889 | 3381 | 3313 | 2880 | 3039 | 2011,64 | 870 | 3153 | 3,627 | 1,859 | 29,201 | 1,90E-187 | 3,04E-186 | 0,0034721 | 0,002694 | 0,0031243 | 0,0031243 | TRUE | 0,0407859 | No |
| orf19.705 | 1553 | 1231 | 1459 | 816 | 1233 | 883 | 1028 | 1410 | 1414 | 1245 | 1381 | 1226 | 1097 | 959 | 1053 | 1076 | 1181,16 | 1316 | 1046 | 0,794 | -0,333 | -5,071 | 3,95E-07 | 6,67E-07 | 0,0031437 | 0,003155 | 0,0032871 | 0,0032871 | TRUE | 0,0365078 | No |
| orf19.7050 | 9265 | 8639 | 8712 | 5089 | 11417 | 8501 | 8832 | 12129 | 8435 | 8737 | 8245 | 7644 | 10155 | 9230 | 9045 | 9252 | 8843,05 | 8265 | 9420 | 1,139 | 0,188 | 3,573 | 0,000353 | 0,000513 | 0,0028038 | 0,002186 | 0,0025493 | 0,0025493 | TRUE | 0,0337476 | No |
| orf19.7051 | 98 | 123 | 142 | 70 | 201 | 161 | 149 | 218 | 89 | 124 | 134 | 105 | 179 | 175 | 153 | 166 | 140,7 | 113 | 168 | 1,481 | 0,567 | 4,11 | 3,96E-05 | 6,08E-05 | 0,0096708 | 0,011425 | 0,0107793 | 0,0107793 | TRUE | 0,2399466 | No |
| orf19.7052 | 786 | 782 | 817 | 454 | 1524 | 1291 | 1452 | 1695 | 716 | 791 | 773 | 682 | 1356 | 1402 | 1487 | 1293 | 1062,37 | 740 | 1384 | 1,863 | 0,898 | 13,677 | 1,39E-42 | 5,68E-42 | 0,0026868 | 0,00328 | 0,003074 | 0,003074 | TRUE | 0,0300912 | No |
| orf19.7053 | 41900 | 40319 | 43666 | 27996 | 18160 | 16271 | 16287 | 22392 | 38148 | 40778 | 41326 | 42054 | 16153 | 17666 | 16680 | 17081 | 28735,8 | 40576 | 16895 | 0,416 | -1,264 | -29,248 | 4,83E-188 | 7,73E-187 | 0,0015389 | 0,002083 | 0,0017535 | 0,0017535 | TRUE | 0,0199239 | No |
| orf19.7054 | 54 | 43 | 43 | 38 | 10 | 12 | 14 | 27 | 49 | 43 | 41 | 57 | 9 | 13 | 14 | 21 | 30,91 | 48 | 14 | 0,305 | -1,712 | -5,978 | 2,26E-09 | 4,18E-09 | 0,0098261 | 0,044772 | 0,035155 | 0,035155 | TRUE | 0,5445115 | No |
| orf19.7055 | 0 | 5 | 2 | 0 | 27 | 21 | 20 | 24 | 0 | 5 | 2 | 0 | 24 | 23 | 20 | 18 | 11,57 | 2 | 21 | 11,662 | 3,544 | 5,543 | 2,97E-08 | 5,23E-08 | 1,26E-08 | 0,116217 | 0,0828481 | 0,0828481 | TRUE | 1,2382898 | No |
| orf19.7056 | 316 | 257 | 295 | 199 | 130 | 116 | 140 | 140 | 288 | 260 | 279 | 299 | 116 | 126 | 143 | 107 | 202,19 | 282 | 123 | 0,435 | -1,201 | -10,696 | 1,06E-26 | 3,15E-26 | 0,0020891 | 0,008571 | 0,0063941 | 0,0063941 | TRUE | 0,1275211 | No |
| orf19.7057 | 6279 | 5103 | 5628 | 3570 | 8114 | 6418 | 7072 | 9276 | 5717 | 5161 | 5326 | 5363 | 7217 | 6968 | 7243 | 7076 | 6258,84 | 5392 | 7126 | 1,321 | 0,402 | 9,982 | 1,82E-23 | 5,03E-23 | 0,0008846 | 0,002248 | 0,001392 | 0,001392 | TRUE | 0,0206401 | No |
| orf19.7058 | 1344 | 1142 | 1214 | 863 | 1709 | 1509 | 1454 | 1992 | 1224 | 1155 | 1149 | 1296 | 1520 | 1638 | 1489 | 1520 | 1373,88 | 1206 | 1542 | 1,281 | 0,357 | 6,414 | 1,41E-10 | 2,73E-10 | 0,0015736 | 0,002999 | 0,002224 | 0,002224 | TRUE | 0,0271834 | No |
| orf19.7059 | 872 | 740 | 826 | 510 | 1444 | 1205 | 1266 | 1737 | 794 | 748 | 782 | 766 | 1284 | 1308 | 1297 | 1325 | 1038,05 | 772 | 1304 | 1,686 | 0,754 | 14,921 | 2,41E-50 | 1,11E-49 | 1,00E-08 | 0,00331 | 0,0014027 | 0,0014027 | TRUE | 0,0057843 | No |
| orf19.706 | 21533 | 17923 | 19676 | 12211 | 20917 | 16657 | 18959 | 25366 | 19605 | 18127 | 18622 | 18343 | 18605 | 18085 | 19416 | 19349 | 18769 | 18674 | 18864 | 1,01 | 0,015 | 0,367 | 0,713983 | 0,747396 | 0,000986 | 0,002107 | 0,001452 | 0,001452 | TRUE | 0,0138626 | No |
| orf19.7060 | 1566 | 1322 | 1503 | 908 | 451 | 348 | 451 | 521 | 1426 | 1337 | 1422 | 1364 | 401 | 378 | 462 | 397 | 898,44 | 1387 | 410 | 0,295 | -1,762 | -26,663 | 1,27E-156 | 1,73E-155 | 0,0014116 | 0,003507 | 0,0026683 | 0,0026683 | TRUE | 0,080441 | No |
| orf19.7061 | 150 | 107 | 141 | 84 | 96 | 60 | 58 | 96 | 137 | 108 | 133 | 126 | 85 | 65 | 59 | 73 | 98,45 | 126 | 70 | 0,564 | -0,827 | -5,357 | 8,44E-08 | 1,46E-07 | 0,0060002 | 0,015455 | 0,012083 | 0,012083 | TRUE | 0,1751829 | No |
| orf19.7062 | 14493 | 12405 | 13546 | 8571 | 22285 | 18333 | 20038 | 27502 | 13195 | 12546 | 12820 | 12875 | 19822 | 19905 | 20521 | 20979 | 16582,9 | 12859 | 20307 | 1,579 | 0,659 | 19,765 | 5,98E-87 | 4,59E-86 | 0,0004912 | 0,002117 | 0,0010035 | 0,0010035 | TRUE | 0,0061355 | No |
| orf19.7063 | 709 | 580 | 624 | 429 | 862 | 696 | 714 | 1051 | 646 | 587 | 591 | 644 | 767 | 756 | 731 | 802 | 690,3 | 617 | 764 | 1,242 | 0,313 | 4,92 | 8,64E-07 | 1,44E-06 | 0,0006799 | 0,003951 | 0,002402 | 0,002402 | TRUE | 0,0140522 | No |
| orf19.7064 | 7770 | 6871 | 7364 | 4869 | 8533 | 6858 | 7372 | 10053 | 7074 | 6949 | 6969 | 7314 | 7590 | 7446 | 7550 | 7669 | 7320,12 | 7076 | 7564 | 1,07 | 0,097 | 3,068 | 0,002152 | 0,002962 | 0,0001725 | 0,002217 | 0,0008237 | 0,0008237 | TRUE | 0,0058554 | No |
| orf19.7065 | 746 | 654 | 647 | 378 | 523 | 349 | 366 | 530 | 679 | 661 | 612 | 568 | 465 | 379 | 375 | 404 | 518 | 630 | 406 | 0,643 | -0,637 | -7,336 | 2,20E-13 | 4,65E-13 | 0,0057585 | 0,004587 | 0,0052307 | 0,0052307 | TRUE | 0,1124328 | No |

| | | | | | | | | | | | | | | | | | | | | | | | | | | | | | | | |
|---|---|---|---|---|---|---|---|---|---|---|---|---|---|---|---|---|---|---|---|---|---|---|---|---|---|---|---|---|---|---|---|
| orf19.7067 | 2841 | 2562 | 2777 | 1720 | 3938 | 3263 | 3372 | 4615 | 2587 | 2591 | 2628 | 2584 | 3503 | 3543 | 3453 | 3520 | 3051,1 | 2598 | 3505 | 1,349 | 0,432 | 12,569 | 3,14E-36 | 1,15E-35 | 1,00E-08 | 0,00247 | 0,0007945 | 0,0007945 | TRUE | 0,0011478 | No |
| orf19.7068 | 486 | 389 | 438 | 293 | 863 | 736 | 726 | 1058 | 442 | 393 | 415 | 440 | 768 | 799 | 744 | 807 | 600,98 | 422 | 780 | 1,848 | 0,886 | 13,158 | 1,53E-39 | 5,93E-39 | 1,00E-08 | 0,004235 | 0,0024892 | 0,0024892 | TRUE | 0,0251108 | No |
| orf19.7069 | 3156 | 2370 | 2554 | 1822 | 5925 | 4534 | 5131 | 7278 | 2873 | 2397 | 2417 | 2737 | 5270 | 4923 | 5255 | 5552 | 3927,97 | 2606 | 5250 | 2,016 | 1,011 | 15,553 | 1,52E-54 | 7,61E-54 | 0,0049409 | 0,002373 | 0,0037645 | 0,0037645 | TRUE | 0,0600759 | No |
| orf19.707 | 1045 | 889 | 956 | 619 | 452 | 416 | 410 | 568 | 951 | 899 | 905 | 930 | 402 | 452 | 420 | 433 | 674 | 921 | 427 | 0,463 | -1,111 | -17,698 | 4,31E-70 | 2,71E-69 | 1,00E-08 | 0,003997 | 0,002113 | 0,002113 | TRUE | 0,0178354 | No |
| orf19.7070 | 6 | 8 | 10 | 4 | 15 | 9 | 3 | 12 | 5 | 8 | 9 | 6 | 13 | 10 | 3 | 9 | 8,05 | 7 | 9 | 1,218 | 0,285 | 0,555 | 0,579115 | 0,619544 | 1,00E-08 | 0,166222 | 0,1276163 | 0,1276163 | TRUE | 0,426729 | No |
| orf19.7071 | 11 | 10 | 5 | 8 | 6 | 13 | 9 | 7 | 10 | 10 | 5 | 12 | 5 | 14 | 9 | 5 | 8,86 | 9 | 8 | 0,916 | -0,126 | -0,254 | 0,799685 | 0,826121 | 0,0721225 | 0,151121 | 0,1231063 | 0,1231063 | TRUE | 0,5645987 | No |
| orf19.7072 | 7753 | 6593 | 7525 | 4759 | 5277 | 4597 | 4718 | 6410 | 7059 | 6668 | 7122 | 7149 | 4694 | 4991 | 4832 | 4890 | 5925,45 | 7000 | 4852 | 0,693 | -0,529 | -13,812 | 2,15E-43 | 8,92E-43 | 0,0006067 | 0,00226 | 0,0012334 | 0,0012334 | TRUE | 0,0124546 | No |
| orf19.7073 | 229 | 251 | 231 | 153 | 390 | 262 | 300 | 370 | 208 | 254 | 219 | 230 | 347 | 284 | 307 | 282 | 266,45 | 228 | 305 | 1,343 | 0,425 | 4,141 | 3,46E-05 | 5,32E-05 | 0,0052819 | 0,006995 | 0,0062411 | 0,0062411 | TRUE | 0,0990029 | No |
| orf19.7074 | 867 | 796 | 804 | 538 | 800 | 666 | 655 | 945 | 789 | 805 | 761 | 808 | 712 | 723 | 671 | 721 | 748,73 | 791 | 707 | 0,895 | -0,16 | -2,775 | 0,005528 | 0,007389 | 1,00E-08 | 0,003801 | 0,001858 | 0,001858 | TRUE | 0,0131552 | No |
| orf19.7076 | 5436 | 4698 | 5131 | 2995 | 5417 | 4106 | 4355 | 6145 | 4949 | 4752 | 4856 | 4499 | 4818 | 4458 | 4460 | 4687 | 4684,94 | 4764 | 4606 | 0,966 | -0,049 | -1,103 | 0,269857 | 0,305946 | 0,001292 | 0,002319 | 0,0017083 | 0,0017083 | TRUE | 0,0163241 | No |
| orf19.7077 | 2406 | 2040 | 2245 | 1513 | 4800 | 3980 | 4214 | 6046 | 2191 | 2063 | 2125 | 2273 | 4269 | 4321 | 4316 | 4612 | 3271,19 | 2163 | 4380 | 2,028 | 1,02 | 22,377 | 6,62E-111 | 6,28E-110 | 0,0010424 | 0,002441 | 0,00164 | 0,00164 | TRUE | 0,0159301 | No |
| orf19.7078 | 565 | 443 | 464 | 275 | 165 | 106 | 117 | 170 | 514 | 448 | 439 | 413 | 147 | 115 | 120 | 130 | 290,75 | 454 | 128 | 0,282 | -1,826 | -17,002 | 7,90E-65 | 4,63E-64 | 0,0058322 | 0,00658 | 0,0062725 | 0,0062725 | TRUE | 0,1013324 | No |
| orf19.7079 | 165 | 140 | 187 | 128 | 331 | 264 | 294 | 442 | 150 | 142 | 177 | 192 | 294 | 287 | 301 | 337 | 235,05 | 165 | 305 | 1,864 | 0,898 | 8,114 | 4,90E-16 | 1,11E-15 | 0,0058958 | 0,007657 | 0,0069882 | 0,0069882 | TRUE | 0,1020509 | No |
| orf19.708 | 399 | 360 | 426 | 226 | 175 | 143 | 153 | 204 | 363 | 364 | 403 | 339 | 156 | 155 | 157 | 156 | 261,66 | 367 | 156 | 0,422 | -1,243 | -13,115 | 2,71E-39 | 1,04E-38 | 1,00E-08 | 0,007086 | 0,0041964 | 0,0041964 | TRUE | 0,0491192 | No |
| orf19.7080 | 788 | 682 | 644 | 579 | 839 | 722 | 744 | 1099 | 717 | 690 | 609 | 870 | 746 | 784 | 762 | 838 | 752,11 | 722 | 782 | 1,09 | 0,124 | 1,342 | 0,179469 | 0,208666 | 0,0107309 | 0,003793 | 0,006869 | 0,006869 | TRUE | 0,2098164 | No |
| orf19.7081 | 18219 | 14872 | 16153 | 11126 | 9381 | 8126 | 8531 | 11557 | 16588 | 15041 | 15287 | 16713 | 8344 | 8823 | 8737 | 8816 | 12293,6 | 15907 | 8680 | 0,546 | -0,874 | -19,19 | 4,48E-82 | 3,26E-81 | 0,0017032 | 0,002144 | 0,0019036 | 0,0019036 | TRUE | 0,0164017 | No |
| orf19.7082 | 114 | 97 | 107 | 78 | 77 | 58 | 75 | 101 | 104 | 98 | 101 | 117 | 68 | 63 | 77 | 77 | 88,21 | 105 | 71 | 0,685 | -0,545 | -3,65 | 0,000262 | 0,000384 | 1,00E-08 | 0,017013 | 0,0097944 | 0,0097944 | TRUE | 0,0502695 | No |
| orf19.7083 | 78 | 82 | 61 | 65 | 84 | 44 | 61 | 87 | 71 | 83 | 58 | 98 | 75 | 48 | 62 | 66 | 70,08 | 78 | 63 | 0,828 | -0,273 | -1,394 | 0,163406 | 0,191421 | 0,0250578 | 0,020887 | 0,0224536 | 0,0224536 | TRUE | 0,1908058 | No |
| orf19.7084 | 19530 | 16783 | 18836 | 11159 | 4754 | 3835 | 4561 | 5936 | 17781 | 16974 | 17827 | 16763 | 4229 | 4164 | 4671 | 4528 | 10867 | 17336 | 4398 | 0,254 | -1,979 | -42,725 | 0 | 0 | 0,0018167 | 0,002159 | 0,0019233 | 0,0019233 | TRUE | 0,0208866 | No |
| orf19.7085 | 13245 | 11224 | 12349 | 8677 | 1133 | 937 | 957 | 1335 | 12059 | 11352 | 11687 | 13034 | 1008 | 1017 | 980 | 1018 | 6519,48 | 12033 | 1006 | 0,084 | -3,58 | -68,339 | 0 | 0 | 0,0020524 | 0,00224 | 0,0021261 | 0,0021261 | TRUE | 0,0377589 | No |
| orf19.7086 | 7339 | 6452 | 6870 | 4375 | 800 | 690 | 700 | 910 | 6682 | 6526 | 6502 | 6572 | 712 | 749 | 717 | 694 | 3644,11 | 6570 | 718 | 0,109 | -3,197 | -75,208 | 0 | 0 | 1,00E-08 | 0,0024 | 0,001004 | 0,001004 | TRUE | 0,0095601 | No |
| orf19.7088 | 3252 | 2623 | 2987 | 1920 | 2890 | 2434 | 2593 | 3519 | 2961 | 2653 | 2827 | 2884 | 2571 | 2643 | 2656 | 2684 | 2734,73 | 2831 | 2638 | 0,932 | -0,101 | -2,265 | 0,023505 | 0,029969 | 0,0009208 | 0,00252 | 0,0015598 | 0,0015598 | TRUE | 0,0222045 | No |
| orf19.7089 | 1800 | 1515 | 1735 | 1065 | 2946 | 2549 | 2807 | 3717 | 1639 | 1532 | 1642 | 1600 | 2620 | 2768 | 2875 | 2835 | 2188,86 | 1603 | 2774 | 1,73 | 0,791 | 16,634 | 3,97E-62 | 2,24E-61 | 0,0009817 | 0,002641 | 0,001666 | 0,001666 | TRUE | 0,0172165 | No |
| orf19.709 | 4441 | 3740 | 3988 | 2446 | 4142 | 3208 | 3348 | 4538 | 4043 | 3783 | 3774 | 3674 | 3684 | 3483 | 3429 | 3462 | 3666,51 | 3818 | 3514 | 0,92 | -0,121 | -2,731 | 0,006309 | 0,008403 | 0,0010271 | 0,002397 | 0,0016033 | 0,0016033 | TRUE | 0,0193574 | No |
| orf19.7091 | 999 | 744 | 874 | 660 | 931 | 850 | 794 | 1156 | 910 | 752 | 827 | 991 | 828 | 923 | 813 | 882 | 865,82 | 870 | 862 | 0,993 | -0,011 | -0,13 | 0,896857 | 0,911586 | 0,0072462 | 0,003563 | 0,0053395 | 0,0053395 | TRUE | 0,0972958 | No |
| orf19.7092 | 1665 | 1359 | 1464 | 972 | 1686 | 1359 | 1451 | 1945 | 1516 | 1374 | 1386 | 1460 | 1500 | 1475 | 1486 | 1484 | 1460,11 | 1434 | 1486 | 1,037 | 0,052 | 1,074 | 0,282759 | 0,319325 | 0,0004376 | 0,002942 | 0,0015755 | 0,0015755 | TRUE | 0,0196782 | No |
| orf19.7093 | 333 | 282 | 316 | 190 | 503 | 387 | 419 | 561 | 303 | 285 | 299 | 285 | 447 | 420 | 429 | 428 | 362,19 | 293 | 431 | 1,468 | 0,554 | 7,241 | 4,46E-13 | 9,38E-13 | 1,00E-08 | 0,005684 | 0,0027268 | 0,0027268 | TRUE | 0,0075513 | No |
| orf19.7094 | 12 | 16 | 15 | 10 | 118 | 75 | 85 | 122 | 11 | 16 | 14 | 15 | 105 | 81 | 87 | 93 | 52,85 | 14 | 92 | 6,591 | 2,72 | 10,841 | 2,20E-27 | 6,66E-27 | 1,00E-08 | 0,027031 | 0,0175332 | 0,0175332 | TRUE | 0,121386 | No |
| orf19.7095 | 2618 | 2072 | 2335 | 1401 | 2214 | 1734 | 2078 | 2901 | 2384 | 2096 | 2210 | 2105 | 1969 | 1883 | 2128 | 2213 | 2123,32 | 2199 | 2048 | 0,932 | -0,102 | -1,607 | 0,107992 | 0,129221 | 0,0038945 | 0,002659 | 0,00342 | 0,00342 | TRUE | 0,0395107 | No |
| orf19.7096 | 3187 | 2767 | 2979 | 1850 | 1970 | 1611 | 1796 | 2427 | 2902 | 2799 | 2819 | 2779 | 1752 | 1749 | 1839 | 1851 | 2311,31 | 2825 | 1798 | 0,637 | -0,652 | -15,767 | 5,23E-56 | 2,68E-55 | 0,0001731 | 0,002609 | 0,0011908 | 0,0011908 | TRUE | 0,0051497 | No |
| orf19.7097 | 2445 | 1923 | 2161 | 1366 | 2999 | 2224 | 2494 | 3402 | 2226 | 1945 | 2045 | 2052 | 2668 | 2415 | 2554 | 2595 | 2312,44 | 2067 | 2558 | 1,237 | 0,307 | 5,778 | 7,55E-09 | 1,37E-08 | 0,0018771 | 0,002608 | 0,0022742 | 0,0022742 | TRUE | 0,0337852 | No |
| orf19.7098 | 1507 | 1291 | 1447 | 869 | 2842 | 2120 | 2466 | 3381 | 1372 | 1306 | 1369 | 1305 | 2528 | 2302 | 2525 | 2579 | 1910,84 | 1338 | 2484 | 1,855 | 0,892 | 17,542 | 6,89E-69 | 4,25E-68 | 0,0012713 | 0,002728 | 0,0018921 | 0,0018921 | TRUE | 0,028753 | No |
| orf19.71 | 1203 | 1035 | 1156 | 762 | 651 | 621 | 539 | 778 | 1095 | 1047 | 1094 | 1145 | 579 | 674 | 552 | 593 | 847,44 | 1095 | 600 | 0,547 | -0,869 | -12,968 | 1,85E-38 | 7,03E-38 | 0,0023601 | 0,003596 | 0,0030415 | 0,0030415 | TRUE | 0,0770283 | No |
| orf19.7100 | 190 | 195 | 194 | 133 | 687 | 549 | 593 | 769 | 173 | 197 | 184 | 200 | 611 | 596 | 607 | 587 | 394,33 | 188 | 600 | 3,203 | 1,679 | 20,542 | 9,08E-94 | 7,33E-93 | 1,00E-08 | 0,005387 | 0,0028091 | 0,0028091 | TRUE | 0,0345944 | No |
| orf19.7101 | 445 | 431 | 438 | 252 | 467 | 412 | 432 | 625 | 405 | 436 | 415 | 379 | 415 | 447 | 442 | 477 | 427 | 409 | 445 | 1,088 | 0,122 | 1,584 | 0,113266 | 0,135144 | 0,0010865 | 0,005131 | 0,0033183 | 0,0033183 | TRUE | 0,0288606 | No |
| orf19.7102 | 377 | 304 | 310 | 235 | 624 | 486 | 566 | 744 | 343 | 307 | 293 | 353 | 555 | 528 | 580 | 568 | 440,87 | 324 | 558 | 1,728 | 0,789 | 10 | 1,53E-23 | 4,23E-23 | 0,0010344 | 0,005033 | 0,0034843 | 0,0034843 | TRUE | 0,0476499 | No |
| orf19.7103 | 1256 | 1056 | 1156 | 736 | 1070 | 812 | 911 | 1271 | 1144 | 1068 | 1094 | 1106 | 952 | 882 | 933 | 970 | 1018,38 | 1103 | 934 | 0,848 | -0,238 | -4,425 | 9,66E-06 | 1,53E-05 | 1,00E-08 | 0,003334 | 0,0017889 | 0,0017889 | TRUE | 0,016272 | No |
| orf19.7104 | 712 | 534 | 581 | 416 | 708 | 601 | 645 | 859 | 648 | 540 | 550 | 625 | 630 | 653 | 661 | 655 | 620,15 | 591 | 650 | 1,101 | 0,139 | 1,91 | 0,056142 | 0,069255 | 0,0026208 | 0,004167 | 0,003445 | 0,003445 | TRUE | 0,0575289 | No |
| orf19.7105 | 848 | 766 | 907 | 628 | 394 | 430 | 414 | 582 | 772 | 775 | 858 | 943 | 350 | 467 | 424 | 444 | 629,22 | 837 | 421 | 0,504 | -0,988 | -10,442 | 1,60E-25 | 4,64E-25 | 0,0094961 | 0,004136 | 0,0068486 | 0,0068486 | TRUE | 0,1509891 | No |
| orf19.7106 | 590 | 467 | 561 | 342 | 967 | 820 | 877 | 1228 | 537 | 472 | 531 | 514 | 860 | 890 | 898 | 937 | 704,93 | 514 | 896 | 1,745 | 0,803 | 12,469 | 1,11E-35 | 4,00E-35 | 0,0007353 | 0,003911 | 0,0024246 | 0,0024246 | TRUE | 0,0362782 | No |
| orf19.7107 | 6998 | 5489 | 6247 | 4540 | 10126 | 8157 | 8533 | 12073 | 6371 | 5552 | 5912 | 6820 | 9007 | 8856 | 8739 | 9209 | 7558,27 | 6164 | 8953 | 1,453 | 0,54 | 9,109 | 8,31E-20 | 2,10E-19 | 0,004074 | 0,002212 | 0,0032299 | 0,0032299 | TRUE | 0,0613373 | No |
| orf19.7108 | 2356 | 1904 | 1896 | 1429 | 3797 | 3093 | 3309 | 4579 | 2145 | 1926 | 1794 | 2147 | 3377 | 3358 | 3389 | 3493 | 2703,61 | 2003 | 3404 | 1,702 | 0,767 | 12,947 | 2,43E-38 | 9,23E-38 | 0,0033797 | 0,002526 | 0,0029646 | 0,0029646 | TRUE | 0,0602924 | No |
| orf19.7109 | 770 | 590 | 648 | 479 | 1207 | 955 | 986 | 1345 | 701 | 597 | 613 | 720 | 1074 | 1037 | 1010 | 1026 | 847,1 | 658 | 1037 | 1,581 | 0,661 | 9,693 | 3,23E-22 | 8,61E-22 | 0,0027133 | 0,003596 | 0,0031966 | 0,0031966 | TRUE | 0,0446872 | No |
| orf19.711 | 23492 | 19694 | 21152 | 14136 | 9379 | 7811 | 8161 | 11439 | 21388 | 19918 | 20018 | 21235 | 8342 | 8481 | 8358 | 8726 | 14558,3 | 20640 | 8477 | 0,411 | -1,284 | -33,976 | 5,06E-253 | 1,07E-251 | 0,0008199 | 0,002128 | 0,0012896 | 0,0012896 | TRUE | 0,0074251 | No |
| orf19.7110 | 1562 | 1283 | 1452 | 1093 | 1371 | 1207 | 1184 | 1804 | 1422 | 1298 | 1374 | 1642 | 1219 | 1310 | 1213 | 1376 | 1356,8 | 1434 | 1280 | 0,895 | -0,16 | -2,158 | 0,03094 | 0,039073 | 0,0061956 | 0,003011 | 0,0045492 | 0,0045492 | TRUE | 0,1081699 | No |
| orf19.7111 | 1238 | 933 | 1075 | 696 | 1379 | 1033 | 1150 | 1595 | 1127 | 944 | 1017 | 1046 | 1227 | 1122 | 1178 | 1217 | 1109,53 | 1034 | 1186 | 1,148 | 0,199 | 3,158 | 0,001588 | 0,002212 | 0,0026037 | 0,003228 | 0,0029044 | 0,0029044 | TRUE | 0,047783 | No |
| orf19.7111.1 | 4858 | 3765 | 4106 | 3008 | 3225 | 2490 | 2673 | 3981 | 4423 | 3808 | 3886 | 4518 | 2869 | 2703 | 2737 | 3037 | 3497,7 | 4159 | 2836 | 0,683 | -0,551 | -8,413 | 4,00E-17 | 9,39E-17 | 0,0048495 | 0,002415 | 0,0038179 | 0,0038179 | TRUE | 0,0401494 | No |
| orf19.7112 | 382 | 357 | 340 | 235 | 659 | 527 | 527 | 749 | 348 | 361 | 322 | 353 | 586 | 572 | 540 | 571 | 456,63 | 346 | 567 | 1,644 | 0,717 | 9,906 | 3,93E-23 | 1,07E-22 | 1,00E-08 | 0,00493 | 0,0026665 | 0,0026665 | TRUE | 0,026947 | No |
| orf19.7114 | 1733 | 1464 | 1472 | 1278 | 1258 | 1339 | 1182 | 1865 | 1578 | 1481 | 1393 | 1920 | 1119 | 1454 | 1211 | 1423 | 1447,16 | 1593 | 1302 | 0,818 | -0,289 | -2,795 | 0,005191 | 0,006957 | 0,0173707 | 0,00295 | 0,0095783 | 0,0095783 | TRUE | 0,223641 | No |
| orf19.7115 | 2911 | 2417 | 2780 | 2002 | 2516 | 2025 | 2288 | 3201 | 2650 | 2445 | 2631 | 3007 | 2238 | 2199 | 2343 | 2442 | 2494,33 | 2683 | 2306 | 0,861 | -0,216 | -3,379 | 0,000727 | 0,001036 | 0,0044292 | 0,002567 | 0,0035343 | 0,0035343 | TRUE | 0,076959 | No |
| orf19.7116 | 2166 | 1637 | 1901 | 1097 | 3861 | 2822 | 3713 | 4643 | 1972 | 1656 | 1799 | 1648 | 3434 | 3064 | 3803 | 3542 | 2614,64 | 1769 | 3461 | 1,955 | 0,967 | 12,899 | 4,54E-38 | 1,71E-37 | 0,0070606 | 0,002542 | 0,0049567 | 0,0049567 | TRUE | 0,0730928 | No |
| orf19.7118 | 1157 | 897 | 936 | 574 | 2103 | 1649 | 1942 | 2497 | 1053 | 907 | 886 | 862 | 1871 | 1790 | 1989 | 1905 | 1407,9 | 927 | 1889 | 2,032 | 1,023 | 15,319 | 5,67E-53 | 2,76E-52 | 0,0041909 | 0,002975 | 0,0034554 | 0,0034554 | TRUE | 0,1031007 | No |
| orf19.7119 | 7380 | 5735 | 6526 | 3678 | 4960 | 3746 | 4806 | 6713 | 6719 | 5800 | 6176 | 5525 | 4412 | 4067 | 4922 | 5121 | 5342,79 | 6055 | 4630 | 0,765 | -0,387 | -4,954 | 7,27E-07 | 1,22E-06 | 0,0087577 | 0,002284 | 0,0056792 | 0,0056792 | TRUE | 0,0813203 | No |
| orf19.712 | 476 | 534 | 521 | 341 | 762 | 676 | 703 | 873 | 433 | 540 | 493 | 512 | 678 | 734 | 720 | 666 | 597,05 | 494 | 700 | 1,415 | 0,501 | 6,532 | 6,47E-11 | 1,26E-10 | 0,0034845 | 0,00425 | 0,0038962 | 0,0038962 | TRUE | 0,0866909 | No |
| orf19.7121 | 881 | 646 | 733 | 483 | 1330 | 996 | 1171 | 1623 | 802 | 653 | 694 | 726 | 1183 | 1081 | 1199 | 1238 | 947,05 | 719 | 1175 | 1,636 | 0,711 | 9,869 | 5,65E-23 | 1,54E-22 | 0,0039597 | 0,003432 | 0,0038314 | 0,0038314 | TRUE | 0,0777088 | No |
| orf19.7123 | 3697 | 2947 | 3330 | 2656 | 2331 | 1903 | 1967 | 2840 | 3366 | 2981 | 3152 | 3990 | 2073 | 2066 | 2014 | 2166 | 2726,02 | 3372 | 2080 | 0,618 | -0,695 | -8,889 | 6,19E-19 | 1,53E-18 | 0,0084962 | 0,002522 | 0,0054858 | 0,0054858 | TRUE | 0,1809015 | No |
| orf19.7124 | 16015 | 12375 | 13844 | 8237 | 5251 | 3935 | 4585 | 6253 | 14581 | 12516 | 13102 | 12373 | 4671 | 4272 | 4696 | 4770 | 8872,59 | 13143 | 4602 | 0,35 | -1,514 | -25,618 | 9,65E-145 | 1,22E-143 | 0,0039351 | 0,002186 | 0,0032122 | 0,0032122 | TRUE | 0,0665364 | No |
| orf19.7125 | 1421 | 1136 | 1285 | 738 | 1553 | 1289 | 1573 | 2011 | 1294 | 1149 | 1216 | 1109 | 1381 | 1399 | 1611 | 1534 | 1336,65 | 1192 | 1481 | 1,241 | 0,311 | 4,583 | 4,58E-06 | 7,40E-06 | 0,004062 | 0,003025 | 0,00366 | 0,00366 | TRUE | 0,0406649 | No |
| orf19.7127 | 26 | 19 | 28 | 16 | 89 | 68 | 69 | 104 | 24 | 19 | 26 | 24 | 79 | 74 | 71 | 79 | 49,55 | 23 | 76 | 3,251 | 1,701 | 7,893 | 2,96E-15 | 6,59E-15 | 1,00E-08 | 0,028696 | 0,0158014 | 0,0158014 | TRUE | 0,0887459 | No |
| orf19.7127.1 | 58 | 47 | 40 | 35 | 249 | 181 | 284 | 358 | 53 | 48 | 38 | 53 | 221 | 197 | 291 | 273 | 146,59 | 48 | 246 | 5,187 | 2,375 | 13,785 | 3,13E-43 | 1,29E-42 | 0,0214916 | 0,011049 | 0,0153783 | 0,0153783 | TRUE | 0,0815132 | No |
| orf19.7128 | 436 | 333 | 356 | 259 | 625 | 461 | 526 | 813 | 397 | 337 | 337 | 389 | 556 | 501 | 539 | 620 | 459,38 | 365 | 554 | 1,524 | 0,608 | 6,784 | 1,17E-11 | 2,35E-11 | 0,0059629 | 0,004913 | 0,0054038 | 0,0054038 | TRUE | 0,0794543 | No |
| orf19.713 | 3530 | 3025 | 3351 | 2260 | 1190 | 881 | 903 | 1259 | 3214 | 3059 | 3171 | 3395 | 1058 | 957 | 925 | 960 | 2092,48 | 3210 | 975 | 0,304 | -1,717 | -31,353 | 8,81E-216 | 1,60E-214 | 0,0018131 | 0,002668 | 0,0022325 | 0,0022325 | TRUE | 0,0400769 | No |
| orf19.7130 | 41 | 42 | 33 | 30 | 115 | 62 | 94 | 137 | 37 | 42 | 31 | 45 | 102 | 67 | 96 | 105 | 65,81 | 39 | 92 | 2,413 | 1,271 | 6,291 | 3,16E-10 | 6,02E-10 | 0,0175451 | 0,02211 | 0,0203828 | 0,0203828 | TRUE | 0,3033091 | No |
| orf19.7131 | 582 | 443 | 543 | 407 | 943 | 802 | 955 | 1327 | 530 | 448 | 514 | 611 | 839 | 871 | 978 | 1012 | 725,38 | 526 | 925 | 1,767 | 0,822 | 8,876 | 6,91E-19 | 1,70E-18 | 0,0102192 | 0,003858 | 0,0066931 | 0,0066931 | TRUE | 0,1260682 | No |
| orf19.7136 | 1671 | 1298 | 1486 | 935 | 5068 | 4059 | 4963 | 6627 | 1521 | 1313 | 1406 | 1405 | 4508 | 4407 | 5083 | 5055 | 3087,21 | 1411 | 4763 | 3,374 | 1,755 | 27,28 | 7,32E-164 | 1,05E-162 | 0,0040071 | 0,002465 | 0,0034928 | 0,0034928 | TRUE | 0,0345497 | No |
| orf19.7139 | 657 | 473 | 497 | 369 | 1042 | 904 | 882 | 1353 | 598 | 478 | 470 | 554 | 927 | 981 | 903 | 1032 | 743,11 | 525 | 961 | 1,832 | 0,874 | 10,438 | 1,67E-25 | 4,84E-25 | 0,0067039 | 0,003815 | 0,0052173 | 0,0052173 | TRUE | 0,1125818 | No |
| orf19.714 | 497 | 499 | 510 | 341 | 655 | 557 | 587 | 774 | 452 | 505 | 483 | 512 | 583 | 605 | 601 | 590 | 541,38 | 488 | 595 | 1,222 | 0,289 | 4,345 | 1,39E-05 | 2,20E-05 | 1,00E-08 | 0,004477 | 0,0023741 | 0,0023741 | TRUE | 0,0292336 | No |

| | | | | | | | | | | | | | | | | | | | | | | | | | | | | | | | |
|---|---|---|---|---|---|---|---|---|---|---|---|---|---|---|---|---|---|---|---|---|---|---|---|---|---|---|---|---|---|---|---|
| orf19.7140 | 1777 | 1422 | 1599 | 1073 | 1471 | 1120 | 1222 | 1697 | 1618 | 1438 | 1513 | 1612 | 1308 | 1216 | 1251 | 1294 | 1406,45 | 1545 | 1267 | 0,822 | -0,283 | -5,198 | 2,01E-07 | 3,44E-07 | 0,0014204 | 0,002976 | 0,0021345 | 0,0021345 | TRUE | 0,0263058 | No |
| orf19.7141 | 3192 | 2519 | 2783 | 1726 | 2420 | 1873 | 2329 | 3042 | 2906 | 2548 | 2634 | 2593 | 2153 | 2034 | 2385 | 2320 | 2446,52 | 2670 | 2223 | 0,832 | -0,265 | -4,237 | 2,27E-05 | 3,53E-05 | 0,0038599 | 0,002577 | 0,0033361 | 0,0033361 | TRUE | 0,0437623 | No |
| orf19.7144 | 4068 | 3368 | 3790 | 2468 | 4546 | 3631 | 4311 | 5704 | 3704 | 3406 | 3587 | 3707 | 4044 | 3942 | 4415 | 4351 | 3894,52 | 3601 | 4188 | 1,164 | 0,219 | 4,334 | 1,46E-05 | 2,31E-05 | 0,0019406 | 0,002376 | 0,0021817 | 0,0021817 | TRUE | 0,0186849 | No |
| orf19.7148 | 201 | 162 | 167 | 111 | 275 | 197 | 219 | 313 | 183 | 164 | 158 | 167 | 245 | 214 | 224 | 239 | 199,15 | 168 | 230 | 1,374 | 0,458 | 4,43 | 9,42E-06 | 1,50E-05 | 1,00E-08 | 0,00867 | 0,0050983 | 0,0050983 | TRUE | 0,0444137 | No |
| orf19.7149 | 921 | 776 | 818 | 597 | 3767 | 2964 | 3357 | 4503 | 839 | 785 | 774 | 897 | 3351 | 3218 | 3438 | 3435 | 2091,99 | 824 | 3360 | 4,097 | 2,035 | 37,943 | 0 | 0 | 0,0011968 | 0,002668 | 0,001969 | 0,001969 | TRUE | 0,0362486 | No |
| orf19.715 | 1894 | 1897 | 1908 | 1257 | 524 | 458 | 373 | 554 | 1724 | 1919 | 1806 | 1888 | 466 | 497 | 382 | 423 | 1138,11 | 1834 | 442 | 0,241 | -2,054 | -27,612 | 7,93E-168 | 1,16E-166 | 0,0050067 | 0,003198 | 0,0039613 | 0,0039613 | TRUE | 0,0903521 | No |
| orf19.7150 | 12441 | 10435 | 12253 | 8258 | 5269 | 4605 | 4941 | 6744 | 11327 | 10554 | 11596 | 12405 | 4687 | 5000 | 5060 | 5144 | 8221,63 | 11470 | 4973 | 0,434 | -1,205 | -22,432 | 1,91E-111 | 1,82E-110 | 0,0028309 | 0,002198 | 0,0026323 | 0,0026323 | TRUE | 0,036257 | No |
| orf19.7151 | 3 | 5 | 8 | 3 | 9 | 5 | 7 | 6 | 3 | 5 | 8 | 5 | 8 | 5 | 7 | 5 | 5,63 | 5 | 6 | 1,258 | 0,331 | 0,57 | 0,568509 | 0,609219 | 1,00E-08 | 0,236643 | 0,1421645 | 0,1421645 | TRUE | 0,2931328 | No |
| orf19.7152 | 8504 | 6615 | 7470 | 5170 | 8558 | 6966 | 7567 | 11012 | 7743 | 6690 | 7070 | 7766 | 7612 | 7563 | 7750 | 8400 | 7574,19 | 7317 | 7831 | 1,071 | 0,099 | 1,708 | 0,087595 | 0,105761 | 0,0036515 | 0,002211 | 0,0030666 | 0,0030666 | TRUE | 0,0406155 | No |
| orf19.7153 | 4767 | 4695 | 4888 | 3131 | 5583 | 4798 | 4944 | 6256 | 4340 | 4748 | 4626 | 4703 | 4966 | 5209 | 5063 | 4772 | 4803,57 | 4604 | 5002 | 1,087 | 0,12 | 2,698 | 0,006974 | 0,009259 | 0,0011863 | 0,002312 | 0,0016792 | 0,0016792 | TRUE | 0,018569 | No |
| orf19.7154 | 10386 | 9263 | 10313 | 5981 | 11288 | 8825 | 10030 | 12926 | 9456 | 9369 | 9760 | 8984 | 10040 | 9581 | 10272 | 9860 | 9665,39 | 9392 | 9938 | 1,058 | 0,081 | 2,1 | 0,035707 | 0,044845 | 0,0008715 | 0,002174 | 0,0013218 | 0,0013218 | TRUE | 0,0101248 | No |
| orf19.7156 | 1801 | 1614 | 1753 | 1117 | 34981 | 29255 | 31989 | 44016 | 1640 | 1632 | 1659 | 1678 | 31114 | 31763 | 32761 | 33576 | 16977,9 | 1652 | 32304 | 19,564 | 4,29 | 107,739 | 0 | 0 | 0,0007301 | 0,002115 | 0,0011834 | 0,0011834 | TRUE | 0,0086598 | No |
| orf19.7157 | 143 | 138 | 143 | 111 | 316 | 238 | 257 | 346 | 130 | 140 | 135 | 167 | 281 | 258 | 263 | 264 | 204,81 | 143 | 266 | 1,888 | 0,917 | 8,706 | 3,15E-18 | 7,62E-18 | 3,64E-05 | 0,008487 | 0,0051392 | 0,0051392 | TRUE | 0,1004574 | No |
| orf19.7158 | 1533 | 1302 | 1457 | 896 | 865 | 804 | 839 | 1201 | 1396 | 1317 | 1379 | 1346 | 769 | 873 | 859 | 916 | 1106,89 | 1360 | 854 | 0,629 | -0,67 | -11,027 | 2,84E-28 | 8,76E-28 | 0,0019986 | 0,00323 | 0,0026029 | 0,0026029 | TRUE | 0,0553519 | No |
| orf19.7159 | 935 | 822 | 977 | 601 | 1666 | 1507 | 1441 | 1961 | 851 | 831 | 925 | 903 | 1482 | 1636 | 1476 | 1496 | 1199,97 | 878 | 1522 | 1,735 | 0,795 | 13,69 | 1,17E-42 | 4,78E-42 | 0,0015698 | 0,003138 | 0,0023213 | 0,0023213 | TRUE | 0,0294618 | No |
| orf19.716 | 1044 | 1115 | 1084 | 575 | 261 | 203 | 219 | 290 | 951 | 1128 | 1026 | 864 | 232 | 220 | 224 | 221 | 608,24 | 992 | 224 | 0,226 | -2,147 | -23,734 | 1,61E-124 | 1,71E-123 | 0,0062854 | 0,004209 | 0,0052374 | 0,0052374 | TRUE | 0,0974288 | No |
| orf19.7160 | 3004 | 2454 | 2738 | 1813 | 3521 | 2944 | 2913 | 4147 | 2735 | 2482 | 2591 | 2723 | 3132 | 3196 | 2983 | 3163 | 2875,81 | 2633 | 3118 | 1,185 | 0,245 | 5,336 | 9,50E-08 | 1,64E-07 | 0,0010249 | 0,002496 | 0,0016746 | 0,0016746 | TRUE | 0,0181337 | No |
| orf19.7161 | 9260 | 7413 | 8109 | 4743 | 9159 | 6720 | 7985 | 10664 | 8431 | 7497 | 7674 | 7125 | 8147 | 7296 | 8178 | 8135 | 7810,3 | 7682 | 7939 | 1,033 | 0,047 | 0,803 | 0,421826 | 0,462927 | 0,0035913 | 0,002206 | 0,0031623 | 0,0031623 | TRUE | 0,0527537 | No |
| orf19.7163 | 170 | 127 | 126 | 88 | 382 | 275 | 322 | 467 | 155 | 128 | 119 | 132 | 340 | 299 | 330 | 356 | 232,38 | 134 | 331 | 2,478 | 1,309 | 11,948 | 6,66E-33 | 2,27E-32 | 0,0036387 | 0,007722 | 0,0061504 | 0,0061504 | TRUE | 0,1333182 | No |
| orf19.7164 | 1511 | 1300 | 1494 | 943 | 1116 | 941 | 961 | 1362 | 1376 | 1315 | 1414 | 1417 | 993 | 1022 | 984 | 1039 | 1194,8 | 1380 | 1010 | 0,732 | -0,45 | -8,957 | 3,35E-19 | 8,32E-19 | 1,00E-08 | 0,003143 | 0,0015686 | 0,0015686 | TRUE | 0,012282 | No |
| orf19.7165 | 548 | 493 | 469 | 327 | 371 | 268 | 301 | 390 | 499 | 499 | 444 | 491 | 330 | 291 | 308 | 297 | 394,92 | 483 | 306 | 0,636 | -0,654 | -8,247 | 1,63E-16 | 3,76E-16 | 1,00E-08 | 0,005382 | 0,003406 | 0,003406 | TRUE | 0,037237 | No |
| orf19.7166 | 10195 | 8453 | 9545 | 5909 | 7645 | 5985 | 6947 | 9331 | 9282 | 8549 | 9033 | 8876 | 6800 | 6498 | 7115 | 7118 | 7908,94 | 8935 | 6883 | 0,77 | -0,376 | -8,644 | 5,41E-18 | 1,30E-17 | 0,0013463 | 0,002204 | 0,0016929 | 0,0016929 | TRUE | 0,0170694 | No |
| orf19.7167 | 51 | 30 | 31 | 29 | 28 | 25 | 37 | 42 | 46 | 30 | 29 | 44 | 25 | 27 | 38 | 32 | 33,96 | 37 | 30 | 0,82 | -0,287 | -1,126 | 0,260061 | 0,295769 | 0,0180799 | 0,040939 | 0,0326405 | 0,0326405 | TRUE | 0,2020434 | No |
| orf19.717 | 9116 | 8064 | 8737 | 5169 | 6304 | 4856 | 5065 | 7210 | 8300 | 8156 | 8269 | 7765 | 5607 | 5272 | 5187 | 5500 | 6756,95 | 8122 | 5392 | 0,664 | -0,591 | -14,702 | 6,30E-49 | 2,85E-48 | 0,0009089 | 0,002233 | 0,0014008 | 0,0014008 | TRUE | 0,0103736 | No |
| orf19.7170 | 18 | 6 | 11 | 15 | 11 | 9 | 8 | 7 | 16 | 6 | 10 | 23 | 10 | 10 | 8 | 5 | 11,06 | 14 | 8 | 0,606 | -0,722 | -1,548 | 0,121544 | 0,144641 | 0,0962382 | 0,121466 | 0,1122465 | 0,1122465 | TRUE | 0,1568 | No |
| orf19.7173 | 404 | 313 | 371 | 235 | 860 | 721 | 705 | 933 | 368 | 317 | 351 | 353 | 765 | 783 | 722 | 712 | 546,25 | 347 | 746 | 2,145 | 1,101 | 15,155 | 7,05E-52 | 3,37E-51 | 0,0009531 | 0,004455 | 0,0029054 | 0,0029054 | TRUE | 0,0421066 | No |
| orf19.7175 | 1062 | 858 | 1049 | 613 | 1334 | 1055 | 1185 | 1645 | 967 | 868 | 993 | 921 | 1187 | 1145 | 1214 | 1255 | 1068,59 | 937 | 1200 | 1,28 | 0,357 | 6,057 | 1,39E-09 | 2,59E-09 | 0,0015041 | 0,003273 | 0,002366 | 0,002366 | TRUE | 0,031482 | No |
| orf19.7176 | 797 | 752 | 815 | 511 | 1172 | 930 | 1081 | 1419 | 726 | 761 | 771 | 768 | 1042 | 1010 | 1107 | 1082 | 908,35 | 756 | 1060 | 1,404 | 0,49 | 8,744 | 2,24E-18 | 5,44E-18 | 1,00E-08 | 0,003491 | 0,0018644 | 0,0018644 | TRUE | 0,0116347 | No |
| orf19.7177 | 5185 | 4989 | 5360 | 3372 | 2628 | 2315 | 2231 | 3125 | 4721 | 5046 | 5073 | 5065 | 2338 | 2513 | 2285 | 2384 | 3678,02 | 4976 | 2380 | 0,478 | -1,064 | -23,825 | 1,82E-125 | 1,96E-124 | 0,0010181 | 0,002396 | 0,0016103 | 0,0016103 | TRUE | 0,0166133 | No |
| orf19.7178 | 7268 | 6044 | 6510 | 4373 | 6096 | 4678 | 5143 | 7040 | 6617 | 6113 | 6161 | 6569 | 5422 | 5079 | 5267 | 5370 | 5824,82 | 6365 | 5284 | 0,831 | -0,268 | -6,352 | 2,13E-10 | 4,08E-10 | 0,0010452 | 0,002264 | 0,0015302 | 0,0015302 | TRUE | 0,0090765 | No |
| orf19.7179 | 1606 | 1307 | 1479 | 933 | 1018 | 815 | 1030 | 1268 | 1462 | 1322 | 1400 | 1402 | 905 | 885 | 1055 | 967 | 1174,72 | 1396 | 953 | 0,683 | -0,551 | -8,628 | 6,25E-18 | 1,50E-17 | 0,0028774 | 0,003162 | 0,0030407 | 0,0030407 | TRUE | 0,0592662 | No |
| orf19.718 | 6031 | 6082 | 6381 | 3709 | 6261 | 5134 | 5538 | 6782 | 5491 | 6151 | 6039 | 5572 | 5569 | 5574 | 5672 | 5173 | 5655,1 | 5813 | 5497 | 0,945 | -0,082 | -1,613 | 0,106843 | 0,128031 | 0,0021867 | 0,002271 | 0,0022756 | 0,0022756 | TRUE | 0,0194506 | No |
| orf19.7181 | 878 | 807 | 881 | 605 | 1151 | 1024 | 1008 | 1421 | 799 | 816 | 834 | 909 | 1024 | 1112 | 1032 | 1084 | 951,25 | 840 | 1063 | 1,27 | 0,345 | 5,881 | 4,09E-09 | 7,47E-09 | 0,0010128 | 0,003426 | 0,0022305 | 0,0022305 | TRUE | 0,0318737 | No |
| orf19.7182 | 1881 | 1629 | 1705 | 963 | 2654 | 1989 | 2154 | 2868 | 1713 | 1648 | 1614 | 1447 | 2361 | 2159 | 2206 | 2188 | 1916,77 | 1606 | 2228 | 1,385 | 0,47 | 8,213 | 2,15E-16 | 4,93E-16 | 0,0025984 | 0,002726 | 0,0026003 | 0,0026003 | TRUE | 0,0488663 | No |
| orf19.7183 | 1297 | 1039 | 1114 | 739 | 581 | 451 | 472 | 624 | 1181 | 1051 | 1054 | 1110 | 517 | 490 | 483 | 476 | 795,24 | 1099 | 492 | 0,447 | -1,161 | -18,103 | 3,00E-73 | 1,96E-72 | 0,0014127 | 0,003698 | 0,0025146 | 0,0025146 | TRUE | 0,0325074 | No |
| orf19.7184 | 2812 | 2509 | 2696 | 1952 | 980 | 775 | 720 | 975 | 2560 | 2538 | 2552 | 2932 | 872 | 841 | 737 | 744 | 1721,97 | 2646 | 798 | 0,302 | -1,727 | -24,452 | 4,82E-132 | 5,49E-131 | 0,0051738 | 0,002804 | 0,0040009 | 0,0040009 | TRUE | 0,0623315 | No |
| orf19.7185 | 3018 | 2526 | 2602 | 1645 | 2641 | 2020 | 2275 | 3146 | 2748 | 2555 | 2463 | 2471 | 2349 | 2193 | 2330 | 2400 | 2438,51 | 2559 | 2318 | 0,906 | -0,143 | -2,827 | 0,0047 | 0,006311 | 0,0015181 | 0,002579 | 0,0020461 | 0,0020461 | TRUE | 0,0306198 | No |
| orf19.7186 | 246 | 223 | 272 | 147 | 244 | 197 | 181 | 281 | 224 | 226 | 257 | 221 | 217 | 214 | 185 | 214 | 219,8 | 232 | 208 | 0,894 | -0,161 | -1,596 | 0,110495 | 0,132035 | 1,00E-08 | 0,008047 | 0,005237 | 0,005237 | TRUE | 0,0633701 | No |
| orf19.7187 | 6229 | 5563 | 5660 | 3662 | 6242 | 4833 | 4896 | 6745 | 5671 | 5626 | 5357 | 5501 | 5552 | 5247 | 5014 | 5145 | 5389,23 | 5539 | 5240 | 0,946 | -0,08 | -1,891 | 0,058598 | 0,072144 | 0,0010332 | 0,002282 | 0,0015365 | 0,0015365 | TRUE | 0,0197862 | No |
| orf19.7188 | 2064 | 1674 | 1782 | 1568 | 3576 | 2971 | 2560 | 4153 | 1879 | 1693 | 1687 | 2355 | 3181 | 3226 | 2622 | 3168 | 2476,28 | 1904 | 3049 | 1,605 | 0,682 | 6,818 | 9,21E-12 | 1,86E-11 | 0,0174503 | 0,00257 | 0,0091794 | 0,0091794 | TRUE | 0,3011328 | No |
| orf19.719 | 1294 | 1295 | 1304 | 755 | 801 | 733 | 727 | 923 | 1178 | 1310 | 1234 | 1134 | 712 | 796 | 745 | 704 | 976,63 | 1214 | 739 | 0,607 | -0,721 | -11,238 | 2,65E-29 | 8,38E-29 | 0,0023587 | 0,00339 | 0,0028718 | 0,0028718 | TRUE | 0,0332727 | No |
| orf19.7190 | 584 | 523 | 533 | 346 | 1290 | 1020 | 1089 | 1351 | 532 | 529 | 504 | 520 | 1147 | 1107 | 1115 | 1031 | 810,69 | 521 | 1100 | 2,109 | 1,076 | 17,616 | 1,85E-69 | 1,15E-68 | 1,00E-08 | 0,003666 | 0,0021357 | 0,0021357 | TRUE | 0,0233343 | No |
| orf19.7192 | 18 | 15 | 22 | 18 | 46 | 36 | 32 | 33 | 16 | 15 | 21 | 27 | 41 | 39 | 33 | 25 | 27,17 | 20 | 34 | 1,766 | 0,82 | 2,839 | 0,004526 | 0,006083 | 0,0190547 | 0,050656 | 0,0391741 | 0,0391741 | TRUE | 0,2874453 | No |
| orf19.7193 | 8825 | 7760 | 8425 | 5315 | 7038 | 5858 | 6359 | 8516 | 8035 | 7848 | 7973 | 7984 | 6260 | 6360 | 6512 | 6496 | 7183,68 | 7960 | 6407 | 0,805 | -0,313 | -10,336 | 4,86E-25 | 1,40E-24 | 9,00E-05 | 0,002221 | 0,0007401 | 0,0007401 | TRUE | 0,0029719 | No |
| orf19.7194 | 1541 | 1590 | 1584 | 973 | 1413 | 1061 | 1091 | 1406 | 1403 | 1608 | 1499 | 1462 | 1257 | 1152 | 1117 | 1073 | 1321,31 | 1493 | 1150 | 0,77 | -0,378 | -5,892 | 3,83E-09 | 7,01E-09 | 0,0032999 | 0,003037 | 0,0031778 | 0,0031778 | TRUE | 0,0485363 | No |
| orf19.7195 | 1952 | 1715 | 1796 | 1040 | 3228 | 2564 | 2878 | 3905 | 1777 | 1735 | 1700 | 1562 | 2871 | 2784 | 2947 | 2979 | 2294,37 | 1694 | 2895 | 1,706 | 0,771 | 15,704 | 1,42E-55 | 7,24E-55 | 0,0013361 | 0,002613 | 0,0018348 | 0,0018348 | TRUE | 0,0295385 | No |
| orf19.7196 | 14158 | 12463 | 13205 | 9155 | 29494 | 26233 | 24918 | 35612 | 12890 | 12605 | 12497 | 13752 | 26234 | 28482 | 25519 | 27165 | 19893,1 | 12936 | 26850 | 2,076 | 1,054 | 22,558 | 1,12E-112 | 1,08E-111 | 0,0020258 | 0,002103 | 0,0020378 | 0,0020378 | TRUE | 0,0217257 | No |
| orf19.7197 | 8747 | 9018 | 9558 | 5369 | 11207 | 9055 | 9822 | 12220 | 7964 | 9121 | 9046 | 8065 | 9968 | 9831 | 10059 | 9322 | 9171,92 | 8549 | 9795 | 1,145 | 0,196 | 3,612 | 0,000304 | 0,000444 | 0,0030422 | 0,002181 | 0,0027129 | 0,0027129 | TRUE | 0,0264553 | No |
| orf19.7198 | 1169 | 1116 | 1104 | 674 | 1887 | 1423 | 1569 | 2051 | 1064 | 1129 | 1045 | 1012 | 1678 | 1545 | 1607 | 1565 | 1330,64 | 1062 | 1599 | 1,502 | 0,587 | 11,122 | 9,85E-29 | 3,06E-28 | 0,0008484 | 0,00303 | 0,0018811 | 0,0018811 | TRUE | 0,0205985 | No |
| orf19.7199 | 474 | 533 | 496 | 249 | 859 | 694 | 714 | 878 | 432 | 539 | 469 | 374 | 764 | 753 | 731 | 670 | 591,58 | 454 | 730 | 1,6 | 0,678 | 7,143 | 9,10E-13 | 1,89E-12 | 0,0101761 | 0,00427 | 0,006837 | 0,006837 | TRUE | 0,1816447 | No |
| orf19.720 | 235 | 208 | 244 | 132 | 479 | 390 | 384 | 481 | 214 | 210 | 231 | 198 | 426 | 423 | 393 | 367 | 307,9 | 213 | 402 | 1,874 | 0,906 | 9,925 | 3,23E-23 | 8,84E-23 | 1,00E-08 | 0,006327 | 0,0043581 | 0,0043581 | TRUE | 0,044515 | No |
| orf19.7200 | 377 | 426 | 423 | 270 | 1079 | 918 | 861 | 1097 | 343 | 431 | 400 | 406 | 960 | 997 | 882 | 837 | 656,88 | 395 | 919 | 2,328 | 1,219 | 14,459 | 2,20E-47 | 9,70E-47 | 0,0059826 | 0,004048 | 0,0049585 | 0,0049585 | TRUE | 0,096427 | No |
| orf19.7201 | 3327 | 3383 | 3621 | 2103 | 2217 | 1902 | 1892 | 2453 | 3029 | 3422 | 3427 | 3159 | 1972 | 2065 | 1938 | 1871 | 2610,3 | 3259 | 1962 | 0,601 | -0,734 | -13,443 | 3,38E-41 | 1,34E-40 | 0,0021461 | 0,002543 | 0,0024581 | 0,0024581 | TRUE | 0,0286246 | No |
| orf19.7202 | 298 | 325 | 356 | 173 | 439 | 328 | 380 | 419 | 271 | 329 | 337 | 260 | 390 | 356 | 389 | 320 | 331,52 | 299 | 364 | 1,209 | 0,274 | 2,597 | 0,009395 | 0,012344 | 0,0096285 | 0,006022 | 0,0076126 | 0,0076126 | TRUE | 0,0828056 | No |
| orf19.7203 | 2864 | 2780 | 2863 | 1564 | 2366 | 1801 | 1925 | 2514 | 2608 | 2812 | 2710 | 2349 | 2104 | 1955 | 1971 | 1918 | 2303,4 | 2620 | 1987 | 0,758 | -0,401 | -6,739 | 1,59E-11 | 3,19E-11 | 0,0031173 | 0,002611 | 0,0029528 | 0,0029528 | TRUE | 0,0553591 | No |
| orf19.7204 | 113 | 104 | 104 | 54 | 72 | 39 | 62 | 60 | 103 | 105 | 98 | 81 | 64 | 42 | 63 | 46 | 75,41 | 97 | 54 | 0,552 | -0,857 | -4,81 | 1,51E-06 | 2,49E-06 | 0,0096989 | 0,019555 | 0,0163917 | 0,0163917 | TRUE | 0,1481812 | No |
| orf19.7205 | 1336 | 1225 | 1301 | 832 | 1678 | 1429 | 1426 | 2128 | 1216 | 1239 | 1231 | 1250 | 1493 | 1551 | 1460 | 1623 | 1383,01 | 1234 | 1532 | 1,243 | 0,314 | 6,229 | 4,68E-10 | 8,86E-10 | 0,0007126 | 0,002992 | 0,0016987 | 0,0016987 | TRUE | 0,0206066 | No |
| orf19.7206 | 161 | 121 | 147 | 118 | 130 | 100 | 90 | 155 | 147 | 122 | 139 | 177 | 116 | 109 | 92 | 118 | 127,49 | 146 | 109 | 0,752 | -0,411 | -2,921 | 0,003492 | 0,004723 | 0,0088567 | 0,012398 | 0,0110749 | 0,0110749 | TRUE | 0,1715076 | No |
| orf19.7207 | 4751 | 4363 | 4696 | 2915 | 4183 | 3524 | 3617 | 4957 | 4326 | 4413 | 4444 | 4379 | 3721 | 3826 | 3704 | 3781 | 4074,21 | 4390 | 3758 | 0,856 | -0,224 | -6,855 | 7,13E-12 | 1,44E-11 | 1,00E-08 | 0,002361 | 0,0007825 | 0,0007825 | TRUE | 0,001707 | No |
| orf19.7208 | 16 | 8 | 10 | 3 | 21 | 16 | 21 | 26 | 15 | 8 | 9 | 5 | 19 | 17 | 22 | 20 | 14,25 | 9 | 20 | 2,03 | 1,022 | 2,672 | 0,00755 | 0,009992 | 0,0004008 | 0,094723 | 0,0607341 | 0,0607341 | TRUE | 0,5388694 | No |
| orf19.7209 | 95 | 81 | 89 | 47 | 319 | 255 | 270 | 292 | 86 | 82 | 84 | 71 | 284 | 277 | 277 | 223 | 172,89 | 81 | 265 | 3,24 | 1,696 | 12,945 | 2,51E-38 | 9,52E-38 | 1,00E-08 | 0,009678 | 0,0081994 | 0,0081994 | TRUE | 0,1405591 | No |
| orf19.721 | 2236 | 2360 | 2505 | 1459 | 2295 | 1969 | 2014 | 2575 | 2036 | 2387 | 2371 | 2192 | 2041 | 2138 | 2063 | 1964 | 2148,88 | 2246 | 2052 | 0,913 | -0,132 | -2,246 | 0,024675 | 0,031404 | 0,0027106 | 0,002652 | 0,0028419 | 0,0028419 | TRUE | 0,0504215 | No |
| orf19.7210 | 2910 | 2389 | 2588 | 1939 | 2392 | 2043 | 2043 | 2958 | 2649 | 2416 | 2449 | 2913 | 2128 | 2218 | 2092 | 2256 | 2390,26 | 2607 | 2174 | 0,835 | -0,26 | -4,188 | 2,82E-05 | 4,36E-05 | 0,0038954 | 0,00259 | 0,0032738 | 0,0032738 | TRUE | 0,0723053 | No |
| orf19.7212 | 701 | 603 | 665 | 433 | 396 | 347 | 373 | 498 | 638 | 610 | 629 | 650 | 352 | 377 | 382 | 380 | 502,34 | 632 | 373 | 0,59 | -0,761 | -11,252 | 2,26E-29 | 7,14E-29 | 1,00E-08 | 0,004667 | 0,0022913 | 0,0022913 | TRUE | 0,0162841 | No |
| orf19.7213 | 2203 | 1974 | 2179 | 1360 | 2461 | 2132 | 2278 | 2962 | 2006 | 1996 | 2062 | 2043 | 2189 | 2315 | 2333 | 2259 | 2150,44 | 2027 | 2274 | 1,122 | 0,166 | 4,021 | 5,79E-05 | 8,78E-05 | 1,00E-08 | 0,002651 | 0,0011577 | 0,0011577 | TRUE | 0,0076495 | No |
| orf19.7214 | 2781 | 2356 | 2598 | 1519 | 1585 | 1275 | 1419 | 1941 | 2532 | 2383 | 2459 | 2282 | 1410 | 1384 | 1453 | 1481 | 1922,92 | 2414 | 1432 | 0,593 | -0,755 | -15,592 | 8,30E-55 | 4,18E-54 | 0,0009181 | 0,002724 | 0,0017001 | 0,0017001 | TRUE | 0,0151885 | No |
| orf19.7215 | 7056 | 6549 | 6851 | 4570 | 10993 | 9779 | 9450 | 13211 | 6424 | 6624 | 6484 | 6865 | 9778 | 10617 | 9678 | 10077 | 8318,4 | 6599 | 10038 | 1,522 | 0,606 | 14,536 | 7,15E-48 | 3,18E-47 | 0,001157 | 0,002196 | 0,0015391 | 0,0015391 | TRUE | 0,0182638 | No |

| | | | | | | | | | | | | | | | | | | | | | | | | | | | | | | | |
|---|---|---|---|---|---|---|---|---|---|---|---|---|---|---|---|---|---|---|---|---|---|---|---|---|---|---|---|---|---|---|---|
| orf19.7215.3 | 2742 | 2200 | 2368 | 1428 | 1114 | 795 | 857 | 1286 | 2496 | 2225 | 2241 | 2145 | 991 | 863 | 878 | 981 | 1602,55 | 2277 | 928 | 0,408 | -1,294 | -19,539 | 5,07E-85 | 3,83E-84 | 0,0038361 | 0,002861 | 0,0034699 | 0,0034699 | TRUE | 0,0516544 | No |
| orf19.7216 | 2814 | 2425 | 2668 | 1563 | 1865 | 1463 | 1625 | 2139 | 2562 | 2453 | 2525 | 2348 | 1659 | 1588 | 1664 | 1632 | 2053,83 | 2472 | 1636 | 0,661 | -0,598 | -13,452 | 2,99E-41 | 1,19E-40 | 0,0004208 | 0,00268 | 0,001392 | 0,001392 | TRUE | 0,012627 | No |
| orf19.7217 | 22091 | 16748 | 18556 | 13687 | 28981 | 25681 | 25669 | 37958 | 20113 | 16939 | 17562 | 20560 | 25778 | 27882 | 26288 | 28955 | 23009,6 | 18794 | 27226 | 1,449 | 0,535 | 7,99 | 1,35E-15 | 3,03E-15 | 0,0060117 | 0,002094 | 0,0042613 | 0,0042613 | TRUE | 0,0539841 | No |
| orf19.7218 | 99 | 54 | 60 | 76 | 108 | 89 | 103 | 142 | 90 | 55 | 57 | 114 | 96 | 97 | 105 | 108 | 90,27 | 79 | 102 | 1,309 | 0,388 | 1,899 | 0,05758 | 0,070932 | 0,0483664 | 0,01667 | 0,0286225 | 0,0286225 | TRUE | 0,4990012 | No |
| orf19.7219 | 12335 | 10440 | 11780 | 7694 | 17761 | 14526 | 15681 | 22098 | 11230 | 10559 | 11149 | 11558 | 15798 | 15771 | 16059 | 16857 | 13622,6 | 11124 | 16121 | 1,45 | 0,536 | 13,398 | 6,18E-41 | 2,44E-40 | 0,0010929 | 0,002134 | 0,0014586 | 0,0014586 | TRUE | 0,0142704 | No |
| orf19.7221 | 2214 | 1928 | 2126 | 1260 | 3158 | 2558 | 2832 | 3907 | 2016 | 1950 | 2012 | 1893 | 2809 | 2777 | 2900 | 2980 | 2417,17 | 1968 | 2866 | 1,456 | 0,542 | 12,479 | 9,67E-36 | 3,50E-35 | 0,0005495 | 0,002584 | 0,001373 | 0,001373 | TRUE | 0,0089735 | No |
| orf19.7222 | 3209 | 2487 | 2751 | 1715 | 1826 | 1272 | 1600 | 2075 | 2922 | 2515 | 2604 | 2576 | 1624 | 1381 | 1639 | 1583 | 2105,43 | 2654 | 1557 | 0,587 | -0,77 | -11,514 | 1,12E-30 | 3,67E-30 | 0,0046175 | 0,002664 | 0,0037878 | 0,0037878 | TRUE | 0,0677424 | No |
| orf19.7223 | 2492 | 2380 | 2410 | 1660 | 2239 | 1843 | 1990 | 2541 | 2269 | 2407 | 2281 | 2494 | 1992 | 2001 | 2038 | 1938 | 2177,4 | 2363 | 1992 | 0,844 | -0,244 | -5,341 | 9,23E-08 | 1,59E-07 | 0,00063 | 0,002644 | 0,001545 | 0,001545 | TRUE | 0,0155816 | No |
| orf19.7224 | 2945 | 2524 | 2728 | 1688 | 3737 | 2990 | 3208 | 4481 | 2681 | 2553 | 2582 | 2536 | 3324 | 3246 | 3285 | 3418 | 2953,16 | 2588 | 3318 | 1,282 | 0,358 | 9,219 | 3,01E-20 | 7,68E-20 | 0,0002544 | 0,002484 | 0,0011022 | 0,0011022 | TRUE | 0,0073703 | No |
| orf19.7225 | 85 | 82 | 104 | 74 | 193 | 194 | 204 | 238 | 77 | 83 | 98 | 111 | 172 | 211 | 209 | 182 | 142,83 | 92 | 194 | 2,108 | 1,076 | 7,789 | 6,78E-15 | 1,50E-14 | 0,0081808 | 0,011285 | 0,0100737 | 0,0100737 | TRUE | 0,1353588 | No |
| orf19.7227 | 2577 | 2082 | 2284 | 1559 | 254 | 204 | 210 | 294 | 2346 | 2106 | 2162 | 2342 | 226 | 221 | 215 | 224 | 1230,27 | 2239 | 222 | 0,099 | -3,334 | -47,434 | 0 | 0 | 1,00E-08 | 0,003111 | 0,0024093 | 0,0024093 | TRUE | 0,0193462 | No |
| orf19.7228 | 1167 | 1147 | 1117 | 756 | 820 | 631 | 663 | 971 | 1063 | 1160 | 1057 | 1136 | 729 | 685 | 679 | 741 | 906,19 | 1104 | 708 | 0,644 | -0,635 | -10,662 | 1,54E-26 | 4,55E-26 | 0,0010602 | 0,003495 | 0,0022621 | 0,0022621 | TRUE | 0,0153894 | No |
| orf19.7229 | 6771 | 5778 | 6212 | 4225 | 4381 | 3628 | 3865 | 5452 | 6165 | 5844 | 5879 | 6347 | 3897 | 3939 | 3958 | 4159 | 5023,39 | 6059 | 3988 | 0,659 | -0,602 | -14,353 | 1,02E-46 | 4,45E-46 | 0,0009338 | 0,0023 | 0,0014827 | 0,0014827 | TRUE | 0,0119693 | No |
| orf19.723 | 7405 | 6761 | 7347 | 5297 | 6357 | 5820 | 5917 | 8161 | 6742 | 6838 | 6953 | 7957 | 5654 | 6319 | 6060 | 6225 | 6593,56 | 7122 | 6064 | 0,852 | -0,231 | -3,892 | 9,93E-05 | 0,000149 | 0,0040926 | 0,002237 | 0,0032337 | 0,0032337 | TRUE | 0,0746736 | No |
| orf19.7231 | 1200 | 1038 | 1119 | 716 | 1777 | 1535 | 1721 | 2188 | 1093 | 1050 | 1059 | 1076 | 1581 | 1667 | 1763 | 1669 | 1369,46 | 1070 | 1670 | 1,561 | 0,642 | 12,646 | 1,18E-36 | 4,36E-36 | 0,0006687 | 0,003002 | 0,0016967 | 0,0016967 | TRUE | 0,0165778 | No |
| orf19.7232 | 515 | 430 | 447 | 284 | 694 | 590 | 662 | 804 | 469 | 435 | 423 | 427 | 617 | 641 | 678 | 613 | 537,82 | 438 | 637 | 1,449 | 0,535 | 7,7 | 1,36E-14 | 2,99E-14 | 1,00E-08 | 0,004493 | 0,0026894 | 0,0026894 | TRUE | 0,027401 | No |
| orf19.7234 | 2746 | 2135 | 2472 | 1617 | 3328 | 2768 | 3000 | 4040 | 2500 | 2159 | 2340 | 2429 | 2960 | 3005 | 3072 | 3082 | 2693,44 | 2357 | 3030 | 1,286 | 0,363 | 7,195 | 6,26E-13 | 1,31E-12 | 0,0016295 | 0,002527 | 0,0020603 | 0,0020603 | TRUE | 0,039154 | No |
| orf19.7235 | 5037 | 4383 | 4809 | 2862 | 2845 | 2279 | 2613 | 3665 | 4586 | 4433 | 4551 | 4299 | 2531 | 2474 | 2676 | 2796 | 3543,25 | 4467 | 2619 | 0,586 | -0,77 | -15,987 | 1,58E-57 | 8,28E-57 | 0,0015546 | 0,00241 | 0,0019293 | 0,0019293 | TRUE | 0,0257606 | No |
| orf19.7236 | 6701 | 5229 | 5720 | 3568 | 10590 | 7492 | 9160 | 12613 | 6101 | 5289 | 5413 | 5360 | 9419 | 8134 | 9381 | 9621 | 7339,84 | 5541 | 9139 | 1,649 | 0,722 | 11,308 | 1,20E-29 | 3,81E-29 | 0,0048744 | 0,002217 | 0,0037657 | 0,0037657 | TRUE | 0,0666855 | No |
| orf19.7237 | 451 | 374 | 378 | 295 | 1062 | 887 | 920 | 1300 | 411 | 378 | 358 | 443 | 945 | 963 | 942 | 992 | 678,91 | 398 | 960 | 2,432 | 1,282 | 18,379 | 1,92E-75 | 1,28E-74 | 0,0008292 | 0,003983 | 0,0028349 | 0,0028349 | TRUE | 0,0562832 | No |
| orf19.7238 | 20168 | 16087 | 17492 | 10768 | 27830 | 21217 | 24522 | 32665 | 18362 | 16270 | 16555 | 16175 | 24754 | 23036 | 25114 | 24917 | 20647,8 | 16840 | 24455 | 1,452 | 0,538 | 10,739 | 6,68E-27 | 2,00E-26 | 0,0025596 | 0,002101 | 0,0023603 | 0,0023603 | TRUE | 0,0454424 | No |
| orf19.7239 | 23539 | 19510 | 22024 | 14145 | 12376 | 10188 | 11423 | 15236 | 21431 | 19732 | 20844 | 21248 | 11008 | 11061 | 11699 | 11622 | 16080,7 | 20814 | 11348 | 0,545 | -0,875 | -21,897 | 2,80E-106 | 2,55E-105 | 0,0010911 | 0,002119 | 0,0014668 | 0,0014668 | TRUE | 0,0150151 | No |
| orf19.7242 | 4252 | 3574 | 3883 | 2448 | 4199 | 3490 | 3839 | 5407 | 3871 | 3615 | 3675 | 3677 | 3735 | 3789 | 3932 | 4125 | 3802,29 | 3710 | 3895 | 1,05 | 0,071 | 1,569 | 0,116627 | 0,139023 | 0,0011 | 0,002384 | 0,0016768 | 0,0016768 | TRUE | 0,0196877 | No |
| orf19.7243 | 747 | 705 | 690 | 473 | 1196 | 979 | 1113 | 1433 | 680 | 713 | 653 | 711 | 1064 | 1063 | 1140 | 1093 | 889,55 | 689 | 1090 | 1,584 | 0,664 | 11,671 | 1,80E-31 | 5,99E-31 | 1,00E-08 | 0,003522 | 0,0019023 | 0,0019023 | TRUE | 0,0150751 | No |
| orf19.7244 | 1028 | 793 | 883 | 623 | 1323 | 960 | 1072 | 1544 | 936 | 802 | 836 | 936 | 1177 | 1042 | 1098 | 1178 | 1000,53 | 878 | 1124 | 1,284 | 0,361 | 5,238 | 1,62E-07 | 2,77E-07 | 0,0034559 | 0,003357 | 0,0035293 | 0,0035293 | TRUE | 0,0395702 | No |
| orf19.7245 | 2640 | 2138 | 2387 | 1630 | 2162 | 1693 | 1837 | 2639 | 2404 | 2162 | 2259 | 2449 | 1923 | 1838 | 1881 | 2013 | 2116,14 | 2318 | 1914 | 0,827 | -0,274 | -5,163 | 2,43E-07 | 4,13E-07 | 0,001739 | 0,002661 | 0,002229 | 0,002229 | TRUE | 0,0247114 | No |
| orf19.7247 | 14525 | 12081 | 13291 | 8649 | 16193 | 13250 | 14801 | 20252 | 13224 | 12219 | 12579 | 12992 | 14403 | 14386 | 15158 | 15448 | 13801,2 | 12754 | 14849 | 1,164 | 0,22 | 5,392 | 6,97E-08 | 1,21E-07 | 0,0011691 | 0,002133 | 0,001521 | 0,001521 | TRUE | 0,0097803 | No |
| orf19.725 | 208 | 248 | 234 | 187 | 250 | 232 | 212 | 243 | 189 | 251 | 221 | 281 | 222 | 252 | 217 | 185 | 227,41 | 236 | 219 | 0,934 | -0,099 | -0,758 | 0,448544 | 0,490728 | 0,0172481 | 0,007846 | 0,0119182 | 0,0119182 | TRUE | 0,1980513 | No |
| orf19.7250 | 2314 | 1742 | 1857 | 1330 | 2815 | 1931 | 2425 | 3354 | 2107 | 1762 | 1757 | 1998 | 2504 | 2097 | 2484 | 2558 | 2158,29 | 1906 | 2411 | 1,266 | 0,34 | 4,385 | 1,16E-05 | 1,84E-05 | 0,0074341 | 0,002649 | 0,0053027 | 0,0053027 | TRUE | 0,0922717 | No |
| orf19.7251 | 768 | 651 | 698 | 573 | 1291 | 1105 | 1300 | 1783 | 699 | 658 | 661 | 861 | 1148 | 1200 | 1331 | 1360 | 989,81 | 720 | 1260 | 1,759 | 0,815 | 9,222 | 2,91E-20 | 7,42E-20 | 0,0097689 | 0,003372 | 0,0063679 | 0,0063679 | TRUE | 0,190007 | No |
| orf19.7252 | 9 | 3 | 7 | 5 | 3 | 2 | 1 | 3 | 8 | 3 | 7 | 8 | 3 | 2 | 1 | 2 | 4,19 | 6 | 2 | 0,327 | -1,612 | -2,229 | 0,025793 | 0,032768 | 1,00E-08 | 0,317345 | 0,1926029 | 0,1926029 | TRUE | 0,3126284 | No |
| orf19.7254 | 1609 | 1424 | 1581 | 1057 | 1443 | 1277 | 1315 | 1996 | 1465 | 1440 | 1496 | 1588 | 1284 | 1386 | 1347 | 1523 | 1441,06 | 1497 | 1385 | 0,927 | -0,109 | -1,78 | 0,075128 | 0,091366 | 0,0026696 | 0,002954 | 0,0029362 | 0,0029362 | TRUE | 0,0565106 | No |
| orf19.7255 | 2065 | 1308 | 1466 | 915 | 2671 | 1798 | 2236 | 3310 | 1880 | 1323 | 1387 | 1374 | 2376 | 1952 | 2290 | 2525 | 1888,46 | 1491 | 2286 | 1,532 | 0,616 | 4,19 | 2,79E-05 | 4,32E-05 | 0,0201817 | 0,002737 | 0,0104133 | 0,0201817 | TRUE | 0,3739812 | No |
| orf19.7256 | 1394 | 961 | 1019 | 549 | 1877 | 1124 | 1420 | 2130 | 1269 | 972 | 964 | 825 | 1670 | 1220 | 1454 | 1625 | 1249,89 | 1008 | 1492 | 1,48 | 0,566 | 3,419 | 0,000629 | 0,000899 | 0,0254418 | 0,003094 | 0,0129816 | 0,0254418 | TRUE | 0,3667424 | No |
| orf19.7257 | 2537 | 1847 | 2275 | 1628 | 2290 | 2002 | 2158 | 3097 | 2310 | 1868 | 2153 | 2446 | 2037 | 2174 | 2210 | 2362 | 2194,93 | 2194 | 2196 | 1,002 | 0,003 | 0,036 | 0,97133 | 0,974693 | 0,0079383 | 0,002639 | 0,005365 | 0,005365 | TRUE | 0,1211409 | No |
| orf19.7258 | 1715 | 1278 | 1428 | 927 | 2713 | 2080 | 2405 | 3405 | 1561 | 1293 | 1351 | 1393 | 2413 | 2258 | 2463 | 2597 | 1916,22 | 1400 | 2433 | 1,739 | 0,798 | 12,06 | 1,72E-33 | 5,96E-33 | 0,0043112 | 0,002726 | 0,0036261 | 0,0036261 | TRUE | 0,0760613 | No |
| orf19.726 | 2216 | 2143 | 2196 | 1294 | 1810 | 1560 | 1644 | 2098 | 2018 | 2167 | 2078 | 1944 | 1610 | 1694 | 1684 | 1600 | 1849,35 | 2052 | 1647 | 0,801 | -0,32 | -6,6 | 4,12E-11 | 8,10E-11 | 0,0007703 | 0,002751 | 0,0017126 | 0,0017126 | TRUE | 0,0170511 | No |
| orf19.7260 | 422 | 323 | 396 | 280 | 473 | 358 | 348 | 611 | 384 | 327 | 375 | 421 | 421 | 389 | 356 | 466 | 392,27 | 377 | 408 | 1,09 | 0,125 | 1,237 | 0,216254 | 0,249041 | 0,0092905 | 0,005405 | 0,0071736 | 0,0071736 | TRUE | 0,1110039 | No |
| orf19.7261 | 2947 | 2187 | 2579 | 1669 | 3428 | 2796 | 3297 | 4470 | 2683 | 2212 | 2441 | 2507 | 3049 | 3036 | 3377 | 3410 | 2839,25 | 2461 | 3218 | 1,308 | 0,387 | 5,903 | 3,58E-09 | 6,57E-09 | 0,0046481 | 0,002502 | 0,0037706 | 0,0037706 | TRUE | 0,056913 | No |
| orf19.7263 | 2535 | 1966 | 2188 | 1497 | 4230 | 3434 | 3780 | 5662 | 2308 | 1988 | 2071 | 2249 | 3762 | 3728 | 3871 | 4319 | 3037,11 | 2154 | 3920 | 1,822 | 0,865 | 13,531 | 1,02E-41 | 4,09E-41 | 0,0043489 | 0,002472 | 0,0035564 | 0,0035564 | TRUE | 0,0578412 | No |
| orf19.7264 | 5749 | 4015 | 4693 | 2899 | 7725 | 6056 | 7316 | 9920 | 5234 | 4061 | 4441 | 4355 | 6871 | 6575 | 7493 | 7567 | 5824,63 | 4523 | 7126 | 1,575 | 0,656 | 8,693 | 3,52E-18 | 8,50E-18 | 0,0081944 | 0,002264 | 0,0052793 | 0,0052793 | TRUE | 0,1377579 | No |
| orf19.7265 | 978 | 768 | 834 | 563 | 1496 | 1214 | 1369 | 1983 | 890 | 777 | 789 | 846 | 1331 | 1318 | 1402 | 1513 | 1108,2 | 826 | 1391 | 1,687 | 0,754 | 11,356 | 6,96E-30 | 2,23E-29 | 0,0032706 | 0,003229 | 0,0032511 | 0,0032511 | TRUE | 0,0433623 | No |
| orf19.7266 | 630 | 509 | 576 | 316 | 982 | 693 | 898 | 1185 | 574 | 515 | 545 | 475 | 873 | 752 | 920 | 904 | 694,71 | 527 | 862 | 1,632 | 0,706 | 8,742 | 2,30E-18 | 5,58E-18 | 0,0055705 | 0,003939 | 0,0047113 | 0,0047113 | TRUE | 0,084703 | No |
| orf19.7267 | 1024 | 743 | 948 | 714 | 1557 | 1286 | 1407 | 1991 | 932 | 751 | 897 | 1073 | 1385 | 1396 | 1441 | 1519 | 1174,29 | 913 | 1435 | 1,576 | 0,657 | 7,579 | 3,49E-14 | 7,55E-14 | 0,0101229 | 0,003162 | 0,0062922 | 0,0062922 | TRUE | 0,1682317 | No |
| orf19.7269 | 6009 | 4459 | 5104 | 3643 | 6826 | 5369 | 5986 | 8677 | 5471 | 4510 | 4830 | 5472 | 6072 | 5829 | 6130 | 6619 | 5616,7 | 5071 | 6162 | 1,216 | 0,282 | 4,185 | 2,85E-05 | 4,41E-05 | 0,0056999 | 0,002272 | 0,0041819 | 0,0041819 | TRUE | 0,0676727 | No |
| orf19.727 | 46 | 41 | 46 | 39 | 125 | 78 | 93 | 123 | 42 | 41 | 44 | 59 | 111 | 85 | 95 | 94 | 71,3 | 46 | 96 | 2,118 | 1,083 | 6,052 | 1,43E-09 | 2,66E-09 | 0,0024268 | 0,020564 | 0,0142195 | 0,0142195 | TRUE | 0,1742866 | No |
| orf19.7270 | 3741 | 2872 | 3242 | 1766 | 6992 | 5126 | 6433 | 8818 | 3406 | 2905 | 3068 | 2653 | 6219 | 5565 | 6588 | 6726 | 4641,38 | 3008 | 6274 | 2,084 | 1,06 | 13,596 | 4,23E-42 | 1,72E-41 | 0,0086002 | 0,002322 | 0,0055805 | 0,0055805 | TRUE | 0,0965274 | No |
| orf19.7271 | 2637 | 2290 | 2499 | 1450 | 2362 | 1851 | 1924 | 2655 | 2401 | 2316 | 2365 | 2178 | 2101 | 2010 | 1970 | 2025 | 2170,81 | 2315 | 2026 | 0,874 | -0,194 | -4,257 | 2,07E-05 | 3,23E-05 | 0,000637 | 0,002646 | 0,0015325 | 0,0015325 | TRUE | 0,0175286 | No |
| orf19.7272 | 1411 | 1155 | 1205 | 829 | 1673 | 1280 | 1402 | 1993 | 1285 | 1168 | 1140 | 1245 | 1488 | 1390 | 1436 | 1520 | 1334,05 | 1210 | 1458 | 1,208 | 0,272 | 4,863 | 1,16E-06 | 1,92E-06 | 0,00249 | 0,003027 | 0,0022467 | 0,0022467 | TRUE | 0,0231821 | No |
| orf19.7274 | 162 | 116 | 135 | 98 | 334 | 270 | 325 | 416 | 147 | 117 | 128 | 147 | 297 | 293 | 333 | 317 | 222,52 | 135 | 310 | 2,308 | 1,207 | 11,285 | 1,56E-29 | 4,94E-29 | 0,0008214 | 0,007973 | 0,005521 | 0,005521 | TRUE | 0,0792997 | No |
| orf19.7275 | 7 | 6 | 11 | 7 | 46 | 41 | 37 | 58 | 6 | 6 | 10 | 11 | 41 | 45 | 38 | 44 | 25,12 | 8 | 42 | 5,132 | 2,36 | 7,052 | 1,76E-12 | 3,62E-12 | 1,00E-08 | 0,054632 | 0,0317529 | 0,0317529 | TRUE | 0,1245818 | No |
| orf19.7276.1 | 136 | 132 | 112 | 78 | 244 | 157 | 220 | 283 | 124 | 134 | 106 | 117 | 217 | 170 | 225 | 216 | 163,65 | 120 | 207 | 1,728 | 0,789 | 6,236 | 4,50E-10 | 8,52E-10 | 0,0064967 | 0,010109 | 0,0086886 | 0,0086886 | TRUE | 0,1445825 | No |
| orf19.7277 | 10 | 0 | 8 | 7 | 39 | 25 | 29 | 50 | 9 | 0 | 8 | 11 | 35 | 27 | 30 | 38 | 19,61 | 7 | 32 | 4,931 | 2,302 | 5,621 | 1,90E-08 | 3,38E-08 | 0,0241141 | 0,069408 | 0,0665742 | 0,0665742 | TRUE | 1,217546 | No |
| orf19.7278 | 0 | 0 | 0 | 0 | 0 | 0 | 0 | 0 | 0 | 0 | 0 | 0 | 0 | 0 | 0 | 0 | 0 | 0 | 0 | NA | NA | NA | NA | NA | NA | NA | NA | NA | NA | NA | NA |
| orf19.7279 | 136 | 120 | 120 | 112 | 487 | 378 | 383 | 525 | 124 | 121 | 114 | 168 | 433 | 410 | 392 | 400 | 270,41 | 132 | 409 | 3,161 | 1,66 | 15,338 | 4,25E-53 | 2,07E-52 | 0,0031728 | 0,006922 | 0,0060004 | 0,0060004 | TRUE | 0,2766466 | No |
| orf19.7279.1 | 2912 | 2468 | 2654 | 1710 | 5538 | 4236 | 4315 | 5848 | 2651 | 2496 | 2512 | 2569 | 4926 | 4599 | 4419 | 4461 | 3579,1 | 2557 | 4601 | 1,8 | 0,848 | 18,034 | 1,05E-72 | 6,80E-72 | 0,0014363 | 0,002406 | 0,0018089 | 0,0018089 | TRUE | 0,027734 | No |
| orf19.728 | 658 | 752 | 786 | 418 | 975 | 785 | 804 | 1023 | 599 | 761 | 744 | 628 | 867 | 852 | 823 | 780 | 756,84 | 683 | 830 | 1,214 | 0,28 | 3,394 | 0,000688 | 0,000981 | 0,0063584 | 0,003782 | 0,0051747 | 0,0051747 | TRUE | 0,088208 | No |
| orf19.7280 | 4 | 2 | 2 | 3 | 12 | 2 | 3 | 5 | 4 | 2 | 2 | 5 | 11 | 2 | 3 | 4 | 3,97 | 3 | 5 | 1,705 | 0,769 | 1,02 | 0,307495 | 0,345949 | 0,1807268 | 0,334412 | 0,2712068 | 0,2712068 | TRUE | 1,3573605 | No |
| orf19.7281 | 2115 | 1813 | 1971 | 1252 | 8157 | 6659 | 6973 | 9506 | 1926 | 1834 | 1865 | 1881 | 7255 | 7230 | 7141 | 7251 | 4547,88 | 1876 | 7219 | 3,847 | 1,944 | 55,977 | 0 | 0 | 1,00E-08 | 0,002327 | 0,000808 | 0,000808 | TRUE | 0,0041783 | No |
| orf19.7282 | 2788 | 2166 | 2466 | 1625 | 7850 | 6340 | 7203 | 9851 | 2538 | 2191 | 2334 | 2441 | 6982 | 6883 | 7377 | 7514 | 4782,61 | 2376 | 7189 | 3,027 | 1,598 | 29,991 | 1,30E-197 | 2,19E-196 | 0,00249 | 0,002313 | 0,0024341 | 0,0024341 | TRUE | 0,0338369 | No |
| orf19.7283 | 20935 | 16851 | 18239 | 12063 | 2852 | 2114 | 2504 | 3274 | 19060 | 17043 | 17262 | 18121 | 2537 | 2295 | 2564 | 2497 | 10172,4 | 17872 | 2473 | 0,138 | -2,852 | -56,088 | 0 | 0 | 0,0022312 | 0,002167 | 0,0022653 | 0,0022653 | TRUE | 0,0277797 | No |
| orf19.7284 | 13003 | 10369 | 11428 | 6671 | 5476 | 4099 | 4802 | 6313 | 11839 | 10487 | 10816 | 10021 | 4871 | 4450 | 4918 | 4816 | 7777,11 | 10791 | 4764 | 0,441 | -1,18 | -20,926 | 3,07E-97 | 2,61E-96 | 0,0033508 | 0,002207 | 0,0029056 | 0,0029056 | TRUE | 0,0523835 | No |
| orf19.7285 | 2522 | 2080 | 2310 | 1700 | 3710 | 3212 | 3236 | 4620 | 2296 | 2104 | 2186 | 2554 | 3300 | 3487 | 3314 | 3524 | 2845,66 | 2285 | 3406 | 1,494 | 0,579 | 9,714 | 2,64E-22 | 7,05E-22 | 0,0037603 | 0,002501 | 0,0030359 | 0,0030359 | TRUE | 0,0719621 | No |
| orf19.7286 | 2845 | 2646 | 2858 | 1885 | 6223 | 4820 | 4814 | 6476 | 2590 | 2676 | 2705 | 2832 | 5535 | 5233 | 4930 | 4940 | 3930,15 | 2701 | 5160 | 1,912 | 0,935 | 18,565 | 6,11E-77 | 4,15E-76 | 0,0018568 | 0,002373 | 0,0021466 | 0,0021466 | TRUE | 0,0295525 | No |
| orf19.7288 | 616 | 507 | 560 | 389 | 29490 | 24463 | 25914 | 36007 | 561 | 513 | 530 | 584 | 26230 | 26560 | 26539 | 27466 | 13623 | 547 | 26699 | 49,006 | 5,615 | 117,856 | 0 | 0 | 0,0003905 | 0,002134 | 0,0011871 | 0,0011871 | TRUE | 0,0193757 | No |
| orf19.729 | 1109 | 1030 | 1042 | 595 | 1362 | 993 | 1079 | 1403 | 1010 | 1042 | 986 | 894 | 1211 | 1078 | 1105 | 1070 | 1049,53 | 983 | 1116 | 1,133 | 0,18 | 2,813 | 0,004904 | 0,006583 | 0,0025249 | 0,003296 | 0,0029578 | 0,0029578 | TRUE | 0,0402716 | No |
| orf19.7290 | 1321 | 1161 | 1169 | 720 | 2366 | 1649 | 2068 | 2602 | 1203 | 1174 | 1106 | 1082 | 2104 | 1790 | 2118 | 1985 | 1570,3 | 1141 | 1999 | 1,75 | 0,807 | 12,555 | 3,72E-36 | 1,36E-35 | 0,0036031 | 0,002878 | 0,0032668 | 0,0032668 | TRUE | 0,0583822 | No |

| | | | | | | | | | | | | | | | | | | | | | | | | | | | | | | | |
|---|---|---|---|---|---|---|---|---|---|---|---|---|---|---|---|---|---|---|---|---|---|---|---|---|---|---|---|---|---|---|---|
| orf19.7291 | 4096 | 3213 | 3769 | 2474 | 5464 | 4479 | 5160 | 6796 | 3729 | 3250 | 3567 | 3716 | 4860 | 4863 | 5284 | 5184 | 4306,72 | 3566 | 5048 | 1,416 | 0,502 | 9,336 | 1,00E-20 | 2,59E-20 | 0,0025808 | 0,002344 | 0,0025353 | 0,0025353 | TRUE | 0,0435193 | No |
| orf19.729.1 | 1528 | 1315 | 1444 | 1011 | 269 | 221 | 232 | 302 | 1391 | 1330 | 1367 | 1519 | 239 | 240 | 238 | 230 | 819,2 | 1402 | 237 | 0,169 | -2,564 | -36,019 | 4,23E-284 | 1,02E-282 | 1,00E-08 | 0,00365 | 0,0025158 | 0,0025158 | TRUE | 0,0346253 | No |
| orf19.7292 | 2406 | 1944 | 2107 | 1329 | 3040 | 2271 | 2732 | 3663 | 2191 | 1966 | 1994 | 1996 | 2704 | 2466 | 2798 | 2794 | 2363,61 | 2037 | 2690 | 1,321 | 0,401 | 7,145 | 8,97E-13 | 1,87E-12 | 0,0023954 | 0,002596 | 0,0025916 | 0,0025916 | TRUE | 0,0376404 | No |
| orf19.7293 | 446 | 413 | 437 | 278 | 779 | 659 | 729 | 949 | 406 | 418 | 414 | 418 | 693 | 715 | 747 | 724 | 566,73 | 414 | 720 | 1,74 | 0,799 | 12,415 | 2,16E-35 | 7,76E-35 | 1,00E-08 | 0,004368 | 0,0020448 | 0,0020448 | TRUE | 0,0075986 | No |
| orf19.7295 | 846 | 719 | 858 | 610 | 1261 | 1144 | 1157 | 1578 | 770 | 727 | 812 | 916 | 1122 | 1242 | 1185 | 1204 | 997,26 | 806 | 1188 | 1,479 | 0,564 | 8,04 | 8,96E-16 | 2,02E-15 | 0,0040153 | 0,003362 | 0,003669 | 0,003669 | TRUE | 0,0895999 | No |
| orf19.7296 | 7941 | 6914 | 7200 | 4954 | 3883 | 3120 | 3257 | 4468 | 7230 | 6993 | 6814 | 7442 | 3454 | 3387 | 3336 | 3408 | 5257,95 | 7120 | 3396 | 0,477 | -1,067 | -27,272 | 9,17E-164 | 1,31E-162 | 0,0006979 | 0,002288 | 0,001256 | 0,001256 | TRUE | 0,0108552 | No |
| orf19.7297 | 2203 | 1940 | 2009 | 1207 | 1376 | 1063 | 1063 | 1390 | 2006 | 1962 | 1901 | 1813 | 1224 | 1154 | 1089 | 1060 | 1526,16 | 1920 | 1132 | 0,588 | -0,765 | -13,157 | 1,55E-39 | 5,98E-39 | 0,0020843 | 0,002903 | 0,0025568 | 0,0025568 | TRUE | 0,0373469 | No |
| orf19.7298 | 1607 | 1624 | 1714 | 1002 | 2872 | 2464 | 2718 | 3534 | 1463 | 1643 | 1622 | 1505 | 2555 | 2675 | 2784 | 2696 | 2117,75 | 1558 | 2678 | 1,717 | 0,78 | 14,979 | 1,00E-50 | 4,69E-50 | 0,0015264 | 0,002661 | 0,0020815 | 0,0020815 | TRUE | 0,0222871 | No |
| orf19.7299.1 | 6 | 7 | 6 | 5 | 10 | 6 | 7 | 4 | 5 | 7 | 6 | 8 | 9 | 7 | 7 | 3 | 6,42 | 6 | 6 | 0,995 | -0,007 | -0,013 | 0,989497 | 0,990274 | 1,00E-08 | 0,207796 | 0,127651 | 0,127651 | TRUE | 0,4407849 | No |
| orf19.73 | 5017 | 5057 | 5243 | 3071 | 2439 | 2086 | 2031 | 2614 | 4568 | 5115 | 4962 | 4613 | 2169 | 2265 | 2080 | 1994 | 3470,72 | 4814 | 2127 | 0,441 | -1,18 | -21,396 | 1,44E-101 | 1,27E-100 | 0,0025807 | 0,002418 | 0,0025886 | 0,0025886 | TRUE | 0,0226683 | No |
| orf19.730 | 2085 | 1697 | 1934 | 1258 | 421 | 324 | 378 | 481 | 1898 | 1716 | 1830 | 1890 | 374 | 352 | 387 | 367 | 1101,87 | 1834 | 370 | 0,202 | -2,308 | -37,319 | 8,09E-305 | 2,12E-303 | 1,00E-08 | 0,003236 | 0,0021214 | 0,0021214 | TRUE | 0,0226315 | No |
| orf19.7300 | 2112 | 2121 | 2280 | 1400 | 1379 | 1162 | 1084 | 1500 | 1923 | 2145 | 2158 | 2103 | 1227 | 1262 | 1110 | 1144 | 1633,93 | 2082 | 1186 | 0,569 | -0,813 | -13,818 | 1,98E-43 | 8,23E-43 | 0,0022441 | 0,002845 | 0,0026677 | 0,0026677 | TRUE | 0,0333326 | No |
| orf19.7301 | 198 | 165 | 157 | 109 | 452 | 293 | 369 | 491 | 180 | 167 | 149 | 164 | 402 | 318 | 378 | 375 | 266,51 | 165 | 368 | 2,236 | 1,161 | 11,069 | 1,77E-28 | 5,45E-28 | 0,0047924 | 0,006994 | 0,0060593 | 0,0060593 | TRUE | 0,0907668 | No |
| orf19.7303 | 10 | 11 | 9 | 3 | 7 | 12 | 5 | 7 | 9 | 11 | 9 | 5 | 6 | 13 | 5 | 5 | 7,87 | 8 | 7 | 0,856 | -0,225 | -0,442 | 0,658711 | 0,695475 | 0,0241141 | 0,169865 | 0,1219142 | 0,1219142 | TRUE | 0,6394606 | No |
| orf19.7304 | 1088 | 1166 | 1251 | 719 | 374 | 338 | 338 | 411 | 991 | 1179 | 1184 | 1080 | 333 | 367 | 346 | 314 | 724,15 | 1108 | 340 | 0,306 | -1,71 | -21,419 | 8,88E-102 | 7,85E-101 | 0,0046657 | 0,003861 | 0,0042601 | 0,0042601 | TRUE | 0,0650283 | No |
| orf19.7305 | 504 | 480 | 435 | 230 | 602 | 422 | 502 | 657 | 459 | 485 | 412 | 345 | 535 | 458 | 514 | 501 | 463,8 | 425 | 502 | 1,175 | 0,233 | 2,401 | 0,016372 | 0,021158 | 0,0096206 | 0,004885 | 0,0068207 | 0,0068207 | TRUE | 0,1605479 | No |
| orf19.7306 | 681 | 622 | 663 | 407 | 896 | 661 | 704 | 997 | 620 | 629 | 627 | 611 | 797 | 718 | 721 | 761 | 685,51 | 622 | 749 | 1,205 | 0,269 | 4,388 | 1,14E-05 | 1,81E-05 | 1,00E-08 | 0,003964 | 0,0021314 | 0,0021314 | TRUE | 0,0216615 | No |
| orf19.7307 | 2994 | 2678 | 2864 | 2183 | 3579 | 3244 | 2838 | 4065 | 2726 | 2709 | 2711 | 3279 | 3183 | 3522 | 2906 | 3101 | 3017,12 | 2856 | 3178 | 1,114 | 0,156 | 2,09 | 0,036659 | 0,045995 | 0,007708 | 0,002475 | 0,005002 | 0,005002 | TRUE | 0,1175231 | No |
| orf19.7308 | 4789 | 3920 | 4311 | 2869 | 4982 | 4016 | 4466 | 6089 | 4360 | 3965 | 4080 | 4310 | 4431 | 4360 | 4574 | 4645 | 4340,57 | 4179 | 4502 | 1,078 | 0,109 | 2,466 | 0,013655 | 0,017748 | 0,0011169 | 0,002341 | 0,001629 | 0,001629 | TRUE | 0,0145057 | No |
| orf19.731 | 2220 | 1870 | 2056 | 1372 | 2384 | 2017 | 2243 | 2910 | 2021 | 1891 | 1946 | 2061 | 2120 | 2190 | 2297 | 2220 | 2093,32 | 1980 | 2207 | 1,116 | 0,158 | 3,412 | 0,000646 | 0,000923 | 0,0006459 | 0,002668 | 0,0015715 | 0,0015715 | TRUE | 0,0107738 | No |
| orf19.7310 | 8335 | 7769 | 8074 | 5074 | 1603 | 1360 | 1443 | 1957 | 7589 | 7858 | 7641 | 7622 | 1426 | 1477 | 1478 | 1493 | 4572,81 | 7678 | 1468 | 0,191 | -2,387 | -65,133 | 0 | 0 | 1,00E-08 | 0,002326 | 0,0009015 | 0,0009015 | TRUE | 0,0047405 | No |
| orf19.7311 | 754 | 712 | 757 | 538 | 466 | 466 | 437 | 528 | 686 | 720 | 716 | 808 | 414 | 506 | 448 | 403 | 587,74 | 732 | 443 | 0,604 | -0,728 | -8,747 | 2,19E-18 | 5,33E-18 | 0,005601 | 0,004285 | 0,0048498 | 0,0048498 | TRUE | 0,1041408 | No |
| orf19.7312 | 2362 | 2122 | 2121 | 1380 | 4112 | 3326 | 3439 | 4866 | 2151 | 2146 | 2007 | 2073 | 3657 | 3611 | 3522 | 3712 | 2859,92 | 2094 | 3626 | 1,731 | 0,792 | 19,272 | 9,26E-83 | 6,82E-82 | 0,0003949 | 0,002499 | 0,0012357 | 0,0012357 | TRUE | 0,0102085 | No |
| orf19.7313 | 3293 | 2965 | 3225 | 2156 | 3449 | 2840 | 2887 | 3978 | 2998 | 2999 | 3052 | 3239 | 3068 | 3083 | 2957 | 3034 | 3053,76 | 3072 | 3036 | 0,99 | -0,015 | -0,374 | 0,708173 | 0,742045 | 0,0004248 | 0,00247 | 0,0012269 | 0,0012269 | TRUE | 0,0149123 | No |
| orf19.7314 | 4952 | 4213 | 4436 | 3023 | 2873 | 2328 | 2334 | 3422 | 4509 | 4261 | 4198 | 4541 | 2555 | 2528 | 2390 | 2610 | 3449,07 | 4377 | 2521 | 0,577 | -0,795 | -17,507 | 1,28E-68 | 7,84E-68 | 0,0010523 | 0,00242 | 0,0016694 | 0,0016694 | TRUE | 0,0146426 | No |
| orf19.7316 | 1517 | 1263 | 1532 | 884 | 1520 | 1225 | 1335 | 1831 | 1381 | 1277 | 1450 | 1328 | 1352 | 1330 | 1367 | 1397 | 1360,28 | 1359 | 1362 | 1,001 | 0,002 | 0,031 | 0,974922 | 0,977375 | 0,0009772 | 0,003008 | 0,0019121 | 0,0019121 | TRUE | 0,0249954 | No |
| orf19.7317 | 1787 | 1563 | 1757 | 1004 | 1583 | 1317 | 1398 | 1787 | 1627 | 1581 | 1663 | 1508 | 1408 | 1430 | 1432 | 1363 | 1501,45 | 1595 | 1408 | 0,881 | -0,184 | -3,815 | 0,000136 | 0,000202 | 0,0003831 | 0,002917 | 0,0015533 | 0,0015533 | TRUE | 0,0142896 | No |
| orf19.7319 | 1590 | 1427 | 1570 | 957 | 1271 | 1103 | 1143 | 1553 | 1448 | 1443 | 1486 | 1438 | 1131 | 1198 | 1171 | 1185 | 1312,2 | 1454 | 1171 | 0,805 | -0,313 | -6,792 | 1,11E-11 | 2,23E-11 | 1,00E-08 | 0,003044 | 0,0012752 | 0,0012752 | TRUE | 0,0065641 | No |
| orf19.732 | 4191 | 3278 | 3569 | 2526 | 5967 | 4889 | 5107 | 7299 | 3816 | 3315 | 3378 | 3794 | 5307 | 5308 | 5230 | 5568 | 4464,6 | 3576 | 5353 | 1,498 | 0,583 | 10,699 | 1,03E-26 | 3,06E-26 | 0,0028377 | 0,002333 | 0,0026163 | 0,0026163 | TRUE | 0,0291644 | No |
| orf19.7320 | 105 | 81 | 94 | 77 | 71 | 65 | 78 | 74 | 96 | 82 | 89 | 116 | 63 | 71 | 80 | 56 | 81,53 | 96 | 68 | 0,708 | -0,497 | -2,946 | 0,003214 | 0,004365 | 0,0085947 | 0,018241 | 0,0147705 | 0,0147705 | TRUE | 0,1521623 | No |
| orf19.7321 | 621 | 553 | 670 | 369 | 609 | 433 | 534 | 703 | 565 | 559 | 634 | 554 | 542 | 470 | 547 | 536 | 551 | 578 | 524 | 0,905 | -0,143 | -1,914 | 0,05565 | 0,068687 | 0,0025478 | 0,004434 | 0,0035706 | 0,0035706 | TRUE | 0,0521383 | No |
| orf19.7322 | 5513 | 4619 | 4705 | 3432 | 6176 | 4606 | 4618 | 6596 | 5019 | 4672 | 4453 | 5155 | 5493 | 5001 | 4729 | 5031 | 4944,29 | 4825 | 5064 | 1,05 | 0,071 | 1,182 | 0,237269 | 0,271317 | 0,0039009 | 0,002304 | 0,0032323 | 0,0032323 | TRUE | 0,0398469 | No |
| orf19.7323 | 5907 | 5677 | 5986 | 3727 | 3747 | 2840 | 3254 | 4214 | 5378 | 5742 | 5665 | 5599 | 3333 | 3083 | 3333 | 3214 | 4418,35 | 5596 | 3241 | 0,579 | -0,788 | -19,169 | 6,78E-82 | 4,92E-81 | 0,0007382 | 0,002336 | 0,0013804 | 0,0013804 | TRUE | 0,0125244 | No |
| orf19.7324 | 386 | 317 | 330 | 226 | 217 | 162 | 187 | 281 | 351 | 321 | 312 | 339 | 193 | 176 | 192 | 214 | 262,33 | 331 | 194 | 0,588 | -0,765 | -8,068 | 7,13E-16 | 1,61E-15 | 1,00E-08 | 0,007073 | 0,0046254 | 0,0046254 | TRUE | 0,0616287 | No |
| orf19.7325 | 4315 | 3563 | 3964 | 2487 | 3789 | 2912 | 3212 | 4413 | 3929 | 3604 | 3752 | 3736 | 3370 | 3162 | 3289 | 3366 | 3525,9 | 3755 | 3297 | 0,878 | -0,187 | -4,447 | 8,73E-06 | 1,39E-05 | 0,0006896 | 0,002412 | 0,0014197 | 0,0014197 | TRUE | 0,0122197 | No |
| orf19.7326 | 2863 | 2642 | 2776 | 1943 | 2610 | 2389 | 2282 | 3074 | 2607 | 2672 | 2627 | 2919 | 2322 | 2594 | 2337 | 2345 | 2552,74 | 2706 | 2400 | 0,887 | -0,172 | -3,193 | 0,001407 | 0,001967 | 0,0023656 | 0,002554 | 0,0024051 | 0,0024051 | TRUE | 0,0353424 | No |
| orf19.7327 | 5006 | 4125 | 4486 | 3409 | 10522 | 8654 | 8627 | 12021 | 4558 | 4172 | 4246 | 5121 | 9359 | 9396 | 8835 | 9170 | 6856,98 | 4524 | 9190 | 2,033 | 1,024 | 16,696 | 1,39E-62 | 7,88E-62 | 0,0047198 | 0,00223 | 0,0034419 | 0,0034419 | TRUE | 0,0936651 | No |
| orf19.7328 | 730 | 708 | 680 | 485 | 1765 | 1356 | 1394 | 1936 | 665 | 716 | 644 | 729 | 1570 | 1472 | 1428 | 1477 | 1087,42 | 688 | 1487 | 2,168 | 1,116 | 18,865 | 2,21E-79 | 1,56E-78 | 0,0011341 | 0,003252 | 0,0022676 | 0,0022676 | TRUE | 0,0223419 | No |
| orf19.7329 | 5413 | 4499 | 4563 | 3312 | 2790 | 2264 | 2361 | 3502 | 4928 | 4550 | 4318 | 4975 | 2482 | 2458 | 2418 | 2671 | 3600,15 | 4693 | 2507 | 0,535 | -0,903 | -15,984 | 1,65E-57 | 8,65E-57 | 0,0028361 | 0,002404 | 0,0027629 | 0,0027629 | TRUE | 0,0353942 | No |
| orf19.733 | 2122 | 1678 | 1940 | 1338 | 1121 | 897 | 1003 | 1423 | 1932 | 1697 | 1836 | 2010 | 997 | 974 | 1027 | 1085 | 1444,84 | 1869 | 1021 | 0,548 | -0,869 | -13,846 | 1,35E-43 | 5,64E-43 | 0,0031106 | 0,002951 | 0,0030364 | 0,0030364 | TRUE | 0,0459589 | No |
| orf19.7330 | 32 | 43 | 41 | 28 | 33 | 33 | 34 | 30 | 29 | 43 | 39 | 42 | 29 | 36 | 35 | 23 | 34,55 | 38 | 31 | 0,797 | -0,327 | -1,321 | 0,186413 | 0,216502 | 1,00E-08 | 0,040275 | 0,0293464 | 0,0293464 | TRUE | 0,2446338 | No |
| orf19.7331 | 198 | 180 | 237 | 141 | 122 | 116 | 99 | 164 | 180 | 182 | 224 | 212 | 109 | 126 | 101 | 125 | 157,42 | 200 | 115 | 0,58 | -0,787 | -6,237 | 4,45E-10 | 8,42E-10 | 0,0057541 | 0,010428 | 0,0085542 | 0,0085542 | TRUE | 0,0861628 | No |
| orf19.7332 | 35068 | 32443 | 34566 | 20422 | 44035 | 36649 | 40005 | 51965 | 31928 | 32813 | 32714 | 30677 | 39168 | 39791 | 40970 | 39639 | 35962,4 | 32033 | 39892 | 1,245 | 0,316 | 9,425 | 4,29E-21 | 1,12E-20 | 0,0006213 | 0,002074 | 0,0010533 | 0,0010533 | TRUE | 0,0098161 | No |
| orf19.7335 | 5784 | 4940 | 5316 | 3115 | 4745 | 3649 | 3991 | 5265 | 5266 | 4996 | 5031 | 4679 | 4221 | 3962 | 4087 | 4016 | 4532,31 | 4993 | 4072 | 0,815 | -0,296 | -6,625 | 3,48E-11 | 6,86E-11 | 0,0012468 | 0,002328 | 0,0016896 | 0,0016896 | TRUE | 0,0208944 | No |
| orf19.7336 | 5 | 16 | 13 | 3 | 5 | 5 | 3 | 5 | 5 | 16 | 12 | 5 | 4 | 5 | 3 | 4 | 6,79 | 10 | 4 | 0,437 | -1,196 | -2,013 | 0,044084 | 0,054868 | 1,00E-08 | 0,196635 | 0,1711004 | 0,1711004 | TRUE | 0,1946724 | No |
| orf19.7337 | 2352 | 2429 | 2416 | 1497 | 2578 | 2287 | 2160 | 2900 | 2141 | 2457 | 2287 | 2249 | 2293 | 2483 | 2212 | 2212 | 2291,71 | 2284 | 2300 | 1,007 | 0,01 | 0,171 | 0,8641 | 0,884278 | 0,0027074 | 0,002613 | 0,0026657 | 0,0026657 | TRUE | 0,0340667 | No |
| orf19.7339 | 510 | 456 | 472 | 285 | 262 | 203 | 238 | 292 | 464 | 461 | 447 | 428 | 233 | 220 | 244 | 223 | 340,03 | 450 | 230 | 0,509 | -0,973 | -12,036 | 2,29E-33 | 7,88E-33 | 1,00E-08 | 0,005922 | 0,0030694 | 0,0030694 | TRUE | 0,0167588 | No |
| orf19.734 | 14428 | 12173 | 13318 | 8421 | 1448 | 1293 | 1283 | 1802 | 13136 | 12312 | 12604 | 12650 | 1288 | 1404 | 1314 | 1375 | 7010,26 | 12676 | 1345 | 0,106 | -3,237 | -76,834 | 0 | 0 | 0,0006032 | 0,002225 | 0,0013147 | 0,0013147 | TRUE | 0,0100501 | No |
| orf19.7340 | 13 | 26 | 19 | 26 | 12 | 17 | 13 | 24 | 12 | 26 | 18 | 39 | 11 | 18 | 13 | 18 | 19,49 | 24 | 15 | 0,658 | -0,604 | -1,587 | 0,112578 | 0,134423 | 0,1093641 | 0,069814 | 0,085408 | 0,085408 | TRUE | 0,2715827 | No |
| orf19.7341 | 4699 | 3927 | 4271 | 2649 | 2749 | 2211 | 2419 | 3250 | 4278 | 3972 | 4042 | 3979 | 2445 | 2401 | 2477 | 2479 | 3259,19 | 4068 | 2450 | 0,602 | -0,732 | -18,213 | 4,08E-74 | 2,68E-73 | 0,0005202 | 0,002442 | 0,0012267 | 0,0012267 | TRUE | 0,0151267 | No |
| orf19.7341.1 | 30955 | 32246 | 32201 | 17266 | 16473 | 12164 | 13244 | 16730 | 28183 | 32613 | 30475 | 25936 | 14652 | 13207 | 13564 | 12762 | 21424,1 | 29302 | 13546 | 0,462 | -1,113 | -16,107 | 2,28E-58 | 1,22E-57 | 0,0065405 | 0,002099 | 0,0045373 | 0,0045373 | TRUE | 0,0729493 | No |
| orf19.7342 | 795 | 813 | 882 | 568 | 359 | 353 | 295 | 415 | 724 | 822 | 835 | 853 | 319 | 383 | 302 | 317 | 569,41 | 808 | 330 | 0,408 | -1,292 | -15,148 | 7,86E-52 | 3,75E-51 | 0,0056894 | 0,004357 | 0,0049076 | 0,0049076 | TRUE | 0,1242072 | No |
| orf19.7343 | 1816 | 1808 | 1946 | 994 | 1913 | 1624 | 1563 | 2158 | 1653 | 1829 | 1842 | 1493 | 1702 | 1763 | 1601 | 1646 | 1691,06 | 1704 | 1678 | 0,982 | -0,026 | -0,376 | 0,706845 | 0,741019 | 0,0048183 | 0,002818 | 0,0038512 | 0,0038512 | TRUE | 0,078573 | No |
| orf19.7344 | 1231 | 1462 | 1522 | 872 | 702 | 612 | 644 | 771 | 1121 | 1479 | 1440 | 1310 | 624 | 664 | 660 | 588 | 985,78 | 1338 | 634 | 0,474 | -1,078 | -12,432 | 1,76E-35 | 6,36E-35 | 0,0087018 | 0,003377 | 0,0060842 | 0,0060842 | TRUE | 0,1479844 | No |
| orf19.7345 | 1240 | 1343 | 1306 | 747 | 499 | 388 | 373 | 544 | 1129 | 1358 | 1236 | 1122 | 444 | 421 | 382 | 415 | 813,43 | 1211 | 416 | 0,343 | -1,544 | -19,716 | 1,56E-86 | 1,19E-85 | 0,0049893 | 0,003661 | 0,0043235 | 0,0043235 | TRUE | 0,0799591 | No |
| orf19.7347 | 794 | 737 | 841 | 587 | 1497 | 1239 | 1314 | 1761 | 723 | 745 | 796 | 882 | 1332 | 1345 | 1346 | 1343 | 1063,97 | 786 | 1342 | 1,714 | 0,777 | 12,61 | 1,85E-36 | 6,82E-36 | 0,0017485 | 0,003279 | 0,0026145 | 0,0026145 | TRUE | 0,0691797 | No |
| orf19.7348 | 1 | 2 | 2 | 4 | 2 | 0 | 1 | 7 | 1 | 2 | 2 | 6 | 2 | 0 | 1 | 5 | 2,37 | 3 | 2 | 0,848 | -0,238 | -0,237 | 0,812714 | 0,837409 | 0,5132847 | 0,558908 | 0,542344 | 0,542344 | TRUE | 1,4274884 | No |
| orf19.7349 | 6439 | 5910 | 6496 | 4123 | 7148 | 6234 | 6325 | 8377 | 5862 | 5977 | 6148 | 6193 | 6358 | 6768 | 6478 | 6390 | 6271,87 | 6045 | 6498 | 1,075 | 0,104 | 2,84 | 0,004506 | 0,00606 | 0,0005223 | 0,002248 | 0,0011383 | 0,0011383 | TRUE | 0,0094076 | No |
| orf19.7350 | 24412 | 18996 | 21408 | 11558 | 21193 | 15217 | 19024 | 25511 | 22226 | 19212 | 20261 | 17362 | 18851 | 16521 | 19483 | 19460 | 19172 | 19765 | 18579 | 0,94 | -0,089 | -1,195 | 0,232136 | 0,265876 | 0,0080623 | 0,002106 | 0,0053372 | 0,0053372 | TRUE | 0,0859899 | No |
| orf19.7353 | 746 | 709 | 739 | 472 | 1624 | 1400 | 1428 | 1989 | 679 | 717 | 699 | 709 | 1444 | 1520 | 1462 | 1517 | 1093,61 | 701 | 1486 | 2,122 | 1,085 | 20,957 | 1,63E-97 | 1,39E-96 | 1,00E-08 | 0,003245 | 0,0014987 | 0,0014987 | TRUE | 0,0056673 | No |
| orf19.7354 | 1131 | 1243 | 1305 | 672 | 2006 | 1832 | 1939 | 2253 | 1030 | 1257 | 1235 | 1009 | 1784 | 1989 | 1986 | 1719 | 1501,14 | 1133 | 1870 | 1,646 | 0,719 | 8,784 | 1,58E-18 | 3,85E-18 | 0,0085358 | 0,002917 | 0,0057155 | 0,0057155 | TRUE | 0,0644199 | No |
| orf19.7355 | 651 | 699 | 660 | 453 | 845 | 719 | 756 | 937 | 593 | 707 | 625 | 680 | 752 | 781 | 774 | 715 | 703,25 | 651 | 756 | 1,161 | 0,216 | 3,089 | 0,002011 | 0,002775 | 0,0025227 | 0,003915 | 0,003244 | 0,003244 | TRUE | 0,0449694 | No |
| orf19.7356 | 414 | 418 | 445 | 273 | 1027 | 650 | 727 | 1085 | 377 | 423 | 421 | 410 | 913 | 706 | 745 | 828 | 602,79 | 408 | 798 | 1,961 | 0,972 | 10,959 | 5,98E-28 | 1,83E-27 | 0,0070821 | 0,004228 | 0,005647 | 0,005647 | TRUE | 0,1122479 | No |
| orf19.7357 | 1907 | 1816 | 1927 | 1257 | 4764 | 4148 | 4125 | 5181 | 1736 | 1837 | 1824 | 1888 | 4237 | 4504 | 4225 | 3952 | 3025,31 | 1821 | 4230 | 2,324 | 1,216 | 23,959 | 7,44E-127 | 8,07E-126 | 0,0017936 | 0,002474 | 0,0020685 | 0,0020685 | TRUE | 0,0241401 | No |
| orf19.7358 | 881 | 1070 | 1104 | 549 | 755 | 661 | 640 | 792 | 802 | 1082 | 1045 | 825 | 672 | 718 | 655 | 604 | 800,33 | 938 | 662 | 0,704 | -0,506 | -5,041 | 4,62E-07 | 7,79E-07 | 0,0128024 | 0,003688 | 0,0084117 | 0,0084117 | TRUE | 0,1204862 | No |
| orf19.7359 | 2808 | 2569 | 2752 | 1798 | 2422 | 2229 | 2274 | 3072 | 2557 | 2598 | 2605 | 2701 | 2154 | 2420 | 2329 | 2343 | 2463,35 | 2615 | 2312 | 0,884 | -0,177 | -3,807 | 0,000141 | 0,000209 | 0,0009087 | 0,002573 | 0,0016762 | 0,0016762 | TRUE | 0,0256232 | No |
| orf19.736 | 1859 | 1875 | 1984 | 1221 | 877 | 813 | 790 | 939 | 1693 | 1896 | 1878 | 1834 | 780 | 883 | 809 | 716 | 1311,1 | 1825 | 797 | 0,436 | -1,198 | -17,678 | 6,16E-70 | 3,86E-69 | 0,0038235 | 0,003045 | 0,0035299 | 0,0035299 | TRUE | 0,0611262 | No |

| | | | | | | | | | | | | | | | | | | | | | | | | | | | | | | | |
|---|---|---|---|---|---|---|---|---|---|---|---|---|---|---|---|---|---|---|---|---|---|---|---|---|---|---|---|---|---|---|---|
| orf19.7360 | 498 | 588 | 575 | 313 | 404 | 347 | 343 | 430 | 453 | 595 | 544 | 470 | 359 | 377 | 351 | 328 | 434,73 | 516 | 354 | 0,684 | -0,548 | -5,739 | 9,54E-09 | 1,72E-08 | 0,0078717 | 0,005076 | 0,0063969 | 0,0063969 | TRUE | 0,1245641 | No |
| orf19.7361 | 3184 | 3080 | 3278 | 1946 | 2812 | 2238 | 2360 | 2954 | 2899 | 3115 | 3102 | 2923 | 2501 | 2430 | 2417 | 2253 | 2705,1 | 3010 | 2400 | 0,797 | -0,328 | -6,818 | 9,24E-12 | 1,87E-11 | 0,0013849 | 0,002525 | 0,0018468 | 0,0018468 | TRUE | 0,0214988 | No |
| orf19.7362 | 1051 | 1066 | 1091 | 726 | 1828 | 1560 | 1502 | 2140 | 957 | 1078 | 1033 | 1091 | 1626 | 1694 | 1538 | 1632 | 1331,06 | 1040 | 1622 | 1,564 | 0,646 | 11,414 | 3,57E-30 | 1,15E-29 | 0,001418 | 0,003029 | 0,0022659 | 0,0022659 | TRUE | 0,0354471 | No |
| orf19.7363 | 10888 | 10306 | 11175 | 6791 | 10811 | 9453 | 9686 | 13373 | 9913 | 10423 | 10576 | 10201 | 9616 | 10263 | 9920 | 10201 | 10139,2 | 10278 | 10000 | 0,973 | -0,04 | -1,073 | 0,283218 | 0,319729 | 0,0007008 | 0,002167 | 0,0012158 | 0,0012158 | TRUE | 0,0082136 | No |
| orf19.7364 | 36 | 31 | 28 | 20 | 28 | 26 | 41 | 60 | 33 | 31 | 26 | 30 | 25 | 28 | 42 | 46 | 32,7 | 30 | 35 | 1,174 | 0,232 | 0,881 | 0,37858 | 0,418856 | 0,0252699 | 0,04244 | 0,0357205 | 0,0357205 | TRUE | 0,1765854 | No |
| orf19.7365 | 1185 | 1239 | 1310 | 827 | 155 | 109 | 147 | 161 | 1079 | 1253 | 1240 | 1242 | 138 | 118 | 151 | 123 | 667,96 | 1204 | 132 | 0,11 | -3,185 | -34,683 | 1,40E-263 | 3,11E-262 | 0,004311 | 0,004015 | 0,0041366 | 0,0041366 | TRUE | 0,0810948 | No |
| orf19.7366 | 2973 | 2787 | 3005 | 1482 | 2498 | 1840 | 2101 | 2728 | 2707 | 2819 | 2844 | 2226 | 2222 | 1998 | 2152 | 2081 | 2380,99 | 2649 | 2113 | 0,797 | -0,328 | -4,47 | 7,81E-06 | 1,25E-05 | 0,0063782 | 0,002592 | 0,0047478 | 0,0047478 | TRUE | 0,1349451 | No |
| orf19.7367 | 2879 | 3039 | 3096 | 1890 | 2643 | 2416 | 2280 | 3127 | 2621 | 3074 | 2930 | 2839 | 2351 | 2623 | 2335 | 2385 | 2644,78 | 2866 | 2424 | 0,845 | -0,242 | -4,086 | 4,39E-05 | 6,72E-05 | 0,0032247 | 0,002536 | 0,0029979 | 0,0029979 | TRUE | 0,0414012 | No |
| orf19.7368 | 1624 | 1425 | 1577 | 1046 | 2807 | 2400 | 2551 | 3399 | 1479 | 1441 | 1492 | 1571 | 2497 | 2606 | 2613 | 2593 | 2036,42 | 1496 | 2577 | 1,726 | 0,787 | 18,049 | 8,03E-73 | 5,21E-72 | 0,0001847 | 0,002686 | 0,0012861 | 0,0012861 | TRUE | 0,0120267 | No |
| orf19.7369 | 613 | 705 | 663 | 415 | 630 | 536 | 564 | 708 | 558 | 713 | 627 | 623 | 560 | 582 | 578 | 540 | 597,75 | 630 | 565 | 0,896 | -0,159 | -2,027 | 0,042634 | 0,053167 | 0,0042311 | 0,004247 | 0,0042347 | 0,0042347 | TRUE | 0,095153 | No |
| orf19.7370 | 3240 | 3039 | 3196 | 1952 | 2706 | 2164 | 2154 | 2917 | 2950 | 3074 | 3025 | 2932 | 2407 | 2350 | 2206 | 2225 | 2645,99 | 2995 | 2297 | 0,766 | -0,384 | -8,749 | 2,15E-18 | 5,22E-18 | 0,0006232 | 0,002536 | 0,0014644 | 0,0014644 | TRUE | 0,0129709 | No |
| orf19.7371 | 2356 | 2379 | 2527 | 1546 | 2862 | 2403 | 2495 | 3298 | 2145 | 2406 | 2392 | 2322 | 2546 | 2609 | 2555 | 2516 | 2436,33 | 2316 | 2556 | 1,104 | 0,142 | 3,027 | 0,002467 | 0,00338 | 0,0010955 | 0,002579 | 0,0017087 | 0,0017087 | TRUE | 0,0315955 | No |
| orf19.7372 | 1580 | 1420 | 1548 | 1001 | 1941 | 1659 | 1647 | 2247 | 1439 | 1436 | 1465 | 1504 | 1726 | 1801 | 1687 | 1714 | 1596,48 | 1461 | 1732 | 1,187 | 0,247 | 5,534 | 3,14E-08 | 5,53E-08 | 1,00E-08 | 0,002864 | 0,0012737 | 0,0012737 | TRUE | 0,0082473 | No |
| orf19.7374 | 8162 | 7571 | 8377 | 5243 | 5303 | 4554 | 4576 | 6425 | 7431 | 7657 | 7928 | 7876 | 4717 | 4944 | 4686 | 4901 | 6267,62 | 7723 | 4812 | 0,623 | -0,682 | -18,112 | 2,57E-73 | 1,68E-72 | 0,0005699 | 0,002248 | 0,0011948 | 0,0011948 | TRUE | 0,0080308 | No |
| orf19.7375 | 361 | 328 | 307 | 201 | 298 | 209 | 226 | 315 | 329 | 332 | 291 | 302 | 265 | 227 | 231 | 240 | 277,08 | 314 | 241 | 0,769 | -0,378 | -4,112 | 3,93E-05 | 6,03E-05 | 1,00E-08 | 0,006805 | 0,004485 | 0,004485 | TRUE | 0,0507008 | No |
| orf19.7376 | 494 | 540 | 518 | 346 | 574 | 506 | 486 | 626 | 450 | 546 | 490 | 520 | 511 | 549 | 498 | 478 | 505,13 | 502 | 509 | 1,015 | 0,022 | 0,275 | 0,783616 | 0,813343 | 0,003339 | 0,004652 | 0,0040406 | 0,0040406 | TRUE | 0,0595008 | No |
| orf19.7377 | 256 | 241 | 216 | 153 | 85 | 90 | 104 | 139 | 233 | 244 | 204 | 230 | 76 | 98 | 107 | 106 | 162,12 | 228 | 97 | 0,424 | -1,239 | -9,663 | 4,32E-22 | 1,15E-21 | 0,0056305 | 0,010185 | 0,008602 | 0,008602 | TRUE | 0,2194405 | No |
| orf19.7378 | 3691 | 3379 | 3597 | 2264 | 3110 | 2567 | 2863 | 3840 | 3360 | 3417 | 3404 | 3401 | 2766 | 2787 | 2932 | 2929 | 3124,71 | 3396 | 2854 | 0,841 | -0,25 | -6,603 | 4,03E-11 | 7,93E-11 | 0,0001998 | 0,00246 | 0,0010588 | 0,0010588 | TRUE | 0,0053666 | No |
| orf19.7379 | 6918 | 6502 | 6870 | 4523 | 13431 | 11891 | 12020 | 16426 | 6299 | 6576 | 6502 | 6794 | 11946 | 12910 | 12310 | 12530 | 9483,42 | 6543 | 12424 | 1,9 | 0,926 | 23,824 | 1,87E-125 | 2,01E-124 | 0,0008615 | 0,002176 | 0,0013305 | 0,0013305 | TRUE | 0,0084048 | No |
| orf19.738 | 4165 | 3847 | 4302 | 2598 | 3321 | 2737 | 2890 | 3910 | 3792 | 3891 | 4071 | 3903 | 2954 | 2972 | 2960 | 2983 | 3440,6 | 3914 | 2967 | 0,758 | -0,4 | -10,79 | 3,83E-27 | 1,15E-26 | 0,0002553 | 0,002421 | 0,0010227 | 0,0010227 | TRUE | 0,0093034 | No |
| orf19.7380 | 233 | 193 | 227 | 145 | 579 | 499 | 523 | 762 | 212 | 195 | 215 | 218 | 515 | 542 | 536 | 581 | 376,7 | 210 | 544 | 2,595 | 1,376 | 16,433 | 1,10E-60 | 6,10E-60 | 1,00E-08 | 0,005544 | 0,0033364 | 0,0033364 | TRUE | 0,0280814 | No |
| orf19.7381 | 10097 | 10064 | 10954 | 6822 | 3023 | 2765 | 2657 | 3519 | 9193 | 10179 | 10367 | 10248 | 2689 | 3002 | 2721 | 2684 | 6385,32 | 9997 | 2774 | 0,277 | -1,85 | -34,427 | 1,01E-259 | 2,20E-258 | 0,0026774 | 0,002244 | 0,0025509 | 0,0025509 | TRUE | 0,0365647 | No |
| orf19.7382 | 11169 | 9580 | 10161 | 6894 | 13756 | 11525 | 11598 | 16089 | 10169 | 9689 | 9616 | 10356 | 12236 | 12513 | 11878 | 12273 | 11091,2 | 9958 | 12225 | 1,228 | 0,296 | 7,994 | 1,31E-15 | 2,95E-15 | 0,0007665 | 0,002156 | 0,0012285 | 0,0012285 | TRUE | 0,0085768 | No |
| orf19.7383 | 2855 | 2672 | 2733 | 1778 | 3569 | 2865 | 2979 | 3922 | 2599 | 2702 | 2587 | 2671 | 3175 | 3111 | 3051 | 2992 | 2860,86 | 2640 | 3082 | 1,168 | 0,224 | 5,722 | 1,05E-08 | 1,89E-08 | 0,0002751 | 0,002499 | 0,0011134 | 0,0011134 | TRUE | 0,0051565 | No |
| orf19.7384 | 24913 | 22133 | 23988 | 13916 | 37109 | 29415 | 33159 | 43765 | 22682 | 22385 | 22702 | 20904 | 33007 | 31936 | 33959 | 33384 | 27620,1 | 22168 | 33072 | 1,491 | 0,577 | 15,012 | 6,15E-51 | 2,89E-50 | 0,001026 | 0,002085 | 0,0013796 | 0,0013796 | TRUE | 0,0178066 | No |
| orf19.7385 | 414 | 316 | 370 | 223 | 1146 | 973 | 1119 | 1449 | 377 | 320 | 350 | 335 | 1019 | 1056 | 1146 | 1105 | 713,59 | 346 | 1082 | 3,123 | 1,643 | 22,861 | 1,14E-115 | 1,13E-114 | 0,0019151 | 0,003888 | 0,0029869 | 0,0029869 | TRUE | 0,0462572 | No |
| orf19.7386 | 2722 | 2540 | 2682 | 1853 | 1514 | 1350 | 1320 | 1659 | 2478 | 2569 | 2538 | 2784 | 1347 | 1466 | 1352 | 1265 | 1974,84 | 2592 | 1358 | 0,524 | -0,933 | -16,378 | 2,76E-60 | 1,51E-59 | 0,0024054 | 0,002706 | 0,0025656 | 0,0025656 | TRUE | 0,0340269 | No |
| orf19.7387 | 10073 | 9012 | 9484 | 5945 | 7417 | 6279 | 6586 | 8773 | 9171 | 9115 | 8976 | 8930 | 6597 | 6817 | 6745 | 6692 | 7880,4 | 9048 | 6713 | 0,742 | -0,431 | -14,805 | 1,36E-49 | 6,22E-49 | 8,67E-07 | 0,002205 | 0,0006856 | 0,0006856 | TRUE | 0,0016338 | No |
| orf19.7388 | 10961 | 9606 | 10507 | 6582 | 12372 | 10269 | 11228 | 15153 | 9980 | 9715 | 9944 | 9887 | 11005 | 11149 | 11499 | 11559 | 10592,2 | 9882 | 11303 | 1,144 | 0,194 | 6,246 | 4,22E-10 | 8,01E-10 | 0,0002262 | 0,002162 | 0,0008301 | 0,0008301 | TRUE | 0,0039175 | No |
| orf19.7389 | 707 | 647 | 700 | 451 | 1940 | 1721 | 1751 | 2340 | 644 | 654 | 662 | 677 | 1726 | 1869 | 1793 | 1785 | 1226,29 | 659 | 1793 | 2,722 | 1,445 | 27,663 | 1,92E-168 | 2,84E-167 | 1,00E-08 | 0,003114 | 0,0015475 | 0,0015475 | TRUE | 0,0093431 | No |
| orf19.7391 | 4073 | 3494 | 3880 | 2489 | 2119 | 1722 | 1836 | 2392 | 3708 | 3534 | 3672 | 3739 | 1885 | 1870 | 1880 | 1825 | 2764,05 | 3663 | 1865 | 0,509 | -0,974 | -25,111 | 3,79E-139 | 4,61E-138 | 1,00E-08 | 0,002515 | 0,0010474 | 0,0010474 | TRUE | 0,006902 | No |
| orf19.7392 | 39678 | 32854 | 36561 | 24303 | 55890 | 46481 | 49740 | 71880 | 36125 | 33228 | 34602 | 36507 | 49712 | 50465 | 50940 | 54831 | 43301,3 | 35116 | 51487 | 1,466 | 0,552 | 12,223 | 2,34E-34 | 8,26E-34 | 0,0018699 | 0,002068 | 0,0019368 | 0,0019368 | TRUE | 0,0234626 | No |
| orf19.7393 | 3289 | 3002 | 3243 | 2161 | 4666 | 4099 | 4350 | 5761 | 2994 | 3036 | 3069 | 3246 | 4150 | 4450 | 4455 | 4395 | 3724,52 | 3086 | 4362 | 1,415 | 0,501 | 11,786 | 4,59E-32 | 1,54E-31 | 0,0008207 | 0,002392 | 0,0014508 | 0,0014508 | TRUE | 0,0137003 | No |
| orf19.7394 | 10600 | 9323 | 10321 | 6443 | 16465 | 14199 | 14951 | 19964 | 9651 | 9429 | 9768 | 9678 | 14645 | 15416 | 15312 | 15229 | 12391 | 9632 | 15150 | 1,573 | 0,653 | 20,998 | 6,89E-98 | 5,89E-97 | 0,0002584 | 0,002144 | 0,0008432 | 0,0008432 | TRUE | 0,0061815 | No |
| orf19.7395 | 0 | 2 | 2 | 3 | 6 | 3 | 5 | 14 | 0 | 2 | 2 | 5 | 5 | 3 | 5 | 11 | 4,1 | 2 | 6 | 3,251 | 1,701 | 2,111 | 0,034747 | 0,043673 | 0,1578397 | 0,324074 | 0,2710572 | 0,2710572 | TRUE | 0,7867741 | No |
| orf19.7396 | 674 | 624 | 689 | 464 | 606 | 510 | 556 | 681 | 614 | 631 | 652 | 697 | 539 | 554 | 569 | 519 | 596,93 | 648 | 545 | 0,843 | -0,247 | -3,722 | 0,000198 | 0,000291 | 0,0004593 | 0,00425 | 0,002544 | 0,002544 | TRUE | 0,025654 | No |
| orf19.7397 | 1396 | 1312 | 1473 | 859 | 1099 | 981 | 995 | 1344 | 1271 | 1327 | 1394 | 1290 | 978 | 1065 | 1019 | 1025 | 1171,15 | 1320 | 1022 | 0,773 | -0,372 | -7,005 | 2,47E-12 | 5,08E-12 | 0,0006732 | 0,003165 | 0,0018466 | 0,0018466 | TRUE | 0,0175271 | No |
| orf19.7397.1 | 37 | 37 | 52 | 70 | 42 | 40 | 30 | 43 | 34 | 37 | 49 | 105 | 37 | 43 | 31 | 33 | 46,22 | 56 | 36 | 0,653 | -0,615 | -1,934 | 0,053129 | 0,065652 | 0,1335663 | 0,030616 | 0,0744656 | 0,0744656 | TRUE | 2,3927518 | No |
| orf19.7397.3 | 4 | 4 | 5 | 0 | 5 | 0 | 2 | 3 | 4 | 4 | 5 | 0 | 4 | 0 | 2 | 2 | 2,65 | 3 | 2 | 0,692 | -0,532 | -0,588 | 0,556577 | 0,59794 | 0,0679177 | 0,500445 | 0,4147079 | 0,4147079 | TRUE | 0,3403824 | No |
| orf19.7398 | 0 | 0 | 0 | 0 | 0 | 0 | 0 | 0 | 0 | 0 | 0 | 0 | 0 | 0 | 0 | 0 | 0 | 0 | 0 | NA | NA | NA | NA | NA | NA | NA | NA | NA | NA | NA | NA |
| orf19.7398.1 | 2 | 1 | 0 | 0 | 2 | 0 | 3 | 2 | 2 | 1 | 0 | 0 | 2 | 0 | 3 | 2 | 1,15 | 1 | 2 | 2,108 | 1,076 | 0,78 | 0,43558 | 0,477281 | 1,00E-08 | 1,149578 | 0,8401775 | 0,8401775 | TRUE | 0,2886241 | No |
| orf19.74 | 1188 | 1462 | 1366 | 843 | 882 | 832 | 768 | 942 | 1082 | 1479 | 1293 | 1266 | 785 | 903 | 787 | 719 | 1039,04 | 1280 | 798 | 0,623 | -0,683 | -7,252 | 4,10E-13 | 8,64E-13 | 0,0113332 | 0,003308 | 0,0075007 | 0,0075007 | TRUE | 0,134297 | No |
| orf19.740 | 45082 | 42743 | 46629 | 28225 | 8854 | 7478 | 7930 | 10486 | 41045 | 43230 | 44130 | 42398 | 7875 | 8119 | 8121 | 7999 | 25364,8 | 42701 | 8028 | 0,188 | -2,411 | -71,551 | 0 | 0 | 0,000479 | 0,002089 | 0,00102 | 0,00102 | TRUE | 0,0084087 | No |
| orf19.7400 | 1611 | 1342 | 1483 | 1002 | 1902 | 1609 | 1806 | 2406 | 1467 | 1357 | 1404 | 1505 | 1692 | 1747 | 1850 | 1835 | 1607,04 | 1433 | 1781 | 1,245 | 0,316 | 6,041 | 1,53E-09 | 2,85E-09 | 0,0010926 | 0,002859 | 0,0019825 | 0,0019825 | TRUE | 0,0149936 | No |
| orf19.7401 | 1744 | 1560 | 1758 | 1077 | 2500 | 2150 | 2322 | 3117 | 1588 | 1578 | 1664 | 1618 | 2224 | 2334 | 2378 | 2378 | 1970,11 | 1612 | 2328 | 1,445 | 0,531 | 11,979 | 4,58E-33 | 1,57E-32 | 0,0004015 | 0,002708 | 0,0013512 | 0,0013512 | TRUE | 0,0113406 | No |
| orf19.7402 | 1897 | 1649 | 1688 | 1083 | 2387 | 1952 | 2081 | 2621 | 1727 | 1668 | 1598 | 1627 | 2123 | 2119 | 2131 | 1999 | 1874,04 | 1655 | 2093 | 1,263 | 0,337 | 7,301 | 2,86E-13 | 6,05E-13 | 0,0006009 | 0,002742 | 0,0014995 | 0,0014995 | TRUE | 0,0115979 | No |
| orf19.7403 | 2844 | 2535 | 2761 | 1654 | 2041 | 1614 | 1818 | 2400 | 2589 | 2564 | 2613 | 2485 | 1815 | 1752 | 1862 | 1831 | 2188,9 | 2563 | 1815 | 0,708 | -0,499 | -12,215 | 2,59E-34 | 9,12E-34 | 1,00E-08 | 0,002641 | 0,001133 | 0,001133 | TRUE | 0,006154 | No |
| orf19.7404 | 2683 | 2087 | 2234 | 1589 | 2806 | 2325 | 2472 | 3545 | 2443 | 2111 | 2114 | 2387 | 2496 | 2524 | 2532 | 2704 | 2413,84 | 2264 | 2564 | 1,134 | 0,181 | 3,056 | 0,002246 | 0,003085 | 0,0031512 | 0,002584 | 0,0029569 | 0,0029569 | TRUE | 0,0364454 | No |
| orf19.7405 | 1399 | 1229 | 1264 | 841 | 1262 | 1039 | 1062 | 1481 | 1274 | 1243 | 1196 | 1263 | 1123 | 1128 | 1088 | 1130 | 1180,53 | 1244 | 1117 | 0,899 | -0,154 | -3,191 | 0,001417 | 0,00198 | 1,00E-08 | 0,003156 | 0,0013918 | 0,0013918 | TRUE | 0,0084499 | No |
| orf19.7406 | 1254 | 1145 | 1307 | 739 | 1340 | 1103 | 1061 | 1562 | 1142 | 1158 | 1237 | 1110 | 1192 | 1198 | 1087 | 1192 | 1164,29 | 1162 | 1167 | 1,003 | 0,005 | 0,087 | 0,930429 | 0,940312 | 0,0012181 | 0,003172 | 0,0021595 | 0,0021595 | TRUE | 0,025025 | No |
| orf19.7409 | 832 | 698 | 725 | 492 | 760 | 619 | 632 | 902 | 758 | 706 | 686 | 739 | 676 | 672 | 647 | 688 | 696,5 | 722 | 671 | 0,93 | -0,104 | -1,716 | 0,086073 | 0,104003 | 1,00E-08 | 0,003934 | 0,0020891 | 0,0020891 | TRUE | 0,0148878 | No |
| orf19.7409.1 | 374 | 340 | 331 | 221 | 347 | 269 | 278 | 388 | 341 | 344 | 313 | 332 | 309 | 292 | 285 | 296 | 313,88 | 332 | 296 | 0,889 | -0,169 | -2,074 | 0,038038 | 0,047622 | 1,00E-08 | 0,006246 | 0,0031964 | 0,0031964 | TRUE | 0,0189578 | No |
| orf19.741 | 8 | 11 | 11 | 5 | 9 | 7 | 4 | 11 | 7 | 11 | 10 | 8 | 8 | 8 | 4 | 8 | 8,05 | 9 | 7 | 0,775 | -0,367 | -0,768 | 0,442466 | 0,484327 | 1,00E-08 | 0,166078 | 0,0954399 | 0,0954399 | TRUE | 0,1964237 | No |
| orf19.7410 | 4638 | 4449 | 4637 | 2964 | 5761 | 5224 | 5206 | 7005 | 4223 | 4500 | 4389 | 4452 | 5124 | 5672 | 5332 | 5343 | 4879,3 | 4391 | 5368 | 1,223 | 0,29 | 6,804 | 1,02E-11 | 2,05E-11 | 0,0009924 | 0,002308 | 0,0015331 | 0,0015331 | TRUE | 0,0174597 | No |
| orf19.7411 | 665 | 497 | 579 | 379 | 696 | 626 | 645 | 834 | 605 | 503 | 548 | 569 | 619 | 680 | 661 | 636 | 602,61 | 556 | 649 | 1,165 | 0,22 | 3,075 | 0,002107 | 0,002902 | 0,0021272 | 0,004229 | 0,0032526 | 0,0032526 | TRUE | 0,0512262 | No |
| orf19.7412 | 4149 | 3729 | 4087 | 2410 | 4337 | 3344 | 3831 | 5172 | 3777 | 3771 | 3868 | 3620 | 3858 | 3631 | 3923 | 3945 | 3799,26 | 3759 | 3839 | 1,021 | 0,03 | 0,718 | 0,472672 | 0,514387 | 0,0007251 | 0,002385 | 0,0013974 | 0,0013974 | TRUE | 0,0159135 | No |
| orf19.7413 | 777 | 758 | 745 | 542 | 900 | 727 | 791 | 1060 | 707 | 767 | 705 | 814 | 801 | 789 | 810 | 809 | 775,23 | 748 | 802 | 1,077 | 0,107 | 1,73 | 0,083709 | 0,101242 | 0,0007931 | 0,003741 | 0,0023537 | 0,0023537 | TRUE | 0,0361994 | No |
| orf19.7414 | 2323 | 2073 | 2245 | 1470 | 5230 | 4723 | 4928 | 6979 | 2115 | 2097 | 2125 | 2208 | 4652 | 5128 | 5047 | 5324 | 3586,85 | 2136 | 5038 | 2,36 | 1,239 | 25,034 | 2,63E-138 | 3,16E-137 | 0,0017531 | 0,002405 | 0,002007 | 0,002007 | TRUE | 0,0327262 | No |
| orf19.7417 | 3 | 1 | 1 | 1 | 0 | 2 | 1 | 0 | 3 | 1 | 1 | 2 | 0 | 2 | 1 | 0 | 1,17 | 2 | 1 | 0,47 | -1,09 | -0,808 | 0,41914 | 0,460534 | 1,00E-08 | 1,127859 | 0,7725772 | 0,7725772 | TRUE | 0,4795566 | No |
| orf19.7419 | 48 | 37 | 39 | 18 | 83 | 55 | 67 | 90 | 44 | 37 | 37 | 27 | 74 | 60 | 69 | 69 | 51,99 | 36 | 68 | 1,837 | 0,877 | 4,4 | 1,08E-05 | 1,71E-05 | 1,00E-08 | 0,027448 | 0,0169443 | 0,0169443 | TRUE | 0,1378019 | No |
| orf19.742 | 3265 | 2797 | 2989 | 1939 | 1881 | 1590 | 1601 | 2231 | 2973 | 2829 | 2829 | 2913 | 1673 | 1726 | 1640 | 1702 | 2285,48 | 2886 | 1685 | 0,584 | -0,776 | -18,939 | 5,44E-80 | 3,85E-79 | 1,00E-08 | 0,002615 | 0,0011477 | 0,0011477 | TRUE | 0,0053903 | No |
| orf19.7420 | 1 | 1 | 1 | 1 | 2 | 3 | 4 | 4 | 1 | 1 | 1 | 2 | 2 | 3 | 4 | 3 | 2,07 | 1 | 3 | 2,855 | 1,513 | 1,493 | 0,135335 | 0,159919 | 1,00E-08 | 0,640422 | 0,3327421 | 0,3327421 | TRUE | 0,1169338 | No |
| orf19.7421 | 2425 | 1898 | 2094 | 1500 | 1633 | 1254 | 1534 | 2073 | 2208 | 1920 | 1982 | 2253 | 1453 | 1361 | 1571 | 1581 | 1791,1 | 2091 | 1492 | 0,715 | -0,485 | -7,07 | 1,55E-12 | 3,20E-12 | 0,0047912 | 0,002775 | 0,0039445 | 0,0039445 | TRUE | 0,0405976 | No |
| orf19.7422 | 4606 | 3435 | 3945 | 2529 | 5021 | 3881 | 4880 | 6506 | 4194 | 3474 | 3734 | 3799 | 4466 | 4214 | 4998 | 4963 | 4230,06 | 3800 | 4660 | 1,226 | 0,294 | 4,21 | 2,55E-05 | 3,96E-05 | 0,0061426 | 0,002349 | 0,0044544 | 0,0044544 | TRUE | 0,0600438 | No |
| orf19.7424 | 1 | 0 | 2 | 1 | 2 | 1 | 1 | 2 | 1 | 0 | 2 | 2 | 2 | 1 | 1 | 2 | 1,21 | 1 | 2 | 1,297 | 0,375 | 0,308 | 0,758346 | 0,789781 | 1,00E-08 | 1,089296 | 0,5844741 | 0,5844741 | TRUE | 0,2350004 | No |
| orf19.7425 | 914 | 870 | 929 | 594 | 849 | 732 | 779 | 1001 | 832 | 880 | 879 | 892 | 755 | 795 | 798 | 764 | 824,35 | 871 | 778 | 0,893 | -0,163 | -2,925 | 0,003446 | 0,004664 | 1,00E-08 | 0,003639 | 0,0017513 | 0,0017513 | TRUE | 0,0112251 | No |
| orf19.7426 | 919 | 788 | 911 | 521 | 950 | 752 | 789 | 1134 | 837 | 797 | 862 | 783 | 845 | 816 | 808 | 865 | 826,63 | 820 | 834 | 1,016 | 0,023 | 0,396 | 0,692077 | 0,727331 | 1,00E-08 | 0,003635 | 0,0019949 | 0,0019949 | TRUE | 0,014322 | No |

| | | | | | | | | | | | | | | | | | | | | | | | | | | | | | | | |
|---|---|---|---|---|---|---|---|---|---|---|---|---|---|---|---|---|---|---|---|---|---|---|---|---|---|---|---|---|---|---|---|
| orf19.7427 | 6879 | 6008 | 6528 | 4271 | 7126 | 5889 | 6360 | 8166 | 6263 | 6076 | 6178 | 6416 | 6338 | 6394 | 6513 | 6229 | 6301,01 | 6233 | 6368 | 1,022 | 0,031 | 0,939 | 0,347608 | 0,387654 | 0,000226 | 0,002247 | 0,0009108 | 0,0009108 | TRUE | 0,0044394 | No |
| orf19.7428 | 3791 | 3264 | 3658 | 1990 | 5221 | 4146 | 4419 | 6206 | 3452 | 3301 | 3462 | 2989 | 4644 | 4501 | 4526 | 4734 | 3951,11 | 3301 | 4601 | 1,392 | 0,477 | 9,406 | 5,17E-21 | 1,34E-20 | 0,0021587 | 0,002371 | 0,002207 | 0,002207 | TRUE | 0,0465923 | No |
| orf19.7433 | 6234 | 5462 | 6141 | 3799 | 5426 | 4636 | 5020 | 6925 | 5676 | 5524 | 5812 | 5707 | 4826 | 5033 | 5141 | 5282 | 5375,23 | 5680 | 5070 | 0,893 | -0,163 | -4,126 | 3,69E-05 | 5,68E-05 | 0,0006893 | 0,002283 | 0,0013174 | 0,0013174 | TRUE | 0,0130889 | No |
| orf19.7434 | 1040 | 1065 | 1070 | 627 | 446 | 361 | 430 | 522 | 947 | 1077 | 1013 | 942 | 397 | 392 | 440 | 398 | 700,72 | 995 | 407 | 0,408 | -1,293 | -18,403 | 1,25E-75 | 8,34E-75 | 0,002117 | 0,003922 | 0,0030519 | 0,0030519 | TRUE | 0,0371239 | No |
| orf19.7436 | 9072 | 8106 | 9347 | 6214 | 4216 | 4025 | 4239 | 5798 | 8260 | 8198 | 8846 | 9334 | 3750 | 4370 | 4341 | 4423 | 6440,32 | 8660 | 4221 | 0,488 | -1,036 | -16,573 | 1,10E-61 | 6,13E-61 | 0,0044819 | 0,002242 | 0,0035833 | 0,0035833 | TRUE | 0,069096 | No |
| orf19.7436.1 | 607 | 514 | 532 | 393 | 482 | 446 | 507 | 667 | 553 | 520 | 503 | 590 | 429 | 484 | 519 | 509 | 513,42 | 542 | 485 | 0,899 | -0,154 | -1,905 | 0,056766 | 0,069956 | 0,0040157 | 0,00461 | 0,0043366 | 0,0043366 | TRUE | 0,0737193 | No |
| orf19.7437 | 777 | 643 | 708 | 468 | 1086 | 883 | 956 | 1282 | 707 | 650 | 670 | 703 | 966 | 959 | 979 | 978 | 826,56 | 682 | 970 | 1,423 | 0,509 | 9,18 | 4,32E-20 | 1,10E-19 | 1,00E-08 | 0,003635 | 0,0016913 | 0,0016913 | TRUE | 0,0120468 | No |
| orf19.7438 | 5535 | 4887 | 5339 | 3549 | 4827 | 4025 | 4203 | 5868 | 5039 | 4943 | 5053 | 5331 | 4293 | 4370 | 4304 | 4476 | 4726,27 | 5092 | 4361 | 0,857 | -0,222 | -5,939 | 2,86E-09 | 5,27E-09 | 0,0004399 | 0,002317 | 0,0011271 | 0,0011271 | TRUE | 0,0115117 | No |
| orf19.744 | 4888 | 4289 | 4834 | 3059 | 6412 | 5637 | 5554 | 7551 | 4450 | 4338 | 4575 | 4595 | 5703 | 6120 | 5688 | 5760 | 5153,71 | 4490 | 5818 | 1,296 | 0,374 | 9,433 | 3,97E-21 | 1,04E-20 | 0,0007067 | 0,002293 | 0,0013098 | 0,0013098 | TRUE | 0,0146631 | No |
| orf19.7440 | 44 | 66 | 58 | 38 | 174 | 128 | 167 | 209 | 40 | 67 | 55 | 57 | 155 | 139 | 171 | 159 | 105,37 | 55 | 156 | 2,876 | 1,524 | 9,478 | 2,59E-21 | 6,77E-21 | 0,0057088 | 0,014573 | 0,0120542 | 0,0120542 | TRUE | 0,3164766 | No |
| orf19.7441 | 1527 | 1532 | 1600 | 1046 | 2793 | 2371 | 2444 | 3130 | 1390 | 1549 | 1514 | 1571 | 2484 | 2574 | 2503 | 2388 | 1996,79 | 1506 | 2487 | 1,653 | 0,725 | 14,271 | 3,33E-46 | 1,44E-45 | 0,0012081 | 0,002699 | 0,001934 | 0,001934 | TRUE | 0,0335108 | No |
| orf19.7442 | 1214 | 1101 | 1176 | 694 | 2992 | 2224 | 2413 | 3225 | 1105 | 1114 | 1113 | 1042 | 2661 | 2415 | 2471 | 2460 | 1797,69 | 1094 | 2502 | 2,284 | 1,192 | 23,78 | 5,36E-125 | 5,72E-124 | 0,0009625 | 0,002772 | 0,0017347 | 0,0017347 | TRUE | 0,0225036 | No |
| orf19.7443 | 477 | 580 | 575 | 360 | 405 | 358 | 365 | 383 | 434 | 587 | 544 | 541 | 360 | 389 | 374 | 292 | 440,09 | 526 | 354 | 0,671 | -0,576 | -5,254 | 1,49E-07 | 2,55E-07 | 0,0139585 | 0,005039 | 0,0092014 | 0,0092014 | TRUE | 0,1696112 | No |
| orf19.7444 | 784 | 788 | 786 | 483 | 402 | 375 | 379 | 480 | 714 | 797 | 744 | 726 | 358 | 407 | 388 | 366 | 562,4 | 745 | 380 | 0,508 | -0,977 | -13,789 | 2,98E-43 | 1,23E-42 | 0,0010349 | 0,004386 | 0,0028699 | 0,0028699 | TRUE | 0,0265648 | No |
| orf19.7445 | 8828 | 8407 | 9260 | 5898 | 4427 | 3957 | 4149 | 5567 | 8038 | 8503 | 8764 | 8860 | 3938 | 4296 | 4249 | 4247 | 6361,66 | 8541 | 4182 | 0,49 | -1,03 | -22,529 | 2,14E-112 | 2,05E-111 | 0,0014661 | 0,002245 | 0,0018317 | 0,0018317 | TRUE | 0,019391 | No |
| orf19.7446 | 214 | 208 | 211 | 126 | 610 | 543 | 514 | 676 | 195 | 210 | 200 | 189 | 543 | 590 | 526 | 516 | 371,04 | 198 | 544 | 2,726 | 1,447 | 16,933 | 2,55E-64 | 1,48E-63 | 1,00E-08 | 0,005597 | 0,0034808 | 0,0034808 | TRUE | 0,036299 | No |
| orf19.7447 | 39 | 28 | 39 | 18 | 92 | 94 | 90 | 127 | 36 | 28 | 37 | 27 | 82 | 102 | 92 | 97 | 62,59 | 32 | 93 | 2,875 | 1,524 | 7,891 | 2,99E-15 | 6,66E-15 | 1,00E-08 | 0,023143 | 0,014434 | 0,014434 | TRUE | 0,0715298 | No |
| orf19.7448 | 1618 | 1380 | 1466 | 961 | 2635 | 2076 | 2006 | 2870 | 1473 | 1396 | 1387 | 1444 | 2344 | 2254 | 2054 | 2189 | 1817,65 | 1425 | 2210 | 1,552 | 0,634 | 12,061 | 1,69E-33 | 5,85E-33 | 0,0015402 | 0,002764 | 0,0020641 | 0,0020641 | TRUE | 0,0267779 | No |
| orf19.7449 | 2298 | 2246 | 2315 | 1452 | 1630 | 1368 | 1452 | 1782 | 2092 | 2272 | 2191 | 2181 | 1450 | 1485 | 1487 | 1359 | 1814,67 | 2184 | 1445 | 0,661 | -0,597 | -12,179 | 4,02E-34 | 1,40E-33 | 0,0009627 | 0,002765 | 0,0017374 | 0,0017374 | TRUE | 0,0203737 | No |
| orf19.745 | 6994 | 6542 | 6985 | 4995 | 3744 | 3594 | 3424 | 4614 | 6368 | 6617 | 6611 | 7503 | 3330 | 3902 | 3507 | 3520 | 5169,59 | 6775 | 3565 | 0,526 | -0,926 | -14,754 | 2,90E-49 | 1,32E-48 | 0,0047178 | 0,002293 | 0,0035728 | 0,0035728 | TRUE | 0,0629981 | No |
| orf19.7450 | 1042 | 1020 | 1028 | 606 | 1389 | 1160 | 1213 | 1641 | 949 | 1032 | 973 | 910 | 1235 | 1259 | 1242 | 1252 | 1106,56 | 966 | 1247 | 1,288 | 0,366 | 6,987 | 2,81E-12 | 5,76E-12 | 1,00E-08 | 0,003231 | 0,0017016 | 0,0017016 | TRUE | 0,0243344 | No |
| orf19.7451 | 6407 | 5970 | 6432 | 4274 | 2374 | 1999 | 2093 | 2834 | 5833 | 6038 | 6087 | 6420 | 2112 | 2170 | 2143 | 2162 | 4120,76 | 6094 | 2147 | 0,353 | -1,504 | -36,632 | 8,96E-294 | 2,26E-292 | 0,0007165 | 0,002358 | 0,0013131 | 0,0013131 | TRUE | 0,015109 | No |
| orf19.7452 | 113 | 125 | 120 | 65 | 91 | 75 | 94 | 116 | 103 | 126 | 114 | 98 | 81 | 81 | 96 | 88 | 98,45 | 110 | 86 | 0,784 | -0,351 | -2,429 | 0,015153 | 0,019642 | 1,00E-08 | 0,015454 | 0,009853 | 0,009853 | TRUE | 0,0959026 | No |
| orf19.7453 | 65 | 56 | 70 | 42 | 52 | 43 | 49 | 63 | 59 | 57 | 66 | 63 | 46 | 47 | 50 | 48 | 54,54 | 61 | 48 | 0,781 | -0,357 | -1,959 | 0,050065 | 0,06201 | 1,00E-08 | 0,026257 | 0,013485 | 0,013485 | TRUE | 0,0303822 | No |
| orf19.7454 | 1238 | 1094 | 1165 | 759 | 1090 | 1025 | 999 | 1426 | 1127 | 1106 | 1103 | 1140 | 970 | 1113 | 1023 | 1088 | 1083,7 | 1119 | 1048 | 0,937 | -0,094 | -1,651 | 0,098798 | 0,118748 | 0,0011677 | 0,003256 | 0,0021705 | 0,0021705 | TRUE | 0,031281 | No |
| orf19.7455 | 343 | 300 | 351 | 206 | 19 | 10 | 13 | 13 | 312 | 303 | 332 | 309 | 17 | 11 | 13 | 10 | 163,54 | 314 | 13 | 0,04 | -4,63 | -21,551 | 5,13E-103 | 4,61E-102 | 1,00E-08 | 0,010114 | 0,0062501 | 0,0062501 | TRUE | 0,2343678 | No |
| orf19.7456 | 56 | 52 | 38 | 31 | 49 | 39 | 39 | 47 | 51 | 53 | 36 | 47 | 44 | 42 | 40 | 36 | 43,48 | 47 | 40 | 0,867 | -0,206 | -0,97 | 0,331882 | 0,37164 | 1,00E-08 | 0,03242 | 0,0204549 | 0,0204549 | TRUE | 0,2097999 | No |
| orf19.7457 | 1015 | 852 | 940 | 544 | 691 | 512 | 568 | 739 | 924 | 862 | 890 | 817 | 615 | 556 | 582 | 564 | 726,07 | 873 | 579 | 0,661 | -0,596 | -9,385 | 6,28E-21 | 1,63E-20 | 0,0009 | 0,003856 | 0,0024576 | 0,0024576 | TRUE | 0,0206839 | No |
| orf19.7458 | 393 | 322 | 328 | 209 | 283 | 246 | 255 | 344 | 358 | 326 | 310 | 314 | 252 | 267 | 261 | 262 | 293,78 | 327 | 260 | 0,795 | -0,332 | -3,85 | 0,000118 | 0,000176 | 1,00E-08 | 0,006534 | 0,0037146 | 0,0037146 | TRUE | 0,0491964 | No |
| orf19.7459 | 16260 | 15324 | 16376 | 11299 | 9853 | 8893 | 8701 | 11607 | 14804 | 15499 | 15498 | 16973 | 8764 | 9655 | 8911 | 8854 | 12369,8 | 15694 | 9046 | 0,577 | -0,795 | -15,63 | 4,53E-55 | 2,29E-54 | 0,0026089 | 0,002144 | 0,0023967 | 0,0023967 | TRUE | 0,0364825 | No |
| orf19.746 | 2492 | 2880 | 2943 | 1722 | 2468 | 2130 | 2256 | 2701 | 2269 | 2913 | 2785 | 2587 | 2195 | 2313 | 2310 | 2060 | 2429,03 | 2638 | 2220 | 0,841 | -0,25 | -3,355 | 0,000795 | 0,001129 | 0,0065495 | 0,002581 | 0,0049193 | 0,0049193 | TRUE | 0,1103584 | No |
| orf19.7460 | 2177 | 1798 | 1961 | 1235 | 2485 | 1954 | 1988 | 2782 | 1982 | 1818 | 1856 | 1855 | 2210 | 2121 | 2036 | 2122 | 2000,19 | 1878 | 2122 | 1,13 | 0,176 | 3,747 | 0,000179 | 0,000265 | 0,0008237 | 0,002697 | 0,0016158 | 0,0016158 | TRUE | 0,0176826 | No |
| orf19.7463 | 510 | 431 | 518 | 337 | 245 | 232 | 227 | 294 | 464 | 436 | 490 | 506 | 218 | 252 | 232 | 224 | 352,91 | 474 | 232 | 0,488 | -1,034 | -12,119 | 8,35E-34 | 2,91E-33 | 1,00E-08 | 0,00578 | 0,0038427 | 0,0038427 | TRUE | 0,0357429 | No |
| orf19.7464 | 16 | 8 | 11 | 3 | 31 | 22 | 20 | 37 | 15 | 8 | 10 | 5 | 28 | 24 | 20 | 28 | 17,22 | 10 | 25 | 2,568 | 1,361 | 3,749 | 0,000178 | 0,000263 | 1,00E-08 | 0,07876 | 0,0538905 | 0,0538905 | TRUE | 0,4687184 | No |
| orf19.7466 | 6592 | 5967 | 6412 | 3819 | 56276 | 46929 | 50893 | 68287 | 6002 | 6035 | 6068 | 5737 | 50056 | 50952 | 52121 | 52090 | 28632,5 | 5960 | 51305 | 8,601 | 3,105 | 95,982 | 0 | 0 | 0,0003865 | 0,002083 | 0,0009052 | 0,0009052 | TRUE | 0,0072922 | No |
| orf19.7468 | 3510 | 3095 | 3249 | 2171 | 1314 | 1166 | 1153 | 1637 | 3196 | 3130 | 3075 | 3261 | 1169 | 1266 | 1181 | 1249 | 2190,79 | 3166 | 1216 | 0,384 | -1,379 | -30,421 | 2,89E-203 | 5,02E-202 | 0,0003587 | 0,00264 | 0,0014199 | 0,0014199 | TRUE | 0,0084857 | No |
| orf19.7469 | 30259 | 25123 | 27164 | 15885 | 21101 | 16317 | 18009 | 24626 | 27550 | 25409 | 25708 | 23862 | 18769 | 17716 | 18443 | 18785 | 22030,2 | 25632 | 18428 | 0,719 | -0,476 | -10,137 | 3,79E-24 | 1,07E-23 | 0,0020481 | 0,002097 | 0,0020741 | 0,0020741 | TRUE | 0,0310597 | No |
| orf19.747 | 3951 | 3509 | 3896 | 2307 | 1923 | 1581 | 1567 | 2423 | 3597 | 3549 | 3687 | 3465 | 1710 | 1717 | 1605 | 1848 | 2647,37 | 3574 | 1720 | 0,481 | -1,055 | -21,293 | 1,31E-100 | 1,15E-99 | 0,0014423 | 0,002536 | 0,001933 | 0,001933 | TRUE | 0,0317225 | No |
| orf19.7472 | 1645 | 1446 | 1651 | 994 | 909 | 821 | 828 | 1058 | 1498 | 1462 | 1563 | 1493 | 809 | 891 | 848 | 807 | 1171,35 | 1504 | 839 | 0,556 | -0,846 | -15,888 | 7,73E-57 | 4,01E-56 | 0,0004825 | 0,003165 | 0,0018066 | 0,0018066 | TRUE | 0,0204604 | No |
| orf19.7473 | 1862 | 1861 | 2018 | 1295 | 318 | 309 | 334 | 385 | 1695 | 1882 | 1910 | 1945 | 283 | 335 | 342 | 294 | 1085,84 | 1858 | 314 | 0,168 | -2,572 | -34,355 | 1,18E-258 | 2,57E-257 | 0,0037804 | 0,003254 | 0,0035989 | 0,0035989 | TRUE | 0,0475704 | No |
| orf19.7475 | 940 | 1016 | 1033 | 670 | 529 | 514 | 505 | 628 | 856 | 1028 | 978 | 1006 | 471 | 558 | 517 | 479 | 736,54 | 967 | 506 | 0,523 | -0,935 | -11,927 | 8,60E-33 | 2,93E-32 | 0,0050319 | 0,003831 | 0,0044182 | 0,0044182 | TRUE | 0,0739915 | No |
| orf19.7477 | 5167 | 4813 | 5093 | 2718 | 5314 | 3946 | 4477 | 5876 | 4704 | 4868 | 4820 | 4083 | 4727 | 4284 | 4585 | 4482 | 4569,16 | 4619 | 4520 | 0,978 | -0,033 | -0,551 | 0,581528 | 0,62202 | 0,0033404 | 0,002326 | 0,0031328 | 0,0031328 | TRUE | 0,0720965 | No |
| orf19.7478 | 3094 | 2933 | 3107 | 1952 | 2705 | 2272 | 2318 | 3067 | 2817 | 2966 | 2940 | 2932 | 2406 | 2467 | 2374 | 2340 | 2655,29 | 2914 | 2397 | 0,822 | -0,282 | -7,196 | 6,22E-13 | 1,30E-12 | 0,0001941 | 0,002535 | 0,0010984 | 0,0010984 | TRUE | 0,0064198 | No |
| orf19.7479 | 5064 | 4775 | 5134 | 3062 | 9404 | 7853 | 7991 | 10622 | 4611 | 4829 | 4859 | 4600 | 8365 | 8526 | 8184 | 8103 | 6509,44 | 4725 | 8294 | 1,754 | 0,811 | 22,312 | 2,84E-110 | 2,68E-109 | 0,0004698 | 0,00224 | 0,0010983 | 0,0010983 | TRUE | 0,0044778 | No |
| orf19.748 | 3843 | 3840 | 4160 | 2340 | 3672 | 3116 | 3205 | 4117 | 3499 | 3884 | 3937 | 3515 | 3266 | 3383 | 3282 | 3140 | 3488,35 | 3709 | 3268 | 0,88 | -0,184 | -3,56 | 0,000371 | 0,000538 | 0,0020711 | 0,002416 | 0,0022739 | 0,0022739 | TRUE | 0,0210153 | No |
| orf19.7480 | 6 | 5 | 2 | 4 | 12 | 9 | 11 | 14 | 5 | 5 | 2 | 6 | 11 | 10 | 11 | 11 | 7,6 | 4 | 11 | 2,358 | 1,238 | 2,377 | 0,017473 | 0,022534 | 1,00E-08 | 0,175821 | 0,0987945 | 0,0987945 | TRUE | 0,3404749 | No |
| orf19.7481 | 2810 | 2452 | 2714 | 1891 | 3460 | 2859 | 2687 | 4093 | 2558 | 2480 | 2569 | 2841 | 3078 | 3104 | 2752 | 3122 | 2812,89 | 2612 | 3014 | 1,156 | 0,209 | 3,631 | 0,000283 | 0,000414 | 0,0030414 | 0,002507 | 0,0028162 | 0,0028162 | TRUE | 0,0412772 | No |
| orf19.7482 | 38 | 62 | 44 | 20 | 19 | 14 | 17 | 26 | 35 | 63 | 42 | 30 | 17 | 15 | 17 | 20 | 29,79 | 42 | 17 | 0,409 | -1,289 | -4,379 | 1,19E-05 | 1,89E-05 | 0,04043 | 0,046378 | 0,0439473 | 0,0439473 | TRUE | 0,715212 | No |
| orf19.7483 | 6166 | 6127 | 6502 | 3869 | 1453 | 1243 | 1234 | 1660 | 5614 | 6197 | 6154 | 5812 | 1292 | 1350 | 1264 | 1266 | 3618,51 | 5944 | 1293 | 0,217 | -2,202 | -45,493 | 0 | 0 | 0,0014034 | 0,002402 | 0,0017974 | 0,0017974 | TRUE | 0,0176036 | No |
| orf19.7484 | 1973 | 1606 | 1766 | 1127 | 2989 | 2288 | 2370 | 3362 | 1796 | 1624 | 1671 | 1693 | 2659 | 2484 | 2427 | 2565 | 2114,93 | 1696 | 2534 | 1,494 | 0,579 | 11,696 | 1,33E-31 | 4,45E-31 | 0,0012549 | 0,002662 | 0,001852 | 0,001852 | TRUE | 0,0201883 | No |
| orf19.7485 | 3583 | 3114 | 3342 | 1922 | 3547 | 2645 | 2969 | 3934 | 3262 | 3149 | 3163 | 2887 | 3155 | 2872 | 3041 | 3001 | 3066,24 | 3115 | 3017 | 0,967 | -0,048 | -0,973 | 0,330787 | 0,370479 | 0,001638 | 0,002468 | 0,002 | 0,002 | TRUE | 0,0278522 | No |
| orf19.7486 | 947 | 941 | 976 | 602 | 1671 | 1312 | 1354 | 1819 | 862 | 952 | 924 | 904 | 1486 | 1424 | 1387 | 1388 | 1165,86 | 910 | 1421 | 1,561 | 0,643 | 12,043 | 2,12E-33 | 7,30E-33 | 0,000591 | 0,00317 | 0,0018175 | 0,0018175 | TRUE | 0,0168134 | No |
| orf19.7487 | 13 | 16 | 14 | 7 | 18 | 22 | 18 | 32 | 12 | 16 | 13 | 11 | 16 | 24 | 18 | 24 | 16,82 | 13 | 20 | 1,584 | 0,664 | 1,946 | 0,051617 | 0,063845 | 1,00E-08 | 0,080596 | 0,0486276 | 0,0486276 | TRUE | 0,1434636 | No |
| orf19.7488 | 7342 | 6906 | 6739 | 4565 | 8610 | 6554 | 6765 | 9005 | 6685 | 6985 | 6378 | 6857 | 7658 | 7116 | 6928 | 6869 | 6934,49 | 6726 | 7143 | 1,062 | 0,087 | 1,835 | 0,06655 | 0,081395 | 0,0018362 | 0,002228 | 0,0020167 | 0,0020167 | TRUE | 0,0290019 | No |
| orf19.7489 | 2466 | 2133 | 2424 | 1515 | 4128 | 3617 | 3820 | 5124 | 2245 | 2157 | 2294 | 2276 | 3672 | 3927 | 3912 | 3909 | 3048,99 | 2243 | 3855 | 1,719 | 0,781 | 18,668 | 8,97E-78 | 6,17E-77 | 0,0006033 | 0,00247 | 0,0013213 | 0,0013213 | TRUE | 0,0125611 | No |
| orf19.7489.3 | 1259 | 1048 | 1087 | 613 | 829 | 571 | 651 | 770 | 1146 | 1060 | 1029 | 921 | 737 | 620 | 667 | 587 | 845,9 | 1039 | 653 | 0,627 | -0,674 | -7,997 | 1,27E-15 | 2,87E-15 | 0,007323 | 0,003599 | 0,005589 | 0,005589 | TRUE | 0,0922086 | No |
| orf19.7490 | 298 | 279 | 303 | 162 | 396 | 384 | 391 | 487 | 271 | 282 | 287 | 243 | 352 | 417 | 400 | 371 | 328,08 | 271 | 385 | 1,411 | 0,497 | 5,602 | 2,12E-08 | 3,77E-08 | 0,002038 | 0,006063 | 0,0043844 | 0,0043844 | TRUE | 0,0418065 | No |
| orf19.7491 | 72 | 68 | 82 | 47 | 91 | 82 | 85 | 127 | 66 | 69 | 78 | 71 | 81 | 89 | 87 | 97 | 79,55 | 71 | 88 | 1,255 | 0,328 | 2,11 | 0,034883 | 0,043828 | 1,00E-08 | 0,018642 | 0,0104156 | 0,0104156 | TRUE | 0,0464065 | No |
| orf19.7492 | 2033 | 1863 | 1958 | 1220 | 2293 | 1880 | 2132 | 2735 | 1851 | 1884 | 1853 | 1833 | 2040 | 2041 | 2183 | 2086 | 1971,41 | 1855 | 2088 | 1,125 | 0,17 | 4,006 | 6,18E-05 | 9,36E-05 | 1,00E-08 | 0,002707 | 0,0012062 | 0,0012062 | TRUE | 0,011164 | No |
| orf19.7494 | 215 | 214 | 240 | 141 | 494 | 393 | 375 | 554 | 196 | 216 | 227 | 212 | 439 | 427 | 384 | 423 | 315,48 | 213 | 418 | 1,968 | 0,976 | 11,025 | 2,90E-28 | 8,91E-28 | 1,00E-08 | 0,006224 | 0,0039136 | 0,0039136 | TRUE | 0,0369595 | No |
| orf19.7495 | 173 | 177 | 180 | 85 | 630 | 491 | 542 | 743 | 158 | 179 | 170 | 128 | 560 | 533 | 555 | 567 | 356,23 | 159 | 554 | 3,452 | 1,787 | 19,463 | 2,27E-84 | 1,70E-83 | 1,00E-08 | 0,005745 | 0,0039473 | 0,0039473 | TRUE | 0,1315604 | No |
| orf19.7497 | 1791 | 1629 | 1824 | 1011 | 2324 | 1886 | 2120 | 2632 | 1631 | 1648 | 1726 | 1519 | 2067 | 2048 | 2171 | 2008 | 1852,1 | 1631 | 2074 | 1,268 | 0,343 | 6,784 | 1,17E-11 | 2,35E-11 | 0,0012109 | 0,00275 | 0,0018971 | 0,0018971 | TRUE | 0,0232064 | No |
| orf19.7498 | 3053 | 2794 | 3039 | 1982 | 1751 | 1455 | 1451 | 1996 | 2780 | 2826 | 2876 | 2977 | 1557 | 1580 | 1486 | 1523 | 2200,58 | 2865 | 1536 | 0,537 | -0,897 | -20,916 | 3,82E-97 | 3,23E-96 | 0,0002613 | 0,002637 | 0,0012759 | 0,0012759 | TRUE | 0,0078038 | No |
| orf19.7499 | 2746 | 2220 | 2476 | 1642 | 3127 | 2607 | 2677 | 3749 | 2500 | 2245 | 2343 | 2467 | 2781 | 2830 | 2742 | 2860 | 2596,06 | 2389 | 2803 | 1,174 | 0,232 | 5,085 | 3,68E-07 | 6,23E-07 | 0,0008578 | 0,002546 | 0,0016051 | 0,0016051 | TRUE | 0,0199641 | No |
| orf19.7499.1 | 564 | 486 | 514 | 338 | 637 | 472 | 501 | 715 | 513 | 492 | 486 | 508 | 567 | 512 | 513 | 545 | 517,1 | 500 | 534 | 1,072 | 0,1 | 1,474 | 0,140382 | 0,165483 | 1,00E-08 | 0,004592 | 0,0024699 | 0,0024699 | TRUE | 0,0188077 | No |
| orf19.750 | 234 | 160 | 177 | 143 | 326 | 375 | 327 | 482 | 213 | 162 | 168 | 215 | 290 | 407 | 335 | 368 | 269,61 | 190 | 350 | 1,854 | 0,891 | 6,975 | 3,06E-12 | 6,25E-12 | 0,0174313 | 0,006937 | 0,0114845 | 0,0114845 | TRUE | 0,1543937 | No |
| orf19.7500 | 5575 | 5261 | 5527 | 3518 | 4898 | 4323 | 4517 | 5628 | 5076 | 5321 | 5231 | 5285 | 4357 | 4694 | 4626 | 4293 | 4860,17 | 5228 | 4492 | 0,859 | -0,219 | -5,251 | 1,52E-07 | 2,60E-07 | 0,0008995 | 0,002309 | 0,0014688 | 0,0014688 | TRUE | 0,0110427 | No |

| | | | | | | | | | | | | | | | | | | | | | | | | | | | | | | | |
|---|---|---|---|---|---|---|---|---|---|---|---|---|---|---|---|---|---|---|---|---|---|---|---|---|---|---|---|---|---|---|---|
| orf19.7501 | 28628 | 24688 | 26588 | 15327 | 28136 | 21366 | 24426 | 32858 | 26065 | 24969 | 25163 | 23024 | 25026 | 23198 | 25015 | 25064 | 24690,5 | 24805 | 24576 | 0,991 | -0,014 | -0,292 | 0,770284 | 0,800418 | 0,0017292 | 0,002091 | 0,0020458 | 0,0020458 | TRUE | 0,0283861 | No |
| orf19.7502 | 9371 | 11307 | 11023 | 6778 | 15738 | 15143 | 14011 | 17340 | 8532 | 11436 | 10432 | 10182 | 13998 | 16441 | 14349 | 13227 | 12324,7 | 10146 | 14504 | 1,43 | 0,516 | 6,049 | 1,45E-09 | 2,71E-09 | 0,0114217 | 0,002144 | 0,0068932 | 0,0068932 | TRUE | 0,1407183 | No |
| orf19.7503 | 1135 | 1070 | 1123 | 789 | 1268 | 1147 | 1086 | 1546 | 1033 | 1082 | 1063 | 1185 | 1128 | 1245 | 1112 | 1179 | 1128,53 | 1091 | 1166 | 1,072 | 0,1 | 1,669 | 0,0952 | 0,114639 | 0,0019732 | 0,003208 | 0,002561 | 0,002561 | TRUE | 0,0365137 | No |
| orf19.7504 | 19456 | 18081 | 19554 | 12525 | 16368 | 13452 | 13892 | 19095 | 17714 | 18287 | 18506 | 18815 | 14559 | 14605 | 14227 | 14566 | 16409,8 | 18330 | 14489 | 0,791 | -0,339 | -11,038 | 2,50E-28 | 7,70E-28 | 0,0003124 | 0,002118 | 0,0008435 | 0,0008435 | TRUE | 0,0063058 | No |
| orf19.7506 | 2010 | 1888 | 2010 | 1168 | 2610 | 2072 | 2196 | 2869 | 1830 | 1910 | 1902 | 1755 | 2322 | 2250 | 2249 | 2188 | 2050,62 | 1849 | 2252 | 1,216 | 0,282 | 6,307 | 2,85E-10 | 5,44E-10 | 0,0004048 | 0,002681 | 0,0014205 | 0,0014205 | TRUE | 0,0126695 | No |
| orf19.7507 | 1100 | 893 | 961 | 621 | 1098 | 801 | 939 | 1309 | 1002 | 903 | 909 | 933 | 977 | 870 | 962 | 999 | 944,19 | 937 | 952 | 1,017 | 0,024 | 0,387 | 0,698399 | 0,733008 | 0,0019649 | 0,003436 | 0,0026854 | 0,0026854 | TRUE | 0,0386265 | No |
| orf19.7509.1 | 2937 | 2151 | 2217 | 1565 | 5682 | 3926 | 4453 | 6323 | 2674 | 2176 | 2098 | 2351 | 5054 | 4263 | 4560 | 4823 | 3499,84 | 2325 | 4675 | 2,011 | 1,008 | 13,05 | 6,40E-39 | 2,44E-38 | 0,008284 | 0,002414 | 0,0054012 | 0,0054012 | TRUE | 0,1268947 | No |
| orf19.7509.2 | 2736 | 2060 | 2188 | 1100 | 3178 | 2036 | 2750 | 3509 | 2491 | 2083 | 2071 | 1652 | 2827 | 2211 | 2816 | 2677 | 2353,49 | 2074 | 2633 | 1,268 | 0,343 | 2,405 | 0,016192 | 0,020934 | 0,0190989 | 0,002598 | 0,0105093 | 0,0190989 | TRUE | 0,2233938 | No |
| orf19.751 | 38 | 32 | 40 | 15 | 60 | 45 | 53 | 70 | 35 | 32 | 38 | 23 | 53 | 49 | 54 | 53 | 42,16 | 32 | 52 | 1,617 | 0,693 | 3,218 | 0,001293 | 0,001812 | 1,00E-08 | 0,033373 | 0,0193574 | 0,0193574 | TRUE | 0,1686586 | No |
| orf19.7510 | 4325 | 3399 | 4002 | 2645 | 3653 | 3190 | 3441 | 4941 | 3938 | 3438 | 3788 | 3973 | 3249 | 3463 | 3524 | 3769 | 3642,74 | 3784 | 3501 | 0,926 | -0,111 | -1,854 | 0,06372 | 0,078178 | 0,0035857 | 0,0024 | 0,0031728 | 0,0031728 | TRUE | 0,0463351 | No |
| orf19.7511 | 5344 | 4792 | 5180 | 3184 | 4231 | 3477 | 3874 | 5045 | 4865 | 4847 | 4902 | 4783 | 3763 | 3775 | 3967 | 3848 | 4343,95 | 4849 | 3838 | 0,791 | -0,338 | -9,91 | 3,76E-23 | 1,03E-22 | 4,07E-05 | 0,002341 | 0,0008829 | 0,0008829 | TRUE | 0,0060421 | No |
| orf19.7512 | 1485 | 1222 | 1324 | 879 | 1303 | 1033 | 1078 | 1567 | 1352 | 1236 | 1253 | 1320 | 1159 | 1122 | 1104 | 1195 | 1217,66 | 1290 | 1145 | 0,889 | -0,17 | -3,199 | 0,001377 | 0,001925 | 0,0008208 | 0,003122 | 0,0018876 | 0,0018876 | TRUE | 0,0138511 | No |
| orf19.7513 | 1876 | 1443 | 1585 | 970 | 5144 | 4182 | 4465 | 6291 | 1708 | 1459 | 1500 | 1457 | 4575 | 4540 | 4573 | 4799 | 3076,51 | 1531 | 4622 | 3,014 | 1,592 | 28,753 | 8,34E-182 | 1,30E-180 | 0,0028001 | 0,002466 | 0,0024906 | 0,0024906 | TRUE | 0,0750424 | No |
| orf19.7514 | 40286 | 32257 | 35024 | 23935 | 85879 | 70149 | 74942 | 1E+05 | 36679 | 32625 | 33147 | 35954 | 76387 | 76162 | 76750 | 81723 | 56178,3 | 34601 | 77756 | 2,247 | 1,168 | 24,452 | 4,83E-132 | 5,50E-131 | 0,0022495 | 0,002061 | 0,0021716 | 0,0021716 | TRUE | 0,0201326 | No |
| orf19.7516 | 2078 | 1697 | 1914 | 1288 | 2484 | 2196 | 2420 | 3237 | 1892 | 1716 | 1811 | 1935 | 2209 | 2384 | 2478 | 2469 | 2111,97 | 1838 | 2385 | 1,299 | 0,377 | 6,81 | 9,78E-12 | 1,98E-11 | 0,0020937 | 0,002662 | 0,0024532 | 0,0024532 | TRUE | 0,0302727 | No |
| orf19.7517 | 273 | 224 | 209 | 143 | 310 | 278 | 301 | 437 | 249 | 227 | 198 | 215 | 276 | 302 | 308 | 333 | 263,36 | 222 | 305 | 1,372 | 0,457 | 4,509 | 6,50E-06 | 1,04E-05 | 0,0044014 | 0,007053 | 0,00591 | 0,00591 | TRUE | 0,0786089 | No |
| orf19.7518 | 1308 | 1104 | 1149 | 836 | 2827 | 2400 | 2614 | 3600 | 1191 | 1117 | 1087 | 1256 | 2515 | 2606 | 2677 | 2746 | 1899,26 | 1163 | 2636 | 2,273 | 1,185 | 21,517 | 1,07E-102 | 9,56E-102 | 0,0020085 | 0,002733 | 0,0022705 | 0,0022705 | TRUE | 0,0310021 | No |
| orf19.7519 | 2354 | 1799 | 1963 | 1330 | 1802 | 1361 | 1590 | 2070 | 2143 | 1819 | 1858 | 1998 | 1603 | 1478 | 1628 | 1579 | 1763,28 | 1954 | 1572 | 0,805 | -0,314 | -5,129 | 2,92E-07 | 4,95E-07 | 0,0029554 | 0,002786 | 0,003018 | 0,003018 | TRUE | 0,0535111 | No |
| orf19.752 | 1849 | 1675 | 1835 | 1136 | 2368 | 2069 | 2089 | 2975 | 1683 | 1694 | 1737 | 1706 | 2106 | 2246 | 2139 | 2269 | 1947,75 | 1705 | 2190 | 1,285 | 0,362 | 8,174 | 2,99E-16 | 6,84E-16 | 0,0004201 | 0,002715 | 0,0013509 | 0,0013509 | TRUE | 0,0083519 | No |
| orf19.7520 | 7323 | 5773 | 6383 | 4397 | 83468 | 68154 | 71775 | 1E+05 | 6667 | 5839 | 6041 | 6605 | 74242 | 73996 | 73507 | 78749 | 40705,8 | 6288 | 75124 | 11,951 | 3,579 | 71,507 | 0 | 0 | 0,0025489 | 0,002069 | 0,0023144 | 0,0023144 | TRUE | 0,028233 | No |
| orf19.7521 | 1009 | 819 | 980 | 628 | 1963 | 1669 | 1851 | 2440 | 919 | 828 | 927 | 943 | 1746 | 1812 | 1896 | 1861 | 1366,6 | 904 | 1829 | 2,024 | 1,018 | 18,334 | 4,40E-75 | 2,91E-74 | 0,001167 | 0,003004 | 0,0021084 | 0,0021084 | TRUE | 0,038585 | No |
| orf19.7522 | 793 | 669 | 703 | 471 | 1923 | 1563 | 1709 | 2282 | 722 | 677 | 665 | 708 | 1710 | 1697 | 1750 | 1741 | 1208,73 | 693 | 1724 | 2,492 | 1,317 | 25,884 | 1,01E-147 | 1,31E-146 | 1,00E-08 | 0,00313 | 0,0014451 | 0,0014451 | TRUE | 0,0113382 | No |
| orf19.7523 | 3170 | 2637 | 3015 | 1830 | 4059 | 3113 | 3676 | 4780 | 2886 | 2667 | 2853 | 2749 | 3610 | 3380 | 3765 | 3646 | 3194,59 | 2789 | 3600 | 1,291 | 0,368 | 7,842 | 4,43E-15 | 9,81E-15 | 0,0012208 | 0,002451 | 0,0017924 | 0,0017924 | TRUE | 0,020158 | No |
| orf19.7527 | 2967 | 2497 | 2714 | 1793 | 2007 | 1753 | 1955 | 2496 | 2701 | 2525 | 2569 | 2693 | 1785 | 1903 | 2002 | 1904 | 2260,41 | 2622 | 1898 | 0,724 | -0,466 | -9,616 | 6,85E-22 | 1,81E-21 | 0,001059 | 0,002621 | 0,0018007 | 0,0018007 | TRUE | 0,0197136 | No |
| orf19.7529 | 18782 | 15228 | 17404 | 10995 | 9398 | 7472 | 8339 | 11512 | 17100 | 15401 | 16471 | 16516 | 8359 | 8113 | 8540 | 8781 | 12410,3 | 16372 | 8448 | 0,516 | -0,954 | -22,174 | 6,16E-109 | 5,75E-108 | 0,0013755 | 0,002143 | 0,0016908 | 0,0016908 | TRUE | 0,0195971 | No |
| orf19.753 | 1363 | 1332 | 1514 | 854 | 2056 | 1697 | 1719 | 2238 | 1241 | 1347 | 1433 | 1283 | 1829 | 1842 | 1760 | 1707 | 1555,34 | 1326 | 1784 | 1,344 | 0,427 | 7,6 | 2,96E-14 | 6,41E-14 | 0,0017324 | 0,002886 | 0,0023608 | 0,0023608 | TRUE | 0,0366598 | No |
| orf19.7531 | 3731 | 3036 | 3158 | 2298 | 2260 | 1835 | 1922 | 2736 | 3397 | 3071 | 2989 | 3452 | 2010 | 1992 | 1968 | 2087 | 2620,77 | 3227 | 2014 | 0,625 | -0,678 | -12,186 | 3,70E-34 | 1,29E-33 | 0,0023589 | 0,002541 | 0,0025732 | 0,0025732 | TRUE | 0,0301742 | No |
| orf19.7534 | 18264 | 14646 | 16874 | 11800 | 23332 | 20406 | 22366 | 30669 | 16629 | 14813 | 15970 | 17725 | 20753 | 22155 | 22906 | 23395 | 19293,1 | 16284 | 22302 | 1,37 | 0,454 | 7,73 | 1,08E-14 | 2,36E-14 | 0,0040653 | 0,002105 | 0,0032602 | 0,0032602 | TRUE | 0,0452156 | No |
| orf19.7537 | 151 | 98 | 146 | 103 | 498 | 375 | 436 | 612 | 137 | 99 | 138 | 155 | 443 | 407 | 447 | 467 | 286,62 | 132 | 441 | 3,362 | 1,749 | 15,695 | 1,65E-55 | 8,39E-55 | 0,0072578 | 0,006646 | 0,0068115 | 0,0068115 | TRUE | 0,294068 | No |
| orf19.7538 | 902 | 792 | 925 | 552 | 1900 | 1452 | 1748 | 2339 | 821 | 801 | 875 | 829 | 1690 | 1576 | 1790 | 1784 | 1270,96 | 832 | 1710 | 2,057 | 1,041 | 17,626 | 1,54E-69 | 9,63E-69 | 0,0018773 | 0,003076 | 0,0024288 | 0,0024288 | TRUE | 0,032433 | No |
| orf19.7539 | 629 | 442 | 602 | 410 | 2694 | 2223 | 2462 | 3571 | 573 | 447 | 570 | 616 | 2396 | 2414 | 2521 | 2724 | 1532,56 | 552 | 2514 | 4,574 | 2,193 | 26,493 | 1,17E-154 | 1,57E-153 | 0,0090066 | 0,002899 | 0,0054183 | 0,0054183 | TRUE | 0,1907434 | No |
| orf19.7539.1 | 1549 | 1253 | 1334 | 913 | 3195 | 2641 | 2815 | 4041 | 1410 | 1267 | 1263 | 1371 | 2842 | 2867 | 2883 | 3083 | 2123,28 | 1328 | 2919 | 2,201 | 1,138 | 21,55 | 5,34E-103 | 4,79E-102 | 0,0015594 | 0,002659 | 0,0021119 | 0,0021119 | TRUE | 0,0232489 | No |
| orf19.754 | 5302 | 4624 | 4932 | 3056 | 8661 | 6943 | 7551 | 9967 | 4827 | 4677 | 4668 | 4591 | 7704 | 7538 | 7733 | 7603 | 6167,52 | 4691 | 7644 | 1,629 | 0,704 | 22,111 | 2,49E-108 | 2,32E-107 | 6,29E-05 | 0,002251 | 0,0007969 | 0,0007969 | TRUE | 0,0047632 | No |
| orf19.7544 | 279 | 213 | 260 | 142 | 153 | 114 | 128 | 200 | 254 | 215 | 246 | 213 | 136 | 124 | 131 | 153 | 184,04 | 232 | 136 | 0,585 | -0,772 | -6,784 | 1,17E-11 | 2,34E-11 | 0,0030533 | 0,009215 | 0,0067415 | 0,0067415 | TRUE | 0,0785618 | No |
| orf19.7545 | 36 | 41 | 41 | 30 | 304 | 259 | 279 | 335 | 33 | 41 | 39 | 45 | 270 | 281 | 286 | 256 | 156,37 | 40 | 273 | 6,999 | 2,807 | 18,708 | 4,22E-78 | 2,93E-77 | 1,00E-08 | 0,010485 | 0,0063594 | 0,0063594 | TRUE | 0,108516 | No |
| orf19.7546 | 7478 | 7132 | 7512 | 4854 | 6480 | 5729 | 5528 | 7447 | 6808 | 7213 | 7109 | 7291 | 5764 | 6220 | 5661 | 5681 | 6468,55 | 7105 | 5832 | 0,821 | -0,285 | -6,622 | 3,55E-11 | 7,01E-11 | 0,0012518 | 0,002241 | 0,0016236 | 0,0016236 | TRUE | 0,0242256 | No |
| orf19.7547 | 3309 | 2993 | 3012 | 1910 | 3646 | 2918 | 2906 | 4115 | 3013 | 3027 | 2851 | 2869 | 3243 | 3168 | 2976 | 3139 | 3035,71 | 2940 | 3132 | 1,065 | 0,091 | 2,091 | 0,036545 | 0,045862 | 0,000705 | 0,002472 | 0,0014725 | 0,0014725 | TRUE | 0,0133405 | No |
| orf19.7548 | 278 | 245 | 246 | 197 | 264 | 270 | 229 | 346 | 253 | 248 | 233 | 296 | 235 | 293 | 235 | 264 | 257,01 | 258 | 257 | 1,002 | 0,003 | 0,032 | 0,974574 | 0,977179 | 0,0063445 | 0,007177 | 0,0068391 | 0,0068391 | TRUE | 0,0953651 | No |
| orf19.7549 | 216 | 208 | 211 | 135 | 536 | 427 | 405 | 548 | 197 | 210 | 200 | 203 | 477 | 464 | 415 | 418 | 322,83 | 202 | 444 | 2,19 | 1,131 | 12,658 | 1,02E-36 | 3,76E-36 | 1,00E-08 | 0,006129 | 0,0039826 | 0,0039826 | TRUE | 0,0316766 | No |
| orf19.755 | 1047 | 939 | 969 | 537 | 1122 | 816 | 922 | 1192 | 953 | 950 | 917 | 807 | 998 | 886 | 944 | 909 | 920,52 | 907 | 934 | 1,027 | 0,039 | 0,584 | 0,559459 | 0,600733 | 0,0027369 | 0,003472 | 0,0031379 | 0,0031379 | TRUE | 0,0581856 | No |
| orf19.7550 | 652 | 556 | 640 | 397 | 114 | 84 | 104 | 120 | 594 | 562 | 606 | 596 | 101 | 91 | 107 | 92 | 343,58 | 590 | 98 | 0,165 | -2,596 | -26,84 | 1,10E-158 | 1,52E-157 | 1,00E-08 | 0,005882 | 0,0033182 | 0,0033182 | TRUE | 0,0334277 | No |
| orf19.7551 | 5826 | 5170 | 5529 | 3329 | 4906 | 3937 | 4193 | 5743 | 5304 | 5229 | 5233 | 5001 | 4364 | 4274 | 4294 | 4381 | 4759,97 | 5192 | 4328 | 0,833 | -0,263 | -7,732 | 1,06E-14 | 2,33E-14 | 0,0001124 | 0,002315 | 0,0009016 | 0,0009016 | TRUE | 0,0069565 | No |
| orf19.7552 | 16602 | 14540 | 16299 | 8957 | 20568 | 15943 | 18571 | 24019 | 15115 | 14706 | 15426 | 13455 | 18295 | 17310 | 19019 | 18322 | 16455,8 | 14676 | 18236 | 1,242 | 0,313 | 6,293 | 3,12E-10 | 5,95E-10 | 0,0023997 | 0,002117 | 0,0023141 | 0,0023141 | TRUE | 0,037871 | No |
| orf19.7553 | 232 | 245 | 285 | 140 | 182 | 131 | 168 | 186 | 211 | 248 | 270 | 210 | 162 | 142 | 172 | 142 | 194,64 | 235 | 154 | 0,655 | -0,611 | -5,139 | 2,77E-07 | 4,70E-07 | 0,0074843 | 0,008824 | 0,008267 | 0,008267 | TRUE | 0,1137072 | No |
| orf19.7554 | 860 | 834 | 969 | 563 | 767 | 711 | 656 | 912 | 783 | 844 | 917 | 846 | 682 | 772 | 672 | 696 | 776,37 | 848 | 706 | 0,832 | -0,266 | -3,839 | 0,000124 | 0,000184 | 0,0028653 | 0,003739 | 0,0033056 | 0,0033056 | TRUE | 0,0452544 | No |
| orf19.7555 | 38 | 19 | 34 | 25 | 28 | 25 | 29 | 28 | 35 | 19 | 32 | 38 | 25 | 27 | 30 | 21 | 28,33 | 31 | 26 | 0,836 | -0,259 | -0,947 | 0,343854 | 0,383765 | 1,00E-08 | 0,048663 | 0,0363269 | 0,0363269 | TRUE | 0,4417166 | No |
| orf19.7556 | 439 | 428 | 470 | 267 | 350 | 276 | 311 | 366 | 400 | 433 | 445 | 401 | 311 | 300 | 319 | 279 | 360,89 | 420 | 302 | 0,717 | -0,479 | -5,887 | 3,93E-09 | 7,19E-09 | 1,00E-08 | 0,005697 | 0,0035445 | 0,0035445 | TRUE | 0,0334332 | No |
| orf19.7557 | 20 | 17 | 12 | 6 | 27 | 15 | 15 | 25 | 18 | 17 | 11 | 9 | 24 | 16 | 15 | 19 | 16,31 | 14 | 18 | 1,319 | 0,399 | 1,138 | 0,254971 | 0,290413 | 1,00E-08 | 0,083014 | 0,0561466 | 0,0561466 | TRUE | 0,2030953 | No |
| orf19.7558 | 4315 | 3892 | 4308 | 2736 | 3935 | 3553 | 3887 | 5068 | 3929 | 3936 | 4077 | 4110 | 3500 | 3858 | 3981 | 3866 | 3907,04 | 4013 | 3801 | 0,947 | -0,078 | -1,65 | 0,098994 | 0,118961 | 0,0014358 | 0,002375 | 0,0018926 | 0,0018926 | TRUE | 0,034817 | No |
| orf19.756 | 10 | 8 | 8 | 8 | 48 | 33 | 29 | 60 | 9 | 8 | 8 | 12 | 43 | 36 | 30 | 46 | 23,85 | 9 | 39 | 4,318 | 2,11 | 6,249 | 4,13E-10 | 7,83E-10 | 1,00E-08 | 0,057432 | 0,038584 | 0,038584 | TRUE | 0,1570015 | No |
| orf19.7561 | 27732 | 22638 | 25991 | 17695 | 8428 | 7211 | 8014 | 11029 | 25249 | 22896 | 24598 | 26581 | 7496 | 7829 | 8207 | 8413 | 16408,7 | 24831 | 7986 | 0,322 | -1,636 | -30,216 | 1,48E-200 | 2,53E-199 | 0,0030863 | 0,002118 | 0,0027369 | 0,0027369 | TRUE | 0,0336707 | No |
| orf19.7563 | 734 | 652 | 723 | 488 | 605 | 480 | 527 | 743 | 668 | 659 | 684 | 733 | 538 | 521 | 540 | 567 | 613,85 | 686 | 542 | 0,793 | -0,335 | -5,236 | 1,64E-07 | 2,80E-07 | 1,00E-08 | 0,004189 | 0,0022923 | 0,0022923 | TRUE | 0,0211442 | No |
| orf19.7564 | 2143 | 2059 | 2305 | 1340 | 4060 | 3327 | 3497 | 4543 | 1951 | 2082 | 2181 | 2013 | 3611 | 3612 | 3581 | 3465 | 2812,27 | 2057 | 3567 | 1,733 | 0,793 | 17,616 | 1,87E-69 | 1,16E-68 | 0,0008333 | 0,002507 | 0,001555 | 0,001555 | TRUE | 0,0208574 | No |
| orf19.7565 | 2321 | 2056 | 2172 | 1353 | 5254 | 4499 | 4776 | 6424 | 2113 | 2079 | 2056 | 2032 | 4673 | 4885 | 4891 | 4900 | 3453,76 | 2070 | 4837 | 2,335 | 1,223 | 32,575 | 9,20E-233 | 1,80E-231 | 1,00E-08 | 0,002419 | 0,0009984 | 0,0009984 | TRUE | 0,0064271 | No |
| orf19.7566 | 33293 | 28720 | 31850 | 20347 | 64238 | 54594 | 57611 | 81039 | 30312 | 29047 | 30143 | 30564 | 57138 | 59274 | 59001 | 61817 | 44662 | 30016 | 59308 | 1,976 | 0,983 | 28,032 | 6,69E-173 | 1,01E-171 | 0,000741 | 0,002067 | 0,0011545 | 0,0011545 | TRUE | 0,0099774 | No |
| orf19.7567 | 4165 | 3736 | 4026 | 2721 | 1028 | 955 | 961 | 1212 | 3792 | 3779 | 3810 | 4087 | 914 | 1037 | 984 | 925 | 2416,02 | 3867 | 965 | 0,249 | -2,004 | -38,162 | 0 | 0 | 0,0016059 | 0,002584 | 0,0020244 | 0,0020244 | TRUE | 0,029146 | No |
| orf19.7568 | 102 | 89 | 96 | 47 | 147 | 128 | 127 | 181 | 93 | 90 | 91 | 71 | 131 | 139 | 130 | 138 | 110,27 | 86 | 134 | 1,543 | 0,626 | 4,648 | 3,35E-06 | 5,44E-06 | 1,00E-08 | 0,014016 | 0,0078652 | 0,0078652 | TRUE | 0,0983081 | No |
| orf19.7569 | 24272 | 21260 | 22155 | 15441 | 38916 | 33685 | 32382 | 45995 | 22099 | 21502 | 20968 | 23195 | 34615 | 36573 | 33163 | 35085 | 28399,9 | 21941 | 34859 | 1,589 | 0,668 | 15,124 | 1,13E-51 | 5,39E-51 | 0,0017104 | 0,002084 | 0,0018372 | 0,0018372 | TRUE | 0,0179409 | No |
| orf19.757 | 2388 | 2221 | 2423 | 1454 | 1559 | 1486 | 1477 | 1875 | 2174 | 2246 | 2293 | 2184 | 1387 | 1613 | 1513 | 1430 | 1855,09 | 2224 | 1486 | 0,667 | -0,585 | -10,8 | 3,44E-27 | 1,04E-26 | 0,0018245 | 0,002749 | 0,0022577 | 0,0022577 | TRUE | 0,0394132 | No |
| orf19.7570 | 15452 | 14110 | 15778 | 8905 | 10721 | 8577 | 9076 | 12366 | 14068 | 14271 | 14932 | 13377 | 9536 | 9312 | 9295 | 9433 | 11778,1 | 14162 | 9394 | 0,663 | -0,593 | -15,077 | 2,30E-51 | 1,09E-50 | 0,0009768 | 0,002149 | 0,0013962 | 0,0013962 | TRUE | 0,0166959 | No |
| orf19.7571 | 8293 | 7668 | 8056 | 5354 | 8830 | 7221 | 7070 | 9584 | 7550 | 7755 | 7624 | 8043 | 7854 | 7840 | 7241 | 7311 | 7652,24 | 7743 | 7562 | 0,977 | -0,034 | -0,804 | 0,421129 | 0,462241 | 0,0011752 | 0,00221 | 0,001568 | 0,001568 | TRUE | 0,0098048 | No |
| orf19.757.1 | 677 | 556 | 584 | 481 | 621 | 577 | 569 | 723 | 616 | 562 | 553 | 723 | 552 | 626 | 583 | 552 | 595,88 | 614 | 578 | 0,946 | -0,08 | -0,92 | 0,357731 | 0,397721 | 0,0076841 | 0,004254 | 0,0056335 | 0,0056335 | TRUE | 0,1523498 | No |
| orf19.7572 | 1742 | 1606 | 1762 | 1049 | 2420 | 1980 | 2229 | 2858 | 1586 | 1624 | 1668 | 1576 | 2153 | 2150 | 2283 | 2180 | 1902,35 | 1614 | 2192 | 1,357 | 0,44 | 10,054 | 8,87E-24 | 2,47E-23 | 0,0002177 | 0,002731 | 0,0012952 | 0,0012952 | TRUE | 0,0092964 | No |
| orf19.7574 | 3034 | 2838 | 3122 | 1975 | 3124 | 2628 | 2615 | 3690 | 2762 | 2870 | 2955 | 2967 | 2779 | 2853 | 2678 | 2815 | 2834,87 | 2888 | 2781 | 0,964 | -0,053 | -1,278 | 0,201306 | 0,23295 | 0,0005425 | 0,002503 | 0,0013196 | 0,0013196 | TRUE | 0,010787 | No |
| orf19.7576 | 1261 | 1012 | 1127 | 662 | 911 | 678 | 703 | 923 | 1148 | 1024 | 1067 | 994 | 810 | 736 | 720 | 704 | 900,39 | 1058 | 742 | 0,7 | -0,514 | -7,661 | 1,85E-14 | 4,03E-14 | 0,0028786 | 0,003504 | 0,0031865 | 0,0031865 | TRUE | 0,0459988 | No |
| orf19.7577 | 3839 | 3440 | 3641 | 2435 | 2908 | 2624 | 2716 | 3538 | 3495 | 3479 | 3446 | 3658 | 2587 | 2849 | 2782 | 2699 | 3124,24 | 3520 | 2729 | 0,776 | -0,367 | -8,454 | 2,82E-17 | 6,65E-17 | 0,0007842 | 0,00246 | 0,0014826 | 0,0014826 | TRUE | 0,0149007 | No |
| orf19.7578 | 93 | 75 | 71 | 63 | 94 | 70 | 68 | 124 | 85 | 76 | 67 | 95 | 84 | 76 | 70 | 95 | 80,77 | 81 | 81 | 1,02 | 0,029 | 0,174 | 0,861921 | 0,882832 | 0,0069718 | 0,018391 | 0,014166 | 0,014166 | TRUE | 0,1347256 | No |

| | | | | | | | | | | | | | | | | | | | | | | | | | | | | | | | |
|---|---|---|---|---|---|---|---|---|---|---|---|---|---|---|---|---|---|---|---|---|---|---|---|---|---|---|---|---|---|---|---|
| orf19.7579 | 791 | 651 | 717 | 517 | 663 | 571 | 591 | 789 | 720 | 658 | 679 | 777 | 590 | 620 | 605 | 602 | 656,32 | 708 | 604 | 0,855 | -0,225 | -3,404 | 0,000664 | 0,000948 | 0,0008839 | 0,00405 | 0,0026694 | 0,0026694 | TRUE | 0,0426612 | No |
| orf19.758 | 1645 | 1355 | 1517 | 957 | 1336 | 1216 | 1160 | 1646 | 1498 | 1370 | 1436 | 1438 | 1188 | 1320 | 1188 | 1256 | 1336,69 | 1436 | 1238 | 0,862 | -0,214 | -3,974 | 7,06E-05 | 0,000107 | 0,001176 | 0,003025 | 0,002039 | 0,002039 | TRUE | 0,0226299 | No |
| orf19.7580 | 660 | 522 | 501 | 290 | 314 | 224 | 271 | 350 | 601 | 528 | 474 | 436 | 279 | 243 | 278 | 267 | 388,21 | 510 | 267 | 0,522 | -0,939 | -9,125 | 7,20E-20 | 1,82E-19 | 0,0095749 | 0,00544 | 0,0073599 | 0,0073599 | TRUE | 0,1713404 | No |
| orf19.7581 | 7330 | 6267 | 6980 | 4019 | 3165 | 2435 | 2823 | 3586 | 6674 | 6338 | 6606 | 6037 | 2815 | 2644 | 2891 | 2735 | 4592,58 | 6414 | 2771 | 0,432 | -1,211 | -26,06 | 1,04E-149 | 1,34E-148 | 0,0014107 | 0,002325 | 0,0018226 | 0,0018226 | TRUE | 0,0184566 | No |
| orf19.7583 | 1847 | 1794 | 1901 | 1146 | 2874 | 2547 | 2557 | 3327 | 1682 | 1814 | 1799 | 1721 | 2556 | 2765 | 2619 | 2538 | 2186,86 | 1754 | 2620 | 1,492 | 0,577 | 12,15 | 5,72E-34 | 1,99E-33 | 0,0009867 | 0,002641 | 0,0016806 | 0,0016806 | TRUE | 0,0167056 | No |
| orf19.7585 | 1523 | 1271 | 1408 | 864 | 19585 | 16173 | 16443 | 23270 | 1387 | 1285 | 1333 | 1298 | 17420 | 17559 | 16840 | 17751 | 9359,05 | 1326 | 17392 | 13,103 | 3,712 | 92,183 | 0 | 0 | 0,0004613 | 0,002178 | 0,001127 | 0,001127 | TRUE | 0,012104 | No |
| orf19.7586 | 630 | 563 | 608 | 429 | 339 | 310 | 309 | 437 | 574 | 569 | 575 | 644 | 302 | 337 | 316 | 333 | 456,34 | 590 | 322 | 0,547 | -0,87 | -11,548 | 7,55E-31 | 2,48E-30 | 1,00E-08 | 0,004932 | 0,0030944 | 0,0030944 | TRUE | 0,0378859 | No |
| orf19.7588 | 721 | 721 | 760 | 415 | 952 | 785 | 893 | 1009 | 656 | 729 | 719 | 623 | 847 | 852 | 915 | 770 | 763,95 | 682 | 846 | 1,235 | 0,305 | 4,158 | 3,21E-05 | 4,95E-05 | 0,0038586 | 0,003766 | 0,0038116 | 0,0038116 | TRUE | 0,0459328 | No |
| orf19.7589 | 789 | 712 | 710 | 435 | 616 | 503 | 527 | 745 | 718 | 720 | 672 | 653 | 548 | 546 | 540 | 568 | 620,74 | 691 | 550 | 0,796 | -0,33 | -5,231 | 1,69E-07 | 2,89E-07 | 1,00E-08 | 0,004165 | 0,0022033 | 0,0022033 | TRUE | 0,013235 | No |
| orf19.759 | 1321 | 1182 | 1236 | 797 | 1768 | 1573 | 1527 | 2142 | 1203 | 1195 | 1170 | 1197 | 1573 | 1708 | 1564 | 1634 | 1405,42 | 1191 | 1620 | 1,359 | 0,443 | 9,125 | 7,20E-20 | 1,82E-19 | 1,00E-08 | 0,002977 | 0,0015235 | 0,0015235 | TRUE | 0,0153246 | No |
| orf19.7590 | 4896 | 4438 | 4748 | 2802 | 7847 | 6642 | 6757 | 9175 | 4458 | 4489 | 4494 | 4209 | 6980 | 7211 | 6920 | 6999 | 5719,82 | 4412 | 7028 | 1,591 | 0,67 | 18,783 | 1,05E-78 | 7,34E-78 | 0,0003952 | 0,002268 | 0,0010327 | 0,0010327 | TRUE | 0,0108404 | No |
| orf19.7592 | 35265 | 33549 | 36495 | 22061 | 44375 | 39453 | 40331 | 52888 | 32107 | 33931 | 34539 | 33139 | 39470 | 42835 | 41304 | 40343 | 37208,7 | 33429 | 40988 | 1,226 | 0,294 | 7,606 | 2,83E-14 | 6,12E-14 | 0,0010764 | 0,002073 | 0,0014079 | 0,0014079 | TRUE | 0,0112514 | No |
| orf19.7593 | 2854 | 2709 | 2849 | 1744 | 2385 | 2008 | 2005 | 2698 | 2598 | 2740 | 2696 | 2620 | 2121 | 2180 | 2053 | 2058 | 2383,42 | 2664 | 2103 | 0,789 | -0,342 | -8,253 | 1,55E-16 | 3,58E-16 | 0,0002769 | 0,002591 | 0,0012243 | 0,0012243 | TRUE | 0,0070304 | No |
| orf19.7594 | 707 | 701 | 683 | 430 | 283 | 254 | 272 | 351 | 644 | 709 | 646 | 646 | 252 | 276 | 279 | 268 | 464,85 | 661 | 269 | 0,405 | -1,303 | -17,38 | 1,18E-67 | 7,15E-67 | 1,00E-08 | 0,004879 | 0,0028553 | 0,0028553 | TRUE | 0,028506 | No |
| orf19.7595 | 258 | 245 | 292 | 166 | 665 | 567 | 591 | 704 | 235 | 248 | 276 | 249 | 591 | 616 | 605 | 537 | 419,72 | 252 | 587 | 2,324 | 1,217 | 14,562 | 4,90E-48 | 2,18E-47 | 0,0021107 | 0,005184 | 0,0037936 | 0,0037936 | TRUE | 0,0516566 | No |
| orf19.7596 | 164 | 142 | 151 | 89 | 1366 | 1113 | 1120 | 1584 | 149 | 144 | 143 | 134 | 1215 | 1208 | 1147 | 1208 | 668,53 | 142 | 1194 | 8,348 | 3,061 | 38,134 | 0 | 0 | 1,00E-08 | 0,004013 | 0,0021068 | 0,0021068 | TRUE | 0,0123695 | No |
| orf19.7597 | 4 | 1 | 6 | 4 | 16 | 14 | 9 | 22 | 4 | 1 | 6 | 6 | 14 | 15 | 9 | 17 | 8,97 | 4 | 14 | 3,512 | 1,812 | 3,412 | 0,000646 | 0,000923 | 1,00E-08 | 0,149283 | 0,1041065 | 0,1041065 | TRUE | 0,4428141 | No |
| orf19.7598 | 5774 | 4958 | 5529 | 3549 | 5404 | 4238 | 4764 | 6415 | 5257 | 5014 | 5233 | 5331 | 4807 | 4601 | 4879 | 4893 | 5001,96 | 5209 | 4795 | 0,921 | -0,118 | -3,153 | 0,001617 | 0,00225 | 0,0004689 | 0,002301 | 0,0011545 | 0,0011545 | TRUE | 0,008823 | No |
| orf19.7599 | 13512 | 11795 | 12869 | 8855 | 14789 | 12637 | 13242 | 17769 | 12302 | 11929 | 12179 | 13302 | 13154 | 13720 | 13561 | 13554 | 12962,9 | 12428 | 13497 | 1,087 | 0,12 | 2,936 | 0,003329 | 0,004515 | 0,0012324 | 0,002139 | 0,0015189 | 0,0015189 | TRUE | 0,026817 | No |
| orf19.76 | 17714 | 16596 | 17809 | 10622 | 19979 | 16672 | 17735 | 22613 | 16128 | 16785 | 16855 | 15956 | 17771 | 18101 | 18163 | 17249 | 17125,9 | 16431 | 17821 | 1,084 | 0,117 | 3,413 | 0,000642 | 0,000917 | 0,0005816 | 0,002114 | 0,0010642 | 0,0010642 | TRUE | 0,0057532 | No |
| orf19.760 | 781 | 619 | 689 | 420 | 692 | 481 | 556 | 741 | 711 | 626 | 652 | 631 | 616 | 522 | 569 | 565 | 611,56 | 655 | 568 | 0,867 | -0,206 | -2,866 | 0,004151 | 0,005589 | 0,0023851 | 0,004197 | 0,003339 | 0,003339 | TRUE | 0,0408467 | No |
| orf19.7600 | 11623 | 9376 | 10329 | 6532 | 11255 | 8746 | 9956 | 13972 | 10582 | 9483 | 9775 | 9812 | 10011 | 9496 | 10196 | 10658 | 10001,7 | 9913 | 10090 | 1,018 | 0,026 | 0,529 | 0,597129 | 0,63721 | 0,0019234 | 0,002169 | 0,0021596 | 0,0021596 | TRUE | 0,0254931 | No |
| orf19.7601 | 2451 | 2138 | 2331 | 1580 | 2354 | 2062 | 2013 | 2902 | 2232 | 2162 | 2206 | 2373 | 2094 | 2239 | 2062 | 2214 | 2197,65 | 2243 | 2152 | 0,961 | -0,058 | -1,206 | 0,227755 | 0,261375 | 0,0010451 | 0,002638 | 0,0017503 | 0,0017503 | TRUE | 0,0169437 | No |
| orf19.7602 | 1674 | 1510 | 1530 | 953 | 549 | 450 | 481 | 608 | 1524 | 1527 | 1448 | 1432 | 488 | 489 | 493 | 464 | 983,02 | 1483 | 484 | 0,325 | -1,621 | -28,66 | 1,19E-180 | 1,85E-179 | 1,00E-08 | 0,003381 | 0,0017454 | 0,0017454 | TRUE | 0,0096571 | No |
| orf19.7603 | 1382 | 1144 | 1277 | 719 | 1466 | 993 | 1163 | 1399 | 1258 | 1157 | 1209 | 1080 | 1304 | 1078 | 1191 | 1067 | 1168,03 | 1176 | 1160 | 0,985 | -0,022 | -0,299 | 0,765157 | 0,795738 | 0,0057688 | 0,003168 | 0,0045035 | 0,0045035 | TRUE | 0,0849976 | No |
| orf19.7604 | 849 | 730 | 811 | 554 | 915 | 798 | 852 | 1155 | 773 | 738 | 768 | 832 | 814 | 866 | 873 | 881 | 818,11 | 778 | 858 | 1,107 | 0,147 | 2,48 | 0,013141 | 0,017086 | 0,0004109 | 0,003652 | 0,0021233 | 0,0021233 | TRUE | 0,0224119 | No |
| orf19.7605 | 10849 | 8330 | 9551 | 6309 | 6293 | 4965 | 5286 | 7585 | 9878 | 8425 | 9039 | 9477 | 5597 | 5391 | 5414 | 5786 | 7375,77 | 9205 | 5547 | 0,603 | -0,73 | -13,798 | 2,62E-43 | 1,09E-42 | 0,002646 | 0,002216 | 0,0025475 | 0,0025475 | TRUE | 0,0399048 | No |
| orf19.7606 | 25 | 47 | 32 | 29 | 48 | 46 | 41 | 73 | 23 | 48 | 30 | 44 | 43 | 50 | 42 | 56 | 41,81 | 36 | 48 | 1,345 | 0,428 | 1,727 | 0,084122 | 0,101703 | 0,0347764 | 0,033634 | 0,0339927 | 0,0339927 | TRUE | 0,3796733 | No |
| orf19.7608 | 4 | 3 | 3 | 3 | 11 | 9 | 6 | 14 | 4 | 3 | 3 | 5 | 10 | 10 | 6 | 11 | 6,3 | 4 | 9 | 2,667 | 1,415 | 2,445 | 0,01448 | 0,018793 | 1,00E-08 | 0,211713 | 0,1165944 | 0,1165944 | TRUE | 0,1470214 | No |
| orf19.7609 | 20 | 12 | 11 | 11 | 26 | 18 | 12 | 39 | 18 | 12 | 10 | 17 | 23 | 20 | 12 | 30 | 17,75 | 14 | 21 | 1,508 | 0,593 | 1,637 | 0,101578 | 0,12202 | 0,0511103 | 0,076466 | 0,0667903 | 0,0667903 | TRUE | 0,1905457 | No |
| orf19.761 | 4390 | 3520 | 4209 | 2864 | 2258 | 1961 | 2182 | 2969 | 3997 | 3560 | 3983 | 4302 | 2008 | 2129 | 2235 | 2265 | 3059,95 | 3960 | 2159 | 0,546 | -0,874 | -13,964 | 2,59E-44 | 1,10E-43 | 0,0039333 | 0,002469 | 0,0034099 | 0,0034099 | TRUE | 0,0562711 | No |
| orf19.7610 | 3386 | 3101 | 3416 | 2127 | 4196 | 3675 | 3903 | 5139 | 3083 | 3136 | 3233 | 3195 | 3732 | 3990 | 3997 | 3920 | 3535,83 | 3162 | 3910 | 1,237 | 0,306 | 7,782 | 7,14E-15 | 1,57E-14 | 0,0004932 | 0,002411 | 0,0011985 | 0,0011985 | TRUE | 0,0114287 | No |
| orf19.7611 | 6296 | 6450 | 6126 | 4224 | 3342 | 2714 | 2392 | 3477 | 5732 | 6523 | 5798 | 6345 | 2973 | 2947 | 2450 | 2652 | 4427,47 | 6100 | 2756 | 0,452 | -1,146 | -16,684 | 1,72E-62 | 9,74E-62 | 0,0058601 | 0,002335 | 0,0042757 | 0,0042757 | TRUE | 0,0668967 | No |
| orf19.7612 | 22724 | 20731 | 21695 | 13760 | 3277 | 2706 | 2730 | 3778 | 20689 | 20967 | 20532 | 20670 | 2915 | 2938 | 2796 | 2882 | 11798,6 | 20714 | 2883 | 0,139 | -2,845 | -92,349 | 0 | 0 | 3,37E-05 | 0,002149 | 0,0007236 | 0,0007236 | TRUE | 0,0047632 | No |
| orf19.7613 | 5032 | 4132 | 4557 | 2812 | 5156 | 4001 | 4541 | 6154 | 4581 | 4179 | 4313 | 4224 | 4586 | 4344 | 4651 | 4694 | 4446,54 | 4324 | 4569 | 1,056 | 0,079 | 1,789 | 0,073617 | 0,089614 | 0,001153 | 0,002334 | 0,0016469 | 0,0016469 | TRUE | 0,019882 | No |
| orf19.7614 | 4 | 3 | 5 | 2 | 25 | 18 | 13 | 17 | 4 | 3 | 5 | 3 | 22 | 20 | 13 | 13 | 10,31 | 4 | 17 | 4,632 | 2,212 | 4,276 | 1,90E-05 | 2,97E-05 | 1,00E-08 | 0,130175 | 0,0849694 | 0,0849694 | TRUE | 0,1531679 | No |
| orf19.7615 | 527 | 441 | 436 | 276 | 836 | 703 | 736 | 872 | 480 | 446 | 413 | 415 | 744 | 763 | 754 | 665 | 584,86 | 438 | 732 | 1,661 | 0,732 | 9,677 | 3,79E-22 | 1,01E-21 | 0,0029326 | 0,004296 | 0,0036438 | 0,0036438 | TRUE | 0,0494685 | No |
| orf19.7617 | 2015 | 1565 | 1748 | 1112 | 1653 | 1317 | 1406 | 1984 | 1835 | 1583 | 1654 | 1670 | 1470 | 1430 | 1440 | 1513 | 1574,46 | 1686 | 1463 | 0,868 | -0,204 | -3,706 | 0,000211 | 0,00031 | 0,0017892 | 0,002876 | 0,0022672 | 0,0022672 | TRUE | 0,0448013 | No |
| orf19.7618 | 8933 | 7456 | 7980 | 5337 | 8621 | 6521 | 7244 | 10181 | 8133 | 7541 | 7552 | 8017 | 7668 | 7080 | 7419 | 7766 | 7647,06 | 7811 | 7483 | 0,959 | -0,061 | -1,377 | 0,16837 | 0,196766 | 0,0014432 | 0,00221 | 0,0017604 | 0,0017604 | TRUE | 0,0159647 | No |
| orf19.7619 | 9827 | 8924 | 9659 | 5677 | 8353 | 6656 | 7415 | 9439 | 8947 | 9026 | 9141 | 8528 | 7430 | 7227 | 7594 | 7200 | 8136,52 | 8910 | 7363 | 0,826 | -0,276 | -7,586 | 3,29E-14 | 7,11E-14 | 0,0006011 | 0,002199 | 0,0011484 | 0,0011484 | TRUE | 0,0098325 | No |
| orf19.762 | 32 | 30 | 21 | 23 | 56 | 40 | 55 | 68 | 29 | 30 | 20 | 35 | 50 | 43 | 56 | 52 | 39,42 | 28 | 50 | 1,799 | 0,847 | 3,664 | 0,000248 | 0,000365 | 1,00E-08 | 0,03555 | 0,0231664 | 0,0231664 | TRUE | 0,288229 | No |
| orf19.7620 | 1062 | 866 | 925 | 550 | 708 | 485 | 543 | 805 | 967 | 876 | 875 | 826 | 630 | 527 | 556 | 614 | 733,86 | 886 | 582 | 0,657 | -0,607 | -8,123 | 4,56E-16 | 1,04E-15 | 0,003769 | 0,003837 | 0,0039552 | 0,0039552 | TRUE | 0,0461966 | No |
| orf19.7621 | 417 | 376 | 392 | 233 | 793 | 671 | 637 | 859 | 380 | 380 | 371 | 350 | 705 | 729 | 652 | 655 | 527,8 | 370 | 685 | 1,844 | 0,883 | 12,403 | 2,52E-35 | 9,04E-35 | 1,00E-08 | 0,00454 | 0,0027436 | 0,0027436 | TRUE | 0,020117 | No |
| orf19.7622 | 1214 | 1059 | 1123 | 669 | 584 | 442 | 467 | 612 | 1105 | 1071 | 1063 | 1005 | 519 | 480 | 478 | 467 | 773,57 | 1061 | 486 | 0,457 | -1,129 | -18,299 | 8,48E-75 | 5,61E-74 | 1,00E-08 | 0,003745 | 0,0021938 | 0,0021938 | TRUE | 0,0260301 | No |
| orf19.7624 | 9807 | 8206 | 9061 | 5570 | 9464 | 7284 | 8449 | 11131 | 8929 | 8299 | 8575 | 8367 | 8418 | 7908 | 8653 | 8491 | 8455,1 | 8542 | 8368 | 0,979 | -0,03 | -0,725 | 0,468487 | 0,510357 | 0,0011018 | 0,002193 | 0,0015325 | 0,0015325 | TRUE | 0,0164621 | No |
| orf19.7625 | 1893 | 1733 | 1803 | 1165 | 3104 | 2359 | 2615 | 3493 | 1723 | 1753 | 1706 | 1750 | 2761 | 2561 | 2678 | 2664 | 2199,67 | 1733 | 2666 | 1,54 | 0,623 | 14,915 | 2,64E-50 | 1,22E-49 | 1,00E-08 | 0,002638 | 0,0011886 | 0,0011886 | TRUE | 0,0082005 | No |
| orf19.7626 | 3779 | 3203 | 3297 | 2139 | 5127 | 3964 | 4173 | 5736 | 3441 | 3239 | 3120 | 3213 | 4560 | 4304 | 4274 | 4375 | 3815,85 | 3253 | 4378 | 1,346 | 0,428 | 9,846 | 7,15E-23 | 1,94E-22 | 0,0009337 | 0,002383 | 0,0015439 | 0,0015439 | TRUE | 0,0189167 | No |
| orf19.7627 | 953 | 885 | 879 | 654 | 791 | 638 | 639 | 911 | 868 | 895 | 832 | 982 | 704 | 693 | 654 | 695 | 790,33 | 894 | 686 | 0,771 | -0,375 | -5,846 | 5,03E-09 | 9,18E-09 | 0,0015508 | 0,003708 | 0,0026778 | 0,0026778 | TRUE | 0,0466282 | No |
| orf19.7629 | 530 | 505 | 538 | 353 | 481 | 411 | 420 | 623 | 483 | 511 | 509 | 530 | 428 | 446 | 430 | 475 | 476,52 | 508 | 445 | 0,879 | -0,186 | -2,628 | 0,008593 | 0,011321 | 1,00E-08 | 0,004809 | 0,0027209 | 0,0027209 | TRUE | 0,0272209 | No |
| orf19.763 | 3687 | 3331 | 3395 | 1769 | 3093 | 2124 | 2426 | 3242 | 3357 | 3369 | 3213 | 2657 | 2751 | 2306 | 2485 | 2473 | 2826,37 | 3149 | 2504 | 0,794 | -0,332 | -4,273 | 1,93E-05 | 3,01E-05 | 0,0078943 | 0,002504 | 0,0054446 | 0,0054446 | TRUE | 0,1305946 | No |
| orf19.7631 | 126 | 110 | 144 | 88 | 222 | 151 | 175 | 263 | 115 | 111 | 136 | 132 | 197 | 164 | 179 | 201 | 154,46 | 124 | 185 | 1,511 | 0,595 | 4,841 | 1,29E-06 | 2,14E-06 | 0,0030533 | 0,010589 | 0,0076828 | 0,0076828 | TRUE | 0,061785 | No |
| orf19.7632 | 666 | 550 | 625 | 388 | 384 | 329 | 364 | 482 | 606 | 556 | 592 | 583 | 342 | 357 | 373 | 368 | 472,02 | 584 | 360 | 0,615 | -0,7 | -9,972 | 2,02E-23 | 5,59E-23 | 1,00E-08 | 0,004836 | 0,0025264 | 0,0025264 | TRUE | 0,0139753 | No |
| orf19.7634 | 97 | 88 | 106 | 50 | 93 | 66 | 72 | 113 | 88 | 89 | 100 | 75 | 83 | 72 | 74 | 86 | 83,38 | 88 | 79 | 0,888 | -0,171 | -1,108 | 0,268002 | 0,303951 | 1,00E-08 | 0,01788 | 0,0110921 | 0,0110921 | TRUE | 0,0794185 | No |
| orf19.7635 | 13785 | 11738 | 13103 | 8562 | 16894 | 13455 | 15565 | 21219 | 12551 | 11872 | 12401 | 12861 | 15027 | 14608 | 15941 | 16186 | 13930,8 | 12421 | 15440 | 1,243 | 0,314 | 7,06 | 1,66E-12 | 3,43E-12 | 0,0016244 | 0,002132 | 0,0018301 | 0,0018301 | TRUE | 0,0160297 | No |
| orf19.7637 | 1879 | 1650 | 1748 | 1033 | 1806 | 1489 | 1646 | 2143 | 1711 | 1669 | 1654 | 1552 | 1606 | 1617 | 1686 | 1635 | 1641,13 | 1646 | 1636 | 0,991 | -0,013 | -0,271 | 0,786431 | 0,815601 | 0,0002599 | 0,002842 | 0,001463 | 0,001463 | TRUE | 0,0159876 | No |
| orf19.7638 | 2796 | 2197 | 2387 | 1727 | 1857 | 1582 | 1786 | 2355 | 2546 | 2222 | 2259 | 2594 | 1652 | 1718 | 1829 | 1796 | 2076,98 | 2405 | 1749 | 0,728 | -0,458 | -7,326 | 2,37E-13 | 5,01E-13 | 0,0035622 | 0,002673 | 0,0032682 | 0,0032682 | TRUE | 0,0321856 | No |
| orf19.764 | 376 | 415 | 439 | 272 | 383 | 327 | 393 | 453 | 342 | 420 | 415 | 409 | 341 | 355 | 402 | 346 | 378,73 | 396 | 361 | 0,911 | -0,135 | -1,484 | 0,137888 | 0,162681 | 0,0050644 | 0,005525 | 0,0053157 | 0,0053157 | TRUE | 0,1047777 | No |
| orf19.7642 | 645 | 577 | 598 | 400 | 885 | 754 | 769 | 1026 | 587 | 584 | 566 | 601 | 787 | 819 | 788 | 783 | 689,21 | 584 | 794 | 1,36 | 0,443 | 7,618 | 2,58E-14 | 5,59E-14 | 1,00E-08 | 0,003954 | 0,0017519 | 0,0017519 | TRUE | 0,0053276 | No |
| orf19.7643 | 1641 | 1380 | 1514 | 996 | 2061 | 1776 | 1848 | 2688 | 1494 | 1396 | 1433 | 1496 | 1833 | 1928 | 1893 | 2050 | 1690,41 | 1455 | 1926 | 1,326 | 0,407 | 7,908 | 2,61E-15 | 5,84E-15 | 0,0012659 | 0,002818 | 0,0019287 | 0,0019287 | TRUE | 0,0239774 | No |
| orf19.7644 | 397 | 338 | 350 | 254 | 492 | 369 | 461 | 636 | 361 | 342 | 331 | 382 | 438 | 401 | 472 | 485 | 401,45 | 354 | 449 | 1,275 | 0,351 | 4,136 | 3,54E-05 | 5,45E-05 | 0,003113 | 0,005328 | 0,0043425 | 0,0043425 | TRUE | 0,0576548 | No |
| orf19.7645 | 1703 | 1423 | 1510 | 971 | 2842 | 2225 | 2410 | 3503 | 1551 | 1439 | 1429 | 1459 | 2528 | 2416 | 2468 | 2672 | 1995,16 | 1470 | 2521 | 1,716 | 0,779 | 15,498 | 3,58E-54 | 1,78E-53 | 0,0012824 | 0,002699 | 0,0018769 | 0,0018769 | TRUE | 0,020288 | No |
| orf19.7646 | 443 | 348 | 424 | 241 | 344 | 232 | 300 | 411 | 403 | 352 | 401 | 362 | 306 | 252 | 307 | 314 | 337,15 | 380 | 295 | 0,777 | -0,365 | -3,967 | 7,29E-05 | 0,00011 | 0,0039964 | 0,005955 | 0,0051375 | 0,0051375 | TRUE | 0,1007283 | No |
| orf19.7648 | 575 | 496 | 542 | 392 | 597 | 457 | 480 | 767 | 524 | 502 | 513 | 589 | 531 | 496 | 492 | 585 | 528,85 | 532 | 526 | 0,996 | -0,006 | -0,077 | 0,938292 | 0,947507 | 0,0038939 | 0,004535 | 0,0042357 | 0,0042357 | TRUE | 0,0704936 | No |
| orf19.7650 | 5963 | 4866 | 5582 | 3385 | 5605 | 4457 | 5159 | 6894 | 5429 | 4921 | 5283 | 5085 | 4985 | 4839 | 5283 | 5259 | 5135,62 | 5180 | 5092 | 0,983 | -0,025 | -0,533 | 0,593695 | 0,633758 | 0,0014474 | 0,002294 | 0,0018916 | 0,0018916 | TRUE | 0,0140729 | No |
| orf19.7652 | 9060 | 7461 | 8625 | 5245 | 7990 | 6453 | 7100 | 9725 | 8249 | 7546 | 8163 | 7879 | 7107 | 7006 | 7271 | 7418 | 7579,87 | 7959 | 7200 | 0,905 | -0,145 | -3,587 | 0,000335 | 0,000487 | 0,0009377 | 0,002211 | 0,0014265 | 0,0014265 | TRUE | 0,0151862 | No |
| orf19.7654 | 5214 | 4071 | 4576 | 3111 | 2518 | 2106 | 2234 | 3155 | 4747 | 4117 | 4331 | 4673 | 2240 | 2287 | 2288 | 2407 | 3386,16 | 4467 | 2306 | 0,516 | -0,953 | -17,696 | 4,49E-70 | 2,82E-69 | 0,0022258 | 0,002427 | 0,0024644 | 0,0024644 | TRUE | 0,0339308 | No |
| orf19.7655 | 4447 | 3755 | 4085 | 2527 | 7718 | 6182 | 7095 | 9813 | 4049 | 3798 | 3866 | 3796 | 6865 | 6712 | 7266 | 7485 | 5479,64 | 3877 | 7082 | 1,826 | 0,869 | 18,741 | 2,31E-78 | 1,61E-77 | 0,0014701 | 0,002278 | 0,0018594 | 0,0018594 | TRUE | 0,0181831 | No |

| | | | | | | | | | | | | | | | | | | | | | | | | | | | | | | | |
|---|---|---|---|---|---|---|---|---|---|---|---|---|---|---|---|---|---|---|---|---|---|---|---|---|---|---|---|---|---|---|---|
| orf19.7657 | 3291 | 2509 | 2725 | 1852 | 2853 | 2207 | 2308 | 3298 | 2996 | 2538 | 2579 | 2782 | 2538 | 2396 | 2364 | 2516 | 2588,52 | 2724 | 2454 | 0,901 | -0,15 | -2,55 | 0,010763 | 0,014088 | 0,003096 | 0,002547 | 0,0029401 | 0,0029401 | TRUE | 0,0571376 | No |
| orf19.7658 | 1920 | 1623 | 1793 | 1198 | 2030 | 1644 | 1680 | 2464 | 1748 | 1641 | 1697 | 1800 | 1806 | 1785 | 1721 | 1880 | 1759,59 | 1722 | 1798 | 1,046 | 0,065 | 1,345 | 0,178599 | 0,207806 | 0,0008416 | 0,002788 | 0,0016986 | 0,0016986 | TRUE | 0,0118387 | No |
| orf19.7660 | 630 | 556 | 539 | 417 | 502 | 422 | 404 | 624 | 574 | 562 | 510 | 626 | 447 | 458 | 414 | 476 | 508,36 | 568 | 449 | 0,794 | -0,333 | -4,236 | 2,27E-05 | 3,54E-05 | 0,0031272 | 0,004636 | 0,0039512 | 0,0039512 | TRUE | 0,0554444 | No |
| orf19.7661 | 1002 | 890 | 924 | 597 | 1555 | 1250 | 1372 | 1916 | 912 | 900 | 874 | 897 | 1383 | 1357 | 1405 | 1462 | 1148,83 | 896 | 1402 | 1,566 | 0,647 | 12,832 | 1,09E-37 | 4,08E-37 | 1,00E-08 | 0,003187 | 0,0015127 | 0,0015127 | TRUE | 0,0105097 | No |
| orf19.7662 | 3772 | 3143 | 3445 | 2066 | 3723 | 2794 | 3245 | 4361 | 3434 | 3179 | 3260 | 3103 | 3311 | 3034 | 3323 | 3327 | 3246,47 | 3244 | 3249 | 1,001 | 0,001 | 0,028 | 0,978053 | 0,979897 | 0,0016042 | 0,002444 | 0,0019477 | 0,0019477 | TRUE | 0,0236811 | No |
| orf19.7663 | 756 | 588 | 718 | 420 | 514 | 410 | 481 | 559 | 688 | 595 | 680 | 631 | 457 | 445 | 493 | 426 | 551,85 | 648 | 455 | 0,7 | -0,514 | -6,861 | 6,85E-12 | 1,39E-11 | 0,0025548 | 0,004431 | 0,0035446 | 0,0035446 | TRUE | 0,0392212 | No |
| orf19.7664 | 1677 | 1495 | 1650 | 1137 | 3263 | 2403 | 2741 | 3567 | 1527 | 1512 | 1562 | 1708 | 2902 | 2609 | 2807 | 2721 | 2168,47 | 1577 | 2760 | 1,754 | 0,811 | 15,189 | 4,16E-52 | 1,99E-51 | 0,0019765 | 0,002646 | 0,0022255 | 0,0022255 | TRUE | 0,034333 | No |
| orf19.7665 | 1224 | 910 | 986 | 677 | 1881 | 1338 | 1441 | 2100 | 1114 | 920 | 933 | 1017 | 1673 | 1453 | 1476 | 1602 | 1273,54 | 996 | 1551 | 1,558 | 0,64 | 8,76 | 1,95E-18 | 4,74E-18 | 0,0052269 | 0,003074 | 0,0042786 | 0,0042786 | TRUE | 0,0808479 | No |
| orf19.7666 | 18 | 13 | 16 | 11 | 43 | 57 | 29 | 56 | 16 | 13 | 15 | 17 | 38 | 62 | 30 | 43 | 29,22 | 15 | 43 | 2,819 | 1,495 | 4,917 | 8,78E-07 | 1,46E-06 | 1,00E-08 | 0,047247 | 0,0430607 | 0,0430607 | TRUE | 0,3205208 | No |
| orf19.7667 | 1528 | 1313 | 1358 | 990 | 1476 | 1254 | 1374 | 1848 | 1391 | 1328 | 1285 | 1487 | 1313 | 1361 | 1407 | 1410 | 1372,83 | 1373 | 1373 | 1,002 | 0,004 | 0,063 | 0,949495 | 0,955792 | 0,0014843 | 0,002999 | 0,0022924 | 0,0022924 | TRUE | 0,0341223 | No |
| orf19.7668 | 416 | 306 | 351 | 218 | 406 | 341 | 403 | 544 | 379 | 309 | 332 | 327 | 361 | 370 | 413 | 415 | 363,37 | 337 | 390 | 1,155 | 0,208 | 2,329 | 0,019871 | 0,025478 | 0,0038965 | 0,005672 | 0,004878 | 0,004878 | TRUE | 0,0851952 | No |
| orf19.767 | 1189 | 1295 | 935 | 708 | 1758 | 1930 | 1302 | 2214 | 1083 | 1310 | 885 | 1064 | 1564 | 2095 | 1333 | 1689 | 1377,76 | 1086 | 1670 | 1,539 | 0,622 | 3,467 | 0,000525 | 0,000756 | 0,0301454 | 0,002996 | 0,0149283 | 0,0301454 | TRUE | 0,3269194 | No |
| orf19.7670 | 1873 | 1434 | 1706 | 1074 | 1395 | 1171 | 1389 | 1909 | 1705 | 1450 | 1615 | 1613 | 1241 | 1271 | 1423 | 1456 | 1471,8 | 1596 | 1348 | 0,845 | -0,243 | -3,503 | 0,00046 | 0,000664 | 0,0044765 | 0,002935 | 0,0039426 | 0,0039426 | TRUE | 0,0463437 | No |
| orf19.7672 | 2284 | 1796 | 1983 | 1400 | 2642 | 2053 | 2280 | 3389 | 2079 | 1816 | 1877 | 2103 | 2350 | 2229 | 2335 | 2585 | 2171,85 | 1969 | 2375 | 1,208 | 0,273 | 4,26 | 2,05E-05 | 3,19E-05 | 0,0040451 | 0,002645 | 0,0034654 | 0,0034654 | TRUE | 0,0439136 | No |
| orf19.7673 | 425 | 307 | 338 | 196 | 745 | 457 | 583 | 846 | 387 | 310 | 320 | 294 | 663 | 496 | 597 | 645 | 464,12 | 328 | 600 | 1,826 | 0,869 | 8,078 | 6,56E-16 | 1,49E-15 | 0,01323 | 0,004883 | 0,0087015 | 0,0087015 | TRUE | 0,1716688 | No |
| orf19.7675 | 751 | 545 | 602 | 453 | 1238 | 911 | 1061 | 1519 | 684 | 551 | 570 | 680 | 1101 | 989 | 1087 | 1159 | 852,59 | 621 | 1084 | 1,75 | 0,807 | 9,835 | 7,92E-23 | 2,14E-22 | 0,0068895 | 0,003586 | 0,0051734 | 0,0051734 | TRUE | 0,0673578 | No |
| orf19.7676 | 19956 | 15522 | 16642 | 12942 | 5834 | 4735 | 4881 | 7243 | 18169 | 15699 | 15750 | 19441 | 5189 | 5141 | 4999 | 5525 | 11239,1 | 17265 | 5214 | 0,302 | -1,727 | -24,757 | 2,62E-135 | 3,06E-134 | 0,0065064 | 0,002155 | 0,0045543 | 0,0045543 | TRUE | 0,0876302 | No |
| orf19.7678 | 5979 | 4593 | 4954 | 3449 | 9778 | 7088 | 7929 | 11268 | 5444 | 4645 | 4689 | 5181 | 8697 | 7696 | 8120 | 8595 | 6633,36 | 4990 | 8277 | 1,659 | 0,731 | 11,895 | 1,26E-32 | 4,30E-32 | 0,0043694 | 0,002236 | 0,0034605 | 0,0034605 | TRUE | 0,0467467 | No |
| orf19.768 | 559 | 546 | 578 | 316 | 1087 | 894 | 987 | 1304 | 509 | 552 | 547 | 475 | 967 | 971 | 1011 | 995 | 753,23 | 521 | 986 | 1,885 | 0,915 | 14,694 | 7,09E-49 | 3,20E-48 | 1,00E-08 | 0,003791 | 0,0022258 | 0,0022258 | TRUE | 0,0333054 | No |
| orf19.7680 | 523 | 396 | 447 | 293 | 534 | 401 | 522 | 668 | 476 | 401 | 423 | 440 | 475 | 435 | 535 | 510 | 461,79 | 435 | 489 | 1,124 | 0,169 | 1,993 | 0,046242 | 0,05742 | 0,0045043 | 0,004898 | 0,0047174 | 0,0047174 | TRUE | 0,0590565 | No |
| orf19.769 | 91 | 76 | 69 | 42 | 109 | 100 | 104 | 133 | 83 | 77 | 65 | 63 | 97 | 109 | 107 | 101 | 87,7 | 72 | 104 | 1,42 | 0,506 | 3,343 | 0,000828 | 0,001175 | 1,00E-08 | 0,0171 | 0,0101748 | 0,0101748 | TRUE | 0,097526 | No |
| orf19.771 | 662 | 641 | 599 | 386 | 726 | 614 | 656 | 793 | 603 | 648 | 567 | 580 | 646 | 667 | 672 | 605 | 623,36 | 600 | 648 | 1,077 | 0,107 | 1,566 | 0,117417 | 0,139886 | 0,0014157 | 0,004156 | 0,0028517 | 0,0028517 | TRUE | 0,0358354 | No |
| orf19.77.1 | 3 | 7 | 10 | 3 | 6 | 8 | 6 | 8 | 3 | 7 | 9 | 5 | 5 | 9 | 6 | 6 | 6,26 | 6 | 6 | 1,084 | 0,116 | 0,208 | 0,835114 | 0,858132 | 1,00E-08 | 0,213179 | 0,1396033 | 0,1396033 | TRUE | 0,3160716 | No |
| orf19.772 | 2789 | 2453 | 2566 | 1584 | 4502 | 3457 | 3628 | 5039 | 2539 | 2481 | 2428 | 2379 | 4004 | 3753 | 3716 | 3844 | 3143,15 | 2457 | 3829 | 1,558 | 0,639 | 15,259 | 1,44E-52 | 6,95E-52 | 0,0006449 | 0,002457 | 0,0013443 | 0,0013443 | TRUE | 0,0115262 | No |
| orf19.772.1 | 2162 | 1971 | 1981 | 1374 | 1458 | 1345 | 1230 | 1857 | 1968 | 1993 | 1875 | 2064 | 1297 | 1460 | 1260 | 1417 | 1666,75 | 1975 | 1358 | 0,688 | -0,539 | -9,241 | 2,44E-20 | 6,23E-20 | 0,0024207 | 0,00283 | 0,0026449 | 0,0026449 | TRUE | 0,0290476 | No |
| orf19.773 | 3965 | 3905 | 3989 | 2551 | 3619 | 2795 | 3028 | 3737 | 3610 | 3949 | 3775 | 3832 | 3219 | 3035 | 3101 | 2851 | 3421,49 | 3792 | 3052 | 0,805 | -0,313 | -6,384 | 1,73E-10 | 3,32E-10 | 0,0015728 | 0,002423 | 0,0020198 | 0,0020198 | TRUE | 0,0246683 | No |
| orf19.775 | 3400 | 3349 | 3393 | 2121 | 1833 | 1604 | 1651 | 2225 | 3096 | 3387 | 3211 | 3186 | 1630 | 1741 | 1691 | 1697 | 2454,99 | 3220 | 1690 | 0,525 | -0,931 | -20,771 | 7,85E-96 | 6,46E-95 | 0,0007564 | 0,002575 | 0,0014837 | 0,0014837 | TRUE | 0,0151498 | No |
| orf19.776 | 117 | 110 | 110 | 65 | 176 | 153 | 191 | 247 | 107 | 111 | 104 | 98 | 157 | 166 | 196 | 188 | 140,78 | 105 | 177 | 1,679 | 0,748 | 5,922 | 3,19E-09 | 5,85E-09 | 1,00E-08 | 0,011421 | 0,0076138 | 0,0076138 | TRUE | 0,0668801 | No |
| orf19.778 | 10178 | 8599 | 8989 | 6013 | 5400 | 4332 | 4650 | 6200 | 9267 | 8697 | 8507 | 9032 | 4803 | 4703 | 4762 | 4729 | 6812,68 | 8876 | 4749 | 0,535 | -0,902 | -23,925 | 1,68E-126 | 1,82E-125 | 0,0006991 | 0,002231 | 0,0012053 | 0,0012053 | TRUE | 0,0110921 | No |
| orf19.779 | 849 | 735 | 796 | 451 | 785 | 656 | 773 | 912 | 773 | 743 | 753 | 677 | 698 | 712 | 792 | 696 | 730,62 | 736 | 724 | 0,979 | -0,031 | -0,461 | 0,644835 | 0,682408 | 0,0017172 | 0,003845 | 0,0029223 | 0,0029223 | TRUE | 0,045276 | No |
| orf19.780 | 2115 | 1923 | 2251 | 1285 | 13286 | 11627 | 12052 | 16098 | 1926 | 1945 | 2130 | 1930 | 11817 | 12624 | 12343 | 12280 | 7124,35 | 1983 | 12266 | 6,181 | 2,628 | 59,19 | 0 | 0 | 0,0012203 | 0,002222 | 0,0015834 | 0,0015834 | TRUE | 0,0313365 | No |
| orf19.781 | 170 | 158 | 157 | 113 | 463 | 417 | 401 | 539 | 155 | 160 | 149 | 170 | 412 | 453 | 411 | 411 | 289,91 | 158 | 422 | 2,675 | 1,42 | 15,368 | 2,70E-53 | 1,33E-52 | 1,00E-08 | 0,006594 | 0,0037147 | 0,0037147 | TRUE | 0,0269202 | No |
| orf19.782 | 4160 | 3853 | 4334 | 2745 | 4355 | 3853 | 3898 | 5150 | 3788 | 3897 | 4102 | 4123 | 3874 | 4183 | 3992 | 3928 | 3985,87 | 3978 | 3994 | 1,005 | 0,007 | 0,149 | 0,881623 | 0,8988 | 0,0009545 | 0,002368 | 0,0016108 | 0,0016108 | TRUE | 0,0127503 | No |
| orf19.783 | 3769 | 3518 | 3611 | 2497 | 3125 | 2653 | 2808 | 3911 | 3432 | 3558 | 3417 | 3751 | 2780 | 2880 | 2876 | 2983 | 3209,63 | 3540 | 2880 | 0,815 | -0,295 | -6,649 | 2,95E-11 | 5,85E-11 | 0,00095 | 0,002449 | 0,0015813 | 0,0015813 | TRUE | 0,0185479 | No |
| orf19.787 | 17 | 24 | 30 | 21 | 34 | 34 | 35 | 44 | 15 | 24 | 28 | 32 | 30 | 37 | 36 | 34 | 29,53 | 25 | 34 | 1,392 | 0,478 | 1,793 | 0,072965 | 0,088889 | 1,00E-08 | 0,046769 | 0,0326554 | 0,0326554 | TRUE | 0,4493928 | No |
| orf19.787.1 | 6930 | 6727 | 6584 | 4589 | 4268 | 3426 | 3172 | 4393 | 6309 | 6804 | 6231 | 6893 | 3796 | 3720 | 3249 | 3351 | 5044,14 | 6559 | 3529 | 0,538 | -0,894 | -14,882 | 4,30E-50 | 1,98E-49 | 0,0039513 | 0,002299 | 0,0032516 | 0,0032516 | TRUE | 0,0343144 | No |
| orf19.789 | 5776 | 5511 | 5687 | 3568 | 6144 | 5083 | 5121 | 6869 | 5259 | 5574 | 5382 | 5360 | 5465 | 5519 | 5245 | 5240 | 5380,3 | 5394 | 5367 | 0,995 | -0,007 | -0,203 | 0,83927 | 0,862124 | 0,0004931 | 0,002283 | 0,0011075 | 0,0011075 | TRUE | 0,0062754 | No |
| orf19.789.1 | 837 | 734 | 771 | 427 | 363 | 265 | 309 | 398 | 762 | 742 | 730 | 641 | 323 | 288 | 316 | 304 | 513,27 | 719 | 308 | 0,427 | -1,229 | -15,945 | 3,07E-57 | 1,60E-56 | 0,001961 | 0,004611 | 0,003446 | 0,003446 | TRUE | 0,0544386 | No |
| orf19.791 | 0 | 0 | 0 | 0 | 0 | 0 | 0 | 0 | 0 | 0 | 0 | 0 | 0 | 0 | 0 | 0 | 0 | 0 | 0 | NA | NA | NA | NA | NA | NA | NA | NA | NA | NA | NA | NA |
| orf19.792 | 2084 | 2296 | 2190 | 1822 | 5229 | 4271 | 4429 | 6021 | 1897 | 2322 | 2073 | 2737 | 4651 | 4637 | 4536 | 4593 | 3430,75 | 2257 | 4604 | 2,043 | 1,031 | 11,754 | 6,76E-32 | 2,27E-31 | 0,0126382 | 0,002422 | 0,0070498 | 0,0070498 | TRUE | 0,2420854 | No |
| orf19.793 | 1452 | 1588 | 1708 | 1013 | 1391 | 1304 | 1197 | 1542 | 1322 | 1606 | 1616 | 1522 | 1237 | 1416 | 1226 | 1176 | 1390,17 | 1516 | 1264 | 0,833 | -0,264 | -3,418 | 0,000631 | 0,000902 | 0,0065163 | 0,002987 | 0,0050035 | 0,0050035 | TRUE | 0,0928561 | No |
| orf19.794 | 1193 | 1182 | 1234 | 828 | 1009 | 950 | 897 | 1197 | 1086 | 1195 | 1168 | 1244 | 897 | 1031 | 919 | 913 | 1056,74 | 1173 | 940 | 0,802 | -0,319 | -5,004 | 5,62E-07 | 9,44E-07 | 0,0024463 | 0,003287 | 0,0029374 | 0,0029374 | TRUE | 0,0490979 | No |
| orf19.795 | 2424 | 2347 | 2415 | 1365 | 1609 | 1288 | 1487 | 1812 | 2207 | 2374 | 2286 | 2050 | 1431 | 1398 | 1523 | 1382 | 1831,42 | 2229 | 1434 | 0,642 | -0,64 | -11,379 | 5,29E-30 | 1,70E-29 | 0,0020929 | 0,002758 | 0,0024655 | 0,0024655 | TRUE | 0,0328125 | No |
| orf19.796 | 1238 | 1188 | 1213 | 784 | 1121 | 939 | 997 | 1281 | 1127 | 1202 | 1148 | 1178 | 997 | 1019 | 1021 | 977 | 1083,65 | 1164 | 1004 | 0,863 | -0,213 | -4,263 | 2,02E-05 | 3,15E-05 | 1,00E-08 | 0,003256 | 0,0014788 | 0,0014788 | TRUE | 0,0063384 | No |
| orf19.797 | 2262 | 1902 | 2027 | 1331 | 2181 | 1755 | 1865 | 2618 | 2059 | 1924 | 1918 | 1999 | 1940 | 1905 | 1910 | 1997 | 1956,66 | 1975 | 1938 | 0,982 | -0,026 | -0,593 | 0,553024 | 0,594624 | 0,0003828 | 0,002712 | 0,0013536 | 0,0013536 | TRUE | 0,0108931 | No |
| orf19.798 | 2777 | 2479 | 2548 | 1631 | 3378 | 2857 | 2968 | 3942 | 2528 | 2507 | 2411 | 2450 | 3005 | 3102 | 3040 | 3007 | 2756,27 | 2474 | 3038 | 1,227 | 0,295 | 7,89 | 3,02E-15 | 6,73E-15 | 1,00E-08 | 0,002516 | 0,0009736 | 0,0009736 | TRUE | 0,0038872 | No |
| orf19.799 | 184 | 131 | 181 | 131 | 459 | 392 | 385 | 503 | 168 | 132 | 171 | 197 | 408 | 426 | 394 | 384 | 284,99 | 167 | 403 | 2,431 | 1,282 | 12,316 | 7,42E-35 | 2,65E-34 | 0,0044908 | 0,006672 | 0,0060183 | 0,0060183 | TRUE | 0,2015411 | No |
| orf19.8 | 116 | 124 | 117 | 88 | 156 | 137 | 127 | 177 | 106 | 125 | 111 | 132 | 139 | 149 | 130 | 135 | 128,32 | 118 | 138 | 1,174 | 0,232 | 1,833 | 0,066755 | 0,081619 | 1,00E-08 | 0,012332 | 0,0074336 | 0,0074336 | TRUE | 0,0549649 | No |
| orf19.801 | 6166 | 5090 | 5890 | 3765 | 5541 | 4682 | 5089 | 6741 | 5614 | 5148 | 5574 | 5656 | 4929 | 5083 | 5212 | 5142 | 5294,7 | 5498 | 5092 | 0,926 | -0,11 | -2,639 | 0,008314 | 0,010965 | 0,0009471 | 0,002287 | 0,0014931 | 0,0014931 | TRUE | 0,0225261 | No |
| orf19.802 | 1813 | 1467 | 1656 | 1126 | 1827 | 1493 | 1603 | 2114 | 1651 | 1484 | 1567 | 1691 | 1625 | 1621 | 1642 | 1613 | 1611,67 | 1598 | 1625 | 1,018 | 0,026 | 0,516 | 0,60585 | 0,645436 | 0,0008653 | 0,002857 | 0,0018151 | 0,0018151 | TRUE | 0,0277868 | No |
| orf19.804 | 462 | 410 | 464 | 263 | 1403 | 1131 | 1323 | 1666 | 421 | 415 | 439 | 395 | 1248 | 1228 | 1355 | 1271 | 846,39 | 418 | 1276 | 3,045 | 1,607 | 25,518 | 1,26E-143 | 1,57E-142 | 1,00E-08 | 0,003598 | 0,0021673 | 0,0021673 | TRUE | 0,0204385 | No |
| orf19.804.1 | 6019 | 4147 | 4227 | 2955 | 4850 | 3356 | 3445 | 5365 | 5480 | 4194 | 4000 | 4439 | 4314 | 3644 | 3528 | 4092 | 4211,48 | 4528 | 3894 | 0,86 | -0,217 | -2,324 | 0,020136 | 0,025802 | 0,0148099 | 0,002351 | 0,0081706 | 0,0081706 | TRUE | 0,2456269 | No |
| orf19.805 | 904 | 743 | 840 | 510 | 971 | 786 | 855 | 1124 | 823 | 751 | 795 | 766 | 864 | 853 | 876 | 857 | 823,21 | 784 | 862 | 1,098 | 0,135 | 2,435 | 0,014883 | 0,019301 | 1,00E-08 | 0,003642 | 0,0017452 | 0,0017452 | TRUE | 0,0140866 | No |
| orf19.806 | 279 | 200 | 252 | 146 | 276 | 208 | 243 | 291 | 254 | 202 | 238 | 219 | 245 | 226 | 249 | 222 | 232,03 | 228 | 236 | 1,026 | 0,037 | 0,368 | 0,713014 | 0,746627 | 0,0024908 | 0,00773 | 0,0056087 | 0,0056087 | TRUE | 0,0726155 | No |
| orf19.807 | 2730 | 2289 | 2475 | 1478 | 2254 | 1863 | 2126 | 2751 | 2486 | 2315 | 2342 | 2220 | 2005 | 2023 | 2177 | 2098 | 2208,32 | 2341 | 2076 | 0,886 | -0,175 | -3,558 | 0,000373 | 0,000541 | 0,001216 | 0,002635 | 0,0018758 | 0,0018758 | TRUE | 0,0211585 | No |
| orf19.808 | 3821 | 3470 | 3855 | 2349 | 3642 | 3101 | 3373 | 4468 | 3479 | 3510 | 3648 | 3529 | 3239 | 3367 | 3454 | 3408 | 3454,28 | 3542 | 3367 | 0,951 | -0,073 | -1,912 | 0,055865 | 0,06894 | 0,0003467 | 0,002419 | 0,0011092 | 0,0011092 | TRUE | 0,0080448 | No |
| orf19.809 | 10894 | 9526 | 10956 | 6318 | 12063 | 10132 | 10694 | 14607 | 9919 | 9635 | 10369 | 9491 | 10730 | 11001 | 10952 | 11142 | 10404,6 | 9854 | 10956 | 1,112 | 0,153 | 4,096 | 4,20E-05 | 6,43E-05 | 0,0007652 | 0,002164 | 0,0012341 | 0,0012341 | TRUE | 0,0152052 | No |
| orf19.81 | 2468 | 3131 | 3019 | 1723 | 1797 | 1562 | 1556 | 1805 | 2247 | 3167 | 2857 | 2588 | 1598 | 1696 | 1594 | 1377 | 2140,47 | 2715 | 1566 | 0,576 | -0,795 | -8,331 | 8,00E-17 | 1,86E-16 | 0,0134156 | 0,002654 | 0,0082402 | 0,0082402 | TRUE | 0,1535917 | No |
| orf19.810 | 713 | 615 | 699 | 426 | 846 | 711 | 783 | 1067 | 649 | 622 | 662 | 640 | 752 | 772 | 802 | 814 | 714,11 | 643 | 785 | 1,221 | 0,288 | 4,853 | 1,22E-06 | 2,01E-06 | 1,00E-08 | 0,003887 | 0,0019486 | 0,0019486 | TRUE | 0,0098331 | No |
| orf19.811 | 739 | 649 | 725 | 412 | 785 | 612 | 703 | 941 | 673 | 656 | 686 | 619 | 698 | 664 | 720 | 718 | 679,34 | 658 | 700 | 1,061 | 0,085 | 1,386 | 0,165875 | 0,194135 | 1,00E-08 | 0,003982 | 0,0021504 | 0,0021504 | TRUE | 0,0162568 | No |
| orf19.812 | 42 | 20 | 42 | 18 | 67 | 50 | 62 | 64 | 38 | 20 | 40 | 27 | 60 | 54 | 63 | 49 | 43,93 | 31 | 56 | 1,778 | 0,83 | 3,544 | 0,000395 | 0,000572 | 0,0175422 | 0,032107 | 0,0276388 | 0,0276388 | TRUE | 0,425937 | No |
| orf19.813 | 37436 | 34497 | 36870 | 21246 | 47469 | 37362 | 42063 | 54673 | 34084 | 34890 | 34894 | 31915 | 42222 | 40565 | 43078 | 41705 | 37919,1 | 33946 | 41892 | 1,234 | 0,303 | 7,72 | 1,16E-14 | 2,56E-14 | 0,0011373 | 0,002072 | 0,0014554 | 0,0014554 | TRUE | 0,0196925 | No |
| orf19.814 | 4982 | 4400 | 4903 | 2967 | 2953 | 2386 | 2605 | 3610 | 4536 | 4450 | 4640 | 4457 | 2627 | 2591 | 2668 | 2754 | 3590,24 | 4521 | 2660 | 0,588 | -0,765 | -20,407 | 1,46E-92 | 1,17E-91 | 0,00026 | 0,002405 | 0,0010546 | 0,0010546 | TRUE | 0,0072045 | No |
| orf19.815 | 11540 | 11105 | 11905 | 7181 | 11266 | 9370 | 9654 | 12875 | 10507 | 11232 | 11267 | 10787 | 10021 | 10173 | 9887 | 9821 | 10461,8 | 10948 | 9976 | 0,911 | -0,134 | -3,816 | 0,000136 | 0,000202 | 0,0005544 | 0,002163 | 0,0010968 | 0,0010968 | TRUE | 0,0092596 | No |
| orf19.816 | 709 | 634 | 696 | 416 | 359 | 299 | 301 | 418 | 646 | 641 | 659 | 625 | 319 | 325 | 308 | 319 | 480,18 | 643 | 318 | 0,494 | -1,018 | -14,912 | 2,75E-50 | 1,27E-49 | 1,00E-08 | 0,004788 | 0,0021798 | 0,0021798 | TRUE | 0,0042141 | No |
| orf19.819 | 904 | 773 | 838 | 559 | 1343 | 929 | 1006 | 1331 | 823 | 782 | 793 | 840 | 1195 | 1009 | 1030 | 1015 | 935,8 | 810 | 1062 | 1,314 | 0,394 | 5,832 | 5,48E-09 | 9,99E-09 | 0,0031372 | 0,003449 | 0,003285 | 0,003285 | TRUE | 0,0853604 | No |
| orf19.820 | 1352 | 1142 | 1205 | 742 | 1365 | 997 | 1192 | 1521 | 1231 | 1155 | 1140 | 1115 | 1214 | 1082 | 1221 | 1160 | 1164,82 | 1160 | 1169 | 1,007 | 0,01 | 0,171 | 0,863959 | 0,884278 | 0,0015117 | 0,003171 | 0,0023107 | 0,0023107 | TRUE | 0,0284283 | No |

| | | | | | | | | | | | | | | | | | | | | | | | | | | | | | | | |
|---|---|---|---|---|---|---|---|---|---|---|---|---|---|---|---|---|---|---|---|---|---|---|---|---|---|---|---|---|---|---|---|
| orf19.822 | 3E+05 | 2E+05 | 2E+05 | 1E+05 | 28221 | 20859 | 21674 | 30402 | 2E+05 | 2E+05 | 2E+05 | 2E+05 | 25102 | 22647 | 22197 | 23191 | 124526 | 225767 | 23284 | 0,103 | -3,277 | -58,449 | 0 | 0 | 0,0037329 | 0,002048 | 0,0029986 | 0,0029986 | TRUE | 0,0481334 | No |
| orf19.823 | 988 | 1262 | 1242 | 1826 | 177 | 296 | 112 | 195 | 900 | 1276 | 1175 | 2743 | 157 | 321 | 115 | 149 | 854,57 | 1524 | 186 | 0,122 | -3,038 | -6,339 | 2,31E-10 | 4,42E-10 | 0,2177788 | 0,003583 | 0,0787746 | 0,2177788 | TRUE | 1,0256197 | No |
| orf19.825 | 2018 | 1730 | 1857 | 1217 | 2025 | 1550 | 1758 | 2362 | 1837 | 1750 | 1757 | 1828 | 1801 | 1683 | 1800 | 1802 | 1782,35 | 1793 | 1772 | 0,989 | -0,016 | -0,348 | 0,727782 | 0,7603 | 0,0002952 | 0,002778 | 0,0013936 | 0,0013936 | TRUE | 0,0131198 | No |
| orf19.826 | 820 | 781 | 852 | 480 | 407 | 354 | 351 | 474 | 747 | 790 | 806 | 721 | 362 | 384 | 359 | 362 | 566,41 | 766 | 367 | 0,477 | -1,067 | -15,657 | 2,98E-55 | 1,51E-54 | 1,00E-08 | 0,004369 | 0,0024901 | 0,0024901 | TRUE | 0,0157576 | No |
| orf19.827 | 1449 | 1207 | 1374 | 942 | 1853 | 1543 | 1554 | 2093 | 1319 | 1221 | 1300 | 1415 | 1648 | 1675 | 1591 | 1597 | 1470,86 | 1314 | 1628 | 1,242 | 0,312 | 5,907 | 3,49E-09 | 6,40E-09 | 0,0012409 | 0,002935 | 0,0019846 | 0,0019846 | TRUE | 0,0286483 | No |
| orf19.827.1 | 11957 | 9751 | 9817 | 6475 | 15685 | 10974 | 11319 | 16454 | 10886 | 9862 | 9291 | 9726 | 13951 | 11915 | 11592 | 12551 | 11221,9 | 9941 | 12502 | 1,258 | 0,331 | 5,035 | 4,78E-07 | 8,04E-07 | 0,0056803 | 0,002155 | 0,0040519 | 0,0040519 | TRUE | 0,0744762 | No |
| orf19.8278 | 2106 | 2043 | 2095 | 1289 | 2024 | 1582 | 1730 | 2154 | 1917 | 2066 | 1983 | 1936 | 1800 | 1718 | 1772 | 1643 | 1854,43 | 1976 | 1733 | 0,876 | -0,19 | -3,943 | 8,06E-05 | 0,000121 | 0,0009131 | 0,002749 | 0,0016925 | 0,0016925 | TRUE | 0,0157347 | No |
| orf19.828 | 2450 | 2219 | 2289 | 1472 | 1962 | 1546 | 1645 | 2164 | 2231 | 2244 | 2166 | 2211 | 1745 | 1679 | 1685 | 1651 | 1951,43 | 2213 | 1690 | 0,763 | -0,389 | -9,455 | 3,24E-21 | 8,45E-21 | 1,00E-08 | 0,002714 | 0,0011099 | 0,0011099 | TRUE | 0,0061021 | No |
| orf19.829 | 5573 | 5088 | 5406 | 3553 | 6462 | 5562 | 5917 | 7973 | 5074 | 5146 | 5116 | 5337 | 5748 | 6039 | 6060 | 6082 | 5575,19 | 5168 | 5982 | 1,158 | 0,212 | 5,917 | 3,28E-09 | 6,02E-09 | 0,0003582 | 0,002274 | 0,0010484 | 0,0010484 | TRUE | 0,0085672 | No |
| orf19.831 | 2854 | 2710 | 2919 | 1813 | 1551 | 1314 | 1337 | 1832 | 2598 | 2741 | 2763 | 2723 | 1380 | 1427 | 1369 | 1397 | 2049,78 | 2706 | 1393 | 0,515 | -0,958 | -22,523 | 2,47E-112 | 2,37E-111 | 1,00E-08 | 0,002681 | 0,0012022 | 0,0012022 | TRUE | 0,0092157 | No |
| orf19.832 | 1183 | 1155 | 1277 | 711 | 833 | 805 | 814 | 1042 | 1077 | 1168 | 1209 | 1068 | 741 | 874 | 834 | 795 | 970,66 | 1130 | 811 | 0,715 | -0,483 | -7,181 | 6,91E-13 | 1,44E-12 | 0,0032004 | 0,003398 | 0,0032982 | 0,0032982 | TRUE | 0,0398234 | No |
| orf19.834 | 3889 | 3854 | 4106 | 2434 | 3029 | 2382 | 2719 | 3328 | 3541 | 3898 | 3886 | 3656 | 2694 | 2586 | 2785 | 2539 | 3198,06 | 3745 | 2651 | 0,707 | -0,499 | -10,16 | 3,00E-24 | 8,45E-24 | 0,0015403 | 0,00245 | 0,0019992 | 0,0019992 | TRUE | 0,0171757 | No |
| orf19.835 | 1152 | 1124 | 1186 | 719 | 2004 | 1639 | 1824 | 2427 | 1049 | 1137 | 1122 | 1080 | 1782 | 1779 | 1868 | 1851 | 1458,69 | 1097 | 1820 | 1,659 | 0,73 | 15,098 | 1,66E-51 | 7,90E-51 | 0,0002442 | 0,002943 | 0,0015027 | 0,0015027 | TRUE | 0,0118251 | No |
| orf19.836 | 209 | 214 | 197 | 133 | 32 | 34 | 43 | 30 | 190 | 216 | 186 | 200 | 28 | 37 | 44 | 23 | 115,66 | 198 | 33 | 0,163 | -2,613 | -15,086 | 2,00E-51 | 9,50E-51 | 0,0048996 | 0,013459 | 0,0117679 | 0,0117679 | TRUE | 0,380069 | No |
| orf19.836.1 | 339 | 249 | 258 | 118 | 366 | 239 | 281 | 351 | 309 | 252 | 244 | 177 | 326 | 259 | 288 | 268 | 265,31 | 246 | 285 | 1,152 | 0,205 | 1,493 | 0,135481 | 0,160032 | 0,024302 | 0,007016 | 0,0142314 | 0,0142314 | TRUE | 0,3040352 | No |
| orf19.837 | 1214 | 1133 | 1126 | 611 | 1815 | 1261 | 1505 | 1883 | 1105 | 1146 | 1066 | 918 | 1614 | 1369 | 1541 | 1436 | 1274,48 | 1059 | 1490 | 1,404 | 0,49 | 6,551 | 5,70E-11 | 1,12E-10 | 0,0060068 | 0,003074 | 0,0045408 | 0,0045408 | TRUE | 0,0880466 | No |
| orf19.837.1 | 919 | 856 | 945 | 499 | 2264 | 1688 | 1815 | 2479 | 837 | 866 | 894 | 750 | 2014 | 1833 | 1859 | 1891 | 1367,83 | 837 | 1899 | 2,262 | 1,178 | 19,562 | 3,26E-85 | 2,47E-84 | 0,0021534 | 0,003003 | 0,0025946 | 0,0025946 | TRUE | 0,0503891 | No |
| orf19.838.1 | 19928 | 16350 | 17684 | 10478 | 33600 | 24498 | 26365 | 37553 | 18144 | 16536 | 16736 | 15740 | 29886 | 26598 | 27001 | 28646 | 22410,8 | 16789 | 28033 | 1,669 | 0,739 | 13,753 | 4,87E-43 | 2,00E-42 | 0,0031826 | 0,002096 | 0,002728 | 0,002728 | TRUE | 0,0361542 | No |
| orf19.839 | 3240 | 2873 | 3104 | 1856 | 3700 | 2906 | 3297 | 4368 | 2950 | 2906 | 2938 | 2788 | 3291 | 3155 | 3377 | 3332 | 3091,99 | 2896 | 3289 | 1,135 | 0,183 | 4,595 | 4,32E-06 | 6,99E-06 | 0,0003158 | 0,002464 | 0,0011889 | 0,0011889 | TRUE | 0,0087714 | No |
| orf19.84 | 5341 | 4742 | 5175 | 3394 | 1357 | 1273 | 1191 | 1744 | 4863 | 4796 | 4898 | 5098 | 1207 | 1382 | 1220 | 1330 | 3099,25 | 4914 | 1285 | 0,262 | -1,935 | -37,957 | 0 | 0 | 0,0019002 | 0,002463 | 0,002022 | 0,002022 | TRUE | 0,0293262 | No |
| orf19.840 | 18678 | 14805 | 15522 | 11308 | 27442 | 21230 | 21819 | 31684 | 17006 | 14974 | 14690 | 16986 | 24409 | 23050 | 22345 | 24169 | 19703,6 | 15914 | 23493 | 1,477 | 0,562 | 9,737 | 2,11E-22 | 5,64E-22 | 0,0038864 | 0,002104 | 0,0031498 | 0,0031498 | TRUE | 0,0328407 | No |
| orf19.841 | 1848 | 1658 | 1763 | 1135 | 2065 | 1763 | 1909 | 2361 | 1683 | 1677 | 1669 | 1705 | 1837 | 1914 | 1955 | 1801 | 1779,98 | 1684 | 1877 | 1,114 | 0,156 | 3,491 | 0,000482 | 0,000695 | 1,00E-08 | 0,002779 | 0,0013532 | 0,0013532 | TRUE | 0,0096528 | No |
| orf19.842 | 16568 | 15340 | 15962 | 9439 | 5709 | 4174 | 4466 | 5759 | 15084 | 15515 | 15107 | 14179 | 5078 | 4532 | 4574 | 4393 | 9807,65 | 14971 | 4644 | 0,31 | -1,689 | -32,869 | 6,06E-237 | 1,20E-235 | 0,0025992 | 0,002172 | 0,0023996 | 0,0023996 | TRUE | 0,0485898 | No |
| orf19.843 | 1277 | 1212 | 1236 | 723 | 1141 | 997 | 1129 | 1421 | 1163 | 1226 | 1170 | 1086 | 1015 | 1082 | 1156 | 1084 | 1122,73 | 1161 | 1084 | 0,932 | -0,102 | -1,748 | 0,080398 | 0,097422 | 0,0014514 | 0,003214 | 0,0023898 | 0,0023898 | TRUE | 0,0233246 | No |
| orf19.844 | 10225 | 9579 | 10630 | 6246 | 2335 | 1916 | 2144 | 2813 | 9309 | 9688 | 10060 | 9382 | 2077 | 2080 | 2196 | 2146 | 5867,38 | 9610 | 2125 | 0,221 | -2,178 | -52,881 | 0 | 0 | 0,0008376 | 0,002262 | 0,0013524 | 0,0013524 | TRUE | 0,0122623 | No |
| orf19.846 | 6823 | 6156 | 6779 | 3942 | 4849 | 4015 | 4479 | 6067 | 6212 | 6226 | 6416 | 5922 | 4313 | 4359 | 4587 | 4628 | 5332,83 | 6194 | 4472 | 0,722 | -0,471 | -11,382 | 5,15E-30 | 1,66E-29 | 0,0009151 | 0,002285 | 0,0014496 | 0,0014496 | TRUE | 0,0102435 | No |
| orf19.847 | 4618 | 4469 | 4563 | 2951 | 5740 | 4648 | 4579 | 6044 | 4204 | 4520 | 4318 | 4433 | 5106 | 5046 | 4689 | 4610 | 4615,95 | 4369 | 4863 | 1,113 | 0,155 | 3,291 | 0,000998 | 0,001408 | 0,001517 | 0,002323 | 0,0019013 | 0,0019013 | TRUE | 0,0152409 | No |
| orf19.848 | 0 | 1 | 0 | 0 | 3 | 3 | 1 | 0 | 0 | 1 | 0 | 0 | 3 | 3 | 1 | 0 | 1 | 0 | 2 | 5,245 | 2,391 | 1,485 | 0,137461 | 0,16222 | 1,00E-08 | 1,329491 | 1,2153354 | 1,2153354 | TRUE | 0,2158702 | No |
| orf19.849 | 84 | 103 | 88 | 64 | 497 | 436 | 459 | 527 | 76 | 104 | 83 | 96 | 442 | 473 | 470 | 402 | 268,45 | 90 | 447 | 4,992 | 2,32 | 19,868 | 7,67E-88 | 5,94E-87 | 0,0046453 | 0,006958 | 0,0061017 | 0,0061017 | TRUE | 0,1263717 | No |
| orf19.85 | 2040 | 2072 | 2058 | 1183 | 1106 | 778 | 826 | 1144 | 1857 | 2096 | 1948 | 1777 | 984 | 845 | 846 | 873 | 1403,09 | 1920 | 887 | 0,462 | -1,114 | -16,315 | 7,78E-60 | 4,24E-59 | 0,0041972 | 0,002979 | 0,0036729 | 0,0036729 | TRUE | 0,0650446 | No |
| orf19.850 | 1828 | 1711 | 1925 | 1125 | 1364 | 1083 | 1122 | 1452 | 1664 | 1730 | 1822 | 1690 | 1213 | 1176 | 1149 | 1108 | 1444,04 | 1726 | 1162 | 0,672 | -0,574 | -11,1 | 1,25E-28 | 3,87E-28 | 0,000969 | 0,002952 | 0,0018514 | 0,0018514 | TRUE | 0,0171975 | No |
| orf19.851 | 3812 | 3786 | 4011 | 2491 | 833 | 784 | 803 | 1034 | 3471 | 3829 | 3796 | 3742 | 741 | 851 | 822 | 789 | 2255,12 | 3710 | 801 | 0,216 | -2,214 | -39,677 | 0 | 0 | 0,0017062 | 0,002623 | 0,0022598 | 0,0022598 | TRUE | 0,029862 | No |
| orf19.852 | 875 | 783 | 854 | 640 | 849 | 782 | 683 | 1046 | 797 | 792 | 808 | 961 | 755 | 849 | 699 | 798 | 807,47 | 840 | 775 | 0,927 | -0,11 | -1,374 | 0,169434 | 0,197792 | 0,0064142 | 0,003673 | 0,0048559 | 0,0048559 | TRUE | 0,1042422 | No |
| orf19.853 | 105 | 86 | 102 | 58 | 86 | 85 | 80 | 111 | 96 | 87 | 97 | 87 | 76 | 92 | 82 | 85 | 87,7 | 92 | 84 | 0,91 | -0,135 | -0,924 | 0,355331 | 0,395191 | 1,00E-08 | 0,017099 | 0,0092468 | 0,0092468 | TRUE | 0,0386057 | No |
| orf19.854 | 3364 | 2845 | 3273 | 1984 | 3838 | 2985 | 3193 | 4279 | 3063 | 2877 | 3098 | 2980 | 3414 | 3241 | 3270 | 3264 | 3150,85 | 3004 | 3297 | 1,097 | 0,134 | 3,296 | 0,000982 | 0,001387 | 0,0005166 | 0,002456 | 0,0012602 | 0,0012602 | TRUE | 0,0102871 | No |
| orf19.855 | 42 | 28 | 29 | 16 | 33 | 18 | 23 | 30 | 38 | 28 | 27 | 24 | 29 | 20 | 24 | 23 | 26,67 | 29 | 24 | 0,801 | -0,32 | -1,178 | 0,238746 | 0,272956 | 1,00E-08 | 0,051565 | 0,0337434 | 0,0337434 | TRUE | 0,2819555 | No |
| orf19.856 | 1225 | 1129 | 1222 | 738 | 2440 | 1954 | 2148 | 2867 | 1115 | 1142 | 1157 | 1109 | 2170 | 2121 | 2200 | 2187 | 1650,11 | 1131 | 2170 | 1,918 | 0,939 | 21,627 | 1,01E-103 | 9,11E-103 | 1,00E-08 | 0,002838 | 0,0011246 | 0,0011246 | TRUE | 0,0028969 | No |
| orf19.857 | 1140 | 981 | 1060 | 671 | 3394 | 2834 | 2988 | 4332 | 1038 | 992 | 1003 | 1008 | 3019 | 3077 | 3060 | 3304 | 2062,7 | 1010 | 3115 | 3,084 | 1,625 | 33,292 | 5,09E-243 | 1,03E-241 | 0,0009096 | 0,002677 | 0,0016072 | 0,0016072 | TRUE | 0,0210635 | No |
| orf19.86 | 3072 | 2647 | 2765 | 1596 | 1850 | 1329 | 1499 | 1956 | 2797 | 2677 | 2617 | 2397 | 1646 | 1443 | 1535 | 1492 | 2075,5 | 2622 | 1529 | 0,582 | -0,78 | -13,091 | 3,69E-39 | 1,41E-38 | 0,0029445 | 0,002673 | 0,0028951 | 0,0028951 | TRUE | 0,0383096 | No |
| orf19.860 | 462 | 383 | 387 | 281 | 1173 | 1022 | 1116 | 1499 | 421 | 387 | 366 | 422 | 1043 | 1110 | 1143 | 1143 | 754,46 | 399 | 1110 | 2,789 | 1,48 | 21,809 | 1,90E-105 | 1,72E-104 | 0,0013154 | 0,003788 | 0,0026582 | 0,0026582 | TRUE | 0,0368014 | No |
| orf19.861 | 751 | 596 | 589 | 434 | 630 | 558 | 583 | 844 | 684 | 603 | 557 | 652 | 560 | 606 | 597 | 644 | 612,87 | 624 | 602 | 0,966 | -0,05 | -0,636 | 0,524617 | 0,566469 | 0,0042368 | 0,004192 | 0,0042141 | 0,0042141 | TRUE | 0,0634536 | No |
| orf19.862 | 380 | 337 | 340 | 229 | 449 | 404 | 395 | 580 | 346 | 341 | 322 | 344 | 399 | 439 | 405 | 442 | 379,69 | 338 | 421 | 1,248 | 0,319 | 4,117 | 3,84E-05 | 5,91E-05 | 1,00E-08 | 0,005516 | 0,0030846 | 0,0030846 | TRUE | 0,015073 | No |
| orf19.863 | 2798 | 2557 | 2854 | 1599 | 2848 | 2172 | 2358 | 3216 | 2547 | 2586 | 2701 | 2402 | 2533 | 2358 | 2415 | 2453 | 2499,51 | 2559 | 2440 | 0,952 | -0,07 | -1,493 | 0,135366 | 0,159926 | 0,0010827 | 0,002566 | 0,0017284 | 0,0017284 | TRUE | 0,0191351 | No |
| orf19.864 | 2896 | 2773 | 3084 | 1874 | 1188 | 1000 | 1092 | 1426 | 2637 | 2805 | 2919 | 2815 | 1057 | 1086 | 1118 | 1088 | 1940,45 | 2794 | 1087 | 0,389 | -1,362 | -28,096 | 1,08E-173 | 1,64E-172 | 0,0008858 | 0,002718 | 0,0016333 | 0,0016333 | TRUE | 0,0181775 | No |
| orf19.866 | 1467 | 1376 | 1473 | 895 | 6844 | 5922 | 6254 | 8271 | 1336 | 1392 | 1394 | 1344 | 6088 | 6430 | 6405 | 6309 | 3837,13 | 1366 | 6308 | 4,613 | 2,206 | 54,509 | 0 | 0 | 1,00E-08 | 0,002381 | 0,0011066 | 0,0011066 | TRUE | 0,0067536 | No |
| orf19.867 | 971 | 828 | 917 | 526 | 1117 | 892 | 1045 | 1274 | 884 | 837 | 868 | 790 | 994 | 968 | 1070 | 972 | 922,94 | 845 | 1001 | 1,181 | 0,24 | 4,045 | 5,22E-05 | 7,94E-05 | 0,0010225 | 0,003468 | 0,0022691 | 0,0022691 | TRUE | 0,0251681 | No |
| orf19.868 | 29071 | 25846 | 28612 | 19574 | 30413 | 27937 | 28935 | 38634 | 26468 | 26140 | 27079 | 29403 | 27051 | 30332 | 29633 | 29470 | 28197,1 | 27272 | 29122 | 1,068 | 0,095 | 1,88 | 0,060047 | 0,073843 | 0,002616 | 0,002084 | 0,0024071 | 0,0024071 | TRUE | 0,0336631 | No |
| orf19.87 | 212 | 196 | 186 | 112 | 393 | 269 | 285 | 438 | 193 | 198 | 176 | 168 | 350 | 292 | 292 | 334 | 250,39 | 184 | 317 | 1,721 | 0,783 | 7,636 | 2,24E-14 | 4,88E-14 | 0,003607 | 0,007313 | 0,0057423 | 0,0057423 | TRUE | 0,0531653 | No |
| orf19.871 | 5993 | 5496 | 5894 | 3703 | 4108 | 3638 | 3697 | 5094 | 5456 | 5559 | 5578 | 5562 | 3654 | 3950 | 3786 | 3886 | 4678,92 | 5539 | 3819 | 0,689 | -0,536 | -14,641 | 1,55E-48 | 6,95E-48 | 0,0003258 | 0,002319 | 0,0010697 | 0,0010697 | TRUE | 0,0104046 | No |
| orf19.872 | 1356 | 1165 | 1305 | 913 | 1704 | 1434 | 1526 | 2095 | 1235 | 1178 | 1235 | 1371 | 1516 | 1557 | 1563 | 1598 | 1406,61 | 1255 | 1558 | 1,246 | 0,317 | 5,901 | 3,62E-09 | 6,64E-09 | 0,0011177 | 0,002976 | 0,0020458 | 0,0020458 | TRUE | 0,0418202 | No |
| orf19.873 | 140 | 131 | 147 | 77 | 147 | 140 | 133 | 180 | 127 | 132 | 139 | 116 | 131 | 152 | 136 | 137 | 133,88 | 128 | 139 | 1,072 | 0,1 | 0,824 | 0,410042 | 0,451315 | 1,00E-08 | 0,011904 | 0,0067148 | 0,0067148 | TRUE | 0,0371668 | No |
| orf19.873.1 | 4854 | 4656 | 4764 | 2960 | 6552 | 5367 | 5224 | 7012 | 4419 | 4709 | 4509 | 4446 | 5828 | 5827 | 5350 | 5349 | 5054,65 | 4521 | 5588 | 1,236 | 0,305 | 6,714 | 1,90E-11 | 3,79E-11 | 0,0013719 | 0,002298 | 0,0017838 | 0,0017838 | TRUE | 0,0103737 | No |
| orf19.874 | 2007 | 1636 | 1886 | 1177 | 890 | 721 | 822 | 1094 | 1827 | 1655 | 1785 | 1768 | 792 | 783 | 842 | 835 | 1285,71 | 1759 | 813 | 0,462 | -1,113 | -20,711 | 2,73E-95 | 2,23E-94 | 0,0008963 | 0,003064 | 0,001894 | 0,001894 | TRUE | 0,0197936 | No |
| orf19.875 | 1649 | 1588 | 1635 | 988 | 1646 | 1317 | 1376 | 1947 | 1501 | 1606 | 1547 | 1484 | 1464 | 1430 | 1409 | 1485 | 1490,91 | 1534 | 1447 | 0,943 | -0,085 | -1,81 | 0,070286 | 0,085788 | 0,0002177 | 0,002923 | 0,0014523 | 0,0014523 | TRUE | 0,0116884 | No |
| orf19.876 | 1326 | 1025 | 1168 | 975 | 2815 | 2505 | 2501 | 3476 | 1207 | 1037 | 1105 | 1465 | 2504 | 2720 | 2561 | 2652 | 1906,3 | 1204 | 2609 | 2,174 | 1,121 | 12,981 | 1,57E-38 | 5,97E-38 | 0,0120176 | 0,00273 | 0,0065284 | 0,0065284 | TRUE | 0,2433137 | No |
| orf19.878 | 3098 | 2417 | 2554 | 1727 | 1847 | 1468 | 1581 | 2191 | 2821 | 2445 | 2417 | 2594 | 1643 | 1594 | 1619 | 1671 | 2100,45 | 2569 | 1632 | 0,635 | -0,654 | -11,606 | 3,82E-31 | 1,26E-30 | 0,0024768 | 0,002666 | 0,0025553 | 0,0025553 | TRUE | 0,054781 | No |
| orf19.879 | 971 | 879 | 992 | 658 | 1432 | 1239 | 1311 | 1632 | 884 | 889 | 939 | 988 | 1274 | 1345 | 1343 | 1245 | 1113,35 | 925 | 1302 | 1,409 | 0,495 | 8,795 | 1,43E-18 | 3,48E-18 | 0,0009119 | 0,003224 | 0,0021019 | 0,0021019 | TRUE | 0,0220303 | No |
| orf19.88 | 6833 | 5704 | 5885 | 3957 | 13317 | 10827 | 11714 | 16533 | 6221 | 5769 | 5570 | 5944 | 11845 | 11755 | 11997 | 12612 | 8964,01 | 5876 | 12052 | 2,051 | 1,037 | 23,51 | 3,24E-122 | 3,36E-121 | 0,0014501 | 0,002184 | 0,0017368 | 0,0017368 | TRUE | 0,0196585 | No |
| orf19.880 | 3606 | 3163 | 3394 | 2377 | 2465 | 2040 | 2183 | 3159 | 3283 | 3199 | 3212 | 3571 | 2193 | 2215 | 2236 | 2410 | 2789,72 | 3316 | 2264 | 0,684 | -0,549 | -10,722 | 8,06E-27 | 2,41E-26 | 0,0018359 | 0,002511 | 0,0021445 | 0,0021445 | TRUE | 0,0308104 | No |
| orf19.881 | 1011 | 892 | 925 | 640 | 1089 | 916 | 1036 | 1361 | 920 | 902 | 875 | 961 | 969 | 995 | 1061 | 1038 | 965,22 | 914 | 1016 | 1,113 | 0,154 | 2,749 | 0,005981 | 0,007976 | 0,0005132 | 0,003406 | 0,0019633 | 0,0019633 | TRUE | 0,0120294 | No |
| orf19.882 | 11494 | 10177 | 11057 | 6781 | 2681 | 2224 | 2458 | 3274 | 10465 | 10293 | 10464 | 10186 | 2385 | 2415 | 2517 | 2497 | 6402,8 | 10352 | 2454 | 0,237 | -2,077 | -60,685 | 0 | 0 | 9,07E-05 | 0,002243 | 0,0008826 | 0,0008826 | TRUE | 0,0044944 | No |
| orf19.886 | 6730 | 5711 | 6645 | 4130 | 4630 | 4096 | 4166 | 5680 | 6127 | 5776 | 6289 | 6204 | 4118 | 4447 | 4267 | 4333 | 5195,11 | 6099 | 4291 | 0,704 | -0,507 | -12,111 | 9,24E-34 | 3,21E-33 | 0,000929 | 0,002291 | 0,0014873 | 0,0014873 | TRUE | 0,0156343 | No |
| orf19.889 | 428 | 399 | 399 | 295 | 969 | 849 | 958 | 1224 | 390 | 404 | 378 | 443 | 862 | 922 | 981 | 934 | 664,05 | 404 | 925 | 2,303 | 1,204 | 17,014 | 6,51E-65 | 3,82E-64 | 0,0015233 | 0,004026 | 0,0029697 | 0,0029697 | TRUE | 0,0418423 | No |
| orf19.89 | 1715 | 1387 | 1494 | 1020 | 2957 | 2485 | 2831 | 3778 | 1561 | 1403 | 1414 | 1532 | 2630 | 2698 | 2899 | 2882 | 2127,47 | 1478 | 2777 | 1,881 | 0,912 | 16,566 | 1,23E-61 | 6,89E-61 | 0,001971 | 0,002658 | 0,0023759 | 0,0023759 | TRUE | 0,019302 | No |
| orf19.891 | 5248 | 4880 | 5190 | 3338 | 7618 | 6558 | 6830 | 8910 | 4778 | 4936 | 4912 | 5014 | 6776 | 7120 | 6995 | 6797 | 5915,91 | 4910 | 6922 | 1,41 | 0,496 | 14,381 | 6,83E-47 | 3,00E-46 | 0,0002517 | 0,00226 | 0,000964 | 0,000964 | TRUE | 0,0044961 | No |
| orf19.893 | 40 | 44 | 40 | 36 | 171 | 154 | 118 | 188 | 36 | 45 | 38 | 54 | 152 | 167 | 121 | 143 | 94,55 | 43 | 146 | 3,441 | 1,783 | 10,201 | 1,96E-24 | 5,56E-24 | 0,0091733 | 0,016008 | 0,0135999 | 0,0135999 | TRUE | 0,1546617 | No |
| orf19.894 | 598 | 494 | 522 | 324 | 987 | 740 | 814 | 1091 | 544 | 500 | 494 | 487 | 878 | 803 | 834 | 832 | 671,5 | 506 | 837 | 1,65 | 0,722 | 11,273 | 1,78E-29 | 5,63E-29 | 1,00E-08 | 0,004004 | 0,0023332 | 0,0023332 | TRUE | 0,031951 | No |

| | | | | | | | | | | | | | | | | | | | | | | | | | | | | | | | |
|---|---|---|---|---|---|---|---|---|---|---|---|---|---|---|---|---|---|---|---|---|---|---|---|---|---|---|---|---|---|---|---|
| orf19.895 | 5031 | 4450 | 4769 | 3210 | 6258 | 5532 | 5701 | 7574 | 4581 | 4501 | 4513 | 4822 | 5566 | 6006 | 5839 | 5778 | 5200,64 | 4604 | 5797 | 1,26 | 0,333 | 8,376 | 5,49E-17 | 1,29E-16 | 0,0007578 | 0,002291 | 0,0013224 | 0,0013224 | TRUE | 0,0116573 | No |
| orf19.896 | 4679 | 4308 | 4739 | 2811 | 5159 | 4430 | 4574 | 6222 | 4260 | 4357 | 4485 | 4223 | 4589 | 4810 | 4684 | 4746 | 4519,22 | 4331 | 4707 | 1,086 | 0,119 | 3,289 | 0,001006 | 0,00142 | 0,0002705 | 0,002329 | 0,0010374 | 0,0010374 | TRUE | 0,0070371 | No |
| orf19.897 | 2370 | 1987 | 2142 | 1511 | 615 | 564 | 546 | 753 | 2158 | 2010 | 2027 | 2270 | 547 | 612 | 559 | 574 | 1344,67 | 2116 | 573 | 0,271 | -1,884 | -31,012 | 3,77E-211 | 6,73E-210 | 0,0019648 | 0,003019 | 0,0024734 | 0,0024734 | TRUE | 0,0269703 | No |
| orf19.898 | 1106 | 860 | 934 | 682 | 1469 | 1277 | 1260 | 1808 | 1007 | 870 | 884 | 1024 | 1307 | 1386 | 1290 | 1379 | 1143,48 | 946 | 1340 | 1,42 | 0,506 | 7,802 | 6,08E-15 | 1,34E-14 | 0,0031071 | 0,003192 | 0,0031183 | 0,0031183 | TRUE | 0,0346021 | No |
| orf19.899 | 680 | 620 | 578 | 406 | 930 | 847 | 803 | 1166 | 619 | 627 | 547 | 610 | 827 | 920 | 822 | 889 | 732,71 | 601 | 864 | 1,44 | 0,526 | 7,766 | 8,12E-15 | 1,79E-14 | 0,0020515 | 0,00384 | 0,0029778 | 0,0029778 | TRUE | 0,0454296 | No |
| orf19.9 | 128 | 119 | 108 | 73 | 107 | 97 | 112 | 119 | 117 | 120 | 102 | 110 | 95 | 105 | 115 | 91 | 106,84 | 112 | 102 | 0,899 | -0,154 | -1,109 | 0,267601 | 0,303551 | 1,00E-08 | 0,014401 | 0,0090637 | 0,0090637 | TRUE | 0,0758669 | No |
| orf19.90 | 1234 | 1148 | 1274 | 751 | 1277 | 1094 | 1130 | 1543 | 1124 | 1161 | 1206 | 1128 | 1136 | 1188 | 1157 | 1177 | 1159,54 | 1155 | 1164 | 1,007 | 0,011 | 0,218 | 0,827543 | 0,851039 | 1,00E-08 | 0,003176 | 0,0014826 | 0,0014826 | TRUE | 0,0110601 | No |
| orf19.900 | 5177 | 4131 | 4581 | 3143 | 4305 | 3473 | 3920 | 5220 | 4713 | 4178 | 4336 | 4721 | 3829 | 3771 | 4015 | 3982 | 4193,07 | 4487 | 3899 | 0,87 | -0,202 | -4,011 | 6,05E-05 | 9,17E-05 | 0,0019949 | 0,002352 | 0,0021889 | 0,0021889 | TRUE | 0,0261495 | No |
| orf19.903 | 13840 | 11224 | 11723 | 8984 | 5766 | 4393 | 4576 | 6741 | 12601 | 11352 | 11095 | 13495 | 5129 | 4770 | 4686 | 5142 | 8533,69 | 12136 | 4932 | 0,407 | -1,298 | -19,992 | 6,51E-89 | 5,07E-88 | 0,00523 | 0,002192 | 0,0039127 | 0,0039127 | TRUE | 0,0689544 | No |
| orf19.904 | 1222 | 1124 | 1165 | 747 | 496 | 371 | 415 | 548 | 1113 | 1137 | 1103 | 1122 | 441 | 403 | 425 | 418 | 770,13 | 1119 | 422 | 0,377 | -1,405 | -23,96 | 7,32E-127 | 7,95E-126 | 1,00E-08 | 0,003752 | 0,001723 | 0,001723 | TRUE | 0,0112073 | No |
| orf19.905 | 2095 | 1813 | 1982 | 1259 | 2575 | 2157 | 2143 | 3008 | 1907 | 1834 | 1876 | 1891 | 2290 | 2342 | 2195 | 2295 | 2078,7 | 1877 | 2280 | 1,215 | 0,281 | 6,799 | 1,05E-11 | 2,12E-11 | 1,00E-08 | 0,002673 | 0,0011504 | 0,0011504 | TRUE | 0,0074661 | No |
| orf19.906 | 5158 | 4870 | 5354 | 3189 | 3763 | 3283 | 3261 | 4463 | 4696 | 4925 | 5067 | 4790 | 3347 | 3564 | 3340 | 3404 | 4141,83 | 4870 | 3414 | 0,701 | -0,513 | -12,538 | 4,64E-36 | 1,69E-35 | 0,0006873 | 0,002356 | 0,0013607 | 0,0013607 | TRUE | 0,0103918 | No |
| orf19.908 | 4697 | 4526 | 4748 | 2804 | 5830 | 4519 | 4692 | 6012 | 4276 | 4578 | 4494 | 4212 | 5186 | 4906 | 4805 | 4586 | 4630,35 | 4390 | 4871 | 1,109 | 0,149 | 3,06 | 0,002216 | 0,003048 | 0,0017988 | 0,002322 | 0,0020615 | 0,0020615 | TRUE | 0,0234021 | No |
| orf19.909 | 12697 | 11384 | 12262 | 7703 | 14452 | 12156 | 12669 | 17256 | 11560 | 11514 | 11605 | 11571 | 12855 | 13198 | 12975 | 13163 | 12305 | 11562 | 13048 | 1,128 | 0,174 | 6,712 | 1,92E-11 | 3,83E-11 | 1,00E-08 | 0,002144 | 0,0005653 | 0,0005653 | TRUE | 0,0012236 | No |
| orf19.909.1 | 887 | 755 | 739 | 567 | 1225 | 912 | 969 | 1309 | 808 | 764 | 699 | 852 | 1090 | 990 | 992 | 999 | 899,12 | 781 | 1018 | 1,308 | 0,388 | 5,674 | 1,39E-08 | 2,49E-08 | 0,0030726 | 0,003506 | 0,0033285 | 0,0033285 | TRUE | 0,0588238 | No |
| orf19.91 | 1132 | 1030 | 1110 | 664 | 1171 | 942 | 970 | 1308 | 1031 | 1042 | 1051 | 997 | 1042 | 1023 | 993 | 998 | 1021,97 | 1030 | 1014 | 0,983 | -0,025 | -0,502 | 0,615436 | 0,654555 | 1,00E-08 | 0,00333 | 0,0014369 | 0,0014369 | TRUE | 0,004583 | No |
| orf19.910 | 4263 | 4033 | 4210 | 2780 | 5114 | 4520 | 4537 | 6020 | 3881 | 4079 | 3984 | 4176 | 4549 | 4907 | 4646 | 4592 | 4351,92 | 4030 | 4674 | 1,16 | 0,214 | 5,217 | 1,82E-07 | 3,10E-07 | 0,0007761 | 0,002341 | 0,0013868 | 0,0013868 | TRUE | 0,0135881 | No |
| orf19.913.2 | 2776 | 2468 | 2610 | 1578 | 5824 | 4266 | 5149 | 6484 | 2527 | 2496 | 2470 | 2370 | 5180 | 4632 | 5273 | 4946 | 3736,91 | 2466 | 5008 | 2,029 | 1,021 | 20,566 | 5,51E-94 | 4,47E-93 | 0,0016071 | 0,002391 | 0,0020548 | 0,0020548 | TRUE | 0,0306044 | No |
| orf19.915 | 1236 | 1418 | 1373 | 946 | 662 | 613 | 530 | 768 | 1125 | 1434 | 1299 | 1421 | 589 | 666 | 543 | 586 | 957,87 | 1320 | 596 | 0,452 | -1,147 | -13,224 | 6,37E-40 | 2,47E-39 | 0,0087139 | 0,003416 | 0,0060255 | 0,0060255 | TRUE | 0,120693 | No |
| orf19.916 | 3922 | 4065 | 4182 | 2162 | 1789 | 1358 | 1513 | 1979 | 3571 | 4111 | 3958 | 3248 | 1591 | 1474 | 1550 | 1510 | 2626,56 | 3722 | 1531 | 0,411 | -1,283 | -18,206 | 4,66E-74 | 3,06E-73 | 0,0054031 | 0,00254 | 0,0043159 | 0,0043159 | TRUE | 0,0869071 | No |
| orf19.918 | 3911 | 3299 | 3748 | 2269 | 3848 | 3279 | 3543 | 4846 | 3561 | 3337 | 3547 | 3408 | 3423 | 3560 | 3628 | 3697 | 3520,09 | 3463 | 3577 | 1,032 | 0,046 | 1,098 | 0,27206 | 0,308116 | 0,0007958 | 0,002412 | 0,0014048 | 0,0014048 | TRUE | 0,0104773 | No |
| orf19.92 | 2071 | 1863 | 2096 | 1236 | 2800 | 2330 | 2613 | 3454 | 1886 | 1884 | 1984 | 1857 | 2491 | 2530 | 2676 | 2635 | 2242,64 | 1903 | 2583 | 1,357 | 0,44 | 9,928 | 3,14E-23 | 8,61E-23 | 0,0005822 | 0,002626 | 0,001422 | 0,001422 | TRUE | 0,0101633 | No |
| orf19.920 | 2694 | 2665 | 2678 | 1653 | 2859 | 2364 | 2559 | 3198 | 2453 | 2695 | 2534 | 2483 | 2543 | 2567 | 2621 | 2439 | 2541,94 | 2541 | 2542 | 1 | -0,001 | -0,015 | 0,988051 | 0,989448 | 0,0009762 | 0,002557 | 0,0016168 | 0,0016168 | TRUE | 0,0204189 | No |
| orf19.921 | 5304 | 5323 | 5801 | 3505 | 2685 | 2386 | 2429 | 3124 | 4829 | 5384 | 5490 | 5265 | 2388 | 2591 | 2488 | 2383 | 3852,16 | 5242 | 2462 | 0,47 | -1,091 | -21,196 | 1,04E-99 | 9,06E-99 | 0,0019279 | 0,00238 | 0,0022502 | 0,0022502 | TRUE | 0,0350156 | No |
| orf19.922 | 2653 | 2655 | 2782 | 1663 | 5179 | 4686 | 4482 | 6006 | 2415 | 2685 | 2633 | 2498 | 4607 | 5088 | 4590 | 4581 | 3637,19 | 2558 | 4716 | 1,843 | 0,882 | 16,92 | 3,18E-64 | 1,84E-63 | 0,0021417 | 0,0024 | 0,0022987 | 0,0022987 | TRUE | 0,0338285 | No |
| orf19.923 | 775 | 674 | 758 | 515 | 708 | 669 | 564 | 835 | 706 | 682 | 717 | 774 | 630 | 726 | 578 | 637 | 681,11 | 720 | 643 | 0,894 | -0,161 | -2,126 | 0,033475 | 0,042133 | 0,0041172 | 0,003976 | 0,0040335 | 0,0040335 | TRUE | 0,0857108 | No |
| orf19.925 | 527 | 652 | 670 | 354 | 959 | 884 | 819 | 1018 | 480 | 659 | 634 | 532 | 853 | 960 | 839 | 777 | 716,65 | 576 | 857 | 1,484 | 0,57 | 5,658 | 1,54E-08 | 2,74E-08 | 0,0128227 | 0,00388 | 0,0082518 | 0,0082518 | TRUE | 0,1591777 | No |
| orf19.926 | 1310 | 1151 | 1244 | 787 | 1848 | 1513 | 1690 | 2184 | 1193 | 1164 | 1177 | 1182 | 1644 | 1643 | 1731 | 1666 | 1424,94 | 1179 | 1671 | 1,417 | 0,503 | 11,135 | 8,45E-29 | 2,63E-28 | 1,00E-08 | 0,002964 | 0,0012232 | 0,0012232 | TRUE | 0,0067808 | No |
| orf19.927 | 215 | 242 | 219 | 144 | 118 | 111 | 101 | 160 | 196 | 245 | 207 | 216 | 105 | 121 | 103 | 122 | 164,38 | 216 | 113 | 0,523 | -0,934 | -7,739 | 1,00E-14 | 2,21E-14 | 1,00E-08 | 0,010073 | 0,007393 | 0,007393 | TRUE | 0,0935493 | No |
| orf19.928 | 1209 | 1051 | 1168 | 814 | 885 | 746 | 814 | 1030 | 1101 | 1063 | 1105 | 1223 | 787 | 810 | 834 | 786 | 963,54 | 1123 | 804 | 0,718 | -0,479 | -8,183 | 2,76E-16 | 6,32E-16 | 0,0008699 | 0,003408 | 0,0022257 | 0,0022257 | TRUE | 0,038045 | No |
| orf19.929 | 102 | 102 | 111 | 73 | 43 | 36 | 51 | 53 | 93 | 103 | 105 | 110 | 38 | 39 | 52 | 40 | 72,59 | 103 | 42 | 0,414 | -1,272 | -7,381 | 1,57E-13 | 3,34E-13 | 1,00E-08 | 0,020235 | 0,0123435 | 0,0123435 | TRUE | 0,1705665 | No |
| orf19.93 | 5490 | 4324 | 4780 | 3957 | 3892 | 3269 | 3331 | 4678 | 4998 | 4373 | 4524 | 5944 | 3462 | 3549 | 3411 | 3568 | 4228,8 | 4960 | 3498 | 0,706 | -0,503 | -6,191 | 5,97E-10 | 1,12E-09 | 0,01009 | 0,002349 | 0,00609 | 0,00609 | TRUE | 0,2145103 | No |
| orf19.930 | 64742 | 53137 | 56479 | 37722 | 88802 | 70714 | 72098 | 1E+05 | 58945 | 53742 | 53452 | 56664 | 78987 | 76776 | 73837 | 81831 | 66779,3 | 55701 | 77858 | 1,398 | 0,483 | 10,468 | 1,21E-25 | 3,53E-25 | 0,002018 | 0,002057 | 0,0020315 | 0,0020315 | TRUE | 0,0188921 | No |
| orf19.931 | 1163 | 1141 | 1110 | 719 | 1116 | 1046 | 949 | 1338 | 1059 | 1154 | 1051 | 1080 | 993 | 1136 | 972 | 1021 | 1058,03 | 1086 | 1030 | 0,948 | -0,077 | -1,23 | 0,21888 | 0,251961 | 0,0021959 | 0,003286 | 0,0028099 | 0,0028099 | TRUE | 0,0542455 | No |
| orf19.932 | 6300 | 5670 | 6128 | 3893 | 2500 | 2188 | 2330 | 2966 | 5736 | 5735 | 5800 | 5848 | 2224 | 2376 | 2386 | 2262 | 4045,74 | 5780 | 2312 | 0,4 | -1,323 | -35,167 | 6,47E-271 | 1,48E-269 | 0,0001417 | 0,002364 | 0,0010632 | 0,0010632 | TRUE | 0,0078414 | No |
| orf19.933 | 4326 | 3750 | 4215 | 2403 | 3590 | 2850 | 3028 | 4048 | 3939 | 3793 | 3989 | 3610 | 3193 | 3094 | 3101 | 3088 | 3475,82 | 3833 | 3119 | 0,813 | -0,299 | -7,085 | 1,40E-12 | 2,88E-12 | 0,0007266 | 0,002417 | 0,0014191 | 0,0014191 | TRUE | 0,0174912 | No |
| orf19.934 | 832 | 764 | 793 | 509 | 1288 | 900 | 1141 | 1527 | 758 | 773 | 751 | 765 | 1146 | 977 | 1169 | 1165 | 937,68 | 762 | 1114 | 1,466 | 0,551 | 8,279 | 1,25E-16 | 2,89E-16 | 0,002847 | 0,003446 | 0,0031358 | 0,0031358 | TRUE | 0,0789849 | No |
| orf19.935 | 24 | 12 | 15 | 24 | 1836 | 1692 | 1620 | 2299 | 22 | 12 | 14 | 36 | 1633 | 1837 | 1659 | 1754 | 870,89 | 21 | 1721 | 87,11 | 6,445 | 36,304 | 1,38E-288 | 3,41E-287 | 0,0029539 | 0,003554 | 0,0033735 | 0,0033735 | TRUE | 0,775043 | No |
| orf19.936 | 8 | 9 | 7 | 4 | 26 | 20 | 24 | 32 | 7 | 9 | 7 | 6 | 23 | 22 | 25 | 24 | 15,36 | 7 | 24 | 3,204 | 1,68 | 4,428 | 9,52E-06 | 1,51E-05 | 1,00E-08 | 0,088061 | 0,0462882 | 0,0462882 | TRUE | 0,0770383 | No |
| orf19.937 | 7 | 11 | 4 | 2 | 0 | 0 | 0 | 0 | 6 | 11 | 4 | 3 | 0 | 0 | 0 | 0 | 3,04 | 6 | 0 | 0,028 | -5,168 | -4,178 | 2,94E-05 | 4,55E-05 | 0,2202387 | 0,43714 | 0,3827454 | 0,3827454 | TRUE | 0,3398331 | No |
| orf19.938 | 358 | 316 | 362 | 228 | 58 | 41 | 54 | 76 | 326 | 320 | 343 | 342 | 52 | 45 | 55 | 58 | 192,5 | 333 | 52 | 0,159 | -2,655 | -20,891 | 6,53E-97 | 5,51E-96 | 1,00E-08 | 0,008899 | 0,0051193 | 0,0051193 | TRUE | 0,0729315 | No |
| orf19.939 | 7891 | 7226 | 7977 | 4994 | 7990 | 6888 | 6985 | 9363 | 7184 | 7308 | 7550 | 7502 | 7107 | 7478 | 7154 | 7142 | 7303,13 | 7386 | 7220 | 0,978 | -0,033 | -0,951 | 0,3418 | 0,381808 | 0,0003706 | 0,002218 | 0,0009976 | 0,0009976 | TRUE | 0,0069579 | No |
| orf19.94 | 4167 | 3740 | 3905 | 2130 | 2270 | 1762 | 1946 | 2640 | 3794 | 3783 | 3696 | 3200 | 2019 | 1913 | 1993 | 2014 | 2801,34 | 3618 | 1985 | 0,548 | -0,868 | -14,963 | 1,28E-50 | 5,98E-50 | 0,0028307 | 0,002509 | 0,0028452 | 0,0028452 | TRUE | 0,0707471 | No |
| orf19.940 | 3815 | 4108 | 4219 | 2433 | 1796 | 1537 | 1482 | 1885 | 3473 | 4155 | 3993 | 3655 | 1597 | 1669 | 1518 | 1438 | 2687,22 | 3819 | 1556 | 0,407 | -1,297 | -19,281 | 7,79E-83 | 5,76E-82 | 0,0047602 | 0,002529 | 0,0039053 | 0,0039053 | TRUE | 0,0462839 | No |
| orf19.941 | 24587 | 23159 | 23541 | 14967 | 29660 | 23347 | 23433 | 32350 | 22385 | 23423 | 22279 | 22483 | 26382 | 25348 | 23998 | 24677 | 23872 | 22642 | 25101 | 1,109 | 0,149 | 3,854 | 0,000116 | 0,000174 | 0,0010338 | 0,002092 | 0,0013883 | 0,0013883 | TRUE | 0,0144677 | No |
| orf19.942 | 96 | 86 | 107 | 66 | 172 | 142 | 148 | 197 | 87 | 87 | 101 | 99 | 153 | 154 | 152 | 150 | 122,97 | 94 | 152 | 1,63 | 0,705 | 5,568 | 2,58E-08 | 4,57E-08 | 1,00E-08 | 0,012779 | 0,0066519 | 0,0066519 | TRUE | 0,03605 | No |
| orf19.943 | 3300 | 3148 | 3234 | 2042 | 4936 | 4362 | 4283 | 5815 | 3005 | 3184 | 3061 | 3067 | 4390 | 4736 | 4386 | 4436 | 3783,11 | 3079 | 4487 | 1,457 | 0,543 | 13,148 | 1,75E-39 | 6,75E-39 | 0,0006298 | 0,002386 | 0,0013579 | 0,0013579 | TRUE | 0,0166115 | No |
| orf19.944 | 3359 | 3230 | 3471 | 2090 | 5529 | 4465 | 4768 | 6010 | 3058 | 3267 | 3285 | 3140 | 4918 | 4848 | 4883 | 4584 | 3997,83 | 3188 | 4808 | 1,508 | 0,592 | 14,097 | 3,96E-45 | 1,70E-44 | 0,0008744 | 0,002367 | 0,0014294 | 0,0014294 | TRUE | 0,0122963 | No |
| orf19.945 | 1239 | 1035 | 1109 | 706 | 1358 | 984 | 1096 | 1449 | 1128 | 1047 | 1050 | 1061 | 1208 | 1068 | 1122 | 1105 | 1098,62 | 1072 | 1126 | 1,051 | 0,072 | 1,268 | 0,204844 | 0,23683 | 0,0012076 | 0,003239 | 0,0021656 | 0,0021656 | TRUE | 0,0290034 | No |
| orf19.946 | 1787 | 1564 | 1554 | 970 | 3822 | 2915 | 2950 | 3929 | 1627 | 1582 | 1471 | 1457 | 3400 | 3165 | 3021 | 2997 | 2339,91 | 1534 | 3146 | 2,048 | 1,034 | 17,974 | 3,14E-72 | 2,02E-71 | 0,0025549 | 0,002602 | 0,0026787 | 0,0026787 | TRUE | 0,0363042 | No |
| orf19.947 | 1690 | 1376 | 1491 | 875 | 978 | 738 | 837 | 1177 | 1539 | 1392 | 1411 | 1314 | 870 | 801 | 857 | 898 | 1135,25 | 1414 | 856 | 0,605 | -0,724 | -11,651 | 2,26E-31 | 7,50E-31 | 0,0024024 | 0,003201 | 0,002784 | 0,002784 | TRUE | 0,0428269 | No |
| orf19.948 | 1 | 1 | 1 | 4 | 2 | 0 | 2 | 2 | 1 | 1 | 1 | 6 | 2 | 0 | 2 | 2 | 1,78 | 2 | 2 | 0,678 | -0,561 | -0,506 | 0,612912 | 0,65234 | 0,2489739 | 0,744708 | 0,5746745 | 0,5746745 | TRUE | 0,8389212 | No |
| orf19.949 | 918 | 901 | 955 | 556 | 549 | 483 | 534 | 680 | 836 | 911 | 904 | 835 | 488 | 524 | 547 | 519 | 695,55 | 872 | 520 | 0,595 | -0,75 | -11,661 | 2,02E-31 | 6,72E-31 | 0,0008003 | 0,003936 | 0,0024541 | 0,0024541 | TRUE | 0,0193201 | No |
| orf19.95 | 81 | 75 | 100 | 54 | 263 | 173 | 217 | 316 | 74 | 76 | 95 | 81 | 234 | 188 | 222 | 241 | 151,3 | 82 | 221 | 2,729 | 1,448 | 10,717 | 8,46E-27 | 2,53E-26 | 0,0059742 | 0,010768 | 0,0088972 | 0,0088972 | TRUE | 0,1287385 | No |
| orf19.951 | 9358 | 9665 | 10208 | 6020 | 1618 | 1274 | 1366 | 1740 | 8520 | 9775 | 9661 | 9043 | 1439 | 1383 | 1399 | 1327 | 5318,47 | 9250 | 1387 | 0,15 | -2,738 | -50,345 | 0 | 0 | 0,002554 | 0,002285 | 0,0024452 | 0,0024452 | TRUE | 0,0349813 | No |
| orf19.952 | 635 | 567 | 679 | 442 | 505 | 475 | 458 | 659 | 578 | 573 | 643 | 664 | 449 | 516 | 469 | 503 | 549,35 | 614 | 484 | 0,79 | -0,34 | -4,482 | 7,38E-06 | 1,18E-05 | 0,0028117 | 0,004442 | 0,0036911 | 0,0036911 | TRUE | 0,0301135 | No |
| orf19.953.1 | 1617 | 1375 | 1494 | 1010 | 2380 | 1789 | 1761 | 2592 | 1472 | 1391 | 1414 | 1517 | 2117 | 1942 | 1803 | 1977 | 1704,25 | 1448 | 1960 | 1,355 | 0,439 | 7,601 | 2,93E-14 | 6,35E-14 | 0,0024073 | 0,002812 | 0,0025883 | 0,0025883 | TRUE | 0,0351583 | No |
| orf19.954 | 2724 | 2284 | 2516 | 1543 | 2752 | 2140 | 2452 | 3259 | 2480 | 2310 | 2381 | 2318 | 2448 | 2323 | 2511 | 2486 | 2407,19 | 2372 | 2442 | 1,029 | 0,041 | 0,928 | 0,353325 | 0,393234 | 0,0007013 | 0,002586 | 0,0014794 | 0,0014794 | TRUE | 0,0125119 | No |
| orf19.955 | 2170 | 1918 | 1944 | 1298 | 1942 | 1486 | 1598 | 2153 | 1976 | 1940 | 1840 | 1950 | 1727 | 1613 | 1637 | 1642 | 1790,6 | 1926 | 1655 | 0,86 | -0,218 | -4,773 | 1,81E-06 | 2,98E-06 | 0,0004375 | 0,002775 | 0,001441 | 0,001441 | TRUE | 0,0115415 | No |
| orf19.956 | 991 | 923 | 1028 | 611 | 1111 | 913 | 1008 | 1302 | 902 | 934 | 973 | 918 | 988 | 991 | 1032 | 993 | 966,43 | 932 | 1001 | 1,074 | 0,103 | 1,961 | 0,049936 | 0,061861 | 1,00E-08 | 0,003404 | 0,0015917 | 0,0015917 | TRUE | 0,0112688 | No |
| orf19.96 | 11272 | 11367 | 11968 | 6858 | 17315 | 14260 | 15240 | 19916 | 10263 | 11497 | 11327 | 10302 | 15401 | 15482 | 15608 | 15192 | 13133,9 | 10847 | 15421 | 1,421 | 0,507 | 11,143 | 7,79E-29 | 2,43E-28 | 0,0017875 | 0,002138 | 0,0019102 | 0,0019102 | TRUE | 0,0198237 | No |
| orf19.961.2 | 1284 | 1148 | 1106 | 664 | 1233 | 864 | 966 | 1335 | 1169 | 1161 | 1047 | 997 | 1097 | 938 | 989 | 1018 | 1052,09 | 1094 | 1010 | 0,923 | -0,116 | -1,662 | 0,096438 | 0,116042 | 0,0039058 | 0,003293 | 0,0036905 | 0,0036905 | TRUE | 0,0391877 | No |
| orf19.962 | 4907 | 3809 | 4174 | 2530 | 5407 | 3842 | 4502 | 6183 | 4468 | 3852 | 3950 | 3800 | 4809 | 4171 | 4611 | 4716 | 4297,32 | 4018 | 4577 | 1,139 | 0,188 | 2,986 | 0,002828 | 0,003857 | 0,0044972 | 0,002344 | 0,0035546 | 0,0035546 | TRUE | 0,0698161 | No |
| orf19.964 | 4202 | 3877 | 4125 | 2474 | 4641 | 3710 | 4200 | 5469 | 3826 | 3921 | 3904 | 3716 | 4128 | 4028 | 4301 | 4172 | 3999,55 | 3842 | 4157 | 1,081 | 0,113 | 3,002 | 0,002684 | 0,003666 | 0,000333 | 0,002367 | 0,0011062 | 0,0011062 | TRUE | 0,006428 | No |
| orf19.966 | 3866 | 2964 | 3206 | 2233 | 3543 | 2746 | 3161 | 4275 | 3520 | 2998 | 3034 | 3354 | 3151 | 2981 | 3237 | 3261 | 3192,14 | 3226 | 3158 | 0,979 | -0,03 | -0,509 | 0,611007 | 0,650496 | 0,0031762 | 0,002451 | 0,0030804 | 0,0030804 | TRUE | 0,0471818 | No |
| orf19.967 | 2165 | 1721 | 1887 | 1308 | 2481 | 2090 | 2042 | 2918 | 1971 | 1741 | 1786 | 1965 | 2207 | 2269 | 2091 | 2226 | 2031,94 | 1866 | 2198 | 1,18 | 0,238 | 4,374 | 1,22E-05 | 1,92E-05 | 0,0019937 | 0,002687 | 0,0023466 | 0,0023466 | TRUE | 0,02447 | No |
| orf19.968 | 16506 | 12580 | 13337 | 9678 | 20733 | 15673 | 16511 | 24479 | 15028 | 12723 | 12622 | 14538 | 18441 | 17017 | 16909 | 18673 | 15743,9 | 13728 | 17760 | 1,294 | 0,372 | 5,775 | 7,69E-09 | 1,39E-08 | 0,0053226 | 0,002121 | 0,0039163 | 0,0039163 | TRUE | 0,0501527 | No |

| | | | | | | | | | | | | | | | | | | | | | | | | | | | | | | | |
|---|---|---|---|---|---|---|---|---|---|---|---|---|---|---|---|---|---|---|---|---|---|---|---|---|---|---|---|---|---|---|---|
| orf19.969 | 12714 | 10275 | 11122 | 7444 | 15240 | 12814 | 13280 | 18527 | 11576 | 10392 | 10526 | 11182 | 13555 | 13912 | 13600 | 14133 | 12359,6 | 10919 | 13800 | 1,264 | 0,338 | 7,923 | 2,32E-15 | 5,20E-15 | 0,0013891 | 0,002144 | 0,0016656 | 0,0016656 | TRUE | 0,0204389 | No |
| orf19.97 | 48269 | 43659 | 46590 | 27529 | 80415 | 67616 | 70270 | 96878 | 43947 | 44156 | 44093 | 41353 | 71527 | 73412 | 71965 | 73899 | 58044,1 | 43387 | 72701 | 1,675 | 0,745 | 22,677 | 7,56E-114 | 7,37E-113 | 0,0005879 | 0,00206 | 0,0010167 | 0,0010167 | TRUE | 0,0120901 | No |
| orf19.970 | 764 | 664 | 694 | 463 | 401 | 379 | 381 | 444 | 696 | 672 | 657 | 695 | 357 | 411 | 390 | 339 | 527,06 | 680 | 374 | 0,549 | -0,866 | -11,622 | 3,19E-31 | 1,05E-30 | 0,0014567 | 0,004543 | 0,0032955 | 0,0032955 | TRUE | 0,0504578 | No |
| orf19.971 | 547 | 485 | 533 | 336 | 858 | 719 | 786 | 1080 | 498 | 491 | 504 | 505 | 763 | 781 | 805 | 824 | 646,29 | 500 | 793 | 1,59 | 0,669 | 10,911 | 1,02E-27 | 3,11E-27 | 1,00E-08 | 0,004081 | 0,0019536 | 0,0019536 | TRUE | 0,0088563 | No |
| orf19.972 | 338 | 247 | 270 | 224 | 444 | 394 | 366 | 599 | 308 | 250 | 256 | 336 | 395 | 428 | 375 | 457 | 350,5 | 288 | 414 | 1,449 | 0,535 | 5,061 | 4,17E-07 | 7,04E-07 | 0,010493 | 0,005806 | 0,0077371 | 0,0077371 | TRUE | 0,1272181 | No |
| orf19.974 | 638 | 545 | 659 | 353 | 590 | 437 | 499 | 683 | 581 | 551 | 624 | 530 | 525 | 474 | 511 | 521 | 539,66 | 572 | 508 | 0,887 | -0,173 | -2,402 | 0,016288 | 0,021054 | 0,0014961 | 0,004485 | 0,0031427 | 0,0031427 | TRUE | 0,04416 | No |
| orf19.976 | 12047 | 10710 | 11657 | 7107 | 8329 | 6753 | 7622 | 9877 | 10968 | 10832 | 11032 | 10676 | 7408 | 7332 | 7806 | 7534 | 9198,61 | 10877 | 7520 | 0,691 | -0,533 | -16,232 | 2,97E-59 | 1,61E-58 | 0,0003381 | 0,002181 | 0,0009229 | 0,0009229 | TRUE | 0,007854 | No |
| orf19.978 | 4798 | 4030 | 4590 | 2919 | 6838 | 5533 | 6212 | 8230 | 4368 | 4076 | 4344 | 4385 | 6082 | 6007 | 6362 | 6278 | 5237,8 | 4293 | 6182 | 1,441 | 0,527 | 13,357 | 1,09E-40 | 4,26E-40 | 0,0006718 | 0,002289 | 0,0012915 | 0,0012915 | TRUE | 0,0142722 | No |
| orf19.979 | 6339 | 5623 | 5955 | 4092 | 40706 | 35221 | 35633 | 50305 | 5771 | 5687 | 5636 | 6147 | 36207 | 38240 | 36493 | 38373 | 21569,2 | 5810 | 37328 | 6,43 | 2,685 | 67,091 | 0 | 0 | 0,0011232 | 0,002098 | 0,0014325 | 0,0014325 | TRUE | 0,0177545 | No |
| orf19.980 | 7867 | 6686 | 7536 | 4950 | 6024 | 5333 | 5902 | 8099 | 7163 | 6762 | 7132 | 7436 | 5358 | 5790 | 6044 | 6178 | 6482,91 | 7123 | 5842 | 0,821 | -0,285 | -5,502 | 3,75E-08 | 6,58E-08 | 0,0024538 | 0,002241 | 0,0024281 | 0,0024281 | TRUE | 0,0383336 | No |
| orf19.984 | 87 | 75 | 87 | 65 | 148 | 103 | 128 | 180 | 79 | 76 | 82 | 98 | 132 | 112 | 131 | 137 | 105,86 | 84 | 128 | 1,549 | 0,632 | 4,468 | 7,88E-06 | 1,26E-05 | 1,00E-08 | 0,014515 | 0,0091381 | 0,0091381 | TRUE | 0,0860853 | No |
| orf19.985 | 2473 | 2321 | 2626 | 1637 | 2322 | 1983 | 2135 | 2912 | 2252 | 2347 | 2485 | 2459 | 2065 | 2153 | 2187 | 2221 | 2271,18 | 2386 | 2156 | 0,905 | -0,144 | -3,025 | 0,002483 | 0,003401 | 0,0011296 | 0,002619 | 0,0017447 | 0,0017447 | TRUE | 0,0179267 | No |
| orf19.986 | 38637 | 32894 | 35259 | 22100 | 38780 | 30852 | 32160 | 44896 | 35177 | 33269 | 33369 | 33198 | 34494 | 33497 | 32936 | 34247 | 33773,3 | 33753 | 33794 | 1,001 | 0,002 | 0,05 | 0,959972 | 0,964662 | 0,0005011 | 0,002076 | 0,001021 | 0,001021 | TRUE | 0,0099262 | No |
| orf19.988 | 2580 | 2203 | 2371 | 1621 | 1740 | 1378 | 1471 | 1978 | 2349 | 2228 | 2244 | 2435 | 1548 | 1496 | 1506 | 1509 | 1914,39 | 2314 | 1515 | 0,656 | -0,609 | -13,421 | 4,57E-41 | 1,81E-40 | 0,0004617 | 0,002727 | 0,0014362 | 0,0014362 | TRUE | 0,0136016 | No |
| orf19.989 | 4212 | 3871 | 4153 | 2976 | 3453 | 2930 | 2920 | 3894 | 3835 | 3915 | 3930 | 4470 | 3071 | 3181 | 2990 | 2970 | 3545,52 | 4038 | 3053 | 0,757 | -0,402 | -7,411 | 1,25E-13 | 2,67E-13 | 0,0027144 | 0,00241 | 0,0025339 | 0,0025339 | TRUE | 0,0611629 | No |
| orf19.99 | 1124 | 908 | 926 | 690 | 1086 | 829 | 922 | 1322 | 1023 | 918 | 876 | 1036 | 966 | 900 | 944 | 1008 | 959,16 | 963 | 954 | 0,994 | -0,009 | -0,132 | 0,895346 | 0,910195 | 0,0033602 | 0,003414 | 0,0033868 | 0,0033868 | TRUE | 0,045045 | No |
| orf19.991 | 3328 | 2850 | 3137 | 1928 | 2655 | 2031 | 2299 | 3120 | 3030 | 2882 | 2969 | 2896 | 2362 | 2205 | 2354 | 2380 | 2634,82 | 2944 | 2325 | 0,79 | -0,34 | -8,118 | 4,73E-16 | 1,08E-15 | 0,0004566 | 0,002538 | 0,0013042 | 0,0013042 | TRUE | 0,0141647 | No |
| orf19.992 | 3853 | 3421 | 3667 | 2197 | 3462 | 2860 | 2958 | 4056 | 3508 | 3460 | 3470 | 3300 | 3079 | 3105 | 3029 | 3094 | 3255,81 | 3434 | 3077 | 0,895 | -0,16 | -4,406 | 1,05E-05 | 1,67E-05 | 3,49E-06 | 0,002443 | 0,0009623 | 0,0009623 | TRUE | 0,0076775 | No |
| orf19.993 | 241 | 207 | 216 | 131 | 282 | 209 | 200 | 291 | 219 | 209 | 204 | 197 | 251 | 227 | 205 | 222 | 216,82 | 207 | 226 | 1,088 | 0,121 | 1,205 | 0,228193 | 0,261783 | 1,00E-08 | 0,00813 | 0,0050803 | 0,0050803 | TRUE | 0,0610983 | No |
| orf19.994 | 3 | 1 | 2 | 3 | 5 | 2 | 4 | 2 | 3 | 1 | 2 | 5 | 4 | 2 | 4 | 2 | 2,8 | 3 | 3 | 1,254 | 0,327 | 0,392 | 0,695064 | 0,72974 | 1,00E-08 | 0,474181 | 0,293346 | 0,293346 | TRUE | 0,2226914 | No |
| orf19.996 | 118 | 135 | 112 | 73 | 153 | 121 | 148 | 191 | 107 | 137 | 106 | 110 | 136 | 131 | 152 | 146 | 128,04 | 115 | 141 | 1,229 | 0,297 | 2,284 | 0,022379 | 0,028568 | 1,00E-08 | 0,012354 | 0,0083412 | 0,0083412 | TRUE | 0,1676064 | No |
| orf19.997 | 824 | 796 | 695 | 453 | 552 | 398 | 350 | 559 | 750 | 805 | 658 | 680 | 491 | 432 | 358 | 426 | 575,18 | 723 | 427 | 0,59 | -0,76 | -7,872 | 3,49E-15 | 7,75E-15 | 0,0098932 | 0,004334 | 0,0071196 | 0,0071196 | TRUE | 0,1319187 | No |
| orf19.998 | 175 | 183 | 157 | 122 | 227 | 209 | 183 | 272 | 159 | 185 | 149 | 183 | 202 | 227 | 187 | 207 | 187,5 | 169 | 206 | 1,224 | 0,292 | 2,595 | 0,009447 | 0,012408 | 0,0027222 | 0,009082 | 0,0067046 | 0,0067046 | TRUE | 0,0723372 | No |
| orf19.999 | 6022 | 4984 | 5414 | 3877 | 2170 | 2003 | 1877 | 2848 | 5483 | 5041 | 5124 | 5824 | 1930 | 2175 | 1922 | 2172 | 3708,86 | 5368 | 2050 | 0,382 | -1,388 | -22,184 | 4,90E-109 | 4,59E-108 | 0,0042214 | 0,002393 | 0,0034302 | 0,0034302 | TRUE | 0,0389925 | No |
| RDN18 | 80311 | 73414 | 80379 | 46846 | 37122 | 32062 | 38925 | 47508 | 73120 | 74250 | 76071 | 70370 | 33019 | 34810 | 39864 | 36239 | 54718,1 | 73453 | 35983 | 0,49 | -1,03 | -18,287 | 1,04E-74 | 6,89E-74 | 0,0037562 | 0,002061 | 0,0030252 | 0,0030252 | TRUE | 0,0645176 | No |
| RDN25 | 73724 | 71932 | 77944 | 51278 | 50721 | 50151 | 62143 | 73916 | 67123 | 72752 | 73767 | 77028 | 45115 | 54450 | 63642 | 56384 | 63782,4 | 72668 | 54898 | 0,755 | -0,405 | -4,781 | 1,74E-06 | 2,87E-06 | 0,0112423 | 0,002058 | 0,0068638 | 0,0068638 | TRUE | 0,1763755 | No |
| RDN5 | 276 | 705 | 672 | 479 | 127 | 243 | 193 | 206 | 251 | 713 | 636 | 720 | 113 | 264 | 198 | 157 | 381,43 | 580 | 183 | 0,315 | -1,666 | -4,017 | 5,89E-05 | 8,94E-05 | 0,1616833 | 0,0055 | 0,0653078 | 0,1616833 | TRUE | 1,7211831 | No |
| RDN58 | 2598 | 2571 | 2991 | 2394 | 950 | 889 | 948 | 1196 | 2365 | 2600 | 2831 | 3596 | 845 | 965 | 971 | 912 | 1885,74 | 2848 | 923 | 0,324 | -1,624 | -15,331 | 4,77E-53 | 2,33E-52 | 0,0186053 | 0,002738 | 0,0100813 | 0,0100813 | TRUE | 0,3740543 | No |
| RPR1 | 169 | 131 | 143 | 95 | 242 | 161 | 188 | 279 | 154 | 132 | 135 | 143 | 215 | 175 | 193 | 213 | 169,98 | 141 | 199 | 1,415 | 0,5 | 4,326 | 1,52E-05 | 2,39E-05 | 1,00E-08 | 0,009809 | 0,0067334 | 0,0067334 | TRUE | 0,0681625 | No |
| SCR1 | 501 | 410 | 394 | 243 | 191 | 114 | 156 | 161 | 456 | 415 | 373 | 365 | 170 | 124 | 160 | 123 | 273,12 | 402 | 144 | 0,357 | -1,486 | -12,209 | 2,78E-34 | 9,80E-34 | 0,0134515 | 0,006874 | 0,0096548 | 0,0096548 | TRUE | 0,1616971 | No |
| snR10a | 588 | 563 | 540 | 408 | 301 | 234 | 225 | 303 | 535 | 569 | 511 | 613 | 268 | 254 | 230 | 231 | 401,51 | 557 | 246 | 0,443 | -1,176 | -13,519 | 1,20E-41 | 4,82E-41 | 0,0031865 | 0,005327 | 0,0044016 | 0,0044016 | TRUE | 0,0470329 | No |
| snR10b | 0 | 0 | 1 | 1 | 0 | 0 | 0 | 3 | 0 | 0 | 1 | 2 | 0 | 0 | 0 | 2 | 0,59 | 1 | 0 | 1,135 | 0,182 | 0,089 | 0,928847 | 0,938862 | 1,00E-08 | 2,232984 | 2,3472609 | 2,3472609 | TRUE | 1,4530292 | No |
| snR10c | 1360 | 1192 | 1128 | 847 | 383 | 291 | 270 | 391 | 1238 | 1206 | 1068 | 1272 | 341 | 316 | 277 | 298 | 751,88 | 1196 | 308 | 0,258 | -1,955 | -24,228 | 1,12E-129 | 1,23E-128 | 0,0048671 | 0,003794 | 0,0042887 | 0,0042887 | TRUE | 0,0637233 | No |
| snR128 | 2875 | 2539 | 2784 | 1733 | 2524 | 2188 | 2049 | 2782 | 2618 | 2568 | 2635 | 2603 | 2245 | 2376 | 2098 | 2122 | 2408,09 | 2606 | 2210 | 0,848 | -0,239 | -4,963 | 6,94E-07 | 1,16E-06 | 0,0011669 | 0,002586 | 0,0018008 | 0,0018008 | TRUE | 0,029767 | No |
| snR13 | 890 | 688 | 701 | 526 | 841 | 562 | 666 | 921 | 810 | 696 | 663 | 790 | 748 | 610 | 682 | 703 | 712,82 | 740 | 686 | 0,929 | -0,106 | -1,264 | 0,206179 | 0,238243 | 0,0068267 | 0,00389 | 0,0053154 | 0,0053154 | TRUE | 0,0628843 | No |
| snR161a | 130 | 76 | 88 | 65 | 115 | 59 | 89 | 122 | 118 | 77 | 83 | 98 | 102 | 64 | 91 | 93 | 90,84 | 94 | 88 | 0,936 | -0,095 | -0,523 | 0,601141 | 0,640955 | 0,0268798 | 0,016579 | 0,0207242 | 0,0207242 | TRUE | 0,3280848 | No |
| snR17A | 3673 | 3282 | 3188 | 2409 | 5399 | 5089 | 3728 | 5991 | 3344 | 3319 | 3017 | 3619 | 4802 | 5525 | 3818 | 4570 | 4001,84 | 3325 | 4679 | 1,408 | 0,493 | 5,301 | 1,15E-07 | 1,99E-07 | 0,0136951 | 0,002367 | 0,0080595 | 0,0080595 | TRUE | 0,179879 | No |
| snR18 | 0 | 0 | 0 | 0 | 0 | 0 | 0 | 0 | 0 | 0 | 0 | 0 | 0 | 0 | 0 | 0 | 0 | 0 | 0 | NA | NA | NA | NA | NA | NA | NA | NA | NA | NA | NA | NA |
| snR189a | 3 | 3 | 1 | 1 | 2 | 0 | 1 | 2 | 3 | 3 | 1 | 2 | 2 | 0 | 1 | 2 | 1,57 | 2 | 1 | 0,542 | -0,884 | -0,805 | 0,420588 | 0,461727 | 1,00E-08 | 0,844591 | 0,4877849 | 0,4877849 | TRUE | 0,1883467 | No |
| snR189b | 118 | 99 | 92 | 88 | 38 | 32 | 39 | 50 | 107 | 100 | 87 | 132 | 34 | 35 | 40 | 38 | 71,68 | 106 | 37 | 0,347 | -1,525 | -8,143 | 3,85E-16 | 8,78E-16 | 1,00E-08 | 0,020466 | 0,0159536 | 0,0159536 | TRUE | 0,2063115 | No |
| snR189c | 183 | 205 | 203 | 144 | 121 | 91 | 86 | 141 | 167 | 207 | 192 | 216 | 108 | 99 | 88 | 108 | 148,05 | 196 | 101 | 0,519 | -0,946 | -7,368 | 1,73E-13 | 3,67E-13 | 0,0046228 | 0,010959 | 0,0084674 | 0,0084674 | TRUE | 0,1132086 | No |
| snR189d | 0 | 1 | 1 | 0 | 0 | 0 | 0 | 0 | 0 | 1 | 1 | 0 | 0 | 0 | 0 | 0 | 0,24 | 0 | 0 | 0,321 | -1,64 | -0,58 | 0,562213 | 0,603284 | 1,00E-08 | 5,399916 | 5,7781006 | 5,7781006 | TRUE | 0,0501011 | No |
| snR190 | 1277 | 1862 | 1795 | 872 | 1325 | 1328 | 1342 | 1410 | 1163 | 1883 | 1699 | 1310 | 1179 | 1442 | 1374 | 1076 | 1390,61 | 1514 | 1268 | 0,837 | -0,257 | -1,382 | 0,166836 | 0,195116 | 0,0324252 | 0,002987 | 0,0162457 | 0,0324252 | TRUE | 0,1783651 | No |
| snR24 | 0 | 0 | 0 | 0 | 0 | 0 | 0 | 0 | 0 | 0 | 0 | 0 | 0 | 0 | 0 | 0 | 0 | 0 | 0 | NA | NA | NA | NA | NA | NA | NA | NA | NA | NA | NA | NA |
| snR31a | 44 | 29 | 34 | 25 | 47 | 28 | 24 | 61 | 40 | 29 | 32 | 38 | 42 | 30 | 25 | 47 | 35,3 | 35 | 36 | 1,046 | 0,065 | 0,253 | 0,800225 | 0,826545 | 0,0263147 | 0,039454 | 0,0344905 | 0,0344905 | TRUE | 0,207362 | No |
| snR32a | 37 | 39 | 30 | 10 | 46 | 26 | 25 | 40 | 34 | 39 | 28 | 15 | 41 | 28 | 26 | 31 | 30,23 | 29 | 32 | 1,055 | 0,077 | 0,271 | 0,786246 | 0,815542 | 0,0429677 | 0,045742 | 0,0448951 | 0,0448951 | TRUE | 0,4362261 | No |
| snR33a | 53991 | 35181 | 38180 | 32840 | 7570 | 5482 | 5406 | 8185 | 49157 | 35582 | 36134 | 49331 | 6733 | 5952 | 5536 | 6244 | 24333,6 | 42551 | 6116 | 0,144 | -2,798 | -19,465 | 2,20E-84 | 1,64E-83 | 0,0197713 | 0,002091 | 0,0104426 | 0,0197713 | TRUE | 0,0435782 | No |
| snR35a | 15 | 19 | 25 | 15 | 40 | 26 | 42 | 41 | 14 | 19 | 24 | 23 | 36 | 28 | 43 | 31 | 27,15 | 20 | 34 | 1,764 | 0,819 | 2,908 | 0,003641 | 0,004923 | 1,00E-08 | 0,050701 | 0,0354105 | 0,0354105 | TRUE | 0,2727102 | No |
| snR35c | 36 | 17 | 38 | 19 | 28 | 23 | 23 | 37 | 33 | 17 | 36 | 29 | 25 | 25 | 24 | 28 | 27,02 | 29 | 26 | 0,888 | -0,172 | -0,615 | 0,538644 | 0,580239 | 1,00E-08 | 0,050933 | 0,0381888 | 0,0381888 | TRUE | 0,4989415 | No |
| snR39b | 0 | 0 | 0 | 0 | 0 | 0 | 0 | 0 | 0 | 0 | 0 | 0 | 0 | 0 | 0 | 0 | 0 | 0 | 0 | NA | NA | NA | NA | NA | NA | NA | NA | NA | NA | NA | NA |
| snR3a | 87 | 83 | 88 | 44 | 103 | 75 | 69 | 107 | 79 | 84 | 83 | 66 | 92 | 81 | 71 | 82 | 79,73 | 78 | 82 | 1,034 | 0,049 | 0,307 | 0,758511 | 0,789781 | 1,00E-08 | 0,018605 | 0,011431 | 0,011431 | TRUE | 0,0715195 | No |
| snR40 | 30 | 36 | 25 | 23 | 41 | 36 | 26 | 48 | 27 | 36 | 24 | 35 | 36 | 39 | 27 | 37 | 32,59 | 30 | 35 | 1,153 | 0,205 | 0,822 | 0,410882 | 0,452161 | 1,00E-08 | 0,042569 | 0,0287209 | 0,0287209 | TRUE | 0,1624306 | No |
| snR41 | 1747 | 1609 | 1700 | 1006 | 1288 | 1139 | 1136 | 1515 | 1591 | 1627 | 1609 | 1511 | 1146 | 1237 | 1163 | 1156 | 1379,91 | 1584 | 1176 | 0,74 | -0,435 | -8,894 | 5,87E-19 | 1,45E-18 | 0,0001564 | 0,002994 | 0,0015567 | 0,0015567 | TRUE | 0,0140766 | No |
| snR41b | 602 | 568 | 520 | 495 | 628 | 488 | 519 | 689 | 548 | 574 | 492 | 744 | 559 | 530 | 532 | 526 | 562,97 | 590 | 537 | 0,915 | -0,128 | -1,228 | 0,219331 | 0,252434 | 0,0137796 | 0,004383 | 0,0086568 | 0,0086568 | TRUE | 0,3377594 | No |
| snR42a | 157 | 131 | 117 | 125 | 147 | 130 | 134 | 200 | 143 | 132 | 111 | 188 | 131 | 141 | 137 | 153 | 141,95 | 144 | 140 | 0,992 | -0,011 | -0,075 | 0,94002 | 0,949101 | 0,0171236 | 0,011343 | 0,0136702 | 0,0136702 | TRUE | 0,3754161 | No |
| snR43a | 3 | 4 | 1 | 3 | 3 | 4 | 5 | 9 | 3 | 4 | 1 | 5 | 3 | 4 | 5 | 7 | 3,9 | 3 | 5 | 1,627 | 0,702 | 0,979 | 0,327674 | 0,367315 | 1,00E-08 | 0,340461 | 0,2169354 | 0,2169354 | TRUE | 0,3195776 | No |
| snR43b | 29 | 25 | 17 | 19 | 21 | 24 | 19 | 35 | 26 | 25 | 16 | 29 | 19 | 26 | 19 | 27 | 23,4 | 24 | 23 | 0,956 | -0,064 | -0,219 | 0,826897 | 0,850511 | 1,00E-08 | 0,058487 | 0,0403375 | 0,0403375 | TRUE | 0,3255337 | No |
| snR44a | 20 | 22 | 24 | 18 | 25 | 30 | 38 | 35 | 18 | 22 | 23 | 27 | 22 | 33 | 39 | 27 | 26,33 | 22 | 30 | 1,342 | 0,424 | 1,503 | 0,132712 | 0,157052 | 1,00E-08 | 0,052209 | 0,036869 | 0,036869 | TRUE | 0,2346526 | No |
| snR44b | 62 | 65 | 64 | 42 | 66 | 64 | 61 | 66 | 56 | 66 | 61 | 63 | 59 | 69 | 62 | 50 | 60,86 | 62 | 60 | 0,975 | -0,036 | -0,202 | 0,839892 | 0,862485 | 1,00E-08 | 0,023744 | 0,0144069 | 0,0144069 | TRUE | 0,1241838 | No |
| snR44c | 12 | 18 | 17 | 17 | 26 | 37 | 20 | 20 | 11 | 18 | 16 | 26 | 23 | 40 | 20 | 15 | 21,22 | 18 | 24 | 1,416 | 0,502 | 1,373 | 0,16974 | 0,198076 | 0,0993375 | 0,064277 | 0,0783784 | 0,0783784 | TRUE | 0,9460571 | No |
| snR44d | 382 | 414 | 339 | 254 | 572 | 415 | 424 | 563 | 348 | 419 | 321 | 382 | 509 | 451 | 434 | 429 | 411,49 | 368 | 456 | 1,244 | 0,315 | 3,311 | 0,00093 | 0,001315 | 0,007344 | 0,005247 | 0,0062067 | 0,0062067 | TRUE | 0,1097379 | No |
| snR46a | 0 | 0 | 0 | 0 | 1 | 0 | 0 | 2 | 0 | 0 | 0 | 0 | 1 | 0 | 0 | 2 | 0,3 | 0 | 1 | 3,171 | 1,665 | 0,634 | 0,526375 | 0,567982 | 1,00E-08 | 4,37786 | 4,8842113 | 4,8842113 | TRUE | 0,25714 | No |
| snR47 | 706 | 490 | 535 | 317 | 763 | 469 | 481 | 704 | 643 | 496 | 506 | 476 | 679 | 509 | 493 | 537 | 542,3 | 530 | 554 | 1,044 | 0,062 | 0,535 | 0,5927 | 0,632903 | 0,0187508 | 0,004473 | 0,0111126 | 0,0111126 | TRUE | 0,2688462 | No |
| snR48 | 1252 | 1381 | 1302 | 581 | 1062 | 797 | 850 | 1047 | 1140 | 1397 | 1232 | 873 | 945 | 865 | 871 | 799 | 1015,09 | 1160 | 870 | 0,748 | -0,419 | -3,754 | 0,000174 | 0,000258 | 0,0188566 | 0,003338 | 0,0109878 | 0,0109878 | TRUE | 0,3116526 | No |
| snR49a | 0 | 0 | 0 | 0 | 0 | 0 | 0 | 0 | 0 | 0 | 0 | 0 | 0 | 0 | 0 | 0 | 0 | 0 | 0 | NA | NA | NA | NA | NA | NA | NA | NA | NA | NA | NA | NA |
| snR50 | 2 | 5 | 5 | 2 | 3 | 5 | 6 | 8 | 2 | 5 | 5 | 3 | 3 | 5 | 6 | 6 | 4,37 | 4 | 5 | 1,383 | 0,468 | 0,707 | 0,479698 | 0,521589 | 1,00E-08 | 0,304338 | 0,1839925 | 0,1839925 | TRUE | 0,2417818 | No |
| snR52 | 11 | 8 | 11 | 4 | 23 | 19 | 21 | 20 | 10 | 8 | 10 | 6 | 20 | 21 | 22 | 15 | 14,05 | 8 | 20 | 2,185 | 1,128 | 2,966 | 0,003016 | 0,004105 | 1,00E-08 | 0,096079 | 0,0547489 | 0,0547489 | TRUE | 0,1444379 | No |

| | | | | | | | | | | | | | | | | | | | | | | | | | | | | | | | |
|---|---|---|---|---|---|---|---|---|---|---|---|---|---|---|---|---|---|---|---|---|---|---|---|---|---|---|---|---|---|---|---|
| snR54 | 0 | 0 | 0 | 0 | 0 | 0 | 0 | 0 | 0 | 0 | 0 | 0 | 0 | 0 | 0 | 0 | 0 | 0 | 0 | NA | NA | NA | NA | NA | NA | NA | NA | NA | NA | NA | NA |
| snR55 | 13 | 7 | 13 | 11 | 10 | 4 | 4 | 7 | 12 | 7 | 12 | 17 | 9 | 4 | 4 | 5 | 8,8 | 12 | 6 | 0,489 | -1,032 | -2,11 | 0,034865 | 0,043813 | 1,00E-08 | 0,152115 | 0,103073 | 0,103073 | TRUE | 0,2520695 | No |
| snR56 | 570 | 388 | 434 | 344 | 441 | 378 | 288 | 530 | 519 | 392 | 411 | 517 | 392 | 410 | 295 | 404 | 417,6 | 460 | 375 | 0,819 | -0,288 | -2,373 | 0,017649 | 0,022744 | 0,0196767 | 0,0052 | 0,0117008 | 0,0117008 | TRUE | 0,2345975 | No |
| snR57 | 99 | 126 | 129 | 35 | 59 | 67 | 70 | 74 | 90 | 127 | 122 | 53 | 52 | 73 | 72 | 56 | 80,7 | 98 | 63 | 0,636 | -0,653 | -2,811 | 0,004935 | 0,006623 | 0,0702182 | 0,018406 | 0,0389701 | 0,0389701 | TRUE | 0,4121449 | No |
| snR57b | 833 | 853 | 790 | 348 | 777 | 511 | 659 | 794 | 758 | 863 | 748 | 523 | 691 | 555 | 675 | 606 | 677,25 | 723 | 632 | 0,871 | -0,199 | -1,619 | 0,105389 | 0,126408 | 0,0245064 | 0,003988 | 0,0130344 | 0,0130344 | TRUE | 0,3864316 | No |
| snR58 | 287 | 234 | 196 | 152 | 258 | 242 | 167 | 283 | 261 | 237 | 185 | 228 | 229 | 263 | 171 | 216 | 223,87 | 228 | 220 | 0,963 | -0,054 | -0,392 | 0,69517 | 0,72974 | 0,0209776 | 0,007938 | 0,0135835 | 0,0135835 | TRUE | 0,2349512 | No |
| snR5a | 362 | 285 | 254 | 229 | 234 | 148 | 136 | 230 | 330 | 288 | 240 | 344 | 208 | 161 | 139 | 175 | 235,72 | 300 | 171 | 0,572 | -0,805 | -5,826 | 5,68E-09 | 1,03E-08 | 0,0220073 | 0,007641 | 0,0138298 | 0,0138298 | TRUE | 0,2313052 | No |
| snR5b | 29 | 9 | 20 | 11 | 24 | 13 | 10 | 17 | 26 | 9 | 19 | 17 | 21 | 14 | 10 | 13 | 16,2 | 18 | 14 | 0,823 | -0,282 | -0,727 | 0,467368 | 0,509224 | 0,0810461 | 0,083562 | 0,0826041 | 0,0826041 | TRUE | 0,2180677 | No |
| snR5c | 6 | 6 | 7 | 0 | 11 | 9 | 17 | 10 | 5 | 6 | 7 | 0 | 10 | 10 | 17 | 8 | 7,84 | 4 | 11 | 2,282 | 1,19 | 2,159 | 0,030868 | 0,03899 | 0,0669194 | 0,170451 | 0,1397663 | 0,1397663 | TRUE | 0,5249909 | No |
| snR5d | 215 | 195 | 175 | 171 | 287 | 198 | 189 | 301 | 196 | 197 | 166 | 257 | 255 | 215 | 194 | 230 | 213,61 | 204 | 224 | 1,108 | 0,148 | 1,125 | 0,260736 | 0,296397 | 0,0176508 | 0,008221 | 0,0119685 | 0,0119685 | TRUE | 0,2880767 | No |
| snR60 | 0 | 0 | 3 | 2 | 0 | 1 | 0 | 1 | 0 | 0 | 3 | 3 | 0 | 1 | 0 | 1 | 0,96 | 2 | 0 | 0,328 | -1,609 | -0,964 | 0,334812 | 0,374724 | 1,00E-08 | 1,375916 | 1,3422968 | 1,3422968 | TRUE | 0,073339 | No |
| snR61 | 427 | 505 | 463 | 348 | 369 | 397 | 261 | 363 | 389 | 511 | 438 | 523 | 328 | 431 | 267 | 277 | 395,49 | 465 | 326 | 0,701 | -0,513 | -3,633 | 0,00028 | 0,00041 | 0,03225 | 0,005377 | 0,0165777 | 0,0165777 | TRUE | 0,5238181 | No |
| snR62 | 200 | 223 | 176 | 126 | 141 | 143 | 107 | 113 | 182 | 226 | 167 | 189 | 125 | 155 | 110 | 86 | 154,99 | 191 | 119 | 0,62 | -0,689 | -4,208 | 2,58E-05 | 4,01E-05 | 0,030706 | 0,01056 | 0,0189875 | 0,0189875 | TRUE | 0,4081932 | No |
| snR63 | 226 | 182 | 156 | 80 | 228 | 154 | 197 | 204 | 206 | 184 | 148 | 120 | 203 | 167 | 202 | 156 | 173,13 | 164 | 182 | 1,095 | 0,131 | 0,84 | 0,400951 | 0,441691 | 0,0287426 | 0,009667 | 0,0175482 | 0,0175482 | TRUE | 0,2326765 | No |
| snR64 | 18 | 12 | 7 | 13 | 10 | 4 | 11 | 10 | 16 | 12 | 7 | 20 | 9 | 4 | 11 | 8 | 10,85 | 14 | 8 | 0,602 | -0,733 | -1,602 | 0,109135 | 0,130483 | 0,0628457 | 0,123775 | 0,1031118 | 0,1031118 | TRUE | 0,2659626 | No |
| snR65 | 20 | 7 | 16 | 6 | 14 | 7 | 8 | 16 | 18 | 7 | 15 | 9 | 12 | 8 | 8 | 12 | 11,24 | 12 | 10 | 0,812 | -0,3 | -0,693 | 0,488275 | 0,530011 | 0,0404248 | 0,119596 | 0,0918669 | 0,0918669 | TRUE | 0,20977 | No |
| snR66 | 82 | 93 | 132 | 113 | 246 | 241 | 171 | 274 | 75 | 94 | 125 | 170 | 219 | 262 | 175 | 209 | 166 | 116 | 216 | 1,889 | 0,918 | 4,594 | 4,35E-06 | 7,04E-06 | 0,0645108 | 0,009995 | 0,0314669 | 0,0314669 | TRUE | 0,7729167 | No |
| snR67 | 2 | 2 | 8 | 5 | 4 | 4 | 3 | 8 | 2 | 2 | 8 | 8 | 4 | 4 | 3 | 6 | 4,5 | 5 | 4 | 0,944 | -0,083 | -0,121 | 0,903329 | 0,917191 | 1,00E-08 | 0,295581 | 0,2252232 | 0,2252232 | TRUE | 0,1520435 | No |
| snR68 | 36 | 57 | 40 | 46 | 28 | 16 | 22 | 25 | 33 | 58 | 38 | 69 | 25 | 17 | 23 | 19 | 35,16 | 50 | 21 | 0,433 | -1,207 | -4,227 | 2,37E-05 | 3,69E-05 | 0,0530108 | 0,039611 | 0,0450865 | 0,0450865 | TRUE | 0,1346505 | No |
| snR69 | 488 | 389 | 422 | 372 | 350 | 267 | 294 | 380 | 444 | 393 | 399 | 559 | 311 | 290 | 301 | 290 | 373,51 | 449 | 298 | 0,668 | -0,582 | -5,453 | 4,95E-08 | 8,64E-08 | 0,0114513 | 0,005574 | 0,0081597 | 0,0081597 | TRUE | 0,2844394 | No |
| snR70 | 1546 | 1457 | 1509 | 921 | 1136 | 914 | 970 | 1235 | 1408 | 1474 | 1428 | 1383 | 1010 | 992 | 993 | 942 | 1203,88 | 1423 | 984 | 0,691 | -0,534 | -10,763 | 5,15E-27 | 1,54E-26 | 1,00E-08 | 0,003134 | 0,0015141 | 0,0015141 | TRUE | 0,0110486 | No |
| snR71 | 11 | 13 | 14 | 6 | 9 | 6 | 9 | 7 | 10 | 13 | 13 | 9 | 8 | 7 | 9 | 5 | 9,31 | 11 | 7 | 0,628 | -0,672 | -1,505 | 0,132315 | 0,15667 | 1,00E-08 | 0,143885 | 0,0804872 | 0,0804872 | TRUE | 0,1247833 | No |
| snR72 | 5 | 11 | 7 | 8 | 11 | 14 | 6 | 14 | 5 | 11 | 7 | 12 | 10 | 15 | 6 | 11 | 9,52 | 9 | 10 | 1,252 | 0,324 | 0,683 | 0,49453 | 0,535979 | 0,0360402 | 0,140855 | 0,1070574 | 0,1070574 | TRUE | 0,2955076 | No |
| snR73 | 419 | 407 | 397 | 299 | 328 | 229 | 186 | 393 | 381 | 412 | 376 | 449 | 292 | 249 | 190 | 300 | 331,08 | 404 | 258 | 0,641 | -0,641 | -5,137 | 2,79E-07 | 4,74E-07 | 0,0196003 | 0,006027 | 0,0118132 | 0,0118132 | TRUE | 0,3389391 | No |
| snR74 | 82 | 172 | 112 | 106 | 51 | 59 | 47 | 50 | 75 | 174 | 106 | 159 | 45 | 64 | 48 | 38 | 88,69 | 128 | 49 | 0,381 | -1,394 | -5,535 | 3,12E-08 | 5,50E-08 | 0,0865722 | 0,016931 | 0,0470756 | 0,0470756 | TRUE | 0,141905 | No |
| snR75 | 163 | 226 | 207 | 171 | 96 | 75 | 88 | 132 | 148 | 229 | 196 | 257 | 85 | 81 | 90 | 101 | 148,42 | 208 | 89 | 0,436 | -1,199 | -7,307 | 2,73E-13 | 5,78E-13 | 0,0269343 | 0,010937 | 0,0180262 | 0,0180262 | TRUE | 0,1979465 | No |
| snR76 | 409 | 399 | 420 | 228 | 213 | 152 | 196 | 243 | 372 | 404 | 397 | 342 | 189 | 165 | 201 | 185 | 282,06 | 379 | 185 | 0,487 | -1,037 | -10,974 | 5,10E-28 | 1,56E-27 | 0,0011574 | 0,00672 | 0,0046573 | 0,0046573 | TRUE | 0,051071 | No |
| snR77 | 259 | 321 | 356 | 231 | 186 | 164 | 131 | 197 | 236 | 325 | 337 | 347 | 165 | 178 | 134 | 150 | 234,04 | 311 | 157 | 0,506 | -0,983 | -7,342 | 2,10E-13 | 4,45E-13 | 0,0185862 | 0,007681 | 0,0124944 | 0,0124944 | TRUE | 0,3141903 | No |
| snR77b | 103 | 237 | 220 | 127 | 112 | 122 | 106 | 101 | 94 | 240 | 208 | 191 | 100 | 132 | 109 | 77 | 143,77 | 183 | 104 | 0,57 | -0,812 | -2,501 | 0,0124 | 0,016152 | 0,0938994 | 0,011225 | 0,0464501 | 0,0938994 | TRUE | 0,7235966 | No |
| snR78 | 51 | 96 | 100 | 32 | 30 | 27 | 32 | 27 | 46 | 97 | 95 | 48 | 27 | 29 | 33 | 21 | 49,45 | 72 | 28 | 0,377 | -1,409 | -4,94 | 7,82E-07 | 1,31E-06 | 0,0862804 | 0,028751 | 0,0534726 | 0,0534726 | TRUE | 0,0685516 | No |
| snR79 | 118 | 170 | 154 | 126 | 143 | 119 | 124 | 140 | 107 | 172 | 146 | 189 | 127 | 129 | 127 | 107 | 138,07 | 154 | 122 | 0,803 | -0,316 | -1,955 | 0,050578 | 0,062584 | 0,0261203 | 0,011605 | 0,0177665 | 0,0177665 | TRUE | 0,3967117 | No |
| snR87 | 280 | 271 | 268 | 168 | 204 | 138 | 138 | 188 | 255 | 274 | 254 | 252 | 181 | 150 | 141 | 143 | 206,38 | 259 | 154 | 0,595 | -0,748 | -6,964 | 3,30E-12 | 6,73E-12 | 0,00176 | 0,008438 | 0,0060041 | 0,0060041 | TRUE | 0,1586112 | No |
| snR8a | 235 | 234 | 215 | 133 | 322 | 228 | 256 | 314 | 214 | 237 | 203 | 200 | 286 | 248 | 262 | 240 | 236,19 | 214 | 259 | 1,209 | 0,274 | 2,723 | 0,006462 | 0,008596 | 0,0023069 | 0,00763 | 0,0054198 | 0,0054198 | TRUE | 0,0600917 | No |
| snR8b | 18 | 28 | 23 | 22 | 45 | 30 | 35 | 42 | 16 | 28 | 22 | 33 | 40 | 33 | 36 | 32 | 30 | 25 | 35 | 1,447 | 0,533 | 2 | 0,045499 | 0,056552 | 0,0027221 | 0,04607 | 0,03292 | 0,03292 | TRUE | 0,3339483 | No |
| snR8c | 101 | 122 | 113 | 87 | 94 | 79 | 91 | 110 | 92 | 123 | 107 | 131 | 84 | 86 | 93 | 84 | 99,93 | 113 | 87 | 0,771 | -0,375 | -2,519 | 0,011768 | 0,015357 | 1,00E-08 | 0,015256 | 0,0111249 | 0,0111249 | TRUE | 0,1689962 | No |
| SNRNAU1 | 163 | 137 | 121 | 68 | 111 | 71 | 87 | 144 | 148 | 139 | 115 | 102 | 99 | 77 | 89 | 110 | 109,8 | 126 | 94 | 0,743 | -0,428 | -2,712 | 0,006696 | 0,008898 | 0,0158881 | 0,014068 | 0,0147801 | 0,0147801 | TRUE | 0,1485803 | No |
| SNRNAU2 | 39 | 37 | 34 | 22 | 28 | 29 | 13 | 26 | 36 | 37 | 32 | 33 | 25 | 31 | 13 | 20 | 28,46 | 34 | 22 | 0,643 | -0,636 | -2,309 | 0,020935 | 0,026788 | 0,0057412 | 0,04845 | 0,0365565 | 0,0365565 | TRUE | 0,2933673 | No |
| SNRNAU4 | 151 | 162 | 180 | 78 | 124 | 81 | 113 | 131 | 137 | 164 | 170 | 117 | 110 | 88 | 116 | 100 | 125,34 | 147 | 104 | 0,698 | -0,519 | -3,554 | 0,00038 | 0,000551 | 0,0118902 | 0,012576 | 0,0123575 | 0,0123575 | TRUE | 0,1573632 | No |
| SNRNAU5 | 20 | 20 | 19 | 16 | 8 | 3 | 5 | 10 | 18 | 20 | 18 | 24 | 7 | 3 | 5 | 8 | 12,95 | 20 | 6 | 0,299 | -1,744 | -4,19 | 2,79E-05 | 4,33E-05 | 1,00E-08 | 0,104069 | 0,0614566 | 0,0614566 | TRUE | 0,1687902 | No |
| SNRNAU6 | 31 | 21 | 35 | 17 | 10 | 13 | 10 | 15 | 28 | 21 | 33 | 26 | 9 | 14 | 10 | 11 | 19,1 | 27 | 11 | 0,409 | -1,29 | -3,894 | 9,88E-05 | 0,000148 | 1,00E-08 | 0,071192 | 0,0439698 | 0,0439698 | TRUE | 0,1572697 | No |
| tA(AGC)1 | 0 | 0 | 0 | 1 | 0 | 0 | 0 | 0 | 0 | 0 | 0 | 2 | 0 | 0 | 0 | 0 | 0,19 | 0 | 0 | 0,528 | -0,92 | -0,26 | 0,794572 | 0,821771 | 1,00E-08 | 7,03723 | 10 | 10 | TRUE | 0,6271619 | No |
| tA(AGC)2 | 0 | 0 | 0 | 0 | 0 | 0 | 1 | 0 | 0 | 0 | 0 | 0 | 0 | 0 | 1 | 0 | 0,13 | 0 | 0 | 1,436 | 0,523 | 0,148 | 0,882445 | 0,8988 | 1,00E-08 | 10,32106 | 10 | 10 | TRUE | 0,417491 | No |
| tA(AGC)3 | 0 | 0 | 0 | 0 | 0 | 0 | 0 | 0 | 0 | 0 | 0 | 0 | 0 | 0 | 0 | 0 | 0 | 0 | 0 | NA | NA | NA | NA | NA | NA | NA | NA | NA | NA | NA | NA |
| tA(AGC)4 | 0 | 0 | 1 | 0 | 0 | 0 | 1 | 0 | 0 | 0 | 1 | 0 | 0 | 0 | 1 | 0 | 0,25 | 0 | 0 | 0,871 | -0,199 | -0,07 | 0,944114 | 0,951878 | 1,00E-08 | 5,365037 | 5,7273304 | 5,7273304 | TRUE | 0,4174904 | No |
| tA(AGC)5 | 0 | 0 | 0 | 0 | 1 | 2 | 0 | 1 | 0 | 0 | 0 | 0 | 1 | 2 | 0 | 1 | 0,48 | 0 | 1 | 4,808 | 2,265 | 1,118 | 0,263455 | 0,29938 | 1,00E-08 | 2,765833 | 2,4486035 | 2,4486035 | TRUE | 0,1871684 | No |
| tA(AGC)7 | 0 | 0 | 0 | 1 | 0 | 0 | 0 | 1 | 0 | 0 | 0 | 2 | 0 | 0 | 0 | 1 | 0,28 | 0 | 0 | 0,871 | -0,199 | -0,073 | 0,94161 | 0,950026 | 1,00E-08 | 4,667879 | 5,0775645 | 5,0775645 | TRUE | 0,6271667 | No |
| tA(UGC)1 | 1 | 0 | 0 | 0 | 0 | 0 | 0 | 0 | 1 | 0 | 0 | 0 | 0 | 0 | 0 | 0 | 0,11 | 0 | 0 | 0,528 | -0,92 | -0,26 | 0,794572 | 0,821771 | 1,00E-08 | 11,60937 | 10 | 10 | TRUE | 0,2548005 | No |
| tA(UGC)2 | 0 | 0 | 0 | 0 | 0 | 0 | 0 | 0 | 0 | 0 | 0 | 0 | 0 | 0 | 0 | 0 | 0 | 0 | 0 | NA | NA | NA | NA | NA | NA | NA | NA | NA | NA | NA | NA |
| tC(GCA)1 | 0 | 0 | 0 | 0 | 0 | 0 | 0 | 0 | 0 | 0 | 0 | 0 | 0 | 0 | 0 | 0 | 0 | 0 | 0 | NA | NA | NA | NA | NA | NA | NA | NA | NA | NA | NA | NA |
| tC(GCA)2 | 0 | 0 | 0 | 0 | 0 | 0 | 0 | 0 | 0 | 0 | 0 | 0 | 0 | 0 | 0 | 0 | 0 | 0 | 0 | NA | NA | NA | NA | NA | NA | NA | NA | NA | NA | NA | NA |
| tD(GUC)2 | 0 | 0 | 1 | 0 | 0 | 0 | 0 | 0 | 0 | 0 | 1 | 0 | 0 | 0 | 0 | 0 | 0,12 | 0 | 0 | 0,528 | -0,92 | -0,26 | 0,794572 | 0,821771 | 1,00E-08 | 11,16843 | 10 | 10 | TRUE | 0,2762646 | No |
| tD(GUC)3 | 0 | 0 | 0 | 0 | 0 | 0 | 0 | 0 | 0 | 0 | 0 | 0 | 0 | 0 | 0 | 0 | 0 | 0 | 0 | NA | NA | NA | NA | NA | NA | NA | NA | NA | NA | NA | NA |
| tD(GUC)4 | 0 | 0 | 0 | 1 | 0 | 0 | 0 | 1 | 0 | 0 | 0 | 2 | 0 | 0 | 0 | 1 | 0,28 | 0 | 0 | 0,871 | -0,199 | -0,073 | 0,94161 | 0,950026 | 1,00E-08 | 4,667879 | 5,0775645 | 5,0775645 | TRUE | 0,6271667 | No |
| tD(GUC)5 | 1 | 1 | 0 | 0 | 0 | 0 | 0 | 0 | 1 | 1 | 0 | 0 | 0 | 0 | 0 | 0 | 0,24 | 0 | 0 | 0,325 | -1,623 | -0,571 | 0,568181 | 0,60897 | 1,00E-08 | 5,500893 | 5,8463886 | 5,8463886 | TRUE | 0,0523072 | No |
| tD(GUC)6 | 0 | 0 | 0 | 0 | 2 | 1 | 0 | 0 | 0 | 0 | 0 | 0 | 2 | 1 | 0 | 0 | 0,36 | 0 | 1 | 3,618 | 1,855 | 0,737 | 0,461417 | 0,50317 | 1,00E-08 | 3,691127 | 4,4383727 | 4,4383727 | TRUE | 0,1863969 | No |
| tD(GUC)7 | 0 | 0 | 0 | 0 | 0 | 0 | 0 | 0 | 0 | 0 | 0 | 0 | 0 | 0 | 0 | 0 | 0 | 0 | 0 | NA | NA | NA | NA | NA | NA | NA | NA | NA | NA | NA | NA |
| tE(CUC)1 | 1 | 0 | 0 | 0 | 0 | 0 | 0 | 0 | 1 | 0 | 0 | 0 | 0 | 0 | 0 | 0 | 0,11 | 0 | 0 | 0,528 | -0,92 | -0,26 | 0,794572 | 0,821771 | 1,00E-08 | 11,60937 | 10 | 10 | TRUE | 0,2548005 | No |
| TER1 | 3391 | 2952 | 3071 | 2110 | 715 | 586 | 551 | 775 | 3087 | 2986 | 2906 | 3170 | 636 | 636 | 564 | 591 | 1822,08 | 3037 | 607 | 0,2 | -2,323 | -42,071 | 0 | 0 | 0,001106 | 0,002762 | 0,001981 | 0,001981 | TRUE | 0,0242852 | No |
| tE(UUC)1 | 8 | 4 | 6 | 3 | 6 | 4 | 3 | 7 | 7 | 4 | 6 | 5 | 5 | 4 | 3 | 5 | 4,95 | 6 | 4 | 0,838 | -0,254 | -0,425 | 0,670892 | 0,707399 | 1,00E-08 | 0,268865 | 0,1448695 | 0,1448695 | TRUE | 0,1394591 | No |
| tE(UUC)2 | 0 | 0 | 0 | 0 | 0 | 0 | 0 | 0 | 0 | 0 | 0 | 0 | 0 | 0 | 0 | 0 | 0 | 0 | 0 | NA | NA | NA | NA | NA | NA | NA | NA | NA | NA | NA | NA |
| tE(UUC)3 | 1 | 0 | 0 | 1 | 1 | 0 | 0 | 1 | 1 | 0 | 0 | 2 | 1 | 0 | 0 | 1 | 0,51 | 1 | 0 | 0,849 | -0,236 | -0,121 | 0,903796 | 0,917374 | 1,00E-08 | 2,601855 | 1,7988787 | 1,7988787 | TRUE | 0,2360405 | No |
| tE(UUC)4 | 0 | 0 | 0 | 0 | 0 | 0 | 0 | 0 | 0 | 0 | 0 | 0 | 0 | 0 | 0 | 0 | 0 | 0 | 0 | NA | NA | NA | NA | NA | NA | NA | NA | NA | NA | NA | NA |
| tE(UUC)5 | 0 | 0 | 0 | 0 | 1 | 0 | 0 | 0 | 0 | 0 | 0 | 0 | 1 | 0 | 0 | 0 | 0,11 | 0 | 0 | 1,436 | 0,523 | 0,148 | 0,882445 | 0,8988 | 1,00E-08 | 11,88327 | 10 | 10 | TRUE | 0,3214251 | No |
| tE(UUC)6 | 2 | 1 | 0 | 2 | 1 | 1 | 4 | 2 | 2 | 1 | 0 | 3 | 1 | 1 | 4 | 2 | 1,68 | 2 | 2 | 1,375 | 0,459 | 0,413 | 0,679813 | 0,715504 | 1,00E-08 | 0,7887 | 0,5512623 | 0,5512623 | TRUE | 0,4343725 | No |
| tE(UUC)7 | 7 | 7 | 4 | 1 | 6 | 7 | 7 | 11 | 6 | 7 | 4 | 2 | 5 | 8 | 7 | 8 | 5,9 | 5 | 7 | 1,459 | 0,545 | 0,958 | 0,337929 | 0,378014 | 1,00E-08 | 0,225758 | 0,1371475 | 0,1371475 | TRUE | 0,2153088 | No |
| tF(GAA)1 | 3 | 3 | 2 | 3 | 3 | 1 | 2 | 5 | 3 | 3 | 2 | 5 | 3 | 1 | 2 | 4 | 2,72 | 3 | 2 | 0,842 | -0,247 | -0,303 | 0,761802 | 0,792508 | 1,00E-08 | 0,487224 | 0,2713591 | 0,2713591 | TRUE | 0,2232729 | No |
| tF(GAA)2 | 0 | 0 | 1 | 0 | 0 | 0 | 0 | 0 | 0 | 0 | 1 | 0 | 0 | 0 | 0 | 0 | 0,12 | 0 | 0 | 0,528 | -0,92 | -0,26 | 0,794572 | 0,821771 | 1,00E-08 | 11,16843 | 10 | 10 | TRUE | 0,2762646 | No |

| | | | | | | | | | | | | | | | | | | | | | | | | | | | | | | | |
|---|---|---|---|---|---|---|---|---|---|---|---|---|---|---|---|---|---|---|---|---|---|---|---|---|---|---|---|---|---|---|---|
| tF(GAA)3 | 1 | 2 | 0 | 0 | 1 | 1 | 0 | 1 | 1 | 2 | 0 | 0 | 1 | 1 | 0 | 1 | 0,71 | 1 | 1 | 0,897 | -0,157 | -0,096 | 0,923256 | 0,93395 | 1,00E-08 | 1,86547 | 1,188179 | 1,188179 | TRUE | 0,459264 | No |
| tF(GAA)4 | 0 | 0 | 0 | 0 | 0 | 0 | 0 | 0 | 0 | 0 | 0 | 0 | 0 | 0 | 0 | 0 | 0 | 0 | 0 | NA | NA | NA | NA | NA | NA | NA | NA | NA | NA | NA | NA |
| tF(GAA)5 | 0 | 0 | 0 | 0 | 0 | 0 | 0 | 0 | 0 | 0 | 0 | 0 | 0 | 0 | 0 | 0 | 0 | 0 | 0 | NA | NA | NA | NA | NA | NA | NA | NA | NA | NA | NA | NA |
| tG(CCC)1 | 2 | 1 | 0 | 0 | 0 | 0 | 0 | 0 | 2 | 1 | 0 | 0 | 0 | 0 | 0 | 0 | 0,35 | 1 | 0 | 0,234 | -2,098 | -0,837 | 0,402518 | 0,44334 | 1,00E-08 | 3,733262 | 4,3038296 | 4,3038296 | TRUE | 0,2355702 | No |
| tG(GCC)1 | 0 | 0 | 0 | 0 | 0 | 0 | 0 | 0 | 0 | 0 | 0 | 0 | 0 | 0 | 0 | 0 | 0 | 0 | 0 | NA | NA | NA | NA | NA | NA | NA | NA | NA | NA | NA | NA |
| tG(GCC)2 | 0 | 0 | 0 | 1 | 0 | 0 | 0 | 0 | 0 | 0 | 0 | 2 | 0 | 0 | 0 | 0 | 0,19 | 0 | 0 | 0,528 | -0,92 | -0,26 | 0,794572 | 0,821771 | 1,00E-08 | 7,03723 | 10 | 10 | TRUE | 0,6271619 | No |
| tG(GCC)3 | 0 | 0 | 0 | 0 | 0 | 0 | 0 | 0 | 0 | 0 | 0 | 0 | 0 | 0 | 0 | 0 | 0 | 0 | 0 | NA | NA | NA | NA | NA | NA | NA | NA | NA | NA | NA | NA |
| tG(GCC)4 | 0 | 1 | 2 | 0 | 0 | 0 | 0 | 1 | 0 | 1 | 2 | 0 | 0 | 0 | 0 | 1 | 0,46 | 1 | 0 | 0,375 | -1,413 | -0,671 | 0,502368 | 0,544104 | 1,00E-08 | 2,883936 | 2,5517509 | 2,5517509 | TRUE | 0,4097428 | No |
| tG(GCC)5 | 0 | 0 | 1 | 0 | 1 | 1 | 0 | 0 | 0 | 0 | 1 | 0 | 1 | 1 | 0 | 0 | 0,37 | 0 | 0 | 1,471 | 0,557 | 0,246 | 0,805702 | 0,831394 | 1,00E-08 | 3,619228 | 2,9960624 | 2,9960624 | TRUE | 0,2762674 | No |
| tG(GCC)6 | 3 | 0 | 0 | 0 | 2 | 1 | 1 | 7 | 3 | 0 | 0 | 0 | 2 | 1 | 1 | 5 | 1,49 | 1 | 2 | 3,167 | 1,663 | 1,225 | 0,220662 | 0,253874 | 0,8148391 | 0,885661 | 0,8656455 | 0,8656455 | TRUE | 1,4514671 | No |
| tG(UCC)1 | 0 | 0 | 0 | 0 | 0 | 0 | 0 | 0 | 0 | 0 | 0 | 0 | 0 | 0 | 0 | 0 | 0 | 0 | 0 | NA | NA | NA | NA | NA | NA | NA | NA | NA | NA | NA | NA |
| tG(UCC)2 | 0 | 2 | 0 | 0 | 0 | 0 | 0 | 0 | 0 | 2 | 0 | 0 | 0 | 0 | 0 | 0 | 0,25 | 0 | 0 | 0,314 | -1,673 | -0,501 | 0,616545 | 0,655407 | 10 | 5,226504 | 8,8150756 | 8,8150756 | TRUE | 0,8523697 | No |
| tH(GUG)1 | 2 | 0 | 0 | 0 | 2 | 1 | 0 | 1 | 2 | 0 | 0 | 0 | 2 | 1 | 0 | 1 | 0,68 | 0 | 1 | 1,762 | 0,817 | 0,463 | 0,643459 | 0,681178 | 1,00E-08 | 1,941692 | 1,5430885 | 1,5430885 | TRUE | 0,8262211 | No |
| tH(GUG)3 | 10 | 4 | 9 | 2 | 3 | 8 | 7 | 14 | 9 | 4 | 9 | 3 | 3 | 9 | 7 | 11 | 6,73 | 6 | 8 | 1,148 | 0,2 | 0,348 | 0,727861 | 0,7603 | 0,1131576 | 0,198196 | 0,1688647 | 0,1688647 | TRUE | 0,3892047 | No |
| tI(AAU)1 | 0 | 0 | 0 | 0 | 0 | 0 | 0 | 0 | 0 | 0 | 0 | 0 | 0 | 0 | 0 | 0 | 0 | 0 | 0 | NA | NA | NA | NA | NA | NA | NA | NA | NA | NA | NA | NA |
| tI(AAU)2 | 0 | 0 | 0 | 0 | 0 | 0 | 0 | 0 | 0 | 0 | 0 | 0 | 0 | 0 | 0 | 0 | 0 | 0 | 0 | NA | NA | NA | NA | NA | NA | NA | NA | NA | NA | NA | NA |
| tI(AAU)3 | 0 | 0 | 0 | 0 | 5 | 5 | 2 | 5 | 0 | 0 | 0 | 0 | 4 | 5 | 2 | 4 | 1,97 | 0 | 4 | 19,975 | 4,32 | 3,364 | 0,000767 | 0,001091 | 1,00E-08 | 0,673521 | 0,4654557 | 0,4654557 | TRUE | 0,1608302 | No |
| tI(AAU)4 | 0 | 0 | 0 | 0 | 0 | 0 | 0 | 1 | 0 | 0 | 0 | 0 | 0 | 0 | 0 | 1 | 0,1 | 0 | 0 | 1,436 | 0,523 | 0,148 | 0,882445 | 0,8988 | 1,00E-08 | 13,85609 | 10 | 10 | TRUE | 0,2341787 | No |
| tI(AAU)5 | 0 | 0 | 0 | 0 | 0 | 0 | 0 | 0 | 0 | 0 | 0 | 0 | 0 | 0 | 0 | 0 | 0 | 0 | 0 | NA | NA | NA | NA | NA | NA | NA | NA | NA | NA | NA | NA |
| tI(UAU)1 | 0 | 0 | 1 | 0 | 1 | 0 | 1 | 0 | 0 | 0 | 1 | 0 | 1 | 0 | 1 | 0 | 0,36 | 0 | 0 | 1,471 | 0,557 | 0,245 | 0,806545 | 0,831994 | 1,00E-08 | 3,697131 | 3,0443473 | 3,0443473 | TRUE | 0,2724658 | No |
| tK(CUU)1 | 3 | 6 | 2 | 3 | 6 | 9 | 3 | 8 | 3 | 6 | 2 | 5 | 5 | 10 | 3 | 6 | 4,94 | 4 | 6 | 1,621 | 0,697 | 1,072 | 0,283535 | 0,320031 | 1,00E-08 | 0,269702 | 0,1859709 | 0,1859709 | TRUE | 0,3023659 | No |
| tK(CUU)2 | 1 | 1 | 5 | 3 | 9 | 2 | 1 | 3 | 1 | 1 | 5 | 5 | 8 | 2 | 1 | 2 | 3,08 | 3 | 3 | 1,264 | 0,338 | 0,381 | 0,703534 | 0,738034 | 0,4110444 | 0,430766 | 0,4229796 | 0,4229796 | TRUE | 0,7286598 | No |
| tK(UUU)1 | 0 | 0 | 0 | 0 | 0 | 0 | 0 | 0 | 0 | 0 | 0 | 0 | 0 | 0 | 0 | 0 | 0 | 0 | 0 | NA | NA | NA | NA | NA | NA | NA | NA | NA | NA | NA | NA |
| tK(UUU)2 | 0 | 0 | 0 | 0 | 0 | 0 | 0 | 0 | 0 | 0 | 0 | 0 | 0 | 0 | 0 | 0 | 0 | 0 | 0 | NA | NA | NA | NA | NA | NA | NA | NA | NA | NA | NA | NA |
| tK(UUU)3 | 1 | 0 | 0 | 0 | 1 | 0 | 0 | 0 | 1 | 0 | 0 | 0 | 1 | 0 | 0 | 0 | 0,22 | 0 | 0 | 0,871 | -0,199 | -0,068 | 0,945712 | 0,952989 | 1,00E-08 | 5,873379 | 6,1895949 | 6,1895949 | TRUE | 0,3214243 | No |
| tK(UUU)4 | 0 | 0 | 0 | 0 | 0 | 0 | 0 | 0 | 0 | 0 | 0 | 0 | 0 | 0 | 0 | 0 | 0 | 0 | 0 | NA | NA | NA | NA | NA | NA | NA | NA | NA | NA | NA | NA |
| tK(UUU)5 | 0 | 0 | 0 | 0 | 0 | 0 | 0 | 1 | 0 | 0 | 0 | 0 | 0 | 0 | 0 | 1 | 0,1 | 0 | 0 | 1,436 | 0,523 | 0,148 | 0,882445 | 0,8988 | 1,00E-08 | 13,85609 | 10 | 10 | TRUE | 0,2341787 | No |
| tL(AAG)1 | 3 | 3 | 2 | 4 | 5 | 3 | 2 | 5 | 3 | 3 | 2 | 6 | 4 | 3 | 2 | 4 | 3,4 | 4 | 3 | 1,058 | 0,081 | 0,11 | 0,912615 | 0,925242 | 1,00E-08 | 0,390083 | 0,2226611 | 0,2226611 | TRUE | 0,2272489 | No |
| tL(AAG)2 | 0 | 0 | 1 | 1 | 0 | 0 | 2 | 1 | 0 | 0 | 1 | 2 | 0 | 0 | 2 | 1 | 0,66 | 1 | 1 | 1,291 | 0,368 | 0,202 | 0,83979 | 0,862485 | 1,00E-08 | 2,011301 | 1,6084171 | 1,6084171 | TRUE | 0,5133243 | No |
| tL(CAA)1 | 0 | 0 | 2 | 0 | 0 | 0 | 0 | 0 | 0 | 0 | 2 | 0 | 0 | 0 | 0 | 0 | 0,24 | 0 | 0 | 0,328 | -1,608 | -0,471 | 0,637838 | 0,675901 | 1,00E-08 | 5,585235 | 9,2797791 | 9,2797791 | TRUE | 0,8327704 | No |
| tL(CAA)2 | 0 | 0 | 1 | 0 | 1 | 0 | 0 | 0 | 0 | 0 | 1 | 0 | 1 | 0 | 0 | 0 | 0,23 | 0 | 0 | 0,871 | -0,199 | -0,069 | 0,945365 | 0,952989 | 1,00E-08 | 5,758401 | 6,0858471 | 6,0858471 | TRUE | 0,3214244 | No |
| tL(CAA)3 | 0 | 1 | 0 | 0 | 0 | 0 | 0 | 1 | 0 | 1 | 0 | 0 | 0 | 0 | 0 | 1 | 0,22 | 0 | 0 | 0,871 | -0,199 | -0,068 | 0,945963 | 0,952989 | 1,00E-08 | 5,95851 | 6,2661203 | 6,2661203 | TRUE | 0,3156264 | No |
| tL(CAA)4 | 71 | 48 | 74 | 40 | 77 | 66 | 77 | 122 | 65 | 49 | 70 | 60 | 68 | 72 | 79 | 93 | 69,42 | 61 | 78 | 1,285 | 0,362 | 2,007 | 0,044784 | 0,055707 | 0,0091968 | 0,021065 | 0,016505 | 0,016505 | TRUE | 0,1797691 | No |
| tL(CAA)5 | 0 | 0 | 0 | 0 | 0 | 0 | 0 | 0 | 0 | 0 | 0 | 0 | 0 | 0 | 0 | 0 | 0 | 0 | 0 | NA | NA | NA | NA | NA | NA | NA | NA | NA | NA | NA | NA |
| tL(CAA)6 | 2 | 0 | 1 | 0 | 1 | 1 | 0 | 0 | 2 | 0 | 1 | 0 | 1 | 1 | 0 | 0 | 0,59 | 1 | 0 | 0,64 | -0,644 | -0,348 | 0,728063 | 0,760387 | 1,00E-08 | 2,230387 | 1,6422148 | 1,6422148 | TRUE | 0,4201697 | No |
| tL(UAA)1 | 0 | 0 | 0 | 0 | 0 | 0 | 0 | 0 | 0 | 0 | 0 | 0 | 0 | 0 | 0 | 0 | 0 | 0 | 0 | NA | NA | NA | NA | NA | NA | NA | NA | NA | NA | NA | NA |
| tL(UAA)2 | 0 | 0 | 1 | 0 | 0 | 0 | 0 | 0 | 0 | 0 | 1 | 0 | 0 | 0 | 0 | 0 | 0,12 | 0 | 0 | 0,528 | -0,92 | -0,26 | 0,794572 | 0,821771 | 1,00E-08 | 11,16843 | 10 | 10 | TRUE | 0,2762646 | No |
| tL(UAA)3 | 1 | 0 | 1 | 0 | 1 | 0 | 0 | 0 | 1 | 0 | 1 | 0 | 1 | 0 | 0 | 0 | 0,34 | 0 | 0 | 0,544 | -0,878 | -0,384 | 0,701276 | 0,735786 | 1,00E-08 | 3,850067 | 3,099213 | 3,099213 | TRUE | 0,2922827 | No |
| tL(UAA)4 | 1 | 0 | 0 | 1 | 4 | 0 | 2 | 3 | 1 | 0 | 0 | 2 | 4 | 0 | 2 | 2 | 1,29 | 1 | 2 | 3,772 | 1,915 | 1,355 | 0,175272 | 0,204159 | 1,00E-08 | 1,027341 | 0,7677647 | 0,7677647 | TRUE | 0,2364685 | No |
| tL(UAA)5 | 0 | 0 | 0 | 0 | 0 | 1 | 0 | 0 | 0 | 0 | 0 | 0 | 0 | 1 | 0 | 0 | 0,14 | 0 | 0 | 1,436 | 0,523 | 0,148 | 0,882445 | 0,8988 | 1,00E-08 | 9,73564 | 10 | 10 | TRUE | 0,4622362 | No |
| tM(CAU)1 | 0 | 0 | 0 | 0 | 0 | 0 | 0 | 0 | 0 | 0 | 0 | 0 | 0 | 0 | 0 | 0 | 0 | 0 | 0 | NA | NA | NA | NA | NA | NA | NA | NA | NA | NA | NA | NA |
| tM(CAU)2 | 1 | 1 | 0 | 0 | 0 | 0 | 0 | 1 | 1 | 1 | 0 | 0 | 0 | 0 | 0 | 1 | 0,34 | 0 | 0 | 0,535 | -0,904 | -0,393 | 0,694404 | 0,729295 | 1,00E-08 | 3,938471 | 3,1566505 | 3,1566505 | TRUE | 0,2041508 | No |
| tM(CAU)3 | 0 | 0 | 0 | 1 | 0 | 0 | 0 | 0 | 0 | 0 | 0 | 2 | 0 | 0 | 0 | 0 | 0,19 | 0 | 0 | 0,528 | -0,92 | -0,26 | 0,794572 | 0,821771 | 1,00E-08 | 7,03723 | 10 | 10 | TRUE | 0,6271619 | No |
| tM(CAU)4 | 0 | 0 | 0 | 0 | 0 | 2 | 0 | 0 | 0 | 0 | 0 | 0 | 0 | 2 | 0 | 0 | 0,27 | 0 | 0 | 2,582 | 1,368 | 0,416 | 0,677143 | 0,713165 | 10 | 4,868838 | 8,4753903 | 8,4753903 | TRUE | 1,0614604 | No |
| tN(GUU)1 | 2 | 0 | 0 | 0 | 0 | 0 | 0 | 0 | 2 | 0 | 0 | 0 | 0 | 0 | 0 | 0 | 0,23 | 0 | 0 | 0,337 | -1,57 | -0,453 | 0,650347 | 0,68767 | 1,00E-08 | 5,805701 | 9,5796318 | 9,5796318 | TRUE | 0,8211616 | No |
| tN(GUU)2 | 0 | 0 | 0 | 0 | 0 | 0 | 0 | 0 | 0 | 0 | 0 | 0 | 0 | 0 | 0 | 0 | 0 | 0 | 0 | NA | NA | NA | NA | NA | NA | NA | NA | NA | NA | NA | NA |
| tN(GUU)3 | 0 | 1 | 0 | 0 | 1 | 0 | 1 | 0 | 0 | 1 | 0 | 0 | 1 | 0 | 1 | 0 | 0,37 | 0 | 0 | 1,471 | 0,557 | 0,246 | 0,805573 | 0,831394 | 1,00E-08 | 3,615037 | 2,9887284 | 2,9887284 | TRUE | 0,3156284 | No |
| tN(GUU)4 | 0 | 1 | 0 | 0 | 0 | 0 | 0 | 0 | 0 | 1 | 0 | 0 | 0 | 0 | 0 | 0 | 0,13 | 0 | 0 | 0,528 | -0,92 | -0,26 | 0,794572 | 0,821771 | 1,00E-08 | 10,45097 | 10 | 10 | TRUE | 0,3156251 | No |
| tP(AGG)1 | 1 | 0 | 2 | 1 | 2 | 1 | 1 | 4 | 1 | 0 | 2 | 2 | 2 | 1 | 1 | 3 | 1,41 | 1 | 2 | 1,692 | 0,758 | 0,648 | 0,516939 | 0,559031 | 1,00E-08 | 0,941793 | 0,5520396 | 0,5520396 | TRUE | 0,2788799 | No |
| tP(UGG)1 | 0 | 0 | 1 | 0 | 0 | 0 | 0 | 0 | 0 | 0 | 1 | 0 | 0 | 0 | 0 | 0 | 0,12 | 0 | 0 | 0,528 | -0,92 | -0,26 | 0,794572 | 0,821771 | 1,00E-08 | 11,16843 | 10 | 10 | TRUE | 0,2762646 | No |
| tP(UGG)2 | 2 | 0 | 1 | 0 | 0 | 0 | 0 | 1 | 2 | 0 | 1 | 0 | 0 | 0 | 0 | 1 | 0,44 | 1 | 0 | 0,386 | -1,372 | -0,647 | 0,51789 | 0,559869 | 1,00E-08 | 2,995687 | 2,5970863 | 2,5970863 | TRUE | 0,4205281 | No |
| tP(UGG)3 | 4 | 4 | 2 | 3 | 1 | 2 | 2 | 2 | 4 | 4 | 2 | 5 | 1 | 2 | 2 | 2 | 2,59 | 4 | 2 | 0,471 | -1,086 | -1,259 | 0,208209 | 0,240459 | 1,00E-08 | 0,512038 | 0,2761102 | 0,2761102 | TRUE | 0,1612491 | No |
| tP(UGG)4 | 0 | 0 | 0 | 0 | 1 | 0 | 0 | 0 | 0 | 0 | 0 | 0 | 1 | 0 | 0 | 0 | 0,11 | 0 | 0 | 1,436 | 0,523 | 0,148 | 0,882445 | 0,8988 | 1,00E-08 | 11,88327 | 10 | 10 | TRUE | 0,3214251 | No |
| tP(UGG)5 | 0 | 0 | 0 | 1 | 0 | 1 | 0 | 1 | 0 | 0 | 0 | 2 | 0 | 1 | 0 | 1 | 0,42 | 0 | 0 | 1,407 | 0,493 | 0,226 | 0,821268 | 0,845267 | 1,00E-08 | 3,15601 | 2,618715 | 2,618715 | TRUE | 0,6271695 | No |
| tQ(CUG)1 | 0 | 0 | 0 | 0 | 0 | 2 | 0 | 0 | 0 | 0 | 0 | 0 | 0 | 2 | 0 | 0 | 0,27 | 0 | 0 | 2,582 | 1,368 | 0,416 | 0,677143 | 0,713165 | 10 | 4,868838 | 8,4753903 | 8,4753903 | TRUE | 1,0614604 | No |
| tQ(UUG)1 | 0 | 0 | 2 | 1 | 0 | 0 | 0 | 0 | 0 | 0 | 2 | 2 | 0 | 0 | 0 | 0 | 0,42 | 1 | 0 | 0,206 | -2,276 | -0,949 | 0,342405 | 0,382416 | 1,00E-08 | 3,114864 | 3,8722757 | 3,8722757 | TRUE | 0,0832387 | No |
| tQ(UUG)3 | 0 | 2 | 0 | 1 | 1 | 1 | 2 | 1 | 0 | 2 | 0 | 2 | 1 | 1 | 2 | 1 | 1,04 | 1 | 1 | 1,432 | 0,518 | 0,372 | 0,710044 | 0,743883 | 1,00E-08 | 1,273573 | 0,8209074 | 0,8209074 | TRUE | 0,3936781 | No |
| tQ(UUG)4 | 0 | 0 | 0 | 0 | 0 | 0 | 0 | 0 | 0 | 0 | 0 | 0 | 0 | 0 | 0 | 0 | 0 | 0 | 0 | NA | NA | NA | NA | NA | NA | NA | NA | NA | NA | NA | NA |
| tQ(UUG)5 | 0 | 1 | 1 | 0 | 1 | 0 | 0 | 0 | 0 | 1 | 1 | 0 | 1 | 0 | 0 | 0 | 0,36 | 0 | 0 | 0,529 | -0,918 | -0,405 | 0,685591 | 0,720871 | 1,00E-08 | 3,713655 | 3,0229463 | 3,0229463 | TRUE | 0,2825643 | No |
| tR(ACG)1 | 0 | 0 | 1 | 0 | 0 | 0 | 2 | 0 | 0 | 0 | 1 | 0 | 0 | 0 | 2 | 0 | 0,37 | 0 | 0 | 1,535 | 0,619 | 0,242 | 0,809004 | 0,834126 | 1,00E-08 | 3,530977 | 4,3807 | 4,3807 | TRUE | 0,9804015 | No |
| tR(ACG)2 | 0 | 0 | 0 | 0 | 1 | 1 | 2 | 2 | 0 | 0 | 0 | 0 | 1 | 1 | 2 | 2 | 0,69 | 0 | 2 | 7,045 | 2,817 | 1,709 | 0,087477 | 0,105639 | 1,00E-08 | 1,906501 | 1,2734633 | 1,2734633 | TRUE | 0,0616775 | No |
| tR(CCG)1 | 6 | 6 | 8 | 9 | 10 | 6 | 5 | 4 | 5 | 6 | 8 | 14 | 9 | 7 | 5 | 3 | 7,03 | 8 | 6 | 0,743 | -0,429 | -0,785 | 0,432334 | 0,473887 | 1,00E-08 | 0,19007 | 0,1389008 | 0,1389008 | TRUE | 0,3735526 | No |
| tR(CCU)1 | 53 | 78 | 59 | 50 | 35 | 23 | 15 | 36 | 48 | 79 | 56 | 75 | 31 | 25 | 15 | 27 | 44,63 | 64 | 24 | 0,39 | -1,359 | -5,375 | 7,67E-08 | 1,33E-07 | 0,0389893 | 0,031638 | 0,0343095 | 0,0343095 | TRUE | 0,2068957 | No |
| tR(UCU)1 | 3 | 1 | 1 | 0 | 0 | 0 | 0 | 0 | 3 | 1 | 1 | 0 | 0 | 0 | 0 | 0 | 0,59 | 1 | 0 | 0,141 | -2,831 | -1,487 | 0,136955 | 0,161653 | 1,00E-08 | 2,255734 | 2,0398045 | 2,0398045 | TRUE | 0,2261017 | No |
| tR(UCU)2 | 0 | 0 | 0 | 0 | 0 | 0 | 0 | 0 | 0 | 0 | 0 | 0 | 0 | 0 | 0 | 0 | 0 | 0 | 0 | NA | NA | NA | NA | NA | NA | NA | NA | NA | NA | NA | NA |
| tR(UCU)3 | 0 | 0 | 0 | 0 | 0 | 1 | 0 | 0 | 0 | 0 | 0 | 0 | 0 | 1 | 0 | 0 | 0,14 | 0 | 0 | 1,436 | 0,523 | 0,148 | 0,882445 | 0,8988 | 1,00E-08 | 9,73564 | 10 | 10 | TRUE | 0,4622362 | No |
| tR(UCU)4 | 4 | 2 | 6 | 3 | 3 | 5 | 7 | 11 | 4 | 2 | 6 | 5 | 3 | 5 | 7 | 8 | 4,94 | 4 | 6 | 1,508 | 0,593 | 0,927 | 0,353721 | 0,393607 | 1,00E-08 | 0,269538 | 0,1796121 | 0,1796121 | TRUE | 0,3698062 | No |

| | | | | | | | | | | | | | | | | | | | | | | | | | | | | | | | |
|---|---|---|---|---|---|---|---|---|---|---|---|---|---|---|---|---|---|---|---|---|---|---|---|---|---|---|---|---|---|---|---|
| tR(UCU)5 | 0 | 0 | 0 | 0 | 0 | 0 | 0 | 0 | 0 | 0 | 0 | 0 | 0 | 0 | 0 | 0 | 0 | 0 | 0 | NA | NA | NA | NA | NA | NA | NA | NA | NA | NA | NA | NA |
| tS(AGA)1 | 2 | 4 | 2 | 1 | 1 | 0 | 1 | 5 | 2 | 4 | 2 | 2 | 1 | 0 | 1 | 4 | 1,87 | 2 | 2 | 0,652 | -0,617 | -0,592 | 0,553888 | 0,595353 | 0,1349565 | 0,707081 | 0,5108199 | 0,5108199 | TRUE | 1,1011914 | No |
| tS(AGA)2 | 0 | 0 | 0 | 0 | 0 | 0 | 0 | 0 | 0 | 0 | 0 | 0 | 0 | 0 | 0 | 0 | 0 | 0 | 0 | NA | NA | NA | NA | NA | NA | NA | NA | NA | NA | NA | NA |
| tS(AGA)3 | 1 | 0 | 1 | 0 | 0 | 2 | 3 | 3 | 1 | 0 | 1 | 0 | 0 | 2 | 3 | 2 | 1,17 | 0 | 2 | 3,553 | 1,829 | 1,265 | 0,205775 | 0,23782 | 1,00E-08 | 1,127594 | 0,8278224 | 0,8278224 | TRUE | 0,4437123 | No |
| tS(AGA)4 | 2 | 0 | 0 | 1 | 2 | 0 | 1 | 0 | 2 | 0 | 0 | 2 | 2 | 0 | 1 | 0 | 0,77 | 1 | 1 | 0,868 | -0,204 | -0,118 | 0,905864 | 0,919275 | 1,00E-08 | 1,727101 | 1,5173113 | 1,5173113 | TRUE | 0,3129416 | No |
| tS(CAG)1 | 0 | 0 | 1 | 0 | 1 | 0 | 1 | 0 | 0 | 0 | 1 | 0 | 1 | 0 | 1 | 0 | 0,36 | 0 | 0 | 1,471 | 0,557 | 0,245 | 0,806545 | 0,831994 | 1,00E-08 | 3,697131 | 3,0443473 | 3,0443473 | TRUE | 0,2724658 | No |
| tS(CGA)1 | 0 | 0 | 1 | 0 | 1 | 0 | 0 | 4 | 0 | 0 | 1 | 0 | 1 | 0 | 0 | 3 | 0,61 | 0 | 1 | 3,177 | 1,668 | 0,846 | 0,397647 | 0,438505 | 2,7400784 | 2,164457 | 2,2821572 | 2,2821572 | TRUE | 1,1994928 | No |
| tS(GCU)1 | 10 | 13 | 8 | 5 | 55 | 25 | 25 | 32 | 9 | 13 | 8 | 8 | 49 | 27 | 26 | 24 | 20,43 | 10 | 32 | 3,352 | 1,745 | 4,616 | 3,90E-06 | 6,33E-06 | 0,0654335 | 0,066708 | 0,0660226 | 0,0660226 | TRUE | 1,0096645 | No |
| tS(GCU)2 | 8 | 6 | 5 | 7 | 32 | 14 | 13 | 33 | 7 | 6 | 5 | 11 | 28 | 15 | 13 | 25 | 13,84 | 7 | 20 | 3,003 | 1,587 | 3,645 | 0,000268 | 0,000392 | 1,00E-08 | 0,09746 | 0,0827946 | 0,0827946 | TRUE | 0,0974751 | No |
| tS(UGA)1 | 0 | 0 | 0 | 0 | 0 | 1 | 0 | 0 | 0 | 0 | 0 | 0 | 0 | 1 | 0 | 0 | 0,14 | 0 | 0 | 1,436 | 0,523 | 0,148 | 0,882445 | 0,8988 | 1,00E-08 | 9,73564 | 10 | 10 | TRUE | 0,4622362 | No |
| tS(UGA)2 | 0 | 0 | 1 | 0 | 0 | 0 | 0 | 1 | 0 | 0 | 1 | 0 | 0 | 0 | 0 | 1 | 0,21 | 0 | 0 | 0,871 | -0,199 | -0,067 | 0,946611 | 0,953341 | 1,00E-08 | 6,184973 | 6,4683949 | 6,4683949 | TRUE | 0,2762655 | No |
| tS(UGA)3 | 0 | 0 | 0 | 0 | 0 | 0 | 0 | 1 | 0 | 0 | 0 | 0 | 0 | 0 | 0 | 1 | 0,1 | 0 | 0 | 1,436 | 0,523 | 0,148 | 0,882445 | 0,8988 | 1,00E-08 | 13,85609 | 10 | 10 | TRUE | 0,2341787 | No |
| tT(AGU)1 | 0 | 0 | 0 | 0 | 0 | 0 | 0 | 0 | 0 | 0 | 0 | 0 | 0 | 0 | 0 | 0 | 0 | 0 | 0 | NA | NA | NA | NA | NA | NA | NA | NA | NA | NA | NA | NA |
| tT(AGU)3 | 0 | 0 | 0 | 0 | 0 | 0 | 0 | 1 | 0 | 0 | 0 | 0 | 0 | 0 | 0 | 1 | 0,1 | 0 | 0 | 1,436 | 0,523 | 0,148 | 0,882445 | 0,8988 | 1,00E-08 | 13,85609 | 10 | 10 | TRUE | 0,2341787 | No |
| tT(AGU)4 | 0 | 0 | 0 | 0 | 1 | 0 | 0 | 0 | 0 | 0 | 0 | 0 | 1 | 0 | 0 | 0 | 0,11 | 0 | 0 | 1,436 | 0,523 | 0,148 | 0,882445 | 0,8988 | 1,00E-08 | 11,88327 | 10 | 10 | TRUE | 0,3214251 | No |
| tT(AGU)6 | 1 | 0 | 0 | 0 | 0 | 0 | 0 | 0 | 1 | 0 | 0 | 0 | 0 | 0 | 0 | 0 | 0,11 | 0 | 0 | 0,528 | -0,92 | -0,26 | 0,794572 | 0,821771 | 1,00E-08 | 11,60937 | 10 | 10 | TRUE | 0,2548005 | No |
| tT(CGU)1 | 0 | 0 | 0 | 0 | 0 | 1 | 0 | 0 | 0 | 0 | 0 | 0 | 0 | 1 | 0 | 0 | 0,14 | 0 | 0 | 1,436 | 0,523 | 0,148 | 0,882445 | 0,8988 | 1,00E-08 | 9,73564 | 10 | 10 | TRUE | 0,4622362 | No |
| tT(UGU)1 | 0 | 0 | 0 | 0 | 0 | 0 | 0 | 0 | 0 | 0 | 0 | 0 | 0 | 0 | 0 | 0 | 0 | 0 | 0 | NA | NA | NA | NA | NA | NA | NA | NA | NA | NA | NA | NA |
| tT(UGU)2 | 0 | 0 | 0 | 0 | 0 | 0 | 0 | 0 | 0 | 0 | 0 | 0 | 0 | 0 | 0 | 0 | 0 | 0 | 0 | NA | NA | NA | NA | NA | NA | NA | NA | NA | NA | NA | NA |
| tV(AAC)1 | 1 | 0 | 0 | 0 | 1 | 0 | 0 | 1 | 1 | 0 | 0 | 0 | 1 | 0 | 0 | 1 | 0,32 | 0 | 0 | 1,397 | 0,482 | 0,207 | 0,836221 | 0,859132 | 1,00E-08 | 4,125751 | 3,2724409 | 3,2724409 | TRUE | 0,2548029 | No |
| tV(AAC)2 | 0 | 0 | 0 | 0 | 0 | 0 | 0 | 0 | 0 | 0 | 0 | 0 | 0 | 0 | 0 | 0 | 0 | 0 | 0 | NA | NA | NA | NA | NA | NA | NA | NA | NA | NA | NA | NA |
| tV(AAC)3 | 0 | 0 | 0 | 0 | 0 | 0 | 0 | 0 | 0 | 0 | 0 | 0 | 0 | 0 | 0 | 0 | 0 | 0 | 0 | NA | NA | NA | NA | NA | NA | NA | NA | NA | NA | NA | NA |
| tV(AAC)4 | 0 | 0 | 0 | 0 | 0 | 1 | 0 | 0 | 0 | 0 | 0 | 0 | 0 | 1 | 0 | 0 | 0,14 | 0 | 0 | 1,436 | 0,523 | 0,148 | 0,882445 | 0,8988 | 1,00E-08 | 9,73564 | 10 | 10 | TRUE | 0,4622362 | No |
| tV(AAC)5 | 2 | 1 | 0 | 1 | 0 | 2 | 1 | 0 | 2 | 1 | 0 | 2 | 0 | 2 | 1 | 0 | 0,94 | 1 | 1 | 0,697 | -0,521 | -0,343 | 0,731923 | 0,764168 | 1,00E-08 | 1,405481 | 1,0927178 | 1,0927178 | TRUE | 0,4780112 | No |
| tV(AAC)6 | 1 | 2 | 0 | 0 | 2 | 1 | 1 | 1 | 1 | 2 | 0 | 0 | 2 | 1 | 1 | 1 | 0,95 | 1 | 1 | 1,502 | 0,587 | 0,417 | 0,676793 | 0,713131 | 1,00E-08 | 1,395341 | 0,81389 | 0,81389 | TRUE | 0,4553501 | No |
| tV(CAC)1 | 1 | 0 | 2 | 1 | 2 | 0 | 0 | 2 | 1 | 0 | 2 | 2 | 2 | 0 | 0 | 2 | 0,95 | 1 | 1 | 0,827 | -0,275 | -0,189 | 0,850175 | 0,871922 | 1,00E-08 | 1,390736 | 0,9962997 | 0,9962997 | TRUE | 0,2117241 | No |
| tV(UAC)1 | 8 | 9 | 9 | 10 | 10 | 15 | 16 | 20 | 7 | 9 | 9 | 15 | 9 | 16 | 16 | 15 | 12,09 | 10 | 14 | 1,467 | 0,553 | 1,345 | 0,178679 | 0,207861 | 1,00E-08 | 0,111269 | 0,0743621 | 0,0743621 | TRUE | 0,308263 | No |
| tW(CCA)1 | 0 | 0 | 1 | 1 | 0 | 0 | 0 | 0 | 0 | 0 | 1 | 2 | 0 | 0 | 0 | 0 | 0,31 | 1 | 0 | 0,313 | -1,675 | -0,631 | 0,527957 | 0,569497 | 1,00E-08 | 4,31802 | 4,8682685 | 4,8682685 | TRUE | 0,1642732 | No |
| tW(CCA)2 | 1 | 0 | 0 | 0 | 1 | 0 | 1 | 0 | 1 | 0 | 0 | 0 | 1 | 0 | 1 | 0 | 0,35 | 0 | 0 | 1,472 | 0,557 | 0,244 | 0,807092 | 0,832424 | 1,00E-08 | 3,744171 | 3,0760699 | 3,0760699 | TRUE | 0,247888 | No |
| tY(GUA)1 | 0 | 1 | 0 | 0 | 0 | 0 | 0 | 1 | 0 | 1 | 0 | 0 | 0 | 0 | 0 | 1 | 0,22 | 0 | 0 | 0,871 | -0,199 | -0,068 | 0,945963 | 0,952989 | 1,00E-08 | 5,95851 | 6,2661203 | 6,2661203 | TRUE | 0,3156264 | No |
| tY(GUA)2 | 1 | 0 | 0 | 0 | 2 | 0 | 1 | 1 | 1 | 0 | 0 | 0 | 2 | 0 | 1 | 1 | 0,56 | 0 | 1 | 2,79 | 1,48 | 0,815 | 0,414884 | 0,456251 | 1,00E-08 | 2,362896 | 1,6516297 | 1,6516297 | TRUE | 0,213387 | No |
| tY(GUA)3 | 8 | 4 | 5 | 5 | 3 | 9 | 9 | 11 | 7 | 4 | 5 | 8 | 3 | 10 | 9 | 8 | 6,7 | 6 | 8 | 1,281 | 0,357 | 0,642 | 0,52107 | 0,562925 | 1,00E-08 | 0,199127 | 0,1427338 | 0,1427338 | TRUE | 0,5952673 | No |
| tY(GUA)4 | 0 | 0 | 0 | 0 | 0 | 0 | 0 | 0 | 0 | 0 | 0 | 0 | 0 | 0 | 0 | 0 | 0 | 0 | 0 | NA | NA | NA | NA | NA | NA | NA | NA | NA | NA | NA | NA |

$P_{TET}$-*ZCF26* doxycycline-treated complete data

| | WT dox | | | | ZCF26 | | | | norm. WT dox | | | | norm. ZCF26 | | | | | | | | | | | | | | | | | | |
|---|---|---|---|---|---|---|---|---|---|---|---|---|---|---|---|---|---|---|---|---|---|---|---|---|---|---|---|---|---|---|---|
| Gene ID | 1 | 2 | 3 | 4 | 1 | 2 | 3 | 4 | 1 | 2 | 3 | 4 | 1 | 2 | 3 | 4 | baseMean | WT | ZCF26 | FC | log2FC | stat | pvalue | padj | dispGeneEst | dispFit | dispMAP | dispersion | betaConv | maxCooks | outlier |
| CaalfMp01 | 20 | 33 | 22 | 23 | 14 | 13 | 13 | 17 | 19 | 35 | 22 | 36 | 11 | 15 | 11 | 14 | 20,32 | 28 | 13 | 0,46 | -1,121 | -3,253 | 0,00114 | 0,0015311 | 1,00E-08 | 0,082881 | 0,0584311 | 0,0584311 | TRUE | 0,1065319 | No |
| CaalfMp02 | 1 | 3 | 4 | 3 | 5 | 1 | 2 | 1 | 1 | 3 | 4 | 5 | 4 | 1 | 2 | 1 | 2,55 | 3 | 2 | 0,63 | -0,668 | -0,728 | 0,4663604 | 0,5028377 | 1,00E-08 | 0,6449878 | 0,3993878 | 0,3993878 | TRUE | 0,5978211 | No |
| CaalfMp03 | 8 | 6 | 7 | 5 | 13 | 7 | 9 | 2 | 8 | 6 | 7 | 8 | 10 | 8 | 8 | 2 | 7,05 | 7 | 7 | 0,961 | -0,057 | -0,1 | 0,9204514 | 0,9275743 | 1,00E-08 | 0,2351392 | 0,1718531 | 0,1718531 | TRUE | 0,8733482 | No |
| CaalfMp04 | 0 | 0 | 0 | 0 | 0 | 0 | 0 | 0 | 0 | 0 | 0 | 0 | 0 | 0 | 0 | 0 | 0 | 0 | 0 | NA | NA | NA | NA | NA | NA | NA | NA | NA | NA | NA | NA |
| CaalfMp05 | 23 | 40 | 27 | 32 | 14 | 11 | 14 | 12 | 22 | 42 | 27 | 50 | 11 | 13 | 12 | 10 | 23,3 | 35 | 12 | 0,326 | -1,616 | -4,56 | 5,10E-06 | 7,67E-06 | 1,00E-08 | 0,0725611 | 0,0649908 | 0,0649908 | TRUE | 0,0974586 | No |
| CaalfMp06 | 5 | 19 | 13 | 13 | 9 | 2 | 7 | 7 | 5 | 20 | 13 | 20 | 7 | 2 | 6 | 6 | 9,9 | 14 | 5 | 0,38 | -1,397 | -2,547 | 0,010872 | 0,0137702 | 0,1665639 | 0,167971 | 0,167441 | 0,167441 | TRUE | 0,4556861 | No |
| CaalfMp07 | 0 | 0 | 1 | 0 | 1 | 1 | 0 | 0 | 0 | 0 | 1 | 0 | 1 | 1 | 0 | 0 | 0,37 | 0 | 0 | 1,323 | 0,404 | 0,17 | 0,8650426 | 0,8826091 | 1,00E-08 | 4,4908238 | 3,5016363 | 3,5016363 | TRUE | 0,275731 | No |
| CaalfMp08 | 21 | 21 | 20 | 17 | 26 | 14 | 13 | 6 | 20 | 22 | 20 | 27 | 21 | 16 | 11 | 5 | 17,66 | 22 | 13 | 0,596 | -0,746 | -1,866 | 0,0620692 | 0,0739062 | 0,0929134 | 0,0950621 | 0,094153 | 0,094153 | TRUE | 0,5049154 | No |
| CaalfMp08.1 | 0 | 0 | 0 | 0 | 0 | 0 | 0 | 0 | 0 | 0 | 0 | 0 | 0 | 0 | 0 | 0 | 0 | 0 | 0 | NA | NA | NA | NA | NA | NA | NA | NA | NA | NA | NA | NA |
| CaalfMp08.2 | 0 | 0 | 0 | 0 | 0 | 0 | 0 | 0 | 0 | 0 | 0 | 0 | 0 | 0 | 0 | 0 | 0 | 0 | 0 | NA | NA | NA | NA | NA | NA | NA | NA | NA | NA | NA | NA |
| CaalfMp08.3 | 0 | 0 | 0 | 0 | 0 | 0 | 0 | 0 | 0 | 0 | 0 | 0 | 0 | 0 | 0 | 0 | 0 | 0 | 0 | NA | NA | NA | NA | NA | NA | NA | NA | NA | NA | NA | NA |
| CaalfMp08.4 | 0 | 0 | 0 | 0 | 0 | 0 | 0 | 0 | 0 | 0 | 0 | 0 | 0 | 0 | 0 | 0 | 0 | 0 | 0 | NA | NA | NA | NA | NA | NA | NA | NA | NA | NA | NA | NA |
| CaalfMp09 | 5 | 5 | 3 | 2 | 3 | 4 | 3 | 0 | 5 | 5 | 3 | 3 | 2 | 5 | 3 | 0 | 3,21 | 4 | 2 | 0,558 | -0,841 | -1,009 | 0,3131475 | 0,3462903 | 1,00E-08 | 0,5141897 | 0,3353967 | 0,3353967 | TRUE | 0,554102 | No |
| CaalfMp10 | 2 | 1 | 1 | 0 | 0 | 3 | 0 | 0 | 2 | 1 | 1 | 0 | 0 | 3 | 0 | 0 | 0,92 | 1 | 1 | 0,744 | -0,427 | -0,246 | 0,8060289 | 0,8279573 | 1,7676618 | 1,7859613 | 1,7809602 | 1,7809602 | TRUE | 1,4604747 | No |
| CaalfMp11 | 80 | 76 | 97 | 82 | 57 | 64 | 47 | 41 | 76 | 80 | 96 | 128 | 46 | 73 | 40 | 33 | 71,58 | 95 | 48 | 0,501 | -0,997 | -4,031 | 5,54E-05 | 7,94E-05 | 0,0664712 | 0,0250308 | 0,0432961 | 0,0432961 | TRUE | 0,8846031 | No |
| CaalfMp11.1 | 0 | 0 | 0 | 0 | 0 | 0 | 0 | 0 | 0 | 0 | 0 | 0 | 0 | 0 | 0 | 0 | 0 | 0 | 0 | NA | NA | NA | NA | NA | NA | NA | NA | NA | NA | NA | NA |
| CaalfMp12 | 0 | 0 | 0 | 0 | 0 | 0 | 1 | 0 | 0 | 0 | 0 | 0 | 0 | 0 | 1 | 0 | 0,11 | 0 | 0 | 1,314 | 0,394 | 0,111 | 0,9112982 | 0,9205309 | 1,00E-08 | 15,473589 | 10 | 10 | TRUE | 0,3207401 | No |
| CaalfMp13 | 5 | 6 | 10 | 6 | 5 | 8 | 5 | 2 | 5 | 6 | 10 | 9 | 4 | 9 | 4 | 2 | 6,17 | 8 | 5 | 0,608 | -0,718 | -1,168 | 0,2427149 | 0,2724778 | 0,0689502 | 0,2681815 | 0,1910399 | 0,1910399 | TRUE | 0,6147008 | No |
| CaalfMp14 | 3 | 3 | 5 | 3 | 5 | 5 | 10 | 4 | 3 | 3 | 5 | 5 | 4 | 6 | 8 | 3 | 4,64 | 4 | 5 | 1,378 | 0,462 | 0,697 | 0,4855966 | 0,5219121 | 1,00E-08 | 0,3560869 | 0,1946588 | 0,1946588 | TRUE | 0,4073629 | No |
| CaalfMp15 | 0 | 0 | 0 | 0 | 0 | 0 | 0 | 0 | 0 | 0 | 0 | 0 | 0 | 0 | 0 | 0 | 0 | 0 | 0 | NA | NA | NA | NA | NA | NA | NA | NA | NA | NA | NA | NA |
| CaalfMr16 | 1586 | 2242 | 2176 | 1673 | 5945 | 5815 | 3375 | 2104 | 1512 | 2371 | 2150 | 2621 | 4764 | 6612 | 2865 | 1711 | 3075,64 | 2164 | 3988 | 1,843 | 0,882 | 2,098 | 0,0359177 | 0,0437053 | 0,169644 | 0,0026251 | 0,0704374 | 0,169644 | TRUE | 0,183731 | No |
| CaalfMr17 | 399 | 590 | 654 | 470 | 383 | 403 | 310 | 323 | 380 | 624 | 646 | 736 | 307 | 458 | 263 | 263 | 459,7 | 596 | 323 | 0,541 | -0,887 | -3,221 | 0,0012783 | 0,0017096 | 0,0705678 | 0,005663 | 0,0349539 | 0,0705678 | TRUE | 0,8262222 | No |
| CaalfMt18 | 0 | 0 | 0 | 0 | 0 | 0 | 0 | 0 | 0 | 0 | 0 | 0 | 0 | 0 | 0 | 0 | 0 | 0 | 0 | NA | NA | NA | NA | NA | NA | NA | NA | NA | NA | NA | NA |
| CaalfMt19 | 0 | 0 | 0 | 0 | 0 | 0 | 0 | 0 | 0 | 0 | 0 | 0 | 0 | 0 | 0 | 0 | 0 | 0 | 0 | NA | NA | NA | NA | NA | NA | NA | NA | NA | NA | NA | NA |
| CaalfMt20 | 0 | 0 | 0 | 0 | 0 | 0 | 0 | 0 | 0 | 0 | 0 | 0 | 0 | 0 | 0 | 0 | 0 | 0 | 0 | NA | NA | NA | NA | NA | NA | NA | NA | NA | NA | NA | NA |
| CaalfMt21 | 0 | 0 | 0 | 0 | 0 | 0 | 0 | 0 | 0 | 0 | 0 | 0 | 0 | 0 | 0 | 0 | 0 | 0 | 0 | NA | NA | NA | NA | NA | NA | NA | NA | NA | NA | NA | NA |
| CaalfMt22 | 0 | 0 | 0 | 0 | 0 | 0 | 0 | 0 | 0 | 0 | 0 | 0 | 0 | 0 | 0 | 0 | 0 | 0 | 0 | NA | NA | NA | NA | NA | NA | NA | NA | NA | NA | NA | NA |
| CaalfMt23 | 0 | 0 | 0 | 0 | 0 | 0 | 0 | 0 | 0 | 0 | 0 | 0 | 0 | 0 | 0 | 0 | 0 | 0 | 0 | NA | NA | NA | NA | NA | NA | NA | NA | NA | NA | NA | NA |
| CaalfMt24 | 0 | 0 | 0 | 0 | 0 | 0 | 0 | 0 | 0 | 0 | 0 | 0 | 0 | 0 | 0 | 0 | 0 | 0 | 0 | NA | NA | NA | NA | NA | NA | NA | NA | NA | NA | NA | NA |
| CaalfMt25 | 0 | 0 | 0 | 0 | 0 | 0 | 0 | 0 | 0 | 0 | 0 | 0 | 0 | 0 | 0 | 0 | 0 | 0 | 0 | NA | NA | NA | NA | NA | NA | NA | NA | NA | NA | NA | NA |
| CaalfMt26 | 0 | 0 | 1 | 0 | 1 | 0 | 0 | 0 | 0 | 0 | 1 | 0 | 1 | 0 | 0 | 0 | 0,22 | 0 | 0 | 0,797 | -0,328 | -0,104 | 0,9169674 | 0,9246474 | 1,00E-08 | 7,3435845 | 7,4919227 | 7,4919227 | TRUE | 0,2859076 | No |
| CaalfMt27 | 2 | 0 | 1 | 0 | 2 | 0 | 0 | 0 | 2 | 0 | 1 | 0 | 2 | 0 | 0 | 0 | 0,56 | 1 | 0 | 0,57 | -0,81 | -0,378 | 0,7054299 | 0,7357549 | 1,00E-08 | 2,9228276 | 2,7283473 | 2,7283473 | TRUE | 0,871649 | No |
| CaalfMt28 | 0 | 0 | 1 | 0 | 0 | 0 | 0 | 0 | 0 | 0 | 1 | 0 | 0 | 0 | 0 | 0 | 0,12 | 0 | 0 | 0,483 | -1,049 | -0,297 | 0,7665832 | 0,7911318 | 1,00E-08 | 13,298132 | 10 | 10 | TRUE | 0,2757288 | No |
| CaalfMt29 | 0 | 0 | 0 | 0 | 0 | 0 | 0 | 0 | 0 | 0 | 0 | 0 | 0 | 0 | 0 | 0 | 0 | 0 | 0 | NA | NA | NA | NA | NA | NA | NA | NA | NA | NA | NA | NA |
| CaalfMt30 | 0 | 0 | 0 | 0 | 0 | 0 | 0 | 0 | 0 | 0 | 0 | 0 | 0 | 0 | 0 | 0 | 0 | 0 | 0 | NA | NA | NA | NA | NA | NA | NA | NA | NA | NA | NA | NA |
| CaalfMt31 | 0 | 0 | 0 | 0 | 0 | 0 | 0 | 0 | 0 | 0 | 0 | 0 | 0 | 0 | 0 | 0 | 0 | 0 | 0 | NA | NA | NA | NA | NA | NA | NA | NA | NA | NA | NA | NA |
| CaalfMt32 | 0 | 0 | 0 | 0 | 0 | 0 | 0 | 0 | 0 | 0 | 0 | 0 | 0 | 0 | 0 | 0 | 0 | 0 | 0 | NA | NA | NA | NA | NA | NA | NA | NA | NA | NA | NA | NA |
| CaalfMt33 | 0 | 0 | 0 | 0 | 0 | 1 | 0 | 0 | 0 | 0 | 0 | 0 | 0 | 1 | 0 | 0 | 0,14 | 0 | 0 | 1,314 | 0,394 | 0,111 | 0,9112982 | 0,9205309 | 1,00E-08 | 11,553708 | 10 | 10 | TRUE | 0,5388425 | No |
| CaalfMt34 | 0 | 0 | 0 | 0 | 0 | 0 | 0 | 0 | 0 | 0 | 0 | 0 | 0 | 0 | 0 | 0 | 0 | 0 | 0 | NA | NA | NA | NA | NA | NA | NA | NA | NA | NA | NA | NA |
| CaalfMt35 | 0 | 0 | 0 | 0 | 0 | 0 | 0 | 0 | 0 | 0 | 0 | 0 | 0 | 0 | 0 | 0 | 0 | 0 | 0 | NA | NA | NA | NA | NA | NA | NA | NA | NA | NA | NA | NA |
| CaalfMt36 | 0 | 0 | 0 | 0 | 0 | 0 | 0 | 0 | 0 | 0 | 0 | 0 | 0 | 0 | 0 | 0 | 0 | 0 | 0 | NA | NA | NA | NA | NA | NA | NA | NA | NA | NA | NA | NA |
| CaalfMt37 | 0 | 0 | 0 | 0 | 0 | 0 | 0 | 0 | 0 | 0 | 0 | 0 | 0 | 0 | 0 | 0 | 0 | 0 | 0 | NA | NA | NA | NA | NA | NA | NA | NA | NA | NA | NA | NA |
| CaalfMt38 | 0 | 1 | 1 | 1 | 0 | 0 | 0 | 1 | 0 | 1 | 1 | 2 | 0 | 0 | 0 | 1 | 0,55 | 1 | 0 | 0,311 | -1,684 | -0,871 | 0,3836564 | 0,418982 | 1,00E-08 | 2,9705529 | 1,9450255 | 1,9450255 | TRUE | 0,2944903 | No |
| CaalfMt39 | 0 | 0 | 0 | 0 | 1 | 0 | 0 | 0 | 0 | 0 | 0 | 0 | 1 | 0 | 0 | 0 | 0,1 | 0 | 0 | 1,314 | 0,394 | 0,111 | 0,9112982 | 0,9205309 | 1,00E-08 | 16,395074 | 10 | 10 | TRUE | 0,2859086 | No |
| CaalfMt40 | 0 | 0 | 0 | 0 | 0 | 0 | 0 | 0 | 0 | 0 | 0 | 0 | 0 | 0 | 0 | 0 | 0 | 0 | 0 | NA | NA | NA | NA | NA | NA | NA | NA | NA | NA | NA | NA |
| CaalfMt41 | 0 | 0 | 0 | 0 | 0 | 1 | 0 | 0 | 0 | 0 | 0 | 0 | 0 | 1 | 0 | 0 | 0,14 | 0 | 0 | 1,314 | 0,394 | 0,111 | 0,9112982 | 0,9205309 | 1,00E-08 | 11,553708 | 10 | 10 | TRUE | 0,5388425 | No |
| CaalfMt42 | 0 | 0 | 0 | 0 | 0 | 0 | 0 | 0 | 0 | 0 | 0 | 0 | 0 | 0 | 0 | 0 | 0 | 0 | 0 | NA | NA | NA | NA | NA | NA | NA | NA | NA | NA | NA | NA |
| CaalfMt43 | 0 | 0 | 0 | 0 | 0 | 0 | 0 | 0 | 0 | 0 | 0 | 0 | 0 | 0 | 0 | 0 | 0 | 0 | 0 | NA | NA | NA | NA | NA | NA | NA | NA | NA | NA | NA | NA |
| CaalfMt44 | 0 | 0 | 0 | 0 | 0 | 0 | 0 | 0 | 0 | 0 | 0 | 0 | 0 | 0 | 0 | 0 | 0 | 0 | 0 | NA | NA | NA | NA | NA | NA | NA | NA | NA | NA | NA | NA |
| CaalfMt45 | 0 | 0 | 0 | 0 | 0 | 0 | 0 | 0 | 0 | 0 | 0 | 0 | 0 | 0 | 0 | 0 | 0 | 0 | 0 | NA | NA | NA | NA | NA | NA | NA | NA | NA | NA | NA | NA |
| CaalfMt46 | 0 | 0 | 0 | 0 | 0 | 0 | 0 | 0 | 0 | 0 | 0 | 0 | 0 | 0 | 0 | 0 | 0 | 0 | 0 | NA | NA | NA | NA | NA | NA | NA | NA | NA | NA | NA | NA |
| CaalfMt47 | 0 | 0 | 0 | 0 | 0 | 0 | 0 | 0 | 0 | 0 | 0 | 0 | 0 | 0 | 0 | 0 | 0 | 0 | 0 | NA | NA | NA | NA | NA | NA | NA | NA | NA | NA | NA | NA |
| CD39 | 0 | 0 | 0 | 0 | 0 | 0 | 0 | 0 | 0 | 0 | 0 | 0 | 0 | 0 | 0 | 0 | 0 | 0 | 0 | NA | NA | NA | NA | NA | NA | NA | NA | NA | NA | NA | NA |
| ITS1 | 192 | 224 | 222 | 233 | 86 | 71 | 70 | 61 | 183 | 237 | 219 | 365 | 69 | 81 | 59 | 50 | 157,86 | 251 | 65 | 0,258 | -1,956 | -9,217 | 3,06E-20 | 6,90E-20 | 0,0579441 | 0,0124922 | 0,0340271 | 0,0340271 | TRUE | 0,5665395 | No |
| ITS2 | 444 | 314 | 315 | 222 | 132 | 52 | 102 | 108 | 423 | 332 | 311 | 348 | 106 | 59 | 87 | 88 | 219,2 | 354 | 85 | 0,242 | -2,045 | -13,327 | 1,61E-40 | 5,12E-40 | 0,0255302 | 0,0095817 | 0,0158308 | 0,0158308 | TRUE | 0,3393675 | No |
| LSU-C2809 | 108 | 116 | 123 | 93 | 89 | 62 | 76 | 58 | 103 | 123 | 122 | 146 | 71 | 70 | 65 | 47 | 93,29 | 124 | 63 | 0,516 | -0,954 | -5,48 | 4,26E-08 | 6,95E-08 | 0,014362 | 0,0196913 | 0,0174433 | 0,0174433 | TRUE | 0,28723 | No |
| LSU-G1431 | 11 | 18 | 17 | 9 | 6 | 5 | 2 | 1 | 10 | 19 | 17 | 14 | 5 | 6 | 2 | 1 | 9,18 | 15 | 4 | 0,207 | -2,27 | -3,979 | 6,93E-05 | 9,89E-05 | 0,0264438 | 0,1810025 | 0,1340231 | 0,1340231 | TRUE | 0,4301126 | No |
| orf19.10 | 54 | 62 | 48 | 40 | 99 | 69 | 97 | 90 | 51 | 66 | 47 | 63 | 79 | 78 | 82 | 73 | 67,55 | 57 | 78 | 1,391 | 0,476 | 2,695 | 0,0070466 | 0,00902 | 1,00E-08 | 0,0263961 | 0,0144553 | 0,0144553 | TRUE | 0,1078563 | No |
| orf19.100 | 2501 | 2105 | 2433 | 1417 | 1022 | 680 | 872 | 1044 | 2384 | 2226 | 2404 | 2220 | 819 | 773 | 740 | 849 | 1551,86 | 2308 | 795 | 0,345 | -1,537 | -26,467 | 2,31E-154 | 2,39E-153 | 0,0018644 | 0,0031493 | 0,0024347 | 0,0024347 | TRUE | 0,0273133 | No |
| orf19.1002 | 38 | 55 | 61 | 34 | 150 | 140 | 161 | 181 | 36 | 58 | 60 | 53 | 120 | 159 | 137 | 147 | 96,39 | 52 | 141 | 2,712 | 1,439 | 8,008 | 1,17E-15 | 2,39E-15 | 0,013882 | 0,0191245 | 0,0170954 | 0,0170954 | TRUE | 0,3951631 | No |
| orf19.1005 | 97 | 115 | 104 | 76 | 136 | 112 | 137 | 181 | 92 | 122 | 103 | 119 | 109 | 127 | 116 | 147 | 116,96 | 109 | 125 | 1,153 | 0,206 | 1,363 | 0,1729472 | 0,1973138 | 0,0096917 | 0,0161293 | 0,0131431 | 0,0131431 | TRUE | 0,1604432 | No |
| orf19.1007 | 5607 | 5297 | 5561 | 3619 | 3233 | 2238 | 3118 | 3194 | 5346 | 5601 | 5494 | 5669 | 2591 | 2545 | 2647 | 2597 | 4061,15 | 5528 | 2595 | 0,47 | -1,089 | -30,248 | 5,53E-201 | 7,59E-200 | 0,0002456 | 0,0024956 | 0,0009718 | 0,0009718 | TRUE | 0,0060372 | No |
| orf19.1008 | 57 | 54 | 43 | 41 | 19 | 19 | 23 | 30 | 54 | 57 | 42 | 64 | 15 | 22 | 20 | 24 | 37,36 | 54 | 20 | 0,372 | -1,426 | -5,6 | 2,14E-08 | 3,53E-08 | 1,00E-08 | 0,0460367 | 0,0298844 | 0,0298844 | TRUE | 0,1902421 | No |
| orf19.1009 | 100 | 63 | 92 | 75 | 82 | 71 | 97 | 101 | 95 | 67 | 91 | 117 | 66 | 81 | 82 | 82 | 85,16 | 92 | 78 | 0,847 | -0,24 | -1,32 | 0,1868563 | 0,2123473 | 0,0178681 | 0,0213725 | 0,019877 | 0,019877 | TRUE | 0,2244591 | No |
| orf19.101 | 1027 | 926 | 1006 | 703 | 597 | 486 | 652 | 660 | 979 | 979 | 994 | 1101 | 478 | 553 | 554 | 537 | 771,82 | 1013 | 530 | 0,524 | -0,933 | -13,39 | 6,93E-41 | 2,22E-40 | 0,0023453 | 0,0042186 | 0,0032607 | 0,0032607 | TRUE | 0,0530225 | No |
| orf19.1010 | 1853 | 1986 | 1805 | 1295 | 1464 | 1185 | 1405 | 1261 | 1767 | 2100 | 1783 | 2029 | 1173 | 1347 | 1193 | 1025 | 1552,14 | 1920 | 1184 | 0,617 | -0,697 | -8,095 | 5,72E-16 | 1,18E-15 | 0,0091859 | 0,0031491 | 0,0064495 | 0,0064495 | TRUE | 0,0999633 | No |
| orf19.1011 | 2041 | 1971 | 2098 | 1314 | 2321 | 1712 | 2132 | 2387 | 1946 | 2084 | 2073 | 2058 | 1860 | 1947 | 1810 | 1941 | 1964,8 | 2040 | 1890 | 0,926 | -0,111 | -2,401 | 0,0163296 | 0,0204272 | 0,0006618 | 0,0029269 | 0,0015375 | 0,0015375 | TRUE | 0,0123625 | No |
| orf19.1012 | 523 | 442 | 470 | 241 | 1068 | 660 | 923 | 961 | 499 | 467 | 464 | 378 | 856 | 750 | 784 | 781 | 622,39 | 452 | 793 | 1,744 | 0,803 | 9,641 | 5,36E-22 | 1,25E-21 | 0,0050102 | 0,0047294 | 0,0048584 | 0,0048584 | TRUE | 0,1211874 | No |
| orf19.102 | 766 | 742 | 784 | 457 | 1203 | 895 | 1141 | 1187 | 730 | 785 | 775 | 716 | 964 | 1018 | 969 | 965 | 865,1 | 752 | 979 | 1,299 | 0,377 | 6,66 | 2,74E-11 | 4,97E-11 | 1,00E-08 | 0,0039892 | 0,0018826 | 0,0018826 | TRUE | 0,0099594 | No |
| orf19.1026 | 4451 | 4136 | 4317 | 2865 | 4868 | 3648 | 4746 | 4932 | 4244 | 4373 | 4265 | 4488 | 3901 | 4148 | 4029 | 4010 | 4182,25 | 4342 | 4022 | 0,927 | -0,11 | -2,97 | 0,002977 | 0,0038999 | 0,0003343 | 0,0024839 | 0,0010774 | 0,0010774 | TRUE | 0,005803 | No |

| | | | | | | | | | | | | | | | | | | | | | | | | | | | | | | | |
|---|---|---|---|---|---|---|---|---|---|---|---|---|---|---|---|---|---|---|---|---|---|---|---|---|---|---|---|---|---|---|---|
| orf19.1026.1 | 353 | 299 | 296 | 186 | 797 | 518 | 673 | 617 | 337 | 316 | 292 | 291 | 639 | 589 | 571 | 502 | 442,15 | 309 | 575 | 1,855 | 0,891 | 9,505 | 2,00E-21 | 4,63E-21 | 0,0059144 | 0,0058048 | 0,0058621 | 0,0058621 | TRUE | 0,0900923 | No |
| orf19.1027 | 4527 | 4448 | 4524 | 2647 | 2479 | 1602 | 2411 | 2503 | 4316 | 4703 | 4469 | 4147 | 1986 | 1822 | 2047 | 2035 | 3190,65 | 4409 | 1972 | 0,448 | -1,16 | -21,319 | 7,52E-101 | 4,96E-100 | 0,0023373 | 0,0026059 | 0,0024877 | 0,0024877 | TRUE | 0,0306963 | No |
| orf19.1028 | 2052 | 2048 | 2180 | 1512 | 1620 | 1256 | 1564 | 1547 | 1956 | 2166 | 2154 | 2369 | 1298 | 1428 | 1328 | 1258 | 1744,51 | 2161 | 1328 | 0,615 | -0,702 | -10,737 | 6,81E-27 | 1,75E-26 | 0,0037234 | 0,0030325 | 0,0035027 | 0,0035027 | TRUE | 0,0494568 | No |
| orf19.1029 | 4948 | 5425 | 5372 | 3297 | 3717 | 2836 | 3409 | 3261 | 4717 | 5736 | 5307 | 5165 | 2978 | 3225 | 2894 | 2652 | 4084,3 | 5231 | 2937 | 0,561 | -0,833 | -11,7 | 1,27E-31 | 3,53E-31 | 0,0061115 | 0,0024933 | 0,0046144 | 0,0046144 | TRUE | 0,054023 | No |
| orf19.103 | 153 | 151 | 165 | 119 | 191 | 166 | 172 | 192 | 146 | 160 | 163 | 186 | 153 | 189 | 146 | 156 | 162,36 | 164 | 161 | 0,985 | -0,022 | -0,173 | 0,8625295 | 0,8806082 | 0,0044424 | 0,0122039 | 0,0088574 | 0,0088574 | TRUE | 0,1281053 | No |
| orf19.1030 | 26080 | 23650 | 25185 | 14868 | 34514 | 22795 | 33673 | 35554 | 24864 | 25007 | 24880 | 23292 | 27655 | 25920 | 28588 | 28909 | 26139,41 | 24511 | 27768 | 1,133 | 0,18 | 4,051 | 5,10E-05 | 7,32E-05 | 0,0016698 | 0,0021541 | 0,0018562 | 0,0018562 | TRUE | 0,0244647 | No |
| orf19.1031 | 1617 | 1427 | 1672 | 1007 | 3848 | 2865 | 3480 | 3739 | 1542 | 1509 | 1652 | 1578 | 3083 | 3258 | 2955 | 3040 | 2326,94 | 1570 | 3084 | 1,963 | 0,973 | 19,85 | 1,11E-87 | 6,48E-87 | 0,0011723 | 0,0027969 | 0,0018053 | 0,0018053 | TRUE | 0,0168607 | No |
| orf19.1032 | 4068 | 3561 | 3902 | 2678 | 13151 | 9225 | 12490 | 13352 | 3878 | 3765 | 3855 | 4195 | 10538 | 10490 | 10604 | 10856 | 7272,66 | 3923 | 10622 | 2,711 | 1,439 | 36,086 | 3,72E-285 | 7,71E-284 | 0,0007671 | 0,002317 | 0,0013411 | 0,0013411 | TRUE | 0,0247398 | No |
| orf19.1033 | 4691 | 4344 | 4561 | 2484 | 7904 | 5603 | 7292 | 7778 | 4472 | 4593 | 4506 | 3891 | 6333 | 6371 | 6191 | 6324 | 5335,28 | 4366 | 6305 | 1,442 | 0,528 | 10,11 | 5,01E-24 | 1,22E-23 | 0,0025112 | 0,002399 | 0,0024238 | 0,0024238 | TRUE | 0,0625154 | No |
| orf19.1034 | 2002 | 1872 | 1942 | 1091 | 1510 | 917 | 1324 | 1310 | 1909 | 1979 | 1919 | 1709 | 1210 | 1043 | 1124 | 1065 | 1494,7 | 1879 | 1110 | 0,591 | -0,759 | -11,769 | 5,64E-32 | 1,58E-31 | 0,0032585 | 0,0031898 | 0,0032977 | 0,0032977 | TRUE | 0,0441566 | No |
| orf19.1035 | 3933 | 3545 | 3802 | 2767 | 2967 | 2248 | 2745 | 3054 | 3750 | 3748 | 3756 | 4335 | 2377 | 2556 | 2330 | 2483 | 3167 | 3897 | 2436 | 0,626 | -0,676 | -11,612 | 3,58E-31 | 9,86E-31 | 0,0032326 | 0,0026097 | 0,0029291 | 0,0029291 | TRUE | 0,0671724 | No |
| orf19.1036 | 1184 | 1100 | 1068 | 666 | 2291 | 1568 | 2212 | 2344 | 1129 | 1163 | 1055 | 1043 | 1836 | 1783 | 1878 | 1906 | 1474,11 | 1098 | 1851 | 1,684 | 0,752 | 14,425 | 3,58E-47 | 1,27E-46 | 0,0008373 | 0,0032051 | 0,0018584 | 0,0018584 | TRUE | 0,0187706 | No |
| orf19.1037 | 93 | 97 | 93 | 52 | 269 | 201 | 248 | 270 | 89 | 103 | 92 | 81 | 216 | 229 | 211 | 220 | 154,84 | 91 | 219 | 2,38 | 1,251 | 10,242 | 1,28E-24 | 3,15E-24 | 1,00E-08 | 0,0126949 | 0,0062364 | 0,0062364 | TRUE | 0,0638539 | No |
| orf19.1039 | 259 | 390 | 313 | 181 | 477 | 307 | 410 | 290 | 247 | 412 | 309 | 284 | 382 | 349 | 348 | 236 | 320,91 | 313 | 329 | 1,049 | 0,069 | 0,411 | 0,6813273 | 0,7128298 | 0,0426852 | 0,0072078 | 0,0241099 | 0,0241099 | TRUE | 0,5211283 | No |
| orf19.104 | 252 | 258 | 310 | 182 | 329 | 260 | 288 | 347 | 240 | 273 | 306 | 285 | 264 | 296 | 245 | 282 | 273,79 | 276 | 272 | 0,983 | -0,025 | -0,241 | 0,8096831 | 0,8314431 | 0,0050864 | 0,0080882 | 0,0066504 | 0,0066504 | TRUE | 0,0929531 | No |
| orf19.1040 | 343 | 305 | 333 | 189 | 329 | 247 | 275 | 327 | 327 | 322 | 329 | 296 | 264 | 281 | 233 | 266 | 289,8 | 318 | 261 | 0,814 | -0,297 | -3,274 | 0,0010606 | 0,001429 | 1,00E-08 | 0,0077569 | 0,0044275 | 0,0044275 | TRUE | 0,0567372 | No |
| orf19.1041 | 428 | 512 | 467 | 354 | 880 | 690 | 791 | 781 | 408 | 541 | 461 | 555 | 705 | 785 | 672 | 635 | 595,2 | 491 | 699 | 1,427 | 0,513 | 4,979 | 6,39E-07 | 9,96E-07 | 0,0117511 | 0,0048498 | 0,0083855 | 0,0083855 | TRUE | 0,154938 | No |
| orf19.1042 | 16601 | 14194 | 14985 | 10665 | 14639 | 11473 | 13793 | 15101 | 15827 | 15008 | 14804 | 16707 | 11730 | 13046 | 11710 | 12279 | 13888,88 | 15586 | 12191 | 0,782 | -0,354 | -6,791 | 1,12E-11 | 2,05E-11 | 0,0027795 | 0,0022095 | 0,0025437 | 0,0025437 | TRUE | 0,0283507 | No |
| orf19.1043 | 690 | 794 | 858 | 434 | 1052 | 776 | 982 | 823 | 658 | 840 | 848 | 680 | 843 | 882 | 834 | 669 | 781,64 | 756 | 807 | 1,065 | 0,091 | 0,847 | 0,3969647 | 0,432701 | 0,0150705 | 0,0041919 | 0,0097167 | 0,0097167 | TRUE | 0,159401 | No |
| orf19.1045 | 2228 | 2289 | 2250 | 1022 | 2896 | 1889 | 2645 | 2350 | 2124 | 2420 | 2223 | 1601 | 2320 | 2148 | 2246 | 1911 | 2124,14 | 2092 | 2156 | 1,029 | 0,042 | 0,392 | 0,6948353 | 0,7260101 | 0,0179689 | 0,0028643 | 0,0103314 | 0,0103314 | TRUE | 0,2939122 | No |
| orf19.1047 | 8238 | 8502 | 8740 | 5130 | 13475 | 9823 | 12257 | 13247 | 7854 | 8990 | 8634 | 8036 | 10797 | 11170 | 10406 | 10771 | 9582,31 | 8378 | 10786 | 1,287 | 0,364 | 7,34 | 2,13E-13 | 4,11E-13 | 0,00225 | 0,0022626 | 0,0022525 | 0,0022525 | TRUE | 0,0294913 | No |
| orf19.1048 | 564 | 522 | 546 | 326 | 131 | 114 | 126 | 152 | 538 | 552 | 539 | 511 | 105 | 130 | 107 | 124 | 325,61 | 535 | 116 | 0,216 | -2,212 | -22,983 | 6,84E-117 | 5,28E-116 | 1,00E-08 | 0,0071338 | 0,0040172 | 0,0040172 | TRUE | 0,049975 | No |
| orf19.1049 | 71 | 101 | 91 | 59 | 151 | 123 | 135 | 140 | 68 | 107 | 90 | 92 | 121 | 140 | 115 | 114 | 105,76 | 89 | 122 | 1,372 | 0,456 | 2,854 | 0,0043153 | 0,005599 | 0,0103996 | 0,0176155 | 0,0146031 | 0,0146031 | TRUE | 0,2792828 | No |
| orf19.105 | 1075 | 747 | 801 | 537 | 2327 | 1455 | 2252 | 2277 | 1025 | 790 | 791 | 841 | 1865 | 1654 | 1912 | 1851 | 1341,21 | 862 | 1820 | 2,112 | 1,078 | 12,604 | 2,01E-36 | 6,00E-36 | 0,0091526 | 0,0033155 | 0,0061302 | 0,0061302 | TRUE | 0,2003656 | No |
| orf19.1050 | 176 | 207 | 215 | 110 | 255 | 187 | 231 | 225 | 168 | 219 | 212 | 172 | 204 | 213 | 196 | 183 | 195,93 | 193 | 199 | 1,026 | 0,037 | 0,314 | 0,7537579 | 0,7815615 | 0,0057426 | 0,0104714 | 0,0083545 | 0,0083545 | TRUE | 0,0975512 | No |
| orf19.1051 | 3102 | 2566 | 2657 | 2028 | 3505 | 2531 | 3243 | 3668 | 2957 | 2713 | 2625 | 3177 | 2808 | 2878 | 2753 | 2982 | 2861,83 | 2868 | 2855 | 0,997 | -0,005 | -0,073 | 0,9420635 | 0,9483053 | 0,0034131 | 0,002665 | 0,0033523 | 0,0033523 | TRUE | 0,0611642 | No |
| orf19.1052 | 1519 | 1302 | 1421 | 1077 | 1270 | 945 | 1203 | 1218 | 1448 | 1377 | 1404 | 1687 | 1018 | 1075 | 1021 | 990 | 1252,47 | 1479 | 1026 | 0,695 | -0,525 | -7,659 | 1,87E-14 | 3,73E-14 | 0,004091 | 0,0034022 | 0,0036888 | 0,0036888 | TRUE | 0,1010368 | No |
| orf19.1053 | 1174 | 1122 | 1222 | 701 | 1382 | 1063 | 1289 | 1433 | 1119 | 1186 | 1207 | 1098 | 1107 | 1209 | 1094 | 1165 | 1148,33 | 1152 | 1144 | 0,99 | -0,015 | -0,266 | 0,790259 | 0,8133295 | 0,0011336 | 0,0035211 | 0,0021951 | 0,0021951 | TRUE | 0,0162597 | No |
| orf19.1054 | 413 | 407 | 451 | 317 | 525 | 479 | 505 | 437 | 394 | 430 | 446 | 497 | 421 | 545 | 429 | 355 | 439,46 | 442 | 438 | 0,99 | -0,014 | -0,119 | 0,905372 | 0,9186164 | 0,0178055 | 0,0058275 | 0,0117597 | 0,0117597 | TRUE | 0,3041159 | No |
| orf19.1055 | 2136 | 2050 | 2055 | 1365 | 2380 | 1741 | 2166 | 2421 | 2036 | 2168 | 2030 | 2138 | 1907 | 1980 | 1839 | 1969 | 2008,33 | 2093 | 1924 | 0,919 | -0,121 | -2,637 | 0,0083613 | 0,0106664 | 0,0006559 | 0,0029088 | 0,0015319 | 0,0015319 | TRUE | 0,0106183 | No |
| orf19.1057 | 1437 | 1302 | 1294 | 859 | 1369 | 927 | 1276 | 1457 | 1370 | 1377 | 1278 | 1346 | 1097 | 1054 | 1083 | 1185 | 1223,72 | 1343 | 1105 | 0,824 | -0,28 | -5,073 | 3,92E-07 | 6,17E-07 | 0,0010919 | 0,003433 | 0,0020935 | 0,0020935 | TRUE | 0,0288005 | No |
| orf19.1058 | 3888 | 3456 | 3695 | 2232 | 4770 | 3041 | 4607 | 4643 | 3707 | 3654 | 3650 | 3497 | 3822 | 3458 | 3911 | 3775 | 3684,3 | 3627 | 3742 | 1,032 | 0,045 | 0,96 | 0,3371052 | 0,3709378 | 0,0013423 | 0,0025369 | 0,0018363 | 0,0018363 | TRUE | 0,0306931 | No |
| orf19.1059 | 3262 | 2761 | 2890 | 1699 | 8316 | 4987 | 7579 | 8098 | 3110 | 2919 | 2855 | 2662 | 6663 | 5671 | 6435 | 6584 | 4612,38 | 2886 | 6338 | 2,194 | 1,134 | 17,829 | 4,18E-71 | 2,03E-70 | 0,0042936 | 0,0024473 | 0,0036186 | 0,0036186 | TRUE | 0,0607186 | No |
| orf19.1060 | 108 | 88 | 98 | 56 | 305 | 184 | 204 | 150 | 103 | 93 | 97 | 88 | 244 | 209 | 173 | 122 | 141,17 | 95 | 187 | 1,959 | 0,97 | 4,977 | 6,45E-07 | 1,01E-06 | 0,0425477 | 0,0137223 | 0,0282126 | 0,0282126 | TRUE | 0,1903248 | No |
| orf19.1061 | 2149 | 1731 | 1842 | 953 | 1171 | 686 | 1034 | 1152 | 2049 | 1830 | 1820 | 1493 | 938 | 780 | 878 | 937 | 1340,58 | 1798 | 883 | 0,491 | -1,026 | -11,1 | 1,26E-28 | 3,32E-28 | 0,0105343 | 0,003316 | 0,0073894 | 0,0073894 | TRUE | 0,1516251 | No |
| orf19.1062 | 92 | 93 | 78 | 47 | 63 | 27 | 36 | 64 | 88 | 98 | 77 | 74 | 50 | 31 | 31 | 52 | 62,56 | 84 | 41 | 0,49 | -1,03 | -4,724 | 2,31E-06 | 3,52E-06 | 0,0281562 | 0,0283347 | 0,0282701 | 0,0282701 | TRUE | 0,2636824 | No |
| orf19.1063 | 409 | 484 | 453 | 284 | 620 | 495 | 588 | 483 | 390 | 512 | 448 | 445 | 497 | 563 | 499 | 393 | 468,21 | 449 | 488 | 1,086 | 0,119 | 1,037 | 0,2995451 | 0,3324169 | 0,015235 | 0,005598 | 0,0104713 | 0,0104713 | TRUE | 0,2042495 | No |
| orf19.1064 | 4444 | 4108 | 4375 | 2750 | 21852 | 16347 | 20583 | 23127 | 4237 | 4344 | 4322 | 4308 | 17509 | 18588 | 17475 | 18804 | 11198,42 | 4303 | 18094 | 4,205 | 2,072 | 54,289 | 0 | 0 | 0,0008288 | 0,0022379 | 0,0012436 | 0,0012436 | TRUE | 0,0086426 | No |
| orf19.1065 | 1289 | 1045 | 1228 | 786 | 2361 | 1690 | 2343 | 2651 | 1229 | 1105 | 1213 | 1231 | 1892 | 1922 | 1989 | 2156 | 1592,07 | 1194 | 1990 | 1,667 | 0,737 | 12,311 | 7,94E-35 | 2,32E-34 | 0,0024837 | 0,0031226 | 0,0027493 | 0,0027493 | TRUE | 0,0387238 | No |
| orf19.1066 | 177 | 164 | 210 | 89 | 363 | 218 | 293 | 290 | 169 | 173 | 207 | 139 | 291 | 248 | 249 | 236 | 214,04 | 172 | 256 | 1,474 | 0,559 | 4,478 | 7,55E-06 | 1,13E-05 | 0,0103512 | 0,0097622 | 0,0100256 | 0,0100256 | TRUE | 0,1985365 | No |
| orf19.1067 | 1172 | 1077 | 1095 | 633 | 729 | 499 | 713 | 734 | 1117 | 1139 | 1082 | 992 | 584 | 567 | 605 | 597 | 835,4 | 1082 | 588 | 0,543 | -0,882 | -14,258 | 4,02E-46 | 1,41E-45 | 0,000863 | 0,0040567 | 0,0024002 | 0,0024002 | TRUE | 0,0334447 | No |
| orf19.1069 | 44639 | 39590 | 43620 | 27657 | 27764 | 19643 | 26139 | 27563 | 42558 | 41861 | 43092 | 43326 | 22246 | 22336 | 22192 | 22411 | 32503,01 | 42709 | 22296 | 0,522 | -0,938 | -38,975 | 0 | 0 | 7,99E-05 | 0,0021418 | 0,0005227 | 0,0005227 | TRUE | 0,0021864 | No |
| orf19.107 | 7797 | 7215 | 7799 | 4674 | 6657 | 4587 | 6370 | 6760 | 7434 | 7629 | 7705 | 7322 | 5334 | 5216 | 5408 | 5497 | 6442,98 | 7522 | 5364 | 0,713 | -0,488 | -14,375 | 7,39E-47 | 2,61E-46 | 0,0003052 | 0,0023461 | 0,0009487 | 0,0009487 | TRUE | 0,0040298 | No |
| orf19.1070 | 338 | 308 | 366 | 198 | 503 | 377 | 481 | 455 | 322 | 326 | 362 | 310 | 403 | 429 | 408 | 370 | 366,21 | 330 | 402 | 1,214 | 0,279 | 3,306 | 0,0009469 | 0,0012792 | 0,0012974 | 0,0065747 | 0,0040661 | 0,0040661 | TRUE | 0,0485321 | No |
| orf19.1075 | 116 | 114 | 110 | 86 | 166 | 123 | 139 | 127 | 111 | 121 | 109 | 135 | 133 | 140 | 118 | 103 | 121,08 | 119 | 124 | 1,045 | 0,063 | 0,444 | 0,657004 | 0,68919 | 0,005687 | 0,0156513 | 0,0112542 | 0,0112542 | TRUE | 0,1306271 | No |
| orf19.1077 | 2593 | 2494 | 2552 | 1720 | 1548 | 1135 | 1451 | 1530 | 2472 | 2637 | 2521 | 2694 | 1240 | 1291 | 1232 | 1244 | 1916,47 | 2581 | 1252 | 0,485 | -1,043 | -22,546 | 1,47E-112 | 1,09E-111 | 0,0005471 | 0,002948 | 0,0014814 | 0,0014814 | TRUE | 0,0098056 | No |
| orf19.1078 | 2823 | 2775 | 2704 | 1813 | 2495 | 1866 | 2324 | 2479 | 2691 | 2934 | 2671 | 2840 | 1999 | 2122 | 1973 | 2016 | 2405,85 | 2784 | 2028 | 0,728 | -0,458 | -9,64 | 5,42E-22 | 1,26E-21 | 0,000961 | 0,0027737 | 0,0017426 | 0,0017426 | TRUE | 0,0167305 | No |
| orf19.1080 | 18 | 18 | 13 | 16 | 42 | 29 | 34 | 39 | 17 | 19 | 13 | 25 | 34 | 33 | 29 | 32 | 25,16 | 18 | 32 | 1,76 | 0,816 | 2,809 | 0,0049672 | 0,0064214 | 1,00E-08 | 0,067341 | 0,0361902 | 0,0361902 | TRUE | 0,2211017 | No |
| orf19.1082 | 1029 | 1053 | 1009 | 606 | 802 | 575 | 791 | 801 | 981 | 1113 | 997 | 949 | 643 | 654 | 672 | 651 | 832,48 | 1010 | 655 | 0,647 | -0,628 | -9,803 | 1,10E-22 | 2,59E-22 | 0,0015643 | 0,0040636 | 0,0027012 | 0,0027012 | TRUE | 0,0563967 | No |
| orf19.1082.1 | 6626 | 5220 | 5190 | 4152 | 10672 | 7201 | 9589 | 10641 | 6317 | 5519 | 5127 | 6504 | 8551 | 8188 | 8141 | 8652 | 7125,09 | 5867 | 8383 | 1,43 | 0,516 | 7,272 | 3,55E-13 | 6,82E-13 | 0,006457 | 0,0023217 | 0,0046807 | 0,0046807 | TRUE | 0,0881544 | No |
| orf19.1083 | 682 | 600 | 640 | 415 | 1345 | 909 | 1268 | 1348 | 650 | 634 | 632 | 650 | 1078 | 1034 | 1077 | 1096 | 856,36 | 642 | 1071 | 1,672 | 0,741 | 13,542 | 8,84E-42 | 2,88E-41 | 1,00E-08 | 0,0040086 | 0,001592 | 0,001592 | TRUE | 0,0060149 | No |
| orf19.1084 | 8396 | 8020 | 8317 | 5363 | 6925 | 5237 | 6087 | 6859 | 8005 | 8480 | 8216 | 8401 | 5549 | 5955 | 5168 | 5577 | 6918,91 | 8276 | 5562 | 0,672 | -0,574 | -12,128 | 7,54E-34 | 2,16E-33 | 0,001798 | 0,0023286 | 0,002 | 0,002 | TRUE | 0,0277018 | No |
| orf19.1085 | 966 | 883 | 971 | 627 | 2489 | 1633 | 2322 | 2396 | 921 | 934 | 959 | 982 | 1994 | 1857 | 1971 | 1948 | 1445,86 | 949 | 1942 | 2,053 | 1,037 | 21,205 | 8,61E-100 | 5,63E-99 | 1,00E-08 | 0,0032269 | 0,0014789 | 0,0014789 | TRUE | 0,0100292 | No |
| orf19.1086 | 2821 | 2880 | 3071 | 2000 | 2833 | 2117 | 2729 | 2923 | 2690 | 3045 | 3034 | 3133 | 2270 | 2407 | 2317 | 2377 | 2659,07 | 2976 | 2343 | 0,788 | -0,344 | -6,368 | 1,91E-10 | 3,39E-10 | 0,0019965 | 0,0027088 | 0,0024198 | 0,0024198 | TRUE | 0,0517622 | No |
| orf19.1087 | 1773 | 1951 | 2091 | 1119 | 1538 | 1102 | 1423 | 1385 | 1690 | 2063 | 2066 | 1753 | 1232 | 1253 | 1208 | 1126 | 1548,96 | 1893 | 1205 | 0,636 | -0,654 | -8,645 | 5,39E-18 | 1,16E-17 | 0,005713 | 0,0031513 | 0,0048191 | 0,0048191 | TRUE | 0,0656089 | No |
| orf19.1089 | 1271 | 1531 | 1466 | 870 | 4297 | 3770 | 3597 | 3081 | 1212 | 1619 | 1448 | 1363 | 3443 | 4287 | 3054 | 2505 | 2366,33 | 1410 | 3322 | 2,355 | 1,236 | 6,713 | 1,91E-11 | 3,49E-11 | 0,0320215 | 0,0027851 | 0,0165083 | 0,0320215 | TRUE | 0,1908753 | No |
| orf19.109 | 1541 | 1287 | 1426 | 870 | 1479 | 983 | 1377 | 1451 | 1469 | 1361 | 1409 | 1363 | 1185 | 1118 | 1169 | 1180 | 1281,67 | 1400 | 1163 | 0,83 | -0,268 | -5,407 | 6,40E-08 | 1,04E-07 | 1,00E-08 | 0,0033723 | 0,0015844 | 0,0015844 | TRUE | 0,0135657 | No |
| orf19.1090 | 257 | 213 | 220 | 170 | 269 | 174 | 250 | 303 | 245 | 225 | 217 | 266 | 216 | 198 | 212 | 246 | 228,24 | 238 | 218 | 0,922 | -0,118 | -1,093 | 0,2742829 | 0,3055516 | 0,0036445 | 0,0092851 | 0,0067242 | 0,0067242 | TRUE | 0,0863803 | No |
| orf19.1091 | 6769 | 6749 | 7154 | 3907 | 7366 | 5041 | 6527 | 6518 | 6454 | 7136 | 7067 | 6121 | 5902 | 5732 | 5541 | 5300 | 6156,64 | 6694 | 5619 | 0,839 | -0,254 | -4,35 | 1,36E-05 | 2,01E-05 | 0,0035265 | 0,002358 | 0,0030987 | 0,0030987 | TRUE | 0,0397238 | No |
| orf19.1092 | 910 | 1083 | 1107 | 656 | 774 | 602 | 647 | 592 | 868 | 1145 | 1094 | 1028 | 620 | 685 | 549 | 481 | 808,67 | 1034 | 584 | 0,564 | -0,826 | -7,399 | 1,37E-13 | 2,67E-13 | 0,016344 | 0,0041217 | 0,0106621 | 0,0106621 | TRUE | 0,1658811 | No |
| orf19.1093 | 8049 | 6936 | 7919 | 5419 | 7194 | 4987 | 7165 | 7393 | 7674 | 7334 | 7823 | 8489 | 5764 | 5671 | 6083 | 6011 | 6856,18 | 7830 | 5882 | 0,752 | -0,412 | -8,183 | 2,77E-16 | 5,74E-16 | 0,0022749 | 0,0023308 | 0,0022814 | 0,0022814 | TRUE | 0,0383171 | No |
| orf19.1095 | 2590 | 2383 | 2527 | 1772 | 2304 | 1781 | 2160 | 2456 | 2469 | 2520 | 2496 | 2776 | 1846 | 2025 | 1834 | 1997 | 2245,43 | 2565 | 1926 | 0,751 | -0,413 | -7,456 | 8,95E-14 | 1,74E-13 | 0,0022606 | 0,0028225 | 0,0024894 | 0,0024894 | TRUE | 0,0349988 | No |
| orf19.1096 | 1151 | 1313 | 1370 | 772 | 1307 | 1013 | 1209 | 1216 | 1097 | 1388 | 1353 | 1209 | 1047 | 1152 | 1026 | 989 | 1157,85 | 1262 | 1054 | 0,834 | -0,263 | -3,233 | 0,0012247 | 0,0016404 | 0,0067083 | 0,0035093 | 0,0054657 | 0,0054657 | TRUE | 0,0970007 | No |
| orf19.1097 | 2298 | 1849 | 2067 | 1588 | 4969 | 3454 | 4396 | 4885 | 2191 | 1955 | 2042 | 2488 | 3982 | 3928 | 3732 | 3972 | 3036,11 | 2169 | 3904 | 1,803 | 0,85 | 12,47 | 1,09E-35 | 3,21E-35 | 0,005775 | 0,0026321 | 0,0040918 | 0,0040918 | TRUE | 0,1132089 | No |
| orf19.11 | 480 | 348 | 412 | 230 | 215 | 114 | 206 | 200 | 458 | 368 | 407 | 360 | 172 | 130 | 175 | 163 | 279,04 | 398 | 160 | 0,402 | -1,313 | -11,294 | 1,40E-29 | 3,75E-29 | 0,0098017 | 0,0079754 | 0,0088031 | 0,0088031 | TRUE | 0,1506414 | No |
| orf19.1105 | 44 | 47 | 64 | 34 | 32 | 32 | 25 | 30 | 42 | 50 | 63 | 53 | 26 | 36 | 21 | 24 | 39,47 | 52 | 27 | 0,511 | -0,97 | -3,917 | 8,95E-05 | 0,0001271 | 0,013613 | 0,0436874 | 0,0312979 | 0,0312979 | TRUE | 0,2980389 | No |
| orf19.1105.2 | 69796 | 53337 | 58691 | 41323 | 77826 | 49476 | 73853 | 81161 | 66543 | 56397 | 57981 | 64735 | 62360 | 56259 | 62701 | 65992 | 61620,85 | 61414 | 61828 | 1,007 | 0,01 | 0,149 | 0,8812685 | 0,8974424 | 0,0046496 | 0,0021179 | 0,0040868 | 0,0040868 | TRUE | 0,0450089 | No |
| orf19.1105.3 | 13 | 8 | 7 | 13 | 8 | 8 | 9 | 13 | 12 | 8 | 7 | 20 | 6 | 9 | 8 | 11 | 10,23 | 12 | 8 | 0,723 | -0,467 | -0,993 | 0,3208798 | 0,3541221 | 0,0370715 | 0,1625672 | 0,111952 | 0,111952 | TRUE | 0,5303889 | No |
| orf19.1106 | 277 | 302 | 300 | 195 | 140 | 118 | 153 | 146 | 264 | 319 | 296 | 305 | 112 | 134 | 130 | 119 | 210,03 | 296 | 124 | 0,417 | -1,263 | -11,333 | 9,01E-30 | 2,42E-29 | 1,00E-08 | 0,0099088 | 0,0064202 | 0,0064202 | TRUE | 0,0646752 | No |
| orf19.1107 | 4921 | 4095 | 4409 | 3251 | 5965 | 4431 | 5891 | 6335 | 4692 | 4330 | 4356 | 5093 | 4780 | 5038 | 5001 | 5151 | 4805,07 | 4618 | 4992 | 1,082 | 0,114 | 2,023 | 0,0430289 | 0,0520112 | 0,0031993 | 0,002433 | 0,0028339 | 0,0028339 | TRUE | 0,0566979 | No |
| orf19.1108 | 3164 | 2859 | 2984 | 2078 | 3224 | 2613 | 3032 | 3174 | 3017 | 3023 | 2948 | 3255 | 2583 | 2971 | 2574 | 2581 | 2869,03 | 3061 | 2677 | 0,875 | -0,193 | -3,325 | 0,0008846 | 0,0011982 | 0,0031712 | 0,0026636 | 0,0028905 | 0,0028905 | TRUE | 0,0645595 | No |
| orf19.1109 | 2217 | 2192 | 2289 | 1250 | 2092 | 1464 | 1980 | 1867 | 2114 | 2318 | 2261 | 1958 | 1676 | 1665 | 1681 | 1518 | 1898,87 | 2163 | 1635 | 0,755 | -0,406 | -6,513 | 7,37E-11 | 1,32E-10 | 0,0032091 | 0,002956 | 0,0031986 | 0,0031986 | TRUE | 0,0460458 | No |
| orf19.111 | 25665 | 23125 | 24607 | 15595 | 19165 | 13690 | 17738 | 19552 | 24469 | 24452 | 24309 | 24430 | 15356 | 15567 | 15059 | 15898 | 19942,57 | 24415 | 15470 | 0,634 | -0,658 | -23,833 | 1,53E-125 | 1,28E-124 | 0,0001942 | 0,0021736 | 0,0006808 | 0,0006808 | TRUE | 0,0042997 | No |

| | | | | | | | | | | | | | | | | | | | | | | | | | | | | | | | |
|---|---|---|---|---|---|---|---|---|---|---|---|---|---|---|---|---|---|---|---|---|---|---|---|---|---|---|---|---|---|---|---|
| orf19.1110 | 426 | 377 | 409 | 270 | 818 | 618 | 756 | 673 | 406 | 399 | 404 | 423 | 655 | 703 | 642 | 547 | 522,38 | 408 | 637 | 1,56 | 0,642 | 7,495 | 6,61E-14 | 1,29E-13 | 0,0047211 | 0,0052344 | 0,0049584 | 0,0049584 | TRUE | 0,1076228 | No |
| orf19.1111 | 756 | 720 | 837 | 557 | 1009 | 752 | 974 | 972 | 721 | 761 | 827 | 873 | 808 | 855 | 827 | 790 | 807,79 | 796 | 820 | 1,034 | 0,048 | 0,698 | 0,4854791 | 0,5218737 | 0,0025028 | 0,0041239 | 0,0033019 | 0,0033019 | TRUE | 0,0479644 | No |
| orf19.1112 | 1208 | 1156 | 1244 | 727 | 680 | 518 | 632 | 629 | 1152 | 1222 | 1229 | 1139 | 545 | 589 | 537 | 511 | 865,47 | 1186 | 546 | 0,458 | -1,125 | -18,117 | 2,33E-73 | 1,17E-72 | 0,0007489 | 0,0039884 | 0,0024128 | 0,0024128 | TRUE | 0,0304458 | No |
| orf19.1113 | 535 | 444 | 511 | 279 | 1413 | 971 | 1329 | 1393 | 510 | 469 | 505 | 437 | 1132 | 1104 | 1128 | 1133 | 802,34 | 480 | 1124 | 2,328 | 1,219 | 19,72 | 1,44E-86 | 8,36E-86 | 1,00E-08 | 0,0041377 | 0,0021196 | 0,0021196 | TRUE | 0,0345167 | No |
| orf19.1114 | 5176 | 4457 | 4583 | 2909 | 5856 | 3498 | 5682 | 5814 | 4935 | 4713 | 4528 | 4557 | 4692 | 3978 | 4824 | 4727 | 4619,16 | 4683 | 4555 | 0,973 | -0,039 | -0,626 | 0,531223 | 0,5671297 | 0,0037378 | 0,0024467 | 0,0035901 | 0,0035901 | TRUE | 0,0874875 | No |
| orf19.1115 | 2402 | 2249 | 2357 | 1372 | 3351 | 2352 | 3141 | 3368 | 2290 | 2378 | 2328 | 2149 | 2685 | 2674 | 2667 | 2739 | 2488,82 | 2286 | 2691 | 1,175 | 0,233 | 5,518 | 3,44E-08 | 5,62E-08 | 0,0002683 | 0,002751 | 0,0012966 | 0,0012966 | TRUE | 0,0176131 | No |
| orf19.1116 | 192 | 165 | 175 | 110 | 215 | 164 | 231 | 261 | 183 | 174 | 173 | 172 | 172 | 186 | 196 | 212 | 183,73 | 176 | 192 | 1,091 | 0,126 | 1,139 | 0,2546927 | 0,2851218 | 1,00E-08 | 0,0110279 | 0,0061796 | 0,0061796 | TRUE | 0,0599927 | No |
| orf19.1117 | 299 | 264 | 271 | 152 | 167 | 140 | 171 | 169 | 285 | 279 | 268 | 238 | 134 | 159 | 145 | 137 | 205,71 | 268 | 144 | 0,533 | -0,909 | -8,451 | 2,89E-17 | 6,11E-17 | 1,00E-08 | 0,0100731 | 0,0059232 | 0,0059232 | TRUE | 0,0498206 | No |
| orf19.1119 | 2020 | 1955 | 2126 | 1329 | 1768 | 1380 | 1711 | 1863 | 1926 | 2067 | 2100 | 2082 | 1417 | 1569 | 1453 | 1515 | 1766,06 | 2044 | 1488 | 0,728 | -0,458 | -8,916 | 4,85E-19 | 1,07E-18 | 0,0012401 | 0,003021 | 0,0019606 | 0,0019606 | TRUE | 0,0189974 | No |
| orf19.1120 | 465 | 393 | 460 | 280 | 940 | 622 | 899 | 894 | 443 | 416 | 454 | 439 | 753 | 707 | 763 | 727 | 587,82 | 438 | 738 | 1,685 | 0,753 | 11,413 | 3,59E-30 | 9,72E-30 | 1,00E-08 | 0,0048845 | 0,0023052 | 0,0023052 | TRUE | 0,014639 | No |
| orf19.1121 | 4245 | 4242 | 4295 | 2583 | 3264 | 2482 | 3049 | 3040 | 4047 | 4485 | 4243 | 4046 | 2615 | 2822 | 2589 | 2472 | 3415 | 4205 | 2624 | 0,623 | -0,682 | -12,723 | 4,41E-37 | 1,34E-36 | 0,0023574 | 0,0025721 | 0,0024539 | 0,0024539 | TRUE | 0,0302697 | No |
| orf19.1122 | 60 | 77 | 79 | 53 | 44 | 29 | 33 | 36 | 57 | 81 | 78 | 83 | 35 | 33 | 28 | 29 | 53,15 | 75 | 31 | 0,421 | -1,248 | -5,887 | 3,93E-09 | 6,62E-09 | 1,00E-08 | 0,0329822 | 0,0214731 | 0,0214731 | TRUE | 0,2509684 | No |
| orf19.1123 | 4 | 0 | 3 | 1 | 1 | 4 | 0 | 1 | 4 | 0 | 3 | 2 | 1 | 5 | 0 | 1 | 1,81 | 2 | 2 | 0,677 | -0,563 | -0,459 | 0,6461522 | 0,678567 | 0,804324 | 0,9075592 | 0,8695334 | 0,8695334 | TRUE | 0,8232373 | No |
| orf19.1124 | 317 | 256 | 260 | 161 | 544 | 388 | 470 | 520 | 302 | 271 | 257 | 252 | 436 | 441 | 399 | 423 | 347,61 | 270 | 425 | 1,562 | 0,643 | 7,418 | 1,19E-13 | 2,32E-13 | 0,0008154 | 0,0068146 | 0,0041208 | 0,0041208 | TRUE | 0,0734708 | No |
| orf19.1124.2 | 374 | 215 | 270 | 183 | 776 | 434 | 707 | 720 | 357 | 227 | 267 | 287 | 622 | 493 | 600 | 585 | 429,78 | 284 | 575 | 2,024 | 1,017 | 8,158 | 3,42E-16 | 7,06E-16 | 0,0194251 | 0,0059116 | 0,012176 | 0,012176 | TRUE | 0,3496234 | No |
| orf19.113 | 2155 | 2036 | 2074 | 1309 | 1154 | 843 | 1000 | 1050 | 2055 | 2153 | 2049 | 2051 | 925 | 959 | 849 | 854 | 1486,61 | 2077 | 897 | 0,431 | -1,215 | -22,686 | 6,12E-114 | 4,59E-113 | 0,0010446 | 0,0031957 | 0,0019839 | 0,0019839 | TRUE | 0,0236236 | No |
| orf19.1133 | 957 | 908 | 946 | 699 | 1169 | 889 | 1076 | 1286 | 912 | 960 | 935 | 1095 | 937 | 1011 | 914 | 1046 | 976,1 | 976 | 977 | 1,004 | 0,006 | 0,083 | 0,9336973 | 0,9404771 | 0,004099 | 0,0037734 | 0,003948 | 0,003948 | TRUE | 0,0743317 | No |
| orf19.1135 | 411 | 507 | 497 | 265 | 318 | 227 | 294 | 312 | 392 | 536 | 491 | 415 | 255 | 258 | 250 | 254 | 356,28 | 458 | 254 | 0,553 | -0,855 | -8,057 | 7,80E-16 | 1,60E-15 | 0,0088915 | 0,0066997 | 0,0078303 | 0,0078303 | TRUE | 0,1504574 | No |
| orf19.1136 | 197 | 202 | 173 | 113 | 200 | 106 | 154 | 145 | 188 | 214 | 171 | 177 | 160 | 121 | 131 | 118 | 159,85 | 188 | 132 | 0,708 | -0,499 | -3,715 | 0,0002029 | 0,0002831 | 0,0093387 | 0,0123631 | 0,0109553 | 0,0109553 | TRUE | 0,2208701 | No |
| orf19.1137 | 1646 | 1506 | 1529 | 1031 | 2268 | 1806 | 2108 | 2146 | 1569 | 1592 | 1511 | 1615 | 1817 | 2054 | 1790 | 1745 | 1711,6 | 1572 | 1852 | 1,177 | 0,235 | 3,98 | 6,88E-05 | 9,81E-05 | 0,0024059 | 0,0030506 | 0,0027524 | 0,0027524 | TRUE | 0,0629323 | No |
| orf19.1139 | 7250 | 6628 | 7452 | 4614 | 6605 | 4549 | 6263 | 6776 | 6912 | 7008 | 7362 | 7228 | 5292 | 5173 | 5317 | 5510 | 6225,27 | 7128 | 5323 | 0,747 | -0,42 | -11,377 | 5,47E-30 | 1,47E-29 | 0,0005369 | 0,002355 | 0,0011484 | 0,0011484 | TRUE | 0,0068093 | No |
| orf19.114 | 153 | 187 | 188 | 107 | 531 | 385 | 556 | 566 | 146 | 198 | 186 | 168 | 425 | 438 | 472 | 460 | 311,56 | 174 | 449 | 2,576 | 1,365 | 13,42 | 4,59E-41 | 1,48E-40 | 0,0032484 | 0,0073613 | 0,0057489 | 0,0057489 | TRUE | 0,1488546 | No |
| orf19.1140 | 743 | 743 | 768 | 459 | 2106 | 1540 | 1990 | 1906 | 708 | 786 | 759 | 719 | 1687 | 1751 | 1690 | 1550 | 1206,2 | 743 | 1670 | 2,243 | 1,166 | 19,349 | 2,07E-83 | 1,16E-82 | 0,001708 | 0,0034525 | 0,0024638 | 0,0024638 | TRUE | 0,0284364 | No |
| orf19.1142 | 1322 | 1274 | 1378 | 853 | 1926 | 1507 | 1848 | 2006 | 1260 | 1347 | 1361 | 1336 | 1543 | 1714 | 1569 | 1631 | 1470,24 | 1326 | 1614 | 1,216 | 0,282 | 5,373 | 7,76E-08 | 1,25E-07 | 0,0010712 | 0,003208 | 0,0019539 | 0,0019539 | TRUE | 0,0195728 | No |
| orf19.1143 | 429 | 394 | 422 | 221 | 1098 | 664 | 1066 | 1069 | 409 | 417 | 417 | 346 | 880 | 755 | 905 | 869 | 624,72 | 397 | 852 | 2,138 | 1,096 | 13,336 | 1,42E-40 | 4,54E-40 | 0,0042894 | 0,0047195 | 0,0045612 | 0,0045612 | TRUE | 0,0715145 | No |
| orf19.1144 | 730 | 677 | 738 | 427 | 628 | 434 | 618 | 692 | 696 | 716 | 729 | 669 | 503 | 493 | 525 | 563 | 611,73 | 702 | 521 | 0,741 | -0,431 | -6,403 | 1,52E-10 | 2,71E-10 | 0,0006481 | 0,0047753 | 0,0027111 | 0,0027111 | TRUE | 0,0346441 | No |
| orf19.1146 | 54 | 42 | 49 | 38 | 71 | 49 | 59 | 74 | 51 | 44 | 48 | 60 | 57 | 56 | 50 | 60 | 53,34 | 51 | 56 | 1,105 | 0,143 | 0,745 | 0,4564977 | 0,4930453 | 1,00E-08 | 0,0328748 | 0,0166267 | 0,0166267 | TRUE | 0,0723859 | No |
| orf19.1148 | 4 | 3 | 7 | 6 | 7 | 8 | 10 | 12 | 4 | 3 | 7 | 9 | 6 | 9 | 8 | 10 | 7,03 | 6 | 8 | 1,463 | 0,549 | 1,001 | 0,3168706 | 0,3501214 | 1,00E-08 | 0,2355937 | 0,1358926 | 0,1358926 | TRUE | 0,226529 | No |
| orf19.1149 | 2892 | 2328 | 2534 | 1631 | 3179 | 2178 | 2911 | 3145 | 2757 | 2462 | 2503 | 2555 | 2547 | 2477 | 2471 | 2557 | 2541,2 | 2569 | 2513 | 0,978 | -0,032 | -0,687 | 0,4920063 | 0,5280007 | 0,0009378 | 0,0027374 | 0,0016989 | 0,0016989 | TRUE | 0,0307509 | No |
| orf19.115 | 129 | 149 | 110 | 59 | 399 | 286 | 368 | 347 | 123 | 158 | 109 | 92 | 320 | 325 | 312 | 282 | 215,14 | 120 | 310 | 2,546 | 1,348 | 10,034 | 1,08E-23 | 2,61E-23 | 0,0141054 | 0,0097231 | 0,0113035 | 0,0113035 | TRUE | 0,4162414 | No |
| orf19.1150 | 5702 | 5076 | 5481 | 3883 | 5659 | 4149 | 5674 | 5706 | 5436 | 5367 | 5415 | 6083 | 4534 | 4718 | 4817 | 4640 | 5126,25 | 5575 | 4677 | 0,84 | -0,252 | -5,212 | 1,87E-07 | 2,98E-07 | 0,0019217 | 0,0024116 | 0,0020502 | 0,0020502 | TRUE | 0,0442985 | No |
| orf19.1150.1 | 200 | 184 | 194 | 149 | 547 | 384 | 522 | 539 | 191 | 195 | 192 | 233 | 438 | 437 | 443 | 438 | 320,83 | 203 | 439 | 2,19 | 1,131 | 12,831 | 1,10E-37 | 3,35E-37 | 1,00E-08 | 0,0072089 | 0,0036748 | 0,0036748 | TRUE | 0,0884463 | No |
| orf19.1152 | 55367 | 49905 | 53037 | 32479 | 10852 | 7477 | 9985 | 10224 | 52786 | 52768 | 52396 | 50880 | 8695 | 8502 | 8477 | 8313 | 30352,25 | 52208 | 8497 | 0,163 | -2,619 | -90,892 | 0 | 0 | 0,0002283 | 0,0021454 | 0,0007343 | 0,0007343 | TRUE | 0,00355 | No |
| orf19.1153 | 7748 | 6907 | 7682 | 4561 | 1935 | 1314 | 1794 | 1979 | 7387 | 7303 | 7589 | 7145 | 1550 | 1494 | 1523 | 1609 | 4450,14 | 7356 | 1544 | 0,21 | -2,251 | -56,316 | 0 | 0 | 0,0004401 | 0,0024602 | 0,0011695 | 0,0011695 | TRUE | 0,0097097 | No |
| orf19.1154 | 2071 | 1698 | 1779 | 1223 | 7174 | 5153 | 6660 | 7354 | 1974 | 1795 | 1757 | 1916 | 5748 | 5859 | 5654 | 5980 | 3835,6 | 1860 | 5810 | 3,124 | 1,644 | 35,067 | 2,16E-269 | 4,01E-268 | 0,0013069 | 0,0025193 | 0,0017303 | 0,0017303 | TRUE | 0,0224231 | No |
| orf19.1155 | 274 | 256 | 256 | 177 | 1100 | 740 | 1051 | 1124 | 261 | 271 | 253 | 277 | 881 | 841 | 892 | 914 | 573,89 | 266 | 882 | 3,338 | 1,739 | 24,125 | 1,36E-128 | 1,17E-127 | 1,00E-08 | 0,0049523 | 0,0023873 | 0,0023873 | TRUE | 0,0112654 | No |
| orf19.1156 | 208 | 189 | 220 | 136 | 250 | 195 | 264 | 291 | 198 | 200 | 217 | 213 | 200 | 222 | 224 | 237 | 213,92 | 207 | 221 | 1,067 | 0,093 | 0,915 | 0,3600123 | 0,3947145 | 1,00E-08 | 0,0097667 | 0,0052374 | 0,0052374 | TRUE | 0,0465087 | No |
| orf19.1158 | 458 | 423 | 417 | 335 | 381 | 298 | 344 | 430 | 437 | 447 | 412 | 525 | 305 | 339 | 292 | 350 | 388,31 | 455 | 322 | 0,709 | -0,496 | -5,166 | 2,39E-07 | 3,79E-07 | 0,0057741 | 0,0063196 | 0,0062172 | 0,0062172 | TRUE | 0,1089581 | No |
| orf19.1159 | 3963 | 3282 | 3733 | 2139 | 5839 | 3356 | 5563 | 5878 | 3778 | 3470 | 3688 | 3351 | 4679 | 3816 | 4723 | 4779 | 4035,54 | 3572 | 4499 | 1,26 | 0,333 | 4,542 | 5,56E-06 | 8,34E-06 | 0,0063361 | 0,0024981 | 0,0049054 | 0,0049054 | TRUE | 0,1258524 | No |
| orf19.1160 | 1247 | 1013 | 1183 | 776 | 1626 | 1174 | 1433 | 1493 | 1189 | 1071 | 1169 | 1216 | 1303 | 1335 | 1217 | 1214 | 1214,09 | 1161 | 1267 | 1,092 | 0,126 | 2,145 | 0,0319429 | 0,039025 | 0,0017511 | 0,0034437 | 0,0024975 | 0,0024975 | TRUE | 0,0326477 | No |
| orf19.1161 | 3654 | 3059 | 3446 | 2090 | 4449 | 2949 | 4001 | 4430 | 3484 | 3235 | 3404 | 3274 | 3565 | 3353 | 3397 | 3602 | 3414,2 | 3349 | 3479 | 1,039 | 0,055 | 1,262 | 0,2070636 | 0,2345593 | 0,0008229 | 0,0025722 | 0,0015128 | 0,0015128 | TRUE | 0,0090333 | No |
| orf19.1162 | 2002 | 1668 | 1764 | 971 | 2240 | 1441 | 2159 | 2275 | 1909 | 1764 | 1743 | 1521 | 1795 | 1639 | 1833 | 1850 | 1756,54 | 1734 | 1779 | 1,025 | 0,035 | 0,5 | 0,6172452 | 0,6509076 | 0,0050121 | 0,003026 | 0,0041139 | 0,0041139 | TRUE | 0,0776619 | No |
| orf19.1162.1 | 502 | 447 | 444 | 278 | 631 | 437 | 598 | 633 | 479 | 473 | 439 | 436 | 506 | 497 | 508 | 515 | 481,28 | 457 | 506 | 1,107 | 0,147 | 2,127 | 0,0334489 | 0,0407946 | 1,00E-08 | 0,0055028 | 0,002487 | 0,002487 | TRUE | 0,0127865 | No |
| orf19.1163 | 1732 | 1526 | 1683 | 1209 | 1596 | 1130 | 1524 | 1522 | 1651 | 1614 | 1663 | 1894 | 1279 | 1285 | 1294 | 1238 | 1489,57 | 1706 | 1274 | 0,749 | -0,417 | -7,241 | 4,44E-13 | 8,52E-13 | 0,0020712 | 0,0031935 | 0,0025078 | 0,0025078 | TRUE | 0,0617155 | No |
| orf19.1164 | 9610 | 7379 | 8223 | 5485 | 21637 | 13288 | 20689 | 21265 | 9162 | 7802 | 8124 | 8593 | 17337 | 15110 | 17565 | 17290 | 12622,83 | 8420 | 16826 | 1,999 | 0,999 | 15,797 | 3,26E-56 | 1,30E-55 | 0,0046856 | 0,0022214 | 0,0037481 | 0,0037481 | TRUE | 0,0574511 | No |
| orf19.1166 | 6185 | 5158 | 5779 | 3889 | 6175 | 4105 | 5833 | 6337 | 5897 | 5454 | 5709 | 6092 | 4948 | 4668 | 4952 | 5153 | 5359,06 | 5788 | 4930 | 0,853 | -0,23 | -4,873 | 1,10E-06 | 1,70E-06 | 0,0016722 | 0,0023977 | 0,0019546 | 0,0019546 | TRUE | 0,0183181 | No |
| orf19.1167 | 898 | 786 | 808 | 550 | 811 | 571 | 809 | 868 | 856 | 831 | 798 | 862 | 650 | 649 | 687 | 706 | 754,85 | 837 | 673 | 0,806 | -0,311 | -5,169 | 2,35E-07 | 3,73E-07 | 1,00E-08 | 0,0042664 | 0,0021477 | 0,0021477 | TRUE | 0,0130266 | No |
| orf19.1168 | 9210 | 7383 | 8350 | 5658 | 5039 | 3430 | 4869 | 4930 | 8781 | 7807 | 8249 | 8864 | 4038 | 3900 | 4134 | 4009 | 6222,5 | 8425 | 4020 | 0,477 | -1,067 | -22,081 | 4,82E-108 | 3,42E-107 | 0,0017605 | 0,0023551 | 0,0020632 | 0,0020632 | TRUE | 0,029819 | No |
| orf19.1169 | 6683 | 5818 | 6486 | 3468 | 3040 | 2098 | 3024 | 3169 | 6372 | 6152 | 6408 | 5433 | 2436 | 2386 | 2567 | 2577 | 4291,15 | 6091 | 2492 | 0,409 | -1,29 | -22,065 | 6,78E-108 | 4,79E-107 | 0,0031231 | 0,0024739 | 0,0030125 | 0,0030125 | TRUE | 0,0629899 | No |
| orf19.117 | 4006 | 3732 | 4081 | 2527 | 3446 | 2496 | 3396 | 3373 | 3819 | 3946 | 4032 | 3959 | 2761 | 2838 | 2883 | 2743 | 3372,61 | 3939 | 2806 | 0,712 | -0,489 | -13,042 | 7,05E-39 | 2,19E-38 | 0,0002797 | 0,0025781 | 0,0010495 | 0,0010495 | TRUE | 0,005331 | No |
| orf19.1170 | 1777 | 1564 | 1681 | 1061 | 2145 | 1540 | 1922 | 2053 | 1694 | 1654 | 1661 | 1662 | 1719 | 1751 | 1632 | 1669 | 1680,2 | 1668 | 1693 | 1,014 | 0,02 | 0,459 | 0,6462391 | 0,678567 | 1,00E-08 | 0,0030685 | 0,0012371 | 0,0012371 | TRUE | 0,0070041 | No |
| orf19.1171 | 30 | 31 | 29 | 22 | 32 | 18 | 31 | 36 | 29 | 33 | 29 | 34 | 26 | 20 | 26 | 29 | 28,27 | 31 | 25 | 0,83 | -0,269 | -1,032 | 0,3020926 | 0,3350109 | 1,00E-08 | 0,0601623 | 0,0296843 | 0,0296843 | TRUE | 0,0856793 | No |
| orf19.1177 | 479 | 363 | 445 | 251 | 1124 | 787 | 1081 | 1162 | 457 | 384 | 440 | 393 | 901 | 895 | 918 | 945 | 666,43 | 418 | 915 | 2,178 | 1,123 | 16,007 | 1,14E-57 | 4,63E-57 | 0,0007317 | 0,004555 | 0,0029081 | 0,0029081 | TRUE | 0,0449709 | No |
| orf19.1178 | 130 | 104 | 123 | 84 | 214 | 157 | 202 | 227 | 124 | 110 | 122 | 132 | 171 | 179 | 171 | 185 | 149,13 | 122 | 176 | 1,456 | 0,542 | 4,594 | 4,36E-06 | 6,57E-06 | 1,00E-08 | 0,0131008 | 0,0062416 | 0,0062416 | TRUE | 0,041404 | No |
| orf19.1179 | 1708 | 1222 | 1268 | 1048 | 3350 | 2442 | 3109 | 3390 | 1628 | 1292 | 1253 | 1642 | 2684 | 2777 | 2640 | 2756 | 2083,98 | 1454 | 2714 | 1,87 | 0,903 | 10,563 | 4,44E-26 | 1,12E-25 | 0,0100133 | 0,0028791 | 0,0064621 | 0,0064621 | TRUE | 0,1050803 | No |
| orf19.118 | 18331 | 17865 | 18334 | 11007 | 51181 | 37314 | 46386 | 46771 | 17477 | 18890 | 18112 | 17243 | 41010 | 42429 | 39381 | 38029 | 29071,49 | 17930 | 40212 | 2,242 | 1,165 | 25,228 | 1,96E-140 | 1,85E-139 | 0,0019269 | 0,0021478 | 0,0020061 | 0,0020061 | TRUE | 0,0168243 | No |
| orf19.1180 | 296 | 209 | 281 | 190 | 865 | 518 | 817 | 852 | 282 | 221 | 278 | 298 | 693 | 589 | 694 | 693 | 468,37 | 270 | 667 | 2,487 | 1,315 | 13,491 | 1,76E-41 | 5,70E-41 | 0,0072231 | 0,0055969 | 0,0063655 | 0,0063655 | TRUE | 0,1655541 | No |
| orf19.1181 | 93 | 69 | 91 | 50 | 336 | 245 | 310 | 355 | 89 | 73 | 90 | 78 | 269 | 279 | 263 | 289 | 178,69 | 82 | 275 | 3,314 | 1,729 | 14,204 | 8,63E-46 | 3,00E-45 | 1,00E-08 | 0,0112799 | 0,0059212 | 0,0059212 | TRUE | 0,0643711 | No |
| orf19.1182 | 707 | 608 | 642 | 405 | 1026 | 627 | 999 | 1017 | 674 | 643 | 634 | 634 | 822 | 713 | 848 | 827 | 724,47 | 646 | 802 | 1,243 | 0,314 | 4,546 | 5,47E-06 | 8,20E-06 | 0,0021 | 0,0043576 | 0,003166 | 0,003166 | TRUE | 0,0620868 | No |
| orf19.1183 | 2597 | 2139 | 2513 | 1473 | 4527 | 2741 | 4196 | 4748 | 2476 | 2262 | 2483 | 2308 | 3627 | 3117 | 3562 | 3861 | 2961,86 | 2382 | 3542 | 1,487 | 0,572 | 8,579 | 9,54E-18 | 2,04E-17 | 0,0045297 | 0,0026456 | 0,0039136 | 0,0039136 | TRUE | 0,0780224 | No |
| orf19.1185 | 4333 | 3438 | 3895 | 2614 | 5078 | 3539 | 4901 | 5249 | 4131 | 3635 | 3848 | 4095 | 4069 | 4024 | 4161 | 4268 | 4028,88 | 3927 | 4130 | 1,052 | 0,074 | 1,5 | 0,1336931 | 0,1540997 | 0,0017522 | 0,0024988 | 0,0020689 | 0,0020689 | TRUE | 0,0305966 | No |
| orf19.1186 | 949 | 727 | 807 | 481 | 1601 | 1072 | 1565 | 1587 | 905 | 769 | 797 | 754 | 1283 | 1219 | 1329 | 1290 | 1043,14 | 806 | 1280 | 1,585 | 0,664 | 10,06 | 8,34E-24 | 2,02E-23 | 0,0028693 | 0,0036653 | 0,0031431 | 0,0031431 | TRUE | 0,0831404 | No |
| orf19.1187 | 36280 | 33438 | 37018 | 23905 | 14300 | 11034 | 13843 | 15232 | 34589 | 35356 | 36570 | 37448 | 11458 | 12547 | 11753 | 12385 | 24013,35 | 35991 | 12036 | 0,334 | -1,58 | -37,005 | 9,67E-300 | 2,14E-298 | 0,001464 | 0,0021597 | 0,0016995 | 0,0016995 | TRUE | 0,0127859 | No |
| orf19.1189 | 27688 | 22718 | 25701 | 17458 | 5125 | 3611 | 5131 | 5427 | 26398 | 24021 | 25390 | 27349 | 4107 | 4106 | 4356 | 4413 | 15017,42 | 25790 | 4246 | 0,165 | -2,602 | -52,885 | 0 | 0 | 0,0020727 | 0,0022006 | 0,0021988 | 0,0021988 | TRUE | 0,0260584 | No |
| orf19.119 | 255 | 248 | 254 | 171 | 745 | 604 | 692 | 554 | 243 | 262 | 251 | 268 | 597 | 687 | 588 | 450 | 418,23 | 256 | 580 | 2,267 | 1,18 | 9,863 | 6,02E-23 | 1,43E-22 | 0,0151648 | 0,0060171 | 0,0107863 | 0,0107863 | TRUE | 0,2400209 | No |
| orf19.1190 | 1669 | 1433 | 1656 | 971 | 2800 | 1977 | 2644 | 2774 | 1591 | 1515 | 1636 | 1521 | 2244 | 2248 | 2245 | 2256 | 1906,92 | 1566 | 2248 | 1,433 | 0,519 | 12,133 | 7,06E-34 | 2,02E-33 | 1,00E-08 | 0,0029523 | 0,0012045 | 0,0012045 | TRUE | 0,0110228 | No |
| orf19.1191 | 4888 | 4380 | 4864 | 2929 | 3675 | 2515 | 3476 | 3766 | 4660 | 4631 | 4805 | 4588 | 2945 | 2860 | 2951 | 3062 | 3812,84 | 4671 | 2954 | 0,633 | -0,661 | -17,794 | 7,88E-71 | 3,81E-70 | 0,0002286 | 0,0025219 | 0,0010521 | 0,0010521 | TRUE | 0,0073253 | No |
| orf19.1192 | 745 | 680 | 680 | 471 | 1366 | 917 | 1234 | 1424 | 710 | 719 | 672 | 738 | 1095 | 1043 | 1048 | 1158 | 897,71 | 710 | 1086 | 1,534 | 0,618 | 10,192 | 2,16E-24 | 5,28E-24 | 0,0010509 | 0,0039203 | 0,0023249 | 0,0023249 | TRUE | 0,0242901 | No |
| orf19.1193 | 124718 | 104779 | 116396 | 73853 | 54443 | 38048 | 51742 | 56075 | 118905 | 110791 | 114988 | 115695 | 43624 | 43264 | 43929 | 45594 | 79598,65 | 115095 | 44103 | 0,383 | -1,384 | -41,287 | 0 | 0 | 0,0006727 | 0,0021119 | 0,0010643 | 0,0010643 | TRUE | 0,0077769 | No |
| orf19.1195 | 1529 | 1366 | 1531 | 922 | 3028 | 1981 | 2922 | 3147 | 1458 | 1444 | 1512 | 1444 | 2426 | 2253 | 2481 | 2559 | 1947,17 | 1464 | 2430 | 1,659 | 0,731 | 14,257 | 4,07E-46 | 1,42E-45 | 0,0013338 | 0,0029345 | 0,0019556 | 0,0019556 | TRUE | 0,0279029 | No |
| orf19.1196 | 1441 | 1320 | 1443 | 915 | 1699 | 1092 | 1579 | 1663 | 1374 | 1396 | 1426 | 1433 | 1361 | 1242 | 1341 | 1352 | 1365,54 | 1407 | 1324 | 0,943 | -0,085 | -1,721 | 0,0853283 | 0,1002714 | 0,0002442 | 0,0032937 | 0,0015812 | 0,0015812 | TRUE | 0,0193722 | No |
| orf19.1198 | 1540 | 1364 | 1439 | 940 | 846 | 608 | 819 | 868 | 1468 | 1442 | 1422 | 1473 | 678 | 691 | 695 | 706 | 1071,87 | 1451 | 692 | 0,478 | -1,066 | -21,399 | 1,36E-101 | 8,98E-101 | 1,00E-08 | 0,0036231 | 0,0013559 | 0,0013559 | TRUE | 0,002652 | No |
| orf19.1199 | 22316 | 19379 | 21051 | 14824 | 41584 | 31540 | 39692 | 41348 | 21276 | 20491 | 20796 | 23223 | 33320 | 35864 | 33698 | 33620 | 27785,97 | 21446 | 34126 | 1,591 | 0,67 | 14,158 | 1,67E-45 | 5,76E-45 | 0,0021751 | 0,0021504 | 0,0021147 | 0,0021147 | TRUE | 0,0376986 | No |

| | | | | | | | | | | | | | | | | | | | | | | | | | | | | | | | |
|---|---|---|---|---|---|---|---|---|---|---|---|---|---|---|---|---|---|---|---|---|---|---|---|---|---|---|---|---|---|---|---|
| orf19.120 | 197 | 233 | 206 | 138 | 466 | 329 | 391 | 378 | 188 | 246 | 204 | 216 | 373 | 374 | 332 | 307 | 280,09 | 214 | 346 | 1,625 | 0,7 | 6,345 | 2,22E-10 | 3,92E-10 | 0,0075973 | 0,0079534 | 0,0077829 | 0,0077829 | TRUE | 0,1258313 | No |
| orf19.1200 | 2386 | 2034 | 2188 | 1489 | 2973 | 1995 | 2847 | 2915 | 2275 | 2151 | 2162 | 2333 | 2382 | 2268 | 2417 | 2370 | 2294,7 | 2230 | 2359 | 1,06 | 0,084 | 1,892 | 0,0584647 | 0,0697706 | 0,0005725 | 0,0028068 | 0,0014603 | 0,0014603 | TRUE | 0,0104804 | No |
| orf19.1201 | 2793 | 2281 | 2559 | 1640 | 2041 | 1498 | 1944 | 2114 | 2663 | 2412 | 2528 | 2569 | 1635 | 1703 | 1650 | 1719 | 2110 | 2543 | 1677 | 0,659 | -0,601 | -13,031 | 8,16E-39 | 2,54E-38 | 0,0007616 | 0,0028694 | 0,0015555 | 0,0015555 | TRUE | 0,0149648 | No |
| orf19.1202 | 127 | 142 | 146 | 113 | 110 | 67 | 97 | 106 | 121 | 150 | 144 | 177 | 88 | 76 | 82 | 86 | 115,67 | 148 | 83 | 0,569 | -0,813 | -5,502 | 3,75E-08 | 6,12E-08 | 0,006172 | 0,0162862 | 0,0118073 | 0,0118073 | TRUE | 0,1614208 | No |
| orf19.1203 | 976 | 831 | 874 | 523 | 1640 | 1158 | 1548 | 1641 | 931 | 879 | 863 | 819 | 1314 | 1317 | 1314 | 1334 | 1096,41 | 873 | 1320 | 1,507 | 0,592 | 11,213 | 3,50E-29 | 9,31E-29 | 1,00E-08 | 0,0035888 | 0,0016976 | 0,0016976 | TRUE | 0,0235785 | No |
| orf19.1203.1 | 380 | 314 | 313 | 242 | 660 | 442 | 534 | 635 | 362 | 332 | 309 | 379 | 529 | 503 | 453 | 516 | 422,97 | 346 | 500 | 1,454 | 0,54 | 6,181 | 6,39E-10 | 1,11E-09 | 0,0035738 | 0,0059732 | 0,0048039 | 0,0048039 | TRUE | 0,0582776 | No |
| orf19.1204 | 1244 | 1075 | 1232 | 763 | 1066 | 696 | 991 | 1047 | 1186 | 1137 | 1217 | 1195 | 854 | 791 | 841 | 851 | 1009,16 | 1184 | 834 | 0,706 | -0,502 | -9,44 | 3,73E-21 | 8,55E-21 | 1,00E-08 | 0,0037183 | 0,0017049 | 0,0017049 | TRUE | 0,0129667 | No |
| orf19.1207 | 9 | 3 | 9 | 4 | 5 | 1 | 3 | 2 | 9 | 3 | 9 | 6 | 4 | 1 | 3 | 2 | 4,53 | 7 | 2 | 0,353 | -1,503 | -2,107 | 0,0351355 | 0,0427699 | 1,00E-08 | 0,3646761 | 0,2210239 | 0,2210239 | TRUE | 0,3347922 | No |
| orf19.1208 | 20 | 20 | 27 | 19 | 135 | 72 | 118 | 121 | 19 | 21 | 27 | 30 | 108 | 82 | 100 | 98 | 60,66 | 24 | 97 | 4,113 | 2,04 | 9,3 | 1,40E-20 | 3,18E-20 | 1,00E-08 | 0,0291596 | 0,0183177 | 0,0183177 | TRUE | 0,1281201 | No |
| orf19.121 | 339 | 379 | 347 | 174 | 528 | 343 | 467 | 371 | 323 | 401 | 343 | 273 | 423 | 390 | 396 | 302 | 356,32 | 335 | 378 | 1,123 | 0,168 | 1,305 | 0,192016 | 0,217978 | 0,0197713 | 0,0066992 | 0,0130443 | 0,0130443 | TRUE | 0,2186385 | No |
| orf19.1210 | 4505 | 4400 | 4617 | 2773 | 2356 | 1641 | 2138 | 2125 | 4295 | 4652 | 4561 | 4344 | 1888 | 1866 | 1815 | 1728 | 3143,68 | 4463 | 1824 | 0,408 | -1,292 | -27,739 | 2,37E-169 | 2,70E-168 | 0,0011777 | 0,0026136 | 0,0017115 | 0,0017115 | TRUE | 0,0156416 | No |
| orf19.1212 | 804 | 711 | 753 | 557 | 3662 | 2754 | 3463 | 3599 | 767 | 752 | 744 | 873 | 2934 | 3132 | 2940 | 2926 | 1883,37 | 784 | 2983 | 3,824 | 1,935 | 34,298 | 8,49E-258 | 1,51E-256 | 0,0013833 | 0,0029631 | 0,0021888 | 0,0021888 | TRUE | 0,0593073 | No |
| orf19.1214 | 6763 | 6081 | 6547 | 3806 | 10239 | 6956 | 9851 | 9964 | 6448 | 6430 | 6468 | 5962 | 8204 | 7910 | 8363 | 8102 | 7235,84 | 6327 | 8145 | 1,286 | 0,363 | 9,385 | 6,27E-21 | 1,43E-20 | 0,0008266 | 0,0023182 | 0,0012958 | 0,0012958 | TRUE | 0,0175035 | No |
| orf19.1215 | 11 | 14 | 3 | 12 | 5 | 8 | 4 | 6 | 10 | 15 | 3 | 19 | 4 | 9 | 3 | 5 | 8,55 | 12 | 5 | 0,454 | -1,139 | -1,925 | 0,0542881 | 0,0649929 | 0,2129283 | 0,1940411 | 0,2022068 | 0,2022068 | TRUE | 0,2567638 | No |
| orf19.1217 | 763 | 821 | 858 | 466 | 1262 | 995 | 1143 | 1190 | 727 | 868 | 848 | 730 | 1011 | 1131 | 970 | 968 | 906,72 | 793 | 1020 | 1,282 | 0,358 | 4,508 | 6,54E-06 | 9,78E-06 | 0,005813 | 0,0039021 | 0,0049236 | 0,0049236 | TRUE | 0,0619081 | No |
| orf19.1219 | 309 | 313 | 320 | 157 | 550 | 373 | 481 | 574 | 295 | 331 | 316 | 246 | 441 | 424 | 408 | 467 | 365,94 | 297 | 435 | 1,456 | 0,542 | 5,723 | 1,05E-08 | 1,74E-08 | 0,0045578 | 0,006578 | 0,0057054 | 0,0057054 | TRUE | 0,1224276 | No |
| orf19.122 | 707 | 678 | 716 | 445 | 630 | 432 | 616 | 614 | 674 | 717 | 707 | 697 | 505 | 491 | 523 | 499 | 601,71 | 699 | 504 | 0,723 | -0,468 | -7,522 | 5,40E-14 | 1,06E-13 | 1,00E-08 | 0,00482 | 0,0020434 | 0,0020434 | TRUE | 0,007506 | No |
| orf19.1220 | 12106 | 10859 | 12070 | 7778 | 7019 | 4715 | 6137 | 6442 | 11542 | 11482 | 11924 | 12185 | 5624 | 5361 | 5210 | 5238 | 8570,77 | 11783 | 5358 | 0,455 | -1,136 | -29,231 | 7,94E-188 | 1,01E-186 | 0,0008366 | 0,0022829 | 0,0013202 | 0,0013202 | TRUE | 0,0138607 | No |
| orf19.1221 | 900 | 862 | 925 | 540 | 1342 | 997 | 1148 | 1184 | 858 | 911 | 914 | 846 | 1075 | 1134 | 975 | 963 | 959,45 | 882 | 1037 | 1,172 | 0,229 | 3,369 | 0,0007544 | 0,0010268 | 0,0029911 | 0,0038026 | 0,0033535 | 0,0033535 | TRUE | 0,0449912 | No |
| orf19.1223 | 958 | 1086 | 1084 | 642 | 688 | 547 | 657 | 631 | 913 | 1148 | 1071 | 1006 | 551 | 622 | 558 | 513 | 797,8 | 1034 | 561 | 0,541 | -0,886 | -10,506 | 8,13E-26 | 2,04E-25 | 0,0066701 | 0,0041493 | 0,0054894 | 0,0054894 | TRUE | 0,0781091 | No |
| orf19.1224 | 5623 | 5785 | 5913 | 4173 | 17520 | 13486 | 16378 | 16143 | 5361 | 6117 | 5841 | 6537 | 14038 | 15335 | 13905 | 13126 | 10032,53 | 5964 | 14101 | 2,365 | 1,242 | 18,847 | 3,08E-79 | 1,67E-78 | 0,0053732 | 0,0022549 | 0,0040459 | 0,0040459 | TRUE | 0,0566971 | No |
| orf19.1225 | 1597 | 1639 | 1637 | 937 | 3168 | 2205 | 2768 | 2657 | 1523 | 1733 | 1617 | 1468 | 2538 | 2507 | 2350 | 2160 | 1987,1 | 1585 | 2389 | 1,505 | 0,59 | 8,594 | 8,40E-18 | 1,80E-17 | 0,0046224 | 0,0029176 | 0,0039789 | 0,0039789 | TRUE | 0,0508163 | No |
| orf19.1227 | 133 | 120 | 129 | 76 | 44 | 35 | 60 | 64 | 127 | 127 | 127 | 119 | 35 | 40 | 51 | 52 | 84,78 | 125 | 44 | 0,357 | -1,488 | -8,813 | 1,22E-18 | 2,65E-18 | 1,00E-08 | 0,0214586 | 0,0129769 | 0,0129769 | TRUE | 0,1892621 | No |
| orf19.1228 | 3775 | 3579 | 3831 | 2404 | 1928 | 1324 | 1834 | 1798 | 3599 | 3784 | 3785 | 3766 | 1545 | 1506 | 1557 | 1462 | 2625,43 | 3734 | 1518 | 0,407 | -1,298 | -31,265 | 1,41E-214 | 2,06E-213 | 0,0003392 | 0,0027167 | 0,0012112 | 0,0012112 | TRUE | 0,0077437 | No |
| orf19.1229 | 1634 | 2169 | 2077 | 1214 | 1810 | 1400 | 1477 | 1299 | 1558 | 2293 | 2052 | 1902 | 1450 | 1592 | 1254 | 1056 | 1644,67 | 1951 | 1338 | 0,686 | -0,545 | -3,222 | 0,0012723 | 0,0017023 | 0,0268358 | 0,0030896 | 0,0149611 | 0,0268358 | TRUE | 0,2430975 | No |
| orf19.123 | 414 | 456 | 492 | 228 | 671 | 433 | 626 | 526 | 395 | 482 | 486 | 357 | 538 | 492 | 531 | 428 | 463,66 | 430 | 497 | 1,152 | 0,204 | 1,831 | 0,0670872 | 0,079777 | 0,0134624 | 0,0056325 | 0,0097511 | 0,0097511 | TRUE | 0,1334042 | No |
| orf19.1232 | 3432 | 3901 | 3708 | 2047 | 7355 | 5330 | 6417 | 5821 | 3272 | 4125 | 3663 | 3207 | 5893 | 6061 | 5448 | 4733 | 4550,23 | 3567 | 5534 | 1,551 | 0,633 | 6,949 | 3,69E-12 | 6,87E-12 | 0,0124752 | 0,0024521 | 0,0077306 | 0,0077306 | TRUE | 0,1349167 | No |
| orf19.1233 | 9158 | 9322 | 9884 | 5906 | 14880 | 11256 | 14116 | 14181 | 8731 | 9857 | 9764 | 9252 | 11923 | 12799 | 11984 | 11531 | 10730,18 | 9401 | 12059 | 1,282 | 0,359 | 7,108 | 1,18E-12 | 2,22E-12 | 0,0023781 | 0,0022443 | 0,0023525 | 0,0023525 | TRUE | 0,0285471 | No |
| orf19.1234 | 39 | 42 | 53 | 33 | 163 | 103 | 161 | 159 | 37 | 44 | 52 | 52 | 131 | 117 | 137 | 129 | 87,42 | 46 | 128 | 2,8 | 1,485 | 8,811 | 1,24E-18 | 2,69E-18 | 1,00E-08 | 0,0208735 | 0,0117729 | 0,0117729 | TRUE | 0,1590151 | No |
| orf19.1235 | 2594 | 2448 | 2579 | 1626 | 2862 | 2185 | 2677 | 2927 | 2473 | 2588 | 2548 | 2547 | 2293 | 2485 | 2273 | 2380 | 2448,38 | 2539 | 2358 | 0,928 | -0,108 | -2,483 | 0,0130361 | 0,0164362 | 0,0005946 | 0,0027619 | 0,0014075 | 0,0014075 | TRUE | 0,0150574 | No |
| orf19.1236 | 2785 | 2663 | 2654 | 1989 | 2491 | 2139 | 2414 | 2533 | 2655 | 2816 | 2622 | 3116 | 1996 | 2432 | 2049 | 2060 | 2468,25 | 2802 | 2134 | 0,762 | -0,393 | -5,239 | 1,61E-07 | 2,57E-07 | 0,0071061 | 0,0027565 | 0,0049811 | 0,0049811 | TRUE | 0,1048715 | No |
| orf19.1237 | 823 | 778 | 772 | 528 | 684 | 508 | 707 | 679 | 785 | 823 | 763 | 827 | 548 | 578 | 600 | 552 | 684,39 | 800 | 570 | 0,713 | -0,487 | -7,741 | 9,88E-15 | 1,98E-14 | 1,00E-08 | 0,0044904 | 0,002324 | 0,002324 | TRUE | 0,0166053 | No |
| orf19.1238 | 235 | 230 | 226 | 144 | 516 | 335 | 479 | 483 | 224 | 243 | 223 | 226 | 413 | 381 | 407 | 393 | 313,73 | 229 | 398 | 1,742 | 0,801 | 9,417 | 4,65E-21 | 1,06E-20 | 1,00E-08 | 0,0073247 | 0,0033983 | 0,0033983 | TRUE | 0,0198848 | No |
| orf19.1239 | 10 | 3 | 6 | 4 | 8 | 4 | 10 | 7 | 10 | 3 | 6 | 6 | 6 | 5 | 8 | 6 | 6,25 | 6 | 6 | 1,012 | 0,018 | 0,031 | 0,975009 | 0,9776135 | 1,00E-08 | 0,2645867 | 0,1480859 | 0,1480859 | TRUE | 0,3540534 | No |
| orf19.124 | 8265 | 8415 | 8613 | 5433 | 8822 | 6587 | 8534 | 8308 | 7880 | 8898 | 8509 | 8511 | 7069 | 7490 | 7245 | 6755 | 7794,61 | 8450 | 7140 | 0,845 | -0,243 | -4,976 | 6,49E-07 | 1,01E-06 | 0,0020092 | 0,0023019 | 0,0021636 | 0,0021636 | TRUE | 0,0254876 | No |
| orf19.1240 | 9978 | 9008 | 9729 | 6093 | 4221 | 3056 | 3782 | 4197 | 9513 | 9525 | 9611 | 9545 | 3382 | 3475 | 3211 | 3413 | 6459,33 | 9548 | 3370 | 0,353 | -1,503 | -43,633 | 0 | 0 | 0,0003048 | 0,0023455 | 0,000948 | 0,000948 | TRUE | 0,0123207 | No |
| orf19.1244 | 2597 | 2492 | 2626 | 1542 | 2538 | 1769 | 2350 | 2567 | 2476 | 2635 | 2594 | 2416 | 2034 | 2012 | 1995 | 2087 | 2281,04 | 2530 | 2032 | 0,802 | -0,318 | -7,255 | 4,02E-13 | 7,72E-13 | 0,0004143 | 0,0028111 | 0,0013995 | 0,0013995 | TRUE | 0,0101582 | No |
| orf19.1246 | 4766 | 4092 | 4683 | 3028 | 5660 | 3947 | 5309 | 5750 | 4544 | 4327 | 4626 | 4744 | 4535 | 4488 | 4507 | 4675 | 4555,8 | 4560 | 4551 | 0,999 | -0,001 | -0,038 | 0,9696251 | 0,9726737 | 0,0005637 | 0,0024517 | 0,001259 | 0,001259 | TRUE | 0,0143615 | No |
| orf19.1247 | 266 | 232 | 249 | 144 | 641 | 396 | 609 | 662 | 254 | 245 | 246 | 226 | 514 | 450 | 517 | 538 | 373,71 | 243 | 505 | 2,077 | 1,055 | 12,036 | 2,30E-33 | 6,55E-33 | 1,00E-08 | 0,0064848 | 0,0042058 | 0,0042058 | TRUE | 0,0564946 | No |
| orf19.1248 | 4050 | 3417 | 3707 | 2565 | 5346 | 3681 | 5089 | 5279 | 3861 | 3613 | 3662 | 4018 | 4284 | 4186 | 4321 | 4292 | 4029,59 | 3788 | 4271 | 1,129 | 0,175 | 4,101 | 4,12E-05 | 5,93E-05 | 0,0009794 | 0,0024987 | 0,0014873 | 0,0014873 | TRUE | 0,0189337 | No |
| orf19.1249 | 738 | 587 | 640 | 414 | 2101 | 1458 | 2021 | 2165 | 704 | 621 | 632 | 649 | 1683 | 1658 | 1716 | 1760 | 1177,83 | 652 | 1704 | 2,616 | 1,387 | 24,546 | 4,74E-133 | 4,27E-132 | 0,0005104 | 0,0034853 | 0,0019489 | 0,0019489 | TRUE | 0,038109 | No |
| orf19.125 | 9690 | 9205 | 9602 | 5680 | 6657 | 4953 | 6264 | 6625 | 9238 | 9733 | 9486 | 8898 | 5334 | 5632 | 5318 | 5387 | 7378,29 | 9339 | 5418 | 0,58 | -0,787 | -19,791 | 3,59E-87 | 2,09E-86 | 0,0008969 | 0,0023138 | 0,0013746 | 0,0013746 | TRUE | 0,0119241 | No |
| orf19.1250 | 3919 | 3250 | 3574 | 1957 | 8599 | 5324 | 8627 | 8682 | 3736 | 3436 | 3531 | 3066 | 6890 | 6054 | 7324 | 7059 | 5137,11 | 3442 | 6832 | 1,983 | 0,988 | 13,825 | 1,80E-43 | 6,04E-43 | 0,0062538 | 0,0024109 | 0,0046758 | 0,0046758 | TRUE | 0,0711844 | No |
| orf19.1251 | 1912 | 1739 | 1863 | 1230 | 2051 | 1457 | 1882 | 2062 | 1823 | 1839 | 1840 | 1927 | 1643 | 1657 | 1598 | 1677 | 1750,44 | 1857 | 1644 | 0,886 | -0,174 | -4,081 | 4,49E-05 | 6,46E-05 | 1,00E-08 | 0,0030293 | 0,0011777 | 0,0011777 | TRUE | 0,0067734 | No |
| orf19.1252 | 20845 | 17709 | 19667 | 12803 | 13099 | 9311 | 12699 | 13620 | 19873 | 18725 | 19429 | 20057 | 10496 | 10587 | 10781 | 11074 | 15127,91 | 19521 | 10734 | 0,55 | -0,862 | -24,502 | 1,41E-132 | 1,25E-131 | 0,0006482 | 0,0021998 | 0,0011195 | 0,0011195 | TRUE | 0,0092222 | No |
| orf19.1253 | 2165 | 1835 | 2106 | 1387 | 1616 | 1174 | 1517 | 1686 | 2064 | 1940 | 2081 | 2173 | 1295 | 1335 | 1288 | 1371 | 1693,29 | 2064 | 1322 | 0,641 | -0,641 | -12,846 | 9,09E-38 | 2,78E-37 | 0,0008946 | 0,0030609 | 0,0017826 | 0,0017826 | TRUE | 0,0195451 | No |
| orf19.1254 | 1002 | 837 | 839 | 532 | 2013 | 1425 | 1962 | 2259 | 955 | 885 | 829 | 833 | 1613 | 1620 | 1666 | 1837 | 1279,8 | 876 | 1684 | 1,921 | 0,942 | 14,177 | 1,27E-45 | 4,40E-45 | 0,003031 | 0,0033742 | 0,0033295 | 0,0033295 | TRUE | 0,0461679 | No |
| orf19.1255 | 61 | 42 | 58 | 43 | 515 | 355 | 413 | 518 | 58 | 44 | 57 | 67 | 413 | 404 | 351 | 421 | 226,92 | 56 | 397 | 7,072 | 2,822 | 20,417 | 1,19E-92 | 7,30E-92 | 0,0046609 | 0,0093269 | 0,00737 | 0,00737 | TRUE | 0,174391 | No |
| orf19.1257 | 12 | 9 | 10 | 9 | 20 | 22 | 21 | 30 | 11 | 10 | 10 | 14 | 16 | 25 | 18 | 24 | 16,02 | 11 | 21 | 1,87 | 0,903 | 2,458 | 0,0139885 | 0,0175987 | 1,00E-08 | 0,1045515 | 0,0579789 | 0,0579789 | TRUE | 0,1535167 | No |
| orf19.1258 | 92 | 91 | 70 | 52 | 183 | 141 | 176 | 171 | 88 | 96 | 69 | 81 | 147 | 160 | 149 | 139 | 116,25 | 84 | 149 | 1,774 | 0,827 | 5,831 | 5,52E-09 | 9,25E-09 | 1,00E-08 | 0,0162156 | 0,0096896 | 0,0096896 | TRUE | 0,1448713 | No |
| orf19.1259 | 3926 | 3647 | 3837 | 2424 | 4384 | 3095 | 4187 | 4546 | 3743 | 3856 | 3791 | 3797 | 3513 | 3519 | 3555 | 3696 | 3683,79 | 3797 | 3571 | 0,941 | -0,088 | -2,504 | 0,0122948 | 0,0155352 | 4,98E-07 | 0,002537 | 0,0009136 | 0,0009136 | TRUE | 0,006951 | No |
| orf19.1260 | 681 | 614 | 598 | 379 | 1010 | 703 | 928 | 997 | 649 | 649 | 591 | 594 | 809 | 799 | 788 | 811 | 711,27 | 621 | 802 | 1,289 | 0,366 | 6,022 | 1,72E-09 | 2,93E-09 | 1,00E-08 | 0,0043997 | 0,0021006 | 0,0021006 | TRUE | 0,0148651 | No |
| orf19.1261 | 3116 | 2822 | 3221 | 1903 | 2137 | 1477 | 2116 | 2169 | 2971 | 2984 | 3182 | 2981 | 1712 | 1679 | 1796 | 1764 | 2383,72 | 3030 | 1738 | 0,574 | -0,801 | -18,18 | 7,45E-74 | 3,77E-73 | 0,0006142 | 0,0027801 | 0,0014231 | 0,0014231 | TRUE | 0,0143598 | No |
| orf19.1263 | 2360 | 2188 | 2438 | 1560 | 2753 | 1973 | 2573 | 2897 | 2250 | 2314 | 2409 | 2444 | 2206 | 2243 | 2184 | 2356 | 2300,66 | 2354 | 2247 | 0,956 | -0,065 | -1,416 | 0,1567411 | 0,1796273 | 0,0008615 | 0,0028049 | 0,0015971 | 0,0015971 | TRUE | 0,0131228 | No |
| orf19.1264 | 619 | 561 | 575 | 404 | 749 | 585 | 750 | 815 | 590 | 593 | 568 | 633 | 600 | 665 | 637 | 663 | 618,63 | 596 | 641 | 1,078 | 0,108 | 1,627 | 0,1037482 | 0,1209385 | 1,00E-08 | 0,0047454 | 0,0025917 | 0,0025917 | TRUE | 0,0226737 | No |
| orf19.1265 | 351 | 354 | 319 | 208 | 596 | 416 | 563 | 593 | 335 | 374 | 315 | 326 | 478 | 473 | 478 | 482 | 407,58 | 338 | 478 | 1,414 | 0,5 | 6,438 | 1,21E-10 | 2,16E-10 | 1,00E-08 | 0,0061197 | 0,0032142 | 0,0032142 | TRUE | 0,0628968 | No |
| orf19.1266 | 1542 | 1499 | 1509 | 802 | 998 | 731 | 961 | 995 | 1470 | 1585 | 1491 | 1256 | 800 | 831 | 816 | 809 | 1132,26 | 1450 | 814 | 0,559 | -0,838 | -12,028 | 2,52E-33 | 7,19E-33 | 0,0038781 | 0,0035414 | 0,0037243 | 0,0037243 | TRUE | 0,0902205 | No |
| orf19.1267 | 7180 | 6286 | 7137 | 4305 | 7096 | 4883 | 6643 | 6931 | 6845 | 6647 | 7051 | 6744 | 5686 | 5552 | 5640 | 5636 | 6225,05 | 6822 | 5628 | 0,825 | -0,278 | -8,582 | 9,31E-18 | 1,99E-17 | 0,0002194 | 0,002355 | 0,0008424 | 0,0008424 | TRUE | 0,0063453 | No |
| orf19.1267.1 | 1501 | 1396 | 1383 | 871 | 1330 | 939 | 1125 | 1158 | 1431 | 1476 | 1366 | 1364 | 1066 | 1068 | 955 | 942 | 1208,5 | 1409 | 1008 | 0,714 | -0,487 | -7,979 | 1,47E-15 | 3,00E-15 | 0,0021707 | 0,0034499 | 0,0027339 | 0,0027339 | TRUE | 0,0234973 | No |
| orf19.1268 | 190 | 156 | 158 | 138 | 186 | 148 | 132 | 155 | 181 | 165 | 156 | 216 | 149 | 168 | 112 | 126 | 159,22 | 180 | 139 | 0,775 | -0,368 | -2,45 | 0,0142736 | 0,0179292 | 0,0183904 | 0,0124032 | 0,0153168 | 0,0153168 | TRUE | 0,1924643 | No |
| orf19.1270 | 1179 | 1091 | 1183 | 738 | 1200 | 864 | 1199 | 1241 | 1124 | 1154 | 1169 | 1156 | 962 | 982 | 1018 | 1009 | 1071,68 | 1151 | 993 | 0,863 | -0,212 | -4,269 | 1,96E-05 | 2,87E-05 | 1,00E-08 | 0,0036234 | 0,0014368 | 0,0014368 | TRUE | 0,0058294 | No |
| orf19.1272 | 1009 | 930 | 922 | 621 | 2306 | 1622 | 2109 | 2368 | 962 | 983 | 911 | 973 | 1848 | 1844 | 1791 | 1925 | 1404,63 | 957 | 1852 | 1,937 | 0,954 | 19,05 | 6,61E-81 | 3,64E-80 | 1,00E-08 | 0,0032602 | 0,0015814 | 0,0015814 | TRUE | 0,0133548 | No |
| orf19.1273 | 178 | 125 | 122 | 82 | 357 | 234 | 362 | 374 | 170 | 132 | 121 | 128 | 286 | 266 | 307 | 304 | 214,3 | 138 | 291 | 2,105 | 1,074 | 8,895 | 5,85E-19 | 1,28E-18 | 0,0065569 | 0,0097529 | 0,008439 | 0,008439 | TRUE | 0,2748285 | No |
| orf19.1274 | 12 | 8 | 15 | 13 | 95 | 79 | 83 | 83 | 11 | 8 | 15 | 20 | 76 | 90 | 70 | 67 | 44,87 | 14 | 76 | 5,693 | 2,509 | 9,046 | 1,48E-19 | 3,30E-19 | 0,0036271 | 0,0386809 | 0,0263878 | 0,0263878 | TRUE | 0,2604642 | No |
| orf19.1275 | 282 | 239 | 258 | 185 | 1511 | 1131 | 1474 | 1583 | 269 | 253 | 255 | 290 | 1211 | 1286 | 1251 | 1287 | 762,7 | 267 | 1259 | 4,747 | 2,247 | 32,438 | 8,06E-231 | 1,24E-229 | 1,00E-08 | 0,004244 | 0,0021613 | 0,0021613 | TRUE | 0,028985 | No |
| orf19.1276 | 201 | 223 | 225 | 136 | 287 | 224 | 352 | 287 | 192 | 236 | 222 | 213 | 230 | 255 | 299 | 233 | 234,95 | 216 | 254 | 1,178 | 0,237 | 2,035 | 0,0418361 | 0,0506559 | 0,0081897 | 0,0090795 | 0,0086375 | 0,0086375 | TRUE | 0,1634217 | No |
| orf19.1277 | 15782 | 13278 | 14984 | 10449 | 3351 | 2573 | 3148 | 3403 | 15046 | 14040 | 14803 | 16369 | 2685 | 2926 | 2673 | 2767 | 8913,54 | 15064 | 2763 | 0,183 | -2,447 | -45,442 | 0 | 0 | 0,0026753 | 0,0022755 | 0,002587 | 0,002587 | TRUE | 0,0411104 | No |
| orf19.1279 | 4956 | 4818 | 5094 | 3458 | 7991 | 6132 | 7665 | 7824 | 4725 | 5094 | 5032 | 5417 | 6403 | 6973 | 6508 | 6362 | 5814,22 | 5067 | 6562 | 1,295 | 0,373 | 7,384 | 1,54E-13 | 2,98E-13 | 0,002252 | 0,0023737 | 0,0022778 | 0,0022778 | TRUE | 0,0255169 | No |
| orf19.1280 | 17903 | 17965 | 17399 | 9922 | 21442 | 15116 | 19649 | 19537 | 17069 | 18996 | 17189 | 15543 | 17181 | 17188 | 16682 | 15885 | 16966,58 | 17199 | 16734 | 0,973 | -0,04 | -0,679 | 0,4974313 | 0,5332845 | 0,0036855 | 0,0021881 | 0,0032613 | 0,0032613 | TRUE | 0,0604106 | No |
| orf19.1281 | 4474 | 3926 | 4186 | 2621 | 2352 | 1517 | 2237 | 2258 | 4265 | 4151 | 4135 | 4106 | 1885 | 1725 | 1899 | 1836 | 3000,34 | 4164 | 1836 | 0,441 | -1,18 | -28,067 | 2,46E-173 | 2,86E-172 | 0,0005412 | 0,0026385 | 0,0013192 | 0,0013192 | TRUE | 0,0186347 | No |
| orf19.1282 | 1174 | 1140 | 1195 | 669 | 1273 | 867 | 1269 | 1231 | 1119 | 1205 | 1181 | 1048 | 1020 | 986 | 1077 | 1001 | 1079,68 | 1138 | 1021 | 0,895 | -0,159 | -2,681 | 0,0073352 | 0,0093819 | 0,0014351 | 0,003612 | 0,0024649 | 0,0024649 | TRUE | 0,0300664 | No |
| orf19.1283 | 669 | 661 | 667 | 467 | 1019 | 797 | 942 | 1010 | 638 | 699 | 659 | 732 | 816 | 906 | 800 | 821 | 758,87 | 682 | 836 | 1,228 | 0,296 | 4,396 | 1,10E-05 | 1,63E-05 | 0,0017328 | 0,0042549 | 0,0030024 | 0,0030024 | TRUE | 0,0355869 | No |
| orf19.1285 | 5777 | 5533 | 5973 | 3978 | 2413 | 1877 | 2384 | 2501 | 5508 | 5850 | 5901 | 6232 | 1933 | 2134 | 2024 | 2034 | 3952,01 | 5873 | 2031 | 0,346 | -1,532 | -30,751 | 1,20E-207 | 1,68E-206 | 0,0016487 | 0,0025067 | 0,0020665 | 0,0020665 | TRUE | 0,0214158 | No |

| | | | | | | | | | | | | | | | | | | | | | | | | | | | | | | |
|---|---|---|---|---|---|---|---|---|---|---|---|---|---|---|---|---|---|---|---|---|---|---|---|---|---|---|---|---|---|---|
| orf19.1286 | 144 | 88 | 116 | 67 | 55 | 43 | 59 | 74 | 137 | 93 | 115 | 105 | 44 | 49 | 50 | 60 | 81,64 | 112 | 51 | 0,45 | -1,151 | -6,33 | 2,45E-10 | 4,32E-10 | 0,0131306 | 0,0222029 | 0,0180419 | 0,0180419 | TRUE | 0,233265 | No |
| orf19.1287 | 10769 | 10359 | 10676 | 6717 | 7556 | 5493 | 6928 | 7106 | 10267 | 10953 | 10547 | 10523 | 6054 | 6246 | 5882 | 5778 | 8281,25 | 10572 | 5990 | 0,566 | -0,82 | -21,507 | 1,34E-102 | 8,93E-102 | 0,0007729 | 0,0022895 | 0,0012688 | 0,0012688 | TRUE | 0,0098243 | No |
| orf19.1288 | 6296 | 5354 | 5631 | 3891 | 64810 | 44689 | 59613 | 68681 | 6003 | 5661 | 5563 | 6095 | 51930 | 50815 | 50611 | 55844 | 29065,41 | 5830 | 52300 | 8,974 | 3,166 | 67,416 | 0 | 0 | 0,0019357 | 0,0021478 | 0,0020128 | 0,0020128 | TRUE | 0,0255141 | No |
| orf19.1289 | 2030 | 1898 | 2044 | 1165 | 2128 | 1453 | 1828 | 1993 | 1935 | 2007 | 2019 | 1825 | 1705 | 1652 | 1552 | 1620 | 1789,54 | 1946 | 1632 | 0,837 | -0,256 | -5,028 | 4,96E-07 | 7,76E-07 | 0,0010655 | 0,0030088 | 0,0019352 | 0,0019352 | TRUE | 0,0193998 | No |
| orf19.1290 | 4791 | 4396 | 4762 | 2926 | 971 | 693 | 895 | 902 | 4568 | 4648 | 4704 | 4584 | 778 | 788 | 760 | 733 | 2695,42 | 4626 | 765 | 0,165 | -2,599 | -60,654 | 0 | 0 | 1,00E-08 | 0,0027004 | 0,001059 | 0,001059 | TRUE | 0,0093012 | No |
| orf19.1291 | 2356 | 2071 | 2223 | 1355 | 1681 | 1179 | 1645 | 1667 | 2246 | 2190 | 2196 | 2123 | 1347 | 1341 | 1397 | 1355 | 1774,3 | 2189 | 1360 | 0,621 | -0,688 | -16,22 | 3,64E-59 | 1,52E-58 | 1,00E-08 | 0,0030167 | 0,0011429 | 0,0011429 | TRUE | 0,0044434 | No |
| orf19.1295 | 5607 | 5834 | 5999 | 3443 | 11329 | 7953 | 9874 | 10597 | 5346 | 6169 | 5926 | 5394 | 9078 | 9043 | 8383 | 8616 | 7244,34 | 5709 | 8780 | 1,537 | 0,62 | 11,271 | 1,82E-29 | 4,87E-29 | 0,0030537 | 0,0023179 | 0,0027612 | 0,0027612 | TRUE | 0,0358723 | No |
| orf19.1296 | 1436 | 1533 | 1586 | 901 | 1328 | 1030 | 1249 | 1187 | 1369 | 1621 | 1567 | 1411 | 1064 | 1171 | 1060 | 965 | 1278,64 | 1492 | 1065 | 0,712 | -0,489 | -6,534 | 6,42E-11 | 1,15E-10 | 0,0053327 | 0,0033754 | 0,0045836 | 0,0045836 | TRUE | 0,0516055 | No |
| orf19.1297 | 924 | 949 | 985 | 612 | 724 | 514 | 684 | 689 | 881 | 1003 | 973 | 959 | 580 | 584 | 581 | 560 | 765,21 | 954 | 576 | 0,604 | -0,727 | -11,648 | 2,34E-31 | 6,46E-31 | 1,00E-08 | 0,004237 | 0,0023764 | 0,0023764 | TRUE | 0,0341711 | No |
| orf19.1298 | 3162 | 3341 | 3549 | 1888 | 2853 | 2112 | 2664 | 2599 | 3015 | 3533 | 3506 | 2958 | 2286 | 2402 | 2262 | 2113 | 2759,19 | 3253 | 2266 | 0,696 | -0,523 | -7,506 | 6,11E-14 | 1,20E-13 | 0,0053788 | 0,0026863 | 0,0042999 | 0,0042999 | TRUE | 0,0437558 | No |
| orf19.1299 | 1890 | 2400 | 2371 | 1527 | 2500 | 1688 | 2172 | 1963 | 1802 | 2538 | 2342 | 2392 | 2003 | 1919 | 1844 | 1596 | 2054,6 | 2268 | 1840 | 0,812 | -0,301 | -2,989 | 0,0027972 | 0,0036726 | 0,01396 | 0,0028904 | 0,0092484 | 0,0092484 | TRUE | 0,2349239 | No |
| orf19.130 | 406 | 444 | 465 | 277 | 859 | 685 | 771 | 792 | 387 | 469 | 459 | 434 | 688 | 779 | 655 | 644 | 564,45 | 437 | 692 | 1,578 | 0,658 | 7,562 | 3,97E-14 | 7,80E-14 | 0,0056648 | 0,0050001 | 0,0053403 | 0,0053403 | TRUE | 0,0813977 | No |
| orf19.1300 | 2655 | 2469 | 2605 | 1538 | 2303 | 1653 | 2094 | 2249 | 2531 | 2611 | 2573 | 2409 | 1845 | 1880 | 1778 | 1829 | 2182,02 | 2531 | 1833 | 0,723 | -0,468 | -10,91 | 1,03E-27 | 2,68E-27 | 0,0002373 | 0,0028438 | 0,0013045 | 0,0013045 | TRUE | 0,0114588 | No |
| orf19.1301 | 433 | 482 | 448 | 319 | 913 | 650 | 814 | 715 | 413 | 510 | 443 | 500 | 732 | 739 | 691 | 581 | 575,99 | 466 | 686 | 1,474 | 0,559 | 5,752 | 8,80E-09 | 1,46E-08 | 0,0093418 | 0,0049419 | 0,0072177 | 0,0072177 | TRUE | 0,1266785 | No |
| orf19.1302 | 44 | 34 | 27 | 23 | 17 | 13 | 21 | 23 | 42 | 36 | 27 | 36 | 14 | 15 | 18 | 19 | 25,69 | 35 | 16 | 0,464 | -1,109 | -3,81 | 0,0001389 | 0,0001952 | 1,00E-08 | 0,0659978 | 0,0381578 | 0,0381578 | TRUE | 0,2071938 | No |
| orf19.1303 | 1904 | 1746 | 1887 | 1149 | 1871 | 1227 | 1672 | 1733 | 1815 | 1846 | 1864 | 1800 | 1499 | 1395 | 1420 | 1409 | 1631,07 | 1831 | 1431 | 0,781 | -0,356 | -7,961 | 1,71E-15 | 3,47E-15 | 1,00E-08 | 0,0030979 | 0,0013015 | 0,0013015 | TRUE | 0,0128264 | No |
| orf19.1304 | 5115 | 4919 | 5232 | 2790 | 4798 | 3095 | 4464 | 4453 | 4877 | 5201 | 5169 | 4371 | 3844 | 3519 | 3790 | 3621 | 4298,96 | 4904 | 3694 | 0,753 | -0,41 | -6,856 | 7,10E-12 | 1,31E-11 | 0,003585 | 0,0024732 | 0,0031965 | 0,0031965 | TRUE | 0,0634663 | No |
| orf19.1305 | 1202 | 1419 | 1458 | 887 | 1562 | 1154 | 1344 | 1175 | 1146 | 1500 | 1440 | 1390 | 1252 | 1312 | 1141 | 955 | 1267,06 | 1369 | 1165 | 0,851 | -0,233 | -2,238 | 0,025197 | 0,031039 | 0,0144361 | 0,0033871 | 0,0096484 | 0,0096484 | TRUE | 0,1776398 | No |
| orf19.1306 | 2372 | 2332 | 2436 | 1514 | 3249 | 2214 | 3045 | 3275 | 2261 | 2466 | 2407 | 2372 | 2603 | 2518 | 2585 | 2663 | 2484,31 | 2376 | 2592 | 1,092 | 0,126 | 2,95 | 0,0031726 | 0,0041476 | 0,0005221 | 0,0027522 | 0,0013553 | 0,0013553 | TRUE | 0,0136792 | No |
| orf19.1307 | 764 | 711 | 698 | 501 | 337 | 247 | 300 | 333 | 728 | 752 | 690 | 785 | 270 | 281 | 255 | 271 | 503,87 | 739 | 269 | 0,365 | -1,455 | -19,433 | 4,09E-84 | 2,31E-83 | 1,00E-08 | 0,0053499 | 0,0029702 | 0,0029702 | TRUE | 0,0236143 | No |
| orf19.1308 | 4137 | 3649 | 3881 | 2514 | 2793 | 2013 | 2576 | 2734 | 3944 | 3858 | 3834 | 3938 | 2238 | 2289 | 2187 | 2223 | 3063,98 | 3894 | 2234 | 0,574 | -0,802 | -22,478 | 6,78E-112 | 4,99E-111 | 1,00E-08 | 0,0026272 | 0,0008776 | 0,0008776 | TRUE | 0,0030238 | No |
| orf19.1309 | 747 | 542 | 604 | 380 | 267 | 162 | 244 | 287 | 712 | 573 | 597 | 595 | 214 | 184 | 207 | 233 | 414,49 | 619 | 210 | 0,339 | -1,559 | -15,638 | 4,00E-55 | 1,58E-54 | 0,0069916 | 0,0060525 | 0,0065185 | 0,0065185 | TRUE | 0,1245145 | No |
| orf19.131 | 368 | 331 | 334 | 206 | 575 | 370 | 513 | 524 | 351 | 350 | 330 | 323 | 461 | 421 | 436 | 426 | 387,07 | 338 | 436 | 1,286 | 0,363 | 4,711 | 2,47E-06 | 3,76E-06 | 1,00E-08 | 0,0063332 | 0,0030378 | 0,0030378 | TRUE | 0,0178521 | No |
| orf19.1310 | 745 | 673 | 819 | 493 | 867 | 616 | 817 | 835 | 710 | 712 | 809 | 772 | 695 | 700 | 694 | 679 | 721,38 | 751 | 692 | 0,922 | -0,117 | -1,83 | 0,0671945 | 0,0798748 | 0,0008375 | 0,0043674 | 0,0025173 | 0,0025173 | TRUE | 0,0360787 | No |
| orf19.1311 | 14 | 13 | 12 | 1 | 19 | 8 | 7 | 12 | 13 | 14 | 12 | 2 | 15 | 9 | 6 | 10 | 10,07 | 10 | 10 | 0,956 | -0,065 | -0,127 | 0,8987981 | 0,9129625 | 0,1376085 | 0,1651894 | 0,155783 | 0,155783 | TRUE | 0,6702354 | No |
| orf19.131.2 | 3045 | 2581 | 2569 | 1887 | 3724 | 2862 | 3414 | 3865 | 2903 | 2729 | 2538 | 2956 | 2984 | 3254 | 2898 | 3143 | 2925,69 | 2782 | 3070 | 1,104 | 0,143 | 2,407 | 0,0160906 | 0,02014 | 0,003201 | 0,0026525 | 0,0030358 | 0,0030358 | TRUE | 0,0425879 | No |
| orf19.1313 | 602 | 517 | 550 | 394 | 1391 | 1134 | 1329 | 1453 | 574 | 547 | 543 | 617 | 1115 | 1289 | 1128 | 1181 | 874,37 | 570 | 1178 | 2,07 | 1,05 | 15,142 | 8,60E-52 | 3,25E-51 | 0,0026244 | 0,0039691 | 0,0032507 | 0,0032507 | TRUE | 0,0459377 | No |
| orf19.1314 | 4584 | 4462 | 4678 | 2902 | 3764 | 2830 | 3687 | 3647 | 4370 | 4718 | 4621 | 4546 | 3016 | 3218 | 3130 | 2965 | 3823,18 | 4564 | 3082 | 0,675 | -0,567 | -13,244 | 4,89E-40 | 1,55E-39 | 0,0008366 | 0,0025207 | 0,0014922 | 0,0014922 | TRUE | 0,0103125 | No |
| orf19.1317 | 1246 | 1141 | 1211 | 802 | 2846 | 2071 | 2745 | 2971 | 1188 | 1206 | 1196 | 1256 | 2280 | 2355 | 2330 | 2416 | 1778,58 | 1212 | 2345 | 1,939 | 0,955 | 21,514 | 1,14E-102 | 7,64E-102 | 1,00E-08 | 0,0030144 | 0,0012405 | 0,0012405 | TRUE | 0,006278 | No |
| orf19.1318 | 1137 | 1183 | 1195 | 668 | 1233 | 920 | 1209 | 1103 | 1084 | 1251 | 1181 | 1046 | 988 | 1046 | 1026 | 897 | 1064,91 | 1140 | 989 | 0,865 | -0,209 | -2,87 | 0,0041073 | 0,00534 | 0,0042908 | 0,0036331 | 0,0041359 | 0,0041359 | TRUE | 0,0501479 | No |
| orf19.132 | 3268 | 3043 | 3222 | 2075 | 2514 | 1943 | 2393 | 2466 | 3116 | 3218 | 3183 | 3251 | 2014 | 2209 | 2032 | 2005 | 2628,43 | 3192 | 2065 | 0,646 | -0,629 | -14,269 | 3,44E-46 | 1,21E-45 | 0,0007301 | 0,002716 | 0,0014751 | 0,0014751 | TRUE | 0,0252697 | No |
| orf19.1321 | 13694 | 11416 | 13298 | 8965 | 888 | 615 | 901 | 911 | 13056 | 12071 | 13137 | 14044 | 712 | 699 | 765 | 741 | 6903,07 | 13077 | 729 | 0,056 | -4,163 | -72,717 | 0 | 0 | 0,0026024 | 0,0023291 | 0,0024936 | 0,0024936 | TRUE | 0,0327304 | No |
| orf19.1323 | 1728 | 1630 | 1735 | 1078 | 1934 | 1395 | 1841 | 2024 | 1647 | 1724 | 1714 | 1689 | 1550 | 1586 | 1563 | 1646 | 1639,79 | 1694 | 1586 | 0,937 | -0,094 | -2,13 | 0,0331341 | 0,0404261 | 1,00E-08 | 0,0030926 | 0,0012688 | 0,0012688 | TRUE | 0,0080984 | No |
| orf19.1324 | 1809 | 1568 | 1755 | 1084 | 1916 | 1350 | 1822 | 1988 | 1725 | 1658 | 1734 | 1698 | 1535 | 1535 | 1547 | 1616 | 1631,02 | 1704 | 1558 | 0,915 | -0,128 | -2,926 | 0,003429 | 0,0044782 | 1,00E-08 | 0,003098 | 0,0012341 | 0,0012341 | TRUE | 0,0078006 | No |
| orf19.1325 | 369 | 307 | 353 | 204 | 279 | 180 | 261 | 263 | 352 | 325 | 349 | 320 | 224 | 205 | 222 | 214 | 276,05 | 336 | 216 | 0,642 | -0,64 | -7,216 | 5,35E-13 | 1,02E-12 | 1,00E-08 | 0,0080392 | 0,003846 | 0,003846 | TRUE | 0,0115797 | No |
| orf19.1326 | 267 | 212 | 204 | 136 | 1459 | 1035 | 1422 | 1493 | 255 | 224 | 202 | 213 | 1169 | 1177 | 1207 | 1214 | 707,56 | 224 | 1192 | 5,316 | 2,41 | 31,742 | 4,13E-221 | 6,14E-220 | 1,00E-08 | 0,0044118 | 0,0026886 | 0,0026886 | TRUE | 0,1082133 | No |
| orf19.1327 | 159 | 138 | 150 | 109 | 232 | 178 | 213 | 220 | 152 | 146 | 148 | 171 | 186 | 202 | 181 | 179 | 170,56 | 154 | 187 | 1,218 | 0,284 | 2,551 | 0,0107388 | 0,0136123 | 1,00E-08 | 0,011718 | 0,0058781 | 0,0058781 | TRUE | 0,0424851 | No |
| orf19.133 | 0 | 0 | 0 | 0 | 0 | 2 | 0 | 2 | 0 | 0 | 0 | 0 | 0 | 2 | 0 | 2 | 0,49 | 0 | 1 | 4,651 | 2,218 | 0,898 | 0,3692642 | 0,4042331 | 0,0028521 | 3,369778 | 4,3936766 | 4,3936766 | TRUE | 0,0907559 | No |
| orf19.1330 | 204 | 164 | 209 | 120 | 145 | 105 | 138 | 136 | 194 | 173 | 206 | 188 | 116 | 119 | 117 | 111 | 153,21 | 190 | 116 | 0,606 | -0,723 | -6,162 | 7,19E-10 | 1,24E-09 | 1,00E-08 | 0,012808 | 0,0064485 | 0,0064485 | TRUE | 0,0427411 | No |
| orf19.1331 | 5106 | 5566 | 5505 | 3189 | 1204 | 975 | 1114 | 1069 | 4868 | 5885 | 5438 | 4996 | 965 | 1109 | 946 | 869 | 3134,49 | 5297 | 972 | 0,183 | -2,449 | -30,723 | 2,78E-207 | 3,90E-206 | 0,0082842 | 0,0026151 | 0,0055345 | 0,0055345 | TRUE | 0,1029754 | No |
| orf19.1332 | 966 | 826 | 936 | 623 | 878 | 708 | 905 | 927 | 921 | 873 | 925 | 976 | 704 | 805 | 768 | 754 | 840,71 | 924 | 758 | 0,82 | -0,286 | -4,569 | 4,90E-06 | 7,38E-06 | 0,0012234 | 0,0040443 | 0,0025604 | 0,0025604 | TRUE | 0,0278602 | No |
| orf19.1333 | 987 | 898 | 1009 | 651 | 1725 | 1231 | 1629 | 1750 | 941 | 950 | 997 | 1020 | 1382 | 1400 | 1383 | 1423 | 1186,88 | 977 | 1397 | 1,433 | 0,519 | 10,484 | 1,03E-25 | 2,57E-25 | 1,00E-08 | 0,0034747 | 0,0014633 | 0,0014633 | TRUE | 0,0088545 | No |
| orf19.1334 | 783 | 731 | 882 | 484 | 292 | 223 | 258 | 282 | 747 | 773 | 871 | 758 | 234 | 254 | 219 | 229 | 510,61 | 787 | 234 | 0,296 | -1,757 | -21,194 | 1,09E-99 | 7,10E-99 | 0,0027252 | 0,0053069 | 0,0039767 | 0,0039767 | TRUE | 0,0634758 | No |
| orf19.1335 | 7704 | 7521 | 8033 | 4862 | 11482 | 8274 | 10558 | 11322 | 7345 | 7953 | 7936 | 7617 | 9200 | 9408 | 8964 | 9206 | 8453,49 | 7713 | 9194 | 1,192 | 0,253 | 6,706 | 2,00E-11 | 3,64E-11 | 0,000742 | 0,0022855 | 0,0012482 | 0,0012482 | TRUE | 0,01298 | No |
| orf19.1336 | 4453 | 3706 | 3812 | 2386 | 4154 | 2849 | 3709 | 4148 | 4245 | 3919 | 3766 | 3738 | 3328 | 3240 | 3149 | 3373 | 3594,68 | 3917 | 3272 | 0,835 | -0,26 | -5,161 | 2,46E-07 | 3,90E-07 | 0,0018546 | 0,002548 | 0,0021615 | 0,0021615 | TRUE | 0,0393347 | No |
| orf19.1336.2 | 1645 | 1464 | 1435 | 728 | 927 | 576 | 817 | 808 | 1568 | 1548 | 1418 | 1140 | 743 | 655 | 694 | 657 | 1052,85 | 1418 | 687 | 0,484 | -1,048 | -11,108 | 1,15E-28 | 3,03E-28 | 0,011104 | 0,0036508 | 0,0075064 | 0,0075064 | TRUE | 0,1996207 | No |
| orf19.1338 | 1371 | 1212 | 1338 | 748 | 1728 | 1174 | 1591 | 1724 | 1307 | 1282 | 1322 | 1172 | 1385 | 1335 | 1351 | 1402 | 1319,29 | 1271 | 1368 | 1,074 | 0,103 | 1,99 | 0,046593 | 0,056181 | 0,000469 | 0,0033358 | 0,0018106 | 0,0018106 | TRUE | 0,0287571 | No |
| orf19.1339 | 10661 | 9007 | 9824 | 6833 | 25664 | 19800 | 24128 | 26976 | 10164 | 9524 | 9705 | 10704 | 20564 | 22514 | 20485 | 21934 | 15699,27 | 10024 | 21374 | 2,133 | 1,093 | 21,92 | 1,69E-106 | 1,17E-105 | 0,0024029 | 0,0021959 | 0,0023103 | 0,0023103 | TRUE | 0,0250128 | No |
| orf19.134 | 0 | 1 | 0 | 0 | 1 | 0 | 0 | 1 | 0 | 1 | 0 | 0 | 1 | 0 | 0 | 1 | 0,33 | 0 | 0 | 1,288 | 0,365 | 0,15 | 0,8805006 | 0,8969467 | 1,00E-08 | 4,9184476 | 3,6995225 | 3,6995225 | TRUE | 0,3160008 | No |
| orf19.1340 | 142 | 118 | 153 | 76 | 422 | 331 | 421 | 423 | 135 | 125 | 151 | 119 | 338 | 376 | 357 | 344 | 243,28 | 132 | 354 | 2,644 | 1,403 | 13,376 | 8,38E-41 | 2,69E-40 | 1,00E-08 | 0,0088403 | 0,0051437 | 0,0051437 | TRUE | 0,0910831 | No |
| orf19.1341 | 445 | 391 | 401 | 252 | 819 | 585 | 769 | 819 | 424 | 413 | 396 | 395 | 656 | 665 | 653 | 666 | 533,61 | 407 | 660 | 1,618 | 0,694 | 10,546 | 5,32E-26 | 1,34E-25 | 1,00E-08 | 0,0051683 | 0,0021178 | 0,0021178 | TRUE | 0,0096738 | No |
| orf19.1342 | 6082 | 5453 | 5850 | 3576 | 14041 | 9646 | 13542 | 14548 | 5799 | 5766 | 5779 | 5602 | 11251 | 10968 | 11497 | 11829 | 8561,33 | 5736 | 11386 | 1,984 | 0,989 | 27,969 | 3,89E-172 | 4,48E-171 | 0,0004658 | 0,0022831 | 0,0010627 | 0,0010627 | TRUE | 0,0083767 | No |
| orf19.1343 | 809 | 813 | 822 | 473 | 957 | 721 | 920 | 1012 | 771 | 860 | 812 | 741 | 767 | 820 | 781 | 823 | 796,82 | 796 | 798 | 0,999 | -0,001 | -0,021 | 0,9831366 | 0,984989 | 0,0010333 | 0,0041518 | 0,0026048 | 0,0026048 | TRUE | 0,0334278 | No |
| orf19.1344 | 1589 | 1830 | 1816 | 1063 | 4950 | 3565 | 4470 | 3535 | 1515 | 1935 | 1794 | 1665 | 3966 | 4054 | 3795 | 2874 | 2699,82 | 1727 | 3672 | 2,125 | 1,088 | 10,41 | 2,24E-25 | 5,58E-25 | 0,0157476 | 0,0026994 | 0,0100391 | 0,0100391 | TRUE | 0,2614895 | No |
| orf19.1345 | 47 | 28 | 38 | 15 | 127 | 105 | 126 | 130 | 45 | 30 | 38 | 23 | 102 | 119 | 107 | 106 | 71,16 | 34 | 108 | 3,114 | 1,639 | 8,362 | 6,19E-17 | 1,30E-16 | 1,00E-08 | 0,0251644 | 0,0167917 | 0,0167917 | TRUE | 0,3277344 | No |
| orf19.1348 | 1 | 0 | 3 | 3 | 0 | 2 | 4 | 0 | 1 | 0 | 3 | 5 | 0 | 2 | 3 | 0 | 1,79 | 2 | 1 | 0,67 | -0,579 | -0,436 | 0,6628087 | 0,6950505 | 1,470397 | 0,9214791 | 1,0969484 | 1,0969484 | TRUE | 0,3324218 | No |
| orf19.1349 | 95 | 60 | 88 | 62 | 105 | 70 | 87 | 110 | 91 | 63 | 87 | 97 | 84 | 80 | 74 | 89 | 83,14 | 84 | 82 | 0,974 | -0,038 | -0,223 | 0,8237703 | 0,8445499 | 0,0052969 | 0,0218403 | 0,0152007 | 0,0152007 | TRUE | 0,2649199 | No |
| orf19.135 | 1168 | 1183 | 1288 | 664 | 1810 | 1321 | 1667 | 1617 | 1114 | 1251 | 1272 | 1040 | 1450 | 1502 | 1415 | 1315 | 1294,94 | 1169 | 1420 | 1,211 | 0,277 | 3,817 | 0,0001349 | 0,0001898 | 0,0050274 | 0,0033592 | 0,0042532 | 0,0042532 | TRUE | 0,0606462 | No |
| orf19.1350 | 7818 | 7918 | 8155 | 5370 | 4316 | 3247 | 4202 | 4114 | 7454 | 8372 | 8056 | 8412 | 3458 | 3692 | 3567 | 3345 | 5794,7 | 8074 | 3516 | 0,435 | -1,2 | -23,49 | 5,17E-122 | 4,19E-121 | 0,0021258 | 0,0023746 | 0,0023075 | 0,0023075 | TRUE | 0,0328929 | No |
| orf19.1351 | 430 | 395 | 453 | 262 | 499 | 319 | 515 | 469 | 410 | 418 | 448 | 410 | 400 | 363 | 437 | 381 | 408,34 | 422 | 395 | 0,939 | -0,091 | -1,11 | 0,2668776 | 0,2977179 | 0,001768 | 0,0061122 | 0,0039831 | 0,0039831 | TRUE | 0,0584865 | No |
| orf19.1352 | 4902 | 4264 | 4794 | 2897 | 1962 | 1220 | 1819 | 1988 | 4674 | 4509 | 4736 | 4538 | 1572 | 1387 | 1544 | 1616 | 3072,07 | 4614 | 1530 | 0,332 | -1,59 | -31,377 | 4,22E-216 | 6,19E-215 | 0,0018193 | 0,0026257 | 0,0020533 | 0,0020533 | TRUE | 0,0447018 | No |
| orf19.1353 | 244 | 173 | 205 | 176 | 76 | 36 | 37 | 33 | 233 | 183 | 203 | 276 | 61 | 41 | 31 | 27 | 131,73 | 224 | 40 | 0,18 | -2,475 | -11,286 | 1,54E-29 | 4,11E-29 | 0,0613713 | 0,0145552 | 0,0325233 | 0,0325233 | TRUE | 0,726712 | No |
| orf19.1354 | 147029 | 144734 | 147807 | 82571 | 8425 | 6453 | 7890 | 7868 | 140176 | 153038 | 146019 | 129352 | 6751 | 7338 | 6699 | 6397 | 74471,21 | 142146 | 6796 | 0,048 | -4,387 | -73,912 | 0 | 0 | 0,0040553 | 0,0021133 | 0,003315 | 0,003315 | TRUE | 0,0449514 | No |
| orf19.1355 | 3259 | 3300 | 3399 | 2180 | 4152 | 3012 | 3742 | 3857 | 3107 | 3489 | 3358 | 3415 | 3327 | 3425 | 3177 | 3136 | 3304,28 | 3342 | 3266 | 0,977 | -0,033 | -0,662 | 0,5078748 | 0,5439323 | 0,0016254 | 0,0025882 | 0,0020965 | 0,0020965 | TRUE | 0,0279652 | No |
| orf19.1356 | 552 | 572 | 613 | 388 | 527 | 410 | 461 | 480 | 526 | 605 | 606 | 608 | 422 | 466 | 391 | 390 | 501,83 | 586 | 417 | 0,711 | -0,491 | -5,914 | 3,33E-09 | 5,63E-09 | 0,0038217 | 0,0053631 | 0,0045903 | 0,0045903 | TRUE | 0,0652549 | No |
| orf19.1357 | 25327 | 23935 | 26421 | 15406 | 26450 | 18077 | 24809 | 25518 | 24147 | 25308 | 26101 | 24134 | 21194 | 20555 | 21063 | 20749 | 22906,33 | 24922 | 20890 | 0,838 | -0,255 | -7,129 | 1,01E-12 | 1,91E-12 | 0,0007795 | 0,002163 | 0,0011827 | 0,0011827 | TRUE | 0,0124473 | No |
| orf19.1358 | 19307 | 16929 | 18143 | 12938 | 19531 | 15471 | 18436 | 20537 | 18407 | 17900 | 17924 | 20268 | 15650 | 17592 | 15652 | 16699 | 17511,4 | 18625 | 16398 | 0,881 | -0,184 | -3,327 | 0,000877 | 0,0011882 | 0,0033637 | 0,002185 | 0,0028661 | 0,0028661 | TRUE | 0,0428373 | No |
| orf19.1359 | 1236 | 1324 | 1410 | 945 | 2117 | 1503 | 1900 | 1928 | 1178 | 1400 | 1393 | 1480 | 1696 | 1709 | 1613 | 1568 | 1504,72 | 1363 | 1646 | 1,21 | 0,275 | 3,932 | 8,44E-05 | 0,00012 | 0,004447 | 0,0031824 | 0,0040186 | 0,0040186 | TRUE | 0,1020542 | No |
| orf19.136 | 3774 | 3525 | 3872 | 2522 | 3723 | 2751 | 3442 | 3662 | 3598 | 3727 | 3825 | 3951 | 2983 | 3128 | 2922 | 2978 | 3389,05 | 3775 | 3003 | 0,796 | -0,33 | -7,665 | 1,79E-14 | 3,56E-14 | 0,0008143 | 0,0025758 | 0,0014785 | 0,0014785 | TRUE | 0,0122249 | No |
| orf19.1360 | 533 | 489 | 543 | 334 | 709 | 474 | 686 | 648 | 508 | 517 | 536 | 523 | 568 | 539 | 582 | 527 | 537,66 | 521 | 554 | 1,064 | 0,09 | 1,331 | 0,1831963 | 0,2085594 | 1,00E-08 | 0,0051451 | 0,0024822 | 0,0024822 | TRUE | 0,0141102 | No |
| orf19.1360.1 | 274 | 204 | 196 | 176 | 360 | 284 | 321 | 315 | 261 | 216 | 194 | 276 | 288 | 323 | 273 | 256 | 260,79 | 237 | 285 | 1,208 | 0,273 | 2,212 | 0,0270009 | 0,0331842 | 0,0130113 | 0,0083872 | 0,0106654 | 0,0106654 | TRUE | 0,1682433 | No |
| orf19.1361 | 4349 | 4166 | 4281 | 2433 | 5101 | 3527 | 4678 | 5157 | 4146 | 4405 | 4229 | 3811 | 4087 | 4011 | 3972 | 4193 | 4106,81 | 4148 | 4066 | 0,979 | -0,03 | -0,623 | 0,5335594 | 0,569243 | 0,0016836 | 0,0024911 | 0,0020262 | 0,0020262 | TRUE | 0,0345166 | No |
| orf19.1362 | 7737 | 7457 | 7946 | 4596 | 5442 | 3871 | 4964 | 5158 | 7376 | 7885 | 7850 | 7200 | 4361 | 4402 | 4214 | 4194 | 5935,19 | 7578 | 4293 | 0,566 | -0,821 | -19,296 | 5,76E-83 | 3,24E-82 | 0,0010702 | 0,0023679 | 0,0015596 | 0,0015596 | TRUE | 0,013257 | No |

| | | | | | | | | | | | | | | | | | | | | | | | | | | | | | | |
|---|---|---|---|---|---|---|---|---|---|---|---|---|---|---|---|---|---|---|---|---|---|---|---|---|---|---|---|---|---|---|
| orf19.1363 | 34139 | 32582 | 33618 | 22204 | 13542 | 10044 | 12358 | 12362 | 32548 | 34451 | 33211 | 34784 | 10851 | 11421 | 10492 | 10051 | 22226,18 | 33748 | 10704 | 0,317 | -1,657 | -36,171 | 1,72E-286 | 3,64E-285 | 0,0018641 | 0,0021652 | 0,0019574 | 0,0019574 | TRUE | 0,0246137 | No |
| orf19.1364 | 2693 | 2579 | 2780 | 1850 | 1389 | 974 | 1294 | 1417 | 2567 | 2727 | 2746 | 2898 | 1113 | 1108 | 1099 | 1152 | 1926,27 | 2734 | 1118 | 0,409 | -1,288 | -25,751 | 3,15E-146 | 3,11E-145 | 0,0010678 | 0,0029437 | 0,0017978 | 0,0017978 | TRUE | 0,0207073 | No |
| orf19.1365 | 1250 | 1059 | 1103 | 704 | 687 | 536 | 635 | 729 | 1192 | 1120 | 1090 | 1103 | 550 | 609 | 539 | 593 | 849,48 | 1126 | 573 | 0,507 | -0,979 | -15,572 | 1,12E-54 | 4,41E-54 | 0,0011506 | 0,0040241 | 0,0025152 | 0,0025152 | TRUE | 0,0194647 | No |
| orf19.1366 | 94 | 64 | 81 | 50 | 31 | 30 | 31 | 31 | 90 | 68 | 80 | 78 | 25 | 34 | 26 | 25 | 53,26 | 79 | 28 | 0,345 | -1,535 | -7,293 | 3,02E-13 | 5,82E-13 | 1,00E-08 | 0,0329168 | 0,0191884 | 0,0191884 | TRUE | 0,1312589 | No |
| orf19.1367 | 162 | 186 | 177 | 96 | 136 | 126 | 132 | 111 | 154 | 197 | 175 | 150 | 109 | 143 | 112 | 90 | 141,37 | 169 | 114 | 0,665 | -0,589 | -3,841 | 0,0001225 | 0,0001727 | 0,0168114 | 0,0137057 | 0,0152875 | 0,0152875 | TRUE | 0,2806461 | No |
| orf19.1367.1 | 1521 | 1200 | 1250 | 639 | 2408 | 1491 | 2223 | 2193 | 1450 | 1269 | 1235 | 1001 | 1929 | 1695 | 1887 | 1783 | 1531,27 | 1239 | 1824 | 1,469 | 0,555 | 5,992 | 2,07E-09 | 3,52E-09 | 0,0118573 | 0,0031635 | 0,0075333 | 0,0075333 | TRUE | 0,1897644 | No |
| orf19.1368 | 2545 | 2250 | 2518 | 1694 | 1143 | 822 | 1092 | 1189 | 2426 | 2379 | 2488 | 2654 | 916 | 935 | 927 | 967 | 1711,4 | 2487 | 936 | 0,377 | -1,407 | -27,681 | 1,19E-168 | 1,34E-167 | 0,000905 | 0,0030507 | 0,0017784 | 0,0017784 | TRUE | 0,0229127 | No |
| orf19.1369 | 457 | 422 | 505 | 245 | 450 | 283 | 375 | 465 | 436 | 446 | 499 | 384 | 361 | 322 | 318 | 378 | 392,93 | 441 | 345 | 0,779 | -0,359 | -3,73 | 0,0001912 | 0,0002669 | 0,0064409 | 0,0062699 | 0,0063531 | 0,0063531 | TRUE | 0,0886934 | No |
| orf19.137 | 3162 | 3713 | 3615 | 2285 | 3305 | 2678 | 3085 | 2765 | 3015 | 3926 | 3571 | 3580 | 2648 | 3045 | 2619 | 2248 | 3081,52 | 3523 | 2640 | 0,749 | -0,417 | -4,437 | 9,12E-06 | 1,35E-05 | 0,0128912 | 0,0026241 | 0,0081402 | 0,0081402 | TRUE | 0,1281047 | No |
| orf19.1370 | 5 | 14 | 5 | 4 | 1 | 1 | 1 | 2 | 5 | 15 | 5 | 6 | 1 | 1 | 1 | 2 | 4,4 | 8 | 1 | 0,143 | -2,805 | -3,192 | 0,0014137 | 0,001888 | 1,00E-08 | 0,3753628 | 0,2671873 | 0,2671873 | TRUE | 1,1030784 | No |
| orf19.1371 | 571 | 538 | 541 | 344 | 643 | 436 | 613 | 629 | 544 | 569 | 534 | 539 | 515 | 496 | 520 | 511 | 528,68 | 546 | 510 | 0,935 | -0,097 | -1,5 | 0,1335136 | 0,1539206 | 1,00E-08 | 0,0051969 | 0,0021443 | 0,0021443 | TRUE | 0,008946 | No |
| orf19.1372 | 431 | 442 | 420 | 281 | 772 | 535 | 738 | 748 | 411 | 467 | 415 | 440 | 619 | 608 | 627 | 608 | 524,38 | 433 | 616 | 1,423 | 0,509 | 7,364 | 1,79E-13 | 3,46E-13 | 1,00E-08 | 0,0052224 | 0,0025759 | 0,0025759 | TRUE | 0,0354404 | No |
| orf19.1373 | 170 | 141 | 163 | 106 | 432 | 339 | 428 | 429 | 162 | 149 | 161 | 166 | 346 | 385 | 363 | 349 | 260,26 | 160 | 361 | 2,262 | 1,177 | 12,233 | 2,08E-34 | 6,03E-34 | 1,00E-08 | 0,0084 | 0,0041726 | 0,0041726 | TRUE | 0,0217464 | No |
| orf19.1374 | 381 | 377 | 337 | 232 | 1079 | 690 | 983 | 982 | 363 | 399 | 333 | 363 | 865 | 785 | 835 | 798 | 592,55 | 364 | 821 | 2,254 | 1,173 | 15,859 | 1,21E-56 | 4,85E-56 | 0,001347 | 0,0048622 | 0,0031762 | 0,0031762 | TRUE | 0,0480023 | No |
| orf19.1375 | 27705 | 23957 | 25968 | 16330 | 14104 | 9814 | 13402 | 14679 | 26414 | 25331 | 25654 | 25582 | 11301 | 11159 | 11378 | 11935 | 18594,39 | 25745 | 11443 | 0,445 | -1,17 | -35,465 | 1,72E-275 | 3,34E-274 | 0,000522 | 0,0021796 | 0,0009838 | 0,0009838 | TRUE | 0,0102848 | No |
| orf19.1376 | 2669 | 2541 | 2555 | 1746 | 4140 | 2911 | 3800 | 3823 | 2545 | 2687 | 2524 | 2735 | 3317 | 3310 | 3226 | 3108 | 2931,58 | 2623 | 3240 | 1,237 | 0,307 | 6,951 | 3,63E-12 | 6,76E-12 | 0,0007799 | 0,0026514 | 0,0015161 | 0,0015161 | TRUE | 0,0093764 | No |
| orf19.1377 | 2005 | 1824 | 1801 | 1227 | 1198 | 893 | 1155 | 1244 | 1912 | 1929 | 1779 | 1922 | 960 | 1015 | 981 | 1011 | 1438,63 | 1886 | 992 | 0,526 | -0,927 | -18,597 | 3,37E-77 | 1,77E-76 | 0,0005359 | 0,0032326 | 0,0016362 | 0,0016362 | TRUE | 0,0179791 | No |
| orf19.1378 | 10508 | 8709 | 9980 | 6277 | 13953 | 9243 | 13234 | 14460 | 10018 | 9209 | 9859 | 9833 | 11180 | 10510 | 11236 | 11757 | 10450,34 | 9730 | 11171 | 1,148 | 0,2 | 4,429 | 9,45E-06 | 1,40E-05 | 0,001593 | 0,0022484 | 0,0018521 | 0,0018521 | TRUE | 0,0191665 | No |
| orf19.138 | 100 | 83 | 90 | 74 | 93 | 48 | 69 | 58 | 95 | 88 | 89 | 116 | 75 | 55 | 59 | 47 | 77,85 | 97 | 59 | 0,612 | -0,708 | -3,86 | 0,0001132 | 0,0001598 | 0,0131113 | 0,0231827 | 0,0189066 | 0,0189066 | TRUE | 0,3154779 | No |
| orf19.1381 | 2848 | 2370 | 2455 | 1564 | 1134 | 768 | 1118 | 1197 | 2715 | 2506 | 2425 | 2450 | 909 | 873 | 949 | 973 | 1725,13 | 2524 | 926 | 0,367 | -1,446 | -25,587 | 2,12E-144 | 2,06E-143 | 0,0018766 | 0,003043 | 0,0023609 | 0,0023609 | TRUE | 0,0322902 | No |
| orf19.1382 | 574 | 493 | 494 | 316 | 487 | 379 | 517 | 517 | 547 | 521 | 488 | 495 | 390 | 431 | 439 | 420 | 466,51 | 513 | 420 | 0,817 | -0,292 | -3,902 | 9,55E-05 | 0,0001354 | 1,00E-08 | 0,0056108 | 0,0032309 | 0,0032309 | TRUE | 0,0279287 | No |
| orf19.1383 | 748 | 722 | 752 | 474 | 1190 | 824 | 1160 | 1237 | 713 | 763 | 743 | 743 | 954 | 937 | 985 | 1006 | 855,39 | 740 | 970 | 1,312 | 0,392 | 6,996 | 2,63E-12 | 4,91E-12 | 1,00E-08 | 0,0040108 | 0,0018007 | 0,0018007 | TRUE | 0,0080495 | No |
| orf19.1386 | 2390 | 2177 | 2332 | 1439 | 1887 | 1155 | 1743 | 1765 | 2279 | 2302 | 2304 | 2254 | 1512 | 1313 | 1480 | 1435 | 1859,85 | 2285 | 1435 | 0,629 | -0,669 | -13,239 | 5,24E-40 | 1,66E-39 | 0,0011356 | 0,0029741 | 0,0018931 | 0,0018931 | TRUE | 0,0366178 | No |
| orf19.1387 | 4628 | 3768 | 4093 | 2744 | 3587 | 2446 | 3598 | 3795 | 4412 | 3984 | 4043 | 4299 | 2874 | 2781 | 3055 | 3086 | 3566,81 | 4184 | 2949 | 0,705 | -0,504 | -9,735 | 2,13E-22 | 5,01E-22 | 0,002022 | 0,0025516 | 0,0022853 | 0,0022853 | TRUE | 0,0172013 | No |
| orf19.1388 | 5760 | 5207 | 5575 | 3128 | 8839 | 5530 | 8709 | 8774 | 5492 | 5506 | 5508 | 4900 | 7082 | 6288 | 7394 | 7134 | 6162,95 | 5352 | 6974 | 1,303 | 0,382 | 6,402 | 1,53E-10 | 2,72E-10 | 0,0037037 | 0,0023577 | 0,0032462 | 0,0032462 | TRUE | 0,0530258 | No |
| orf19.1389 | 3758 | 3363 | 3496 | 2040 | 4914 | 3033 | 4866 | 4964 | 3583 | 3556 | 3454 | 3196 | 3937 | 3449 | 4131 | 4036 | 3667,74 | 3447 | 3888 | 1,128 | 0,174 | 2,767 | 0,0056631 | 0,0072973 | 0,0039939 | 0,0025389 | 0,003501 | 0,003501 | TRUE | 0,0692533 | No |
| orf19.139 | 1031 | 970 | 1069 | 682 | 2863 | 2100 | 2588 | 2821 | 983 | 1026 | 1056 | 1068 | 2294 | 2388 | 2197 | 2294 | 1663,24 | 1033 | 2293 | 2,222 | 1,152 | 23,205 | 4,01E-119 | 3,16E-118 | 0,0005986 | 0,0030785 | 0,001628 | 0,001628 | TRUE | 0,0133597 | No |
| orf19.1390 | 586 | 507 | 539 | 382 | 888 | 614 | 788 | 923 | 559 | 536 | 532 | 598 | 712 | 698 | 669 | 750 | 631,86 | 556 | 707 | 1,276 | 0,352 | 5,206 | 1,92E-07 | 3,07E-07 | 0,0009063 | 0,0046898 | 0,0027481 | 0,0027481 | TRUE | 0,025354 | No |
| orf19.1391 | 115 | 100 | 86 | 70 | 163 | 101 | 129 | 147 | 110 | 106 | 85 | 110 | 131 | 115 | 110 | 120 | 110,56 | 103 | 119 | 1,164 | 0,219 | 1,541 | 0,1234162 | 0,1426665 | 1,00E-08 | 0,0169418 | 0,0102078 | 0,0102078 | TRUE | 0,1396013 | No |
| orf19.1392 | 430 | 390 | 441 | 259 | 993 | 680 | 928 | 1055 | 410 | 412 | 436 | 406 | 796 | 773 | 788 | 858 | 609,79 | 416 | 804 | 1,931 | 0,95 | 14,035 | 9,56E-45 | 3,26E-44 | 1,00E-08 | 0,0047839 | 0,0025039 | 0,0025039 | TRUE | 0,0249636 | No |
| orf19.1393 | 3387 | 3010 | 3417 | 2147 | 1895 | 1357 | 1812 | 1894 | 3229 | 3183 | 3376 | 3363 | 1518 | 1543 | 1538 | 1540 | 2411,34 | 3288 | 1535 | 0,467 | -1,098 | -27,166 | 1,63E-162 | 1,79E-161 | 1,00E-08 | 0,0027722 | 0,0011078 | 0,0011078 | TRUE | 0,0055706 | No |
| orf19.1394 | 1485 | 1325 | 1348 | 656 | 2181 | 1297 | 2137 | 2256 | 1416 | 1401 | 1332 | 1028 | 1748 | 1475 | 1814 | 1834 | 1505,9 | 1294 | 1718 | 1,325 | 0,406 | 4,084 | 4,43E-05 | 6,38E-05 | 0,0135932 | 0,0031816 | 0,0088153 | 0,0088153 | TRUE | 0,2199552 | No |
| orf19.1395 | 356 | 269 | 296 | 193 | 204 | 145 | 204 | 198 | 339 | 284 | 292 | 302 | 163 | 165 | 173 | 161 | 235,14 | 304 | 166 | 0,543 | -0,881 | -8,757 | 2,00E-18 | 4,33E-18 | 1,00E-08 | 0,0090739 | 0,0052023 | 0,0052023 | TRUE | 0,0729454 | No |
| orf19.1396 | 2146 | 1796 | 1962 | 1300 | 1678 | 1186 | 1596 | 1734 | 2046 | 1899 | 1938 | 2037 | 1345 | 1349 | 1355 | 1410 | 1672,23 | 1980 | 1365 | 0,69 | -0,536 | -11,518 | 1,07E-30 | 2,92E-30 | 0,0003974 | 0,0030731 | 0,0014657 | 0,0014657 | TRUE | 0,0091209 | No |
| orf19.1397 | 1110 | 925 | 1019 | 688 | 1064 | 792 | 913 | 1097 | 1058 | 978 | 1007 | 1078 | 853 | 901 | 775 | 892 | 942,63 | 1030 | 855 | 0,83 | -0,268 | -4,153 | 3,29E-05 | 4,76E-05 | 0,0021998 | 0,0038331 | 0,0029443 | 0,0029443 | TRUE | 0,0475496 | No |
| orf19.1400 | 1463 | 1455 | 1542 | 825 | 1255 | 947 | 1141 | 1134 | 1395 | 1538 | 1523 | 1292 | 1006 | 1077 | 969 | 922 | 1215,28 | 1437 | 994 | 0,689 | -0,538 | -7,584 | 3,35E-14 | 6,60E-14 | 0,0042916 | 0,0034423 | 0,0039819 | 0,0039819 | TRUE | 0,0513054 | No |
| orf19.1401 | 1009 | 880 | 998 | 713 | 2261 | 1757 | 2200 | 2354 | 962 | 930 | 986 | 1117 | 1812 | 1998 | 1868 | 1914 | 1448,34 | 999 | 1898 | 1,906 | 0,931 | 14,993 | 8,20E-51 | 3,05E-50 | 0,0026009 | 0,0032249 | 0,0028987 | 0,0028987 | TRUE | 0,0677203 | No |
| orf19.1402 | 5678 | 4683 | 5075 | 3636 | 9158 | 6670 | 8790 | 9633 | 5413 | 4952 | 5014 | 5696 | 7338 | 7584 | 7463 | 7833 | 6411,54 | 5269 | 7554 | 1,435 | 0,521 | 10,231 | 1,45E-24 | 3,55E-24 | 0,002391 | 0,0023474 | 0,0023242 | 0,0023242 | TRUE | 0,0351322 | No |
| orf19.1403 | 5164 | 4503 | 4983 | 3275 | 4296 | 2970 | 4089 | 4186 | 4923 | 4761 | 4923 | 5130 | 3442 | 3377 | 3472 | 3404 | 4179,06 | 4934 | 3424 | 0,695 | -0,526 | -14,604 | 2,65E-48 | 9,55E-48 | 0,000214 | 0,0024842 | 0,0010004 | 0,0010004 | TRUE | 0,0080943 | No |
| orf19.1404 | 7218 | 5914 | 6527 | 4326 | 6125 | 4207 | 5974 | 6636 | 6882 | 6253 | 6448 | 6777 | 4908 | 4784 | 5072 | 5396 | 5814,87 | 6590 | 5040 | 0,765 | -0,386 | -7,699 | 1,37E-14 | 2,74E-14 | 0,0021218 | 0,0023736 | 0,0022426 | 0,0022426 | TRUE | 0,0276512 | No |
| orf19.1405 | 722 | 655 | 633 | 435 | 354 | 248 | 376 | 342 | 688 | 693 | 625 | 681 | 284 | 282 | 319 | 278 | 481,33 | 672 | 291 | 0,433 | -1,207 | -15,653 | 3,15E-55 | 1,25E-54 | 1,00E-08 | 0,0055024 | 0,0033422 | 0,0033422 | TRUE | 0,0508672 | No |
| orf19.1406 | 787 | 819 | 804 | 465 | 1215 | 937 | 1118 | 992 | 750 | 866 | 794 | 728 | 974 | 1065 | 949 | 807 | 866,72 | 784 | 949 | 1,206 | 0,27 | 3,047 | 0,0023134 | 0,0030544 | 0,0078155 | 0,0039857 | 0,0063449 | 0,0063449 | TRUE | 0,122581 | No |
| orf19.1409 | 929 | 791 | 901 | 580 | 937 | 674 | 844 | 1009 | 886 | 836 | 890 | 909 | 751 | 766 | 717 | 820 | 821,87 | 880 | 764 | 0,868 | -0,204 | -3,274 | 0,0010622 | 0,0014308 | 0,0011473 | 0,0040891 | 0,0024912 | 0,0024912 | TRUE | 0,03146 | No |
| orf19.1409.1 | 9708 | 8118 | 8666 | 5476 | 25745 | 18106 | 23248 | 25284 | 9256 | 8584 | 8561 | 8578 | 20629 | 20588 | 19737 | 20558 | 14561,44 | 8745 | 20378 | 2,33 | 1,22 | 32,448 | 5,78E-231 | 8,96E-230 | 0,0008706 | 0,002204 | 0,0012716 | 0,0012716 | TRUE | 0,0191838 | No |
| orf19.1409.2 | 327 | 302 | 301 | 209 | 257 | 159 | 215 | 240 | 312 | 319 | 297 | 327 | 206 | 181 | 183 | 195 | 252,53 | 314 | 191 | 0,612 | -0,709 | -7,577 | 3,53E-14 | 6,96E-14 | 1,00E-08 | 0,008593 | 0,0043013 | 0,0043013 | TRUE | 0,0306641 | No |
| orf19.1409.3 | 19 | 14 | 18 | 9 | 70 | 34 | 73 | 46 | 18 | 15 | 18 | 14 | 56 | 39 | 62 | 37 | 32,37 | 16 | 48 | 2,978 | 1,574 | 5,318 | 1,05E-07 | 1,68E-07 | 1,00E-08 | 0,0528205 | 0,0410995 | 0,0410995 | TRUE | 0,1178663 | No |
| orf19.1411 | 1185 | 1100 | 1151 | 770 | 1254 | 1003 | 1226 | 1267 | 1130 | 1163 | 1137 | 1206 | 1005 | 1141 | 1041 | 1030 | 1106,57 | 1159 | 1054 | 0,909 | -0,138 | -2,445 | 0,0145003 | 0,0181995 | 0,0008335 | 0,0035751 | 0,0021312 | 0,0021312 | TRUE | 0,0339301 | No |
| orf19.1412 | 2297 | 2042 | 2225 | 1329 | 2009 | 1334 | 2062 | 2105 | 2190 | 2159 | 2198 | 2082 | 1610 | 1517 | 1751 | 1712 | 1902,25 | 2157 | 1648 | 0,764 | -0,389 | -7,261 | 3,85E-13 | 7,40E-13 | 0,0016284 | 0,0029544 | 0,0022194 | 0,0022194 | TRUE | 0,0323065 | No |
| orf19.1413 | 1225 | 1027 | 1099 | 702 | 1210 | 819 | 1182 | 1193 | 1168 | 1086 | 1086 | 1100 | 970 | 931 | 1004 | 970 | 1039,2 | 1110 | 969 | 0,873 | -0,196 | -3,688 | 0,0002263 | 0,0003151 | 1,00E-08 | 0,0036712 | 0,0017455 | 0,0017455 | TRUE | 0,0159598 | No |
| orf19.1414 | 2293 | 1824 | 2049 | 1220 | 2614 | 1753 | 2721 | 2852 | 2186 | 1929 | 2024 | 1911 | 2095 | 1993 | 2310 | 2319 | 2095,89 | 2012 | 2179 | 1,082 | 0,114 | 1,744 | 0,081222 | 0,0957292 | 0,0040717 | 0,0028747 | 0,0036442 | 0,0036442 | TRUE | 0,0412771 | No |
| orf19.1415 | 16264 | 14770 | 15800 | 10398 | 44578 | 32851 | 41373 | 46438 | 15506 | 15617 | 15609 | 16289 | 35719 | 37355 | 35125 | 37759 | 26122,37 | 15755 | 36490 | 2,316 | 1,212 | 33,525 | 2,06E-246 | 3,45E-245 | 0,000802 | 0,0021541 | 0,0012073 | 0,0012073 | TRUE | 0,0077575 | No |
| orf19.1416 | 2652 | 2190 | 2213 | 1487 | 2681 | 1778 | 2627 | 2845 | 2528 | 2316 | 2186 | 2329 | 2148 | 2022 | 2230 | 2313 | 2259,16 | 2340 | 2178 | 0,931 | -0,102 | -1,725 | 0,0846174 | 0,0994921 | 0,0028579 | 0,0028181 | 0,0029409 | 0,0029409 | TRUE | 0,0368727 | No |
| orf19.1417 | 1139 | 988 | 1012 | 600 | 523 | 332 | 465 | 458 | 1086 | 1045 | 1000 | 940 | 419 | 378 | 395 | 372 | 704,26 | 1018 | 391 | 0,384 | -1,382 | -19,556 | 3,67E-85 | 2,09E-84 | 0,001866 | 0,0044227 | 0,0031054 | 0,0031054 | TRUE | 0,0283319 | No |
| orf19.1418 | 684 | 644 | 685 | 425 | 1195 | 935 | 1217 | 1237 | 652 | 681 | 677 | 666 | 958 | 1063 | 1033 | 1006 | 841,91 | 669 | 1015 | 1,515 | 0,6 | 10,244 | 1,26E-24 | 3,09E-24 | 1,00E-08 | 0,0040415 | 0,0020141 | 0,0020141 | TRUE | 0,0175701 | No |
| orf19.1420 | 124 | 109 | 121 | 78 | 301 | 191 | 291 | 306 | 118 | 115 | 120 | 122 | 241 | 217 | 247 | 249 | 178,68 | 119 | 238 | 2,017 | 1,012 | 9,038 | 1,60E-19 | 3,55E-19 | 1,00E-08 | 0,0112804 | 0,0055098 | 0,0055098 | TRUE | 0,0356337 | No |
| orf19.1421 | 162 | 120 | 143 | 109 | 583 | 374 | 538 | 574 | 154 | 127 | 141 | 171 | 467 | 425 | 457 | 467 | 301,16 | 148 | 454 | 3,092 | 1,628 | 16,334 | 5,61E-60 | 2,36E-59 | 1,19E-08 | 0,0075433 | 0,004786 | 0,004786 | TRUE | 0,0933686 | No |
| orf19.1422 | 8541 | 7897 | 8494 | 5312 | 5628 | 4252 | 5326 | 5923 | 8143 | 8350 | 8391 | 8322 | 4510 | 4835 | 4522 | 4816 | 6486 | 8302 | 4671 | 0,562 | -0,83 | -22,242 | 1,36E-109 | 9,86E-109 | 0,0005991 | 0,0023444 | 0,0011746 | 0,0011746 | TRUE | 0,0065894 | No |
| orf19.1424 | 901 | 819 | 868 | 529 | 956 | 708 | 950 | 1051 | 859 | 866 | 858 | 829 | 766 | 805 | 807 | 855 | 830,42 | 853 | 808 | 0,946 | -0,079 | -1,379 | 0,1678601 | 0,1917935 | 1,00E-08 | 0,0040685 | 0,0019823 | 0,0019823 | TRUE | 0,0189317 | No |
| orf19.1426 | 6851 | 6139 | 6695 | 4436 | 4754 | 3375 | 4555 | 4950 | 6532 | 6491 | 6614 | 6949 | 3809 | 3838 | 3867 | 4025 | 5265,63 | 6646 | 3885 | 0,585 | -0,774 | -20,498 | 2,25E-93 | 1,39E-92 | 0,0005261 | 0,0024031 | 0,0011677 | 0,0011677 | TRUE | 0,0109322 | No |
| orf19.1427 | 296 | 269 | 295 | 210 | 653 | 482 | 555 | 652 | 282 | 284 | 291 | 329 | 523 | 548 | 471 | 530 | 407,46 | 296 | 518 | 1,754 | 0,811 | 9,648 | 5,02E-22 | 1,17E-21 | 0,0017748 | 0,0061209 | 0,0040305 | 0,0040305 | TRUE | 0,0489632 | No |
| orf19.1428 | 168 | 156 | 168 | 87 | 428 | 258 | 373 | 417 | 160 | 165 | 166 | 136 | 343 | 293 | 317 | 339 | 239,93 | 157 | 323 | 2,049 | 1,035 | 9,949 | 2,54E-23 | 6,09E-23 | 0,0010865 | 0,0089346 | 0,0054928 | 0,0054928 | TRUE | 0,0611178 | No |
| orf19.1429 | 85 | 76 | 85 | 46 | 412 | 250 | 400 | 384 | 81 | 80 | 84 | 72 | 330 | 284 | 340 | 312 | 197,96 | 79 | 316 | 3,971 | 1,99 | 16,154 | 1,06E-58 | 4,40E-58 | 1,00E-08 | 0,0103855 | 0,0062309 | 0,0062309 | TRUE | 0,0485325 | No |
| orf19.1430 | 76 | 69 | 93 | 53 | 27 | 25 | 28 | 21 | 72 | 73 | 92 | 83 | 22 | 28 | 24 | 17 | 51,4 | 80 | 23 | 0,28 | -1,835 | -8,191 | 2,60E-16 | 5,38E-16 | 1,00E-08 | 0,0340328 | 0,0214097 | 0,0214097 | TRUE | 0,1818148 | No |
| orf19.1433 | 2255 | 2007 | 2046 | 1300 | 1907 | 1326 | 1690 | 1923 | 2150 | 2122 | 2021 | 2037 | 1528 | 1508 | 1435 | 1564 | 1795,5 | 2082 | 1509 | 0,724 | -0,466 | -9,907 | 3,88E-23 | 9,26E-23 | 0,0006081 | 0,0030057 | 0,0015576 | 0,0015576 | TRUE | 0,0134918 | No |
| orf19.1433.1 | 4 | 4 | 5 | 9 | 1 | 0 | 2 | 2 | 4 | 4 | 5 | 14 | 1 | 0 | 2 | 2 | 3,9 | 7 | 1 | 0,168 | -2,576 | -2,797 | 0,0051584 | 0,0066618 | 1,00E-08 | 0,4229975 | 0,3199374 | 0,3199374 | TRUE | 0,8874377 | No |
| orf19.1434 | 1169 | 1044 | 1184 | 733 | 1330 | 938 | 1290 | 1378 | 1115 | 1104 | 1170 | 1148 | 1066 | 1067 | 1095 | 1120 | 1110,54 | 1134 | 1087 | 0,959 | -0,06 | -1,198 | 0,2310735 | 0,2602329 | 1,00E-08 | 0,0035698 | 0,0014902 | 0,0014902 | TRUE | 0,0059913 | No |
| orf19.1435 | 11795 | 8193 | 9107 | 5622 | 25961 | 16492 | 24415 | 29224 | 11245 | 8663 | 8997 | 8807 | 20802 | 18753 | 20728 | 23762 | 15219,64 | 9428 | 21011 | 2,228 | 1,156 | 12,971 | 1,78E-38 | 5,49E-38 | 0,0130866 | 0,0021992 | 0,0075491 | 0,0075491 | TRUE | 0,206309 | No |
| orf19.1438 | 378 | 356 | 390 | 239 | 223 | 173 | 229 | 197 | 360 | 376 | 385 | 374 | 179 | 197 | 194 | 160 | 278,31 | 374 | 182 | 0,486 | -1,041 | -11,04 | 2,46E-28 | 6,45E-28 | 1,00E-08 | 0,0079908 | 0,0045933 | 0,0045933 | TRUE | 0,0764182 | No |
| orf19.1439 | 1628 | 1784 | 1863 | 1137 | 1495 | 1147 | 1305 | 1300 | 1552 | 1886 | 1840 | 1781 | 1198 | 1304 | 1108 | 1057 | 1465,9 | 1765 | 1167 | 0,66 | -0,598 | -7,497 | 6,53E-14 | 1,28E-13 | 0,0069125 | 0,0032113 | 0,0054158 | 0,0054158 | TRUE | 0,0816874 | No |
| orf19.144 | 2494 | 2821 | 2959 | 1674 | 2718 | 1941 | 2457 | 2248 | 2378 | 2983 | 2923 | 2622 | 2178 | 2207 | 2086 | 1828 | 2400,62 | 2726 | 2075 | 0,761 | -0,395 | -4,831 | 1,36E-06 | 2,09E-06 | 0,0081998 | 0,0027752 | 0,005998 | 0,005998 | TRUE | 0,0919309 | No |
| orf19.1440 | 960 | 1021 | 1009 | 632 | 2009 | 1483 | 1847 | 1862 | 915 | 1080 | 997 | 990 | 1610 | 1686 | 1568 | 1514 | 1294,98 | 996 | 1594 | 1,601 | 0,679 | 10,987 | 4,42E-28 | 1,15E-27 | 0,0022279 | 0,0033592 | 0,0028243 | 0,0028243 | TRUE | 0,0402583 | No |
| orf19.1440.1 | 9 | 23 | 9 | 9 | 58 | 37 | 44 | 33 | 9 | 24 | 9 | 14 | 46 | 42 | 37 | 27 | 26,08 | 14 | 38 | 2,752 | 1,461 | 4,073 | 4,64E-05 | 6,66E-05 | 0,0819873 | 0,0650525 | 0,0715364 | 0,0715364 | TRUE | 1,1453259 | No |
| orf19.1440.2 | 240 | 212 | 213 | 101 | 703 | 363 | 634 | 663 | 229 | 224 | 210 | 158 | 563 | 413 | 538 | 539 | 359,38 | 205 | 513 | 2,484 | 1,312 | 10,477 | 1,11E-25 | 2,77E-25 | 0,0166917 | 0,00666 | 0,0114913 | 0,0114913 | TRUE | 0,2152931 | No |

| | | | | | | | | | | | | | | | | | | | | | | | | | | | | | | | |
|---|---|---|---|---|---|---|---|---|---|---|---|---|---|---|---|---|---|---|---|---|---|---|---|---|---|---|---|---|---|---|---|
| orf19.1441 | 4737 | 4366 | 4800 | 2814 | 4083 | 2802 | 4036 | 3868 | 4516 | 4616 | 4742 | 4408 | 3272 | 3186 | 3427 | 3145 | 3914,03 | 4570 | 3258 | 0,712 | -0,489 | -11,462 | 2,05E-30 | 5,56E-30 | 0,0008649 | 0,0025108 | 0,0014893 | 0,0014893 | TRUE | 0,0149717 | No |
| orf19.1442 | 441 | 374 | 372 | 265 | 447 | 327 | 426 | 501 | 420 | 395 | 368 | 415 | 358 | 372 | 362 | 407 | 387,2 | 400 | 375 | 0,94 | -0,09 | -1,093 | 0,2745995 | 0,3058508 | 0,0013996 | 0,0063318 | 0,0039115 | 0,0039115 | TRUE | 0,0419031 | No |
| orf19.1444 | 2408 | 2138 | 2475 | 1734 | 3006 | 2286 | 2938 | 3212 | 2296 | 2261 | 2445 | 2716 | 2409 | 2599 | 2494 | 2612 | 2478,99 | 2430 | 2528 | 1,042 | 0,06 | 0,967 | 0,3335365 | 0,3672643 | 0,0036805 | 0,0027536 | 0,0032643 | 0,0032643 | TRUE | 0,0729884 | No |
| orf19.1445 | 85 | 74 | 78 | 44 | 36 | 10 | 19 | 20 | 81 | 78 | 77 | 69 | 29 | 11 | 16 | 16 | 47,23 | 76 | 18 | 0,242 | -2,047 | -8,118 | 4,75E-16 | 9,78E-16 | 0,0021636 | 0,0368517 | 0,0298188 | 0,0298188 | TRUE | 0,9185599 | No |
| orf19.1446 | 3728 | 3390 | 3747 | 2246 | 1257 | 983 | 1262 | 1221 | 3554 | 3584 | 3702 | 3518 | 1007 | 1118 | 1071 | 993 | 2318,51 | 3590 | 1047 | 0,291 | -1,781 | -36,749 | 1,19E-295 | 2,61E-294 | 0,0007392 | 0,0027995 | 0,0016711 | 0,0016711 | TRUE | 0,0226253 | No |
| orf19.1447 | 288 | 237 | 293 | 193 | 489 | 371 | 484 | 536 | 275 | 251 | 289 | 302 | 392 | 422 | 411 | 436 | 347,17 | 279 | 415 | 1,492 | 0,577 | 6,721 | 1,81E-11 | 3,30E-11 | 0,0005778 | 0,0068206 | 0,003996 | 0,003996 | TRUE | 0,0520653 | No |
| orf19.1448 | 3386 | 2972 | 3084 | 1870 | 8551 | 5865 | 7967 | 8320 | 3228 | 3143 | 3047 | 2929 | 6852 | 6669 | 6764 | 6765 | 4924,56 | 3087 | 6762 | 2,188 | 1,13 | 29,462 | 8,96E-191 | 1,17E-189 | 0,000391 | 0,0024247 | 0,0011634 | 0,0011634 | TRUE | 0,0130198 | No |
| orf19.1448.1 | 542 | 536 | 495 | 380 | 821 | 593 | 761 | 810 | 517 | 567 | 489 | 595 | 658 | 674 | 646 | 659 | 600,58 | 542 | 659 | 1,222 | 0,289 | 4,019 | 5,84E-05 | 8,35E-05 | 0,0014488 | 0,0048252 | 0,0032489 | 0,0032489 | TRUE | 0,049998 | No |
| orf19.1449 | 221 | 238 | 260 | 138 | 837 | 536 | 780 | 816 | 211 | 252 | 257 | 216 | 671 | 609 | 662 | 663 | 442,66 | 234 | 651 | 2,78 | 1,475 | 17,235 | 1,46E-66 | 6,70E-66 | 0,0012414 | 0,0058005 | 0,0039782 | 0,0039782 | TRUE | 0,060484 | No |
| orf19.145 | 1372 | 1362 | 1420 | 861 | 1565 | 1196 | 1426 | 1449 | 1308 | 1440 | 1403 | 1349 | 1254 | 1360 | 1211 | 1178 | 1312,83 | 1375 | 1251 | 0,908 | -0,139 | -2,346 | 0,0189569 | 0,0235705 | 0,0018844 | 0,0033419 | 0,0026179 | 0,0026179 | TRUE | 0,039281 | No |
| orf19.1450 | 13 | 12 | 19 | 9 | 3 | 2 | 6 | 7 | 12 | 13 | 19 | 14 | 2 | 2 | 5 | 6 | 9,18 | 14 | 4 | 0,272 | -1,881 | -3,647 | 0,0002657 | 0,0003691 | 1,00E-08 | 0,1810078 | 0,1044161 | 0,1044161 | TRUE | 0,1965874 | No |
| orf19.1451 | 822 | 760 | 750 | 537 | 1295 | 936 | 1225 | 1273 | 784 | 804 | 741 | 841 | 1038 | 1064 | 1040 | 1035 | 918,31 | 792 | 1044 | 1,322 | 0,402 | 7,209 | 5,65E-13 | 1,08E-12 | 1,00E-08 | 0,0038792 | 0,0018541 | 0,0018541 | TRUE | 0,0225306 | No |
| orf19.1453 | 14603 | 11721 | 13429 | 7928 | 15343 | 9626 | 14014 | 15139 | 13922 | 12393 | 13267 | 12420 | 12294 | 10946 | 11898 | 12309 | 12431,11 | 13000 | 11862 | 0,912 | -0,132 | -2,458 | 0,0139743 | 0,0175843 | 0,0029508 | 0,0022234 | 0,002702 | 0,002702 | TRUE | 0,0328133 | No |
| orf19.1456 | 20 | 6 | 2 | 4 | 12 | 3 | 4 | 5 | 19 | 6 | 2 | 6 | 10 | 3 | 3 | 4 | 6,77 | 8 | 5 | 0,608 | -0,717 | -1,025 | 0,3055087 | 0,3386227 | 0,3961856 | 0,2446979 | 0,3185776 | 0,3185776 | TRUE | 1,0658656 | No |
| orf19.1457 | 1954 | 1750 | 1922 | 1222 | 2492 | 1768 | 2487 | 2511 | 1863 | 1850 | 1899 | 1914 | 1997 | 2010 | 2111 | 2042 | 1960,84 | 1882 | 2040 | 1,085 | 0,118 | 2,856 | 0,0042906 | 0,0055703 | 1,00E-08 | 0,0029286 | 0,0011203 | 0,0011203 | TRUE | 0,0068022 | No |
| orf19.1458 | 446 | 437 | 417 | 277 | 634 | 546 | 660 | 669 | 425 | 462 | 412 | 434 | 508 | 621 | 560 | 544 | 495,79 | 433 | 558 | 1,286 | 0,362 | 4,491 | 7,10E-06 | 1,06E-05 | 0,0029883 | 0,005403 | 0,0041597 | 0,0041597 | TRUE | 0,0622428 | No |
| orf19.1459 | 554 | 513 | 595 | 387 | 1132 | 748 | 1040 | 1124 | 528 | 542 | 588 | 606 | 907 | 851 | 883 | 914 | 727,39 | 566 | 889 | 1,577 | 0,657 | 10,171 | 2,68E-24 | 6,54E-24 | 0,0005353 | 0,0043485 | 0,0025156 | 0,0025156 | TRUE | 0,0243077 | No |
| orf19.1460 | 2833 | 2316 | 2719 | 1620 | 7015 | 4822 | 6703 | 7278 | 2701 | 2449 | 2686 | 2538 | 5621 | 5483 | 5691 | 5918 | 4135,78 | 2594 | 5678 | 2,188 | 1,13 | 24,676 | 1,94E-134 | 1,76E-133 | 0,0012305 | 0,0024883 | 0,0017174 | 0,0017174 | TRUE | 0,0179328 | No |
| orf19.1461 | 1663 | 1352 | 1600 | 1061 | 1547 | 1108 | 1511 | 1543 | 1585 | 1430 | 1581 | 1662 | 1240 | 1260 | 1283 | 1255 | 1411,84 | 1564 | 1260 | 0,806 | -0,311 | -5,719 | 1,07E-08 | 1,78E-08 | 0,0012703 | 0,0032542 | 0,0021256 | 0,0021256 | TRUE | 0,040377 | No |
| orf19.1462 | 2175 | 1851 | 2029 | 1372 | 7548 | 5040 | 6952 | 7678 | 2074 | 1957 | 2004 | 2149 | 6048 | 5731 | 5902 | 6243 | 4013,59 | 2046 | 5981 | 2,928 | 1,55 | 34,239 | 6,39E-257 | 1,13E-255 | 0,0010615 | 0,0025004 | 0,0016179 | 0,0016179 | TRUE | 0,0125596 | No |
| orf19.1464 | 5196 | 4681 | 5211 | 2968 | 1702 | 1212 | 1699 | 1657 | 4954 | 4950 | 5148 | 4650 | 1364 | 1378 | 1442 | 1347 | 3154,07 | 4926 | 1383 | 0,281 | -1,834 | -39,673 | 0 | 0 | 0,0010209 | 0,0026118 | 0,0016146 | 0,0016146 | TRUE | 0,0164669 | No |
| orf19.1465 | 85 | 76 | 73 | 48 | 144 | 73 | 106 | 134 | 81 | 80 | 72 | 75 | 115 | 83 | 90 | 109 | 88,26 | 77 | 99 | 1,293 | 0,371 | 2,28 | 0,0226353 | 0,0279752 | 1,00E-08 | 0,0206951 | 0,0137573 | 0,0137573 | TRUE | 0,1183689 | No |
| orf19.1466 | 1958 | 1709 | 1790 | 951 | 2630 | 1713 | 2401 | 2509 | 1867 | 1807 | 1768 | 1490 | 2107 | 1948 | 2038 | 2040 | 1883,2 | 1733 | 2033 | 1,171 | 0,228 | 3,407 | 0,0006563 | 0,0008962 | 0,0044546 | 0,0029631 | 0,0037388 | 0,0037388 | TRUE | 0,1007939 | No |
| orf19.1467 | 8783 | 7302 | 7533 | 5809 | 12287 | 8775 | 11703 | 12254 | 8374 | 7721 | 7442 | 9100 | 9845 | 9978 | 9936 | 9964 | 9044,9 | 8159 | 9931 | 1,218 | 0,284 | 4,822 | 1,42E-06 | 2,18E-06 | 0,0039698 | 0,0022728 | 0,0032238 | 0,0032238 | TRUE | 0,0724511 | No |
| orf19.1468 | 4421 | 3907 | 4124 | 2494 | 3430 | 2285 | 3294 | 3507 | 4215 | 4131 | 4074 | 3907 | 2748 | 2598 | 2797 | 2852 | 3415,24 | 4082 | 2749 | 0,673 | -0,57 | -13,09 | 3,76E-39 | 1,17E-38 | 0,0008853 | 0,002572 | 0,0015234 | 0,0015234 | TRUE | 0,0156951 | No |
| orf19.147 | 13321 | 13905 | 15000 | 9272 | 1697 | 1353 | 1606 | 1450 | 12700 | 14703 | 14819 | 14525 | 1360 | 1538 | 1363 | 1179 | 7773,41 | 14187 | 1360 | 0,096 | -3,384 | -44,336 | 0 | 0 | 0,0077615 | 0,0023025 | 0,0052287 | 0,0052287 | TRUE | 0,0971987 | No |
| orf19.1470 | 10651 | 8522 | 9081 | 8039 | 32529 | 25283 | 30347 | 33772 | 10155 | 9011 | 8971 | 12594 | 26065 | 28749 | 25764 | 27460 | 18596,02 | 10183 | 27010 | 2,653 | 1,408 | 15,088 | 1,95E-51 | 7,32E-51 | 0,015116 | 0,0021796 | 0,0082943 | 0,0082943 | TRUE | 0,3089196 | No |
| orf19.1471 | 5897 | 5129 | 5138 | 3238 | 12848 | 8147 | 11704 | 12664 | 5622 | 5423 | 5076 | 5072 | 10295 | 9264 | 9937 | 10297 | 7623,26 | 5298 | 9948 | 1,877 | 0,909 | 17,826 | 4,42E-71 | 2,15E-70 | 0,0023151 | 0,0023067 | 0,0023435 | 0,0023435 | TRUE | 0,0259456 | No |
| orf19.1473 | 356 | 357 | 380 | 202 | 485 | 345 | 444 | 503 | 339 | 377 | 375 | 316 | 389 | 392 | 377 | 409 | 371,95 | 352 | 392 | 1,107 | 0,147 | 1,782 | 0,0747688 | 0,088401 | 0,0004969 | 0,0065056 | 0,0038139 | 0,0038139 | TRUE | 0,0427467 | No |
| orf19.1474 | 5234 | 4465 | 4970 | 3282 | 6359 | 4650 | 6101 | 6701 | 4990 | 4721 | 4910 | 5141 | 5095 | 5287 | 5180 | 5449 | 5096,7 | 4940 | 5253 | 1,064 | 0,089 | 2,214 | 0,0268056 | 0,0329633 | 0,0007863 | 0,0024134 | 0,0013581 | 0,0013581 | TRUE | 0,0107934 | No |
| orf19.1476 | 33 | 29 | 32 | 20 | 38 | 29 | 32 | 39 | 31 | 31 | 32 | 31 | 30 | 33 | 27 | 32 | 30,92 | 31 | 30 | 0,975 | -0,037 | -0,15 | 0,8808916 | 0,8972018 | 1,00E-08 | 0,0551903 | 0,024847 | 0,024847 | TRUE | 0,0404984 | No |
| orf19.1477 | 1899 | 1910 | 1981 | 1264 | 784 | 528 | 666 | 615 | 1810 | 2020 | 1957 | 1980 | 628 | 600 | 565 | 500 | 1257,66 | 1942 | 573 | 0,295 | -1,76 | -24,375 | 3,15E-131 | 2,78E-130 | 0,0042475 | 0,0033968 | 0,0039386 | 0,0039386 | TRUE | 0,0907123 | No |
| orf19.1478 | 1181 | 1104 | 1173 | 738 | 2064 | 1445 | 1890 | 2113 | 1126 | 1167 | 1159 | 1156 | 1654 | 1643 | 1605 | 1718 | 1403,48 | 1152 | 1655 | 1,437 | 0,523 | 11,188 | 4,68E-29 | 1,24E-28 | 1,00E-08 | 0,0032612 | 0,0013467 | 0,0013467 | TRUE | 0,0082383 | No |
| orf19.1479 | 3460 | 2979 | 3222 | 2238 | 2837 | 2089 | 2744 | 2963 | 3299 | 3150 | 3183 | 3506 | 2273 | 2375 | 2330 | 2409 | 2815,63 | 3284 | 2347 | 0,715 | -0,483 | -10,483 | 1,03E-25 | 2,58E-25 | 0,001028 | 0,0026744 | 0,0016789 | 0,0016789 | TRUE | 0,0234467 | No |
| orf19.1480 | 1974 | 1675 | 1770 | 1287 | 5896 | 4129 | 5530 | 5974 | 1882 | 1771 | 1749 | 2016 | 4724 | 4695 | 4695 | 4857 | 3298,69 | 1854 | 4743 | 2,564 | 1,358 | 27,887 | 3,85E-171 | 4,41E-170 | 0,0013167 | 0,002589 | 0,0018801 | 0,0018801 | TRUE | 0,0378222 | No |
| orf19.1481 | 7286 | 6788 | 7489 | 4549 | 2260 | 1616 | 2218 | 2292 | 6946 | 7177 | 7398 | 7126 | 1811 | 1838 | 1883 | 1864 | 4505,45 | 7162 | 1849 | 0,258 | -1,954 | -52,218 | 0 | 0 | 0,0003684 | 0,0024557 | 0,0010243 | 0,0010243 | TRUE | 0,0061864 | No |
| orf19.1482 | 1737 | 1310 | 1469 | 792 | 607 | 406 | 576 | 574 | 1656 | 1385 | 1451 | 1241 | 486 | 462 | 489 | 467 | 954,61 | 1433 | 476 | 0,332 | -1,593 | -18,774 | 1,24E-78 | 6,63E-78 | 0,0070595 | 0,0038112 | 0,0055828 | 0,0055828 | TRUE | 0,1319274 | No |
| orf19.1483 | 1290 | 1339 | 1284 | 737 | 1168 | 745 | 952 | 1049 | 1230 | 1416 | 1268 | 1155 | 936 | 847 | 808 | 853 | 1064,11 | 1267 | 861 | 0,679 | -0,559 | -7,641 | 2,15E-14 | 4,28E-14 | 0,0043984 | 0,0036343 | 0,0041768 | 0,0041768 | TRUE | 0,0741328 | No |
| orf19.1484 | 1039 | 1252 | 1224 | 773 | 551 | 428 | 450 | 433 | 991 | 1324 | 1209 | 1211 | 442 | 487 | 382 | 352 | 799,61 | 1184 | 416 | 0,35 | -1,513 | -13,806 | 2,33E-43 | 7,79E-43 | 0,0152752 | 0,0041447 | 0,0099765 | 0,0099765 | TRUE | 0,1475841 | No |
| orf19.1485 | 923 | 822 | 823 | 465 | 1619 | 1035 | 1459 | 1467 | 880 | 869 | 813 | 728 | 1297 | 1177 | 1239 | 1193 | 1024,53 | 822 | 1226 | 1,486 | 0,571 | 8,538 | 1,37E-17 | 2,91E-17 | 0,0031661 | 0,0036939 | 0,0032562 | 0,0032562 | TRUE | 0,0618659 | No |
| orf19.1486 | 2013 | 2099 | 2271 | 1619 | 704 | 571 | 734 | 663 | 1919 | 2219 | 2244 | 2536 | 564 | 649 | 623 | 539 | 1411,75 | 2230 | 594 | 0,266 | -1,909 | -21,646 | 6,65E-104 | 4,49E-103 | 0,0086652 | 0,0032543 | 0,0064624 | 0,0064624 | TRUE | 0,1002104 | No |
| orf19.1488 | 460 | 504 | 478 | 317 | 291 | 233 | 255 | 303 | 439 | 533 | 472 | 497 | 233 | 265 | 216 | 246 | 362,66 | 485 | 240 | 0,494 | -1,016 | -10,833 | 2,41E-27 | 6,24E-27 | 0,0041712 | 0,0066187 | 0,0054281 | 0,0054281 | TRUE | 0,054727 | No |
| orf19.149 | 17 | 12 | 11 | 15 | 7 | 8 | 3 | 6 | 16 | 13 | 11 | 23 | 6 | 9 | 3 | 5 | 10,67 | 16 | 6 | 0,347 | -1,526 | -3,11 | 0,0018697 | 0,0024814 | 0,0379079 | 0,1559129 | 0,112402 | 0,112402 | TRUE | 0,3342698 | No |
| orf19.1490 | 3578 | 3311 | 3619 | 2607 | 2298 | 1673 | 2179 | 2356 | 3411 | 3501 | 3575 | 4084 | 1841 | 1902 | 1850 | 1916 | 2760,09 | 3643 | 1877 | 0,516 | -0,955 | -16,19 | 5,91E-59 | 2,45E-58 | 0,0030859 | 0,0026862 | 0,002945 | 0,002945 | TRUE | 0,0780278 | No |
| orf19.1490.1 | 43 | 36 | 30 | 15 | 59 | 37 | 63 | 58 | 41 | 38 | 30 | 23 | 47 | 42 | 53 | 47 | 40,27 | 33 | 47 | 1,417 | 0,502 | 2,162 | 0,0305835 | 0,0374502 | 1,00E-08 | 0,0428598 | 0,0259657 | 0,0259657 | TRUE | 0,1792189 | No |
| orf19.1491 | 2251 | 2271 | 2313 | 1388 | 1562 | 1186 | 1476 | 1403 | 2146 | 2401 | 2285 | 2174 | 1252 | 1349 | 1253 | 1141 | 1750,11 | 2252 | 1249 | 0,553 | -0,853 | -13,943 | 3,48E-44 | 1,18E-43 | 0,0028205 | 0,0030295 | 0,0029885 | 0,0029885 | TRUE | 0,0407583 | No |
| orf19.1492 | 856 | 992 | 930 | 686 | 734 | 564 | 622 | 586 | 816 | 1049 | 919 | 1075 | 588 | 641 | 528 | 476 | 761,55 | 965 | 558 | 0,579 | -0,788 | -7,381 | 1,57E-13 | 3,04E-13 | 0,0141082 | 0,0042473 | 0,0095663 | 0,0095663 | TRUE | 0,1297081 | No |
| orf19.1493 | 3457 | 3088 | 3362 | 1999 | 5445 | 3862 | 5197 | 5433 | 3296 | 3265 | 3321 | 3132 | 4363 | 4391 | 4412 | 4418 | 3824,76 | 3254 | 4396 | 1,349 | 0,432 | 12,689 | 6,82E-37 | 2,05E-36 | 1,00E-08 | 0,0025206 | 0,0008406 | 0,0008406 | TRUE | 0,0069204 | No |
| orf19.1494 | 9007 | 7641 | 8335 | 5984 | 13766 | 10559 | 13084 | 13632 | 8587 | 8079 | 8234 | 9374 | 11030 | 12007 | 11108 | 11084 | 9938,03 | 8568 | 11307 | 1,32 | 0,401 | 7,498 | 6,49E-14 | 1,27E-13 | 0,0029884 | 0,0022565 | 0,002637 | 0,002637 | TRUE | 0,0480221 | No |
| orf19.1495 | 622 | 747 | 718 | 457 | 1151 | 905 | 929 | 828 | 593 | 790 | 709 | 716 | 922 | 1029 | 789 | 673 | 777,67 | 702 | 853 | 1,215 | 0,281 | 2,268 | 0,0233158 | 0,028805 | 0,0215154 | 0,0042026 | 0,0133714 | 0,0133714 | TRUE | 0,1876453 | No |
| orf19.1496 | 462 | 510 | 489 | 284 | 855 | 626 | 722 | 772 | 440 | 539 | 483 | 445 | 685 | 712 | 613 | 628 | 568,16 | 477 | 660 | 1,378 | 0,463 | 5,502 | 3,75E-08 | 6,12E-08 | 0,0047087 | 0,0049811 | 0,0049484 | 0,0049484 | TRUE | 0,0902455 | No |
| orf19.1497 | 489 | 462 | 509 | 294 | 865 | 669 | 832 | 842 | 466 | 489 | 503 | 461 | 693 | 761 | 706 | 685 | 595,37 | 480 | 711 | 1,477 | 0,563 | 8,359 | 6,35E-17 | 1,33E-16 | 1,00E-08 | 0,0048491 | 0,0025674 | 0,0025674 | TRUE | 0,0238916 | No |
| orf19.1499 | 2920 | 2793 | 3238 | 2021 | 4238 | 3213 | 4200 | 4226 | 2784 | 2953 | 3199 | 3166 | 3396 | 3653 | 3566 | 3436 | 3269,15 | 3026 | 3513 | 1,162 | 0,216 | 4,052 | 5,08E-05 | 7,29E-05 | 0,0022237 | 0,0025935 | 0,0024174 | 0,0024174 | TRUE | 0,0357733 | No |
| orf19.150 | 5012 | 4113 | 4425 | 2809 | 3477 | 2282 | 3307 | 3294 | 4778 | 4349 | 4371 | 4400 | 2786 | 2595 | 2808 | 2678 | 3595,77 | 4474 | 2717 | 0,607 | -0,72 | -15,318 | 5,81E-53 | 2,23E-52 | 0,0013032 | 0,0025479 | 0,0018294 | 0,0018294 | TRUE | 0,02598 | No |
| orf19.1500 | 945 | 925 | 987 | 613 | 856 | 687 | 853 | 874 | 901 | 978 | 975 | 960 | 686 | 781 | 724 | 711 | 839,54 | 954 | 726 | 0,759 | -0,397 | -6,457 | 1,07E-10 | 1,91E-10 | 0,0009878 | 0,004047 | 0,0024237 | 0,0024237 | TRUE | 0,029078 | No |
| orf19.1502 | 483 | 561 | 566 | 346 | 189 | 164 | 186 | 151 | 460 | 593 | 559 | 542 | 151 | 186 | 158 | 123 | 346,68 | 538 | 154 | 0,285 | -1,809 | -14,82 | 1,08E-49 | 3,98E-49 | 0,0142024 | 0,0068273 | 0,0103582 | 0,0103582 | TRUE | 0,209381 | No |
| orf19.1504 | 2559 | 2611 | 2706 | 1516 | 4394 | 3267 | 4168 | 4228 | 2440 | 2761 | 2673 | 2375 | 3521 | 3715 | 3539 | 3438 | 3057,59 | 2562 | 3553 | 1,385 | 0,47 | 8,474 | 2,37E-17 | 5,03E-17 | 0,0026759 | 0,0026283 | 0,0026036 | 0,0026036 | TRUE | 0,032599 | No |
| orf19.1505 | 2297 | 2270 | 2269 | 1430 | 4182 | 3145 | 3941 | 3764 | 2190 | 2400 | 2242 | 2240 | 3351 | 3576 | 3346 | 3060 | 2800,67 | 2268 | 3333 | 1,469 | 0,555 | 9,955 | 2,40E-23 | 5,76E-23 | 0,0023258 | 0,0026775 | 0,0025981 | 0,0025981 | TRUE | 0,0370473 | No |
| orf19.1506 | 0 | 0 | 0 | 0 | 0 | 0 | 0 | 0 | 0 | 0 | 0 | 0 | 0 | 0 | 0 | 0 | 0 | 0 | 0 | NA | NA | NA | NA | NA | NA | NA | NA | NA | NA | NA | NA |
| orf19.1507 | 255 | 281 | 303 | 147 | 261 | 216 | 232 | 253 | 243 | 297 | 299 | 230 | 209 | 246 | 197 | 206 | 240,91 | 267 | 214 | 0,795 | -0,331 | -2,835 | 0,0045894 | 0,0059414 | 0,0089746 | 0,0089067 | 0,00893 | 0,00893 | TRUE | 0,0964533 | No |
| orf19.1508 | 26 | 20 | 19 | 15 | 17 | 25 | 26 | 22 | 25 | 21 | 19 | 23 | 14 | 28 | 22 | 18 | 21,28 | 22 | 20 | 0,918 | -0,124 | -0,387 | 0,6984533 | 0,7291935 | 1,00E-08 | 0,0792597 | 0,051208 | 0,051208 | TRUE | 0,3017035 | No |
| orf19.1509 | 9211 | 9071 | 9649 | 5932 | 3834 | 2871 | 3723 | 3719 | 8782 | 9591 | 9532 | 9293 | 3072 | 3265 | 3161 | 3024 | 6214,95 | 9300 | 3130 | 0,336 | -1,572 | -36,281 | 3,19E-288 | 6,79E-287 | 0,0010736 | 0,0023555 | 0,0015985 | 0,0015985 | TRUE | 0,0174847 | No |
| orf19.151 | 1172 | 1129 | 1268 | 797 | 816 | 571 | 734 | 737 | 1117 | 1194 | 1253 | 1249 | 654 | 649 | 623 | 599 | 917,24 | 1203 | 631 | 0,525 | -0,929 | -15,096 | 1,71E-51 | 6,42E-51 | 0,0013265 | 0,0038813 | 0,0024645 | 0,0024645 | TRUE | 0,0286918 | No |
| orf19.1510 | 1222 | 1151 | 1251 | 742 | 1481 | 1108 | 1436 | 1509 | 1165 | 1217 | 1236 | 1162 | 1187 | 1260 | 1219 | 1227 | 1209,13 | 1195 | 1223 | 1,022 | 0,031 | 0,626 | 0,5314156 | 0,5672404 | 1,00E-08 | 0,0034492 | 0,0014871 | 0,0014871 | TRUE | 0,0064417 | No |
| orf19.1513 | 6521 | 6281 | 6545 | 3767 | 1980 | 1390 | 1749 | 1941 | 6217 | 6641 | 6466 | 5901 | 1587 | 1581 | 1485 | 1578 | 3931,96 | 6306 | 1558 | 0,247 | -2,019 | -41,203 | 0 | 0 | 0,0015866 | 0,0025089 | 0,0019289 | 0,0019289 | TRUE | 0,0220055 | No |
| orf19.1514 | 2396 | 2069 | 2312 | 1342 | 3386 | 2413 | 3351 | 3428 | 2284 | 2188 | 2284 | 2102 | 2713 | 2744 | 2845 | 2787 | 2493,45 | 2214 | 2772 | 1,25 | 0,322 | 7,536 | 4,85E-14 | 9,52E-14 | 0,0004536 | 0,0027498 | 0,0013343 | 0,0013343 | TRUE | 0,0127044 | No |
| orf19.1515 | 259 | 257 | 237 | 151 | 312 | 223 | 313 | 311 | 247 | 272 | 234 | 237 | 250 | 254 | 266 | 253 | 251,44 | 248 | 256 | 1,032 | 0,045 | 0,482 | 0,6300115 | 0,6628365 | 1,00E-08 | 0,0086212 | 0,0042954 | 0,0042954 | TRUE | 0,0492818 | No |
| orf19.1516 | 1185 | 1164 | 1198 | 653 | 3092 | 2141 | 2918 | 3079 | 1130 | 1231 | 1184 | 1023 | 2478 | 2435 | 2477 | 2504 | 1807,49 | 1142 | 2474 | 2,159 | 1,111 | 20,24 | 4,35E-91 | 2,63E-90 | 0,0013945 | 0,0029997 | 0,0022202 | 0,0022202 | TRUE | 0,0512492 | No |
| orf19.1517 | 2073 | 1709 | 1802 | 1143 | 4283 | 2796 | 4266 | 4416 | 1976 | 1807 | 1780 | 1791 | 3432 | 3179 | 3622 | 3591 | 2647,23 | 1838 | 3456 | 1,879 | 0,91 | 16,034 | 7,40E-58 | 3,02E-57 | 0,0024246 | 0,0027115 | 0,0026594 | 0,0026594 | TRUE | 0,0344157 | No |
| orf19.1519 | 1148 | 1003 | 1108 | 713 | 1564 | 1243 | 1637 | 1630 | 1094 | 1061 | 1095 | 1117 | 1253 | 1413 | 1390 | 1325 | 1218,54 | 1092 | 1345 | 1,232 | 0,301 | 5,473 | 4,43E-08 | 7,22E-08 | 0,0010741 | 0,0034387 | 0,0020562 | 0,0020562 | TRUE | 0,0259986 | No |
| orf19.1522 | 72 | 73 | 73 | 42 | 89 | 60 | 101 | 106 | 69 | 77 | 72 | 66 | 71 | 68 | 86 | 86 | 74,4 | 71 | 78 | 1,098 | 0,136 | 0,811 | 0,4170929 | 0,4531704 | 1,00E-08 | 0,0241591 | 0,0132661 | 0,0132661 | TRUE | 0,0545232 | No |
| orf19.1523 | 1491 | 1198 | 1390 | 898 | 941 | 667 | 851 | 930 | 1422 | 1267 | 1373 | 1407 | 754 | 758 | 722 | 756 | 1057,41 | 1367 | 748 | 0,547 | -0,87 | -15,347 | 3,70E-53 | 1,42E-52 | 0,0008914 | 0,003644 | 0,0020785 | 0,0020785 | TRUE | 0,029805 | No |
| orf19.1524 | 5 | 6 | 5 | 4 | 5 | 11 | 13 | 14 | 5 | 6 | 5 | 6 | 4 | 13 | 11 | 11 | 7,66 | 6 | 10 | 1,743 | 0,802 | 1,471 | 0,1413687 | 0,1625065 | 1,00E-08 | 0,2165385 | 0,1385067 | 0,1385067 | TRUE | 0,6592285 | No |

| | | | | | | | | | | | | | | | | | | | | | | | | | | | | | | | |
|---|---|---|---|---|---|---|---|---|---|---|---|---|---|---|---|---|---|---|---|---|---|---|---|---|---|---|---|---|---|---|---|
| orf19.1525 | 1543 | 1524 | 1580 | 1082 | 1833 | 1343 | 1722 | 1797 | 1471 | 1611 | 1561 | 1695 | 1469 | 1527 | 1462 | 1461 | 1532,17 | 1584 | 1480 | 0,936 | -0,096 | -1,823 | 0,0682512 | 0,0810404 | 0,0011088 | 0,0031629 | 0,0020146 | 0,0020146 | TRUE | 0,028252 | No |
| orf19.1526 | 3719 | 3285 | 3732 | 2354 | 3428 | 2452 | 3204 | 3451 | 3546 | 3473 | 3687 | 3688 | 2747 | 2788 | 2720 | 2806 | 3181,84 | 3598 | 2765 | 0,769 | -0,379 | -10,029 | 1,13E-23 | 2,73E-23 | 0,0002363 | 0,0026073 | 0,0010553 | 0,0010553 | TRUE | 0,0065938 | No |
| orf19.1527 | 1130 | 1019 | 1074 | 723 | 1388 | 969 | 1190 | 1167 | 1077 | 1077 | 1061 | 1133 | 1112 | 1102 | 1010 | 949 | 1065,2 | 1087 | 1043 | 0,96 | -0,059 | -0,925 | 0,3551869 | 0,3898929 | 0,0024186 | 0,0036327 | 0,0029484 | 0,0029484 | TRUE | 0,0453213 | No |
| orf19.1528 | 3731 | 3727 | 3950 | 2253 | 3844 | 2724 | 3615 | 3729 | 3557 | 3941 | 3902 | 3529 | 3080 | 3097 | 3069 | 3032 | 3401,03 | 3732 | 3070 | 0,822 | -0,283 | -5,972 | 2,35E-09 | 3,98E-09 | 0,0013701 | 0,002574 | 0,0018642 | 0,0018642 | TRUE | 0,0169775 | No |
| orf19.1529 | 17 | 9 | 12 | 14 | 6 | 10 | 6 | 13 | 16 | 10 | 12 | 22 | 5 | 11 | 5 | 11 | 11,42 | 15 | 8 | 0,537 | -0,898 | -1,932 | 0,0533725 | 0,0639688 | 0,0609333 | 0,1458761 | 0,1122303 | 0,1122303 | TRUE | 0,222889 | No |
| orf19.1530 | 1 | 5 | 7 | 0 | 1 | 1 | 1 | 0 | 1 | 5 | 7 | 0 | 1 | 1 | 1 | 0 | 1,99 | 3 | 1 | 0,198 | -2,334 | -1,824 | 0,0681992 | 0,0809936 | 1,00E-08 | 0,8259806 | 0,7483157 | 0,7483157 | TRUE | 0,0805983 | No |
| orf19.1531 | 3032 | 3448 | 3332 | 2018 | 3194 | 2484 | 2885 | 2823 | 2891 | 3646 | 3292 | 3161 | 2559 | 2825 | 2449 | 2295 | 2889,75 | 3248 | 2532 | 0,779 | -0,36 | -4,551 | 5,34E-06 | 8,03E-06 | 0,0080764 | 0,0026595 | 0,0056542 | 0,0056542 | TRUE | 0,0834871 | No |
| orf19.1532 | 466 | 494 | 499 | 355 | 899 | 594 | 804 | 707 | 444 | 522 | 493 | 556 | 720 | 675 | 683 | 575 | 583,62 | 504 | 663 | 1,321 | 0,401 | 4,4 | 1,08E-05 | 1,60E-05 | 0,007403 | 0,0049046 | 0,0061997 | 0,0061997 | TRUE | 0,0982052 | No |
| orf19.1533 | 2622 | 2660 | 2720 | 1558 | 3137 | 2397 | 3052 | 2852 | 2500 | 2813 | 2687 | 2441 | 2514 | 2726 | 2591 | 2319 | 2573,68 | 2610 | 2538 | 0,971 | -0,043 | -0,675 | 0,4994217 | 0,5353284 | 0,003593 | 0,0027292 | 0,0034532 | 0,0034532 | TRUE | 0,0408724 | No |
| orf19.1534 | 776 | 818 | 882 | 470 | 689 | 487 | 621 | 568 | 740 | 865 | 871 | 736 | 552 | 554 | 527 | 462 | 663,41 | 803 | 524 | 0,65 | -0,621 | -7,226 | 4,98E-13 | 9,54E-13 | 0,0064203 | 0,0045662 | 0,0055347 | 0,0055347 | TRUE | 0,0761445 | No |
| orf19.1535 | 144 | 113 | 136 | 66 | 102 | 79 | 74 | 73 | 137 | 119 | 134 | 103 | 82 | 90 | 63 | 59 | 98,53 | 123 | 74 | 0,586 | -0,772 | -4,54 | 5,62E-06 | 8,42E-06 | 0,0149256 | 0,0187548 | 0,0171559 | 0,0171559 | TRUE | 0,1803922 | No |
| orf19.1536 | 1257 | 1403 | 1423 | 1030 | 2607 | 1950 | 2291 | 2398 | 1198 | 1483 | 1406 | 1614 | 2089 | 2217 | 1945 | 1950 | 1737,79 | 1425 | 2050 | 1,441 | 0,527 | 6,318 | 2,65E-10 | 4,67E-10 | 0,0086516 | 0,0030361 | 0,0060666 | 0,0060666 | TRUE | 0,1397046 | No |
| orf19.1538 | 247 | 223 | 226 | 143 | 378 | 235 | 330 | 327 | 235 | 236 | 223 | 224 | 303 | 267 | 280 | 266 | 254,34 | 230 | 279 | 1,215 | 0,281 | 3,021 | 0,0025187 | 0,0033186 | 1,00E-08 | 0,0085468 | 0,0043205 | 0,0043205 | TRUE | 0,0390993 | No |
| orf19.1539 | 277 | 318 | 302 | 212 | 103 | 62 | 87 | 79 | 264 | 336 | 298 | 332 | 83 | 70 | 74 | 64 | 190,24 | 308 | 73 | 0,238 | -2,072 | -15,612 | 6,01E-55 | 2,37E-54 | 0,0069775 | 0,010722 | 0,0089967 | 0,0089967 | TRUE | 0,1070689 | No |
| orf19.154 | 2489 | 2664 | 2685 | 1616 | 2043 | 1477 | 1840 | 1941 | 2373 | 2817 | 2653 | 2532 | 1637 | 1679 | 1562 | 1578 | 2103,84 | 2594 | 1614 | 0,622 | -0,685 | -11,618 | 3,35E-31 | 9,23E-31 | 0,0025142 | 0,0028717 | 0,0028439 | 0,0028439 | TRUE | 0,041356 | No |
| orf19.1541 | 30 | 40 | 40 | 23 | 59 | 49 | 79 | 49 | 29 | 42 | 40 | 36 | 47 | 56 | 67 | 40 | 44,54 | 37 | 52 | 1,431 | 0,517 | 2,154 | 0,0312595 | 0,0382194 | 0,0228754 | 0,0389519 | 0,0315545 | 0,0315545 | TRUE | 0,2765115 | No |
| orf19.1542 | 4469 | 3948 | 4337 | 2562 | 3456 | 2225 | 3287 | 3420 | 4261 | 4175 | 4285 | 4014 | 2769 | 2530 | 2791 | 2781 | 3450,49 | 4184 | 2718 | 0,65 | -0,622 | -13,81 | 2,21E-43 | 7,38E-43 | 0,0010899 | 0,0025671 | 0,0016485 | 0,0016485 | TRUE | 0,0250337 | No |
| orf19.1543 | 1017 | 916 | 966 | 637 | 1901 | 1435 | 1867 | 1951 | 970 | 969 | 954 | 998 | 1523 | 1632 | 1585 | 1586 | 1277,09 | 973 | 1582 | 1,627 | 0,702 | 14,509 | 1,07E-47 | 3,82E-47 | 1,00E-08 | 0,0033769 | 0,001392 | 0,001392 | TRUE | 0,0075275 | No |
| orf19.1544 | 312 | 271 | 294 | 219 | 1355 | 910 | 1310 | 1327 | 297 | 287 | 290 | 343 | 1086 | 1035 | 1112 | 1079 | 691,15 | 304 | 1078 | 3,573 | 1,837 | 26,092 | 4,49E-150 | 4,52E-149 | 1,00E-08 | 0,0044669 | 0,0025056 | 0,0025056 | TRUE | 0,0647223 | No |
| orf19.1545 | 1507 | 1230 | 1313 | 855 | 1291 | 859 | 1191 | 1313 | 1437 | 1301 | 1297 | 1339 | 1034 | 977 | 1011 | 1068 | 1182,98 | 1344 | 1022 | 0,762 | -0,393 | -7,049 | 1,81E-12 | 3,39E-12 | 0,0011207 | 0,0034792 | 0,0021279 | 0,0021279 | TRUE | 0,0280191 | No |
| orf19.1546 | 1567 | 1420 | 1459 | 895 | 1592 | 1020 | 1517 | 1607 | 1494 | 1501 | 1441 | 1402 | 1276 | 1160 | 1288 | 1307 | 1358,61 | 1460 | 1258 | 0,862 | -0,214 | -4,001 | 6,32E-05 | 9,03E-05 | 0,0009197 | 0,0032998 | 0,0020204 | 0,0020204 | TRUE | 0,030513 | No |
| orf19.1547 | 352 | 333 | 340 | 227 | 502 | 392 | 496 | 573 | 336 | 352 | 336 | 356 | 402 | 446 | 421 | 466 | 389,27 | 345 | 434 | 1,259 | 0,333 | 4,142 | 3,44E-05 | 4,97E-05 | 1,00E-08 | 0,0063092 | 0,0035467 | 0,0035467 | TRUE | 0,0316145 | No |
| orf19.1548 | 1059 | 814 | 898 | 519 | 1254 | 859 | 1275 | 1318 | 1010 | 861 | 887 | 813 | 1005 | 977 | 1082 | 1072 | 963,28 | 893 | 1034 | 1,156 | 0,209 | 2,871 | 0,0040891 | 0,0053185 | 0,004248 | 0,0037958 | 0,0040183 | 0,0040183 | TRUE | 0,0941114 | No |
| orf19.1549 | 2575 | 2496 | 2742 | 1400 | 5921 | 3945 | 5548 | 5718 | 2455 | 2639 | 2709 | 2193 | 4744 | 4486 | 4710 | 4649 | 3573,23 | 2499 | 4647 | 1,857 | 0,893 | 14,459 | 2,20E-47 | 7,82E-47 | 0,0041416 | 0,0025508 | 0,0033406 | 0,0033406 | TRUE | 0,0780142 | No |
| orf19.155 | 2285 | 2371 | 2402 | 1551 | 1684 | 1383 | 1534 | 1521 | 2179 | 2507 | 2373 | 2430 | 1349 | 1573 | 1302 | 1237 | 1868,65 | 2372 | 1365 | 0,575 | -0,799 | -10,593 | 3,21E-26 | 8,17E-26 | 0,0066763 | 0,0029699 | 0,0048926 | 0,0048926 | TRUE | 0,12203 | No |
| orf19.1551 | 10 | 13 | 12 | 4 | 12 | 9 | 7 | 13 | 10 | 14 | 12 | 6 | 10 | 10 | 6 | 11 | 9,72 | 10 | 9 | 0,858 | -0,221 | -0,49 | 0,6241122 | 0,6572801 | 1,00E-08 | 0,1710046 | 0,093879 | 0,093879 | TRUE | 0,2047911 | No |
| orf19.1552 | 3135 | 2407 | 2418 | 1645 | 7591 | 5028 | 7045 | 7897 | 2989 | 2545 | 2389 | 2577 | 6082 | 5717 | 5981 | 6421 | 4337,71 | 2625 | 6050 | 2,305 | 1,205 | 17,775 | 1,10E-70 | 5,28E-70 | 0,0055559 | 0,0024698 | 0,0041215 | 0,0041215 | TRUE | 0,1079602 | No |
| orf19.1553 | 6783 | 5957 | 6498 | 4109 | 9210 | 6599 | 9009 | 9365 | 6467 | 6299 | 6419 | 6437 | 7380 | 7504 | 7649 | 7615 | 6971,08 | 6406 | 7537 | 1,177 | 0,235 | 7,938 | 2,06E-15 | 4,18E-15 | 9,20E-05 | 0,0023268 | 0,0006924 | 0,0006924 | TRUE | 0,0024726 | No |
| orf19.1555 | 2182 | 2111 | 2191 | 1361 | 2439 | 1725 | 2241 | 2316 | 2080 | 2232 | 2165 | 2132 | 1954 | 1961 | 1903 | 1883 | 2038,81 | 2152 | 1925 | 0,894 | -0,161 | -3,818 | 0,0001346 | 0,0001894 | 1,00E-08 | 0,0028966 | 0,0012228 | 0,0012228 | TRUE | 0,0077638 | No |
| orf19.1557 | 149 | 106 | 141 | 95 | 296 | 228 | 262 | 296 | 142 | 112 | 139 | 149 | 237 | 259 | 222 | 241 | 187,73 | 136 | 240 | 1,773 | 0,826 | 7,106 | 1,19E-12 | 2,25E-12 | 0,0011067 | 0,0108376 | 0,0069973 | 0,0069973 | TRUE | 0,1404186 | No |
| orf19.1559 | 5067 | 4228 | 4786 | 3069 | 8215 | 5543 | 7864 | 8397 | 4831 | 4471 | 4728 | 4808 | 6582 | 6303 | 6676 | 6828 | 5653,33 | 4710 | 6597 | 1,402 | 0,487 | 11,761 | 6,24E-32 | 1,74E-31 | 0,000932 | 0,0023817 | 0,0014603 | 0,0014603 | TRUE | 0,0143079 | No |
| orf19.156 | 3098 | 2892 | 3073 | 1769 | 2620 | 1892 | 2604 | 2656 | 2954 | 3058 | 3036 | 2771 | 2099 | 2151 | 2211 | 2160 | 2554,96 | 2955 | 2155 | 0,728 | -0,457 | -10,3 | 7,07E-25 | 1,75E-24 | 0,0006608 | 0,0027339 | 0,0014949 | 0,0014949 | TRUE | 0,0194862 | No |
| orf19.1560 | 5786 | 5000 | 5449 | 3380 | 7474 | 5181 | 7278 | 7853 | 5516 | 5287 | 5383 | 5295 | 5989 | 5891 | 6179 | 6385 | 5740,68 | 5370 | 6111 | 1,138 | 0,186 | 4,884 | 1,04E-06 | 1,60E-06 | 0,0006025 | 0,0023773 | 0,0012208 | 0,0012208 | TRUE | 0,0111688 | No |
| orf19.1562 | 32160 | 29870 | 31589 | 22185 | 1367 | 1010 | 1297 | 1301 | 30661 | 31584 | 31207 | 34754 | 1095 | 1148 | 1101 | 1058 | 16576,07 | 32052 | 1100 | 0,034 | -4,865 | -93,92 | 0 | 0 | 0,0015938 | 0,0021903 | 0,0021539 | 0,0021539 | TRUE | 0,0392306 | No |
| orf19.1563 | 2834 | 2517 | 2683 | 1762 | 6380 | 4345 | 5967 | 6170 | 2702 | 2661 | 2651 | 2760 | 5112 | 4941 | 5066 | 5017 | 3863,71 | 2694 | 5034 | 1,871 | 0,904 | 26,509 | 7,58E-155 | 7,90E-154 | 1,00E-08 | 0,0025162 | 0,0008193 | 0,0008193 | TRUE | 0,0029602 | No |
| orf19.1564 | 1483 | 1381 | 1438 | 1016 | 3195 | 2305 | 2873 | 3013 | 1414 | 1460 | 1421 | 1592 | 2560 | 2621 | 2439 | 2450 | 1994,55 | 1472 | 2518 | 1,715 | 0,778 | 15,164 | 6,14E-52 | 2,32E-51 | 0,0014682 | 0,0029145 | 0,001967 | 0,001967 | TRUE | 0,0327206 | No |
| orf19.1565 | 984 | 875 | 921 | 589 | 2759 | 1972 | 2709 | 2956 | 938 | 925 | 910 | 923 | 2211 | 2242 | 2300 | 2404 | 1606,55 | 924 | 2289 | 2,478 | 1,309 | 26,632 | 2,90E-156 | 3,06E-155 | 1,00E-08 | 0,0031133 | 0,0015208 | 0,0015208 | TRUE | 0,0140006 | No |
| orf19.1566 | 18399 | 16883 | 17981 | 11461 | 20490 | 14321 | 19279 | 21295 | 17541 | 17852 | 17764 | 17954 | 16418 | 16284 | 16368 | 17315 | 17186,98 | 17778 | 16596 | 0,934 | -0,099 | -3,167 | 0,0015404 | 0,0020538 | 0,0003934 | 0,0021868 | 0,0008785 | 0,0008785 | TRUE | 0,0104484 | No |
| orf19.1567 | 3284 | 3156 | 3346 | 1915 | 2213 | 1523 | 2083 | 2336 | 3131 | 3337 | 3306 | 3000 | 1773 | 1732 | 1768 | 1899 | 2493,29 | 3194 | 1793 | 0,561 | -0,833 | -16,409 | 1,66E-60 | 7,02E-60 | 0,0014701 | 0,0027498 | 0,0020498 | 0,0020498 | TRUE | 0,0194305 | No |
| orf19.1569 | 10173 | 9891 | 10562 | 6839 | 9778 | 7503 | 9230 | 10053 | 9699 | 10458 | 10434 | 10714 | 7835 | 8532 | 7836 | 8174 | 9210,24 | 10326 | 8094 | 0,784 | -0,351 | -7,78 | 7,23E-15 | 1,45E-14 | 0,0015918 | 0,0022695 | 0,0018484 | 0,0018484 | TRUE | 0,0205874 | No |
| orf19.1570 | 1001 | 889 | 925 | 526 | 1207 | 798 | 1207 | 1281 | 954 | 940 | 914 | 824 | 967 | 907 | 1025 | 1042 | 946,63 | 908 | 985 | 1,083 | 0,115 | 1,73 | 0,0836586 | 0,0984192 | 0,0025727 | 0,0038258 | 0,0031976 | 0,0031976 | TRUE | 0,040916 | No |
| orf19.1571 | 1 | 3 | 2 | 1 | 1 | 0 | 1 | 1 | 1 | 3 | 2 | 2 | 1 | 0 | 1 | 1 | 1,27 | 2 | 1 | 0,338 | -1,567 | -1,216 | 0,2240209 | 0,2527812 | 1,00E-08 | 1,2985989 | 0,6208687 | 0,6208687 | TRUE | 0,1664991 | No |
| orf19.1573 | 2338 | 2099 | 2171 | 1376 | 3891 | 2763 | 3490 | 3892 | 2229 | 2219 | 2145 | 2156 | 3118 | 3142 | 2963 | 3165 | 2641,98 | 2187 | 3097 | 1,415 | 0,501 | 12,322 | 6,90E-35 | 2,01E-34 | 0,0003622 | 0,0027127 | 0,0011849 | 0,0011849 | TRUE | 0,0104041 | No |
| orf19.1574 | 2188 | 1819 | 2106 | 1194 | 1799 | 1066 | 1524 | 1801 | 2086 | 1923 | 2081 | 1870 | 1441 | 1212 | 1294 | 1464 | 1671,53 | 1990 | 1353 | 0,68 | -0,556 | -8,002 | 1,23E-15 | 2,50E-15 | 0,0046608 | 0,0030736 | 0,0040267 | 0,0040267 | TRUE | 0,0567914 | No |
| orf19.1575 | 11725 | 10114 | 10546 | 6747 | 14193 | 9753 | 13119 | 14337 | 11179 | 10694 | 10418 | 10570 | 11372 | 11090 | 11138 | 11657 | 11014,82 | 10715 | 11314 | 1,056 | 0,078 | 2,218 | 0,0265762 | 0,0326938 | 0,0006133 | 0,0022403 | 0,0011093 | 0,0011093 | TRUE | 0,0105209 | No |
| orf19.1576 | 1574 | 1379 | 1547 | 967 | 1173 | 910 | 1116 | 1134 | 1501 | 1458 | 1528 | 1515 | 940 | 1035 | 947 | 922 | 1230,76 | 1500 | 961 | 0,639 | -0,646 | -12,314 | 7,62E-35 | 2,22E-34 | 0,0004361 | 0,0034253 | 0,0018006 | 0,0018006 | TRUE | 0,0291928 | No |
| orf19.1577 | 69 | 69 | 65 | 53 | 70 | 40 | 60 | 57 | 66 | 73 | 64 | 83 | 56 | 45 | 51 | 46 | 60,61 | 72 | 50 | 0,705 | -0,504 | -2,742 | 0,0061037 | 0,0078492 | 1,00E-08 | 0,029183 | 0,0157179 | 0,0157179 | TRUE | 0,0796573 | No |
| orf19.1578 | 12235 | 12656 | 12802 | 7283 | 19987 | 14087 | 17641 | 18892 | 11665 | 13382 | 12647 | 11409 | 16015 | 16018 | 14977 | 15361 | 13934,32 | 12276 | 15593 | 1,27 | 0,345 | 6,3 | 2,97E-10 | 5,23E-10 | 0,0031956 | 0,0022091 | 0,0028009 | 0,0028009 | TRUE | 0,0449489 | No |
| orf19.158 | 5391 | 6152 | 6296 | 3163 | 6779 | 4870 | 6120 | 5530 | 5140 | 6505 | 6220 | 4955 | 5432 | 5538 | 5196 | 4496 | 5435,16 | 5705 | 5166 | 0,905 | -0,144 | -1,556 | 0,119736 | 0,1386131 | 0,0130156 | 0,0023934 | 0,0080421 | 0,0080421 | TRUE | 0,1084961 | No |
| orf19.1580 | 293 | 308 | 303 | 201 | 107 | 74 | 81 | 102 | 279 | 326 | 299 | 315 | 86 | 84 | 69 | 83 | 192,6 | 305 | 80 | 0,264 | -1,921 | -15,98 | 1,76E-57 | 7,14E-57 | 1,00E-08 | 0,0106162 | 0,0065174 | 0,0065174 | TRUE | 0,0949283 | No |
| orf19.1581 | 21 | 12 | 18 | 7 | 5 | 4 | 13 | 11 | 20 | 13 | 18 | 11 | 4 | 5 | 11 | 9 | 11,25 | 16 | 7 | 0,463 | -1,112 | -2,416 | 0,0156783 | 0,0196394 | 0,0338098 | 0,1480487 | 0,1059702 | 0,1059702 | TRUE | 0,3077679 | No |
| orf19.1582 | 89 | 55 | 84 | 51 | 84 | 70 | 87 | 91 | 85 | 58 | 83 | 80 | 67 | 80 | 74 | 74 | 75,08 | 76 | 74 | 0,961 | -0,058 | -0,336 | 0,7368668 | 0,76554 | 1,00E-08 | 0,0239599 | 0,0151946 | 0,0151946 | TRUE | 0,2475752 | No |
| orf19.1584 | 17607 | 14400 | 15754 | 9476 | 12801 | 8407 | 12280 | 13048 | 16786 | 15226 | 15563 | 14845 | 10257 | 9560 | 10426 | 10609 | 12909,02 | 15605 | 10213 | 0,654 | -0,612 | -12,264 | 1,42E-34 | 4,11E-34 | 0,0023509 | 0,0022185 | 0,0023097 | 0,0023097 | TRUE | 0,0318516 | No |
| orf19.1585 | 1219 | 1034 | 1158 | 797 | 2293 | 1606 | 2144 | 2393 | 1162 | 1093 | 1144 | 1249 | 1837 | 1826 | 1820 | 1946 | 1509,69 | 1162 | 1857 | 1,603 | 0,681 | 12,909 | 4,01E-38 | 1,23E-37 | 0,0011243 | 0,0031789 | 0,001947 | 0,001947 | TRUE | 0,0263168 | No |
| orf19.1586 | 149 | 117 | 111 | 73 | 251 | 181 | 220 | 266 | 142 | 124 | 110 | 114 | 201 | 206 | 187 | 216 | 162,47 | 122 | 202 | 1,646 | 0,719 | 5,876 | 4,19E-09 | 7,05E-09 | 1,00E-08 | 0,012197 | 0,0076337 | 0,0076337 | TRUE | 0,1280201 | No |
| orf19.1587 | 102 | 110 | 99 | 79 | 271 | 190 | 257 | 282 | 97 | 116 | 98 | 124 | 217 | 216 | 218 | 229 | 164,47 | 109 | 220 | 2,047 | 1,034 | 8,643 | 5,48E-18 | 1,18E-17 | 1,00E-08 | 0,012074 | 0,006552 | 0,006552 | TRUE | 0,0644669 | No |
| orf19.1588 | 1124 | 940 | 1089 | 598 | 1477 | 972 | 1404 | 1404 | 1072 | 994 | 1076 | 937 | 1183 | 1105 | 1192 | 1142 | 1087,56 | 1020 | 1156 | 1,131 | 0,177 | 2,984 | 0,0028474 | 0,0037355 | 0,001518 | 0,003601 | 0,002453 | 0,002453 | TRUE | 0,0314686 | No |
| orf19.1589 | 1835 | 1628 | 1789 | 1134 | 3100 | 2241 | 2868 | 3105 | 1749 | 1721 | 1767 | 1776 | 2484 | 2548 | 2435 | 2525 | 2125,81 | 1753 | 2498 | 1,425 | 0,51 | 12,887 | 5,32E-38 | 1,63E-37 | 1,00E-08 | 0,0028636 | 0,0010085 | 0,0010085 | TRUE | 0,0034778 | No |
| orf19.1589.1 | 172 | 139 | 155 | 75 | 916 | 543 | 950 | 1078 | 164 | 147 | 153 | 117 | 734 | 617 | 807 | 877 | 452,01 | 145 | 759 | 5,176 | 2,372 | 18,834 | 4,00E-79 | 2,16E-78 | 0,0166947 | 0,0057238 | 0,0108313 | 0,0108313 | TRUE | 0,1318666 | No |
| orf19.1590 | 695 | 744 | 813 | 452 | 812 | 612 | 809 | 740 | 663 | 787 | 803 | 708 | 651 | 696 | 687 | 602 | 699,45 | 740 | 659 | 0,888 | -0,171 | -2,141 | 0,0323016 | 0,0394481 | 0,0049044 | 0,0044387 | 0,0046868 | 0,0046868 | TRUE | 0,0639396 | No |
| orf19.1591 | 1521 | 1327 | 1370 | 1009 | 6100 | 4549 | 5720 | 6486 | 1450 | 1403 | 1353 | 1581 | 4888 | 5173 | 4856 | 5274 | 3247,21 | 1447 | 5048 | 3,497 | 1,806 | 33,074 | 6,98E-240 | 1,13E-238 | 0,0025238 | 0,0025969 | 0,002386 | 0,002386 | TRUE | 0,0423913 | No |
| orf19.1592 | 1555 | 1505 | 1551 | 1165 | 2532 | 1801 | 2289 | 2214 | 1483 | 1591 | 1532 | 1825 | 2029 | 2048 | 1943 | 1800 | 1781,43 | 1608 | 1955 | 1,219 | 0,285 | 4,053 | 5,05E-05 | 7,25E-05 | 0,0053492 | 0,003013 | 0,0041727 | 0,0041727 | TRUE | 0,0944538 | No |
| orf19.1595 | 659 | 778 | 778 | 447 | 981 | 776 | 948 | 904 | 628 | 823 | 769 | 700 | 786 | 882 | 805 | 735 | 766,01 | 730 | 802 | 1,097 | 0,134 | 1,494 | 0,1352525 | 0,1558408 | 0,0078224 | 0,0042347 | 0,0063436 | 0,0063436 | TRUE | 0,1104661 | No |
| orf19.1596 | 447 | 419 | 491 | 318 | 782 | 629 | 717 | 824 | 426 | 443 | 485 | 498 | 627 | 715 | 609 | 670 | 559,12 | 463 | 655 | 1,417 | 0,503 | 6,322 | 2,58E-10 | 4,55E-10 | 0,0033357 | 0,0050279 | 0,0041711 | 0,0041711 | TRUE | 0,0420614 | No |
| orf19.1597 | 1744 | 1683 | 1667 | 1141 | 2346 | 1783 | 2162 | 2394 | 1663 | 1780 | 1647 | 1787 | 1880 | 2027 | 1836 | 1947 | 1820,73 | 1719 | 1922 | 1,119 | 0,162 | 3,149 | 0,0016368 | 0,0021809 | 0,0011548 | 0,0029931 | 0,0019801 | 0,0019801 | TRUE | 0,0155668 | No |
| orf19.1598 | 649 | 489 | 570 | 408 | 728 | 521 | 674 | 711 | 619 | 517 | 563 | 639 | 583 | 592 | 572 | 578 | 583,02 | 584 | 581 | 0,997 | -0,004 | -0,05 | 0,9604955 | 0,9644252 | 0,0022019 | 0,0049075 | 0,0036638 | 0,0036638 | TRUE | 0,0699539 | No |
| orf19.1599 | 13 | 15 | 8 | 5 | 12 | 6 | 9 | 7 | 12 | 16 | 8 | 8 | 10 | 7 | 8 | 6 | 9,22 | 11 | 8 | 0,671 | -0,576 | -1,225 | 0,2207593 | 0,2492771 | 1,00E-08 | 0,1801692 | 0,1030952 | 0,1030952 | TRUE | 0,3090144 | No |
| orf19.1600 | 1483 | 1348 | 1386 | 983 | 1527 | 1139 | 1486 | 1558 | 1414 | 1425 | 1369 | 1540 | 1224 | 1295 | 1262 | 1267 | 1349,43 | 1437 | 1262 | 0,88 | -0,185 | -3,611 | 0,0003051 | 0,0004227 | 0,0005595 | 0,003308 | 0,0017722 | 0,0017722 | TRUE | 0,0248837 | No |
| orf19.1601 | 27325 | 20869 | 22874 | 15894 | 38788 | 26145 | 36232 | 40372 | 26051 | 22066 | 22597 | 24899 | 31080 | 29729 | 30761 | 32826 | 27501,24 | 23903 | 31099 | 1,301 | 0,38 | 6,518 | 7,11E-11 | 1,28E-10 | 0,0039248 | 0,002151 | 0,0032234 | 0,0032234 | TRUE | 0,0450487 | No |
| orf19.1604 | 2610 | 2321 | 2521 | 1608 | 2572 | 1788 | 2401 | 2643 | 2488 | 2454 | 2491 | 2519 | 2061 | 2033 | 2038 | 2149 | 2279,18 | 2488 | 2070 | 0,833 | -0,264 | -6,726 | 1,74E-11 | 3,19E-11 | 1,00E-08 | 0,0028117 | 0,0010363 | 0,0010363 | TRUE | 0,0080838 | No |
| orf19.1605 | 90 | 81 | 90 | 58 | 58 | 43 | 69 | 59 | 86 | 86 | 89 | 91 | 46 | 49 | 59 | 48 | 69,14 | 88 | 50 | 0,576 | -0,795 | -4,675 | 2,94E-06 | 4,47E-06 | 1,00E-08 | 0,0258377 | 0,0126457 | 0,0126457 | TRUE | 0,1048002 | No |
| orf19.1606 | 1299 | 1369 | 1365 | 817 | 1235 | 881 | 1248 | 1240 | 1238 | 1448 | 1348 | 1280 | 990 | 1002 | 1060 | 1008 | 1171,69 | 1328 | 1015 | 0,763 | -0,389 | -6,418 | 1,38E-10 | 2,47E-10 | 0,0020524 | 0,0034926 | 0,0026715 | 0,0026715 | TRUE | 0,0441685 | No |
| orf19.1607 | 11431 | 11329 | 12154 | 6959 | 8068 | 6108 | 7568 | 8272 | 10898 | 11979 | 12007 | 10902 | 6465 | 6945 | 6425 | 6726 | 9043,37 | 11446 | 6640 | 0,58 | -0,786 | -16,197 | 5,30E-59 | 2,20E-58 | 0,0020326 | 0,0022728 | 0,0021468 | 0,0021468 | TRUE | 0,013301 | No |

| | | | | | | | | | | | | | | | | | | | | | | | | | | | | | | | |
|---|---|---|---|---|---|---|---|---|---|---|---|---|---|---|---|---|---|---|---|---|---|---|---|---|---|---|---|---|---|---|---|
| orf19.1608 | 750 | 712 | 749 | 434 | 1099 | 830 | 1088 | 1103 | 715 | 753 | 740 | 680 | 881 | 944 | 924 | 897 | 816,58 | 722 | 912 | 1,258 | 0,331 | 5,69 | 1,27E-08 | 2,11E-08 | 1,00E-08 | 0,004102 | 0,0019804 | 0,0019804 | TRUE | 0,0149948 | No |
| orf19.1609 | 10109 | 9374 | 10243 | 6100 | 13835 | 9632 | 13333 | 13148 | 9638 | 9912 | 10119 | 9556 | 11086 | 10952 | 11320 | 10691 | 10409,13 | 9806 | 11012 | 1,123 | 0,167 | 4,876 | 1,08E-06 | 1,67E-06 | 0,0005026 | 0,002249 | 0,0010274 | 0,0010274 | TRUE | 0,0056621 | No |
| orf19.1610 | 116 | 97 | 109 | 55 | 151 | 118 | 146 | 153 | 111 | 103 | 108 | 86 | 121 | 134 | 124 | 124 | 113,82 | 102 | 126 | 1,222 | 0,289 | 2,15 | 0,031525 | 0,0385292 | 1,00E-08 | 0,0165173 | 0,0083698 | 0,0083698 | TRUE | 0,0758502 | No |
| orf19.1611 | 328 | 358 | 337 | 215 | 171 | 112 | 155 | 114 | 313 | 379 | 333 | 337 | 137 | 127 | 132 | 93 | 231,21 | 340 | 122 | 0,359 | -1,48 | -11,723 | 9,69E-32 | 2,70E-31 | 0,0110817 | 0,0091928 | 0,0099943 | 0,0099943 | TRUE | 0,2923186 | No |
| orf19.1613 | 5363 | 4999 | 5488 | 3636 | 15711 | 12109 | 14805 | 15397 | 5113 | 5286 | 5422 | 5696 | 12589 | 13769 | 12569 | 12519 | 9120,36 | 5379 | 12862 | 2,392 | 1,258 | 26,273 | 3,86E-152 | 3,93E-151 | 0,0019797 | 0,0022713 | 0,0020635 | 0,0020635 | TRUE | 0,0273669 | No |
| orf19.1614 | 13144 | 12209 | 13313 | 7459 | 10392 | 6793 | 9640 | 9583 | 12531 | 12909 | 13152 | 11685 | 8327 | 7724 | 8184 | 7792 | 10288,13 | 12569 | 8007 | 0,637 | -0,651 | -13,868 | 9,91E-44 | 3,33E-43 | 0,0018463 | 0,0022509 | 0,0020155 | 0,0020155 | TRUE | 0,026962 | No |
| orf19.1616 | 2455 | 2233 | 2377 | 1584 | 2096 | 1585 | 1957 | 2132 | 2341 | 2361 | 2348 | 2481 | 1679 | 1802 | 1661 | 1734 | 2051,01 | 2383 | 1719 | 0,722 | -0,471 | -10,543 | 5,47E-26 | 1,38E-25 | 0,0003968 | 0,0028918 | 0,0014175 | 0,0014175 | TRUE | 0,0119064 | No |
| orf19.1617 | 497 | 527 | 521 | 296 | 903 | 624 | 820 | 889 | 474 | 557 | 515 | 464 | 724 | 710 | 696 | 723 | 607,7 | 502 | 713 | 1,416 | 0,501 | 7,108 | 1,18E-12 | 2,23E-12 | 0,0009492 | 0,0047931 | 0,0030445 | 0,0030445 | TRUE | 0,061887 | No |
| orf19.1618 | 8029 | 8034 | 8233 | 4951 | 6737 | 5074 | 6105 | 6446 | 7655 | 8495 | 8133 | 7756 | 5398 | 5770 | 5183 | 5241 | 6703,91 | 8010 | 5398 | 0,674 | -0,57 | -11,5 | 1,32E-30 | 3,61E-30 | 0,0021337 | 0,0023362 | 0,0022077 | 0,0022077 | TRUE | 0,0257149 | No |
| orf19.1618.1 | 528 | 543 | 515 | 343 | 677 | 403 | 603 | 591 | 503 | 574 | 509 | 537 | 542 | 458 | 512 | 481 | 514,6 | 531 | 498 | 0,942 | -0,087 | -1,103 | 0,2698152 | 0,3007846 | 0,0027409 | 0,0052819 | 0,0039849 | 0,0039849 | TRUE | 0,0420166 | No |
| orf19.1619 | 4620 | 4301 | 4516 | 2801 | 3371 | 2328 | 3228 | 3532 | 4405 | 4548 | 4461 | 4388 | 2701 | 2647 | 2741 | 2872 | 3595,3 | 4450 | 2740 | 0,616 | -0,699 | -18,04 | 9,48E-73 | 4,72E-72 | 0,000349 | 0,002548 | 0,0011526 | 0,0011526 | TRUE | 0,0128178 | No |
| orf19.1620 | 7 | 13 | 13 | 6 | 7 | 4 | 6 | 3 | 7 | 14 | 13 | 9 | 6 | 5 | 5 | 2 | 7,54 | 11 | 4 | 0,413 | -1,276 | -2,361 | 0,0182093 | 0,0226807 | 1,00E-08 | 0,2197342 | 0,1277532 | 0,1277532 | TRUE | 0,2732377 | No |
| orf19.1621 | 745 | 689 | 725 | 483 | 1220 | 935 | 1101 | 1190 | 710 | 729 | 716 | 757 | 978 | 1063 | 935 | 968 | 856,84 | 728 | 986 | 1,354 | 0,438 | 7,24 | 4,49E-13 | 8,61E-13 | 0,0008303 | 0,0040075 | 0,002286 | 0,002286 | TRUE | 0,0309523 | No |
| orf19.1622 | 2920 | 2744 | 3050 | 1790 | 1745 | 1270 | 1621 | 1655 | 2784 | 2901 | 3013 | 2804 | 1398 | 1444 | 1376 | 1346 | 2133,35 | 2876 | 1391 | 0,483 | -1,05 | -22,849 | 1,48E-115 | 1,13E-114 | 0,0006736 | 0,0028609 | 0,0015104 | 0,0015104 | TRUE | 0,0128473 | No |
| orf19.1623 | 10715 | 9628 | 10333 | 6261 | 6743 | 4669 | 6456 | 6816 | 10216 | 10180 | 10208 | 9808 | 5403 | 5309 | 5481 | 5542 | 7768,43 | 10103 | 5434 | 0,538 | -0,895 | -28,528 | 5,32E-179 | 6,32E-178 | 0,0001737 | 0,0023026 | 0,000807 | 0,000807 | TRUE | 0,0045022 | No |
| orf19.1624 | 1069 | 1218 | 1133 | 703 | 2158 | 1635 | 1914 | 1951 | 1019 | 1288 | 1119 | 1101 | 1729 | 1859 | 1625 | 1586 | 1415,91 | 1132 | 1700 | 1,501 | 0,586 | 7,535 | 4,87E-14 | 9,55E-14 | 0,0065586 | 0,0032509 | 0,0050374 | 0,0050374 | TRUE | 0,1049054 | No |
| orf19.1624.1 | 28 | 37 | 25 | 25 | 62 | 39 | 66 | 63 | 27 | 39 | 25 | 39 | 50 | 44 | 56 | 51 | 41,37 | 32 | 50 | 1,584 | 0,663 | 2,841 | 0,004504 | 0,0058368 | 1,00E-08 | 0,0417791 | 0,0260869 | 0,0260869 | TRUE | 0,1783745 | No |
| orf19.1625 | 2435 | 2137 | 2200 | 1443 | 4408 | 2866 | 4256 | 4405 | 2322 | 2260 | 2173 | 2261 | 3532 | 3259 | 3613 | 3582 | 2875,12 | 2254 | 3496 | 1,553 | 0,635 | 13,576 | 5,53E-42 | 1,81E-41 | 0,0011706 | 0,0026624 | 0,0017212 | 0,0017212 | TRUE | 0,0245408 | No |
| orf19.1626 | 2469 | 2107 | 2412 | 1416 | 6274 | 4213 | 6015 | 6417 | 2354 | 2228 | 2383 | 2218 | 5027 | 4791 | 5107 | 5218 | 3665,62 | 2296 | 5036 | 2,192 | 1,132 | 25,483 | 3,02E-143 | 2,92E-142 | 0,0009058 | 0,0025392 | 0,0015619 | 0,0015619 | TRUE | 0,0127132 | No |
| orf19.1628 | 1150 | 1005 | 1053 | 631 | 2471 | 1726 | 2428 | 2547 | 1096 | 1063 | 1040 | 988 | 1980 | 1963 | 2061 | 2071 | 1532,84 | 1047 | 2019 | 1,924 | 0,944 | 19,121 | 1,68E-81 | 9,30E-81 | 0,0003449 | 0,0031624 | 0,0015865 | 0,0015865 | TRUE | 0,0143811 | No |
| orf19.163 | 2298 | 2164 | 2302 | 1443 | 2435 | 1816 | 2348 | 2266 | 2191 | 2288 | 2274 | 2261 | 1951 | 2065 | 1993 | 1842 | 2108,21 | 2254 | 1963 | 0,87 | -0,2 | -4,266 | 1,99E-05 | 2,91E-05 | 0,0008619 | 0,0028701 | 0,0016388 | 0,0016388 | TRUE | 0,0207744 | No |
| orf19.1630 | 3281 | 2978 | 3368 | 2246 | 2102 | 1433 | 2075 | 2111 | 3128 | 3149 | 3327 | 3518 | 1684 | 1629 | 1762 | 1716 | 2489,31 | 3280 | 1698 | 0,519 | -0,947 | -18,583 | 4,39E-77 | 2,30E-76 | 0,0014797 | 0,0027509 | 0,0020612 | 0,0020612 | TRUE | 0,0274065 | No |
| orf19.1631 | 2288 | 2192 | 2248 | 1314 | 16512 | 12058 | 15406 | 15722 | 2181 | 2318 | 2221 | 2058 | 13231 | 13711 | 13080 | 12783 | 7697,89 | 2194 | 13201 | 6,005 | 2,586 | 59,001 | 0 | 0 | 0,0012724 | 0,0023046 | 0,0015553 | 0,0015553 | TRUE | 0,0189816 | No |
| orf19.1632 | 404 | 419 | 435 | 241 | 2036 | 1485 | 1871 | 1883 | 385 | 443 | 430 | 378 | 1631 | 1689 | 1588 | 1531 | 1009,37 | 409 | 1610 | 3,921 | 1,971 | 29,409 | 4,19E-190 | 5,43E-189 | 0,001543 | 0,0037179 | 0,0026776 | 0,0026776 | TRUE | 0,0348274 | No |
| orf19.1633 | 10624 | 10652 | 11253 | 6640 | 14279 | 10479 | 13357 | 13411 | 10129 | 11263 | 11117 | 10402 | 11441 | 11916 | 11340 | 10904 | 11064,02 | 10728 | 11400 | 1,062 | 0,087 | 1,87 | 0,0614977 | 0,0732668 | 0,0018497 | 0,0022397 | 0,0020034 | 0,0020034 | TRUE | 0,0175912 | No |
| orf19.1634 | 3381 | 3035 | 3201 | 1969 | 4063 | 2700 | 3822 | 4023 | 3223 | 3209 | 3162 | 3085 | 3256 | 3070 | 3245 | 3271 | 3190,13 | 3170 | 3210 | 1,013 | 0,018 | 0,478 | 0,6326875 | 0,6652132 | 0,0002067 | 0,002606 | 0,0010943 | 0,0010943 | TRUE | 0,009956 | No |
| orf19.1635 | 17038 | 14160 | 14512 | 10374 | 51754 | 37111 | 49271 | 54678 | 16244 | 14972 | 14336 | 16251 | 41469 | 42199 | 41831 | 44458 | 28970,13 | 15451 | 42489 | 2,75 | 1,46 | 29,679 | 1,45E-193 | 1,94E-192 | 0,0023563 | 0,002148 | 0,0022768 | 0,0022768 | TRUE | 0,0289321 | No |
| orf19.1636 | 1930 | 1569 | 1694 | 1150 | 2196 | 1549 | 2197 | 2248 | 1840 | 1659 | 1674 | 1802 | 1760 | 1761 | 1865 | 1828 | 1773,52 | 1744 | 1804 | 1,035 | 0,05 | 0,986 | 0,3242437 | 0,3577107 | 0,0011816 | 0,0030171 | 0,0019273 | 0,0019273 | TRUE | 0,0184407 | No |
| orf19.1637 | 627 | 475 | 532 | 317 | 1302 | 793 | 1289 | 1246 | 598 | 502 | 526 | 497 | 1043 | 902 | 1094 | 1013 | 771,83 | 531 | 1013 | 1,907 | 0,931 | 11,411 | 3,69E-30 | 9,99E-30 | 0,0051672 | 0,0042186 | 0,0048972 | 0,0048972 | TRUE | 0,0878126 | No |
| orf19.1638 | 729 | 664 | 761 | 493 | 1338 | 1065 | 1147 | 1403 | 695 | 702 | 752 | 772 | 1072 | 1211 | 974 | 1141 | 914,86 | 730 | 1100 | 1,506 | 0,591 | 7,789 | 6,76E-15 | 1,36E-14 | 0,0047296 | 0,003886 | 0,0043488 | 0,0043488 | TRUE | 0,0707877 | No |
| orf19.1639 | 104 | 66 | 82 | 46 | 339 | 282 | 340 | 344 | 99 | 70 | 81 | 72 | 272 | 321 | 289 | 280 | 185,33 | 80 | 290 | 3,56 | 1,832 | 13,911 | 5,43E-44 | 1,83E-43 | 0,0037201 | 0,0109505 | 0,0082792 | 0,0082792 | TRUE | 0,2424799 | No |
| orf19.164 | 1170 | 1198 | 1195 | 734 | 1641 | 1173 | 1490 | 1532 | 1115 | 1267 | 1181 | 1150 | 1315 | 1334 | 1265 | 1246 | 1233,99 | 1178 | 1290 | 1,094 | 0,129 | 2,325 | 0,0200565 | 0,0248892 | 0,001147 | 0,0034218 | 0,0021413 | 0,0021413 | TRUE | 0,031374 | No |
| orf19.1641 | 38 | 37 | 50 | 32 | 460 | 300 | 428 | 427 | 36 | 39 | 49 | 50 | 369 | 341 | 363 | 347 | 199,39 | 44 | 355 | 8,233 | 3,041 | 21,2 | 9,56E-100 | 6,25E-99 | 1,00E-08 | 0,0103258 | 0,0057338 | 0,0057338 | TRUE | 0,113359 | No |
| orf19.1642 | 7872 | 6899 | 7351 | 4468 | 8399 | 5500 | 8435 | 8507 | 7505 | 7295 | 7262 | 6999 | 6730 | 6254 | 7161 | 6917 | 7015,44 | 7265 | 6766 | 0,931 | -0,103 | -2,156 | 0,0310567 | 0,0380151 | 0,0018286 | 0,0023253 | 0,0020382 | 0,0020382 | TRUE | 0,0310778 | No |
| orf19.1643 | 2767 | 2427 | 2667 | 1841 | 1755 | 1293 | 1793 | 1806 | 2638 | 2566 | 2635 | 2884 | 1406 | 1470 | 1522 | 1468 | 2073,78 | 2681 | 1466 | 0,548 | -0,868 | -17,165 | 4,89E-66 | 2,22E-65 | 0,0011741 | 0,002883 | 0,0019405 | 0,0019405 | TRUE | 0,0294993 | No |
| orf19.1646 | 12545 | 10706 | 12504 | 7437 | 14315 | 9545 | 13605 | 14599 | 11960 | 11320 | 12353 | 11650 | 11470 | 10854 | 11551 | 11870 | 11628,56 | 11821 | 11436 | 0,968 | -0,048 | -1,135 | 0,2562578 | 0,2865722 | 0,0012705 | 0,0022325 | 0,0016052 | 0,0016052 | TRUE | 0,0142038 | No |
| orf19.1647 | 31 | 33 | 27 | 21 | 63 | 45 | 83 | 75 | 30 | 35 | 27 | 33 | 50 | 51 | 70 | 61 | 44,64 | 31 | 58 | 1,896 | 0,923 | 4,069 | 4,72E-05 | 6,79E-05 | 1,00E-08 | 0,0388725 | 0,0237613 | 0,0237613 | TRUE | 0,177919 | No |
| orf19.1648 | 443 | 420 | 453 | 268 | 846 | 621 | 759 | 841 | 422 | 444 | 448 | 420 | 678 | 706 | 644 | 684 | 555,75 | 434 | 678 | 1,56 | 0,642 | 9,582 | 9,51E-22 | 2,20E-21 | 1,00E-08 | 0,0050457 | 0,0023659 | 0,0023659 | TRUE | 0,0131207 | No |
| orf19.1649 | 3494 | 3137 | 3161 | 2089 | 5484 | 3955 | 5189 | 5488 | 3331 | 3317 | 3123 | 3273 | 4394 | 4497 | 4405 | 4462 | 3850,31 | 3261 | 4440 | 1,361 | 0,445 | 12,334 | 5,96E-35 | 1,74E-34 | 0,0001329 | 0,0025177 | 0,0009778 | 0,0009778 | TRUE | 0,0103861 | No |
| orf19.1650 | 995 | 738 | 710 | 482 | 2243 | 1366 | 1999 | 2375 | 949 | 780 | 701 | 755 | 1797 | 1553 | 1697 | 1931 | 1270,53 | 796 | 1744 | 2,189 | 1,131 | 11,944 | 6,96E-33 | 1,97E-32 | 0,0119422 | 0,0033836 | 0,0076407 | 0,0076407 | TRUE | 0,2024181 | No |
| orf19.1651 | 14 | 22 | 16 | 9 | 39 | 26 | 55 | 27 | 13 | 23 | 16 | 14 | 31 | 30 | 47 | 22 | 24,5 | 16 | 32 | 1,939 | 0,955 | 2,835 | 0,0045814 | 0,0059323 | 0,05369 | 0,0691155 | 0,0619208 | 0,0619208 | TRUE | 0,4573656 | No |
| orf19.1652 | 11107 | 9726 | 10252 | 7484 | 80882 | 60408 | 75601 | 85131 | 10589 | 10284 | 10128 | 11724 | 64808 | 68689 | 64185 | 69220 | 38703,46 | 10681 | 66726 | 6,25 | 2,644 | 50,6 | 0 | 0 | 0,0029627 | 0,0021337 | 0,0025631 | 0,0025631 | TRUE | 0,0520094 | No |
| orf19.1653 | 2277 | 2200 | 2223 | 1227 | 977 | 608 | 913 | 860 | 2171 | 2326 | 2196 | 1922 | 783 | 691 | 775 | 699 | 1445,49 | 2154 | 737 | 0,342 | -1,547 | -21,971 | 5,41E-107 | 3,77E-106 | 0,0040232 | 0,0032272 | 0,0038955 | 0,0038955 | TRUE | 0,0601078 | No |
| orf19.1654 | 1520 | 1294 | 1379 | 1024 | 4003 | 2957 | 3727 | 3940 | 1449 | 1368 | 1362 | 1604 | 3207 | 3362 | 3164 | 3204 | 2340,19 | 1446 | 3234 | 2,243 | 1,165 | 21,009 | 5,37E-98 | 3,45E-97 | 0,0021109 | 0,0027929 | 0,0024259 | 0,0024259 | TRUE | 0,0593353 | No |
| orf19.1655 | 2404 | 2002 | 2213 | 1558 | 27977 | 20060 | 26261 | 29787 | 2292 | 2117 | 2186 | 2441 | 22417 | 22810 | 22295 | 24220 | 12597,26 | 2259 | 22936 | 10,17 | 3,346 | 66,549 | 0 | 0 | 0,0023647 | 0,0022216 | 0,0021591 | 0,0021591 | TRUE | 0,0329333 | No |
| orf19.1655.3 | 3569 | 3178 | 3256 | 2046 | 2662 | 1758 | 2336 | 2450 | 3403 | 3360 | 3217 | 3205 | 2133 | 1999 | 1983 | 1992 | 2661,51 | 3296 | 2027 | 0,615 | -0,702 | -16,063 | 4,62E-58 | 1,90E-57 | 0,0007053 | 0,0027082 | 0,0014427 | 0,0014427 | TRUE | 0,0154986 | No |
| orf19.1656 | 1069 | 1042 | 1055 | 714 | 2478 | 1827 | 2438 | 2604 | 1019 | 1102 | 1042 | 1119 | 1986 | 2077 | 2070 | 2117 | 1566,49 | 1070 | 2062 | 1,931 | 0,949 | 19,097 | 2,66E-81 | 1,47E-80 | 0,0005041 | 0,0031394 | 0,0016328 | 0,0016328 | TRUE | 0,0125099 | No |
| orf19.1658 | 11833 | 10175 | 11442 | 7319 | 10264 | 6727 | 9567 | 10234 | 11281 | 10759 | 11304 | 11466 | 8224 | 7649 | 8122 | 8321 | 9640,81 | 11202 | 8079 | 0,722 | -0,471 | -12,051 | 1,91E-33 | 5,45E-33 | 0,0009134 | 0,0022616 | 0,0013599 | 0,0013599 | TRUE | 0,0153713 | No |
| orf19.1659 | 661 | 751 | 795 | 477 | 1303 | 1002 | 1199 | 1169 | 630 | 794 | 785 | 747 | 1044 | 1139 | 1018 | 951 | 888,6 | 739 | 1038 | 1,404 | 0,489 | 5,812 | 6,18E-09 | 1,03E-08 | 0,0071073 | 0,003939 | 0,00561 | 0,00561 | TRUE | 0,1228649 | No |
| orf19.166 | 2737 | 3004 | 3160 | 1966 | 2249 | 1900 | 2082 | 2116 | 2609 | 3176 | 3122 | 3080 | 1802 | 2160 | 1768 | 1721 | 2429,76 | 2997 | 1863 | 0,621 | -0,687 | -8,302 | 1,02E-16 | 2,13E-16 | 0,009062 | 0,002767 | 0,0061497 | 0,0061497 | TRUE | 0,1374973 | No |
| orf19.1661 | 5015 | 4533 | 4883 | 2904 | 8860 | 6491 | 8801 | 9164 | 4781 | 4793 | 4824 | 4549 | 7099 | 7381 | 7472 | 7451 | 6043,86 | 4737 | 7351 | 1,55 | 0,633 | 17,788 | 8,68E-71 | 4,19E-70 | 0,0003987 | 0,0023629 | 0,0010348 | 0,0010348 | TRUE | 0,0080479 | No |
| orf19.1662 | 3147 | 2860 | 3085 | 1904 | 2742 | 1958 | 2536 | 2628 | 3000 | 3024 | 3048 | 2983 | 2197 | 2226 | 2153 | 2137 | 2596,02 | 3014 | 2178 | 0,722 | -0,47 | -12,9 | 4,51E-38 | 1,39E-37 | 1,00E-08 | 0,0027237 | 0,0008814 | 0,0008814 | TRUE | 0,0024544 | No |
| orf19.1663 | 2194 | 2211 | 2411 | 1324 | 4304 | 3162 | 4139 | 4026 | 2092 | 2338 | 2382 | 2074 | 3449 | 3595 | 3514 | 3274 | 2839,65 | 2222 | 3458 | 1,555 | 0,636 | 11,054 | 2,10E-28 | 5,51E-28 | 0,0029177 | 0,0026695 | 0,0028009 | 0,0028009 | TRUE | 0,0287191 | No |
| orf19.1664 | 363 | 326 | 298 | 238 | 489 | 349 | 416 | 506 | 346 | 345 | 294 | 373 | 392 | 397 | 353 | 411 | 363,91 | 340 | 388 | 1,149 | 0,2 | 2,182 | 0,029079 | 0,0356765 | 0,0037685 | 0,0066031 | 0,0052509 | 0,0052509 | TRUE | 0,0932633 | No |
| orf19.1665 | 9438 | 8913 | 9468 | 6098 | 19958 | 15113 | 19009 | 19917 | 8998 | 9424 | 9353 | 9553 | 15992 | 17185 | 16139 | 16194 | 12854,8 | 9332 | 16378 | 1,755 | 0,812 | 22,013 | 2,18E-107 | 1,53E-106 | 0,0007655 | 0,002219 | 0,0012178 | 0,0012178 | TRUE | 0,0133344 | No |
| orf19.1666 | 1296 | 1056 | 1256 | 859 | 1296 | 911 | 1283 | 1244 | 1236 | 1117 | 1241 | 1346 | 1038 | 1036 | 1089 | 1011 | 1139,22 | 1235 | 1044 | 0,847 | -0,239 | -3,83 | 0,0001282 | 0,0001805 | 0,0021869 | 0,0035325 | 0,0028542 | 0,0028542 | TRUE | 0,0488855 | No |
| orf19.1667 | 1016 | 989 | 1066 | 598 | 1069 | 791 | 949 | 1032 | 969 | 1046 | 1053 | 937 | 857 | 899 | 806 | 839 | 925,64 | 1001 | 850 | 0,846 | -0,24 | -3,902 | 9,56E-05 | 0,0001355 | 0,0012473 | 0,0038651 | 0,002562 | 0,002562 | TRUE | 0,0194648 | No |
| orf19.1667.1 | 599 | 476 | 474 | 244 | 874 | 455 | 794 | 780 | 571 | 503 | 468 | 382 | 700 | 517 | 674 | 634 | 556,36 | 481 | 631 | 1,309 | 0,388 | 3,221 | 0,0012782 | 0,0017096 | 0,0190774 | 0,0050424 | 0,0120894 | 0,0120894 | TRUE | 0,2033968 | No |
| orf19.1668 | 849 | 814 | 861 | 534 | 745 | 535 | 674 | 736 | 809 | 861 | 851 | 837 | 597 | 608 | 572 | 598 | 716,65 | 840 | 594 | 0,707 | -0,5 | -8,561 | 1,12E-17 | 2,39E-17 | 1,00E-08 | 0,0043824 | 0,0018543 | 0,0018543 | TRUE | 0,00758 | No |
| orf19.1669 | 6266 | 5727 | 6336 | 3908 | 5097 | 3609 | 5007 | 5117 | 5974 | 6056 | 6259 | 6122 | 4084 | 4104 | 4251 | 4161 | 5126,29 | 6103 | 4150 | 0,68 | -0,556 | -16,736 | 7,15E-63 | 3,13E-62 | 0,0001949 | 0,0024116 | 0,0008602 | 0,0008602 | TRUE | 0,00381 | No |
| orf19.1670 | 1433 | 1482 | 1608 | 860 | 889 | 642 | 803 | 816 | 1366 | 1567 | 1589 | 1347 | 712 | 730 | 682 | 663 | 1082,07 | 1467 | 697 | 0,474 | -1,078 | -15,169 | 5,63E-52 | 2,14E-51 | 0,0039902 | 0,0036086 | 0,0038203 | 0,0038203 | TRUE | 0,0373049 | No |
| orf19.1671 | 8775 | 8378 | 8615 | 5593 | 6307 | 4570 | 5805 | 6329 | 8366 | 8859 | 8511 | 8762 | 5054 | 5197 | 4928 | 5146 | 6852,73 | 8624 | 5081 | 0,589 | -0,763 | -21,699 | 2,08E-104 | 1,41E-103 | 0,0004039 | 0,0023309 | 0,0010345 | 0,0010345 | TRUE | 0,0050254 | No |
| orf19.1671.1 | 10 | 7 | 4 | 2 | 2 | 2 | 8 | 0 | 10 | 7 | 4 | 3 | 2 | 2 | 7 | 0 | 4,34 | 6 | 3 | 0,435 | -1,201 | -1,448 | 0,14773 | 0,1696662 | 0,4370148 | 0,3807487 | 0,4015184 | 0,4015184 | TRUE | 1,0003451 | No |
| orf19.1672 | 3731 | 3582 | 3875 | 2278 | 6521 | 4557 | 5668 | 6109 | 3557 | 3788 | 3828 | 3569 | 5225 | 5182 | 4812 | 4967 | 4365,93 | 3686 | 5046 | 1,368 | 0,453 | 10,114 | 4,81E-24 | 1,17E-23 | 0,0011634 | 0,0024674 | 0,0016812 | 0,0016812 | TRUE | 0,0119583 | No |
| orf19.1673 | 2705 | 2787 | 2733 | 1535 | 1623 | 1087 | 1539 | 1393 | 2579 | 2947 | 2700 | 2405 | 1300 | 1236 | 1307 | 1133 | 1950,77 | 2658 | 1244 | 0,468 | -1,096 | -15,462 | 6,21E-54 | 2,42E-53 | 0,0049944 | 0,0029329 | 0,0042578 | 0,0042578 | TRUE | 0,0646699 | No |
| orf19.1674 | 2799 | 2830 | 2917 | 1787 | 2257 | 1745 | 2073 | 2145 | 2669 | 2992 | 2882 | 2799 | 1808 | 1984 | 1760 | 1744 | 2329,85 | 2836 | 1824 | 0,643 | -0,638 | -11,374 | 5,66E-30 | 1,53E-29 | 0,0023816 | 0,002796 | 0,0025786 | 0,0025786 | TRUE | 0,0404441 | No |
| orf19.1675 | 1765 | 1510 | 1707 | 989 | 2917 | 1904 | 2772 | 3035 | 1683 | 1597 | 1686 | 1549 | 2337 | 2165 | 2353 | 2468 | 1979,82 | 1629 | 2331 | 1,43 | 0,516 | 9,608 | 7,39E-22 | 1,72E-21 | 0,0015928 | 0,0029206 | 0,0022351 | 0,0022351 | TRUE | 0,0266985 | No |
| orf19.1676 | 1555 | 1546 | 1587 | 953 | 1628 | 1261 | 1457 | 1499 | 1483 | 1635 | 1568 | 1493 | 1304 | 1434 | 1237 | 1219 | 1421,51 | 1545 | 1298 | 0,839 | -0,253 | -4,02 | 5,83E-05 | 8,34E-05 | 0,0029318 | 0,0032463 | 0,0031037 | 0,0031037 | TRUE | 0,0564184 | No |
| orf19.1677 | 2522 | 2048 | 2404 | 1529 | 3917 | 2799 | 3941 | 4128 | 2404 | 2166 | 2375 | 2395 | 3139 | 3183 | 3346 | 3356 | 2795,47 | 2335 | 3256 | 1,395 | 0,48 | 9,887 | 4,76E-23 | 1,13E-22 | 0,001282 | 0,0026786 | 0,001889 | 0,001889 | TRUE | 0,0293422 | No |
| orf19.1678 | 1147 | 972 | 1070 | 713 | 1867 | 1402 | 1929 | 1968 | 1094 | 1028 | 1057 | 1117 | 1496 | 1594 | 1638 | 1600 | 1327,92 | 1074 | 1582 | 1,475 | 0,561 | 10,728 | 7,51E-27 | 1,93E-26 | 0,0007401 | 0,0033277 | 0,0018174 | 0,0018174 | TRUE | 0,0167592 | No |
| orf19.1679 | 64 | 39 | 51 | 37 | 172 | 99 | 132 | 165 | 61 | 41 | 50 | 58 | 138 | 113 | 112 | 134 | 88,4 | 52 | 124 | 2,374 | 1,247 | 7,201 | 6,00E-13 | 1,14E-12 | 0,0060426 | 0,0206643 | 0,0146405 | 0,0146405 | TRUE | 0,1777541 | No |
| orf19.168 | 6503 | 5851 | 6045 | 3561 | 4485 | 2947 | 4359 | 4327 | 6200 | 6187 | 5972 | 5578 | 3594 | 3351 | 3701 | 3518 | 4762,59 | 5984 | 3541 | 0,592 | -0,757 | -15,656 | 3,00E-55 | 1,19E-54 | 0,0017374 | 0,002436 | 0,0020269 | 0,0020269 | TRUE | 0,0245396 | No |
| orf19.1680 | 2017 | 1821 | 1929 | 1198 | 4893 | 3466 | 4470 | 4960 | 1923 | 1925 | 1906 | 1877 | 3921 | 3941 | 3795 | 4033 | 2915,08 | 1908 | 3922 | 2,055 | 1,039 | 26,953 | 5,25E-160 | 5,70E-159 | 1,00E-08 | 0,0026545 | 0,0010193 | 0,0010193 | TRUE | 0,0059215 | No |

| | | | | | | | | | | | | | | | | | | | | | | | | | | | | | | | |
|---|---|---|---|---|---|---|---|---|---|---|---|---|---|---|---|---|---|---|---|---|---|---|---|---|---|---|---|---|---|---|---|
| orf19.1681 | 241 | 264 | 267 | 191 | 312 | 209 | 290 | 347 | 230 | 279 | 264 | 299 | 250 | 238 | 246 | 282 | 260,99 | 268 | 254 | 0,955 | -0,066 | -0,626 | 0,5315195 | 0,5672564 | 0,0048929 | 0,0083824 | 0,0067506 | 0,0067506 | TRUE | 0,1064933 | No |
| orf19.1682 | 5701 | 4614 | 5189 | 3426 | 12145 | 8643 | 11813 | 12933 | 5435 | 4879 | 5126 | 5367 | 9731 | 9828 | 10029 | 10516 | 7613,94 | 5202 | 10026 | 1,928 | 0,947 | 20,697 | 3,65E-95 | 2,28E-94 | 0,001594 | 0,0023069 | 0,0018583 | 0,0018583 | TRUE | 0,0214057 | No |
| orf19.1683 | 1890 | 1694 | 1851 | 1167 | 4467 | 2952 | 4261 | 4518 | 1802 | 1791 | 1829 | 1828 | 3579 | 3357 | 3618 | 3674 | 2684,62 | 1812 | 3557 | 1,964 | 0,974 | 22,589 | 5,58E-113 | 4,15E-112 | 0,0006135 | 0,0027029 | 0,0013504 | 0,0013504 | TRUE | 0,0167094 | No |
| orf19.1684 | 701 | 669 | 754 | 486 | 712 | 557 | 697 | 666 | 668 | 707 | 745 | 761 | 571 | 633 | 592 | 542 | 652,38 | 720 | 584 | 0,811 | -0,302 | -4,23 | 2,33E-05 | 3,40E-05 | 0,0021882 | 0,0046081 | 0,003362 | 0,003362 | TRUE | 0,034699 | No |
| orf19.1685 | 433 | 383 | 435 | 301 | 550 | 367 | 495 | 508 | 413 | 405 | 430 | 472 | 441 | 417 | 420 | 413 | 426,3 | 430 | 423 | 0,989 | -0,016 | -0,208 | 0,8351806 | 0,8551488 | 1,00E-08 | 0,0059428 | 0,0031236 | 0,0031236 | TRUE | 0,0409167 | No |
| orf19.1686 | 680 | 586 | 735 | 387 | 962 | 647 | 908 | 989 | 648 | 620 | 726 | 606 | 771 | 736 | 771 | 804 | 710,23 | 650 | 770 | 1,183 | 0,242 | 3,456 | 0,0005483 | 0,0007517 | 0,0021717 | 0,0044031 | 0,0032744 | 0,0032744 | TRUE | 0,0742484 | No |
| orf19.1687 | 13647 | 12223 | 13389 | 8458 | 17413 | 12476 | 16454 | 17708 | 13011 | 12924 | 13227 | 13250 | 13953 | 14186 | 13969 | 14398 | 13614,84 | 13103 | 14126 | 1,078 | 0,109 | 4,039 | 5,37E-05 | 7,70E-05 | 8,76E-05 | 0,0022119 | 0,0006208 | 0,0006208 | TRUE | 0,0020883 | No |
| orf19.169 | 1686 | 1865 | 1977 | 1049 | 4924 | 3652 | 4299 | 4049 | 1607 | 1972 | 1953 | 1643 | 3945 | 4153 | 3650 | 3292 | 2777 | 1794 | 3760 | 2,094 | 1,066 | 12,253 | 1,61E-34 | 4,68E-34 | 0,010327 | 0,0026825 | 0,0068389 | 0,0068389 | TRUE | 0,0856382 | No |
| orf19.1690 | 3664 | 3064 | 3341 | 2157 | 6690 | 4618 | 6325 | 6962 | 3493 | 3240 | 3301 | 3379 | 5361 | 5251 | 5370 | 5661 | 4381,86 | 3353 | 5411 | 1,614 | 0,691 | 16,709 | 1,13E-62 | 4,95E-62 | 0,0007351 | 0,002466 | 0,0013903 | 0,0013903 | TRUE | 0,0118844 | No |
| orf19.1691 | 406 | 373 | 376 | 268 | 183 | 128 | 144 | 167 | 387 | 394 | 371 | 420 | 147 | 146 | 122 | 136 | 265,37 | 393 | 138 | 0,35 | -1,514 | -15,336 | 4,41E-53 | 1,69E-52 | 1,00E-08 | 0,0082784 | 0,0046908 | 0,0046908 | TRUE | 0,0603123 | No |
| orf19.1693 | 815 | 737 | 821 | 488 | 2225 | 1663 | 1969 | 2215 | 777 | 779 | 811 | 764 | 1783 | 1891 | 1672 | 1801 | 1284,79 | 783 | 1787 | 2,277 | 1,187 | 21,167 | 1,91E-99 | 1,25E-98 | 0,0011479 | 0,0033692 | 0,0020574 | 0,0020574 | TRUE | 0,0225402 | No |
| orf19.1697 | 3724 | 3316 | 3435 | 2043 | 5309 | 3222 | 5063 | 5391 | 3550 | 3506 | 3393 | 3200 | 4254 | 3664 | 4298 | 4383 | 3781,26 | 3412 | 4150 | 1,216 | 0,282 | 4,547 | 5,45E-06 | 8,18E-06 | 0,0038243 | 0,0025255 | 0,0034239 | 0,0034239 | TRUE | 0,0744656 | No |
| orf19.1698 | 764 | 671 | 776 | 508 | 1379 | 993 | 1347 | 1526 | 728 | 709 | 767 | 796 | 1105 | 1129 | 1144 | 1241 | 952,35 | 750 | 1155 | 1,544 | 0,626 | 10,065 | 7,86E-24 | 1,90E-23 | 0,0015892 | 0,0038153 | 0,0025836 | 0,0025836 | TRUE | 0,0311288 | No |
| orf19.1699 | 870 | 882 | 891 | 563 | 1959 | 1379 | 1897 | 1912 | 829 | 933 | 880 | 882 | 1570 | 1568 | 1611 | 1555 | 1228,4 | 881 | 1576 | 1,79 | 0,84 | 16,21 | 4,28E-59 | 1,79E-58 | 8,05E-05 | 0,0034279 | 0,0016591 | 0,0016591 | TRUE | 0,0203205 | No |
| orf19.17 | 738 | 850 | 827 | 469 | 798 | 670 | 704 | 614 | 704 | 899 | 817 | 735 | 639 | 762 | 598 | 499 | 706,53 | 789 | 624 | 0,79 | -0,34 | -2,877 | 0,0040118 | 0,0052211 | 0,0194384 | 0,0044152 | 0,0119868 | 0,0119868 | TRUE | 0,2506637 | No |
| orf19.1700 | 10363 | 9784 | 9502 | 6754 | 33930 | 26423 | 30795 | 33275 | 9880 | 10345 | 9387 | 10581 | 27187 | 30045 | 26145 | 27056 | 18828,24 | 10048 | 27608 | 2,748 | 1,458 | 26,77 | 7,20E-158 | 7,78E-157 | 0,0031637 | 0,0021785 | 0,0027791 | 0,0027791 | TRUE | 0,0431178 | No |
| orf19.1701 | 1628 | 1564 | 1584 | 1105 | 2603 | 2067 | 2404 | 2587 | 1552 | 1654 | 1565 | 1731 | 2086 | 2350 | 2041 | 2103 | 1885,28 | 1626 | 2145 | 1,321 | 0,401 | 6,76 | 1,38E-11 | 2,54E-11 | 0,002713 | 0,0029622 | 0,002827 | 0,002827 | TRUE | 0,0485912 | No |
| orf19.1702 | 1072 | 969 | 942 | 687 | 2274 | 1647 | 2085 | 2254 | 1022 | 1025 | 931 | 1076 | 1822 | 1873 | 1770 | 1833 | 1418,9 | 1014 | 1824 | 1,805 | 0,852 | 15,92 | 4,60E-57 | 1,85E-56 | 0,0007707 | 0,0032484 | 0,0019483 | 0,0019483 | TRUE | 0,0366673 | No |
| orf19.1704 | 6422 | 5352 | 5903 | 3504 | 37594 | 25599 | 36092 | 38823 | 6123 | 5659 | 5832 | 5489 | 30123 | 29108 | 30642 | 31567 | 18067,83 | 5776 | 30360 | 5,253 | 2,393 | 54,659 | 0 | 0 | 0,001529 | 0,0021822 | 0,001729 | 0,001729 | TRUE | 0,0200264 | No |
| orf19.1706 | 324 | 414 | 398 | 245 | 627 | 508 | 489 | 425 | 309 | 438 | 393 | 384 | 502 | 578 | 415 | 346 | 420,55 | 381 | 460 | 1,207 | 0,271 | 1,811 | 0,0701898 | 0,0832183 | 0,0329336 | 0,0059955 | 0,0190313 | 0,0190313 | TRUE | 0,1828945 | No |
| orf19.1707 | 20 | 48 | 34 | 12 | 72 | 79 | 54 | 31 | 19 | 51 | 34 | 19 | 58 | 90 | 46 | 25 | 42,6 | 31 | 55 | 1,763 | 0,818 | 2,087 | 0,0368476 | 0,0448027 | 0,2143193 | 0,0406356 | 0,1210138 | 0,1210138 | TRUE | 0,2454798 | No |
| orf19.1708 | 3571 | 3682 | 3816 | 2282 | 3491 | 2237 | 3319 | 2891 | 3405 | 3893 | 3770 | 3575 | 2797 | 2544 | 2818 | 2351 | 3143,99 | 3661 | 2628 | 0,718 | -0,478 | -7,084 | 1,40E-12 | 2,64E-12 | 0,0048828 | 0,0026135 | 0,0040559 | 0,0040559 | TRUE | 0,0620912 | No |
| orf19.1709 | 1862 | 1714 | 1679 | 921 | 6192 | 3955 | 5687 | 6130 | 1775 | 1812 | 1659 | 1443 | 4961 | 4497 | 4828 | 4984 | 3245,03 | 1672 | 4818 | 2,875 | 1,524 | 22,377 | 6,63E-111 | 4,85E-110 | 0,0054993 | 0,0025973 | 0,0040235 | 0,0040235 | TRUE | 0,0963519 | No |
| orf19.171 | 49630 | 46105 | 49665 | 30828 | 62792 | 43043 | 58075 | 63699 | 47317 | 48750 | 49064 | 48294 | 50313 | 48944 | 49305 | 51793 | 49222,63 | 48356 | 50089 | 1,036 | 0,051 | 1,692 | 0,0905561 | 0,1061423 | 0,0004212 | 0,0021246 | 0,0008471 | 0,0008471 | TRUE | 0,0064332 | No |
| orf19.1710 | 1819 | 1910 | 1883 | 1190 | 4704 | 3648 | 4328 | 4113 | 1734 | 2020 | 1860 | 1864 | 3769 | 4148 | 3674 | 3344 | 2801,78 | 1870 | 3734 | 1,997 | 0,998 | 14,176 | 1,29E-45 | 4,46E-45 | 0,0053398 | 0,0026773 | 0,0043342 | 0,0043342 | TRUE | 0,0669511 | No |
| orf19.1711 | 1458 | 1620 | 1653 | 803 | 1234 | 857 | 1160 | 1006 | 1390 | 1713 | 1633 | 1258 | 989 | 974 | 985 | 818 | 1220 | 1498 | 942 | 0,627 | -0,673 | -6,819 | 9,15E-12 | 1,69E-11 | 0,0126181 | 0,0034371 | 0,0085027 | 0,0085027 | TRUE | 0,1346265 | No |
| orf19.1712 | 929 | 1013 | 967 | 658 | 1910 | 1564 | 1776 | 1784 | 886 | 1071 | 955 | 1031 | 1530 | 1778 | 1508 | 1451 | 1276,27 | 986 | 1567 | 1,59 | 0,669 | 8,402 | 4,38E-17 | 9,23E-17 | 0,0066556 | 0,0033778 | 0,0052286 | 0,0052286 | TRUE | 0,0971969 | No |
| orf19.1714 | 96 | 125 | 114 | 111 | 69 | 68 | 73 | 81 | 92 | 132 | 113 | 174 | 55 | 77 | 62 | 66 | 96,33 | 128 | 65 | 0,514 | -0,961 | -4,792 | 1,65E-06 | 2,53E-06 | 0,0365735 | 0,0191356 | 0,0272829 | 0,0272829 | TRUE | 0,256973 | No |
| orf19.1715 | 740 | 715 | 814 | 515 | 832 | 571 | 763 | 830 | 706 | 756 | 804 | 807 | 667 | 649 | 648 | 675 | 713,88 | 768 | 660 | 0,861 | -0,215 | -3,366 | 0,0007613 | 0,0010361 | 0,0008283 | 0,0043912 | 0,0025271 | 0,0025271 | TRUE | 0,0373673 | No |
| orf19.1716 | 2608 | 2200 | 2285 | 1392 | 2671 | 1769 | 2638 | 2682 | 2486 | 2326 | 2257 | 2181 | 2140 | 2012 | 2240 | 2181 | 2227,84 | 2312 | 2143 | 0,926 | -0,111 | -2,039 | 0,0414377 | 0,0502116 | 0,0020043 | 0,0028283 | 0,002378 | 0,002378 | TRUE | 0,031328 | No |
| orf19.1717 | 166 | 177 | 184 | 134 | 314 | 235 | 273 | 288 | 158 | 187 | 182 | 210 | 252 | 267 | 232 | 234 | 215,23 | 184 | 246 | 1,344 | 0,427 | 3,892 | 9,95E-05 | 0,000141 | 0,0025586 | 0,0097197 | 0,006673 | 0,006673 | TRUE | 0,0997833 | No |
| orf19.1718 | 898 | 982 | 1004 | 584 | 605 | 453 | 532 | 554 | 856 | 1038 | 992 | 915 | 485 | 515 | 452 | 450 | 712,9 | 950 | 476 | 0,499 | -1,002 | -12,722 | 4,48E-37 | 1,36E-36 | 0,0044614 | 0,0043944 | 0,0044302 | 0,0044302 | TRUE | 0,0567936 | No |
| orf19.1719 | 2259 | 2591 | 2404 | 1616 | 2897 | 2349 | 2642 | 2691 | 2154 | 2740 | 2375 | 2532 | 2321 | 2671 | 2243 | 2188 | 2402,9 | 2450 | 2356 | 0,961 | -0,057 | -0,693 | 0,4884268 | 0,5246888 | 0,0080641 | 0,0027746 | 0,0060321 | 0,0060321 | TRUE | 0,0968439 | No |
| orf19.172 | 4746 | 4901 | 4703 | 3027 | 7980 | 5854 | 6860 | 7075 | 4525 | 5182 | 4646 | 4742 | 6394 | 6657 | 5824 | 5753 | 5465,31 | 4774 | 6157 | 1,289 | 0,367 | 5,976 | 2,29E-09 | 3,89E-09 | 0,0041024 | 0,0023917 | 0,0034275 | 0,0034275 | TRUE | 0,0408137 | No |
| orf19.1720 | 4388 | 4671 | 4879 | 2981 | 9653 | 7172 | 8784 | 8632 | 4183 | 4939 | 4820 | 4670 | 7735 | 8155 | 7458 | 7019 | 6122,31 | 4653 | 7592 | 1,631 | 0,706 | 11,262 | 2,01E-29 | 5,38E-29 | 0,0042915 | 0,0023595 | 0,0035956 | 0,0035956 | TRUE | 0,0571416 | No |
| orf19.1721 | 21670 | 22389 | 22725 | 13009 | 47255 | 33212 | 43143 | 45762 | 20660 | 23674 | 22450 | 20379 | 37864 | 37765 | 36628 | 37209 | 29578,65 | 21791 | 37366 | 1,715 | 0,778 | 15,389 | 1,93E-53 | 7,47E-53 | 0,002613 | 0,0021468 | 0,0024166 | 0,0024166 | TRUE | 0,0413886 | No |
| orf19.1722 | 72 | 48 | 54 | 55 | 55 | 48 | 45 | 58 | 69 | 51 | 53 | 86 | 44 | 55 | 38 | 47 | 55,37 | 65 | 46 | 0,716 | -0,481 | -2,21 | 0,0270729 | 0,0332664 | 0,0205323 | 0,0317473 | 0,0270276 | 0,0270276 | TRUE | 0,3332034 | No |
| orf19.1723 | 1457 | 1343 | 1434 | 965 | 1054 | 786 | 1084 | 1158 | 1389 | 1420 | 1417 | 1512 | 845 | 894 | 920 | 942 | 1167,21 | 1434 | 900 | 0,629 | -0,67 | -12,157 | 5,25E-34 | 1,51E-33 | 0,0008336 | 0,003498 | 0,0020255 | 0,0020255 | TRUE | 0,0219342 | No |
| orf19.1724 | 0 | 1 | 1 | 1 | 4 | 2 | 1 | 3 | 0 | 1 | 1 | 2 | 3 | 2 | 1 | 2 | 1,55 | 1 | 2 | 2,603 | 1,38 | 1,139 | 0,2547749 | 0,2851545 | 1,00E-08 | 1,0631483 | 0,5514211 | 0,5514211 | TRUE | 0,2269532 | No |
| orf19.1725 | 1347 | 1075 | 1355 | 965 | 1256 | 993 | 1353 | 1393 | 1284 | 1137 | 1339 | 1512 | 1006 | 1129 | 1149 | 1133 | 1211,01 | 1318 | 1104 | 0,839 | -0,253 | -3,056 | 0,0022395 | 0,0029599 | 0,0075714 | 0,0034471 | 0,0057218 | 0,0057218 | TRUE | 0,1114301 | No |
| orf19.1727 | 14135 | 13832 | 14788 | 9072 | 6936 | 5256 | 6438 | 6916 | 13476 | 14626 | 14609 | 14212 | 5558 | 5977 | 5466 | 5623 | 9943,26 | 14231 | 5656 | 0,397 | -1,332 | -30,921 | 6,26E-210 | 8,87E-209 | 0,00133 | 0,0022564 | 0,001663 | 0,001663 | TRUE | 0,0173767 | No |
| orf19.1728 | 509 | 500 | 543 | 328 | 962 | 718 | 826 | 884 | 485 | 529 | 536 | 514 | 771 | 816 | 701 | 719 | 633,94 | 516 | 752 | 1,455 | 0,541 | 7,496 | 6,60E-14 | 1,29E-13 | 0,0021053 | 0,0046813 | 0,0033147 | 0,0033147 | TRUE | 0,0368824 | No |
| orf19.1729 | 1054 | 1274 | 1251 | 700 | 1144 | 924 | 1092 | 995 | 1005 | 1347 | 1236 | 1097 | 917 | 1051 | 927 | 809 | 1048,49 | 1171 | 926 | 0,789 | -0,341 | -3,436 | 0,0005896 | 0,0008068 | 0,0127162 | 0,0036573 | 0,0085157 | 0,0085157 | TRUE | 0,1232884 | No |
| orf19.173 | 24307 | 22932 | 25099 | 15373 | 7517 | 5153 | 7046 | 7315 | 23174 | 24248 | 24795 | 24083 | 6023 | 5859 | 5982 | 5948 | 15014,03 | 24075 | 5953 | 0,247 | -2,016 | -62,215 | 0 | 0 | 0,0004453 | 0,0022006 | 0,0009095 | 0,0009095 | TRUE | 0,0078679 | No |
| orf19.1730 | 1097 | 920 | 1021 | 584 | 1068 | 773 | 1077 | 1119 | 1046 | 973 | 1009 | 915 | 856 | 879 | 914 | 910 | 937,64 | 986 | 890 | 0,9 | -0,152 | -2,584 | 0,0097522 | 0,0123987 | 0,0008353 | 0,0038424 | 0,0022413 | 0,0022413 | TRUE | 0,0243194 | No |
| orf19.1734 | 2399 | 2498 | 2449 | 1590 | 4205 | 3194 | 3753 | 4053 | 2287 | 2641 | 2419 | 2491 | 3369 | 3632 | 3186 | 3295 | 2915,21 | 2460 | 3370 | 1,37 | 0,454 | 7,849 | 4,21E-15 | 8,50E-15 | 0,0029137 | 0,0026545 | 0,002858 | 0,002858 | TRUE | 0,0323363 | No |
| orf19.1735 | 485 | 479 | 473 | 269 | 408 | 248 | 331 | 304 | 462 | 506 | 467 | 421 | 327 | 282 | 281 | 247 | 374,33 | 464 | 284 | 0,611 | -0,711 | -7,239 | 4,51E-13 | 8,65E-13 | 0,0064997 | 0,0064775 | 0,0064916 | 0,0064916 | TRUE | 0,1228653 | No |
| orf19.1736 | 156 | 107 | 126 | 93 | 188 | 137 | 207 | 186 | 149 | 113 | 124 | 146 | 151 | 156 | 176 | 151 | 145,68 | 133 | 158 | 1,195 | 0,257 | 1,98 | 0,047706 | 0,0574905 | 0,0035032 | 0,013362 | 0,009123 | 0,009123 | TRUE | 0,1036832 | No |
| orf19.1737 | 21 | 29 | 29 | 15 | 24 | 18 | 26 | 16 | 20 | 31 | 29 | 23 | 19 | 20 | 22 | 13 | 22,2 | 26 | 18 | 0,722 | -0,47 | -1,522 | 0,1281029 | 0,1479769 | 1,00E-08 | 0,076045 | 0,0460613 | 0,0460613 | TRUE | 0,2589431 | No |
| orf19.1738 | 8865 | 7398 | 8135 | 4875 | 7775 | 5284 | 7316 | 8408 | 8452 | 7822 | 8037 | 7637 | 6230 | 6008 | 6211 | 6837 | 7154,23 | 7987 | 6322 | 0,791 | -0,338 | -6,627 | 3,43E-11 | 6,20E-11 | 0,0023737 | 0,0023208 | 0,0023504 | 0,0023504 | TRUE | 0,0368359 | No |
| orf19.1738.1 | 374 | 318 | 299 | 180 | 527 | 342 | 530 | 495 | 357 | 336 | 295 | 282 | 422 | 389 | 450 | 402 | 366,72 | 318 | 416 | 1,305 | 0,384 | 4,113 | 3,91E-05 | 5,65E-05 | 0,0043292 | 0,0065685 | 0,0055394 | 0,0055394 | TRUE | 0,0785749 | No |
| orf19.1741 | 3 | 4 | 8 | 1 | 3 | 0 | 6 | 4 | 3 | 4 | 8 | 2 | 2 | 0 | 5 | 3 | 3,41 | 4 | 2 | 0,653 | -0,616 | -0,738 | 0,4606669 | 0,4971276 | 0,1865921 | 0,4830654 | 0,3771973 | 0,3771973 | TRUE | 0,4879298 | No |
| orf19.1742 | 6501 | 7783 | 6691 | 4831 | 7047 | 5824 | 5784 | 6456 | 6198 | 8230 | 6610 | 7568 | 5647 | 6622 | 4911 | 5249 | 6379,32 | 7152 | 5607 | 0,784 | -0,351 | -3,507 | 0,0004539 | 0,0006238 | 0,0167816 | 0,0023487 | 0,0094703 | 0,0094703 | TRUE | 0,180455 | No |
| orf19.1743 | 3455 | 2959 | 3085 | 1981 | 10744 | 7672 | 9926 | 11292 | 3294 | 3129 | 3048 | 3103 | 8609 | 8724 | 8427 | 9181 | 5939,37 | 3144 | 8735 | 2,778 | 1,474 | 34,548 | 1,50E-261 | 2,70E-260 | 0,0009566 | 0,0023677 | 0,0015178 | 0,0015178 | TRUE | 0,0145909 | No |
| orf19.1744 | 2570 | 2766 | 2846 | 1631 | 1829 | 1291 | 1676 | 1557 | 2450 | 2925 | 2812 | 2555 | 1466 | 1468 | 1423 | 1266 | 2045,49 | 2686 | 1406 | 0,523 | -0,935 | -13,234 | 5,61E-40 | 1,77E-39 | 0,0049406 | 0,002894 | 0,0042677 | 0,0042677 | TRUE | 0,0546446 | No |
| orf19.1745 | 726 | 749 | 774 | 524 | 505 | 463 | 543 | 563 | 692 | 792 | 765 | 821 | 405 | 526 | 461 | 458 | 614,94 | 768 | 462 | 0,602 | -0,732 | -8,407 | 4,20E-17 | 8,86E-17 | 0,0060908 | 0,0047613 | 0,005582 | 0,005582 | TRUE | 0,0945208 | No |
| orf19.1746 | 96 | 96 | 109 | 69 | 119 | 111 | 130 | 104 | 92 | 102 | 108 | 108 | 95 | 126 | 110 | 85 | 103,16 | 102 | 104 | 1,015 | 0,021 | 0,134 | 0,8931751 | 0,9079736 | 0,0093306 | 0,0180068 | 0,014113 | 0,014113 | TRUE | 0,1811568 | No |
| orf19.1747 | 102 | 102 | 93 | 57 | 106 | 81 | 96 | 116 | 97 | 108 | 92 | 89 | 85 | 92 | 82 | 94 | 92,39 | 96 | 88 | 0,909 | -0,138 | -0,935 | 0,3499105 | 0,3844316 | 1,00E-08 | 0,0198625 | 0,0100599 | 0,0100599 | TRUE | 0,058363 | No |
| orf19.1748 | 168 | 151 | 173 | 126 | 390 | 322 | 371 | 440 | 160 | 160 | 171 | 197 | 312 | 366 | 315 | 358 | 254,94 | 172 | 338 | 1,976 | 0,983 | 9,256 | 2,13E-20 | 4,83E-20 | 0,0035315 | 0,0085317 | 0,0062366 | 0,0062366 | TRUE | 0,0825979 | No |
| orf19.175 | 167 | 166 | 203 | 105 | 258 | 187 | 254 | 244 | 159 | 176 | 201 | 164 | 207 | 213 | 216 | 198 | 191,65 | 175 | 208 | 1,186 | 0,247 | 2,247 | 0,0246612 | 0,0303966 | 1,00E-08 | 0,0106586 | 0,0062452 | 0,0062452 | TRUE | 0,11007 | No |
| orf19.1750 | 23658 | 22443 | 24658 | 14735 | 17373 | 11623 | 15929 | 15456 | 22555 | 23731 | 24360 | 23083 | 13920 | 13216 | 13524 | 12567 | 18369,58 | 23432 | 13307 | 0,568 | -0,816 | -19,208 | 3,18E-82 | 1,77E-81 | 0,0014013 | 0,0021807 | 0,0016775 | 0,0016775 | TRUE | 0,017262 | No |
| orf19.1751 | 12327 | 12083 | 12800 | 8129 | 12919 | 9347 | 11785 | 12646 | 11752 | 12776 | 12645 | 12735 | 10352 | 10628 | 10005 | 10282 | 11397,03 | 12477 | 10317 | 0,827 | -0,274 | -6,997 | 2,62E-12 | 4,89E-12 | 0,0009487 | 0,0022353 | 0,0013862 | 0,0013862 | TRUE | 0,0188911 | No |
| orf19.1753 | 8119 | 7681 | 8093 | 4812 | 8427 | 6447 | 8211 | 8495 | 7741 | 8122 | 7995 | 7538 | 6752 | 7331 | 6971 | 6907 | 7419,64 | 7849 | 6990 | 0,89 | -0,168 | -4,174 | 2,99E-05 | 4,34E-05 | 0,0009851 | 0,0023126 | 0,0014242 | 0,0014242 | TRUE | 0,0128595 | No |
| orf19.1754 | 1930 | 1775 | 1772 | 1146 | 3452 | 2450 | 3162 | 3320 | 1840 | 1877 | 1751 | 1795 | 2766 | 2786 | 2685 | 2699 | 2274,82 | 1816 | 2734 | 1,505 | 0,59 | 14,304 | 2,07E-46 | 7,27E-46 | 0,0001032 | 0,0028131 | 0,0011594 | 0,0011594 | TRUE | 0,0077048 | No |
| orf19.1755 | 3383 | 3287 | 3501 | 2094 | 2289 | 1704 | 2182 | 2205 | 3225 | 3476 | 3459 | 3280 | 1834 | 1938 | 1853 | 1793 | 2607,13 | 3360 | 1854 | 0,551 | -0,859 | -19,032 | 9,28E-81 | 5,11E-80 | 0,0008505 | 0,0027211 | 0,0015496 | 0,0015496 | TRUE | 0,0104164 | No |
| orf19.1756 | 7614 | 7039 | 7851 | 4780 | 19611 | 14410 | 18079 | 19049 | 7259 | 7443 | 7756 | 7488 | 15714 | 16385 | 15349 | 15489 | 11610,37 | 7486 | 15734 | 2,101 | 1,071 | 29,503 | 2,63E-191 | 3,45E-190 | 0,0006704 | 0,0022327 | 0,0011627 | 0,0011627 | TRUE | 0,0093843 | No |
| orf19.1757 | 3194 | 3068 | 3278 | 1939 | 6754 | 4849 | 6507 | 6753 | 3045 | 3244 | 3238 | 3038 | 5412 | 5514 | 5524 | 5491 | 4313,23 | 3141 | 5485 | 1,745 | 0,803 | 21,521 | 9,94E-103 | 6,66E-102 | 0,0002776 | 0,0024719 | 0,0010763 | 0,0010763 | TRUE | 0,005803 | No |
| orf19.1759 | 2705 | 2442 | 2502 | 1677 | 2965 | 2032 | 2732 | 2724 | 2579 | 2582 | 2472 | 2627 | 2376 | 2311 | 2319 | 2215 | 2435,07 | 2565 | 2305 | 0,899 | -0,153 | -3,666 | 0,0002468 | 0,0003433 | 0,0003943 | 0,0027656 | 0,0012601 | 0,0012601 | TRUE | 0,0087485 | No |
| orf19.176 | 2920 | 2632 | 2665 | 1796 | 389 | 305 | 385 | 423 | 2784 | 2783 | 2633 | 2814 | 312 | 347 | 327 | 344 | 1542,81 | 2754 | 332 | 0,12 | -3,053 | -52,464 | 0 | 0 | 1,00E-08 | 0,0031555 | 0,0017087 | 0,0017087 | TRUE | 0,020942 | No |
| orf19.1760 | 11030 | 10671 | 11070 | 7002 | 5429 | 4064 | 5054 | 5433 | 10516 | 11283 | 10936 | 10969 | 4350 | 4621 | 4291 | 4418 | 7672,98 | 10926 | 4420 | 0,404 | -1,306 | -34,033 | 7,19E-254 | 1,25E-252 | 0,0007331 | 0,0023053 | 0,0012621 | 0,0012621 | TRUE | 0,0110356 | No |
| orf19.1761 | 273 | 300 | 253 | 169 | 382 | 275 | 366 | 328 | 260 | 317 | 250 | 265 | 306 | 313 | 311 | 267 | 286,05 | 273 | 299 | 1,094 | 0,129 | 1,259 | 0,208125 | 0,2357029 | 0,0051695 | 0,0078312 | 0,0065544 | 0,0065544 | TRUE | 0,1372624 | No |
| orf19.1762 | 1517 | 1333 | 1568 | 882 | 1610 | 1112 | 1462 | 1562 | 1446 | 1409 | 1549 | 1382 | 1290 | 1264 | 1241 | 1270 | 1356,54 | 1446 | 1266 | 0,874 | -0,194 | -3,805 | 0,000142 | 0,0001993 | 0,0006209 | 0,0033016 | 0,0017642 | 0,0017642 | TRUE | 0,0276568 | No |
| orf19.1763 | 1395 | 1483 | 1516 | 938 | 1437 | 1156 | 1365 | 1335 | 1330 | 1568 | 1498 | 1469 | 1151 | 1314 | 1159 | 1085 | 1321,93 | 1466 | 1177 | 0,802 | -0,318 | -4,428 | 9,49E-06 | 1,41E-05 | 0,004696 | 0,0033333 | 0,0041915 | 0,0041915 | TRUE | 0,0706749 | No |

| | | | | | | | | | | | | | | | | | | | | | | | | | | | | | | | |
|---|---|---|---|---|---|---|---|---|---|---|---|---|---|---|---|---|---|---|---|---|---|---|---|---|---|---|---|---|---|---|---|
| orf19.1764 | 5674 | 5573 | 5979 | 3800 | 3986 | 2880 | 3717 | 3492 | 5410 | 5893 | 5907 | 5953 | 3194 | 3275 | 3156 | 2839 | 4453,2 | 5791 | 3116 | 0,538 | -0,894 | -16,446 | 8,91E-61 | 3,79E-60 | 0,0025206 | 0,00246 | 0,0025984 | 0,0025984 | TRUE | 0,0437773 | No |
| orf19.1765 | 880 | 822 | 921 | 461 | 1062 | 810 | 1008 | 963 | 839 | 869 | 910 | 722 | 851 | 921 | 856 | 783 | 843,87 | 835 | 853 | 1,016 | 0,023 | 0,297 | 0,7665333 | 0,7911318 | 0,0052847 | 0,004037 | 0,004677 | 0,004677 | TRUE | 0,0885383 | No |
| orf19.1766 | 138 | 136 | 127 | 69 | 213 | 206 | 194 | 221 | 132 | 144 | 125 | 108 | 171 | 234 | 165 | 180 | 157,28 | 127 | 188 | 1,455 | 0,541 | 3,771 | 0,0001628 | 0,000228 | 0,0135412 | 0,0125307 | 0,0129956 | 0,0129956 | TRUE | 0,2846464 | No |
| orf19.1767 | 485 | 622 | 652 | 350 | 501 | 343 | 464 | 422 | 462 | 658 | 644 | 548 | 401 | 390 | 394 | 343 | 480,13 | 578 | 382 | 0,66 | -0,598 | -5,348 | 8,91E-08 | 1,44E-07 | 0,0140251 | 0,005511 | 0,0098826 | 0,0098826 | TRUE | 0,2242949 | No |
| orf19.1768 | 293 | 334 | 368 | 219 | 257 | 194 | 240 | 235 | 279 | 353 | 364 | 343 | 206 | 221 | 204 | 191 | 270,06 | 335 | 206 | 0,613 | -0,706 | -6,637 | 3,21E-11 | 5,82E-11 | 0,005848 | 0,008171 | 0,0070192 | 0,0070192 | TRUE | 0,1522757 | No |
| orf19.1769 | 886 | 852 | 915 | 507 | 1502 | 982 | 1394 | 1436 | 845 | 901 | 904 | 794 | 1204 | 1117 | 1183 | 1168 | 1014,37 | 861 | 1168 | 1,354 | 0,437 | 7,526 | 5,23E-14 | 1,03E-13 | 0,0007475 | 0,0037099 | 0,0022047 | 0,0022047 | TRUE | 0,0274662 | No |
| orf19.177 | 496 | 532 | 509 | 372 | 649 | 461 | 608 | 660 | 473 | 563 | 503 | 583 | 520 | 524 | 516 | 537 | 527,26 | 530 | 524 | 0,994 | -0,009 | -0,12 | 0,9042311 | 0,9176047 | 0,0022516 | 0,0052053 | 0,0038704 | 0,0038704 | TRUE | 0,0629579 | No |
| orf19.1770 | 8255 | 6835 | 6983 | 4786 | 5902 | 3849 | 5378 | 6037 | 7870 | 7227 | 6899 | 7498 | 4729 | 4377 | 4566 | 4909 | 6009,22 | 7374 | 4645 | 0,63 | -0,666 | -12,597 | 2,19E-36 | 6,52E-36 | 0,002523 | 0,0023645 | 0,0025116 | 0,0025116 | TRUE | 0,0256084 | No |
| orf19.1771 | 1740 | 1803 | 1813 | 1138 | 1932 | 1315 | 1685 | 1724 | 1659 | 1906 | 1791 | 1783 | 1548 | 1495 | 1431 | 1402 | 1626,85 | 1785 | 1469 | 0,823 | -0,281 | -4,919 | 8,71E-07 | 1,35E-06 | 0,0021047 | 0,0031005 | 0,0025112 | 0,0025112 | TRUE | 0,0285564 | No |
| orf19.1772 | 3054 | 3485 | 3620 | 2194 | 3249 | 2530 | 3012 | 2821 | 2912 | 3685 | 3576 | 3437 | 2603 | 2877 | 2557 | 2294 | 2992,62 | 3402 | 2583 | 0,759 | -0,398 | -4,803 | 1,57E-06 | 2,40E-06 | 0,0088893 | 0,0026399 | 0,0062625 | 0,0062625 | TRUE | 0,1161377 | No |
| orf19.1773 | 2507 | 2391 | 2437 | 1586 | 2345 | 1728 | 2315 | 2117 | 2390 | 2528 | 2408 | 2485 | 1879 | 1965 | 1965 | 1721 | 2167,63 | 2453 | 1882 | 0,767 | -0,382 | -7,213 | 5,48E-13 | 1,05E-12 | 0,0015897 | 0,0028487 | 0,0022269 | 0,0022269 | TRUE | 0,040826 | No |
| orf19.1774 | 501 | 499 | 472 | 343 | 719 | 525 | 675 | 740 | 478 | 528 | 466 | 537 | 576 | 597 | 573 | 602 | 544,59 | 502 | 587 | 1,173 | 0,23 | 3,268 | 0,0010846 | 0,0014597 | 0,0003334 | 0,0051062 | 0,0028894 | 0,0028894 | TRUE | 0,0265518 | No |
| orf19.1776 | 311 | 305 | 334 | 188 | 470 | 354 | 425 | 424 | 297 | 322 | 330 | 295 | 377 | 403 | 361 | 345 | 341,02 | 311 | 372 | 1,188 | 0,249 | 2,892 | 0,00383 | 0,0049907 | 1,00E-08 | 0,0069059 | 0,0041102 | 0,0041102 | TRUE | 0,0335591 | No |
| orf19.1777 | 11727 | 12362 | 12818 | 7314 | 4708 | 3540 | 4171 | 4368 | 11180 | 13071 | 12663 | 11458 | 3772 | 4025 | 3541 | 3552 | 7907,86 | 12093 | 3722 | 0,308 | -1,701 | -27,024 | 7,62E-161 | 8,33E-160 | 0,004515 | 0,0022989 | 0,0036371 | 0,0036371 | TRUE | 0,0362599 | No |
| orf19.1778 | 93 | 97 | 84 | 58 | 46 | 33 | 36 | 32 | 89 | 103 | 83 | 91 | 37 | 38 | 31 | 26 | 62,01 | 92 | 33 | 0,357 | -1,488 | -7,612 | 2,69E-14 | 5,32E-14 | 1,00E-08 | 0,0285715 | 0,0168949 | 0,0168949 | TRUE | 0,1497231 | No |
| orf19.1779 | 20322 | 17443 | 19167 | 13593 | 51781 | 40701 | 49363 | 51801 | 19375 | 18444 | 18935 | 21294 | 41491 | 46281 | 41909 | 42119 | 31230,92 | 19512 | 42950 | 2,202 | 1,139 | 20,863 | 1,16E-96 | 7,31E-96 | 0,0033492 | 0,0021439 | 0,0028219 | 0,0028219 | TRUE | 0,0459151 | No |
| orf19.178 | 959 | 1077 | 1094 | 670 | 1467 | 1201 | 1408 | 1352 | 914 | 1139 | 1081 | 1050 | 1175 | 1366 | 1195 | 1099 | 1127,41 | 1046 | 1209 | 1,155 | 0,208 | 2,498 | 0,0124814 | 0,0157617 | 0,0072716 | 0,0035476 | 0,0057356 | 0,0057356 | TRUE | 0,0891449 | No |
| orf19.1780 | 24 | 34 | 47 | 27 | 56 | 25 | 46 | 28 | 23 | 36 | 46 | 42 | 45 | 28 | 39 | 23 | 35,33 | 37 | 34 | 0,926 | -0,111 | -0,374 | 0,7086559 | 0,7386366 | 0,0645069 | 0,0485581 | 0,0558413 | 0,0558413 | TRUE | 0,3168086 | No |
| orf19.1782 | 535 | 472 | 474 | 352 | 1304 | 891 | 1121 | 1203 | 510 | 499 | 468 | 551 | 1045 | 1013 | 952 | 978 | 752,09 | 507 | 997 | 1,974 | 0,981 | 14,76 | 2,66E-49 | 9,71E-49 | 0,0011452 | 0,0042744 | 0,0026884 | 0,0026884 | TRUE | 0,0334523 | No |
| orf19.1782.1 | 26 | 19 | 29 | 10 | 26 | 21 | 46 | 32 | 25 | 20 | 29 | 16 | 21 | 24 | 39 | 26 | 24,87 | 22 | 28 | 1,212 | 0,277 | 0,886 | 0,3754276 | 0,4104169 | 0,0346046 | 0,0681048 | 0,0533342 | 0,0533342 | TRUE | 0,5702867 | No |
| orf19.1783 | 12123 | 12235 | 12792 | 7193 | 17299 | 11982 | 15345 | 16130 | 11558 | 12937 | 12637 | 11268 | 13861 | 13625 | 13028 | 13115 | 12753,66 | 12100 | 13407 | 1,108 | 0,148 | 2,873 | 0,0040637 | 0,0052865 | 0,0025906 | 0,00222 | 0,002454 | 0,002454 | TRUE | 0,0264137 | No |
| orf19.1785 | 29607 | 27220 | 29944 | 20314 | 2807 | 2153 | 2667 | 2648 | 28227 | 28782 | 29582 | 31823 | 2249 | 2448 | 2264 | 2153 | 15941,03 | 29604 | 2278 | 0,077 | -3,701 | -70,442 | 0 | 0 | 0,0025735 | 0,0021943 | 0,0024356 | 0,0024356 | TRUE | 0,0310197 | No |
| orf19.1789 | 273 | 394 | 338 | 182 | 566 | 423 | 505 | 418 | 260 | 417 | 334 | 285 | 454 | 481 | 429 | 340 | 374,88 | 324 | 426 | 1,311 | 0,39 | 2,68 | 0,0073676 | 0,0094214 | 0,0293014 | 0,0064711 | 0,0175945 | 0,0175945 | TRUE | 0,4223004 | No |
| orf19.1789.1 | 1763 | 1697 | 1743 | 1186 | 3922 | 2977 | 3544 | 3712 | 1681 | 1794 | 1722 | 1858 | 3143 | 3385 | 3009 | 3018 | 2451,22 | 1764 | 3139 | 1,781 | 0,833 | 15,418 | 1,24E-53 | 4,80E-53 | 0,0019556 | 0,0027611 | 0,0023388 | 0,0023388 | TRUE | 0,033005 | No |
| orf19.1791 | 6886 | 6867 | 7214 | 4031 | 6927 | 4496 | 6305 | 5905 | 6565 | 7261 | 7127 | 6315 | 5550 | 5112 | 5353 | 4801 | 6010,57 | 6817 | 5204 | 0,763 | -0,39 | -6,465 | 1,01E-10 | 1,81E-10 | 0,0038818 | 0,0023644 | 0,0033242 | 0,0033242 | TRUE | 0,0334939 | No |
| orf19.1792 | 488 | 513 | 583 | 320 | 553 | 481 | 556 | 520 | 465 | 542 | 576 | 501 | 443 | 547 | 472 | 423 | 496,23 | 521 | 471 | 0,901 | -0,151 | -1,55 | 0,1210488 | 0,1400567 | 0,008241 | 0,0054 | 0,0070713 | 0,0070713 | TRUE | 0,1289072 | No |
| orf19.1793 | 1556 | 1416 | 1537 | 927 | 1122 | 836 | 1024 | 1139 | 1483 | 1497 | 1518 | 1452 | 899 | 951 | 869 | 926 | 1199,56 | 1488 | 911 | 0,611 | -0,71 | -14,237 | 5,38E-46 | 1,88E-45 | 1,00E-08 | 0,00346 | 0,0015244 | 0,0015244 | TRUE | 0,0112437 | No |
| orf19.1794 | 183 | 200 | 175 | 83 | 267 | 197 | 246 | 228 | 174 | 211 | 173 | 130 | 214 | 224 | 209 | 185 | 190,13 | 172 | 208 | 1,195 | 0,257 | 1,926 | 0,0541178 | 0,0648012 | 0,0131867 | 0,010727 | 0,0117636 | 0,0117636 | TRUE | 0,2353932 | No |
| orf19.1795 | 2354 | 2365 | 2381 | 1364 | 1789 | 1435 | 1798 | 1853 | 2244 | 2501 | 2352 | 2137 | 1433 | 1632 | 1526 | 1507 | 1916,54 | 2308 | 1524 | 0,659 | -0,602 | -9,891 | 4,53E-23 | 1,08E-22 | 0,0029816 | 0,002948 | 0,0030145 | 0,0030145 | TRUE | 0,0376157 | No |
| orf19.1795.1 | 268 | 266 | 270 | 184 | 285 | 239 | 240 | 251 | 256 | 281 | 267 | 288 | 228 | 272 | 204 | 204 | 249,97 | 273 | 227 | 0,829 | -0,27 | -2,455 | 0,0140955 | 0,0177298 | 0,0059986 | 0,0086598 | 0,0075692 | 0,0075692 | TRUE | 0,1770367 | No |
| orf19.1796 | 269 | 268 | 293 | 153 | 590 | 384 | 483 | 479 | 256 | 283 | 289 | 240 | 473 | 437 | 410 | 389 | 347,24 | 267 | 427 | 1,591 | 0,67 | 7,012 | 2,35E-12 | 4,40E-12 | 0,0044244 | 0,0068197 | 0,0056378 | 0,0056378 | TRUE | 0,063604 | No |
| orf19.1797 | 6377 | 6646 | 6891 | 4306 | 6255 | 4720 | 5768 | 5563 | 6080 | 7027 | 6808 | 6746 | 5012 | 5367 | 4897 | 4523 | 5807,45 | 6665 | 4950 | 0,742 | -0,43 | -6,957 | 3,47E-12 | 6,46E-12 | 0,0041107 | 0,002374 | 0,003487 | 0,003487 | TRUE | 0,0431105 | No |
| orf19.1798 | 724 | 759 | 839 | 463 | 584 | 446 | 545 | 530 | 690 | 803 | 829 | 725 | 468 | 507 | 463 | 431 | 614,46 | 762 | 467 | 0,611 | -0,71 | -8,803 | 1,34E-18 | 2,91E-18 | 0,0043575 | 0,0047634 | 0,0045505 | 0,0045505 | TRUE | 0,0516523 | No |
| orf19.1799 | 15248 | 13446 | 14503 | 8524 | 12618 | 8213 | 11591 | 12033 | 14537 | 14217 | 14328 | 13353 | 10110 | 9339 | 9841 | 9784 | 11938,71 | 14109 | 9768 | 0,692 | -0,531 | -13,155 | 1,59E-39 | 4,99E-39 | 0,0011022 | 0,0022288 | 0,0014768 | 0,0014768 | TRUE | 0,0155834 | No |
| orf19.18 | 32285 | 27614 | 28971 | 18080 | 49879 | 35139 | 45715 | 50648 | 30780 | 29198 | 28621 | 28323 | 39967 | 39956 | 38812 | 41182 | 34604,84 | 29230 | 39979 | 1,368 | 0,452 | 12,144 | 6,15E-34 | 1,77E-33 | 0,0009553 | 0,0021387 | 0,001298 | 0,001298 | TRUE | 0,0156474 | No |
| orf19.1800 | 10424 | 9833 | 10849 | 6225 | 3056 | 2214 | 2925 | 2801 | 9938 | 10397 | 10718 | 9752 | 2449 | 2518 | 2483 | 2277 | 6316,49 | 10201 | 2432 | 0,238 | -2,07 | -43,657 | 0 | 0 | 0,0015448 | 0,0023512 | 0,0019197 | 0,0019197 | TRUE | 0,0223873 | No |
| orf19.1801 | 1830 | 1796 | 1931 | 1080 | 2723 | 1946 | 2503 | 2512 | 1745 | 1899 | 1908 | 1692 | 2182 | 2213 | 2125 | 2042 | 1975,68 | 1811 | 2140 | 1,18 | 0,239 | 4,45 | 8,60E-06 | 1,28E-05 | 0,0017441 | 0,0029223 | 0,0022401 | 0,0022401 | TRUE | 0,0214515 | No |
| orf19.1802 | 6960 | 6655 | 6950 | 3922 | 6856 | 4755 | 6670 | 6961 | 6636 | 7037 | 6866 | 6144 | 5494 | 5407 | 5663 | 5660 | 6113,19 | 6671 | 5556 | 0,832 | -0,265 | -5,614 | 1,98E-08 | 3,27E-08 | 0,0017296 | 0,0023599 | 0,0019691 | 0,0019691 | TRUE | 0,0333191 | No |
| orf19.1803 | 163 | 192 | 163 | 130 | 158 | 108 | 141 | 138 | 155 | 203 | 161 | 204 | 127 | 123 | 120 | 112 | 150,55 | 181 | 120 | 0,67 | -0,578 | -4,381 | 1,18E-05 | 1,75E-05 | 0,0062819 | 0,0129971 | 0,0098955 | 0,0098955 | TRUE | 0,0978971 | No |
| orf19.1804 | 4051 | 3892 | 4245 | 2461 | 3273 | 2224 | 3113 | 3094 | 3862 | 4115 | 4194 | 3855 | 2623 | 2529 | 2643 | 2516 | 3292,07 | 4006 | 2578 | 0,643 | -0,637 | -14,444 | 2,74E-47 | 9,74E-47 | 0,0008285 | 0,00259 | 0,0015531 | 0,0015531 | TRUE | 0,0121081 | No |
| orf19.1805 | 3067 | 2917 | 3158 | 2082 | 5772 | 4247 | 5537 | 5658 | 2924 | 3084 | 3120 | 3262 | 4625 | 4829 | 4701 | 4600 | 3893,17 | 3098 | 4689 | 1,515 | 0,6 | 14,159 | 1,64E-45 | 5,65E-45 | 0,000885 | 0,002513 | 0,0014452 | 0,0014452 | TRUE | 0,0173963 | No |
| orf19.1806 | 45 | 38 | 43 | 22 | 33 | 24 | 25 | 24 | 43 | 40 | 42 | 34 | 26 | 27 | 21 | 20 | 31,81 | 40 | 24 | 0,582 | -0,782 | -3,116 | 0,0018314 | 0,002432 | 1,00E-08 | 0,0537035 | 0,0276608 | 0,0276608 | TRUE | 0,0918604 | No |
| orf19.1808 | 349 | 389 | 434 | 248 | 423 | 325 | 323 | 365 | 333 | 411 | 429 | 389 | 339 | 370 | 274 | 297 | 355,1 | 390 | 320 | 0,818 | -0,291 | -2,574 | 0,0100581 | 0,0127774 | 0,0121111 | 0,006715 | 0,0093987 | 0,0093987 | TRUE | 0,1208865 | No |
| orf19.1813 | 3086 | 2757 | 2898 | 1770 | 5235 | 3891 | 4786 | 5094 | 2942 | 2915 | 2863 | 2773 | 4195 | 4424 | 4063 | 4142 | 3539,67 | 2873 | 4206 | 1,462 | 0,548 | 13,17 | 1,30E-39 | 4,09E-39 | 0,0006279 | 0,0025551 | 0,001358 | 0,001358 | TRUE | 0,0143507 | No |
| orf19.1814 | 2185 | 2220 | 2239 | 1202 | 2153 | 1690 | 1975 | 2139 | 2083 | 2347 | 2212 | 1883 | 1725 | 1922 | 1677 | 1739 | 1948,53 | 2131 | 1766 | 0,827 | -0,275 | -3,866 | 0,0001108 | 0,0001565 | 0,005457 | 0,0029339 | 0,0043303 | 0,0043303 | TRUE | 0,070736 | No |
| orf19.1815 | 12663 | 10467 | 10748 | 6838 | 16335 | 10872 | 15279 | 16911 | 12073 | 11068 | 10618 | 10712 | 13089 | 12362 | 12972 | 13750 | 12080,46 | 11118 | 13043 | 1,173 | 0,23 | 4,469 | 7,86E-06 | 1,17E-05 | 0,0026317 | 0,0022272 | 0,0024672 | 0,0024672 | TRUE | 0,041149 | No |
| orf19.1816 | 7074 | 6242 | 6875 | 4597 | 5159 | 3378 | 4614 | 4816 | 6744 | 6600 | 6792 | 7201 | 4134 | 3841 | 3917 | 3916 | 5393,21 | 6834 | 3952 | 0,579 | -0,789 | -18,98 | 2,50E-80 | 1,37E-79 | 0,0009564 | 0,0023957 | 0,001463 | 0,001463 | TRUE | 0,0153391 | No |
| orf19.182 | 2718 | 2414 | 2571 | 1495 | 2691 | 1872 | 2555 | 2641 | 2591 | 2552 | 2540 | 2342 | 2156 | 2129 | 2169 | 2147 | 2328,39 | 2506 | 2150 | 0,856 | -0,224 | -5,174 | 2,29E-07 | 3,65E-07 | 0,0004499 | 0,0027964 | 0,0013622 | 0,0013622 | TRUE | 0,0214163 | No |
| orf19.1821 | 0 | 0 | 2 | 1 | 1 | 0 | 0 | 1 | 0 | 0 | 2 | 2 | 1 | 0 | 0 | 1 | 0,64 | 1 | 0 | 0,512 | -0,966 | -0,508 | 0,6114025 | 0,6451727 | 1,00E-08 | 2,5492919 | 1,8648468 | 1,8648468 | TRUE | 0,1763341 | No |
| orf19.1822 | 77 | 65 | 63 | 47 | 41 | 30 | 43 | 33 | 73 | 69 | 62 | 74 | 33 | 34 | 37 | 27 | 51,04 | 70 | 33 | 0,469 | -1,094 | -5,388 | 7,13E-08 | 1,15E-07 | 1,00E-08 | 0,034261 | 0,017838 | 0,017838 | TRUE | 0,1127449 | No |
| orf19.1823 | 167 | 180 | 180 | 122 | 160 | 139 | 140 | 162 | 159 | 190 | 178 | 191 | 128 | 158 | 119 | 132 | 156,92 | 180 | 134 | 0,746 | -0,424 | -3,361 | 0,0007775 | 0,0010569 | 0,0037733 | 0,0125549 | 0,0087864 | 0,0087864 | TRUE | 0,1303744 | No |
| orf19.1824 | 0 | 1 | 0 | 0 | 1 | 0 | 0 | 0 | 0 | 1 | 0 | 0 | 1 | 0 | 0 | 0 | 0,23 | 0 | 0 | 0,797 | -0,328 | -0,106 | 0,9159347 | 0,9244826 | 1,00E-08 | 7,0691906 | 7,259296 | 7,259296 | TRUE | 0,3159987 | No |
| orf19.1825 | 62 | 49 | 66 | 47 | 110 | 76 | 104 | 123 | 59 | 52 | 65 | 74 | 88 | 86 | 88 | 100 | 76,58 | 62 | 90 | 1,471 | 0,556 | 3,322 | 0,0008928 | 0,0012086 | 1,00E-08 | 0,0235325 | 0,0130401 | 0,0130401 | TRUE | 0,1067727 | No |
| orf19.1826 | 20168 | 20208 | 21038 | 13363 | 6125 | 4798 | 5447 | 5778 | 19228 | 21367 | 20784 | 20934 | 4908 | 5456 | 4624 | 4698 | 12749,86 | 20578 | 4922 | 0,239 | -2,065 | -35,728 | 1,44E-279 | 2,89E-278 | 0,0037771 | 0,0022201 | 0,0030896 | 0,0030896 | TRUE | 0,0641796 | No |
| orf19.1827 | 17 | 18 | 17 | 8 | 8 | 10 | 17 | 6 | 16 | 19 | 17 | 13 | 6 | 11 | 14 | 5 | 12,71 | 16 | 9 | 0,561 | -0,833 | -1,944 | 0,0519252 | 0,0623512 | 0,0281157 | 0,131299 | 0,0934543 | 0,0934543 | TRUE | 0,5420805 | No |
| orf19.1828 | 453 | 400 | 462 | 226 | 781 | 581 | 772 | 721 | 432 | 423 | 456 | 354 | 626 | 661 | 655 | 586 | 524,17 | 416 | 632 | 1,507 | 0,591 | 7,152 | 8,54E-13 | 1,62E-12 | 0,0036872 | 0,0052236 | 0,0045203 | 0,0045203 | TRUE | 0,0980436 | No |
| orf19.183 | 1367 | 1214 | 1175 | 803 | 1798 | 1273 | 1726 | 1853 | 1303 | 1284 | 1161 | 1258 | 1441 | 1448 | 1465 | 1507 | 1358,24 | 1252 | 1465 | 1,171 | 0,228 | 4,392 | 1,13E-05 | 1,66E-05 | 0,0007277 | 0,0033001 | 0,0018369 | 0,0018369 | TRUE | 0,0304277 | No |
| orf19.1830 | 3333 | 3492 | 3543 | 2149 | 1553 | 994 | 1488 | 1453 | 3178 | 3692 | 3500 | 3367 | 1244 | 1130 | 1263 | 1181 | 2319,51 | 3434 | 1204 | 0,351 | -1,51 | -25,476 | 3,65E-143 | 3,52E-142 | 0,0028663 | 0,0027991 | 0,0028378 | 0,0028378 | TRUE | 0,0317222 | No |
| orf19.1831 | 1 | 3 | 0 | 2 | 2 | 0 | 0 | 3 | 1 | 3 | 0 | 3 | 2 | 0 | 0 | 2 | 1,41 | 2 | 1 | 0,605 | -0,725 | -0,534 | 0,5934792 | 0,6278168 | 0,7234712 | 1,1644547 | 1,0305029 | 1,0305029 | TRUE | 0,3386585 | No |
| orf19.1832 | 2551 | 2236 | 2561 | 1485 | 3906 | 2742 | 3667 | 3780 | 2432 | 2364 | 2530 | 2326 | 3130 | 3118 | 3113 | 3074 | 2760,9 | 2413 | 3109 | 1,286 | 0,363 | 9,143 | 6,09E-20 | 1,36E-19 | 0,0002254 | 0,002686 | 0,0011399 | 0,0011399 | TRUE | 0,0129084 | No |
| orf19.1833 | 18265 | 17982 | 18976 | 10723 | 33411 | 23597 | 31324 | 31743 | 17414 | 19014 | 18746 | 16798 | 26771 | 26832 | 26594 | 25810 | 22247,41 | 17993 | 26502 | 1,473 | 0,558 | 12,276 | 1,22E-34 | 3,55E-34 | 0,0018328 | 0,0021651 | 0,0019389 | 0,0019389 | TRUE | 0,024127 | No |
| orf19.1834 | 299 | 341 | 385 | 227 | 373 | 344 | 403 | 389 | 285 | 361 | 380 | 356 | 299 | 391 | 342 | 316 | 341,26 | 346 | 337 | 0,974 | -0,038 | -0,331 | 0,7404095 | 0,7689702 | 0,0118341 | 0,0069026 | 0,0093481 | 0,0093481 | TRUE | 0,1675009 | No |
| orf19.1835 | 4228 | 4317 | 4578 | 2829 | 2266 | 1751 | 2163 | 2203 | 4031 | 4565 | 4523 | 4432 | 1816 | 1991 | 1836 | 1791 | 3123,05 | 4388 | 1858 | 0,423 | -1,24 | -22,716 | 3,15E-114 | 2,37E-113 | 0,0022149 | 0,002617 | 0,0024938 | 0,0024938 | TRUE | 0,0372377 | No |
| orf19.1836 | 152 | 152 | 139 | 85 | 508 | 297 | 424 | 414 | 145 | 161 | 137 | 133 | 407 | 338 | 360 | 337 | 252,18 | 144 | 360 | 2,497 | 1,32 | 12,027 | 2,56E-33 | 7,31E-33 | 0,0042893 | 0,008602 | 0,0064818 | 0,0064818 | TRUE | 0,0902707 | No |
| orf19.1837 | 5851 | 5002 | 5690 | 3193 | 3476 | 2110 | 3300 | 3464 | 5578 | 5289 | 5621 | 5002 | 2785 | 2399 | 2802 | 2817 | 4036,65 | 5372 | 2701 | 0,503 | -0,992 | -16,161 | 9,55E-59 | 3,96E-58 | 0,0038429 | 0,002498 | 0,0033474 | 0,0033474 | TRUE | 0,0667589 | No |
| orf19.1839 | 13989 | 13377 | 14033 | 7808 | 21191 | 14622 | 19245 | 21136 | 13337 | 14144 | 13863 | 12232 | 16980 | 16627 | 16339 | 17186 | 15088,4 | 13394 | 16783 | 1,253 | 0,325 | 6,721 | 1,80E-11 | 3,29E-11 | 0,0021465 | 0,0022001 | 0,0021756 | 0,0021756 | TRUE | 0,0410886 | No |
| orf19.184 | 567 | 579 | 661 | 340 | 652 | 514 | 642 | 580 | 541 | 612 | 653 | 533 | 522 | 584 | 545 | 472 | 557,75 | 585 | 531 | 0,904 | -0,145 | -1,609 | 0,1076113 | 0,1251672 | 0,0069577 | 0,0050351 | 0,0060193 | 0,0060193 | TRUE | 0,0735548 | No |
| orf19.1840 | 2296 | 2068 | 2174 | 1340 | 3484 | 2536 | 3399 | 3706 | 2189 | 2187 | 2148 | 2099 | 2792 | 2884 | 2886 | 3013 | 2524,61 | 2156 | 2894 | 1,341 | 0,424 | 10,245 | 1,24E-24 | 3,06E-24 | 0,0004071 | 0,0027416 | 0,0012262 | 0,0012262 | TRUE | 0,0097296 | No |
| orf19.1841 | 51 | 87 | 80 | 40 | 30 | 29 | 34 | 36 | 49 | 92 | 79 | 63 | 24 | 33 | 29 | 29 | 49,68 | 71 | 29 | 0,405 | -1,305 | -5,309 | 1,10E-07 | 1,77E-07 | 0,0338646 | 0,0351392 | 0,034541 | 0,034541 | TRUE | 0,1259707 | No |
| orf19.1842 | 3217 | 3298 | 3462 | 1969 | 2889 | 2019 | 2507 | 2650 | 3067 | 3487 | 3420 | 3085 | 2315 | 2296 | 2128 | 2155 | 2744,09 | 3265 | 2224 | 0,68 | -0,555 | -9,685 | 3,49E-22 | 8,18E-22 | 0,0026882 | 0,0026896 | 0,0027848 | 0,0027848 | TRUE | 0,025446 | No |
| orf19.1843 | 209 | 265 | 252 | 174 | 602 | 445 | 506 | 526 | 199 | 280 | 249 | 273 | 482 | 506 | 430 | 428 | 355,83 | 250 | 462 | 1,85 | 0,888 | 7,995 | 1,30E-15 | 2,64E-15 | 0,0108235 | 0,0067056 | 0,0086185 | 0,0086185 | TRUE | 0,2216823 | No |
| orf19.1844 | 11 | 17 | 16 | 13 | 12 | 11 | 18 | 14 | 10 | 18 | 16 | 20 | 10 | 13 | 15 | 11 | 14,18 | 16 | 12 | 0,766 | -0,384 | -1,009 | 0,3127411 | 0,345918 | 1,00E-08 | 0,1178989 | 0,0674066 | 0,0674066 | TRUE | 0,2916308 | No |
| orf19.1847 | 1025 | 872 | 1033 | 685 | 2095 | 1666 | 2101 | 2152 | 977 | 922 | 1021 | 1073 | 1679 | 1894 | 1784 | 1750 | 1387,43 | 998 | 1777 | 1,782 | 0,834 | 13,738 | 6,01E-43 | 1,99E-42 | 0,002302 | 0,0032747 | 0,0027208 | 0,0027208 | TRUE | 0,0306924 | No |

| | | | | | | | | | | | | | | | | | | | | | | | | | | | | | | | |
|---|---|---|---|---|---|---|---|---|---|---|---|---|---|---|---|---|---|---|---|---|---|---|---|---|---|---|---|---|---|---|---|
| orf19.1848.1 | 503 | 388 | 342 | 267 | 276 | 172 | 206 | 220 | 480 | 410 | 338 | 418 | 221 | 196 | 175 | 179 | 302,06 | 412 | 193 | 0,468 | -1,095 | -8,978 | 2,75E-19 | 6,08E-19 | 0,013611 | 0,007527 | 0,0105931 | 0,0105931 | TRUE | 0,1739081 | No |
| orf19.1849 | 5328 | 5114 | 5436 | 3209 | 5532 | 3967 | 5102 | 5399 | 5080 | 5407 | 5370 | 5027 | 4433 | 4511 | 4332 | 4390 | 4818,67 | 5221 | 4416 | 0,845 | -0,243 | -6,306 | 2,87E-10 | 5,04E-10 | 0,000568 | 0,002432 | 0,0012135 | 0,0012135 | TRUE | 0,0071507 | No |
| orf19.185 | 3020 | 2755 | 2755 | 1614 | 2934 | 1945 | 2506 | 2807 | 2879 | 2913 | 2722 | 2528 | 2351 | 2212 | 2128 | 2282 | 2501,86 | 2760 | 2243 | 0,812 | -0,301 | -5,389 | 7,08E-08 | 1,15E-07 | 0,0024006 | 0,0027476 | 0,0025932 | 0,0025932 | TRUE | 0,0366965 | No |
| orf19.1850 | 1002 | 908 | 981 | 601 | 1230 | 852 | 1100 | 1102 | 955 | 960 | 969 | 941 | 986 | 969 | 934 | 896 | 951,29 | 956 | 946 | 0,988 | -0,018 | -0,326 | 0,7442421 | 0,7724478 | 1,00E-08 | 0,0038173 | 0,001777 | 0,001777 | TRUE | 0,0156948 | No |
| orf19.1852 | 1066 | 868 | 960 | 628 | 1355 | 896 | 1339 | 1365 | 1016 | 918 | 948 | 984 | 1086 | 1019 | 1137 | 1110 | 1027,19 | 966 | 1088 | 1,127 | 0,173 | 2,975 | 0,0029292 | 0,003838 | 0,0010628 | 0,0036897 | 0,0022501 | 0,0022501 | TRUE | 0,0201887 | No |
| orf19.1853 | 1338 | 1030 | 1032 | 648 | 682 | 442 | 593 | 649 | 1276 | 1089 | 1020 | 1015 | 546 | 503 | 503 | 528 | 809,95 | 1100 | 520 | 0,472 | -1,082 | -13,149 | 1,73E-39 | 5,42E-39 | 0,0059665 | 0,0041184 | 0,0051324 | 0,0051324 | TRUE | 0,141509 | No |
| orf19.1854 | 3489 | 3062 | 3215 | 2048 | 1482 | 993 | 1269 | 1390 | 3326 | 3238 | 3176 | 3208 | 1187 | 1129 | 1077 | 1130 | 2184,08 | 3237 | 1131 | 0,349 | -1,517 | -33,753 | 9,69E-250 | 1,65E-248 | 0,000336 | 0,002843 | 0,0013718 | 0,0013718 | TRUE | 0,0143103 | No |
| orf19.1855 | 335 | 345 | 362 | 234 | 381 | 232 | 326 | 354 | 319 | 365 | 358 | 367 | 305 | 264 | 277 | 288 | 317,76 | 352 | 284 | 0,809 | -0,305 | -3,46 | 0,0005403 | 0,000741 | 1,00E-08 | 0,0072584 | 0,0043192 | 0,0043192 | TRUE | 0,0476093 | No |
| orf19.1856 | 2920 | 2761 | 2893 | 1716 | 2425 | 1495 | 2227 | 2118 | 2784 | 2919 | 2858 | 2688 | 1943 | 1700 | 1891 | 1722 | 2313,18 | 2812 | 1814 | 0,645 | -0,632 | -11,382 | 5,13E-30 | 1,38E-29 | 0,002251 | 0,0028011 | 0,0025172 | 0,0025172 | TRUE | 0,0279224 | No |
| orf19.1857 | 763 | 661 | 773 | 439 | 1961 | 1235 | 1941 | 2001 | 727 | 699 | 764 | 688 | 1571 | 1404 | 1648 | 1627 | 1141,03 | 720 | 1562 | 2,171 | 1,118 | 16,803 | 2,32E-63 | 1,02E-62 | 0,002918 | 0,0035302 | 0,0031882 | 0,0031882 | TRUE | 0,0536282 | No |
| orf19.1860 | 5006 | 3788 | 4333 | 3096 | 11945 | 8489 | 11327 | 12420 | 4773 | 4005 | 4281 | 4850 | 9571 | 9653 | 9617 | 10099 | 7105,98 | 4477 | 9735 | 2,176 | 1,121 | 18,49 | 2,48E-76 | 1,29E-75 | 0,0041535 | 0,0023223 | 0,0033613 | 0,0033613 | TRUE | 0,0612405 | No |
| orf19.1860.1 | 233 | 242 | 235 | 137 | 525 | 376 | 567 | 507 | 222 | 256 | 232 | 215 | 421 | 428 | 481 | 412 | 333,33 | 231 | 436 | 1,878 | 0,909 | 9,794 | 1,19E-22 | 2,82E-22 | 0,0025763 | 0,0070171 | 0,0048458 | 0,0048458 | TRUE | 0,0602944 | No |
| orf19.1861 | 15241 | 13677 | 14933 | 9177 | 2444 | 1765 | 2373 | 2495 | 14531 | 14462 | 14752 | 14376 | 1958 | 2007 | 2015 | 2029 | 8266,2 | 14530 | 2002 | 0,138 | -2,86 | -90,807 | 0 | 0 | 1,00E-08 | 0,0022899 | 0,0006911 | 0,0006911 | TRUE | 0,0027913 | No |
| orf19.1862 | 7477 | 6116 | 6479 | 3788 | 6475 | 4128 | 6025 | 6230 | 7129 | 6467 | 6401 | 5934 | 5188 | 4694 | 5115 | 5066 | 5749,13 | 6483 | 5016 | 0,773 | -0,371 | -6,32 | 2,62E-10 | 4,62E-10 | 0,003605 | 0,0023769 | 0,003126 | 0,003126 | TRUE | 0,0550221 | No |
| orf19.1863 | 22 | 21 | 17 | 12 | 42 | 27 | 15 | 40 | 21 | 22 | 17 | 19 | 34 | 31 | 13 | 33 | 23,55 | 20 | 28 | 1,387 | 0,472 | 1,376 | 0,1687928 | 0,1927809 | 0,0647106 | 0,0718163 | 0,0685779 | 0,0685779 | TRUE | 0,9192195 | No |
| orf19.1864 | 3094 | 2645 | 3026 | 2047 | 5425 | 3801 | 5299 | 5694 | 2950 | 2797 | 2989 | 3207 | 4347 | 4322 | 4499 | 4630 | 3717,53 | 2986 | 4450 | 1,493 | 0,578 | 11,858 | 1,95E-32 | 5,49E-32 | 0,0017516 | 0,0025329 | 0,0019901 | 0,0019901 | TRUE | 0,0283123 | No |
| orf19.1865 | 948 | 845 | 882 | 541 | 1384 | 946 | 1376 | 1363 | 904 | 893 | 871 | 848 | 1109 | 1076 | 1168 | 1108 | 997,16 | 879 | 1115 | 1,268 | 0,342 | 6,395 | 1,60E-10 | 2,85E-10 | 1,00E-08 | 0,0037379 | 0,0017123 | 0,0017123 | TRUE | 0,0121741 | No |
| orf19.1866 | 436 | 317 | 340 | 236 | 1487 | 914 | 1419 | 1432 | 416 | 335 | 336 | 370 | 1191 | 1039 | 1205 | 1164 | 757,04 | 364 | 1150 | 3,161 | 1,66 | 19,924 | 2,52E-88 | 1,48E-87 | 0,0051895 | 0,0042601 | 0,0047437 | 0,0047437 | TRUE | 0,1153888 | No |
| orf19.1867 | 619 | 529 | 551 | 394 | 704 | 594 | 761 | 811 | 590 | 559 | 544 | 617 | 564 | 675 | 646 | 659 | 607,01 | 578 | 636 | 1,102 | 0,14 | 1,831 | 0,0671232 | 0,079805 | 0,0032302 | 0,0047962 | 0,003973 | 0,003973 | TRUE | 0,0708423 | No |
| orf19.1868 | 7428 | 6458 | 7075 | 4361 | 1608 | 1062 | 1505 | 1650 | 7082 | 6829 | 6989 | 6832 | 1288 | 1208 | 1278 | 1342 | 4105,86 | 6933 | 1279 | 0,185 | -2,436 | -58,833 | 0 | 0 | 0,0003103 | 0,0024912 | 0,0012177 | 0,0012177 | TRUE | 0,0153239 | No |
| orf19.1871 | 3420 | 2924 | 3239 | 2041 | 3349 | 2356 | 3148 | 3467 | 3261 | 3092 | 3200 | 3197 | 2683 | 2679 | 2673 | 2819 | 2950,45 | 3188 | 2714 | 0,852 | -0,232 | -5,974 | 2,32E-09 | 3,93E-09 | 0,0002899 | 0,0026478 | 0,001106 | 0,001106 | TRUE | 0,0085067 | No |
| orf19.1872 | 3379 | 2693 | 2876 | 2111 | 8928 | 6160 | 8575 | 9090 | 3222 | 2848 | 2841 | 3307 | 7154 | 7004 | 7280 | 7391 | 5130,83 | 3054 | 7207 | 2,362 | 1,24 | 22,206 | 3,03E-109 | 2,18E-108 | 0,0031713 | 0,0024113 | 0,0027494 | 0,0027494 | TRUE | 0,0358603 | No |
| orf19.1873 | 1242 | 1056 | 1152 | 807 | 4570 | 3437 | 4367 | 4412 | 1184 | 1117 | 1138 | 1264 | 3662 | 3908 | 3708 | 3587 | 2445,99 | 1176 | 3716 | 3,168 | 1,663 | 32,556 | 1,72E-232 | 2,70E-231 | 0,0014803 | 0,0027625 | 0,0019082 | 0,0019082 | TRUE | 0,0268841 | No |
| orf19.1874 | 4 | 2 | 3 | 0 | 9 | 4 | 13 | 11 | 4 | 2 | 3 | 0 | 7 | 5 | 11 | 9 | 5,08 | 2 | 8 | 3,337 | 1,739 | 2,454 | 0,0141471 | 0,0177842 | 1,00E-08 | 0,3253577 | 0,1964403 | 0,1964403 | TRUE | 0,187086 | No |
| orf19.1876 | 1881 | 1508 | 1691 | 1213 | 1912 | 1380 | 1944 | 2030 | 1793 | 1595 | 1671 | 1900 | 1532 | 1569 | 1650 | 1651 | 1670,11 | 1740 | 1600 | 0,922 | -0,117 | -1,919 | 0,0549524 | 0,0657389 | 0,0028962 | 0,0030744 | 0,002991 | 0,002991 | TRUE | 0,0432433 | No |
| orf19.1877 | 632 | 583 | 644 | 402 | 294 | 249 | 338 | 341 | 603 | 616 | 636 | 630 | 236 | 283 | 287 | 277 | 445,99 | 621 | 271 | 0,435 | -1,201 | -14,956 | 1,42E-50 | 5,29E-50 | 0,0006099 | 0,0057728 | 0,0036401 | 0,0036401 | TRUE | 0,0917714 | No |
| orf19.1878 | 1323 | 1072 | 1233 | 786 | 1220 | 739 | 1197 | 1207 | 1261 | 1134 | 1218 | 1231 | 978 | 840 | 1016 | 981 | 1082,47 | 1211 | 954 | 0,789 | -0,341 | -5,045 | 4,53E-07 | 7,11E-07 | 0,0030901 | 0,0036081 | 0,0034592 | 0,0034592 | TRUE | 0,0719426 | No |
| orf19.188 | 2284 | 2117 | 2236 | 1384 | 1797 | 1225 | 1674 | 1694 | 2178 | 2238 | 2209 | 2168 | 1440 | 1393 | 1421 | 1377 | 1803,06 | 2198 | 1408 | 0,64 | -0,643 | -15,564 | 1,28E-54 | 5,00E-54 | 1,00E-08 | 0,0030019 | 0,0010672 | 0,0010672 | TRUE | 0,0029382 | No |
| orf19.1880 | 3800 | 3321 | 3640 | 2206 | 5364 | 3594 | 5127 | 5403 | 3623 | 3512 | 3596 | 3456 | 4298 | 4087 | 4353 | 4393 | 3914,61 | 3547 | 4283 | 1,207 | 0,272 | 7,05 | 1,79E-12 | 3,36E-12 | 0,000398 | 0,0025107 | 0,0011653 | 0,0011653 | TRUE | 0,0110862 | No |
| orf19.1881 | 1402 | 1276 | 1230 | 835 | 939 | 643 | 847 | 931 | 1337 | 1349 | 1215 | 1308 | 752 | 731 | 719 | 757 | 1021,09 | 1302 | 740 | 0,569 | -0,814 | -14,481 | 1,59E-47 | 5,67E-47 | 1,00E-08 | 0,0036993 | 0,0020043 | 0,0020043 | TRUE | 0,0258143 | No |
| orf19.1883 | 1956 | 1663 | 1853 | 1157 | 1738 | 1206 | 1603 | 1810 | 1865 | 1758 | 1831 | 1813 | 1393 | 1371 | 1361 | 1472 | 1607,86 | 1817 | 1399 | 0,77 | -0,376 | -8,024 | 1,03E-15 | 2,10E-15 | 0,0003835 | 0,0031125 | 0,0014838 | 0,0014838 | TRUE | 0,015055 | No |
| orf19.1885 | 3025 | 2545 | 2838 | 1638 | 2711 | 1853 | 2773 | 2762 | 2884 | 2691 | 2804 | 2566 | 2172 | 2107 | 2354 | 2246 | 2478 | 2736 | 2220 | 0,811 | -0,303 | -5,741 | 9,44E-09 | 1,57E-08 | 0,001875 | 0,0027539 | 0,0022639 | 0,0022639 | TRUE | 0,0200516 | No |
| orf19.1886 | 6333 | 5347 | 5799 | 3325 | 8201 | 5397 | 7894 | 8286 | 6038 | 5654 | 5729 | 5209 | 6571 | 6137 | 6702 | 6737 | 6097,08 | 5658 | 6537 | 1,155 | 0,208 | 3,973 | 7,09E-05 | 0,0001011 | 0,0025002 | 0,0023606 | 0,0024576 | 0,0024576 | TRUE | 0,0337126 | No |
| orf19.1887 | 1359 | 1146 | 1254 | 768 | 1471 | 976 | 1385 | 1548 | 1296 | 1212 | 1239 | 1203 | 1179 | 1110 | 1176 | 1259 | 1209,04 | 1238 | 1181 | 0,954 | -0,067 | -1,217 | 0,2235805 | 0,2523735 | 0,0010799 | 0,0034493 | 0,0021048 | 0,0021048 | TRUE | 0,0236861 | No |
| orf19.1888 | 1198 | 1028 | 1091 | 703 | 1046 | 689 | 1049 | 1070 | 1142 | 1087 | 1078 | 1101 | 838 | 783 | 891 | 870 | 973,8 | 1102 | 846 | 0,769 | -0,38 | -6,622 | 3,53E-11 | 6,39E-11 | 0,0006669 | 0,0037774 | 0,002123 | 0,002123 | TRUE | 0,026383 | No |
| orf19.1889 | 528 | 476 | 442 | 316 | 1246 | 845 | 1232 | 1369 | 503 | 503 | 437 | 495 | 998 | 961 | 1046 | 1113 | 757,09 | 484 | 1030 | 2,129 | 1,09 | 14,871 | 5,12E-50 | 1,89E-49 | 0,0029563 | 0,00426 | 0,003573 | 0,003573 | TRUE | 0,0549746 | No |
| orf19.1890 | 4080 | 3460 | 3845 | 2680 | 2895 | 2107 | 2767 | 3016 | 3890 | 3659 | 3798 | 4198 | 2320 | 2396 | 2349 | 2452 | 3132,78 | 3886 | 2379 | 0,613 | -0,706 | -14,29 | 2,54E-46 | 8,91E-46 | 0,001577 | 0,0026154 | 0,0020134 | 0,0020134 | TRUE | 0,0338318 | No |
| orf19.1891 | 12297 | 9732 | 10461 | 6787 | 20898 | 13933 | 19973 | 21790 | 11724 | 10290 | 10334 | 10632 | 16745 | 15843 | 16957 | 17717 | 13780,41 | 10745 | 16816 | 1,565 | 0,646 | 12,198 | 3,19E-34 | 9,20E-34 | 0,0028929 | 0,0022104 | 0,0026162 | 0,0026162 | TRUE | 0,0463569 | No |
| orf19.1893 | 1426 | 1209 | 1309 | 803 | 1766 | 1210 | 1718 | 1767 | 1360 | 1278 | 1293 | 1258 | 1415 | 1376 | 1459 | 1437 | 1359,41 | 1297 | 1422 | 1,095 | 0,131 | 2,721 | 0,0064997 | 0,0083383 | 1,00E-08 | 0,0032991 | 0,0014747 | 0,0014747 | TRUE | 0,0129332 | No |
| orf19.1896 | 22154 | 18958 | 20649 | 13991 | 29284 | 21121 | 28055 | 30728 | 21121 | 20046 | 20399 | 21918 | 23464 | 24016 | 23819 | 24985 | 22471,04 | 20871 | 24071 | 1,154 | 0,206 | 5,284 | 1,26E-07 | 2,02E-07 | 0,0010945 | 0,0021643 | 0,0014157 | 0,0014157 | TRUE | 0,0137477 | No |
| orf19.1897 | 3151 | 2661 | 2890 | 1827 | 2259 | 1432 | 2168 | 2321 | 3004 | 2814 | 2855 | 2862 | 1810 | 1628 | 1841 | 1887 | 2337,64 | 2884 | 1792 | 0,622 | -0,685 | -12,974 | 1,73E-38 | 5,35E-38 | 0,001813 | 0,0027937 | 0,0022329 | 0,0022329 | TRUE | 0,0431488 | No |
| orf19.190 | 502 | 570 | 499 | 294 | 484 | 297 | 415 | 325 | 479 | 603 | 493 | 461 | 388 | 338 | 352 | 264 | 422,12 | 509 | 336 | 0,659 | -0,602 | -4,93 | 8,21E-07 | 1,27E-06 | 0,0179282 | 0,0059809 | 0,0118975 | 0,0118975 | TRUE | 0,2430771 | No |
| orf19.1900 | 3124 | 2732 | 2941 | 1864 | 3097 | 2023 | 3033 | 3096 | 2978 | 2889 | 2905 | 2920 | 2482 | 2300 | 2575 | 2517 | 2695,86 | 2923 | 2468 | 0,845 | -0,242 | -5,447 | 5,14E-08 | 8,35E-08 | 0,0007444 | 0,0027003 | 0,0015296 | 0,0015296 | TRUE | 0,0241684 | No |
| orf19.1901 | 9793 | 8687 | 9435 | 6000 | 12809 | 8993 | 11953 | 13122 | 9337 | 9185 | 9321 | 9399 | 10263 | 10226 | 10148 | 10669 | 9818,62 | 9310 | 10326 | 1,109 | 0,15 | 5,006 | 5,56E-07 | 8,69E-07 | 0,0001662 | 0,0022585 | 0,0007559 | 0,0007559 | TRUE | 0,0061678 | No |
| orf19.1902 | 10423 | 9557 | 10461 | 6067 | 9730 | 6937 | 9310 | 9733 | 9937 | 10105 | 10334 | 9504 | 7796 | 7888 | 7904 | 7914 | 8922,96 | 9970 | 7876 | 0,789 | -0,341 | -9,885 | 4,85E-23 | 1,16E-22 | 0,0004908 | 0,0022753 | 0,0010302 | 0,0010302 | TRUE | 0,0116327 | No |
| orf19.1906 | 78 | 81 | 68 | 55 | 136 | 92 | 108 | 117 | 74 | 86 | 67 | 86 | 109 | 105 | 92 | 95 | 89,22 | 78 | 100 | 1,286 | 0,363 | 2,332 | 0,0197169 | 0,0244821 | 1,00E-08 | 0,0204942 | 0,0116324 | 0,0116324 | TRUE | 0,0871674 | No |
| orf19.1907 | 4181 | 3427 | 4085 | 2558 | 8750 | 5682 | 7998 | 8800 | 3986 | 3624 | 4036 | 4007 | 7011 | 6461 | 6790 | 7155 | 5383,77 | 3913 | 6854 | 1,753 | 0,809 | 16,287 | 1,22E-59 | 5,12E-59 | 0,0019342 | 0,0023963 | 0,0021627 | 0,0021627 | TRUE | 0,0304701 | No |
| orf19.191 | 4539 | 4218 | 4559 | 2940 | 2605 | 1965 | 2573 | 2638 | 4327 | 4460 | 4504 | 4606 | 2087 | 2234 | 2184 | 2145 | 3318,51 | 4474 | 2162 | 0,483 | -1,049 | -26,479 | 1,71E-154 | 1,77E-153 | 0,00039 | 0,0025861 | 0,0011744 | 0,0011744 | TRUE | 0,0067099 | No |
| orf19.1910 | 648 | 477 | 585 | 381 | 1317 | 791 | 1193 | 1276 | 618 | 504 | 578 | 597 | 1055 | 899 | 1013 | 1038 | 787,75 | 574 | 1001 | 1,747 | 0,805 | 10,273 | 9,33E-25 | 2,30E-24 | 0,0047592 | 0,0041756 | 0,0044751 | 0,0044751 | TRUE | 0,0803861 | No |
| orf19.1911 | 4663 | 3646 | 3960 | 2731 | 7697 | 5123 | 7376 | 8102 | 4446 | 3855 | 3912 | 4278 | 6167 | 5825 | 6262 | 6588 | 5166,72 | 4123 | 6210 | 1,507 | 0,592 | 10,197 | 2,04E-24 | 4,98E-24 | 0,003403 | 0,0024091 | 0,0030266 | 0,0030266 | TRUE | 0,0348989 | No |
| orf19.1913 | 875 | 582 | 681 | 570 | 1021 | 691 | 927 | 1074 | 834 | 615 | 673 | 893 | 818 | 786 | 787 | 873 | 784,93 | 754 | 816 | 1,086 | 0,119 | 1,124 | 0,2610535 | 0,2915775 | 0,0147031 | 0,0041831 | 0,0094198 | 0,0094198 | TRUE | 0,179575 | No |
| orf19.1914 | 543 | 417 | 458 | 335 | 1546 | 1047 | 1379 | 1499 | 518 | 441 | 452 | 525 | 1239 | 1191 | 1171 | 1219 | 844,35 | 484 | 1205 | 2,499 | 1,321 | 19,545 | 4,59E-85 | 2,61E-84 | 0,0013206 | 0,0040359 | 0,0028584 | 0,0028584 | TRUE | 0,040438 | No |
| orf19.1915 | 6484 | 4823 | 5370 | 3078 | 15905 | 9133 | 14809 | 15323 | 6182 | 5100 | 5305 | 4822 | 12744 | 10385 | 12573 | 12459 | 8696,19 | 5352 | 12040 | 2,249 | 1,169 | 14,075 | 5,44E-45 | 1,86E-44 | 0,010092 | 0,0022801 | 0,0064877 | 0,0064877 | TRUE | 0,1331658 | No |
| orf19.1917 | 3139 | 2434 | 2834 | 1814 | 4023 | 2630 | 3812 | 4040 | 2993 | 2574 | 2800 | 2842 | 3224 | 2991 | 3236 | 3285 | 2992,89 | 2802 | 3184 | 1,137 | 0,185 | 3,39 | 0,0006994 | 0,000954 | 0,0023188 | 0,0026399 | 0,0025213 | 0,0025213 | TRUE | 0,0371243 | No |
| orf19.192 | 1765 | 1785 | 1882 | 1271 | 1582 | 1136 | 1393 | 1339 | 1683 | 1887 | 1859 | 1991 | 1268 | 1292 | 1183 | 1089 | 1531,4 | 1855 | 1208 | 0,651 | -0,619 | -8,849 | 8,85E-19 | 1,93E-18 | 0,0043805 | 0,0031634 | 0,004019 | 0,004019 | TRUE | 0,053431 | No |
| orf19.1920 | 32 | 12 | 18 | 10 | 4 | 6 | 9 | 3 | 31 | 13 | 18 | 16 | 3 | 7 | 8 | 2 | 12,09 | 20 | 5 | 0,254 | -1,976 | -3,918 | 8,93E-05 | 0,0001268 | 0,1073056 | 0,1378521 | 0,1251259 | 0,1251259 | TRUE | 0,8986598 | No |
| orf19.1923 | 2683 | 2023 | 2429 | 1775 | 3623 | 2288 | 3366 | 3472 | 2558 | 2139 | 2400 | 2781 | 2903 | 2602 | 2858 | 2823 | 2632,84 | 2470 | 2796 | 1,134 | 0,182 | 2,453 | 0,0141805 | 0,0178227 | 0,0066151 | 0,0027149 | 0,0048796 | 0,0048796 | TRUE | 0,0978355 | No |
| orf19.1925 | 37 | 37 | 32 | 28 | 91 | 57 | 93 | 88 | 35 | 39 | 32 | 44 | 73 | 65 | 79 | 72 | 54,76 | 38 | 72 | 1,955 | 0,967 | 4,876 | 1,08E-06 | 1,67E-06 | 1,00E-08 | 0,0320726 | 0,0166115 | 0,0166115 | TRUE | 0,0795697 | No |
| orf19.1926 | 247 | 234 | 217 | 189 | 918 | 715 | 760 | 795 | 235 | 247 | 214 | 296 | 736 | 813 | 645 | 646 | 479,2 | 248 | 710 | 2,878 | 1,525 | 14,006 | 1,44E-44 | 4,89E-44 | 0,012549 | 0,0055176 | 0,0084802 | 0,0084802 | TRUE | 0,1586946 | No |
| orf19.1927 | 1726 | 1644 | 1713 | 1062 | 1798 | 1190 | 1692 | 1718 | 1646 | 1738 | 1692 | 1664 | 1441 | 1353 | 1436 | 1397 | 1545,88 | 1685 | 1407 | 0,836 | -0,259 | -5,691 | 1,26E-08 | 2,09E-08 | 1,00E-08 | 0,0031534 | 0,0013425 | 0,0013425 | TRUE | 0,0072676 | No |
| orf19.193 | 433 | 509 | 497 | 287 | 280 | 206 | 233 | 204 | 413 | 538 | 491 | 450 | 224 | 234 | 198 | 166 | 339,24 | 473 | 206 | 0,433 | -1,207 | -10,047 | 9,45E-24 | 2,28E-23 | 0,0144051 | 0,0069313 | 0,0105001 | 0,0105001 | TRUE | 0,1917761 | No |
| orf19.1930 | 147 | 141 | 144 | 86 | 17 | 26 | 15 | 24 | 140 | 149 | 142 | 135 | 14 | 30 | 13 | 20 | 80,21 | 142 | 19 | 0,129 | -2,957 | -13,678 | 1,38E-42 | 4,55E-42 | 1,00E-08 | 0,0225621 | 0,0166822 | 0,0166822 | TRUE | 0,6153218 | No |
| orf19.1932 | 1051 | 867 | 910 | 655 | 643 | 456 | 616 | 697 | 1002 | 917 | 899 | 1026 | 515 | 519 | 523 | 567 | 745,91 | 961 | 531 | 0,554 | -0,852 | -12,526 | 5,41E-36 | 1,60E-35 | 0,0019081 | 0,0042925 | 0,0030206 | 0,0030206 | TRUE | 0,0252086 | No |
| orf19.1933 | 1685 | 1569 | 1660 | 997 | 1733 | 1221 | 1636 | 1750 | 1606 | 1659 | 1640 | 1562 | 1389 | 1388 | 1389 | 1423 | 1507,02 | 1617 | 1397 | 0,863 | -0,212 | -4,836 | 1,32E-06 | 2,04E-06 | 1,00E-08 | 0,0031808 | 0,0011835 | 0,0011835 | TRUE | 0,0054103 | No |
| orf19.1934 | 2698 | 2438 | 2704 | 1821 | 2753 | 1940 | 2855 | 3064 | 2572 | 2578 | 2671 | 2853 | 2206 | 2206 | 2424 | 2491 | 2500,15 | 2668 | 2332 | 0,875 | -0,192 | -3,359 | 0,000781 | 0,0010613 | 0,0026789 | 0,002748 | 0,0027426 | 0,0027426 | TRUE | 0,0258752 | No |
| orf19.1935 | 11 | 13 | 10 | 7 | 10 | 9 | 9 | 12 | 10 | 14 | 10 | 11 | 8 | 10 | 8 | 10 | 10,09 | 11 | 9 | 0,786 | -0,347 | -0,809 | 0,4187762 | 0,4546896 | 1,00E-08 | 0,1648116 | 0,0770347 | 0,0770347 | TRUE | 0,0848575 | No |
| orf19.1936 | 8020 | 6740 | 7596 | 4880 | 5097 | 3726 | 5090 | 5433 | 7646 | 7127 | 7504 | 7645 | 4084 | 4237 | 4321 | 4418 | 5872,7 | 7480 | 4265 | 0,57 | -0,81 | -19,906 | 3,64E-88 | 2,14E-87 | 0,0008408 | 0,0023709 | 0,0014111 | 0,0014111 | TRUE | 0,012432 | No |
| orf19.1938 | 1076 | 963 | 1037 | 687 | 1740 | 1100 | 1657 | 1748 | 1026 | 1018 | 1024 | 1076 | 1394 | 1251 | 1407 | 1421 | 1202,23 | 1036 | 1368 | 1,325 | 0,406 | 7,175 | 7,21E-13 | 1,37E-12 | 0,0012269 | 0,003457 | 0,0021994 | 0,0021994 | TRUE | 0,0376523 | No |
| orf19.1939 | 20 | 18 | 19 | 8 | 24 | 15 | 31 | 34 | 19 | 19 | 19 | 13 | 19 | 17 | 26 | 28 | 19,96 | 18 | 22 | 1,289 | 0,366 | 1,135 | 0,2565578 | 0,2868574 | 1,00E-08 | 0,0843643 | 0,0488166 | 0,0488166 | TRUE | 0,1455584 | No |
| orf19.194 | 1356 | 1648 | 1508 | 982 | 1669 | 1264 | 1468 | 1263 | 1293 | 1743 | 1490 | 1538 | 1337 | 1437 | 1246 | 1027 | 1388,92 | 1516 | 1262 | 0,832 | -0,265 | -2,488 | 0,0128358 | 0,0161899 | 0,0160715 | 0,0032734 | 0,0101735 | 0,0101735 | TRUE | 0,190308 | No |
| orf19.1940 | 2851 | 2350 | 2646 | 1669 | 2719 | 1859 | 2573 | 2721 | 2718 | 2485 | 2614 | 2615 | 2179 | 2114 | 2184 | 2212 | 2390,12 | 2608 | 2172 | 0,833 | -0,263 | -6,164 | 7,09E-10 | 1,22E-09 | 0,0004733 | 0,0027782 | 0,0013286 | 0,0013286 | TRUE | 0,0126693 | No |
| orf19.1941 | 935 | 794 | 849 | 493 | 686 | 521 | 665 | 643 | 891 | 840 | 839 | 772 | 550 | 592 | 565 | 523 | 696,44 | 836 | 558 | 0,664 | -0,591 | -8,771 | 1,77E-18 | 3,83E-18 | 0,0013823 | 0,0044488 | 0,0028914 | 0,0028914 | TRUE | 0,0269908 | No |
| orf19.1942 | 3049 | 2818 | 3009 | 1970 | 4623 | 3341 | 4313 | 4571 | 2907 | 2980 | 2973 | 3086 | 3704 | 3799 | 3662 | 3717 | 3353,37 | 2986 | 3720 | 1,247 | 0,318 | 8,859 | 8,05E-19 | 1,76E-18 | 2,51E-05 | 0,0025809 | 0,0009313 | 0,0009313 | TRUE | 0,0055412 | No |

| | | | | | | | | | | | | | | | | | | | | | | | | | | | | | | | |
|---|---|---|---|---|---|---|---|---|---|---|---|---|---|---|---|---|---|---|---|---|---|---|---|---|---|---|---|---|---|---|---|
| orf19.1943 | 431 | 446 | 430 | 296 | 623 | 495 | 564 | 582 | 411 | 472 | 425 | 464 | 499 | 563 | 479 | 473 | 473,14 | 443 | 504 | 1,137 | 0,185 | 2,242 | 0,0249586 | 0,0307512 | 0,0032493 | 0,0055615 | 0,0044078 | 0,0044078 | TRUE | 0,0680259 | No |
| orf19.1944 | 20649 | 17719 | 20142 | 12906 | 3790 | 2716 | 3668 | 3896 | 19687 | 18736 | 19898 | 20218 | 3037 | 3088 | 3114 | 3168 | 11368,2 | 19635 | 3102 | 0,158 | -2,662 | -72,486 | 0 | 0 | 0,0006538 | 0,0022357 | 0,0011225 | 0,0011225 | TRUE | 0,0116282 | No |
| orf19.1945 | 19365 | 16882 | 18378 | 11219 | 12278 | 8321 | 11285 | 12160 | 18462 | 17851 | 18156 | 17575 | 9838 | 9462 | 9581 | 9887 | 13851,47 | 18011 | 9692 | 0,538 | -0,894 | -28,607 | 5,50E-180 | 6,60E-179 | 0,0003419 | 0,0022098 | 0,0008606 | 0,0008606 | TRUE | 0,003485 | No |
| orf19.1946 | 2556 | 2191 | 2292 | 1374 | 2777 | 1968 | 2569 | 2818 | 2437 | 2317 | 2264 | 2152 | 2225 | 2238 | 2181 | 2291 | 2263,2 | 2292 | 2234 | 0,973 | -0,04 | -0,845 | 0,3980535 | 0,4338137 | 0,0009439 | 0,0028168 | 0,0016998 | 0,0016998 | TRUE | 0,0217917 | No |
| orf19.1948 | 1151 | 939 | 1037 | 733 | 1174 | 765 | 1199 | 1251 | 1097 | 993 | 1024 | 1148 | 941 | 870 | 1018 | 1017 | 1013,58 | 1066 | 962 | 0,905 | -0,144 | -2,051 | 0,0402622 | 0,0488242 | 0,003582 | 0,0037112 | 0,0037294 | 0,0037294 | TRUE | 0,0462056 | No |
| orf19.1949 | 4098 | 3521 | 3831 | 2389 | 5414 | 3714 | 5202 | 5812 | 3907 | 3723 | 3785 | 3742 | 4338 | 4223 | 4416 | 4726 | 4107,57 | 3789 | 4426 | 1,168 | 0,224 | 5,032 | 4,86E-07 | 7,62E-07 | 0,0010918 | 0,002491 | 0,0016571 | 0,0016571 | TRUE | 0,0254726 | No |
| orf19.1950 | 583 | 442 | 497 | 389 | 407 | 273 | 344 | 451 | 556 | 467 | 491 | 609 | 326 | 310 | 292 | 367 | 427,36 | 531 | 324 | 0,613 | -0,707 | -6,869 | 6,44E-12 | 1,19E-11 | 0,0095094 | 0,0059333 | 0,0077171 | 0,0077171 | TRUE | 0,1044914 | No |
| orf19.1953 | 3206 | 2721 | 2961 | 1953 | 2887 | 1940 | 2791 | 2901 | 3057 | 2877 | 2925 | 3059 | 2313 | 2206 | 2370 | 2359 | 2645,74 | 2980 | 2312 | 0,777 | -0,364 | -8,504 | 1,83E-17 | 3,89E-17 | 0,0004939 | 0,0027119 | 0,0013779 | 0,0013779 | TRUE | 0,0108468 | No |
| orf19.1954 | 4939 | 4418 | 4921 | 2826 | 5071 | 3355 | 4939 | 5170 | 4709 | 4671 | 4861 | 4427 | 4063 | 3815 | 4193 | 4204 | 4367,99 | 4667 | 4069 | 0,872 | -0,198 | -4,284 | 1,83E-05 | 2,69E-05 | 0,0014248 | 0,0024672 | 0,0018258 | 0,0018258 | TRUE | 0,0208368 | No |
| orf19.1955 | 887 | 766 | 802 | 541 | 942 | 600 | 861 | 867 | 846 | 810 | 792 | 848 | 755 | 682 | 731 | 705 | 771,05 | 824 | 718 | 0,874 | -0,194 | -3,246 | 0,0011711 | 0,001571 | 1,00E-08 | 0,0042207 | 0,0021319 | 0,0021319 | TRUE | 0,0139821 | No |
| orf19.1956 | 3764 | 3379 | 3537 | 2316 | 4688 | 3236 | 4386 | 4523 | 3589 | 3573 | 3494 | 3628 | 3756 | 3680 | 3724 | 3678 | 3640,14 | 3571 | 3710 | 1,039 | 0,056 | 1,694 | 0,0902411 | 0,1058119 | 1,00E-08 | 0,0025423 | 0,0007671 | 0,0007671 | TRUE | 0,0025748 | No |
| orf19.1957 | 10981 | 8991 | 9929 | 5810 | 10621 | 6667 | 10011 | 10763 | 10469 | 9507 | 9809 | 9102 | 8510 | 7581 | 8499 | 8751 | 9028,57 | 9722 | 8335 | 0,857 | -0,222 | -3,855 | 0,0001159 | 0,0001635 | 0,0035276 | 0,0022731 | 0,0030754 | 0,0030754 | TRUE | 0,0449392 | No |
| orf19.1958 | 13 | 13 | 17 | 13 | 27 | 18 | 25 | 21 | 12 | 14 | 17 | 20 | 22 | 20 | 21 | 17 | 17,96 | 16 | 20 | 1,294 | 0,372 | 1,123 | 0,2612612 | 0,2917584 | 1,00E-08 | 0,0934977 | 0,0471742 | 0,0471742 | TRUE | 0,1084813 | No |
| orf19.1959 | 1320 | 1259 | 1266 | 894 | 4092 | 2844 | 4031 | 4104 | 1258 | 1331 | 1251 | 1400 | 3279 | 3234 | 3422 | 3337 | 2314,1 | 1310 | 3318 | 2,541 | 1,345 | 28,123 | 5,16E-174 | 6,01E-173 | 0,0007061 | 0,0028008 | 0,0016333 | 0,0016333 | TRUE | 0,02285 | No |
| orf19.1960 | 6339 | 5973 | 6402 | 4019 | 4072 | 3003 | 3710 | 4222 | 6044 | 6316 | 6325 | 6296 | 3263 | 3415 | 3150 | 3433 | 4779,99 | 6245 | 3315 | 0,531 | -0,914 | -22,476 | 7,05E-112 | 5,18E-111 | 0,000742 | 0,0024348 | 0,0013626 | 0,0013626 | TRUE | 0,0137314 | No |
| orf19.1961 | 9144 | 8859 | 9615 | 5557 | 7190 | 5183 | 6796 | 7215 | 8718 | 9367 | 9499 | 8705 | 5761 | 5894 | 5770 | 5866 | 7447,51 | 9072 | 5823 | 0,642 | -0,64 | -15,807 | 2,80E-56 | 1,12E-55 | 0,00095 | 0,0023117 | 0,0014375 | 0,0014375 | TRUE | 0,0122709 | No |
| orf19.1963 | 5059 | 4545 | 5046 | 2765 | 12223 | 8145 | 12144 | 12993 | 4823 | 4806 | 4985 | 4332 | 9794 | 9262 | 10310 | 10565 | 7359,47 | 4736 | 9983 | 2,106 | 1,075 | 19,024 | 1,08E-80 | 5,93E-80 | 0,003236 | 0,0023144 | 0,0029017 | 0,0029017 | TRUE | 0,0390752 | No |
| orf19.1964 | 2680 | 2592 | 2703 | 1789 | 5889 | 4429 | 5616 | 5927 | 2555 | 2741 | 2670 | 2803 | 4719 | 5036 | 4768 | 4819 | 3763,84 | 2692 | 4836 | 1,798 | 0,846 | 19,87 | 7,38E-88 | 4,33E-87 | 0,0008154 | 0,0025275 | 0,0014396 | 0,0014396 | TRUE | 0,0144483 | No |
| orf19.1966 | 4567 | 3888 | 4219 | 2598 | 5710 | 3594 | 5406 | 5632 | 4354 | 4111 | 4168 | 4070 | 4575 | 4087 | 4590 | 4579 | 4316,76 | 4176 | 4458 | 1,068 | 0,095 | 1,942 | 0,0521097 | 0,0625611 | 0,0016831 | 0,0024716 | 0,0020417 | 0,0020417 | TRUE | 0,0373046 | No |
| orf19.1967 | 1565 | 1301 | 1449 | 789 | 2608 | 1613 | 2410 | 2586 | 1492 | 1376 | 1431 | 1236 | 2090 | 1834 | 2046 | 2103 | 1700,97 | 1384 | 2018 | 1,456 | 0,542 | 8,063 | 7,47E-16 | 1,53E-15 | 0,0042322 | 0,0030566 | 0,0037132 | 0,0037132 | TRUE | 0,0575425 | No |
| orf19.1968 | 106 | 95 | 117 | 78 | 320 | 231 | 270 | 271 | 101 | 100 | 116 | 122 | 256 | 263 | 229 | 220 | 175,99 | 110 | 242 | 2,216 | 1,148 | 9,385 | 6,30E-21 | 1,44E-20 | 1,00E-08 | 0,0114207 | 0,0074178 | 0,0074178 | TRUE | 0,0423722 | No |
| orf19.1968.1 | 245 | 233 | 210 | 137 | 437 | 293 | 387 | 388 | 234 | 246 | 207 | 215 | 350 | 333 | 329 | 315 | 278,67 | 226 | 332 | 1,468 | 0,554 | 5,972 | 2,35E-09 | 3,98E-09 | 1,00E-08 | 0,0079831 | 0,004459 | 0,004459 | TRUE | 0,0422945 | No |
| orf19.1969 | 50 | 31 | 40 | 34 | 56 | 33 | 47 | 57 | 48 | 33 | 40 | 53 | 45 | 38 | 40 | 46 | 42,73 | 44 | 42 | 0,989 | -0,016 | -0,072 | 0,9424351 | 0,9485297 | 1,00E-08 | 0,0405128 | 0,0244608 | 0,0244608 | TRUE | 0,1982696 | No |
| orf19.1970 | 2499 | 2112 | 2370 | 1632 | 2325 | 1620 | 2161 | 2369 | 2383 | 2233 | 2341 | 2557 | 1863 | 1842 | 1835 | 1926 | 2122,45 | 2378 | 1866 | 0,786 | -0,347 | -6,99 | 2,75E-12 | 5,13E-12 | 0,0012089 | 0,0028649 | 0,0018889 | 0,0018889 | TRUE | 0,0284474 | No |
| orf19.1971 | 12433 | 10958 | 11911 | 7586 | 15739 | 11007 | 14409 | 15429 | 11854 | 11587 | 11767 | 11884 | 12611 | 12516 | 12233 | 12545 | 12124,58 | 11773 | 12476 | 1,06 | 0,084 | 3,152 | 0,0016228 | 0,0021627 | 8,17E-05 | 0,0022267 | 0,000597 | 0,000597 | TRUE | 0,0021275 | No |
| orf19.1972 | 4432 | 3654 | 3903 | 2581 | 6276 | 4158 | 5882 | 6381 | 4225 | 3864 | 3856 | 4043 | 5029 | 4728 | 4994 | 5188 | 4490,89 | 3997 | 4985 | 1,248 | 0,319 | 6,908 | 4,91E-12 | 9,11E-12 | 0,0013934 | 0,0024569 | 0,0018206 | 0,0018206 | TRUE | 0,0187434 | No |
| orf19.1973 | 1248 | 950 | 1152 | 692 | 1561 | 998 | 1495 | 1543 | 1190 | 1005 | 1138 | 1084 | 1251 | 1135 | 1269 | 1255 | 1165,74 | 1104 | 1228 | 1,112 | 0,153 | 2,343 | 0,0191372 | 0,0237808 | 0,0027112 | 0,0034997 | 0,0032156 | 0,0032156 | TRUE | 0,0462609 | No |
| orf19.1974 | 4953 | 4094 | 4594 | 3278 | 5326 | 3940 | 5117 | 5420 | 4722 | 4329 | 4538 | 5135 | 4268 | 4480 | 4344 | 4407 | 4527,95 | 4681 | 4375 | 0,935 | -0,097 | -1,815 | 0,0695859 | 0,0825945 | 0,0025983 | 0,0024539 | 0,002494 | 0,002494 | TRUE | 0,050145 | No |
| orf19.1975 | 588 | 558 | 534 | 298 | 1099 | 762 | 973 | 1004 | 561 | 590 | 528 | 467 | 881 | 866 | 826 | 816 | 691,81 | 536 | 847 | 1,572 | 0,652 | 8,876 | 6,91E-19 | 1,51E-18 | 0,0026258 | 0,0044646 | 0,003618 | 0,003618 | TRUE | 0,0746881 | No |
| orf19.1976 | 451 | 432 | 469 | 270 | 538 | 398 | 465 | 508 | 430 | 457 | 463 | 423 | 431 | 453 | 395 | 413 | 433,07 | 443 | 423 | 0,95 | -0,074 | -0,971 | 0,3316527 | 0,3654423 | 1,00E-08 | 0,0058826 | 0,0032328 | 0,0032328 | TRUE | 0,0230894 | No |
| orf19.1977 | 1387 | 1203 | 1265 | 925 | 3119 | 2264 | 2890 | 3170 | 1322 | 1272 | 1250 | 1449 | 2499 | 2574 | 2454 | 2578 | 1924,72 | 1323 | 2526 | 1,915 | 0,937 | 17,865 | 2,19E-71 | 1,07E-70 | 0,0012555 | 0,0029443 | 0,0020393 | 0,0020393 | TRUE | 0,0438817 | No |
| orf19.1978 | 955 | 873 | 937 | 575 | 2253 | 1520 | 2034 | 2202 | 910 | 923 | 926 | 901 | 1805 | 1728 | 1727 | 1790 | 1338,87 | 915 | 1762 | 1,926 | 0,946 | 20,03 | 3,01E-89 | 1,79E-88 | 1,00E-08 | 0,0033176 | 0,0012782 | 0,0012782 | TRUE | 0,0032047 | No |
| orf19.1979 | 22144 | 20492 | 21615 | 12990 | 6723 | 4619 | 6204 | 6462 | 21112 | 21668 | 21354 | 20350 | 5387 | 5252 | 5267 | 5254 | 13205,41 | 21121 | 5290 | 0,25 | -1,997 | -62,28 | 0 | 0 | 0,0003802 | 0,0022156 | 0,0008767 | 0,0008767 | TRUE | 0,0072242 | No |
| orf19.198 | 5538 | 4894 | 5401 | 3618 | 17617 | 13185 | 17164 | 18944 | 5280 | 5175 | 5336 | 5668 | 14116 | 14993 | 14572 | 15403 | 10067,76 | 5365 | 14771 | 2,755 | 1,462 | 34,118 | 4,02E-255 | 7,01E-254 | 0,0013188 | 0,0022544 | 0,0016288 | 0,0016288 | TRUE | 0,0168928 | No |
| orf19.1980 | 111 | 107 | 110 | 54 | 61 | 37 | 64 | 65 | 106 | 113 | 109 | 85 | 49 | 42 | 54 | 53 | 76,3 | 103 | 50 | 0,48 | -1,06 | -6,203 | 5,53E-10 | 9,60E-10 | 1,00E-08 | 0,0236115 | 0,0137506 | 0,0137506 | TRUE | 0,1078705 | No |
| orf19.1981 | 2014 | 1736 | 1842 | 1073 | 1097 | 675 | 1065 | 1069 | 1920 | 1836 | 1820 | 1681 | 879 | 768 | 904 | 869 | 1334,53 | 1814 | 855 | 0,472 | -1,084 | -17,021 | 5,76E-65 | 2,59E-64 | 0,0028184 | 0,0033216 | 0,003067 | 0,003067 | TRUE | 0,0523364 | No |
| orf19.1985 | 643 | 584 | 680 | 414 | 6355 | 4963 | 5804 | 6541 | 613 | 618 | 672 | 649 | 5092 | 5643 | 4928 | 5318 | 2941,54 | 638 | 5245 | 8,228 | 3,041 | 50,532 | 0 | 0 | 0,0024187 | 0,0026495 | 0,0025139 | 0,0025139 | TRUE | 0,0312908 | No |
| orf19.1986 | 2656 | 2283 | 2489 | 1519 | 6834 | 4650 | 6743 | 7211 | 2532 | 2414 | 2459 | 2380 | 5476 | 5287 | 5725 | 5863 | 4017,01 | 2446 | 5588 | 2,283 | 1,191 | 26,125 | 1,90E-150 | 1,92E-149 | 0,0012468 | 0,0025 | 0,0016864 | 0,0016864 | TRUE | 0,0155891 | No |
| orf19.1988 | 1378 | 1139 | 1245 | 858 | 1459 | 1113 | 1451 | 1481 | 1314 | 1204 | 1230 | 1344 | 1169 | 1266 | 1232 | 1204 | 1245,36 | 1273 | 1218 | 0,957 | -0,063 | -1,156 | 0,2477871 | 0,277781 | 0,0008619 | 0,0034097 | 0,0020573 | 0,0020573 | TRUE | 0,0154063 | No |
| orf19.1989 | 4210 | 3733 | 4028 | 2580 | 7063 | 4904 | 6759 | 6790 | 4014 | 3947 | 3979 | 4042 | 5659 | 5576 | 5738 | 5521 | 4809,61 | 3996 | 5624 | 1,408 | 0,494 | 15,569 | 1,19E-54 | 4,67E-54 | 1,00E-08 | 0,0024327 | 0,0007457 | 0,0007457 | TRUE | 0,0023128 | No |
| orf19.199 | 2665 | 2790 | 2955 | 1650 | 2537 | 1758 | 2159 | 2189 | 2541 | 2950 | 2919 | 2585 | 2033 | 1999 | 1833 | 1780 | 2329,95 | 2749 | 1911 | 0,695 | -0,526 | -7,787 | 6,87E-15 | 1,38E-14 | 0,0045839 | 0,002796 | 0,0039368 | 0,0039368 | TRUE | 0,0327731 | No |
| orf19.1990 | 3063 | 3345 | 3462 | 2059 | 2446 | 1799 | 2276 | 2257 | 2920 | 3537 | 3420 | 3226 | 1960 | 2046 | 1932 | 1835 | 2609,48 | 3276 | 1943 | 0,593 | -0,754 | -11,709 | 1,14E-31 | 3,17E-31 | 0,0038347 | 0,0027205 | 0,0035804 | 0,0035804 | TRUE | 0,0665078 | No |
| orf19.1991 | 3881 | 3731 | 3837 | 2448 | 4035 | 2943 | 3680 | 3907 | 3700 | 3945 | 3791 | 3835 | 3233 | 3346 | 3124 | 3177 | 3518,92 | 3818 | 3220 | 0,843 | -0,246 | -6,232 | 4,59E-10 | 8,02E-10 | 0,0004108 | 0,0025579 | 0,0012099 | 0,0012099 | TRUE | 0,008057 | No |
| orf19.1992 | 3360 | 3456 | 3692 | 2316 | 3181 | 2366 | 3006 | 3051 | 3203 | 3654 | 3647 | 3628 | 2549 | 2690 | 2552 | 2481 | 3050,65 | 3533 | 2568 | 0,727 | -0,46 | -8,567 | 1,06E-17 | 2,27E-17 | 0,0020802 | 0,0026295 | 0,0024391 | 0,0024391 | TRUE | 0,0491315 | No |
| orf19.1993 | 1265 | 1651 | 1531 | 935 | 1904 | 1493 | 1590 | 1259 | 1206 | 1746 | 1512 | 1465 | 1526 | 1698 | 1350 | 1024 | 1440,73 | 1482 | 1400 | 0,944 | -0,084 | -0,444 | 0,6572926 | 0,6893794 | 0,0333941 | 0,0032309 | 0,017382 | 0,0333941 | TRUE | 0,3948917 | No |
| orf19.1994 | 1492 | 1642 | 1744 | 1157 | 1883 | 1464 | 1718 | 1591 | 1422 | 1736 | 1723 | 1813 | 1509 | 1665 | 1459 | 1294 | 1577,47 | 1674 | 1482 | 0,885 | -0,176 | -1,976 | 0,0481824 | 0,0580207 | 0,0097566 | 0,0031321 | 0,0069463 | 0,0069463 | TRUE | 0,1247539 | No |
| orf19.1995 | 544 | 543 | 525 | 302 | 747 | 580 | 658 | 686 | 519 | 574 | 519 | 473 | 599 | 660 | 559 | 558 | 557,38 | 521 | 594 | 1,133 | 0,18 | 2,213 | 0,0269299 | 0,0331098 | 0,0039976 | 0,005037 | 0,0045542 | 0,0045542 | TRUE | 0,0613616 | No |
| orf19.1996 | 7192 | 6588 | 7097 | 4291 | 7157 | 5255 | 6853 | 7140 | 6857 | 6966 | 7011 | 6722 | 5735 | 5975 | 5818 | 5806 | 6361,22 | 6889 | 5834 | 0,846 | -0,241 | -7,677 | 1,63E-14 | 3,25E-14 | 0,0001151 | 0,0023494 | 0,0007891 | 0,0007891 | TRUE | 0,003143 | No |
| orf19.1998 | 367 | 319 | 337 | 221 | 346 | 259 | 349 | 346 | 350 | 337 | 333 | 346 | 277 | 295 | 296 | 281 | 314,46 | 342 | 287 | 0,841 | -0,25 | -3,082 | 0,0020551 | 0,0027207 | 1,00E-08 | 0,0073126 | 0,0031585 | 0,0031585 | TRUE | 0,0065208 | No |
| orf19.1999 | 1911 | 1601 | 1723 | 1055 | 877 | 585 | 847 | 958 | 1822 | 1693 | 1702 | 1653 | 703 | 665 | 719 | 779 | 1216,95 | 1718 | 716 | 0,418 | -1,26 | -20,755 | 1,12E-95 | 7,00E-95 | 0,0017668 | 0,0034405 | 0,0025901 | 0,0025901 | TRUE | 0,0410715 | No |
| orf19.20 | 2552 | 2444 | 2486 | 1629 | 2381 | 1721 | 2151 | 2433 | 2433 | 2584 | 2456 | 2552 | 1908 | 1957 | 1826 | 1978 | 2211,79 | 2506 | 1917 | 0,765 | -0,386 | -8,621 | 6,61E-18 | 1,42E-17 | 0,0006291 | 0,0028336 | 0,0014665 | 0,0014665 | TRUE | 0,0125104 | No |
| orf19.200 | 1072 | 998 | 1118 | 599 | 2661 | 1733 | 2694 | 2614 | 1022 | 1055 | 1104 | 938 | 2132 | 1971 | 2287 | 2125 | 1579,44 | 1030 | 2129 | 2,063 | 1,045 | 16,35 | 4,32E-60 | 1,82E-59 | 0,0031844 | 0,0031308 | 0,0031651 | 0,0031651 | TRUE | 0,0382997 | No |
| orf19.2000 | 3654 | 3792 | 4004 | 2245 | 3027 | 2158 | 2728 | 2961 | 3484 | 4010 | 3956 | 3517 | 2425 | 2454 | 2316 | 2408 | 3071,08 | 3742 | 2401 | 0,641 | -0,641 | -11,31 | 1,17E-29 | 3,14E-29 | 0,0026451 | 0,0026259 | 0,0027499 | 0,0027499 | TRUE | 0,0281681 | No |
| orf19.2001 | 607 | 678 | 702 | 388 | 596 | 435 | 511 | 485 | 579 | 717 | 694 | 608 | 478 | 495 | 434 | 394 | 549,66 | 650 | 450 | 0,691 | -0,533 | -5,58 | 2,40E-08 | 3,95E-08 | 0,0086008 | 0,0050784 | 0,0068884 | 0,0068884 | TRUE | 0,0822842 | No |
| orf19.2002 | 3030 | 3061 | 3031 | 2009 | 4592 | 3456 | 4265 | 4282 | 2889 | 3237 | 2994 | 3147 | 3679 | 3930 | 3621 | 3482 | 3372,35 | 3067 | 3678 | 1,199 | 0,262 | 4,959 | 7,09E-07 | 1,10E-06 | 0,0021777 | 0,0025782 | 0,0023814 | 0,0023814 | TRUE | 0,0252461 | No |
| orf19.2003 | 8360 | 7676 | 8230 | 4854 | 9069 | 6370 | 8535 | 8840 | 7970 | 8116 | 8130 | 7604 | 7267 | 7243 | 7246 | 7188 | 7595,65 | 7955 | 7236 | 0,909 | -0,138 | -4,209 | 2,57E-05 | 3,73E-05 | 0,0003096 | 0,0023074 | 0,0008963 | 0,0008963 | TRUE | 0,0102165 | No |
| orf19.2005 | 18724 | 16446 | 17945 | 11023 | 9214 | 6314 | 8496 | 9305 | 17851 | 17390 | 17728 | 17268 | 7383 | 7180 | 7213 | 7566 | 12447,3 | 17559 | 7336 | 0,418 | -1,259 | -40,636 | 0 | 0 | 0,0002871 | 0,0022232 | 0,0008291 | 0,0008291 | TRUE | 0,0054858 | No |
| orf19.2006 | 908 | 867 | 935 | 566 | 1478 | 1039 | 1374 | 1438 | 866 | 917 | 924 | 887 | 1184 | 1181 | 1167 | 1169 | 1036,78 | 898 | 1175 | 1,308 | 0,387 | 7,701 | 1,35E-14 | 2,69E-14 | 1,00E-08 | 0,0036749 | 0,0014281 | 0,0014281 | TRUE | 0,008167 | No |
| orf19.2006.1 | 2255 | 1972 | 1955 | 1188 | 1057 | 760 | 943 | 998 | 2150 | 2085 | 1931 | 1861 | 847 | 864 | 801 | 811 | 1418,83 | 2007 | 831 | 0,413 | -1,276 | -21,138 | 3,54E-99 | 2,30E-98 | 0,0022633 | 0,0032485 | 0,0026802 | 0,0026802 | TRUE | 0,0278907 | No |
| orf19.2007 | 536 | 526 | 510 | 317 | 983 | 705 | 906 | 960 | 511 | 556 | 504 | 497 | 788 | 802 | 769 | 781 | 650,84 | 517 | 785 | 1,515 | 0,599 | 9,546 | 1,34E-21 | 3,11E-21 | 1,00E-08 | 0,004614 | 0,002139 | 0,002139 | TRUE | 0,0305366 | No |
| orf19.2008 | 1670 | 1670 | 1786 | 1042 | 2280 | 1748 | 2018 | 2233 | 1592 | 1766 | 1764 | 1632 | 1827 | 1988 | 1713 | 1816 | 1762,27 | 1688 | 1836 | 1,086 | 0,118 | 1,983 | 0,0473881 | 0,0571181 | 0,0025612 | 0,003023 | 0,002845 | 0,002845 | TRUE | 0,0359869 | No |
| orf19.2009 | 1934 | 1752 | 1864 | 1136 | 1871 | 1416 | 1778 | 1801 | 1844 | 1853 | 1841 | 1780 | 1499 | 1610 | 1510 | 1464 | 1675,08 | 1830 | 1521 | 0,829 | -0,27 | -5,845 | 5,06E-09 | 8,49E-09 | 0,0002121 | 0,0030715 | 0,0014475 | 0,0014475 | TRUE | 0,0175112 | No |
| orf19.201 | 262 | 213 | 254 | 130 | 321 | 213 | 304 | 346 | 250 | 225 | 251 | 204 | 257 | 242 | 258 | 281 | 246,05 | 232 | 260 | 1,112 | 0,153 | 1,525 | 0,1271703 | 0,1469528 | 0,0014203 | 0,0087643 | 0,005592 | 0,005592 | TRUE | 0,061252 | No |
| orf19.2010 | 14 | 20 | 23 | 9 | 18 | 19 | 11 | 27 | 13 | 21 | 23 | 14 | 14 | 22 | 9 | 22 | 17,33 | 18 | 17 | 0,929 | -0,107 | -0,283 | 0,7768891 | 0,8007316 | 0,0541314 | 0,0968375 | 0,0786681 | 0,0786681 | TRUE | 0,4908212 | No |
| orf19.2012 | 3759 | 3403 | 3540 | 2238 | 3983 | 2784 | 3804 | 3898 | 3584 | 3598 | 3497 | 3506 | 3191 | 3166 | 3230 | 3169 | 3367,66 | 3546 | 3189 | 0,899 | -0,154 | -4,602 | 4,19E-06 | 6,32E-06 | 1,00E-08 | 0,0025788 | 0,0007703 | 0,0007703 | TRUE | 0,0011824 | No |
| orf19.2013 | 10555 | 8738 | 9585 | 6144 | 10892 | 7313 | 10561 | 11437 | 10063 | 9239 | 9469 | 9625 | 8727 | 8316 | 8966 | 9299 | 9213,12 | 9599 | 8827 | 0,92 | -0,121 | -2,664 | 0,0077219 | 0,0098616 | 0,0016098 | 0,0022695 | 0,0018601 | 0,0018601 | TRUE | 0,0183386 | No |
| orf19.2014 | 6808 | 5818 | 6304 | 4158 | 6522 | 4687 | 6471 | 6911 | 6491 | 6152 | 6228 | 6514 | 5226 | 5330 | 5494 | 5619 | 5881,57 | 6346 | 5417 | 0,854 | -0,228 | -5,841 | 5,17E-09 | 8,68E-09 | 0,0007058 | 0,0023704 | 0,0012872 | 0,0012872 | TRUE | 0,0077905 | No |
| orf19.2016 | 219 | 247 | 207 | 138 | 261 | 195 | 242 | 248 | 209 | 261 | 204 | 216 | 209 | 222 | 205 | 202 | 216,08 | 222 | 210 | 0,939 | -0,091 | -0,838 | 0,4017546 | 0,4373991 | 0,0028183 | 0,00969 | 0,0065757 | 0,0065757 | TRUE | 0,1548261 | No |
| orf19.2017 | 6121 | 5487 | 5798 | 3879 | 9541 | 7063 | 8877 | 9417 | 5836 | 5802 | 5728 | 6077 | 7645 | 8031 | 7537 | 7657 | 6788,96 | 5861 | 7718 | 1,317 | 0,398 | 11,05 | 2,20E-28 | 5,78E-28 | 0,0005238 | 0,0023331 | 0,0010891 | 0,0010891 | TRUE | 0,0089264 | No |
| orf19.2018 | 1651 | 1361 | 1460 | 1108 | 3645 | 2418 | 3359 | 3584 | 1574 | 1439 | 1442 | 1736 | 2921 | 2749 | 2852 | 2914 | 2203,41 | 1548 | 2859 | 1,853 | 0,89 | 14,511 | 1,04E-47 | 3,71E-47 | 0,0032995 | 0,0028364 | 0,0030874 | 0,0030874 | TRUE | 0,0745624 | No |
| orf19.2018.1 | 116 | 106 | 122 | 83 | 135 | 104 | 120 | 131 | 111 | 112 | 121 | 130 | 108 | 118 | 102 | 107 | 113,51 | 118 | 109 | 0,921 | -0,118 | -0,89 | 0,373552 | 0,4086465 | 1,00E-08 | 0,0165566 | 0,0081724 | 0,0081724 | TRUE | 0,0339695 | No |
| orf19.2018.2 | 18 | 12 | 9 | 7 | 9 | 12 | 9 | 11 | 17 | 13 | 9 | 11 | 7 | 14 | 8 | 9 | 10,89 | 12 | 10 | 0,733 | -0,448 | -1,026 | 0,3048121 | 0,3379093 | 1,00E-08 | 0,1528144 | 0,090441 | 0,090441 | TRUE | 0,2976191 | No |
| orf19.2019 | 2861 | 2363 | 2529 | 1850 | 2705 | 1967 | 2622 | 2792 | 2728 | 2499 | 2498 | 2898 | 2167 | 2237 | 2226 | 2270 | 2440,39 | 2656 | 2225 | 0,839 | -0,253 | -4,602 | 4,18E-06 | 6,31E-06 | 0,0023134 | 0,0027641 | 0,0024931 | 0,0024931 | TRUE | 0,0432119 | No |

| | | | | | | | | | | | | | | | | | | | | | | | | | | | | | | | |
|---|---|---|---|---|---|---|---|---|---|---|---|---|---|---|---|---|---|---|---|---|---|---|---|---|---|---|---|---|---|---|---|
| orf19.202 | 5247 | 4506 | 4771 | 3194 | 7248 | 5056 | 6911 | 7456 | 5002 | 4765 | 4713 | 5004 | 5808 | 5749 | 5867 | 6062 | 5371,3 | 4871 | 5872 | 1,206 | 0,271 | 7,232 | 4,77E-13 | 9,13E-13 | 0,0005172 | 0,002397 | 0,001152 | 0,001152 | TRUE | 0,0058749 | No |
| orf19.2020 | 11044 | 9528 | 10493 | 6820 | 3061 | 2158 | 2578 | 3240 | 10529 | 10075 | 10366 | 10684 | 2453 | 2454 | 2189 | 2634 | 6422,95 | 10414 | 2432 | 0,234 | -2,098 | -38,461 | 0 | 0 | 0,0028219 | 0,0023469 | 0,0026195 | 0,0026195 | TRUE | 0,0559419 | No |
| orf19.2021 | 30152 | 25220 | 27159 | 16389 | 50653 | 33363 | 46634 | 52942 | 28747 | 26667 | 26831 | 25674 | 40587 | 37937 | 39592 | 43047 | 33635,11 | 26980 | 40291 | 1,493 | 0,578 | 11,597 | 4,27E-31 | 1,17E-30 | 0,0024993 | 0,0021401 | 0,0023586 | 0,0023586 | TRUE | 0,0259843 | No |
| orf19.2023 | 27155 | 21870 | 24432 | 15567 | 5800 | 3992 | 5624 | 6237 | 25889 | 23125 | 24137 | 24387 | 4647 | 4539 | 4775 | 5071 | 14571,23 | 24384 | 4758 | 0,195 | -2,357 | -48,486 | 0 | 0 | 0,0020962 | 0,002204 | 0,0021531 | 0,0021531 | TRUE | 0,024013 | No |
| orf19.2024 | 39 | 32 | 26 | 17 | 13 | 11 | 11 | 11 | 37 | 34 | 26 | 27 | 10 | 13 | 9 | 9 | 20,57 | 31 | 10 | 0,328 | -1,607 | -4,828 | 1,38E-06 | 2,12E-06 | 1,00E-08 | 0,08192 | 0,0449691 | 0,0449691 | TRUE | 0,1375224 | No |
| orf19.2025 | 1 | 4 | 3 | 0 | 3 | 1 | 1 | 3 | 1 | 4 | 3 | 0 | 2 | 1 | 1 | 2 | 1,87 | 2 | 2 | 0,816 | -0,294 | -0,281 | 0,7789295 | 0,802705 | 1,00E-08 | 0,8791889 | 0,5357049 | 0,5357049 | TRUE | 0,1812228 | No |
| orf19.2026 | 569 | 540 | 540 | 329 | 830 | 612 | 857 | 888 | 542 | 571 | 533 | 515 | 665 | 696 | 728 | 722 | 621,61 | 540 | 703 | 1,297 | 0,375 | 5,745 | 9,22E-09 | 1,53E-08 | 1,00E-08 | 0,0047326 | 0,0024367 | 0,0024367 | TRUE | 0,0164726 | No |
| orf19.2028 | 4196 | 3596 | 3849 | 2568 | 6733 | 4265 | 6376 | 6455 | 4000 | 3802 | 3802 | 4023 | 5395 | 4850 | 5413 | 5249 | 4566,81 | 3907 | 5227 | 1,339 | 0,421 | 9,002 | 2,22E-19 | 4,92E-19 | 0,0014379 | 0,0024508 | 0,001872 | 0,001872 | TRUE | 0,0280714 | No |
| orf19.2029 | 517 | 522 | 549 | 330 | 1029 | 691 | 972 | 1000 | 493 | 552 | 542 | 517 | 825 | 786 | 825 | 813 | 669,09 | 526 | 812 | 1,545 | 0,627 | 9,963 | 2,21E-23 | 5,32E-23 | 1,00E-08 | 0,0045452 | 0,0021996 | 0,0021996 | TRUE | 0,0241616 | No |
| orf19.203 | 3379 | 2955 | 3321 | 2300 | 1104 | 728 | 1043 | 1066 | 3222 | 3125 | 3281 | 3603 | 885 | 828 | 886 | 867 | 2086,83 | 3308 | 866 | 0,263 | -1,93 | -34,294 | 9,61E-258 | 1,71E-256 | 0,0019957 | 0,0028781 | 0,0023546 | 0,0023546 | TRUE | 0,0418834 | No |
| orf19.2030 | 11158 | 8570 | 9276 | 6447 | 3251 | 2143 | 2978 | 3303 | 10638 | 9062 | 9164 | 10100 | 2605 | 2437 | 2528 | 2686 | 6152,34 | 9741 | 2564 | 0,263 | -1,925 | -32,129 | 1,72E-226 | 2,62E-225 | 0,0035667 | 0,0023582 | 0,0032154 | 0,0032154 | TRUE | 0,0475832 | No |
| orf19.2031 | 2101 | 1922 | 2084 | 1208 | 1118 | 798 | 1123 | 1064 | 2003 | 2032 | 2059 | 1892 | 896 | 907 | 953 | 865 | 1451,04 | 1996 | 905 | 0,453 | -1,143 | -22,158 | 8,73E-109 | 6,25E-108 | 0,0006462 | 0,0032228 | 0,001783 | 0,001783 | TRUE | 0,0158742 | No |
| orf19.2032 | 8 | 4 | 9 | 3 | 4 | 4 | 3 | 3 | 8 | 4 | 9 | 5 | 3 | 5 | 3 | 2 | 4,77 | 6 | 3 | 0,482 | -1,054 | -1,617 | 0,1058215 | 0,1232427 | 1,00E-08 | 0,3460611 | 0,1794269 | 0,1794269 | TRUE | 0,1767761 | No |
| orf19.2033 | 6 | 2 | 11 | 8 | 3 | 4 | 3 | 4 | 6 | 2 | 11 | 13 | 2 | 5 | 3 | 3 | 5,5 | 8 | 3 | 0,41 | -1,285 | -1,89 | 0,0587813 | 0,0701353 | 1,00E-08 | 0,300714 | 0,2285665 | 0,2285665 | TRUE | 0,1689665 | No |
| orf19.2034 | 16 | 14 | 17 | 5 | 21 | 7 | 13 | 19 | 15 | 15 | 17 | 8 | 17 | 8 | 11 | 15 | 13,24 | 14 | 13 | 0,93 | -0,105 | -0,261 | 0,7938428 | 0,8167544 | 1,00E-08 | 0,1260591 | 0,0802052 | 0,0802052 | TRUE | 0,2173399 | No |
| orf19.2035 | 141 | 128 | 111 | 89 | 241 | 169 | 236 | 240 | 134 | 135 | 110 | 139 | 193 | 192 | 200 | 195 | 162,45 | 130 | 195 | 1,513 | 0,598 | 5,127 | 2,94E-07 | 4,65E-07 | 1,00E-08 | 0,0121982 | 0,0064522 | 0,0064522 | TRUE | 0,1180175 | No |
| orf19.2036 | 487 | 421 | 459 | 309 | 1276 | 957 | 1295 | 1363 | 464 | 445 | 453 | 484 | 1022 | 1088 | 1099 | 1108 | 770,66 | 462 | 1079 | 2,343 | 1,229 | 19,758 | 6,83E-87 | 3,98E-86 | 1,00E-08 | 0,0042218 | 0,0020904 | 0,0020904 | TRUE | 0,015905 | No |
| orf19.2037 | 545 | 464 | 474 | 306 | 1369 | 978 | 1363 | 1355 | 520 | 491 | 468 | 479 | 1097 | 1112 | 1157 | 1102 | 803,22 | 490 | 1117 | 2,279 | 1,188 | 19,68 | 3,19E-86 | 1,84E-85 | 1,00E-08 | 0,0041354 | 0,0019643 | 0,0019643 | TRUE | 0,0218604 | No |
| orf19.2038 | 576 | 615 | 609 | 427 | 1034 | 774 | 926 | 923 | 549 | 650 | 602 | 669 | 829 | 880 | 786 | 750 | 714,41 | 618 | 811 | 1,317 | 0,397 | 5,047 | 4,48E-07 | 7,03E-07 | 0,004246 | 0,0043895 | 0,0044695 | 0,0044695 | TRUE | 0,0669257 | No |
| orf19.2039 | 2359 | 1901 | 2235 | 1324 | 3112 | 2102 | 3008 | 3282 | 2249 | 2010 | 2208 | 2074 | 2494 | 2390 | 2554 | 2669 | 2330,91 | 2135 | 2527 | 1,183 | 0,242 | 4,525 | 6,05E-06 | 9,05E-06 | 0,0018602 | 0,0027957 | 0,002313 | 0,002313 | TRUE | 0,019832 | No |
| orf19.204 | 880 | 746 | 862 | 532 | 631 | 433 | 575 | 564 | 839 | 789 | 852 | 833 | 506 | 492 | 488 | 459 | 657,19 | 828 | 486 | 0,587 | -0,769 | -12,152 | 5,61E-34 | 1,61E-33 | 1,00E-08 | 0,0045897 | 0,0022554 | 0,0022554 | TRUE | 0,0181146 | No |
| orf19.2040 | 2568 | 2253 | 2474 | 1488 | 2992 | 2130 | 2850 | 3037 | 2448 | 2382 | 2444 | 2331 | 2397 | 2422 | 2420 | 2469 | 2414,26 | 2401 | 2427 | 1,01 | 0,014 | 0,359 | 0,7197439 | 0,7492146 | 1,00E-08 | 0,0027714 | 0,0009718 | 0,0009718 | TRUE | 0,004188 | No |
| orf19.2041 | 1001 | 985 | 1039 | 582 | 1214 | 850 | 1202 | 1280 | 954 | 1042 | 1026 | 912 | 973 | 967 | 1020 | 1041 | 991,82 | 984 | 1000 | 1,015 | 0,022 | 0,363 | 0,7169322 | 0,7464096 | 0,0012442 | 0,0037467 | 0,0024448 | 0,0024448 | TRUE | 0,0249619 | No |
| orf19.2042 | 1669 | 1369 | 1496 | 876 | 2145 | 1422 | 2074 | 2205 | 1591 | 1448 | 1478 | 1372 | 1719 | 1617 | 1761 | 1793 | 1597,29 | 1472 | 1722 | 1,168 | 0,225 | 3,909 | 9,27E-05 | 0,0001315 | 0,0020534 | 0,0031192 | 0,0025283 | 0,0025283 | TRUE | 0,035863 | No |
| orf19.2043 | 899 | 777 | 857 | 567 | 997 | 729 | 1008 | 1057 | 857 | 822 | 847 | 888 | 799 | 829 | 856 | 859 | 844,57 | 854 | 836 | 0,981 | -0,028 | -0,488 | 0,6259024 | 0,6588392 | 1,00E-08 | 0,0040353 | 0,00187 | 0,00187 | TRUE | 0,0113205 | No |
| orf19.2044 | 488 | 489 | 471 | 321 | 709 | 561 | 665 | 756 | 465 | 517 | 465 | 503 | 568 | 638 | 565 | 615 | 541,97 | 488 | 596 | 1,223 | 0,291 | 3,954 | 7,68E-05 | 0,0001094 | 0,001525 | 0,0051208 | 0,0032951 | 0,0032951 | TRUE | 0,0237963 | No |
| orf19.2045 | 5803 | 5492 | 5705 | 3795 | 9132 | 6357 | 8621 | 8969 | 5533 | 5807 | 5636 | 5945 | 7317 | 7228 | 7319 | 7293 | 6509,78 | 5730 | 7289 | 1,273 | 0,349 | 10,377 | 3,17E-25 | 7,88E-25 | 0,0003168 | 0,0023435 | 0,0009242 | 0,0009242 | TRUE | 0,0073213 | No |
| orf19.2046 | 798 | 645 | 749 | 438 | 890 | 583 | 817 | 892 | 761 | 682 | 740 | 686 | 713 | 663 | 694 | 725 | 707,98 | 717 | 699 | 0,973 | -0,039 | -0,609 | 0,5427589 | 0,5784775 | 0,0007322 | 0,0044104 | 0,0025273 | 0,0025273 | TRUE | 0,0201125 | No |
| orf19.2047 | 1643 | 1426 | 1521 | 931 | 374 | 241 | 319 | 357 | 1566 | 1508 | 1503 | 1458 | 300 | 274 | 271 | 290 | 896,27 | 1509 | 284 | 0,188 | -2,41 | -37,795 | 0 | 0 | 1,00E-08 | 0,0039232 | 0,0019705 | 0,0019705 | TRUE | 0,0168278 | No |
| orf19.2048 | 15558 | 12582 | 13136 | 9718 | 980 | 769 | 924 | 1130 | 14833 | 13304 | 12977 | 15224 | 785 | 874 | 784 | 919 | 7462,57 | 14084 | 840 | 0,06 | -4,067 | -57,618 | 0 | 0 | 0,0054399 | 0,0023113 | 0,0042138 | 0,0042138 | TRUE | 0,048966 | No |
| orf19.2049 | 681 | 658 | 623 | 377 | 251 | 147 | 195 | 210 | 649 | 696 | 615 | 591 | 201 | 167 | 166 | 171 | 406,96 | 638 | 176 | 0,276 | -1,855 | -19,951 | 1,48E-88 | 8,73E-88 | 0,0034472 | 0,0061259 | 0,0049067 | 0,0049067 | TRUE | 0,1047345 | No |
| orf19.2050 | 7142 | 6226 | 6871 | 4275 | 1605 | 1124 | 1525 | 1672 | 6809 | 6583 | 6788 | 6697 | 1286 | 1278 | 1295 | 1359 | 4011,95 | 6719 | 1304 | 0,194 | -2,364 | -61,89 | 0 | 0 | 1,00E-08 | 0,0025005 | 0,0009759 | 0,0009759 | TRUE | 0,0099509 | No |
| orf19.2051 | 406 | 387 | 437 | 292 | 1337 | 911 | 1245 | 1373 | 387 | 409 | 432 | 457 | 1071 | 1036 | 1057 | 1116 | 745,75 | 421 | 1070 | 2,555 | 1,353 | 20,352 | 4,45E-92 | 2,71E-91 | 0,0004625 | 0,004293 | 0,0024988 | 0,0024988 | TRUE | 0,0352332 | No |
| orf19.2054 | 2150 | 1905 | 2224 | 1356 | 1890 | 1337 | 1947 | 1933 | 2050 | 2014 | 2197 | 2124 | 1514 | 1520 | 1653 | 1572 | 1830,6 | 2096 | 1565 | 0,747 | -0,421 | -8,432 | 3,38E-17 | 7,15E-17 | 0,001083 | 0,0029882 | 0,0018383 | 0,0018383 | TRUE | 0,0174735 | No |
| orf19.2055 | 2806 | 2307 | 2498 | 1493 | 3773 | 2296 | 3618 | 3684 | 2675 | 2439 | 2468 | 2339 | 3023 | 2611 | 3072 | 2995 | 2702,79 | 2480 | 2925 | 1,179 | 0,238 | 3,786 | 0,0001533 | 0,000215 | 0,0037509 | 0,0026988 | 0,0034092 | 0,0034092 | TRUE | 0,0621266 | No |
| orf19.2057 | 6606 | 5900 | 6590 | 4203 | 7363 | 5480 | 7219 | 7958 | 6298 | 6239 | 6510 | 6584 | 5900 | 6231 | 6129 | 6471 | 6295,21 | 6408 | 6183 | 0,965 | -0,051 | -1,274 | 0,2026148 | 0,229683 | 0,0008557 | 0,0023521 | 0,0013902 | 0,0013902 | TRUE | 0,0122357 | No |
| orf19.2059 | 1442 | 1292 | 1403 | 957 | 4565 | 3254 | 4460 | 4588 | 1375 | 1366 | 1386 | 1499 | 3658 | 3700 | 3787 | 3730 | 2562,63 | 1406 | 3719 | 2,652 | 1,407 | 33,08 | 5,72E-240 | 9,31E-239 | 5,81E-05 | 0,002732 | 0,0012178 | 0,0012178 | TRUE | 0,0203607 | No |
| orf19.206 | 1439 | 1228 | 1302 | 945 | 5630 | 4054 | 5341 | 5801 | 1372 | 1298 | 1286 | 1480 | 4511 | 4610 | 4534 | 4717 | 2976,15 | 1359 | 4593 | 3,39 | 1,761 | 35,965 | 3,00E-283 | 6,17E-282 | 0,0010284 | 0,002643 | 0,0017911 | 0,0017911 | TRUE | 0,0382314 | No |
| orf19.2060 | 54 | 36 | 41 | 33 | 91 | 87 | 76 | 104 | 51 | 38 | 41 | 52 | 73 | 99 | 65 | 85 | 62,83 | 46 | 80 | 1,766 | 0,82 | 4,005 | 6,19E-05 | 8,85E-05 | 0,0149233 | 0,0282218 | 0,0223599 | 0,0223599 | TRUE | 0,2056202 | No |
| orf19.2061 | 7 | 4 | 1 | 1 | 5 | 0 | 2 | 4 | 7 | 4 | 1 | 2 | 4 | 0 | 2 | 3 | 2,8 | 4 | 2 | 0,674 | -0,569 | -0,61 | 0,5418028 | 0,5775549 | 0,3088179 | 0,588108 | 0,4832222 | 0,4832222 | TRUE | 0,3819285 | No |
| orf19.2062 | 340 | 282 | 279 | 250 | 548 | 405 | 498 | 598 | 324 | 298 | 276 | 392 | 439 | 461 | 423 | 486 | 387,28 | 322 | 452 | 1,413 | 0,499 | 4,835 | 1,33E-06 | 2,05E-06 | 0,0088628 | 0,0063309 | 0,007484 | 0,007484 | TRUE | 0,2056104 | No |
| orf19.2063 | 3845 | 3511 | 3841 | 2453 | 3515 | 2471 | 3183 | 3554 | 3666 | 3712 | 3795 | 3843 | 2816 | 2810 | 2702 | 2890 | 3279,23 | 3754 | 2804 | 0,748 | -0,42 | -10,969 | 5,40E-28 | 1,41E-27 | 0,0003089 | 0,002592 | 0,0010961 | 0,0010961 | TRUE | 0,007492 | No |
| orf19.2064 | 2696 | 2179 | 2351 | 1439 | 1443 | 934 | 1404 | 1595 | 2570 | 2304 | 2323 | 2254 | 1156 | 1062 | 1192 | 1297 | 1769,79 | 2363 | 1177 | 0,498 | -1,005 | -14,862 | 5,82E-50 | 2,15E-49 | 0,0043396 | 0,003019 | 0,003771 | 0,003771 | TRUE | 0,0566475 | No |
| orf19.2065 | 111 | 88 | 134 | 54 | 193 | 99 | 149 | 167 | 106 | 93 | 132 | 85 | 155 | 113 | 127 | 136 | 118,17 | 104 | 133 | 1,267 | 0,341 | 2,098 | 0,0359469 | 0,0437325 | 0,0170878 | 0,0159858 | 0,0166957 | 0,0166957 | TRUE | 0,3337195 | No |
| orf19.2066 | 1060 | 944 | 987 | 654 | 2609 | 1817 | 2560 | 2664 | 1011 | 998 | 975 | 1025 | 2091 | 2066 | 2173 | 2166 | 1563,06 | 1002 | 2124 | 2,123 | 1,086 | 23,483 | 6,03E-122 | 4,87E-121 | 1,00E-08 | 0,0031417 | 0,0012864 | 0,0012864 | TRUE | 0,0039852 | No |
| orf19.2066.1 | 2408 | 1878 | 1897 | 1279 | 4436 | 2851 | 4017 | 4284 | 2296 | 1986 | 1874 | 2004 | 3554 | 3242 | 3410 | 3483 | 2731,15 | 2040 | 3422 | 1,678 | 0,746 | 11,732 | 8,72E-32 | 2,43E-31 | 0,0041395 | 0,0026925 | 0,0034802 | 0,0034802 | TRUE | 0,0889547 | No |
| orf19.2067 | 4618 | 4025 | 4257 | 2472 | 2251 | 1469 | 2199 | 2181 | 4403 | 4256 | 4206 | 3873 | 1804 | 1670 | 1867 | 1773 | 2981,38 | 4184 | 1778 | 0,425 | -1,234 | -23,324 | 2,55E-120 | 2,03E-119 | 0,0019614 | 0,002642 | 0,0023041 | 0,0023041 | TRUE | 0,0293337 | No |
| orf19.2068 | 544 | 494 | 531 | 320 | 1691 | 1322 | 1706 | 1662 | 519 | 522 | 525 | 501 | 1355 | 1503 | 1448 | 1351 | 965,6 | 517 | 1414 | 2,73 | 1,449 | 23,488 | 5,45E-122 | 4,42E-121 | 1,00E-08 | 0,0037917 | 0,002259 | 0,002259 | TRUE | 0,0203703 | No |
| orf19.2069 | 2614 | 2210 | 2411 | 1584 | 3338 | 2354 | 3182 | 3456 | 2492 | 2337 | 2382 | 2481 | 2675 | 2677 | 2702 | 2810 | 2569,39 | 2423 | 2716 | 1,122 | 0,166 | 4,013 | 5,99E-05 | 8,56E-05 | 0,0003767 | 0,0027303 | 0,0012406 | 0,0012406 | TRUE | 0,0069481 | No |
| orf19.207 | 2516 | 2078 | 2292 | 1647 | 3911 | 2865 | 3582 | 4135 | 2399 | 2197 | 2264 | 2580 | 3134 | 3258 | 3041 | 3362 | 2779,39 | 2360 | 3199 | 1,357 | 0,441 | 7,623 | 2,47E-14 | 4,90E-14 | 0,003036 | 0,002682 | 0,0028296 | 0,0028296 | TRUE | 0,0449997 | No |
| orf19.2070 | 3312 | 3040 | 3241 | 2039 | 3273 | 2259 | 3165 | 3447 | 3158 | 3214 | 3202 | 3194 | 2623 | 2569 | 2687 | 2803 | 2931,14 | 3192 | 2670 | 0,837 | -0,256 | -6,392 | 1,64E-10 | 2,91E-10 | 0,0004026 | 0,0026514 | 0,0012015 | 0,0012015 | TRUE | 0,0136305 | No |
| orf19.2071 | 1 | 1 | 3 | 1 | 4 | 2 | 5 | 3 | 1 | 1 | 3 | 2 | 3 | 2 | 4 | 2 | 2,34 | 2 | 3 | 1,866 | 0,9 | 0,972 | 0,3309209 | 0,3647618 | 1,00E-08 | 0,7043385 | 0,3396371 | 0,3396371 | TRUE | 0,2549146 | No |
| orf19.2072 | 6 | 9 | 10 | 17 | 42 | 37 | 33 | 51 | 6 | 10 | 10 | 27 | 34 | 42 | 28 | 41 | 24,62 | 13 | 36 | 2,989 | 1,58 | 4,119 | 3,80E-05 | 5,48E-05 | 0,1136463 | 0,0687819 | 0,0828691 | 0,0828691 | TRUE | 1,3875125 | No |
| orf19.2073 | 10479 | 8744 | 9716 | 6082 | 5447 | 3755 | 5174 | 5523 | 9991 | 9246 | 9598 | 9528 | 4365 | 4270 | 4393 | 4491 | 6985,03 | 9591 | 4380 | 0,457 | -1,131 | -30,756 | 1,01E-207 | 1,42E-206 | 0,0005066 | 0,0023263 | 0,001137 | 0,001137 | TRUE | 0,009809 | No |
| orf19.2074 | 254 | 194 | 204 | 122 | 105 | 69 | 101 | 104 | 242 | 205 | 202 | 191 | 84 | 78 | 86 | 85 | 146,61 | 210 | 83 | 0,395 | -1,339 | -10,336 | 4,87E-25 | 1,21E-24 | 1,00E-08 | 0,0132907 | 0,0081302 | 0,0081302 | TRUE | 0,1207534 | No |
| orf19.2075 | 3332 | 2879 | 3019 | 1780 | 4002 | 2652 | 3910 | 4103 | 3177 | 3044 | 2982 | 2788 | 3207 | 3016 | 3320 | 3336 | 3108,72 | 2998 | 3220 | 1,073 | 0,102 | 1,947 | 0,051556 | 0,0619312 | 0,0020721 | 0,0026194 | 0,0023031 | 0,0023031 | TRUE | 0,0253903 | No |
| orf19.2076 | 184 | 199 | 193 | 119 | 338 | 270 | 337 | 369 | 175 | 210 | 191 | 186 | 271 | 307 | 286 | 300 | 240,86 | 190 | 291 | 1,523 | 0,607 | 6,099 | 1,07E-09 | 1,83E-09 | 1,00E-08 | 0,008908 | 0,0050516 | 0,0050516 | TRUE | 0,0544471 | No |
| orf19.2077 | 3291 | 2852 | 3063 | 2069 | 2756 | 1882 | 2728 | 2850 | 3138 | 3016 | 3026 | 3241 | 2208 | 2140 | 2316 | 2317 | 2675,26 | 3105 | 2245 | 0,724 | -0,465 | -10,323 | 5,54E-25 | 1,37E-24 | 0,0008222 | 0,002705 | 0,0015701 | 0,0015701 | TRUE | 0,0114235 | No |
| orf19.2078 | 1352 | 1238 | 1309 | 740 | 1086 | 733 | 1029 | 1059 | 1289 | 1309 | 1293 | 1159 | 870 | 833 | 874 | 861 | 1061,1 | 1262 | 860 | 0,679 | -0,558 | -10,056 | 8,62E-24 | 2,09E-23 | 0,0004328 | 0,0036386 | 0,0019909 | 0,0019909 | TRUE | 0,0319831 | No |
| orf19.2079 | 4523 | 3582 | 3630 | 2955 | 1397 | 1000 | 1191 | 1484 | 4312 | 3788 | 3586 | 4629 | 1119 | 1137 | 1011 | 1207 | 2598,65 | 4079 | 1118 | 0,274 | -1,866 | -22,254 | 1,03E-109 | 7,49E-109 | 0,0087187 | 0,0027231 | 0,0062121 | 0,0062121 | TRUE | 0,0986744 | No |
| orf19.2081 | 2518 | 2298 | 2486 | 1523 | 2407 | 1780 | 2356 | 2610 | 2401 | 2430 | 2456 | 2386 | 1929 | 2024 | 2000 | 2122 | 2218,42 | 2418 | 2019 | 0,834 | -0,261 | -5,998 | 2,00E-09 | 3,40E-09 | 0,0005022 | 0,0028314 | 0,0013647 | 0,0013647 | TRUE | 0,0150346 | No |
| orf19.2082 | 4 | 4 | 1 | 3 | 3 | 4 | 5 | 0 | 4 | 4 | 1 | 5 | 2 | 5 | 4 | 0 | 3,12 | 4 | 3 | 0,811 | -0,302 | -0,341 | 0,7330371 | 0,7617972 | 1,00E-08 | 0,5290227 | 0,4250749 | 0,4250749 | TRUE | 0,7019186 | No |
| orf19.2084 | 2196 | 1800 | 2087 | 1323 | 2401 | 1620 | 2298 | 2497 | 2094 | 1903 | 2062 | 2073 | 1924 | 1842 | 1951 | 2030 | 1984,81 | 2033 | 1937 | 0,954 | -0,068 | -1,362 | 0,1730742 | 0,1973881 | 0,0012289 | 0,0029185 | 0,0019028 | 0,0019028 | TRUE | 0,0226512 | No |
| orf19.2087 | 902 | 761 | 833 | 570 | 949 | 701 | 932 | 985 | 860 | 805 | 823 | 893 | 760 | 797 | 791 | 801 | 816,27 | 845 | 787 | 0,933 | -0,099 | -1,723 | 0,084974 | 0,099893 | 1,00E-08 | 0,0041028 | 0,0019662 | 0,0019662 | TRUE | 0,0145547 | No |
| orf19.2088 | 695 | 516 | 620 | 382 | 1070 | 663 | 1031 | 1091 | 663 | 546 | 613 | 598 | 857 | 754 | 875 | 887 | 724,1 | 605 | 843 | 1,395 | 0,481 | 6,172 | 6,74E-10 | 1,17E-09 | 0,0043949 | 0,0043588 | 0,00437 | 0,00437 | TRUE | 0,0573124 | No |
| orf19.2089 | 3928 | 3486 | 3597 | 2257 | 2980 | 2020 | 2781 | 2980 | 3745 | 3686 | 3553 | 3536 | 2388 | 2297 | 2361 | 2423 | 2998,62 | 3630 | 2367 | 0,652 | -0,617 | -15,733 | 8,98E-56 | 3,57E-55 | 0,0003053 | 0,0026388 | 0,0011326 | 0,0011326 | TRUE | 0,0055996 | No |
| orf19.209 | 350 | 304 | 269 | 218 | 290 | 219 | 260 | 256 | 334 | 321 | 266 | 342 | 232 | 249 | 221 | 208 | 271,58 | 316 | 228 | 0,721 | -0,471 | -4,472 | 7,76E-06 | 1,16E-05 | 0,0055641 | 0,008137 | 0,0069105 | 0,0069105 | TRUE | 0,1332286 | No |
| orf19.2090 | 8730 | 7576 | 8205 | 4715 | 12592 | 8113 | 11837 | 12666 | 8323 | 8011 | 8106 | 7386 | 10090 | 9225 | 10050 | 10299 | 8936,11 | 7956 | 9916 | 1,246 | 0,317 | 6,327 | 2,50E-10 | 4,41E-10 | 0,0022636 | 0,002275 | 0,0022978 | 0,0022978 | TRUE | 0,0277661 | No |
| orf19.2091 | 3481 | 3031 | 3331 | 2161 | 8705 | 6203 | 8850 | 9072 | 3319 | 3205 | 3291 | 3385 | 6975 | 7053 | 7514 | 7376 | 5264,77 | 3300 | 7230 | 2,192 | 1,132 | 28,492 | 1,47E-178 | 1,74E-177 | 0,000754 | 0,0024031 | 0,0012846 | 0,0012846 | TRUE | 0,0085509 | No |
| orf19.2092 | 2851 | 2409 | 2570 | 1741 | 4559 | 3161 | 4506 | 4744 | 2718 | 2547 | 2539 | 2727 | 3653 | 3594 | 3826 | 3857 | 3182,73 | 2633 | 3732 | 1,419 | 0,505 | 11,252 | 2,25E-29 | 6,00E-29 | 0,0009402 | 0,0026072 | 0,0016016 | 0,0016016 | TRUE | 0,0072879 | No |
| orf19.2093 | 3738 | 3094 | 3453 | 2388 | 4588 | 3341 | 4302 | 4605 | 3564 | 3272 | 3411 | 3741 | 3676 | 3799 | 3652 | 3744 | 3607,42 | 3497 | 3718 | 1,064 | 0,09 | 1,913 | 0,0556913 | 0,0665979 | 0,0014555 | 0,0025464 | 0,001832 | 0,001832 | TRUE | 0,0252586 | No |
| orf19.2094 | 6578 | 6162 | 6594 | 4214 | 1502 | 1214 | 1468 | 1563 | 6271 | 6516 | 6514 | 6601 | 1204 | 1380 | 1246 | 1271 | 3875,47 | 6476 | 1275 | 0,197 | -2,347 | -50,041 | 0 | 0 | 0,0010047 | 0,0025149 | 0,0016744 | 0,0016744 | TRUE | 0,0343731 | No |
| orf19.2095 | 13608 | 11724 | 12599 | 8669 | 7222 | 5020 | 6600 | 7256 | 12974 | 12397 | 12447 | 13580 | 5787 | 5708 | 5603 | 5900 | 9299,45 | 12850 | 5750 | 0,448 | -1,16 | -28,757 | 7,30E-182 | 8,88E-181 | 0,0009952 | 0,0022678 | 0,0014398 | 0,0014398 | TRUE | 0,0175434 | No |

| | | | | | | | | | | | | | | | | | | | | | | | | | | | | | | | |
|---|---|---|---|---|---|---|---|---|---|---|---|---|---|---|---|---|---|---|---|---|---|---|---|---|---|---|---|---|---|---|---|
| orf19.2097 | 701 | 615 | 703 | 402 | 1767 | 1238 | 1741 | 1898 | 668 | 650 | 694 | 630 | 1416 | 1408 | 1478 | 1543 | 1060,97 | 660 | 1461 | 2,207 | 1,142 | 19,623 | 9,93E-86 | 5,70E-85 | 0,0009193 | 0,0036388 | 0,0021055 | 0,0021055 | TRUE | 0,0174371 | No |
| orf19.2098 | 1986 | 1579 | 1793 | 1247 | 2161 | 1410 | 2119 | 2118 | 1893 | 1670 | 1771 | 1953 | 1732 | 1603 | 1799 | 1722 | 1767,98 | 1822 | 1714 | 0,943 | -0,085 | -1,409 | 0,1586993 | 0,1816757 | 0,0027912 | 0,00302 | 0,0029361 | 0,0029361 | TRUE | 0,0379768 | No |
| orf19.2099 | 1887 | 1552 | 1750 | 1128 | 2029 | 1388 | 1901 | 2086 | 1799 | 1641 | 1729 | 1767 | 1626 | 1578 | 1614 | 1696 | 1681,26 | 1734 | 1628 | 0,94 | -0,089 | -1,847 | 0,0646816 | 0,0769594 | 0,0006568 | 0,0030679 | 0,0016345 | 0,0016345 | TRUE | 0,0160196 | No |
| orf19.21 | 630 | 532 | 573 | 428 | 381 | 291 | 345 | 368 | 601 | 563 | 566 | 670 | 305 | 331 | 293 | 299 | 453,5 | 600 | 307 | 0,513 | -0,964 | -11,767 | 5,81E-32 | 1,62E-31 | 0,0022485 | 0,0057118 | 0,0040513 | 0,0040513 | TRUE | 0,0644215 | No |
| orf19.210 | 146 | 152 | 128 | 97 | 126 | 94 | 137 | 159 | 139 | 161 | 126 | 152 | 101 | 107 | 116 | 129 | 128,97 | 144 | 113 | 0,788 | -0,343 | -2,506 | 0,0122049 | 0,015434 | 0,0045016 | 0,0148221 | 0,0102189 | 0,0102189 | TRUE | 0,0963856 | No |
| orf19.2101 | 1135 | 920 | 1009 | 631 | 1473 | 962 | 1366 | 1424 | 1082 | 973 | 997 | 988 | 1180 | 1094 | 1160 | 1158 | 1078,99 | 1010 | 1148 | 1,136 | 0,184 | 3,318 | 0,0009068 | 0,0012267 | 0,0007151 | 0,003613 | 0,0020215 | 0,0020215 | TRUE | 0,0291341 | No |
| orf19.2102 | 4914 | 4087 | 4480 | 2726 | 3589 | 2315 | 3509 | 3551 | 4685 | 4321 | 4426 | 4270 | 2876 | 2632 | 2979 | 2887 | 3634,66 | 4426 | 2844 | 0,643 | -0,638 | -12,693 | 6,48E-37 | 1,95E-36 | 0,0018434 | 0,002543 | 0,0021405 | 0,0021405 | TRUE | 0,0292833 | No |
| orf19.2103 | 25 | 23 | 32 | 16 | 33 | 29 | 41 | 43 | 24 | 24 | 32 | 25 | 26 | 33 | 35 | 35 | 29,25 | 26 | 32 | 1,227 | 0,295 | 1,13 | 0,2585716 | 0,2889066 | 1,00E-08 | 0,0582195 | 0,0303986 | 0,0303986 | TRUE | 0,131807 | No |
| orf19.2104 | 593 | 507 | 492 | 348 | 832 | 549 | 817 | 859 | 565 | 536 | 486 | 545 | 667 | 624 | 694 | 698 | 601,96 | 533 | 671 | 1,261 | 0,335 | 4,685 | 2,80E-06 | 4,26E-06 | 0,0015792 | 0,0048189 | 0,00319 | 0,00319 | TRUE | 0,0441905 | No |
| orf19.2105 | 1134 | 966 | 1052 | 779 | 1758 | 1298 | 1693 | 1683 | 1081 | 1021 | 1039 | 1220 | 1409 | 1476 | 1437 | 1368 | 1256,57 | 1090 | 1422 | 1,308 | 0,388 | 6,28 | 3,39E-10 | 5,94E-10 | 0,0023103 | 0,0033979 | 0,0028291 | 0,0028291 | TRUE | 0,0690771 | No |
| orf19.2106 | 1498 | 1128 | 1307 | 805 | 2364 | 1486 | 2241 | 2361 | 1428 | 1193 | 1291 | 1261 | 1894 | 1690 | 1903 | 1920 | 1572,43 | 1293 | 1852 | 1,432 | 0,518 | 7,774 | 7,60E-15 | 1,53E-14 | 0,0037636 | 0,0031355 | 0,0035796 | 0,0035796 | TRUE | 0,0612273 | No |
| orf19.2107 | 869 | 683 | 739 | 520 | 2782 | 1809 | 2814 | 2900 | 828 | 722 | 730 | 815 | 2229 | 2057 | 2389 | 2358 | 1516,07 | 774 | 2258 | 2,924 | 1,548 | 22,547 | 1,45E-112 | 1,07E-111 | 0,003649 | 0,0031743 | 0,0036026 | 0,0036026 | TRUE | 0,042369 | No |
| orf19.2107.1 | 110649 | 78158 | 83372 | 44334 | 10052 | 5055 | 9164 | 9961 | 105492 | 82642 | 82364 | 69452 | 8054 | 5748 | 7780 | 8099 | 46203,88 | 84988 | 7420 | 0,087 | -3,518 | -20,986 | 8,75E-98 | 5,60E-97 | 0,0269281 | 0,0021268 | 0,0140254 | 0,0269281 | TRUE | 0,1743244 | No |
| orf19.2108 | 4061 | 3383 | 3667 | 2304 | 1244 | 791 | 1127 | 1271 | 3872 | 3577 | 3623 | 3609 | 997 | 899 | 957 | 1033 | 2320,91 | 3670 | 972 | 0,265 | -1,915 | -36,087 | 3,59E-285 | 7,46E-284 | 0,0015609 | 0,0027987 | 0,0020923 | 0,0020923 | TRUE | 0,0274915 | No |
| orf19.211 | 1916 | 1627 | 1802 | 1232 | 327 | 226 | 337 | 352 | 1827 | 1720 | 1780 | 1930 | 262 | 257 | 286 | 286 | 1043,57 | 1814 | 273 | 0,151 | -2,728 | -40,388 | 0 | 0 | 2,91E-07 | 0,0036646 | 0,0024463 | 0,0024463 | TRUE | 0,0204681 | No |
| orf19.2110 | 250 | 200 | 213 | 118 | 380 | 245 | 346 | 345 | 238 | 211 | 210 | 185 | 304 | 279 | 294 | 281 | 250,31 | 211 | 290 | 1,361 | 0,444 | 4,5 | 6,78E-06 | 1,01E-05 | 0,0003447 | 0,0086509 | 0,0051976 | 0,0051976 | TRUE | 0,0812036 | No |
| orf19.2111 | 3763 | 2907 | 3351 | 2192 | 3710 | 2486 | 3705 | 3833 | 3588 | 3074 | 3310 | 3434 | 2973 | 2827 | 3146 | 3117 | 3183,42 | 3352 | 3016 | 0,9 | -0,152 | -2,641 | 0,0082545 | 0,0105344 | 0,0028608 | 0,002607 | 0,0028446 | 0,0028446 | TRUE | 0,0381913 | No |
| orf19.2111.2 | 6678 | 5169 | 5233 | 3337 | 20674 | 14553 | 19265 | 21537 | 6367 | 5466 | 5170 | 5228 | 16565 | 16548 | 16356 | 17512 | 11151,34 | 5558 | 16745 | 3,012 | 1,591 | 24,761 | 2,35E-135 | 2,14E-134 | 0,0053003 | 0,0022385 | 0,0038361 | 0,0038361 | TRUE | 0,118125 | No |
| orf19.2112 | 1015 | 782 | 957 | 671 | 1575 | 1111 | 1636 | 1712 | 968 | 827 | 945 | 1051 | 1262 | 1263 | 1389 | 1392 | 1137,18 | 948 | 1326 | 1,404 | 0,489 | 6,601 | 4,08E-11 | 7,36E-11 | 0,0051122 | 0,0035351 | 0,0043416 | 0,0043416 | TRUE | 0,0871352 | No |
| orf19.2113 | 2533 | 2180 | 2340 | 1552 | 4050 | 2725 | 3632 | 3918 | 2415 | 2305 | 2312 | 2431 | 3245 | 3099 | 3084 | 3186 | 2759,5 | 2366 | 3154 | 1,334 | 0,416 | 10,365 | 3,60E-25 | 8,93E-25 | 0,0003118 | 0,0026863 | 0,0011691 | 0,0011691 | TRUE | 0,004734 | No |
| orf19.2114 | 201 | 157 | 183 | 154 | 832 | 646 | 856 | 864 | 192 | 166 | 181 | 241 | 667 | 735 | 727 | 703 | 451,27 | 195 | 708 | 3,675 | 1,878 | 19,562 | 3,27E-85 | 1,86E-84 | 0,0044354 | 0,0057297 | 0,0053204 | 0,0053204 | TRUE | 0,2194925 | No |
| orf19.2115 | 2784 | 2207 | 2510 | 1548 | 7946 | 5038 | 7900 | 8265 | 2654 | 2334 | 2480 | 2425 | 6367 | 5729 | 6707 | 6720 | 4426,92 | 2473 | 6381 | 2,58 | 1,367 | 22,088 | 4,09E-108 | 2,91E-107 | 0,0037635 | 0,0024622 | 0,0033817 | 0,0033817 | TRUE | 0,0571032 | No |
| orf19.2116 | 2491 | 1970 | 2162 | 1317 | 3141 | 1865 | 2894 | 3132 | 2375 | 2083 | 2136 | 2063 | 2517 | 2121 | 2457 | 2547 | 2287,25 | 2164 | 2410 | 1,114 | 0,155 | 2,249 | 0,0244903 | 0,0301976 | 0,004949 | 0,0028091 | 0,0041408 | 0,0041408 | TRUE | 0,0774541 | No |
| orf19.2117 | 3896 | 3145 | 3511 | 2333 | 3706 | 2410 | 3407 | 3651 | 3714 | 3325 | 3469 | 3655 | 2970 | 2740 | 2893 | 2969 | 3216,77 | 3541 | 2893 | 0,818 | -0,29 | -5,906 | 3,51E-09 | 5,93E-09 | 0,001535 | 0,0026017 | 0,0020049 | 0,0020049 | TRUE | 0,0204884 | No |
| orf19.2119 | 8972 | 7230 | 8601 | 6332 | 7746 | 5523 | 7457 | 7981 | 8554 | 7645 | 8497 | 9919 | 6207 | 6280 | 6331 | 6489 | 7490,26 | 8654 | 6327 | 0,731 | -0,451 | -6,628 | 3,40E-11 | 6,15E-11 | 0,0059044 | 0,0023105 | 0,0043141 | 0,0043141 | TRUE | 0,117056 | No |
| orf19.212 | 1767 | 1562 | 1724 | 1180 | 851 | 596 | 820 | 857 | 1685 | 1652 | 1703 | 1849 | 682 | 678 | 696 | 697 | 1205,07 | 1722 | 688 | 0,401 | -1,319 | -24,283 | 2,96E-130 | 2,57E-129 | 0,0006481 | 0,0034538 | 0,0018611 | 0,0018611 | TRUE | 0,0266977 | No |
| orf19.2123 | 1 | 0 | 2 | 0 | 4 | 2 | 4 | 8 | 1 | 0 | 2 | 0 | 3 | 2 | 3 | 7 | 2,29 | 1 | 4 | 4,841 | 2,275 | 2,008 | 0,0446792 | 0,0539753 | 1,00E-08 | 0,7195021 | 0,4316979 | 0,4316979 | TRUE | 0,4299827 | No |
| orf19.2124 | 2947 | 2637 | 2731 | 1792 | 7830 | 5259 | 7139 | 7984 | 2810 | 2788 | 2698 | 2807 | 6274 | 5980 | 6061 | 6492 | 4488,73 | 2776 | 6202 | 2,236 | 1,161 | 28,627 | 3,08E-180 | 3,71E-179 | 0,0007497 | 0,0024571 | 0,0013039 | 0,0013039 | TRUE | 0,0121214 | No |
| orf19.2125 | 5095 | 3726 | 3907 | 3136 | 1529 | 1040 | 1496 | 1515 | 4858 | 3940 | 3860 | 4913 | 1225 | 1183 | 1270 | 1232 | 2809,93 | 4393 | 1228 | 0,28 | -1,838 | -22,275 | 6,44E-110 | 4,67E-109 | 0,00816 | 0,0026756 | 0,0060484 | 0,0060484 | TRUE | 0,0586742 | No |
| orf19.2127 | 449 | 376 | 422 | 266 | 491 | 324 | 482 | 572 | 428 | 398 | 417 | 417 | 393 | 368 | 409 | 465 | 411,92 | 415 | 409 | 0,989 | -0,017 | -0,193 | 0,8470774 | 0,8660793 | 0,0033293 | 0,0060772 | 0,0046847 | 0,0046847 | TRUE | 0,0998255 | No |
| orf19.2128 | 755 | 563 | 595 | 444 | 1987 | 1253 | 1825 | 1954 | 720 | 595 | 588 | 696 | 1592 | 1425 | 1549 | 1589 | 1094,2 | 650 | 1539 | 2,375 | 1,248 | 16,504 | 3,42E-61 | 1,47E-60 | 0,0052931 | 0,0035918 | 0,0043326 | 0,0043326 | TRUE | 0,0699913 | No |
| orf19.2129 | 706 | 667 | 642 | 477 | 2037 | 1338 | 1933 | 1999 | 673 | 705 | 634 | 747 | 1632 | 1521 | 1641 | 1625 | 1147,49 | 690 | 1605 | 2,338 | 1,225 | 20,4 | 1,68E-92 | 1,03E-91 | 0,0012007 | 0,0035221 | 0,0023717 | 0,0023717 | TRUE | 0,0344876 | No |
| orf19.213 | 2749 | 2736 | 2776 | 1906 | 3516 | 2831 | 3248 | 3283 | 2621 | 2893 | 2742 | 2986 | 2817 | 3219 | 2758 | 2669 | 2838,18 | 2810 | 2866 | 1,02 | 0,028 | 0,425 | 0,6704943 | 0,7024174 | 0,0042821 | 0,0026698 | 0,0038472 | 0,0038472 | TRUE | 0,0820788 | No |
| orf19.2131 | 3005 | 2622 | 2809 | 1697 | 2538 | 1751 | 2425 | 2628 | 2865 | 2772 | 2775 | 2658 | 2034 | 1991 | 2059 | 2137 | 2411,39 | 2768 | 2055 | 0,742 | -0,43 | -10,088 | 6,25E-24 | 1,52E-23 | 0,000346 | 0,0027722 | 0,0013267 | 0,0013267 | TRUE | 0,008737 | No |
| orf19.2132 | 1510 | 1166 | 1277 | 787 | 1414 | 853 | 1375 | 1416 | 1440 | 1233 | 1262 | 1233 | 1133 | 970 | 1167 | 1151 | 1198,57 | 1292 | 1105 | 0,856 | -0,224 | -2,986 | 0,0028264 | 0,0037094 | 0,0052665 | 0,0034612 | 0,0045726 | 0,0045726 | TRUE | 0,0779222 | No |
| orf19.2133 | 57 | 49 | 56 | 41 | 42 | 36 | 40 | 68 | 54 | 52 | 55 | 64 | 34 | 41 | 34 | 55 | 48,69 | 56 | 41 | 0,731 | -0,452 | -2,038 | 0,0415147 | 0,0502953 | 0,0139476 | 0,0358107 | 0,0263889 | 0,0263889 | TRUE | 0,4845153 | No |
| orf19.2135 | 1967 | 1721 | 1805 | 1156 | 2067 | 1414 | 2078 | 2110 | 1875 | 1820 | 1783 | 1811 | 1656 | 1608 | 1764 | 1716 | 1754,14 | 1822 | 1686 | 0,926 | -0,111 | -2,391 | 0,0168138 | 0,0210041 | 0,0004961 | 0,0030273 | 0,001508 | 0,001508 | TRUE | 0,011595 | No |
| orf19.2137 | 814 | 671 | 708 | 498 | 894 | 615 | 861 | 905 | 776 | 709 | 699 | 780 | 716 | 699 | 731 | 736 | 730,95 | 741 | 720 | 0,975 | -0,037 | -0,592 | 0,553762 | 0,5888278 | 0,000365 | 0,0043375 | 0,002323 | 0,002323 | TRUE | 0,0173731 | No |
| orf19.2138 | 13730 | 11723 | 12719 | 8135 | 17669 | 12510 | 16168 | 18852 | 13090 | 12396 | 12565 | 12744 | 14158 | 14225 | 13727 | 15328 | 13529,05 | 12699 | 14360 | 1,131 | 0,177 | 4,229 | 2,35E-05 | 3,42E-05 | 0,0013052 | 0,0022126 | 0,0016145 | 0,0016145 | TRUE | 0,0253805 | No |
| orf19.2143 | 4455 | 3651 | 4245 | 2821 | 7081 | 4943 | 6887 | 7276 | 4247 | 3860 | 4194 | 4419 | 5674 | 5621 | 5847 | 5916 | 4972,29 | 4180 | 5764 | 1,38 | 0,465 | 9,959 | 2,30E-23 | 5,52E-23 | 0,0015842 | 0,0024215 | 0,0018799 | 0,0018799 | TRUE | 0,0321912 | No |
| orf19.2146 | 943 | 758 | 834 | 570 | 1329 | 869 | 1239 | 1298 | 899 | 801 | 824 | 893 | 1065 | 988 | 1052 | 1055 | 947,21 | 854 | 1040 | 1,221 | 0,288 | 4,85 | 1,23E-06 | 1,90E-06 | 0,0007928 | 0,0038247 | 0,0022887 | 0,0022887 | TRUE | 0,0204815 | No |
| orf19.2149 | 854 | 646 | 750 | 490 | 2488 | 1578 | 2360 | 2507 | 814 | 683 | 741 | 768 | 1994 | 1794 | 2004 | 2038 | 1354,47 | 752 | 1958 | 2,608 | 1,383 | 20,784 | 6,02E-96 | 3,79E-95 | 0,0030165 | 0,0033035 | 0,0032748 | 0,0032748 | TRUE | 0,0457896 | No |
| orf19.215 | 10232 | 8792 | 9481 | 6411 | 3066 | 2160 | 3099 | 3181 | 9755 | 9296 | 9366 | 10043 | 2457 | 2456 | 2631 | 2586 | 6073,92 | 9615 | 2532 | 0,264 | -1,924 | -45,116 | 0 | 0 | 0,0009566 | 0,0023616 | 0,0015109 | 0,0015109 | TRUE | 0,0106563 | No |
| orf19.2150 | 1079 | 842 | 907 | 639 | 3872 | 2464 | 3634 | 4055 | 1029 | 890 | 896 | 1001 | 3103 | 2802 | 3085 | 3297 | 2012,84 | 954 | 3072 | 3,225 | 1,689 | 24,949 | 2,21E-137 | 2,04E-136 | 0,0040738 | 0,002907 | 0,003667 | 0,003667 | TRUE | 0,0416404 | No |
| orf19.2151 | 12561 | 10014 | 11528 | 7117 | 6165 | 3873 | 5999 | 6379 | 11976 | 10589 | 11389 | 11149 | 4940 | 4404 | 5093 | 5187 | 8090,68 | 11276 | 4906 | 0,435 | -1,2 | -20,473 | 3,78E-93 | 2,33E-92 | 0,0036454 | 0,0022942 | 0,0031594 | 0,0031594 | TRUE | 0,0569358 | No |
| orf19.2154 | 2297 | 1716 | 2033 | 1220 | 3332 | 2133 | 3272 | 3224 | 2190 | 1814 | 2008 | 1911 | 2670 | 2425 | 2778 | 2621 | 2302,32 | 1981 | 2624 | 1,324 | 0,405 | 6,169 | 6,88E-10 | 1,19E-09 | 0,0042598 | 0,0028044 | 0,0036813 | 0,0036813 | TRUE | 0,0621728 | No |
| orf19.2156 | 2096 | 1576 | 1804 | 1201 | 3112 | 1885 | 3001 | 2992 | 1998 | 1666 | 1782 | 1881 | 2494 | 2143 | 2548 | 2433 | 2118,24 | 1832 | 2404 | 1,314 | 0,393 | 5,572 | 2,51E-08 | 4,12E-08 | 0,0051473 | 0,0028664 | 0,0042943 | 0,0042943 | TRUE | 0,0632293 | No |
| orf19.2157 | 5435 | 4302 | 4715 | 3257 | 3187 | 2151 | 2947 | 3266 | 5182 | 4549 | 4658 | 5102 | 2554 | 2446 | 2502 | 2656 | 3705,98 | 4873 | 2540 | 0,522 | -0,939 | -17,381 | 1,15E-67 | 5,33E-67 | 0,0023171 | 0,0025343 | 0,0025121 | 0,0025121 | TRUE | 0,0243985 | No |
| orf19.2158 | 272 | 212 | 237 | 166 | 958 | 664 | 891 | 912 | 259 | 224 | 234 | 260 | 768 | 755 | 756 | 742 | 499,79 | 244 | 755 | 3,1 | 1,632 | 21,576 | 3,01E-103 | 2,02E-102 | 1,00E-08 | 0,0053765 | 0,0026264 | 0,0026264 | TRUE | 0,0337011 | No |
| orf19.216 | 5434 | 4831 | 5353 | 3051 | 1965 | 1336 | 1979 | 1974 | 5181 | 5108 | 5288 | 4780 | 1574 | 1519 | 1680 | 1605 | 3341,95 | 5089 | 1594 | 0,313 | -1,674 | -34,63 | 8,84E-263 | 1,59E-261 | 0,0012656 | 0,0025826 | 0,0018537 | 0,0018537 | TRUE | 0,0195922 | No |
| orf19.2160 | 44 | 14 | 27 | 21 | 50 | 35 | 60 | 55 | 42 | 15 | 27 | 33 | 40 | 40 | 51 | 45 | 36,48 | 29 | 44 | 1,514 | 0,598 | 2,108 | 0,0350218 | 0,0426454 | 0,0492907 | 0,0470988 | 0,0479037 | 0,0479037 | TRUE | 0,4674138 | No |
| orf19.216.1 | 2472 | 1853 | 1767 | 1275 | 3416 | 2281 | 3089 | 3496 | 2357 | 1959 | 1746 | 1997 | 2737 | 2594 | 2623 | 2843 | 2356,88 | 2015 | 2699 | 1,34 | 0,422 | 5,275 | 1,33E-07 | 2,13E-07 | 0,00823 | 0,0027879 | 0,0056936 | 0,0056936 | TRUE | 0,1615703 | No |
| orf19.2163 | 1074 | 945 | 1083 | 605 | 5446 | 3385 | 5054 | 5297 | 1024 | 999 | 1070 | 948 | 4364 | 3849 | 4291 | 4307 | 2606,42 | 1010 | 4203 | 4,153 | 2,054 | 35,658 | 1,78E-278 | 3,56E-277 | 0,0023865 | 0,0027212 | 0,002526 | 0,002526 | TRUE | 0,0382331 | No |
| orf19.2164.1 | 0 | 0 | 1 | 2 | 0 | 1 | 1 | 0 | 0 | 0 | 1 | 3 | 0 | 1 | 1 | 0 | 0,76 | 1 | 0 | 0,485 | -1,043 | -0,559 | 0,5760014 | 0,610137 | 0,9502532 | 2,1529049 | 1,8134026 | 1,8134026 | TRUE | 0,564892 | No |
| orf19.2165 | 1430 | 1650 | 1664 | 938 | 2942 | 2211 | 2840 | 2394 | 1363 | 1745 | 1644 | 1469 | 2357 | 2514 | 2411 | 1947 | 1931,31 | 1555 | 2307 | 1,482 | 0,568 | 6,165 | 7,07E-10 | 1,22E-09 | 0,0111695 | 0,0029414 | 0,007594 | 0,007594 | TRUE | 0,1350704 | No |
| orf19.2166 | 1339 | 1573 | 1587 | 980 | 2287 | 1819 | 1961 | 1809 | 1277 | 1663 | 1568 | 1535 | 1833 | 2068 | 1665 | 1471 | 1634,94 | 1511 | 1759 | 1,164 | 0,219 | 2,108 | 0,0350266 | 0,0426454 | 0,0155566 | 0,0030955 | 0,0097449 | 0,0097449 | TRUE | 0,1669282 | No |
| orf19.2167 | 9837 | 9265 | 9602 | 5517 | 12078 | 8260 | 11505 | 11820 | 9379 | 9797 | 9486 | 8643 | 9678 | 9392 | 9768 | 9611 | 9469,03 | 9326 | 9612 | 1,03 | 0,043 | 0,999 | 0,3178646 | 0,3510981 | 0,0013711 | 0,0022647 | 0,0016675 | 0,0016675 | TRUE | 0,0288874 | No |
| orf19.2168 | 459 | 435 | 466 | 270 | 914 | 695 | 813 | 839 | 438 | 460 | 460 | 423 | 732 | 790 | 690 | 682 | 584,49 | 445 | 724 | 1,619 | 0,695 | 9,544 | 1,37E-21 | 3,18E-21 | 0,0016818 | 0,0049004 | 0,0032188 | 0,0032188 | TRUE | 0,0420736 | No |
| orf19.2168.3 | 2647 | 2487 | 2522 | 1765 | 6845 | 5517 | 6346 | 6476 | 2524 | 2630 | 2491 | 2765 | 5485 | 6273 | 5388 | 5266 | 4102,64 | 2602 | 5603 | 2,154 | 1,107 | 17,806 | 6,38E-71 | 3,09E-70 | 0,0040592 | 0,0024915 | 0,0034152 | 0,0034152 | TRUE | 0,0782982 | No |
| orf19.2169 | 12 | 7 | 13 | 14 | 34 | 20 | 20 | 34 | 11 | 7 | 13 | 22 | 27 | 23 | 17 | 28 | 18,53 | 13 | 24 | 1,836 | 0,876 | 2,376 | 0,0175124 | 0,0218341 | 0,0315715 | 0,0907069 | 0,0687839 | 0,0687839 | TRUE | 0,4862845 | No |
| orf19.217 | 138 | 131 | 121 | 67 | 116 | 81 | 92 | 91 | 132 | 139 | 120 | 105 | 93 | 92 | 78 | 74 | 103,97 | 124 | 84 | 0,675 | -0,567 | -3,788 | 0,0001518 | 0,0002129 | 0,0030533 | 0,0178839 | 0,0117582 | 0,0117582 | TRUE | 0,0806914 | No |
| orf19.2170 | 6274 | 5988 | 6388 | 4084 | 9019 | 6874 | 8295 | 9214 | 5982 | 6332 | 6311 | 6398 | 7227 | 7816 | 7042 | 7492 | 6824,87 | 6256 | 7394 | 1,182 | 0,241 | 5,533 | 3,14E-08 | 5,15E-08 | 0,0012818 | 0,0023319 | 0,0016761 | 0,0016761 | TRUE | 0,0177102 | No |
| orf19.2172 | 11777 | 11357 | 11125 | 6229 | 4273 | 3000 | 3837 | 4064 | 11228 | 12009 | 10990 | 9758 | 3424 | 3411 | 3258 | 3304 | 7172,79 | 10996 | 3349 | 0,304 | -1,716 | -28,609 | 5,21E-180 | 6,26E-179 | 0,0036346 | 0,0023202 | 0,0032704 | 0,0032704 | TRUE | 0,0693043 | No |
| orf19.2173 | 13142 | 11478 | 12638 | 7630 | 1741 | 1248 | 1621 | 1860 | 12529 | 12137 | 12485 | 11953 | 1395 | 1419 | 1376 | 1512 | 6850,83 | 12276 | 1426 | 0,116 | -3,106 | -74,967 | 0 | 0 | 0,0006481 | 0,0023309 | 0,00129 | 0,00129 | TRUE | 0,0211278 | No |
| orf19.2174 | 575 | 484 | 516 | 290 | 916 | 574 | 903 | 912 | 548 | 512 | 510 | 454 | 734 | 653 | 767 | 742 | 614,86 | 506 | 724 | 1,427 | 0,512 | 6,675 | 2,47E-11 | 4,50E-11 | 0,0030175 | 0,0047617 | 0,0039424 | 0,0039424 | TRUE | 0,0481055 | No |
| orf19.2175 | 21385 | 18393 | 20029 | 12354 | 2541 | 1810 | 2451 | 2563 | 20388 | 19448 | 19787 | 19353 | 2036 | 2058 | 2081 | 2084 | 10904,44 | 19744 | 2065 | 0,105 | -3,258 | -95,755 | 0 | 0 | 1,00E-08 | 0,0022419 | 0,000867 | 0,000867 | TRUE | 0,0059374 | No |
| orf19.2176 | 1796 | 1396 | 1636 | 980 | 1525 | 933 | 1385 | 1485 | 1712 | 1476 | 1616 | 1535 | 1222 | 1061 | 1176 | 1207 | 1375,75 | 1585 | 1166 | 0,736 | -0,441 | -6,81 | 9,75E-12 | 1,79E-11 | 0,003057 | 0,0032847 | 0,0032975 | 0,0032975 | TRUE | 0,0422471 | No |
| orf19.2177 | 520 | 436 | 495 | 388 | 922 | 700 | 946 | 989 | 496 | 461 | 489 | 608 | 739 | 796 | 803 | 804 | 649,46 | 514 | 786 | 1,539 | 0,622 | 7,571 | 3,71E-14 | 7,32E-14 | 0,0050566 | 0,0046194 | 0,0048212 | 0,0048212 | TRUE | 0,1569371 | No |
| orf19.2178 | 3490 | 3007 | 3359 | 2511 | 4593 | 3618 | 4447 | 4725 | 3327 | 3180 | 3318 | 3934 | 3680 | 4114 | 3775 | 3842 | 3646,3 | 3440 | 3853 | 1,121 | 0,165 | 2,454 | 0,0141184 | 0,0177551 | 0,0055771 | 0,0025416 | 0,004063 | 0,004063 | TRUE | 0,1103399 | No |
| orf19.2178.1 | 836 | 666 | 660 | 399 | 891 | 509 | 782 | 819 | 797 | 704 | 652 | 625 | 714 | 579 | 664 | 666 | 675,11 | 694 | 656 | 0,943 | -0,084 | -0,94 | 0,3470439 | 0,3814639 | 0,0076531 | 0,0045233 | 0,0062558 | 0,0062558 | TRUE | 0,1183746 | No |
| orf19.2179 | 11681 | 9583 | 10435 | 6393 | 14918 | 9781 | 14046 | 14955 | 11137 | 10133 | 10309 | 10015 | 11953 | 11122 | 11925 | 12160 | 11094,15 | 10398 | 11790 | 1,134 | 0,181 | 3,909 | 9,26E-05 | 0,0001314 | 0,0018201 | 0,0022393 | 0,0019685 | 0,0019685 | TRUE | 0,0280962 | No |
| orf19.2179.2 | 15850 | 12553 | 12815 | 8071 | 39558 | 25423 | 37049 | 39599 | 15111 | 13273 | 12660 | 12644 | 31697 | 28908 | 31454 | 32198 | 22243,16 | 13422 | 31064 | 2,314 | 1,21 | 19,475 | 1,79E-84 | 1,02E-83 | 0,0048071 | 0,0021651 | 0,0036552 | 0,0036552 | TRUE | 0,0880857 | No |
| orf19.218 | 28 | 20 | 24 | 15 | 61 | 46 | 58 | 62 | 27 | 21 | 24 | 23 | 49 | 52 | 49 | 50 | 36,99 | 24 | 50 | 2,103 | 1,073 | 4,549 | 5,38E-06 | 8,08E-06 | 1,00E-08 | 0,0464839 | 0,0212742 | 0,0212742 | TRUE | 0,0476616 | No |

| | | | | | | | | | | | | | | | | | | | | | | | | | | | | | | | |
|---|---|---|---|---|---|---|---|---|---|---|---|---|---|---|---|---|---|---|---|---|---|---|---|---|---|---|---|---|---|---|---|
| orf19.2180 | 3070 | 2811 | 3147 | 2115 | 2057 | 1448 | 1929 | 2002 | 2927 | 2972 | 3109 | 3313 | 1648 | 1647 | 1638 | 1628 | 2360,2 | 3080 | 1640 | 0,533 | -0,907 | -18,718 | 3,52E-78 | 1,88E-77 | 0,0011881 | 0,0027869 | 0,0018001 | 0,0018001 | TRUE | 0,029543 | No |
| orf19.2181 | 23 | 27 | 19 | 12 | 52 | 29 | 38 | 41 | 22 | 29 | 19 | 19 | 42 | 33 | 32 | 33 | 28,54 | 22 | 35 | 1,587 | 0,666 | 2,454 | 0,0141439 | 0,0177837 | 1,00E-08 | 0,0596297 | 0,0330164 | 0,0330164 | TRUE | 0,2216152 | No |
| orf19.2182 | 7587 | 7146 | 7412 | 4787 | 7661 | 5599 | 6960 | 7902 | 7233 | 7556 | 7322 | 7499 | 6139 | 6367 | 5909 | 6425 | 6806,25 | 7402 | 6210 | 0,839 | -0,253 | -6,566 | 5,16E-11 | 9,29E-11 | 0,0007287 | 0,0023325 | 0,0012825 | 0,0012825 | TRUE | 0,0130181 | No |
| orf19.2183 | 24152 | 20980 | 22497 | 15119 | 46626 | 32778 | 44923 | 48900 | 23026 | 22184 | 22225 | 23685 | 37360 | 37272 | 38139 | 39760 | 30456,38 | 22780 | 38133 | 1,674 | 0,744 | 20,208 | 8,35E-91 | 5,03E-90 | 0,0009059 | 0,0021452 | 0,0012642 | 0,0012642 | TRUE | 0,0101267 | No |
| orf19.2184 | 234 | 182 | 201 | 135 | 721 | 470 | 668 | 686 | 223 | 192 | 199 | 211 | 578 | 534 | 567 | 558 | 382,83 | 206 | 559 | 2,714 | 1,441 | 17,194 | 2,93E-66 | 1,34E-65 | 1,00E-08 | 0,0063801 | 0,0032537 | 0,0032537 | TRUE | 0,0387368 | No |
| orf19.2185 | 4931 | 4107 | 4640 | 2764 | 7417 | 4954 | 7171 | 7546 | 4701 | 4343 | 4584 | 4330 | 5943 | 5633 | 6088 | 6136 | 5219,7 | 4490 | 5950 | 1,325 | 0,406 | 9,085 | 1,04E-19 | 2,32E-19 | 0,0012974 | 0,0024058 | 0,0017166 | 0,0017166 | TRUE | 0,0153324 | No |
| orf19.2186 | 1090 | 917 | 1026 | 668 | 1230 | 776 | 1094 | 1178 | 1039 | 970 | 1014 | 1046 | 986 | 882 | 929 | 958 | 977,93 | 1017 | 939 | 0,925 | -0,112 | -1,98 | 0,0477367 | 0,0575166 | 0,0006087 | 0,0037702 | 0,0020624 | 0,0020624 | TRUE | 0,0178578 | No |
| orf19.2187 | 835 | 707 | 859 | 562 | 1913 | 1402 | 1792 | 1939 | 796 | 748 | 849 | 880 | 1533 | 1594 | 1521 | 1577 | 1187,21 | 818 | 1556 | 1,907 | 0,931 | 16,058 | 5,02E-58 | 2,06E-57 | 0,0009786 | 0,0034743 | 0,002256 | 0,002256 | TRUE | 0,0390406 | No |
| orf19.2190 | 1007 | 797 | 951 | 612 | 992 | 738 | 900 | 1121 | 960 | 843 | 939 | 959 | 795 | 839 | 764 | 911 | 876,33 | 925 | 827 | 0,895 | -0,161 | -2,215 | 0,0267286 | 0,0328749 | 0,0038355 | 0,0039649 | 0,0038939 | 0,0038939 | TRUE | 0,0580836 | No |
| orf19.2191 | 1661 | 1359 | 1447 | 944 | 2532 | 1735 | 2426 | 2626 | 1584 | 1437 | 1429 | 1479 | 2029 | 1973 | 2060 | 2135 | 1765,68 | 1482 | 2049 | 1,383 | 0,468 | 9,214 | 3,13E-20 | 7,07E-20 | 0,0010885 | 0,0030212 | 0,0018757 | 0,0018757 | TRUE | 0,0272716 | No |
| orf19.2191.1 | 251 | 199 | 194 | 172 | 657 | 350 | 526 | 644 | 239 | 210 | 192 | 269 | 526 | 398 | 447 | 524 | 350,68 | 228 | 474 | 2,098 | 1,069 | 8,689 | 3,67E-18 | 7,90E-18 | 0,0155605 | 0,0067733 | 0,0111095 | 0,0111095 | TRUE | 0,1431885 | No |
| orf19.2192 | 81189 | 72951 | 78116 | 50388 | 122181 | 88544 | 113653 | 127304 | 77405 | 77136 | 77171 | 78935 | 97900 | 100682 | 96491 | 103510 | 88653,98 | 77662 | 99646 | 1,283 | 0,36 | 11,451 | 2,34E-30 | 6,34E-30 | 0,0005326 | 0,0021098 | 0,000936 | 0,000936 | TRUE | 0,0083694 | No |
| orf19.2193 | 9165 | 7676 | 8212 | 5366 | 10083 | 7133 | 9642 | 10449 | 8738 | 8116 | 8113 | 8406 | 8079 | 8111 | 8186 | 8496 | 8280,65 | 8343 | 8218 | 0,985 | -0,022 | -0,573 | 0,5664175 | 0,6007829 | 0,0007199 | 0,0022896 | 0,0012494 | 0,0012494 | TRUE | 0,012733 | No |
| orf19.2194 | 984 | 903 | 928 | 583 | 1164 | 744 | 1034 | 1022 | 938 | 955 | 917 | 913 | 933 | 846 | 878 | 831 | 901,32 | 931 | 872 | 0,937 | -0,094 | -1,623 | 0,1045974 | 0,1218775 | 0,0006611 | 0,003913 | 0,0021059 | 0,0021059 | TRUE | 0,0272234 | No |
| orf19.2195 | 7 | 9 | 3 | 5 | 3 | 5 | 3 | 3 | 7 | 10 | 3 | 8 | 2 | 6 | 3 | 2 | 5,01 | 7 | 3 | 0,473 | -1,079 | -1,62 | 0,1051679 | 0,1225039 | 1,00E-08 | 0,3299628 | 0,2017343 | 0,2017343 | TRUE | 0,3998074 | No |
| orf19.2196 | 983 | 939 | 1045 | 589 | 508 | 404 | 534 | 536 | 937 | 993 | 1032 | 923 | 407 | 459 | 453 | 436 | 705,09 | 971 | 439 | 0,45 | -1,151 | -16,966 | 1,46E-64 | 6,54E-64 | 0,0013308 | 0,0044199 | 0,0028254 | 0,0028254 | TRUE | 0,0286627 | No |
| orf19.2197 | 65 | 38 | 45 | 41 | 59 | 47 | 54 | 57 | 62 | 40 | 44 | 64 | 47 | 53 | 46 | 46 | 50,47 | 52 | 48 | 0,92 | -0,12 | -0,565 | 0,5721194 | 0,6064275 | 0,0064397 | 0,0346247 | 0,0235039 | 0,0235039 | TRUE | 0,1696061 | No |
| orf19.2198 | 3973 | 3634 | 3934 | 2495 | 4798 | 3520 | 4567 | 5093 | 3788 | 3842 | 3886 | 3909 | 3844 | 4003 | 3877 | 4141 | 3911,35 | 3856 | 3966 | 1,029 | 0,041 | 1,071 | 0,2841504 | 0,3162128 | 0,0004916 | 0,0025111 | 0,0011447 | 0,0011447 | TRUE | 0,0110649 | No |
| orf19.2199 | 3640 | 3281 | 3538 | 2438 | 5912 | 4326 | 5600 | 5952 | 3470 | 3469 | 3495 | 3819 | 4737 | 4919 | 4754 | 4840 | 4188,02 | 3563 | 4812 | 1,352 | 0,435 | 10,426 | 1,88E-25 | 4,70E-25 | 0,0009768 | 0,0024833 | 0,0014237 | 0,0014237 | TRUE | 0,0265271 | No |
| orf19.22 | 3427 | 3108 | 3220 | 2059 | 2410 | 1787 | 2247 | 2480 | 3267 | 3286 | 3181 | 3226 | 1931 | 2032 | 1908 | 2016 | 2605,92 | 3240 | 1972 | 0,608 | -0,718 | -18,093 | 3,63E-73 | 1,81E-72 | 0,0001376 | 0,0027213 | 0,0011105 | 0,0011105 | TRUE | 0,0057153 | No |
| orf19.220 | 14471 | 12613 | 13578 | 9559 | 13285 | 10187 | 12366 | 12838 | 13797 | 13337 | 13414 | 14975 | 10645 | 11584 | 10499 | 10439 | 12335,91 | 13881 | 10792 | 0,778 | -0,363 | -7,118 | 1,09E-12 | 2,06E-12 | 0,0026059 | 0,0022244 | 0,0024164 | 0,0024164 | TRUE | 0,0340231 | No |
| orf19.2200 | 1503 | 1372 | 1509 | 924 | 2539 | 1798 | 2379 | 2486 | 1433 | 1451 | 1491 | 1447 | 2034 | 2044 | 2020 | 2021 | 1742,74 | 1456 | 2030 | 1,394 | 0,479 | 11,739 | 8,06E-32 | 2,25E-31 | 1,00E-08 | 0,0030334 | 0,0009978 | 0,0009978 | TRUE | 0,0034352 | No |
| orf19.2201 | 1142 | 1015 | 999 | 620 | 1329 | 961 | 1250 | 1293 | 1089 | 1073 | 987 | 971 | 1065 | 1093 | 1061 | 1051 | 1048,8 | 1030 | 1068 | 1,033 | 0,047 | 0,851 | 0,3944946 | 0,4303762 | 0,0005362 | 0,0036568 | 0,0020226 | 0,0020226 | TRUE | 0,0177526 | No |
| orf19.2202 | 128 | 122 | 125 | 86 | 463 | 317 | 505 | 489 | 122 | 129 | 123 | 135 | 371 | 360 | 429 | 398 | 258,38 | 127 | 390 | 3,077 | 1,622 | 15,276 | 1,11E-52 | 4,24E-52 | 1,00E-08 | 0,0084459 | 0,0052988 | 0,0052988 | TRUE | 0,0531711 | No |
| orf19.2203 | 3 | 5 | 5 | 3 | 7 | 4 | 6 | 2 | 3 | 5 | 5 | 5 | 6 | 5 | 5 | 2 | 4,33 | 4 | 4 | 0,952 | -0,071 | -0,105 | 0,9164127 | 0,9245264 | 1,00E-08 | 0,381025 | 0,2067693 | 0,2067693 | TRUE | 0,3859413 | No |
| orf19.2203.1 | 59 | 56 | 55 | 43 | 26 | 20 | 31 | 25 | 56 | 59 | 54 | 67 | 21 | 23 | 26 | 20 | 40,92 | 59 | 22 | 0,383 | -1,386 | -6,034 | 1,60E-09 | 2,73E-09 | 1,00E-08 | 0,0422135 | 0,0214813 | 0,0214813 | TRUE | 0,0856771 | No |
| orf19.2204 | 6978 | 5840 | 6374 | 4202 | 5232 | 3663 | 5116 | 5389 | 6653 | 6175 | 6297 | 6583 | 4192 | 4165 | 4343 | 4382 | 5348,75 | 6427 | 4270 | 0,665 | -0,589 | -14,919 | 2,48E-50 | 9,17E-50 | 0,0006872 | 0,0023982 | 0,0013039 | 0,0013039 | TRUE | 0,0085031 | No |
| orf19.2204.2 | 1005 | 815 | 890 | 645 | 832 | 581 | 808 | 828 | 958 | 862 | 879 | 1010 | 667 | 661 | 686 | 673 | 799,51 | 927 | 672 | 0,727 | -0,461 | -7,106 | 1,19E-12 | 2,25E-12 | 0,0015292 | 0,0041449 | 0,0027686 | 0,0027686 | TRUE | 0,0380756 | No |
| orf19.2208 | 7226 | 6154 | 6634 | 4384 | 5732 | 3994 | 5394 | 5662 | 6889 | 6507 | 6554 | 6868 | 4593 | 4542 | 4579 | 4604 | 5641,93 | 6704 | 4580 | 0,683 | -0,549 | -16,066 | 4,42E-58 | 1,81E-57 | 0,0003205 | 0,0023823 | 0,0009407 | 0,0009407 | TRUE | 0,0047663 | No |
| orf19.2209 | 1356 | 1086 | 1289 | 743 | 2529 | 1733 | 2329 | 2510 | 1293 | 1148 | 1273 | 1164 | 2026 | 1971 | 1977 | 2041 | 1611,7 | 1220 | 2004 | 1,64 | 0,714 | 13,69 | 1,16E-42 | 3,85E-42 | 0,0011152 | 0,00311 | 0,0019267 | 0,0019267 | TRUE | 0,0206087 | No |
| orf19.2210 | 2050 | 1824 | 1985 | 1351 | 1388 | 987 | 1294 | 1423 | 1954 | 1929 | 1961 | 2116 | 1112 | 1122 | 1099 | 1157 | 1556,33 | 1990 | 1122 | 0,565 | -0,823 | -16,968 | 1,41E-64 | 6,29E-64 | 0,0004266 | 0,0031463 | 0,0015799 | 0,0015799 | TRUE | 0,0199391 | No |
| orf19.2211 | 11 | 13 | 9 | 9 | 8 | 6 | 8 | 14 | 10 | 14 | 9 | 14 | 6 | 7 | 7 | 11 | 9,83 | 12 | 8 | 0,678 | -0,56 | -1,24 | 0,2148486 | 0,2428608 | 1,00E-08 | 0,1691396 | 0,0915785 | 0,0915785 | TRUE | 0,3274707 | No |
| orf19.2212 | 35 | 23 | 42 | 15 | 21 | 20 | 18 | 26 | 33 | 24 | 41 | 23 | 17 | 23 | 15 | 21 | 24,83 | 30 | 19 | 0,607 | -0,721 | -2,376 | 0,0174793 | 0,0218013 | 1,00E-08 | 0,0682072 | 0,0468903 | 0,0468903 | TRUE | 0,378983 | No |
| orf19.2213 | 125 | 129 | 152 | 88 | 235 | 186 | 242 | 285 | 119 | 136 | 150 | 138 | 188 | 211 | 205 | 232 | 172,57 | 136 | 209 | 1,541 | 0,624 | 5,157 | 2,51E-07 | 3,97E-07 | 1,00E-08 | 0,0116056 | 0,0078283 | 0,0078283 | TRUE | 0,0800324 | No |
| orf19.2214 | 2492 | 2131 | 2307 | 1546 | 2704 | 1903 | 2532 | 2732 | 2376 | 2253 | 2279 | 2422 | 2167 | 2164 | 2150 | 2221 | 2253,96 | 2332 | 2176 | 0,934 | -0,099 | -2,392 | 0,0167688 | 0,0209561 | 0,0001565 | 0,0028197 | 0,0011921 | 0,0011921 | TRUE | 0,0071603 | No |
| orf19.2215 | 5108 | 4387 | 4811 | 3150 | 3343 | 2398 | 3288 | 3331 | 4870 | 4639 | 4753 | 4935 | 2679 | 2727 | 2791 | 2708 | 3762,67 | 4799 | 2726 | 0,568 | -0,815 | -22,099 | 3,20E-108 | 2,28E-107 | 0,0002762 | 0,0025276 | 0,0010252 | 0,0010252 | TRUE | 0,0061052 | No |
| orf19.2216 | 238 | 229 | 225 | 145 | 693 | 501 | 649 | 721 | 227 | 242 | 222 | 227 | 555 | 570 | 551 | 586 | 397,58 | 230 | 566 | 2,462 | 1,3 | 16,417 | 1,44E-60 | 6,10E-60 | 1,00E-08 | 0,006221 | 0,0028167 | 0,0028167 | TRUE | 0,0155527 | No |
| orf19.2219 | 3019 | 2689 | 3026 | 1971 | 2603 | 1951 | 2501 | 2720 | 2878 | 2843 | 2989 | 3088 | 2086 | 2218 | 2123 | 2212 | 2554,72 | 2950 | 2160 | 0,733 | -0,449 | -10,129 | 4,10E-24 | 9,98E-24 | 0,0006374 | 0,002734 | 0,0014872 | 0,0014872 | TRUE | 0,0111235 | No |
| orf19.2220 | 285 | 266 | 226 | 161 | 185 | 108 | 149 | 179 | 272 | 281 | 223 | 252 | 148 | 123 | 127 | 146 | 196,44 | 257 | 136 | 0,53 | -0,916 | -7,757 | 8,69E-15 | 1,74E-14 | 0,0050349 | 0,0104494 | 0,0079444 | 0,0079444 | TRUE | 0,0950962 | No |
| orf19.2222 | 2727 | 2514 | 2842 | 1696 | 2404 | 1697 | 2424 | 2444 | 2600 | 2658 | 2808 | 2657 | 1926 | 1930 | 2058 | 1987 | 2327,96 | 2681 | 1975 | 0,737 | -0,44 | -9,927 | 3,19E-23 | 7,63E-23 | 0,0006593 | 0,0027966 | 0,0014536 | 0,0014536 | TRUE | 0,0128333 | No |
| orf19.2227 | 146 | 158 | 131 | 88 | 159 | 119 | 140 | 157 | 139 | 167 | 129 | 138 | 127 | 135 | 119 | 128 | 135,35 | 143 | 127 | 0,885 | -0,176 | -1,369 | 0,1710152 | 0,1952143 | 1,00E-08 | 0,0142225 | 0,0083943 | 0,0083943 | TRUE | 0,132504 | No |
| orf19.2228 | 195 | 226 | 203 | 146 | 253 | 221 | 204 | 208 | 186 | 239 | 201 | 229 | 203 | 251 | 173 | 169 | 206,31 | 214 | 199 | 0,93 | -0,104 | -0,737 | 0,4613168 | 0,4976607 | 0,018813 | 0,0100497 | 0,0142492 | 0,0142492 | TRUE | 0,3165388 | No |
| orf19.2229 | 1259 | 1083 | 1175 | 727 | 2000 | 1299 | 1877 | 1932 | 1200 | 1145 | 1161 | 1139 | 1603 | 1477 | 1594 | 1571 | 1361,15 | 1161 | 1561 | 1,344 | 0,427 | 8,679 | 3,98E-18 | 8,56E-18 | 1,00E-08 | 0,0032975 | 0,0015568 | 0,0015568 | TRUE | 0,0146704 | No |
| orf19.223 | 10082 | 9243 | 9935 | 6231 | 8872 | 6136 | 8371 | 8859 | 9612 | 9773 | 9815 | 9761 | 7109 | 6977 | 7107 | 7203 | 8419,71 | 9740 | 7099 | 0,729 | -0,456 | -16,646 | 3,21E-62 | 1,39E-61 | 1,00E-08 | 0,0022863 | 0,0005994 | 0,0005994 | TRUE | 0,0015582 | No |
| orf19.2230 | 550 | 490 | 534 | 290 | 821 | 596 | 751 | 751 | 524 | 518 | 528 | 454 | 658 | 678 | 638 | 611 | 576,01 | 506 | 646 | 1,268 | 0,343 | 4,861 | 1,17E-06 | 1,80E-06 | 0,0008092 | 0,0049417 | 0,0029893 | 0,0029893 | TRUE | 0,0458015 | No |
| orf19.2231 | 2191 | 2044 | 2094 | 1567 | 1837 | 1397 | 1686 | 1762 | 2089 | 2161 | 2069 | 2455 | 1472 | 1589 | 1431 | 1433 | 1837,27 | 2194 | 1481 | 0,676 | -0,565 | -8,818 | 1,16E-18 | 2,54E-18 | 0,0038147 | 0,0029849 | 0,003382 | 0,003382 | TRUE | 0,0741404 | No |
| orf19.2232 | 12829 | 10735 | 10913 | 6754 | 33848 | 21696 | 30838 | 35340 | 12231 | 11351 | 10781 | 10581 | 27121 | 24670 | 26181 | 28735 | 18956,42 | 11236 | 26677 | 2,374 | 1,247 | 20,962 | 1,47E-97 | 9,36E-97 | 0,0041194 | 0,0021779 | 0,0033344 | 0,0033344 | TRUE | 0,0435946 | No |
| orf19.2233 | 4739 | 4026 | 4291 | 2560 | 5412 | 3844 | 5189 | 5201 | 4518 | 4257 | 4239 | 4010 | 4336 | 4371 | 4405 | 4229 | 4295,8 | 4256 | 4335 | 1,017 | 0,025 | 0,578 | 0,5633157 | 0,5978905 | 0,0009965 | 0,0024735 | 0,0015431 | 0,0015431 | TRUE | 0,0209482 | No |
| orf19.2236 | 4447 | 4008 | 4436 | 2735 | 3799 | 2696 | 3647 | 3944 | 4240 | 4238 | 4382 | 4285 | 3044 | 3066 | 3096 | 3207 | 3694,66 | 4286 | 3103 | 0,724 | -0,466 | -13,126 | 2,33E-39 | 7,30E-39 | 0,0001557 | 0,0025357 | 0,0009331 | 0,0009331 | TRUE | 0,0062953 | No |
| orf19.2237 | 76 | 56 | 67 | 41 | 40 | 26 | 28 | 32 | 72 | 59 | 66 | 64 | 32 | 30 | 24 | 26 | 46,69 | 65 | 28 | 0,423 | -1,24 | -5,796 | 6,80E-09 | 1,14E-08 | 1,00E-08 | 0,0372597 | 0,0194992 | 0,0194992 | TRUE | 0,0838774 | No |
| orf19.2237.1 | 2362 | 2450 | 2452 | 1556 | 2706 | 1867 | 2419 | 2466 | 2252 | 2591 | 2422 | 2438 | 2168 | 2123 | 2054 | 2005 | 2256,55 | 2426 | 2088 | 0,861 | -0,216 | -4,09 | 4,31E-05 | 6,21E-05 | 0,0016364 | 0,0028189 | 0,0022385 | 0,0022385 | TRUE | 0,0292496 | No |
| orf19.2238 | 1040 | 1095 | 1113 | 658 | 700 | 526 | 705 | 693 | 992 | 1158 | 1100 | 1031 | 561 | 598 | 599 | 563 | 825,09 | 1070 | 580 | 0,541 | -0,885 | -13,238 | 5,30E-40 | 1,68E-39 | 0,0021265 | 0,0040813 | 0,0029991 | 0,0029991 | TRUE | 0,0369824 | No |
| orf19.2241 | 5701 | 4703 | 5235 | 3228 | 2576 | 1803 | 2646 | 2788 | 5435 | 4973 | 5172 | 5057 | 2064 | 2050 | 2246 | 2267 | 3658,03 | 5159 | 2157 | 0,418 | -1,258 | -25,056 | 1,49E-138 | 1,39E-137 | 0,0017606 | 0,0025401 | 0,0021034 | 0,0021034 | TRUE | 0,0160457 | No |
| orf19.2242 | 10748 | 9808 | 10517 | 6796 | 11187 | 7735 | 10490 | 11227 | 10247 | 10371 | 10390 | 10646 | 8964 | 8795 | 8906 | 9129 | 9680,96 | 10414 | 8948 | 0,86 | -0,218 | -7,56 | 4,03E-14 | 7,93E-14 | 0,0001116 | 0,0022609 | 0,0006937 | 0,0006937 | TRUE | 0,0026365 | No |
| orf19.2244 | 805 | 711 | 788 | 549 | 4855 | 3598 | 4565 | 5038 | 767 | 752 | 778 | 860 | 3890 | 4091 | 3876 | 4096 | 2388,91 | 789 | 3988 | 5,073 | 2,343 | 45,37 | 0 | 0 | 0,000854 | 0,0027786 | 0,001738 | 0,001738 | TRUE | 0,0360651 | No |
| orf19.2245 | 642 | 597 | 603 | 365 | 2003 | 1612 | 2097 | 2119 | 612 | 631 | 596 | 572 | 1605 | 1833 | 1780 | 1723 | 1169,01 | 603 | 1735 | 2,87 | 1,521 | 24,426 | 9,10E-132 | 8,07E-131 | 0,0018303 | 0,0034958 | 0,0025405 | 0,0025405 | TRUE | 0,031227 | No |
| orf19.2246 | 1003 | 876 | 896 | 505 | 1981 | 1281 | 1975 | 1952 | 956 | 926 | 885 | 791 | 1587 | 1457 | 1677 | 1587 | 1233,33 | 890 | 1577 | 1,768 | 0,822 | 12,123 | 7,99E-34 | 2,29E-33 | 0,0037453 | 0,0034226 | 0,0035013 | 0,0035013 | TRUE | 0,0588891 | No |
| orf19.2247 | 102 | 96 | 90 | 81 | 83 | 46 | 82 | 74 | 97 | 102 | 89 | 127 | 67 | 52 | 70 | 60 | 82,89 | 104 | 62 | 0,61 | -0,713 | -4,202 | 2,65E-05 | 3,85E-05 | 0,0050587 | 0,0218985 | 0,0150968 | 0,0150968 | TRUE | 0,1777352 | No |
| orf19.2248 | 1205 | 1128 | 1244 | 804 | 1412 | 1066 | 1437 | 1443 | 1149 | 1193 | 1229 | 1260 | 1131 | 1212 | 1220 | 1173 | 1195,86 | 1208 | 1184 | 0,982 | -0,027 | -0,505 | 0,6135524 | 0,6473343 | 0,0005448 | 0,0034643 | 0,001815 | 0,001815 | TRUE | 0,0130875 | No |
| orf19.2249 | 361 | 273 | 328 | 202 | 1258 | 835 | 1089 | 1240 | 344 | 289 | 324 | 316 | 1008 | 949 | 925 | 1008 | 645,45 | 318 | 972 | 3,052 | 1,61 | 21,821 | 1,48E-105 | 1,01E-104 | 0,0013096 | 0,0046351 | 0,0030214 | 0,0030214 | TRUE | 0,0481856 | No |
| orf19.225 | 3602 | 3276 | 3568 | 2043 | 4456 | 3079 | 4479 | 4680 | 3434 | 3464 | 3525 | 3200 | 3570 | 3501 | 3803 | 3805 | 3537,86 | 3406 | 3670 | 1,077 | 0,107 | 2,257 | 0,0240118 | 0,0296477 | 0,001412 | 0,0025554 | 0,0018611 | 0,0018611 | TRUE | 0,0188332 | No |
| orf19.2250 | 4273 | 3530 | 3599 | 2541 | 8439 | 6187 | 7579 | 9010 | 4074 | 3733 | 3555 | 3981 | 6762 | 7035 | 6435 | 7326 | 5362,51 | 3836 | 6890 | 1,797 | 0,845 | 14,824 | 1,02E-49 | 3,75E-49 | 0,0031475 | 0,0023975 | 0,0029112 | 0,0029112 | TRUE | 0,0297828 | No |
| orf19.2251 | 9027 | 7569 | 7943 | 5390 | 7957 | 5873 | 7434 | 7897 | 8606 | 8003 | 7847 | 8444 | 6376 | 6678 | 6311 | 6421 | 7335,81 | 8225 | 6446 | 0,784 | -0,352 | -8,474 | 2,38E-17 | 5,04E-17 | 0,0010758 | 0,0023151 | 0,0015151 | 0,0015151 | TRUE | 0,0122905 | No |
| orf19.2252 | 5 | 8 | 3 | 3 | 5 | 0 | 4 | 3 | 5 | 8 | 3 | 5 | 4 | 0 | 3 | 2 | 3,84 | 5 | 2 | 0,492 | -1,023 | -1,337 | 0,1811593 | 0,2063508 | 1,00E-08 | 0,429518 | 0,2822734 | 0,2822734 | TRUE | 0,3664075 | No |
| orf19.2253 | 273 | 201 | 231 | 152 | 343 | 236 | 285 | 324 | 260 | 213 | 228 | 238 | 275 | 268 | 242 | 263 | 248,47 | 235 | 262 | 1,116 | 0,158 | 1,595 | 0,1107193 | 0,1286091 | 1,00E-08 | 0,0086994 | 0,0053306 | 0,0053306 | TRUE | 0,0666211 | No |
| orf19.2256 | 4175 | 3294 | 3645 | 2261 | 5867 | 3739 | 5815 | 5844 | 3980 | 3483 | 3601 | 3542 | 4701 | 4252 | 4937 | 4752 | 4155,95 | 3652 | 4660 | 1,276 | 0,352 | 5,92 | 3,23E-09 | 5,45E-09 | 0,0035094 | 0,0024864 | 0,0031427 | 0,0031427 | TRUE | 0,045356 | No |
| orf19.2257 | 1604 | 1403 | 1507 | 898 | 4682 | 3464 | 4598 | 4796 | 1529 | 1483 | 1489 | 1407 | 3752 | 3939 | 3904 | 3900 | 2675,25 | 1477 | 3874 | 2,616 | 1,387 | 33,078 | 6,25E-240 | 1,01E-238 | 0,000206 | 0,002705 | 0,0011954 | 0,0011954 | TRUE | 0,0106386 | No |
| orf19.2258 | 1115 | 1015 | 1138 | 602 | 1487 | 1005 | 1511 | 1538 | 1063 | 1073 | 1124 | 943 | 1191 | 1143 | 1283 | 1251 | 1133,9 | 1051 | 1217 | 1,155 | 0,208 | 3,231 | 0,0012345 | 0,0016528 | 0,0027895 | 0,0035393 | 0,0030963 | 0,0030963 | TRUE | 0,0509622 | No |
| orf19.2259 | 1951 | 1698 | 1774 | 1044 | 1903 | 1313 | 1810 | 1925 | 1860 | 1795 | 1753 | 1635 | 1525 | 1493 | 1537 | 1565 | 1645,4 | 1761 | 1530 | 0,867 | -0,205 | -4,105 | 4,04E-05 | 5,83E-05 | 0,0008184 | 0,0030892 | 0,001794 | 0,001794 | TRUE | 0,0249526 | No |
| orf19.2260 | 2216 | 1976 | 2275 | 1293 | 3546 | 2487 | 3360 | 3603 | 2113 | 2089 | 2247 | 2026 | 2841 | 2828 | 2853 | 2930 | 2490,82 | 2119 | 2863 | 1,349 | 0,432 | 9,97 | 2,06E-23 | 4,94E-23 | 0,0005505 | 0,0027505 | 0,0013846 | 0,0013846 | TRUE | 0,0203636 | No |
| orf19.2261 | 1930 | 1745 | 1848 | 1167 | 1609 | 1077 | 1609 | 1640 | 1840 | 1845 | 1826 | 1828 | 1289 | 1225 | 1366 | 1333 | 1569,05 | 1835 | 1303 | 0,711 | -0,491 | -10,347 | 4,31E-25 | 1,07E-24 | 0,0002399 | 0,0031377 | 0,0015181 | 0,0015181 | TRUE | 0,0181538 | No |
| orf19.2262 | 2957 | 2610 | 2879 | 1698 | 1356 | 906 | 1312 | 1406 | 2819 | 2760 | 2844 | 2660 | 1087 | 1030 | 1114 | 1143 | 1932,12 | 2771 | 1094 | 0,395 | -1,341 | -28,003 | 1,47E-172 | 1,70E-171 | 0,0005542 | 0,0029411 | 0,0015905 | 0,0015905 | TRUE | 0,0165161 | No |
| orf19.2263 | 875 | 789 | 838 | 539 | 1270 | 872 | 1183 | 1268 | 834 | 834 | 828 | 844 | 1018 | 992 | 1004 | 1031 | 923,16 | 835 | 1011 | 1,212 | 0,278 | 5,429 | 5,66E-08 | 9,20E-08 | 1,00E-08 | 0,0038699 | 0,001403 | 0,001403 | TRUE | 0,0020786 | No |

| | | | | | | | | | | | | | | | | | | | | | | | | | | | | | | | |
|---|---|---|---|---|---|---|---|---|---|---|---|---|---|---|---|---|---|---|---|---|---|---|---|---|---|---|---|---|---|---|---|
| orf19.2264 | 5565 | 4724 | 5115 | 3406 | 5085 | 3731 | 4800 | 5177 | 5306 | 4995 | 5053 | 5336 | 4074 | 4242 | 4075 | 4209 | 4661,37 | 5172 | 4150 | 0,803 | -0,317 | -8,318 | 8,96E-17 | 1,87E-16 | 0,000498 | 0,0024435 | 0,0011805 | 0,0011805 | TRUE | 0,0064308 | No |
| orf19.2265 | 2284 | 2001 | 2199 | 1487 | 3679 | 2577 | 3478 | 3657 | 2178 | 2116 | 2172 | 2329 | 2948 | 2930 | 2953 | 2973 | 2574,96 | 2199 | 2951 | 1,345 | 0,428 | 10,499 | 8,72E-26 | 2,19E-25 | 1,00E-08 | 0,0027289 | 0,0011854 | 0,0011854 | TRUE | 0,0172401 | No |
| orf19.2266 | 1425 | 1305 | 1294 | 866 | 2926 | 1921 | 2771 | 2929 | 1359 | 1380 | 1278 | 1357 | 2345 | 2184 | 2353 | 2382 | 1829,55 | 1344 | 2316 | 1,726 | 0,788 | 16,175 | 7,55E-59 | 3,13E-58 | 0,0007707 | 0,0029887 | 0,0016666 | 0,0016666 | TRUE | 0,0167889 | No |
| orf19.2267 | 2681 | 2058 | 2194 | 1545 | 4655 | 3233 | 4408 | 4798 | 2556 | 2176 | 2167 | 2420 | 3730 | 3676 | 3742 | 3901 | 3046,21 | 2330 | 3762 | 1,616 | 0,692 | 11,793 | 4,26E-32 | 1,19E-31 | 0,0032258 | 0,0026303 | 0,0029478 | 0,0029478 | TRUE | 0,0543027 | No |
| orf19.2268 | 7054 | 6247 | 6832 | 4505 | 11524 | 8214 | 11341 | 11990 | 6725 | 6605 | 6749 | 7057 | 9234 | 9340 | 9628 | 9749 | 8136,09 | 6784 | 9488 | 1,399 | 0,485 | 13,602 | 3,92E-42 | 1,29E-41 | 0,0005571 | 0,0022931 | 0,0010902 | 0,0010902 | TRUE | 0,0085091 | No |
| orf19.2269 | 689 | 699 | 654 | 409 | 1261 | 855 | 1026 | 1223 | 657 | 739 | 646 | 641 | 1010 | 972 | 871 | 994 | 816,36 | 671 | 962 | 1,433 | 0,519 | 7,209 | 5,65E-13 | 1,08E-12 | 0,0033119 | 0,0041025 | 0,0036754 | 0,0036754 | TRUE | 0,0560639 | No |
| orf19.227 | 2901 | 2558 | 2673 | 1582 | 5254 | 3376 | 4414 | 4339 | 2766 | 2705 | 2641 | 2478 | 4210 | 3839 | 3747 | 3528 | 3239,21 | 2648 | 3831 | 1,446 | 0,532 | 8,797 | 1,41E-18 | 3,07E-18 | 0,0034193 | 0,0025982 | 0,0031812 | 0,0031812 | TRUE | 0,054689 | No |
| orf19.2270 | 8753 | 7895 | 8530 | 5228 | 6411 | 4569 | 5977 | 6470 | 8345 | 8348 | 8427 | 8190 | 5137 | 5195 | 5074 | 5261 | 6747,16 | 8328 | 5167 | 0,62 | -0,689 | -23,384 | 6,20E-121 | 4,98E-120 | 1,00E-08 | 0,0023346 | 0,0006799 | 0,0006799 | TRUE | 0,0018986 | No |
| orf19.2272 | 2942 | 2668 | 3017 | 1954 | 1675 | 1293 | 1698 | 1762 | 2805 | 2821 | 2981 | 3061 | 1342 | 1470 | 1442 | 1433 | 2169,27 | 2917 | 1422 | 0,487 | -1,037 | -20,958 | 1,58E-97 | 1,01E-96 | 0,0011785 | 0,0028482 | 0,0018414 | 0,0018414 | TRUE | 0,0174355 | No |
| orf19.2274 | 49 | 53 | 57 | 36 | 18 | 13 | 15 | 13 | 47 | 56 | 56 | 56 | 14 | 15 | 13 | 11 | 33,5 | 54 | 13 | 0,243 | -2,042 | -7,514 | 5,74E-14 | 1,12E-13 | 1,00E-08 | 0,0511081 | 0,0265484 | 0,0265484 | TRUE | 0,0863104 | No |
| orf19.2275 | 1519 | 1271 | 1361 | 931 | 2254 | 1384 | 1860 | 2204 | 1448 | 1344 | 1345 | 1458 | 1806 | 1574 | 1579 | 1792 | 1543,26 | 1399 | 1688 | 1,209 | 0,274 | 4,262 | 2,03E-05 | 2,97E-05 | 0,0030535 | 0,0031552 | 0,0032885 | 0,0032885 | TRUE | 0,02687 | No |
| orf19.2276 | 1020 | 824 | 846 | 475 | 1859 | 1219 | 1633 | 1833 | 972 | 871 | 836 | 744 | 1490 | 1386 | 1386 | 1490 | 1147,01 | 856 | 1438 | 1,675 | 0,744 | 9,959 | 2,30E-23 | 5,52E-23 | 0,0057045 | 0,0035227 | 0,0043908 | 0,0043908 | TRUE | 0,0999229 | No |
| orf19.2277 | 2187 | 1956 | 2155 | 1416 | 2402 | 1794 | 2374 | 2545 | 2085 | 2068 | 2129 | 2218 | 1925 | 2040 | 2016 | 2069 | 2068,74 | 2125 | 2012 | 0,948 | -0,077 | -1,737 | 0,0823892 | 0,0970331 | 0,0004689 | 0,002885 | 0,0014078 | 0,0014078 | TRUE | 0,0107505 | No |
| orf19.2278 | 2482 | 2255 | 2357 | 1588 | 2764 | 2070 | 2538 | 2659 | 2366 | 2384 | 2328 | 2488 | 2215 | 2354 | 2155 | 2162 | 2306,52 | 2392 | 2222 | 0,929 | -0,106 | -2,352 | 0,018677 | 0,0232452 | 0,0006889 | 0,0028031 | 0,0015301 | 0,0015301 | TRUE | 0,0184844 | No |
| orf19.2280 | 727 | 653 | 732 | 458 | 1491 | 1083 | 1485 | 1551 | 693 | 690 | 723 | 717 | 1195 | 1231 | 1261 | 1261 | 971,53 | 706 | 1237 | 1,753 | 0,81 | 15,171 | 5,48E-52 | 2,08E-51 | 1,00E-08 | 0,0037813 | 0,0015877 | 0,0015877 | TRUE | 0,0067681 | No |
| orf19.2281 | 125 | 99 | 107 | 60 | 339 | 227 | 335 | 338 | 119 | 105 | 106 | 94 | 272 | 258 | 284 | 275 | 189,07 | 106 | 272 | 2,552 | 1,352 | 11,901 | 1,16E-32 | 3,28E-32 | 1,00E-08 | 0,0107755 | 0,0055587 | 0,0055587 | TRUE | 0,0714323 | No |
| orf19.2282 | 1179 | 1030 | 1047 | 618 | 541 | 361 | 506 | 499 | 1124 | 1089 | 1034 | 968 | 433 | 410 | 430 | 406 | 736,86 | 1054 | 420 | 0,398 | -1,331 | -19,617 | 1,10E-85 | 6,31E-85 | 0,001508 | 0,0043195 | 0,0028267 | 0,0028267 | TRUE | 0,0319356 | No |
| orf19.2283 | 1933 | 1515 | 1650 | 1115 | 3140 | 1953 | 2883 | 3091 | 1843 | 1602 | 1630 | 1747 | 2516 | 2221 | 2448 | 2513 | 2064,9 | 1706 | 2424 | 1,423 | 0,509 | 8,279 | 1,24E-16 | 2,59E-16 | 0,0031078 | 0,0028864 | 0,0031152 | 0,0031152 | TRUE | 0,0376514 | No |
| orf19.2284 | 1792 | 1466 | 1613 | 1136 | 1027 | 669 | 920 | 965 | 1708 | 1550 | 1593 | 1780 | 823 | 761 | 781 | 785 | 1222,63 | 1658 | 788 | 0,476 | -1,071 | -17,959 | 4,07E-72 | 2,01E-71 | 0,0017991 | 0,0034342 | 0,002506 | 0,002506 | TRUE | 0,0269229 | No |
| orf19.2285 | 111 | 108 | 112 | 70 | 75 | 62 | 60 | 66 | 106 | 114 | 111 | 110 | 60 | 70 | 51 | 54 | 84,44 | 110 | 59 | 0,53 | -0,917 | -5,76 | 8,43E-09 | 1,40E-08 | 1,00E-08 | 0,0215357 | 0,0116296 | 0,0116296 | TRUE | 0,1290984 | No |
| orf19.2286 | 5580 | 4521 | 4802 | 3309 | 7823 | 5392 | 7411 | 8360 | 5320 | 4780 | 4744 | 5184 | 6268 | 6131 | 6292 | 6797 | 5689,61 | 5007 | 6372 | 1,273 | 0,349 | 6,649 | 2,95E-11 | 5,36E-11 | 0,0024739 | 0,0023799 | 0,002456 | 0,002456 | TRUE | 0,0247295 | No |
| orf19.2287 | 1802 | 1401 | 1536 | 1042 | 3181 | 2254 | 2973 | 3122 | 1718 | 1481 | 1517 | 1632 | 2549 | 2563 | 2524 | 2538 | 2065,44 | 1587 | 2544 | 1,603 | 0,681 | 12,934 | 2,90E-38 | 8,95E-38 | 0,0016684 | 0,0028862 | 0,0021303 | 0,0021303 | TRUE | 0,0401202 | No |
| orf19.2288 | 5162 | 4275 | 4932 | 2885 | 9603 | 6193 | 9390 | 10233 | 4921 | 4520 | 4872 | 4520 | 7695 | 7042 | 7972 | 8320 | 6232,83 | 4708 | 7757 | 1,647 | 0,72 | 12,526 | 5,39E-36 | 1,60E-35 | 0,003284 | 0,0023547 | 0,0029971 | 0,0029971 | TRUE | 0,0465713 | No |
| orf19.2289 | 1312 | 1199 | 1189 | 842 | 1969 | 1348 | 1877 | 2045 | 1251 | 1268 | 1175 | 1319 | 1578 | 1533 | 1594 | 1663 | 1422,39 | 1253 | 1592 | 1,274 | 0,349 | 6,689 | 2,24E-11 | 4,08E-11 | 0,0007616 | 0,0032456 | 0,0018872 | 0,0018872 | TRUE | 0,0214418 | No |
| orf19.229 | 1953 | 1910 | 1896 | 1086 | 1802 | 1303 | 1599 | 1609 | 1862 | 2020 | 1873 | 1701 | 1444 | 1482 | 1358 | 1308 | 1630,9 | 1864 | 1398 | 0,748 | -0,419 | -6,629 | 3,37E-11 | 6,10E-11 | 0,0032034 | 0,003098 | 0,0032051 | 0,0032051 | TRUE | 0,0388061 | No |
| orf19.2290 | 6307 | 5542 | 6064 | 3802 | 3397 | 2435 | 3164 | 3493 | 6013 | 5860 | 5991 | 5956 | 2722 | 2769 | 2686 | 2840 | 4354,6 | 5955 | 2754 | 0,462 | -1,113 | -32,537 | 3,20E-232 | 4,99E-231 | 1,65E-06 | 0,0024683 | 0,0008661 | 0,0008661 | TRUE | 0,0056156 | No |
| orf19.2296 | 101341 | 82301 | 92018 | 62502 | 27901 | 19967 | 24885 | 27399 | 96618 | 87023 | 90905 | 97913 | 22356 | 22704 | 21127 | 22278 | 57615,52 | 93115 | 22116 | 0,238 | -2,074 | -44,914 | 0 | 0 | 0,001928 | 0,0021198 | 0,0020223 | 0,0020223 | TRUE | 0,0237669 | No |
| orf19.2297 | 228 | 198 | 233 | 145 | 463 | 324 | 508 | 542 | 217 | 209 | 230 | 227 | 371 | 368 | 431 | 441 | 311,93 | 221 | 403 | 1,828 | 0,87 | 8,922 | 4,58E-19 | 1,01E-18 | 0,0037532 | 0,0073549 | 0,0055055 | 0,0055055 | TRUE | 0,0467576 | No |
| orf19.2298 | 1555 | 1439 | 1412 | 1060 | 2618 | 1900 | 2580 | 2724 | 1483 | 1522 | 1395 | 1661 | 2098 | 2160 | 2190 | 2215 | 1840,38 | 1515 | 2166 | 1,434 | 0,52 | 9,366 | 7,51E-21 | 1,71E-20 | 0,0018948 | 0,0029834 | 0,0023791 | 0,0023791 | TRUE | 0,0461906 | No |
| orf19.2299 | 905 | 682 | 779 | 492 | 3490 | 2066 | 3138 | 3369 | 863 | 721 | 770 | 771 | 2796 | 2349 | 2664 | 2739 | 1709,18 | 781 | 2637 | 3,376 | 1,755 | 24,19 | 2,87E-129 | 2,47E-128 | 0,0046283 | 0,0030519 | 0,0041677 | 0,0041677 | TRUE | 0,0642173 | No |
| orf19.23 | 3901 | 3437 | 3770 | 2275 | 6049 | 4175 | 5582 | 5890 | 3719 | 3634 | 3724 | 3564 | 4847 | 4747 | 4739 | 4789 | 4220,52 | 3660 | 4780 | 1,305 | 0,384 | 11,674 | 1,72E-31 | 4,76E-31 | 1,00E-08 | 0,0024803 | 0,0007922 | 0,0007922 | TRUE | 0,0034615 | No |
| orf19.230 | 670 | 811 | 691 | 405 | 682 | 631 | 626 | 516 | 639 | 858 | 683 | 634 | 546 | 718 | 531 | 420 | 628,55 | 704 | 554 | 0,785 | -0,349 | -2,444 | 0,0145425 | 0,0182489 | 0,0336667 | 0,0047035 | 0,0179889 | 0,0179889 | TRUE | 0,4541279 | No |
| orf19.2301 | 2001 | 1714 | 1891 | 1101 | 2197 | 1559 | 2032 | 2115 | 1908 | 1812 | 1868 | 1725 | 1760 | 1773 | 1725 | 1720 | 1786,37 | 1828 | 1744 | 0,952 | -0,071 | -1,561 | 0,1185318 | 0,137244 | 0,0003057 | 0,0030104 | 0,0014387 | 0,0014387 | TRUE | 0,0156324 | No |
| orf19.2302 | 945 | 861 | 1002 | 616 | 2051 | 1586 | 2049 | 2233 | 901 | 910 | 990 | 965 | 1643 | 1803 | 1740 | 1816 | 1346,04 | 942 | 1750 | 1,86 | 0,895 | 15,866 | 1,08E-56 | 4,33E-56 | 0,0014042 | 0,0033111 | 0,0022062 | 0,0022062 | TRUE | 0,0209285 | No |
| orf19.2303 | 233 | 225 | 211 | 153 | 610 | 415 | 575 | 588 | 222 | 238 | 208 | 240 | 489 | 472 | 488 | 478 | 354,39 | 227 | 482 | 2,132 | 1,093 | 13,243 | 4,93E-40 | 1,56E-39 | 1,00E-08 | 0,0067243 | 0,0031535 | 0,0031535 | TRUE | 0,0342078 | No |
| orf19.2304 | 778 | 709 | 710 | 544 | 1192 | 822 | 1078 | 1268 | 742 | 750 | 701 | 852 | 955 | 935 | 915 | 1031 | 860,13 | 761 | 959 | 1,266 | 0,34 | 4,831 | 1,36E-06 | 2,09E-06 | 0,0033829 | 0,0040002 | 0,0035515 | 0,0035515 | TRUE | 0,0682664 | No |
| orf19.2305 | 499 | 398 | 399 | 239 | 922 | 585 | 805 | 927 | 476 | 421 | 394 | 374 | 739 | 665 | 683 | 754 | 563,29 | 416 | 710 | 1,701 | 0,766 | 9,109 | 8,35E-20 | 1,87E-19 | 0,0048382 | 0,0050061 | 0,0048228 | 0,0048228 | TRUE | 0,1092876 | No |
| orf19.2306 | 702 | 625 | 666 | 341 | 1756 | 1193 | 1731 | 1685 | 669 | 661 | 658 | 534 | 1407 | 1357 | 1470 | 1370 | 1015,69 | 630 | 1401 | 2,209 | 1,143 | 16,41 | 1,62E-60 | 6,88E-60 | 0,0031516 | 0,0037078 | 0,0034588 | 0,0034588 | TRUE | 0,1066221 | No |
| orf19.2307 | 2298 | 1945 | 2264 | 1328 | 2463 | 1839 | 2462 | 2658 | 2191 | 2057 | 2237 | 2080 | 1974 | 2091 | 2090 | 2161 | 2110,07 | 2141 | 2079 | 0,97 | -0,044 | -0,909 | 0,3635507 | 0,3982519 | 0,0011003 | 0,0028694 | 0,0017818 | 0,0017818 | TRUE | 0,0145904 | No |
| orf19.2308 | 30929 | 28640 | 30843 | 19118 | 12502 | 9033 | 12117 | 11756 | 29487 | 30283 | 30470 | 29949 | 10017 | 10271 | 10287 | 9559 | 20040,6 | 30047 | 10034 | 0,334 | -1,583 | -46,588 | 0 | 0 | 0,0005881 | 0,0021732 | 0,0010452 | 0,0010452 | TRUE | 0,0124862 | No |
| orf19.2309 | 5332 | 4940 | 5377 | 3548 | 6564 | 5042 | 6205 | 6877 | 5083 | 5223 | 5312 | 5558 | 5260 | 5733 | 5268 | 5592 | 5378,68 | 5294 | 5463 | 1,032 | 0,046 | 1,014 | 0,3107577 | 0,3440599 | 0,0014056 | 0,0023965 | 0,0017693 | 0,0017693 | TRUE | 0,0132147 | No |
| orf19.2309.2 | 30325 | 24404 | 25995 | 17536 | 76429 | 49834 | 72300 | 78335 | 28912 | 25804 | 25681 | 27471 | 61240 | 56666 | 61382 | 63694 | 43856,22 | 26967 | 60746 | 2,253 | 1,172 | 22,697 | 4,80E-114 | 3,61E-113 | 0,0027522 | 0,0021287 | 0,0025321 | 0,0025321 | TRUE | 0,0290142 | No |
| orf19.231 | 725 | 739 | 773 | 418 | 1366 | 929 | 1200 | 1282 | 691 | 781 | 764 | 655 | 1095 | 1056 | 1019 | 1042 | 887,89 | 723 | 1053 | 1,452 | 0,538 | 8,201 | 2,38E-16 | 4,94E-16 | 0,0019295 | 0,0039405 | 0,0029397 | 0,0029397 | TRUE | 0,040328 | No |
| orf19.2310 | 1102 | 1055 | 1135 | 720 | 623 | 442 | 629 | 630 | 1051 | 1116 | 1121 | 1128 | 499 | 503 | 534 | 512 | 807,93 | 1104 | 512 | 0,465 | -1,106 | -18,89 | 1,37E-79 | 7,48E-79 | 1,00E-08 | 0,0041235 | 0,001915 | 0,001915 | TRUE | 0,0132483 | No |
| orf19.2310.1 | 9479 | 7163 | 7316 | 5824 | 25297 | 16499 | 22053 | 24897 | 9037 | 7574 | 7228 | 9124 | 20270 | 18761 | 18723 | 20244 | 13869,93 | 8241 | 19500 | 2,367 | 1,243 | 16,445 | 9,10E-61 | 3,87E-60 | 0,007972 | 0,0022097 | 0,0053981 | 0,0053981 | TRUE | 0,0838321 | No |
| orf19.2311 | 14631 | 12221 | 13426 | 9909 | 25635 | 19988 | 24170 | 26272 | 13949 | 12922 | 13264 | 15523 | 20541 | 22728 | 20520 | 21362 | 17601,05 | 13914 | 21288 | 1,53 | 0,614 | 10,076 | 7,07E-24 | 1,72E-23 | 0,0045484 | 0,0021846 | 0,0035036 | 0,0035036 | TRUE | 0,0734272 | No |
| orf19.2312 | 69 | 46 | 65 | 47 | 134 | 109 | 141 | 153 | 66 | 49 | 64 | 74 | 107 | 124 | 120 | 124 | 90,96 | 63 | 119 | 1,898 | 0,924 | 5,745 | 9,19E-09 | 1,53E-08 | 1,00E-08 | 0,0201419 | 0,0122217 | 0,0122217 | TRUE | 0,2025704 | No |
| orf19.2313 | 1326 | 1173 | 1215 | 884 | 1899 | 1313 | 1787 | 1896 | 1264 | 1240 | 1200 | 1385 | 1522 | 1493 | 1517 | 1542 | 1395,38 | 1272 | 1518 | 1,198 | 0,26 | 4,965 | 6,87E-07 | 1,07E-06 | 0,0006591 | 0,003268 | 0,001899 | 0,001899 | TRUE | 0,0376101 | No |
| orf19.2314 | 7229 | 6271 | 6724 | 4081 | 7826 | 5050 | 7843 | 7777 | 6892 | 6631 | 6643 | 6393 | 6271 | 5742 | 6659 | 6323 | 6444,23 | 6640 | 6249 | 0,941 | -0,087 | -1,756 | 0,0791255 | 0,0933444 | 0,0021018 | 0,0023461 | 0,0022271 | 0,0022271 | TRUE | 0,0357049 | No |
| orf19.2315 | 3416 | 2831 | 3249 | 2193 | 2036 | 1441 | 1961 | 2099 | 3257 | 2993 | 3210 | 3435 | 1631 | 1639 | 1665 | 1707 | 2442,11 | 3224 | 1660 | 0,516 | -0,955 | -19,133 | 1,35E-81 | 7,46E-81 | 0,0014427 | 0,0027636 | 0,0019506 | 0,0019506 | TRUE | 0,027861 | No |
| orf19.2317 | 101 | 77 | 67 | 54 | 39 | 24 | 47 | 33 | 96 | 81 | 66 | 85 | 31 | 27 | 40 | 27 | 56,72 | 82 | 31 | 0,383 | -1,384 | -6,478 | 9,32E-11 | 1,67E-10 | 0,0121088 | 0,0310383 | 0,0228633 | 0,0228633 | TRUE | 0,2499423 | No |
| orf19.2318.1 | 1527 | 1269 | 1357 | 922 | 2468 | 1552 | 2339 | 2584 | 1456 | 1342 | 1341 | 1444 | 1978 | 1765 | 1986 | 2101 | 1676,47 | 1396 | 1958 | 1,405 | 0,491 | 7,886 | 3,11E-15 | 6,31E-15 | 0,0027669 | 0,0030707 | 0,003083 | 0,003083 | TRUE | 0,0511135 | No |
| orf19.2319 | 7428 | 6436 | 7092 | 3844 | 9960 | 6127 | 9726 | 9836 | 7082 | 6805 | 7006 | 6022 | 7981 | 6967 | 8257 | 7998 | 7264,71 | 6729 | 7801 | 1,159 | 0,213 | 3,224 | 0,0012637 | 0,0016912 | 0,0050888 | 0,0023173 | 0,0040451 | 0,0040451 | TRUE | 0,0627261 | No |
| orf19.232 | 384 | 395 | 418 | 277 | 330 | 254 | 298 | 374 | 366 | 418 | 413 | 434 | 264 | 289 | 253 | 304 | 342,62 | 408 | 278 | 0,682 | -0,551 | -5,986 | 2,15E-09 | 3,66E-09 | 0,0033295 | 0,0068834 | 0,0051632 | 0,0051632 | TRUE | 0,0565573 | No |
| orf19.2320 | 10101 | 8433 | 9526 | 4994 | 10670 | 6588 | 10494 | 10671 | 9630 | 8917 | 9411 | 7823 | 8550 | 7491 | 8909 | 8677 | 8675,98 | 8945 | 8407 | 0,94 | -0,09 | -1,242 | 0,214324 | 0,2423106 | 0,0066905 | 0,0022805 | 0,0049208 | 0,0049208 | TRUE | 0,0861583 | No |
| orf19.2322.3 | 1122 | 930 | 1025 | 613 | 2738 | 1769 | 2732 | 2809 | 1070 | 983 | 1013 | 960 | 2194 | 2012 | 2319 | 2284 | 1604,35 | 1006 | 2202 | 2,186 | 1,128 | 18,632 | 1,77E-77 | 9,34E-77 | 0,0025028 | 0,0031147 | 0,0027615 | 0,0027615 | TRUE | 0,0396488 | No |
| orf19.2324 | 1697 | 1393 | 1507 | 1115 | 3011 | 2151 | 2958 | 3176 | 1618 | 1473 | 1489 | 1747 | 2413 | 2446 | 2511 | 2582 | 2034,82 | 1582 | 2488 | 1,577 | 0,657 | 11,174 | 5,47E-29 | 1,45E-28 | 0,0030311 | 0,0028982 | 0,0027859 | 0,0027859 | TRUE | 0,0548197 | No |
| orf19.2325 | 952 | 795 | 927 | 607 | 815 | 615 | 797 | 816 | 908 | 841 | 916 | 951 | 653 | 699 | 677 | 663 | 788,43 | 904 | 673 | 0,745 | -0,424 | -7,039 | 1,93E-12 | 3,62E-12 | 1,11E-05 | 0,0041738 | 0,0022053 | 0,0022053 | TRUE | 0,0260009 | No |
| orf19.2326 | 2812 | 2444 | 2731 | 1548 | 1814 | 1243 | 1842 | 1934 | 2681 | 2584 | 2698 | 2425 | 1454 | 1413 | 1564 | 1573 | 2048,93 | 2597 | 1501 | 0,578 | -0,792 | -14,509 | 1,07E-47 | 3,81E-47 | 0,0018655 | 0,0028926 | 0,0023451 | 0,0023451 | TRUE | 0,0226461 | No |
| orf19.2328 | 2428 | 2099 | 2363 | 1393 | 1540 | 1039 | 1508 | 1511 | 2315 | 2219 | 2334 | 2182 | 1234 | 1181 | 1280 | 1229 | 1746,89 | 2262 | 1231 | 0,544 | -0,878 | -18,784 | 1,02E-78 | 5,47E-78 | 0,0004022 | 0,0030312 | 0,0014899 | 0,0014899 | TRUE | 0,0085146 | No |
| orf19.2329.1 | 15691 | 13976 | 13706 | 8625 | 41666 | 28892 | 38258 | 42150 | 14960 | 14778 | 13540 | 13512 | 33386 | 32853 | 32481 | 34272 | 23722,59 | 14198 | 33248 | 2,341 | 1,227 | 27,597 | 1,21E-167 | 1,36E-166 | 0,0017114 | 0,0021605 | 0,0018469 | 0,0018469 | TRUE | 0,0160203 | No |
| orf19.2330 | 5895 | 5040 | 5533 | 3286 | 7573 | 5165 | 7153 | 7832 | 5620 | 5329 | 5466 | 5148 | 6068 | 5873 | 6073 | 6368 | 5743,16 | 5391 | 6096 | 1,13 | 0,177 | 4,231 | 2,33E-05 | 3,39E-05 | 0,0010232 | 0,0023772 | 0,0014952 | 0,0014952 | TRUE | 0,0110828 | No |
| orf19.2331 | 1727 | 1573 | 1774 | 1026 | 1714 | 1277 | 1622 | 1777 | 1647 | 1663 | 1753 | 1607 | 1373 | 1452 | 1377 | 1445 | 1539,62 | 1668 | 1412 | 0,845 | -0,243 | -5,037 | 4,73E-07 | 7,41E-07 | 0,0004602 | 0,0031577 | 0,0015783 | 0,0015783 | TRUE | 0,0144217 | No |
| orf19.233.1 | 2106 | 1818 | 1766 | 1177 | 4323 | 2930 | 4043 | 4248 | 2008 | 1922 | 1745 | 1844 | 3464 | 3332 | 3432 | 3454 | 2650,09 | 1880 | 3420 | 1,819 | 0,863 | 17,515 | 1,11E-68 | 5,20E-68 | 0,0013891 | 0,0027108 | 0,0019032 | 0,0019032 | TRUE | 0,0302378 | No |
| orf19.2332 | 1 | 2 | 0 | 1 | 0 | 0 | 1 | 0 | 1 | 2 | 0 | 2 | 0 | 0 | 1 | 0 | 0,69 | 1 | 0 | 0,238 | -2,071 | -1,138 | 0,2553132 | 0,285666 | 1,00E-08 | 2,3974189 | 1,6888892 | 1,6888892 | TRUE | 0,3078457 | No |
| orf19.2333 | 9513 | 8095 | 9249 | 5565 | 5267 | 3698 | 5160 | 5497 | 9070 | 8559 | 9137 | 8718 | 4220 | 4205 | 4381 | 4470 | 6594,96 | 8871 | 4319 | 0,487 | -1,038 | -26,765 | 8,32E-158 | 8,95E-157 | 0,0006993 | 0,0023402 | 0,0012785 | 0,0012785 | TRUE | 0,0070056 | No |
| orf19.2334 | 943 | 959 | 883 | 526 | 1151 | 901 | 1041 | 1038 | 899 | 1014 | 872 | 824 | 922 | 1025 | 884 | 844 | 910,5 | 902 | 919 | 1,015 | 0,021 | 0,263 | 0,7921734 | 0,8151683 | 0,0060445 | 0,0038946 | 0,0050332 | 0,0050332 | TRUE | 0,0818971 | No |
| orf19.2335 | 1251 | 1210 | 1298 | 769 | 2377 | 1744 | 2217 | 2465 | 1193 | 1279 | 1282 | 1205 | 1905 | 1983 | 1882 | 2004 | 1591,66 | 1240 | 1944 | 1,566 | 0,647 | 13,228 | 6,05E-40 | 1,91E-39 | 0,000555 | 0,0031228 | 0,0016147 | 0,0016147 | TRUE | 0,009035 | No |
| orf19.2336 | 229 | 195 | 190 | 136 | 241 | 188 | 230 | 279 | 218 | 206 | 188 | 213 | 193 | 214 | 195 | 227 | 206,78 | 206 | 207 | 1,005 | 0,007 | 0,069 | 0,9448166 | 0,9504769 | 1,00E-08 | 0,0100315 | 0,0059673 | 0,0059673 | TRUE | 0,0493047 | No |
| orf19.2337 | 14599 | 12810 | 13661 | 7770 | 19247 | 12707 | 17952 | 19771 | 13919 | 13545 | 13496 | 12172 | 15422 | 14449 | 15241 | 16076 | 14289,92 | 13283 | 15297 | 1,151 | 0,203 | 3,983 | 6,80E-05 | 9,71E-05 | 0,0025186 | 0,0022062 | 0,0024318 | 0,0024318 | TRUE | 0,0382263 | No |
| orf19.234 | 727 | 653 | 719 | 469 | 1369 | 1068 | 1242 | 1239 | 693 | 690 | 710 | 735 | 1097 | 1214 | 1054 | 1007 | 900,23 | 707 | 1093 | 1,545 | 0,628 | 9,17 | 4,73E-20 | 1,06E-19 | 0,0028132 | 0,0039152 | 0,0032999 | 0,0032999 | TRUE | 0,0632615 | No |
| orf19.2340 | 21992 | 17998 | 19671 | 12859 | 24072 | 16710 | 22611 | 25126 | 20967 | 19031 | 19433 | 20144 | 19288 | 19001 | 19197 | 20430 | 19686,31 | 19894 | 19479 | 0,979 | -0,03 | -0,72 | 0,4715112 | 0,5079711 | 0,001336 | 0,0021747 | 0,0016488 | 0,0016488 | TRUE | 0,0163097 | No |
| orf19.2341 | 5624 | 4982 | 5407 | 3320 | 2418 | 1711 | 2275 | 2314 | 5362 | 5268 | 5342 | 5201 | 1937 | 1946 | 1931 | 1882 | 3608,53 | 5293 | 1924 | 0,363 | -1,461 | -42,191 | 0 | 0 | 1,00E-08 | 0,0025463 | 0,0008137 | 0,0008137 | TRUE | 0,0027471 | No |

| | | | | | | | | | | | | | | | | | | | | | | | | | | | | | | | |
|---|---|---|---|---|---|---|---|---|---|---|---|---|---|---|---|---|---|---|---|---|---|---|---|---|---|---|---|---|---|---|---|
| orf19.2342 | 1949 | 1623 | 1750 | 992 | 1813 | 1168 | 1797 | 1874 | 1858 | 1716 | 1729 | 1554 | 1453 | 1328 | 1526 | 1524 | 1585,92 | 1714 | 1458 | 0,85 | -0,235 | -3,556 | 0,0003764 | 0,0005194 | 0,0038201 | 0,0031266 | 0,0035674 | 0,0035674 | TRUE | 0,0447589 | No |
| orf19.2343 | 270 | 267 | 253 | 195 | 521 | 341 | 453 | 569 | 257 | 282 | 250 | 305 | 417 | 388 | 385 | 463 | 343,45 | 274 | 413 | 1,52 | 0,605 | 6,246 | 4,20E-10 | 7,35E-10 | 0,0047959 | 0,0068719 | 0,0058601 | 0,0058601 | TRUE | 0,0769843 | No |
| orf19.2343.1 | 369 | 312 | 345 | 241 | 1257 | 861 | 1302 | 1239 | 352 | 330 | 341 | 378 | 1007 | 979 | 1105 | 1007 | 687,39 | 350 | 1024 | 2,942 | 1,557 | 21,709 | 1,70E-104 | 1,15E-103 | 0,0014822 | 0,0044799 | 0,0029046 | 0,0029046 | TRUE | 0,0337294 | No |
| orf19.2344 | 2352 | 1820 | 1963 | 1502 | 10098 | 7042 | 9493 | 10359 | 2242 | 1924 | 1939 | 2353 | 8091 | 8007 | 8060 | 8423 | 5130 | 2114 | 8145 | 3,858 | 1,948 | 30,287 | 1,72E-201 | 2,38E-200 | 0,0050896 | 0,0024113 | 0,0036504 | 0,0036504 | TRUE | 0,0659492 | No |
| orf19.2346 | 989 | 891 | 962 | 692 | 3284 | 2186 | 3076 | 3226 | 943 | 942 | 950 | 1084 | 2631 | 2486 | 2612 | 2623 | 1783,88 | 980 | 2588 | 2,654 | 1,408 | 26,205 | 2,34E-151 | 2,37E-150 | 0,0010253 | 0,0030117 | 0,0020262 | 0,0020262 | TRUE | 0,053294 | No |
| orf19.2347 | 2887 | 2579 | 2664 | 1745 | 5843 | 4229 | 5557 | 6199 | 2752 | 2727 | 2632 | 2734 | 4682 | 4809 | 4718 | 5040 | 3761,71 | 2711 | 4812 | 1,775 | 0,828 | 20,696 | 3,77E-95 | 2,35E-94 | 0,0006137 | 0,0025278 | 0,0012372 | 0,0012372 | TRUE | 0,0126851 | No |
| orf19.2350 | 528 | 413 | 437 | 277 | 293 | 215 | 298 | 284 | 503 | 437 | 432 | 434 | 235 | 244 | 253 | 231 | 346,11 | 452 | 241 | 0,532 | -0,911 | -10,383 | 2,96E-25 | 7,36E-25 | 1,00E-08 | 0,0068351 | 0,0043078 | 0,0043078 | TRUE | 0,0736866 | No |
| orf19.2351 | 1290 | 1159 | 1164 | 827 | 711 | 512 | 686 | 713 | 1230 | 1225 | 1150 | 1296 | 570 | 582 | 582 | 580 | 901,86 | 1225 | 578 | 0,473 | -1,08 | -18,67 | 8,75E-78 | 4,64E-77 | 1,00E-08 | 0,0039119 | 0,0019842 | 0,0019842 | TRUE | 0,0205482 | No |
| orf19.2352 | 437 | 357 | 383 | 208 | 1240 | 790 | 1246 | 1267 | 417 | 377 | 378 | 326 | 994 | 898 | 1058 | 1030 | 684,78 | 374 | 995 | 2,643 | 1,402 | 17,057 | 3,09E-65 | 1,39E-64 | 0,0049473 | 0,004489 | 0,0045554 | 0,0045554 | TRUE | 0,0732306 | No |
| orf19.2356 | 28074 | 25541 | 28171 | 17897 | 6962 | 5075 | 6619 | 6743 | 26766 | 27006 | 27830 | 28037 | 5578 | 5771 | 5619 | 5483 | 16511,27 | 27410 | 5613 | 0,205 | -2,288 | -71,32 | 0 | 0 | 0,0003394 | 0,0021907 | 0,0008884 | 0,0008884 | TRUE | 0,0042097 | No |
| orf19.236 | 10652 | 9144 | 9387 | 5525 | 35477 | 22979 | 33379 | 35711 | 10156 | 9669 | 9273 | 8655 | 28427 | 26129 | 28339 | 29036 | 18710,47 | 9438 | 27983 | 2,964 | 1,567 | 28,58 | 1,17E-179 | 1,40E-178 | 0,003202 | 0,002179 | 0,0028137 | 0,0028137 | TRUE | 0,0374822 | No |
| orf19.2360 | 8826 | 8153 | 8552 | 5580 | 43611 | 32677 | 38543 | 44000 | 8415 | 8621 | 8449 | 8741 | 34944 | 37157 | 32723 | 35776 | 21853,16 | 8556 | 35150 | 4,109 | 2,039 | 46,523 | 0 | 0 | 0,0014592 | 0,0021664 | 0,0017659 | 0,0017659 | TRUE | 0,0264638 | No |
| orf19.2361 | 3823 | 3544 | 3918 | 2484 | 1572 | 1172 | 1557 | 1610 | 3645 | 3747 | 3871 | 3891 | 1260 | 1333 | 1322 | 1309 | 2547,16 | 3788 | 1306 | 0,345 | -1,537 | -35,762 | 4,32E-280 | 8,71E-279 | 0,0004413 | 0,0027359 | 0,0012828 | 0,0012828 | TRUE | 0,0080826 | No |
| orf19.2362 | 3635 | 3188 | 3316 | 2188 | 3327 | 2327 | 3323 | 3204 | 3466 | 3371 | 3276 | 3428 | 2666 | 2646 | 2821 | 2605 | 3034,78 | 3385 | 2684 | 0,793 | -0,334 | -8,026 | 1,01E-15 | 2,06E-15 | 0,0006202 | 0,0026323 | 0,0013291 | 0,0013291 | TRUE | 0,0144538 | No |
| orf19.2363 | 3989 | 3693 | 4079 | 2430 | 3059 | 2074 | 2856 | 3077 | 3803 | 3905 | 4030 | 3807 | 2451 | 2358 | 2425 | 2502 | 3160,05 | 3886 | 2434 | 0,626 | -0,675 | -17,209 | 2,28E-66 | 1,04E-65 | 0,0003612 | 0,0026109 | 0,0011477 | 0,0011477 | TRUE | 0,0076718 | No |
| orf19.2364 | 1588 | 1419 | 1502 | 1014 | 5410 | 3758 | 4985 | 5568 | 1514 | 1500 | 1484 | 1588 | 4335 | 4273 | 4232 | 4527 | 2931,79 | 1522 | 4342 | 2,859 | 1,516 | 35,299 | 6,17E-273 | 1,18E-271 | 0,0005359 | 0,0026513 | 0,0012985 | 0,0012985 | TRUE | 0,0101909 | No |
| orf19.2365 | 2473 | 2329 | 2400 | 1404 | 3255 | 2157 | 2816 | 3095 | 2358 | 2463 | 2371 | 2199 | 2608 | 2453 | 2391 | 2517 | 2419,86 | 2348 | 2492 | 1,06 | 0,084 | 1,736 | 0,0826143 | 0,0972803 | 0,0012266 | 0,0027698 | 0,0018561 | 0,0018561 | TRUE | 0,0200571 | No |
| orf19.2366 | 1 | 3 | 2 | 2 | 3 | 0 | 0 | 1 | 1 | 3 | 2 | 3 | 2 | 0 | 0 | 1 | 1,56 | 2 | 1 | 0,378 | -1,404 | -1,141 | 0,2538586 | 0,2842379 | 1,00E-08 | 1,0570162 | 0,6859464 | 0,6859464 | TRUE | 0,8317292 | No |
| orf19.2367 | 1423 | 1439 | 1467 | 672 | 699 | 425 | 670 | 612 | 1357 | 1522 | 1449 | 1053 | 560 | 483 | 569 | 498 | 936,25 | 1345 | 528 | 0,392 | -1,352 | -12,805 | 1,53E-37 | 4,66E-37 | 0,0149728 | 0,003845 | 0,0094413 | 0,0094413 | TRUE | 0,2460679 | No |
| orf19.2368 | 2116 | 2133 | 2214 | 1125 | 2164 | 1380 | 2010 | 1937 | 2017 | 2255 | 2187 | 1762 | 1734 | 1569 | 1706 | 1575 | 1850,87 | 2055 | 1646 | 0,8 | -0,322 | -4,269 | 1,96E-05 | 2,87E-05 | 0,0063514 | 0,0029784 | 0,0049249 | 0,0049249 | TRUE | 0,1063115 | No |
| orf19.2369 | 621 | 631 | 642 | 410 | 894 | 654 | 747 | 839 | 592 | 667 | 634 | 642 | 716 | 744 | 634 | 682 | 664,02 | 634 | 694 | 1,094 | 0,13 | 1,83 | 0,0672333 | 0,0799061 | 0,0022295 | 0,004564 | 0,0033303 | 0,0033303 | TRUE | 0,0398662 | No |
| orf19.2369.1 | 1469 | 1332 | 1377 | 702 | 944 | 555 | 887 | 822 | 1401 | 1408 | 1360 | 1100 | 756 | 631 | 753 | 668 | 1009,74 | 1317 | 702 | 0,533 | -0,909 | -10,171 | 2,66E-24 | 6,49E-24 | 0,008696 | 0,0037173 | 0,0066044 | 0,0066044 | TRUE | 0,1403018 | No |
| orf19.2370 | 342 | 346 | 367 | 204 | 296 | 210 | 300 | 279 | 326 | 366 | 363 | 320 | 237 | 239 | 255 | 227 | 291,45 | 344 | 240 | 0,695 | -0,525 | -5,817 | 5,98E-09 | 1,00E-08 | 1,00E-08 | 0,0077249 | 0,0043523 | 0,0043523 | TRUE | 0,0218544 | No |
| orf19.2371 | 2275 | 1898 | 1992 | 1259 | 2205 | 1461 | 2033 | 2104 | 2169 | 2007 | 1968 | 1972 | 1767 | 1661 | 1726 | 1711 | 1872,61 | 2029 | 1716 | 0,846 | -0,242 | -4,997 | 5,81E-07 | 9,09E-07 | 0,0009058 | 0,0029681 | 0,0017159 | 0,0017159 | TRUE | 0,0270229 | No |
| orf19.2372 | 2856 | 2500 | 2645 | 1648 | 4004 | 2701 | 3532 | 3872 | 2723 | 2643 | 2613 | 2582 | 3208 | 3071 | 2999 | 3148 | 2873,44 | 2640 | 3106 | 1,176 | 0,234 | 5,821 | 5,84E-09 | 9,78E-09 | 0,0004113 | 0,0026627 | 0,0011948 | 0,0011948 | TRUE | 0,0068895 | No |
| orf19.2376 | 65 | 67 | 69 | 45 | 100 | 68 | 101 | 97 | 62 | 71 | 68 | 70 | 80 | 77 | 86 | 79 | 74,19 | 68 | 80 | 1,192 | 0,253 | 1,576 | 0,1150145 | 0,133244 | 1,00E-08 | 0,0242215 | 0,0111092 | 0,0111092 | TRUE | 0,0335155 | No |
| orf19.2378 | 468 | 417 | 430 | 305 | 1000 | 659 | 882 | 888 | 446 | 441 | 425 | 478 | 801 | 749 | 749 | 722 | 601,4 | 448 | 755 | 1,695 | 0,761 | 11,161 | 6,32E-29 | 1,67E-28 | 1,00E-08 | 0,0048214 | 0,002626 | 0,002626 | TRUE | 0,0208938 | No |
| orf19.2379 | 4079 | 3420 | 3826 | 2559 | 4219 | 2857 | 4102 | 4278 | 3889 | 3616 | 3780 | 4009 | 3381 | 3249 | 3483 | 3478 | 3610,48 | 3824 | 3398 | 0,89 | -0,169 | -3,757 | 0,0001717 | 0,0002402 | 0,0011044 | 0,002546 | 0,0016535 | 0,0016535 | TRUE | 0,0160494 | No |
| orf19.238 | 7210 | 6274 | 6632 | 3910 | 2644 | 1755 | 2570 | 2609 | 6874 | 6634 | 6552 | 6125 | 2119 | 1996 | 2182 | 2121 | 4325,3 | 6546 | 2104 | 0,321 | -1,637 | -34,27 | 2,17E-257 | 3,84E-256 | 0,0014086 | 0,0024709 | 0,0018934 | 0,0018934 | TRUE | 0,0221289 | No |
| orf19.2381 | 802 | 635 | 690 | 500 | 313 | 242 | 325 | 366 | 765 | 671 | 682 | 783 | 251 | 275 | 276 | 298 | 500,06 | 725 | 275 | 0,38 | -1,397 | -16,498 | 3,81E-61 | 1,63E-60 | 0,0036078 | 0,0053747 | 0,0044884 | 0,0044884 | TRUE | 0,0425793 | No |
| orf19.2382 | 453 | 438 | 455 | 278 | 1512 | 1104 | 1457 | 1556 | 432 | 463 | 449 | 436 | 1212 | 1255 | 1237 | 1265 | 843,63 | 445 | 1242 | 2,788 | 1,479 | 25,239 | 1,51E-140 | 1,43E-139 | 1,00E-08 | 0,0040375 | 0,0016926 | 0,0016926 | TRUE | 0,0087602 | No |
| orf19.2383 | 133 | 130 | 127 | 112 | 450 | 295 | 419 | 471 | 127 | 137 | 125 | 175 | 361 | 335 | 356 | 383 | 249,99 | 141 | 359 | 2,582 | 1,368 | 12,29 | 1,03E-34 | 3,00E-34 | 0,0029204 | 0,0086593 | 0,006661 | 0,006661 | TRUE | 0,2149613 | No |
| orf19.2384 | 2831 | 2343 | 2477 | 1600 | 2297 | 1525 | 2230 | 2277 | 2699 | 2477 | 2447 | 2506 | 1841 | 1734 | 1893 | 1851 | 2181,16 | 2532 | 1830 | 0,723 | -0,468 | -9,503 | 2,04E-21 | 4,70E-21 | 0,0012441 | 0,002844 | 0,0018645 | 0,0018645 | TRUE | 0,0247378 | No |
| orf19.2385 | 2872 | 2557 | 2624 | 1650 | 3521 | 2406 | 3417 | 3709 | 2738 | 2704 | 2592 | 2585 | 2821 | 2736 | 2901 | 3016 | 2761,6 | 2655 | 2868 | 1,08 | 0,112 | 2,477 | 0,013247 | 0,0166955 | 0,000808 | 0,0026858 | 0,0015842 | 0,0015842 | TRUE | 0,0145388 | No |
| orf19.2386 | 5500 | 4865 | 5232 | 3364 | 4894 | 3364 | 4872 | 4810 | 5244 | 5144 | 5169 | 5270 | 3921 | 3825 | 4136 | 3911 | 4577,53 | 5207 | 3948 | 0,759 | -0,398 | -10,918 | 9,45E-28 | 2,46E-27 | 0,0003315 | 0,00245 | 0,0010558 | 0,0010558 | TRUE | 0,012484 | No |
| orf19.2387 | 4580 | 4030 | 4499 | 2747 | 4284 | 3056 | 3992 | 4437 | 4367 | 4261 | 4445 | 4303 | 3433 | 3475 | 3389 | 3608 | 3910,02 | 4344 | 3476 | 0,8 | -0,322 | -8,787 | 1,54E-18 | 3,34E-18 | 0,0003167 | 0,0025112 | 0,0010304 | 0,0010304 | TRUE | 0,0081439 | No |
| orf19.2389 | 1166 | 1001 | 1010 | 640 | 1187 | 752 | 1199 | 1175 | 1112 | 1058 | 998 | 1003 | 951 | 855 | 1018 | 955 | 993,75 | 1043 | 945 | 0,907 | -0,141 | -2,13 | 0,0331892 | 0,0404857 | 0,0027771 | 0,0037435 | 0,0032179 | 0,0032179 | TRUE | 0,0456176 | No |
| orf19.239 | 4165 | 3995 | 4302 | 2513 | 3828 | 2698 | 3481 | 3915 | 3971 | 4224 | 4250 | 3937 | 3067 | 3068 | 2955 | 3183 | 3581,94 | 4096 | 3068 | 0,749 | -0,417 | -9,562 | 1,15E-21 | 2,67E-21 | 0,0008862 | 0,0025497 | 0,0015458 | 0,0015458 | TRUE | 0,0078977 | No |
| orf19.2391 | 899 | 727 | 828 | 601 | 1790 | 1223 | 1730 | 1781 | 857 | 769 | 818 | 941 | 1434 | 1391 | 1469 | 1448 | 1140,89 | 846 | 1436 | 1,703 | 0,768 | 12,348 | 5,02E-35 | 1,47E-34 | 0,001907 | 0,0035304 | 0,0027426 | 0,0027426 | TRUE | 0,0595116 | No |
| orf19.2392 | 0 | 0 | 0 | 0 | 0 | 0 | 0 | 0 | 0 | 0 | 0 | 0 | 0 | 0 | 0 | 0 | 0 | 0 | 0 | NA | NA | NA | NA | NA | NA | NA | NA | NA | NA | NA | NA |
| orf19.2393 | 6773 | 5616 | 6314 | 3769 | 6564 | 4564 | 6406 | 6651 | 6457 | 5938 | 6238 | 5904 | 5260 | 5190 | 5439 | 5408 | 5729,16 | 6134 | 5324 | 0,868 | -0,205 | -4,999 | 5,75E-07 | 9,00E-07 | 0,0009293 | 0,0023779 | 0,0014395 | 0,0014395 | TRUE | 0,0154203 | No |
| orf19.2394 | 28 | 31 | 35 | 32 | 23 | 15 | 27 | 21 | 27 | 33 | 35 | 50 | 18 | 17 | 23 | 17 | 27,46 | 36 | 19 | 0,536 | -0,9 | -3,106 | 0,0018946 | 0,002514 | 1,00E-08 | 0,0618883 | 0,0413498 | 0,0413498 | TRUE | 0,3583923 | No |
| orf19.2395 | 14 | 13 | 17 | 12 | 13 | 17 | 13 | 9 | 13 | 14 | 17 | 19 | 10 | 19 | 11 | 7 | 13,85 | 16 | 12 | 0,756 | -0,403 | -1,009 | 0,3131862 | 0,3462903 | 0,0152344 | 0,1206529 | 0,0792439 | 0,0792439 | TRUE | 0,5552533 | No |
| orf19.2396 | 643 | 568 | 628 | 396 | 2543 | 1844 | 2304 | 2450 | 613 | 601 | 620 | 620 | 2038 | 2097 | 1956 | 1992 | 1317,12 | 614 | 2021 | 3,293 | 1,719 | 33,024 | 3,73E-239 | 6,00E-238 | 1,00E-08 | 0,0033379 | 0,0014739 | 0,0014739 | TRUE | 0,0073082 | No |
| orf19.2397 | 229 | 208 | 176 | 138 | 320 | 203 | 302 | 278 | 218 | 220 | 174 | 216 | 256 | 231 | 256 | 226 | 224,75 | 207 | 242 | 1,175 | 0,233 | 2,139 | 0,0324739 | 0,039651 | 0,0037392 | 0,0093968 | 0,006837 | 0,006837 | TRUE | 0,1360102 | No |
| orf19.2397.3 | 6566 | 6653 | 6954 | 3814 | 2508 | 1692 | 2386 | 2300 | 6260 | 7035 | 6870 | 5975 | 2010 | 1924 | 2026 | 1870 | 4246,1 | 6535 | 1958 | 0,299 | -1,74 | -29,221 | 1,06E-187 | 1,35E-186 | 0,003191 | 0,002478 | 0,00309 | 0,00309 | TRUE | 0,039657 | No |
| orf19.2398 | 19972 | 17786 | 19466 | 12883 | 2044 | 1518 | 1990 | 2009 | 19041 | 18806 | 19231 | 20182 | 1638 | 1726 | 1690 | 1634 | 10493,37 | 19315 | 1672 | 0,086 | -3,531 | -89,839 | 0 | 0 | 0,0006704 | 0,0022478 | 0,0011876 | 0,0011876 | TRUE | 0,0109747 | No |
| orf19.2399 | 463 | 489 | 520 | 253 | 553 | 413 | 557 | 563 | 441 | 517 | 514 | 396 | 443 | 470 | 473 | 458 | 463,99 | 467 | 461 | 0,981 | -0,027 | -0,306 | 0,7593748 | 0,7871297 | 0,0050729 | 0,00563 | 0,005458 | 0,005458 | TRUE | 0,1034543 | No |
| orf19.24 | 4698 | 4031 | 4267 | 2773 | 2845 | 2041 | 2681 | 2931 | 4479 | 4262 | 4215 | 4344 | 2280 | 2321 | 2276 | 2383 | 3320,06 | 4325 | 2315 | 0,535 | -0,902 | -23,365 | 9,67E-121 | 7,76E-120 | 0,0003638 | 0,0025858 | 0,0011074 | 0,0011074 | TRUE | 0,0073595 | No |
| orf19.240 | 1857 | 1820 | 1766 | 908 | 1857 | 1229 | 1769 | 1774 | 1770 | 1924 | 1745 | 1422 | 1488 | 1397 | 1502 | 1442 | 1586,46 | 1715 | 1457 | 0,848 | -0,238 | -2,989 | 0,0028015 | 0,0036776 | 0,007237 | 0,0031262 | 0,0054416 | 0,0054416 | TRUE | 0,151387 | No |
| orf19.2400 | 4499 | 4240 | 4511 | 2486 | 4140 | 2747 | 4034 | 4087 | 4289 | 4483 | 4456 | 3894 | 3317 | 3124 | 3425 | 3323 | 3789,04 | 4280 | 3297 | 0,77 | -0,378 | -7,073 | 1,52E-12 | 2,85E-12 | 0,002382 | 0,0025246 | 0,0024686 | 0,0024686 | TRUE | 0,0431337 | No |
| orf19.2401 | 306 | 330 | 336 | 205 | 725 | 441 | 591 | 633 | 292 | 349 | 332 | 321 | 581 | 501 | 502 | 515 | 424,07 | 324 | 525 | 1,625 | 0,701 | 8,089 | 6,03E-16 | 1,24E-15 | 0,0033222 | 0,005963 | 0,004626 | 0,004626 | TRUE | 0,0627794 | No |
| orf19.2402 | 3380 | 2821 | 2988 | 2034 | 2626 | 1643 | 2340 | 2416 | 3222 | 2983 | 2952 | 3186 | 2104 | 1868 | 1987 | 1964 | 2533,38 | 3086 | 1981 | 0,643 | -0,638 | -12,352 | 4,77E-35 | 1,40E-34 | 0,0016479 | 0,0027394 | 0,0021506 | 0,0021506 | TRUE | 0,0213072 | No |
| orf19.2404 | 3935 | 3729 | 4122 | 2436 | 3082 | 2249 | 3031 | 3376 | 3752 | 3943 | 4072 | 3816 | 2470 | 2557 | 2573 | 2745 | 3241 | 3896 | 2586 | 0,664 | -0,591 | -12,532 | 4,96E-36 | 1,47E-35 | 0,0012277 | 0,0025979 | 0,0018195 | 0,0018195 | TRUE | 0,0211871 | No |
| orf19.2406 | 2487 | 2323 | 2451 | 1420 | 2169 | 1513 | 2052 | 2195 | 2371 | 2456 | 2421 | 2225 | 1738 | 1720 | 1742 | 1785 | 2057,31 | 2368 | 1746 | 0,736 | -0,441 | -9,878 | 5,16E-23 | 1,23E-22 | 0,0003872 | 0,0028894 | 0,0014253 | 0,0014253 | TRUE | 0,0182405 | No |
| orf19.2407 | 4061 | 3657 | 3741 | 2374 | 6520 | 4720 | 6116 | 7076 | 3872 | 3867 | 3696 | 3719 | 5224 | 5367 | 5192 | 5753 | 4586,32 | 3788 | 5384 | 1,421 | 0,507 | 11,324 | 9,99E-30 | 2,68E-29 | 0,0011518 | 0,0024493 | 0,0016919 | 0,0016919 | TRUE | 0,0263488 | No |
| orf19.2408 | 1691 | 1617 | 1604 | 1030 | 678 | 559 | 734 | 776 | 1612 | 1710 | 1585 | 1614 | 543 | 636 | 623 | 631 | 1119,14 | 1630 | 608 | 0,373 | -1,425 | -22,828 | 2,41E-115 | 1,83E-114 | 0,0016333 | 0,0035584 | 0,00266 | 0,00266 | TRUE | 0,0636426 | No |
| orf19.2410 | 2360 | 2205 | 2397 | 1452 | 2637 | 1748 | 2517 | 2600 | 2250 | 2332 | 2368 | 2275 | 2113 | 1988 | 2137 | 2114 | 2196,96 | 2306 | 2088 | 0,906 | -0,142 | -3,344 | 0,0008247 | 0,0011197 | 0,0002975 | 0,0028386 | 0,0012847 | 0,0012847 | TRUE | 0,0118027 | No |
| orf19.2411 | 1476 | 1352 | 1429 | 866 | 1178 | 868 | 1033 | 1180 | 1407 | 1430 | 1412 | 1357 | 944 | 987 | 877 | 959 | 1171,56 | 1402 | 942 | 0,671 | -0,576 | -10,851 | 1,98E-27 | 5,13E-27 | 0,0004024 | 0,0034927 | 0,0018354 | 0,0018354 | TRUE | 0,0256666 | No |
| orf19.2412 | 383 | 345 | 369 | 189 | 560 | 414 | 502 | 485 | 365 | 365 | 365 | 296 | 449 | 471 | 426 | 394 | 391,32 | 348 | 435 | 1,241 | 0,311 | 3,428 | 0,0006071 | 0,0008302 | 0,0042537 | 0,0062871 | 0,005296 | 0,005296 | TRUE | 0,0944195 | No |
| orf19.2414 | 9629 | 8265 | 8602 | 5697 | 1928 | 1399 | 1804 | 2004 | 9180 | 8739 | 8498 | 8925 | 1545 | 1591 | 1532 | 1629 | 5204,84 | 8836 | 1574 | 0,178 | -2,489 | -59,697 | 0 | 0 | 0,000764 | 0,0024067 | 0,0013219 | 0,0013219 | TRUE | 0,0086708 | No |
| orf19.2415 | 602 | 492 | 570 | 331 | 1415 | 980 | 1422 | 1565 | 574 | 520 | 563 | 519 | 1134 | 1114 | 1207 | 1272 | 862,96 | 544 | 1182 | 2,169 | 1,117 | 16,44 | 9,89E-61 | 4,20E-60 | 0,0022362 | 0,0039939 | 0,0030304 | 0,0030304 | TRUE | 0,0321926 | No |
| orf19.2416.1 | 401 | 308 | 322 | 243 | 453 | 263 | 442 | 531 | 382 | 326 | 318 | 381 | 363 | 299 | 375 | 432 | 359,47 | 352 | 367 | 1,05 | 0,07 | 0,608 | 0,5432107 | 0,5787656 | 0,0131754 | 0,0066588 | 0,0098772 | 0,0098772 | TRUE | 0,1679023 | No |
| orf19.2417 | 811 | 720 | 810 | 504 | 2070 | 1485 | 1949 | 2128 | 773 | 761 | 800 | 790 | 1659 | 1689 | 1655 | 1730 | 1232,05 | 781 | 1683 | 2,156 | 1,108 | 22,44 | 1,60E-111 | 1,17E-110 | 1,00E-08 | 0,0034239 | 0,0013626 | 0,0013626 | TRUE | 0,0045281 | No |
| orf19.2418 | 903 | 656 | 723 | 495 | 1082 | 650 | 1032 | 1003 | 861 | 694 | 714 | 775 | 867 | 739 | 876 | 816 | 792,75 | 761 | 824 | 1,085 | 0,117 | 1,41 | 0,1586025 | 0,1815974 | 0,0060868 | 0,0041624 | 0,0053824 | 0,0053824 | TRUE | 0,0981784 | No |
| orf19.2419 | 1732 | 1461 | 1542 | 985 | 2450 | 1738 | 2307 | 2564 | 1651 | 1545 | 1523 | 1543 | 1963 | 1976 | 1959 | 2085 | 1780,66 | 1566 | 1996 | 1,274 | 0,349 | 7,386 | 1,51E-13 | 2,93E-13 | 0,0006299 | 0,0030134 | 0,0015667 | 0,0015667 | TRUE | 0,0172854 | No |
| orf19.242 | 156 | 154 | 165 | 99 | 260 | 197 | 228 | 256 | 149 | 163 | 163 | 155 | 208 | 224 | 194 | 208 | 182,96 | 158 | 208 | 1,321 | 0,402 | 3,734 | 0,0001881 | 0,0002626 | 1,00E-08 | 0,0110652 | 0,005433 | 0,005433 | TRUE | 0,025168 | No |
| orf19.2422 | 3373 | 2930 | 3210 | 1947 | 6177 | 4330 | 5899 | 6565 | 3216 | 3098 | 3171 | 3050 | 4949 | 4924 | 5008 | 5338 | 4094,3 | 3134 | 5055 | 1,612 | 0,689 | 16,675 | 1,98E-62 | 8,62E-62 | 0,0008104 | 0,0024923 | 0,0013716 | 0,0013716 | TRUE | 0,0175049 | No |
| orf19.242.2 | 153 | 147 | 153 | 68 | 376 | 238 | 388 | 327 | 146 | 155 | 151 | 107 | 301 | 271 | 329 | 266 | 215,77 | 140 | 292 | 2,066 | 1,047 | 8,36 | 6,29E-17 | 1,32E-16 | 0,0094505 | 0,0097007 | 0,0095872 | 0,0095872 | TRUE | 0,2027348 | No |
| orf19.2423 | 2322 | 1941 | 2221 | 1539 | 1821 | 1336 | 1744 | 1893 | 2214 | 2052 | 2194 | 2411 | 1459 | 1519 | 1481 | 1539 | 1858,66 | 2218 | 1500 | 0,677 | -0,562 | -10,28 | 8,66E-25 | 2,13E-24 | 0,0017582 | 0,0029747 | 0,0023223 | 0,0023223 | TRUE | 0,038697 | No |
| orf19.2425 | 263 | 238 | 252 | 155 | 208 | 136 | 216 | 215 | 251 | 252 | 249 | 243 | 167 | 155 | 183 | 175 | 209,21 | 249 | 170 | 0,685 | -0,546 | -5,456 | 4,88E-08 | 7,95E-08 | 1,00E-08 | 0,0099394 | 0,0047631 | 0,0047631 | TRUE | 0,0329359 | No |
| orf19.2426 | 494 | 434 | 440 | 316 | 421 | 318 | 454 | 532 | 471 | 459 | 435 | 495 | 337 | 362 | 385 | 433 | 422,07 | 465 | 379 | 0,818 | -0,289 | -3,195 | 0,0013979 | 0,0018673 | 0,0049785 | 0,0059814 | 0,0054726 | 0,0054726 | TRUE | 0,1074086 | No |
| orf19.2429 | 35 | 29 | 32 | 17 | 45 | 43 | 45 | 38 | 33 | 31 | 32 | 27 | 36 | 49 | 38 | 31 | 34,54 | 31 | 38 | 1,236 | 0,306 | 1,244 | 0,2135388 | 0,2415085 | 1,00E-08 | 0,0496256 | 0,0283963 | 0,0283963 | TRUE | 0,2104288 | No |

| | | | | | | | | | | | | | | | | | | | | | | | | | | | | | | |
|---|---|---|---|---|---|---|---|---|---|---|---|---|---|---|---|---|---|---|---|---|---|---|---|---|---|---|---|---|---|---|
| orf19.243 | 1195 | 974 | 1087 | 678 | 1352 | 923 | 1280 | 1394 | 1139 | 1030 | 1074 | 1062 | 1083 | 1050 | 1087 | 1133 | 1082,27 | 1076 | 1088 | 1,011 | 0,016 | 0,293 | 0,7696637 | 0,7941824 | 0,0005247 | 0,0036084 | 0,0018913 | 0,0018913 | TRUE | 0,0197386 | No |
| orf19.2430 | 9 | 8 | 4 | 1 | 10 | 8 | 13 | 4 | 9 | 8 | 4 | 2 | 8 | 9 | 11 | 3 | 6,74 | 6 | 8 | 1,331 | 0,413 | 0,683 | 0,4943386 | 0,5302362 | 0,1237946 | 0,2455326 | 0,1980495 | 0,1980495 | TRUE | 0,4558921 | No |
| orf19.2431 | 32 | 30 | 36 | 24 | 21 | 28 | 29 | 25 | 31 | 32 | 36 | 38 | 17 | 32 | 25 | 20 | 28,63 | 34 | 24 | 0,684 | -0,547 | -1,988 | 0,0467837 | 0,0564003 | 1,00E-08 | 0,0594491 | 0,0368657 | 0,0368657 | TRUE | 0,294133 | No |
| orf19.2432 | 6562 | 6629 | 7026 | 4325 | 3332 | 2368 | 3143 | 3198 | 6256 | 7009 | 6941 | 6775 | 2670 | 2693 | 2668 | 2600 | 4701,62 | 6745 | 2658 | 0,394 | -1,344 | -29,666 | 2,12E-193 | 2,82E-192 | 0,0013483 | 0,0024405 | 0,0017185 | 0,0017185 | TRUE | 0,0297271 | No |
| orf19.2433 | 150 | 181 | 174 | 94 | 91 | 77 | 87 | 77 | 143 | 191 | 172 | 147 | 73 | 88 | 74 | 63 | 118,81 | 163 | 74 | 0,45 | -1,152 | -7,495 | 6,63E-14 | 1,29E-13 | 0,0098787 | 0,0159107 | 0,0132406 | 0,0132406 | TRUE | 0,1401366 | No |
| orf19.2434 | 1819 | 1857 | 1904 | 1177 | 2971 | 2181 | 2788 | 2766 | 1734 | 1964 | 1881 | 1844 | 2381 | 2480 | 2367 | 2249 | 2112,39 | 1856 | 2369 | 1,276 | 0,352 | 6,727 | 1,73E-11 | 3,16E-11 | 0,0014716 | 0,0028686 | 0,0021336 | 0,0021336 | TRUE | 0,0248122 | No |
| orf19.2435 | 10574 | 10403 | 10638 | 6488 | 22975 | 17536 | 21357 | 22999 | 10081 | 11000 | 10509 | 10164 | 18409 | 19940 | 18132 | 18700 | 14616,97 | 10438 | 18795 | 1,8 | 0,848 | 19,19 | 4,51E-82 | 2,51E-81 | 0,0016014 | 0,0022036 | 0,0017984 | 0,0017984 | TRUE | 0,0204354 | No |
| orf19.2436 | 8296 | 7833 | 8404 | 5187 | 3535 | 2559 | 3360 | 3552 | 7909 | 8282 | 8302 | 8126 | 2832 | 2910 | 2853 | 2888 | 5512,86 | 8155 | 2871 | 0,352 | -1,507 | -44,927 | 0 | 0 | 0,0002291 | 0,0023891 | 0,0008553 | 0,0008553 | TRUE | 0,0052162 | No |
| orf19.2437 | 742 | 870 | 841 | 596 | 1415 | 1226 | 1245 | 1098 | 707 | 920 | 831 | 934 | 1134 | 1394 | 1057 | 893 | 983,68 | 848 | 1120 | 1,32 | 0,401 | 3,236 | 0,0012106 | 0,0016229 | 0,0235783 | 0,0037604 | 0,0136742 | 0,0136742 | TRUE | 0,322188 | No |
| orf19.2438 | 2572 | 2316 | 2340 | 1327 | 2084 | 1463 | 2047 | 1898 | 2452 | 2449 | 2312 | 2079 | 1670 | 1664 | 1738 | 1543 | 1988,26 | 2323 | 1654 | 0,711 | -0,493 | -7,847 | 4,26E-15 | 8,60E-15 | 0,0033573 | 0,0029171 | 0,0032751 | 0,0032751 | TRUE | 0,0572722 | No |
| orf19.2439 | 1668 | 1518 | 1481 | 900 | 3334 | 2157 | 2898 | 2754 | 1590 | 1605 | 1463 | 1410 | 2671 | 2453 | 2460 | 2239 | 1986,52 | 1517 | 2456 | 1,617 | 0,693 | 10,543 | 5,46E-26 | 1,38E-25 | 0,0038622 | 0,0029178 | 0,0035962 | 0,0035962 | TRUE | 0,0435572 | No |
| orf19.2439.1 | 2202 | 1831 | 1829 | 1080 | 3967 | 2230 | 3337 | 3826 | 2099 | 1936 | 1807 | 1692 | 3179 | 2536 | 2833 | 3111 | 2399,07 | 1884 | 2915 | 1,546 | 0,629 | 7,598 | 3,01E-14 | 5,95E-14 | 0,0088078 | 0,0027757 | 0,0061301 | 0,0061301 | TRUE | 0,0917614 | No |
| orf19.244 | 79 | 66 | 58 | 42 | 134 | 89 | 121 | 128 | 75 | 70 | 57 | 66 | 107 | 101 | 103 | 104 | 85,45 | 67 | 104 | 1,548 | 0,63 | 4,056 | 5,00E-05 | 7,18E-05 | 1,00E-08 | 0,0213068 | 0,010652 | 0,010652 | TRUE | 0,0985737 | No |
| orf19.2440 | 965 | 1037 | 1029 | 615 | 1220 | 914 | 1144 | 1094 | 920 | 1096 | 1017 | 963 | 978 | 1039 | 971 | 890 | 984,27 | 999 | 970 | 0,969 | -0,046 | -0,647 | 0,5175134 | 0,5538834 | 0,0038927 | 0,0037594 | 0,0038326 | 0,0038326 | TRUE | 0,0521104 | No |
| orf19.2442 | 408 | 474 | 492 | 296 | 379 | 248 | 375 | 283 | 389 | 501 | 486 | 464 | 304 | 282 | 318 | 230 | 371,76 | 460 | 284 | 0,617 | -0,697 | -6,033 | 1,61E-09 | 2,75E-09 | 0,013563 | 0,0065078 | 0,010024 | 0,010024 | TRUE | 0,1908713 | No |
| orf19.2443 | 1099 | 1112 | 1115 | 677 | 1729 | 1336 | 1545 | 1629 | 1048 | 1176 | 1102 | 1061 | 1385 | 1519 | 1312 | 1325 | 1240,8 | 1097 | 1385 | 1,261 | 0,334 | 5,233 | 1,67E-07 | 2,67E-07 | 0,0026418 | 0,0034145 | 0,003087 | 0,003087 | TRUE | 0,0487235 | No |
| orf19.2444 | 3217 | 3034 | 3106 | 1945 | 5596 | 3895 | 4919 | 5073 | 3067 | 3208 | 3068 | 3047 | 4484 | 4429 | 4176 | 4125 | 3700,55 | 3098 | 4304 | 1,389 | 0,474 | 10,977 | 4,92E-28 | 1,29E-27 | 0,0009634 | 0,002535 | 0,0015017 | 0,0015017 | TRUE | 0,0100666 | No |
| orf19.2445 | 92 | 79 | 84 | 68 | 301 | 208 | 246 | 245 | 88 | 84 | 83 | 107 | 241 | 237 | 209 | 199 | 155,81 | 90 | 222 | 2,48 | 1,31 | 9,873 | 5,46E-23 | 1,30E-22 | 0,0038286 | 0,0126289 | 0,0086592 | 0,0086592 | TRUE | 0,1041226 | No |
| orf19.2446 | 678 | 715 | 682 | 473 | 794 | 636 | 797 | 777 | 646 | 756 | 674 | 741 | 636 | 723 | 677 | 632 | 685,62 | 704 | 667 | 0,947 | -0,078 | -1,047 | 0,2951929 | 0,3278152 | 0,003288 | 0,004486 | 0,0038586 | 0,0038586 | TRUE | 0,0371769 | No |
| orf19.2447 | 1484 | 1404 | 1483 | 907 | 3361 | 2386 | 3110 | 2939 | 1415 | 1485 | 1465 | 1421 | 2693 | 2713 | 2640 | 2390 | 2027,69 | 1446 | 2609 | 1,802 | 0,85 | 15,901 | 6,28E-57 | 2,53E-56 | 0,001742 | 0,002901 | 0,0021833 | 0,0021833 | TRUE | 0,039402 | No |
| orf19.2449 | 10 | 7 | 12 | 3 | 4 | 5 | 6 | 6 | 10 | 7 | 12 | 5 | 3 | 6 | 5 | 5 | 6,54 | 8 | 5 | 0,543 | -0,882 | -1,59 | 0,1118687 | 0,1298116 | 1,00E-08 | 0,252988 | 0,1351901 | 0,1351901 | TRUE | 0,1499226 | No |
| orf19.245 | 541 | 562 | 600 | 377 | 1033 | 758 | 989 | 953 | 516 | 594 | 593 | 591 | 828 | 862 | 840 | 775 | 699,69 | 574 | 826 | 1,442 | 0,528 | 7,637 | 2,21E-14 | 4,39E-14 | 0,0017871 | 0,0044379 | 0,0030727 | 0,0030727 | TRUE | 0,0574651 | No |
| orf19.2451 | 568 | 590 | 571 | 409 | 12254 | 10390 | 10969 | 11047 | 542 | 624 | 564 | 641 | 9819 | 11814 | 9313 | 8982 | 5287,28 | 593 | 9982 | 16,885 | 4,078 | 45,734 | 0 | 0 | 0,0100767 | 0,0024018 | 0,0066382 | 0,0066382 | TRUE | 0,1860867 | No |
| orf19.2452 | 974 | 1041 | 1057 | 629 | 6112 | 4426 | 5517 | 5449 | 929 | 1101 | 1044 | 985 | 4897 | 5033 | 4684 | 4431 | 2887,94 | 1015 | 4761 | 4,689 | 2,229 | 35,504 | 4,28E-276 | 8,35E-275 | 0,0034073 | 0,0026598 | 0,0031385 | 0,0031385 | TRUE | 0,0422534 | No |
| orf19.2454 | 12521 | 12228 | 12522 | 8083 | 13270 | 9742 | 12012 | 12671 | 11937 | 12930 | 12371 | 12662 | 10633 | 11078 | 10198 | 10303 | 11513,91 | 12475 | 10553 | 0,846 | -0,241 | -5,896 | 3,72E-09 | 6,26E-09 | 0,001167 | 0,0022339 | 0,0015237 | 0,0015237 | TRUE | 0,0135075 | No |
| orf19.2455 | 3630 | 3562 | 3736 | 2227 | 4944 | 3643 | 4537 | 4636 | 3461 | 3766 | 3691 | 3489 | 3961 | 4142 | 3852 | 3770 | 3766,5 | 3602 | 3931 | 1,091 | 0,125 | 2,679 | 0,0073789 | 0,009434 | 0,0013555 | 0,0025272 | 0,0018207 | 0,0018207 | TRUE | 0,015452 | No |
| orf19.2457 | 1179 | 1160 | 1173 | 742 | 701 | 439 | 614 | 680 | 1124 | 1227 | 1159 | 1162 | 562 | 499 | 521 | 553 | 850,86 | 1168 | 534 | 0,458 | -1,126 | -18,361 | 2,67E-75 | 1,38E-74 | 0,0006923 | 0,004021 | 0,0022973 | 0,0022973 | TRUE | 0,0198971 | No |
| orf19.2458 | 21292 | 19620 | 21289 | 14171 | 7723 | 5642 | 7075 | 7482 | 20300 | 20746 | 21032 | 22200 | 6188 | 6415 | 6007 | 6084 | 13621,29 | 21070 | 6174 | 0,293 | -1,771 | -44,603 | 0 | 0 | 0,0010164 | 0,0022118 | 0,0014152 | 0,0014152 | TRUE | 0,01576 | No |
| orf19.2459 | 5436 | 4999 | 5332 | 3531 | 4485 | 3108 | 4138 | 4393 | 5183 | 5286 | 5268 | 5531 | 3594 | 3534 | 3513 | 3572 | 4435,04 | 5317 | 3553 | 0,669 | -0,58 | -16,706 | 1,18E-62 | 5,16E-62 | 0,0001372 | 0,0024615 | 0,0009253 | 0,0009253 | TRUE | 0,0084164 | No |
| orf19.246 | 481 | 428 | 457 | 287 | 1262 | 865 | 1131 | 1213 | 459 | 453 | 451 | 450 | 1011 | 984 | 960 | 986 | 719,19 | 453 | 985 | 2,174 | 1,12 | 18,777 | 1,17E-78 | 6,29E-78 | 1,00E-08 | 0,0043743 | 0,0017399 | 0,0017399 | TRUE | 0,0039366 | No |
| orf19.2460 | 13 | 9 | 7 | 6 | 11 | 15 | 8 | 8 | 12 | 10 | 7 | 9 | 9 | 17 | 7 | 7 | 9,67 | 10 | 10 | 0,995 | -0,007 | -0,014 | 0,9890541 | 0,990451 | 0,0465584 | 0,1718142 | 0,1183497 | 0,1183497 | TRUE | 0,7212484 | No |
| orf19.2461 | 1161 | 1029 | 1132 | 678 | 1662 | 1085 | 1598 | 1637 | 1107 | 1088 | 1118 | 1062 | 1332 | 1234 | 1357 | 1331 | 1203,57 | 1094 | 1314 | 1,201 | 0,264 | 5,149 | 2,62E-07 | 4,14E-07 | 1,00E-08 | 0,0034555 | 0,0016712 | 0,0016712 | TRUE | 0,0184221 | No |
| orf19.2462 | 2895 | 2573 | 2811 | 1655 | 2680 | 1843 | 2464 | 2643 | 2760 | 2721 | 2777 | 2593 | 2147 | 2096 | 2092 | 2149 | 2416,79 | 2713 | 2121 | 0,781 | -0,357 | -8,94 | 3,88E-19 | 8,56E-19 | 6,52E-05 | 0,0027707 | 0,0011101 | 0,0011101 | TRUE | 0,0096769 | No |
| orf19.2463 | 939 | 942 | 979 | 523 | 849 | 543 | 793 | 823 | 895 | 996 | 967 | 819 | 680 | 617 | 673 | 669 | 789,74 | 919 | 660 | 0,717 | -0,48 | -6,719 | 1,83E-11 | 3,34E-11 | 0,0029128 | 0,0041703 | 0,0036204 | 0,0036204 | TRUE | 0,0568939 | No |
| orf19.2465 | 3805 | 3248 | 3540 | 2109 | 3219 | 2077 | 3142 | 3238 | 3628 | 3434 | 3497 | 3304 | 2579 | 2362 | 2668 | 2633 | 3013,05 | 3466 | 2560 | 0,739 | -0,437 | -8,636 | 5,81E-18 | 1,25E-17 | 0,0017397 | 0,0026362 | 0,0021188 | 0,0021188 | TRUE | 0,0317703 | No |
| orf19.2467 | 3007 | 2443 | 2694 | 1692 | 2115 | 1390 | 1957 | 2140 | 2867 | 2583 | 2661 | 2651 | 1695 | 1581 | 1661 | 1740 | 2179,85 | 2690 | 1669 | 0,621 | -0,688 | -13,657 | 1,83E-42 | 6,04E-42 | 0,0013931 | 0,0028445 | 0,0019605 | 0,0019605 | TRUE | 0,0243841 | No |
| orf19.2468 | 8 | 8 | 3 | 3 | 20 | 18 | 22 | 30 | 8 | 8 | 3 | 5 | 16 | 20 | 19 | 24 | 12,91 | 6 | 20 | 3,305 | 1,725 | 3,86 | 0,0001135 | 0,0001602 | 1,00E-08 | 0,129231 | 0,077232 | 0,077232 | TRUE | 0,320359 | No |
| orf19.2469 | 161 | 146 | 149 | 94 | 713 | 522 | 759 | 815 | 153 | 154 | 147 | 147 | 571 | 594 | 644 | 663 | 384,28 | 150 | 618 | 4,101 | 2,036 | 21,538 | 6,88E-103 | 4,61E-102 | 1,00E-08 | 0,0063639 | 0,0041795 | 0,0041795 | TRUE | 0,0326879 | No |
| orf19.247 | 202 | 203 | 188 | 98 | 228 | 138 | 192 | 187 | 193 | 215 | 186 | 154 | 183 | 157 | 163 | 152 | 175,14 | 187 | 164 | 0,872 | -0,198 | -1,607 | 0,10801 | 0,1256082 | 0,0054311 | 0,0114659 | 0,0088601 | 0,0088601 | TRUE | 0,1219309 | No |
| orf19.2471 | 898 | 804 | 853 | 549 | 1696 | 1196 | 1546 | 1675 | 856 | 850 | 843 | 860 | 1359 | 1360 | 1313 | 1362 | 1100,3 | 852 | 1348 | 1,583 | 0,662 | 13,563 | 6,59E-42 | 2,16E-41 | 1,00E-08 | 0,0035835 | 0,0013045 | 0,0013045 | TRUE | 0,0039247 | No |
| orf19.2472 | 814 | 695 | 734 | 502 | 977 | 673 | 886 | 958 | 776 | 735 | 725 | 786 | 783 | 765 | 752 | 779 | 762,72 | 756 | 770 | 1,021 | 0,03 | 0,519 | 0,6038578 | 0,638161 | 1,00E-08 | 0,004244 | 0,0018646 | 0,0018646 | TRUE | 0,0087928 | No |
| orf19.2472.1 | 1720 | 1359 | 1361 | 911 | 1238 | 715 | 1097 | 1163 | 1640 | 1437 | 1345 | 1427 | 992 | 813 | 931 | 946 | 1191,31 | 1462 | 920 | 0,63 | -0,666 | -8,542 | 1,32E-17 | 2,81E-17 | 0,0059292 | 0,0034695 | 0,0049662 | 0,0049662 | TRUE | 0,0828892 | No |
| orf19.2473 | 3305 | 3002 | 3155 | 2074 | 2805 | 1957 | 2530 | 2783 | 3151 | 3174 | 3117 | 3249 | 2248 | 2225 | 2148 | 2263 | 2696,84 | 3173 | 2221 | 0,701 | -0,513 | -13,536 | 9,53E-42 | 3,10E-41 | 1,00E-08 | 0,0027001 | 0,0010029 | 0,0010029 | TRUE | 0,0061074 | No |
| orf19.2474 | 1256 | 1113 | 1202 | 813 | 2852 | 2001 | 2831 | 2975 | 1197 | 1177 | 1187 | 1274 | 2285 | 2275 | 2404 | 2419 | 1777,3 | 1209 | 2346 | 1,946 | 0,96 | 20,169 | 1,82E-90 | 1,09E-89 | 0,0005039 | 0,0030151 | 0,0015222 | 0,0015222 | TRUE | 0,0134481 | No |
| orf19.2475 | 5 | 6 | 8 | 2 | 2 | 4 | 2 | 2 | 5 | 6 | 8 | 3 | 2 | 5 | 2 | 2 | 3,95 | 6 | 3 | 0,4 | -1,324 | -1,786 | 0,0741446 | 0,0877116 | 1,00E-08 | 0,4174655 | 0,2324804 | 0,2324804 | TRUE | 0,2858324 | No |
| orf19.2476 | 1644 | 1575 | 1583 | 1012 | 2089 | 1541 | 1879 | 2037 | 1567 | 1665 | 1564 | 1585 | 1674 | 1752 | 1595 | 1656 | 1632,45 | 1595 | 1669 | 1,045 | 0,064 | 1,329 | 0,1838315 | 0,2092452 | 0,0005777 | 0,0030971 | 0,0016074 | 0,0016074 | TRUE | 0,0125786 | No |
| orf19.2478 | 94 | 75 | 77 | 34 | 47 | 33 | 38 | 40 | 90 | 79 | 76 | 53 | 38 | 38 | 32 | 33 | 54,78 | 74 | 35 | 0,462 | -1,115 | -5,308 | 1,11E-07 | 1,78E-07 | 1,00E-08 | 0,0320651 | 0,0222243 | 0,0222243 | TRUE | 0,2517577 | No |
| orf19.2478.1 | 22386 | 18853 | 19644 | 14457 | 47754 | 34149 | 45706 | 50226 | 21343 | 19935 | 19406 | 22648 | 38264 | 38830 | 38804 | 40839 | 30008,56 | 20833 | 39184 | 1,881 | 0,912 | 17,601 | 2,42E-69 | 1,15E-68 | 0,002825 | 0,002146 | 0,0025395 | 0,0025395 | TRUE | 0,0417565 | No |
| orf19.2479 | 161 | 155 | 163 | 98 | 728 | 587 | 760 | 807 | 153 | 164 | 161 | 154 | 583 | 667 | 645 | 656 | 398,02 | 158 | 638 | 4,028 | 2,01 | 22,103 | 2,99E-108 | 2,13E-107 | 1,00E-08 | 0,0062165 | 0,0037362 | 0,0037362 | TRUE | 0,0405329 | No |
| orf19.248 | 1031 | 1087 | 1111 | 678 | 1980 | 1555 | 1856 | 1700 | 983 | 1149 | 1098 | 1062 | 1587 | 1768 | 1576 | 1382 | 1325,59 | 1073 | 1578 | 1,469 | 0,555 | 7,06 | 1,67E-12 | 3,13E-12 | 0,006068 | 0,0033299 | 0,0051305 | 0,0051305 | TRUE | 0,0845879 | No |
| orf19.2480.1 | 17088 | 14277 | 15811 | 9393 | 6448 | 4137 | 6046 | 6228 | 16292 | 15096 | 15620 | 14715 | 5167 | 4704 | 5133 | 5064 | 10223,73 | 15431 | 5017 | 0,325 | -1,621 | -35,086 | 1,12E-269 | 2,09E-268 | 0,001705 | 0,0022519 | 0,0019237 | 0,0019237 | TRUE | 0,0209795 | No |
| orf19.2481 | 541 | 462 | 499 | 379 | 140 | 96 | 145 | 163 | 516 | 489 | 493 | 594 | 112 | 109 | 123 | 133 | 320,99 | 523 | 119 | 0,23 | -2,121 | -20,17 | 1,81E-90 | 1,09E-89 | 0,0041353 | 0,0072063 | 0,0058276 | 0,0058276 | TRUE | 0,0866873 | No |
| orf19.2483 | 2182 | 1678 | 1694 | 1469 | 1165 | 949 | 1120 | 1136 | 2080 | 1774 | 1674 | 2301 | 933 | 1079 | 951 | 924 | 1464,56 | 1957 | 972 | 0,497 | -1,01 | -10,513 | 7,53E-26 | 1,89E-25 | 0,0125163 | 0,0032124 | 0,0081099 | 0,0081099 | TRUE | 0,1642759 | No |
| orf19.2484 | 2262 | 1830 | 1975 | 1572 | 3918 | 3039 | 3746 | 4029 | 2157 | 1935 | 1951 | 2463 | 3139 | 3456 | 3180 | 3276 | 2694,57 | 2126 | 3263 | 1,537 | 0,62 | 8,412 | 4,03E-17 | 8,50E-17 | 0,0070957 | 0,0027006 | 0,0048142 | 0,0048142 | TRUE | 0,1317611 | No |
| orf19.2485 | 3296 | 3290 | 3556 | 2191 | 4213 | 3149 | 3889 | 4360 | 3142 | 3479 | 3513 | 3432 | 3376 | 3581 | 3302 | 3545 | 3421,22 | 3392 | 3451 | 1,017 | 0,025 | 0,504 | 0,6144997 | 0,6482266 | 0,00177 | 0,0025712 | 0,0020676 | 0,0020676 | TRUE | 0,0306647 | No |
| orf19.2487 | 2634 | 2332 | 2555 | 1545 | 3775 | 2815 | 3552 | 3670 | 2511 | 2466 | 2524 | 2420 | 3025 | 3201 | 3016 | 2984 | 2768,36 | 2480 | 3056 | 1,23 | 0,299 | 7,454 | 9,08E-14 | 1,77E-13 | 0,0003119 | 0,0026844 | 0,0011709 | 0,0011709 | TRUE | 0,0117032 | No |
| orf19.2488 | 4821 | 4260 | 4675 | 2668 | 6508 | 4313 | 6379 | 6519 | 4596 | 4504 | 4618 | 4180 | 5215 | 4904 | 5416 | 5301 | 4841,75 | 4474 | 5209 | 1,163 | 0,218 | 4,616 | 3,92E-06 | 5,92E-06 | 0,0016352 | 0,0024304 | 0,0019375 | 0,0019375 | TRUE | 0,0229258 | No |
| orf19.2489 | 17737 | 16166 | 17228 | 11717 | 38061 | 29567 | 34544 | 39492 | 16910 | 17093 | 17020 | 18355 | 30497 | 33620 | 29328 | 32111 | 24366,84 | 17344 | 31389 | 1,81 | 0,856 | 17,079 | 2,14E-65 | 9,69E-65 | 0,0024878 | 0,0021587 | 0,0023664 | 0,0023664 | TRUE | 0,0279605 | No |
| orf19.2492 | 38 | 22 | 24 | 23 | 69 | 58 | 67 | 86 | 36 | 23 | 24 | 36 | 55 | 66 | 57 | 70 | 45,91 | 30 | 62 | 2,099 | 1,07 | 4,635 | 3,58E-06 | 5,42E-06 | 0,0041097 | 0,0378549 | 0,0251086 | 0,0251086 | TRUE | 0,1868873 | No |
| orf19.2494 | 534 | 428 | 521 | 317 | 1037 | 814 | 1073 | 1077 | 509 | 453 | 515 | 497 | 831 | 926 | 911 | 876 | 689,52 | 494 | 886 | 1,793 | 0,842 | 12,342 | 5,39E-35 | 1,58E-34 | 0,0013104 | 0,0044725 | 0,0028374 | 0,0028374 | TRUE | 0,0377318 | No |
| orf19.2495 | 663 | 564 | 657 | 368 | 1102 | 772 | 1038 | 1048 | 632 | 596 | 649 | 576 | 883 | 878 | 881 | 852 | 743,53 | 613 | 874 | 1,419 | 0,504 | 8,393 | 4,72E-17 | 9,94E-17 | 1,00E-08 | 0,0042995 | 0,0020508 | 0,0020508 | TRUE | 0,0172248 | No |
| orf19.2496 | 2143 | 1778 | 1878 | 1371 | 12815 | 9165 | 11885 | 12826 | 2043 | 1880 | 1855 | 2148 | 10268 | 10421 | 10090 | 10429 | 6141,87 | 1982 | 10302 | 5,21 | 2,381 | 48,822 | 0 | 0 | 0,001635 | 0,0023586 | 0,0019565 | 0,0019565 | TRUE | 0,0352317 | No |
| orf19.2498 | 11072 | 9143 | 10296 | 6861 | 5657 | 3884 | 5731 | 5978 | 10556 | 9668 | 10171 | 10748 | 4533 | 4416 | 4866 | 4861 | 7477,33 | 10286 | 4669 | 0,454 | -1,139 | -23,049 | 1,49E-117 | 1,16E-116 | 0,0020727 | 0,0023109 | 0,0021936 | 0,0021936 | TRUE | 0,0199964 | No |
| orf19.2499 | 688 | 569 | 625 | 357 | 1213 | 860 | 1109 | 1192 | 656 | 602 | 617 | 559 | 972 | 978 | 942 | 969 | 786,86 | 608 | 965 | 1,578 | 0,658 | 10,762 | 5,18E-27 | 1,33E-26 | 1,00E-08 | 0,0041779 | 0,0022167 | 0,0022167 | TRUE | 0,0319306 | No |
| orf19.25 | 3180 | 2748 | 2978 | 1896 | 2566 | 1919 | 2600 | 2735 | 3032 | 2906 | 2942 | 2970 | 2056 | 2182 | 2207 | 2224 | 2564,87 | 2962 | 2167 | 0,731 | -0,451 | -10,81 | 3,08E-27 | 7,94E-27 | 0,0004483 | 0,0027314 | 0,0012771 | 0,0012771 | TRUE | 0,0150247 | No |
| orf19.250 | 3790 | 4159 | 4322 | 2654 | 6936 | 5297 | 6453 | 6251 | 3613 | 4398 | 4270 | 4158 | 5558 | 6023 | 5479 | 5083 | 4822,54 | 4110 | 5536 | 1,347 | 0,43 | 6,186 | 6,16E-10 | 1,07E-09 | 0,0056712 | 0,0024317 | 0,0044144 | 0,0044144 | TRUE | 0,0815963 | No |
| orf19.2500 | 524 | 495 | 494 | 324 | 1029 | 674 | 965 | 948 | 500 | 523 | 488 | 508 | 825 | 766 | 819 | 771 | 649,95 | 505 | 795 | 1,579 | 0,659 | 10,367 | 3,49E-25 | 8,67E-25 | 1,00E-08 | 0,0046175 | 0,002209 | 0,002209 | TRUE | 0,0079301 | No |
| orf19.2501 | 9940 | 9948 | 10472 | 6720 | 7486 | 5887 | 6762 | 7255 | 9477 | 10519 | 10345 | 10527 | 5998 | 6694 | 5741 | 5899 | 8150,04 | 10217 | 6083 | 0,595 | -0,748 | -13,196 | 9,28E-40 | 2,92E-39 | 0,0034191 | 0,0022927 | 0,0029614 | 0,0029614 | TRUE | 0,0551195 | No |
| orf19.2503 | 16 | 24 | 14 | 9 | 22 | 13 | 15 | 22 | 15 | 25 | 14 | 14 | 18 | 15 | 13 | 18 | 16,45 | 17 | 16 | 0,918 | -0,124 | -0,349 | 0,7274034 | 0,7566941 | 1,00E-08 | 0,1019075 | 0,0614051 | 0,0614051 | TRUE | 0,511931 | No |
| orf19.2504 | 14899 | 14026 | 14711 | 8875 | 18115 | 12886 | 16814 | 17636 | 14205 | 14831 | 14533 | 13903 | 14515 | 14653 | 14275 | 14340 | 14406,73 | 14368 | 14446 | 1,005 | 0,007 | 0,233 | 0,8159608 | 0,8373506 | 0,0003312 | 0,0022052 | 0,0008624 | 0,0008624 | TRUE | 0,005704 | No |
| orf19.2505 | 21 | 13 | 17 | 11 | 14 | 11 | 19 | 17 | 20 | 14 | 17 | 17 | 11 | 13 | 16 | 14 | 15,18 | 17 | 14 | 0,792 | -0,337 | -0,951 | 0,3414075 | 0,3754776 | 1,00E-08 | 0,1102243 | 0,0545495 | 0,0545495 | TRUE | 0,0900295 | No |
| orf19.2506 | 782 | 710 | 778 | 478 | 2120 | 1431 | 2178 | 2178 | 746 | 751 | 769 | 749 | 1699 | 1627 | 1849 | 1771 | 1244,95 | 754 | 1736 | 2,306 | 1,206 | 21,078 | 1,27E-98 | 8,20E-98 | 0,0012843 | 0,0034101 | 0,002142 | 0,002142 | TRUE | 0,0227849 | No |
| orf19.2507 | 1462 | 1582 | 1610 | 943 | 1865 | 1351 | 1743 | 1673 | 1394 | 1673 | 1591 | 1477 | 1494 | 1536 | 1480 | 1360 | 1500,64 | 1534 | 1468 | 0,956 | -0,065 | -0,971 | 0,331619 | 0,3654423 | 0,0036437 | 0,0031854 | 0,0036102 | 0,0036102 | TRUE | 0,047922 | No |

| | | | | | | | | | | | | | | | | | | | | | | | | | | | | | | | |
|---|---|---|---|---|---|---|---|---|---|---|---|---|---|---|---|---|---|---|---|---|---|---|---|---|---|---|---|---|---|---|---|
| orf19.2508 | 154 | 173 | 159 | 106 | 135 | 91 | 133 | 118 | 147 | 183 | 157 | 166 | 108 | 103 | 113 | 96 | 134,17 | 163 | 105 | 0,646 | -0,631 | -4,884 | 1,04E-06 | 1,60E-06 | 1,00E-08 | 0,0143284 | 0,0083698 | 0,0083698 | TRUE | 0,0756994 | No |
| orf19.2509 | 0 | 0 | 0 | 0 | 0 | 0 | 0 | 0 | 0 | 0 | 0 | 0 | 0 | 0 | 0 | 0 | 0 | 0 | 0 | NA | NA | NA | NA | NA | NA | NA | NA | NA | NA | NA | NA |
| orf19.2509.1 | 88 | 89 | 71 | 55 | 706 | 474 | 588 | 598 | 84 | 94 | 70 | 86 | 566 | 539 | 499 | 486 | 303,05 | 84 | 522 | 6,274 | 2,649 | 22,616 | 3,04E-113 | 2,28E-112 | 0,0033728 | 0,0075092 | 0,0056721 | 0,0056721 | TRUE | 0,121614 | No |
| orf19.251 | 11165 | 9856 | 9932 | 5974 | 4459 | 3060 | 4051 | 4397 | 10645 | 10421 | 9812 | 9359 | 3573 | 3479 | 3439 | 3575 | 6787,92 | 10059 | 3516 | 0,349 | -1,517 | -31,91 | 1,92E-223 | 2,88E-222 | 0,0016221 | 0,0023332 | 0,0019866 | 0,0019866 | TRUE | 0,0262863 | No |
| orf19.2510 | 113 | 172 | 192 | 83 | 520 | 426 | 457 | 377 | 108 | 182 | 190 | 130 | 417 | 484 | 388 | 307 | 275,61 | 152 | 399 | 2,608 | 1,383 | 7,761 | 8,42E-15 | 1,69E-14 | 0,0474194 | 0,0080486 | 0,0256754 | 0,0256754 | TRUE | 0,2338531 | No |
| orf19.2511.1 | 1647 | 1315 | 1294 | 748 | 2143 | 1264 | 1865 | 1951 | 1570 | 1390 | 1278 | 1172 | 1717 | 1437 | 1583 | 1586 | 1466,87 | 1352 | 1581 | 1,167 | 0,223 | 2,574 | 0,0100606 | 0,012778 | 0,0096254 | 0,0032106 | 0,0065195 | 0,0065195 | TRUE | 0,1414822 | No |
| orf19.2511.2 | 911 | 1060 | 1015 | 559 | 1127 | 886 | 1081 | 899 | 869 | 1121 | 1003 | 876 | 903 | 1007 | 918 | 731 | 928,38 | 967 | 890 | 0,918 | -0,123 | -1,162 | 0,2451142 | 0,2750261 | 0,014629 | 0,0038599 | 0,0096517 | 0,0096517 | TRUE | 0,1738528 | No |
| orf19.2512 | 170 | 174 | 196 | 128 | 113 | 77 | 103 | 120 | 162 | 184 | 194 | 201 | 91 | 88 | 87 | 98 | 137,92 | 185 | 91 | 0,494 | -1,018 | -7,938 | 2,05E-15 | 4,17E-15 | 1,00E-08 | 0,0139964 | 0,0078768 | 0,0078768 | TRUE | 0,0794357 | No |
| orf19.2513 | 431 | 611 | 587 | 365 | 342 | 291 | 329 | 249 | 411 | 646 | 580 | 572 | 274 | 331 | 279 | 202 | 411,92 | 552 | 272 | 0,491 | -1,026 | -6,691 | 2,21E-11 | 4,03E-11 | 0,0352901 | 0,0060772 | 0,019905 | 0,019905 | TRUE | 0,3557282 | No |
| orf19.2514 | 1130 | 1128 | 1199 | 666 | 1044 | 806 | 946 | 870 | 1077 | 1193 | 1184 | 1043 | 837 | 916 | 803 | 707 | 970,18 | 1124 | 816 | 0,723 | -0,467 | -5,72 | 1,07E-08 | 1,77E-08 | 0,0064683 | 0,0037837 | 0,0053549 | 0,0053549 | TRUE | 0,0961487 | No |
| orf19.2515 | 11265 | 11190 | 11752 | 6926 | 5246 | 3936 | 5078 | 5407 | 10740 | 11832 | 11610 | 10850 | 4203 | 4476 | 4311 | 4396 | 7802,31 | 11258 | 4346 | 0,386 | -1,374 | -31,065 | 7,08E-212 | 1,02E-210 | 0,0013158 | 0,0023017 | 0,0017249 | 0,0017249 | TRUE | 0,0143944 | No |
| orf19.2516 | 497 | 538 | 540 | 316 | 935 | 628 | 889 | 757 | 474 | 569 | 533 | 495 | 749 | 714 | 755 | 616 | 613,09 | 518 | 708 | 1,367 | 0,451 | 5,17 | 2,35E-07 | 3,73E-07 | 0,0063049 | 0,0047693 | 0,0055815 | 0,0055815 | TRUE | 0,0952835 | No |
| orf19.2517 | 281 | 264 | 256 | 174 | 303 | 201 | 263 | 302 | 268 | 279 | 253 | 273 | 243 | 229 | 223 | 246 | 251,59 | 268 | 235 | 0,879 | -0,186 | -2,028 | 0,0425945 | 0,0515057 | 1,00E-08 | 0,0086174 | 0,0041146 | 0,0041146 | TRUE | 0,0173774 | No |
| orf19.2518 | 3974 | 3679 | 3902 | 2350 | 3485 | 2494 | 3428 | 3589 | 3789 | 3890 | 3855 | 3681 | 2792 | 2836 | 2910 | 2918 | 3333,99 | 3804 | 2864 | 0,753 | -0,41 | -11,056 | 2,05E-28 | 5,39E-28 | 0,0001819 | 0,0025838 | 0,0010182 | 0,0010182 | TRUE | 0,0052035 | No |
| orf19.2519 | 522 | 446 | 466 | 328 | 195 | 106 | 154 | 172 | 498 | 472 | 460 | 514 | 156 | 121 | 131 | 140 | 311,35 | 486 | 137 | 0,284 | -1,816 | -18,548 | 8,51E-77 | 4,45E-76 | 0,0008898 | 0,0073647 | 0,0048254 | 0,0048254 | TRUE | 0,0971651 | No |
| orf19.252 | 445 | 515 | 518 | 236 | 762 | 499 | 723 | 643 | 424 | 545 | 512 | 370 | 611 | 567 | 614 | 523 | 520,61 | 463 | 579 | 1,246 | 0,317 | 2,852 | 0,0043462 | 0,0056368 | 0,0151222 | 0,0052451 | 0,0099057 | 0,0099057 | TRUE | 0,189223 | No |
| orf19.2520 | 2829 | 2561 | 2757 | 1755 | 2644 | 1904 | 2665 | 2780 | 2697 | 2708 | 2724 | 2749 | 2119 | 2165 | 2263 | 2260 | 2460,57 | 2720 | 2202 | 0,81 | -0,304 | -7,581 | 3,42E-14 | 6,74E-14 | 0,0001941 | 0,0027586 | 0,0011332 | 0,0011332 | TRUE | 0,0083293 | No |
| orf19.2521 | 430 | 392 | 445 | 257 | 1560 | 1078 | 1521 | 1673 | 410 | 414 | 440 | 403 | 1250 | 1226 | 1291 | 1360 | 849,26 | 417 | 1282 | 3,073 | 1,62 | 25,078 | 8,70E-139 | 8,12E-138 | 1,00E-08 | 0,0040246 | 0,002323 | 0,002323 | TRUE | 0,0206247 | No |
| orf19.2524 | 7296 | 5970 | 6383 | 4267 | 4189 | 3031 | 4023 | 4456 | 6956 | 6313 | 6306 | 6684 | 3357 | 3447 | 3416 | 3623 | 5012,56 | 6565 | 3461 | 0,527 | -0,924 | -19,922 | 2,60E-88 | 1,53E-87 | 0,0014081 | 0,0024188 | 0,0018488 | 0,0018488 | TRUE | 0,0202007 | No |
| orf19.2525 | 2070 | 1662 | 1791 | 1222 | 5207 | 3479 | 4929 | 5316 | 1974 | 1757 | 1769 | 1914 | 4172 | 3956 | 4185 | 4322 | 3006,23 | 1854 | 4159 | 2,246 | 1,167 | 22,614 | 3,12E-113 | 2,33E-112 | 0,0017197 | 0,0026374 | 0,0021469 | 0,0021469 | TRUE | 0,0248077 | No |
| orf19.2527 | 4143 | 3418 | 3829 | 2335 | 4419 | 2858 | 4250 | 4567 | 3950 | 3614 | 3783 | 3658 | 3541 | 3250 | 3608 | 3713 | 3639,61 | 3751 | 3528 | 0,941 | -0,088 | -1,708 | 0,0875693 | 0,1027735 | 0,0020234 | 0,0025424 | 0,0022807 | 0,0022807 | TRUE | 0,0333422 | No |
| orf19.2528 | 3618 | 2957 | 3330 | 2146 | 3396 | 2263 | 3166 | 3599 | 3449 | 3127 | 3290 | 3362 | 2721 | 2573 | 2688 | 2926 | 3017,02 | 3307 | 2727 | 0,825 | -0,277 | -5,363 | 8,17E-08 | 1,32E-07 | 0,0018642 | 0,0026355 | 0,0022265 | 0,0022265 | TRUE | 0,0294419 | No |
| orf19.2529 | 1302 | 1190 | 1246 | 786 | 1285 | 945 | 1248 | 1378 | 1241 | 1258 | 1231 | 1231 | 1030 | 1075 | 1060 | 1120 | 1155,75 | 1240 | 1071 | 0,863 | -0,212 | -4,275 | 1,91E-05 | 2,80E-05 | 1,00E-08 | 0,0035119 | 0,0015015 | 0,0015015 | TRUE | 0,0123101 | No |
| orf19.2529.1 | 17392 | 12949 | 13002 | 7510 | 5219 | 2710 | 4750 | 5116 | 16581 | 13692 | 12845 | 11765 | 4182 | 3082 | 4033 | 4160 | 8792,34 | 13721 | 3864 | 0,282 | -1,828 | -12,514 | 6,29E-36 | 1,86E-35 | 0,0203478 | 0,002278 | 0,0112774 | 0,0203478 | TRUE | 0,2411228 | No |
| orf19.2530 | 8 | 9 | 8 | 4 | 8 | 6 | 13 | 9 | 8 | 10 | 8 | 6 | 6 | 7 | 11 | 7 | 7,86 | 8 | 8 | 1,001 | 0,002 | 0,004 | 0,9966703 | 0,9972954 | 1,00E-08 | 0,2109169 | 0,1040179 | 0,1040179 | TRUE | 0,2429537 | No |
| orf19.2531 | 11271 | 10210 | 10723 | 7177 | 10779 | 8471 | 10043 | 10651 | 10746 | 10796 | 10593 | 11243 | 8637 | 9632 | 8526 | 8660 | 9854,23 | 10844 | 8864 | 0,817 | -0,291 | -6,256 | 3,95E-10 | 6,92E-10 | 0,0018864 | 0,0022579 | 0,0019776 | 0,0019776 | TRUE | 0,0410996 | No |
| orf19.2532 | 1132 | 994 | 1138 | 679 | 1265 | 899 | 1219 | 1187 | 1079 | 1051 | 1124 | 1064 | 1014 | 1022 | 1035 | 965 | 1044,26 | 1080 | 1009 | 0,933 | -0,099 | -1,902 | 0,0571183 | 0,0682531 | 1,00E-08 | 0,0036636 | 0,0016622 | 0,0016622 | TRUE | 0,0106131 | No |
| orf19.2533.1 | 1152 | 997 | 1061 | 706 | 2963 | 1887 | 2581 | 2824 | 1098 | 1054 | 1048 | 1106 | 2374 | 2146 | 2191 | 2296 | 1664,24 | 1076 | 2252 | 2,095 | 1,067 | 20,697 | 3,66E-95 | 2,28E-94 | 0,0010328 | 0,0030779 | 0,0018334 | 0,0018334 | TRUE | 0,0162545 | No |
| orf19.2534 | 3945 | 3523 | 3920 | 2587 | 3152 | 2411 | 2989 | 3055 | 3761 | 3725 | 3873 | 4053 | 2526 | 2742 | 2538 | 2484 | 3212,54 | 3853 | 2572 | 0,668 | -0,583 | -12,474 | 1,04E-35 | 3,07E-35 | 0,001301 | 0,0026024 | 0,0017777 | 0,0017777 | TRUE | 0,02286 | No |
| orf19.2537 | 22 | 26 | 30 | 19 | 27 | 18 | 21 | 24 | 21 | 27 | 30 | 30 | 22 | 20 | 18 | 20 | 23,41 | 27 | 20 | 0,742 | -0,431 | -1,501 | 0,1332365 | 0,1536458 | 1,00E-08 | 0,0722157 | 0,0359969 | 0,0359969 | TRUE | 0,1578321 | No |
| orf19.2538 | 22341 | 21422 | 23143 | 13111 | 36821 | 24846 | 34056 | 34511 | 21300 | 22651 | 22863 | 20539 | 29504 | 28252 | 28913 | 28061 | 25260,35 | 21838 | 28682 | 1,313 | 0,393 | 9,203 | 3,48E-20 | 7,84E-20 | 0,0014986 | 0,0021563 | 0,0017111 | 0,0017111 | TRUE | 0,0193917 | No |
| orf19.2539 | 1 | 1 | 2 | 3 | 2 | 3 | 3 | 1 | 1 | 1 | 2 | 5 | 2 | 3 | 3 | 1 | 2,13 | 2 | 2 | 1,009 | 0,013 | 0,013 | 0,9895711 | 0,9907741 | 1,00E-08 | 0,7720284 | 0,4416186 | 0,4416186 | TRUE | 0,3486065 | No |
| orf19.254 | 7588 | 6946 | 7546 | 5258 | 7437 | 5734 | 6984 | 7865 | 7234 | 7345 | 7455 | 8237 | 5959 | 6520 | 5929 | 6395 | 6884,25 | 7568 | 6201 | 0,82 | -0,287 | -5,382 | 7,37E-08 | 1,19E-07 | 0,0027913 | 0,0023298 | 0,0025815 | 0,0025815 | TRUE | 0,0424123 | No |
| orf19.2540 | 1289 | 1168 | 1259 | 675 | 1739 | 1178 | 1688 | 1718 | 1229 | 1235 | 1244 | 1057 | 1393 | 1339 | 1433 | 1397 | 1291 | 1191 | 1390 | 1,164 | 0,219 | 3,706 | 0,0002102 | 0,000293 | 0,0018167 | 0,0033631 | 0,0025632 | 0,0025632 | TRUE | 0,0610368 | No |
| orf19.2541 | 726 | 769 | 790 | 431 | 952 | 671 | 816 | 820 | 692 | 813 | 780 | 675 | 763 | 763 | 693 | 667 | 730,78 | 740 | 722 | 0,972 | -0,042 | -0,527 | 0,5983976 | 0,6326003 | 0,0049066 | 0,0043381 | 0,0046366 | 0,0046366 | TRUE | 0,0509407 | No |
| orf19.2542 | 526 | 535 | 538 | 339 | 498 | 324 | 483 | 408 | 501 | 566 | 531 | 531 | 399 | 368 | 410 | 332 | 454,87 | 532 | 377 | 0,709 | -0,496 | -5,852 | 4,85E-09 | 8,15E-09 | 0,0036152 | 0,0057009 | 0,0046525 | 0,0046525 | TRUE | 0,0814998 | No |
| orf19.2544 | 396 | 358 | 351 | 234 | 682 | 428 | 642 | 677 | 378 | 379 | 347 | 367 | 546 | 487 | 545 | 550 | 449,76 | 368 | 532 | 1,452 | 0,538 | 7,1 | 1,25E-12 | 2,36E-12 | 1,00E-08 | 0,0057419 | 0,0031661 | 0,0031661 | TRUE | 0,0350532 | No |
| orf19.2545 | 5566 | 4620 | 5235 | 3245 | 7126 | 4836 | 6932 | 7171 | 5307 | 4885 | 5172 | 5083 | 5710 | 5499 | 5885 | 5831 | 5421,45 | 5112 | 5731 | 1,121 | 0,165 | 4,101 | 4,11E-05 | 5,93E-05 | 0,0007978 | 0,0023941 | 0,0013698 | 0,0013698 | TRUE | 0,0111491 | No |
| orf19.2546 | 2366 | 2240 | 2432 | 1491 | 2799 | 1968 | 2618 | 2735 | 2256 | 2369 | 2403 | 2336 | 2243 | 2238 | 2223 | 2224 | 2286,2 | 2341 | 2232 | 0,954 | -0,069 | -1,757 | 0,0789658 | 0,0931904 | 1,00E-08 | 0,0028095 | 0,0010278 | 0,0010278 | TRUE | 0,0078409 | No |
| orf19.2547 | 3365 | 3428 | 3570 | 2131 | 2253 | 1673 | 2164 | 2206 | 3208 | 3625 | 3527 | 3338 | 1805 | 1902 | 1837 | 1794 | 2629,56 | 3424 | 1834 | 0,535 | -0,902 | -18,019 | 1,37E-72 | 6,83E-72 | 0,0015655 | 0,0027157 | 0,0019962 | 0,0019962 | TRUE | 0,0230417 | No |
| orf19.2549 | 3335 | 2969 | 3134 | 1878 | 4267 | 2982 | 4327 | 4511 | 3180 | 3139 | 3096 | 2942 | 3419 | 3391 | 3674 | 3668 | 3313,54 | 3089 | 3538 | 1,144 | 0,195 | 4,255 | 2,09E-05 | 3,05E-05 | 0,001108 | 0,0025868 | 0,0017027 | 0,0017027 | TRUE | 0,0116577 | No |
| orf19.255 | 11443 | 11114 | 11718 | 7165 | 12163 | 8778 | 11065 | 11548 | 10910 | 11752 | 11576 | 11224 | 9746 | 9981 | 9394 | 9390 | 10496,62 | 11366 | 9628 | 0,847 | -0,24 | -6,253 | 4,02E-10 | 7,04E-10 | 0,0008656 | 0,0022477 | 0,0013158 | 0,0013158 | TRUE | 0,0091125 | No |
| orf19.2551 | 7293 | 6391 | 6606 | 4724 | 64331 | 46073 | 60007 | 68724 | 6953 | 6758 | 6526 | 7400 | 51547 | 52389 | 50946 | 55879 | 29799,73 | 6909 | 52690 | 7,631 | 2,932 | 60,876 | 0 | 0 | 0,0022097 | 0,0021464 | 0,0021378 | 0,0021378 | TRUE | 0,0271955 | No |
| orf19.2552 | 943 | 906 | 930 | 601 | 112 | 84 | 114 | 114 | 899 | 958 | 919 | 941 | 90 | 96 | 97 | 93 | 511,5 | 929 | 94 | 0,101 | -3,312 | -36,428 | 1,51E-290 | 3,25E-289 | 1,00E-08 | 0,0053012 | 0,0026019 | 0,0026019 | TRUE | 0,0085621 | No |
| orf19.2553 | 329 | 348 | 357 | 200 | 41 | 30 | 26 | 35 | 314 | 368 | 353 | 313 | 33 | 34 | 22 | 28 | 183,14 | 337 | 29 | 0,086 | -3,533 | -21,905 | 2,32E-106 | 1,60E-105 | 1,00E-08 | 0,0110566 | 0,0081474 | 0,0081474 | TRUE | 0,2002543 | No |
| orf19.2555 | 1849 | 1680 | 1703 | 1148 | 3490 | 2747 | 3351 | 3701 | 1763 | 1776 | 1682 | 1798 | 2796 | 3124 | 2845 | 3009 | 2349,29 | 1755 | 2944 | 1,677 | 0,746 | 14,916 | 2,59E-50 | 9,57E-50 | 0,0014064 | 0,0027902 | 0,0019305 | 0,0019305 | TRUE | 0,0198808 | No |
| orf19.2557 | 2302 | 1944 | 2108 | 1303 | 2536 | 1724 | 2549 | 2637 | 2195 | 2056 | 2083 | 2041 | 2032 | 1960 | 2164 | 2144 | 2084,32 | 2094 | 2075 | 0,991 | -0,013 | -0,259 | 0,795314 | 0,8177407 | 0,0011477 | 0,002879 | 0,0018263 | 0,0018263 | TRUE | 0,0158864 | No |
| orf19.2558 | 881 | 821 | 880 | 560 | 819 | 618 | 728 | 733 | 840 | 868 | 869 | 877 | 656 | 703 | 618 | 596 | 753,46 | 864 | 643 | 0,743 | -0,428 | -6,59 | 4,40E-11 | 7,94E-11 | 0,0011933 | 0,0042704 | 0,0026998 | 0,0026998 | TRUE | 0,0417288 | No |
| orf19.2559 | 4400 | 3795 | 4130 | 2542 | 3976 | 2882 | 3842 | 4108 | 4195 | 4013 | 4080 | 3982 | 3186 | 3277 | 3262 | 3340 | 3666,86 | 4068 | 3266 | 0,802 | -0,318 | -8,795 | 1,43E-18 | 3,10E-18 | 0,0002119 | 0,002539 | 0,0009768 | 0,0009768 | TRUE | 0,0055039 | No |
| orf19.2560 | 7977 | 6872 | 7414 | 4871 | 17312 | 12204 | 15974 | 18235 | 7605 | 7266 | 7324 | 7631 | 13872 | 13877 | 13562 | 14827 | 10745,47 | 7456 | 14034 | 1,883 | 0,913 | 23,131 | 2,23E-118 | 1,75E-117 | 0,0009448 | 0,0022441 | 0,001388 | 0,001388 | TRUE | 0,0177753 | No |
| orf19.2563 | 1618 | 1406 | 1619 | 1016 | 1707 | 1248 | 1679 | 1768 | 1543 | 1487 | 1599 | 1592 | 1368 | 1419 | 1425 | 1438 | 1483,77 | 1555 | 1412 | 0,909 | -0,138 | -2,951 | 0,0031649 | 0,0041384 | 1,00E-08 | 0,0031979 | 0,0014228 | 0,0014228 | TRUE | 0,0105967 | No |
| orf19.2564 | 2040 | 1739 | 1959 | 1200 | 2289 | 1375 | 2150 | 2156 | 1945 | 1839 | 1935 | 1880 | 1834 | 1563 | 1825 | 1753 | 1821,86 | 1900 | 1744 | 0,919 | -0,122 | -2,11 | 0,0348388 | 0,0424492 | 0,0022067 | 0,0029925 | 0,0026495 | 0,0026495 | TRUE | 0,0559425 | No |
| orf19.2565 | 2550 | 2343 | 2507 | 1446 | 1266 | 854 | 1191 | 1261 | 2431 | 2477 | 2477 | 2265 | 1014 | 971 | 1011 | 1025 | 1709,06 | 2412 | 1005 | 0,417 | -1,264 | -26,092 | 4,52E-150 | 4,55E-149 | 0,0005139 | 0,003052 | 0,0015762 | 0,0015762 | TRUE | 0,018655 | No |
| orf19.2568 | 6834 | 5240 | 6051 | 4055 | 4582 | 2833 | 4384 | 4589 | 6515 | 5541 | 5978 | 6352 | 3671 | 3221 | 3722 | 3731 | 4841,55 | 6096 | 3586 | 0,589 | -0,764 | -11,838 | 2,49E-32 | 7,01E-32 | 0,0045645 | 0,0024304 | 0,0037881 | 0,0037881 | TRUE | 0,056143 | No |
| orf19.257 | 4 | 3 | 4 | 1 | 2 | 1 | 0 | 0 | 4 | 3 | 4 | 2 | 2 | 1 | 0 | 0 | 1,91 | 3 | 1 | 0,208 | -2,264 | -1,914 | 0,0556105 | 0,0665137 | 1,00E-08 | 0,8637853 | 0,5050052 | 0,5050052 | TRUE | 0,398854 | No |
| orf19.2570 | 1982 | 1728 | 1912 | 1149 | 4423 | 2897 | 4131 | 4479 | 1890 | 1827 | 1889 | 1800 | 3544 | 3294 | 3507 | 3642 | 2674,1 | 1852 | 3497 | 1,888 | 0,917 | 20,187 | 1,26E-90 | 7,60E-90 | 0,0008816 | 0,0027053 | 0,0015503 | 0,0015503 | TRUE | 0,0178196 | No |
| orf19.2571 | 4179 | 3852 | 4319 | 2484 | 5384 | 3847 | 5084 | 5249 | 3984 | 4073 | 4267 | 3891 | 4314 | 4374 | 4316 | 4268 | 4186 | 4054 | 4318 | 1,064 | 0,09 | 2,322 | 0,0202242 | 0,0250876 | 0,0005127 | 0,0024835 | 0,0011874 | 0,0011874 | TRUE | 0,0153119 | No |
| orf19.2573 | 6129 | 5396 | 5837 | 3713 | 12163 | 9031 | 11872 | 13082 | 5843 | 5706 | 5766 | 5817 | 9746 | 10269 | 10079 | 10637 | 7982,88 | 5783 | 10183 | 1,761 | 0,816 | 22,205 | 3,06E-109 | 2,20E-108 | 0,0005418 | 0,002297 | 0,0011563 | 0,0011563 | TRUE | 0,0111831 | No |
| orf19.2574 | 3906 | 3302 | 3693 | 2232 | 3902 | 2702 | 3836 | 3974 | 3724 | 3491 | 3648 | 3497 | 3127 | 3072 | 3257 | 3231 | 3380,91 | 3590 | 3172 | 0,883 | -0,179 | -4,419 | 9,91E-06 | 1,47E-05 | 0,0004943 | 0,0025769 | 0,0012818 | 0,0012818 | TRUE | 0,0077482 | No |
| orf19.2575 | 4112 | 3627 | 3907 | 2502 | 5191 | 3594 | 4848 | 5334 | 3920 | 3835 | 3860 | 3920 | 4159 | 4087 | 4116 | 4337 | 4029,22 | 3884 | 4175 | 1,076 | 0,105 | 2,977 | 0,0029148 | 0,0038207 | 0,0002307 | 0,0024988 | 0,000945 | 0,000945 | TRUE | 0,0084323 | No |
| orf19.2579 | 1081 | 948 | 1129 | 679 | 1374 | 1049 | 1370 | 1486 | 1031 | 1002 | 1115 | 1064 | 1101 | 1193 | 1163 | 1208 | 1109,65 | 1053 | 1166 | 1,107 | 0,147 | 2,589 | 0,0096366 | 0,0122614 | 0,0011109 | 0,0035709 | 0,0021757 | 0,0021757 | TRUE | 0,0205204 | No |
| orf19.258 | 428 | 397 | 410 | 283 | 662 | 502 | 591 | 545 | 408 | 420 | 405 | 443 | 530 | 571 | 502 | 443 | 465,29 | 419 | 512 | 1,221 | 0,288 | 3,281 | 0,0010337 | 0,0013934 | 0,0047424 | 0,00562 | 0,0051647 | 0,0051647 | TRUE | 0,096208 | No |
| orf19.2580 | 827 | 744 | 843 | 449 | 905 | 618 | 904 | 909 | 788 | 787 | 833 | 703 | 725 | 703 | 767 | 739 | 755,72 | 778 | 734 | 0,94 | -0,089 | -1,358 | 0,1743568 | 0,1987799 | 0,0013287 | 0,0042639 | 0,0027613 | 0,0027613 | TRUE | 0,0424291 | No |
| orf19.2581 | 95 | 80 | 100 | 62 | 150 | 132 | 161 | 205 | 91 | 85 | 99 | 97 | 120 | 150 | 137 | 167 | 118,09 | 93 | 144 | 1,547 | 0,629 | 4,26 | 2,04E-05 | 2,98E-05 | 0,0072231 | 0,0159948 | 0,0118241 | 0,0118241 | TRUE | 0,1377642 | No |
| orf19.2582 | 1939 | 1850 | 1916 | 1106 | 3708 | 2629 | 3455 | 3792 | 1849 | 1956 | 1893 | 1733 | 2971 | 2989 | 2933 | 3083 | 2425,91 | 1858 | 2994 | 1,609 | 0,686 | 14,928 | 2,17E-50 | 8,06E-50 | 0,0007078 | 0,0027681 | 0,0015768 | 0,0015768 | TRUE | 0,0219974 | No |
| orf19.2583 | 9 | 7 | 13 | 4 | 27 | 17 | 14 | 19 | 9 | 7 | 13 | 6 | 22 | 19 | 12 | 15 | 12,92 | 9 | 17 | 1,901 | 0,927 | 2,218 | 0,0265718 | 0,0326938 | 1,00E-08 | 0,1291358 | 0,0795209 | 0,0795209 | TRUE | 0,3057667 | No |
| orf19.2585 | 9 | 12 | 17 | 19 | 14 | 19 | 12 | 14 | 9 | 13 | 17 | 30 | 11 | 22 | 10 | 11 | 15,28 | 17 | 14 | 0,815 | -0,296 | -0,675 | 0,4995235 | 0,5353476 | 0,1258312 | 0,1095618 | 0,1166985 | 0,1166985 | TRUE | 0,5980899 | No |
| orf19.2587 | 6 | 7 | 5 | 6 | 46 | 22 | 36 | 42 | 6 | 7 | 5 | 9 | 37 | 25 | 31 | 34 | 19,26 | 7 | 32 | 4,8 | 2,263 | 5,792 | 6,95E-09 | 1,16E-08 | 1,00E-08 | 0,0873578 | 0,0493866 | 0,0493866 | TRUE | 0,1162672 | No |
| orf19.259 | 1562 | 1439 | 1487 | 905 | 1697 | 1068 | 1692 | 1749 | 1489 | 1522 | 1469 | 1418 | 1360 | 1214 | 1436 | 1422 | 1416,28 | 1474 | 1358 | 0,922 | -0,117 | -1,938 | 0,0525783 | 0,0630644 | 0,0022838 | 0,0032506 | 0,0028095 | 0,0028095 | TRUE | 0,0577617 | No |
| orf19.2590 | 13 | 4 | 4 | 10 | 65 | 55 | 59 | 76 | 12 | 4 | 4 | 16 | 52 | 63 | 50 | 62 | 32,84 | 9 | 57 | 6,561 | 2,714 | 7,819 | 5,33E-15 | 1,07E-14 | 0,0040683 | 0,0520828 | 0,0436021 | 0,0436021 | TRUE | 0,4850802 | No |
| orf19.2591 | 51 | 33 | 47 | 29 | 92 | 77 | 96 | 88 | 49 | 35 | 46 | 45 | 74 | 88 | 82 | 72 | 61,21 | 44 | 79 | 1,784 | 0,835 | 4,405 | 1,06E-05 | 1,57E-05 | 1,00E-08 | 0,0289139 | 0,0161702 | 0,0161702 | TRUE | 0,1524291 | No |
| orf19.2593 | 94 | 84 | 93 | 69 | 241 | 161 | 233 | 271 | 90 | 89 | 92 | 108 | 193 | 183 | 198 | 220 | 146,59 | 95 | 198 | 2,124 | 1,087 | 8,463 | 2,61E-17 | 5,53E-17 | 1,00E-08 | 0,0132917 | 0,0076729 | 0,0076729 | TRUE | 0,0649566 | No |
| orf19.2594 | 4970 | 4331 | 4753 | 2970 | 7580 | 5388 | 7384 | 7743 | 4738 | 4579 | 4696 | 4653 | 6074 | 6127 | 6269 | 6296 | 5428,88 | 4666 | 6192 | 1,327 | 0,408 | 12,84 | 9,80E-38 | 2,99E-37 | 0,0001226 | 0,0023937 | 0,0007759 | 0,0007759 | TRUE | 0,0020805 | No |

| | | | | | | | | | | | | | | | | | | | | | | | | | | | | | | |
|---|---|---|---|---|---|---|---|---|---|---|---|---|---|---|---|---|---|---|---|---|---|---|---|---|---|---|---|---|---|---|
| orf19.2597 | 708 | 627 | 721 | 431 | 1207 | 846 | 1165 | 1245 | 675 | 663 | 712 | 675 | 967 | 962 | 989 | 1012 | 831,99 | 681 | 982 | 1,442 | 0,528 | 9,431 | 4,07E-21 | 9,32E-21 | 1,00E-08 | 0,0040647 | 0,0017389 | 0,0017389 | TRUE | 0,0117966 | No |
| orf19.2598 | 2272 | 2080 | 2118 | 1285 | 4517 | 2912 | 4347 | 4593 | 2166 | 2199 | 2092 | 2013 | 3619 | 3311 | 3691 | 3735 | 2853,32 | 2118 | 3589 | 1,694 | 0,76 | 14,758 | 2,72E-49 | 9,94E-49 | 0,0016833 | 0,0026667 | 0,0021594 | 0,0021594 | TRUE | 0,0320656 | No |
| orf19.2599 | 2339 | 1889 | 2144 | 1456 | 13318 | 9603 | 12470 | 14128 | 2230 | 1997 | 2118 | 2281 | 10671 | 10919 | 10587 | 11487 | 6536,44 | 2156 | 10916 | 5,068 | 2,341 | 46,922 | 0 | 0 | 0,00203 | 0,0023425 | 0,0020888 | 0,0020888 | TRUE | 0,029721 | No |
| orf19.26 | 2122 | 1983 | 2055 | 1365 | 1327 | 920 | 1273 | 1272 | 2023 | 2097 | 2030 | 2138 | 1063 | 1046 | 1081 | 1034 | 1564,1 | 2072 | 1056 | 0,51 | -0,97 | -21,468 | 3,09E-102 | 2,06E-101 | 1,00E-08 | 0,003141 | 0,0012681 | 0,0012681 | TRUE | 0,0050445 | No |
| orf19.260 | 6578 | 6064 | 6574 | 3882 | 3067 | 2188 | 2726 | 3091 | 6271 | 6412 | 6494 | 6081 | 2457 | 2488 | 2314 | 2513 | 4379,03 | 6314 | 2443 | 0,387 | -1,371 | -33,112 | 2,00E-240 | 3,27E-239 | 0,0006937 | 0,0024662 | 0,001374 | 0,001374 | TRUE | 0,0154972 | No |
| orf19.2600 | 1203 | 1164 | 1234 | 807 | 940 | 687 | 866 | 875 | 1147 | 1231 | 1219 | 1264 | 753 | 781 | 735 | 711 | 980,26 | 1215 | 745 | 0,613 | -0,705 | -12,435 | 1,69E-35 | 4,96E-35 | 0,0005567 | 0,0037663 | 0,0020255 | 0,0020255 | TRUE | 0,0176449 | No |
| orf19.2601 | 10697 | 9158 | 9865 | 6861 | 7875 | 6003 | 7316 | 8279 | 10198 | 9683 | 9746 | 10748 | 6310 | 6826 | 6211 | 6732 | 8306,81 | 10094 | 6520 | 0,646 | -0,63 | -12,873 | 6,42E-38 | 1,97E-37 | 0,0021129 | 0,0022889 | 0,0021792 | 0,0021792 | TRUE | 0,0228222 | No |
| orf19.2602 | 994 | 801 | 915 | 609 | 2471 | 1741 | 2362 | 2563 | 948 | 847 | 904 | 954 | 1980 | 1980 | 2005 | 2084 | 1462,69 | 913 | 2012 | 2,208 | 1,142 | 21,831 | 1,17E-105 | 8,04E-105 | 0,0005503 | 0,0032138 | 0,0017941 | 0,0017941 | TRUE | 0,028378 | No |
| orf19.2604 | 6324 | 5295 | 5911 | 3611 | 4999 | 3413 | 4808 | 5165 | 6029 | 5599 | 5840 | 5657 | 4006 | 3881 | 4082 | 4200 | 4911,55 | 5781 | 4042 | 0,699 | -0,516 | -12,553 | 3,83E-36 | 1,14E-35 | 0,0008347 | 0,0024256 | 0,001416 | 0,001416 | TRUE | 0,0103195 | No |
| orf19.2605 | 2641 | 2409 | 2564 | 1618 | 2996 | 2086 | 2880 | 3123 | 2518 | 2547 | 2533 | 2535 | 2401 | 2372 | 2445 | 2539 | 2486,22 | 2533 | 2439 | 0,964 | -0,054 | -1,37 | 0,1707695 | 0,1950036 | 1,00E-08 | 0,0027517 | 0,0010648 | 0,0010648 | TRUE | 0,0093581 | No |
| orf19.2606 | 2151 | 1989 | 2107 | 1415 | 2397 | 1638 | 2255 | 2315 | 2051 | 2103 | 2082 | 2217 | 1921 | 1863 | 1914 | 1882 | 2004,01 | 2113 | 1895 | 0,899 | -0,154 | -3,655 | 0,0002575 | 0,0003579 | 2,37E-05 | 0,0029106 | 0,0012001 | 0,0012001 | TRUE | 0,0118007 | No |
| orf19.2607 | 939 | 773 | 895 | 633 | 1379 | 985 | 1383 | 1388 | 895 | 817 | 884 | 992 | 1105 | 1120 | 1174 | 1129 | 1014,51 | 897 | 1132 | 1,266 | 0,341 | 5,453 | 4,94E-08 | 8,04E-08 | 0,001767 | 0,0037097 | 0,0027248 | 0,0027248 | TRUE | 0,0525996 | No |
| orf19.2608 | 150148 | 120586 | 134822 | 82223 | 28649 | 18183 | 26996 | 28427 | 143150 | 127504 | 133191 | 128807 | 22956 | 20676 | 22919 | 23114 | 77789,63 | 133163 | 22416 | 0,168 | -2,57 | -49,981 | 0 | 0 | 0,0027288 | 0,0021124 | 0,0025171 | 0,0025171 | TRUE | 0,0333044 | No |
| orf19.2609 | 9664 | 8023 | 9319 | 5670 | 5910 | 3797 | 5856 | 6272 | 9214 | 8483 | 9206 | 8882 | 4736 | 4318 | 4972 | 5100 | 6863,75 | 8946 | 4782 | 0,535 | -0,903 | -16,149 | 1,15E-58 | 4,76E-58 | 0,0031176 | 0,0023305 | 0,0028491 | 0,0028491 | TRUE | 0,0510203 | No |
| orf19.261 | 1738 | 1831 | 1810 | 1080 | 2924 | 2173 | 2548 | 2710 | 1657 | 1936 | 1788 | 1692 | 2343 | 2471 | 2163 | 2203 | 2031,7 | 1768 | 2295 | 1,296 | 0,375 | 5,879 | 4,14E-09 | 6,95E-09 | 0,0036909 | 0,0028994 | 0,0033852 | 0,0033852 | TRUE | 0,0495686 | No |
| orf19.2610 | 1320 | 1044 | 1093 | 766 | 1324 | 919 | 1294 | 1367 | 1258 | 1104 | 1080 | 1200 | 1061 | 1045 | 1099 | 1112 | 1119,76 | 1160 | 1079 | 0,931 | -0,103 | -1,673 | 0,0942985 | 0,1104071 | 0,0021408 | 0,0035576 | 0,0027613 | 0,0027613 | TRUE | 0,0422851 | No |
| orf19.2611 | 2317 | 1938 | 2134 | 1281 | 4778 | 3187 | 4554 | 4967 | 2209 | 2049 | 2108 | 2007 | 3828 | 3624 | 3866 | 4039 | 2966,31 | 2093 | 3839 | 1,833 | 0,874 | 17,82 | 4,98E-71 | 2,42E-70 | 0,0013679 | 0,0026448 | 0,0019253 | 0,0019253 | TRUE | 0,0169484 | No |
| orf19.2612 | 2433 | 1736 | 2067 | 1535 | 1342 | 986 | 1311 | 1376 | 2320 | 1836 | 2042 | 2405 | 1075 | 1121 | 1113 | 1119 | 1628,77 | 2151 | 1107 | 0,515 | -0,957 | -12,036 | 2,29E-33 | 6,53E-33 | 0,0066324 | 0,0030993 | 0,0053996 | 0,0053996 | TRUE | 0,1174227 | No |
| orf19.2613 | 16506 | 13696 | 15090 | 9402 | 11111 | 7397 | 10922 | 11337 | 15737 | 14482 | 14907 | 14729 | 8903 | 8411 | 9273 | 9218 | 11957,43 | 14964 | 8951 | 0,598 | -0,741 | -16,849 | 1,07E-63 | 4,72E-63 | 0,0015169 | 0,0022286 | 0,0017708 | 0,0017708 | TRUE | 0,0198632 | No |
| orf19.2614 | 2263 | 1910 | 2120 | 1425 | 2976 | 1973 | 2924 | 3007 | 2158 | 2020 | 2094 | 2232 | 2385 | 2243 | 2482 | 2445 | 2257,41 | 2126 | 2389 | 1,126 | 0,171 | 3,434 | 0,0005943 | 0,0008129 | 0,0012812 | 0,0028186 | 0,0019286 | 0,0019286 | TRUE | 0,0194292 | No |
| orf19.2615 | 743 | 626 | 743 | 405 | 746 | 520 | 729 | 787 | 708 | 662 | 734 | 634 | 598 | 591 | 619 | 640 | 648,33 | 684 | 612 | 0,891 | -0,166 | -2,461 | 0,0138539 | 0,0174397 | 0,0010219 | 0,0046238 | 0,0028095 | 0,0028095 | TRUE | 0,027012 | No |
| orf19.2616 | 4883 | 4392 | 4796 | 2839 | 3943 | 2600 | 3622 | 4018 | 4655 | 4644 | 4738 | 4447 | 3159 | 2956 | 3075 | 3267 | 3867,85 | 4621 | 3114 | 0,674 | -0,569 | -13,321 | 1,75E-40 | 5,57E-40 | 0,0008873 | 0,0025158 | 0,0014876 | 0,0014876 | TRUE | 0,0135918 | No |
| orf19.2618 | 9361 | 7726 | 8456 | 5278 | 6377 | 4066 | 5873 | 6169 | 8925 | 8169 | 8354 | 8268 | 5110 | 4623 | 4986 | 5016 | 6681,4 | 8429 | 4934 | 0,586 | -0,772 | -16,937 | 2,41E-64 | 1,07E-63 | 0,0015309 | 0,002337 | 0,0018393 | 0,0018393 | TRUE | 0,0213404 | No |
| orf19.2619 | 423 | 416 | 429 | 310 | 390 | 353 | 389 | 441 | 403 | 440 | 424 | 486 | 312 | 401 | 330 | 359 | 394,41 | 438 | 350 | 0,801 | -0,321 | -3,363 | 0,0007709 | 0,0010485 | 0,0058822 | 0,0062542 | 0,0061636 | 0,0061636 | TRUE | 0,1002682 | No |
| orf19.262 | 95 | 93 | 121 | 66 | 263 | 191 | 219 | 244 | 91 | 98 | 120 | 103 | 211 | 217 | 186 | 198 | 153,01 | 103 | 203 | 1,97 | 0,978 | 7,652 | 1,98E-14 | 3,93E-14 | 1,00E-08 | 0,012822 | 0,0080818 | 0,0080818 | TRUE | 0,1309567 | No |
| orf19.2620 | 262 | 203 | 255 | 184 | 90 | 75 | 103 | 109 | 250 | 215 | 252 | 288 | 72 | 85 | 87 | 89 | 167,26 | 251 | 83 | 0,333 | -1,586 | -11,992 | 3,91E-33 | 1,11E-32 | 0,0058089 | 0,0119078 | 0,0092141 | 0,0092141 | TRUE | 0,1033475 | No |
| orf19.2621 | 795 | 833 | 787 | 502 | 1180 | 853 | 1124 | 1125 | 758 | 881 | 777 | 786 | 946 | 970 | 954 | 915 | 873,38 | 800 | 946 | 1,181 | 0,24 | 3,861 | 0,000113 | 0,0001596 | 0,0011005 | 0,0039712 | 0,0025367 | 0,0025367 | TRUE | 0,0558179 | No |
| orf19.2622 | 2251 | 2154 | 2175 | 1297 | 5486 | 3778 | 5219 | 5421 | 2146 | 2278 | 2149 | 2032 | 4396 | 4296 | 4431 | 4408 | 3266,82 | 2151 | 4383 | 2,035 | 1,025 | 24,504 | 1,34E-132 | 1,20E-131 | 0,0003835 | 0,0025939 | 0,0013167 | 0,0013167 | TRUE | 0,0184467 | No |
| orf19.2623 | 15021 | 13342 | 14429 | 9492 | 3611 | 2615 | 3401 | 3671 | 14321 | 14107 | 14254 | 14870 | 2893 | 2973 | 2887 | 2985 | 8661,47 | 14388 | 2934 | 0,204 | -2,293 | -68,649 | 0 | 0 | 0,0002957 | 0,0022808 | 0,0008805 | 0,0008805 | TRUE | 0,006018 | No |
| orf19.2624 | 5 | 14 | 6 | 3 | 8 | 8 | 7 | 8 | 5 | 15 | 6 | 5 | 6 | 9 | 6 | 7 | 7,27 | 8 | 7 | 0,905 | -0,144 | -0,26 | 0,7949012 | 0,8175797 | 0,0599635 | 0,2279692 | 0,1559834 | 0,1559834 | TRUE | 1,1464427 | No |
| orf19.2625 | 1 | 2 | 2 | 0 | 0 | 0 | 0 | 0 | 1 | 2 | 2 | 0 | 0 | 0 | 0 | 0 | 0,63 | 1 | 0 | 0,126 | -2,988 | -1,545 | 0,122339 | 0,1414469 | 1,00E-08 | 2,6062467 | 2,1602021 | 2,1602021 | TRUE | 0,1502336 | No |
| orf19.2629 | 1181 | 1149 | 1264 | 736 | 1806 | 1201 | 1604 | 1702 | 1126 | 1215 | 1249 | 1153 | 1447 | 1366 | 1362 | 1384 | 1287,62 | 1186 | 1390 | 1,172 | 0,229 | 4,34 | 1,42E-05 | 2,09E-05 | 0,0007357 | 0,0033664 | 0,0018678 | 0,0018678 | TRUE | 0,0159894 | No |
| orf19.2630 | 649 | 655 | 590 | 372 | 896 | 587 | 878 | 905 | 619 | 693 | 583 | 583 | 718 | 667 | 745 | 736 | 667,95 | 620 | 716 | 1,157 | 0,21 | 2,874 | 0,0040555 | 0,005277 | 0,0026582 | 0,0045494 | 0,0036227 | 0,0036227 | TRUE | 0,0748781 | No |
| orf19.2631 | 1311 | 1287 | 1251 | 815 | 1765 | 1281 | 1577 | 1738 | 1250 | 1361 | 1236 | 1277 | 1414 | 1457 | 1339 | 1413 | 1343,28 | 1281 | 1406 | 1,097 | 0,134 | 2,552 | 0,0107143 | 0,0135839 | 0,0008106 | 0,0033136 | 0,0018779 | 0,0018779 | TRUE | 0,0221387 | No |
| orf19.263.1 | 7503 | 5863 | 6582 | 3739 | 3572 | 1989 | 3246 | 3446 | 7153 | 6199 | 6502 | 5857 | 2862 | 2262 | 2756 | 2802 | 4549,25 | 6428 | 2670 | 0,416 | -1,267 | -15,762 | 5,65E-56 | 2,25E-55 | 0,0086539 | 0,0024522 | 0,00595 | 0,00595 | TRUE | 0,1268161 | No |
| orf19.2633 | 3499 | 2988 | 3282 | 2188 | 5509 | 3734 | 5197 | 5360 | 3336 | 3159 | 3242 | 3428 | 4414 | 4246 | 4412 | 4358 | 3824,47 | 3291 | 4358 | 1,326 | 0,407 | 10,583 | 3,59E-26 | 9,11E-26 | 0,0004293 | 0,0025206 | 0,0011451 | 0,0011451 | TRUE | 0,008649 | No |
| orf19.2633.1 | 236 | 217 | 230 | 140 | 150 | 131 | 159 | 138 | 225 | 229 | 227 | 219 | 120 | 149 | 135 | 112 | 177,17 | 225 | 129 | 0,569 | -0,815 | -7,06 | 1,66E-12 | 3,12E-12 | 1,00E-08 | 0,0113588 | 0,0068403 | 0,0068403 | TRUE | 0,0965358 | No |
| orf19.2637 | 1217 | 1136 | 1164 | 694 | 983 | 700 | 930 | 987 | 1160 | 1201 | 1150 | 1087 | 788 | 796 | 790 | 803 | 971,78 | 1150 | 794 | 0,689 | -0,537 | -10,138 | 3,75E-24 | 9,13E-24 | 1,00E-08 | 0,0037809 | 0,0016498 | 0,0016498 | TRUE | 0,0136741 | No |
| orf19.2638 | 467 | 443 | 460 | 300 | 152 | 94 | 119 | 106 | 445 | 468 | 454 | 470 | 122 | 107 | 101 | 86 | 281,74 | 459 | 104 | 0,226 | -2,144 | -20,119 | 5,06E-90 | 3,03E-89 | 1,00E-08 | 0,0079189 | 0,0054073 | 0,0054073 | TRUE | 0,153645 | No |
| orf19.2638.1 | 2999 | 2620 | 2603 | 1476 | 1417 | 856 | 1260 | 1297 | 2859 | 2770 | 2572 | 2312 | 1135 | 973 | 1070 | 1055 | 1843,3 | 2628 | 1058 | 0,403 | -1,313 | -17,934 | 6,42E-72 | 3,16E-71 | 0,0053577 | 0,002982 | 0,0045089 | 0,0045089 | TRUE | 0,0764577 | No |
| orf19.2639 | 1774 | 1532 | 1725 | 1026 | 1409 | 975 | 1315 | 1448 | 1691 | 1620 | 1704 | 1607 | 1129 | 1109 | 1116 | 1177 | 1394,26 | 1656 | 1133 | 0,684 | -0,548 | -11,506 | 1,23E-30 | 3,35E-30 | 1,00E-08 | 0,0032689 | 0,0014487 | 0,0014487 | TRUE | 0,0085653 | No |
| orf19.2639.1 | 1255 | 1147 | 1199 | 625 | 2349 | 1286 | 2173 | 2309 | 1197 | 1213 | 1184 | 979 | 1882 | 1462 | 1845 | 1877 | 1454,96 | 1143 | 1766 | 1,544 | 0,626 | 6,84 | 7,92E-12 | 1,46E-11 | 0,0103126 | 0,0032198 | 0,0073095 | 0,0073095 | TRUE | 0,1592721 | No |
| orf19.264 | 881 | 940 | 1004 | 603 | 804 | 650 | 730 | 833 | 840 | 994 | 992 | 945 | 644 | 739 | 620 | 677 | 806,35 | 943 | 670 | 0,71 | -0,494 | -6,408 | 1,47E-10 | 2,62E-10 | 0,0047308 | 0,0041275 | 0,0044502 | 0,0044502 | TRUE | 0,0675453 | No |
| orf19.2640 | 1693 | 1574 | 1560 | 913 | 3088 | 2172 | 2991 | 3190 | 1614 | 1664 | 1541 | 1430 | 2474 | 2470 | 2539 | 2594 | 2040,87 | 1562 | 2519 | 1,609 | 0,686 | 13,208 | 7,89E-40 | 2,49E-39 | 0,0013732 | 0,0028958 | 0,0020543 | 0,0020543 | TRUE | 0,0349491 | No |
| orf19.2641 | 135 | 114 | 125 | 70 | 132 | 91 | 130 | 115 | 129 | 121 | 123 | 110 | 106 | 103 | 110 | 94 | 111,94 | 121 | 103 | 0,851 | -0,232 | -1,734 | 0,0829571 | 0,0976299 | 1,00E-08 | 0,0167591 | 0,0083675 | 0,0083675 | TRUE | 0,0444887 | No |
| orf19.2642 | 35 | 49 | 51 | 28 | 116 | 61 | 110 | 109 | 33 | 52 | 50 | 44 | 93 | 69 | 93 | 89 | 65,47 | 45 | 86 | 1,932 | 0,95 | 4,817 | 1,46E-06 | 2,24E-06 | 0,0079669 | 0,0271703 | 0,0198023 | 0,0198023 | TRUE | 0,2722368 | No |
| orf19.2643 | 5342 | 4441 | 5065 | 3075 | 8550 | 5669 | 8158 | 8651 | 5093 | 4696 | 5004 | 4817 | 6851 | 6446 | 6926 | 7034 | 5858,37 | 4902 | 6814 | 1,39 | 0,475 | 11,006 | 3,58E-28 | 9,38E-28 | 0,0011453 | 0,0023715 | 0,0016084 | 0,0016084 | TRUE | 0,015817 | No |
| orf19.2644 | 2837 | 2685 | 2665 | 1956 | 10356 | 8345 | 9894 | 10600 | 2705 | 2839 | 2633 | 3064 | 8298 | 9489 | 8400 | 8619 | 5755,81 | 2810 | 8702 | 3,1 | 1,632 | 27,237 | 2,35E-163 | 2,62E-162 | 0,0039029 | 0,0023765 | 0,0031963 | 0,0031963 | TRUE | 0,0450145 | No |
| orf19.2645 | 40 | 50 | 35 | 27 | 24 | 21 | 17 | 25 | 38 | 53 | 35 | 42 | 19 | 24 | 14 | 20 | 30,72 | 42 | 19 | 0,461 | -1,118 | -4,086 | 4,39E-05 | 6,31E-05 | 1,00E-08 | 0,0555413 | 0,0354625 | 0,0354625 | TRUE | 0,2453019 | No |
| orf19.2646 | 1871 | 2001 | 2141 | 1261 | 2029 | 1387 | 1810 | 1903 | 1784 | 2116 | 2115 | 1975 | 1626 | 1577 | 1537 | 1547 | 1784,63 | 1998 | 1572 | 0,787 | -0,346 | -5,633 | 1,77E-08 | 2,92E-08 | 0,00308 | 0,0030113 | 0,0030507 | 0,0030507 | TRUE | 0,0654949 | No |
| orf19.2647 | 131 | 141 | 135 | 82 | 284 | 205 | 291 | 248 | 125 | 149 | 133 | 128 | 228 | 233 | 247 | 202 | 180,65 | 134 | 228 | 1,693 | 0,76 | 6,484 | 8,96E-11 | 1,61E-10 | 1,00E-08 | 0,0111803 | 0,0070764 | 0,0070764 | TRUE | 0,0676082 | No |
| orf19.2649 | 267 | 273 | 334 | 169 | 263 | 198 | 240 | 216 | 255 | 289 | 330 | 265 | 211 | 225 | 204 | 176 | 244,15 | 285 | 204 | 0,713 | -0,489 | -4,241 | 2,23E-05 | 3,25E-05 | 0,008337 | 0,0088163 | 0,0085859 | 0,0085859 | TRUE | 0,1350836 | No |
| orf19.265 | 5820 | 5882 | 5786 | 3606 | 520 | 395 | 476 | 425 | 5549 | 6219 | 5716 | 5649 | 417 | 449 | 404 | 346 | 3093,59 | 5783 | 404 | 0,07 | -3,845 | -54,387 | 0 | 0 | 0,0053043 | 0,002622 | 0,0035959 | 0,0035959 | TRUE | 0,1118794 | No |
| orf19.2650 | 957 | 1128 | 1099 | 679 | 1489 | 1110 | 1369 | 1303 | 912 | 1193 | 1086 | 1064 | 1193 | 1262 | 1162 | 1059 | 1116,44 | 1064 | 1169 | 1,099 | 0,136 | 1,623 | 0,104611 | 0,1218775 | 0,0072965 | 0,0035619 | 0,0058421 | 0,0058421 | TRUE | 0,1138654 | No |
| orf19.2650.1 | 494 | 369 | 465 | 206 | 463 | 306 | 462 | 458 | 471 | 390 | 459 | 323 | 371 | 348 | 392 | 372 | 390,85 | 411 | 371 | 0,898 | -0,155 | -1,414 | 0,1572242 | 0,1801163 | 0,0117623 | 0,0062921 | 0,0090184 | 0,0090184 | TRUE | 0,2125082 | No |
| orf19.2651 | 3548 | 3566 | 3520 | 2213 | 9563 | 6869 | 8634 | 9008 | 3383 | 3771 | 3477 | 3467 | 7663 | 7811 | 7330 | 7324 | 5528,16 | 3524 | 7532 | 2,137 | 1,095 | 24,168 | 4,83E-129 | 4,14E-128 | 0,0014039 | 0,0023883 | 0,0017531 | 0,0017531 | TRUE | 0,0273136 | No |
| orf19.2652 | 95 | 106 | 118 | 62 | 125 | 109 | 136 | 99 | 91 | 112 | 117 | 97 | 100 | 124 | 115 | 80 | 104,55 | 104 | 105 | 1,001 | 0,002 | 0,01 | 0,9920722 | 0,9930059 | 0,0158821 | 0,0177955 | 0,0169162 | 0,0169162 | TRUE | 0,2571354 | No |
| orf19.2653 | 2485 | 2546 | 2559 | 1619 | 1691 | 1356 | 1656 | 1744 | 2369 | 2692 | 2528 | 2536 | 1355 | 1542 | 1406 | 1418 | 1980,8 | 2531 | 1430 | 0,564 | -0,825 | -14,529 | 7,94E-48 | 2,85E-47 | 0,0021652 | 0,0029202 | 0,0025627 | 0,0025627 | TRUE | 0,0316647 | No |
| orf19.2654 | 3420 | 3528 | 3554 | 2110 | 2801 | 2067 | 2495 | 2510 | 3261 | 3730 | 3511 | 3305 | 2244 | 2350 | 2118 | 2041 | 2820,16 | 3452 | 2188 | 0,633 | -0,659 | -10,889 | 1,30E-27 | 3,36E-27 | 0,0033987 | 0,0026735 | 0,0031491 | 0,0031491 | TRUE | 0,0359695 | No |
| orf19.2655 | 24787 | 23324 | 24963 | 14754 | 2120 | 1478 | 1993 | 2169 | 23632 | 24662 | 24661 | 23113 | 1699 | 1681 | 1692 | 1764 | 12862,86 | 24017 | 1709 | 0,071 | -3,813 | -97,651 | 0 | 0 | 0,0007035 | 0,0022189 | 0,0011796 | 0,0011796 | TRUE | 0,0077372 | No |
| orf19.2657 | 22 | 32 | 28 | 20 | 59 | 42 | 48 | 62 | 21 | 34 | 28 | 31 | 47 | 48 | 41 | 50 | 37,5 | 28 | 46 | 1,653 | 0,725 | 3,027 | 0,0024732 | 0,00326 | 1,00E-08 | 0,0458754 | 0,025759 | 0,025759 | TRUE | 0,2285677 | No |
| orf19.2660 | 3036 | 3255 | 3287 | 2209 | 2615 | 2000 | 2508 | 2423 | 2894 | 3442 | 3247 | 3461 | 2095 | 2274 | 2129 | 1970 | 2689,12 | 3261 | 2117 | 0,649 | -0,623 | -9,351 | 8,64E-21 | 1,97E-20 | 0,0044684 | 0,0027019 | 0,0038777 | 0,0038777 | TRUE | 0,0703784 | No |
| orf19.2661 | 1402 | 1053 | 1151 | 699 | 912 | 539 | 888 | 939 | 1337 | 1113 | 1137 | 1095 | 731 | 613 | 754 | 763 | 942,9 | 1170 | 715 | 0,611 | -0,71 | -8,219 | 2,05E-16 | 4,26E-16 | 0,0078186 | 0,0038326 | 0,0060537 | 0,0060537 | TRUE | 0,1113404 | No |
| orf19.2662 | 212 | 180 | 194 | 114 | 177 | 88 | 164 | 199 | 202 | 190 | 192 | 179 | 142 | 100 | 139 | 162 | 163,2 | 191 | 136 | 0,716 | -0,483 | -3,514 | 0,0004411 | 0,0006066 | 0,0115829 | 0,0121518 | 0,0119323 | 0,0119323 | TRUE | 0,2855025 | No |
| orf19.2663 | 794 | 668 | 815 | 465 | 688 | 502 | 651 | 626 | 757 | 706 | 805 | 728 | 551 | 571 | 553 | 509 | 647,59 | 749 | 546 | 0,727 | -0,461 | -6,74 | 1,58E-11 | 2,90E-11 | 0,0013615 | 0,0046267 | 0,0029234 | 0,0029234 | TRUE | 0,0308149 | No |
| orf19.2664 | 2568 | 2423 | 2558 | 1618 | 1827 | 1353 | 1689 | 1812 | 2448 | 2562 | 2527 | 2535 | 1464 | 1538 | 1434 | 1473 | 1997,72 | 2518 | 1477 | 0,586 | -0,77 | -18,204 | 4,82E-74 | 2,45E-73 | 1,00E-08 | 0,0029132 | 0,0011939 | 0,0011939 | TRUE | 0,0085216 | No |
| orf19.2665 | 39545 | 35601 | 38783 | 23638 | 6125 | 4103 | 5802 | 6017 | 37702 | 37644 | 38314 | 37030 | 4908 | 4665 | 4926 | 4892 | 21260,14 | 37672 | 4848 | 0,129 | -2,957 | -97,001 | 0 | 0 | 0,0002603 | 0,0021685 | 0,0007857 | 0,0007857 | TRUE | 0,0074266 | No |
| orf19.2667 | 11398 | 9230 | 10163 | 5968 | 13072 | 8094 | 12645 | 13193 | 10867 | 9760 | 10040 | 9349 | 10474 | 9204 | 10736 | 10727 | 10144,52 | 10004 | 10285 | 1,028 | 0,04 | 0,641 | 0,5212074 | 0,5575564 | 0,0040584 | 0,0022531 | 0,0036165 | 0,0036165 | TRUE | 0,0608518 | No |
| orf19.2668 | 1099 | 946 | 996 | 707 | 2842 | 2194 | 2762 | 2872 | 1048 | 1000 | 984 | 1108 | 2277 | 2495 | 2345 | 2335 | 1698,96 | 1035 | 2363 | 2,288 | 1,194 | 22,358 | 1,00E-110 | 7,30E-110 | 0,0012822 | 0,0030577 | 0,0020057 | 0,0020057 | TRUE | 0,0231928 | No |
| orf19.2669 | 1653 | 1511 | 1691 | 1034 | 3596 | 2475 | 3135 | 3629 | 1576 | 1598 | 1671 | 1620 | 2881 | 2814 | 2662 | 2951 | 2221,5 | 1616 | 2827 | 1,75 | 0,807 | 16,855 | 9,67E-64 | 4,28E-63 | 0,0010406 | 0,0028304 | 0,0016968 | 0,0016968 | TRUE | 0,0192461 | No |
| orf19.267 | 3106 | 2788 | 2964 | 1951 | 4021 | 2941 | 3954 | 4328 | 2961 | 2948 | 2928 | 3056 | 3222 | 3344 | 3357 | 3519 | 3166,97 | 2973 | 3360 | 1,131 | 0,178 | 4,34 | 1,42E-05 | 2,09E-05 | 0,0006023 | 0,0026097 | 0,0012875 | 0,0012875 | TRUE | 0,0126 | No |
| orf19.2670 | 1383 | 1240 | 1299 | 815 | 953 | 681 | 935 | 1018 | 1319 | 1311 | 1283 | 1277 | 764 | 774 | 794 | 828 | 1043,65 | 1298 | 790 | 0,609 | -0,716 | -13,864 | 1,05E-43 | 3,54E-43 | 1,00E-08 | 0,0036645 | 0,0015663 | 0,0015663 | TRUE | 0,0128946 | No |
| orf19.2671 | 545 | 441 | 457 | 363 | 583 | 422 | 590 | 639 | 520 | 466 | 451 | 569 | 467 | 480 | 501 | 520 | 496,69 | 502 | 492 | 0,986 | -0,021 | -0,245 | 0,8065207 | 0,828329 | 0,0039501 | 0,005397 | 0,0047482 | 0,0047482 | TRUE | 0,0823465 | No |

| | | | | | | | | | | | | | | | | | | | | | | | | | | | | | | | |
|---|---|---|---|---|---|---|---|---|---|---|---|---|---|---|---|---|---|---|---|---|---|---|---|---|---|---|---|---|---|---|---|
| orf19.2672 | 2730 | 2450 | 2582 | 1696 | 2934 | 2022 | 2687 | 2920 | 2603 | 2591 | 2551 | 2657 | 2351 | 2299 | 2281 | 2374 | 2463,32 | 2600 | 2326 | 0,896 | -0,159 | -4,212 | 2,53E-05 | 3,69E-05 | 1,00E-08 | 0,0027578 | 0,0009589 | 0,0009589 | TRUE | 0,0023482 | No |
| orf19.2673 | 474 | 463 | 457 | 248 | 1717 | 1043 | 1660 | 1702 | 452 | 490 | 451 | 389 | 1376 | 1186 | 1409 | 1384 | 892,05 | 446 | 1339 | 2,995 | 1,583 | 19,691 | 2,60E-86 | 1,50E-85 | 0,0053811 | 0,0039319 | 0,0046181 | 0,0046181 | TRUE | 0,0719649 | No |
| orf19.2674 | 413 | 292 | 342 | 240 | 444 | 348 | 469 | 496 | 394 | 309 | 338 | 376 | 356 | 396 | 398 | 403 | 371,16 | 354 | 388 | 1,097 | 0,134 | 1,437 | 0,1506056 | 0,1728443 | 0,0046776 | 0,006515 | 0,0056236 | 0,0056236 | TRUE | 0,0859979 | No |
| orf19.2675 | 1887 | 1698 | 1844 | 1167 | 3287 | 2416 | 3091 | 3325 | 1799 | 1795 | 1822 | 1828 | 2634 | 2747 | 2624 | 2704 | 2244,14 | 1811 | 2677 | 1,478 | 0,564 | 14,42 | 3,89E-47 | 1,38E-46 | 1,00E-08 | 0,0028229 | 0,0009912 | 0,0009912 | TRUE | 0,0035174 | No |
| orf19.2676 | 1979 | 1749 | 1845 | 1267 | 2180 | 1702 | 2013 | 2084 | 1887 | 1849 | 1823 | 1985 | 1747 | 1935 | 1709 | 1694 | 1828,65 | 1886 | 1771 | 0,939 | -0,091 | -1,639 | 0,1011622 | 0,1180534 | 0,001959 | 0,0029892 | 0,00238 | 0,00238 | TRUE | 0,0448206 | No |
| orf19.2677 | 363 | 327 | 376 | 262 | 733 | 646 | 761 | 870 | 346 | 346 | 371 | 410 | 587 | 735 | 646 | 707 | 518,64 | 368 | 669 | 1,821 | 0,864 | 9,484 | 2,45E-21 | 5,63E-21 | 0,0062723 | 0,0052571 | 0,0057792 | 0,0057792 | TRUE | 0,0809188 | No |
| orf19.2678 | 726 | 573 | 659 | 438 | 581 | 389 | 544 | 626 | 692 | 606 | 651 | 686 | 466 | 442 | 462 | 509 | 564,24 | 659 | 470 | 0,715 | -0,484 | -6,577 | 4,79E-11 | 8,63E-11 | 0,0018511 | 0,0050012 | 0,0034001 | 0,0034001 | TRUE | 0,0375755 | No |
| orf19.268 | 566 | 461 | 530 | 333 | 1382 | 908 | 1320 | 1383 | 540 | 487 | 524 | 522 | 1107 | 1032 | 1121 | 1125 | 807,17 | 518 | 1096 | 2,118 | 1,083 | 17,521 | 9,99E-69 | 4,69E-68 | 1,00E-08 | 0,0041254 | 0,002185 | 0,002185 | TRUE | 0,0195072 | No |
| orf19.2680 | 1992 | 1714 | 1821 | 1211 | 2808 | 1934 | 2609 | 2712 | 1899 | 1812 | 1799 | 1897 | 2250 | 2199 | 2215 | 2205 | 2034,6 | 1852 | 2217 | 1,199 | 0,261 | 6,404 | 1,51E-10 | 2,69E-10 | 1,00E-08 | 0,0028983 | 0,0010945 | 0,0010945 | TRUE | 0,0045542 | No |
| orf19.2681 | 106 | 86 | 106 | 63 | 141 | 99 | 111 | 131 | 101 | 91 | 105 | 99 | 113 | 113 | 94 | 107 | 102,71 | 99 | 107 | 1,075 | 0,105 | 0,744 | 0,4566029 | 0,4930755 | 1,00E-08 | 0,0180765 | 0,0092763 | 0,0092763 | TRUE | 0,0629453 | No |
| orf19.2682 | 2891 | 2683 | 2910 | 1583 | 2424 | 1623 | 2341 | 2414 | 2756 | 2837 | 2875 | 2480 | 1942 | 1845 | 1987 | 1963 | 2335,74 | 2737 | 1934 | 0,706 | -0,502 | -9,24 | 2,46E-20 | 5,56E-20 | 0,0020522 | 0,0027942 | 0,0023983 | 0,0023983 | TRUE | 0,0455773 | No |
| orf19.2684 | 1866 | 1854 | 1941 | 1128 | 830 | 634 | 800 | 886 | 1779 | 1960 | 1918 | 1767 | 665 | 721 | 679 | 720 | 1276,2 | 1856 | 696 | 0,374 | -1,417 | -24,332 | 9,03E-131 | 7,89E-130 | 0,0015068 | 0,0033778 | 0,0023133 | 0,0023133 | TRUE | 0,0170891 | No |
| orf19.2685 | 321 | 307 | 299 | 238 | 97 | 91 | 108 | 89 | 306 | 325 | 295 | 373 | 78 | 103 | 92 | 72 | 205,52 | 325 | 86 | 0,265 | -1,918 | -14,634 | 1,71E-48 | 6,18E-48 | 0,009034 | 0,0100804 | 0,0095548 | 0,0095548 | TRUE | 0,1477788 | No |
| orf19.2686 | 2340 | 2121 | 2202 | 1516 | 2200 | 1617 | 1946 | 2135 | 2231 | 2243 | 2175 | 2375 | 1763 | 1839 | 1652 | 1736 | 2001,68 | 2256 | 1748 | 0,775 | -0,368 | -7,507 | 6,07E-14 | 1,19E-13 | 0,0009953 | 0,0029115 | 0,0017995 | 0,0017995 | TRUE | 0,0161254 | No |
| orf19.2687.1 | 4727 | 4066 | 4203 | 2461 | 7124 | 4179 | 6632 | 6758 | 4507 | 4299 | 4152 | 3855 | 5708 | 4752 | 5631 | 5495 | 4799,88 | 4203 | 5396 | 1,284 | 0,36 | 5,361 | 8,29E-08 | 1,34E-07 | 0,0051433 | 0,0024334 | 0,0041175 | 0,0041175 | TRUE | 0,0779641 | No |
| orf19.2688 | 15612 | 15136 | 16085 | 10202 | 12245 | 9072 | 11117 | 12202 | 14884 | 16004 | 15890 | 15982 | 9812 | 10316 | 9438 | 9921 | 12781,01 | 15690 | 9872 | 0,629 | -0,669 | -16,389 | 2,31E-60 | 9,74E-60 | 0,00115 | 0,0022197 | 0,0015174 | 0,0015174 | TRUE | 0,0147435 | No |
| orf19.269 | 4359 | 3994 | 4151 | 2791 | 10531 | 7285 | 10016 | 11100 | 4156 | 4223 | 4101 | 4372 | 8438 | 8284 | 8504 | 9025 | 6387,85 | 4213 | 8563 | 2,034 | 1,025 | 25,271 | 6,60E-141 | 6,28E-140 | 0,0008586 | 0,0023483 | 0,0013926 | 0,0013926 | TRUE | 0,0162324 | No |
| orf19.2690 | 9652 | 8819 | 9507 | 6244 | 7168 | 5434 | 6722 | 7296 | 9202 | 9325 | 9392 | 9782 | 5744 | 6179 | 5707 | 5932 | 7657,8 | 9425 | 5890 | 0,625 | -0,678 | -17,49 | 1,71E-68 | 8,00E-68 | 0,0008423 | 0,0023057 | 0,0013082 | 0,0013082 | TRUE | 0,0129092 | No |
| orf19.2691 | 1018 | 972 | 979 | 731 | 1241 | 870 | 1090 | 1298 | 971 | 1028 | 967 | 1145 | 994 | 989 | 925 | 1055 | 1009,38 | 1028 | 991 | 0,968 | -0,047 | -0,686 | 0,4927742 | 0,5287358 | 0,0033856 | 0,0037179 | 0,0035216 | 0,0035216 | TRUE | 0,0646657 | No |
| orf19.2693 | 2146 | 1846 | 1969 | 1151 | 3902 | 2575 | 3657 | 3935 | 2046 | 1952 | 1945 | 1803 | 3127 | 2928 | 3105 | 3200 | 2513,13 | 1936 | 3090 | 1,593 | 0,672 | 13,469 | 2,37E-41 | 7,66E-41 | 0,0014721 | 0,0027446 | 0,0019555 | 0,0019555 | TRUE | 0,0237335 | No |
| orf19.2694 | 6547 | 5720 | 6138 | 3697 | 8600 | 6040 | 8382 | 8830 | 6242 | 6048 | 6064 | 5792 | 6891 | 6868 | 7116 | 7180 | 6525,03 | 6036 | 7014 | 1,161 | 0,216 | 5,967 | 2,42E-09 | 4,11E-09 | 0,0005178 | 0,0023429 | 0,0010974 | 0,0010974 | TRUE | 0,0086056 | No |
| orf19.2695 | 3680 | 3381 | 3602 | 2171 | 4757 | 3414 | 4454 | 4888 | 3508 | 3575 | 3558 | 3401 | 3812 | 3882 | 3781 | 3974 | 3686,55 | 3510 | 3862 | 1,099 | 0,136 | 3,76 | 0,0001702 | 0,0002381 | 0,0002118 | 0,0025367 | 0,0009885 | 0,0009885 | TRUE | 0,0048769 | No |
| orf19.2697 | 1148 | 1087 | 1141 | 687 | 1725 | 1126 | 1482 | 1624 | 1094 | 1149 | 1127 | 1076 | 1382 | 1280 | 1258 | 1320 | 1211,06 | 1112 | 1310 | 1,178 | 0,236 | 4,503 | 6,70E-06 | 1,00E-05 | 0,0006123 | 0,003447 | 0,0018028 | 0,0018028 | TRUE | 0,0168245 | No |
| orf19.2698 | 539 | 487 | 534 | 304 | 611 | 438 | 527 | 596 | 514 | 515 | 528 | 476 | 490 | 498 | 447 | 485 | 494,03 | 508 | 480 | 0,941 | -0,088 | -1,252 | 0,2105784 | 0,2382871 | 1,00E-08 | 0,0054148 | 0,0027663 | 0,0027663 | TRUE | 0,0245314 | No |
| orf19.2699 | 3394 | 2739 | 3107 | 1907 | 6838 | 4534 | 6710 | 7139 | 3236 | 2896 | 3069 | 2987 | 5479 | 5156 | 5697 | 5805 | 4290,61 | 3047 | 5534 | 1,816 | 0,861 | 16,563 | 1,28E-61 | 5,53E-61 | 0,0021206 | 0,002474 | 0,0023273 | 0,0023273 | TRUE | 0,0253831 | No |
| orf19.27 | 1896 | 1604 | 1809 | 1189 | 1233 | 917 | 1165 | 1256 | 1808 | 1696 | 1787 | 1863 | 988 | 1043 | 989 | 1021 | 1399,3 | 1788 | 1010 | 0,565 | -0,824 | -16,611 | 5,75E-62 | 2,49E-61 | 0,0003561 | 0,0032647 | 0,0016043 | 0,0016043 | TRUE | 0,0144016 | No |
| orf19.270 | 413 | 316 | 317 | 230 | 447 | 291 | 444 | 488 | 394 | 334 | 313 | 360 | 358 | 331 | 377 | 397 | 358,02 | 350 | 366 | 1,047 | 0,066 | 0,691 | 0,4894051 | 0,5256512 | 0,0051578 | 0,0066773 | 0,0059309 | 0,0059309 | TRUE | 0,088472 | No |
| orf19.2701 | 2932 | 2679 | 2535 | 1733 | 5658 | 3748 | 5059 | 5403 | 2795 | 2833 | 2504 | 2715 | 4534 | 4262 | 4295 | 4393 | 3541,36 | 2712 | 4371 | 1,612 | 0,689 | 14,313 | 1,81E-46 | 6,36E-46 | 0,001456 | 0,0025549 | 0,0019164 | 0,0019164 | TRUE | 0,0334055 | No |
| orf19.2703 | 5572 | 4691 | 5270 | 3273 | 5119 | 3487 | 5047 | 5200 | 5312 | 4960 | 5206 | 5127 | 4102 | 3965 | 4285 | 4228 | 4648,22 | 5151 | 4145 | 0,805 | -0,313 | -7,769 | 7,91E-15 | 1,59E-14 | 0,0007138 | 0,0024445 | 0,0013439 | 0,0013439 | TRUE | 0,0099949 | No |
| orf19.2704 | 2 | 4 | 3 | 2 | 2 | 2 | 0 | 1 | 2 | 4 | 3 | 3 | 2 | 2 | 0 | 1 | 2,12 | 3 | 1 | 0,369 | -1,436 | -1,406 | 0,1597125 | 0,1828028 | 1,00E-08 | 0,7782711 | 0,4238569 | 0,4238569 | TRUE | 0,2930824 | No |
| orf19.2706 | 608 | 553 | 570 | 400 | 351 | 235 | 282 | 303 | 580 | 585 | 563 | 627 | 281 | 267 | 239 | 246 | 423,55 | 589 | 258 | 0,44 | -1,184 | -14,654 | 1,27E-48 | 4,61E-48 | 0,0005778 | 0,0059678 | 0,0035918 | 0,0035918 | TRUE | 0,0440941 | No |
| orf19.2707.1 | 2067 | 1418 | 1569 | 1108 | 5648 | 3141 | 5486 | 5336 | 1971 | 1499 | 1550 | 1736 | 4526 | 3572 | 4658 | 4339 | 2981,15 | 1689 | 4274 | 2,531 | 1,34 | 13,836 | 1,55E-43 | 5,19E-43 | 0,0139035 | 0,002642 | 0,0085682 | 0,0085682 | TRUE | 0,1555376 | No |
| orf19.2708 | 2343 | 2066 | 2185 | 1443 | 2753 | 1977 | 2556 | 2710 | 2234 | 2185 | 2159 | 2261 | 2206 | 2248 | 2170 | 2203 | 2208,11 | 2210 | 2207 | 0,999 | -0,001 | -0,026 | 0,9789089 | 0,9810613 | 1,00E-08 | 0,0028349 | 0,0009866 | 0,0009866 | TRUE | 0,0028984 | No |
| orf19.2709 | 12457 | 10702 | 11615 | 7308 | 17027 | 11372 | 16598 | 17650 | 11876 | 11316 | 11475 | 11448 | 13643 | 12931 | 14092 | 14351 | 12641,54 | 11529 | 13754 | 1,193 | 0,255 | 6,255 | 3,97E-10 | 6,96E-10 | 0,0011359 | 0,0022212 | 0,0015126 | 0,0015126 | TRUE | 0,0196351 | No |
| orf19.271 | 71 | 63 | 84 | 49 | 37 | 20 | 22 | 38 | 68 | 67 | 83 | 77 | 30 | 23 | 19 | 31 | 49,5 | 74 | 26 | 0,35 | -1,515 | -6,774 | 1,25E-11 | 2,30E-11 | 1,00E-08 | 0,0352598 | 0,0231138 | 0,0231138 | TRUE | 0,2366737 | No |
| orf19.2710 | 4553 | 4036 | 4442 | 2258 | 2200 | 1451 | 2265 | 2202 | 4341 | 4268 | 4388 | 3537 | 1763 | 1650 | 1923 | 1790 | 2957,5 | 4134 | 1782 | 0,431 | -1,215 | -16,496 | 3,91E-61 | 1,67E-60 | 0,0061426 | 0,0026464 | 0,004822 | 0,004822 | TRUE | 0,1120209 | No |
| orf19.2711 | 7602 | 6535 | 7165 | 4133 | 7585 | 5020 | 7087 | 7709 | 7248 | 6910 | 7078 | 6475 | 6078 | 5708 | 6017 | 6268 | 6472,67 | 6928 | 6018 | 0,868 | -0,204 | -4,362 | 1,29E-05 | 1,90E-05 | 0,0016877 | 0,0023449 | 0,0019379 | 0,0019379 | TRUE | 0,0229522 | No |
| orf19.2711.1 | 197 | 179 | 177 | 116 | 551 | 363 | 485 | 540 | 188 | 189 | 175 | 182 | 442 | 413 | 412 | 439 | 304,85 | 184 | 426 | 2,326 | 1,218 | 13,702 | 9,90E-43 | 3,28E-42 | 1,00E-08 | 0,0074773 | 0,0035161 | 0,0035161 | TRUE | 0,0122813 | No |
| orf19.2712 | 10876 | 9470 | 10241 | 6073 | 10022 | 6834 | 9731 | 10074 | 10369 | 10013 | 10117 | 9514 | 8030 | 7771 | 8262 | 8191 | 9033,39 | 10003 | 8064 | 0,806 | -0,311 | -8,077 | 6,62E-16 | 1,36E-15 | 0,0008592 | 0,002273 | 0,0013157 | 0,0013157 | TRUE | 0,0128749 | No |
| orf19.2713 | 59 | 55 | 58 | 49 | 112 | 83 | 93 | 108 | 56 | 58 | 57 | 77 | 90 | 94 | 79 | 88 | 74,92 | 62 | 88 | 1,431 | 0,517 | 3,054 | 0,0022604 | 0,0029856 | 1,00E-08 | 0,0240068 | 0,013323 | 0,013323 | TRUE | 0,1574819 | No |
| orf19.2715 | 5595 | 4991 | 5479 | 3214 | 4349 | 2956 | 4192 | 4405 | 5334 | 5277 | 5413 | 5035 | 3485 | 3361 | 3559 | 3582 | 4380,73 | 5265 | 3497 | 0,664 | -0,591 | -15,102 | 1,58E-51 | 5,92E-51 | 0,0005655 | 0,0024661 | 0,0012348 | 0,0012348 | TRUE | 0,0099474 | No |
| orf19.2717 | 16771 | 15058 | 15637 | 8737 | 17315 | 10626 | 16280 | 16939 | 15989 | 15922 | 15448 | 13687 | 13874 | 12083 | 13822 | 13773 | 14324,69 | 15262 | 13388 | 0,877 | -0,189 | -3,023 | 0,0025006 | 0,0032954 | 0,0045132 | 0,0022059 | 0,0036891 | 0,0036891 | TRUE | 0,058519 | No |
| orf19.272 | 2394 | 2254 | 2424 | 1579 | 19622 | 14677 | 18548 | 19385 | 2282 | 2383 | 2395 | 2474 | 15723 | 16689 | 15747 | 15762 | 9181,83 | 2384 | 15980 | 6,712 | 2,747 | 69,242 | 0 | 0 | 0,0006579 | 0,0022701 | 0,0012472 | 0,0012472 | TRUE | 0,0107943 | No |
| orf19.2720 | 6055 | 5245 | 5753 | 4136 | 7609 | 5903 | 7161 | 7843 | 5773 | 5546 | 5683 | 6479 | 6097 | 6712 | 6080 | 6377 | 6093,41 | 5870 | 6316 | 1,077 | 0,106 | 1,882 | 0,0598748 | 0,0713999 | 0,0034036 | 0,0023607 | 0,0029052 | 0,0029052 | TRUE | 0,0580675 | No |
| orf19.2721 | 3544 | 3235 | 3305 | 2169 | 2513 | 1780 | 2345 | 2483 | 3379 | 3421 | 3265 | 3398 | 2014 | 2024 | 1991 | 2019 | 2688,72 | 3366 | 2012 | 0,598 | -0,742 | -20,17 | 1,81E-90 | 1,09E-89 | 1,00E-08 | 0,0027019 | 0,0009107 | 0,0009107 | TRUE | 0,0050858 | No |
| orf19.2722 | 723 | 668 | 696 | 398 | 263 | 202 | 268 | 257 | 689 | 706 | 688 | 623 | 211 | 230 | 228 | 209 | 447,95 | 676 | 220 | 0,322 | -1,634 | -20,587 | 3,57E-94 | 2,21E-93 | 1,00E-08 | 0,0057566 | 0,0031916 | 0,0031916 | TRUE | 0,0283482 | No |
| orf19.2723 | 2769 | 2725 | 2840 | 1980 | 2352 | 1577 | 2222 | 2262 | 2640 | 2881 | 2806 | 3102 | 1885 | 1793 | 1886 | 1839 | 2354,02 | 2857 | 1851 | 0,649 | -0,623 | -11,554 | 7,01E-31 | 1,92E-30 | 0,001943 | 0,0027888 | 0,0023563 | 0,0023563 | TRUE | 0,0383123 | No |
| orf19.2724 | 58125 | 58917 | 61853 | 41939 | 24543 | 19747 | 23112 | 23099 | 55416 | 62297 | 61105 | 65700 | 19666 | 22454 | 19622 | 18782 | 40630,14 | 61130 | 20131 | 0,329 | -1,603 | -24,434 | 7,52E-132 | 6,69E-131 | 0,0055684 | 0,0021317 | 0,0041019 | 0,0041019 | TRUE | 0,0736985 | No |
| orf19.2725 | 6 | 12 | 5 | 10 | 12 | 6 | 5 | 6 | 6 | 13 | 5 | 16 | 10 | 7 | 4 | 5 | 8,07 | 10 | 6 | 0,678 | -0,561 | -1,017 | 0,3089558 | 0,3422651 | 0,1080723 | 0,2055009 | 0,1633094 | 0,1633094 | TRUE | 0,1596204 | No |
| orf19.2726 | 4309 | 3895 | 4249 | 2632 | 5033 | 3730 | 4745 | 4838 | 4108 | 4118 | 4198 | 4123 | 4033 | 4241 | 4028 | 3934 | 4097,97 | 4137 | 4059 | 0,98 | -0,028 | -0,783 | 0,433474 | 0,4696091 | 0,0002351 | 0,0024919 | 0,0010247 | 0,0010247 | TRUE | 0,0106347 | No |
| orf19.2727 | 4548 | 4225 | 4265 | 2855 | 5899 | 4505 | 5635 | 5849 | 4336 | 4467 | 4213 | 4473 | 4727 | 5123 | 4784 | 4756 | 4609,82 | 4372 | 4848 | 1,109 | 0,149 | 3,602 | 0,0003153 | 0,0004364 | 0,0008364 | 0,0024475 | 0,0014142 | 0,0014142 | TRUE | 0,0173068 | No |
| orf19.2728 | 881 | 889 | 918 | 506 | 1986 | 1408 | 1863 | 1872 | 840 | 940 | 907 | 793 | 1591 | 1601 | 1582 | 1522 | 1221,96 | 870 | 1574 | 1,803 | 0,851 | 14,458 | 2,24E-47 | 7,98E-47 | 0,0013411 | 0,0034349 | 0,0023982 | 0,0023982 | TRUE | 0,0362028 | No |
| orf19.273 | 1362 | 1316 | 1394 | 939 | 1542 | 1184 | 1434 | 1418 | 1299 | 1392 | 1377 | 1471 | 1236 | 1346 | 1217 | 1153 | 1311,31 | 1385 | 1238 | 0,894 | -0,161 | -2,615 | 0,0089309 | 0,0113816 | 0,0023905 | 0,0033434 | 0,0028662 | 0,0028662 | TRUE | 0,0394943 | No |
| orf19.2730 | 2418 | 1715 | 2096 | 1483 | 736 | 494 | 707 | 740 | 2305 | 1813 | 2071 | 2323 | 590 | 562 | 600 | 602 | 1358,24 | 2128 | 588 | 0,277 | -1,853 | -22,901 | 4,50E-116 | 3,46E-115 | 0,0068323 | 0,0033001 | 0,0052599 | 0,0052599 | TRUE | 0,1202315 | No |
| orf19.2731 | 147 | 122 | 117 | 57 | 390 | 200 | 370 | 350 | 140 | 129 | 116 | 89 | 312 | 227 | 314 | 285 | 201,58 | 118 | 284 | 2,382 | 1,252 | 8,587 | 8,92E-18 | 1,91E-17 | 0,0192463 | 0,0102364 | 0,0141847 | 0,0141847 | TRUE | 0,2153236 | No |
| orf19.2733 | 829 | 765 | 795 | 491 | 776 | 537 | 692 | 697 | 790 | 809 | 785 | 769 | 622 | 611 | 588 | 567 | 692,56 | 788 | 597 | 0,755 | -0,405 | -6,711 | 1,93E-11 | 3,52E-11 | 1,00E-08 | 0,0044621 | 0,0020342 | 0,0020342 | TRUE | 0,0140329 | No |
| orf19.2734 | 114 | 103 | 100 | 68 | 216 | 160 | 221 | 219 | 109 | 109 | 99 | 107 | 173 | 182 | 188 | 178 | 142,95 | 106 | 180 | 1,704 | 0,769 | 6,396 | 1,60E-10 | 2,84E-10 | 1,00E-08 | 0,013577 | 0,0061633 | 0,0061633 | TRUE | 0,0215016 | No |
| orf19.2735 | 11850 | 11019 | 11883 | 7323 | 9582 | 6989 | 9399 | 9541 | 11298 | 11651 | 11739 | 11472 | 7678 | 7947 | 7980 | 7758 | 9690,3 | 11540 | 7841 | 0,679 | -0,558 | -18,493 | 2,34E-76 | 1,22E-75 | 0,0001754 | 0,0022607 | 0,0007682 | 0,0007682 | TRUE | 0,0025319 | No |
| orf19.2736 | 903 | 801 | 882 | 557 | 1595 | 1123 | 1451 | 1533 | 861 | 847 | 871 | 873 | 1278 | 1277 | 1232 | 1246 | 1060,64 | 863 | 1258 | 1,458 | 0,544 | 11,012 | 3,34E-28 | 8,75E-28 | 1,00E-08 | 0,0036393 | 0,0013456 | 0,0013456 | TRUE | 0,0024052 | No |
| orf19.2737 | 8395 | 6930 | 7605 | 5080 | 12354 | 8727 | 11623 | 12818 | 8004 | 7328 | 7513 | 7958 | 9899 | 9923 | 9868 | 10422 | 8864,36 | 7701 | 10028 | 1,303 | 0,381 | 9,256 | 2,12E-20 | 4,80E-20 | 0,0011273 | 0,0022765 | 0,0015135 | 0,0015135 | TRUE | 0,0129841 | No |
| orf19.2738 | 3229 | 2865 | 3144 | 2100 | 12203 | 8894 | 11305 | 12730 | 3079 | 3029 | 3106 | 3290 | 9778 | 10113 | 9598 | 10351 | 6542,93 | 3126 | 9960 | 3,19 | 1,674 | 40,214 | 0 | 0 | 0,0009537 | 0,0023422 | 0,0014379 | 0,0014379 | TRUE | 0,0140013 | No |
| orf19.2739 | 307 | 289 | 311 | 181 | 792 | 548 | 763 | 784 | 293 | 306 | 307 | 284 | 635 | 623 | 648 | 637 | 466,5 | 298 | 636 | 2,134 | 1,094 | 15,284 | 9,74E-53 | 3,73E-52 | 1,00E-08 | 0,0056109 | 0,0023516 | 0,0023516 | TRUE | 0,0082119 | No |
| orf19.274 | 499 | 452 | 526 | 269 | 697 | 448 | 598 | 667 | 476 | 478 | 520 | 421 | 558 | 509 | 508 | 542 | 501,58 | 474 | 529 | 1,114 | 0,155 | 2,001 | 0,0453682 | 0,0547766 | 0,0020688 | 0,0053647 | 0,0037651 | 0,0037651 | TRUE | 0,0537812 | No |
| orf19.2740 | 659 | 605 | 668 | 416 | 2256 | 1639 | 2127 | 2337 | 628 | 640 | 660 | 652 | 1808 | 1864 | 1806 | 1900 | 1244,62 | 645 | 1844 | 2,861 | 1,517 | 29,427 | 2,50E-190 | 3,24E-189 | 1,00E-08 | 0,0034105 | 0,001445 | 0,001445 | TRUE | 0,0053099 | No |
| orf19.2742 | 135 | 94 | 108 | 49 | 246 | 128 | 223 | 216 | 129 | 99 | 107 | 77 | 197 | 146 | 189 | 176 | 139,9 | 103 | 177 | 1,703 | 0,768 | 4,903 | 9,46E-07 | 1,46E-06 | 0,0183121 | 0,0138279 | 0,0157008 | 0,0157008 | TRUE | 0,2770026 | No |
| orf19.2743 | 1197 | 922 | 1006 | 773 | 455 | 288 | 411 | 430 | 1141 | 975 | 994 | 1211 | 365 | 327 | 349 | 350 | 713,94 | 1080 | 348 | 0,323 | -1,63 | -19,287 | 6,88E-83 | 3,86E-82 | 0,0056315 | 0,0043911 | 0,0050546 | 0,0050546 | TRUE | 0,0736276 | No |
| orf19.2745 | 20 | 17 | 21 | 11 | 7 | 5 | 3 | 12 | 19 | 18 | 21 | 17 | 6 | 6 | 3 | 10 | 12,33 | 19 | 6 | 0,315 | -1,666 | -3,771 | 0,0001626 | 0,0002277 | 1,00E-08 | 0,1352819 | 0,0830931 | 0,0830931 | TRUE | 0,5725479 | No |
| orf19.2746 | 2523 | 2607 | 2658 | 1418 | 3406 | 2467 | 3041 | 3066 | 2405 | 2757 | 2626 | 2221 | 2729 | 2805 | 2582 | 2493 | 2577,29 | 2502 | 2652 | 1,058 | 0,082 | 1,18 | 0,2378648 | 0,2673152 | 0,0053732 | 0,0027283 | 0,0042152 | 0,0042152 | TRUE | 0,0661614 | No |
| orf19.2747 | 923 | 1046 | 1125 | 739 | 947 | 743 | 810 | 849 | 880 | 1106 | 1111 | 1158 | 759 | 845 | 688 | 690 | 904,59 | 1064 | 746 | 0,701 | -0,513 | -5,373 | 7,76E-08 | 1,25E-07 | 0,01028 | 0,0039064 | 0,0076212 | 0,0076212 | TRUE | 0,1651329 | No |
| orf19.2748 | 1392 | 1582 | 1567 | 949 | 737 | 552 | 624 | 648 | 1327 | 1673 | 1548 | 1487 | 591 | 628 | 530 | 527 | 1038,68 | 1509 | 569 | 0,376 | -1,41 | -16,811 | 2,02E-63 | 8,90E-63 | 0,0072856 | 0,003672 | 0,005594 | 0,005594 | TRUE | 0,0816441 | No |
| orf19.2749 | 5668 | 6031 | 6434 | 3841 | 868 | 672 | 894 | 840 | 5404 | 6377 | 6356 | 6017 | 696 | 764 | 759 | 683 | 3381,97 | 6038 | 726 | 0,12 | -3,059 | -45,858 | 0 | 0 | 0,0043063 | 0,0025768 | 0,0035643 | 0,0035643 | TRUE | 0,0619166 | No |
| orf19.275 | 463 | 362 | 381 | 227 | 794 | 530 | 783 | 774 | 441 | 383 | 376 | 356 | 636 | 603 | 665 | 629 | 511,14 | 389 | 633 | 1,621 | 0,697 | 8,839 | 9,67E-19 | 2,11E-18 | 0,0021442 | 0,0053035 | 0,0038405 | 0,0038405 | TRUE | 0,0975768 | No |
| orf19.2751 | 1022 | 1118 | 1151 | 637 | 1494 | 1164 | 1436 | 1371 | 974 | 1182 | 1137 | 998 | 1197 | 1324 | 1219 | 1115 | 1143,26 | 1073 | 1214 | 1,129 | 0,175 | 2,253 | 0,0242711 | 0,0299621 | 0,0058824 | 0,0035274 | 0,0048846 | 0,0048846 | TRUE | 0,0560088 | No |

| | | | | | | | | | | | | | | | | | | | | | | | | | | | | | | | |
|---|---|---|---|---|---|---|---|---|---|---|---|---|---|---|---|---|---|---|---|---|---|---|---|---|---|---|---|---|---|---|---|
| orf19.2752 | 2150 | 2164 | 2305 | 1391 | 6211 | 4540 | 5708 | 5848 | 2050 | 2288 | 2277 | 2179 | 4977 | 5162 | 4846 | 4755 | 3566,78 | 2198 | 4935 | 2,244 | 1,166 | 23,937 | 1,25E-126 | 1,06E-125 | 0,0014691 | 0,0025516 | 0,0019321 | 0,0019321 | TRUE | 0,0265942 | No |
| orf19.2753 | 2106 | 2040 | 2206 | 1375 | 3643 | 2834 | 3554 | 3739 | 2008 | 2157 | 2179 | 2154 | 2919 | 3223 | 3017 | 3040 | 2587,16 | 2124 | 3050 | 1,435 | 0,521 | 10,943 | 7,22E-28 | 1,88E-27 | 0,0010138 | 0,0027259 | 0,0017676 | 0,0017676 | TRUE | 0,0173042 | No |
| orf19.2754 | 527 | 494 | 473 | 375 | 1152 | 762 | 1047 | 1094 | 502 | 522 | 467 | 587 | 923 | 866 | 889 | 890 | 705,93 | 520 | 892 | 1,728 | 0,789 | 11,11 | 1,12E-28 | 2,95E-28 | 0,0017922 | 0,0044171 | 0,0032562 | 0,0032562 | TRUE | 0,0765678 | No |
| orf19.2755 | 5445 | 5081 | 5411 | 3402 | 5597 | 4330 | 5264 | 5494 | 5191 | 5373 | 5346 | 5329 | 4485 | 4924 | 4469 | 4467 | 4947,91 | 5310 | 4586 | 0,863 | -0,212 | -5,007 | 5,53E-07 | 8,65E-07 | 0,0010499 | 0,0024231 | 0,0015218 | 0,0015218 | TRUE | 0,029052 | No |
| orf19.2756 | 193 | 310 | 271 | 129 | 528 | 439 | 387 | 251 | 184 | 328 | 268 | 202 | 423 | 499 | 329 | 204 | 304,56 | 246 | 364 | 1,48 | 0,566 | 1,718 | 0,0858794 | 0,100883 | 0,100667 | 0,0074823 | 0,0488591 | 0,100667 | TRUE | 0,2465812 | No |
| orf19.2757 | 670 | 696 | 678 | 518 | 1150 | 1046 | 1089 | 1073 | 639 | 736 | 670 | 811 | 921 | 1189 | 925 | 872 | 845,48 | 714 | 977 | 1,37 | 0,454 | 4,337 | 1,45E-05 | 2,13E-05 | 0,0149412 | 0,0040333 | 0,0092637 | 0,0092637 | TRUE | 0,2501159 | No |
| orf19.2758 | 414 | 335 | 360 | 269 | 179 | 131 | 151 | 177 | 395 | 354 | 356 | 421 | 143 | 149 | 128 | 144 | 261,31 | 382 | 141 | 0,371 | -1,432 | -13,962 | 2,66E-44 | 9,02E-44 | 1,00E-08 | 0,0083746 | 0,0054706 | 0,0054706 | TRUE | 0,0484209 | No |
| orf19.276 | 897 | 858 | 862 | 538 | 806 | 654 | 766 | 752 | 855 | 907 | 852 | 843 | 646 | 744 | 650 | 611 | 763,51 | 864 | 663 | 0,764 | -0,388 | -5,576 | 2,46E-08 | 4,05E-08 | 0,0024407 | 0,0042418 | 0,0033235 | 0,0033235 | TRUE | 0,0739686 | No |
| orf19.2760 | 1950 | 2103 | 2125 | 1189 | 2857 | 2025 | 2514 | 2812 | 1859 | 2224 | 2099 | 1863 | 2289 | 2303 | 2134 | 2286 | 2132,17 | 2011 | 2253 | 1,119 | 0,162 | 2,511 | 0,0120362 | 0,0152267 | 0,0040375 | 0,0028613 | 0,0035359 | 0,0035359 | TRUE | 0,0609515 | No |
| orf19.2761 | 253 | 291 | 290 | 185 | 829 | 671 | 817 | 818 | 241 | 308 | 286 | 290 | 664 | 763 | 694 | 665 | 488,9 | 281 | 696 | 2,479 | 1,31 | 14,762 | 2,56E-49 | 9,35E-49 | 0,0043144 | 0,0054497 | 0,0049223 | 0,0049223 | TRUE | 0,1129232 | No |
| orf19.2762 | 15534 | 13772 | 14189 | 9355 | 14700 | 10442 | 13092 | 13995 | 14810 | 14562 | 14017 | 14655 | 11779 | 11873 | 11115 | 11379 | 13023,89 | 14511 | 11536 | 0,795 | -0,331 | -9,345 | 9,20E-21 | 2,10E-20 | 0,0006474 | 0,0022173 | 0,0011277 | 0,0011277 | TRUE | 0,0073946 | No |
| orf19.2763 | 139 | 148 | 131 | 59 | 331 | 208 | 245 | 255 | 133 | 156 | 129 | 92 | 265 | 237 | 208 | 207 | 178,49 | 128 | 229 | 1,775 | 0,828 | 5,643 | 1,67E-08 | 2,76E-08 | 0,0186383 | 0,01129 | 0,0143911 | 0,0143911 | TRUE | 0,2751785 | No |
| orf19.2765 | 92176 | 76457 | 85080 | 71014 | 106255 | 82043 | 98992 | 109075 | 87880 | 80844 | 84051 | 111247 | 85139 | 93290 | 84044 | 88688 | 89397,89 | 91006 | 87790 | 0,965 | -0,052 | -0,598 | 0,5496657 | 0,585057 | 0,0114987 | 0,0021096 | 0,0072016 | 0,0072016 | TRUE | 0,2745732 | No |
| orf19.2766 | 9 | 12 | 12 | 4 | 0 | 1 | 4 | 3 | 9 | 13 | 12 | 6 | 0 | 1 | 3 | 2 | 5,8 | 10 | 2 | 0,176 | -2,503 | -3,461 | 0,0005381 | 0,0007382 | 1,00E-08 | 0,2854074 | 0,1946521 | 0,1946521 | TRUE | 0,4774382 | No |
| orf19.2767 | 27267 | 22772 | 24868 | 18897 | 38958 | 27483 | 35500 | 38113 | 25996 | 24079 | 24567 | 29603 | 31216 | 31251 | 30139 | 30990 | 28480,06 | 26061 | 30899 | 1,186 | 0,246 | 4,026 | 5,66E-05 | 8,11E-05 | 0,0046861 | 0,0021489 | 0,003548 | 0,003548 | TRUE | 0,1020219 | No |
| orf19.2768 | 40960 | 40990 | 42868 | 23944 | 4385 | 3108 | 4106 | 3939 | 39051 | 43342 | 42350 | 37510 | 3514 | 3534 | 3486 | 3203 | 21998,52 | 40563 | 3434 | 0,085 | -3,563 | -63,698 | 0 | 0 | 0,0030957 | 0,0021659 | 0,0028609 | 0,0028609 | TRUE | 0,0313213 | No |
| orf19.2769 | 16305 | 13916 | 14030 | 8185 | 5413 | 3265 | 4971 | 5153 | 15545 | 14714 | 13860 | 12822 | 4337 | 3713 | 4220 | 4190 | 9175,27 | 14235 | 4115 | 0,289 | -1,79 | -26,719 | 2,81E-157 | 3,01E-156 | 0,0053931 | 0,0022702 | 0,0041645 | 0,0041645 | TRUE | 0,054206 | No |
| orf19.277 | 955 | 846 | 867 | 624 | 1185 | 829 | 1093 | 1195 | 910 | 895 | 857 | 978 | 950 | 943 | 928 | 972 | 928,85 | 910 | 948 | 1,046 | 0,064 | 1,136 | 0,2558431 | 0,2862088 | 0,000204 | 0,0038589 | 0,001988 | 0,001988 | TRUE | 0,0257924 | No |
| orf19.2770 | 5078 | 4615 | 4754 | 3049 | 527 | 329 | 536 | 566 | 4841 | 4880 | 4697 | 4776 | 422 | 374 | 455 | 460 | 2613,21 | 4798 | 428 | 0,09 | -3,48 | -61,2 | 0 | 0 | 0,0003402 | 0,0027196 | 0,001954 | 0,001954 | TRUE | 0,0711451 | No |
| orf19.2770.1 | 7699 | 6124 | 6291 | 4266 | 2385 | 1581 | 2014 | 2268 | 7340 | 6475 | 6215 | 6683 | 1911 | 1798 | 1710 | 1844 | 4247,01 | 6678 | 1816 | 0,272 | -1,879 | -31,5 | 8,56E-218 | 1,26E-216 | 0,0032091 | 0,0024779 | 0,0030858 | 0,0030858 | TRUE | 0,0550753 | No |
| orf19.2771 | 1508 | 1440 | 1549 | 919 | 2068 | 1573 | 2004 | 2109 | 1438 | 1523 | 1530 | 1440 | 1657 | 1789 | 1701 | 1715 | 1599,02 | 1483 | 1716 | 1,155 | 0,208 | 4,364 | 1,28E-05 | 1,89E-05 | 0,0004193 | 0,0031181 | 0,0015398 | 0,0015398 | TRUE | 0,0093374 | No |
| orf19.2772 | 6301 | 5902 | 6326 | 3990 | 1379 | 1008 | 1241 | 1456 | 6007 | 6241 | 6249 | 6251 | 1105 | 1146 | 1054 | 1184 | 3654,57 | 6187 | 1122 | 0,181 | -2,463 | -54,868 | 0 | 0 | 0,0007275 | 0,0025406 | 0,001447 | 0,001447 | TRUE | 0,0206744 | No |
| orf19.2774 | 6012 | 5104 | 5308 | 3173 | 4957 | 3116 | 4766 | 4979 | 5732 | 5397 | 5244 | 4971 | 3972 | 3543 | 4046 | 4048 | 4619,11 | 5336 | 3902 | 0,731 | -0,451 | -7,716 | 1,20E-14 | 2,39E-14 | 0,0033991 | 0,0024467 | 0,0030663 | 0,0030663 | TRUE | 0,0458392 | No |
| orf19.2775 | 4236 | 3852 | 4125 | 2520 | 6496 | 4755 | 6125 | 6222 | 4039 | 4073 | 4075 | 3948 | 5205 | 5407 | 5200 | 5059 | 4625,69 | 4034 | 5218 | 1,292 | 0,37 | 10,517 | 7,23E-26 | 1,82E-25 | 0,0002787 | 0,0024462 | 0,0009627 | 0,0009627 | TRUE | 0,0069835 | No |
| orf19.2778 | 822 | 1053 | 924 | 618 | 1174 | 953 | 956 | 898 | 784 | 1113 | 913 | 968 | 941 | 1084 | 812 | 730 | 918,02 | 944 | 892 | 0,943 | -0,084 | -0,675 | 0,4997373 | 0,5354868 | 0,0242739 | 0,0038798 | 0,0138577 | 0,0138577 | TRUE | 0,2460739 | No |
| orf19.2779 | 2 | 1 | 2 | 1 | 4 | 0 | 6 | 1 | 2 | 1 | 2 | 2 | 3 | 0 | 5 | 1 | 1,95 | 2 | 2 | 1,432 | 0,518 | 0,479 | 0,6319704 | 0,6645687 | 1,00E-08 | 0,8430864 | 0,5915538 | 0,5915538 | TRUE | 0,2670436 | No |
| orf19.278 | 1349 | 1157 | 1295 | 908 | 4056 | 3083 | 3825 | 4106 | 1286 | 1223 | 1279 | 1422 | 3250 | 3506 | 3247 | 3339 | 2319,11 | 1302 | 3336 | 2,566 | 1,36 | 25,543 | 6,49E-144 | 6,30E-143 | 0,0016315 | 0,0027993 | 0,0021545 | 0,0021545 | TRUE | 0,0409555 | No |
| orf19.2781 | 1796 | 1995 | 2022 | 1200 | 1692 | 1353 | 1478 | 1416 | 1712 | 2109 | 1998 | 1880 | 1356 | 1538 | 1255 | 1151 | 1624,94 | 1925 | 1325 | 0,688 | -0,54 | -5,987 | 2,13E-09 | 3,63E-09 | 0,0107907 | 0,0031017 | 0,0071924 | 0,0071924 | TRUE | 0,1381649 | No |
| orf19.2782 | 2141 | 2019 | 2099 | 1433 | 3802 | 2868 | 3507 | 3545 | 2041 | 2135 | 2074 | 2245 | 3046 | 3261 | 2977 | 2882 | 2582,75 | 2124 | 3042 | 1,433 | 0,519 | 10,01 | 1,37E-23 | 3,31E-23 | 0,0017577 | 0,002727 | 0,0021732 | 0,0021732 | TRUE | 0,027761 | No |
| orf19.2783 | 1 | 3 | 6 | 1 | 3 | 3 | 5 | 4 | 1 | 3 | 6 | 2 | 2 | 3 | 4 | 3 | 3,12 | 3 | 3 | 1,115 | 0,157 | 0,195 | 0,8457271 | 0,8649759 | 1,00E-08 | 0,5289347 | 0,305435 | 0,305435 | TRUE | 0,6556711 | No |
| orf19.2784 | 22 | 31 | 26 | 33 | 97 | 90 | 91 | 77 | 21 | 33 | 26 | 52 | 78 | 102 | 77 | 63 | 56,38 | 33 | 80 | 2,507 | 1,326 | 5,088 | 3,63E-07 | 5,71E-07 | 0,0590708 | 0,0312119 | 0,0420211 | 0,0420211 | TRUE | 0,6967671 | No |
| orf19.2785 | 2304 | 2291 | 2085 | 1316 | 5492 | 4012 | 4308 | 4587 | 2197 | 2422 | 2060 | 2062 | 4401 | 4562 | 3657 | 3730 | 3136,27 | 2185 | 4088 | 1,869 | 0,902 | 10,986 | 4,46E-28 | 1,17E-27 | 0,0089295 | 0,0026148 | 0,0061136 | 0,0061136 | TRUE | 0,0736976 | No |
| orf19.2786 | 625 | 683 | 759 | 416 | 1266 | 889 | 1048 | 1013 | 596 | 722 | 750 | 652 | 1014 | 1011 | 890 | 824 | 807,28 | 680 | 935 | 1,373 | 0,457 | 4,953 | 7,31E-07 | 1,14E-06 | 0,0093584 | 0,0041251 | 0,0068743 | 0,0068743 | TRUE | 0,087103 | No |
| orf19.2787 | 9 | 11 | 11 | 7 | 1 | 1 | 4 | 3 | 9 | 12 | 11 | 11 | 1 | 1 | 3 | 2 | 6,23 | 11 | 2 | 0,189 | -2,405 | -3,636 | 0,0002773 | 0,000385 | 1,00E-08 | 0,2657551 | 0,1431152 | 0,1431152 | TRUE | 0,2574342 | No |
| orf19.2788 | 601 | 557 | 627 | 398 | 997 | 699 | 937 | 1026 | 573 | 589 | 619 | 623 | 799 | 795 | 796 | 834 | 703,54 | 601 | 806 | 1,344 | 0,426 | 7,083 | 1,41E-12 | 2,66E-12 | 1,00E-08 | 0,0044251 | 0,001996 | 0,001996 | TRUE | 0,0122362 | No |
| orf19.2789 | 1925 | 1914 | 1891 | 1257 | 1234 | 934 | 1198 | 1292 | 1835 | 2024 | 1868 | 1969 | 989 | 1062 | 1017 | 1051 | 1476,85 | 1924 | 1030 | 0,535 | -0,902 | -17,242 | 1,29E-66 | 5,92E-66 | 0,0010037 | 0,003203 | 0,0019014 | 0,0019014 | TRUE | 0,0156875 | No |
| orf19.279 | 45 | 26 | 28 | 19 | 52 | 30 | 67 | 57 | 43 | 27 | 28 | 30 | 42 | 34 | 57 | 46 | 38,35 | 32 | 45 | 1,4 | 0,485 | 1,903 | 0,0570672 | 0,0682049 | 0,0225077 | 0,044901 | 0,0352396 | 0,0352396 | TRUE | 0,4043423 | No |
| orf19.2790 | 1718 | 1429 | 1545 | 965 | 2263 | 1587 | 2166 | 2321 | 1638 | 1511 | 1526 | 1512 | 1813 | 1805 | 1839 | 1887 | 1691,36 | 1547 | 1836 | 1,186 | 0,246 | 5,307 | 1,12E-07 | 1,79E-07 | 0,000354 | 0,003062 | 0,0014606 | 0,0014606 | TRUE | 0,019888 | No |
| orf19.2791 | 5422 | 4782 | 5346 | 3286 | 3569 | 2578 | 3500 | 3871 | 5169 | 5056 | 5281 | 5148 | 2860 | 2931 | 2971 | 3147 | 4070,6 | 5164 | 2977 | 0,577 | -0,794 | -19,463 | 2,24E-84 | 1,27E-83 | 0,0006519 | 0,0024946 | 0,0013392 | 0,0013392 | TRUE | 0,0183475 | No |
| orf19.2792 | 978 | 818 | 955 | 599 | 681 | 560 | 627 | 646 | 932 | 865 | 943 | 938 | 546 | 637 | 532 | 525 | 739,9 | 920 | 560 | 0,607 | -0,72 | -9,953 | 2,44E-23 | 5,84E-23 | 0,0031773 | 0,0043104 | 0,0036194 | 0,0036194 | TRUE | 0,0922832 | No |
| orf19.2794 | 1123 | 1018 | 1037 | 624 | 2121 | 1511 | 1954 | 1906 | 1071 | 1076 | 1024 | 978 | 1699 | 1718 | 1659 | 1550 | 1346,92 | 1037 | 1656 | 1,593 | 0,671 | 12,133 | 7,03E-34 | 2,02E-33 | 0,0012527 | 0,0033103 | 0,0021306 | 0,0021306 | TRUE | 0,0231183 | No |
| orf19.2795 | 4382 | 4566 | 4507 | 2704 | 9594 | 6835 | 8939 | 8574 | 4178 | 4828 | 4452 | 4236 | 7687 | 7772 | 7589 | 6971 | 5964,28 | 4424 | 7505 | 1,696 | 0,762 | 13,549 | 7,98E-42 | 2,61E-41 | 0,0031475 | 0,0023666 | 0,0028512 | 0,0028512 | TRUE | 0,046294 | No |
| orf19.2796 | 522 | 422 | 482 | 310 | 263 | 228 | 239 | 255 | 498 | 446 | 476 | 486 | 211 | 259 | 203 | 207 | 348,24 | 476 | 220 | 0,459 | -1,123 | -12,084 | 1,29E-33 | 3,68E-33 | 0,0026985 | 0,0068061 | 0,0050822 | 0,0050822 | TRUE | 0,1405373 | No |
| orf19.2797 | 89 | 82 | 73 | 54 | 72 | 45 | 78 | 70 | 85 | 87 | 72 | 85 | 58 | 51 | 66 | 57 | 70,03 | 82 | 58 | 0,711 | -0,491 | -2,883 | 0,003941 | 0,0051321 | 1,00E-08 | 0,0255359 | 0,0134666 | 0,0134666 | TRUE | 0,0794001 | No |
| orf19.2798 | 187 | 155 | 168 | 116 | 511 | 387 | 537 | 537 | 178 | 164 | 166 | 182 | 409 | 440 | 456 | 437 | 303,99 | 172 | 436 | 2,531 | 1,34 | 14,632 | 1,77E-48 | 6,38E-48 | 1,00E-08 | 0,0074925 | 0,0038013 | 0,0038013 | TRUE | 0,0198769 | No |
| orf19.2799 | 1657 | 1367 | 1576 | 971 | 1679 | 1176 | 1550 | 1695 | 1580 | 1445 | 1557 | 1521 | 1345 | 1337 | 1316 | 1378 | 1434,99 | 1526 | 1344 | 0,881 | -0,183 | -3,809 | 0,0001394 | 0,0001958 | 1,00E-08 | 0,0032355 | 0,0015237 | 0,0015237 | TRUE | 0,0159483 | No |
| orf19.28 | 8061 | 7261 | 7685 | 4231 | 4265 | 2653 | 4179 | 4414 | 7685 | 7678 | 7592 | 6628 | 3417 | 3017 | 3548 | 3589 | 5394,26 | 7396 | 3393 | 0,459 | -1,124 | -16,857 | 9,30E-64 | 4,12E-63 | 0,0050867 | 0,0023957 | 0,0040633 | 0,0040633 | TRUE | 0,0665206 | No |
| orf19.280 | 35 | 30 | 43 | 25 | 50 | 36 | 63 | 42 | 33 | 32 | 42 | 39 | 40 | 41 | 53 | 34 | 39,42 | 36 | 42 | 1,153 | 0,205 | 0,864 | 0,3874376 | 0,4230389 | 1,00E-08 | 0,0437416 | 0,0282232 | 0,0282232 | TRUE | 0,2768394 | No |
| orf19.2800 | 0 | 0 | 0 | 0 | 0 | 0 | 0 | 0 | 0 | 0 | 0 | 0 | 0 | 0 | 0 | 0 | 0 | 0 | 0 | NA | NA | NA | NA | NA | NA | NA | NA | NA | NA | NA | NA |
| orf19.2803 | 2875 | 2482 | 2689 | 1703 | 1728 | 1053 | 1610 | 1717 | 2741 | 2624 | 2656 | 2668 | 1385 | 1197 | 1367 | 1396 | 2004,33 | 2672 | 1336 | 0,501 | -0,997 | -18,494 | 2,29E-76 | 1,19E-75 | 0,0018623 | 0,0029105 | 0,0022473 | 0,0022473 | TRUE | 0,0551396 | No |
| orf19.2804 | 5 | 1 | 6 | 1 | 6 | 6 | 2 | 6 | 5 | 1 | 6 | 2 | 5 | 7 | 2 | 5 | 3,94 | 4 | 5 | 1,296 | 0,374 | 0,493 | 0,6222979 | 0,6558023 | 0,0852605 | 0,4187513 | 0,2940583 | 0,2940583 | TRUE | 0,2774162 | No |
| orf19.2805 | 288 | 291 | 321 | 194 | 783 | 658 | 668 | 670 | 275 | 308 | 317 | 304 | 627 | 748 | 567 | 545 | 461,35 | 301 | 622 | 2,064 | 1,045 | 9,753 | 1,79E-22 | 4,23E-22 | 0,0110003 | 0,0056502 | 0,0084397 | 0,0084397 | TRUE | 0,2118683 | No |
| orf19.2808 | 754 | 832 | 853 | 493 | 1239 | 880 | 1146 | 1138 | 719 | 880 | 843 | 772 | 993 | 1001 | 973 | 925 | 888,16 | 804 | 973 | 1,21 | 0,275 | 3,89 | 0,0001003 | 0,0001419 | 0,0030872 | 0,0039399 | 0,0036269 | 0,0036269 | TRUE | 0,0650156 | No |
| orf19.2809 | 1565 | 1381 | 1406 | 988 | 3029 | 2280 | 2898 | 3234 | 1492 | 1460 | 1389 | 1548 | 2427 | 2593 | 2460 | 2630 | 1999,82 | 1472 | 2528 | 1,719 | 0,782 | 15,489 | 4,11E-54 | 1,60E-53 | 0,001059 | 0,0029123 | 0,001886 | 0,001886 | TRUE | 0,017533 | No |
| orf19.281 | 205 | 183 | 176 | 134 | 323 | 226 | 305 | 325 | 195 | 193 | 174 | 210 | 259 | 257 | 259 | 264 | 226,47 | 193 | 260 | 1,352 | 0,436 | 4,476 | 7,59E-06 | 1,13E-05 | 1,00E-08 | 0,0093414 | 0,0044854 | 0,0044854 | TRUE | 0,0497133 | No |
| orf19.2810 | 387 | 320 | 361 | 241 | 877 | 639 | 757 | 842 | 369 | 338 | 357 | 378 | 703 | 727 | 643 | 685 | 524,77 | 360 | 690 | 1,914 | 0,937 | 12,798 | 1,68E-37 | 5,09E-37 | 1,00E-08 | 0,0052201 | 0,002946 | 0,002946 | TRUE | 0,0243231 | No |
| orf19.2812 | 357 | 297 | 351 | 205 | 1493 | 1036 | 1422 | 1502 | 340 | 314 | 347 | 321 | 1196 | 1178 | 1207 | 1221 | 765,65 | 330 | 1200 | 3,624 | 1,857 | 29,152 | 7,78E-187 | 9,81E-186 | 1,00E-08 | 0,0042358 | 0,0018535 | 0,0018535 | TRUE | 0,0150337 | No |
| orf19.2813 | 3 | 2 | 0 | 0 | 2 | 1 | 1 | 0 | 3 | 2 | 0 | 0 | 2 | 1 | 1 | 0 | 1,07 | 1 | 1 | 0,681 | -0,555 | -0,38 | 0,7043018 | 0,7346983 | 1,00E-08 | 1,5359443 | 1,1230488 | 1,1230488 | TRUE | 0,2187399 | No |
| orf19.2818 | 1591 | 1498 | 1656 | 1037 | 1417 | 1070 | 1336 | 1389 | 1517 | 1584 | 1636 | 1625 | 1135 | 1217 | 1134 | 1129 | 1372,13 | 1590 | 1154 | 0,725 | -0,464 | -9,258 | 2,07E-20 | 4,70E-20 | 0,0004559 | 0,0032879 | 0,0016671 | 0,0016671 | TRUE | 0,0148013 | No |
| orf19.2819 | 138 | 181 | 180 | 107 | 246 | 216 | 202 | 177 | 132 | 191 | 178 | 168 | 197 | 246 | 171 | 144 | 178,32 | 167 | 190 | 1,13 | 0,177 | 1,045 | 0,2960091 | 0,3286643 | 0,0329943 | 0,0112991 | 0,0217622 | 0,0217622 | TRUE | 0,3536463 | No |
| orf19.2820 | 4432 | 3959 | 4294 | 2769 | 4395 | 2972 | 4112 | 4330 | 4225 | 4186 | 4242 | 4338 | 3522 | 3379 | 3491 | 3521 | 3863,03 | 4248 | 3478 | 0,82 | -0,287 | -8,492 | 2,03E-17 | 4,31E-17 | 1,00E-08 | 0,0025163 | 0,0008339 | 0,0008339 | TRUE | 0,0041772 | No |
| orf19.2821 | 2194 | 2140 | 2187 | 1407 | 4125 | 3024 | 3903 | 4033 | 2092 | 2263 | 2161 | 2204 | 3305 | 3439 | 3314 | 3279 | 2756,98 | 2180 | 3334 | 1,53 | 0,613 | 14,996 | 7,81E-51 | 2,91E-50 | 0,000269 | 0,0026868 | 0,0012148 | 0,0012148 | TRUE | 0,0093941 | No |
| orf19.2822 | 255 | 258 | 239 | 154 | 179 | 143 | 166 | 183 | 243 | 273 | 236 | 241 | 143 | 163 | 141 | 149 | 198,63 | 248 | 149 | 0,597 | -0,744 | -7,003 | 2,50E-12 | 4,67E-12 | 1,00E-08 | 0,0103574 | 0,0055876 | 0,0055876 | TRUE | 0,0501688 | No |
| orf19.2823 | 3512 | 3236 | 3505 | 2598 | 3238 | 2774 | 3172 | 3498 | 3348 | 3422 | 3463 | 4070 | 2595 | 3154 | 2693 | 2844 | 3198,56 | 3576 | 2822 | 0,789 | -0,341 | -4,494 | 6,99E-06 | 1,04E-05 | 0,0076649 | 0,0026046 | 0,005219 | 0,005219 | TRUE | 0,1030058 | No |
| orf19.2825 | 7112 | 6561 | 6923 | 4081 | 3318 | 2297 | 3003 | 2875 | 6781 | 6937 | 6839 | 6393 | 2659 | 2612 | 2550 | 2338 | 4638,5 | 6738 | 2540 | 0,377 | -1,409 | -28,35 | 8,36E-177 | 9,82E-176 | 0,0018234 | 0,0024453 | 0,002112 | 0,002112 | TRUE | 0,0353087 | No |
| orf19.2826 | 2118 | 1840 | 2048 | 1246 | 1762 | 1143 | 1777 | 1828 | 2019 | 1946 | 2023 | 1952 | 1412 | 1300 | 1509 | 1486 | 1705,82 | 1985 | 1427 | 0,719 | -0,475 | -8,723 | 2,70E-18 | 5,84E-18 | 0,0016008 | 0,0030538 | 0,0022511 | 0,0022511 | TRUE | 0,0405258 | No |
| orf19.2827 | 296 | 348 | 374 | 194 | 442 | 302 | 390 | 389 | 282 | 368 | 369 | 304 | 354 | 343 | 331 | 316 | 333,57 | 331 | 336 | 1,014 | 0,019 | 0,19 | 0,8496084 | 0,8681107 | 0,0072285 | 0,0070136 | 0,0071179 | 0,0071179 | TRUE | 0,1252317 | No |
| orf19.2828 | 451 | 427 | 460 | 241 | 524 | 343 | 550 | 477 | 430 | 451 | 454 | 378 | 420 | 390 | 467 | 388 | 422,27 | 428 | 416 | 0,968 | -0,046 | -0,517 | 0,6054008 | 0,639474 | 0,0045964 | 0,0059796 | 0,0053063 | 0,0053063 | TRUE | 0,0788804 | No |
| orf19.2829 | 1627 | 1535 | 1548 | 907 | 2126 | 1398 | 2079 | 1906 | 1551 | 1623 | 1529 | 1421 | 1704 | 1590 | 1765 | 1550 | 1591,54 | 1531 | 1652 | 1,078 | 0,108 | 1,795 | 0,0727278 | 0,0860835 | 0,0024736 | 0,0031229 | 0,0028355 | 0,0028355 | TRUE | 0,0258282 | No |
| orf19.2830 | 10887 | 10294 | 10555 | 6740 | 8574 | 6245 | 8239 | 8291 | 10380 | 10885 | 10427 | 10559 | 6870 | 7101 | 6995 | 6741 | 8744,7 | 10563 | 6927 | 0,656 | -0,609 | -18,593 | 3,65E-77 | 1,92E-76 | 0,0003252 | 0,002279 | 0,0009124 | 0,0009124 | TRUE | 0,0052369 | No |
| orf19.2831 | 2771 | 2448 | 2656 | 1440 | 4788 | 2926 | 4586 | 4473 | 2642 | 2588 | 2624 | 2256 | 3836 | 3327 | 3893 | 3637 | 3100,51 | 2528 | 3673 | 1,452 | 0,538 | 8,181 | 2,80E-16 | 5,80E-16 | 0,0045846 | 0,0026208 | 0,0038103 | 0,0038103 | TRUE | 0,0606352 | No |
| orf19.2832 | 4022 | 3903 | 4244 | 2561 | 1332 | 994 | 1300 | 1407 | 3835 | 4127 | 4193 | 4012 | 1067 | 1130 | 1104 | 1144 | 2576,42 | 4042 | 1111 | 0,275 | -1,863 | -39,331 | 0 | 0 | 0,0009572 | 0,0027286 | 0,0016145 | 0,0016145 | TRUE | 0,0150889 | No |
| orf19.2833 | 88 | 84 | 100 | 75 | 39 | 37 | 36 | 34 | 84 | 89 | 99 | 117 | 31 | 42 | 31 | 28 | 65,07 | 97 | 33 | 0,337 | -1,57 | -7,803 | 6,04E-15 | 1,22E-14 | 0,0070695 | 0,0273256 | 0,019396 | 0,019396 | TRUE | 0,2041744 | No |

| | | | | | | | | | | | | | | | | | | | | | | | | | | | | | | | |
|---|---|---|---|---|---|---|---|---|---|---|---|---|---|---|---|---|---|---|---|---|---|---|---|---|---|---|---|---|---|---|---|
| orf19.2834 | 3465 | 3230 | 3535 | 2148 | 1883 | 1284 | 1748 | 1787 | 3304 | 3415 | 3492 | 3365 | 1509 | 1460 | 1484 | 1453 | 2435,24 | 3394 | 1476 | 0,435 | -1,201 | -29,955 | 3,76E-197 | 5,07E-196 | 1,00E-08 | 0,0027655 | 0,001075 | 0,001075 | TRUE | 0,0048015 | No |
| orf19.2835 | 591 | 618 | 639 | 359 | 1070 | 870 | 974 | 1068 | 563 | 653 | 631 | 562 | 857 | 989 | 827 | 868 | 744,06 | 602 | 885 | 1,464 | 0,55 | 7,025 | 2,14E-12 | 4,00E-12 | 0,004399 | 0,0042979 | 0,0044498 | 0,0044498 | TRUE | 0,070616 | No |
| orf19.2836 | 262 | 178 | 205 | 143 | 505 | 306 | 448 | 467 | 250 | 188 | 203 | 224 | 405 | 348 | 380 | 380 | 297,15 | 216 | 378 | 1,754 | 0,81 | 7,88 | 3,27E-15 | 6,61E-15 | 0,0047474 | 0,0076168 | 0,0063891 | 0,0063891 | TRUE | 0,1369375 | No |
| orf19.2837 | 204 | 174 | 176 | 112 | 259 | 204 | 245 | 266 | 194 | 184 | 174 | 175 | 208 | 232 | 208 | 216 | 198,95 | 182 | 216 | 1,181 | 0,24 | 2,329 | 0,0198338 | 0,0246225 | 1,00E-08 | 0,0103442 | 0,0050492 | 0,0050492 | TRUE | 0,0248799 | No |
| orf19.2838 | 2453 | 2390 | 2564 | 1499 | 342 | 286 | 318 | 302 | 2339 | 2527 | 2533 | 2348 | 274 | 325 | 270 | 246 | 1357,73 | 2437 | 279 | 0,114 | -3,138 | -42,369 | 0 | 0 | 0,0036321 | 0,0033006 | 0,003431 | 0,003431 | TRUE | 0,1255042 | No |
| orf19.2839 | 4738 | 4177 | 4483 | 2617 | 459 | 293 | 440 | 454 | 4517 | 4417 | 4429 | 4100 | 368 | 333 | 374 | 369 | 2363,24 | 4366 | 361 | 0,083 | -3,593 | -61,758 | 0 | 0 | 0,0011694 | 0,002786 | 0,0018947 | 0,0018947 | TRUE | 0,0260794 | No |
| orf19.284 | 117 | 84 | 82 | 72 | 192 | 102 | 166 | 207 | 112 | 89 | 81 | 113 | 154 | 116 | 141 | 168 | 121,65 | 99 | 145 | 1,485 | 0,571 | 3,556 | 0,0003768 | 0,0005199 | 0,0163877 | 0,0155877 | 0,0159425 | 0,0159425 | TRUE | 0,1696117 | No |
| orf19.2841 | 8829 | 9009 | 9147 | 5089 | 2417 | 1669 | 2249 | 2202 | 8417 | 9526 | 9036 | 7972 | 1937 | 1898 | 1909 | 1790 | 5310,78 | 8738 | 1884 | 0,215 | -2,215 | -37,186 | 1,16E-302 | 2,60E-301 | 0,0032043 | 0,0024004 | 0,0031045 | 0,0031045 | TRUE | 0,0448397 | No |
| orf19.2842 | 9774 | 8734 | 9780 | 6401 | 6915 | 4965 | 6813 | 6977 | 9318 | 9235 | 9662 | 10028 | 5541 | 5646 | 5784 | 5673 | 7610,8 | 9561 | 5661 | 0,592 | -0,755 | -20,074 | 1,25E-89 | 7,47E-89 | 0,0006848 | 0,002307 | 0,0012218 | 0,0012218 | TRUE | 0,0127658 | No |
| orf19.2843 | 4508 | 3923 | 3958 | 2346 | 10013 | 6064 | 9152 | 9565 | 4298 | 4148 | 3910 | 3675 | 8023 | 6895 | 7770 | 7777 | 5812,12 | 4008 | 7616 | 1,899 | 0,926 | 14,925 | 2,27E-50 | 8,39E-50 | 0,0041821 | 0,0023738 | 0,0034947 | 0,0034947 | TRUE | 0,0491695 | No |
| orf19.2844 | 724 | 754 | 789 | 436 | 960 | 686 | 903 | 913 | 690 | 797 | 779 | 683 | 769 | 780 | 767 | 742 | 751,03 | 737 | 764 | 1,034 | 0,048 | 0,714 | 0,4750908 | 0,5114821 | 0,0016677 | 0,0042775 | 0,0029843 | 0,0029843 | TRUE | 0,0342215 | No |
| orf19.2846 | 342 | 291 | 361 | 215 | 580 | 381 | 510 | 511 | 326 | 308 | 357 | 337 | 465 | 433 | 433 | 415 | 384,21 | 332 | 436 | 1,317 | 0,397 | 4,899 | 9,62E-07 | 1,49E-06 | 1,00E-08 | 0,0063648 | 0,0036031 | 0,0036031 | TRUE | 0,0322315 | No |
| orf19.2847 | 3999 | 4431 | 4452 | 2800 | 3301 | 2697 | 3027 | 2988 | 3813 | 4685 | 4398 | 4386 | 2645 | 3067 | 2570 | 2430 | 3499,19 | 4320 | 2678 | 0,619 | -0,691 | -8,699 | 3,34E-18 | 7,20E-18 | 0,0084241 | 0,0025605 | 0,0057615 | 0,0057615 | TRUE | 0,1144585 | No |
| orf19.2848 | 2954 | 2498 | 2742 | 1786 | 2605 | 1791 | 2411 | 2616 | 2816 | 2641 | 2709 | 2798 | 2087 | 2037 | 2047 | 2127 | 2407,77 | 2741 | 2074 | 0,757 | -0,401 | -9,813 | 9,95E-23 | 2,35E-22 | 0,0002501 | 0,0027732 | 0,0011811 | 0,0011811 | TRUE | 0,0073197 | No |
| orf19.2849 | 24746 | 19196 | 20827 | 12641 | 20438 | 12706 | 18850 | 20717 | 23593 | 20297 | 20575 | 19803 | 16376 | 14448 | 16004 | 16845 | 18492,57 | 21067 | 15918 | 0,756 | -0,404 | -6,214 | 5,18E-10 | 9,00E-10 | 0,0053651 | 0,0021801 | 0,0040131 | 0,0040131 | TRUE | 0,0798711 | No |
| orf19.285 | 6130 | 5272 | 5735 | 3081 | 2044 | 1208 | 1884 | 1913 | 5844 | 5574 | 5666 | 4827 | 1638 | 1374 | 1600 | 1555 | 3509,66 | 5478 | 1542 | 0,282 | -1,829 | -25,7 | 1,17E-145 | 1,14E-144 | 0,0056725 | 0,0025591 | 0,0044697 | 0,0044697 | TRUE | 0,0766041 | No |
| orf19.2850 | 6961 | 6192 | 6785 | 4059 | 2633 | 1763 | 2436 | 2630 | 6637 | 6547 | 6703 | 6359 | 2110 | 2005 | 2068 | 2138 | 4320,8 | 6562 | 2080 | 0,317 | -1,657 | -44,043 | 0 | 0 | 0,0002267 | 0,0024713 | 0,0010585 | 0,0010585 | TRUE | 0,0067144 | No |
| orf19.2851 | 857 | 884 | 908 | 602 | 1270 | 981 | 1204 | 1302 | 817 | 935 | 897 | 943 | 1018 | 1115 | 1022 | 1059 | 975,72 | 898 | 1054 | 1,175 | 0,232 | 3,701 | 0,0002151 | 0,0002998 | 0,0018303 | 0,003774 | 0,0027272 | 0,0027272 | TRUE | 0,0455238 | No |
| orf19.2852 | 3546 | 3193 | 3264 | 2106 | 3388 | 2453 | 3178 | 3296 | 3381 | 3376 | 3225 | 3299 | 2715 | 2789 | 2698 | 2680 | 3020,33 | 3320 | 2720 | 0,819 | -0,288 | -7,825 | 5,10E-15 | 1,03E-14 | 1,00E-08 | 0,0026349 | 0,0009708 | 0,0009708 | TRUE | 0,0048876 | No |
| orf19.2853 | 1457 | 1446 | 1509 | 1029 | 488 | 410 | 467 | 476 | 1389 | 1529 | 1491 | 1612 | 391 | 466 | 396 | 387 | 957,69 | 1505 | 410 | 0,272 | -1,879 | -25,444 | 8,17E-143 | 7,84E-142 | 0,0040089 | 0,0038057 | 0,0037727 | 0,0037727 | TRUE | 0,0888524 | No |
| orf19.2857 | 8865 | 7774 | 8404 | 5119 | 8847 | 5896 | 8178 | 8954 | 8452 | 8220 | 8302 | 8019 | 7089 | 6704 | 6943 | 7280 | 7626,26 | 8248 | 7004 | 0,849 | -0,236 | -6,346 | 2,21E-10 | 3,91E-10 | 0,0006651 | 0,0023066 | 0,0011949 | 0,0011949 | TRUE | 0,0098906 | No |
| orf19.2859 | 7255 | 6151 | 6816 | 4134 | 12642 | 8194 | 11783 | 12415 | 6917 | 6504 | 6734 | 6476 | 10130 | 9317 | 10004 | 10095 | 8271,97 | 6658 | 9886 | 1,485 | 0,57 | 13,82 | 1,94E-43 | 6,49E-43 | 0,0010701 | 0,0022898 | 0,0015054 | 0,0015054 | TRUE | 0,0180703 | No |
| orf19.286 | 80 | 67 | 74 | 55 | 68 | 56 | 57 | 71 | 76 | 71 | 73 | 86 | 54 | 64 | 48 | 58 | 66,33 | 76 | 56 | 0,733 | -0,448 | -2,563 | 0,0103921 | 0,0131885 | 1,00E-08 | 0,0268436 | 0,014054 | 0,014054 | TRUE | 0,073502 | No |
| orf19.2862 | 4231 | 3440 | 3887 | 2436 | 3909 | 2427 | 3663 | 3950 | 4034 | 3637 | 3840 | 3816 | 3132 | 2760 | 3110 | 3212 | 3442,6 | 3832 | 3054 | 0,797 | -0,327 | -5,851 | 4,89E-09 | 8,21E-09 | 0,0026615 | 0,0025682 | 0,0027004 | 0,0027004 | TRUE | 0,0495453 | No |
| orf19.2863 | 447 | 365 | 409 | 250 | 620 | 455 | 586 | 676 | 426 | 386 | 404 | 392 | 497 | 517 | 498 | 550 | 458,64 | 402 | 516 | 1,279 | 0,356 | 4,795 | 1,63E-06 | 2,49E-06 | 1,00E-08 | 0,0056712 | 0,0030303 | 0,0030303 | TRUE | 0,0250892 | No |
| orf19.2863.1 | 593 | 458 | 545 | 315 | 895 | 573 | 864 | 902 | 565 | 484 | 538 | 493 | 717 | 652 | 734 | 733 | 614,64 | 520 | 709 | 1,361 | 0,444 | 6,025 | 1,69E-09 | 2,88E-09 | 0,0022675 | 0,0047626 | 0,0035184 | 0,0035184 | TRUE | 0,0406013 | No |
| orf19.2864.1 | 20529 | 16209 | 17035 | 10274 | 51043 | 29880 | 48779 | 50891 | 19572 | 17139 | 16829 | 16095 | 40899 | 33976 | 41413 | 41379 | 28412,85 | 17409 | 39417 | 2,264 | 1,179 | 15,583 | 9,43E-55 | 3,71E-54 | 0,0079447 | 0,0021491 | 0,0054551 | 0,0054551 | TRUE | 0,1056429 | No |
| orf19.2866 | 109 | 74 | 80 | 59 | 79 | 61 | 67 | 95 | 104 | 78 | 79 | 92 | 63 | 69 | 57 | 77 | 77,55 | 88 | 66 | 0,754 | -0,407 | -2,333 | 0,0196424 | 0,0243991 | 0,0072231 | 0,023263 | 0,0161674 | 0,0161674 | TRUE | 0,1526919 | No |
| orf19.2867 | 4169 | 3762 | 4061 | 2608 | 2563 | 1794 | 2510 | 2579 | 3975 | 3978 | 4012 | 4086 | 2054 | 2040 | 2131 | 2097 | 3046,44 | 4013 | 2080 | 0,519 | -0,946 | -26,442 | 4,51E-154 | 4,65E-153 | 1,00E-08 | 0,0026302 | 0,0008752 | 0,0008752 | TRUE | 0,0032181 | No |
| orf19.2868 | 3 | 4 | 4 | 2 | 0 | 1 | 3 | 2 | 3 | 4 | 4 | 3 | 0 | 1 | 3 | 2 | 2,44 | 4 | 2 | 0,373 | -1,424 | -1,505 | 0,1322646 | 0,152591 | 1,00E-08 | 0,6762229 | 0,3679736 | 0,3679736 | TRUE | 0,3850097 | No |
| orf19.2869 | 5 | 2 | 5 | 0 | 1 | 4 | 1 | 6 | 5 | 2 | 5 | 0 | 1 | 5 | 1 | 5 | 2,86 | 3 | 3 | 0,873 | -0,196 | -0,208 | 0,8353322 | 0,8551668 | 0,4006095 | 0,5757213 | 0,5093227 | 0,5093227 | TRUE | 0,2003908 | No |
| orf19.287 | 4030 | 3267 | 3560 | 2283 | 8944 | 5756 | 8578 | 9054 | 3842 | 3454 | 3517 | 3576 | 7167 | 6545 | 7283 | 7362 | 5343,26 | 3597 | 7089 | 1,971 | 0,979 | 18,898 | 1,19E-79 | 6,47E-79 | 0,0022167 | 0,0023986 | 0,0023565 | 0,0023565 | TRUE | 0,0321182 | No |
| orf19.2870 | 2828 | 2341 | 2453 | 1299 | 1213 | 704 | 997 | 1117 | 2696 | 2475 | 2423 | 2035 | 972 | 801 | 846 | 908 | 1644,61 | 2407 | 882 | 0,366 | -1,449 | -16,65 | 3,02E-62 | 1,31E-61 | 0,00907 | 0,0030896 | 0,0065355 | 0,0065355 | TRUE | 0,1273913 | No |
| orf19.2871 | 7784 | 6696 | 7059 | 4763 | 21785 | 14932 | 20704 | 23039 | 7421 | 7080 | 6974 | 7461 | 17456 | 16979 | 17578 | 18733 | 12460,22 | 7234 | 17686 | 2,446 | 1,29 | 30,269 | 2,92E-201 | 4,01E-200 | 0,0013227 | 0,0022231 | 0,0016421 | 0,0016421 | TRUE | 0,0194503 | No |
| orf19.2873 | 2513 | 2349 | 2581 | 1551 | 4342 | 3125 | 3950 | 4395 | 2396 | 2484 | 2550 | 2430 | 3479 | 3553 | 3354 | 3574 | 2977,35 | 2465 | 3490 | 1,415 | 0,501 | 12,254 | 1,60E-34 | 4,64E-34 | 0,0004941 | 0,0026427 | 0,0012485 | 0,0012485 | TRUE | 0,0084744 | No |
| orf19.2875 | 1788 | 1654 | 1710 | 961 | 2831 | 1796 | 2576 | 2641 | 1705 | 1749 | 1689 | 1505 | 2268 | 2042 | 2187 | 2147 | 1911,67 | 1662 | 2161 | 1,298 | 0,377 | 6,646 | 3,01E-11 | 5,46E-11 | 0,0023251 | 0,0029502 | 0,0025383 | 0,0025383 | TRUE | 0,0442966 | No |
| orf19.2876 | 3371 | 2850 | 3096 | 1994 | 1119 | 772 | 1114 | 1145 | 3214 | 3014 | 3059 | 3124 | 897 | 878 | 946 | 931 | 2007,61 | 3103 | 913 | 0,295 | -1,763 | -37,908 | 0 | 0 | 0,000411 | 0,0029091 | 0,0014089 | 0,0014089 | TRUE | 0,0075278 | No |
| orf19.2877 | 28896 | 24558 | 26727 | 18340 | 17496 | 12468 | 16708 | 18779 | 27549 | 25967 | 26404 | 28731 | 14019 | 14177 | 14185 | 15269 | 20787,62 | 27163 | 14412 | 0,531 | -0,914 | -20,286 | 1,69E-91 | 1,03E-90 | 0,0017703 | 0,0021703 | 0,0018984 | 0,0018984 | TRUE | 0,0196592 | No |
| orf19.2878 | 81 | 72 | 95 | 48 | 41 | 22 | 49 | 41 | 77 | 76 | 94 | 75 | 33 | 25 | 42 | 33 | 56,9 | 80 | 33 | 0,415 | -1,27 | -6,229 | 4,69E-10 | 8,17E-10 | 1,00E-08 | 0,0309468 | 0,0197618 | 0,0197618 | TRUE | 0,2056004 | No |
| orf19.2879 | 230 | 193 | 211 | 154 | 214 | 134 | 169 | 227 | 219 | 204 | 208 | 241 | 171 | 152 | 143 | 185 | 190,62 | 218 | 163 | 0,752 | -0,411 | -3,544 | 0,0003942 | 0,0005429 | 0,0036936 | 0,0107049 | 0,0075964 | 0,0075964 | TRUE | 0,0879261 | No |
| orf19.288 | 2035 | 1854 | 2040 | 1341 | 8499 | 6276 | 8146 | 8760 | 1940 | 1960 | 2015 | 2101 | 6810 | 7136 | 6916 | 7123 | 4500,2 | 2004 | 6996 | 3,497 | 1,806 | 45,478 | 0 | 0 | 0,0004753 | 0,0024561 | 0,00117 | 0,00117 | TRUE | 0,0112744 | No |
| orf19.2881 | 8320 | 8986 | 8936 | 5199 | 7235 | 5496 | 6510 | 6715 | 7932 | 9502 | 8828 | 8145 | 5797 | 6249 | 5527 | 5460 | 7179,97 | 8602 | 5758 | 0,669 | -0,58 | -8,87 | 7,34E-19 | 1,61E-18 | 0,0051631 | 0,00232 | 0,0039581 | 0,0039581 | TRUE | 0,0606863 | No |
| orf19.2882 | 2708 | 2496 | 2670 | 1610 | 5300 | 3801 | 5150 | 5537 | 2582 | 2639 | 2638 | 2522 | 4247 | 4322 | 4372 | 4502 | 3478,01 | 2595 | 4361 | 1,679 | 0,748 | 19,865 | 8,10E-88 | 4,75E-87 | 0,0002882 | 0,0025634 | 0,0010413 | 0,0010413 | TRUE | 0,0059165 | No |
| orf19.2883 | 1281 | 1229 | 1243 | 886 | 3312 | 2594 | 3089 | 3123 | 1221 | 1300 | 1228 | 1388 | 2654 | 2950 | 2623 | 2539 | 1987,75 | 1284 | 2692 | 2,099 | 1,07 | 17,168 | 4,61E-66 | 2,10E-65 | 0,0032726 | 0,0029173 | 0,0031219 | 0,0031219 | TRUE | 0,0494467 | No |
| orf19.2884 | 4127 | 4181 | 4262 | 2427 | 5509 | 3886 | 4958 | 5010 | 3935 | 4421 | 4210 | 3802 | 4414 | 4419 | 4209 | 4074 | 4185,48 | 4092 | 4279 | 1,045 | 0,063 | 1,151 | 0,2497148 | 0,2798469 | 0,0027532 | 0,0024836 | 0,0026516 | 0,0026516 | TRUE | 0,0353873 | No |
| orf19.2885 | 54 | 63 | 52 | 33 | 23 | 23 | 24 | 24 | 51 | 67 | 51 | 52 | 18 | 26 | 20 | 20 | 38,2 | 55 | 21 | 0,377 | -1,409 | -5,791 | 7,00E-09 | 1,17E-08 | 1,00E-08 | 0,0450678 | 0,0254603 | 0,0254603 | TRUE | 0,1614495 | No |
| orf19.2886 | 964 | 1134 | 1136 | 773 | 1287 | 1079 | 1213 | 1194 | 919 | 1199 | 1122 | 1211 | 1031 | 1227 | 1030 | 971 | 1088,77 | 1113 | 1065 | 0,957 | -0,063 | -0,657 | 0,5110868 | 0,5471887 | 0,0121198 | 0,0035993 | 0,0080131 | 0,0080131 | TRUE | 0,1675609 | No |
| orf19.2887 | 4514 | 3934 | 4215 | 2808 | 6046 | 4110 | 5692 | 6089 | 4304 | 4160 | 4164 | 4399 | 4844 | 4673 | 4832 | 4951 | 4540,95 | 4257 | 4825 | 1,135 | 0,183 | 4,981 | 6,34E-07 | 9,89E-07 | 0,000358 | 0,0024529 | 0,0010655 | 0,0010655 | TRUE | 0,005699 | No |
| orf19.2888 | 170 | 223 | 197 | 98 | 497 | 293 | 406 | 421 | 162 | 236 | 195 | 154 | 398 | 333 | 345 | 342 | 270,55 | 187 | 354 | 1,893 | 0,921 | 7,206 | 5,78E-13 | 1,10E-12 | 0,0156446 | 0,00816 | 0,011436 | 0,011436 | TRUE | 0,3344601 | No |
| orf19.2889 | 861 | 785 | 831 | 502 | 769 | 488 | 678 | 672 | 821 | 830 | 821 | 786 | 616 | 555 | 576 | 546 | 693,92 | 814 | 573 | 0,704 | -0,507 | -8,083 | 6,34E-16 | 1,30E-15 | 1,00E-08 | 0,0044574 | 0,002318 | 0,002318 | TRUE | 0,0311747 | No |
| orf19.2890 | 1012 | 986 | 919 | 623 | 1573 | 1220 | 1380 | 1429 | 965 | 1043 | 908 | 976 | 1260 | 1387 | 1172 | 1162 | 1109,05 | 973 | 1245 | 1,279 | 0,355 | 4,981 | 6,32E-07 | 9,87E-07 | 0,0039127 | 0,0035717 | 0,0039306 | 0,0039306 | TRUE | 0,0678918 | No |
| orf19.2891 | 1006 | 1074 | 1112 | 722 | 1814 | 1448 | 1673 | 1667 | 959 | 1136 | 1099 | 1131 | 1454 | 1647 | 1420 | 1355 | 1275,02 | 1081 | 1469 | 1,359 | 0,443 | 5,834 | 5,42E-09 | 9,09E-09 | 0,0055058 | 0,003379 | 0,0046992 | 0,0046992 | TRUE | 0,0774308 | No |
| orf19.2892 | 3694 | 4010 | 4122 | 2655 | 2427 | 2005 | 2347 | 2390 | 3522 | 4240 | 4072 | 4159 | 1945 | 2280 | 1993 | 1943 | 3019,21 | 3998 | 2040 | 0,51 | -0,971 | -13,546 | 8,33E-42 | 2,72E-41 | 0,0059417 | 0,0026351 | 0,0045793 | 0,0045793 | TRUE | 0,0792167 | No |
| orf19.2893 | 903 | 1121 | 1176 | 642 | 996 | 821 | 844 | 776 | 861 | 1185 | 1162 | 1006 | 798 | 934 | 717 | 631 | 911,61 | 1054 | 770 | 0,73 | -0,455 | -3,696 | 0,0002188 | 0,0003048 | 0,0227703 | 0,0038924 | 0,0134309 | 0,0134309 | TRUE | 0,2380607 | No |
| orf19.2895 | 696 | 590 | 645 | 387 | 1364 | 922 | 1221 | 1263 | 664 | 624 | 637 | 606 | 1093 | 1048 | 1037 | 1027 | 841,97 | 633 | 1051 | 1,657 | 0,729 | 12,694 | 6,40E-37 | 1,93E-36 | 1,00E-08 | 0,0040414 | 0,0018621 | 0,0018621 | TRUE | 0,0128338 | No |
| orf19.2896 | 1669 | 1318 | 1522 | 956 | 3700 | 2528 | 3548 | 3739 | 1591 | 1394 | 1504 | 1498 | 2965 | 2875 | 3012 | 3040 | 2234,71 | 1497 | 2973 | 1,986 | 0,99 | 20,122 | 4,75E-90 | 2,85E-89 | 0,0010875 | 0,002826 | 0,0018 | 0,0018 | TRUE | 0,0269375 | No |
| orf19.2897 | 112 | 80 | 105 | 69 | 137 | 89 | 96 | 127 | 107 | 85 | 104 | 108 | 110 | 101 | 82 | 103 | 99,87 | 101 | 99 | 0,984 | -0,023 | -0,151 | 0,8798432 | 0,8965633 | 0,0051185 | 0,0185322 | 0,0126818 | 0,0126818 | TRUE | 0,1470815 | No |
| orf19.2898 | 1419 | 1360 | 1449 | 910 | 1695 | 1129 | 1592 | 1658 | 1353 | 1438 | 1431 | 1426 | 1358 | 1284 | 1352 | 1348 | 1373,7 | 1412 | 1336 | 0,947 | -0,078 | -1,65 | 0,0989252 | 0,1155909 | 1,00E-08 | 0,0032865 | 0,001415 | 0,001415 | TRUE | 0,0102197 | No |
| orf19.2899 | 112 | 90 | 121 | 75 | 72 | 50 | 66 | 87 | 107 | 95 | 120 | 117 | 58 | 57 | 56 | 71 | 85,04 | 110 | 60 | 0,553 | -0,855 | -5,273 | 1,34E-07 | 2,15E-07 | 1,00E-08 | 0,0213995 | 0,0127997 | 0,0127997 | TRUE | 0,128863 | No |
| orf19.29 | 5432 | 5613 | 5711 | 3754 | 2591 | 1960 | 2382 | 2224 | 5179 | 5935 | 5642 | 5881 | 2076 | 2229 | 2022 | 1808 | 3846,51 | 5659 | 2034 | 0,359 | -1,477 | -21,901 | 2,56E-106 | 1,77E-105 | 0,0050647 | 0,0025181 | 0,0040496 | 0,0040496 | TRUE | 0,0677959 | No |
| orf19.290 | 1005 | 913 | 1046 | 578 | 1398 | 996 | 1268 | 1364 | 958 | 965 | 1033 | 905 | 1120 | 1133 | 1077 | 1109 | 1037,58 | 965 | 1110 | 1,146 | 0,196 | 3,552 | 0,0003817 | 0,0005263 | 0,00042 | 0,0036737 | 0,0019426 | 0,0019426 | TRUE | 0,0264954 | No |
| orf19.2901 | 2995 | 2899 | 3018 | 2073 | 1998 | 1521 | 1895 | 1983 | 2855 | 3065 | 2981 | 3247 | 1601 | 1730 | 1609 | 1612 | 2337,67 | 3037 | 1638 | 0,539 | -0,89 | -17,239 | 1,36E-66 | 6,23E-66 | 0,0015387 | 0,0027936 | 0,0021031 | 0,0021031 | TRUE | 0,0249514 | No |
| orf19.2903 | 195 | 270 | 237 | 138 | 644 | 530 | 568 | 628 | 186 | 285 | 234 | 216 | 516 | 603 | 482 | 511 | 379,16 | 230 | 528 | 2,286 | 1,193 | 9,799 | 1,14E-22 | 2,70E-22 | 0,017114 | 0,0064217 | 0,0109465 | 0,0109465 | TRUE | 0,2919843 | No |
| orf19.2905 | 16 | 18 | 15 | 4 | 12 | 8 | 12 | 7 | 15 | 19 | 15 | 6 | 10 | 9 | 10 | 6 | 11,25 | 14 | 9 | 0,607 | -0,72 | -1,636 | 0,1018465 | 0,1188085 | 1,00E-08 | 0,1480977 | 0,0953297 | 0,0953297 | TRUE | 0,3540087 | No |
| orf19.2906 | 161 | 123 | 158 | 111 | 190 | 130 | 167 | 156 | 153 | 130 | 156 | 174 | 152 | 148 | 142 | 127 | 147,78 | 153 | 142 | 0,931 | -0,103 | -0,81 | 0,4178472 | 0,4538354 | 0,0024268 | 0,0132019 | 0,00867 | 0,00867 | TRUE | 0,1068381 | No |
| orf19.2907 | 4 | 6 | 6 | 7 | 4 | 7 | 5 | 5 | 4 | 6 | 6 | 11 | 3 | 8 | 4 | 4 | 5,82 | 7 | 5 | 0,727 | -0,46 | -0,763 | 0,4457471 | 0,4822503 | 1,00E-08 | 0,2844097 | 0,1699112 | 0,1699112 | TRUE | 0,2956724 | No |
| orf19.2907.1 | 353 | 258 | 294 | 150 | 356 | 183 | 313 | 329 | 337 | 273 | 290 | 235 | 285 | 208 | 266 | 268 | 270,17 | 284 | 257 | 0,903 | -0,147 | -1,169 | 0,2422153 | 0,2720126 | 0,0153978 | 0,0081686 | 0,0115599 | 0,0115599 | TRUE | 0,1766836 | No |
| orf19.2908 | 353 | 318 | 333 | 244 | 586 | 439 | 637 | 666 | 337 | 336 | 329 | 382 | 470 | 499 | 541 | 542 | 429,38 | 346 | 513 | 1,49 | 0,576 | 6,93 | 4,22E-12 | 7,84E-12 | 0,0023536 | 0,0059151 | 0,0041344 | 0,0041344 | TRUE | 0,0470418 | No |
| orf19.2909 | 942 | 877 | 919 | 582 | 1390 | 1134 | 1298 | 1437 | 898 | 927 | 908 | 912 | 1114 | 1289 | 1102 | 1168 | 1039,83 | 911 | 1168 | 1,28 | 0,356 | 5,742 | 9,37E-09 | 1,56E-08 | 0,0019468 | 0,0036703 | 0,0026941 | 0,0026941 | TRUE | 0,0550917 | No |
| orf19.291 | 1536 | 1329 | 1452 | 908 | 3025 | 1968 | 2798 | 3073 | 1464 | 1405 | 1434 | 1422 | 2424 | 2238 | 2375 | 2499 | 1907,79 | 1431 | 2384 | 1,666 | 0,737 | 15,225 | 2,41E-52 | 9,20E-52 | 0,0008751 | 0,0029519 | 0,0016684 | 0,0016684 | TRUE | 0,0196513 | No |
| orf19.2910 | 130 | 98 | 107 | 81 | 280 | 198 | 262 | 267 | 124 | 104 | 106 | 127 | 224 | 225 | 222 | 217 | 168,65 | 115 | 222 | 1,942 | 0,958 | 8,282 | 1,21E-16 | 2,52E-16 | 1,00E-08 | 0,0118269 | 0,0059783 | 0,0059783 | TRUE | 0,0421379 | No |
| orf19.2911 | 2162 | 1930 | 2137 | 1251 | 1538 | 1124 | 1475 | 1593 | 2061 | 2041 | 2111 | 1960 | 1232 | 1278 | 1252 | 1295 | 1653,86 | 2043 | 1264 | 0,618 | -0,695 | -15,407 | 1,47E-53 | 5,72E-53 | 1,00E-08 | 0,0030841 | 0,001326 | 0,001326 | TRUE | 0,008159 | No |
| orf19.2914 | 688 | 542 | 637 | 376 | 481 | 349 | 449 | 500 | 656 | 573 | 629 | 589 | 385 | 397 | 381 | 407 | 502,17 | 612 | 392 | 0,64 | -0,644 | -8,862 | 7,89E-19 | 1,73E-18 | 1,00E-08 | 0,0053609 | 0,0030263 | 0,0030263 | TRUE | 0,0286282 | No |

| | | | | | | | | | | | | | | | | | | | | | | | | | | | | | | | |
|---|---|---|---|---|---|---|---|---|---|---|---|---|---|---|---|---|---|---|---|---|---|---|---|---|---|---|---|---|---|---|---|
| orf19.2915 | 3275 | 3220 | 3381 | 2047 | 2495 | 1726 | 2300 | 2509 | 3122 | 3405 | 3340 | 3207 | 1999 | 1963 | 1953 | 2040 | 2628,56 | 3268 | 1989 | 0,608 | -0,717 | -16,529 | 2,27E-61 | 9,79E-61 | 0,0006796 | 0,0027159 | 0,0014088 | 0,0014088 | TRUE | 0,0117481 | No |
| orf19.2916 | 63 | 79 | 85 | 32 | 47 | 29 | 37 | 55 | 60 | 84 | 84 | 50 | 38 | 33 | 31 | 45 | 53,06 | 70 | 37 | 0,526 | -0,928 | -4,078 | 4,55E-05 | 6,54E-05 | 0,0251935 | 0,0330366 | 0,0296347 | 0,0296347 | TRUE | 0,1495663 | No |
| orf19.2917 | 15512 | 13669 | 14351 | 8884 | 24560 | 16868 | 24291 | 25457 | 14789 | 14453 | 14177 | 13917 | 19679 | 19180 | 20623 | 20699 | 17189,81 | 14334 | 20045 | 1,398 | 0,484 | 12,683 | 7,32E-37 | 2,20E-36 | 0,0009419 | 0,0021868 | 0,0013352 | 0,0013352 | TRUE | 0,0102352 | No |
| orf19.2917.1 | 715 | 550 | 585 | 455 | 234 | 172 | 243 | 243 | 682 | 582 | 578 | 713 | 187 | 196 | 206 | 198 | 417,61 | 639 | 197 | 0,309 | -1,694 | -17,68 | 6,02E-70 | 2,88E-69 | 0,0053472 | 0,0060229 | 0,0056731 | 0,0056731 | TRUE | 0,0648189 | No |
| orf19.2919 | 599 | 558 | 585 | 355 | 625 | 422 | 592 | 650 | 571 | 590 | 578 | 556 | 501 | 480 | 503 | 529 | 538,36 | 574 | 503 | 0,877 | -0,19 | -2,886 | 0,0038967 | 0,0050756 | 1,00E-08 | 0,0051411 | 0,0022927 | 0,0022927 | TRUE | 0,0137367 | No |
| orf19.292 | 106 | 77 | 70 | 60 | 282 | 156 | 233 | 239 | 101 | 81 | 69 | 94 | 226 | 177 | 198 | 194 | 142,64 | 86 | 199 | 2,317 | 1,212 | 8,352 | 6,69E-17 | 1,40E-16 | 0,0086192 | 0,0136022 | 0,0115215 | 0,0115215 | TRUE | 0,1852827 | No |
| orf19.2920 | 2723 | 2173 | 2371 | 1479 | 4427 | 3045 | 4277 | 4607 | 2596 | 2298 | 2342 | 2317 | 3547 | 3462 | 3631 | 3746 | 2992,47 | 2388 | 3596 | 1,505 | 0,59 | 11,561 | 6,51E-31 | 1,78E-30 | 0,0018137 | 0,00264 | 0,0021403 | 0,0021403 | TRUE | 0,0427603 | No |
| orf19.2921 | 105 | 121 | 114 | 80 | 504 | 377 | 522 | 537 | 100 | 128 | 113 | 125 | 404 | 429 | 443 | 437 | 272,29 | 116 | 428 | 3,702 | 1,888 | 17,953 | 4,56E-72 | 2,25E-71 | 1,00E-08 | 0,0081212 | 0,0047862 | 0,0047862 | TRUE | 0,0981508 | No |
| orf19.2922 | 31 | 14 | 17 | 12 | 25 | 20 | 17 | 16 | 30 | 15 | 17 | 19 | 20 | 23 | 14 | 13 | 18,77 | 20 | 18 | 0,864 | -0,212 | -0,597 | 0,5502907 | 0,5855269 | 0,0379132 | 0,0895618 | 0,0670563 | 0,0670563 | TRUE | 0,6278718 | No |
| orf19.2923 | 288 | 260 | 245 | 170 | 419 | 288 | 458 | 387 | 275 | 275 | 242 | 266 | 336 | 327 | 389 | 315 | 303,07 | 264 | 342 | 1,293 | 0,371 | 3,813 | 0,0001373 | 0,000193 | 0,0038827 | 0,0075088 | 0,0056824 | 0,0056824 | TRUE | 0,1012441 | No |
| orf19.2925 | 289 | 316 | 315 | 174 | 336 | 213 | 315 | 288 | 276 | 334 | 311 | 273 | 269 | 242 | 267 | 234 | 275,81 | 298 | 253 | 0,847 | -0,239 | -2,384 | 0,0171228 | 0,0213733 | 0,0037094 | 0,0080443 | 0,0060052 | 0,0060052 | TRUE | 0,0721594 | No |
| orf19.2926 | 2136 | 2073 | 2309 | 1310 | 3198 | 2184 | 3026 | 2874 | 2036 | 2192 | 2281 | 2052 | 2562 | 2483 | 2569 | 2337 | 2314,18 | 2140 | 2488 | 1,161 | 0,216 | 4,018 | 5,87E-05 | 8,39E-05 | 0,0018554 | 0,0028008 | 0,0023301 | 0,0023301 | TRUE | 0,0240582 | No |
| orf19.2927 | 1507 | 1968 | 1961 | 1166 | 1051 | 789 | 803 | 773 | 1437 | 2081 | 1937 | 1827 | 842 | 897 | 682 | 629 | 1291,39 | 1820 | 762 | 0,418 | -1,257 | -10,08 | 6,75E-24 | 1,64E-23 | 0,0242994 | 0,0033627 | 0,0140387 | 0,0140387 | TRUE | 0,2460589 | No |
| orf19.2928 | 778 | 857 | 909 | 521 | 895 | 656 | 750 | 661 | 742 | 906 | 898 | 816 | 717 | 746 | 637 | 537 | 749,92 | 840 | 659 | 0,783 | -0,353 | -3,39 | 0,0006978 | 0,0009521 | 0,0135744 | 0,0042807 | 0,0090396 | 0,0090396 | TRUE | 0,1855827 | No |
| orf19.2929 | 9400 | 9250 | 9451 | 6113 | 24093 | 18838 | 21106 | 23818 | 8962 | 9781 | 9337 | 9576 | 19305 | 21421 | 17919 | 19366 | 14458,29 | 9414 | 19503 | 2,072 | 1,051 | 18,794 | 8,48E-79 | 4,56E-78 | 0,0033292 | 0,0022048 | 0,0029204 | 0,0029204 | TRUE | 0,0534287 | No |
| orf19.2930 | 4404 | 4052 | 4384 | 2995 | 5408 | 4202 | 5022 | 5367 | 4199 | 4284 | 4331 | 4692 | 4333 | 4778 | 4264 | 4364 | 4405,61 | 4376 | 4435 | 1,014 | 0,02 | 0,38 | 0,7036188 | 0,7341059 | 0,0020558 | 0,002464 | 0,0023195 | 0,0023195 | TRUE | 0,0324204 | No |
| orf19.2933 | 1767 | 1719 | 1840 | 1082 | 1190 | 765 | 1043 | 1096 | 1685 | 1818 | 1818 | 1695 | 954 | 870 | 886 | 891 | 1326,88 | 1754 | 900 | 0,513 | -0,962 | -17,808 | 6,16E-71 | 2,99E-70 | 0,0009898 | 0,0033287 | 0,0019868 | 0,0019868 | TRUE | 0,0195918 | No |
| orf19.2934 | 6255 | 6512 | 6561 | 4012 | 6498 | 4785 | 6052 | 5911 | 5963 | 6886 | 6482 | 6285 | 5207 | 5441 | 5138 | 4806 | 5775,96 | 6404 | 5148 | 0,804 | -0,315 | -5,733 | 9,88E-09 | 1,64E-08 | 0,0028765 | 0,0023755 | 0,002733 | 0,002733 | TRUE | 0,0314939 | No |
| orf19.2935 | 26374 | 21971 | 23633 | 14688 | 66253 | 44361 | 63148 | 70168 | 25145 | 23232 | 23347 | 23010 | 53087 | 50442 | 53612 | 57053 | 38615,97 | 23684 | 53548 | 2,261 | 1,177 | 24,923 | 4,16E-137 | 3,82E-136 | 0,0021105 | 0,0021338 | 0,0021102 | 0,0021102 | TRUE | 0,0237858 | No |
| orf19.2936 | 311 | 324 | 340 | 177 | 679 | 445 | 673 | 540 | 297 | 343 | 336 | 277 | 544 | 506 | 571 | 439 | 414,1 | 313 | 515 | 1,64 | 0,713 | 6,904 | 5,07E-12 | 9,40E-12 | 0,0091089 | 0,0060563 | 0,0075979 | 0,0075979 | TRUE | 0,1201242 | No |
| orf19.2937 | 1402 | 1385 | 1392 | 783 | 2941 | 2069 | 2468 | 2271 | 1337 | 1464 | 1375 | 1227 | 2357 | 2353 | 2095 | 1847 | 1756,74 | 1351 | 2163 | 1,599 | 0,677 | 8,07 | 7,03E-16 | 1,44E-15 | 0,0082074 | 0,0030259 | 0,0061301 | 0,0061301 | TRUE | 0,1182221 | No |
| orf19.2938 | 1498 | 1581 | 1616 | 993 | 2162 | 1599 | 2054 | 2089 | 1428 | 1672 | 1596 | 1556 | 1732 | 1818 | 1744 | 1699 | 1655,61 | 1563 | 1748 | 1,118 | 0,161 | 2,868 | 0,004125 | 0,0053619 | 0,0017349 | 0,003083 | 0,0024139 | 0,0024139 | TRUE | 0,0430246 | No |
| orf19.2939 | 457 | 375 | 407 | 230 | 880 | 600 | 730 | 771 | 436 | 397 | 402 | 360 | 705 | 682 | 620 | 627 | 528,58 | 399 | 658 | 1,643 | 0,716 | 9,023 | 1,83E-19 | 4,05E-19 | 0,0027605 | 0,0051975 | 0,0039727 | 0,0039727 | TRUE | 0,0447258 | No |
| orf19.2940 | 523 | 485 | 556 | 315 | 678 | 445 | 640 | 586 | 499 | 513 | 549 | 493 | 543 | 506 | 543 | 476 | 515,41 | 514 | 517 | 1,006 | 0,009 | 0,117 | 0,9068756 | 0,9197876 | 0,0015729 | 0,0052769 | 0,003369 | 0,003369 | TRUE | 0,0354492 | No |
| orf19.2941 | 3 | 7 | 5 | 3 | 6 | 1 | 5 | 11 | 3 | 7 | 5 | 5 | 5 | 1 | 4 | 9 | 4,88 | 5 | 5 | 0,991 | -0,013 | -0,019 | 0,9850072 | 0,9865533 | 0,099068 | 0,3385924 | 0,2415 | 0,2415 | TRUE | 0,3855228 | No |
| orf19.2942 | 43201 | 38206 | 41933 | 25897 | 47553 | 33397 | 45615 | 48017 | 41187 | 40398 | 41426 | 40569 | 38103 | 37975 | 38727 | 39042 | 39678,49 | 40895 | 38462 | 0,94 | -0,089 | -3,61 | 0,0003066 | 0,0004246 | 0,0001252 | 0,0021327 | 0,0005523 | 0,0005523 | TRUE | 0,001266 | No |
| orf19.2943.5 | 2 | 2 | 1 | 3 | 1 | 2 | 4 | 0 | 2 | 2 | 1 | 5 | 1 | 2 | 3 | 0 | 2,02 | 2 | 2 | 0,685 | -0,547 | -0,511 | 0,6090948 | 0,6429502 | 1,00E-08 | 0,8138835 | 0,5738072 | 0,5738072 | TRUE | 0,4339837 | No |
| orf19.2945 | 56 | 54 | 52 | 17 | 127 | 66 | 109 | 90 | 53 | 57 | 51 | 27 | 102 | 75 | 93 | 73 | 66,38 | 47 | 86 | 1,784 | 0,835 | 3,874 | 0,0001071 | 0,0001514 | 0,0294426 | 0,0268272 | 0,0277918 | 0,0277918 | TRUE | 0,4682038 | No |
| orf19.2946 | 245 | 257 | 250 | 166 | 361 | 281 | 321 | 367 | 234 | 272 | 247 | 260 | 289 | 320 | 273 | 298 | 274,01 | 253 | 295 | 1,165 | 0,221 | 2,35 | 0,0187932 | 0,0233806 | 1,00E-08 | 0,0080835 | 0,004748 | 0,004748 | TRUE | 0,0321907 | No |
| orf19.2947 | 4150 | 3605 | 3792 | 2438 | 1419 | 992 | 1414 | 1562 | 3957 | 3812 | 3746 | 3819 | 1137 | 1128 | 1200 | 1270 | 2508,67 | 3834 | 1184 | 0,309 | -1,694 | -34,807 | 1,88E-265 | 3,44E-264 | 0,0010775 | 0,0027458 | 0,00175 | 0,00175 | TRUE | 0,0293051 | No |
| orf19.2948 | 1707 | 1436 | 1615 | 1078 | 604 | 415 | 562 | 640 | 1627 | 1518 | 1595 | 1689 | 484 | 472 | 477 | 520 | 1047,93 | 1607 | 488 | 0,305 | -1,715 | -28,682 | 6,49E-181 | 7,83E-180 | 0,00088 | 0,0036581 | 0,0021732 | 0,0021732 | TRUE | 0,0234916 | No |
| orf19.2949 | 1273 | 1268 | 1302 | 805 | 1179 | 803 | 1106 | 1156 | 1214 | 1341 | 1286 | 1261 | 945 | 913 | 939 | 940 | 1104,81 | 1276 | 934 | 0,733 | -0,448 | -8,612 | 7,17E-18 | 1,54E-17 | 1,00E-08 | 0,0035774 | 0,0016851 | 0,0016851 | TRUE | 0,0147561 | No |
| orf19.295 | 158 | 119 | 141 | 88 | 199 | 139 | 180 | 192 | 151 | 126 | 139 | 138 | 159 | 158 | 153 | 156 | 147,51 | 138 | 156 | 1,13 | 0,176 | 1,502 | 0,133023 | 0,1534381 | 1,00E-08 | 0,0132223 | 0,0062845 | 0,0062845 | TRUE | 0,0417297 | No |
| orf19.2951 | 699 | 637 | 691 | 468 | 2053 | 1587 | 1970 | 2174 | 666 | 674 | 683 | 733 | 1645 | 1805 | 1673 | 1768 | 1205,69 | 689 | 1723 | 2,506 | 1,326 | 23,179 | 7,38E-119 | 5,80E-118 | 0,0010066 | 0,0034531 | 0,0020669 | 0,0020669 | TRUE | 0,0185403 | No |
| orf19.2952 | 568 | 493 | 566 | 375 | 1100 | 736 | 1113 | 1076 | 542 | 521 | 559 | 587 | 881 | 837 | 945 | 875 | 718,44 | 552 | 884 | 1,608 | 0,685 | 10,419 | 2,03E-25 | 5,06E-25 | 0,001021 | 0,0043766 | 0,0026322 | 0,0026322 | TRUE | 0,0250338 | No |
| orf19.2953 | 2016 | 1753 | 1780 | 1150 | 6498 | 4452 | 6245 | 6603 | 1922 | 1854 | 1758 | 1802 | 5207 | 5062 | 5302 | 5369 | 3534,43 | 1834 | 5235 | 2,853 | 1,513 | 35,51 | 3,40E-276 | 6,65E-275 | 0,0005696 | 0,0025558 | 0,0013513 | 0,0013513 | TRUE | 0,0133242 | No |
| orf19.2954 | 4402 | 3651 | 3861 | 2482 | 9693 | 6556 | 9151 | 9799 | 4197 | 3860 | 3814 | 3888 | 7767 | 7455 | 7769 | 7968 | 5839,74 | 3940 | 7740 | 1,964 | 0,974 | 22,854 | 1,35E-115 | 1,03E-114 | 0,001094 | 0,0023724 | 0,001543 | 0,001543 | TRUE | 0,0241823 | No |
| orf19.2956 | 1360 | 1200 | 1286 | 876 | 2223 | 1579 | 2186 | 2361 | 1297 | 1269 | 1270 | 1372 | 1781 | 1795 | 1856 | 1920 | 1570,07 | 1302 | 1838 | 1,415 | 0,501 | 10,233 | 1,41E-24 | 3,47E-24 | 0,0005678 | 0,003137 | 0,0016276 | 0,0016276 | TRUE | 0,0138266 | No |
| orf19.2957 | 19 | 29 | 25 | 15 | 33 | 23 | 47 | 32 | 18 | 31 | 25 | 23 | 26 | 26 | 40 | 26 | 26,94 | 24 | 30 | 1,227 | 0,295 | 1,016 | 0,3094635 | 0,3427086 | 1,00E-08 | 0,0630461 | 0,0430516 | 0,0430516 | TRUE | 0,3953382 | No |
| orf19.2958 | 399 | 328 | 386 | 240 | 576 | 374 | 502 | 519 | 380 | 347 | 381 | 376 | 462 | 425 | 426 | 422 | 402,44 | 371 | 434 | 1,17 | 0,226 | 2,956 | 0,0031183 | 0,0040808 | 1,00E-08 | 0,0061711 | 0,0030966 | 0,0030966 | TRUE | 0,0232247 | No |
| orf19.2959 | 7 | 7 | 8 | 3 | 2 | 7 | 2 | 3 | 7 | 7 | 8 | 5 | 2 | 8 | 2 | 2 | 5,05 | 7 | 4 | 0,477 | -1,069 | -1,572 | 0,1159878 | 0,1343227 | 0,0191667 | 0,3274002 | 0,2233633 | 0,2233633 | TRUE | 0,9699907 | No |
| orf19.2959.1 | 2842 | 2460 | 2334 | 1436 | 1851 | 1065 | 1579 | 1484 | 2710 | 2601 | 2306 | 2250 | 1483 | 1211 | 1341 | 1207 | 1888,42 | 2467 | 1310 | 0,531 | -0,913 | -11,125 | 9,52E-29 | 2,51E-28 | 0,0083222 | 0,0029607 | 0,0058941 | 0,0058941 | TRUE | 0,0957931 | No |
| orf19.296 | 53 | 48 | 53 | 39 | 62 | 35 | 44 | 47 | 51 | 51 | 52 | 61 | 50 | 40 | 37 | 38 | 47,47 | 54 | 41 | 0,777 | -0,365 | -1,77 | 0,0767166 | 0,0906366 | 1,00E-08 | 0,0366771 | 0,0195422 | 0,0195422 | TRUE | 0,1655776 | No |
| orf19.2960 | 3469 | 3054 | 3227 | 2007 | 8520 | 5664 | 8020 | 8854 | 3307 | 3229 | 3188 | 3144 | 6827 | 6440 | 6809 | 7199 | 5018 | 3217 | 6819 | 2,119 | 1,084 | 24,933 | 3,25E-137 | 2,98E-136 | 0,0009601 | 0,0024185 | 0,001573 | 0,001573 | TRUE | 0,0171905 | No |
| orf19.2961 | 720 | 663 | 719 | 420 | 1656 | 1186 | 1562 | 1714 | 686 | 701 | 710 | 658 | 1327 | 1349 | 1326 | 1394 | 1018,88 | 689 | 1349 | 1,953 | 0,966 | 18,144 | 1,42E-73 | 7,15E-73 | 1,00E-08 | 0,0037028 | 0,0015827 | 0,0015827 | TRUE | 0,0087519 | No |
| orf19.2962 | 916 | 899 | 972 | 572 | 80 | 62 | 81 | 72 | 873 | 951 | 960 | 896 | 64 | 70 | 69 | 59 | 492,76 | 920 | 66 | 0,071 | -3,821 | -36,135 | 6,31E-286 | 1,32E-284 | 1,00E-08 | 0,0054233 | 0,0033378 | 0,0033378 | TRUE | 0,0484793 | No |
| orf19.2963 | 156 | 140 | 156 | 73 | 311 | 194 | 277 | 273 | 149 | 148 | 154 | 114 | 249 | 221 | 235 | 222 | 186,52 | 141 | 232 | 1,624 | 0,7 | 6,093 | 1,11E-09 | 1,90E-09 | 0,0005151 | 0,010894 | 0,0068377 | 0,0068377 | TRUE | 0,1276792 | No |
| orf19.2964 | 5263 | 4679 | 4934 | 3174 | 3663 | 2524 | 3506 | 3924 | 5018 | 4947 | 4874 | 4972 | 2935 | 2870 | 2977 | 3191 | 3972,99 | 4953 | 2993 | 0,605 | -0,726 | -17,369 | 1,42E-67 | 6,59E-67 | 0,0007911 | 0,0025045 | 0,0014132 | 0,0014132 | TRUE | 0,0242034 | No |
| orf19.2965 | 2193 | 2016 | 2040 | 1416 | 6026 | 4238 | 5789 | 6250 | 2091 | 2132 | 2015 | 2218 | 4828 | 4819 | 4915 | 5082 | 3512,52 | 2114 | 4911 | 2,328 | 1,219 | 28,993 | 7,99E-185 | 9,94E-184 | 0,0006564 | 0,0025587 | 0,0013398 | 0,0013398 | TRUE | 0,0120228 | No |
| orf19.2965.1 | 443 | 410 | 434 | 249 | 814 | 621 | 747 | 806 | 422 | 434 | 429 | 390 | 652 | 706 | 634 | 655 | 540,33 | 419 | 662 | 1,573 | 0,653 | 9,373 | 7,06E-21 | 1,61E-20 | 1,00E-08 | 0,00513 | 0,0026608 | 0,0026608 | TRUE | 0,0217278 | No |
| orf19.2966 | 268 | 265 | 241 | 135 | 764 | 605 | 737 | 811 | 256 | 280 | 238 | 211 | 612 | 688 | 626 | 659 | 446,32 | 246 | 646 | 2,602 | 1,38 | 15,642 | 3,74E-55 | 1,48E-54 | 0,0027605 | 0,0057701 | 0,0045304 | 0,0045304 | TRUE | 0,0881859 | No |
| orf19.2967 | 8542 | 7044 | 7680 | 4656 | 17975 | 11432 | 17587 | 18194 | 8144 | 7448 | 7587 | 7294 | 14403 | 12999 | 14931 | 14793 | 10949,98 | 7618 | 14282 | 1,874 | 0,906 | 16,653 | 2,88E-62 | 1,25E-61 | 0,002962 | 0,0022412 | 0,0027406 | 0,0027406 | TRUE | 0,0444269 | No |
| orf19.2968 | 352 | 318 | 331 | 191 | 423 | 268 | 387 | 417 | 336 | 336 | 327 | 299 | 339 | 305 | 329 | 339 | 326,17 | 324 | 328 | 1,009 | 0,012 | 0,148 | 0,8821611 | 0,8982081 | 1,00E-08 | 0,0071252 | 0,0035765 | 0,0035765 | TRUE | 0,0249475 | No |
| orf19.2969 | 22894 | 19524 | 21950 | 13394 | 11980 | 8034 | 11602 | 12053 | 21827 | 20644 | 21685 | 20982 | 9599 | 9135 | 9850 | 9800 | 15440,37 | 21284 | 9596 | 0,451 | -1,149 | -30,931 | 4,52E-210 | 6,43E-209 | 0,0008271 | 0,0021976 | 0,0012524 | 0,0012524 | TRUE | 0,0125305 | No |
| orf19.297 | 900 | 618 | 696 | 432 | 1122 | 680 | 1016 | 1155 | 858 | 653 | 688 | 677 | 899 | 773 | 863 | 939 | 793,72 | 719 | 868 | 1,207 | 0,272 | 2,889 | 0,0038675 | 0,0050384 | 0,0101402 | 0,0041599 | 0,0072015 | 0,0072015 | TRUE | 0,2064809 | No |
| orf19.2970 | 2833 | 2542 | 2639 | 1669 | 6288 | 4399 | 5958 | 6513 | 2701 | 2688 | 2607 | 2615 | 5038 | 5002 | 5058 | 5296 | 3875,61 | 2653 | 5098 | 1,922 | 0,942 | 25,303 | 2,99E-141 | 2,84E-140 | 0,0003353 | 0,0025149 | 0,0010327 | 0,0010327 | TRUE | 0,0083466 | No |
| orf19.2971 | 739 | 781 | 815 | 511 | 1022 | 779 | 996 | 1046 | 705 | 826 | 805 | 801 | 819 | 886 | 846 | 850 | 817,1 | 784 | 850 | 1,085 | 0,118 | 1,803 | 0,0713193 | 0,0844633 | 0,0017192 | 0,0041007 | 0,0028373 | 0,0028373 | TRUE | 0,0589805 | No |
| orf19.2972 | 11936 | 11120 | 11897 | 7501 | 6498 | 4989 | 6161 | 6645 | 11380 | 11758 | 11753 | 11751 | 5207 | 5673 | 5231 | 5403 | 8519,35 | 11660 | 5378 | 0,461 | -1,117 | -29,813 | 2,64E-195 | 3,55E-194 | 0,0007467 | 0,002284 | 0,0012168 | 0,0012168 | TRUE | 0,0160808 | No |
| orf19.2973 | 7081 | 6572 | 7268 | 4692 | 4799 | 3650 | 4681 | 4799 | 6751 | 6949 | 7180 | 7350 | 3845 | 4150 | 3974 | 3902 | 5512,78 | 7058 | 3968 | 0,562 | -0,831 | -19,835 | 1,48E-87 | 8,65E-87 | 0,0009863 | 0,0023891 | 0,0014922 | 0,0014922 | TRUE | 0,0112877 | No |
| orf19.2974 | 1466 | 1459 | 1523 | 850 | 2056 | 1300 | 1945 | 2005 | 1398 | 1543 | 1505 | 1332 | 1647 | 1478 | 1651 | 1630 | 1522,96 | 1444 | 1602 | 1,108 | 0,149 | 2,427 | 0,0152084 | 0,0190621 | 0,0027148 | 0,0031694 | 0,0029278 | 0,0029278 | TRUE | 0,0310817 | No |
| orf19.2975 | 1002 | 909 | 932 | 565 | 1903 | 1293 | 1795 | 1866 | 955 | 961 | 921 | 885 | 1525 | 1470 | 1524 | 1517 | 1219,82 | 930 | 1509 | 1,619 | 0,695 | 14,105 | 3,54E-45 | 1,21E-44 | 1,00E-08 | 0,0034373 | 0,0014364 | 0,0014364 | TRUE | 0,0106799 | No |
| orf19.2977 | 5616 | 4918 | 5560 | 3655 | 5513 | 3977 | 5398 | 5562 | 5354 | 5200 | 5493 | 5726 | 4417 | 4522 | 4583 | 4522 | 4977,23 | 5443 | 4511 | 0,829 | -0,27 | -6,875 | 6,21E-12 | 1,15E-11 | 0,0006856 | 0,0024212 | 0,0012768 | 0,0012768 | TRUE | 0,0140944 | No |
| orf19.2978 | 1714 | 1373 | 1509 | 937 | 2751 | 1744 | 2617 | 2697 | 1634 | 1452 | 1491 | 1468 | 2204 | 1983 | 2222 | 2193 | 1830,83 | 1511 | 2150 | 1,423 | 0,509 | 8,857 | 8,24E-19 | 1,80E-18 | 0,0021394 | 0,0029881 | 0,0025901 | 0,0025901 | TRUE | 0,0373834 | No |
| orf19.298 | 41 | 31 | 41 | 25 | 38 | 20 | 27 | 43 | 39 | 33 | 41 | 39 | 30 | 23 | 23 | 35 | 32,83 | 38 | 28 | 0,74 | -0,434 | -1,726 | 0,0844105 | 0,0992671 | 1,00E-08 | 0,052109 | 0,0302591 | 0,0302591 | TRUE | 0,2137322 | No |
| orf19.2980 | 1748 | 1491 | 1590 | 1055 | 1657 | 1138 | 1557 | 1736 | 1667 | 1577 | 1571 | 1653 | 1328 | 1294 | 1322 | 1412 | 1477,71 | 1617 | 1339 | 0,829 | -0,27 | -5,485 | 4,14E-08 | 6,75E-08 | 0,0005506 | 0,0032024 | 0,0016458 | 0,0016458 | TRUE | 0,016336 | No |
| orf19.2982 | 300 | 305 | 303 | 180 | 743 | 521 | 715 | 670 | 286 | 322 | 299 | 282 | 595 | 592 | 607 | 545 | 441,18 | 297 | 585 | 1,961 | 0,972 | 12,348 | 5,02E-35 | 1,47E-34 | 1,00E-08 | 0,0058129 | 0,0033151 | 0,0033151 | TRUE | 0,035801 | No |
| orf19.2983 | 1205 | 1038 | 1141 | 802 | 2673 | 1853 | 2486 | 2666 | 1149 | 1098 | 1127 | 1256 | 2142 | 2107 | 2111 | 2168 | 1644,64 | 1158 | 2132 | 1,848 | 0,886 | 17,882 | 1,62E-71 | 7,95E-71 | 0,0001731 | 0,0030896 | 0,0016639 | 0,0016639 | TRUE | 0,034296 | No |
| orf19.2984 | 1743 | 1468 | 1625 | 991 | 1349 | 906 | 1366 | 1451 | 1662 | 1552 | 1605 | 1552 | 1081 | 1030 | 1160 | 1180 | 1352,8 | 1593 | 1113 | 0,699 | -0,517 | -9,081 | 1,07E-19 | 2,39E-19 | 0,0014762 | 0,003305 | 0,0023562 | 0,0023562 | TRUE | 0,0276865 | No |
| orf19.2985 | 1329 | 1076 | 1191 | 775 | 1367 | 869 | 1361 | 1376 | 1267 | 1138 | 1177 | 1214 | 1095 | 988 | 1155 | 1119 | 1144,15 | 1199 | 1089 | 0,91 | -0,136 | -2,171 | 0,0299164 | 0,0366615 | 0,0023348 | 0,0035263 | 0,0028664 | 0,0028664 | TRUE | 0,0439438 | No |
| orf19.2986 | 547 | 411 | 447 | 335 | 466 | 328 | 420 | 515 | 522 | 435 | 442 | 525 | 373 | 373 | 357 | 419 | 430,52 | 481 | 380 | 0,794 | -0,333 | -3,642 | 0,0002707 | 0,0003759 | 0,005434 | 0,0059051 | 0,0056673 | 0,0056673 | TRUE | 0,055186 | No |
| orf19.2987 | 4280 | 3590 | 3913 | 2598 | 3119 | 2176 | 2901 | 3184 | 4081 | 3796 | 3866 | 4070 | 2499 | 2474 | 2463 | 2589 | 3229,67 | 3953 | 2506 | 0,635 | -0,656 | -15,824 | 2,14E-56 | 8,54E-56 | 0,0006492 | 0,0025997 | 0,001331 | 0,001331 | TRUE | 0,0086765 | No |
| orf19.2988 | 2356 | 2005 | 2120 | 1461 | 986 | 692 | 921 | 973 | 2246 | 2120 | 2094 | 2289 | 790 | 787 | 782 | 791 | 1487,41 | 2187 | 788 | 0,36 | -1,472 | -29,153 | 7,67E-187 | 9,69E-186 | 1,00E-08 | 0,0031951 | 0,0016253 | 0,0016253 | TRUE | 0,0106876 | No |
| orf19.2989 | 4404 | 3676 | 4081 | 2654 | 1495 | 1048 | 1505 | 1512 | 4199 | 3887 | 4032 | 4158 | 1198 | 1192 | 1278 | 1229 | 2646,45 | 4069 | 1224 | 0,301 | -1,732 | -38,088 | 0 | 0 | 0,0007389 | 0,0027117 | 0,0014819 | 0,0014819 | TRUE | 0,0110899 | No |

| | | | | | | | | | | | | | | | | | | | | | | | | | | | | | | | |
|---|---|---|---|---|---|---|---|---|---|---|---|---|---|---|---|---|---|---|---|---|---|---|---|---|---|---|---|---|---|---|---|
| orf19.299 | 2983 | 2681 | 2891 | 2089 | 3181 | 2455 | 3083 | 3225 | 2844 | 2835 | 2856 | 3273 | 2549 | 2792 | 2617 | 2622 | 2798,43 | 2952 | 2645 | 0,897 | -0,157 | -2,774 | 0,0055426 | 0,0071477 | 0,0029117 | 0,002678 | 0,0027073 | 0,0027073 | TRUE | 0,0619797 | No |
| orf19.2990 | 2176 | 1913 | 2027 | 1418 | 3442 | 2526 | 3321 | 3558 | 2075 | 2023 | 2002 | 2221 | 2758 | 2872 | 2820 | 2893 | 2458 | 2080 | 2836 | 1,366 | 0,45 | 10,042 | 9,93E-24 | 2,40E-23 | 0,0006248 | 0,0027593 | 0,0014962 | 0,0014962 | TRUE | 0,0227716 | No |
| orf19.2991 | 123 | 93 | 104 | 90 | 395 | 345 | 453 | 454 | 117 | 98 | 103 | 141 | 317 | 392 | 385 | 369 | 240,23 | 115 | 366 | 3,218 | 1,686 | 13,518 | 1,23E-41 | 4,00E-41 | 0,008625 | 0,0089259 | 0,00881 | 0,00881 | TRUE | 0,1794198 | No |
| orf19.2992 | 6796 | 5684 | 5906 | 4283 | 23336 | 17149 | 20313 | 21801 | 6479 | 6010 | 5835 | 6710 | 18698 | 19500 | 17246 | 17726 | 12275,48 | 6258 | 18292 | 2,924 | 1,548 | 27,309 | 3,35E-164 | 3,74E-163 | 0,0034712 | 0,002225 | 0,0029719 | 0,0029719 | TRUE | 0,0280795 | No |
| orf19.2994 | 21751 | 17635 | 19237 | 12741 | 63953 | 42106 | 60997 | 63242 | 20737 | 18647 | 19004 | 19959 | 51244 | 47878 | 51786 | 51422 | 35084,72 | 19587 | 50582 | 2,583 | 1,369 | 30,504 | 2,32E-204 | 3,22E-203 | 0,0017644 | 0,0021381 | 0,001897 | 0,001897 | TRUE | 0,0193184 | No |
| orf19.2994.1 | 12694 | 10396 | 10736 | 7298 | 35555 | 23475 | 32585 | 36454 | 12102 | 10992 | 10606 | 11433 | 28489 | 26693 | 27665 | 29641 | 19702,64 | 11283 | 28122 | 2,493 | 1,318 | 26,017 | 3,19E-149 | 3,17E-148 | 0,0025244 | 0,0021746 | 0,0023981 | 0,0023981 | TRUE | 0,0295525 | No |
| orf19.2995 | 1175 | 947 | 1102 | 722 | 1286 | 974 | 1241 | 1408 | 1120 | 1001 | 1089 | 1131 | 1030 | 1108 | 1054 | 1145 | 1084,71 | 1085 | 1084 | 0,999 | -0,001 | -0,013 | 0,9896872 | 0,9907741 | 0,0017425 | 0,003605 | 0,0025635 | 0,0025635 | TRUE | 0,0326612 | No |
| orf19.2996 | 1800 | 1702 | 1918 | 1109 | 2402 | 1676 | 2229 | 2402 | 1716 | 1800 | 1895 | 1737 | 1925 | 1906 | 1892 | 1953 | 1852,97 | 1787 | 1919 | 1,073 | 0,102 | 2,206 | 0,0273755 | 0,0336317 | 0,000506 | 0,0029774 | 0,0015001 | 0,0015001 | TRUE | 0,0206088 | No |
| orf19.2998 | 2590 | 2077 | 2170 | 1361 | 5292 | 3486 | 5163 | 5341 | 2469 | 2196 | 2144 | 2132 | 4240 | 3964 | 4383 | 4343 | 3233,95 | 2235 | 4232 | 1,892 | 0,92 | 15,954 | 2,68E-57 | 1,08E-56 | 0,003053 | 0,002599 | 0,0028368 | 0,0028368 | TRUE | 0,0613777 | No |
| orf19.30 | 2565 | 2389 | 2494 | 1575 | 8693 | 6445 | 7724 | 8667 | 2445 | 2526 | 2464 | 2467 | 6965 | 7329 | 6558 | 7047 | 4725,18 | 2476 | 6975 | 2,816 | 1,494 | 34,147 | 1,46E-255 | 2,56E-254 | 0,0011037 | 0,0024388 | 0,0015465 | 0,0015465 | TRUE | 0,019755 | No |
| orf19.300 | 1720 | 1360 | 1463 | 998 | 4478 | 3092 | 4473 | 4666 | 1640 | 1438 | 1445 | 1563 | 3588 | 3516 | 3798 | 3794 | 2597,75 | 1522 | 3674 | 2,416 | 1,273 | 23,113 | 3,45E-118 | 2,70E-117 | 0,0021636 | 0,0027233 | 0,0024203 | 0,0024203 | TRUE | 0,0355463 | No |
| orf19.3000 | 715 | 605 | 672 | 469 | 1241 | 913 | 1228 | 1290 | 682 | 640 | 664 | 735 | 994 | 1038 | 1043 | 1049 | 855,5 | 680 | 1031 | 1,521 | 0,605 | 10,255 | 1,12E-24 | 2,76E-24 | 0,0001032 | 0,0040105 | 0,0020888 | 0,0020888 | TRUE | 0,0288814 | No |
| orf19.3001 | 216 | 199 | 247 | 140 | 228 | 171 | 220 | 234 | 206 | 210 | 244 | 219 | 183 | 194 | 187 | 190 | 204,23 | 220 | 188 | 0,856 | -0,224 | -2,181 | 0,0292014 | 0,0358128 | 1,00E-08 | 0,0101307 | 0,0052539 | 0,0052539 | TRUE | 0,0658783 | No |
| orf19.3002 | 14698 | 12055 | 12543 | 8012 | 43372 | 30666 | 40743 | 46769 | 14013 | 12747 | 12391 | 12551 | 34753 | 34870 | 34591 | 38028 | 24242,89 | 12926 | 35560 | 2,751 | 1,46 | 28,74 | 1,21E-181 | 1,46E-180 | 0,0026408 | 0,002159 | 0,0024225 | 0,0024225 | TRUE | 0,0394663 | No |
| orf19.3003 | 2850 | 2656 | 2765 | 1658 | 2401 | 1524 | 2164 | 2346 | 2717 | 2808 | 2732 | 2597 | 1924 | 1733 | 1837 | 1908 | 2282 | 2714 | 1850 | 0,682 | -0,552 | -11,533 | 8,99E-31 | 2,46E-30 | 0,0010125 | 0,0028108 | 0,0017505 | 0,0017505 | TRUE | 0,0208295 | No |
| orf19.3003.1 | 14572 | 12058 | 12889 | 9273 | 37351 | 27309 | 35924 | 38267 | 13893 | 12750 | 12733 | 14527 | 29928 | 31053 | 30499 | 31115 | 22062,19 | 13476 | 30649 | 2,275 | 1,186 | 24,571 | 2,59E-133 | 2,33E-132 | 0,0022555 | 0,0021657 | 0,0021815 | 0,0021815 | TRUE | 0,0332283 | No |
| orf19.3004 | 6796 | 6407 | 6991 | 4045 | 4679 | 2901 | 4361 | 4515 | 6479 | 6775 | 6906 | 6337 | 3749 | 3299 | 3702 | 3671 | 5114,8 | 6624 | 3605 | 0,544 | -0,877 | -17,266 | 8,47E-67 | 3,90E-66 | 0,0021439 | 0,0024123 | 0,0022691 | 0,0022691 | TRUE | 0,0388044 | No |
| orf19.3006 | 3130 | 2872 | 3002 | 1994 | 3234 | 2316 | 3082 | 3326 | 2984 | 3037 | 2966 | 3124 | 2591 | 2634 | 2617 | 2704 | 2832,01 | 3028 | 2636 | 0,872 | -0,198 | -5,276 | 1,32E-07 | 2,12E-07 | 1,00E-08 | 0,002671 | 0,0009968 | 0,0009968 | TRUE | 0,0050315 | No |
| orf19.3007 | 759 | 605 | 664 | 423 | 638 | 385 | 623 | 611 | 724 | 640 | 656 | 663 | 511 | 438 | 529 | 497 | 582,08 | 671 | 494 | 0,738 | -0,439 | -5,761 | 8,36E-09 | 1,39E-08 | 0,0027526 | 0,004912 | 0,00383 | 0,00383 | TRUE | 0,0614729 | No |
| orf19.3007.2 | 912 | 802 | 859 | 441 | 1753 | 1108 | 1711 | 1766 | 869 | 848 | 849 | 691 | 1405 | 1260 | 1453 | 1436 | 1101,26 | 814 | 1388 | 1,699 | 0,765 | 9,985 | 1,78E-23 | 4,28E-23 | 0,0058825 | 0,0035822 | 0,0046285 | 0,0046285 | TRUE | 0,110968 | No |
| orf19.3008 | 1449 | 1221 | 1343 | 805 | 1759 | 1173 | 1708 | 1785 | 1381 | 1291 | 1327 | 1261 | 1409 | 1334 | 1450 | 1451 | 1363,13 | 1315 | 1411 | 1,072 | 0,101 | 1,949 | 0,0512917 | 0,061637 | 0,0007002 | 0,0032958 | 0,0018275 | 0,0018275 | TRUE | 0,0152456 | No |
| orf19.3009 | 655 | 672 | 730 | 470 | 681 | 487 | 613 | 645 | 624 | 711 | 721 | 736 | 546 | 554 | 520 | 524 | 617,1 | 698 | 536 | 0,769 | -0,379 | -5,357 | 8,44E-08 | 1,36E-07 | 0,0016943 | 0,004752 | 0,0031554 | 0,0031554 | TRUE | 0,0625157 | No |
| orf19.301 | 2355 | 1843 | 2078 | 1646 | 760 | 555 | 732 | 805 | 2245 | 1949 | 2053 | 2579 | 609 | 631 | 621 | 655 | 1417,68 | 2206 | 629 | 0,286 | -1,808 | -21,669 | 4,04E-104 | 2,73E-103 | 0,0077645 | 0,0032494 | 0,0057215 | 0,0057215 | TRUE | 0,1508827 | No |
| orf19.3010 | 602 | 468 | 500 | 277 | 758 | 506 | 736 | 756 | 574 | 495 | 494 | 434 | 607 | 575 | 625 | 615 | 552,37 | 499 | 606 | 1,208 | 0,272 | 3,292 | 0,000994 | 0,0013403 | 0,004331 | 0,0050637 | 0,0047144 | 0,0047144 | TRUE | 0,1190121 | No |
| orf19.3010.1 | 9256 | 7967 | 8283 | 6021 | 17071 | 13361 | 15803 | 17870 | 8825 | 8424 | 8183 | 9432 | 13679 | 15193 | 13417 | 14530 | 11460,2 | 8716 | 14205 | 1,63 | 0,705 | 12,487 | 8,80E-36 | 2,60E-35 | 0,0034739 | 0,0022346 | 0,0029663 | 0,0029663 | TRUE | 0,0367856 | No |
| orf19.3012 | 1251 | 1127 | 1248 | 866 | 1274 | 933 | 1117 | 1326 | 1193 | 1192 | 1233 | 1357 | 1021 | 1061 | 948 | 1078 | 1135,26 | 1244 | 1027 | 0,828 | -0,273 | -4,32 | 1,56E-05 | 2,29E-05 | 0,0023536 | 0,0035376 | 0,0029433 | 0,0029433 | TRUE | 0,0410489 | No |
| orf19.3013 | 1270 | 1113 | 1134 | 715 | 1989 | 1372 | 2071 | 2045 | 1211 | 1177 | 1120 | 1120 | 1594 | 1560 | 1758 | 1663 | 1400,36 | 1157 | 1644 | 1,42 | 0,506 | 9,032 | 1,68E-19 | 3,74E-19 | 0,0015414 | 0,0032638 | 0,002256 | 0,002256 | TRUE | 0,0265171 | No |
| orf19.3014 | 9551 | 8316 | 8868 | 6199 | 11803 | 8570 | 11215 | 12102 | 9106 | 8793 | 8761 | 9711 | 9457 | 9745 | 9521 | 9840 | 9366,83 | 9093 | 9641 | 1,061 | 0,085 | 2,068 | 0,0386465 | 0,0469185 | 0,0012082 | 0,0022666 | 0,0015245 | 0,0015245 | TRUE | 0,0248831 | No |
| orf19.3015 | 11540 | 9842 | 10909 | 7214 | 18354 | 13482 | 17643 | 19618 | 11002 | 10407 | 10777 | 11301 | 14707 | 15330 | 14979 | 15951 | 13056,74 | 10872 | 15242 | 1,402 | 0,488 | 12,109 | 9,48E-34 | 2,71E-33 | 0,0011185 | 0,002217 | 0,0014777 | 0,0014777 | TRUE | 0,0121088 | No |
| orf19.3016 | 1434 | 1138 | 1319 | 717 | 3486 | 2053 | 3306 | 3445 | 1367 | 1203 | 1303 | 1123 | 2793 | 2334 | 2807 | 2801 | 1966,54 | 1249 | 2684 | 2,146 | 1,101 | 14,234 | 5,60E-46 | 1,95E-45 | 0,0067035 | 0,0029262 | 0,0051326 | 0,0051326 | TRUE | 0,0915763 | No |
| orf19.3018 | 1072 | 961 | 1032 | 645 | 1644 | 1118 | 1587 | 1644 | 1022 | 1016 | 1020 | 1010 | 1317 | 1271 | 1347 | 1337 | 1167,59 | 1017 | 1318 | 1,297 | 0,375 | 7,827 | 4,99E-15 | 1,01E-14 | 1,00E-08 | 0,0034975 | 0,0013174 | 0,0013174 | TRUE | 0,0062694 | No |
| orf19.3019 | 414 | 359 | 385 | 223 | 1604 | 1152 | 1580 | 1621 | 395 | 380 | 380 | 349 | 1285 | 1310 | 1341 | 1318 | 844,82 | 376 | 1314 | 3,476 | 1,797 | 29,369 | 1,38E-189 | 1,78E-188 | 1,00E-08 | 0,0040348 | 0,0017881 | 0,0017881 | TRUE | 0,020121 | No |
| orf19.302 | 344 | 306 | 314 | 185 | 582 | 374 | 560 | 530 | 328 | 324 | 310 | 290 | 466 | 425 | 475 | 431 | 381,19 | 313 | 449 | 1,433 | 0,519 | 6,396 | 1,60E-10 | 2,84E-10 | 1,00E-08 | 0,0063986 | 0,003555 | 0,003555 | TRUE | 0,0221885 | No |
| orf19.3021 | 2105 | 1788 | 1903 | 1197 | 857 | 615 | 846 | 897 | 2007 | 1891 | 1880 | 1875 | 687 | 699 | 718 | 729 | 1310,78 | 1913 | 708 | 0,37 | -1,434 | -28,037 | 5,71E-173 | 6,63E-172 | 1,00E-08 | 0,0033439 | 0,0015902 | 0,0015902 | TRUE | 0,0136696 | No |
| orf19.3022 | 3917 | 3129 | 3530 | 2247 | 4512 | 2987 | 4276 | 4624 | 3734 | 3309 | 3487 | 3520 | 3615 | 3396 | 3630 | 3760 | 3556,52 | 3512 | 3600 | 1,025 | 0,036 | 0,725 | 0,4681587 | 0,504615 | 0,0016905 | 0,0025529 | 0,0021052 | 0,0021052 | TRUE | 0,0227018 | No |
| orf19.3023 | 1908 | 1522 | 1653 | 988 | 1866 | 1213 | 1869 | 2012 | 1819 | 1609 | 1633 | 1548 | 1495 | 1379 | 1587 | 1636 | 1588,29 | 1652 | 1524 | 0,922 | -0,117 | -1,697 | 0,0897148 | 0,1052335 | 0,0044032 | 0,003125 | 0,0039132 | 0,0039132 | TRUE | 0,0564048 | No |
| orf19.3026 | 3352 | 2780 | 3176 | 2058 | 3371 | 2222 | 3264 | 3413 | 3196 | 2939 | 3138 | 3224 | 2701 | 2527 | 2771 | 2775 | 2908,84 | 3124 | 2694 | 0,863 | -0,212 | -4,404 | 1,06E-05 | 1,57E-05 | 0,0013146 | 0,0026557 | 0,0018838 | 0,0018838 | TRUE | 0,0202391 | No |
| orf19.3027 | 5752 | 5138 | 5548 | 3635 | 2488 | 1857 | 2375 | 2677 | 5484 | 5433 | 5481 | 5694 | 1994 | 2112 | 2016 | 2177 | 3798,77 | 5523 | 2075 | 0,376 | -1,412 | -34 | 2,19E-253 | 3,79E-252 | 0,000623 | 0,0025235 | 0,0013397 | 0,0013397 | TRUE | 0,0140463 | No |
| orf19.3029 | 3221 | 3017 | 3105 | 2089 | 10259 | 7738 | 10009 | 10857 | 3071 | 3190 | 3067 | 3273 | 8220 | 8799 | 8498 | 8828 | 5868,17 | 3150 | 8586 | 2,728 | 1,448 | 35,56 | 5,80E-277 | 1,14E-275 | 0,0007641 | 0,0023711 | 0,001361 | 0,001361 | TRUE | 0,0101217 | No |
| orf19.3030 | 2078 | 1852 | 2059 | 1314 | 2219 | 1475 | 2146 | 2159 | 1981 | 1958 | 2034 | 2058 | 1778 | 1677 | 1822 | 1755 | 1883,07 | 2008 | 1758 | 0,877 | -0,189 | -4,272 | 1,94E-05 | 2,84E-05 | 0,0002406 | 0,0029632 | 0,0013482 | 0,0013482 | TRUE | 0,0107396 | No |
| orf19.3031 | 1859 | 1452 | 1532 | 934 | 3520 | 2122 | 3176 | 3461 | 1772 | 1535 | 1513 | 1463 | 2820 | 2413 | 2696 | 2814 | 2128,53 | 1571 | 2686 | 1,708 | 0,773 | 10,749 | 6,02E-27 | 1,55E-26 | 0,0056823 | 0,0028627 | 0,0044377 | 0,0044377 | TRUE | 0,0910393 | No |
| orf19.3034 | 10893 | 9671 | 10661 | 6499 | 20413 | 14529 | 19549 | 21037 | 10385 | 10226 | 10532 | 10181 | 16356 | 16521 | 16597 | 17105 | 13487,93 | 10331 | 16645 | 1,611 | 0,688 | 23,469 | 8,53E-122 | 6,89E-121 | 0,0002037 | 0,002213 | 0,0007434 | 0,0007434 | TRUE | 0,0042755 | No |
| orf19.3035 | 5029 | 3987 | 4634 | 2806 | 4633 | 2915 | 4156 | 4567 | 4795 | 4216 | 4578 | 4396 | 3712 | 3315 | 3528 | 3713 | 4031,6 | 4496 | 3567 | 0,794 | -0,334 | -6,101 | 1,05E-09 | 1,80E-09 | 0,0025968 | 0,0024985 | 0,00262 | 0,00262 | TRUE | 0,0269972 | No |
| orf19.3037 | 25733 | 21208 | 23228 | 15940 | 65191 | 45761 | 61743 | 67266 | 24534 | 22425 | 22947 | 24971 | 52236 | 52034 | 52420 | 54694 | 38282,45 | 23719 | 52846 | 2,228 | 1,156 | 26,9 | 2,17E-159 | 2,35E-158 | 0,001569 | 0,0021342 | 0,0017419 | 0,0017419 | TRUE | 0,0164923 | No |
| orf19.3038 | 70134 | 61043 | 66769 | 41592 | 9817 | 6738 | 8973 | 9906 | 66865 | 64545 | 65961 | 65156 | 7866 | 7662 | 7618 | 8055 | 36716,04 | 65632 | 7800 | 0,119 | -3,073 | -100,527 | 0 | 0 | 0,0003544 | 0,002136 | 0,0008314 | 0,0008314 | TRUE | 0,005926 | No |
| orf19.304 | 1662 | 1502 | 1561 | 959 | 3479 | 2438 | 3274 | 3365 | 1585 | 1588 | 1542 | 1502 | 2788 | 2772 | 2780 | 2736 | 2161,59 | 1554 | 2769 | 1,779 | 0,831 | 20,95 | 1,89E-97 | 1,21E-96 | 1,00E-08 | 0,0028509 | 0,0009896 | 0,0009896 | TRUE | 0,0051583 | No |
| orf19.3040 | 6462 | 5643 | 6012 | 3867 | 6549 | 4628 | 6294 | 6817 | 6161 | 5967 | 5939 | 6058 | 5248 | 5262 | 5344 | 5543 | 5690,14 | 6031 | 5349 | 0,887 | -0,173 | -5,09 | 3,57E-07 | 5,63E-07 | 0,0003138 | 0,0023798 | 0,0009295 | 0,0009295 | TRUE | 0,0073503 | No |
| orf19.3041 | 7375 | 6559 | 7263 | 4379 | 7357 | 5023 | 7004 | 7649 | 7031 | 6935 | 7175 | 6860 | 5895 | 5712 | 5946 | 6219 | 6471,74 | 7000 | 5943 | 0,849 | -0,236 | -6,333 | 2,41E-10 | 4,25E-10 | 0,0006052 | 0,002345 | 0,0011795 | 0,0011795 | TRUE | 0,0119723 | No |
| orf19.3042 | 207 | 206 | 203 | 164 | 168 | 140 | 186 | 168 | 197 | 218 | 201 | 257 | 135 | 159 | 158 | 137 | 182,62 | 218 | 147 | 0,677 | -0,562 | -4,623 | 3,79E-06 | 5,73E-06 | 0,0053769 | 0,0110822 | 0,0085545 | 0,0085545 | TRUE | 0,1319881 | No |
| orf19.3043 | 415 | 374 | 385 | 289 | 894 | 642 | 877 | 879 | 396 | 395 | 380 | 453 | 716 | 730 | 745 | 715 | 566,23 | 406 | 726 | 1,8 | 0,848 | 12,202 | 3,02E-34 | 8,72E-34 | 1,00E-08 | 0,004991 | 0,0026454 | 0,0026454 | TRUE | 0,0562836 | No |
| orf19.3045 | 2070 | 1896 | 2045 | 1304 | 4906 | 3463 | 4980 | 5314 | 1974 | 2005 | 2020 | 2043 | 3931 | 3938 | 4228 | 4321 | 3057,36 | 2010 | 4104 | 2,043 | 1,031 | 22,212 | 2,61E-109 | 1,88E-108 | 0,0011731 | 0,0026283 | 0,0016801 | 0,0016801 | TRUE | 0,0154246 | No |
| orf19.3047 | 5448 | 5084 | 5647 | 3470 | 1344 | 1055 | 1338 | 1398 | 5194 | 5376 | 5579 | 5436 | 1077 | 1200 | 1136 | 1137 | 3266,7 | 5396 | 1138 | 0,21 | -2,248 | -49,054 | 0 | 0 | 0,000697 | 0,0025939 | 0,0015203 | 0,0015203 | TRUE | 0,0153935 | No |
| orf19.3048 | 161 | 133 | 169 | 117 | 349 | 198 | 328 | 380 | 153 | 141 | 167 | 183 | 280 | 225 | 278 | 309 | 217,08 | 161 | 273 | 1,712 | 0,776 | 6,253 | 4,03E-10 | 7,05E-10 | 0,0096383 | 0,009655 | 0,0096499 | 0,0096499 | TRUE | 0,1440803 | No |
| orf19.3049 | 66 | 50 | 44 | 40 | 53 | 26 | 56 | 44 | 63 | 53 | 43 | 63 | 42 | 30 | 48 | 36 | 47,16 | 56 | 39 | 0,71 | -0,494 | -2,206 | 0,027382 | 0,0336333 | 0,0121006 | 0,0369074 | 0,0267487 | 0,0267487 | TRUE | 0,1885205 | No |
| orf19.305 | 419 | 403 | 433 | 256 | 783 | 597 | 714 | 730 | 399 | 426 | 428 | 401 | 627 | 679 | 606 | 594 | 520,05 | 414 | 626 | 1,509 | 0,594 | 8,178 | 2,88E-16 | 5,95E-16 | 1,00E-08 | 0,0052485 | 0,0030055 | 0,0030055 | TRUE | 0,0342164 | No |
| orf19.3050 | 3777 | 3618 | 3968 | 2156 | 2402 | 1682 | 2274 | 2448 | 3601 | 3826 | 3920 | 3377 | 1925 | 1913 | 1931 | 1990 | 2810,29 | 3681 | 1940 | 0,526 | -0,926 | -17,59 | 2,94E-69 | 1,39E-68 | 0,0018072 | 0,0026755 | 0,002276 | 0,002276 | TRUE | 0,0356159 | No |
| orf19.3051 | 1054 | 947 | 975 | 623 | 6013 | 4295 | 5657 | 5674 | 1005 | 1001 | 963 | 976 | 4818 | 4884 | 4803 | 4614 | 2882,94 | 986 | 4780 | 4,842 | 2,276 | 52,621 | 0 | 0 | 1,00E-08 | 0,0026608 | 0,0011371 | 0,0011371 | TRUE | 0,0067219 | No |
| orf19.3052 | 8718 | 7438 | 7966 | 4471 | 9854 | 6331 | 9276 | 9544 | 8312 | 7865 | 7870 | 7004 | 7896 | 7199 | 7875 | 7760 | 7722,53 | 7763 | 7682 | 0,989 | -0,016 | -0,276 | 0,7825434 | 0,8060387 | 0,002824 | 0,0023039 | 0,0029043 | 0,0029043 | TRUE | 0,0518048 | No |
| orf19.3053 | 1732 | 1474 | 1508 | 1156 | 1357 | 968 | 1264 | 1388 | 1651 | 1559 | 1490 | 1811 | 1087 | 1101 | 1073 | 1129 | 1362,53 | 1628 | 1098 | 0,676 | -0,565 | -8,892 | 5,99E-19 | 1,32E-18 | 0,0028595 | 0,0032963 | 0,0031261 | 0,0031261 | TRUE | 0,0645013 | No |
| orf19.3054 | 7752 | 6709 | 7329 | 4716 | 9014 | 6810 | 9114 | 9648 | 7391 | 7094 | 7240 | 7388 | 7223 | 7744 | 7738 | 7845 | 7457,7 | 7278 | 7638 | 1,049 | 0,069 | 1,828 | 0,06762 | 0,0803507 | 0,0006855 | 0,0023114 | 0,0012521 | 0,0012521 | TRUE | 0,016503 | No |
| orf19.3055 | 1848 | 1666 | 1752 | 1145 | 1610 | 1187 | 1535 | 1661 | 1762 | 1762 | 1731 | 1794 | 1290 | 1350 | 1303 | 1351 | 1542,69 | 1762 | 1324 | 0,751 | -0,413 | -9,461 | 3,06E-21 | 7,03E-21 | 1,00E-08 | 0,0031556 | 0,0011725 | 0,0011725 | TRUE | 0,0035349 | No |
| orf19.3057 | 742 | 578 | 690 | 366 | 1050 | 764 | 1002 | 1041 | 707 | 611 | 682 | 573 | 841 | 869 | 851 | 846 | 747,6 | 643 | 852 | 1,317 | 0,397 | 5,639 | 1,71E-08 | 2,82E-08 | 0,0023742 | 0,0042875 | 0,0033692 | 0,0033692 | TRUE | 0,0548419 | No |
| orf19.3058 | 1142 | 970 | 1024 | 702 | 1414 | 1005 | 1337 | 1520 | 1089 | 1026 | 1012 | 1100 | 1133 | 1143 | 1135 | 1236 | 1109,07 | 1057 | 1162 | 1,102 | 0,14 | 2,497 | 0,0125276 | 0,0158106 | 0,0009538 | 0,0035717 | 0,0020874 | 0,0020874 | TRUE | 0,0229054 | No |
| orf19.3059 | 32410 | 26084 | 28912 | 18704 | 16110 | 10422 | 15618 | 16423 | 30899 | 27581 | 28562 | 29301 | 12908 | 11851 | 13260 | 13353 | 20964,42 | 29086 | 12843 | 0,442 | -1,179 | -23,235 | 2,01E-119 | 1,59E-118 | 0,0025205 | 0,0021696 | 0,0024197 | 0,0024197 | TRUE | 0,0328337 | No |
| orf19.3059.1 | 0 | 4 | 1 | 2 | 5 | 3 | 4 | 4 | 0 | 4 | 1 | 3 | 4 | 3 | 3 | 3 | 2,8 | 2 | 3 | 1,776 | 0,829 | 0,927 | 0,3541118 | 0,3887796 | 1,00E-08 | 0,5880539 | 0,3616114 | 0,3616114 | TRUE | 0,3683794 | No |
| orf19.306 | 211 | 191 | 198 | 133 | 270 | 156 | 217 | 217 | 201 | 202 | 196 | 208 | 216 | 177 | 184 | 176 | 195,19 | 202 | 188 | 0,939 | -0,091 | -0,839 | 0,4011914 | 0,4369351 | 1,00E-08 | 0,0105033 | 0,0062026 | 0,0062026 | TRUE | 0,1126894 | No |
| orf19.3060 | 963 | 829 | 850 | 519 | 2345 | 1732 | 2135 | 2378 | 918 | 877 | 840 | 813 | 1879 | 1969 | 1813 | 1934 | 1380,25 | 862 | 1899 | 2,196 | 1,135 | 20,659 | 8,02E-95 | 5,00E-94 | 0,0009269 | 0,0032808 | 0,0020139 | 0,0020139 | TRUE | 0,0233259 | No |
| orf19.3061 | 402 | 300 | 331 | 203 | 767 | 478 | 759 | 705 | 383 | 317 | 327 | 318 | 615 | 544 | 644 | 573 | 465,15 | 336 | 594 | 1,762 | 0,818 | 9,223 | 2,89E-20 | 6,53E-20 | 0,004641 | 0,0056211 | 0,0051365 | 0,0051365 | TRUE | 0,1057618 | No |
| orf19.3061.1 | 7120 | 6400 | 6402 | 4724 | 5971 | 4448 | 5397 | 5683 | 6788 | 6767 | 6325 | 7400 | 4784 | 5058 | 4582 | 4621 | 5790,67 | 6820 | 4761 | 0,698 | -0,518 | -9,492 | 2,27E-21 | 5,22E-21 | 0,0028939 | 0,0023748 | 0,0026842 | 0,0026842 | TRUE | 0,0391875 | No |
| orf19.3062 | 996 | 1070 | 1073 | 689 | 2162 | 1610 | 1902 | 2264 | 950 | 1131 | 1060 | 1079 | 1732 | 1831 | 1615 | 1841 | 1404,88 | 1055 | 1755 | 1,664 | 0,735 | 10,922 | 9,04E-28 | 2,35E-27 | 0,0034196 | 0,00326 | 0,0035585 | 0,0035585 | TRUE | 0,0566484 | No |
| orf19.3063 | 786 | 626 | 662 | 451 | 1292 | 886 | 1285 | 1350 | 749 | 662 | 654 | 707 | 1035 | 1007 | 1091 | 1098 | 875,39 | 693 | 1058 | 1,528 | 0,612 | 9,565 | 1,13E-21 | 2,61E-21 | 0,0015726 | 0,0039669 | 0,0027019 | 0,0027019 | TRUE | 0,0396244 | No |
| orf19.3064 | 3040 | 2479 | 2638 | 1653 | 3594 | 2383 | 3530 | 3611 | 2898 | 2621 | 2606 | 2590 | 2880 | 2710 | 2997 | 2936 | 2779,71 | 2679 | 2881 | 1,075 | 0,104 | 1,993 | 0,0462566 | 0,055807 | 0,0019443 | 0,002682 | 0,0022742 | 0,0022742 | TRUE | 0,0377683 | No |
| orf19.3064.1 | 465 | 369 | 398 | 292 | 813 | 566 | 798 | 841 | 443 | 390 | 393 | 457 | 651 | 644 | 677 | 684 | 542,56 | 421 | 664 | 1,584 | 0,664 | 9,09 | 9,90E-20 | 2,21E-19 | 0,0006296 | 0,0051175 | 0,0031211 | 0,0031211 | TRUE | 0,0318378 | No |

| | | | | | | | | | | | | | | | | | | | | | | | | | | | | | | | |
|---|---|---|---|---|---|---|---|---|---|---|---|---|---|---|---|---|---|---|---|---|---|---|---|---|---|---|---|---|---|---|---|
| orf19.3065 | 1035 | 935 | 1024 | 648 | 4905 | 3564 | 4842 | 5130 | 987 | 989 | 1012 | 1015 | 3930 | 4053 | 4111 | 4171 | 2533,37 | 1001 | 4066 | 4,067 | 2,024 | 46,523 | 0 | 0 | 1,00E-08 | 0,0027394 | 0,0011499 | 0,0011499 | TRUE | 0,0063676 | No |
| orf19.3066 | 2177 | 1998 | 2210 | 1464 | 2426 | 1852 | 2239 | 2568 | 2076 | 2113 | 2183 | 2293 | 1944 | 2106 | 1901 | 2088 | 2087,95 | 2166 | 2010 | 0,928 | -0,107 | -2,029 | 0,0425041 | 0,0514061 | 0,001642 | 0,0028777 | 0,0021967 | 0,0021967 | TRUE | 0,0175824 | No |
| orf19.307 | 2587 | 2168 | 2440 | 1674 | 2149 | 1593 | 2110 | 2186 | 2466 | 2292 | 2410 | 2622 | 1722 | 1811 | 1791 | 1777 | 2111,73 | 2448 | 1775 | 0,726 | -0,462 | -9,296 | 1,45E-20 | 3,30E-20 | 0,0011536 | 0,0028688 | 0,0018868 | 0,0018868 | TRUE | 0,0257756 | No |
| orf19.3071 | 7979 | 6977 | 7846 | 5231 | 1940 | 1469 | 1867 | 2198 | 7607 | 7377 | 7751 | 8195 | 1554 | 1670 | 1585 | 1787 | 4690,91 | 7732 | 1649 | 0,213 | -2,229 | -41,017 | 0 | 0 | 0,0024973 | 0,0024413 | 0,0024918 | 0,0024918 | TRUE | 0,0396319 | No |
| orf19.3073 | 0 | 0 | 0 | 0 | 0 | 0 | 1 | 0 | 0 | 0 | 0 | 0 | 0 | 0 | 1 | 0 | 0,11 | 0 | 0 | 1,314 | 0,394 | 0,111 | 0,9112982 | 0,9205309 | 1,00E-08 | 15,473589 | 10 | 10 | TRUE | 0,3207401 | No |
| orf19.3074 | 131 | 105 | 135 | 84 | 288 | 178 | 289 | 292 | 125 | 111 | 133 | 132 | 231 | 202 | 245 | 237 | 177,1 | 125 | 229 | 1,839 | 0,879 | 7,482 | 7,30E-14 | 1,42E-13 | 1,00E-08 | 0,0113622 | 0,0068517 | 0,0068517 | TRUE | 0,060071 | No |
| orf19.3076 | 859 | 680 | 728 | 410 | 2769 | 1386 | 2599 | 2493 | 819 | 719 | 719 | 642 | 2219 | 1576 | 2207 | 2027 | 1365,97 | 725 | 2007 | 2,765 | 1,467 | 13,764 | 4,20E-43 | 1,40E-42 | 0,0164721 | 0,0032933 | 0,0099186 | 0,0099186 | TRUE | 0,2494228 | No |
| orf19.3077 | 1978 | 1515 | 1807 | 1255 | 3134 | 1948 | 2975 | 3229 | 1886 | 1602 | 1785 | 1966 | 2511 | 2215 | 2526 | 2625 | 2139,55 | 1810 | 2469 | 1,367 | 0,451 | 6,23 | 4,66E-10 | 8,11E-10 | 0,0056963 | 0,0028587 | 0,0045359 | 0,0045359 | TRUE | 0,0720363 | No |
| orf19.308 | 1505 | 1358 | 1386 | 999 | 2757 | 1880 | 2624 | 2835 | 1435 | 1436 | 1369 | 1565 | 2209 | 2138 | 2228 | 2305 | 1835,59 | 1451 | 2220 | 1,535 | 0,618 | 12,078 | 1,38E-33 | 3,95E-33 | 0,0010992 | 0,0029858 | 0,0019239 | 0,0019239 | TRUE | 0,030054 | No |
| orf19.3080 | 1504 | 1116 | 1390 | 976 | 2289 | 1488 | 1976 | 2372 | 1434 | 1180 | 1373 | 1529 | 1834 | 1692 | 1678 | 1929 | 1581,06 | 1379 | 1783 | 1,295 | 0,373 | 4,718 | 2,38E-06 | 3,63E-06 | 0,0071158 | 0,0031298 | 0,0053582 | 0,0053582 | TRUE | 0,1132637 | No |
| orf19.3083 | 2265 | 1788 | 2188 | 1480 | 570 | 427 | 563 | 570 | 2159 | 1891 | 2162 | 2318 | 457 | 486 | 478 | 463 | 1301,72 | 2132 | 471 | 0,221 | -2,179 | -30,937 | 3,84E-210 | 5,46E-209 | 0,0036997 | 0,0033526 | 0,0035505 | 0,0035505 | TRUE | 0,0703884 | No |
| orf19.3084 | 2281 | 1752 | 1942 | 1114 | 592 | 342 | 588 | 604 | 2175 | 1853 | 1919 | 1745 | 474 | 389 | 499 | 491 | 1193,05 | 1923 | 463 | 0,241 | -2,05 | -23,286 | 6,13E-120 | 4,88E-119 | 0,0087156 | 0,0034675 | 0,0061887 | 0,0061887 | TRUE | 0,1262603 | No |
| orf19.3086 | 2221 | 1752 | 1873 | 1160 | 3791 | 2382 | 3657 | 3780 | 2117 | 1853 | 1850 | 1817 | 3038 | 2709 | 3105 | 3074 | 2445,25 | 1909 | 2982 | 1,561 | 0,642 | 10,003 | 1,48E-23 | 3,57E-23 | 0,0040026 | 0,0027627 | 0,0035125 | 0,0035125 | TRUE | 0,0663247 | No |
| orf19.3087 | 17946 | 13740 | 14644 | 10516 | 58710 | 39926 | 56940 | 61398 | 17110 | 14528 | 14467 | 16474 | 47043 | 45399 | 48342 | 49922 | 31660,62 | 15645 | 47676 | 3,048 | 1,608 | 26,38 | 2,31E-153 | 2,37E-152 | 0,0044781 | 0,0021431 | 0,0035234 | 0,0035234 | TRUE | 0,0490197 | No |
| orf19.3087.1 | 195 | 125 | 132 | 90 | 447 | 253 | 396 | 433 | 186 | 132 | 130 | 141 | 358 | 288 | 336 | 352 | 240,45 | 147 | 334 | 2,26 | 1,177 | 9,135 | 6,52E-20 | 1,46E-19 | 0,0137555 | 0,0089197 | 0,0108093 | 0,0108093 | TRUE | 0,3515874 | No |
| orf19.3087.2 | 417 | 348 | 341 | 281 | 622 | 518 | 627 | 689 | 398 | 368 | 337 | 440 | 498 | 589 | 532 | 560 | 465,32 | 386 | 545 | 1,418 | 0,504 | 5,522 | 3,36E-08 | 5,50E-08 | 0,0060846 | 0,0056198 | 0,0057283 | 0,0057283 | TRUE | 0,0896943 | No |
| orf19.3088 | 2957 | 2450 | 2577 | 1644 | 3736 | 2435 | 3474 | 3649 | 2819 | 2591 | 2546 | 2575 | 2994 | 2769 | 2949 | 2967 | 2776,22 | 2633 | 2920 | 1,109 | 0,149 | 3,097 | 0,0019524 | 0,002588 | 0,0012704 | 0,0026827 | 0,0018622 | 0,0018622 | TRUE | 0,0284628 | No |
| orf19.3089 | 7642 | 6313 | 6747 | 3775 | 4195 | 2784 | 3905 | 4225 | 7286 | 6675 | 6665 | 5914 | 3361 | 3166 | 3315 | 3435 | 4977,23 | 6635 | 3319 | 0,5 | -1 | -16,341 | 5,06E-60 | 2,13E-59 | 0,003869 | 0,0024212 | 0,0033756 | 0,0033756 | TRUE | 0,0640707 | No |
| orf19.309 | 375 | 315 | 362 | 242 | 212 | 163 | 190 | 237 | 358 | 333 | 358 | 379 | 170 | 185 | 161 | 193 | 267,07 | 357 | 177 | 0,497 | -1,008 | -10,488 | 9,82E-26 | 2,46E-25 | 1,00E-08 | 0,0082391 | 0,0047807 | 0,0047807 | TRUE | 0,0421222 | No |
| orf19.3091 | 4879 | 4260 | 4653 | 2808 | 7107 | 4841 | 6599 | 7316 | 4652 | 4504 | 4597 | 4399 | 5695 | 5505 | 5603 | 5949 | 5112,75 | 4538 | 5688 | 1,253 | 0,325 | 8,416 | 3,90E-17 | 8,23E-17 | 0,000595 | 0,0024124 | 0,0012321 | 0,0012321 | TRUE | 0,0116856 | No |
| orf19.3093 | 2796 | 2320 | 2559 | 1712 | 3832 | 2595 | 3592 | 4079 | 2666 | 2453 | 2528 | 2682 | 3070 | 2951 | 3050 | 3317 | 2839,53 | 2582 | 3097 | 1,201 | 0,264 | 5,17 | 2,33E-07 | 3,71E-07 | 0,0017005 | 0,0026695 | 0,002138 | 0,002138 | TRUE | 0,0278868 | No |
| orf19.3097 | 14516 | 11428 | 12973 | 7962 | 25928 | 16040 | 24497 | 26736 | 13839 | 12084 | 12816 | 12473 | 20775 | 18239 | 20798 | 21739 | 16595,39 | 12803 | 20388 | 1,592 | 0,671 | 10,945 | 7,01E-28 | 1,83E-27 | 0,0043557 | 0,0021902 | 0,0035472 | 0,0035472 | TRUE | 0,0614181 | No |
| orf19.3098 | 1577 | 1434 | 1503 | 972 | 3064 | 2222 | 2808 | 3017 | 1503 | 1516 | 1485 | 1523 | 2455 | 2527 | 2384 | 2453 | 1980,76 | 1507 | 2455 | 1,629 | 0,704 | 17,076 | 2,22E-65 | 1,00E-64 | 1,00E-08 | 0,0029202 | 0,0010787 | 0,0010787 | TRUE | 0,004496 | No |
| orf19.3099 | 1047 | 869 | 972 | 652 | 1491 | 1049 | 1437 | 1476 | 998 | 919 | 960 | 1021 | 1195 | 1193 | 1220 | 1200 | 1088,29 | 974 | 1202 | 1,236 | 0,305 | 5,893 | 3,80E-09 | 6,39E-09 | 1,00E-08 | 0,0036 | 0,0016298 | 0,0016298 | TRUE | 0,0173579 | No |
| orf19.31 | 210 | 157 | 153 | 112 | 627 | 460 | 585 | 692 | 200 | 166 | 151 | 175 | 502 | 523 | 497 | 563 | 347,2 | 173 | 521 | 3,008 | 1,589 | 15,978 | 1,83E-57 | 7,42E-57 | 0,0035856 | 0,0068203 | 0,0054258 | 0,0054258 | TRUE | 0,1358275 | No |
| orf19.310 | 41 | 25 | 34 | 21 | 186 | 165 | 188 | 250 | 39 | 26 | 34 | 33 | 149 | 188 | 160 | 203 | 103,94 | 33 | 175 | 5,272 | 2,398 | 12,557 | 3,66E-36 | 1,09E-35 | 1,00E-08 | 0,0178873 | 0,0157467 | 0,0157467 | TRUE | 0,1050308 | No |
| orf19.3100 | 265 | 190 | 241 | 150 | 427 | 325 | 388 | 454 | 253 | 201 | 238 | 235 | 342 | 370 | 329 | 369 | 292,11 | 232 | 352 | 1,519 | 0,603 | 6,264 | 3,74E-10 | 6,56E-10 | 0,0021688 | 0,0077121 | 0,0052319 | 0,0052319 | TRUE | 0,0926487 | No |
| orf19.3102 | 1733 | 1469 | 1581 | 989 | 896 | 582 | 922 | 918 | 1652 | 1553 | 1562 | 1549 | 718 | 662 | 783 | 746 | 1153,2 | 1579 | 727 | 0,461 | -1,116 | -18,691 | 5,89E-78 | 3,13E-77 | 0,0016074 | 0,0035151 | 0,0024576 | 0,0024576 | TRUE | 0,0393364 | No |
| orf19.3103 | 6955 | 5874 | 6537 | 3873 | 12044 | 7964 | 11009 | 12014 | 6631 | 6211 | 6458 | 6067 | 9651 | 9056 | 9347 | 9769 | 7898,56 | 6342 | 9456 | 1,49 | 0,576 | 13,677 | 1,40E-42 | 4,62E-42 | 0,0011832 | 0,0022992 | 0,0015655 | 0,0015655 | TRUE | 0,0114903 | No |
| orf19.3104 | 1316 | 1161 | 1297 | 706 | 2062 | 1406 | 2029 | 2053 | 1255 | 1228 | 1281 | 1106 | 1652 | 1599 | 1723 | 1669 | 1439,06 | 1218 | 1661 | 1,36 | 0,444 | 8,075 | 6,76E-16 | 1,39E-15 | 0,001249 | 0,0032322 | 0,0021759 | 0,0021759 | TRUE | 0,0402066 | No |
| orf19.3105 | 950 | 787 | 890 | 595 | 2269 | 1671 | 2267 | 2367 | 906 | 832 | 879 | 932 | 1818 | 1900 | 1925 | 1925 | 1389,58 | 887 | 1892 | 2,136 | 1,095 | 21,125 | 4,65E-99 | 3,02E-98 | 0,0004072 | 0,0032729 | 0,0017122 | 0,0017122 | TRUE | 0,0204229 | No |
| orf19.3106 | 1358 | 1124 | 1185 | 878 | 3394 | 2382 | 3312 | 3493 | 1295 | 1188 | 1171 | 1375 | 2720 | 2709 | 2812 | 2840 | 2013,67 | 1257 | 2770 | 2,21 | 1,144 | 20,548 | 8,04E-94 | 4,98E-93 | 0,0018898 | 0,0029067 | 0,0023664 | 0,0023664 | TRUE | 0,0429556 | No |
| orf19.3107 | 97 | 102 | 94 | 45 | 309 | 245 | 304 | 304 | 92 | 108 | 93 | 70 | 248 | 279 | 258 | 247 | 174,39 | 91 | 258 | 2,792 | 1,481 | 11,737 | 8,21E-32 | 2,29E-31 | 1,00E-08 | 0,0115062 | 0,0075345 | 0,0075345 | TRUE | 0,1511219 | No |
| orf19.3108 | 407 | 371 | 367 | 198 | 888 | 567 | 893 | 939 | 388 | 392 | 363 | 310 | 712 | 645 | 758 | 763 | 541,37 | 363 | 720 | 1,972 | 0,979 | 11,218 | 3,33E-29 | 8,87E-29 | 0,0055614 | 0,0051241 | 0,0051686 | 0,0051686 | TRUE | 0,0921014 | No |
| orf19.3109 | 27 | 26 | 42 | 20 | 32 | 21 | 27 | 33 | 26 | 27 | 41 | 31 | 26 | 24 | 23 | 27 | 28,17 | 31 | 25 | 0,789 | -0,342 | -1,267 | 0,2052109 | 0,2325845 | 1,00E-08 | 0,0603843 | 0,0346117 | 0,0346117 | TRUE | 0,3440149 | No |
| orf19.3110 | 3812 | 3332 | 3537 | 2156 | 5529 | 3702 | 5152 | 5707 | 3634 | 3523 | 3494 | 3377 | 4430 | 4210 | 4374 | 4640 | 3960,41 | 3507 | 4414 | 1,258 | 0,331 | 7,665 | 1,79E-14 | 3,55E-14 | 0,0009236 | 0,0025059 | 0,0015298 | 0,0015298 | TRUE | 0,0145808 | No |
| orf19.3111 | 51 | 38 | 50 | 37 | 54 | 46 | 69 | 61 | 49 | 40 | 49 | 58 | 43 | 52 | 59 | 50 | 49,99 | 49 | 51 | 1,046 | 0,065 | 0,319 | 0,7499932 | 0,7779112 | 1,00E-08 | 0,0349363 | 0,0202866 | 0,0202866 | TRUE | 0,1127533 | No |
| orf19.3112 | 513 | 486 | 535 | 348 | 497 | 368 | 506 | 527 | 489 | 514 | 529 | 545 | 398 | 418 | 430 | 429 | 468,93 | 519 | 419 | 0,809 | -0,307 | -4,301 | 1,70E-05 | 2,49E-05 | 1,00E-08 | 0,0055927 | 0,0027379 | 0,0027379 | TRUE | 0,018269 | No |
| orf19.3113 | 501 | 413 | 471 | 304 | 605 | 458 | 617 | 661 | 478 | 437 | 465 | 476 | 485 | 521 | 524 | 537 | 490,34 | 464 | 517 | 1,114 | 0,156 | 2,195 | 0,0281396 | 0,0345372 | 1,00E-08 | 0,0054398 | 0,0027792 | 0,0027792 | TRUE | 0,0215971 | No |
| orf19.3114 | 884 | 757 | 796 | 497 | 1227 | 776 | 1156 | 1193 | 843 | 800 | 786 | 779 | 983 | 882 | 981 | 970 | 878,15 | 802 | 954 | 1,19 | 0,251 | 4,209 | 2,57E-05 | 3,74E-05 | 0,0007559 | 0,003961 | 0,0022571 | 0,0022571 | TRUE | 0,0282385 | No |
| orf19.3115 | 838 | 778 | 800 | 496 | 1107 | 724 | 1048 | 1144 | 799 | 823 | 790 | 777 | 887 | 823 | 890 | 930 | 839,89 | 797 | 882 | 1,108 | 0,148 | 2,521 | 0,0117134 | 0,0148241 | 1,00E-08 | 0,0040462 | 0,0021016 | 0,0021016 | TRUE | 0,0222567 | No |
| orf19.3116 | 1793 | 1444 | 1593 | 787 | 1119 | 689 | 1064 | 1106 | 1709 | 1527 | 1574 | 1233 | 897 | 783 | 903 | 899 | 1190,7 | 1511 | 870 | 0,576 | -0,797 | -8,7 | 3,32E-18 | 7,16E-18 | 0,0098578 | 0,0034702 | 0,0071683 | 0,0071683 | TRUE | 0,1762784 | No |
| orf19.3117 | 318 | 240 | 215 | 165 | 38 | 37 | 32 | 39 | 303 | 254 | 212 | 258 | 30 | 42 | 27 | 32 | 144,9 | 257 | 33 | 0,126 | -2,988 | -16,841 | 1,23E-63 | 5,41E-63 | 0,0155745 | 0,0134223 | 0,0143555 | 0,0143555 | TRUE | 0,1338644 | No |
| orf19.3120 | 40 | 29 | 41 | 26 | 10 | 4 | 9 | 20 | 38 | 31 | 41 | 41 | 8 | 5 | 8 | 16 | 23,31 | 38 | 9 | 0,25 | -2 | -5,664 | 1,48E-08 | 2,45E-08 | 0,0171177 | 0,0725222 | 0,0572996 | 0,0572996 | TRUE | 1,0450777 | No |
| orf19.3121 | 40 | 35 | 40 | 25 | 70 | 48 | 72 | 61 | 38 | 37 | 40 | 39 | 56 | 55 | 61 | 50 | 46,9 | 38 | 56 | 1,441 | 0,527 | 2,581 | 0,0098424 | 0,0125084 | 1,00E-08 | 0,0370981 | 0,0174995 | 0,0174995 | TRUE | 0,0471577 | No |
| orf19.3122 | 1464 | 1229 | 1344 | 875 | 2153 | 1509 | 2033 | 2118 | 1396 | 1300 | 1328 | 1371 | 1725 | 1716 | 1726 | 1722 | 1535,36 | 1349 | 1722 | 1,278 | 0,354 | 7,97 | 1,58E-15 | 3,22E-15 | 1,00E-08 | 0,0031607 | 0,0012177 | 0,0012177 | TRUE | 0,0076622 | No |
| orf19.3122.2 | 2489 | 1882 | 1914 | 1233 | 2667 | 1565 | 2449 | 2585 | 2373 | 1990 | 1891 | 1932 | 2137 | 1780 | 2079 | 2102 | 2035,37 | 2046 | 2024 | 0,989 | -0,016 | -0,19 | 0,8493661 | 0,8680326 | 0,0085178 | 0,002898 | 0,0059466 | 0,0059466 | TRUE | 0,1416777 | No |
| orf19.3123 | 9063 | 7261 | 7977 | 5172 | 7685 | 5275 | 7402 | 7996 | 8641 | 7678 | 7881 | 8102 | 6158 | 5998 | 6284 | 6502 | 7155,33 | 8076 | 6236 | 0,772 | -0,373 | -7,98 | 1,46E-15 | 2,97E-15 | 0,0017113 | 0,0023207 | 0,0019542 | 0,0019542 | TRUE | 0,0276028 | No |
| orf19.3123.2 | 308 | 242 | 253 | 182 | 684 | 427 | 613 | 660 | 294 | 256 | 250 | 285 | 548 | 486 | 520 | 537 | 396,91 | 271 | 523 | 1,935 | 0,953 | 11,261 | 2,05E-29 | 5,47E-29 | 0,0012156 | 0,006228 | 0,0039607 | 0,0039607 | TRUE | 0,0423832 | No |
| orf19.3124 | 6407 | 5283 | 5760 | 3975 | 8283 | 5975 | 8059 | 8508 | 6108 | 5586 | 5690 | 6227 | 6637 | 6794 | 6842 | 6918 | 6350,35 | 5903 | 6798 | 1,152 | 0,204 | 4,663 | 3,11E-06 | 4,73E-06 | 0,0013421 | 0,0023498 | 0,0016847 | 0,0016847 | TRUE | 0,0159739 | No |
| orf19.3125 | 540 | 491 | 508 | 315 | 619 | 472 | 640 | 632 | 515 | 519 | 502 | 493 | 496 | 537 | 543 | 514 | 514,91 | 507 | 522 | 1,027 | 0,039 | 0,574 | 0,5660464 | 0,6004891 | 1,00E-08 | 0,00528 | 0,0024241 | 0,0024241 | TRUE | 0,0141303 | No |
| orf19.3126 | 5261 | 4631 | 5049 | 3184 | 7973 | 5692 | 7842 | 8475 | 5016 | 4897 | 4988 | 4988 | 6389 | 6472 | 6658 | 6891 | 5787,25 | 4972 | 6602 | 1,328 | 0,409 | 11,29 | 1,47E-29 | 3,92E-29 | 0,0005289 | 0,002375 | 0,0010817 | 0,0010817 | TRUE | 0,0106802 | No |
| orf19.3127 | 2096 | 1679 | 2029 | 1234 | 821 | 561 | 809 | 901 | 1998 | 1775 | 2004 | 1933 | 658 | 638 | 687 | 733 | 1303,3 | 1928 | 679 | 0,353 | -1,504 | -23,815 | 2,34E-125 | 1,96E-124 | 0,0025056 | 0,0033511 | 0,0028813 | 0,0028813 | TRUE | 0,0351038 | No |
| orf19.3128 | 1328 | 1207 | 1334 | 866 | 2097 | 1545 | 2070 | 2194 | 1266 | 1276 | 1318 | 1357 | 1680 | 1757 | 1757 | 1784 | 1524,41 | 1304 | 1744 | 1,34 | 0,422 | 9,038 | 1,60E-19 | 3,55E-19 | 1,00E-08 | 0,0031684 | 0,0014069 | 0,0014069 | TRUE | 0,0077453 | No |
| orf19.3129 | 5692 | 4711 | 5196 | 3219 | 5392 | 3800 | 5134 | 5650 | 5427 | 4981 | 5133 | 5043 | 4320 | 4321 | 4359 | 4594 | 4772,25 | 5146 | 4398 | 0,855 | -0,227 | -5,42 | 5,97E-08 | 9,70E-08 | 0,0009008 | 0,0024353 | 0,0014704 | 0,0014704 | TRUE | 0,0167566 | No |
| orf19.313 | 7 | 8 | 4 | 5 | 57 | 33 | 53 | 56 | 7 | 8 | 4 | 8 | 46 | 38 | 45 | 46 | 25,08 | 7 | 44 | 6,601 | 2,723 | 7,405 | 1,31E-13 | 2,54E-13 | 1,00E-08 | 0,0675567 | 0,0361194 | 0,0361194 | TRUE | 0,220939 | No |
| orf19.3130 | 6433 | 5721 | 5947 | 3831 | 5770 | 4110 | 5473 | 5830 | 6133 | 6049 | 5875 | 6001 | 4623 | 4673 | 4647 | 4740 | 5342,83 | 6014 | 4671 | 0,777 | -0,365 | -11,697 | 1,32E-31 | 3,66E-31 | 1,00E-08 | 0,0023986 | 0,0007453 | 0,0007453 | TRUE | 0,0031329 | No |
| orf19.3131 | 2124 | 1851 | 1877 | 1326 | 1705 | 1302 | 1561 | 1709 | 2025 | 1957 | 1854 | 2077 | 1366 | 1480 | 1325 | 1390 | 1684,41 | 1978 | 1390 | 0,703 | -0,509 | -9,386 | 6,24E-21 | 1,43E-20 | 0,001483 | 0,003066 | 0,0022162 | 0,0022162 | TRUE | 0,0217905 | No |
| orf19.3132 | 2856 | 2593 | 2639 | 1758 | 2165 | 1555 | 2053 | 2153 | 2723 | 2742 | 2607 | 2754 | 1735 | 1768 | 1743 | 1751 | 2227,78 | 2706 | 1749 | 0,647 | -0,629 | -15,957 | 2,56E-57 | 1,04E-56 | 1,00E-08 | 0,0028283 | 0,0010306 | 0,0010306 | TRUE | 0,0076158 | No |
| orf19.3133 | 20477 | 17515 | 18699 | 12784 | 26485 | 19903 | 25740 | 28366 | 19523 | 18520 | 18473 | 20027 | 21222 | 22632 | 21853 | 23064 | 20664,09 | 19136 | 22193 | 1,16 | 0,214 | 5,063 | 4,12E-07 | 6,47E-07 | 0,0014176 | 0,0021707 | 0,0016667 | 0,0016667 | TRUE | 0,0118692 | No |
| orf19.3134 | 4 | 4 | 5 | 5 | 3 | 2 | 4 | 3 | 4 | 4 | 5 | 8 | 2 | 2 | 3 | 2 | 3,92 | 5 | 2 | 0,523 | -0,935 | -1,324 | 0,1856289 | 0,2111028 | 1,00E-08 | 0,4213638 | 0,2010261 | 0,2010261 | TRUE | 0,1421168 | No |
| orf19.3135 | 2420 | 2263 | 2402 | 1513 | 2335 | 1674 | 2194 | 2336 | 2307 | 2393 | 2373 | 2370 | 1871 | 1903 | 1863 | 1899 | 2122,47 | 2361 | 1884 | 0,798 | -0,325 | -8,484 | 2,18E-17 | 4,62E-17 | 1,00E-08 | 0,0028649 | 0,0009375 | 0,0009375 | TRUE | 0,0030182 | No |
| orf19.3136 | 6056 | 5488 | 5859 | 3512 | 3682 | 2597 | 3641 | 3806 | 5774 | 5803 | 5788 | 5502 | 2950 | 2953 | 3091 | 3095 | 4369,45 | 5717 | 3022 | 0,529 | -0,92 | -24,6 | 1,26E-133 | 1,13E-132 | 0,0003433 | 0,002467 | 0,0010967 | 0,0010967 | TRUE | 0,007352 | No |
| orf19.3138 | 24277 | 17839 | 20810 | 13803 | 44000 | 28424 | 41751 | 44147 | 23145 | 18862 | 20558 | 21623 | 35256 | 32321 | 35446 | 35896 | 27888,52 | 21047 | 34730 | 1,65 | 0,723 | 11,562 | 6,41E-31 | 1,75E-30 | 0,0047151 | 0,0021502 | 0,0037132 | 0,0037132 | TRUE | 0,0598795 | No |
| orf19.3139 | 3333 | 2802 | 2991 | 1875 | 3432 | 2321 | 3470 | 3609 | 3178 | 2963 | 2955 | 2937 | 2750 | 2639 | 2946 | 2934 | 2912,77 | 3008 | 2817 | 0,937 | -0,094 | -1,868 | 0,0617928 | 0,0735908 | 0,0016512 | 0,002655 | 0,0021035 | 0,0021035 | TRUE | 0,0211986 | No |
| orf19.314 | 922 | 850 | 954 | 592 | 1041 | 712 | 933 | 1034 | 879 | 899 | 942 | 927 | 834 | 810 | 792 | 841 | 865,53 | 912 | 819 | 0,9 | -0,153 | -2,766 | 0,0056734 | 0,0073091 | 1,00E-08 | 0,0039883 | 0,0017723 | 0,0017723 | TRUE | 0,007489 | No |
| orf19.3140 | 23 | 19 | 18 | 18 | 7 | 7 | 14 | 16 | 22 | 20 | 18 | 28 | 6 | 8 | 12 | 13 | 15,81 | 22 | 10 | 0,446 | -1,166 | -3,056 | 0,0022403 | 0,0029604 | 1,00E-08 | 0,105958 | 0,0680978 | 0,0680978 | TRUE | 0,3523686 | No |
| orf19.3140.1 | 4355 | 3321 | 3793 | 2510 | 5458 | 3526 | 5302 | 5420 | 4152 | 3512 | 3747 | 3932 | 4373 | 4009 | 4501 | 4407 | 4079,23 | 3836 | 4322 | 1,128 | 0,173 | 2,923 | 0,0034638 | 0,0045228 | 0,0034365 | 0,0024938 | 0,0031198 | 0,0031198 | TRUE | 0,0397175 | No |
| orf19.3141 | 5012 | 4268 | 4808 | 3249 | 5513 | 3792 | 5292 | 5688 | 4778 | 4513 | 4750 | 5090 | 4417 | 4312 | 4493 | 4625 | 4622,23 | 4783 | 4462 | 0,934 | -0,099 | -2,177 | 0,0294504 | 0,0361043 | 0,0013721 | 0,0024465 | 0,0017499 | 0,0017499 | TRUE | 0,0217004 | No |
| orf19.3142 | 646 | 610 | 614 | 442 | 396 | 285 | 345 | 346 | 616 | 645 | 607 | 692 | 317 | 324 | 293 | 281 | 471,94 | 640 | 304 | 0,475 | -1,073 | -13,628 | 2,71E-42 | 8,92E-42 | 0,0013901 | 0,0055704 | 0,0036034 | 0,0036034 | TRUE | 0,0315041 | No |
| orf19.3147 | 571 | 795 | 858 | 425 | 702 | 549 | 639 | 624 | 544 | 841 | 848 | 666 | 562 | 624 | 543 | 507 | 641,88 | 725 | 559 | 0,77 | -0,376 | -2,935 | 0,0033355 | 0,0043579 | 0,0240353 | 0,0046492 | 0,0142006 | 0,0142006 | TRUE | 0,3429588 | No |
| orf19.3148 | 731 | 682 | 788 | 429 | 520 | 327 | 465 | 460 | 697 | 721 | 778 | 672 | 417 | 372 | 395 | 374 | 553,23 | 717 | 390 | 0,543 | -0,882 | -11,908 | 1,07E-32 | 3,04E-32 | 0,0016908 | 0,0050591 | 0,0033485 | 0,0033485 | TRUE | 0,0395988 | No |
| orf19.3149 | 7794 | 7369 | 7374 | 4847 | 7394 | 5572 | 6706 | 7142 | 7431 | 7792 | 7285 | 7593 | 5925 | 6336 | 5693 | 5807 | 6732,67 | 7525 | 5940 | 0,789 | -0,342 | -7,832 | 4,79E-15 | 9,65E-15 | 0,0013303 | 0,0023352 | 0,001677 | 0,001677 | TRUE | 0,0240041 | No |

| | | | | | | | | | | | | | | | | | | | | | | | | | | | | | | | |
|---|---|---|---|---|---|---|---|---|---|---|---|---|---|---|---|---|---|---|---|---|---|---|---|---|---|---|---|---|---|---|---|
| orf19.315 | 93 | 66 | 66 | 59 | 111 | 80 | 122 | 115 | 89 | 70 | 65 | 92 | 89 | 91 | 104 | 94 | 86,63 | 79 | 94 | 1,204 | 0,267 | 1,64 | 0,1009855 | 0,1179119 | 0,0020271 | 0,0210435 | 0,0136699 | 0,0136699 | TRUE | 0,1320231 | No |
| orf19.3150 | 2578 | 2455 | 2396 | 1612 | 1262 | 962 | 1245 | 1247 | 2458 | 2596 | 2367 | 2525 | 1011 | 1094 | 1057 | 1014 | 1765,25 | 2486 | 1044 | 0,42 | -1,253 | -24,975 | 1,13E-137 | 1,04E-136 | 0,0009072 | 0,0030214 | 0,0017635 | 0,0017635 | TRUE | 0,0129837 | No |
| orf19.3151 | 421 | 450 | 402 | 259 | 637 | 497 | 552 | 544 | 401 | 476 | 397 | 406 | 510 | 565 | 469 | 442 | 458,32 | 420 | 496 | 1,179 | 0,238 | 2,481 | 0,0131161 | 0,0165338 | 0,0074272 | 0,0056737 | 0,0065665 | 0,0065665 | TRUE | 0,0948355 | No |
| orf19.3152 | 1455 | 1354 | 1331 | 860 | 1481 | 1137 | 1352 | 1491 | 1387 | 1432 | 1315 | 1347 | 1187 | 1293 | 1148 | 1212 | 1290,09 | 1370 | 1210 | 0,881 | -0,182 | -3,331 | 0,0008645 | 0,0011718 | 0,0011165 | 0,003364 | 0,0020915 | 0,0020915 | TRUE | 0,0238327 | No |
| orf19.3153 | 6349 | 6114 | 6861 | 3927 | 4576 | 3224 | 4292 | 4502 | 6053 | 6465 | 6778 | 6152 | 3667 | 3666 | 3644 | 3661 | 5010,6 | 6362 | 3660 | 0,575 | -0,798 | -18,335 | 4,32E-75 | 2,21E-74 | 0,0010482 | 0,0024419 | 0,0016103 | 0,0016103 | TRUE | 0,0239234 | No |
| orf19.3154 | 1949 | 1996 | 2060 | 1221 | 1895 | 1444 | 1771 | 1851 | 1858 | 2111 | 2035 | 1913 | 1518 | 1642 | 1504 | 1505 | 1760,69 | 1979 | 1542 | 0,778 | -0,362 | -6,479 | 9,24E-11 | 1,66E-10 | 0,001847 | 0,0030238 | 0,0024242 | 0,0024242 | TRUE | 0,0242909 | No |
| orf19.3156 | 1810 | 1628 | 1779 | 1091 | 831 | 566 | 845 | 810 | 1726 | 1721 | 1757 | 1709 | 666 | 644 | 717 | 659 | 1199,89 | 1728 | 672 | 0,389 | -1,363 | -26,304 | 1,73E-152 | 1,77E-151 | 1,00E-08 | 0,0034597 | 0,0015918 | 0,0015918 | TRUE | 0,0252636 | No |
| orf19.3158 | 2542 | 2333 | 2520 | 1614 | 1299 | 964 | 1295 | 1340 | 2424 | 2467 | 2490 | 2528 | 1041 | 1096 | 1099 | 1090 | 1779,29 | 2477 | 1082 | 0,437 | -1,195 | -27,542 | 5,52E-167 | 6,21E-166 | 1,00E-08 | 0,0030141 | 0,00117 | 0,00117 | TRUE | 0,00803 | No |
| orf19.3159 | 3380 | 3547 | 3504 | 2001 | 7868 | 6177 | 7255 | 7312 | 3222 | 3751 | 3462 | 3135 | 6304 | 7024 | 6159 | 5945 | 4875,29 | 3392 | 6358 | 1,873 | 0,905 | 13,278 | 3,10E-40 | 9,86E-40 | 0,0057094 | 0,0024281 | 0,004227 | 0,004227 | TRUE | 0,0612837 | No |
| orf19.3159.4 | 0 | 2 | 1 | 0 | 3 | 4 | 3 | 1 | 0 | 2 | 1 | 0 | 2 | 5 | 3 | 1 | 1,68 | 1 | 3 | 3,095 | 1,63 | 1,313 | 0,1891089 | 0,2148307 | 1,00E-08 | 0,9812429 | 0,6211496 | 0,6211496 | TRUE | 0,3712821 | No |
| orf19.316 | 1212 | 939 | 1062 | 724 | 2235 | 1485 | 2063 | 2287 | 1156 | 993 | 1049 | 1134 | 1791 | 1689 | 1751 | 1860 | 1427,77 | 1083 | 1773 | 1,64 | 0,713 | 11,839 | 2,46E-32 | 6,92E-32 | 0,0022618 | 0,0032413 | 0,0027158 | 0,0027158 | TRUE | 0,0377209 | No |
| orf19.3160 | 2415 | 1873 | 1853 | 1233 | 3157 | 1897 | 3036 | 3194 | 2302 | 1980 | 1831 | 1932 | 2530 | 2157 | 2578 | 2597 | 2238,29 | 2011 | 2466 | 1,226 | 0,294 | 3,652 | 0,00026 | 0,0003613 | 0,0080774 | 0,0028248 | 0,005753 | 0,005753 | TRUE | 0,1168595 | No |
| orf19.3161 | 6650 | 5573 | 5989 | 3475 | 8712 | 5304 | 8668 | 8928 | 6340 | 5893 | 5917 | 5444 | 6981 | 6031 | 7359 | 7259 | 6402,92 | 5898 | 6908 | 1,171 | 0,228 | 3,332 | 0,0008627 | 0,0011698 | 0,005563 | 0,0023477 | 0,0043262 | 0,0043262 | TRUE | 0,0883181 | No |
| orf19.3163 | 831 | 704 | 790 | 484 | 963 | 689 | 934 | 984 | 792 | 744 | 780 | 758 | 772 | 783 | 793 | 800 | 777,93 | 768 | 787 | 1,023 | 0,032 | 0,58 | 0,5621062 | 0,596706 | 1,00E-08 | 0,0042019 | 0,0016893 | 0,0016893 | TRUE | 0,0060533 | No |
| orf19.3166 | 619 | 578 | 599 | 385 | 954 | 751 | 912 | 1021 | 590 | 611 | 592 | 603 | 764 | 854 | 774 | 830 | 702,38 | 599 | 806 | 1,344 | 0,426 | 6,736 | 1,62E-11 | 2,97E-11 | 1,00E-08 | 0,0044289 | 0,0023544 | 0,0023544 | TRUE | 0,0177367 | No |
| orf19.3167 | 3675 | 2737 | 3090 | 2226 | 3874 | 2696 | 3618 | 3869 | 3504 | 2894 | 3053 | 3487 | 3104 | 3066 | 3072 | 3146 | 3165,6 | 3234 | 3097 | 0,958 | -0,062 | -0,967 | 0,3335357 | 0,3672643 | 0,0041947 | 0,00261 | 0,0035831 | 0,0035831 | TRUE | 0,0609609 | No |
| orf19.3168 | 6312 | 5632 | 6024 | 3595 | 6555 | 4328 | 6621 | 6886 | 6018 | 5955 | 5951 | 5632 | 5252 | 4921 | 5621 | 5599 | 5618,71 | 5889 | 5348 | 0,908 | -0,139 | -2,758 | 0,0058142 | 0,007486 | 0,0021153 | 0,0023835 | 0,002255 | 0,002255 | TRUE | 0,0345534 | No |
| orf19.3169 | 7882 | 6592 | 7370 | 4604 | 5647 | 3703 | 5425 | 5619 | 7515 | 6970 | 7281 | 7212 | 4525 | 4211 | 4606 | 4569 | 5861,02 | 7244 | 4478 | 0,618 | -0,694 | -16,435 | 1,08E-60 | 4,56E-60 | 0,0010539 | 0,0023714 | 0,0015326 | 0,0015326 | TRUE | 0,0189922 | No |
| orf19.317 | 1105 | 914 | 1027 | 634 | 1107 | 749 | 1101 | 1227 | 1053 | 966 | 1015 | 993 | 887 | 852 | 935 | 998 | 962,35 | 1007 | 918 | 0,912 | -0,133 | -2,088 | 0,0367749 | 0,0447229 | 0,0020512 | 0,0037974 | 0,0028314 | 0,0028314 | TRUE | 0,0412495 | No |
| orf19.3170 | 2908 | 2698 | 2999 | 1721 | 5130 | 3449 | 5137 | 5211 | 2772 | 2853 | 2963 | 2696 | 4111 | 3922 | 4361 | 4237 | 3489,34 | 2821 | 4158 | 1,473 | 0,559 | 11,647 | 2,39E-31 | 6,59E-31 | 0,0014398 | 0,0025618 | 0,0019047 | 0,0019047 | TRUE | 0,0172458 | No |
| orf19.3171 | 9767 | 8394 | 8929 | 6676 | 12713 | 9568 | 11997 | 13589 | 9312 | 8876 | 8821 | 10458 | 10187 | 10880 | 10185 | 11049 | 9970,93 | 9367 | 10575 | 1,13 | 0,176 | 2,967 | 0,0030024 | 0,0039316 | 0,0040835 | 0,0022559 | 0,0032657 | 0,0032657 | TRUE | 0,0741997 | No |
| orf19.3172 | 13 | 7 | 16 | 14 | 13 | 21 | 14 | 26 | 12 | 7 | 16 | 22 | 10 | 24 | 12 | 21 | 15,61 | 14 | 17 | 1,184 | 0,244 | 0,576 | 0,5644926 | 0,5990399 | 0,1026369 | 0,1072945 | 0,1054872 | 0,1054872 | TRUE | 0,2206648 | No |
| orf19.3173 | 613 | 494 | 531 | 295 | 1655 | 996 | 1619 | 1735 | 584 | 522 | 525 | 462 | 1326 | 1133 | 1375 | 1411 | 917,17 | 523 | 1311 | 2,498 | 1,321 | 15,23 | 2,22E-52 | 8,47E-52 | 0,0072356 | 0,0038815 | 0,0058101 | 0,0058101 | TRUE | 0,0977235 | No |
| orf19.3174 | 1202 | 1102 | 1231 | 811 | 1973 | 1460 | 1958 | 2063 | 1146 | 1165 | 1216 | 1270 | 1581 | 1660 | 1662 | 1677 | 1422,32 | 1199 | 1645 | 1,375 | 0,459 | 9,11 | 8,21E-20 | 1,84E-19 | 0,0004727 | 0,0032457 | 0,0016984 | 0,0016984 | TRUE | 0,0166349 | No |
| orf19.3175 | 6107 | 5247 | 5637 | 3617 | 6355 | 4122 | 5934 | 6448 | 5822 | 5548 | 5569 | 5666 | 5092 | 4687 | 5038 | 5243 | 5333,17 | 5651 | 5015 | 0,888 | -0,172 | -3,994 | 6,50E-05 | 9,27E-05 | 0,0010936 | 0,0023991 | 0,0015825 | 0,0015825 | TRUE | 0,0229784 | No |
| orf19.3176 | 2033 | 1734 | 1920 | 1278 | 1288 | 1038 | 1320 | 1446 | 1938 | 1833 | 1897 | 2002 | 1032 | 1180 | 1121 | 1176 | 1522,41 | 1918 | 1127 | 0,588 | -0,767 | -13,523 | 1,15E-41 | 3,74E-41 | 0,0016162 | 0,0031698 | 0,0023976 | 0,0023976 | TRUE | 0,0397145 | No |
| orf19.3177 | 509 | 444 | 477 | 258 | 2206 | 1379 | 2072 | 2234 | 485 | 469 | 471 | 404 | 1768 | 1568 | 1759 | 1816 | 1092,67 | 457 | 1728 | 3,759 | 1,91 | 26,701 | 4,54E-157 | 4,84E-156 | 0,0033067 | 0,0035939 | 0,0034414 | 0,0034414 | TRUE | 0,0592835 | No |
| orf19.3178 | 2878 | 2583 | 2843 | 1554 | 2304 | 1507 | 2128 | 2269 | 2744 | 2731 | 2809 | 2434 | 1846 | 1714 | 1807 | 1845 | 2241,17 | 2680 | 1803 | 0,672 | -0,573 | -10,619 | 2,43E-26 | 6,19E-26 | 0,0019169 | 0,0028239 | 0,0023389 | 0,0023389 | TRUE | 0,0431551 | No |
| orf19.318 | 1006 | 908 | 944 | 576 | 1426 | 971 | 1329 | 1528 | 959 | 960 | 933 | 902 | 1143 | 1104 | 1128 | 1242 | 1046,45 | 938 | 1154 | 1,229 | 0,298 | 5,212 | 1,87E-07 | 2,98E-07 | 0,0010206 | 0,0036603 | 0,0021545 | 0,0021545 | TRUE | 0,0321182 | No |
| orf19.3180 | 1346 | 1152 | 1249 | 778 | 1398 | 980 | 1284 | 1410 | 1283 | 1218 | 1234 | 1219 | 1120 | 1114 | 1090 | 1146 | 1178,14 | 1238 | 1118 | 0,902 | -0,149 | -3,083 | 0,0020466 | 0,0027099 | 1,00E-08 | 0,0034849 | 0,0013947 | 0,0013947 | TRUE | 0,007462 | No |
| orf19.3181.1 | 424 | 318 | 357 | 209 | 629 | 431 | 575 | 590 | 404 | 336 | 353 | 327 | 504 | 490 | 488 | 480 | 422,82 | 355 | 490 | 1,374 | 0,459 | 5,631 | 1,79E-08 | 2,96E-08 | 0,0013695 | 0,0059745 | 0,0039 | 0,0039 | TRUE | 0,1028592 | No |
| orf19.3182 | 13072 | 10295 | 11098 | 7595 | 25026 | 16897 | 23897 | 24981 | 12463 | 10886 | 10964 | 11898 | 20053 | 19213 | 20288 | 20312 | 15759,57 | 11553 | 19966 | 1,729 | 0,79 | 15,813 | 2,53E-56 | 1,01E-55 | 0,0024142 | 0,0021955 | 0,002324 | 0,002324 | TRUE | 0,0348883 | No |
| orf19.3183 | 1268 | 1043 | 1181 | 903 | 1466 | 1036 | 1323 | 1333 | 1209 | 1103 | 1167 | 1415 | 1175 | 1178 | 1123 | 1084 | 1181,6 | 1224 | 1140 | 0,934 | -0,098 | -1,315 | 0,1883443 | 0,2140002 | 0,005722 | 0,0034808 | 0,0044704 | 0,0044704 | TRUE | 0,1240119 | No |
| orf19.3184 | 2536 | 2111 | 2241 | 1405 | 1598 | 1109 | 1635 | 1538 | 2418 | 2232 | 2214 | 2201 | 1280 | 1261 | 1388 | 1251 | 1780,62 | 2266 | 1295 | 0,571 | -0,808 | -15,016 | 5,75E-51 | 2,15E-50 | 0,0016222 | 0,0030134 | 0,0021867 | 0,0021867 | TRUE | 0,0285242 | No |
| orf19.3185 | 4943 | 4496 | 4768 | 3135 | 6007 | 4392 | 5653 | 6001 | 4713 | 4754 | 4710 | 4911 | 4813 | 4994 | 4799 | 4879 | 4821,77 | 4772 | 4871 | 1,021 | 0,03 | 0,918 | 0,3585107 | 0,3932708 | 8,76E-05 | 0,0024318 | 0,0008406 | 0,0008406 | TRUE | 0,0043316 | No |
| orf19.3186 | 0 | 1 | 3 | 0 | 1 | 0 | 1 | 3 | 0 | 1 | 3 | 0 | 1 | 0 | 1 | 2 | 1,01 | 1 | 1 | 1,003 | 0,005 | 0,003 | 0,9974339 | 0,9977466 | 0,4598656 | 1,6215965 | 1,2148668 | 1,2148668 | TRUE | 0,8192836 | No |
| orf19.3187 | 3880 | 3734 | 3933 | 2465 | 3863 | 2940 | 3739 | 3912 | 3699 | 3948 | 3885 | 3862 | 3095 | 3343 | 3174 | 3181 | 3523,49 | 3848 | 3198 | 0,831 | -0,268 | -6,601 | 4,08E-11 | 7,36E-11 | 0,0005269 | 0,0025573 | 0,0012931 | 0,0012931 | TRUE | 0,0107754 | No |
| orf19.3188 | 1955 | 1880 | 1967 | 1265 | 2014 | 1526 | 1898 | 1971 | 1864 | 1988 | 1943 | 1982 | 1614 | 1735 | 1611 | 1603 | 1792,45 | 1944 | 1641 | 0,844 | -0,245 | -5,241 | 1,60E-07 | 2,55E-07 | 0,0005417 | 0,0030073 | 0,0015441 | 0,0015441 | TRUE | 0,016947 | No |
| orf19.319 | 192 | 158 | 181 | 99 | 344 | 265 | 324 | 356 | 183 | 167 | 179 | 155 | 276 | 301 | 275 | 289 | 228,19 | 171 | 285 | 1,655 | 0,727 | 7,312 | 2,64E-13 | 5,09E-13 | 1,00E-08 | 0,0092866 | 0,004691 | 0,004691 | TRUE | 0,0317747 | No |
| orf19.3190 | 7022 | 6486 | 6792 | 4490 | 2759 | 2043 | 2559 | 2748 | 6695 | 6858 | 6710 | 7034 | 2211 | 2323 | 2173 | 2234 | 4529,66 | 6824 | 2235 | 0,327 | -1,611 | -43,019 | 0 | 0 | 0,0002807 | 0,0024538 | 0,0010637 | 0,0010637 | TRUE | 0,0078918 | No |
| orf19.3192 | 6510 | 6062 | 6419 | 4334 | 3932 | 2928 | 3669 | 4118 | 6207 | 6410 | 6341 | 6789 | 3151 | 3329 | 3115 | 3348 | 4836,31 | 6437 | 3236 | 0,503 | -0,992 | -22,608 | 3,58E-113 | 2,67E-112 | 0,0011324 | 0,0024308 | 0,0016218 | 0,0016218 | TRUE | 0,0160218 | No |
| orf19.3193 | 1620 | 1598 | 1642 | 1104 | 1811 | 1456 | 1763 | 1849 | 1544 | 1690 | 1622 | 1729 | 1451 | 1656 | 1497 | 1503 | 1586,59 | 1646 | 1527 | 0,927 | -0,109 | -1,882 | 0,0597768 | 0,0712964 | 0,0020721 | 0,0031261 | 0,0025691 | 0,0025691 | TRUE | 0,0371396 | No |
| orf19.3195 | 10526 | 9494 | 10692 | 6442 | 9541 | 6586 | 9032 | 9687 | 10035 | 10039 | 10563 | 10092 | 7645 | 7489 | 7668 | 7876 | 8925,86 | 10182 | 7670 | 0,753 | -0,409 | -12,285 | 1,09E-34 | 3,16E-34 | 0,0003767 | 0,0022752 | 0,0009491 | 0,0009491 | TRUE | 0,0078662 | No |
| orf19.3196 | 259 | 278 | 265 | 205 | 186 | 199 | 162 | 194 | 247 | 294 | 262 | 321 | 149 | 226 | 138 | 158 | 224,3 | 281 | 168 | 0,595 | -0,75 | -4,982 | 6,29E-07 | 9,82E-07 | 0,0259825 | 0,0094114 | 0,0170443 | 0,0170443 | TRUE | 0,552227 | No |
| orf19.3197 | 1435 | 1273 | 1349 | 825 | 2152 | 1505 | 2026 | 2103 | 1368 | 1346 | 1333 | 1292 | 1724 | 1711 | 1720 | 1710 | 1525,61 | 1335 | 1716 | 1,284 | 0,361 | 8,386 | 5,02E-17 | 1,05E-16 | 1,00E-08 | 0,0031675 | 0,0010968 | 0,0010968 | TRUE | 0,0046492 | No |
| orf19.3198 | 2695 | 2190 | 2390 | 1464 | 3055 | 1997 | 3022 | 3136 | 2569 | 2316 | 2361 | 2293 | 2448 | 2271 | 2566 | 2550 | 2421,72 | 2385 | 2459 | 1,031 | 0,044 | 0,78 | 0,4351238 | 0,4712364 | 0,0023506 | 0,0027693 | 0,0025941 | 0,0025941 | TRUE | 0,0334719 | No |
| orf19.3199 | 1411 | 1256 | 1344 | 784 | 1442 | 1006 | 1304 | 1402 | 1345 | 1328 | 1328 | 1228 | 1155 | 1144 | 1107 | 1140 | 1221,95 | 1307 | 1136 | 0,867 | -0,206 | -4,143 | 3,43E-05 | 4,96E-05 | 1,00E-08 | 0,003435 | 0,0015457 | 0,0015457 | TRUE | 0,0172648 | No |
| orf19.320 | 2226 | 1882 | 2062 | 1541 | 1396 | 974 | 1382 | 1432 | 2122 | 1990 | 2037 | 2414 | 1119 | 1108 | 1173 | 1164 | 1640,89 | 2141 | 1141 | 0,534 | -0,904 | -13,99 | 1,79E-44 | 6,07E-44 | 0,0034621 | 0,0030919 | 0,003358 | 0,003358 | TRUE | 0,0846775 | No |
| orf19.3200 | 103 | 124 | 106 | 92 | 151 | 128 | 150 | 149 | 98 | 131 | 105 | 144 | 121 | 146 | 127 | 121 | 124,15 | 120 | 129 | 1,085 | 0,118 | 0,795 | 0,4265263 | 0,4626864 | 0,0097724 | 0,0153165 | 0,0129839 | 0,0129839 | TRUE | 0,1568629 | No |
| orf19.3201 | 1140 | 967 | 1064 | 704 | 1221 | 851 | 1116 | 1123 | 1087 | 1022 | 1051 | 1103 | 978 | 968 | 947 | 913 | 1008,74 | 1066 | 952 | 0,894 | -0,162 | -3,051 | 0,0022793 | 0,0030099 | 1,00E-08 | 0,003719 | 0,0017244 | 0,0017244 | TRUE | 0,0092149 | No |
| orf19.3202 | 870 | 1042 | 978 | 628 | 598 | 505 | 595 | 573 | 829 | 1102 | 966 | 984 | 479 | 574 | 505 | 466 | 738,21 | 970 | 506 | 0,521 | -0,941 | -9,924 | 3,26E-23 | 7,79E-23 | 0,0091476 | 0,0043155 | 0,0071736 | 0,0071736 | TRUE | 0,1176847 | No |
| orf19.3203 | 464 | 541 | 587 | 320 | 702 | 539 | 729 | 676 | 442 | 572 | 580 | 501 | 562 | 613 | 619 | 550 | 554,95 | 524 | 586 | 1,117 | 0,159 | 1,713 | 0,0866441 | 0,1017438 | 0,0078366 | 0,00505 | 0,0064738 | 0,0064738 | TRUE | 0,1384497 | No |
| orf19.3204 | 1935 | 2394 | 2341 | 1262 | 1564 | 1141 | 1482 | 1340 | 1845 | 2531 | 2313 | 1977 | 1253 | 1297 | 1258 | 1090 | 1695,53 | 2166 | 1224 | 0,565 | -0,824 | -8,601 | 7,92E-18 | 1,70E-17 | 0,0124164 | 0,0030597 | 0,0081972 | 0,0081972 | TRUE | 0,1558734 | No |
| orf19.3205 | 4323 | 3781 | 4093 | 2300 | 3271 | 2122 | 3163 | 3182 | 4122 | 3998 | 4043 | 3603 | 2621 | 2413 | 2685 | 2587 | 3259,06 | 3942 | 2576 | 0,653 | -0,614 | -11,394 | 4,49E-30 | 1,21E-29 | 0,0023162 | 0,0025951 | 0,0024721 | 0,0024721 | TRUE | 0,0389582 | No |
| orf19.3206 | 5872 | 5819 | 6254 | 3661 | 8117 | 6007 | 7681 | 7760 | 5598 | 6153 | 6178 | 5735 | 6504 | 6830 | 6521 | 6310 | 6228,73 | 5916 | 6541 | 1,105 | 0,144 | 3,138 | 0,0016992 | 0,0022622 | 0,0015585 | 0,0023549 | 0,0018626 | 0,0018626 | TRUE | 0,0165318 | No |
| orf19.3207 | 135 | 137 | 160 | 90 | 197 | 153 | 156 | 167 | 129 | 145 | 158 | 141 | 158 | 174 | 132 | 136 | 146,58 | 143 | 150 | 1,043 | 0,06 | 0,457 | 0,6474575 | 0,6796523 | 0,0058068 | 0,0132923 | 0,0098968 | 0,0098968 | TRUE | 0,1084056 | No |
| orf19.3208 | 843 | 844 | 973 | 515 | 453 | 367 | 481 | 467 | 804 | 892 | 961 | 807 | 363 | 417 | 408 | 380 | 629,06 | 866 | 392 | 0,451 | -1,149 | -14,121 | 2,82E-45 | 9,69E-45 | 0,0044396 | 0,0047014 | 0,004564 | 0,004564 | TRUE | 0,0662486 | No |
| orf19.3209 | 23 | 18 | 22 | 15 | 43 | 13 | 33 | 32 | 22 | 19 | 22 | 23 | 34 | 15 | 28 | 26 | 23,68 | 22 | 26 | 1,221 | 0,288 | 0,927 | 0,3540924 | 0,3887796 | 1,00E-08 | 0,071419 | 0,049835 | 0,049835 | TRUE | 0,3761374 | No |
| orf19.321 | 1413 | 1290 | 1354 | 985 | 3909 | 2730 | 3781 | 4027 | 1347 | 1364 | 1338 | 1543 | 3132 | 3104 | 3210 | 3274 | 2289,08 | 1398 | 3180 | 2,283 | 1,191 | 22,952 | 1,39E-116 | 1,07E-115 | 0,0013776 | 0,0028086 | 0,0020427 | 0,0020427 | TRUE | 0,0528298 | No |
| orf19.3210 | 23 | 35 | 36 | 18 | 39 | 36 | 44 | 37 | 22 | 37 | 36 | 28 | 31 | 41 | 37 | 30 | 32,79 | 31 | 35 | 1,128 | 0,174 | 0,671 | 0,5024366 | 0,5381984 | 1,00E-08 | 0,0521639 | 0,0338611 | 0,0338611 | TRUE | 0,293114 | No |
| orf19.3211 | 1266 | 1187 | 1365 | 808 | 1810 | 1337 | 1638 | 1730 | 1207 | 1255 | 1348 | 1266 | 1450 | 1520 | 1391 | 1407 | 1355,53 | 1269 | 1442 | 1,135 | 0,183 | 3,378 | 0,0007295 | 0,000994 | 0,0011382 | 0,0033025 | 0,002058 | 0,002058 | TRUE | 0,0227072 | No |
| orf19.3212 | 1980 | 2007 | 2057 | 1261 | 1298 | 909 | 1142 | 1115 | 1888 | 2122 | 2032 | 1975 | 1040 | 1034 | 970 | 907 | 1495,9 | 2004 | 988 | 0,492 | -1,023 | -16,829 | 1,50E-63 | 6,59E-63 | 0,0025287 | 0,0031889 | 0,0028127 | 0,0028127 | TRUE | 0,0371438 | No |
| orf19.3213 | 2120 | 2136 | 2317 | 1444 | 2875 | 2027 | 2764 | 2758 | 2021 | 2259 | 2289 | 2262 | 2304 | 2305 | 2347 | 2243 | 2253,56 | 2208 | 2300 | 1,042 | 0,06 | 1,197 | 0,231376 | 0,2605277 | 0,0013935 | 0,0028199 | 0,0019509 | 0,0019509 | TRUE | 0,040722 | No |
| orf19.3214 | 424 | 477 | 504 | 297 | 915 | 732 | 846 | 703 | 404 | 504 | 498 | 465 | 733 | 832 | 718 | 572 | 590,89 | 468 | 714 | 1,524 | 0,607 | 5,394 | 6,88E-08 | 1,12E-07 | 0,0155957 | 0,00487 | 0,0103448 | 0,0103448 | TRUE | 0,215386 | No |
| orf19.3215 | 2122 | 2040 | 2101 | 1237 | 2059 | 1375 | 2032 | 1884 | 2023 | 2157 | 2076 | 1938 | 1650 | 1563 | 1725 | 1532 | 1832,99 | 2048 | 1618 | 0,789 | -0,342 | -6,238 | 4,44E-10 | 7,75E-10 | 0,0017292 | 0,002987 | 0,0023284 | 0,0023284 | TRUE | 0,0243168 | No |
| orf19.3216 | 1142 | 1277 | 1381 | 894 | 998 | 781 | 892 | 980 | 1089 | 1350 | 1364 | 1400 | 800 | 888 | 757 | 797 | 1055,71 | 1301 | 810 | 0,623 | -0,682 | -8,041 | 8,95E-16 | 1,83E-15 | 0,0078966 | 0,0036465 | 0,0059311 | 0,0059311 | TRUE | 0,1480078 | No |
| orf19.3218 | 766 | 781 | 804 | 466 | 855 | 650 | 772 | 758 | 730 | 826 | 794 | 730 | 685 | 739 | 655 | 616 | 722,04 | 770 | 674 | 0,872 | -0,197 | -2,671 | 0,0075672 | 0,0096707 | 0,0033654 | 0,0043652 | 0,00384 | 0,00384 | TRUE | 0,0466856 | No |
| orf19.3219 | 250 | 372 | 366 | 200 | 353 | 262 | 332 | 307 | 238 | 393 | 362 | 313 | 283 | 298 | 282 | 250 | 302,35 | 326 | 278 | 0,85 | -0,234 | -1,692 | 0,0907148 | 0,1063088 | 0,0227583 | 0,0075217 | 0,01501 | 0,01501 | TRUE | 0,3976881 | No |
| orf19.322 | 9 | 6 | 6 | 9 | 5 | 2 | 1 | 8 | 9 | 6 | 6 | 14 | 4 | 2 | 1 | 7 | 6,07 | 9 | 4 | 0,409 | -1,292 | -1,992 | 0,0464019 | 0,0559611 | 0,1006377 | 0,2724443 | 0,2101967 | 0,2101967 | TRUE | 0,6678035 | No |
| orf19.3220 | 7007 | 6462 | 6648 | 3854 | 9050 | 5889 | 8311 | 8377 | 6680 | 6833 | 6568 | 6037 | 7252 | 6696 | 7056 | 6811 | 6741,67 | 6530 | 6954 | 1,064 | 0,09 | 1,893 | 0,0583173 | 0,0696076 | 0,0018425 | 0,0023348 | 0,0020238 | 0,0020238 | TRUE | 0,0303873 | No |
| orf19.3221 | 11925 | 11127 | 12024 | 7275 | 6174 | 4611 | 5830 | 6321 | 11369 | 11765 | 11879 | 11397 | 4947 | 5243 | 4950 | 5140 | 8336,15 | 11602 | 5070 | 0,437 | -1,195 | -34,055 | 3,40E-254 | 5,91E-253 | 0,0004758 | 0,0022882 | 0,0010462 | 0,0010462 | TRUE | 0,0062136 | No |
| orf19.3222 | 811 | 824 | 930 | 521 | 1252 | 928 | 1029 | 1177 | 773 | 871 | 919 | 816 | 1003 | 1055 | 874 | 957 | 908,56 | 845 | 972 | 1,149 | 0,2 | 2,617 | 0,00888 | 0,0113214 | 0,0049913 | 0,0038984 | 0,004491 | 0,004491 | TRUE | 0,0557377 | No |
| orf19.3223 | 5584 | 5052 | 5083 | 3465 | 13296 | 10169 | 12036 | 13421 | 5324 | 5342 | 5022 | 5428 | 10654 | 11563 | 10219 | 10913 | 8057,88 | 5279 | 10837 | 2,053 | 1,038 | 22,062 | 7,28E-108 | 5,14E-107 | 0,0017241 | 0,002295 | 0,0019774 | 0,0019774 | TRUE | 0,0246137 | No |

| | | | | | | | | | | | | | | | | | | | | | | | | | | | | | | | |
|---|---|---|---|---|---|---|---|---|---|---|---|---|---|---|---|---|---|---|---|---|---|---|---|---|---|---|---|---|---|---|---|
| orf19.3223.1 | 1620 | 1469 | 1384 | 934 | 4700 | 3038 | 4271 | 4717 | 1544 | 1553 | 1367 | 1463 | 3766 | 3454 | 3626 | 3835 | 2576,26 | 1482 | 3670 | 2,477 | 1,309 | 23,56 | 9,88E-123 | 8,08E-122 | 0,0021271 | 0,0027286 | 0,0024617 | 0,0024617 | TRUE | 0,0346918 | No |
| orf19.3225 | 815 | 814 | 888 | 500 | 2621 | 1987 | 2445 | 2490 | 777 | 861 | 877 | 783 | 2100 | 2259 | 2076 | 2025 | 1469,77 | 824 | 2115 | 2,559 | 1,355 | 22,554 | 1,23E-112 | 9,16E-112 | 0,001996 | 0,0032084 | 0,0025777 | 0,0025777 | TRUE | 0,0246671 | No |
| orf19.3226 | 965 | 988 | 996 | 647 | 1928 | 1476 | 1715 | 1824 | 920 | 1045 | 984 | 1014 | 1545 | 1678 | 1456 | 1483 | 1265,57 | 991 | 1540 | 1,555 | 0,637 | 10,037 | 1,05E-23 | 2,54E-23 | 0,002721 | 0,0033886 | 0,0030114 | 0,0030114 | TRUE | 0,0418491 | No |
| orf19.3227 | 2375 | 2526 | 2660 | 1402 | 3022 | 2185 | 2796 | 2648 | 2264 | 2671 | 2628 | 2196 | 2421 | 2485 | 2374 | 2153 | 2399,03 | 2440 | 2358 | 0,965 | -0,051 | -0,689 | 0,4906169 | 0,5265982 | 0,0058908 | 0,0027757 | 0,0048194 | 0,0048194 | TRUE | 0,0524613 | No |
| orf19.3228 | 1208 | 1410 | 1465 | 853 | 2311 | 1875 | 2153 | 2009 | 1152 | 1491 | 1447 | 1336 | 1852 | 2132 | 1828 | 1634 | 1608,92 | 1356 | 1862 | 1,371 | 0,455 | 4,883 | 1,04E-06 | 1,61E-06 | 0,0114495 | 0,0031118 | 0,007697 | 0,007697 | TRUE | 0,1277242 | No |
| orf19.3229 | 4 | 8 | 2 | 0 | 8 | 6 | 5 | 4 | 4 | 8 | 2 | 0 | 6 | 7 | 4 | 3 | 4,37 | 4 | 5 | 1,379 | 0,463 | 0,62 | 0,5349821 | 0,5706655 | 0,1510799 | 0,3776149 | 0,2989305 | 0,2989305 | TRUE | 0,7449791 | No |
| orf19.323 | 2120 | 1786 | 1879 | 1158 | 1849 | 1238 | 1657 | 1877 | 2021 | 1888 | 1856 | 1814 | 1482 | 1408 | 1407 | 1526 | 1675,28 | 1895 | 1456 | 0,768 | -0,381 | -7,272 | 3,54E-13 | 6,81E-13 | 0,0013296 | 0,0030714 | 0,0020383 | 0,0020383 | TRUE | 0,0247771 | No |
| orf19.3230 | 6073 | 5393 | 6068 | 3870 | 5468 | 3951 | 5228 | 5523 | 5790 | 5702 | 5995 | 6063 | 4381 | 4493 | 4439 | 4491 | 5169,1 | 5888 | 4451 | 0,756 | -0,403 | -11,693 | 1,38E-31 | 3,81E-31 | 0,00022 | 0,0024089 | 0,0009442 | 0,0009442 | TRUE | 0,0053741 | No |
| orf19.3231 | 476 | 584 | 547 | 331 | 683 | 534 | 607 | 565 | 454 | 618 | 540 | 519 | 547 | 607 | 515 | 459 | 532,43 | 533 | 532 | 0,998 | -0,003 | -0,03 | 0,9758553 | 0,9783084 | 0,0129686 | 0,0051751 | 0,0091357 | 0,0091357 | TRUE | 0,1376881 | No |
| orf19.3232 | 438 | 422 | 411 | 290 | 325 | 280 | 322 | 347 | 418 | 446 | 406 | 454 | 260 | 318 | 273 | 282 | 357,31 | 431 | 283 | 0,657 | -0,606 | -6,968 | 3,22E-12 | 6,00E-12 | 0,0016272 | 0,0066865 | 0,0043928 | 0,0043928 | TRUE | 0,0688712 | No |
| orf19.3233 | 2864 | 2531 | 2750 | 1820 | 2873 | 2033 | 2736 | 2932 | 2731 | 2676 | 2717 | 2851 | 2302 | 2312 | 2323 | 2384 | 2536,9 | 2744 | 2330 | 0,85 | -0,234 | -6,033 | 1,61E-09 | 2,75E-09 | 1,00E-08 | 0,0027385 | 0,0010437 | 0,0010437 | TRUE | 0,0075619 | No |
| orf19.3234 | 255 | 243 | 248 | 151 | 451 | 311 | 425 | 406 | 243 | 257 | 245 | 237 | 361 | 354 | 361 | 330 | 298,44 | 246 | 352 | 1,429 | 0,515 | 6,001 | 1,96E-09 | 3,34E-09 | 1,00E-08 | 0,0075928 | 0,0035507 | 0,0035507 | TRUE | 0,0204929 | No |
| orf19.3234.1 | 749 | 644 | 629 | 363 | 348 | 282 | 358 | 350 | 714 | 681 | 621 | 569 | 279 | 321 | 304 | 285 | 471,64 | 646 | 297 | 0,457 | -1,131 | -12,698 | 6,04E-37 | 1,82E-36 | 0,0049105 | 0,0055725 | 0,0052415 | 0,0052415 | TRUE | 0,0684893 | No |
| orf19.3235 | 673 | 728 | 743 | 462 | 1095 | 735 | 1002 | 904 | 642 | 770 | 734 | 724 | 877 | 836 | 851 | 735 | 771,01 | 718 | 825 | 1,15 | 0,202 | 2,582 | 0,0098146 | 0,0124755 | 0,0048456 | 0,0042208 | 0,0045586 | 0,0045586 | TRUE | 0,0658111 | No |
| orf19.3237 | 2265 | 2207 | 2163 | 1261 | 3531 | 2475 | 3360 | 3430 | 2159 | 2334 | 2137 | 1975 | 2829 | 2814 | 2853 | 2789 | 2486,31 | 2151 | 2821 | 1,309 | 0,389 | 7,625 | 2,45E-14 | 4,85E-14 | 0,0018196 | 0,0027517 | 0,0020751 | 0,0020751 | TRUE | 0,0385105 | No |
| orf19.3238 | 5 | 6 | 6 | 5 | 13 | 7 | 10 | 9 | 5 | 6 | 6 | 8 | 10 | 8 | 8 | 7 | 7,38 | 6 | 8 | 1,407 | 0,493 | 0,97 | 0,3322567 | 0,3660445 | 1,00E-08 | 0,2245135 | 0,1049144 | 0,1049144 | TRUE | 0,0838215 | No |
| orf19.3239 | 952 | 761 | 878 | 512 | 1754 | 1245 | 1780 | 1830 | 908 | 805 | 867 | 802 | 1405 | 1416 | 1511 | 1488 | 1150,25 | 846 | 1455 | 1,717 | 0,78 | 13,326 | 1,64E-40 | 5,23E-40 | 0,001243 | 0,0035187 | 0,0023219 | 0,0023219 | TRUE | 0,0296833 | No |
| orf19.3240 | 562 | 476 | 609 | 408 | 819 | 641 | 807 | 818 | 536 | 503 | 602 | 639 | 656 | 729 | 685 | 665 | 626,91 | 570 | 684 | 1,203 | 0,267 | 3,299 | 0,0009718 | 0,0013121 | 0,0048237 | 0,0047103 | 0,0046274 | 0,0046274 | TRUE | 0,0705119 | No |
| orf19.3241 | 1411 | 1265 | 1350 | 889 | 1801 | 1207 | 1646 | 1830 | 1345 | 1338 | 1334 | 1393 | 1443 | 1372 | 1397 | 1488 | 1388,76 | 1352 | 1425 | 1,056 | 0,079 | 1,653 | 0,0982402 | 0,1148336 | 1,00E-08 | 0,0032736 | 0,0014675 | 0,0014675 | TRUE | 0,0105745 | No |
| orf19.3242 | 1619 | 1311 | 1354 | 904 | 2152 | 1375 | 2025 | 2108 | 1544 | 1386 | 1338 | 1416 | 1724 | 1563 | 1719 | 1714 | 1550,58 | 1421 | 1680 | 1,183 | 0,243 | 4,121 | 3,77E-05 | 5,45E-05 | 0,0021257 | 0,0031502 | 0,0026728 | 0,0026728 | TRUE | 0,0429129 | No |
| orf19.3243 | 3528 | 3020 | 3361 | 2058 | 5701 | 3888 | 5648 | 5975 | 3364 | 3193 | 3320 | 3224 | 4568 | 4421 | 4795 | 4858 | 3967,95 | 3275 | 4660 | 1,423 | 0,509 | 11,695 | 1,36E-31 | 3,76E-31 | 0,0008963 | 0,0025051 | 0,0015506 | 0,0015506 | TRUE | 0,014149 | No |
| orf19.3244 | 389 | 336 | 339 | 214 | 869 | 525 | 794 | 862 | 371 | 355 | 335 | 335 | 696 | 597 | 674 | 701 | 508,07 | 349 | 667 | 1,911 | 0,935 | 11,807 | 3,57E-32 | 1,00E-31 | 0,0022302 | 0,0053229 | 0,0037522 | 0,0037522 | TRUE | 0,0547434 | No |
| orf19.3245 | 1839 | 1731 | 1774 | 1123 | 1140 | 795 | 1158 | 1155 | 1753 | 1830 | 1753 | 1759 | 913 | 904 | 983 | 939 | 1354,39 | 1774 | 935 | 0,527 | -0,923 | -18,889 | 1,41E-79 | 7,63E-79 | 1,00E-08 | 0,0033036 | 0,0014996 | 0,0014996 | TRUE | 0,0144039 | No |
| orf19.3247 | 547 | 667 | 626 | 396 | 639 | 454 | 508 | 503 | 522 | 705 | 618 | 620 | 512 | 516 | 431 | 409 | 541,76 | 616 | 467 | 0,757 | -0,401 | -3,751 | 0,0001761 | 0,0002461 | 0,0129143 | 0,005122 | 0,0091018 | 0,0091018 | TRUE | 0,1315506 | No |
| orf19.3248 | 394 | 473 | 401 | 277 | 320 | 261 | 289 | 284 | 376 | 500 | 396 | 434 | 256 | 297 | 245 | 231 | 341,92 | 426 | 257 | 0,602 | -0,732 | -6,55 | 5,74E-11 | 1,03E-10 | 0,0109752 | 0,0068934 | 0,0089245 | 0,0089245 | TRUE | 0,162941 | No |
| orf19.3249 | 1883 | 1982 | 1977 | 1148 | 4713 | 3170 | 4062 | 4168 | 1795 | 2096 | 1953 | 1798 | 3776 | 3605 | 3449 | 3389 | 2732,63 | 1910 | 3555 | 1,859 | 0,894 | 14,723 | 4,57E-49 | 1,66E-48 | 0,0034846 | 0,0026921 | 0,0031233 | 0,0031233 | TRUE | 0,0513837 | No |
| orf19.325 | 3062 | 2807 | 3029 | 1921 | 3201 | 2245 | 3072 | 3224 | 2919 | 2968 | 2992 | 3009 | 2565 | 2553 | 2608 | 2621 | 2779,53 | 2972 | 2587 | 0,871 | -0,199 | -5,646 | 1,64E-08 | 2,71E-08 | 1,00E-08 | 0,002682 | 0,0008343 | 0,0008343 | TRUE | 0,0017975 | No |
| orf19.3250 | 1316 | 1340 | 1391 | 783 | 4238 | 3046 | 3886 | 4025 | 1255 | 1417 | 1374 | 1227 | 3396 | 3464 | 3299 | 3273 | 2337,95 | 1318 | 3358 | 2,542 | 1,346 | 25,161 | 1,07E-139 | 1,00E-138 | 0,0016848 | 0,0027936 | 0,0021893 | 0,0021893 | TRUE | 0,0298362 | No |
| orf19.3251 | 964 | 965 | 958 | 641 | 871 | 619 | 742 | 690 | 919 | 1020 | 946 | 1004 | 698 | 704 | 630 | 561 | 810,35 | 972 | 648 | 0,666 | -0,586 | -7,384 | 1,54E-13 | 2,98E-13 | 0,0053723 | 0,0041175 | 0,0047793 | 0,0047793 | TRUE | 0,0988351 | No |
| orf19.3252 | 2365 | 2300 | 2501 | 1574 | 1525 | 1092 | 1450 | 1513 | 2255 | 2432 | 2471 | 2466 | 1222 | 1242 | 1231 | 1230 | 1818,52 | 2406 | 1231 | 0,512 | -0,966 | -20,53 | 1,16E-93 | 7,18E-93 | 0,0006498 | 0,0029942 | 0,0015275 | 0,0015275 | TRUE | 0,0224899 | No |
| orf19.3254 | 573 | 536 | 593 | 355 | 978 | 710 | 853 | 907 | 546 | 567 | 586 | 556 | 784 | 807 | 724 | 737 | 663,46 | 564 | 763 | 1,351 | 0,434 | 6,704 | 2,03E-11 | 3,70E-11 | 1,00E-08 | 0,0045661 | 0,0024511 | 0,0024511 | TRUE | 0,0164457 | No |
| orf19.3255 | 495 | 570 | 525 | 262 | 596 | 383 | 520 | 456 | 472 | 603 | 519 | 410 | 478 | 436 | 441 | 371 | 466,13 | 501 | 432 | 0,858 | -0,221 | -1,906 | 0,0567129 | 0,0677941 | 0,015517 | 0,0056137 | 0,0107134 | 0,0107134 | TRUE | 0,2137852 | No |
| orf19.3256 | 7104 | 6884 | 7177 | 4526 | 5591 | 4162 | 5229 | 5586 | 6773 | 7279 | 7090 | 7090 | 4480 | 4733 | 4439 | 4542 | 5803,26 | 7058 | 4548 | 0,644 | -0,634 | -16,642 | 3,48E-62 | 1,51E-61 | 0,0006083 | 0,0023742 | 0,0012171 | 0,0012171 | TRUE | 0,0092581 | No |
| orf19.3259 | 492 | 438 | 478 | 265 | 501 | 367 | 471 | 518 | 469 | 463 | 472 | 415 | 401 | 417 | 400 | 421 | 432,42 | 455 | 410 | 0,897 | -0,158 | -2,144 | 0,0320534 | 0,0391526 | 1,00E-08 | 0,0058883 | 0,002877 | 0,002877 | TRUE | 0,0327658 | No |
| orf19.326 | 372 | 302 | 304 | 205 | 358 | 226 | 355 | 386 | 355 | 319 | 300 | 321 | 287 | 257 | 301 | 314 | 306,82 | 324 | 290 | 0,897 | -0,157 | -1,665 | 0,0959239 | 0,1122484 | 0,002796 | 0,0074427 | 0,0052528 | 0,0052528 | TRUE | 0,0583864 | No |
| orf19.3260 | 3857 | 3425 | 3825 | 2324 | 5595 | 3715 | 5033 | 5591 | 3677 | 3622 | 3779 | 3641 | 4483 | 4224 | 4273 | 4546 | 4030,57 | 3680 | 4382 | 1,191 | 0,252 | 6,383 | 1,74E-10 | 3,08E-10 | 0,0006238 | 0,0024986 | 0,0012443 | 0,0012443 | TRUE | 0,0077459 | No |
| orf19.3261 | 102 | 70 | 86 | 50 | 120 | 82 | 127 | 97 | 97 | 74 | 85 | 78 | 96 | 93 | 108 | 79 | 88,83 | 84 | 94 | 1,118 | 0,161 | 0,987 | 0,3234858 | 0,3569364 | 0,0064397 | 0,020575 | 0,0141463 | 0,0141463 | TRUE | 0,1256635 | No |
| orf19.3263 | 27 | 26 | 31 | 19 | 24 | 19 | 31 | 33 | 26 | 27 | 31 | 30 | 19 | 22 | 26 | 27 | 25,95 | 28 | 24 | 0,832 | -0,266 | -0,974 | 0,3298632 | 0,3636588 | 1,00E-08 | 0,0653603 | 0,032822 | 0,032822 | TRUE | 0,1110326 | No |
| orf19.3264 | 340 | 381 | 377 | 237 | 102 | 94 | 84 | 72 | 324 | 403 | 372 | 371 | 82 | 107 | 71 | 59 | 223,65 | 368 | 80 | 0,214 | -2,224 | -15,096 | 1,72E-51 | 6,46E-51 | 0,0213246 | 0,0094327 | 0,0136576 | 0,0136576 | TRUE | 0,4357498 | No |
| orf19.3264.1 | 6478 | 4812 | 5025 | 2697 | 6894 | 3645 | 5488 | 6067 | 6176 | 5088 | 4964 | 4225 | 5524 | 4145 | 4659 | 4933 | 4964,3 | 5113 | 4815 | 0,941 | -0,087 | -0,82 | 0,4124912 | 0,4483997 | 0,0178182 | 0,002422 | 0,0106304 | 0,0106304 | TRUE | 0,2388573 | No |
| orf19.3265 | 1512 | 1421 | 1519 | 878 | 2606 | 1838 | 2429 | 2633 | 1442 | 1503 | 1501 | 1375 | 2088 | 2090 | 2062 | 2141 | 1775,16 | 1455 | 2095 | 1,437 | 0,523 | 11,623 | 3,14E-31 | 8,65E-31 | 3,89E-05 | 0,0030162 | 0,0013482 | 0,0013482 | TRUE | 0,0140127 | No |
| orf19.3265.1 | 293 | 249 | 271 | 175 | 528 | 348 | 510 | 504 | 279 | 263 | 268 | 274 | 423 | 396 | 433 | 410 | 343,26 | 271 | 416 | 1,535 | 0,618 | 7,676 | 1,64E-14 | 3,27E-14 | 1,00E-08 | 0,0068746 | 0,0031033 | 0,0031033 | TRUE | 0,0104198 | No |
| orf19.3266 | 188 | 157 | 202 | 164 | 768 | 615 | 698 | 841 | 179 | 166 | 200 | 257 | 615 | 699 | 593 | 684 | 424,1 | 200 | 648 | 3,268 | 1,708 | 14,833 | 8,90E-50 | 3,28E-49 | 0,0141175 | 0,0059628 | 0,0092197 | 0,0092197 | TRUE | 0,3268646 | No |
| orf19.3267 | 83 | 74 | 89 | 65 | 262 | 233 | 266 | 282 | 79 | 78 | 88 | 102 | 210 | 265 | 226 | 229 | 159,64 | 87 | 232 | 2,699 | 1,432 | 10,713 | 8,89E-27 | 2,28E-26 | 0,004091 | 0,0123763 | 0,00877 | 0,00877 | TRUE | 0,0940257 | No |
| orf19.3268 | 11881 | 9287 | 9628 | 7778 | 38936 | 28030 | 36325 | 39932 | 11327 | 9820 | 9512 | 12185 | 31198 | 31873 | 30840 | 32469 | 21152,81 | 10711 | 31595 | 2,951 | 1,561 | 21,755 | 6,20E-105 | 4,21E-104 | 0,007129 | 0,0021689 | 0,0048804 | 0,0048804 | TRUE | 0,1040031 | No |
| orf19.3269 | 445 | 371 | 431 | 262 | 539 | 392 | 510 | 540 | 424 | 392 | 426 | 410 | 432 | 446 | 433 | 439 | 425,31 | 413 | 438 | 1,057 | 0,08 | 1,117 | 0,2641684 | 0,2948502 | 1,00E-08 | 0,0059518 | 0,002544 | 0,002544 | TRUE | 0,0145073 | No |
| orf19.327 | 544 | 496 | 526 | 342 | 926 | 631 | 893 | 902 | 519 | 524 | 520 | 536 | 742 | 718 | 758 | 733 | 631,19 | 525 | 738 | 1,409 | 0,495 | 8,172 | 3,04E-16 | 6,29E-16 | 1,00E-08 | 0,0046925 | 0,0018569 | 0,0018569 | TRUE | 0,0040081 | No |
| orf19.3271 | 2 | 5 | 2 | 1 | 1 | 4 | 2 | 0 | 2 | 5 | 2 | 2 | 1 | 5 | 2 | 0 | 2,22 | 3 | 2 | 0,606 | -0,722 | -0,691 | 0,4897874 | 0,5259733 | 0,3130088 | 0,7407044 | 0,5738421 | 0,5738421 | TRUE | 0,7699969 | No |
| orf19.3272 | 1023 | 891 | 950 | 714 | 1222 | 924 | 1211 | 1284 | 975 | 942 | 939 | 1119 | 979 | 1051 | 1028 | 1044 | 1009,56 | 994 | 1026 | 1,036 | 0,05 | 0,778 | 0,4366456 | 0,4727241 | 0,0024135 | 0,0037176 | 0,0030407 | 0,0030407 | TRUE | 0,0768277 | No |
| orf19.3273 | 317 | 272 | 260 | 190 | 260 | 158 | 262 | 235 | 302 | 288 | 257 | 298 | 208 | 180 | 222 | 191 | 243,23 | 286 | 200 | 0,704 | -0,507 | -4,904 | 9,40E-07 | 1,46E-06 | 0,0027376 | 0,0088417 | 0,0060729 | 0,0060729 | TRUE | 0,0589764 | No |
| orf19.3275 | 2624 | 2125 | 2246 | 1380 | 1489 | 912 | 1353 | 1433 | 2502 | 2247 | 2219 | 2162 | 1193 | 1037 | 1149 | 1165 | 1709,16 | 2282 | 1136 | 0,498 | -1,006 | -15,935 | 3,63E-57 | 1,46E-56 | 0,0031112 | 0,0030519 | 0,0031864 | 0,0031864 | TRUE | 0,0514717 | No |
| orf19.3276 | 8846 | 7907 | 8591 | 5343 | 11329 | 8135 | 10428 | 11320 | 8434 | 8361 | 8487 | 8370 | 9078 | 9250 | 8853 | 9204 | 8754,62 | 8413 | 9096 | 1,081 | 0,112 | 3,887 | 0,0001015 | 0,0001436 | 0,0001258 | 0,0022788 | 0,0006852 | 0,0006852 | TRUE | 0,0039566 | No |
| orf19.3277 | 26 | 17 | 24 | 20 | 8 | 4 | 9 | 2 | 25 | 18 | 24 | 31 | 6 | 5 | 8 | 2 | 14,75 | 24 | 5 | 0,209 | -2,255 | -5,035 | 4,78E-07 | 7,49E-07 | 1,00E-08 | 0,1133785 | 0,0820489 | 0,0820489 | TRUE | 0,4319131 | No |
| orf19.3278 | 26585 | 22972 | 24811 | 16191 | 7067 | 5020 | 6596 | 7332 | 25346 | 24290 | 24511 | 25364 | 5663 | 5708 | 5600 | 5962 | 15305,41 | 24878 | 5733 | 0,23 | -2,117 | -62,175 | 0 | 0 | 0,0004919 | 0,0021986 | 0,0010131 | 0,0010131 | TRUE | 0,0089476 | No |
| orf19.3279 | 2485 | 2175 | 2412 | 1536 | 2779 | 1979 | 2645 | 2944 | 2369 | 2300 | 2383 | 2406 | 2227 | 2250 | 2246 | 2394 | 2321,8 | 2364 | 2279 | 0,965 | -0,052 | -1,233 | 0,2176884 | 0,2459401 | 0,0004051 | 0,0027984 | 0,0012769 | 0,0012769 | TRUE | 0,0142905 | No |
| orf19.328 | 533 | 518 | 559 | 313 | 681 | 473 | 619 | 639 | 508 | 548 | 552 | 490 | 546 | 538 | 526 | 520 | 528,38 | 524 | 532 | 1,011 | 0,016 | 0,233 | 0,8154719 | 0,8369834 | 1,00E-08 | 0,0051987 | 0,0025812 | 0,0025812 | TRUE | 0,0183856 | No |
| orf19.3280 | 2 | 1 | 2 | 1 | 1 | 2 | 4 | 0 | 2 | 1 | 2 | 2 | 1 | 2 | 3 | 0 | 1,62 | 2 | 2 | 0,966 | -0,05 | -0,044 | 0,9651653 | 0,9686568 | 1,00E-08 | 1,0142099 | 0,6210354 | 0,6210354 | TRUE | 0,3207652 | No |
| orf19.3281 | 829 | 745 | 752 | 543 | 255 | 183 | 251 | 268 | 790 | 788 | 743 | 851 | 204 | 208 | 213 | 218 | 501,88 | 793 | 211 | 0,267 | -1,906 | -24,54 | 5,49E-133 | 4,94E-132 | 1,00E-08 | 0,0053628 | 0,0029798 | 0,0029798 | TRUE | 0,0250606 | No |
| orf19.3282 | 1894 | 1646 | 1888 | 1054 | 1154 | 855 | 1037 | 1115 | 1806 | 1740 | 1865 | 1651 | 925 | 972 | 880 | 907 | 1343,3 | 1766 | 921 | 0,52 | -0,944 | -16,838 | 1,28E-63 | 5,66E-63 | 0,0013495 | 0,0033136 | 0,0022118 | 0,0022118 | TRUE | 0,0210295 | No |
| orf19.3283 | 95 | 93 | 104 | 57 | 227 | 164 | 204 | 247 | 91 | 98 | 103 | 89 | 182 | 186 | 173 | 201 | 140,42 | 95 | 186 | 1,942 | 0,957 | 7,581 | 3,43E-14 | 6,76E-14 | 1,00E-08 | 0,0137842 | 0,0071089 | 0,0071089 | TRUE | 0,035775 | No |
| orf19.3284 | 0 | 1 | 0 | 1 | 0 | 0 | 1 | 3 | 0 | 1 | 0 | 2 | 0 | 0 | 1 | 2 | 0,74 | 1 | 1 | 1,469 | 0,555 | 0,301 | 0,7632643 | 0,7905193 | 1,00E-08 | 2,2238037 | 1,834958 | 1,834958 | TRUE | 0,8314359 | No |
| orf19.3285 | 151 | 124 | 110 | 77 | 306 | 181 | 285 | 315 | 144 | 131 | 109 | 121 | 245 | 206 | 242 | 256 | 181,68 | 126 | 237 | 1,883 | 0,913 | 7,362 | 1,81E-13 | 3,51E-13 | 0,0049806 | 0,0111285 | 0,0084869 | 0,0084869 | TRUE | 0,1023011 | No |
| orf19.3286 | 845 | 771 | 787 | 490 | 1281 | 893 | 1264 | 1256 | 806 | 815 | 777 | 768 | 1026 | 1015 | 1073 | 1021 | 912,77 | 792 | 1034 | 1,305 | 0,384 | 7,145 | 9,03E-13 | 1,71E-12 | 1,00E-08 | 0,0038901 | 0,0016407 | 0,0016407 | TRUE | 0,0078317 | No |
| orf19.3287 | 8810 | 7443 | 8325 | 4770 | 7407 | 4793 | 7139 | 7146 | 8399 | 7870 | 8224 | 7472 | 5935 | 5450 | 6061 | 5810 | 6902,83 | 7991 | 5814 | 0,727 | -0,459 | -9,213 | 3,19E-20 | 7,19E-20 | 0,0021451 | 0,0023291 | 0,0022388 | 0,0022388 | TRUE | 0,022821 | No |
| orf19.3288 | 9764 | 8266 | 9233 | 5676 | 13248 | 8925 | 12661 | 13567 | 9309 | 8740 | 9121 | 8892 | 10615 | 10149 | 10749 | 11031 | 9825,8 | 9016 | 10636 | 1,18 | 0,239 | 6,205 | 5,47E-10 | 9,50E-10 | 0,0008494 | 0,0022584 | 0,0013145 | 0,0013145 | TRUE | 0,0114516 | No |
| orf19.3289 | 620 | 531 | 566 | 322 | 752 | 482 | 742 | 759 | 591 | 561 | 559 | 504 | 603 | 548 | 630 | 617 | 576,74 | 554 | 600 | 1,08 | 0,111 | 1,518 | 0,1289547 | 0,14888 | 0,0016162 | 0,0049382 | 0,0033621 | 0,0033621 | TRUE | 0,0361107 | No |
| orf19.329 | 3430 | 2897 | 3175 | 2006 | 3108 | 2097 | 2959 | 3122 | 3270 | 3063 | 3137 | 3143 | 2490 | 2384 | 2512 | 2538 | 2817,24 | 3153 | 2481 | 0,787 | -0,345 | -8,538 | 1,36E-17 | 2,90E-17 | 0,0003831 | 0,0026741 | 0,0012089 | 0,0012089 | TRUE | 0,007953 | No |
| orf19.3290 | 2577 | 2078 | 2241 | 1441 | 5170 | 3534 | 4915 | 5054 | 2457 | 2197 | 2214 | 2257 | 4143 | 4018 | 4173 | 4109 | 3196,08 | 2281 | 4111 | 1,801 | 0,849 | 18,715 | 3,72E-78 | 1,99E-77 | 0,0010551 | 0,002605 | 0,001621 | 0,001621 | TRUE | 0,0340864 | No |
| orf19.3291 | 10237 | 8696 | 9281 | 5296 | 18863 | 12659 | 18153 | 19983 | 9760 | 9195 | 9169 | 8296 | 15114 | 14394 | 15412 | 16248 | 12198,59 | 9105 | 15292 | 1,679 | 0,748 | 13,355 | 1,11E-40 | 3,55E-40 | 0,0033425 | 0,0022259 | 0,0029187 | 0,0029187 | TRUE | 0,0429434 | No |
| orf19.3292 | 2618 | 2151 | 2270 | 1634 | 1959 | 1353 | 1885 | 1924 | 2496 | 2274 | 2243 | 2560 | 1570 | 1538 | 1600 | 1564 | 1980,7 | 2393 | 1568 | 0,656 | -0,608 | -11,269 | 1,87E-29 | 4,98E-29 | 0,0016131 | 0,0029202 | 0,0022703 | 0,0022703 | TRUE | 0,0247864 | No |
| orf19.3293 | 875 | 783 | 847 | 485 | 1944 | 1443 | 1918 | 2000 | 834 | 828 | 837 | 760 | 1558 | 1641 | 1628 | 1626 | 1213,97 | 815 | 1613 | 1,972 | 0,98 | 18,961 | 3,61E-80 | 1,98E-79 | 5,95E-05 | 0,0034438 | 0,0016052 | 0,0016052 | TRUE | 0,0202105 | No |
| orf19.3294 | 15847 | 12436 | 13316 | 8048 | 13654 | 8462 | 13149 | 13589 | 15108 | 13149 | 13155 | 12608 | 10941 | 9622 | 11163 | 11049 | 12099,46 | 13505 | 10694 | 0,792 | -0,337 | -5,097 | 3,44E-07 | 5,43E-07 | 0,0054692 | 0,002227 | 0,0041097 | 0,0041097 | TRUE | 0,0783035 | No |
| orf19.3295 | 1248 | 1127 | 1172 | 731 | 795 | 565 | 768 | 772 | 1190 | 1192 | 1158 | 1145 | 637 | 642 | 652 | 628 | 905,46 | 1171 | 640 | 0,546 | -0,874 | -16,581 | 9,53E-62 | 4,12E-61 | 1,00E-08 | 0,0039046 | 0,0014948 | 0,0014948 | TRUE | 0,0022771 | No |

| | | | | | | | | | | | | | | | | | | | | | | | | | | | | | | | |
|---|---|---|---|---|---|---|---|---|---|---|---|---|---|---|---|---|---|---|---|---|---|---|---|---|---|---|---|---|---|---|---|
| orf19.3296 | 197 | 164 | 171 | 100 | 108 | 81 | 91 | 139 | 188 | 173 | 169 | 157 | 87 | 92 | 77 | 113 | 131,97 | 172 | 92 | 0,535 | -0,901 | -6,344 | 2,24E-10 | 3,95E-10 | 0,0068878 | 0,0145331 | 0,0113095 | 0,0113095 | TRUE | 0,2459196 | No |
| orf19.3297 | 2134 | 1984 | 2038 | 1426 | 2065 | 1487 | 1981 | 2167 | 2035 | 2098 | 2013 | 2234 | 1655 | 1691 | 1682 | 1762 | 1896,12 | 2095 | 1698 | 0,812 | -0,301 | -6,24 | 4,39E-10 | 7,67E-10 | 0,0007776 | 0,0029572 | 0,0016969 | 0,0016969 | TRUE | 0,0221093 | No |
| orf19.3298 | 2271 | 2093 | 2338 | 1349 | 2432 | 1667 | 2212 | 2321 | 2165 | 2213 | 2310 | 2113 | 1949 | 1896 | 1878 | 1887 | 2051,33 | 2200 | 1902 | 0,864 | -0,212 | -4,855 | 1,20E-06 | 1,86E-06 | 0,0003375 | 0,0028917 | 0,0013332 | 0,0013332 | TRUE | 0,0136543 | No |
| orf19.33 | 4209 | 3677 | 3783 | 2480 | 2767 | 1910 | 2526 | 2606 | 4013 | 3888 | 3737 | 3885 | 2217 | 2172 | 2145 | 2119 | 3021,94 | 3881 | 2163 | 0,557 | -0,843 | -21,332 | 5,77E-101 | 3,81E-100 | 0,0003868 | 0,0026346 | 0,0011493 | 0,0011493 | TRUE | 0,0078954 | No |
| orf19.3300 | 17731 | 15674 | 16858 | 10315 | 20684 | 14362 | 19477 | 21171 | 16905 | 16573 | 16654 | 16159 | 16573 | 16331 | 16536 | 17214 | 16618,16 | 16573 | 16664 | 1,005 | 0,008 | 0,25 | 0,8024252 | 0,8247866 | 0,0003227 | 0,0021901 | 0,0008466 | 0,0008466 | TRUE | 0,0060713 | No |
| orf19.3301 | 29617 | 26467 | 29114 | 17610 | 13963 | 10164 | 13626 | 14040 | 28237 | 27986 | 28762 | 27587 | 11188 | 11557 | 11568 | 11416 | 19787,6 | 28143 | 11432 | 0,406 | -1,3 | -46,435 | 0 | 0 | 0,0001906 | 0,0021743 | 0,0006937 | 0,0006937 | TRUE | 0,0026835 | No |
| orf19.330.1 | 444 | 340 | 381 | 269 | 1082 | 700 | 1095 | 1150 | 423 | 360 | 376 | 421 | 867 | 796 | 930 | 935 | 638,53 | 395 | 882 | 2,24 | 1,164 | 14,333 | 1,37E-46 | 4,83E-46 | 0,0041491 | 0,0046627 | 0,0044034 | 0,0044034 | TRUE | 0,0484558 | No |
| orf19.3302 | 23393 | 22188 | 23519 | 14726 | 1741 | 1322 | 1642 | 1788 | 22303 | 23461 | 23235 | 23069 | 1395 | 1503 | 1394 | 1454 | 12226,68 | 23017 | 1436 | 0,062 | -4,004 | -103,685 | 0 | 0 | 0,0004137 | 0,0022256 | 0,001097 | 0,001097 | TRUE | 0,0107006 | No |
| orf19.3303 | 3710 | 3263 | 3506 | 2095 | 2488 | 1728 | 2277 | 2515 | 3537 | 3450 | 3464 | 3282 | 1994 | 1965 | 1933 | 2045 | 2708,67 | 3433 | 1984 | 0,577 | -0,792 | -19,261 | 1,14E-82 | 6,36E-82 | 0,0003728 | 0,0026974 | 0,001236 | 0,001236 | TRUE | 0,0098677 | No |
| orf19.3304 | 6525 | 5849 | 6098 | 3329 | 5393 | 3543 | 5346 | 5594 | 6221 | 6185 | 6024 | 5215 | 4321 | 4029 | 4539 | 4548 | 5135,24 | 5911 | 4359 | 0,737 | -0,44 | -6,882 | 5,90E-12 | 1,09E-11 | 0,004488 | 0,002411 | 0,0037266 | 0,0037266 | TRUE | 0,0750474 | No |
| orf19.3305 | 2598 | 2352 | 2493 | 1549 | 2486 | 1739 | 2360 | 2531 | 2477 | 2487 | 2463 | 2427 | 1992 | 1977 | 2004 | 2058 | 2235,53 | 2464 | 2008 | 0,815 | -0,295 | -7,799 | 6,25E-15 | 1,26E-14 | 1,00E-08 | 0,0028257 | 0,0009264 | 0,0009264 | TRUE | 0,0034927 | No |
| orf19.3306 | 20 | 20 | 21 | 14 | 118 | 101 | 100 | 107 | 19 | 21 | 21 | 22 | 95 | 115 | 85 | 87 | 58,02 | 21 | 96 | 4,6 | 2,202 | 9,612 | 7,09E-22 | 1,65E-21 | 1,00E-08 | 0,0303884 | 0,0188426 | 0,0188426 | TRUE | 0,1657867 | No |
| orf19.3307 | 261 | 242 | 276 | 177 | 748 | 565 | 736 | 730 | 249 | 256 | 273 | 277 | 599 | 642 | 625 | 594 | 439,36 | 264 | 615 | 2,338 | 1,225 | 15,861 | 1,18E-56 | 4,73E-56 | 1,00E-08 | 0,0058283 | 0,0028872 | 0,0028872 | TRUE | 0,0166483 | No |
| orf19.3308 | 506 | 470 | 528 | 321 | 1353 | 1027 | 1251 | 1336 | 482 | 497 | 522 | 503 | 1084 | 1168 | 1062 | 1086 | 800,52 | 501 | 1100 | 2,194 | 1,133 | 18,381 | 1,87E-75 | 9,65E-75 | 1,00E-08 | 0,0041423 | 0,0021311 | 0,0021311 | TRUE | 0,0191136 | No |
| orf19.3309 | 3325 | 2736 | 2885 | 1842 | 5630 | 3726 | 5356 | 5731 | 3170 | 2893 | 2850 | 2886 | 4511 | 4237 | 4547 | 4660 | 3719,22 | 2950 | 4489 | 1,522 | 0,606 | 12,295 | 9,68E-35 | 2,82E-34 | 0,0016854 | 0,0025327 | 0,0020392 | 0,0020392 | TRUE | 0,031577 | No |
| orf19.331 | 1157 | 922 | 1006 | 710 | 1766 | 1301 | 1694 | 1792 | 1103 | 975 | 994 | 1112 | 1415 | 1479 | 1438 | 1457 | 1246,72 | 1046 | 1447 | 1,386 | 0,471 | 8,272 | 1,32E-16 | 2,74E-16 | 0,0014302 | 0,0034083 | 0,0022636 | 0,0022636 | TRUE | 0,0245334 | No |
| orf19.3310 | 942 | 785 | 920 | 599 | 639 | 452 | 632 | 658 | 898 | 830 | 909 | 938 | 512 | 514 | 537 | 535 | 709,12 | 894 | 524 | 0,588 | -0,766 | -12,206 | 2,90E-34 | 8,39E-34 | 1,00E-08 | 0,0044067 | 0,0023072 | 0,0023072 | TRUE | 0,0272699 | No |
| orf19.3311 | 189 | 161 | 164 | 92 | 929 | 579 | 885 | 995 | 180 | 170 | 162 | 144 | 744 | 658 | 751 | 809 | 452,46 | 164 | 740 | 4,486 | 2,165 | 22,183 | 5,01E-109 | 3,59E-108 | 0,0046059 | 0,0057201 | 0,0051851 | 0,0051851 | TRUE | 0,0628191 | No |
| orf19.3312 | 545 | 517 | 524 | 403 | 1633 | 1212 | 1598 | 1786 | 520 | 547 | 518 | 631 | 1308 | 1378 | 1357 | 1452 | 963,84 | 554 | 1374 | 2,497 | 1,32 | 18,943 | 5,06E-80 | 2,76E-79 | 0,0028004 | 0,0037948 | 0,0033194 | 0,0033194 | TRUE | 0,0897463 | No |
| orf19.3314 | 244 | 217 | 229 | 173 | 207 | 158 | 177 | 195 | 233 | 229 | 226 | 271 | 166 | 180 | 150 | 159 | 201,71 | 240 | 164 | 0,684 | -0,548 | -5,077 | 3,84E-07 | 6,04E-07 | 1,00E-08 | 0,0102312 | 0,0061038 | 0,0061038 | TRUE | 0,0695665 | No |
| orf19.3315 | 2353 | 2155 | 2460 | 1559 | 578 | 425 | 553 | 575 | 2243 | 2279 | 2430 | 2442 | 463 | 483 | 469 | 468 | 1409,74 | 2348 | 471 | 0,2 | -2,319 | -41,384 | 0 | 0 | 1,00E-08 | 0,003256 | 0,0018292 | 0,0018292 | TRUE | 0,0112003 | No |
| orf19.3317 | 289 | 318 | 286 | 217 | 518 | 365 | 432 | 437 | 276 | 336 | 283 | 340 | 415 | 415 | 367 | 355 | 348,31 | 309 | 388 | 1,262 | 0,336 | 3,376 | 0,0007356 | 0,0010022 | 0,0062617 | 0,0068053 | 0,0065432 | 0,0065432 | TRUE | 0,0594394 | No |
| orf19.3318 | 313 | 316 | 327 | 187 | 1555 | 1071 | 1372 | 1415 | 298 | 334 | 323 | 293 | 1246 | 1218 | 1165 | 1151 | 753,46 | 312 | 1195 | 3,815 | 1,932 | 27,911 | 1,96E-171 | 2,25E-170 | 1,00E-08 | 0,0042704 | 0,0024501 | 0,0024501 | TRUE | 0,024284 | No |
| orf19.3319 | 1453 | 1316 | 1316 | 967 | 2848 | 2282 | 2784 | 3108 | 1385 | 1392 | 1300 | 1515 | 2282 | 2595 | 2364 | 2527 | 1919,91 | 1398 | 2442 | 1,75 | 0,807 | 13,214 | 7,26E-40 | 2,29E-39 | 0,0029923 | 0,0029465 | 0,0029944 | 0,0029944 | TRUE | 0,0351239 | No |
| orf19.332 | 4369 | 3789 | 4280 | 2607 | 4230 | 2946 | 4159 | 4370 | 4165 | 4006 | 4228 | 4084 | 3389 | 3350 | 3531 | 3553 | 3788,43 | 4121 | 3456 | 0,839 | -0,254 | -6,614 | 3,74E-11 | 6,75E-11 | 0,0004478 | 0,0025247 | 0,0011476 | 0,0011476 | TRUE | 0,0049218 | No |
| orf19.3320 | 25 | 11 | 20 | 20 | 34 | 26 | 26 | 34 | 24 | 12 | 20 | 31 | 27 | 30 | 22 | 28 | 24,14 | 22 | 27 | 1,254 | 0,327 | 1,043 | 0,2971129 | 0,3298325 | 0,0152071 | 0,0701206 | 0,0511539 | 0,0511539 | TRUE | 0,2947689 | No |
| orf19.3321 | 5751 | 4933 | 5504 | 3324 | 8186 | 6054 | 8140 | 8580 | 5483 | 5216 | 5437 | 5207 | 6559 | 6884 | 6911 | 6976 | 6084,25 | 5336 | 6832 | 1,28 | 0,356 | 9,611 | 7,19E-22 | 1,67E-21 | 0,0005173 | 0,0023611 | 0,0011448 | 0,0011448 | TRUE | 0,0089935 | No |
| orf19.332.1 | 1695 | 1366 | 1391 | 963 | 3195 | 2150 | 2972 | 3116 | 1616 | 1444 | 1374 | 1509 | 2560 | 2445 | 2523 | 2534 | 2000,6 | 1486 | 2516 | 1,694 | 0,76 | 13,956 | 2,88E-44 | 9,75E-44 | 0,00187 | 0,002912 | 0,0022936 | 0,0022936 | TRUE | 0,0450666 | No |
| orf19.3322 | 579 | 389 | 469 | 355 | 785 | 500 | 718 | 783 | 552 | 411 | 463 | 556 | 629 | 569 | 610 | 637 | 553,32 | 496 | 611 | 1,237 | 0,307 | 3,188 | 0,0014338 | 0,0019141 | 0,0089763 | 0,0050586 | 0,0070397 | 0,0070397 | TRUE | 0,1527248 | No |
| orf19.3323 | 704 | 725 | 703 | 482 | 1084 | 747 | 964 | 985 | 671 | 767 | 694 | 755 | 869 | 849 | 818 | 801 | 778,08 | 722 | 834 | 1,158 | 0,212 | 3,281 | 0,0010337 | 0,0013934 | 0,0013497 | 0,0042015 | 0,0027015 | 0,0027015 | TRUE | 0,0271901 | No |
| orf19.3324 | 21151 | 18138 | 19788 | 11826 | 35171 | 23916 | 34836 | 37348 | 20165 | 19179 | 19549 | 18526 | 28182 | 27195 | 29576 | 30367 | 24092,23 | 19355 | 28830 | 1,489 | 0,575 | 12,717 | 4,75E-37 | 1,44E-36 | 0,0017768 | 0,0021594 | 0,0019176 | 0,0019176 | TRUE | 0,0177744 | No |
| orf19.3325 | 1907 | 1657 | 1718 | 1132 | 3958 | 2700 | 3732 | 4018 | 1818 | 1752 | 1697 | 1773 | 3171 | 3070 | 3168 | 3267 | 2464,72 | 1760 | 3169 | 1,802 | 0,849 | 20,224 | 6,03E-91 | 3,64E-90 | 0,0003217 | 0,0027574 | 0,0012344 | 0,0012344 | TRUE | 0,0074542 | No |
| orf19.3325.3 | 8301 | 6301 | 6284 | 3776 | 26507 | 15517 | 24394 | 26759 | 7914 | 6663 | 6208 | 5915 | 21239 | 17644 | 20710 | 21758 | 13506,43 | 6675 | 20338 | 3,046 | 1,607 | 17,973 | 3,15E-72 | 1,56E-71 | 0,0127051 | 0,0022128 | 0,0075734 | 0,0075734 | TRUE | 0,1910722 | No |
| orf19.3327 | 5071 | 4450 | 4798 | 3124 | 6913 | 5088 | 6634 | 7282 | 4835 | 4705 | 4740 | 4894 | 5539 | 5786 | 5632 | 5921 | 5256,47 | 4794 | 5720 | 1,194 | 0,255 | 7,089 | 1,35E-12 | 2,54E-12 | 0,0004481 | 0,0024036 | 0,0010489 | 0,0010489 | TRUE | 0,0070652 | No |
| orf19.3328 | 2513 | 2197 | 2410 | 1584 | 1911 | 1352 | 1774 | 2023 | 2396 | 2323 | 2381 | 2481 | 1531 | 1537 | 1506 | 1645 | 1975,1 | 2395 | 1555 | 0,65 | -0,622 | -13,477 | 2,14E-41 | 6,94E-41 | 0,0005914 | 0,0029226 | 0,0015206 | 0,0015206 | TRUE | 0,0189283 | No |
| orf19.3329 | 861 | 640 | 724 | 491 | 4573 | 3246 | 4423 | 4727 | 821 | 677 | 715 | 769 | 3664 | 3691 | 3755 | 3844 | 2241,98 | 746 | 3738 | 5,018 | 2,327 | 39,85 | 0 | 0 | 0,0019289 | 0,0028236 | 0,0024035 | 0,0024035 | TRUE | 0,0615496 | No |
| orf19.333 | 560 | 489 | 514 | 302 | 1370 | 934 | 1266 | 1373 | 534 | 517 | 508 | 473 | 1098 | 1062 | 1075 | 1116 | 797,85 | 508 | 1088 | 2,134 | 1,093 | 18,327 | 4,99E-75 | 2,55E-74 | 1,00E-08 | 0,0041492 | 0,0019161 | 0,0019161 | TRUE | 0,0199204 | No |
| orf19.3330 | 2407 | 1909 | 2264 | 1529 | 1986 | 1277 | 1860 | 1992 | 2295 | 2019 | 2237 | 2395 | 1591 | 1452 | 1579 | 1620 | 1898,43 | 2236 | 1560 | 0,699 | -0,516 | -8,427 | 3,55E-17 | 7,50E-17 | 0,0029446 | 0,0029562 | 0,0030655 | 0,0030655 | TRUE | 0,0519394 | No |
| orf19.3330.3 | 155 | 160 | 131 | 108 | 104 | 71 | 88 | 108 | 148 | 169 | 129 | 169 | 83 | 81 | 75 | 88 | 117,77 | 154 | 82 | 0,534 | -0,905 | -6,331 | 2,44E-10 | 4,31E-10 | 1,00E-08 | 0,016033 | 0,0105251 | 0,0105251 | TRUE | 0,1239378 | No |
| orf19.3331 | 23800 | 21332 | 23257 | 15506 | 11327 | 8403 | 10985 | 11635 | 22691 | 22556 | 22976 | 24291 | 9076 | 9555 | 9326 | 9460 | 16241,36 | 23128 | 9354 | 0,404 | -1,306 | -36,169 | 1,85E-286 | 3,90E-285 | 0,0007372 | 0,0021924 | 0,0011798 | 0,0011798 | TRUE | 0,013812 | No |
| orf19.3332 | 2421 | 2041 | 2272 | 1462 | 2191 | 1550 | 2087 | 2159 | 2308 | 2158 | 2245 | 2290 | 1756 | 1762 | 1772 | 1755 | 2005,81 | 2250 | 1761 | 0,783 | -0,353 | -8,584 | 9,20E-18 | 1,97E-17 | 1,00E-08 | 0,0029099 | 0,0011205 | 0,0011205 | TRUE | 0,0093395 | No |
| orf19.3333 | 7684 | 6479 | 7065 | 5050 | 9716 | 6800 | 9289 | 9596 | 7326 | 6851 | 6980 | 7911 | 7785 | 7732 | 7886 | 7802 | 7534,17 | 7267 | 7801 | 1,074 | 0,104 | 2,175 | 0,0296081 | 0,0362907 | 0,0019844 | 0,0023092 | 0,0020407 | 0,0020407 | TRUE | 0,0423124 | No |
| orf19.3334 | 20239 | 15571 | 16619 | 10620 | 68789 | 44639 | 64487 | 71619 | 19296 | 16464 | 16418 | 16637 | 55119 | 50759 | 54749 | 58233 | 35959,29 | 17204 | 54715 | 3,18 | 1,669 | 26,925 | 1,13E-159 | 1,23E-158 | 0,0048196 | 0,0021369 | 0,0036514 | 0,0036514 | TRUE | 0,0822808 | No |
| orf19.3335 | 13244 | 11188 | 11997 | 8482 | 20123 | 14016 | 18992 | 20074 | 12627 | 11830 | 11852 | 13288 | 16124 | 15937 | 16124 | 16322 | 14262,96 | 12399 | 16127 | 1,301 | 0,38 | 8,775 | 1,71E-18 | 3,72E-18 | 0,00154 | 0,0022064 | 0,0017272 | 0,0017272 | TRUE | 0,0279052 | No |
| orf19.3336 | 6 | 7 | 6 | 2 | 8 | 4 | 2 | 10 | 6 | 7 | 6 | 3 | 6 | 5 | 2 | 8 | 5,37 | 6 | 5 | 0,925 | -0,113 | -0,179 | 0,857828 | 0,8760884 | 1,00E-08 | 0,3077751 | 0,1986264 | 0,1986264 | TRUE | 0,4573605 | No |
| orf19.3337 | 28155 | 24424 | 27176 | 18157 | 7176 | 5248 | 7001 | 7413 | 26843 | 25825 | 26847 | 28444 | 5750 | 5967 | 5944 | 6027 | 16455,99 | 26990 | 5922 | 0,219 | -2,188 | -56,638 | 0 | 0 | 0,000872 | 0,0021911 | 0,0013372 | 0,0013372 | TRUE | 0,0159169 | No |
| orf19.334 | 6384 | 6225 | 6353 | 4352 | 6703 | 4816 | 6098 | 5940 | 6086 | 6582 | 6276 | 6818 | 5371 | 5476 | 5177 | 4830 | 5827,06 | 6440 | 5214 | 0,81 | -0,305 | -5,764 | 8,22E-09 | 1,37E-08 | 0,002519 | 0,0023731 | 0,0025083 | 0,0025083 | TRUE | 0,0300788 | No |
| orf19.3340 | 2726 | 2290 | 2416 | 1834 | 3529 | 2655 | 3410 | 3629 | 2599 | 2421 | 2387 | 2873 | 2828 | 3019 | 2895 | 2951 | 2746,58 | 2570 | 2923 | 1,139 | 0,188 | 3,12 | 0,001807 | 0,0024012 | 0,0036426 | 0,0026891 | 0,0031054 | 0,0031054 | TRUE | 0,0727644 | No |
| orf19.3341 | 9456 | 8392 | 8889 | 5851 | 15565 | 11400 | 14628 | 16311 | 9015 | 8873 | 8781 | 9166 | 12472 | 12963 | 12419 | 13262 | 10869,03 | 8959 | 12779 | 1,427 | 0,513 | 14,669 | 1,02E-48 | 3,71E-48 | 0,000551 | 0,0022423 | 0,0010752 | 0,0010752 | TRUE | 0,0080434 | No |
| orf19.3342 | 2859 | 2643 | 2751 | 1700 | 2620 | 1929 | 2445 | 2737 | 2726 | 2795 | 2718 | 2663 | 2099 | 2193 | 2076 | 2225 | 2436,91 | 2726 | 2148 | 0,788 | -0,344 | -8,279 | 1,24E-16 | 2,59E-16 | 0,0003493 | 0,002765 | 0,0012485 | 0,0012485 | TRUE | 0,0075021 | No |
| orf19.3344 | 2486 | 2210 | 2375 | 1503 | 2593 | 1797 | 2551 | 2776 | 2370 | 2337 | 2346 | 2355 | 2078 | 2043 | 2166 | 2257 | 2243,96 | 2352 | 2136 | 0,909 | -0,138 | -3,102 | 0,0019236 | 0,0025514 | 0,0006216 | 0,002823 | 0,0014473 | 0,0014473 | TRUE | 0,0177978 | No |
| orf19.3345 | 3220 | 2805 | 3003 | 1943 | 3010 | 1989 | 2844 | 3017 | 3070 | 2966 | 2967 | 3044 | 2412 | 2262 | 2415 | 2453 | 2698,44 | 3012 | 2386 | 0,793 | -0,335 | -8,197 | 2,47E-16 | 5,12E-16 | 0,0002842 | 0,0026997 | 0,0012259 | 0,0012259 | TRUE | 0,0138433 | No |
| orf19.3347 | 1078 | 983 | 1075 | 640 | 1216 | 858 | 1242 | 1216 | 1028 | 1039 | 1062 | 1003 | 974 | 976 | 1054 | 989 | 1015,61 | 1033 | 998 | 0,966 | -0,05 | -0,95 | 0,3420838 | 0,3761566 | 1,00E-08 | 0,0037079 | 0,001719 | 0,001719 | TRUE | 0,0174332 | No |
| orf19.3348 | 2046 | 1669 | 1664 | 1056 | 2448 | 1601 | 2388 | 2500 | 1951 | 1765 | 1644 | 1654 | 1962 | 1820 | 2027 | 2033 | 1856,96 | 1754 | 1960 | 1,118 | 0,16 | 2,469 | 0,0135589 | 0,0170785 | 0,0038492 | 0,0029755 | 0,0035028 | 0,0035028 | TRUE | 0,0706273 | No |
| orf19.3349 | 6569 | 5799 | 6459 | 3756 | 8881 | 5767 | 8272 | 8989 | 6263 | 6132 | 6381 | 5884 | 7116 | 6558 | 7023 | 7309 | 6583,11 | 6165 | 7002 | 1,135 | 0,183 | 4,088 | 4,35E-05 | 6,27E-05 | 0,0014317 | 0,0023407 | 0,0017764 | 0,0017764 | TRUE | 0,0218302 | No |
| orf19.335 | 1305 | 1536 | 1503 | 999 | 473 | 380 | 367 | 334 | 1244 | 1624 | 1485 | 1565 | 379 | 432 | 312 | 272 | 914,05 | 1480 | 349 | 0,235 | -2,089 | -16,623 | 4,72E-62 | 2,04E-61 | 0,0248539 | 0,0038876 | 0,0135054 | 0,0135054 | TRUE | 0,2842496 | No |
| orf19.3350 | 3404 | 3188 | 3288 | 2186 | 3476 | 2780 | 3326 | 3519 | 3245 | 3371 | 3248 | 3424 | 2785 | 3161 | 2824 | 2861 | 3115,04 | 3322 | 2908 | 0,875 | -0,193 | -3,893 | 9,90E-05 | 0,0001402 | 0,0016785 | 0,0026184 | 0,0020307 | 0,0020307 | TRUE | 0,0404004 | No |
| orf19.3351 | 1945 | 2047 | 2105 | 1246 | 1424 | 1073 | 1310 | 1293 | 1854 | 2164 | 2080 | 1952 | 1141 | 1220 | 1112 | 1051 | 1571,86 | 2012 | 1131 | 0,561 | -0,833 | -12,8 | 1,64E-37 | 4,98E-37 | 0,0033144 | 0,0031358 | 0,0033958 | 0,0033958 | TRUE | 0,0354854 | No |
| orf19.3352 | 749 | 723 | 754 | 409 | 1092 | 855 | 1044 | 1071 | 714 | 764 | 745 | 641 | 875 | 972 | 886 | 871 | 808,57 | 716 | 901 | 1,252 | 0,324 | 4,731 | 2,24E-06 | 3,41E-06 | 0,0023292 | 0,0041219 | 0,0032266 | 0,0032266 | TRUE | 0,0510274 | No |
| orf19.3353 | 3046 | 2741 | 2832 | 1647 | 3776 | 2537 | 3646 | 3849 | 2904 | 2898 | 2798 | 2580 | 3026 | 2885 | 3095 | 3130 | 2914,45 | 2795 | 3034 | 1,084 | 0,117 | 2,35 | 0,0187711 | 0,0233577 | 0,0016464 | 0,0026546 | 0,0020254 | 0,0020254 | TRUE | 0,0303512 | No |
| orf19.3354 | 9353 | 7345 | 7884 | 4287 | 30962 | 18825 | 28656 | 31809 | 8917 | 7766 | 7789 | 6716 | 24809 | 21406 | 24329 | 25864 | 15949,4 | 7797 | 24102 | 3,09 | 1,628 | 20,009 | 4,56E-89 | 2,70E-88 | 0,0096683 | 0,0021942 | 0,0062663 | 0,0062663 | TRUE | 0,1142404 | No |
| orf19.3355 | 2775 | 2693 | 2808 | 1810 | 1599 | 1135 | 1422 | 1591 | 2646 | 2848 | 2774 | 2835 | 1281 | 1291 | 1207 | 1294 | 2021,93 | 2776 | 1268 | 0,457 | -1,129 | -24,31 | 1,55E-130 | 1,35E-129 | 0,0006352 | 0,0029033 | 0,0015181 | 0,0015181 | TRUE | 0,0127893 | No |
| orf19.3356 | 468 | 590 | 518 | 293 | 307 | 208 | 253 | 282 | 446 | 624 | 512 | 459 | 246 | 237 | 215 | 229 | 370,92 | 510 | 232 | 0,453 | -1,142 | -10,074 | 7,22E-24 | 1,75E-23 | 0,0119161 | 0,0065178 | 0,0093025 | 0,0093025 | TRUE | 0,26343 | No |
| orf19.3357 | 1014 | 885 | 990 | 556 | 1316 | 877 | 1287 | 1201 | 967 | 936 | 978 | 871 | 1054 | 997 | 1093 | 977 | 984,05 | 938 | 1030 | 1,096 | 0,132 | 2,169 | 0,0300625 | 0,0368335 | 0,0015193 | 0,0037598 | 0,0025388 | 0,0025388 | TRUE | 0,0240604 | No |
| orf19.3358 | 2489 | 1927 | 2081 | 1468 | 8373 | 5453 | 7661 | 8653 | 2373 | 2038 | 2056 | 2300 | 6709 | 6201 | 6504 | 7036 | 4401,94 | 2192 | 6612 | 3,02 | 1,594 | 25,776 | 1,66E-146 | 1,65E-145 | 0,0040153 | 0,0024643 | 0,0033498 | 0,0033498 | TRUE | 0,0396679 | No |
| orf19.3359 | 1665 | 1692 | 1727 | 938 | 2059 | 1403 | 1992 | 1903 | 1587 | 1789 | 1706 | 1469 | 1650 | 1595 | 1691 | 1547 | 1629,46 | 1638 | 1621 | 0,988 | -0,018 | -0,274 | 0,7843218 | 0,8077401 | 0,0031867 | 0,0030989 | 0,0033531 | 0,0033531 | TRUE | 0,0534896 | No |
| orf19.336 | 2640 | 2257 | 2385 | 1426 | 1861 | 1204 | 1910 | 1799 | 2517 | 2386 | 2356 | 2234 | 1491 | 1369 | 1622 | 1463 | 1929,76 | 2373 | 1486 | 0,626 | -0,675 | -11,202 | 4,00E-29 | 1,06E-28 | 0,0029318 | 0,0029421 | 0,0029537 | 0,0029537 | TRUE | 0,0450709 | No |
| orf19.3360 | 9339 | 9626 | 9941 | 6252 | 2212 | 1639 | 2127 | 1985 | 8904 | 10178 | 9821 | 9794 | 1772 | 1864 | 1806 | 1614 | 5719,1 | 9674 | 1764 | 0,182 | -2,456 | -42,741 | 0 | 0 | 0,0029616 | 0,0023784 | 0,0028602 | 0,0028602 | TRUE | 0,0399761 | No |
| orf19.3362 | 253 | 229 | 242 | 161 | 351 | 194 | 298 | 316 | 241 | 242 | 239 | 252 | 281 | 221 | 253 | 257 | 248,3 | 244 | 253 | 1,045 | 0,063 | 0,637 | 0,523957 | 0,5603099 | 1,00E-08 | 0,0087038 | 0,0053562 | 0,0053562 | TRUE | 0,0733675 | No |
| orf19.3363 | 30621 | 27690 | 29468 | 17871 | 4361 | 3128 | 4032 | 4653 | 29194 | 29279 | 29112 | 27996 | 3494 | 3557 | 3423 | 3783 | 16229,7 | 28895 | 3564 | 0,123 | -3,019 | -76,46 | 0 | 0 | 0,0009635 | 0,0021924 | 0,0013527 | 0,0013527 | TRUE | 0,0212984 | No |
| orf19.3364 | 834 | 790 | 864 | 488 | 159 | 114 | 159 | 159 | 795 | 835 | 854 | 764 | 127 | 130 | 135 | 129 | 471,22 | 812 | 130 | 0,16 | -2,642 | -30,647 | 2,91E-206 | 4,07E-205 | 1,00E-08 | 0,0055756 | 0,0030492 | 0,0030492 | TRUE | 0,0161087 | No |
| orf19.3365 | 3069 | 2566 | 2773 | 1751 | 1692 | 1210 | 1613 | 1701 | 2926 | 2713 | 2739 | 2743 | 1356 | 1376 | 1369 | 1383 | 2075,73 | 2780 | 1371 | 0,493 | -1,021 | -23,444 | 1,54E-121 | 1,24E-120 | 1,00E-08 | 0,0028823 | 0,001294 | 0,001294 | TRUE | 0,015711 | No |
| orf19.3366 | 3792 | 3066 | 3231 | 2094 | 6908 | 4672 | 6753 | 6998 | 3615 | 3242 | 3192 | 3280 | 5535 | 5312 | 5733 | 5690 | 4450,05 | 3332 | 5568 | 1,671 | 0,74 | 14,885 | 4,12E-50 | 1,52E-49 | 0,0019245 | 0,0024602 | 0,0021262 | 0,0021262 | TRUE | 0,040825 | No |

| | | | | | | | | | | | | | | | | | | | | | | | | | | | | | | | |
|---|---|---|---|---|---|---|---|---|---|---|---|---|---|---|---|---|---|---|---|---|---|---|---|---|---|---|---|---|---|---|---|
| orf19.3366.1 | 1480 | 1317 | 1366 | 896 | 3095 | 2042 | 2694 | 2970 | 1411 | 1393 | 1349 | 1404 | 2480 | 2322 | 2287 | 2415 | 1882,58 | 1389 | 2376 | 1,712 | 0,776 | 16,754 | 5,30E-63 | 2,32E-62 | 0,000583 | 0,0029634 | 0,0014665 | 0,0014665 | TRUE | 0,0106696 | No |
| orf19.3367 | 4334 | 4123 | 4139 | 2605 | 3951 | 2631 | 3572 | 3580 | 4132 | 4360 | 4089 | 4081 | 3166 | 2992 | 3033 | 2911 | 3595,3 | 4166 | 3026 | 0,726 | -0,461 | -10,883 | 1,40E-27 | 3,62E-27 | 0,0007367 | 0,002548 | 0,0014439 | 0,0014439 | TRUE | 0,0121825 | No |
| orf19.3368 | 319 | 333 | 307 | 241 | 107 | 77 | 107 | 87 | 304 | 352 | 303 | 378 | 86 | 88 | 91 | 71 | 208,99 | 334 | 84 | 0,251 | -1,994 | -15,716 | 1,17E-55 | 4,64E-55 | 0,0065973 | 0,0099476 | 0,0084495 | 0,0084495 | TRUE | 0,1133744 | No |
| orf19.3369 | 22127 | 23586 | 24885 | 15033 | 4248 | 3231 | 3761 | 3620 | 21096 | 24939 | 24584 | 23550 | 3404 | 3674 | 3193 | 2943 | 13422,89 | 23542 | 3304 | 0,14 | -2,834 | -39,112 | 0 | 0 | 0,0068988 | 0,0022136 | 0,0048842 | 0,0048842 | TRUE | 0,0683406 | No |
| orf19.337 | 3055 | 2932 | 2842 | 1826 | 2245 | 1488 | 1937 | 2155 | 2913 | 3100 | 2808 | 2861 | 1799 | 1692 | 1645 | 1752 | 2321,07 | 2920 | 1722 | 0,59 | -0,762 | -15,425 | 1,11E-53 | 4,32E-53 | 0,0013235 | 0,0027987 | 0,00189 | 0,00189 | TRUE | 0,021124 | No |
| orf19.3370 | 12294 | 11725 | 12505 | 7878 | 10408 | 7410 | 9708 | 9898 | 11721 | 12398 | 12354 | 12341 | 8340 | 8426 | 8242 | 8048 | 10233,66 | 12204 | 8264 | 0,677 | -0,562 | -16,862 | 8,57E-64 | 3,80E-63 | 0,0004069 | 0,0022517 | 0,000968 | 0,000968 | TRUE | 0,0088007 | No |
| orf19.3371 | 2231 | 2097 | 2287 | 1256 | 1471 | 1018 | 1359 | 1522 | 2127 | 2217 | 2259 | 1968 | 1179 | 1158 | 1154 | 1238 | 1662,35 | 2143 | 1182 | 0,551 | -0,86 | -15,563 | 1,29E-54 | 5,06E-54 | 0,001724 | 0,003079 | 0,0022935 | 0,0022935 | TRUE | 0,0338584 | No |
| orf19.3373 | 14416 | 12283 | 13125 | 9620 | 1140 | 894 | 1031 | 1136 | 13744 | 12988 | 12966 | 15070 | 913 | 1017 | 875 | 924 | 7312,16 | 13692 | 932 | 0,068 | -3,878 | -61,275 | 0 | 0 | 0,0039032 | 0,0023158 | 0,003327 | 0,003327 | TRUE | 0,0556332 | No |
| orf19.3374 | 16501 | 13556 | 14242 | 10402 | 536 | 371 | 527 | 550 | 15732 | 14334 | 14070 | 16295 | 429 | 422 | 447 | 447 | 7772,08 | 15108 | 436 | 0,029 | -5,111 | -80,094 | 0 | 0 | 0,0032301 | 0,0023025 | 0,0028551 | 0,0028551 | TRUE | 0,0339104 | No |
| orf19.3375 | 2 | 3 | 1 | 0 | 2 | 5 | 0 | 3 | 2 | 3 | 1 | 0 | 2 | 6 | 0 | 2 | 1,97 | 2 | 2 | 1,473 | 0,558 | 0,489 | 0,6247377 | 0,6578302 | 0,4979066 | 0,833746 | 0,7105906 | 0,7105906 | TRUE | 0,392011 | No |
| orf19.3376 | 135 | 149 | 134 | 89 | 46 | 30 | 54 | 55 | 129 | 158 | 132 | 139 | 37 | 34 | 46 | 45 | 89,95 | 140 | 40 | 0,292 | -1,777 | -10,525 | 6,60E-26 | 1,66E-25 | 1,00E-08 | 0,0203449 | 0,0124487 | 0,0124487 | TRUE | 0,082592 | No |
| orf19.3378 | 94 | 70 | 68 | 44 | 8 | 5 | 2 | 4 | 90 | 74 | 67 | 69 | 6 | 6 | 2 | 3 | 39,6 | 75 | 4 | 0,056 | -4,164 | -10,784 | 4,10E-27 | 1,06E-26 | 1,00E-08 | 0,0435552 | 0,0304541 | 0,0304541 | TRUE | 0,3448418 | No |
| orf19.338 | 3947 | 3291 | 3542 | 2355 | 10153 | 7088 | 9608 | 10338 | 3763 | 3480 | 3499 | 3689 | 8135 | 8060 | 8157 | 8406 | 5898,65 | 3608 | 8190 | 2,271 | 1,183 | 30,639 | 3,68E-206 | 5,13E-205 | 0,0006562 | 0,0023696 | 0,0012218 | 0,0012218 | TRUE | 0,010988 | No |
| orf19.3380 | 23 | 18 | 13 | 11 | 10 | 9 | 8 | 10 | 22 | 19 | 13 | 17 | 8 | 10 | 7 | 8 | 13,03 | 18 | 8 | 0,461 | -1,117 | -2,793 | 0,0052144 | 0,0067314 | 1,00E-08 | 0,128143 | 0,068118 | 0,068118 | TRUE | 0,1985672 | No |
| orf19.339 | 8216 | 7279 | 7632 | 5111 | 12478 | 9208 | 11872 | 13325 | 7833 | 7697 | 7540 | 8007 | 9998 | 10470 | 10079 | 10834 | 9057,3 | 7769 | 10345 | 1,332 | 0,414 | 10,611 | 2,65E-26 | 6,75E-26 | 0,0008714 | 0,0022726 | 0,0013436 | 0,0013436 | TRUE | 0,0125348 | No |
| orf19.3391 | 1260 | 1086 | 1165 | 810 | 4870 | 3501 | 4691 | 5160 | 1201 | 1148 | 1151 | 1269 | 3902 | 3981 | 3983 | 4196 | 2603,84 | 1192 | 4016 | 3,376 | 1,756 | 36,79 | 2,69E-296 | 5,91E-295 | 0,0008914 | 0,0027218 | 0,0016042 | 0,0016042 | TRUE | 0,0193318 | No |
| orf19.3392 | 670 | 584 | 628 | 409 | 1188 | 826 | 1140 | 1178 | 639 | 618 | 620 | 641 | 952 | 939 | 968 | 958 | 791,78 | 630 | 954 | 1,518 | 0,602 | 10,93 | 8,27E-28 | 2,15E-27 | 1,00E-08 | 0,004165 | 0,0015632 | 0,0015632 | TRUE | 0,001808 | No |
| orf19.3393 | 10852 | 10150 | 10835 | 6735 | 12938 | 9381 | 12413 | 13170 | 10346 | 10732 | 10704 | 10551 | 10367 | 10667 | 10539 | 10708 | 10576,77 | 10583 | 10570 | 0,999 | -0,002 | -0,067 | 0,9469324 | 0,9524552 | 0,000167 | 0,0022465 | 0,0006973 | 0,0006973 | TRUE | 0,0028835 | No |
| orf19.3394 | 483 | 445 | 462 | 350 | 858 | 629 | 755 | 792 | 460 | 471 | 456 | 548 | 687 | 715 | 641 | 644 | 577,93 | 484 | 672 | 1,395 | 0,48 | 6,306 | 2,87E-10 | 5,04E-10 | 0,0024229 | 0,0049323 | 0,0037402 | 0,0037402 | TRUE | 0,0799598 | No |
| orf19.3395 | 2124 | 1850 | 1972 | 1211 | 4080 | 2771 | 3899 | 4025 | 2025 | 1956 | 1948 | 1897 | 3269 | 3151 | 3310 | 3273 | 2603,67 | 1956 | 3251 | 1,66 | 0,731 | 18,376 | 2,03E-75 | 1,05E-74 | 0,0001032 | 0,0027219 | 0,0010969 | 0,0010969 | TRUE | 0,0067752 | No |
| orf19.3396 | 1314 | 1025 | 1258 | 711 | 1556 | 976 | 1401 | 1525 | 1253 | 1084 | 1243 | 1114 | 1247 | 1110 | 1189 | 1240 | 1184,89 | 1174 | 1196 | 1,019 | 0,027 | 0,415 | 0,6781448 | 0,7099659 | 0,0032376 | 0,003477 | 0,0033444 | 0,0033444 | TRUE | 0,03775 | No |
| orf19.3399 | 1922 | 1540 | 1770 | 1095 | 2127 | 1280 | 2077 | 2015 | 1832 | 1628 | 1749 | 1715 | 1704 | 1455 | 1763 | 1638 | 1685,78 | 1731 | 1640 | 0,948 | -0,077 | -1,16 | 0,2458534 | 0,2757585 | 0,003705 | 0,0030653 | 0,0035766 | 0,0035766 | TRUE | 0,0667931 | No |
| orf19.34 | 383 | 334 | 354 | 206 | 135 | 82 | 102 | 104 | 365 | 353 | 350 | 323 | 108 | 93 | 87 | 85 | 220,41 | 348 | 93 | 0,267 | -1,904 | -16,864 | 8,33E-64 | 3,69E-63 | 1,00E-08 | 0,0095405 | 0,0058684 | 0,0058684 | TRUE | 0,1308389 | No |
| orf19.340 | 581 | 551 | 588 | 324 | 331 | 224 | 285 | 300 | 554 | 583 | 581 | 508 | 265 | 255 | 242 | 244 | 403,85 | 556 | 252 | 0,45 | -1,151 | -14,264 | 3,64E-46 | 1,28E-45 | 1,00E-08 | 0,0061569 | 0,0034774 | 0,0034774 | TRUE | 0,0343326 | No |
| orf19.3400 | 1551 | 1301 | 1400 | 750 | 2444 | 1587 | 2388 | 2578 | 1479 | 1376 | 1383 | 1175 | 1958 | 1805 | 2027 | 2096 | 1662,35 | 1353 | 1972 | 1,454 | 0,54 | 7,43 | 1,08E-13 | 2,11E-13 | 0,0056264 | 0,003079 | 0,0044355 | 0,0044355 | TRUE | 0,0879769 | No |
| orf19.3401 | 2327 | 1948 | 2170 | 1365 | 3105 | 2134 | 2946 | 3025 | 2219 | 2060 | 2144 | 2138 | 2488 | 2427 | 2501 | 2460 | 2304,46 | 2140 | 2469 | 1,154 | 0,206 | 5,109 | 3,23E-07 | 5,10E-07 | 1,00E-08 | 0,0028038 | 0,0011203 | 0,0011203 | TRUE | 0,0081229 | No |
| orf19.3402 | 2872 | 2334 | 2584 | 1850 | 2791 | 1925 | 2647 | 2796 | 2738 | 2468 | 2553 | 2898 | 2236 | 2189 | 2247 | 2273 | 2450,36 | 2664 | 2236 | 0,841 | -0,25 | -4,607 | 4,09E-06 | 6,17E-06 | 0,0021861 | 0,0027613 | 0,0024177 | 0,0024177 | TRUE | 0,0398695 | No |
| orf19.3404 | 7 | 15 | 12 | 4 | 7 | 7 | 6 | 8 | 7 | 16 | 12 | 6 | 6 | 8 | 5 | 7 | 8,23 | 10 | 6 | 0,603 | -0,729 | -1,432 | 0,1522556 | 0,1746438 | 1,00E-08 | 0,201646 | 0,1234666 | 0,1234666 | TRUE | 0,3411527 | No |
| orf19.3405 | 470 | 424 | 477 | 343 | 1374 | 1095 | 1315 | 1468 | 448 | 448 | 471 | 537 | 1101 | 1245 | 1116 | 1194 | 820,14 | 476 | 1164 | 2,456 | 1,296 | 17,722 | 2,84E-70 | 1,36E-69 | 0,0030311 | 0,0040933 | 0,0035711 | 0,0035711 | TRUE | 0,0736721 | No |
| orf19.3406 | 1231 | 1091 | 1167 | 754 | 562 | 391 | 653 | 617 | 1174 | 1154 | 1153 | 1181 | 450 | 445 | 554 | 502 | 826,53 | 1166 | 488 | 0,419 | -1,253 | -17,299 | 4,80E-67 | 2,22E-66 | 0,0031664 | 0,0040778 | 0,0036443 | 0,0036443 | TRUE | 0,0989342 | No |
| orf19.3407 | 1384 | 1128 | 1301 | 874 | 1153 | 691 | 1047 | 1130 | 1319 | 1193 | 1285 | 1369 | 924 | 786 | 889 | 919 | 1085,49 | 1292 | 880 | 0,683 | -0,55 | -8,18 | 2,84E-16 | 5,88E-16 | 0,0030568 | 0,0036039 | 0,0033968 | 0,0033968 | TRUE | 0,0572337 | No |
| orf19.3409 | 1570 | 1330 | 1476 | 948 | 2612 | 1779 | 2432 | 2651 | 1497 | 1406 | 1458 | 1485 | 2093 | 2023 | 2065 | 2156 | 1772,81 | 1462 | 2084 | 1,427 | 0,513 | 11,485 | 1,58E-30 | 4,29E-30 | 1,00E-08 | 0,0030174 | 0,0013208 | 0,0013208 | TRUE | 0,0079365 | No |
| orf19.341 | 376 | 291 | 346 | 213 | 1138 | 872 | 1086 | 1190 | 358 | 308 | 342 | 334 | 912 | 992 | 922 | 968 | 641,83 | 336 | 948 | 2,82 | 1,496 | 21,052 | 2,18E-98 | 1,40E-97 | 0,0006481 | 0,0046494 | 0,0027195 | 0,0027195 | TRUE | 0,0380979 | No |
| orf19.3411 | 204 | 166 | 179 | 102 | 782 | 581 | 693 | 817 | 194 | 176 | 177 | 160 | 627 | 661 | 588 | 664 | 405,82 | 177 | 635 | 3,566 | 1,834 | 20,416 | 1,21E-92 | 7,42E-92 | 1,00E-08 | 0,0061372 | 0,0039274 | 0,0039274 | TRUE | 0,0496585 | No |
| orf19.3412 | 5005 | 4056 | 4652 | 2881 | 3382 | 2257 | 3163 | 3420 | 4772 | 4289 | 4596 | 4513 | 2710 | 2566 | 2685 | 2781 | 3613,98 | 4542 | 2686 | 0,591 | -0,758 | -16,494 | 4,06E-61 | 1,74E-60 | 0,0011308 | 0,0025456 | 0,0017358 | 0,0017358 | TRUE | 0,0176259 | No |
| orf19.3413 | 3 | 7 | 0 | 1 | 3 | 4 | 5 | 2 | 3 | 7 | 0 | 2 | 2 | 5 | 4 | 2 | 3,08 | 3 | 3 | 1,054 | 0,076 | 0,084 | 0,9332641 | 0,9401892 | 0,3576062 | 0,5349241 | 0,4666898 | 0,4666898 | TRUE | 0,9130226 | No |
| orf19.3414 | 9053 | 8538 | 9260 | 5360 | 6606 | 4625 | 6061 | 6254 | 8631 | 9028 | 9148 | 8397 | 5293 | 5259 | 5146 | 5085 | 6998,34 | 8801 | 5196 | 0,59 | -0,761 | -19,591 | 1,85E-85 | 1,06E-84 | 0,000754 | 0,0023259 | 0,0012995 | 0,0012995 | TRUE | 0,0112504 | No |
| orf19.3415 | 11953 | 10233 | 11804 | 7584 | 7937 | 5437 | 7380 | 8164 | 11396 | 10820 | 11661 | 11881 | 6360 | 6182 | 6266 | 6638 | 8900,46 | 11440 | 6362 | 0,556 | -0,846 | -20,382 | 2,43E-92 | 1,48E-91 | 0,0011237 | 0,0022758 | 0,0015349 | 0,0015349 | TRUE | 0,0162273 | No |
| orf19.3415.1 | 15953 | 13458 | 13657 | 8936 | 38994 | 24369 | 35225 | 38634 | 15209 | 14230 | 13492 | 13999 | 31245 | 27710 | 29906 | 31413 | 22150,46 | 14232 | 30068 | 2,113 | 1,079 | 20,473 | 3,73E-93 | 2,30E-92 | 0,0028315 | 0,0021654 | 0,0026143 | 0,0026143 | TRUE | 0,0340485 | No |
| orf19.3417 | 114 | 114 | 128 | 86 | 264 | 191 | 263 | 286 | 109 | 121 | 126 | 135 | 212 | 217 | 223 | 233 | 171,87 | 123 | 221 | 1,818 | 0,862 | 7,535 | 4,89E-14 | 9,59E-14 | 1,00E-08 | 0,0116445 | 0,0059915 | 0,0059915 | TRUE | 0,0622087 | No |
| orf19.3418 | 132 | 120 | 133 | 57 | 207 | 145 | 198 | 249 | 126 | 127 | 131 | 89 | 166 | 165 | 168 | 202 | 146,84 | 118 | 175 | 1,465 | 0,551 | 3,981 | 6,86E-05 | 9,78E-05 | 0,0082829 | 0,0132729 | 0,0112121 | 0,0112121 | TRUE | 0,2083376 | No |
| orf19.3419 | 913 | 845 | 859 | 585 | 755 | 509 | 730 | 836 | 870 | 893 | 849 | 916 | 605 | 579 | 620 | 680 | 751,53 | 882 | 621 | 0,706 | -0,502 | -7,656 | 1,92E-14 | 3,82E-14 | 0,0013763 | 0,004276 | 0,002774 | 0,002774 | TRUE | 0,0488133 | No |
| orf19.342 | 35 | 33 | 24 | 23 | 101 | 84 | 109 | 117 | 33 | 35 | 24 | 36 | 81 | 96 | 93 | 95 | 61,51 | 32 | 91 | 2,869 | 1,521 | 7,57 | 3,73E-14 | 7,35E-14 | 1,00E-08 | 0,0287824 | 0,0163443 | 0,0163443 | TRUE | 0,225131 | No |
| orf19.3420 | 0 | 0 | 0 | 0 | 0 | 1 | 1 | 0 | 0 | 0 | 0 | 0 | 0 | 1 | 1 | 0 | 0,25 | 0 | 0 | 2,249 | 1,17 | 0,384 | 0,701175 | 0,7317955 | 1,00E-08 | 6,6157187 | 7,0103865 | 7,0103865 | TRUE | 0,1184769 | No |
| orf19.3421.1 | 1902 | 1763 | 1771 | 1216 | 1321 | 926 | 1213 | 1292 | 1813 | 1864 | 1750 | 1905 | 1058 | 1053 | 1030 | 1051 | 1440,47 | 1833 | 1048 | 0,573 | -0,805 | -16,961 | 1,59E-64 | 7,10E-64 | 1,00E-08 | 0,0032311 | 0,0014286 | 0,0014286 | TRUE | 0,011336 | No |
| orf19.3422 | 824 | 771 | 868 | 502 | 2330 | 1727 | 2107 | 2367 | 786 | 815 | 858 | 786 | 1867 | 1964 | 1789 | 1925 | 1348,61 | 811 | 1886 | 2,321 | 1,214 | 22,18 | 5,41E-109 | 3,88E-108 | 0,0009412 | 0,0033088 | 0,0019534 | 0,0019534 | TRUE | 0,0182091 | No |
| orf19.3423 | 10311 | 8900 | 10229 | 6156 | 18514 | 12595 | 18183 | 18869 | 9830 | 9411 | 10105 | 9644 | 14835 | 14322 | 15437 | 15342 | 12365,75 | 9748 | 14984 | 1,537 | 0,62 | 15,994 | 1,40E-57 | 5,70E-57 | 0,0009317 | 0,0022241 | 0,0013571 | 0,0013571 | TRUE | 0,0107065 | No |
| orf19.3425 | 2533 | 2377 | 2564 | 1634 | 2921 | 2189 | 2883 | 2948 | 2415 | 2513 | 2533 | 2560 | 2341 | 2489 | 2448 | 2397 | 2461,91 | 2505 | 2419 | 0,966 | -0,051 | -1,239 | 0,2154994 | 0,2435101 | 0,0002808 | 0,0027582 | 0,0011926 | 0,0011926 | TRUE | 0,0073631 | No |
| orf19.3426 | 13165 | 11876 | 12008 | 9136 | 33444 | 25698 | 31012 | 33479 | 12551 | 12557 | 11863 | 14312 | 26798 | 29221 | 26329 | 27222 | 20106,61 | 12821 | 27392 | 2,137 | 1,096 | 18,354 | 3,09E-75 | 1,58E-74 | 0,0043319 | 0,0021729 | 0,003364 | 0,003364 | TRUE | 0,0742656 | No |
| orf19.3427 | 224 | 213 | 243 | 188 | 173 | 151 | 168 | 175 | 214 | 225 | 240 | 295 | 139 | 172 | 143 | 142 | 196,07 | 244 | 149 | 0,613 | -0,705 | -5,611 | 2,02E-08 | 3,33E-08 | 0,0089336 | 0,0104652 | 0,0098288 | 0,0098288 | TRUE | 0,1909403 | No |
| orf19.3428 | 9425 | 8804 | 9674 | 5936 | 4300 | 3068 | 4092 | 4297 | 8986 | 9309 | 9557 | 9299 | 3445 | 3489 | 3474 | 3494 | 6381,61 | 9288 | 3476 | 0,374 | -1,418 | -42,879 | 0 | 0 | 0,0002879 | 0,0023486 | 0,0008612 | 0,0008612 | TRUE | 0,0060394 | No |
| orf19.3429 | 298 | 318 | 317 | 216 | 1069 | 811 | 945 | 1025 | 284 | 336 | 313 | 338 | 857 | 922 | 802 | 833 | 585,79 | 318 | 854 | 2,693 | 1,429 | 18,042 | 9,17E-73 | 4,57E-72 | 0,0024982 | 0,0048941 | 0,0037392 | 0,0037392 | TRUE | 0,0615062 | No |
| orf19.3430 | 142 | 137 | 145 | 84 | 160 | 86 | 105 | 131 | 135 | 145 | 143 | 132 | 128 | 98 | 89 | 107 | 122,09 | 139 | 106 | 0,761 | -0,394 | -2,774 | 0,0055437 | 0,0071477 | 0,005733 | 0,0155394 | 0,0111892 | 0,0111892 | TRUE | 0,225784 | No |
| orf19.3431 | 1615 | 1689 | 1692 | 977 | 1588 | 1133 | 1428 | 1608 | 1540 | 1786 | 1672 | 1531 | 1272 | 1288 | 1212 | 1307 | 1451,03 | 1632 | 1270 | 0,777 | -0,363 | -5,985 | 2,16E-09 | 3,67E-09 | 0,0023475 | 0,0032228 | 0,0028429 | 0,0028429 | TRUE | 0,0484715 | No |
| orf19.3432 | 354 | 335 | 350 | 192 | 250 | 186 | 216 | 230 | 338 | 354 | 346 | 301 | 200 | 211 | 183 | 187 | 265,06 | 335 | 195 | 0,58 | -0,785 | -8,256 | 1,51E-16 | 3,14E-16 | 1,00E-08 | 0,0082857 | 0,004747 | 0,004747 | TRUE | 0,0429914 | No |
| orf19.3432.1 | 28 | 32 | 28 | 31 | 41 | 41 | 28 | 30 | 27 | 34 | 28 | 49 | 33 | 47 | 24 | 24 | 33,05 | 34 | 32 | 0,94 | -0,089 | -0,301 | 0,7632574 | 0,7905193 | 0,0561607 | 0,051772 | 0,0536914 | 0,0536914 | TRUE | 0,5803999 | No |
| orf19.3433 | 568 | 570 | 554 | 365 | 219 | 161 | 214 | 220 | 542 | 603 | 547 | 572 | 175 | 183 | 182 | 179 | 372,8 | 566 | 180 | 0,318 | -1,654 | -19,738 | 1,01E-86 | 5,87E-86 | 1,00E-08 | 0,0064955 | 0,0032822 | 0,0032822 | TRUE | 0,0241686 | No |
| orf19.3434 | 602 | 651 | 679 | 425 | 1105 | 819 | 937 | 973 | 574 | 688 | 671 | 666 | 885 | 931 | 796 | 791 | 750,27 | 650 | 851 | 1,309 | 0,389 | 4,839 | 1,30E-06 | 2,00E-06 | 0,0052823 | 0,0042797 | 0,004812 | 0,004812 | TRUE | 0,0766848 | No |
| orf19.3435 | 210 | 168 | 214 | 154 | 328 | 261 | 345 | 352 | 200 | 178 | 211 | 241 | 263 | 297 | 293 | 286 | 246,15 | 208 | 285 | 1,38 | 0,464 | 4,422 | 9,78E-06 | 1,45E-05 | 0,0028243 | 0,0087615 | 0,0063121 | 0,0063121 | TRUE | 0,1039227 | No |
| orf19.3436 | 69 | 67 | 76 | 30 | 175 | 113 | 141 | 145 | 66 | 71 | 75 | 47 | 140 | 128 | 120 | 118 | 95,63 | 65 | 126 | 1,925 | 0,945 | 5,883 | 4,03E-09 | 6,78E-09 | 0,0022829 | 0,0192609 | 0,0127785 | 0,0127785 | TRUE | 0,2037575 | No |
| orf19.3437 | 88 | 89 | 89 | 69 | 437 | 319 | 383 | 364 | 84 | 94 | 88 | 108 | 350 | 363 | 325 | 296 | 213,5 | 94 | 334 | 3,601 | 1,848 | 15,001 | 7,19E-51 | 2,68E-50 | 0,0042374 | 0,0097815 | 0,0072331 | 0,0072331 | TRUE | 0,0782646 | No |
| orf19.3438 | 1087 | 975 | 1120 | 571 | 2532 | 1489 | 2211 | 2395 | 1036 | 1031 | 1106 | 895 | 2029 | 1693 | 1877 | 1947 | 1451,83 | 1017 | 1886 | 1,851 | 0,889 | 11,913 | 1,01E-32 | 2,85E-32 | 0,0056668 | 0,0032222 | 0,004553 | 0,004553 | TRUE | 0,0718841 | No |
| orf19.3439 | 3312 | 3561 | 3734 | 2208 | 1048 | 770 | 1040 | 833 | 3158 | 3765 | 3689 | 3459 | 840 | 876 | 883 | 677 | 2168,29 | 3518 | 819 | 0,233 | -2,104 | -24,601 | 1,23E-133 | 1,11E-132 | 0,0089177 | 0,0028485 | 0,0063208 | 0,0063208 | TRUE | 0,1642667 | No |
| orf19.344 | 939 | 864 | 852 | 630 | 951 | 712 | 971 | 920 | 895 | 914 | 842 | 987 | 762 | 810 | 824 | 748 | 847,68 | 910 | 786 | 0,866 | -0,207 | -3,175 | 0,0014961 | 0,0019955 | 0,0018937 | 0,0040282 | 0,0028969 | 0,0028969 | TRUE | 0,0349996 | No |
| orf19.3440 | 118 | 133 | 119 | 81 | 64 | 31 | 50 | 61 | 113 | 141 | 118 | 127 | 51 | 35 | 42 | 50 | 84,52 | 125 | 44 | 0,364 | -1,46 | -8,501 | 1,87E-17 | 3,98E-17 | 1,00E-08 | 0,0215174 | 0,0139689 | 0,0139689 | TRUE | 0,1280625 | No |
| orf19.3441 | 3189 | 2984 | 3111 | 2135 | 2350 | 1714 | 2126 | 2240 | 3040 | 3155 | 3073 | 3345 | 1883 | 1949 | 1805 | 1821 | 2508,97 | 3153 | 1864 | 0,592 | -0,757 | -16,121 | 1,83E-58 | 7,52E-58 | 0,000983 | 0,0027457 | 0,0017002 | 0,0017002 | TRUE | 0,0190087 | No |
| orf19.3442 | 3612 | 3387 | 3625 | 2371 | 10927 | 7941 | 10144 | 10779 | 3444 | 3581 | 3581 | 3714 | 8755 | 9030 | 8612 | 8764 | 6185,26 | 3580 | 8790 | 2,458 | 1,297 | 36,112 | 1,49E-285 | 3,11E-284 | 0,0004103 | 0,0023567 | 0,0010311 | 0,0010311 | TRUE | 0,0080518 | No |
| orf19.3443 | 2896 | 2805 | 2867 | 1845 | 2757 | 1925 | 2624 | 2682 | 2761 | 2966 | 2832 | 2890 | 2209 | 2189 | 2228 | 2181 | 2532,01 | 2862 | 2202 | 0,77 | -0,378 | -9,573 | 1,04E-21 | 2,42E-21 | 1,00E-08 | 0,0027397 | 0,0010959 | 0,0010959 | TRUE | 0,0076381 | No |
| orf19.3444 | 509 | 349 | 373 | 289 | 112 | 89 | 130 | 126 | 485 | 369 | 368 | 453 | 90 | 101 | 110 | 102 | 259,91 | 419 | 101 | 0,241 | -2,052 | -16,009 | 1,10E-57 | 4,49E-57 | 0,0116316 | 0,0084085 | 0,010021 | 0,010021 | TRUE | 0,1275693 | No |
| orf19.3445 | 807 | 855 | 853 | 581 | 1849 | 1371 | 1580 | 1698 | 769 | 904 | 843 | 910 | 1482 | 1559 | 1341 | 1381 | 1148,61 | 856 | 1441 | 1,684 | 0,752 | 10,511 | 7,69E-26 | 1,93E-25 | 0,0040323 | 0,0035208 | 0,0039485 | 0,0039485 | TRUE | 0,0574253 | No |
| orf19.3446 | 667 | 692 | 703 | 348 | 552 | 350 | 485 | 434 | 636 | 732 | 694 | 545 | 442 | 398 | 412 | 353 | 526,53 | 652 | 401 | 0,614 | -0,704 | -7 | 2,57E-12 | 4,79E-12 | 0,010291 | 0,0052097 | 0,0077521 | 0,0077521 | TRUE | 0,1298116 | No |
| orf19.3447 | 304 | 383 | 331 | 244 | 1264 | 974 | 1131 | 1041 | 290 | 405 | 327 | 382 | 1013 | 1108 | 960 | 846 | 666,38 | 351 | 982 | 2,805 | 1,488 | 13,465 | 2,52E-41 | 8,15E-41 | 0,0146722 | 0,0045552 | 0,00966 | 0,00966 | TRUE | 0,163067 | No |
| orf19.3448 | 1776 | 2920 | 2753 | 1405 | 651 | 516 | 620 | 400 | 1693 | 3088 | 2720 | 2201 | 522 | 587 | 526 | 325 | 1457,68 | 2426 | 490 | 0,202 | -2,308 | -8,914 | 4,94E-19 | 1,09E-18 | 0,063278 | 0,0032177 | 0,0288808 | 0,063278 | TRUE | 0,1243341 | No |

| | | | | | | | | | | | | | | | | | | | | | | | | | | | | | | | |
|---|---|---|---|---|---|---|---|---|---|---|---|---|---|---|---|---|---|---|---|---|---|---|---|---|---|---|---|---|---|---|---|
| orf19.3449 | 2699 | 2425 | 2747 | 1853 | 2146 | 1451 | 2027 | 1977 | 2573 | 2564 | 2714 | 2903 | 1720 | 1650 | 1721 | 1607 | 2181,47 | 2688 | 1674 | 0,624 | -0,681 | -12,91 | 3,97E-38 | 1,22E-37 | 0,0016424 | 0,0028439 | 0,0021922 | 0,0021922 | TRUE | 0,0328542 | No |
| orf19.3449.2 | 909 | 962 | 936 | 629 | 1240 | 933 | 1234 | 1169 | 867 | 1017 | 925 | 985 | 994 | 1061 | 1048 | 951 | 980,81 | 948 | 1014 | 1,069 | 0,096 | 1,449 | 0,147292 | 0,1691935 | 0,0027527 | 0,0037653 | 0,0032095 | 0,0032095 | TRUE | 0,0417335 | No |
| orf19.345 | 203 | 194 | 208 | 132 | 303 | 213 | 291 | 308 | 194 | 205 | 205 | 207 | 243 | 242 | 247 | 250 | 224,18 | 203 | 246 | 1,214 | 0,28 | 2,967 | 0,0030057 | 0,0039351 | 1,00E-08 | 0,0094154 | 0,0039911 | 0,0039911 | TRUE | 0,0110488 | No |
| orf19.3450.1 | 356 | 337 | 368 | 221 | 499 | 333 | 517 | 468 | 339 | 356 | 364 | 346 | 400 | 379 | 439 | 381 | 375,43 | 351 | 400 | 1,138 | 0,186 | 2,281 | 0,0225451 | 0,027869 | 1,00E-08 | 0,0064647 | 0,0036965 | 0,0036965 | TRUE | 0,0515598 | No |
| orf19.3453 | 164 | 166 | 169 | 112 | 664 | 469 | 593 | 625 | 156 | 176 | 167 | 175 | 532 | 533 | 503 | 508 | 343,91 | 168 | 519 | 3,09 | 1,628 | 18,505 | 1,89E-76 | 9,83E-76 | 1,00E-08 | 0,0068655 | 0,0032702 | 0,0032702 | TRUE | 0,0272822 | No |
| orf19.3455 | 2098 | 2156 | 2258 | 1292 | 1038 | 864 | 1065 | 951 | 2000 | 2280 | 2231 | 2024 | 832 | 982 | 904 | 773 | 1503,27 | 2134 | 873 | 0,408 | -1,293 | -16,729 | 8,02E-63 | 3,51E-62 | 0,0065233 | 0,0031835 | 0,0049592 | 0,0049592 | TRUE | 0,0814989 | No |
| orf19.3456 | 866 | 870 | 900 | 591 | 487 | 347 | 453 | 435 | 826 | 920 | 889 | 926 | 390 | 395 | 385 | 354 | 635,45 | 890 | 381 | 0,428 | -1,224 | -17,591 | 2,87E-69 | 1,36E-68 | 0,0010865 | 0,0046751 | 0,0028501 | 0,0028501 | TRUE | 0,0293815 | No |
| orf19.3457 | 542 | 539 | 525 | 314 | 1050 | 735 | 949 | 971 | 517 | 570 | 519 | 492 | 841 | 836 | 806 | 790 | 671,19 | 524 | 818 | 1,556 | 0,637 | 9,784 | 1,32E-22 | 3,11E-22 | 0,0003201 | 0,0045375 | 0,002468 | 0,002468 | TRUE | 0,039181 | No |
| orf19.3458 | 1614 | 1531 | 1625 | 843 | 1650 | 1202 | 1509 | 1359 | 1539 | 1619 | 1605 | 1321 | 1322 | 1367 | 1281 | 1105 | 1394,82 | 1521 | 1269 | 0,832 | -0,265 | -3,26 | 0,0011133 | 0,0014974 | 0,0072367 | 0,0032684 | 0,0056314 | 0,0056314 | TRUE | 0,0917661 | No |
| orf19.3459 | 303 | 307 | 315 | 205 | 566 | 401 | 563 | 577 | 289 | 325 | 311 | 321 | 454 | 456 | 478 | 469 | 387,81 | 312 | 464 | 1,495 | 0,58 | 7,517 | 5,59E-14 | 1,10E-13 | 1,00E-08 | 0,0063251 | 0,0029666 | 0,0029666 | TRUE | 0,0291063 | No |
| orf19.346 | 3617 | 3121 | 3398 | 2043 | 5130 | 3398 | 4795 | 5145 | 3448 | 3300 | 3357 | 3200 | 4111 | 3864 | 4071 | 4183 | 3691,81 | 3326 | 4057 | 1,219 | 0,286 | 6,881 | 5,95E-12 | 1,10E-11 | 0,0006933 | 0,002536 | 0,0013784 | 0,0013784 | TRUE | 0,0120754 | No |
| orf19.3460 | 416 | 461 | 474 | 271 | 349 | 242 | 346 | 303 | 397 | 487 | 468 | 425 | 280 | 275 | 294 | 246 | 358,98 | 444 | 274 | 0,615 | -0,7 | -7,395 | 1,41E-13 | 2,74E-13 | 0,0047751 | 0,0066651 | 0,0057193 | 0,0057193 | TRUE | 0,0666225 | No |
| orf19.3461 | 11 | 10 | 18 | 11 | 83 | 61 | 75 | 81 | 10 | 11 | 18 | 17 | 67 | 69 | 64 | 66 | 40,18 | 14 | 66 | 4,812 | 2,267 | 8,406 | 4,23E-17 | 8,91E-17 | 1,00E-08 | 0,04295 | 0,0229871 | 0,0229871 | TRUE | 0,195509 | No |
| orf19.3462 | 1679 | 1582 | 1494 | 993 | 3213 | 2351 | 2894 | 3178 | 1601 | 1673 | 1476 | 1556 | 2574 | 2673 | 2457 | 2584 | 2074,23 | 1576 | 2572 | 1,631 | 0,706 | 13,764 | 4,18E-43 | 1,39E-42 | 0,0012822 | 0,0028829 | 0,001993 | 0,001993 | TRUE | 0,0235658 | No |
| orf19.3463 | 15181 | 14783 | 15533 | 9084 | 13187 | 9062 | 11863 | 12773 | 14473 | 15631 | 15345 | 14231 | 10566 | 10304 | 10072 | 10386 | 12626,03 | 14920 | 10332 | 0,692 | -0,53 | -13,16 | 1,48E-39 | 4,66E-39 | 0,0011072 | 0,0022213 | 0,0014794 | 0,0014794 | TRUE | 0,0125387 | No |
| orf19.3465 | 20221 | 17666 | 17905 | 11984 | 61806 | 45188 | 57751 | 64083 | 19279 | 18680 | 17688 | 18774 | 49523 | 51383 | 49030 | 52106 | 34557,78 | 18605 | 50510 | 2,715 | 1,441 | 37,7 | 0 | 0 | 0,0010129 | 0,0021388 | 0,0013642 | 0,0013642 | TRUE | 0,0135742 | No |
| orf19.3466 | 1273 | 1332 | 1372 | 821 | 1402 | 1060 | 1250 | 1212 | 1214 | 1408 | 1355 | 1286 | 1123 | 1205 | 1061 | 985 | 1204,88 | 1316 | 1094 | 0,83 | -0,269 | -3,695 | 0,0002197 | 0,000306 | 0,0045856 | 0,003454 | 0,0042474 | 0,0042474 | TRUE | 0,0540046 | No |
| orf19.3467 | 3079 | 3107 | 3430 | 1839 | 4630 | 3238 | 4127 | 4244 | 2935 | 3285 | 3389 | 2881 | 3710 | 3682 | 3504 | 3451 | 3354,57 | 3122 | 3587 | 1,147 | 0,198 | 3,321 | 0,0008969 | 0,0012138 | 0,0035011 | 0,0025807 | 0,0031233 | 0,0031233 | TRUE | 0,0400174 | No |
| orf19.3468 | 3958 | 4238 | 4200 | 2689 | 3490 | 2634 | 3098 | 2972 | 3774 | 4481 | 4149 | 4212 | 2796 | 2995 | 2630 | 2417 | 3431,82 | 4154 | 2710 | 0,652 | -0,617 | -8,561 | 1,12E-17 | 2,38E-17 | 0,0061732 | 0,0025697 | 0,0046885 | 0,0046885 | TRUE | 0,0645427 | No |
| orf19.3469 | 3772 | 3481 | 3908 | 2989 | 2950 | 2442 | 2830 | 3118 | 3596 | 3681 | 3861 | 4682 | 2364 | 2777 | 2403 | 2535 | 3237,31 | 3955 | 2520 | 0,637 | -0,65 | -7,766 | 8,13E-15 | 1,63E-14 | 0,0102124 | 0,0025985 | 0,0064045 | 0,0064045 | TRUE | 0,1834898 | No |
| orf19.347 | 4986 | 4248 | 4594 | 3217 | 1253 | 946 | 1249 | 1278 | 4754 | 4492 | 4538 | 5040 | 1004 | 1076 | 1060 | 1039 | 2875,32 | 4706 | 1045 | 0,222 | -2,171 | -42,256 | 0 | 0 | 0,0015667 | 0,0026623 | 0,0019879 | 0,0019879 | TRUE | 0,026618 | No |
| orf19.3470 | 5746 | 5429 | 5881 | 3561 | 6885 | 4780 | 6316 | 6739 | 5478 | 5740 | 5810 | 5578 | 5517 | 5435 | 5362 | 5479 | 5550,1 | 5652 | 5448 | 0,964 | -0,053 | -1,577 | 0,1147144 | 0,1329445 | 0,0002637 | 0,0023871 | 0,0009045 | 0,0009045 | TRUE | 0,0055466 | No |
| orf19.3471 | 211 | 166 | 190 | 105 | 300 | 165 | 282 | 238 | 201 | 176 | 188 | 164 | 240 | 188 | 239 | 194 | 198,73 | 182 | 215 | 1,179 | 0,237 | 1,939 | 0,0525206 | 0,063007 | 0,0080888 | 0,0103534 | 0,0092766 | 0,0092766 | TRUE | 0,0739143 | No |
| orf19.3473 | 498 | 461 | 508 | 366 | 612 | 472 | 632 | 631 | 475 | 487 | 502 | 573 | 490 | 537 | 537 | 513 | 514,27 | 509 | 519 | 1,024 | 0,034 | 0,445 | 0,6562744 | 0,6885378 | 0,0017819 | 0,005284 | 0,0036239 | 0,0036239 | TRUE | 0,0716464 | No |
| orf19.3474 | 1928 | 1942 | 2126 | 1201 | 508 | 327 | 445 | 466 | 1838 | 2053 | 2100 | 1881 | 407 | 372 | 378 | 379 | 1176,11 | 1968 | 384 | 0,195 | -2,357 | -34,942 | 1,69E-267 | 3,12E-266 | 0,0024967 | 0,0034873 | 0,0029261 | 0,0029261 | TRUE | 0,0255874 | No |
| orf19.3475 | 1546 | 1442 | 1516 | 1079 | 676 | 534 | 631 | 696 | 1474 | 1525 | 1498 | 1690 | 542 | 607 | 536 | 566 | 1054,64 | 1547 | 563 | 0,364 | -1,459 | -22,372 | 7,40E-111 | 5,40E-110 | 0,0020557 | 0,0036481 | 0,0029241 | 0,0029241 | TRUE | 0,0436597 | No |
| orf19.3476 | 12550 | 11587 | 11980 | 7398 | 8916 | 6132 | 8569 | 8934 | 11965 | 12252 | 11835 | 11589 | 7144 | 6973 | 7275 | 7264 | 9537,17 | 11910 | 7164 | 0,601 | -0,733 | -22,833 | 2,18E-115 | 1,66E-114 | 0,000312 | 0,0022634 | 0,0008811 | 0,0008811 | TRUE | 0,0045136 | No |
| orf19.3477 | 8942 | 8989 | 9557 | 5114 | 10662 | 7046 | 9838 | 9859 | 8525 | 9505 | 9441 | 8011 | 8543 | 8012 | 8352 | 8016 | 8550,82 | 8870 | 8231 | 0,928 | -0,108 | -1,855 | 0,0636071 | 0,0757091 | 0,0037095 | 0,0022833 | 0,0031675 | 0,0031675 | TRUE | 0,0510687 | No |
| orf19.3478 | 8103 | 7344 | 7716 | 4105 | 7817 | 4877 | 7084 | 7122 | 7725 | 7765 | 7623 | 6431 | 6264 | 5546 | 6014 | 5791 | 6644,79 | 7386 | 5904 | 0,799 | -0,324 | -4,957 | 7,17E-07 | 1,12E-06 | 0,0048993 | 0,0023384 | 0,0039469 | 0,0039469 | TRUE | 0,0910502 | No |
| orf19.348 | 1110 | 934 | 1107 | 631 | 2860 | 2049 | 2810 | 2926 | 1058 | 988 | 1094 | 988 | 2292 | 2330 | 2386 | 2379 | 1689,29 | 1032 | 2347 | 2,269 | 1,182 | 24,002 | 2,64E-127 | 2,24E-126 | 0,0003359 | 0,0030632 | 0,0015968 | 0,0015968 | TRUE | 0,0192979 | No |
| orf19.3480 | 3799 | 3650 | 3723 | 2551 | 4249 | 3392 | 3776 | 3978 | 3622 | 3859 | 3678 | 3996 | 3405 | 3857 | 3206 | 3234 | 3607,19 | 3789 | 3426 | 0,904 | -0,146 | -2,285 | 0,0223034 | 0,0275863 | 0,0045077 | 0,0025465 | 0,0036239 | 0,0036239 | TRUE | 0,085994 | No |
| orf19.3481 | 6129 | 5967 | 6303 | 4220 | 10481 | 8265 | 9764 | 10660 | 5843 | 6309 | 6227 | 6611 | 8398 | 9398 | 8290 | 8668 | 7467,96 | 6248 | 8688 | 1,391 | 0,476 | 8,949 | 3,58E-19 | 7,91E-19 | 0,002746 | 0,0023111 | 0,0025773 | 0,0025773 | TRUE | 0,0365887 | No |
| orf19.3482 | 3365 | 3394 | 3530 | 2048 | 3017 | 2143 | 2794 | 2754 | 3208 | 3589 | 3487 | 3208 | 2417 | 2437 | 2372 | 2239 | 2869,76 | 3373 | 2366 | 0,701 | -0,513 | -9,779 | 1,39E-22 | 3,27E-22 | 0,0018961 | 0,0026634 | 0,0022845 | 0,0022845 | TRUE | 0,0224013 | No |
| orf19.3483 | 10181 | 11277 | 11144 | 6422 | 6063 | 4499 | 5225 | 4919 | 9706 | 11924 | 11009 | 10060 | 4858 | 5116 | 4436 | 4000 | 7638,7 | 10675 | 4602 | 0,431 | -1,214 | -14,703 | 6,18E-49 | 2,24E-48 | 0,009827 | 0,0023062 | 0,0064032 | 0,0064032 | TRUE | 0,094969 | No |
| orf19.3487 | 614 | 489 | 585 | 350 | 786 | 512 | 748 | 769 | 585 | 517 | 578 | 548 | 630 | 582 | 635 | 625 | 587,62 | 557 | 618 | 1,11 | 0,15 | 2,202 | 0,0276553 | 0,0339558 | 0,0004593 | 0,0048854 | 0,0027281 | 0,0027281 | TRUE | 0,029722 | No |
| orf19.349 | 1427 | 1304 | 1405 | 1001 | 1945 | 1481 | 1793 | 1805 | 1360 | 1379 | 1388 | 1568 | 1558 | 1684 | 1522 | 1468 | 1490,98 | 1424 | 1558 | 1,096 | 0,132 | 2,109 | 0,0349114 | 0,0425295 | 0,0031562 | 0,0031925 | 0,0030786 | 0,0030786 | TRUE | 0,0517086 | No |
| orf19.3490 | 39 | 28 | 29 | 18 | 105 | 52 | 111 | 100 | 37 | 30 | 29 | 28 | 84 | 59 | 94 | 81 | 55,31 | 31 | 80 | 2,576 | 1,365 | 6,165 | 7,04E-10 | 1,22E-09 | 1,00E-08 | 0,031779 | 0,023689 | 0,023689 | TRUE | 0,2474981 | No |
| orf19.3494 | 75 | 69 | 63 | 26 | 86 | 69 | 97 | 89 | 72 | 73 | 62 | 41 | 69 | 78 | 82 | 72 | 68,69 | 62 | 75 | 1,197 | 0,26 | 1,38 | 0,1675188 | 0,1914629 | 0,0065854 | 0,0259945 | 0,0191979 | 0,0191979 | TRUE | 0,3225558 | No |
| orf19.3496 | 2746 | 2782 | 2770 | 1811 | 6443 | 4807 | 5430 | 6259 | 2618 | 2942 | 2736 | 2837 | 5163 | 5466 | 4610 | 5089 | 3932,62 | 2783 | 5082 | 1,826 | 0,869 | 14,591 | 3,22E-48 | 1,16E-47 | 0,0032968 | 0,0025088 | 0,0031143 | 0,0031143 | TRUE | 0,047654 | No |
| orf19.3498 | 339 | 323 | 312 | 221 | 596 | 420 | 541 | 584 | 323 | 342 | 308 | 346 | 478 | 478 | 459 | 475 | 401,06 | 330 | 472 | 1,437 | 0,523 | 6,931 | 4,17E-12 | 7,74E-12 | 1,00E-08 | 0,0061852 | 0,0028283 | 0,0028283 | TRUE | 0,0222184 | No |
| orf19.3499 | 2073 | 1863 | 1968 | 1327 | 895 | 660 | 802 | 886 | 1976 | 1970 | 1944 | 2079 | 717 | 750 | 681 | 720 | 1354,77 | 1992 | 717 | 0,36 | -1,474 | -28,85 | 5,09E-183 | 6,26E-182 | 1,00E-08 | 0,0033032 | 0,001603 | 0,001603 | TRUE | 0,0135625 | No |
| orf19.35 | 18923 | 20636 | 21636 | 11880 | 8622 | 6287 | 8024 | 7691 | 18041 | 21820 | 21374 | 18611 | 6909 | 7149 | 6812 | 6254 | 13371,16 | 19962 | 6781 | 0,34 | -1,558 | -22,59 | 5,47E-113 | 4,08E-112 | 0,0060155 | 0,0022141 | 0,004476 | 0,004476 | TRUE | 0,0516168 | No |
| orf19.350 | 4484 | 3876 | 4153 | 2598 | 4426 | 2945 | 4225 | 4410 | 4275 | 4098 | 4103 | 4070 | 3546 | 3349 | 3587 | 3586 | 3826,75 | 4136 | 3517 | 0,85 | -0,234 | -6,018 | 1,77E-09 | 3,02E-09 | 0,0004275 | 0,0025203 | 0,0011852 | 0,0011852 | TRUE | 0,0120177 | No |
| orf19.3501 | 1665 | 1511 | 1653 | 1052 | 2046 | 1505 | 1943 | 2100 | 1587 | 1598 | 1633 | 1648 | 1639 | 1711 | 1650 | 1708 | 1646,74 | 1616 | 1677 | 1,038 | 0,053 | 1,242 | 0,2140884 | 0,2420872 | 1,00E-08 | 0,0030883 | 0,0011613 | 0,0011613 | TRUE | 0,0027789 | No |
| orf19.3503 | 244 | 146 | 149 | 96 | 110 | 80 | 63 | 121 | 233 | 154 | 147 | 150 | 88 | 91 | 53 | 98 | 126,95 | 171 | 82 | 0,481 | -1,055 | -5,137 | 2,80E-07 | 4,42E-07 | 0,0511689 | 0,0150251 | 0,0318246 | 0,0318246 | TRUE | 0,6428903 | No |
| orf19.3504 | 18030 | 14785 | 15592 | 9821 | 34905 | 22196 | 33225 | 35197 | 17190 | 15633 | 15403 | 15385 | 27968 | 25239 | 28208 | 28619 | 21705,63 | 15903 | 27508 | 1,73 | 0,791 | 14,799 | 1,49E-49 | 5,46E-49 | 0,0029743 | 0,0021669 | 0,0026897 | 0,0026897 | TRUE | 0,0376512 | No |
| orf19.3505 | 8757 | 7587 | 8261 | 5476 | 3904 | 2649 | 3540 | 3942 | 8349 | 8022 | 8161 | 8578 | 3128 | 3012 | 3005 | 3205 | 5682,71 | 8278 | 3088 | 0,373 | -1,421 | -36,447 | 7,77E-291 | 1,68E-289 | 0,0006157 | 0,0023802 | 0,0012479 | 0,0012479 | TRUE | 0,0080218 | No |
| orf19.3506 | 242 | 216 | 199 | 119 | 338 | 253 | 365 | 378 | 231 | 228 | 197 | 186 | 271 | 288 | 310 | 307 | 252,23 | 210 | 294 | 1,388 | 0,473 | 4,617 | 3,89E-06 | 5,88E-06 | 0,002628 | 0,0086007 | 0,0059438 | 0,0059438 | TRUE | 0,051018 | No |
| orf19.3507 | 2570 | 2105 | 2141 | 1403 | 7741 | 5206 | 7385 | 8239 | 2450 | 2226 | 2115 | 2198 | 6203 | 5920 | 6270 | 6699 | 4260,03 | 2247 | 6273 | 2,79 | 1,481 | 26,07 | 7,91E-150 | 7,93E-149 | 0,0029474 | 0,0024767 | 0,0027753 | 0,0027753 | TRUE | 0,046127 | No |
| orf19.3508 | 482 | 405 | 414 | 306 | 1085 | 718 | 970 | 1014 | 460 | 428 | 409 | 479 | 869 | 816 | 824 | 824 | 638,74 | 444 | 833 | 1,886 | 0,915 | 13,467 | 2,45E-41 | 7,92E-41 | 0,0003299 | 0,0046618 | 0,0026375 | 0,0026375 | TRUE | 0,0328403 | No |
| orf19.351 | 268 | 206 | 218 | 123 | 169 | 152 | 169 | 183 | 256 | 218 | 215 | 193 | 135 | 173 | 143 | 149 | 185,24 | 220 | 150 | 0,674 | -0,569 | -4,65 | 3,32E-06 | 5,04E-06 | 0,0063823 | 0,010955 | 0,0088917 | 0,0088917 | TRUE | 0,1287706 | No |
| orf19.35.1 | 15 | 5 | 8 | 4 | 14 | 12 | 12 | 12 | 14 | 5 | 8 | 6 | 11 | 14 | 10 | 10 | 9,82 | 8 | 11 | 1,287 | 0,364 | 0,785 | 0,4326279 | 0,468772 | 1,00E-08 | 0,1692802 | 0,1024994 | 0,1024994 | TRUE | 0,73378 | No |
| orf19.3512 | 350 | 306 | 334 | 197 | 426 | 283 | 403 | 380 | 334 | 324 | 330 | 309 | 341 | 322 | 342 | 309 | 326,26 | 324 | 328 | 1,012 | 0,017 | 0,209 | 0,8344774 | 0,8547031 | 1,00E-08 | 0,0071238 | 0,0034272 | 0,0034272 | TRUE | 0,0202773 | No |
| orf19.3515 | 1651 | 1467 | 1567 | 904 | 900 | 551 | 840 | 855 | 1574 | 1551 | 1548 | 1416 | 721 | 627 | 713 | 695 | 1105,68 | 1522 | 689 | 0,453 | -1,143 | -18,711 | 4,04E-78 | 2,15E-77 | 0,0016557 | 0,0035762 | 0,0025675 | 0,0025675 | TRUE | 0,0399925 | No |
| orf19.3516 | 1382 | 1080 | 1140 | 681 | 2742 | 1707 | 2514 | 2765 | 1318 | 1142 | 1126 | 1067 | 2197 | 1941 | 2134 | 2248 | 1646,66 | 1163 | 2130 | 1,829 | 0,871 | 12,082 | 1,32E-33 | 3,77E-33 | 0,0054097 | 0,0030884 | 0,004297 | 0,004297 | TRUE | 0,0972535 | No |
| orf19.3517 | 1195 | 1111 | 1155 | 613 | 1111 | 797 | 1057 | 1146 | 1139 | 1175 | 1141 | 960 | 890 | 906 | 897 | 932 | 1005,13 | 1104 | 906 | 0,818 | -0,289 | -4,381 | 1,18E-05 | 1,75E-05 | 0,0027283 | 0,0037248 | 0,0031925 | 0,0031925 | TRUE | 0,0822347 | No |
| orf19.3518 | 2873 | 2570 | 2657 | 1717 | 5794 | 4173 | 5675 | 6075 | 2739 | 2717 | 2625 | 2690 | 4643 | 4745 | 4818 | 4940 | 3739,55 | 2693 | 4786 | 1,778 | 0,83 | 22,13 | 1,64E-108 | 1,17E-107 | 0,0003513 | 0,0025303 | 0,0010485 | 0,0010485 | TRUE | 0,0057747 | No |
| orf19.3519 | 534 | 482 | 539 | 362 | 1169 | 794 | 1102 | 1122 | 509 | 510 | 532 | 567 | 937 | 903 | 936 | 912 | 725,72 | 530 | 922 | 1,749 | 0,806 | 13,335 | 1,46E-40 | 4,65E-40 | 1,00E-08 | 0,0043537 | 0,0019709 | 0,0019709 | TRUE | 0,0215998 | No |
| orf19.352 | 1831 | 1739 | 1804 | 1169 | 1730 | 1334 | 1651 | 1661 | 1746 | 1839 | 1782 | 1831 | 1386 | 1517 | 1402 | 1351 | 1606,65 | 1800 | 1414 | 0,785 | -0,349 | -6,962 | 3,35E-12 | 6,25E-12 | 0,0007492 | 0,0031132 | 0,0017886 | 0,0017886 | TRUE | 0,0271331 | No |
| orf19.3521 | 451 | 390 | 401 | 286 | 542 | 309 | 456 | 535 | 430 | 412 | 396 | 448 | 434 | 351 | 387 | 435 | 411,79 | 422 | 402 | 0,959 | -0,06 | -0,685 | 0,4933378 | 0,5292516 | 0,0039766 | 0,0060785 | 0,0050501 | 0,0050501 | TRUE | 0,075724 | No |
| orf19.3522 | 2 | 1 | 2 | 1 | 0 | 0 | 2 | 4 | 2 | 1 | 2 | 2 | 0 | 0 | 2 | 3 | 1,43 | 2 | 1 | 0,784 | -0,351 | -0,284 | 0,7762349 | 0,8001865 | 1,00E-08 | 1,1485825 | 0,777813 | 0,777813 | TRUE | 0,835558 | No |
| orf19.3523 | 2441 | 2073 | 2270 | 1392 | 3318 | 2218 | 3226 | 3386 | 2327 | 2192 | 2243 | 2181 | 2659 | 2522 | 2739 | 2753 | 2451,88 | 2236 | 2668 | 1,193 | 0,255 | 5,622 | 1,89E-08 | 3,11E-08 | 0,0008247 | 0,0027609 | 0,0015578 | 0,0015578 | TRUE | 0,015693 | No |
| orf19.3526 | 8574 | 7359 | 8023 | 5125 | 6698 | 4844 | 6372 | 6770 | 8174 | 7781 | 7926 | 8029 | 5367 | 5508 | 5410 | 5505 | 6712,45 | 7978 | 5448 | 0,683 | -0,551 | -17,628 | 1,51E-69 | 7,22E-69 | 0,0001795 | 0,0023359 | 0,0007841 | 0,0007841 | TRUE | 0,0035239 | No |
| orf19.3527 | 4427 | 3683 | 3989 | 2647 | 12214 | 8136 | 11798 | 12242 | 4221 | 3894 | 3941 | 4147 | 9787 | 9251 | 10016 | 9954 | 6901,36 | 4051 | 9752 | 2,409 | 1,268 | 29,431 | 2,18E-190 | 2,84E-189 | 0,0011606 | 0,0023292 | 0,0015985 | 0,0015985 | TRUE | 0,0143861 | No |
| orf19.3528 | 746 | 695 | 738 | 402 | 1324 | 776 | 1153 | 1095 | 711 | 735 | 729 | 630 | 1061 | 882 | 979 | 890 | 827,18 | 701 | 953 | 1,357 | 0,44 | 5,643 | 1,67E-08 | 2,77E-08 | 0,0046553 | 0,0040762 | 0,0045724 | 0,0045724 | TRUE | 0,0701327 | No |
| orf19.3529 | 231 | 221 | 249 | 152 | 110 | 90 | 97 | 103 | 220 | 234 | 246 | 238 | 88 | 102 | 82 | 84 | 161,82 | 234 | 89 | 0,378 | -1,404 | -11,653 | 2,23E-31 | 6,15E-31 | 1,00E-08 | 0,0122375 | 0,0065409 | 0,0065409 | TRUE | 0,0795882 | No |
| orf19.353 | 1340 | 1123 | 1271 | 736 | 2500 | 1687 | 2448 | 2541 | 1278 | 1187 | 1256 | 1153 | 2003 | 1918 | 2078 | 2066 | 1617,43 | 1218 | 2016 | 1,652 | 0,725 | 14,026 | 1,08E-44 | 3,66E-44 | 0,000919 | 0,0031064 | 0,0018813 | 0,0018813 | TRUE | 0,0138333 | No |
| orf19.3530 | 3933 | 4027 | 4170 | 2384 | 2627 | 1908 | 2476 | 2422 | 3750 | 4258 | 4120 | 3735 | 2105 | 2170 | 2102 | 1969 | 3025,99 | 3966 | 2086 | 0,526 | -0,928 | -16,463 | 6,75E-61 | 2,88E-60 | 0,0025416 | 0,0026339 | 0,0026943 | 0,0026943 | TRUE | 0,0298952 | No |
| orf19.3531 | 538 | 499 | 568 | 339 | 660 | 472 | 610 | 650 | 513 | 528 | 561 | 531 | 529 | 537 | 518 | 529 | 530,59 | 533 | 528 | 0,99 | -0,015 | -0,23 | 0,8180061 | 0,8391795 | 1,00E-08 | 0,0051858 | 0,0022364 | 0,0022364 | TRUE | 0,0159891 | No |
| orf19.3532 | 2157 | 1954 | 2084 | 1352 | 2306 | 1701 | 2329 | 2327 | 2056 | 2066 | 2059 | 2118 | 1848 | 1934 | 1977 | 1892 | 1993,83 | 2075 | 1913 | 0,922 | -0,117 | -2,802 | 0,0050754 | 0,0065586 | 1,00E-08 | 0,0029148 | 0,0011627 | 0,0011627 | TRUE | 0,0065618 | No |
| orf19.3533 | 940 | 753 | 846 | 540 | 2454 | 1610 | 2340 | 2415 | 896 | 796 | 836 | 846 | 1966 | 1831 | 1987 | 1964 | 1390,18 | 844 | 1937 | 2,297 | 1,2 | 21,9 | 2,58E-106 | 1,78E-105 | 0,0009437 | 0,0032724 | 0,0019884 | 0,0019884 | TRUE | 0,023028 | No |
| orf19.3534 | 703 | 623 | 686 | 421 | 1784 | 1216 | 1751 | 1802 | 670 | 659 | 678 | 660 | 1429 | 1383 | 1487 | 1465 | 1053,77 | 667 | 1441 | 2,162 | 1,112 | 20,915 | 3,88E-97 | 2,46E-96 | 1,00E-08 | 0,0036494 | 0,0015718 | 0,0015718 | TRUE | 0,0082342 | No |
| orf19.3535 | 4274 | 3367 | 3892 | 2412 | 3475 | 2398 | 3466 | 3718 | 4075 | 3560 | 3845 | 3779 | 2784 | 2727 | 2943 | 3023 | 3341,91 | 3815 | 2869 | 0,752 | -0,411 | -7,63 | 2,34E-14 | 4,64E-14 | 0,0022801 | 0,0025826 | 0,002478 | 0,002478 | TRUE | 0,0262003 | No |

| | | | | | | | | | | | | | | | | | | | | | | | | | | | | | | | |
|---|---|---|---|---|---|---|---|---|---|---|---|---|---|---|---|---|---|---|---|---|---|---|---|---|---|---|---|---|---|---|---|
| orf19.3536 | 1091 | 1031 | 1148 | 714 | 2663 | 2062 | 2566 | 2690 | 1040 | 1090 | 1134 | 1119 | 2134 | 2345 | 2179 | 2187 | 1653,39 | 1096 | 2211 | 2,018 | 1,013 | 19,622 | 1,01E-85 | 5,79E-85 | 0,0009763 | 0,0030843 | 0,0018447 | 0,0018447 | TRUE | 0,0191567 | No |
| orf19.3537 | 2762 | 2208 | 2140 | 1683 | 518 | 324 | 466 | 479 | 2633 | 2335 | 2114 | 2637 | 415 | 368 | 396 | 389 | 1410,89 | 2430 | 392 | 0,162 | -2,628 | -32,048 | 2,36E-225 | 3,55E-224 | 0,0065895 | 0,003255 | 0,0050885 | 0,0050885 | TRUE | 0,0934915 | No |
| orf19.3538 | 5906 | 5193 | 5902 | 3730 | 1720 | 1213 | 1726 | 1757 | 5631 | 5491 | 5831 | 5843 | 1378 | 1379 | 1465 | 1429 | 3555,87 | 5699 | 1413 | 0,248 | -2,011 | -47,838 | 0 | 0 | 0,000563 | 0,002553 | 0,0012822 | 0,0012822 | TRUE | 0,0075493 | No |
| orf19.3539 | 2769 | 2420 | 2535 | 1680 | 2901 | 1986 | 2740 | 3006 | 2640 | 2559 | 2504 | 2632 | 2324 | 2258 | 2326 | 2444 | 2461,01 | 2584 | 2338 | 0,906 | -0,142 | -3,362 | 0,000774 | 0,0010524 | 0,0004659 | 0,0027584 | 0,0013125 | 0,0013125 | TRUE | 0,0113319 | No |
| orf19.354 | 1130 | 966 | 1085 | 695 | 1513 | 1082 | 1403 | 1589 | 1077 | 1021 | 1072 | 1089 | 1212 | 1230 | 1191 | 1292 | 1148,15 | 1065 | 1231 | 1,157 | 0,211 | 4,079 | 4,52E-05 | 6,50E-05 | 1,00E-08 | 0,0035213 | 0,0016785 | 0,0016785 | TRUE | 0,0137808 | No |
| orf19.3540 | 7946 | 6403 | 6933 | 4422 | 12553 | 8158 | 12296 | 12959 | 7576 | 6770 | 6849 | 6927 | 10058 | 9276 | 10439 | 10537 | 8554,17 | 7030 | 10078 | 1,433 | 0,52 | 9,655 | 4,69E-22 | 1,10E-21 | 0,002831 | 0,0022832 | 0,0026566 | 0,0026566 | TRUE | 0,0347093 | No |
| orf19.3541 | 11370 | 9837 | 10769 | 6355 | 20201 | 14243 | 19894 | 21287 | 10840 | 10401 | 10639 | 9955 | 16186 | 16196 | 16890 | 17308 | 13552 | 10459 | 16645 | 1,591 | 0,67 | 16,711 | 1,09E-62 | 4,77E-62 | 0,0011069 | 0,0022124 | 0,0014625 | 0,0014625 | TRUE | 0,0125057 | No |
| orf19.3542 | 6455 | 5412 | 5729 | 3709 | 6460 | 4488 | 6040 | 6835 | 6154 | 5723 | 5660 | 5810 | 5176 | 5103 | 5128 | 5558 | 5538,95 | 5837 | 5241 | 0,898 | -0,155 | -3,493 | 0,0004784 | 0,0006569 | 0,001298 | 0,0023877 | 0,0017121 | 0,0017121 | TRUE | 0,0202818 | No |
| orf19.3543 | 2 | 0 | 1 | 2 | 0 | 1 | 2 | 0 | 2 | 0 | 1 | 3 | 0 | 1 | 2 | 0 | 1,11 | 2 | 1 | 0,474 | -1,078 | -0,72 | 0,4717549 | 0,5081479 | 1,00E-08 | 1,4841415 | 1,1159911 | 1,1159911 | TRUE | 0,2456801 | No |
| orf19.3544 | 292 | 317 | 319 | 196 | 396 | 276 | 364 | 376 | 278 | 335 | 315 | 307 | 317 | 314 | 309 | 306 | 310,21 | 309 | 312 | 1,009 | 0,013 | 0,145 | 0,8846932 | 0,9002116 | 1,00E-08 | 0,0073842 | 0,0039921 | 0,0039921 | TRUE | 0,0562693 | No |
| orf19.3545 | 1592 | 1480 | 1473 | 975 | 1995 | 1451 | 1874 | 2087 | 1518 | 1565 | 1455 | 1527 | 1599 | 1650 | 1591 | 1697 | 1575,21 | 1516 | 1634 | 1,078 | 0,108 | 2,302 | 0,0213256 | 0,0264077 | 0,0003784 | 0,0031336 | 0,0014833 | 0,0014833 | TRUE | 0,0093195 | No |
| orf19.3546 | 2610 | 2213 | 2320 | 1375 | 2355 | 1751 | 2315 | 2335 | 2488 | 2340 | 2292 | 2154 | 1887 | 1991 | 1965 | 1899 | 2127,04 | 2318 | 1936 | 0,833 | -0,264 | -5,12 | 3,05E-07 | 4,82E-07 | 0,0014799 | 0,0028632 | 0,0020745 | 0,0020745 | TRUE | 0,0294799 | No |
| orf19.3547 | 14660 | 13512 | 14167 | 8585 | 22210 | 15098 | 20413 | 22181 | 13977 | 14287 | 13996 | 13449 | 17796 | 17168 | 17331 | 18035 | 15754,79 | 13927 | 17582 | 1,262 | 0,336 | 10,257 | 1,10E-24 | 2,71E-24 | 0,0004904 | 0,0021955 | 0,0009646 | 0,0009646 | TRUE | 0,006334 | No |
| orf19.3548 | 2531 | 2099 | 2335 | 1413 | 3775 | 2479 | 3558 | 3959 | 2413 | 2219 | 2307 | 2214 | 3025 | 2819 | 3021 | 3219 | 2654,52 | 2288 | 3021 | 1,32 | 0,4 | 7,654 | 1,95E-14 | 3,87E-14 | 0,001786 | 0,0027098 | 0,0022348 | 0,0022348 | TRUE | 0,0238449 | No |
| orf19.3548.1 | 82538 | 59849 | 60697 | 46945 | 82737 | 61904 | 72948 | 88956 | 78691 | 63283 | 59963 | 73542 | 66295 | 70390 | 61933 | 72330 | 68303,25 | 68870 | 67737 | 0,984 | -0,024 | -0,289 | 0,772484 | 0,7967463 | 0,0090276 | 0,0021153 | 0,0065576 | 0,0065576 | TRUE | 0,1130528 | No |
| orf19.3549 | 1433 | 1191 | 1257 | 790 | 1316 | 915 | 1260 | 1367 | 1366 | 1259 | 1242 | 1238 | 1054 | 1040 | 1070 | 1112 | 1172,63 | 1276 | 1069 | 0,837 | -0,257 | -4,761 | 1,93E-06 | 2,95E-06 | 0,0007659 | 0,0034915 | 0,0019332 | 0,0019332 | TRUE | 0,028231 | No |
| orf19.355 | 99 | 88 | 81 | 62 | 103 | 64 | 104 | 107 | 94 | 93 | 80 | 97 | 83 | 73 | 88 | 87 | 86,9 | 91 | 83 | 0,915 | -0,129 | -0,844 | 0,3988881 | 0,434649 | 1,00E-08 | 0,0209859 | 0,010893 | 0,010893 | TRUE | 0,068254 | No |
| orf19.3550 | 1176 | 1035 | 1117 | 694 | 362 | 251 | 359 | 395 | 1121 | 1094 | 1103 | 1087 | 290 | 285 | 305 | 321 | 700,96 | 1101 | 300 | 0,273 | -1,872 | -28,945 | 3,23E-184 | 4,00E-183 | 1,00E-08 | 0,0044336 | 0,0020349 | 0,0020349 | TRUE | 0,0254121 | No |
| orf19.3551 | 311 | 243 | 257 | 140 | 64 | 35 | 69 | 61 | 297 | 257 | 254 | 219 | 51 | 40 | 59 | 50 | 153,24 | 257 | 50 | 0,195 | -2,36 | -15,579 | 1,02E-54 | 4,00E-54 | 0,0085301 | 0,0128059 | 0,0110303 | 0,0110303 | TRUE | 0,1101494 | No |
| orf19.3552 | 2964 | 2904 | 3115 | 1866 | 3206 | 2462 | 3088 | 3205 | 2826 | 3071 | 3077 | 2923 | 2569 | 2800 | 2622 | 2606 | 2811,63 | 2974 | 2649 | 0,89 | -0,169 | -3,585 | 0,0003375 | 0,0004665 | 0,0011156 | 0,0026752 | 0,0017643 | 0,0017643 | TRUE | 0,017047 | No |
| orf19.3553 | 9901 | 9452 | 9631 | 5028 | 14466 | 9191 | 13607 | 13714 | 9440 | 9994 | 9515 | 7877 | 11591 | 10451 | 11552 | 11151 | 10196,28 | 9206 | 11186 | 1,215 | 0,281 | 4,045 | 5,24E-05 | 7,51E-05 | 0,0059781 | 0,0022523 | 0,0045202 | 0,0045202 | TRUE | 0,1141154 | No |
| orf19.3554 | 3454 | 3385 | 3386 | 2089 | 4600 | 3520 | 4035 | 4395 | 3293 | 3579 | 3345 | 3273 | 3686 | 4003 | 3426 | 3574 | 3522,18 | 3372 | 3672 | 1,088 | 0,121 | 2,186 | 0,0288241 | 0,0353705 | 0,0027198 | 0,0025574 | 0,0026753 | 0,0026753 | TRUE | 0,0437402 | No |
| orf19.3555 | 10230 | 8880 | 9872 | 5584 | 2503 | 1705 | 2474 | 2555 | 9753 | 9389 | 9753 | 8748 | 2006 | 1939 | 2100 | 2077 | 5720,64 | 9411 | 2030 | 0,216 | -2,212 | -45,665 | 0 | 0 | 0,001522 | 0,0023783 | 0,0019742 | 0,0019742 | TRUE | 0,0268777 | No |
| orf19.3556 | 3688 | 3937 | 4117 | 2720 | 5272 | 4237 | 4763 | 5099 | 3516 | 4163 | 4067 | 4261 | 4224 | 4818 | 4044 | 4146 | 4154,89 | 4002 | 4308 | 1,077 | 0,107 | 1,477 | 0,1395785 | 0,1605644 | 0,0064078 | 0,0024865 | 0,0047504 | 0,0047504 | TRUE | 0,0819611 | No |
| orf19.3557 | 281 | 246 | 254 | 137 | 268 | 171 | 232 | 247 | 268 | 260 | 251 | 215 | 215 | 194 | 197 | 201 | 225,07 | 248 | 202 | 0,808 | -0,308 | -3,036 | 0,0023974 | 0,0031633 | 1,00E-08 | 0,0093864 | 0,005442 | 0,005442 | TRUE | 0,0743002 | No |
| orf19.3558 | 749 | 848 | 788 | 481 | 1245 | 869 | 1044 | 949 | 714 | 897 | 778 | 754 | 998 | 988 | 886 | 772 | 848,3 | 786 | 911 | 1,158 | 0,212 | 2,212 | 0,0269631 | 0,0331442 | 0,010146 | 0,0040268 | 0,0075924 | 0,0075924 | TRUE | 0,1282699 | No |
| orf19.3559 | 6055 | 5602 | 5562 | 3670 | 4753 | 3323 | 4390 | 4778 | 5773 | 5923 | 5495 | 5749 | 3808 | 3779 | 3727 | 3885 | 4767,4 | 5735 | 3800 | 0,663 | -0,594 | -16,076 | 3,79E-58 | 1,56E-57 | 0,0004766 | 0,0024357 | 0,0010932 | 0,0010932 | TRUE | 0,0099869 | No |
| orf19.356 | 208 | 195 | 193 | 105 | 371 | 302 | 343 | 349 | 198 | 206 | 191 | 164 | 297 | 343 | 291 | 284 | 246,91 | 190 | 304 | 1,584 | 0,663 | 6,32 | 2,62E-10 | 4,61E-10 | 0,0027521 | 0,008741 | 0,0061872 | 0,0061872 | TRUE | 0,0793422 | No |
| orf19.3561 | 2966 | 2920 | 3228 | 1978 | 2033 | 1506 | 1903 | 2020 | 2828 | 3088 | 3189 | 3099 | 1629 | 1712 | 1616 | 1642 | 2350,3 | 3051 | 1650 | 0,541 | -0,887 | -17,902 | 1,15E-71 | 5,62E-71 | 0,0013693 | 0,0027899 | 0,0019026 | 0,0019026 | TRUE | 0,0304248 | No |
| orf19.3563 | 3398 | 3019 | 3348 | 2390 | 1739 | 1320 | 1704 | 1751 | 3240 | 3192 | 3308 | 3744 | 1393 | 1501 | 1447 | 1424 | 2406,02 | 3371 | 1441 | 0,428 | -1,225 | -21,028 | 3,67E-98 | 2,36E-97 | 0,0026589 | 0,0027737 | 0,0027801 | 0,0027801 | TRUE | 0,0648186 | No |
| orf19.3564 | 8036 | 7118 | 7665 | 4263 | 8810 | 6141 | 8460 | 8984 | 7661 | 7526 | 7572 | 6678 | 7059 | 6983 | 7182 | 7305 | 7245,97 | 7359 | 7132 | 0,968 | -0,046 | -0,958 | 0,3380063 | 0,3718011 | 0,0019462 | 0,0023179 | 0,0020983 | 0,0020983 | TRUE | 0,0461152 | No |
| orf19.3565 | 480 | 427 | 457 | 302 | 470 | 336 | 453 | 472 | 458 | 451 | 451 | 473 | 377 | 382 | 385 | 384 | 420,09 | 458 | 382 | 0,834 | -0,262 | -3,714 | 0,0002041 | 0,0002848 | 1,00E-08 | 0,0059997 | 0,0023838 | 0,0023838 | TRUE | 0,004351 | No |
| orf19.3567 | 2209 | 2099 | 2057 | 1385 | 697 | 506 | 689 | 689 | 2106 | 2219 | 2032 | 2170 | 558 | 575 | 585 | 560 | 1350,79 | 2132 | 570 | 0,267 | -1,903 | -35,624 | 5,89E-278 | 1,18E-276 | 1,00E-08 | 0,0033068 | 0,0016963 | 0,0016963 | TRUE | 0,0122819 | No |
| orf19.3568 | 1758 | 1674 | 1869 | 1115 | 1063 | 733 | 936 | 1010 | 1676 | 1770 | 1846 | 1747 | 852 | 833 | 795 | 821 | 1292,54 | 1760 | 825 | 0,469 | -1,093 | -20,816 | 3,10E-96 | 1,96E-95 | 0,0006889 | 0,0033616 | 0,0017871 | 0,0017871 | TRUE | 0,0139643 | No |
| orf19.3569 | 2642 | 2606 | 2672 | 1556 | 2269 | 1607 | 2091 | 2130 | 2519 | 2756 | 2640 | 2438 | 1818 | 1827 | 1775 | 1732 | 2188,02 | 2588 | 1788 | 0,69 | -0,535 | -10,83 | 2,48E-27 | 6,41E-27 | 0,0011168 | 0,0028417 | 0,00188 | 0,00188 | TRUE | 0,022708 | No |
| orf19.357 | 1917 | 1532 | 1748 | 1066 | 2772 | 1716 | 2702 | 2675 | 1828 | 1620 | 1727 | 1670 | 2221 | 1951 | 2294 | 2175 | 1935,72 | 1711 | 2160 | 1,263 | 0,337 | 5,46 | 4,77E-08 | 7,76E-08 | 0,0029979 | 0,0029395 | 0,0031143 | 0,0031143 | TRUE | 0,0496768 | No |
| orf19.3571 | 255 | 224 | 222 | 180 | 91 | 75 | 89 | 80 | 243 | 237 | 219 | 282 | 73 | 85 | 76 | 65 | 160,01 | 245 | 75 | 0,304 | -1,717 | -12,865 | 7,08E-38 | 2,17E-37 | 0,0038521 | 0,0123526 | 0,0088168 | 0,0088168 | TRUE | 0,0952758 | No |
| orf19.3572 | 322 | 376 | 347 | 205 | 212 | 179 | 178 | 186 | 307 | 398 | 343 | 321 | 170 | 204 | 151 | 151 | 255,53 | 342 | 169 | 0,491 | -1,028 | -8,421 | 3,72E-17 | 7,85E-17 | 0,0115478 | 0,0085166 | 0,0100025 | 0,0100025 | TRUE | 0,18371 | No |
| orf19.3572.3 | 11419 | 9967 | 10117 | 6395 | 33985 | 22752 | 32402 | 34387 | 10887 | 10539 | 9995 | 10018 | 27231 | 25871 | 27509 | 27960 | 18751,21 | 10360 | 27143 | 2,62 | 1,389 | 33,213 | 7,01E-242 | 1,15E-240 | 0,0013296 | 0,0021788 | 0,00161 | 0,00161 | TRUE | 0,0144427 | No |
| orf19.3573 | 1537 | 1469 | 1540 | 916 | 11432 | 8200 | 10530 | 10992 | 1465 | 1553 | 1521 | 1435 | 9160 | 9324 | 8940 | 8938 | 5292,09 | 1494 | 9090 | 6,075 | 2,603 | 65,447 | 0 | 0 | 0,0003332 | 0,0024015 | 0,0010959 | 0,0010959 | TRUE | 0,0083141 | No |
| orf19.3574 | 628 | 577 | 589 | 411 | 979 | 625 | 907 | 919 | 599 | 610 | 582 | 644 | 784 | 711 | 770 | 747 | 680,87 | 609 | 753 | 1,243 | 0,314 | 4,967 | 6,80E-07 | 1,06E-06 | 1,00E-08 | 0,0045028 | 0,0023244 | 0,0023244 | TRUE | 0,0155106 | No |
| orf19.3575 | 4633 | 3803 | 4074 | 2877 | 5669 | 3998 | 5327 | 5978 | 4417 | 4021 | 4025 | 4507 | 4542 | 4546 | 4523 | 4861 | 4430,22 | 4242 | 4618 | 1,089 | 0,124 | 2,423 | 0,0153738 | 0,0192655 | 0,0021376 | 0,0024619 | 0,0022652 | 0,0022652 | TRUE | 0,0205304 | No |
| orf19.3577 | 1807 | 1458 | 1520 | 1020 | 3260 | 2046 | 3084 | 3428 | 1723 | 1542 | 1502 | 1598 | 2612 | 2326 | 2618 | 2787 | 2088,52 | 1591 | 2586 | 1,627 | 0,702 | 10,679 | 1,28E-26 | 3,27E-26 | 0,0038874 | 0,0028774 | 0,0036205 | 0,0036205 | TRUE | 0,0540333 | No |
| orf19.3577.1 | 97 | 86 | 84 | 69 | 720 | 466 | 606 | 640 | 92 | 91 | 83 | 108 | 577 | 530 | 514 | 520 | 314,52 | 94 | 535 | 5,79 | 2,534 | 23,314 | 3,22E-120 | 2,57E-119 | 1,00E-08 | 0,0073116 | 0,004528 | 0,004528 | TRUE | 0,073439 | No |
| orf19.3578 | 721 | 579 | 633 | 502 | 1183 | 790 | 1067 | 1150 | 687 | 612 | 625 | 786 | 948 | 898 | 906 | 935 | 799,81 | 678 | 922 | 1,367 | 0,451 | 5,898 | 3,68E-09 | 6,20E-09 | 0,0044731 | 0,0041441 | 0,0043031 | 0,0043031 | TRUE | 0,122124 | No |
| orf19.3579 | 5120 | 4222 | 4582 | 3379 | 9965 | 7411 | 9112 | 10046 | 4881 | 4464 | 4527 | 5293 | 7985 | 8427 | 7736 | 8168 | 6435,2 | 4791 | 8079 | 1,687 | 0,755 | 13,129 | 2,23E-39 | 6,99E-39 | 0,0036053 | 0,0023464 | 0,0030015 | 0,0030015 | TRUE | 0,0588844 | No |
| orf19.3581 | 466 | 455 | 487 | 309 | 1091 | 781 | 1034 | 1033 | 444 | 481 | 481 | 484 | 874 | 888 | 878 | 840 | 671,33 | 472 | 870 | 1,843 | 0,882 | 14,123 | 2,73E-45 | 9,39E-45 | 1,00E-08 | 0,004537 | 0,0020531 | 0,0020531 | TRUE | 0,0205569 | No |
| orf19.3582 | 1188 | 1011 | 1047 | 599 | 2271 | 1511 | 2125 | 2287 | 1133 | 1069 | 1034 | 938 | 1820 | 1718 | 1804 | 1860 | 1421,98 | 1044 | 1800 | 1,72 | 0,783 | 13,041 | 7,13E-39 | 2,22E-38 | 0,002148 | 0,0032459 | 0,0026731 | 0,0026731 | TRUE | 0,0487637 | No |
| orf19.3583 | 3846 | 3414 | 3707 | 2183 | 2353 | 1419 | 2297 | 2350 | 3667 | 3610 | 3662 | 3420 | 1885 | 1614 | 1950 | 1911 | 2714,8 | 3590 | 1840 | 0,513 | -0,963 | -15,882 | 8,39E-57 | 3,37E-56 | 0,0034455 | 0,0026961 | 0,0031306 | 0,0031306 | TRUE | 0,0797146 | No |
| orf19.3585 | 988 | 913 | 889 | 531 | 2036 | 1397 | 1777 | 1756 | 942 | 965 | 878 | 832 | 1631 | 1589 | 1509 | 1428 | 1221,72 | 904 | 1539 | 1,697 | 0,763 | 11,699 | 1,28E-31 | 3,56E-31 | 0,002795 | 0,0034352 | 0,0031738 | 0,0031738 | TRUE | 0,0305877 | No |
| orf19.3586 | 2153 | 1827 | 2092 | 1295 | 4423 | 2939 | 4300 | 4503 | 2053 | 1932 | 2067 | 2029 | 3544 | 3342 | 3651 | 3661 | 2784,73 | 2020 | 3550 | 1,758 | 0,814 | 17,654 | 9,45E-70 | 4,51E-69 | 0,001023 | 0,0026809 | 0,0016378 | 0,0016378 | TRUE | 0,0181641 | No |
| orf19.3589 | 99 | 78 | 90 | 59 | 101 | 94 | 124 | 139 | 94 | 82 | 89 | 92 | 81 | 107 | 105 | 113 | 95,54 | 89 | 102 | 1,132 | 0,179 | 1,148 | 0,2511218 | 0,2813711 | 1,00E-08 | 0,019277 | 0,0127688 | 0,0127688 | TRUE | 0,2014175 | No |
| orf19.359 | 28 | 18 | 31 | 22 | 181 | 151 | 161 | 198 | 27 | 19 | 31 | 34 | 145 | 172 | 137 | 161 | 90,65 | 28 | 154 | 5,618 | 2,49 | 12,603 | 2,04E-36 | 6,10E-36 | 0,0045127 | 0,0202031 | 0,0142972 | 0,0142972 | TRUE | 0,2722486 | No |
| orf19.3590 | 2096 | 1812 | 1956 | 1446 | 12312 | 9106 | 11480 | 13066 | 1998 | 1916 | 1932 | 2265 | 9865 | 10354 | 9746 | 10624 | 6087,73 | 2028 | 10147 | 5,015 | 2,326 | 40,79 | 0 | 0 | 0,0036275 | 0,002361 | 0,002801 | 0,002801 | TRUE | 0,0703616 | No |
| orf19.3591 | 1787 | 1663 | 1750 | 1123 | 4585 | 3367 | 4454 | 4548 | 1704 | 1758 | 1729 | 1759 | 3674 | 3829 | 3781 | 3698 | 2741,5 | 1738 | 3746 | 2,156 | 1,109 | 29,073 | 7,93E-186 | 9,98E-185 | 1,00E-08 | 0,0026902 | 0,0009545 | 0,0009545 | TRUE | 0,0025894 | No |
| orf19.3592 | 1408 | 1407 | 1444 | 928 | 1285 | 968 | 1227 | 1302 | 1342 | 1488 | 1427 | 1454 | 1030 | 1101 | 1042 | 1059 | 1242,64 | 1428 | 1058 | 0,741 | -0,433 | -8,241 | 1,70E-16 | 3,54E-16 | 0,0006592 | 0,0034126 | 0,0018323 | 0,0018323 | TRUE | 0,0204202 | No |
| orf19.3593 | 6056 | 5226 | 5400 | 3536 | 7859 | 5508 | 7655 | 8158 | 5774 | 5526 | 5335 | 5539 | 6297 | 6263 | 6499 | 6633 | 5983,27 | 5544 | 6423 | 1,159 | 0,213 | 5,509 | 3,60E-08 | 5,89E-08 | 0,0006709 | 0,0023657 | 0,0012601 | 0,0012601 | TRUE | 0,0098857 | No |
| orf19.3594 | 632 | 565 | 551 | 363 | 455 | 297 | 380 | 399 | 603 | 597 | 544 | 569 | 365 | 338 | 323 | 324 | 457,79 | 578 | 338 | 0,583 | -0,778 | -10,22 | 1,61E-24 | 3,94E-24 | 1,00E-08 | 0,0056779 | 0,0032774 | 0,0032774 | TRUE | 0,0366826 | No |
| orf19.3599 | 15362 | 13806 | 15036 | 9075 | 21593 | 14810 | 20210 | 21655 | 14646 | 14598 | 14854 | 14216 | 17302 | 16840 | 17158 | 17608 | 15902,83 | 14578 | 17227 | 1,181 | 0,241 | 8,241 | 1,71E-16 | 3,55E-16 | 0,0002529 | 0,0021945 | 0,0007539 | 0,0007539 | TRUE | 0,003303 | No |
| orf19.36 | 276 | 326 | 253 | 158 | 475 | 290 | 358 | 365 | 263 | 345 | 250 | 248 | 381 | 330 | 304 | 297 | 302,05 | 276 | 328 | 1,184 | 0,243 | 1,948 | 0,0514602 | 0,0618278 | 0,0161778 | 0,0075272 | 0,0116145 | 0,0116145 | TRUE | 0,3124361 | No |
| orf19.360 | 3 | 3 | 0 | 5 | 35 | 14 | 36 | 28 | 3 | 3 | 0 | 8 | 28 | 16 | 31 | 23 | 13,89 | 4 | 24 | 7,896 | 2,981 | 5,316 | 1,06E-07 | 1,71E-07 | 0,0588922 | 0,1202571 | 0,1057343 | 0,1057343 | TRUE | 0,6302373 | No |
| orf19.3600 | 121 | 136 | 141 | 94 | 108 | 87 | 98 | 95 | 115 | 144 | 139 | 147 | 87 | 99 | 83 | 77 | 111,45 | 136 | 86 | 0,633 | -0,659 | -4,566 | 4,98E-06 | 7,49E-06 | 1,00E-08 | 0,0168231 | 0,0107049 | 0,0107049 | TRUE | 0,1194061 | No |
| orf19.3601 | 1260 | 1346 | 1335 | 788 | 848 | 667 | 759 | 782 | 1201 | 1423 | 1319 | 1234 | 679 | 758 | 644 | 636 | 986,99 | 1294 | 679 | 0,523 | -0,934 | -12,355 | 4,60E-35 | 1,35E-34 | 0,0048164 | 0,0037548 | 0,0043914 | 0,0043914 | TRUE | 0,067893 | No |
| orf19.3603 | 1338 | 1250 | 1356 | 858 | 1270 | 879 | 1213 | 1177 | 1276 | 1322 | 1340 | 1344 | 1018 | 1000 | 1030 | 957 | 1160,63 | 1320 | 1001 | 0,759 | -0,398 | -7,934 | 2,12E-15 | 4,32E-15 | 1,00E-08 | 0,0035059 | 0,0015489 | 0,0015489 | TRUE | 0,0110914 | No |
| orf19.3604 | 881 | 875 | 898 | 478 | 1517 | 960 | 1478 | 1423 | 840 | 925 | 887 | 749 | 1216 | 1092 | 1255 | 1157 | 1015,01 | 850 | 1180 | 1,384 | 0,469 | 6,468 | 9,94E-11 | 1,78E-10 | 0,0044764 | 0,0037089 | 0,004017 | 0,004017 | TRUE | 0,0682261 | No |
| orf19.3605 | 1349 | 1561 | 1492 | 820 | 2107 | 1409 | 1695 | 1403 | 1286 | 1651 | 1474 | 1285 | 1688 | 1602 | 1439 | 1141 | 1445,68 | 1424 | 1468 | 1,03 | 0,042 | 0,366 | 0,7141279 | 0,7437326 | 0,0212465 | 0,003227 | 0,0121364 | 0,0121364 | TRUE | 0,2731142 | No |
| orf19.3606 | 5398 | 4769 | 5342 | 3079 | 4166 | 2521 | 3883 | 3857 | 5146 | 5043 | 5277 | 4823 | 3338 | 2867 | 3297 | 3136 | 4115,91 | 5072 | 3160 | 0,623 | -0,683 | -12,482 | 9,37E-36 | 2,77E-35 | 0,0026608 | 0,0024902 | 0,0026201 | 0,0026201 | TRUE | 0,0460036 | No |
| orf19.3607 | 538 | 587 | 578 | 318 | 356 | 298 | 349 | 284 | 513 | 621 | 571 | 498 | 285 | 339 | 296 | 231 | 419,26 | 551 | 288 | 0,52 | -0,944 | -8,268 | 1,36E-16 | 2,84E-16 | 0,0136029 | 0,0060075 | 0,0099316 | 0,0099316 | TRUE | 0,2041031 | No |
| orf19.3608 | 611 | 727 | 782 | 413 | 1553 | 1081 | 1400 | 1293 | 583 | 769 | 773 | 647 | 1244 | 1229 | 1189 | 1051 | 935,53 | 693 | 1178 | 1,699 | 0,765 | 8,073 | 6,85E-16 | 1,40E-15 | 0,010418 | 0,0038463 | 0,0074237 | 0,0074237 | TRUE | 0,143975 | No |
| orf19.3609 | 8219 | 7901 | 8259 | 5478 | 7749 | 5895 | 7062 | 7574 | 7836 | 8354 | 8159 | 8582 | 6209 | 6703 | 5996 | 6158 | 7249,64 | 8233 | 6266 | 0,761 | -0,394 | -8,47 | 2,45E-17 | 5,18E-17 | 0,001733 | 0,0023178 | 0,0019351 | 0,0019351 | TRUE | 0,0263708 | No |
| orf19.36.1 | 196 | 118 | 120 | 91 | 347 | 186 | 287 | 316 | 187 | 125 | 119 | 143 | 278 | 211 | 244 | 257 | 195,36 | 144 | 248 | 1,73 | 0,791 | 5,303 | 1,14E-07 | 1,83E-07 | 0,0221999 | 0,0104958 | 0,0156492 | 0,0156492 | TRUE | 0,4769963 | No |
| orf19.3610 | 1407 | 1484 | 1533 | 878 | 1161 | 764 | 1066 | 1130 | 1341 | 1569 | 1514 | 1375 | 930 | 869 | 905 | 919 | 1177,91 | 1450 | 906 | 0,625 | -0,679 | -10,734 | 7,05E-27 | 1,81E-26 | 0,0025674 | 0,0034852 | 0,0029623 | 0,0029623 | TRUE | 0,0367132 | No |
| orf19.3611 | 3032 | 2681 | 2717 | 1710 | 4978 | 3606 | 4626 | 4856 | 2891 | 2835 | 2684 | 2679 | 3989 | 4100 | 3927 | 3948 | 3381,67 | 2772 | 3991 | 1,438 | 0,524 | 12,9 | 4,50E-38 | 1,38E-37 | 0,0005257 | 0,0025768 | 0,0012693 | 0,0012693 | TRUE | 0,0102428 | No |

| | | | | | | | | | | | | | | | | | | | | | | | | | | | | | | | |
|---|---|---|---|---|---|---|---|---|---|---|---|---|---|---|---|---|---|---|---|---|---|---|---|---|---|---|---|---|---|---|---|
| orf19.3612 | 896 | 862 | 847 | 598 | 1240 | 854 | 1129 | 1260 | 854 | 911 | 837 | 937 | 994 | 971 | 959 | 1025 | 935,86 | 885 | 987 | 1,119 | 0,162 | 2,816 | 0,0048641 | 0,0062932 | 0,0005665 | 0,0038457 | 0,0021103 | 0,0021103 | TRUE | 0,0162653 | No |
| orf19.3613 | 2823 | 2887 | 3141 | 1666 | 2763 | 1888 | 2622 | 2583 | 2691 | 3053 | 3103 | 2610 | 2214 | 2147 | 2226 | 2100 | 2518 | 2864 | 2172 | 0,758 | -0,401 | -6,42 | 1,36E-10 | 2,43E-10 | 0,0036305 | 0,0027433 | 0,0033368 | 0,0033368 | TRUE | 0,0413422 | No |
| orf19.3615 | 1254 | 1325 | 1316 | 844 | 946 | 638 | 877 | 824 | 1196 | 1401 | 1300 | 1322 | 758 | 725 | 745 | 670 | 1014,61 | 1305 | 724 | 0,556 | -0,848 | -12,731 | 3,98E-37 | 1,21E-36 | 0,0028176 | 0,0037095 | 0,0032094 | 0,0032094 | TRUE | 0,0397691 | No |
| orf19.3616 | 1593 | 1687 | 1678 | 937 | 1392 | 1090 | 1311 | 1209 | 1519 | 1784 | 1658 | 1468 | 1115 | 1239 | 1113 | 983 | 1359,87 | 1607 | 1112 | 0,691 | -0,534 | -6,586 | 4,52E-11 | 8,16E-11 | 0,0073471 | 0,0032987 | 0,0055501 | 0,0055501 | TRUE | 0,0741511 | No |
| orf19.3617 | 823 | 1036 | 1074 | 595 | 890 | 759 | 844 | 661 | 785 | 1095 | 1061 | 932 | 713 | 863 | 717 | 537 | 837,92 | 968 | 708 | 0,73 | -0,455 | -3,423 | 0,0006184 | 0,0008451 | 0,0267188 | 0,0040508 | 0,0157405 | 0,0157405 | TRUE | 0,3128862 | No |
| orf19.3618 | 2010 | 1675 | 1842 | 1420 | 4012 | 3163 | 3508 | 4039 | 1916 | 1771 | 1820 | 2225 | 3215 | 3597 | 2978 | 3284 | 2600,67 | 1933 | 3268 | 1,693 | 0,76 | 9,773 | 1,47E-22 | 3,47E-22 | 0,0080387 | 0,0027226 | 0,0053752 | 0,0053752 | TRUE | 0,119733 | No |
| orf19.362 | 70 | 55 | 57 | 44 | 280 | 140 | 261 | 274 | 67 | 58 | 56 | 69 | 224 | 159 | 222 | 223 | 134,76 | 62 | 207 | 3,344 | 1,742 | 10,869 | 1,62E-27 | 4,19E-27 | 0,0127323 | 0,0142755 | 0,013563 | 0,013563 | TRUE | 0,242362 | No |
| orf19.3621 | 747 | 688 | 755 | 504 | 52 | 31 | 38 | 48 | 712 | 727 | 746 | 790 | 42 | 35 | 32 | 39 | 390,41 | 744 | 37 | 0,05 | -4,317 | -33,066 | 9,24E-240 | 1,49E-238 | 1,00E-08 | 0,0062969 | 0,0037484 | 0,0037484 | TRUE | 0,0669834 | No |
| orf19.3622 | 1240 | 1275 | 1280 | 757 | 1656 | 1124 | 1428 | 1656 | 1182 | 1348 | 1265 | 1186 | 1327 | 1278 | 1212 | 1346 | 1268,08 | 1245 | 1291 | 1,036 | 0,051 | 0,843 | 0,3994421 | 0,4351784 | 0,0020043 | 0,0033861 | 0,0027362 | 0,0027362 | TRUE | 0,037206 | No |
| orf19.3623 | 1211 | 1152 | 1326 | 809 | 1082 | 838 | 1064 | 961 | 1155 | 1218 | 1310 | 1267 | 867 | 953 | 903 | 781 | 1056,82 | 1238 | 876 | 0,707 | -0,5 | -7,083 | 1,41E-12 | 2,65E-12 | 0,0039531 | 0,0036449 | 0,0038134 | 0,0038134 | TRUE | 0,0638205 | No |
| orf19.3624 | 4243 | 4595 | 4628 | 2674 | 4653 | 3598 | 4284 | 4250 | 4045 | 4859 | 4572 | 4189 | 3728 | 4091 | 3637 | 3456 | 4072,15 | 4416 | 3728 | 0,844 | -0,245 | -3,547 | 0,0003897 | 0,0005368 | 0,0057386 | 0,0024945 | 0,0043487 | 0,0043487 | TRUE | 0,0555022 | No |
| orf19.3625 | 492 | 493 | 521 | 265 | 277 | 199 | 253 | 257 | 469 | 521 | 515 | 415 | 222 | 226 | 215 | 209 | 349,02 | 480 | 218 | 0,451 | -1,147 | -12,31 | 7,98E-35 | 2,33E-34 | 0,0033338 | 0,0067956 | 0,0051324 | 0,0051324 | TRUE | 0,0824813 | No |
| orf19.3626 | 1586 | 1742 | 1679 | 1029 | 1609 | 1287 | 1478 | 1445 | 1512 | 1842 | 1659 | 1612 | 1289 | 1463 | 1255 | 1175 | 1475,89 | 1656 | 1296 | 0,781 | -0,356 | -4,525 | 6,04E-06 | 9,04E-06 | 0,0069725 | 0,0032038 | 0,0052664 | 0,0052664 | TRUE | 0,0887012 | No |
| orf19.3626.1 | 46 | 61 | 51 | 24 | 38 | 31 | 19 | 18 | 44 | 64 | 50 | 38 | 30 | 35 | 16 | 15 | 36,6 | 49 | 24 | 0,482 | -1,053 | -3,42 | 0,0006251 | 0,0008541 | 0,0805275 | 0,0469519 | 0,0609328 | 0,0609328 | TRUE | 0,4055202 | No |
| orf19.3627 | 1186 | 1125 | 1240 | 647 | 2906 | 1995 | 2668 | 2442 | 1131 | 1190 | 1225 | 1014 | 2328 | 2268 | 2265 | 1986 | 1675,81 | 1140 | 2212 | 1,936 | 0,953 | 13,453 | 2,95E-41 | 9,53E-41 | 0,0048316 | 0,003071 | 0,0041215 | 0,0041215 | TRUE | 0,0608412 | No |
| orf19.3628 | 6245 | 5797 | 6014 | 3660 | 4531 | 3143 | 4068 | 4431 | 5954 | 6130 | 5941 | 5734 | 3631 | 3574 | 3454 | 3603 | 4752,42 | 5940 | 3566 | 0,6 | -0,737 | -20,263 | 2,72E-91 | 1,65E-90 | 0,0003113 | 0,0024368 | 0,0010507 | 0,0010507 | TRUE | 0,0062423 | No |
| orf19.3629 | 1706 | 1740 | 1741 | 1081 | 1338 | 1051 | 1204 | 1301 | 1626 | 1840 | 1720 | 1693 | 1072 | 1195 | 1022 | 1058 | 1403,36 | 1720 | 1087 | 0,631 | -0,665 | -10,668 | 1,43E-26 | 3,65E-26 | 0,0027328 | 0,0032613 | 0,0029904 | 0,0029904 | TRUE | 0,0508823 | No |
| orf19.3630 | 7191 | 7372 | 7583 | 4734 | 8557 | 6258 | 8045 | 7791 | 6856 | 7795 | 7491 | 7416 | 6856 | 7116 | 6830 | 6335 | 7086,94 | 7390 | 6784 | 0,918 | -0,123 | -2,389 | 0,0168815 | 0,0210804 | 0,0023725 | 0,002323 | 0,0024217 | 0,0024217 | TRUE | 0,0292767 | No |
| orf19.3631 | 229 | 247 | 267 | 165 | 572 | 418 | 502 | 430 | 218 | 261 | 264 | 258 | 458 | 475 | 426 | 350 | 338,9 | 250 | 427 | 1,707 | 0,771 | 6,829 | 8,55E-12 | 1,58E-11 | 0,0108937 | 0,0069361 | 0,0089521 | 0,0089521 | TRUE | 0,1778012 | No |
| orf19.3633 | 360 | 366 | 352 | 228 | 446 | 345 | 420 | 437 | 343 | 387 | 348 | 357 | 357 | 392 | 357 | 355 | 362,09 | 359 | 365 | 1,016 | 0,024 | 0,291 | 0,7710238 | 0,7954572 | 1,00E-08 | 0,0066258 | 0,003501 | 0,003501 | TRUE | 0,0339373 | No |
| orf19.3635 | 1546 | 1394 | 1479 | 931 | 2280 | 1724 | 2071 | 2304 | 1474 | 1474 | 1461 | 1458 | 1827 | 1960 | 1758 | 1873 | 1660,8 | 1467 | 1854 | 1,263 | 0,337 | 7,034 | 2,01E-12 | 3,76E-12 | 0,0006027 | 0,0030799 | 0,0015761 | 0,0015761 | TRUE | 0,0166742 | No |
| orf19.3637 | 0 | 1 | 0 | 1 | 0 | 0 | 1 | 0 | 0 | 1 | 0 | 2 | 0 | 0 | 1 | 0 | 0,43 | 1 | 0 | 0,463 | -1,11 | -0,485 | 0,6273352 | 0,6601296 | 1,00E-08 | 3,7842745 | 3,1331281 | 3,1331281 | TRUE | 0,2837405 | No |
| orf19.3638 | 41 | 42 | 42 | 23 | 25 | 19 | 17 | 24 | 39 | 44 | 41 | 36 | 20 | 22 | 14 | 20 | 29,58 | 40 | 19 | 0,465 | -1,106 | -4,2 | 2,66E-05 | 3,87E-05 | 1,00E-08 | 0,0576067 | 0,0291907 | 0,0291907 | TRUE | 0,1470259 | No |
| orf19.3639 | 879 | 825 | 943 | 561 | 798 | 609 | 768 | 818 | 838 | 872 | 932 | 879 | 639 | 692 | 652 | 665 | 771,23 | 880 | 662 | 0,751 | -0,413 | -6,834 | 8,28E-12 | 1,53E-11 | 1,00E-08 | 0,0042202 | 0,0021887 | 0,0021887 | TRUE | 0,0199072 | No |
| orf19.364 | 1595 | 1317 | 1392 | 913 | 4999 | 3305 | 4976 | 5469 | 1521 | 1393 | 1375 | 1430 | 4006 | 3758 | 4225 | 4447 | 2769,21 | 1430 | 4109 | 2,875 | 1,523 | 24,853 | 2,38E-136 | 2,17E-135 | 0,0029811 | 0,0026842 | 0,0031058 | 0,0031058 | TRUE | 0,0394547 | No |
| orf19.3642 | 9544 | 8061 | 8767 | 5478 | 21096 | 14439 | 19895 | 21397 | 9099 | 8523 | 8661 | 8582 | 16904 | 16418 | 16891 | 17398 | 12809,48 | 8716 | 16903 | 1,939 | 0,955 | 27,192 | 8,18E-163 | 9,04E-162 | 0,0006136 | 0,0022195 | 0,0010941 | 0,0010941 | TRUE | 0,0108582 | No |
| orf19.3643 | 557 | 460 | 535 | 362 | 514 | 335 | 480 | 503 | 531 | 486 | 529 | 567 | 412 | 381 | 408 | 409 | 465,29 | 528 | 402 | 0,765 | -0,386 | -5,244 | 1,57E-07 | 2,51E-07 | 1,00E-08 | 0,00562 | 0,0030337 | 0,0030337 | TRUE | 0,0322258 | No |
| orf19.3644 | 4364 | 3774 | 4106 | 2778 | 2816 | 1988 | 2649 | 2980 | 4161 | 3991 | 4056 | 4352 | 2256 | 2261 | 2249 | 2423 | 3218,54 | 4140 | 2297 | 0,556 | -0,848 | -18,885 | 1,52E-79 | 8,27E-79 | 0,0009433 | 0,0026014 | 0,0016057 | 0,0016057 | TRUE | 0,0168268 | No |
| orf19.3646 | 41323 | 33937 | 35671 | 23957 | 37589 | 25253 | 34614 | 37057 | 39397 | 35884 | 35240 | 37530 | 30119 | 28715 | 29387 | 30131 | 33300,31 | 37013 | 29588 | 0,799 | -0,323 | -7,618 | 2,59E-14 | 5,12E-14 | 0,0014837 | 0,0021406 | 0,0016962 | 0,0016962 | TRUE | 0,0231645 | No |
| orf19.3647 | 537 | 490 | 546 | 335 | 1280 | 973 | 1216 | 1346 | 512 | 518 | 539 | 525 | 1026 | 1106 | 1032 | 1094 | 794,14 | 524 | 1064 | 2,032 | 1,023 | 16,966 | 1,47E-64 | 6,55E-64 | 1,00E-08 | 0,0041588 | 0,0020062 | 0,0020062 | TRUE | 0,0076809 | No |
| orf19.3648 | 425 | 335 | 360 | 291 | 686 | 412 | 680 | 679 | 405 | 354 | 356 | 456 | 550 | 468 | 577 | 552 | 464,81 | 393 | 537 | 1,377 | 0,461 | 4,74 | 2,14E-06 | 3,27E-06 | 0,0085035 | 0,0056237 | 0,0068352 | 0,0068352 | TRUE | 0,117543 | No |
| orf19.3649 | 2262 | 2236 | 2263 | 1504 | 4106 | 3039 | 3734 | 4058 | 2157 | 2364 | 2236 | 2356 | 3290 | 3456 | 3170 | 3300 | 2790,99 | 2278 | 3304 | 1,451 | 0,537 | 11,404 | 3,97E-30 | 1,07E-29 | 0,0010995 | 0,0026796 | 0,0017482 | 0,0017482 | TRUE | 0,0159409 | No |
| orf19.3651 | 20804 | 16377 | 17502 | 11579 | 6231 | 4519 | 5899 | 6674 | 19834 | 17317 | 17290 | 18139 | 4993 | 5139 | 5008 | 5427 | 11643,31 | 18145 | 5142 | 0,283 | -1,819 | -34,236 | 7,02E-257 | 1,23E-255 | 0,0027595 | 0,0022323 | 0,0025948 | 0,0025948 | TRUE | 0,0483607 | No |
| orf19.3653 | 5454 | 4726 | 5023 | 3349 | 14744 | 10680 | 13999 | 15312 | 5200 | 4997 | 4962 | 5246 | 11814 | 12144 | 11885 | 12450 | 8587,36 | 5101 | 12073 | 2,368 | 1,244 | 35,108 | 5,13E-270 | 9,66E-269 | 0,0004899 | 0,0022825 | 0,0010574 | 0,0010574 | TRUE | 0,0054696 | No |
| orf19.3655 | 41 | 44 | 39 | 29 | 41 | 44 | 58 | 68 | 39 | 47 | 39 | 45 | 33 | 50 | 49 | 55 | 44,62 | 42 | 47 | 1,108 | 0,147 | 0,646 | 0,518015 | 0,5543273 | 1,00E-08 | 0,0388859 | 0,0272345 | 0,0272345 | TRUE | 0,3679107 | No |
| orf19.3656 | 4086 | 3935 | 4167 | 2923 | 3957 | 2930 | 3730 | 3994 | 3896 | 4161 | 4117 | 4579 | 3171 | 3332 | 3167 | 3248 | 3708,57 | 4188 | 3230 | 0,772 | -0,374 | -7,163 | 7,89E-13 | 1,50E-12 | 0,0022459 | 0,002534 | 0,0023407 | 0,0023407 | TRUE | 0,0463527 | No |
| orf19.3658 | 237 | 218 | 204 | 143 | 566 | 426 | 536 | 530 | 226 | 231 | 202 | 224 | 454 | 484 | 455 | 431 | 338,24 | 221 | 456 | 2,067 | 1,047 | 12,073 | 1,46E-33 | 4,18E-33 | 1,00E-08 | 0,0069455 | 0,0037234 | 0,0037234 | TRUE | 0,0402559 | No |
| orf19.3659 | 1756 | 1592 | 1751 | 1030 | 1003 | 688 | 925 | 1036 | 1674 | 1683 | 1730 | 1614 | 804 | 782 | 785 | 842 | 1239,32 | 1675 | 803 | 0,479 | -1,061 | -20,924 | 3,22E-97 | 2,04E-96 | 1,00E-08 | 0,0034161 | 0,0015782 | 0,0015782 | TRUE | 0,012966 | No |
| orf19.366 | 1549 | 1091 | 1257 | 650 | 3641 | 1959 | 3371 | 3513 | 1477 | 1154 | 1242 | 1018 | 2917 | 2228 | 2862 | 2856 | 1969,23 | 1223 | 2716 | 2,218 | 1,149 | 10,517 | 7,20E-26 | 1,81E-25 | 0,0188765 | 0,0029251 | 0,0108482 | 0,0108482 | TRUE | 0,23555 | No |
| orf19.3660 | 2320 | 2008 | 2273 | 1338 | 2323 | 1487 | 2092 | 2194 | 2212 | 2123 | 2246 | 2096 | 1861 | 1691 | 1776 | 1784 | 1973,61 | 2169 | 1778 | 0,82 | -0,287 | -6,105 | 1,03E-09 | 1,76E-09 | 0,0006216 | 0,0029232 | 0,0016109 | 0,0016109 | TRUE | 0,0123669 | No |
| orf19.3661 | 4994 | 4921 | 5164 | 3355 | 3375 | 2598 | 3078 | 3405 | 4761 | 5203 | 5102 | 5256 | 2704 | 2954 | 2613 | 2769 | 3920,27 | 5080 | 2760 | 0,543 | -0,881 | -17,363 | 1,56E-67 | 7,24E-67 | 0,0019498 | 0,0025101 | 0,0021972 | 0,0021972 | TRUE | 0,0262976 | No |
| orf19.3663 | 7744 | 7396 | 7846 | 4857 | 9368 | 6484 | 8721 | 9126 | 7383 | 7820 | 7751 | 7609 | 7506 | 7373 | 7404 | 7420 | 7533,36 | 7641 | 7426 | 0,972 | -0,041 | -1,306 | 0,1914202 | 0,2173789 | 0,0001719 | 0,0023092 | 0,000812 | 0,000812 | TRUE | 0,0065628 | No |
| orf19.3663.1 | 873 | 754 | 784 | 504 | 802 | 559 | 670 | 723 | 832 | 797 | 775 | 790 | 643 | 636 | 569 | 588 | 703,57 | 798 | 609 | 0,761 | -0,394 | -6,091 | 1,12E-09 | 1,93E-09 | 0,0008896 | 0,0044249 | 0,0025773 | 0,0025773 | TRUE | 0,0230769 | No |
| orf19.3664 | 10653 | 9513 | 9835 | 6309 | 498 | 362 | 490 | 485 | 10156 | 10059 | 9716 | 9883 | 399 | 412 | 416 | 394 | 5179,47 | 9954 | 405 | 0,041 | -4,62 | -96,54 | 0 | 0 | 1,00E-08 | 0,0024083 | 0,0010478 | 0,0010478 | TRUE | 0,0042464 | No |
| orf19.3665 | 1607 | 1467 | 1502 | 959 | 2226 | 1635 | 2123 | 2209 | 1532 | 1551 | 1484 | 1502 | 1784 | 1859 | 1802 | 1796 | 1663,84 | 1517 | 1810 | 1,192 | 0,253 | 5,945 | 2,77E-09 | 4,69E-09 | 1,00E-08 | 0,0030781 | 0,0011263 | 0,0011263 | TRUE | 0,0036828 | No |
| orf19.3666 | 635 | 626 | 661 | 385 | 1637 | 1149 | 1536 | 1663 | 605 | 662 | 653 | 603 | 1312 | 1307 | 1304 | 1352 | 974,74 | 631 | 1319 | 2,087 | 1,061 | 19,084 | 3,46E-81 | 1,91E-80 | 1,00E-08 | 0,0037757 | 0,0017489 | 0,0017489 | TRUE | 0,0124355 | No |
| orf19.3667 | 56 | 49 | 46 | 33 | 38 | 32 | 47 | 34 | 53 | 52 | 45 | 52 | 30 | 36 | 40 | 28 | 42,09 | 50 | 34 | 0,662 | -0,596 | -2,704 | 0,0068428 | 0,0087678 | 1,00E-08 | 0,0411001 | 0,0223075 | 0,0223075 | TRUE | 0,1330476 | No |
| orf19.3668 | 4258 | 3577 | 3770 | 2602 | 2940 | 1979 | 2640 | 2996 | 4060 | 3782 | 3724 | 4076 | 2356 | 2250 | 2241 | 2436 | 3115,72 | 3910 | 2321 | 0,594 | -0,751 | -15,331 | 4,75E-53 | 1,82E-52 | 0,001438 | 0,0026183 | 0,0019687 | 0,0019687 | TRUE | 0,0136236 | No |
| orf19.3669 | 17247 | 15782 | 17267 | 11387 | 3508 | 2616 | 3247 | 3552 | 16443 | 16687 | 17058 | 17838 | 2811 | 2975 | 2757 | 2888 | 9932,18 | 17006 | 2858 | 0,168 | -2,574 | -62,802 | 0 | 0 | 0,0009203 | 0,0022566 | 0,0014234 | 0,0014234 | TRUE | 0,0130412 | No |
| orf19.367 | 4440 | 3473 | 4148 | 2873 | 7691 | 5541 | 7627 | 8287 | 4233 | 3672 | 4098 | 4501 | 6163 | 6301 | 6475 | 6738 | 5272,56 | 4126 | 6419 | 1,557 | 0,639 | 10,517 | 7,19E-26 | 1,81E-25 | 0,0039922 | 0,0024027 | 0,0033366 | 0,0033366 | TRUE | 0,0665677 | No |
| orf19.3670 | 6097 | 5715 | 6019 | 3658 | 1618 | 1112 | 1456 | 1545 | 5813 | 6043 | 5946 | 5730 | 1296 | 1264 | 1236 | 1256 | 3573,21 | 5883 | 1263 | 0,215 | -2,22 | -56,449 | 0 | 0 | 1,00E-08 | 0,0025508 | 0,0010366 | 0,0010366 | TRUE | 0,0040228 | No |
| orf19.3671 | 1 | 1 | 0 | 2 | 6 | 4 | 12 | 3 | 1 | 1 | 0 | 3 | 5 | 5 | 10 | 2 | 3,39 | 1 | 6 | 4,827 | 2,271 | 2,306 | 0,0211203 | 0,0261686 | 0,1958775 | 0,486302 | 0,365504 | 0,365504 | TRUE | 0,6204155 | No |
| orf19.3672 | 6920 | 6218 | 6519 | 4217 | 1850 | 1322 | 1720 | 1854 | 6597 | 6575 | 6440 | 6606 | 1482 | 1503 | 1460 | 1507 | 4021,48 | 6554 | 1488 | 0,227 | -2,139 | -61,095 | 0 | 0 | 1,00E-08 | 0,0024996 | 0,0007919 | 0,0007919 | TRUE | 0,0019572 | No |
| orf19.3673 | 182 | 139 | 163 | 104 | 276 | 200 | 249 | 282 | 174 | 147 | 161 | 163 | 221 | 227 | 211 | 229 | 191,71 | 161 | 222 | 1,378 | 0,463 | 4,389 | 1,14E-05 | 1,68E-05 | 1,00E-08 | 0,0106557 | 0,0052258 | 0,0052258 | TRUE | 0,0390233 | No |
| orf19.3674 | 2682 | 2505 | 2592 | 1716 | 1421 | 1072 | 1353 | 1352 | 2557 | 2649 | 2561 | 2688 | 1139 | 1219 | 1149 | 1099 | 1882,52 | 2614 | 1152 | 0,44 | -1,184 | -25,265 | 7,77E-141 | 7,38E-140 | 0,0004149 | 0,0029635 | 0,0015052 | 0,0015052 | TRUE | 0,0171717 | No |
| orf19.3675 | 7745 | 7553 | 7704 | 5487 | 2715 | 2177 | 2497 | 2594 | 7384 | 7986 | 7611 | 8596 | 2175 | 2475 | 2120 | 2109 | 5057,11 | 7894 | 2220 | 0,281 | -1,831 | -28,325 | 1,69E-176 | 1,98E-175 | 0,0048926 | 0,002416 | 0,0037405 | 0,0037405 | TRUE | 0,070904 | No |
| orf19.3676 | 3237 | 3287 | 3270 | 2101 | 3343 | 2331 | 3033 | 3165 | 3086 | 3476 | 3230 | 3291 | 2679 | 2651 | 2575 | 2573 | 2945,14 | 3271 | 2620 | 0,801 | -0,32 | -6,959 | 3,42E-12 | 6,37E-12 | 0,0011415 | 0,0026488 | 0,0016883 | 0,0016883 | TRUE | 0,0222825 | No |
| orf19.3677 | 302 | 270 | 355 | 202 | 274 | 249 | 281 | 288 | 288 | 285 | 351 | 316 | 220 | 283 | 239 | 234 | 277 | 310 | 244 | 0,783 | -0,353 | -3,262 | 0,0011045 | 0,001486 | 0,0070936 | 0,0080188 | 0,0075749 | 0,0075749 | TRUE | 0,1197958 | No |
| orf19.3678 | 2926 | 2855 | 3033 | 1925 | 1256 | 926 | 1324 | 1343 | 2790 | 3019 | 2996 | 3016 | 1006 | 1053 | 1124 | 1092 | 2011,97 | 2955 | 1069 | 0,362 | -1,466 | -28,399 | 2,07E-177 | 2,44E-176 | 0,0012903 | 0,0029073 | 0,0019511 | 0,0019511 | TRUE | 0,0197366 | No |
| orf19.3679 | 1802 | 1721 | 1785 | 1095 | 1112 | 773 | 995 | 975 | 1718 | 1820 | 1763 | 1715 | 891 | 879 | 845 | 793 | 1303,01 | 1754 | 852 | 0,485 | -1,044 | -19,402 | 7,41E-84 | 4,17E-83 | 0,0007748 | 0,0033514 | 0,0019359 | 0,0019359 | TRUE | 0,0269273 | No |
| orf19.3680 | 388 | 428 | 425 | 261 | 1178 | 898 | 1058 | 1194 | 370 | 453 | 420 | 409 | 944 | 1021 | 898 | 971 | 685,66 | 413 | 958 | 2,321 | 1,215 | 16,082 | 3,42E-58 | 1,41E-57 | 0,002534 | 0,0044859 | 0,0036569 | 0,0036569 | TRUE | 0,0626981 | No |
| orf19.3681 | 2377 | 2202 | 2398 | 1691 | 3922 | 2990 | 3403 | 3989 | 2266 | 2328 | 2369 | 2649 | 3143 | 3400 | 2889 | 3243 | 2785,96 | 2403 | 3169 | 1,32 | 0,401 | 6,241 | 4,35E-10 | 7,60E-10 | 0,0042821 | 0,0026806 | 0,0035792 | 0,0035792 | TRUE | 0,0549903 | No |
| orf19.3682 | 1634 | 1943 | 2008 | 1208 | 1460 | 1243 | 1286 | 1122 | 1558 | 2054 | 1984 | 1892 | 1170 | 1413 | 1092 | 912 | 1509,47 | 1872 | 1147 | 0,612 | -0,708 | -5,99 | 2,10E-09 | 3,57E-09 | 0,0223788 | 0,003179 | 0,0127446 | 0,0127446 | TRUE | 0,2891095 | No |
| orf19.3683 | 1814 | 1819 | 1977 | 1047 | 2855 | 2030 | 2714 | 2770 | 1729 | 1923 | 1953 | 1640 | 2288 | 2308 | 2304 | 2252 | 2049,81 | 1811 | 2288 | 1,261 | 0,334 | 5,701 | 1,19E-08 | 1,98E-08 | 0,0028889 | 0,0028923 | 0,0027926 | 0,0027926 | TRUE | 0,0451055 | No |
| orf19.3684 | 2405 | 2012 | 2137 | 1482 | 14831 | 11082 | 13966 | 14853 | 2293 | 2127 | 2111 | 2322 | 11884 | 12601 | 11857 | 12077 | 7159 | 2213 | 12105 | 5,475 | 2,453 | 55,429 | 0 | 0 | 0,0012711 | 0,0023206 | 0,0015892 | 0,0015892 | TRUE | 0,0118926 | No |
| orf19.3685 | 620 | 650 | 696 | 392 | 753 | 550 | 644 | 661 | 591 | 687 | 688 | 614 | 603 | 625 | 547 | 537 | 611,63 | 645 | 578 | 0,894 | -0,162 | -2,029 | 0,042421 | 0,0513153 | 0,0041494 | 0,0047758 | 0,0044495 | 0,0044495 | TRUE | 0,0413379 | No |
| orf19.3686 | 780 | 778 | 730 | 487 | 1413 | 1083 | 1193 | 1229 | 744 | 823 | 721 | 763 | 1132 | 1231 | 1013 | 999 | 928,27 | 763 | 1094 | 1,433 | 0,519 | 6,567 | 5,14E-11 | 9,25E-11 | 0,0056686 | 0,0038601 | 0,0048496 | 0,0048496 | TRUE | 0,0825033 | No |
| orf19.3687 | 238 | 225 | 227 | 144 | 371 | 285 | 405 | 314 | 227 | 238 | 224 | 226 | 297 | 324 | 344 | 255 | 266,89 | 229 | 305 | 1,331 | 0,413 | 3,811 | 0,0001385 | 0,0001947 | 0,0065236 | 0,0082432 | 0,0073613 | 0,0073613 | TRUE | 0,142186 | No |
| orf19.3688 | 296 | 278 | 303 | 135 | 483 | 304 | 448 | 466 | 282 | 294 | 299 | 211 | 387 | 346 | 380 | 379 | 322,36 | 272 | 373 | 1,362 | 0,446 | 4,343 | 1,40E-05 | 2,07E-05 | 0,0064126 | 0,0071846 | 0,0068788 | 0,0068788 | TRUE | 0,2041915 | No |
| orf19.3689 | 3880 | 3766 | 4007 | 2278 | 4051 | 2919 | 3606 | 3773 | 3699 | 3982 | 3959 | 3569 | 3246 | 3319 | 3061 | 3068 | 3487,85 | 3802 | 3174 | 0,834 | -0,262 | -5,207 | 1,91E-07 | 3,05E-07 | 0,0018514 | 0,002562 | 0,0021501 | 0,0021501 | TRUE | 0,0197254 | No |
| orf19.3690.2 | 6008 | 5918 | 5344 | 3462 | 21713 | 15792 | 19253 | 19037 | 5728 | 6258 | 5279 | 5423 | 17398 | 17957 | 16346 | 15479 | 11233,48 | 5672 | 16795 | 2,96 | 1,566 | 24,528 | 7,48E-133 | 6,70E-132 | 0,0049092 | 0,0022374 | 0,0037882 | 0,0037882 | TRUE | 0,0590157 | No |
| orf19.3691 | 941 | 966 | 1014 | 496 | 1208 | 774 | 1117 | 1085 | 897 | 1021 | 1002 | 777 | 968 | 880 | 948 | 882 | 921,99 | 924 | 920 | 0,992 | -0,011 | -0,133 | 0,8938656 | 0,9085309 | 0,0071725 | 0,0038721 | 0,00556 | 0,00556 | TRUE | 0,1254324 | No |
| orf19.3693 | 33 | 32 | 27 | 13 | 15 | 10 | 10 | 14 | 31 | 34 | 27 | 20 | 12 | 11 | 8 | 11 | 19,45 | 28 | 10 | 0,38 | -1,397 | -4,154 | 3,27E-05 | 4,74E-05 | 1,00E-08 | 0,086508 | 0,0479303 | 0,0479303 | TRUE | 0,1450043 | No |

| | | | | | | | | | | | | | | | | | | | | | | | | | | | | | | | |
|---|---|---|---|---|---|---|---|---|---|---|---|---|---|---|---|---|---|---|---|---|---|---|---|---|---|---|---|---|---|---|---|
| orf19.3694 | 411 | 408 | 451 | 227 | 186 | 127 | 165 | 128 | 392 | 431 | 446 | 356 | 149 | 144 | 140 | 104 | 270,25 | 406 | 134 | 0,329 | -1,603 | -13,074 | 4,67E-39 | 1,46E-38 | 0,011814 | 0,0081667 | 0,0097403 | 0,0097403 | TRUE | 0,2565675 | No |
| orf19.3695 | 2220 | 2456 | 2459 | 1161 | 2665 | 1825 | 2268 | 2106 | 2117 | 2597 | 2429 | 1819 | 2135 | 2075 | 1926 | 1712 | 2101,24 | 2240 | 1962 | 0,875 | -0,193 | -1,881 | 0,059991 | 0,0715252 | 0,0157912 | 0,0028727 | 0,0096476 | 0,0096476 | TRUE | 0,1891622 | No |
| orf19.3696 | 1753 | 1579 | 1540 | 890 | 2671 | 2003 | 2211 | 2377 | 1671 | 1670 | 1521 | 1394 | 2140 | 2278 | 1877 | 1933 | 1810,52 | 1564 | 2057 | 1,312 | 0,392 | 5,052 | 4,38E-07 | 6,87E-07 | 0,0069531 | 0,0029981 | 0,0052017 | 0,0052017 | TRUE | 0,0618158 | No |
| orf19.3697 | 243 | 314 | 333 | 191 | 781 | 665 | 687 | 599 | 232 | 332 | 329 | 299 | 626 | 756 | 583 | 487 | 455,52 | 298 | 613 | 2,056 | 1,04 | 7,405 | 1,31E-13 | 2,55E-13 | 0,0276707 | 0,0056958 | 0,0163068 | 0,0163068 | TRUE | 0,1870432 | No |
| orf19.3698 | 338 | 436 | 437 | 232 | 174 | 112 | 153 | 121 | 322 | 461 | 432 | 363 | 139 | 127 | 130 | 98 | 259,18 | 394 | 124 | 0,313 | -1,675 | -11,852 | 2,09E-32 | 5,89E-32 | 0,0203768 | 0,0084262 | 0,0141547 | 0,0141547 | TRUE | 0,131103 | No |
| orf19.3699 | 522 | 615 | 657 | 359 | 332 | 296 | 319 | 250 | 498 | 650 | 649 | 562 | 266 | 337 | 271 | 203 | 429,51 | 590 | 269 | 0,454 | -1,138 | -8,36 | 6,30E-17 | 1,32E-16 | 0,0247229 | 0,005914 | 0,0151745 | 0,0151745 | TRUE | 0,3106291 | No |
| orf19.3700 | 3232 | 3644 | 3679 | 2364 | 4520 | 3799 | 4243 | 4132 | 3081 | 3853 | 3635 | 3703 | 3622 | 4320 | 3602 | 3360 | 3646,98 | 3568 | 3726 | 1,044 | 0,062 | 0,73 | 0,4654085 | 0,5019049 | 0,0094253 | 0,0025415 | 0,0066961 | 0,0066961 | TRUE | 0,1389591 | No |
| orf19.3701 | 1029 | 1056 | 1074 | 624 | 1291 | 913 | 1084 | 1119 | 981 | 1117 | 1061 | 978 | 1034 | 1038 | 920 | 910 | 1004,87 | 1034 | 976 | 0,942 | -0,087 | -1,239 | 0,2152484 | 0,2432695 | 0,0033665 | 0,0037252 | 0,0037075 | 0,0037075 | TRUE | 0,0343147 | No |
| orf19.3704 | 7712 | 7132 | 7592 | 4801 | 7505 | 5384 | 7165 | 7701 | 7353 | 7541 | 7500 | 7521 | 6014 | 6122 | 6083 | 6262 | 6799,41 | 7479 | 6120 | 0,818 | -0,289 | -9,695 | 3,17E-22 | 7,44E-22 | 9,20E-05 | 0,0023328 | 0,000705 | 0,000705 | TRUE | 0,0030451 | No |
| orf19.3704.1 | 1256 | 928 | 949 | 588 | 1404 | 770 | 1287 | 1454 | 1197 | 981 | 938 | 921 | 1125 | 876 | 1093 | 1182 | 1039,1 | 1009 | 1069 | 1,059 | 0,083 | 0,795 | 0,4265764 | 0,4626864 | 0,0149195 | 0,0036714 | 0,0093887 | 0,0093887 | TRUE | 0,1907354 | No |
| orf19.3705 | 2870 | 2492 | 2680 | 1966 | 5942 | 4271 | 5742 | 6149 | 2736 | 2635 | 2648 | 3080 | 4761 | 4857 | 4875 | 5000 | 3823,87 | 2775 | 4873 | 1,76 | 0,815 | 15,476 | 5,04E-54 | 1,96E-53 | 0,0026456 | 0,0025207 | 0,0023699 | 0,0023699 | TRUE | 0,0629154 | No |
| orf19.3706 | 3454 | 3379 | 3557 | 1954 | 2720 | 1925 | 2475 | 2655 | 3293 | 3573 | 3514 | 3061 | 2179 | 2189 | 2101 | 2159 | 2758,66 | 3360 | 2157 | 0,641 | -0,641 | -11,954 | 6,18E-33 | 1,75E-32 | 0,0020627 | 0,0026865 | 0,0023901 | 0,0023901 | TRUE | 0,0413723 | No |
| orf19.3707 | 29758 | 26245 | 28293 | 17328 | 6234 | 4460 | 5857 | 6310 | 28371 | 27751 | 27951 | 27145 | 4995 | 5071 | 4973 | 5131 | 16423,45 | 27804 | 5042 | 0,181 | -2,463 | -83,96 | 0 | 0 | 0,0002113 | 0,0021913 | 0,0007179 | 0,0007179 | TRUE | 0,0030589 | No |
| orf19.3708 | 15 | 13 | 17 | 5 | 6 | 5 | 9 | 15 | 14 | 14 | 17 | 8 | 5 | 6 | 8 | 12 | 10,38 | 13 | 8 | 0,57 | -0,811 | -1,751 | 0,0799679 | 0,0942859 | 1,00E-08 | 0,1603387 | 0,1065387 | 0,1065387 | TRUE | 0,5766424 | No |
| orf19.371 | 1115 | 911 | 947 | 695 | 2239 | 1542 | 2241 | 2299 | 1063 | 963 | 936 | 1089 | 1794 | 1753 | 1903 | 1869 | 1421,24 | 1013 | 1830 | 1,812 | 0,858 | 14,228 | 6,17E-46 | 2,15E-45 | 0,0023895 | 0,0032465 | 0,0026887 | 0,0026887 | TRUE | 0,031614 | No |
| orf19.3710 | 3704 | 3854 | 3906 | 2143 | 2337 | 1692 | 2095 | 1979 | 3531 | 4075 | 3859 | 3357 | 1873 | 1924 | 1779 | 1609 | 2750,83 | 3706 | 1796 | 0,484 | -1,047 | -14,266 | 3,59E-46 | 1,26E-45 | 0,0062022 | 0,0026882 | 0,004768 | 0,004768 | TRUE | 0,0598265 | No |
| orf19.3711 | 59 | 50 | 61 | 29 | 197 | 138 | 192 | 204 | 56 | 53 | 60 | 45 | 158 | 157 | 163 | 166 | 107,31 | 54 | 161 | 2,962 | 1,566 | 10,509 | 7,83E-26 | 1,97E-25 | 1,00E-08 | 0,0173922 | 0,0083879 | 0,0083879 | TRUE | 0,058382 | No |
| orf19.3712 | 15372 | 16687 | 16874 | 9718 | 5409 | 3987 | 4821 | 4476 | 14656 | 17644 | 16670 | 15224 | 4334 | 4534 | 4093 | 3639 | 10099,21 | 16048 | 4150 | 0,258 | -1,952 | -25,815 | 5,96E-147 | 5,91E-146 | 0,0076976 | 0,0022539 | 0,0053487 | 0,0053487 | TRUE | 0,0838405 | No |
| orf19.3713 | 4025 | 5711 | 4514 | 3380 | 1737 | 1398 | 1223 | 1290 | 3837 | 6039 | 4459 | 5295 | 1392 | 1590 | 1038 | 1049 | 3087,39 | 4908 | 1267 | 0,258 | -1,954 | -9,367 | 7,45E-21 | 1,70E-20 | 0,0413329 | 0,0026231 | 0,0198214 | 0,0413329 | TRUE | 0,1164692 | No |
| orf19.3714 | 868 | 818 | 897 | 547 | 1237 | 838 | 1189 | 1149 | 828 | 865 | 886 | 857 | 991 | 953 | 1009 | 934 | 915,41 | 859 | 972 | 1,132 | 0,179 | 3,235 | 0,001217 | 0,0016304 | 1,00E-08 | 0,0038849 | 0,0018353 | 0,0018353 | TRUE | 0,008806 | No |
| orf19.3715 | 1176 | 921 | 997 | 582 | 1175 | 704 | 1051 | 1006 | 1121 | 974 | 985 | 912 | 941 | 801 | 892 | 818 | 930,5 | 998 | 863 | 0,864 | -0,211 | -2,704 | 0,0068612 | 0,0087896 | 0,0053688 | 0,0038558 | 0,0047757 | 0,0047757 | TRUE | 0,0835793 | No |
| orf19.3719 | 14 | 9 | 12 | 8 | 9 | 2 | 8 | 8 | 13 | 10 | 12 | 13 | 7 | 2 | 7 | 7 | 8,75 | 12 | 6 | 0,497 | -1,009 | -2,089 | 0,0367075 | 0,0446494 | 1,00E-08 | 0,1896477 | 0,1012047 | 0,1012047 | TRUE | 0,2773204 | No |
| orf19.372 | 717 | 524 | 617 | 394 | 589 | 381 | 515 | 651 | 684 | 554 | 610 | 617 | 472 | 433 | 437 | 529 | 542,02 | 616 | 468 | 0,76 | -0,395 | -4,421 | 9,81E-06 | 1,46E-05 | 0,0064333 | 0,0051205 | 0,0058035 | 0,0058035 | TRUE | 0,0923633 | No |
| orf19.3720 | 4107 | 3550 | 3899 | 2297 | 3279 | 2137 | 3099 | 3465 | 3916 | 3754 | 3852 | 3598 | 2627 | 2430 | 2631 | 2817 | 3203,15 | 3780 | 2626 | 0,695 | -0,525 | -10,078 | 6,95E-24 | 1,68E-23 | 0,0020514 | 0,0026039 | 0,0022877 | 0,0022877 | TRUE | 0,0296903 | No |
| orf19.3721 | 251 | 222 | 239 | 182 | 222 | 206 | 264 | 264 | 239 | 235 | 236 | 285 | 178 | 234 | 204 | 215 | 228,22 | 249 | 208 | 0,836 | -0,258 | -2,295 | 0,0217423 | 0,0269184 | 0,0058148 | 0,0092855 | 0,0077155 | 0,0077155 | TRUE | 0,1054769 | No |
| orf19.3722 | 3306 | 2793 | 2978 | 2003 | 4503 | 3091 | 4170 | 4702 | 3152 | 2953 | 2942 | 3138 | 3608 | 3515 | 3540 | 3823 | 3333,91 | 3046 | 3622 | 1,19 | 0,251 | 5,529 | 3,21E-08 | 5,26E-08 | 0,0010465 | 0,0025838 | 0,0016685 | 0,0016685 | TRUE | 0,0172294 | No |
| orf19.3724 | 3361 | 3008 | 3138 | 2012 | 5144 | 3305 | 4995 | 5156 | 3204 | 3181 | 3100 | 3152 | 4122 | 3758 | 4241 | 4192 | 3618,72 | 3159 | 4078 | 1,292 | 0,369 | 7,962 | 1,69E-15 | 3,44E-15 | 0,0011676 | 0,002545 | 0,0017785 | 0,0017785 | TRUE | 0,0329628 | No |
| orf19.3726 | 895 | 816 | 917 | 523 | 2808 | 1857 | 2665 | 2749 | 853 | 863 | 906 | 819 | 2250 | 2112 | 2263 | 2235 | 1537,58 | 860 | 2215 | 2,57 | 1,362 | 26,812 | 2,35E-158 | 2,54E-157 | 0,0004218 | 0,0031591 | 0,0016281 | 0,0016281 | TRUE | 0,0150549 | No |
| orf19.3727 | 115 | 112 | 108 | 71 | 120 | 66 | 91 | 101 | 110 | 118 | 107 | 111 | 96 | 75 | 77 | 82 | 97,07 | 112 | 82 | 0,745 | -0,425 | -2,9 | 0,0037319 | 0,0048649 | 1,00E-08 | 0,0190058 | 0,0102394 | 0,0102394 | TRUE | 0,1222697 | No |
| orf19.3728 | 2122 | 2011 | 2183 | 1454 | 2759 | 1947 | 2623 | 2761 | 2023 | 2126 | 2157 | 2278 | 2211 | 2214 | 2227 | 2245 | 2185,04 | 2146 | 2224 | 1,039 | 0,055 | 1,215 | 0,2244787 | 0,2531635 | 0,0006071 | 0,0028427 | 0,0014803 | 0,0014803 | TRUE | 0,0188278 | No |
| orf19.3730 | 413 | 432 | 411 | 277 | 1163 | 834 | 1076 | 959 | 394 | 457 | 406 | 434 | 932 | 948 | 914 | 780 | 658 | 423 | 894 | 2,116 | 1,081 | 12,971 | 1,78E-38 | 5,51E-38 | 0,0050624 | 0,0045866 | 0,0048415 | 0,0048415 | TRUE | 0,0890896 | No |
| orf19.3732 | 4912 | 4720 | 4657 | 3050 | 6557 | 5157 | 6016 | 6189 | 4683 | 4991 | 4601 | 4778 | 5254 | 5864 | 5108 | 5032 | 5038,78 | 4763 | 5314 | 1,115 | 0,157 | 2,868 | 0,0041372 | 0,0053766 | 0,0028841 | 0,0024171 | 0,0026911 | 0,0026911 | TRUE | 0,0582293 | No |
| orf19.3733 | 1623 | 1452 | 1525 | 996 | 1974 | 1529 | 1799 | 2016 | 1547 | 1535 | 1507 | 1560 | 1582 | 1739 | 1527 | 1639 | 1579,55 | 1537 | 1622 | 1,054 | 0,076 | 1,474 | 0,1406005 | 0,1616817 | 0,0010543 | 0,0031308 | 0,0019039 | 0,0019039 | TRUE | 0,026755 | No |
| orf19.3734 | 3928 | 3761 | 3996 | 2289 | 1322 | 855 | 1210 | 1264 | 3745 | 3977 | 3948 | 3586 | 1059 | 972 | 1027 | 1028 | 2417,72 | 3814 | 1022 | 0,268 | -1,9 | -37,255 | 8,87E-304 | 1,99E-302 | 0,0013456 | 0,0027704 | 0,0019132 | 0,0019132 | TRUE | 0,0186024 | No |
| orf19.3735 | 12 | 11 | 9 | 18 | 48 | 48 | 47 | 52 | 11 | 12 | 9 | 28 | 38 | 55 | 40 | 42 | 29,42 | 15 | 44 | 3,07 | 1,618 | 4,905 | 9,33E-07 | 1,45E-06 | 0,048215 | 0,0578942 | 0,0550936 | 0,0550936 | TRUE | 1,0261386 | No |
| orf19.3736 | 6 | 9 | 11 | 3 | 14 | 4 | 21 | 14 | 6 | 10 | 11 | 5 | 11 | 5 | 18 | 11 | 9,47 | 8 | 11 | 1,455 | 0,541 | 1,06 | 0,2892862 | 0,3216476 | 0,0904763 | 0,175421 | 0,1406501 | 0,1406501 | TRUE | 0,1752873 | No |
| orf19.3737 | 4730 | 4397 | 4693 | 3076 | 2514 | 1860 | 2486 | 2622 | 4510 | 4649 | 4636 | 4819 | 2014 | 2115 | 2111 | 2132 | 3373,21 | 4654 | 2093 | 0,45 | -1,152 | -29,234 | 7,27E-188 | 9,28E-187 | 0,0003831 | 0,002578 | 0,0011567 | 0,0011567 | TRUE | 0,0080124 | No |
| orf19.3738 | 34 | 37 | 29 | 18 | 74 | 38 | 70 | 42 | 32 | 39 | 29 | 28 | 59 | 43 | 59 | 34 | 40,56 | 32 | 49 | 1,524 | 0,608 | 2,371 | 0,0177432 | 0,0221131 | 0,0301247 | 0,0425736 | 0,0367613 | 0,0367613 | TRUE | 0,3289819 | No |
| orf19.374 | 6878 | 5462 | 6148 | 3701 | 6365 | 4080 | 5965 | 6258 | 6557 | 5775 | 6074 | 5798 | 5100 | 4639 | 5064 | 5088 | 5512,04 | 6051 | 4973 | 0,822 | -0,283 | -5,335 | 9,58E-08 | 1,54E-07 | 0,0025681 | 0,0023892 | 0,0025232 | 0,0025232 | TRUE | 0,0390239 | No |
| orf19.3740 | 370 | 281 | 289 | 262 | 546 | 395 | 473 | 489 | 353 | 297 | 286 | 410 | 437 | 449 | 402 | 398 | 378,96 | 336 | 422 | 1,26 | 0,333 | 3,034 | 0,0024138 | 0,0031837 | 0,0116231 | 0,006424 | 0,0088366 | 0,0088366 | TRUE | 0,2190657 | No |
| orf19.3742 | 808 | 655 | 702 | 485 | 649 | 436 | 574 | 651 | 770 | 693 | 694 | 760 | 520 | 496 | 487 | 529 | 618,58 | 729 | 508 | 0,699 | -0,518 | -7,611 | 2,72E-14 | 5,37E-14 | 0,0009819 | 0,0047456 | 0,0027944 | 0,0027944 | TRUE | 0,0198818 | No |
| orf19.3743 | 510 | 414 | 514 | 370 | 941 | 703 | 892 | 937 | 486 | 438 | 508 | 580 | 754 | 799 | 757 | 762 | 635,49 | 503 | 768 | 1,535 | 0,618 | 7,847 | 4,27E-15 | 8,61E-15 | 0,0037425 | 0,004675 | 0,0042582 | 0,0042582 | TRUE | 0,1062432 | No |
| orf19.3744 | 485 | 473 | 524 | 315 | 1042 | 716 | 921 | 1032 | 462 | 500 | 518 | 493 | 835 | 814 | 782 | 839 | 655,47 | 493 | 818 | 1,658 | 0,729 | 11,288 | 1,50E-29 | 4,03E-29 | 1,00E-08 | 0,0045962 | 0,0023337 | 0,0023337 | TRUE | 0,0235162 | No |
| orf19.3746 | 3145 | 3152 | 3323 | 2078 | 1955 | 1472 | 1756 | 1945 | 2998 | 3333 | 3283 | 3255 | 1566 | 1674 | 1491 | 1581 | 2397,74 | 3217 | 1578 | 0,49 | -1,029 | -19,632 | 8,26E-86 | 4,74E-85 | 0,0015813 | 0,0027761 | 0,0021771 | 0,0021771 | TRUE | 0,0262544 | No |
| orf19.3748 | 59 | 51 | 76 | 34 | 35 | 23 | 18 | 28 | 56 | 54 | 75 | 53 | 28 | 26 | 15 | 23 | 41,35 | 60 | 23 | 0,383 | -1,383 | -5,519 | 3,40E-08 | 5,56E-08 | 0,0163402 | 0,0418028 | 0,031623 | 0,031623 | TRUE | 0,337979 | No |
| orf19.3749 | 15278 | 14480 | 15053 | 9386 | 7727 | 5589 | 6925 | 7703 | 14566 | 15311 | 14871 | 14704 | 6191 | 6355 | 5879 | 6263 | 10517,56 | 14863 | 6172 | 0,415 | -1,268 | -35,019 | 1,15E-268 | 2,14E-267 | 0,0006439 | 0,0022474 | 0,0011493 | 0,0011493 | TRUE | 0,0124838 | No |
| orf19.3751 | 2854 | 2557 | 2802 | 1745 | 1094 | 767 | 1024 | 1121 | 2721 | 2704 | 2768 | 2734 | 877 | 872 | 869 | 911 | 1807 | 2732 | 882 | 0,323 | -1,629 | -37,842 | 0 | 0 | 1,00E-08 | 0,0029999 | 0,0010711 | 0,0010711 | TRUE | 0,0061322 | No |
| orf19.3752 | 6264 | 5149 | 5566 | 3660 | 4318 | 2897 | 4120 | 4568 | 5972 | 5444 | 5499 | 5734 | 3460 | 3294 | 3498 | 3714 | 4576,86 | 5662 | 3492 | 0,617 | -0,697 | -14,166 | 1,48E-45 | 5,10E-45 | 0,0018273 | 0,00245 | 0,0020949 | 0,0020949 | TRUE | 0,0224504 | No |
| orf19.3753 | 9456 | 8391 | 9476 | 6216 | 4374 | 3466 | 4251 | 4753 | 9015 | 8872 | 9361 | 9738 | 3505 | 3941 | 3609 | 3865 | 6488,3 | 9246 | 3730 | 0,403 | -1,31 | -26,115 | 2,45E-150 | 2,48E-149 | 0,0021565 | 0,0023443 | 0,0022359 | 0,0022359 | TRUE | 0,0201893 | No |
| orf19.3755 | 2450 | 2307 | 2254 | 1392 | 2305 | 1600 | 2199 | 2301 | 2336 | 2439 | 2227 | 2181 | 1847 | 1819 | 1867 | 1871 | 2073,34 | 2296 | 1851 | 0,806 | -0,312 | -6,653 | 2,88E-11 | 5,23E-11 | 0,0007033 | 0,0028832 | 0,001624 | 0,001624 | TRUE | 0,0211772 | No |
| orf19.3756 | 13268 | 12851 | 13770 | 8438 | 9509 | 7217 | 9014 | 9269 | 12650 | 13588 | 13603 | 13219 | 7619 | 8206 | 7653 | 7537 | 10509,38 | 13265 | 7754 | 0,584 | -0,775 | -18,663 | 9,86E-78 | 5,21E-77 | 0,0012091 | 0,0022475 | 0,0015567 | 0,0015567 | TRUE | 0,0185355 | No |
| orf19.3757 | 1652 | 1673 | 1621 | 1262 | 5137 | 3776 | 4359 | 4557 | 1575 | 1769 | 1601 | 1977 | 4116 | 4294 | 3701 | 3705 | 2842,28 | 1730 | 3954 | 2,289 | 1,195 | 15,341 | 4,05E-53 | 1,56E-52 | 0,008036 | 0,002669 | 0,0053836 | 0,0053836 | TRUE | 0,1064123 | No |
| orf19.3758 | 4932 | 4944 | 5002 | 3324 | 2050 | 1506 | 1924 | 1899 | 4702 | 5228 | 4942 | 5207 | 1643 | 1712 | 1633 | 1544 | 3326,39 | 5020 | 1633 | 0,325 | -1,62 | -31,753 | 2,87E-221 | 4,27E-220 | 0,0015797 | 0,0025849 | 0,0021145 | 0,0021145 | TRUE | 0,0223343 | No |
| orf19.3759 | 6728 | 6236 | 6900 | 4342 | 3451 | 2647 | 3352 | 3364 | 6414 | 6594 | 6817 | 6802 | 2765 | 3010 | 2846 | 2735 | 4747,86 | 6657 | 2839 | 0,426 | -1,23 | -28,7 | 3,82E-181 | 4,62E-180 | 0,0010161 | 0,0024371 | 0,0015225 | 0,0015225 | TRUE | 0,0191016 | No |
| orf19.376 | 1601 | 1269 | 1512 | 882 | 2754 | 1773 | 2642 | 2926 | 1526 | 1342 | 1494 | 1382 | 2207 | 2016 | 2243 | 2379 | 1823,57 | 1436 | 2211 | 1,539 | 0,622 | 9,688 | 3,39E-22 | 7,96E-22 | 0,0034137 | 0,0029917 | 0,0033697 | 0,0033697 | TRUE | 0,0414782 | No |
| orf19.3760 | 111 | 95 | 88 | 52 | 49 | 28 | 46 | 40 | 106 | 100 | 87 | 81 | 39 | 32 | 39 | 33 | 64,67 | 94 | 36 | 0,38 | -1,395 | -7,378 | 1,61E-13 | 3,12E-13 | 1,00E-08 | 0,0274806 | 0,0160211 | 0,0160211 | TRUE | 0,0744855 | No |
| orf19.3761 | 3014 | 2889 | 3052 | 1760 | 2304 | 1475 | 2086 | 2229 | 2874 | 3055 | 3015 | 2757 | 1846 | 1677 | 1771 | 1812 | 2350,9 | 2925 | 1776 | 0,607 | -0,719 | -14,369 | 8,11E-47 | 2,87E-46 | 0,0013295 | 0,0027897 | 0,0019626 | 0,0019626 | TRUE | 0,0169081 | No |
| orf19.3762 | 713 | 732 | 654 | 473 | 692 | 506 | 588 | 571 | 680 | 774 | 646 | 741 | 554 | 575 | 499 | 464 | 616,77 | 710 | 523 | 0,737 | -0,441 | -5,06 | 4,19E-07 | 6,58E-07 | 0,0064662 | 0,0047534 | 0,005643 | 0,005643 | TRUE | 0,0685621 | No |
| orf19.3763 | 961 | 985 | 1041 | 598 | 686 | 562 | 720 | 728 | 916 | 1042 | 1028 | 937 | 550 | 639 | 611 | 592 | 789,36 | 981 | 598 | 0,608 | -0,718 | -10,187 | 2,27E-24 | 5,55E-24 | 0,0028193 | 0,0041713 | 0,0034495 | 0,0034495 | TRUE | 0,0353908 | No |
| orf19.3764 | 656 | 780 | 790 | 422 | 699 | 479 | 577 | 569 | 625 | 825 | 780 | 661 | 560 | 545 | 490 | 463 | 618,62 | 723 | 514 | 0,71 | -0,494 | -4,909 | 9,14E-07 | 1,42E-06 | 0,0111938 | 0,0047454 | 0,0080745 | 0,0080745 | TRUE | 0,1062444 | No |
| orf19.3765 | 1131 | 1151 | 1179 | 682 | 2498 | 1900 | 2185 | 2371 | 1078 | 1217 | 1165 | 1068 | 2002 | 2160 | 1855 | 1928 | 1559,18 | 1132 | 1986 | 1,751 | 0,808 | 12,547 | 4,14E-36 | 1,23E-35 | 0,00322 | 0,0031443 | 0,0032611 | 0,0032611 | TRUE | 0,0408329 | No |
| orf19.3767 | 740 | 657 | 659 | 384 | 1468 | 1051 | 1300 | 1416 | 706 | 695 | 651 | 602 | 1176 | 1195 | 1104 | 1151 | 909,9 | 664 | 1156 | 1,736 | 0,795 | 12,712 | 5,10E-37 | 1,54E-36 | 0,0011184 | 0,0038958 | 0,0025339 | 0,0025339 | TRUE | 0,0387995 | No |
| orf19.3769 | 1209 | 1129 | 1135 | 715 | 1513 | 1219 | 1559 | 1565 | 1153 | 1194 | 1121 | 1120 | 1212 | 1386 | 1324 | 1272 | 1222,79 | 1147 | 1298 | 1,13 | 0,176 | 3,112 | 0,0018567 | 0,0024652 | 0,0013566 | 0,003434 | 0,0022408 | 0,0022408 | TRUE | 0,0241626 | No |
| orf19.377 | 143 | 125 | 91 | 80 | 152 | 88 | 137 | 159 | 136 | 132 | 90 | 125 | 122 | 100 | 116 | 129 | 118,9 | 121 | 117 | 0,972 | -0,041 | -0,26 | 0,7948868 | 0,8175797 | 0,0141198 | 0,0159007 | 0,0152004 | 0,0152004 | TRUE | 0,3251023 | No |
| orf19.3770 | 7402 | 7204 | 7481 | 4199 | 2272 | 1565 | 2062 | 2179 | 7057 | 7617 | 7391 | 6578 | 1820 | 1780 | 1751 | 1772 | 4470,65 | 7161 | 1781 | 0,249 | -2,009 | -39,078 | 0 | 0 | 0,0020622 | 0,0024585 | 0,0022074 | 0,0022074 | TRUE | 0,0355954 | No |
| orf19.3773 | 2334 | 2196 | 2490 | 1560 | 2200 | 1509 | 2008 | 2043 | 2225 | 2322 | 2460 | 2444 | 1763 | 1716 | 1705 | 1661 | 2036,94 | 2363 | 1711 | 0,725 | -0,464 | -9,646 | 5,11E-22 | 1,19E-21 | 0,0009932 | 0,0028973 | 0,0017238 | 0,0017238 | TRUE | 0,0191438 | No |
| orf19.3774 | 1724 | 1535 | 1621 | 981 | 1796 | 1265 | 1714 | 1915 | 1644 | 1623 | 1601 | 1537 | 1439 | 1438 | 1455 | 1557 | 1536,83 | 1601 | 1472 | 0,919 | -0,122 | -2,522 | 0,011679 | 0,0147835 | 0,0005223 | 0,0031596 | 0,0016043 | 0,0016043 | TRUE | 0,0185052 | No |
| orf19.3774.1 | 93 | 87 | 108 | 68 | 119 | 94 | 99 | 118 | 89 | 92 | 107 | 107 | 95 | 107 | 84 | 96 | 97,01 | 99 | 96 | 0,971 | -0,042 | -0,285 | 0,7755753 | 0,7996358 | 1,00E-08 | 0,0190157 | 0,0108319 | 0,0108319 | TRUE | 0,0643925 | No |
| orf19.3775 | 4353 | 3920 | 4215 | 2584 | 4655 | 3201 | 4227 | 4690 | 4150 | 4145 | 4164 | 4048 | 3730 | 3640 | 3589 | 3813 | 3909,86 | 4127 | 3693 | 0,895 | -0,161 | -4,515 | 6,33E-06 | 9,46E-06 | 0,0002101 | 0,0025112 | 0,0009561 | 0,0009561 | TRUE | 0,0059445 | No |
| orf19.3777 | 293 | 503 | 488 | 193 | 318 | 273 | 291 | 165 | 279 | 532 | 482 | 302 | 255 | 310 | 247 | 134 | 317,76 | 399 | 236 | 0,592 | -0,756 | -2,246 | 0,024696 | 0,0304336 | 0,1054797 | 0,0072584 | 0,0502284 | 0,1054797 | TRUE | 0,2175969 | No |
| orf19.3778 | 8931 | 7877 | 8274 | 5299 | 9225 | 6173 | 8567 | 9446 | 8515 | 8329 | 8174 | 8301 | 7392 | 7019 | 7273 | 7681 | 7835,45 | 8330 | 7341 | 0,882 | -0,182 | -4,884 | 1,04E-06 | 1,61E-06 | 0,0006703 | 0,0023008 | 0,0012023 | 0,0012023 | TRUE | 0,0118089 | No |
| orf19.3780 | 354 | 361 | 349 | 235 | 1010 | 790 | 869 | 918 | 338 | 382 | 345 | 368 | 809 | 898 | 738 | 746 | 577,99 | 358 | 798 | 2,229 | 1,156 | 13,657 | 1,82E-42 | 6,01E-42 | 0,0045967 | 0,004932 | 0,0047548 | 0,0047548 | TRUE | 0,0808791 | No |
| orf19.3781 | 1433 | 1333 | 1449 | 978 | 1523 | 1082 | 1368 | 1425 | 1366 | 1409 | 1431 | 1532 | 1220 | 1230 | 1161 | 1159 | 1313,75 | 1434 | 1192 | 0,833 | -0,264 | -5,017 | 5,24E-07 | 8,21E-07 | 0,0006821 | 0,0033411 | 0,0018946 | 0,0018946 | TRUE | 0,0224604 | No |

| | | | | | | | | | | | | | | | | | | | | | | | | | | | | | | | |
|---|---|---|---|---|---|---|---|---|---|---|---|---|---|---|---|---|---|---|---|---|---|---|---|---|---|---|---|---|---|---|---|
| orf19.3782 | 549 | 490 | 559 | 369 | 1041 | 733 | 1000 | 1061 | 523 | 518 | 552 | 578 | 834 | 833 | 849 | 863 | 693,89 | 543 | 845 | 1,562 | 0,643 | 10,538 | 5,78E-26 | 1,46E-25 | 1,00E-08 | 0,0044575 | 0,0020217 | 0,0020217 | TRUE | 0,0180425 | No |
| orf19.3782.2 | 1261 | 1009 | 1120 | 559 | 3447 | 1994 | 3225 | 3409 | 1202 | 1067 | 1106 | 876 | 2762 | 2267 | 2738 | 2772 | 1848,81 | 1063 | 2635 | 2,473 | 1,307 | 14,09 | 4,37E-45 | 1,50E-44 | 0,0117729 | 0,0029794 | 0,007559 | 0,007559 | TRUE | 0,1580666 | No |
| orf19.3783 | 149 | 138 | 175 | 89 | 397 | 245 | 367 | 393 | 142 | 146 | 173 | 139 | 318 | 279 | 312 | 320 | 228,51 | 150 | 307 | 2,042 | 1,03 | 9,617 | 6,80E-22 | 1,58E-21 | 0,0009094 | 0,0092765 | 0,0058692 | 0,0058692 | TRUE | 0,1150854 | No |
| orf19.3784 | 280 | 296 | 325 | 164 | 519 | 373 | 438 | 454 | 267 | 313 | 321 | 257 | 416 | 424 | 372 | 369 | 342,36 | 290 | 395 | 1,357 | 0,441 | 4,53 | 5,90E-06 | 8,84E-06 | 0,0048664 | 0,0068871 | 0,0060395 | 0,0060395 | TRUE | 0,0603963 | No |
| orf19.3785 | 1005 | 921 | 918 | 636 | 1403 | 940 | 1291 | 1394 | 958 | 974 | 907 | 996 | 1124 | 1069 | 1096 | 1133 | 1032,22 | 959 | 1106 | 1,157 | 0,21 | 3,945 | 7,98E-05 | 0,0001134 | 1,00E-08 | 0,0036819 | 0,0017273 | 0,0017273 | TRUE | 0,0160898 | No |
| orf19.3787 | 1233 | 977 | 1036 | 727 | 2893 | 1973 | 2828 | 3029 | 1176 | 1033 | 1023 | 1139 | 2318 | 2243 | 2401 | 2463 | 1724,54 | 1093 | 2356 | 2,16 | 1,111 | 18,788 | 9,52E-79 | 5,12E-78 | 0,0023546 | 0,0030434 | 0,0026522 | 0,0026522 | TRUE | 0,0345816 | No |
| orf19.3788 | 893 | 733 | 808 | 480 | 1417 | 928 | 1313 | 1406 | 851 | 775 | 798 | 752 | 1135 | 1055 | 1115 | 1143 | 953,15 | 794 | 1112 | 1,398 | 0,484 | 8,252 | 1,56E-16 | 3,25E-16 | 0,0006386 | 0,0038139 | 0,0021932 | 0,0021932 | TRUE | 0,0284808 | No |
| orf19.3788.1 | 19120 | 15439 | 15834 | 11064 | 23449 | 16525 | 21314 | 23944 | 18229 | 16325 | 15642 | 17332 | 18789 | 18790 | 18095 | 19469 | 17834,02 | 16882 | 18786 | 1,113 | 0,154 | 3,005 | 0,0026599 | 0,0035002 | 0,0026298 | 0,0021833 | 0,002475 | 0,002475 | TRUE | 0,0356189 | No |
| orf19.3789 | 7622 | 6689 | 6565 | 4638 | 27057 | 19878 | 23669 | 27681 | 7267 | 7073 | 6486 | 7266 | 21680 | 22603 | 20095 | 22507 | 14372 | 7023 | 21721 | 3,094 | 1,629 | 30,843 | 7,04E-209 | 9,93E-208 | 0,0026865 | 0,0022055 | 0,0025798 | 0,0025798 | TRUE | 0,0326133 | No |
| orf19.3791 | 565 | 463 | 522 | 327 | 1008 | 708 | 1028 | 1089 | 539 | 490 | 516 | 512 | 808 | 805 | 873 | 885 | 678,39 | 514 | 843 | 1,64 | 0,714 | 10,784 | 4,08E-27 | 1,05E-26 | 1,00E-08 | 0,0045116 | 0,002589 | 0,002589 | TRUE | 0,0139988 | No |
| orf19.3792 | 4616 | 4119 | 4475 | 2872 | 4671 | 3231 | 4243 | 4810 | 4401 | 4355 | 4421 | 4499 | 3743 | 3674 | 3602 | 3911 | 4075,77 | 4419 | 3732 | 0,845 | -0,243 | -6,356 | 2,07E-10 | 3,66E-10 | 0,0004925 | 0,0024941 | 0,0011521 | 0,0011521 | TRUE | 0,0128139 | No |
| orf19.3793 | 1609 | 1591 | 1677 | 1057 | 1034 | 733 | 911 | 1002 | 1534 | 1682 | 1657 | 1656 | 829 | 833 | 773 | 815 | 1222,38 | 1632 | 812 | 0,498 | -1,006 | -18,677 | 7,59E-78 | 4,03E-77 | 0,0007659 | 0,0034345 | 0,0018931 | 0,0018931 | TRUE | 0,0207998 | No |
| orf19.3794 | 446 | 447 | 484 | 317 | 748 | 513 | 698 | 734 | 425 | 473 | 478 | 497 | 599 | 583 | 593 | 597 | 530,59 | 468 | 593 | 1,272 | 0,347 | 5,013 | 5,37E-07 | 8,40E-07 | 1,00E-08 | 0,0051858 | 0,0026622 | 0,0026622 | TRUE | 0,0465799 | No |
| orf19.3795 | 3281 | 2867 | 3182 | 2048 | 6761 | 4704 | 6487 | 6949 | 3128 | 3031 | 3144 | 3208 | 5417 | 5349 | 5507 | 5650 | 4304,41 | 3128 | 5481 | 1,754 | 0,811 | 22,232 | 1,69E-109 | 1,22E-108 | 0,0003077 | 0,0024727 | 0,0010154 | 0,0010154 | TRUE | 0,005292 | No |
| orf19.3796 | 1726 | 1643 | 1716 | 1167 | 3662 | 2744 | 3515 | 3642 | 1646 | 1737 | 1695 | 1828 | 2934 | 3120 | 2984 | 2961 | 2363,27 | 1726 | 3000 | 1,74 | 0,799 | 17,492 | 1,66E-68 | 7,78E-68 | 0,0007768 | 0,002786 | 0,001531 | 0,001531 | TRUE | 0,0170263 | No |
| orf19.3797 | 1969 | 1877 | 1854 | 1155 | 2267 | 1518 | 1993 | 2134 | 1877 | 1985 | 1832 | 1809 | 1816 | 1726 | 1692 | 1735 | 1809,08 | 1876 | 1742 | 0,928 | -0,107 | -2,237 | 0,0252632 | 0,0311145 | 0,0007463 | 0,0029989 | 0,0016478 | 0,0016478 | TRUE | 0,0184098 | No |
| orf19.3798 | 5657 | 4686 | 5077 | 2970 | 5218 | 3370 | 5058 | 5110 | 5393 | 4955 | 5016 | 4653 | 4181 | 3832 | 4294 | 4155 | 4559,83 | 5004 | 4116 | 0,822 | -0,282 | -5,18 | 2,21E-07 | 3,52E-07 | 0,002715 | 0,0024514 | 0,0026346 | 0,0026346 | TRUE | 0,0334695 | No |
| orf19.3799 | 3643 | 3440 | 3721 | 2526 | 7833 | 6051 | 7437 | 7794 | 3473 | 3637 | 3676 | 3957 | 6276 | 6881 | 6314 | 6337 | 5068,98 | 3686 | 6452 | 1,752 | 0,809 | 16,024 | 8,66E-58 | 3,53E-57 | 0,0021909 | 0,0024152 | 0,0022247 | 0,0022247 | TRUE | 0,0285469 | No |
| orf19.3801 | 92 | 66 | 98 | 42 | 62 | 54 | 66 | 68 | 88 | 70 | 97 | 66 | 50 | 61 | 56 | 55 | 67,81 | 80 | 56 | 0,685 | -0,547 | -2,959 | 0,0030873 | 0,004041 | 0,0064397 | 0,026303 | 0,0178111 | 0,0178111 | TRUE | 0,1813449 | No |
| orf19.3802 | 601 | 519 | 646 | 382 | 1420 | 1060 | 1253 | 1415 | 573 | 549 | 638 | 598 | 1138 | 1205 | 1064 | 1151 | 864,48 | 590 | 1140 | 1,931 | 0,95 | 14,16 | 1,63E-45 | 5,61E-45 | 0,002064 | 0,0039906 | 0,0029765 | 0,0029765 | TRUE | 0,039596 | No |
| orf19.3803 | 1660 | 1484 | 1654 | 1036 | 1738 | 1315 | 1639 | 1715 | 1583 | 1569 | 1634 | 1623 | 1393 | 1495 | 1392 | 1394 | 1510,32 | 1602 | 1418 | 0,884 | -0,177 | -3,817 | 0,0001352 | 0,0001901 | 1,00E-08 | 0,0031784 | 0,0014024 | 0,0014024 | TRUE | 0,0147668 | No |
| orf19.3804 | 1827 | 1733 | 1759 | 988 | 1588 | 1202 | 1504 | 1632 | 1742 | 1832 | 1738 | 1548 | 1272 | 1367 | 1277 | 1327 | 1512,85 | 1715 | 1311 | 0,762 | -0,392 | -6,713 | 1,91E-11 | 3,49E-11 | 0,0021232 | 0,0031766 | 0,0026056 | 0,0026056 | TRUE | 0,0475542 | No |
| orf19.3806 | 2227 | 1906 | 2095 | 1285 | 1705 | 1148 | 1684 | 1723 | 2123 | 2015 | 2070 | 2013 | 1366 | 1305 | 1430 | 1401 | 1715,43 | 2055 | 1376 | 0,67 | -0,579 | -12,404 | 2,47E-35 | 7,27E-35 | 0,0003761 | 0,0030484 | 0,0014935 | 0,0014935 | TRUE | 0,013009 | No |
| orf19.3809 | 3798 | 3160 | 3604 | 2272 | 3451 | 2345 | 3378 | 3622 | 3621 | 3341 | 3560 | 3559 | 2765 | 2666 | 2868 | 2945 | 3165,81 | 3520 | 2811 | 0,799 | -0,324 | -6,993 | 2,69E-12 | 5,01E-12 | 0,0011146 | 0,0026099 | 0,0017369 | 0,0017369 | TRUE | 0,0145092 | No |
| orf19.3810 | 1757 | 1552 | 1686 | 1048 | 2110 | 1557 | 1976 | 2207 | 1675 | 1641 | 1666 | 1642 | 1691 | 1770 | 1678 | 1795 | 1694,59 | 1656 | 1734 | 1,046 | 0,065 | 1,458 | 0,1449054 | 0,1665392 | 1,00E-08 | 0,0030602 | 0,0012928 | 0,0012928 | TRUE | 0,0073445 | No |
| orf19.3811 | 978 | 858 | 931 | 543 | 1747 | 1206 | 1706 | 1751 | 932 | 907 | 920 | 851 | 1400 | 1371 | 1448 | 1424 | 1156,66 | 902 | 1411 | 1,559 | 0,641 | 12,566 | 3,26E-36 | 9,71E-36 | 1,00E-08 | 0,0035108 | 0,0015657 | 0,0015657 | TRUE | 0,0149399 | No |
| orf19.3812 | 13382 | 11210 | 12271 | 8501 | 26999 | 18907 | 25902 | 27647 | 12758 | 11853 | 12123 | 13317 | 21634 | 21499 | 21991 | 22480 | 17206,76 | 12513 | 21901 | 1,751 | 0,808 | 18,889 | 1,40E-79 | 7,61E-79 | 0,0014948 | 0,0021867 | 0,0016928 | 0,0016928 | TRUE | 0,0224475 | No |
| orf19.3813 | 1342 | 1058 | 1230 | 782 | 1892 | 1223 | 1769 | 1883 | 1279 | 1119 | 1215 | 1225 | 1516 | 1391 | 1502 | 1531 | 1347,24 | 1210 | 1485 | 1,229 | 0,297 | 5,193 | 2,07E-07 | 3,30E-07 | 0,0016694 | 0,00331 | 0,0023845 | 0,0023845 | TRUE | 0,0316389 | No |
| orf19.3815 | 7932 | 6947 | 7849 | 5236 | 3150 | 2242 | 2960 | 3280 | 7562 | 7346 | 7754 | 8202 | 2524 | 2549 | 2513 | 2667 | 5139,72 | 7716 | 2563 | 0,332 | -1,589 | -35,336 | 1,66E-273 | 3,18E-272 | 0,001151 | 0,0024107 | 0,0016942 | 0,0016942 | TRUE | 0,0213187 | No |
| orf19.3817 | 717 | 568 | 652 | 432 | 1029 | 689 | 959 | 947 | 684 | 601 | 644 | 677 | 825 | 783 | 814 | 770 | 724,65 | 652 | 798 | 1,227 | 0,296 | 4,641 | 3,47E-06 | 5,25E-06 | 0,0006608 | 0,0043571 | 0,0024762 | 0,0024762 | TRUE | 0,032694 | No |
| orf19.3818 | 295 | 245 | 306 | 191 | 949 | 664 | 897 | 1000 | 281 | 259 | 302 | 299 | 760 | 755 | 762 | 813 | 528,99 | 285 | 772 | 2,714 | 1,44 | 19,179 | 5,55E-82 | 3,09E-81 | 1,00E-08 | 0,0051951 | 0,0028868 | 0,0028868 | TRUE | 0,0436539 | No |
| orf19.3819 | 98 | 77 | 93 | 61 | 259 | 196 | 255 | 294 | 93 | 81 | 92 | 96 | 208 | 223 | 216 | 239 | 156,03 | 90 | 222 | 2,451 | 1,294 | 10,424 | 1,93E-25 | 4,81E-25 | 1,00E-08 | 0,0126144 | 0,0066365 | 0,0066365 | TRUE | 0,0441437 | No |
| orf19.382 | 10247 | 7690 | 8246 | 6829 | 23831 | 16863 | 21950 | 26438 | 9769 | 8131 | 8146 | 10698 | 19095 | 19175 | 18635 | 21497 | 14393,35 | 9186 | 19600 | 2,134 | 1,094 | 12,722 | 4,45E-37 | 1,35E-36 | 0,0115666 | 0,0022054 | 0,0070181 | 0,0070181 | TRUE | 0,1488956 | No |
| orf19.3820 | 43 | 34 | 38 | 23 | 59 | 31 | 46 | 59 | 41 | 36 | 38 | 36 | 47 | 35 | 39 | 48 | 40,01 | 38 | 42 | 1,131 | 0,177 | 0,796 | 0,42605 | 0,4622725 | 1,00E-08 | 0,04313 | 0,0224596 | 0,0224596 | TRUE | 0,0829938 | No |
| orf19.3821 | 550 | 476 | 496 | 300 | 1359 | 926 | 1403 | 1373 | 524 | 503 | 490 | 470 | 1089 | 1053 | 1191 | 1116 | 804,63 | 497 | 1112 | 2,234 | 1,16 | 17,708 | 3,65E-70 | 1,75E-69 | 0,0012989 | 0,0041319 | 0,0025959 | 0,0025959 | TRUE | 0,0271621 | No |
| orf19.3822 | 17957 | 14290 | 15634 | 10440 | 12679 | 8935 | 12107 | 13289 | 17120 | 15110 | 15445 | 16355 | 10159 | 10160 | 10279 | 10805 | 13179,11 | 16008 | 10351 | 0,647 | -0,629 | -13,323 | 1,71E-40 | 5,44E-40 | 0,0019505 | 0,0022159 | 0,0020617 | 0,0020617 | TRUE | 0,027084 | No |
| orf19.3823 | 3874 | 3202 | 3586 | 2208 | 4831 | 3089 | 4551 | 4809 | 3693 | 3386 | 3543 | 3459 | 3871 | 3512 | 3864 | 3910 | 3654,76 | 3520 | 3789 | 1,077 | 0,107 | 2,2 | 0,0278132 | 0,0341431 | 0,0015397 | 0,0025405 | 0,0019786 | 0,0019786 | TRUE | 0,0285875 | No |
| orf19.3825 | 1059 | 948 | 1043 | 692 | 2480 | 1841 | 2380 | 2627 | 1010 | 1002 | 1030 | 1084 | 1987 | 2093 | 2021 | 2136 | 1545,45 | 1032 | 2059 | 2 | 1 | 20,233 | 5,00E-91 | 3,02E-90 | 0,0005099 | 0,0031537 | 0,0015857 | 0,0015857 | TRUE | 0,0119285 | No |
| orf19.3826 | 5473 | 4596 | 4910 | 3452 | 4154 | 2918 | 4059 | 4186 | 5218 | 4860 | 4851 | 5408 | 3328 | 3318 | 3446 | 3404 | 4229,02 | 5084 | 3374 | 0,664 | -0,59 | -12,886 | 5,43E-38 | 1,66E-37 | 0,0013035 | 0,0024795 | 0,0017707 | 0,0017707 | TRUE | 0,0214232 | No |
| orf19.3827 | 2671 | 2191 | 2503 | 1599 | 3113 | 2214 | 3117 | 3232 | 2547 | 2317 | 2473 | 2505 | 2494 | 2518 | 2646 | 2628 | 2515,87 | 2460 | 2572 | 1,046 | 0,065 | 1,416 | 0,1568141 | 0,1796788 | 0,0008984 | 0,0027439 | 0,0015922 | 0,0015922 | TRUE | 0,0190243 | No |
| orf19.3828 | 10 | 12 | 12 | 7 | 26 | 18 | 28 | 26 | 10 | 13 | 12 | 11 | 21 | 20 | 24 | 21 | 16,41 | 12 | 22 | 1,918 | 0,939 | 2,694 | 0,0070645 | 0,0090411 | 1,00E-08 | 0,1021645 | 0,0469249 | 0,0469249 | TRUE | 0,0494687 | No |
| orf19.3829 | 3002 | 2551 | 2704 | 1736 | 3261 | 2376 | 2961 | 3401 | 2862 | 2697 | 2671 | 2720 | 2613 | 2702 | 2514 | 2765 | 2693,02 | 2738 | 2648 | 0,967 | -0,048 | -1,056 | 0,2908029 | 0,3232776 | 0,0010408 | 0,002701 | 0,0016463 | 0,0016463 | TRUE | 0,0143338 | No |
| orf19.3831 | 7764 | 6118 | 6643 | 4026 | 7495 | 4350 | 7045 | 7031 | 7402 | 6469 | 6563 | 6307 | 6006 | 4946 | 5981 | 5717 | 6173,83 | 6685 | 5662 | 0,847 | -0,239 | -3,38 | 0,0007241 | 0,0009871 | 0,0062841 | 0,0023572 | 0,0046569 | 0,0046569 | TRUE | 0,0875858 | No |
| orf19.3833 | 4956 | 4323 | 4882 | 3053 | 3617 | 2473 | 3402 | 3603 | 4725 | 4571 | 4823 | 4783 | 2898 | 2812 | 2888 | 2930 | 3803,72 | 4726 | 2882 | 0,61 | -0,712 | -19,973 | 9,52E-89 | 5,63E-88 | 0,0001759 | 0,0025229 | 0,0009474 | 0,0009474 | TRUE | 0,0059606 | No |
| orf19.3836 | 2599 | 2209 | 2409 | 1592 | 3437 | 2323 | 3366 | 3518 | 2478 | 2336 | 2380 | 2494 | 2754 | 2641 | 2858 | 2860 | 2600,13 | 2422 | 2778 | 1,149 | 0,2 | 4,489 | 7,14E-06 | 1,06E-05 | 0,0006815 | 0,0027227 | 0,0015149 | 0,0015149 | TRUE | 0,0127224 | No |
| orf19.3838 | 14581 | 12035 | 12630 | 8657 | 25074 | 16493 | 22244 | 26434 | 13901 | 12725 | 12477 | 13562 | 20091 | 18754 | 18885 | 21493 | 16486,16 | 13166 | 19806 | 1,505 | 0,589 | 10,645 | 1,84E-26 | 4,70E-26 | 0,0033036 | 0,0021909 | 0,0028789 | 0,0028789 | TRUE | 0,0403043 | No |
| orf19.3839 | 2254 | 2082 | 2253 | 1386 | 3733 | 2438 | 3512 | 3915 | 2149 | 2201 | 2226 | 2171 | 2991 | 2772 | 2982 | 3183 | 2584,46 | 2187 | 2982 | 1,365 | 0,448 | 9,117 | 7,69E-20 | 1,72E-19 | 0,0014358 | 0,0027266 | 0,0019164 | 0,0019164 | TRUE | 0,026272 | No |
| orf19.384 | 1991 | 1665 | 1842 | 1240 | 2362 | 1751 | 2243 | 2455 | 1898 | 1761 | 1820 | 1943 | 1893 | 1991 | 1904 | 1996 | 1900,63 | 1856 | 1946 | 1,05 | 0,07 | 1,477 | 0,1396954 | 0,1606698 | 0,0006539 | 0,0029552 | 0,0016324 | 0,0016324 | TRUE | 0,01401 | No |
| orf19.3840 | 1750 | 1486 | 1655 | 1033 | 2889 | 1891 | 2752 | 2817 | 1668 | 1571 | 1635 | 1618 | 2315 | 2150 | 2336 | 2290 | 1948,12 | 1623 | 2273 | 1,401 | 0,487 | 10,636 | 2,04E-26 | 5,19E-26 | 0,0005205 | 0,0029341 | 0,001468 | 0,001468 | TRUE | 0,0151045 | No |
| orf19.3841 | 7339 | 6369 | 7080 | 4522 | 1874 | 1437 | 1777 | 1996 | 6997 | 6734 | 6994 | 7084 | 1502 | 1634 | 1509 | 1623 | 4259,61 | 6952 | 1567 | 0,225 | -2,151 | -50,185 | 0 | 0 | 0,0007538 | 0,0024767 | 0,0013977 | 0,0013977 | TRUE | 0,0095477 | No |
| orf19.3843 | 1542 | 1382 | 1494 | 843 | 3540 | 2343 | 3356 | 3560 | 1470 | 1461 | 1476 | 1321 | 2837 | 2664 | 2849 | 2895 | 2121,56 | 1432 | 2811 | 1,959 | 0,97 | 19,286 | 6,99E-83 | 3,92E-82 | 0,0012971 | 0,0028652 | 0,0018799 | 0,0018799 | TRUE | 0,029205 | No |
| orf19.3844 | 1038 | 968 | 977 | 651 | 756 | 520 | 717 | 716 | 990 | 1024 | 965 | 1020 | 606 | 591 | 609 | 582 | 798,27 | 1000 | 597 | 0,598 | -0,742 | -13,195 | 9,40E-40 | 2,96E-39 | 1,00E-08 | 0,0041481 | 0,0017269 | 0,0017269 | TRUE | 0,0065148 | No |
| orf19.3845 | 2399 | 1990 | 2398 | 1534 | 1279 | 889 | 1242 | 1281 | 2287 | 2104 | 2369 | 2403 | 1025 | 1011 | 1054 | 1042 | 1661,9 | 2291 | 1033 | 0,451 | -1,147 | -21,207 | 8,16E-100 | 5,35E-99 | 0,0015006 | 0,0030792 | 0,002132 | 0,002132 | TRUE | 0,036499 | No |
| orf19.3846 | 1735 | 1571 | 1630 | 1012 | 5536 | 3861 | 5248 | 5621 | 1654 | 1661 | 1610 | 1585 | 4436 | 4390 | 4456 | 4570 | 3045,37 | 1628 | 4463 | 2,739 | 1,454 | 38,074 | 0 | 0 | 1,00E-08 | 0,0026304 | 0,0009568 | 0,0009568 | TRUE | 0,0032216 | No |
| orf19.3848 | 573 | 517 | 528 | 309 | 814 | 478 | 695 | 740 | 546 | 547 | 522 | 484 | 652 | 544 | 590 | 602 | 560,77 | 525 | 597 | 1,137 | 0,185 | 2,436 | 0,0148657 | 0,0186398 | 0,0024525 | 0,0050192 | 0,0037186 | 0,0037186 | TRUE | 0,0460163 | No |
| orf19.385 | 7046 | 6168 | 6720 | 4617 | 8731 | 6349 | 8096 | 8992 | 6718 | 6522 | 6639 | 7233 | 6996 | 7219 | 6873 | 7311 | 6938,89 | 6778 | 7100 | 1,048 | 0,068 | 1,589 | 0,112099 | 0,1300552 | 0,00128 | 0,0023279 | 0,0016107 | 0,0016107 | TRUE | 0,0240503 | No |
| orf19.3851 | 735 | 627 | 702 | 460 | 1741 | 1234 | 1766 | 1875 | 701 | 663 | 694 | 721 | 1395 | 1403 | 1499 | 1525 | 1074,99 | 695 | 1456 | 2,1 | 1,071 | 18,58 | 4,63E-77 | 2,43E-76 | 0,0009371 | 0,0036186 | 0,0020759 | 0,0020759 | TRUE | 0,0124315 | No |
| orf19.3852 | 181 | 186 | 200 | 122 | 90 | 67 | 74 | 94 | 173 | 197 | 198 | 191 | 72 | 76 | 63 | 76 | 130,69 | 190 | 72 | 0,379 | -1,4 | -10,495 | 9,06E-26 | 2,27E-25 | 1,00E-08 | 0,014655 | 0,0079428 | 0,0079428 | TRUE | 0,06924 | No |
| orf19.3854 | 8583 | 7519 | 8436 | 5254 | 5580 | 3888 | 5481 | 5773 | 8183 | 7950 | 8334 | 8231 | 4471 | 4421 | 4653 | 4694 | 6367,18 | 8174 | 4560 | 0,558 | -0,842 | -23,761 | 8,37E-125 | 6,99E-124 | 0,000401 | 0,0023491 | 0,0010375 | 0,0010375 | TRUE | 0,004896 | No |
| orf19.3856 | 654 | 646 | 647 | 338 | 1416 | 881 | 1282 | 1356 | 624 | 683 | 639 | 529 | 1135 | 1002 | 1088 | 1103 | 850,32 | 619 | 1082 | 1,742 | 0,801 | 10,534 | 6,02E-26 | 1,52E-25 | 0,0044619 | 0,0040222 | 0,0042326 | 0,0042326 | TRUE | 0,0960687 | No |
| orf19.3858 | 2774 | 2401 | 2560 | 1454 | 2378 | 1405 | 2300 | 2193 | 2645 | 2539 | 2529 | 2278 | 1905 | 1598 | 1953 | 1783 | 2153,64 | 2498 | 1810 | 0,725 | -0,465 | -6,654 | 2,85E-11 | 5,18E-11 | 0,0051376 | 0,0028537 | 0,0042144 | 0,0042144 | TRUE | 0,072887 | No |
| orf19.3859 | 1562 | 1419 | 1511 | 1010 | 2858 | 2140 | 2673 | 2917 | 1489 | 1500 | 1493 | 1582 | 2290 | 2433 | 2269 | 2372 | 1928,64 | 1516 | 2341 | 1,546 | 0,628 | 13,94 | 3,62E-44 | 1,22E-43 | 0,0003754 | 0,0029426 | 0,0013899 | 0,0013899 | TRUE | 0,0090918 | No |
| orf19.386 | 1943 | 1654 | 1725 | 1236 | 5187 | 3807 | 4814 | 5054 | 1852 | 1749 | 1704 | 1936 | 4156 | 4329 | 4087 | 4109 | 2990,41 | 1810 | 4170 | 2,307 | 1,206 | 25,019 | 3,80E-138 | 3,52E-137 | 0,0012104 | 0,0026403 | 0,0018135 | 0,0018135 | TRUE | 0,0238869 | No |
| orf19.3861 | 7648 | 6665 | 7311 | 4313 | 3015 | 2087 | 2925 | 3057 | 7292 | 7047 | 7223 | 6757 | 2416 | 2373 | 2483 | 2486 | 4759,49 | 7080 | 2440 | 0,345 | -1,537 | -39,121 | 0 | 0 | 0,000606 | 0,0024363 | 0,0012204 | 0,0012204 | TRUE | 0,0110212 | No |
| orf19.3862 | 3247 | 2623 | 2902 | 1923 | 3217 | 2150 | 3165 | 3325 | 3096 | 2773 | 2867 | 3012 | 2578 | 2445 | 2687 | 2704 | 2770,2 | 2937 | 2604 | 0,887 | -0,173 | -3,34 | 0,0008382 | 0,0011372 | 0,0017789 | 0,002684 | 0,0022044 | 0,0022044 | TRUE | 0,0196207 | No |
| orf19.3863 | 97 | 107 | 119 | 68 | 104 | 82 | 88 | 105 | 92 | 113 | 118 | 107 | 83 | 93 | 75 | 85 | 95,8 | 108 | 84 | 0,781 | -0,357 | -2,376 | 0,017493 | 0,0218141 | 1,00E-08 | 0,0192309 | 0,0111477 | 0,0111477 | TRUE | 0,0990581 | No |
| orf19.3865 | 2295 | 2101 | 2282 | 1473 | 2144 | 1549 | 2023 | 2025 | 2188 | 2222 | 2254 | 2308 | 1718 | 1761 | 1718 | 1647 | 1976,85 | 2243 | 1711 | 0,763 | -0,39 | -9,205 | 3,41E-20 | 7,69E-20 | 1,00E-08 | 0,0029218 | 0,0012146 | 0,0012146 | TRUE | 0,0078073 | No |
| orf19.3867 | 8785 | 8491 | 9122 | 5490 | 10548 | 7524 | 9841 | 9844 | 8376 | 8978 | 9012 | 8600 | 8452 | 8555 | 8355 | 8004 | 8541,51 | 8742 | 8342 | 0,954 | -0,068 | -1,735 | 0,0827744 | 0,0974508 | 0,0008726 | 0,0022835 | 0,0013552 | 0,0013552 | TRUE | 0,009991 | No |
| orf19.3868 | 1 | 1 | 1 | 1 | 0 | 0 | 2 | 0 | 1 | 1 | 1 | 2 | 0 | 0 | 2 | 0 | 0,78 | 1 | 0 | 0,399 | -1,324 | -0,791 | 0,4291202 | 0,4651293 | 1,00E-08 | 2,0992936 | 1,2560029 | 1,2560029 | TRUE | 0,8858642 | No |
| orf19.3869 | 16240 | 15531 | 16956 | 11323 | 13001 | 9546 | 12173 | 12854 | 15483 | 16422 | 16751 | 17738 | 10417 | 10855 | 10335 | 10452 | 13556,56 | 16598 | 10515 | 0,634 | -0,658 | -14,529 | 7,89E-48 | 2,83E-47 | 0,0017109 | 0,0022124 | 0,001896 | 0,001896 | TRUE | 0,0258154 | No |
| orf19.387 | 2184 | 1978 | 2263 | 1419 | 4375 | 3321 | 4018 | 4347 | 2082 | 2091 | 2236 | 2223 | 3506 | 3776 | 3411 | 3535 | 2857,49 | 2158 | 3557 | 1,648 | 0,721 | 15,053 | 3,32E-51 | 1,24E-50 | 0,0011888 | 0,0026659 | 0,001817 | 0,001817 | TRUE | 0,02031 | No |
| orf19.3870 | 6592 | 6038 | 6279 | 4048 | 11570 | 8494 | 10959 | 12017 | 6285 | 6384 | 6203 | 6341 | 9271 | 9658 | 9304 | 9771 | 7902,24 | 6303 | 9501 | 1,507 | 0,592 | 18,074 | 5,12E-73 | 2,55E-72 | 0,0003552 | 0,0022991 | 0,0008938 | 0,0008938 | TRUE | 0,0045846 | No |
| orf19.3871 | 31 | 29 | 43 | 21 | 107 | 73 | 98 | 96 | 30 | 31 | 42 | 33 | 86 | 83 | 83 | 78 | 58,2 | 34 | 82 | 2,427 | 1,279 | 6,461 | 1,04E-10 | 1,86E-10 | 1,00E-08 | 0,0303028 | 0,0159356 | 0,0159356 | TRUE | 0,2290623 | No |

| | | | | | | | | | | | | | | | | | | | | | | | | | | | | | | | |
|---|---|---|---|---|---|---|---|---|---|---|---|---|---|---|---|---|---|---|---|---|---|---|---|---|---|---|---|---|---|---|---|
| orf19.3872 | 211 | 184 | 217 | 125 | 389 | 212 | 306 | 372 | 201 | 195 | 214 | 196 | 312 | 241 | 260 | 302 | 240,12 | 202 | 279 | 1,387 | 0,472 | 4,289 | 1,79E-05 | 2,63E-05 | 0,0055117 | 0,0089292 | 0,0072457 | 0,0072457 | TRUE | 0,0850014 | No |
| orf19.3873 | 1406 | 1184 | 1287 | 822 | 1766 | 1299 | 1735 | 1869 | 1340 | 1252 | 1271 | 1288 | 1415 | 1477 | 1473 | 1520 | 1379,54 | 1288 | 1471 | 1,142 | 0,191 | 3,949 | 7,85E-05 | 0,0001117 | 1,00E-08 | 0,0032815 | 0,0015163 | 0,0015163 | TRUE | 0,009856 | No |
| orf19.3874 | 4173 | 3962 | 4226 | 2632 | 4355 | 3369 | 4228 | 4646 | 3979 | 4189 | 4175 | 4123 | 3490 | 3831 | 3590 | 3778 | 3894,18 | 4116 | 3672 | 0,892 | -0,165 | -3,843 | 0,0001217 | 0,0001716 | 0,0008754 | 0,0025129 | 0,0015219 | 0,0015219 | TRUE | 0,0136959 | No |
| orf19.3876 | 79 | 63 | 91 | 48 | 311 | 231 | 270 | 295 | 75 | 67 | 90 | 75 | 249 | 263 | 229 | 240 | 161 | 77 | 245 | 3,18 | 1,669 | 12,811 | 1,42E-37 | 4,33E-37 | 1,00E-08 | 0,0122896 | 0,0072761 | 0,0072761 | TRUE | 0,1348004 | No |
| orf19.3877 | 486 | 513 | 573 | 324 | 707 | 512 | 629 | 695 | 463 | 542 | 566 | 508 | 566 | 582 | 534 | 565 | 540,9 | 520 | 562 | 1,08 | 0,111 | 1,438 | 0,1505037 | 0,1727584 | 0,0025318 | 0,0051268 | 0,0038163 | 0,0038163 | TRUE | 0,0694863 | No |
| orf19.3878 | 1015 | 962 | 1011 | 575 | 1084 | 780 | 1079 | 1143 | 968 | 1017 | 999 | 901 | 869 | 887 | 916 | 929 | 935,67 | 971 | 900 | 0,925 | -0,113 | -1,964 | 0,0495009 | 0,0595522 | 0,0004286 | 0,0038461 | 0,0020979 | 0,0020979 | TRUE | 0,0243026 | No |
| orf19.3879 | 69 | 69 | 72 | 64 | 356 | 245 | 311 | 384 | 66 | 73 | 71 | 100 | 285 | 279 | 264 | 312 | 181,28 | 78 | 285 | 3,757 | 1,91 | 14,185 | 1,13E-45 | 3,92E-45 | 0,0039484 | 0,0111486 | 0,0085276 | 0,0085276 | TRUE | 0,2621173 | No |
| orf19.388 | 2042 | 1646 | 1855 | 1246 | 1354 | 922 | 1379 | 1323 | 1947 | 1740 | 1833 | 1952 | 1085 | 1048 | 1171 | 1076 | 1481,45 | 1868 | 1095 | 0,587 | -0,768 | -13,343 | 1,31E-40 | 4,17E-40 | 0,0019419 | 0,0031996 | 0,0024753 | 0,0024753 | TRUE | 0,0260971 | No |
| orf19.3881 | 1512 | 1140 | 1272 | 845 | 1386 | 881 | 1344 | 1381 | 1442 | 1205 | 1257 | 1324 | 1111 | 1002 | 1141 | 1123 | 1200,45 | 1307 | 1094 | 0,838 | -0,255 | -3,707 | 0,0002099 | 0,0002926 | 0,0035759 | 0,003459 | 0,0036932 | 0,0036932 | TRUE | 0,060884 | No |
| orf19.3884 | 338 | 270 | 299 | 202 | 1461 | 938 | 1406 | 1391 | 322 | 285 | 295 | 316 | 1171 | 1067 | 1194 | 1131 | 722,69 | 304 | 1141 | 3,751 | 1,907 | 26,524 | 5,17E-155 | 5,41E-154 | 0,0011685 | 0,0043632 | 0,0027466 | 0,0027466 | TRUE | 0,0213863 | No |
| orf19.3885 | 173 | 182 | 156 | 118 | 163 | 121 | 119 | 151 | 165 | 192 | 154 | 185 | 131 | 138 | 101 | 123 | 148,54 | 174 | 123 | 0,707 | -0,501 | -3,737 | 0,000186 | 0,0002598 | 0,0068707 | 0,0131446 | 0,0103681 | 0,0103681 | TRUE | 0,1515829 | No |
| orf19.3886 | 2852 | 2453 | 2654 | 1779 | 1405 | 1052 | 1408 | 1440 | 2719 | 2594 | 2622 | 2787 | 1126 | 1196 | 1195 | 1171 | 1926,23 | 2680 | 1172 | 0,437 | -1,193 | -26,027 | 2,48E-149 | 2,47E-148 | 0,0004129 | 0,0029437 | 0,0014271 | 0,0014271 | TRUE | 0,0087325 | No |
| orf19.3887 | 2821 | 2414 | 2691 | 1398 | 2514 | 1542 | 2232 | 2455 | 2690 | 2552 | 2658 | 2190 | 2014 | 1753 | 1895 | 1996 | 2218,68 | 2522 | 1914 | 0,758 | -0,399 | -5,597 | 2,19E-08 | 3,60E-08 | 0,0054746 | 0,0028313 | 0,0044235 | 0,0044235 | TRUE | 0,0916853 | No |
| orf19.3888 | 4151 | 3525 | 3802 | 2528 | 1663 | 1240 | 1566 | 1678 | 3958 | 3727 | 3756 | 3960 | 1333 | 1410 | 1330 | 1364 | 2604,68 | 3850 | 1359 | 0,353 | -1,503 | -34,521 | 3,82E-261 | 6,84E-260 | 0,0005319 | 0,0027216 | 0,0013454 | 0,0013454 | TRUE | 0,0070077 | No |
| orf19.3888.2 | 1 | 1 | 3 | 0 | 0 | 0 | 9 | 2 | 1 | 1 | 3 | 0 | 0 | 0 | 8 | 2 | 1,78 | 1 | 2 | 1,803 | 0,851 | 0,606 | 0,5444579 | 0,5799976 | 1,8467695 | 0,9244034 | 1,2715296 | 1,2715296 | TRUE | 2,8673964 | No |
| orf19.389 | 168 | 147 | 136 | 104 | 249 | 181 | 243 | 260 | 160 | 155 | 134 | 163 | 200 | 206 | 206 | 211 | 179,49 | 153 | 206 | 1,348 | 0,431 | 3,945 | 7,99E-05 | 0,0001136 | 1,00E-08 | 0,0112389 | 0,0056519 | 0,0056519 | TRUE | 0,0765758 | No |
| orf19.3893 | 156 | 129 | 162 | 112 | 42 | 24 | 33 | 45 | 149 | 136 | 160 | 175 | 34 | 27 | 28 | 37 | 93,27 | 155 | 32 | 0,205 | -2,286 | -12,88 | 5,81E-38 | 1,78E-37 | 1,00E-08 | 0,0196947 | 0,0126239 | 0,0126239 | TRUE | 0,0911475 | No |
| orf19.3894 | 3210 | 2833 | 2999 | 1850 | 2239 | 1491 | 2227 | 2305 | 3060 | 2996 | 2963 | 2898 | 1794 | 1695 | 1891 | 1874 | 2396,39 | 2979 | 1814 | 0,609 | -0,715 | -15,396 | 1,75E-53 | 6,79E-53 | 0,0008806 | 0,0027764 | 0,0016379 | 0,0016379 | TRUE | 0,021761 | No |
| orf19.3895 | 825 | 659 | 728 | 566 | 903 | 715 | 849 | 915 | 787 | 697 | 719 | 887 | 724 | 813 | 721 | 744 | 761,32 | 772 | 750 | 0,974 | -0,038 | -0,473 | 0,6359651 | 0,6683289 | 0,0056412 | 0,0042479 | 0,0049394 | 0,0049394 | TRUE | 0,1068475 | No |
| orf19.3897 | 24 | 15 | 31 | 10 | 12 | 3 | 12 | 11 | 23 | 16 | 31 | 16 | 10 | 3 | 10 | 9 | 14,65 | 22 | 8 | 0,383 | -1,386 | -3,323 | 0,0008907 | 0,001206 | 0,0451479 | 0,1141751 | 0,0871948 | 0,0871948 | TRUE | 0,4715857 | No |
| orf19.3898 | 298 | 267 | 258 | 158 | 699 | 465 | 712 | 710 | 284 | 282 | 255 | 248 | 560 | 529 | 604 | 577 | 417,43 | 267 | 568 | 2,119 | 1,083 | 13,096 | 3,45E-39 | 1,08E-38 | 0,0010873 | 0,0060246 | 0,0037084 | 0,0037084 | TRUE | 0,0228145 | No |
| orf19.3899 | 324 | 289 | 287 | 164 | 547 | 370 | 586 | 609 | 309 | 306 | 284 | 257 | 438 | 421 | 498 | 495 | 375,83 | 289 | 463 | 1,598 | 0,676 | 7,329 | 2,33E-13 | 4,49E-13 | 0,0040326 | 0,00646 | 0,0052852 | 0,0052852 | TRUE | 0,0503259 | No |
| orf19.390 | 1875 | 1591 | 1858 | 1088 | 3607 | 2584 | 3587 | 3733 | 1788 | 1682 | 1836 | 1704 | 2890 | 2938 | 3045 | 3035 | 2364,86 | 1752 | 2977 | 1,697 | 0,763 | 16,972 | 1,33E-64 | 5,98E-64 | 0,0005514 | 0,0027856 | 0,0014709 | 0,0014709 | TRUE | 0,0124119 | No |
| orf19.3900 | 9397 | 7973 | 9040 | 5511 | 4248 | 2893 | 4072 | 4327 | 8959 | 8430 | 8931 | 8633 | 3404 | 3290 | 3457 | 3518 | 6077,77 | 8738 | 3417 | 0,391 | -1,354 | -35,613 | 8,71E-278 | 1,73E-276 | 0,0005501 | 0,0023614 | 0,001193 | 0,001193 | TRUE | 0,0073154 | No |
| orf19.3901 | 897 | 741 | 822 | 569 | 1003 | 703 | 869 | 1074 | 855 | 784 | 812 | 891 | 804 | 799 | 738 | 873 | 819,53 | 836 | 804 | 0,964 | -0,053 | -0,765 | 0,44437 | 0,4808419 | 0,0028126 | 0,0040948 | 0,0034002 | 0,0034002 | TRUE | 0,0420753 | No |
| orf19.3902 | 36 | 48 | 44 | 38 | 8 | 7 | 6 | 6 | 34 | 51 | 43 | 60 | 6 | 8 | 5 | 5 | 26,55 | 47 | 6 | 0,129 | -2,957 | -7,988 | 1,38E-15 | 2,80E-15 | 1,00E-08 | 0,0639288 | 0,0455633 | 0,0455633 | TRUE | 0,0814141 | No |
| orf19.3903 | 1 | 1 | 1 | 0 | 1 | 3 | 1 | 1 | 1 | 1 | 1 | 0 | 1 | 3 | 1 | 1 | 1,11 | 1 | 2 | 1,742 | 0,801 | 0,591 | 0,5548432 | 0,5896826 | 1,00E-08 | 1,4824001 | 0,7548079 | 0,7548079 | TRUE | 0,4471695 | No |
| orf19.3904 | 10 | 19 | 20 | 6 | 0 | 0 | 1 | 0 | 10 | 20 | 20 | 9 | 0 | 0 | 1 | 0 | 7,45 | 15 | 0 | 0,018 | -5,78 | -5,132 | 2,86E-07 | 4,52E-07 | 1,00E-08 | 0,2223685 | 0,180924 | 0,180924 | TRUE | 0,2613074 | No |
| orf19.3905 | 2 | 7 | 8 | 2 | 3 | 1 | 4 | 5 | 2 | 7 | 8 | 3 | 2 | 1 | 3 | 4 | 3,92 | 5 | 2 | 0,546 | -0,872 | -1,158 | 0,246684 | 0,2766415 | 1,00E-08 | 0,4211164 | 0,2779058 | 0,2779058 | TRUE | 0,1579483 | No |
| orf19.3906 | 5 | 9 | 15 | 9 | 7 | 2 | 1 | 2 | 5 | 10 | 15 | 14 | 6 | 2 | 1 | 2 | 6,69 | 11 | 3 | 0,246 | -2,023 | -2,984 | 0,0028474 | 0,0037355 | 0,1754657 | 0,2473384 | 0,2196808 | 0,2196808 | TRUE | 0,7045511 | No |
| orf19.3907 | 1 | 0 | 0 | 0 | 1 | 0 | 0 | 0 | 1 | 0 | 0 | 0 | 1 | 0 | 0 | 0 | 0,22 | 0 | 0 | 0,797 | -0,328 | -0,104 | 0,9174923 | 0,9248844 | 1,00E-08 | 7,4879881 | 7,6135352 | 7,6135352 | TRUE | 0,2859076 | No |
| orf19.3908 | 23 | 20 | 26 | 19 | 21 | 13 | 23 | 23 | 22 | 21 | 26 | 30 | 17 | 15 | 20 | 19 | 21,05 | 25 | 18 | 0,722 | -0,47 | -1,555 | 0,1198979 | 0,1387754 | 1,00E-08 | 0,0801088 | 0,0395837 | 0,0395837 | TRUE | 0,0774938 | No |
| orf19.391 | 1017 | 937 | 1019 | 718 | 2782 | 1921 | 2492 | 2768 | 970 | 991 | 1007 | 1125 | 2229 | 2184 | 2116 | 2251 | 1608,96 | 1023 | 2195 | 2,155 | 1,108 | 20,431 | 8,80E-93 | 5,39E-92 | 0,0011054 | 0,0031118 | 0,0020678 | 0,0020678 | TRUE | 0,047064 | No |
| orf19.3910 | 328 | 274 | 310 | 206 | 494 | 387 | 480 | 510 | 313 | 290 | 306 | 323 | 396 | 440 | 408 | 415 | 361,18 | 308 | 415 | 1,347 | 0,43 | 5,376 | 7,61E-08 | 1,23E-07 | 1,00E-08 | 0,0066372 | 0,0032395 | 0,0032395 | TRUE | 0,0178773 | No |
| orf19.3911 | 7965 | 6358 | 6457 | 4885 | 32960 | 23350 | 30264 | 35115 | 7594 | 6723 | 6379 | 7653 | 26410 | 26551 | 25694 | 28552 | 16944,36 | 7087 | 26802 | 3,783 | 1,92 | 30,226 | 1,09E-200 | 1,48E-199 | 0,0049444 | 0,0021882 | 0,0037785 | 0,0037785 | TRUE | 0,0554176 | No |
| orf19.3912 | 3820 | 3473 | 3842 | 2564 | 3273 | 2400 | 3273 | 3534 | 3642 | 3672 | 3796 | 4017 | 2623 | 2729 | 2779 | 2873 | 3266,28 | 3782 | 2751 | 0,728 | -0,457 | -9,682 | 3,60E-22 | 8,44E-22 | 0,0013154 | 0,002594 | 0,0018319 | 0,0018319 | TRUE | 0,020202 | No |
| orf19.3914 | 8283 | 7991 | 8517 | 5103 | 6573 | 4825 | 6447 | 6445 | 7897 | 8449 | 8414 | 7994 | 5267 | 5486 | 5473 | 5240 | 6777,7 | 8188 | 5366 | 0,655 | -0,61 | -15,913 | 5,16E-57 | 2,08E-56 | 0,0006943 | 0,0023335 | 0,0012602 | 0,0012602 | TRUE | 0,007305 | No |
| orf19.3915 | 4783 | 4081 | 4368 | 2734 | 8288 | 6061 | 8107 | 8763 | 4560 | 4315 | 4315 | 4283 | 6641 | 6892 | 6883 | 7125 | 5626,77 | 4368 | 6885 | 1,575 | 0,656 | 17,044 | 3,89E-65 | 1,75E-64 | 0,0005971 | 0,0023831 | 0,0012272 | 0,0012272 | TRUE | 0,0108859 | No |
| orf19.3916 | 1058 | 1069 | 1104 | 700 | 2021 | 1389 | 1815 | 1781 | 1009 | 1130 | 1091 | 1097 | 1619 | 1579 | 1541 | 1448 | 1314,26 | 1082 | 1547 | 1,431 | 0,517 | 8,991 | 2,45E-19 | 5,41E-19 | 0,0016646 | 0,0033406 | 0,0023684 | 0,0023684 | TRUE | 0,0260919 | No |
| orf19.3919 | 1483 | 1374 | 1464 | 960 | 1369 | 1089 | 1299 | 1319 | 1414 | 1453 | 1446 | 1504 | 1097 | 1238 | 1103 | 1072 | 1290,93 | 1454 | 1128 | 0,775 | -0,368 | -6,5 | 8,03E-11 | 1,44E-10 | 0,001486 | 0,0033632 | 0,0023019 | 0,0023019 | TRUE | 0,0490207 | No |
| orf19.3919.1 | 792 | 635 | 662 | 457 | 1252 | 772 | 1232 | 1194 | 755 | 671 | 654 | 716 | 1003 | 878 | 1046 | 971 | 836,78 | 699 | 974 | 1,397 | 0,482 | 6,604 | 4,00E-11 | 7,22E-11 | 0,0036794 | 0,0040534 | 0,003853 | 0,003853 | TRUE | 0,0502263 | No |
| orf19.392 | 731 | 588 | 714 | 433 | 1373 | 967 | 1349 | 1407 | 697 | 622 | 705 | 678 | 1100 | 1100 | 1145 | 1144 | 898,92 | 676 | 1122 | 1,661 | 0,732 | 12,457 | 1,28E-35 | 3,77E-35 | 0,0002367 | 0,0039178 | 0,0020919 | 0,0020919 | TRUE | 0,0358276 | No |
| orf19.3920 | 639 | 529 | 570 | 368 | 1967 | 1398 | 1889 | 2166 | 609 | 559 | 563 | 576 | 1576 | 1590 | 1604 | 1761 | 1104,85 | 577 | 1633 | 2,829 | 1,5 | 24,226 | 1,17E-129 | 1,01E-128 | 0,0016163 | 0,0035774 | 0,0024418 | 0,0024418 | TRUE | 0,0345664 | No |
| orf19.3921 | 1590 | 1209 | 1418 | 895 | 2750 | 1861 | 2766 | 2875 | 1516 | 1278 | 1401 | 1402 | 2203 | 2116 | 2348 | 2338 | 1825,35 | 1399 | 2251 | 1,609 | 0,686 | 11,201 | 4,03E-29 | 1,07E-28 | 0,002861 | 0,0029908 | 0,0030052 | 0,0030052 | TRUE | 0,0420115 | No |
| orf19.3922 | 4759 | 3962 | 4227 | 2873 | 2781 | 1927 | 2710 | 2832 | 4537 | 4189 | 4176 | 4501 | 2228 | 2191 | 2301 | 2303 | 3303,26 | 4351 | 2256 | 0,519 | -0,946 | -21,112 | 6,20E-99 | 4,02E-98 | 0,0010562 | 0,0025883 | 0,0016019 | 0,0016019 | TRUE | 0,0107749 | No |
| orf19.3923 | 420 | 374 | 371 | 297 | 127 | 74 | 127 | 141 | 400 | 395 | 367 | 465 | 102 | 84 | 108 | 115 | 254,5 | 407 | 102 | 0,254 | -1,978 | -16,799 | 2,49E-63 | 1,09E-62 | 0,0062945 | 0,0085427 | 0,0075995 | 0,0075995 | TRUE | 0,1153295 | No |
| orf19.3924 | 26 | 27 | 36 | 12 | 184 | 143 | 170 | 196 | 25 | 29 | 36 | 19 | 147 | 163 | 144 | 159 | 90,18 | 27 | 153 | 5,559 | 2,475 | 12,938 | 2,75E-38 | 8,49E-38 | 1,00E-08 | 0,0202984 | 0,0121796 | 0,0121796 | TRUE | 0,2858677 | No |
| orf19.3925 | 8 | 12 | 10 | 5 | 42 | 30 | 36 | 41 | 8 | 13 | 10 | 8 | 34 | 34 | 31 | 33 | 21,21 | 10 | 33 | 3,43 | 1,778 | 5,248 | 1,54E-07 | 2,46E-07 | 1,00E-08 | 0,0794965 | 0,0390709 | 0,0390709 | TRUE | 0,1637722 | No |
| orf19.3926 | 51 | 33 | 29 | 29 | 79 | 62 | 71 | 104 | 49 | 35 | 29 | 45 | 63 | 70 | 60 | 85 | 54,53 | 40 | 70 | 1,78 | 0,832 | 3,755 | 0,0001734 | 0,0002424 | 0,0188122 | 0,0322016 | 0,0265321 | 0,0265321 | TRUE | 0,2709038 | No |
| orf19.3928 | 652 | 516 | 602 | 356 | 815 | 524 | 792 | 892 | 622 | 546 | 595 | 558 | 653 | 596 | 672 | 725 | 620,77 | 580 | 662 | 1,141 | 0,19 | 2,462 | 0,0138159 | 0,0173952 | 0,0035245 | 0,0047362 | 0,0041035 | 0,0041035 | TRUE | 0,0494004 | No |
| orf19.3929 | 778 | 667 | 694 | 480 | 2165 | 1591 | 2184 | 2261 | 742 | 705 | 686 | 752 | 1735 | 1809 | 1854 | 1838 | 1265,13 | 721 | 1809 | 2,514 | 1,33 | 24,856 | 2,21E-136 | 2,02E-135 | 0,0004289 | 0,0033891 | 0,0017258 | 0,0017258 | TRUE | 0,013248 | No |
| orf19.393 | 1059 | 863 | 908 | 651 | 1386 | 952 | 1342 | 1501 | 1010 | 913 | 897 | 1020 | 1111 | 1083 | 1139 | 1220 | 1048,98 | 960 | 1138 | 1,189 | 0,25 | 3,869 | 0,0001092 | 0,0001544 | 0,0023067 | 0,0036565 | 0,0030249 | 0,0030249 | TRUE | 0,0286781 | No |
| orf19.3930 | 3271 | 2821 | 3061 | 2032 | 4341 | 2945 | 4212 | 4381 | 3119 | 2983 | 3024 | 3183 | 3478 | 3349 | 3576 | 3562 | 3284,22 | 3077 | 3491 | 1,136 | 0,184 | 4,551 | 5,33E-06 | 8,01E-06 | 0,0004739 | 0,0025912 | 0,0012609 | 0,0012609 | TRUE | 0,0087684 | No |
| orf19.3931 | 1858 | 1574 | 1686 | 1158 | 8578 | 6407 | 8468 | 8955 | 1771 | 1664 | 1666 | 1814 | 6873 | 7285 | 7189 | 7281 | 4443,07 | 1729 | 7157 | 4,146 | 2,052 | 47,135 | 0 | 0 | 0,0008516 | 0,0024608 | 0,0014318 | 0,0014318 | TRUE | 0,0116965 | No |
| orf19.3931.2 | 12 | 9 | 15 | 16 | 9 | 7 | 7 | 8 | 11 | 10 | 15 | 25 | 7 | 8 | 6 | 7 | 11,06 | 15 | 7 | 0,465 | -1,103 | -2,406 | 0,0161441 | 0,020203 | 1,00E-08 | 0,1505802 | 0,0993164 | 0,0993164 | TRUE | 0,5771214 | No |
| orf19.3932 | 11765 | 9706 | 10734 | 6896 | 3459 | 2430 | 3287 | 3655 | 11217 | 10263 | 10604 | 10803 | 2772 | 2763 | 2791 | 2972 | 6772,99 | 10722 | 2824 | 0,264 | -1,924 | -45,03 | 0 | 0 | 0,0010066 | 0,0023337 | 0,0015426 | 0,0015426 | TRUE | 0,0152571 | No |
| orf19.3932.1 | 1946 | 1361 | 1406 | 1212 | 409 | 299 | 314 | 357 | 1855 | 1439 | 1389 | 1899 | 328 | 340 | 267 | 290 | 975,83 | 1646 | 306 | 0,186 | -2,427 | -20,954 | 1,74E-97 | 1,11E-96 | 0,0186268 | 0,0037739 | 0,0110829 | 0,0110829 | TRUE | 0,0522427 | No |
| orf19.3934 | 13578 | 11100 | 12103 | 7436 | 7250 | 4823 | 6966 | 7498 | 12945 | 11737 | 11957 | 11649 | 5809 | 5484 | 5914 | 6097 | 8948,95 | 12072 | 5826 | 0,483 | -1,051 | -21,71 | 1,65E-104 | 1,12E-103 | 0,0020234 | 0,0022748 | 0,0021273 | 0,0021273 | TRUE | 0,029155 | No |
| orf19.3935 | 4 | 3 | 2 | 2 | 12 | 6 | 6 | 1 | 4 | 3 | 2 | 3 | 10 | 7 | 5 | 1 | 4,3 | 3 | 6 | 1,84 | 0,88 | 1,136 | 0,2560909 | 0,2864358 | 1,00E-08 | 0,3834917 | 0,3119407 | 0,3119407 | TRUE | 0,2740112 | No |
| orf19.3936 | 2512 | 2332 | 2419 | 1497 | 4417 | 3168 | 4190 | 4574 | 2395 | 2466 | 2390 | 2345 | 3539 | 3602 | 3557 | 3719 | 3001,69 | 2399 | 3604 | 1,502 | 0,586 | 15,478 | 4,90E-54 | 1,91E-53 | 0,0001941 | 0,0026383 | 0,0010209 | 0,0010209 | TRUE | 0,0057512 | No |
| orf19.3937 | 621 | 604 | 634 | 406 | 1012 | 751 | 987 | 1076 | 592 | 639 | 626 | 636 | 811 | 854 | 838 | 875 | 733,84 | 623 | 844 | 1,356 | 0,44 | 7,329 | 2,31E-13 | 4,46E-13 | 1,00E-08 | 0,0043287 | 0,0020322 | 0,0020322 | TRUE | 0,0142988 | No |
| orf19.3938 | 2352 | 2134 | 2281 | 1782 | 2146 | 1689 | 2059 | 1985 | 2242 | 2256 | 2253 | 2792 | 1720 | 1921 | 1748 | 1614 | 2068,25 | 2386 | 1751 | 0,734 | -0,445 | -5,582 | 2,38E-08 | 3,92E-08 | 0,0084047 | 0,0028851 | 0,0056252 | 0,0056252 | TRUE | 0,1539599 | No |
| orf19.3939 | 652 | 613 | 676 | 435 | 1162 | 848 | 1130 | 1133 | 622 | 648 | 668 | 681 | 931 | 964 | 959 | 921 | 799,37 | 655 | 944 | 1,444 | 0,53 | 9,228 | 2,76E-20 | 6,24E-20 | 1,00E-08 | 0,0041453 | 0,0018435 | 0,0018435 | TRUE | 0,0142025 | No |
| orf19.394 | 1906 | 1538 | 1738 | 1173 | 1588 | 1130 | 1497 | 1703 | 1817 | 1626 | 1717 | 1838 | 1272 | 1285 | 1271 | 1385 | 1526,37 | 1750 | 1303 | 0,746 | -0,423 | -7,521 | 5,42E-14 | 1,06E-13 | 0,0018013 | 0,003167 | 0,0023714 | 0,0023714 | TRUE | 0,0270692 | No |
| orf19.3940 | 80 | 74 | 70 | 53 | 196 | 159 | 209 | 239 | 76 | 78 | 69 | 83 | 157 | 181 | 177 | 194 | 127,04 | 76 | 177 | 2,326 | 1,218 | 8,75 | 2,13E-18 | 4,61E-18 | 1,00E-08 | 0,0150157 | 0,0088028 | 0,0088028 | TRUE | 0,0692566 | No |
| orf19.3940.1 | 74206 | 42907 | 49512 | 40372 | 25582 | 18020 | 21495 | 26394 | 70747 | 45369 | 48913 | 63245 | 20498 | 20490 | 18249 | 21461 | 38621,58 | 57068 | 20174 | 0,354 | -1,5 | -9,452 | 3,31E-21 | 7,61E-21 | 0,0241708 | 0,0021338 | 0,0126833 | 0,0241708 | TRUE | 0,1233602 | No |
| orf19.3941 | 18056 | 15293 | 16580 | 10269 | 28235 | 19052 | 26681 | 29248 | 17214 | 16170 | 16379 | 16087 | 22624 | 21664 | 22652 | 23781 | 19571,56 | 16462 | 22680 | 1,378 | 0,462 | 11,582 | 5,09E-31 | 1,40E-30 | 0,0011501 | 0,0021752 | 0,0014756 | 0,0014756 | TRUE | 0,0130785 | No |
| orf19.3942 | 386 | 357 | 361 | 206 | 743 | 484 | 657 | 712 | 368 | 377 | 357 | 323 | 595 | 550 | 558 | 579 | 463,41 | 356 | 570 | 1,596 | 0,674 | 9,072 | 1,17E-19 | 2,61E-19 | 1,00E-08 | 0,0056344 | 0,0029649 | 0,0029649 | TRUE | 0,0361531 | No |
| orf19.3942.1 | 15321 | 12228 | 12211 | 7621 | 37667 | 23450 | 33925 | 37522 | 14607 | 12930 | 12063 | 11939 | 30181 | 26665 | 28802 | 30509 | 20961,98 | 12885 | 29039 | 2,253 | 1,172 | 17,087 | 1,84E-65 | 8,34E-65 | 0,006291 | 0,0021696 | 0,0044619 | 0,0044619 | TRUE | 0,0992753 | No |
| orf19.3944 | 10661 | 9403 | 10300 | 6634 | 9170 | 6331 | 8606 | 9270 | 10164 | 9942 | 10175 | 10393 | 7348 | 7199 | 7306 | 7537 | 8758,12 | 10168 | 7348 | 0,723 | -0,468 | -15,247 | 1,72E-52 | 6,55E-52 | 0,0001917 | 0,0022788 | 0,0007884 | 0,0007884 | TRUE | 0,0037018 | No |
| orf19.3945 | 4851 | 4673 | 4942 | 3022 | 4072 | 3001 | 3798 | 3948 | 4625 | 4941 | 4882 | 4734 | 3263 | 3412 | 3224 | 3210 | 4036,52 | 4796 | 3277 | 0,683 | -0,55 | -13,984 | 1,96E-44 | 6,64E-44 | 0,0005038 | 0,002498 | 0,0012328 | 0,0012328 | TRUE | 0,0089123 | No |
| orf19.3946 | 461 | 417 | 420 | 312 | 780 | 570 | 752 | 814 | 440 | 441 | 415 | 489 | 625 | 648 | 638 | 662 | 544,7 | 446 | 643 | 1,45 | 0,536 | 7,724 | 1,13E-14 | 2,26E-14 | 1,00E-08 | 0,0051056 | 0,002671 | 0,002671 | TRUE | 0,0397059 | No |
| orf19.3947 | 1341 | 1088 | 1106 | 688 | 1350 | 872 | 1321 | 1366 | 1278 | 1150 | 1093 | 1078 | 1082 | 992 | 1122 | 1111 | 1113,1 | 1150 | 1077 | 0,936 | -0,096 | -1,404 | 0,160309 | 0,1834526 | 0,0034567 | 0,0035664 | 0,0035978 | 0,0035978 | TRUE | 0,0695882 | No |

| | | | | | | | | | | | | | | | | | | | | | | | | | | | | | | | |
|---|---|---|---|---|---|---|---|---|---|---|---|---|---|---|---|---|---|---|---|---|---|---|---|---|---|---|---|---|---|---|---|
| orf19.3949 | 4641 | 3976 | 4353 | 2756 | 3033 | 1923 | 2839 | 2874 | 4425 | 4204 | 4300 | 4317 | 2430 | 2187 | 2410 | 2337 | 3326,32 | 4312 | 2341 | 0,544 | -0,88 | -20,045 | 2,23E-89 | 1,32E-88 | 0,0008939 | 0,0025849 | 0,0015276 | 0,0015276 | TRUE | 0,0225971 | No |
| orf19.395 | 14978 | 11641 | 12454 | 8741 | 6045 | 4258 | 5635 | 6396 | 14280 | 12309 | 12303 | 13693 | 4844 | 4842 | 4784 | 5201 | 9031,93 | 13146 | 4918 | 0,374 | -1,418 | -24,618 | 8,14E-134 | 7,36E-133 | 0,0034549 | 0,0022731 | 0,0030551 | 0,0030551 | TRUE | 0,0416469 | No |
| orf19.3950 | 337 | 301 | 336 | 207 | 890 | 680 | 850 | 897 | 321 | 318 | 332 | 324 | 713 | 773 | 722 | 729 | 529,14 | 324 | 734 | 2,263 | 1,178 | 16,905 | 4,11E-64 | 1,83E-63 | 1,00E-08 | 0,0051942 | 0,0023424 | 0,0023424 | TRUE | 0,0137645 | No |
| orf19.3951 | 1364 | 1142 | 1326 | 807 | 2505 | 1715 | 2324 | 2405 | 1300 | 1208 | 1310 | 1264 | 2007 | 1950 | 1973 | 1955 | 1621 | 1270 | 1971 | 1,551 | 0,633 | 13,927 | 4,36E-44 | 1,47E-43 | 1,00E-08 | 0,0031042 | 0,001319 | 0,001319 | TRUE | 0,01426 | No |
| orf19.3952 | 2684 | 2275 | 2537 | 1576 | 4084 | 2859 | 3975 | 4175 | 2559 | 2406 | 2506 | 2469 | 3272 | 3251 | 3375 | 3395 | 2904,05 | 2485 | 3323 | 1,337 | 0,419 | 10,767 | 4,91E-27 | 1,27E-26 | 0,0002354 | 0,0026567 | 0,0010953 | 0,0010953 | TRUE | 0,0059415 | No |
| orf19.3954 | 1285 | 1301 | 1324 | 804 | 1767 | 1311 | 1702 | 1811 | 1225 | 1376 | 1308 | 1260 | 1416 | 1491 | 1445 | 1473 | 1374,04 | 1292 | 1456 | 1,126 | 0,171 | 3,308 | 0,0009389 | 0,001269 | 0,0007049 | 0,0032862 | 0,0018216 | 0,0018216 | TRUE | 0,0231688 | No |
| orf19.3954.1 | 39 | 38 | 52 | 32 | 158 | 110 | 145 | 127 | 37 | 40 | 51 | 50 | 127 | 125 | 123 | 103 | 82,11 | 44 | 120 | 2,692 | 1,429 | 8,068 | 7,15E-16 | 1,47E-15 | 1,00E-08 | 0,0220867 | 0,013858 | 0,013858 | TRUE | 0,1107796 | No |
| orf19.3955 | 4608 | 4169 | 4278 | 3020 | 10986 | 8236 | 10410 | 11779 | 4393 | 4408 | 4226 | 4731 | 8803 | 9365 | 8838 | 9577 | 6792,75 | 4440 | 9146 | 2,062 | 1,044 | 22,049 | 9,70E-108 | 6,83E-107 | 0,0017965 | 0,002333 | 0,0019766 | 0,0019766 | TRUE | 0,0227753 | No |
| orf19.3956 | 1275 | 1107 | 1163 | 810 | 1357 | 912 | 1359 | 1439 | 1216 | 1171 | 1149 | 1269 | 1087 | 1037 | 1154 | 1170 | 1156,51 | 1201 | 1112 | 0,929 | -0,107 | -1,825 | 0,0680699 | 0,0808552 | 0,0013525 | 0,003511 | 0,0024027 | 0,0024027 | TRUE | 0,0229737 | No |
| orf19.3957 | 2248 | 1887 | 2067 | 1230 | 3009 | 2018 | 2801 | 2998 | 2143 | 1995 | 2042 | 1927 | 2411 | 2295 | 2378 | 2438 | 2203,59 | 2027 | 2380 | 1,173 | 0,23 | 4,993 | 5,95E-07 | 9,29E-07 | 0,0007364 | 0,0028364 | 0,0015789 | 0,0015789 | TRUE | 0,0181882 | No |
| orf19.3959 | 4464 | 3652 | 3919 | 2647 | 4283 | 2913 | 4081 | 4260 | 4256 | 3862 | 3872 | 4147 | 3432 | 3312 | 3465 | 3464 | 3726,06 | 4034 | 3418 | 0,848 | -0,238 | -5,313 | 1,08E-07 | 1,73E-07 | 0,0010721 | 0,0025319 | 0,0016558 | 0,0016558 | TRUE | 0,0176228 | No |
| orf19.396 | 314 | 286 | 310 | 185 | 1006 | 657 | 1061 | 1010 | 299 | 302 | 306 | 290 | 806 | 747 | 901 | 821 | 559,12 | 299 | 819 | 2,733 | 1,451 | 18,202 | 4,97E-74 | 2,52E-73 | 0,0025866 | 0,0050278 | 0,0036967 | 0,0036967 | TRUE | 0,0536026 | No |
| orf19.3960 | 2927 | 2767 | 2918 | 1773 | 2799 | 1844 | 2840 | 2902 | 2791 | 2926 | 2883 | 2777 | 2243 | 2097 | 2411 | 2360 | 2560,85 | 2844 | 2278 | 0,801 | -0,32 | -6,225 | 4,80E-10 | 8,36E-10 | 0,0016698 | 0,0027324 | 0,0021365 | 0,0021365 | TRUE | 0,033063 | No |
| orf19.3962 | 16540 | 14753 | 16086 | 10102 | 20484 | 14747 | 20133 | 21749 | 15769 | 15599 | 15891 | 15825 | 16413 | 16769 | 17093 | 17684 | 16380,51 | 15771 | 16990 | 1,077 | 0,107 | 3,315 | 0,0009179 | 0,0012415 | 0,0004545 | 0,0021915 | 0,0009474 | 0,0009474 | TRUE | 0,0093215 | No |
| orf19.3963 | 1465 | 1343 | 1417 | 1012 | 1410 | 1046 | 1306 | 1355 | 1397 | 1420 | 1400 | 1585 | 1130 | 1189 | 1109 | 1102 | 1291,46 | 1450 | 1132 | 0,782 | -0,355 | -6,264 | 3,75E-10 | 6,58E-10 | 0,0015156 | 0,0033626 | 0,0022932 | 0,0022932 | TRUE | 0,0428491 | No |
| orf19.3964 | 716 | 580 | 638 | 397 | 888 | 608 | 933 | 940 | 683 | 613 | 630 | 622 | 712 | 691 | 792 | 764 | 688,43 | 637 | 740 | 1,161 | 0,216 | 3,126 | 0,001773 | 0,0023579 | 0,0018696 | 0,0044763 | 0,0030903 | 0,0030903 | TRUE | 0,0288891 | No |
| orf19.3965 | 406 | 381 | 370 | 252 | 449 | 356 | 418 | 455 | 387 | 403 | 366 | 395 | 360 | 405 | 355 | 370 | 379,96 | 388 | 372 | 0,959 | -0,06 | -0,756 | 0,4496412 | 0,4861335 | 1,00E-08 | 0,0064126 | 0,0034156 | 0,0034156 | TRUE | 0,0353471 | No |
| orf19.3966 | 25 | 32 | 26 | 13 | 29 | 16 | 25 | 25 | 24 | 34 | 26 | 20 | 23 | 18 | 21 | 20 | 23,34 | 26 | 20 | 0,797 | -0,328 | -1,127 | 0,2595634 | 0,289964 | 1,00E-08 | 0,0724445 | 0,0386451 | 0,0386451 | TRUE | 0,2475448 | No |
| orf19.3967 | 33248 | 29380 | 31722 | 19913 | 14690 | 10464 | 13621 | 15083 | 31698 | 31066 | 31338 | 31195 | 11771 | 11899 | 11564 | 12264 | 21599,3 | 31324 | 11874 | 0,379 | -1,399 | -48,321 | 0 | 0 | 0,000274 | 0,0021673 | 0,00075 | 0,00075 | TRUE | 0,0060562 | No |
| orf19.3968 | 6 | 14 | 9 | 8 | 5 | 0 | 3 | 1 | 6 | 15 | 9 | 13 | 4 | 0 | 3 | 1 | 6,16 | 11 | 2 | 0,187 | -2,415 | -3,358 | 0,0007858 | 0,0010676 | 0,1052108 | 0,2684525 | 0,2147782 | 0,2147782 | TRUE | 0,3792608 | No |
| orf19.3969 | 6451 | 5738 | 6325 | 3964 | 1977 | 1497 | 1861 | 2197 | 6150 | 6067 | 6249 | 6210 | 1584 | 1702 | 1580 | 1786 | 3916,07 | 6169 | 1663 | 0,27 | -1,892 | -40,26 | 0 | 0 | 0,001373 | 0,0025106 | 0,0017598 | 0,0017598 | TRUE | 0,0315934 | No |
| orf19.397 | 1138 | 984 | 1031 | 796 | 1584 | 1256 | 1613 | 1823 | 1085 | 1040 | 1019 | 1247 | 1269 | 1428 | 1369 | 1482 | 1242,5 | 1098 | 1387 | 1,267 | 0,342 | 4,618 | 3,87E-06 | 5,85E-06 | 0,0054979 | 0,0034127 | 0,0044156 | 0,0044156 | TRUE | 0,0928656 | No |
| orf19.3970 | 1983 | 1706 | 1635 | 1011 | 1605 | 1136 | 1601 | 1660 | 1891 | 1804 | 1615 | 1584 | 1286 | 1292 | 1359 | 1350 | 1522,53 | 1724 | 1322 | 0,766 | -0,385 | -5,979 | 2,24E-09 | 3,81E-09 | 0,0032075 | 0,0031697 | 0,0033134 | 0,0033134 | TRUE | 0,0519456 | No |
| orf19.3971 | 367 | 329 | 355 | 232 | 282 | 231 | 235 | 206 | 350 | 348 | 351 | 363 | 226 | 263 | 200 | 167 | 283,44 | 353 | 214 | 0,603 | -0,729 | -6,063 | 1,33E-09 | 2,28E-09 | 0,013582 | 0,007884 | 0,0102007 | 0,0102007 | TRUE | 0,2399005 | No |
| orf19.3972 | 687 | 781 | 836 | 567 | 720 | 568 | 611 | 501 | 655 | 826 | 826 | 888 | 577 | 646 | 519 | 407 | 667,97 | 799 | 537 | 0,672 | -0,573 | -4,416 | 1,00E-05 | 1,49E-05 | 0,0253214 | 0,0045493 | 0,0146096 | 0,0146096 | TRUE | 0,3143542 | No |
| orf19.3973 | 1606 | 1351 | 1375 | 966 | 857 | 546 | 820 | 808 | 1531 | 1429 | 1358 | 1513 | 687 | 621 | 696 | 657 | 1061,5 | 1458 | 665 | 0,457 | -1,128 | -18,161 | 1,05E-73 | 5,27E-73 | 0,0018473 | 0,0036381 | 0,0026499 | 0,0026499 | TRUE | 0,0260518 | No |
| orf19.3974 | 34189 | 31460 | 33836 | 22477 | 45174 | 34874 | 41815 | 47732 | 32596 | 33265 | 33427 | 35211 | 36197 | 39655 | 35501 | 38811 | 35582,7 | 33625 | 37541 | 1,117 | 0,159 | 3,454 | 0,0005516 | 0,000756 | 0,0019397 | 0,0021374 | 0,0020069 | 0,0020069 | TRUE | 0,0175512 | No |
| orf19.3976 | 423 | 386 | 443 | 240 | 483 | 357 | 479 | 434 | 403 | 408 | 438 | 376 | 387 | 406 | 407 | 353 | 397,19 | 406 | 388 | 0,95 | -0,073 | -0,892 | 0,3725648 | 0,4077064 | 0,0015264 | 0,006225 | 0,0039628 | 0,0039628 | TRUE | 0,0445446 | No |
| orf19.3977 | 3682 | 3088 | 3297 | 1965 | 4301 | 2902 | 4204 | 4369 | 3510 | 3265 | 3257 | 3078 | 3446 | 3300 | 3569 | 3552 | 3372,33 | 3278 | 3467 | 1,057 | 0,08 | 1,616 | 0,1060154 | 0,1234458 | 0,0017096 | 0,0025782 | 0,002046 | 0,002046 | TRUE | 0,0278787 | No |
| orf19.3978 | 5622 | 5602 | 5736 | 2606 | 5206 | 3262 | 4792 | 4586 | 5360 | 5923 | 5667 | 4082 | 4171 | 3709 | 4068 | 3729 | 4588,79 | 5258 | 3919 | 0,745 | -0,425 | -4,403 | 1,07E-05 | 1,58E-05 | 0,0135753 | 0,0024491 | 0,0087133 | 0,0087133 | TRUE | 0,2727869 | No |
| orf19.398 | 124 | 113 | 127 | 96 | 615 | 403 | 520 | 559 | 118 | 119 | 125 | 150 | 493 | 458 | 441 | 455 | 295,07 | 128 | 462 | 3,642 | 1,865 | 18,331 | 4,71E-75 | 2,41E-74 | 0,0006899 | 0,0076557 | 0,0045996 | 0,0045996 | TRUE | 0,100602 | No |
| orf19.3980 | 985 | 924 | 1079 | 635 | 3311 | 2349 | 3130 | 3472 | 939 | 977 | 1066 | 995 | 2653 | 2671 | 2657 | 2823 | 1847,66 | 994 | 2701 | 2,718 | 1,442 | 27,658 | 2,25E-168 | 2,55E-167 | 0,0009696 | 0,0029799 | 0,0018826 | 0,0018826 | TRUE | 0,0305946 | No |
| orf19.3981 | 412 | 344 | 390 | 260 | 199 | 126 | 187 | 191 | 393 | 364 | 385 | 407 | 159 | 143 | 159 | 155 | 270,74 | 387 | 154 | 0,4 | -1,321 | -14,196 | 9,68E-46 | 3,36E-45 | 1,00E-08 | 0,0081558 | 0,0039968 | 0,0039968 | TRUE | 0,02008 | No |
| orf19.3982 | 234 | 195 | 219 | 124 | 621 | 511 | 557 | 604 | 223 | 206 | 216 | 194 | 498 | 581 | 473 | 491 | 360,32 | 210 | 511 | 2,414 | 1,271 | 13,308 | 2,07E-40 | 6,60E-40 | 0,003842 | 0,0066481 | 0,0052427 | 0,0052427 | TRUE | 0,0940344 | No |
| orf19.3983 | 992 | 951 | 974 | 642 | 3049 | 2329 | 2872 | 2894 | 946 | 1006 | 962 | 1006 | 2443 | 2648 | 2438 | 2353 | 1725,26 | 980 | 2470 | 2,523 | 1,335 | 24,844 | 2,98E-136 | 2,72E-135 | 0,001338 | 0,003043 | 0,0020188 | 0,0020188 | TRUE | 0,0273223 | No |
| orf19.3984 | 11600 | 12418 | 12609 | 7504 | 2893 | 2217 | 2572 | 2601 | 11059 | 13130 | 12456 | 11755 | 2318 | 2521 | 2184 | 2115 | 7192,4 | 12100 | 2284 | 0,189 | -2,406 | -35,764 | 4,05E-280 | 8,21E-279 | 0,0054398 | 0,0023196 | 0,0041063 | 0,0041063 | TRUE | 0,0575681 | No |
| orf19.3986 | 1116 | 1270 | 1269 | 737 | 781 | 629 | 677 | 672 | 1064 | 1343 | 1254 | 1155 | 626 | 715 | 575 | 546 | 909,66 | 1204 | 616 | 0,51 | -0,971 | -10,257 | 1,10E-24 | 2,72E-24 | 0,0106522 | 0,0038963 | 0,007414 | 0,007414 | TRUE | 0,1338599 | No |
| orf19.3988 | 9452 | 8183 | 8926 | 5710 | 527 | 351 | 481 | 575 | 9011 | 8652 | 8818 | 8945 | 422 | 399 | 408 | 468 | 4640,54 | 8856 | 424 | 0,048 | -4,379 | -85,088 | 0 | 0 | 0,0002935 | 0,0024451 | 0,0014369 | 0,0014369 | TRUE | 0,0560185 | No |
| orf19.399 | 7580 | 6709 | 7297 | 4881 | 4171 | 2906 | 3852 | 4415 | 7227 | 7094 | 7209 | 7646 | 3342 | 3304 | 3270 | 3590 | 5335,29 | 7294 | 3376 | 0,463 | -1,11 | -25,408 | 2,05E-142 | 1,96E-141 | 0,0011857 | 0,002399 | 0,001623 | 0,001623 | TRUE | 0,0222695 | No |
| orf19.3990 | 668 | 743 | 719 | 424 | 1384 | 898 | 1158 | 1013 | 637 | 786 | 710 | 664 | 1109 | 1021 | 983 | 824 | 841,74 | 699 | 984 | 1,406 | 0,492 | 5,089 | 3,60E-07 | 5,66E-07 | 0,0110032 | 0,0040419 | 0,00771 | 0,00771 | TRUE | 0,1468329 | No |
| orf19.3991 | 2627 | 2320 | 2628 | 1517 | 2481 | 1757 | 2432 | 2602 | 2505 | 2453 | 2596 | 2376 | 1988 | 1998 | 2065 | 2116 | 2262,07 | 2482 | 2042 | 0,822 | -0,283 | -6,33 | 2,44E-10 | 4,31E-10 | 0,0006456 | 0,0028171 | 0,0014803 | 0,0014803 | TRUE | 0,0114207 | No |
| orf19.3992 | 67 | 40 | 46 | 41 | 139 | 77 | 119 | 134 | 64 | 42 | 45 | 64 | 111 | 88 | 101 | 109 | 78,1 | 54 | 102 | 1,919 | 0,941 | 5,128 | 2,93E-07 | 4,63E-07 | 0,0081008 | 0,0231156 | 0,0175631 | 0,0175631 | TRUE | 0,1707279 | No |
| orf19.3994 | 464 | 421 | 426 | 307 | 745 | 560 | 679 | 687 | 442 | 445 | 421 | 481 | 597 | 637 | 576 | 559 | 519,76 | 447 | 592 | 1,326 | 0,407 | 5,568 | 2,58E-08 | 4,24E-08 | 0,0009248 | 0,0052502 | 0,0031346 | 0,0031346 | TRUE | 0,0276651 | No |
| orf19.3995 | 382 | 379 | 363 | 197 | 879 | 625 | 794 | 850 | 364 | 401 | 359 | 309 | 704 | 711 | 674 | 691 | 526,55 | 358 | 695 | 1,928 | 0,947 | 12,308 | 8,23E-35 | 2,40E-34 | 0,0004593 | 0,0052095 | 0,0034926 | 0,0034926 | TRUE | 0,0785357 | No |
| orf19.3996 | 103 | 83 | 102 | 58 | 403 | 315 | 338 | 397 | 98 | 88 | 101 | 91 | 323 | 358 | 287 | 323 | 208,56 | 94 | 323 | 3,398 | 1,765 | 14,637 | 1,62E-48 | 5,88E-48 | 1,00E-08 | 0,009964 | 0,0067138 | 0,0067138 | TRUE | 0,063046 | No |
| orf19.3997 | 75794 | 57465 | 64327 | 42099 | 15638 | 10124 | 14904 | 16331 | 72261 | 60762 | 63549 | 65950 | 12530 | 11512 | 12653 | 13279 | 39062,12 | 65630 | 12494 | 0,19 | -2,393 | -39,298 | 0 | 0 | 0,0044424 | 0,0021333 | 0,0035183 | 0,0035183 | TRUE | 0,0567549 | No |
| orf19.3998 | 158 | 173 | 160 | 122 | 217 | 161 | 203 | 176 | 151 | 183 | 158 | 191 | 174 | 183 | 172 | 143 | 169,39 | 171 | 168 | 0,989 | -0,016 | -0,13 | 0,896386 | 0,9108024 | 0,0060126 | 0,0117841 | 0,0091777 | 0,0091777 | TRUE | 0,1121067 | No |
| orf19.3999 | 966 | 909 | 976 | 612 | 1343 | 905 | 1257 | 1370 | 921 | 961 | 964 | 959 | 1076 | 1029 | 1067 | 1114 | 1011,42 | 951 | 1072 | 1,128 | 0,174 | 3,342 | 0,0008304 | 0,0011272 | 1,00E-08 | 0,0037146 | 0,0016089 | 0,0016089 | TRUE | 0,0084325 | No |
| orf19.400 | 2649 | 2312 | 2435 | 1652 | 2638 | 1898 | 2664 | 2753 | 2526 | 2445 | 2406 | 2588 | 2114 | 2158 | 2262 | 2238 | 2341,98 | 2491 | 2193 | 0,882 | -0,182 | -4,139 | 3,49E-05 | 5,05E-05 | 0,000594 | 0,0027924 | 0,0014262 | 0,0014262 | TRUE | 0,007576 | No |
| orf19.4000 | 15040 | 13746 | 15383 | 9817 | 9301 | 6846 | 8851 | 9808 | 14339 | 14535 | 15197 | 15379 | 7453 | 7785 | 7514 | 7975 | 11271,99 | 14862 | 7682 | 0,517 | -0,952 | -24,334 | 8,52E-131 | 7,46E-130 | 0,0009416 | 0,0022369 | 0,0013738 | 0,0013738 | TRUE | 0,0082131 | No |
| orf19.4001 | 377 | 366 | 393 | 233 | 290 | 176 | 275 | 251 | 359 | 387 | 388 | 365 | 232 | 200 | 233 | 204 | 296,22 | 375 | 217 | 0,581 | -0,782 | -8,626 | 6,33E-18 | 1,36E-17 | 1,00E-08 | 0,0076342 | 0,0043713 | 0,0043713 | TRUE | 0,0274253 | No |
| orf19.4002 | 462 | 395 | 481 | 247 | 650 | 474 | 664 | 664 | 440 | 418 | 475 | 387 | 521 | 539 | 547 | 540 | 483,34 | 430 | 537 | 1,241 | 0,311 | 4,128 | 3,66E-05 | 5,28E-05 | 0,0006643 | 0,0054883 | 0,0033355 | 0,0033355 | TRUE | 0,0563124 | No |
| orf19.4003 | 778 | 748 | 744 | 483 | 719 | 586 | 703 | 750 | 742 | 791 | 735 | 757 | 576 | 666 | 597 | 610 | 684,18 | 756 | 612 | 0,808 | -0,308 | -4,697 | 2,64E-06 | 4,02E-06 | 0,0008622 | 0,0044911 | 0,0026482 | 0,0026482 | TRUE | 0,0379557 | No |
| orf19.4004 | 5834 | 4850 | 5512 | 3351 | 8985 | 5907 | 8568 | 9281 | 5562 | 5128 | 5445 | 5250 | 7199 | 6717 | 7274 | 7546 | 6265,24 | 5346 | 7184 | 1,344 | 0,426 | 9,151 | 5,64E-20 | 1,27E-19 | 0,0015989 | 0,0023533 | 0,0019168 | 0,0019168 | TRUE | 0,0230331 | No |
| orf19.4005 | 407 | 378 | 409 | 282 | 904 | 659 | 929 | 926 | 388 | 400 | 404 | 442 | 724 | 749 | 789 | 753 | 581,11 | 408 | 754 | 1,855 | 0,891 | 13,025 | 8,82E-39 | 2,74E-38 | 1,00E-08 | 0,0049167 | 0,0025341 | 0,0025341 | TRUE | 0,0283625 | No |
| orf19.4006 | 1278 | 1143 | 1126 | 802 | 938 | 716 | 893 | 996 | 1218 | 1209 | 1112 | 1256 | 752 | 814 | 758 | 810 | 991,19 | 1199 | 784 | 0,654 | -0,613 | -10,291 | 7,72E-25 | 1,90E-24 | 0,0012386 | 0,0037478 | 0,0023678 | 0,0023678 | TRUE | 0,0288777 | No |
| orf19.4007 | 259 | 200 | 231 | 164 | 655 | 504 | 642 | 687 | 247 | 211 | 228 | 257 | 525 | 573 | 545 | 559 | 393,14 | 236 | 550 | 2,341 | 1,227 | 14,639 | 1,59E-48 | 5,76E-48 | 1,00E-08 | 0,0062677 | 0,0035693 | 0,0035693 | TRUE | 0,0518657 | No |
| orf19.4009 | 1113 | 983 | 984 | 618 | 1903 | 1227 | 1727 | 1877 | 1061 | 1039 | 972 | 968 | 1525 | 1395 | 1466 | 1526 | 1244,15 | 1010 | 1478 | 1,462 | 0,548 | 9,885 | 4,84E-23 | 1,15E-22 | 0,0010618 | 0,003411 | 0,0020989 | 0,0020989 | TRUE | 0,0161724 | No |
| orf19.401 | 3427 | 2824 | 3035 | 2541 | 7636 | 5945 | 7236 | 7865 | 3267 | 2986 | 2998 | 3981 | 6119 | 6760 | 6143 | 6395 | 4831,13 | 3308 | 6354 | 1,923 | 0,943 | 11,365 | 6,24E-30 | 1,68E-29 | 0,0107483 | 0,0024311 | 0,0063772 | 0,0063772 | TRUE | 0,2231118 | No |
| orf19.4010 | 1985 | 1807 | 1893 | 1426 | 2201 | 1705 | 2073 | 2182 | 1892 | 1911 | 1870 | 2234 | 1764 | 1939 | 1760 | 1774 | 1892,96 | 1977 | 1809 | 0,917 | -0,126 | -1,941 | 0,0522891 | 0,0627636 | 0,0042351 | 0,0029587 | 0,0034981 | 0,0034981 | TRUE | 0,0878656 | No |
| orf19.4011 | 46 | 34 | 53 | 29 | 93 | 63 | 90 | 100 | 44 | 36 | 52 | 45 | 75 | 72 | 76 | 81 | 60,18 | 44 | 76 | 1,714 | 0,777 | 4,138 | 3,50E-05 | 5,06E-05 | 1,00E-08 | 0,0293728 | 0,0155251 | 0,0155251 | TRUE | 0,1331884 | No |
| orf19.4012 | 9852 | 10704 | 11112 | 6808 | 6016 | 4596 | 5739 | 5257 | 9393 | 11318 | 10978 | 10665 | 4820 | 5226 | 4872 | 4274 | 7693,37 | 10588 | 4798 | 0,453 | -1,142 | -16,039 | 6,82E-58 | 2,79E-57 | 0,006303 | 0,0023047 | 0,0047259 | 0,0047259 | TRUE | 0,0710353 | No |
| orf19.4013 | 522 | 460 | 504 | 302 | 1037 | 702 | 935 | 1027 | 498 | 486 | 498 | 473 | 831 | 798 | 794 | 835 | 651,64 | 489 | 814 | 1,664 | 0,735 | 11,974 | 4,84E-33 | 1,37E-32 | 1,00E-08 | 0,0046109 | 0,0019325 | 0,0019325 | TRUE | 0,0043639 | No |
| orf19.4014 | 1087 | 980 | 1063 | 699 | 1246 | 871 | 1214 | 1224 | 1036 | 1036 | 1050 | 1095 | 998 | 990 | 1031 | 995 | 1029,05 | 1054 | 1004 | 0,954 | -0,068 | -1,359 | 0,1742079 | 0,1986456 | 1,00E-08 | 0,0036868 | 0,0014337 | 0,0014337 | TRUE | 0,0068046 | No |
| orf19.4015 | 918 | 826 | 917 | 590 | 1411 | 981 | 1347 | 1445 | 875 | 873 | 906 | 924 | 1131 | 1115 | 1144 | 1175 | 1017,92 | 894 | 1141 | 1,278 | 0,354 | 6,917 | 4,60E-12 | 8,54E-12 | 1,00E-08 | 0,0037043 | 0,0015036 | 0,0015036 | TRUE | 0,0049201 | No |
| orf19.4016 | 6671 | 5375 | 5913 | 4318 | 19085 | 14441 | 18762 | 20526 | 6360 | 5683 | 5841 | 6764 | 15292 | 16421 | 15929 | 16690 | 11122,59 | 6162 | 16083 | 2,611 | 1,385 | 23,886 | 4,27E-126 | 3,61E-125 | 0,0038002 | 0,0022389 | 0,0031093 | 0,0031093 | TRUE | 0,051595 | No |
| orf19.4017 | 417 | 338 | 376 | 267 | 506 | 345 | 520 | 575 | 398 | 357 | 371 | 418 | 405 | 392 | 441 | 468 | 406,43 | 386 | 426 | 1,111 | 0,151 | 1,756 | 0,0790359 | 0,0932559 | 0,0030921 | 0,0061311 | 0,0046385 | 0,0046385 | TRUE | 0,0486872 | No |
| orf19.4018 | 1836 | 1551 | 1788 | 1125 | 2535 | 1808 | 2434 | 2624 | 1750 | 1640 | 1766 | 1762 | 2031 | 2056 | 2066 | 2134 | 1900,78 | 1730 | 2072 | 1,198 | 0,261 | 5,893 | 3,79E-09 | 6,37E-09 | 0,0002743 | 0,0029551 | 0,0013448 | 0,0013448 | TRUE | 0,0150116 | No |
| orf19.4019 | 53 | 48 | 60 | 30 | 153 | 100 | 144 | 170 | 51 | 51 | 59 | 47 | 123 | 114 | 122 | 138 | 88,04 | 52 | 124 | 2,384 | 1,253 | 7,709 | 1,27E-14 | 2,54E-14 | 1,00E-08 | 0,0207403 | 0,0111918 | 0,0111918 | TRUE | 0,0775035 | No |
| orf19.4021 | 1228 | 907 | 1086 | 651 | 2668 | 1688 | 2537 | 2735 | 1171 | 959 | 1073 | 1020 | 2138 | 1919 | 2154 | 2224 | 1582,18 | 1056 | 2109 | 1,996 | 0,997 | 14,166 | 1,48E-45 | 5,10E-45 | 0,0046146 | 0,003129 | 0,0040156 | 0,0040156 | TRUE | 0,0665059 | No |
| orf19.4022 | 1077 | 845 | 930 | 606 | 1728 | 1097 | 1647 | 1753 | 1027 | 893 | 919 | 949 | 1385 | 1247 | 1398 | 1425 | 1155,5 | 947 | 1364 | 1,441 | 0,527 | 8,197 | 2,47E-16 | 5,12E-16 | 0,0026898 | 0,0035122 | 0,0030566 | 0,0030566 | TRUE | 0,0409558 | No |
| orf19.4023 | 1273 | 1005 | 1033 | 756 | 1937 | 1383 | 1837 | 1947 | 1214 | 1063 | 1021 | 1184 | 1552 | 1573 | 1560 | 1583 | 1343,56 | 1120 | 1567 | 1,401 | 0,486 | 7,928 | 2,23E-15 | 4,53E-15 | 0,0024836 | 0,0033133 | 0,0028203 | 0,0028203 | TRUE | 0,0442893 | No |
| orf19.4024 | 1087 | 967 | 1055 | 647 | 1087 | 732 | 1080 | 1166 | 1036 | 1022 | 1042 | 1014 | 871 | 832 | 917 | 948 | 960,37 | 1028 | 892 | 0,868 | -0,204 | -3,573 | 0,0003527 | 0,0004873 | 0,0006262 | 0,0038009 | 0,0020846 | 0,0020846 | TRUE | 0,0220555 | No |

| | | | | | | | | | | | | | | | | | | | | | | | | | | | | | | | |
|---|---|---|---|---|---|---|---|---|---|---|---|---|---|---|---|---|---|---|---|---|---|---|---|---|---|---|---|---|---|---|---|
| orf19.4025 | 4929 | 3940 | 4086 | 2573 | 5568 | 3625 | 5333 | 5566 | 4699 | 4166 | 4037 | 4031 | 4461 | 4122 | 4528 | 4526 | 4321,18 | 4233 | 4409 | 1,041 | 0,058 | 0,999 | 0,3176341 | 0,3509043 | 0,0034071 | 0,0024712 | 0,0030525 | 0,0030525 | TRUE | 0,0676804 | No |
| orf19.4026 | 1179 | 863 | 981 | 521 | 3042 | 1900 | 2807 | 3083 | 1124 | 913 | 969 | 816 | 2437 | 2160 | 2383 | 2507 | 1663,71 | 956 | 2372 | 2,476 | 1,308 | 15,183 | 4,58E-52 | 1,74E-51 | 0,010117 | 0,0030782 | 0,0063501 | 0,0063501 | TRUE | 0,1687586 | No |
| orf19.4028 | 2811 | 2373 | 2570 | 1619 | 3266 | 2225 | 3212 | 3296 | 2680 | 2509 | 2539 | 2536 | 2617 | 2530 | 2727 | 2680 | 2602,28 | 2566 | 2638 | 1,028 | 0,04 | 0,94 | 0,3472114 | 0,3815319 | 0,0005943 | 0,0027222 | 0,0013783 | 0,0013783 | TRUE | 0,0112917 | No |
| orf19.4029 | 17983 | 14420 | 16288 | 10090 | 20739 | 13597 | 20411 | 21344 | 17145 | 15247 | 16091 | 15807 | 16618 | 15461 | 17329 | 17355 | 16381,48 | 16072 | 16691 | 1,038 | 0,054 | 1,066 | 0,2865718 | 0,3187961 | 0,002561 | 0,0021915 | 0,0024506 | 0,0024506 | TRUE | 0,0299147 | No |
| orf19.403 | 26 | 23 | 19 | 17 | 36 | 22 | 30 | 26 | 25 | 24 | 19 | 27 | 29 | 25 | 25 | 21 | 24,37 | 24 | 25 | 1,072 | 0,101 | 0,356 | 0,7215679 | 0,7509908 | 1,00E-08 | 0,0694578 | 0,0353143 | 0,0353143 | TRUE | 0,1199694 | No |
| orf19.4030 | 813 | 672 | 774 | 466 | 1047 | 667 | 976 | 1094 | 775 | 711 | 765 | 730 | 839 | 758 | 829 | 890 | 786,98 | 745 | 829 | 1,113 | 0,155 | 2,353 | 0,0186305 | 0,0231918 | 0,0017409 | 0,0041776 | 0,0028751 | 0,0028751 | TRUE | 0,0360349 | No |
| orf19.4031 | 4283 | 3484 | 3914 | 2647 | 4725 | 3214 | 4536 | 4987 | 4083 | 3684 | 3867 | 4147 | 3786 | 3655 | 3851 | 4055 | 3890,89 | 3945 | 3837 | 0,973 | -0,039 | -0,76 | 0,4475163 | 0,4840823 | 0,002014 | 0,0025133 | 0,0022534 | 0,0022534 | TRUE | 0,0241271 | No |
| orf19.4032 | 5117 | 4407 | 4531 | 2936 | 6202 | 4310 | 5976 | 6240 | 4879 | 4660 | 4476 | 4599 | 4969 | 4901 | 5074 | 5074 | 4828,95 | 4654 | 5004 | 1,075 | 0,105 | 2,744 | 0,0060699 | 0,0078073 | 0,0005213 | 0,0024313 | 0,0011894 | 0,0011894 | TRUE | 0,0133502 | No |
| orf19.4033 | 587 | 459 | 619 | 335 | 2092 | 1416 | 1983 | 2215 | 560 | 485 | 612 | 525 | 1676 | 1610 | 1684 | 1801 | 1119,03 | 546 | 1693 | 3,099 | 1,632 | 22,553 | 1,26E-112 | 9,37E-112 | 0,0043066 | 0,0035585 | 0,0037369 | 0,0037369 | TRUE | 0,0807766 | No |
| orf19.4035 | 9465 | 7588 | 8482 | 5375 | 2572 | 1534 | 2175 | 2656 | 9024 | 8023 | 8379 | 8420 | 2061 | 1744 | 1847 | 2160 | 5207,27 | 8462 | 1953 | 0,231 | -2,114 | -31,245 | 2,61E-214 | 3,80E-213 | 0,0055548 | 0,0024066 | 0,0041017 | 0,0041017 | TRUE | 0,0612899 | No |
| orf19.4036 | 762 | 643 | 732 | 470 | 2523 | 1669 | 2547 | 2574 | 726 | 680 | 723 | 736 | 2022 | 1898 | 2162 | 2093 | 1380,06 | 716 | 2044 | 2,858 | 1,515 | 25,708 | 9,63E-146 | 9,45E-145 | 0,0016548 | 0,003281 | 0,002334 | 0,002334 | TRUE | 0,0266797 | No |
| orf19.4040 | 3025 | 2327 | 2603 | 1809 | 14508 | 9882 | 13641 | 14845 | 2884 | 2461 | 2572 | 2834 | 11625 | 11237 | 11581 | 12070 | 7157,88 | 2688 | 11628 | 4,33 | 2,114 | 38,616 | 0 | 0 | 0,0030123 | 0,0023207 | 0,0026311 | 0,0026311 | TRUE | 0,0392878 | No |
| orf19.4041 | 793 | 618 | 691 | 460 | 4596 | 2965 | 4166 | 4570 | 756 | 653 | 683 | 721 | 3683 | 3371 | 3537 | 3716 | 2139,95 | 703 | 3577 | 5,091 | 2,348 | 40,06 | 0 | 0 | 0,0018531 | 0,0028585 | 0,0023779 | 0,0023779 | TRUE | 0,0341343 | No |
| orf19.4043 | 3422 | 2750 | 3215 | 2187 | 2807 | 2029 | 2728 | 2988 | 3263 | 2908 | 3176 | 3426 | 2249 | 2307 | 2316 | 2430 | 2759,3 | 3193 | 2326 | 0,729 | -0,456 | -8,26 | 1,46E-16 | 3,03E-16 | 0,002332 | 0,0026863 | 0,0025577 | 0,0025577 | TRUE | 0,0437922 | No |
| orf19.4044 | 535 | 430 | 456 | 364 | 1210 | 675 | 1138 | 1211 | 510 | 455 | 450 | 570 | 970 | 768 | 966 | 985 | 709,17 | 496 | 922 | 1,868 | 0,901 | 9,141 | 6,21E-20 | 1,39E-19 | 0,0106245 | 0,0044065 | 0,0077105 | 0,0077105 | TRUE | 0,1463794 | No |
| orf19.4045 | 6662 | 5359 | 5995 | 3656 | 7597 | 4913 | 7462 | 7616 | 6351 | 5666 | 5922 | 5727 | 6087 | 5587 | 6335 | 6193 | 5983,66 | 5916 | 6050 | 1,023 | 0,032 | 0,605 | 0,544882 | 0,5802557 | 0,0023871 | 0,0023657 | 0,0025395 | 0,0025395 | TRUE | 0,0319965 | No |
| orf19.4046 | 1070 | 804 | 824 | 575 | 980 | 576 | 906 | 920 | 1020 | 850 | 814 | 901 | 785 | 655 | 769 | 748 | 817,81 | 896 | 739 | 0,826 | -0,275 | -3,264 | 0,0010975 | 0,0014768 | 0,0064944 | 0,004099 | 0,0056075 | 0,0056075 | TRUE | 0,108158 | No |
| orf19.4048 | 11704 | 9201 | 10285 | 6809 | 8541 | 5457 | 8268 | 8605 | 11158 | 9729 | 10161 | 10667 | 6844 | 6205 | 7019 | 6997 | 8597,45 | 10429 | 6766 | 0,649 | -0,624 | -11,133 | 8,66E-29 | 2,29E-28 | 0,0031961 | 0,0022823 | 0,0028939 | 0,0028939 | TRUE | 0,0375832 | No |
| orf19.405 | 974 | 818 | 911 | 595 | 3121 | 2195 | 2989 | 3291 | 929 | 865 | 900 | 932 | 2501 | 2496 | 2538 | 2676 | 1729,48 | 906 | 2553 | 2,82 | 1,496 | 30,142 | 1,37E-199 | 1,86E-198 | 0,0005944 | 0,0030406 | 0,0015723 | 0,0015723 | TRUE | 0,0130408 | No |
| orf19.4051 | 4386 | 3332 | 3890 | 2249 | 10915 | 7061 | 9941 | 11866 | 4182 | 3523 | 3843 | 3523 | 8746 | 8029 | 8440 | 9648 | 6241,72 | 3768 | 8716 | 2,312 | 1,209 | 17,17 | 4,42E-66 | 2,01E-65 | 0,0063395 | 0,0023543 | 0,0045636 | 0,0045636 | TRUE | 0,0669006 | No |
| orf19.4054 | 560 | 399 | 521 | 310 | 1241 | 797 | 1244 | 1235 | 534 | 422 | 515 | 486 | 994 | 906 | 1056 | 1004 | 739,64 | 489 | 990 | 2,025 | 1,018 | 12,509 | 6,71E-36 | 1,99E-35 | 0,0049481 | 0,0043112 | 0,0047587 | 0,0047587 | TRUE | 0,1043052 | No |
| orf19.4055 | 243 | 172 | 178 | 117 | 655 | 409 | 561 | 607 | 232 | 182 | 176 | 183 | 525 | 465 | 476 | 494 | 341,55 | 193 | 490 | 2,529 | 1,338 | 13,314 | 1,93E-40 | 6,13E-40 | 0,0045652 | 0,0068985 | 0,0059171 | 0,0059171 | TRUE | 0,2121408 | No |
| orf19.4056 | 26344 | 22994 | 25417 | 17877 | 9952 | 7186 | 9438 | 9792 | 25116 | 24313 | 25110 | 28005 | 7974 | 8171 | 8013 | 7962 | 16833,04 | 25636 | 8030 | 0,313 | -1,674 | -35,577 | 3,21E-277 | 6,34E-276 | 0,0019693 | 0,0021888 | 0,0020505 | 0,0020505 | TRUE | 0,0470173 | No |
| orf19.4057 | 14 | 11 | 10 | 7 | 14 | 9 | 11 | 14 | 13 | 12 | 10 | 11 | 11 | 10 | 9 | 11 | 11 | 12 | 10 | 0,919 | -0,122 | -0,299 | 0,7649993 | 0,7911318 | 1,00E-08 | 0,1513596 | 0,0688216 | 0,0688216 | TRUE | 0,0544397 | No |
| orf19.4059 | 2198 | 1932 | 2172 | 1262 | 2391 | 1579 | 2285 | 2240 | 2096 | 2043 | 2146 | 1977 | 1916 | 1795 | 1940 | 1821 | 1966,71 | 2066 | 1868 | 0,904 | -0,146 | -3,091 | 0,0019975 | 0,002646 | 0,0006451 | 0,0029261 | 0,0016231 | 0,0016231 | TRUE | 0,0090949 | No |
| orf19.406 | 2738 | 2393 | 2712 | 1778 | 3257 | 2455 | 2963 | 3271 | 2610 | 2530 | 2679 | 2785 | 2610 | 2792 | 2516 | 2660 | 2647,71 | 2651 | 2644 | 0,998 | -0,003 | -0,07 | 0,9438127 | 0,949759 | 0,0011397 | 0,0027114 | 0,0018496 | 0,0018496 | TRUE | 0,0163997 | No |
| orf19.4060 | 6784 | 5642 | 5990 | 4353 | 16477 | 11683 | 15766 | 17392 | 6468 | 5966 | 5918 | 6819 | 13203 | 13285 | 13385 | 14141 | 9898,01 | 6293 | 13504 | 2,147 | 1,103 | 21,251 | 3,23E-100 | 2,12E-99 | 0,0026943 | 0,0022572 | 0,002463 | 0,002463 | TRUE | 0,0376996 | No |
| orf19.4061 | 612 | 591 | 574 | 425 | 690 | 457 | 602 | 597 | 583 | 625 | 567 | 666 | 553 | 520 | 511 | 485 | 563,78 | 610 | 517 | 0,851 | -0,233 | -3,121 | 0,0018051 | 0,0023992 | 0,0021746 | 0,0050036 | 0,0035675 | 0,0035675 | TRUE | 0,0385999 | No |
| orf19.4062 | 3227 | 2851 | 2980 | 1732 | 3106 | 2095 | 3078 | 3266 | 3077 | 3015 | 2944 | 2713 | 2489 | 2382 | 2613 | 2656 | 2736,02 | 2937 | 2535 | 0,863 | -0,213 | -3,972 | 7,13E-05 | 0,0001017 | 0,0021675 | 0,0026914 | 0,0024041 | 0,0024041 | TRUE | 0,0301847 | No |
| orf19.4063 | 243 | 196 | 228 | 150 | 211 | 158 | 228 | 240 | 232 | 207 | 225 | 235 | 169 | 180 | 194 | 195 | 204,57 | 225 | 184 | 0,822 | -0,283 | -2,729 | 0,0063453 | 0,0081484 | 1,00E-08 | 0,0101173 | 0,0053982 | 0,0053982 | TRUE | 0,0379408 | No |
| orf19.4064 | 715 | 670 | 714 | 415 | 1514 | 1121 | 1385 | 1361 | 682 | 708 | 705 | 650 | 1213 | 1275 | 1176 | 1107 | 939,49 | 686 | 1193 | 1,731 | 0,792 | 12,614 | 1,76E-36 | 5,28E-36 | 0,0016089 | 0,0038389 | 0,0025961 | 0,0025961 | TRUE | 0,0284052 | No |
| orf19.4066 | 1124 | 1439 | 1401 | 873 | 3823 | 2918 | 3445 | 3271 | 1072 | 1522 | 1384 | 1368 | 3063 | 3318 | 2925 | 2660 | 2163,82 | 1336 | 2992 | 2,239 | 1,163 | 11,872 | 1,65E-32 | 4,64E-32 | 0,013542 | 0,0028501 | 0,0086461 | 0,0086461 | TRUE | 0,218241 | No |
| orf19.4067 | 3 | 6 | 3 | 0 | 4 | 1 | 1 | 1 | 3 | 6 | 3 | 0 | 3 | 1 | 1 | 1 | 2,27 | 3 | 2 | 0,482 | -1,053 | -1,04 | 0,2982619 | 0,3310504 | 1,00E-08 | 0,7249033 | 0,5165733 | 0,5165733 | TRUE | 0,1835019 | No |
| orf19.4068 | 561 | 524 | 563 | 362 | 994 | 734 | 1021 | 1154 | 535 | 554 | 556 | 567 | 796 | 835 | 867 | 938 | 706,05 | 553 | 859 | 1,556 | 0,638 | 9,197 | 3,69E-20 | 8,30E-20 | 0,002065 | 0,0044167 | 0,0030884 | 0,0030884 | TRUE | 0,047434 | No |
| orf19.4069 | 55 | 67 | 54 | 46 | 118 | 104 | 126 | 126 | 52 | 71 | 53 | 72 | 95 | 118 | 107 | 102 | 83,86 | 62 | 106 | 1,712 | 0,776 | 4,574 | 4,78E-06 | 7,20E-06 | 0,0025821 | 0,0216692 | 0,0143075 | 0,0143075 | TRUE | 0,0987594 | No |
| orf19.407 | 4134 | 3390 | 3744 | 2269 | 5636 | 3483 | 5332 | 5508 | 3941 | 3584 | 3699 | 3555 | 4516 | 3960 | 4527 | 4479 | 4032,61 | 3695 | 4370 | 1,183 | 0,242 | 4,341 | 1,42E-05 | 2,09E-05 | 0,0027646 | 0,0024984 | 0,0027373 | 0,0027373 | TRUE | 0,0476296 | No |
| orf19.4070 | 258 | 234 | 222 | 146 | 392 | 306 | 373 | 501 | 246 | 247 | 219 | 229 | 314 | 348 | 317 | 407 | 290,94 | 235 | 346 | 1,471 | 0,557 | 5,143 | 2,70E-07 | 4,28E-07 | 0,0074395 | 0,0077347 | 0,0075814 | 0,0075814 | TRUE | 0,168151 | No |
| orf19.4072 | 3314 | 2573 | 2971 | 2006 | 3910 | 2712 | 3933 | 4326 | 3160 | 2721 | 2935 | 3143 | 3133 | 3084 | 3339 | 3517 | 3128,88 | 2990 | 3268 | 1,094 | 0,13 | 2,087 | 0,0369063 | 0,0448656 | 0,0038161 | 0,002616 | 0,0033925 | 0,0033925 | TRUE | 0,0446357 | No |
| orf19.4076 | 8342 | 7612 | 8011 | 5007 | 34971 | 24856 | 32575 | 36455 | 7953 | 8049 | 7914 | 7844 | 28021 | 28264 | 27656 | 29641 | 18167,74 | 7940 | 28396 | 3,576 | 1,838 | 55,747 | 0 | 0 | 0,000417 | 0,0021817 | 0,0009576 | 0,0009576 | TRUE | 0,0107315 | No |
| orf19.4077 | 12385 | 11968 | 12832 | 7787 | 10960 | 7692 | 10135 | 10472 | 11808 | 12655 | 12677 | 12199 | 8782 | 8746 | 8605 | 8515 | 10498,21 | 12335 | 8662 | 0,702 | -0,51 | -14,602 | 2,75E-48 | 9,89E-48 | 0,0005371 | 0,0022477 | 0,0010748 | 0,0010748 | TRUE | 0,0103547 | No |
| orf19.4078 | 3819 | 3672 | 3739 | 2374 | 3125 | 2244 | 2900 | 2910 | 3641 | 3883 | 3694 | 3719 | 2504 | 2552 | 2462 | 2366 | 3102,53 | 3734 | 2471 | 0,661 | -0,596 | -14,442 | 2,81E-47 | 9,97E-47 | 0,0005814 | 0,0026205 | 0,0013062 | 0,0013062 | TRUE | 0,0099856 | No |
| orf19.4079 | 1517 | 1427 | 1537 | 1010 | 1891 | 1289 | 1796 | 1755 | 1446 | 1509 | 1518 | 1582 | 1515 | 1466 | 1525 | 1427 | 1498,56 | 1514 | 1483 | 0,982 | -0,027 | -0,549 | 0,5827434 | 0,6168689 | 0,0004622 | 0,0031869 | 0,0015965 | 0,0015965 | TRUE | 0,010978 | No |
| orf19.4080 | 781 | 859 | 993 | 523 | 838 | 623 | 671 | 637 | 745 | 908 | 981 | 819 | 671 | 708 | 570 | 518 | 740,08 | 863 | 617 | 0,713 | -0,488 | -4,418 | 9,98E-06 | 1,48E-05 | 0,0155181 | 0,0043098 | 0,0103285 | 0,0103285 | TRUE | 0,1388639 | No |
| orf19.4081 | 3 | 0 | 0 | 0 | 0 | 4 | 1 | 3 | 3 | 0 | 0 | 0 | 0 | 5 | 1 | 2 | 1,34 | 1 | 2 | 2,469 | 1,304 | 0,82 | 0,4122152 | 0,4481761 | 2,0888332 | 1,2300479 | 1,4735665 | 1,4735665 | TRUE | 0,8292292 | No |
| orf19.4082 | 7723 | 6095 | 6396 | 4586 | 6664 | 4688 | 6392 | 6991 | 7363 | 6445 | 6319 | 7184 | 5340 | 5331 | 5427 | 5684 | 6136,51 | 6828 | 5446 | 0,798 | -0,326 | -5,772 | 7,82E-09 | 1,31E-08 | 0,0031696 | 0,0023588 | 0,0028956 | 0,0028956 | TRUE | 0,0348156 | No |
| orf19.4084 | 4276 | 3680 | 4179 | 3006 | 3615 | 2668 | 3375 | 3666 | 4077 | 3891 | 4128 | 4709 | 2897 | 3034 | 2865 | 2981 | 3572,73 | 4201 | 2944 | 0,701 | -0,512 | -8,682 | 3,88E-18 | 8,36E-18 | 0,0034974 | 0,0025508 | 0,0030493 | 0,0030493 | TRUE | 0,078097 | No |
| orf19.4085 | 345 | 379 | 342 | 186 | 338 | 274 | 300 | 278 | 329 | 401 | 338 | 291 | 271 | 312 | 255 | 226 | 302,75 | 340 | 266 | 0,777 | -0,364 | -3,036 | 0,0024 | 0,0031661 | 0,0134762 | 0,0075145 | 0,0104762 | 0,0104762 | TRUE | 0,1631907 | No |
| orf19.4086 | 716 | 613 | 557 | 411 | 551 | 388 | 495 | 597 | 683 | 648 | 550 | 644 | 442 | 441 | 420 | 485 | 539,16 | 631 | 447 | 0,709 | -0,496 | -5,916 | 3,29E-09 | 5,56E-09 | 0,0046135 | 0,0051366 | 0,0048616 | 0,0048616 | TRUE | 0,0923762 | No |
| orf19.4088 | 852 | 768 | 777 | 548 | 1587 | 1209 | 1604 | 1707 | 812 | 812 | 768 | 858 | 1272 | 1375 | 1362 | 1388 | 1080,82 | 812 | 1349 | 1,664 | 0,734 | 12,952 | 2,28E-38 | 7,03E-38 | 0,0008412 | 0,0036104 | 0,0020657 | 0,0020657 | TRUE | 0,0185302 | No |
| orf19.4089 | 4572 | 3903 | 4305 | 2609 | 5777 | 3631 | 5599 | 5791 | 4359 | 4127 | 4253 | 4087 | 4629 | 4129 | 4754 | 4709 | 4380,72 | 4206 | 4555 | 1,083 | 0,115 | 2,222 | 0,0262941 | 0,0323593 | 0,0021376 | 0,0024661 | 0,0023455 | 0,0023455 | TRUE | 0,0472895 | No |
| orf19.409 | 53612 | 43511 | 47000 | 30771 | 14642 | 10263 | 13905 | 15442 | 51113 | 46007 | 46432 | 48204 | 11732 | 11670 | 11805 | 12556 | 29939,98 | 47939 | 11941 | 0,249 | -2,005 | -45,022 | 0 | 0 | 0,0017017 | 0,0021461 | 0,0018565 | 0,0018565 | TRUE | 0,0244233 | No |
| orf19.4090 | 132 | 120 | 135 | 102 | 329 | 227 | 308 | 351 | 126 | 127 | 133 | 160 | 264 | 258 | 261 | 285 | 201,81 | 136 | 267 | 1,982 | 0,987 | 8,984 | 2,60E-19 | 5,76E-19 | 1,00E-08 | 0,010227 | 0,0057954 | 0,0057954 | TRUE | 0,103666 | No |
| orf19.4090.1 | 523 | 385 | 422 | 270 | 964 | 600 | 879 | 934 | 499 | 407 | 417 | 423 | 772 | 682 | 746 | 759 | 588,24 | 436 | 740 | 1,695 | 0,761 | 9,462 | 3,02E-21 | 6,93E-21 | 0,0036641 | 0,0048825 | 0,0043286 | 0,0043286 | TRUE | 0,1139926 | No |
| orf19.4091 | 767 | 631 | 754 | 427 | 658 | 411 | 696 | 767 | 731 | 667 | 745 | 669 | 527 | 467 | 591 | 624 | 627,67 | 703 | 552 | 0,786 | -0,347 | -3,82 | 0,0001337 | 0,0001882 | 0,0074114 | 0,0047071 | 0,0063015 | 0,0063015 | TRUE | 0,1177034 | No |
| orf19.4092 | 846 | 754 | 743 | 482 | 943 | 650 | 892 | 954 | 807 | 797 | 734 | 755 | 756 | 739 | 757 | 776 | 765,08 | 773 | 757 | 0,979 | -0,031 | -0,533 | 0,59426 | 0,6285385 | 1,00E-08 | 0,0042373 | 0,0019934 | 0,0019934 | TRUE | 0,0157162 | No |
| orf19.4093 | 18677 | 16417 | 18002 | 10855 | 33364 | 21965 | 32511 | 33680 | 17806 | 17359 | 17784 | 17005 | 26734 | 24976 | 27602 | 27385 | 22081,39 | 17488 | 26674 | 1,525 | 0,609 | 15,103 | 1,54E-51 | 5,78E-51 | 0,0011723 | 0,0021656 | 0,0015127 | 0,0015127 | TRUE | 0,0223503 | No |
| orf19.4094 | 60 | 65 | 59 | 55 | 11 | 11 | 16 | 14 | 57 | 69 | 58 | 86 | 9 | 13 | 14 | 11 | 39,58 | 68 | 12 | 0,172 | -2,538 | -8,941 | 3,86E-19 | 8,52E-19 | 1,00E-08 | 0,0435707 | 0,0304806 | 0,0304806 | TRUE | 0,2387387 | No |
| orf19.4095 | 6 | 7 | 1 | 7 | 3 | 7 | 6 | 7 | 6 | 7 | 1 | 11 | 2 | 8 | 5 | 6 | 5,78 | 6 | 5 | 0,861 | -0,215 | -0,321 | 0,7478771 | 0,7760943 | 0,1958067 | 0,2862509 | 0,2514116 | 0,2514116 | TRUE | 0,3075518 | No |
| orf19.4096 | 1983 | 1653 | 1886 | 1175 | 2028 | 1325 | 1861 | 2089 | 1891 | 1748 | 1863 | 1841 | 1625 | 1507 | 1580 | 1699 | 1719,06 | 1836 | 1603 | 0,874 | -0,194 | -3,758 | 0,0001711 | 0,0002394 | 0,0012526 | 0,0030464 | 0,0019841 | 0,0019841 | TRUE | 0,019534 | No |
| orf19.4097 | 939 | 961 | 942 | 579 | 1424 | 1084 | 1388 | 1571 | 895 | 1016 | 931 | 907 | 1141 | 1233 | 1178 | 1277 | 1072,3 | 937 | 1207 | 1,287 | 0,364 | 5,875 | 4,24E-09 | 7,12E-09 | 0,0020029 | 0,0036225 | 0,002715 | 0,002715 | TRUE | 0,0388477 | No |
| orf19.4099 | 14727 | 12743 | 13838 | 8735 | 42608 | 29627 | 39945 | 43764 | 14041 | 13474 | 13671 | 13684 | 34141 | 33689 | 33913 | 35584 | 24024,47 | 13718 | 34332 | 2,503 | 1,324 | 42,661 | 0 | 0 | 0,0003902 | 0,0021596 | 0,0008703 | 0,0008703 | TRUE | 0,0074047 | No |
| orf19.4101 | 265 | 169 | 207 | 134 | 341 | 223 | 301 | 324 | 253 | 179 | 204 | 210 | 273 | 254 | 256 | 263 | 236,44 | 212 | 262 | 1,235 | 0,305 | 2,727 | 0,0063947 | 0,0082085 | 0,0059464 | 0,0090354 | 0,0076374 | 0,0076374 | TRUE | 0,2057562 | No |
| orf19.4102 | 5541 | 4865 | 5162 | 3595 | 5370 | 3410 | 5088 | 5247 | 5283 | 5144 | 5100 | 5632 | 4303 | 3877 | 4320 | 4266 | 4740,56 | 5290 | 4192 | 0,793 | -0,334 | -6,688 | 2,26E-11 | 4,12E-11 | 0,0020157 | 0,0024376 | 0,0021795 | 0,0021795 | TRUE | 0,0302279 | No |
| orf19.410.3 | 1153 | 931 | 1045 | 717 | 1019 | 807 | 917 | 1048 | 1099 | 984 | 1032 | 1123 | 816 | 918 | 779 | 852 | 950,5 | 1060 | 841 | 0,793 | -0,334 | -4,905 | 9,34E-07 | 1,45E-06 | 0,0028342 | 0,0038187 | 0,0033835 | 0,0033835 | TRUE | 0,0417163 | No |
| orf19.4104 | 148 | 131 | 137 | 74 | 160 | 96 | 132 | 165 | 141 | 139 | 135 | 116 | 128 | 109 | 112 | 134 | 126,81 | 133 | 121 | 0,908 | -0,139 | -1,051 | 0,2934549 | 0,3260553 | 1,00E-08 | 0,015039 | 0,0088825 | 0,0088825 | TRUE | 0,0568647 | No |
| orf19.4105 | 769 | 666 | 698 | 442 | 698 | 499 | 734 | 724 | 733 | 704 | 690 | 692 | 559 | 567 | 623 | 589 | 644,74 | 705 | 584 | 0,829 | -0,27 | -4,236 | 2,28E-05 | 3,32E-05 | 1,00E-08 | 0,0046379 | 0,0023537 | 0,0023537 | TRUE | 0,0235452 | No |
| orf19.4106 | 1269 | 1164 | 1303 | 794 | 1600 | 1147 | 1502 | 1686 | 1210 | 1231 | 1287 | 1244 | 1282 | 1304 | 1275 | 1371 | 1275,51 | 1243 | 1308 | 1,053 | 0,074 | 1,489 | 0,136375 | 0,1569924 | 1,00E-08 | 0,0033785 | 0,0015777 | 0,0015777 | TRUE | 0,0131291 | No |
| orf19.4107 | 1744 | 1442 | 1621 | 952 | 2302 | 1527 | 2273 | 2334 | 1663 | 1525 | 1601 | 1491 | 1845 | 1736 | 1930 | 1898 | 1711,07 | 1570 | 1852 | 1,179 | 0,237 | 4,4 | 1,08E-05 | 1,60E-05 | 0,0014683 | 0,0030509 | 0,0021908 | 0,0021908 | TRUE | 0,020314 | No |
| orf19.4109 | 1223 | 1071 | 1108 | 754 | 1460 | 1047 | 1408 | 1583 | 1166 | 1132 | 1095 | 1181 | 1170 | 1191 | 1195 | 1287 | 1177,14 | 1144 | 1211 | 1,06 | 0,085 | 1,573 | 0,1156931 | 0,1340057 | 0,0007764 | 0,0034861 | 0,0019225 | 0,0019225 | TRUE | 0,0224054 | No |
| orf19.411 | 19940 | 17605 | 18706 | 10985 | 15577 | 9946 | 14244 | 14380 | 19011 | 18615 | 18480 | 17209 | 12481 | 11309 | 12093 | 11692 | 15111,29 | 18329 | 11894 | 0,649 | -0,624 | -13,792 | 2,84E-43 | 9,48E-43 | 0,0017164 | 0,0021999 | 0,0018978 | 0,0018978 | TRUE | 0,0204488 | No |
| orf19.4110 | 1805 | 1613 | 1806 | 1209 | 2673 | 1854 | 2545 | 2810 | 1721 | 1706 | 1784 | 1894 | 2142 | 2108 | 2161 | 2285 | 1975 | 1776 | 2174 | 1,227 | 0,295 | 5,954 | 2,61E-09 | 4,43E-09 | 0,0011081 | 0,0029226 | 0,0018353 | 0,0018353 | TRUE | 0,0217962 | No |
| orf19.4112 | 1806 | 1475 | 1580 | 1034 | 3200 | 2173 | 3073 | 3328 | 1722 | 1560 | 1561 | 1620 | 2564 | 2471 | 2609 | 2706 | 2101,51 | 1616 | 2588 | 1,602 | 0,68 | 13,452 | 2,98E-41 | 9,62E-41 | 0,0011557 | 0,0028726 | 0,0019336 | 0,0019336 | TRUE | 0,0252578 | No |
| orf19.4114 | 2387 | 2152 | 2370 | 1423 | 7976 | 5739 | 7633 | 8259 | 2276 | 2275 | 2341 | 2229 | 6391 | 6526 | 6480 | 6715 | 4404,27 | 2280 | 6528 | 2,86 | 1,516 | 41,932 | 0 | 0 | 0,0002399 | 0,0024641 | 0,0009411 | 0,0009411 | TRUE | 0,0046478 | No |

| | | | | | | | | | | | | | | | | | | | | | | | | | | | | | | | |
|---|---|---|---|---|---|---|---|---|---|---|---|---|---|---|---|---|---|---|---|---|---|---|---|---|---|---|---|---|---|---|---|
| orf19.4115 | 16 | 8 | 8 | 4 | 25 | 24 | 31 | 30 | 15 | 8 | 8 | 6 | 20 | 27 | 26 | 24 | 16,99 | 9 | 24 | 2,503 | 1,324 | 3,534 | 0,0004087 | 0,0005624 | 1,00E-08 | 0,0987333 | 0,0595592 | 0,0595592 | TRUE | 0,6072513 | No |
| orf19.4116 | 716 | 589 | 641 | 421 | 580 | 428 | 548 | 625 | 683 | 623 | 633 | 660 | 465 | 487 | 465 | 508 | 565,38 | 650 | 481 | 0,741 | -0,433 | -6,403 | 1,52E-10 | 2,71E-10 | 1,00E-08 | 0,0049954 | 0,0026003 | 0,0026003 | TRUE | 0,018042 | No |
| orf19.4117 | 1557 | 1450 | 1581 | 898 | 809 | 658 | 744 | 789 | 1484 | 1533 | 1562 | 1407 | 648 | 748 | 632 | 642 | 1081,99 | 1496 | 668 | 0,444 | -1,171 | -17,911 | 9,65E-72 | 4,75E-71 | 0,0025314 | 0,0036088 | 0,0030607 | 0,0030607 | TRUE | 0,0724855 | No |
| orf19.4118 | 28153 | 26406 | 27868 | 17511 | 48122 | 35243 | 44836 | 47657 | 26841 | 27921 | 27531 | 27432 | 38559 | 40074 | 38066 | 38750 | 33146,65 | 27431 | 38862 | 1,417 | 0,502 | 17,259 | 9,58E-67 | 4,40E-66 | 0,0003349 | 0,0021408 | 0,0007821 | 0,0007821 | TRUE | 0,0053617 | No |
| orf19.4119 | 5369 | 5083 | 5394 | 3191 | 3431 | 2315 | 3230 | 3387 | 5119 | 5375 | 5329 | 4999 | 2749 | 2632 | 2742 | 2754 | 3962,34 | 5206 | 2719 | 0,522 | -0,937 | -23,84 | 1,28E-125 | 1,08E-124 | 0,0004345 | 0,0025057 | 0,0012102 | 0,0012102 | TRUE | 0,0081472 | No |
| orf19.412 | 2540 | 2879 | 2684 | 1470 | 4049 | 3166 | 3469 | 3165 | 2422 | 3044 | 2652 | 2303 | 3244 | 3600 | 2945 | 2573 | 2847,89 | 2605 | 3090 | 1,185 | 0,245 | 2,36 | 0,0182796 | 0,0227638 | 0,0173623 | 0,0026678 | 0,0100251 | 0,0100251 | TRUE | 0,156029 | No |
| orf19.4120 | 6174 | 5406 | 5653 | 3595 | 5730 | 3935 | 5646 | 6118 | 5886 | 5716 | 5585 | 5632 | 4591 | 4474 | 4793 | 4975 | 5206,56 | 5705 | 4708 | 0,826 | -0,277 | -6,407 | 1,48E-10 | 2,64E-10 | 0,0011134 | 0,0024066 | 0,0015954 | 0,0015954 | TRUE | 0,0176746 | No |
| orf19.4121 | 1907 | 1599 | 1766 | 1102 | 2376 | 1718 | 2273 | 2382 | 1818 | 1691 | 1745 | 1726 | 1904 | 1954 | 1930 | 1937 | 1837,97 | 1745 | 1931 | 1,105 | 0,145 | 3,408 | 0,000654 | 0,0008932 | 1,00E-08 | 0,0029846 | 0,0011769 | 0,0011769 | TRUE | 0,010182 | No |
| orf19.4122 | 1979 | 1839 | 1890 | 1267 | 4048 | 2716 | 3859 | 4038 | 1887 | 1945 | 1867 | 1985 | 3244 | 3088 | 3276 | 3283 | 2571,83 | 1921 | 3223 | 1,681 | 0,75 | 17,958 | 4,15E-72 | 2,05E-71 | 0,0003382 | 0,0027297 | 0,0012418 | 0,0012418 | TRUE | 0,0091294 | No |
| orf19.4123 | 3394 | 2973 | 3285 | 2003 | 4157 | 2777 | 3782 | 3934 | 3236 | 3144 | 3245 | 3138 | 3331 | 3158 | 3211 | 3199 | 3207,58 | 3191 | 3225 | 1,01 | 0,015 | 0,409 | 0,682892 | 0,7143498 | 0,0001543 | 0,0026032 | 0,0009788 | 0,0009788 | TRUE | 0,0060774 | No |
| orf19.4125 | 3310 | 3100 | 3241 | 2025 | 3137 | 2270 | 2924 | 3043 | 3156 | 3278 | 3202 | 3172 | 2514 | 2581 | 2482 | 2474 | 2857,39 | 3202 | 2513 | 0,784 | -0,351 | -9,615 | 6,93E-22 | 1,61E-21 | 1,00E-08 | 0,0026659 | 0,0009246 | 0,0009246 | TRUE | 0,0037757 | No |
| orf19.4127 | 10413 | 8902 | 9841 | 5774 | 1636 | 981 | 1578 | 1493 | 9928 | 9413 | 9722 | 9045 | 1311 | 1115 | 1340 | 1214 | 5385,96 | 9527 | 1245 | 0,131 | -2,934 | -49,947 | 0 | 0 | 0,0035478 | 0,0023961 | 0,0028963 | 0,0028963 | TRUE | 0,0555397 | No |
| orf19.4128 | 1045 | 955 | 1083 | 594 | 4492 | 3354 | 4173 | 4365 | 996 | 1010 | 1070 | 931 | 3599 | 3814 | 3543 | 3549 | 2313,96 | 1002 | 3626 | 3,608 | 1,851 | 35,768 | 3,53E-280 | 7,17E-279 | 0,0011561 | 0,0028008 | 0,001891 | 0,001891 | TRUE | 0,0246882 | No |
| orf19.413 | 4678 | 4447 | 4784 | 2895 | 805 | 603 | 735 | 817 | 4460 | 4702 | 4726 | 4535 | 645 | 686 | 624 | 664 | 2630,3 | 4606 | 655 | 0,142 | -2,817 | -58,631 | 0 | 0 | 0,0005205 | 0,0027155 | 0,0014143 | 0,0014143 | TRUE | 0,0114827 | No |
| orf19.4131 | 975 | 1049 | 1024 | 602 | 1836 | 1382 | 1607 | 1710 | 930 | 1109 | 1012 | 943 | 1471 | 1571 | 1364 | 1390 | 1223,84 | 998 | 1449 | 1,449 | 0,535 | 7,561 | 4,00E-14 | 7,87E-14 | 0,0043282 | 0,0034329 | 0,0039363 | 0,0039363 | TRUE | 0,0669997 | No |
| orf19.413.1 | 4665 | 4937 | 4999 | 2167 | 2967 | 2017 | 2620 | 2429 | 4448 | 5220 | 4939 | 3395 | 2377 | 2294 | 2224 | 1975 | 3358,92 | 4500 | 2218 | 0,492 | -1,022 | -9,313 | 1,24E-20 | 2,82E-20 | 0,0185357 | 0,0025801 | 0,0112432 | 0,0112432 | TRUE | 0,3291477 | No |
| orf19.4132 | 2179 | 1969 | 1975 | 1178 | 2665 | 1681 | 2329 | 2459 | 2077 | 2082 | 1951 | 1845 | 2135 | 1911 | 1977 | 1999 | 1997,43 | 1989 | 2006 | 1,008 | 0,011 | 0,197 | 0,8435718 | 0,863325 | 0,0018393 | 0,0029133 | 0,0024362 | 0,0024362 | TRUE | 0,0263574 | No |
| orf19.4133 | 2890 | 2478 | 2771 | 1673 | 3334 | 2264 | 3402 | 3482 | 2755 | 2620 | 2737 | 2621 | 2671 | 2574 | 2888 | 2831 | 2712,39 | 2683 | 2741 | 1,022 | 0,031 | 0,636 | 0,5248507 | 0,5610776 | 0,0012635 | 0,0026966 | 0,0018923 | 0,0018923 | TRUE | 0,019547 | No |
| orf19.4134 | 912 | 880 | 937 | 587 | 897 | 629 | 756 | 795 | 869 | 930 | 926 | 920 | 719 | 715 | 642 | 646 | 795,93 | 911 | 680 | 0,746 | -0,422 | -6,67 | 2,55E-11 | 4,65E-11 | 0,001187 | 0,0041542 | 0,0025736 | 0,0025736 | TRUE | 0,0188618 | No |
| orf19.4135 | 946 | 863 | 814 | 562 | 898 | 649 | 828 | 846 | 902 | 913 | 804 | 880 | 720 | 738 | 703 | 688 | 793,42 | 875 | 712 | 0,814 | -0,298 | -4,793 | 1,64E-06 | 2,52E-06 | 0,0009488 | 0,0041607 | 0,0024351 | 0,0024351 | TRUE | 0,0374506 | No |
| orf19.4136 | 385 | 397 | 415 | 231 | 568 | 426 | 572 | 586 | 367 | 420 | 410 | 362 | 455 | 484 | 486 | 476 | 432,54 | 390 | 475 | 1,216 | 0,282 | 3,651 | 0,0002609 | 0,0003625 | 0,0001849 | 0,0058872 | 0,0033445 | 0,0033445 | TRUE | 0,0298056 | No |
| orf19.414 | 6693 | 5998 | 6511 | 3678 | 6312 | 4294 | 6062 | 6498 | 6381 | 6342 | 6432 | 5762 | 5058 | 4883 | 5147 | 5283 | 5660,95 | 6229 | 5093 | 0,817 | -0,291 | -6,292 | 3,13E-10 | 5,49E-10 | 0,0015754 | 0,0023813 | 0,0018817 | 0,0018817 | TRUE | 0,0300503 | No |
| orf19.4142 | 681 | 597 | 578 | 403 | 947 | 638 | 892 | 930 | 649 | 631 | 571 | 631 | 759 | 725 | 757 | 756 | 685,07 | 620 | 749 | 1,209 | 0,274 | 4,354 | 1,34E-05 | 1,97E-05 | 1,00E-08 | 0,004488 | 0,002318 | 0,002318 | TRUE | 0,0367582 | No |
| orf19.4143 | 1748 | 1455 | 1615 | 981 | 1761 | 1065 | 1636 | 1657 | 1667 | 1538 | 1595 | 1537 | 1411 | 1211 | 1389 | 1347 | 1461,94 | 1584 | 1340 | 0,846 | -0,241 | -4,083 | 4,45E-05 | 6,40E-05 | 0,0020699 | 0,0032144 | 0,0026603 | 0,0026603 | TRUE | 0,0473987 | No |
| orf19.4144 | 3381 | 2954 | 3185 | 2139 | 1820 | 1311 | 1682 | 1780 | 3223 | 3123 | 3146 | 3351 | 1458 | 1491 | 1428 | 1447 | 2333,57 | 3211 | 1456 | 0,454 | -1,14 | -27,213 | 4,58E-163 | 5,08E-162 | 0,0002177 | 0,0027949 | 0,0012041 | 0,0012041 | TRUE | 0,0095524 | No |
| orf19.4145 | 2624 | 2462 | 2615 | 1556 | 2575 | 1833 | 2258 | 2529 | 2502 | 2603 | 2583 | 2438 | 2063 | 2084 | 1917 | 2056 | 2280,85 | 2532 | 2030 | 0,801 | -0,32 | -7,089 | 1,35E-12 | 2,54E-12 | 0,0007008 | 0,0028111 | 0,0015164 | 0,0015164 | TRUE | 0,0172078 | No |
| orf19.4146 | 1416 | 1204 | 1192 | 617 | 1759 | 1071 | 1625 | 1760 | 1350 | 1273 | 1178 | 967 | 1409 | 1218 | 1380 | 1431 | 1275,64 | 1192 | 1360 | 1,139 | 0,187 | 2,013 | 0,044132 | 0,0533245 | 0,0111716 | 0,0033784 | 0,0075056 | 0,0075056 | TRUE | 0,183409 | No |
| orf19.4147 | 2664 | 2430 | 2554 | 1686 | 5278 | 3909 | 4879 | 5482 | 2540 | 2569 | 2523 | 2641 | 4229 | 4445 | 4142 | 4457 | 3443,4 | 2568 | 4318 | 1,682 | 0,75 | 18,206 | 4,59E-74 | 2,33E-73 | 0,0006651 | 0,0025681 | 0,0013089 | 0,0013089 | TRUE | 0,0091801 | No |
| orf19.4148 | 557 | 522 | 558 | 327 | 448 | 289 | 388 | 442 | 531 | 552 | 551 | 512 | 359 | 329 | 329 | 359 | 440,36 | 536 | 344 | 0,641 | -0,641 | -8,713 | 2,95E-18 | 6,37E-18 | 1,00E-08 | 0,0058198 | 0,0028661 | 0,0028661 | TRUE | 0,0108435 | No |
| orf19.4149 | 17 | 16 | 11 | 14 | 24 | 22 | 25 | 10 | 16 | 17 | 11 | 22 | 19 | 25 | 21 | 8 | 17,44 | 16 | 18 | 1,123 | 0,167 | 0,423 | 0,672055 | 0,7039368 | 0,0831105 | 0,0962326 | 0,0905755 | 0,0905755 | TRUE | 0,8468877 | No |
| orf19.4149.1 | 14018 | 11936 | 12490 | 8860 | 44696 | 31907 | 41929 | 45241 | 13365 | 12621 | 12339 | 13880 | 35814 | 36281 | 35598 | 36785 | 24585,2 | 13051 | 36120 | 2,769 | 1,469 | 34,704 | 6,75E-264 | 1,23E-262 | 0,0014826 | 0,0021581 | 0,0016659 | 0,0016659 | TRUE | 0,0219172 | No |
| orf19.415 | 1405 | 1155 | 1247 | 853 | 2750 | 1810 | 2587 | 2778 | 1340 | 1221 | 1232 | 1336 | 2203 | 2058 | 2196 | 2259 | 1730,72 | 1282 | 2179 | 1,703 | 0,768 | 14,55 | 5,87E-48 | 2,11E-47 | 0,0012039 | 0,00304 | 0,0020321 | 0,0020321 | TRUE | 0,0161747 | No |
| orf19.4150 | 7482 | 6617 | 6985 | 4566 | 6116 | 4376 | 5857 | 6056 | 7133 | 6997 | 6901 | 7153 | 4901 | 4976 | 4973 | 4924 | 5994,56 | 7046 | 4944 | 0,702 | -0,511 | -17,136 | 7,99E-66 | 3,63E-65 | 1,00E-08 | 0,0023652 | 0,0006837 | 0,0006837 | TRUE | 0,0023809 | No |
| orf19.4151 | 111 | 107 | 103 | 66 | 95 | 77 | 107 | 99 | 106 | 113 | 102 | 103 | 76 | 88 | 91 | 80 | 94,89 | 106 | 84 | 0,787 | -0,345 | -2,389 | 0,0168747 | 0,0210761 | 1,00E-08 | 0,0193943 | 0,009487 | 0,009487 | TRUE | 0,0386432 | No |
| orf19.4152 | 59685 | 52446 | 56865 | 37633 | 118070 | 85210 | 109486 | 127257 | 56903 | 55455 | 56177 | 58954 | 94606 | 96891 | 92953 | 103472 | 76926,53 | 56872 | 96980 | 1,705 | 0,77 | 18,451 | 5,11E-76 | 2,65E-75 | 0,0014674 | 0,0021126 | 0,001659 | 0,001659 | TRUE | 0,0249059 | No |
| orf19.4153 | 898 | 1074 | 1080 | 581 | 614 | 552 | 528 | 537 | 856 | 1136 | 1067 | 910 | 492 | 628 | 448 | 437 | 746,68 | 992 | 501 | 0,503 | -0,99 | -8,224 | 1,96E-16 | 4,08E-16 | 0,0219962 | 0,0042902 | 0,0124584 | 0,0124584 | TRUE | 0,325825 | No |
| orf19.4157 | 403 | 331 | 348 | 237 | 1791 | 1300 | 1620 | 1790 | 384 | 350 | 344 | 371 | 1435 | 1478 | 1375 | 1455 | 899,17 | 362 | 1436 | 3,965 | 1,987 | 31,194 | 1,30E-213 | 1,89E-212 | 1,00E-08 | 0,0039173 | 0,0020519 | 0,0020519 | TRUE | 0,0229652 | No |
| orf19.4159 | 1746 | 1500 | 1655 | 1075 | 2097 | 1453 | 2004 | 2124 | 1665 | 1586 | 1635 | 1684 | 1680 | 1652 | 1701 | 1727 | 1666,32 | 1642 | 1690 | 1,031 | 0,043 | 1,003 | 0,3160218 | 0,3493046 | 1,00E-08 | 0,0030766 | 0,0011942 | 0,0011942 | TRUE | 0,0063646 | No |
| orf19.416 | 126 | 104 | 134 | 80 | 264 | 172 | 235 | 215 | 120 | 110 | 132 | 125 | 212 | 196 | 200 | 175 | 158,66 | 122 | 196 | 1,603 | 0,681 | 5,602 | 2,12E-08 | 3,50E-08 | 1,00E-08 | 0,0124402 | 0,0073487 | 0,0073487 | TRUE | 0,058989 | No |
| orf19.4160 | 2536 | 2238 | 2407 | 1712 | 2621 | 1993 | 2604 | 2690 | 2418 | 2366 | 2378 | 2682 | 2100 | 2266 | 2211 | 2187 | 2326,05 | 2461 | 2191 | 0,892 | -0,166 | -3,214 | 0,0013074 | 0,0017475 | 0,001753 | 0,0027972 | 0,0021167 | 0,0021167 | TRUE | 0,0413574 | No |
| orf19.4161 | 727 | 698 | 682 | 460 | 1024 | 652 | 1012 | 989 | 693 | 738 | 674 | 721 | 821 | 741 | 859 | 804 | 756,34 | 706 | 806 | 1,145 | 0,195 | 2,991 | 0,0027776 | 0,0036499 | 0,0013854 | 0,0042621 | 0,0027437 | 0,0027437 | TRUE | 0,032059 | No |
| orf19.4162 | 295 | 307 | 342 | 185 | 450 | 325 | 367 | 427 | 281 | 325 | 338 | 290 | 361 | 370 | 312 | 347 | 327,8 | 308 | 348 | 1,122 | 0,166 | 1,762 | 0,0780587 | 0,092171 | 0,0036135 | 0,0071001 | 0,0054269 | 0,0054269 | TRUE | 0,0553159 | No |
| orf19.4163 | 1234 | 1208 | 1325 | 830 | 2605 | 1975 | 2376 | 2278 | 1176 | 1277 | 1309 | 1300 | 2087 | 2246 | 2017 | 1852 | 1658,19 | 1266 | 2050 | 1,62 | 0,696 | 10,486 | 1,00E-25 | 2,51E-25 | 0,0039237 | 0,0030815 | 0,0035665 | 0,0035665 | TRUE | 0,051517 | No |
| orf19.4164 | 1547 | 1559 | 1556 | 922 | 2176 | 1405 | 2064 | 2053 | 1475 | 1648 | 1537 | 1444 | 1744 | 1598 | 1752 | 1669 | 1608,46 | 1526 | 1691 | 1,107 | 0,147 | 2,617 | 0,0088608 | 0,0112991 | 0,0018307 | 0,0031121 | 0,0024037 | 0,0024037 | TRUE | 0,0348052 | No |
| orf19.4166 | 5157 | 4319 | 4909 | 3594 | 5320 | 3968 | 4897 | 5503 | 4917 | 4567 | 4850 | 5630 | 4263 | 4512 | 4158 | 4474 | 4671,25 | 4991 | 4352 | 0,873 | -0,197 | -3,119 | 0,0018136 | 0,0024095 | 0,004582 | 0,0024428 | 0,003602 | 0,003602 | TRUE | 0,0885707 | No |
| orf19.4167 | 800 | 731 | 781 | 443 | 2111 | 1289 | 2117 | 2104 | 763 | 773 | 772 | 694 | 1691 | 1466 | 1797 | 1711 | 1208,31 | 750 | 1666 | 2,218 | 1,149 | 16,063 | 4,65E-58 | 1,90E-57 | 0,0042648 | 0,0034501 | 0,0039018 | 0,0039018 | TRUE | 0,0764361 | No |
| orf19.4168 | 1753 | 1343 | 1471 | 895 | 1693 | 1080 | 1638 | 1676 | 1671 | 1420 | 1453 | 1402 | 1357 | 1228 | 1391 | 1363 | 1410,58 | 1486 | 1335 | 0,897 | -0,156 | -2,292 | 0,0219231 | 0,0271317 | 0,004057 | 0,0032553 | 0,0037606 | 0,0037606 | TRUE | 0,0859375 | No |
| orf19.4169 | 87 | 76 | 71 | 52 | 174 | 109 | 181 | 158 | 83 | 80 | 70 | 81 | 139 | 124 | 154 | 128 | 107,55 | 78 | 136 | 1,741 | 0,8 | 5,507 | 3,65E-08 | 5,96E-08 | 1,00E-08 | 0,0173575 | 0,0099084 | 0,0099084 | TRUE | 0,0762864 | No |
| orf19.417 | 3621 | 3201 | 3499 | 2264 | 2522 | 1795 | 2483 | 2546 | 3452 | 3385 | 3457 | 3547 | 2021 | 2041 | 2108 | 2070 | 2760,04 | 3460 | 2060 | 0,596 | -0,747 | -20,052 | 1,96E-89 | 1,17E-88 | 1,00E-08 | 0,0026862 | 0,0009533 | 0,0009533 | TRUE | 0,0031199 | No |
| orf19.4170 | 1398 | 1276 | 1297 | 824 | 384 | 243 | 326 | 400 | 1333 | 1349 | 1281 | 1291 | 308 | 276 | 277 | 325 | 805,03 | 1314 | 296 | 0,226 | -2,143 | -31,55 | 1,79E-218 | 2,66E-217 | 0,0001941 | 0,0041308 | 0,0025089 | 0,0025089 | TRUE | 0,0488086 | No |
| orf19.4171 | 354 | 340 | 403 | 188 | 537 | 383 | 542 | 546 | 338 | 360 | 398 | 295 | 430 | 436 | 460 | 444 | 394,94 | 348 | 442 | 1,266 | 0,34 | 3,781 | 0,0001565 | 0,0002193 | 0,0037383 | 0,0062486 | 0,0051519 | 0,0051519 | TRUE | 0,1078045 | No |
| orf19.4172 | 1343 | 1353 | 1282 | 731 | 1521 | 1161 | 1527 | 1544 | 1280 | 1431 | 1266 | 1145 | 1219 | 1320 | 1296 | 1255 | 1276,68 | 1280 | 1272 | 0,991 | -0,013 | -0,195 | 0,845071 | 0,8644434 | 0,0033381 | 0,0033774 | 0,003513 | 0,003513 | TRUE | 0,0731134 | No |
| orf19.4173 | 1257 | 929 | 1087 | 609 | 1449 | 867 | 1366 | 1580 | 1198 | 982 | 1074 | 954 | 1161 | 986 | 1160 | 1285 | 1099,99 | 1052 | 1148 | 1,09 | 0,125 | 1,369 | 0,1708771 | 0,1950916 | 0,0099896 | 0,0035839 | 0,007046 | 0,007046 | TRUE | 0,1056171 | No |
| orf19.4174 | 2658 | 2222 | 2304 | 1676 | 984 | 687 | 975 | 1045 | 2534 | 2349 | 2276 | 2626 | 788 | 781 | 828 | 850 | 1629,05 | 2446 | 812 | 0,333 | -1,589 | -26,259 | 5,70E-152 | 5,79E-151 | 0,0024715 | 0,0030992 | 0,0027344 | 0,0027344 | TRUE | 0,0277962 | No |
| orf19.4175 | 4573 | 3997 | 4360 | 2816 | 4821 | 3347 | 4532 | 4784 | 4360 | 4226 | 4307 | 4411 | 3863 | 3806 | 3848 | 3890 | 4088,89 | 4326 | 3852 | 0,891 | -0,167 | -5,143 | 2,71E-07 | 4,28E-07 | 1,00E-08 | 0,0024928 | 0,0007617 | 0,0007617 | TRUE | 0,0028995 | No |
| orf19.4176 | 1335 | 1201 | 1245 | 743 | 1889 | 1236 | 1823 | 1998 | 1273 | 1270 | 1230 | 1164 | 1514 | 1405 | 1548 | 1625 | 1378,49 | 1234 | 1523 | 1,233 | 0,302 | 5,223 | 1,76E-07 | 2,80E-07 | 0,0016512 | 0,0032824 | 0,0024726 | 0,0024726 | TRUE | 0,0307377 | No |
| orf19.4177 | 2146 | 1951 | 2048 | 1394 | 2469 | 1789 | 2380 | 2496 | 2046 | 2063 | 2023 | 2184 | 1978 | 2034 | 2021 | 2029 | 2047,33 | 2079 | 2016 | 0,971 | -0,042 | -1,013 | 0,3108708 | 0,3440879 | 2,40E-05 | 0,0028933 | 0,0011754 | 0,0011754 | TRUE | 0,0123962 | No |
| orf19.4179 | 56 | 40 | 47 | 31 | 48 | 24 | 29 | 39 | 53 | 42 | 46 | 49 | 38 | 27 | 25 | 32 | 39,1 | 48 | 30 | 0,644 | -0,635 | -2,723 | 0,0064605 | 0,0082896 | 1,00E-08 | 0,0440888 | 0,0260055 | 0,0260055 | TRUE | 0,2345255 | No |
| orf19.418 | 3209 | 3003 | 3226 | 1865 | 2959 | 2120 | 2820 | 3223 | 3059 | 3175 | 3187 | 2922 | 2371 | 2411 | 2394 | 2621 | 2767,46 | 3086 | 2449 | 0,793 | -0,334 | -6,795 | 1,09E-11 | 2,00E-11 | 0,001427 | 0,0026846 | 0,0019615 | 0,0019615 | TRUE | 0,027501 | No |
| orf19.4180 | 1146 | 1014 | 1015 | 686 | 1811 | 1278 | 1665 | 1947 | 1093 | 1072 | 1003 | 1075 | 1451 | 1453 | 1414 | 1583 | 1267,89 | 1061 | 1475 | 1,392 | 0,477 | 8,474 | 2,36E-17 | 5,01E-17 | 0,0013944 | 0,0033863 | 0,0022152 | 0,0022152 | TRUE | 0,02997 | No |
| orf19.4181 | 569 | 421 | 452 | 309 | 894 | 599 | 763 | 831 | 542 | 445 | 447 | 484 | 716 | 681 | 648 | 676 | 579,89 | 480 | 680 | 1,418 | 0,504 | 6,409 | 1,46E-10 | 2,60E-10 | 0,0033094 | 0,0049227 | 0,0041109 | 0,0041109 | TRUE | 0,0994903 | No |
| orf19.4182 | 1777 | 1580 | 1576 | 1069 | 2098 | 1422 | 1962 | 1965 | 1694 | 1671 | 1557 | 1675 | 1681 | 1617 | 1666 | 1598 | 1644,74 | 1649 | 1640 | 0,996 | -0,006 | -0,136 | 0,8919456 | 0,9070608 | 0,0004572 | 0,0030896 | 0,0015193 | 0,0015193 | TRUE | 0,0178979 | No |
| orf19.4183 | 4016 | 3370 | 3730 | 2399 | 2825 | 1939 | 2665 | 2955 | 3829 | 3563 | 3685 | 3758 | 2264 | 2205 | 2263 | 2403 | 2996,11 | 3709 | 2284 | 0,616 | -0,699 | -16,061 | 4,81E-58 | 1,97E-57 | 0,0006743 | 0,0026393 | 0,0014696 | 0,0014696 | TRUE | 0,0151266 | No |
| orf19.4184 | 2389 | 2104 | 2475 | 1717 | 1994 | 1518 | 1918 | 2049 | 2278 | 2225 | 2445 | 2690 | 1598 | 1726 | 1628 | 1666 | 2031,93 | 2410 | 1654 | 0,688 | -0,54 | -8,558 | 1,14E-17 | 2,44E-17 | 0,0036496 | 0,0028993 | 0,0033251 | 0,0033251 | TRUE | 0,0709067 | No |
| orf19.4185 | 230 | 237 | 222 | 133 | 406 | 301 | 382 | 423 | 219 | 251 | 219 | 208 | 325 | 342 | 324 | 344 | 279,17 | 224 | 334 | 1,482 | 0,568 | 6,23 | 4,65E-10 | 8,11E-10 | 1,00E-08 | 0,0079726 | 0,0041727 | 0,0041727 | TRUE | 0,0668836 | No |
| orf19.4186 | 1179 | 1043 | 1211 | 728 | 1642 | 1171 | 1630 | 1738 | 1124 | 1103 | 1196 | 1140 | 1316 | 1332 | 1384 | 1413 | 1250,99 | 1141 | 1361 | 1,193 | 0,255 | 4,96 | 7,05E-07 | 1,10E-06 | 0,0004826 | 0,0034038 | 0,0017175 | 0,0017175 | TRUE | 0,0137399 | No |
| orf19.4187 | 1380 | 1159 | 1379 | 803 | 1338 | 992 | 1219 | 1443 | 1316 | 1225 | 1362 | 1258 | 1072 | 1128 | 1035 | 1173 | 1196,22 | 1290 | 1102 | 0,853 | -0,23 | -3,85 | 0,0001179 | 0,0001663 | 0,0019019 | 0,0034639 | 0,0025717 | 0,0025717 | TRUE | 0,0239283 | No |
| orf19.4188 | 6954 | 6801 | 6959 | 4124 | 9385 | 6821 | 8350 | 8854 | 6630 | 7191 | 6875 | 6460 | 7520 | 7756 | 7089 | 7199 | 7090,08 | 6789 | 7391 | 1,088 | 0,122 | 2,602 | 0,009279 | 0,0118182 | 0,0017527 | 0,0023229 | 0,0019571 | 0,0019571 | TRUE | 0,0193213 | No |
| orf19.4189 | 22 | 15 | 15 | 11 | 12 | 10 | 6 | 18 | 21 | 16 | 15 | 17 | 10 | 11 | 5 | 15 | 13,7 | 17 | 10 | 0,589 | -0,764 | -1,898 | 0,0577447 | 0,068963 | 1,00E-08 | 0,1219364 | 0,0795354 | 0,0795354 | TRUE | 0,4651617 | No |
| orf19.419 | 7407 | 6750 | 6900 | 4020 | 1248 | 924 | 1118 | 1216 | 7062 | 7137 | 6817 | 6298 | 1000 | 1051 | 949 | 989 | 3912,71 | 6828 | 997 | 0,146 | -2,778 | -51,421 | 0 | 0 | 0,0021156 | 0,0025109 | 0,0022728 | 0,0022728 | TRUE | 0,0324804 | No |
| orf19.4190 | 3479 | 2861 | 3195 | 2078 | 4366 | 2799 | 3937 | 4190 | 3317 | 3025 | 3156 | 3255 | 3498 | 3183 | 3342 | 3407 | 3273,01 | 3188 | 3358 | 1,054 | 0,076 | 1,633 | 0,1024227 | 0,119437 | 0,0011743 | 0,0025929 | 0,0017615 | 0,0017615 | TRUE | 0,0145889 | No |
| orf19.4191 | 11630 | 10606 | 10870 | 6369 | 16150 | 10049 | 15456 | 16384 | 11088 | 11215 | 10739 | 9977 | 12941 | 11427 | 13122 | 13322 | 11728,67 | 10755 | 12703 | 1,181 | 0,24 | 4,161 | 3,18E-05 | 4,60E-05 | 0,0035391 | 0,0022313 | 0,0031106 | 0,0031106 | TRUE | 0,0555941 | No |
| orf19.4191.1 | 1438 | 1479 | 1591 | 929 | 753 | 529 | 752 | 752 | 1371 | 1564 | 1572 | 1455 | 603 | 602 | 638 | 611 | 1052,09 | 1490 | 614 | 0,412 | -1,28 | -20,367 | 3,25E-92 | 1,98E-91 | 0,0019896 | 0,0036519 | 0,0026891 | 0,0026891 | TRUE | 0,0375478 | No |

| | | | | | | | | | | | | | | | | | | | | | | | | | | | | | | | |
|---|---|---|---|---|---|---|---|---|---|---|---|---|---|---|---|---|---|---|---|---|---|---|---|---|---|---|---|---|---|---|---|
| orf19.4192 | 3207 | 2828 | 3000 | 2033 | 1455 | 1148 | 1396 | 1576 | 3058 | 2990 | 2964 | 3185 | 1166 | 1305 | 1185 | 1281 | 2141,77 | 3049 | 1234 | 0,405 | -1,305 | -25,471 | 4,10E-143 | 3,94E-142 | 0,0013372 | 0,0028579 | 0,0019741 | 0,0019741 | TRUE | 0,0169559 | No |
| orf19.4192.1 | 864 | 655 | 734 | 434 | 1386 | 914 | 1381 | 1414 | 824 | 693 | 725 | 680 | 1111 | 1039 | 1172 | 1150 | 924,17 | 730 | 1118 | 1,528 | 0,612 | 8,493 | 2,01E-17 | 4,28E-17 | 0,0038286 | 0,0038679 | 0,003812 | 0,003812 | TRUE | 0,0901563 | No |
| orf19.4193 | 552 | 501 | 512 | 371 | 789 | 576 | 758 | 824 | 526 | 530 | 506 | 581 | 632 | 655 | 644 | 670 | 592,96 | 536 | 650 | 1,219 | 0,285 | 4,331 | 1,48E-05 | 2,18E-05 | 1,00E-08 | 0,0048603 | 0,0024381 | 0,0024381 | TRUE | 0,0317578 | No |
| orf19.4193.1 | 18375 | 14749 | 15131 | 10531 | 38178 | 25705 | 34165 | 39545 | 17519 | 15595 | 14948 | 16497 | 30591 | 29229 | 29006 | 32154 | 23192,35 | 16140 | 30245 | 1,874 | 0,906 | 16,168 | 8,49E-59 | 3,52E-58 | 0,0034868 | 0,0021621 | 0,0029684 | 0,0029684 | TRUE | 0,0407819 | No |
| orf19.4194 | 634 | 567 | 610 | 393 | 712 | 613 | 684 | 721 | 604 | 600 | 603 | 616 | 571 | 697 | 581 | 586 | 607,09 | 606 | 609 | 1,002 | 0,003 | 0,042 | 0,9663996 | 0,969743 | 0,002882 | 0,0047958 | 0,0038091 | 0,0038091 | TRUE | 0,1048514 | No |
| orf19.4195 | 264 | 221 | 241 | 137 | 279 | 214 | 261 | 227 | 252 | 234 | 238 | 215 | 224 | 243 | 222 | 185 | 226,39 | 235 | 218 | 0,923 | -0,115 | -1,055 | 0,2915273 | 0,3240264 | 0,0042624 | 0,0093438 | 0,0070057 | 0,0070057 | TRUE | 0,122989 | No |
| orf19.4195.1 | 2467 | 2025 | 2161 | 1220 | 6652 | 3778 | 6377 | 6409 | 2352 | 2141 | 2135 | 1911 | 5330 | 4296 | 5414 | 5211 | 3598,8 | 2135 | 5063 | 2,37 | 1,245 | 15,387 | 2,01E-53 | 7,77E-53 | 0,0083452 | 0,0025475 | 0,0059337 | 0,0059337 | TRUE | 0,1256816 | No |
| orf19.4196 | 46 | 23 | 38 | 26 | 38 | 18 | 32 | 31 | 44 | 24 | 38 | 41 | 30 | 20 | 27 | 25 | 31,22 | 37 | 26 | 0,713 | -0,488 | -1,81 | 0,0702743 | 0,0833031 | 1,00E-08 | 0,0546877 | 0,0374174 | 0,0374174 | TRUE | 0,3751103 | No |
| orf19.4197 | 2265 | 1711 | 1969 | 1269 | 5106 | 3466 | 5046 | 5411 | 2159 | 1809 | 1945 | 1988 | 4091 | 3941 | 4284 | 4400 | 3077,23 | 1975 | 4179 | 2,116 | 1,081 | 17,986 | 2,52E-72 | 1,25E-71 | 0,0033586 | 0,0026248 | 0,0030779 | 0,0030779 | TRUE | 0,0494452 | No |
| orf19.4199 | 10509 | 9793 | 10586 | 7065 | 6639 | 5109 | 6442 | 6633 | 10019 | 10355 | 10458 | 11068 | 5320 | 5809 | 5469 | 5393 | 7986,41 | 10475 | 5498 | 0,525 | -0,93 | -21,028 | 3,66E-98 | 2,35E-97 | 0,0014618 | 0,0022969 | 0,0017435 | 0,0017435 | TRUE | 0,0173732 | No |
| orf19.420 | 32 | 23 | 12 | 13 | 13 | 11 | 8 | 10 | 31 | 24 | 12 | 20 | 10 | 13 | 7 | 8 | 15,61 | 22 | 10 | 0,428 | -1,224 | -3,009 | 0,0026197 | 0,0034481 | 1,00E-08 | 0,1072609 | 0,0857583 | 0,0857583 | TRUE | 0,2002529 | No |
| orf19.4200 | 22 | 26 | 35 | 36 | 41 | 26 | 37 | 41 | 21 | 27 | 35 | 56 | 33 | 30 | 31 | 33 | 33,33 | 35 | 32 | 0,941 | -0,088 | -0,302 | 0,7623517 | 0,7898305 | 0,0516985 | 0,0513598 | 0,0514834 | 0,0514834 | TRUE | 0,6987491 | No |
| orf19.4201 | 2689 | 2275 | 2637 | 1770 | 3106 | 2237 | 2865 | 3119 | 2564 | 2406 | 2605 | 2773 | 2489 | 2544 | 2432 | 2536 | 2543,49 | 2587 | 2500 | 0,968 | -0,047 | -0,967 | 0,3337534 | 0,3674159 | 0,0013883 | 0,0027368 | 0,001903 | 0,001903 | TRUE | 0,0265108 | No |
| orf19.4203 | 3341 | 3449 | 3659 | 2388 | 2808 | 2322 | 2631 | 2682 | 3185 | 3647 | 3615 | 3741 | 2250 | 2640 | 2234 | 2181 | 2936,57 | 3547 | 2326 | 0,656 | -0,609 | -8,528 | 1,49E-17 | 3,18E-17 | 0,0060835 | 0,0026504 | 0,0045499 | 0,0045499 | TRUE | 0,0982377 | No |
| orf19.4204 | 3618 | 3217 | 3350 | 2150 | 4060 | 2687 | 4091 | 4273 | 3449 | 3402 | 3309 | 3368 | 3253 | 3055 | 3473 | 3474 | 3348,08 | 3382 | 3314 | 0,981 | -0,028 | -0,574 | 0,5658826 | 0,6004152 | 0,0014961 | 0,0025817 | 0,002047 | 0,002047 | TRUE | 0,0324602 | No |
| orf19.4205.1 | 886 | 684 | 759 | 359 | 845 | 434 | 783 | 795 | 845 | 723 | 750 | 562 | 677 | 493 | 665 | 646 | 670,24 | 720 | 620 | 0,861 | -0,217 | -1,758 | 0,0787079 | 0,0929033 | 0,0223071 | 0,004541 | 0,0130813 | 0,0130813 | TRUE | 0,241687 | No |
| orf19.4206 | 603 | 572 | 623 | 364 | 532 | 378 | 493 | 548 | 575 | 605 | 615 | 570 | 426 | 430 | 419 | 446 | 510,7 | 591 | 430 | 0,726 | -0,462 | -6,846 | 7,58E-12 | 1,40E-11 | 1,00E-08 | 0,0053063 | 0,0023861 | 0,0023861 | TRUE | 0,008952 | No |
| orf19.4207 | 235 | 195 | 220 | 160 | 493 | 324 | 489 | 511 | 224 | 206 | 217 | 251 | 395 | 368 | 415 | 415 | 311,54 | 224 | 398 | 1,79 | 0,84 | 9,23 | 2,70E-20 | 6,10E-20 | 0,0004533 | 0,0073616 | 0,0043346 | 0,0043346 | TRUE | 0,0526279 | No |
| orf19.4208 | 1178 | 960 | 1082 | 686 | 1587 | 1044 | 1498 | 1516 | 1123 | 1015 | 1069 | 1075 | 1272 | 1187 | 1272 | 1233 | 1155,62 | 1070 | 1241 | 1,16 | 0,214 | 4,026 | 5,67E-05 | 8,12E-05 | 0,0004795 | 0,0035121 | 0,0018326 | 0,0018326 | TRUE | 0,0151946 | No |
| orf19.4209 | 1391 | 1410 | 1495 | 852 | 1156 | 805 | 1065 | 1082 | 1326 | 1491 | 1477 | 1335 | 926 | 915 | 904 | 880 | 1156,78 | 1407 | 906 | 0,643 | -0,637 | -10,915 | 9,73E-28 | 2,53E-27 | 0,0015277 | 0,0035107 | 0,002375 | 0,002375 | TRUE | 0,0201314 | No |
| orf19.4210 | 1055 | 974 | 1005 | 649 | 2032 | 1442 | 1886 | 2019 | 1006 | 1030 | 993 | 1017 | 1628 | 1640 | 1601 | 1642 | 1319,5 | 1012 | 1628 | 1,61 | 0,687 | 15,091 | 1,86E-51 | 6,96E-51 | 1,00E-08 | 0,0033356 | 0,0011644 | 0,0011644 | TRUE | 0,0020098 | No |
| orf19.4211 | 1934 | 1835 | 1865 | 1212 | 14532 | 10655 | 13706 | 14937 | 1844 | 1940 | 1842 | 1899 | 11644 | 12116 | 11636 | 12145 | 6883,32 | 1881 | 11885 | 6,321 | 2,66 | 70,369 | 0 | 0 | 0,0004185 | 0,0023298 | 0,0010368 | 0,0010368 | TRUE | 0,0058215 | No |
| orf19.4212 | 370 | 318 | 344 | 206 | 2282 | 1610 | 2145 | 2301 | 353 | 336 | 340 | 323 | 1829 | 1831 | 1821 | 1871 | 1087,85 | 338 | 1838 | 5,419 | 2,438 | 41,32 | 0 | 0 | 1,00E-08 | 0,0036006 | 0,0014673 | 0,0014673 | TRUE | 0,0099414 | No |
| orf19.4213 | 975 | 818 | 1000 | 585 | 1749 | 1237 | 1631 | 1813 | 930 | 865 | 988 | 916 | 1401 | 1407 | 1385 | 1474 | 1170,71 | 925 | 1417 | 1,531 | 0,615 | 11,063 | 1,90E-28 | 4,99E-28 | 0,0008368 | 0,0034938 | 0,0020432 | 0,0020432 | TRUE | 0,0266187 | No |
| orf19.4214 | 42 | 38 | 44 | 26 | 17 | 14 | 12 | 21 | 40 | 40 | 43 | 41 | 14 | 16 | 10 | 17 | 27,65 | 41 | 14 | 0,344 | -1,541 | -5,46 | 4,75E-08 | 7,74E-08 | 1,00E-08 | 0,0614663 | 0,0315209 | 0,0315209 | TRUE | 0,1819455 | No |
| orf19.4215 | 16848 | 14225 | 15404 | 9726 | 26519 | 18466 | 25411 | 27357 | 16063 | 15041 | 15218 | 15236 | 21249 | 20998 | 21574 | 22244 | 18452,75 | 15390 | 21516 | 1,398 | 0,483 | 13,831 | 1,66E-43 | 5,54E-43 | 0,0006818 | 0,0021803 | 0,0011159 | 0,0011159 | TRUE | 0,0107363 | No |
| orf19.4216 | 2318 | 1924 | 1917 | 1335 | 3263 | 1997 | 2921 | 3115 | 2210 | 2034 | 1894 | 2091 | 2615 | 2271 | 2480 | 2533 | 2265,94 | 2057 | 2475 | 1,204 | 0,268 | 4,363 | 1,28E-05 | 1,89E-05 | 0,0031998 | 0,0028159 | 0,0031614 | 0,0031614 | TRUE | 0,0362168 | No |
| orf19.4217 | 7 | 3 | 3 | 0 | 5 | 4 | 2 | 0 | 7 | 3 | 3 | 0 | 4 | 5 | 2 | 0 | 2,88 | 3 | 3 | 0,747 | -0,42 | -0,429 | 0,6681405 | 0,7000664 | 0,5934059 | 0,5716457 | 0,5786531 | 0,5786531 | TRUE | 0,2423348 | No |
| orf19.4219 | 6798 | 5688 | 6356 | 3968 | 8738 | 5949 | 8040 | 8902 | 6481 | 6014 | 6279 | 6216 | 7002 | 6765 | 6826 | 7238 | 6602,61 | 6248 | 6958 | 1,114 | 0,155 | 4,008 | 6,12E-05 | 8,74E-05 | 0,0007369 | 0,00234 | 0,0012879 | 0,0012879 | TRUE | 0,0090293 | No |
| orf19.422 | 947 | 797 | 899 | 652 | 901 | 654 | 835 | 976 | 903 | 843 | 888 | 1021 | 722 | 744 | 709 | 794 | 827,9 | 914 | 742 | 0,815 | -0,295 | -4,185 | 2,85E-05 | 4,14E-05 | 0,0032337 | 0,0040745 | 0,0035669 | 0,0035669 | TRUE | 0,0673596 | No |
| orf19.4220 | 1520 | 1306 | 1424 | 959 | 1833 | 1181 | 1635 | 1792 | 1449 | 1381 | 1407 | 1502 | 1469 | 1343 | 1388 | 1457 | 1424,5 | 1435 | 1414 | 0,988 | -0,017 | -0,331 | 0,7409035 | 0,769358 | 0,0007574 | 0,0032439 | 0,0018241 | 0,0018241 | TRUE | 0,0129677 | No |
| orf19.4221 | 2397 | 2052 | 2278 | 1491 | 2344 | 1707 | 2251 | 2500 | 2285 | 2170 | 2250 | 2336 | 1878 | 1941 | 1911 | 2033 | 2100,53 | 2260 | 1941 | 0,859 | -0,219 | -4,847 | 1,25E-06 | 1,93E-06 | 0,000606 | 0,0028729 | 0,0014756 | 0,0014756 | TRUE | 0,0128119 | No |
| orf19.4222 | 2136 | 2065 | 2138 | 1438 | 1446 | 1041 | 1352 | 1508 | 2036 | 2183 | 2112 | 2253 | 1159 | 1184 | 1148 | 1226 | 1662,64 | 2146 | 1179 | 0,55 | -0,862 | -17,326 | 2,99E-67 | 1,38E-66 | 0,0008215 | 0,0030788 | 0,0017353 | 0,0017353 | TRUE | 0,0144147 | No |
| orf19.4223 | 13098 | 10504 | 11788 | 7254 | 18510 | 11632 | 17478 | 18850 | 12488 | 11107 | 11645 | 11364 | 14832 | 13227 | 14839 | 15327 | 13103,39 | 11651 | 14556 | 1,249 | 0,321 | 5,797 | 6,77E-09 | 1,13E-08 | 0,0032015 | 0,0022166 | 0,0028709 | 0,0028709 | TRUE | 0,0459737 | No |
| orf19.4225 | 5405 | 4861 | 5354 | 3318 | 5442 | 3647 | 5097 | 5670 | 5153 | 5140 | 5289 | 5198 | 4361 | 4147 | 4327 | 4610 | 4778,14 | 5195 | 4361 | 0,84 | -0,252 | -6,219 | 5,01E-10 | 8,72E-10 | 0,000759 | 0,0024349 | 0,0013603 | 0,0013603 | TRUE | 0,0180082 | No |
| orf19.4225.1 | 4465 | 3724 | 4043 | 2785 | 3162 | 2272 | 3058 | 3049 | 4257 | 3938 | 3994 | 4363 | 2534 | 2583 | 2596 | 2479 | 3342,99 | 4138 | 2548 | 0,616 | -0,699 | -15,38 | 2,24E-53 | 8,66E-53 | 0,0010183 | 0,0025824 | 0,0016699 | 0,0016699 | TRUE | 0,0153955 | No |
| orf19.4227 | 94 | 76 | 98 | 67 | 32 | 25 | 20 | 40 | 90 | 80 | 97 | 105 | 26 | 28 | 17 | 33 | 59,42 | 93 | 26 | 0,279 | -1,841 | -8,487 | 2,12E-17 | 4,49E-17 | 0,0057145 | 0,0297255 | 0,0219229 | 0,0219229 | TRUE | 0,374561 | No |
| orf19.4228 | 1165 | 982 | 936 | 665 | 3136 | 1923 | 2819 | 3038 | 1111 | 1038 | 925 | 1042 | 2513 | 2187 | 2393 | 2470 | 1709,8 | 1029 | 2391 | 2,326 | 1,218 | 18,168 | 9,33E-74 | 4,70E-73 | 0,0037104 | 0,0030516 | 0,0035809 | 0,0035809 | TRUE | 0,0582458 | No |
| orf19.4229 | 850 | 731 | 812 | 495 | 2295 | 1590 | 2213 | 2239 | 810 | 773 | 802 | 775 | 1839 | 1808 | 1879 | 1821 | 1313,4 | 790 | 1837 | 2,321 | 1,215 | 25,214 | 2,78E-140 | 2,62E-139 | 1,00E-08 | 0,0033414 | 0,0012828 | 0,0012828 | TRUE | 0,0035646 | No |
| orf19.423 | 1197 | 990 | 1053 | 873 | 1593 | 1312 | 1512 | 1679 | 1141 | 1047 | 1040 | 1368 | 1276 | 1492 | 1284 | 1365 | 1251,63 | 1149 | 1354 | 1,181 | 0,24 | 2,72 | 0,0065379 | 0,0083856 | 0,0105142 | 0,0034031 | 0,0066695 | 0,0066695 | TRUE | 0,1862387 | No |
| orf19.4230 | 6338 | 5515 | 5827 | 3761 | 7111 | 4668 | 6618 | 6957 | 6043 | 5831 | 5757 | 5892 | 5698 | 5308 | 5619 | 5657 | 5725,43 | 5881 | 5570 | 0,948 | -0,077 | -2,104 | 0,0354119 | 0,0430981 | 0,0004897 | 0,0023781 | 0,0011268 | 0,0011268 | TRUE | 0,0118584 | No |
| orf19.4231 | 608 | 540 | 599 | 420 | 472 | 319 | 466 | 444 | 580 | 571 | 592 | 658 | 378 | 363 | 396 | 361 | 487,24 | 600 | 374 | 0,627 | -0,674 | -9,066 | 1,23E-19 | 2,75E-19 | 0,0006481 | 0,0054611 | 0,0031743 | 0,0031743 | TRUE | 0,0425284 | No |
| orf19.4232 | 2001 | 1896 | 1994 | 1188 | 2595 | 1844 | 2501 | 2628 | 1908 | 2005 | 1970 | 1861 | 2079 | 2097 | 2123 | 2137 | 2022,46 | 1936 | 2109 | 1,088 | 0,122 | 2,928 | 0,0034134 | 0,0044587 | 1,00E-08 | 0,0029031 | 0,0011639 | 0,0011639 | TRUE | 0,0070999 | No |
| orf19.4233 | 4160 | 3747 | 3954 | 2433 | 8897 | 6264 | 8413 | 9134 | 3966 | 3962 | 3906 | 3811 | 7129 | 7123 | 7143 | 7427 | 5558,34 | 3911 | 7206 | 1,841 | 0,88 | 26,412 | 9,96E-154 | 1,02E-152 | 0,0002133 | 0,0023867 | 0,0008614 | 0,0008614 | TRUE | 0,0052955 | No |
| orf19.4234 | 5399 | 4839 | 5320 | 3373 | 4755 | 3662 | 4545 | 4691 | 5147 | 5117 | 5256 | 5284 | 3810 | 4164 | 3859 | 3814 | 4556,33 | 5201 | 3912 | 0,752 | -0,412 | -10,222 | 1,58E-24 | 3,87E-24 | 0,0007639 | 0,0024516 | 0,0013355 | 0,0013355 | TRUE | 0,0221201 | No |
| orf19.4235 | 2049 | 1933 | 2172 | 1449 | 1497 | 1051 | 1382 | 1513 | 1953 | 2044 | 2146 | 2270 | 1200 | 1195 | 1173 | 1230 | 1651,4 | 2103 | 1200 | 0,571 | -0,807 | -14,625 | 1,94E-48 | 7,00E-48 | 0,0017291 | 0,0030855 | 0,0022851 | 0,0022851 | TRUE | 0,0320745 | No |
| orf19.4236 | 1352 | 1245 | 1246 | 771 | 2764 | 1972 | 2566 | 2732 | 1289 | 1316 | 1231 | 1208 | 2215 | 2242 | 2179 | 2221 | 1737,64 | 1261 | 2214 | 1,752 | 0,809 | 18,04 | 9,45E-73 | 4,70E-72 | 1,00E-08 | 0,0030362 | 0,001288 | 0,001288 | TRUE | 0,0100125 | No |
| orf19.4239 | 11 | 16 | 19 | 15 | 28 | 20 | 17 | 19 | 10 | 17 | 19 | 23 | 22 | 23 | 14 | 15 | 18,09 | 17 | 18 | 1,094 | 0,13 | 0,367 | 0,7133554 | 0,7430494 | 0,0191667 | 0,0928462 | 0,0636978 | 0,0636978 | TRUE | 0,3903899 | No |
| orf19.424 | 1760 | 1415 | 1473 | 1008 | 2572 | 1760 | 2380 | 2605 | 1678 | 1496 | 1455 | 1579 | 2061 | 2001 | 2021 | 2118 | 1801,16 | 1552 | 2050 | 1,322 | 0,402 | 7,446 | 9,62E-14 | 1,87E-13 | 0,0015749 | 0,0030029 | 0,002224 | 0,002224 | TRUE | 0,0386396 | No |
| orf19.4240 | 709 | 622 | 663 | 453 | 875 | 602 | 760 | 866 | 676 | 658 | 655 | 710 | 701 | 685 | 645 | 704 | 679,16 | 675 | 684 | 1,016 | 0,023 | 0,375 | 0,7074022 | 0,7374503 | 1,00E-08 | 0,0045088 | 0,0022183 | 0,0022183 | TRUE | 0,0177612 | No |
| orf19.4241 | 780 | 707 | 744 | 545 | 887 | 660 | 927 | 993 | 744 | 748 | 735 | 854 | 711 | 750 | 787 | 807 | 766,95 | 770 | 764 | 0,996 | -0,005 | -0,078 | 0,9376607 | 0,9440223 | 0,0022998 | 0,0042321 | 0,0033347 | 0,0033347 | TRUE | 0,0563517 | No |
| orf19.4242 | 1901 | 1708 | 1824 | 1217 | 4013 | 2670 | 3617 | 4023 | 1812 | 1806 | 1802 | 1906 | 3216 | 3036 | 3071 | 3271 | 2490,03 | 1832 | 3148 | 1,722 | 0,784 | 17,816 | 5,28E-71 | 2,56E-70 | 0,000632 | 0,0027507 | 0,0014121 | 0,0014121 | TRUE | 0,0083476 | No |
| orf19.4243 | 215 | 198 | 181 | 105 | 340 | 248 | 307 | 409 | 205 | 209 | 179 | 164 | 272 | 282 | 261 | 333 | 238,16 | 189 | 287 | 1,507 | 0,592 | 5,091 | 3,56E-07 | 5,60E-07 | 0,0079103 | 0,0089854 | 0,0084707 | 0,0084707 | TRUE | 0,1357171 | No |
| orf19.4244 | 83 | 113 | 115 | 58 | 193 | 95 | 149 | 169 | 79 | 119 | 114 | 91 | 155 | 108 | 127 | 137 | 116,21 | 101 | 132 | 1,309 | 0,388 | 2,324 | 0,0201459 | 0,0249953 | 0,0199955 | 0,0162203 | 0,0179185 | 0,0179185 | TRUE | 0,2334254 | No |
| orf19.4245 | 1241 | 1045 | 1139 | 769 | 538 | 348 | 513 | 558 | 1183 | 1105 | 1125 | 1205 | 431 | 396 | 436 | 454 | 791,76 | 1154 | 429 | 0,373 | -1,422 | -22,246 | 1,24E-109 | 9,00E-109 | 0,0006044 | 0,004165 | 0,0024033 | 0,0024033 | TRUE | 0,0276891 | No |
| orf19.4246 | 18310 | 15829 | 17288 | 10706 | 19250 | 13075 | 18536 | 19294 | 17457 | 16737 | 17079 | 16772 | 15424 | 14867 | 15737 | 15688 | 16220,12 | 17011 | 15429 | 0,907 | -0,141 | -4,38 | 1,19E-05 | 1,75E-05 | 0,0004449 | 0,0021925 | 0,0009299 | 0,0009299 | TRUE | 0,007232 | No |
| orf19.4247 | 726 | 585 | 654 | 382 | 1019 | 690 | 976 | 972 | 692 | 619 | 646 | 598 | 816 | 785 | 829 | 790 | 721,91 | 639 | 805 | 1,256 | 0,329 | 5,182 | 2,20E-07 | 3,49E-07 | 0,000486 | 0,0043657 | 0,002452 | 0,002452 | TRUE | 0,0378835 | No |
| orf19.4248 | 1263 | 1090 | 1164 | 714 | 2670 | 1875 | 2608 | 2936 | 1204 | 1153 | 1150 | 1119 | 2139 | 2132 | 2214 | 2387 | 1687,25 | 1156 | 2218 | 1,917 | 0,939 | 17,366 | 1,49E-67 | 6,93E-67 | 0,0015124 | 0,0030644 | 0,0021198 | 0,0021198 | TRUE | 0,0322933 | No |
| orf19.425 | 326 | 274 | 313 | 201 | 638 | 451 | 564 | 580 | 311 | 290 | 309 | 315 | 511 | 513 | 479 | 472 | 399,89 | 306 | 494 | 1,612 | 0,689 | 8,916 | 4,82E-19 | 1,06E-18 | 1,00E-08 | 0,0061972 | 0,0030147 | 0,0030147 | TRUE | 0,0148171 | No |
| orf19.4250 | 156 | 121 | 113 | 89 | 282 | 209 | 267 | 246 | 149 | 128 | 112 | 139 | 226 | 238 | 227 | 200 | 177,26 | 132 | 223 | 1,688 | 0,755 | 6,214 | 5,15E-10 | 8,96E-10 | 0,0033512 | 0,0113542 | 0,0079412 | 0,0079412 | TRUE | 0,1194717 | No |
| orf19.4251 | 193 | 189 | 198 | 149 | 390 | 295 | 373 | 414 | 184 | 200 | 196 | 233 | 312 | 335 | 317 | 337 | 264,26 | 203 | 325 | 1,614 | 0,691 | 7,164 | 7,83E-13 | 1,49E-12 | 1,19E-08 | 0,0083044 | 0,0047963 | 0,0047963 | TRUE | 0,0865665 | No |
| orf19.4252 | 3116 | 2755 | 2905 | 1929 | 2129 | 1590 | 2154 | 2261 | 2971 | 2913 | 2870 | 3022 | 1706 | 1808 | 1829 | 1838 | 2369,58 | 2944 | 1795 | 0,61 | -0,713 | -16,731 | 7,76E-63 | 3,39E-62 | 0,0003941 | 0,0027842 | 0,0013036 | 0,0013036 | TRUE | 0,0141471 | No |
| orf19.4253 | 524 | 555 | 555 | 307 | 1616 | 1063 | 1469 | 1480 | 500 | 587 | 548 | 481 | 1295 | 1209 | 1247 | 1203 | 883,72 | 529 | 1238 | 2,335 | 1,223 | 17,905 | 1,07E-71 | 5,26E-71 | 0,0020555 | 0,0039492 | 0,0030672 | 0,0030672 | TRUE | 0,061605 | No |
| orf19.4255 | 434 | 339 | 381 | 292 | 131 | 136 | 154 | 149 | 414 | 358 | 376 | 457 | 105 | 155 | 131 | 121 | 264,69 | 401 | 128 | 0,317 | -1,657 | -13,358 | 1,07E-40 | 3,41E-40 | 0,0122813 | 0,0082943 | 0,0098618 | 0,0098618 | TRUE | 0,1822013 | No |
| orf19.4257 | 3042 | 2815 | 3159 | 1848 | 2362 | 1701 | 2196 | 2421 | 2900 | 2977 | 3121 | 2895 | 1893 | 1934 | 1864 | 1969 | 2444,02 | 2973 | 1915 | 0,644 | -0,636 | -14,729 | 4,17E-49 | 1,52E-48 | 0,0005617 | 0,0027631 | 0,0013671 | 0,0013671 | TRUE | 0,0138532 | No |
| orf19.4258 | 3397 | 2832 | 3266 | 2035 | 3985 | 2558 | 3780 | 3938 | 3239 | 2994 | 3226 | 3188 | 3193 | 2909 | 3209 | 3202 | 3145,06 | 3162 | 3128 | 0,99 | -0,014 | -0,299 | 0,7650939 | 0,7911318 | 0,0014368 | 0,0026133 | 0,0018795 | 0,0018795 | TRUE | 0,0261487 | No |
| orf19.426 | 1107 | 1018 | 1119 | 666 | 1340 | 920 | 1333 | 1424 | 1055 | 1076 | 1105 | 1043 | 1074 | 1046 | 1132 | 1158 | 1086,25 | 1070 | 1102 | 1,03 | 0,043 | 0,794 | 0,4269787 | 0,463044 | 0,0005655 | 0,0036028 | 0,001898 | 0,001898 | TRUE | 0,0137643 | No |
| orf19.4261 | 4059 | 4250 | 4304 | 2625 | 7718 | 5665 | 7084 | 7201 | 3870 | 4494 | 4252 | 4112 | 6184 | 6442 | 6014 | 5855 | 5152,88 | 4182 | 6124 | 1,464 | 0,55 | 10,349 | 4,21E-25 | 1,05E-24 | 0,0025196 | 0,0024099 | 0,0025031 | 0,0025031 | TRUE | 0,0316274 | No |
| orf19.4262 | 986 | 840 | 953 | 684 | 1704 | 1208 | 1459 | 1745 | 940 | 888 | 941 | 1072 | 1365 | 1374 | 1239 | 1419 | 1154,72 | 960 | 1349 | 1,409 | 0,495 | 7,243 | 4,38E-13 | 8,42E-13 | 0,0037663 | 0,0035132 | 0,0035648 | 0,0035648 | TRUE | 0,0655062 | No |
| orf19.4263 | 592 | 610 | 724 | 418 | 1314 | 908 | 1152 | 1272 | 564 | 645 | 715 | 655 | 1053 | 1032 | 978 | 1034 | 834,64 | 645 | 1024 | 1,59 | 0,669 | 9,149 | 5,75E-20 | 1,29E-19 | 0,0034305 | 0,0040585 | 0,0038271 | 0,0038271 | TRUE | 0,0894605 | No |
| orf19.4264 | 473 | 491 | 495 | 322 | 646 | 482 | 618 | 629 | 451 | 519 | 489 | 504 | 518 | 548 | 525 | 511 | 508,17 | 491 | 526 | 1,071 | 0,099 | 1,398 | 0,1620749 | 0,1854402 | 1,00E-08 | 0,0053223 | 0,0028528 | 0,0028528 | TRUE | 0,0374659 | No |
| orf19.4265 | 4080 | 4514 | 4450 | 2716 | 3990 | 3007 | 3601 | 3475 | 3890 | 4773 | 4396 | 4255 | 3197 | 3419 | 3057 | 2826 | 3726,6 | 4328 | 3125 | 0,722 | -0,471 | -6,495 | 8,29E-11 | 1,49E-10 | 0,0063668 | 0,0025319 | 0,0047721 | 0,0047721 | TRUE | 0,0586224 | No |
| orf19.4266 | 860 | 943 | 896 | 592 | 989 | 822 | 895 | 864 | 820 | 997 | 885 | 927 | 792 | 935 | 760 | 703 | 852,39 | 907 | 798 | 0,878 | -0,188 | -2,053 | 0,0401086 | 0,0486564 | 0,009465 | 0,0040175 | 0,0069012 | 0,0069012 | TRUE | 0,1539229 | No |

| | | | | | | | | | | | | | | | | | | | | | | | | | | | | | | | |
|---|---|---|---|---|---|---|---|---|---|---|---|---|---|---|---|---|---|---|---|---|---|---|---|---|---|---|---|---|---|---|---|
| orf19.4268 | 12445 | 11899 | 12805 | 6819 | 18145 | 12645 | 16969 | 18055 | 11865 | 12582 | 12650 | 10682 | 14539 | 14379 | 14407 | 14680 | 13222,96 | 11945 | 14501 | 1,214 | 0,279 | 5,239 | 1,62E-07 | 2,58E-07 | 0,0028681 | 0,0022155 | 0,0026503 | 0,0026503 | TRUE | 0,0609839 | No |
| orf19.4269 | 2088 | 2070 | 2149 | 1339 | 1840 | 1302 | 1679 | 1698 | 1991 | 2189 | 2123 | 2098 | 1474 | 1480 | 1425 | 1381 | 1770,12 | 2100 | 1440 | 0,686 | -0,545 | -11,197 | 4,21E-29 | 1,12E-28 | 0,0008329 | 0,0030188 | 0,0016937 | 0,0016937 | TRUE | 0,0157441 | No |
| orf19.427 | 2706 | 2368 | 2671 | 1513 | 2942 | 2171 | 2730 | 2976 | 2580 | 2504 | 2639 | 2370 | 2357 | 2469 | 2318 | 2420 | 2457,02 | 2523 | 2391 | 0,946 | -0,081 | -1,751 | 0,0799235 | 0,0942509 | 0,0009022 | 0,0027595 | 0,0016329 | 0,0016329 | TRUE | 0,0185634 | No |
| orf19.4270 | 1055 | 1112 | 1134 | 710 | 632 | 536 | 584 | 613 | 1006 | 1176 | 1120 | 1112 | 506 | 609 | 496 | 498 | 815,54 | 1104 | 527 | 0,477 | -1,069 | -13,391 | 6,85E-41 | 2,20E-40 | 0,0056013 | 0,0041046 | 0,0047604 | 0,0047604 | TRUE | 0,119485 | No |
| orf19.4271 | 1913 | 1795 | 1869 | 864 | 2621 | 1394 | 2377 | 2176 | 1824 | 1898 | 1846 | 1354 | 2100 | 1585 | 2018 | 1769 | 1799,29 | 1730 | 1868 | 1,079 | 0,109 | 0,998 | 0,3181661 | 0,3513703 | 0,0191008 | 0,0030038 | 0,010916 | 0,010916 | TRUE | 0,2515626 | No |
| orf19.4273 | 5223 | 6084 | 5950 | 3470 | 5065 | 3568 | 4476 | 4068 | 4980 | 6433 | 5878 | 5436 | 4058 | 4057 | 3800 | 3308 | 4743,74 | 5682 | 3806 | 0,67 | -0,578 | -6,837 | 8,08E-12 | 1,49E-11 | 0,0099952 | 0,0024374 | 0,0066603 | 0,0066603 | TRUE | 0,0969413 | No |
| orf19.4274 | 36042 | 40494 | 41408 | 26188 | 27247 | 21564 | 24679 | 23227 | 34362 | 42817 | 40907 | 41025 | 21832 | 24520 | 20952 | 18886 | 30662,76 | 39778 | 21548 | 0,542 | -0,885 | -10,657 | 1,61E-26 | 4,12E-26 | 0,0101845 | 0,0021448 | 0,0065837 | 0,0065837 | TRUE | 0,1054297 | No |
| orf19.4275 | 835 | 776 | 851 | 496 | 753 | 580 | 738 | 803 | 796 | 821 | 841 | 777 | 603 | 660 | 627 | 653 | 722,08 | 809 | 636 | 0,784 | -0,352 | -5,797 | 6,74E-09 | 1,13E-08 | 1,00E-08 | 0,0043651 | 0,0021378 | 0,0021378 | TRUE | 0,0141416 | No |
| orf19.4278 | 4571 | 4079 | 4295 | 2659 | 5561 | 3810 | 5162 | 5328 | 4358 | 4313 | 4243 | 4165 | 4456 | 4332 | 4383 | 4332 | 4322,8 | 4270 | 4376 | 1,024 | 0,035 | 1,049 | 0,2940071 | 0,326612 | 1,00E-08 | 0,0024711 | 0,0008066 | 0,0008066 | TRUE | 0,0030039 | No |
| orf19.4279 | 21197 | 19906 | 21241 | 12835 | 3395 | 2525 | 3268 | 3578 | 20209 | 21048 | 20984 | 20107 | 2720 | 2871 | 2775 | 2909 | 11702,9 | 20587 | 2819 | 0,137 | -2,869 | -78,236 | 0 | 0 | 0,0004973 | 0,0022316 | 0,0011059 | 0,0011059 | TRUE | 0,0068736 | No |
| orf19.428 | 1758 | 1659 | 1708 | 1038 | 742 | 545 | 708 | 800 | 1676 | 1754 | 1687 | 1626 | 595 | 620 | 601 | 650 | 1151,19 | 1686 | 616 | 0,365 | -1,453 | -26,762 | 8,99E-158 | 9,66E-157 | 1,00E-08 | 0,0035176 | 0,0017758 | 0,0017758 | TRUE | 0,0176191 | No |
| orf19.4280 | 8023 | 7385 | 7803 | 4774 | 895 | 690 | 920 | 892 | 7649 | 7809 | 7709 | 7479 | 717 | 785 | 781 | 725 | 4206,65 | 7662 | 752 | 0,098 | -3,352 | -75,424 | 0 | 0 | 0,0001946 | 0,0024816 | 0,0012307 | 0,0012307 | TRUE | 0,0116227 | No |
| orf19.4282 | 4752 | 4237 | 4647 | 2752 | 11207 | 7607 | 10472 | 11607 | 4531 | 4480 | 4591 | 4311 | 8980 | 8650 | 8891 | 9438 | 6733,82 | 4478 | 8990 | 2,006 | 1,005 | 25,206 | 3,46E-140 | 3,26E-139 | 0,0008094 | 0,0023351 | 0,0013501 | 0,0013501 | TRUE | 0,0137856 | No |
| orf19.4283 | 5800 | 5222 | 5522 | 3320 | 7572 | 5481 | 7420 | 7480 | 5530 | 5522 | 5455 | 5201 | 6067 | 6232 | 6300 | 6082 | 5798,57 | 5427 | 6170 | 1,136 | 0,184 | 5,29 | 1,22E-07 | 1,96E-07 | 0,0003438 | 0,0023744 | 0,0009824 | 0,0009824 | TRUE | 0,0089396 | No |
| orf19.4284 | 545 | 515 | 559 | 303 | 983 | 704 | 956 | 1005 | 520 | 545 | 552 | 475 | 788 | 801 | 812 | 817 | 663,5 | 523 | 804 | 1,531 | 0,615 | 9,602 | 7,87E-22 | 1,83E-21 | 1,00E-08 | 0,0045659 | 0,0023202 | 0,0023202 | TRUE | 0,0361882 | No |
| orf19.4286 | 4476 | 4169 | 4432 | 2616 | 2104 | 1470 | 2069 | 2040 | 4267 | 4408 | 4378 | 4098 | 1686 | 1672 | 1757 | 1659 | 2990,59 | 4288 | 1694 | 0,395 | -1,342 | -32,02 | 5,74E-225 | 8,63E-224 | 0,0005041 | 0,0026403 | 0,0012907 | 0,0012907 | TRUE | 0,0100944 | No |
| orf19.4287 | 808 | 629 | 729 | 432 | 3094 | 2045 | 3049 | 3154 | 770 | 665 | 720 | 677 | 2479 | 2325 | 2589 | 2565 | 1598,74 | 708 | 2490 | 3,509 | 1,811 | 29,414 | 3,59E-190 | 4,65E-189 | 0,0022115 | 0,0031183 | 0,0026716 | 0,0026716 | TRUE | 0,0429253 | No |
| orf19.4288 | 2319 | 2049 | 2223 | 1550 | 4375 | 3120 | 4020 | 4506 | 2211 | 2167 | 2196 | 2428 | 3506 | 3548 | 3413 | 3664 | 2891,47 | 2250 | 3533 | 1,573 | 0,653 | 13,808 | 2,29E-43 | 7,66E-43 | 0,0013684 | 0,0026591 | 0,0017732 | 0,0017732 | TRUE | 0,0314584 | No |
| orf19.429 | 3300 | 3023 | 3207 | 1923 | 4968 | 3419 | 4646 | 5163 | 3146 | 3196 | 3168 | 3012 | 3981 | 3888 | 3944 | 4198 | 3566,77 | 3130 | 4003 | 1,278 | 0,354 | 8,716 | 2,89E-18 | 6,25E-18 | 0,0005343 | 0,0025516 | 0,0012898 | 0,0012898 | TRUE | 0,0132419 | No |
| orf19.4290 | 4759 | 3993 | 4653 | 2749 | 5326 | 3731 | 5290 | 5657 | 4537 | 4222 | 4597 | 4306 | 4268 | 4242 | 4491 | 4600 | 4407,92 | 4416 | 4400 | 0,996 | -0,005 | -0,117 | 0,9069587 | 0,9197876 | 0,0013894 | 0,0024638 | 0,0017698 | 0,0017698 | TRUE | 0,0113925 | No |
| orf19.4292 | 1419 | 1210 | 1231 | 832 | 1316 | 937 | 1280 | 1397 | 1353 | 1279 | 1216 | 1303 | 1054 | 1065 | 1087 | 1136 | 1186,79 | 1288 | 1086 | 0,844 | -0,245 | -4,545 | 5,50E-06 | 8,26E-06 | 0,0008352 | 0,0034748 | 0,0019553 | 0,0019553 | TRUE | 0,0179223 | No |
| orf19.4293 | 3879 | 3097 | 3442 | 2222 | 2860 | 1972 | 2820 | 2894 | 3698 | 3275 | 3400 | 3481 | 2292 | 2242 | 2394 | 2353 | 2891,92 | 3464 | 2320 | 0,67 | -0,577 | -11,929 | 8,34E-33 | 2,36E-32 | 0,0012562 | 0,002659 | 0,001895 | 0,001895 | TRUE | 0,0262607 | No |
| orf19.4294 | 2734 | 2541 | 2822 | 1642 | 2697 | 1820 | 2563 | 2563 | 2607 | 2687 | 2788 | 2572 | 2161 | 2070 | 2176 | 2084 | 2393 | 2664 | 2123 | 0,797 | -0,328 | -7,577 | 3,55E-14 | 6,99E-14 | 0,0005351 | 0,0027774 | 0,0013787 | 0,0013787 | TRUE | 0,0121416 | No |
| orf19.4295 | 1366 | 1213 | 1382 | 778 | 1963 | 1452 | 1860 | 1965 | 1302 | 1283 | 1365 | 1219 | 1573 | 1651 | 1579 | 1598 | 1446,23 | 1292 | 1600 | 1,234 | 0,304 | 6,17 | 6,85E-10 | 1,18E-09 | 0,0002872 | 0,0032266 | 0,001618 | 0,001618 | TRUE | 0,0170817 | No |
| orf19.4297 | 2319 | 2366 | 2443 | 1542 | 1936 | 1456 | 1835 | 1929 | 2211 | 2502 | 2413 | 2416 | 1551 | 1656 | 1558 | 1568 | 1984,37 | 2386 | 1583 | 0,664 | -0,592 | -11,556 | 6,92E-31 | 1,89E-30 | 0,0014087 | 0,0029187 | 0,0019995 | 0,0019995 | TRUE | 0,0305593 | No |
| orf19.4299 | 1788 | 1584 | 1745 | 1145 | 916 | 619 | 910 | 924 | 1705 | 1675 | 1724 | 1794 | 734 | 704 | 773 | 751 | 1232,36 | 1724 | 740 | 0,431 | -1,215 | -23,563 | 9,27E-123 | 7,59E-122 | 1,00E-08 | 0,0034236 | 0,0016286 | 0,0016286 | TRUE | 0,0115755 | No |
| orf19.430 | 2152 | 1795 | 1993 | 1257 | 872 | 647 | 875 | 957 | 2052 | 1898 | 1969 | 1969 | 699 | 736 | 743 | 778 | 1355,39 | 1972 | 739 | 0,375 | -1,417 | -26,636 | 2,58E-156 | 2,73E-155 | 0,000687 | 0,0033027 | 0,0018278 | 0,0018278 | TRUE | 0,0171468 | No |
| orf19.4301 | 435 | 367 | 414 | 248 | 358 | 246 | 295 | 367 | 415 | 388 | 409 | 389 | 287 | 280 | 250 | 298 | 339,46 | 400 | 279 | 0,696 | -0,523 | -6,156 | 7,48E-10 | 1,29E-09 | 1,00E-08 | 0,006928 | 0,0039308 | 0,0039308 | TRUE | 0,055947 | No |
| orf19.4304 | 36040 | 32669 | 34838 | 21350 | 25566 | 18067 | 23561 | 24820 | 34360 | 34543 | 34417 | 33446 | 20485 | 20544 | 20003 | 20181 | 27247,43 | 34192 | 20303 | 0,594 | -0,752 | -29,366 | 1,51E-189 | 1,94E-188 | 0,0001351 | 0,0021515 | 0,0005917 | 0,0005917 | TRUE | 0,0025856 | No |
| orf19.4305 | 154 | 144 | 149 | 94 | 164 | 100 | 162 | 143 | 147 | 152 | 147 | 147 | 131 | 114 | 138 | 116 | 136,56 | 148 | 125 | 0,843 | -0,246 | -2,001 | 0,0453953 | 0,054799 | 1,00E-08 | 0,0141148 | 0,0072193 | 0,0072193 | TRUE | 0,0485406 | No |
| orf19.4305.1 | 1232 | 1018 | 1050 | 615 | 927 | 614 | 853 | 991 | 1175 | 1076 | 1037 | 963 | 743 | 698 | 724 | 806 | 902,83 | 1063 | 743 | 0,698 | -0,519 | -7,102 | 1,23E-12 | 2,31E-12 | 0,0038028 | 0,0039099 | 0,0039975 | 0,0039975 | TRUE | 0,0601049 | No |
| orf19.4306 | 1329 | 1154 | 1247 | 846 | 2316 | 1752 | 2233 | 2362 | 1267 | 1220 | 1232 | 1325 | 1856 | 1992 | 1896 | 1921 | 1588,59 | 1261 | 1916 | 1,521 | 0,605 | 12,671 | 8,51E-37 | 2,55E-36 | 0,0002713 | 0,0031248 | 0,0015118 | 0,0015118 | TRUE | 0,0121433 | No |
| orf19.4307 | 3301 | 2955 | 2888 | 1883 | 2886 | 1903 | 2659 | 2817 | 3147 | 3125 | 2853 | 2950 | 2312 | 2164 | 2257 | 2290 | 2637,36 | 3019 | 2256 | 0,748 | -0,42 | -8,901 | 5,51E-19 | 1,21E-18 | 0,0011737 | 0,0027138 | 0,0017509 | 0,0017509 | TRUE | 0,0174929 | No |
| orf19.4308 | 397 | 353 | 373 | 261 | 1219 | 841 | 1126 | 1351 | 378 | 373 | 368 | 409 | 977 | 956 | 956 | 1098 | 689,58 | 382 | 997 | 2,619 | 1,389 | 18,774 | 1,24E-78 | 6,66E-78 | 0,0024306 | 0,0044723 | 0,0033447 | 0,0033447 | TRUE | 0,057435 | No |
| orf19.4309 | 3683 | 3306 | 3350 | 2226 | 601 | 449 | 599 | 576 | 3511 | 3496 | 3309 | 3487 | 482 | 511 | 509 | 468 | 1971,58 | 3451 | 492 | 0,142 | -2,812 | -54,254 | 0 | 0 | 1,00E-08 | 0,0029241 | 0,001512 | 0,001512 | TRUE | 0,0126377 | No |
| orf19.431 | 7604 | 6593 | 7260 | 4616 | 3758 | 2723 | 3588 | 3960 | 7250 | 6971 | 7172 | 7231 | 3011 | 3096 | 3046 | 3220 | 5124,72 | 7156 | 3093 | 0,432 | -1,21 | -33,54 | 1,24E-246 | 2,08E-245 | 0,0002854 | 0,0024117 | 0,0010262 | 0,0010262 | TRUE | 0,0095504 | No |
| orf19.4310 | 1 | 0 | 1 | 1 | 0 | 0 | 0 | 1 | 1 | 0 | 1 | 2 | 0 | 0 | 0 | 1 | 0,54 | 1 | 0 | 0,317 | -1,655 | -0,854 | 0,3932019 | 0,4291126 | 1,00E-08 | 3,0419876 | 1,9522131 | 1,9522131 | TRUE | 0,2944903 | No |
| orf19.4311 | 2458 | 2068 | 2212 | 1327 | 8884 | 5741 | 8144 | 8554 | 2343 | 2187 | 2185 | 2079 | 7118 | 6528 | 6914 | 6955 | 4538,76 | 2198 | 6879 | 3,125 | 1,644 | 34,678 | 1,71E-263 | 3,10E-262 | 0,0014897 | 0,002453 | 0,0018373 | 0,0018373 | TRUE | 0,0240355 | No |
| orf19.4312 | 913 | 771 | 847 | 487 | 1734 | 1114 | 1668 | 1631 | 870 | 815 | 837 | 763 | 1389 | 1267 | 1416 | 1326 | 1085,47 | 821 | 1350 | 1,64 | 0,713 | 11,803 | 3,78E-32 | 1,06E-31 | 0,0014303 | 0,0036039 | 0,0024973 | 0,0024973 | TRUE | 0,0235896 | No |
| orf19.4315 | 2033 | 2201 | 2308 | 1282 | 2139 | 1658 | 1942 | 1818 | 1938 | 2327 | 2280 | 2008 | 1714 | 1885 | 1649 | 1478 | 1910,01 | 2138 | 1682 | 0,785 | -0,349 | -4,225 | 2,39E-05 | 3,48E-05 | 0,0084095 | 0,0029509 | 0,0060122 | 0,0060122 | TRUE | 0,0803002 | No |
| orf19.4316 | 700 | 694 | 722 | 461 | 2175 | 1720 | 1983 | 2037 | 667 | 734 | 713 | 722 | 1743 | 1956 | 1684 | 1656 | 1234,38 | 709 | 1760 | 2,481 | 1,311 | 19,461 | 2,36E-84 | 1,33E-83 | 0,0032264 | 0,0034214 | 0,0033099 | 0,0033099 | TRUE | 0,0657677 | No |
| orf19.4317 | 2151 | 2085 | 2080 | 1282 | 836 | 603 | 679 | 723 | 2051 | 2205 | 2055 | 2008 | 670 | 686 | 576 | 588 | 1354,8 | 2080 | 630 | 0,302 | -1,727 | -26,046 | 1,50E-149 | 1,50E-148 | 0,0033927 | 0,0033032 | 0,0032389 | 0,0032389 | TRUE | 0,0382856 | No |
| orf19.4318 | 17510 | 16362 | 18446 | 11314 | 6289 | 4569 | 6091 | 6265 | 16694 | 17301 | 18223 | 17724 | 5039 | 5195 | 5171 | 5094 | 11305,17 | 17486 | 5125 | 0,293 | -1,771 | -47,881 | 0 | 0 | 0,0006172 | 0,0022365 | 0,0011939 | 0,0011939 | TRUE | 0,0114816 | No |
| orf19.432 | 2978 | 2591 | 2827 | 1748 | 3503 | 2451 | 3518 | 3686 | 2839 | 2740 | 2793 | 2738 | 2807 | 2787 | 2987 | 2997 | 2835,96 | 2778 | 2894 | 1,042 | 0,06 | 1,414 | 0,1573799 | 0,1802622 | 0,0006175 | 0,0026702 | 0,0013462 | 0,0013462 | TRUE | 0,0071824 | No |
| orf19.4320 | 11 | 5 | 6 | 3 | 4 | 9 | 6 | 4 | 10 | 5 | 6 | 5 | 3 | 10 | 5 | 3 | 6,02 | 6 | 5 | 0,781 | -0,357 | -0,585 | 0,5582619 | 0,5930197 | 0,072699 | 0,2746826 | 0,1891758 | 0,1891758 | TRUE | 0,638239 | No |
| orf19.4321 | 34 | 10 | 20 | 20 | 35 | 22 | 28 | 34 | 32 | 11 | 20 | 31 | 28 | 25 | 24 | 28 | 24,82 | 24 | 26 | 1,125 | 0,17 | 0,512 | 0,6085915 | 0,6425898 | 0,0587385 | 0,0682451 | 0,0645653 | 0,0645653 | TRUE | 0,5420086 | No |
| orf19.4322 | 7371 | 7081 | 7496 | 4323 | 8348 | 5927 | 7668 | 8152 | 7027 | 7487 | 7405 | 6772 | 6689 | 6740 | 6510 | 6628 | 6907,41 | 7173 | 6642 | 0,925 | -0,112 | -2,754 | 0,0058872 | 0,0075738 | 0,0010011 | 0,002329 | 0,0014433 | 0,0014433 | TRUE | 0,0165722 | No |
| orf19.4323 | 1469 | 1444 | 1402 | 781 | 564 | 401 | 527 | 487 | 1401 | 1527 | 1385 | 1223 | 452 | 456 | 447 | 396 | 910,9 | 1384 | 438 | 0,315 | -1,665 | -21,154 | 2,56E-99 | 1,67E-98 | 0,005086 | 0,0038938 | 0,0045216 | 0,0045216 | TRUE | 0,0681598 | No |
| orf19.4324 | 498 | 438 | 438 | 291 | 560 | 336 | 494 | 478 | 475 | 463 | 433 | 456 | 449 | 382 | 419 | 389 | 433,17 | 457 | 410 | 0,899 | -0,153 | -1,937 | 0,0527779 | 0,0632918 | 0,0015496 | 0,0058818 | 0,0037185 | 0,0037185 | TRUE | 0,0487202 | No |
| orf19.4325 | 746 | 814 | 838 | 496 | 753 | 551 | 622 | 638 | 711 | 861 | 828 | 777 | 603 | 627 | 528 | 519 | 681,69 | 794 | 569 | 0,715 | -0,483 | -5,651 | 1,60E-08 | 2,64E-08 | 0,0064587 | 0,0044999 | 0,005528 | 0,005528 | TRUE | 0,0622876 | No |
| orf19.4326 | 1519 | 1563 | 1636 | 957 | 1991 | 1397 | 1873 | 1826 | 1448 | 1653 | 1616 | 1499 | 1595 | 1589 | 1590 | 1485 | 1559,38 | 1554 | 1565 | 1,006 | 0,009 | 0,151 | 0,8800316 | 0,8966122 | 0,0018866 | 0,0031442 | 0,002412 | 0,002412 | TRUE | 0,0274753 | No |
| orf19.4328 | 9486 | 9638 | 10104 | 6236 | 7555 | 5949 | 6948 | 7252 | 9044 | 10191 | 9982 | 9769 | 6054 | 6765 | 5899 | 5897 | 7949,9 | 9746 | 6154 | 0,631 | -0,664 | -11,722 | 9,87E-32 | 2,74E-31 | 0,0033976 | 0,0022978 | 0,0029518 | 0,0029518 | TRUE | 0,0539154 | No |
| orf19.433 | 212 | 169 | 199 | 120 | 493 | 336 | 464 | 517 | 202 | 179 | 197 | 188 | 395 | 382 | 394 | 420 | 294,6 | 192 | 398 | 2,077 | 1,055 | 11,701 | 1,26E-31 | 3,50E-31 | 1,00E-08 | 0,0076646 | 0,0037868 | 0,0037868 | TRUE | 0,0238733 | No |
| orf19.4330 | 485 | 362 | 418 | 238 | 506 | 307 | 444 | 468 | 462 | 383 | 413 | 373 | 405 | 349 | 377 | 381 | 392,87 | 408 | 378 | 0,925 | -0,112 | -1,257 | 0,2086229 | 0,2361999 | 0,0038968 | 0,0062705 | 0,0051113 | 0,0051113 | TRUE | 0,096108 | No |
| orf19.4332 | 822 | 713 | 782 | 494 | 972 | 642 | 901 | 938 | 784 | 754 | 773 | 774 | 779 | 730 | 765 | 763 | 765,06 | 771 | 759 | 0,986 | -0,021 | -0,367 | 0,7133336 | 0,7430494 | 1,00E-08 | 0,0042374 | 0,0016935 | 0,0016935 | TRUE | 0,0070734 | No |
| orf19.4334 | 94 | 71 | 87 | 68 | 110 | 115 | 130 | 134 | 90 | 75 | 86 | 107 | 88 | 131 | 110 | 109 | 99,42 | 90 | 110 | 1,23 | 0,299 | 1,812 | 0,0699704 | 0,0830172 | 0,012147 | 0,0186054 | 0,0157686 | 0,0157686 | TRUE | 0,1812492 | No |
| orf19.4335 | 278 | 296 | 240 | 175 | 3036 | 2140 | 2821 | 2971 | 265 | 313 | 237 | 274 | 2433 | 2433 | 2395 | 2416 | 1345,75 | 272 | 2419 | 8,904 | 3,154 | 45,957 | 0 | 0 | 1,00E-08 | 0,0033113 | 0,0022972 | 0,0022972 | TRUE | 0,123259 | No |
| orf19.4336 | 12170 | 10109 | 10141 | 7408 | 43509 | 31396 | 40771 | 45534 | 11603 | 10689 | 10018 | 11605 | 34863 | 35700 | 34614 | 37023 | 23264,45 | 10979 | 35550 | 3,239 | 1,695 | 32,209 | 1,34E-227 | 2,04E-226 | 0,0028599 | 0,0021619 | 0,0025982 | 0,0025982 | TRUE | 0,0425678 | No |
| orf19.4337 | 3537 | 3275 | 3482 | 2133 | 2995 | 2258 | 2926 | 3001 | 3372 | 3463 | 3440 | 3341 | 2400 | 2568 | 2484 | 2440 | 2938,5 | 3404 | 2473 | 0,725 | -0,463 | -12,101 | 1,04E-33 | 2,97E-33 | 9,64E-05 | 0,00265 | 0,0010597 | 0,0010597 | TRUE | 0,0075632 | No |
| orf19.4339 | 3345 | 2920 | 3231 | 2192 | 4316 | 3048 | 4183 | 4629 | 3189 | 3088 | 3192 | 3434 | 3458 | 3466 | 3551 | 3764 | 3392,72 | 3226 | 3560 | 1,105 | 0,144 | 3,04 | 0,0023637 | 0,0031201 | 0,0013669 | 0,0025752 | 0,001863 | 0,001863 | TRUE | 0,0215803 | No |
| orf19.434 | 546 | 500 | 559 | 326 | 613 | 408 | 598 | 579 | 521 | 529 | 552 | 511 | 491 | 464 | 508 | 471 | 505,72 | 528 | 484 | 0,915 | -0,128 | -1,872 | 0,0611432 | 0,072858 | 1,00E-08 | 0,0053379 | 0,002516 | 0,002516 | TRUE | 0,0132102 | No |
| orf19.4340 | 1014 | 929 | 925 | 551 | 1357 | 894 | 1313 | 1286 | 967 | 982 | 914 | 863 | 1087 | 1017 | 1115 | 1046 | 998,78 | 932 | 1066 | 1,142 | 0,192 | 3,226 | 0,0012541 | 0,0016787 | 0,0011843 | 0,0037352 | 0,0023852 | 0,0023852 | TRUE | 0,0251285 | No |
| orf19.4340.1 | 1132 | 1056 | 1083 | 566 | 1196 | 692 | 1103 | 996 | 1079 | 1117 | 1070 | 887 | 958 | 787 | 936 | 810 | 955,48 | 1038 | 873 | 0,84 | -0,252 | -2,861 | 0,004221 | 0,0054834 | 0,0084797 | 0,0038097 | 0,0063986 | 0,0063986 | TRUE | 0,1072924 | No |
| orf19.4341 | 369 | 413 | 403 | 261 | 888 | 612 | 834 | 832 | 352 | 437 | 398 | 409 | 712 | 696 | 708 | 676 | 548,43 | 399 | 698 | 1,754 | 0,811 | 10,771 | 4,73E-27 | 1,22E-26 | 0,0013082 | 0,0050851 | 0,0034005 | 0,0034005 | TRUE | 0,0790743 | No |
| orf19.4342 | 2662 | 2589 | 2835 | 1674 | 2084 | 1436 | 1940 | 1995 | 2538 | 2738 | 2801 | 2622 | 1670 | 1633 | 1647 | 1622 | 2158,81 | 2675 | 1643 | 0,614 | -0,703 | -15,405 | 1,52E-53 | 5,88E-53 | 0,0007453 | 0,0028518 | 0,0015195 | 0,0015195 | TRUE | 0,015545 | No |
| orf19.4346 | 1880 | 1623 | 1763 | 1198 | 3150 | 2189 | 2833 | 3012 | 1792 | 1716 | 1742 | 1877 | 2524 | 2489 | 2405 | 2449 | 2124,28 | 1782 | 2467 | 1,387 | 0,472 | 10,783 | 4,15E-27 | 1,07E-26 | 0,0003614 | 0,0028642 | 0,0013423 | 0,0013423 | TRUE | 0,0136984 | No |
| orf19.4347 | 8411 | 7558 | 8546 | 5481 | 2525 | 1772 | 2349 | 2500 | 8019 | 7992 | 8443 | 8586 | 2023 | 2015 | 1994 | 2033 | 5138,09 | 8260 | 2016 | 0,244 | -2,034 | -51,739 | 0 | 0 | 0,0006342 | 0,0024108 | 0,0011939 | 0,0011939 | TRUE | 0,008309 | No |
| orf19.4348 | 843 | 992 | 1054 | 584 | 1198 | 882 | 1086 | 1041 | 804 | 1049 | 1041 | 915 | 960 | 1003 | 922 | 846 | 942,5 | 952 | 933 | 0,979 | -0,031 | -0,347 | 0,7288544 | 0,7580798 | 0,0093129 | 0,0038333 | 0,0067255 | 0,0067255 | TRUE | 0,1376798 | No |
| orf19.4349 | 732 | 687 | 786 | 407 | 700 | 492 | 653 | 725 | 698 | 726 | 776 | 638 | 561 | 559 | 554 | 589 | 637,83 | 710 | 566 | 0,795 | -0,331 | -4,668 | 3,04E-06 | 4,62E-06 | 0,0018553 | 0,0046655 | 0,0032502 | 0,0032502 | TRUE | 0,0476186 | No |
| orf19.4353 | 2075 | 2111 | 2270 | 1345 | 1180 | 948 | 1159 | 1297 | 1978 | 2232 | 2243 | 2107 | 946 | 1078 | 984 | 1055 | 1577,75 | 2140 | 1016 | 0,474 | -1,077 | -17,337 | 2,49E-67 | 1,15E-66 | 0,0029086 | 0,0031319 | 0,003004 | 0,003004 | TRUE | 0,0327388 | No |
| orf19.4354 | 3915 | 3629 | 3917 | 2299 | 5613 | 3869 | 5193 | 5844 | 3733 | 3837 | 3870 | 3602 | 4498 | 4399 | 4409 | 4752 | 4137,3 | 3760 | 4514 | 1,2 | 0,263 | 6,235 | 4,51E-10 | 7,88E-10 | 0,0008542 | 0,0024881 | 0,0014593 | 0,0014593 | TRUE | 0,0153973 | No |
| orf19.4355 | 1715 | 1749 | 1803 | 1096 | 1401 | 1034 | 1319 | 1336 | 1635 | 1849 | 1781 | 1717 | 1123 | 1176 | 1120 | 1086 | 1435,88 | 1746 | 1126 | 0,644 | -0,634 | -11,678 | 1,65E-31 | 4,55E-31 | 0,0013249 | 0,0032348 | 0,0021116 | 0,0021116 | TRUE | 0,0234936 | No |
| orf19.4356 | 188 | 219 | 196 | 136 | 1035 | 727 | 904 | 964 | 179 | 232 | 194 | 213 | 829 | 827 | 767 | 784 | 503,1 | 204 | 802 | 3,94 | 1,978 | 22,948 | 1,56E-116 | 1,20E-115 | 0,0014177 | 0,0053549 | 0,0038467 | 0,0038467 | TRUE | 0,0991999 | No |

| | | | | | | | | | | | | | | | | | | | | | | | | | | | | | | | |
|---|---|---|---|---|---|---|---|---|---|---|---|---|---|---|---|---|---|---|---|---|---|---|---|---|---|---|---|---|---|---|---|
| orf19.4357 | 1706 | 1619 | 1652 | 1019 | 2927 | 2064 | 2646 | 2773 | 1626 | 1712 | 1632 | 1596 | 2345 | 2347 | 2246 | 2255 | 1970,02 | 1642 | 2298 | 1,398 | 0,484 | 11,162 | 6,22E-29 | 1,65E-28 | 0,0001941 | 0,0029247 | 0,0012672 | 0,0012672 | TRUE | 0,0099226 | No |
| orf19.4358 | 1458 | 1217 | 1269 | 766 | 987 | 597 | 950 | 1023 | 1390 | 1287 | 1254 | 1200 | 791 | 679 | 807 | 832 | 1029,82 | 1283 | 777 | 0,606 | -0,722 | -9,885 | 4,85E-23 | 1,15E-22 | 0,0043527 | 0,0036856 | 0,0041094 | 0,0041094 | TRUE | 0,0804039 | No |
| orf19.4361 | 75 | 49 | 50 | 34 | 25 | 12 | 24 | 23 | 72 | 52 | 49 | 53 | 20 | 14 | 20 | 19 | 37,34 | 56 | 18 | 0,325 | -1,624 | -6,31 | 2,78E-10 | 4,90E-10 | 1,00E-08 | 0,0460619 | 0,0296053 | 0,0296053 | TRUE | 0,2947739 | No |
| orf19.4362 | 2260 | 2206 | 2436 | 1335 | 2561 | 1797 | 2563 | 2577 | 2155 | 2333 | 2407 | 2091 | 2052 | 2043 | 2176 | 2095 | 2168,98 | 2246 | 2092 | 0,93 | -0,104 | -1,927 | 0,0540023 | 0,0646751 | 0,0019688 | 0,0028483 | 0,0023568 | 0,0023568 | TRUE | 0,027914 | No |
| orf19.4363 | 8027 | 7439 | 8204 | 4695 | 9526 | 6151 | 9303 | 9737 | 7653 | 7866 | 8105 | 7355 | 7633 | 6994 | 7898 | 7917 | 7677,61 | 7745 | 7610 | 0,983 | -0,025 | -0,498 | 0,6181625 | 0,6517672 | 0,0020424 | 0,0023051 | 0,0023257 | 0,0023257 | TRUE | 0,0358142 | No |
| orf19.4365 | 1836 | 1874 | 1941 | 1259 | 1741 | 1346 | 1738 | 1917 | 1750 | 1982 | 1918 | 1972 | 1395 | 1531 | 1476 | 1559 | 1697,69 | 1906 | 1490 | 0,782 | -0,354 | -6,125 | 9,05E-10 | 1,56E-09 | 0,0023013 | 0,0030584 | 0,002612 | 0,002612 | TRUE | 0,0372803 | No |
| orf19.4366 | 44 | 33 | 42 | 34 | 25 | 9 | 21 | 18 | 42 | 35 | 41 | 53 | 20 | 10 | 18 | 15 | 29,29 | 43 | 16 | 0,376 | -1,413 | -4,945 | 7,61E-07 | 1,18E-06 | 1,00E-08 | 0,0581463 | 0,0373411 | 0,0373411 | TRUE | 0,1998982 | No |
| orf19.4367 | 617 | 594 | 603 | 395 | 1796 | 1291 | 1615 | 1793 | 588 | 628 | 596 | 619 | 1439 | 1468 | 1371 | 1458 | 1020,86 | 608 | 1434 | 2,362 | 1,24 | 22,373 | 7,25E-111 | 5,29E-110 | 1,00E-08 | 0,0036996 | 0,001721 | 0,001721 | TRUE | 0,0105768 | No |
| orf19.4368 | 1220 | 1074 | 1216 | 826 | 2718 | 1992 | 2556 | 2772 | 1163 | 1136 | 1201 | 1294 | 2178 | 2265 | 2170 | 2254 | 1707,61 | 1198 | 2217 | 1,855 | 0,891 | 17,759 | 1,46E-70 | 7,00E-70 | 0,0005936 | 0,0030528 | 0,0017471 | 0,0017471 | TRUE | 0,0300772 | No |
| orf19.4369 | 2076 | 1987 | 2095 | 1332 | 1704 | 1146 | 1632 | 1724 | 1979 | 2101 | 2070 | 2087 | 1365 | 1303 | 1386 | 1402 | 1711,55 | 2059 | 1364 | 0,664 | -0,592 | -13,002 | 1,20E-38 | 3,72E-38 | 0,0002442 | 0,0030506 | 0,0013888 | 0,0013888 | TRUE | 0,0099301 | No |
| orf19.437 | 4873 | 4353 | 4774 | 2780 | 13352 | 8874 | 12771 | 13929 | 4646 | 4603 | 4716 | 4355 | 10699 | 10091 | 10843 | 11326 | 7659,64 | 4580 | 10740 | 2,344 | 1,229 | 26,957 | 4,70E-160 | 5,12E-159 | 0,0015034 | 0,0023056 | 0,0018308 | 0,0018308 | TRUE | 0,0199817 | No |
| orf19.4370 | 2691 | 2490 | 2641 | 1615 | 1674 | 1233 | 1536 | 1689 | 2566 | 2633 | 2609 | 2530 | 1341 | 1402 | 1304 | 1373 | 1969,78 | 2584 | 1355 | 0,524 | -0,933 | -21,937 | 1,16E-106 | 8,04E-106 | 1,00E-08 | 0,0029248 | 0,0011922 | 0,0011922 | TRUE | 0,0076464 | No |
| orf19.4371 | 2963 | 2517 | 2659 | 1830 | 4842 | 3388 | 4555 | 4888 | 2825 | 2661 | 2627 | 2867 | 3880 | 3852 | 3867 | 3974 | 3319,22 | 2745 | 3893 | 1,42 | 0,506 | 12,262 | 1,46E-34 | 4,22E-34 | 0,0006188 | 0,0025859 | 0,0013157 | 0,0013157 | TRUE | 0,0102099 | No |
| orf19.4372 | 5333 | 5067 | 5410 | 3454 | 1311 | 949 | 1178 | 1253 | 5084 | 5358 | 5345 | 5411 | 1050 | 1079 | 1000 | 1019 | 3168,26 | 5300 | 1037 | 0,196 | -2,354 | -53,905 | 0 | 0 | 0,0004977 | 0,0026095 | 0,0012954 | 0,0012954 | TRUE | 0,0092601 | No |
| orf19.4373 | 1735 | 1617 | 1577 | 1130 | 1933 | 1404 | 1872 | 1928 | 1654 | 1710 | 1558 | 1770 | 1549 | 1596 | 1589 | 1568 | 1624,29 | 1673 | 1576 | 0,943 | -0,084 | -1,684 | 0,0921041 | 0,1078576 | 0,0007254 | 0,0031021 | 0,0017718 | 0,0017718 | TRUE | 0,0259573 | No |
| orf19.4374 | 598 | 599 | 632 | 354 | 1063 | 716 | 999 | 1034 | 570 | 633 | 624 | 555 | 852 | 814 | 848 | 841 | 717,15 | 596 | 839 | 1,405 | 0,491 | 7,766 | 8,13E-15 | 1,63E-14 | 1,00E-08 | 0,0043808 | 0,0023678 | 0,0023678 | TRUE | 0,0206047 | No |
| orf19.4375 | 2839 | 2354 | 2562 | 1656 | 3581 | 2419 | 3473 | 3631 | 2707 | 2489 | 2531 | 2594 | 2869 | 2751 | 2949 | 2952 | 2730,23 | 2580 | 2880 | 1,117 | 0,159 | 3,583 | 0,0003398 | 0,0004696 | 0,0007222 | 0,0026927 | 0,0015292 | 0,0015292 | TRUE | 0,0139876 | No |
| orf19.4375.1 | 10056 | 7792 | 8021 | 6341 | 20335 | 14022 | 17430 | 21226 | 9587 | 8239 | 7924 | 9934 | 16294 | 15944 | 14798 | 17259 | 12497,34 | 8921 | 16074 | 1,802 | 0,85 | 11,204 | 3,90E-29 | 1,04E-28 | 0,0080141 | 0,0022227 | 0,0054372 | 0,0054372 | TRUE | 0,0706503 | No |
| orf19.4376 | 384 | 371 | 379 | 256 | 381 | 278 | 372 | 379 | 366 | 392 | 374 | 401 | 305 | 316 | 316 | 308 | 347,4 | 383 | 311 | 0,814 | -0,297 | -3,799 | 0,0001451 | 0,0002035 | 1,00E-08 | 0,0068175 | 0,0029918 | 0,0029918 | TRUE | 0,0106884 | No |
| orf19.4377 | 2574 | 2458 | 2626 | 1636 | 1400 | 965 | 1407 | 1390 | 2454 | 2599 | 2594 | 2563 | 1122 | 1097 | 1195 | 1130 | 1844,25 | 2552 | 1136 | 0,446 | -1,166 | -25,14 | 1,82E-139 | 1,70E-138 | 0,000414 | 0,0029816 | 0,0014547 | 0,0014547 | TRUE | 0,0144448 | No |
| orf19.4378 | 715 | 561 | 600 | 357 | 1271 | 764 | 1244 | 1217 | 682 | 593 | 593 | 559 | 1018 | 869 | 1056 | 990 | 794,96 | 607 | 983 | 1,618 | 0,694 | 8,561 | 1,12E-17 | 2,40E-17 | 0,0053553 | 0,0041567 | 0,0049408 | 0,0049408 | TRUE | 0,0831281 | No |
| orf19.4379 | 1701 | 1283 | 1587 | 1019 | 1654 | 1095 | 1524 | 1789 | 1622 | 1357 | 1568 | 1596 | 1325 | 1245 | 1294 | 1455 | 1432,67 | 1536 | 1330 | 0,867 | -0,206 | -2,909 | 0,0036311 | 0,0047363 | 0,0044553 | 0,0032373 | 0,0041104 | 0,0041104 | TRUE | 0,0750457 | No |
| orf19.438 | 720 | 765 | 741 | 541 | 1056 | 718 | 978 | 1063 | 686 | 809 | 732 | 848 | 846 | 816 | 830 | 864 | 804,01 | 769 | 839 | 1,097 | 0,133 | 1,873 | 0,0610365 | 0,0727445 | 0,0031351 | 0,0041334 | 0,0036062 | 0,0036062 | TRUE | 0,0621991 | No |
| orf19.4380.1 | 1803 | 1455 | 1626 | 1059 | 1594 | 1029 | 1481 | 1725 | 1719 | 1538 | 1606 | 1659 | 1277 | 1170 | 1257 | 1403 | 1453,75 | 1630 | 1277 | 0,784 | -0,351 | -5,508 | 3,64E-08 | 5,94E-08 | 0,0029987 | 0,0032207 | 0,0032005 | 0,0032005 | TRUE | 0,0529632 | No |
| orf19.4381 | 2444 | 2317 | 2488 | 1717 | 3719 | 2722 | 3294 | 3566 | 2330 | 2450 | 2458 | 2690 | 2980 | 3095 | 2797 | 2899 | 2712,36 | 2482 | 2943 | 1,187 | 0,248 | 4,605 | 4,12E-06 | 6,22E-06 | 0,0022615 | 0,0026966 | 0,0023943 | 0,0023943 | TRUE | 0,0363284 | No |
| orf19.4382 | 1278 | 1228 | 1202 | 720 | 2121 | 1473 | 1881 | 1878 | 1218 | 1298 | 1187 | 1128 | 1699 | 1675 | 1597 | 1527 | 1416,33 | 1208 | 1624 | 1,342 | 0,424 | 7,234 | 4,68E-13 | 8,97E-13 | 0,0019292 | 0,0032505 | 0,0025612 | 0,0025612 | TRUE | 0,0296862 | No |
| orf19.4383 | 151 | 149 | 162 | 128 | 209 | 175 | 235 | 219 | 144 | 158 | 160 | 201 | 167 | 199 | 200 | 178 | 175,76 | 166 | 186 | 1,133 | 0,18 | 1,442 | 0,1493737 | 0,1714961 | 0,0062646 | 0,0114328 | 0,0091975 | 0,0091975 | TRUE | 0,1756563 | No |
| orf19.4384 | 28 | 11 | 28 | 18 | 49 | 26 | 51 | 38 | 27 | 12 | 28 | 28 | 39 | 30 | 43 | 31 | 29,65 | 24 | 36 | 1,535 | 0,618 | 2,068 | 0,0386319 | 0,0469096 | 0,0359845 | 0,0574655 | 0,0495136 | 0,0495136 | TRUE | 0,6898631 | No |
| orf19.4384.1 | 33 | 20 | 22 | 12 | 55 | 22 | 57 | 52 | 31 | 21 | 22 | 19 | 44 | 25 | 48 | 42 | 31,61 | 23 | 40 | 1,709 | 0,773 | 2,692 | 0,0071105 | 0,0090981 | 0,031084 | 0,0540294 | 0,0445568 | 0,0445568 | TRUE | 0,4083086 | No |
| orf19.4388 | 1920 | 1551 | 1835 | 1084 | 2193 | 1516 | 2148 | 2187 | 1831 | 1640 | 1813 | 1698 | 1757 | 1724 | 1824 | 1778 | 1758,04 | 1746 | 1771 | 1,014 | 0,02 | 0,391 | 0,6954653 | 0,7263116 | 0,0010498 | 0,0030252 | 0,0018579 | 0,0018579 | TRUE | 0,0212691 | No |
| orf19.439 | 2395 | 1919 | 2177 | 1469 | 2838 | 1954 | 2684 | 2814 | 2283 | 2029 | 2151 | 2301 | 2274 | 2222 | 2279 | 2288 | 2228,38 | 2191 | 2266 | 1,035 | 0,05 | 1,021 | 0,3072513 | 0,3404359 | 0,0011725 | 0,0028281 | 0,0018602 | 0,0018602 | TRUE | 0,0299611 | No |
| orf19.4390 | 624 | 592 | 606 | 377 | 835 | 603 | 765 | 874 | 595 | 626 | 599 | 591 | 669 | 686 | 649 | 711 | 640,62 | 603 | 679 | 1,125 | 0,17 | 2,733 | 0,0062807 | 0,0080703 | 1,00E-08 | 0,0046543 | 0,0021547 | 0,0021547 | TRUE | 0,0127488 | No |
| orf19.4391 | 65 | 52 | 50 | 39 | 37 | 36 | 25 | 41 | 62 | 55 | 49 | 61 | 30 | 41 | 21 | 33 | 44,07 | 57 | 31 | 0,546 | -0,874 | -3,757 | 0,0001717 | 0,0002402 | 0,0075424 | 0,039345 | 0,0277288 | 0,0277288 | TRUE | 0,3375239 | No |
| orf19.4392 | 407 | 365 | 381 | 225 | 571 | 415 | 565 | 592 | 388 | 386 | 376 | 352 | 458 | 472 | 480 | 481 | 424,16 | 376 | 473 | 1,253 | 0,326 | 4,497 | 6,90E-06 | 1,03E-05 | 1,00E-08 | 0,0059622 | 0,0026186 | 0,0026186 | TRUE | 0,0157334 | No |
| orf19.4393 | 12760 | 10656 | 11509 | 8198 | 29574 | 21026 | 28537 | 31196 | 12165 | 11267 | 11370 | 12843 | 23697 | 23908 | 24228 | 25365 | 18105,43 | 11911 | 24300 | 2,041 | 1,029 | 21,095 | 8,89E-99 | 5,74E-98 | 0,0023106 | 0,002182 | 0,0022208 | 0,0022208 | TRUE | 0,0333333 | No |
| orf19.4394 | 17 | 11 | 18 | 19 | 28 | 25 | 28 | 26 | 16 | 12 | 18 | 30 | 22 | 28 | 24 | 21 | 21,4 | 19 | 24 | 1,304 | 0,383 | 1,17 | 0,2420587 | 0,2718846 | 0,0051185 | 0,0788333 | 0,053727 | 0,053727 | TRUE | 0,5171492 | No |
| orf19.4395 | 1295 | 1164 | 1246 | 823 | 3869 | 2729 | 3832 | 4052 | 1235 | 1231 | 1231 | 1289 | 3100 | 3103 | 3253 | 3295 | 2217,11 | 1246 | 3188 | 2,563 | 1,358 | 30,45 | 1,21E-203 | 1,68E-202 | 0,0004523 | 0,0028318 | 0,0013191 | 0,0013191 | TRUE | 0,0062365 | No |
| orf19.4396 | 3583 | 3083 | 3534 | 2146 | 8019 | 6005 | 7608 | 8626 | 3416 | 3260 | 3491 | 3362 | 6425 | 6828 | 6459 | 7014 | 5031,94 | 3382 | 6682 | 1,975 | 0,982 | 22,355 | 1,07E-110 | 7,78E-110 | 0,001039 | 0,0024176 | 0,0016183 | 0,0016183 | TRUE | 0,013932 | No |
| orf19.4398 | 2151 | 1853 | 1975 | 1378 | 2490 | 1795 | 2413 | 2581 | 2051 | 1959 | 1951 | 2159 | 1995 | 2041 | 2049 | 2099 | 2037,92 | 2030 | 2046 | 1,01 | 0,014 | 0,3 | 0,7644167 | 0,7911318 | 0,0006722 | 0,002897 | 0,0015756 | 0,0015756 | TRUE | 0,0198969 | No |
| orf19.4399 | 9811 | 8410 | 9100 | 5659 | 9771 | 6551 | 9507 | 10165 | 9354 | 8893 | 8990 | 8865 | 7829 | 7449 | 8071 | 8265 | 8464,51 | 9026 | 7904 | 0,876 | -0,191 | -4,599 | 4,24E-06 | 6,40E-06 | 0,0011434 | 0,0022853 | 0,0015419 | 0,0015419 | TRUE | 0,0179708 | No |
| orf19.440 | 499 | 430 | 457 | 341 | 808 | 548 | 826 | 788 | 476 | 455 | 451 | 534 | 647 | 623 | 701 | 641 | 566,08 | 479 | 653 | 1,372 | 0,456 | 6,133 | 8,62E-10 | 1,48E-09 | 0,0017869 | 0,0049918 | 0,003452 | 0,003452 | TRUE | 0,0590773 | No |
| orf19.4400 | 471 | 358 | 422 | 287 | 622 | 387 | 553 | 604 | 449 | 379 | 417 | 450 | 498 | 440 | 469 | 491 | 449,14 | 424 | 474 | 1,126 | 0,171 | 2,109 | 0,0349427 | 0,0425596 | 0,0021278 | 0,0057469 | 0,0040223 | 0,0040223 | TRUE | 0,0592917 | No |
| orf19.4401 | 3461 | 2937 | 3128 | 2080 | 3969 | 2807 | 3706 | 4127 | 3300 | 3106 | 3090 | 3258 | 3180 | 3192 | 3146 | 3356 | 3203,48 | 3188 | 3218 | 1,01 | 0,014 | 0,344 | 0,7311332 | 0,7602022 | 0,0006901 | 0,0026038 | 0,0013607 | 0,0013607 | TRUE | 0,0102139 | No |
| orf19.4403 | 1007 | 1012 | 1077 | 615 | 1046 | 849 | 1025 | 1043 | 960 | 1070 | 1064 | 963 | 838 | 965 | 870 | 848 | 947,42 | 1014 | 880 | 0,865 | -0,209 | -3,125 | 0,001778 | 0,0023636 | 0,0025688 | 0,0038243 | 0,0032312 | 0,0032312 | TRUE | 0,0472915 | No |
| orf19.4404 | 15 | 15 | 16 | 8 | 15 | 23 | 14 | 9 | 14 | 16 | 16 | 13 | 12 | 26 | 12 | 7 | 14,48 | 15 | 14 | 0,949 | -0,076 | -0,179 | 0,8577955 | 0,8760884 | 0,0856533 | 0,1154469 | 0,1022839 | 0,1022839 | TRUE | 1,1846434 | No |
| orf19.4405 | 36 | 33 | 30 | 31 | 92 | 72 | 89 | 69 | 34 | 35 | 30 | 49 | 74 | 82 | 76 | 56 | 54,33 | 37 | 72 | 1,98 | 0,986 | 4,533 | 5,81E-06 | 8,70E-06 | 0,0116173 | 0,0323103 | 0,0237443 | 0,0237443 | TRUE | 0,2283179 | No |
| orf19.4406 | 426 | 397 | 366 | 332 | 1404 | 1043 | 1249 | 1332 | 406 | 420 | 362 | 520 | 1125 | 1186 | 1060 | 1083 | 770,25 | 427 | 1114 | 2,627 | 1,394 | 15,216 | 2,77E-52 | 1,05E-51 | 0,0089945 | 0,0042229 | 0,0063282 | 0,0063282 | TRUE | 0,2210673 | No |
| orf19.4407 | 3110 | 2827 | 3012 | 1886 | 5044 | 3619 | 4774 | 5073 | 2965 | 2989 | 2976 | 2955 | 4042 | 4115 | 4053 | 4125 | 3527,38 | 2971 | 4084 | 1,374 | 0,458 | 14,144 | 2,02E-45 | 6,95E-45 | 1,00E-08 | 0,0025568 | 0,0007105 | 0,0007105 | TRUE | 0,0006001 | No |
| orf19.4409 | 2378 | 2019 | 2249 | 1525 | 2144 | 1497 | 2087 | 2265 | 2267 | 2135 | 2222 | 2389 | 1718 | 1702 | 1772 | 1842 | 2005,81 | 2253 | 1758 | 0,782 | -0,355 | -7,159 | 8,15E-13 | 1,55E-12 | 0,0010709 | 0,0029099 | 0,0018561 | 0,0018561 | TRUE | 0,0182845 | No |
| orf19.441 | 9876 | 7984 | 8973 | 5326 | 8630 | 5707 | 8366 | 8738 | 9416 | 8442 | 8864 | 8343 | 6915 | 6489 | 7103 | 7105 | 7834,7 | 8766 | 6903 | 0,787 | -0,345 | -6,874 | 6,26E-12 | 1,16E-11 | 0,0022595 | 0,0023008 | 0,0022899 | 0,0022899 | TRUE | 0,0304957 | No |
| orf19.4410 | 1201 | 1088 | 1140 | 704 | 1420 | 1046 | 1318 | 1458 | 1145 | 1150 | 1126 | 1103 | 1138 | 1189 | 1119 | 1185 | 1144,52 | 1131 | 1158 | 1,022 | 0,031 | 0,631 | 0,5278291 | 0,5638838 | 1,00E-08 | 0,0035259 | 0,0014598 | 0,0014598 | TRUE | 0,0060972 | No |
| orf19.4411 | 1246 | 1099 | 1242 | 755 | 1245 | 855 | 1228 | 1219 | 1188 | 1162 | 1227 | 1183 | 998 | 972 | 1043 | 991 | 1095,4 | 1190 | 1001 | 0,841 | -0,249 | -4,949 | 7,45E-07 | 1,16E-06 | 1,00E-08 | 0,0035902 | 0,0015179 | 0,0015179 | TRUE | 0,0093757 | No |
| orf19.4412 | 1220 | 1032 | 1192 | 679 | 1969 | 1350 | 1957 | 2043 | 1163 | 1091 | 1178 | 1064 | 1578 | 1535 | 1661 | 1661 | 1366,38 | 1124 | 1609 | 1,429 | 0,515 | 9,518 | 1,77E-21 | 4,08E-21 | 0,0009627 | 0,0032929 | 0,0020387 | 0,0020387 | TRUE | 0,0137319 | No |
| orf19.4413 | 9297 | 7643 | 8344 | 5335 | 5458 | 3564 | 5313 | 5488 | 8864 | 8082 | 8243 | 8358 | 4373 | 4053 | 4511 | 4462 | 6368,09 | 8387 | 4350 | 0,519 | -0,946 | -20,098 | 7,65E-90 | 4,57E-89 | 0,0017005 | 0,0023491 | 0,00196 | 0,00196 | TRUE | 0,0250789 | No |
| orf19.4414 | 929 | 792 | 849 | 594 | 969 | 640 | 903 | 969 | 886 | 837 | 839 | 931 | 776 | 728 | 767 | 788 | 818,89 | 873 | 765 | 0,879 | -0,185 | -3,117 | 0,001828 | 0,0024281 | 0,0002398 | 0,0040963 | 0,0021753 | 0,0021753 | TRUE | 0,0200207 | No |
| orf19.4415 | 1534 | 1327 | 1496 | 1008 | 696 | 485 | 719 | 702 | 1463 | 1403 | 1478 | 1579 | 558 | 551 | 610 | 571 | 1026,63 | 1481 | 572 | 0,388 | -1,367 | -22,754 | 1,32E-114 | 9,93E-114 | 0,0010882 | 0,0036906 | 0,0023066 | 0,0023066 | TRUE | 0,0234395 | No |
| orf19.4416 | 1284 | 1103 | 1271 | 769 | 1275 | 915 | 1065 | 1231 | 1224 | 1166 | 1256 | 1205 | 1022 | 1040 | 904 | 1001 | 1102,24 | 1213 | 992 | 0,817 | -0,292 | -4,95 | 7,43E-07 | 1,16E-06 | 0,0015152 | 0,0035809 | 0,002427 | 0,002427 | TRUE | 0,0426624 | No |
| orf19.4418 | 1114 | 963 | 1123 | 704 | 1176 | 822 | 1105 | 1238 | 1062 | 1018 | 1109 | 1103 | 942 | 935 | 938 | 1007 | 1014,29 | 1073 | 956 | 0,892 | -0,165 | -3 | 0,0026988 | 0,00355 | 0,0004823 | 0,00371 | 0,0019295 | 0,0019295 | TRUE | 0,0159913 | No |
| orf19.4418.1 | 326 | 246 | 281 | 237 | 268 | 188 | 215 | 266 | 311 | 260 | 278 | 371 | 215 | 214 | 183 | 216 | 255,89 | 305 | 207 | 0,682 | -0,551 | -4,604 | 4,15E-06 | 6,26E-06 | 0,0111608 | 0,0085077 | 0,0097606 | 0,0097606 | TRUE | 0,2118419 | No |
| orf19.442 | 85 | 52 | 58 | 45 | 80 | 57 | 57 | 84 | 81 | 55 | 57 | 70 | 64 | 65 | 48 | 68 | 63,68 | 66 | 61 | 0,931 | -0,102 | -0,518 | 0,6042648 | 0,6384855 | 0,0140251 | 0,0278758 | 0,0217886 | 0,0217886 | TRUE | 0,2428362 | No |
| orf19.4420 | 1769 | 1612 | 1751 | 1121 | 2532 | 1898 | 2485 | 2724 | 1687 | 1704 | 1730 | 1756 | 2029 | 2158 | 2110 | 2215 | 1923,58 | 1719 | 2128 | 1,238 | 0,308 | 6,847 | 7,56E-12 | 1,40E-11 | 0,0005071 | 0,0029448 | 0,0014117 | 0,0014117 | TRUE | 0,0122817 | No |
| orf19.4423 | 1507 | 1412 | 1548 | 925 | 2540 | 1747 | 2394 | 2530 | 1437 | 1493 | 1529 | 1449 | 2035 | 1986 | 2032 | 2057 | 1752,43 | 1477 | 2028 | 1,373 | 0,457 | 10,631 | 2,13E-26 | 5,44E-26 | 1,00E-08 | 0,0030282 | 0,001175 | 0,001175 | TRUE | 0,0071105 | No |
| orf19.4424 | 43 | 42 | 63 | 37 | 44 | 39 | 44 | 52 | 41 | 44 | 62 | 58 | 35 | 44 | 37 | 42 | 45,61 | 51 | 40 | 0,776 | -0,365 | -1,653 | 0,0982411 | 0,1148336 | 1,00E-08 | 0,0380935 | 0,0247552 | 0,0247552 | TRUE | 0,1819631 | No |
| orf19.4426 | 1976 | 1999 | 2118 | 1219 | 2425 | 1886 | 2276 | 2420 | 1884 | 2114 | 2092 | 1910 | 1943 | 2145 | 1932 | 1968 | 1998,41 | 2000 | 1997 | 0,997 | -0,005 | -0,079 | 0,9367509 | 0,9432551 | 0,0024401 | 0,0029129 | 0,0026357 | 0,0026357 | TRUE | 0,0288635 | No |
| orf19.4427 | 6930 | 5924 | 6384 | 3598 | 6908 | 4657 | 6651 | 6830 | 6607 | 6264 | 6307 | 5636 | 5535 | 5295 | 5647 | 5553 | 5855,61 | 6204 | 5508 | 0,887 | -0,173 | -3,37 | 0,000752 | 0,0010238 | 0,0023004 | 0,0023717 | 0,0023469 | 0,0023469 | TRUE | 0,0448699 | No |
| orf19.4428 | 1415 | 1355 | 1442 | 872 | 1139 | 823 | 1080 | 1075 | 1349 | 1433 | 1425 | 1366 | 913 | 936 | 917 | 874 | 1151,48 | 1393 | 910 | 0,652 | -0,617 | -12,162 | 4,95E-34 | 1,42E-33 | 1,00E-08 | 0,0035172 | 0,0015766 | 0,0015766 | TRUE | 0,0085004 | No |
| orf19.443 | 863 | 769 | 804 | 493 | 1912 | 1337 | 1802 | 1839 | 823 | 813 | 794 | 772 | 1532 | 1520 | 1530 | 1495 | 1160 | 800 | 1519 | 1,894 | 0,921 | 18,801 | 7,40E-79 | 3,99E-78 | 1,00E-08 | 0,0035067 | 0,0013174 | 0,0013174 | TRUE | 0,0054751 | No |
| orf19.4430 | 4595 | 4147 | 4472 | 2718 | 8859 | 6341 | 8388 | 8934 | 4381 | 4385 | 4418 | 4258 | 7098 | 7210 | 7121 | 7264 | 5766,99 | 4360 | 7173 | 1,644 | 0,717 | 23,695 | 4,06E-124 | 3,37E-123 | 1,00E-08 | 0,002376 | 0,0006883 | 0,0006883 | TRUE | 0,0027466 | No |
| orf19.4432 | 18400 | 16196 | 17266 | 10464 | 2373 | 1732 | 2290 | 2481 | 17542 | 17125 | 17057 | 16392 | 1901 | 1969 | 1944 | 2017 | 9493,7 | 17029 | 1958 | 0,115 | -3,121 | -82,777 | 0 | 0 | 0,0005595 | 0,0022642 | 0,0011043 | 0,0011043 | TRUE | 0,0075913 | No |
| orf19.4433 | 314 | 248 | 282 | 208 | 719 | 509 | 677 | 741 | 299 | 262 | 279 | 326 | 576 | 579 | 575 | 603 | 437,27 | 292 | 583 | 2,013 | 1,009 | 12,733 | 3,88E-37 | 1,18E-36 | 1,00E-08 | 0,0058461 | 0,0033459 | 0,0033459 | TRUE | 0,0557192 | No |
| orf19.4435 | 4059 | 3940 | 4004 | 2733 | 1338 | 1005 | 1243 | 1305 | 3870 | 4166 | 3956 | 4281 | 1072 | 1143 | 1055 | 1061 | 2575,51 | 4068 | 1083 | 0,266 | -1,91 | -38,008 | 0 | 0 | 0,0013216 | 0,0027288 | 0,0018744 | 0,0018744 | TRUE | 0,0153477 | No |
| orf19.4436 | 412 | 392 | 394 | 271 | 106 | 87 | 109 | 104 | 393 | 414 | 389 | 425 | 85 | 99 | 93 | 85 | 247,75 | 405 | 90 | 0,222 | -2,172 | -20,444 | 6,75E-93 | 4,14E-92 | 1,00E-08 | 0,0087185 | 0,0045132 | 0,0045132 | TRUE | 0,0334562 | No |
| orf19.4437 | 2715 | 2615 | 2826 | 1727 | 4310 | 2934 | 4021 | 4313 | 2588 | 2765 | 2792 | 2705 | 3453 | 3336 | 3414 | 3507 | 3070,14 | 2712 | 3428 | 1,264 | 0,338 | 8,392 | 4,80E-17 | 1,01E-16 | 0,0004384 | 0,0026261 | 0,0012223 | 0,0012223 | TRUE | 0,0123101 | No |

| | | | | | | | | | | | | | | | | | | | | | | | | | | | | | | | |
|---|---|---|---|---|---|---|---|---|---|---|---|---|---|---|---|---|---|---|---|---|---|---|---|---|---|---|---|---|---|---|---|
| orf19.4438 | 911 | 811 | 931 | 654 | 552 | 399 | 572 | 615 | 869 | 858 | 920 | 1025 | 442 | 454 | 486 | 500 | 694 | 918 | 470 | 0,515 | -0,958 | -12,673 | 8,32E-37 | 2,50E-36 | 0,0034317 | 0,0044571 | 0,0039253 | 0,0039253 | TRUE | 0,0665703 | No |
| orf19.4439 | 17 | 11 | 11 | 5 | 23 | 16 | 24 | 22 | 16 | 12 | 11 | 8 | 18 | 18 | 20 | 18 | 15,18 | 12 | 18 | 1,571 | 0,652 | 1,798 | 0,0721029 | 0,0853596 | 1,00E-08 | 0,1102668 | 0,0557254 | 0,0557254 | TRUE | 0,2801264 | No |
| orf19.4439.1 | 425 | 345 | 358 | 287 | 588 | 439 | 569 | 633 | 405 | 365 | 354 | 450 | 471 | 499 | 483 | 515 | 442,67 | 394 | 492 | 1,258 | 0,331 | 3,902 | 9,55E-05 | 0,0001354 | 0,0031045 | 0,0058004 | 0,004575 | 0,004575 | TRUE | 0,0902821 | No |
| orf19.444 | 468 | 394 | 419 | 265 | 735 | 460 | 847 | 719 | 740 | 446 | 417 | 414 | 415 | 589 | 523 | 610 | 602 | 502 | 423 | 581 | 1,375 | 0,46 | 6,135 | 8,50E-10 | 1,46E-09 | 0,0012473 | 0,005362 | 0,0033111 | 0,0033111 | TRUE | 0,0482157 | No |
| orf19.4440 | 893 | 739 | 832 | 509 | 1074 | 847 | 1027 | 1170 | 851 | 781 | 822 | 797 | 861 | 963 | 872 | 951 | 862,38 | 813 | 912 | 1,119 | 0,162 | 2,586 | 0,0097102 | 0,0123476 | 0,0014292 | 0,0039952 | 0,0025808 | 0,0025808 | TRUE | 0,0172172 | No |
| orf19.4441 | 417 | 376 | 436 | 264 | 1150 | 768 | 1090 | 1195 | 398 | 398 | 431 | 414 | 921 | 873 | 925 | 972 | 666,4 | 410 | 923 | 2,256 | 1,174 | 17,619 | 1,76E-69 | 8,35E-69 | 1,00E-08 | 0,0045551 | 0,0024182 | 0,0024182 | TRUE | 0,0153634 | No |
| orf19.4442 | 1075 | 955 | 1039 | 701 | 1501 | 989 | 1360 | 1451 | 1025 | 1010 | 1026 | 1098 | 1203 | 1125 | 1155 | 1180 | 1102,63 | 1040 | 1166 | 1,125 | 0,17 | 3,277 | 0,0010505 | 0,0014157 | 1,00E-08 | 0,0035804 | 0,0016494 | 0,0016494 | TRUE | 0,0145922 | No |
| orf19.4443 | 1474 | 1230 | 1377 | 787 | 5753 | 3433 | 5358 | 5592 | 1405 | 1301 | 1360 | 1233 | 4610 | 3904 | 4549 | 4547 | 2863,52 | 1325 | 4402 | 3,319 | 1,731 | 26,531 | 4,21E-155 | 4,42E-154 | 0,0038119 | 0,0026647 | 0,0035607 | 0,0035607 | TRUE | 0,0697478 | No |
| orf19.4444 | 982 | 913 | 924 | 593 | 2247 | 1523 | 2230 | 2334 | 936 | 965 | 913 | 929 | 1800 | 1732 | 1893 | 1898 | 1383,33 | 936 | 1831 | 1,958 | 0,969 | 18,559 | 6,94E-77 | 3,63E-76 | 0,0007496 | 0,0032782 | 0,0017778 | 0,0017778 | TRUE | 0,0150649 | No |
| orf19.4445 | 23556 | 21849 | 22926 | 15876 | 16239 | 13095 | 15218 | 16200 | 22458 | 23103 | 22649 | 24871 | 13012 | 14890 | 12920 | 13172 | 18384,27 | 23270 | 13498 | 0,58 | -0,786 | -14,225 | 6,39E-46 | 2,22E-45 | 0,0034326 | 0,0021806 | 0,0028739 | 0,0028739 | TRUE | 0,0585957 | No |
| orf19.4446 | 4959 | 4347 | 4732 | 2975 | 5751 | 3859 | 5730 | 5915 | 4728 | 4596 | 4675 | 4660 | 4608 | 4388 | 4865 | 4809 | 4666,24 | 4665 | 4668 | 1,001 | 0,002 | 0,037 | 0,9702611 | 0,9731588 | 0,0009427 | 0,0024431 | 0,0014534 | 0,0014534 | TRUE | 0,0191632 | No |
| orf19.4447 | 1600 | 1246 | 1444 | 880 | 4521 | 2919 | 4250 | 4682 | 1525 | 1317 | 1427 | 1379 | 3623 | 3319 | 3608 | 3807 | 2500,61 | 1412 | 3589 | 2,541 | 1,345 | 22,514 | 3,00E-112 | 2,22E-111 | 0,0028511 | 0,0027479 | 0,0029057 | 0,0029057 | TRUE | 0,0363588 | No |
| orf19.4448 | 2370 | 2010 | 2175 | 1492 | 2213 | 1700 | 2027 | 2430 | 2260 | 2125 | 2149 | 2337 | 1773 | 1933 | 1721 | 1976 | 2034,23 | 2218 | 1851 | 0,835 | -0,26 | -4,459 | 8,25E-06 | 1,23E-05 | 0,0025373 | 0,0028984 | 0,002779 | 0,002779 | TRUE | 0,0270354 | No |
| orf19.4449 | 3557 | 2996 | 3294 | 2121 | 2591 | 1815 | 2506 | 2706 | 3391 | 3168 | 3254 | 3323 | 2076 | 2064 | 2128 | 2200 | 2700,46 | 3284 | 2117 | 0,645 | -0,633 | -15,104 | 1,53E-51 | 5,75E-51 | 0,0005101 | 0,0026993 | 0,001302 | 0,001302 | TRUE | 0,0086176 | No |
| orf19.445 | 3621 | 3191 | 3452 | 2040 | 3988 | 2731 | 3939 | 4124 | 3452 | 3374 | 3410 | 3196 | 3195 | 3105 | 3344 | 3353 | 3303,82 | 3358 | 3249 | 0,967 | -0,048 | -1,109 | 0,2672889 | 0,2981245 | 0,0008458 | 0,0025883 | 0,0015074 | 0,0015074 | TRUE | 0,011987 | No |
| orf19.4450 | 493 | 397 | 446 | 295 | 1257 | 898 | 1206 | 1347 | 470 | 420 | 441 | 462 | 1007 | 1021 | 1024 | 1095 | 742,5 | 448 | 1037 | 2,317 | 1,212 | 18,675 | 7,90E-78 | 4,19E-77 | 1,00E-08 | 0,0043026 | 0,0023709 | 0,0023709 | TRUE | 0,0213137 | No |
| orf19.4450.1 | 4231 | 3527 | 3514 | 2520 | 391 | 283 | 356 | 396 | 4034 | 3729 | 3471 | 3948 | 313 | 322 | 302 | 322 | 2055,21 | 3796 | 315 | 0,083 | -3,592 | -53,093 | 0 | 0 | 0,0027934 | 0,0028902 | 0,0028339 | 0,0028339 | TRUE | 0,0407421 | No |
| orf19.4450.2 | 580 | 458 | 436 | 344 | 871 | 593 | 767 | 803 | 553 | 484 | 431 | 539 | 698 | 674 | 651 | 653 | 585,4 | 502 | 669 | 1,337 | 0,419 | 5,072 | 3,93E-07 | 6,18E-07 | 0,0044033 | 0,0048961 | 0,0047525 | 0,0047525 | TRUE | 0,1101242 | No |
| orf19.4451 | 6857 | 6024 | 6743 | 3823 | 9716 | 6346 | 9249 | 10151 | 6537 | 6370 | 6661 | 5989 | 7785 | 7216 | 7852 | 8254 | 7083,08 | 6389 | 7777 | 1,217 | 0,283 | 5,52 | 3,38E-08 | 5,54E-08 | 0,0023674 | 0,0023231 | 0,0023808 | 0,0023808 | TRUE | 0,0284248 | No |
| orf19.4455 | 3509 | 2887 | 3357 | 2192 | 4255 | 3125 | 4090 | 4338 | 3345 | 3053 | 3316 | 3434 | 3409 | 3553 | 3472 | 3527 | 3388,85 | 3287 | 3490 | 1,062 | 0,087 | 1,956 | 0,0504734 | 0,0606994 | 0,0010193 | 0,0025758 | 0,0016181 | 0,0016181 | TRUE | 0,0280439 | No |
| orf19.4456 | 37945 | 31914 | 34616 | 21680 | 28844 | 19542 | 27572 | 29331 | 36176 | 33745 | 34197 | 33963 | 23112 | 22221 | 23408 | 23849 | 28833,99 | 34520 | 23148 | 0,671 | -0,577 | -15,56 | 1,37E-54 | 5,36E-54 | 0,0009192 | 0,0021482 | 0,0012834 | 0,0012834 | TRUE | 0,0128259 | No |
| orf19.4457 | 2268 | 2179 | 2362 | 1324 | 2549 | 1833 | 2373 | 2613 | 2162 | 2304 | 2333 | 2074 | 2042 | 2084 | 2015 | 2125 | 2142,48 | 2218 | 2066 | 0,93 | -0,105 | -2,125 | 0,0335535 | 0,0409144 | 0,0011564 | 0,0028576 | 0,0018609 | 0,0018609 | TRUE | 0,02119 | No |
| orf19.4459 | 2428 | 2452 | 2565 | 1583 | 4674 | 3552 | 4236 | 4135 | 2315 | 2593 | 2534 | 2480 | 3745 | 4039 | 3596 | 3362 | 3082,99 | 2480 | 3686 | 1,485 | 0,571 | 9,177 | 4,45E-20 | 1,00E-19 | 0,003669 | 0,0026238 | 0,0033667 | 0,0033667 | TRUE | 0,0498155 | No |
| orf19.446 | 868 | 851 | 868 | 543 | 1217 | 804 | 1114 | 1141 | 828 | 900 | 858 | 851 | 975 | 914 | 946 | 928 | 899,8 | 859 | 941 | 1,096 | 0,133 | 2,405 | 0,0161944 | 0,020262 | 1,00E-08 | 0,003916 | 0,0017936 | 0,0017936 | TRUE | 0,012752 | No |
| orf19.4461 | 0 | 0 | 1 | 0 | 3 | 2 | 3 | 2 | 0 | 0 | 1 | 0 | 2 | 2 | 3 | 2 | 1,23 | 0 | 2 | 6,52 | 2,705 | 1,923 | 0,0545114 | 0,065248 | 1,00E-08 | 1,3370975 | 0,6996316 | 0,6996316 | TRUE | 0,2757322 | No |
| orf19.446.1 | 533 | 467 | 462 | 294 | 2168 | 1537 | 1863 | 2078 | 508 | 494 | 456 | 461 | 1737 | 1748 | 1582 | 1690 | 1084,39 | 480 | 1689 | 3,511 | 1,812 | 28,818 | 1,29E-182 | 1,58E-181 | 0,0013449 | 0,0036054 | 0,002372 | 0,002372 | TRUE | 0,0222522 | No |
| orf19.446.2 | 470 | 431 | 397 | 312 | 1566 | 1141 | 1439 | 1467 | 448 | 456 | 392 | 489 | 1255 | 1297 | 1222 | 1193 | 843,94 | 446 | 1242 | 2,795 | 1,483 | 21,302 | 1,09E-100 | 7,18E-100 | 0,0017851 | 0,0040368 | 0,0030355 | 0,0030355 | TRUE | 0,0782958 | No |
| orf19.4463 | 9 | 4 | 8 | 3 | 23 | 15 | 22 | 21 | 9 | 4 | 8 | 5 | 18 | 17 | 19 | 17 | 12,08 | 6 | 18 | 2,734 | 1,451 | 3,366 | 0,0007618 | 0,0010365 | 1,00E-08 | 0,1379944 | 0,0685758 | 0,0685758 | TRUE | 0,1437501 | No |
| orf19.4464 | 804 | 740 | 726 | 439 | 882 | 675 | 824 | 735 | 767 | 782 | 717 | 688 | 707 | 768 | 700 | 598 | 715,67 | 738 | 693 | 0,935 | -0,097 | -1,186 | 0,2357685 | 0,2650993 | 0,0056768 | 0,0043855 | 0,0050642 | 0,0050642 | TRUE | 0,1027392 | No |
| orf19.4465 | 1520 | 1582 | 1634 | 1039 | 1274 | 939 | 1164 | 1090 | 1449 | 1673 | 1614 | 1628 | 1021 | 1068 | 988 | 886 | 1290,86 | 1591 | 991 | 0,622 | -0,684 | -9,823 | 8,96E-23 | 2,12E-22 | 0,0042404 | 0,0033632 | 0,0038516 | 0,0038516 | TRUE | 0,0610427 | No |
| orf19.4466 | 371 | 373 | 339 | 249 | 682 | 462 | 614 | 663 | 354 | 394 | 335 | 390 | 546 | 525 | 521 | 539 | 450,66 | 368 | 533 | 1,454 | 0,54 | 7,076 | 1,49E-12 | 2,80E-12 | 1,88E-08 | 0,0057346 | 0,0032379 | 0,0032379 | TRUE | 0,0433733 | No |
| orf19.4467 | 648 | 709 | 653 | 433 | 1097 | 771 | 991 | 874 | 618 | 750 | 645 | 678 | 879 | 877 | 841 | 711 | 749,82 | 673 | 827 | 1,229 | 0,298 | 3,42 | 0,0006254 | 0,0008544 | 0,0073071 | 0,004281 | 0,0058953 | 0,0058953 | TRUE | 0,1087704 | No |
| orf19.4468 | 2860 | 2540 | 2581 | 1404 | 7265 | 4501 | 6334 | 6145 | 2727 | 2686 | 2550 | 2199 | 5821 | 5118 | 5378 | 4996 | 3934,37 | 2540 | 5328 | 2,095 | 1,067 | 14,813 | 1,20E-49 | 4,40E-49 | 0,0064053 | 0,0025086 | 0,0046752 | 0,0046752 | TRUE | 0,095148 | No |
| orf19.4470 | 298 | 330 | 299 | 171 | 384 | 238 | 282 | 326 | 284 | 349 | 295 | 268 | 308 | 271 | 239 | 265 | 284,89 | 299 | 271 | 0,903 | -0,147 | -1,334 | 0,1820961 | 0,2073438 | 0,0085323 | 0,0078546 | 0,0081845 | 0,0081845 | TRUE | 0,1406123 | No |
| orf19.4471 | 907 | 899 | 974 | 571 | 1216 | 944 | 1109 | 1188 | 865 | 951 | 962 | 895 | 974 | 1073 | 942 | 966 | 953,41 | 918 | 989 | 1,075 | 0,104 | 1,646 | 0,0997286 | 0,1164869 | 0,0018729 | 0,0038134 | 0,0027556 | 0,0027556 | TRUE | 0,0371241 | No |
| orf19.4472 | 41 | 34 | 34 | 29 | 108 | 94 | 133 | 124 | 39 | 36 | 34 | 45 | 87 | 107 | 113 | 101 | 70,15 | 38 | 102 | 2,67 | 1,417 | 7,488 | 6,97E-14 | 1,36E-13 | 1,00E-08 | 0,0254959 | 0,0154023 | 0,0154023 | TRUE | 0,1083676 | No |
| orf19.4473 | 396 | 469 | 389 | 260 | 455 | 327 | 420 | 396 | 378 | 496 | 384 | 407 | 365 | 372 | 357 | 322 | 385 | 416 | 354 | 0,849 | -0,236 | -2,345 | 0,0190393 | 0,0236683 | 0,0078434 | 0,0063559 | 0,0071095 | 0,0071095 | TRUE | 0,1977095 | No |
| orf19.4474 | 2004 | 2266 | 2273 | 1275 | 2594 | 1674 | 2302 | 2191 | 1911 | 2396 | 2246 | 1997 | 2078 | 1903 | 1954 | 1781 | 2033,42 | 2138 | 1929 | 0,902 | -0,148 | -1,929 | 0,0537407 | 0,0643738 | 0,0068337 | 0,0028987 | 0,0051742 | 0,0051742 | TRUE | 0,0804733 | No |
| orf19.4475 | 2123 | 2460 | 2509 | 1427 | 2163 | 1737 | 1991 | 1855 | 2024 | 2601 | 2479 | 2235 | 1733 | 1975 | 1690 | 1508 | 2030,78 | 2335 | 1726 | 0,739 | -0,437 | -4,763 | 1,91E-06 | 2,92E-06 | 0,0115762 | 0,0028998 | 0,0075729 | 0,0075729 | TRUE | 0,1113947 | No |
| orf19.4476 | 510 | 431 | 444 | 352 | 255 | 230 | 240 | 288 | 486 | 456 | 439 | 551 | 204 | 262 | 204 | 234 | 354,47 | 483 | 226 | 0,468 | -1,097 | -10,41 | 2,24E-25 | 5,58E-25 | 0,0084477 | 0,0067232 | 0,0075101 | 0,0075101 | TRUE | 0,112614 | No |
| orf19.4477 | 956 | 825 | 795 | 503 | 483 | 318 | 407 | 488 | 911 | 872 | 785 | 788 | 387 | 362 | 346 | 397 | 606,01 | 839 | 373 | 0,444 | -1,172 | -14,971 | 1,14E-50 | 4,26E-50 | 0,0033055 | 0,0048006 | 0,0040207 | 0,0040207 | TRUE | 0,040707 | No |
| orf19.4478 | 282 | 233 | 277 | 176 | 911 | 578 | 771 | 835 | 269 | 246 | 274 | 276 | 730 | 657 | 655 | 679 | 473,16 | 266 | 680 | 2,562 | 1,357 | 17,251 | 1,11E-66 | 5,08E-66 | 1,00E-08 | 0,0055613 | 0,0031968 | 0,0031968 | TRUE | 0,0295243 | No |
| orf19.4479 | 14988 | 13470 | 14449 | 8285 | 16567 | 11660 | 15715 | 16638 | 14289 | 14243 | 14274 | 12979 | 13275 | 13258 | 13342 | 13528 | 13648,6 | 13946 | 13351 | 0,957 | -0,064 | -1,626 | 0,1040499 | 0,121268 | 0,0010155 | 0,0022116 | 0,0013964 | 0,0013964 | TRUE | 0,0260941 | No |
| orf19.448 | 2735 | 2327 | 2446 | 1532 | 3429 | 2209 | 3292 | 3593 | 2608 | 2461 | 2416 | 2400 | 2748 | 2512 | 2795 | 2921 | 2607,52 | 2471 | 2744 | 1,111 | 0,151 | 2,769 | 0,0056184 | 0,0072412 | 0,0021659 | 0,002721 | 0,0024798 | 0,0024798 | TRUE | 0,0380937 | No |
| orf19.4481 | 10 | 8 | 10 | 6 | 78 | 45 | 66 | 90 | 10 | 8 | 10 | 9 | 62 | 51 | 56 | 73 | 35,02 | 9 | 60 | 6,543 | 2,71 | 8,56 | 1,13E-17 | 2,41E-17 | 1,00E-08 | 0,0489774 | 0,0297491 | 0,0297491 | TRUE | 0,1744311 | No |
| orf19.4482 | 358 | 357 | 352 | 203 | 512 | 381 | 523 | 480 | 341 | 377 | 348 | 318 | 410 | 433 | 444 | 390 | 382,79 | 346 | 419 | 1,206 | 0,27 | 3,256 | 0,0011319 | 0,0015208 | 0,0012563 | 0,0063806 | 0,0039501 | 0,0039501 | TRUE | 0,0405407 | No |
| orf19.4488 | 11215 | 10307 | 11046 | 6873 | 9314 | 6371 | 8770 | 8400 | 10692 | 10898 | 10912 | 10767 | 7463 | 7244 | 7446 | 6830 | 9031,64 | 10817 | 7246 | 0,67 | -0,578 | -15,41 | 1,40E-53 | 5,42E-53 | 0,00072 | 0,0022731 | 0,0012383 | 0,0012383 | TRUE | 0,0184987 | No |
| orf19.449 | 38067 | 35063 | 37919 | 24321 | 2225 | 1645 | 2192 | 2299 | 36293 | 37075 | 37460 | 38100 | 1783 | 1871 | 1861 | 1869 | 19538,95 | 37232 | 1846 | 0,05 | -4,335 | -125,584 | 0 | 0 | 0,0003301 | 0,0021753 | 0,0008875 | 0,0008875 | TRUE | 0,0066481 | No |
| orf19.4490 | 14942 | 12863 | 12830 | 9295 | 41787 | 30159 | 38159 | 42142 | 14246 | 13601 | 12675 | 14561 | 33483 | 34293 | 32397 | 34265 | 23690,13 | 13771 | 33610 | 2,441 | 1,288 | 26,964 | 3,86E-160 | 4,21E-159 | 0,0021117 | 0,0021606 | 0,0021364 | 0,0021364 | TRUE | 0,035202 | No |
| orf19.4490.2 | 2873 | 2472 | 2541 | 1731 | 7127 | 4853 | 6270 | 7195 | 2739 | 2614 | 2510 | 2712 | 5711 | 5518 | 5323 | 5850 | 4122,16 | 2644 | 5600 | 2,12 | 1,084 | 23,279 | 7,27E-120 | 5,78E-119 | 0,0012422 | 0,0024896 | 0,0017893 | 0,0017893 | TRUE | 0,0143392 | No |
| orf19.4491 | 3871 | 4162 | 4377 | 2259 | 5378 | 3793 | 4998 | 4325 | 3691 | 4401 | 4324 | 3539 | 4309 | 4313 | 4243 | 3517 | 4042,05 | 3989 | 4096 | 1,026 | 0,037 | 0,435 | 0,6634531 | 0,6954977 | 0,0105427 | 0,0024975 | 0,0067917 | 0,0067917 | TRUE | 0,1107755 | No |
| orf19.4492 | 14470 | 12945 | 13404 | 8915 | 15860 | 11400 | 15286 | 15424 | 13796 | 13688 | 13242 | 13966 | 12708 | 12963 | 12978 | 12541 | 13235,11 | 13673 | 12798 | 0,936 | -0,095 | -3,125 | 0,0017762 | 0,0023618 | 0,0002907 | 0,0022153 | 0,0008158 | 0,0008158 | TRUE | 0,0055368 | No |
| orf19.4494 | 888 | 918 | 917 | 597 | 1120 | 809 | 1089 | 1093 | 847 | 971 | 906 | 935 | 897 | 920 | 925 | 889 | 911,13 | 915 | 908 | 0,993 | -0,009 | -0,16 | 0,8726042 | 0,8893288 | 0,0007561 | 0,0038933 | 0,0022018 | 0,0022018 | TRUE | 0,0316297 | No |
| orf19.4495 | 5836 | 5236 | 5610 | 3384 | 16352 | 11568 | 15380 | 16782 | 5564 | 5536 | 5542 | 5301 | 13102 | 13154 | 13058 | 13645 | 9362,87 | 5486 | 13240 | 2,412 | 1,27 | 39,034 | 0 | 0 | 0,0002954 | 0,0022666 | 0,0008803 | 0,0008803 | TRUE | 0,0058238 | No |
| orf19.4496 | 2247 | 2178 | 2312 | 1421 | 2124 | 1580 | 2016 | 2098 | 2142 | 2303 | 2284 | 2226 | 1702 | 1797 | 1712 | 1706 | 1983,91 | 2239 | 1729 | 0,772 | -0,374 | -8,465 | 2,57E-17 | 5,43E-17 | 0,0003839 | 0,0029189 | 0,001366 | 0,001366 | TRUE | 0,0109677 | No |
| orf19.4497 | 1167 | 1199 | 1086 | 607 | 903 | 588 | 779 | 718 | 1113 | 1268 | 1073 | 951 | 724 | 669 | 661 | 584 | 880,19 | 1101 | 660 | 0,598 | -0,743 | -8,071 | 6,96E-16 | 1,43E-15 | 0,0092356 | 0,0039567 | 0,0069423 | 0,0069423 | TRUE | 0,1227132 | No |
| orf19.4498 | 388 | 463 | 480 | 247 | 659 | 496 | 629 | 554 | 370 | 490 | 474 | 387 | 528 | 564 | 534 | 450 | 474,64 | 430 | 519 | 1,203 | 0,266 | 2,492 | 0,0127124 | 0,0160375 | 0,0121027 | 0,0055505 | 0,0088147 | 0,0088147 | TRUE | 0,1124875 | No |
| orf19.4499 | 2846 | 2643 | 2772 | 1721 | 2373 | 1720 | 2337 | 2501 | 2713 | 2795 | 2738 | 2696 | 1901 | 1956 | 1984 | 2034 | 2352,17 | 2736 | 1969 | 0,719 | -0,475 | -11,856 | 2,01E-32 | 5,67E-32 | 1,00E-08 | 0,0027893 | 0,0011093 | 0,0011093 | TRUE | 0,0068071 | No |
| orf19.450 | 51 | 51 | 59 | 58 | 29 | 7 | 22 | 21 | 49 | 54 | 58 | 91 | 23 | 8 | 19 | 17 | 39,83 | 63 | 17 | 0,276 | -1,86 | -6,138 | 8,36E-10 | 1,44E-09 | 0,0686277 | 0,0433134 | 0,0530974 | 0,0530974 | TRUE | 0,6282744 | No |
| orf19.4502 | 1347 | 1212 | 1311 | 722 | 2537 | 1817 | 2343 | 2619 | 1284 | 1282 | 1295 | 1131 | 2033 | 2066 | 1989 | 2129 | 1651,2 | 1248 | 2054 | 1,641 | 0,714 | 13,422 | 4,47E-41 | 1,44E-40 | 0,0011625 | 0,0030857 | 0,0020532 | 0,0020532 | TRUE | 0,0419909 | No |
| orf19.4503 | 967 | 720 | 864 | 581 | 2546 | 1689 | 2525 | 2518 | 922 | 761 | 854 | 910 | 2040 | 1921 | 2144 | 2047 | 1449,83 | 862 | 2038 | 2,369 | 1,244 | 18,822 | 5,02E-79 | 2,71E-78 | 0,0035233 | 0,0032238 | 0,0033248 | 0,0033248 | TRUE | 0,074025 | No |
| orf19.4504 | 510 | 443 | 489 | 273 | 1418 | 898 | 1308 | 1459 | 486 | 468 | 483 | 428 | 1136 | 1021 | 1110 | 1186 | 789,94 | 466 | 1113 | 2,381 | 1,252 | 17,899 | 1,19E-71 | 5,85E-71 | 0,0021364 | 0,0041698 | 0,0031013 | 0,0031013 | TRUE | 0,0353021 | No |
| orf19.4505 | 162 | 138 | 129 | 99 | 84 | 62 | 87 | 95 | 154 | 146 | 127 | 155 | 67 | 70 | 74 | 77 | 108,98 | 146 | 72 | 0,498 | -1,007 | -7,104 | 1,21E-12 | 2,28E-12 | 1,00E-08 | 0,0171579 | 0,0092997 | 0,0092997 | TRUE | 0,0784147 | No |
| orf19.4506 | 3545 | 3177 | 3357 | 1939 | 3889 | 2755 | 3417 | 3822 | 3380 | 3359 | 3316 | 3038 | 3116 | 3133 | 2901 | 3108 | 3168,81 | 3273 | 3064 | 0,935 | -0,097 | -2,033 | 0,0420837 | 0,050946 | 0,0014008 | 0,0026094 | 0,0018842 | 0,0018842 | TRUE | 0,0268045 | No |
| orf19.4507 | 79 | 91 | 70 | 48 | 312 | 217 | 316 | 341 | 75 | 96 | 69 | 75 | 250 | 247 | 268 | 277 | 169,77 | 79 | 260 | 3,302 | 1,723 | 13,241 | 5,07E-40 | 1,61E-39 | 0,0005151 | 0,0117625 | 0,007534 | 0,007534 | TRUE | 0,2078301 | No |
| orf19.4508 | 31 | 43 | 29 | 26 | 97 | 58 | 97 | 105 | 30 | 45 | 29 | 41 | 78 | 66 | 82 | 85 | 56,98 | 36 | 78 | 2,193 | 1,133 | 5,382 | 7,37E-08 | 1,19E-07 | 0,0038342 | 0,030909 | 0,0212457 | 0,0212457 | TRUE | 0,2537995 | No |
| orf19.4509 | 8 | 9 | 10 | 4 | 19 | 20 | 19 | 20 | 8 | 10 | 10 | 6 | 15 | 23 | 16 | 17 | 12,96 | 8 | 18 | 2,058 | 1,041 | 2,577 | 0,009964 | 0,0126604 | 1,00E-08 | 0,128821 | 0,0656604 | 0,0656604 | TRUE | 0,1651163 | No |
| orf19.451 | 18265 | 17169 | 18307 | 12512 | 12792 | 9666 | 11558 | 12506 | 17414 | 18154 | 18086 | 19601 | 10250 | 10991 | 9813 | 10169 | 14309,53 | 18314 | 10306 | 0,563 | -0,829 | -16,852 | 1,02E-63 | 4,50E-63 | 0,0023162 | 0,002206 | 0,002253 | 0,002253 | TRUE | 0,0271365 | No |
| orf19.4510 | 375 | 364 | 385 | 215 | 329 | 267 | 317 | 372 | 358 | 385 | 380 | 337 | 264 | 304 | 269 | 302 | 324,8 | 365 | 285 | 0,776 | -0,366 | -4,121 | 3,78E-05 | 5,45E-05 | 0,0012902 | 0,0071464 | 0,0044702 | 0,0044702 | TRUE | 0,0289492 | No |
| orf19.4511 | 684 | 604 | 754 | 444 | 659 | 494 | 661 | 667 | 652 | 639 | 745 | 696 | 528 | 562 | 561 | 542 | 615,56 | 683 | 548 | 0,803 | -0,317 | -4,533 | 5,81E-06 | 8,70E-06 | 0,001523 | 0,0047586 | 0,0030578 | 0,0030578 | TRUE | 0,0482803 | No |
| orf19.4513 | 7353 | 7474 | 8078 | 4696 | 7943 | 5919 | 7591 | 7504 | 7010 | 7903 | 7980 | 7357 | 6364 | 6730 | 6445 | 6101 | 6986,38 | 7562 | 6410 | 0,847 | -0,239 | -4,579 | 4,66E-06 | 7,02E-06 | 0,0024821 | 0,0023263 | 0,0024719 | 0,0024719 | TRUE | 0,0300865 | No |
| orf19.4515 | 202 | 168 | 207 | 114 | 313 | 236 | 322 | 302 | 193 | 178 | 204 | 179 | 251 | 268 | 273 | 246 | 223,92 | 188 | 260 | 1,371 | 0,455 | 4,543 | 5,54E-06 | 8,31E-06 | 1,00E-08 | 0,0094237 | 0,0049685 | 0,0049685 | TRUE | 0,0365755 | No |
| orf19.4516 | 1258 | 1102 | 1265 | 691 | 1649 | 1183 | 1552 | 1607 | 1199 | 1165 | 1250 | 1082 | 1321 | 1345 | 1318 | 1307 | 1248,44 | 1174 | 1323 | 1,122 | 0,167 | 3,134 | 0,001726 | 0,0022969 | 0,0005255 | 0,0034064 | 0,0019021 | 0,0019021 | TRUE | 0,0289265 | No |

| | | | | | | | | | | | | | | | | | | | | | | | | | | | | | | | |
|---|---|---|---|---|---|---|---|---|---|---|---|---|---|---|---|---|---|---|---|---|---|---|---|---|---|---|---|---|---|---|---|
| orf19.4517 | 2900 | 3679 | 3368 | 2042 | 4042 | 3568 | 3695 | 2921 | 2765 | 3890 | 3327 | 3199 | 3239 | 4057 | 3137 | 2375 | 3248,63 | 3295 | 3202 | 0,972 | -0,042 | -0,224 | 0,8228535 | 0,8437454 | 0,0330436 | 0,0025967 | 0,0166966 | 0,0330436 | TRUE | 0,2704647 | No |
| orf19.4518 | 1121 | 982 | 1107 | 670 | 1477 | 1032 | 1412 | 1528 | 1069 | 1038 | 1094 | 1050 | 1183 | 1173 | 1199 | 1242 | 1131,06 | 1063 | 1199 | 1,129 | 0,175 | 3,535 | 0,0004084 | 0,0005621 | 1,00E-08 | 0,0035429 | 0,0014479 | 0,0014479 | TRUE | 0,0070909 | No |
| orf19.4519 | 6786 | 6707 | 6934 | 4591 | 5046 | 3880 | 4776 | 4984 | 6470 | 7092 | 6850 | 7192 | 4043 | 4412 | 4055 | 4052 | 5520,76 | 6901 | 4140 | 0,6 | -0,737 | -15,353 | 3,36E-53 | 1,30E-52 | 0,0017811 | 0,0023887 | 0,0020236 | 0,0020236 | TRUE | 0,0231212 | No |
| orf19.4520 | 165 | 163 | 169 | 84 | 203 | 147 | 193 | 199 | 157 | 172 | 167 | 132 | 163 | 167 | 164 | 162 | 160,46 | 157 | 164 | 1,034 | 0,048 | 0,412 | 0,6801754 | 0,7119749 | 1,00E-08 | 0,0123238 | 0,006624 | 0,006624 | TRUE | 0,0945694 | No |
| orf19.4521 | 458 | 436 | 486 | 276 | 666 | 446 | 604 | 610 | 437 | 461 | 480 | 432 | 534 | 507 | 513 | 496 | 482,47 | 452 | 512 | 1,13 | 0,177 | 2,506 | 0,0122133 | 0,0154414 | 1,00E-08 | 0,0054944 | 0,0026735 | 0,0026735 | TRUE | 0,0199814 | No |
| orf19.4522 | 962 | 757 | 809 | 596 | 800 | 602 | 803 | 850 | 917 | 800 | 799 | 934 | 641 | 685 | 682 | 691 | 768,61 | 862 | 675 | 0,783 | -0,352 | -4,981 | 6,33E-07 | 9,88E-07 | 0,0028302 | 0,0042275 | 0,0034828 | 0,0034828 | TRUE | 0,0324137 | No |
| orf19.4523 | 322 | 292 | 304 | 203 | 550 | 439 | 531 | 551 | 307 | 309 | 300 | 318 | 441 | 499 | 451 | 448 | 384,1 | 308 | 460 | 1,489 | 0,574 | 7,267 | 3,68E-13 | 7,07E-13 | 1,00E-08 | 0,006366 | 0,0032109 | 0,0032109 | TRUE | 0,0352494 | No |
| orf19.4524 | 918 | 886 | 986 | 580 | 1651 | 1166 | 1583 | 1589 | 875 | 937 | 974 | 909 | 1323 | 1326 | 1344 | 1292 | 1122,43 | 924 | 1321 | 1,429 | 0,515 | 9,827 | 8,61E-23 | 2,04E-22 | 1,00E-08 | 0,0035541 | 0,0016987 | 0,0016987 | TRUE | 0,0172027 | No |
| orf19.4525 | 2829 | 2566 | 2690 | 1725 | 3014 | 2291 | 2866 | 2897 | 2697 | 2713 | 2657 | 2702 | 2415 | 2605 | 2433 | 2356 | 2572,37 | 2692 | 2452 | 0,91 | -0,136 | -3,185 | 0,0014457 | 0,0019295 | 0,0004763 | 0,0027296 | 0,0013678 | 0,0013678 | TRUE | 0,0202531 | No |
| orf19.4526 | 39009 | 35560 | 36958 | 23720 | 4238 | 2906 | 3692 | 3970 | 37191 | 37600 | 36511 | 37159 | 3396 | 3304 | 3134 | 3228 | 20190,42 | 37115 | 3266 | 0,088 | -3,507 | -103,342 | 0 | 0 | 0,0004306 | 0,0021726 | 0,0009539 | 0,0009539 | TRUE | 0,0091749 | No |
| orf19.4527 | 9545 | 7787 | 8528 | 5721 | 4785 | 3378 | 4493 | 4933 | 9100 | 8234 | 8425 | 8962 | 3834 | 3841 | 3815 | 4011 | 6277,71 | 8680 | 3875 | 0,447 | -1,163 | -26,503 | 8,87E-155 | 9,23E-154 | 0,0011795 | 0,0023528 | 0,001669 | 0,001669 | TRUE | 0,0146656 | No |
| orf19.4528 | 1355 | 1300 | 1306 | 892 | 2075 | 1558 | 1828 | 1984 | 1292 | 1375 | 1290 | 1397 | 1663 | 1772 | 1552 | 1613 | 1494,17 | 1338 | 1650 | 1,233 | 0,302 | 5,373 | 7,74E-08 | 1,25E-07 | 0,0015473 | 0,0031902 | 0,0023503 | 0,0023503 | TRUE | 0,0283039 | No |
| orf19.4528.1 | 3 | 2 | 2 | 4 | 2 | 3 | 5 | 1 | 3 | 2 | 2 | 6 | 2 | 3 | 4 | 1 | 2,91 | 3 | 2 | 0,786 | -0,348 | -0,405 | 0,6852885 | 0,7165597 | 1,00E-08 | 0,5661053 | 0,3489796 | 0,3489796 | TRUE | 0,2955258 | No |
| orf19.4529 | 910 | 819 | 828 | 580 | 1341 | 1001 | 1188 | 1346 | 868 | 866 | 818 | 909 | 1075 | 1138 | 1009 | 1094 | 971,99 | 865 | 1079 | 1,249 | 0,32 | 5,394 | 6,90E-08 | 1,12E-07 | 0,0011144 | 0,0037805 | 0,0023228 | 0,0023228 | TRUE | 0,0229568 | No |
| orf19.4530.1 | 8342 | 6990 | 7902 | 5093 | 4084 | 2810 | 4108 | 4059 | 7953 | 7391 | 7806 | 7978 | 3272 | 3195 | 3488 | 3300 | 5548,1 | 7782 | 3314 | 0,426 | -1,231 | -28,85 | 5,14E-183 | 6,31E-182 | 0,0010233 | 0,0023872 | 0,0015407 | 0,0015407 | TRUE | 0,0151454 | No |
| orf19.4531 | 26415 | 24471 | 26708 | 15803 | 4756 | 3355 | 4380 | 4914 | 25184 | 25875 | 26385 | 24756 | 3811 | 3815 | 3719 | 3996 | 14692,5 | 25550 | 3835 | 0,15 | -2,736 | -73,593 | 0 | 0 | 0,0006569 | 0,002203 | 0,001189 | 0,001189 | TRUE | 0,0098806 | No |
| orf19.4532 | 2466 | 2173 | 2272 | 1497 | 9402 | 6923 | 8909 | 9966 | 2351 | 2298 | 2245 | 2345 | 7534 | 7872 | 7564 | 8103 | 5038,88 | 2310 | 7768 | 3,365 | 1,75 | 43,055 | 0 | 0 | 0,0007492 | 0,0024171 | 0,0012863 | 0,0012863 | TRUE | 0,0104934 | No |
| orf19.4533 | 961 | 853 | 956 | 644 | 1825 | 1330 | 1706 | 1785 | 916 | 902 | 944 | 1009 | 1462 | 1512 | 1448 | 1451 | 1205,73 | 943 | 1468 | 1,562 | 0,643 | 12,46 | 1,24E-35 | 3,65E-35 | 1,00E-08 | 0,003453 | 0,0016583 | 0,0016583 | TRUE | 0,0224712 | No |
| orf19.4534 | 11102 | 9086 | 10092 | 6261 | 4115 | 2827 | 4131 | 4328 | 10585 | 9607 | 9970 | 9808 | 3297 | 3215 | 3507 | 3519 | 6688,51 | 9992 | 3384 | 0,339 | -1,561 | -33,282 | 7,03E-243 | 1,16E-241 | 0,0016402 | 0,0023368 | 0,0019256 | 0,0019256 | TRUE | 0,0196232 | No |
| orf19.4535 | 520 | 436 | 475 | 270 | 454 | 335 | 388 | 414 | 496 | 461 | 469 | 423 | 364 | 381 | 329 | 337 | 407,47 | 462 | 353 | 0,758 | -0,399 | -4,877 | 1,07E-06 | 1,66E-06 | 0,0015198 | 0,0061208 | 0,0039687 | 0,0039687 | TRUE | 0,0321603 | No |
| orf19.4536 | 7762 | 6685 | 7329 | 4656 | 19233 | 13911 | 18446 | 20756 | 7400 | 7069 | 7240 | 7294 | 15411 | 15818 | 15661 | 16877 | 11596,14 | 7251 | 15942 | 2,199 | 1,137 | 29,269 | 2,57E-188 | 3,29E-187 | 0,0008685 | 0,0022329 | 0,0013427 | 0,0013427 | TRUE | 0,0191931 | No |
| orf19.4537 | 3559 | 2940 | 3216 | 2012 | 3737 | 2519 | 3789 | 3977 | 3393 | 3109 | 3177 | 3152 | 2994 | 2864 | 3217 | 3234 | 3142,5 | 3208 | 3077 | 0,96 | -0,06 | -1,132 | 0,2574977 | 0,2877571 | 0,0020493 | 0,0026138 | 0,0023368 | 0,0023368 | TRUE | 0,0255448 | No |
| orf19.4538 | 3732 | 2967 | 3206 | 1869 | 3357 | 2181 | 3136 | 3362 | 3558 | 3137 | 3167 | 2928 | 2690 | 2480 | 2662 | 2734 | 2919,54 | 3198 | 2642 | 0,826 | -0,277 | -4,467 | 7,94E-06 | 1,18E-05 | 0,0038094 | 0,0026537 | 0,0033355 | 0,0033355 | TRUE | 0,0703462 | No |
| orf19.4539 | 1829 | 1469 | 1699 | 959 | 1809 | 1079 | 1773 | 1835 | 1744 | 1553 | 1678 | 1502 | 1449 | 1227 | 1505 | 1492 | 1518,94 | 1619 | 1418 | 0,876 | -0,191 | -2,561 | 0,0104343 | 0,0132395 | 0,0056631 | 0,0031722 | 0,0046879 | 0,0046879 | TRUE | 0,0959355 | No |
| orf19.454 | 7286 | 6316 | 7187 | 4852 | 6176 | 4582 | 5810 | 6361 | 6946 | 6678 | 7100 | 7601 | 4949 | 5210 | 4933 | 5172 | 6073,67 | 7081 | 5066 | 0,716 | -0,483 | -10,331 | 5,11E-25 | 1,26E-24 | 0,0016858 | 0,0023616 | 0,0019277 | 0,0019277 | TRUE | 0,0288966 | No |
| orf19.4540 | 6306 | 5452 | 5987 | 3765 | 3042 | 2206 | 2885 | 3148 | 6012 | 5765 | 5915 | 5898 | 2437 | 2508 | 2449 | 2560 | 4193,05 | 5898 | 2488 | 0,422 | -1,245 | -35,261 | 2,31E-272 | 4,41E-271 | 0,0001642 | 0,0024829 | 0,0009223 | 0,0009223 | TRUE | 0,0047277 | No |
| orf19.4542 | 894 | 859 | 923 | 568 | 1418 | 997 | 1368 | 1415 | 852 | 908 | 912 | 890 | 1136 | 1134 | 1161 | 1151 | 1018,01 | 890 | 1146 | 1,287 | 0,364 | 7,145 | 9,01E-13 | 1,71E-12 | 1,00E-08 | 0,0037041 | 0,0014745 | 0,0014745 | TRUE | 0,0110976 | No |
| orf19.4543 | 2325 | 1920 | 2186 | 1403 | 1966 | 1343 | 1924 | 2037 | 2217 | 2030 | 2160 | 2198 | 1575 | 1527 | 1633 | 1656 | 1874,55 | 2151 | 1598 | 0,744 | -0,427 | -8,792 | 1,47E-18 | 3,20E-18 | 0,0008922 | 0,0029672 | 0,001724 | 0,001724 | TRUE | 0,0175707 | No |
| orf19.4544 | 1125 | 1002 | 1010 | 639 | 1271 | 822 | 1106 | 1187 | 1073 | 1059 | 998 | 1001 | 1018 | 935 | 939 | 965 | 998,51 | 1033 | 964 | 0,934 | -0,099 | -1,778 | 0,0753397 | 0,0890594 | 0,0005809 | 0,0037356 | 0,0019864 | 0,0019864 | TRUE | 0,0174577 | No |
| orf19.4545 | 2386 | 2102 | 2341 | 1504 | 3483 | 2438 | 3196 | 3534 | 2275 | 2223 | 2313 | 2356 | 2791 | 2772 | 2713 | 2873 | 2539,51 | 2292 | 2787 | 1,218 | 0,284 | 7,12 | 1,08E-12 | 2,04E-12 | 0,0002044 | 0,0027378 | 0,0011241 | 0,0011241 | TRUE | 0,005374 | No |
| orf19.4546 | 6584 | 5836 | 6460 | 3871 | 6741 | 4471 | 6504 | 6787 | 6277 | 6171 | 6382 | 6064 | 5401 | 5084 | 5522 | 5518 | 5802,45 | 6224 | 5381 | 0,865 | -0,21 | -5,345 | 9,05E-08 | 1,46E-07 | 0,0007438 | 0,0023742 | 0,0013028 | 0,0013028 | TRUE | 0,0163661 | No |
| orf19.4548 | 321 | 293 | 339 | 206 | 617 | 415 | 596 | 569 | 306 | 310 | 335 | 323 | 494 | 472 | 506 | 463 | 401,05 | 318 | 484 | 1,522 | 0,606 | 7,851 | 4,13E-15 | 8,35E-15 | 1,00E-08 | 0,0061853 | 0,0030417 | 0,0030417 | TRUE | 0,0159643 | No |
| orf19.4549 | 525 | 461 | 513 | 310 | 777 | 601 | 756 | 792 | 501 | 487 | 507 | 486 | 623 | 683 | 642 | 644 | 571,52 | 495 | 648 | 1,305 | 0,384 | 5,918 | 3,25E-09 | 5,49E-09 | 1,00E-08 | 0,0049641 | 0,0022227 | 0,0022227 | TRUE | 0,0144954 | No |
| orf19.4550 | 1022 | 926 | 974 | 634 | 2867 | 2120 | 2666 | 2918 | 974 | 979 | 962 | 993 | 2297 | 2411 | 2263 | 2373 | 1656,6 | 977 | 2336 | 2,391 | 1,258 | 27,188 | 8,94E-163 | 9,84E-162 | 1,00E-08 | 0,0030824 | 0,001292 | 0,001292 | TRUE | 0,0052715 | No |
| orf19.4551 | 2484 | 2417 | 2567 | 1702 | 4498 | 3433 | 4159 | 4248 | 2368 | 2556 | 2536 | 2666 | 3604 | 3904 | 3531 | 3454 | 3077,36 | 2532 | 3623 | 1,432 | 0,518 | 9,681 | 3,64E-22 | 8,52E-22 | 0,0022084 | 0,0026248 | 0,002403 | 0,002403 | TRUE | 0,0322575 | No |
| orf19.4553 | 6165 | 5595 | 6243 | 4119 | 2901 | 2198 | 2848 | 3089 | 5878 | 5916 | 6167 | 6453 | 2324 | 2499 | 2418 | 2512 | 4270,9 | 6104 | 2438 | 0,4 | -1,323 | -28,911 | 8,64E-184 | 1,07E-182 | 0,0011965 | 0,0024757 | 0,0017361 | 0,0017361 | TRUE | 0,0174493 | No |
| orf19.4555 | 4196 | 3375 | 3868 | 2814 | 14223 | 10457 | 12813 | 14750 | 4000 | 3569 | 3821 | 4408 | 11396 | 11891 | 10878 | 11993 | 7744,62 | 3950 | 11540 | 2,924 | 1,548 | 24,639 | 4,83E-134 | 4,38E-133 | 0,0047604 | 0,0023033 | 0,0036104 | 0,0036104 | TRUE | 0,072183 | No |
| orf19.4557 | 576 | 539 | 535 | 320 | 270 | 188 | 278 | 278 | 549 | 570 | 529 | 501 | 216 | 214 | 236 | 226 | 380,14 | 537 | 223 | 0,415 | -1,27 | -15,517 | 2,67E-54 | 1,05E-53 | 1,00E-08 | 0,0064105 | 0,0034034 | 0,0034034 | TRUE | 0,019891 | No |
| orf19.4560 | 4915 | 4294 | 4683 | 2990 | 8470 | 6134 | 8145 | 8633 | 4686 | 4540 | 4626 | 4684 | 6787 | 6975 | 6915 | 7019 | 5779,1 | 4634 | 6924 | 1,494 | 0,579 | 18,778 | 1,13E-78 | 6,09E-78 | 1,00E-08 | 0,0023754 | 0,0007285 | 0,0007285 | TRUE | 0,0022748 | No |
| orf19.4563 | 5851 | 4982 | 5502 | 3469 | 4919 | 3510 | 4855 | 5012 | 5578 | 5268 | 5435 | 5434 | 3941 | 3991 | 4122 | 4075 | 4730,71 | 5429 | 4032 | 0,743 | -0,429 | -12,244 | 1,81E-34 | 5,24E-34 | 0,0002997 | 0,0024384 | 0,0009644 | 0,0009644 | TRUE | 0,0050016 | No |
| orf19.4565 | 5431 | 4383 | 4582 | 3334 | 11550 | 8370 | 10425 | 11962 | 5178 | 4634 | 4527 | 5223 | 9255 | 9517 | 8851 | 9726 | 7113,87 | 4890 | 9337 | 1,91 | 0,934 | 16,467 | 6,33E-61 | 2,71E-60 | 0,0033342 | 0,0023221 | 0,0029251 | 0,0029251 | TRUE | 0,0306916 | No |
| orf19.4566 | 27 | 23 | 26 | 13 | 26 | 9 | 21 | 23 | 26 | 24 | 26 | 20 | 21 | 10 | 18 | 19 | 20,46 | 24 | 17 | 0,709 | -0,496 | -1,577 | 0,1148456 | 0,1330724 | 1,00E-08 | 0,0823266 | 0,0460097 | 0,0460097 | TRUE | 0,2751295 | No |
| orf19.4567 | 41 | 41 | 36 | 23 | 60 | 46 | 59 | 55 | 39 | 43 | 36 | 36 | 48 | 52 | 50 | 45 | 43,65 | 38 | 49 | 1,259 | 0,333 | 1,586 | 0,1127542 | 0,1307678 | 1,00E-08 | 0,0397032 | 0,0187095 | 0,0187095 | TRUE | 0,0518821 | No |
| orf19.4568 | 274 | 233 | 261 | 164 | 333 | 257 | 357 | 381 | 261 | 246 | 258 | 257 | 267 | 292 | 303 | 310 | 274,29 | 256 | 293 | 1,146 | 0,197 | 2,164 | 0,0304542 | 0,0372991 | 1,00E-08 | 0,0080774 | 0,0042203 | 0,0042203 | TRUE | 0,0444318 | No |
| orf19.4569 | 66 | 48 | 54 | 36 | 845 | 682 | 812 | 836 | 63 | 51 | 53 | 56 | 677 | 775 | 689 | 680 | 380,64 | 56 | 705 | 12,587 | 3,654 | 29,184 | 3,09E-187 | 3,91E-186 | 1,00E-08 | 0,0064048 | 0,0045575 | 0,0045575 | TRUE | 0,0744146 | No |
| orf19.457 | 3486 | 3493 | 3749 | 2089 | 3395 | 2427 | 3050 | 3468 | 3324 | 3693 | 3704 | 3273 | 2720 | 2760 | 2589 | 2820 | 3110,3 | 3498 | 2722 | 0,778 | -0,363 | -6,623 | 3,52E-11 | 6,37E-11 | 0,0024099 | 0,0026192 | 0,0025599 | 0,0025599 | TRUE | 0,0218263 | No |
| orf19.4570 | 140 | 129 | 135 | 91 | 795 | 547 | 761 | 781 | 133 | 136 | 133 | 143 | 637 | 622 | 646 | 635 | 385,74 | 136 | 635 | 4,674 | 2,225 | 25,144 | 1,64E-139 | 1,53E-138 | 1,00E-08 | 0,0063478 | 0,0027488 | 0,0027488 | TRUE | 0,0067067 | No |
| orf19.4571 | 86 | 65 | 82 | 42 | 2085 | 1561 | 2032 | 2182 | 82 | 69 | 81 | 66 | 1671 | 1775 | 1725 | 1774 | 326,16 | 74 | 1736 | 23,042 | 4,526 | 44,923 | 0 | 0 | 1,00E-08 | 0,0039049 | 0,0021813 | 0,0021813 | TRUE | 0,0410372 | No |
| orf19.4573 | 239 | 168 | 220 | 165 | 21821 | 15509 | 20814 | 22198 | 228 | 178 | 217 | 258 | 17485 | 17635 | 17671 | 18049 | 8965,13 | 220 | 17710 | 81,307 | 6,345 | 96,513 | 0 | 0 | 0,0001649 | 0,0022744 | 0,0015891 | 0,0015891 | TRUE | 0,178534 | No |
| orf19.4574 | 1416 | 1270 | 1372 | 857 | 1407 | 905 | 1291 | 1376 | 1350 | 1343 | 1355 | 1343 | 1127 | 1029 | 1096 | 1119 | 1220,27 | 1348 | 1093 | 0,812 | -0,3 | -6,111 | 9,91E-10 | 1,70E-09 | 1,00E-08 | 0,0034368 | 0,0014933 | 0,0014933 | TRUE | 0,0167953 | No |
| orf19.4575 | 999 | 952 | 1030 | 683 | 2159 | 1452 | 2090 | 2166 | 952 | 1007 | 1018 | 1070 | 1730 | 1651 | 1774 | 1761 | 1370,39 | 1012 | 1729 | 1,715 | 0,778 | 14,901 | 3,27E-50 | 1,21E-49 | 0,0006011 | 0,0032894 | 0,001808 | 0,001808 | TRUE | 0,0187665 | No |
| orf19.4577 | 381 | 326 | 342 | 195 | 635 | 375 | 624 | 657 | 363 | 345 | 338 | 305 | 509 | 426 | 530 | 534 | 418,81 | 338 | 500 | 1,477 | 0,563 | 6,104 | 1,03E-09 | 1,77E-09 | 0,0052331 | 0,0060117 | 0,0056302 | 0,0056302 | TRUE | 0,1057341 | No |
| orf19.4577.3 | 1602 | 1355 | 1379 | 922 | 3268 | 2512 | 2881 | 3132 | 1527 | 1433 | 1362 | 1444 | 2619 | 2856 | 2446 | 2547 | 2029,28 | 1442 | 2617 | 1,814 | 0,859 | 14,329 | 1,44E-46 | 5,07E-46 | 0,0026103 | 0,0029004 | 0,0028882 | 0,0028882 | TRUE | 0,0447889 | No |
| orf19.4578 | 1655 | 1451 | 1495 | 887 | 1736 | 1155 | 1732 | 1777 | 1578 | 1534 | 1477 | 1390 | 1391 | 1313 | 1470 | 1445 | 1449,78 | 1495 | 1405 | 0,939 | -0,091 | -1,586 | 0,1128475 | 0,1308523 | 0,0017829 | 0,0032238 | 0,0024468 | 0,0024468 | TRUE | 0,024471 | No |
| orf19.4579 | 476 | 417 | 452 | 290 | 1220 | 927 | 1087 | 1205 | 454 | 441 | 447 | 454 | 978 | 1054 | 923 | 980 | 716,23 | 449 | 984 | 2,189 | 1,13 | 17,193 | 2,99E-66 | 1,36E-65 | 1,00E-08 | 0,0043837 | 0,0024522 | 0,0024522 | TRUE | 0,0258442 | No |
| orf19.458 | 1700 | 1432 | 1565 | 970 | 3024 | 2073 | 3035 | 3182 | 1621 | 1514 | 1546 | 1520 | 2423 | 2357 | 2577 | 2587 | 2018,09 | 1550 | 2486 | 1,603 | 0,681 | 13,697 | 1,06E-42 | 3,50E-42 | 0,0011574 | 0,0029049 | 0,001833 | 0,001833 | TRUE | 0,0140863 | No |
| orf19.4580 | 1319 | 1236 | 1238 | 705 | 1859 | 1117 | 1684 | 1712 | 1258 | 1307 | 1223 | 1104 | 1490 | 1270 | 1430 | 1392 | 1309,16 | 1223 | 1396 | 1,14 | 0,189 | 2,811 | 0,0049421 | 0,0063901 | 0,0036655 | 0,0033454 | 0,0035501 | 0,0035501 | TRUE | 0,0465319 | No |
| orf19.4581 | 194 | 202 | 205 | 118 | 351 | 294 | 301 | 295 | 185 | 214 | 203 | 185 | 281 | 334 | 256 | 240 | 237,11 | 197 | 278 | 1,405 | 0,49 | 4,08 | 4,51E-05 | 6,48E-05 | 0,00983 | 0,0090159 | 0,0094158 | 0,0094158 | TRUE | 0,1969127 | No |
| orf19.458.1 | 294 | 201 | 233 | 141 | 371 | 228 | 328 | 313 | 280 | 213 | 230 | 221 | 297 | 259 | 278 | 254 | 254,17 | 236 | 272 | 1,151 | 0,203 | 1,846 | 0,0649398 | 0,0772522 | 0,0065563 | 0,008551 | 0,0076013 | 0,0076013 | TRUE | 0,1868471 | No |
| orf19.4582 | 1948 | 1685 | 1812 | 1136 | 3621 | 2399 | 3370 | 3606 | 1857 | 1782 | 1790 | 1780 | 2901 | 2728 | 2861 | 2932 | 2328,87 | 1802 | 2856 | 1,585 | 0,664 | 15,698 | 1,56E-55 | 6,21E-55 | 0,0003429 | 0,0027963 | 0,0012522 | 0,0012522 | TRUE | 0,0104386 | No |
| orf19.4583 | 167 | 159 | 161 | 98 | 937 | 635 | 936 | 940 | 159 | 168 | 159 | 154 | 751 | 722 | 795 | 764 | 458,97 | 160 | 758 | 4,731 | 2,242 | 26,439 | 4,82E-154 | 4,97E-153 | 1,00E-08 | 0,0056687 | 0,0028662 | 0,0028662 | TRUE | 0,0124198 | No |
| orf19.4584 | 383 | 359 | 357 | 182 | 424 | 308 | 364 | 469 | 365 | 380 | 353 | 285 | 340 | 350 | 309 | 381 | 345,36 | 346 | 345 | 0,992 | -0,011 | -0,11 | 0,9124293 | 0,9213818 | 0,0074688 | 0,0068455 | 0,0071344 | 0,0071344 | TRUE | 0,1344438 | No |
| orf19.4585 | 10863 | 9315 | 10334 | 6040 | 6690 | 4155 | 6390 | 6603 | 10357 | 9849 | 10209 | 9462 | 5361 | 4725 | 5425 | 5369 | 7594,52 | 9969 | 5220 | 0,524 | -0,933 | -17,654 | 9,41E-70 | 4,49E-69 | 0,002624 | 0,0023075 | 0,0025416 | 0,0025416 | TRUE | 0,0488438 | No |
| orf19.4587 | 4524 | 4038 | 4209 | 2528 | 10094 | 7002 | 9571 | 10129 | 4313 | 4270 | 4158 | 3960 | 8088 | 7962 | 8126 | 8236 | 6139,08 | 4175 | 8103 | 1,939 | 0,955 | 26,045 | 1,53E-149 | 1,53E-148 | 0,0004192 | 0,0023587 | 0,0011012 | 0,0011012 | TRUE | 0,0135407 | No |
| orf19.4589 | 1129 | 1007 | 1102 | 652 | 628 | 479 | 640 | 634 | 1076 | 1065 | 1089 | 1021 | 503 | 545 | 543 | 516 | 794,74 | 1063 | 527 | 0,494 | -1,018 | -17,406 | 7,40E-68 | 3,45E-67 | 1,00E-08 | 0,0041572 | 0,0019092 | 0,0019092 | TRUE | 0,0107741 | No |
| orf19.459 | 2550 | 2306 | 2492 | 1436 | 4119 | 2667 | 3698 | 4103 | 2431 | 2438 | 2462 | 2250 | 3300 | 3033 | 3140 | 3336 | 2798,71 | 2395 | 3202 | 1,336 | 0,418 | 8,621 | 6,62E-18 | 1,42E-17 | 0,0013297 | 0,0026779 | 0,0018803 | 0,0018803 | TRUE | 0,0187085 | No |
| orf19.4590 | 418 | 424 | 455 | 327 | 235 | 192 | 219 | 256 | 399 | 448 | 449 | 512 | 188 | 218 | 186 | 208 | 326,16 | 452 | 200 | 0,444 | -1,173 | -11,71 | 1,13E-31 | 3,13E-31 | 0,0050904 | 0,0071253 | 0,006153 | 0,006153 | TRUE | 0,0823696 | No |
| orf19.4591 | 2955 | 2653 | 2862 | 1929 | 35487 | 27771 | 33973 | 37294 | 2817 | 2805 | 2827 | 3022 | 28435 | 31578 | 28843 | 30324 | 16331,39 | 2868 | 29795 | 10,4 | 3,378 | 72,889 | 0 | 0 | 0,0015574 | 0,0021918 | 0,0018517 | 0,0018517 | TRUE | 0,0198018 | No |
| orf19.4592 | 2153 | 2011 | 2093 | 1331 | 5043 | 3742 | 4909 | 5362 | 2053 | 2126 | 2068 | 2085 | 4041 | 4255 | 4168 | 4360 | 3144,39 | 2083 | 4206 | 2,019 | 1,014 | 25,043 | 2,07E-138 | 1,93E-137 | 0,0005043 | 0,0026134 | 0,0011978 | 0,0011978 | TRUE | 0,0086781 | No |
| orf19.4593 | 10418 | 9199 | 10176 | 6298 | 2923 | 1983 | 2752 | 2977 | 9932 | 9727 | 10053 | 9866 | 2342 | 2255 | 2336 | 2421 | 6116,53 | 9894 | 2338 | 0,237 | -2,08 | -60,074 | 0 | 0 | 0,0001611 | 0,0023597 | 0,0009034 | 0,0009034 | TRUE | 0,0067252 | No |
| orf19.4593.1 | 334 | 306 | 310 | 193 | 572 | 374 | 504 | 562 | 318 | 324 | 306 | 302 | 458 | 425 | 428 | 457 | 377,38 | 312 | 442 | 1,414 | 0,499 | 6,443 | 1,17E-10 | 2,09E-10 | 1,00E-08 | 0,0064421 | 0,0029866 | 0,0029866 | TRUE | 0,0070393 | No |
| orf19.4594 | 929 | 818 | 924 | 514 | 2342 | 1447 | 2279 | 2407 | 886 | 865 | 913 | 805 | 1877 | 1645 | 1935 | 1957 | 1360,32 | 867 | 1854 | 2,135 | 1,094 | 16,101 | 2,50E-58 | 1,03E-57 | 0,0037442 | 0,0032983 | 0,0035465 | 0,0035465 | TRUE | 0,0666693 | No |
| orf19.4595 | 2087 | 2132 | 2130 | 1564 | 1440 | 1279 | 1406 | 1441 | 1990 | 2254 | 2104 | 2450 | 1154 | 1454 | 1194 | 1172 | 1721,49 | 2200 | 1244 | 0,565 | -0,823 | -9,665 | 4,23E-22 | 9,91E-22 | 0,0097386 | 0,0030451 | 0,0063549 | 0,0063549 | TRUE | 0,1527034 | No |

| | | | | | | | | | | | | | | | | | | | | | | | | | | | | | | | |
|---|---|---|---|---|---|---|---|---|---|---|---|---|---|---|---|---|---|---|---|---|---|---|---|---|---|---|---|---|---|---|---|
| orf19.4596 | 1822 | 1512 | 1664 | 1097 | 1615 | 1135 | 1631 | 1645 | 1737 | 1599 | 1644 | 1719 | 1294 | 1291 | 1385 | 1338 | 1500,64 | 1675 | 1327 | 0,793 | -0,334 | -6,721 | 1,81E-11 | 3,31E-11 | 0,0006604 | 0,0031854 | 0,0017042 | 0,0017042 | TRUE | 0,0112552 | No |
| orf19.4597 | 2418 | 1982 | 2207 | 1448 | 1566 | 1078 | 1471 | 1579 | 2305 | 2096 | 2180 | 2268 | 1255 | 1226 | 1249 | 1284 | 1732,88 | 2212 | 1254 | 0,567 | -0,818 | -17,124 | 9,78E-66 | 4,44E-65 | 0,0006509 | 0,0030388 | 0,0015829 | 0,0015829 | TRUE | 0,015425 | No |
| orf19.4599 | 238 | 242 | 234 | 179 | 93 | 70 | 92 | 95 | 227 | 256 | 231 | 280 | 75 | 80 | 78 | 77 | 162,98 | 248 | 78 | 0,313 | -1,677 | -13,269 | 3,49E-40 | 1,11E-39 | 1,00E-08 | 0,0121655 | 0,0073322 | 0,0073322 | TRUE | 0,0697088 | No |
| orf19.460 | 1593 | 1358 | 1528 | 1067 | 1026 | 800 | 1024 | 1074 | 1519 | 1436 | 1510 | 1672 | 822 | 910 | 869 | 873 | 1201,26 | 1534 | 868 | 0,567 | -0,819 | -13,587 | 4,80E-42 | 1,57E-41 | 0,0017329 | 0,0034581 | 0,0026073 | 0,0026073 | TRUE | 0,0401974 | No |
| orf19.4600 | 2641 | 2135 | 2400 | 1407 | 2909 | 1849 | 2869 | 2959 | 2518 | 2257 | 2371 | 2204 | 2331 | 2102 | 2436 | 2406 | 2328,2 | 2338 | 2319 | 0,992 | -0,012 | -0,197 | 0,8440621 | 0,8636883 | 0,0034239 | 0,0027965 | 0,003161 | 0,003161 | TRUE | 0,046297 | No |
| orf19.4600.1 | 662 | 530 | 533 | 372 | 1712 | 1062 | 1598 | 1748 | 631 | 560 | 527 | 583 | 1372 | 1208 | 1357 | 1421 | 957,28 | 575 | 1340 | 2,332 | 1,221 | 16,456 | 7,63E-61 | 3,25E-60 | 0,0041346 | 0,0038065 | 0,0039806 | 0,0039806 | TRUE | 0,0551437 | No |
| orf19.4601 | 552 | 480 | 558 | 366 | 1041 | 714 | 1015 | 1102 | 526 | 508 | 551 | 573 | 834 | 812 | 862 | 896 | 695,27 | 540 | 851 | 1,583 | 0,663 | 10,185 | 2,30E-24 | 5,62E-24 | 0,0006251 | 0,0044528 | 0,0025065 | 0,0025065 | TRUE | 0,0177409 | No |
| orf19.4602 | 2671 | 2047 | 2192 | 1783 | 5524 | 4018 | 5311 | 5794 | 2547 | 2164 | 2165 | 2793 | 4426 | 4569 | 4509 | 4711 | 3485,59 | 2417 | 4554 | 1,886 | 0,916 | 11,948 | 6,69E-33 | 1,90E-32 | 0,0080626 | 0,0025623 | 0,0053095 | 0,0053095 | TRUE | 0,1282894 | No |
| orf19.4603 | 2547 | 2344 | 2541 | 1482 | 1963 | 1178 | 1821 | 1885 | 2428 | 2478 | 2510 | 2322 | 1573 | 1339 | 1546 | 1533 | 1966,22 | 2434 | 1498 | 0,616 | -0,7 | -12,2 | 3,10E-34 | 8,94E-34 | 0,0024268 | 0,0029263 | 0,0026288 | 0,0026288 | TRUE | 0,0578255 | No |
| orf19.4605 | 1126 | 1013 | 1121 | 708 | 2618 | 1755 | 2406 | 2569 | 1074 | 1071 | 1107 | 1109 | 2098 | 1996 | 2043 | 2089 | 1573,26 | 1090 | 2056 | 1,889 | 0,917 | 20,363 | 3,59E-92 | 2,19E-91 | 1,00E-08 | 0,0031349 | 0,0012199 | 0,0012199 | TRUE | 0,004456 | No |
| orf19.4606 | 1023 | 942 | 1024 | 623 | 1932 | 1348 | 1797 | 2020 | 975 | 996 | 1012 | 976 | 1548 | 1533 | 1526 | 1642 | 1275,99 | 990 | 1562 | 1,578 | 0,658 | 13,213 | 7,39E-40 | 2,33E-39 | 1,00E-08 | 0,0033781 | 0,001534 | 0,001534 | TRUE | 0,0147726 | No |
| orf19.4607 | 263 | 251 | 303 | 133 | 316 | 261 | 321 | 320 | 251 | 265 | 299 | 208 | 253 | 297 | 273 | 260 | 263,32 | 256 | 271 | 1,047 | 0,066 | 0,596 | 0,5511385 | 0,5862335 | 0,0080691 | 0,0083268 | 0,0080787 | 0,0080787 | TRUE | 0,1442928 | No |
| orf19.4608 | 244 | 240 | 262 | 158 | 302 | 206 | 313 | 350 | 233 | 254 | 259 | 248 | 242 | 234 | 266 | 285 | 252,41 | 248 | 257 | 1,037 | 0,053 | 0,532 | 0,5945039 | 0,6286923 | 1,00E-08 | 0,0085962 | 0,0053801 | 0,0053801 | TRUE | 0,0613339 | No |
| orf19.4609 | 1641 | 1387 | 1465 | 1077 | 1982 | 1500 | 1863 | 1960 | 1565 | 1467 | 1447 | 1687 | 1588 | 1706 | 1582 | 1594 | 1579,33 | 1542 | 1618 | 1,051 | 0,072 | 1,229 | 0,2189414 | 0,2472681 | 0,0023656 | 0,0031309 | 0,0026077 | 0,0026077 | TRUE | 0,0447474 | No |
| orf19.4610 | 344 | 334 | 341 | 183 | 723 | 497 | 665 | 726 | 328 | 353 | 337 | 287 | 579 | 565 | 565 | 590 | 450,5 | 326 | 575 | 1,752 | 0,809 | 10,6 | 2,99E-26 | 7,60E-26 | 1,00E-08 | 0,0057359 | 0,0031211 | 0,0031211 | TRUE | 0,0588394 | No |
| orf19.4611 | 4824 | 3868 | 4228 | 2772 | 5633 | 3914 | 5364 | 5779 | 4599 | 4090 | 4177 | 4342 | 4514 | 4451 | 4554 | 4699 | 4428,18 | 4302 | 4554 | 1,059 | 0,082 | 1,808 | 0,0705304 | 0,0835755 | 0,0013224 | 0,0024621 | 0,0017658 | 0,0017658 | TRUE | 0,0272653 | No |
| orf19.4612 | 66 | 47 | 65 | 39 | 85 | 63 | 86 | 93 | 63 | 50 | 64 | 61 | 68 | 72 | 73 | 76 | 65,79 | 60 | 72 | 1,212 | 0,277 | 1,597 | 0,1103048 | 0,1282068 | 1,00E-08 | 0,0270487 | 0,0134431 | 0,0134431 | TRUE | 0,1096296 | No |
| orf19.4614 | 1481 | 1438 | 1441 | 862 | 1433 | 1088 | 1352 | 1407 | 1412 | 1521 | 1424 | 1350 | 1148 | 1237 | 1148 | 1144 | 1297,96 | 1427 | 1169 | 0,817 | -0,291 | -5,38 | 7,43E-08 | 1,20E-07 | 0,0009282 | 0,0033563 | 0,002033 | 0,002033 | TRUE | 0,0231889 | No |
| orf19.4615 | 2359 | 2136 | 2192 | 1213 | 2620 | 1672 | 2329 | 2513 | 2249 | 2259 | 2165 | 1900 | 2099 | 1901 | 1977 | 2043 | 2074,31 | 2143 | 2005 | 0,934 | -0,098 | -1,594 | 0,1108746 | 0,1287284 | 0,0033134 | 0,0028828 | 0,0031408 | 0,0031408 | TRUE | 0,0662498 | No |
| orf19.4616 | 489 | 443 | 543 | 353 | 1023 | 778 | 899 | 992 | 466 | 468 | 536 | 553 | 820 | 885 | 763 | 807 | 662,28 | 506 | 819 | 1,621 | 0,697 | 8,98 | 2,70E-19 | 5,98E-19 | 0,0037422 | 0,0045704 | 0,0041305 | 0,0041305 | TRUE | 0,0392417 | No |
| orf19.4617 | 1383 | 1431 | 1331 | 836 | 745 | 507 | 727 | 680 | 1319 | 1513 | 1315 | 1310 | 597 | 577 | 617 | 553 | 974,97 | 1364 | 586 | 0,429 | -1,22 | -17,902 | 1,14E-71 | 5,61E-71 | 0,002893 | 0,0037753 | 0,0032832 | 0,0032832 | TRUE | 0,0657571 | No |
| orf19.4618 | 10564 | 8608 | 9382 | 6692 | 5280 | 4039 | 5181 | 5820 | 10072 | 9102 | 9269 | 10483 | 4231 | 4593 | 4399 | 4732 | 7109,96 | 9732 | 4489 | 0,461 | -1,116 | -19,667 | 4,15E-86 | 2,39E-85 | 0,0032542 | 0,0023222 | 0,002936 | 0,002936 | TRUE | 0,0325628 | No |
| orf19.4620 | 388 | 396 | 361 | 236 | 691 | 441 | 580 | 622 | 370 | 419 | 357 | 370 | 554 | 501 | 492 | 506 | 446,03 | 379 | 513 | 1,356 | 0,439 | 5,499 | 3,82E-08 | 6,24E-08 | 0,0017627 | 0,0057724 | 0,0037874 | 0,0037874 | TRUE | 0,0598597 | No |
| orf19.4621 | 998 | 1071 | 1093 | 670 | 2096 | 1554 | 1936 | 2064 | 951 | 1132 | 1080 | 1050 | 1679 | 1767 | 1644 | 1678 | 1372,71 | 1053 | 1692 | 1,606 | 0,684 | 11,4 | 4,18E-30 | 1,13E-29 | 0,0020627 | 0,0032874 | 0,0026556 | 0,0026556 | TRUE | 0,0543352 | No |
| orf19.4622 | 8890 | 7445 | 8149 | 5254 | 7803 | 5137 | 7427 | 7792 | 8476 | 7872 | 8050 | 8231 | 6252 | 5841 | 6305 | 6336 | 7170,45 | 8157 | 6184 | 0,758 | -0,399 | -9,741 | 2,02E-22 | 4,74E-22 | 0,0010071 | 0,0023203 | 0,0014696 | 0,0014696 | TRUE | 0,0165472 | No |
| orf19.4623 | 3540 | 2850 | 3449 | 2585 | 3031 | 2224 | 3011 | 3011 | 3375 | 3014 | 3407 | 4050 | 2429 | 2529 | 2556 | 2448 | 2975,93 | 3462 | 2490 | 0,72 | -0,473 | -6,193 | 5,92E-10 | 1,03E-09 | 0,0075936 | 0,002643 | 0,0052703 | 0,0052703 | TRUE | 0,1554124 | No |
| orf19.4623.3 | 8847 | 8470 | 8249 | 4941 | 6228 | 3946 | 5640 | 6151 | 8435 | 8956 | 8149 | 7740 | 4990 | 4487 | 4788 | 5001 | 6568,39 | 8320 | 4816 | 0,579 | -0,788 | -14,395 | 5,53E-47 | 1,96E-46 | 0,0029144 | 0,0023413 | 0,0027212 | 0,0027212 | TRUE | 0,0322005 | No |
| orf19.4624 | 422 | 378 | 351 | 207 | 1222 | 950 | 1097 | 1166 | 402 | 400 | 347 | 324 | 979 | 1080 | 931 | 948 | 676,48 | 368 | 984 | 2,656 | 1,409 | 16,979 | 1,17E-64 | 5,23E-64 | 0,0048012 | 0,0045184 | 0,0046507 | 0,0046507 | TRUE | 0,0612101 | No |
| orf19.4625 | 3173 | 3345 | 3356 | 1820 | 3172 | 2151 | 2688 | 2642 | 3025 | 3537 | 3315 | 2851 | 2542 | 2446 | 2282 | 2148 | 2768,3 | 3182 | 2354 | 0,739 | -0,436 | -5,805 | 6,43E-09 | 1,08E-08 | 0,0068217 | 0,0026844 | 0,0050492 | 0,0050492 | TRUE | 0,0680348 | No |
| orf19.4626 | 2090 | 2037 | 2088 | 1100 | 2280 | 1511 | 2024 | 2069 | 1993 | 2154 | 2063 | 1723 | 1827 | 1718 | 1718 | 1682 | 1859,76 | 1983 | 1736 | 0,874 | -0,194 | -2,905 | 0,0036712 | 0,0047867 | 0,0043018 | 0,0029741 | 0,0037526 | 0,0037526 | TRUE | 0,0885713 | No |
| orf19.4627 | 1438 | 1575 | 1697 | 980 | 1873 | 1400 | 1685 | 1665 | 1371 | 1665 | 1676 | 1535 | 1501 | 1592 | 1431 | 1354 | 1515,64 | 1562 | 1470 | 0,94 | -0,089 | -1,188 | 0,2347932 | 0,2640493 | 0,0056692 | 0,0031746 | 0,0047817 | 0,0047817 | TRUE | 0,0848327 | No |
| orf19.4628 | 1657 | 1523 | 1697 | 970 | 1195 | 831 | 1069 | 1189 | 1580 | 1610 | 1676 | 1520 | 958 | 945 | 908 | 967 | 1270,37 | 1596 | 944 | 0,591 | -0,76 | -14,773 | 2,19E-49 | 8,01E-49 | 0,0004457 | 0,0033837 | 0,0017176 | 0,0017176 | TRUE | 0,0136128 | No |
| orf19.4629 | 2561 | 2841 | 2799 | 1800 | 3158 | 2434 | 2781 | 2670 | 2442 | 3004 | 2765 | 2820 | 2530 | 2768 | 2361 | 2171 | 2607,59 | 2758 | 2458 | 0,891 | -0,167 | -2,052 | 0,0401482 | 0,0486952 | 0,0084158 | 0,0027209 | 0,0059472 | 0,0059472 | TRUE | 0,0862404 | No |
| orf19.4630 | 11614 | 10251 | 10895 | 7145 | 4069 | 3110 | 3919 | 4175 | 11073 | 10839 | 10763 | 11193 | 3260 | 3536 | 3327 | 3395 | 7173,34 | 10967 | 3380 | 0,308 | -1,699 | -46,674 | 0 | 0 | 0,0005177 | 0,0023202 | 0,0010888 | 0,0010888 | TRUE | 0,0112743 | No |
| orf19.4631 | 3360 | 3463 | 3468 | 2520 | 971 | 833 | 896 | 848 | 3203 | 3662 | 3426 | 3948 | 778 | 947 | 761 | 690 | 2176,79 | 3560 | 794 | 0,223 | -2,167 | -23,651 | 1,16E-123 | 9,57E-123 | 0,0124497 | 0,0028456 | 0,0073414 | 0,0073414 | TRUE | 0,1940748 | No |
| orf19.4632 | 11607 | 10271 | 10175 | 7688 | 32173 | 25230 | 28369 | 32128 | 11066 | 10860 | 10052 | 12044 | 25779 | 28689 | 24085 | 26123 | 18587,28 | 11006 | 26169 | 2,378 | 1,25 | 19,232 | 2,01E-82 | 1,12E-81 | 0,0053218 | 0,0021796 | 0,0039903 | 0,0039903 | TRUE | 0,051335 | No |
| orf19.4633 | 1580 | 1408 | 1566 | 960 | 2958 | 2153 | 2765 | 2961 | 1506 | 1489 | 1547 | 1504 | 2370 | 2448 | 2347 | 2408 | 1952,43 | 1512 | 2393 | 1,582 | 0,662 | 16,202 | 4,87E-59 | 2,03E-58 | 1,00E-08 | 0,0029322 | 0,0010458 | 0,0010458 | TRUE | 0,0031944 | No |
| orf19.4634 | 4596 | 4248 | 4624 | 2807 | 3561 | 2380 | 3266 | 3226 | 4382 | 4492 | 4568 | 4397 | 2853 | 2706 | 2773 | 2623 | 3599,3 | 4460 | 2739 | 0,614 | -0,703 | -17,62 | 1,74E-69 | 8,26E-69 | 0,0005495 | 0,0025475 | 0,001241 | 0,001241 | TRUE | 0,0102899 | No |
| orf19.4635 | 7645 | 7972 | 7982 | 4619 | 13133 | 9715 | 12023 | 12091 | 7289 | 8429 | 7885 | 7236 | 10523 | 11047 | 10207 | 9831 | 9056 | 7710 | 10402 | 1,349 | 0,432 | 7,418 | 1,19E-13 | 2,31E-13 | 0,003738 | 0,0022726 | 0,0031343 | 0,0031343 | TRUE | 0,048182 | No |
| orf19.4636 | 1 | 5 | 2 | 0 | 5 | 1 | 4 | 4 | 1 | 5 | 2 | 0 | 4 | 1 | 3 | 3 | 2,5 | 2 | 3 | 1,403 | 0,489 | 0,519 | 0,6037256 | 0,6381271 | 0,0191667 | 0,658594 | 0,4431244 | 0,4431244 | TRUE | 0,9385398 | No |
| orf19.4637 | 445 | 463 | 489 | 285 | 568 | 382 | 561 | 549 | 424 | 490 | 483 | 446 | 455 | 434 | 476 | 446 | 456,94 | 461 | 453 | 0,983 | -0,024 | -0,319 | 0,7499937 | 0,7779112 | 0,0008154 | 0,0056845 | 0,0033415 | 0,0033415 | TRUE | 0,0378533 | No |
| orf19.4639 | 738 | 697 | 753 | 435 | 1267 | 1069 | 1178 | 1067 | 704 | 737 | 744 | 681 | 1015 | 1216 | 1000 | 868 | 870,55 | 716 | 1025 | 1,426 | 0,512 | 5,534 | 3,13E-08 | 5,14E-08 | 0,0090638 | 0,0039773 | 0,0069985 | 0,0069985 | TRUE | 0,1830015 | No |
| orf19.4640 | 11180 | 9625 | 10689 | 6320 | 15287 | 10515 | 14414 | 15425 | 10659 | 10177 | 10560 | 9901 | 12249 | 11957 | 12237 | 12542 | 11285,18 | 10324 | 12246 | 1,186 | 0,246 | 6,924 | 4,39E-12 | 8,15E-12 | 0,0006478 | 0,0022368 | 0,0011191 | 0,0011191 | TRUE | 0,0090215 | No |
| orf19.4641 | 2845 | 3015 | 2946 | 1904 | 1964 | 1371 | 1803 | 1987 | 2712 | 3188 | 2910 | 2983 | 1574 | 1559 | 1531 | 1616 | 2259,06 | 2948 | 1570 | 0,533 | -0,908 | -16,445 | 9,17E-61 | 3,90E-60 | 0,0022271 | 0,0028181 | 0,0024549 | 0,0024549 | TRUE | 0,0373795 | No |
| orf19.4642 | 891 | 1006 | 992 | 504 | 1062 | 838 | 988 | 861 | 849 | 1064 | 980 | 790 | 851 | 953 | 839 | 700 | 878,18 | 921 | 836 | 0,905 | -0,144 | -1,343 | 0,1792499 | 0,2042489 | 0,0152958 | 0,0039609 | 0,0098709 | 0,0098709 | TRUE | 0,1432257 | No |
| orf19.4643 | 1842 | 1777 | 1910 | 1030 | 1127 | 787 | 1071 | 1187 | 1756 | 1879 | 1887 | 1614 | 903 | 895 | 909 | 965 | 1350,99 | 1784 | 918 | 0,514 | -0,96 | -15,605 | 6,73E-55 | 2,66E-54 | 0,0024712 | 0,0033066 | 0,0028353 | 0,0028353 | TRUE | 0,0460143 | No |
| orf19.4645 | 5027 | 5704 | 5826 | 3310 | 1814 | 1353 | 1564 | 1518 | 4793 | 6031 | 5756 | 5185 | 1454 | 1538 | 1328 | 1234 | 3414,86 | 5441 | 1388 | 0,255 | -1,972 | -23,51 | 3,19E-122 | 2,60E-121 | 0,0093338 | 0,0025721 | 0,0063302 | 0,0063302 | TRUE | 0,0793863 | No |
| orf19.4646 | 315 | 251 | 295 | 260 | 96 | 73 | 78 | 87 | 300 | 265 | 291 | 407 | 77 | 83 | 66 | 71 | 195,17 | 316 | 74 | 0,235 | -2,087 | -13,682 | 1,30E-42 | 4,30E-42 | 0,0189359 | 0,010504 | 0,0145239 | 0,0145239 | TRUE | 0,3861615 | No |
| orf19.4647 | 323 | 236 | 289 | 176 | 300 | 202 | 273 | 276 | 308 | 250 | 286 | 276 | 240 | 230 | 232 | 224 | 255,62 | 280 | 232 | 0,827 | -0,275 | -2,855 | 0,0043057 | 0,0055877 | 1,00E-08 | 0,0085145 | 0,0049692 | 0,0049692 | TRUE | 0,0634385 | No |
| orf19.4649 | 740 | 887 | 801 | 563 | 1075 | 798 | 924 | 900 | 706 | 938 | 791 | 882 | 861 | 907 | 784 | 732 | 825,21 | 829 | 821 | 0,991 | -0,013 | -0,134 | 0,8931189 | 0,9079736 | 0,0111057 | 0,004081 | 0,0077302 | 0,0077302 | TRUE | 0,1227077 | No |
| orf19.465 | 36 | 18 | 28 | 32 | 12 | 10 | 13 | 16 | 34 | 19 | 28 | 50 | 10 | 11 | 11 | 13 | 22,02 | 33 | 11 | 0,35 | -1,516 | -4,201 | 2,65E-05 | 3,86E-05 | 0,0575173 | 0,0766476 | 0,0680606 | 0,0680606 | TRUE | 0,1892995 | No |
| orf19.4650 | 1129 | 969 | 1088 | 649 | 3739 | 2664 | 3466 | 3739 | 1076 | 1025 | 1075 | 1017 | 2996 | 3029 | 2943 | 3040 | 2025,06 | 1048 | 3002 | 2,858 | 1,515 | 34,927 | 2,91E-267 | 5,36E-266 | 1,00E-08 | 0,0029021 | 0,0011269 | 0,0011269 | TRUE | 0,0041013 | No |
| orf19.4651 | 1459 | 1492 | 1505 | 992 | 5131 | 3778 | 4898 | 5112 | 1391 | 1578 | 1487 | 1554 | 4111 | 4296 | 4158 | 4157 | 2841,45 | 1502 | 4180 | 2,787 | 1,479 | 31,183 | 1,83E-213 | 2,65E-212 | 0,0009128 | 0,0026691 | 0,0016777 | 0,0016777 | TRUE | 0,031224 | No |
| orf19.4652 | 1 | 0 | 1 | 0 | 1 | 0 | 1 | 0 | 1 | 0 | 1 | 0 | 1 | 0 | 1 | 0 | 0,45 | 0 | 0 | 0,812 | -0,3 | -0,146 | 0,8841534 | 0,8998059 | 1,00E-08 | 3,6593361 | 2,2019723 | 2,2019723 | TRUE | 0,1208452 | No |
| orf19.4653 | 12 | 8 | 14 | 14 | 4 | 6 | 8 | 8 | 11 | 8 | 14 | 22 | 3 | 7 | 7 | 7 | 9,87 | 14 | 6 | 0,426 | -1,231 | -2,505 | 0,0122387 | 0,0154675 | 1,00E-08 | 0,1683885 | 0,1137902 | 0,1137902 | TRUE | 0,3409633 | No |
| orf19.4654 | 28 | 15 | 23 | 14 | 12 | 14 | 8 | 16 | 27 | 16 | 23 | 22 | 10 | 16 | 7 | 13 | 16,57 | 22 | 12 | 0,51 | -0,971 | -2,615 | 0,0089268 | 0,0113788 | 1,00E-08 | 0,1011913 | 0,0669988 | 0,0669988 | TRUE | 0,3049213 | No |
| orf19.4655 | 412 | 470 | 473 | 306 | 535 | 424 | 495 | 479 | 393 | 497 | 467 | 479 | 429 | 482 | 420 | 389 | 444,62 | 459 | 430 | 0,937 | -0,095 | -0,997 | 0,3188928 | 0,352051 | 0,0069516 | 0,0057841 | 0,0063715 | 0,0063715 | TRUE | 0,1155536 | No |
| orf19.4656 | 195 | 250 | 236 | 136 | 414 | 320 | 368 | 309 | 186 | 264 | 233 | 213 | 332 | 364 | 312 | 251 | 269,47 | 224 | 315 | 1,401 | 0,486 | 3,592 | 0,0003287 | 0,0004544 | 0,0194155 | 0,0081845 | 0,0136763 | 0,0136763 | TRUE | 0,2146215 | No |
| orf19.4657 | 11544 | 11204 | 12027 | 6889 | 6946 | 4824 | 6759 | 6741 | 11006 | 11847 | 11882 | 10792 | 5566 | 5485 | 5738 | 5481 | 8474,58 | 11382 | 5568 | 0,489 | -1,032 | -23,833 | 1,52E-125 | 1,28E-124 | 0,001284 | 0,002285 | 0,0016707 | 0,0016707 | TRUE | 0,0145149 | No |
| orf19.4658 | 3247 | 3301 | 3505 | 1850 | 4690 | 3303 | 4412 | 4137 | 3096 | 3490 | 3463 | 2898 | 3758 | 3756 | 3746 | 3364 | 3446,26 | 3237 | 3656 | 1,128 | 0,174 | 2,6 | 0,0093355 | 0,0118878 | 0,0050069 | 0,0025677 | 0,004013 | 0,004013 | TRUE | 0,0580707 | No |
| orf19.4659 | 3284 | 3059 | 3296 | 2038 | 5088 | 3687 | 4871 | 4860 | 3131 | 3235 | 3256 | 3193 | 4077 | 4192 | 4135 | 3952 | 3646,33 | 3204 | 4089 | 1,276 | 0,351 | 9,612 | 7,09E-22 | 1,65E-21 | 0,0002534 | 0,0025416 | 0,000999 | 0,000999 | TRUE | 0,0062711 | No |
| orf19.4660 | 20674 | 17816 | 18599 | 11675 | 56123 | 39348 | 53229 | 59032 | 19710 | 18838 | 18374 | 18290 | 44970 | 44742 | 45191 | 47999 | 32264,24 | 18803 | 45726 | 2,432 | 1,282 | 32,92 | 1,15E-237 | 1,84E-236 | 0,001107 | 0,0021422 | 0,0014168 | 0,0014168 | TRUE | 0,0137389 | No |
| orf19.4662 | 1378 | 1111 | 1245 | 909 | 1685 | 1209 | 1603 | 1747 | 1314 | 1175 | 1230 | 1424 | 1350 | 1375 | 1361 | 1420 | 1331,09 | 1286 | 1376 | 1,074 | 0,103 | 1,653 | 0,0982338 | 0,1148336 | 0,002813 | 0,0033248 | 0,0029268 | 0,0029268 | TRUE | 0,0573239 | No |
| orf19.4664 | 3626 | 3316 | 3423 | 1968 | 530 | 337 | 490 | 525 | 3457 | 3506 | 3382 | 3083 | 425 | 383 | 416 | 427 | 1884,82 | 3357 | 413 | 0,123 | -3,022 | -48,666 | 0 | 0 | 0,002054 | 0,0029624 | 0,0024592 | 0,0024592 | TRUE | 0,0348671 | No |
| orf19.4665 | 2799 | 2139 | 2429 | 1335 | 1232 | 743 | 1210 | 1229 | 2669 | 2262 | 2400 | 2091 | 987 | 845 | 1027 | 999 | 1659,98 | 2356 | 964 | 0,41 | -1,288 | -15,598 | 7,50E-55 | 2,96E-54 | 0,0079676 | 0,0030804 | 0,0058414 | 0,0058414 | TRUE | 0,0973083 | No |
| orf19.4666 | 3481 | 3208 | 3108 | 1984 | 2958 | 2076 | 2795 | 2805 | 3319 | 3392 | 3070 | 3108 | 2370 | 2361 | 2373 | 2281 | 2784,21 | 3222 | 2346 | 0,728 | -0,459 | -10,024 | 1,19E-23 | 2,88E-23 | 0,001048 | 0,002681 | 0,0016476 | 0,0016476 | TRUE | 0,015184 | No |
| orf19.4668 | 1407 | 1270 | 1440 | 919 | 1044 | 780 | 973 | 1084 | 1341 | 1343 | 1423 | 1440 | 837 | 887 | 826 | 881 | 1122,18 | 1387 | 858 | 0,619 | -0,692 | -12,939 | 2,71E-38 | 8,38E-38 | 0,0004182 | 0,0035544 | 0,0018246 | 0,0018246 | TRUE | 0,0072407 | No |
| orf19.4669 | 485 | 582 | 534 | 334 | 2135 | 1648 | 2048 | 1829 | 462 | 615 | 528 | 523 | 1711 | 1874 | 1739 | 1487 | 1117,39 | 532 | 1703 | 3,199 | 1,677 | 18,143 | 1,46E-73 | 7,34E-73 | 0,0097921 | 0,0035607 | 0,0068891 | 0,0068891 | TRUE | 0,1335766 | No |
| orf19.467 | 11616 | 10141 | 11279 | 7755 | 1484 | 1118 | 1423 | 1544 | 11075 | 10723 | 11143 | 12149 | 1189 | 1271 | 1208 | 1255 | 6251,57 | 11272 | 1231 | 0,109 | -3,195 | -63,489 | 0 | 0 | 0,0018238 | 0,0023539 | 0,0020198 | 0,0020198 | TRUE | 0,0328948 | No |
| orf19.4670 | 5949 | 5515 | 6067 | 3678 | 2785 | 1993 | 2672 | 2775 | 5672 | 5831 | 5994 | 5762 | 2232 | 2266 | 2269 | 2256 | 4035,15 | 5815 | 2256 | 0,388 | -1,367 | -38,839 | 0 | 0 | 1,00E-08 | 0,0024982 | 0,0008941 | 0,0008941 | TRUE | 0,0053914 | No |
| orf19.4672 | 3 | 4 | 6 | 4 | 1 | 3 | 9 | 1 | 3 | 4 | 6 | 6 | 1 | 3 | 8 | 1 | 3,99 | 5 | 3 | 0,659 | -0,601 | -0,742 | 0,4581564 | 0,4945857 | 0,313077 | 0,4132107 | 0,3702095 | 0,3702095 | TRUE | 1,5320165 | No |
| orf19.4673 | 1537 | 1419 | 1564 | 941 | 237 | 207 | 248 | 254 | 1465 | 1500 | 1545 | 1474 | 190 | 235 | 211 | 207 | 853,42 | 1496 | 211 | 0,14 | -2,839 | -40,177 | 0 | 0 | 2,19E-05 | 0,0040152 | 0,0022989 | 0,0022989 | TRUE | 0,0584701 | No |
| orf19.4674.1 | 310 | 276 | 255 | 149 | 814 | 579 | 727 | 773 | 296 | 292 | 252 | 233 | 652 | 658 | 617 | 629 | 453,63 | 268 | 639 | 2,365 | 1,242 | 14,852 | 6,75E-50 | 2,49E-49 | 0,000949 | 0,0057107 | 0,003948 | 0,003948 | TRUE | 0,0664983 | No |

| | | | | | | | | | | | | | | | | | | | | | | | | | | | | | | | |
|---|---|---|---|---|---|---|---|---|---|---|---|---|---|---|---|---|---|---|---|---|---|---|---|---|---|---|---|---|---|---|---|
| orf19.4675 | 709 | 600 | 712 | 390 | 731 | 527 | 680 | 734 | 676 | 634 | 703 | 611 | 586 | 599 | 577 | 597 | 622,98 | 656 | 590 | 0,895 | -0,16 | -2,434 | 0,0149211 | 0,0187056 | 1,00E-08 | 0,0047269 | 0,002527 | 0,002527 | TRUE | 0,0270036 | No |
| orf19.4676 | 370 | 364 | 370 | 178 | 703 | 458 | 633 | 654 | 353 | 385 | 366 | 279 | 563 | 521 | 537 | 532 | 441,91 | 346 | 538 | 1,547 | 0,629 | 7,04 | 1,93E-12 | 3,61E-12 | 0,004375 | 0,0058068 | 0,0052378 | 0,0052378 | TRUE | 0,1582544 | No |
| orf19.4677 | 1119 | 1133 | 1149 | 709 | 1532 | 988 | 1354 | 1372 | 1067 | 1198 | 1135 | 1111 | 1228 | 1123 | 1150 | 1116 | 1140,84 | 1128 | 1154 | 1,024 | 0,034 | 0,592 | 0,553707 | 0,5888278 | 0,0013805 | 0,0035305 | 0,0023045 | 0,0023045 | TRUE | 0,0226859 | No |
| orf19.4678 | 1182 | 1196 | 1231 | 703 | 1626 | 1062 | 1488 | 1548 | 1127 | 1265 | 1216 | 1101 | 1303 | 1208 | 1263 | 1259 | 1217,67 | 1177 | 1258 | 1,068 | 0,094 | 1,636 | 0,1018868 | 0,1188338 | 0,0014847 | 0,0034397 | 0,0023737 | 0,0023737 | TRUE | 0,0292923 | No |
| orf19.4679 | 23209 | 19566 | 21758 | 14503 | 36089 | 25414 | 34280 | 36826 | 22127 | 20689 | 21495 | 22720 | 28917 | 28898 | 29104 | 29943 | 25486,51 | 21758 | 29216 | 1,343 | 0,426 | 11,568 | 5,96E-31 | 1,63E-30 | 0,0008901 | 0,0021557 | 0,0012585 | 0,0012585 | TRUE | 0,0133627 | No |
| orf19.4680 | 435 | 408 | 437 | 279 | 469 | 315 | 420 | 475 | 415 | 431 | 432 | 437 | 376 | 358 | 357 | 386 | 398,96 | 429 | 369 | 0,863 | -0,212 | -2,864 | 0,0041813 | 0,0054328 | 1,00E-08 | 0,0062067 | 0,0027479 | 0,0027479 | TRUE | 0,0112791 | No |
| orf19.4681 | 7015 | 6617 | 7222 | 4585 | 9711 | 6996 | 9165 | 10096 | 6688 | 6997 | 7135 | 7183 | 7781 | 7955 | 7781 | 8209 | 7466,03 | 7001 | 7932 | 1,133 | 0,181 | 4,84 | 1,30E-06 | 2,00E-06 | 0,0006522 | 0,0023112 | 0,0012022 | 0,0012022 | TRUE | 0,0111845 | No |
| orf19.4682 | 273 | 288 | 317 | 200 | 744 | 565 | 754 | 788 | 260 | 305 | 313 | 313 | 596 | 642 | 640 | 641 | 463,84 | 298 | 630 | 2,122 | 1,085 | 13,53 | 1,04E-41 | 3,39E-41 | 0,0009677 | 0,0056311 | 0,0035906 | 0,0035906 | TRUE | 0,0883917 | No |
| orf19.4683 | 5045 | 4886 | 5143 | 3165 | 3851 | 2798 | 3435 | 3831 | 4810 | 5166 | 5081 | 4958 | 3086 | 3182 | 2916 | 3115 | 4039,21 | 5004 | 3075 | 0,614 | -0,703 | -16,61 | 5,94E-62 | 2,57E-61 | 0,000804 | 0,0024978 | 0,0014635 | 0,0014635 | TRUE | 0,0146232 | No |
| orf19.4684.2 | 4860 | 3794 | 3905 | 2553 | 17591 | 9741 | 15494 | 15954 | 4633 | 4012 | 3858 | 3999 | 14095 | 11076 | 13154 | 12972 | 8475,04 | 4126 | 12824 | 3,108 | 1,636 | 21,1 | 7,97E-99 | 5,15E-98 | 0,0081629 | 0,002285 | 0,0056044 | 0,0056044 | TRUE | 0,1026376 | No |
| orf19.4685 | 11 | 10 | 18 | 6 | 29 | 25 | 20 | 24 | 10 | 11 | 18 | 9 | 23 | 28 | 17 | 20 | 17,05 | 12 | 22 | 1,784 | 0,835 | 2,291 | 0,0219842 | 0,0272022 | 1,00E-08 | 0,0983899 | 0,061911 | 0,061911 | TRUE | 0,4234752 | No |
| orf19.4686 | 3317 | 3273 | 3465 | 2046 | 4683 | 3389 | 4399 | 4306 | 3162 | 3461 | 3423 | 3205 | 3752 | 3854 | 3735 | 3501 | 3511,66 | 3313 | 3710 | 1,119 | 0,162 | 3,372 | 0,0007461 | 0,0010162 | 0,0014624 | 0,0025588 | 0,001935 | 0,001935 | TRUE | 0,017661 | No |
| orf19.4688 | 1842 | 1626 | 1834 | 1243 | 10161 | 7736 | 9579 | 9318 | 1756 | 1719 | 1812 | 1947 | 8142 | 8797 | 8133 | 7576 | 4985,21 | 1808 | 8162 | 4,519 | 2,176 | 38,124 | 0 | 0 | 0,0029444 | 0,0024206 | 0,0027619 | 0,0027619 | TRUE | 0,0331689 | No |
| orf19.4689 | 936 | 781 | 897 | 590 | 706 | 543 | 746 | 697 | 892 | 826 | 886 | 924 | 566 | 617 | 633 | 567 | 738,98 | 882 | 596 | 0,675 | -0,566 | -8,591 | 8,61E-18 | 1,84E-17 | 0,0013717 | 0,0043131 | 0,0027833 | 0,0027833 | TRUE | 0,0229713 | No |
| orf19.469 | 3398 | 2984 | 3327 | 2128 | 1774 | 1196 | 1685 | 1894 | 3240 | 3155 | 3287 | 3334 | 1421 | 1360 | 1431 | 1540 | 2345,9 | 3254 | 1438 | 0,443 | -1,176 | -24,683 | 1,61E-134 | 1,46E-133 | 0,0009611 | 0,0027912 | 0,001695 | 0,001695 | TRUE | 0,0276532 | No |
| orf19.4690 | 181 | 158 | 171 | 96 | 190 | 169 | 182 | 189 | 173 | 167 | 169 | 150 | 152 | 192 | 155 | 154 | 163,94 | 165 | 163 | 0,98 | -0,03 | -0,249 | 0,8035016 | 0,8254941 | 0,0017197 | 0,0121063 | 0,0076369 | 0,0076369 | TRUE | 0,1358135 | No |
| orf19.4691 | 183 | 170 | 192 | 107 | 69 | 43 | 56 | 48 | 174 | 180 | 190 | 168 | 55 | 49 | 48 | 39 | 112,78 | 178 | 48 | 0,267 | -1,904 | -12,587 | 2,49E-36 | 7,44E-36 | 1,00E-08 | 0,0166491 | 0,0095349 | 0,0095349 | TRUE | 0,1402323 | No |
| orf19.4697 | 982 | 930 | 1015 | 564 | 6330 | 4294 | 5612 | 5883 | 936 | 983 | 1003 | 884 | 5072 | 4883 | 4765 | 4783 | 2913,57 | 952 | 4876 | 5,108 | 2,353 | 47,802 | 0 | 0 | 0,0008535 | 0,0026548 | 0,0016489 | 0,0016489 | TRUE | 0,023059 | No |
| orf19.4698 | 2517 | 2627 | 2843 | 1456 | 851 | 575 | 780 | 709 | 2400 | 2778 | 2809 | 2281 | 682 | 654 | 662 | 576 | 1605,17 | 2567 | 644 | 0,25 | -1,997 | -24,427 | 8,87E-132 | 7,87E-131 | 0,0075529 | 0,0031142 | 0,0055076 | 0,0055076 | TRUE | 0,0657569 | No |
| orf19.4699 | 2994 | 3125 | 3198 | 1923 | 2431 | 1696 | 2118 | 2115 | 2854 | 3304 | 3159 | 3012 | 1948 | 1929 | 1798 | 1720 | 2465,6 | 3082 | 1849 | 0,599 | -0,739 | -12,178 | 4,06E-34 | 1,17E-33 | 0,0031141 | 0,0027572 | 0,0031074 | 0,0031074 | TRUE | 0,0311796 | No |
| orf19.470 | 2629 | 2213 | 2484 | 1560 | 1690 | 1117 | 1642 | 1674 | 2506 | 2340 | 2454 | 2444 | 1354 | 1270 | 1394 | 1361 | 1890,46 | 2436 | 1345 | 0,553 | -0,855 | -18,338 | 4,15E-75 | 2,12E-74 | 0,0005277 | 0,0029598 | 0,0015268 | 0,0015268 | TRUE | 0,0154422 | No |
| orf19.4701 | 344 | 445 | 403 | 266 | 464 | 364 | 394 | 369 | 328 | 471 | 398 | 417 | 372 | 414 | 335 | 300 | 379,19 | 404 | 355 | 0,88 | -0,185 | -1,49 | 0,1361319 | 0,1567408 | 0,0179787 | 0,0064213 | 0,0121559 | 0,0121559 | TRUE | 0,1900245 | No |
| orf19.4702 | 117 | 93 | 116 | 78 | 273 | 163 | 224 | 216 | 112 | 98 | 115 | 122 | 219 | 185 | 190 | 176 | 152,07 | 112 | 192 | 1,734 | 0,794 | 6,209 | 5,34E-10 | 9,28E-10 | 1,00E-08 | 0,0128882 | 0,00839 | 0,00839 | TRUE | 0,097617 | No |
| orf19.4703 | 410 | 367 | 379 | 224 | 818 | 593 | 779 | 880 | 391 | 388 | 374 | 351 | 655 | 674 | 661 | 716 | 526,36 | 376 | 676 | 1,793 | 0,842 | 11,919 | 9,46E-33 | 2,68E-32 | 1,00E-08 | 0,0052106 | 0,0026581 | 0,0026581 | TRUE | 0,0186277 | No |
| orf19.4704 | 7364 | 6644 | 7070 | 4433 | 7873 | 5345 | 7057 | 8160 | 7021 | 7025 | 6984 | 6945 | 6308 | 6078 | 5991 | 6635 | 6623,42 | 6994 | 6253 | 0,894 | -0,161 | -4,029 | 5,60E-05 | 8,02E-05 | 0,0008698 | 0,0023392 | 0,0013855 | 0,0013855 | TRUE | 0,0207503 | No |
| orf19.4705 | 1562 | 1414 | 1510 | 1013 | 2491 | 1763 | 2291 | 2531 | 1489 | 1495 | 1492 | 1587 | 1996 | 2005 | 1945 | 2058 | 1758,33 | 1516 | 2001 | 1,323 | 0,404 | 9,128 | 6,96E-20 | 1,56E-19 | 1,00E-08 | 0,0030251 | 0,0012833 | 0,0012833 | TRUE | 0,0104797 | No |
| orf19.4706 | 1293 | 1163 | 1224 | 815 | 1139 | 809 | 1123 | 1129 | 1233 | 1230 | 1209 | 1277 | 913 | 920 | 953 | 918 | 1081,54 | 1237 | 926 | 0,75 | -0,416 | -8,435 | 3,32E-17 | 7,01E-17 | 1,00E-08 | 0,0036094 | 0,0013982 | 0,0013982 | TRUE | 0,0049011 | No |
| orf19.4707 | 181 | 132 | 167 | 105 | 105 | 76 | 95 | 125 | 173 | 140 | 165 | 164 | 84 | 86 | 81 | 102 | 124,31 | 160 | 88 | 0,551 | -0,861 | -6,259 | 3,87E-10 | 6,79E-10 | 1,00E-08 | 0,0152998 | 0,0096416 | 0,0096416 | TRUE | 0,1107884 | No |
| orf19.4711 | 3544 | 3118 | 3431 | 2066 | 2194 | 1510 | 2098 | 2427 | 3379 | 3297 | 3390 | 3236 | 1758 | 1717 | 1781 | 1973 | 2566,41 | 3326 | 1807 | 0,544 | -0,879 | -17,259 | 9,54E-67 | 4,39E-66 | 0,0016491 | 0,002731 | 0,002075 | 0,002075 | TRUE | 0,0467938 | No |
| orf19.4712 | 25 | 17 | 15 | 15 | 47 | 32 | 57 | 76 | 24 | 18 | 15 | 23 | 38 | 36 | 48 | 62 | 33,05 | 20 | 46 | 2,333 | 1,222 | 4,219 | 2,45E-05 | 3,57E-05 | 0,0322587 | 0,0517779 | 0,0428898 | 0,0428898 | TRUE | 0,4535647 | No |
| orf19.4713 | 392 | 412 | 410 | 297 | 632 | 492 | 640 | 679 | 374 | 436 | 405 | 465 | 506 | 559 | 543 | 552 | 480,12 | 420 | 540 | 1,293 | 0,37 | 4,592 | 4,39E-06 | 6,62E-06 | 0,0025315 | 0,005511 | 0,0040837 | 0,0040837 | TRUE | 0,06423 | No |
| orf19.4715 | 1818 | 1685 | 1831 | 1117 | 2046 | 1447 | 1938 | 2030 | 1733 | 1782 | 1809 | 1750 | 1639 | 1645 | 1645 | 1651 | 1706,79 | 1768 | 1645 | 0,93 | -0,105 | -2,558 | 0,0105367 | 0,0133667 | 1,00E-08 | 0,0030533 | 0,0010228 | 0,0010228 | TRUE | 0,0030033 | No |
| orf19.4716 | 16048 | 13178 | 14286 | 9270 | 66175 | 44415 | 62823 | 67619 | 15300 | 13934 | 14113 | 14522 | 53024 | 50504 | 53336 | 54981 | 33714,3 | 14467 | 52961 | 3,661 | 1,872 | 44,365 | 0 | 0 | 0,0014319 | 0,00214 | 0,0016633 | 0,0016633 | TRUE | 0,0185867 | No |
| orf19.4718 | 10656 | 9119 | 9850 | 6451 | 14188 | 9757 | 13574 | 14921 | 10159 | 9642 | 9731 | 10106 | 11368 | 11095 | 11524 | 12132 | 10719,71 | 9910 | 11530 | 1,164 | 0,219 | 5,6 | 2,15E-08 | 3,54E-08 | 0,0009468 | 0,0022444 | 0,0013727 | 0,0013727 | TRUE | 0,0151556 | No |
| orf19.4719 | 798 | 758 | 784 | 431 | 2035 | 1415 | 1906 | 2053 | 761 | 801 | 775 | 675 | 1631 | 1609 | 1618 | 1669 | 1192,38 | 753 | 1632 | 2,158 | 1,109 | 19,717 | 1,55E-86 | 8,98E-86 | 7,32E-05 | 0,0034683 | 0,0020274 | 0,0020274 | TRUE | 0,0475787 | No |
| orf19.4720 | 3542 | 3073 | 3343 | 2143 | 2659 | 1758 | 2595 | 2657 | 3377 | 3249 | 3303 | 3357 | 2131 | 1999 | 2203 | 2160 | 2722,38 | 3322 | 2123 | 0,64 | -0,644 | -15,198 | 3,63E-52 | 1,38E-51 | 0,0004947 | 0,0026944 | 0,0013427 | 0,0013427 | TRUE | 0,0175692 | No |
| orf19.4721 | 4739 | 4327 | 4485 | 2515 | 3316 | 2194 | 3177 | 3416 | 4518 | 4575 | 4431 | 3940 | 2657 | 2495 | 2697 | 2778 | 3511,32 | 4366 | 2657 | 0,608 | -0,717 | -12,678 | 7,87E-37 | 2,36E-36 | 0,0028086 | 0,0025589 | 0,0027775 | 0,0027775 | TRUE | 0,0506849 | No |
| orf19.4722 | 2758 | 2324 | 2631 | 1528 | 935 | 626 | 879 | 952 | 2629 | 2457 | 2599 | 2394 | 749 | 712 | 746 | 774 | 1632,63 | 2520 | 745 | 0,296 | -1,757 | -33,202 | 1,00E-241 | 1,64E-240 | 0,0009956 | 0,003097 | 0,0018695 | 0,0018695 | TRUE | 0,0128257 | No |
| orf19.4723 | 1040 | 864 | 932 | 593 | 1719 | 1321 | 1700 | 1833 | 992 | 914 | 921 | 929 | 1377 | 1502 | 1443 | 1490 | 1196 | 939 | 1453 | 1,546 | 0,628 | 11,572 | 5,69E-31 | 1,56E-30 | 0,0007933 | 0,0034641 | 0,0019276 | 0,0019276 | TRUE | 0,0183945 | No |
| orf19.4724 | 1179 | 1017 | 1145 | 657 | 2330 | 1537 | 2216 | 2367 | 1124 | 1075 | 1131 | 1029 | 1867 | 1748 | 1881 | 1925 | 1472,55 | 1090 | 1855 | 1,7 | 0,765 | 14,518 | 9,33E-48 | 3,34E-47 | 0,0008102 | 0,0032063 | 0,0019138 | 0,0019138 | TRUE | 0,01738 | No |
| orf19.4725 | 1147 | 1030 | 1033 | 753 | 2493 | 1838 | 2375 | 2565 | 1094 | 1089 | 1021 | 1180 | 1998 | 2090 | 2016 | 2086 | 1571,53 | 1096 | 2048 | 1,874 | 0,906 | 17,458 | 2,97E-68 | 1,39E-67 | 0,0007141 | 0,0031361 | 0,001855 | 0,001855 | TRUE | 0,0277914 | No |
| orf19.4726 | 1102 | 1020 | 1109 | 609 | 784 | 549 | 762 | 820 | 1051 | 1079 | 1096 | 954 | 628 | 624 | 647 | 667 | 843,11 | 1045 | 642 | 0,613 | -0,707 | -11,456 | 2,19E-30 | 5,95E-30 | 0,0009155 | 0,0040387 | 0,0024244 | 0,0024244 | TRUE | 0,0357262 | No |
| orf19.4727 | 967 | 855 | 848 | 540 | 1605 | 1047 | 1481 | 1640 | 922 | 904 | 838 | 846 | 1286 | 1191 | 1257 | 1333 | 1072,13 | 878 | 1267 | 1,444 | 0,53 | 9,003 | 2,20E-19 | 4,88E-19 | 0,0012873 | 0,0036227 | 0,0023329 | 0,0023329 | TRUE | 0,0185485 | No |
| orf19.4728 | 4659 | 4068 | 4648 | 2625 | 3906 | 2614 | 3543 | 3896 | 4442 | 4301 | 4592 | 4112 | 3130 | 2972 | 3008 | 3168 | 3715,65 | 4362 | 3070 | 0,703 | -0,508 | -11,127 | 9,23E-29 | 2,44E-28 | 0,0011963 | 0,0025332 | 0,0017243 | 0,0017243 | TRUE | 0,0171806 | No |
| orf19.473 | 1877 | 1494 | 1689 | 1054 | 1964 | 1361 | 1919 | 2040 | 1790 | 1580 | 1669 | 1651 | 1574 | 1548 | 1629 | 1659 | 1637,27 | 1672 | 1602 | 0,958 | -0,062 | -1,182 | 0,2372021 | 0,2666173 | 0,001273 | 0,0030941 | 0,002015 | 0,002015 | TRUE | 0,0281489 | No |
| orf19.4730 | 2151 | 1831 | 2053 | 1225 | 2538 | 1575 | 2256 | 2494 | 2051 | 1936 | 2028 | 1919 | 2034 | 1791 | 1915 | 2028 | 1962,72 | 1984 | 1942 | 0,979 | -0,03 | -0,563 | 0,5734194 | 0,6077046 | 0,0017689 | 0,0029278 | 0,0022397 | 0,0022397 | TRUE | 0,0316523 | No |
| orf19.4731 | 1040 | 934 | 1113 | 619 | 2072 | 1582 | 1983 | 2108 | 992 | 988 | 1100 | 970 | 1660 | 1799 | 1684 | 1714 | 1363,13 | 1012 | 1714 | 1,689 | 0,756 | 13,555 | 7,35E-42 | 2,40E-41 | 0,0012807 | 0,0032958 | 0,0021764 | 0,0021764 | TRUE | 0,041559 | No |
| orf19.4732 | 2160 | 1889 | 2080 | 1402 | 2590 | 1960 | 2472 | 2671 | 2059 | 1997 | 2055 | 2196 | 2075 | 2229 | 2099 | 2172 | 2110,29 | 2077 | 2144 | 1,033 | 0,047 | 1,013 | 0,3111052 | 0,3442876 | 0,0007277 | 0,0028693 | 0,0015804 | 0,0015804 | TRUE | 0,0163706 | No |
| orf19.4733 | 1265 | 1132 | 1201 | 621 | 2936 | 1883 | 2878 | 2990 | 1206 | 1197 | 1186 | 973 | 2353 | 2141 | 2443 | 2431 | 1741,32 | 1140 | 2342 | 2,048 | 1,034 | 14,26 | 3,88E-46 | 1,36E-45 | 0,0057451 | 0,0030342 | 0,0043676 | 0,0043676 | TRUE | 0,1078019 | No |
| orf19.4734 | 768 | 639 | 717 | 489 | 1200 | 875 | 1171 | 1264 | 732 | 676 | 708 | 766 | 962 | 995 | 994 | 1028 | 857,58 | 720 | 995 | 1,384 | 0,469 | 8,007 | 1,17E-15 | 2,40E-15 | 0,0001768 | 0,0040059 | 0,0020646 | 0,0020646 | TRUE | 0,0199762 | No |
| orf19.4735 | 1643 | 1432 | 1570 | 1052 | 1921 | 1351 | 1779 | 2055 | 1566 | 1514 | 1551 | 1648 | 1539 | 1536 | 1510 | 1671 | 1567,04 | 1570 | 1564 | 0,998 | -0,003 | -0,049 | 0,9611417 | 0,9649222 | 0,0010652 | 0,0031391 | 0,0019135 | 0,0019135 | TRUE | 0,0261133 | No |
| orf19.4736 | 1177 | 1053 | 1164 | 729 | 3065 | 2162 | 2881 | 3236 | 1122 | 1113 | 1150 | 1142 | 2456 | 2458 | 2446 | 2631 | 1814,86 | 1132 | 2498 | 2,208 | 1,143 | 24,422 | 1,00E-131 | 8,85E-131 | 1,00E-08 | 0,002996 | 0,0014299 | 0,0014299 | TRUE | 0,0160032 | No |
| orf19.4737 | 1674 | 1540 | 1589 | 1033 | 308 | 238 | 301 | 336 | 1596 | 1628 | 1570 | 1618 | 247 | 271 | 256 | 273 | 932,32 | 1603 | 262 | 0,163 | -2,618 | -41,64 | 0 | 0 | 1,00E-08 | 0,0038524 | 0,0017466 | 0,0017466 | TRUE | 0,0172486 | No |
| orf19.4738 | 1356 | 1173 | 1314 | 811 | 1342 | 1011 | 1263 | 1376 | 1293 | 1240 | 1298 | 1270 | 1075 | 1150 | 1072 | 1119 | 1189,71 | 1275 | 1104 | 0,864 | -0,21 | -4,259 | 2,05E-05 | 3,00E-05 | 1,00E-08 | 0,0034714 | 0,0015013 | 0,0015013 | TRUE | 0,008435 | No |
| orf19.4739 | 21910 | 18915 | 20381 | 13354 | 22466 | 16401 | 21548 | 23051 | 20889 | 20000 | 20134 | 20920 | 18001 | 18649 | 18294 | 18743 | 19453,86 | 20486 | 18422 | 0,899 | -0,153 | -4,966 | 6,84E-07 | 1,07E-06 | 0,000384 | 0,0021757 | 0,0008612 | 0,0008612 | TRUE | 0,0031067 | No |
| orf19.474 | 1350 | 1332 | 1308 | 829 | 2359 | 1723 | 2324 | 2600 | 1287 | 1408 | 1292 | 1299 | 1890 | 1959 | 1973 | 2114 | 1652,86 | 1322 | 1984 | 1,501 | 0,586 | 10,815 | 2,93E-27 | 7,58E-27 | 0,0015553 | 0,0030847 | 0,0021707 | 0,0021707 | TRUE | 0,0240894 | No |
| orf19.4740 | 577 | 485 | 557 | 354 | 510 | 373 | 505 | 510 | 550 | 513 | 550 | 555 | 409 | 424 | 429 | 415 | 480,5 | 542 | 419 | 0,773 | -0,371 | -5,417 | 6,06E-08 | 9,83E-08 | 1,00E-08 | 0,0055084 | 0,0024063 | 0,0024063 | TRUE | 0,0155308 | No |
| orf19.4741 | 18 | 13 | 16 | 7 | 38 | 27 | 23 | 31 | 17 | 14 | 16 | 11 | 30 | 31 | 20 | 25 | 20,45 | 14 | 26 | 1,795 | 0,844 | 2,593 | 0,0095144 | 0,0121108 | 1,00E-08 | 0,0823987 | 0,0471081 | 0,0471081 | TRUE | 0,213846 | No |
| orf19.4742 | 1149 | 973 | 1053 | 690 | 1252 | 966 | 1148 | 1298 | 1095 | 1029 | 1040 | 1081 | 1003 | 1098 | 975 | 1055 | 1047,14 | 1061 | 1033 | 0,972 | -0,041 | -0,713 | 0,4756728 | 0,5120223 | 0,0009392 | 0,0036593 | 0,0021484 | 0,0021484 | TRUE | 0,0201708 | No |
| orf19.4743 | 443 | 456 | 488 | 289 | 586 | 374 | 554 | 572 | 422 | 482 | 482 | 453 | 470 | 425 | 470 | 465 | 458,7 | 460 | 458 | 0,997 | -0,004 | -0,056 | 0,9554196 | 0,9597816 | 0,0008154 | 0,0056708 | 0,0033577 | 0,0033577 | TRUE | 0,0392752 | No |
| orf19.4744 | 2931 | 2607 | 2733 | 1804 | 2316 | 1695 | 2139 | 2339 | 2794 | 2757 | 2700 | 2826 | 1856 | 1927 | 1816 | 1902 | 2322,24 | 2769 | 1875 | 0,677 | -0,562 | -14,004 | 1,48E-44 | 5,04E-44 | 1,00E-08 | 0,0027983 | 0,0011078 | 0,0011078 | TRUE | 0,0054074 | No |
| orf19.4746 | 9692 | 8253 | 9329 | 5456 | 11323 | 7852 | 11129 | 11543 | 9240 | 8727 | 9216 | 8547 | 9073 | 8928 | 9448 | 9386 | 9070,66 | 8932 | 9209 | 1,031 | 0,043 | 1,094 | 0,2741512 | 0,3054582 | 0,0009829 | 0,0022723 | 0,0014049 | 0,0014049 | TRUE | 0,0100017 | No |
| orf19.4747 | 339 | 290 | 343 | 171 | 627 | 470 | 517 | 626 | 323 | 307 | 339 | 268 | 502 | 534 | 439 | 509 | 402,67 | 309 | 496 | 1,593 | 0,672 | 7,216 | 5,37E-13 | 1,03E-12 | 0,0047105 | 0,0061689 | 0,0056187 | 0,0056187 | TRUE | 0,0755281 | No |
| orf19.4748 | 341 | 391 | 355 | 268 | 325 | 234 | 255 | 238 | 325 | 413 | 351 | 420 | 260 | 266 | 216 | 194 | 305,7 | 377 | 234 | 0,621 | -0,687 | -5,527 | 3,25E-08 | 5,32E-08 | 0,015644 | 0,0074623 | 0,0114347 | 0,0114347 | TRUE | 0,1557996 | No |
| orf19.4749 | 969 | 862 | 982 | 656 | 1340 | 1005 | 1328 | 1407 | 924 | 911 | 970 | 1028 | 1074 | 1143 | 1127 | 1144 | 1040,13 | 958 | 1122 | 1,174 | 0,231 | 4,107 | 4,00E-05 | 5,78E-05 | 0,0006566 | 0,0036698 | 0,0020545 | 0,0020545 | TRUE | 0,0245376 | No |
| orf19.475 | 4586 | 4108 | 4425 | 2932 | 6204 | 4621 | 5976 | 6133 | 4372 | 4344 | 4371 | 4593 | 4971 | 5254 | 5074 | 4987 | 4745,81 | 4420 | 5072 | 1,148 | 0,199 | 5,45 | 5,04E-08 | 8,20E-08 | 0,0003916 | 0,0024372 | 0,0010637 | 0,0010637 | TRUE | 0,0078563 | No |
| orf19.4750 | 839 | 883 | 954 | 545 | 808 | 589 | 671 | 664 | 800 | 934 | 942 | 854 | 647 | 670 | 570 | 540 | 744,57 | 882 | 607 | 0,686 | -0,544 | -6,357 | 2,06E-10 | 3,65E-10 | 0,0068669 | 0,0042965 | 0,0056474 | 0,0056474 | TRUE | 0,0654498 | No |
| orf19.4751 | 2558 | 2528 | 2564 | 1548 | 2462 | 1766 | 2154 | 2336 | 2439 | 2673 | 2533 | 2425 | 1973 | 2008 | 1829 | 1899 | 2222,35 | 2518 | 1927 | 0,765 | -0,387 | -7,711 | 1,25E-14 | 2,49E-14 | 0,0012761 | 0,0028301 | 0,0019649 | 0,0019649 | TRUE | 0,0209913 | No |
| orf19.4751.1 | 1268 | 1008 | 996 | 702 | 2831 | 1877 | 2573 | 2740 | 1209 | 1066 | 984 | 1100 | 2268 | 2134 | 2184 | 2228 | 1646,68 | 1090 | 2204 | 2,023 | 1,017 | 16,278 | 1,41E-59 | 5,92E-59 | 0,0030858 | 0,0030884 | 0,0030267 | 0,0030267 | TRUE | 0,0697035 | No |
| orf19.4752 | 1643 | 1398 | 1548 | 979 | 1907 | 1343 | 1667 | 1867 | 1566 | 1478 | 1529 | 1534 | 1528 | 1527 | 1415 | 1518 | 1512 | 1527 | 1497 | 0,98 | -0,029 | -0,605 | 0,5453085 | 0,580613 | 0,0004139 | 0,0031772 | 0,0015348 | 0,0015348 | TRUE | 0,0165906 | No |
| orf19.4753 | 8483 | 7877 | 8677 | 4862 | 3380 | 2363 | 3238 | 3358 | 8088 | 8329 | 8572 | 7617 | 2708 | 2687 | 2749 | 2730 | 5434,98 | 8152 | 2718 | 0,333 | -1,585 | -36,21 | 4,20E-287 | 8,89E-286 | 0,0012034 | 0,0023934 | 0,0016061 | 0,0016061 | TRUE | 0,0230542 | No |
| orf19.4754 | 1540 | 1661 | 1687 | 978 | 4076 | 3250 | 3795 | 3579 | 1468 | 1756 | 1667 | 1532 | 3266 | 3696 | 3222 | 2910 | 2439,59 | 1606 | 3274 | 2,037 | 1,026 | 13,035 | 7,77E-39 | 2,41E-38 | 0,0075258 | 0,0027643 | 0,0054641 | 0,0054641 | TRUE | 0,0905687 | No |
| orf19.4755 | 3667 | 3409 | 3733 | 2299 | 4505 | 3138 | 4317 | 4570 | 3496 | 3605 | 3688 | 3602 | 3610 | 3568 | 3665 | 3716 | 3618,61 | 3598 | 3640 | 1,012 | 0,017 | 0,491 | 0,6235066 | 0,6567507 | 0,0001427 | 0,002545 | 0,0009355 | 0,0009355 | TRUE | 0,0046577 | No |

| | | | | | | | | | | | | | | | | | | | | | | | | | | | | | | | |
|---|---|---|---|---|---|---|---|---|---|---|---|---|---|---|---|---|---|---|---|---|---|---|---|---|---|---|---|---|---|---|---|
| orf19.4756 | 1106 | 1097 | 1072 | 689 | 1063 | 712 | 960 | 945 | 1054 | 1160 | 1059 | 1079 | 852 | 810 | 815 | 768 | 949,69 | 1088 | 811 | 0,746 | -0,424 | -7,202 | 5,94E-13 | 1,13E-12 | 0,0010073 | 0,0038202 | 0,0022552 | 0,0022552 | TRUE | 0,0244368 | No |
| orf19.4757 | 7216 | 6665 | 7218 | 4237 | 6575 | 4639 | 6195 | 6383 | 6880 | 7047 | 7131 | 6637 | 5268 | 5275 | 5260 | 5190 | 6086,01 | 6924 | 5248 | 0,757 | -0,401 | -11,746 | 7,38E-32 | 2,06E-31 | 0,0002985 | 0,0023611 | 0,0009518 | 0,0009518 | TRUE | 0,0089198 | No |
| orf19.4758 | 3194 | 3180 | 3266 | 2118 | 5672 | 4444 | 5292 | 5157 | 3045 | 3362 | 3226 | 3318 | 4545 | 5053 | 4493 | 4193 | 3904,51 | 3238 | 4571 | 1,411 | 0,497 | 8,216 | 2,10E-16 | 4,35E-16 | 0,0036375 | 0,0025118 | 0,0032451 | 0,0032451 | TRUE | 0,060598 | No |
| orf19.4759 | 4186 | 3699 | 3632 | 2978 | 7923 | 5844 | 6887 | 7728 | 3991 | 3911 | 3588 | 4665 | 6348 | 6645 | 5847 | 6284 | 5159,96 | 4039 | 6281 | 1,557 | 0,638 | 8,669 | 4,37E-18 | 9,40E-18 | 0,0073783 | 0,0024095 | 0,004998 | 0,004998 | TRUE | 0,1299854 | No |
| orf19.4760 | 4783 | 5144 | 5289 | 2714 | 6372 | 4456 | 5955 | 5560 | 4560 | 5439 | 5225 | 4252 | 5106 | 5067 | 5056 | 4521 | 4903,13 | 4869 | 4938 | 1,013 | 0,019 | 0,249 | 0,8030743 | 0,8251879 | 0,0079654 | 0,0024261 | 0,0054778 | 0,0054778 | TRUE | 0,0870503 | No |
| orf19.4761 | 1440 | 1461 | 1529 | 915 | 2077 | 1433 | 1971 | 1978 | 1373 | 1545 | 1511 | 1433 | 1664 | 1629 | 1673 | 1608 | 1554,62 | 1466 | 1644 | 1,122 | 0,165 | 3,254 | 0,0011376 | 0,0015281 | 0,0008725 | 0,0031474 | 0,001828 | 0,001828 | TRUE | 0,0238174 | No |
| orf19.4763 | 203 | 183 | 217 | 143 | 549 | 428 | 452 | 439 | 194 | 193 | 214 | 224 | 440 | 487 | 384 | 357 | 311,59 | 206 | 417 | 2,022 | 1,016 | 8,867 | 7,55E-19 | 1,65E-18 | 0,0101926 | 0,0073608 | 0,0088022 | 0,0088022 | TRUE | 0,1392694 | No |
| orf19.4764 | 1214 | 1403 | 1364 | 841 | 1508 | 1197 | 1296 | 1265 | 1157 | 1483 | 1348 | 1317 | 1208 | 1361 | 1100 | 1029 | 1250,52 | 1326 | 1174 | 0,885 | -0,177 | -1,869 | 0,0616406 | 0,0734233 | 0,0116044 | 0,0034043 | 0,0077985 | 0,0077985 | TRUE | 0,133878 | No |
| orf19.4765 | 159 | 146 | 132 | 116 | 123 | 127 | 122 | 126 | 152 | 154 | 130 | 182 | 99 | 144 | 104 | 102 | 133,39 | 154 | 112 | 0,725 | -0,463 | -2,996 | 0,0027349 | 0,0035952 | 0,0164246 | 0,0144007 | 0,0152979 | 0,0152979 | TRUE | 0,337918 | No |
| orf19.4766 | 3187 | 3210 | 3423 | 2020 | 2251 | 1635 | 2123 | 2085 | 3038 | 3394 | 3382 | 3164 | 1804 | 1859 | 1802 | 1695 | 2517,4 | 3244 | 1790 | 0,551 | -0,859 | -16,462 | 6,84E-61 | 2,92E-60 | 0,0018473 | 0,0027435 | 0,0021928 | 0,0021928 | TRUE | 0,0232883 | No |
| orf19.4767 | 511 | 450 | 526 | 319 | 875 | 552 | 797 | 854 | 487 | 476 | 520 | 500 | 701 | 628 | 677 | 694 | 585,27 | 496 | 675 | 1,365 | 0,449 | 6,64 | 3,14E-11 | 5,70E-11 | 1,00E-08 | 0,0048966 | 0,002603 | 0,002603 | TRUE | 0,0239242 | No |
| orf19.4768 | 659 | 808 | 803 | 432 | 962 | 777 | 917 | 864 | 628 | 854 | 793 | 677 | 771 | 884 | 779 | 703 | 761,01 | 738 | 784 | 1,059 | 0,083 | 0,815 | 0,4149244 | 0,4509679 | 0,0129568 | 0,0042488 | 0,0086925 | 0,0086925 | TRUE | 0,133197 | No |
| orf19.4769 | 3245 | 3134 | 3270 | 2001 | 2611 | 2060 | 2467 | 2488 | 3094 | 3314 | 3230 | 3135 | 2092 | 2342 | 2094 | 2023 | 2665,58 | 3193 | 2138 | 0,669 | -0,581 | -10,954 | 6,33E-28 | 1,65E-27 | 0,0021091 | 0,0027072 | 0,002314 | 0,002314 | TRUE | 0,0482714 | No |
| orf19.477 | 1123 | 1069 | 1134 | 685 | 2075 | 1399 | 1837 | 1923 | 1071 | 1130 | 1120 | 1073 | 1663 | 1591 | 1560 | 1564 | 1346,37 | 1098 | 1594 | 1,45 | 0,536 | 10,958 | 6,07E-28 | 1,58E-27 | 1,00E-08 | 0,0033108 | 0,0015101 | 0,0015101 | TRUE | 0,0105475 | No |
| orf19.4771 | 5948 | 5715 | 6135 | 3878 | 1518 | 1167 | 1362 | 1511 | 5671 | 6043 | 6061 | 6075 | 1216 | 1327 | 1156 | 1229 | 3597,22 | 5962 | 1232 | 0,206 | -2,277 | -46,032 | 0 | 0 | 0,0015286 | 0,0025477 | 0,001892 | 0,001892 | TRUE | 0,0299888 | No |
| orf19.4772 | 1200 | 1128 | 1179 | 813 | 457 | 389 | 365 | 468 | 1144 | 1193 | 1165 | 1274 | 366 | 442 | 310 | 381 | 784,26 | 1194 | 375 | 0,313 | -1,676 | -18,636 | 1,64E-77 | 8,64E-77 | 0,0083931 | 0,0041849 | 0,0060944 | 0,0060944 | TRUE | 0,1566361 | No |
| orf19.4773 | 7716 | 6976 | 7287 | 4438 | 7190 | 4983 | 6735 | 7084 | 7356 | 7376 | 7199 | 6952 | 5761 | 5666 | 5718 | 5760 | 6473,63 | 7221 | 5726 | 0,793 | -0,335 | -10,389 | 2,78E-25 | 6,93E-25 | 0,0001898 | 0,0023449 | 0,0008446 | 0,0008446 | TRUE | 0,0071977 | No |
| orf19.4774 | 1072 | 938 | 992 | 700 | 914 | 705 | 937 | 977 | 1022 | 992 | 980 | 1097 | 732 | 802 | 796 | 794 | 901,79 | 1023 | 781 | 0,765 | -0,386 | -6,455 | 1,08E-10 | 1,93E-10 | 0,0009042 | 0,003912 | 0,0023188 | 0,0023188 | TRUE | 0,0249207 | No |
| orf19.4775 | 10028 | 8998 | 9702 | 6360 | 8707 | 6150 | 8418 | 8851 | 9561 | 9514 | 9585 | 9963 | 6977 | 6993 | 7147 | 7197 | 8367,01 | 9656 | 7078 | 0,734 | -0,447 | -14,524 | 8,57E-48 | 3,07E-47 | 0,0001876 | 0,0022875 | 0,0007884 | 0,0007884 | TRUE | 0,0053525 | No |
| orf19.4776 | 1721 | 1652 | 1711 | 1129 | 1707 | 1287 | 1594 | 1772 | 1641 | 1747 | 1690 | 1769 | 1368 | 1463 | 1353 | 1441 | 1558,98 | 1712 | 1406 | 0,822 | -0,283 | -5,748 | 9,04E-09 | 1,50E-08 | 0,0006399 | 0,0031445 | 0,0016784 | 0,0016784 | TRUE | 0,0095072 | No |
| orf19.4777 | 3414 | 3062 | 3315 | 1916 | 2014 | 1321 | 1858 | 1998 | 3255 | 3238 | 3275 | 3002 | 1614 | 1502 | 1577 | 1625 | 2385,85 | 3192 | 1580 | 0,495 | -1,015 | -21,76 | 5,55E-105 | 3,78E-104 | 0,0007884 | 0,0027795 | 0,0016323 | 0,0016323 | TRUE | 0,0182991 | No |
| orf19.4778 | 422 | 367 | 423 | 237 | 493 | 343 | 538 | 519 | 402 | 388 | 418 | 371 | 395 | 390 | 457 | 422 | 405,42 | 395 | 416 | 1,052 | 0,073 | 0,889 | 0,3741443 | 0,4092243 | 0,0016322 | 0,0061412 | 0,0039256 | 0,0039256 | TRUE | 0,0507137 | No |
| orf19.4779 | 7221 | 6279 | 6822 | 4356 | 3186 | 2206 | 2985 | 3087 | 6884 | 6639 | 6739 | 6824 | 2553 | 2508 | 2534 | 2510 | 4649,08 | 6772 | 2526 | 0,373 | -1,422 | -44,004 | 0 | 0 | 1,00E-08 | 0,0024444 | 0,0007431 | 0,0007431 | TRUE | 0,0021452 | No |
| orf19.478 | 1565 | 1347 | 1466 | 986 | 1120 | 877 | 1111 | 1146 | 1492 | 1424 | 1448 | 1545 | 897 | 997 | 943 | 932 | 1209,87 | 1477 | 942 | 0,638 | -0,649 | -12,194 | 3,36E-34 | 9,68E-34 | 0,000448 | 0,0034484 | 0,0018657 | 0,0018657 | TRUE | 0,0168172 | No |
| orf19.4780 | 1804 | 1482 | 1721 | 1090 | 3128 | 2194 | 2781 | 3045 | 1720 | 1567 | 1700 | 1708 | 2506 | 2495 | 2361 | 2476 | 2066,59 | 1674 | 2460 | 1,47 | 0,556 | 11,84 | 2,41E-32 | 6,79E-32 | 0,0006139 | 0,0028858 | 0,0015974 | 0,0015974 | TRUE | 0,0226656 | No |
| orf19.4781 | 433 | 336 | 407 | 314 | 788 | 648 | 740 | 822 | 413 | 355 | 402 | 492 | 631 | 737 | 628 | 668 | 540,87 | 416 | 666 | 1,61 | 0,687 | 7,214 | 5,43E-13 | 1,04E-12 | 0,0084302 | 0,005127 | 0,0066812 | 0,0066812 | TRUE | 0,1549364 | No |
| orf19.4783 | 115 | 103 | 134 | 77 | 25 | 12 | 28 | 32 | 110 | 109 | 132 | 121 | 20 | 14 | 24 | 26 | 69,38 | 118 | 21 | 0,18 | -2,47 | -11,724 | 9,62E-32 | 2,68E-31 | 0,0017197 | 0,0257575 | 0,017041 | 0,017041 | TRUE | 0,2277738 | No |
| orf19.4784 | 1763 | 1507 | 1579 | 1077 | 506 | 384 | 444 | 528 | 1681 | 1593 | 1560 | 1687 | 405 | 437 | 377 | 429 | 1021,22 | 1630 | 412 | 0,252 | -1,986 | -31,368 | 5,58E-216 | 8,17E-215 | 0,0011317 | 0,0036991 | 0,0024214 | 0,0024214 | TRUE | 0,0382155 | No |
| orf19.4785 | 1951 | 1572 | 1798 | 1090 | 1942 | 1258 | 1912 | 1956 | 1860 | 1662 | 1776 | 1708 | 1556 | 1430 | 1623 | 1590 | 1650,78 | 1752 | 1550 | 0,885 | -0,176 | -3,096 | 0,0019645 | 0,0026034 | 0,0018808 | 0,0030859 | 0,0024905 | 0,0024905 | TRUE | 0,0307971 | No |
| orf19.4786 | 588 | 575 | 589 | 371 | 621 | 407 | 515 | 547 | 561 | 608 | 582 | 581 | 498 | 463 | 437 | 445 | 521,75 | 583 | 461 | 0,791 | -0,339 | -4,78 | 1,75E-06 | 2,68E-06 | 1,00E-08 | 0,0052382 | 0,0029068 | 0,0029068 | TRUE | 0,0366725 | No |
| orf19.4787 | 1885 | 1624 | 1875 | 1191 | 1236 | 854 | 1164 | 1369 | 1797 | 1717 | 1852 | 1866 | 990 | 971 | 988 | 1113 | 1411,9 | 1808 | 1016 | 0,563 | -0,83 | -14,284 | 2,76E-46 | 9,67E-46 | 0,0017525 | 0,0032542 | 0,0024883 | 0,0024883 | TRUE | 0,0509475 | No |
| orf19.4788 | 13751 | 11957 | 13040 | 8093 | 8989 | 6271 | 8190 | 9299 | 13110 | 12643 | 12882 | 12678 | 7203 | 7131 | 6953 | 7561 | 10020,13 | 12828 | 7212 | 0,562 | -0,831 | -23,169 | 9,43E-119 | 7,40E-118 | 0,0006228 | 0,0022551 | 0,0011293 | 0,0011293 | TRUE | 0,0131091 | No |
| orf19.4789 | 62 | 60 | 76 | 50 | 25 | 12 | 19 | 26 | 59 | 63 | 75 | 78 | 20 | 14 | 16 | 21 | 43,36 | 69 | 18 | 0,262 | -1,931 | -7,92 | 2,38E-15 | 4,82E-15 | 1,00E-08 | 0,0399548 | 0,0242663 | 0,0242663 | TRUE | 0,0948645 | No |
| orf19.4791 | 1776 | 1962 | 2039 | 1151 | 1100 | 799 | 1042 | 1036 | 1693 | 2075 | 2014 | 1803 | 881 | 909 | 885 | 842 | 1387,77 | 1896 | 879 | 0,463 | -1,11 | -15,672 | 2,34E-55 | 9,27E-55 | 0,0045688 | 0,0032744 | 0,0040151 | 0,0040151 | TRUE | 0,0657009 | No |
| orf19.4792 | 17765 | 15331 | 17092 | 11488 | 3669 | 2675 | 3539 | 3806 | 16937 | 16211 | 16885 | 17997 | 2940 | 3042 | 3005 | 3095 | 10013,78 | 17008 | 3020 | 0,178 | -2,493 | -59,933 | 0 | 0 | 0,0009132 | 0,0022552 | 0,0014809 | 0,0014809 | TRUE | 0,0184472 | No |
| orf19.479.2 | 449 | 384 | 417 | 256 | 608 | 426 | 495 | 570 | 428 | 406 | 412 | 401 | 487 | 484 | 420 | 463 | 437,8 | 412 | 464 | 1,124 | 0,168 | 2,182 | 0,0290986 | 0,0356936 | 1,00E-08 | 0,0058416 | 0,0033778 | 0,0033778 | TRUE | 0,0474294 | No |
| orf19.4793 | 5251 | 4956 | 5108 | 3095 | 5252 | 3513 | 4733 | 4821 | 5006 | 5240 | 5046 | 4848 | 4208 | 3995 | 4018 | 3920 | 4535,3 | 5035 | 4035 | 0,801 | -0,32 | -7,999 | 1,25E-15 | 2,56E-15 | 0,0006769 | 0,0024533 | 0,0013143 | 0,0013143 | TRUE | 0,0103504 | No |
| orf19.4795 | 2047 | 2018 | 2077 | 1296 | 1738 | 1177 | 1561 | 1653 | 1952 | 2134 | 2052 | 2030 | 1393 | 1338 | 1325 | 1344 | 1695,97 | 2042 | 1350 | 0,661 | -0,596 | -12,802 | 1,60E-37 | 4,86E-37 | 0,0004645 | 0,0030594 | 0,0014772 | 0,0014772 | TRUE | 0,0114919 | No |
| orf19.4796 | 2698 | 2322 | 2497 | 1495 | 5570 | 3637 | 5372 | 5583 | 2572 | 2455 | 2467 | 2342 | 4463 | 4136 | 4561 | 4540 | 3441,91 | 2459 | 4425 | 1,798 | 0,846 | 17,81 | 5,95E-71 | 2,89E-70 | 0,0013051 | 0,0025683 | 0,0018397 | 0,0018397 | TRUE | 0,0229318 | No |
| orf19.4798 | 4024 | 3799 | 3874 | 2680 | 6458 | 4702 | 5933 | 6430 | 3836 | 4017 | 3827 | 4198 | 5175 | 5347 | 5037 | 5228 | 4583,18 | 3970 | 5197 | 1,31 | 0,39 | 9,262 | 2,00E-20 | 4,54E-20 | 0,0009881 | 0,0024495 | 0,001474 | 0,001474 | TRUE | 0,0172799 | No |
| orf19.4799 | 5708 | 5207 | 5770 | 3627 | 4707 | 3340 | 4553 | 4945 | 5442 | 5506 | 5700 | 5682 | 3772 | 3798 | 3865 | 4021 | 4723,19 | 5582 | 3864 | 0,693 | -0,53 | -14,257 | 4,05E-46 | 1,41E-45 | 0,0003906 | 0,0024389 | 0,0011104 | 0,0011104 | TRUE | 0,0092384 | No |
| orf19.48 | 4019 | 4635 | 4748 | 3005 | 2105 | 1641 | 1888 | 1859 | 3832 | 4901 | 4691 | 4707 | 1687 | 1866 | 1603 | 1512 | 3099,72 | 4533 | 1667 | 0,368 | -1,444 | -17,213 | 2,13E-66 | 9,72E-66 | 0,0090839 | 0,002621 | 0,0063676 | 0,0063676 | TRUE | 0,1331535 | No |
| orf19.480 | 4638 | 3914 | 4286 | 2741 | 4337 | 3012 | 4293 | 4451 | 4422 | 4139 | 4234 | 4294 | 3475 | 3425 | 3645 | 3619 | 3906,54 | 4272 | 3541 | 0,829 | -0,27 | -6,824 | 8,83E-12 | 1,63E-11 | 0,0005017 | 0,0025116 | 0,0012477 | 0,0012477 | TRUE | 0,0071009 | No |
| orf19.4800 | 2528 | 2253 | 2426 | 1470 | 1007 | 740 | 981 | 1023 | 2410 | 2382 | 2397 | 2303 | 807 | 841 | 833 | 832 | 1600,61 | 2373 | 828 | 0,348 | -1,521 | -33,742 | 1,42E-249 | 2,40E-248 | 1,00E-08 | 0,0031171 | 0,0011776 | 0,0011776 | TRUE | 0,0042975 | No |
| orf19.4801 | 634 | 598 | 605 | 363 | 1284 | 934 | 1303 | 1415 | 604 | 632 | 598 | 569 | 1029 | 1062 | 1106 | 1151 | 843,84 | 601 | 1087 | 1,806 | 0,853 | 13,625 | 2,86E-42 | 9,41E-42 | 0,0011184 | 0,004037 | 0,0024213 | 0,0024213 | TRUE | 0,0191477 | No |
| orf19.4802 | 5052 | 4436 | 4921 | 2801 | 1675 | 1067 | 1553 | 1550 | 4817 | 4691 | 4861 | 4388 | 1342 | 1213 | 1318 | 1260 | 2986,33 | 4689 | 1283 | 0,274 | -1,869 | -36,852 | 2,67E-297 | 5,90E-296 | 0,0013993 | 0,0026411 | 0,0020017 | 0,0020017 | TRUE | 0,0218266 | No |
| orf19.4803 | 14 | 11 | 14 | 9 | 6 | 5 | 6 | 8 | 13 | 12 | 14 | 14 | 5 | 6 | 5 | 7 | 9,38 | 13 | 6 | 0,418 | -1,258 | -2,728 | 0,0063652 | 0,0081723 | 1,00E-08 | 0,1772275 | 0,0815041 | 0,0815041 | TRUE | 0,0420209 | No |
| orf19.4804 | 69 | 68 | 82 | 34 | 44 | 34 | 43 | 48 | 66 | 72 | 81 | 53 | 35 | 39 | 37 | 39 | 52,68 | 68 | 38 | 0,543 | -0,882 | -4,362 | 1,29E-05 | 1,90E-05 | 1,00E-08 | 0,0332611 | 0,0191979 | 0,0191979 | TRUE | 0,1431794 | No |
| orf19.4805 | 5039 | 4495 | 4949 | 3108 | 10765 | 7721 | 9894 | 11004 | 4804 | 4753 | 4889 | 4869 | 8626 | 8779 | 8400 | 8947 | 6758,43 | 4829 | 8688 | 1,799 | 0,848 | 25,092 | 6,06E-139 | 5,66E-138 | 0,000369 | 0,0023342 | 0,0009275 | 0,0009275 | TRUE | 0,0061298 | No |
| orf19.4807 | 1352 | 1273 | 1380 | 860 | 3411 | 2277 | 3270 | 3380 | 1289 | 1346 | 1363 | 1347 | 2733 | 2589 | 2776 | 2748 | 2024,04 | 1336 | 2712 | 2,032 | 1,023 | 22,862 | 1,12E-115 | 8,53E-115 | 0,0003412 | 0,0029025 | 0,0013377 | 0,0013377 | TRUE | 0,0106331 | No |
| orf19.4808 | 4085 | 4000 | 3968 | 2509 | 4199 | 3099 | 3747 | 4229 | 3895 | 4229 | 3920 | 3930 | 3365 | 3524 | 3181 | 3439 | 3685,34 | 3994 | 3377 | 0,845 | -0,243 | -5,174 | 2,30E-07 | 3,65E-07 | 0,001351 | 0,0025368 | 0,0018398 | 0,0018398 | TRUE | 0,0194653 | No |
| orf19.4809 | 1293 | 1139 | 1123 | 736 | 1712 | 1146 | 1670 | 1734 | 1233 | 1204 | 1109 | 1153 | 1372 | 1303 | 1418 | 1410 | 1275,26 | 1175 | 1376 | 1,171 | 0,228 | 4,196 | 2,72E-05 | 3,95E-05 | 0,0009992 | 0,0033788 | 0,0020357 | 0,0020357 | TRUE | 0,0184948 | No |
| orf19.481 | 3700 | 2997 | 3298 | 2215 | 6445 | 4488 | 6378 | 6552 | 3528 | 3169 | 3258 | 3470 | 5164 | 5103 | 5415 | 5327 | 4304,28 | 3356 | 5252 | 1,566 | 0,647 | 14,144 | 2,03E-45 | 6,98E-45 | 0,0013363 | 0,0024727 | 0,0017572 | 0,0017572 | TRUE | 0,0171859 | No |
| orf19.4811 | 2435 | 2068 | 2112 | 1599 | 3033 | 2216 | 3029 | 3250 | 2322 | 2187 | 2086 | 2505 | 2430 | 2520 | 2572 | 2643 | 2407,97 | 2275 | 2541 | 1,119 | 0,162 | 2,719 | 0,0065519 | 0,0084019 | 0,0032316 | 0,0027731 | 0,0029904 | 0,0029904 | TRUE | 0,0530544 | No |
| orf19.4812 | 476 | 386 | 382 | 341 | 1156 | 813 | 1042 | 1098 | 454 | 408 | 377 | 534 | 926 | 924 | 885 | 893 | 675,21 | 443 | 907 | 2,06 | 1,042 | 11,638 | 2,64E-31 | 7,29E-31 | 0,0077337 | 0,004523 | 0,0059348 | 0,0059348 | TRUE | 0,1936043 | No |
| orf19.4813 | 22379 | 18635 | 19951 | 12995 | 45036 | 31269 | 42732 | 47013 | 21336 | 19704 | 19710 | 20357 | 36086 | 35556 | 36279 | 38226 | 28406,78 | 20277 | 36537 | 1,802 | 0,85 | 21,389 | 1,68E-101 | 1,11E-100 | 0,0011797 | 0,0021491 | 0,0014755 | 0,0014755 | TRUE | 0,0152857 | No |
| orf19.4814 | 82 | 70 | 65 | 39 | 117 | 62 | 108 | 107 | 78 | 74 | 64 | 61 | 94 | 70 | 92 | 87 | 77,56 | 69 | 86 | 1,235 | 0,304 | 1,806 | 0,070922 | 0,0840084 | 1,00E-08 | 0,0232619 | 0,0140761 | 0,0140761 | TRUE | 0,1167455 | No |
| orf19.4815 | 10958 | 9181 | 10036 | 6101 | 12672 | 8407 | 12201 | 13166 | 10447 | 9708 | 9915 | 9558 | 10154 | 9560 | 10359 | 10705 | 10050,53 | 9907 | 10194 | 1,029 | 0,041 | 0,893 | 0,3716519 | 0,4067772 | 0,0017487 | 0,0022546 | 0,0019411 | 0,0019411 | TRUE | 0,0212233 | No |
| orf19.4816 | 2145 | 1845 | 1983 | 1301 | 1924 | 1429 | 1874 | 1979 | 2045 | 1951 | 1959 | 2038 | 1542 | 1625 | 1591 | 1609 | 1794,96 | 1998 | 1592 | 0,797 | -0,328 | -7,616 | 2,61E-14 | 5,17E-14 | 1,00E-08 | 0,003006 | 0,0012222 | 0,0012222 | TRUE | 0,0055204 | No |
| orf19.4817 | 562 | 551 | 560 | 384 | 1010 | 753 | 957 | 995 | 536 | 583 | 553 | 602 | 809 | 856 | 812 | 809 | 695,03 | 568 | 822 | 1,45 | 0,536 | 8,604 | 7,69E-18 | 1,65E-17 | 1,00E-08 | 0,0044536 | 0,0021922 | 0,0021922 | TRUE | 0,0174129 | No |
| orf19.4818 | 3642 | 3003 | 3281 | 2218 | 4498 | 3181 | 4578 | 4684 | 3472 | 3175 | 3241 | 3475 | 3604 | 3617 | 3887 | 3809 | 3534,99 | 3341 | 3729 | 1,117 | 0,16 | 3,36 | 0,0007782 | 0,0010576 | 0,0014148 | 0,0025558 | 0,0018828 | 0,0018828 | TRUE | 0,0134662 | No |
| orf19.4819 | 211 | 257 | 196 | 126 | 252 | 215 | 242 | 233 | 201 | 272 | 194 | 197 | 202 | 244 | 205 | 189 | 213,15 | 216 | 210 | 0,969 | -0,046 | -0,346 | 0,7295013 | 0,758629 | 0,0148087 | 0,0097942 | 0,0121862 | 0,0121862 | TRUE | 0,3349268 | No |
| orf19.482 | 5045 | 4295 | 4796 | 2717 | 6989 | 4671 | 6731 | 7123 | 4810 | 4541 | 4738 | 4256 | 5600 | 5311 | 5715 | 5792 | 5095,41 | 4586 | 5604 | 1,221 | 0,288 | 5,9 | 3,64E-09 | 6,13E-09 | 0,001899 | 0,0024135 | 0,0020894 | 0,0020894 | TRUE | 0,027432 | No |
| orf19.4820 | 2598 | 2283 | 2503 | 1544 | 2834 | 2013 | 2577 | 2896 | 2477 | 2414 | 2473 | 2419 | 2271 | 2289 | 2188 | 2355 | 2360,59 | 2446 | 2276 | 0,93 | -0,105 | -2,588 | 0,0096496 | 0,0122755 | 1,00E-08 | 0,0027868 | 0,0011446 | 0,0011446 | TRUE | 0,0083508 | No |
| orf19.4821 | 15 | 19 | 14 | 8 | 15 | 6 | 11 | 10 | 14 | 20 | 14 | 13 | 12 | 7 | 9 | 8 | 12,13 | 15 | 9 | 0,601 | -0,735 | -1,812 | 0,0700357 | 0,0830511 | 1,00E-08 | 0,1374145 | 0,0733279 | 0,0733279 | TRUE | 0,2094221 | No |
| orf19.4822 | 5 | 5 | 7 | 3 | 1 | 3 | 4 | 0 | 5 | 5 | 7 | 5 | 1 | 3 | 3 | 0 | 3,66 | 6 | 2 | 0,333 | -1,588 | -1,933 | 0,0532545 | 0,0638393 | 1,00E-08 | 0,4507395 | 0,3007972 | 0,3007972 | TRUE | 0,4982088 | No |
| orf19.4823 | 282 | 248 | 260 | 178 | 391 | 242 | 320 | 344 | 269 | 262 | 257 | 279 | 313 | 275 | 272 | 280 | 275,83 | 267 | 285 | 1,073 | 0,101 | 1,115 | 0,2646445 | 0,29533 | 1,00E-08 | 0,0080439 | 0,004271 | 0,004271 | TRUE | 0,0533333 | No |
| orf19.4824 | 1610 | 1504 | 1591 | 1027 | 2434 | 1792 | 2419 | 2544 | 1535 | 1590 | 1572 | 1609 | 1950 | 2038 | 2054 | 2069 | 1802,01 | 1576 | 2028 | 1,287 | 0,364 | 8,405 | 4,29E-17 | 9,04E-17 | 1,00E-08 | 0,0030024 | 0,0012277 | 0,0012277 | TRUE | 0,0082932 | No |
| orf19.4825 | 1407 | 1184 | 1251 | 853 | 1240 | 936 | 1137 | 1169 | 1341 | 1252 | 1236 | 1336 | 994 | 1064 | 965 | 951 | 1142,4 | 1291 | 994 | 0,769 | -0,379 | -6,651 | 2,92E-11 | 5,30E-11 | 0,0012026 | 0,0035285 | 0,0022411 | 0,0022411 | TRUE | 0,0255728 | No |
| orf19.4826 | 14758 | 11921 | 12683 | 8864 | 27189 | 18552 | 25728 | 28438 | 14070 | 12605 | 12530 | 13886 | 21786 | 21095 | 21843 | 23123 | 17617,18 | 13273 | 21962 | 1,655 | 0,727 | 14,295 | 2,35E-46 | 8,24E-46 | 0,002562 | 0,0021845 | 0,0024208 | 0,0024208 | TRUE | 0,0203218 | No |
| orf19.4827 | 11444 | 9282 | 9855 | 6333 | 19294 | 12539 | 18270 | 20003 | 10911 | 9815 | 9736 | 9921 | 15460 | 14258 | 15511 | 16264 | 12734,39 | 10096 | 15373 | 1,523 | 0,607 | 11,468 | 1,91E-30 | 5,19E-30 | 0,0028234 | 0,0022202 | 0,0026039 | 0,0026039 | TRUE | 0,0364143 | No |
| orf19.4828 | 12077 | 10530 | 11767 | 6704 | 3194 | 2135 | 3009 | 3276 | 11514 | 11134 | 11625 | 10502 | 2559 | 2428 | 2555 | 2664 | 6872,55 | 11194 | 2552 | 0,228 | -2,133 | -46,104 | 0 | 0 | 0,0014202 | 0,0023302 | 0,0018303 | 0,0018303 | TRUE | 0,0207222 | No |
| orf19.4829 | 1904 | 1548 | 1683 | 1150 | 3243 | 2229 | 3218 | 3365 | 1815 | 1637 | 1663 | 1802 | 2599 | 2535 | 2732 | 2736 | 2189,69 | 1729 | 2650 | 1,535 | 0,618 | 11,929 | 8,32E-33 | 2,36E-32 | 0,0015048 | 0,0028411 | 0,0020844 | 0,0020844 | TRUE | 0,0154674 | No |

| | | | | | | | | | | | | | | | | | | | | | | | | | | | | | | | |
|---|---|---|---|---|---|---|---|---|---|---|---|---|---|---|---|---|---|---|---|---|---|---|---|---|---|---|---|---|---|---|---|
| orf19.4830 | 510 | 406 | 381 | 289 | 740 | 464 | 674 | 785 | 486 | 429 | 376 | 453 | 593 | 528 | 572 | 638 | 509,46 | 436 | 583 | 1,34 | 0,422 | 4,593 | 4,36E-06 | 6,58E-06 | 0,006771 | 0,0053141 | 0,0060498 | 0,0060498 | TRUE | 0,1038373 | No |
| orf19.4831 | 3253 | 2724 | 2989 | 1867 | 12437 | 8573 | 11875 | 13261 | 3101 | 2880 | 2953 | 2925 | 9965 | 9748 | 10082 | 10782 | 6554,65 | 2965 | 10144 | 3,421 | 1,774 | 39,569 | 0 | 0 | 0,0012034 | 0,0023418 | 0,001697 | 0,001697 | TRUE | 0,0219734 | No |
| orf19.4833 | 11923 | 9778 | 10533 | 7209 | 57227 | 39412 | 54828 | 61427 | 11367 | 10339 | 10406 | 11293 | 45854 | 44815 | 46549 | 49946 | 28821,16 | 10851 | 46791 | 4,313 | 2,109 | 42,764 | 0 | 0 | 0,0023566 | 0,0021482 | 0,0022743 | 0,0022743 | TRUE | 0,0252559 | No |
| orf19.4834 | 26 | 31 | 24 | 16 | 49 | 36 | 43 | 41 | 25 | 33 | 24 | 25 | 39 | 41 | 37 | 33 | 32,05 | 27 | 38 | 1,405 | 0,49 | 1,966 | 0,0493198 | 0,0593455 | 1,00E-08 | 0,0533242 | 0,026947 | 0,026947 | TRUE | 0,157409 | No |
| orf19.4835 | 13975 | 11804 | 12981 | 8113 | 14783 | 10005 | 14539 | 15285 | 13324 | 12481 | 12824 | 12709 | 11845 | 11377 | 12344 | 12428 | 12416,48 | 12834 | 11998 | 0,935 | -0,097 | -2,409 | 0,0160082 | 0,0200409 | 0,0011033 | 0,0022235 | 0,0014783 | 0,0014783 | TRUE | 0,014685 | No |
| orf19.4836 | 2272 | 1798 | 1930 | 1347 | 6865 | 4733 | 6592 | 6892 | 2166 | 1901 | 1907 | 2110 | 5501 | 5382 | 5597 | 5604 | 3770,88 | 2021 | 5521 | 2,734 | 1,451 | 28,585 | 1,04E-179 | 1,25E-178 | 0,0020293 | 0,0025267 | 0,0021153 | 0,0021153 | TRUE | 0,0306514 | No |
| orf19.4837 | 514 | 383 | 466 | 349 | 226 | 154 | 231 | 238 | 490 | 405 | 460 | 547 | 181 | 175 | 196 | 194 | 330,99 | 476 | 186 | 0,394 | -1,342 | -12,844 | 9,35E-38 | 2,85E-37 | 0,0067807 | 0,0070518 | 0,0069176 | 0,0069176 | TRUE | 0,1117729 | No |
| orf19.4837.1 | 666 | 577 | 612 | 319 | 389 | 208 | 337 | 292 | 635 | 610 | 605 | 500 | 312 | 237 | 286 | 237 | 427,64 | 588 | 268 | 0,456 | -1,133 | -10,437 | 1,68E-25 | 4,19E-25 | 0,0112406 | 0,0059307 | 0,0087004 | 0,0087004 | TRUE | 0,1392114 | No |
| orf19.4839 | 1240 | 933 | 1082 | 738 | 3331 | 2348 | 3319 | 3326 | 1182 | 987 | 1069 | 1156 | 2669 | 2670 | 2818 | 2704 | 1906,86 | 1098 | 2715 | 2,475 | 1,307 | 22,07 | 6,09E-108 | 4,31E-107 | 0,002762 | 0,0029523 | 0,002693 | 0,002693 | TRUE | 0,0568315 | No |
| orf19.484 | 6909 | 5819 | 6356 | 3942 | 5395 | 3654 | 5243 | 5615 | 6587 | 6153 | 6279 | 6175 | 4323 | 4155 | 4451 | 4566 | 5336,11 | 6298 | 4374 | 0,695 | -0,526 | -12,359 | 4,37E-35 | 1,28E-34 | 0,0010584 | 0,002399 | 0,0015476 | 0,0015476 | TRUE | 0,0133667 | No |
| orf19.4841 | 3360 | 2828 | 3135 | 1863 | 1731 | 1187 | 1796 | 1806 | 3203 | 2990 | 3097 | 2918 | 1387 | 1350 | 1525 | 1468 | 2242,4 | 3052 | 1432 | 0,469 | -1,091 | -20,654 | 9,08E-95 | 5,64E-94 | 0,0016711 | 0,0028235 | 0,0021849 | 0,0021849 | TRUE | 0,0224572 | No |
| orf19.4842 | 1438 | 1104 | 1183 | 940 | 682 | 466 | 684 | 713 | 1371 | 1167 | 1169 | 1473 | 546 | 530 | 581 | 580 | 927,05 | 1295 | 559 | 0,433 | -1,207 | -14,351 | 1,05E-46 | 3,71E-46 | 0,007019 | 0,0038624 | 0,00556 | 0,00556 | TRUE | 0,0965747 | No |
| orf19.4843 | 6186 | 5757 | 6168 | 4012 | 1476 | 1149 | 1398 | 1582 | 5898 | 6087 | 6093 | 6285 | 1183 | 1307 | 1187 | 1286 | 3665,72 | 6091 | 1241 | 0,204 | -2,297 | -49,438 | 0 | 0 | 0,0010627 | 0,0025392 | 0,0016201 | 0,0016201 | TRUE | 0,0141562 | No |
| orf19.4844 | 3863 | 3033 | 3467 | 2089 | 2814 | 1802 | 2810 | 2817 | 3683 | 3207 | 3425 | 3273 | 2255 | 2049 | 2386 | 2290 | 2820,94 | 3397 | 2245 | 0,661 | -0,597 | -9,826 | 8,71E-23 | 2,06E-22 | 0,0034403 | 0,0026733 | 0,0031825 | 0,0031825 | TRUE | 0,0404843 | No |
| orf19.4845 | 499 | 497 | 512 | 270 | 852 | 649 | 768 | 804 | 476 | 526 | 506 | 423 | 683 | 738 | 652 | 654 | 582,06 | 483 | 682 | 1,404 | 0,489 | 6,241 | 4,35E-10 | 7,61E-10 | 0,0031731 | 0,0049122 | 0,00409 | 0,00409 | TRUE | 0,067371 | No |
| orf19.4846 | 1795 | 1459 | 1644 | 1030 | 2824 | 1933 | 2732 | 2924 | 1711 | 1543 | 1624 | 1614 | 2263 | 2198 | 2319 | 2377 | 1956,18 | 1623 | 2289 | 1,411 | 0,496 | 10,203 | 1,92E-24 | 4,69E-24 | 0,0009343 | 0,0029306 | 0,0017325 | 0,0017325 | TRUE | 0,0171536 | No |
| orf19.4847 | 50 | 31 | 35 | 22 | 45 | 22 | 33 | 55 | 48 | 33 | 35 | 34 | 36 | 25 | 28 | 45 | 35,41 | 38 | 34 | 0,898 | -0,155 | -0,589 | 0,5556751 | 0,5903701 | 0,0246712 | 0,0484565 | 0,0380564 | 0,0380564 | TRUE | 0,3877591 | No |
| orf19.4848 | 2511 | 2226 | 2367 | 1497 | 2764 | 1907 | 2577 | 2864 | 2394 | 2354 | 2338 | 2345 | 2215 | 2168 | 2188 | 2329 | 2291,36 | 2358 | 2225 | 0,944 | -0,083 | -2,037 | 0,0416176 | 0,0504104 | 1,00E-08 | 0,0028078 | 0,0011605 | 0,0011605 | TRUE | 0,0121281 | No |
| orf19.4849 | 7368 | 6896 | 7254 | 4868 | 5097 | 3695 | 4867 | 4947 | 7025 | 7292 | 7166 | 7626 | 4084 | 4202 | 4132 | 4022 | 5693,57 | 7277 | 4110 | 0,565 | -0,824 | -21,932 | 1,28E-106 | 8,90E-106 | 0,0005391 | 0,0023797 | 0,0011679 | 0,0011679 | TRUE | 0,0121978 | No |
| orf19.485 | 893 | 794 | 891 | 530 | 1113 | 683 | 1037 | 1043 | 851 | 840 | 880 | 830 | 892 | 777 | 880 | 848 | 849,79 | 850 | 849 | 1 | 0 | -0,005 | 0,9957857 | 0,9965666 | 0,0009702 | 0,0040234 | 0,002407 | 0,002407 | TRUE | 0,036128 | No |
| orf19.4850 | 10183 | 9749 | 10635 | 6082 | 4595 | 3357 | 4257 | 4368 | 9708 | 10308 | 10506 | 9528 | 3682 | 3817 | 3614 | 3552 | 6839,46 | 10012 | 3666 | 0,366 | -1,451 | -32,466 | 3,19E-231 | 4,95E-230 | 0,0013291 | 0,0023313 | 0,0017389 | 0,0017389 | TRUE | 0,0134641 | No |
| orf19.4851 | 1782 | 1578 | 1693 | 1194 | 3976 | 2744 | 3934 | 3945 | 1699 | 1669 | 1673 | 1870 | 3186 | 3120 | 3340 | 3208 | 2470,51 | 1728 | 3214 | 1,865 | 0,899 | 18,613 | 2,53E-77 | 1,33E-76 | 0,0010514 | 0,0027559 | 0,0017771 | 0,0017771 | TRUE | 0,0337462 | No |
| orf19.4853 | 169 | 145 | 167 | 118 | 138 | 106 | 147 | 160 | 161 | 153 | 165 | 185 | 111 | 121 | 125 | 130 | 143,79 | 166 | 122 | 0,736 | -0,442 | -3,628 | 0,0002861 | 0,0003969 | 1,00E-08 | 0,0135104 | 0,0072454 | 0,0072454 | TRUE | 0,0483891 | No |
| orf19.4855 | 930 | 730 | 829 | 518 | 1070 | 731 | 1045 | 1003 | 887 | 772 | 819 | 811 | 857 | 831 | 887 | 816 | 835,04 | 822 | 848 | 1,03 | 0,043 | 0,69 | 0,4900358 | 0,526063 | 0,0012496 | 0,0040575 | 0,002537 | 0,002537 | TRUE | 0,0352098 | No |
| orf19.4856 | 21 | 9 | 21 | 8 | 42 | 29 | 32 | 44 | 20 | 10 | 21 | 13 | 34 | 33 | 27 | 36 | 24,05 | 16 | 32 | 2,024 | 1,017 | 3,248 | 0,0011607 | 0,0015573 | 1,00E-08 | 0,0703656 | 0,0457041 | 0,0457041 | TRUE | 0,3624561 | No |
| orf19.4857 | 2683 | 2392 | 2679 | 1914 | 3366 | 2511 | 3186 | 3414 | 2558 | 2529 | 2647 | 2998 | 2697 | 2855 | 2705 | 2776 | 2720,66 | 2683 | 2758 | 1,03 | 0,042 | 0,734 | 0,4628961 | 0,49928 | 0,0028614 | 0,0026948 | 0,0027876 | 0,0027876 | TRUE | 0,0722469 | No |
| orf19.4858 | 634 | 564 | 614 | 366 | 1479 | 1001 | 1424 | 1442 | 604 | 596 | 607 | 573 | 1185 | 1138 | 1209 | 1172 | 885,69 | 595 | 1176 | 1,973 | 0,981 | 17,715 | 3,23E-70 | 1,55E-69 | 1,00E-08 | 0,0039451 | 0,0016298 | 0,0016298 | TRUE | 0,0057713 | No |
| orf19.486 | 592 | 508 | 553 | 320 | 160 | 98 | 132 | 139 | 564 | 537 | 546 | 501 | 128 | 111 | 112 | 113 | 326,74 | 537 | 116 | 0,216 | -2,21 | -23,085 | 6,56E-118 | 5,11E-117 | 1,00E-08 | 0,0071164 | 0,0039511 | 0,0039511 | TRUE | 0,0535636 | No |
| orf19.4861 | 15 | 27 | 32 | 26 | 36 | 30 | 36 | 56 | 14 | 29 | 32 | 41 | 29 | 34 | 31 | 46 | 31,78 | 29 | 35 | 1,234 | 0,303 | 0,982 | 0,3259019 | 0,359478 | 0,0661353 | 0,0537541 | 0,0589699 | 0,0589699 | TRUE | 0,5501598 | No |
| orf19.4861.1 | 8 | 10 | 15 | 1 | 14 | 18 | 11 | 9 | 8 | 11 | 15 | 2 | 11 | 20 | 9 | 7 | 10,37 | 9 | 12 | 1,323 | 0,404 | 0,759 | 0,4477882 | 0,4842944 | 0,1905771 | 0,160484 | 0,1715084 | 0,1715084 | TRUE | 0,3395126 | No |
| orf19.4862 | 1066 | 1006 | 1144 | 749 | 1486 | 964 | 1450 | 1408 | 1016 | 1064 | 1130 | 1173 | 1191 | 1096 | 1231 | 1145 | 1130,78 | 1096 | 1166 | 1,067 | 0,094 | 1,51 | 0,1310541 | 0,1512217 | 0,0020847 | 0,0035433 | 0,0028138 | 0,0028138 | TRUE | 0,0289886 | No |
| orf19.4862.2 | 2195 | 1999 | 1972 | 1292 | 1767 | 1111 | 1690 | 1707 | 2093 | 2114 | 1948 | 2024 | 1416 | 1263 | 1435 | 1388 | 1710,05 | 2045 | 1376 | 0,674 | -0,57 | -10,585 | 3,50E-26 | 8,89E-26 | 0,0014353 | 0,0030514 | 0,0021843 | 0,0021843 | TRUE | 0,033955 | No |
| orf19.4863 | 1858 | 1699 | 1870 | 1156 | 2241 | 1693 | 2267 | 2402 | 1771 | 1796 | 1847 | 1811 | 1796 | 1925 | 1925 | 1953 | 1853,08 | 1806 | 1900 | 1,051 | 0,072 | 1,59 | 0,1118301 | 0,1297904 | 0,0004898 | 0,0029773 | 0,0014283 | 0,0014283 | TRUE | 0,0169932 | No |
| orf19.4864 | 993 | 914 | 956 | 562 | 1345 | 967 | 1293 | 1264 | 947 | 966 | 944 | 880 | 1078 | 1100 | 1098 | 1028 | 1005,1 | 934 | 1076 | 1,147 | 0,198 | 3,67 | 0,0002423 | 0,0003371 | 1,00E-08 | 0,0037249 | 0,0017935 | 0,0017935 | TRUE | 0,0152808 | No |
| orf19.4865 | 1019 | 1125 | 1138 | 646 | 1792 | 1359 | 1685 | 1613 | 972 | 1190 | 1124 | 1012 | 1436 | 1545 | 1431 | 1312 | 1252,57 | 1074 | 1431 | 1,33 | 0,411 | 5,446 | 5,14E-08 | 8,36E-08 | 0,0055057 | 0,0034021 | 0,0046311 | 0,0046311 | TRUE | 0,0625455 | No |
| orf19.4866 | 4864 | 4501 | 5017 | 3522 | 3432 | 2725 | 3273 | 3393 | 4637 | 4759 | 4956 | 5517 | 2750 | 3099 | 2779 | 2759 | 3907,05 | 4967 | 2847 | 0,573 | -0,803 | -12,695 | 6,28E-37 | 1,89E-36 | 0,0044553 | 0,0025115 | 0,0035723 | 0,0035723 | TRUE | 0,0661505 | No |
| orf19.4867 | 11634 | 10837 | 11881 | 7471 | 966 | 746 | 937 | 1088 | 11092 | 11459 | 11737 | 11704 | 774 | 848 | 796 | 885 | 6161,75 | 11498 | 826 | 0,072 | -3,801 | -77,927 | 0 | 0 | 0,00132 | 0,0023577 | 0,0016955 | 0,0016955 | TRUE | 0,0301625 | No |
| orf19.4869 | 3752 | 3035 | 3591 | 2069 | 4341 | 2979 | 4340 | 4396 | 3577 | 3209 | 3548 | 3241 | 3478 | 3387 | 3685 | 3574 | 3462,47 | 3394 | 3531 | 1,04 | 0,056 | 1,106 | 0,2685362 | 0,299411 | 0,0019271 | 0,0025655 | 0,0022059 | 0,0022059 | TRUE | 0,0170823 | No |
| orf19.487 | 1847 | 1627 | 1827 | 1228 | 782 | 600 | 819 | 792 | 1761 | 1720 | 1805 | 1924 | 627 | 682 | 695 | 644 | 1232,25 | 1802 | 662 | 0,367 | -1,444 | -24,713 | 7,85E-135 | 7,14E-134 | 0,0013488 | 0,0034237 | 0,0022936 | 0,0022936 | TRUE | 0,0227427 | No |
| orf19.4870 | 13486 | 12435 | 13531 | 8673 | 14280 | 10804 | 13739 | 14394 | 12857 | 13148 | 13367 | 13587 | 11442 | 12285 | 11664 | 11704 | 12506,91 | 13240 | 11774 | 0,889 | -0,169 | -4,82 | 1,44E-06 | 2,21E-06 | 0,0006367 | 0,0022226 | 0,0011048 | 0,0011048 | TRUE | 0,0102835 | No |
| orf19.4871 | 18055 | 17150 | 18083 | 11063 | 10127 | 7612 | 9538 | 10192 | 17213 | 18134 | 17864 | 17331 | 8114 | 8656 | 8098 | 8287 | 12962,16 | 17636 | 8289 | 0,47 | -1,09 | -30,55 | 5,59E-205 | 7,78E-204 | 0,0006767 | 0,0022179 | 0,0011364 | 0,0011364 | TRUE | 0,0106033 | No |
| orf19.4872 | 139 | 154 | 163 | 112 | 96 | 96 | 89 | 95 | 133 | 163 | 161 | 175 | 77 | 109 | 76 | 77 | 121,34 | 158 | 85 | 0,534 | -0,906 | -5,841 | 5,19E-09 | 8,70E-09 | 0,0123303 | 0,0156226 | 0,01423 | 0,01423 | TRUE | 0,3150724 | No |
| orf19.4873 | 12611 | 11711 | 11972 | 7825 | 1916 | 1394 | 1800 | 1910 | 12023 | 12383 | 11827 | 12258 | 1535 | 1585 | 1528 | 1553 | 6836,64 | 12123 | 1550 | 0,128 | -2,968 | -83,776 | 0 | 0 | 1,00E-08 | 0,0023314 | 0,0008711 | 0,0008711 | TRUE | 0,0033231 | No |
| orf19.4874 | 5444 | 5082 | 5469 | 3197 | 4393 | 3262 | 4111 | 4414 | 5190 | 5374 | 5403 | 5008 | 3520 | 3709 | 3490 | 3589 | 4410,42 | 5244 | 3577 | 0,681 | -0,554 | -13,84 | 1,46E-43 | 4,88E-43 | 0,0006654 | 0,0024636 | 0,0013037 | 0,0013037 | TRUE | 0,0105088 | No |
| orf19.4875 | 923 | 757 | 857 | 453 | 686 | 427 | 633 | 761 | 880 | 800 | 847 | 710 | 550 | 486 | 537 | 619 | 678,51 | 809 | 548 | 0,676 | -0,564 | -6,39 | 1,66E-10 | 2,95E-10 | 0,0071445 | 0,0045111 | 0,0059705 | 0,0059705 | TRUE | 0,0893501 | No |
| orf19.4878 | 1139 | 921 | 1094 | 658 | 1410 | 962 | 1392 | 1446 | 1086 | 974 | 1081 | 1031 | 1130 | 1094 | 1182 | 1176 | 1094,07 | 1043 | 1146 | 1,098 | 0,135 | 2,391 | 0,0168099 | 0,0210034 | 0,000968 | 0,003592 | 0,0021451 | 0,0021451 | TRUE | 0,0251786 | No |
| orf19.4879.2 | 3664 | 3277 | 3239 | 2201 | 4308 | 3239 | 4012 | 4207 | 3493 | 3465 | 3200 | 3448 | 3452 | 3683 | 3406 | 3421 | 3445,98 | 3402 | 3490 | 1,026 | 0,037 | 0,809 | 0,4187717 | 0,4546896 | 0,0011008 | 0,0025677 | 0,0016913 | 0,0016913 | TRUE | 0,0199103 | No |
| orf19.488 | 4631 | 4238 | 4526 | 2871 | 4515 | 3175 | 4364 | 4743 | 4415 | 4481 | 4471 | 4498 | 3618 | 3610 | 3705 | 3857 | 4081,83 | 4466 | 3698 | 0,828 | -0,272 | -7,545 | 4,52E-14 | 8,87E-14 | 0,0002864 | 0,0024935 | 0,0009985 | 0,0009985 | TRUE | 0,0103568 | No |
| orf19.4880 | 263 | 207 | 252 | 126 | 704 | 485 | 718 | 692 | 251 | 219 | 249 | 197 | 564 | 551 | 610 | 563 | 400,47 | 229 | 572 | 2,477 | 1,308 | 14,81 | 1,27E-49 | 4,65E-49 | 0,0015429 | 0,0061912 | 0,0043046 | 0,0043046 | TRUE | 0,0742574 | No |
| orf19.4882 | 5917 | 4942 | 5415 | 2990 | 4356 | 2734 | 4199 | 4441 | 5641 | 5226 | 5350 | 4684 | 3490 | 3109 | 3565 | 3611 | 4334,41 | 5225 | 3444 | 0,659 | -0,602 | -9,144 | 6,00E-20 | 1,35E-19 | 0,0048242 | 0,0024701 | 0,0039217 | 0,0039217 | TRUE | 0,0579521 | No |
| orf19.4883 | 3483 | 3258 | 3473 | 2209 | 1985 | 1424 | 1795 | 1957 | 3321 | 3445 | 3431 | 3461 | 1591 | 1619 | 1524 | 1591 | 2497,75 | 3414 | 1581 | 0,463 | -1,111 | -27,913 | 1,85E-171 | 2,12E-170 | 1,00E-08 | 0,0027486 | 0,0010733 | 0,0010733 | TRUE | 0,0071817 | No |
| orf19.4884 | 111 | 120 | 118 | 89 | 169 | 122 | 155 | 165 | 106 | 127 | 117 | 139 | 135 | 139 | 132 | 134 | 128,58 | 122 | 135 | 1,114 | 0,156 | 1,216 | 0,2241154 | 0,2528431 | 1,00E-08 | 0,0148613 | 0,0079647 | 0,0079647 | TRUE | 0,0838492 | No |
| orf19.4885 | 15225 | 12508 | 13116 | 9831 | 23824 | 16959 | 22606 | 24467 | 14515 | 13226 | 12957 | 15401 | 19089 | 19284 | 19192 | 19894 | 16694,87 | 14025 | 19365 | 1,381 | 0,466 | 8,387 | 5,00E-17 | 1,05E-16 | 0,0034072 | 0,0021896 | 0,0029018 | 0,0029018 | TRUE | 0,0527959 | No |
| orf19.4886 | 86 | 82 | 80 | 42 | 84 | 70 | 77 | 52 | 82 | 87 | 79 | 66 | 67 | 80 | 65 | 42 | 71,01 | 78 | 64 | 0,8 | -0,321 | -1,594 | 0,1108395 | 0,1287109 | 0,0240982 | 0,0252134 | 0,0247817 | 0,0247817 | TRUE | 0,4849174 | No |
| orf19.4887 | 13227 | 12112 | 12871 | 8651 | 2128 | 1678 | 1928 | 2115 | 12611 | 12807 | 12715 | 13552 | 1705 | 1908 | 1637 | 1720 | 7331,84 | 12921 | 1742 | 0,135 | -2,892 | -57,2 | 0 | 0 | 0,0023372 | 0,0023152 | 0,0021554 | 0,0021554 | TRUE | 0,0470249 | No |
| orf19.4888 | 3555 | 3353 | 3399 | 2097 | 1953 | 1250 | 1776 | 1714 | 3389 | 3545 | 3358 | 3285 | 1565 | 1421 | 1508 | 1394 | 2433,17 | 3394 | 1472 | 0,434 | -1,205 | -24,044 | 9,61E-128 | 8,20E-127 | 0,0013048 | 0,0027661 | 0,0019432 | 0,0019432 | TRUE | 0,0222726 | No |
| orf19.4889 | 5898 | 5561 | 5937 | 3608 | 3586 | 2737 | 3302 | 3575 | 5623 | 5880 | 5865 | 5652 | 2873 | 3112 | 2803 | 2907 | 4339,53 | 5755 | 2924 | 0,507 | -0,979 | -23,02 | 2,94E-117 | 2,27E-116 | 0,0009667 | 0,0024696 | 0,0014842 | 0,0014842 | TRUE | 0,0218238 | No |
| orf19.489 | 553 | 498 | 535 | 345 | 2243 | 1575 | 2217 | 2171 | 527 | 527 | 529 | 540 | 1797 | 1791 | 1882 | 1765 | 1169,8 | 531 | 1809 | 3,413 | 1,771 | 32,736 | 4,86E-235 | 7,66E-234 | 1,00E-08 | 0,0034949 | 0,0015147 | 0,0015147 | TRUE | 0,0091665 | No |
| orf19.4890 | 1905 | 1875 | 1924 | 1301 | 1983 | 1577 | 1916 | 2082 | 1816 | 1983 | 1901 | 2038 | 1589 | 1793 | 1627 | 1693 | 1804,91 | 1934 | 1676 | 0,866 | -0,207 | -3,701 | 0,0002145 | 0,000299 | 0,0019393 | 0,003001 | 0,0024409 | 0,0024409 | TRUE | 0,0258127 | No |
| orf19.4892 | 12078 | 11737 | 12118 | 7205 | 6236 | 4651 | 6072 | 6064 | 11515 | 12410 | 11971 | 11287 | 4997 | 5289 | 5155 | 4931 | 8444,37 | 11796 | 5093 | 0,431 | -1,213 | -28,528 | 5,33E-179 | 6,32E-178 | 0,0012061 | 0,0022857 | 0,0015999 | 0,0015999 | TRUE | 0,0149873 | No |
| orf19.4893 | 2906 | 2713 | 2949 | 1773 | 1696 | 1176 | 1470 | 1646 | 2771 | 2869 | 2913 | 2777 | 1359 | 1337 | 1248 | 1338 | 2076,57 | 2832 | 1320 | 0,466 | -1,102 | -24,357 | 4,88E-131 | 4,28E-130 | 0,0004527 | 0,002882 | 0,001428 | 0,001428 | TRUE | 0,0169103 | No |
| orf19.4894 | 530 | 460 | 456 | 365 | 539 | 364 | 489 | 547 | 505 | 486 | 450 | 572 | 432 | 414 | 415 | 445 | 464,96 | 503 | 426 | 0,852 | -0,231 | -2,814 | 0,0048856 | 0,0063197 | 0,0028168 | 0,0056226 | 0,0042839 | 0,0042839 | TRUE | 0,0843132 | No |
| orf19.4895 | 1604 | 1318 | 1542 | 893 | 2114 | 1404 | 2099 | 2132 | 1529 | 1394 | 1523 | 1399 | 1694 | 1596 | 1782 | 1734 | 1581,38 | 1461 | 1702 | 1,164 | 0,219 | 3,949 | 7,86E-05 | 0,0001117 | 0,0015447 | 0,0031296 | 0,0022961 | 0,0022961 | TRUE | 0,0197405 | No |
| orf19.4896 | 4036 | 3590 | 3811 | 2489 | 5521 | 3621 | 5110 | 5274 | 3848 | 3796 | 3765 | 3899 | 4424 | 4117 | 4338 | 4288 | 4059,47 | 3827 | 4292 | 1,123 | 0,167 | 4,55 | 5,37E-06 | 8,07E-06 | 0,0003277 | 0,0024957 | 0,001042 | 0,001042 | TRUE | 0,008734 | No |
| orf19.4897 | 682 | 627 | 707 | 410 | 817 | 549 | 748 | 803 | 650 | 663 | 698 | 642 | 655 | 624 | 635 | 653 | 652,6 | 663 | 642 | 0,967 | -0,049 | -0,804 | 0,4211383 | 0,4571765 | 1,00E-08 | 0,0046072 | 0,0020245 | 0,0020245 | TRUE | 0,015293 | No |
| orf19.4898 | 2257 | 1904 | 2154 | 1244 | 725 | 465 | 626 | 679 | 2152 | 2013 | 2128 | 1949 | 581 | 529 | 531 | 552 | 1304,38 | 2060 | 548 | 0,266 | -1,91 | -32,881 | 4,09E-237 | 6,48E-236 | 0,0012008 | 0,00335 | 0,0021559 | 0,0021559 | TRUE | 0,0193468 | No |
| orf19.4899 | 1082 | 863 | 936 | 660 | 357 | 228 | 303 | 387 | 1032 | 913 | 925 | 1034 | 286 | 259 | 257 | 315 | 627,49 | 976 | 279 | 0,287 | -1,799 | -21,495 | 1,72E-102 | 1,15E-101 | 0,0043924 | 0,0047079 | 0,0045561 | 0,0045561 | TRUE | 0,0839067 | No |
| orf19.490 | 1169 | 1014 | 1066 | 675 | 2963 | 1998 | 2700 | 2954 | 1115 | 1072 | 1053 | 1057 | 2374 | 2272 | 2292 | 2402 | 1704,68 | 1074 | 2335 | 2,173 | 1,12 | 24,218 | 1,44E-129 | 1,24E-128 | 1,00E-08 | 0,0030545 | 0,0013411 | 0,0013411 | TRUE | 0,0080881 | No |
| orf19.4900 | 1156 | 1224 | 1241 | 755 | 904 | 670 | 859 | 824 | 1102 | 1294 | 1226 | 1183 | 724 | 762 | 729 | 670 | 961,32 | 1201 | 721 | 0,6 | -0,738 | -11,002 | 3,75E-28 | 9,81E-28 | 0,0027719 | 0,0037992 | 0,0032296 | 0,0032296 | TRUE | 0,0394075 | No |
| orf19.4901 | 700 | 636 | 649 | 455 | 680 | 500 | 649 | 607 | 667 | 672 | 641 | 713 | 545 | 569 | 551 | 494 | 606,47 | 673 | 540 | 0,802 | -0,319 | -4,627 | 3,71E-06 | 5,61E-06 | 0,0010093 | 0,0047986 | 0,0028973 | 0,0028973 | TRUE | 0,0395148 | No |
| orf19.4903 | 847 | 994 | 936 | 589 | 2009 | 1543 | 1731 | 1439 | 808 | 1051 | 925 | 923 | 1610 | 1755 | 1470 | 1170 | 1213,73 | 927 | 1501 | 1,619 | 0,695 | 6,084 | 1,17E-09 | 2,01E-09 | 0,018405 | 0,0034441 | 0,0116409 | 0,0116409 | TRUE | 0,2671924 | No |
| orf19.4904 | 979 | 967 | 1003 | 668 | 1143 | 922 | 1059 | 1160 | 933 | 1022 | 991 | 1046 | 916 | 1048 | 899 | 943 | 974,96 | 998 | 952 | 0,953 | -0,069 | -1,06 | 0,2890655 | 0,3215141 | 0,0023116 | 0,0037753 | 0,0030682 | 0,0030682 | TRUE | 0,0526019 | No |
| orf19.4905 | 1129 | 1021 | 1181 | 679 | 1191 | 788 | 1106 | 1138 | 1076 | 1080 | 1167 | 1064 | 954 | 896 | 939 | 925 | 1012,62 | 1097 | 928 | 0,847 | -0,24 | -4,422 | 9,78E-06 | 1,45E-05 | 1,00E-08 | 0,0037127 | 0,0018435 | 0,0018435 | TRUE | 0,0229084 | No |

| | | | | | | | | | | | | | | | | | | | | | | | | | | | | | | | |
|---|---|---|---|---|---|---|---|---|---|---|---|---|---|---|---|---|---|---|---|---|---|---|---|---|---|---|---|---|---|---|---|
| orf19.4906 | 7828 | 6904 | 7642 | 4890 | 6975 | 4808 | 6641 | 6976 | 7463 | 7300 | 7550 | 7660 | 5589 | 5467 | 5638 | 5672 | 6542,45 | 7493 | 5592 | 0,747 | -0,421 | -13,372 | 8,84E-41 | 2,83E-40 | 0,0001264 | 0,0023422 | 0,0007983 | 0,0007983 | TRUE | 0,003641 | No |
| orf19.4907 | 12038 | 11315 | 11786 | 7531 | 936 | 666 | 817 | 982 | 11477 | 11964 | 11643 | 11798 | 750 | 757 | 694 | 798 | 6235,21 | 11720 | 750 | 0,064 | -3,966 | -86,122 | 0 | 0 | 0,0004568 | 0,0023546 | 0,0013953 | 0,0013953 | TRUE | 0,0314925 | No |
| orf19.4908 | 117 | 89 | 109 | 74 | 14 | 8 | 12 | 19 | 112 | 94 | 108 | 116 | 11 | 9 | 10 | 15 | 59,4 | 108 | 11 | 0,109 | -3,199 | -12,706 | 5,49E-37 | 1,66E-36 | 1,00E-08 | 0,0297321 | 0,0177061 | 0,0177061 | TRUE | 0,2411154 | No |
| orf19.4909 | 2528 | 2258 | 2474 | 1572 | 3139 | 2177 | 3026 | 3123 | 2410 | 2388 | 2444 | 2463 | 2515 | 2475 | 2569 | 2539 | 2475,43 | 2426 | 2524 | 1,042 | 0,059 | 1,598 | 0,1099609 | 0,1278303 | 1,00E-08 | 0,0027546 | 0,0008952 | 0,0008952 | TRUE | 0,0019354 | No |
| orf19.4909.1 | 10814 | 8262 | 8737 | 5351 | 33443 | 20874 | 31289 | 33725 | 10310 | 8736 | 8631 | 8383 | 26797 | 23736 | 26564 | 27422 | 17572,29 | 9015 | 26130 | 2,898 | 1,535 | 22,022 | 1,76E-107 | 1,24E-106 | 0,0065826 | 0,0021847 | 0,0045879 | 0,0045879 | TRUE | 0,1146667 | No |
| orf19.491 | 3241 | 2838 | 3188 | 1882 | 5961 | 4110 | 5678 | 6206 | 3090 | 3001 | 3149 | 2948 | 4776 | 4673 | 4821 | 5046 | 3938,12 | 3047 | 4829 | 1,584 | 0,663 | 16,271 | 1,59E-59 | 6,67E-59 | 0,0006079 | 0,0025082 | 0,0013196 | 0,0013196 | TRUE | 0,0112017 | No |
| orf19.4910 | 178 | 164 | 153 | 101 | 86 | 52 | 73 | 62 | 170 | 173 | 151 | 158 | 69 | 59 | 62 | 50 | 111,61 | 163 | 60 | 0,368 | -1,441 | -9,815 | 9,66E-23 | 2,29E-22 | 1,00E-08 | 0,0168018 | 0,0098845 | 0,0098845 | TRUE | 0,1191335 | No |
| orf19.4911 | 478 | 426 | 472 | 301 | 227 | 121 | 207 | 227 | 456 | 450 | 466 | 472 | 182 | 138 | 176 | 185 | 315,47 | 461 | 170 | 0,372 | -1,427 | -15,005 | 6,81E-51 | 2,54E-50 | 1,00E-08 | 0,0072959 | 0,0048706 | 0,0048706 | TRUE | 0,1489473 | No |
| orf19.4912 | 2198 | 1832 | 2055 | 1327 | 2404 | 1606 | 2226 | 2528 | 2096 | 1937 | 2030 | 2079 | 1926 | 1826 | 1890 | 2056 | 1979,93 | 2036 | 1924 | 0,947 | -0,079 | -1,55 | 0,1210875 | 0,140076 | 0,0013831 | 0,0029206 | 0,0019917 | 0,0019917 | TRUE | 0,0255432 | No |
| orf19.4913 | 1526 | 1596 | 1654 | 979 | 1254 | 984 | 1129 | 1151 | 1455 | 1688 | 1634 | 1534 | 1005 | 1119 | 959 | 936 | 1291,02 | 1578 | 1005 | 0,635 | -0,654 | -9,275 | 1,78E-20 | 4,04E-20 | 0,0043633 | 0,0033631 | 0,0039724 | 0,0039724 | TRUE | 0,0669353 | No |
| orf19.4914 | 206 | 171 | 176 | 111 | 344 | 216 | 271 | 313 | 196 | 181 | 174 | 174 | 276 | 246 | 230 | 254 | 216,35 | 181 | 252 | 1,385 | 0,47 | 4,491 | 7,09E-06 | 1,06E-05 | 1,00E-08 | 0,0096804 | 0,0056874 | 0,0056874 | TRUE | 0,0494841 | No |
| orf19.4914.1 | 42186 | 39346 | 39854 | 21332 | 7777 | 4379 | 7118 | 7238 | 40220 | 41603 | 39372 | 33418 | 6231 | 4979 | 6043 | 5885 | 22219 | 38653 | 5784 | 0,15 | -2,74 | -34,274 | 1,93E-257 | 3,42E-256 | 0,0088352 | 0,0021652 | 0,0060484 | 0,0060484 | TRUE | 0,1064163 | No |
| orf19.4916 | 843 | 774 | 866 | 535 | 732 | 563 | 620 | 742 | 804 | 818 | 856 | 838 | 587 | 640 | 526 | 603 | 709,02 | 829 | 589 | 0,71 | -0,495 | -7,145 | 8,98E-13 | 1,70E-12 | 0,0019907 | 0,004407 | 0,0031743 | 0,0031743 | TRUE | 0,0603322 | No |
| orf19.4917 | 93 | 83 | 103 | 51 | 107 | 86 | 93 | 105 | 89 | 88 | 102 | 80 | 86 | 98 | 79 | 85 | 88,24 | 90 | 87 | 0,961 | -0,058 | -0,377 | 0,706054 | 0,7362854 | 1,00E-08 | 0,0206982 | 0,0112048 | 0,0112048 | TRUE | 0,0809542 | No |
| orf19.4918 | 253 | 238 | 301 | 209 | 482 | 382 | 488 | 527 | 241 | 252 | 297 | 327 | 386 | 434 | 414 | 429 | 347,63 | 279 | 416 | 1,498 | 0,583 | 5,762 | 8,33E-09 | 1,39E-08 | 0,0066379 | 0,0068144 | 0,0067365 | 0,0067365 | TRUE | 0,1269652 | No |
| orf19.4919 | 376 | 295 | 320 | 226 | 1110 | 757 | 1014 | 1155 | 358 | 312 | 316 | 354 | 889 | 861 | 861 | 939 | 611,35 | 335 | 888 | 2,657 | 1,41 | 19,233 | 1,94E-82 | 1,09E-81 | 0,0010488 | 0,004777 | 0,0029892 | 0,0029892 | TRUE | 0,0309961 | No |
| orf19.492 | 7265 | 6248 | 6879 | 4822 | 14742 | 10805 | 14178 | 15511 | 6926 | 6606 | 6796 | 7554 | 11812 | 12286 | 12037 | 12612 | 9578,77 | 6970 | 12187 | 1,75 | 0,807 | 17,157 | 5,54E-66 | 2,52E-65 | 0,0019795 | 0,0022627 | 0,0020072 | 0,0020072 | TRUE | 0,037631 | No |
| orf19.4921 | 1125 | 1002 | 1019 | 657 | 1392 | 1027 | 1213 | 1435 | 1073 | 1059 | 1007 | 1029 | 1115 | 1168 | 1030 | 1167 | 1080,97 | 1042 | 1120 | 1,074 | 0,102 | 1,75 | 0,0800337 | 0,094346 | 0,0013954 | 0,0036102 | 0,0023476 | 0,0023476 | TRUE | 0,0354815 | No |
| orf19.4921.1 | 977 | 919 | 865 | 471 | 617 | 418 | 495 | 390 | 931 | 972 | 855 | 738 | 494 | 475 | 420 | 317 | 650,33 | 874 | 426 | 0,487 | -1,038 | -8,138 | 4,03E-16 | 8,30E-16 | 0,0240992 | 0,004616 | 0,0139327 | 0,0139327 | TRUE | 0,3548919 | No |
| orf19.4922 | 764 | 760 | 799 | 461 | 1063 | 772 | 998 | 1026 | 728 | 804 | 789 | 722 | 852 | 878 | 847 | 834 | 806,83 | 761 | 853 | 1,118 | 0,161 | 2,71 | 0,0067303 | 0,0086272 | 6,55E-05 | 0,0041263 | 0,0021154 | 0,0021154 | TRUE | 0,0163789 | No |
| orf19.4923 | 239 | 195 | 229 | 139 | 188 | 147 | 180 | 180 | 228 | 206 | 226 | 218 | 151 | 167 | 153 | 146 | 186,87 | 220 | 154 | 0,699 | -0,516 | -4,891 | 1,00E-06 | 1,55E-06 | 1,00E-08 | 0,0108774 | 0,0052558 | 0,0052558 | TRUE | 0,0289456 | No |
| orf19.4923.1 | 33 | 28 | 22 | 15 | 22 | 19 | 25 | 27 | 31 | 30 | 22 | 23 | 18 | 22 | 21 | 22 | 23,59 | 26 | 21 | 0,768 | -0,381 | -1,324 | 0,1856667 | 0,2111081 | 1,00E-08 | 0,071696 | 0,0370685 | 0,0370685 | TRUE | 0,1131585 | No |
| orf19.4924 | 2550 | 2534 | 2645 | 1596 | 1794 | 1298 | 1740 | 1848 | 2431 | 2679 | 2613 | 2500 | 1437 | 1476 | 1477 | 1503 | 2014,63 | 2556 | 1473 | 0,576 | -0,795 | -17,002 | 7,93E-65 | 3,56E-64 | 0,000779 | 0,0029063 | 0,0015783 | 0,0015783 | TRUE | 0,0140871 | No |
| orf19.4927 | 3915 | 3605 | 3949 | 2350 | 2540 | 1747 | 2355 | 2537 | 3733 | 3812 | 3901 | 3681 | 2035 | 1986 | 1999 | 2063 | 2901,37 | 3782 | 2021 | 0,534 | -0,904 | -23,667 | 7,93E-124 | 6,53E-123 | 1,00E-08 | 0,0026572 | 0,0010325 | 0,0010325 | TRUE | 0,0055312 | No |
| orf19.4928 | 4362 | 4309 | 4628 | 2668 | 3059 | 2130 | 2871 | 3019 | 4159 | 4556 | 4572 | 4180 | 2451 | 2422 | 2437 | 2455 | 3403,97 | 4367 | 2441 | 0,559 | -0,839 | -18,459 | 4,39E-76 | 2,27E-75 | 0,001183 | 0,0025736 | 0,001674 | 0,001674 | TRUE | 0,0132884 | No |
| orf19.4929 | 3113 | 2998 | 3193 | 1845 | 2124 | 1543 | 1999 | 2198 | 2968 | 3170 | 3154 | 2890 | 1702 | 1755 | 1697 | 1787 | 2390,42 | 3046 | 1735 | 0,569 | -0,813 | -17,453 | 3,26E-68 | 1,53E-67 | 0,0009475 | 0,0027781 | 0,0016433 | 0,0016433 | TRUE | 0,013234 | No |
| orf19.493 | 25663 | 20052 | 20963 | 12759 | 65196 | 39935 | 62356 | 66569 | 24467 | 21202 | 20709 | 19988 | 52240 | 45410 | 52940 | 54127 | 36385,34 | 21592 | 51179 | 2,37 | 1,245 | 17,327 | 2,96E-67 | 1,37E-66 | 0,0071036 | 0,0021364 | 0,0049258 | 0,0049258 | TRUE | 0,0985062 | No |
| orf19.4930 | 502 | 464 | 481 | 342 | 624 | 444 | 579 | 629 | 479 | 491 | 475 | 536 | 500 | 505 | 492 | 511 | 498,5 | 495 | 502 | 1,019 | 0,026 | 0,385 | 0,7001432 | 0,7308382 | 1,00E-08 | 0,0053849 | 0,0025133 | 0,0025133 | TRUE | 0,0296379 | No |
| orf19.4931 | 4585 | 4225 | 4546 | 2897 | 6629 | 4810 | 6326 | 6763 | 4371 | 4467 | 4491 | 4538 | 5312 | 5469 | 5371 | 5499 | 4939,85 | 4467 | 5413 | 1,212 | 0,278 | 8,666 | 4,48E-18 | 9,63E-18 | 1,00E-08 | 0,0024237 | 0,0007765 | 0,0007765 | TRUE | 0,0025819 | No |
| orf19.4931.1 | 11232 | 9633 | 9728 | 6462 | 32027 | 21838 | 29736 | 32008 | 10708 | 10186 | 9610 | 10123 | 25662 | 24832 | 25246 | 26026 | 17799,12 | 10157 | 25442 | 2,505 | 1,325 | 33,447 | 2,82E-245 | 4,69E-244 | 0,0010915 | 0,0021835 | 0,0014336 | 0,0014336 | TRUE | 0,0166353 | No |
| orf19.4932 | 4548 | 3951 | 4094 | 2596 | 8051 | 5826 | 7951 | 8586 | 4336 | 4178 | 4044 | 4067 | 6451 | 6625 | 6750 | 6981 | 5429,04 | 4156 | 6702 | 1,612 | 0,689 | 16,892 | 5,13E-64 | 2,28E-63 | 0,0008081 | 0,0023937 | 0,001394 | 0,001394 | TRUE | 0,0105551 | No |
| orf19.4933 | 5888 | 5212 | 5547 | 3822 | 19866 | 15336 | 19299 | 20505 | 5614 | 5511 | 5480 | 5987 | 15918 | 17438 | 16385 | 16673 | 11125,7 | 5648 | 16604 | 2,942 | 1,557 | 35,864 | 1,11E-281 | 2,27E-280 | 0,0014327 | 0,0022389 | 0,0016824 | 0,0016824 | TRUE | 0,0191644 | No |
| orf19.4934 | 25 | 21 | 12 | 14 | 253 | 134 | 204 | 166 | 24 | 22 | 12 | 22 | 203 | 152 | 173 | 135 | 92,89 | 20 | 166 | 8,386 | 3,068 | 13,002 | 1,19E-38 | 3,69E-38 | 0,0274711 | 0,0197679 | 0,0228211 | 0,0228211 | TRUE | 0,3057674 | No |
| orf19.4936 | 96 | 87 | 101 | 59 | 58 | 43 | 51 | 55 | 92 | 92 | 100 | 92 | 46 | 49 | 43 | 45 | 69,89 | 94 | 46 | 0,486 | -1,041 | -6,168 | 6,92E-10 | 1,20E-09 | 1,00E-08 | 0,0255844 | 0,0116935 | 0,0116935 | TRUE | 0,018246 | No |
| orf19.4936.1 | 17231 | 15923 | 16570 | 9585 | 2122 | 1398 | 1898 | 1915 | 16428 | 16837 | 16370 | 15015 | 1700 | 1590 | 1611 | 1557 | 8888,48 | 16162 | 1614 | 0,1 | -3,323 | -68,095 | 0 | 0 | 0,0014691 | 0,002276 | 0,0019762 | 0,0019762 | TRUE | 0,0275293 | No |
| orf19.4937 | 8210 | 7445 | 7933 | 4811 | 8343 | 5717 | 7709 | 8403 | 7827 | 7872 | 7837 | 7537 | 6685 | 6501 | 6545 | 6832 | 7204,54 | 7768 | 6641 | 0,855 | -0,227 | -6,922 | 4,46E-12 | 8,29E-12 | 0,0002695 | 0,0023192 | 0,0008892 | 0,0008892 | TRUE | 0,0046555 | No |
| orf19.4939 | 382 | 436 | 419 | 275 | 314 | 239 | 253 | 296 | 364 | 461 | 414 | 431 | 252 | 272 | 215 | 241 | 331,1 | 418 | 245 | 0,586 | -0,772 | -7,566 | 3,86E-14 | 7,59E-14 | 0,0065844 | 0,0070503 | 0,0068239 | 0,0068239 | TRUE | 0,0899784 | No |
| orf19.494 | 7920 | 6339 | 7163 | 4856 | 8267 | 5439 | 8158 | 8097 | 7551 | 6703 | 7076 | 7607 | 6624 | 6185 | 6926 | 6584 | 6906,95 | 7234 | 6580 | 0,91 | -0,136 | -2,571 | 0,010131 | 0,0128623 | 0,0026184 | 0,002329 | 0,0025438 | 0,0025438 | TRUE | 0,0298753 | No |
| orf19.4940 | 11721 | 9842 | 10929 | 7392 | 11272 | 8174 | 10651 | 11479 | 11175 | 10407 | 10797 | 11580 | 9032 | 9295 | 9043 | 9334 | 10082,6 | 10990 | 9176 | 0,835 | -0,26 | -6,454 | 1,09E-10 | 1,94E-10 | 0,0010684 | 0,0022541 | 0,0014559 | 0,0014559 | TRUE | 0,0155698 | No |
| orf19.4941 | 28783 | 25073 | 28101 | 17479 | 6017 | 4067 | 5807 | 5886 | 27441 | 26512 | 27761 | 27382 | 4821 | 4625 | 4930 | 4786 | 16032,21 | 27274 | 4790 | 0,176 | -2,509 | -76,521 | 0 | 0 | 0,0003773 | 0,0021937 | 0,0009183 | 0,0009183 | TRUE | 0,0063183 | No |
| orf19.4942 | 41 | 46 | 40 | 34 | 25 | 22 | 25 | 19 | 39 | 49 | 40 | 53 | 20 | 25 | 21 | 15 | 32,78 | 45 | 20 | 0,452 | -1,144 | -4,389 | 1,14E-05 | 1,69E-05 | 1,00E-08 | 0,052182 | 0,0307975 | 0,0307975 | TRUE | 0,1672069 | No |
| orf19.4943 | 8732 | 7984 | 8528 | 5467 | 1203 | 909 | 1126 | 1246 | 8325 | 8442 | 8425 | 8564 | 964 | 1034 | 956 | 1013 | 4715,37 | 8439 | 992 | 0,117 | -3,091 | -77,95 | 0 | 0 | 1,00E-08 | 0,0024395 | 0,0009939 | 0,0009939 | TRUE | 0,0085374 | No |
| orf19.4945 | 547 | 516 | 525 | 276 | 1177 | 890 | 1187 | 1253 | 522 | 546 | 519 | 432 | 943 | 1012 | 1008 | 1019 | 749,98 | 505 | 996 | 1,96 | 0,971 | 13,582 | 5,12E-42 | 1,68E-41 | 0,0021618 | 0,0042806 | 0,0033553 | 0,0033553 | TRUE | 0,0897038 | No |
| orf19.4946 | 6 | 5 | 5 | 5 | 14 | 9 | 12 | 12 | 6 | 5 | 5 | 8 | 11 | 10 | 10 | 10 | 8,15 | 6 | 10 | 1,782 | 0,834 | 1,701 | 0,0889201 | 0,1043397 | 1,00E-08 | 0,2036254 | 0,0922957 | 0,0922957 | TRUE | 0,05913 | No |
| orf19.4947 | 7612 | 6886 | 6472 | 4453 | 4057 | 3148 | 3762 | 3734 | 7257 | 7281 | 6394 | 6976 | 3251 | 3580 | 3194 | 3036 | 5121,03 | 6977 | 3265 | 0,468 | -1,096 | -18,057 | 6,95E-73 | 3,46E-72 | 0,0038987 | 0,0024119 | 0,0033232 | 0,0033232 | TRUE | 0,0501387 | No |
| orf19.4948 | 609 | 614 | 638 | 323 | 1366 | 986 | 1366 | 1361 | 581 | 649 | 630 | 506 | 1095 | 1121 | 1160 | 1107 | 856,02 | 592 | 1121 | 1,884 | 0,914 | 12,702 | 5,77E-37 | 1,74E-36 | 0,0031509 | 0,0040093 | 0,0036298 | 0,0036298 | TRUE | 0,0944374 | No |
| orf19.4949 | 1 | 4 | 1 | 5 | 4 | 5 | 4 | 2 | 1 | 4 | 1 | 8 | 3 | 6 | 3 | 2 | 3,49 | 4 | 4 | 1,048 | 0,067 | 0,08 | 0,9359555 | 0,9426029 | 0,2278505 | 0,4726137 | 0,3744765 | 0,3744765 | TRUE | 0,5531869 | No |
| orf19.4950 | 6524 | 6414 | 6792 | 4498 | 4134 | 3201 | 3884 | 4056 | 6220 | 6782 | 6710 | 7046 | 3312 | 3640 | 3297 | 3298 | 5038,23 | 6690 | 3387 | 0,506 | -0,982 | -19,012 | 1,35E-80 | 7,41E-80 | 0,0022736 | 0,0024172 | 0,0023465 | 0,0023465 | TRUE | 0,0299458 | No |
| orf19.4951 | 2724 | 2847 | 2993 | 1878 | 1475 | 1109 | 1373 | 1413 | 2597 | 3010 | 2957 | 2942 | 1182 | 1261 | 1166 | 1149 | 2032,96 | 2876 | 1190 | 0,413 | -1,275 | -21,541 | 6,37E-103 | 4,28E-102 | 0,0027199 | 0,0028989 | 0,0027916 | 0,0027916 | TRUE | 0,0532603 | No |
| orf19.4952 | 6275 | 5791 | 6262 | 3664 | 6312 | 4515 | 5671 | 5956 | 5983 | 6123 | 6186 | 5740 | 5058 | 5134 | 4815 | 4843 | 5485,12 | 6008 | 4962 | 0,825 | -0,277 | -6,937 | 4,00E-12 | 7,43E-12 | 0,0008029 | 0,0023906 | 0,00135 | 0,00135 | TRUE | 0,0104772 | No |
| orf19.4952.1 | 1304 | 1078 | 1146 | 632 | 1440 | 859 | 1350 | 1253 | 1243 | 1140 | 1132 | 990 | 1154 | 977 | 1146 | 1019 | 1100,1 | 1126 | 1074 | 0,952 | -0,071 | -0,885 | 0,376018 | 0,4109215 | 0,006394 | 0,0035838 | 0,005239 | 0,005239 | TRUE | 0,0738412 | No |
| orf19.4953 | 1588 | 1808 | 1646 | 1046 | 1438 | 1063 | 1266 | 1209 | 1514 | 1912 | 1626 | 1639 | 1152 | 1209 | 1075 | 983 | 1388,65 | 1673 | 1105 | 0,66 | -0,6 | -7,183 | 6,84E-13 | 1,30E-12 | 0,008031 | 0,0032736 | 0,0059533 | 0,0059533 | TRUE | 0,1132009 | No |
| orf19.4954 | 835 | 788 | 820 | 536 | 1611 | 1333 | 1534 | 1529 | 796 | 833 | 810 | 840 | 1291 | 1516 | 1302 | 1243 | 1078,9 | 820 | 1338 | 1,631 | 0,706 | 10,155 | 3,13E-24 | 7,64E-24 | 0,0036222 | 0,0036131 | 0,0036148 | 0,0036148 | TRUE | 0,0924438 | No |
| orf19.4955 | 46 | 66 | 69 | 47 | 49 | 33 | 55 | 42 | 44 | 70 | 68 | 74 | 39 | 38 | 47 | 34 | 51,63 | 64 | 40 | 0,623 | -0,682 | -3,077 | 0,0020893 | 0,0027653 | 0,0184865 | 0,0338905 | 0,0270993 | 0,0270993 | TRUE | 0,4223638 | No |
| orf19.4956 | 8033 | 7704 | 7989 | 4956 | 12317 | 8745 | 11374 | 11845 | 7659 | 8146 | 7892 | 7764 | 9869 | 9944 | 9656 | 9631 | 8820,19 | 7865 | 9775 | 1,243 | 0,313 | 9,607 | 7,46E-22 | 1,73E-21 | 0,00033 | 0,0022774 | 0,0009038 | 0,0009038 | TRUE | 0,0071217 | No |
| orf19.4958 | 995 | 1088 | 1181 | 666 | 811 | 645 | 753 | 706 | 949 | 1150 | 1167 | 1043 | 650 | 733 | 639 | 574 | 863,21 | 1077 | 649 | 0,601 | -0,734 | -8,297 | 1,07E-16 | 2,22E-16 | 0,0078924 | 0,0039934 | 0,0063031 | 0,0063031 | TRUE | 0,0859418 | No |
| orf19.4959 | 6207 | 5421 | 5885 | 3898 | 2520 | 1853 | 2459 | 2641 | 5918 | 5732 | 5814 | 6106 | 2019 | 2107 | 2088 | 2147 | 3991,41 | 5892 | 2090 | 0,355 | -1,495 | -38,455 | 0 | 0 | 0,0004195 | 0,0025026 | 0,0011413 | 0,0011413 | TRUE | 0,0068827 | No |
| orf19.496 | 551 | 498 | 587 | 360 | 951 | 674 | 845 | 1004 | 525 | 527 | 580 | 564 | 762 | 766 | 717 | 816 | 657,24 | 549 | 765 | 1,396 | 0,482 | 7,043 | 1,88E-12 | 3,52E-12 | 0,0014068 | 0,0045895 | 0,0028904 | 0,0028904 | TRUE | 0,0247971 | No |
| orf19.4960 | 11514 | 10913 | 11334 | 7025 | 7565 | 5209 | 7041 | 7201 | 10977 | 11539 | 11197 | 11005 | 6062 | 5923 | 5978 | 5855 | 8567 | 11180 | 5954 | 0,533 | -0,909 | -28,802 | 2,01E-182 | 2,46E-181 | 0,00028 | 0,0022829 | 0,0008309 | 0,0008309 | TRUE | 0,0057517 | No |
| orf19.4961 | 39259 | 35803 | 38734 | 25804 | 19785 | 14539 | 18564 | 19301 | 37429 | 37857 | 38266 | 40423 | 15853 | 16532 | 15761 | 15694 | 27226,85 | 38494 | 15960 | 0,415 | -1,27 | -35,135 | 1,94E-270 | 3,67E-269 | 0,0008327 | 0,0021516 | 0,0012128 | 0,0012128 | TRUE | 0,0138267 | No |
| orf19.4962 | 1159 | 1266 | 1347 | 814 | 1121 | 797 | 1049 | 987 | 1105 | 1339 | 1331 | 1275 | 898 | 906 | 891 | 803 | 1068,39 | 1262 | 874 | 0,693 | -0,53 | -7,196 | 6,22E-13 | 1,18E-12 | 0,0047442 | 0,0036281 | 0,0042495 | 0,0042495 | TRUE | 0,0881351 | No |
| orf19.4963 | 1926 | 2286 | 2214 | 1145 | 3233 | 2274 | 2904 | 2606 | 1836 | 2417 | 2187 | 1794 | 2591 | 2586 | 2465 | 2119 | 2249,37 | 2058 | 2440 | 1,184 | 0,244 | 2,514 | 0,011936 | 0,0151028 | 0,0139465 | 0,0028212 | 0,0085851 | 0,0085851 | TRUE | 0,1659083 | No |
| orf19.4964 | 1876 | 1723 | 1846 | 1046 | 1088 | 747 | 982 | 985 | 1789 | 1822 | 1824 | 1639 | 872 | 849 | 834 | 801 | 1303,56 | 1768 | 839 | 0,473 | -1,079 | -19,603 | 1,46E-85 | 8,36E-85 | 0,0010288 | 0,0033508 | 0,0020597 | 0,0020597 | TRUE | 0,0265953 | No |
| orf19.4965 | 1215 | 1400 | 1350 | 769 | 828 | 620 | 778 | 790 | 1158 | 1480 | 1334 | 1205 | 663 | 705 | 661 | 642 | 981,04 | 1294 | 668 | 0,515 | -0,957 | -11,915 | 9,86E-33 | 2,79E-32 | 0,0061825 | 0,0037649 | 0,0050919 | 0,0050919 | TRUE | 0,112955 | No |
| orf19.4966 | 5207 | 5879 | 5875 | 3746 | 5510 | 4309 | 4991 | 4804 | 4964 | 6216 | 5804 | 5868 | 4415 | 4900 | 4237 | 3906 | 5038,88 | 5713 | 4364 | 0,764 | -0,389 | -4,879 | 1,07E-06 | 1,65E-06 | 0,00856 | 0,0024171 | 0,0058967 | 0,0058967 | TRUE | 0,0956769 | No |
| orf19.4967 | 2609 | 2352 | 2310 | 1396 | 2724 | 1846 | 2515 | 2728 | 2487 | 2487 | 2282 | 2187 | 2183 | 2099 | 2135 | 2218 | 2259,8 | 2361 | 2159 | 0,913 | -0,131 | -2,492 | 0,0126984 | 0,016023 | 0,0017472 | 0,0028179 | 0,0021964 | 0,0021964 | TRUE | 0,0276376 | No |
| orf19.4969 | 2959 | 3033 | 3107 | 1813 | 3303 | 2298 | 2850 | 3106 | 2821 | 3207 | 3069 | 2840 | 2647 | 2613 | 2420 | 2525 | 2767,8 | 2984 | 2551 | 0,854 | -0,227 | -4,167 | 3,08E-05 | 4,47E-05 | 0,0022574 | 0,0026845 | 0,002495 | 0,002495 | TRUE | 0,0305839 | No |
| orf19.497 | 2315 | 1901 | 2228 | 1353 | 2557 | 1578 | 2368 | 2421 | 2207 | 2010 | 2201 | 2120 | 2049 | 1794 | 2010 | 1969 | 2044,98 | 2134 | 1956 | 0,917 | -0,125 | -2,271 | 0,0231739 | 0,0286353 | 0,001927 | 0,0028942 | 0,0024276 | 0,0024276 | TRUE | 0,03562 | No |
| orf19.4970 | 2740 | 2574 | 2731 | 1583 | 2101 | 1537 | 1936 | 2076 | 2612 | 2722 | 2698 | 2480 | 1683 | 1748 | 1644 | 1688 | 2159,33 | 2628 | 1691 | 0,642 | -0,639 | -14,051 | 7,58E-45 | 2,58E-44 | 0,0005396 | 0,0028517 | 0,0015096 | 0,0015096 | TRUE | 0,0159121 | No |
| orf19.4972 | 258 | 263 | 275 | 168 | 36 | 27 | 39 | 52 | 246 | 278 | 272 | 263 | 29 | 31 | 33 | 42 | 149,23 | 265 | 34 | 0,128 | -2,964 | -19,057 | 5,76E-81 | 3,18E-80 | 1,00E-08 | 0,0130936 | 0,0080672 | 0,0080672 | TRUE | 0,236124 | No |
| orf19.4975 | 5466 | 4689 | 5034 | 3614 | 441 | 310 | 431 | 460 | 5211 | 4958 | 4973 | 5662 | 353 | 352 | 366 | 374 | 2781,21 | 5201 | 361 | 0,07 | -3,844 | -60,744 | 0 | 0 | 0,0023871 | 0,0026816 | 0,0025089 | 0,0025089 | TRUE | 0,0419955 | No |
| orf19.4976 | 2 | 5 | 7 | 0 | 17 | 6 | 7 | 2 | 2 | 5 | 7 | 0 | 14 | 7 | 6 | 2 | 5,27 | 4 | 7 | 1,903 | 0,928 | 1,166 | 0,2435646 | 0,2733836 | 0,5101345 | 0,3139268 | 0,3932194 | 0,3932194 | TRUE | 0,2984277 | No |
| orf19.4979 | 526 | 511 | 534 | 249 | 430 | 279 | 392 | 410 | 501 | 540 | 528 | 390 | 345 | 317 | 333 | 333 | 410,92 | 490 | 332 | 0,675 | -0,568 | -5,76 | 8,39E-09 | 1,40E-08 | 0,0077057 | 0,0060869 | 0,0068495 | 0,0068495 | TRUE | 0,1919453 | No |

| | | | | | | | | | | | | | | | | | | | | | | | | | | | | | | | |
|---|---|---|---|---|---|---|---|---|---|---|---|---|---|---|---|---|---|---|---|---|---|---|---|---|---|---|---|---|---|---|---|
| orf19.498 | 3763 | 3372 | 3682 | 2370 | 3756 | 2793 | 3737 | 4033 | 3588 | 3565 | 3637 | 3713 | 3010 | 3176 | 3173 | 3279 | 3392,58 | 3626 | 3160 | 0,872 | -0,198 | -4,964 | 6,91E-07 | 1,08E-06 | 0,0005357 | 0,0025752 | 0,0012319 | 0,0012319 | TRUE | 0,0127656 | No |
| orf19.4980 | 61165 | 56134 | 59582 | 36289 | 10693 | 7994 | 10141 | 11418 | 58314 | 59355 | 58861 | 56849 | 8568 | 9090 | 8610 | 9284 | 33616,29 | 58345 | 8888 | 0,152 | -2,715 | -73,432 | 0 | 0 | 0,0008824 | 0,0021401 | 0,0012532 | 0,0012532 | TRUE | 0,0112097 | No |
| orf19.4981 | 688 | 804 | 798 | 480 | 1203 | 957 | 1068 | 1066 | 656 | 850 | 788 | 752 | 964 | 1088 | 907 | 867 | 859 | 762 | 956 | 1,254 | 0,327 | 3,55 | 0,0003856 | 0,0005313 | 0,0092093 | 0,0040027 | 0,0069388 | 0,0069388 | TRUE | 0,1083791 | No |
| orf19.4982 | 1212 | 1257 | 1341 | 732 | 551 | 423 | 541 | 546 | 1156 | 1329 | 1325 | 1147 | 442 | 481 | 459 | 444 | 847,73 | 1239 | 456 | 0,367 | -1,445 | -19,884 | 5,64E-88 | 3,31E-87 | 0,0033241 | 0,0040281 | 0,0036411 | 0,0036411 | TRUE | 0,0281287 | No |
| orf19.4983 | 1643 | 1579 | 1663 | 988 | 1879 | 1280 | 1764 | 1774 | 1566 | 1670 | 1643 | 1548 | 1506 | 1455 | 1498 | 1442 | 1540,97 | 1607 | 1475 | 0,917 | -0,124 | -2,667 | 0,0076622 | 0,0097883 | 0,0001941 | 0,0031568 | 0,0014345 | 0,0014345 | TRUE | 0,0081491 | No |
| orf19.4984 | 2986 | 2802 | 3027 | 1676 | 4684 | 3316 | 4140 | 4428 | 2847 | 2963 | 2990 | 2626 | 3753 | 3771 | 3515 | 3600 | 3258,06 | 2856 | 3660 | 1,279 | 0,355 | 7,069 | 1,56E-12 | 2,93E-12 | 0,0018394 | 0,0025952 | 0,0021038 | 0,0021038 | TRUE | 0,0335954 | No |
| orf19.4985 | 594 | 657 | 644 | 317 | 1481 | 1063 | 1369 | 1245 | 566 | 695 | 636 | 497 | 1187 | 1209 | 1162 | 1012 | 870,48 | 598 | 1142 | 1,901 | 0,927 | 9,646 | 5,11E-22 | 1,19E-21 | 0,0116084 | 0,0039775 | 0,0075411 | 0,0075411 | TRUE | 0,1384803 | No |
| orf19.4987 | 3956 | 3551 | 3934 | 2449 | 4038 | 2711 | 3920 | 3959 | 3772 | 3755 | 3886 | 3836 | 3236 | 3083 | 3328 | 3219 | 3514,32 | 3812 | 3216 | 0,844 | -0,244 | -6,459 | 1,06E-10 | 1,89E-10 | 0,0002396 | 0,0025585 | 0,0010851 | 0,0010851 | TRUE | 0,0089832 | No |
| orf19.4988 | 1659 | 1646 | 1677 | 1023 | 1116 | 773 | 990 | 1066 | 1582 | 1740 | 1657 | 1603 | 894 | 879 | 841 | 867 | 1257,73 | 1646 | 870 | 0,528 | -0,92 | -17,506 | 1,30E-68 | 6,10E-68 | 0,0006501 | 0,0033967 | 0,001799 | 0,001799 | TRUE | 0,018443 | No |
| orf19.499 | 1866 | 1576 | 1729 | 1004 | 1607 | 1070 | 1565 | 1652 | 1779 | 1666 | 1708 | 1573 | 1288 | 1217 | 1329 | 1343 | 1487,82 | 1682 | 1294 | 0,769 | -0,379 | -6,918 | 4,57E-12 | 8,48E-12 | 0,0013601 | 0,0031948 | 0,0022043 | 0,0022043 | TRUE | 0,0208677 | No |
| orf19.4991 | 974 | 937 | 1036 | 692 | 3553 | 2688 | 3199 | 3501 | 929 | 991 | 1023 | 1084 | 2847 | 3056 | 2716 | 2847 | 1936,61 | 1007 | 2866 | 2,854 | 1,513 | 25,751 | 3,12E-146 | 3,09E-145 | 0,0023881 | 0,0029391 | 0,0025994 | 0,0025994 | TRUE | 0,0330027 | No |
| orf19.4993 | 2522 | 2523 | 2644 | 1485 | 3602 | 2543 | 3350 | 3514 | 2404 | 2668 | 2612 | 2326 | 2886 | 2892 | 2844 | 2857 | 2686,21 | 2502 | 2870 | 1,145 | 0,195 | 3,903 | 9,49E-05 | 0,0001346 | 0,0016605 | 0,0027025 | 0,002028 | 0,002028 | TRUE | 0,0251553 | No |
| orf19.4995 | 53 | 49 | 49 | 30 | 82 | 61 | 97 | 89 | 51 | 52 | 48 | 47 | 66 | 69 | 82 | 72 | 60,94 | 50 | 72 | 1,462 | 0,548 | 3,012 | 0,0025956 | 0,0034178 | 1,00E-08 | 0,0290337 | 0,0144749 | 0,0144749 | TRUE | 0,0822379 | No |
| orf19.4996 | 30 | 42 | 39 | 26 | 66 | 57 | 55 | 54 | 29 | 44 | 39 | 41 | 53 | 65 | 47 | 44 | 45,07 | 38 | 52 | 1,367 | 0,451 | 1,975 | 0,0482751 | 0,0581105 | 0,0096245 | 0,0385203 | 0,0266673 | 0,0266673 | TRUE | 0,2313186 | No |
| orf19.4997 | 2097 | 1907 | 2072 | 1305 | 2518 | 1820 | 2348 | 2486 | 1999 | 2016 | 2047 | 2044 | 2018 | 2070 | 1993 | 2021 | 2026,11 | 2026 | 2026 | 0,999 | -0,001 | -0,029 | 0,9768407 | 0,9791424 | 1,00E-08 | 0,0029017 | 0,000949 | 0,000949 | TRUE | 0,0023784 | No |
| orf19.4998 | 3675 | 3554 | 3875 | 2293 | 2601 | 1806 | 2547 | 2534 | 3504 | 3758 | 3828 | 3592 | 2084 | 2054 | 2162 | 2060 | 2880,29 | 3670 | 2090 | 0,569 | -0,812 | -18,465 | 3,97E-76 | 2,06E-75 | 0,0008455 | 0,0026613 | 0,001492 | 0,001492 | TRUE | 0,0120909 | No |
| orf19.50 | 1863 | 1929 | 1919 | 1129 | 1658 | 1172 | 1521 | 1496 | 1776 | 2040 | 1896 | 1769 | 1329 | 1333 | 1291 | 1216 | 1581,15 | 1870 | 1292 | 0,69 | -0,535 | -8,937 | 4,01E-19 | 8,83E-19 | 0,0023093 | 0,0031297 | 0,0027972 | 0,0027972 | TRUE | 0,0450217 | No |
| orf19.500 | 6365 | 5734 | 6214 | 3698 | 7991 | 5332 | 7619 | 8063 | 6068 | 6063 | 6139 | 5793 | 6403 | 6063 | 6468 | 6556 | 6194,21 | 6016 | 6372 | 1,059 | 0,083 | 2,168 | 0,0301364 | 0,0369169 | 0,0006787 | 0,0023564 | 0,0012391 | 0,0012391 | TRUE | 0,0127001 | No |
| orf19.5000 | 4355 | 3924 | 4197 | 2534 | 17330 | 11804 | 16367 | 17245 | 4152 | 4149 | 4146 | 3970 | 13886 | 13422 | 13896 | 14022 | 8955,33 | 4104 | 13806 | 3,361 | 1,749 | 53,954 | 0 | 0 | 0,0002106 | 0,0022746 | 0,0008399 | 0,0008399 | TRUE | 0,0053933 | No |
| orf19.5001 | 2765 | 2430 | 2654 | 1619 | 2501 | 1764 | 2328 | 2419 | 2636 | 2569 | 2622 | 2536 | 2004 | 2006 | 1976 | 1967 | 2289,61 | 2591 | 1988 | 0,767 | -0,384 | -10,2 | 1,99E-24 | 4,86E-24 | 1,00E-08 | 0,0028084 | 0,0009171 | 0,0009171 | TRUE | 0,002182 | No |
| orf19.5003 | 1762 | 1798 | 1819 | 988 | 1506 | 997 | 1350 | 1381 | 1680 | 1901 | 1797 | 1548 | 1207 | 1134 | 1146 | 1123 | 1441,9 | 1732 | 1152 | 0,665 | -0,59 | -8,943 | 3,78E-19 | 8,35E-19 | 0,0034288 | 0,00323 | 0,0034602 | 0,0034602 | TRUE | 0,0571867 | No |
| orf19.5004 | 373 | 350 | 368 | 194 | 297 | 198 | 274 | 318 | 356 | 370 | 364 | 304 | 238 | 225 | 233 | 259 | 293,43 | 348 | 239 | 0,683 | -0,551 | -5,9 | 3,65E-09 | 6,14E-09 | 0,0012582 | 0,0076868 | 0,004907 | 0,004907 | TRUE | 0,0691491 | No |
| orf19.5005 | 3667 | 3324 | 3587 | 2231 | 19411 | 13649 | 17846 | 19451 | 3496 | 3515 | 3544 | 3495 | 15553 | 15520 | 15151 | 15816 | 9511,21 | 3512 | 15510 | 4,415 | 2,142 | 70,305 | 0 | 0 | 1,00E-08 | 0,0022639 | 0,0007032 | 0,0007032 | TRUE | 0,0029953 | No |
| orf19.5006 | 2633 | 2420 | 2428 | 1608 | 4906 | 3372 | 4342 | 4664 | 2510 | 2559 | 2399 | 2519 | 3931 | 3834 | 3686 | 3792 | 3153,84 | 2497 | 3811 | 1,527 | 0,611 | 15,194 | 3,86E-52 | 1,47E-51 | 0,0004643 | 0,0026119 | 0,0012087 | 0,0012087 | TRUE | 0,0088185 | No |
| orf19.5006.1 | 4834 | 4694 | 4879 | 2610 | 5508 | 3772 | 5009 | 5033 | 4609 | 4963 | 4820 | 4089 | 4413 | 4289 | 4253 | 4092 | 4441,02 | 4620 | 4262 | 0,922 | -0,118 | -1,976 | 0,0481818 | 0,0580207 | 0,0036189 | 0,002461 | 0,0031913 | 0,0031913 | TRUE | 0,0707704 | No |
| orf19.5007 | 5516 | 4594 | 4905 | 3441 | 12112 | 8570 | 11370 | 12626 | 5259 | 4858 | 4846 | 5391 | 9705 | 9745 | 9653 | 10266 | 7465,22 | 5088 | 9842 | 1,936 | 0,953 | 20,625 | 1,64E-94 | 1,02E-93 | 0,0017177 | 0,0023112 | 0,0018937 | 0,0018937 | TRUE | 0,0186884 | No |
| orf19.5008.1 | 245 | 218 | 238 | 159 | 446 | 315 | 388 | 409 | 234 | 231 | 235 | 249 | 357 | 358 | 329 | 333 | 290,73 | 237 | 344 | 1,455 | 0,541 | 6,22 | 4,98E-10 | 8,66E-10 | 1,00E-08 | 0,0077389 | 0,003627 | 0,003627 | TRUE | 0,0099636 | No |
| orf19.5009 | 1782 | 1729 | 1799 | 1030 | 3609 | 2385 | 3385 | 3554 | 1699 | 1828 | 1777 | 1614 | 2892 | 2712 | 2874 | 2890 | 2285,66 | 1730 | 2842 | 1,641 | 0,715 | 14,487 | 1,47E-47 | 5,25E-47 | 0,001319 | 0,0028096 | 0,0018555 | 0,0018555 | TRUE | 0,0219447 | No |
| orf19.501 | 16976 | 14359 | 15808 | 9554 | 29537 | 19387 | 28292 | 30090 | 16185 | 15183 | 15617 | 14967 | 23667 | 22045 | 24020 | 24466 | 19518,62 | 15488 | 23550 | 1,52 | 0,604 | 13,873 | 9,23E-44 | 3,11E-43 | 0,0015345 | 0,0021754 | 0,0017683 | 0,0017683 | TRUE | 0,0225214 | No |
| orf19.5010 | 3305 | 3009 | 3145 | 1814 | 3942 | 2952 | 3885 | 4061 | 3151 | 3182 | 3107 | 2842 | 3159 | 3357 | 3298 | 3302 | 3174,61 | 3070 | 3279 | 1,066 | 0,092 | 2,005 | 0,0449351 | 0,0542742 | 0,0011934 | 0,0026085 | 0,0017076 | 0,0017076 | TRUE | 0,0283597 | No |
| orf19.5011 | 922 | 891 | 928 | 574 | 1068 | 790 | 937 | 986 | 879 | 942 | 917 | 899 | 856 | 898 | 796 | 802 | 873,55 | 909 | 838 | 0,92 | -0,121 | -2,002 | 0,0453378 | 0,0547503 | 0,0009564 | 0,0039709 | 0,0023497 | 0,0023497 | TRUE | 0,0257988 | No |
| orf19.5012 | 185 | 193 | 160 | 98 | 543 | 295 | 454 | 466 | 176 | 204 | 158 | 154 | 435 | 335 | 385 | 379 | 278,36 | 173 | 384 | 2,214 | 1,146 | 9,875 | 5,32E-23 | 1,27E-22 | 0,0088585 | 0,0079897 | 0,0085618 | 0,0085618 | TRUE | 0,1527328 | No |
| orf19.5013 | 277 | 286 | 300 | 248 | 622 | 452 | 532 | 586 | 264 | 302 | 296 | 389 | 498 | 514 | 452 | 476 | 398,98 | 313 | 485 | 1,564 | 0,645 | 6,115 | 9,67E-10 | 1,66E-09 | 0,0101337 | 0,0062065 | 0,0079561 | 0,0079561 | TRUE | 0,2626811 | No |
| orf19.5014 | 203 | 172 | 179 | 118 | 687 | 440 | 584 | 589 | 194 | 182 | 177 | 185 | 550 | 500 | 496 | 479 | 345,33 | 184 | 506 | 2,749 | 1,459 | 16,112 | 2,10E-58 | 8,64E-58 | 1,00E-08 | 0,0068459 | 0,0039767 | 0,0039767 | TRUE | 0,042279 | No |
| orf19.5015 | 1362 | 1312 | 1292 | 798 | 2375 | 1727 | 2100 | 2450 | 1299 | 1387 | 1276 | 1250 | 1903 | 1964 | 1783 | 1992 | 1606,75 | 1303 | 1910 | 1,464 | 0,55 | 10,005 | 1,44E-23 | 3,47E-23 | 0,0016227 | 0,0031132 | 0,0022371 | 0,0022371 | TRUE | 0,024658 | No |
| orf19.5016 | 1137 | 1023 | 1016 | 765 | 791 | 544 | 683 | 669 | 1084 | 1082 | 1004 | 1198 | 634 | 619 | 580 | 544 | 843 | 1092 | 594 | 0,545 | -0,876 | -11,995 | 3,77E-33 | 1,07E-32 | 0,0036848 | 0,004039 | 0,0038515 | 0,0038515 | TRUE | 0,0473812 | No |
| orf19.5017 | 70 | 63 | 61 | 42 | 217 | 173 | 201 | 226 | 67 | 67 | 60 | 66 | 174 | 197 | 171 | 184 | 123,05 | 65 | 182 | 2,791 | 1,481 | 10,591 | 3,27E-26 | 8,32E-26 | 1,00E-08 | 0,0154345 | 0,0077536 | 0,0077536 | TRUE | 0,0316767 | No |
| orf19.5019 | 2799 | 2295 | 2572 | 1660 | 1782 | 1264 | 1760 | 1807 | 2669 | 2427 | 2541 | 2600 | 1428 | 1437 | 1494 | 1469 | 2008,15 | 2559 | 1457 | 0,57 | -0,812 | -17,627 | 1,54E-69 | 7,33E-69 | 0,0006632 | 0,0029089 | 0,0015112 | 0,0015112 | TRUE | 0,0150009 | No |
| orf19.5020 | 6710 | 5853 | 6368 | 4469 | 6213 | 4400 | 6125 | 6319 | 6397 | 6189 | 6291 | 7001 | 4978 | 5003 | 5200 | 5138 | 5774,69 | 6470 | 5080 | 0,786 | -0,348 | -7,614 | 2,66E-14 | 5,26E-14 | 0,0015715 | 0,0023756 | 0,0018253 | 0,0018253 | TRUE | 0,0360956 | No |
| orf19.5021 | 5121 | 4428 | 4828 | 3118 | 10617 | 7683 | 10462 | 10793 | 4882 | 4682 | 4770 | 4885 | 8507 | 8736 | 8882 | 8776 | 6764,97 | 4805 | 8725 | 1,817 | 0,861 | 26,603 | 6,29E-156 | 6,63E-155 | 0,000225 | 0,002334 | 0,0008379 | 0,0008379 | TRUE | 0,0035917 | No |
| orf19.5022 | 4509 | 3939 | 4295 | 2809 | 2572 | 1901 | 2541 | 2781 | 4299 | 4165 | 4243 | 4400 | 2061 | 2162 | 2157 | 2261 | 3218,54 | 4277 | 2160 | 0,505 | -0,985 | -23,541 | 1,54E-122 | 1,26E-121 | 0,0005207 | 0,0026014 | 0,0013414 | 0,0013414 | TRUE | 0,0125 | No |
| orf19.5023 | 205 | 168 | 167 | 117 | 192 | 140 | 185 | 226 | 195 | 178 | 165 | 183 | 154 | 159 | 157 | 184 | 171,9 | 180 | 164 | 0,908 | -0,14 | -1,211 | 0,2257493 | 0,2545514 | 1,00E-08 | 0,0116427 | 0,0069326 | 0,0069326 | TRUE | 0,0798461 | No |
| orf19.5024 | 4833 | 4186 | 4469 | 2966 | 12039 | 8245 | 11177 | 12567 | 4608 | 4426 | 4415 | 4646 | 9647 | 9375 | 9489 | 10218 | 7103,06 | 4524 | 9682 | 2,142 | 1,099 | 27,084 | 1,51E-161 | 1,65E-160 | 0,0008843 | 0,0023224 | 0,0014095 | 0,0014095 | TRUE | 0,0170283 | No |
| orf19.5025 | 14259 | 13157 | 13528 | 8291 | 43165 | 29696 | 39724 | 42978 | 13594 | 13912 | 13364 | 12988 | 34587 | 33767 | 33725 | 34945 | 23860,45 | 13464 | 34256 | 2,543 | 1,347 | 41,753 | 0 | 0 | 0,0005052 | 0,0021601 | 0,0009444 | 0,0009444 | TRUE | 0,0067121 | No |
| orf19.5026 | 19789 | 18176 | 19551 | 12735 | 15707 | 11672 | 14415 | 16045 | 18867 | 19219 | 19315 | 19950 | 12586 | 13272 | 12238 | 13046 | 16061,52 | 19338 | 12786 | 0,661 | -0,597 | -16,122 | 1,79E-58 | 7,38E-58 | 0,0008427 | 0,0021935 | 0,0012524 | 0,0012524 | TRUE | 0,0101383 | No |
| orf19.5027 | 3227 | 3039 | 3103 | 1829 | 6943 | 4919 | 6818 | 7264 | 3077 | 3213 | 3065 | 2865 | 5563 | 5593 | 5788 | 5906 | 4384 | 3055 | 5712 | 1,868 | 0,901 | 20,472 | 3,85E-93 | 2,37E-92 | 0,0011738 | 0,0024658 | 0,0015991 | 0,0015991 | TRUE | 0,0196516 | No |
| orf19.5029 | 2356 | 2207 | 2459 | 1489 | 1891 | 1466 | 1835 | 1795 | 2246 | 2334 | 2429 | 2333 | 1515 | 1667 | 1558 | 1460 | 1942,66 | 2336 | 1550 | 0,663 | -0,594 | -11,392 | 4,59E-30 | 1,24E-29 | 0,001403 | 0,0029365 | 0,0020772 | 0,0020772 | TRUE | 0,0295383 | No |
| orf19.5030 | 1773 | 1568 | 1688 | 949 | 657 | 406 | 617 | 705 | 1690 | 1658 | 1668 | 1487 | 526 | 462 | 524 | 573 | 1073,47 | 1626 | 521 | 0,321 | -1,639 | -23,02 | 2,93E-117 | 2,27E-116 | 0,0038261 | 0,0036208 | 0,0036673 | 0,0036673 | TRUE | 0,0631455 | No |
| orf19.5031 | 10198 | 8895 | 9985 | 6676 | 4153 | 3182 | 4103 | 4449 | 9723 | 9405 | 9864 | 10458 | 3328 | 3618 | 3483 | 3617 | 6687,17 | 9862 | 3512 | 0,356 | -1,49 | -32,339 | 1,96E-229 | 3,02E-228 | 0,0015154 | 0,0023368 | 0,0018534 | 0,0018534 | TRUE | 0,019748 | No |
| orf19.5032 | 33778 | 27708 | 29922 | 20112 | 93652 | 61655 | 89032 | 97063 | 32204 | 29298 | 29560 | 31507 | 75041 | 70107 | 75588 | 78921 | 52778,15 | 30642 | 74914 | 2,445 | 1,29 | 26,737 | 1,76E-157 | 1,88E-156 | 0,0022405 | 0,0021224 | 0,0022117 | 0,0022117 | TRUE | 0,022832 | No |
| orf19.5033 | 189 | 169 | 159 | 100 | 312 | 229 | 300 | 323 | 180 | 179 | 157 | 157 | 250 | 260 | 255 | 263 | 212,54 | 168 | 257 | 1,521 | 0,605 | 5,985 | 2,17E-09 | 3,68E-09 | 1,00E-08 | 0,0098163 | 0,004781 | 0,004781 | TRUE | 0,0264806 | No |
| orf19.5034 | 1087 | 1093 | 1144 | 787 | 1784 | 1319 | 1749 | 1764 | 1036 | 1156 | 1130 | 1233 | 1429 | 1500 | 1485 | 1434 | 1300,45 | 1139 | 1462 | 1,287 | 0,364 | 6,276 | 3,47E-10 | 6,08E-10 | 0,0017864 | 0,0033538 | 0,0024365 | 0,0024365 | TRUE | 0,0445521 | No |
| orf19.5035 | 1539 | 1335 | 1496 | 913 | 2834 | 1824 | 2758 | 2861 | 1467 | 1412 | 1478 | 1430 | 2271 | 2074 | 2342 | 2326 | 1849,96 | 1447 | 2253 | 1,558 | 0,64 | 12,353 | 4,67E-35 | 1,37E-34 | 0,0013439 | 0,0029788 | 0,0019907 | 0,0019907 | TRUE | 0,0330888 | No |
| orf19.5037 | 6654 | 7434 | 7287 | 4734 | 4743 | 3788 | 4408 | 4108 | 6344 | 7861 | 7199 | 7416 | 3800 | 4307 | 3742 | 3340 | 5501,2 | 7205 | 3797 | 0,527 | -0,924 | -11,419 | 3,35E-30 | 9,06E-30 | 0,0090912 | 0,0023897 | 0,0060979 | 0,0060979 | TRUE | 0,0984675 | No |
| orf19.5038 | 4898 | 5528 | 5744 | 3301 | 4815 | 3882 | 4290 | 4254 | 4670 | 5845 | 5675 | 5171 | 3858 | 4414 | 3642 | 3459 | 4591,75 | 5340 | 3843 | 0,719 | -0,475 | -5,589 | 2,29E-08 | 3,77E-08 | 0,0104428 | 0,0024489 | 0,0067238 | 0,0067238 | TRUE | 0,1206733 | No |
| orf19.5039 | 3020 | 2804 | 2676 | 1682 | 2785 | 1879 | 2473 | 2660 | 2879 | 2965 | 2644 | 2635 | 2232 | 2137 | 2100 | 2163 | 2469,15 | 2781 | 2158 | 0,775 | -0,367 | -7,143 | 9,16E-13 | 1,74E-12 | 0,0015877 | 0,0027562 | 0,0021257 | 0,0021257 | TRUE | 0,0238726 | No |
| orf19.504 | 3426 | 3010 | 3318 | 2183 | 3741 | 2607 | 3511 | 3814 | 3266 | 3183 | 3278 | 3420 | 2998 | 2964 | 2981 | 3101 | 3148,82 | 3287 | 3011 | 0,917 | -0,124 | -3,232 | 0,0012279 | 0,0016444 | 0,0002329 | 0,0026127 | 0,0011033 | 0,0011033 | TRUE | 0,0082311 | No |
| orf19.5040 | 4089 | 4330 | 4573 | 2453 | 2881 | 1983 | 2729 | 2548 | 3898 | 4578 | 4518 | 3843 | 2308 | 2255 | 2317 | 2072 | 3223,66 | 4209 | 2238 | 0,531 | -0,912 | -13,238 | 5,26E-40 | 1,66E-39 | 0,0051943 | 0,0026006 | 0,0042273 | 0,0042273 | TRUE | 0,0421837 | No |
| orf19.5041 | 225 | 202 | 176 | 125 | 95 | 78 | 102 | 115 | 215 | 214 | 174 | 196 | 76 | 89 | 87 | 94 | 142,84 | 200 | 86 | 0,431 | -1,213 | -9,173 | 4,60E-20 | 1,03E-19 | 1,00E-08 | 0,0135861 | 0,0088228 | 0,0088228 | TRUE | 0,088848 | No |
| orf19.5042 | 59 | 87 | 77 | 45 | 39 | 33 | 36 | 29 | 56 | 92 | 76 | 70 | 31 | 38 | 31 | 24 | 52,22 | 74 | 31 | 0,413 | -1,275 | -5,586 | 2,32E-08 | 3,82E-08 | 0,0206142 | 0,0335362 | 0,0278255 | 0,0278255 | TRUE | 0,146969 | No |
| orf19.5043 | 203 | 195 | 209 | 112 | 168 | 123 | 209 | 169 | 194 | 206 | 206 | 175 | 135 | 140 | 177 | 137 | 171,37 | 195 | 147 | 0,751 | -0,413 | -3,314 | 0,0009207 | 0,001245 | 0,0057769 | 0,0116722 | 0,009009 | 0,009009 | TRUE | 0,2077835 | No |
| orf19.5045 | 8709 | 9723 | 10201 | 5931 | 2326 | 1790 | 2035 | 1982 | 8303 | 10281 | 10078 | 9291 | 1864 | 2035 | 1728 | 1612 | 5648,9 | 9488 | 1810 | 0,191 | -2,392 | -29,303 | 9,59E-189 | 1,23E-187 | 0,0091212 | 0,0023819 | 0,0060925 | 0,0060925 | TRUE | 0,0869223 | No |
| orf19.5046 | 270 | 284 | 293 | 180 | 656 | 442 | 589 | 601 | 257 | 300 | 289 | 282 | 526 | 503 | 500 | 489 | 393,26 | 282 | 504 | 1,788 | 0,839 | 10,503 | 8,37E-26 | 2,10E-25 | 1,00E-08 | 0,0062664 | 0,003264 | 0,003264 | TRUE | 0,0448707 | No |
| orf19.5049 | 5125 | 4417 | 4635 | 2984 | 5780 | 3838 | 5491 | 5596 | 4886 | 4670 | 4579 | 4675 | 4631 | 4364 | 4662 | 4550 | 4627,19 | 4702 | 4552 | 0,968 | -0,047 | -1,208 | 0,2270356 | 0,2559132 | 0,0005262 | 0,0024461 | 0,0012084 | 0,0012084 | TRUE | 0,0090347 | No |
| orf19.505 | 3397 | 2905 | 3202 | 2081 | 2683 | 1990 | 2575 | 2924 | 3239 | 3072 | 3163 | 3260 | 2150 | 2263 | 2186 | 2377 | 2713,74 | 3184 | 2244 | 0,705 | -0,504 | -11,019 | 3,10E-28 | 8,13E-28 | 0,000858 | 0,0026963 | 0,0016332 | 0,0016332 | TRUE | 0,020156 | No |
| orf19.5050 | 2426 | 2217 | 2349 | 1467 | 2608 | 1833 | 2560 | 2734 | 2313 | 2344 | 2321 | 2298 | 2090 | 2084 | 2173 | 2223 | 2230,79 | 2319 | 2142 | 0,924 | -0,114 | -2,798 | 0,0051478 | 0,0066495 | 1,00E-08 | 0,0028273 | 0,0011349 | 0,0011349 | TRUE | 0,0078253 | No |
| orf19.5051 | 2815 | 2485 | 2680 | 1768 | 1884 | 1353 | 1804 | 1875 | 2684 | 2628 | 2648 | 2770 | 1510 | 1538 | 1532 | 1525 | 2104,11 | 2682 | 1526 | 0,569 | -0,812 | -20,482 | 3,09E-93 | 1,91E-92 | 1,00E-08 | 0,0028716 | 0,0010092 | 0,0010092 | TRUE | 0,0052161 | No |
| orf19.5052 | 646 | 559 | 551 | 410 | 939 | 717 | 859 | 1019 | 616 | 591 | 544 | 642 | 752 | 815 | 729 | 829 | 689,89 | 598 | 781 | 1,308 | 0,388 | 5,319 | 1,04E-07 | 1,68E-07 | 0,0027971 | 0,0044712 | 0,0035953 | 0,0035953 | TRUE | 0,0441418 | No |
| orf19.5053 | 1292 | 1213 | 1340 | 844 | 367 | 265 | 353 | 315 | 1232 | 1283 | 1324 | 1322 | 294 | 301 | 300 | 256 | 788,94 | 1290 | 288 | 0,223 | -2,166 | -31,303 | 4,19E-215 | 6,12E-214 | 0,0007445 | 0,0041724 | 0,0026159 | 0,0026159 | TRUE | 0,0656023 | No |
| orf19.5054 | 2484 | 2270 | 2239 | 1322 | 2472 | 1760 | 2256 | 2570 | 2368 | 2400 | 2212 | 2071 | 1981 | 2001 | 1915 | 2090 | 2129,8 | 2263 | 1997 | 0,881 | -0,183 | -3,256 | 0,0011305 | 0,0015193 | 0,0022794 | 0,0028622 | 0,0025531 | 0,0025531 | TRUE | 0,036773 | No |
| orf19.5056 | 162 | 131 | 131 | 91 | 313 | 241 | 296 | 315 | 154 | 139 | 129 | 143 | 251 | 274 | 251 | 256 | 199,65 | 141 | 258 | 1,823 | 0,867 | 8,14 | 3,94E-16 | 8,13E-16 | 1,00E-08 | 0,0103152 | 0,0052199 | 0,0052199 | TRUE | 0,0484389 | No |
| orf19.5057 | 18 | 22 | 14 | 8 | 40 | 27 | 22 | 43 | 17 | 23 | 14 | 13 | 32 | 31 | 19 | 35 | 22,9 | 17 | 29 | 1,722 | 0,784 | 2,384 | 0,0171029 | 0,0213527 | 0,0309411 | 0,0737981 | 0,0555617 | 0,0555617 | TRUE | 0,4240098 | No |
| orf19.5058 | 21958 | 19774 | 21530 | 12778 | 5956 | 4276 | 5829 | 5928 | 20935 | 20909 | 21270 | 20017 | 4772 | 4862 | 4949 | 4820 | 12816,69 | 20783 | 4851 | 0,233 | -2,1 | -64,623 | 0 | 0 | 0,0003896 | 0,0022194 | 0,0008945 | 0,0008945 | TRUE | 0,0073532 | No |

| | | | | | | | | | | | | | | | | | | | | | | | | | | | | | | | |
|---|---|---|---|---|---|---|---|---|---|---|---|---|---|---|---|---|---|---|---|---|---|---|---|---|---|---|---|---|---|---|---|
| orf19.5059 | 9309 | 9485 | 10118 | 5908 | 14852 | 11264 | 13921 | 14030 | 8875 | 10029 | 9996 | 9255 | 11900 | 12808 | 11819 | 11408 | 10761,3 | 9539 | 11984 | 1,256 | 0,329 | 6,158 | 7,38E-10 | 1,27E-09 | 0,0028648 | 0,0022439 | 0,0026428 | 0,0026428 | TRUE | 0,0272197 | No |
| orf19.506 | 9252 | 7847 | 8248 | 5234 | 18460 | 12729 | 17803 | 19087 | 8821 | 8297 | 8148 | 8199 | 14791 | 14474 | 15115 | 15520 | 11670,66 | 8366 | 14975 | 1,79 | 0,84 | 21,265 | 2,38E-100 | 1,57E-99 | 0,001005 | 0,002232 | 0,0014002 | 0,0014002 | TRUE | 0,0165637 | No |
| orf19.5061 | 8545 | 7739 | 8296 | 5017 | 17620 | 12553 | 16325 | 17763 | 8147 | 8183 | 8196 | 7859 | 14118 | 14274 | 13860 | 14443 | 11135 | 8096 | 14174 | 1,75 | 0,807 | 26,765 | 8,21E-158 | 8,85E-157 | 0,0002168 | 0,0022387 | 0,0007721 | 0,0007721 | TRUE | 0,004461 | No |
| orf19.5063 | 4013 | 3818 | 3650 | 2698 | 7665 | 6074 | 6674 | 6350 | 3826 | 4037 | 3606 | 4227 | 6142 | 6907 | 5666 | 5163 | 4946,65 | 3924 | 5970 | 1,522 | 0,606 | 7,288 | 3,14E-13 | 6,05E-13 | 0,0097655 | 0,0024232 | 0,0064122 | 0,0064122 | TRUE | 0,1355528 | No |
| orf19.5064 | 4917 | 4810 | 4772 | 2908 | 4082 | 2939 | 3706 | 3708 | 4688 | 5086 | 4714 | 4556 | 3271 | 3342 | 3146 | 3015 | 3977,21 | 4761 | 3194 | 0,67 | -0,577 | -11,64 | 2,59E-31 | 7,15E-31 | 0,0018083 | 0,0025041 | 0,0021045 | 0,0021045 | TRUE | 0,0257163 | No |
| orf19.5064.1 | 725 | 577 | 568 | 340 | 1857 | 1212 | 1554 | 1828 | 691 | 610 | 561 | 533 | 1488 | 1378 | 1319 | 1486 | 1008,36 | 599 | 1418 | 2,36 | 1,239 | 15,289 | 9,08E-53 | 3,48E-52 | 0,0068722 | 0,0037196 | 0,0050536 | 0,0050536 | TRUE | 0,129126 | No |
| orf19.5065 | 457 | 596 | 544 | 318 | 471 | 434 | 415 | 360 | 436 | 630 | 537 | 498 | 377 | 493 | 352 | 293 | 452,18 | 525 | 379 | 0,719 | -0,476 | -3,216 | 0,0013004 | 0,0017385 | 0,0336766 | 0,0057224 | 0,0187582 | 0,0187582 | TRUE | 0,4606762 | No |
| orf19.5066 | 4123 | 3803 | 3849 | 2223 | 4647 | 3385 | 4493 | 4447 | 3931 | 4021 | 3802 | 3482 | 3724 | 3849 | 3815 | 3616 | 3779,98 | 3809 | 3751 | 0,983 | -0,024 | -0,486 | 0,627195 | 0,6600909 | 0,0017903 | 0,0025256 | 0,0021634 | 0,0021634 | TRUE | 0,0385546 | No |
| orf19.5067 | 3528 | 3413 | 3382 | 1884 | 3062 | 1958 | 2852 | 2689 | 3364 | 3609 | 3341 | 2951 | 2453 | 2226 | 2421 | 2186 | 2819,07 | 3316 | 2322 | 0,7 | -0,515 | -7,799 | 6,24E-15 | 1,26E-14 | 0,0045602 | 0,0026737 | 0,0038294 | 0,0038294 | TRUE | 0,0642312 | No |
| orf19.5068 | 3256 | 3356 | 3443 | 2040 | 2568 | 1841 | 2383 | 2416 | 3104 | 3549 | 3401 | 3196 | 2058 | 2093 | 2023 | 1964 | 2673,57 | 3312 | 2034 | 0,614 | -0,704 | -13,548 | 8,09E-42 | 2,64E-41 | 0,0016671 | 0,0027054 | 0,0022046 | 0,0022046 | TRUE | 0,0281107 | No |
| orf19.5069 | 10880 | 10053 | 10907 | 6945 | 2917 | 2091 | 2922 | 2955 | 10373 | 10630 | 10775 | 10880 | 2337 | 2378 | 2481 | 2403 | 6531,99 | 10664 | 2400 | 0,225 | -2,151 | -60,8 | 0 | 0 | 0,0003161 | 0,0023426 | 0,0009635 | 0,0009635 | TRUE | 0,0064134 | No |
| orf19.5070 | 2148 | 1979 | 2207 | 1497 | 2130 | 1632 | 2108 | 2088 | 2048 | 2093 | 2180 | 2345 | 1707 | 1856 | 1790 | 1698 | 1964,47 | 2166 | 1763 | 0,814 | -0,296 | -5,373 | 7,76E-08 | 1,25E-07 | 0,0020457 | 0,0029271 | 0,0024084 | 0,0024084 | TRUE | 0,0348716 | No |
| orf19.5071 | 218 | 190 | 241 | 161 | 565 | 357 | 453 | 560 | 208 | 201 | 238 | 252 | 453 | 406 | 385 | 455 | 324,7 | 225 | 425 | 1,903 | 0,928 | 9,166 | 4,92E-20 | 1,11E-19 | 0,0053366 | 0,0071479 | 0,0062941 | 0,0062941 | TRUE | 0,060289 | No |
| orf19.5073 | 733 | 629 | 674 | 383 | 1649 | 995 | 1592 | 1653 | 699 | 665 | 666 | 600 | 1321 | 1131 | 1352 | 1344 | 972,26 | 658 | 1287 | 1,954 | 0,967 | 13,081 | 4,24E-39 | 1,32E-38 | 0,0042795 | 0,00378 | 0,0040468 | 0,0040468 | TRUE | 0,0762672 | No |
| orf19.5074 | 984 | 913 | 896 | 590 | 1697 | 1191 | 1705 | 1790 | 938 | 965 | 885 | 924 | 1360 | 1354 | 1448 | 1455 | 1166,24 | 928 | 1404 | 1,514 | 0,598 | 11,061 | 1,94E-28 | 5,09E-28 | 0,0006762 | 0,0034991 | 0,0018881 | 0,0018881 | TRUE | 0,0126157 | No |
| orf19.5076 | 1550 | 1368 | 1384 | 924 | 3830 | 2578 | 3677 | 3890 | 1478 | 1446 | 1367 | 1447 | 3069 | 2931 | 3122 | 3163 | 2253 | 1434 | 3071 | 2,143 | 1,099 | 23,946 | 1,02E-126 | 8,62E-126 | 0,0006391 | 0,00282 | 0,0014878 | 0,0014878 | TRUE | 0,0128166 | No |
| orf19.5076.1 | 902 | 832 | 884 | 713 | 1452 | 1188 | 1376 | 1509 | 860 | 880 | 873 | 1117 | 1163 | 1351 | 1168 | 1227 | 1079,93 | 932 | 1227 | 1,32 | 0,401 | 4,601 | 4,21E-06 | 6,35E-06 | 0,0089063 | 0,0036117 | 0,0063144 | 0,0063144 | TRUE | 0,198045 | No |
| orf19.5077 | 3045 | 2617 | 2909 | 1806 | 7939 | 5570 | 7583 | 7905 | 2903 | 2767 | 2874 | 2829 | 6361 | 6334 | 6438 | 6428 | 4616,7 | 2843 | 6390 | 2,247 | 1,168 | 35,538 | 1,26E-276 | 2,47E-275 | 1,00E-08 | 0,0024469 | 0,0007699 | 0,0007699 | TRUE | 0,0042089 | No |
| orf19.5078 | 792 | 657 | 713 | 417 | 779 | 517 | 766 | 807 | 755 | 695 | 704 | 653 | 624 | 588 | 650 | 656 | 665,75 | 702 | 630 | 0,896 | -0,159 | -2,349 | 0,0188 | 0,0233846 | 0,0012295 | 0,0045575 | 0,0028797 | 0,0028797 | TRUE | 0,0309502 | No |
| orf19.5079 | 81762 | 70731 | 78184 | 51925 | 6652 | 4702 | 6334 | 6538 | 77951 | 74789 | 77238 | 81343 | 5330 | 5347 | 5378 | 5316 | 41586,52 | 77830 | 5343 | 0,069 | -3,865 | -112,058 | 0 | 0 | 0,000483 | 0,0021308 | 0,0010516 | 0,0010516 | TRUE | 0,0112518 | No |
| orf19.5079.1 | 2261 | 1808 | 2037 | 1200 | 5648 | 3464 | 5297 | 5371 | 2156 | 1912 | 2012 | 1880 | 4526 | 3939 | 4497 | 4367 | 3161,04 | 1990 | 4332 | 2,175 | 1,121 | 18,461 | 4,22E-76 | 2,19E-75 | 0,0034134 | 0,0026107 | 0,0031568 | 0,0031568 | TRUE | 0,0447145 | No |
| orf19.508 | 2538 | 2303 | 2476 | 1642 | 540 | 411 | 521 | 517 | 2420 | 2435 | 2446 | 2572 | 433 | 467 | 442 | 420 | 1454,49 | 2468 | 440 | 0,178 | -2,488 | -45,811 | 0 | 0 | 1,00E-08 | 0,0032201 | 0,0015939 | 0,0015939 | TRUE | 0,0166934 | No |
| orf19.5081 | 10991 | 9044 | 9989 | 6339 | 30103 | 19741 | 28715 | 30332 | 10479 | 9563 | 9868 | 9930 | 24121 | 22447 | 24379 | 24663 | 16931,23 | 9960 | 23902 | 2,4 | 1,263 | 28,878 | 2,23E-183 | 2,74E-182 | 0,0015043 | 0,0021883 | 0,0017619 | 0,0017619 | TRUE | 0,0204471 | No |
| orf19.5083 | 3301 | 2642 | 3015 | 1933 | 8008 | 5269 | 7622 | 8182 | 3147 | 2794 | 2979 | 3028 | 6417 | 5991 | 6471 | 6653 | 4684,89 | 2987 | 6383 | 2,138 | 1,096 | 21,985 | 4,04E-107 | 2,82E-106 | 0,0018449 | 0,0024417 | 0,0021286 | 0,0021286 | TRUE | 0,0234277 | No |
| orf19.5085 | 4489 | 4023 | 4373 | 2870 | 10036 | 7220 | 9364 | 10215 | 4280 | 4254 | 4320 | 4496 | 8042 | 8210 | 7950 | 8306 | 6232,1 | 4338 | 8127 | 1,875 | 0,907 | 26,681 | 7,84E-157 | 8,34E-156 | 0,0002765 | 0,0023547 | 0,000925 | 0,000925 | TRUE | 0,0067876 | No |
| orf19.5087 | 3086 | 2535 | 2895 | 1682 | 1410 | 938 | 1397 | 1505 | 2942 | 2680 | 2860 | 2635 | 1130 | 1067 | 1186 | 1224 | 1965,46 | 2779 | 1152 | 0,414 | -1,271 | -21,852 | 7,41E-106 | 5,10E-105 | 0,0023252 | 0,0029267 | 0,0026569 | 0,0026569 | TRUE | 0,0276958 | No |
| orf19.5089 | 427 | 438 | 476 | 309 | 527 | 357 | 441 | 521 | 407 | 463 | 470 | 484 | 422 | 406 | 374 | 424 | 431,35 | 456 | 406 | 0,894 | -0,161 | -1,964 | 0,0495852 | 0,0596425 | 0,0023202 | 0,0058977 | 0,0041206 | 0,0041206 | TRUE | 0,063845 | No |
| orf19.5090 | 566 | 471 | 515 | 362 | 547 | 367 | 530 | 625 | 540 | 498 | 509 | 567 | 438 | 417 | 450 | 508 | 490,91 | 528 | 453 | 0,862 | -0,214 | -2,63 | 0,0085378 | 0,0108894 | 0,0031927 | 0,0054359 | 0,0043092 | 0,0043092 | TRUE | 0,0781562 | No |
| orf19.5092 | 19 | 19 | 16 | 9 | 12 | 14 | 15 | 28 | 18 | 20 | 16 | 14 | 10 | 16 | 13 | 23 | 16,14 | 17 | 16 | 0,888 | -0,172 | -0,462 | 0,6437905 | 0,6762184 | 1,00E-08 | 0,1037995 | 0,0713771 | 0,0713771 | TRUE | 0,6220749 | No |
| orf19.5093 | 3272 | 2830 | 3066 | 2003 | 3053 | 2047 | 2846 | 3067 | 3119 | 2992 | 3029 | 3138 | 2446 | 2328 | 2416 | 2494 | 2745,31 | 3070 | 2421 | 0,79 | -0,341 | -8,548 | 1,25E-17 | 2,66E-17 | 0,0002591 | 0,0026894 | 0,0011563 | 0,0011563 | TRUE | 0,0076644 | No |
| orf19.5094 | 1565 | 1401 | 1487 | 890 | 353 | 263 | 367 | 311 | 1492 | 1481 | 1469 | 1394 | 283 | 299 | 312 | 253 | 872,88 | 1459 | 287 | 0,196 | -2,353 | -33,742 | 1,41E-249 | 2,40E-248 | 0,0007237 | 0,0039723 | 0,0027315 | 0,0027315 | TRUE | 0,074871 | No |
| orf19.5095 | 5115 | 4358 | 4944 | 2903 | 7658 | 5099 | 7370 | 8061 | 4877 | 4608 | 4884 | 4548 | 6136 | 5798 | 6257 | 6554 | 5457,77 | 4729 | 6186 | 1,308 | 0,387 | 8,091 | 5,93E-16 | 1,22E-15 | 0,0017264 | 0,0023921 | 0,0020084 | 0,0020084 | TRUE | 0,0213973 | No |
| orf19.5097 | 740 | 683 | 731 | 424 | 2537 | 1760 | 2381 | 2641 | 706 | 722 | 722 | 664 | 2033 | 2001 | 2021 | 2147 | 1377,13 | 704 | 2050 | 2,908 | 1,54 | 29,317 | 6,23E-189 | 7,99E-188 | 1,00E-08 | 0,0032835 | 0,0016411 | 0,0016411 | TRUE | 0,0135737 | No |
| orf19.5098 | 1832 | 1465 | 1633 | 1049 | 4158 | 2607 | 3944 | 4173 | 1747 | 1549 | 1613 | 1643 | 3332 | 2964 | 3348 | 3393 | 2448,72 | 1638 | 3259 | 1,991 | 0,993 | 17,053 | 3,34E-65 | 1,51E-64 | 0,0025044 | 0,0027618 | 0,0027778 | 0,0027778 | TRUE | 0,0439964 | No |
| orf19.51 | 2375 | 2738 | 2647 | 1457 | 3216 | 2419 | 2864 | 2885 | 2264 | 2895 | 2615 | 2282 | 2577 | 2751 | 2432 | 2346 | 2520,21 | 2514 | 2526 | 1,004 | 0,005 | 0,066 | 0,9473883 | 0,9527635 | 0,0089786 | 0,0027428 | 0,0061113 | 0,0061113 | TRUE | 0,1260814 | No |
| orf19.510 | 44 | 41 | 42 | 19 | 19 | 19 | 12 | 13 | 42 | 43 | 41 | 30 | 15 | 22 | 10 | 11 | 26,77 | 39 | 14 | 0,356 | -1,49 | -4,867 | 1,13E-06 | 1,75E-06 | 1,19E-08 | 0,0634292 | 0,0443612 | 0,0443612 | TRUE | 0,4123632 | No |
| orf19.5100 | 3541 | 3111 | 3497 | 2178 | 7360 | 5148 | 6728 | 7418 | 3376 | 3289 | 3455 | 3412 | 5897 | 5854 | 5712 | 6032 | 4628,35 | 3383 | 5874 | 1,737 | 0,796 | 22,462 | 9,78E-112 | 7,18E-111 | 0,0002926 | 0,002446 | 0,0009645 | 0,0009645 | TRUE | 0,0042834 | No |
| orf19.5101 | 4760 | 4154 | 4725 | 2954 | 4919 | 3388 | 4597 | 4904 | 4538 | 4392 | 4668 | 4628 | 3941 | 3852 | 3903 | 3987 | 4238,76 | 4556 | 3921 | 0,861 | -0,216 | -6,118 | 9,48E-10 | 1,63E-09 | 0,0002288 | 0,0024786 | 0,0009576 | 0,0009576 | TRUE | 0,0072224 | No |
| orf19.5102 | 1683 | 1473 | 1590 | 971 | 1546 | 1047 | 1453 | 1476 | 1605 | 1558 | 1571 | 1521 | 1239 | 1191 | 1234 | 1200 | 1389,62 | 1564 | 1216 | 0,777 | -0,364 | -8,038 | 9,13E-16 | 1,87E-15 | 1,00E-08 | 0,0032728 | 0,0012434 | 0,0012434 | TRUE | 0,0037543 | No |
| orf19.5103 | 569 | 470 | 517 | 356 | 662 | 446 | 671 | 679 | 542 | 497 | 511 | 558 | 530 | 507 | 570 | 552 | 533,4 | 527 | 540 | 1,029 | 0,041 | 0,573 | 0,5669334 | 0,6012302 | 0,0005151 | 0,0051694 | 0,0029274 | 0,0029274 | TRUE | 0,0174632 | No |
| orf19.5104 | 1753 | 1438 | 1462 | 1260 | 3751 | 2784 | 3563 | 3812 | 1671 | 1521 | 1444 | 1974 | 3006 | 3166 | 3025 | 3100 | 2363,21 | 1652 | 3074 | 1,864 | 0,899 | 11,005 | 3,62E-28 | 9,47E-28 | 0,0096975 | 0,0027861 | 0,0059175 | 0,0059175 | TRUE | 0,1981918 | No |
| orf19.5105 | 3309 | 2854 | 3186 | 2153 | 3567 | 2620 | 3353 | 3664 | 3155 | 3018 | 3147 | 3373 | 2858 | 2979 | 2847 | 2979 | 3044,49 | 3173 | 2916 | 0,92 | -0,121 | -2,709 | 0,0067422 | 0,0086406 | 0,0009536 | 0,0026306 | 0,0015715 | 0,0015715 | TRUE | 0,0202558 | No |
| orf19.5106 | 16350 | 14711 | 16180 | 9220 | 15106 | 10026 | 14135 | 14918 | 15588 | 15555 | 15984 | 14444 | 12104 | 11400 | 12001 | 12130 | 13650,71 | 15393 | 11909 | 0,774 | -0,37 | -8,953 | 3,47E-19 | 7,68E-19 | 0,0012416 | 0,0022116 | 0,0015709 | 0,0015709 | TRUE | 0,0207066 | No |
| orf19.5107 | 5907 | 4916 | 5484 | 3731 | 4710 | 3277 | 4618 | 4976 | 5632 | 5198 | 5418 | 5845 | 3774 | 3726 | 3921 | 4046 | 4694,88 | 5523 | 3867 | 0,701 | -0,513 | -10,702 | 9,98E-27 | 2,56E-26 | 0,0016812 | 0,002441 | 0,0019896 | 0,0019896 | TRUE | 0,0189986 | No |
| orf19.511 | 2728 | 2347 | 2506 | 1613 | 2106 | 1330 | 2067 | 2191 | 2601 | 2482 | 2476 | 2527 | 1687 | 1512 | 1755 | 1781 | 2102,65 | 2522 | 1684 | 0,669 | -0,58 | -10,399 | 2,50E-25 | 6,24E-25 | 0,0021886 | 0,0028722 | 0,0024998 | 0,0024998 | TRUE | 0,0540531 | No |
| orf19.5110 | 6078 | 5183 | 5939 | 3857 | 7295 | 4984 | 7189 | 7470 | 5795 | 5480 | 5867 | 6042 | 5845 | 5667 | 6103 | 6074 | 5859,28 | 5796 | 5922 | 1,023 | 0,032 | 0,746 | 0,4557856 | 0,4925263 | 0,0011739 | 0,0023715 | 0,0016202 | 0,0016202 | TRUE | 0,0163432 | No |
| orf19.5112 | 5727 | 4746 | 5123 | 3670 | 15657 | 11252 | 14417 | 16653 | 5460 | 5018 | 5061 | 5749 | 12546 | 12795 | 12240 | 13540 | 9051,14 | 5322 | 12780 | 2,403 | 1,265 | 23,65 | 1,18E-123 | 9,68E-123 | 0,0028963 | 0,0022727 | 0,0026062 | 0,0026062 | TRUE | 0,0345154 | No |
| orf19.5113 | 4356 | 3074 | 3294 | 2418 | 1954 | 1273 | 1709 | 2037 | 4153 | 3250 | 3254 | 3788 | 1566 | 1448 | 1451 | 1656 | 2570,73 | 3611 | 1530 | 0,424 | -1,238 | -14,822 | 1,06E-49 | 3,89E-49 | 0,0089779 | 0,00273 | 0,0062527 | 0,0062527 | TRUE | 0,1260983 | No |
| orf19.5114 | 1064 | 894 | 926 | 509 | 815 | 461 | 766 | 837 | 1014 | 945 | 915 | 797 | 653 | 524 | 650 | 681 | 772,5 | 918 | 627 | 0,683 | -0,55 | -5,902 | 3,59E-09 | 6,06E-09 | 0,0092901 | 0,0042167 | 0,0070165 | 0,0070165 | TRUE | 0,1361445 | No |
| orf19.5114.1 | 22816 | 16304 | 17523 | 9896 | 3346 | 1943 | 3037 | 3272 | 21753 | 17239 | 17311 | 15503 | 2681 | 2209 | 2578 | 2660 | 10241,87 | 17952 | 2532 | 0,141 | -2,825 | -29,818 | 2,26E-195 | 3,04E-194 | 0,0145157 | 0,0022516 | 0,0084163 | 0,0084163 | TRUE | 0,2488598 | No |
| orf19.5117 | 156093 | 130019 | 143837 | 95503 | 209061 | 145622 | 200445 | 218229 | 148818 | 137479 | 142097 | 149611 | 167515 | 165585 | 170177 | 177441 | 157340,26 | 144501 | 170180 | 1,178 | 0,236 | 5,936 | 2,92E-09 | 4,94E-09 | 0,0012541 | 0,0021017 | 0,0015124 | 0,0015124 | TRUE | 0,0131174 | No |
| orf19.5118 | 48886 | 42179 | 46387 | 31412 | 8722 | 6184 | 8307 | 9059 | 46608 | 44599 | 45826 | 49209 | 6989 | 7032 | 7053 | 7366 | 26834,99 | 46560 | 7110 | 0,153 | -2,711 | -68,124 | 0 | 0 | 0,0010639 | 0,0021525 | 0,0014463 | 0,0014463 | TRUE | 0,0178462 | No |
| orf19.512 | 9834 | 8762 | 9776 | 6272 | 16874 | 11934 | 15561 | 17259 | 9376 | 9265 | 9658 | 9825 | 13521 | 13570 | 13211 | 14033 | 11557,33 | 9531 | 13584 | 1,426 | 0,512 | 14,814 | 1,18E-49 | 4,34E-49 | 0,0005555 | 0,0022333 | 0,0010539 | 0,0010539 | TRUE | 0,0061222 | No |
| orf19.5121 | 54 | 68 | 69 | 24 | 78 | 56 | 71 | 67 | 51 | 72 | 68 | 38 | 62 | 64 | 60 | 54 | 58,76 | 57 | 60 | 1,034 | 0,048 | 0,231 | 0,8176395 | 0,8389383 | 0,0146524 | 0,0300338 | 0,0242568 | 0,0242568 | TRUE | 0,3152155 | No |
| orf19.5124 | 1514 | 1207 | 1404 | 1028 | 2315 | 1712 | 2295 | 2566 | 1443 | 1276 | 1387 | 1610 | 1855 | 1947 | 1948 | 2086 | 1694,2 | 1429 | 1959 | 1,374 | 0,458 | 6,569 | 5,06E-11 | 9,12E-11 | 0,0051292 | 0,0030604 | 0,0040497 | 0,0040497 | TRUE | 0,0817823 | No |
| orf19.5125 | 6690 | 6414 | 6442 | 4394 | 4959 | 3900 | 4544 | 4774 | 6378 | 6782 | 6364 | 6883 | 3974 | 4435 | 3858 | 3882 | 5319,43 | 6602 | 4037 | 0,611 | -0,71 | -13,177 | 1,19E-39 | 3,73E-39 | 0,0028019 | 0,0023999 | 0,0025924 | 0,0025924 | TRUE | 0,0525005 | No |
| orf19.5126 | 2656 | 2442 | 2515 | 1882 | 2869 | 2150 | 2720 | 2867 | 2532 | 2582 | 2485 | 2948 | 2299 | 2445 | 2309 | 2331 | 2491,39 | 2637 | 2346 | 0,891 | -0,166 | -2,871 | 0,0040969 | 0,0053276 | 0,0030878 | 0,0027503 | 0,0028225 | 0,0028225 | TRUE | 0,0730394 | No |
| orf19.5129 | 423 | 408 | 448 | 260 | 878 | 740 | 856 | 840 | 403 | 431 | 443 | 407 | 704 | 841 | 727 | 683 | 579,91 | 421 | 739 | 1,747 | 0,805 | 9,817 | 9,50E-23 | 2,25E-22 | 0,0041771 | 0,0049226 | 0,0045259 | 0,0045259 | TRUE | 0,0985533 | No |
| orf19.513 | 0 | 2 | 4 | 1 | 8 | 2 | 6 | 4 | 0 | 2 | 4 | 2 | 6 | 2 | 5 | 3 | 3,08 | 2 | 4 | 2,251 | 1,171 | 1,326 | 0,184774 | 0,210243 | 1,00E-08 | 0,5346662 | 0,3556238 | 0,3556238 | TRUE | 0,4497107 | No |
| orf19.5130 | 3746 | 3192 | 3354 | 2272 | 8922 | 6625 | 8675 | 9632 | 3571 | 3375 | 3313 | 3559 | 7149 | 7533 | 7365 | 7832 | 5462,26 | 3454 | 7470 | 2,163 | 1,113 | 25,153 | 1,30E-139 | 1,22E-138 | 0,0011712 | 0,0023919 | 0,0016576 | 0,0016576 | TRUE | 0,013183 | No |
| orf19.5131 | 5690 | 5102 | 5669 | 3449 | 1463 | 1097 | 1377 | 1495 | 5425 | 5395 | 5600 | 5403 | 1172 | 1247 | 1169 | 1216 | 3328,41 | 5456 | 1201 | 0,22 | -2,186 | -54,681 | 0 | 0 | 1,00E-08 | 0,0025846 | 0,0010596 | 0,0010596 | TRUE | 0,0072504 | No |
| orf19.5132 | 112 | 126 | 116 | 72 | 121 | 91 | 84 | 83 | 107 | 133 | 115 | 113 | 97 | 103 | 71 | 67 | 100,83 | 117 | 84 | 0,721 | -0,472 | -2,772 | 0,0055746 | 0,0071862 | 0,0167629 | 0,0183754 | 0,0177465 | 0,0177465 | TRUE | 0,1881867 | No |
| orf19.5133 | 4115 | 3688 | 4043 | 2614 | 3515 | 2508 | 3182 | 3641 | 3923 | 3900 | 3994 | 4095 | 2816 | 2852 | 2702 | 2960 | 3405,27 | 3978 | 2832 | 0,712 | -0,489 | -11,887 | 1,38E-32 | 3,90E-32 | 0,00055 | 0,0025734 | 0,0013262 | 0,0013262 | TRUE | 0,0119956 | No |
| orf19.5134 | 2455 | 2342 | 2499 | 1475 | 2280 | 1616 | 2139 | 2222 | 2341 | 2476 | 2469 | 2311 | 1827 | 1838 | 1816 | 1807 | 2110,44 | 2399 | 1822 | 0,758 | -0,399 | -9,492 | 2,27E-21 | 5,23E-21 | 0,0001731 | 0,0028693 | 0,0012171 | 0,0012171 | TRUE | 0,006625 | No |
| orf19.5136 | 1439 | 1312 | 1395 | 824 | 1409 | 929 | 1331 | 1507 | 1372 | 1387 | 1378 | 1291 | 1129 | 1056 | 1130 | 1225 | 1246,11 | 1357 | 1135 | 0,837 | -0,257 | -4,511 | 6,47E-06 | 9,66E-06 | 0,0013244 | 0,0034089 | 0,0023225 | 0,0023225 | TRUE | 0,0343377 | No |
| orf19.5137 | 12 | 19 | 14 | 9 | 22 | 19 | 21 | 16 | 11 | 20 | 14 | 14 | 18 | 22 | 18 | 13 | 16,19 | 15 | 18 | 1,169 | 0,225 | 0,632 | 0,5274353 | 0,5636518 | 1,00E-08 | 0,1034969 | 0,0590685 | 0,0590685 | TRUE | 0,2617129 | No |
| orf19.5137.1 | 1841 | 1524 | 1549 | 1120 | 1631 | 1304 | 1429 | 1583 | 1755 | 1611 | 1530 | 1755 | 1307 | 1483 | 1213 | 1287 | 1492,68 | 1663 | 1322 | 0,795 | -0,331 | -4,622 | 3,79E-06 | 5,74E-06 | 0,0050628 | 0,0031913 | 0,004252 | 0,004252 | TRUE | 0,0771534 | No |
| orf19.5138 | 336 | 350 | 342 | 206 | 861 | 636 | 848 | 858 | 320 | 370 | 338 | 323 | 690 | 723 | 720 | 698 | 522,71 | 338 | 708 | 2,09 | 1,064 | 14,806 | 1,35E-49 | 4,94E-49 | 1,00E-08 | 0,0052324 | 0,0026815 | 0,0026815 | TRUE | 0,047738 | No |
| orf19.5139 | 2678 | 2458 | 2651 | 1496 | 2244 | 1525 | 2057 | 2210 | 2553 | 2599 | 2619 | 2344 | 1798 | 1734 | 1746 | 1797 | 2148,77 | 2529 | 1769 | 0,699 | -0,518 | -10,988 | 4,38E-28 | 1,14E-27 | 0,0007884 | 0,0028554 | 0,0016557 | 0,0016557 | TRUE | 0,0269745 | No |
| orf19.514 | 861 | 806 | 796 | 514 | 1423 | 1001 | 1302 | 1267 | 821 | 852 | 786 | 805 | 1140 | 1138 | 1105 | 1030 | 959,84 | 816 | 1103 | 1,351 | 0,434 | 7,441 | 1,00E-13 | 1,95E-13 | 0,0009131 | 0,0038019 | 0,0021707 | 0,0021707 | TRUE | 0,0246115 | No |
| orf19.5140 | 1597 | 1337 | 1486 | 903 | 70 | 77 | 93 | 95 | 1523 | 1414 | 1468 | 1415 | 56 | 88 | 79 | 77 | 764,84 | 1455 | 75 | 0,051 | -4,297 | -43,039 | 0 | 0 | 0,0009136 | 0,004238 | 0,003213 | 0,003213 | TRUE | 0,2943515 | No |
| orf19.5141 | 15 | 25 | 20 | 18 | 40 | 29 | 31 | 39 | 14 | 26 | 20 | 28 | 32 | 33 | 26 | 32 | 26,47 | 22 | 31 | 1,415 | 0,5 | 1,737 | 0,082302 | 0,0969662 | 1,00E-08 | 0,0641238 | 0,0395156 | 0,0395156 | TRUE | 0,3563789 | No |
| orf19.5142 | 81 | 86 | 82 | 52 | 320 | 209 | 299 | 344 | 77 | 91 | 81 | 81 | 256 | 238 | 254 | 280 | 169,78 | 82 | 257 | 3,117 | 1,64 | 13,137 | 2,01E-39 | 6,30E-39 | 1,00E-08 | 0,011762 | 0,0065253 | 0,0065253 | TRUE | 0,0450549 | No |

| | | | | | | | | | | | | | | | | | | | | | | | | | | | | | | | |
|---|---|---|---|---|---|---|---|---|---|---|---|---|---|---|---|---|---|---|---|---|---|---|---|---|---|---|---|---|---|---|---|
| orf19.5143 | 3716 | 2939 | 3394 | 2010 | 6559 | 4270 | 6497 | 6727 | 3543 | 3108 | 3353 | 3149 | 5256 | 4855 | 5516 | 5470 | 4281,08 | 3288 | 5274 | 1,603 | 0,681 | 11,809 | 3,50E-32 | 9,82E-32 | 0,0031694 | 0,0024748 | 0,0029397 | 0,0029397 | TRUE | 0,0342981 | No |
| orf19.5144 | 182 | 142 | 166 | 112 | 188 | 128 | 157 | 162 | 174 | 150 | 164 | 175 | 151 | 146 | 133 | 132 | 153,04 | 166 | 140 | 0,847 | -0,239 | -2,029 | 0,0424123 | 0,0513146 | 1,00E-08 | 0,01282 | 0,0067717 | 0,0067717 | TRUE | 0,0425772 | No |
| orf19.5145 | 482 | 429 | 468 | 289 | 387 | 264 | 371 | 407 | 460 | 454 | 462 | 453 | 310 | 300 | 315 | 331 | 385,55 | 457 | 314 | 0,688 | -0,54 | -7,219 | 5,23E-13 | 1,00E-12 | 1,00E-08 | 0,0063499 | 0,0027315 | 0,0027315 | TRUE | 0,0151247 | No |
| orf19.5147 | 1647 | 1644 | 1602 | 1001 | 2486 | 1863 | 2343 | 2553 | 1570 | 1738 | 1583 | 1568 | 1992 | 2118 | 1989 | 2076 | 1829,34 | 1615 | 2044 | 1,264 | 0,338 | 6,697 | 2,13E-11 | 3,88E-11 | 0,0010577 | 0,0029888 | 0,0018838 | 0,0018838 | TRUE | 0,0323896 | No |
| orf19.5148 | 4093 | 3575 | 3891 | 2390 | 3514 | 2376 | 3278 | 3502 | 3902 | 3780 | 3844 | 3744 | 2816 | 2702 | 2783 | 2847 | 3302,28 | 3818 | 2787 | 0,73 | -0,454 | -12,538 | 4,61E-36 | 1,37E-35 | 7,32E-05 | 0,0025885 | 0,000951 | 0,000951 | TRUE | 0,0048186 | No |
| orf19.515 | 489 | 393 | 450 | 282 | 621 | 346 | 548 | 595 | 466 | 416 | 445 | 442 | 498 | 393 | 465 | 484 | 451,02 | 442 | 460 | 1,044 | 0,062 | 0,718 | 0,4730355 | 0,5094413 | 0,0041078 | 0,0057317 | 0,0049283 | 0,0049283 | TRUE | 0,1013082 | No |
| orf19.5155 | 340 | 282 | 292 | 203 | 652 | 505 | 578 | 667 | 324 | 298 | 288 | 318 | 522 | 574 | 491 | 542 | 419,82 | 307 | 532 | 1,733 | 0,793 | 9,658 | 4,56E-22 | 1,07E-21 | 0,001582 | 0,0060023 | 0,0038146 | 0,0038146 | TRUE | 0,0322053 | No |
| orf19.5156 | 708 | 652 | 604 | 442 | 2056 | 1548 | 1928 | 2095 | 675 | 689 | 597 | 692 | 1647 | 1760 | 1637 | 1703 | 1175,18 | 663 | 1687 | 2,547 | 1,349 | 22,313 | 2,76E-110 | 2,00E-109 | 0,0014798 | 0,0034884 | 0,0023995 | 0,0023995 | TRUE | 0,0564916 | No |
| orf19.5157 | 260 | 266 | 262 | 161 | 575 | 432 | 512 | 569 | 248 | 281 | 259 | 252 | 461 | 491 | 435 | 463 | 361,19 | 260 | 462 | 1,773 | 0,826 | 9,906 | 3,91E-23 | 9,34E-23 | 1,00E-08 | 0,0066372 | 0,0035791 | 0,0035791 | TRUE | 0,0344011 | No |
| orf19.5158 | 1254 | 1225 | 1186 | 810 | 1061 | 851 | 921 | 942 | 1196 | 1295 | 1172 | 1269 | 850 | 968 | 782 | 766 | 1037,13 | 1233 | 842 | 0,681 | -0,553 | -7,188 | 6,58E-13 | 1,25E-12 | 0,0057445 | 0,0036744 | 0,0047004 | 0,0047004 | TRUE | 0,1156499 | No |
| orf19.5159 | 3966 | 3631 | 3816 | 2338 | 1704 | 1193 | 1589 | 1639 | 3781 | 3839 | 3770 | 3663 | 1365 | 1357 | 1349 | 1333 | 2557,07 | 3763 | 1351 | 0,359 | -1,479 | -38,432 | 0 | 0 | 1,00E-08 | 0,0027334 | 0,0009435 | 0,0009435 | TRUE | 0,0035767 | No |
| orf19.516 | 299 | 300 | 339 | 178 | 637 | 441 | 586 | 590 | 285 | 317 | 335 | 279 | 510 | 501 | 498 | 480 | 400,64 | 304 | 497 | 1,629 | 0,704 | 8,675 | 4,13E-18 | 8,89E-18 | 3,09E-05 | 0,0061895 | 0,0035953 | 0,0035953 | TRUE | 0,0536392 | No |
| orf19.5160 | 2369 | 2014 | 2272 | 1312 | 3510 | 2538 | 3381 | 3417 | 2259 | 2130 | 2245 | 2055 | 2812 | 2886 | 2870 | 2778 | 2504,4 | 2172 | 2836 | 1,303 | 0,382 | 8,756 | 2,02E-18 | 4,38E-18 | 0,0006099 | 0,0027469 | 0,0014123 | 0,0014123 | TRUE | 0,0143124 | No |
| orf19.5161 | 1797 | 1459 | 1456 | 1054 | 1816 | 1355 | 1775 | 1912 | 1713 | 1543 | 1438 | 1651 | 1455 | 1541 | 1507 | 1555 | 1550,37 | 1586 | 1514 | 0,955 | -0,066 | -1,086 | 0,2773665 | 0,3088788 | 0,0024877 | 0,0031503 | 0,0029274 | 0,0029274 | TRUE | 0,048917 | No |
| orf19.5162 | 1214 | 1168 | 1243 | 754 | 1399 | 949 | 1293 | 1434 | 1157 | 1235 | 1228 | 1181 | 1121 | 1079 | 1098 | 1166 | 1158,17 | 1200 | 1116 | 0,93 | -0,104 | -2,031 | 0,042277 | 0,0511703 | 1,00E-08 | 0,0035089 | 0,0016717 | 0,0016717 | TRUE | 0,0110054 | No |
| orf19.5163 | 1342 | 1263 | 1325 | 825 | 1906 | 1368 | 1847 | 1919 | 1279 | 1335 | 1309 | 1292 | 1527 | 1556 | 1568 | 1560 | 1428,44 | 1304 | 1553 | 1,19 | 0,251 | 5,724 | 1,04E-08 | 1,73E-08 | 1,00E-08 | 0,0032407 | 0,0011368 | 0,0011368 | TRUE | 0,0032296 | No |
| orf19.5164 | 358 | 341 | 351 | 271 | 2179 | 1580 | 1999 | 2104 | 341 | 361 | 347 | 425 | 1746 | 1797 | 1697 | 1711 | 1052,95 | 368 | 1738 | 4,762 | 2,252 | 33,786 | 3,17E-250 | 5,41E-249 | 0,000421 | 0,0036506 | 0,0024814 | 0,0024814 | TRUE | 0,0979634 | No |
| orf19.5165 | 1155 | 1057 | 1172 | 650 | 1180 | 779 | 1103 | 1164 | 1101 | 1118 | 1158 | 1018 | 946 | 886 | 936 | 946 | 1013,63 | 1099 | 928 | 0,844 | -0,245 | -4,338 | 1,44E-05 | 2,11E-05 | 0,0005845 | 0,0037111 | 0,0020741 | 0,0020741 | TRUE | 0,0253157 | No |
| orf19.5166 | 1661 | 1379 | 1626 | 1060 | 1955 | 1413 | 2043 | 2007 | 1584 | 1458 | 1606 | 1661 | 1566 | 1607 | 1734 | 1632 | 1606,02 | 1577 | 1635 | 1,038 | 0,054 | 0,969 | 0,3325575 | 0,3663127 | 0,0016053 | 0,0031136 | 0,0023332 | 0,0023332 | TRUE | 0,0309743 | No |
| orf19.5167 | 5764 | 4837 | 5481 | 3292 | 5737 | 3817 | 5617 | 5884 | 5495 | 5115 | 5415 | 5157 | 4597 | 4340 | 4769 | 4784 | 4958,99 | 5296 | 4622 | 0,873 | -0,196 | -4,319 | 1,56E-05 | 2,30E-05 | 0,0013577 | 0,0024224 | 0,0017736 | 0,0017736 | TRUE | 0,0200309 | No |
| orf19.5168 | 4005 | 3439 | 3767 | 2363 | 4709 | 3095 | 4458 | 4783 | 3818 | 3636 | 3721 | 3702 | 3773 | 3519 | 3785 | 3889 | 3730,52 | 3719 | 3742 | 1,006 | 0,009 | 0,222 | 0,8240776 | 0,8447291 | 0,0008228 | 0,0025314 | 0,0014218 | 0,0014218 | TRUE | 0,0187149 | No |
| orf19.5169 | 201 | 179 | 175 | 131 | 308 | 259 | 280 | 332 | 192 | 189 | 173 | 205 | 247 | 295 | 238 | 270 | 226 | 190 | 262 | 1,383 | 0,468 | 4,368 | 1,26E-05 | 1,85E-05 | 0,0029334 | 0,0093565 | 0,0063793 | 0,0063793 | TRUE | 0,0695353 | No |
| orf19.517 | 1288 | 1097 | 1225 | 870 | 1998 | 1531 | 1884 | 1887 | 1228 | 1160 | 1210 | 1363 | 1601 | 1741 | 1600 | 1534 | 1429,58 | 1240 | 1619 | 1,307 | 0,387 | 6,237 | 4,47E-10 | 7,80E-10 | 0,0029395 | 0,0032398 | 0,0029594 | 0,0029594 | TRUE | 0,0483532 | No |
| orf19.5170 | 6910 | 6510 | 6886 | 4192 | 10618 | 7342 | 9683 | 10793 | 6588 | 6884 | 6803 | 6567 | 8508 | 8349 | 8221 | 8776 | 7586,77 | 6710 | 8464 | 1,261 | 0,335 | 9,41 | 4,98E-21 | 1,14E-20 | 0,0004952 | 0,0023077 | 0,0010777 | 0,0010777 | TRUE | 0,0076064 | No |
| orf19.5171 | 1216 | 1195 | 1325 | 876 | 1694 | 1331 | 1629 | 1819 | 1159 | 1264 | 1309 | 1372 | 1357 | 1513 | 1383 | 1479 | 1354,63 | 1276 | 1433 | 1,125 | 0,17 | 2,664 | 0,0077227 | 0,0098616 | 0,0028464 | 0,0033034 | 0,0031354 | 0,0031354 | TRUE | 0,0463496 | No |
| orf19.5172 | 13 | 7 | 8 | 4 | 4 | 0 | 3 | 5 | 12 | 7 | 8 | 6 | 3 | 0 | 3 | 4 | 5,47 | 8 | 2 | 0,3 | -1,737 | -2,608 | 0,0091196 | 0,0116176 | 1,00E-08 | 0,3021007 | 0,1898014 | 0,1898014 | TRUE | 0,3549311 | No |
| orf19.5174 | 925 | 672 | 694 | 452 | 1440 | 853 | 1340 | 1514 | 882 | 711 | 686 | 708 | 1154 | 970 | 1138 | 1231 | 934,82 | 747 | 1123 | 1,504 | 0,589 | 6,224 | 4,85E-10 | 8,45E-10 | 0,0107231 | 0,0038477 | 0,0074443 | 0,0074443 | TRUE | 0,1817682 | No |
| orf19.5175 | 3734 | 3341 | 3649 | 2526 | 7412 | 5608 | 6732 | 7490 | 3560 | 3533 | 3605 | 3957 | 5939 | 6377 | 5715 | 6090 | 4847 | 3664 | 6030 | 1,647 | 0,72 | 14,182 | 1,18E-45 | 4,10E-45 | 0,0022254 | 0,00243 | 0,0022485 | 0,0022485 | TRUE | 0,0337777 | No |
| orf19.5177 | 2857 | 2534 | 2720 | 1662 | 1009 | 766 | 971 | 1107 | 2724 | 2679 | 2687 | 2604 | 808 | 871 | 824 | 900 | 1762,24 | 2674 | 851 | 0,318 | -1,654 | -33,698 | 6,14E-249 | 1,03E-247 | 0,0003714 | 0,003023 | 0,0015779 | 0,0015779 | TRUE | 0,0197178 | No |
| orf19.5178 | 3075 | 2641 | 2897 | 1801 | 6151 | 4510 | 5990 | 6569 | 2932 | 2793 | 2862 | 2821 | 4929 | 5128 | 5085 | 5341 | 3986,39 | 2852 | 5121 | 1,795 | 0,844 | 21,179 | 1,50E-99 | 9,77E-99 | 0,000633 | 0,0025032 | 0,0012399 | 0,0012399 | TRUE | 0,0104575 | No |
| orf19.5179 | 23 | 19 | 29 | 25 | 37 | 24 | 21 | 26 | 22 | 20 | 29 | 39 | 30 | 27 | 18 | 21 | 25,72 | 28 | 24 | 0,889 | -0,17 | -0,554 | 0,579408 | 0,6135417 | 0,029562 | 0,0659358 | 0,0509257 | 0,0509257 | TRUE | 0,3622423 | No |
| orf19.518 | 7857 | 7052 | 7651 | 4481 | 11683 | 8196 | 11455 | 12531 | 7491 | 7457 | 7558 | 7020 | 9361 | 9320 | 9725 | 10189 | 8515,07 | 7382 | 9649 | 1,307 | 0,386 | 9,054 | 1,37E-19 | 3,06E-19 | 0,0012653 | 0,0022841 | 0,001622 | 0,001622 | TRUE | 0,017423 | No |
| orf19.5180 | 1165 | 862 | 1007 | 680 | 1441 | 900 | 1394 | 1461 | 1111 | 911 | 995 | 1065 | 1155 | 1023 | 1183 | 1188 | 1078,96 | 1020 | 1137 | 1,117 | 0,159 | 2,134 | 0,0328396 | 0,0400822 | 0,0047074 | 0,003613 | 0,0043884 | 0,0043884 | TRUE | 0,0626486 | No |
| orf19.5181 | 9089 | 7678 | 8503 | 4947 | 11940 | 7590 | 11343 | 12007 | 8665 | 8119 | 8400 | 7750 | 9567 | 8631 | 9630 | 9763 | 8815,56 | 8234 | 9398 | 1,141 | 0,191 | 3,679 | 0,0002338 | 0,0003254 | 0,0025013 | 0,0022775 | 0,002461 | 0,002461 | TRUE | 0,0365289 | No |
| orf19.5182 | 3613 | 3139 | 3584 | 2115 | 6031 | 4246 | 5766 | 6391 | 3445 | 3319 | 3541 | 3313 | 4832 | 4828 | 4895 | 5196 | 4171,25 | 3404 | 4938 | 1,45 | 0,536 | 12,719 | 4,64E-37 | 1,40E-36 | 0,0008113 | 0,0024849 | 0,0014486 | 0,0014486 | TRUE | 0,0152836 | No |
| orf19.5184 | 4994 | 3980 | 4524 | 2704 | 5978 | 3716 | 5884 | 6049 | 4761 | 4208 | 4469 | 4236 | 4790 | 4225 | 4995 | 4918 | 4575,52 | 4418 | 4732 | 1,071 | 0,099 | 1,587 | 0,1124089 | 0,130391 | 0,0041045 | 0,0024501 | 0,0035102 | 0,0035102 | TRUE | 0,0624384 | No |
| orf19.5188 | 217 | 212 | 214 | 137 | 247 | 203 | 230 | 269 | 207 | 224 | 211 | 215 | 198 | 231 | 195 | 219 | 212,48 | 214 | 211 | 0,981 | -0,028 | -0,277 | 0,7818525 | 0,805457 | 1,00E-08 | 0,0098187 | 0,0052778 | 0,0052778 | TRUE | 0,0399865 | No |
| orf19.519 | 1112 | 1039 | 1080 | 610 | 1436 | 1091 | 1456 | 1471 | 1060 | 1099 | 1067 | 956 | 1151 | 1241 | 1236 | 1196 | 1125,59 | 1046 | 1206 | 1,149 | 0,201 | 3,512 | 0,0004449 | 0,0006116 | 0,0011791 | 0,00355 | 0,0022273 | 0,0022273 | TRUE | 0,0346572 | No |
| orf19.5190 | 1 | 2 | 1 | 1 | 2 | 7 | 3 | 3 | 1 | 2 | 1 | 2 | 2 | 8 | 3 | 2 | 2,52 | 2 | 4 | 2,529 | 1,339 | 1,354 | 0,1757606 | 0,2003445 | 1,00E-08 | 0,6532836 | 0,4105814 | 0,4105814 | TRUE | 0,8419987 | No |
| orf19.5191 | 41 | 32 | 25 | 23 | 114 | 86 | 110 | 113 | 39 | 34 | 25 | 36 | 91 | 98 | 93 | 92 | 63,51 | 34 | 94 | 2,811 | 1,491 | 7,654 | 1,94E-14 | 3,86E-14 | 1,00E-08 | 0,0279451 | 0,0150163 | 0,0150163 | TRUE | 0,2407941 | No |
| orf19.5194.1 | 876 | 703 | 791 | 491 | 1041 | 776 | 996 | 1116 | 835 | 743 | 781 | 769 | 834 | 882 | 846 | 907 | 824,83 | 782 | 867 | 1,107 | 0,147 | 2,398 | 0,0164669 | 0,0205909 | 0,0008969 | 0,0040819 | 0,0023555 | 0,0023555 | TRUE | 0,026137 | No |
| orf19.5195 | 2321 | 1915 | 2105 | 1367 | 3502 | 2423 | 3384 | 3624 | 2213 | 2025 | 2080 | 2141 | 2806 | 2755 | 2873 | 2947 | 2479,95 | 2115 | 2845 | 1,346 | 0,429 | 9,585 | 9,21E-22 | 2,14E-21 | 0,0007375 | 0,0027534 | 0,0015 | 0,0015 | TRUE | 0,0127573 | No |
| orf19.5196 | 2115 | 1632 | 1841 | 1094 | 2653 | 1662 | 2556 | 2580 | 2016 | 1726 | 1819 | 1714 | 2126 | 1890 | 2170 | 2098 | 1944,75 | 1819 | 2071 | 1,138 | 0,187 | 2,84 | 0,0045102 | 0,0058424 | 0,0040734 | 0,0029356 | 0,0036219 | 0,0036219 | TRUE | 0,0657131 | No |
| orf19.5197 | 9023 | 7635 | 8235 | 5573 | 11432 | 8336 | 10807 | 11943 | 8602 | 8073 | 8135 | 8730 | 9160 | 9479 | 9175 | 9711 | 8883,26 | 8385 | 9381 | 1,119 | 0,162 | 4,042 | 5,29E-05 | 7,59E-05 | 0,0010194 | 0,0022761 | 0,0014349 | 0,0014349 | TRUE | 0,00905 | No |
| orf19.5198 | 24411 | 21059 | 22778 | 13004 | 25979 | 16512 | 25543 | 26215 | 23273 | 22267 | 22503 | 20371 | 20816 | 18776 | 21686 | 21315 | 21375,94 | 22104 | 20648 | 0,934 | -0,098 | -1,737 | 0,0823866 | 0,0970331 | 0,0034678 | 0,0021681 | 0,003033 | 0,003033 | TRUE | 0,0454483 | No |
| orf19.52 | 2521 | 2346 | 2439 | 1556 | 2929 | 2193 | 2770 | 2815 | 2404 | 2481 | 2410 | 2438 | 2347 | 2494 | 2352 | 2289 | 2401,54 | 2433 | 2370 | 0,973 | -0,039 | -0,929 | 0,352681 | 0,3874088 | 0,0003417 | 0,002775 | 0,0012534 | 0,0012534 | TRUE | 0,0140301 | No |
| orf19.520 | 1483 | 1169 | 1360 | 947 | 2026 | 1511 | 2015 | 2095 | 1414 | 1236 | 1344 | 1484 | 1623 | 1718 | 1711 | 1703 | 1529,09 | 1370 | 1689 | 1,235 | 0,305 | 5,141 | 2,73E-07 | 4,33E-07 | 0,0024328 | 0,0031651 | 0,0027053 | 0,0027053 | TRUE | 0,0509913 | No |
| orf19.5200 | 3239 | 2810 | 3164 | 1967 | 5126 | 3721 | 4870 | 5234 | 3088 | 2971 | 3126 | 3081 | 4107 | 4231 | 4135 | 4256 | 3624,4 | 3066 | 4182 | 1,364 | 0,447 | 12,573 | 2,96E-36 | 8,82E-36 | 0,0001226 | 0,0025443 | 0,0009256 | 0,0009256 | TRUE | 0,0054869 | No |
| orf19.5201 | 2906 | 2511 | 2676 | 1745 | 2695 | 2134 | 2542 | 2803 | 2771 | 2655 | 2644 | 2734 | 2159 | 2427 | 2158 | 2279 | 2478,27 | 2701 | 2256 | 0,834 | -0,261 | -5,319 | 1,05E-07 | 1,68E-07 | 0,0013052 | 0,0027538 | 0,0019059 | 0,0019059 | TRUE | 0,0300884 | No |
| orf19.5201.1 | 7393 | 6652 | 6500 | 3999 | 7103 | 4351 | 6685 | 6641 | 7048 | 7034 | 6421 | 6265 | 5691 | 4947 | 5676 | 5400 | 6060,29 | 6692 | 5428 | 0,811 | -0,302 | -5,095 | 3,49E-07 | 5,51E-07 | 0,0036864 | 0,0023622 | 0,0032063 | 0,0032063 | TRUE | 0,042809 | No |
| orf19.5203 | 507 | 418 | 497 | 298 | 1297 | 944 | 1287 | 1338 | 483 | 442 | 491 | 467 | 1039 | 1073 | 1093 | 1088 | 772,05 | 471 | 1073 | 2,277 | 1,187 | 19,461 | 2,34E-84 | 1,32E-83 | 1,00E-08 | 0,004218 | 0,0019755 | 0,0019755 | TRUE | 0,0214478 | No |
| orf19.5204 | 2145 | 1739 | 1910 | 1462 | 1236 | 877 | 1196 | 1289 | 2045 | 1839 | 1887 | 2290 | 990 | 997 | 1015 | 1048 | 1514,01 | 2015 | 1012 | 0,504 | -0,99 | -14,095 | 4,10E-45 | 1,40E-44 | 0,0044307 | 0,0031758 | 0,0040122 | 0,0040122 | TRUE | 0,097138 | No |
| orf19.5205 | 727 | 696 | 758 | 551 | 643 | 552 | 621 | 621 | 693 | 736 | 749 | 863 | 515 | 628 | 527 | 505 | 652,01 | 760 | 544 | 0,716 | -0,483 | -5,415 | 6,14E-08 | 9,96E-08 | 0,0078914 | 0,0046095 | 0,0060696 | 0,0060696 | TRUE | 0,1196072 | No |
| orf19.5206 | 1856 | 1730 | 1828 | 1255 | 2453 | 1882 | 2374 | 2550 | 1769 | 1829 | 1806 | 1966 | 1966 | 2140 | 2016 | 2073 | 1945,64 | 1842 | 2049 | 1,113 | 0,155 | 3,132 | 0,0017388 | 0,0023135 | 0,0010715 | 0,0029352 | 0,0018217 | 0,0018217 | TRUE | 0,0223618 | No |
| orf19.5207 | 4564 | 3931 | 4190 | 2407 | 4345 | 2945 | 4228 | 4469 | 4351 | 4157 | 4139 | 3771 | 3482 | 3349 | 3590 | 3634 | 3808,92 | 4104 | 3514 | 0,855 | -0,225 | -4,408 | 1,04E-05 | 1,54E-05 | 0,0020451 | 0,0025223 | 0,0022459 | 0,0022459 | TRUE | 0,0349022 | No |
| orf19.5209 | 364 | 406 | 409 | 233 | 610 | 495 | 592 | 588 | 347 | 429 | 404 | 365 | 489 | 563 | 503 | 478 | 447,22 | 386 | 508 | 1,31 | 0,39 | 4,382 | 1,18E-05 | 1,74E-05 | 0,0047593 | 0,0057627 | 0,0052704 | 0,0052704 | TRUE | 0,064779 | No |
| orf19.5210 | 395 | 315 | 393 | 291 | 1017 | 800 | 1022 | 987 | 377 | 333 | 388 | 456 | 815 | 910 | 868 | 803 | 618,57 | 388 | 849 | 2,197 | 1,135 | 12,954 | 2,23E-38 | 6,87E-38 | 0,0061955 | 0,0047456 | 0,0053957 | 0,0053957 | TRUE | 0,1343959 | No |
| orf19.5210.1 | 59 | 59 | 71 | 65 | 82 | 68 | 63 | 56 | 56 | 62 | 70 | 102 | 66 | 77 | 53 | 46 | 66,58 | 72 | 60 | 0,84 | -0,252 | -1,109 | 0,2674958 | 0,2983032 | 0,0436334 | 0,0267515 | 0,0342998 | 0,0342998 | TRUE | 0,5317586 | No |
| orf19.5211 | 628 | 489 | 505 | 377 | 2781 | 1936 | 2622 | 2911 | 599 | 517 | 499 | 591 | 2228 | 2201 | 2226 | 2367 | 1403,5 | 552 | 2256 | 4,104 | 2,037 | 31,155 | 4,33E-213 | 6,25E-212 | 0,0024572 | 0,0032611 | 0,0028893 | 0,0028893 | TRUE | 0,0496753 | No |
| orf19.5212 | 11 | 17 | 17 | 13 | 43 | 47 | 35 | 52 | 10 | 18 | 17 | 20 | 34 | 53 | 30 | 42 | 28,19 | 16 | 40 | 2,462 | 1,3 | 4,155 | 3,26E-05 | 4,72E-05 | 0,0319452 | 0,0603365 | 0,0482337 | 0,0482337 | TRUE | 0,2426374 | No |
| orf19.5213 | 2701 | 2517 | 2825 | 1812 | 3032 | 2189 | 2917 | 3203 | 2575 | 2661 | 2791 | 2839 | 2429 | 2489 | 2477 | 2604 | 2608,17 | 2716 | 2500 | 0,921 | -0,118 | -2,557 | 0,010564 | 0,0133987 | 0,0009104 | 0,0027208 | 0,0016657 | 0,0016657 | TRUE | 0,0151048 | No |
| orf19.5213.1 | 7334 | 5872 | 6094 | 3668 | 11172 | 6959 | 10058 | 11199 | 6992 | 6209 | 6020 | 5746 | 8952 | 7913 | 8539 | 9106 | 7434,67 | 6242 | 8628 | 1,382 | 0,467 | 7,049 | 1,80E-12 | 3,37E-12 | 0,0053986 | 0,0023121 | 0,0040658 | 0,0040658 | TRUE | 0,080202 | No |
| orf19.5213.2 | 4602 | 3548 | 3554 | 2991 | 6326 | 4230 | 5472 | 5902 | 4388 | 3752 | 3511 | 4686 | 5069 | 4810 | 4646 | 4799 | 4457,37 | 4084 | 4831 | 1,184 | 0,243 | 2,992 | 0,0027735 | 0,0036452 | 0,0092805 | 0,0024596 | 0,0061279 | 0,0061279 | TRUE | 0,1175797 | No |
| orf19.5214 | 979 | 892 | 1021 | 611 | 960 | 693 | 898 | 953 | 933 | 943 | 1009 | 957 | 769 | 788 | 762 | 775 | 867,11 | 960 | 774 | 0,805 | -0,313 | -5,746 | 9,13E-09 | 1,52E-08 | 1,00E-08 | 0,0039848 | 0,0016983 | 0,0016983 | TRUE | 0,014623 | No |
| orf19.5215 | 2342 | 1940 | 2077 | 1465 | 2738 | 1855 | 2717 | 2797 | 2233 | 2051 | 2052 | 2295 | 2194 | 2109 | 2307 | 2274 | 2189,39 | 2158 | 2221 | 1,031 | 0,044 | 0,83 | 0,4064376 | 0,4423466 | 0,0018643 | 0,0028412 | 0,0022811 | 0,0022811 | TRUE | 0,0205381 | No |
| orf19.5216 | 677 | 661 | 633 | 427 | 1042 | 739 | 931 | 1036 | 645 | 699 | 625 | 669 | 835 | 840 | 790 | 842 | 743,33 | 660 | 827 | 1,255 | 0,328 | 5,361 | 8,25E-08 | 1,33E-07 | 1,00E-08 | 0,0043001 | 0,0021959 | 0,0021959 | TRUE | 0,0205937 | No |
| orf19.5217 | 494 | 443 | 447 | 301 | 2158 | 1450 | 2039 | 2076 | 471 | 468 | 442 | 472 | 1729 | 1649 | 1731 | 1688 | 1081,19 | 463 | 1699 | 3,676 | 1,878 | 33,424 | 6,15E-245 | 1,02E-243 | 1,00E-08 | 0,0036099 | 0,0015683 | 0,0015683 | TRUE | 0,0121938 | No |
| orf19.5219 | 4002 | 3935 | 4095 | 2418 | 4461 | 3232 | 3826 | 4193 | 3815 | 4161 | 4045 | 3788 | 3574 | 3675 | 3248 | 3409 | 3714,59 | 3952 | 3476 | 0,879 | -0,186 | -3,573 | 0,0003529 | 0,0004874 | 0,0021404 | 0,0025333 | 0,0023379 | 0,0023379 | TRUE | 0,0237814 | No |
| orf19.522 | 6315 | 5637 | 5977 | 3673 | 8715 | 6173 | 8251 | 8874 | 6021 | 5960 | 5905 | 5754 | 6983 | 7019 | 7005 | 7215 | 6482,82 | 5910 | 7056 | 1,193 | 0,255 | 8,191 | 2,59E-16 | 5,36E-16 | 9,57E-05 | 0,0023446 | 0,0007707 | 0,0007707 | TRUE | 0,0035711 | No |
| orf19.5220 | 3641 | 3213 | 3570 | 2191 | 2832 | 1952 | 2785 | 2695 | 3471 | 3397 | 3527 | 3432 | 2269 | 2220 | 2364 | 2191 | 2859,04 | 3457 | 2261 | 0,654 | -0,612 | -15,343 | 3,93E-53 | 1,51E-52 | 0,0003218 | 0,0026656 | 0,0011696 | 0,0011696 | TRUE | 0,0115917 | No |
| orf19.5221 | 1654 | 1556 | 1734 | 1039 | 1092 | 767 | 1043 | 1106 | 1577 | 1645 | 1713 | 1628 | 875 | 872 | 886 | 899 | 1261,85 | 1641 | 883 | 0,538 | -0,894 | -18,077 | 4,85E-73 | 2,42E-72 | 1,00E-08 | 0,0033925 | 0,0015001 | 0,0015001 | TRUE | 0,0112579 | No |
| orf19.5223 | 29 | 28 | 29 | 15 | 41 | 24 | 25 | 24 | 28 | 30 | 29 | 23 | 33 | 27 | 21 | 20 | 26,29 | 28 | 25 | 0,913 | -0,131 | -0,463 | 0,6434511 | 0,6759733 | 1,00E-08 | 0,0645556 | 0,0386213 | 0,0386213 | TRUE | 0,3130189 | No |
| orf19.5224 | 2674 | 2587 | 2788 | 1771 | 4436 | 3263 | 4200 | 4308 | 2549 | 2735 | 2754 | 2774 | 3554 | 3710 | 3566 | 3503 | 3143,35 | 2703 | 3583 | 1,326 | 0,407 | 9,615 | 6,94E-22 | 1,62E-21 | 0,000614 | 0,0026136 | 0,0013889 | 0,0013889 | TRUE | 0,0184529 | No |

| | | | | | | | | | | | | | | | | | | | | | | | | | | | | | | | |
|---|---|---|---|---|---|---|---|---|---|---|---|---|---|---|---|---|---|---|---|---|---|---|---|---|---|---|---|---|---|---|---|
| orf19.5225 | 556 | 561 | 519 | 346 | 746 | 618 | 692 | 831 | 530 | 593 | 513 | 542 | 598 | 703 | 588 | 676 | 592,71 | 544 | 641 | 1,175 | 0,233 | 2,86 | 0,0042428 | 0,0055094 | 0,0044623 | 0,0048614 | 0,0046514 | 0,0046514 | TRUE | 0,0466389 | No |
| orf19.5225.2 | 14223 | 11965 | 12432 | 9059 | 37149 | 27179 | 35361 | 37617 | 13560 | 12651 | 12282 | 14191 | 29766 | 30905 | 30021 | 30586 | 21745,44 | 13171 | 30320 | 2,303 | 1,203 | 24,965 | 1,47E-137 | 1,35E-136 | 0,002239 | 0,0021668 | 0,0021744 | 0,0021744 | TRUE | 0,0327675 | No |
| orf19.5226 | 2211 | 2020 | 2267 | 1319 | 4536 | 3167 | 4308 | 4545 | 2108 | 2136 | 2240 | 2066 | 3635 | 3601 | 3657 | 3696 | 2892,3 | 2138 | 3647 | 1,705 | 0,769 | 19,699 | 2,18E-86 | 1,26E-85 | 3,47E-05 | 0,002659 | 0,0010801 | 0,0010801 | TRUE | 0,0127036 | No |
| orf19.5227 | 1789 | 1614 | 1781 | 993 | 1105 | 880 | 1026 | 1037 | 1706 | 1707 | 1759 | 1556 | 885 | 1001 | 871 | 843 | 1290,94 | 1682 | 900 | 0,533 | -0,907 | -14,023 | 1,12E-44 | 3,82E-44 | 0,0031905 | 0,0033631 | 0,003189 | 0,003189 | TRUE | 0,0634421 | No |
| orf19.5228 | 6173 | 5548 | 6109 | 3945 | 3147 | 2410 | 3026 | 3091 | 5885 | 5866 | 6035 | 6180 | 2522 | 2740 | 2569 | 2513 | 4288,89 | 5992 | 2586 | 0,431 | -1,213 | -29,554 | 5,86E-192 | 7,73E-191 | 0,0007663 | 0,0024741 | 0,0013508 | 0,0013508 | TRUE | 0,018603 | No |
| orf19.5229 | 9449 | 9124 | 9410 | 5797 | 10612 | 7957 | 9509 | 10044 | 9009 | 9647 | 9296 | 9081 | 8503 | 9048 | 8073 | 8167 | 8853,04 | 9258 | 8448 | 0,912 | -0,133 | -2,907 | 0,0036546 | 0,004766 | 0,0016889 | 0,0022767 | 0,0018911 | 0,0018911 | TRUE | 0,0275601 | No |
| orf19.5230 | 5030 | 4569 | 4987 | 2798 | 4240 | 2953 | 4169 | 4248 | 4796 | 4831 | 4927 | 4383 | 3397 | 3358 | 3539 | 3454 | 4085,66 | 4734 | 3437 | 0,725 | -0,463 | -10,265 | 1,01E-24 | 2,49E-24 | 0,0012175 | 0,0024931 | 0,0017059 | 0,0017059 | TRUE | 0,0288569 | No |
| orf19.5231 | 38 | 49 | 39 | 18 | 129 | 80 | 109 | 104 | 36 | 52 | 39 | 28 | 103 | 91 | 93 | 85 | 65,78 | 39 | 93 | 2,367 | 1,243 | 6,322 | 2,58E-10 | 4,54E-10 | 0,0021636 | 0,0270537 | 0,0181856 | 0,0181856 | TRUE | 0,3575836 | No |
| orf19.5231.2 | 1311 | 1117 | 1193 | 685 | 4534 | 2699 | 4310 | 4318 | 1250 | 1181 | 1179 | 1073 | 3633 | 3069 | 3659 | 3511 | 2319,34 | 1171 | 3468 | 2,958 | 1,565 | 22,945 | 1,67E-116 | 1,28E-115 | 0,0042956 | 0,0027992 | 0,0038569 | 0,0038569 | TRUE | 0,071649 | No |
| orf19.5232 | 5019 | 5436 | 5175 | 3463 | 6591 | 5231 | 5674 | 5274 | 4785 | 5748 | 5112 | 5425 | 5281 | 5948 | 4817 | 4288 | 5175,64 | 5268 | 5084 | 0,965 | -0,052 | -0,568 | 0,5697901 | 0,6041593 | 0,0108419 | 0,0024085 | 0,0076951 | 0,0076951 | TRUE | 0,1588866 | No |
| orf19.5233 | 0 | 0 | 1 | 1 | 1 | 2 | 0 | 1 | 0 | 0 | 1 | 2 | 1 | 2 | 0 | 1 | 0,81 | 1 | 1 | 1,649 | 0,722 | 0,431 | 0,6665011 | 0,6984633 | 1,00E-08 | 2,0407715 | 1,3016672 | 1,3016672 | TRUE | 0,307739 | No |
| orf19.5234 | 1540 | 1373 | 1507 | 894 | 2801 | 2087 | 2689 | 2721 | 1468 | 1452 | 1489 | 1400 | 2244 | 2373 | 2283 | 2212 | 1865,26 | 1452 | 2278 | 1,565 | 0,646 | 14,498 | 1,25E-47 | 4,44E-47 | 0,0002395 | 0,0029715 | 0,0013254 | 0,0013254 | TRUE | 0,0089976 | No |
| orf19.5235 | 797 | 1044 | 1025 | 537 | 1052 | 883 | 923 | 703 | 760 | 1104 | 1013 | 841 | 843 | 1004 | 784 | 572 | 864,98 | 930 | 801 | 0,861 | -0,217 | -1,078 | 0,2808321 | 0,312629 | 0,0375811 | 0,0039895 | 0,020551 | 0,0375811 | TRUE | 0,3940356 | No |
| orf19.5236 | 484 | 528 | 537 | 330 | 459 | 337 | 447 | 419 | 461 | 558 | 531 | 517 | 368 | 383 | 380 | 341 | 442,3 | 517 | 368 | 0,711 | -0,492 | -5,954 | 2,62E-09 | 4,44E-09 | 0,0027539 | 0,0058035 | 0,0042493 | 0,0042493 | TRUE | 0,0657799 | No |
| orf19.5237 | 679 | 592 | 645 | 378 | 525 | 322 | 488 | 471 | 647 | 626 | 637 | 592 | 421 | 366 | 414 | 383 | 510,85 | 626 | 396 | 0,633 | -0,66 | -9,125 | 7,20E-20 | 1,61E-19 | 0,0003073 | 0,0053054 | 0,0030035 | 0,0030035 | TRUE | 0,0262386 | No |
| orf19.5238 | 3024 | 2814 | 2967 | 1979 | 2472 | 1795 | 2214 | 2501 | 2883 | 2975 | 2931 | 3100 | 1981 | 2041 | 1880 | 2034 | 2478,11 | 2972 | 1984 | 0,668 | -0,582 | -13,026 | 8,75E-39 | 2,72E-38 | 0,000634 | 0,0027538 | 0,0015053 | 0,0015053 | TRUE | 0,0151868 | No |
| orf19.5239 | 535 | 513 | 513 | 327 | 709 | 498 | 728 | 663 | 510 | 542 | 507 | 512 | 568 | 566 | 618 | 539 | 545,38 | 518 | 573 | 1,106 | 0,145 | 2,076 | 0,0378942 | 0,0460314 | 1,00E-08 | 0,0051018 | 0,0028561 | 0,0028561 | TRUE | 0,0342847 | No |
| orf19.524 | 1362 | 1123 | 1324 | 757 | 612 | 366 | 534 | 584 | 1299 | 1187 | 1308 | 1186 | 490 | 416 | 453 | 475 | 851,82 | 1245 | 458 | 0,369 | -1,438 | -20,944 | 2,15E-97 | 1,37E-96 | 0,0022305 | 0,0040188 | 0,0031083 | 0,0031083 | TRUE | 0,0406039 | No |
| orf19.5241 | 7932 | 6839 | 7505 | 4650 | 3188 | 2190 | 3084 | 3177 | 7562 | 7231 | 7414 | 7284 | 2554 | 2490 | 2618 | 2583 | 4967,32 | 7373 | 2561 | 0,348 | -1,525 | -43,904 | 0 | 0 | 0,0002055 | 0,0024218 | 0,0009077 | 0,0009077 | TRUE | 0,0039427 | No |
| orf19.5242 | 75 | 97 | 81 | 42 | 184 | 156 | 197 | 150 | 72 | 103 | 80 | 66 | 147 | 177 | 167 | 122 | 116,74 | 80 | 153 | 1,901 | 0,927 | 5,376 | 7,60E-08 | 1,23E-07 | 0,0217307 | 0,0161559 | 0,0186543 | 0,0186543 | TRUE | 0,3278942 | No |
| orf19.5243 | 1473 | 1538 | 1596 | 854 | 2018 | 1422 | 1906 | 1933 | 1404 | 1626 | 1577 | 1338 | 1617 | 1617 | 1618 | 1572 | 1546,12 | 1486 | 1606 | 1,078 | 0,109 | 1,698 | 0,0895759 | 0,1050899 | 0,0035012 | 0,0031532 | 0,0032883 | 0,0032883 | TRUE | 0,0498601 | No |
| orf19.5244 | 450 | 517 | 552 | 323 | 1020 | 760 | 898 | 929 | 429 | 547 | 545 | 506 | 817 | 864 | 762 | 755 | 653,28 | 507 | 800 | 1,578 | 0,658 | 7,614 | 2,65E-14 | 5,24E-14 | 0,0063686 | 0,0046046 | 0,0054972 | 0,0054972 | TRUE | 0,1337205 | No |
| orf19.5245 | 1781 | 1677 | 1871 | 1222 | 607 | 513 | 570 | 626 | 1698 | 1773 | 1848 | 1914 | 486 | 583 | 484 | 509 | 1162,07 | 1808 | 516 | 0,284 | -1,814 | -26,261 | 5,39E-152 | 5,48E-151 | 0,0032974 | 0,0035042 | 0,0034016 | 0,0034016 | TRUE | 0,0835505 | No |
| orf19.5246 | 327 | 285 | 328 | 220 | 313 | 250 | 300 | 321 | 312 | 301 | 324 | 345 | 251 | 284 | 255 | 261 | 291,57 | 320 | 263 | 0,82 | -0,286 | -3,236 | 0,0012126 | 0,0016249 | 1,00E-08 | 0,0077225 | 0,0040444 | 0,0040444 | TRUE | 0,0302267 | No |
| orf19.5247 | 1863 | 1666 | 1678 | 1053 | 1674 | 1127 | 1555 | 1544 | 1776 | 1762 | 1658 | 1650 | 1341 | 1282 | 1320 | 1255 | 1505,43 | 1712 | 1300 | 0,759 | -0,398 | -8,097 | 5,64E-16 | 1,16E-15 | 0,0005791 | 0,0031819 | 0,0016513 | 0,0016513 | TRUE | 0,0079616 | No |
| orf19.5248 | 4183 | 3305 | 3756 | 2379 | 1963 | 1355 | 2060 | 2049 | 3988 | 3495 | 3711 | 3727 | 1573 | 1541 | 1749 | 1666 | 2681,08 | 3730 | 1632 | 0,438 | -1,192 | -20,881 | 7,94E-97 | 5,02E-96 | 0,0025461 | 0,0027037 | 0,0027027 | 0,0027027 | TRUE | 0,0279184 | No |
| orf19.5249 | 778 | 612 | 677 | 432 | 451 | 295 | 352 | 442 | 742 | 647 | 669 | 677 | 361 | 335 | 299 | 359 | 511,18 | 684 | 338 | 0,496 | -1,013 | -12,289 | 1,04E-34 | 3,03E-34 | 0,0034618 | 0,0053032 | 0,0043858 | 0,0043858 | TRUE | 0,0758506 | No |
| orf19.5250 | 1216 | 1095 | 1135 | 751 | 2561 | 1841 | 2421 | 2577 | 1159 | 1158 | 1121 | 1176 | 2052 | 2093 | 2055 | 2095 | 1613,89 | 1154 | 2074 | 1,799 | 0,847 | 19,65 | 5,81E-86 | 3,34E-85 | 1,00E-08 | 0,0031086 | 0,0010864 | 0,0010864 | TRUE | 0,0043885 | No |
| orf19.5251 | 5538 | 4921 | 5500 | 3773 | 3873 | 2928 | 3638 | 3941 | 5280 | 5203 | 5433 | 5911 | 3103 | 3329 | 3089 | 3204 | 4319,14 | 5457 | 3181 | 0,583 | -0,778 | -15,565 | 1,26E-54 | 4,93E-54 | 0,0019702 | 0,0024714 | 0,0021539 | 0,0021539 | TRUE | 0,0369569 | No |
| orf19.5253 | 869 | 815 | 905 | 529 | 1530 | 1046 | 1347 | 1483 | 828 | 862 | 894 | 829 | 1226 | 1189 | 1144 | 1206 | 1022,22 | 853 | 1191 | 1,395 | 0,48 | 8,899 | 5,63E-19 | 1,24E-18 | 1,00E-08 | 0,0036975 | 0,0017629 | 0,0017629 | TRUE | 0,0127461 | No |
| orf19.5254 | 429 | 423 | 439 | 312 | 452 | 370 | 410 | 388 | 409 | 447 | 434 | 489 | 362 | 421 | 348 | 315 | 403,15 | 445 | 362 | 0,813 | -0,299 | -3,068 | 0,0021518 | 0,0028463 | 0,0068746 | 0,006164 | 0,0065898 | 0,0065898 | TRUE | 0,1290261 | No |
| orf19.5255 | 3542 | 3163 | 3491 | 2224 | 4686 | 3461 | 4342 | 4784 | 3377 | 3344 | 3449 | 3484 | 3755 | 3935 | 3686 | 3890 | 3615,07 | 3414 | 3816 | 1,118 | 0,161 | 4,236 | 2,27E-05 | 3,32E-05 | 0,0003801 | 0,0025455 | 0,0011076 | 0,0011076 | TRUE | 0,0063802 | No |
| orf19.5256 | 72 | 57 | 78 | 43 | 94 | 55 | 81 | 66 | 69 | 60 | 77 | 67 | 75 | 63 | 69 | 54 | 66,7 | 68 | 65 | 0,952 | -0,071 | -0,392 | 0,6953652 | 0,7263116 | 1,00E-08 | 0,0267064 | 0,0166272 | 0,0166272 | TRUE | 0,1412427 | No |
| orf19.5257 | 979 | 834 | 905 | 607 | 1073 | 785 | 1015 | 1086 | 933 | 882 | 894 | 951 | 860 | 893 | 862 | 883 | 894,66 | 915 | 874 | 0,956 | -0,064 | -1,195 | 0,2319647 | 0,2610984 | 1,00E-08 | 0,0039265 | 0,0016613 | 0,0016613 | TRUE | 0,0069205 | No |
| orf19.5258 | 1249 | 1257 | 1330 | 791 | 584 | 470 | 586 | 603 | 1191 | 1329 | 1314 | 1239 | 468 | 534 | 498 | 490 | 882,89 | 1268 | 498 | 0,391 | -1,354 | -20,887 | 7,01E-97 | 4,44E-96 | 0,0016077 | 0,003951 | 0,0026956 | 0,0026956 | TRUE | 0,0262639 | No |
| orf19.5259 | 664 | 619 | 667 | 433 | 307 | 223 | 235 | 306 | 633 | 655 | 659 | 678 | 246 | 254 | 200 | 249 | 446,59 | 656 | 237 | 0,361 | -1,47 | -17,397 | 8,74E-68 | 4,07E-67 | 0,0015168 | 0,0057678 | 0,0041224 | 0,0041224 | TRUE | 0,1295911 | No |
| orf19.526 | 10843 | 9646 | 9999 | 5503 | 18620 | 12127 | 17533 | 18239 | 10338 | 10199 | 9878 | 8621 | 14920 | 13789 | 14885 | 14830 | 12182,56 | 9759 | 14606 | 1,496 | 0,581 | 9,878 | 5,19E-23 | 1,23E-22 | 0,0038015 | 0,0022261 | 0,0032372 | 0,0032372 | TRUE | 0,0742312 | No |
| orf19.5260 | 6356 | 5629 | 6119 | 4134 | 9753 | 7151 | 9508 | 10067 | 6060 | 5952 | 6045 | 6476 | 7815 | 8131 | 8072 | 8185 | 7092,08 | 6133 | 8051 | 1,314 | 0,394 | 10,441 | 1,61E-25 | 4,02E-25 | 0,0007412 | 0,0023228 | 0,0012174 | 0,0012174 | TRUE | 0,0164424 | No |
| orf19.5262 | 43 | 21 | 24 | 14 | 90 | 61 | 93 | 89 | 41 | 22 | 24 | 22 | 72 | 69 | 79 | 72 | 50,21 | 27 | 73 | 2,646 | 1,404 | 6,058 | 1,38E-09 | 2,35E-09 | 1,00E-08 | 0,0347952 | 0,0254072 | 0,0254072 | TRUE | 0,8041147 | No |
| orf19.5263 | 7556 | 6613 | 7013 | 4695 | 19578 | 14417 | 18186 | 20058 | 7204 | 6992 | 6928 | 7355 | 15687 | 16393 | 15440 | 16309 | 11538,63 | 7120 | 15957 | 2,242 | 1,165 | 32,053 | 1,98E-225 | 2,99E-224 | 0,0006722 | 0,0022336 | 0,001161 | 0,001161 | TRUE | 0,0058269 | No |
| orf19.5264 | 3 | 11 | 2 | 7 | 7 | 6 | 6 | 9 | 3 | 12 | 2 | 11 | 6 | 7 | 5 | 7 | 6,53 | 7 | 6 | 0,938 | -0,092 | -0,147 | 0,8835155 | 0,8993001 | 0,1734511 | 0,253357 | 0,2205388 | 0,2205388 | TRUE | 0,1582194 | No |
| orf19.5265 | 3359 | 3061 | 3214 | 1910 | 1939 | 1365 | 1874 | 1929 | 3202 | 3237 | 3175 | 2992 | 1554 | 1552 | 1591 | 1568 | 2358,95 | 3152 | 1566 | 0,496 | -1,01 | -24,226 | 1,18E-129 | 1,02E-128 | 0,0002177 | 0,0027873 | 0,0012071 | 0,0012071 | TRUE | 0,0128526 | No |
| orf19.5266 | 64 | 45 | 47 | 22 | 25 | 11 | 12 | 18 | 61 | 48 | 46 | 34 | 20 | 13 | 10 | 15 | 30,86 | 47 | 14 | 0,302 | -1,729 | -5,769 | 7,96E-09 | 1,33E-08 | 0,0259839 | 0,0553009 | 0,0438188 | 0,0438188 | TRUE | 0,2053285 | No |
| orf19.5267 | 138 | 120 | 122 | 95 | 47 | 43 | 55 | 39 | 132 | 127 | 121 | 149 | 38 | 49 | 47 | 32 | 86,59 | 132 | 42 | 0,311 | -1,685 | -9,626 | 6,22E-22 | 1,45E-21 | 0,001439 | 0,021052 | 0,0142828 | 0,0142828 | TRUE | 0,201717 | No |
| orf19.5268 | 169 | 139 | 163 | 122 | 248 | 181 | 223 | 200 | 161 | 147 | 161 | 191 | 199 | 206 | 189 | 163 | 177,09 | 165 | 189 | 1,151 | 0,203 | 1,67 | 0,0949916 | 0,1111778 | 0,0047931 | 0,0113629 | 0,0084547 | 0,0084547 | TRUE | 0,1006586 | No |
| orf19.5269 | 658 | 545 | 599 | 353 | 1294 | 1000 | 1386 | 1392 | 627 | 576 | 592 | 553 | 1037 | 1137 | 1177 | 1132 | 853,85 | 587 | 1121 | 1,902 | 0,927 | 14,154 | 1,77E-45 | 6,09E-45 | 0,0017529 | 0,0040142 | 0,0027747 | 0,0027747 | TRUE | 0,0313846 | No |
| orf19.5269.1 | 374 | 382 | 320 | 215 | 856 | 556 | 768 | 759 | 357 | 404 | 316 | 337 | 686 | 632 | 652 | 617 | 500,09 | 354 | 647 | 1,829 | 0,871 | 10,368 | 3,47E-25 | 8,63E-25 | 0,0037787 | 0,0053745 | 0,0045074 | 0,0045074 | TRUE | 0,1078407 | No |
| orf19.527 | 1662 | 1487 | 1582 | 1111 | 1909 | 1504 | 1866 | 2011 | 1585 | 1572 | 1563 | 1740 | 1530 | 1710 | 1584 | 1635 | 1614,92 | 1615 | 1615 | 1,001 | 0,001 | 0,026 | 0,9791741 | 0,981173 | 0,001499 | 0,003108 | 0,0022575 | 0,0022575 | TRUE | 0,0300686 | No |
| orf19.5270 | 182 | 170 | 196 | 113 | 697 | 470 | 600 | 609 | 174 | 180 | 194 | 177 | 558 | 534 | 509 | 495 | 352,68 | 181 | 524 | 2,892 | 1,532 | 17,116 | 1,13E-65 | 5,13E-65 | 1,00E-08 | 0,0067468 | 0,0037815 | 0,0037815 | TRUE | 0,0258666 | No |
| orf19.5271 | 213 | 232 | 253 | 141 | 466 | 360 | 432 | 448 | 203 | 245 | 250 | 221 | 373 | 409 | 367 | 364 | 304,12 | 230 | 378 | 1,641 | 0,715 | 7,547 | 4,45E-14 | 8,75E-14 | 0,001783 | 0,0074901 | 0,0050122 | 0,0050122 | TRUE | 0,0786829 | No |
| orf19.5274 | 3597 | 3374 | 3837 | 2271 | 3296 | 2139 | 3076 | 3329 | 3429 | 3568 | 3791 | 3558 | 2641 | 2432 | 2612 | 2707 | 3092,09 | 3586 | 2598 | 0,725 | -0,464 | -9,501 | 2,09E-21 | 4,81E-21 | 0,0014537 | 0,0026223 | 0,0019623 | 0,0019623 | TRUE | 0,0214615 | No |
| orf19.5275 | 488 | 517 | 510 | 333 | 301 | 155 | 255 | 263 | 465 | 547 | 504 | 522 | 241 | 176 | 216 | 214 | 360,65 | 510 | 212 | 0,419 | -1,255 | -12,653 | 1,08E-36 | 3,24E-36 | 0,0057957 | 0,0066439 | 0,0062492 | 0,0062492 | TRUE | 0,1241661 | No |
| orf19.5276 | 805 | 866 | 867 | 538 | 617 | 437 | 515 | 441 | 767 | 916 | 857 | 843 | 494 | 497 | 437 | 359 | 646,2 | 846 | 447 | 0,528 | -0,923 | -9,075 | 1,14E-19 | 2,53E-19 | 0,0118939 | 0,0046321 | 0,0082591 | 0,0082591 | TRUE | 0,2103885 | No |
| orf19.5277 | 1814 | 1676 | 1873 | 1058 | 1404 | 929 | 1363 | 1289 | 1729 | 1772 | 1850 | 1657 | 1125 | 1056 | 1157 | 1048 | 1424,5 | 1752 | 1096 | 0,625 | -0,677 | -12,226 | 2,27E-34 | 6,57E-34 | 0,0014469 | 0,0032439 | 0,0022186 | 0,0022186 | TRUE | 0,0170352 | No |
| orf19.5278 | 1940 | 1784 | 1824 | 1071 | 4017 | 2715 | 3667 | 3674 | 1850 | 1886 | 1802 | 1678 | 3219 | 3087 | 3113 | 2987 | 2452,77 | 1804 | 3102 | 1,716 | 0,779 | 16,257 | 1,99E-59 | 8,35E-59 | 0,0011553 | 0,0027607 | 0,0017498 | 0,0017498 | TRUE | 0,0239597 | No |
| orf19.5279 | 2962 | 2606 | 2608 | 1482 | 2169 | 1235 | 1968 | 1990 | 2824 | 2756 | 2576 | 2322 | 1738 | 1404 | 1671 | 1618 | 2113,59 | 2620 | 1608 | 0,614 | -0,704 | -9,092 | 9,74E-20 | 2,18E-19 | 0,0070209 | 0,0028681 | 0,0052681 | 0,0052681 | TRUE | 0,0851068 | No |
| orf19.528 | 3213 | 3089 | 3182 | 1957 | 5364 | 3845 | 5007 | 5595 | 3063 | 3266 | 3144 | 3066 | 4298 | 4372 | 4251 | 4549 | 3751,13 | 3135 | 4368 | 1,393 | 0,478 | 11,793 | 4,23E-32 | 1,18E-31 | 0,0005465 | 0,002529 | 0,0012955 | 0,0012955 | TRUE | 0,0097701 | No |
| orf19.5280 | 68553 | 57925 | 63100 | 41383 | 74644 | 52947 | 70488 | 77505 | 65358 | 61248 | 62337 | 64829 | 59810 | 60206 | 59844 | 63019 | 62081,28 | 63443 | 60720 | 0,957 | -0,063 | -1,812 | 0,0700307 | 0,0830511 | 0,000783 | 0,0021177 | 0,0011543 | 0,0011543 | TRUE | 0,0079754 | No |
| orf19.5281 | 1603 | 1477 | 1600 | 1002 | 4456 | 3333 | 4196 | 4669 | 1528 | 1562 | 1581 | 1570 | 3570 | 3790 | 3562 | 3796 | 2619,93 | 1560 | 3680 | 2,359 | 1,238 | 28,537 | 4,06E-179 | 4,84E-178 | 0,0005943 | 0,002718 | 0,0013261 | 0,0013261 | TRUE | 0,0058206 | No |
| orf19.5282 | 9071 | 7863 | 8643 | 5760 | 3867 | 2846 | 3669 | 4049 | 8648 | 8314 | 8538 | 9023 | 3099 | 3236 | 3115 | 3292 | 5908,25 | 8631 | 3186 | 0,369 | -1,438 | -35,831 | 3,64E-281 | 7,42E-280 | 0,0007376 | 0,0023692 | 0,0013404 | 0,0013404 | TRUE | 0,0110415 | No |
| orf19.5284 | 63 | 58 | 60 | 51 | 93 | 90 | 86 | 128 | 60 | 61 | 59 | 80 | 75 | 102 | 73 | 104 | 76,81 | 65 | 88 | 1,369 | 0,453 | 2,395 | 0,0166164 | 0,0207697 | 0,0173354 | 0,0234665 | 0,0206584 | 0,0206584 | TRUE | 0,1246594 | No |
| orf19.5285 | 1366 | 1089 | 1071 | 848 | 932 | 677 | 857 | 915 | 1302 | 1151 | 1058 | 1328 | 747 | 770 | 728 | 744 | 978,56 | 1210 | 747 | 0,618 | -0,694 | -9,109 | 8,30E-20 | 1,85E-19 | 0,0051092 | 0,0037692 | 0,0045077 | 0,0045077 | TRUE | 0,0869347 | No |
| orf19.5286 | 3661 | 2881 | 3100 | 2443 | 2591 | 1836 | 2499 | 2560 | 3490 | 3046 | 3063 | 3827 | 2076 | 2088 | 2122 | 2082 | 2724,15 | 3356 | 2092 | 0,624 | -0,681 | -9,592 | 8,64E-22 | 2,01E-21 | 0,0057739 | 0,002694 | 0,0044544 | 0,0044544 | TRUE | 0,1051941 | No |
| orf19.5287 | 128 | 103 | 116 | 98 | 250 | 165 | 255 | 280 | 122 | 109 | 115 | 154 | 200 | 188 | 216 | 228 | 166,4 | 125 | 208 | 1,692 | 0,759 | 5,904 | 3,54E-09 | 5,98E-09 | 0,0050633 | 0,0119588 | 0,0091949 | 0,0091949 | TRUE | 0,190241 | No |
| orf19.5288 | 175079 | 144775 | 154847 | 104159 | 12876 | 9093 | 12136 | 13434 | 166919 | 153081 | 152974 | 163171 | 10317 | 10340 | 10303 | 10923 | 84753,52 | 159036 | 10471 | 0,066 | -3,925 | -94,213 | 0 | 0 | 0,0013457 | 0,0021107 | 0,0016211 | 0,0016211 | TRUE | 0,0136688 | No |
| orf19.5288.1 | 796 | 765 | 759 | 441 | 1258 | 842 | 1124 | 1327 | 759 | 809 | 750 | 691 | 1008 | 957 | 954 | 1079 | 875,89 | 752 | 1000 | 1,327 | 0,408 | 6,089 | 1,13E-09 | 1,94E-09 | 0,0022193 | 0,0039658 | 0,0031179 | 0,0031179 | TRUE | 0,0345897 | No |
| orf19.5289 | 5 | 3 | 11 | 4 | 13 | 20 | 16 | 16 | 5 | 3 | 11 | 6 | 10 | 23 | 14 | 13 | 10,6 | 6 | 15 | 2,339 | 1,226 | 2,451 | 0,0142286 | 0,0178761 | 0,0687025 | 0,156944 | 0,1214253 | 0,1214253 | TRUE | 0,670497 | No |
| orf19.529 | 2659 | 2208 | 2530 | 1680 | 1746 | 1304 | 1681 | 1886 | 2535 | 2335 | 2499 | 2632 | 1399 | 1483 | 1427 | 1533 | 1980,43 | 2500 | 1460 | 0,585 | -0,775 | -14,766 | 2,44E-49 | 8,92E-49 | 0,001566 | 0,0029203 | 0,0021108 | 0,0021108 | TRUE | 0,0239387 | No |
| orf19.5290 | 0 | 0 | 0 | 0 | 0 | 0 | 0 | 0 | 0 | 0 | 0 | 0 | 0 | 0 | 0 | 0 | 0 | 0 | 0 | NA | NA | NA | NA | NA | NA | NA | NA | NA | NA | NA | NA |
| orf19.5291 | 919 | 1049 | 1032 | 703 | 1848 | 1422 | 1696 | 1560 | 876 | 1109 | 1020 | 1101 | 1481 | 1617 | 1440 | 1268 | 1239,02 | 1026 | 1452 | 1,415 | 0,501 | 5,589 | 2,29E-08 | 3,76E-08 | 0,0093341 | 0,0034164 | 0,0068542 | 0,0068542 | TRUE | 0,1187274 | No |
| orf19.529.1 | 64 | 77 | 97 | 58 | 90 | 60 | 80 | 90 | 61 | 81 | 96 | 91 | 72 | 68 | 68 | 73 | 76,32 | 82 | 70 | 0,862 | -0,215 | -1,217 | 0,2234447 | 0,2522648 | 0,0072231 | 0,0236047 | 0,0167191 | 0,0167191 | TRUE | 0,3112114 | No |
| orf19.5292 | 914 | 945 | 873 | 538 | 2036 | 1604 | 1854 | 1934 | 871 | 999 | 862 | 843 | 1631 | 1824 | 1574 | 1573 | 1272,21 | 894 | 1650 | 1,842 | 0,881 | 12,324 | 6,74E-35 | 1,97E-34 | 0,0046115 | 0,0033819 | 0,0040121 | 0,0040121 | TRUE | 0,0750709 | No |
| orf19.5293 | 2404 | 2684 | 2742 | 1651 | 5008 | 3836 | 4571 | 4204 | 2292 | 2838 | 2709 | 2586 | 4013 | 4362 | 3881 | 3418 | 3262,35 | 2606 | 3918 | 1,503 | 0,588 | 7,219 | 5,24E-13 | 1,00E-12 | 0,0085456 | 0,0025946 | 0,0060395 | 0,0060395 | TRUE | 0,0901277 | No |

| | | | | | | | | | | | | | | | | | | | | | | | | | | | | | | | |
|---|---|---|---|---|---|---|---|---|---|---|---|---|---|---|---|---|---|---|---|---|---|---|---|---|---|---|---|---|---|---|---|
| orf19.5294 | 9527 | 8395 | 8787 | 6011 | 18829 | 14013 | 17297 | 18924 | 9083 | 8877 | 8681 | 9417 | 15087 | 15934 | 14685 | 15387 | 12143,77 | 9014 | 15273 | 1,695 | 0,761 | 19,008 | 1,46E-80 | 8,02E-80 | 0,0010842 | 0,0022265 | 0,0014482 | 0,0014482 | TRUE | 0,0106976 | No |
| orf19.5295 | 1104 | 1055 | 1023 | 670 | 2269 | 1735 | 2261 | 2253 | 1053 | 1116 | 1011 | 1050 | 1818 | 1973 | 1920 | 1832 | 1471,34 | 1058 | 1886 | 1,782 | 0,834 | 15,887 | 7,86E-57 | 3,16E-56 | 0,0009124 | 0,0032072 | 0,0018776 | 0,0018776 | TRUE | 0,0172072 | No |
| orf19.5296 | 1687 | 1473 | 1589 | 915 | 1755 | 1226 | 1611 | 1634 | 1608 | 1558 | 1570 | 1433 | 1406 | 1394 | 1368 | 1329 | 1458,21 | 1542 | 1374 | 0,888 | -0,171 | -3,379 | 0,0007284 | 0,0009927 | 0,0006013 | 0,0032173 | 0,0017596 | 0,0017596 | TRUE | 0,0241792 | No |
| orf19.5297 | 1103 | 1141 | 1159 | 712 | 1223 | 844 | 1100 | 1176 | 1052 | 1206 | 1145 | 1115 | 980 | 960 | 934 | 956 | 1043,52 | 1130 | 958 | 0,848 | -0,238 | -4,175 | 2,98E-05 | 4,32E-05 | 0,0010037 | 0,0036647 | 0,0021682 | 0,0021682 | TRUE | 0,028158 | No |
| orf19.5299 | 3697 | 3624 | 3606 | 1957 | 4185 | 2616 | 4011 | 3976 | 3525 | 3832 | 3562 | 3066 | 3353 | 2975 | 3405 | 3233 | 3368,86 | 3496 | 3242 | 0,927 | -0,11 | -1,59 | 0,1117554 | 0,1297273 | 0,0054926 | 0,0025787 | 0,0042889 | 0,0042889 | TRUE | 0,0808619 | No |
| orf19.5299.1 | 1261 | 1184 | 1348 | 588 | 651 | 431 | 621 | 598 | 1202 | 1252 | 1332 | 921 | 522 | 490 | 527 | 486 | 841,52 | 1177 | 506 | 0,429 | -1,22 | -12,206 | 2,90E-34 | 8,37E-34 | 0,0122375 | 0,0040424 | 0,0082349 | 0,0082349 | TRUE | 0,2426381 | No |
| orf19.53 | 942 | 815 | 957 | 532 | 1414 | 902 | 1375 | 1359 | 898 | 862 | 945 | 833 | 1133 | 1026 | 1167 | 1105 | 996,21 | 884 | 1108 | 1,251 | 0,323 | 5,179 | 2,23E-07 | 3,54E-07 | 0,0016262 | 0,0037394 | 0,0026989 | 0,0026989 | TRUE | 0,027883 | No |
| orf19.530 | 284 | 314 | 326 | 190 | 320 | 285 | 319 | 255 | 271 | 332 | 322 | 298 | 256 | 324 | 271 | 207 | 285,14 | 306 | 264 | 0,862 | -0,214 | -1,661 | 0,09674 | 0,1131619 | 0,0172125 | 0,0078495 | 0,0124151 | 0,0124151 | TRUE | 0,2424795 | No |
| orf19.5300 | 2077 | 1980 | 2055 | 1318 | 3690 | 2629 | 3372 | 3629 | 1980 | 2094 | 2030 | 2065 | 2957 | 2989 | 2863 | 2951 | 2491,04 | 2042 | 2940 | 1,44 | 0,526 | 13,422 | 4,51E-41 | 1,45E-40 | 1,00E-08 | 0,0027504 | 0,0010496 | 0,0010496 | TRUE | 0,0052983 | No |
| orf19.5302 | 127 | 104 | 111 | 80 | 505 | 347 | 467 | 451 | 121 | 110 | 110 | 125 | 405 | 395 | 396 | 367 | 253,55 | 116 | 391 | 3,368 | 1,752 | 16,903 | 4,29E-64 | 1,91E-63 | 1,00E-08 | 0,0085668 | 0,0043942 | 0,0043942 | TRUE | 0,0206272 | No |
| orf19.5303 | 1 | 4 | 1 | 0 | 1 | 0 | 0 | 0 | 1 | 4 | 1 | 0 | 1 | 0 | 0 | 0 | 0,87 | 2 | 0 | 0,17 | -2,553 | -1,472 | 0,1409008 | 0,1619978 | 0,763433 | 1,8860715 | 1,5345337 | 1,5345337 | TRUE | 0,9031723 | No |
| orf19.5304 | 0 | 0 | 0 | 0 | 0 | 0 | 0 | 0 | 0 | 0 | 0 | 0 | 0 | 0 | 0 | 0 | 0 | 0 | 0 | NA | NA | NA | NA | NA | NA | NA | NA | NA | NA | NA | NA |
| orf19.5305 | 3624 | 2964 | 3130 | 2256 | 1735 | 1120 | 1523 | 1605 | 3455 | 3134 | 3092 | 3534 | 1390 | 1274 | 1293 | 1305 | 2309,65 | 3304 | 1316 | 0,399 | -1,326 | -22,824 | 2,65E-115 | 2,01E-114 | 0,0026882 | 0,0028022 | 0,0027333 | 0,0027333 | TRUE | 0,0255517 | No |
| orf19.5306 | 0 | 0 | 0 | 0 | 2 | 0 | 0 | 0 | 0 | 0 | 0 | 0 | 2 | 0 | 0 | 0 | 0,2 | 0 | 0 | 2,103 | 1,073 | 0,304 | 0,7610817 | 0,7887709 | 1,00E-08 | 8,1985829 | 10 | 10 | TRUE | 0,8681513 | No |
| orf19.5307 | 1372 | 1154 | 1221 | 781 | 2787 | 1898 | 2642 | 2739 | 1308 | 1220 | 1206 | 1223 | 2233 | 2158 | 2243 | 2227 | 1727,43 | 1239 | 2215 | 1,787 | 0,837 | 18,351 | 3,25E-75 | 1,67E-74 | 1,17E-05 | 0,0030418 | 0,0013461 | 0,0013461 | TRUE | 0,0178948 | No |
| orf19.5308 | 92 | 81 | 82 | 48 | 102 | 95 | 127 | 108 | 88 | 86 | 81 | 75 | 82 | 108 | 108 | 88 | 89,37 | 82 | 96 | 1,158 | 0,211 | 1,326 | 0,1848525 | 0,2102948 | 1,00E-08 | 0,0204635 | 0,0129974 | 0,0129974 | TRUE | 0,1074421 | No |
| orf19.5311 | 2132 | 1880 | 2105 | 1252 | 1544 | 1083 | 1494 | 1704 | 2033 | 1988 | 2080 | 1961 | 1237 | 1231 | 1268 | 1386 | 1647,99 | 2016 | 1280 | 0,635 | -0,654 | -12,546 | 4,19E-36 | 1,25E-35 | 0,0010515 | 0,0030876 | 0,0019824 | 0,0019824 | TRUE | 0,037327 | No |
| orf19.5312 | 2645 | 2323 | 2580 | 1684 | 4061 | 3100 | 4025 | 4005 | 2522 | 2456 | 2549 | 2638 | 3254 | 3525 | 3417 | 3256 | 2952,18 | 2541 | 3363 | 1,324 | 0,405 | 9,193 | 3,81E-20 | 8,58E-20 | 0,0007514 | 0,0026474 | 0,001506 | 0,001506 | TRUE | 0,0123072 | No |
| orf19.5316 | 165 | 147 | 128 | 88 | 445 | 280 | 397 | 379 | 157 | 155 | 126 | 138 | 357 | 318 | 337 | 308 | 237,15 | 144 | 330 | 2,284 | 1,192 | 11,021 | 3,02E-28 | 7,90E-28 | 0,0022916 | 0,0090147 | 0,0060292 | 0,0060292 | TRUE | 0,083783 | No |
| orf19.5318 | 1453 | 1342 | 1474 | 937 | 1268 | 967 | 1261 | 1350 | 1385 | 1419 | 1456 | 1468 | 1016 | 1100 | 1071 | 1098 | 1251,52 | 1432 | 1071 | 0,748 | -0,419 | -8,341 | 7,36E-17 | 1,54E-16 | 1,00E-08 | 0,0034032 | 0,0016137 | 0,0016137 | TRUE | 0,0147756 | No |
| orf19.532 | 723 | 605 | 558 | 386 | 2265 | 1653 | 1951 | 2206 | 689 | 640 | 551 | 605 | 1815 | 1880 | 1656 | 1794 | 1203,69 | 621 | 1786 | 2,871 | 1,521 | 20,942 | 2,24E-97 | 1,42E-96 | 0,0043172 | 0,0034553 | 0,0039153 | 0,0039153 | TRUE | 0,0737668 | No |
| orf19.5320 | 271 | 227 | 274 | 184 | 224 | 178 | 246 | 257 | 258 | 240 | 271 | 288 | 179 | 202 | 209 | 209 | 232,13 | 264 | 200 | 0,759 | -0,398 | -3,918 | 8,94E-05 | 0,000127 | 1,00E-08 | 0,0091645 | 0,005578 | 0,005578 | TRUE | 0,0562011 | No |
| orf19.5321 | 2304 | 1929 | 2211 | 1375 | 3738 | 2587 | 3516 | 3818 | 2197 | 2040 | 2184 | 2154 | 2995 | 2942 | 2985 | 3104 | 2575,1 | 2144 | 3006 | 1,403 | 0,488 | 11,66 | 2,04E-31 | 5,65E-31 | 0,0004036 | 0,0027289 | 0,0012744 | 0,0012744 | TRUE | 0,0133686 | No |
| orf19.5322 | 776 | 604 | 696 | 388 | 2372 | 1584 | 2264 | 2374 | 740 | 639 | 688 | 608 | 1901 | 1801 | 1922 | 1930 | 1278,51 | 669 | 1888 | 2,814 | 1,492 | 23,628 | 1,98E-123 | 1,62E-122 | 0,0020902 | 0,0033755 | 0,0027609 | 0,0027609 | TRUE | 0,0611901 | No |
| orf19.5323 | 1022 | 784 | 868 | 609 | 3847 | 2557 | 3644 | 3936 | 974 | 829 | 858 | 954 | 3082 | 2908 | 3094 | 3200 | 1987,37 | 904 | 3071 | 3,405 | 1,767 | 29,054 | 1,39E-185 | 1,73E-184 | 0,0029354 | 0,0029174 | 0,0027864 | 0,0027864 | TRUE | 0,03733 | No |
| orf19.5325 | 3061 | 2351 | 2700 | 1573 | 2167 | 1268 | 2148 | 2241 | 2918 | 2486 | 2667 | 2464 | 1736 | 1442 | 1824 | 1822 | 2169,97 | 2634 | 1706 | 0,648 | -0,626 | -7,705 | 1,31E-14 | 2,61E-14 | 0,0081589 | 0,0028479 | 0,0058645 | 0,0058645 | TRUE | 0,1279034 | No |
| orf19.5326 | 2953 | 2411 | 2639 | 1916 | 2631 | 1725 | 2495 | 2726 | 2815 | 2549 | 2607 | 3002 | 2108 | 1961 | 2118 | 2216 | 2422,21 | 2743 | 2101 | 0,767 | -0,382 | -6,208 | 5,37E-10 | 9,34E-10 | 0,0035272 | 0,0027691 | 0,0032199 | 0,0032199 | TRUE | 0,0464947 | No |
| orf19.5328 | 1885 | 1605 | 1722 | 1045 | 6946 | 4701 | 6193 | 6612 | 1797 | 1697 | 1701 | 1637 | 5566 | 5345 | 5258 | 5376 | 3547,19 | 1708 | 5386 | 3,148 | 1,654 | 39,257 | 0 | 0 | 0,0005266 | 0,0025542 | 0,0012947 | 0,0012947 | TRUE | 0,0150826 | No |
| orf19.533 | 0 | 0 | 0 | 1 | 2 | 1 | 0 | 0 | 0 | 0 | 0 | 2 | 2 | 1 | 0 | 0 | 0,54 | 0 | 1 | 1,994 | 0,995 | 0,464 | 0,6423937 | 0,6749735 | 1,00E-08 | 3,0524072 | 2,7543076 | 2,7543076 | TRUE | 0,6024198 | No |
| orf19.5334 | 3949 | 3043 | 3312 | 2275 | 2686 | 1799 | 2653 | 2651 | 3765 | 3218 | 3272 | 3564 | 2152 | 2046 | 2252 | 2156 | 2803,02 | 3455 | 2152 | 0,623 | -0,682 | -11,51 | 1,17E-30 | 3,19E-30 | 0,0030302 | 0,002677 | 0,0030028 | 0,0030028 | TRUE | 0,0459755 | No |
| orf19.5335 | 2468 | 2020 | 2303 | 1477 | 3018 | 1939 | 2773 | 3077 | 2353 | 2136 | 2275 | 2314 | 2418 | 2205 | 2354 | 2502 | 2319,63 | 2270 | 2370 | 1,045 | 0,064 | 1,215 | 0,2244226 | 0,2531448 | 0,001695 | 0,0027991 | 0,0022315 | 0,0022315 | TRUE | 0,0256001 | No |
| orf19.5337 | 416 | 316 | 335 | 300 | 511 | 341 | 475 | 450 | 397 | 334 | 331 | 470 | 409 | 388 | 403 | 366 | 387,25 | 383 | 392 | 1,029 | 0,042 | 0,383 | 0,7015934 | 0,7321124 | 0,0108218 | 0,0063312 | 0,0087588 | 0,0087588 | TRUE | 0,2391746 | No |
| orf19.5338 | 1709 | 1466 | 1610 | 1124 | 1642 | 1038 | 1527 | 1772 | 1629 | 1550 | 1591 | 1761 | 1316 | 1180 | 1296 | 1441 | 1470,5 | 1633 | 1308 | 0,804 | -0,316 | -4,695 | 2,67E-06 | 4,06E-06 | 0,0039706 | 0,0032078 | 0,0036504 | 0,0036504 | TRUE | 0,0558206 | No |
| orf19.5339 | 2825 | 1961 | 2175 | 1511 | 1723 | 1066 | 1598 | 1606 | 2693 | 2074 | 2149 | 2367 | 1381 | 1212 | 1357 | 1306 | 1817,23 | 2321 | 1314 | 0,567 | -0,82 | -10,017 | 1,28E-23 | 3,08E-23 | 0,0079881 | 0,0029948 | 0,0058493 | 0,0058493 | TRUE | 0,1447722 | No |
| orf19.5340 | 5155 | 4336 | 4802 | 2916 | 2755 | 1828 | 2628 | 2947 | 4915 | 4585 | 4744 | 4568 | 2208 | 2079 | 2231 | 2396 | 3465,62 | 4703 | 2228 | 0,474 | -1,077 | -21,287 | 1,51E-100 | 9,96E-100 | 0,0018651 | 0,002565 | 0,0021374 | 0,0021374 | TRUE | 0,0311738 | No |
| orf19.5341 | 7571 | 5535 | 5844 | 4185 | 28414 | 19218 | 26619 | 31039 | 7218 | 5853 | 5773 | 6556 | 22767 | 21853 | 22599 | 25238 | 14732,13 | 6350 | 23114 | 3,64 | 1,864 | 25,34 | 1,15E-141 | 1,10E-140 | 0,0075774 | 0,0022027 | 0,0050907 | 0,0050907 | TRUE | 0,1045216 | No |
| orf19.5342 | 1211 | 1098 | 1182 | 841 | 1852 | 1321 | 1783 | 1875 | 1155 | 1161 | 1168 | 1317 | 1484 | 1502 | 1514 | 1525 | 1353,14 | 1200 | 1506 | 1,26 | 0,333 | 6,285 | 3,28E-10 | 5,76E-10 | 0,0006141 | 0,0033047 | 0,0019299 | 0,0019299 | TRUE | 0,0458703 | No |
| orf19.5342.1 | 1495 | 1066 | 1269 | 800 | 2272 | 1392 | 2242 | 2218 | 1425 | 1127 | 1254 | 1253 | 1820 | 1583 | 1903 | 1803 | 1521,2 | 1265 | 1777 | 1,406 | 0,491 | 6,247 | 4,19E-10 | 7,33E-10 | 0,0067514 | 0,0031706 | 0,0052425 | 0,0052425 | TRUE | 0,0903919 | No |
| orf19.5342.2 | 190 | 123 | 157 | 101 | 652 | 348 | 677 | 653 | 181 | 130 | 155 | 158 | 522 | 396 | 575 | 531 | 331,05 | 156 | 506 | 3,245 | 1,698 | 12,775 | 2,27E-37 | 6,87E-37 | 0,0184893 | 0,007051 | 0,0125015 | 0,0125015 | TRUE | 0,2405204 | No |
| orf19.5343 | 323 | 288 | 345 | 231 | 156 | 119 | 156 | 172 | 308 | 305 | 341 | 362 | 125 | 135 | 132 | 140 | 230,97 | 329 | 133 | 0,406 | -1,299 | -12,403 | 2,53E-35 | 7,43E-35 | 1,00E-08 | 0,0091999 | 0,0054865 | 0,0054865 | TRUE | 0,0438339 | No |
| orf19.5345 | 3409 | 2671 | 2985 | 2179 | 5137 | 3873 | 5213 | 5629 | 3250 | 2824 | 2949 | 3414 | 4116 | 4404 | 4426 | 4577 | 3744,95 | 3109 | 4381 | 1,41 | 0,496 | 7,824 | 5,11E-15 | 1,03E-14 | 0,00438 | 0,0025297 | 0,0035754 | 0,0035754 | TRUE | 0,0507301 | No |
| orf19.5346 | 1446 | 1213 | 1423 | 821 | 1162 | 715 | 1161 | 1185 | 1379 | 1283 | 1406 | 1286 | 931 | 813 | 986 | 964 | 1130,8 | 1338 | 924 | 0,691 | -0,534 | -7,869 | 3,58E-15 | 7,24E-15 | 0,0033633 | 0,0035433 | 0,0035195 | 0,0035195 | TRUE | 0,0722926 | No |
| orf19.5348 | 11880 | 9871 | 11245 | 7228 | 1484 | 1006 | 1395 | 1533 | 11326 | 10437 | 11109 | 11323 | 1189 | 1144 | 1184 | 1246 | 6119,94 | 11049 | 1191 | 0,108 | -3,212 | -70,631 | 0 | 0 | 0,0011044 | 0,0023596 | 0,0015609 | 0,0015609 | TRUE | 0,0170316 | No |
| orf19.535 | 58 | 38 | 39 | 18 | 669 | 425 | 543 | 655 | 55 | 40 | 39 | 28 | 536 | 483 | 461 | 533 | 271,89 | 40 | 503 | 12,077 | 3,594 | 23,994 | 3,22E-127 | 2,74E-126 | 0,005861 | 0,0081302 | 0,0074204 | 0,0074204 | TRUE | 0,4565635 | No |
| orf19.5350 | 907 | 749 | 869 | 549 | 1941 | 1496 | 1888 | 2039 | 865 | 792 | 858 | 860 | 1555 | 1701 | 1603 | 1658 | 1236,55 | 844 | 1629 | 1,93 | 0,949 | 17,322 | 3,22E-67 | 1,49E-66 | 0,0008056 | 0,0034191 | 0,0019423 | 0,0019423 | TRUE | 0,0208745 | No |
| orf19.5351 | 5765 | 4410 | 4902 | 2748 | 16297 | 9425 | 15563 | 15798 | 5496 | 4663 | 4843 | 4305 | 13058 | 10717 | 13213 | 12845 | 8642,56 | 4827 | 12458 | 2,58 | 1,367 | 16,689 | 1,58E-62 | 6,90E-62 | 0,0095719 | 0,0022813 | 0,006297 | 0,006297 | TRUE | 0,1079314 | No |
| orf19.5352 | 2701 | 2219 | 2469 | 1659 | 2719 | 1899 | 2523 | 2604 | 2575 | 2346 | 2439 | 2599 | 2179 | 2159 | 2142 | 2117 | 2319,6 | 2490 | 2149 | 0,864 | -0,211 | -4,659 | 3,18E-06 | 4,83E-06 | 0,0006339 | 0,0027991 | 0,0015344 | 0,0015344 | TRUE | 0,0181592 | No |
| orf19.5353 | 13 | 11 | 14 | 6 | 36 | 21 | 23 | 44 | 12 | 12 | 14 | 9 | 29 | 24 | 20 | 36 | 19,41 | 12 | 27 | 2,264 | 1,179 | 3,363 | 0,0007702 | 0,0010477 | 1,00E-08 | 0,0866803 | 0,0552355 | 0,0552355 | TRUE | 0,218637 | No |
| orf19.5356 | 6200 | 5202 | 5729 | 3921 | 9975 | 6917 | 9154 | 10327 | 5911 | 5500 | 5660 | 6142 | 7993 | 7865 | 7772 | 8397 | 6905,02 | 5803 | 8007 | 1,381 | 0,465 | 10,244 | 1,26E-24 | 3,09E-24 | 0,0015762 | 0,0023291 | 0,0018286 | 0,0018286 | TRUE | 0,0181424 | No |
| orf19.5357 | 2319 | 1975 | 2249 | 1587 | 3117 | 2176 | 2899 | 3269 | 2211 | 2088 | 2222 | 2486 | 2498 | 2474 | 2461 | 2658 | 2387,28 | 2252 | 2523 | 1,123 | 0,167 | 2,915 | 0,0035615 | 0,0046484 | 0,0028637 | 0,0027791 | 0,0027219 | 0,0027219 | TRUE | 0,0558843 | No |
| orf19.5358 | 1585 | 1302 | 1351 | 1011 | 2865 | 1978 | 2772 | 3018 | 1511 | 1377 | 1335 | 1584 | 2296 | 2249 | 2353 | 2454 | 1894,8 | 1452 | 2338 | 1,615 | 0,691 | 11,381 | 5,18E-30 | 1,40E-29 | 0,0031557 | 0,0029578 | 0,002962 | 0,002962 | TRUE | 0,0415395 | No |
| orf19.536 | 2944 | 2502 | 2835 | 1698 | 3517 | 2306 | 3329 | 3506 | 2807 | 2646 | 2801 | 2660 | 2818 | 2622 | 2826 | 2851 | 2753,78 | 2728 | 2779 | 1,019 | 0,027 | 0,602 | 0,5469746 | 0,5822898 | 0,000752 | 0,0026875 | 0,0015344 | 0,0015344 | TRUE | 0,0167771 | No |
| orf19.5360 | 1095 | 799 | 843 | 612 | 2042 | 1175 | 1827 | 2042 | 1044 | 845 | 833 | 959 | 1636 | 1336 | 1551 | 1660 | 1233,01 | 920 | 1546 | 1,683 | 0,751 | 8,423 | 3,68E-17 | 7,76E-17 | 0,0093279 | 0,0034229 | 0,0067262 | 0,0067262 | TRUE | 0,1030975 | No |
| orf19.5362 | 261 | 180 | 225 | 159 | 387 | 245 | 338 | 367 | 249 | 190 | 222 | 249 | 310 | 279 | 287 | 298 | 260,57 | 228 | 294 | 1,296 | 0,374 | 3,618 | 0,0002973 | 0,0004121 | 0,0033865 | 0,0083925 | 0,006265 | 0,006265 | TRUE | 0,1347092 | No |
| orf19.5363 | 1149 | 881 | 967 | 642 | 1511 | 902 | 1351 | 1468 | 1095 | 932 | 955 | 1006 | 1211 | 1026 | 1147 | 1194 | 1070,63 | 997 | 1144 | 1,149 | 0,201 | 2,778 | 0,0054713 | 0,0070573 | 0,0040604 | 0,0036249 | 0,004055 | 0,004055 | TRUE | 0,0558633 | No |
| orf19.5364 | 1712 | 1361 | 1529 | 997 | 4286 | 2742 | 4264 | 4177 | 1632 | 1439 | 1511 | 1562 | 3434 | 3118 | 3620 | 3396 | 2464,03 | 1536 | 3392 | 2,21 | 1,144 | 19,389 | 9,47E-84 | 5,33E-83 | 0,0026481 | 0,0027576 | 0,002845 | 0,002845 | TRUE | 0,0351405 | No |
| orf19.5365 | 841 | 782 | 861 | 562 | 3346 | 2482 | 3206 | 3672 | 802 | 827 | 851 | 880 | 2681 | 2822 | 2722 | 2986 | 1821,32 | 840 | 2803 | 3,344 | 1,742 | 31,534 | 3,00E-218 | 4,43E-217 | 0,0015104 | 0,0029928 | 0,0021019 | 0,0021019 | TRUE | 0,0240507 | No |
| orf19.5365.1 | 743 | 559 | 556 | 424 | 1241 | 880 | 1013 | 1272 | 708 | 591 | 549 | 664 | 994 | 1001 | 860 | 1034 | 800,28 | 628 | 972 | 1,549 | 0,631 | 7,049 | 1,80E-12 | 3,37E-12 | 0,0078539 | 0,0041429 | 0,0063492 | 0,0063492 | TRUE | 0,0941891 | No |
| orf19.5367 | 559 | 493 | 517 | 341 | 1575 | 1060 | 1530 | 1537 | 533 | 521 | 511 | 534 | 1262 | 1205 | 1299 | 1250 | 889,4 | 525 | 1254 | 2,394 | 1,259 | 21,999 | 2,93E-107 | 2,05E-106 | 1,00E-08 | 0,0039374 | 0,0017299 | 0,0017299 | TRUE | 0,0075453 | No |
| orf19.5368 | 828 | 657 | 732 | 617 | 1912 | 1292 | 1770 | 1786 | 789 | 695 | 723 | 967 | 1532 | 1469 | 1503 | 1452 | 1141,23 | 794 | 1489 | 1,886 | 0,915 | 10,53 | 6,27E-26 | 1,58E-25 | 0,0091267 | 0,00353 | 0,0062368 | 0,0062368 | TRUE | 0,2367103 | No |
| orf19.5369 | 1293 | 1064 | 1173 | 901 | 5982 | 4102 | 5694 | 6246 | 1233 | 1125 | 1159 | 1411 | 4793 | 4664 | 4834 | 5079 | 3037,3 | 1232 | 4842 | 3,945 | 1,98 | 30,372 | 1,29E-202 | 1,79E-201 | 0,0043816 | 0,0026319 | 0,0035301 | 0,0035301 | TRUE | 0,1062025 | No |
| orf19.537 | 260 | 190 | 239 | 147 | 456 | 289 | 450 | 439 | 248 | 201 | 236 | 230 | 365 | 329 | 382 | 357 | 293,52 | 229 | 358 | 1,568 | 0,649 | 6,792 | 1,10E-11 | 2,03E-11 | 0,0017726 | 0,0076851 | 0,0050768 | 0,0050768 | TRUE | 0,0785358 | No |
| orf19.5370 | 746 | 603 | 637 | 429 | 1453 | 895 | 1403 | 1444 | 711 | 638 | 629 | 672 | 1164 | 1018 | 1191 | 1174 | 899,67 | 662 | 1137 | 1,719 | 0,781 | 11,142 | 7,86E-29 | 2,08E-28 | 0,0031175 | 0,0039163 | 0,0034843 | 0,0034843 | TRUE | 0,0564746 | No |
| orf19.5376 | 648 | 605 | 674 | 404 | 592 | 419 | 568 | 569 | 618 | 640 | 666 | 633 | 474 | 476 | 482 | 463 | 556,49 | 639 | 474 | 0,741 | -0,432 | -6,743 | 1,55E-11 | 2,84E-11 | 1,00E-08 | 0,0050417 | 0,0021335 | 0,0021335 | TRUE | 0,0101152 | No |
| orf19.5377 | 2526 | 2198 | 2396 | 1439 | 1844 | 1269 | 1699 | 1923 | 2408 | 2324 | 2367 | 2254 | 1478 | 1443 | 1442 | 1564 | 1910,03 | 2338 | 1482 | 0,633 | -0,659 | -14,244 | 4,91E-46 | 1,71E-45 | 0,0005446 | 0,0029509 | 0,0015138 | 0,0015138 | TRUE | 0,0170436 | No |
| orf19.5378 | 2644 | 2321 | 2545 | 1377 | 4292 | 2945 | 4012 | 4187 | 2521 | 2454 | 2514 | 2157 | 3439 | 3349 | 3406 | 3404 | 2905,59 | 2412 | 3400 | 1,407 | 0,492 | 9,479 | 2,57E-21 | 5,91E-21 | 0,002139 | 0,0026564 | 0,0022261 | 0,0022261 | TRUE | 0,0568571 | No |
| orf19.5379 | 1985 | 1766 | 1751 | 1284 | 1526 | 1154 | 1421 | 1562 | 1892 | 1867 | 1730 | 2011 | 1223 | 1312 | 1206 | 1270 | 1564,06 | 1875 | 1253 | 0,669 | -0,581 | -10,22 | 1,61E-24 | 3,95E-24 | 0,0017416 | 0,003141 | 0,0024418 | 0,0024418 | TRUE | 0,0330204 | No |
| orf19.538 | 314 | 279 | 297 | 234 | 363 | 288 | 348 | 404 | 299 | 295 | 293 | 367 | 291 | 327 | 295 | 328 | 312,08 | 314 | 310 | 0,996 | -0,005 | -0,055 | 0,9562306 | 0,960445 | 0,0036987 | 0,0073524 | 0,0057014 | 0,0057014 | TRUE | 0,1238904 | No |
| orf19.5380 | 761 | 734 | 767 | 432 | 1199 | 805 | 1010 | 1141 | 726 | 776 | 758 | 677 | 961 | 915 | 857 | 928 | 824,68 | 734 | 915 | 1,244 | 0,315 | 4,913 | 8,95E-07 | 1,39E-06 | 0,001428 | 0,0040822 | 0,002686 | 0,002686 | TRUE | 0,0277274 | No |
| orf19.5381 | 1352 | 1333 | 1412 | 971 | 1295 | 946 | 1209 | 1234 | 1289 | 1409 | 1395 | 1521 | 1038 | 1076 | 1026 | 1003 | 1219,7 | 1404 | 1036 | 0,739 | -0,436 | -7,38 | 1,58E-13 | 3,07E-13 | 0,0015952 | 0,0034374 | 0,0025158 | 0,0025158 | TRUE | 0,0366037 | No |
| orf19.5382 | 0 | 0 | 0 | 0 | 0 | 0 | 0 | 0 | 0 | 0 | 0 | 0 | 0 | 0 | 0 | 0 | 0 | 0 | 0 | NA | NA | NA | NA | NA | NA | NA | NA | NA | NA | NA | NA |
| orf19.5383 | 4131 | 3571 | 3894 | 2657 | 6335 | 4552 | 5717 | 6544 | 3938 | 3776 | 3847 | 4162 | 5076 | 5176 | 4854 | 5321 | 4518,78 | 3931 | 5107 | 1,3 | 0,379 | 8,325 | 8,44E-17 | 1,76E-16 | 0,0013525 | 0,0024546 | 0,001757 | 0,001757 | TRUE | 0,0180862 | No |
| orf19.5384 | 969 | 910 | 962 | 626 | 1763 | 1132 | 1600 | 1654 | 924 | 962 | 950 | 981 | 1413 | 1287 | 1358 | 1345 | 1152,52 | 954 | 1351 | 1,419 | 0,505 | 9,704 | 2,91E-22 | 6,83E-22 | 1,00E-08 | 0,0035159 | 0,0016854 | 0,0016854 | TRUE | 0,0114373 | No |

| | | | | | | | | | | | | | | | | | | | | | | | | | | | | | | | |
|---|---|---|---|---|---|---|---|---|---|---|---|---|---|---|---|---|---|---|---|---|---|---|---|---|---|---|---|---|---|---|---|
| orf19.5387 | 777 | 834 | 806 | 527 | 997 | 794 | 923 | 1049 | 741 | 882 | 796 | 826 | 799 | 903 | 784 | 853 | 822,84 | 811 | 835 | 1,029 | 0,041 | 0,562 | 0,5739637 | 0,6081804 | 0,0035403 | 0,0040867 | 0,003793 | 0,003793 | TRUE | 0,0432981 | No |
| orf19.5388 | 1108 | 958 | 1065 | 669 | 1306 | 858 | 1289 | 1310 | 1056 | 1013 | 1052 | 1048 | 1046 | 976 | 1094 | 1065 | 1043,88 | 1042 | 1045 | 1,005 | 0,007 | 0,121 | 0,9033546 | 0,9168611 | 0,000365 | 0,0036642 | 0,0018681 | 0,0018681 | TRUE | 0,0220598 | No |
| orf19.5389 | 1226 | 1116 | 1192 | 866 | 609 | 515 | 628 | 623 | 1169 | 1180 | 1178 | 1357 | 488 | 586 | 533 | 507 | 874,55 | 1221 | 528 | 0,433 | -1,209 | -16,195 | 5,44E-59 | 2,26E-58 | 0,0041799 | 0,0039687 | 0,0040401 | 0,0040401 | TRUE | 0,0625263 | No |
| orf19.539 | 831 | 739 | 780 | 491 | 1440 | 1042 | 1373 | 1623 | 792 | 781 | 771 | 769 | 1154 | 1185 | 1166 | 1320 | 992,18 | 778 | 1206 | 1,549 | 0,632 | 10,3 | 7,06E-25 | 1,74E-24 | 0,0016705 | 0,0037461 | 0,0025204 | 0,0025204 | TRUE | 0,0498164 | No |
| orf19.5390 | 0 | 0 | 0 | 0 | 0 | 0 | 0 | 0 | 0 | 0 | 0 | 0 | 0 | 0 | 0 | 0 | 0 | 0 | 0 | NA | NA | NA | NA | NA | NA | NA | NA | NA | NA | NA | NA |
| orf19.5391 | 1707 | 1737 | 1818 | 1133 | 2127 | 1585 | 1871 | 2151 | 1627 | 1837 | 1796 | 1775 | 1704 | 1802 | 1588 | 1749 | 1734,88 | 1759 | 1711 | 0,973 | -0,04 | -0,695 | 0,4871438 | 0,5233987 | 0,0022871 | 0,0030377 | 0,0025959 | 0,0025959 | TRUE | 0,0318424 | No |
| orf19.5392 | 54 | 34 | 35 | 22 | 31 | 24 | 20 | 35 | 51 | 36 | 35 | 34 | 25 | 27 | 17 | 28 | 31,76 | 39 | 24 | 0,617 | -0,697 | -2,573 | 0,0100698 | 0,0127871 | 0,0170878 | 0,0537961 | 0,0379996 | 0,0379996 | TRUE | 0,3626176 | No |
| orf19.5393 | 5202 | 4989 | 5375 | 3109 | 9019 | 6372 | 8242 | 8943 | 4960 | 5275 | 5310 | 4870 | 7227 | 7246 | 6997 | 7272 | 6144,54 | 5104 | 7186 | 1,407 | 0,492 | 12,346 | 5,11E-35 | 1,50E-34 | 0,0008735 | 0,0023585 | 0,0013551 | 0,0013551 | TRUE | 0,0108835 | No |
| orf19.5394.1 | 1696 | 1395 | 1450 | 998 | 1093 | 759 | 1040 | 1106 | 1617 | 1475 | 1432 | 1563 | 876 | 863 | 883 | 899 | 1201,12 | 1522 | 880 | 0,579 | -0,788 | -14,199 | 9,34E-46 | 3,24E-45 | 0,0011183 | 0,0034583 | 0,0020839 | 0,0020839 | TRUE | 0,0233103 | No |
| orf19.5395 | 2957 | 2706 | 2877 | 1838 | 1661 | 1198 | 1527 | 1667 | 2819 | 2861 | 2842 | 2879 | 1331 | 1362 | 1296 | 1355 | 2093,37 | 2850 | 1336 | 0,469 | -1,093 | -27,466 | 4,51E-166 | 5,05E-165 | 1,00E-08 | 0,0028756 | 0,0009907 | 0,0009907 | TRUE | 0,0048462 | No |
| orf19.5397 | 324 | 312 | 327 | 202 | 511 | 361 | 480 | 498 | 309 | 330 | 323 | 316 | 409 | 410 | 408 | 405 | 363,83 | 320 | 408 | 1,277 | 0,352 | 4,645 | 3,41E-06 | 5,16E-06 | 1,00E-08 | 0,0066041 | 0,0026919 | 0,0026919 | TRUE | 0,0067201 | No |
| orf19.5399 | 31 | 31 | 15 | 18 | 69 | 37 | 78 | 75 | 30 | 33 | 15 | 28 | 55 | 42 | 66 | 61 | 41,24 | 26 | 56 | 2,156 | 1,109 | 4,179 | 2,93E-05 | 4,26E-05 | 0,0325389 | 0,0419054 | 0,0384184 | 0,0384184 | TRUE | 0,5984555 | No |
| orf19.54 | 302 | 250 | 282 | 203 | 1873 | 1547 | 1789 | 1835 | 288 | 264 | 279 | 318 | 1501 | 1759 | 1519 | 1492 | 927,45 | 287 | 1568 | 5,48 | 2,454 | 30,034 | 3,53E-198 | 4,81E-197 | 0,0044699 | 0,0038616 | 0,0041738 | 0,0041738 | TRUE | 0,0790089 | No |
| orf19.540 | 393 | 303 | 385 | 215 | 1022 | 677 | 959 | 1039 | 375 | 320 | 380 | 337 | 819 | 770 | 814 | 845 | 582,49 | 353 | 812 | 2,294 | 1,198 | 16,071 | 4,07E-58 | 1,67E-57 | 0,0010779 | 0,00491 | 0,0032114 | 0,0032114 | TRUE | 0,0496767 | No |
| orf19.5402 | 0 | 0 | 0 | 0 | 0 | 0 | 0 | 0 | 0 | 0 | 0 | 0 | 0 | 0 | 0 | 0 | 0 | 0 | 0 | NA | NA | NA | NA | NA | NA | NA | NA | NA | NA | NA | NA |
| orf19.5404.1 | 1129 | 993 | 1111 | 744 | 2231 | 1569 | 2105 | 2355 | 1076 | 1050 | 1098 | 1166 | 1788 | 1784 | 1787 | 1915 | 1457,89 | 1098 | 1818 | 1,662 | 0,733 | 14,193 | 1,02E-45 | 3,52E-45 | 0,0006656 | 0,0032175 | 0,0018021 | 0,0018021 | TRUE | 0,0181099 | No |
| orf19.5406 | 4036 | 3538 | 3864 | 2249 | 3201 | 2310 | 3033 | 3289 | 3848 | 3741 | 3817 | 3523 | 2565 | 2627 | 2575 | 2674 | 3171,27 | 3732 | 2610 | 0,698 | -0,518 | -12,512 | 6,42E-36 | 1,90E-35 | 0,000524 | 0,002609 | 0,0013236 | 0,0013236 | TRUE | 0,0161072 | No |
| orf19.5407 | 14021 | 11945 | 12862 | 7879 | 16052 | 10691 | 15402 | 16569 | 13368 | 12630 | 12706 | 12343 | 12862 | 12157 | 13076 | 13472 | 12826,77 | 12762 | 12892 | 1,01 | 0,015 | 0,341 | 0,7331448 | 0,7617972 | 0,0012304 | 0,0022193 | 0,0016743 | 0,0016743 | TRUE | 0,0178288 | No |
| orf19.5408 | 7561 | 6412 | 7223 | 4628 | 6426 | 4490 | 6224 | 6579 | 7209 | 6780 | 7136 | 7250 | 5149 | 5106 | 5284 | 5349 | 6157,77 | 7094 | 5222 | 0,737 | -0,441 | -12,131 | 7,25E-34 | 2,08E-33 | 0,0004733 | 0,0023579 | 0,0011052 | 0,0011052 | TRUE | 0,0108647 | No |
| orf19.5409 | 501 | 418 | 490 | 308 | 492 | 369 | 511 | 527 | 478 | 442 | 484 | 482 | 394 | 420 | 434 | 429 | 445,29 | 472 | 419 | 0,889 | -0,17 | -2,32 | 0,0203234 | 0,0252008 | 1,00E-08 | 0,0057785 | 0,0028927 | 0,0028927 | TRUE | 0,0211361 | No |
| orf19.541 | 2366 | 2025 | 2264 | 1499 | 4795 | 3599 | 4751 | 5152 | 2256 | 2141 | 2237 | 2348 | 3842 | 4092 | 4034 | 4189 | 3142,36 | 2246 | 4039 | 1,801 | 0,849 | 18,762 | 1,53E-78 | 8,19E-78 | 0,0009084 | 0,0026138 | 0,0016024 | 0,0016024 | TRUE | 0,0133834 | No |
| orf19.5410 | 1293 | 1044 | 1253 | 759 | 1287 | 859 | 1184 | 1172 | 1233 | 1104 | 1238 | 1189 | 1031 | 977 | 1005 | 953 | 1091,21 | 1191 | 992 | 0,833 | -0,264 | -4,617 | 3,89E-06 | 5,88E-06 | 0,0011748 | 0,0035959 | 0,0022293 | 0,0022293 | TRUE | 0,0302736 | No |
| orf19.5411 | 967 | 799 | 963 | 639 | 992 | 701 | 948 | 1000 | 922 | 845 | 951 | 1001 | 795 | 797 | 805 | 813 | 866,13 | 930 | 802 | 0,865 | -0,209 | -3,382 | 0,000719 | 0,0009803 | 0,0009765 | 0,003987 | 0,0024915 | 0,0024915 | TRUE | 0,0443035 | No |
| orf19.5412 | 1549 | 1305 | 1470 | 941 | 1597 | 1216 | 1564 | 1663 | 1477 | 1380 | 1452 | 1474 | 1280 | 1383 | 1328 | 1352 | 1390,67 | 1446 | 1336 | 0,923 | -0,115 | -2,354 | 0,0185839 | 0,0231384 | 0,0003073 | 0,0032719 | 0,0015709 | 0,0015709 | TRUE | 0,0114892 | No |
| orf19.5413 | 1467 | 1196 | 1324 | 859 | 1445 | 972 | 1376 | 1433 | 1399 | 1265 | 1308 | 1346 | 1158 | 1105 | 1168 | 1165 | 1239,17 | 1330 | 1149 | 0,865 | -0,209 | -4,045 | 5,24E-05 | 7,51E-05 | 0,0004671 | 0,0034163 | 0,0017431 | 0,0017431 | TRUE | 0,0162687 | No |
| orf19.5416 | 7673 | 6941 | 7685 | 4787 | 3859 | 2722 | 3687 | 3966 | 7315 | 7339 | 7592 | 7499 | 3092 | 3095 | 3130 | 3225 | 5286 | 7436 | 3136 | 0,422 | -1,245 | -37,305 | 1,38E-304 | 3,10E-303 | 0,0001712 | 0,0024019 | 0,0008519 | 0,0008519 | TRUE | 0,0045508 | No |
| orf19.5417 | 13564 | 10673 | 11939 | 7925 | 14083 | 9260 | 13602 | 14489 | 12932 | 11285 | 11795 | 12415 | 11284 | 10529 | 11548 | 11781 | 11696,18 | 12107 | 11286 | 0,932 | -0,101 | -1,902 | 0,0572168 | 0,0683581 | 0,0028053 | 0,0022317 | 0,0026227 | 0,0026227 | TRUE | 0,0260472 | No |
| orf19.5418 | 1564 | 1341 | 1413 | 937 | 3019 | 2075 | 2953 | 3033 | 1491 | 1418 | 1396 | 1468 | 2419 | 2359 | 2507 | 2466 | 1940,56 | 1443 | 2438 | 1,691 | 0,758 | 17,052 | 3,37E-65 | 1,52E-64 | 0,0002543 | 0,0029374 | 0,0013256 | 0,0013256 | TRUE | 0,006857 | No |
| orf19.5419 | 4880 | 4043 | 4373 | 2986 | 13873 | 9681 | 13539 | 13976 | 4653 | 4275 | 4320 | 4678 | 11116 | 11008 | 11495 | 11364 | 7863,49 | 4482 | 11246 | 2,511 | 1,328 | 32,094 | 5,28E-226 | 8,01E-225 | 0,001121 | 0,0023001 | 0,0014797 | 0,0014797 | TRUE | 0,0115952 | No |
| orf19.542 | 4983 | 4098 | 4554 | 2720 | 1946 | 1393 | 1844 | 2102 | 4751 | 4333 | 4499 | 4261 | 1559 | 1584 | 1566 | 1709 | 3032,72 | 4461 | 1604 | 0,36 | -1,476 | -28,793 | 2,66E-182 | 3,24E-181 | 0,0015888 | 0,0026327 | 0,0021185 | 0,0021185 | TRUE | 0,0239026 | No |
| orf19.5420 | 5162 | 4070 | 4609 | 3058 | 5803 | 4053 | 5811 | 5939 | 4921 | 4304 | 4553 | 4791 | 4650 | 4609 | 4934 | 4829 | 4698,7 | 4642 | 4756 | 1,025 | 0,035 | 0,707 | 0,4793644 | 0,515735 | 0,0018403 | 0,0024407 | 0,0021792 | 0,0021792 | TRUE | 0,0295665 | No |
| orf19.5422 | 56 | 43 | 66 | 46 | 107 | 78 | 100 | 104 | 53 | 45 | 65 | 72 | 86 | 89 | 85 | 85 | 72,5 | 59 | 86 | 1,473 | 0,559 | 3,182 | 0,0014643 | 0,0019539 | 1,00E-08 | 0,0247378 | 0,0148862 | 0,0148862 | TRUE | 0,1946017 | No |
| orf19.542.2 | 1253 | 1019 | 1130 | 741 | 1205 | 805 | 1072 | 1229 | 1195 | 1077 | 1116 | 1161 | 966 | 915 | 910 | 999 | 1042,44 | 1137 | 948 | 0,834 | -0,261 | -4,511 | 6,45E-06 | 9,65E-06 | 0,0011515 | 0,0036663 | 0,0022586 | 0,0022586 | TRUE | 0,016304 | No |
| orf19.5423 | 42 | 49 | 35 | 32 | 75 | 55 | 63 | 98 | 40 | 52 | 35 | 50 | 60 | 63 | 53 | 80 | 54,05 | 44 | 64 | 1,465 | 0,551 | 2,551 | 0,0107467 | 0,0136196 | 0,0157667 | 0,0324712 | 0,0250493 | 0,0250493 | TRUE | 0,266215 | No |
| orf19.5424 | 3 | 0 | 3 | 0 | 2 | 1 | 4 | 1 | 3 | 0 | 3 | 0 | 2 | 1 | 3 | 1 | 1,6 | 2 | 2 | 1,124 | 0,169 | 0,143 | 0,8862838 | 0,9016863 | 0,1697919 | 1,0304823 | 0,7288569 | 0,7288569 | TRUE | 0,1504594 | No |
| orf19.5425 | 6491 | 5568 | 6298 | 3975 | 7725 | 5328 | 7158 | 7763 | 6188 | 5887 | 6222 | 6227 | 6190 | 6058 | 6077 | 6312 | 6145,28 | 6131 | 6159 | 1,005 | 0,007 | 0,209 | 0,8348313 | 0,8549283 | 0,0002987 | 0,0023585 | 0,0009767 | 0,0009767 | TRUE | 0,0088168 | No |
| orf19.5426 | 1874 | 1631 | 1751 | 1118 | 2469 | 1649 | 2412 | 2632 | 1787 | 1725 | 1730 | 1751 | 1978 | 1875 | 2048 | 2140 | 1879,21 | 1748 | 2010 | 1,151 | 0,203 | 4,009 | 6,10E-05 | 8,72E-05 | 0,0012475 | 0,002965 | 0,0019205 | 0,0019205 | TRUE | 0,02354 | No |
| orf19.5428 | 134 | 150 | 122 | 89 | 297 | 196 | 274 | 307 | 128 | 159 | 121 | 139 | 238 | 223 | 233 | 250 | 186,18 | 137 | 236 | 1,735 | 0,795 | 6,883 | 5,84E-12 | 1,08E-11 | 0,0006091 | 0,0109104 | 0,0068219 | 0,0068219 | TRUE | 0,1327398 | No |
| orf19.5429 | 2080 | 1836 | 2032 | 1303 | 863 | 645 | 819 | 928 | 1983 | 1941 | 2007 | 2041 | 691 | 733 | 695 | 755 | 1355,98 | 1993 | 718 | 0,361 | -1,472 | -29,057 | 1,26E-185 | 1,57E-184 | 1,00E-08 | 0,0033021 | 0,0015613 | 0,0015613 | TRUE | 0,0147828 | No |
| orf19.543 | 774 | 689 | 713 | 465 | 595 | 410 | 590 | 632 | 738 | 729 | 704 | 728 | 477 | 466 | 501 | 514 | 607,13 | 725 | 490 | 0,677 | -0,564 | -8,876 | 6,93E-19 | 1,52E-18 | 1,00E-08 | 0,0047957 | 0,002195 | 0,002195 | TRUE | 0,0132693 | No |
| orf19.5430 | 5163 | 4137 | 4540 | 2727 | 7734 | 4654 | 7531 | 7521 | 4922 | 4374 | 4485 | 4272 | 6197 | 5292 | 6394 | 6115 | 5256,49 | 4513 | 6000 | 1,329 | 0,41 | 6,193 | 5,92E-10 | 1,03E-09 | 0,0050313 | 0,0024036 | 0,0040208 | 0,0040208 | TRUE | 0,0761059 | No |
| orf19.5431 | 851 | 781 | 854 | 629 | 725 | 530 | 728 | 753 | 811 | 826 | 844 | 985 | 581 | 603 | 618 | 612 | 735,01 | 866 | 604 | 0,699 | -0,516 | -7,203 | 5,87E-13 | 1,12E-12 | 0,0027221 | 0,0043251 | 0,0035316 | 0,0035316 | TRUE | 0,0912469 | No |
| orf19.5432 | 1224 | 1102 | 1183 | 780 | 1787 | 1328 | 1882 | 1918 | 1167 | 1165 | 1169 | 1222 | 1432 | 1510 | 1598 | 1560 | 1352,75 | 1181 | 1525 | 1,293 | 0,371 | 7,096 | 1,29E-12 | 2,43E-12 | 0,0008852 | 0,003305 | 0,0018569 | 0,0018569 | TRUE | 0,0212853 | No |
| orf19.5433 | 556 | 474 | 475 | 300 | 768 | 434 | 719 | 759 | 530 | 501 | 469 | 470 | 615 | 493 | 610 | 617 | 538,37 | 492 | 584 | 1,187 | 0,247 | 2,852 | 0,0043383 | 0,0056278 | 0,0055097 | 0,0051411 | 0,0053275 | 0,0053275 | TRUE | 0,1195952 | No |
| orf19.5436 | 12961 | 11348 | 12340 | 7960 | 6602 | 4665 | 6351 | 6970 | 12357 | 11999 | 12191 | 12470 | 5290 | 5305 | 5392 | 5667 | 8833,78 | 12254 | 5414 | 0,442 | -1,178 | -33,805 | 1,66E-250 | 2,84E-249 | 0,0004856 | 0,0022771 | 0,0010378 | 0,0010378 | TRUE | 0,012304 | No |
| orf19.5437 | 86361 | 67816 | 73595 | 49806 | 199241 | 135333 | 193204 | 207360 | 82336 | 71707 | 72705 | 78024 | 159646 | 153886 | 164029 | 168603 | 118867 | 76193 | 161541 | 2,12 | 1,084 | 20,937 | 2,45E-97 | 1,56E-96 | 0,0028491 | 0,0021051 | 0,0025664 | 0,0025664 | TRUE | 0,036168 | No |
| orf19.5438 | 1702 | 1341 | 1437 | 933 | 5159 | 3148 | 4930 | 4968 | 1623 | 1418 | 1420 | 1462 | 4134 | 3580 | 4186 | 4039 | 2732,52 | 1481 | 3985 | 2,691 | 1,428 | 21,98 | 4,50E-107 | 3,14E-106 | 0,0039373 | 0,0026922 | 0,0035631 | 0,0035631 | TRUE | 0,0560414 | No |
| orf19.5439 | 462 | 454 | 461 | 314 | 1277 | 882 | 1193 | 1189 | 440 | 480 | 455 | 492 | 1023 | 1003 | 1013 | 967 | 734,2 | 467 | 1002 | 2,153 | 1,106 | 17,792 | 8,17E-71 | 3,95E-70 | 1,00E-08 | 0,0043276 | 0,0020688 | 0,0020688 | TRUE | 0,0171431 | No |
| orf19.544 | 888 | 793 | 802 | 546 | 989 | 767 | 991 | 1099 | 847 | 838 | 792 | 855 | 792 | 872 | 841 | 894 | 841,54 | 833 | 850 | 1,02 | 0,029 | 0,48 | 0,6313981 | 0,6640763 | 0,0009902 | 0,0040424 | 0,0023584 | 0,0023584 | TRUE | 0,0256461 | No |
| orf19.5440 | 17491 | 13971 | 15724 | 9461 | 13431 | 9009 | 13042 | 13815 | 16676 | 14773 | 15534 | 14821 | 10762 | 10244 | 11073 | 11233 | 13139,34 | 15451 | 10828 | 0,701 | -0,513 | -10,296 | 7,33E-25 | 1,81E-24 | 0,0023547 | 0,0022162 | 0,002307 | 0,002307 | TRUE | 0,0349715 | No |
| orf19.5441 | 1554 | 1406 | 1593 | 986 | 2339 | 1587 | 2128 | 2356 | 1482 | 1487 | 1574 | 1545 | 1874 | 1805 | 1807 | 1916 | 1685,96 | 1522 | 1850 | 1,217 | 0,284 | 6,167 | 6,97E-10 | 1,20E-09 | 0,0003353 | 0,0030652 | 0,0014238 | 0,0014238 | TRUE | 0,0071119 | No |
| orf19.544.1 | 7242 | 5679 | 6078 | 3804 | 6551 | 4513 | 6265 | 6740 | 6904 | 6005 | 6004 | 5959 | 5249 | 5132 | 5319 | 5480 | 5756,62 | 6218 | 5295 | 0,851 | -0,232 | -4,283 | 1,85E-05 | 2,70E-05 | 0,0028577 | 0,0023765 | 0,0026495 | 0,0026495 | TRUE | 0,0680107 | No |
| orf19.5442 | 2757 | 2331 | 2642 | 1564 | 1159 | 741 | 1110 | 1248 | 2629 | 2465 | 2610 | 2450 | 929 | 843 | 942 | 1015 | 1735,22 | 2538 | 932 | 0,368 | -1,443 | -24,237 | 9,18E-130 | 7,94E-129 | 0,002577 | 0,0030375 | 0,0027054 | 0,0027054 | TRUE | 0,0462597 | No |
| orf19.5443 | 6086 | 5043 | 5542 | 3193 | 2743 | 1801 | 2638 | 2839 | 5802 | 5332 | 5475 | 5002 | 2198 | 2048 | 2240 | 2308 | 3800,68 | 5403 | 2198 | 0,407 | -1,297 | -23,14 | 1,82E-118 | 1,43E-117 | 0,0027169 | 0,0025233 | 0,0027105 | 0,0027105 | TRUE | 0,0301842 | No |
| orf19.5444 | 3661 | 3043 | 3342 | 2071 | 6803 | 4673 | 6694 | 7013 | 3490 | 3218 | 3302 | 3244 | 5451 | 5314 | 5683 | 5702 | 4425,9 | 3314 | 5538 | 1,671 | 0,74 | 17,236 | 1,42E-66 | 6,52E-66 | 0,0009436 | 0,0024623 | 0,0015216 | 0,0015216 | TRUE | 0,0161074 | No |
| orf19.5445 | 2890 | 2208 | 2669 | 1741 | 5565 | 3822 | 5539 | 5859 | 2755 | 2335 | 2637 | 2727 | 4459 | 4346 | 4703 | 4764 | 3590,7 | 2614 | 4568 | 1,749 | 0,806 | 13,686 | 1,24E-42 | 4,08E-42 | 0,0032559 | 0,0025485 | 0,0030213 | 0,0030213 | TRUE | 0,0630717 | No |
| orf19.5446 | 0 | 2 | 2 | 4 | 4 | 2 | 5 | 4 | 0 | 2 | 2 | 6 | 3 | 2 | 4 | 3 | 2,92 | 2 | 3 | 1,393 | 0,478 | 0,539 | 0,590058 | 0,6243011 | 1,26E-08 | 0,5650281 | 0,382763 | 0,382763 | TRUE | 0,5723716 | No |
| orf19.5447 | 3376 | 2733 | 2974 | 2074 | 5974 | 4166 | 5434 | 6428 | 3219 | 2890 | 2938 | 3249 | 4787 | 4737 | 4613 | 5227 | 3957,43 | 3074 | 4841 | 1,576 | 0,656 | 11,553 | 7,16E-31 | 1,96E-30 | 0,0030027 | 0,0025062 | 0,0028246 | 0,0028246 | TRUE | 0,0353081 | No |
| orf19.5449 | 60 | 51 | 54 | 44 | 633 | 417 | 575 | 598 | 57 | 54 | 53 | 69 | 507 | 474 | 488 | 486 | 273,65 | 58 | 489 | 8,514 | 3,09 | 25,037 | 2,44E-138 | 2,26E-137 | 1,00E-08 | 0,0080913 | 0,0041203 | 0,0041203 | TRUE | 0,0827094 | No |
| orf19.5450 | 2658 | 1985 | 2180 | 1502 | 3760 | 2412 | 3672 | 3691 | 2534 | 2099 | 2154 | 2353 | 3013 | 2743 | 3118 | 3001 | 2626,71 | 2285 | 2969 | 1,3 | 0,379 | 5,663 | 1,49E-08 | 2,47E-08 | 0,0047108 | 0,0027164 | 0,003894 | 0,003894 | TRUE | 0,067688 | No |
| orf19.5451 | 681 | 529 | 652 | 574 | 171 | 126 | 145 | 171 | 649 | 559 | 644 | 899 | 137 | 143 | 123 | 139 | 411,8 | 688 | 136 | 0,198 | -2,339 | -16,968 | 1,41E-64 | 6,31E-64 | 0,0223131 | 0,0060785 | 0,0141353 | 0,0141353 | TRUE | 0,4794284 | No |
| orf19.5454 | 603 | 469 | 520 | 363 | 1446 | 944 | 1391 | 1442 | 575 | 496 | 514 | 569 | 1159 | 1073 | 1181 | 1172 | 842,33 | 538 | 1146 | 2,136 | 1,095 | 16,33 | 6,07E-60 | 2,55E-59 | 0,0017941 | 0,0040405 | 0,0028881 | 0,0028881 | TRUE | 0,0326222 | No |
| orf19.5455 | 2806 | 2170 | 2244 | 1369 | 2100 | 1304 | 2045 | 2150 | 2675 | 2295 | 2217 | 2145 | 1683 | 1483 | 1736 | 1748 | 1997,62 | 2333 | 1662 | 0,713 | -0,489 | -6,298 | 3,01E-10 | 5,28E-10 | 0,007188 | 0,0029132 | 0,0052768 | 0,0052768 | TRUE | 0,119417 | No |
| orf19.5457 | 2483 | 1887 | 2130 | 1363 | 3784 | 2321 | 3714 | 3669 | 2367 | 1995 | 2104 | 2135 | 3032 | 2639 | 3153 | 2983 | 2551,2 | 2150 | 2952 | 1,373 | 0,457 | 6,692 | 2,20E-11 | 4,01E-11 | 0,0048984 | 0,0027349 | 0,0040728 | 0,0040728 | TRUE | 0,0605332 | No |
| orf19.5459 | 6804 | 4955 | 5986 | 3946 | 5052 | 3246 | 4776 | 5025 | 6487 | 5239 | 5914 | 6182 | 4048 | 3691 | 4055 | 4086 | 4962,63 | 5956 | 3970 | 0,667 | -0,584 | -8,87 | 7,33E-19 | 1,61E-18 | 0,0048116 | 0,0024221 | 0,0039633 | 0,0039633 | TRUE | 0,080283 | No |
| orf19.5461 | 51 | 23 | 28 | 24 | 24 | 26 | 20 | 28 | 49 | 24 | 28 | 38 | 19 | 30 | 17 | 23 | 28,34 | 35 | 22 | 0,634 | -0,658 | -2,138 | 0,0325265 | 0,0397076 | 0,0473085 | 0,0600214 | 0,0543011 | 0,0543011 | TRUE | 0,4054349 | No |
| orf19.5462 | 5 | 1 | 2 | 0 | 2 | 1 | 1 | 1 | 5 | 1 | 2 | 0 | 2 | 1 | 1 | 1 | 1,53 | 2 | 1 | 0,532 | -0,911 | -0,78 | 0,4351047 | 0,4712364 | 1,00E-08 | 1,0785846 | 0,6368288 | 0,6368288 | TRUE | 0,524369 | No |
| orf19.5463 | 1533 | 1236 | 1466 | 816 | 2778 | 1729 | 2676 | 2756 | 1462 | 1307 | 1448 | 1278 | 2226 | 1966 | 2272 | 2241 | 1774,98 | 1374 | 2176 | 1,583 | 0,662 | 10,139 | 3,70E-24 | 9,03E-24 | 0,0036898 | 0,0030163 | 0,0034821 | 0,0034821 | TRUE | 0,0496408 | No |
| orf19.5464 | 103 | 76 | 103 | 65 | 218 | 142 | 177 | 221 | 98 | 80 | 102 | 102 | 175 | 161 | 150 | 180 | 131,03 | 96 | 166 | 1,75 | 0,807 | 5,992 | 2,07E-09 | 3,53E-09 | 1,00E-08 | 0,0146219 | 0,0089235 | 0,0089235 | TRUE | 0,1121811 | No |
| orf19.5465 | 1283 | 1053 | 1248 | 782 | 2703 | 1775 | 2573 | 2784 | 1223 | 1113 | 1233 | 1225 | 2166 | 2018 | 2184 | 2264 | 1678,36 | 1198 | 2158 | 1,803 | 0,85 | 15,432 | 9,99E-54 | 3,88E-53 | 0,0016349 | 0,0030696 | 0,0022388 | 0,0022388 | TRUE | 0,0280489 | No |
| orf19.5466 | 12074 | 9100 | 9664 | 6308 | 42722 | 24497 | 37677 | 40867 | 11511 | 9622 | 9547 | 9882 | 34232 | 27855 | 31988 | 33229 | 20983,23 | 10140 | 31826 | 3,138 | 1,65 | 21,907 | 2,20E-106 | 1,53E-105 | 0,0079483 | 0,0021695 | 0,0053805 | 0,0053805 | TRUE | 0,1016572 | No |
| orf19.5467 | 12 | 12 | 17 | 9 | 49 | 32 | 54 | 44 | 11 | 13 | 17 | 14 | 39 | 36 | 46 | 36 | 26,54 | 14 | 39 | 2,87 | 1,521 | 5,109 | 3,23E-07 | 5,10E-07 | 1,00E-08 | 0,0639642 | 0,033446 | 0,033446 | TRUE | 0,113547 | No |
| orf19.5468 | 94 | 80 | 77 | 44 | 374 | 188 | 295 | 295 | 90 | 85 | 76 | 69 | 300 | 214 | 250 | 240 | 165,37 | 80 | 251 | 3,128 | 1,645 | 11,272 | 1,80E-29 | 4,82E-29 | 0,0114911 | 0,0120199 | 0,011766 | 0,011766 | TRUE | 0,1945592 | No |

| | | | | | | | | | | | | | | | | | | | | | | | | | | | | | | | |
|---|---|---|---|---|---|---|---|---|---|---|---|---|---|---|---|---|---|---|---|---|---|---|---|---|---|---|---|---|---|---|---|
| orf19.5469 | 363 | 321 | 300 | 197 | 1646 | 1037 | 1452 | 1414 | 346 | 339 | 296 | 309 | 1319 | 1179 | 1233 | 1150 | 771,37 | 322 | 1220 | 3,775 | 1,917 | 25,024 | 3,37E-138 | 3,13E-137 | 0,002904 | 0,0042198 | 0,003542 | 0,003542 | TRUE | 0,039759 | No |
| orf19.547 | 1083 | 887 | 995 | 602 | 2038 | 1346 | 1872 | 1937 | 1033 | 938 | 983 | 943 | 1633 | 1531 | 1589 | 1575 | 1278,03 | 974 | 1582 | 1,622 | 0,698 | 13,457 | 2,82E-41 | 9,10E-41 | 0,0003705 | 0,003376 | 0,0017265 | 0,0017265 | TRUE | 0,0201927 | No |
| orf19.5472 | 2 | 3 | 4 | 2 | 9 | 4 | 8 | 9 | 2 | 3 | 4 | 3 | 7 | 5 | 7 | 7 | 4,75 | 3 | 6 | 2,162 | 1,112 | 1,69 | 0,0910154 | 0,1066219 | 1,00E-08 | 0,3474529 | 0,1648403 | 0,1648403 | TRUE | 0,1017988 | No |
| orf19.5474 | 359 | 275 | 274 | 141 | 859 | 501 | 787 | 789 | 342 | 291 | 271 | 221 | 688 | 570 | 668 | 642 | 461,54 | 281 | 642 | 2,268 | 1,181 | 10,522 | 6,84E-26 | 1,72E-25 | 0,014248 | 0,0056488 | 0,0094285 | 0,0094285 | TRUE | 0,2392578 | No |
| orf19.5475 | 15 | 14 | 22 | 20 | 74 | 45 | 94 | 76 | 14 | 15 | 22 | 31 | 59 | 51 | 80 | 62 | 41,78 | 20 | 63 | 3,187 | 1,672 | 6,065 | 1,32E-09 | 2,25E-09 | 0,0318673 | 0,0413909 | 0,0379217 | 0,0379217 | TRUE | 0,4435191 | No |
| orf19.5479 | 4 | 3 | 1 | 0 | 10 | 7 | 13 | 9 | 4 | 3 | 1 | 0 | 8 | 8 | 11 | 7 | 5,29 | 2 | 8 | 3,994 | 1,998 | 2,802 | 0,0050799 | 0,006563 | 1,00E-08 | 0,3126114 | 0,1792199 | 0,1792199 | TRUE | 0,3308939 | No |
| orf19.548 | 579 | 553 | 587 | 364 | 896 | 641 | 813 | 903 | 552 | 585 | 580 | 570 | 718 | 729 | 690 | 734 | 644,77 | 572 | 718 | 1,255 | 0,328 | 5,38 | 7,46E-08 | 1,21E-07 | 1,00E-08 | 0,0046378 | 0,0019751 | 0,0019751 | TRUE | 0,0080833 | No |
| orf19.5480 | 1810 | 1497 | 1718 | 1057 | 3276 | 2224 | 3151 | 3457 | 1726 | 1583 | 1697 | 1656 | 2625 | 2529 | 2675 | 2811 | 2162,69 | 1666 | 2660 | 1,597 | 0,676 | 13,565 | 6,49E-42 | 2,12E-41 | 0,0012615 | 0,0028505 | 0,0018781 | 0,0018781 | TRUE | 0,0177396 | No |
| orf19.5483 | 1524 | 1363 | 1598 | 922 | 1595 | 1082 | 1593 | 1674 | 1453 | 1441 | 1579 | 1444 | 1278 | 1230 | 1352 | 1361 | 1392,39 | 1479 | 1305 | 0,883 | -0,18 | -3,26 | 0,0011135 | 0,0014974 | 0,0014204 | 0,0032705 | 0,0022033 | 0,0022033 | TRUE | 0,0253833 | No |
| orf19.5484 | 2515 | 2300 | 2363 | 1492 | 2408 | 1739 | 2253 | 2432 | 2398 | 2432 | 2334 | 2337 | 1929 | 1977 | 1913 | 1977 | 2162,32 | 2375 | 1949 | 0,82 | -0,286 | -7,28 | 3,34E-13 | 6,43E-13 | 1,00E-08 | 0,0028506 | 0,0010214 | 0,0010214 | TRUE | 0,0030831 | No |
| orf19.5485 | 605 | 527 | 551 | 316 | 1078 | 759 | 1062 | 1147 | 577 | 557 | 544 | 495 | 864 | 863 | 902 | 933 | 716,81 | 543 | 890 | 1,632 | 0,707 | 10,838 | 2,28E-27 | 5,89E-27 | 0,0007911 | 0,0043819 | 0,0025581 | 0,0025581 | TRUE | 0,034601 | No |
| orf19.5486 | 4142 | 3734 | 4158 | 2319 | 4804 | 3343 | 4594 | 4973 | 3949 | 3948 | 4108 | 3633 | 3849 | 3801 | 3900 | 4044 | 3904,02 | 3910 | 3898 | 0,996 | -0,005 | -0,118 | 0,9057787 | 0,9188829 | 0,0011635 | 0,0025118 | 0,0017483 | 0,0017483 | TRUE | 0,0260641 | No |
| orf19.5486.1 | 2202 | 1891 | 1791 | 1056 | 1424 | 920 | 1197 | 1310 | 2099 | 1999 | 1769 | 1654 | 1141 | 1046 | 1016 | 1065 | 1473,88 | 1880 | 1067 | 0,567 | -0,819 | -10,577 | 3,82E-26 | 9,68E-26 | 0,0061823 | 0,0032053 | 0,0050435 | 0,0050435 | TRUE | 0,0755551 | No |
| orf19.5487 | 4707 | 4159 | 4413 | 2724 | 5381 | 3750 | 5250 | 5739 | 4488 | 4398 | 4360 | 4267 | 4312 | 4264 | 4457 | 4666 | 4401,43 | 4378 | 4425 | 1,011 | 0,015 | 0,37 | 0,7115656 | 0,7414272 | 0,0007076 | 0,0024643 | 0,0013853 | 0,0013853 | TRUE | 0,0165614 | No |
| orf19.5488 | 321 | 276 | 298 | 200 | 1041 | 594 | 952 | 976 | 306 | 292 | 294 | 313 | 834 | 675 | 808 | 794 | 539,62 | 301 | 778 | 2,591 | 1,374 | 16,212 | 4,18E-59 | 1,74E-58 | 0,0038765 | 0,005134 | 0,0044658 | 0,0044658 | TRUE | 0,0876987 | No |
| orf19.5489 | 137 | 150 | 151 | 98 | 301 | 241 | 280 | 291 | 131 | 159 | 149 | 154 | 241 | 274 | 238 | 237 | 197,68 | 148 | 248 | 1,672 | 0,742 | 6,704 | 2,03E-11 | 3,70E-11 | 1,00E-08 | 0,010397 | 0,0061765 | 0,0061765 | TRUE | 0,0718083 | No |
| orf19.549 | 2955 | 2689 | 2748 | 2061 | 4169 | 3026 | 3997 | 4095 | 2817 | 2843 | 2715 | 3229 | 3341 | 3441 | 3393 | 3330 | 3138,55 | 2901 | 3376 | 1,166 | 0,221 | 4,075 | 4,61E-05 | 6,62E-05 | 0,0027065 | 0,0026144 | 0,0025048 | 0,0025048 | TRUE | 0,0667865 | No |
| orf19.5491 | 66 | 107 | 84 | 64 | 100 | 67 | 62 | 73 | 63 | 113 | 83 | 100 | 80 | 76 | 53 | 59 | 78,45 | 90 | 67 | 0,749 | -0,416 | -1,954 | 0,0506883 | 0,0609463 | 0,0393562 | 0,0230202 | 0,030659 | 0,030659 | TRUE | 0,2643912 | No |
| orf19.5491.1 | 3179 | 2682 | 2762 | 2170 | 6843 | 5070 | 6617 | 6763 | 3031 | 2836 | 2729 | 3399 | 5483 | 5765 | 5618 | 5499 | 4294,95 | 2999 | 5591 | 1,867 | 0,901 | 14,263 | 3,73E-46 | 1,30E-45 | 0,00477 | 0,0024736 | 0,003563 | 0,003563 | TRUE | 0,0946999 | No |
| orf19.5492 | 2253 | 1846 | 1926 | 1120 | 3196 | 2034 | 3012 | 3084 | 2148 | 1952 | 1903 | 1755 | 2561 | 2313 | 2557 | 2508 | 2211,95 | 1940 | 2485 | 1,28 | 0,356 | 5,561 | 2,68E-08 | 4,40E-08 | 0,0040071 | 0,0028336 | 0,003459 | 0,003459 | TRUE | 0,0636635 | No |
| orf19.5493 | 9408 | 8328 | 8640 | 5398 | 20295 | 13308 | 18441 | 21573 | 8970 | 8806 | 8536 | 8456 | 16262 | 15132 | 15656 | 17541 | 12419,81 | 8692 | 16148 | 1,858 | 0,894 | 17,87 | 2,02E-71 | 9,86E-71 | 0,0023193 | 0,0022235 | 0,0023092 | 0,0023092 | TRUE | 0,0413039 | No |
| orf19.5495 | 2099 | 1920 | 2177 | 1340 | 3108 | 2165 | 2971 | 3242 | 2001 | 2030 | 2151 | 2099 | 2490 | 2462 | 2522 | 2636 | 2298,97 | 2070 | 2528 | 1,222 | 0,289 | 6,56 | 5,39E-11 | 9,71E-11 | 0,0005903 | 0,0028055 | 0,001419 | 0,001419 | TRUE | 0,0102335 | No |
| orf19.5496 | 1897 | 1673 | 1959 | 1312 | 2544 | 1932 | 2574 | 2556 | 1809 | 1769 | 1935 | 2055 | 2038 | 2197 | 2185 | 2078 | 2008,38 | 1892 | 2124 | 1,125 | 0,169 | 2,997 | 0,0027293 | 0,0035894 | 0,0023737 | 0,0029088 | 0,0025571 | 0,0025571 | TRUE | 0,0378386 | No |
| orf19.5498 | 282 | 260 | 299 | 173 | 382 | 273 | 398 | 365 | 269 | 275 | 295 | 271 | 306 | 310 | 338 | 297 | 295,17 | 278 | 313 | 1,125 | 0,17 | 1,94 | 0,0524109 | 0,0628872 | 1,00E-08 | 0,0076538 | 0,0039875 | 0,0039875 | TRUE | 0,0347739 | No |
| orf19.5499 | 160 | 182 | 179 | 98 | 52 | 34 | 37 | 34 | 153 | 192 | 177 | 154 | 42 | 39 | 31 | 28 | 101,84 | 169 | 35 | 0,205 | -2,287 | -12,917 | 3,61E-38 | 1,11E-37 | 0,0073399 | 0,0182135 | 0,0139963 | 0,0139963 | TRUE | 0,1592563 | No |
| orf19.55 | 3222 | 3467 | 3505 | 2029 | 8868 | 6294 | 7871 | 7143 | 3072 | 3666 | 3463 | 3179 | 7106 | 7157 | 6682 | 5808 | 5016,47 | 3345 | 6688 | 1,999 | 0,999 | 13,143 | 1,86E-39 | 5,83E-39 | 0,0073207 | 0,0024186 | 0,005316 | 0,005316 | TRUE | 0,096138 | No |
| orf19.550 | 23973 | 20045 | 22250 | 14692 | 4761 | 3271 | 4601 | 4756 | 22856 | 21195 | 21981 | 23016 | 3815 | 3719 | 3906 | 3867 | 13044,37 | 22262 | 3827 | 0,172 | -2,54 | -65,444 | 0 | 0 | 0,000738 | 0,0022172 | 0,0013043 | 0,0013043 | TRUE | 0,0127367 | No |
| orf19.5500 | 17716 | 15895 | 17628 | 9014 | 13166 | 8112 | 12869 | 13291 | 16890 | 16807 | 17415 | 14121 | 10550 | 9224 | 10926 | 10807 | 13342,39 | 16308 | 10377 | 0,636 | -0,652 | -8,929 | 4,30E-19 | 9,47E-19 | 0,0068843 | 0,0022143 | 0,0050512 | 0,0050512 | TRUE | 0,0991525 | No |
| orf19.5501 | 1088 | 1050 | 1074 | 704 | 825 | 539 | 762 | 736 | 1037 | 1110 | 1061 | 1103 | 661 | 613 | 647 | 598 | 853,84 | 1078 | 630 | 0,586 | -0,772 | -13,1 | 3,29E-39 | 1,03E-38 | 1,00E-08 | 0,0040142 | 0,0021086 | 0,0021086 | TRUE | 0,0146842 | No |
| orf19.5502 | 336 | 397 | 420 | 334 | 488 | 487 | 428 | 414 | 320 | 420 | 415 | 523 | 391 | 554 | 363 | 337 | 415,38 | 420 | 411 | 0,981 | -0,028 | -0,171 | 0,8640225 | 0,8818502 | 0,042164 | 0,0060441 | 0,0229688 | 0,0229688 | TRUE | 0,5133512 | No |
| orf19.5503 | 51 | 43 | 45 | 27 | 23 | 20 | 14 | 14 | 49 | 45 | 44 | 42 | 18 | 23 | 12 | 11 | 30,66 | 45 | 16 | 0,348 | -1,521 | -5,433 | 5,56E-08 | 9,03E-08 | 1,00E-08 | 0,0556426 | 0,0348586 | 0,0348586 | TRUE | 0,2958369 | No |
| orf19.5504 | 2270 | 2688 | 2726 | 1554 | 1434 | 1069 | 1313 | 1316 | 2164 | 2842 | 2693 | 2434 | 1149 | 1216 | 1115 | 1070 | 1835,4 | 2533 | 1138 | 0,449 | -1,157 | -14,092 | 4,22E-45 | 1,45E-44 | 0,0075379 | 0,0029859 | 0,0058537 | 0,0058537 | TRUE | 0,1194024 | No |
| orf19.5505 | 2317 | 2231 | 2531 | 1360 | 2315 | 1736 | 2135 | 2278 | 2209 | 2359 | 2500 | 2131 | 1855 | 1974 | 1813 | 1852 | 2086,58 | 2300 | 1874 | 0,813 | -0,299 | -5,152 | 2,58E-07 | 4,08E-07 | 0,002589 | 0,0028782 | 0,002741 | 0,002741 | TRUE | 0,0418987 | No |
| orf19.5506 | 1084 | 1194 | 1105 | 713 | 1076 | 818 | 985 | 966 | 1033 | 1263 | 1092 | 1117 | 862 | 930 | 836 | 785 | 989,82 | 1126 | 853 | 0,757 | -0,402 | -5,288 | 1,24E-07 | 1,98E-07 | 0,0048552 | 0,0037501 | 0,0045135 | 0,0045135 | TRUE | 0,0816438 | No |
| orf19.5507 | 9368 | 9190 | 9617 | 5654 | 7560 | 5444 | 6861 | 7222 | 8931 | 9717 | 9501 | 8857 | 6058 | 6190 | 5825 | 5872 | 7618,96 | 9252 | 5986 | 0,647 | -0,629 | -14,623 | 2,00E-48 | 7,22E-48 | 0,0012609 | 0,0023068 | 0,0016403 | 0,0016403 | TRUE | 0,0139713 | No |
| orf19.5508 | 527 | 497 | 506 | 349 | 542 | 386 | 442 | 480 | 502 | 526 | 500 | 547 | 434 | 439 | 375 | 390 | 464,16 | 519 | 410 | 0,79 | -0,339 | -4,347 | 1,38E-05 | 2,03E-05 | 0,0016108 | 0,0056286 | 0,0036733 | 0,0036733 | TRUE | 0,0369296 | No |
| orf19.551 | 18926 | 17083 | 18210 | 11867 | 5807 | 4175 | 5383 | 5783 | 18044 | 18063 | 17990 | 18590 | 4653 | 4747 | 4570 | 4702 | 11419,96 | 18172 | 4668 | 0,257 | -1,961 | -67,016 | 0 | 0 | 0,0001211 | 0,0022351 | 0,0006954 | 0,0006954 | TRUE | 0,0028535 | No |
| orf19.5510 | 2157 | 2075 | 2314 | 1261 | 4585 | 3162 | 4201 | 4254 | 2056 | 2194 | 2286 | 1975 | 3674 | 3595 | 3567 | 3459 | 2850,85 | 2128 | 3574 | 1,677 | 0,746 | 14,513 | 1,01E-47 | 3,61E-47 | 0,0019345 | 0,0026672 | 0,0021453 | 0,0021453 | TRUE | 0,030178 | No |
| orf19.5513 | 2189 | 1966 | 2118 | 1531 | 1570 | 1196 | 1424 | 1575 | 2087 | 2079 | 2092 | 2398 | 1258 | 1360 | 1209 | 1281 | 1720,51 | 2164 | 1277 | 0,591 | -0,76 | -12,406 | 2,42E-35 | 7,12E-35 | 0,0029918 | 0,0030456 | 0,0029904 | 0,0029904 | TRUE | 0,0607105 | No |
| orf19.5514 | 793 | 748 | 765 | 488 | 741 | 529 | 706 | 801 | 756 | 791 | 756 | 764 | 594 | 602 | 599 | 651 | 689,14 | 767 | 612 | 0,798 | -0,325 | -5,336 | 9,48E-08 | 1,53E-07 | 1,00E-08 | 0,0044738 | 0,0021145 | 0,0021145 | TRUE | 0,0237589 | No |
| orf19.5515 | 2828 | 2756 | 2798 | 1789 | 3749 | 2616 | 3412 | 3745 | 2696 | 2914 | 2764 | 2803 | 3004 | 2975 | 2897 | 3045 | 2887,18 | 2794 | 2980 | 1,067 | 0,094 | 2,31 | 0,0208832 | 0,0258849 | 0,0004295 | 0,00266 | 0,0012222 | 0,0012222 | TRUE | 0,0105667 | No |
| orf19.5516 | 1274 | 1387 | 1474 | 884 | 2233 | 1728 | 2085 | 2033 | 1215 | 1467 | 1456 | 1385 | 1789 | 1965 | 1770 | 1653 | 1587,44 | 1381 | 1794 | 1,299 | 0,378 | 5,147 | 2,65E-07 | 4,19E-07 | 0,0052732 | 0,0031256 | 0,0045086 | 0,0045086 | TRUE | 0,0818072 | No |
| orf19.5517 | 2123 | 1805 | 1901 | 1279 | 7481 | 5690 | 7091 | 7474 | 2024 | 1909 | 1878 | 2004 | 5994 | 6470 | 6020 | 6077 | 4046,99 | 1954 | 6140 | 3,144 | 1,653 | 37,118 | 1,43E-301 | 3,17E-300 | 0,0008706 | 0,002497 | 0,0015429 | 0,0015429 | TRUE | 0,0155898 | No |
| orf19.5518 | 2492 | 2431 | 2613 | 1700 | 1425 | 1166 | 1365 | 1394 | 2376 | 2570 | 2581 | 2663 | 1142 | 1326 | 1159 | 1133 | 1868,86 | 2548 | 1190 | 0,467 | -1,1 | -17,837 | 3,65E-71 | 1,78E-70 | 0,0032566 | 0,0029698 | 0,0030538 | 0,0030538 | TRUE | 0,0674242 | No |
| orf19.5519 | 1282 | 1161 | 1245 | 725 | 1207 | 937 | 1161 | 1266 | 1222 | 1228 | 1230 | 1136 | 967 | 1065 | 986 | 1029 | 1107,9 | 1204 | 1012 | 0,838 | -0,256 | -4,694 | 2,68E-06 | 4,09E-06 | 0,0004574 | 0,0035733 | 0,0019428 | 0,0019428 | TRUE | 0,0152175 | No |
| orf19.552 | 4 | 2 | 1 | 2 | 4 | 2 | 4 | 2 | 4 | 2 | 1 | 3 | 3 | 2 | 3 | 2 | 2,57 | 2 | 2 | 1,062 | 0,087 | 0,101 | 0,9193289 | 0,9265894 | 1,00E-08 | 0,6412514 | 0,3197895 | 0,3197895 | TRUE | 0,206426 | No |
| orf19.5520 | 3 | 2 | 2 | 2 | 1 | 3 | 0 | 1 | 3 | 2 | 2 | 3 | 1 | 3 | 0 | 1 | 1,89 | 2 | 1 | 0,464 | -1,107 | -1,024 | 0,3058233 | 0,3389125 | 1,00E-08 | 0,8714283 | 0,514178 | 0,514178 | TRUE | 0,6428949 | No |
| orf19.5521 | 4509 | 3933 | 4260 | 2598 | 3319 | 2253 | 3292 | 3212 | 4299 | 4159 | 4208 | 4070 | 2659 | 2562 | 2795 | 2612 | 3420,46 | 4184 | 2657 | 0,635 | -0,655 | -15,975 | 1,89E-57 | 7,69E-57 | 0,0005623 | 0,0025713 | 0,0013127 | 0,0013127 | TRUE | 0,0148005 | No |
| orf19.5522 | 505 | 527 | 508 | 257 | 1280 | 833 | 1137 | 1118 | 481 | 557 | 502 | 403 | 1026 | 947 | 965 | 909 | 723,79 | 486 | 962 | 1,97 | 0,978 | 11,405 | 3,97E-30 | 1,07E-29 | 0,0067389 | 0,0043598 | 0,0054476 | 0,0054476 | TRUE | 0,1327594 | No |
| orf19.5523 | 443 | 499 | 416 | 320 | 779 | 570 | 701 | 760 | 422 | 528 | 411 | 501 | 624 | 648 | 595 | 618 | 543,46 | 466 | 621 | 1,339 | 0,421 | 4,816 | 1,47E-06 | 2,25E-06 | 0,0057815 | 0,0051125 | 0,0054048 | 0,0054048 | TRUE | 0,1026985 | No |
| orf19.5524 | 49 | 58 | 56 | 37 | 18 | 9 | 14 | 19 | 47 | 61 | 55 | 58 | 14 | 10 | 12 | 15 | 34,17 | 55 | 13 | 0,239 | -2,066 | -7,568 | 3,78E-14 | 7,44E-14 | 1,00E-08 | 0,0501493 | 0,0277555 | 0,0277555 | TRUE | 0,1008634 | No |
| orf19.5525 | 1285 | 1065 | 1175 | 796 | 710 | 500 | 646 | 735 | 1225 | 1126 | 1161 | 1247 | 569 | 569 | 548 | 598 | 880,31 | 1190 | 571 | 0,481 | -1,057 | -17,614 | 1,94E-69 | 9,20E-69 | 0,0007269 | 0,0039564 | 0,0022053 | 0,0022053 | TRUE | 0,0152597 | No |
| orf19.5526 | 130 | 115 | 119 | 90 | 213 | 132 | 216 | 210 | 124 | 122 | 118 | 141 | 171 | 150 | 183 | 171 | 147,37 | 126 | 169 | 1,353 | 0,436 | 3,562 | 0,0003687 | 0,000509 | 1,00E-08 | 0,0132324 | 0,0073388 | 0,0073388 | TRUE | 0,0514111 | No |
| orf19.5527 | 335 | 327 | 326 | 238 | 430 | 336 | 455 | 518 | 319 | 346 | 322 | 373 | 345 | 382 | 386 | 421 | 361,77 | 340 | 384 | 1,133 | 0,181 | 1,995 | 0,0460725 | 0,0555955 | 0,0034668 | 0,0066299 | 0,0050548 | 0,0050548 | TRUE | 0,0576218 | No |
| orf19.5528 | 1002 | 886 | 972 | 624 | 422 | 291 | 401 | 404 | 955 | 937 | 960 | 978 | 338 | 331 | 340 | 328 | 645,98 | 958 | 334 | 0,35 | -1,515 | -24,197 | 2,39E-129 | 2,06E-128 | 1,00E-08 | 0,004633 | 0,0018549 | 0,0018549 | TRUE | 0,0024191 | No |
| orf19.553 | 11 | 16 | 18 | 9 | 1 | 1 | 2 | 2 | 10 | 17 | 18 | 14 | 1 | 1 | 2 | 2 | 8,07 | 15 | 2 | 0,089 | -3,487 | -4,884 | 1,04E-06 | 1,61E-06 | 1,00E-08 | 0,2055833 | 0,1176853 | 0,1176853 | TRUE | 0,2021995 | No |
| orf19.5530 | 6582 | 5652 | 6443 | 4001 | 5769 | 4111 | 5393 | 5915 | 6275 | 5976 | 6365 | 6268 | 4623 | 4675 | 4579 | 4809 | 5446,19 | 6221 | 4672 | 0,751 | -0,413 | -11,531 | 9,22E-31 | 2,52E-30 | 0,0004455 | 0,0023928 | 0,0010472 | 0,0010472 | TRUE | 0,0086966 | No |
| orf19.5531 | 2360 | 2050 | 2154 | 1444 | 1193 | 839 | 1125 | 1273 | 2250 | 2168 | 2128 | 2262 | 956 | 954 | 955 | 1035 | 1588,47 | 2202 | 975 | 0,443 | -1,173 | -23,71 | 2,83E-124 | 2,35E-123 | 0,0005429 | 0,0031249 | 0,0016386 | 0,0016386 | TRUE | 0,0212653 | No |
| orf19.5532 | 182 | 155 | 143 | 94 | 183 | 149 | 170 | 143 | 174 | 164 | 141 | 147 | 147 | 169 | 144 | 116 | 150,32 | 156 | 144 | 0,914 | -0,13 | -0,942 | 0,3463728 | 0,3808072 | 0,0098327 | 0,0130137 | 0,0115533 | 0,0115533 | TRUE | 0,183843 | No |
| orf19.5533 | 1586 | 1421 | 1579 | 961 | 1274 | 880 | 1275 | 1319 | 1512 | 1503 | 1560 | 1505 | 1021 | 1001 | 1082 | 1072 | 1282,05 | 1520 | 1044 | 0,687 | -0,541 | -11,066 | 1,84E-28 | 4,83E-28 | 1,00E-08 | 0,003372 | 0,0014981 | 0,0014981 | TRUE | 0,008452 | No |
| orf19.5534 | 1593 | 1341 | 1370 | 935 | 2012 | 1405 | 1903 | 2202 | 1519 | 1418 | 1353 | 1465 | 1612 | 1598 | 1616 | 1790 | 1546,34 | 1439 | 1654 | 1,151 | 0,202 | 3,494 | 0,0004764 | 0,0006543 | 0,0021525 | 0,0031531 | 0,0025631 | 0,0025631 | TRUE | 0,0378548 | No |
| orf19.5535 | 1008 | 930 | 1012 | 699 | 2424 | 1750 | 2359 | 2439 | 961 | 983 | 1000 | 1095 | 1942 | 1990 | 2003 | 1983 | 1494,66 | 1010 | 1980 | 1,968 | 0,977 | 19,457 | 2,56E-84 | 1,45E-83 | 1,00E-08 | 0,0031898 | 0,0016376 | 0,0016376 | TRUE | 0,0331407 | No |
| orf19.5536 | 521 | 451 | 504 | 334 | 231 | 196 | 226 | 252 | 497 | 477 | 498 | 523 | 185 | 223 | 192 | 205 | 349,93 | 499 | 201 | 0,402 | -1,314 | -15,266 | 1,28E-52 | 4,91E-52 | 1,00E-08 | 0,0067833 | 0,0037665 | 0,0037665 | TRUE | 0,0496175 | No |
| orf19.5537 | 8943 | 7389 | 8145 | 5499 | 4760 | 3130 | 4725 | 4902 | 8526 | 7813 | 8046 | 8614 | 3814 | 3559 | 4012 | 3986 | 6046,32 | 8250 | 3843 | 0,466 | -1,101 | -21,429 | 7,23E-102 | 4,81E-101 | 0,0022969 | 0,0023628 | 0,0023516 | 0,0023516 | TRUE | 0,0292636 | No |
| orf19.5539 | 644 | 622 | 694 | 420 | 791 | 603 | 770 | 761 | 614 | 658 | 686 | 658 | 634 | 686 | 654 | 619 | 650,9 | 654 | 648 | 0,99 | -0,015 | -0,226 | 0,8215206 | 0,8425141 | 1,00E-08 | 0,0046138 | 0,0025388 | 0,0025388 | TRUE | 0,0217256 | No |
| orf19.554 | 652 | 580 | 610 | 408 | 203 | 143 | 158 | 173 | 622 | 613 | 603 | 639 | 163 | 163 | 134 | 141 | 384,59 | 619 | 150 | 0,242 | -2,048 | -23,21 | 3,58E-119 | 2,83E-118 | 1,00E-08 | 0,0063605 | 0,0036043 | 0,0036043 | TRUE | 0,0538823 | No |
| orf19.5541 | 4160 | 3858 | 4315 | 2692 | 5662 | 4061 | 5250 | 5522 | 3966 | 4079 | 4263 | 4217 | 4537 | 4618 | 4457 | 4490 | 4328,39 | 4131 | 4526 | 1,096 | 0,132 | 3,555 | 0,0003784 | 0,000522 | 0,0003435 | 0,0024706 | 0,0010858 | 0,0010858 | TRUE | 0,0090896 | No |
| orf19.5542 | 67 | 55 | 82 | 58 | 137 | 99 | 128 | 136 | 64 | 58 | 81 | 91 | 110 | 113 | 109 | 111 | 91,94 | 74 | 111 | 1,522 | 0,606 | 3,768 | 0,0001648 | 0,0002307 | 1,00E-08 | 0,0199502 | 0,013104 | 0,013104 | TRUE | 0,1689001 | No |
| orf19.5543 | 1582 | 1472 | 1642 | 932 | 1455 | 994 | 1388 | 1477 | 1508 | 1556 | 1622 | 1460 | 1166 | 1130 | 1178 | 1201 | 1352,79 | 1536 | 1169 | 0,76 | -0,396 | -7,811 | 5,69E-15 | 1,15E-14 | 0,0005172 | 0,003305 | 0,0017213 | 0,0017213 | TRUE | 0,016868 | No |
| orf19.5544 | 7138 | 6360 | 6793 | 3988 | 6900 | 4629 | 6622 | 7167 | 6805 | 6725 | 6711 | 6247 | 5529 | 5264 | 5622 | 5827 | 6091,29 | 6622 | 5560 | 0,839 | -0,252 | -5,654 | 1,57E-08 | 2,60E-08 | 0,0013959 | 0,0023608 | 0,0017489 | 0,0017489 | TRUE | 0,0170895 | No |
| orf19.5547 | 2470 | 2066 | 2220 | 1274 | 4542 | 3011 | 4390 | 4640 | 2355 | 2185 | 2193 | 1996 | 3639 | 3424 | 3727 | 3773 | 2911,42 | 2182 | 3641 | 1,666 | 0,737 | 13,174 | 1,23E-39 | 3,87E-39 | 0,0026806 | 0,0026552 | 0,0026208 | 0,0026208 | TRUE | 0,0373984 | No |
| orf19.5548 | 720 | 690 | 710 | 457 | 868 | 610 | 852 | 865 | 686 | 730 | 701 | 716 | 696 | 694 | 723 | 703 | 706,14 | 708 | 704 | 0,995 | -0,007 | -0,122 | 0,9027244 | 0,9165131 | 1,00E-08 | 0,0044164 | 0,0017899 | 0,0017899 | TRUE | 0,0054633 | No |
| orf19.5549 | 28 | 32 | 38 | 25 | 33 | 18 | 37 | 38 | 27 | 34 | 38 | 39 | 26 | 20 | 31 | 31 | 30,81 | 34 | 27 | 0,811 | -0,303 | -1,162 | 0,2452835 | 0,2751678 | 1,00E-08 | 0,0553879 | 0,0326417 | 0,0326417 | TRUE | 0,1716667 | No |

| | | | | | | | | | | | | | | | | | | | | | | | | | | | | | | | |
|---|---|---|---|---|---|---|---|---|---|---|---|---|---|---|---|---|---|---|---|---|---|---|---|---|---|---|---|---|---|---|---|
| orf19.555 | 2 | 1 | 2 | 3 | 1 | 2 | 4 | 3 | 2 | 1 | 2 | 5 | 1 | 2 | 3 | 2 | 2,32 | 2 | 2 | 0,974 | -0,038 | -0,041 | 0,967449 | 0,9706434 | 1,00E-08 | 0,7101766 | 0,3863537 | 0,3863537 | TRUE | 0,2503167 | No |
| orf19.5550 | 9470 | 9208 | 9334 | 4899 | 13073 | 8272 | 12042 | 12306 | 9029 | 9736 | 9221 | 7675 | 10475 | 9406 | 10224 | 10006 | 9471,39 | 8915 | 10028 | 1,124 | 0,169 | 2,474 | 0,0133429 | 0,016813 | 0,0057601 | 0,0022646 | 0,0043813 | 0,0043813 | TRUE | 0,1058467 | No |
| orf19.5551 | 6529 | 5747 | 6293 | 3782 | 1975 | 1298 | 1894 | 2027 | 6225 | 6077 | 6217 | 5925 | 1583 | 1476 | 1608 | 1648 | 3844,7 | 6111 | 1579 | 0,259 | -1,951 | -45,036 | 0 | 0 | 0,0008235 | 0,0025183 | 0,0014278 | 0,0014278 | TRUE | 0,0213758 | No |
| orf19.5552 | 4808 | 4416 | 4769 | 2861 | 1165 | 817 | 1080 | 1211 | 4584 | 4669 | 4711 | 4482 | 933 | 929 | 917 | 985 | 2776,32 | 4612 | 941 | 0,204 | -2,293 | -53,27 | 0 | 0 | 1,00E-08 | 0,0026827 | 0,0011843 | 0,0011843 | TRUE | 0,0121801 | No |
| orf19.5553 | 1033 | 969 | 1052 | 603 | 937 | 589 | 913 | 978 | 985 | 1025 | 1039 | 945 | 751 | 670 | 775 | 795 | 873,03 | 998 | 748 | 0,75 | -0,416 | -6,297 | 3,03E-10 | 5,32E-10 | 0,0020604 | 0,003972 | 0,0030296 | 0,0030296 | TRUE | 0,0537627 | No |
| orf19.5555 | 2266 | 2032 | 2197 | 1396 | 2340 | 1663 | 2193 | 2367 | 2160 | 2149 | 2170 | 2187 | 1875 | 1891 | 1862 | 1925 | 2027,34 | 2166 | 1888 | 0,872 | -0,198 | -5,153 | 2,57E-07 | 4,07E-07 | 1,00E-08 | 0,0029012 | 0,0009225 | 0,0009225 | TRUE | 0,0021639 | No |
| orf19.5556 | 1 | 2 | 1 | 0 | 7 | 2 | 2 | 3 | 1 | 2 | 1 | 0 | 6 | 2 | 2 | 2 | 2,01 | 1 | 3 | 2,82 | 1,496 | 1,385 | 0,1659253 | 0,1897096 | 1,00E-08 | 0,8191416 | 0,4587959 | 0,4587959 | TRUE | 0,5608243 | No |
| orf19.5557 | 630 | 685 | 640 | 444 | 3945 | 3007 | 3697 | 3964 | 601 | 724 | 632 | 696 | 3161 | 3419 | 3139 | 3223 | 1949,35 | 663 | 3236 | 4,891 | 2,29 | 36,294 | 1,99E-288 | 4,24E-287 | 0,0027389 | 0,0029336 | 0,0028385 | 0,0028385 | TRUE | 0,0505041 | No |
| orf19.5558 | 1625 | 1262 | 1508 | 1082 | 794 | 541 | 726 | 857 | 1549 | 1334 | 1490 | 1695 | 636 | 615 | 616 | 697 | 1079,13 | 1517 | 641 | 0,424 | -1,239 | -15,881 | 8,60E-57 | 3,45E-56 | 0,0057585 | 0,0036128 | 0,0047759 | 0,0047759 | TRUE | 0,0784529 | No |
| orf19.5559 | 621 | 555 | 559 | 350 | 1098 | 793 | 1022 | 1161 | 592 | 587 | 552 | 548 | 880 | 902 | 868 | 944 | 734,08 | 570 | 898 | 1,573 | 0,654 | 10,581 | 3,65E-26 | 9,25E-26 | 1,00E-08 | 0,004328 | 0,0021911 | 0,0021911 | TRUE | 0,0147654 | No |
| orf19.556 | 995 | 1094 | 991 | 651 | 3007 | 2288 | 2685 | 2545 | 949 | 1157 | 979 | 1020 | 2409 | 2602 | 2280 | 2069 | 1683,02 | 1026 | 2340 | 2,28 | 1,189 | 14,547 | 6,13E-48 | 2,20E-47 | 0,0076477 | 0,0030668 | 0,0056749 | 0,0056749 | TRUE | 0,0899912 | No |
| orf19.5561 | 3914 | 3929 | 4145 | 2547 | 5515 | 4230 | 5154 | 5391 | 3732 | 4154 | 4095 | 3990 | 4419 | 4810 | 4376 | 4383 | 4244,86 | 3993 | 4497 | 1,126 | 0,171 | 3,439 | 0,0005845 | 0,0008001 | 0,0018648 | 0,0024781 | 0,0021351 | 0,0021351 | TRUE | 0,0261533 | No |
| orf19.5563 | 1649 | 1344 | 1471 | 896 | 5230 | 3437 | 5067 | 5172 | 1572 | 1421 | 1453 | 1404 | 4191 | 3908 | 4302 | 4205 | 2807,01 | 1462 | 4152 | 2,835 | 1,504 | 29,227 | 8,74E-188 | 1,11E-186 | 0,0015645 | 0,0026762 | 0,00205 | 0,00205 | TRUE | 0,0315131 | No |
| orf19.5564 | 437 | 422 | 427 | 267 | 1812 | 1357 | 1836 | 1782 | 417 | 446 | 422 | 418 | 1452 | 1543 | 1559 | 1449 | 963,2 | 426 | 1501 | 3,521 | 1,816 | 29,587 | 2,21E-192 | 2,92E-191 | 1,00E-08 | 0,0037959 | 0,002013 | 0,002013 | TRUE | 0,0124066 | No |
| orf19.5565 | 298 | 302 | 264 | 199 | 727 | 450 | 671 | 674 | 284 | 319 | 261 | 312 | 583 | 512 | 570 | 548 | 423,49 | 294 | 553 | 1,892 | 0,92 | 10,718 | 8,38E-27 | 2,15E-26 | 0,0022541 | 0,0059684 | 0,0043591 | 0,0043591 | TRUE | 0,0672571 | No |
| orf19.5566 | 1986 | 1748 | 1947 | 1198 | 2211 | 1480 | 2033 | 2140 | 1893 | 1848 | 1923 | 1877 | 1772 | 1683 | 1726 | 1740 | 1807,81 | 1885 | 1730 | 0,918 | -0,123 | -2,976 | 0,0029249 | 0,0038332 | 1,00E-08 | 0,0029995 | 0,0010961 | 0,0010961 | TRUE | 0,0037571 | No |
| orf19.5567 | 2333 | 2020 | 2251 | 1339 | 1924 | 1268 | 1923 | 2012 | 2224 | 2136 | 2224 | 2098 | 1542 | 1442 | 1633 | 1636 | 1866,7 | 2170 | 1563 | 0,72 | -0,473 | -9,003 | 2,19E-19 | 4,86E-19 | 0,0014258 | 0,0029709 | 0,0021057 | 0,0021057 | TRUE | 0,0308746 | No |
| orf19.5568 | 363 | 348 | 435 | 243 | 374 | 232 | 328 | 353 | 346 | 368 | 430 | 381 | 300 | 264 | 278 | 287 | 331,68 | 381 | 282 | 0,742 | -0,43 | -4,654 | 3,26E-06 | 4,95E-06 | 0,003234 | 0,0070416 | 0,0051529 | 0,0051529 | TRUE | 0,0916326 | No |
| orf19.5569 | 10795 | 9605 | 10421 | 6263 | 5510 | 3739 | 5302 | 5453 | 10292 | 10156 | 10295 | 9811 | 4415 | 4252 | 4501 | 4434 | 7269,5 | 10138 | 4400 | 0,434 | -1,204 | -35,436 | 4,84E-275 | 9,39E-274 | 0,0003245 | 0,0023171 | 0,0009516 | 0,0009516 | TRUE | 0,0060167 | No |
| orf19.557 | 51 | 58 | 49 | 28 | 47 | 31 | 49 | 33 | 49 | 61 | 48 | 44 | 38 | 35 | 42 | 27 | 42,95 | 50 | 36 | 0,695 | -0,525 | -2,321 | 0,0203093 | 0,0251882 | 1,00E-08 | 0,0403235 | 0,0256318 | 0,0256318 | TRUE | 0,2189057 | No |
| orf19.5572 | 412 | 378 | 398 | 228 | 1575 | 1122 | 1452 | 1551 | 393 | 400 | 393 | 357 | 1262 | 1276 | 1233 | 1261 | 821,81 | 386 | 1258 | 3,245 | 1,698 | 27,992 | 2,05E-172 | 2,36E-171 | 1,00E-08 | 0,0040892 | 0,0017464 | 0,0017464 | TRUE | 0,0217194 | No |
| orf19.5573 | 266 | 264 | 294 | 167 | 146 | 115 | 125 | 154 | 254 | 279 | 290 | 262 | 117 | 131 | 106 | 125 | 195,49 | 271 | 120 | 0,44 | -1,186 | -10,654 | 1,67E-26 | 4,27E-26 | 1,00E-08 | 0,0104903 | 0,0061168 | 0,0061168 | TRUE | 0,0604166 | No |
| orf19.5574 | 1238 | 1139 | 1203 | 731 | 1281 | 924 | 1188 | 1269 | 1180 | 1204 | 1188 | 1145 | 1026 | 1051 | 1009 | 1032 | 1104,47 | 1179 | 1030 | 0,871 | -0,199 | -4,129 | 3,65E-05 | 5,27E-05 | 1,00E-08 | 0,0035779 | 0,0013287 | 0,0013287 | TRUE | 0,0038871 | No |
| orf19.5575 | 1616 | 1734 | 1692 | 1163 | 1580 | 1248 | 1491 | 1530 | 1541 | 1833 | 1672 | 1822 | 1266 | 1419 | 1266 | 1244 | 1507,82 | 1717 | 1299 | 0,757 | -0,402 | -5,819 | 5,91E-09 | 9,89E-09 | 0,0043019 | 0,0031802 | 0,0039195 | 0,0039195 | TRUE | 0,0585957 | No |
| orf19.5576 | 4774 | 4731 | 4807 | 3119 | 9929 | 7246 | 9140 | 9595 | 4551 | 5002 | 4749 | 4886 | 7956 | 8239 | 7760 | 7802 | 6368,19 | 4797 | 7939 | 1,655 | 0,727 | 17,689 | 5,05E-70 | 2,42E-69 | 0,0009675 | 0,0023491 | 0,0014488 | 0,0014488 | TRUE | 0,0148017 | No |
| orf19.5578 | 464 | 430 | 477 | 321 | 990 | 690 | 883 | 960 | 442 | 455 | 471 | 503 | 793 | 785 | 750 | 781 | 622,4 | 468 | 777 | 1,668 | 0,738 | 11,359 | 6,69E-30 | 1,80E-29 | 1,00E-08 | 0,0047293 | 0,0022856 | 0,0022856 | TRUE | 0,0241624 | No |
| orf19.5579 | 306 | 330 | 322 | 222 | 529 | 389 | 428 | 489 | 292 | 349 | 318 | 348 | 424 | 442 | 363 | 398 | 366,72 | 327 | 407 | 1,248 | 0,319 | 3,424 | 0,0006172 | 0,0008437 | 0,004456 | 0,0065686 | 0,0055351 | 0,0055351 | TRUE | 0,0613607 | No |
| orf19.558 | 6477 | 5626 | 6252 | 3885 | 7834 | 5446 | 7699 | 8155 | 6175 | 5949 | 6176 | 6086 | 6277 | 6193 | 6536 | 6631 | 6252,91 | 6096 | 6409 | 1,052 | 0,072 | 2 | 0,0455263 | 0,0549468 | 0,0004673 | 0,0023539 | 0,0010983 | 0,0010983 | TRUE | 0,0066138 | No |
| orf19.5580 | 707 | 626 | 696 | 365 | 1378 | 918 | 1280 | 1337 | 674 | 662 | 688 | 572 | 1104 | 1044 | 1087 | 1087 | 864,65 | 649 | 1080 | 1,657 | 0,729 | 11,521 | 1,03E-30 | 2,82E-30 | 0,0007243 | 0,0039902 | 0,0025714 | 0,0025714 | TRUE | 0,0631779 | No |
| orf19.5584 | 611 | 615 | 629 | 371 | 759 | 500 | 779 | 766 | 583 | 650 | 621 | 581 | 608 | 569 | 661 | 623 | 612,04 | 609 | 615 | 1,011 | 0,016 | 0,219 | 0,8266108 | 0,8467817 | 0,0016878 | 0,004774 | 0,003206 | 0,003206 | TRUE | 0,0294597 | No |
| orf19.5585 | 920 | 792 | 833 | 591 | 367 | 305 | 342 | 447 | 877 | 837 | 823 | 926 | 294 | 347 | 290 | 363 | 594,75 | 866 | 324 | 0,374 | -1,42 | -16,653 | 2,90E-62 | 1,26E-61 | 0,0051173 | 0,0048519 | 0,0049508 | 0,0049508 | TRUE | 0,086343 | No |
| orf19.5586 | 801 | 656 | 727 | 497 | 994 | 700 | 893 | 954 | 764 | 694 | 718 | 779 | 796 | 796 | 758 | 776 | 760,05 | 739 | 782 | 1,06 | 0,084 | 1,395 | 0,1631588 | 0,1866134 | 1,00E-08 | 0,0042516 | 0,0021649 | 0,0021649 | TRUE | 0,0193112 | No |
| orf19.5587 | 931 | 894 | 889 | 580 | 1209 | 893 | 1139 | 1281 | 888 | 945 | 878 | 909 | 969 | 1015 | 967 | 1042 | 951,56 | 905 | 998 | 1,103 | 0,142 | 2,554 | 0,0106592 | 0,0135168 | 1,00E-08 | 0,0038168 | 0,0018906 | 0,0018906 | TRUE | 0,0114423 | No |
| orf19.5588 | 86 | 72 | 63 | 43 | 31 | 32 | 24 | 31 | 82 | 76 | 62 | 67 | 25 | 36 | 20 | 25 | 49,32 | 72 | 26 | 0,364 | -1,457 | -6,451 | 1,11E-10 | 1,98E-10 | 0,0059069 | 0,0353846 | 0,024342 | 0,024342 | TRUE | 0,3105278 | No |
| orf19.5589 | 217 | 211 | 243 | 136 | 528 | 503 | 563 | 577 | 207 | 223 | 240 | 213 | 423 | 572 | 478 | 469 | 353,16 | 221 | 486 | 2,189 | 1,131 | 10,533 | 6,10E-26 | 1,54E-25 | 0,0084303 | 0,0067405 | 0,0076098 | 0,0076098 | TRUE | 0,1591221 | No |
| orf19.559 | 2942 | 2859 | 3148 | 1886 | 4270 | 3335 | 4058 | 4374 | 2805 | 3023 | 3110 | 2955 | 3421 | 3792 | 3445 | 3556 | 3263,46 | 2973 | 3554 | 1,195 | 0,257 | 5,136 | 2,80E-07 | 4,43E-07 | 0,0016651 | 0,0025944 | 0,0020794 | 0,0020794 | TRUE | 0,0241258 | No |
| orf19.5591 | 2676 | 2392 | 2503 | 1782 | 7606 | 6264 | 6962 | 7503 | 2551 | 2529 | 2473 | 2792 | 6094 | 7123 | 5911 | 6101 | 4446,68 | 2586 | 6307 | 2,44 | 1,287 | 19,622 | 1,00E-85 | 5,73E-85 | 0,0049334 | 0,0024605 | 0,0038426 | 0,0038426 | TRUE | 0,0916754 | No |
| orf19.5592 | 20 | 19 | 23 | 8 | 26 | 14 | 35 | 25 | 19 | 20 | 23 | 13 | 21 | 16 | 30 | 20 | 20,15 | 19 | 22 | 1,154 | 0,207 | 0,63 | 0,5285005 | 0,5645065 | 1,00E-08 | 0,0835719 | 0,0533387 | 0,0533387 | TRUE | 0,3755098 | No |
| orf19.5595 | 750 | 667 | 729 | 486 | 1190 | 914 | 1069 | 1198 | 715 | 705 | 720 | 761 | 954 | 1039 | 908 | 974 | 847,04 | 725 | 969 | 1,336 | 0,418 | 6,79 | 1,12E-11 | 2,05E-11 | 0,0010151 | 0,0040297 | 0,002401 | 0,002401 | TRUE | 0,0267197 | No |
| orf19.5596 | 1744 | 1591 | 1619 | 1175 | 1231 | 923 | 1079 | 1222 | 1663 | 1682 | 1599 | 1841 | 986 | 1050 | 916 | 994 | 1341,34 | 1696 | 986 | 0,582 | -0,781 | -12,781 | 2,09E-37 | 6,35E-37 | 0,0022983 | 0,0033154 | 0,0027991 | 0,0027991 | TRUE | 0,0368319 | No |
| orf19.5597 | 3895 | 3773 | 4036 | 2318 | 3550 | 2615 | 3438 | 3476 | 3713 | 3989 | 3987 | 3631 | 2845 | 2973 | 2919 | 2826 | 3360,57 | 3830 | 2891 | 0,754 | -0,408 | -9,038 | 1,59E-19 | 3,54E-19 | 0,0010455 | 0,0025799 | 0,0016533 | 0,0016533 | TRUE | 0,0139563 | No |
| orf19.5597.1 | 1238 | 1097 | 1089 | 900 | 2809 | 2171 | 2509 | 2742 | 1180 | 1160 | 1076 | 1410 | 2251 | 2469 | 2130 | 2230 | 1738,12 | 1206 | 2270 | 1,886 | 0,915 | 11,598 | 4,20E-31 | 1,15E-30 | 0,0081467 | 0,0030359 | 0,0053167 | 0,0053167 | TRUE | 0,1455 | No |
| orf19.5599 | 3638 | 3169 | 3542 | 2221 | 3184 | 2320 | 3086 | 3312 | 3468 | 3351 | 3499 | 3479 | 2551 | 2638 | 2620 | 2693 | 3037,5 | 3449 | 2626 | 0,761 | -0,394 | -10,554 | 4,84E-26 | 1,23E-25 | 1,00E-08 | 0,0026318 | 0,0010031 | 0,0010031 | TRUE | 0,0045842 | No |
| orf19.56 | 2318 | 2141 | 2291 | 1440 | 1517 | 1136 | 1379 | 1432 | 2210 | 2264 | 2263 | 2256 | 1216 | 1292 | 1171 | 1164 | 1729,41 | 2248 | 1211 | 0,538 | -0,895 | -19,067 | 4,76E-81 | 2,63E-80 | 0,0002352 | 0,0030407 | 0,0014985 | 0,0014985 | TRUE | 0,022404 | No |
| orf19.5601 | 1288 | 1141 | 1272 | 879 | 1749 | 1188 | 1730 | 1770 | 1228 | 1206 | 1257 | 1377 | 1401 | 1351 | 1469 | 1439 | 1341,03 | 1267 | 1415 | 1,121 | 0,165 | 2,968 | 0,0029938 | 0,003921 | 0,0012557 | 0,0033156 | 0,0021956 | 0,0021956 | TRUE | 0,0366308 | No |
| orf19.5602 | 2277 | 2055 | 2197 | 1428 | 1457 | 1072 | 1447 | 1564 | 2171 | 2173 | 2170 | 2237 | 1167 | 1219 | 1228 | 1272 | 1704,73 | 2188 | 1222 | 0,559 | -0,84 | -18,713 | 3,90E-78 | 2,08E-77 | 1,00E-08 | 0,0030544 | 0,0013112 | 0,0013112 | TRUE | 0,0114518 | No |
| orf19.5604 | 1404 | 1236 | 1370 | 847 | 835 | 577 | 767 | 812 | 1339 | 1307 | 1353 | 1327 | 669 | 656 | 651 | 660 | 995,29 | 1332 | 659 | 0,495 | -1,014 | -20,02 | 3,66E-89 | 2,17E-88 | 1,00E-08 | 0,0037409 | 0,0013701 | 0,0013701 | TRUE | 0,0019786 | No |
| orf19.5605 | 14609 | 12284 | 14029 | 9124 | 8171 | 5688 | 7875 | 8385 | 13928 | 12989 | 13859 | 14293 | 6547 | 6468 | 6686 | 6818 | 10198,5 | 13767 | 6630 | 0,482 | -1,054 | -26,642 | 2,20E-156 | 2,33E-155 | 0,0009288 | 0,0022523 | 0,0013939 | 0,0013939 | TRUE | 0,0177248 | No |
| orf19.5606 | 55 | 53 | 55 | 43 | 50 | 59 | 62 | 71 | 52 | 56 | 54 | 67 | 40 | 67 | 53 | 58 | 55,96 | 57 | 54 | 0,946 | -0,081 | -0,391 | 0,6959052 | 0,7266522 | 0,0108714 | 0,0314311 | 0,0227492 | 0,0227492 | TRUE | 0,2877403 | No |
| orf19.5607 | 10441 | 9627 | 10041 | 6245 | 5209 | 3500 | 4909 | 5305 | 9954 | 10179 | 9920 | 9783 | 4174 | 3980 | 4168 | 4313 | 7058,9 | 9959 | 4159 | 0,418 | -1,259 | -35,591 | 1,93E-277 | 3,83E-276 | 0,0004512 | 0,0023239 | 0,0010381 | 0,0010381 | TRUE | 0,0097544 | No |
| orf19.5608 | 3616 | 3465 | 3508 | 1902 | 3152 | 2103 | 2941 | 2875 | 3447 | 3664 | 3466 | 2980 | 2526 | 2391 | 2497 | 2338 | 2913,48 | 3389 | 2438 | 0,719 | -0,477 | -7,595 | 3,07E-14 | 6,07E-14 | 0,0038486 | 0,0026548 | 0,0034361 | 0,0034361 | TRUE | 0,0773008 | No |
| orf19.5609 | 2229 | 1848 | 2033 | 1299 | 2649 | 1991 | 2794 | 2810 | 2125 | 1954 | 2008 | 2035 | 2123 | 2264 | 2372 | 2285 | 2145,74 | 2030 | 2261 | 1,113 | 0,154 | 3,1 | 0,001937 | 0,0025681 | 0,0013303 | 0,0028565 | 0,0019108 | 0,0019108 | TRUE | 0,021154 | No |
| orf19.5610 | 3599 | 3539 | 3692 | 2163 | 958 | 741 | 906 | 856 | 3431 | 3742 | 3647 | 3388 | 768 | 843 | 769 | 696 | 2160,56 | 3552 | 769 | 0,216 | -2,212 | -35,756 | 5,26E-280 | 1,06E-278 | 0,0031174 | 0,0028512 | 0,0029347 | 0,0029347 | TRUE | 0,0484933 | No |
| orf19.5611 | 236 | 236 | 240 | 118 | 42 | 28 | 28 | 33 | 225 | 250 | 237 | 185 | 34 | 32 | 24 | 27 | 126,57 | 224 | 29 | 0,128 | -2,962 | -16,785 | 3,14E-63 | 1,38E-62 | 1,00E-08 | 0,0150632 | 0,0121034 | 0,0121034 | TRUE | 0,1284051 | No |
| orf19.5612 | 1036 | 1070 | 1138 | 602 | 1954 | 1297 | 1776 | 1646 | 988 | 1131 | 1124 | 943 | 1566 | 1475 | 1508 | 1338 | 1259,13 | 1046 | 1472 | 1,403 | 0,489 | 6,579 | 4,72E-11 | 8,51E-11 | 0,0052288 | 0,0033953 | 0,0044592 | 0,0044592 | TRUE | 0,0482834 | No |
| orf19.5614 | 7951 | 6518 | 7279 | 4557 | 2452 | 1745 | 2464 | 2582 | 7580 | 6892 | 7191 | 7139 | 1965 | 1984 | 2092 | 2099 | 4617,8 | 7200 | 2035 | 0,283 | -1,823 | -40,769 | 0 | 0 | 0,0011727 | 0,0024468 | 0,0016218 | 0,0016218 | TRUE | 0,0156686 | No |
| orf19.5615 | 1827 | 1525 | 1615 | 1061 | 3202 | 2197 | 2862 | 3099 | 1742 | 1612 | 1595 | 1662 | 2566 | 2498 | 2430 | 2520 | 2078,17 | 1653 | 2504 | 1,515 | 0,599 | 13,231 | 5,81E-40 | 1,83E-39 | 0,0005201 | 0,0028814 | 0,0014494 | 0,0014494 | TRUE | 0,0171819 | No |
| orf19.5616 | 42 | 40 | 39 | 35 | 98 | 73 | 107 | 112 | 40 | 42 | 39 | 55 | 79 | 83 | 91 | 91 | 64,89 | 44 | 86 | 1,991 | 0,994 | 5,366 | 8,06E-08 | 1,30E-07 | 1,00E-08 | 0,0273934 | 0,0148921 | 0,0148921 | TRUE | 0,1482712 | No |
| orf19.5617 | 4949 | 4286 | 4572 | 2695 | 3804 | 2589 | 3742 | 3600 | 4718 | 4532 | 4517 | 4222 | 3048 | 2944 | 3177 | 2927 | 3760,61 | 4497 | 3024 | 0,672 | -0,574 | -12,186 | 3,68E-34 | 1,06E-33 | 0,0013968 | 0,0025279 | 0,0018542 | 0,0018542 | TRUE | 0,0197416 | No |
| orf19.5618 | 1455 | 1489 | 1539 | 980 | 1170 | 893 | 1056 | 1060 | 1387 | 1574 | 1520 | 1535 | 937 | 1015 | 897 | 862 | 1216,07 | 1504 | 928 | 0,616 | -0,699 | -10,715 | 8,61E-27 | 2,21E-26 | 0,0028352 | 0,0034415 | 0,0032258 | 0,0032258 | TRUE | 0,0453476 | No |
| orf19.5619 | 1628 | 1463 | 1536 | 966 | 1993 | 1483 | 1884 | 1911 | 1552 | 1547 | 1517 | 1513 | 1597 | 1686 | 1600 | 1554 | 1570,79 | 1532 | 1609 | 1,048 | 0,068 | 1,501 | 0,1332512 | 0,1536458 | 1,00E-08 | 0,0031366 | 0,0013253 | 0,0013253 | TRUE | 0,0116577 | No |
| orf19.562 | 698 | 617 | 730 | 452 | 1552 | 1147 | 1522 | 1576 | 665 | 652 | 721 | 708 | 1244 | 1304 | 1292 | 1281 | 983,57 | 686 | 1280 | 1,866 | 0,9 | 16,19 | 5,95E-59 | 2,47E-58 | 1,00E-08 | 0,0037606 | 0,0018046 | 0,0018046 | TRUE | 0,015701 | No |
| orf19.5620 | 795 | 777 | 751 | 485 | 449 | 290 | 383 | 362 | 758 | 822 | 742 | 760 | 360 | 330 | 325 | 294 | 548,78 | 770 | 327 | 0,425 | -1,235 | -15,963 | 2,31E-57 | 9,35E-57 | 0,0021373 | 0,0050832 | 0,0036606 | 0,0036606 | TRUE | 0,0563548 | No |
| orf19.5621 | 681 | 730 | 763 | 434 | 1107 | 894 | 1027 | 963 | 649 | 772 | 754 | 680 | 887 | 1017 | 872 | 783 | 801,66 | 714 | 890 | 1,243 | 0,314 | 3,566 | 0,000363 | 0,0005013 | 0,0075366 | 0,0041394 | 0,0061678 | 0,0061678 | TRUE | 0,1060168 | No |
| orf19.5622 | 3019 | 2963 | 3198 | 1763 | 3466 | 2325 | 3010 | 3301 | 2878 | 3133 | 3159 | 2762 | 2777 | 2644 | 2555 | 2684 | 2824,11 | 2983 | 2665 | 0,893 | -0,164 | -3,032 | 0,0024323 | 0,0032073 | 0,0022475 | 0,0026727 | 0,0024525 | 0,0024525 | TRUE | 0,0285535 | No |
| orf19.5623 | 1560 | 1568 | 1648 | 999 | 1932 | 1442 | 1862 | 1893 | 1487 | 1658 | 1628 | 1565 | 1548 | 1640 | 1581 | 1539 | 1580,76 | 1584 | 1577 | 0,994 | -0,008 | -0,163 | 0,8707102 | 0,887682 | 0,000933 | 0,00313 | 0,0018398 | 0,0018398 | TRUE | 0,0222585 | No |
| orf19.5625 | 703 | 763 | 767 | 438 | 544 | 422 | 492 | 478 | 670 | 807 | 758 | 686 | 436 | 480 | 418 | 389 | 580,37 | 730 | 431 | 0,587 | -0,768 | -8,889 | 6,14E-19 | 1,35E-18 | 0,0057672 | 0,0049203 | 0,0053523 | 0,0053523 | TRUE | 0,0638763 | No |
| orf19.5626 | 27896 | 26829 | 28764 | 19222 | 2625 | 1925 | 2383 | 2462 | 26596 | 28368 | 28416 | 30112 | 2103 | 2189 | 2023 | 2002 | 15226,21 | 28373 | 2079 | 0,073 | -3,771 | -77,302 | 0 | 0 | 0,0017746 | 0,0021991 | 0,0020511 | 0,0020511 | TRUE | 0,0217922 | No |
| orf19.5627 | 10552 | 10280 | 11084 | 6798 | 10386 | 7750 | 10435 | 10465 | 10060 | 10870 | 10950 | 10649 | 8322 | 8812 | 8859 | 8509 | 9629,02 | 10632 | 8626 | 0,811 | -0,302 | -7,489 | 6,94E-14 | 1,35E-13 | 0,0010123 | 0,0022618 | 0,0014557 | 0,0014557 | TRUE | 0,0163157 | No |
| orf19.5628 | 1875 | 1727 | 1822 | 1156 | 3325 | 2323 | 3132 | 3143 | 1788 | 1826 | 1800 | 1811 | 2664 | 2641 | 2659 | 2556 | 2218,11 | 1806 | 2630 | 1,456 | 0,542 | 13,945 | 3,36E-44 | 1,14E-43 | 1,00E-08 | 0,0028315 | 0,0009726 | 0,0009726 | TRUE | 0,0045704 | No |
| orf19.5629 | 2815 | 2579 | 2543 | 1579 | 6723 | 4685 | 6035 | 6031 | 2684 | 2727 | 2512 | 2474 | 5387 | 5327 | 5124 | 4904 | 3892,28 | 2599 | 5186 | 1,993 | 0,995 | 20,214 | 7,33E-91 | 4,42E-90 | 0,0016618 | 0,0025131 | 0,002023 | 0,002023 | TRUE | 0,0164323 | No |
| orf19.563 | 8292 | 6797 | 7622 | 4362 | 10642 | 6504 | 10332 | 10624 | 7906 | 7187 | 7530 | 6833 | 8527 | 7396 | 8772 | 8638 | 7848,57 | 7364 | 8333 | 1,131 | 0,178 | 2,792 | 0,0052319 | 0,0067525 | 0,0046624 | 0,0023005 | 0,003783 | 0,003783 | TRUE | 0,0695357 | No |
| orf19.5630 | 1680 | 1761 | 1795 | 1096 | 1725 | 1280 | 1669 | 1586 | 1602 | 1862 | 1773 | 1717 | 1382 | 1455 | 1417 | 1290 | 1562,27 | 1738 | 1386 | 0,797 | -0,328 | -5,388 | 7,11E-08 | 1,15E-07 | 0,0027444 | 0,0031423 | 0,0029114 | 0,0029114 | TRUE | 0,0356008 | No |

| | | | | | | | | | | | | | | | | | | | | | | | | | | | | | | | |
|---|---|---|---|---|---|---|---|---|---|---|---|---|---|---|---|---|---|---|---|---|---|---|---|---|---|---|---|---|---|---|---|
| orf19.5633 | 6258 | 5481 | 5717 | 3714 | 4931 | 3345 | 4627 | 4816 | 5966 | 5795 | 5648 | 5818 | 3951 | 3804 | 3928 | 3916 | 4853,33 | 5807 | 3900 | 0,672 | -0,574 | -16,914 | 3,56E-64 | 1,58E-63 | 0,0002146 | 0,0024296 | 0,0008936 | 0,0008936 | TRUE | 0,004372 | No |
| orf19.5634 | 4214 | 3498 | 3830 | 2331 | 1540 | 1036 | 1487 | 1541 | 4018 | 3699 | 3784 | 3652 | 1234 | 1178 | 1262 | 1253 | 2509,88 | 3788 | 1232 | 0,325 | -1,62 | -33,963 | 7,96E-253 | 1,37E-251 | 0,0010246 | 0,0027455 | 0,0016749 | 0,0016749 | TRUE | 0,0205597 | No |
| orf19.5635 | 3366 | 2570 | 2830 | 2270 | 1360 | 1014 | 1258 | 1407 | 3209 | 2717 | 2796 | 3556 | 1090 | 1153 | 1068 | 1144 | 2091,65 | 3070 | 1114 | 0,363 | -1,462 | -17,875 | 1,86E-71 | 9,11E-71 | 0,007923 | 0,0028763 | 0,0058366 | 0,0058366 | TRUE | 0,1348542 | No |
| orf19.5636 | 23644 | 18043 | 19957 | 15914 | 7458 | 5365 | 6943 | 7741 | 22542 | 19078 | 19716 | 24930 | 5976 | 6100 | 5895 | 6294 | 13816,38 | 21566 | 6066 | 0,281 | -1,83 | -24,008 | 2,29E-127 | 1,95E-126 | 0,0081117 | 0,0022101 | 0,0054804 | 0,0054804 | TRUE | 0,1344342 | No |
| orf19.5639 | 5998 | 5118 | 5728 | 3708 | 5828 | 4086 | 5613 | 6131 | 5718 | 5412 | 5659 | 5809 | 4670 | 4646 | 4765 | 4985 | 5208,01 | 5650 | 4766 | 0,844 | -0,244 | -6,158 | 7,38E-10 | 1,27E-09 | 0,0007207 | 0,0024065 | 0,0013169 | 0,0013169 | TRUE | 0,0117075 | No |
| orf19.564 | 435 | 401 | 492 | 268 | 327 | 250 | 352 | 355 | 415 | 424 | 486 | 420 | 262 | 284 | 299 | 289 | 359,8 | 436 | 284 | 0,649 | -0,625 | -7,167 | 7,66E-13 | 1,46E-12 | 0,0022065 | 0,0066547 | 0,0044429 | 0,0044429 | TRUE | 0,0719138 | No |
| orf19.5640 | 13546 | 10993 | 12318 | 8335 | 16397 | 11095 | 16010 | 16800 | 12915 | 11624 | 12169 | 13057 | 13138 | 12616 | 13592 | 13660 | 12846,43 | 12441 | 13252 | 1,066 | 0,092 | 1,921 | 0,0547036 | 0,0654535 | 0,0020096 | 0,0022191 | 0,0021016 | 0,0021016 | TRUE | 0,0239126 | No |
| orf19.5641 | 17672 | 13823 | 15433 | 9742 | 23718 | 15043 | 22810 | 24928 | 16848 | 14616 | 15246 | 15261 | 19005 | 17105 | 19366 | 20269 | 17214,54 | 15493 | 18936 | 1,222 | 0,29 | 4,776 | 1,79E-06 | 2,74E-06 | 0,0042822 | 0,0021867 | 0,0034721 | 0,0034721 | TRUE | 0,0516585 | No |
| orf19.5642 | 7754 | 6023 | 6412 | 4436 | 10332 | 6458 | 9446 | 10350 | 7393 | 6369 | 6334 | 6949 | 8279 | 7343 | 8020 | 8416 | 7387,76 | 6761 | 8014 | 1,186 | 0,246 | 3,949 | 7,85E-05 | 0,0001117 | 0,0043941 | 0,0023135 | 0,003581 | 0,003581 | TRUE | 0,0489704 | No |
| orf19.5643 | 980 | 835 | 943 | 600 | 1500 | 1004 | 1402 | 1601 | 934 | 883 | 932 | 940 | 1202 | 1142 | 1190 | 1302 | 1065,55 | 922 | 1209 | 1,313 | 0,393 | 6,774 | 1,26E-11 | 2,31E-11 | 0,0012289 | 0,0036322 | 0,0022564 | 0,0022564 | TRUE | 0,0323616 | No |
| orf19.5644 | 274 | 251 | 275 | 187 | 422 | 353 | 392 | 416 | 261 | 265 | 272 | 293 | 338 | 401 | 333 | 338 | 312,73 | 273 | 352 | 1,293 | 0,37 | 3,999 | 6,37E-05 | 9,09E-05 | 0,0022015 | 0,0073415 | 0,0049082 | 0,0049082 | TRUE | 0,090487 | No |
| orf19.5645 | 3024 | 2384 | 2672 | 1892 | 18607 | 13028 | 17699 | 19329 | 2883 | 2521 | 2640 | 2964 | 14909 | 14814 | 15026 | 15716 | 8934,17 | 2752 | 15116 | 5,5 | 2,459 | 46,169 | 0 | 0 | 0,0029008 | 0,0022751 | 0,0024905 | 0,0024905 | TRUE | 0,038482 | No |
| orf19.5646 | 407 | 311 | 342 | 233 | 1531 | 1018 | 1412 | 1510 | 388 | 329 | 338 | 365 | 1227 | 1158 | 1199 | 1228 | 778,83 | 355 | 1203 | 3,393 | 1,763 | 25,974 | 9,75E-149 | 9,67E-148 | 0,0001731 | 0,0041995 | 0,0024827 | 0,0024827 | TRUE | 0,0530021 | No |
| orf19.5647 | 5038 | 4306 | 4677 | 2995 | 11289 | 8387 | 10663 | 12389 | 4803 | 4553 | 4620 | 4692 | 9046 | 9537 | 9053 | 10073 | 7047,14 | 4667 | 9427 | 2,02 | 1,014 | 22,403 | 3,68E-111 | 2,70E-110 | 0,0013907 | 0,0023243 | 0,0018002 | 0,0018002 | TRUE | 0,0262971 | No |
| orf19.5648 | 531 | 454 | 444 | 279 | 2770 | 1608 | 2528 | 2718 | 506 | 480 | 439 | 437 | 2220 | 1828 | 2146 | 2210 | 1283,28 | 466 | 2101 | 4,506 | 2,172 | 27,265 | 1,12E-163 | 1,24E-162 | 0,0058313 | 0,0033707 | 0,0046802 | 0,0046802 | TRUE | 0,0901324 | No |
| orf19.5649 | 45 | 42 | 30 | 27 | 96 | 69 | 100 | 82 | 43 | 44 | 30 | 42 | 77 | 78 | 85 | 67 | 58,28 | 40 | 77 | 1,935 | 0,952 | 4,763 | 1,91E-06 | 2,92E-06 | 1,00E-08 | 0,0302663 | 0,0185512 | 0,0185512 | TRUE | 0,2443646 | No |
| orf19.5650 | 6629 | 5221 | 5722 | 3631 | 9857 | 6693 | 9566 | 10418 | 6320 | 5521 | 5653 | 5688 | 7898 | 7611 | 8121 | 8471 | 6910,32 | 5796 | 8025 | 1,385 | 0,47 | 8,824 | 1,11E-18 | 2,41E-18 | 0,0027252 | 0,0023289 | 0,0025666 | 0,0025666 | TRUE | 0,0459335 | No |
| orf19.5651 | 7528 | 6128 | 6905 | 4362 | 5076 | 3412 | 4864 | 5083 | 7177 | 6480 | 6821 | 6833 | 4067 | 3880 | 4130 | 4133 | 5440,12 | 6828 | 4052 | 0,594 | -0,752 | -17,589 | 2,98E-69 | 1,41E-68 | 0,0010388 | 0,0023931 | 0,0015634 | 0,0015634 | TRUE | 0,0148042 | No |
| orf19.5653 | 13174 | 10753 | 11642 | 7883 | 31618 | 21056 | 29572 | 33158 | 12560 | 11370 | 11501 | 12349 | 25335 | 23943 | 25106 | 26961 | 18640,57 | 11945 | 25336 | 2,121 | 1,085 | 21,838 | 1,01E-105 | 6,92E-105 | 0,0023699 | 0,0021794 | 0,0023068 | 0,0023068 | TRUE | 0,0228206 | No |
| orf19.5654 | 2799 | 2492 | 2610 | 1758 | 4946 | 3590 | 4688 | 5066 | 2669 | 2635 | 2578 | 2754 | 3963 | 4082 | 3980 | 4119 | 3347,55 | 2659 | 4036 | 1,519 | 0,604 | 16,106 | 2,30E-58 | 9,48E-58 | 0,0001445 | 0,0025818 | 0,0010263 | 0,0010263 | TRUE | 0,0062804 | No |
| orf19.5655 | 135 | 166 | 123 | 74 | 208 | 138 | 208 | 216 | 129 | 176 | 122 | 116 | 167 | 157 | 177 | 176 | 152,18 | 136 | 169 | 1,244 | 0,315 | 2,252 | 0,0243213 | 0,0300124 | 0,0109057 | 0,0128802 | 0,0120343 | 0,0120343 | TRUE | 0,4073181 | No |
| orf19.5657 | 3759 | 3125 | 3593 | 2251 | 2814 | 1896 | 2506 | 2890 | 3584 | 3304 | 3550 | 3526 | 2255 | 2156 | 2128 | 2350 | 2856,51 | 3491 | 2222 | 0,637 | -0,651 | -13,562 | 6,69E-42 | 2,19E-41 | 0,0012037 | 0,0026661 | 0,0018494 | 0,0018494 | TRUE | 0,0183545 | No |
| orf19.5658 | 1039 | 805 | 850 | 584 | 2991 | 2045 | 2887 | 3205 | 991 | 851 | 840 | 915 | 2397 | 2325 | 2451 | 2606 | 1671,91 | 899 | 2445 | 2,721 | 1,444 | 22,563 | 9,94E-113 | 7,39E-112 | 0,0031507 | 0,0030733 | 0,0031251 | 0,0031251 | TRUE | 0,060607 | No |
| orf19.5660 | 828 | 695 | 715 | 495 | 1875 | 1298 | 1866 | 1966 | 789 | 735 | 706 | 775 | 1502 | 1476 | 1584 | 1599 | 1145,9 | 751 | 1540 | 2,053 | 1,038 | 18,002 | 1,86E-72 | 9,26E-72 | 0,0010321 | 0,0035241 | 0,0021575 | 0,0021575 | TRUE | 0,0203519 | No |
| orf19.5660.1 | 2592 | 2101 | 2156 | 1572 | 8033 | 5263 | 7563 | 7995 | 2471 | 2222 | 2130 | 2463 | 6437 | 5985 | 6421 | 6501 | 4328,51 | 2322 | 6336 | 2,733 | 1,45 | 25,602 | 1,44E-144 | 1,41E-143 | 0,0030584 | 0,0024706 | 0,002768 | 0,002768 | TRUE | 0,0377992 | No |
| orf19.5661 | 2061 | 1649 | 1765 | 1210 | 4125 | 3078 | 4027 | 4715 | 1965 | 1744 | 1744 | 1896 | 3305 | 3500 | 3419 | 3834 | 2675,7 | 1837 | 3514 | 1,914 | 0,937 | 15,225 | 2,42E-52 | 9,22E-52 | 0,0033785 | 0,0027049 | 0,0032006 | 0,0032006 | TRUE | 0,0460925 | No |
| orf19.5662 | 1436 | 1225 | 1328 | 834 | 1484 | 998 | 1461 | 1500 | 1369 | 1295 | 1312 | 1307 | 1189 | 1135 | 1240 | 1220 | 1258,34 | 1321 | 1196 | 0,906 | -0,142 | -2,839 | 0,0045276 | 0,0058638 | 1,00E-08 | 0,0033961 | 0,0016138 | 0,0016138 | TRUE | 0,0129904 | No |
| orf19.5663 | 1534 | 1349 | 1391 | 856 | 1634 | 1082 | 1474 | 1590 | 1463 | 1426 | 1374 | 1341 | 1309 | 1230 | 1251 | 1293 | 1335,99 | 1401 | 1271 | 0,906 | -0,142 | -2,848 | 0,004402 | 0,005708 | 0,0003288 | 0,0033203 | 0,0016265 | 0,0016265 | TRUE | 0,0105726 | No |
| orf19.5664 | 144 | 159 | 170 | 86 | 152 | 113 | 149 | 168 | 137 | 168 | 168 | 135 | 122 | 128 | 127 | 137 | 140,18 | 152 | 128 | 0,84 | -0,251 | -1,971 | 0,048696 | 0,0586061 | 1,00E-08 | 0,0138039 | 0,0084825 | 0,0084825 | TRUE | 0,0555903 | No |
| orf19.5665 | 546 | 452 | 498 | 322 | 1701 | 1122 | 1652 | 1659 | 521 | 478 | 492 | 504 | 1363 | 1276 | 1403 | 1349 | 923,14 | 499 | 1348 | 2,705 | 1,436 | 23,837 | 1,37E-125 | 1,15E-124 | 1,00E-08 | 0,0038699 | 0,0020351 | 0,0020351 | TRUE | 0,0143974 | No |
| orf19.5666 | 899 | 729 | 768 | 436 | 2714 | 1538 | 2707 | 2590 | 857 | 771 | 759 | 683 | 2175 | 1749 | 2298 | 2106 | 1424,66 | 768 | 2082 | 2,708 | 1,437 | 15,928 | 4,05E-57 | 1,63E-56 | 0,0095458 | 0,0032438 | 0,0068733 | 0,0068733 | TRUE | 0,1378404 | No |
| orf19.5667 | 6904 | 5628 | 6325 | 3781 | 5999 | 3768 | 5730 | 6158 | 6582 | 5951 | 6249 | 5923 | 4807 | 4285 | 4865 | 5007 | 5458,49 | 6176 | 4741 | 0,768 | -0,381 | -6,684 | 2,32E-11 | 4,22E-11 | 0,0032117 | 0,0023921 | 0,0029413 | 0,0029413 | TRUE | 0,0502873 | No |
| orf19.5669 | 730 | 592 | 649 | 391 | 1807 | 1167 | 1639 | 1843 | 696 | 626 | 641 | 613 | 1448 | 1327 | 1392 | 1499 | 1030,07 | 644 | 1416 | 2,196 | 1,135 | 17,871 | 1,99E-71 | 9,72E-71 | 0,0018561 | 0,0036853 | 0,0026902 | 0,0026902 | TRUE | 0,0360965 | No |
| orf19.567 | 927 | 1030 | 1070 | 670 | 575 | 423 | 538 | 479 | 884 | 1089 | 1057 | 1050 | 461 | 481 | 457 | 389 | 733,44 | 1020 | 447 | 0,438 | -1,191 | -13,614 | 3,30E-42 | 1,08E-41 | 0,0071356 | 0,0043299 | 0,0057985 | 0,0057985 | TRUE | 0,0997357 | No |
| orf19.5671 | 5360 | 4573 | 4938 | 3464 | 2232 | 1532 | 2017 | 2380 | 5110 | 4835 | 4878 | 5427 | 1788 | 1742 | 1712 | 1935 | 3428,55 | 5062 | 1794 | 0,355 | -1,495 | -27,196 | 7,25E-163 | 8,02E-162 | 0,0024467 | 0,0025702 | 0,0025401 | 0,0025401 | TRUE | 0,0341598 | No |
| orf19.5672 | 42 | 51 | 43 | 40 | 343 | 257 | 332 | 346 | 40 | 54 | 42 | 63 | 275 | 292 | 282 | 281 | 166,17 | 50 | 282 | 5,81 | 2,539 | 17,618 | 1,81E-69 | 8,58E-69 | 1,00E-08 | 0,011972 | 0,0069877 | 0,0069877 | TRUE | 0,1676927 | No |
| orf19.5673 | 2127 | 1841 | 2087 | 1499 | 4018 | 3083 | 3859 | 4119 | 2028 | 1947 | 2062 | 2348 | 3220 | 3506 | 3276 | 3349 | 2716,89 | 2096 | 3338 | 1,595 | 0,674 | 11,291 | 1,45E-29 | 3,89E-29 | 0,0036525 | 0,0026956 | 0,0030149 | 0,0030149 | TRUE | 0,0747773 | No |
| orf19.5674 | 460 | 332 | 410 | 308 | 293 | 254 | 278 | 321 | 439 | 351 | 405 | 482 | 235 | 289 | 236 | 261 | 337,22 | 419 | 255 | 0,609 | -0,716 | -6,483 | 8,98E-11 | 1,61E-10 | 0,0103664 | 0,0069602 | 0,0086066 | 0,0086066 | TRUE | 0,1366937 | No |
| orf19.5675 | 5733 | 4972 | 5471 | 3400 | 4794 | 3433 | 4718 | 5005 | 5466 | 5257 | 5405 | 5326 | 3841 | 3904 | 4006 | 4070 | 4659,27 | 5364 | 3955 | 0,737 | -0,44 | -12,541 | 4,44E-36 | 1,32E-35 | 0,0002851 | 0,0024437 | 0,0009619 | 0,0009619 | TRUE | 0,0047836 | No |
| orf19.5676 | 2752 | 2123 | 2554 | 1503 | 3661 | 2279 | 3443 | 3582 | 2624 | 2245 | 2523 | 2355 | 2933 | 2591 | 2923 | 2913 | 2638,33 | 2437 | 2840 | 1,166 | 0,221 | 3,551 | 0,0003842 | 0,0005297 | 0,0036151 | 0,0027136 | 0,0033319 | 0,0033319 | TRUE | 0,0411724 | No |
| orf19.5677 | 80 | 81 | 86 | 47 | 125 | 84 | 110 | 98 | 76 | 86 | 85 | 74 | 100 | 96 | 93 | 80 | 86,16 | 80 | 92 | 1,144 | 0,194 | 1,253 | 0,2103849 | 0,2381103 | 1,00E-08 | 0,0211485 | 0,0113802 | 0,0113802 | TRUE | 0,0882116 | No |
| orf19.5678 | 1741 | 1309 | 1566 | 981 | 1833 | 1173 | 1800 | 1907 | 1660 | 1384 | 1547 | 1537 | 1469 | 1334 | 1528 | 1551 | 1501,14 | 1532 | 1470 | 0,961 | -0,058 | -0,849 | 0,3961348 | 0,431944 | 0,0040351 | 0,0031851 | 0,0038329 | 0,0038329 | TRUE | 0,0520811 | No |
| orf19.568 | 3187 | 3160 | 3311 | 1847 | 4538 | 3254 | 4231 | 4298 | 3038 | 3341 | 3271 | 2893 | 3636 | 3700 | 3592 | 3495 | 3370,9 | 3136 | 3606 | 1,148 | 0,199 | 3,876 | 0,000106 | 0,0001498 | 0,0020704 | 0,0025784 | 0,0022368 | 0,0022368 | TRUE | 0,0309963 | No |
| orf19.5680 | 1929 | 1591 | 1725 | 1075 | 2384 | 1680 | 2431 | 2502 | 1839 | 1682 | 1704 | 1684 | 1910 | 1910 | 2064 | 2034 | 1853,55 | 1727 | 1980 | 1,145 | 0,196 | 3,841 | 0,0001226 | 0,0001727 | 0,0012705 | 0,0029771 | 0,0019482 | 0,0019482 | TRUE | 0,0237029 | No |
| orf19.5681 | 1139 | 965 | 969 | 642 | 1565 | 1000 | 1516 | 1476 | 1086 | 1020 | 957 | 1006 | 1254 | 1137 | 1287 | 1200 | 1118,45 | 1017 | 1220 | 1,2 | 0,262 | 4,297 | 1,73E-05 | 2,53E-05 | 0,0019645 | 0,0035593 | 0,0026651 | 0,0026651 | TRUE | 0,0262117 | No |
| orf19.5682 | 2944 | 2365 | 2582 | 1812 | 6684 | 4697 | 6405 | 6829 | 2807 | 2501 | 2551 | 2839 | 5356 | 5341 | 5438 | 5553 | 4047,99 | 2674 | 5422 | 2,03 | 1,021 | 20,762 | 9,55E-96 | 6,00E-95 | 0,0018843 | 0,0024969 | 0,0020304 | 0,0020304 | TRUE | 0,022898 | No |
| orf19.5683 | 1761 | 1460 | 1616 | 923 | 3222 | 1984 | 3189 | 3140 | 1679 | 1544 | 1596 | 1446 | 2582 | 2256 | 2707 | 2553 | 2045,42 | 1566 | 2524 | 1,61 | 0,687 | 10,339 | 4,68E-25 | 1,16E-24 | 0,0041203 | 0,002894 | 0,0037087 | 0,0037087 | TRUE | 0,0606291 | No |
| orf19.5684 | 2145 | 1556 | 1731 | 1068 | 3362 | 1906 | 3151 | 3181 | 2045 | 1645 | 1710 | 1673 | 2694 | 2167 | 2675 | 2586 | 2149,53 | 1768 | 2530 | 1,431 | 0,517 | 6,031 | 1,63E-09 | 2,79E-09 | 0,0096655 | 0,0028551 | 0,0065605 | 0,0065605 | TRUE | 0,135841 | No |
| orf19.5684.1 | 110 | 95 | 90 | 78 | 373 | 227 | 318 | 385 | 105 | 100 | 89 | 122 | 299 | 258 | 270 | 313 | 194,56 | 104 | 285 | 2,775 | 1,472 | 11,818 | 3,14E-32 | 8,80E-32 | 0,0045307 | 0,0105305 | 0,007941 | 0,007941 | TRUE | 0,1004303 | No |
| orf19.5685 | 4357 | 3552 | 3982 | 2430 | 14052 | 8755 | 13288 | 14441 | 4154 | 3756 | 3934 | 3807 | 11259 | 9955 | 11281 | 11742 | 7486,04 | 3913 | 11059 | 2,826 | 1,499 | 26,086 | 5,24E-150 | 5,26E-149 | 0,0032159 | 0,0023106 | 0,0029865 | 0,0029865 | TRUE | 0,0548395 | No |
| orf19.5686 | 5857 | 4399 | 4676 | 2874 | 831 | 560 | 819 | 787 | 5584 | 4651 | 4619 | 4502 | 666 | 637 | 695 | 640 | 2749,37 | 4839 | 660 | 0,136 | -2,875 | -38,647 | 0 | 0 | 0,0059161 | 0,0026885 | 0,004523 | 0,004523 | TRUE | 0,1316747 | No |
| orf19.5688 | 2757 | 2157 | 2306 | 1372 | 1930 | 1107 | 1859 | 1860 | 2629 | 2281 | 2278 | 2149 | 1546 | 1259 | 1578 | 1512 | 1904,07 | 2334 | 1474 | 0,632 | -0,663 | -8,155 | 3,50E-16 | 7,21E-16 | 0,0080448 | 0,0029536 | 0,0057993 | 0,0057993 | TRUE | 0,1130276 | No |
| orf19.5689 | 1096 | 845 | 957 | 619 | 2292 | 1598 | 2179 | 2328 | 1045 | 893 | 945 | 970 | 1837 | 1817 | 1850 | 1893 | 1406,24 | 963 | 1849 | 1,919 | 0,94 | 17,06 | 2,94E-65 | 1,33E-64 | 0,001235 | 0,0032589 | 0,0020951 | 0,0020951 | TRUE | 0,0422213 | No |
| orf19.5691 | 742 | 597 | 670 | 367 | 1059 | 617 | 962 | 995 | 707 | 631 | 662 | 575 | 849 | 702 | 817 | 809 | 718,92 | 644 | 794 | 1,231 | 0,3 | 3,707 | 0,0002094 | 0,000292 | 0,005095 | 0,0043751 | 0,0048446 | 0,0048446 | TRUE | 0,0689057 | No |
| orf19.5692 | 484 | 398 | 413 | 320 | 899 | 570 | 839 | 834 | 461 | 421 | 408 | 501 | 720 | 648 | 712 | 678 | 568,81 | 448 | 690 | 1,55 | 0,632 | 8,156 | 3,45E-16 | 7,13E-16 | 0,0025855 | 0,0049778 | 0,0038618 | 0,0038618 | TRUE | 0,0630258 | No |
| orf19.5693 | 1036 | 792 | 922 | 560 | 3371 | 2121 | 3309 | 3325 | 988 | 837 | 911 | 877 | 2701 | 2412 | 2809 | 2704 | 1779,87 | 903 | 2656 | 2,939 | 1,555 | 23,325 | 2,45E-120 | 1,96E-119 | 0,0035679 | 0,0030138 | 0,0034788 | 0,0034788 | TRUE | 0,0490057 | No |
| orf19.5694 | 153 | 129 | 171 | 118 | 603 | 421 | 582 | 684 | 146 | 136 | 169 | 185 | 483 | 479 | 494 | 556 | 331,03 | 159 | 503 | 3,195 | 1,676 | 15,94 | 3,37E-57 | 1,36E-56 | 0,0050072 | 0,0070513 | 0,0061937 | 0,0061937 | TRUE | 0,0976441 | No |
| orf19.5698 | 2260 | 1775 | 2001 | 1481 | 2704 | 1817 | 2468 | 2801 | 2155 | 1877 | 1977 | 2320 | 2167 | 2066 | 2095 | 2277 | 2116,74 | 2082 | 2151 | 1,035 | 0,05 | 0,742 | 0,4583878 | 0,4947518 | 0,0043408 | 0,002867 | 0,0038802 | 0,0038802 | TRUE | 0,0680265 | No |
| orf19.57 | 693 | 625 | 649 | 405 | 782 | 542 | 723 | 801 | 661 | 661 | 641 | 634 | 627 | 616 | 614 | 651 | 638,15 | 649 | 627 | 0,965 | -0,051 | -0,847 | 0,3969629 | 0,432701 | 1,00E-08 | 0,0046642 | 0,0019217 | 0,0019217 | TRUE | 0,008287 | No |
| orf19.570 | 194 | 166 | 190 | 111 | 141 | 109 | 145 | 163 | 185 | 176 | 188 | 174 | 113 | 124 | 123 | 133 | 151,83 | 181 | 123 | 0,68 | -0,556 | -4,792 | 1,65E-06 | 2,53E-06 | 1,00E-08 | 0,0129054 | 0,0062364 | 0,0062364 | TRUE | 0,0353347 | No |
| orf19.5700 | 57 | 49 | 59 | 33 | 217 | 117 | 235 | 212 | 54 | 52 | 58 | 52 | 174 | 133 | 200 | 172 | 111,87 | 54 | 170 | 3,144 | 1,653 | 9,826 | 8,70E-23 | 2,06E-22 | 1,00E-08 | 0,0167684 | 0,0142989 | 0,0142989 | TRUE | 0,1928919 | No |
| orf19.5701 | 26 | 25 | 30 | 13 | 46 | 44 | 39 | 42 | 25 | 26 | 30 | 20 | 37 | 50 | 33 | 34 | 31,92 | 25 | 38 | 1,489 | 0,575 | 2,203 | 0,0275614 | 0,0338471 | 1,00E-08 | 0,0535261 | 0,0319596 | 0,0319596 | TRUE | 0,2584367 | No |
| orf19.5702 | 1671 | 1657 | 1690 | 1033 | 1845 | 1338 | 1717 | 1842 | 1593 | 1752 | 1670 | 1618 | 1478 | 1521 | 1458 | 1498 | 1573,53 | 1658 | 1489 | 0,897 | -0,157 | -3,298 | 0,0009746 | 0,0013157 | 0,0004102 | 0,0031347 | 0,0015357 | 0,0015357 | TRUE | 0,0177298 | No |
| orf19.5704 | 4242 | 3745 | 4335 | 2622 | 3193 | 2220 | 2878 | 3292 | 4044 | 3960 | 4283 | 4108 | 2558 | 2524 | 2443 | 2677 | 3324,64 | 4099 | 2550 | 0,622 | -0,684 | -15,458 | 6,69E-54 | 2,60E-53 | 0,0008538 | 0,0025851 | 0,0015669 | 0,0015669 | TRUE | 0,0136919 | No |
| orf19.5705 | 938 | 889 | 960 | 583 | 2066 | 1548 | 1990 | 2247 | 894 | 940 | 948 | 913 | 1655 | 1760 | 1690 | 1827 | 1328,52 | 924 | 1733 | 1,875 | 0,907 | 16,95 | 1,94E-64 | 8,64E-64 | 0,0009288 | 0,0033272 | 0,0018833 | 0,0018833 | TRUE | 0,0168783 | No |
| orf19.5710 | 5921 | 5328 | 5784 | 3667 | 4843 | 3465 | 4495 | 5012 | 5645 | 5634 | 5714 | 5745 | 3881 | 3940 | 3816 | 4075 | 4806,17 | 5684 | 3928 | 0,691 | -0,533 | -15,504 | 3,27E-54 | 1,28E-53 | 0,0002294 | 0,0024329 | 0,0009215 | 0,0009215 | TRUE | 0,0080105 | No |
| orf19.5711 | 958 | 929 | 1002 | 663 | 4292 | 3032 | 3938 | 4348 | 913 | 982 | 990 | 1039 | 3439 | 3448 | 3343 | 3535 | 2211,19 | 981 | 3441 | 3,519 | 1,815 | 37,327 | 5,94E-305 | 1,34E-303 | 0,000524 | 0,0028338 | 0,0015691 | 0,0015691 | TRUE | 0,0264261 | No |
| orf19.5713 | 6403 | 5970 | 6462 | 4157 | 3290 | 2336 | 3029 | 3238 | 6105 | 6313 | 6384 | 6512 | 2636 | 2656 | 2572 | 2633 | 4476,24 | 6328 | 2624 | 0,415 | -1,269 | -35,408 | 1,28E-274 | 2,47E-273 | 0,0002949 | 0,0024581 | 0,0009749 | 0,0009749 | TRUE | 0,0069805 | No |
| orf19.5714 | 842 | 713 | 777 | 476 | 984 | 756 | 908 | 1074 | 803 | 754 | 768 | 746 | 788 | 860 | 771 | 873 | 795,27 | 768 | 823 | 1,07 | 0,097 | 1,519 | 0,1286923 | 0,148604 | 0,001461 | 0,0041559 | 0,0026763 | 0,0026763 | TRUE | 0,0218013 | No |
| orf19.5716 | 74 | 80 | 60 | 51 | 52 | 33 | 43 | 48 | 71 | 85 | 59 | 80 | 42 | 38 | 37 | 39 | 56,13 | 74 | 39 | 0,53 | -0,917 | -4,639 | 3,51E-06 | 5,32E-06 | 1,00E-08 | 0,0313434 | 0,0183927 | 0,0183927 | TRUE | 0,1639048 | No |
| orf19.5717 | 2 | 11 | 5 | 4 | 12 | 8 | 12 | 6 | 2 | 12 | 5 | 6 | 10 | 9 | 10 | 5 | 7,32 | 6 | 8 | 1,377 | 0,461 | 0,793 | 0,4276321 | 0,4636739 | 0,1098169 | 0,2265402 | 0,1803268 | 0,1803268 | TRUE | 0,6725049 | No |
| orf19.5718 | 541 | 547 | 553 | 375 | 613 | 433 | 527 | 536 | 516 | 578 | 546 | 587 | 491 | 492 | 447 | 436 | 511,84 | 557 | 466 | 0,839 | -0,253 | -3,341 | 0,0008341 | 0,0011319 | 0,0018212 | 0,0052991 | 0,0035481 | 0,0035481 | TRUE | 0,0298515 | No |
| orf19.5719 | 1905 | 1592 | 1728 | 1074 | 3423 | 2301 | 3252 | 3568 | 1816 | 1683 | 1707 | 1682 | 2743 | 2616 | 2761 | 2901 | 2238,8 | 1722 | 2755 | 1,599 | 0,677 | 13,885 | 7,85E-44 | 2,64E-43 | 0,0011542 | 0,0028247 | 0,001798 | 0,001798 | TRUE | 0,0168553 | No |

| | | | | | | | | | | | | | | | | | | | | | | | | | | | | | | | |
|---|---|---|---|---|---|---|---|---|---|---|---|---|---|---|---|---|---|---|---|---|---|---|---|---|---|---|---|---|---|---|---|
| orf19.5720 | 4265 | 3693 | 3969 | 2277 | 4458 | 3101 | 4147 | 4575 | 4066 | 3905 | 3921 | 3567 | 3572 | 3526 | 3521 | 3720 | 3724,75 | 3865 | 3585 | 0,926 | -0,11 | -2,332 | 0,0196753 | 0,0244352 | 0,0014734 | 0,0025321 | 0,0018715 | 0,0018715 | TRUE | 0,0310066 | No |
| orf19.5722 | 1100 | 872 | 938 | 611 | 1726 | 1125 | 1629 | 1727 | 1049 | 922 | 927 | 957 | 1383 | 1279 | 1383 | 1404 | 1163 | 964 | 1362 | 1,414 | 0,5 | 8,36 | 6,27E-17 | 1,32E-16 | 0,0015366 | 0,0035031 | 0,0025263 | 0,0025263 | TRUE | 0,0451508 | No |
| orf19.5723 | 3726 | 3173 | 3397 | 2376 | 15721 | 11837 | 14802 | 16496 | 3552 | 3355 | 3356 | 3722 | 12597 | 13460 | 12567 | 13413 | 8252,7 | 3496 | 13009 | 3,724 | 1,897 | 40,515 | 0 | 0 | 0,0017575 | 0,0022902 | 0,0019097 | 0,0019097 | TRUE | 0,0216928 | No |
| orf19.5724 | 8 | 5 | 11 | 4 | 7 | 6 | 7 | 6 | 8 | 5 | 11 | 6 | 6 | 7 | 6 | 5 | 6,66 | 8 | 6 | 0,758 | -0,4 | -0,75 | 0,4534592 | 0,4901784 | 1,00E-08 | 0,2485301 | 0,1236387 | 0,1236387 | TRUE | 0,2671914 | No |
| orf19.5725 | 379 | 391 | 477 | 294 | 580 | 439 | 548 | 585 | 361 | 413 | 471 | 461 | 465 | 499 | 465 | 476 | 451,42 | 426 | 476 | 1,119 | 0,162 | 1,809 | 0,0704431 | 0,0834876 | 0,0051259 | 0,0057284 | 0,0054336 | 0,0054336 | TRUE | 0,1294462 | No |
| orf19.5727 | 1374 | 1123 | 1189 | 705 | 1225 | 856 | 1145 | 1251 | 1310 | 1187 | 1175 | 1104 | 982 | 973 | 972 | 1017 | 1090,08 | 1194 | 986 | 0,824 | -0,28 | -4,629 | 3,68E-06 | 5,57E-06 | 0,0017664 | 0,0035975 | 0,0025805 | 0,0025805 | TRUE | 0,0518093 | No |
| orf19.5728 | 269 | 262 | 243 | 173 | 2044 | 1451 | 1945 | 2060 | 256 | 277 | 240 | 271 | 1638 | 1650 | 1651 | 1675 | 957,32 | 261 | 1654 | 6,356 | 2,668 | 40,697 | 0 | 0 | 1,00E-08 | 0,0038064 | 0,00173 | 0,00173 | TRUE | 0,0346743 | No |
| orf19.5729 | 418 | 383 | 404 | 277 | 674 | 552 | 651 | 624 | 399 | 405 | 399 | 434 | 540 | 628 | 553 | 507 | 483,04 | 409 | 557 | 1,361 | 0,445 | 5,434 | 5,51E-08 | 8,96E-08 | 0,0030847 | 0,0054904 | 0,004254 | 0,004254 | TRUE | 0,0799036 | No |
| orf19.5730 | 2136 | 1966 | 2092 | 1379 | 1902 | 1407 | 1786 | 2015 | 2036 | 2079 | 2067 | 2160 | 1524 | 1600 | 1516 | 1638 | 1827,6 | 2086 | 1570 | 0,753 | -0,409 | -8,891 | 6,07E-19 | 1,33E-18 | 0,0004645 | 0,0029897 | 0,0014762 | 0,0014762 | TRUE | 0,0111877 | No |
| orf19.5731 | 2012 | 1847 | 1969 | 1175 | 2328 | 1520 | 2237 | 2402 | 1918 | 1953 | 1945 | 1841 | 1865 | 1728 | 1899 | 1953 | 1887,88 | 1914 | 1861 | 0,973 | -0,04 | -0,804 | 0,4215851 | 0,4575059 | 0,0009767 | 0,002961 | 0,0018505 | 0,0018505 | TRUE | 0,0264416 | No |
| orf19.5732 | 29322 | 26514 | 28239 | 16621 | 25168 | 16501 | 23679 | 25190 | 27955 | 28035 | 27897 | 26038 | 20166 | 18763 | 20103 | 20482 | 23680,06 | 27481 | 19878 | 0,723 | -0,467 | -11,344 | 7,97E-30 | 2,14E-29 | 0,0012922 | 0,0021606 | 0,0015871 | 0,0015871 | TRUE | 0,0173405 | No |
| orf19.5734 | 3364 | 2896 | 3246 | 2088 | 2103 | 1482 | 2023 | 2111 | 3207 | 3062 | 3207 | 3271 | 1685 | 1685 | 1718 | 1716 | 2443,91 | 3187 | 1701 | 0,534 | -0,904 | -22,818 | 3,02E-115 | 2,29E-114 | 1,00E-08 | 0,0027631 | 0,0010699 | 0,0010699 | TRUE | 0,0083878 | No |
| orf19.5735 | 1926 | 1627 | 1746 | 982 | 2394 | 1673 | 2370 | 2556 | 1836 | 1720 | 1725 | 1538 | 1918 | 1902 | 2012 | 2078 | 1841,35 | 1705 | 1978 | 1,158 | 0,211 | 3,56 | 0,0003702 | 0,000511 | 0,0027617 | 0,002983 | 0,002828 | 0,002828 | TRUE | 0,0482381 | No |
| orf19.5735.3 | 3054 | 2483 | 2673 | 1687 | 346 | 244 | 341 | 360 | 2912 | 2625 | 2641 | 2643 | 277 | 277 | 290 | 293 | 1494,68 | 2705 | 284 | 0,105 | -3,25 | -50,326 | 0 | 0 | 1,00E-08 | 0,0031898 | 0,0022362 | 0,0022362 | TRUE | 0,0329104 | No |
| orf19.5736 | 4827 | 4138 | 4711 | 3143 | 5465 | 3927 | 4948 | 5650 | 4602 | 4375 | 4654 | 4924 | 4379 | 4465 | 4201 | 4594 | 4524,28 | 4639 | 4410 | 0,951 | -0,072 | -1,526 | 0,1271253 | 0,1469275 | 0,0015829 | 0,0024542 | 0,0019174 | 0,0019174 | TRUE | 0,0199045 | No |
| orf19.5741 | 51254 | 43385 | 48418 | 33794 | 133886 | 98982 | 125287 | 138574 | 48865 | 45874 | 47832 | 52940 | 107279 | 112551 | 106368 | 112674 | 79298,03 | 48878 | 109718 | 2,245 | 1,167 | 24,237 | 9,13E-130 | 7,91E-129 | 0,002307 | 0,002112 | 0,0022108 | 0,0022108 | TRUE | 0,0381931 | No |
| orf19.5742 | 1742 | 1472 | 1669 | 1051 | 2312 | 1811 | 2165 | 2453 | 1661 | 1556 | 1649 | 1646 | 1853 | 2059 | 1838 | 1995 | 1782,12 | 1628 | 1936 | 1,188 | 0,249 | 4,692 | 2,71E-06 | 4,12E-06 | 0,0015311 | 0,0030126 | 0,0021228 | 0,0021228 | TRUE | 0,0211537 | No |
| orf19.5746 | 8669 | 8205 | 8664 | 5168 | 14231 | 10328 | 13409 | 14463 | 8265 | 8676 | 8559 | 8096 | 11403 | 11744 | 11384 | 11760 | 9985,82 | 8399 | 11573 | 1,377 | 0,462 | 13,416 | 4,87E-41 | 1,57E-40 | 0,0005242 | 0,0022557 | 0,0010315 | 0,0010315 | TRUE | 0,0068643 | No |
| orf19.5747 | 2620 | 2536 | 2663 | 1764 | 3375 | 2534 | 3175 | 3139 | 2498 | 2681 | 2631 | 2763 | 2704 | 2881 | 2696 | 2552 | 2675,89 | 2643 | 2708 | 1,025 | 0,035 | 0,698 | 0,4851844 | 0,5216448 | 0,0016157 | 0,0027049 | 0,0021009 | 0,0021009 | TRUE | 0,0216856 | No |
| orf19.5748 | 17 | 18 | 20 | 12 | 102 | 66 | 89 | 95 | 16 | 19 | 20 | 19 | 82 | 75 | 76 | 77 | 47,92 | 18 | 78 | 4,212 | 2,075 | 8,832 | 1,03E-18 | 2,25E-18 | 1,00E-08 | 0,036353 | 0,0172217 | 0,0172217 | TRUE | 0,0415784 | No |
| orf19.5749 | 4462 | 4053 | 4311 | 2718 | 1422 | 1009 | 1324 | 1452 | 4254 | 4286 | 4259 | 4258 | 1139 | 1147 | 1124 | 1181 | 2705,97 | 4264 | 1148 | 0,269 | -1,893 | -49,522 | 0 | 0 | 1,00E-08 | 0,0026981 | 0,0008838 | 0,0008838 | TRUE | 0,0047328 | No |
| orf19.575 | 9303 | 7467 | 8420 | 5451 | 8895 | 6108 | 8284 | 9464 | 8869 | 7895 | 8318 | 8539 | 7127 | 6945 | 7033 | 7695 | 7802,89 | 8405 | 7200 | 0,857 | -0,223 | -4,524 | 6,08E-06 | 9,09E-06 | 0,0021129 | 0,0023017 | 0,0022048 | 0,0022048 | TRUE | 0,0262522 | No |
| orf19.5750 | 4209 | 3815 | 4109 | 2438 | 14031 | 9990 | 13366 | 14087 | 4013 | 4034 | 4059 | 3819 | 11243 | 11360 | 11348 | 11454 | 7666,15 | 3981 | 11351 | 2,847 | 1,51 | 47,708 | 0 | 0 | 1,38E-05 | 0,0023055 | 0,0007812 | 0,0007812 | TRUE | 0,0082356 | No |
| orf19.5751 | 4615 | 4492 | 4752 | 3051 | 2853 | 2010 | 2582 | 2594 | 4400 | 4750 | 4695 | 4780 | 2286 | 2286 | 2192 | 2109 | 3437,07 | 4656 | 2218 | 0,476 | -1,07 | -23,438 | 1,76E-121 | 1,42E-120 | 0,000992 | 0,002569 | 0,0016772 | 0,0016772 | TRUE | 0,0169934 | No |
| orf19.5752 | 2129 | 2086 | 2203 | 1253 | 2608 | 1735 | 2440 | 2558 | 2030 | 2206 | 2176 | 1963 | 2090 | 1973 | 2072 | 2080 | 2073,59 | 2094 | 2054 | 0,98 | -0,029 | -0,582 | 0,5608885 | 0,5955124 | 0,0012488 | 0,0028831 | 0,0019165 | 0,0019165 | TRUE | 0,019435 | No |
| orf19.5753 | 5390 | 4940 | 5312 | 3376 | 29878 | 21700 | 27096 | 29438 | 5139 | 5223 | 5248 | 5289 | 23940 | 24675 | 23004 | 23936 | 14556,77 | 5225 | 23889 | 4,573 | 2,193 | 65,693 | 0 | 0 | 0,0004765 | 0,0022041 | 0,0009439 | 0,0009439 | TRUE | 0,0075977 | No |
| orf19.5754 | 10340 | 10691 | 11027 | 6505 | 5160 | 3880 | 4624 | 4767 | 9858 | 11304 | 10894 | 10190 | 4135 | 4412 | 3926 | 3876 | 7324,35 | 10562 | 4087 | 0,387 | -1,37 | -23,733 | 1,64E-124 | 1,36E-123 | 0,0034876 | 0,0023155 | 0,0030402 | 0,0030402 | TRUE | 0,0342567 | No |
| orf19.5755 | 3292 | 3064 | 3368 | 1981 | 709 | 490 | 684 | 641 | 3139 | 3240 | 3327 | 3103 | 568 | 557 | 581 | 521 | 1879,52 | 3202 | 557 | 0,174 | -2,525 | -47,945 | 0 | 0 | 0,0006631 | 0,0029649 | 0,0016886 | 0,0016886 | TRUE | 0,0234206 | No |
| orf19.5757 | 1916 | 1884 | 2030 | 1243 | 1562 | 1243 | 1525 | 1440 | 1827 | 1992 | 2005 | 1947 | 1252 | 1413 | 1295 | 1171 | 1612,75 | 1943 | 1283 | 0,659 | -0,601 | -9,541 | 1,41E-21 | 3,27E-21 | 0,0031243 | 0,0031094 | 0,0031734 | 0,0031734 | TRUE | 0,0541279 | No |
| orf19.5758 | 7575 | 7146 | 7461 | 4425 | 9863 | 7367 | 8949 | 9516 | 7222 | 7556 | 7371 | 6932 | 7903 | 8377 | 7598 | 7737 | 7586,96 | 7270 | 7904 | 1,086 | 0,119 | 2,726 | 0,0064094 | 0,0082258 | 0,0014026 | 0,0023077 | 0,0017108 | 0,0017108 | TRUE | 0,0195216 | No |
| orf19.5759 | 4075 | 3883 | 4187 | 2456 | 8450 | 6181 | 7931 | 8391 | 3885 | 4106 | 4136 | 3847 | 6771 | 7028 | 6733 | 6823 | 5416,23 | 3994 | 6839 | 1,711 | 0,775 | 20,315 | 9,40E-92 | 5,71E-91 | 0,0006114 | 0,0023944 | 0,0011899 | 0,0011899 | TRUE | 0,0070388 | No |
| orf19.576 | 270 | 223 | 240 | 165 | 398 | 282 | 356 | 360 | 257 | 236 | 237 | 258 | 319 | 321 | 302 | 293 | 277,91 | 247 | 309 | 1,25 | 0,322 | 3,612 | 0,0003037 | 0,0004208 | 1,00E-08 | 0,0079992 | 0,0039063 | 0,0039063 | TRUE | 0,0141063 | No |
| orf19.5760 | 5180 | 3905 | 4443 | 3473 | 1236 | 929 | 1127 | 1327 | 4939 | 4129 | 4389 | 5441 | 990 | 1056 | 957 | 1079 | 2872,5 | 4724 | 1020 | 0,216 | -2,21 | -26,592 | 8,49E-156 | 8,94E-155 | 0,0084113 | 0,0026629 | 0,0060778 | 0,0060778 | TRUE | 0,123676 | No |
| orf19.5762 | 106 | 63 | 71 | 59 | 144 | 87 | 122 | 146 | 101 | 67 | 70 | 92 | 115 | 99 | 104 | 119 | 95,86 | 82 | 109 | 1,331 | 0,413 | 2,471 | 0,0134834 | 0,0169867 | 0,0113211 | 0,0192203 | 0,0159376 | 0,0159376 | TRUE | 0,2544147 | No |
| orf19.5763 | 195 | 136 | 143 | 98 | 169 | 128 | 165 | 192 | 186 | 144 | 141 | 154 | 135 | 146 | 140 | 156 | 150,21 | 156 | 144 | 0,922 | -0,117 | -0,912 | 0,3615172 | 0,3962296 | 0,0046923 | 0,0130221 | 0,0091881 | 0,0091881 | TRUE | 0,1914321 | No |
| orf19.5764 | 1259 | 1007 | 1085 | 669 | 1989 | 1300 | 1901 | 2057 | 1200 | 1065 | 1072 | 1048 | 1594 | 1478 | 1614 | 1673 | 1342,93 | 1096 | 1590 | 1,449 | 0,535 | 8,699 | 3,33E-18 | 7,19E-18 | 0,002329 | 0,0033139 | 0,0028393 | 0,0028393 | TRUE | 0,0505117 | No |
| orf19.5765 | 1644 | 1503 | 1666 | 1005 | 2352 | 1675 | 2232 | 2472 | 1567 | 1589 | 1646 | 1574 | 1885 | 1905 | 1895 | 2010 | 1758,87 | 1594 | 1924 | 1,206 | 0,271 | 6,054 | 1,42E-09 | 2,42E-09 | 1,00E-08 | 0,0030248 | 0,0013348 | 0,0013348 | TRUE | 0,011425 | No |
| orf19.5767 | 730 | 677 | 703 | 410 | 1005 | 648 | 957 | 955 | 696 | 716 | 694 | 642 | 805 | 737 | 812 | 777 | 734,96 | 687 | 783 | 1,138 | 0,186 | 3 | 0,0026977 | 0,0035493 | 0,0001884 | 0,0043253 | 0,0023121 | 0,0023121 | TRUE | 0,0190409 | No |
| orf19.5768 | 7483 | 6481 | 6947 | 4334 | 5719 | 3687 | 5153 | 5694 | 7134 | 6853 | 6863 | 6789 | 4582 | 4192 | 4375 | 4630 | 5677,38 | 6910 | 4445 | 0,644 | -0,636 | -15,176 | 5,07E-52 | 1,92E-51 | 0,0010258 | 0,0023805 | 0,0015043 | 0,0015043 | TRUE | 0,0172672 | No |
| orf19.577 | 4089 | 3534 | 3854 | 2133 | 6024 | 3784 | 5680 | 6127 | 3898 | 3737 | 3807 | 3341 | 4827 | 4303 | 4822 | 4982 | 4214,72 | 3696 | 4734 | 1,28 | 0,356 | 5,821 | 5,84E-09 | 9,78E-09 | 0,0038464 | 0,0024808 | 0,0033462 | 0,0033462 | TRUE | 0,0489703 | No |
| orf19.5770 | 7355 | 6628 | 7127 | 4140 | 7532 | 5077 | 6946 | 7559 | 7012 | 7008 | 7041 | 6486 | 6035 | 5773 | 5897 | 6146 | 6424,79 | 6887 | 5963 | 0,865 | -0,208 | -5,178 | 2,24E-07 | 3,57E-07 | 0,000919 | 0,0023468 | 0,0013999 | 0,0013999 | TRUE | 0,0179997 | No |
| orf19.5771 | 8586 | 7005 | 7871 | 5502 | 8345 | 5814 | 7976 | 8495 | 8186 | 7407 | 7776 | 8619 | 6687 | 6611 | 6772 | 6907 | 7370,52 | 7997 | 6744 | 0,844 | -0,245 | -4,966 | 6,84E-07 | 1,07E-06 | 0,0021319 | 0,002314 | 0,0022 | 0,0022 | TRUE | 0,0327003 | No |
| orf19.5772 | 1302 | 1125 | 1213 | 759 | 1319 | 966 | 1320 | 1417 | 1241 | 1190 | 1198 | 1189 | 1057 | 1098 | 1121 | 1152 | 1155,79 | 1204 | 1107 | 0,918 | -0,123 | -2,416 | 0,0157133 | 0,0196794 | 1,00E-08 | 0,0035119 | 0,0016088 | 0,0016088 | TRUE | 0,0119451 | No |
| orf19.5773 | 1921 | 1831 | 1916 | 1247 | 3617 | 2618 | 3433 | 3789 | 1831 | 1936 | 1893 | 1953 | 2898 | 2977 | 2915 | 3081 | 2435,54 | 1903 | 2968 | 1,561 | 0,642 | 15,128 | 1,06E-51 | 3,99E-51 | 0,0004363 | 0,0027654 | 0,0012855 | 0,0012855 | TRUE | 0,0082844 | No |
| orf19.5775 | 47 | 57 | 50 | 22 | 130 | 85 | 119 | 152 | 45 | 60 | 49 | 34 | 104 | 97 | 101 | 124 | 76,8 | 47 | 106 | 2,226 | 1,154 | 6,229 | 4,69E-10 | 8,17E-10 | 0,0070926 | 0,023471 | 0,01718 | 0,01718 | TRUE | 0,2498912 | No |
| orf19.5776 | 4457 | 4037 | 4286 | 2571 | 5799 | 3947 | 5175 | 5641 | 4249 | 4269 | 4234 | 4028 | 4647 | 4488 | 4394 | 4587 | 4361,82 | 4195 | 4529 | 1,079 | 0,109 | 2,956 | 0,0031197 | 0,0040817 | 0,000355 | 0,0024677 | 0,0010823 | 0,0010823 | TRUE | 0,0081103 | No |
| orf19.5777 | 286 | 213 | 300 | 188 | 560 | 407 | 527 | 500 | 273 | 225 | 296 | 295 | 449 | 463 | 447 | 407 | 356,78 | 272 | 442 | 1,624 | 0,7 | 7,23 | 4,82E-13 | 9,24E-13 | 0,0051185 | 0,0066933 | 0,0059205 | 0,0059205 | TRUE | 0,1527346 | No |
| orf19.5779 | 4492 | 3859 | 4266 | 2823 | 4645 | 3386 | 4397 | 4669 | 4283 | 4080 | 4214 | 4422 | 3722 | 3850 | 3733 | 3796 | 4012,66 | 4250 | 3775 | 0,889 | -0,17 | -4,542 | 5,57E-06 | 8,34E-06 | 0,0003526 | 0,0025005 | 0,001095 | 0,001095 | TRUE | 0,0086223 | No |
| orf19.578 | 3785 | 3431 | 3620 | 2075 | 2991 | 2067 | 2947 | 3193 | 3609 | 3628 | 3576 | 3251 | 2397 | 2350 | 2502 | 2596 | 2988,55 | 3516 | 2461 | 0,7 | -0,515 | -10,063 | 8,09E-24 | 1,96E-23 | 0,0018225 | 0,0026407 | 0,0021771 | 0,0021771 | TRUE | 0,0297231 | No |
| orf19.5780 | 1378 | 1218 | 1336 | 896 | 1934 | 1357 | 1874 | 2024 | 1314 | 1288 | 1320 | 1404 | 1550 | 1543 | 1591 | 1646 | 1456,82 | 1332 | 1582 | 1,192 | 0,253 | 5,216 | 1,82E-07 | 2,91E-07 | 0,0002347 | 0,0032183 | 0,0015564 | 0,0015564 | TRUE | 0,0140056 | No |
| orf19.5782 | 8804 | 7776 | 8659 | 4841 | 5657 | 3784 | 5394 | 5796 | 8394 | 8222 | 8554 | 7584 | 4533 | 4303 | 4579 | 4713 | 6360,18 | 8188 | 4532 | 0,553 | -0,854 | -17,729 | 2,49E-70 | 1,19E-69 | 0,0018305 | 0,0023494 | 0,0020598 | 0,0020598 | TRUE | 0,0294388 | No |
| orf19.5783 | 1112 | 909 | 1003 | 652 | 2187 | 1467 | 2121 | 2192 | 1060 | 961 | 991 | 1021 | 1752 | 1668 | 1801 | 1782 | 1379,64 | 1008 | 1751 | 1,738 | 0,797 | 15,35 | 3,55E-53 | 1,37E-52 | 0,0006078 | 0,0032814 | 0,0017813 | 0,0017813 | TRUE | 0,0158993 | No |
| orf19.5784 | 786 | 710 | 707 | 463 | 1512 | 1064 | 1418 | 1558 | 749 | 751 | 698 | 725 | 1212 | 1210 | 1204 | 1267 | 976,99 | 731 | 1223 | 1,673 | 0,743 | 13,763 | 4,24E-43 | 1,41E-42 | 1,00E-08 | 0,0037719 | 0,0016675 | 0,0016675 | TRUE | 0,0118737 | No |
| orf19.5785 | 1876 | 1758 | 1884 | 1175 | 705 | 534 | 658 | 646 | 1789 | 1859 | 1861 | 1841 | 565 | 607 | 559 | 525 | 1200,67 | 1838 | 564 | 0,306 | -1,708 | -30,946 | 2,86E-210 | 4,08E-209 | 7,32E-05 | 0,0034588 | 0,0018274 | 0,0018274 | TRUE | 0,0276914 | No |
| orf19.5788 | 41699 | 33997 | 37247 | 24690 | 89157 | 61436 | 84088 | 96885 | 39755 | 35948 | 36797 | 38678 | 71439 | 69858 | 71390 | 78777 | 55330,25 | 37794 | 72866 | 1,928 | 0,947 | 19 | 1,70E-80 | 9,30E-80 | 0,0025402 | 0,002121 | 0,0023665 | 0,0023665 | TRUE | 0,0365564 | No |
| orf19.5789 | 3918 | 3191 | 3447 | 2275 | 4845 | 3455 | 4605 | 4954 | 3735 | 3374 | 3405 | 3564 | 3882 | 3929 | 3910 | 4028 | 3728,4 | 3520 | 3937 | 1,119 | 0,162 | 3,752 | 0,0001756 | 0,0002455 | 0,0008851 | 0,0025317 | 0,0015133 | 0,0015133 | TRUE | 0,0217804 | No |
| orf19.579 | 1371 | 1244 | 1277 | 834 | 1432 | 1044 | 1354 | 1484 | 1307 | 1315 | 1262 | 1307 | 1147 | 1187 | 1150 | 1207 | 1235,16 | 1298 | 1173 | 0,904 | -0,146 | -3,074 | 0,0021111 | 0,002793 | 1,00E-08 | 0,0034206 | 0,0013544 | 0,0013544 | TRUE | 0,0049515 | No |
| orf19.5791 | 7603 | 6005 | 6648 | 4725 | 17142 | 12465 | 16366 | 17603 | 7249 | 6350 | 6568 | 7402 | 13735 | 14174 | 13895 | 14313 | 10460,57 | 6892 | 14029 | 2,037 | 1,026 | 19,504 | 1,02E-84 | 5,77E-84 | 0,0027907 | 0,0022482 | 0,0025449 | 0,0025449 | TRUE | 0,0341511 | No |
| orf19.5792 | 477 | 457 | 497 | 335 | 1033 | 814 | 930 | 997 | 455 | 483 | 491 | 525 | 828 | 926 | 790 | 811 | 663,41 | 488 | 839 | 1,719 | 0,782 | 10,592 | 3,24E-26 | 8,24E-26 | 0,0026022 | 0,0045662 | 0,0035472 | 0,0035472 | TRUE | 0,0548544 | No |
| orf19.5793 | 9273 | 7706 | 8210 | 4965 | 10839 | 7276 | 10478 | 11208 | 8841 | 8148 | 8111 | 7778 | 8685 | 8273 | 8896 | 9113 | 8480,62 | 8220 | 8742 | 1,063 | 0,089 | 1,802 | 0,0715459 | 0,0847159 | 0,0021556 | 0,0022849 | 0,0022006 | 0,0022006 | TRUE | 0,0317757 | No |
| orf19.5796 | 1222 | 1010 | 1139 | 654 | 1918 | 1315 | 1867 | 2018 | 1165 | 1068 | 1125 | 1025 | 1537 | 1495 | 1585 | 1641 | 1330,09 | 1096 | 1564 | 1,425 | 0,511 | 9,066 | 1,23E-19 | 2,75E-19 | 0,0013682 | 0,0033257 | 0,0022549 | 0,0022549 | TRUE | 0,0215663 | No |
| orf19.5797 | 2083 | 1866 | 1854 | 1217 | 1716 | 1282 | 1747 | 1754 | 1986 | 1973 | 1832 | 1906 | 1375 | 1458 | 1483 | 1426 | 1679,89 | 1924 | 1436 | 0,745 | -0,424 | -8,742 | 2,30E-18 | 4,97E-18 | 0,0007224 | 0,0030687 | 0,0016539 | 0,0016539 | TRUE | 0,0135864 | No |
| orf19.5798 | 2431 | 2190 | 2350 | 1408 | 3740 | 2563 | 3575 | 3908 | 2318 | 2316 | 2322 | 2206 | 2997 | 2914 | 3035 | 3178 | 2660,56 | 2290 | 3031 | 1,323 | 0,404 | 9,434 | 3,95E-21 | 9,05E-21 | 0,0005929 | 0,0027084 | 0,0013643 | 0,0013643 | TRUE | 0,0129408 | No |
| orf19.5799 | 2656 | 2254 | 2404 | 1746 | 6788 | 4885 | 6453 | 7342 | 2532 | 2383 | 2375 | 2735 | 5439 | 5555 | 5479 | 5970 | 4058,46 | 2506 | 5611 | 2,242 | 1,165 | 21,241 | 3,99E-100 | 2,62E-99 | 0,0028299 | 0,0024958 | 0,0025831 | 0,0025831 | TRUE | 0,0431672 | No |
| orf19.58 | 14747 | 13186 | 14392 | 9359 | 13992 | 10198 | 13288 | 14626 | 14060 | 13943 | 14218 | 14661 | 11211 | 11596 | 11281 | 11892 | 12857,84 | 14220 | 11495 | 0,809 | -0,307 | -9,128 | 6,99E-20 | 1,56E-19 | 0,0005113 | 0,002219 | 0,001005 | 0,001005 | TRUE | 0,0067117 | No |
| orf19.580 | 1155 | 1018 | 1056 | 607 | 756 | 494 | 703 | 727 | 1101 | 1076 | 1043 | 951 | 606 | 562 | 597 | 591 | 815,89 | 1043 | 589 | 0,564 | -0,827 | -13,05 | 6,37E-39 | 1,98E-38 | 0,0011329 | 0,0041037 | 0,0025641 | 0,0025641 | TRUE | 0,0371864 | No |
| orf19.5801 | 10033 | 8863 | 9426 | 5789 | 8620 | 6043 | 7956 | 8867 | 9565 | 9372 | 9312 | 9069 | 6907 | 6871 | 6755 | 7210 | 8132,55 | 9330 | 6936 | 0,743 | -0,428 | -12,331 | 6,20E-35 | 1,81E-34 | 0,000464 | 0,0022932 | 0,0010328 | 0,0010328 | TRUE | 0,0087466 | No |
| orf19.5802 | 1579 | 1217 | 1411 | 841 | 2956 | 1855 | 2887 | 2946 | 1505 | 1287 | 1394 | 1317 | 2369 | 2109 | 2451 | 2395 | 1853,49 | 1376 | 2331 | 1,693 | 0,76 | 11,522 | 1,02E-30 | 2,78E-30 | 0,0038537 | 0,0029771 | 0,0035756 | 0,0035756 | TRUE | 0,0493356 | No |
| orf19.5804 | 1701 | 1502 | 1696 | 1081 | 2129 | 1451 | 2007 | 2215 | 1622 | 1588 | 1675 | 1693 | 1706 | 1650 | 1704 | 1801 | 1679,95 | 1644 | 1715 | 1,045 | 0,063 | 1,335 | 0,1817706 | 0,2070103 | 0,0005107 | 0,0030686 | 0,0015444 | 0,0015444 | TRUE | 0,0137912 | No |
| orf19.5805 | 773 | 610 | 726 | 475 | 3157 | 2211 | 3100 | 3250 | 737 | 645 | 717 | 744 | 2530 | 2514 | 2632 | 2643 | 1645,18 | 711 | 2580 | 3,636 | 1,862 | 34,004 | 1,97E-253 | 3,41E-252 | 0,0007345 | 0,0030893 | 0,00192 | 0,00192 | TRUE | 0,0464261 | No |
| orf19.5806 | 60096 | 48214 | 53462 | 36525 | 230190 | 160805 | 218692 | 240353 | 57295 | 50980 | 52815 | 57218 | 184445 | 182850 | 185668 | 195430 | 120837,73 | 54577 | 187098 | 3,428 | 1,777 | 37,649 | 0 | 0 | 0,0021452 | 0,0021049 | 0,002129 | 0,002129 | TRUE | 0,0241175 | No |
| orf19.5808 | 3724 | 3232 | 3539 | 2332 | 3659 | 2346 | 3477 | 3602 | 3550 | 3417 | 3496 | 3653 | 2932 | 2668 | 2952 | 2929 | 3199,68 | 3529 | 2870 | 0,814 | -0,296 | -6,5 | 8,01E-11 | 1,44E-10 | 0,0010843 | 0,0026044 | 0,0016769 | 0,0016769 | TRUE | 0,0262079 | No |
| orf19.5809 | 66 | 55 | 67 | 57 | 495 | 340 | 498 | 513 | 63 | 58 | 66 | 89 | 397 | 387 | 423 | 417 | 237,47 | 69 | 406 | 6,007 | 2,587 | 20,707 | 2,98E-95 | 1,86E-94 | 1,00E-08 | 0,0090056 | 0,0057084 | 0,0057084 | TRUE | 0,2376182 | No |

| | | | | | | | | | | | | | | | | | | | | | | | | | | | | | | | |
|---|---|---|---|---|---|---|---|---|---|---|---|---|---|---|---|---|---|---|---|---|---|---|---|---|---|---|---|---|---|---|---|
| orf19.581 | 14233 | 12148 | 13839 | 9161 | 14133 | 10057 | 13436 | 14316 | 13570 | 12845 | 13672 | 14351 | 11324 | 11436 | 11407 | 11640 | 12530,61 | 13610 | 11452 | 0,842 | -0,249 | -6,38 | 1,77E-10 | 3,15E-10 | 0,0009873 | 0,0022223 | 0,001377 | 0,001377 | TRUE | 0,0174085 | No |
| orf19.5811 | 6038 | 4433 | 5083 | 3639 | 7156 | 4927 | 7078 | 7643 | 5757 | 4687 | 5022 | 5701 | 5734 | 5602 | 6009 | 6214 | 5590,77 | 5292 | 5890 | 1,114 | 0,155 | 2,26 | 0,0238432 | 0,0294452 | 0,0057399 | 0,002385 | 0,0043535 | 0,0043535 | TRUE | 0,0720914 | No |
| orf19.5812 | 5939 | 5284 | 5749 | 3719 | 7494 | 5083 | 7095 | 7712 | 5662 | 5587 | 5679 | 5826 | 6005 | 5780 | 6024 | 6271 | 5854,2 | 5688 | 6020 | 1,059 | 0,083 | 2,265 | 0,0235217 | 0,0290538 | 0,0004816 | 0,0023717 | 0,0011097 | 0,0011097 | TRUE | 0,0095983 | No |
| orf19.5813 | 15116 | 13131 | 14448 | 9098 | 10436 | 6936 | 9994 | 10314 | 14411 | 13884 | 14273 | 14253 | 8362 | 7887 | 8485 | 8386 | 11242,71 | 14205 | 8280 | 0,583 | -0,778 | -22,782 | 6,97E-115 | 5,27E-114 | 0,00052 | 0,0022373 | 0,0010269 | 0,0010269 | TRUE | 0,0121587 | No |
| orf19.5814 | 963 | 863 | 872 | 573 | 1199 | 941 | 1065 | 971 | 918 | 913 | 861 | 898 | 961 | 1070 | 904 | 790 | 914,27 | 898 | 931 | 1,036 | 0,051 | 0,596 | 0,5510863 | 0,5862335 | 0,0074328 | 0,0038871 | 0,005795 | 0,005795 | TRUE | 0,1259813 | No |
| orf19.5814.1 | 574 | 493 | 535 | 310 | 769 | 481 | 811 | 763 | 547 | 521 | 529 | 486 | 616 | 547 | 689 | 620 | 569,34 | 521 | 618 | 1,186 | 0,246 | 3,065 | 0,0021749 | 0,0028763 | 0,003878 | 0,0049751 | 0,00441 | 0,00441 | TRUE | 0,0684729 | No |
| orf19.5815 | 1175 | 1092 | 1127 | 722 | 2549 | 1825 | 2435 | 2602 | 1120 | 1155 | 1113 | 1131 | 2042 | 2075 | 2067 | 2116 | 1602,49 | 1130 | 2075 | 1,837 | 0,877 | 20,216 | 7,07E-91 | 4,26E-90 | 1,00E-08 | 0,0031159 | 0,0011 | 0,0011 | TRUE | 0,002807 | No |
| orf19.5816 | 1102 | 961 | 1098 | 715 | 1702 | 1182 | 1582 | 1728 | 1051 | 1016 | 1085 | 1120 | 1364 | 1344 | 1343 | 1405 | 1215,94 | 1068 | 1364 | 1,28 | 0,357 | 7,079 | 1,45E-12 | 2,73E-12 | 1,00E-08 | 0,0034416 | 0,0015842 | 0,0015842 | TRUE | 0,0122087 | No |
| orf19.5817 | 989 | 950 | 988 | 593 | 1067 | 769 | 1025 | 1040 | 943 | 1005 | 976 | 929 | 855 | 874 | 870 | 846 | 912,21 | 963 | 861 | 0,893 | -0,164 | -3,075 | 0,0021033 | 0,0027834 | 1,00E-08 | 0,0038912 | 0,0016306 | 0,0016306 | TRUE | 0,0097307 | No |
| orf19.5818 | 49848 | 43607 | 47803 | 28523 | 31747 | 22069 | 30081 | 31762 | 47525 | 46109 | 47225 | 44683 | 25438 | 25094 | 25539 | 25826 | 35929,72 | 46386 | 25474 | 0,549 | -0,865 | -28,433 | 7,86E-178 | 9,29E-177 | 0,0004152 | 0,002137 | 0,0008589 | 0,0008589 | TRUE | 0,0074029 | No |
| orf19.5820 | 21 | 8 | 25 | 16 | 35 | 40 | 33 | 41 | 20 | 8 | 25 | 25 | 28 | 45 | 28 | 33 | 26,64 | 20 | 34 | 1,728 | 0,789 | 2,381 | 0,0172525 | 0,021531 | 0,0623662 | 0,063723 | 0,0631959 | 0,0631959 | TRUE | 0,7772142 | No |
| orf19.5821 | 1290 | 1225 | 1298 | 773 | 742 | 642 | 721 | 785 | 1230 | 1295 | 1282 | 1211 | 595 | 730 | 612 | 638 | 949,17 | 1254 | 644 | 0,511 | -0,969 | -14,147 | 1,95E-45 | 6,70E-45 | 0,0028957 | 0,0038211 | 0,003357 | 0,003357 | TRUE | 0,0891084 | No |
| orf19.5823 | 5255 | 4252 | 4555 | 3078 | 8259 | 5486 | 7673 | 8300 | 5010 | 4496 | 4500 | 4822 | 6618 | 6238 | 6514 | 6749 | 5618,32 | 4707 | 6530 | 1,388 | 0,473 | 9,905 | 3,96E-23 | 9,47E-23 | 0,0017553 | 0,0023835 | 0,0020008 | 0,0020008 | TRUE | 0,0237569 | No |
| orf19.5824 | 7128 | 6190 | 6573 | 4555 | 6841 | 4936 | 6396 | 7178 | 6796 | 6545 | 6494 | 7136 | 5481 | 5613 | 5430 | 5836 | 6166,35 | 6743 | 5590 | 0,83 | -0,27 | -6,221 | 4,95E-10 | 8,61E-10 | 0,0012546 | 0,0023575 | 0,0016388 | 0,0016388 | TRUE | 0,018139 | No |
| orf19.5825 | 2571 | 2102 | 2238 | 1429 | 2049 | 1433 | 2032 | 2011 | 2451 | 2223 | 2211 | 2239 | 1642 | 1629 | 1725 | 1635 | 1969,36 | 2281 | 1658 | 0,727 | -0,461 | -9,264 | 1,98E-20 | 4,49E-20 | 0,0011958 | 0,002925 | 0,0018607 | 0,0018607 | TRUE | 0,0318066 | No |
| orf19.5825.1 | 899 | 732 | 737 | 509 | 1223 | 839 | 1196 | 1261 | 857 | 774 | 728 | 797 | 980 | 954 | 1015 | 1025 | 891,41 | 789 | 994 | 1,26 | 0,334 | 5,259 | 1,45E-07 | 2,32E-07 | 0,0016354 | 0,0039332 | 0,0027077 | 0,0027077 | TRUE | 0,0438701 | No |
| orf19.5826 | 16 | 16 | 17 | 10 | 30 | 24 | 42 | 46 | 15 | 17 | 17 | 16 | 24 | 27 | 36 | 37 | 23,63 | 16 | 31 | 1,928 | 0,947 | 3,102 | 0,0019241 | 0,0025515 | 1,00E-08 | 0,0715822 | 0,0410034 | 0,0410034 | TRUE | 0,1770167 | No |
| orf19.5827 | 153 | 154 | 159 | 126 | 110 | 68 | 89 | 99 | 146 | 163 | 157 | 197 | 88 | 77 | 76 | 80 | 123,09 | 166 | 80 | 0,49 | -1,028 | -7,283 | 3,27E-13 | 6,29E-13 | 1,00E-08 | 0,0154308 | 0,0102079 | 0,0102079 | TRUE | 0,1438901 | No |
| orf19.5828 | 8705 | 7485 | 8333 | 5177 | 8959 | 5999 | 8617 | 9141 | 8299 | 7914 | 8232 | 8110 | 7179 | 6821 | 7316 | 7433 | 7663,03 | 8139 | 7187 | 0,883 | -0,179 | -4,726 | 2,28E-06 | 3,48E-06 | 0,0007203 | 0,0023055 | 0,0012464 | 0,0012464 | TRUE | 0,0139714 | No |
| orf19.583 | 12120 | 10405 | 11307 | 7465 | 11411 | 7380 | 10909 | 11564 | 11555 | 11002 | 11170 | 11694 | 9143 | 8392 | 9262 | 9403 | 10202,63 | 11355 | 9050 | 0,797 | -0,327 | -7,36 | 1,84E-13 | 3,56E-13 | 0,0015172 | 0,0022522 | 0,0017936 | 0,0017936 | TRUE | 0,0288659 | No |
| orf19.5830 | 1376 | 1189 | 1405 | 833 | 3098 | 2147 | 2988 | 3279 | 1312 | 1257 | 1388 | 1305 | 2482 | 2441 | 2537 | 2666 | 1923,58 | 1316 | 2532 | 1,924 | 0,944 | 18,809 | 6,33E-79 | 3,42E-78 | 0,0010699 | 0,0029448 | 0,0018181 | 0,0018181 | TRUE | 0,0173532 | No |
| orf19.5831 | 37 | 41 | 42 | 27 | 43 | 34 | 45 | 48 | 35 | 43 | 41 | 42 | 34 | 39 | 38 | 39 | 39,1 | 40 | 38 | 0,928 | -0,108 | -0,491 | 0,623339 | 0,6566825 | 1,00E-08 | 0,0440883 | 0,0206463 | 0,0206463 | TRUE | 0,0660244 | No |
| orf19.5832 | 10298 | 8643 | 9182 | 5574 | 15858 | 10273 | 14719 | 15886 | 9818 | 9139 | 9071 | 8732 | 12707 | 11681 | 12496 | 12917 | 10820,11 | 9190 | 12450 | 1,355 | 0,438 | 9,111 | 8,17E-20 | 1,83E-19 | 0,0020421 | 0,002243 | 0,0021204 | 0,0021204 | TRUE | 0,0259643 | No |
| orf19.5833 | 5304 | 4432 | 4877 | 3085 | 5466 | 3723 | 5188 | 5418 | 5057 | 4686 | 4818 | 4833 | 4380 | 4233 | 4405 | 4405 | 4602,12 | 4848 | 4356 | 0,899 | -0,154 | -4,148 | 3,35E-05 | 4,85E-05 | 0,0003911 | 0,0024481 | 0,0011113 | 0,0011113 | TRUE | 0,010579 | No |
| orf19.5834 | 6435 | 5757 | 6201 | 4021 | 6413 | 4606 | 5858 | 6688 | 6135 | 6087 | 6126 | 6299 | 5139 | 5237 | 4973 | 5438 | 5679,36 | 6162 | 5197 | 0,844 | -0,245 | -6,476 | 9,39E-11 | 1,68E-10 | 0,0005775 | 0,0023804 | 0,0011999 | 0,0011999 | TRUE | 0,0121741 | No |
| orf19.5835 | 8778 | 7343 | 8215 | 5251 | 6618 | 4758 | 6314 | 6778 | 8369 | 7764 | 8116 | 8226 | 5303 | 5410 | 5361 | 5511 | 6757,45 | 8119 | 5396 | 0,665 | -0,589 | -16,513 | 2,95E-61 | 1,27E-60 | 0,0004333 | 0,0023343 | 0,0010704 | 0,0010704 | TRUE | 0,0106546 | No |
| orf19.5837 | 1129 | 936 | 1090 | 643 | 1120 | 725 | 1048 | 1164 | 1076 | 990 | 1077 | 1007 | 897 | 824 | 890 | 946 | 963,52 | 1038 | 889 | 0,858 | -0,221 | -3,633 | 0,0002806 | 0,0003894 | 0,0014586 | 0,0037953 | 0,0025282 | 0,0025282 | TRUE | 0,0267254 | No |
| orf19.5838 | 924 | 791 | 839 | 547 | 1955 | 1296 | 1920 | 1997 | 881 | 836 | 829 | 857 | 1566 | 1474 | 1630 | 1624 | 1212,13 | 851 | 1574 | 1,852 | 0,889 | 16,244 | 2,47E-59 | 1,03E-58 | 0,0008222 | 0,0034458 | 0,0019347 | 0,0019347 | TRUE | 0,0206128 | No |
| orf19.5839 | 634 | 493 | 565 | 393 | 1530 | 1008 | 1438 | 1519 | 604 | 521 | 558 | 616 | 1226 | 1146 | 1221 | 1235 | 890,95 | 575 | 1207 | 2,107 | 1,075 | 16,64 | 3,59E-62 | 1,56E-61 | 0,001247 | 0,0039341 | 0,0026649 | 0,0026649 | TRUE | 0,045628 | No |
| orf19.584 | 7 | 10 | 11 | 7 | 4 | 3 | 7 | 3 | 7 | 11 | 11 | 11 | 3 | 3 | 6 | 2 | 6,76 | 10 | 4 | 0,387 | -1,369 | -2,417 | 0,0156604 | 0,0196209 | 1,00E-08 | 0,2449813 | 0,1322168 | 0,1322168 | TRUE | 0,2990106 | No |
| orf19.5840 | 47 | 42 | 36 | 44 | 47 | 33 | 27 | 31 | 45 | 44 | 36 | 69 | 38 | 38 | 23 | 25 | 39,63 | 48 | 31 | 0,644 | -0,635 | -2,348 | 0,0188957 | 0,023499 | 0,0446997 | 0,0435241 | 0,0439728 | 0,0439728 | TRUE | 0,4973205 | No |
| orf19.5841 | 534 | 435 | 473 | 332 | 1143 | 850 | 1105 | 1226 | 509 | 460 | 467 | 520 | 916 | 967 | 938 | 997 | 721,73 | 489 | 954 | 1,956 | 0,968 | 14,752 | 3,00E-49 | 1,10E-48 | 0,0007239 | 0,0043662 | 0,0025228 | 0,0025228 | TRUE | 0,017856 | No |
| orf19.5842 | 655 | 612 | 682 | 409 | 999 | 643 | 994 | 878 | 624 | 647 | 674 | 641 | 800 | 731 | 844 | 714 | 709,43 | 646 | 772 | 1,195 | 0,258 | 3,618 | 0,0002974 | 0,0004122 | 0,0026221 | 0,0044057 | 0,0034242 | 0,0034242 | TRUE | 0,0463602 | No |
| orf19.5843 | 2802 | 2790 | 2926 | 1856 | 574 | 439 | 513 | 584 | 2671 | 2950 | 2891 | 2908 | 460 | 499 | 436 | 475 | 1661,14 | 2855 | 468 | 0,163 | -2,613 | -44,085 | 0 | 0 | 0,0015206 | 0,0030797 | 0,0022229 | 0,0022229 | TRUE | 0,0239777 | No |
| orf19.5844 | 6 | 3 | 4 | 4 | 6 | 9 | 7 | 9 | 6 | 3 | 4 | 6 | 5 | 10 | 6 | 7 | 5,93 | 5 | 7 | 1,48 | 0,566 | 0,967 | 0,3337894 | 0,3674159 | 1,00E-08 | 0,2791323 | 0,1470255 | 0,1470255 | TRUE | 0,1997545 | No |
| orf19.5845 | 4843 | 4288 | 4752 | 2993 | 5606 | 3924 | 5288 | 5675 | 4617 | 4534 | 4695 | 4689 | 4492 | 4462 | 4489 | 4614 | 4574,02 | 4634 | 4514 | 0,975 | -0,037 | -1,144 | 0,2525754 | 0,28295 | 1,00E-08 | 0,0024502 | 0,0007789 | 0,0007789 | TRUE | 0,0027406 | No |
| orf19.5846 | 3361 | 2888 | 3216 | 2093 | 1700 | 1216 | 1560 | 1808 | 3204 | 3054 | 3177 | 3279 | 1362 | 1383 | 1324 | 1470 | 2281,66 | 3178 | 1385 | 0,436 | -1,198 | -25,331 | 1,46E-141 | 1,39E-140 | 0,0007364 | 0,0028109 | 0,0016467 | 0,0016467 | TRUE | 0,0215517 | No |
| orf19.5847 | 10411 | 8906 | 9811 | 6183 | 12134 | 8502 | 11341 | 12296 | 9926 | 9417 | 9692 | 9686 | 9723 | 9668 | 9628 | 9998 | 9717,19 | 9680 | 9754 | 1,008 | 0,011 | 0,354 | 0,7229989 | 0,7523574 | 0,0002344 | 0,0022602 | 0,0008201 | 0,0008201 | TRUE | 0,0041785 | No |
| orf19.5848 | 6486 | 5569 | 5956 | 4021 | 1108 | 818 | 1056 | 1171 | 6184 | 5889 | 5884 | 6299 | 888 | 930 | 897 | 952 | 3490,24 | 6064 | 917 | 0,151 | -2,726 | -59,135 | 0 | 0 | 0,000791 | 0,0025617 | 0,001461 | 0,001461 | TRUE | 0,0088625 | No |
| orf19.5849 | 3485 | 2757 | 3144 | 2030 | 2343 | 1519 | 2212 | 2343 | 3323 | 2915 | 3106 | 3180 | 1877 | 1727 | 1878 | 1905 | 2488,94 | 3131 | 1847 | 0,59 | -0,76 | -14,205 | 8,50E-46 | 2,95E-45 | 0,0018512 | 0,002751 | 0,0023271 | 0,0023271 | TRUE | 0,026534 | No |
| orf19.585 | 3872 | 3501 | 3660 | 2382 | 3348 | 2552 | 3321 | 3554 | 3692 | 3702 | 3616 | 3732 | 2683 | 2902 | 2820 | 2890 | 3254,3 | 3686 | 2824 | 0,766 | -0,385 | -9,737 | 2,10E-22 | 4,93E-22 | 0,0004341 | 0,0025958 | 0,0011869 | 0,0011869 | TRUE | 0,013949 | No |
| orf19.5850 | 16022 | 14594 | 16116 | 10022 | 26442 | 19411 | 25196 | 26455 | 15275 | 15431 | 15921 | 15700 | 21187 | 22072 | 21391 | 21510 | 18561,08 | 15582 | 21540 | 1,382 | 0,467 | 16,206 | 4,59E-59 | 1,91E-58 | 0,0002496 | 0,0021797 | 0,000741 | 0,000741 | TRUE | 0,0033361 | No |
| orf19.5851 | 771 | 703 | 747 | 513 | 2475 | 1739 | 2416 | 2646 | 735 | 743 | 738 | 804 | 1983 | 1977 | 2051 | 2151 | 1397,9 | 755 | 2040 | 2,713 | 1,44 | 26,47 | 2,14E-154 | 2,21E-153 | 0,0008714 | 0,0032658 | 0,0018806 | 0,0018806 | TRUE | 0,0187893 | No |
| orf19.5852 | 259 | 281 | 271 | 168 | 825 | 646 | 790 | 797 | 247 | 297 | 268 | 263 | 661 | 735 | 671 | 648 | 473,66 | 269 | 679 | 2,521 | 1,334 | 16,207 | 4,49E-59 | 1,87E-58 | 0,0017322 | 0,0055577 | 0,0037712 | 0,0037712 | TRUE | 0,0592299 | No |
| orf19.5854 | 4547 | 4216 | 4487 | 2757 | 5772 | 3999 | 5680 | 5880 | 4335 | 4458 | 4433 | 4319 | 4625 | 4547 | 4822 | 4781 | 4540,02 | 4386 | 4694 | 1,07 | 0,098 | 2,764 | 0,0057086 | 0,007353 | 0,000297 | 0,0024529 | 0,0009819 | 0,0009819 | TRUE | 0,0051464 | No |
| orf19.5854.1 | 126 | 122 | 134 | 90 | 214 | 169 | 201 | 164 | 120 | 129 | 132 | 141 | 171 | 192 | 171 | 133 | 148,77 | 130 | 167 | 1,278 | 0,354 | 2,591 | 0,0095581 | 0,012164 | 0,0084196 | 0,0131281 | 0,010895 | 0,010895 | TRUE | 0,2033304 | No |
| orf19.5855 | 4107 | 3523 | 4013 | 2631 | 2691 | 1915 | 2580 | 2843 | 3916 | 3725 | 3964 | 4122 | 2156 | 2178 | 2190 | 2312 | 3070,32 | 3932 | 2209 | 0,562 | -0,83 | -18,377 | 2,00E-75 | 1,03E-74 | 0,0008925 | 0,002626 | 0,0016145 | 0,0016145 | TRUE | 0,0150616 | No |
| orf19.5856 | 2142 | 2040 | 2143 | 1280 | 1843 | 1343 | 1663 | 1819 | 2042 | 2157 | 2117 | 2005 | 1477 | 1527 | 1412 | 1479 | 1777,03 | 2080 | 1474 | 0,707 | -0,5 | -10,791 | 3,80E-27 | 9,81E-27 | 0,0004228 | 0,0030152 | 0,0014879 | 0,0014879 | TRUE | 0,0095277 | No |
| orf19.5857 | 1231 | 1070 | 1207 | 761 | 1272 | 874 | 1280 | 1306 | 1174 | 1131 | 1192 | 1192 | 1019 | 994 | 1087 | 1062 | 1106,4 | 1172 | 1040 | 0,889 | -0,17 | -3,258 | 0,0011204 | 0,0015061 | 1,00E-08 | 0,0035753 | 0,0017069 | 0,0017069 | TRUE | 0,0105241 | No |
| orf19.5858 | 2808 | 2122 | 2282 | 1372 | 7811 | 4871 | 7311 | 8057 | 2677 | 2244 | 2254 | 2149 | 6259 | 5539 | 6207 | 6551 | 4235,02 | 2331 | 6139 | 2,631 | 1,396 | 18,87 | 2,00E-79 | 1,08E-78 | 0,0071339 | 0,002479 | 0,0049407 | 0,0049407 | TRUE | 0,12216 | No |
| orf19.5859 | 759 | 589 | 695 | 430 | 358 | 303 | 406 | 377 | 724 | 623 | 687 | 674 | 287 | 345 | 345 | 307 | 498,66 | 677 | 321 | 0,472 | -1,082 | -12,894 | 4,89E-38 | 1,50E-37 | 0,0036732 | 0,0053839 | 0,0045442 | 0,0045442 | TRUE | 0,0591233 | No |
| orf19.586 | 1429 | 1282 | 1308 | 772 | 2479 | 1666 | 2438 | 2532 | 1362 | 1356 | 1292 | 1209 | 1986 | 1894 | 2070 | 2059 | 1653,61 | 1305 | 2002 | 1,532 | 0,615 | 11,433 | 2,88E-30 | 7,80E-30 | 0,0014566 | 0,0030842 | 0,0021294 | 0,0021294 | TRUE | 0,0258298 | No |
| orf19.5860 | 134 | 128 | 128 | 77 | 176 | 124 | 179 | 170 | 128 | 135 | 126 | 121 | 141 | 141 | 152 | 138 | 135,3 | 128 | 143 | 1,119 | 0,162 | 1,342 | 0,1795541 | 0,204559 | 1,00E-08 | 0,0142266 | 0,0064873 | 0,0064873 | TRUE | 0,0193759 | No |
| orf19.5861 | 4917 | 4308 | 4513 | 3276 | 7907 | 6048 | 7225 | 7756 | 4688 | 4555 | 4458 | 5132 | 6336 | 6877 | 6134 | 6306 | 5560,82 | 4708 | 6413 | 1,363 | 0,447 | 8,155 | 3,49E-16 | 7,21E-16 | 0,0030118 | 0,0023865 | 0,002692 | 0,002692 | TRUE | 0,0433221 | No |
| orf19.5861.1 | 451 | 397 | 460 | 261 | 810 | 620 | 803 | 830 | 430 | 420 | 454 | 409 | 649 | 705 | 682 | 675 | 552,96 | 428 | 678 | 1,576 | 0,656 | 9,695 | 3,16E-22 | 7,42E-22 | 1,00E-08 | 0,0050606 | 0,0024411 | 0,0024411 | TRUE | 0,019659 | No |
| orf19.5862 | 744 | 635 | 710 | 470 | 854 | 616 | 820 | 907 | 709 | 671 | 701 | 736 | 684 | 700 | 696 | 737 | 704,6 | 704 | 704 | 1,002 | 0,003 | 0,05 | 0,9604445 | 0,9644252 | 1,00E-08 | 0,0044215 | 0,0020746 | 0,0020746 | TRUE | 0,0123474 | No |
| orf19.5863 | 71 | 79 | 73 | 32 | 118 | 128 | 123 | 141 | 68 | 84 | 72 | 50 | 95 | 146 | 104 | 115 | 91,58 | 68 | 115 | 1,65 | 0,722 | 3,863 | 0,0001122 | 0,0001585 | 0,0235248 | 0,0200199 | 0,0215659 | 0,0215659 | TRUE | 0,301561 | No |
| orf19.5864 | 5885 | 5804 | 6192 | 3443 | 4673 | 3102 | 4250 | 4273 | 5611 | 6137 | 6117 | 5394 | 3744 | 3527 | 3608 | 3474 | 4701,58 | 5815 | 3588 | 0,617 | -0,697 | -13,343 | 1,30E-40 | 4,16E-40 | 0,002278 | 0,0024405 | 0,0023986 | 0,0023986 | TRUE | 0,0280371 | No |
| orf19.5865 | 2181 | 2142 | 2415 | 1499 | 2897 | 2157 | 2764 | 2840 | 2079 | 2265 | 2386 | 2348 | 2321 | 2453 | 2347 | 2309 | 2313,51 | 2270 | 2358 | 1,039 | 0,055 | 1,06 | 0,2891864 | 0,3215926 | 0,0016171 | 0,002801 | 0,0021808 | 0,0021808 | TRUE | 0,0397999 | No |
| orf19.5866 | 27 | 24 | 20 | 14 | 11 | 9 | 8 | 8 | 26 | 25 | 20 | 22 | 9 | 10 | 7 | 7 | 15,64 | 23 | 8 | 0,343 | -1,544 | -4,133 | 3,59E-05 | 5,19E-05 | 1,00E-08 | 0,1070447 | 0,0544325 | 0,0544325 | TRUE | 0,073658 | No |
| orf19.5867 | 4279 | 3874 | 4088 | 2645 | 2449 | 1781 | 2357 | 2593 | 4080 | 4096 | 4039 | 4144 | 1962 | 2025 | 2001 | 2108 | 3056,85 | 4090 | 2024 | 0,495 | -1,014 | -26,651 | 1,73E-156 | 1,83E-155 | 9,20E-05 | 0,0026284 | 0,001032 | 0,001032 | TRUE | 0,0099542 | No |
| orf19.5869 | 4730 | 4341 | 4722 | 3001 | 2768 | 1936 | 2687 | 2860 | 4510 | 4590 | 4665 | 4701 | 2218 | 2201 | 2281 | 2325 | 3436,47 | 4616 | 2256 | 0,489 | -1,031 | -27,84 | 1,41E-170 | 1,60E-169 | 0,0001525 | 0,0025691 | 0,0009978 | 0,0009978 | TRUE | 0,0051738 | No |
| orf19.587 | 357 | 310 | 304 | 195 | 597 | 474 | 617 | 600 | 340 | 328 | 300 | 305 | 478 | 539 | 524 | 488 | 412,87 | 318 | 507 | 1,586 | 0,665 | 8,235 | 1,80E-16 | 3,73E-16 | 1,00E-08 | 0,0060681 | 0,0036396 | 0,0036396 | TRUE | 0,0257024 | No |
| orf19.5870 | 484 | 449 | 450 | 266 | 1334 | 897 | 1289 | 1351 | 461 | 475 | 445 | 417 | 1069 | 1020 | 1094 | 1098 | 759,9 | 450 | 1070 | 2,375 | 1,248 | 19,698 | 2,26E-86 | 1,30E-85 | 1,44E-05 | 0,004252 | 0,0022035 | 0,0022035 | TRUE | 0,0219149 | No |
| orf19.5871 | 640 | 539 | 597 | 367 | 1220 | 820 | 1124 | 1213 | 610 | 570 | 590 | 575 | 978 | 932 | 954 | 986 | 774,41 | 586 | 962 | 1,641 | 0,715 | 12,343 | 5,30E-35 | 1,55E-34 | 1,00E-08 | 0,0042115 | 0,0018077 | 0,0018077 | TRUE | 0,0094025 | No |
| orf19.5873 | 539 | 527 | 518 | 328 | 1549 | 1118 | 1488 | 1657 | 514 | 557 | 512 | 514 | 1241 | 1271 | 1263 | 1347 | 902,46 | 524 | 1280 | 2,443 | 1,288 | 21,438 | 5,95E-102 | 3,96E-101 | 1,00E-08 | 0,0039106 | 0,002057 | 0,002057 | TRUE | 0,0216555 | No |
| orf19.5874 | 2 | 1 | 1 | 1 | 11 | 5 | 10 | 6 | 2 | 1 | 1 | 2 | 9 | 6 | 8 | 5 | 4,17 | 2 | 7 | 5,105 | 2,352 | 2,814 | 0,0048948 | 0,0063302 | 1,00E-08 | 0,3955197 | 0,2050411 | 0,2050411 | TRUE | 0,139093 | No |
| orf19.5875 | 254 | 198 | 222 | 151 | 913 | 568 | 883 | 942 | 242 | 209 | 219 | 237 | 732 | 646 | 750 | 766 | 475,05 | 227 | 724 | 3,2 | 1,678 | 19,159 | 8,13E-82 | 4,52E-81 | 0,0030053 | 0,0055475 | 0,0042903 | 0,0042903 | TRUE | 0,0574718 | No |
| orf19.5876 | 21643 | 17430 | 18729 | 11093 | 13800 | 8283 | 13077 | 13777 | 20634 | 18430 | 18502 | 17378 | 11058 | 9419 | 11102 | 11202 | 14715,62 | 18736 | 10695 | 0,571 | -0,809 | -11,968 | 5,24E-33 | 1,49E-32 | 0,005741 | 0,0022029 | 0,0043161 | 0,0043161 | TRUE | 0,0785134 | No |
| orf19.5877 | 13681 | 12497 | 13790 | 8256 | 6264 | 4318 | 6016 | 6281 | 13043 | 13214 | 13623 | 12933 | 5019 | 4910 | 5108 | 5107 | 9119,72 | 13203 | 5036 | 0,381 | -1,39 | -43,006 | 0 | 0 | 0,000253 | 0,0022713 | 0,0008724 | 0,0008724 | TRUE | 0,0056339 | No |
| orf19.5879 | 150 | 110 | 112 | 66 | 245 | 173 | 241 | 274 | 143 | 116 | 111 | 103 | 196 | 197 | 205 | 223 | 161,72 | 118 | 205 | 1,72 | 0,783 | 6,198 | 5,71E-10 | 9,91E-10 | 0,0030141 | 0,0122439 | 0,008471 | 0,008471 | TRUE | 0,2080127 | No |
| orf19.588 | 600 | 556 | 527 | 372 | 1124 | 831 | 935 | 979 | 572 | 588 | 521 | 583 | 901 | 945 | 794 | 796 | 712,34 | 566 | 859 | 1,518 | 0,602 | 7,693 | 1,44E-14 | 2,87E-14 | 0,0043298 | 0,0043962 | 0,0043604 | 0,0043604 | TRUE | 0,05136 | No |
| orf19.5880 | 552 | 437 | 524 | 341 | 1343 | 904 | 1203 | 1326 | 526 | 462 | 518 | 534 | 1076 | 1028 | 1021 | 1078 | 780,47 | 510 | 1051 | 2,065 | 1,046 | 16,422 | 1,33E-60 | 5,65E-60 | 0,0004026 | 0,004195 | 0,0023771 | 0,0023771 | TRUE | 0,0472357 | No |

| | | | | | | | | | | | | | | | | | | | | | | | | | | | | | | | |
|---|---|---|---|---|---|---|---|---|---|---|---|---|---|---|---|---|---|---|---|---|---|---|---|---|---|---|---|---|---|---|---|
| orf19.5884 | 1836 | 1650 | 1800 | 1050 | 4450 | 3077 | 4054 | 4482 | 1750 | 1745 | 1778 | 1645 | 3566 | 3499 | 3442 | 3644 | 2633,6 | 1730 | 3538 | 2,042 | 1,03 | 24,506 | 1,28E-132 | 1,15E-131 | 0,0002988 | 0,0027147 | 0,0012461 | 0,0012461 | TRUE | 0,0114423 | No |
| orf19.5885 | 15771 | 13780 | 14068 | 9321 | 22300 | 15335 | 20037 | 21888 | 15036 | 14571 | 13898 | 14602 | 17868 | 17437 | 17011 | 17797 | 16027,53 | 14527 | 17528 | 1,207 | 0,271 | 7,636 | 2,24E-14 | 4,45E-14 | 0,0006906 | 0,0021937 | 0,0011464 | 0,0011464 | TRUE | 0,0105241 | No |
| orf19.5886 | 5306 | 4320 | 4492 | 3243 | 7857 | 5344 | 7010 | 7736 | 5059 | 4568 | 4438 | 5080 | 6296 | 6077 | 5951 | 6290 | 5469,79 | 4786 | 6154 | 1,287 | 0,363 | 6,893 | 5,48E-12 | 1,02E-11 | 0,0025295 | 0,0023915 | 0,0024804 | 0,0024804 | TRUE | 0,0294362 | No |
| orf19.5887 | 3704 | 3473 | 3611 | 2287 | 3049 | 2206 | 2940 | 3121 | 3531 | 3672 | 3567 | 3583 | 2443 | 2508 | 2496 | 2538 | 3042,36 | 3588 | 2496 | 0,696 | -0,524 | -14,69 | 7,45E-49 | 2,70E-48 | 1,00E-08 | 0,002631 | 0,0008846 | 0,0008846 | TRUE | 0,0031179 | No |
| orf19.589 | 20209 | 16518 | 18226 | 11740 | 12742 | 8162 | 12385 | 12595 | 19267 | 17466 | 18006 | 18391 | 10210 | 9281 | 10515 | 10241 | 14172,02 | 18282 | 10062 | 0,55 | -0,861 | -17,612 | 2,01E-69 | 9,52E-69 | 0,0021881 | 0,0022071 | 0,0022221 | 0,0022221 | TRUE | 0,0329905 | No |
| orf19.5890 | 4013 | 3530 | 3759 | 2387 | 3039 | 2189 | 2876 | 3068 | 3826 | 3733 | 3714 | 3739 | 2435 | 2489 | 2442 | 2495 | 3108,98 | 3753 | 2465 | 0,657 | -0,607 | -17,595 | 2,71E-69 | 1,28E-68 | 1,00E-08 | 0,0026194 | 0,0008119 | 0,0008119 | TRUE | 0,0022083 | No |
| orf19.5892 | 249 | 230 | 257 | 175 | 732 | 470 | 699 | 710 | 237 | 243 | 254 | 274 | 587 | 534 | 593 | 577 | 412,54 | 252 | 573 | 2,289 | 1,195 | 14,783 | 1,88E-49 | 6,87E-49 | 1,00E-08 | 0,0060712 | 0,0032762 | 0,0032762 | TRUE | 0,0300032 | No |
| orf19.5893 | 6832 | 5490 | 5945 | 3696 | 20413 | 12958 | 19489 | 20547 | 6514 | 5805 | 5873 | 5790 | 16356 | 14734 | 16546 | 16707 | 11040,64 | 5996 | 16086 | 2,682 | 1,424 | 25,743 | 3,85E-146 | 3,79E-145 | 0,0031337 | 0,00224 | 0,0028156 | 0,0028156 | TRUE | 0,041688 | No |
| orf19.5894 | 30 | 25 | 27 | 22 | 5 | 12 | 7 | 10 | 29 | 26 | 27 | 34 | 4 | 14 | 6 | 8 | 18,49 | 29 | 8 | 0,265 | -1,917 | -5,025 | 5,04E-07 | 7,89E-07 | 1,26E-08 | 0,0909034 | 0,0621689 | 0,0621689 | TRUE | 0,557821 | No |
| orf19.5896 | 121 | 108 | 102 | 75 | 83 | 62 | 77 | 75 | 115 | 114 | 101 | 117 | 67 | 70 | 65 | 61 | 88,9 | 112 | 66 | 0,588 | -0,766 | -5,093 | 3,52E-07 | 5,55E-07 | 1,00E-08 | 0,020561 | 0,009986 | 0,009986 | TRUE | 0,0476747 | No |
| orf19.5897 | 550 | 492 | 579 | 380 | 834 | 583 | 769 | 873 | 524 | 520 | 572 | 595 | 668 | 663 | 653 | 710 | 613,22 | 553 | 674 | 1,223 | 0,29 | 4,236 | 2,28E-05 | 3,32E-05 | 0,0007009 | 0,0047688 | 0,0028311 | 0,0028311 | TRUE | 0,026138 | No |
| orf19.59 | 11987 | 10379 | 11078 | 7019 | 10860 | 7249 | 10244 | 10397 | 11428 | 10974 | 10944 | 10996 | 8702 | 8243 | 8697 | 8454 | 9804,73 | 11086 | 8524 | 0,769 | -0,379 | -11,391 | 4,64E-30 | 1,25E-29 | 0,0004099 | 0,0022587 | 0,0009593 | 0,0009593 | TRUE | 0,0058617 | No |
| orf19.590 | 1739 | 1356 | 1516 | 1012 | 2732 | 1811 | 2630 | 2805 | 1658 | 1434 | 1498 | 1585 | 2189 | 2059 | 2233 | 2281 | 1867,09 | 1544 | 2190 | 1,42 | 0,506 | 8,8 | 1,37E-18 | 2,98E-18 | 0,0022344 | 0,0029707 | 0,0026105 | 0,0026105 | TRUE | 0,0320916 | No |
| orf19.5901 | 4463 | 3720 | 4137 | 2726 | 3777 | 2677 | 3660 | 3952 | 4255 | 3933 | 4087 | 4270 | 3026 | 3044 | 3107 | 3213 | 3617,11 | 4136 | 3098 | 0,749 | -0,416 | -9,832 | 8,20E-23 | 1,94E-22 | 0,000733 | 0,0025452 | 0,0014397 | 0,0014397 | TRUE | 0,0132901 | No |
| orf19.5902 | 5768 | 5316 | 5793 | 3986 | 2106 | 1552 | 2077 | 2125 | 5499 | 5621 | 5723 | 6244 | 1687 | 1765 | 1763 | 1728 | 3753,85 | 5772 | 1736 | 0,301 | -1,733 | -35,385 | 2,91E-274 | 5,60E-273 | 0,0016119 | 0,0025287 | 0,001947 | 0,001947 | TRUE | 0,0358074 | No |
| orf19.5903 | 3698 | 3340 | 3896 | 2379 | 4830 | 3589 | 4603 | 4836 | 3526 | 3532 | 3849 | 3727 | 3846 | 4081 | 3908 | 3932 | 3800,02 | 3658 | 3942 | 1,077 | 0,108 | 2,499 | 0,0124703 | 0,0157508 | 0,0008979 | 0,0025234 | 0,0015138 | 0,0015138 | TRUE | 0,015764 | No |
| orf19.5904 | 23427 | 19785 | 20655 | 13114 | 54881 | 36488 | 51357 | 57082 | 22335 | 20920 | 20405 | 20544 | 43975 | 41490 | 43602 | 46413 | 32460,49 | 21051 | 43870 | 2,084 | 1,059 | 23,166 | 9,95E-119 | 7,80E-118 | 0,0018911 | 0,0021419 | 0,0019717 | 0,0019717 | TRUE | 0,020729 | No |
| orf19.5905 | 4868 | 4107 | 4387 | 2411 | 7888 | 4697 | 7110 | 7485 | 4641 | 4343 | 4334 | 3777 | 6320 | 5341 | 6036 | 6086 | 5109,8 | 4274 | 5946 | 1,391 | 0,476 | 6,844 | 7,68E-12 | 1,42E-11 | 0,0058097 | 0,0024126 | 0,0044318 | 0,0044318 | TRUE | 0,0727382 | No |
| orf19.5906 | 9784 | 8326 | 8706 | 6151 | 13985 | 9579 | 13032 | 14446 | 9328 | 8804 | 8601 | 9636 | 11206 | 10892 | 11064 | 11746 | 10159,54 | 9092 | 11227 | 1,235 | 0,305 | 6,634 | 3,26E-11 | 5,91E-11 | 0,0017702 | 0,0022529 | 0,001929 | 0,001929 | TRUE | 0,0193401 | No |
| orf19.5908 | 10566 | 9139 | 10326 | 6845 | 1786 | 1280 | 1692 | 1857 | 10074 | 9663 | 10201 | 10723 | 1431 | 1455 | 1436 | 1510 | 5811,75 | 10165 | 1458 | 0,144 | -2,801 | -62,663 | 0 | 0 | 0,0011244 | 0,0023738 | 0,0015568 | 0,0015568 | TRUE | 0,0162444 | No |
| orf19.5910 | 834 | 684 | 757 | 495 | 970 | 703 | 939 | 1064 | 795 | 723 | 748 | 775 | 777 | 799 | 797 | 865 | 785,08 | 760 | 810 | 1,066 | 0,092 | 1,479 | 0,1392532 | 0,160219 | 0,0009398 | 0,0041827 | 0,0024131 | 0,0024131 | TRUE | 0,0263283 | No |
| orf19.5911 | 1957 | 1728 | 1852 | 1318 | 2079 | 1468 | 2093 | 2191 | 1866 | 1827 | 1830 | 2065 | 1666 | 1669 | 1777 | 1781 | 1810,1 | 1897 | 1723 | 0,911 | -0,135 | -2,498 | 0,0124967 | 0,0157747 | 0,0017471 | 0,0029984 | 0,0022301 | 0,0022301 | TRUE | 0,0395663 | No |
| orf19.5912 | 24050 | 21002 | 23064 | 13614 | 26832 | 17584 | 25214 | 26448 | 22929 | 22207 | 22785 | 21327 | 21500 | 19995 | 21407 | 21505 | 21706,72 | 22312 | 21102 | 0,946 | -0,081 | -2,055 | 0,0398844 | 0,0483936 | 0,0010831 | 0,0021669 | 0,0014307 | 0,0014307 | TRUE | 0,0151448 | No |
| orf19.5915 | 280 | 220 | 259 | 174 | 444 | 315 | 402 | 499 | 267 | 233 | 256 | 273 | 356 | 358 | 341 | 406 | 311,12 | 257 | 365 | 1,426 | 0,512 | 5,465 | 4,62E-08 | 7,53E-08 | 0,0025776 | 0,0073686 | 0,0050142 | 0,0050142 | TRUE | 0,0670108 | No |
| orf19.5917 | 2916 | 2522 | 2873 | 1721 | 5223 | 3908 | 4970 | 5317 | 2780 | 2667 | 2838 | 2696 | 4185 | 4444 | 4220 | 4323 | 3519,07 | 2745 | 4293 | 1,562 | 0,643 | 16,244 | 2,46E-59 | 1,03E-58 | 0,000378 | 0,0025579 | 0,0011974 | 0,0011974 | TRUE | 0,0065609 | No |
| orf19.5917.3 | 7939 | 7433 | 7900 | 5492 | 10157 | 7870 | 9129 | 10010 | 7569 | 7859 | 7804 | 8604 | 8139 | 8949 | 7750 | 8139 | 8101,67 | 7959 | 8244 | 1,036 | 0,051 | 0,918 | 0,3585742 | 0,3932729 | 0,0031964 | 0,0022939 | 0,0028529 | 0,0028529 | TRUE | 0,0400649 | No |
| orf19.5918 | 9 | 10 | 7 | 2 | 9 | 7 | 4 | 6 | 9 | 11 | 7 | 3 | 7 | 8 | 3 | 5 | 6,58 | 8 | 6 | 0,77 | -0,377 | -0,671 | 0,502276 | 0,5381167 | 1,00E-08 | 0,2515812 | 0,1514025 | 0,1514025 | TRUE | 0,2276459 | No |
| orf19.5919 | 464 | 553 | 567 | 376 | 676 | 530 | 589 | 554 | 442 | 585 | 560 | 589 | 542 | 603 | 500 | 450 | 533,89 | 544 | 524 | 0,963 | -0,055 | -0,494 | 0,6215539 | 0,6551264 | 0,0143859 | 0,0051667 | 0,0098159 | 0,0098159 | TRUE | 0,1911037 | No |
| orf19.592 | 2635 | 2036 | 2442 | 1536 | 1408 | 856 | 1306 | 1368 | 2512 | 2153 | 2412 | 2406 | 1128 | 973 | 1109 | 1112 | 1725,79 | 2371 | 1080 | 0,457 | -1,131 | -17,365 | 1,53E-67 | 7,08E-67 | 0,003414 | 0,0030427 | 0,0034241 | 0,0034241 | TRUE | 0,0505774 | No |
| orf19.5920 | 2437 | 2071 | 2191 | 1059 | 1741 | 1179 | 1501 | 1606 | 2323 | 2190 | 2165 | 1659 | 1395 | 1341 | 1274 | 1306 | 1706,57 | 2084 | 1329 | 0,636 | -0,652 | -7,307 | 2,74E-13 | 5,28E-13 | 0,0097437 | 0,0030534 | 0,0070379 | 0,0070379 | TRUE | 0,220269 | No |
| orf19.5921 | 1046 | 1063 | 993 | 732 | 2233 | 1810 | 2029 | 2149 | 997 | 1124 | 981 | 1147 | 1789 | 2058 | 1723 | 1747 | 1445,78 | 1062 | 1829 | 1,724 | 0,786 | 10,47 | 1,19E-25 | 2,98E-25 | 0,0058136 | 0,0032269 | 0,0046329 | 0,0046329 | TRUE | 0,0835872 | No |
| orf19.5924 | 802 | 810 | 906 | 500 | 931 | 680 | 849 | 898 | 765 | 856 | 895 | 783 | 746 | 773 | 721 | 730 | 783,7 | 825 | 742 | 0,898 | -0,155 | -2,326 | 0,0200084 | 0,0248344 | 0,0019435 | 0,0041864 | 0,0029912 | 0,0029912 | TRUE | 0,0401419 | No |
| orf19.5925 | 712 | 700 | 730 | 417 | 904 | 654 | 805 | 832 | 679 | 740 | 721 | 653 | 724 | 744 | 683 | 676 | 702,67 | 698 | 707 | 1,009 | 0,012 | 0,19 | 0,849169 | 0,8680326 | 0,0009616 | 0,004428 | 0,002669 | 0,002669 | TRUE | 0,0187894 | No |
| orf19.5926 | 1050 | 963 | 1052 | 640 | 900 | 611 | 859 | 879 | 1001 | 1018 | 1039 | 1003 | 721 | 695 | 729 | 715 | 865,14 | 1015 | 715 | 0,705 | -0,505 | -9,528 | 1,60E-21 | 3,70E-21 | 1,00E-08 | 0,0039891 | 0,0015282 | 0,0015282 | TRUE | 0,0037794 | No |
| orf19.5927 | 13795 | 11278 | 11473 | 8004 | 33899 | 22277 | 30260 | 34627 | 13152 | 11925 | 11334 | 12539 | 27162 | 25331 | 25691 | 28155 | 19411,12 | 12238 | 26585 | 2,173 | 1,119 | 20,466 | 4,31E-93 | 2,65E-92 | 0,0031883 | 0,0021759 | 0,0028114 | 0,0028114 | TRUE | 0,031301 | No |
| orf19.5928 | 7115 | 5470 | 5372 | 4482 | 19574 | 14028 | 16817 | 20209 | 6783 | 5784 | 5307 | 7021 | 15684 | 15951 | 14278 | 16432 | 10905,01 | 6224 | 15586 | 2,505 | 1,325 | 15,931 | 3,85E-57 | 1,55E-56 | 0,0101463 | 0,0022418 | 0,0065257 | 0,0065257 | TRUE | 0,1202682 | No |
| orf19.5929 | 860 | 769 | 803 | 567 | 960 | 746 | 898 | 1018 | 820 | 813 | 793 | 888 | 769 | 848 | 762 | 828 | 815,27 | 828 | 802 | 0,969 | -0,045 | -0,717 | 0,4732402 | 0,5095757 | 0,0011834 | 0,0041052 | 0,0025762 | 0,0025762 | TRUE | 0,0242813 | No |
| orf19.593 | 2883 | 2439 | 2722 | 1724 | 2457 | 1698 | 2289 | 2580 | 2749 | 2579 | 2689 | 2701 | 1969 | 1931 | 1943 | 2098 | 2332,25 | 2680 | 1985 | 0,741 | -0,432 | -9,654 | 4,74E-22 | 1,11E-21 | 0,0006962 | 0,0027953 | 0,0014854 | 0,0014854 | TRUE | 0,0179474 | No |
| orf19.5930 | 1929 | 1481 | 1650 | 1180 | 1727 | 1288 | 1667 | 1778 | 1839 | 1566 | 1630 | 1849 | 1384 | 1465 | 1415 | 1446 | 1574,12 | 1721 | 1428 | 0,83 | -0,269 | -4,273 | 1,93E-05 | 2,82E-05 | 0,0029718 | 0,0031343 | 0,0031545 | 0,0031545 | TRUE | 0,0440954 | No |
| orf19.5931 | 166 | 183 | 165 | 74 | 254 | 181 | 242 | 210 | 158 | 193 | 163 | 116 | 204 | 206 | 205 | 171 | 177,03 | 158 | 196 | 1,232 | 0,301 | 2,15 | 0,031568 | 0,0385744 | 0,0153777 | 0,011366 | 0,0130491 | 0,0130491 | TRUE | 0,2672896 | No |
| orf19.5932 | 2542 | 2238 | 2471 | 1486 | 1405 | 1072 | 1380 | 1523 | 2424 | 2366 | 2441 | 2328 | 1126 | 1219 | 1172 | 1238 | 1789,21 | 2390 | 1189 | 0,497 | -1,01 | -21,442 | 5,38E-102 | 3,59E-101 | 0,0004238 | 0,003009 | 0,0015186 | 0,0015186 | TRUE | 0,0157904 | No |
| orf19.5933 | 72 | 71 | 86 | 45 | 33 | 25 | 30 | 27 | 69 | 75 | 85 | 70 | 26 | 28 | 25 | 22 | 50,18 | 75 | 25 | 0,339 | -1,561 | -7,357 | 1,88E-13 | 3,64E-13 | 1,00E-08 | 0,0348095 | 0,01832 | 0,01832 | TRUE | 0,0809403 | No |
| orf19.5934 | 281 | 275 | 281 | 156 | 1164 | 797 | 1216 | 1230 | 268 | 291 | 278 | 244 | 933 | 906 | 1032 | 1000 | 619,01 | 270 | 968 | 3,568 | 1,835 | 23,326 | 2,42E-120 | 1,94E-119 | 0,0021378 | 0,0047437 | 0,0034315 | 0,0034315 | TRUE | 0,0354282 | No |
| orf19.5935 | 1090 | 970 | 1076 | 708 | 1170 | 888 | 1170 | 1265 | 1039 | 1026 | 1063 | 1109 | 937 | 1010 | 993 | 1029 | 1025,76 | 1059 | 992 | 0,938 | -0,092 | -1,688 | 0,091453 | 0,1071148 | 0,0004593 | 0,003692 | 0,001893 | 0,001893 | TRUE | 0,0172459 | No |
| orf19.5938 | 2342 | 2185 | 2308 | 1478 | 3547 | 2669 | 3417 | 3575 | 2233 | 2310 | 2280 | 2315 | 2842 | 3035 | 2901 | 2907 | 2602,94 | 2284 | 2921 | 1,278 | 0,354 | 9,002 | 2,21E-19 | 4,90E-19 | 0,0001731 | 0,0027221 | 0,0010893 | 0,0010893 | TRUE | 0,0078882 | No |
| orf19.5940 | 1169 | 996 | 1083 | 749 | 1762 | 1262 | 1607 | 1848 | 1115 | 1053 | 1070 | 1173 | 1412 | 1435 | 1364 | 1503 | 1265,59 | 1103 | 1428 | 1,298 | 0,377 | 6,854 | 7,16E-12 | 1,32E-11 | 0,0009269 | 0,0033886 | 0,0020763 | 0,0020763 | TRUE | 0,0194374 | No |
| orf19.5941 | 642 | 546 | 572 | 333 | 1758 | 1126 | 1629 | 1806 | 612 | 577 | 565 | 522 | 1409 | 1280 | 1383 | 1468 | 977,08 | 569 | 1385 | 2,427 | 1,279 | 19,133 | 1,34E-81 | 7,46E-81 | 0,0021042 | 0,0037717 | 0,0029916 | 0,0029916 | TRUE | 0,0310692 | No |
| orf19.5942 | 2792 | 2452 | 2670 | 1548 | 2629 | 1839 | 2474 | 2732 | 2662 | 2593 | 2638 | 2425 | 2107 | 2091 | 2100 | 2221 | 2354,59 | 2580 | 2130 | 0,825 | -0,278 | -6,135 | 8,52E-10 | 1,47E-09 | 0,0007002 | 0,0027886 | 0,0015453 | 0,0015453 | TRUE | 0,0180294 | No |
| orf19.5943 | 1312 | 1289 | 1359 | 768 | 1753 | 1306 | 1757 | 1814 | 1251 | 1363 | 1343 | 1203 | 1405 | 1485 | 1492 | 1475 | 1376,97 | 1290 | 1464 | 1,132 | 0,179 | 3,312 | 0,0009256 | 0,0012513 | 0,0011249 | 0,0032837 | 0,0020749 | 0,0020749 | TRUE | 0,021593 | No |
| orf19.5943.1 | 7773 | 6414 | 6865 | 4467 | 17782 | 12764 | 17772 | 18710 | 7411 | 6782 | 6782 | 6998 | 14248 | 14514 | 15088 | 15213 | 10879,48 | 6993 | 14766 | 2,111 | 1,078 | 25,665 | 2,88E-145 | 2,82E-144 | 0,0012435 | 0,0022422 | 0,0015842 | 0,0015842 | TRUE | 0,0201914 | No |
| orf19.5947 | 2604 | 2335 | 2558 | 1658 | 3597 | 2601 | 3219 | 3612 | 2483 | 2469 | 2527 | 2597 | 2882 | 2958 | 2733 | 2937 | 2698,2 | 2519 | 2878 | 1,143 | 0,193 | 4,575 | 4,76E-06 | 7,17E-06 | 0,0005454 | 0,0026998 | 0,0013243 | 0,0013243 | TRUE | 0,013945 | No |
| orf19.5949 | 6066 | 5619 | 6144 | 3856 | 71785 | 49681 | 63731 | 74569 | 5783 | 5941 | 6070 | 6041 | 57519 | 56492 | 54107 | 60632 | 31573,13 | 5959 | 57188 | 9,599 | 3,263 | 77,195 | 0 | 0 | 0,00123 | 0,0021433 | 0,0016136 | 0,0016136 | TRUE | 0,0201641 | No |
| orf19.5952 | 272 | 249 | 295 | 159 | 889 | 692 | 903 | 1042 | 259 | 263 | 291 | 249 | 712 | 787 | 767 | 847 | 522,03 | 266 | 778 | 2,917 | 1,545 | 18,273 | 1,36E-74 | 6,91E-74 | 0,0032507 | 0,0052365 | 0,0041962 | 0,0041962 | TRUE | 0,0482374 | No |
| orf19.5953 | 7255 | 5963 | 6990 | 4468 | 6849 | 4684 | 6680 | 6934 | 6917 | 6305 | 6905 | 6999 | 5488 | 5326 | 5671 | 5638 | 6156,27 | 6782 | 5531 | 0,816 | -0,293 | -6,622 | 3,54E-11 | 6,40E-11 | 0,001306 | 0,002358 | 0,0017216 | 0,0017216 | TRUE | 0,027436 | No |
| orf19.5954 | 100 | 71 | 63 | 41 | 22 | 28 | 22 | 39 | 95 | 75 | 62 | 64 | 18 | 32 | 19 | 32 | 49,59 | 74 | 25 | 0,331 | -1,595 | -6,231 | 4,64E-10 | 8,09E-10 | 0,0405234 | 0,0351996 | 0,0373748 | 0,0373748 | TRUE | 0,3507843 | No |
| orf19.5956 | 12060 | 9884 | 10840 | 7587 | 7857 | 5253 | 7511 | 7836 | 11498 | 10451 | 10709 | 11885 | 6296 | 5973 | 6377 | 6371 | 8695,03 | 11136 | 6254 | 0,562 | -0,831 | -16,909 | 3,86E-64 | 1,72E-63 | 0,0021066 | 0,0022801 | 0,0022006 | 0,0022006 | TRUE | 0,0246571 | No |
| orf19.5958 | 3771 | 3399 | 3634 | 2015 | 3636 | 2470 | 3269 | 3791 | 3595 | 3594 | 3590 | 3157 | 2913 | 2809 | 2775 | 3082 | 3189,47 | 3484 | 2895 | 0,83 | -0,269 | -4,801 | 1,58E-06 | 2,42E-06 | 0,0026792 | 0,0026061 | 0,0026875 | 0,0026875 | TRUE | 0,0464822 | No |
| orf19.5959 | 13492 | 13161 | 13982 | 7954 | 12568 | 8632 | 11577 | 12044 | 12863 | 13916 | 13813 | 12460 | 10070 | 9815 | 9829 | 9793 | 11570 | 13263 | 9877 | 0,744 | -0,426 | -9,847 | 7,09E-23 | 1,68E-22 | 0,0014211 | 0,0022332 | 0,0017076 | 0,0017076 | TRUE | 0,0198951 | No |
| orf19.5960 | 38765 | 35479 | 38516 | 28931 | 15264 | 11634 | 13766 | 14285 | 36958 | 37515 | 38050 | 45322 | 12231 | 13229 | 11687 | 11615 | 25825,84 | 39461 | 12190 | 0,309 | -1,695 | -24,315 | 1,34E-130 | 1,17E-129 | 0,0067605 | 0,0021549 | 0,0046162 | 0,0046162 | TRUE | 0,1221676 | No |
| orf19.5961 | 4536 | 3928 | 4210 | 2650 | 2034 | 1499 | 1831 | 2000 | 4325 | 4153 | 4159 | 4151 | 1630 | 1704 | 1555 | 1626 | 2912,92 | 4197 | 1629 | 0,388 | -1,368 | -32,92 | 1,13E-237 | 1,80E-236 | 0,0003444 | 0,0026549 | 0,0012489 | 0,0012489 | TRUE | 0,0111702 | No |
| orf19.596.1 | 2835 | 2267 | 2588 | 1696 | 7104 | 4151 | 6397 | 6958 | 2703 | 2397 | 2557 | 2657 | 5692 | 4720 | 5431 | 5658 | 3976,79 | 2578 | 5375 | 2,086 | 1,061 | 16,042 | 6,48E-58 | 2,65E-57 | 0,0045297 | 0,0025042 | 0,0038981 | 0,0038981 | TRUE | 0,0811597 | No |
| orf19.5962 | 2440 | 2397 | 2366 | 1416 | 2690 | 1877 | 2344 | 2499 | 2326 | 2535 | 2337 | 2218 | 2155 | 2134 | 1990 | 2032 | 2216,02 | 2354 | 2078 | 0,882 | -0,182 | -3,467 | 0,0005264 | 0,0007223 | 0,0016961 | 0,0028322 | 0,0021908 | 0,0021908 | TRUE | 0,0319706 | No |
| orf19.596.2 | 549 | 430 | 494 | 276 | 720 | 402 | 664 | 667 | 523 | 455 | 488 | 432 | 577 | 457 | 564 | 542 | 504,82 | 474 | 535 | 1,126 | 0,172 | 1,889 | 0,0589132 | 0,0702796 | 0,0065145 | 0,0053437 | 0,0059271 | 0,0059271 | TRUE | 0,1050702 | No |
| orf19.5963 | 7248 | 6400 | 6908 | 3731 | 4103 | 2692 | 3805 | 3980 | 6910 | 6767 | 6824 | 5845 | 3288 | 3061 | 3230 | 3236 | 4895,23 | 6586 | 3204 | 0,486 | -1,04 | -18,176 | 7,93E-74 | 4,01E-73 | 0,0030229 | 0,0024267 | 0,0029217 | 0,0029217 | TRUE | 0,0685258 | No |
| orf19.5964 | 4698 | 4202 | 4580 | 2603 | 8777 | 6058 | 8569 | 8977 | 4479 | 4443 | 4525 | 4078 | 7033 | 6888 | 7275 | 7299 | 5752,49 | 4381 | 7124 | 1,624 | 0,7 | 16,198 | 5,18E-59 | 2,15E-58 | 0,001234 | 0,0023767 | 0,0016018 | 0,0016018 | TRUE | 0,0250287 | No |
| orf19.5964.2 | 10104 | 9677 | 9414 | 5548 | 26661 | 18401 | 23552 | 23820 | 9633 | 10232 | 9300 | 8691 | 21363 | 20924 | 19996 | 19368 | 14938,3 | 9464 | 20413 | 2,156 | 1,108 | 20,267 | 2,50E-91 | 1,52E-90 | 0,0031809 | 0,0022012 | 0,0027913 | 0,0027913 | TRUE | 0,0362979 | No |
| orf19.5965 | 1689 | 1691 | 1772 | 1060 | 3385 | 2492 | 2993 | 3208 | 1610 | 1788 | 1751 | 1661 | 2712 | 2834 | 2541 | 2608 | 2188,1 | 1702 | 2674 | 1,569 | 0,65 | 12,213 | 2,65E-34 | 7,66E-34 | 0,0016435 | 0,0028417 | 0,0022217 | 0,0022217 | TRUE | 0,0189535 | No |
| orf19.5966 | 12 | 7 | 9 | 8 | 12 | 11 | 7 | 7 | 11 | 7 | 9 | 13 | 10 | 13 | 6 | 6 | 9,25 | 10 | 9 | 0,832 | -0,265 | -0,559 | 0,5758351 | 0,6100621 | 1,00E-08 | 0,1795374 | 0,1050837 | 0,1050837 | TRUE | 0,2638501 | No |
| orf19.5967 | 1233 | 1009 | 1199 | 777 | 1414 | 980 | 1380 | 1451 | 1176 | 1067 | 1184 | 1217 | 1133 | 1114 | 1172 | 1180 | 1155,36 | 1161 | 1150 | 0,992 | -0,011 | -0,202 | 0,8397816 | 0,859584 | 0,0007936 | 0,0035124 | 0,0021037 | 0,0021037 | TRUE | 0,0357481 | No |
| orf19.5968 | 3161 | 2990 | 2907 | 1975 | 3850 | 2629 | 3723 | 3941 | 3014 | 3162 | 2872 | 3094 | 3085 | 2989 | 3161 | 3204 | 3072,57 | 3036 | 3110 | 1,026 | 0,037 | 0,822 | 0,4113158 | 0,4472744 | 0,0008679 | 0,0026257 | 0,0015774 | 0,0015774 | TRUE | 0,0163009 | No |
| orf19.597 | 3467 | 2676 | 3094 | 2125 | 6212 | 4160 | 6029 | 6190 | 3305 | 2830 | 3057 | 3329 | 4977 | 4730 | 5119 | 5033 | 4047,49 | 3130 | 4965 | 1,588 | 0,667 | 11,754 | 6,71E-32 | 1,87E-31 | 0,0030411 | 0,0024969 | 0,0028208 | 0,0028208 | TRUE | 0,0508182 | No |
| orf19.5970 | 696 | 628 | 718 | 373 | 1075 | 758 | 1039 | 1138 | 664 | 664 | 709 | 584 | 861 | 862 | 882 | 925 | 768,99 | 655 | 882 | 1,341 | 0,424 | 6,358 | 2,04E-10 | 3,61E-10 | 0,0015023 | 0,0042264 | 0,0029082 | 0,0029082 | TRUE | 0,0532456 | No |
| orf19.5971 | 1174 | 1046 | 1066 | 681 | 784 | 591 | 787 | 849 | 1119 | 1106 | 1053 | 1067 | 628 | 672 | 668 | 690 | 875,49 | 1086 | 664 | 0,611 | -0,71 | -12,481 | 9,54E-36 | 2,82E-35 | 1,00E-08 | 0,0039667 | 0,0019257 | 0,0019257 | TRUE | 0,017161 | No |

| | | | | | | | | | | | | | | | | | | | | | | | | | | | | | | | |
|---|---|---|---|---|---|---|---|---|---|---|---|---|---|---|---|---|---|---|---|---|---|---|---|---|---|---|---|---|---|---|---|
| orf19.5973 | 3931 | 3704 | 3813 | 2424 | 3031 | 2237 | 2982 | 3163 | 3748 | 3917 | 3767 | 3797 | 2429 | 2544 | 2532 | 2572 | 3163,04 | 3807 | 2519 | 0,662 | -0,596 | -15,793 | 3,46E-56 | 1,38E-55 | 0,0002252 | 0,0026104 | 0,0010431 | 0,0010431 | TRUE | 0,0073086 | No |
| orf19.5974 | 1379 | 1557 | 1646 | 946 | 2324 | 1639 | 2068 | 2098 | 1315 | 1646 | 1626 | 1482 | 1862 | 1864 | 1756 | 1706 | 1657,07 | 1517 | 1797 | 1,184 | 0,243 | 3,336 | 0,0008505 | 0,0011534 | 0,0052784 | 0,0030821 | 0,0044953 | 0,0044953 | TRUE | 0,1013863 | No |
| orf19.5975 | 1678 | 1391 | 1493 | 898 | 4015 | 2596 | 3756 | 3781 | 1600 | 1471 | 1475 | 1407 | 3217 | 2952 | 3189 | 3074 | 2298,05 | 1488 | 3108 | 2,086 | 1,06 | 20,399 | 1,72E-92 | 1,05E-91 | 0,0015681 | 0,0028058 | 0,0020746 | 0,0020746 | TRUE | 0,0312511 | No |
| orf19.5976 | 1505 | 1421 | 1548 | 969 | 1553 | 1105 | 1438 | 1487 | 1435 | 1503 | 1529 | 1518 | 1244 | 1256 | 1221 | 1209 | 1364,43 | 1496 | 1232 | 0,824 | -0,279 | -6,007 | 1,89E-09 | 3,22E-09 | 1,00E-08 | 0,0032946 | 0,0013362 | 0,0013362 | TRUE | 0,0097331 | No |
| orf19.5977 | 1607 | 1323 | 1451 | 912 | 2430 | 1727 | 2224 | 2549 | 1532 | 1399 | 1433 | 1429 | 1947 | 1964 | 1888 | 2073 | 1708,09 | 1448 | 1968 | 1,358 | 0,441 | 8,704 | 3,19E-18 | 6,89E-18 | 0,0010676 | 0,0030525 | 0,0018541 | 0,0018541 | TRUE | 0,0192321 | No |
| orf19.5978 | 264 | 248 | 281 | 209 | 253 | 190 | 249 | 244 | 252 | 262 | 278 | 327 | 203 | 216 | 211 | 198 | 243,44 | 280 | 207 | 0,745 | -0,424 | -4,103 | 4,08E-05 | 5,88E-05 | 0,0026599 | 0,0088359 | 0,0061055 | 0,0061055 | TRUE | 0,1242119 | No |
| orf19.598 | 5 | 6 | 1 | 1 | 3 | 1 | 1 | 2 | 5 | 6 | 1 | 2 | 2 | 1 | 1 | 2 | 2,46 | 4 | 2 | 0,437 | -1,193 | -1,249 | 0,2117171 | 0,2395331 | 1,00E-08 | 0,6694724 | 0,4257372 | 0,4257372 | TRUE | 0,1513897 | No |
| orf19.5980 | 1288 | 1224 | 1287 | 806 | 897 | 636 | 839 | 888 | 1228 | 1294 | 1271 | 1263 | 719 | 723 | 712 | 722 | 991,57 | 1264 | 719 | 0,569 | -0,814 | -16,024 | 8,71E-58 | 3,55E-57 | 1,00E-08 | 0,0037471 | 0,0014153 | 0,0014153 | TRUE | 0,0048546 | No |
| orf19.5982 | 17187 | 13356 | 14325 | 8991 | 46968 | 31052 | 44260 | 49143 | 16386 | 14122 | 14152 | 14085 | 37634 | 35309 | 37577 | 39958 | 26152,8 | 14686 | 37620 | 2,561 | 1,357 | 22,965 | 1,04E-116 | 8,01E-116 | 0,0041883 | 0,0021541 | 0,0033035 | 0,0033035 | TRUE | 0,0745448 | No |
| orf19.5984 | 979 | 839 | 898 | 606 | 1878 | 1184 | 1761 | 1692 | 933 | 887 | 887 | 949 | 1505 | 1346 | 1495 | 1376 | 1172,36 | 914 | 1430 | 1,568 | 0,649 | 11,077 | 1,62E-28 | 4,26E-28 | 0,0015496 | 0,0034918 | 0,0023724 | 0,0023724 | TRUE | 0,0178016 | No |
| orf19.5985 | 1665 | 1386 | 1477 | 944 | 2393 | 1552 | 2358 | 2396 | 1587 | 1466 | 1459 | 1479 | 1917 | 1765 | 2002 | 1948 | 1702,9 | 1498 | 1908 | 1,275 | 0,35 | 6,416 | 1,40E-10 | 2,49E-10 | 0,0016862 | 0,0030555 | 0,0022486 | 0,0022486 | TRUE | 0,0294099 | No |
| orf19.5986 | 764 | 631 | 656 | 545 | 609 | 463 | 540 | 662 | 728 | 667 | 648 | 854 | 488 | 526 | 458 | 538 | 613,58 | 724 | 502 | 0,696 | -0,522 | -5,585 | 2,33E-08 | 3,84E-08 | 0,0086054 | 0,0047672 | 0,0067216 | 0,0067216 | TRUE | 0,1577464 | No |
| orf19.5987 | 3805 | 3187 | 3415 | 2167 | 6370 | 4440 | 6160 | 6572 | 3628 | 3370 | 3374 | 3395 | 5104 | 5049 | 5230 | 5344 | 4311,52 | 3442 | 5182 | 1,505 | 0,59 | 14,604 | 2,64E-48 | 9,52E-48 | 0,0006574 | 0,0024721 | 0,0013175 | 0,0013175 | TRUE | 0,0167146 | No |
| orf19.5989 | 5557 | 4555 | 5195 | 3959 | 8917 | 6645 | 8501 | 8847 | 5298 | 4816 | 5132 | 6202 | 7145 | 7556 | 7217 | 7193 | 6320,02 | 5362 | 7278 | 1,358 | 0,442 | 6,436 | 1,23E-10 | 2,19E-10 | 0,0062628 | 0,0023511 | 0,0043605 | 0,0043605 | TRUE | 0,1331225 | No |
| orf19.599 | 1 | 0 | 0 | 0 | 2 | 0 | 1 | 1 | 1 | 0 | 0 | 0 | 2 | 0 | 1 | 1 | 0,53 | 0 | 1 | 2,483 | 1,312 | 0,69 | 0,4898854 | 0,52599 | 1,00E-08 | 3,1161617 | 1,9400517 | 1,9400517 | TRUE | 0,2511084 | No |
| orf19.5991 | 19515 | 17617 | 19336 | 11538 | 21240 | 14536 | 19620 | 22202 | 18605 | 18628 | 19102 | 18075 | 17019 | 16529 | 16657 | 18052 | 17833,45 | 18602 | 17064 | 0,917 | -0,125 | -3,243 | 0,0011837 | 0,0015875 | 0,0009986 | 0,0021833 | 0,0013611 | 0,0013611 | TRUE | 0,0186355 | No |
| orf19.5992 | 3279 | 2860 | 3159 | 2280 | 940 | 705 | 929 | 982 | 3126 | 3024 | 3121 | 3572 | 753 | 802 | 789 | 798 | 1998,1 | 3211 | 786 | 0,245 | -2,03 | -32,904 | 1,91E-237 | 3,04E-236 | 0,0029103 | 0,002913 | 0,0029103 | 0,0029103 | TRUE | 0,0667973 | No |
| orf19.5994 | 2362 | 2254 | 2221 | 1393 | 3706 | 2500 | 3280 | 3348 | 2252 | 2383 | 2194 | 2182 | 2970 | 2843 | 2785 | 2722 | 2541,35 | 2253 | 2830 | 1,255 | 0,328 | 6,942 | 3,85E-12 | 7,17E-12 | 0,0009974 | 0,0027374 | 0,0017355 | 0,0017355 | TRUE | 0,0184137 | No |
| orf19.5995 | 3682 | 3173 | 3311 | 2256 | 3764 | 2733 | 3518 | 3862 | 3510 | 3355 | 3271 | 3534 | 3016 | 3108 | 2987 | 3140 | 3240,14 | 3418 | 3063 | 0,897 | -0,157 | -3,797 | 0,0001462 | 0,0002051 | 0,0005504 | 0,002598 | 0,0013401 | 0,0013401 | TRUE | 0,0102631 | No |
| orf19.5996.1 | 17196 | 13051 | 13830 | 8931 | 52049 | 33173 | 46618 | 51714 | 16395 | 13800 | 13663 | 13991 | 41705 | 37721 | 39578 | 42048 | 27362,61 | 14462 | 40263 | 2,784 | 1,477 | 23,221 | 2,77E-119 | 2,19E-118 | 0,0052031 | 0,0021513 | 0,0038373 | 0,0038373 | TRUE | 0,0993904 | No |
| orf19.5999 | 1264 | 1134 | 1196 | 712 | 1173 | 782 | 1068 | 1117 | 1205 | 1199 | 1182 | 1115 | 940 | 889 | 907 | 908 | 1043,14 | 1175 | 911 | 0,774 | -0,369 | -7,16 | 8,06E-13 | 1,53E-12 | 1,00E-08 | 0,0036653 | 0,0015917 | 0,0015917 | TRUE | 0,0121069 | No |
| orf19.600 | 3736 | 3053 | 3602 | 2016 | 4276 | 2856 | 4036 | 4481 | 3562 | 3228 | 3558 | 3158 | 3426 | 3248 | 3427 | 3643 | 3406,3 | 3376 | 3436 | 1,017 | 0,024 | 0,435 | 0,6633299 | 0,6954827 | 0,002445 | 0,0025733 | 0,0026886 | 0,0026886 | TRUE | 0,0221144 | No |
| orf19.6000 | 75534 | 71159 | 76660 | 44869 | 57530 | 40800 | 52397 | 58582 | 72013 | 75242 | 75733 | 70290 | 46097 | 46393 | 44485 | 47633 | 59735,7 | 73320 | 46152 | 0,629 | -0,668 | -17,789 | 8,68E-71 | 4,19E-70 | 0,0010029 | 0,0021188 | 0,001337 | 0,001337 | TRUE | 0,0094374 | No |
| orf19.6001 | 612 | 631 | 672 | 378 | 272 | 197 | 281 | 275 | 583 | 667 | 664 | 592 | 218 | 224 | 239 | 224 | 426,35 | 626 | 226 | 0,36 | -1,473 | -17,6 | 2,48E-69 | 1,17E-68 | 1,00E-08 | 0,0059423 | 0,0038693 | 0,0038693 | TRUE | 0,0288303 | No |
| orf19.6002 | 10673 | 10072 | 10401 | 7337 | 30460 | 23306 | 28075 | 29571 | 10176 | 10650 | 10275 | 11494 | 24407 | 26501 | 23836 | 24044 | 17672,72 | 10649 | 24697 | 2,32 | 1,214 | 23,608 | 3,19E-123 | 2,62E-122 | 0,0027258 | 0,0021842 | 0,0024699 | 0,0024699 | TRUE | 0,0343847 | No |
| orf19.6003 | 3127 | 3122 | 3481 | 2140 | 2252 | 1694 | 2051 | 2225 | 2981 | 3301 | 3439 | 3352 | 1804 | 1926 | 1741 | 1809 | 2544,35 | 3268 | 1820 | 0,557 | -0,845 | -15,13 | 1,03E-51 | 3,89E-51 | 0,0021939 | 0,0027366 | 0,0025759 | 0,0025759 | TRUE | 0,0435316 | No |
| orf19.6005 | 348 | 350 | 361 | 229 | 1273 | 1016 | 1244 | 1257 | 332 | 370 | 357 | 359 | 1020 | 1155 | 1056 | 1022 | 708,84 | 354 | 1063 | 3 | 1,585 | 22,058 | 8,02E-108 | 5,66E-107 | 0,0016906 | 0,0044076 | 0,0029635 | 0,0029635 | TRUE | 0,0383178 | No |
| orf19.6007 | 2445 | 2961 | 2822 | 1801 | 8433 | 6381 | 7669 | 6909 | 2331 | 3131 | 2788 | 2821 | 6757 | 7256 | 6511 | 5618 | 4651,58 | 2768 | 6536 | 2,361 | 1,24 | 13,515 | 1,28E-41 | 4,17E-41 | 0,0122522 | 0,0024443 | 0,007811 | 0,007811 | TRUE | 0,1386146 | No |
| orf19.6008 | 312 | 392 | 401 | 249 | 392 | 314 | 366 | 388 | 297 | 414 | 396 | 390 | 314 | 357 | 311 | 315 | 349,44 | 374 | 324 | 0,866 | -0,207 | -1,907 | 0,0564796 | 0,0675279 | 0,0101782 | 0,0067899 | 0,0084735 | 0,0084735 | TRUE | 0,233189 | No |
| orf19.6008.3 | 295 | 265 | 292 | 168 | 534 | 322 | 511 | 504 | 281 | 280 | 288 | 263 | 428 | 366 | 434 | 410 | 343,85 | 278 | 410 | 1,471 | 0,557 | 6,454 | 1,09E-10 | 1,94E-10 | 1,00E-08 | 0,0068664 | 0,0040662 | 0,0040662 | TRUE | 0,0526063 | No |
| orf19.6008.4 | 2460 | 1903 | 1999 | 1306 | 2069 | 1264 | 1919 | 2024 | 2345 | 2012 | 1975 | 2046 | 1658 | 1437 | 1629 | 1646 | 1843,54 | 2094 | 1592 | 0,761 | -0,395 | -5,714 | 1,10E-08 | 1,83E-08 | 0,0046831 | 0,0029819 | 0,0040302 | 0,0040302 | TRUE | 0,0805108 | No |
| orf19.6009 | 1345 | 1278 | 1342 | 846 | 1737 | 1212 | 1728 | 1824 | 1282 | 1351 | 1326 | 1325 | 1392 | 1378 | 1467 | 1483 | 1375,6 | 1321 | 1430 | 1,084 | 0,116 | 2,379 | 0,0173472 | 0,0216407 | 1,00E-08 | 0,0032849 | 0,0015416 | 0,0015416 | TRUE | 0,007537 | No |
| orf19.6010 | 557 | 531 | 549 | 339 | 392 | 251 | 351 | 414 | 531 | 561 | 542 | 531 | 314 | 285 | 298 | 337 | 425,01 | 541 | 308 | 0,571 | -0,808 | -10,486 | 1,00E-25 | 2,51E-25 | 1,00E-08 | 0,0059545 | 0,0032276 | 0,0032276 | TRUE | 0,0429387 | No |
| orf19.6010.1 | 1383 | 1257 | 1241 | 833 | 1624 | 1343 | 1488 | 1644 | 1319 | 1329 | 1226 | 1305 | 1301 | 1527 | 1263 | 1337 | 1325,87 | 1295 | 1357 | 1,047 | 0,066 | 0,997 | 0,318693 | 0,3518913 | 0,0033795 | 0,0033296 | 0,0034286 | 0,0034286 | TRUE | 0,0820633 | No |
| orf19.6011 | 5898 | 5257 | 5762 | 3514 | 9501 | 6871 | 8728 | 9629 | 5623 | 5559 | 5692 | 5505 | 7613 | 7813 | 7410 | 7829 | 6630,51 | 5595 | 7666 | 1,37 | 0,454 | 13,629 | 2,70E-42 | 8,90E-42 | 0,0003237 | 0,0023389 | 0,0009051 | 0,0009051 | TRUE | 0,0061981 | No |
| orf19.6012 | 1435 | 1402 | 1498 | 902 | 1316 | 986 | 1223 | 1379 | 1368 | 1482 | 1480 | 1413 | 1054 | 1121 | 1038 | 1121 | 1259,84 | 1436 | 1084 | 0,754 | -0,407 | -7,637 | 2,23E-14 | 4,42E-14 | 0,0008429 | 0,0033945 | 0,0019291 | 0,0019291 | TRUE | 0,0132873 | No |
| orf19.6013 | 1262 | 1068 | 1205 | 743 | 1061 | 722 | 914 | 985 | 1203 | 1129 | 1190 | 1164 | 850 | 821 | 776 | 801 | 991,85 | 1172 | 812 | 0,693 | -0,53 | -9,742 | 1,99E-22 | 4,69E-22 | 1,00E-08 | 0,0037467 | 0,0018151 | 0,0018151 | TRUE | 0,0128512 | No |
| orf19.6014 | 7947 | 6759 | 7524 | 3949 | 11814 | 7880 | 11676 | 11797 | 7577 | 7147 | 7433 | 6186 | 9466 | 8960 | 9913 | 9592 | 8284,27 | 7086 | 9483 | 1,338 | 0,42 | 6,633 | 3,30E-11 | 5,97E-11 | 0,0046051 | 0,0022895 | 0,0037179 | 0,0037179 | TRUE | 0,08757 | No |
| orf19.6016 | 1869 | 1828 | 1954 | 1200 | 1184 | 950 | 1182 | 1243 | 1782 | 1933 | 1930 | 1880 | 949 | 1080 | 1004 | 1011 | 1446,02 | 1881 | 1011 | 0,536 | -0,899 | -16,406 | 1,73E-60 | 7,32E-60 | 0,0013143 | 0,0032267 | 0,0021388 | 0,0021388 | TRUE | 0,0237735 | No |
| orf19.6017 | 506 | 451 | 494 | 313 | 375 | 292 | 360 | 385 | 482 | 477 | 488 | 490 | 300 | 332 | 306 | 313 | 398,6 | 484 | 313 | 0,644 | -0,634 | -8,556 | 1,17E-17 | 2,50E-17 | 1,00E-08 | 0,0062104 | 0,0026941 | 0,0026941 | TRUE | 0,0169452 | No |
| orf19.6018 | 1476 | 1357 | 1524 | 906 | 3586 | 2433 | 3483 | 3671 | 1407 | 1435 | 1506 | 1419 | 2873 | 2767 | 2957 | 2985 | 2168,59 | 1442 | 2896 | 2,008 | 1,006 | 21,96 | 6,90E-107 | 4,81E-106 | 0,0006102 | 0,0028484 | 0,0014725 | 0,0014725 | TRUE | 0,0111887 | No |
| orf19.6020 | 1226 | 1367 | 1357 | 736 | 1396 | 990 | 1228 | 1147 | 1169 | 1445 | 1341 | 1153 | 1119 | 1126 | 1043 | 933 | 1165,92 | 1277 | 1055 | 0,825 | -0,278 | -3,209 | 0,0013299 | 0,0017771 | 0,0085468 | 0,0034995 | 0,006353 | 0,006353 | TRUE | 0,0942077 | No |
| orf19.6021 | 283 | 247 | 251 | 169 | 768 | 465 | 730 | 798 | 270 | 261 | 248 | 265 | 615 | 529 | 620 | 649 | 432,05 | 261 | 603 | 2,319 | 1,214 | 13,879 | 8,50E-44 | 2,86E-43 | 0,0031236 | 0,0058915 | 0,0044662 | 0,0044662 | TRUE | 0,0753409 | No |
| orf19.6022 | 96 | 83 | 110 | 70 | 52 | 35 | 43 | 51 | 92 | 88 | 109 | 110 | 42 | 40 | 37 | 41 | 69,63 | 100 | 40 | 0,403 | -1,31 | -7,309 | 2,69E-13 | 5,19E-13 | 1,00E-08 | 0,0256711 | 0,0140962 | 0,0140962 | TRUE | 0,058274 | No |
| orf19.6023 | 1500 | 1423 | 1564 | 922 | 3009 | 2222 | 2764 | 2810 | 1430 | 1505 | 1545 | 1444 | 2411 | 2527 | 2347 | 2285 | 1936,65 | 1481 | 2392 | 1,613 | 0,69 | 13,824 | 1,83E-43 | 6,11E-43 | 0,0010932 | 0,0029391 | 0,0018235 | 0,0018235 | TRUE | 0,0165753 | No |
| orf19.6024 | 282 | 308 | 323 | 183 | 316 | 229 | 228 | 219 | 269 | 326 | 319 | 287 | 253 | 260 | 194 | 178 | 260,69 | 300 | 221 | 0,735 | -0,445 | -3,351 | 0,0008056 | 0,0010939 | 0,0179947 | 0,0083895 | 0,0130324 | 0,0130324 | TRUE | 0,1961801 | No |
| orf19.6025 | 36 | 50 | 51 | 27 | 48 | 35 | 59 | 50 | 34 | 53 | 50 | 42 | 38 | 40 | 50 | 41 | 43,61 | 45 | 42 | 0,94 | -0,089 | -0,399 | 0,6897904 | 0,7210932 | 1,00E-08 | 0,0397415 | 0,0253219 | 0,0253219 | TRUE | 0,2326683 | No |
| orf19.6026 | 166 | 188 | 183 | 112 | 685 | 546 | 619 | 644 | 158 | 199 | 181 | 175 | 549 | 621 | 526 | 524 | 366,52 | 178 | 555 | 3,106 | 1,635 | 16,51 | 3,12E-61 | 1,34E-60 | 0,0042462 | 0,006571 | 0,0054845 | 0,0054845 | TRUE | 0,0710055 | No |
| orf19.6027 | 146 | 140 | 152 | 80 | 178 | 111 | 167 | 155 | 139 | 148 | 150 | 125 | 143 | 126 | 142 | 126 | 137,42 | 140 | 134 | 0,95 | -0,074 | -0,597 | 0,5502682 | 0,5855269 | 1,00E-08 | 0,0140393 | 0,0074051 | 0,0074051 | TRUE | 0,0421415 | No |
| orf19.6028 | 332 | 342 | 333 | 214 | 38 | 29 | 36 | 35 | 317 | 362 | 329 | 335 | 30 | 33 | 31 | 28 | 183,1 | 336 | 30 | 0,091 | -3,461 | -22,634 | 2,00E-113 | 1,50E-112 | 1,00E-08 | 0,0110585 | 0,0062677 | 0,0062677 | TRUE | 0,0325932 | No |
| orf19.6029 | 558 | 570 | 574 | 364 | 940 | 582 | 897 | 811 | 532 | 603 | 567 | 570 | 753 | 662 | 762 | 659 | 638,49 | 568 | 709 | 1,251 | 0,323 | 4,261 | 2,04E-05 | 2,98E-05 | 0,0032267 | 0,0046628 | 0,0038915 | 0,0038915 | TRUE | 0,0291727 | No |
| orf19.603 | 12142 | 10075 | 10608 | 7014 | 12799 | 8468 | 12264 | 13508 | 11576 | 10653 | 10480 | 10988 | 10255 | 9629 | 10412 | 10983 | 10622,04 | 10924 | 10320 | 0,945 | -0,082 | -1,637 | 0,101678 | 0,1186336 | 0,0023079 | 0,0022459 | 0,0023035 | 0,0023035 | TRUE | 0,0245963 | No |
| orf19.6030 | 2492 | 2098 | 2276 | 1358 | 1807 | 1030 | 1769 | 1818 | 2376 | 2218 | 2248 | 2127 | 1448 | 1171 | 1502 | 1478 | 1821,16 | 2242 | 1400 | 0,625 | -0,679 | -8,953 | 3,45E-19 | 7,61E-19 | 0,0063263 | 0,0029929 | 0,0049458 | 0,0049458 | TRUE | 0,1404106 | No |
| orf19.6031 | 5005 | 4838 | 5076 | 3124 | 2917 | 2134 | 2806 | 3074 | 4772 | 5116 | 5015 | 4894 | 2337 | 2427 | 2382 | 2499 | 3680,18 | 4949 | 2411 | 0,487 | -1,038 | -25,564 | 3,80E-144 | 3,69E-143 | 0,0006327 | 0,0025374 | 0,0012826 | 0,0012826 | TRUE | 0,0076594 | No |
| orf19.6032 | 4484 | 4487 | 4773 | 3010 | 10127 | 8079 | 9562 | 9617 | 4275 | 4744 | 4715 | 4715 | 8114 | 9187 | 8118 | 7820 | 6461,09 | 4612 | 8310 | 1,801 | 0,849 | 14,496 | 1,28E-47 | 4,56E-47 | 0,0035858 | 0,0023454 | 0,00312 | 0,00312 | TRUE | 0,0611539 | No |
| orf19.6033 | 1539 | 1396 | 1539 | 1004 | 3497 | 2670 | 3232 | 3581 | 1467 | 1476 | 1520 | 1573 | 2802 | 3036 | 2744 | 2912 | 2191,29 | 1509 | 2874 | 1,905 | 0,93 | 19,064 | 5,06E-81 | 2,79E-80 | 0,0010772 | 0,0028406 | 0,0017578 | 0,0017578 | TRUE | 0,0169327 | No |
| orf19.6034 | 2073 | 1876 | 2014 | 1274 | 3548 | 2663 | 3324 | 3657 | 1976 | 1984 | 1990 | 1996 | 2843 | 3028 | 2822 | 2973 | 2451,5 | 1986 | 2916 | 1,468 | 0,554 | 13,432 | 3,94E-41 | 1,27E-40 | 0,0003526 | 0,002761 | 0,001195 | 0,001195 | TRUE | 0,0076093 | No |
| orf19.6035 | 971 | 933 | 919 | 508 | 2357 | 1534 | 2095 | 1939 | 926 | 987 | 908 | 796 | 1889 | 1744 | 1779 | 1577 | 1325,51 | 904 | 1747 | 1,927 | 0,946 | 12,614 | 1,77E-36 | 5,30E-36 | 0,0054925 | 0,00333 | 0,0045293 | 0,0045293 | TRUE | 0,0697652 | No |
| orf19.6036 | 177 | 213 | 215 | 126 | 154 | 123 | 161 | 134 | 169 | 225 | 212 | 197 | 123 | 140 | 137 | 109 | 164,08 | 201 | 127 | 0,631 | -0,664 | -5,03 | 4,91E-07 | 7,70E-07 | 0,0085419 | 0,0120979 | 0,0104388 | 0,0104388 | TRUE | 0,1390656 | No |
| orf19.6037 | 930 | 796 | 900 | 624 | 1584 | 1169 | 1415 | 1600 | 887 | 842 | 889 | 978 | 1269 | 1329 | 1201 | 1301 | 1086,97 | 899 | 1275 | 1,422 | 0,508 | 8,451 | 2,89E-17 | 6,10E-17 | 0,0016681 | 0,0036018 | 0,0024974 | 0,0024974 | TRUE | 0,0361953 | No |
| orf19.6038 | 518 | 395 | 434 | 301 | 838 | 603 | 846 | 891 | 494 | 418 | 429 | 472 | 671 | 686 | 718 | 724 | 576,46 | 453 | 700 | 1,548 | 0,63 | 8,586 | 8,97E-18 | 1,92E-17 | 0,0013667 | 0,0049395 | 0,0032985 | 0,0032985 | TRUE | 0,0494842 | No |
| orf19.6039 | 1325 | 1182 | 1298 | 851 | 1261 | 953 | 1172 | 1408 | 1263 | 1250 | 1282 | 1333 | 1010 | 1084 | 995 | 1145 | 1170,3 | 1282 | 1058 | 0,826 | -0,275 | -4,655 | 3,23E-06 | 4,91E-06 | 0,0017465 | 0,0034943 | 0,0025004 | 0,0025004 | TRUE | 0,0381425 | No |
| orf19.604 | 32 | 33 | 36 | 36 | 25 | 18 | 14 | 25 | 31 | 35 | 36 | 56 | 20 | 20 | 12 | 20 | 28,76 | 40 | 18 | 0,47 | -1,09 | -3,636 | 0,0002774 | 0,000385 | 0,0301885 | 0,0591823 | 0,0473096 | 0,0473096 | TRUE | 0,4691958 | No |
| orf19.6040 | 5253 | 4686 | 5237 | 3037 | 4328 | 2688 | 4269 | 4241 | 5008 | 4955 | 5174 | 4758 | 3468 | 3056 | 3624 | 3448 | 4186,43 | 4974 | 3399 | 0,684 | -0,549 | -9,896 | 4,33E-23 | 1,03E-22 | 0,0027923 | 0,0024835 | 0,0027069 | 0,0027069 | TRUE | 0,0546123 | No |
| orf19.6041 | 11197 | 10288 | 10978 | 7332 | 13640 | 9979 | 12563 | 13795 | 10675 | 10878 | 10845 | 11486 | 10929 | 11347 | 10666 | 11217 | 11005,44 | 10971 | 11040 | 1,007 | 0,009 | 0,256 | 0,7980189 | 0,8203897 | 0,0006907 | 0,0022405 | 0,0012221 | 0,0012221 | TRUE | 0,0118588 | No |
| orf19.6043 | 525 | 426 | 481 | 326 | 686 | 522 | 697 | 743 | 501 | 450 | 475 | 511 | 550 | 594 | 592 | 604 | 534,5 | 484 | 585 | 1,21 | 0,275 | 3,895 | 9,81E-05 | 0,0001389 | 1,00E-08 | 0,0051632 | 0,0028522 | 0,0028522 | TRUE | 0,0251882 | No |
| orf19.6044 | 2775 | 2381 | 2564 | 1874 | 2855 | 2084 | 2698 | 2980 | 2646 | 2518 | 2533 | 2936 | 2288 | 2370 | 2291 | 2423 | 2500,36 | 2658 | 2343 | 0,883 | -0,18 | -3,25 | 0,001154 | 0,0015486 | 0,002514 | 0,0027479 | 0,0025356 | 0,0025356 | TRUE | 0,0567691 | No |
| orf19.6045 | 5504 | 4507 | 5259 | 3024 | 3396 | 2173 | 3463 | 3584 | 5247 | 4766 | 5195 | 4737 | 2721 | 2471 | 2940 | 2914 | 3873,99 | 4986 | 2762 | 0,554 | -0,852 | -13,51 | 1,36E-41 | 4,43E-41 | 0,0042047 | 0,0025151 | 0,0035461 | 0,0035461 | TRUE | 0,0594572 | No |
| orf19.6046 | 984 | 800 | 816 | 565 | 870 | 588 | 857 | 884 | 938 | 846 | 806 | 885 | 697 | 669 | 728 | 719 | 785,92 | 869 | 703 | 0,81 | -0,303 | -4,635 | 3,56E-06 | 5,40E-06 | 0,0016846 | 0,0041804 | 0,0028303 | 0,0028303 | TRUE | 0,0377636 | No |
| orf19.6047 | 8705 | 7055 | 8010 | 5150 | 18704 | 12461 | 17968 | 19479 | 8299 | 7460 | 7913 | 8068 | 14987 | 14169 | 15255 | 15838 | 11498,65 | 7935 | 15062 | 1,899 | 0,925 | 19,446 | 3,13E-84 | 1,77E-83 | 0,0019283 | 0,0022341 | 0,0020723 | 0,0020723 | TRUE | 0,0199787 | No |
| orf19.6048 | 1440 | 1248 | 1314 | 871 | 1275 | 853 | 1177 | 1222 | 1373 | 1320 | 1298 | 1364 | 1022 | 970 | 999 | 994 | 1167,44 | 1339 | 996 | 0,745 | -0,424 | -8,687 | 3,72E-18 | 8,01E-18 | 1,00E-08 | 0,0034977 | 0,001425 | 0,001425 | TRUE | 0,0052031 | No |
| orf19.6049 | 2375 | 1921 | 2124 | 1461 | 1779 | 1227 | 1750 | 1800 | 2264 | 2031 | 2098 | 2289 | 1425 | 1395 | 1486 | 1464 | 1806,57 | 2170 | 1442 | 0,666 | -0,587 | -11,086 | 1,46E-28 | 3,85E-28 | 0,0015284 | 0,0030001 | 0,002125 | 0,002125 | TRUE | 0,0224458 | No |
| orf19.605 | 85 | 79 | 85 | 76 | 193 | 190 | 203 | 227 | 81 | 84 | 84 | 119 | 155 | 216 | 172 | 185 | 136,9 | 92 | 182 | 2,002 | 1,001 | 6,41 | 1,46E-10 | 2,60E-10 | 0,015747 | 0,0140846 | 0,0148018 | 0,0148018 | TRUE | 0,2956802 | No |

| | | | | | | | | | | | | | | | | | | | | | | | | | | | | | | | |
|---|---|---|---|---|---|---|---|---|---|---|---|---|---|---|---|---|---|---|---|---|---|---|---|---|---|---|---|---|---|---|---|
| orf19.6052 | 3722 | 3277 | 3596 | 2358 | 3361 | 2454 | 3294 | 3437 | 3549 | 3465 | 3553 | 3694 | 2693 | 2790 | 2797 | 2795 | 3166,83 | 3565 | 2769 | 0,777 | -0,364 | -9,723 | 2,42E-22 | 5,68E-22 | 0,0001529 | 0,0026097 | 0,0010241 | 0,0010241 | TRUE | 0,0065709 | No |
| orf19.6053 | 2990 | 2660 | 3051 | 1801 | 4606 | 3128 | 4366 | 4706 | 2851 | 2813 | 3014 | 2821 | 3691 | 3557 | 3707 | 3826 | 3284,92 | 2875 | 3695 | 1,285 | 0,362 | 8,7 | 3,33E-18 | 7,19E-18 | 0,0005722 | 0,0025911 | 0,0013473 | 0,0013473 | TRUE | 0,0132639 | No |
| orf19.6054 | 10249 | 8847 | 9859 | 6110 | 7456 | 5193 | 7036 | 7635 | 9771 | 9355 | 9740 | 9572 | 5974 | 5905 | 5974 | 6208 | 7812,25 | 9610 | 6015 | 0,626 | -0,676 | -20,751 | 1,19E-95 | 7,46E-95 | 0,0002605 | 0,0023015 | 0,0008851 | 0,0008851 | TRUE | 0,0057302 | No |
| orf19.6055 | 2592 | 2210 | 2410 | 1590 | 1327 | 968 | 1194 | 1464 | 2471 | 2337 | 2381 | 2491 | 1063 | 1101 | 1014 | 1190 | 1755,97 | 2420 | 1092 | 0,451 | -1,147 | -20,26 | 2,91E-91 | 1,76E-90 | 0,0019072 | 0,0030263 | 0,0024376 | 0,0024376 | TRUE | 0,0460177 | No |
| orf19.6056 | 2443 | 2053 | 2243 | 1248 | 2397 | 1486 | 2231 | 2391 | 2329 | 2171 | 2216 | 1955 | 1921 | 1690 | 1894 | 1944 | 2014,93 | 2168 | 1862 | 0,858 | -0,22 | -3,397 | 0,0006816 | 0,0009301 | 0,0039108 | 0,0029061 | 0,0035344 | 0,0035344 | TRUE | 0,0500056 | No |
| orf19.6057 | 876 | 638 | 783 | 499 | 2262 | 1561 | 2329 | 2303 | 835 | 675 | 774 | 782 | 1812 | 1775 | 1977 | 1873 | 1312,79 | 766 | 1859 | 2,427 | 1,279 | 18,667 | 9,26E-78 | 4,90E-77 | 0,0036271 | 0,003342 | 0,0035374 | 0,0035374 | TRUE | 0,079294 | No |
| orf19.6058 | 1234 | 956 | 1007 | 591 | 1757 | 1056 | 1635 | 1788 | 1176 | 1011 | 995 | 926 | 1408 | 1201 | 1388 | 1454 | 1194,81 | 1027 | 1363 | 1,325 | 0,406 | 4,926 | 8,40E-07 | 1,30E-06 | 0,0075546 | 0,0034655 | 0,00566 | 0,00566 | TRUE | 0,1160848 | No |
| orf19.6059 | 7134 | 5375 | 5420 | 4169 | 4925 | 3558 | 4379 | 5187 | 6801 | 5683 | 5354 | 6531 | 3946 | 4046 | 3718 | 4218 | 5037,2 | 6092 | 3982 | 0,654 | -0,613 | -8,097 | 5,62E-16 | 1,16E-15 | 0,007414 | 0,0024172 | 0,0053063 | 0,0053063 | TRUE | 0,0815428 | No |
| orf19.606 | 5343 | 4866 | 5310 | 3504 | 3537 | 2720 | 3365 | 3695 | 5094 | 5145 | 5246 | 5489 | 2834 | 3102 | 2857 | 3004 | 4096,43 | 5244 | 2949 | 0,562 | -0,83 | -18,721 | 3,31E-78 | 1,77E-77 | 0,0011297 | 0,0024921 | 0,001629 | 0,001629 | TRUE | 0,014157 | No |
| orf19.6060 | 2821 | 2402 | 2661 | 1613 | 4572 | 3074 | 4322 | 4596 | 2690 | 2540 | 2629 | 2527 | 3663 | 3495 | 3669 | 3737 | 3118,77 | 2596 | 3641 | 1,402 | 0,487 | 11,987 | 4,15E-33 | 1,18E-32 | 0,000398 | 0,0026177 | 0,0012486 | 0,0012486 | TRUE | 0,0084661 | No |
| orf19.6061 | 1810 | 1587 | 1735 | 1030 | 2111 | 1499 | 1958 | 2178 | 1726 | 1678 | 1714 | 1614 | 1691 | 1704 | 1662 | 1771 | 1695,06 | 1683 | 1707 | 1,013 | 0,019 | 0,412 | 0,6805721 | 0,7121787 | 0,0001941 | 0,0030599 | 0,0013575 | 0,0013575 | TRUE | 0,0081625 | No |
| orf19.6062 | 5416 | 4374 | 4552 | 2898 | 2817 | 1759 | 2729 | 2827 | 5164 | 4625 | 4497 | 4540 | 2257 | 2000 | 2317 | 2299 | 3462,27 | 4706 | 2218 | 0,472 | -1,084 | -17,399 | 8,40E-68 | 3,92E-67 | 0,0039304 | 0,0025655 | 0,0034099 | 0,0034099 | TRUE | 0,052691 | No |
| orf19.6062.3 | 1198 | 929 | 1033 | 725 | 2949 | 2026 | 2772 | 3125 | 1142 | 982 | 1021 | 1136 | 2363 | 2304 | 2353 | 2541 | 1730,22 | 1070 | 2390 | 2,238 | 1,162 | 18,931 | 6,31E-80 | 3,44E-79 | 0,0029027 | 0,0030402 | 0,0029047 | 0,0029047 | TRUE | 0,0363782 | No |
| orf19.6063 | 7959 | 7195 | 7891 | 4519 | 8506 | 5796 | 7963 | 8666 | 7588 | 7608 | 7796 | 7079 | 6816 | 6591 | 6761 | 7046 | 7160,46 | 7518 | 6804 | 0,905 | -0,145 | -3,541 | 0,0003986 | 0,0005488 | 0,0010291 | 0,0023206 | 0,0014627 | 0,0014627 | TRUE | 0,0181307 | No |
| orf19.6064 | 1716 | 1499 | 1623 | 1091 | 2572 | 1718 | 2422 | 2665 | 1636 | 1585 | 1603 | 1709 | 2061 | 1954 | 2056 | 2167 | 1846,38 | 1633 | 2060 | 1,264 | 0,338 | 6,944 | 3,82E-12 | 7,10E-12 | 0,0007245 | 0,0029805 | 0,001711 | 0,001711 | TRUE | 0,0149116 | No |
| orf19.6065 | 1117 | 1020 | 1061 | 615 | 3974 | 2728 | 3730 | 3943 | 1065 | 1079 | 1048 | 963 | 3184 | 3102 | 3167 | 3206 | 2101,76 | 1039 | 3165 | 3,037 | 1,603 | 35,153 | 1,04E-270 | 1,97E-269 | 1,00E-08 | 0,0028725 | 0,0013186 | 0,0013186 | TRUE | 0,0236441 | No |
| orf19.6066 | 1586 | 1433 | 1488 | 980 | 3713 | 2881 | 3487 | 3851 | 1512 | 1515 | 1470 | 1535 | 2975 | 3276 | 2960 | 3131 | 2296,91 | 1508 | 3086 | 2,046 | 1,033 | 21,428 | 7,37E-102 | 4,89E-101 | 0,0010756 | 0,0028061 | 0,001713 | 0,001713 | TRUE | 0,0201632 | No |
| orf19.6068 | 9829 | 8271 | 8910 | 5979 | 16134 | 11590 | 15465 | 17232 | 9371 | 8746 | 8802 | 9366 | 12928 | 13179 | 13130 | 14011 | 11191,58 | 9071 | 13312 | 1,468 | 0,554 | 13,261 | 3,88E-40 | 1,23E-39 | 0,0012429 | 0,002238 | 0,0015789 | 0,0015789 | TRUE | 0,0153882 | No |
| orf19.607 | 4544 | 3904 | 4436 | 2690 | 4937 | 3138 | 4860 | 5079 | 4332 | 4128 | 4382 | 4214 | 3956 | 3568 | 4126 | 4130 | 4104,56 | 4264 | 3945 | 0,926 | -0,112 | -2,111 | 0,0347477 | 0,0423544 | 0,0022767 | 0,0024913 | 0,0024353 | 0,0024353 | TRUE | 0,0491107 | No |
| orf19.6070 | 9478 | 8431 | 9024 | 6144 | 11644 | 8393 | 10903 | 12156 | 9036 | 8915 | 8915 | 9625 | 9330 | 9544 | 9257 | 9884 | 9313,11 | 9123 | 9504 | 1,042 | 0,06 | 1,515 | 0,1297488 | 0,1497426 | 0,0009881 | 0,0022676 | 0,0013861 | 0,0013861 | TRUE | 0,0162693 | No |
| orf19.6071 | 2626 | 2530 | 2726 | 1726 | 2784 | 2146 | 2500 | 2898 | 2504 | 2675 | 2693 | 2704 | 2231 | 2440 | 2122 | 2356 | 2465,68 | 2644 | 2287 | 0,865 | -0,209 | -3,891 | 9,99E-05 | 0,0001415 | 0,0019559 | 0,0027572 | 0,0023689 | 0,0023689 | TRUE | 0,0284408 | No |
| orf19.6072 | 370 | 333 | 346 | 232 | 1242 | 825 | 1143 | 1218 | 353 | 352 | 342 | 363 | 995 | 938 | 970 | 990 | 663,02 | 352 | 973 | 2,77 | 1,47 | 22,77 | 9,03E-115 | 6,82E-114 | 1,00E-08 | 0,0045677 | 0,0019658 | 0,0019658 | TRUE | 0,0065803 | No |
| orf19.6072.1 | 852 | 823 | 833 | 627 | 618 | 466 | 564 | 543 | 812 | 870 | 823 | 982 | 495 | 530 | 479 | 442 | 679,13 | 872 | 486 | 0,559 | -0,84 | -10,397 | 2,55E-25 | 6,34E-25 | 0,0046855 | 0,0045089 | 0,0047039 | 0,0047039 | TRUE | 0,0792927 | No |
| orf19.6073 | 25710 | 21227 | 23025 | 14898 | 21112 | 14676 | 20147 | 21842 | 24512 | 22445 | 22747 | 23339 | 16916 | 16688 | 17105 | 17760 | 20188,79 | 23261 | 17117 | 0,736 | -0,442 | -11,342 | 8,16E-30 | 2,19E-29 | 0,0010654 | 0,0021726 | 0,0014109 | 0,0014109 | TRUE | 0,0161978 | No |
| orf19.6074 | 5195 | 4332 | 4447 | 2619 | 6604 | 4062 | 6547 | 6932 | 4953 | 4581 | 4393 | 4103 | 5292 | 4619 | 5558 | 5636 | 4891,83 | 4508 | 5276 | 1,17 | 0,227 | 3,105 | 0,0019037 | 0,0025254 | 0,006689 | 0,0024269 | 0,0049216 | 0,0049216 | TRUE | 0,0849689 | No |
| orf19.6075 | 1355 | 1126 | 1171 | 928 | 2042 | 1446 | 1906 | 2030 | 1292 | 1191 | 1157 | 1454 | 1636 | 1644 | 1618 | 1651 | 1455,28 | 1274 | 1637 | 1,29 | 0,367 | 5,451 | 5,00E-08 | 8,13E-08 | 0,0040202 | 0,0032195 | 0,003643 | 0,003643 | TRUE | 0,101113 | No |
| orf19.6076 | 569 | 469 | 532 | 399 | 613 | 453 | 655 | 672 | 542 | 496 | 526 | 625 | 491 | 515 | 556 | 546 | 537,22 | 547 | 527 | 0,968 | -0,046 | -0,566 | 0,5713959 | 0,6057613 | 0,0042149 | 0,0051476 | 0,004598 | 0,004598 | TRUE | 0,0938815 | No |
| orf19.6077 | 2331 | 1919 | 2112 | 1586 | 3322 | 2453 | 3228 | 3499 | 2222 | 2029 | 2086 | 2485 | 2662 | 2789 | 2741 | 2845 | 2482,39 | 2206 | 2759 | 1,253 | 0,326 | 5,216 | 1,83E-07 | 2,92E-07 | 0,0040816 | 0,0027527 | 0,0033264 | 0,0033264 | TRUE | 0,0832032 | No |
| orf19.6078 | 3114 | 2668 | 2897 | 1646 | 1377 | 883 | 1252 | 1452 | 2969 | 2821 | 2862 | 2579 | 1103 | 1004 | 1063 | 1181 | 1947,68 | 2808 | 1088 | 0,388 | -1,368 | -21,888 | 3,41E-106 | 2,35E-105 | 0,0031977 | 0,0029343 | 0,0031385 | 0,0031385 | TRUE | 0,0395697 | No |
| orf19.6078.1 | 0 | 4 | 3 | 3 | 0 | 0 | 0 | 1 | 0 | 4 | 3 | 5 | 0 | 0 | 0 | 1 | 1,59 | 3 | 0 | 0,093 | -3,425 | -2,285 | 0,0223134 | 0,0275934 | 1,00E-08 | 1,0358756 | 0,9648712 | 0,9648712 | TRUE | 0,6505023 | No |
| orf19.6079 | 468 | 366 | 354 | 245 | 322 | 219 | 348 | 385 | 446 | 387 | 350 | 384 | 258 | 249 | 295 | 313 | 335,28 | 392 | 279 | 0,713 | -0,488 | -4,627 | 3,70E-06 | 5,60E-06 | 0,0083136 | 0,0069884 | 0,0076478 | 0,0076478 | TRUE | 0,1072528 | No |
| orf19.6080 | 3012 | 2491 | 2717 | 1655 | 2980 | 1888 | 2872 | 2999 | 2872 | 2634 | 2684 | 2593 | 2388 | 2147 | 2438 | 2438 | 2524,22 | 2696 | 2353 | 0,873 | -0,196 | -3,553 | 0,0003814 | 0,0005259 | 0,0022687 | 0,0027417 | 0,0025221 | 0,0025221 | TRUE | 0,0405697 | No |
| orf19.6081 | 13887 | 11419 | 12710 | 8623 | 23186 | 16449 | 22457 | 24839 | 13240 | 12074 | 12556 | 13508 | 18578 | 18704 | 19066 | 20196 | 15990,4 | 12844 | 19136 | 1,49 | 0,576 | 12,329 | 6,36E-35 | 1,86E-34 | 0,0019403 | 0,002194 | 0,0020262 | 0,0020262 | TRUE | 0,0199093 | No |
| orf19.6082 | 4258 | 3623 | 3947 | 2541 | 6837 | 4785 | 6713 | 7059 | 4060 | 3831 | 3899 | 3981 | 5478 | 5441 | 5699 | 5740 | 4766,06 | 3943 | 5590 | 1,418 | 0,504 | 13,486 | 1,88E-41 | 6,09E-41 | 0,000396 | 0,0024358 | 0,0011189 | 0,0011189 | TRUE | 0,0051692 | No |
| orf19.6083 | 613 | 539 | 564 | 357 | 734 | 483 | 714 | 815 | 584 | 570 | 557 | 559 | 588 | 549 | 606 | 663 | 584,62 | 568 | 602 | 1,061 | 0,086 | 1,185 | 0,235973 | 0,2652826 | 0,0018633 | 0,0048998 | 0,0033058 | 0,0033058 | TRUE | 0,0553747 | No |
| orf19.6084 | 11794 | 10222 | 10864 | 7148 | 3780 | 2658 | 3475 | 3712 | 11244 | 10808 | 10733 | 11198 | 3029 | 3022 | 2950 | 3018 | 7000,35 | 10996 | 3005 | 0,273 | -1,871 | -56,122 | 0 | 0 | 0,0002884 | 0,0023258 | 0,0008678 | 0,0008678 | TRUE | 0,0031986 | No |
| orf19.6085 | 18708 | 15845 | 16961 | 10695 | 53749 | 36911 | 52373 | 55067 | 17836 | 16754 | 16756 | 16754 | 43068 | 41971 | 44464 | 44775 | 30297,26 | 17025 | 43570 | 2,559 | 1,356 | 36,71 | 5,02E-295 | 1,10E-293 | 0,0008941 | 0,0021455 | 0,0012664 | 0,0012664 | TRUE | 0,0126884 | No |
| orf19.6086 | 486 | 386 | 406 | 278 | 3199 | 2181 | 2987 | 3413 | 463 | 408 | 401 | 436 | 2563 | 2480 | 2536 | 2775 | 1507,8 | 427 | 2588 | 6,067 | 2,601 | 39,722 | 0 | 0 | 0,0021574 | 0,0031802 | 0,0026383 | 0,0026383 | TRUE | 0,0437416 | No |
| orf19.609 | 48 | 45 | 45 | 37 | 27 | 21 | 19 | 25 | 46 | 48 | 44 | 58 | 22 | 24 | 16 | 20 | 34,72 | 49 | 20 | 0,42 | -1,251 | -5,019 | 5,20E-07 | 8,14E-07 | 1,00E-08 | 0,0493855 | 0,0263832 | 0,0263832 | TRUE | 0,1226019 | No |
| orf19.6090 | 31343 | 25417 | 27699 | 18183 | 78035 | 51578 | 74836 | 80263 | 29882 | 26875 | 27364 | 28485 | 62527 | 58649 | 63535 | 65261 | 45322,37 | 28152 | 62493 | 2,22 | 1,151 | 24,361 | 4,47E-131 | 3,93E-130 | 0,002096 | 0,0021275 | 0,002116 | 0,002116 | TRUE | 0,0210912 | No |
| orf19.6091 | 4651 | 3889 | 4425 | 2969 | 2996 | 1925 | 2766 | 3027 | 4434 | 4112 | 4371 | 4651 | 2401 | 2189 | 2348 | 2461 | 3371 | 4392 | 2350 | 0,536 | -0,9 | -17,197 | 2,79E-66 | 1,27E-65 | 0,0020746 | 0,0025783 | 0,0023133 | 0,0023133 | TRUE | 0,0247704 | No |
| orf19.6092 | 1430 | 1173 | 1358 | 933 | 3972 | 2697 | 3609 | 4174 | 1363 | 1240 | 1342 | 1462 | 3183 | 3067 | 3064 | 3394 | 2264,26 | 1352 | 3177 | 2,356 | 1,236 | 21,195 | 1,07E-99 | 6,99E-99 | 0,0027411 | 0,0028164 | 0,0027089 | 0,0027089 | TRUE | 0,0362927 | No |
| orf19.6094 | 1082 | 863 | 927 | 545 | 2699 | 1650 | 2720 | 2769 | 1032 | 913 | 916 | 854 | 2163 | 1876 | 2309 | 2251 | 1539,15 | 929 | 2150 | 2,313 | 1,21 | 15,553 | 1,51E-54 | 5,93E-54 | 0,0061813 | 0,003158 | 0,0049945 | 0,0049945 | TRUE | 0,0868894 | No |
| orf19.6096 | 768 | 604 | 663 | 410 | 2659 | 1643 | 2559 | 2505 | 732 | 639 | 655 | 642 | 2131 | 1868 | 2173 | 2037 | 1359,54 | 667 | 2052 | 3,073 | 1,62 | 23,78 | 5,33E-125 | 4,47E-124 | 0,0034787 | 0,003299 | 0,0033975 | 0,0033975 | TRUE | 0,0534581 | No |
| orf19.6099 | 5230 | 3867 | 4519 | 3373 | 10901 | 7367 | 9618 | 11061 | 4986 | 4089 | 4464 | 5284 | 8735 | 8377 | 8166 | 8994 | 6636,79 | 4706 | 8568 | 1,822 | 0,865 | 11,814 | 3,29E-32 | 9,21E-32 | 0,0071166 | 0,0023387 | 0,004982 | 0,004982 | TRUE | 0,0947366 | No |
| orf19.610 | 7007 | 5678 | 6610 | 4667 | 3179 | 2158 | 3017 | 3250 | 6680 | 6004 | 6530 | 7311 | 2547 | 2454 | 2561 | 2643 | 4591,3 | 6631 | 2551 | 0,385 | -1,377 | -23,201 | 4,42E-119 | 3,48E-118 | 0,0033897 | 0,0024489 | 0,0031217 | 0,0031217 | TRUE | 0,0569141 | No |
| orf19.6100 | 689 | 486 | 517 | 380 | 1951 | 1277 | 1881 | 2017 | 657 | 514 | 511 | 595 | 1563 | 1452 | 1597 | 1640 | 1066,14 | 569 | 1563 | 2,75 | 1,459 | 17,692 | 4,86E-70 | 2,33E-69 | 0,0073623 | 0,0036313 | 0,0052597 | 0,0052597 | TRUE | 0,1375684 | No |
| orf19.6102 | 1897 | 1533 | 1858 | 1059 | 2005 | 1156 | 1857 | 2173 | 1809 | 1621 | 1836 | 1659 | 1607 | 1314 | 1577 | 1767 | 1648,56 | 1731 | 1566 | 0,905 | -0,144 | -1,721 | 0,0853257 | 0,1002714 | 0,0082535 | 0,0030872 | 0,0060754 | 0,0060754 | TRUE | 0,1374387 | No |
| orf19.6103 | 353 | 300 | 332 | 223 | 332 | 183 | 297 | 286 | 337 | 317 | 328 | 349 | 266 | 208 | 252 | 233 | 286,24 | 333 | 240 | 0,725 | -0,464 | -4,852 | 1,22E-06 | 1,88E-06 | 0,0019248 | 0,0078275 | 0,0052445 | 0,0052445 | TRUE | 0,0769718 | No |
| orf19.6105 | 1705 | 1139 | 1382 | 1107 | 2711 | 1689 | 2504 | 2748 | 1626 | 1204 | 1365 | 1734 | 2172 | 1921 | 2126 | 2234 | 1797,8 | 1482 | 2113 | 1,428 | 0,514 | 5,169 | 2,35E-07 | 3,73E-07 | 0,0145708 | 0,0030046 | 0,0089086 | 0,0089086 | TRUE | 0,1914154 | No |
| orf19.6109 | 6045 | 4454 | 5209 | 3291 | 8787 | 5557 | 8015 | 8926 | 5763 | 4710 | 5146 | 5156 | 7041 | 6319 | 6805 | 7258 | 6024,54 | 5194 | 6856 | 1,32 | 0,401 | 6,144 | 8,06E-10 | 1,39E-09 | 0,0049762 | 0,0023638 | 0,0039106 | 0,0039106 | TRUE | 0,0672745 | No |
| orf19.6110 | 10 | 5 | 5 | 4 | 16 | 6 | 19 | 22 | 10 | 5 | 5 | 6 | 13 | 7 | 16 | 18 | 9,96 | 6 | 14 | 2,077 | 1,055 | 2,156 | 0,0310709 | 0,0380252 | 1,00E-08 | 0,1669236 | 0,1122629 | 0,1122629 | TRUE | 0,3826903 | No |
| orf19.6112 | 322 | 290 | 304 | 180 | 769 | 480 | 764 | 740 | 307 | 307 | 300 | 282 | 616 | 546 | 649 | 602 | 451,03 | 299 | 603 | 2,015 | 1,011 | 12,525 | 5,46E-36 | 1,62E-35 | 1,00E-08 | 0,0057316 | 0,003657 | 0,003657 | TRUE | 0,044063 | No |
| orf19.6113 | 90 | 64 | 77 | 37 | 239 | 127 | 238 | 260 | 86 | 68 | 76 | 58 | 192 | 144 | 202 | 211 | 129,61 | 72 | 187 | 2,586 | 1,371 | 8,352 | 6,69E-17 | 1,40E-16 | 0,0172077 | 0,0147592 | 0,0158302 | 0,0158302 | TRUE | 0,2364007 | No |
| orf19.6114 | 2421 | 1762 | 2145 | 1200 | 4469 | 2482 | 4416 | 4638 | 2308 | 1863 | 2119 | 1880 | 3581 | 2822 | 3749 | 3771 | 2761,7 | 2042 | 3481 | 1,704 | 0,769 | 8,001 | 1,24E-15 | 2,52E-15 | 0,0131316 | 0,0026858 | 0,0084619 | 0,0084619 | TRUE | 0,1955387 | No |
| orf19.6115 | 4 | 0 | 2 | 0 | 5 | 2 | 3 | 2 | 4 | 0 | 2 | 0 | 4 | 2 | 3 | 2 | 2,03 | 2 | 3 | 1,677 | 0,746 | 0,707 | 0,4795249 | 0,5158207 | 0,0359689 | 0,8107568 | 0,5446743 | 0,5446743 | TRUE | 0,8159706 | No |
| orf19.6116 | 2015 | 1700 | 1939 | 1119 | 305 | 171 | 279 | 300 | 1921 | 1798 | 1916 | 1753 | 244 | 194 | 237 | 244 | 1038,35 | 1847 | 230 | 0,125 | -2,997 | -40,609 | 0 | 0 | 0,0020642 | 0,0036725 | 0,0030087 | 0,0030087 | TRUE | 0,100584 | No |
| orf19.6117 | 8545 | 6833 | 7600 | 5667 | 3469 | 2681 | 3381 | 3520 | 8147 | 7225 | 7508 | 8878 | 2780 | 3049 | 2870 | 2862 | 5414,78 | 7940 | 2890 | 0,364 | -1,458 | -22,477 | 6,90E-112 | 5,08E-111 | 0,0047357 | 0,0023945 | 0,0038141 | 0,0038141 | TRUE | 0,0761896 | No |
| orf19.6118 | 350 | 301 | 279 | 176 | 407 | 292 | 402 | 424 | 334 | 318 | 276 | 276 | 326 | 332 | 341 | 345 | 318,44 | 301 | 336 | 1,113 | 0,154 | 1,711 | 0,0871375 | 0,1023044 | 0,0013286 | 0,0072474 | 0,0046027 | 0,0046027 | TRUE | 0,0631499 | No |
| orf19.6119 | 243 | 191 | 239 | 164 | 289 | 223 | 279 | 259 | 232 | 202 | 236 | 257 | 232 | 254 | 237 | 211 | 232,41 | 232 | 234 | 1,008 | 0,012 | 0,111 | 0,9118597 | 0,9209523 | 0,0026394 | 0,0091561 | 0,0062928 | 0,0062928 | TRUE | 0,0801489 | No |
| orf19.612 | 532 | 501 | 447 | 325 | 772 | 468 | 695 | 761 | 507 | 530 | 442 | 509 | 619 | 532 | 590 | 619 | 543,4 | 497 | 590 | 1,191 | 0,253 | 3,138 | 0,0017007 | 0,0022637 | 0,0035654 | 0,0051128 | 0,0043347 | 0,0043347 | TRUE | 0,0689955 | No |
| orf19.6121 | 8452 | 7340 | 8294 | 5299 | 2296 | 1658 | 2151 | 2358 | 8058 | 7761 | 8194 | 8301 | 1840 | 1885 | 1826 | 1917 | 4972,82 | 8078 | 1867 | 0,231 | -2,113 | -54,554 | 0 | 0 | 0,0004982 | 0,0024215 | 0,0011319 | 0,0011319 | TRUE | 0,0085252 | No |
| orf19.6124 | 1746 | 1630 | 1700 | 1077 | 756 | 546 | 701 | 757 | 1665 | 1724 | 1679 | 1687 | 606 | 621 | 595 | 616 | 1149 | 1689 | 610 | 0,361 | -1,471 | -29,531 | 1,17E-191 | 1,53E-190 | 1,00E-08 | 0,0035203 | 0,0013222 | 0,0013222 | TRUE | 0,0028488 | No |
| orf19.6126 | 5981 | 5076 | 5667 | 3410 | 17187 | 11402 | 15910 | 18413 | 5702 | 5367 | 5598 | 5342 | 13771 | 12965 | 13508 | 14972 | 9653,18 | 5502 | 13804 | 2,508 | 1,327 | 26,41 | 1,06E-153 | 1,09E-152 | 0,0022291 | 0,0022614 | 0,0022888 | 0,0022888 | TRUE | 0,0397082 | No |
| orf19.6127 | 19612 | 15662 | 17550 | 12195 | 26190 | 18176 | 24892 | 27460 | 18698 | 16561 | 17338 | 19104 | 20985 | 20668 | 21133 | 22328 | 19601,77 | 17925 | 21278 | 1,187 | 0,248 | 4,803 | 1,56E-06 | 2,39E-06 | 0,0026889 | 0,002175 | 0,0025027 | 0,0025027 | TRUE | 0,0321098 | No |
| orf19.6129 | 2108 | 1831 | 1970 | 1176 | 3021 | 1852 | 2857 | 3142 | 2010 | 1936 | 1946 | 1842 | 2421 | 2106 | 2426 | 2555 | 2155,14 | 1934 | 2377 | 1,229 | 0,298 | 4,782 | 1,74E-06 | 2,66E-06 | 0,003239 | 0,0028531 | 0,0032466 | 0,0032466 | TRUE | 0,06923 | No |
| orf19.6131 | 781 | 596 | 715 | 513 | 727 | 544 | 687 | 752 | 745 | 630 | 706 | 804 | 583 | 619 | 583 | 611 | 660,07 | 721 | 599 | 0,833 | -0,264 | -3,434 | 0,000594 | 0,0008126 | 0,0035316 | 0,0045787 | 0,0041664 | 0,0041664 | TRUE | 0,0840342 | No |
| orf19.6132 | 1463 | 1224 | 1291 | 744 | 2211 | 1484 | 2096 | 2253 | 1395 | 1294 | 1275 | 1166 | 1772 | 1687 | 1779 | 1832 | 1525,05 | 1282 | 1768 | 1,375 | 0,46 | 7,833 | 4,75E-15 | 9,58E-15 | 0,0023588 | 0,0031679 | 0,0026156 | 0,0026156 | TRUE | 0,0415782 | No |
| orf19.6133 | 4407 | 3780 | 4345 | 2519 | 3557 | 2388 | 3479 | 3643 | 4202 | 3997 | 4292 | 3946 | 2850 | 2715 | 2954 | 2962 | 3489,79 | 4109 | 2870 | 0,698 | -0,518 | -11,226 | 3,05E-29 | 8,12E-29 | 0,001213 | 0,0025618 | 0,0017503 | 0,0017503 | TRUE | 0,0153681 | No |
| orf19.6134 | 805 | 720 | 793 | 506 | 763 | 598 | 747 | 823 | 767 | 761 | 783 | 793 | 611 | 680 | 634 | 669 | 712,45 | 776 | 648 | 0,835 | -0,26 | -4,277 | 1,89E-05 | 2,77E-05 | 1,00E-08 | 0,0043959 | 0,0021376 | 0,0021376 | TRUE | 0,0181122 | No |
| orf19.6135 | 1003 | 840 | 941 | 598 | 1524 | 1099 | 1457 | 1539 | 956 | 888 | 930 | 937 | 1221 | 1250 | 1237 | 1251 | 1083,75 | 928 | 1240 | 1,336 | 0,418 | 8,385 | 5,08E-17 | 1,07E-16 | 1,00E-08 | 0,0036063 | 0,0014257 | 0,0014257 | TRUE | 0,0102005 | No |
| orf19.6135.1 | 981 | 836 | 792 | 465 | 1226 | 720 | 1075 | 1162 | 935 | 884 | 782 | 728 | 982 | 819 | 913 | 945 | 873,58 | 832 | 915 | 1,097 | 0,134 | 1,547 | 0,1218914 | 0,1409549 | 0,0077833 | 0,0039708 | 0,0059967 | 0,0059967 | TRUE | 0,0820842 | No |
| orf19.6136 | 4031 | 3178 | 3607 | 2111 | 3163 | 2303 | 3220 | 3442 | 3843 | 3360 | 3563 | 3307 | 2534 | 2619 | 2734 | 2799 | 3094,93 | 3518 | 2672 | 0,759 | -0,398 | -6,919 | 4,54E-12 | 8,43E-12 | 0,0029181 | 0,0026218 | 0,0028572 | 0,0028572 | TRUE | 0,0472718 | No |

| | | | | | | | | | | | | | | | | | | | | | | | | | | | | | | | |
|---|---|---|---|---|---|---|---|---|---|---|---|---|---|---|---|---|---|---|---|---|---|---|---|---|---|---|---|---|---|---|---|
| orf19.6137 | 2891 | 2544 | 2778 | 1702 | 3272 | 2380 | 3099 | 3338 | 2756 | 2690 | 2744 | 2666 | 2622 | 2706 | 2631 | 2714 | 2691,26 | 2714 | 2668 | 0,982 | -0,026 | -0,711 | 0,4771883 | 0,5135669 | 1,00E-08 | 0,0027014 | 0,000923 | 0,000923 | TRUE | 0,0017561 | No |
| orf19.6138 | 44 | 37 | 67 | 27 | 48 | 41 | 47 | 42 | 42 | 39 | 66 | 42 | 38 | 47 | 40 | 34 | 43,59 | 47 | 40 | 0,829 | -0,27 | -1,139 | 0,2548113 | 0,2851545 | 0,0202626 | 0,0397612 | 0,0310974 | 0,0310974 | TRUE | 0,6020022 | No |
| orf19.6139 | 2189 | 1950 | 2103 | 1322 | 4239 | 2856 | 4211 | 4394 | 2087 | 2062 | 2078 | 2071 | 3397 | 3248 | 3575 | 3573 | 2761,17 | 2074 | 3448 | 1,663 | 0,734 | 16,459 | 7,28E-61 | 3,10E-60 | 0,0008676 | 0,0026859 | 0,0015091 | 0,0015091 | TRUE | 0,0178699 | No |
| orf19.6140 | 889 | 857 | 955 | 620 | 1989 | 1410 | 1774 | 1970 | 848 | 906 | 943 | 971 | 1594 | 1603 | 1506 | 1602 | 1246,67 | 917 | 1576 | 1,723 | 0,785 | 14,052 | 7,51E-45 | 2,56E-44 | 0,0009941 | 0,0034083 | 0,0021005 | 0,0021005 | TRUE | 0,0320557 | No |
| orf19.6141 | 867 | 764 | 847 | 549 | 2364 | 1601 | 2278 | 2410 | 827 | 808 | 837 | 860 | 1894 | 1820 | 1934 | 1960 | 1367,43 | 833 | 1902 | 2,289 | 1,195 | 23,856 | 8,74E-126 | 7,39E-125 | 1,00E-08 | 0,003292 | 0,0015036 | 0,0015036 | TRUE | 0,0094362 | No |
| orf19.6142 | 13 | 6 | 5 | 4 | 12 | 5 | 13 | 7 | 12 | 6 | 5 | 6 | 10 | 6 | 11 | 6 | 7,75 | 7 | 8 | 1,065 | 0,091 | 0,172 | 0,8635245 | 0,8814829 | 1,00E-08 | 0,2140408 | 0,1384902 | 0,1384902 | TRUE | 0,6139881 | No |
| orf19.6143 | 4771 | 3909 | 4233 | 2864 | 9121 | 6409 | 8814 | 9388 | 4549 | 4133 | 4182 | 4487 | 7308 | 7288 | 7483 | 7633 | 5882,84 | 4338 | 7428 | 1,713 | 0,777 | 18,099 | 3,25E-73 | 1,63E-72 | 0,0011719 | 0,0023704 | 0,0015793 | 0,0015793 | TRUE | 0,0138522 | No |
| orf19.6146 | 12158 | 9752 | 10461 | 7078 | 11357 | 7566 | 10846 | 11238 | 11591 | 10312 | 10334 | 11088 | 9100 | 8603 | 9208 | 9138 | 9921,8 | 10831 | 9012 | 0,832 | -0,265 | -5,531 | 3,19E-08 | 5,23E-08 | 0,0019834 | 0,0022568 | 0,0021017 | 0,0021017 | TRUE | 0,0277169 | No |
| orf19.6147 | 729 | 632 | 621 | 317 | 1623 | 813 | 1580 | 1484 | 695 | 668 | 613 | 497 | 1300 | 924 | 1341 | 1207 | 905,79 | 618 | 1193 | 1,926 | 0,946 | 7,776 | 7,46E-15 | 1,50E-14 | 0,0226853 | 0,003904 | 0,0129177 | 0,0129177 | TRUE | 0,2707542 | No |
| orf19.6148 | 352 | 341 | 403 | 245 | 359 | 240 | 347 | 363 | 336 | 361 | 398 | 384 | 288 | 273 | 295 | 295 | 328,55 | 370 | 288 | 0,781 | -0,356 | -4,126 | 3,69E-05 | 5,32E-05 | 1,00E-08 | 0,0070887 | 0,0040936 | 0,0040936 | TRUE | 0,0471585 | No |
| orf19.6151 | 2333 | 1833 | 1963 | 1225 | 2087 | 1307 | 1923 | 2073 | 2224 | 1938 | 1939 | 1919 | 1672 | 1486 | 1633 | 1686 | 1812,16 | 2005 | 1619 | 0,808 | -0,308 | -4,826 | 1,39E-06 | 2,14E-06 | 0,0035145 | 0,0029973 | 0,0033647 | 0,0033647 | TRUE | 0,0668121 | No |
| orf19.6152 | 2487 | 2149 | 2511 | 1445 | 2858 | 1916 | 2832 | 2976 | 2371 | 2272 | 2481 | 2264 | 2290 | 2179 | 2404 | 2420 | 2335,06 | 2347 | 2323 | 0,99 | -0,015 | -0,289 | 0,772523 | 0,7967463 | 0,0016467 | 0,0027944 | 0,0020993 | 0,0020993 | TRUE | 0,0203387 | No |
| orf19.6155 | 4935 | 4521 | 4708 | 2965 | 5156 | 3807 | 4880 | 5075 | 4705 | 4780 | 4651 | 4645 | 4131 | 4329 | 4143 | 4126 | 4438,88 | 4695 | 4182 | 0,89 | -0,168 | -4,996 | 5,87E-07 | 9,17E-07 | 6,20E-05 | 0,0024612 | 0,0008601 | 0,0008601 | TRUE | 0,0064302 | No |
| orf19.6156 | 1496 | 1119 | 1200 | 708 | 1677 | 1121 | 1675 | 1655 | 1426 | 1183 | 1185 | 1109 | 1344 | 1275 | 1422 | 1346 | 1286,28 | 1226 | 1347 | 1,096 | 0,133 | 1,735 | 0,0827935 | 0,0974553 | 0,0063082 | 0,0033678 | 0,004843 | 0,004843 | TRUE | 0,1471504 | No |
| orf19.6160 | 8309 | 7308 | 7834 | 5127 | 11099 | 7666 | 10276 | 11506 | 7922 | 7727 | 7739 | 8032 | 8893 | 8717 | 8724 | 9355 | 8388,75 | 7855 | 8922 | 1,136 | 0,185 | 5,12 | 3,05E-07 | 4,81E-07 | 0,000582 | 0,002287 | 0,0011251 | 0,0011251 | TRUE | 0,0131185 | No |
| orf19.6163 | 1205 | 975 | 1071 | 601 | 939 | 516 | 887 | 937 | 1149 | 1031 | 1058 | 941 | 752 | 587 | 753 | 762 | 879,17 | 1045 | 714 | 0,683 | -0,549 | -6,085 | 1,17E-09 | 2,00E-09 | 0,0087992 | 0,0039588 | 0,0066665 | 0,0066665 | TRUE | 0,1610498 | No |
| orf19.6164 | 17 | 10 | 6 | 2 | 6 | 2 | 2 | 5 | 16 | 11 | 6 | 3 | 5 | 2 | 2 | 4 | 6,09 | 9 | 3 | 0,354 | -1,496 | -2,249 | 0,0244974 | 0,0302005 | 0,1695054 | 0,2718793 | 0,2280098 | 0,2280098 | TRUE | 0,1917827 | No |
| orf19.6165 | 15448 | 12889 | 14048 | 9354 | 20784 | 14197 | 19252 | 22111 | 14728 | 13628 | 13878 | 14654 | 16654 | 16143 | 16345 | 17978 | 15501,03 | 14222 | 16780 | 1,18 | 0,239 | 5,168 | 2,36E-07 | 3,75E-07 | 0,0018771 | 0,0021972 | 0,0019865 | 0,0019865 | TRUE | 0,0283371 | No |
| orf19.6166 | 1062 | 1014 | 1037 | 624 | 1183 | 867 | 1113 | 1221 | 1013 | 1072 | 1024 | 978 | 948 | 986 | 945 | 993 | 994,77 | 1022 | 968 | 0,945 | -0,081 | -1,523 | 0,1277832 | 0,1476343 | 1,00E-08 | 0,0037418 | 0,0017148 | 0,0017148 | TRUE | 0,0128198 | No |
| orf19.6167 | 1000 | 810 | 977 | 656 | 1651 | 1237 | 1687 | 1739 | 953 | 856 | 965 | 1028 | 1323 | 1407 | 1432 | 1414 | 1172,3 | 950 | 1394 | 1,469 | 0,555 | 9,009 | 2,08E-19 | 4,62E-19 | 0,002197 | 0,0034919 | 0,0027357 | 0,0027357 | TRUE | 0,0523548 | No |
| orf19.6168 | 771 | 733 | 777 | 440 | 868 | 627 | 870 | 960 | 735 | 775 | 768 | 689 | 696 | 713 | 739 | 781 | 736,83 | 742 | 732 | 0,985 | -0,022 | -0,343 | 0,7315934 | 0,7605568 | 0,0011976 | 0,0043196 | 0,0026884 | 0,0026884 | TRUE | 0,0245776 | No |
| orf19.6169 | 450 | 443 | 490 | 339 | 1667 | 1196 | 1599 | 1667 | 429 | 468 | 484 | 531 | 1336 | 1360 | 1358 | 1355 | 915,15 | 478 | 1352 | 2,846 | 1,509 | 23,694 | 4,15E-124 | 3,44E-123 | 1,00E-08 | 0,0038854 | 0,0023908 | 0,0023908 | TRUE | 0,0563548 | No |
| orf19.6170 | 907 | 756 | 797 | 494 | 890 | 549 | 808 | 971 | 865 | 799 | 787 | 774 | 713 | 624 | 686 | 790 | 754,78 | 806 | 703 | 0,873 | -0,196 | -2,532 | 0,0113432 | 0,0143642 | 0,0042181 | 0,0042666 | 0,0044291 | 0,0044291 | TRUE | 0,0807631 | No |
| orf19.6171 | 1320 | 1222 | 1212 | 863 | 2695 | 1806 | 2678 | 2866 | 1258 | 1292 | 1197 | 1352 | 2159 | 2054 | 2274 | 2330 | 1739,6 | 1275 | 2204 | 1,734 | 0,794 | 13,764 | 4,17E-43 | 1,39E-42 | 0,0020158 | 0,0030351 | 0,002549 | 0,002549 | TRUE | 0,0246514 | No |
| orf19.6173 | 2791 | 2320 | 2548 | 2083 | 1552 | 1210 | 1494 | 1635 | 2661 | 2453 | 2517 | 3263 | 1244 | 1376 | 1268 | 1329 | 2013,95 | 2724 | 1304 | 0,479 | -1,061 | -12,572 | 3,01E-36 | 8,98E-36 | 0,0095252 | 0,0029065 | 0,0062874 | 0,0062874 | TRUE | 0,2104983 | No |
| orf19.6175 | 2810 | 2401 | 2628 | 1719 | 4134 | 2625 | 3979 | 4066 | 2679 | 2539 | 2596 | 2693 | 3312 | 2985 | 3378 | 3306 | 2936,05 | 2627 | 3245 | 1,237 | 0,307 | 6,232 | 4,61E-10 | 8,05E-10 | 0,0014045 | 0,0026505 | 0,0019773 | 0,0019773 | TRUE | 0,0342901 | No |
| orf19.6176 | 1976 | 1580 | 1684 | 1158 | 4513 | 3081 | 4224 | 4534 | 1884 | 1671 | 1664 | 1814 | 3616 | 3503 | 3586 | 3687 | 2678,06 | 1758 | 3598 | 2,048 | 1,034 | 20,597 | 2,91E-94 | 1,81E-93 | 0,0015738 | 0,0027044 | 0,001978 | 0,001978 | TRUE | 0,0305234 | No |
| orf19.6177 | 599 | 472 | 494 | 371 | 1601 | 988 | 1446 | 1538 | 571 | 499 | 488 | 581 | 1283 | 1123 | 1228 | 1251 | 877,98 | 535 | 1221 | 2,293 | 1,197 | 16,208 | 4,40E-59 | 1,83E-58 | 0,0038401 | 0,0039614 | 0,003821 | 0,003821 | TRUE | 0,0410694 | No |
| orf19.6178 | 3497 | 2647 | 2813 | 2131 | 4696 | 3075 | 4299 | 4634 | 3334 | 2799 | 2779 | 3338 | 3763 | 3497 | 3650 | 3768 | 3365,9 | 3062 | 3670 | 1,199 | 0,262 | 3,804 | 0,0001425 | 0,0002 | 0,0055286 | 0,0025791 | 0,0042562 | 0,0042562 | TRUE | 0,0473965 | No |
| orf19.6179 | 7 | 4 | 6 | 6 | 7 | 4 | 12 | 9 | 7 | 4 | 6 | 9 | 6 | 5 | 10 | 7 | 6,74 | 6 | 7 | 1,093 | 0,128 | 0,234 | 0,8150463 | 0,8366812 | 1,00E-08 | 0,245819 | 0,1355792 | 0,1355792 | TRUE | 0,2987714 | No |
| orf19.6180 | 1063 | 817 | 905 | 571 | 1770 | 1038 | 1648 | 1788 | 1013 | 864 | 894 | 895 | 1418 | 1180 | 1399 | 1454 | 1139,67 | 916 | 1363 | 1,488 | 0,573 | 7,35 | 1,98E-13 | 3,83E-13 | 0,0055206 | 0,003532 | 0,0048975 | 0,0048975 | TRUE | 0,0944449 | No |
| orf19.6181 | 23 | 17 | 22 | 18 | 30 | 24 | 35 | 30 | 22 | 18 | 22 | 28 | 24 | 27 | 30 | 24 | 24,41 | 22 | 26 | 1,189 | 0,25 | 0,881 | 0,3781625 | 0,4131944 | 1,00E-08 | 0,0693579 | 0,034958 | 0,034958 | TRUE | 0,1076206 | No |
| orf19.6182 | 1685 | 1325 | 1459 | 1056 | 2503 | 1778 | 2407 | 2670 | 1606 | 1401 | 1441 | 1654 | 2006 | 2022 | 2044 | 2171 | 1793,12 | 1526 | 2061 | 1,353 | 0,436 | 7,083 | 1,41E-12 | 2,66E-12 | 0,0031801 | 0,0030069 | 0,0030544 | 0,0030544 | TRUE | 0,0359144 | No |
| orf19.6183 | 450 | 319 | 354 | 270 | 966 | 582 | 876 | 843 | 429 | 337 | 350 | 423 | 774 | 662 | 744 | 685 | 550,5 | 385 | 716 | 1,869 | 0,902 | 9,65 | 4,92E-22 | 1,15E-21 | 0,0077063 | 0,0050739 | 0,0063062 | 0,0063062 | TRUE | 0,0794768 | No |
| orf19.6184 | 52 | 38 | 60 | 24 | 105 | 57 | 108 | 91 | 50 | 40 | 59 | 38 | 84 | 65 | 92 | 74 | 62,66 | 47 | 79 | 1,672 | 0,741 | 3,667 | 0,0002454 | 0,0003414 | 0,0124165 | 0,0282958 | 0,0218315 | 0,0218315 | TRUE | 0,2614904 | No |
| orf19.6185 | 0 | 0 | 0 | 0 | 0 | 0 | 0 | 0 | 0 | 0 | 0 | 0 | 0 | 0 | 0 | 0 | 0 | 0 | 0 | NA | NA | NA | NA | NA | NA | NA | NA | NA | NA | NA | NA |
| orf19.6186 | 571 | 458 | 565 | 385 | 656 | 407 | 558 | 587 | 544 | 484 | 558 | 603 | 526 | 463 | 474 | 477 | 516,18 | 547 | 485 | 0,89 | -0,169 | -2,082 | 0,0373553 | 0,0453941 | 0,0034095 | 0,0052722 | 0,0043493 | 0,0043493 | TRUE | 0,0693384 | No |
| orf19.6187 | 1285 | 1022 | 1133 | 737 | 2259 | 1256 | 2077 | 2163 | 1225 | 1081 | 1119 | 1155 | 1810 | 1428 | 1763 | 1759 | 1417,49 | 1145 | 1690 | 1,478 | 0,564 | 7,214 | 5,44E-13 | 1,04E-12 | 0,0066311 | 0,0032496 | 0,0051059 | 0,0051059 | TRUE | 0,1276954 | No |
| orf19.6188 | 9 | 3 | 3 | 1 | 1 | 1 | 2 | 7 | 9 | 3 | 3 | 2 | 1 | 1 | 2 | 6 | 3,2 | 4 | 2 | 0,566 | -0,821 | -0,922 | 0,3565061 | 0,3912064 | 0,3519446 | 0,514967 | 0,4406986 | 0,4406986 | TRUE | 1,2097528 | No |
| orf19.6189 | 2163 | 1688 | 1866 | 1145 | 2132 | 1301 | 2050 | 2172 | 2062 | 1785 | 1843 | 1794 | 1708 | 1479 | 1740 | 1766 | 1772,29 | 1871 | 1673 | 0,895 | -0,161 | -2,303 | 0,0212622 | 0,0263342 | 0,004751 | 0,0030177 | 0,0040961 | 0,0040961 | TRUE | 0,0711735 | No |
| orf19.6190 | 5399 | 4050 | 4371 | 2805 | 11086 | 7185 | 10527 | 11375 | 5147 | 4282 | 4318 | 4394 | 8883 | 8170 | 8937 | 9249 | 6672,66 | 4535 | 8810 | 1,942 | 0,958 | 14,675 | 9,26E-49 | 3,36E-48 | 0,005214 | 0,0023373 | 0,0039147 | 0,0039147 | TRUE | 0,1016394 | No |
| orf19.6191 | 22 | 14 | 18 | 9 | 112 | 56 | 99 | 86 | 21 | 15 | 18 | 14 | 90 | 64 | 84 | 70 | 46,88 | 17 | 77 | 4,491 | 2,167 | 8,461 | 2,65E-17 | 5,62E-17 | 1,00E-08 | 0,0371134 | 0,0250245 | 0,0250245 | TRUE | 0,1040885 | No |
| orf19.6192 | 394 | 267 | 288 | 279 | 722 | 586 | 688 | 777 | 376 | 282 | 285 | 437 | 579 | 666 | 584 | 632 | 480,03 | 345 | 615 | 1,793 | 0,843 | 6,743 | 1,55E-11 | 2,84E-11 | 0,0211422 | 0,0055117 | 0,0126197 | 0,0126197 | TRUE | 0,3323658 | No |
| orf19.6193 | 3691 | 3229 | 3538 | 2219 | 2924 | 1961 | 2622 | 2830 | 3519 | 3414 | 3495 | 3476 | 2343 | 2230 | 2226 | 2301 | 2875,56 | 3476 | 2275 | 0,655 | -0,611 | -16,405 | 1,77E-60 | 7,49E-60 | 1,00E-08 | 0,0026623 | 0,000974 | 0,000974 | TRUE | 0,004963 | No |
| orf19.6194 | 178 | 188 | 175 | 108 | 170 | 132 | 153 | 144 | 170 | 199 | 173 | 169 | 136 | 150 | 130 | 117 | 155,48 | 178 | 133 | 0,746 | -0,423 | -3,449 | 0,0005631 | 0,0007713 | 1,00E-08 | 0,0126514 | 0,0079375 | 0,0079375 | TRUE | 0,0714558 | No |
| orf19.6195 | 6493 | 5972 | 6584 | 3848 | 5638 | 3968 | 5494 | 5807 | 6190 | 6315 | 6504 | 6028 | 4518 | 4512 | 4664 | 4722 | 5431,63 | 6259 | 4604 | 0,735 | -0,444 | -11,837 | 2,52E-32 | 7,09E-32 | 0,0005276 | 0,0023936 | 0,0011622 | 0,0011622 | TRUE | 0,008462 | No |
| orf19.6196 | 1090 | 939 | 992 | 725 | 1867 | 1268 | 1765 | 1813 | 1039 | 993 | 980 | 1136 | 1496 | 1442 | 1498 | 1474 | 1257,28 | 1037 | 1478 | 1,431 | 0,517 | 9,373 | 7,05E-21 | 1,61E-20 | 0,0007515 | 0,0033972 | 0,0020746 | 0,0020746 | TRUE | 0,042964 | No |
| orf19.6197 | 591 | 517 | 549 | 359 | 724 | 536 | 654 | 755 | 563 | 547 | 542 | 562 | 580 | 609 | 555 | 614 | 571,7 | 554 | 590 | 1,065 | 0,09 | 1,378 | 0,1683364 | 0,1922941 | 1,00E-08 | 0,0049632 | 0,0023752 | 0,0023752 | TRUE | 0,0183463 | No |
| orf19.6198.1 | 1742 | 1516 | 1605 | 899 | 3852 | 2287 | 3443 | 3274 | 1661 | 1603 | 1586 | 1408 | 3086 | 2601 | 2923 | 2662 | 2191,24 | 1564 | 2818 | 1,799 | 0,847 | 12,169 | 4,57E-34 | 1,32E-33 | 0,0050085 | 0,0028406 | 0,0041358 | 0,0041358 | TRUE | 0,0509233 | No |
| orf19.6199 | 731 | 604 | 716 | 398 | 1670 | 1196 | 1701 | 1747 | 697 | 639 | 707 | 623 | 1338 | 1360 | 1444 | 1420 | 1028,64 | 666 | 1390 | 2,079 | 1,056 | 17,522 | 9,67E-69 | 4,55E-68 | 0,0011203 | 0,0036875 | 0,0023315 | 0,0023315 | TRUE | 0,0191658 | No |
| orf19.6200 | 27 | 17 | 22 | 16 | 0 | 1 | 1 | 3 | 26 | 18 | 22 | 25 | 0 | 1 | 1 | 2 | 11,87 | 23 | 1 | 0,049 | -4,355 | -5,971 | 2,36E-09 | 4,00E-09 | 1,00E-08 | 0,1404427 | 0,0856191 | 0,0856191 | TRUE | 0,4767147 | No |
| orf19.6202 | 1446 | 1319 | 1317 | 1027 | 2767 | 1955 | 2494 | 2827 | 1379 | 1395 | 1301 | 1609 | 2217 | 2223 | 2117 | 2299 | 1817,42 | 1421 | 2214 | 1,563 | 0,645 | 10,095 | 5,83E-24 | 1,42E-23 | 0,0035867 | 0,0029947 | 0,0033149 | 0,0033149 | TRUE | 0,0887461 | No |
| orf19.6203 | 2545 | 2226 | 2445 | 1549 | 2272 | 1570 | 2071 | 2250 | 2426 | 2354 | 2415 | 2427 | 1820 | 1785 | 1758 | 1829 | 2101,95 | 2406 | 1798 | 0,748 | -0,419 | -10,693 | 1,10E-26 | 2,82E-26 | 1,00E-08 | 0,0028724 | 0,000993 | 0,000993 | TRUE | 0,0028813 | No |
| orf19.6205 | 323 | 286 | 319 | 222 | 254 | 253 | 261 | 279 | 308 | 302 | 315 | 348 | 204 | 288 | 222 | 227 | 276,61 | 318 | 235 | 0,736 | -0,443 | -3,963 | 7,39E-05 | 0,0001053 | 0,008606 | 0,008027 | 0,0082934 | 0,0082934 | TRUE | 0,2332106 | No |
| orf19.6208 | 2809 | 2225 | 2468 | 1794 | 3778 | 2516 | 3679 | 3785 | 2678 | 2353 | 2438 | 2810 | 3027 | 2861 | 3123 | 3078 | 2796,05 | 2570 | 3022 | 1,178 | 0,236 | 3,882 | 0,0001036 | 0,0001465 | 0,0036222 | 0,0026785 | 0,0031895 | 0,0031895 | TRUE | 0,0457792 | No |
| orf19.6209 | 1278 | 1045 | 1179 | 731 | 1614 | 1166 | 1516 | 1700 | 1218 | 1105 | 1165 | 1145 | 1293 | 1326 | 1287 | 1382 | 1240,21 | 1158 | 1322 | 1,14 | 0,19 | 3,607 | 0,0003093 | 0,0004282 | 0,0006614 | 0,0034152 | 0,0018292 | 0,0018292 | TRUE | 0,0154954 | No |
| orf19.6211 | 5670 | 4992 | 5503 | 3288 | 1393 | 932 | 1279 | 1459 | 5406 | 5278 | 5436 | 5151 | 1116 | 1060 | 1086 | 1186 | 3214,94 | 5318 | 1112 | 0,209 | -2,256 | -49,291 | 0 | 0 | 0,0007834 | 0,002602 | 0,0015058 | 0,0015058 | TRUE | 0,0244694 | No |
| orf19.6213 | 7472 | 5913 | 6336 | 4333 | 14404 | 9721 | 13975 | 14486 | 7124 | 6252 | 6259 | 6788 | 11542 | 11054 | 11865 | 11779 | 9082,7 | 6606 | 11560 | 1,75 | 0,808 | 15,882 | 8,42E-57 | 3,38E-56 | 0,002424 | 0,0022721 | 0,0023599 | 0,0023599 | TRUE | 0,0347968 | No |
| orf19.6214 | 1834 | 1572 | 1705 | 1163 | 6342 | 4464 | 5761 | 6481 | 1749 | 1662 | 1684 | 1822 | 5082 | 5076 | 4891 | 5270 | 3404,42 | 1729 | 5080 | 2,943 | 1,557 | 35,082 | 1,26E-269 | 2,36E-268 | 0,0008083 | 0,0025736 | 0,0014786 | 0,0014786 | TRUE | 0,0138848 | No |
| orf19.6217 | 1285 | 1010 | 1223 | 832 | 3957 | 2846 | 3675 | 4170 | 1225 | 1068 | 1208 | 1303 | 3171 | 3236 | 3120 | 3391 | 2215,26 | 1201 | 3230 | 2,694 | 1,43 | 23,732 | 1,69E-124 | 1,40E-123 | 0,0032296 | 0,0028325 | 0,002878 | 0,002878 | TRUE | 0,066391 | No |
| orf19.6219 | 134 | 128 | 93 | 75 | 813 | 554 | 752 | 863 | 128 | 135 | 92 | 117 | 651 | 630 | 638 | 702 | 386,75 | 118 | 655 | 5,557 | 2,474 | 23,239 | 1,83E-119 | 1,45E-118 | 0,0035746 | 0,0063367 | 0,0055017 | 0,0055017 | TRUE | 0,2552174 | No |
| orf19.6220 | 3015 | 2781 | 2881 | 1967 | 3298 | 2303 | 3102 | 3287 | 2874 | 2941 | 2846 | 3081 | 2643 | 2619 | 2634 | 2673 | 2788,77 | 2936 | 2642 | 0,902 | -0,15 | -3,812 | 0,0001376 | 0,0001935 | 0,0001962 | 0,00268 | 0,0011169 | 0,0011169 | TRUE | 0,0123712 | No |
| orf19.6220.3 | 1528 | 1173 | 1331 | 969 | 1699 | 1215 | 1605 | 1707 | 1457 | 1240 | 1315 | 1518 | 1361 | 1382 | 1363 | 1388 | 1377,94 | 1382 | 1374 | 0,995 | -0,007 | -0,105 | 0,9161896 | 0,924513 | 0,0029739 | 0,0032828 | 0,0032981 | 0,0032981 | TRUE | 0,0569312 | No |
| orf19.6220.4 | 14906 | 11699 | 12207 | 8082 | 36602 | 22929 | 33279 | 38156 | 14211 | 12370 | 12059 | 12661 | 29328 | 26072 | 28254 | 31024 | 20747,55 | 12825 | 28670 | 2,235 | 1,161 | 17,874 | 1,87E-71 | 9,14E-71 | 0,005318 | 0,0021704 | 0,0039905 | 0,0039905 | TRUE | 0,0650442 | No |
| orf19.6222 | 106 | 103 | 97 | 67 | 289 | 198 | 268 | 287 | 101 | 109 | 96 | 105 | 232 | 225 | 228 | 233 | 166,04 | 103 | 230 | 2,241 | 1,164 | 10,122 | 4,42E-24 | 1,08E-23 | 1,00E-08 | 0,0119796 | 0,0053517 | 0,0053517 | TRUE | 0,0212333 | No |
| orf19.6222.1 | 3104 | 2416 | 2458 | 1473 | 3561 | 2132 | 3052 | 3620 | 2959 | 2555 | 2428 | 2308 | 2853 | 2424 | 2591 | 2943 | 2632,74 | 2562 | 2703 | 1,054 | 0,076 | 0,917 | 0,3593429 | 0,3940483 | 0,0089012 | 0,0027149 | 0,0063006 | 0,0063006 | TRUE | 0,1326416 | No |
| orf19.6224 | 155 | 132 | 173 | 84 | 250 | 173 | 227 | 226 | 148 | 140 | 171 | 132 | 200 | 197 | 193 | 184 | 170,42 | 148 | 194 | 1,302 | 0,38 | 3,294 | 0,0009873 | 0,0013317 | 1,00E-08 | 0,0117257 | 0,006746 | 0,006746 | TRUE | 0,1207042 | No |
| orf19.6225 | 1549 | 1433 | 1644 | 1067 | 2499 | 1824 | 2383 | 2494 | 1477 | 1515 | 1624 | 1672 | 2002 | 2074 | 2023 | 2028 | 1801,89 | 1572 | 2032 | 1,295 | 0,373 | 7,472 | 7,87E-14 | 1,53E-13 | 0,0010384 | 0,0030025 | 0,0018125 | 0,0018125 | TRUE | 0,020193 | No |
| orf19.6225.1 | 1282 | 1076 | 1107 | 687 | 1159 | 760 | 1112 | 1104 | 1222 | 1138 | 1094 | 1076 | 929 | 864 | 944 | 898 | 1020,55 | 1132 | 909 | 0,802 | -0,319 | -5,345 | 9,06E-08 | 1,46E-07 | 0,0014019 | 0,0037001 | 0,0024303 | 0,0024303 | TRUE | 0,0350714 | No |
| orf19.6227 | 4893 | 4025 | 4614 | 2789 | 4285 | 2760 | 4224 | 4270 | 4665 | 4256 | 4558 | 4369 | 3433 | 3138 | 3586 | 3472 | 3934,76 | 4462 | 3407 | 0,764 | -0,389 | -7,547 | 4,45E-14 | 8,74E-14 | 0,002072 | 0,0025086 | 0,0022881 | 0,0022881 | TRUE | 0,0333711 | No |
| orf19.6229 | 7493 | 6494 | 7055 | 5035 | 65953 | 47343 | 62459 | 69811 | 7144 | 6867 | 6970 | 7888 | 52846 | 53833 | 53027 | 56763 | 30667,19 | 7217 | 54117 | 7,504 | 2,908 | 58,928 | 0 | 0 | 0,0025026 | 0,0021448 | 0,0022523 | 0,0022523 | TRUE | 0,046612 | No |
| orf19.6230 | 4745 | 4696 | 5005 | 2758 | 6145 | 4431 | 5759 | 5736 | 4524 | 4965 | 4944 | 4321 | 4924 | 5038 | 4889 | 4664 | 4783,73 | 4688 | 4879 | 1,04 | 0,056 | 1,053 | 0,2925434 | 0,3250992 | 0,0025481 | 0,0024345 | 0,0025092 | 0,0025092 | TRUE | 0,0327294 | No |
| orf19.6231 | 2310 | 1747 | 1922 | 1478 | 3116 | 2094 | 2860 | 3038 | 2202 | 1847 | 1899 | 2315 | 2497 | 2381 | 2428 | 2470 | 2254,98 | 2066 | 2444 | 1,185 | 0,245 | 3,456 | 0,000549 | 0,0007525 | 0,0057342 | 0,0028194 | 0,0043641 | 0,0043641 | TRUE | 0,0762004 | No |

| | | | | | | | | | | | | | | | | | | | | | | | | | | | | | | | |
|---|---|---|---|---|---|---|---|---|---|---|---|---|---|---|---|---|---|---|---|---|---|---|---|---|---|---|---|---|---|---|---|
| orf19.6232 | 5510 | 4758 | 5183 | 3157 | 4613 | 2978 | 4364 | 4527 | 5253 | 5031 | 5120 | 4946 | 3696 | 3386 | 3705 | 3681 | 4352,31 | 5088 | 3617 | 0,711 | -0,492 | -11,606 | 3,83E-31 | 1,05E-30 | 0,0009275 | 0,0024685 | 0,0014895 | 0,0014895 | TRUE | 0,0215913 | No |
| orf19.6233 | 3894 | 3135 | 3293 | 2039 | 4400 | 2903 | 4300 | 4590 | 3713 | 3315 | 3253 | 3194 | 3526 | 3301 | 3651 | 3732 | 3460,51 | 3369 | 3552 | 1,054 | 0,076 | 1,283 | 0,1993191 | 0,2260674 | 0,0034348 | 0,0025657 | 0,0030897 | 0,0030897 | TRUE | 0,0580858 | No |
| orf19.6234 | 9983 | 8670 | 9419 | 5888 | 10420 | 7413 | 9733 | 10456 | 9518 | 9167 | 9305 | 9224 | 8349 | 8429 | 8263 | 8502 | 8844,7 | 9304 | 8386 | 0,901 | -0,15 | -5,239 | 1,62E-07 | 2,58E-07 | 0,0001114 | 0,0022769 | 0,0006755 | 0,0006755 | TRUE | 0,0029999 | No |
| orf19.6235 | 923 | 714 | 816 | 600 | 639 | 450 | 621 | 612 | 880 | 755 | 806 | 940 | 512 | 512 | 527 | 498 | 678,69 | 845 | 512 | 0,608 | -0,719 | -9,483 | 2,48E-21 | 5,69E-21 | 0,0035009 | 0,0045105 | 0,0039788 | 0,0039788 | TRUE | 0,0605597 | No |
| orf19.6236 | 6354 | 5751 | 5984 | 3488 | 6117 | 3850 | 5863 | 5676 | 6058 | 6081 | 5912 | 5464 | 4901 | 4378 | 4978 | 4615 | 5298,32 | 5879 | 4718 | 0,802 | -0,318 | -5,916 | 3,29E-09 | 5,56E-09 | 0,0026339 | 0,0025766 | 0,0025766 | 0,0025766 | TRUE | 0,0281596 | No |
| orf19.6237 | 2774 | 2491 | 2682 | 1722 | 3346 | 2329 | 3165 | 3412 | 2645 | 2634 | 2650 | 2698 | 2681 | 2648 | 2687 | 2774 | 2677,06 | 2657 | 2698 | 1,017 | 0,024 | 0,648 | 0,5170707 | 0,5535024 | 1,00E-08 | 0,0027046 | 0,0009018 | 0,0009018 | TRUE | 0,0044924 | No |
| orf19.6238 | 337 | 330 | 337 | 202 | 753 | 492 | 731 | 737 | 321 | 349 | 333 | 316 | 603 | 559 | 621 | 599 | 462,78 | 330 | 596 | 1,805 | 0,852 | 11,525 | 9,82E-31 | 2,68E-30 | 1,00E-08 | 0,0056392 | 0,0028186 | 0,0028186 | TRUE | 0,0177999 | No |
| orf19.6239 | 1614 | 1377 | 1477 | 1020 | 1623 | 1155 | 1491 | 1630 | 1539 | 1456 | 1459 | 1598 | 1300 | 1313 | 1266 | 1325 | 1407,1 | 1513 | 1301 | 0,862 | -0,215 | -4,362 | 1,29E-05 | 1,90E-05 | 0,0002628 | 0,0032582 | 0,001618 | 0,001618 | TRUE | 0,0151443 | No |
| orf19.6240 | 665 | 581 | 600 | 394 | 512 | 344 | 497 | 540 | 634 | 614 | 593 | 617 | 410 | 391 | 422 | 439 | 515,09 | 614 | 416 | 0,678 | -0,561 | -8,187 | 2,68E-16 | 5,56E-16 | 1,00E-08 | 0,0052789 | 0,0025326 | 0,0025326 | TRUE | 0,0166529 | No |
| orf19.6242 | 270 | 242 | 275 | 153 | 228 | 164 | 242 | 232 | 257 | 256 | 272 | 240 | 183 | 186 | 205 | 189 | 223,49 | 256 | 191 | 0,743 | -0,429 | -4,399 | 1,09E-05 | 1,61E-05 | 1,00E-08 | 0,0094379 | 0,0046487 | 0,0046487 | TRUE | 0,0302216 | No |
| orf19.6243 | 814 | 775 | 871 | 500 | 1115 | 817 | 1012 | 1130 | 776 | 819 | 860 | 783 | 893 | 929 | 859 | 919 | 854,96 | 810 | 900 | 1,109 | 0,15 | 2,53 | 0,0114118 | 0,0144482 | 0,0005501 | 0,0040117 | 0,0021716 | 0,0021716 | TRUE | 0,0218723 | No |
| orf19.6244 | 1352 | 1173 | 1324 | 905 | 2842 | 2184 | 2730 | 3006 | 1289 | 1240 | 1308 | 1418 | 2277 | 2483 | 2318 | 2444 | 1847,19 | 1314 | 2380 | 1,816 | 0,86 | 15,941 | 3,31E-57 | 1,34E-56 | 0,0017001 | 0,0029801 | 0,0021812 | 0,0021812 | TRUE | 0,0304704 | No |
| orf19.6245 | 14100 | 12871 | 13791 | 8222 | 5073 | 3598 | 4814 | 5195 | 13443 | 13609 | 13624 | 12880 | 4065 | 4091 | 4087 | 4224 | 8752,98 | 13389 | 4117 | 0,307 | -1,702 | -50,564 | 0 | 0 | 0,0003828 | 0,0022789 | 0,0009373 | 0,0009373 | TRUE | 0,0077509 | No |
| orf19.6246 | 1461 | 1196 | 1269 | 759 | 943 | 597 | 815 | 894 | 1393 | 1265 | 1254 | 1189 | 756 | 679 | 692 | 727 | 994,18 | 1275 | 714 | 0,559 | -0,839 | -12,879 | 5,92E-38 | 1,81E-37 | 0,0024149 | 0,0037428 | 0,0030092 | 0,0030092 | TRUE | 0,0471393 | No |
| orf19.6247 | 80 | 63 | 81 | 49 | 96 | 65 | 75 | 84 | 76 | 67 | 80 | 77 | 77 | 74 | 64 | 68 | 72,81 | 75 | 71 | 0,943 | -0,085 | -0,513 | 0,6079854 | 0,6420978 | 1,00E-08 | 0,024642 | 0,012454 | 0,012454 | TRUE | 0,0525623 | No |
| orf19.6247.1 | 4590 | 3806 | 3990 | 1710 | 1652 | 886 | 1368 | 1565 | 4376 | 4024 | 3942 | 2679 | 1324 | 1007 | 1161 | 1272 | 2473,26 | 3755 | 1191 | 0,317 | -1,656 | -9,468 | 2,85E-21 | 6,54E-21 | 0,0288802 | 0,0027551 | 0,0150245 | 0,0288802 | TRUE | 0,4497184 | No |
| orf19.6248 | 45 | 66 | 60 | 32 | 29 | 18 | 31 | 27 | 43 | 70 | 59 | 50 | 23 | 20 | 26 | 22 | 39,26 | 56 | 23 | 0,415 | -1,269 | -5,122 | 3,02E-07 | 4,77E-07 | 1,00E-08 | 0,0439139 | 0,0296219 | 0,0296219 | TRUE | 0,1657401 | No |
| orf19.6249 | 2885 | 2568 | 2819 | 1622 | 618 | 451 | 617 | 660 | 2751 | 2715 | 2785 | 2541 | 495 | 513 | 524 | 537 | 1607,53 | 2698 | 517 | 0,191 | -2,385 | -43,809 | 0 | 0 | 1,00E-08 | 0,0031127 | 0,0017776 | 0,0017776 | TRUE | 0,0172492 | No |
| orf19.6250 | 6215 | 4632 | 5435 | 3645 | 5496 | 3502 | 5163 | 5472 | 5925 | 4898 | 5369 | 5710 | 4404 | 3982 | 4383 | 4449 | 4890,12 | 5476 | 4304 | 0,787 | -0,346 | -5,46 | 4,75E-08 | 7,75E-08 | 0,0043404 | 0,002427 | 0,0036585 | 0,0036585 | TRUE | 0,0617953 | No |
| orf19.6252 | 3544 | 2996 | 3344 | 2096 | 5149 | 3479 | 4974 | 5147 | 3379 | 3168 | 3304 | 3283 | 4126 | 3956 | 4223 | 4185 | 3702,92 | 3284 | 4122 | 1,256 | 0,329 | 8,392 | 4,76E-17 | 1,00E-16 | 0,0003979 | 0,0025347 | 0,0011937 | 0,0011937 | TRUE | 0,008621 | No |
| orf19.6253 | 16846 | 13493 | 14563 | 9070 | 51024 | 34584 | 48525 | 53817 | 16061 | 14267 | 14387 | 14209 | 40884 | 39325 | 41197 | 43758 | 28011,07 | 14731 | 41291 | 2,803 | 1,487 | 28,945 | 3,25E-184 | 4,02E-183 | 0,0027526 | 0,0021499 | 0,0024858 | 0,0024858 | TRUE | 0,0453993 | No |
| orf19.6254 | 3570 | 3022 | 3291 | 2158 | 25234 | 17210 | 24120 | 25751 | 3404 | 3195 | 3251 | 3381 | 20219 | 19569 | 20478 | 20938 | 11804,4 | 3308 | 20301 | 6,141 | 2,618 | 69,075 | 0 | 0 | 0,0006235 | 0,0022304 | 0,0011878 | 0,0011878 | TRUE | 0,0071387 | No |
| orf19.6255 | 1307 | 1219 | 1289 | 751 | 1994 | 1514 | 1948 | 2043 | 1246 | 1289 | 1273 | 1176 | 1598 | 1722 | 1654 | 1661 | 1452,4 | 1246 | 1659 | 1,327 | 0,408 | 8,301 | 1,03E-16 | 2,15E-16 | 0,0003265 | 0,0032218 | 0,0016018 | 0,0016018 | TRUE | 0,0146355 | No |
| orf19.6257 | 4608 | 4220 | 4502 | 2739 | 15896 | 11018 | 14460 | 15936 | 4393 | 4462 | 4448 | 4291 | 12737 | 12528 | 12276 | 12957 | 8511,64 | 4398 | 12624 | 2,869 | 1,52 | 46,161 | 0 | 0 | 0,000326 | 0,0022842 | 0,0008785 | 0,0008785 | TRUE | 0,0042749 | No |
| orf19.6259 | 4470 | 4158 | 4464 | 2610 | 5011 | 3460 | 4668 | 4933 | 4262 | 4397 | 4410 | 4089 | 4015 | 3934 | 3963 | 4011 | 4135,07 | 4290 | 3981 | 0,927 | -0,109 | -2,953 | 0,0031479 | 0,004117 | 0,0003328 | 0,0024883 | 0,0010614 | 0,0010614 | TRUE | 0,0111236 | No |
| orf19.6260 | 3616 | 3299 | 3566 | 2277 | 5340 | 3756 | 4882 | 5408 | 3447 | 3488 | 3523 | 3567 | 4279 | 4271 | 4145 | 4397 | 3889,67 | 3506 | 4273 | 1,219 | 0,286 | 8,093 | 5,82E-16 | 1,20E-15 | 0,0002087 | 0,0025134 | 0,0009351 | 0,0009351 | TRUE | 0,0050738 | No |
| orf19.6261 | 2125 | 2013 | 2199 | 1245 | 1163 | 835 | 1047 | 1085 | 2026 | 2128 | 2172 | 1950 | 932 | 949 | 889 | 882 | 1491,21 | 2069 | 913 | 0,44 | -1,184 | -22,079 | 4,99E-108 | 3,54E-107 | 0,0011224 | 0,0031923 | 0,0019993 | 0,0019993 | TRUE | 0,0165354 | No |
| orf19.6263 | 4770 | 3944 | 4101 | 2890 | 4232 | 2942 | 4004 | 4308 | 4548 | 4170 | 4051 | 4527 | 3391 | 3345 | 3399 | 3503 | 3866,9 | 4324 | 3410 | 0,789 | -0,342 | -7,109 | 1,17E-12 | 2,21E-12 | 0,0015281 | 0,0025159 | 0,001957 | 0,001957 | TRUE | 0,022174 | No |
| orf19.6264 | 425 | 362 | 401 | 278 | 453 | 367 | 441 | 533 | 405 | 383 | 396 | 436 | 363 | 417 | 374 | 433 | 400,96 | 405 | 397 | 0,982 | -0,026 | -0,304 | 0,7614673 | 0,7890424 | 0,0028176 | 0,0061862 | 0,0045049 | 0,0045049 | TRUE | 0,0481808 | No |
| orf19.6264.3 | 5554 | 5495 | 5505 | 2746 | 5299 | 3574 | 5055 | 4490 | 5295 | 5810 | 5438 | 4302 | 4246 | 4064 | 4292 | 3651 | 4637,24 | 5211 | 4063 | 0,779 | -0,36 | -4,231 | 2,33E-05 | 3,39E-05 | 0,0099272 | 0,0024453 | 0,0067376 | 0,0067376 | TRUE | 0,1657624 | No |
| orf19.6264.4 | 7119 | 5447 | 5871 | 3676 | 18913 | 10771 | 17196 | 18128 | 6787 | 5760 | 5800 | 5759 | 15154 | 12248 | 14599 | 14740 | 10105,81 | 6026 | 14185 | 2,354 | 1,235 | 16,364 | 3,47E-60 | 1,47E-59 | 0,0076743 | 0,0022538 | 0,0053461 | 0,0053461 | TRUE | 0,103111 | No |
| orf19.6265 | 8804 | 6569 | 6532 | 4459 | 27383 | 17490 | 24851 | 27756 | 8394 | 6946 | 6453 | 6985 | 21941 | 19888 | 21098 | 22568 | 14284,17 | 7194 | 21374 | 2,971 | 1,571 | 20,969 | 1,26E-97 | 8,08E-97 | 0,0080882 | 0,0022062 | 0,0052916 | 0,0052916 | TRUE | 0,1547292 | No |
| orf19.6265.1 | 16128 | 12924 | 13608 | 8399 | 41670 | 24859 | 38251 | 40798 | 15376 | 13665 | 13443 | 13157 | 33389 | 28267 | 32475 | 33173 | 22868,27 | 13910 | 31826 | 2,288 | 1,194 | 18,215 | 3,95E-74 | 2,01E-73 | 0,0053637 | 0,0021631 | 0,0040731 | 0,0040731 | TRUE | 0,0692766 | No |
| orf19.6266 | 2490 | 2635 | 2700 | 1599 | 2580 | 1979 | 2346 | 2433 | 2374 | 2786 | 2667 | 2505 | 2067 | 2250 | 1992 | 1978 | 2327,5 | 2583 | 2072 | 0,801 | -0,32 | -5,071 | 3,95E-07 | 6,22E-07 | 0,0036993 | 0,0027967 | 0,0033813 | 0,0033813 | TRUE | 0,0394833 | No |
| orf19.6267 | 537 | 500 | 534 | 309 | 78 | 47 | 71 | 73 | 512 | 529 | 528 | 484 | 62 | 53 | 60 | 59 | 285,98 | 513 | 58 | 0,115 | -3,118 | -27,191 | 8,23E-163 | 9,07E-162 | 1,00E-08 | 0,0078326 | 0,0040666 | 0,0040666 | TRUE | 0,0260658 | No |
| orf19.6268 | 1758 | 1647 | 1757 | 1080 | 1536 | 1162 | 1430 | 1451 | 1676 | 1741 | 1736 | 1692 | 1231 | 1321 | 1214 | 1180 | 1473,89 | 1711 | 1236 | 0,721 | -0,472 | -9,509 | 1,93E-21 | 4,45E-21 | 0,0004421 | 0,0032053 | 0,0016735 | 0,0016735 | TRUE | 0,0236935 | No |
| orf19.6269 | 686 | 766 | 804 | 465 | 690 | 466 | 631 | 595 | 654 | 810 | 794 | 728 | 553 | 530 | 536 | 484 | 636,12 | 746 | 526 | 0,704 | -0,507 | -6,163 | 7,12E-10 | 1,23E-09 | 0,0050908 | 0,0046724 | 0,0048966 | 0,0048966 | TRUE | 0,0884454 | No |
| orf19.6271 | 3313 | 3188 | 3529 | 2311 | 8171 | 6131 | 7620 | 7953 | 3159 | 3371 | 3486 | 3620 | 6547 | 6971 | 6469 | 6467 | 5011,34 | 3409 | 6614 | 1,942 | 0,957 | 19,245 | 1,56E-82 | 8,75E-82 | 0,0020164 | 0,0024189 | 0,0021429 | 0,0021429 | TRUE | 0,0300377 | No |
| orf19.6272 | 879 | 1258 | 1200 | 705 | 1326 | 1027 | 1071 | 843 | 838 | 1330 | 1185 | 1104 | 1062 | 1168 | 909 | 685 | 1035,39 | 1114 | 956 | 0,858 | -0,221 | -1,032 | 0,3020176 | 0,334986 | 0,0432486 | 0,0036771 | 0,021828 | 0,0432486 | TRUE | 0,4368849 | No |
| orf19.6273 | 696 | 532 | 570 | 361 | 1062 | 704 | 1070 | 1056 | 664 | 563 | 563 | 566 | 851 | 801 | 908 | 859 | 721,65 | 589 | 855 | 1,45 | 0,536 | 7,265 | 3,73E-13 | 7,16E-13 | 0,003021 | 0,0043665 | 0,0037582 | 0,0037582 | TRUE | 0,090826 | No |
| orf19.6274 | 201 | 164 | 156 | 131 | 53 | 44 | 55 | 47 | 192 | 173 | 154 | 205 | 42 | 50 | 47 | 38 | 112,72 | 181 | 44 | 0,244 | -2,033 | -12,564 | 3,35E-36 | 9,96E-36 | 1,00E-08 | 0,0166571 | 0,0119504 | 0,0119504 | TRUE | 0,1098978 | No |
| orf19.6275 | 1265 | 1314 | 1339 | 721 | 1328 | 817 | 1223 | 1243 | 1206 | 1389 | 1323 | 1129 | 1064 | 929 | 1038 | 1011 | 1136,23 | 1262 | 1010 | 0,8 | -0,322 | -4,393 | 1,12E-05 | 1,65E-05 | 0,004742 | 0,0035363 | 0,0042563 | 0,0042563 | TRUE | 0,0551347 | No |
| orf19.6276 | 1667 | 1613 | 1804 | 1009 | 848 | 618 | 770 | 792 | 1589 | 1706 | 1782 | 1581 | 679 | 703 | 654 | 644 | 1167,2 | 1664 | 670 | 0,402 | -1,316 | -21,809 | 1,89E-105 | 1,30E-104 | 0,0017332 | 0,003498 | 0,0024945 | 0,0024945 | TRUE | 0,0277047 | No |
| orf19.6277 | 50 | 46 | 31 | 26 | 16 | 15 | 20 | 21 | 48 | 49 | 31 | 41 | 13 | 17 | 17 | 17 | 28,95 | 42 | 16 | 0,379 | -1,398 | -4,903 | 9,43E-07 | 1,46E-06 | 1,00E-08 | 0,0588074 | 0,0368646 | 0,0368646 | TRUE | 0,277686 | No |
| orf19.6280 | 5 | 4 | 3 | 2 | 3 | 7 | 0 | 4 | 5 | 4 | 3 | 3 | 2 | 8 | 0 | 3 | 3,59 | 4 | 3 | 0,856 | -0,224 | -0,267 | 0,7897294 | 0,8129155 | 0,2776507 | 0,4596196 | 0,3924782 | 0,3924782 | TRUE | 0,5881996 | No |
| orf19.6282 | 798 | 773 | 761 | 529 | 131 | 112 | 162 | 162 | 761 | 817 | 752 | 829 | 105 | 127 | 138 | 132 | 457,53 | 790 | 126 | 0,159 | -2,655 | -28,305 | 3,01E-176 | 3,51E-175 | 0,001781 | 0,0056799 | 0,0041909 | 0,0041909 | TRUE | 0,1388366 | No |
| orf19.6283 | 3410 | 3145 | 3277 | 2005 | 2662 | 1902 | 2449 | 2618 | 3251 | 3325 | 3237 | 3141 | 2133 | 2163 | 2079 | 2129 | 2682,3 | 3238 | 2126 | 0,656 | -0,609 | -16,121 | 1,81E-58 | 7,44E-58 | 1,00E-08 | 0,0027034 | 0,0009873 | 0,0009873 | TRUE | 0,0045094 | No |
| orf19.6284 | 3713 | 3568 | 3822 | 2265 | 2788 | 1890 | 2587 | 2584 | 3540 | 3773 | 3776 | 3235 | 2234 | 2149 | 2196 | 2101 | 2875,47 | 3581 | 2170 | 0,605 | -0,724 | -13,017 | 9,86E-39 | 3,06E-38 | 0,0024091 | 0,0026623 | 0,0026085 | 0,0026085 | TRUE | 0,0490657 | No |
| orf19.6285 | 6181 | 5510 | 6224 | 3629 | 9661 | 6354 | 9290 | 9969 | 5893 | 5826 | 6149 | 5685 | 7741 | 7225 | 7887 | 8106 | 6813,98 | 5888 | 7740 | 1,314 | 0,394 | 8,657 | 4,85E-18 | 1,04E-17 | 0,0015141 | 0,0023322 | 0,0018395 | 0,0018395 | TRUE | 0,0240811 | No |
| orf19.6286 | 474 | 369 | 431 | 242 | 1327 | 857 | 1306 | 1303 | 452 | 390 | 426 | 379 | 1063 | 974 | 1109 | 1059 | 731,62 | 412 | 1051 | 2,544 | 1,347 | 18,375 | 2,07E-75 | 1,06E-74 | 0,0024506 | 0,0043355 | 0,0033881 | 0,0033881 | TRUE | 0,0501917 | No |
| orf19.6286.2 | 9817 | 8388 | 8621 | 7149 | 23298 | 19618 | 21557 | 23692 | 9359 | 8869 | 8517 | 11199 | 18668 | 22307 | 18302 | 19264 | 14560,73 | 9486 | 19635 | 2,07 | 1,05 | 12,154 | 5,45E-34 | 1,57E-33 | 0,0121261 | 0,002204 | 0,0070879 | 0,0070879 | TRUE | 0,1794361 | No |
| orf19.6287 | 4746 | 4065 | 4354 | 3002 | 9092 | 6611 | 8629 | 9817 | 4525 | 4298 | 4301 | 4703 | 7285 | 7517 | 7326 | 7982 | 5992,22 | 4457 | 7528 | 1,69 | 0,757 | 16,279 | 1,39E-59 | 5,84E-59 | 0,0016227 | 0,0023653 | 0,0018922 | 0,0018922 | TRUE | 0,0203789 | No |
| orf19.6288 | 158 | 127 | 130 | 92 | 37 | 30 | 37 | 31 | 151 | 134 | 128 | 144 | 30 | 34 | 31 | 25 | 84,73 | 139 | 30 | 0,214 | -2,221 | -12,427 | 1,86E-35 | 5,48E-35 | 1,00E-08 | 0,021469 | 0,0118298 | 0,0118298 | TRUE | 0,0887483 | No |
| orf19.6291 | 3531 | 3142 | 3291 | 2054 | 7091 | 4745 | 6737 | 7021 | 3366 | 3322 | 3251 | 3218 | 5682 | 5395 | 5720 | 5709 | 4457,92 | 3289 | 5626 | 1,71 | 0,774 | 21,029 | 3,52E-98 | 2,27E-97 | 0,0003677 | 0,0024596 | 0,0010509 | 0,0010509 | TRUE | 0,0089855 | No |
| orf19.6293 | 832 | 690 | 785 | 445 | 1863 | 1284 | 1783 | 1795 | 793 | 730 | 776 | 697 | 1493 | 1460 | 1514 | 1460 | 1115,19 | 749 | 1482 | 1,971 | 0,979 | 17,862 | 2,32E-71 | 1,13E-70 | 1,00E-08 | 0,0035636 | 0,0018398 | 0,0018398 | TRUE | 0,0215134 | No |
| orf19.6294 | 310 | 387 | 370 | 229 | 871 | 641 | 736 | 786 | 296 | 409 | 366 | 359 | 698 | 729 | 625 | 639 | 514,97 | 358 | 673 | 1,883 | 0,913 | 9,318 | 1,19E-20 | 2,70E-20 | 0,0083861 | 0,0052796 | 0,006989 | 0,006989 | TRUE | 0,1665072 | No |
| orf19.6295 | 3489 | 3209 | 3505 | 2096 | 3846 | 2790 | 3655 | 3888 | 3326 | 3393 | 3463 | 3283 | 3082 | 3172 | 3103 | 3161 | 3248,02 | 3366 | 3130 | 0,929 | -0,107 | -2,979 | 0,0028955 | 0,0037962 | 1,00E-08 | 0,0025968 | 0,0009216 | 0,0009216 | TRUE | 0,0045197 | No |
| orf19.6296 | 325 | 355 | 342 | 189 | 360 | 251 | 322 | 311 | 310 | 375 | 338 | 296 | 288 | 285 | 273 | 253 | 302,41 | 330 | 275 | 0,83 | -0,268 | -2,763 | 0,0057209 | 0,0073674 | 0,0037573 | 0,0075207 | 0,0057471 | 0,0057471 | TRUE | 0,0962743 | No |
| orf19.6297 | 3003 | 2718 | 2874 | 1740 | 3391 | 2549 | 3299 | 3450 | 2863 | 2874 | 2839 | 2726 | 2717 | 2898 | 2801 | 2805 | 2815,45 | 2826 | 2805 | 0,991 | -0,013 | -0,327 | 0,7437217 | 0,7720332 | 0,0001651 | 0,0026745 | 0,0011163 | 0,0011163 | TRUE | 0,0061693 | No |
| orf19.6298 | 3120 | 3013 | 3036 | 1780 | 3532 | 2531 | 3124 | 3329 | 2975 | 3186 | 2999 | 2788 | 2830 | 2878 | 2652 | 2707 | 2876,91 | 2987 | 2767 | 0,925 | -0,113 | -2,228 | 0,0258898 | 0,031874 | 0,0017197 | 0,002662 | 0,0021055 | 0,0021055 | TRUE | 0,0239668 | No |
| orf19.6302 | 31 | 23 | 11 | 7 | 10 | 6 | 5 | 3 | 30 | 24 | 11 | 11 | 8 | 7 | 4 | 2 | 12,15 | 19 | 5 | 0,278 | -1,849 | -3,544 | 0,0003945 | 0,0005433 | 0,1640077 | 0,1371897 | 0,1488014 | 0,1488014 | TRUE | 0,1298465 | No |
| orf19.6304 | 76 | 114 | 99 | 54 | 156 | 134 | 147 | 101 | 72 | 121 | 98 | 85 | 125 | 152 | 125 | 82 | 107,46 | 94 | 121 | 1,283 | 0,359 | 1,741 | 0,0816891 | 0,0962619 | 0,0464688 | 0,0173703 | 0,0311415 | 0,0311415 | TRUE | 0,3996678 | No |
| orf19.6305 | 988 | 1124 | 1084 | 626 | 1948 | 1488 | 1746 | 1823 | 942 | 1188 | 1071 | 981 | 1561 | 1692 | 1482 | 1482 | 1299,93 | 1046 | 1554 | 1,484 | 0,57 | 7,345 | 2,05E-13 | 3,97E-13 | 0,0064509 | 0,0033543 | 0,0049532 | 0,0049532 | TRUE | 0,1019742 | No |
| orf19.6306 | 986 | 914 | 976 | 524 | 575 | 398 | 522 | 564 | 940 | 966 | 964 | 821 | 461 | 453 | 443 | 459 | 688,33 | 923 | 454 | 0,49 | -1,029 | -14,934 | 1,97E-50 | 7,33E-50 | 0,0015225 | 0,0044766 | 0,0029726 | 0,0029726 | TRUE | 0,057973 | No |
| orf19.6307 | 636 | 808 | 797 | 505 | 661 | 473 | 566 | 540 | 606 | 854 | 787 | 791 | 530 | 538 | 481 | 439 | 628,28 | 760 | 497 | 0,654 | -0,613 | -5,724 | 1,04E-08 | 1,73E-08 | 0,0137759 | 0,0047046 | 0,0093644 | 0,0093644 | TRUE | 0,2258495 | No |
| orf19.6308 | 300 | 449 | 453 | 236 | 336 | 260 | 291 | 255 | 286 | 475 | 448 | 370 | 269 | 296 | 247 | 207 | 324,66 | 395 | 255 | 0,644 | -0,634 | -4,098 | 4,17E-05 | 6,01E-05 | 0,0332822 | 0,0071486 | 0,0197998 | 0,0197998 | TRUE | 0,3395597 | No |
| orf19.6309 | 1949 | 1668 | 1987 | 1227 | 1376 | 1031 | 1255 | 1334 | 1858 | 1764 | 1963 | 1922 | 1103 | 1172 | 1065 | 1085 | 1491,5 | 1877 | 1106 | 0,589 | -0,764 | -14,225 | 6,44E-46 | 2,24E-45 | 0,0012437 | 0,0031921 | 0,0020648 | 0,0020648 | TRUE | 0,0200747 | No |
| orf19.6310 | 720 | 863 | 857 | 627 | 524 | 401 | 446 | 487 | 686 | 913 | 847 | 982 | 420 | 456 | 379 | 396 | 634,79 | 857 | 413 | 0,482 | -1,053 | -9,892 | 4,51E-23 | 1,08E-22 | 0,0133887 | 0,0046778 | 0,0091357 | 0,0091357 | TRUE | 0,2167901 | No |
| orf19.6311 | 25777 | 23140 | 23162 | 14463 | 18906 | 12315 | 17220 | 17498 | 24576 | 24468 | 22882 | 22657 | 15149 | 14003 | 14620 | 14228 | 19072,68 | 23646 | 14500 | 0,613 | -0,706 | -16,524 | 2,47E-61 | 1,06E-60 | 0,0014498 | 0,0021774 | 0,0016969 | 0,0016969 | TRUE | 0,0111287 | No |
| orf19.6312 | 13314 | 11277 | 11986 | 8021 | 42161 | 31335 | 38697 | 44063 | 12693 | 11924 | 11841 | 12565 | 33782 | 35631 | 32854 | 35827 | 23389,74 | 12256 | 34524 | 2,817 | 1,494 | 34,987 | 3,56E-268 | 6,58E-267 | 0,0014341 | 0,0021615 | 0,0016932 | 0,0016932 | TRUE | 0,0129904 | No |
| orf19.6313 | 216 | 277 | 265 | 162 | 680 | 508 | 634 | 597 | 206 | 293 | 262 | 254 | 545 | 578 | 538 | 485 | 395,07 | 254 | 536 | 2,116 | 1,082 | 10,293 | 7,60E-25 | 1,87E-24 | 0,008725 | 0,0062472 | 0,0075588 | 0,0075588 | TRUE | 0,1950175 | No |
| orf19.6313.2 | 1899 | 1778 | 1924 | 973 | 2816 | 1802 | 2662 | 2669 | 1810 | 1880 | 1901 | 1524 | 2256 | 2049 | 2260 | 2170 | 1981,39 | 1779 | 2184 | 1,225 | 0,293 | 4,219 | 2,46E-05 | 3,57E-05 | 0,0051363 | 0,0029199 | 0,0041229 | 0,0041229 | TRUE | 0,10542 | No |
| orf19.6314 | 2833 | 2492 | 2519 | 1388 | 5157 | 3374 | 4861 | 4771 | 2701 | 2635 | 2489 | 2174 | 4132 | 3837 | 4127 | 3879 | 3246,72 | 2500 | 3994 | 1,595 | 0,674 | 10,308 | 6,49E-25 | 1,60E-24 | 0,0047419 | 0,002597 | 0,0037662 | 0,0037662 | TRUE | 0,0887377 | No |

| | | | | | | | | | | | | | | | | | | | | | | | | | | | | | | | |
|---|---|---|---|---|---|---|---|---|---|---|---|---|---|---|---|---|---|---|---|---|---|---|---|---|---|---|---|---|---|---|---|
| orf19.6315 | 91 | 74 | 75 | 54 | 191 | 125 | 152 | 160 | 87 | 78 | 74 | 85 | 153 | 142 | 129 | 130 | 109,75 | 81 | 138 | 1,716 | 0,779 | 5,487 | 4,09E-08 | 6,68E-08 | 1,00E-08 | 0,0170515 | 0,0092499 | 0,0092499 | TRUE | 0,0573284 | No |
| orf19.6316 | 381 | 415 | 422 | 228 | 488 | 377 | 489 | 474 | 363 | 439 | 417 | 357 | 391 | 429 | 415 | 385 | 399,55 | 394 | 405 | 1,023 | 0,033 | 0,376 | 0,7072658 | 0,7374286 | 0,0033861 | 0,0062007 | 0,0048609 | 0,0048609 | TRUE | 0,0655587 | No |
| orf19.6316.4 | 3116 | 3229 | 3384 | 1781 | 7070 | 4586 | 6439 | 5650 | 2971 | 3414 | 3343 | 2790 | 5665 | 5215 | 5467 | 4594 | 4182,31 | 3130 | 5235 | 1,672 | 0,741 | 9,396 | 5,66E-21 | 1,29E-20 | 0,0081576 | 0,0024839 | 0,0057126 | 0,0057126 | TRUE | 0,0833898 | No |
| orf19.6317 | 3590 | 3502 | 3677 | 2263 | 12628 | 9379 | 11295 | 12094 | 3423 | 3703 | 3633 | 3545 | 10118 | 10665 | 9589 | 9834 | 6813,68 | 3576 | 10052 | 2,81 | 1,491 | 32,747 | 3,30E-235 | 5,21E-234 | 0,0013761 | 0,0023323 | 0,0017879 | 0,0017879 | TRUE | 0,0203606 | No |
| orf19.6318 | 1259 | 1075 | 983 | 780 | 1468 | 1028 | 1351 | 1553 | 1200 | 1137 | 971 | 1222 | 1176 | 1169 | 1147 | 1263 | 1160,62 | 1132 | 1189 | 1,052 | 0,073 | 0,979 | 0,3275018 | 0,3611178 | 0,0051972 | 0,003506 | 0,0044252 | 0,0044252 | TRUE | 0,1125528 | No |
| orf19.6319 | 7939 | 6679 | 7659 | 4805 | 4019 | 2711 | 3847 | 4334 | 7569 | 7062 | 7566 | 7527 | 3220 | 3083 | 3266 | 3524 | 5352,23 | 7431 | 3273 | 0,441 | -1,182 | -24,326 | 1,05E-130 | 9,14E-130 | 0,0018451 | 0,0023981 | 0,0020549 | 0,0020549 | TRUE | 0,0324431 | No |
| orf19.6321 | 4179 | 3036 | 3266 | 2859 | 1508 | 1247 | 1490 | 1580 | 3984 | 3210 | 3226 | 4479 | 1208 | 1418 | 1265 | 1285 | 2509,46 | 3725 | 1294 | 0,347 | -1,525 | -15,12 | 1,19E-51 | 4,48E-51 | 0,0154589 | 0,0027456 | 0,0092765 | 0,0092765 | TRUE | 0,2032696 | No |
| orf19.6322 | 37147 | 27718 | 31701 | 20491 | 45667 | 28306 | 44047 | 46250 | 35416 | 29308 | 31318 | 32100 | 36592 | 32186 | 37396 | 37606 | 33990,16 | 32036 | 35945 | 1,122 | 0,166 | 2,502 | 0,0123504 | 0,0156025 | 0,0056118 | 0,0021396 | 0,0042084 | 0,0042084 | TRUE | 0,0619282 | No |
| orf19.6323 | 425 | 342 | 376 | 256 | 637 | 430 | 582 | 583 | 405 | 362 | 371 | 401 | 510 | 489 | 494 | 474 | 438,35 | 385 | 492 | 1,281 | 0,357 | 4,811 | 1,50E-06 | 2,30E-06 | 1,00E-08 | 0,0058369 | 0,0029252 | 0,0029252 | TRUE | 0,0187608 | No |
| orf19.6324 | 10438 | 8100 | 9353 | 5383 | 10429 | 6486 | 10014 | 10349 | 9952 | 8565 | 9240 | 8433 | 8356 | 7375 | 8502 | 8415 | 8604,63 | 9048 | 8162 | 0,902 | -0,149 | -2,309 | 0,0209268 | 0,0259339 | 0,0049203 | 0,0022821 | 0,0038647 | 0,0038647 | TRUE | 0,0554439 | No |
| orf19.6325.1 | 153 | 104 | 130 | 113 | 496 | 267 | 490 | 511 | 146 | 110 | 128 | 177 | 397 | 304 | 416 | 415 | 261,73 | 140 | 383 | 2,763 | 1,466 | 10,177 | 2,51E-24 | 6,12E-24 | 0,0239958 | 0,0083646 | 0,0147402 | 0,0147402 | TRUE | 0,2629703 | No |
| orf19.6326 | 880 | 652 | 759 | 473 | 1850 | 1260 | 1743 | 1872 | 839 | 689 | 750 | 741 | 1482 | 1433 | 1480 | 1522 | 1117,02 | 755 | 1479 | 1,958 | 0,969 | 15,274 | 1,13E-52 | 4,33E-52 | 0,002359 | 0,0035612 | 0,0028222 | 0,0028222 | TRUE | 0,0711105 | No |
| orf19.6327 | 932 | 720 | 820 | 577 | 2228 | 1473 | 2082 | 2282 | 889 | 761 | 810 | 904 | 1785 | 1675 | 1768 | 1855 | 1305,89 | 841 | 1771 | 2,112 | 1,078 | 16,89 | 5,31E-64 | 2,35E-63 | 0,0028994 | 0,0033486 | 0,0029932 | 0,0029932 | TRUE | 0,0478586 | No |
| orf19.6328 | 207 | 143 | 172 | 121 | 571 | 330 | 494 | 502 | 197 | 151 | 170 | 190 | 458 | 375 | 419 | 408 | 296,05 | 177 | 415 | 2,355 | 1,236 | 11,598 | 4,24E-31 | 1,16E-30 | 0,0054849 | 0,0076374 | 0,0066723 | 0,0066723 | TRUE | 0,1045194 | No |
| orf19.6329 | 445 | 368 | 439 | 270 | 636 | 441 | 601 | 650 | 424 | 389 | 434 | 423 | 510 | 501 | 510 | 529 | 464,98 | 418 | 512 | 1,228 | 0,297 | 4,205 | 2,61E-05 | 3,80E-05 | 1,00E-08 | 0,0056224 | 0,0025822 | 0,0025822 | TRUE | 0,0252307 | No |
| orf19.633 | 1405 | 1574 | 1661 | 922 | 2063 | 1510 | 1797 | 1648 | 1340 | 1664 | 1641 | 1444 | 1653 | 1717 | 1526 | 1340 | 1540,59 | 1522 | 1559 | 1,023 | 0,033 | 0,365 | 0,7154753 | 0,7450143 | 0,0106632 | 0,003157 | 0,0071386 | 0,0071386 | TRUE | 0,1085965 | No |
| orf19.6336 | 73 | 46 | 52 | 52 | 171 | 116 | 133 | 144 | 70 | 49 | 51 | 81 | 137 | 132 | 113 | 117 | 93,75 | 63 | 125 | 2,015 | 1,011 | 5,76 | 8,41E-09 | 1,40E-08 | 0,0124422 | 0,0196051 | 0,0169899 | 0,0169899 | TRUE | 0,2540898 | No |
| orf19.6337 | 25 | 16 | 28 | 19 | 111 | 64 | 109 | 89 | 24 | 17 | 28 | 30 | 89 | 73 | 93 | 72 | 53,1 | 25 | 82 | 3,377 | 1,756 | 7,647 | 2,05E-14 | 4,07E-14 | 1,00E-08 | 0,0330123 | 0,0223154 | 0,0223154 | TRUE | 0,2561024 | No |
| orf19.6338 | 1 | 0 | 1 | 4 | 6 | 4 | 6 | 6 | 1 | 0 | 1 | 6 | 5 | 5 | 5 | 5 | 3,44 | 2 | 5 | 2,743 | 1,456 | 1,655 | 0,0978717 | 0,1144646 | 1,00E-08 | 0,4791107 | 0,3362787 | 0,3362787 | TRUE | 1,0734296 | No |
| orf19.6339 | 0 | 0 | 0 | 0 | 0 | 0 | 0 | 0 | 0 | 0 | 0 | 0 | 0 | 0 | 0 | 0 | 0 | 0 | 0 | NA | NA | NA | NA | NA | NA | NA | NA | NA | NA | NA | NA |
| orf19.634 | 2 | 3 | 4 | 2 | 2 | 3 | 2 | 2 | 2 | 3 | 4 | 3 | 2 | 3 | 2 | 2 | 2,56 | 3 | 2 | 0,667 | -0,583 | -0,681 | 0,4957959 | 0,5317098 | 1,00E-08 | 0,6427833 | 0,301037 | 0,301037 | TRUE | 0,136385 | No |
| orf19.6340 | 5127 | 6115 | 5563 | 3746 | 4659 | 3403 | 3976 | 3864 | 4888 | 6466 | 5496 | 5868 | 3733 | 3870 | 3376 | 3142 | 4604,74 | 5680 | 3530 | 0,621 | -0,686 | -7,877 | 3,35E-15 | 6,77E-15 | 0,010912 | 0,0024478 | 0,0070637 | 0,0070637 | TRUE | 0,1079721 | No |
| orf19.6341 | 2169 | 2020 | 2251 | 1372 | 2517 | 1852 | 2438 | 2512 | 2068 | 2136 | 2224 | 2149 | 2017 | 2106 | 2070 | 2042 | 2101,49 | 2144 | 2059 | 0,96 | -0,059 | -1,416 | 0,1567387 | 0,1796273 | 1,00E-08 | 0,0028726 | 0,0012027 | 0,0012027 | TRUE | 0,0081298 | No |
| orf19.6342 | 1003 | 879 | 932 | 428 | 1576 | 882 | 1413 | 1400 | 956 | 929 | 921 | 670 | 1263 | 1003 | 1200 | 1138 | 1010,07 | 869 | 1151 | 1,321 | 0,401 | 3,8 | 0,0001446 | 0,0002029 | 0,0150717 | 0,0037168 | 0,0096813 | 0,0096813 | TRUE | 0,2642822 | No |
| orf19.6343 | 10020 | 11251 | 11240 | 6409 | 11202 | 8329 | 10065 | 8770 | 9553 | 11897 | 11104 | 10040 | 8976 | 9471 | 8545 | 7131 | 9589,53 | 10648 | 8531 | 0,801 | -0,32 | -3,61 | 0,0003063 | 0,0004243 | 0,0117283 | 0,0022625 | 0,0074538 | 0,0074538 | TRUE | 0,1493626 | No |
| orf19.6344 | 2584 | 2469 | 2593 | 1532 | 2807 | 1976 | 2651 | 2628 | 2464 | 2611 | 2562 | 2400 | 2249 | 2247 | 2251 | 2137 | 2364,92 | 2509 | 2221 | 0,884 | -0,178 | -4,064 | 4,83E-05 | 6,93E-05 | 0,0004692 | 0,0027856 | 0,001414 | 0,001414 | TRUE | 0,0093509 | No |
| orf19.6345 | 6169 | 6490 | 6728 | 4221 | 10812 | 8417 | 9876 | 9845 | 5881 | 6862 | 6647 | 6612 | 8663 | 9571 | 8385 | 8005 | 7578,34 | 6500 | 8656 | 1,331 | 0,413 | 6,372 | 1,87E-10 | 3,31E-10 | 0,0049504 | 0,0023079 | 0,0038968 | 0,0038968 | TRUE | 0,0614705 | No |
| orf19.6346 | 2357 | 2545 | 2478 | 1708 | 4034 | 3242 | 3610 | 3787 | 2247 | 2691 | 2448 | 2676 | 3232 | 3686 | 3065 | 3079 | 2890,59 | 2516 | 3266 | 1,299 | 0,377 | 5,014 | 5,33E-07 | 8,35E-07 | 0,0069926 | 0,0026593 | 0,0050666 | 0,0050666 | TRUE | 0,0902656 | No |
| orf19.6347 | 3110 | 3203 | 3347 | 2064 | 2910 | 2244 | 2737 | 2562 | 2965 | 3387 | 3307 | 3233 | 2332 | 2552 | 2324 | 2083 | 2772,74 | 3223 | 2323 | 0,72 | -0,474 | -7,155 | 8,39E-13 | 1,59E-12 | 0,0045126 | 0,0026834 | 0,0038406 | 0,0038406 | TRUE | 0,0585435 | No |
| orf19.6348 | 657 | 742 | 735 | 491 | 1737 | 1298 | 1536 | 1467 | 626 | 785 | 726 | 769 | 1392 | 1476 | 1304 | 1193 | 1033,86 | 726 | 1341 | 1,848 | 0,886 | 10,294 | 7,52E-25 | 1,86E-24 | 0,0074778 | 0,0036794 | 0,0060029 | 0,0060029 | TRUE | 0,1054241 | No |
| orf19.6349 | 1797 | 2101 | 2138 | 1150 | 633 | 468 | 574 | 494 | 1713 | 2222 | 2112 | 1802 | 507 | 532 | 487 | 402 | 1222,1 | 1962 | 482 | 0,245 | -2,028 | -19,971 | 9,77E-89 | 5,77E-88 | 0,0128706 | 0,0034348 | 0,0086871 | 0,0086871 | TRUE | 0,1184084 | No |
| orf19.635 | 10 | 7 | 11 | 13 | 2 | 4 | 4 | 2 | 10 | 7 | 11 | 20 | 2 | 5 | 3 | 2 | 7,42 | 12 | 3 | 0,23 | -2,121 | -3,402 | 0,0006686 | 0,0009128 | 1,00E-08 | 0,2234438 | 0,16039 | 0,16039 | TRUE | 0,5619064 | No |
| orf19.6350 | 216 | 165 | 184 | 138 | 63 | 42 | 57 | 49 | 206 | 174 | 182 | 216 | 50 | 48 | 48 | 40 | 120,6 | 194 | 46 | 0,24 | -2,058 | -13,443 | 3,41E-41 | 1,10E-40 | 1,00E-08 | 0,0157053 | 0,0100862 | 0,0100862 | TRUE | 0,088453 | No |
| orf19.6351 | 2970 | 2792 | 3080 | 1684 | 1771 | 1268 | 1770 | 1787 | 2832 | 2952 | 3043 | 2638 | 1419 | 1442 | 1503 | 1453 | 2160,15 | 2866 | 1454 | 0,507 | -0,981 | -18,56 | 6,83E-77 | 3,58E-76 | 0,0017331 | 0,0028514 | 0,0021786 | 0,0021786 | TRUE | 0,0326438 | No |
| orf19.6353 | 988 | 897 | 945 | 606 | 1055 | 806 | 955 | 1066 | 942 | 948 | 934 | 949 | 845 | 916 | 811 | 867 | 901,59 | 943 | 860 | 0,91 | -0,136 | -2,409 | 0,0159905 | 0,0200227 | 1,00E-08 | 0,0039124 | 0,0019372 | 0,0019372 | TRUE | 0,0212966 | No |
| orf19.6354 | 168 | 193 | 184 | 123 | 381 | 301 | 341 | 272 | 160 | 204 | 182 | 193 | 305 | 342 | 290 | 221 | 237,12 | 185 | 290 | 1,567 | 0,648 | 4,681 | 2,86E-06 | 4,35E-06 | 0,0186645 | 0,0090158 | 0,0138216 | 0,0138216 | TRUE | 0,2924466 | No |
| orf19.6355 | 16757 | 14504 | 15782 | 9142 | 15840 | 10810 | 15188 | 16355 | 15976 | 15336 | 15591 | 14321 | 12692 | 12292 | 12895 | 13298 | 14050,19 | 15306 | 12794 | 0,836 | -0,259 | -5,938 | 2,88E-09 | 4,88E-09 | 0,0015081 | 0,0022081 | 0,0017547 | 0,0017547 | TRUE | 0,0225897 | No |
| orf19.6356 | 1039 | 1224 | 1247 | 710 | 1731 | 1292 | 1445 | 1404 | 991 | 1294 | 1232 | 1112 | 1387 | 1469 | 1227 | 1142 | 1231,68 | 1157 | 1306 | 1,128 | 0,173 | 1,783 | 0,07454 | 0,0881467 | 0,012146 | 0,0034243 | 0,0082267 | 0,0082267 | TRUE | 0,1164911 | No |
| orf19.6357 | 492 | 547 | 590 | 367 | 668 | 508 | 622 | 502 | 469 | 578 | 583 | 575 | 535 | 578 | 528 | 408 | 531,8 | 551 | 512 | 0,929 | -0,107 | -0,961 | 0,3363949 | 0,3702201 | 0,0145106 | 0,0051787 | 0,0099108 | 0,0099108 | TRUE | 0,2229448 | No |
| orf19.6358 | 1179 | 1550 | 1379 | 912 | 952 | 780 | 787 | 849 | 1124 | 1639 | 1362 | 1429 | 763 | 887 | 668 | 690 | 1070,28 | 1388 | 752 | 0,541 | -0,886 | -7,793 | 6,54E-15 | 1,32E-14 | 0,0188916 | 0,0036254 | 0,011409 | 0,011409 | TRUE | 0,1676303 | No |
| orf19.6359 | 500 | 464 | 457 | 272 | 315 | 266 | 299 | 306 | 477 | 491 | 451 | 426 | 252 | 302 | 254 | 249 | 362,8 | 461 | 264 | 0,569 | -0,814 | -9,141 | 6,17E-20 | 1,38E-19 | 0,0023083 | 0,0066169 | 0,0047111 | 0,0047111 | TRUE | 0,0942832 | No |
| orf19.6360 | 3006 | 2830 | 2918 | 1796 | 1730 | 1212 | 1642 | 1588 | 2866 | 2992 | 2883 | 2814 | 1386 | 1378 | 1394 | 1291 | 2125,51 | 2889 | 1362 | 0,471 | -1,086 | -24,297 | 2,09E-130 | 1,82E-129 | 0,0004157 | 0,0028638 | 0,0013946 | 0,0013946 | TRUE | 0,0154918 | No |
| orf19.6362 | 6606 | 5978 | 6438 | 4056 | 8949 | 6427 | 8645 | 9195 | 6298 | 6321 | 6360 | 6354 | 7171 | 7308 | 7340 | 7476 | 6828,47 | 6333 | 7324 | 1,156 | 0,21 | 7,202 | 5,94E-13 | 1,13E-12 | 1,00E-08 | 0,0023317 | 0,0006646 | 0,0006646 | TRUE | 0,0024928 | No |
| orf19.6365 | 6681 | 5043 | 5703 | 3976 | 6452 | 4465 | 6380 | 6640 | 6370 | 5332 | 5634 | 6229 | 5170 | 5077 | 5417 | 5399 | 5578,38 | 5891 | 5266 | 0,894 | -0,161 | -2,701 | 0,0069202 | 0,0088635 | 0,0037363 | 0,0023856 | 0,003247 | 0,003247 | TRUE | 0,049824 | No |
| orf19.6366 | 84 | 70 | 64 | 38 | 47 | 34 | 61 | 48 | 80 | 74 | 63 | 60 | 38 | 39 | 52 | 39 | 55,5 | 69 | 42 | 0,6 | -0,736 | -3,662 | 0,0002505 | 0,0003484 | 1,00E-08 | 0,0316755 | 0,0201727 | 0,0201727 | TRUE | 0,2161646 | No |
| orf19.6367 | 43685 | 34842 | 38478 | 27585 | 112061 | 79964 | 107726 | 120169 | 41649 | 36841 | 38013 | 43213 | 89791 | 90926 | 91459 | 97709 | 66200,15 | 39929 | 92471 | 2,316 | 1,212 | 21,698 | 2,14E-104 | 1,45E-103 | 0,0035349 | 0,0021161 | 0,0029774 | 0,0029774 | TRUE | 0,0374146 | No |
| orf19.6369 | 9898 | 8076 | 8840 | 5080 | 12062 | 7281 | 11697 | 12053 | 9437 | 8539 | 8733 | 7958 | 9665 | 8279 | 9931 | 9800 | 9042,78 | 8667 | 9419 | 1,087 | 0,12 | 1,764 | 0,0776616 | 0,0917192 | 0,0056472 | 0,0022729 | 0,0043226 | 0,0043226 | TRUE | 0,0805834 | No |
| orf19.637 | 4718 | 3955 | 4203 | 2885 | 6572 | 4916 | 6399 | 6689 | 4498 | 4182 | 4152 | 4520 | 5266 | 5590 | 5433 | 5439 | 4884,89 | 4338 | 5432 | 1,253 | 0,325 | 7,598 | 3,00E-14 | 5,93E-14 | 0,001061 | 0,0024274 | 0,0015442 | 0,0015442 | TRUE | 0,0101644 | No |
| orf19.6371 | 223 | 179 | 197 | 100 | 446 | 298 | 407 | 462 | 213 | 189 | 195 | 157 | 357 | 339 | 346 | 376 | 271,32 | 188 | 354 | 1,865 | 0,899 | 8,993 | 2,40E-19 | 5,31E-19 | 0,0010986 | 0,0081428 | 0,0054103 | 0,0054103 | TRUE | 0,1061737 | No |
| orf19.6373 | 8204 | 6875 | 7796 | 4820 | 4120 | 2877 | 3981 | 4150 | 7822 | 7269 | 7702 | 7551 | 3301 | 3271 | 3380 | 3374 | 5458,8 | 7586 | 3332 | 0,439 | -1,187 | -32,286 | 1,11E-228 | 1,70E-227 | 0,0005115 | 0,0023921 | 0,0010894 | 0,0010894 | TRUE | 0,0098482 | No |
| orf19.6374 | 630 | 563 | 623 | 340 | 1845 | 1184 | 1876 | 1908 | 601 | 595 | 615 | 533 | 1478 | 1346 | 1593 | 1551 | 1039,1 | 586 | 1492 | 2,54 | 1,345 | 19,008 | 1,46E-80 | 8,02E-80 | 0,0034514 | 0,0036714 | 0,0035535 | 0,0035535 | TRUE | 0,0499758 | No |
| orf19.6375 | 8818 | 7232 | 7436 | 4467 | 31359 | 19089 | 29326 | 31109 | 8407 | 7647 | 7346 | 6998 | 25127 | 21706 | 24898 | 25295 | 15927,87 | 7600 | 24256 | 3,191 | 1,674 | 25,233 | 1,75E-140 | 1,66E-139 | 0,0054825 | 0,0021944 | 0,0041356 | 0,0041356 | TRUE | 0,0626502 | No |
| orf19.6376 | 2530 | 2144 | 2409 | 1445 | 3170 | 2094 | 3032 | 3258 | 2412 | 2267 | 2380 | 2264 | 2540 | 2381 | 2574 | 2649 | 2433,37 | 2331 | 2536 | 1,088 | 0,122 | 2,567 | 0,0102635 | 0,0130279 | 0,0009822 | 0,002766 | 0,0017452 | 0,0017452 | TRUE | 0,0196037 | No |
| orf19.6377 | 1415 | 1177 | 1401 | 800 | 1568 | 1027 | 1493 | 1630 | 1349 | 1245 | 1384 | 1253 | 1256 | 1168 | 1268 | 1325 | 1280,99 | 1308 | 1254 | 0,959 | -0,061 | -1,027 | 0,3043586 | 0,3374652 | 0,0017181 | 0,003373 | 0,002552 | 0,002552 | TRUE | 0,0243346 | No |
| orf19.6378 | 2536 | 1996 | 2176 | 1321 | 4143 | 2479 | 4011 | 4135 | 2418 | 2111 | 2150 | 2069 | 3320 | 2819 | 3405 | 3362 | 2706,68 | 2187 | 3226 | 1,475 | 0,561 | 7,844 | 4,37E-15 | 8,82E-15 | 0,0056853 | 0,0026979 | 0,0045136 | 0,0045136 | TRUE | 0,0864531 | No |
| orf19.6379 | 668 | 653 | 636 | 451 | 564 | 461 | 559 | 606 | 637 | 690 | 628 | 707 | 452 | 524 | 475 | 493 | 575,7 | 666 | 486 | 0,73 | -0,453 | -6,224 | 4,84E-10 | 8,43E-10 | 0,0017231 | 0,0049433 | 0,0033314 | 0,0033314 | TRUE | 0,0299696 | No |
| orf19.638 | 14229 | 12308 | 12655 | 8815 | 6176 | 4479 | 5570 | 6240 | 13566 | 13014 | 12502 | 13809 | 4949 | 5093 | 4729 | 5074 | 9091,92 | 13223 | 4961 | 0,375 | -1,414 | -32,124 | 2,02E-226 | 3,07E-225 | 0,0013618 | 0,0022719 | 0,0017287 | 0,0017287 | TRUE | 0,0165133 | No |
| orf19.6382 | 3383 | 2871 | 3196 | 2202 | 4727 | 3436 | 4454 | 4804 | 3225 | 3036 | 3157 | 3450 | 3788 | 3907 | 3781 | 3906 | 3531,27 | 3217 | 3846 | 1,197 | 0,259 | 5,739 | 9,55E-09 | 1,59E-08 | 0,0012461 | 0,0025562 | 0,0016682 | 0,0016682 | TRUE | 0,0268684 | No |
| orf19.6385 | 16005 | 12914 | 14236 | 10114 | 56724 | 39613 | 52725 | 60513 | 15259 | 13655 | 14064 | 15844 | 45451 | 45044 | 44763 | 49203 | 30410,36 | 14706 | 46115 | 3,137 | 1,649 | 29,959 | 3,33E-197 | 4,50E-196 | 0,0033585 | 0,0021453 | 0,0028634 | 0,0028634 | TRUE | 0,0328829 | No |
| orf19.6387 | 32317 | 26284 | 29280 | 19494 | 2973 | 2046 | 2869 | 3090 | 30811 | 27792 | 28926 | 30538 | 2382 | 2326 | 2436 | 2512 | 15965,49 | 29517 | 2414 | 0,082 | -3,611 | -78,501 | 0 | 0 | 0,0013959 | 0,0021941 | 0,0018284 | 0,0018284 | TRUE | 0,0189598 | No |
| orf19.639 | 164 | 140 | 144 | 108 | 727 | 509 | 685 | 694 | 156 | 148 | 142 | 169 | 583 | 579 | 582 | 564 | 365,37 | 154 | 577 | 3,773 | 1,916 | 21,74 | 8,60E-105 | 5,83E-104 | 1,00E-08 | 0,006585 | 0,0030604 | 0,0030604 | TRUE | 0,0337568 | No |
| orf19.6390 | 1 | 1 | 4 | 3 | 0 | 1 | 2 | 1 | 1 | 1 | 4 | 5 | 0 | 1 | 2 | 1 | 1,79 | 3 | 1 | 0,347 | -1,525 | -1,307 | 0,1913776 | 0,2173692 | 1,00E-08 | 0,9199825 | 0,5946411 | 0,5946411 | TRUE | 0,1502411 | No |
| orf19.6391 | 161 | 143 | 158 | 135 | 51 | 26 | 53 | 62 | 153 | 151 | 156 | 211 | 41 | 30 | 45 | 50 | 104,76 | 168 | 42 | 0,252 | -1,99 | -10,9 | 1,15E-27 | 3,00E-27 | 0,0185108 | 0,0177637 | 0,0180404 | 0,0180404 | TRUE | 0,2757519 | No |
| orf19.639.1 | 690 | 532 | 641 | 412 | 949 | 682 | 879 | 977 | 658 | 563 | 633 | 645 | 760 | 775 | 746 | 794 | 696,95 | 625 | 769 | 1,232 | 0,301 | 4,548 | 5,42E-06 | 8,14E-06 | 0,0009831 | 0,0044471 | 0,0027235 | 0,0027235 | TRUE | 0,0542739 | No |
| orf19.6392 | 452 | 375 | 459 | 270 | 1420 | 901 | 1374 | 1489 | 431 | 397 | 453 | 423 | 1138 | 1025 | 1167 | 1211 | 780,43 | 426 | 1135 | 2,666 | 1,415 | 19,174 | 6,16E-82 | 3,42E-81 | 0,0029346 | 0,0041951 | 0,0035228 | 0,0035228 | TRUE | 0,0487025 | No |
| orf19.6393 | 1392 | 1146 | 1254 | 878 | 1691 | 1085 | 1594 | 1632 | 1327 | 1212 | 1239 | 1375 | 1355 | 1234 | 1353 | 1327 | 1302,76 | 1288 | 1317 | 1,026 | 0,036 | 0,629 | 0,5290723 | 0,5650227 | 0,0016467 | 0,0033516 | 0,0024347 | 0,0024347 | TRUE | 0,0222737 | No |
| orf19.6396 | 2477 | 2253 | 2331 | 1504 | 3666 | 2442 | 3428 | 3751 | 2362 | 2382 | 2303 | 2356 | 2937 | 2777 | 2910 | 3050 | 2634,65 | 2351 | 2918 | 1,243 | 0,314 | 7,427 | 1,11E-13 | 2,17E-13 | 0,00054 | 0,0027145 | 0,0013196 | 0,0013196 | TRUE | 0,0123653 | No |
| orf19.6398 | 410 | 344 | 392 | 256 | 1318 | 981 | 1179 | 1325 | 391 | 364 | 387 | 401 | 1056 | 1115 | 1001 | 1077 | 724,1 | 386 | 1062 | 2,757 | 1,463 | 21,927 | 1,45E-106 | 1,00E-105 | 1,00E-08 | 0,0043588 | 0,0024087 | 0,0024087 | TRUE | 0,0180794 | No |
| orf19.6399 | 577 | 448 | 500 | 314 | 982 | 608 | 903 | 987 | 550 | 474 | 494 | 492 | 787 | 691 | 767 | 803 | 632,13 | 502 | 762 | 1,517 | 0,601 | 8,063 | 7,46E-16 | 1,53E-15 | 0,0026475 | 0,0046887 | 0,0036401 | 0,0036401 | TRUE | 0,0508771 | No |
| orf19.640 | 1010 | 1007 | 1027 | 621 | 1346 | 932 | 1240 | 1208 | 963 | 1065 | 1015 | 973 | 1079 | 1060 | 1053 | 982 | 1023,55 | 1004 | 1044 | 1,038 | 0,054 | 0,94 | 0,3470897 | 0,3814639 | 0,0009955 | 0,0036954 | 0,0021885 | 0,0021885 | TRUE | 0,0201221 | No |
| orf19.6400 | 1539 | 1238 | 1309 | 989 | 2335 | 1656 | 2391 | 2583 | 1467 | 1309 | 1293 | 1549 | 1871 | 1883 | 2030 | 2100 | 1687,87 | 1404 | 1971 | 1,406 | 0,492 | 7,128 | 1,02E-12 | 1,93E-12 | 0,0047351 | 0,003064 | 0,0039461 | 0,0039461 | TRUE | 0,0537882 | No |
| orf19.6402 | 69240 | 54656 | 59709 | 43340 | 121090 | 84391 | 116994 | 123602 | 66013 | 57792 | 58987 | 67894 | 97026 | 95960 | 99327 | 100500 | 80437,45 | 62672 | 98203 | 1,567 | 0,648 | 11,72 | 1,01E-31 | 2,81E-31 | 0,0034243 | 0,0021117 | 0,0029245 | 0,0029245 | TRUE | 0,0384685 | No |
| orf19.6403 | 3182 | 2713 | 3014 | 1637 | 4566 | 3054 | 4341 | 4582 | 3034 | 2869 | 2978 | 2564 | 3659 | 3473 | 3685 | 3726 | 3248,34 | 2861 | 3636 | 1,269 | 0,344 | 6,193 | 5,89E-10 | 1,02E-09 | 0,0027341 | 0,0025967 | 0,0026363 | 0,0026363 | TRUE | 0,056028 | No |

| | | | | | | | | | | | | | | | | | | | | | | | | | | | | | | | |
|---|---|---|---|---|---|---|---|---|---|---|---|---|---|---|---|---|---|---|---|---|---|---|---|---|---|---|---|---|---|---|---|
| orf19.6403.1 | 3499 | 2828 | 2957 | 2316 | 13482 | 8799 | 11998 | 12969 | 3336 | 2990 | 2921 | 3628 | 10803 | 10005 | 10186 | 10545 | 6801,85 | 3219 | 10385 | 3,23 | 1,692 | 25,728 | 5,66E-146 | 5,57E-145 | 0,0055023 | 0,0023327 | 0,0039342 | 0,0039342 | TRUE | 0,0861307 | No |
| orf19.6404 | 2083 | 1659 | 1907 | 1122 | 4041 | 2578 | 4049 | 4049 | 1986 | 1754 | 1884 | 1758 | 3238 | 2931 | 3438 | 3292 | 2535,11 | 1846 | 3225 | 1,746 | 0,804 | 12,983 | 1,53E-38 | 4,72E-38 | 0,0034355 | 0,0027389 | 0,0032432 | 0,0032432 | TRUE | 0,0445534 | No |
| orf19.6405 | 14 | 11 | 9 | 8 | 25 | 19 | 14 | 19 | 13 | 12 | 9 | 13 | 20 | 22 | 12 | 15 | 14,42 | 12 | 17 | 1,482 | 0,567 | 1,484 | 0,1377924 | 0,1585954 | 1,00E-08 | 0,115941 | 0,066169 | 0,066169 | TRUE | 0,2395442 | No |
| orf19.6406 | 2173 | 1686 | 1861 | 1257 | 2126 | 1356 | 2031 | 2089 | 2072 | 1783 | 1838 | 1969 | 1704 | 1542 | 1724 | 1699 | 1791,3 | 1916 | 1667 | 0,871 | -0,199 | -3,251 | 0,0011486 | 0,001542 | 0,0028367 | 0,0030079 | 0,0030225 | 0,0030225 | TRUE | 0,0384089 | No |
| orf19.6407 | 2943 | 2403 | 2666 | 1588 | 4011 | 2540 | 3932 | 4039 | 2806 | 2541 | 2634 | 2488 | 3214 | 2888 | 3338 | 3284 | 2899,07 | 2617 | 3181 | 1,215 | 0,281 | 4,817 | 1,46E-06 | 2,24E-06 | 0,0029703 | 0,0026576 | 0,0029187 | 0,0029187 | TRUE | 0,0455296 | No |
| orf19.6408 | 6033 | 5022 | 5788 | 3760 | 2098 | 1503 | 2182 | 2129 | 5752 | 5310 | 5718 | 5890 | 1681 | 1709 | 1853 | 1731 | 3705,49 | 5668 | 1744 | 0,308 | -1,7 | -34,664 | 2,80E-263 | 5,05E-262 | 0,001409 | 0,0025344 | 0,001954 | 0,001954 | TRUE | 0,0219988 | No |
| orf19.641 | 874 | 904 | 979 | 497 | 1733 | 1184 | 1634 | 1570 | 833 | 956 | 967 | 779 | 1389 | 1346 | 1387 | 1277 | 1116,7 | 884 | 1350 | 1,523 | 0,607 | 8,319 | 8,89E-17 | 1,86E-16 | 0,0050089 | 0,0035616 | 0,0041418 | 0,0041418 | TRUE | 0,0682271 | No |
| orf19.6411 | 1586 | 1304 | 1451 | 985 | 1131 | 705 | 1058 | 1161 | 1512 | 1379 | 1433 | 1543 | 906 | 802 | 898 | 944 | 1177,19 | 1467 | 888 | 0,607 | -0,72 | -11,371 | 5,84E-30 | 1,57E-29 | 0,0024231 | 0,003486 | 0,0029668 | 0,0029668 | TRUE | 0,0470196 | No |
| orf19.6413 | 798 | 764 | 881 | 504 | 1453 | 1031 | 1351 | 1515 | 761 | 808 | 870 | 790 | 1164 | 1172 | 1147 | 1232 | 992,99 | 807 | 1179 | 1,46 | 0,546 | 9,255 | 2,15E-20 | 4,86E-20 | 0,0010119 | 0,0037448 | 0,002266 | 0,002266 | TRUE | 0,0350048 | No |
| orf19.6414 | 1097 | 906 | 1089 | 699 | 2449 | 1772 | 2397 | 2422 | 1046 | 958 | 1076 | 1095 | 1962 | 2015 | 2035 | 1969 | 1519,54 | 1044 | 1995 | 1,915 | 0,937 | 18,128 | 1,93E-73 | 9,69E-73 | 0,0005265 | 0,0031718 | 0,0018044 | 0,0018044 | TRUE | 0,0365119 | No |
| orf19.6414.3 | 3607 | 3151 | 3536 | 2123 | 3630 | 2372 | 3324 | 3488 | 3439 | 3332 | 3493 | 3326 | 2909 | 2697 | 2822 | 2836 | 3106,7 | 3398 | 2816 | 0,829 | -0,271 | -6,772 | 1,27E-11 | 2,34E-11 | 0,000347 | 0,0026198 | 0,0012092 | 0,0012092 | TRUE | 0,0092726 | No |
| orf19.6415.1 | 4106 | 2937 | 3079 | 2163 | 10793 | 6973 | 9436 | 11217 | 3915 | 3106 | 3042 | 3388 | 8648 | 7929 | 8011 | 9120 | 5894,88 | 3363 | 8427 | 2,506 | 1,326 | 16,766 | 4,31E-63 | 1,89E-62 | 0,0089474 | 0,0023698 | 0,0057833 | 0,0057833 | TRUE | 0,1507643 | No |
| orf19.6416 | 4480 | 3712 | 4180 | 2844 | 3259 | 2299 | 2981 | 3375 | 4271 | 3925 | 4129 | 4455 | 2611 | 2614 | 2531 | 2744 | 3410,18 | 4195 | 2625 | 0,626 | -0,675 | -13,717 | 7,99E-43 | 2,65E-42 | 0,0015802 | 0,0025728 | 0,0020211 | 0,0020211 | TRUE | 0,0227051 | No |
| orf19.6417 | 18389 | 16197 | 17757 | 10164 | 19588 | 12927 | 19115 | 20001 | 17532 | 17126 | 17542 | 15922 | 15695 | 14699 | 16229 | 16263 | 16376,08 | 17030 | 15722 | 0,923 | -0,116 | -2,438 | 0,0147543 | 0,0185074 | 0,0020003 | 0,0021915 | 0,0020965 | 0,0020965 | TRUE | 0,0232736 | No |
| orf19.6418 | 5621 | 4717 | 5268 | 3052 | 7690 | 5126 | 7251 | 7663 | 5359 | 4988 | 5204 | 4781 | 6162 | 5829 | 6156 | 6231 | 5588,67 | 5083 | 6094 | 1,198 | 0,261 | 5,793 | 6,91E-09 | 1,15E-08 | 0,0014299 | 0,0023851 | 0,0017637 | 0,0017637 | TRUE | 0,0186786 | No |
| orf19.6419.1 | 828 | 714 | 696 | 399 | 1111 | 699 | 1020 | 1046 | 789 | 755 | 688 | 625 | 890 | 795 | 866 | 850 | 782,32 | 714 | 850 | 1,187 | 0,248 | 3,236 | 0,0012113 | 0,0016235 | 0,0046342 | 0,0041901 | 0,0043089 | 0,0043089 | TRUE | 0,0739041 | No |
| orf19.642 | 911 | 852 | 946 | 547 | 1588 | 1147 | 1453 | 1593 | 869 | 901 | 935 | 857 | 1272 | 1304 | 1234 | 1295 | 1083,3 | 890 | 1276 | 1,431 | 0,517 | 9,856 | 6,46E-23 | 1,53E-22 | 1,00E-08 | 0,0036069 | 0,0016628 | 0,0016628 | TRUE | 0,013618 | No |
| orf19.6420 | 8235 | 6767 | 7567 | 5932 | 1913 | 1441 | 1827 | 1960 | 7851 | 7155 | 7475 | 9293 | 1533 | 1639 | 1551 | 1594 | 4761,36 | 7944 | 1579 | 0,199 | -2,33 | -31,043 | 1,43E-211 | 2,05E-210 | 0,0069905 | 0,0024361 | 0,0050593 | 0,0050593 | TRUE | 0,1579514 | No |
| orf19.6422 | 3310 | 3068 | 3249 | 2117 | 2716 | 1974 | 2704 | 2911 | 3156 | 3244 | 3210 | 3316 | 2176 | 2245 | 2296 | 2367 | 2751,16 | 3232 | 2271 | 0,703 | -0,508 | -12,218 | 2,51E-34 | 7,25E-34 | 0,0004865 | 0,0026881 | 0,0012867 | 0,0012867 | TRUE | 0,0101295 | No |
| orf19.6423 | 2164 | 1874 | 2010 | 1324 | 2794 | 1970 | 2737 | 2878 | 2063 | 1982 | 1986 | 2074 | 2239 | 2240 | 2324 | 2340 | 2155,88 | 2026 | 2286 | 1,129 | 0,176 | 4,259 | 2,06E-05 | 3,00E-05 | 1,00E-08 | 0,0028529 | 0,0011585 | 0,0011585 | TRUE | 0,0031446 | No |
| orf19.6424 | 2645 | 1965 | 2109 | 1545 | 2952 | 1761 | 2699 | 2919 | 2522 | 2078 | 2083 | 2420 | 2365 | 2002 | 2291 | 2373 | 2266,99 | 2276 | 2258 | 0,993 | -0,01 | -0,122 | 0,9031342 | 0,9167832 | 0,0076918 | 0,0028155 | 0,0054865 | 0,0054865 | TRUE | 0,0690592 | No |
| orf19.6425 | 1822 | 1657 | 1741 | 1030 | 3087 | 1934 | 2897 | 3257 | 1737 | 1752 | 1720 | 1614 | 2474 | 2199 | 2460 | 2648 | 2075,39 | 1706 | 2445 | 1,433 | 0,519 | 8,535 | 1,40E-17 | 2,98E-17 | 0,0028796 | 0,0028824 | 0,0030425 | 0,0030425 | TRUE | 0,0539509 | No |
| orf19.643 | 485 | 436 | 437 | 279 | 860 | 603 | 791 | 828 | 462 | 461 | 432 | 437 | 689 | 686 | 672 | 673 | 563,97 | 448 | 680 | 1,515 | 0,599 | 9,318 | 1,19E-20 | 2,70E-20 | 1,00E-08 | 0,0050026 | 0,0020765 | 0,0020765 | TRUE | 0,0083884 | No |
| orf19.6431 | 1197 | 1049 | 1090 | 674 | 1134 | 752 | 1115 | 1214 | 1141 | 1109 | 1077 | 1056 | 909 | 855 | 947 | 987 | 1010,07 | 1096 | 924 | 0,844 | -0,245 | -4,104 | 4,07E-05 | 5,86E-05 | 0,0013391 | 0,0037168 | 0,0024237 | 0,0024237 | TRUE | 0,027972 | No |
| orf19.6432 | 2036 | 1677 | 1915 | 1200 | 3066 | 1966 | 2962 | 3204 | 1941 | 1773 | 1892 | 1880 | 2457 | 2236 | 2515 | 2605 | 2162,27 | 1872 | 2453 | 1,312 | 0,392 | 6,925 | 4,35E-12 | 8,09E-12 | 0,0021424 | 0,0028506 | 0,0025869 | 0,0025869 | TRUE | 0,041713 | No |
| orf19.6434 | 5702 | 4666 | 5110 | 3392 | 10149 | 6851 | 9781 | 10583 | 5436 | 4934 | 5048 | 5314 | 8132 | 7790 | 8304 | 8605 | 6695,4 | 5183 | 8208 | 1,584 | 0,664 | 14,239 | 5,21E-46 | 1,82E-45 | 0,0016512 | 0,0023365 | 0,0019242 | 0,0019242 | TRUE | 0,014124 | No |
| orf19.6435 | 906 | 739 | 762 | 594 | 2265 | 1370 | 2079 | 2128 | 864 | 781 | 753 | 931 | 1815 | 1558 | 1765 | 1730 | 1274,56 | 832 | 1717 | 2,071 | 1,051 | 14,04 | 8,90E-45 | 3,03E-44 | 0,0056319 | 0,0033795 | 0,0044396 | 0,0044396 | TRUE | 0,0681769 | No |
| orf19.6436 | 1507 | 1257 | 1405 | 889 | 1366 | 927 | 1289 | 1352 | 1437 | 1329 | 1388 | 1393 | 1095 | 1054 | 1094 | 1099 | 1236,1 | 1387 | 1086 | 0,783 | -0,352 | -7,241 | 4,46E-13 | 8,55E-13 | 1,00E-08 | 0,0034196 | 0,0014566 | 0,0014566 | TRUE | 0,009797 | No |
| orf19.6437 | 725 | 644 | 678 | 481 | 839 | 556 | 763 | 835 | 691 | 681 | 670 | 754 | 672 | 632 | 648 | 679 | 678,33 | 699 | 658 | 0,946 | -0,08 | -1,276 | 0,2019519 | 0,2289721 | 1,00E-08 | 0,0045118 | 0,0023205 | 0,0023205 | TRUE | 0,0278195 | No |
| orf19.6438 | 2973 | 2538 | 2884 | 1750 | 6726 | 4555 | 6408 | 6837 | 2834 | 2684 | 2849 | 2741 | 5389 | 5179 | 5440 | 5559 | 4084,62 | 2777 | 5392 | 1,941 | 0,957 | 24,048 | 8,82E-128 | 7,53E-127 | 0,0004683 | 0,0024933 | 0,0012355 | 0,0012355 | TRUE | 0,008323 | No |
| orf19.644 | 217 | 195 | 198 | 131 | 103 | 60 | 83 | 89 | 207 | 206 | 196 | 205 | 83 | 68 | 70 | 72 | 138,44 | 204 | 73 | 0,362 | -1,464 | -11,498 | 1,35E-30 | 3,68E-30 | 1,00E-08 | 0,0139517 | 0,006809 | 0,006809 | TRUE | 0,0697786 | No |
| orf19.6440 | 6923 | 5801 | 6557 | 3893 | 2532 | 1714 | 2500 | 2453 | 6600 | 6134 | 6478 | 6099 | 2029 | 1949 | 2122 | 1995 | 4175,66 | 6328 | 2024 | 0,32 | -1,645 | -36,568 | 9,35E-293 | 2,03E-291 | 0,0010034 | 0,0024845 | 0,0016327 | 0,0016327 | TRUE | 0,0130175 | No |
| orf19.6442 | 2875 | 2430 | 2725 | 1680 | 5703 | 3824 | 5133 | 5785 | 2741 | 2569 | 2692 | 2632 | 4570 | 4348 | 4358 | 4704 | 3576,73 | 2658 | 4495 | 1,691 | 0,758 | 17,525 | 9,17E-69 | 4,32E-68 | 0,0009173 | 0,0025503 | 0,0014832 | 0,0014832 | TRUE | 0,0119587 | No |
| orf19.6443 | 852 | 703 | 712 | 591 | 1415 | 1016 | 1381 | 1507 | 812 | 743 | 703 | 926 | 1134 | 1155 | 1172 | 1225 | 983,97 | 796 | 1172 | 1,478 | 0,564 | 7,202 | 5,95E-13 | 1,13E-12 | 0,0058236 | 0,0037599 | 0,0047932 | 0,0047932 | TRUE | 0,1296954 | No |
| orf19.6444 | 1966 | 1613 | 1873 | 1276 | 2206 | 1473 | 2156 | 2250 | 1874 | 1706 | 1850 | 1999 | 1768 | 1675 | 1830 | 1829 | 1816,45 | 1857 | 1776 | 0,958 | -0,062 | -1,082 | 0,27944 | 0,3111336 | 0,0022152 | 0,0029952 | 0,0025575 | 0,0025575 | TRUE | 0,0360762 | No |
| orf19.6445 | 1076 | 850 | 960 | 673 | 5860 | 3705 | 5561 | 6065 | 1026 | 899 | 948 | 1054 | 4695 | 4213 | 4721 | 4931 | 2811,04 | 982 | 4640 | 4,737 | 2,244 | 34,194 | 3,02E-256 | 5,29E-255 | 0,0039505 | 0,0026754 | 0,0034654 | 0,0034654 | TRUE | 0,0461496 | No |
| orf19.6447 | 9477 | 7534 | 8049 | 5073 | 3732 | 2264 | 3474 | 3568 | 9035 | 7966 | 7952 | 7947 | 2990 | 2574 | 2949 | 2901 | 5539,45 | 8225 | 2854 | 0,347 | -1,526 | -24,86 | 2,00E-136 | 1,84E-135 | 0,0040544 | 0,0023877 | 0,003396 | 0,003396 | TRUE | 0,0540697 | No |
| orf19.6448 | 4123 | 3306 | 3813 | 2397 | 2563 | 1706 | 2387 | 2508 | 3931 | 3496 | 3767 | 3755 | 2054 | 1940 | 2027 | 2039 | 2875,97 | 3737 | 2015 | 0,539 | -0,89 | -18,918 | 8,13E-80 | 4,43E-79 | 0,0012332 | 0,0026622 | 0,0017553 | 0,0017553 | TRUE | 0,0234937 | No |
| orf19.6449 | 321 | 237 | 270 | 172 | 584 | 374 | 578 | 592 | 306 | 251 | 267 | 269 | 468 | 425 | 491 | 481 | 369,76 | 273 | 466 | 1,707 | 0,772 | 8,668 | 4,42E-18 | 9,50E-18 | 0,0023772 | 0,0065317 | 0,0046156 | 0,0046156 | TRUE | 0,0804731 | No |
| orf19.6450 | 3 | 1 | 1 | 1 | 1 | 2 | 5 | 2 | 3 | 1 | 1 | 2 | 1 | 2 | 4 | 2 | 1,93 | 2 | 2 | 1,361 | 0,444 | 0,433 | 0,6649849 | 0,6969889 | 1,00E-08 | 0,8539978 | 0,4700836 | 0,4700836 | TRUE | 0,4521583 | No |
| orf19.645.1 | 1580 | 1656 | 1615 | 1087 | 1890 | 1568 | 1749 | 1657 | 1506 | 1751 | 1595 | 1703 | 1514 | 1783 | 1485 | 1347 | 1585,65 | 1639 | 1532 | 0,934 | -0,098 | -1,176 | 0,239437 | 0,2690346 | 0,008358 | 0,0031268 | 0,0060035 | 0,0060035 | TRUE | 0,1428354 | No |
| orf19.6452 | 2712 | 2313 | 2276 | 1507 | 3728 | 2694 | 3287 | 3560 | 2586 | 2446 | 2248 | 2361 | 2987 | 3063 | 2791 | 2895 | 2672,04 | 2410 | 2934 | 1,216 | 0,282 | 5,267 | 1,39E-07 | 2,23E-07 | 0,0020419 | 0,0027058 | 0,0023766 | 0,0023766 | TRUE | 0,0300566 | No |
| orf19.6453 | 1337 | 1413 | 1419 | 897 | 1068 | 801 | 1045 | 922 | 1275 | 1494 | 1402 | 1405 | 856 | 911 | 887 | 750 | 1122,4 | 1394 | 851 | 0,61 | -0,713 | -9,697 | 3,11E-22 | 7,31E-22 | 0,0048552 | 0,0035541 | 0,0042676 | 0,0042676 | TRUE | 0,0776267 | No |
| orf19.6455 | 1716 | 1466 | 1561 | 1013 | 1415 | 951 | 1248 | 1318 | 1636 | 1550 | 1542 | 1587 | 1134 | 1081 | 1060 | 1072 | 1332,69 | 1579 | 1087 | 0,688 | -0,539 | -11,083 | 1,52E-28 | 4,01E-28 | 1,00E-08 | 0,0033233 | 0,0015022 | 0,0015022 | TRUE | 0,0108034 | No |
| orf19.6456 | 964 | 960 | 1077 | 757 | 922 | 761 | 899 | 851 | 919 | 1015 | 1064 | 1186 | 739 | 865 | 763 | 692 | 905,41 | 1046 | 765 | 0,732 | -0,451 | -5,087 | 3,64E-07 | 5,73E-07 | 0,008706 | 0,0039047 | 0,0064139 | 0,0064139 | TRUE | 0,0913511 | No |
| orf19.6457 | 1553 | 1517 | 1677 | 914 | 2038 | 1359 | 1891 | 1964 | 1481 | 1604 | 1657 | 1432 | 1633 | 1545 | 1605 | 1597 | 1569,23 | 1544 | 1595 | 1,032 | 0,046 | 0,824 | 0,4101536 | 0,4462059 | 0,0017165 | 0,0031376 | 0,0023235 | 0,0023235 | TRUE | 0,0294171 | No |
| orf19.6458 | 19 | 40 | 39 | 30 | 32 | 17 | 25 | 19 | 18 | 42 | 39 | 47 | 26 | 19 | 21 | 15 | 28,45 | 36 | 20 | 0,568 | -0,815 | -2,506 | 0,0122268 | 0,0154555 | 0,0699958 | 0,0598085 | 0,0643622 | 0,0643622 | TRUE | 0,8990603 | No |
| orf19.6458.1 | 2135 | 2488 | 2326 | 1107 | 2786 | 1711 | 2461 | 2116 | 2035 | 2631 | 2298 | 1734 | 2232 | 1946 | 2089 | 1721 | 2085,76 | 2174 | 1997 | 0,918 | -0,124 | -1,1 | 0,2715441 | 0,3026062 | 0,0206155 | 0,0028785 | 0,0117643 | 0,0117643 | TRUE | 0,2226634 | No |
| orf19.6459 | 620 | 595 | 647 | 353 | 882 | 720 | 751 | 802 | 591 | 629 | 639 | 553 | 707 | 819 | 638 | 652 | 653,44 | 603 | 704 | 1,162 | 0,216 | 2,452 | 0,0142166 | 0,0178646 | 0,0068904 | 0,004604 | 0,0058936 | 0,0058936 | TRUE | 0,1363499 | No |
| orf19.646 | 149720 | 126823 | 134450 | 78054 | 169844 | 108746 | 157780 | 170532 | 142742 | 134099 | 132824 | 122276 | 136091 | 123654 | 133954 | 138659 | 133037,39 | 132985 | 133090 | 1,001 | 0,001 | 0,02 | 0,9837198 | 0,9854185 | 0,0027445 | 0,0021036 | 0,0028252 | 0,0028252 | TRUE | 0,0359772 | No |
| orf19.6460 | 1909 | 1876 | 2080 | 1249 | 5802 | 4546 | 5353 | 5631 | 1820 | 1984 | 2055 | 1957 | 4649 | 5169 | 4545 | 4579 | 3344,57 | 1954 | 4736 | 2,423 | 1,277 | 22,377 | 6,58E-111 | 4,81E-110 | 0,0026919 | 0,0025822 | 0,0027432 | 0,0027432 | TRUE | 0,0456128 | No |
| orf19.6461 | 1135 | 957 | 1035 | 623 | 1900 | 1233 | 1845 | 1963 | 1082 | 1012 | 1022 | 976 | 1522 | 1402 | 1566 | 1596 | 1272,43 | 1023 | 1522 | 1,486 | 0,572 | 9,783 | 1,34E-22 | 3,15E-22 | 0,001719 | 0,0033817 | 0,0024403 | 0,0024403 | TRUE | 0,0319727 | No |
| orf19.6462 | 1666 | 1394 | 1552 | 1020 | 1694 | 1311 | 1740 | 1708 | 1588 | 1474 | 1533 | 1598 | 1357 | 1491 | 1477 | 1389 | 1488,44 | 1548 | 1428 | 0,922 | -0,117 | -2,225 | 0,0260649 | 0,0320833 | 0,0010812 | 0,0031944 | 0,0019607 | 0,0019607 | TRUE | 0,0136389 | No |
| orf19.6463 | 3383 | 3411 | 3532 | 2002 | 3568 | 2439 | 3253 | 3656 | 3225 | 3607 | 3489 | 3136 | 2859 | 2773 | 2762 | 2973 | 3103,04 | 3364 | 2842 | 0,844 | -0,244 | -4,539 | 5,64E-06 | 8,45E-06 | 0,0023011 | 0,0026204 | 0,0024589 | 0,0024589 | TRUE | 0,028275 | No |
| orf19.6464 | 2484 | 2523 | 2667 | 1635 | 1720 | 1296 | 1555 | 1676 | 2368 | 2668 | 2635 | 2561 | 1378 | 1474 | 1320 | 1363 | 1970,85 | 2558 | 1384 | 0,54 | -0,888 | -16,123 | 1,77E-58 | 7,29E-58 | 0,0019904 | 0,0029244 | 0,002368 | 0,002368 | TRUE | 0,0314349 | No |
| orf19.6465 | 213 | 173 | 202 | 156 | 230 | 184 | 219 | 203 | 203 | 183 | 200 | 244 | 184 | 209 | 186 | 165 | 196,81 | 208 | 186 | 0,9 | -0,151 | -1,278 | 0,2012321 | 0,2281965 | 0,0058482 | 0,0104341 | 0,0083633 | 0,0083633 | TRUE | 0,1288676 | No |
| orf19.6469 | 670 | 623 | 681 | 467 | 652 | 478 | 607 | 695 | 639 | 659 | 673 | 732 | 522 | 544 | 515 | 565 | 606,03 | 676 | 536 | 0,797 | -0,327 | -4,812 | 1,50E-06 | 2,30E-06 | 0,0007316 | 0,0048005 | 0,0027758 | 0,0027758 | TRUE | 0,031859 | No |
| orf19.6470 | 820 | 726 | 808 | 496 | 1439 | 1014 | 1299 | 1279 | 782 | 768 | 798 | 777 | 1153 | 1153 | 1103 | 1040 | 946,69 | 781 | 1112 | 1,422 | 0,508 | 8,864 | 7,69E-19 | 1,68E-18 | 1,00E-08 | 0,0038256 | 0,0020359 | 0,0020359 | TRUE | 0,0234833 | No |
| orf19.6472 | 5306 | 4112 | 4318 | 3078 | 6231 | 4261 | 5778 | 6538 | 5059 | 4348 | 4266 | 4822 | 4993 | 4845 | 4905 | 5316 | 4819,2 | 4624 | 5015 | 1,085 | 0,118 | 1,931 | 0,0535055 | 0,0641162 | 0,0039705 | 0,002432 | 0,0033641 | 0,0033641 | TRUE | 0,0501656 | No |
| orf19.647.3 | 173 | 184 | 187 | 100 | 260 | 158 | 232 | 223 | 165 | 195 | 185 | 157 | 208 | 180 | 197 | 181 | 183,4 | 176 | 192 | 1,09 | 0,125 | 1,102 | 0,2705763 | 0,3015803 | 0,0013669 | 0,0110441 | 0,0068687 | 0,0068687 | TRUE | 0,0558097 | No |
| orf19.6474 | 3666 | 3045 | 3377 | 2001 | 2798 | 1833 | 2615 | 2723 | 3495 | 3220 | 3336 | 3135 | 2242 | 2084 | 2220 | 2214 | 2743,26 | 3296 | 2190 | 0,664 | -0,59 | -12,411 | 2,27E-35 | 6,69E-35 | 0,0011485 | 0,0026898 | 0,0017981 | 0,0017981 | TRUE | 0,0201261 | No |
| orf19.6475 | 32 | 26 | 29 | 20 | 93 | 77 | 95 | 62 | 31 | 27 | 29 | 31 | 75 | 88 | 81 | 50 | 51,39 | 30 | 74 | 2,479 | 1,31 | 5,548 | 2,89E-08 | 4,74E-08 | 1,00E-08 | 0,034041 | 0,0284938 | 0,0284938 | TRUE | 0,4328333 | No |
| orf19.6476 | 2276 | 2351 | 2546 | 1415 | 1647 | 1173 | 1609 | 1616 | 2170 | 2486 | 2515 | 2217 | 1320 | 1334 | 1366 | 1314 | 1840,15 | 2347 | 1334 | 0,568 | -0,817 | -13,757 | 4,61E-43 | 1,53E-42 | 0,0026944 | 0,0029835 | 0,0028107 | 0,0028107 | TRUE | 0,0331665 | No |
| orf19.6477 | 2895 | 3174 | 3291 | 1871 | 4466 | 3414 | 3937 | 3796 | 2760 | 3356 | 3251 | 2931 | 3578 | 3882 | 3342 | 3087 | 3273,49 | 3074 | 3472 | 1,129 | 0,175 | 2,179 | 0,0293183 | 0,0359493 | 0,0084123 | 0,0025929 | 0,0058538 | 0,0058538 | TRUE | 0,0758956 | No |
| orf19.6478 | 13153 | 13445 | 13848 | 7578 | 4637 | 3335 | 4077 | 4317 | 12540 | 14216 | 13681 | 11871 | 3715 | 3792 | 3461 | 3510 | 8348,42 | 13077 | 3620 | 0,277 | -1,854 | -30,281 | 2,03E-201 | 2,80E-200 | 0,0040497 | 0,002288 | 0,0034335 | 0,0034335 | TRUE | 0,0465959 | No |
| orf19.6479 | 906 | 860 | 841 | 497 | 1256 | 973 | 1223 | 1159 | 864 | 909 | 831 | 779 | 1006 | 1106 | 1038 | 942 | 934,5 | 846 | 1023 | 1,206 | 0,27 | 3,928 | 8,57E-05 | 0,0001218 | 0,0029041 | 0,0038483 | 0,0034263 | 0,0034263 | TRUE | 0,0340148 | No |
| orf19.6480 | 508 | 581 | 564 | 321 | 737 | 507 | 624 | 501 | 484 | 614 | 557 | 503 | 591 | 577 | 530 | 407 | 532,86 | 540 | 526 | 0,973 | -0,04 | -0,335 | 0,7379248 | 0,7665144 | 0,0178668 | 0,0051726 | 0,0115632 | 0,0115632 | TRUE | 0,2758586 | No |
| orf19.6481 | 2935 | 2803 | 3029 | 1765 | 2170 | 1495 | 1994 | 2114 | 2798 | 2964 | 2992 | 2765 | 1739 | 1700 | 1693 | 1719 | 2296,23 | 2880 | 1713 | 0,594 | -0,751 | -17,281 | 6,58E-67 | 3,03E-66 | 0,0005031 | 0,0028063 | 0,0013568 | 0,0013568 | TRUE | 0,0083935 | No |
| orf19.6482 | 304 | 349 | 348 | 192 | 275 | 249 | 262 | 243 | 290 | 369 | 344 | 301 | 220 | 283 | 222 | 198 | 278,37 | 326 | 231 | 0,704 | -0,507 | -4,112 | 3,92E-05 | 5,66E-05 | 0,013582 | 0,0079896 | 0,0109117 | 0,0109117 | TRUE | 0,239041 | No |
| orf19.6484 | 229 | 171 | 164 | 134 | 97 | 53 | 79 | 86 | 218 | 181 | 162 | 210 | 78 | 60 | 67 | 70 | 130,76 | 193 | 69 | 0,36 | -1,475 | -9,919 | 3,45E-23 | 8,24E-23 | 0,0087163 | 0,0146482 | 0,0118958 | 0,0118958 | TRUE | 0,1303163 | No |
| orf19.6486 | 31 | 22 | 27 | 26 | 46 | 39 | 49 | 58 | 30 | 23 | 27 | 41 | 37 | 44 | 42 | 47 | 36,27 | 30 | 42 | 1,441 | 0,527 | 2,155 | 0,0311308 | 0,0380766 | 1,00E-08 | 0,0473561 | 0,0281127 | 0,0281127 | TRUE | 0,252153 | No |
| orf19.6487 | 16 | 19 | 19 | 19 | 99 | 86 | 104 | 115 | 15 | 20 | 19 | 30 | 79 | 98 | 88 | 94 | 55,35 | 21 | 90 | 4,422 | 2,145 | 9,212 | 3,19E-20 | 7,20E-20 | 1,00E-08 | 0,0317556 | 0,0197332 | 0,0197332 | TRUE | 0,2765485 | No |
| orf19.6488 | 8 | 7 | 11 | 5 | 1 | 2 | 3 | 1 | 8 | 7 | 11 | 8 | 1 | 2 | 3 | 1 | 5,02 | 8 | 2 | 0,184 | -2,442 | -3,299 | 0,0009687 | 0,0013082 | 1,00E-08 | 0,3291311 | 0,1760321 | 0,1760321 | TRUE | 0,1616939 | No |

| | | | | | | | | | | | | | | | | | | | | | | | | | | | | | | | |
|---|---|---|---|---|---|---|---|---|---|---|---|---|---|---|---|---|---|---|---|---|---|---|---|---|---|---|---|---|---|---|---|
| orf19.6489 | 10048 | 8935 | 9554 | 6141 | 1879 | 1252 | 1675 | 1876 | 9580 | 9448 | 9438 | 9620 | 1506 | 1424 | 1422 | 1525 | 5495,33 | 9522 | 1469 | 0,154 | -2,694 | -71,902 | 0 | 0 | 3,11E-06 | 0,0023901 | 0,0009862 | 0,0009862 | TRUE | 0,0079689 | No |
| orf19.649 | 8118 | 8269 | 8736 | 4802 | 1883 | 1327 | 1721 | 1655 | 7740 | 8743 | 8630 | 7523 | 1509 | 1509 | 1461 | 1346 | 4807,56 | 8159 | 1456 | 0,178 | -2,487 | -39,59 | 0 | 0 | 0,0037242 | 0,0024328 | 0,0034153 | 0,0034153 | TRUE | 0,0330863 | No |
| orf19.6491 | 29 | 41 | 42 | 18 | 29 | 18 | 32 | 19 | 28 | 43 | 41 | 28 | 23 | 20 | 27 | 15 | 28,38 | 35 | 21 | 0,61 | -0,714 | -2,463 | 0,0137737 | 0,0173456 | 0,0224946 | 0,0599527 | 0,0442848 | 0,0442848 | TRUE | 0,1710306 | No |
| orf19.6492 | 1970 | 2270 | 2457 | 1481 | 1128 | 823 | 987 | 911 | 1878 | 2400 | 2427 | 2320 | 904 | 936 | 838 | 741 | 1555,51 | 2256 | 855 | 0,378 | -1,402 | -15,057 | 3,08E-51 | 1,15E-50 | 0,0115676 | 0,0031468 | 0,0075483 | 0,0075483 | TRUE | 0,1565798 | No |
| orf19.6493 | 9 | 9 | 8 | 4 | 7 | 6 | 13 | 9 | 9 | 10 | 8 | 6 | 6 | 7 | 11 | 7 | 7,88 | 8 | 8 | 0,944 | -0,083 | -0,169 | 0,8658449 | 0,8832863 | 1,00E-08 | 0,2104131 | 0,1069668 | 0,1069668 | TRUE | 0,2866975 | No |
| orf19.6494 | 2671 | 2199 | 2470 | 1730 | 1898 | 1267 | 1595 | 1818 | 2547 | 2325 | 2440 | 2710 | 1521 | 1441 | 1354 | 1478 | 1976,97 | 2506 | 1448 | 0,579 | -0,788 | -13,286 | 2,78E-40 | 8,84E-40 | 0,0026046 | 0,0029218 | 0,0028482 | 0,0028482 | TRUE | 0,0346435 | No |
| orf19.6496 | 1691 | 1347 | 1410 | 907 | 953 | 674 | 885 | 963 | 1612 | 1424 | 1393 | 1421 | 764 | 766 | 751 | 783 | 1114,33 | 1462 | 766 | 0,523 | -0,934 | -15,256 | 1,51E-52 | 5,76E-52 | 0,0019811 | 0,0035647 | 0,0026355 | 0,0026355 | TRUE | 0,0593874 | No |
| orf19.6498 | 3065 | 2469 | 2731 | 1748 | 2852 | 1848 | 2711 | 2901 | 2922 | 2611 | 2698 | 2738 | 2285 | 2101 | 2302 | 2359 | 2502,01 | 2742 | 2262 | 0,825 | -0,277 | -5,252 | 1,51E-07 | 2,41E-07 | 0,0018219 | 0,0027475 | 0,0022675 | 0,0022675 | TRUE | 0,0264934 | No |
| orf19.6499 | 1011 | 976 | 1076 | 698 | 1301 | 990 | 1204 | 1360 | 964 | 1032 | 1063 | 1093 | 1042 | 1126 | 1022 | 1106 | 1056,06 | 1038 | 1074 | 1,036 | 0,051 | 0,851 | 0,3949872 | 0,4307891 | 0,0015071 | 0,003646 | 0,0024657 | 0,0024657 | TRUE | 0,0284646 | No |
| orf19.6500 | 3040 | 2487 | 2667 | 1888 | 2203 | 1450 | 1949 | 2175 | 2898 | 2630 | 2635 | 2958 | 1765 | 1649 | 1655 | 1768 | 2244,69 | 2780 | 1709 | 0,616 | -0,699 | -12,647 | 1,17E-36 | 3,50E-36 | 0,0020519 | 0,0028227 | 0,0024734 | 0,0024734 | TRUE | 0,0211448 | No |
| orf19.6501 | 46 | 31 | 38 | 32 | 3 | 2 | 4 | 8 | 44 | 33 | 38 | 50 | 2 | 2 | 3 | 7 | 22,36 | 41 | 4 | 0,092 | -3,449 | -7,887 | 3,08E-15 | 6,25E-15 | 1,00E-08 | 0,0755203 | 0,0514118 | 0,0514118 | TRUE | 0,4469592 | No |
| orf19.6502 | 23 | 38 | 18 | 16 | 21 | 14 | 19 | 14 | 22 | 40 | 18 | 25 | 17 | 16 | 16 | 11 | 20,65 | 26 | 15 | 0,574 | -0,801 | -2,303 | 0,0212565 | 0,0263324 | 0,0458043 | 0,0815949 | 0,0651893 | 0,0651893 | TRUE | 0,8204296 | No |
| orf19.6503 | 1357 | 1340 | 1368 | 1020 | 1395 | 1076 | 1242 | 1321 | 1294 | 1417 | 1351 | 1598 | 1118 | 1224 | 1054 | 1074 | 1266,23 | 1415 | 1118 | 0,791 | -0,339 | -4,568 | 4,93E-06 | 7,42E-06 | 0,0055585 | 0,003388 | 0,0044777 | 0,0044777 | TRUE | 0,0859847 | No |
| orf19.6506 | 3343 | 2800 | 2998 | 1995 | 2171 | 1411 | 1999 | 2119 | 3187 | 2961 | 2962 | 3125 | 1740 | 1604 | 1697 | 1723 | 2374,87 | 3059 | 1691 | 0,554 | -0,853 | -18,356 | 2,95E-75 | 1,51E-74 | 0,0009036 | 0,0027826 | 0,0016253 | 0,0016253 | TRUE | 0,0134269 | No |
| orf19.6507 | 8871 | 7361 | 8156 | 5350 | 10728 | 7676 | 10675 | 11329 | 8458 | 7783 | 8057 | 8381 | 8596 | 8728 | 9063 | 9212 | 8534,78 | 8170 | 8900 | 1,09 | 0,124 | 3,013 | 0,002584 | 0,0034032 | 0,0010828 | 0,0022837 | 0,0015033 | 0,0015033 | TRUE | 0,0124222 | No |
| orf19.6508 | 1083 | 904 | 904 | 618 | 1756 | 1174 | 1635 | 1758 | 1033 | 956 | 893 | 968 | 1407 | 1335 | 1388 | 1429 | 1176,14 | 962 | 1390 | 1,445 | 0,531 | 9,312 | 1,25E-20 | 2,84E-20 | 0,001012 | 0,0034873 | 0,0022228 | 0,0022228 | TRUE | 0,0315127 | No |
| orf19.6509 | 634 | 606 | 612 | 445 | 801 | 632 | 798 | 832 | 604 | 641 | 605 | 697 | 642 | 719 | 677 | 676 | 657,67 | 637 | 678 | 1,069 | 0,096 | 1,397 | 0,162418 | 0,1857994 | 0,001401 | 0,0045878 | 0,0029807 | 0,0029807 | TRUE | 0,0416174 | No |
| orf19.651 | 1197 | 1230 | 1214 | 691 | 1349 | 978 | 1251 | 1220 | 1141 | 1301 | 1199 | 1082 | 1081 | 1112 | 1062 | 992 | 1121,33 | 1181 | 1062 | 0,897 | -0,157 | -2,368 | 0,0178792 | 0,0222783 | 0,0030539 | 0,0035555 | 0,0033367 | 0,0033367 | TRUE | 0,0549233 | No |
| orf19.6510 | 203 | 209 | 210 | 158 | 160 | 118 | 157 | 170 | 194 | 221 | 207 | 248 | 128 | 134 | 133 | 138 | 175,43 | 218 | 133 | 0,619 | -0,693 | -6,038 | 1,56E-09 | 2,66E-09 | 1,00E-08 | 0,0114508 | 0,0067288 | 0,0067288 | TRUE | 0,0789349 | No |
| orf19.6511 | 6903 | 6393 | 6768 | 4266 | 5831 | 4351 | 5446 | 5980 | 6581 | 6760 | 6686 | 6683 | 4672 | 4947 | 4624 | 4862 | 5726,97 | 6678 | 4776 | 0,715 | -0,484 | -13,807 | 2,30E-43 | 7,69E-43 | 0,0003384 | 0,002378 | 0,0010017 | 0,0010017 | TRUE | 0,0067929 | No |
| orf19.6512 | 1283 | 1249 | 1285 | 731 | 518 | 373 | 528 | 523 | 1223 | 1321 | 1269 | 1145 | 415 | 424 | 448 | 425 | 833,9 | 1240 | 428 | 0,345 | -1,536 | -23,519 | 2,63E-122 | 2,14E-121 | 0,00134 | 0,0040602 | 0,0026064 | 0,0026064 | TRUE | 0,0281257 | No |
| orf19.6514 | 39369 | 32758 | 36518 | 22629 | 20332 | 13449 | 19851 | 20267 | 37534 | 34637 | 36076 | 35450 | 16291 | 15293 | 16853 | 16479 | 26076,75 | 35924 | 16229 | 0,452 | -1,146 | -27,484 | 2,71E-166 | 3,04E-165 | 0,0013644 | 0,0021542 | 0,0016277 | 0,0016277 | TRUE | 0,0182937 | No |
| orf19.6515 | 14726 | 12804 | 13549 | 8381 | 13195 | 9143 | 12567 | 14433 | 14040 | 13539 | 13385 | 13129 | 10573 | 10396 | 10669 | 11735 | 12183,32 | 13523 | 10843 | 0,802 | -0,319 | -6,899 | 5,24E-12 | 9,71E-12 | 0,0018491 | 0,002226 | 0,001966 | 0,001966 | TRUE | 0,0376104 | No |
| orf19.6517 | 1026 | 847 | 1002 | 600 | 1849 | 1171 | 1790 | 1918 | 978 | 896 | 990 | 940 | 1482 | 1332 | 1520 | 1560 | 1211,99 | 951 | 1474 | 1,55 | 0,632 | 9,954 | 2,43E-23 | 5,83E-23 | 0,0026247 | 0,003446 | 0,002983 | 0,002983 | TRUE | 0,0479909 | No |
| orf19.6518 | 453 | 330 | 400 | 304 | 1032 | 671 | 1008 | 1046 | 432 | 349 | 395 | 476 | 827 | 763 | 856 | 850 | 618,55 | 413 | 824 | 2,007 | 1,005 | 11,46 | 2,09E-30 | 5,68E-30 | 0,0063746 | 0,0047457 | 0,0054808 | 0,0054808 | TRUE | 0,122984 | No |
| orf19.652 | 553 | 535 | 550 | 310 | 936 | 601 | 877 | 915 | 527 | 566 | 543 | 486 | 750 | 683 | 745 | 744 | 630,48 | 530 | 730 | 1,374 | 0,459 | 6,768 | 1,31E-11 | 2,40E-11 | 0,0005516 | 0,0046955 | 0,0027497 | 0,0027497 | TRUE | 0,0310083 | No |
| orf19.6520 | 61 | 51 | 49 | 41 | 23 | 14 | 19 | 17 | 58 | 54 | 48 | 64 | 18 | 16 | 16 | 14 | 36,13 | 56 | 16 | 0,288 | -1,794 | -7,016 | 2,28E-12 | 4,27E-12 | 1,00E-08 | 0,0475386 | 0,0252226 | 0,0252226 | TRUE | 0,0738292 | No |
| orf19.6522 | 1377 | 1125 | 1219 | 784 | 2238 | 1499 | 2155 | 2292 | 1313 | 1190 | 1204 | 1228 | 1793 | 1704 | 1830 | 1864 | 1515,72 | 1234 | 1798 | 1,458 | 0,544 | 10,376 | 3,20E-25 | 7,96E-25 | 0,0010326 | 0,0031745 | 0,001934 | 0,001934 | TRUE | 0,0239703 | No |
| orf19.6524 | 5095 | 4206 | 4504 | 3204 | 13478 | 9731 | 12900 | 13888 | 4858 | 4447 | 4450 | 5019 | 10800 | 11065 | 10952 | 11292 | 7860,31 | 4694 | 11027 | 2,351 | 1,234 | 26,259 | 5,71E-152 | 5,79E-151 | 0,0018899 | 0,0023002 | 0,0019587 | 0,0019587 | TRUE | 0,0254591 | No |
| orf19.6525 | 1414 | 1314 | 1296 | 842 | 1187 | 778 | 1033 | 1179 | 1348 | 1389 | 1280 | 1319 | 951 | 885 | 877 | 959 | 1126,03 | 1334 | 918 | 0,689 | -0,538 | -9,738 | 2,07E-22 | 4,88E-22 | 0,0008296 | 0,0035494 | 0,0020274 | 0,0020274 | TRUE | 0,0116249 | No |
| orf19.6526 | 1632 | 1357 | 1475 | 1000 | 1888 | 1356 | 1836 | 1778 | 1556 | 1435 | 1457 | 1567 | 1513 | 1542 | 1559 | 1446 | 1509,2 | 1504 | 1515 | 1,008 | 0,012 | 0,229 | 0,8188375 | 0,8398975 | 0,0008576 | 0,0031792 | 0,0018272 | 0,0018272 | TRUE | 0,0116246 | No |
| orf19.6527 | 19284 | 16571 | 18573 | 11379 | 15548 | 10776 | 14931 | 15972 | 18385 | 17522 | 18348 | 17826 | 12458 | 12253 | 12676 | 12987 | 15306,96 | 18020 | 12594 | 0,699 | -0,517 | -15,67 | 2,42E-55 | 9,59E-55 | 0,0004913 | 0,0021985 | 0,0009782 | 0,0009782 | TRUE | 0,0054073 | No |
| orf19.6528 | 960 | 821 | 913 | 563 | 1044 | 688 | 1048 | 1046 | 915 | 868 | 902 | 882 | 837 | 782 | 890 | 850 | 865,8 | 892 | 840 | 0,943 | -0,085 | -1,458 | 0,144929 | 0,1665392 | 0,0004593 | 0,0039877 | 0,0021308 | 0,0021308 | TRUE | 0,0228782 | No |
| orf19.6529 | 17513 | 15009 | 16743 | 10314 | 18013 | 11952 | 17005 | 18057 | 16697 | 15870 | 16540 | 16157 | 14433 | 13590 | 14437 | 14682 | 15300,98 | 16316 | 14286 | 0,876 | -0,192 | -5,316 | 1,06E-07 | 1,70E-07 | 0,0007345 | 0,0021986 | 0,0011813 | 0,0011813 | TRUE | 0,0129545 | No |
| orf19.653 | 1 | 2 | 1 | 1 | 1 | 1 | 1 | 4 | 1 | 2 | 1 | 2 | 1 | 1 | 1 | 3 | 1,46 | 2 | 2 | 1,103 | 0,141 | 0,123 | 0,9018547 | 0,9157757 | 1,00E-08 | 1,128386 | 0,5673572 | 0,5673572 | TRUE | 0,533567 | No |
| orf19.6530 | 3802 | 3413 | 3695 | 2511 | 1833 | 1364 | 1758 | 1861 | 3625 | 3609 | 3650 | 3934 | 1469 | 1551 | 1493 | 1513 | 2605,37 | 3704 | 1506 | 0,407 | -1,297 | -29,065 | 9,98E-186 | 1,25E-184 | 0,0006955 | 0,0027215 | 0,0014646 | 0,0014646 | TRUE | 0,0197841 | No |
| orf19.6531 | 6390 | 5629 | 5928 | 3877 | 13273 | 8774 | 12502 | 13678 | 6092 | 5952 | 5856 | 6074 | 10635 | 9977 | 10614 | 11122 | 8290,21 | 5994 | 10587 | 1,767 | 0,821 | 20,094 | 8,29E-90 | 4,95E-89 | 0,0009791 | 0,0022893 | 0,0014687 | 0,0014687 | TRUE | 0,0181324 | No |
| orf19.6531.1 | 1859 | 1557 | 1436 | 1074 | 2525 | 1772 | 2128 | 2347 | 1772 | 1646 | 1419 | 1682 | 2023 | 2015 | 1807 | 1908 | 1784,12 | 1630 | 1938 | 1,189 | 0,25 | 3,52 | 0,0004308 | 0,0005928 | 0,0048994 | 0,0030116 | 0,0042766 | 0,0042766 | TRUE | 0,0942176 | No |
| orf19.6532 | 601 | 501 | 544 | 356 | 505 | 375 | 464 | 535 | 573 | 530 | 537 | 558 | 405 | 426 | 394 | 435 | 482,23 | 550 | 415 | 0,755 | -0,406 | -5,724 | 1,04E-08 | 1,73E-08 | 1,00E-08 | 0,0054961 | 0,0027309 | 0,0027309 | TRUE | 0,013677 | No |
| orf19.6533 | 739 | 746 | 775 | 497 | 1208 | 797 | 1076 | 1219 | 705 | 789 | 766 | 779 | 968 | 906 | 914 | 991 | 852,05 | 760 | 945 | 1,247 | 0,319 | 5,166 | 2,39E-07 | 3,79E-07 | 0,0011107 | 0,0040183 | 0,0024492 | 0,0024492 | TRUE | 0,0297455 | No |
| orf19.6534 | 7 | 6 | 9 | 9 | 8 | 9 | 6 | 6 | 7 | 6 | 9 | 14 | 6 | 10 | 5 | 5 | 7,83 | 9 | 6 | 0,746 | -0,422 | -0,814 | 0,4159162 | 0,4519689 | 1,00E-08 | 0,2118367 | 0,1266535 | 0,1266535 | TRUE | 0,286825 | No |
| orf19.6534.2 | 1693 | 1433 | 1625 | 927 | 3845 | 2458 | 3839 | 3952 | 1614 | 1515 | 1605 | 1452 | 3081 | 2795 | 3259 | 3213 | 2316,92 | 1546 | 3087 | 1,994 | 0,996 | 16,464 | 6,66E-61 | 2,85E-60 | 0,0028907 | 0,0027999 | 0,0030045 | 0,0030045 | TRUE | 0,0480749 | No |
| orf19.6536 | 432 | 436 | 466 | 263 | 155 | 154 | 202 | 189 | 412 | 461 | 460 | 412 | 124 | 175 | 171 | 154 | 296,22 | 436 | 156 | 0,355 | -1,492 | -13,444 | 3,32E-41 | 1,07E-40 | 0,0073757 | 0,0076342 | 0,0076742 | 0,0076742 | TRUE | 0,2128656 | No |
| orf19.6537 | 188 | 213 | 201 | 116 | 449 | 304 | 384 | 431 | 179 | 225 | 199 | 182 | 360 | 346 | 326 | 350 | 270,83 | 196 | 346 | 1,757 | 0,813 | 8,273 | 1,30E-16 | 2,71E-16 | 0,0009139 | 0,0081537 | 0,0051464 | 0,0051464 | TRUE | 0,1077862 | No |
| orf19.6538 | 843 | 687 | 732 | 476 | 2312 | 1528 | 2056 | 2177 | 804 | 726 | 723 | 746 | 1853 | 1737 | 1746 | 1770 | 1263,08 | 750 | 1776 | 2,369 | 1,244 | 22,707 | 3,81E-114 | 2,87E-113 | 0,000638 | 0,0033912 | 0,0018863 | 0,0018863 | TRUE | 0,0305794 | No |
| orf19.6539 | 6173 | 5162 | 5652 | 3594 | 8799 | 5898 | 8684 | 9032 | 5885 | 5458 | 5584 | 5630 | 7050 | 6707 | 7373 | 7344 | 6378,85 | 5639 | 7118 | 1,263 | 0,336 | 7,697 | 1,39E-14 | 2,77E-14 | 0,0012469 | 0,0023487 | 0,001671 | 0,001671 | TRUE | 0,0181578 | No |
| orf19.654 | 10 | 8 | 8 | 3 | 8 | 6 | 10 | 9 | 10 | 8 | 8 | 5 | 6 | 7 | 8 | 7 | 7,45 | 8 | 7 | 0,927 | -0,109 | -0,219 | 0,82634 | 0,8466403 | 1,00E-08 | 0,2223467 | 0,105104 | 0,105104 | TRUE | 0,1066994 | No |
| orf19.6540 | 26616 | 22236 | 24327 | 15514 | 6699 | 4616 | 6192 | 6900 | 25375 | 23512 | 24033 | 24304 | 5368 | 5249 | 5257 | 5610 | 14838,42 | 24306 | 5371 | 0,221 | -2,178 | -56,603 | 0 | 0 | 0,0008626 | 0,0022019 | 0,0013154 | 0,0013154 | TRUE | 0,0110622 | No |
| orf19.6541 | 19339 | 15612 | 16947 | 10505 | 56023 | 37500 | 54360 | 57762 | 18438 | 16508 | 16742 | 16457 | 44890 | 42641 | 46151 | 46966 | 31098,99 | 17036 | 45162 | 2,651 | 1,406 | 28,757 | 7,30E-182 | 8,88E-181 | 0,0023483 | 0,0021441 | 0,0022547 | 0,0022547 | TRUE | 0,0376966 | No |
| orf19.6544 | 15501 | 13228 | 14532 | 8970 | 4502 | 3140 | 4265 | 4677 | 14779 | 13987 | 14356 | 14052 | 3607 | 3570 | 3621 | 3803 | 8971,91 | 14294 | 3650 | 0,255 | -1,969 | -54,44 | 0 | 0 | 0,0004948 | 0,0022743 | 0,0010945 | 0,0010945 | TRUE | 0,0097396 | No |
| orf19.6547 | 15 | 5 | 6 | 8 | 18 | 9 | 17 | 27 | 14 | 5 | 6 | 13 | 14 | 10 | 14 | 22 | 12,39 | 10 | 15 | 1,641 | 0,715 | 1,579 | 0,1143473 | 0,1325432 | 0,0697041 | 0,1346488 | 0,1084392 | 0,1084392 | TRUE | 0,3835421 | No |
| orf19.6548 | 13083 | 11008 | 11938 | 7706 | 12273 | 8507 | 11491 | 12011 | 12473 | 11640 | 11794 | 12072 | 9834 | 9673 | 9756 | 9766 | 10875,92 | 11995 | 9757 | 0,814 | -0,298 | -9,205 | 3,43E-20 | 7,72E-20 | 0,000364 | 0,0022422 | 0,0009119 | 0,0009119 | TRUE | 0,0090596 | No |
| orf19.655 | 55048 | 47273 | 51091 | 31929 | 55820 | 37701 | 51833 | 57832 | 52482 | 49985 | 50473 | 50018 | 44727 | 42869 | 44006 | 47023 | 47698,05 | 50740 | 44656 | 0,88 | -0,184 | -4,899 | 9,65E-07 | 1,49E-06 | 0,0010144 | 0,0021257 | 0,0013377 | 0,0013377 | TRUE | 0,015599 | No |
| orf19.6550 | 1150 | 971 | 1021 | 592 | 1681 | 1169 | 1509 | 1630 | 1096 | 1027 | 1009 | 927 | 1347 | 1329 | 1281 | 1325 | 1167,73 | 1015 | 1320 | 1,297 | 0,375 | 6,546 | 5,93E-11 | 1,07E-10 | 0,0011318 | 0,0034973 | 0,0022622 | 0,0022622 | TRUE | 0,0350826 | No |
| orf19.6551 | 641 | 693 | 675 | 379 | 1047 | 683 | 953 | 879 | 611 | 733 | 667 | 594 | 839 | 777 | 809 | 715 | 717,98 | 651 | 785 | 1,203 | 0,267 | 3,294 | 0,0009861 | 0,0013303 | 0,0050678 | 0,0043781 | 0,0048612 | 0,0048612 | TRUE | 0,0830739 | No |
| orf19.6551.1 | 163 | 130 | 163 | 109 | 257 | 226 | 219 | 293 | 155 | 137 | 161 | 171 | 206 | 257 | 186 | 238 | 188,97 | 156 | 222 | 1,42 | 0,506 | 3,923 | 8,73E-05 | 0,0001241 | 0,0099415 | 0,0107802 | 0,0103798 | 0,0103798 | TRUE | 0,1307769 | No |
| orf19.6552 | 2351 | 2197 | 2429 | 1601 | 2011 | 1446 | 1840 | 1887 | 2241 | 2323 | 2400 | 2508 | 1611 | 1644 | 1562 | 1534 | 1978,02 | 2368 | 1588 | 0,671 | -0,575 | -11,679 | 1,64E-31 | 4,53E-31 | 0,0009273 | 0,0029214 | 0,0018101 | 0,0018101 | TRUE | 0,0177698 | No |
| orf19.6553 | 1035 | 989 | 997 | 635 | 2731 | 2000 | 2506 | 2540 | 987 | 1046 | 985 | 995 | 2188 | 2274 | 2128 | 2065 | 1583,44 | 1003 | 2164 | 2,155 | 1,108 | 21,761 | 5,45E-105 | 3,71E-104 | 0,0007878 | 0,0031282 | 0,0017245 | 0,0017245 | TRUE | 0,0135691 | No |
| orf19.6554 | 6990 | 7925 | 7525 | 5061 | 1417 | 1258 | 1314 | 1276 | 6664 | 8380 | 7434 | 7928 | 1135 | 1430 | 1116 | 1038 | 4390,64 | 7602 | 1180 | 0,155 | -2,69 | -28,34 | 1,12E-176 | 1,31E-175 | 0,0146952 | 0,0024652 | 0,0082004 | 0,0082004 | TRUE | 0,2403567 | No |
| orf19.6555 | 523 | 572 | 524 | 338 | 205 | 148 | 196 | 193 | 499 | 605 | 518 | 529 | 164 | 168 | 166 | 157 | 350,81 | 538 | 164 | 0,305 | -1,715 | -18,146 | 1,39E-73 | 6,97E-73 | 1,00E-08 | 0,0067716 | 0,0048161 | 0,0048161 | TRUE | 0,0856488 | No |
| orf19.6556 | 11712 | 11539 | 11786 | 7553 | 5321 | 3441 | 4844 | 4816 | 11166 | 12201 | 11643 | 11832 | 4264 | 3913 | 4113 | 3916 | 7880,93 | 11710 | 4052 | 0,346 | -1,531 | -34,743 | 1,74E-264 | 3,18E-263 | 0,0013314 | 0,0022996 | 0,0017059 | 0,0017059 | TRUE | 0,0152967 | No |
| orf19.6557 | 776 | 674 | 795 | 497 | 697 | 476 | 606 | 728 | 740 | 713 | 785 | 779 | 558 | 541 | 514 | 592 | 652,83 | 754 | 551 | 0,733 | -0,449 | -6,588 | 4,46E-11 | 8,05E-11 | 0,0013226 | 0,0046063 | 0,0029004 | 0,0029004 | TRUE | 0,029612 | No |
| orf19.6558 | 3002 | 2967 | 3064 | 1960 | 1551 | 1251 | 1588 | 1708 | 2862 | 3137 | 3027 | 3070 | 1243 | 1422 | 1348 | 1389 | 2187,37 | 3024 | 1350 | 0,446 | -1,164 | -21,566 | 3,77E-103 | 2,54E-102 | 0,0017052 | 0,0028419 | 0,0022799 | 0,0022799 | TRUE | 0,0353837 | No |
| orf19.6559 | 2137 | 1772 | 1911 | 1297 | 1737 | 1272 | 1698 | 1757 | 2037 | 1874 | 1888 | 2032 | 1392 | 1446 | 1442 | 1429 | 1692,4 | 1958 | 1427 | 0,729 | -0,455 | -9,481 | 2,52E-21 | 5,79E-21 | 0,0006497 | 0,0030614 | 0,001614 | 0,001614 | TRUE | 0,0102271 | No |
| orf19.656 | 82 | 60 | 68 | 46 | 89 | 43 | 80 | 88 | 78 | 63 | 67 | 72 | 71 | 49 | 68 | 72 | 67,57 | 70 | 65 | 0,933 | -0,1 | -0,55 | 0,5826376 | 0,6168592 | 1,00E-08 | 0,0263915 | 0,0167862 | 0,0167862 | TRUE | 0,2066291 | No |
| orf19.6561 | 6769 | 5740 | 6275 | 4286 | 13892 | 9894 | 13071 | 14828 | 6454 | 6069 | 6199 | 6714 | 11131 | 11250 | 11097 | 12057 | 8871,45 | 6359 | 11384 | 1,791 | 0,841 | 18,546 | 8,87E-77 | 4,63E-76 | 0,0016375 | 0,0022764 | 0,0018469 | 0,0018469 | TRUE | 0,0194588 | No |
| orf19.6562 | 334 | 234 | 267 | 204 | 642 | 458 | 617 | 633 | 318 | 247 | 264 | 320 | 514 | 521 | 524 | 515 | 402,87 | 287 | 518 | 1,812 | 0,858 | 9,525 | 1,65E-21 | 3,82E-21 | 0,0031428 | 0,0061668 | 0,0049701 | 0,0049701 | TRUE | 0,0967305 | No |
| orf19.6563 | 2348 | 2046 | 2222 | 1481 | 2291 | 1676 | 2195 | 2337 | 2239 | 2163 | 2195 | 2320 | 1836 | 1906 | 1864 | 1900 | 2052,8 | 2229 | 1876 | 0,842 | -0,247 | -5,949 | 2,70E-09 | 4,57E-09 | 1,00E-08 | 0,0028911 | 0,0011702 | 0,0011702 | TRUE | 0,0080709 | No |
| orf19.6563.1 | 3385 | 2616 | 2717 | 1424 | 1927 | 1082 | 1828 | 1898 | 3227 | 2766 | 2684 | 2231 | 1544 | 1230 | 1552 | 1543 | 2097,23 | 2727 | 1467 | 0,538 | -0,894 | -8,577 | 9,77E-18 | 2,09E-17 | 0,0163068 | 0,0028742 | 0,0099332 | 0,0099332 | TRUE | 0,1438197 | No |
| orf19.6565 | 1117 | 946 | 1056 | 664 | 2647 | 1793 | 2363 | 2644 | 1065 | 1000 | 1043 | 1040 | 2121 | 2039 | 2006 | 2150 | 1558,05 | 1037 | 2079 | 2,005 | 1,004 | 20,714 | 2,60E-95 | 1,63E-94 | 1,00E-08 | 0,0031451 | 0,0015004 | 0,0015004 | TRUE | 0,0072096 | No |
| orf19.6566 | 1115 | 979 | 1113 | 685 | 1207 | 966 | 1175 | 1284 | 1063 | 1035 | 1100 | 1073 | 967 | 1098 | 998 | 1044 | 1047,25 | 1068 | 1027 | 0,96 | -0,059 | -1,033 | 0,3015743 | 0,3346107 | 0,0009451 | 0,0036591 | 0,0021485 | 0,0021485 | TRUE | 0,0244997 | No |
| orf19.6568 | 5365 | 5018 | 5270 | 2883 | 6762 | 4454 | 6281 | 6748 | 5115 | 5306 | 5206 | 4516 | 5418 | 5065 | 5333 | 5487 | 5180,7 | 5036 | 5326 | 1,057 | 0,08 | 1,456 | 0,1455264 | 0,1671956 | 0,0028308 | 0,0024082 | 0,0026839 | 0,0026839 | TRUE | 0,0568518 | No |
| orf19.6569 | 449 | 463 | 493 | 244 | 1410 | 1031 | 1191 | 1227 | 428 | 490 | 487 | 382 | 1130 | 1172 | 1011 | 998 | 762,23 | 447 | 1078 | 2,399 | 1,263 | 14,341 | 1,21E-46 | 4,27E-46 | 0,0077714 | 0,0042453 | 0,0057771 | 0,0057771 | TRUE | 0,0937413 | No |

| | | | | | | | | | | | | | | | | | | | | | | | | | | | | | | | |
|---|---|---|---|---|---|---|---|---|---|---|---|---|---|---|---|---|---|---|---|---|---|---|---|---|---|---|---|---|---|---|---|
| orf19.657 | 1246 | 1058 | 1103 | 803 | 4410 | 3167 | 3646 | 4223 | 1188 | 1119 | 1090 | 1258 | 3534 | 3601 | 3095 | 3434 | 2289,77 | 1164 | 3416 | 2,941 | 1,556 | 24,354 | 5,32E-131 | 4,66E-130 | 0,0034811 | 0,0028083 | 0,0033063 | 0,0033063 | TRUE | 0,048685 | No |
| orf19.6570 | 142 | 130 | 154 | 99 | 381 | 247 | 313 | 296 | 135 | 137 | 152 | 155 | 305 | 281 | 266 | 241 | 209,08 | 145 | 273 | 1,89 | 0,918 | 7,988 | 1,37E-15 | 2,79E-15 | 0,004415 | 0,0099443 | 0,0071986 | 0,0071986 | TRUE | 0,076545 | No |
| orf19.6573 | 3758 | 3829 | 3931 | 2303 | 4952 | 3611 | 4480 | 4783 | 3583 | 4049 | 3883 | 3608 | 3968 | 4106 | 3803 | 3889 | 3861,15 | 3781 | 3942 | 1,042 | 0,059 | 1,169 | 0,2423549 | 0,2721215 | 0,0019519 | 0,0025165 | 0,00218 | 0,00218 | TRUE | 0,0276847 | No |
| orf19.6574 | 29 | 33 | 28 | 18 | 92 | 90 | 85 | 93 | 28 | 35 | 28 | 28 | 74 | 102 | 72 | 76 | 55,28 | 30 | 81 | 2,712 | 1,439 | 6,589 | 4,43E-11 | 7,99E-11 | 1,00E-08 | 0,031793 | 0,0215508 | 0,0215508 | TRUE | 0,2656279 | No |
| orf19.6575 | 2 | 1 | 5 | 5 | 1 | 0 | 0 | 0 | 2 | 1 | 5 | 8 | 1 | 0 | 0 | 0 | 2,07 | 4 | 0 | 0,071 | -3,815 | -2,778 | 0,0054655 | 0,0070512 | 1,00E-08 | 0,7963499 | 0,6633922 | 0,6633922 | TRUE | 0,2046112 | No |
| orf19.6577 | 12358 | 11112 | 11530 | 7942 | 4371 | 3261 | 4015 | 4335 | 11782 | 11750 | 11391 | 12442 | 3502 | 3708 | 3409 | 3525 | 7688,44 | 11841 | 3536 | 0,299 | -1,744 | -41,7 | 0 | 0 | 0,0010665 | 0,0023048 | 0,0015055 | 0,0015055 | TRUE | 0,0139456 | No |
| orf19.6578 | 18536 | 17412 | 18835 | 11809 | 3425 | 2429 | 3242 | 3573 | 17672 | 18411 | 18607 | 18499 | 2744 | 2762 | 2752 | 2905 | 10544,21 | 18297 | 2791 | 0,153 | -2,712 | -75,985 | 0 | 0 | 0,0003692 | 0,002247 | 0,0010329 | 0,0010329 | TRUE | 0,0094662 | No |
| orf19.6579 | 715 | 853 | 932 | 487 | 729 | 469 | 634 | 573 | 682 | 902 | 921 | 763 | 584 | 533 | 538 | 466 | 673,61 | 817 | 530 | 0,649 | -0,624 | -5,953 | 2,63E-09 | 4,46E-09 | 0,0132094 | 0,0045288 | 0,0090186 | 0,0090186 | TRUE | 0,1542542 | No |
| orf19.658 | 245 | 234 | 250 | 151 | 531 | 356 | 500 | 555 | 234 | 247 | 247 | 237 | 425 | 405 | 424 | 451 | 333,82 | 241 | 426 | 1,77 | 0,824 | 9,887 | 4,74E-23 | 1,13E-22 | 1,00E-08 | 0,0070098 | 0,0033234 | 0,0033234 | TRUE | 0,017804 | No |
| orf19.6580 | 5318 | 4788 | 5040 | 3317 | 5110 | 3758 | 4742 | 4875 | 5070 | 5063 | 4979 | 5196 | 4094 | 4273 | 4026 | 3964 | 4583,2 | 5077 | 4089 | 0,805 | -0,312 | -8,456 | 2,77E-17 | 5,85E-17 | 0,0003954 | 0,0024495 | 0,0010896 | 0,0010896 | TRUE | 0,0106771 | No |
| orf19.6581 | 875 | 824 | 874 | 489 | 2145 | 1423 | 2013 | 2017 | 834 | 871 | 863 | 766 | 1719 | 1618 | 1709 | 1640 | 1252,6 | 834 | 1672 | 1,999 | 0,999 | 18,295 | 9,06E-75 | 4,62E-74 | 0,0005538 | 0,0034021 | 0,0019282 | 0,0019282 | TRUE | 0,0297199 | No |
| orf19.6582 | 5855 | 5244 | 5503 | 3248 | 5274 | 3449 | 4838 | 5144 | 5582 | 5545 | 5436 | 5088 | 4226 | 3922 | 4107 | 4183 | 4761,16 | 5413 | 4110 | 0,759 | -0,398 | -9,171 | 4,69E-20 | 1,05E-19 | 0,0011136 | 0,0024361 | 0,0015955 | 0,0015955 | TRUE | 0,0189817 | No |
| orf19.6583 | 266 | 291 | 317 | 205 | 579 | 428 | 527 | 497 | 254 | 308 | 313 | 321 | 464 | 487 | 447 | 404 | 374,72 | 299 | 450 | 1,511 | 0,595 | 6,172 | 6,76E-10 | 1,17E-09 | 0,0056204 | 0,006473 | 0,0060628 | 0,0060628 | TRUE | 0,1247904 | No |
| orf19.6584 | 9405 | 8861 | 9341 | 5707 | 12247 | 9117 | 11055 | 12183 | 8967 | 9369 | 9228 | 8940 | 9813 | 10367 | 9386 | 9906 | 9497 | 9126 | 9868 | 1,081 | 0,112 | 2,842 | 0,004478 | 0,0058042 | 0,0009503 | 0,0022642 | 0,0013881 | 0,0013881 | TRUE | 0,0139297 | No |
| orf19.6585 | 660 | 713 | 660 | 421 | 733 | 476 | 582 | 623 | 629 | 754 | 652 | 660 | 587 | 541 | 494 | 507 | 602,99 | 674 | 532 | 0,79 | -0,34 | -4,132 | 3,60E-05 | 5,20E-05 | 0,0048541 | 0,0048142 | 0,0048259 | 0,0048259 | TRUE | 0,0779051 | No |
| orf19.6586 | 3178 | 3100 | 3298 | 2015 | 1914 | 1284 | 1823 | 1899 | 3030 | 3278 | 3258 | 3157 | 1534 | 1460 | 1548 | 1544 | 2350,99 | 3181 | 1522 | 0,479 | -1,063 | -23,778 | 5,65E-125 | 4,73E-124 | 0,0006543 | 0,0027897 | 0,0014475 | 0,0014475 | TRUE | 0,0130626 | No |
| orf19.6587 | 1829 | 1883 | 1905 | 1131 | 1502 | 1103 | 1487 | 1353 | 1744 | 1991 | 1882 | 1772 | 1204 | 1254 | 1262 | 1100 | 1526,1 | 1847 | 1205 | 0,652 | -0,618 | -9,754 | 1,78E-22 | 4,20E-22 | 0,0028757 | 0,0031672 | 0,0031797 | 0,0031797 | TRUE | 0,0420043 | No |
| orf19.6588 | 2563 | 2467 | 2531 | 1472 | 943 | 645 | 909 | 886 | 2444 | 2609 | 2500 | 2306 | 756 | 733 | 772 | 720 | 1604,95 | 2465 | 745 | 0,302 | -1,727 | -31,689 | 2,20E-220 | 3,27E-219 | 0,0012462 | 0,0031143 | 0,0020252 | 0,0020252 | TRUE | 0,0210059 | No |
| orf19.6589 | 6977 | 7157 | 7339 | 4236 | 7583 | 5278 | 7311 | 7118 | 6652 | 7568 | 7250 | 6636 | 6076 | 6002 | 6207 | 5788 | 6522,23 | 7026 | 6018 | 0,856 | -0,224 | -4,363 | 1,28E-05 | 1,89E-05 | 0,0023661 | 0,002343 | 0,0023771 | 0,0023771 | TRUE | 0,0327724 | No |
| orf19.6590 | 123 | 128 | 101 | 68 | 81 | 61 | 83 | 66 | 117 | 135 | 100 | 107 | 65 | 69 | 70 | 54 | 89,66 | 115 | 64 | 0,56 | -0,837 | -5,08 | 3,77E-07 | 5,94E-07 | 0,0064397 | 0,020403 | 0,0142905 | 0,0142905 | TRUE | 0,147224 | No |
| orf19.6591 | 2300 | 2136 | 2236 | 1532 | 1284 | 929 | 1189 | 1204 | 2193 | 2259 | 2209 | 2400 | 1029 | 1056 | 1009 | 979 | 1641,73 | 2265 | 1018 | 0,45 | -1,152 | -23,191 | 5,65E-119 | 4,44E-118 | 0,0006594 | 0,0030914 | 0,0016836 | 0,0016836 | TRUE | 0,017903 | No |
| orf19.6592 | 581 | 552 | 598 | 358 | 181 | 144 | 183 | 180 | 554 | 584 | 591 | 561 | 145 | 164 | 155 | 146 | 362,46 | 572 | 152 | 0,265 | -1,914 | -22,229 | 1,81E-109 | 1,31E-108 | 1,00E-08 | 0,0066212 | 0,0032214 | 0,0032214 | TRUE | 0,0214001 | No |
| orf19.6594 | 24334 | 23595 | 24880 | 14880 | 4942 | 3524 | 4666 | 5094 | 23200 | 24949 | 24579 | 23310 | 3960 | 4007 | 3961 | 4142 | 14013,53 | 24010 | 4018 | 0,167 | -2,579 | -67,294 | 0 | 0 | 0,0007137 | 0,0022084 | 0,0012763 | 0,0012763 | TRUE | 0,0084836 | No |
| orf19.6595 | 65 | 55 | 57 | 35 | 26 | 19 | 28 | 19 | 62 | 58 | 56 | 55 | 21 | 22 | 24 | 15 | 39,12 | 58 | 20 | 0,351 | -1,512 | -6,362 | 1,99E-10 | 3,52E-10 | 1,00E-08 | 0,044067 | 0,0228169 | 0,0228169 | TRUE | 0,1761228 | No |
| orf19.6596 | 1805 | 2004 | 1915 | 1355 | 1998 | 1661 | 1930 | 1792 | 1721 | 2119 | 1892 | 2123 | 1601 | 1889 | 1639 | 1457 | 1804,96 | 1964 | 1646 | 0,838 | -0,254 | -2,898 | 0,0037613 | 0,0049022 | 0,0101079 | 0,0030009 | 0,0068367 | 0,0068367 | TRUE | 0,1163697 | No |
| orf19.6597 | 918 | 1188 | 1174 | 606 | 1723 | 1277 | 1644 | 1414 | 875 | 1256 | 1160 | 949 | 1381 | 1452 | 1396 | 1150 | 1202,33 | 1060 | 1345 | 1,267 | 0,341 | 3,084 | 0,0020418 | 0,0027041 | 0,0178655 | 0,0034569 | 0,0108848 | 0,0108848 | TRUE | 0,1854114 | No |
| orf19.6598 | 3418 | 2989 | 3315 | 2202 | 1409 | 963 | 1391 | 1441 | 3259 | 3160 | 3275 | 3450 | 1129 | 1095 | 1181 | 1172 | 2215,03 | 3286 | 1144 | 0,349 | -1,519 | -32,412 | 1,83E-230 | 2,82E-229 | 0,0006692 | 0,0028325 | 0,0015479 | 0,0015479 | TRUE | 0,0127139 | No |
| orf19.6599.1 | 2579 | 1903 | 1927 | 969 | 4603 | 2304 | 4102 | 4442 | 2459 | 2012 | 1904 | 1518 | 3688 | 2620 | 3483 | 3612 | 2661,89 | 1973 | 3351 | 1,697 | 0,763 | 4,319 | 1,57E-05 | 2,30E-05 | 0,0295909 | 0,0027081 | 0,0157299 | 0,0295909 | TRUE | 0,3322713 | No |
| orf19.660 | 1450 | 1230 | 1385 | 899 | 1260 | 962 | 1156 | 1316 | 1382 | 1301 | 1368 | 1408 | 1010 | 1094 | 981 | 1070 | 1201,82 | 1365 | 1039 | 0,761 | -0,394 | -7,202 | 5,92E-13 | 1,13E-12 | 0,0009096 | 0,0034575 | 0,0020388 | 0,0020388 | TRUE | 0,0162892 | No |
| orf19.6600 | 2029 | 2151 | 2092 | 1165 | 1427 | 976 | 1328 | 1309 | 1934 | 2274 | 2067 | 1825 | 1143 | 1110 | 1127 | 1064 | 1568,2 | 2025 | 1111 | 0,548 | -0,868 | -12,621 | 1,62E-36 | 4,85E-36 | 0,0041002 | 0,0031383 | 0,0038581 | 0,0038581 | TRUE | 0,0823733 | No |
| orf19.6601 | 11864 | 11490 | 11880 | 6829 | 3510 | 2308 | 3321 | 3112 | 11311 | 12149 | 11736 | 10698 | 2812 | 2624 | 2820 | 2530 | 7085,16 | 11474 | 2696 | 0,235 | -2,089 | -38,967 | 0 | 0 | 0,0025618 | 0,002323 | 0,002546 | 0,002546 | TRUE | 0,0249149 | No |
| orf19.6601.1 | 23400 | 22578 | 22798 | 13295 | 4502 | 3027 | 4217 | 4107 | 22309 | 23873 | 22522 | 20827 | 3607 | 3442 | 3580 | 3339 | 12937,65 | 22383 | 3492 | 0,156 | -2,68 | -54,548 | 0 | 0 | 0,0019722 | 0,0022182 | 0,002166 | 0,002166 | TRUE | 0,0265396 | No |
| orf19.6602 | 933 | 1189 | 1122 | 727 | 865 | 654 | 728 | 661 | 890 | 1257 | 1108 | 1139 | 693 | 744 | 618 | 537 | 873,29 | 1098 | 648 | 0,59 | -0,762 | -6,629 | 3,37E-11 | 6,11E-11 | 0,0177577 | 0,0039714 | 0,011493 | 0,011493 | TRUE | 0,2001165 | No |
| orf19.6604 | 5319 | 6054 | 6020 | 3740 | 1556 | 1210 | 1401 | 1299 | 5071 | 6401 | 5947 | 5859 | 1247 | 1376 | 1189 | 1056 | 3518,35 | 5820 | 1217 | 0,209 | -2,259 | -26,516 | 6,29E-155 | 6,57E-154 | 0,0097821 | 0,0025579 | 0,006507 | 0,006507 | TRUE | 0,095559 | No |
| orf19.6605 | 638 | 679 | 703 | 461 | 1092 | 821 | 1034 | 847 | 608 | 718 | 694 | 722 | 875 | 934 | 878 | 689 | 764,75 | 686 | 844 | 1,231 | 0,3 | 3,095 | 0,0019697 | 0,0026098 | 0,0107352 | 0,0042383 | 0,0076469 | 0,0076469 | TRUE | 0,1848663 | No |
| orf19.6606 | 239 | 271 | 230 | 194 | 336 | 256 | 279 | 319 | 228 | 287 | 227 | 304 | 269 | 291 | 237 | 259 | 262,76 | 262 | 264 | 1,016 | 0,022 | 0,19 | 0,8493959 | 0,8680326 | 0,0105061 | 0,0083399 | 0,0093559 | 0,0093559 | TRUE | 0,1166024 | No |
| orf19.6607 | 3111 | 3198 | 2979 | 1500 | 4095 | 2842 | 3726 | 3888 | 2966 | 3381 | 2943 | 2350 | 3281 | 3232 | 3163 | 3161 | 3059,72 | 2910 | 3209 | 1,102 | 0,14 | 1,616 | 0,1060346 | 0,1234458 | 0,01036 | 0,0026279 | 0,0068239 | 0,0068239 | TRUE | 0,198444 | No |
| orf19.6608 | 2937 | 2812 | 3061 | 1752 | 373 | 292 | 360 | 390 | 2800 | 2973 | 3024 | 2745 | 299 | 332 | 306 | 317 | 1599,46 | 2886 | 314 | 0,108 | -3,208 | -51,37 | 0 | 0 | 1,00E-08 | 0,0031178 | 0,0021286 | 0,0021286 | TRUE | 0,0155948 | No |
| orf19.661 | 10545 | 9457 | 9686 | 6395 | 9643 | 6866 | 8954 | 9024 | 10054 | 10000 | 9569 | 10018 | 7727 | 7807 | 7602 | 7337 | 8764,16 | 9910 | 7618 | 0,769 | -0,38 | -10,933 | 8,04E-28 | 2,09E-27 | 0,0004748 | 0,0022786 | 0,0010418 | 0,0010418 | TRUE | 0,0075704 | No |
| orf19.6610 | 334 | 315 | 332 | 227 | 407 | 299 | 334 | 374 | 318 | 333 | 328 | 356 | 326 | 340 | 284 | 304 | 323,65 | 334 | 314 | 0,94 | -0,089 | -1,012 | 0,3115384 | 0,3447072 | 0,0010865 | 0,007165 | 0,0043402 | 0,0043402 | TRUE | 0,0466357 | No |
| orf19.6612 | 1074 | 1177 | 1201 | 674 | 3111 | 2149 | 2867 | 2576 | 1024 | 1245 | 1186 | 1056 | 2493 | 2444 | 2434 | 2095 | 1746,97 | 1128 | 2366 | 2,096 | 1,068 | 13,803 | 2,44E-43 | 8,15E-43 | 0,006445 | 0,0030311 | 0,0050568 | 0,0050568 | TRUE | 0,0732014 | No |
| orf19.6621 | 15333 | 14187 | 15491 | 8977 | 2454 | 1700 | 2288 | 2603 | 14618 | 15001 | 15304 | 14063 | 1966 | 1933 | 1943 | 2116 | 8368,03 | 14746 | 1990 | 0,135 | -2,889 | -64,617 | 0 | 0 | 0,0012067 | 0,0022875 | 0,0016577 | 0,0016577 | TRUE | 0,0226901 | No |
| orf19.6623 | 1682 | 1489 | 1422 | 929 | 2506 | 1712 | 2337 | 2446 | 1604 | 1574 | 1405 | 1455 | 2008 | 1947 | 1984 | 1989 | 1745,72 | 1510 | 1982 | 1,312 | 0,392 | 7,481 | 7,39E-14 | 1,44E-13 | 0,0012781 | 0,0030318 | 0,0020339 | 0,0020339 | TRUE | 0,0286896 | No |
| orf19.6624 | 887 | 749 | 782 | 521 | 1541 | 1109 | 1421 | 1572 | 846 | 792 | 773 | 816 | 1235 | 1261 | 1206 | 1278 | 1025,84 | 807 | 1245 | 1,544 | 0,626 | 11,621 | 3,24E-31 | 8,92E-31 | 1,00E-08 | 0,0036918 | 0,0017392 | 0,0017392 | TRUE | 0,0143919 | No |
| orf19.6625 | 961 | 874 | 921 | 513 | 2922 | 1858 | 2835 | 2934 | 916 | 924 | 910 | 804 | 2341 | 2113 | 2407 | 2386 | 1600,05 | 888 | 2312 | 2,595 | 1,376 | 22,029 | 1,52E-107 | 1,07E-106 | 0,0027238 | 0,0031174 | 0,0029235 | 0,0029235 | TRUE | 0,0430601 | No |
| orf19.6626 | 237 | 186 | 220 | 147 | 880 | 557 | 815 | 837 | 226 | 197 | 217 | 230 | 705 | 633 | 692 | 681 | 447,65 | 218 | 678 | 3,129 | 1,646 | 20 | 5,55E-89 | 3,29E-88 | 1,00E-08 | 0,0057591 | 0,0032838 | 0,0032838 | TRUE | 0,0455907 | No |
| orf19.6627 | 1362 | 1090 | 1110 | 771 | 3071 | 2090 | 2938 | 3195 | 1299 | 1153 | 1097 | 1208 | 2461 | 2377 | 2494 | 2598 | 1835,61 | 1189 | 2482 | 2,089 | 1,063 | 18,001 | 1,90E-72 | 9,42E-72 | 0,0024268 | 0,0029858 | 0,0026955 | 0,0026955 | TRUE | 0,0498664 | No |
| orf19.6628 | 617 | 433 | 463 | 321 | 1047 | 680 | 991 | 1025 | 588 | 458 | 457 | 503 | 839 | 773 | 841 | 833 | 661,66 | 502 | 822 | 1,638 | 0,712 | 8,396 | 4,62E-17 | 9,73E-17 | 0,0061789 | 0,0045728 | 0,0052497 | 0,0052497 | TRUE | 0,1695919 | No |
| orf19.6629 | 462 | 369 | 402 | 265 | 613 | 409 | 572 | 604 | 440 | 390 | 397 | 415 | 491 | 465 | 486 | 491 | 446,99 | 410 | 483 | 1,178 | 0,236 | 3,24 | 0,0011943 | 0,0016014 | 1,00E-08 | 0,0057645 | 0,0028248 | 0,0028248 | TRUE | 0,0311411 | No |
| orf19.663 | 1315 | 1220 | 1321 | 755 | 1328 | 950 | 1279 | 1387 | 1254 | 1290 | 1305 | 1183 | 1064 | 1080 | 1086 | 1128 | 1173,68 | 1258 | 1090 | 0,864 | -0,21 | -4,072 | 4,66E-05 | 6,69E-05 | 0,0002177 | 0,0034902 | 0,0017065 | 0,0017065 | TRUE | 0,0167995 | No |
| orf19.6630 | 364 | 246 | 252 | 142 | 1007 | 646 | 963 | 1048 | 347 | 260 | 249 | 222 | 807 | 735 | 818 | 852 | 536,21 | 270 | 803 | 2,958 | 1,565 | 13,844 | 1,38E-43 | 4,64E-43 | 0,0154963 | 0,0051533 | 0,0096443 | 0,0096443 | TRUE | 0,4270511 | No |
| orf19.6632 | 2119 | 1702 | 1848 | 1252 | 4431 | 3103 | 4347 | 4625 | 2020 | 1800 | 1826 | 1961 | 3550 | 3528 | 3691 | 3761 | 2767,1 | 1902 | 3632 | 1,912 | 0,935 | 18,737 | 2,46E-78 | 1,31E-77 | 0,0015332 | 0,0026846 | 0,0019712 | 0,0019712 | TRUE | 0,0231082 | No |
| orf19.6634 | 1720 | 1502 | 1685 | 1017 | 4605 | 3263 | 4370 | 4788 | 1640 | 1588 | 1665 | 1593 | 3690 | 3710 | 3710 | 3893 | 2686,15 | 1622 | 3751 | 2,312 | 1,209 | 29,52 | 1,61E-191 | 2,12E-190 | 1,00E-08 | 0,0027025 | 0,0011461 | 0,0011461 | TRUE | 0,008089 | No |
| orf19.6635 | 866 | 829 | 886 | 510 | 1231 | 943 | 1203 | 1228 | 826 | 877 | 875 | 799 | 986 | 1072 | 1021 | 998 | 931,86 | 844 | 1019 | 1,204 | 0,267 | 4,678 | 2,89E-06 | 4,40E-06 | 0,0003864 | 0,0038532 | 0,0020386 | 0,0020386 | TRUE | 0,0134009 | No |
| orf19.6636 | 76 | 86 | 84 | 48 | 354 | 243 | 370 | 398 | 72 | 91 | 83 | 75 | 284 | 276 | 314 | 324 | 189,91 | 80 | 300 | 3,721 | 1,896 | 15,006 | 6,67E-51 | 2,49E-50 | 1,00E-08 | 0,010737 | 0,0069617 | 0,0069617 | TRUE | 0,0725203 | No |
| orf19.6637 | 3699 | 3198 | 3517 | 2209 | 3116 | 2203 | 2952 | 3074 | 3527 | 3381 | 3474 | 3461 | 2497 | 2505 | 2506 | 2499 | 2981,31 | 3461 | 2502 | 0,723 | -0,468 | -13,503 | 1,51E-41 | 4,90E-41 | 1,00E-08 | 0,002642 | 0,0008147 | 0,0008147 | TRUE | 0,0030201 | No |
| orf19.6638 | 985 | 880 | 849 | 560 | 894 | 644 | 806 | 787 | 939 | 930 | 839 | 877 | 716 | 732 | 684 | 640 | 794,8 | 896 | 693 | 0,772 | -0,374 | -5,615 | 1,96E-08 | 3,24E-08 | 0,00198 | 0,0041571 | 0,0029859 | 0,0029859 | TRUE | 0,0323178 | No |
| orf19.6639 | 3986 | 3582 | 3768 | 2397 | 1676 | 1148 | 1550 | 1634 | 3800 | 3788 | 3722 | 3755 | 1343 | 1305 | 1316 | 1329 | 2544,75 | 3766 | 1323 | 0,351 | -1,508 | -40,322 | 0 | 0 | 1,00E-08 | 0,0027365 | 0,0008584 | 0,0008584 | TRUE | 0,001216 | No |
| orf19.664 | 85 | 93 | 87 | 63 | 65 | 24 | 53 | 32 | 81 | 98 | 86 | 99 | 52 | 27 | 45 | 26 | 64,3 | 91 | 38 | 0,419 | -1,256 | -5,548 | 2,89E-08 | 4,74E-08 | 0,0371981 | 0,0276262 | 0,0313215 | 0,0313215 | TRUE | 0,5426258 | No |
| orf19.6640 | 9078 | 8090 | 8555 | 5240 | 2618 | 1817 | 2553 | 2677 | 8655 | 8554 | 8452 | 8209 | 2098 | 2066 | 2167 | 2177 | 5297,16 | 8468 | 2127 | 0,251 | -1,993 | -54,62 | 0 | 0 | 0,0002494 | 0,0024012 | 0,0010021 | 0,0010021 | TRUE | 0,0049354 | No |
| orf19.6641 | 1179 | 1032 | 1030 | 733 | 383 | 309 | 398 | 412 | 1124 | 1091 | 1018 | 1148 | 307 | 351 | 338 | 335 | 714,03 | 1095 | 333 | 0,304 | -1,72 | -24,533 | 6,58E-133 | 5,90E-132 | 0,0014096 | 0,0043908 | 0,0028667 | 0,0028667 | TRUE | 0,0325124 | No |
| orf19.6642 | 1921 | 1789 | 1939 | 1360 | 2482 | 1835 | 2342 | 2350 | 1831 | 1892 | 1916 | 2131 | 1989 | 2087 | 1988 | 1911 | 1967,95 | 1942 | 1994 | 1,028 | 0,04 | 0,722 | 0,4700153 | 0,506445 | 0,0021302 | 0,0029256 | 0,0024317 | 0,0024317 | TRUE | 0,0478677 | No |
| orf19.6643 | 3384 | 2846 | 3357 | 2300 | 1734 | 1335 | 1737 | 1856 | 3226 | 3009 | 3316 | 3603 | 1389 | 1518 | 1475 | 1509 | 2380,78 | 3288 | 1473 | 0,448 | -1,158 | -19,201 | 3,61E-82 | 2,01E-81 | 0,0029355 | 0,0027809 | 0,0030181 | 0,0030181 | TRUE | 0,0482767 | No |
| orf19.6644 | 182 | 192 | 245 | 167 | 101 | 93 | 122 | 100 | 174 | 203 | 242 | 262 | 81 | 106 | 104 | 81 | 156,47 | 220 | 93 | 0,422 | -1,243 | -7,877 | 3,35E-15 | 6,79E-15 | 0,0209945 | 0,0125848 | 0,0165324 | 0,0165324 | TRUE | 0,1440593 | No |
| orf19.6645 | 18657 | 15509 | 16704 | 11931 | 6630 | 4606 | 6203 | 6698 | 17787 | 16399 | 16502 | 18691 | 5312 | 5237 | 5266 | 5446 | 11330,13 | 17345 | 5315 | 0,307 | -1,706 | -35,196 | 2,31E-271 | 4,38E-270 | 0,0019882 | 0,0022362 | 0,0021394 | 0,0021394 | TRUE | 0,0330022 | No |
| orf19.6648 | 19444 | 16582 | 18279 | 10578 | 27399 | 17720 | 26555 | 27439 | 18538 | 17533 | 18058 | 16571 | 21954 | 20149 | 22545 | 22311 | 19707,36 | 17675 | 21740 | 1,23 | 0,298 | 6,031 | 1,63E-09 | 2,79E-09 | 0,0023198 | 0,0021746 | 0,0023006 | 0,0023006 | TRUE | 0,02955 | No |
| orf19.6649 | 4787 | 4057 | 4538 | 2736 | 3523 | 2392 | 3415 | 3641 | 4564 | 4290 | 4483 | 4286 | 2823 | 2720 | 2899 | 2960 | 3628,18 | 4406 | 2850 | 0,647 | -0,628 | -14,751 | 3,02E-49 | 1,10E-48 | 0,0007817 | 0,0025438 | 0,0014558 | 0,0014558 | TRUE | 0,0110188 | No |
| orf19.665 | 4785 | 4350 | 4703 | 2777 | 6257 | 4443 | 6030 | 6270 | 4562 | 4600 | 4646 | 4350 | 5014 | 5052 | 5119 | 5098 | 4805,15 | 4540 | 5071 | 1,116 | 0,158 | 4,658 | 3,19E-06 | 4,84E-06 | 0,000107 | 0,002433 | 0,0008954 | 0,0008954 | TRUE | 0,0088008 | No |
| orf19.6650 | 588 | 528 | 577 | 291 | 460 | 298 | 403 | 475 | 561 | 558 | 570 | 456 | 369 | 339 | 342 | 386 | 447,57 | 536 | 359 | 0,667 | -0,584 | -6,667 | 2,61E-11 | 4,75E-11 | 0,0043273 | 0,0057597 | 0,0050807 | 0,0050807 | TRUE | 0,1033292 | No |
| orf19.6652 | 6616 | 5646 | 6209 | 4006 | 7763 | 5320 | 7550 | 7973 | 6308 | 5970 | 6134 | 6276 | 6220 | 6049 | 6410 | 6483 | 6231,18 | 6172 | 6290 | 1,02 | 0,028 | 0,757 | 0,4489116 | 0,485427 | 0,0005801 | 0,0023548 | 0,0011763 | 0,0011763 | TRUE | 0,0078954 | No |
| orf19.6653 | 3067 | 2478 | 2892 | 1589 | 3547 | 2335 | 3582 | 3820 | 2924 | 2620 | 2857 | 2489 | 2842 | 2655 | 3041 | 3106 | 2816,85 | 2722 | 2911 | 1,069 | 0,096 | 1,433 | 0,1518694 | 0,1742321 | 0,0047671 | 0,0026742 | 0,0039221 | 0,0039221 | TRUE | 0,0416664 | No |
| orf19.6654 | 6560 | 5384 | 5778 | 3756 | 10758 | 7235 | 10453 | 10923 | 6254 | 5693 | 5708 | 5884 | 8620 | 8227 | 8875 | 8881 | 7267,77 | 5885 | 8651 | 1,47 | 0,556 | 12,491 | 8,33E-36 | 2,47E-35 | 0,0014264 | 0,0023172 | 0,0017554 | 0,0017554 | TRUE | 0,0223247 | No |
| orf19.6656 | 2558 | 2051 | 2311 | 1471 | 4698 | 3238 | 4622 | 4934 | 2439 | 2169 | 2283 | 2304 | 3764 | 3682 | 3924 | 4012 | 3072,13 | 2299 | 3846 | 1,673 | 0,743 | 15,08 | 2,18E-51 | 8,16E-51 | 0,0014546 | 0,0026257 | 0,0019672 | 0,0019672 | TRUE | 0,021371 | No |

| | | | | | | | | | | | | | | | | | | | | | | | | | | | | | | | |
|---|---|---|---|---|---|---|---|---|---|---|---|---|---|---|---|---|---|---|---|---|---|---|---|---|---|---|---|---|---|---|---|
| orf19.6658 | 5295 | 4428 | 4720 | 3067 | 8861 | 5955 | 8477 | 9158 | 5048 | 4682 | 4663 | 4805 | 7100 | 6771 | 7197 | 7446 | 5964,06 | 4800 | 7128 | 1,486 | 0,571 | 13,06 | 5,54E-39 | 1,73E-38 | 0,0012177 | 0,0023666 | 0,0016561 | 0,0016561 | TRUE | 0,0153103 | No |
| orf19.6659 | 41019 | 34069 | 38302 | 24353 | 42352 | 28821 | 40548 | 43366 | 39107 | 36024 | 37839 | 38150 | 33935 | 32772 | 34425 | 35261 | 35939,14 | 37780 | 34098 | 0,903 | -0,148 | -3,893 | 9,89E-05 | 0,0001401 | 0,00101 | 0,002137 | 0,001357 | 0,001357 | TRUE | 0,0120267 | No |
| orf19.666 | 976 | 891 | 1022 | 627 | 1923 | 1425 | 1771 | 1846 | 931 | 942 | 1010 | 982 | 1541 | 1620 | 1504 | 1501 | 1253,78 | 966 | 1542 | 1,595 | 0,674 | 12,808 | 1,48E-37 | 4,51E-37 | 0,0005271 | 0,0034008 | 0,001786 | 0,001786 | TRUE | 0,0134186 | No |
| orf19.6660 | 17193 | 13334 | 15370 | 10221 | 11623 | 8110 | 11337 | 12191 | 16392 | 14099 | 15184 | 16012 | 9313 | 9222 | 9625 | 9912 | 12469,88 | 15422 | 9518 | 0,617 | -0,696 | -13,376 | 8,31E-41 | 2,67E-40 | 0,0026015 | 0,0022229 | 0,0025169 | 0,0025169 | TRUE | 0,0408586 | No |
| orf19.6661 | 353 | 307 | 353 | 214 | 605 | 410 | 604 | 570 | 337 | 325 | 349 | 335 | 485 | 466 | 513 | 463 | 409,05 | 336 | 482 | 1,433 | 0,519 | 6,821 | 9,02E-12 | 1,66E-11 | 1,00E-08 | 0,0061053 | 0,0029839 | 0,0029839 | TRUE | 0,0221144 | No |
| orf19.6662 | 391 | 306 | 334 | 196 | 1077 | 707 | 1024 | 1112 | 373 | 324 | 330 | 307 | 863 | 804 | 869 | 904 | 596,72 | 334 | 860 | 2,57 | 1,362 | 17,846 | 3,10E-71 | 1,51E-70 | 0,0019252 | 0,0048428 | 0,0034094 | 0,0034094 | TRUE | 0,0746153 | No |
| orf19.6663 | 12813 | 10175 | 10635 | 8442 | 37674 | 26088 | 34140 | 35430 | 12216 | 10759 | 10506 | 13225 | 30187 | 29664 | 28985 | 28808 | 20543,75 | 11676 | 29411 | 2,52 | 1,333 | 19,634 | 7,98E-86 | 4,59E-85 | 0,0061433 | 0,0021712 | 0,0043668 | 0,0043668 | TRUE | 0,0966027 | No |
| orf19.6665 | 4202 | 3566 | 4119 | 2740 | 7544 | 5254 | 7208 | 7801 | 4006 | 3771 | 4069 | 4292 | 6045 | 5974 | 6120 | 6343 | 5077,48 | 4034 | 6120 | 1,519 | 0,603 | 13,24 | 5,18E-40 | 1,64E-39 | 0,0014811 | 0,0024146 | 0,0017779 | 0,0017779 | TRUE | 0,0233539 | No |
| orf19.6667 | 1496 | 1280 | 1378 | 829 | 1274 | 738 | 1154 | 1181 | 1426 | 1353 | 1361 | 1299 | 1021 | 839 | 980 | 960 | 1154,96 | 1360 | 950 | 0,699 | -0,516 | -7,874 | 3,44E-15 | 6,96E-15 | 0,0029241 | 0,0035129 | 0,0032416 | 0,0032416 | TRUE | 0,0689573 | No |
| orf19.6668 | 1493 | 1224 | 1394 | 949 | 1828 | 1212 | 1688 | 1801 | 1423 | 1294 | 1377 | 1487 | 1465 | 1378 | 1433 | 1464 | 1415,23 | 1395 | 1435 | 1,031 | 0,044 | 0,823 | 0,4102633 | 0,4462059 | 0,0011457 | 0,0032515 | 0,0020682 | 0,0020682 | TRUE | 0,0281276 | No |
| orf19.6670 | 269 | 262 | 241 | 168 | 633 | 425 | 602 | 653 | 256 | 277 | 238 | 263 | 507 | 483 | 511 | 531 | 383,41 | 258 | 508 | 1,971 | 0,979 | 12,019 | 2,84E-33 | 8,07E-33 | 1,00E-08 | 0,0063737 | 0,0033365 | 0,0033365 | TRUE | 0,0344707 | No |
| orf19.6671 | 757 | 721 | 780 | 455 | 1385 | 932 | 1410 | 1435 | 722 | 762 | 771 | 713 | 1110 | 1060 | 1197 | 1167 | 937,6 | 742 | 1134 | 1,527 | 0,611 | 10,043 | 9,90E-24 | 2,39E-23 | 0,0012372 | 0,0038424 | 0,0024072 | 0,0024072 | TRUE | 0,0212837 | No |
| orf19.667.1 | 13216 | 10336 | 10359 | 8168 | 35750 | 23660 | 30807 | 34301 | 12600 | 10929 | 10234 | 12796 | 28645 | 26904 | 26155 | 27890 | 19519,04 | 11640 | 27398 | 2,354 | 1,235 | 17,58 | 3,48E-69 | 1,64E-68 | 0,0065463 | 0,0021754 | 0,0046797 | 0,0046797 | TRUE | 0,0809821 | No |
| orf19.6672 | 8119 | 7010 | 7630 | 4685 | 4068 | 2727 | 3838 | 4224 | 7741 | 7412 | 7538 | 7339 | 3260 | 3101 | 3258 | 3435 | 5385,4 | 7508 | 3264 | 0,435 | -1,201 | -29,46 | 9,31E-191 | 1,22E-189 | 0,0008532 | 0,0023962 | 0,0013851 | 0,0013851 | TRUE | 0,0151405 | No |
| orf19.6673 | 531 | 431 | 510 | 344 | 1726 | 1175 | 1640 | 1781 | 506 | 456 | 504 | 539 | 1383 | 1336 | 1392 | 1448 | 945,53 | 501 | 1390 | 2,785 | 1,478 | 23,676 | 6,43E-124 | 5,31E-123 | 0,0006561 | 0,0038278 | 0,0023025 | 0,0023025 | TRUE | 0,0422853 | No |
| orf19.6674 | 344 | 275 | 321 | 141 | 839 | 561 | 778 | 780 | 328 | 291 | 317 | 221 | 672 | 638 | 661 | 634 | 470,21 | 289 | 651 | 2,229 | 1,156 | 11,891 | 1,32E-32 | 3,73E-32 | 0,0080099 | 0,0055832 | 0,0064767 | 0,0064767 | TRUE | 0,2343855 | No |
| orf19.6675 | 839 | 823 | 821 | 574 | 1510 | 1026 | 1414 | 1367 | 800 | 870 | 811 | 899 | 1210 | 1167 | 1200 | 1112 | 1008,62 | 845 | 1172 | 1,391 | 0,477 | 8,059 | 7,72E-16 | 1,58E-15 | 0,0011247 | 0,0037192 | 0,0023118 | 0,0023118 | TRUE | 0,0193666 | No |
| orf19.6676 | 162 | 114 | 138 | 102 | 342 | 178 | 251 | 345 | 154 | 121 | 136 | 160 | 274 | 202 | 213 | 281 | 192,65 | 143 | 242 | 1,711 | 0,775 | 5,413 | 6,19E-08 | 1,00E-07 | 0,017407 | 0,0106142 | 0,0138958 | 0,0138958 | TRUE | 0,1008664 | No |
| orf19.6678 | 367 | 332 | 352 | 246 | 90 | 67 | 85 | 79 | 350 | 351 | 348 | 385 | 72 | 76 | 72 | 64 | 214,84 | 358 | 71 | 0,198 | -2,334 | -20,144 | 3,03E-90 | 1,82E-89 | 1,00E-08 | 0,0097336 | 0,005073 | 0,005073 | TRUE | 0,0420847 | No |
| orf19.6679 | 1407 | 1131 | 1230 | 865 | 1872 | 1305 | 1793 | 1966 | 1341 | 1196 | 1215 | 1355 | 1500 | 1484 | 1522 | 1599 | 1401,52 | 1277 | 1526 | 1,198 | 0,26 | 4,548 | 5,41E-06 | 8,12E-06 | 0,0017051 | 0,0032628 | 0,0024108 | 0,0024108 | TRUE | 0,0214763 | No |
| orf19.668 | 978 | 964 | 976 | 762 | 340 | 203 | 299 | 341 | 932 | 1019 | 964 | 1194 | 272 | 231 | 254 | 277 | 643 | 1027 | 258 | 0,253 | -1,981 | -21,122 | 4,95E-99 | 3,21E-98 | 0,0076921 | 0,0046448 | 0,0061766 | 0,0061766 | TRUE | 0,1335913 | No |
| orf19.6680 | 2747 | 2466 | 2747 | 1778 | 4927 | 3786 | 4800 | 5117 | 2619 | 2607 | 2714 | 2785 | 3948 | 4305 | 4075 | 4161 | 3401,78 | 2681 | 4122 | 1,538 | 0,621 | 14,455 | 2,33E-47 | 8,27E-47 | 0,0007434 | 0,0025739 | 0,0014544 | 0,0014544 | TRUE | 0,0105103 | No |
| orf19.6681 | 2003 | 1889 | 2047 | 1253 | 2068 | 1494 | 1974 | 2142 | 1910 | 1997 | 2022 | 1963 | 1657 | 1699 | 1676 | 1742 | 1833,2 | 1973 | 1694 | 0,858 | -0,221 | -5,156 | 2,52E-07 | 3,99E-07 | 1,00E-08 | 0,0029869 | 0,0012115 | 0,0012115 | TRUE | 0,0061838 | No |
| orf19.6684 | 612 | 556 | 583 | 353 | 586 | 416 | 546 | 601 | 583 | 588 | 576 | 553 | 470 | 473 | 464 | 489 | 524,39 | 575 | 474 | 0,822 | -0,282 | -4,342 | 1,41E-05 | 2,08E-05 | 1,00E-08 | 0,0052224 | 0,002155 | 0,002155 | TRUE | 0,0064001 | No |
| orf19.6685 | 1903 | 1765 | 1831 | 965 | 1549 | 1053 | 1538 | 1454 | 1814 | 1866 | 1809 | 1512 | 1241 | 1197 | 1306 | 1182 | 1490,96 | 1750 | 1232 | 0,702 | -0,51 | -7,345 | 2,05E-13 | 3,97E-13 | 0,0043262 | 0,0031925 | 0,0039428 | 0,0039428 | TRUE | 0,0952966 | No |
| orf19.6686 | 11994 | 11727 | 12382 | 7567 | 10498 | 7825 | 9848 | 10660 | 11435 | 12400 | 12232 | 11854 | 8412 | 8898 | 8361 | 8668 | 10282,39 | 11980 | 8585 | 0,716 | -0,481 | -12,304 | 8,59E-35 | 2,50E-34 | 0,0009267 | 0,002251 | 0,0013699 | 0,0013699 | TRUE | 0,0117053 | No |
| orf19.6687 | 9 | 16 | 7 | 13 | 18 | 25 | 21 | 14 | 9 | 17 | 7 | 20 | 14 | 28 | 18 | 11 | 15,61 | 13 | 18 | 1,387 | 0,472 | 1,077 | 0,2816205 | 0,313452 | 0,1263654 | 0,1073066 | 0,1155525 | 0,1155525 | TRUE | 0,3607912 | No |
| orf19.6688 | 8126 | 7806 | 8399 | 4907 | 7934 | 5608 | 7661 | 7460 | 7747 | 8254 | 8297 | 7687 | 6357 | 6377 | 6504 | 6066 | 7161,19 | 7996 | 6326 | 0,791 | -0,339 | -8,163 | 3,26E-16 | 6,73E-16 | 0,0010606 | 0,0023206 | 0,001512 | 0,001512 | TRUE | 0,0094671 | No |
| orf19.6689 | 4526 | 4341 | 4373 | 2606 | 1553 | 1078 | 1495 | 1507 | 4315 | 4590 | 4320 | 4082 | 1244 | 1226 | 1269 | 1225 | 2784,05 | 4327 | 1241 | 0,287 | -1,802 | -38,054 | 0 | 0 | 0,0010754 | 0,002681 | 0,0016648 | 0,0016648 | TRUE | 0,0201556 | No |
| orf19.669 | 158 | 157 | 134 | 83 | 224 | 152 | 221 | 229 | 151 | 166 | 132 | 130 | 179 | 173 | 188 | 186 | 163,15 | 145 | 182 | 1,25 | 0,322 | 2,739 | 0,0061712 | 0,0079344 | 1,00E-08 | 0,0121551 | 0,0069671 | 0,0069671 | TRUE | 0,0980951 | No |
| orf19.6690 | 2460 | 2267 | 2436 | 1374 | 747 | 577 | 709 | 727 | 2345 | 2397 | 2407 | 2152 | 599 | 656 | 602 | 591 | 1468,64 | 2325 | 612 | 0,262 | -1,932 | -33,294 | 4,74E-243 | 7,84E-242 | 0,0014733 | 0,0032093 | 0,0022607 | 0,0022607 | TRUE | 0,0281699 | No |
| orf19.6691 | 11639 | 11692 | 12095 | 7275 | 10182 | 7387 | 9329 | 9557 | 11097 | 12363 | 11949 | 11397 | 8159 | 8400 | 7920 | 7771 | 9881,75 | 11702 | 8062 | 0,689 | -0,538 | -11,906 | 1,10E-32 | 3,11E-32 | 0,001611 | 0,0022574 | 0,0018564 | 0,0018564 | TRUE | 0,0177615 | No |
| orf19.6692 | 2060 | 2162 | 2237 | 1276 | 2481 | 1871 | 2325 | 2506 | 1964 | 2286 | 2210 | 1999 | 1988 | 2127 | 1974 | 2038 | 2073,23 | 2115 | 2032 | 0,96 | -0,06 | -1,014 | 0,3107916 | 0,3440599 | 0,0027189 | 0,0028832 | 0,0028376 | 0,0028376 | TRUE | 0,0359046 | No |
| orf19.6693 | 1943 | 2074 | 2237 | 1191 | 2331 | 1878 | 2106 | 2275 | 1852 | 2193 | 2210 | 1866 | 1868 | 2135 | 1788 | 1850 | 1970,27 | 2030 | 1910 | 0,939 | -0,09 | -1,158 | 0,2468656 | 0,2767966 | 0,00732 | 0,0029246 | 0,0053158 | 0,0053158 | TRUE | 0,0745626 | No |
| orf19.6694 | 2147 | 1828 | 2078 | 1237 | 1596 | 1063 | 1530 | 1563 | 2047 | 1933 | 2053 | 1938 | 1279 | 1209 | 1299 | 1271 | 1628,49 | 1993 | 1264 | 0,635 | -0,656 | -13,927 | 4,37E-44 | 1,47E-43 | 0,0003622 | 0,0030995 | 0,001498 | 0,001498 | TRUE | 0,0096295 | No |
| orf19.6696 | 969 | 979 | 917 | 504 | 1706 | 1204 | 1529 | 1371 | 924 | 1035 | 906 | 790 | 1367 | 1369 | 1298 | 1115 | 1100,42 | 914 | 1287 | 1,405 | 0,491 | 5,524 | 3,32E-08 | 5,44E-08 | 0,0091263 | 0,0035834 | 0,0066098 | 0,0066098 | TRUE | 0,0990056 | No |
| orf19.6698 | 857 | 688 | 738 | 467 | 1396 | 985 | 1324 | 1441 | 817 | 727 | 729 | 732 | 1119 | 1120 | 1124 | 1172 | 942,44 | 751 | 1134 | 1,507 | 0,591 | 10,12 | 4,50E-24 | 1,09E-23 | 0,0004825 | 0,0038335 | 0,0021412 | 0,0021412 | TRUE | 0,0439208 | No |
| orf19.6699 | 339 | 260 | 314 | 200 | 950 | 694 | 879 | 933 | 323 | 275 | 310 | 313 | 761 | 789 | 746 | 759 | 534,61 | 305 | 764 | 2,5 | 1,322 | 18,181 | 7,27E-74 | 3,68E-73 | 1,00E-08 | 0,0051625 | 0,0026701 | 0,0026701 | TRUE | 0,0531804 | No |
| orf19.670 | 5624 | 4298 | 4510 | 2821 | 7182 | 4138 | 6633 | 7036 | 5362 | 4545 | 4455 | 4419 | 5755 | 4705 | 5631 | 5721 | 5074,19 | 4695 | 5453 | 1,161 | 0,216 | 2,732 | 0,0062993 | 0,0080926 | 0,0085022 | 0,0024149 | 0,0057939 | 0,0057939 | TRUE | 0,1119845 | No |
| orf19.6701 | 3832 | 3334 | 3639 | 2305 | 8015 | 5774 | 7655 | 8476 | 3653 | 3525 | 3595 | 3611 | 6422 | 6566 | 6499 | 6892 | 5095,4 | 3596 | 6595 | 1,834 | 0,875 | 23,682 | 5,47E-124 | 4,52E-123 | 0,0005101 | 0,0024135 | 0,0010864 | 0,0010864 | TRUE | 0,0114184 | No |
| orf19.6702 | 3634 | 3114 | 3228 | 2304 | 6546 | 4762 | 6457 | 7023 | 3465 | 3293 | 3189 | 3609 | 5245 | 5415 | 5482 | 5710 | 4425,98 | 3389 | 5463 | 1,614 | 0,69 | 14,163 | 1,55E-45 | 5,36E-45 | 0,001806 | 0,0024622 | 0,0020342 | 0,0020342 | TRUE | 0,0220522 | No |
| orf19.670.2 | 22344 | 17276 | 18283 | 8704 | 11542 | 6006 | 10640 | 11365 | 21303 | 18267 | 18062 | 13635 | 9248 | 6829 | 9033 | 9241 | 13202,34 | 17817 | 8588 | 0,482 | -1,053 | -6,52 | 7,01E-11 | 1,26E-10 | 0,0249685 | 0,0022156 | 0,0138152 | 0,0249685 | TRUE | 0,2324073 | No |
| orf19.6703 | 978 | 903 | 1051 | 526 | 752 | 513 | 630 | 712 | 932 | 955 | 1038 | 824 | 603 | 583 | 535 | 579 | 756,15 | 937 | 575 | 0,611 | -0,71 | -9,253 | 2,18E-20 | 4,94E-20 | 0,0043129 | 0,0042627 | 0,0042778 | 0,0042778 | TRUE | 0,0714676 | No |
| orf19.6704 | 150 | 132 | 152 | 90 | 207 | 113 | 174 | 196 | 143 | 140 | 150 | 141 | 166 | 128 | 148 | 159 | 146,9 | 144 | 150 | 1,053 | 0,074 | 0,606 | 0,5448322 | 0,5802557 | 1,00E-08 | 0,0132685 | 0,0076544 | 0,0076544 | TRUE | 0,0878226 | No |
| orf19.6705 | 5736 | 5259 | 5666 | 3651 | 4030 | 2984 | 3912 | 4095 | 5469 | 5561 | 5597 | 5719 | 3229 | 3393 | 3321 | 3330 | 4452,43 | 5586 | 3318 | 0,594 | -0,751 | -22,066 | 6,69E-108 | 4,73E-107 | 0,0001201 | 0,00246 | 0,000878 | 0,000878 | TRUE | 0,004012 | No |
| orf19.6706 | 12393 | 11890 | 12588 | 6971 | 5401 | 3562 | 5028 | 5212 | 11815 | 12572 | 12436 | 10920 | 4328 | 4050 | 4269 | 4238 | 8078,54 | 11936 | 4221 | 0,354 | -1,5 | -29,588 | 2,14E-192 | 2,83E-191 | 0,0021783 | 0,0022945 | 0,0023147 | 0,0023147 | TRUE | 0,0394398 | No |
| orf19.6707 | 4011 | 3772 | 3939 | 2328 | 1836 | 1272 | 1685 | 1786 | 3824 | 3988 | 3891 | 3647 | 1471 | 1446 | 1431 | 1452 | 2643,88 | 3838 | 1450 | 0,378 | -1,405 | -32,957 | 3,37E-238 | 5,41E-237 | 0,0004775 | 0,0027123 | 0,001292 | 0,001292 | TRUE | 0,0126084 | No |
| orf19.6709 | 164 | 136 | 165 | 101 | 357 | 202 | 343 | 355 | 156 | 144 | 163 | 158 | 286 | 230 | 291 | 289 | 214,62 | 155 | 274 | 1,771 | 0,825 | 7,32 | 2,49E-13 | 4,80E-13 | 0,0037935 | 0,0097414 | 0,0069828 | 0,0069828 | TRUE | 0,1196508 | No |
| orf19.671 | 2355 | 2282 | 2418 | 1571 | 2859 | 2127 | 2790 | 2927 | 2245 | 2413 | 2389 | 2461 | 2291 | 2419 | 2369 | 2380 | 2370,75 | 2377 | 2365 | 0,996 | -0,006 | -0,147 | 0,8831434 | 0,8990647 | 0,0005819 | 0,0027838 | 0,0014185 | 0,0014185 | TRUE | 0,0173084 | No |
| orf19.6710 | 16102 | 13506 | 14858 | 9156 | 17474 | 11573 | 16778 | 17605 | 15352 | 14281 | 14678 | 14343 | 14001 | 13160 | 14244 | 14315 | 14296,75 | 14664 | 13930 | 0,95 | -0,074 | -1,812 | 0,0699811 | 0,0830172 | 0,0011928 | 0,0022061 | 0,0015317 | 0,0015317 | TRUE | 0,0167719 | No |
| orf19.6712 | 2851 | 2428 | 2708 | 1846 | 1852 | 1257 | 1870 | 2047 | 2718 | 2567 | 2675 | 2892 | 1484 | 1429 | 1588 | 1664 | 2127,23 | 2713 | 1541 | 0,569 | -0,813 | -13,688 | 1,20E-42 | 3,96E-42 | 0,0028837 | 0,0028631 | 0,0028902 | 0,0028902 | TRUE | 0,0349383 | No |
| orf19.6713 | 9617 | 8243 | 9408 | 7386 | 5758 | 4494 | 5542 | 6156 | 9169 | 8716 | 9294 | 11571 | 4614 | 5110 | 4705 | 5005 | 7272,98 | 9688 | 4858 | 0,502 | -0,995 | -12,374 | 3,60E-35 | 1,06E-34 | 0,0097088 | 0,002317 | 0,0060628 | 0,0060628 | TRUE | 0,2077807 | No |
| orf19.6715 | 9 | 8 | 10 | 10 | 19 | 9 | 14 | 10 | 9 | 8 | 10 | 16 | 15 | 10 | 12 | 8 | 11,01 | 11 | 11 | 1,104 | 0,143 | 0,33 | 0,7417116 | 0,7700719 | 1,00E-08 | 0,1512553 | 0,0871784 | 0,0871784 | TRUE | 0,2433664 | No |
| orf19.6716 | 3313 | 2660 | 2906 | 1814 | 2797 | 1812 | 2663 | 2820 | 3159 | 2813 | 2871 | 2842 | 2241 | 2060 | 2261 | 2293 | 2567,39 | 2921 | 2214 | 0,758 | -0,4 | -7,395 | 1,41E-13 | 2,74E-13 | 0,0020885 | 0,0027308 | 0,0024089 | 0,0024089 | TRUE | 0,0372321 | No |
| orf19.6717 | 1531 | 1254 | 1355 | 927 | 2124 | 1435 | 2021 | 2191 | 1460 | 1326 | 1339 | 1452 | 1702 | 1632 | 1716 | 1781 | 1550,92 | 1394 | 1708 | 1,227 | 0,295 | 5,573 | 2,51E-08 | 4,12E-08 | 0,0010984 | 0,0031499 | 0,0020343 | 0,0020343 | TRUE | 0,0136571 | No |
| orf19.6718 | 2965 | 2439 | 2542 | 1694 | 2375 | 1712 | 2163 | 2486 | 2827 | 2579 | 2511 | 2654 | 1903 | 1947 | 1836 | 2021 | 2284,77 | 2643 | 1927 | 0,729 | -0,456 | -8,731 | 2,52E-18 | 5,45E-18 | 0,0017754 | 0,0028099 | 0,0021737 | 0,0021737 | TRUE | 0,0278788 | No |
| orf19.6719 | 1131 | 1060 | 1148 | 658 | 1155 | 734 | 1023 | 1169 | 1078 | 1121 | 1134 | 1031 | 925 | 835 | 869 | 951 | 992,89 | 1091 | 895 | 0,82 | -0,286 | -4,71 | 2,48E-06 | 3,77E-06 | 0,0014898 | 0,0037449 | 0,0025297 | 0,0025297 | TRUE | 0,0226325 | No |
| orf19.672 | 1291 | 1138 | 1306 | 799 | 1473 | 1064 | 1466 | 1563 | 1231 | 1203 | 1290 | 1252 | 1180 | 1210 | 1245 | 1271 | 1235,2 | 1244 | 1226 | 0,986 | -0,02 | -0,398 | 0,6905808 | 0,7218011 | 1,00E-08 | 0,0034205 | 0,001613 | 0,001613 | TRUE | 0,0082873 | No |
| orf19.6720 | 22975 | 18692 | 20027 | 13189 | 11434 | 8128 | 10972 | 11702 | 21904 | 19764 | 19785 | 20661 | 9162 | 9242 | 9315 | 9515 | 14918,58 | 20528 | 9308 | 0,453 | -1,141 | -27,519 | 1,03E-166 | 1,16E-165 | 0,0012328 | 0,0022013 | 0,0015758 | 0,0015758 | TRUE | 0,0251388 | No |
| orf19.6722 | 3421 | 2930 | 3259 | 2042 | 3899 | 2532 | 3641 | 3952 | 3262 | 3098 | 3220 | 3199 | 3124 | 2879 | 3091 | 3213 | 3135,74 | 3195 | 3077 | 0,964 | -0,053 | -1,191 | 0,2334712 | 0,262655 | 0,0008698 | 0,0026149 | 0,0015756 | 0,0015756 | TRUE | 0,0218224 | No |
| orf19.6723 | 1445 | 1348 | 1424 | 940 | 1495 | 1074 | 1452 | 1596 | 1378 | 1425 | 1407 | 1473 | 1198 | 1221 | 1233 | 1298 | 1328,99 | 1421 | 1238 | 0,873 | -0,196 | -3,995 | 6,48E-05 | 9,25E-05 | 1,00E-08 | 0,0033267 | 0,0015664 | 0,0015664 | TRUE | 0,0134347 | No |
| orf19.6724 | 2556 | 1877 | 2189 | 1615 | 7884 | 6090 | 7535 | 8131 | 2437 | 1985 | 2163 | 2530 | 6317 | 6925 | 6397 | 6611 | 4420,58 | 2279 | 6562 | 2,883 | 1,528 | 21,414 | 9,91E-102 | 6,57E-101 | 0,0065003 | 0,0024627 | 0,0045717 | 0,0045717 | TRUE | 0,0910063 | No |
| orf19.6726 | 1833 | 1512 | 1749 | 1095 | 2892 | 1765 | 2816 | 2843 | 1748 | 1599 | 1728 | 1715 | 2317 | 2007 | 2391 | 2312 | 1977,02 | 1698 | 2257 | 1,331 | 0,412 | 6,671 | 2,53E-11 | 4,61E-11 | 0,002941 | 0,0029218 | 0,0031399 | 0,0031399 | TRUE | 0,0650487 | No |
| orf19.6727 | 2860 | 2336 | 2557 | 1612 | 3777 | 2595 | 3483 | 3939 | 2727 | 2470 | 2526 | 2525 | 3026 | 2951 | 2957 | 3203 | 2798,13 | 2562 | 3034 | 1,184 | 0,244 | 5,064 | 4,10E-07 | 6,44E-07 | 0,0012401 | 0,0026781 | 0,0018584 | 0,0018584 | TRUE | 0,0235137 | No |
| orf19.6729 | 1776 | 1583 | 1826 | 1006 | 1342 | 963 | 1179 | 1337 | 1693 | 1674 | 1804 | 1576 | 1075 | 1095 | 1001 | 1087 | 1375,66 | 1687 | 1064 | 0,63 | -0,667 | -11,798 | 3,99E-32 | 1,12E-31 | 0,0015883 | 0,0032848 | 0,0023187 | 0,0023187 | TRUE | 0,026199 | No |
| orf19.673 | 750 | 625 | 634 | 449 | 628 | 435 | 541 | 646 | 715 | 661 | 626 | 703 | 503 | 495 | 459 | 525 | 586 | 676 | 496 | 0,734 | -0,446 | -6,149 | 7,78E-10 | 1,34E-09 | 0,0018846 | 0,0048932 | 0,0033294 | 0,0033294 | TRUE | 0,0304796 | No |
| orf19.6730 | 2576 | 2208 | 2402 | 1501 | 4555 | 2808 | 4355 | 4515 | 2456 | 2335 | 2373 | 2351 | 3650 | 3193 | 3697 | 3671 | 2965,78 | 2379 | 3553 | 1,494 | 0,579 | 10,641 | 1,92E-26 | 4,90E-26 | 0,0020827 | 0,0026449 | 0,0024851 | 0,0024851 | TRUE | 0,0550933 | No |
| orf19.6731 | 1604 | 1348 | 1472 | 975 | 2469 | 1598 | 2324 | 2493 | 1529 | 1425 | 1454 | 1527 | 1978 | 1817 | 1973 | 2027 | 1716,46 | 1484 | 1949 | 1,316 | 0,396 | 7,713 | 1,23E-14 | 2,46E-14 | 0,0011371 | 0,0030478 | 0,0019209 | 0,0019209 | TRUE | 0,0237696 | No |
| orf19.6731.1 | 1048 | 867 | 861 | 462 | 2439 | 1313 | 2266 | 2215 | 999 | 917 | 851 | 724 | 1954 | 1493 | 1924 | 1801 | 1332,8 | 873 | 1793 | 2,051 | 1,036 | 10,092 | 5,99E-24 | 1,45E-23 | 0,0146582 | 0,0033232 | 0,0092338 | 0,0092338 | TRUE | 0,1508415 | No |
| orf19.6732 | 1341 | 1262 | 1344 | 926 | 1439 | 1050 | 1400 | 1443 | 1278 | 1334 | 1328 | 1451 | 1153 | 1194 | 1189 | 1173 | 1262,52 | 1348 | 1177 | 0,876 | -0,192 | -3,68 | 0,0002334 | 0,0003249 | 0,0005246 | 0,0033918 | 0,0018056 | 0,0018056 | TRUE | 0,0282096 | No |
| orf19.6734 | 8968 | 7828 | 9103 | 6171 | 8916 | 6563 | 8497 | 9356 | 8550 | 8277 | 8993 | 9667 | 7144 | 7463 | 7214 | 7607 | 8114,42 | 8872 | 7357 | 0,83 | -0,269 | -5,187 | 2,14E-07 | 3,41E-07 | 0,002603 | 0,0022936 | 0,0024678 | 0,0024678 | TRUE | 0,0436671 | No |
| orf19.6736 | 8969 | 8462 | 9094 | 5878 | 6244 | 5002 | 5974 | 6677 | 8551 | 8947 | 8984 | 9208 | 5003 | 5688 | 5072 | 5429 | 7110,31 | 8922 | 5298 | 0,594 | -0,752 | -15,305 | 7,13E-53 | 2,73E-52 | 0,0020922 | 0,0023222 | 0,0021734 | 0,0021734 | TRUE | 0,029368 | No |
| orf19.6737 | 1027 | 880 | 939 | 672 | 1406 | 1120 | 1344 | 1420 | 979 | 930 | 928 | 1053 | 1127 | 1274 | 1141 | 1155 | 1073,22 | 972 | 1174 | 1,208 | 0,273 | 4,359 | 1,31E-05 | 1,93E-05 | 0,0022011 | 0,0036212 | 0,0028123 | 0,0028123 | TRUE | 0,0370385 | No |

| | | | | | | | | | | | | | | | | | | | | | | | | | | | | | | | |
|---|---|---|---|---|---|---|---|---|---|---|---|---|---|---|---|---|---|---|---|---|---|---|---|---|---|---|---|---|---|---|---|
| orf19.6738 | 3387 | 2802 | 3081 | 1994 | 4280 | 3003 | 4189 | 4159 | 3229 | 2963 | 3044 | 3124 | 3429 | 3415 | 3556 | 3382 | 3267,7 | 3090 | 3446 | 1,115 | 0,157 | 3,823 | 0,000132 | 0,0001858 | 0,0005155 | 0,0025937 | 0,0013163 | 0,0013163 | TRUE | 0,0119136 | No |
| orf19.6739 | 2218 | 1824 | 2071 | 1219 | 1455 | 950 | 1414 | 1559 | 2115 | 1929 | 2046 | 1910 | 1166 | 1080 | 1200 | 1268 | 1589,13 | 2000 | 1178 | 0,59 | -0,762 | -12,668 | 8,85E-37 | 2,66E-36 | 0,0024779 | 0,0031245 | 0,0028161 | 0,0028161 | TRUE | 0,0355924 | No |
| orf19.674 | 2673 | 2369 | 2611 | 1773 | 3243 | 2378 | 3172 | 3414 | 2548 | 2505 | 2579 | 2777 | 2599 | 2704 | 2693 | 2776 | 2647,71 | 2602 | 2693 | 1,036 | 0,051 | 1,134 | 0,256711 | 0,2869784 | 0,0009351 | 0,0027114 | 0,0015955 | 0,0015955 | TRUE | 0,0228783 | No |
| orf19.6740 | 2241 | 1877 | 2159 | 1388 | 3806 | 2646 | 3671 | 3931 | 2137 | 1985 | 2133 | 2174 | 3050 | 3009 | 3117 | 3196 | 2599,98 | 2107 | 3093 | 1,469 | 0,555 | 12,602 | 2,05E-36 | 6,11E-36 | 0,00061 | 0,0027228 | 0,0014528 | 0,0014528 | TRUE | 0,0185963 | No |
| orf19.6741 | 227 | 187 | 194 | 147 | 202 | 151 | 201 | 221 | 216 | 198 | 192 | 230 | 162 | 172 | 171 | 180 | 190 | 209 | 171 | 0,822 | -0,283 | -2,639 | 0,0083081 | 0,0106007 | 1,00E-08 | 0,010733 | 0,0057493 | 0,0057493 | TRUE | 0,0405748 | No |
| orf19.6742 | 1762 | 1537 | 1784 | 1015 | 1802 | 1281 | 1847 | 1897 | 1680 | 1625 | 1762 | 1590 | 1444 | 1457 | 1568 | 1542 | 1583,57 | 1664 | 1503 | 0,902 | -0,149 | -2,84 | 0,0045072 | 0,0058397 | 0,0012079 | 0,0031281 | 0,0020015 | 0,0020015 | TRUE | 0,018995 | No |
| orf19.6744 | 146 | 103 | 150 | 75 | 419 | 327 | 425 | 446 | 139 | 109 | 148 | 117 | 336 | 372 | 361 | 363 | 243,1 | 128 | 358 | 2,761 | 1,465 | 13,261 | 3,90E-40 | 1,24E-39 | 0,0012847 | 0,0088453 | 0,0061821 | 0,0061821 | TRUE | 0,121733 | No |
| orf19.6745 | 8296 | 6371 | 6859 | 4490 | 8109 | 5378 | 7616 | 8273 | 7909 | 6737 | 6776 | 7034 | 6498 | 6115 | 6466 | 6727 | 6782,65 | 7114 | 6452 | 0,907 | -0,141 | -2,45 | 0,0142945 | 0,0179519 | 0,0035181 | 0,0023334 | 0,0030322 | 0,0030322 | TRUE | 0,0699575 | No |
| orf19.6747 | 4698 | 3815 | 4213 | 2714 | 4484 | 3125 | 4495 | 4663 | 4479 | 4034 | 4162 | 4252 | 3593 | 3553 | 3816 | 3791 | 3960,07 | 4232 | 3688 | 0,872 | -0,198 | -4,274 | 1,92E-05 | 2,82E-05 | 0,0013095 | 0,0025059 | 0,0018031 | 0,0018031 | TRUE | 0,0194975 | No |
| orf19.6748 | 5720 | 5076 | 5336 | 3192 | 10357 | 7233 | 10338 | 10961 | 5453 | 5367 | 5271 | 5000 | 8299 | 8225 | 8777 | 8912 | 6913,14 | 5273 | 8553 | 1,621 | 0,697 | 15,987 | 1,57E-57 | 6,38E-57 | 0,0012906 | 0,0023288 | 0,0016663 | 0,0016663 | TRUE | 0,0141209 | No |
| orf19.6749 | 10253 | 8869 | 9814 | 6448 | 21530 | 15417 | 20150 | 23222 | 9775 | 9378 | 9695 | 10101 | 17251 | 17531 | 17107 | 18882 | 13715,03 | 9737 | 17693 | 1,818 | 0,862 | 20,085 | 9,99E-90 | 5,95E-89 | 0,0014122 | 0,002211 | 0,001686 | 0,001686 | TRUE | 0,0251539 | No |
| orf19.675 | 10778 | 8632 | 9305 | 6927 | 2005 | 1413 | 1925 | 2023 | 10276 | 9127 | 9192 | 10852 | 1607 | 1607 | 1634 | 1645 | 5742,42 | 9862 | 1623 | 0,165 | -2,602 | -42,825 | 0 | 0 | 0,0032586 | 0,0023772 | 0,0032142 | 0,0032142 | TRUE | 0,0550241 | No |
| orf19.6751 | 3364 | 2729 | 2949 | 2222 | 3168 | 2434 | 3121 | 3496 | 3207 | 2886 | 2913 | 3481 | 2538 | 2768 | 2650 | 2843 | 2910,67 | 3122 | 2700 | 0,866 | -0,208 | -3,147 | 0,001651 | 0,0021994 | 0,0047915 | 0,0026554 | 0,00386 | 0,00386 | TRUE | 0,0703628 | No |
| orf19.675.1 | 196 | 152 | 169 | 125 | 249 | 188 | 244 | 242 | 187 | 161 | 167 | 196 | 200 | 214 | 207 | 197 | 190,95 | 178 | 204 | 1,154 | 0,207 | 1,93 | 0,0535945 | 0,0642108 | 1,00E-08 | 0,01069 | 0,0056947 | 0,0056947 | TRUE | 0,0413867 | No |
| orf19.6752 | 2759 | 2394 | 2519 | 1659 | 3239 | 2314 | 3100 | 3238 | 2630 | 2531 | 2489 | 2599 | 2595 | 2631 | 2632 | 2633 | 2592,55 | 2562 | 2623 | 1,024 | 0,034 | 0,91 | 0,3628505 | 0,3976213 | 1,00E-08 | 0,0027246 | 0,0009607 | 0,0009607 | TRUE | 0,0047399 | No |
| orf19.6753 | 1225 | 1021 | 1089 | 632 | 2540 | 1454 | 2434 | 2505 | 1168 | 1080 | 1076 | 990 | 2035 | 1653 | 2066 | 2037 | 1513,15 | 1078 | 1948 | 1,805 | 0,852 | 10,751 | 5,85E-27 | 1,50E-26 | 0,0064182 | 0,0031764 | 0,0052812 | 0,0052812 | TRUE | 0,1223781 | No |
| orf19.6754 | 3860 | 3055 | 3480 | 2610 | 4395 | 2920 | 4283 | 4249 | 3680 | 3230 | 3438 | 4089 | 3522 | 3320 | 3636 | 3455 | 3546,25 | 3609 | 3483 | 0,966 | -0,049 | -0,725 | 0,4686131 | 0,5050194 | 0,0052996 | 0,0025543 | 0,0041867 | 0,0041867 | TRUE | 0,0945426 | No |
| orf19.6755 | 544 | 483 | 520 | 358 | 1025 | 736 | 1004 | 1090 | 519 | 511 | 514 | 561 | 821 | 837 | 852 | 886 | 687,6 | 526 | 849 | 1,621 | 0,697 | 11,167 | 5,89E-29 | 1,56E-28 | 1,00E-08 | 0,0044792 | 0,0021501 | 0,0021501 | TRUE | 0,0189938 | No |
| orf19.6756 | 1931 | 1554 | 1721 | 1020 | 1612 | 971 | 1485 | 1539 | 1841 | 1643 | 1700 | 1598 | 1292 | 1104 | 1261 | 1251 | 1461,26 | 1696 | 1227 | 0,724 | -0,466 | -7,176 | 7,18E-13 | 1,37E-12 | 0,0033296 | 0,0032149 | 0,0033609 | 0,0033609 | TRUE | 0,0518144 | No |
| orf19.6757 | 1175 | 1026 | 1075 | 664 | 3059 | 2098 | 2907 | 3102 | 1120 | 1085 | 1062 | 1040 | 2451 | 2386 | 2468 | 2522 | 1766,78 | 1077 | 2457 | 2,278 | 1,188 | 26,035 | 1,97E-149 | 1,96E-148 | 1,00E-08 | 0,0030206 | 0,0013 | 0,0013 | TRUE | 0,0089952 | No |
| orf19.6758 | 946 | 818 | 902 | 531 | 862 | 545 | 793 | 796 | 902 | 865 | 891 | 832 | 691 | 620 | 673 | 647 | 765,08 | 872 | 658 | 0,754 | -0,408 | -6,739 | 1,59E-11 | 2,91E-11 | 1,00E-08 | 0,0042373 | 0,0022034 | 0,0022034 | TRUE | 0,0160584 | No |
| orf19.676 | 773 | 551 | 682 | 415 | 1252 | 964 | 1222 | 1273 | 737 | 583 | 674 | 650 | 1003 | 1096 | 1037 | 1035 | 851,92 | 661 | 1043 | 1,575 | 0,655 | 8,955 | 3,41E-19 | 7,54E-19 | 0,0036967 | 0,0040186 | 0,0038599 | 0,0038599 | TRUE | 0,0789973 | No |
| orf19.6760 | 1688 | 1539 | 1719 | 1034 | 3140 | 2242 | 2988 | 3218 | 1609 | 1627 | 1698 | 1620 | 2516 | 2549 | 2537 | 2617 | 2096,67 | 1638 | 2555 | 1,558 | 0,64 | 15,623 | 5,06E-55 | 2,00E-54 | 1,00E-08 | 0,0028744 | 0,0010949 | 0,0010949 | TRUE | 0,0076018 | No |
| orf19.6763 | 39002 | 34465 | 37600 | 22679 | 8934 | 5773 | 8412 | 8788 | 37184 | 36442 | 37145 | 35528 | 7159 | 6564 | 7142 | 7145 | 21788,73 | 36575 | 7002 | 0,192 | -2,384 | -62,014 | 0 | 0 | 0,0009826 | 0,0021666 | 0,0013408 | 0,0013408 | TRUE | 0,02115 | No |
| orf19.6766 | 10484 | 8734 | 9241 | 5479 | 10607 | 6886 | 10490 | 10542 | 9995 | 9235 | 9129 | 8583 | 8499 | 7830 | 8906 | 8572 | 8843,69 | 9236 | 8452 | 0,915 | -0,128 | -2,294 | 0,0217722 | 0,0269503 | 0,0032528 | 0,0022769 | 0,0028871 | 0,0028871 | TRUE | 0,0375223 | No |
| orf19.6769 | 8081 | 7497 | 8132 | 4722 | 9352 | 6300 | 9131 | 9535 | 7704 | 7927 | 8034 | 7397 | 7493 | 7164 | 7752 | 7753 | 7653,07 | 7766 | 7540 | 0,971 | -0,043 | -1,017 | 0,3093553 | 0,3426482 | 0,001168 | 0,0023058 | 0,0015616 | 0,0015616 | TRUE | 0,0135555 | No |
| orf19.677 | 2922 | 2524 | 2857 | 1722 | 4847 | 3518 | 4827 | 4940 | 2786 | 2669 | 2822 | 2698 | 3884 | 4000 | 4098 | 4017 | 3371,69 | 2744 | 4000 | 1,457 | 0,543 | 14,208 | 8,13E-46 | 2,83E-45 | 0,0003108 | 0,0025782 | 0,0010844 | 0,0010844 | TRUE | 0,0047706 | No |
| orf19.6770 | 8877 | 7978 | 8670 | 6032 | 4060 | 3161 | 4009 | 4184 | 8463 | 8436 | 8565 | 9449 | 3253 | 3594 | 3404 | 3402 | 6070,84 | 8728 | 3413 | 0,391 | -1,354 | -27,1 | 9,82E-162 | 1,08E-160 | 0,0021041 | 0,0023617 | 0,0022036 | 0,0022036 | TRUE | 0,0370051 | No |
| orf19.6771 | 100753 | 80824 | 85226 | 54254 | 94335 | 57729 | 88072 | 94554 | 96057 | 85461 | 84195 | 84992 | 75588 | 65643 | 74773 | 76881 | 80448,8 | 87676 | 73221 | 0,835 | -0,26 | -4,263 | 2,02E-05 | 2,95E-05 | 0,0044822 | 0,0021117 | 0,0035596 | 0,0035596 | TRUE | 0,0594294 | No |
| orf19.6773 | 2549 | 2362 | 2587 | 1526 | 4353 | 3039 | 3841 | 4385 | 2430 | 2498 | 2556 | 2391 | 3488 | 3456 | 3261 | 3565 | 2955,49 | 2469 | 3442 | 1,393 | 0,479 | 10,87 | 1,61E-27 | 4,17E-27 | 0,0008695 | 0,0026468 | 0,0015048 | 0,0015048 | TRUE | 0,015574 | No |
| orf19.6776 | 7797 | 6954 | 7453 | 4957 | 6609 | 4993 | 6459 | 6985 | 7434 | 7353 | 7363 | 7765 | 5296 | 5677 | 5484 | 5679 | 6506,38 | 7479 | 5534 | 0,74 | -0,434 | -11,413 | 3,59E-30 | 9,72E-30 | 0,0006788 | 0,0023436 | 0,0012332 | 0,0012332 | TRUE | 0,0103688 | No |
| orf19.6777 | 85 | 73 | 83 | 64 | 163 | 88 | 118 | 151 | 81 | 77 | 82 | 100 | 131 | 100 | 100 | 123 | 99,26 | 85 | 114 | 1,351 | 0,434 | 2,755 | 0,0058765 | 0,0075617 | 0,0066924 | 0,0186321 | 0,013322 | 0,013322 | TRUE | 0,1084511 | No |
| orf19.6778 | 1181 | 1149 | 1223 | 804 | 2690 | 1955 | 2407 | 2520 | 1126 | 1215 | 1208 | 1260 | 2155 | 2223 | 2044 | 2049 | 1659,94 | 1202 | 2118 | 1,764 | 0,819 | 15,518 | 2,61E-54 | 1,02E-53 | 0,0010475 | 0,0030804 | 0,0019949 | 0,0019949 | TRUE | 0,0223959 | No |
| orf19.6779 | 1438 | 1514 | 1517 | 915 | 2237 | 1659 | 2091 | 2157 | 1371 | 1601 | 1499 | 1433 | 1792 | 1886 | 1775 | 1754 | 1638,98 | 1476 | 1802 | 1,22 | 0,286 | 5,051 | 4,39E-07 | 6,89E-07 | 0,0019194 | 0,0030931 | 0,0024584 | 0,0024584 | TRUE | 0,0397073 | No |
| orf19.6780 | 417 | 391 | 401 | 251 | 953 | 585 | 739 | 763 | 398 | 413 | 396 | 393 | 764 | 665 | 627 | 620 | 534,62 | 400 | 669 | 1,672 | 0,742 | 8,841 | 9,45E-19 | 2,06E-18 | 0,0043177 | 0,0051624 | 0,0046926 | 0,0046926 | TRUE | 0,1111764 | No |
| orf19.6781 | 3003 | 3587 | 3714 | 2160 | 1329 | 1055 | 1297 | 1105 | 2863 | 3793 | 3669 | 3384 | 1065 | 1200 | 1101 | 898 | 2246,6 | 3427 | 1066 | 0,311 | -1,686 | -17,187 | 3,34E-66 | 1,52E-65 | 0,0133664 | 0,0028221 | 0,0086609 | 0,0086609 | TRUE | 0,1509322 | No |
| orf19.6782 | 170 | 162 | 158 | 111 | 891 | 700 | 861 | 835 | 162 | 171 | 156 | 174 | 714 | 796 | 731 | 679 | 447,9 | 166 | 730 | 4,41 | 2,141 | 23,693 | 4,29E-124 | 3,55E-123 | 1,00E-08 | 0,0057571 | 0,0038686 | 0,0038686 | TRUE | 0,0412725 | No |
| orf19.6783 | 3530 | 3429 | 3751 | 2228 | 3725 | 2831 | 3554 | 3531 | 3365 | 3626 | 3706 | 3490 | 2985 | 3219 | 3017 | 2871 | 3284,92 | 3547 | 3023 | 0,852 | -0,232 | -4,677 | 2,91E-06 | 4,43E-06 | 0,001636 | 0,0025911 | 0,0020518 | 0,0020518 | TRUE | 0,0224387 | No |
| orf19.6784 | 213 | 184 | 181 | 129 | 292 | 220 | 268 | 264 | 203 | 195 | 179 | 202 | 234 | 250 | 228 | 215 | 213,11 | 195 | 232 | 1,189 | 0,25 | 2,448 | 0,0143645 | 0,0180361 | 1,00E-08 | 0,0097959 | 0,0051974 | 0,0051974 | TRUE | 0,0344914 | No |
| orf19.6785 | 9560 | 8163 | 8326 | 4930 | 26740 | 17784 | 23964 | 27282 | 9114 | 8631 | 8225 | 7723 | 21426 | 20222 | 20345 | 22183 | 14733,8 | 8423 | 21044 | 2,497 | 1,32 | 23,771 | 6,71E-125 | 5,61E-124 | 0,0033375 | 0,0022027 | 0,0028755 | 0,0028755 | TRUE | 0,0375838 | No |
| orf19.6786 | 419 | 361 | 392 | 254 | 409 | 269 | 445 | 437 | 399 | 382 | 387 | 398 | 328 | 306 | 378 | 355 | 366,63 | 392 | 342 | 0,876 | -0,192 | -2,251 | 0,0243797 | 0,0300729 | 0,0016317 | 0,0065696 | 0,0042366 | 0,0042366 | TRUE | 0,0567268 | No |
| orf19.6787 | 1470 | 1724 | 1634 | 936 | 2983 | 2142 | 2596 | 2171 | 1401 | 1823 | 1614 | 1466 | 2390 | 2436 | 2204 | 1765 | 1887,5 | 1576 | 2199 | 1,394 | 0,479 | 4,574 | 4,79E-06 | 7,22E-06 | 0,0159901 | 0,0029612 | 0,0099811 | 0,0099811 | TRUE | 0,2147101 | No |
| orf19.6788 | 906 | 942 | 939 | 563 | 1083 | 687 | 954 | 956 | 864 | 996 | 928 | 882 | 868 | 781 | 810 | 777 | 863,21 | 918 | 809 | 0,882 | -0,181 | -2,733 | 0,0062695 | 0,0080575 | 0,002275 | 0,0039934 | 0,0030542 | 0,0030542 | TRUE | 0,0401246 | No |
| orf19.6789 | 518 | 512 | 519 | 265 | 505 | 346 | 469 | 456 | 494 | 541 | 513 | 415 | 405 | 393 | 398 | 371 | 441,26 | 491 | 392 | 0,794 | -0,333 | -3,857 | 0,0001146 | 0,0001618 | 0,0037167 | 0,0058122 | 0,004878 | 0,004878 | TRUE | 0,1069984 | No |
| orf19.679 | 759 | 622 | 657 | 450 | 837 | 575 | 700 | 804 | 724 | 658 | 649 | 705 | 671 | 654 | 594 | 654 | 663,48 | 684 | 643 | 0,941 | -0,087 | -1,285 | 0,1987496 | 0,2254616 | 0,0014685 | 0,004566 | 0,0029319 | 0,0029319 | TRUE | 0,0317386 | No |
| orf19.6790 | 554 | 454 | 506 | 315 | 948 | 630 | 867 | 931 | 528 | 480 | 500 | 493 | 760 | 716 | 736 | 757 | 621,33 | 500 | 742 | 1,483 | 0,568 | 8,938 | 3,97E-19 | 8,75E-19 | 1,00E-08 | 0,0047339 | 0,0021721 | 0,0021721 | TRUE | 0,017662 | No |
| orf19.6791 | 17656 | 16147 | 17569 | 8313 | 11735 | 6622 | 10849 | 10970 | 16833 | 17073 | 17357 | 13023 | 9403 | 7530 | 9211 | 8920 | 12418,61 | 16072 | 8766 | 0,545 | -0,875 | -9,449 | 3,43E-21 | 7,87E-21 | 0,0126424 | 0,0022235 | 0,0081464 | 0,0081464 | TRUE | 0,1987046 | No |
| orf19.6792 | 872 | 812 | 876 | 599 | 841 | 629 | 782 | 821 | 831 | 859 | 865 | 938 | 674 | 715 | 664 | 668 | 776,78 | 873 | 680 | 0,78 | -0,358 | -5,902 | 3,59E-09 | 6,05E-09 | 0,0002364 | 0,004205 | 0,002241 | 0,002241 | TRUE | 0,0259275 | No |
| orf19.6793 | 3638 | 3431 | 3608 | 2057 | 3969 | 2547 | 3657 | 3723 | 3468 | 3628 | 3564 | 3222 | 3180 | 2896 | 3105 | 3027 | 3261,42 | 3470 | 3052 | 0,879 | -0,186 | -3,756 | 0,0001724 | 0,000241 | 0,0016952 | 0,0025947 | 0,0020522 | 0,0020522 | TRUE | 0,0266191 | No |
| orf19.6794 | 5076 | 4893 | 4914 | 3020 | 7908 | 6148 | 7478 | 7341 | 4839 | 5174 | 4855 | 4731 | 6336 | 6991 | 6349 | 5969 | 5655,46 | 4900 | 6411 | 1,308 | 0,387 | 7,187 | 6,61E-13 | 1,26E-12 | 0,0027121 | 0,0023816 | 0,0025988 | 0,0025988 | TRUE | 0,0446397 | No |
| orf19.6795 | 457 | 540 | 503 | 293 | 846 | 539 | 709 | 684 | 436 | 571 | 497 | 459 | 678 | 613 | 602 | 556 | 551,43 | 491 | 612 | 1,247 | 0,318 | 3,337 | 0,0008459 | 0,0011474 | 0,0081928 | 0,0050688 | 0,0068399 | 0,0068399 | TRUE | 0,1429647 | No |
| orf19.6796 | 437 | 378 | 423 | 280 | 761 | 548 | 738 | 786 | 417 | 400 | 418 | 439 | 610 | 623 | 627 | 639 | 521,42 | 418 | 625 | 1,498 | 0,583 | 8,737 | 2,39E-18 | 5,18E-18 | 1,00E-08 | 0,0052402 | 0,0022225 | 0,0022225 | TRUE | 0,0098485 | No |
| orf19.6797 | 1328 | 1251 | 1341 | 873 | 1668 | 1290 | 1488 | 1533 | 1266 | 1323 | 1325 | 1368 | 1337 | 1467 | 1263 | 1246 | 1324,3 | 1320 | 1328 | 1,005 | 0,008 | 0,128 | 0,8979741 | 0,9122708 | 0,0024949 | 0,0033311 | 0,0028567 | 0,0028567 | TRUE | 0,0564165 | No |
| orf19.6798 | 27587 | 23611 | 26920 | 17883 | 9844 | 7003 | 9384 | 9937 | 26301 | 24966 | 26594 | 28015 | 7888 | 7963 | 7967 | 8080 | 17221,68 | 26469 | 7974 | 0,301 | -1,731 | -43,224 | 0 | 0 | 0,0010569 | 0,0021866 | 0,0014624 | 0,0014624 | TRUE | 0,0187113 | No |
| orf19.680 | 3636 | 3160 | 3491 | 2119 | 7486 | 4869 | 7002 | 7546 | 3467 | 3341 | 3449 | 3320 | 5998 | 5536 | 5945 | 6136 | 4648,9 | 3394 | 5904 | 1,739 | 0,799 | 18,657 | 1,11E-77 | 5,89E-77 | 0,0008834 | 0,0024445 | 0,0015177 | 0,0015177 | TRUE | 0,020879 | No |
| orf19.6800 | 1513 | 1318 | 1488 | 947 | 1866 | 1304 | 1792 | 1903 | 1442 | 1394 | 1470 | 1484 | 1495 | 1483 | 1521 | 1547 | 1479,54 | 1448 | 1512 | 1,046 | 0,065 | 1,434 | 0,1516169 | 0,1739737 | 1,00E-08 | 0,003201 | 0,0012702 | 0,0012702 | TRUE | 0,007464 | No |
| orf19.6801 | 2730 | 2534 | 2683 | 1673 | 3524 | 2481 | 3541 | 3624 | 2603 | 2679 | 2651 | 2621 | 2824 | 2821 | 3006 | 2947 | 2768,91 | 2638 | 2900 | 1,099 | 0,137 | 3,454 | 0,0005529 | 0,0007575 | 0,0003091 | 0,0026843 | 0,0011336 | 0,0011336 | TRUE | 0,0074604 | No |
| orf19.6802 | 298 | 350 | 344 | 163 | 377 | 251 | 348 | 360 | 284 | 370 | 340 | 255 | 302 | 285 | 295 | 293 | 303,13 | 312 | 294 | 0,937 | -0,094 | -0,852 | 0,394036 | 0,4299493 | 0,0097707 | 0,0075078 | 0,0084935 | 0,0084935 | TRUE | 0,1684227 | No |
| orf19.6803 | 598 | 559 | 570 | 326 | 660 | 504 | 638 | 663 | 570 | 591 | 563 | 511 | 529 | 573 | 542 | 539 | 552,21 | 559 | 546 | 0,971 | -0,042 | -0,616 | 0,5380816 | 0,5738758 | 6,76E-05 | 0,0050646 | 0,0026918 | 0,0026918 | TRUE | 0,0324086 | No |
| orf19.6804 | 1115 | 1430 | 1334 | 845 | 1516 | 1219 | 1259 | 1060 | 1063 | 1512 | 1318 | 1324 | 1215 | 1386 | 1069 | 862 | 1218,54 | 1304 | 1133 | 0,868 | -0,204 | -1,172 | 0,2410637 | 0,2708147 | 0,0281541 | 0,0034387 | 0,0158515 | 0,0281541 | TRUE | 0,3126601 | No |
| orf19.6805 | 912 | 1027 | 1128 | 666 | 715 | 569 | 581 | 606 | 869 | 1086 | 1114 | 1043 | 573 | 647 | 493 | 493 | 789,88 | 1028 | 552 | 0,535 | -0,901 | -8,758 | 1,99E-18 | 4,32E-18 | 0,0128448 | 0,00417 | 0,0088122 | 0,0088122 | TRUE | 0,1525869 | No |
| orf19.6807 | 52 | 43 | 45 | 25 | 80 | 42 | 37 | 56 | 50 | 45 | 44 | 39 | 64 | 48 | 31 | 46 | 45,93 | 44 | 47 | 1,052 | 0,073 | 0,3 | 0,7642986 | 0,7911318 | 0,0302712 | 0,0378364 | 0,0343283 | 0,0343283 | TRUE | 0,3062889 | No |
| orf19.6809 | 774 | 718 | 750 | 411 | 1386 | 950 | 1208 | 1332 | 738 | 759 | 741 | 644 | 1111 | 1080 | 1026 | 1083 | 897,67 | 720 | 1075 | 1,485 | 0,571 | 9,152 | 5,61E-20 | 1,26E-19 | 0,0011009 | 0,0039204 | 0,0025457 | 0,0025457 | TRUE | 0,0512404 | No |
| orf19.681 | 3496 | 2988 | 3377 | 2115 | 3298 | 2411 | 3213 | 3377 | 3333 | 3159 | 3336 | 3313 | 2643 | 2742 | 2728 | 2746 | 2999,96 | 3285 | 2715 | 0,826 | -0,276 | -7,273 | 3,51E-13 | 6,75E-13 | 0,0001978 | 0,0026386 | 0,0010439 | 0,0010439 | TRUE | 0,0082988 | No |
| orf19.6810 | 1508 | 1522 | 1573 | 1061 | 2830 | 2259 | 2572 | 2500 | 1438 | 1609 | 1554 | 1662 | 2268 | 2569 | 2184 | 2033 | 1914,47 | 1566 | 2264 | 1,446 | 0,532 | 7,111 | 1,15E-12 | 2,18E-12 | 0,0061355 | 0,0029489 | 0,0048134 | 0,0048134 | TRUE | 0,0981765 | No |
| orf19.6811 | 5432 | 4757 | 5145 | 2627 | 3067 | 1868 | 2813 | 2810 | 5179 | 5030 | 5083 | 4115 | 2457 | 2124 | 2388 | 2285 | 3582,68 | 4852 | 2314 | 0,477 | -1,069 | -14,206 | 8,39E-46 | 2,91E-45 | 0,0067365 | 0,0025496 | 0,0051276 | 0,0051276 | TRUE | 0,1247209 | No |
| orf19.6812 | 1505 | 1239 | 1336 | 773 | 2958 | 2154 | 2838 | 3036 | 1435 | 1310 | 1320 | 1211 | 2370 | 2449 | 2409 | 2469 | 1871,65 | 1319 | 2424 | 1,832 | 0,874 | 16,399 | 1,96E-60 | 8,27E-60 | 0,0013907 | 0,0029685 | 0,0021165 | 0,0021165 | TRUE | 0,0421291 | No |
| orf19.6813 | 6 | 4 | 9 | 5 | 3 | 3 | 4 | 3 | 6 | 4 | 9 | 8 | 2 | 3 | 3 | 2 | 4,79 | 7 | 2 | 0,435 | -1,2 | -1,834 | 0,066583 | 0,0791922 | 1,00E-08 | 0,3448319 | 0,1724507 | 0,1724507 | TRUE | 0,1529231 | No |
| orf19.6814 | 105363 | 81646 | 89182 | 67659 | 25071 | 18395 | 24052 | 26323 | 100452 | 86330 | 88103 | 105991 | 20089 | 20917 | 20420 | 21403 | 57963,24 | 95219 | 20707 | 0,217 | -2,201 | -33,844 | 4,39E-251 | 7,53E-250 | 0,0053666 | 0,0021196 | 0,0040365 | 0,0040365 | TRUE | 0,0709933 | No |
| orf19.6816 | 762 | 659 | 640 | 467 | 1524 | 1081 | 1313 | 1469 | 726 | 697 | 632 | 732 | 1221 | 1229 | 1115 | 1194 | 943,33 | 697 | 1190 | 1,71 | 0,774 | 12,044 | 2,08E-33 | 5,92E-33 | 0,0018574 | 0,0038318 | 0,0027872 | 0,0027872 | TRUE | 0,0476899 | No |
| orf19.6817 | 2736 | 2468 | 2695 | 1732 | 991 | 728 | 1006 | 1041 | 2608 | 2610 | 2662 | 2713 | 794 | 828 | 854 | 846 | 1739,52 | 2648 | 830 | 0,314 | -1,672 | -36,389 | 6,39E-290 | 1,37E-288 | 1,00E-08 | 0,0030352 | 0,0012791 | 0,0012791 | TRUE | 0,0113241 | No |
| orf19.6818 | 2157 | 1976 | 2162 | 1292 | 5425 | 3746 | 4964 | 5505 | 2056 | 2089 | 2136 | 2024 | 4347 | 4260 | 4214 | 4476 | 3200,33 | 2076 | 4324 | 2,081 | 1,057 | 26,759 | 9,75E-158 | 1,05E-156 | 0,0003643 | 0,0026043 | 0,0011261 | 0,0011261 | TRUE | 0,0068866 | No |
| orf19.68.2 | 389 | 457 | 429 | 309 | 552 | 434 | 511 | 435 | 371 | 483 | 424 | 484 | 442 | 493 | 434 | 354 | 435,66 | 440 | 431 | 0,979 | -0,031 | -0,268 | 0,7890582 | 0,8123556 | 0,0149422 | 0,00586 | 0,0104011 | 0,0104011 | TRUE | 0,1711688 | No |

| | | | | | | | | | | | | | | | | | | | | | | | | | | | | | | | |
|---|---|---|---|---|---|---|---|---|---|---|---|---|---|---|---|---|---|---|---|---|---|---|---|---|---|---|---|---|---|---|---|
| orf19.6820 | 4509 | 3916 | 4405 | 2884 | 2047 | 1471 | 2004 | 2034 | 4299 | 4141 | 4352 | 4518 | 1640 | 1673 | 1701 | 1654 | 2997,16 | 4328 | 1667 | 0,385 | -1,375 | -33,079 | 5,91E-240 | 9,60E-239 | 0,0005058 | 0,0026391 | 0,0012617 | 0,0012617 | TRUE | 0,0100907 | No |
| orf19.6821 | 208 | 177 | 188 | 121 | 363 | 233 | 298 | 317 | 198 | 187 | 186 | 190 | 291 | 265 | 253 | 258 | 228,41 | 190 | 267 | 1,402 | 0,488 | 4,939 | 7,84E-07 | 1,22E-06 | 1,00E-08 | 0,0092796 | 0,0047599 | 0,0047599 | TRUE | 0,0452028 | No |
| orf19.6822 | 1869 | 1680 | 1850 | 966 | 1770 | 1134 | 1522 | 1543 | 1782 | 1776 | 1828 | 1513 | 1418 | 1289 | 1292 | 1255 | 1519,21 | 1725 | 1314 | 0,76 | -0,396 | -5,84 | 5,23E-09 | 8,77E-09 | 0,0040308 | 0,003172 | 0,0037409 | 0,0037409 | TRUE | 0,076965 | No |
| orf19.6824 | 1104 | 1085 | 1100 | 675 | 447 | 346 | 449 | 497 | 1053 | 1147 | 1087 | 1057 | 358 | 393 | 381 | 404 | 735,1 | 1086 | 384 | 0,353 | -1,501 | -22,901 | 4,53E-116 | 3,47E-115 | 1,00E-08 | 0,0043248 | 0,0024514 | 0,0024514 | TRUE | 0,0256657 | No |
| orf19.6826 | 53 | 35 | 37 | 24 | 39 | 27 | 29 | 21 | 51 | 37 | 37 | 38 | 31 | 31 | 25 | 17 | 33,17 | 41 | 26 | 0,634 | -0,658 | -2,46 | 0,0139027 | 0,0174976 | 0,0181433 | 0,0515957 | 0,0377661 | 0,0377661 | TRUE | 0,3776659 | No |
| orf19.6827 | 1158 | 1051 | 1137 | 688 | 1114 | 703 | 1052 | 1001 | 1104 | 1111 | 1123 | 1078 | 893 | 799 | 893 | 814 | 976,93 | 1104 | 850 | 0,77 | -0,377 | -6,54 | 6,15E-11 | 1,11E-10 | 0,0007652 | 0,003772 | 0,002155 | 0,002155 | TRUE | 0,0172571 | No |
| orf19.6828 | 8307 | 7338 | 7699 | 4094 | 9154 | 5681 | 8530 | 8943 | 7920 | 7759 | 7606 | 6413 | 7335 | 6460 | 7242 | 7272 | 7250,78 | 7424 | 7077 | 0,953 | -0,07 | -1,007 | 0,3139447 | 0,3470688 | 0,0057457 | 0,0023177 | 0,004459 | 0,004459 | TRUE | 0,1011203 | No |
| orf19.6828.1 | 679 | 588 | 564 | 276 | 1682 | 968 | 1461 | 1599 | 647 | 622 | 557 | 432 | 1348 | 1101 | 1240 | 1300 | 905,95 | 564 | 1247 | 2,2 | 1,137 | 10,583 | 3,58E-26 | 9,09E-26 | 0,0164273 | 0,0039036 | 0,0097423 | 0,0097423 | TRUE | 0,2653024 | No |
| orf19.6829 | 3725 | 3527 | 3694 | 2282 | 2959 | 2053 | 2712 | 2878 | 3551 | 3729 | 3649 | 3575 | 2371 | 2334 | 2302 | 2340 | 2981,61 | 3626 | 2337 | 0,644 | -0,634 | -17,406 | 7,39E-68 | 3,45E-67 | 1,00E-08 | 0,002642 | 0,0009283 | 0,0009283 | TRUE | 0,0044915 | No |
| orf19.6830 | 456 | 436 | 438 | 268 | 1525 | 1044 | 1266 | 1434 | 435 | 461 | 433 | 420 | 1222 | 1187 | 1075 | 1166 | 799,77 | 437 | 1162 | 2,655 | 1,408 | 20,998 | 6,79E-98 | 4,35E-97 | 0,0014929 | 0,0041442 | 0,0026618 | 0,0026618 | TRUE | 0,0314694 | No |
| orf19.6831 | 11484 | 9887 | 10793 | 6193 | 11483 | 7604 | 11240 | 11722 | 10949 | 10454 | 10662 | 9702 | 9201 | 8646 | 9543 | 9531 | 9836,05 | 10442 | 9230 | 0,884 | -0,178 | -3,6 | 0,0003182 | 0,0004401 | 0,0022053 | 0,0022582 | 0,0022516 | 0,0022516 | TRUE | 0,0273286 | No |
| orf19.6832 | 1084 | 932 | 1014 | 576 | 2097 | 1470 | 1998 | 2132 | 1033 | 985 | 1002 | 902 | 1680 | 1672 | 1696 | 1734 | 1338,08 | 980 | 1696 | 1,722 | 0,784 | 15,213 | 2,92E-52 | 1,11E-51 | 9,20E-05 | 0,0033183 | 0,0017219 | 0,0017219 | TRUE | 0,0294376 | No |
| orf19.6833.2 | 54 | 36 | 42 | 60 | 46 | 48 | 54 | 50 | 51 | 38 | 41 | 94 | 37 | 55 | 46 | 41 | 50,37 | 56 | 45 | 0,803 | -0,317 | -1,132 | 0,2574447 | 0,2877482 | 0,0791269 | 0,0346871 | 0,0547832 | 0,0547832 | TRUE | 1,3743284 | No |
| orf19.6834.10 | 27 | 21 | 24 | 6 | 34 | 34 | 11 | 16 | 26 | 22 | 24 | 9 | 27 | 39 | 9 | 13 | 21,16 | 20 | 22 | 1,056 | 0,079 | 0,175 | 0,8610355 | 0,8792235 | 0,2258471 | 0,079673 | 0,1462717 | 0,1462717 | TRUE | 0,3454402 | No |
| orf19.6837 | 279 | 213 | 250 | 183 | 703 | 491 | 660 | 728 | 266 | 225 | 247 | 287 | 563 | 558 | 560 | 592 | 412,34 | 256 | 568 | 2,233 | 1,159 | 13,917 | 5,02E-44 | 1,69E-43 | 1,00E-08 | 0,0060732 | 0,0036909 | 0,0036909 | TRUE | 0,0706353 | No |
| orf19.6838 | 1739 | 1501 | 1552 | 1057 | 7112 | 5112 | 6443 | 7482 | 1658 | 1587 | 1533 | 1656 | 5699 | 5813 | 5470 | 6084 | 3687,41 | 1608 | 5766 | 3,588 | 1,843 | 37,904 | 0 | 0 | 0,0014373 | 0,0025366 | 0,0018434 | 0,0018434 | TRUE | 0,0169849 | No |
| orf19.684 | 5090 | 4576 | 5043 | 3183 | 3001 | 2185 | 2881 | 3120 | 4853 | 4839 | 4982 | 4986 | 2405 | 2485 | 2446 | 2537 | 3691,45 | 4915 | 2468 | 0,502 | -0,994 | -27,842 | 1,36E-170 | 1,55E-169 | 1,00E-08 | 0,0025361 | 0,0009275 | 0,0009275 | TRUE | 0,0045342 | No |
| orf19.6840 | 996 | 987 | 1005 | 788 | 1180 | 920 | 1156 | 1137 | 950 | 1044 | 993 | 1234 | 946 | 1046 | 981 | 924 | 1014,76 | 1055 | 974 | 0,926 | -0,111 | -1,368 | 0,1712124 | 0,1954045 | 0,0072617 | 0,0037093 | 0,005329 | 0,005329 | TRUE | 0,1466952 | No |
| orf19.6842 | 11970 | 10749 | 12105 | 7345 | 1307 | 973 | 1306 | 1503 | 11412 | 11366 | 11959 | 11506 | 1047 | 1106 | 1109 | 1222 | 6340,91 | 11561 | 1121 | 0,097 | -3,366 | -68,368 | 0 | 0 | 0,0017477 | 0,0023502 | 0,0018811 | 0,0018811 | TRUE | 0,0459826 | No |
| orf19.6843 | 7817 | 6687 | 7302 | 3982 | 3923 | 2401 | 3754 | 3812 | 7453 | 7071 | 7214 | 6238 | 3143 | 2730 | 3187 | 3100 | 5016,9 | 6994 | 3040 | 0,435 | -1,202 | -18,158 | 1,12E-73 | 5,62E-73 | 0,0049124 | 0,0024186 | 0,003981 | 0,003981 | TRUE | 0,0635316 | No |
| orf19.6844 | 30395 | 23819 | 25861 | 18126 | 79640 | 54492 | 76936 | 84029 | 28978 | 25186 | 25548 | 28395 | 63813 | 61962 | 65318 | 68324 | 45940,62 | 27027 | 64854 | 2,4 | 1,263 | 22,852 | 1,41E-115 | 1,08E-114 | 0,0033935 | 0,002127 | 0,002907 | 0,002907 | TRUE | 0,0291118 | No |
| orf19.6845 | 4082 | 3767 | 4111 | 2555 | 3850 | 2834 | 3676 | 3895 | 3892 | 3983 | 4061 | 4003 | 3085 | 3223 | 3121 | 3167 | 3566,75 | 3985 | 3149 | 0,79 | -0,34 | -9,727 | 2,31E-22 | 5,44E-22 | 1,00E-08 | 0,0025516 | 0,0008908 | 0,0008908 | TRUE | 0,0031565 | No |
| orf19.6846 | 3512 | 3252 | 3335 | 2270 | 4346 | 3019 | 4165 | 4353 | 3348 | 3439 | 3295 | 3556 | 3482 | 3433 | 3536 | 3539 | 3453,54 | 3410 | 3498 | 1,027 | 0,039 | 1,033 | 0,3016596 | 0,334647 | 0,000281 | 0,0025667 | 0,0010857 | 0,0010857 | TRUE | 0,0093831 | No |
| orf19.6847 | 1906 | 1678 | 1891 | 1158 | 2720 | 1983 | 2775 | 2904 | 1817 | 1774 | 1868 | 1814 | 2179 | 2255 | 2356 | 2361 | 2053,14 | 1818 | 2288 | 1,258 | 0,331 | 7,303 | 2,82E-13 | 5,43E-13 | 0,0006429 | 0,002891 | 0,0014721 | 0,0014721 | TRUE | 0,0129266 | No |
| orf19.6848 | 848 | 769 | 864 | 538 | 1017 | 761 | 950 | 1025 | 808 | 813 | 854 | 843 | 815 | 865 | 807 | 833 | 829,77 | 830 | 830 | 1 | 0 | 0,004 | 0,9968541 | 0,997323 | 1,00E-08 | 0,00407 | 0,0017684 | 0,0017684 | TRUE | 0,0087934 | No |
| orf19.6849 | 850 | 742 | 818 | 542 | 1195 | 785 | 1068 | 1128 | 810 | 785 | 808 | 849 | 958 | 893 | 907 | 917 | 865,77 | 813 | 919 | 1,133 | 0,18 | 3,251 | 0,0011495 | 0,0015429 | 1,00E-08 | 0,0039877 | 0,0017778 | 0,0017778 | TRUE | 0,0099562 | No |
| orf19.685 | 2513 | 2060 | 2181 | 1597 | 1909 | 1396 | 1918 | 2053 | 2396 | 2178 | 2155 | 2502 | 1530 | 1587 | 1628 | 1669 | 1955,64 | 2308 | 1604 | 0,696 | -0,523 | -8,793 | 1,46E-18 | 3,18E-18 | 0,0026671 | 0,0029309 | 0,0028737 | 0,0028737 | TRUE | 0,0366081 | No |
| orf19.6850 | 940 | 871 | 912 | 540 | 947 | 680 | 877 | 930 | 896 | 921 | 901 | 846 | 759 | 773 | 745 | 756 | 824,61 | 891 | 758 | 0,848 | -0,237 | -4,312 | 1,62E-05 | 2,38E-05 | 1,00E-08 | 0,0040824 | 0,0016891 | 0,0016891 | TRUE | 0,0115217 | No |
| orf19.6851 | 154 | 150 | 183 | 95 | 169 | 146 | 176 | 199 | 147 | 159 | 181 | 149 | 135 | 166 | 149 | 162 | 155,96 | 159 | 153 | 0,959 | -0,06 | -0,488 | 0,6254398 | 0,6584608 | 0,0030533 | 0,0126189 | 0,0082967 | 0,0082967 | TRUE | 0,0966732 | No |
| orf19.685.1 | 1292 | 1309 | 1284 | 916 | 827 | 652 | 740 | 745 | 1232 | 1384 | 1268 | 1435 | 663 | 741 | 628 | 606 | 994,67 | 1330 | 660 | 0,496 | -1,013 | -13,297 | 2,40E-40 | 7,64E-40 | 0,0051185 | 0,003742 | 0,0044663 | 0,0044663 | TRUE | 0,0775085 | No |
| orf19.6852 | 515 | 443 | 433 | 306 | 828 | 575 | 802 | 836 | 491 | 468 | 428 | 479 | 663 | 654 | 681 | 680 | 568,06 | 466 | 670 | 1,437 | 0,524 | 7,747 | 9,42E-15 | 1,89E-14 | 1,00E-08 | 0,0049817 | 0,0025234 | 0,0025234 | TRUE | 0,0392221 | No |
| orf19.6852.1 | 2360 | 2037 | 1879 | 1197 | 1746 | 1186 | 1309 | 1461 | 2250 | 2154 | 1856 | 1875 | 1399 | 1349 | 1111 | 1188 | 1647,77 | 2034 | 1262 | 0,62 | -0,691 | -7,903 | 2,73E-15 | 5,54E-15 | 0,0095934 | 0,0030877 | 0,0067008 | 0,0067008 | TRUE | 0,0777989 | No |
| orf19.6853 | 2720 | 2305 | 2627 | 1625 | 3914 | 2784 | 3745 | 4022 | 2593 | 2437 | 2595 | 2546 | 3136 | 3166 | 3179 | 3270 | 2865,37 | 2543 | 3188 | 1,254 | 0,326 | 8,342 | 7,32E-17 | 1,53E-16 | 0,0002359 | 0,0026643 | 0,001106 | 0,001106 | TRUE | 0,0098585 | No |
| orf19.6854 | 13796 | 11253 | 11982 | 8155 | 34079 | 23697 | 30727 | 35990 | 13153 | 11899 | 11837 | 12775 | 27307 | 26946 | 26087 | 29263 | 19908,31 | 12416 | 27401 | 2,207 | 1,142 | 22,648 | 1,47E-113 | 1,10E-112 | 0,0025016 | 0,0021738 | 0,0023817 | 0,0023817 | TRUE | 0,0256888 | No |
| orf19.6855 | 2219 | 2111 | 2232 | 1462 | 4518 | 3221 | 4338 | 4596 | 2116 | 2232 | 2205 | 2290 | 3620 | 3663 | 3683 | 3737 | 2943,2 | 2211 | 3676 | 1,665 | 0,736 | 18,927 | 6,77E-80 | 3,69E-79 | 0,0001456 | 0,0026491 | 0,0010752 | 0,0010752 | TRUE | 0,0103412 | No |
| orf19.6858 | 1679 | 1551 | 1529 | 944 | 1850 | 1254 | 1749 | 1851 | 1601 | 1640 | 1511 | 1479 | 1482 | 1426 | 1485 | 1505 | 1516,03 | 1558 | 1474 | 0,946 | -0,08 | -1,613 | 0,1068135 | 0,124262 | 0,0005582 | 0,0031743 | 0,0017139 | 0,0017139 | TRUE | 0,0148042 | No |
| orf19.6859 | 57 | 54 | 47 | 47 | 30 | 27 | 33 | 46 | 54 | 57 | 46 | 74 | 24 | 31 | 28 | 37 | 43,96 | 58 | 30 | 0,525 | -0,928 | -3,954 | 7,67E-05 | 0,0001092 | 0,0122691 | 0,0394434 | 0,0283192 | 0,0283192 | TRUE | 0,2168264 | No |
| orf19.686 | 1664 | 1476 | 1645 | 977 | 2146 | 1537 | 2162 | 2404 | 1586 | 1561 | 1625 | 1531 | 1720 | 1748 | 1836 | 1955 | 1695,02 | 1576 | 1815 | 1,151 | 0,203 | 3,78 | 0,0001566 | 0,0002194 | 0,0015807 | 0,0030599 | 0,0021646 | 0,0021646 | TRUE | 0,0333236 | No |
| orf19.6860 | 1165 | 1212 | 1192 | 736 | 1530 | 1283 | 1408 | 1512 | 1111 | 1282 | 1178 | 1153 | 1226 | 1459 | 1195 | 1229 | 1229,05 | 1181 | 1277 | 1,08 | 0,111 | 1,492 | 0,135797 | 0,1564399 | 0,0053649 | 0,0034272 | 0,0044856 | 0,0044856 | TRUE | 0,1061604 | No |
| orf19.6861 | 1176 | 1418 | 1325 | 793 | 507 | 376 | 433 | 412 | 1121 | 1499 | 1309 | 1242 | 406 | 428 | 368 | 335 | 838,52 | 1293 | 384 | 0,296 | -1,754 | -17,525 | 9,17E-69 | 4,32E-68 | 0,0118445 | 0,0040494 | 0,008021 | 0,008021 | TRUE | 0,1400754 | No |
| orf19.6862 | 3476 | 4253 | 4349 | 2460 | 3512 | 2851 | 3380 | 2936 | 3314 | 4497 | 4296 | 3854 | 2814 | 3242 | 2870 | 2387 | 3409,24 | 3990 | 2828 | 0,708 | -0,497 | -4,893 | 9,92E-07 | 1,53E-06 | 0,0159051 | 0,0025729 | 0,0096219 | 0,0096219 | TRUE | 0,1599397 | No |
| orf19.6863 | 4219 | 3950 | 4351 | 2480 | 6521 | 4556 | 5915 | 6122 | 4022 | 4177 | 4298 | 3885 | 5225 | 5181 | 5022 | 4978 | 4598,46 | 4096 | 5102 | 1,244 | 0,315 | 7,508 | 6,01E-14 | 1,18E-13 | 0,0009507 | 0,0024483 | 0,0014683 | 0,0014683 | TRUE | 0,0136605 | No |
| orf19.6864 | 232 | 206 | 267 | 155 | 582 | 430 | 579 | 548 | 221 | 218 | 264 | 243 | 466 | 489 | 492 | 446 | 354,75 | 236 | 473 | 2,002 | 1,001 | 11,255 | 2,19E-29 | 5,85E-29 | 0,001248 | 0,0067196 | 0,004296 | 0,004296 | TRUE | 0,0766667 | No |
| orf19.6866 | 1891 | 1637 | 1827 | 1146 | 1833 | 1280 | 1832 | 1899 | 1803 | 1731 | 1805 | 1795 | 1469 | 1455 | 1555 | 1544 | 1644,7 | 1784 | 1506 | 0,845 | -0,243 | -5,353 | 8,65E-08 | 1,40E-07 | 1,00E-08 | 0,0030896 | 0,0013665 | 0,0013665 | TRUE | 0,0057818 | No |
| orf19.6867 | 658 | 565 | 634 | 298 | 581 | 365 | 563 | 549 | 627 | 597 | 626 | 467 | 466 | 415 | 478 | 446 | 515,36 | 579 | 451 | 0,776 | -0,366 | -3,848 | 0,0001193 | 0,0001682 | 0,0079008 | 0,0052772 | 0,0067525 | 0,0067525 | TRUE | 0,179458 | No |
| orf19.6868 | 695 | 548 | 673 | 469 | 723 | 505 | 639 | 674 | 663 | 579 | 665 | 735 | 579 | 574 | 543 | 548 | 610,71 | 660 | 561 | 0,852 | -0,231 | -3,02 | 0,0025305 | 0,0033335 | 0,0029674 | 0,0047798 | 0,0039697 | 0,0039697 | TRUE | 0,0787736 | No |
| orf19.6869 | 997 | 942 | 1013 | 562 | 1232 | 972 | 1240 | 1222 | 951 | 996 | 1001 | 880 | 987 | 1105 | 1053 | 994 | 995,81 | 957 | 1035 | 1,077 | 0,107 | 1,709 | 0,0875051 | 0,1027171 | 0,0018006 | 0,0037401 | 0,0027277 | 0,0027277 | TRUE | 0,0302694 | No |
| orf19.687 | 219 | 183 | 185 | 140 | 679 | 493 | 656 | 668 | 209 | 193 | 183 | 219 | 544 | 561 | 557 | 543 | 376,14 | 201 | 551 | 2,755 | 1,462 | 17,448 | 3,58E-68 | 1,67E-67 | 1,00E-08 | 0,0064564 | 0,0031597 | 0,0031597 | TRUE | 0,0413039 | No |
| orf19.6871 | 2568 | 2379 | 2612 | 1736 | 3015 | 2197 | 2896 | 3161 | 2448 | 2515 | 2580 | 2720 | 2416 | 2498 | 2459 | 2570 | 2525,83 | 2566 | 2486 | 0,97 | -0,044 | -0,96 | 0,3372032 | 0,3709816 | 0,0008373 | 0,0027413 | 0,001585 | 0,001585 | TRUE | 0,0181722 | No |
| orf19.687.1 | 16456 | 13868 | 14372 | 11335 | 38535 | 30294 | 36773 | 40134 | 15689 | 14664 | 14198 | 17757 | 30877 | 34447 | 31220 | 32633 | 23935,58 | 15577 | 32294 | 2,074 | 1,052 | 15,4 | 1,65E-53 | 6,38E-53 | 0,0063524 | 0,0021599 | 0,0044355 | 0,0044355 | TRUE | 0,1079232 | No |
| orf19.6872 | 2337 | 2180 | 2260 | 1418 | 1548 | 1138 | 1484 | 1562 | 2228 | 2305 | 2233 | 2221 | 1240 | 1294 | 1260 | 1270 | 1756,44 | 2247 | 1266 | 0,563 | -0,829 | -19,806 | 2,62E-87 | 1,53E-86 | 1,00E-08 | 0,0030261 | 0,0010802 | 0,0010802 | TRUE | 0,0037345 | No |
| orf19.6873 | 23255 | 18224 | 19882 | 12889 | 54260 | 35414 | 51046 | 56133 | 22171 | 19270 | 19642 | 20191 | 43477 | 40269 | 43338 | 45641 | 31749,84 | 20318 | 43181 | 2,125 | 1,088 | 19,897 | 4,33E-88 | 2,54E-87 | 0,0032823 | 0,002143 | 0,0028327 | 0,0028327 | TRUE | 0,0463363 | No |
| orf19.6873.1 | 1836 | 1448 | 1394 | 938 | 1726 | 1050 | 1447 | 1555 | 1750 | 1531 | 1377 | 1469 | 1383 | 1194 | 1228 | 1264 | 1399,73 | 1532 | 1267 | 0,827 | -0,274 | -3,502 | 0,0004612 | 0,0006336 | 0,0065119 | 0,0032643 | 0,0051413 | 0,0051413 | TRUE | 0,1136035 | No |
| orf19.6874 | 3680 | 3316 | 3663 | 2503 | 1405 | 1023 | 1296 | 1480 | 3508 | 3506 | 3619 | 3921 | 1126 | 1163 | 1100 | 1203 | 2393,4 | 3638 | 1148 | 0,316 | -1,663 | -31,389 | 2,90E-216 | 4,26E-215 | 0,0017305 | 0,0027773 | 0,0021503 | 0,0021503 | TRUE | 0,0316614 | No |
| orf19.6875 | 1793 | 1700 | 1745 | 1047 | 1323 | 954 | 1252 | 1316 | 1709 | 1798 | 1724 | 1640 | 1060 | 1085 | 1063 | 1070 | 1393,61 | 1718 | 1070 | 0,621 | -0,686 | -14,473 | 1,81E-47 | 6,43E-47 | 1,00E-08 | 0,0032694 | 0,0014166 | 0,0014166 | TRUE | 0,0114079 | No |
| orf19.6877 | 23697 | 21380 | 22949 | 15369 | 19459 | 14404 | 17822 | 20349 | 22593 | 22607 | 22671 | 24076 | 15592 | 16379 | 15131 | 16546 | 19449,25 | 22987 | 15912 | 0,692 | -0,531 | -12,847 | 8,95E-38 | 2,74E-37 | 0,0013064 | 0,0021757 | 0,0015858 | 0,0015858 | TRUE | 0,0133691 | No |
| orf19.6879 | 2211 | 1930 | 2051 | 1402 | 1725 | 1232 | 1559 | 1658 | 2108 | 2041 | 2026 | 2196 | 1382 | 1401 | 1324 | 1348 | 1728,25 | 2093 | 1364 | 0,652 | -0,617 | -13,343 | 1,31E-40 | 4,18E-40 | 0,0003309 | 0,0030413 | 0,0014549 | 0,0014549 | TRUE | 0,0120628 | No |
| orf19.688 | 2193 | 1911 | 2091 | 1440 | 3121 | 2291 | 3152 | 3201 | 2091 | 2021 | 2066 | 2256 | 2501 | 2605 | 2676 | 2603 | 2352,2 | 2108 | 2596 | 1,234 | 0,303 | 6,478 | 9,32E-11 | 1,67E-10 | 0,0009967 | 0,0027893 | 0,0016604 | 0,0016604 | TRUE | 0,0244424 | No |
| orf19.6880 | 604 | 575 | 624 | 433 | 863 | 707 | 855 | 863 | 576 | 608 | 616 | 678 | 691 | 804 | 726 | 702 | 675,2 | 620 | 731 | 1,181 | 0,24 | 3,269 | 0,0010775 | 0,0014509 | 0,0028248 | 0,004523 | 0,0036646 | 0,0036646 | TRUE | 0,0505105 | No |
| orf19.6881 | 8603 | 7504 | 8042 | 5410 | 1356 | 1002 | 1335 | 1376 | 8202 | 7935 | 7945 | 8475 | 1087 | 1139 | 1133 | 1119 | 4629,31 | 8139 | 1120 | 0,138 | -2,862 | -68,611 | 0 | 0 | 0,0005512 | 0,002446 | 0,0012037 | 0,0012037 | TRUE | 0,0090567 | No |
| orf19.6882 | 38975 | 32439 | 36365 | 21310 | 6790 | 4528 | 6578 | 7138 | 37158 | 34300 | 35925 | 33383 | 5441 | 5149 | 5585 | 5804 | 20343,12 | 35192 | 5495 | 0,156 | -2,679 | -54,324 | 0 | 0 | 0,0022491 | 0,002172 | 0,0022386 | 0,0022386 | TRUE | 0,021478 | No |
| orf19.6882.1 | 13889 | 11557 | 11907 | 7298 | 34659 | 22729 | 33151 | 35099 | 13242 | 12220 | 11763 | 11433 | 27771 | 25845 | 28145 | 28539 | 19869,69 | 12164 | 27575 | 2,266 | 1,18 | 22,237 | 1,51E-109 | 1,09E-108 | 0,0029555 | 0,0021739 | 0,0026444 | 0,0026444 | TRUE | 0,0435959 | No |
| orf19.6883 | 2161 | 1884 | 2014 | 1365 | 4561 | 3377 | 4262 | 4558 | 2060 | 1992 | 1990 | 2138 | 3655 | 3840 | 3618 | 3706 | 2874,93 | 2045 | 3705 | 1,813 | 0,859 | 20,656 | 8,70E-95 | 5,42E-94 | 0,0004273 | 0,0026624 | 0,0012639 | 0,0012639 | TRUE | 0,0101132 | No |
| orf19.6884 | 355 | 297 | 358 | 225 | 365 | 288 | 322 | 385 | 338 | 314 | 354 | 352 | 292 | 327 | 273 | 313 | 320,63 | 340 | 301 | 0,887 | -0,173 | -1,941 | 0,0522981 | 0,0627636 | 0,0013848 | 0,0072122 | 0,0045222 | 0,0045222 | TRUE | 0,0446349 | No |
| orf19.6885 | 2705 | 2348 | 2631 | 1655 | 2038 | 1467 | 1935 | 2092 | 2579 | 2483 | 2599 | 2593 | 1633 | 1668 | 1643 | 1701 | 2112,3 | 2564 | 1661 | 0,648 | -0,626 | -15,526 | 2,32E-54 | 9,09E-54 | 1,00E-08 | 0,0028686 | 0,0010715 | 0,0010715 | TRUE | 0,0055298 | No |
| orf19.6886 | 12232 | 11181 | 12189 | 6313 | 17451 | 10833 | 17066 | 17919 | 11662 | 11822 | 12042 | 9890 | 13983 | 12318 | 14489 | 14570 | 12596,94 | 11354 | 13840 | 1,219 | 0,285 | 3,983 | 6,82E-05 | 9,73E-05 | 0,0065461 | 0,0022216 | 0,0048509 | 0,0048509 | TRUE | 0,0913274 | No |
| orf19.6887 | 4539 | 3465 | 3852 | 2234 | 7177 | 4080 | 6801 | 6746 | 4327 | 3664 | 3805 | 3500 | 5751 | 4639 | 5774 | 5485 | 4618,19 | 3824 | 5412 | 1,415 | 0,501 | 6,19 | 6,01E-10 | 1,04E-09 | 0,0089121 | 0,0024468 | 0,0060567 | 0,0060567 | TRUE | 0,1118562 | No |
| orf19.6888 | 15625 | 13967 | 15316 | 10006 | 2517 | 1809 | 2288 | 2431 | 14897 | 14768 | 15131 | 15675 | 2017 | 2057 | 1943 | 1977 | 8557,96 | 15118 | 1998 | 0,132 | -2,92 | -79,059 | 0 | 0 | 0,0004652 | 0,0022831 | 0,0010493 | 0,0010493 | TRUE | 0,0073437 | No |
| orf19.6889 | 2618 | 2321 | 2461 | 1586 | 3120 | 2140 | 2797 | 3159 | 2496 | 2454 | 2431 | 2485 | 2500 | 2433 | 2375 | 2569 | 2467,81 | 2466 | 2469 | 1,002 | 0,002 | 0,058 | 0,9533695 | 0,9580237 | 0,0003492 | 0,0027566 | 0,001204 | 0,001204 | TRUE | 0,0090763 | No |
| orf19.689 | 25 | 19 | 18 | 12 | 21 | 17 | 22 | 41 | 24 | 20 | 18 | 19 | 17 | 19 | 19 | 33 | 21,08 | 20 | 22 | 1,095 | 0,131 | 0,396 | 0,6918329 | 0,7229915 | 0,0321675 | 0,079963 | 0,0577621 | 0,0577621 | TRUE | 0,7875283 | No |
| orf19.6891 | 2047 | 1788 | 1886 | 1259 | 2642 | 1972 | 2473 | 2698 | 1952 | 1891 | 1863 | 1972 | 2117 | 2242 | 2100 | 2194 | 2041,28 | 1920 | 2163 | 1,127 | 0,173 | 3,963 | 7,42E-05 | 0,0001056 | 0,000328 | 0,0028956 | 0,0013254 | 0,0013254 | TRUE | 0,006902 | No |
| orf19.6893 | 1274 | 1116 | 1291 | 809 | 2192 | 1548 | 2156 | 2222 | 1215 | 1180 | 1275 | 1267 | 1756 | 1760 | 1830 | 1807 | 1511,39 | 1234 | 1788 | 1,451 | 0,537 | 11,433 | 2,87E-30 | 7,79E-30 | 1,00E-08 | 0,0031776 | 0,0014134 | 0,0014134 | TRUE | 0,010473 | No |
| orf19.6896 | 18 | 16 | 27 | 9 | 41 | 30 | 38 | 56 | 17 | 17 | 27 | 14 | 33 | 34 | 32 | 46 | 27,45 | 19 | 36 | 1,906 | 0,931 | 3,173 | 0,0015101 | 0,0020138 | 0,0121088 | 0,0619025 | 0,0411156 | 0,0411156 | TRUE | 0,4384828 | No |

| | | | | | | | | | | | | | | | | | | | | | | | | | | | | | | | |
|---|---|---|---|---|---|---|---|---|---|---|---|---|---|---|---|---|---|---|---|---|---|---|---|---|---|---|---|---|---|---|---|
| orf19.6897 | 0 | 0 | 0 | 0 | 0 | 0 | 0 | 0 | 0 | 0 | 0 | 0 | 0 | 0 | 0 | 0 | 0 | 0 | 0 | NA | NA | NA | NA | NA | NA | NA | NA | NA | NA | NA | NA |
| orf19.6898 | 2104 | 1913 | 2111 | 1262 | 1887 | 1272 | 1763 | 1806 | 2006 | 2023 | 2085 | 1977 | 1512 | 1446 | 1497 | 1468 | 1751,84 | 2023 | 1481 | 0,732 | -0,45 | -10,604 | 2,87E-26 | 7,29E-26 | 1,00E-08 | 0,0030285 | 0,0011524 | 0,0011524 | TRUE | 0,0053264 | No |
| orf19.6898.1 | 1379 | 1182 | 1201 | 891 | 2746 | 1932 | 2516 | 2609 | 1315 | 1250 | 1186 | 1396 | 2200 | 2197 | 2136 | 2121 | 1725,18 | 1287 | 2164 | 1,686 | 0,753 | 13,962 | 2,65E-44 | 9,00E-44 | 0,0013833 | 0,003043 | 0,0021523 | 0,0021523 | TRUE | 0,0349092 | No |
| orf19.6899 | 6608 | 5600 | 6041 | 3743 | 4909 | 3218 | 4438 | 4978 | 6300 | 5921 | 5968 | 5864 | 3933 | 3659 | 3768 | 4048 | 4932,61 | 6013 | 3852 | 0,641 | -0,642 | -14,439 | 2,94E-47 | 1,04E-46 | 0,0012482 | 0,0024241 | 0,0016902 | 0,0016902 | TRUE | 0,0141816 | No |
| orf19.69 | 420 | 411 | 452 | 275 | 532 | 419 | 561 | 491 | 400 | 435 | 447 | 431 | 426 | 476 | 476 | 399 | 436,32 | 428 | 444 | 1,037 | 0,052 | 0,623 | 0,5334209 | 0,5691904 | 0,0030272 | 0,0058543 | 0,0044083 | 0,0044083 | TRUE | 0,0557145 | No |
| orf19.690 | 76 | 84 | 76 | 56 | 64 | 44 | 57 | 71 | 72 | 89 | 75 | 88 | 51 | 50 | 48 | 58 | 66,44 | 81 | 52 | 0,645 | -0,632 | -3,602 | 0,0003153 | 0,0004364 | 1,00E-08 | 0,0268038 | 0,0141025 | 0,0141025 | TRUE | 0,0533858 | No |
| orf19.6900 | 1347 | 1247 | 1395 | 790 | 1374 | 944 | 1319 | 1286 | 1284 | 1319 | 1378 | 1238 | 1101 | 1073 | 1120 | 1046 | 1194,79 | 1305 | 1085 | 0,83 | -0,268 | -5,066 | 4,06E-07 | 6,39E-07 | 0,0005904 | 0,0034655 | 0,0018542 | 0,0018542 | TRUE | 0,0172484 | No |
| orf19.6901 | 720 | 646 | 626 | 424 | 949 | 663 | 905 | 936 | 686 | 683 | 618 | 664 | 760 | 754 | 768 | 761 | 711,98 | 663 | 761 | 1,148 | 0,199 | 3,324 | 0,0008878 | 0,0012022 | 1,00E-08 | 0,0043974 | 0,0020238 | 0,0020238 | TRUE | 0,0264593 | No |
| orf19.6902 | 12966 | 11138 | 12346 | 7281 | 16732 | 11090 | 16173 | 17181 | 12362 | 11777 | 12197 | 11406 | 13407 | 12610 | 13731 | 13970 | 12682,41 | 11936 | 13430 | 1,125 | 0,17 | 3,868 | 0,0001096 | 0,0001549 | 0,0015184 | 0,0022207 | 0,0017745 | 0,0017745 | TRUE | 0,0204184 | No |
| orf19.6903 | 4714 | 3893 | 4116 | 2693 | 4428 | 2918 | 4194 | 4364 | 4494 | 4116 | 4066 | 4219 | 3548 | 3318 | 3561 | 3548 | 3858,84 | 4224 | 3494 | 0,827 | -0,273 | -5,959 | 2,54E-09 | 4,30E-09 | 0,0012386 | 0,0025168 | 0,0017572 | 0,0017572 | TRUE | 0,0233547 | No |
| orf19.6904 | 3356 | 2617 | 2904 | 1876 | 2855 | 1950 | 2695 | 3011 | 3200 | 2767 | 2869 | 2939 | 2288 | 2217 | 2288 | 2448 | 2626,96 | 2944 | 2310 | 0,785 | -0,349 | -6,285 | 3,28E-10 | 5,76E-10 | 0,0023476 | 0,0027163 | 0,0025791 | 0,0025791 | TRUE | 0,0429053 | No |
| orf19.6905 | 4254 | 3622 | 3913 | 2791 | 1429 | 1012 | 1296 | 1462 | 4056 | 3830 | 3866 | 4372 | 1145 | 1151 | 1100 | 1189 | 2588,53 | 4031 | 1146 | 0,285 | -1,813 | -33,272 | 9,66E-243 | 1,59E-241 | 0,0020566 | 0,0027256 | 0,0023206 | 0,0023206 | TRUE | 0,037928 | No |
| orf19.6906 | 16576 | 13650 | 14234 | 9265 | 54518 | 35732 | 50848 | 55803 | 15803 | 14433 | 14062 | 14514 | 43684 | 40630 | 43170 | 45373 | 28958,7 | 14703 | 43214 | 2,939 | 1,555 | 31,835 | 2,14E-222 | 3,20E-221 | 0,002305 | 0,002148 | 0,0022445 | 0,0022445 | TRUE | 0,0312717 | No |
| orf19.6907 | 2524 | 2217 | 2488 | 1386 | 2576 | 1693 | 2333 | 2549 | 2406 | 2344 | 2458 | 2171 | 2064 | 1925 | 1981 | 2073 | 2177,77 | 2345 | 2011 | 0,857 | -0,223 | -4,404 | 1,06E-05 | 1,57E-05 | 0,0014143 | 0,0028452 | 0,0020099 | 0,0020099 | TRUE | 0,0278618 | No |
| orf19.6908 | 1112 | 1018 | 1106 | 775 | 910 | 662 | 851 | 875 | 1060 | 1076 | 1093 | 1214 | 729 | 753 | 722 | 711 | 919,89 | 1111 | 729 | 0,658 | -0,604 | -10,118 | 4,61E-24 | 1,12E-23 | 0,0007634 | 0,0038762 | 0,0023052 | 0,0023052 | TRUE | 0,0417146 | No |
| orf19.6909 | 253 | 268 | 252 | 167 | 512 | 290 | 441 | 413 | 241 | 283 | 249 | 262 | 410 | 330 | 374 | 336 | 310,67 | 259 | 362 | 1,406 | 0,492 | 4,878 | 1,07E-06 | 1,66E-06 | 0,0053617 | 0,0073763 | 0,0063626 | 0,0063626 | TRUE | 0,0921485 | No |
| orf19.691 | 172796 | 148559 | 162491 | 99615 | 196560 | 135873 | 186383 | 201344 | 164742 | 157082 | 160526 | 156052 | 157498 | 154500 | 158238 | 163712 | 159043,81 | 159600 | 158487 | 0,993 | -0,01 | -0,317 | 0,7514781 | 0,7793242 | 0,0005185 | 0,0021016 | 0,0009744 | 0,0009744 | TRUE | 0,006038 | No |
| orf19.6910 | 77 | 61 | 62 | 44 | 348 | 257 | 326 | 312 | 73 | 64 | 61 | 69 | 279 | 292 | 277 | 254 | 171,2 | 67 | 276 | 4,106 | 2,038 | 15,656 | 3,04E-55 | 1,20E-54 | 1,00E-08 | 0,0116817 | 0,0063825 | 0,0063825 | TRUE | 0,0458015 | No |
| orf19.6911 | 1 | 1 | 3 | 0 | 3 | 3 | 3 | 7 | 1 | 1 | 3 | 0 | 2 | 3 | 3 | 6 | 2,38 | 1 | 4 | 2,632 | 1,396 | 1,416 | 0,1566823 | 0,1796245 | 1,00E-08 | 0,6923938 | 0,3923626 | 0,3923626 | TRUE | 0,4817453 | No |
| orf19.6912 | 1075 | 1070 | 1116 | 705 | 1138 | 870 | 1068 | 1145 | 1025 | 1131 | 1103 | 1104 | 912 | 989 | 907 | 931 | 1012,76 | 1091 | 935 | 0,856 | -0,224 | -3,962 | 7,43E-05 | 0,0001058 | 0,000719 | 0,0037125 | 0,0020693 | 0,0020693 | TRUE | 0,0210886 | No |
| orf19.6913 | 1222 | 1221 | 1248 | 692 | 909 | 648 | 807 | 859 | 1165 | 1291 | 1233 | 1084 | 728 | 737 | 685 | 698 | 952,73 | 1193 | 712 | 0,595 | -0,749 | -11,498 | 1,35E-30 | 3,69E-30 | 0,0022652 | 0,0038146 | 0,0029771 | 0,0029771 | TRUE | 0,0407576 | No |
| orf19.6915 | 1507 | 1411 | 1513 | 917 | 1778 | 1374 | 1792 | 1809 | 1437 | 1492 | 1495 | 1437 | 1425 | 1562 | 1521 | 1471 | 1479,91 | 1465 | 1495 | 1,019 | 0,027 | 0,547 | 0,5841447 | 0,6181472 | 0,0004388 | 0,0032007 | 0,0015742 | 0,0015742 | TRUE | 0,0120877 | No |
| orf19.6916 | 816 | 785 | 786 | 429 | 849 | 612 | 814 | 819 | 778 | 830 | 776 | 672 | 680 | 696 | 691 | 666 | 723,72 | 764 | 683 | 0,89 | -0,168 | -2,447 | 0,0144225 | 0,0181054 | 0,0018587 | 0,00436 | 0,0031451 | 0,0031451 | TRUE | 0,0673628 | No |
| orf19.6917 | 704 | 802 | 821 | 445 | 649 | 448 | 540 | 461 | 671 | 848 | 811 | 697 | 520 | 509 | 458 | 375 | 611,27 | 757 | 466 | 0,614 | -0,704 | -6,327 | 2,51E-10 | 4,42E-10 | 0,0155284 | 0,0047774 | 0,0101755 | 0,0101755 | TRUE | 0,2056196 | No |
| orf19.6918 | 788 | 1040 | 981 | 554 | 607 | 508 | 561 | 484 | 751 | 1100 | 969 | 868 | 486 | 578 | 476 | 394 | 702,72 | 922 | 484 | 0,523 | -0,935 | -7,391 | 1,46E-13 | 2,83E-13 | 0,0229454 | 0,0044278 | 0,0138189 | 0,0138189 | TRUE | 0,1452037 | No |
| orf19.6919 | 126 | 152 | 142 | 80 | 26 | 19 | 24 | 25 | 120 | 161 | 140 | 125 | 21 | 22 | 20 | 20 | 78,7 | 136 | 21 | 0,152 | -2,723 | -13,219 | 6,78E-40 | 2,14E-39 | 1,00E-08 | 0,0229542 | 0,0152891 | 0,0152891 | TRUE | 0,1461004 | No |
| orf19.692 | 3940 | 3187 | 3665 | 2297 | 6943 | 4560 | 6973 | 7380 | 3756 | 3370 | 3621 | 3598 | 5563 | 5185 | 5920 | 6001 | 4626,79 | 3586 | 5667 | 1,581 | 0,661 | 11,734 | 8,57E-32 | 2,39E-31 | 0,0028663 | 0,0024461 | 0,0028078 | 0,0028078 | TRUE | 0,0394032 | No |
| orf19.69.2 | 17 | 32 | 15 | 13 | 12 | 9 | 9 | 5 | 16 | 34 | 15 | 20 | 10 | 10 | 8 | 4 | 14,6 | 21 | 8 | 0,366 | -1,448 | -3,307 | 0,0009434 | 0,0012748 | 0,0835059 | 0,1145669 | 0,100793 | 0,100793 | TRUE | 0,8970614 | No |
| orf19.6920 | 1485 | 2036 | 1871 | 1111 | 474 | 313 | 423 | 378 | 1416 | 2153 | 1848 | 1740 | 380 | 356 | 359 | 307 | 1069,95 | 1789 | 350 | 0,196 | -2,352 | -20,468 | 4,19E-93 | 2,58E-92 | 0,0184703 | 0,0036258 | 0,0110921 | 0,0110921 | TRUE | 0,0924089 | No |
| orf19.6921 | 1978 | 1912 | 2069 | 1124 | 1208 | 827 | 1116 | 1133 | 1886 | 2022 | 2044 | 1761 | 968 | 940 | 947 | 921 | 1436,16 | 1928 | 944 | 0,489 | -1,032 | -17,727 | 2,60E-70 | 1,25E-69 | 0,0019489 | 0,0032345 | 0,0024927 | 0,0024927 | TRUE | 0,0379287 | No |
| orf19.6922 | 0 | 0 | 0 | 0 | 0 | 0 | 0 | 0 | 0 | 0 | 0 | 0 | 0 | 0 | 0 | 0 | 0 | 0 | 0 | NA | NA | NA | NA | NA | NA | NA | NA | NA | NA | NA | NA |
| orf19.6923 | 2143 | 1832 | 1942 | 1149 | 2838 | 1613 | 2727 | 2736 | 2043 | 1937 | 1919 | 1800 | 2274 | 1834 | 2315 | 2225 | 2043,34 | 1925 | 2162 | 1,123 | 0,168 | 2,252 | 0,0242997 | 0,0299916 | 0,0059421 | 0,0028948 | 0,0048374 | 0,0048374 | TRUE | 0,1232384 | No |
| orf19.6923.1 | 1820 | 1375 | 1383 | 860 | 4243 | 2206 | 3876 | 4131 | 1735 | 1454 | 1366 | 1347 | 3400 | 2508 | 3291 | 3359 | 2307,55 | 1476 | 3140 | 2,127 | 1,089 | 10,554 | 4,87E-26 | 1,23E-25 | 0,0157859 | 0,0028028 | 0,0096946 | 0,0096946 | TRUE | 0,2205562 | No |
| orf19.6924 | 4832 | 3699 | 3959 | 2726 | 6270 | 4132 | 5583 | 6319 | 4607 | 3911 | 3911 | 4270 | 5024 | 4698 | 4740 | 5138 | 4537,48 | 4175 | 4900 | 1,174 | 0,231 | 3,82 | 0,0001332 | 0,0001875 | 0,0038648 | 0,0024531 | 0,0033002 | 0,0033002 | TRUE | 0,0604646 | No |
| orf19.6925 | 6157 | 5565 | 6051 | 4462 | 7700 | 5725 | 7262 | 7556 | 5870 | 5884 | 5978 | 6990 | 6170 | 6510 | 6165 | 6144 | 6213,86 | 6180 | 6247 | 1,012 | 0,017 | 0,279 | 0,7798668 | 0,8035411 | 0,0034678 | 0,0023555 | 0,0031921 | 0,0031921 | TRUE | 0,0929453 | No |
| orf19.6926 | 2978 | 2603 | 2920 | 1802 | 2911 | 2149 | 2759 | 3112 | 2839 | 2752 | 2885 | 2823 | 2333 | 2444 | 2342 | 2530 | 2618,5 | 2825 | 2412 | 0,854 | -0,228 | -5,328 | 9,93E-08 | 1,60E-07 | 0,0006249 | 0,0027183 | 0,0013779 | 0,0013779 | TRUE | 0,013779 | No |
| orf19.6927 | 1413 | 1200 | 1336 | 838 | 1522 | 992 | 1422 | 1560 | 1347 | 1269 | 1320 | 1313 | 1220 | 1128 | 1207 | 1268 | 1258,98 | 1312 | 1206 | 0,92 | -0,12 | -2,285 | 0,0222941 | 0,0275801 | 0,0006549 | 0,0033954 | 0,0018483 | 0,0018483 | TRUE | 0,0209402 | No |
| orf19.6928 | 19667 | 17258 | 18386 | 12115 | 9044 | 6440 | 8657 | 9699 | 18750 | 18248 | 18164 | 18979 | 7247 | 7323 | 7350 | 7886 | 12993,31 | 18535 | 7452 | 0,402 | -1,314 | -34,7 | 7,88E-264 | 1,43E-262 | 0,0008733 | 0,0022176 | 0,0012876 | 0,0012876 | TRUE | 0,0190327 | No |
| orf19.6929 | 1182 | 978 | 1052 | 714 | 1962 | 1248 | 1808 | 1936 | 1127 | 1034 | 1039 | 1119 | 1572 | 1419 | 1535 | 1574 | 1302,39 | 1080 | 1525 | 1,416 | 0,502 | 8,887 | 6,30E-19 | 1,38E-18 | 0,0013884 | 0,003352 | 0,0022458 | 0,0022458 | TRUE | 0,0249178 | No |
| orf19.693 | 169 | 176 | 216 | 119 | 46 | 34 | 46 | 43 | 161 | 186 | 213 | 186 | 37 | 39 | 39 | 35 | 112,07 | 186 | 38 | 0,2 | -2,323 | -14,077 | 5,26E-45 | 1,80E-44 | 1,00E-08 | 0,0167419 | 0,0112758 | 0,0112758 | TRUE | 0,1077961 | No |
| orf19.6930 | 2568 | 2235 | 2450 | 1664 | 2175 | 1597 | 2312 | 2174 | 2448 | 2363 | 2420 | 2607 | 1743 | 1816 | 1963 | 1768 | 2140,99 | 2460 | 1822 | 0,742 | -0,431 | -8,128 | 4,37E-16 | 9,01E-16 | 0,0017225 | 0,0028582 | 0,002228 | 0,002228 | TRUE | 0,0333103 | No |
| orf19.6931 | 2477 | 2190 | 2373 | 1422 | 2900 | 2043 | 2857 | 2911 | 2362 | 2316 | 2344 | 2228 | 2324 | 2323 | 2426 | 2367 | 2336,05 | 2312 | 2360 | 1,019 | 0,028 | 0,695 | 0,4867461 | 0,5230594 | 1,00E-08 | 0,0027941 | 0,0010944 | 0,0010944 | TRUE | 0,0065423 | No |
| orf19.6933 | 988 | 861 | 887 | 646 | 1226 | 831 | 1073 | 1219 | 942 | 910 | 876 | 1012 | 982 | 945 | 911 | 991 | 946,25 | 935 | 957 | 1,027 | 0,039 | 0,639 | 0,5225992 | 0,5589516 | 0,0012432 | 0,0038264 | 0,0024864 | 0,0024864 | TRUE | 0,0320287 | No |
| orf19.6934 | 1421 | 1196 | 1428 | 834 | 1737 | 1105 | 1647 | 1608 | 1355 | 1265 | 1411 | 1307 | 1392 | 1256 | 1398 | 1307 | 1336,33 | 1334 | 1338 | 1,003 | 0,005 | 0,087 | 0,9306051 | 0,9376584 | 0,0016866 | 0,0033199 | 0,002385 | 0,002385 | TRUE | 0,0192249 | No |
| orf19.6936 | 1736 | 1451 | 1728 | 1025 | 1415 | 933 | 1384 | 1435 | 1655 | 1534 | 1707 | 1606 | 1134 | 1061 | 1175 | 1167 | 1379,83 | 1626 | 1134 | 0,698 | -0,518 | -9,433 | 4,00E-21 | 9,16E-21 | 0,0013152 | 0,0032812 | 0,0021594 | 0,0021594 | TRUE | 0,0209686 | No |
| orf19.6937 | 65122 | 55033 | 60206 | 40419 | 83838 | 58137 | 77958 | 86001 | 62087 | 58190 | 59478 | 63319 | 67177 | 66107 | 66186 | 69927 | 64058,82 | 60768 | 67349 | 1,108 | 0,148 | 3,889 | 0,0001006 | 0,0001424 | 0,0010779 | 0,0021169 | 0,0013841 | 0,0013841 | TRUE | 0,009983 | No |
| orf19.6938 | 2529 | 1977 | 2112 | 1392 | 8328 | 5615 | 8029 | 8526 | 2411 | 2090 | 2086 | 2181 | 6673 | 6385 | 6817 | 6932 | 4446,93 | 2192 | 6702 | 3,057 | 1,612 | 29,696 | 8,54E-194 | 1,14E-192 | 0,0026327 | 0,0024605 | 0,0025059 | 0,0025059 | TRUE | 0,0569415 | No |
| orf19.6939 | 4261 | 3633 | 3906 | 2331 | 3896 | 2534 | 3756 | 3943 | 4062 | 3841 | 3859 | 3652 | 3122 | 2881 | 3189 | 3206 | 3476,53 | 3854 | 3100 | 0,804 | -0,314 | -6,33 | 2,45E-10 | 4,32E-10 | 0,0017303 | 0,0025636 | 0,0020778 | 0,0020778 | TRUE | 0,026352 | No |
| orf19.6940 | 28 | 19 | 22 | 9 | 21 | 18 | 22 | 20 | 27 | 20 | 22 | 14 | 17 | 20 | 19 | 16 | 19,36 | 21 | 18 | 0,851 | -0,232 | -0,728 | 0,466431 | 0,5028377 | 1,00E-08 | 0,0869156 | 0,0465021 | 0,0465021 | TRUE | 0,2117585 | No |
| orf19.6941 | 2489 | 2183 | 2492 | 1526 | 3414 | 2516 | 3402 | 3511 | 2373 | 2308 | 2462 | 2391 | 2736 | 2861 | 2888 | 2855 | 2609,15 | 2384 | 2835 | 1,189 | 0,25 | 6,187 | 6,13E-10 | 1,06E-09 | 0,0003102 | 0,0027206 | 0,0011728 | 0,0011728 | TRUE | 0,0069446 | No |
| orf19.6942 | 772 | 675 | 770 | 464 | 1336 | 942 | 1203 | 1297 | 736 | 714 | 761 | 727 | 1070 | 1071 | 1021 | 1055 | 894,36 | 734 | 1054 | 1,434 | 0,52 | 9,635 | 5,68E-22 | 1,33E-21 | 1,00E-08 | 0,0039271 | 0,0016177 | 0,0016177 | TRUE | 0,007288 | No |
| orf19.6943 | 799 | 717 | 747 | 412 | 1097 | 681 | 1063 | 1057 | 762 | 758 | 738 | 645 | 879 | 774 | 902 | 859 | 789,82 | 726 | 854 | 1,174 | 0,231 | 3,209 | 0,0013318 | 0,0017793 | 0,0032685 | 0,0041701 | 0,0036936 | 0,0036936 | TRUE | 0,0575562 | No |
| orf19.6944 | 2050 | 1709 | 1869 | 1156 | 2241 | 1357 | 2213 | 2238 | 1954 | 1807 | 1846 | 1811 | 1796 | 1543 | 1879 | 1820 | 1807,01 | 1854 | 1760 | 0,949 | -0,075 | -1,165 | 0,2441811 | 0,2740273 | 0,0035054 | 0,0029999 | 0,0034354 | 0,0034354 | TRUE | 0,0796002 | No |
| orf19.6947 | 647 | 605 | 673 | 453 | 686 | 505 | 624 | 617 | 617 | 640 | 665 | 710 | 550 | 574 | 530 | 502 | 598,3 | 658 | 539 | 0,82 | -0,286 | -4,016 | 5,91E-05 | 8,45E-05 | 0,0015062 | 0,0048355 | 0,0031722 | 0,0031722 | TRUE | 0,0289139 | No |
| orf19.6948 | 1612 | 1383 | 1507 | 982 | 5886 | 4195 | 5573 | 5936 | 1537 | 1462 | 1489 | 1538 | 4716 | 4770 | 4731 | 4827 | 3133,84 | 1506 | 4761 | 3,163 | 1,661 | 43,968 | 0 | 0 | 1,00E-08 | 0,0026152 | 0,0009057 | 0,0009057 | TRUE | 0,0046545 | No |
| orf19.6949 | 1 | 2 | 2 | 2 | 5 | 4 | 2 | 4 | 1 | 2 | 2 | 3 | 4 | 5 | 2 | 3 | 2,71 | 2 | 4 | 1,705 | 0,77 | 0,89 | 0,3734407 | 0,4085948 | 1,00E-08 | 0,6078998 | 0,3025115 | 0,3025115 | TRUE | 0,1894625 | No |
| orf19.695 | 1440 | 1590 | 1509 | 982 | 1206 | 911 | 1129 | 1063 | 1373 | 1681 | 1491 | 1538 | 966 | 1036 | 959 | 864 | 1238,53 | 1521 | 956 | 0,628 | -0,67 | -8,932 | 4,17E-19 | 9,18E-19 | 0,0050987 | 0,003417 | 0,0045697 | 0,0045697 | TRUE | 0,0624362 | No |
| orf19.6950 | 1693 | 1497 | 1688 | 1152 | 452 | 294 | 418 | 427 | 1614 | 1583 | 1668 | 1805 | 362 | 334 | 355 | 347 | 1008,47 | 1668 | 350 | 0,211 | -2,248 | -34,343 | 1,77E-258 | 3,16E-257 | 0,0015332 | 0,0037194 | 0,0025068 | 0,0025068 | TRUE | 0,0341186 | No |
| orf19.6951 | 3202 | 2864 | 3096 | 1850 | 2178 | 1437 | 2151 | 2205 | 3053 | 3028 | 3059 | 2898 | 1745 | 1634 | 1826 | 1793 | 2379,5 | 3010 | 1750 | 0,582 | -0,782 | -16,88 | 6,36E-64 | 2,82E-63 | 0,000861 | 0,0027813 | 0,0016186 | 0,0016186 | TRUE | 0,022292 | No |
| orf19.6952 | 2076 | 1931 | 2008 | 1193 | 1735 | 1223 | 1699 | 1804 | 1979 | 2042 | 1984 | 1869 | 1390 | 1391 | 1442 | 1467 | 1695,47 | 1968 | 1422 | 0,722 | -0,47 | -10,117 | 4,64E-24 | 1,13E-23 | 0,0003254 | 0,0030597 | 0,0014743 | 0,0014743 | TRUE | 0,0124586 | No |
| orf19.6953 | 3352 | 2806 | 3238 | 2090 | 2252 | 1493 | 2160 | 2241 | 3196 | 2967 | 3199 | 3274 | 1804 | 1698 | 1834 | 1822 | 2474,23 | 3159 | 1790 | 0,567 | -0,818 | -17,311 | 3,92E-67 | 1,81E-66 | 0,0009108 | 0,0027549 | 0,0017144 | 0,0017144 | TRUE | 0,0203575 | No |
| orf19.6955 | 13289 | 11010 | 12433 | 7685 | 16564 | 10595 | 16241 | 16561 | 12670 | 11642 | 12283 | 12039 | 13272 | 12047 | 13789 | 13466 | 12650,85 | 12158 | 13144 | 1,081 | 0,113 | 2,282 | 0,0224751 | 0,027788 | 0,0022002 | 0,0022211 | 0,0022574 | 0,0022574 | TRUE | 0,0382276 | No |
| orf19.6956 | 765 | 686 | 728 | 394 | 862 | 586 | 895 | 949 | 729 | 725 | 719 | 617 | 691 | 666 | 760 | 772 | 709,95 | 698 | 722 | 1,032 | 0,046 | 0,608 | 0,5431673 | 0,5787656 | 0,0036256 | 0,004404 | 0,0040706 | 0,0040706 | TRUE | 0,0626754 | No |
| orf19.6957 | 0 | 0 | 0 | 0 | 0 | 0 | 0 | 0 | 0 | 0 | 0 | 0 | 0 | 0 | 0 | 0 | 0 | 0 | 0 | NA | NA | NA | NA | NA | NA | NA | NA | NA | NA | NA | NA |
| orf19.6958 | 61 | 40 | 57 | 33 | 388 | 281 | 344 | 399 | 58 | 42 | 56 | 52 | 311 | 320 | 292 | 324 | 181,92 | 52 | 312 | 5,954 | 2,574 | 18,642 | 1,48E-77 | 7,81E-77 | 1,00E-08 | 0,0111167 | 0,0063197 | 0,0063197 | TRUE | 0,1437255 | No |
| orf19.696 | 25 | 23 | 24 | 12 | 24 | 13 | 24 | 16 | 24 | 24 | 24 | 19 | 19 | 15 | 20 | 13 | 19,76 | 23 | 17 | 0,74 | -0,435 | -1,384 | 0,1664855 | 0,1903161 | 1,00E-08 | 0,0851936 | 0,0444286 | 0,0444286 | TRUE | 0,1465106 | No |
| orf19.6966 | 1052 | 1299 | 1290 | 783 | 1677 | 1304 | 1466 | 1320 | 1003 | 1374 | 1274 | 1227 | 1344 | 1483 | 1245 | 1073 | 1252,74 | 1220 | 1286 | 1,054 | 0,076 | 0,707 | 0,4798065 | 0,5160366 | 0,0171282 | 0,0034019 | 0,0103428 | 0,0103428 | TRUE | 0,1749816 | No |
| orf19.6967 | 3407 | 3291 | 3669 | 2126 | 4098 | 2760 | 3905 | 4001 | 3248 | 3480 | 3625 | 3330 | 3284 | 3138 | 3315 | 3253 | 3334,2 | 3421 | 3248 | 0,949 | -0,075 | -1,652 | 0,098525 | 0,1151444 | 0,0010636 | 0,0025837 | 0,0016759 | 0,0016759 | TRUE | 0,0199935 | No |
| orf19.6968 | 4307 | 4604 | 4610 | 3037 | 4776 | 3483 | 4183 | 4618 | 4106 | 4868 | 4554 | 4758 | 3827 | 3960 | 3551 | 3755 | 4172,48 | 4572 | 3773 | 0,826 | -0,276 | -4,649 | 3,33E-06 | 5,06E-06 | 0,0034084 | 0,0024848 | 0,0031548 | 0,0031548 | TRUE | 0,057889 | No |
| orf19.697 | 3172 | 3564 | 3619 | 2107 | 2391 | 1794 | 2165 | 1973 | 3024 | 3768 | 3575 | 3301 | 1916 | 2040 | 1838 | 1604 | 2633,34 | 3417 | 1850 | 0,541 | -0,887 | -10,657 | 1,62E-26 | 4,13E-26 | 0,0088794 | 0,0027148 | 0,0062425 | 0,0062425 | TRUE | 0,0969405 | No |
| orf19.6970 | 31 | 36 | 49 | 30 | 62 | 43 | 59 | 47 | 30 | 38 | 48 | 47 | 50 | 49 | 50 | 38 | 43,74 | 41 | 47 | 1,155 | 0,208 | 0,909 | 0,3630874 | 0,3978126 | 1,00E-08 | 0,0396308 | 0,0269381 | 0,0269381 | TRUE | 0,2856736 | No |
| orf19.6971 | 102 | 137 | 107 | 82 | 205 | 166 | 156 | 144 | 97 | 145 | 106 | 128 | 164 | 189 | 132 | 117 | 134,85 | 119 | 150 | 1,265 | 0,339 | 1,901 | 0,0572956 | 0,0684394 | 0,0322934 | 0,0142669 | 0,022872 | 0,022872 | TRUE | 0,2151042 | No |
| orf19.6972 | 838 | 812 | 936 | 434 | 675 | 505 | 586 | 607 | 799 | 859 | 925 | 680 | 541 | 574 | 498 | 494 | 671,03 | 816 | 527 | 0,643 | -0,638 | -6,808 | 9,87E-12 | 1,82E-11 | 0,0086978 | 0,0045381 | 0,0068921 | 0,0068921 | TRUE | 0,136786 | No |

| | | | | | | | | | | | | | | | | | | | | | | | | | | | | | | | |
|---|---|---|---|---|---|---|---|---|---|---|---|---|---|---|---|---|---|---|---|---|---|---|---|---|---|---|---|---|---|---|---|
| orf19.6973 | 12171 | 12133 | 12924 | 7151 | 5995 | 4297 | 5570 | 5539 | 11604 | 12829 | 12768 | 11202 | 4804 | 4886 | 4729 | 4504 | 8415,66 | 12101 | 4731 | 0,391 | -1,356 | -25,244 | 1,30E-140 | 1,24E-139 | 0,0027169 | 0,0022864 | 0,0026291 | 0,0026291 | TRUE | 0,0300817 | No |
| orf19.6975 | 18719 | 14823 | 16028 | 11352 | 57017 | 39729 | 53914 | 60130 | 17847 | 15673 | 15834 | 17784 | 45686 | 45175 | 45773 | 48891 | 31582,92 | 16784 | 46381 | 2,764 | 1,467 | 27,224 | 3,34E-163 | 3,71E-162 | 0,0031352 | 0,0021433 | 0,0027452 | 0,0027452 | TRUE | 0,0242727 | No |
| orf19.6976 | 1776 | 1790 | 1768 | 1032 | 3579 | 2570 | 3125 | 3121 | 1693 | 1893 | 1747 | 1617 | 2868 | 2922 | 2653 | 2538 | 2241,26 | 1738 | 2745 | 1,578 | 0,658 | 10,311 | 6,29E-25 | 1,56E-24 | 0,0037883 | 0,0028239 | 0,0034217 | 0,0034217 | TRUE | 0,0433821 | No |
| orf19.6977 | 257 | 375 | 332 | 174 | 307 | 261 | 260 | 220 | 245 | 397 | 328 | 273 | 246 | 297 | 221 | 179 | 273,06 | 311 | 236 | 0,755 | -0,405 | -2,397 | 0,0165275 | 0,0206626 | 0,0401219 | 0,0081042 | 0,0236393 | 0,0236393 | TRUE | 0,3609601 | No |
| orf19.6978 | 98 | 112 | 106 | 67 | 142 | 108 | 124 | 100 | 93 | 118 | 105 | 105 | 114 | 123 | 105 | 81 | 105,59 | 105 | 106 | 1 | 0 | 0,001 | 0,9993943 | 0,9993943 | 0,0117408 | 0,0176413 | 0,0149505 | 0,0149505 | TRUE | 0,2551041 | No |
| orf19.6979 | 2282 | 2084 | 2277 | 1363 | 4017 | 2920 | 3721 | 4177 | 2176 | 2204 | 2249 | 2135 | 3219 | 3320 | 3159 | 3396 | 2732,29 | 2191 | 3274 | 1,493 | 0,578 | 13,935 | 3,89E-44 | 1,31E-43 | 0,0004877 | 0,0026922 | 0,0012586 | 0,0012586 | TRUE | 0,0080517 | No |
| orf19.698 | 5070 | 5069 | 5631 | 3211 | 3281 | 2525 | 3194 | 3178 | 4834 | 5360 | 5563 | 5030 | 2629 | 2871 | 2712 | 2584 | 3947,81 | 5197 | 2699 | 0,519 | -0,946 | -17,007 | 7,30E-65 | 3,28E-64 | 0,0026973 | 0,0025072 | 0,0026999 | 0,0026999 | TRUE | 0,0277325 | No |
| orf19.6980 | 1564 | 1609 | 1778 | 1139 | 1550 | 1094 | 1439 | 1430 | 1491 | 1701 | 1756 | 1784 | 1242 | 1244 | 1222 | 1163 | 1450,45 | 1683 | 1218 | 0,724 | -0,466 | -7,322 | 2,44E-13 | 4,71E-13 | 0,0031506 | 0,0032233 | 0,0031812 | 0,0031812 | TRUE | 0,0731471 | No |
| orf19.6981 | 931 | 974 | 974 | 565 | 407 | 305 | 359 | 366 | 888 | 1030 | 962 | 885 | 326 | 347 | 305 | 298 | 630,02 | 941 | 319 | 0,337 | -1,568 | -19,885 | 5,53E-88 | 3,25E-87 | 0,0032701 | 0,0046974 | 0,00397 | 0,00397 | TRUE | 0,047806 | No |
| orf19.6982 | 1135 | 1267 | 1175 | 728 | 676 | 497 | 622 | 615 | 1082 | 1340 | 1161 | 1140 | 542 | 565 | 528 | 500 | 857,24 | 1181 | 534 | 0,451 | -1,148 | -14,862 | 5,80E-50 | 2,14E-49 | 0,0047469 | 0,0040066 | 0,0044133 | 0,0044133 | TRUE | 0,1000067 | No |
| orf19.6983 | 5226 | 5546 | 5779 | 3226 | 2219 | 1607 | 2009 | 1947 | 4982 | 5864 | 5709 | 5054 | 1778 | 1827 | 1706 | 1583 | 3562,93 | 5402 | 1724 | 0,319 | -1,65 | -24,402 | 1,63E-131 | 1,44E-130 | 0,0048417 | 0,0025521 | 0,004025 | 0,004025 | TRUE | 0,0403271 | No |
| orf19.6984 | 2520 | 2794 | 2896 | 1709 | 2479 | 1967 | 2390 | 2276 | 2403 | 2954 | 2861 | 2677 | 1986 | 2237 | 2029 | 1851 | 2374,72 | 2724 | 2026 | 0,743 | -0,428 | -5,694 | 1,24E-08 | 2,05E-08 | 0,0065258 | 0,0027827 | 0,0050049 | 0,0050049 | TRUE | 0,0782081 | No |
| orf19.6985 | 1824 | 1919 | 2034 | 1231 | 1110 | 815 | 1017 | 1030 | 1739 | 2029 | 2009 | 1928 | 889 | 927 | 863 | 837 | 1402,87 | 1926 | 879 | 0,456 | -1,133 | -17,934 | 6,41E-72 | 3,16E-71 | 0,0028623 | 0,0032617 | 0,0030303 | 0,0030303 | TRUE | 0,0540186 | No |
| orf19.6986 | 389 | 445 | 438 | 290 | 902 | 665 | 884 | 849 | 371 | 471 | 433 | 454 | 723 | 756 | 751 | 690 | 581,02 | 432 | 730 | 1,694 | 0,76 | 9,439 | 3,76E-21 | 8,61E-21 | 0,003504 | 0,0049172 | 0,0043129 | 0,0043129 | TRUE | 0,1120859 | No |
| orf19.6987 | 7097 | 6899 | 7230 | 4493 | 8328 | 6268 | 7387 | 8079 | 6766 | 7295 | 7143 | 7039 | 6673 | 7127 | 6272 | 6569 | 6860,36 | 7061 | 6660 | 0,943 | -0,085 | -1,808 | 0,0706055 | 0,083649 | 0,0017219 | 0,0023306 | 0,0019602 | 0,0019602 | TRUE | 0,0267486 | No |
| orf19.6988 | 483 | 449 | 523 | 288 | 842 | 622 | 766 | 741 | 460 | 475 | 517 | 451 | 675 | 707 | 650 | 603 | 567,23 | 476 | 659 | 1,379 | 0,464 | 6,119 | 9,41E-10 | 1,62E-09 | 0,002419 | 0,0049859 | 0,0036604 | 0,0036604 | TRUE | 0,0399264 | No |
| orf19.6989 | 947 | 858 | 974 | 585 | 1025 | 798 | 947 | 976 | 903 | 907 | 962 | 916 | 821 | 907 | 804 | 794 | 876,88 | 922 | 832 | 0,899 | -0,153 | -2,507 | 0,0121657 | 0,0153874 | 0,0010817 | 0,0039637 | 0,0024283 | 0,0024283 | TRUE | 0,0413838 | No |
| orf19.699 | 805 | 1362 | 1277 | 641 | 395 | 319 | 324 | 202 | 767 | 1440 | 1262 | 1004 | 317 | 363 | 275 | 164 | 698,99 | 1118 | 280 | 0,25 | -2,001 | -6,559 | 5,43E-11 | 9,77E-11 | 0,0873398 | 0,0044403 | 0,039269 | 0,0873398 | TRUE | 0,1784166 | No |
| orf19.6990 | 1165 | 1159 | 1166 | 720 | 2126 | 1533 | 1873 | 2099 | 1111 | 1225 | 1152 | 1128 | 1704 | 1743 | 1590 | 1707 | 1419,94 | 1154 | 1686 | 1,46 | 0,545 | 10,259 | 1,08E-24 | 2,66E-24 | 0,0010255 | 0,0032476 | 0,0019641 | 0,0019641 | TRUE | 0,0212265 | No |
| orf19.6991 | 4592 | 4115 | 4259 | 2290 | 5017 | 3301 | 4451 | 4848 | 4378 | 4351 | 4207 | 3587 | 4020 | 3754 | 3779 | 3942 | 4002,28 | 4131 | 3874 | 0,937 | -0,094 | -1,491 | 0,1358834 | 0,1565112 | 0,004228 | 0,0025015 | 0,0035808 | 0,0035808 | TRUE | 0,0925808 | No |
| orf19.6992 | 11882 | 10421 | 11239 | 7027 | 5198 | 3835 | 4995 | 5235 | 11328 | 11019 | 11103 | 11008 | 4165 | 4361 | 4241 | 4257 | 7685,17 | 11114 | 4256 | 0,383 | -1,386 | -45,644 | 0 | 0 | 9,73E-05 | 0,0023049 | 0,0007296 | 0,0007296 | TRUE | 0,0031753 | No |
| orf19.6993 | 12993 | 10874 | 11723 | 7632 | 21621 | 14911 | 20326 | 22040 | 12387 | 11498 | 11581 | 11956 | 17324 | 16955 | 17257 | 17921 | 14609,9 | 11856 | 17364 | 1,465 | 0,551 | 15,129 | 1,04E-51 | 3,92E-51 | 0,0007646 | 0,0022037 | 0,0011993 | 0,0011993 | TRUE | 0,0113929 | No |
| orf19.6994 | 5397 | 4502 | 4859 | 3383 | 9921 | 7167 | 9599 | 10516 | 5145 | 4760 | 4800 | 5300 | 7949 | 8150 | 8150 | 8551 | 6600,57 | 5001 | 8200 | 1,641 | 0,714 | 15,51 | 2,98E-54 | 1,16E-53 | 0,0016462 | 0,00234 | 0,0018696 | 0,0018696 | TRUE | 0,0187978 | No |
| orf19.6995 | 406 | 350 | 396 | 270 | 427 | 322 | 395 | 474 | 387 | 370 | 391 | 423 | 342 | 366 | 335 | 385 | 375,05 | 393 | 357 | 0,912 | -0,133 | -1,613 | 0,1066862 | 0,1241365 | 0,0009146 | 0,0064691 | 0,0038383 | 0,0038383 | TRUE | 0,0352437 | No |
| orf19.6996 | 4568 | 4594 | 4642 | 2832 | 3613 | 2779 | 3374 | 3765 | 4355 | 4858 | 4586 | 4436 | 2895 | 3160 | 2865 | 3061 | 3776,97 | 4559 | 2995 | 0,657 | -0,607 | -12,021 | 2,76E-33 | 7,87E-33 | 0,0018952 | 0,002526 | 0,0021762 | 0,0021762 | TRUE | 0,0238601 | No |
| orf19.6997 | 270 | 247 | 240 | 146 | 155 | 112 | 139 | 151 | 257 | 261 | 237 | 229 | 124 | 127 | 118 | 123 | 184,59 | 246 | 123 | 0,498 | -1,007 | -9,292 | 1,52E-20 | 3,45E-20 | 1,00E-08 | 0,010986 | 0,0053938 | 0,0053938 | TRUE | 0,0199823 | No |
| orf19.6998 | 17 | 11 | 25 | 16 | 13 | 7 | 13 | 15 | 16 | 12 | 25 | 25 | 10 | 8 | 11 | 12 | 14,9 | 20 | 10 | 0,549 | -0,866 | -2,228 | 0,0258712 | 0,0318572 | 1,00E-08 | 0,1122762 | 0,0737248 | 0,0737248 | TRUE | 0,2341566 | No |
| orf19.6999 | 3 | 14 | 13 | 5 | 5 | 4 | 5 | 3 | 3 | 15 | 13 | 8 | 4 | 5 | 4 | 2 | 6,7 | 10 | 4 | 0,393 | -1,348 | -2,151 | 0,0314597 | 0,0384567 | 1,00E-08 | 0,2472525 | 0,2009853 | 0,2009853 | TRUE | 0,2446471 | No |
| orf19.700 | 2001 | 1942 | 2045 | 1130 | 4539 | 2929 | 4324 | 4456 | 1908 | 2053 | 2020 | 1770 | 3637 | 3331 | 3671 | 3623 | 2751,67 | 1938 | 3566 | 1,838 | 0,878 | 15,671 | 2,40E-55 | 9,51E-55 | 0,0026312 | 0,002688 | 0,0025975 | 0,0025975 | TRUE | 0,0378329 | No |
| orf19.7001 | 13878 | 12731 | 13760 | 8770 | 10475 | 7262 | 10029 | 10400 | 13231 | 13461 | 13594 | 13739 | 8393 | 8258 | 8515 | 8456 | 10955,81 | 13506 | 8406 | 0,623 | -0,684 | -24,416 | 1,15E-131 | 1,01E-130 | 8,47E-05 | 0,0022411 | 0,0006578 | 0,0006578 | TRUE | 0,0023138 | No |
| orf19.7002 | 11 | 10 | 9 | 4 | 8 | 5 | 10 | 8 | 10 | 11 | 9 | 6 | 6 | 6 | 8 | 7 | 7,91 | 9 | 7 | 0,739 | -0,437 | -0,902 | 0,3671987 | 0,40211 | 1,00E-08 | 0,2095702 | 0,0996837 | 0,0996837 | TRUE | 0,0855196 | No |
| orf19.7006 | 7566 | 6403 | 7318 | 4580 | 2010 | 1339 | 1820 | 2006 | 7213 | 6770 | 7229 | 7175 | 1611 | 1523 | 1545 | 1631 | 4337,17 | 7097 | 1578 | 0,222 | -2,168 | -51,448 | 0 | 0 | 0,0007199 | 0,0024698 | 0,0013435 | 0,0013435 | TRUE | 0,0118432 | No |
| orf19.7007 | 23 | 17 | 21 | 12 | 8 | 4 | 13 | 10 | 22 | 18 | 21 | 19 | 6 | 5 | 11 | 8 | 13,7 | 20 | 8 | 0,384 | -1,38 | -3,473 | 0,0005154 | 0,0007075 | 1,00E-08 | 0,1219662 | 0,0658227 | 0,0658227 | TRUE | 0,3003006 | No |
| orf19.7009 | 110 | 106 | 99 | 66 | 209 | 163 | 198 | 211 | 105 | 112 | 98 | 103 | 167 | 185 | 168 | 172 | 138,83 | 104 | 173 | 1,652 | 0,724 | 5,9 | 3,64E-09 | 6,14E-09 | 1,00E-08 | 0,0139182 | 0,0065758 | 0,0065758 | TRUE | 0,0243169 | No |
| orf19.701 | 31 | 38 | 27 | 19 | 20 | 17 | 24 | 23 | 30 | 40 | 27 | 30 | 16 | 19 | 20 | 19 | 25,08 | 32 | 18 | 0,588 | -0,766 | -2,688 | 0,0071912 | 0,0091995 | 1,00E-08 | 0,0675689 | 0,0364298 | 0,0364298 | TRUE | 0,2356973 | No |
| orf19.7010 | 1810 | 1549 | 1705 | 1130 | 2249 | 1572 | 2112 | 2301 | 1726 | 1638 | 1684 | 1770 | 1802 | 1788 | 1793 | 1871 | 1758,96 | 1704 | 1814 | 1,066 | 0,092 | 2,069 | 0,0385856 | 0,0468623 | 0,0001158 | 0,0030247 | 0,0013113 | 0,0013113 | TRUE | 0,008116 | No |
| orf19.7011 | 13436 | 12240 | 13300 | 8077 | 24671 | 17889 | 23164 | 25455 | 12810 | 12942 | 13139 | 12653 | 19768 | 20341 | 19666 | 20697 | 16502,16 | 12886 | 20118 | 1,561 | 0,642 | 20,886 | 7,12E-97 | 4,50E-96 | 0,0003362 | 0,0021908 | 0,0008424 | 0,0008424 | TRUE | 0,0046492 | No |
| orf19.7012 | 2124 | 1866 | 1799 | 1226 | 2482 | 1638 | 2433 | 2509 | 2025 | 1973 | 1777 | 1921 | 1989 | 1863 | 2066 | 2040 | 1956,61 | 1924 | 1990 | 1,035 | 0,049 | 0,887 | 0,375048 | 0,4101424 | 0,0018867 | 0,0029304 | 0,002441 | 0,002441 | TRUE | 0,0333012 | No |
| orf19.7013 | 235 | 278 | 296 | 203 | 355 | 333 | 336 | 364 | 224 | 294 | 292 | 318 | 284 | 379 | 285 | 296 | 296,59 | 282 | 311 | 1,104 | 0,142 | 1,114 | 0,2652037 | 0,2959022 | 0,0163954 | 0,0076271 | 0,0122191 | 0,0122191 | TRUE | 0,2299294 | No |
| orf19.7014 | 20 | 6 | 9 | 18 | 53 | 32 | 35 | 29 | 19 | 6 | 9 | 28 | 42 | 36 | 30 | 24 | 24,33 | 16 | 33 | 2,187 | 1,129 | 2,846 | 0,0044313 | 0,0057448 | 0,1520156 | 0,0695725 | 0,1005593 | 0,1005593 | TRUE | 0,5956599 | No |
| orf19.7015 | 13605 | 10792 | 11269 | 8168 | 51082 | 34646 | 46025 | 54617 | 12971 | 11411 | 11133 | 12796 | 40931 | 39396 | 39075 | 44409 | 26515,05 | 12078 | 40953 | 3,391 | 1,762 | 28,312 | 2,43E-176 | 2,84E-175 | 0,0047353 | 0,0021532 | 0,0036629 | 0,0036629 | TRUE | 0,0395678 | No |
| orf19.7016 | 2767 | 2553 | 2753 | 1710 | 4255 | 2870 | 4045 | 4260 | 2638 | 2699 | 2720 | 2679 | 3409 | 3263 | 3434 | 3464 | 3038,36 | 2684 | 3392 | 1,265 | 0,339 | 9,079 | 1,10E-19 | 2,44E-19 | 1,00E-08 | 0,0026317 | 0,0009981 | 0,0009981 | TRUE | 0,0075589 | No |
| orf19.7017 | 86 | 70 | 92 | 52 | 26 | 15 | 29 | 24 | 82 | 74 | 91 | 81 | 21 | 17 | 25 | 20 | 51,3 | 82 | 21 | 0,252 | -1,991 | -9,038 | 1,60E-19 | 3,56E-19 | 1,00E-08 | 0,0340987 | 0,0183753 | 0,0183753 | TRUE | 0,1053309 | No |
| orf19.7018 | 16107 | 13885 | 14334 | 10241 | 43339 | 29852 | 41543 | 44229 | 15356 | 14682 | 14161 | 16043 | 34726 | 33944 | 35270 | 35962 | 25018,07 | 15060 | 34976 | 2,323 | 1,216 | 27,447 | 7,52E-166 | 8,40E-165 | 0,0017087 | 0,0021569 | 0,0018356 | 0,0018356 | TRUE | 0,0232491 | No |
| orf19.7019 | 2283 | 2036 | 2273 | 1533 | 3389 | 2161 | 2954 | 3104 | 2177 | 2153 | 2246 | 2402 | 2716 | 2457 | 2508 | 2524 | 2397,62 | 2244 | 2551 | 1,139 | 0,188 | 3,661 | 0,0002514 | 0,0003495 | 0,001673 | 0,0027761 | 0,0021065 | 0,0021065 | TRUE | 0,0250552 | No |
| orf19.702 | 6542 | 5993 | 6461 | 3833 | 7292 | 5097 | 6957 | 7082 | 6237 | 6337 | 6383 | 6005 | 5843 | 5796 | 5906 | 5758 | 6033,1 | 6240 | 5826 | 0,933 | -0,1 | -3,056 | 0,0022441 | 0,0029647 | 0,0002036 | 0,0023634 | 0,0008651 | 0,0008651 | TRUE | 0,0073671 | No |
| orf19.7020 | 7749 | 7253 | 7700 | 4772 | 6921 | 4893 | 6642 | 7014 | 7388 | 7669 | 7607 | 7476 | 5546 | 5564 | 5639 | 5703 | 6573,86 | 7535 | 5613 | 0,745 | -0,425 | -14,08 | 5,05E-45 | 1,72E-44 | 9,53E-05 | 0,002341 | 0,0007197 | 0,0007197 | TRUE | 0,0022289 | No |
| orf19.7021 | 33380 | 30412 | 31617 | 19542 | 12110 | 8599 | 10805 | 12181 | 31824 | 32157 | 31235 | 30614 | 9703 | 9778 | 9173 | 9904 | 20548,52 | 31458 | 9640 | 0,306 | -1,707 | -48,145 | 0 | 0 | 0,0007001 | 0,0021712 | 0,0011426 | 0,0011426 | TRUE | 0,0130342 | No |
| orf19.7022 | 1326 | 1250 | 1287 | 738 | 378 | 289 | 371 | 354 | 1264 | 1322 | 1271 | 1156 | 303 | 329 | 315 | 288 | 780,97 | 1253 | 309 | 0,245 | -2,029 | -28,984 | 1,05E-184 | 1,31E-183 | 0,0014075 | 0,0041937 | 0,0028143 | 0,0028143 | TRUE | 0,0294213 | No |
| orf19.7023 | 272 | 264 | 266 | 176 | 579 | 392 | 562 | 577 | 259 | 279 | 263 | 276 | 464 | 446 | 477 | 469 | 366,62 | 269 | 464 | 1,729 | 0,79 | 10,056 | 8,67E-24 | 2,10E-23 | 1,00E-08 | 0,0065698 | 0,0028999 | 0,0028999 | TRUE | 0,0081016 | No |
| orf19.7024 | 6 | 3 | 4 | 4 | 16 | 9 | 15 | 19 | 6 | 3 | 4 | 6 | 13 | 10 | 13 | 15 | 8,79 | 5 | 13 | 2,752 | 1,461 | 2,879 | 0,0039948 | 0,0052001 | 1,00E-08 | 0,1888086 | 0,0946478 | 0,0946478 | TRUE | 0,0925018 | No |
| orf19.7025 | 2166 | 1981 | 2106 | 1430 | 1247 | 830 | 1135 | 1162 | 2065 | 2095 | 2081 | 2240 | 999 | 944 | 964 | 945 | 1541,47 | 2120 | 963 | 0,455 | -1,135 | -23,28 | 7,05E-120 | 5,61E-119 | 0,0003688 | 0,0031564 | 0,0015553 | 0,0015553 | TRUE | 0,0159961 | No |
| orf19.7027 | 14477 | 12915 | 14901 | 8440 | 1295 | 841 | 1177 | 1289 | 13802 | 13656 | 14721 | 13222 | 1038 | 956 | 999 | 1048 | 7430,25 | 13850 | 1010 | 0,073 | -3,775 | -76,59 | 0 | 0 | 0,0015682 | 0,0023123 | 0,0018519 | 0,0018519 | TRUE | 0,0219326 | No |
| orf19.7029 | 168 | 182 | 178 | 123 | 227 | 133 | 156 | 196 | 160 | 192 | 176 | 193 | 182 | 151 | 132 | 159 | 168,26 | 180 | 156 | 0,872 | -0,198 | -1,55 | 0,1210281 | 0,1400567 | 0,0073405 | 0,0118494 | 0,009734 | 0,009734 | TRUE | 0,1381068 | No |
| orf19.703 | 1682 | 1396 | 1479 | 1009 | 818 | 566 | 782 | 826 | 1604 | 1476 | 1461 | 1581 | 655 | 644 | 664 | 672 | 1094,5 | 1530 | 659 | 0,431 | -1,214 | -21,801 | 2,29E-105 | 1,56E-104 | 1,00E-08 | 0,0035914 | 0,0019348 | 0,0019348 | TRUE | 0,0139703 | No |
| orf19.7030 | 2105 | 1646 | 1944 | 1404 | 4374 | 3758 | 4021 | 4414 | 2007 | 1740 | 1920 | 2199 | 3505 | 4250 | 3414 | 3589 | 2828,16 | 1966 | 3690 | 1,877 | 0,908 | 10,96 | 5,92E-28 | 1,55E-27 | 0,0095061 | 0,0026718 | 0,0061898 | 0,0061898 | TRUE | 0,1262792 | No |
| orf19.7032 | 1045 | 1307 | 1229 | 780 | 1758 | 1402 | 1574 | 1506 | 996 | 1382 | 1214 | 1222 | 1409 | 1594 | 1336 | 1225 | 1297,25 | 1204 | 1391 | 1,155 | 0,208 | 2,065 | 0,0388859 | 0,0472001 | 0,0138529 | 0,003357 | 0,0089728 | 0,0089728 | TRUE | 0,1649752 | No |
| orf19.7033 | 1349 | 1591 | 1550 | 943 | 2178 | 1637 | 1943 | 1835 | 1286 | 1682 | 1531 | 1477 | 1745 | 1861 | 1650 | 1492 | 1590,64 | 1494 | 1687 | 1,128 | 0,174 | 1,992 | 0,0463523 | 0,055912 | 0,0094425 | 0,0031235 | 0,0067178 | 0,0067178 | TRUE | 0,1088077 | No |
| orf19.7034 | 2684 | 2573 | 2920 | 1789 | 2074 | 1397 | 1914 | 2051 | 2559 | 2721 | 2885 | 2803 | 1662 | 1589 | 1625 | 1668 | 2188,72 | 2742 | 1636 | 0,597 | -0,743 | -15,128 | 1,06E-51 | 4,01E-51 | 0,0012346 | 0,0028414 | 0,0018406 | 0,0018406 | TRUE | 0,0253052 | No |
| orf19.7035 | 1186 | 1168 | 1241 | 759 | 1356 | 1056 | 1262 | 1358 | 1131 | 1235 | 1226 | 1189 | 1087 | 1201 | 1071 | 1104 | 1155,46 | 1195 | 1116 | 0,932 | -0,101 | -1,784 | 0,0744035 | 0,0880016 | 0,0012317 | 0,0035123 | 0,0022362 | 0,0022362 | TRUE | 0,0294376 | No |
| orf19.7036 | 3835 | 3811 | 4125 | 2352 | 3935 | 2722 | 3552 | 3562 | 3656 | 4030 | 4075 | 3685 | 3153 | 3095 | 3016 | 2896 | 3450,7 | 3862 | 3040 | 0,787 | -0,346 | -6,731 | 1,68E-11 | 3,08E-11 | 0,0019236 | 0,0025671 | 0,002245 | 0,002245 | TRUE | 0,0169963 | No |
| orf19.7037 | 954 | 904 | 902 | 616 | 1361 | 944 | 1233 | 1240 | 910 | 956 | 891 | 965 | 1091 | 1073 | 1047 | 1008 | 992,56 | 930 | 1055 | 1,135 | 0,183 | 3,329 | 0,000871 | 0,0011803 | 1,00E-08 | 0,0037455 | 0,0018834 | 0,0018834 | TRUE | 0,0109194 | No |
| orf19.7038 | 785 | 913 | 915 | 559 | 1161 | 853 | 1056 | 1051 | 748 | 965 | 904 | 876 | 930 | 970 | 897 | 855 | 883,09 | 873 | 913 | 1,045 | 0,063 | 0,791 | 0,4290537 | 0,4651293 | 0,006021 | 0,0039297 | 0,0050505 | 0,0050505 | TRUE | 0,1160022 | No |
| orf19.704 | 2418 | 1993 | 2135 | 1417 | 2727 | 1859 | 2583 | 2775 | 2305 | 2107 | 2109 | 2220 | 2185 | 2114 | 2193 | 2256 | 2186,23 | 2185 | 2187 | 1,002 | 0,002 | 0,048 | 0,9617854 | 0,9654165 | 0,0008587 | 0,0028423 | 0,0016227 | 0,0016227 | TRUE | 0,0177851 | No |
| orf19.7041 | 4271 | 4055 | 4240 | 2704 | 3969 | 2838 | 3639 | 3760 | 4072 | 4288 | 4189 | 4236 | 3180 | 3227 | 3089 | 3057 | 3667,29 | 4196 | 3138 | 0,748 | -0,419 | -11,304 | 1,26E-29 | 3,38E-29 | 0,0002904 | 0,002539 | 0,0010437 | 0,0010437 | TRUE | 0,0049882 | No |
| orf19.7042 | 1119 | 1105 | 1079 | 712 | 431 | 278 | 365 | 363 | 1067 | 1168 | 1066 | 1115 | 345 | 316 | 310 | 295 | 710,38 | 1104 | 316 | 0,287 | -1,801 | -25,478 | 3,46E-143 | 3,34E-142 | 0,0013395 | 0,0044026 | 0,002883 | 0,002883 | TRUE | 0,0464629 | No |
| orf19.7043 | 38 | 65 | 40 | 20 | 98 | 54 | 67 | 32 | 36 | 69 | 40 | 31 | 79 | 61 | 57 | 26 | 49,83 | 44 | 56 | 1,259 | 0,332 | 1,003 | 0,3161009 | 0,3493314 | 0,1403431 | 0,0350418 | 0,0845877 | 0,0845877 | TRUE | 0,4791289 | No |
| orf19.7043.1 | 739 | 749 | 662 | 450 | 4991 | 3301 | 4293 | 3738 | 705 | 792 | 654 | 705 | 3999 | 3754 | 3645 | 3039 | 2161,53 | 714 | 3609 | 5,056 | 2,338 | 26,709 | 3,67E-157 | 3,91E-156 | 0,0096022 | 0,0028509 | 0,0064451 | 0,0064451 | TRUE | 0,1383544 | No |
| orf19.7044 | 4141 | 3632 | 4062 | 2444 | 3405 | 2466 | 3168 | 3386 | 3948 | 3840 | 4013 | 3829 | 2728 | 2804 | 2690 | 2753 | 3325,63 | 3908 | 2744 | 0,701 | -0,512 | -14,143 | 2,08E-45 | 7,13E-45 | 1,00E-08 | 0,002585 | 0,0009501 | 0,0009501 | TRUE | 0,0040141 | No |
| orf19.7046 | 3637 | 3502 | 3506 | 2123 | 5046 | 3487 | 4667 | 4774 | 3467 | 3703 | 3464 | 3326 | 4043 | 3965 | 3962 | 3882 | 3726,5 | 3490 | 3963 | 1,135 | 0,182 | 4,369 | 1,25E-05 | 1,84E-05 | 0,000785 | 0,0025319 | 0,0013944 | 0,0013944 | TRUE | 0,0203213 | No |
| orf19.7047 | 3249 | 3131 | 3359 | 1943 | 6553 | 4318 | 6433 | 6454 | 3098 | 3311 | 3318 | 3044 | 5251 | 4910 | 5462 | 5248 | 4205,05 | 3193 | 5218 | 1,634 | 0,708 | 14,695 | 6,94E-49 | 2,52E-48 | 0,0016157 | 0,0024817 | 0,0019677 | 0,0019677 | TRUE | 0,0187657 | No |
| orf19.7048.1 | 5817 | 4405 | 4654 | 3228 | 18430 | 11842 | 16948 | 18228 | 5546 | 4658 | 4598 | 5057 | 14767 | 13465 | 14389 | 14821 | 9662,61 | 4965 | 14360 | 2,893 | 1,533 | 24,486 | 2,09E-132 | 1,86E-131 | 0,0046621 | 0,0022612 | 0,0036184 | 0,0036184 | TRUE | 0,077097 | No |
| orf19.7049 | 911 | 902 | 898 | 592 | 6455 | 4492 | 5608 | 5798 | 869 | 954 | 887 | 927 | 5172 | 5108 | 4761 | 4714 | 2924,04 | 909 | 4939 | 5,438 | 2,443 | 45,467 | 0 | 0 | 0,0016814 | 0,0026528 | 0,0020654 | 0,0020654 | TRUE | 0,0142191 | No |

| | | | | | | | | | | | | | | | | | | | | | | | | | | | | | | | |
|---|---|---|---|---|---|---|---|---|---|---|---|---|---|---|---|---|---|---|---|---|---|---|---|---|---|---|---|---|---|---|---|
| orf19.705 | 1553 | 1231 | 1459 | 816 | 1744 | 1179 | 1694 | 1884 | 1481 | 1302 | 1441 | 1278 | 1397 | 1341 | 1438 | 1532 | 1401,25 | 1376 | 1427 | 1,036 | 0,051 | 0,789 | 0,4302882 | 0,466316 | 0,0032778 | 0,003263 | 0,0033574 | 0,0033574 | TRUE | 0,0317885 | No |
| orf19.7050 | 9265 | 8639 | 8712 | 5089 | 11382 | 7326 | 10422 | 11031 | 8833 | 9135 | 8607 | 7972 | 9120 | 8330 | 8848 | 8969 | 8726,82 | 8637 | 8817 | 1,021 | 0,029 | 0,591 | 0,5545733 | 0,589494 | 0,0020345 | 0,0022794 | 0,0022637 | 0,0022637 | TRUE | 0,0320174 | No |
| orf19.7051 | 98 | 123 | 142 | 70 | 337 | 223 | 300 | 265 | 93 | 130 | 140 | 110 | 270 | 254 | 255 | 215 | 183,4 | 118 | 248 | 2,094 | 1,066 | 7,722 | 1,15E-14 | 2,29E-14 | 0,0127659 | 0,0110439 | 0,0117899 | 0,0117899 | TRUE | 0,2342005 | No |
| orf19.7052 | 786 | 782 | 817 | 454 | 1829 | 1339 | 1719 | 1806 | 749 | 827 | 807 | 711 | 1466 | 1523 | 1459 | 1468 | 1126,32 | 774 | 1479 | 1,904 | 0,929 | 16,315 | 7,67E-60 | 3,22E-59 | 0,0004069 | 0,003549 | 0,0020929 | 0,0020929 | TRUE | 0,0290934 | No |
| orf19.7053 | 41900 | 40319 | 43666 | 27996 | 14764 | 11188 | 13785 | 15066 | 39947 | 42632 | 43138 | 43857 | 11830 | 12722 | 11703 | 12250 | 27259,96 | 42394 | 12126 | 0,286 | -1,806 | -42,247 | 0 | 0 | 0,0014611 | 0,0021515 | 0,0017052 | 0,0017052 | TRUE | 0,0185243 | No |
| orf19.7054 | 54 | 43 | 43 | 38 | 8 | 4 | 10 | 16 | 51 | 45 | 42 | 60 | 6 | 5 | 8 | 13 | 28,93 | 50 | 8 | 0,168 | -2,57 | -7,744 | 9,61E-15 | 1,92E-14 | 0,0008154 | 0,0588512 | 0,0411046 | 0,0411046 | TRUE | 0,5708054 | No |
| orf19.7055 | 0 | 5 | 2 | 0 | 14 | 10 | 10 | 18 | 0 | 5 | 2 | 0 | 11 | 11 | 8 | 15 | 6,62 | 2 | 11 | 6,006 | 2,586 | 3,515 | 0,0004405 | 0,0006059 | 1,58E-08 | 0,2500331 | 0,1908359 | 0,1908359 | TRUE | 1,2466369 | No |
| orf19.7056 | 316 | 257 | 295 | 199 | 166 | 106 | 164 | 185 | 301 | 272 | 291 | 312 | 133 | 121 | 139 | 150 | 214,92 | 294 | 136 | 0,465 | -1,104 | -10,415 | 2,13E-25 | 5,30E-25 | 1,00E-08 | 0,0097307 | 0,0056347 | 0,0056347 | TRUE | 0,0548716 | No |
| orf19.7057 | 6279 | 5103 | 5628 | 3570 | 10387 | 7352 | 10120 | 10993 | 5986 | 5396 | 5560 | 5593 | 8323 | 8360 | 8592 | 8938 | 7093,44 | 5634 | 8553 | 1,518 | 0,602 | 13,736 | 6,21E-43 | 2,06E-42 | 0,0013389 | 0,0023227 | 0,0016939 | 0,0016939 | TRUE | 0,0221501 | No |
| orf19.7058 | 1344 | 1142 | 1214 | 863 | 1815 | 1313 | 1787 | 1854 | 1281 | 1208 | 1199 | 1352 | 1454 | 1493 | 1517 | 1507 | 1376,51 | 1260 | 1493 | 1,188 | 0,248 | 4,786 | 1,71E-06 | 2,61E-06 | 0,0006182 | 0,0032841 | 0,0018406 | 0,0018406 | TRUE | 0,0253888 | No |
| orf19.7059 | 872 | 740 | 826 | 510 | 1689 | 1179 | 1625 | 1737 | 831 | 782 | 816 | 799 | 1353 | 1341 | 1380 | 1412 | 1089,34 | 807 | 1372 | 1,698 | 0,764 | 14,966 | 1,23E-50 | 4,55E-50 | 1,00E-08 | 0,0035985 | 0,0014881 | 0,0014881 | TRUE | 0,0055988 | No |
| orf19.706 | 21533 | 17923 | 19676 | 12211 | 21973 | 14449 | 21101 | 22666 | 20529 | 18951 | 19438 | 19129 | 17606 | 16430 | 17915 | 18430 | 18553,54 | 19512 | 17595 | 0,902 | -0,149 | -3,3 | 0,0009681 | 0,0013076 | 0,0017462 | 0,0021798 | 0,0019075 | 0,0019075 | TRUE | 0,024154 | No |
| orf19.7060 | 1566 | 1322 | 1503 | 908 | 482 | 330 | 485 | 538 | 1493 | 1398 | 1485 | 1422 | 386 | 375 | 412 | 437 | 926,1 | 1450 | 402 | 0,278 | -1,845 | -29,026 | 3,09E-185 | 3,85E-184 | 0,0007833 | 0,0038642 | 0,0023916 | 0,0023916 | TRUE | 0,0397365 | No |
| orf19.7061 | 150 | 107 | 141 | 84 | 133 | 105 | 121 | 139 | 143 | 113 | 139 | 132 | 107 | 119 | 103 | 113 | 121,09 | 132 | 110 | 0,834 | -0,261 | -1,954 | 0,0507403 | 0,0609973 | 1,00E-08 | 0,0156503 | 0,0089166 | 0,0089166 | TRUE | 0,0985239 | No |
| orf19.7062 | 14493 | 12405 | 13546 | 8571 | 27198 | 18407 | 25417 | 27803 | 13818 | 13117 | 13382 | 13427 | 21793 | 20930 | 21579 | 22606 | 17581,51 | 13436 | 21727 | 1,617 | 0,694 | 19,818 | 2,08E-87 | 1,22E-86 | 0,0006573 | 0,0021847 | 0,0011137 | 0,0011137 | TRUE | 0,0090862 | No |
| orf19.7063 | 709 | 580 | 624 | 429 | 1436 | 990 | 1354 | 1459 | 676 | 613 | 616 | 672 | 1151 | 1126 | 1150 | 1186 | 898,74 | 644 | 1153 | 1,793 | 0,843 | 14,569 | 4,40E-48 | 1,58E-47 | 3,05E-05 | 0,0039182 | 0,0019581 | 0,0019581 | TRUE | 0,0155805 | No |
| orf19.7064 | 7770 | 6871 | 7364 | 4869 | 10667 | 7200 | 10071 | 11060 | 7408 | 7265 | 7275 | 7628 | 8547 | 8187 | 8550 | 8993 | 7981,6 | 7394 | 8569 | 1,16 | 0,214 | 5,508 | 3,62E-08 | 5,92E-08 | 0,0008334 | 0,002297 | 0,0013187 | 0,0013187 | TRUE | 0,0135275 | No |
| orf19.7065 | 746 | 654 | 647 | 378 | 884 | 514 | 789 | 869 | 711 | 692 | 639 | 592 | 708 | 584 | 670 | 707 | 662,91 | 658 | 667 | 1,013 | 0,018 | 0,22 | 0,8261564 | 0,8465881 | 0,0053808 | 0,0045681 | 0,0049801 | 0,0049801 | TRUE | 0,0774519 | No |
| orf19.7067 | 2841 | 2562 | 2777 | 1720 | 5138 | 3624 | 4998 | 5204 | 2709 | 2709 | 2743 | 2694 | 4117 | 4121 | 4243 | 4231 | 3445,98 | 2714 | 4178 | 1,539 | 0,622 | 18,314 | 6,37E-75 | 3,25E-74 | 1,00E-08 | 0,0025677 | 0,0007955 | 0,0007955 | TRUE | 0,0013337 | No |
| orf19.7068 | 486 | 389 | 438 | 293 | 1146 | 841 | 1151 | 1209 | 463 | 411 | 433 | 459 | 918 | 956 | 977 | 983 | 700,14 | 442 | 958 | 2,174 | 1,12 | 17,248 | 1,15E-66 | 5,29E-66 | 1,00E-08 | 0,0044364 | 0,0023231 | 0,0023231 | TRUE | 0,0247632 | No |
| orf19.7069 | 3156 | 2370 | 2554 | 1822 | 5618 | 3619 | 5422 | 5647 | 3009 | 2506 | 2523 | 2854 | 4502 | 4115 | 4603 | 4592 | 3587,97 | 2723 | 4453 | 1,636 | 0,71 | 10,574 | 3,95E-26 | 1,00E-25 | 0,0051172 | 0,0025489 | 0,0040289 | 0,0040289 | TRUE | 0,0627365 | No |
| orf19.707 | 1045 | 889 | 956 | 619 | 528 | 390 | 518 | 561 | 996 | 940 | 944 | 970 | 423 | 443 | 440 | 456 | 701,61 | 962 | 440 | 0,458 | -1,128 | -18,52 | 1,41E-76 | 7,37E-76 | 1,00E-08 | 0,0044315 | 0,0019697 | 0,0019697 | TRUE | 0,0090489 | No |
| orf19.7070 | 6 | 8 | 10 | 4 | 10 | 7 | 6 | 11 | 6 | 8 | 10 | 6 | 8 | 8 | 5 | 9 | 7,54 | 8 | 8 | 0,98 | -0,029 | -0,057 | 0,9547382 | 0,9592481 | 1,00E-08 | 0,2197963 | 0,1110717 | 0,1110717 | TRUE | 0,1551667 | No |
| orf19.7071 | 11 | 10 | 5 | 8 | 3 | 3 | 3 | 2 | 10 | 11 | 5 | 13 | 2 | 3 | 3 | 2 | 6,07 | 10 | 2 | 0,258 | -1,956 | -3,04 | 0,002367 | 0,0031238 | 1,00E-08 | 0,2728021 | 0,1574226 | 0,1574226 | TRUE | 0,386521 | No |
| orf19.7072 | 7753 | 6593 | 7525 | 4759 | 5150 | 3744 | 5056 | 5159 | 7392 | 6971 | 7434 | 7455 | 4127 | 4257 | 4293 | 4195 | 5765,4 | 7313 | 4218 | 0,577 | -0,794 | -21,842 | 9,21E-106 | 6,33E-105 | 0,0005092 | 0,0023761 | 0,0010857 | 0,0010857 | TRUE | 0,012165 | No |
| orf19.7073 | 229 | 251 | 231 | 153 | 398 | 287 | 358 | 377 | 218 | 265 | 228 | 240 | 319 | 326 | 304 | 307 | 275,92 | 238 | 314 | 1,32 | 0,401 | 4,319 | 1,57E-05 | 2,30E-05 | 1,00E-08 | 0,008042 | 0,0045108 | 0,0045108 | TRUE | 0,0723879 | No |
| orf19.7074 | 867 | 796 | 804 | 538 | 781 | 547 | 760 | 812 | 827 | 842 | 794 | 843 | 626 | 622 | 645 | 660 | 732,32 | 826 | 638 | 0,774 | -0,37 | -6,374 | 1,84E-10 | 3,27E-10 | 1,00E-08 | 0,0043333 | 0,0018553 | 0,0018553 | TRUE | 0,0083731 | No |
| orf19.7076 | 5436 | 4698 | 5131 | 2995 | 6820 | 4374 | 6196 | 6604 | 5183 | 4968 | 5069 | 4692 | 5465 | 4974 | 5260 | 5370 | 5122,41 | 4978 | 5267 | 1,058 | 0,081 | 1,772 | 0,0764344 | 0,09032 | 0,0014438 | 0,0024118 | 0,0018062 | 0,0018062 | TRUE | 0,0174478 | No |
| orf19.7077 | 2406 | 2040 | 2245 | 1513 | 4643 | 3326 | 4569 | 5003 | 2294 | 2157 | 2218 | 2370 | 3720 | 3782 | 3879 | 4068 | 3061,02 | 2260 | 3862 | 1,711 | 0,775 | 16,527 | 2,33E-61 | 1,00E-60 | 0,001143 | 0,0026277 | 0,0017476 | 0,0017476 | TRUE | 0,0158315 | No |
| orf19.7078 | 565 | 443 | 464 | 275 | 338 | 219 | 343 | 338 | 539 | 468 | 458 | 431 | 271 | 249 | 291 | 275 | 372,77 | 474 | 272 | 0,572 | -0,806 | -8,774 | 1,73E-18 | 3,75E-18 | 0,0040625 | 0,0064959 | 0,0052924 | 0,0052924 | TRUE | 0,1000273 | No |
| orf19.7079 | 165 | 140 | 187 | 128 | 399 | 313 | 359 | 374 | 157 | 148 | 185 | 201 | 320 | 356 | 305 | 304 | 246,89 | 173 | 321 | 1,87 | 0,903 | 8,085 | 6,20E-16 | 1,27E-15 | 0,0053374 | 0,0087417 | 0,007334 | 0,007334 | TRUE | 0,1000393 | No |
| orf19.708 | 399 | 360 | 426 | 226 | 228 | 167 | 217 | 247 | 380 | 381 | 421 | 354 | 183 | 190 | 184 | 201 | 286,7 | 384 | 190 | 0,491 | -1,025 | -11,063 | 1,91E-28 | 5,01E-28 | 1,00E-08 | 0,0078182 | 0,0044413 | 0,0044413 | TRUE | 0,0479714 | No |
| orf19.7080 | 788 | 682 | 644 | 579 | 2104 | 1541 | 2031 | 2194 | 751 | 721 | 636 | 907 | 1686 | 1752 | 1724 | 1784 | 1245,25 | 754 | 1736 | 2,314 | 1,211 | 14,173 | 1,35E-45 | 4,66E-45 | 0,0088257 | 0,0034098 | 0,0059994 | 0,0059994 | TRUE | 0,2040825 | No |
| orf19.7081 | 18219 | 14872 | 16153 | 11126 | 10051 | 7125 | 9611 | 10317 | 17370 | 15725 | 15958 | 17429 | 8054 | 8102 | 8160 | 8389 | 12398,24 | 16620 | 8176 | 0,492 | -1,023 | -23,063 | 1,08E-117 | 8,41E-117 | 0,0015252 | 0,0022237 | 0,001802 | 0,001802 | TRUE | 0,0160626 | No |
| orf19.7082 | 114 | 97 | 107 | 78 | 104 | 83 | 120 | 134 | 109 | 103 | 106 | 122 | 83 | 94 | 102 | 109 | 103,46 | 110 | 97 | 0,891 | -0,167 | -1,151 | 0,2497176 | 0,2798469 | 1,00E-08 | 0,0179609 | 0,0105852 | 0,0105852 | TRUE | 0,1016119 | No |
| orf19.7083 | 78 | 82 | 61 | 65 | 136 | 112 | 123 | 127 | 74 | 87 | 60 | 102 | 109 | 127 | 104 | 103 | 95,9 | 81 | 111 | 1,389 | 0,474 | 2,794 | 0,0052104 | 0,0067275 | 0,0125579 | 0,0192129 | 0,0165915 | 0,0165915 | TRUE | 0,2747839 | No |
| orf19.7084 | 19530 | 16783 | 18836 | 11159 | 6019 | 4138 | 5987 | 6039 | 18620 | 17746 | 18608 | 17481 | 4823 | 4705 | 5083 | 4910 | 11497,04 | 18114 | 4880 | 0,269 | -1,892 | -48,439 | 0 | 0 | 0,0008839 | 0,0022341 | 0,0013425 | 0,0013425 | TRUE | 0,0094802 | No |
| orf19.7085 | 13245 | 11224 | 12349 | 8677 | 569 | 437 | 578 | 622 | 12628 | 11868 | 12200 | 13593 | 456 | 497 | 491 | 506 | 6529,69 | 12572 | 488 | 0,039 | -4,69 | -78,583 | 0 | 0 | 0,0025457 | 0,0023427 | 0,0024617 | 0,0024617 | TRUE | 0,0360437 | No |
| orf19.7086 | 7339 | 6452 | 6870 | 4375 | 1334 | 919 | 1171 | 1393 | 6997 | 6822 | 6787 | 6854 | 1069 | 1045 | 994 | 1133 | 3962,55 | 6865 | 1060 | 0,155 | -2,694 | -59,978 | 0 | 0 | 0,0004697 | 0,0025056 | 0,0014351 | 0,0014351 | TRUE | 0,026356 | No |
| orf19.7088 | 3252 | 2623 | 2987 | 1920 | 3309 | 2199 | 3153 | 3330 | 3100 | 2773 | 2951 | 3008 | 2651 | 2500 | 2677 | 2708 | 2796,12 | 2958 | 2634 | 0,891 | -0,166 | -3,466 | 0,0005277 | 0,000724 | 0,0012144 | 0,0026785 | 0,0018489 | 0,0018489 | TRUE | 0,0217924 | No |
| orf19.7089 | 1800 | 1515 | 1735 | 1065 | 4277 | 2996 | 4243 | 4622 | 1716 | 1602 | 1714 | 1668 | 3427 | 3407 | 3602 | 3758 | 2611,82 | 1675 | 3548 | 2,118 | 1,083 | 22,09 | 3,92E-108 | 2,79E-107 | 0,0013309 | 0,0027199 | 0,0018467 | 0,0018467 | TRUE | 0,0193643 | No |
| orf19.709 | 4441 | 3740 | 3988 | 2446 | 5519 | 3886 | 5344 | 5679 | 4234 | 3955 | 3940 | 3832 | 4422 | 4419 | 4537 | 4618 | 4244,46 | 3990 | 4499 | 1,127 | 0,172 | 4,136 | 3,54E-05 | 5,11E-05 | 0,0008389 | 0,0024781 | 0,0014224 | 0,0014224 | TRUE | 0,0209148 | No |
| orf19.7091 | 999 | 744 | 874 | 660 | 1354 | 1000 | 1256 | 1326 | 952 | 787 | 863 | 1034 | 1085 | 1137 | 1066 | 1078 | 1000,37 | 909 | 1092 | 1,204 | 0,267 | 3,445 | 0,0005702 | 0,0007807 | 0,0060294 | 0,0037326 | 0,0047556 | 0,0047556 | TRUE | 0,0963847 | No |
| orf19.7092 | 1665 | 1359 | 1464 | 972 | 3205 | 2086 | 3008 | 3095 | 1587 | 1437 | 1446 | 1523 | 2568 | 2372 | 2554 | 2517 | 2000,46 | 1498 | 2503 | 1,672 | 0,741 | 14,726 | 4,40E-49 | 1,60E-48 | 0,0010452 | 0,002912 | 0,0018807 | 0,0018807 | TRUE | 0,0210094 | No |
| orf19.7093 | 333 | 282 | 316 | 190 | 2622 | 1726 | 2555 | 2680 | 317 | 298 | 312 | 298 | 2101 | 1963 | 2169 | 2179 | 1204,66 | 306 | 2103 | 6,853 | 2,777 | 41,781 | 0 | 0 | 1,00E-08 | 0,0034542 | 0,0022214 | 0,0022214 | TRUE | 0,0233419 | No |
| orf19.7094 | 12 | 16 | 15 | 10 | 24 | 13 | 30 | 32 | 11 | 17 | 15 | 16 | 19 | 15 | 25 | 26 | 18,04 | 15 | 21 | 1,478 | 0,564 | 1,64 | 0,1010583 | 0,1179753 | 1,00E-08 | 0,0930905 | 0,0547996 | 0,0547996 | TRUE | 0,1990438 | No |
| orf19.7095 | 2618 | 2072 | 2335 | 1401 | 2382 | 1433 | 2295 | 2511 | 2496 | 2191 | 2307 | 2195 | 1909 | 1629 | 1948 | 2042 | 2089,57 | 2297 | 1882 | 0,82 | -0,287 | -3,946 | 7,95E-05 | 0,000113 | 0,0057371 | 0,002877 | 0,0045942 | 0,0045942 | TRUE | 0,095974 | No |
| orf19.7096 | 3187 | 2767 | 2979 | 1850 | 2120 | 1402 | 1974 | 2107 | 3038 | 2926 | 2943 | 2898 | 1699 | 1594 | 1676 | 1713 | 2310,91 | 2951 | 1670 | 0,566 | -0,82 | -19,746 | 8,71E-87 | 5,07E-86 | 0,0001594 | 0,0028018 | 0,0011997 | 0,0011997 | TRUE | 0,0105075 | No |
| orf19.7097 | 2445 | 1923 | 2161 | 1366 | 4801 | 3133 | 4601 | 4764 | 2331 | 2033 | 2135 | 2140 | 3847 | 3563 | 3906 | 3874 | 2978,55 | 2160 | 3798 | 1,758 | 0,814 | 15,351 | 3,47E-53 | 1,34E-52 | 0,0020414 | 0,0026425 | 0,0023221 | 0,0023221 | TRUE | 0,0358117 | No |
| orf19.7098 | 1507 | 1291 | 1447 | 869 | 3173 | 1986 | 3232 | 3300 | 1437 | 1365 | 1429 | 1361 | 2542 | 2258 | 2744 | 2683 | 1977,57 | 1398 | 2557 | 1,829 | 0,871 | 13,472 | 2,27E-41 | 7,36E-41 | 0,0038041 | 0,0029215 | 0,0034378 | 0,0034378 | TRUE | 0,0729931 | No |
| orf19.71 | 1203 | 1035 | 1156 | 762 | 1297 | 1023 | 1283 | 1442 | 1147 | 1094 | 1142 | 1194 | 1039 | 1163 | 1089 | 1172 | 1130,16 | 1144 | 1116 | 0,976 | -0,035 | -0,61 | 0,5417796 | 0,5775549 | 0,0014376 | 0,0035441 | 0,0023507 | 0,0023507 | TRUE | 0,0263714 | No |
| orf19.7100 | 190 | 195 | 194 | 133 | 565 | 457 | 566 | 567 | 181 | 206 | 192 | 208 | 453 | 520 | 481 | 461 | 337,66 | 197 | 479 | 2,437 | 1,285 | 14,108 | 3,37E-45 | 1,16E-44 | 1,00E-08 | 0,0069539 | 0,0041909 | 0,0041909 | TRUE | 0,0360863 | No |
| orf19.7101 | 445 | 431 | 438 | 252 | 1232 | 846 | 1104 | 1204 | 424 | 456 | 433 | 395 | 987 | 962 | 937 | 979 | 696,61 | 427 | 966 | 2,255 | 1,173 | 18,387 | 1,66E-75 | 8,59E-75 | 1,00E-08 | 0,0044483 | 0,0021498 | 0,0021498 | TRUE | 0,0230407 | No |
| orf19.7102 | 377 | 304 | 310 | 235 | 1110 | 679 | 1054 | 1067 | 359 | 321 | 306 | 368 | 889 | 772 | 895 | 868 | 597,4 | 338 | 856 | 2,54 | 1,345 | 16,462 | 6,86E-61 | 2,93E-60 | 0,0033262 | 0,0048397 | 0,0042325 | 0,0042325 | TRUE | 0,0488592 | No |
| orf19.7103 | 1256 | 1056 | 1156 | 736 | 1221 | 865 | 1215 | 1312 | 1197 | 1117 | 1142 | 1153 | 978 | 984 | 1032 | 1067 | 1083,66 | 1152 | 1015 | 0,881 | -0,182 | -3,389 | 0,0007011 | 0,0009561 | 0,0005151 | 0,0036064 | 0,0018493 | 0,0018493 | TRUE | 0,0143951 | No |
| orf19.7104 | 712 | 534 | 581 | 416 | 1055 | 692 | 1052 | 1089 | 679 | 565 | 574 | 652 | 845 | 787 | 893 | 885 | 734,99 | 618 | 852 | 1,385 | 0,47 | 6,108 | 1,01E-09 | 1,73E-09 | 0,0039716 | 0,0043252 | 0,0042476 | 0,0042476 | TRUE | 0,0592723 | No |
| orf19.7105 | 848 | 766 | 907 | 628 | 432 | 341 | 402 | 439 | 808 | 810 | 896 | 984 | 346 | 388 | 341 | 357 | 616,3 | 874 | 358 | 0,41 | -1,287 | -15,651 | 3,28E-55 | 1,30E-54 | 0,0044574 | 0,0047554 | 0,004602 | 0,004602 | TRUE | 0,0768779 | No |
| orf19.7106 | 590 | 467 | 561 | 342 | 2043 | 1484 | 2063 | 2154 | 563 | 494 | 554 | 536 | 1637 | 1687 | 1751 | 1751 | 1121,7 | 537 | 1706 | 3,178 | 1,668 | 28,01 | 1,22E-172 | 1,41E-171 | 0,000784 | 0,003555 | 0,0021062 | 0,0021062 | TRUE | 0,0359936 | No |
| orf19.7107 | 6998 | 5489 | 6247 | 4540 | 11529 | 7697 | 10445 | 11110 | 6672 | 5804 | 6171 | 7112 | 9238 | 8752 | 8868 | 9033 | 7706,33 | 6440 | 8973 | 1,394 | 0,479 | 8,014 | 1,11E-15 | 2,27E-15 | 0,0040463 | 0,0023043 | 0,003301 | 0,003301 | TRUE | 0,059036 | No |
| orf19.7108 | 2356 | 1904 | 1896 | 1429 | 5941 | 4038 | 5559 | 5940 | 2246 | 2013 | 1873 | 2239 | 4760 | 4592 | 4720 | 4830 | 3409,05 | 2093 | 4726 | 2,261 | 1,177 | 19,735 | 1,08E-86 | 6,26E-86 | 0,0035473 | 0,0025729 | 0,0030507 | 0,0030507 | TRUE | 0,0613471 | No |
| orf19.7109 | 770 | 590 | 648 | 479 | 1813 | 1231 | 1750 | 1868 | 734 | 624 | 640 | 750 | 1453 | 1400 | 1486 | 1519 | 1075,7 | 687 | 1464 | 2,138 | 1,097 | 16,143 | 1,28E-58 | 5,27E-58 | 0,0029844 | 0,0036176 | 0,0033075 | 0,0033075 | TRUE | 0,0442303 | No |
| orf19.711 | 23492 | 19694 | 21152 | 14136 | 12938 | 8572 | 11896 | 11966 | 22397 | 20824 | 20896 | 22145 | 10367 | 9747 | 10100 | 9730 | 15775,64 | 21566 | 9986 | 0,463 | -1,11 | -27,771 | 9,66E-170 | 1,10E-168 | 0,0010974 | 0,0021954 | 0,001465 | 0,001465 | TRUE | 0,0083707 | No |
| orf19.7110 | 1562 | 1283 | 1452 | 1093 | 1543 | 1257 | 1593 | 1641 | 1489 | 1357 | 1434 | 1712 | 1236 | 1429 | 1352 | 1334 | 1418,11 | 1498 | 1338 | 0,895 | -0,161 | -2,155 | 0,0311574 | 0,0381018 | 0,0060142 | 0,0032491 | 0,0046377 | 0,0046377 | TRUE | 0,1052219 | No |
| orf19.7111 | 1238 | 933 | 1075 | 696 | 1567 | 1010 | 1445 | 1612 | 1180 | 987 | 1062 | 1090 | 1256 | 1148 | 1227 | 1311 | 1157,59 | 1080 | 1236 | 1,145 | 0,195 | 2,917 | 0,0035317 | 0,0046104 | 0,0030888 | 0,0035097 | 0,0034294 | 0,0034294 | TRUE | 0,0498649 | No |
| orf19.7111.1 | 4858 | 3765 | 4106 | 3008 | 5116 | 3235 | 4778 | 4911 | 4632 | 3981 | 4056 | 4712 | 4099 | 3678 | 4056 | 3993 | 4151,07 | 4345 | 3956 | 0,911 | -0,134 | -2,065 | 0,0389671 | 0,0472897 | 0,0046587 | 0,0024868 | 0,0037901 | 0,0037901 | TRUE | 0,038621 | No |
| orf19.7112 | 382 | 357 | 340 | 235 | 592 | 433 | 618 | 663 | 364 | 377 | 336 | 368 | 474 | 492 | 525 | 539 | 434,52 | 361 | 508 | 1,407 | 0,493 | 6,274 | 3,53E-10 | 6,19E-10 | 1,00E-08 | 0,0058699 | 0,0035035 | 0,0035035 | TRUE | 0,0275676 | No |
| orf19.7114 | 1733 | 1464 | 1472 | 1278 | 2132 | 1763 | 1982 | 2268 | 1652 | 1548 | 1454 | 2002 | 1708 | 2005 | 1683 | 1844 | 1737,04 | 1664 | 1810 | 1,089 | 0,123 | 1,322 | 0,1860791 | 0,2115394 | 0,0126712 | 0,0030365 | 0,0077581 | 0,0077581 | TRUE | 0,2179433 | No |
| orf19.7115 | 2911 | 2417 | 2780 | 2002 | 2925 | 1943 | 2608 | 3004 | 2775 | 2556 | 2746 | 3136 | 2344 | 2209 | 2214 | 2443 | 2552,93 | 2803 | 2302 | 0,823 | -0,281 | -4,343 | 1,41E-05 | 2,07E-05 | 0,004397 | 0,0027344 | 0,0036391 | 0,0036391 | TRUE | 0,0744789 | No |
| orf19.7116 | 2166 | 1637 | 1901 | 1097 | 4780 | 2782 | 4966 | 4790 | 2065 | 1731 | 1878 | 1719 | 3830 | 3163 | 4216 | 3895 | 2812,1 | 1848 | 3776 | 2,042 | 1,03 | 11,812 | 3,37E-32 | 9,45E-32 | 0,0101423 | 0,0026752 | 0,0068828 | 0,0068828 | TRUE | 0,1437235 | No |
| orf19.7118 | 1157 | 897 | 936 | 574 | 2743 | 1627 | 2517 | 2567 | 1103 | 948 | 925 | 899 | 2198 | 1850 | 2137 | 2087 | 1518,44 | 969 | 2068 | 2,132 | 1,092 | 14,314 | 1,80E-46 | 6,32E-46 | 0,0061781 | 0,0031726 | 0,0047943 | 0,0047943 | TRUE | 0,1061036 | No |
| orf19.7119 | 7380 | 5735 | 6526 | 3678 | 6416 | 3669 | 6105 | 6393 | 7036 | 6064 | 6447 | 5762 | 5141 | 4172 | 5183 | 5198 | 5625,39 | 6327 | 4924 | 0,778 | -0,362 | -4,508 | 6,56E-06 | 9,80E-06 | 0,0087201 | 0,0023832 | 0,006011 | 0,006011 | TRUE | 0,1276098 | No |
| orf19.712 | 476 | 534 | 521 | 341 | 695 | 557 | 615 | 637 | 454 | 565 | 515 | 534 | 557 | 633 | 522 | 518 | 537,21 | 517 | 558 | 1,078 | 0,109 | 1,2 | 0,230182 | 0,2593205 | 0,0067489 | 0,0051476 | 0,0059681 | 0,0059681 | TRUE | 0,0926864 | No |
| orf19.7121 | 881 | 646 | 733 | 483 | 1931 | 1184 | 1868 | 1839 | 840 | 683 | 724 | 757 | 1547 | 1346 | 1586 | 1495 | 1122,32 | 751 | 1494 | 1,99 | 0,993 | 13,063 | 5,39E-39 | 1,68E-38 | 0,005063 | 0,0035542 | 0,0045012 | 0,0045012 | TRUE | 0,0802607 | No |

| | | | | | | | | | | | | | | | | | | | | | | | | | | | | | | | |
|---|---|---|---|---|---|---|---|---|---|---|---|---|---|---|---|---|---|---|---|---|---|---|---|---|---|---|---|---|---|---|---|
| orf19.7123 | 3697 | 2947 | 3330 | 2656 | 3037 | 2107 | 2852 | 2987 | 3525 | 3116 | 3290 | 4161 | 2433 | 2396 | 2421 | 2429 | 2971,33 | 3523 | 2420 | 0,688 | -0,54 | -6,981 | 2,94E-12 | 5,48E-12 | 0,0079763 | 0,0026439 | 0,0054034 | 0,0054034 | TRUE | 0,176707 | No |
| orf19.7124 | 16015 | 12375 | 13844 | 8237 | 6729 | 4232 | 6466 | 6635 | 15269 | 13085 | 13677 | 12904 | 5392 | 4812 | 5490 | 5395 | 9502,78 | 13734 | 5272 | 0,384 | -1,381 | -21,988 | 3,73E-107 | 2,61E-106 | 0,0046187 | 0,0022641 | 0,003664 | 0,003664 | TRUE | 0,0694332 | No |
| orf19.7125 | 1421 | 1136 | 1285 | 738 | 2819 | 1612 | 2602 | 2529 | 1355 | 1201 | 1269 | 1156 | 2259 | 1833 | 2209 | 2056 | 1667,34 | 1245 | 2089 | 1,676 | 0,745 | 9,927 | 3,16E-23 | 7,57E-23 | 0,0057544 | 0,003076 | 0,004748 | 0,004748 | TRUE | 0,0805612 | No |
| orf19.7127 | 26 | 19 | 28 | 16 | 72 | 69 | 83 | 78 | 25 | 20 | 28 | 25 | 58 | 78 | 70 | 63 | 45,96 | 24 | 67 | 2,751 | 1,46 | 6,319 | 2,63E-10 | 4,63E-10 | 1,00E-08 | 0,0378195 | 0,0219025 | 0,0219025 | TRUE | 0,0946698 | No |
| orf19.7127.1 | 58 | 47 | 40 | 35 | 377 | 246 | 450 | 455 | 55 | 50 | 40 | 55 | 302 | 280 | 382 | 370 | 191,64 | 50 | 334 | 6,748 | 2,754 | 16,8 | 2,44E-63 | 1,07E-62 | 0,0163936 | 0,0106588 | 0,0132667 | 0,0132667 | TRUE | 0,1051163 | No |
| orf19.7128 | 436 | 333 | 356 | 259 | 1067 | 690 | 1022 | 1142 | 416 | 352 | 352 | 406 | 855 | 785 | 868 | 929 | 620,12 | 382 | 859 | 2,261 | 1,177 | 14,342 | 1,19E-46 | 4,21E-46 | 0,0038609 | 0,004739 | 0,0044744 | 0,0044744 | TRUE | 0,0496508 | No |
| orf19.713 | 3530 | 3025 | 3351 | 2260 | 1710 | 1093 | 1534 | 1716 | 3365 | 3199 | 3310 | 3540 | 1370 | 1243 | 1302 | 1395 | 2340,69 | 3354 | 1328 | 0,397 | -1,334 | -25,665 | 2,87E-145 | 2,81E-144 | 0,001546 | 0,0027927 | 0,0020886 | 0,0020886 | TRUE | 0,0208405 | No |
| orf19.7130 | 41 | 42 | 33 | 30 | 193 | 109 | 147 | 181 | 39 | 44 | 33 | 47 | 155 | 124 | 125 | 147 | 89,21 | 41 | 138 | 3,429 | 1,778 | 9,891 | 4,54E-23 | 1,08E-22 | 1,00E-08 | 0,0204968 | 0,014022 | 0,014022 | TRUE | 0,1419439 | No |
| orf19.7131 | 582 | 443 | 543 | 407 | 1800 | 1279 | 1842 | 1964 | 555 | 468 | 536 | 638 | 1442 | 1454 | 1564 | 1597 | 1031,84 | 549 | 1514 | 2,771 | 1,47 | 18,141 | 1,50E-73 | 7,53E-73 | 0,0064782 | 0,0036825 | 0,0049868 | 0,0049868 | TRUE | 0,1204099 | No |
| orf19.7136 | 1671 | 1298 | 1486 | 935 | 9871 | 6295 | 9292 | 9589 | 1593 | 1372 | 1468 | 1465 | 7909 | 7158 | 7889 | 7797 | 4581,41 | 1474 | 7688 | 5,212 | 2,382 | 43,175 | 0 | 0 | 0,0024255 | 0,0024497 | 0,0024834 | 0,0024834 | TRUE | 0,0370078 | No |
| orf19.7139 | 657 | 473 | 497 | 369 | 1567 | 1091 | 1444 | 1533 | 626 | 500 | 491 | 578 | 1256 | 1241 | 1226 | 1246 | 895,52 | 549 | 1242 | 2,266 | 1,18 | 15,457 | 6,81E-54 | 2,65E-53 | 0,0045347 | 0,0039247 | 0,0042139 | 0,0042139 | TRUE | 0,1179458 | No |
| orf19.714 | 497 | 499 | 510 | 341 | 708 | 539 | 638 | 600 | 474 | 528 | 504 | 534 | 567 | 613 | 542 | 488 | 531,15 | 510 | 552 | 1,083 | 0,115 | 1,39 | 0,1644215 | 0,188024 | 0,0042546 | 0,0051825 | 0,0047018 | 0,0047018 | TRUE | 0,0734755 | No |
| orf19.7140 | 1777 | 1422 | 1599 | 1073 | 1838 | 1116 | 1695 | 1834 | 1694 | 1504 | 1580 | 1681 | 1473 | 1269 | 1439 | 1491 | 1516,29 | 1615 | 1418 | 0,88 | -0,184 | -2,855 | 0,0043016 | 0,0055835 | 0,0032387 | 0,0031741 | 0,0033373 | 0,0033373 | TRUE | 0,0575514 | No |
| orf19.7141 | 3192 | 2519 | 2783 | 1726 | 3011 | 1811 | 2887 | 2954 | 3043 | 2664 | 2749 | 2704 | 2413 | 2059 | 2451 | 2402 | 2560,6 | 2790 | 2331 | 0,836 | -0,259 | -3,906 | 9,37E-05 | 0,000133 | 0,0044496 | 0,0027325 | 0,0038153 | 0,0038153 | TRUE | 0,0727612 | No |
| orf19.7144 | 4068 | 3368 | 3790 | 2468 | 6374 | 4174 | 6239 | 6508 | 3878 | 3561 | 3744 | 3866 | 5107 | 4746 | 5297 | 5292 | 4436,51 | 3762 | 5110 | 1,359 | 0,443 | 9,027 | 1,76E-19 | 3,92E-19 | 0,0017361 | 0,0024614 | 0,0020741 | 0,0020741 | TRUE | 0,0274645 | No |
| orf19.7148 | 201 | 162 | 167 | 111 | 255 | 158 | 260 | 274 | 192 | 171 | 165 | 174 | 204 | 180 | 221 | 223 | 191,16 | 176 | 207 | 1,183 | 0,242 | 2,156 | 0,0311179 | 0,0380681 | 1,00E-08 | 0,0106803 | 0,0067811 | 0,0067811 | TRUE | 0,075646 | No |
| orf19.7149 | 921 | 776 | 818 | 597 | 5988 | 4187 | 5700 | 6110 | 878 | 821 | 808 | 935 | 4798 | 4761 | 4839 | 4968 | 2851,03 | 860 | 4842 | 5,65 | 2,498 | 50,231 | 0 | 0 | 0,0004518 | 0,0026672 | 0,0016313 | 0,0016313 | TRUE | 0,034078 | No |
| orf19.715 | 1894 | 1897 | 1908 | 1257 | 838 | 600 | 681 | 749 | 1806 | 2006 | 1885 | 1969 | 671 | 682 | 578 | 609 | 1275,82 | 1916 | 635 | 0,331 | -1,594 | -24,364 | 4,14E-131 | 3,64E-130 | 0,0027775 | 0,0033782 | 0,0031145 | 0,0031145 | TRUE | 0,043072 | No |
| orf19.7150 | 12441 | 10435 | 12253 | 8258 | 5722 | 4122 | 5451 | 5984 | 11861 | 11034 | 12105 | 12937 | 4585 | 4687 | 4628 | 4866 | 8337,7 | 11984 | 4692 | 0,392 | -1,353 | -26,526 | 4,81E-155 | 5,04E-154 | 0,0022841 | 0,0022882 | 0,0023548 | 0,0023548 | TRUE | 0,034745 | No |
| orf19.7151 | 3 | 5 | 8 | 3 | 2 | 5 | 4 | 7 | 3 | 5 | 8 | 5 | 2 | 6 | 3 | 6 | 4,64 | 5 | 4 | 0,774 | -0,37 | -0,549 | 0,583335 | 0,6173927 | 1,00E-08 | 0,3558965 | 0,2196742 | 0,2196742 | TRUE | 0,3675735 | No |
| orf19.7152 | 8504 | 6615 | 7470 | 5170 | 14082 | 9463 | 13538 | 14093 | 8108 | 6995 | 7380 | 8099 | 11284 | 10760 | 11494 | 11459 | 9447,17 | 7646 | 11249 | 1,472 | 0,558 | 10,349 | 4,22E-25 | 1,05E-24 | 0,0029121 | 0,0022651 | 0,0026766 | 0,0026766 | TRUE | 0,0400995 | No |
| orf19.7153 | 4767 | 4695 | 4888 | 3131 | 5115 | 3906 | 4689 | 5112 | 4545 | 4964 | 4829 | 4905 | 4099 | 4441 | 3981 | 4157 | 4490,05 | 4811 | 4170 | 0,867 | -0,207 | -4,381 | 1,18E-05 | 1,74E-05 | 0,0015469 | 0,002457 | 0,0019126 | 0,0019126 | TRUE | 0,0228929 | No |
| orf19.7154 | 10386 | 9263 | 10313 | 5981 | 13347 | 8888 | 12947 | 13663 | 9902 | 9794 | 10188 | 9370 | 10695 | 10106 | 10992 | 11109 | 10269,56 | 9814 | 10726 | 1,093 | 0,128 | 2,989 | 0,0027959 | 0,0036716 | 0,0013411 | 0,0022512 | 0,0016621 | 0,0016621 | TRUE | 0,018218 | No |
| orf19.7156 | 1801 | 1614 | 1753 | 1117 | 22784 | 15881 | 21868 | 23447 | 1717 | 1707 | 1732 | 1750 | 18256 | 18058 | 18566 | 19065 | 10106,26 | 1726 | 18486 | 10,716 | 3,422 | 94,706 | 0 | 0 | 1,00E-08 | 0,0022537 | 0,0009053 | 0,0009053 | TRUE | 0,0054418 | No |
| orf19.7157 | 143 | 138 | 143 | 111 | 411 | 277 | 432 | 413 | 136 | 146 | 141 | 174 | 329 | 315 | 367 | 336 | 243,04 | 149 | 337 | 2,282 | 1,191 | 11,231 | 2,88E-29 | 7,66E-29 | 0,0015063 | 0,0088471 | 0,0057026 | 0,0057026 | TRUE | 0,098884 | No |
| orf19.7158 | 1533 | 1302 | 1457 | 896 | 787 | 515 | 754 | 762 | 1462 | 1377 | 1439 | 1404 | 631 | 586 | 640 | 620 | 1019,65 | 1420 | 619 | 0,436 | -1,196 | -22,092 | 3,76E-108 | 2,68E-107 | 1,00E-08 | 0,0037015 | 0,001704 | 0,001704 | TRUE | 0,0136245 | No |
| orf19.7159 | 935 | 822 | 977 | 601 | 2481 | 1727 | 2347 | 2575 | 891 | 869 | 965 | 941 | 1988 | 1964 | 1993 | 2094 | 1463,16 | 916 | 2010 | 2,195 | 1,134 | 21,79 | 2,87E-105 | 1,96E-104 | 0,000514 | 0,0032134 | 0,0017686 | 0,0017686 | TRUE | 0,0170556 | No |
| orf19.716 | 1044 | 1115 | 1084 | 575 | 572 | 487 | 500 | 483 | 995 | 1179 | 1071 | 901 | 458 | 554 | 424 | 393 | 746,91 | 1036 | 457 | 0,439 | -1,187 | -10,958 | 6,09E-28 | 1,59E-27 | 0,0159487 | 0,0042895 | 0,0097396 | 0,0097396 | TRUE | 0,2237453 | No |
| orf19.7160 | 3004 | 2454 | 2738 | 1813 | 5007 | 3568 | 4788 | 5104 | 2864 | 2595 | 2705 | 2840 | 4012 | 4057 | 4065 | 4150 | 3411 | 2751 | 4071 | 1,481 | 0,566 | 13,395 | 6,47E-41 | 2,08E-40 | 0,0007345 | 0,0025726 | 0,0014024 | 0,0014024 | TRUE | 0,0178048 | No |
| orf19.7161 | 9260 | 7413 | 8109 | 4743 | 10653 | 6637 | 10363 | 11178 | 8828 | 7838 | 8011 | 7430 | 8536 | 7547 | 8798 | 9089 | 8259,69 | 8027 | 8492 | 1,058 | 0,081 | 1,201 | 0,2296126 | 0,2587247 | 0,0056223 | 0,0022901 | 0,0042633 | 0,0042633 | TRUE | 0,0681719 | No |
| orf19.7163 | 170 | 127 | 126 | 88 | 403 | 216 | 320 | 343 | 162 | 134 | 124 | 138 | 323 | 246 | 272 | 279 | 209,72 | 140 | 280 | 2,006 | 1,004 | 8,076 | 6,72E-16 | 1,38E-15 | 0,0085327 | 0,0099202 | 0,0092617 | 0,0092617 | TRUE | 0,1344806 | No |
| orf19.7164 | 1511 | 1300 | 1494 | 943 | 1543 | 1082 | 1417 | 1574 | 1441 | 1375 | 1476 | 1477 | 1236 | 1230 | 1203 | 1280 | 1339,74 | 1442 | 1237 | 0,859 | -0,219 | -4,519 | 6,20E-06 | 9,28E-06 | 1,00E-08 | 0,0033168 | 0,0015149 | 0,0015149 | TRUE | 0,0120071 | No |
| orf19.7165 | 548 | 493 | 469 | 327 | 540 | 379 | 499 | 559 | 522 | 521 | 463 | 512 | 433 | 431 | 424 | 455 | 470,14 | 504 | 436 | 0,864 | -0,211 | -2,91 | 0,0036114 | 0,0047116 | 1,00E-08 | 0,0055836 | 0,0029365 | 0,0029365 | TRUE | 0,038366 | No |
| orf19.7166 | 10195 | 8453 | 9545 | 5909 | 6426 | 4266 | 6108 | 6705 | 9720 | 8938 | 9430 | 9257 | 5149 | 4851 | 5186 | 5452 | 7247,67 | 9336 | 5160 | 0,553 | -0,855 | -18,807 | 6,67E-79 | 3,60E-78 | 0,0015493 | 0,0023178 | 0,0018388 | 0,0018388 | TRUE | 0,0192963 | No |
| orf19.7167 | 51 | 30 | 31 | 29 | 8 | 4 | 4 | 9 | 49 | 32 | 31 | 45 | 6 | 5 | 3 | 7 | 22,26 | 39 | 5 | 0,141 | -2,825 | -7,139 | 9,42E-13 | 1,78E-12 | 1,00E-08 | 0,075855 | 0,0553767 | 0,0553767 | TRUE | 0,1076923 | No |
| orf19.717 | 9116 | 8064 | 8737 | 5169 | 8967 | 5910 | 8353 | 9241 | 8691 | 8527 | 8631 | 8098 | 7185 | 6720 | 7092 | 7514 | 7807,16 | 8487 | 7128 | 0,84 | -0,252 | -5,761 | 8,36E-09 | 1,39E-08 | 0,0013761 | 0,0023016 | 0,0017067 | 0,0017067 | TRUE | 0,0177559 | No |
| orf19.7170 | 18 | 6 | 11 | 15 | 10 | 6 | 18 | 3 | 17 | 6 | 11 | 23 | 8 | 7 | 15 | 2 | 11,3 | 14 | 8 | 0,574 | -0,801 | -1,442 | 0,1493771 | 0,1714961 | 0,2720192 | 0,1473491 | 0,2024017 | 0,2024017 | TRUE | 0,3392651 | No |
| orf19.7173 | 404 | 313 | 371 | 235 | 619 | 471 | 599 | 664 | 385 | 331 | 367 | 368 | 496 | 536 | 509 | 540 | 441,35 | 363 | 520 | 1,432 | 0,518 | 6,753 | 1,45E-11 | 2,66E-11 | 1,00E-08 | 0,0058115 | 0,0032608 | 0,0032608 | TRUE | 0,0415787 | No |
| orf19.7175 | 1062 | 858 | 1049 | 613 | 1887 | 1138 | 1805 | 1862 | 1013 | 907 | 1036 | 960 | 1512 | 1294 | 1532 | 1514 | 1221,1 | 979 | 1463 | 1,495 | 0,58 | 8,326 | 8,33E-17 | 1,74E-16 | 0,0040652 | 0,0034359 | 0,0037838 | 0,0037838 | TRUE | 0,069973 | No |
| orf19.7176 | 797 | 752 | 815 | 511 | 2393 | 1771 | 2436 | 2585 | 760 | 795 | 805 | 801 | 1917 | 2014 | 2068 | 2102 | 1407,74 | 790 | 2025 | 2,565 | 1,359 | 25,623 | 8,49E-145 | 8,28E-144 | 1,00E-08 | 0,0032576 | 0,0017738 | 0,0017738 | TRUE | 0,0161833 | No |
| orf19.7177 | 5185 | 4989 | 5360 | 3372 | 3829 | 2767 | 3428 | 3866 | 4943 | 5275 | 5295 | 5282 | 3068 | 3146 | 2910 | 3143 | 4133,04 | 5199 | 3067 | 0,59 | -0,761 | -17,806 | 6,37E-71 | 3,08E-70 | 0,000843 | 0,0024885 | 0,0015013 | 0,0015013 | TRUE | 0,0144573 | No |
| orf19.7178 | 7268 | 6044 | 6510 | 4373 | 8524 | 5701 | 8195 | 8628 | 6929 | 6391 | 6431 | 6851 | 6830 | 6483 | 6958 | 7015 | 6735,91 | 6650 | 6822 | 1,026 | 0,037 | 0,858 | 0,3907559 | 0,4265891 | 0,0012936 | 0,002335 | 0,0016762 | 0,0016762 | TRUE | 0,0133793 | No |
| orf19.7179 | 1606 | 1307 | 1479 | 933 | 1866 | 1172 | 1915 | 2026 | 1531 | 1382 | 1461 | 1462 | 1495 | 1333 | 1626 | 1647 | 1492,1 | 1459 | 1525 | 1,047 | 0,066 | 0,929 | 0,3528924 | 0,3875742 | 0,0044931 | 0,0031917 | 0,0041308 | 0,0041308 | TRUE | 0,0838585 | No |
| orf19.718 | 6031 | 6082 | 6381 | 3709 | 4755 | 3333 | 4539 | 4355 | 5750 | 6431 | 6304 | 5810 | 3810 | 3790 | 3854 | 3541 | 4911,2 | 6074 | 3749 | 0,617 | -0,697 | -13,764 | 4,23E-43 | 1,40E-42 | 0,002022 | 0,0024256 | 0,0022497 | 0,0022497 | TRUE | 0,0190863 | No |
| orf19.7181 | 878 | 807 | 881 | 605 | 1657 | 1273 | 1595 | 1719 | 837 | 853 | 870 | 948 | 1328 | 1448 | 1354 | 1398 | 1129,45 | 877 | 1382 | 1,579 | 0,659 | 11,61 | 3,66E-31 | 1,01E-30 | 0,0010401 | 0,003545 | 0,0021332 | 0,0021332 | TRUE | 0,030282 | No |
| orf19.7182 | 1881 | 1629 | 1705 | 963 | 3056 | 2098 | 2965 | 3233 | 1793 | 1722 | 1684 | 1509 | 2449 | 2386 | 2517 | 2629 | 2086,13 | 1677 | 2495 | 1,485 | 0,57 | 9,76 | 1,68E-22 | 3,96E-22 | 0,0028131 | 0,0028783 | 0,0027633 | 0,0027633 | TRUE | 0,0508462 | No |
| orf19.7183 | 1297 | 1039 | 1114 | 739 | 802 | 521 | 810 | 800 | 1237 | 1099 | 1101 | 1158 | 643 | 592 | 688 | 650 | 895,82 | 1149 | 643 | 0,561 | -0,833 | -12,598 | 2,18E-36 | 6,50E-36 | 0,0022502 | 0,0039241 | 0,0030203 | 0,0030203 | TRUE | 0,0342031 | No |
| orf19.7184 | 2812 | 2509 | 2696 | 1952 | 1061 | 736 | 972 | 1051 | 2681 | 2653 | 2663 | 3058 | 850 | 837 | 825 | 855 | 1802,75 | 2764 | 842 | 0,305 | -1,712 | -29,39 | 7,29E-190 | 9,41E-189 | 0,0021978 | 0,0030021 | 0,0025253 | 0,0025253 | TRUE | 0,0590811 | No |
| orf19.7185 | 3018 | 2526 | 2602 | 1645 | 3748 | 2479 | 3590 | 3925 | 2877 | 2671 | 2571 | 2577 | 3003 | 2819 | 3048 | 3191 | 2844,63 | 2674 | 3015 | 1,128 | 0,173 | 3,188 | 0,0014326 | 0,0019128 | 0,0022606 | 0,0026685 | 0,0024739 | 0,0024739 | TRUE | 0,0324486 | No |
| orf19.7186 | 246 | 223 | 272 | 147 | 348 | 232 | 315 | 362 | 235 | 236 | 269 | 230 | 279 | 264 | 267 | 294 | 259,22 | 242 | 276 | 1,138 | 0,186 | 1,975 | 0,0482616 | 0,0581052 | 1,00E-08 | 0,0084253 | 0,0046564 | 0,0046564 | TRUE | 0,0622042 | No |
| orf19.7187 | 6229 | 5563 | 5660 | 3662 | 8567 | 5714 | 7704 | 7994 | 5939 | 5882 | 5592 | 5737 | 6864 | 6497 | 6541 | 6500 | 6193,93 | 5788 | 6600 | 1,141 | 0,19 | 5,167 | 2,38E-07 | 3,77E-07 | 0,0005058 | 0,0023564 | 0,0011298 | 0,0011298 | TRUE | 0,0089463 | No |
| orf19.7188 | 2064 | 1674 | 1782 | 1568 | 7898 | 6282 | 7022 | 8489 | 1968 | 1770 | 1760 | 2456 | 6328 | 7143 | 5962 | 6902 | 4286,29 | 1988 | 6584 | 3,316 | 1,729 | 17,76 | 1,45E-70 | 6,98E-70 | 0,0163416 | 0,0024743 | 0,0087576 | 0,0087576 | TRUE | 0,2953737 | No |
| orf19.719 | 1294 | 1295 | 1304 | 755 | 1104 | 807 | 1035 | 1035 | 1234 | 1369 | 1288 | 1183 | 885 | 918 | 879 | 842 | 1074,56 | 1268 | 881 | 0,692 | -0,531 | -8,863 | 7,81E-19 | 1,71E-18 | 0,0015675 | 0,0036193 | 0,0024911 | 0,0024911 | TRUE | 0,0336901 | No |
| orf19.7190 | 584 | 523 | 533 | 346 | 1263 | 901 | 1214 | 1241 | 557 | 553 | 527 | 542 | 1012 | 1025 | 1031 | 1009 | 781,83 | 545 | 1019 | 1,87 | 0,903 | 15,958 | 2,50E-57 | 1,01E-56 | 1,00E-08 | 0,0041914 | 0,0016178 | 0,0016178 | TRUE | 0,0066156 | No |
| orf19.7192 | 18 | 15 | 22 | 18 | 30 | 22 | 24 | 40 | 17 | 16 | 22 | 28 | 24 | 25 | 20 | 33 | 23,11 | 21 | 26 | 1,257 | 0,33 | 1,069 | 0,2849569 | 0,317055 | 1,00E-08 | 0,0731281 | 0,0465727 | 0,0465727 | TRUE | 0,2479313 | No |
| orf19.7193 | 8825 | 7760 | 8425 | 5315 | 7748 | 5173 | 7406 | 7542 | 8414 | 8205 | 8323 | 8326 | 6208 | 5882 | 6288 | 6132 | 7222,33 | 8317 | 6128 | 0,737 | -0,44 | -13,41 | 5,27E-41 | 1,69E-40 | 0,000274 | 0,0023186 | 0,0008936 | 0,0008936 | TRUE | 0,0085349 | No |
| orf19.7194 | 1541 | 1590 | 1584 | 973 | 1555 | 1073 | 1440 | 1329 | 1469 | 1681 | 1565 | 1524 | 1246 | 1220 | 1223 | 1081 | 1376,09 | 1560 | 1192 | 0,764 | -0,389 | -6,094 | 1,10E-09 | 1,89E-09 | 0,0030807 | 0,0032844 | 0,0031678 | 0,0031678 | TRUE | 0,0490145 | No |
| orf19.7195 | 1952 | 1715 | 1796 | 1040 | 3633 | 2178 | 3297 | 3480 | 1861 | 1813 | 1774 | 1629 | 2911 | 2477 | 2799 | 2830 | 2261,78 | 1769 | 2754 | 1,555 | 0,637 | 10,226 | 1,51E-24 | 3,70E-24 | 0,0033815 | 0,0028172 | 0,0032472 | 0,0032472 | TRUE | 0,054439 | No |
| orf19.7196 | 14158 | 12463 | 13205 | 9155 | 27079 | 21462 | 25342 | 27644 | 13498 | 13178 | 13045 | 14342 | 21698 | 24404 | 21515 | 22477 | 18019,69 | 13516 | 22524 | 1,667 | 0,737 | 14,565 | 4,70E-48 | 1,69E-47 | 0,0025707 | 0,0021824 | 0,0023981 | 0,0023981 | TRUE | 0,0385235 | No |
| orf19.7197 | 8747 | 9018 | 9558 | 5369 | 10757 | 7478 | 10001 | 9413 | 8339 | 9535 | 9442 | 8411 | 8619 | 8503 | 8491 | 7654 | 8624,35 | 8932 | 8317 | 0,931 | -0,103 | -1,732 | 0,0833536 | 0,0980784 | 0,003888 | 0,0022817 | 0,0033068 | 0,0033068 | TRUE | 0,0353513 | No |
| orf19.7198 | 1169 | 1116 | 1104 | 674 | 2458 | 1615 | 2158 | 2168 | 1115 | 1180 | 1091 | 1056 | 1970 | 1836 | 1832 | 1763 | 1480,24 | 1110 | 1850 | 1,664 | 0,735 | 13,244 | 4,89E-40 | 1,55E-39 | 0,0015045 | 0,0032005 | 0,0022101 | 0,0022101 | TRUE | 0,0234684 | No |
| orf19.7199 | 474 | 533 | 496 | 249 | 1000 | 639 | 922 | 854 | 452 | 564 | 490 | 390 | 801 | 727 | 783 | 694 | 612,57 | 474 | 751 | 1,579 | 0,659 | 6,658 | 2,77E-11 | 5,04E-11 | 0,0107856 | 0,0047716 | 0,0076347 | 0,0076347 | TRUE | 0,1832093 | No |
| orf19.720 | 235 | 208 | 244 | 132 | 234 | 149 | 194 | 226 | 224 | 220 | 241 | 207 | 187 | 169 | 165 | 184 | 199,65 | 223 | 176 | 0,789 | -0,341 | -3,27 | 0,0010752 | 0,0014481 | 1,00E-08 | 0,0103152 | 0,0054521 | 0,0054521 | TRUE | 0,0324517 | No |
| orf19.7200 | 377 | 426 | 423 | 270 | 824 | 611 | 696 | 675 | 359 | 450 | 418 | 423 | 660 | 695 | 591 | 549 | 518,18 | 412 | 624 | 1,511 | 0,596 | 6,169 | 6,86E-10 | 1,19E-09 | 0,0083924 | 0,0052599 | 0,0068736 | 0,0068736 | TRUE | 0,0923087 | No |
| orf19.7201 | 3327 | 3383 | 3621 | 2103 | 2566 | 1847 | 2458 | 2290 | 3172 | 3577 | 3577 | 3294 | 2056 | 2100 | 2087 | 1862 | 2715,72 | 3405 | 2026 | 0,595 | -0,75 | -12,817 | 1,31E-37 | 4,00E-37 | 0,0027859 | 0,0026959 | 0,0029016 | 0,0029016 | TRUE | 0,0365362 | No |
| orf19.7202 | 298 | 325 | 356 | 173 | 739 | 531 | 641 | 529 | 284 | 344 | 352 | 271 | 592 | 604 | 544 | 430 | 427,59 | 313 | 542 | 1,728 | 0,789 | 6,437 | 1,22E-10 | 2,17E-10 | 0,0178038 | 0,0059312 | 0,0118098 | 0,0118098 | TRUE | 0,2323494 | No |
| orf19.7203 | 2864 | 2780 | 2863 | 1564 | 2633 | 1844 | 2533 | 2450 | 2731 | 2939 | 2828 | 2450 | 2110 | 2097 | 2151 | 1992 | 2412,2 | 2737 | 2088 | 0,761 | -0,393 | -6,692 | 2,20E-11 | 4,00E-11 | 0,0028896 | 0,0027719 | 0,0028965 | 0,0028965 | TRUE | 0,0572052 | No |
| orf19.7204 | 113 | 104 | 104 | 54 | 79 | 39 | 65 | 63 | 108 | 110 | 103 | 85 | 63 | 44 | 55 | 51 | 77,39 | 102 | 53 | 0,528 | -0,92 | -5,361 | 8,27E-08 | 1,33E-07 | 1,00E-08 | 0,0233082 | 0,0145711 | 0,0145711 | TRUE | 0,1341039 | No |
| orf19.7205 | 1336 | 1225 | 1301 | 832 | 1821 | 1338 | 1693 | 1902 | 1274 | 1295 | 1285 | 1303 | 1459 | 1521 | 1437 | 1547 | 1390,26 | 1289 | 1491 | 1,157 | 0,21 | 4,446 | 8,75E-06 | 1,30E-05 | 1,00E-08 | 0,0032723 | 0,0014052 | 0,0014052 | TRUE | 0,0081182 | No |
| orf19.7206 | 161 | 121 | 147 | 118 | 310 | 181 | 259 | 268 | 153 | 128 | 145 | 185 | 248 | 206 | 220 | 218 | 187,94 | 153 | 223 | 1,476 | 0,562 | 4,553 | 5,29E-06 | 7,95E-06 | 0,006285 | 0,0108276 | 0,0089405 | 0,0089405 | TRUE | 0,1651821 | No |
| orf19.7207 | 4751 | 4363 | 4696 | 2915 | 4489 | 3140 | 4184 | 4647 | 4530 | 4613 | 4639 | 4567 | 3597 | 3570 | 3552 | 3778 | 4105,83 | 4587 | 3624 | 0,79 | -0,339 | -9,545 | 1,36E-21 | 3,14E-21 | 0,0002507 | 0,0024912 | 0,0009687 | 0,0009687 | TRUE | 0,0101372 | No |
| orf19.7208 | 16 | 8 | 10 | 3 | 26 | 13 | 19 | 20 | 15 | 8 | 10 | 5 | 21 | 15 | 16 | 16 | 13,29 | 10 | 17 | 1,719 | 0,782 | 1,922 | 0,0546328 | 0,0653811 | 1,00E-08 | 0,1256588 | 0,0769214 | 0,0769214 | TRUE | 0,5441612 | No |
| orf19.7209 | 95 | 81 | 89 | 47 | 348 | 235 | 351 | 383 | 91 | 86 | 88 | 74 | 279 | 267 | 298 | 311 | 186,66 | 85 | 289 | 3,394 | 1,763 | 14,447 | 2,63E-47 | 9,36E-47 | 1,00E-08 | 0,0108877 | 0,0062633 | 0,0062633 | TRUE | 0,0482781 | No |

| | | | | | | | | | | | | | | | | | | | | | | | | | | | | | | | |
|---|---|---|---|---|---|---|---|---|---|---|---|---|---|---|---|---|---|---|---|---|---|---|---|---|---|---|---|---|---|---|---|
| orf19.721 | 2236 | 2360 | 2505 | 1459 | 1805 | 1374 | 1631 | 1660 | 2132 | 2495 | 2475 | 2286 | 1446 | 1562 | 1385 | 1350 | 1891,32 | 2347 | 1436 | 0,611 | -0,711 | -10,671 | 1,39E-26 | 3,56E-26 | 0,0040409 | 0,0029594 | 0,0037078 | 0,0037078 | TRUE | 0,0478273 | No |
| orf19.7210 | 2910 | 2389 | 2588 | 1939 | 2753 | 2010 | 2582 | 2891 | 2774 | 2526 | 2557 | 3038 | 2206 | 2286 | 2192 | 2351 | 2491,11 | 2724 | 2259 | 0,83 | -0,268 | -4,318 | 1,58E-05 | 2,31E-05 | 0,0037707 | 0,0027504 | 0,003299 | 0,003299 | TRUE | 0,0698287 | No |
| orf19.7212 | 701 | 603 | 665 | 433 | 614 | 439 | 546 | 577 | 668 | 638 | 657 | 678 | 492 | 499 | 464 | 469 | 570,63 | 660 | 481 | 0,728 | -0,457 | -7,12 | 1,08E-12 | 2,04E-12 | 1,00E-08 | 0,0049686 | 0,0021894 | 0,0021894 | TRUE | 0,0067653 | No |
| orf19.7213 | 2203 | 1974 | 2179 | 1360 | 3391 | 2325 | 3088 | 3469 | 2100 | 2087 | 2153 | 2131 | 2717 | 2644 | 2622 | 2821 | 2409,24 | 2118 | 2701 | 1,276 | 0,352 | 8,479 | 2,27E-17 | 4,82E-17 | 0,0003806 | 0,0027728 | 0,0012227 | 0,0012227 | TRUE | 0,0109808 | No |
| orf19.7214 | 2781 | 2356 | 2598 | 1519 | 2478 | 1646 | 2466 | 2592 | 2651 | 2491 | 2567 | 2380 | 1986 | 1872 | 2094 | 2108 | 2268,39 | 2522 | 2015 | 0,799 | -0,325 | -6,031 | 1,63E-09 | 2,78E-09 | 0,0019366 | 0,0028151 | 0,002337 | 0,002337 | TRUE | 0,0263641 | No |
| orf19.7215 | 7056 | 6549 | 6851 | 4570 | 14288 | 10378 | 12891 | 14615 | 6727 | 6925 | 6768 | 7159 | 11449 | 11801 | 10944 | 11883 | 9207,03 | 6895 | 11519 | 1,671 | 0,741 | 18,664 | 9,81E-78 | 5,19E-77 | 0,0009358 | 0,0022696 | 0,0013937 | 0,0013937 | TRUE | 0,0138314 | No |
| orf19.7215.3 | 2742 | 2200 | 2368 | 1428 | 1909 | 1153 | 1734 | 1944 | 2614 | 2326 | 2339 | 2237 | 1530 | 1311 | 1472 | 1581 | 1926,29 | 2379 | 1474 | 0,62 | -0,691 | -10,02 | 1,25E-23 | 3,00E-23 | 0,0047671 | 0,0029436 | 0,0040246 | 0,0040246 | TRUE | 0,0637992 | No |
| orf19.7216 | 2814 | 2425 | 2668 | 1563 | 2233 | 1653 | 2176 | 2338 | 2683 | 2564 | 2636 | 2449 | 1789 | 1880 | 1847 | 1901 | 2218,56 | 2583 | 1854 | 0,717 | -0,481 | -10,678 | 1,29E-26 | 3,29E-26 | 0,0006326 | 0,0028314 | 0,0014875 | 0,0014875 | TRUE | 0,0136236 | No |
| orf19.7217 | 22091 | 16748 | 18556 | 13687 | 44672 | 32022 | 41991 | 47641 | 21061 | 17709 | 18332 | 21441 | 35794 | 36412 | 35650 | 38737 | 28142,05 | 19636 | 36648 | 1,867 | 0,9 | 13,865 | 1,03E-43 | 3,47E-43 | 0,0053382 | 0,0021496 | 0,0040117 | 0,0040117 | TRUE | 0,0533871 | No |
| orf19.7218 | 99 | 54 | 60 | 76 | 46 | 29 | 45 | 57 | 94 | 57 | 59 | 119 | 37 | 33 | 38 | 46 | 60,53 | 82 | 38 | 0,475 | -1,073 | -4,204 | 2,63E-05 | 3,82E-05 | 0,0608809 | 0,0292189 | 0,0441043 | 0,0441043 | TRUE | 0,2269158 | No |
| orf19.7219 | 12335 | 10440 | 11780 | 7694 | 22873 | 15877 | 22582 | 25023 | 11760 | 11039 | 11638 | 12053 | 18327 | 18054 | 19172 | 20346 | 15298,6 | 11622 | 18975 | 1,633 | 0,708 | 14,817 | 1,15E-49 | 4,21E-49 | 0,0020553 | 0,0021986 | 0,0021186 | 0,0021186 | TRUE | 0,029009 | No |
| orf19.7221 | 2214 | 1928 | 2126 | 1260 | 2986 | 1997 | 2759 | 3007 | 2111 | 2039 | 2100 | 1974 | 2393 | 2271 | 2342 | 2445 | 2209,29 | 2056 | 2363 | 1,148 | 0,2 | 4,566 | 4,98E-06 | 7,49E-06 | 0,0003608 | 0,0028345 | 0,0013743 | 0,0013743 | TRUE | 0,0078573 | No |
| orf19.7222 | 3209 | 2487 | 2751 | 1715 | 2269 | 1423 | 2171 | 2270 | 3059 | 2630 | 2718 | 2687 | 1818 | 1618 | 1843 | 1846 | 2277,32 | 2774 | 1781 | 0,643 | -0,638 | -10,055 | 8,75E-24 | 2,12E-23 | 0,0037256 | 0,0028123 | 0,0034142 | 0,0034142 | TRUE | 0,059571 | No |
| orf19.7223 | 2492 | 2380 | 2410 | 1660 | 2688 | 1955 | 2488 | 2770 | 2376 | 2517 | 2381 | 2600 | 2154 | 2223 | 2112 | 2252 | 2326,89 | 2468 | 2185 | 0,886 | -0,174 | -3,737 | 0,0001865 | 0,0002605 | 0,0008361 | 0,0027969 | 0,0016455 | 0,0016455 | TRUE | 0,014566 | No |
| orf19.7224 | 2945 | 2524 | 2728 | 1688 | 4831 | 3213 | 4522 | 4939 | 2808 | 2669 | 2695 | 2644 | 3871 | 3653 | 3839 | 4016 | 3274,42 | 2704 | 3845 | 1,422 | 0,508 | 11,792 | 4,30E-32 | 1,20E-31 | 0,0008324 | 0,0025927 | 0,0014559 | 0,0014559 | TRUE | 0,0131736 | No |
| orf19.7225 | 85 | 82 | 104 | 74 | 239 | 196 | 221 | 245 | 81 | 87 | 103 | 116 | 192 | 223 | 188 | 199 | 148,45 | 97 | 200 | 2,093 | 1,066 | 7,873 | 3,46E-15 | 7,00E-15 | 0,0035095 | 0,0131514 | 0,0095264 | 0,0095264 | TRUE | 0,1322078 | No |
| orf19.7227 | 2577 | 2082 | 2284 | 1559 | 275 | 186 | 262 | 300 | 2457 | 2201 | 2256 | 2442 | 220 | 211 | 222 | 244 | 1281,9 | 2339 | 224 | 0,096 | -3,376 | -47,049 | 0 | 0 | 0,0021594 | 0,0033721 | 0,0027317 | 0,0027317 | TRUE | 0,0384831 | No |
| orf19.7228 | 1167 | 1147 | 1117 | 756 | 1209 | 921 | 1139 | 1313 | 1113 | 1213 | 1103 | 1184 | 969 | 1047 | 967 | 1068 | 1082,98 | 1153 | 1013 | 0,879 | -0,187 | -3,141 | 0,0016814 | 0,0022394 | 0,0015883 | 0,0036074 | 0,0024687 | 0,0024687 | TRUE | 0,0171578 | No |
| orf19.7229 | 6771 | 5778 | 6212 | 4225 | 3812 | 2579 | 3677 | 3926 | 6455 | 6110 | 6137 | 6619 | 3054 | 2933 | 3122 | 3192 | 4702,68 | 6330 | 3075 | 0,486 | -1,04 | -23,746 | 1,22E-124 | 1,02E-123 | 0,001093 | 0,0024404 | 0,0016075 | 0,0016075 | TRUE | 0,011364 | No |
| orf19.723 | 7405 | 6761 | 7347 | 5297 | 4843 | 3728 | 4365 | 4755 | 7060 | 7149 | 7258 | 8298 | 3881 | 4239 | 3706 | 3866 | 5682,09 | 7441 | 3923 | 0,527 | -0,923 | -14,927 | 2,20E-50 | 8,17E-50 | 0,0043929 | 0,0023802 | 0,0034855 | 0,0034855 | TRUE | 0,0722194 | No |
| orf19.7231 | 1200 | 1038 | 1119 | 716 | 2542 | 1835 | 2449 | 2469 | 1144 | 1098 | 1105 | 1122 | 2037 | 2087 | 2079 | 2008 | 1584,86 | 1117 | 2053 | 1,836 | 0,877 | 19,844 | 1,23E-87 | 7,21E-87 | 1,00E-08 | 0,0031273 | 0,0011576 | 0,0011576 | TRUE | 0,003547 | No |
| orf19.7232 | 515 | 430 | 447 | 284 | 1050 | 769 | 952 | 1127 | 491 | 455 | 442 | 445 | 841 | 874 | 808 | 916 | 659,07 | 458 | 860 | 1,874 | 0,906 | 13,002 | 1,20E-38 | 3,72E-38 | 0,0015045 | 0,0045825 | 0,0029267 | 0,0029267 | TRUE | 0,0291028 | No |
| orf19.7234 | 2746 | 2135 | 2472 | 1617 | 3733 | 2762 | 3636 | 3920 | 2618 | 2257 | 2442 | 2533 | 2991 | 3141 | 3087 | 3187 | 2782,1 | 2462 | 3102 | 1,26 | 0,333 | 6,391 | 1,64E-10 | 2,92E-10 | 0,0018513 | 0,0026814 | 0,0022332 | 0,0022332 | TRUE | 0,038645 | No |
| orf19.7235 | 5037 | 4383 | 4809 | 2862 | 2998 | 2057 | 2865 | 2946 | 4802 | 4634 | 4751 | 4483 | 2402 | 2339 | 2432 | 2395 | 3530 | 4668 | 2392 | 0,512 | -0,965 | -25,424 | 1,38E-142 | 1,32E-141 | 0,0003027 | 0,0025564 | 0,0010766 | 0,0010766 | TRUE | 0,0080383 | No |
| orf19.7236 | 6701 | 5229 | 5720 | 3568 | 14019 | 8432 | 13144 | 13567 | 6389 | 5529 | 5651 | 5589 | 11233 | 9588 | 11159 | 11031 | 8271,17 | 5790 | 10753 | 1,857 | 0,893 | 13,844 | 1,38E-43 | 4,63E-43 | 0,0048782 | 0,0022898 | 0,0038588 | 0,0038588 | TRUE | 0,0646432 | No |
| orf19.7237 | 451 | 374 | 378 | 295 | 1730 | 1167 | 1625 | 1766 | 430 | 395 | 373 | 462 | 1386 | 1327 | 1380 | 1436 | 898,72 | 415 | 1382 | 3,35 | 1,744 | 25,473 | 3,89E-143 | 3,75E-142 | 0,0013009 | 0,0039182 | 0,0028279 | 0,0028279 | TRUE | 0,0543211 | No |
| orf19.7238 | 20168 | 16087 | 17492 | 10768 | 32167 | 19303 | 30235 | 30399 | 19228 | 17010 | 17280 | 16869 | 25774 | 21949 | 25669 | 24717 | 21062,18 | 17597 | 24527 | 1,394 | 0,479 | 7,753 | 8,97E-15 | 1,80E-14 | 0,0045353 | 0,0021692 | 0,0036181 | 0,0036181 | TRUE | 0,0611458 | No |
| orf19.7239 | 23539 | 19510 | 22024 | 14145 | 18711 | 12336 | 17878 | 19734 | 22442 | 20629 | 21758 | 22159 | 14993 | 14027 | 15178 | 16046 | 18403,93 | 21747 | 15061 | 0,693 | -0,53 | -11,053 | 2,11E-28 | 5,54E-28 | 0,0021065 | 0,0021805 | 0,0021508 | 0,0021508 | TRUE | 0,0259331 | No |
| orf19.7242 | 4252 | 3574 | 3883 | 2448 | 4660 | 3200 | 4301 | 4673 | 4054 | 3779 | 3836 | 3835 | 3734 | 3639 | 3652 | 3800 | 3790,95 | 3876 | 3706 | 0,956 | -0,065 | -1,693 | 0,0904346 | 0,1060193 | 0,0004536 | 0,0025244 | 0,001144 | 0,001144 | TRUE | 0,012045 | No |
| orf19.7243 | 747 | 705 | 690 | 473 | 1828 | 1196 | 1723 | 1814 | 712 | 745 | 682 | 741 | 1465 | 1360 | 1463 | 1475 | 1080,34 | 720 | 1441 | 2,007 | 1,005 | 17,912 | 9,53E-72 | 4,69E-71 | 0,0005013 | 0,0036111 | 0,0019347 | 0,0019347 | TRUE | 0,0159919 | No |
| orf19.7244 | 1028 | 793 | 883 | 623 | 1929 | 1275 | 1811 | 2045 | 980 | 838 | 872 | 976 | 1546 | 1450 | 1538 | 1663 | 1232,83 | 916 | 1549 | 1,694 | 0,76 | 11,199 | 4,12E-29 | 1,09E-28 | 0,0035322 | 0,0034231 | 0,0035206 | 0,0035206 | TRUE | 0,0391027 | No |
| orf19.7245 | 2640 | 2138 | 2387 | 1630 | 2586 | 1845 | 2556 | 2697 | 2517 | 2261 | 2358 | 2553 | 2072 | 2098 | 2170 | 2193 | 2277,77 | 2422 | 2133 | 0,882 | -0,182 | -3,593 | 0,0003263 | 0,0004513 | 0,0013773 | 0,0028121 | 0,0020106 | 0,0020106 | TRUE | 0,0242995 | No |
| orf19.7247 | 14525 | 12081 | 13291 | 8649 | 10126 | 6990 | 9588 | 10273 | 13848 | 12774 | 13130 | 13549 | 8114 | 7948 | 8140 | 8353 | 10732,07 | 13325 | 8139 | 0,611 | -0,711 | -19,35 | 2,03E-83 | 1,14E-82 | 0,0007007 | 0,0022443 | 0,0011992 | 0,0011992 | TRUE | 0,0095294 | No |
| orf19.725 | 208 | 248 | 234 | 187 | 318 | 232 | 306 | 280 | 198 | 262 | 231 | 293 | 255 | 264 | 260 | 228 | 248,84 | 246 | 252 | 1,029 | 0,041 | 0,343 | 0,7318695 | 0,7607198 | 0,0109211 | 0,0086895 | 0,0097229 | 0,0097229 | TRUE | 0,1943603 | No |
| orf19.7250 | 2314 | 1742 | 1857 | 1330 | 2438 | 1401 | 2334 | 2366 | 2206 | 1842 | 1835 | 2084 | 1953 | 1593 | 1982 | 1924 | 1927,26 | 1992 | 1863 | 0,937 | -0,094 | -1,142 | 0,253508 | 0,2838951 | 0,0083192 | 0,0029432 | 0,006053 | 0,006053 | TRUE | 0,1124362 | No |
| orf19.7251 | 768 | 651 | 698 | 573 | 1268 | 862 | 1164 | 1265 | 732 | 688 | 690 | 898 | 1016 | 980 | 988 | 1029 | 877,59 | 752 | 1003 | 1,342 | 0,424 | 5,282 | 1,27E-07 | 2,04E-07 | 0,0060584 | 0,0039622 | 0,0049883 | 0,0049883 | TRUE | 0,1834729 | No |
| orf19.7252 | 9 | 3 | 7 | 5 | 3 | 1 | 1 | 7 | 9 | 3 | 7 | 8 | 2 | 1 | 1 | 6 | 4,57 | 7 | 2 | 0,393 | -1,347 | -1,826 | 0,0678772 | 0,0806413 | 0,1008536 | 0,3611507 | 0,266367 | 0,266367 | TRUE | 0,9737117 | No |
| orf19.7254 | 1609 | 1424 | 1581 | 1057 | 2148 | 1441 | 2008 | 2232 | 1534 | 1506 | 1562 | 1656 | 1721 | 1639 | 1705 | 1815 | 1642,09 | 1564 | 1720 | 1,102 | 0,141 | 2,757 | 0,0058275 | 0,0075016 | 0,0009474 | 0,0030912 | 0,0018788 | 0,0018788 | TRUE | 0,0167425 | No |
| orf19.7255 | 2065 | 1308 | 1466 | 915 | 4560 | 2642 | 4042 | 4402 | 1969 | 1383 | 1448 | 1433 | 3654 | 3004 | 3432 | 3579 | 2487,79 | 1558 | 3417 | 2,191 | 1,132 | 10,85 | 1,99E-27 | 5,16E-27 | 0,01856 | 0,0027513 | 0,00996 | 0,00996 | TRUE | 0,3831146 | No |
| orf19.7256 | 1394 | 961 | 1019 | 549 | 3078 | 1613 | 2884 | 2738 | 1329 | 1016 | 1007 | 860 | 2466 | 1834 | 2449 | 2226 | 1648,38 | 1053 | 2244 | 2,129 | 1,09 | 6,631 | 3,34E-11 | 6,04E-11 | 0,0252392 | 0,0030873 | 0,0134003 | 0,0252392 | TRUE | 0,3738171 | No |
| orf19.7257 | 2537 | 1847 | 2275 | 1628 | 2583 | 1819 | 2561 | 2750 | 2419 | 1953 | 2247 | 2550 | 2070 | 2068 | 2174 | 2236 | 2214,74 | 2292 | 2137 | 0,933 | -0,099 | -1,322 | 0,1861969 | 0,2116356 | 0,0064776 | 0,0028326 | 0,0049794 | 0,0049794 | TRUE | 0,1202545 | No |
| orf19.7258 | 1715 | 1278 | 1428 | 927 | 5799 | 3470 | 5469 | 5702 | 1635 | 1351 | 1411 | 1452 | 4647 | 3946 | 4643 | 4636 | 2965,13 | 1462 | 4468 | 3,055 | 1,611 | 22,138 | 1,35E-108 | 9,66E-108 | 0,0059328 | 0,002645 | 0,0046024 | 0,0046024 | TRUE | 0,078658 | No |
| orf19.726 | 2216 | 2143 | 2196 | 1294 | 1818 | 1286 | 1674 | 1694 | 2113 | 2266 | 2169 | 2027 | 1457 | 1462 | 1421 | 1377 | 1786,6 | 2144 | 1429 | 0,666 | -0,587 | -11,972 | 4,97E-33 | 1,41E-32 | 0,0008579 | 0,0030103 | 0,0017365 | 0,0017365 | TRUE | 0,0174019 | No |
| orf19.7260 | 422 | 323 | 396 | 280 | 661 | 403 | 641 | 656 | 402 | 342 | 391 | 439 | 530 | 458 | 544 | 533 | 454,9 | 394 | 516 | 1,321 | 0,401 | 4,47 | 7,81E-06 | 1,16E-05 | 0,0050927 | 0,0057007 | 0,005439 | 0,005439 | TRUE | 0,089003 | No |
| orf19.7261 | 2947 | 2187 | 2579 | 1669 | 4435 | 2909 | 4332 | 4587 | 2810 | 2312 | 2548 | 2615 | 3554 | 3308 | 3678 | 3730 | 3069,18 | 2571 | 3568 | 1,388 | 0,473 | 7,406 | 1,30E-13 | 2,53E-13 | 0,0041259 | 0,0026262 | 0,003572 | 0,003572 | TRUE | 0,0563834 | No |
| orf19.7263 | 2535 | 1966 | 2188 | 1497 | 5571 | 3552 | 5307 | 5694 | 2417 | 2079 | 2162 | 2345 | 4464 | 4039 | 4506 | 4630 | 3330,07 | 2251 | 4410 | 1,961 | 0,972 | 15,769 | 5,05E-56 | 2,01E-55 | 0,0036857 | 0,0025843 | 0,0032938 | 0,0032938 | TRUE | 0,0383675 | No |
| orf19.7264 | 5749 | 4015 | 4693 | 2899 | 11543 | 7278 | 11163 | 11536 | 5481 | 4245 | 4636 | 4541 | 9249 | 8276 | 9477 | 9380 | 6910,76 | 4726 | 9096 | 1,924 | 0,944 | 12,487 | 8,76E-36 | 2,59E-35 | 0,0079359 | 0,0023289 | 0,0053245 | 0,0053245 | TRUE | 0,1420725 | No |
| orf19.7265 | 978 | 768 | 834 | 563 | 2946 | 1746 | 2737 | 3013 | 932 | 812 | 824 | 882 | 2361 | 1985 | 2324 | 2450 | 1571,23 | 862 | 2280 | 2,646 | 1,404 | 18,699 | 5,05E-78 | 2,69E-77 | 0,0057142 | 0,0031363 | 0,0045644 | 0,0045644 | TRUE | 0,0895331 | No |
| orf19.7266 | 630 | 509 | 576 | 316 | 1808 | 942 | 1718 | 1737 | 601 | 538 | 569 | 495 | 1449 | 1071 | 1459 | 1412 | 949,21 | 551 | 1348 | 2,444 | 1,289 | 12,753 | 2,98E-37 | 9,04E-37 | 0,012865 | 0,003821 | 0,0084613 | 0,0084613 | TRUE | 0,2241596 | No |
| orf19.7267 | 1024 | 743 | 948 | 714 | 1927 | 1247 | 1650 | 1903 | 976 | 786 | 937 | 1119 | 1544 | 1418 | 1401 | 1547 | 1215,89 | 954 | 1478 | 1,553 | 0,635 | 7,051 | 1,78E-12 | 3,33E-12 | 0,0105694 | 0,0034417 | 0,0068975 | 0,0068975 | TRUE | 0,1673164 | No |
| orf19.7269 | 6009 | 4459 | 5104 | 3643 | 8535 | 5530 | 8095 | 8476 | 5729 | 4715 | 5042 | 5707 | 6839 | 6288 | 6873 | 6892 | 6010,54 | 5298 | 6723 | 1,27 | 0,344 | 5,189 | 2,11E-07 | 3,36E-07 | 0,0052126 | 0,0023645 | 0,0040567 | 0,0040567 | TRUE | 0,0670286 | No |
| orf19.727 | 46 | 41 | 46 | 39 | 176 | 141 | 165 | 167 | 44 | 43 | 45 | 61 | 141 | 160 | 140 | 136 | 96,37 | 48 | 144 | 3,026 | 1,598 | 9,74 | 2,04E-22 | 4,79E-22 | 1,00E-08 | 0,0191285 | 0,0110773 | 0,0110773 | TRUE | 0,168694 | No |
| orf19.7270 | 3741 | 2872 | 3242 | 1766 | 11668 | 6100 | 10612 | 10910 | 3567 | 3037 | 3203 | 2767 | 9349 | 6936 | 9010 | 8871 | 5842,33 | 3144 | 8542 | 2,716 | 1,442 | 15,161 | 6,45E-52 | 2,44E-51 | 0,0135044 | 0,0023723 | 0,0084542 | 0,0084542 | TRUE | 0,1948572 | No |
| orf19.7271 | 2637 | 2290 | 2499 | 1450 | 2276 | 1567 | 2155 | 2266 | 2514 | 2421 | 2469 | 2272 | 1824 | 1782 | 1830 | 1842 | 2119,17 | 2419 | 1820 | 0,751 | -0,413 | -9,286 | 1,60E-20 | 3,64E-20 | 0,00041 | 0,0028661 | 0,0014221 | 0,0014221 | TRUE | 0,0185345 | No |
| orf19.7272 | 1411 | 1155 | 1205 | 829 | 1735 | 1092 | 1628 | 1790 | 1345 | 1221 | 1190 | 1299 | 1390 | 1242 | 1382 | 1455 | 1315,64 | 1264 | 1367 | 1,084 | 0,116 | 1,818 | 0,0690185 | 0,0819362 | 0,0027194 | 0,0033393 | 0,0031525 | 0,0031525 | TRUE | 0,0438527 | No |
| orf19.7274 | 162 | 116 | 135 | 98 | 346 | 281 | 352 | 374 | 154 | 123 | 133 | 154 | 277 | 320 | 299 | 304 | 220,46 | 141 | 300 | 2,131 | 1,092 | 9,969 | 2,08E-23 | 5,00E-23 | 0,0011024 | 0,0095388 | 0,0060678 | 0,0060678 | TRUE | 0,0787908 | No |
| orf19.7275 | 7 | 6 | 11 | 7 | 20 | 20 | 26 | 14 | 7 | 6 | 11 | 11 | 16 | 23 | 22 | 11 | 13,38 | 9 | 18 | 2,087 | 1,061 | 2,498 | 0,0124865 | 0,015765 | 1,00E-08 | 0,1247641 | 0,0835638 | 0,0835638 | TRUE | 0,3495874 | No |
| orf19.7276.1 | 136 | 132 | 112 | 78 | 520 | 329 | 564 | 549 | 130 | 140 | 111 | 122 | 417 | 374 | 479 | 446 | 277,26 | 126 | 429 | 3,423 | 1,775 | 15,346 | 3,76E-53 | 1,45E-52 | 0,0066698 | 0,0080132 | 0,0073775 | 0,0073775 | TRUE | 0,0799198 | No |
| orf19.7277 | 10 | 0 | 8 | 7 | 17 | 16 | 14 | 18 | 10 | 0 | 8 | 11 | 14 | 18 | 12 | 15 | 10,84 | 7 | 15 | 2,087 | 1,062 | 2,03 | 0,0423549 | 0,0512548 | 0,1421912 | 0,1535239 | 0,1516636 | 0,1516636 | TRUE | 1,2108228 | No |
| orf19.7278 | 0 | 0 | 0 | 0 | 0 | 0 | 0 | 0 | 0 | 0 | 0 | 0 | 0 | 0 | 0 | 0 | 0 | 0 | 0 | NA | NA | NA | NA | NA | NA | NA | NA | NA | NA | NA | NA |
| orf19.7279 | 136 | 120 | 120 | 112 | 558 | 419 | 528 | 606 | 130 | 127 | 119 | 175 | 447 | 476 | 448 | 493 | 301,89 | 138 | 466 | 3,444 | 1,784 | 16,333 | 5,71E-60 | 2,40E-59 | 0,0038345 | 0,0075301 | 0,0063864 | 0,0063864 | TRUE | 0,2738706 | No |
| orf19.7279.1 | 2912 | 2468 | 2654 | 1710 | 11694 | 8092 | 10880 | 11248 | 2776 | 2610 | 2622 | 2679 | 9370 | 9201 | 9237 | 9146 | 5955,09 | 2672 | 9238 | 3,458 | 1,79 | 52,085 | 0 | 0 | 0,0001361 | 0,002367 | 0,0008756 | 0,0008756 | TRUE | 0,0091402 | No |
| orf19.728 | 658 | 752 | 786 | 418 | 1031 | 732 | 883 | 1019 | 627 | 795 | 776 | 655 | 826 | 832 | 750 | 829 | 761,31 | 713 | 809 | 1,132 | 0,179 | 2,111 | 0,0347634 | 0,0423654 | 0,0064891 | 0,004248 | 0,005559 | 0,005559 | TRUE | 0,0850319 | No |
| orf19.7280 | 4 | 2 | 2 | 3 | 4 | 7 | 5 | 2 | 4 | 2 | 2 | 5 | 3 | 8 | 4 | 2 | 3,7 | 3 | 4 | 1,344 | 0,427 | 0,555 | 0,5789756 | 0,6131856 | 1,00E-08 | 0,4452551 | 0,2788316 | 0,2788316 | TRUE | 0,496919 | No |
| orf19.7281 | 2115 | 1813 | 1971 | 1252 | 8512 | 5984 | 8196 | 8630 | 2016 | 1917 | 1947 | 1961 | 6820 | 6804 | 6958 | 7017 | 4430,26 | 1960 | 6900 | 3,519 | 1,815 | 50,936 | 0 | 0 | 1,00E-08 | 0,0024619 | 0,0008695 | 0,0008695 | TRUE | 0,004879 | No |
| orf19.7282 | 2788 | 2166 | 2466 | 1625 | 10037 | 6395 | 9500 | 9810 | 2658 | 2290 | 2436 | 2546 | 8042 | 7272 | 8065 | 7976 | 5160,77 | 2482 | 7839 | 3,159 | 1,66 | 29,768 | 1,00E-194 | 1,35E-193 | 0,0028205 | 0,0024094 | 0,0027005 | 0,0027005 | TRUE | 0,0333597 | No |
| orf19.7283 | 20935 | 16851 | 18239 | 12063 | 2965 | 1938 | 2889 | 2959 | 19959 | 17818 | 18018 | 18897 | 2376 | 2204 | 2453 | 2406 | 10516,37 | 18673 | 2360 | 0,126 | -2,983 | -58,966 | 0 | 0 | 0,0021107 | 0,0022474 | 0,0022391 | 0,0022391 | TRUE | 0,0265164 | No |
| orf19.7284 | 13003 | 10369 | 11428 | 6671 | 9500 | 6107 | 9197 | 9518 | 12397 | 10964 | 11290 | 10450 | 7612 | 6944 | 7808 | 7739 | 9400,58 | 11275 | 7526 | 0,667 | -0,583 | -9,843 | 7,37E-23 | 1,75E-22 | 0,0039067 | 0,0022659 | 0,003265 | 0,003265 | TRUE | 0,0549355 | No |
| orf19.7285 | 2522 | 2080 | 2310 | 1700 | 3616 | 2618 | 3488 | 3712 | 2404 | 2199 | 2282 | 2663 | 2897 | 2977 | 2961 | 3018 | 2675,35 | 2387 | 2963 | 1,244 | 0,315 | 5,426 | 5,76E-08 | 9,35E-08 | 0,00323 | 0,002705 | 0,0028395 | 0,0028395 | TRUE | 0,0692989 | No |
| orf19.7286 | 2845 | 2646 | 2858 | 1885 | 6012 | 4148 | 5279 | 6086 | 2712 | 2798 | 2823 | 2953 | 4817 | 4717 | 4482 | 4948 | 3781,35 | 2822 | 4741 | 1,682 | 0,75 | 16,486 | 4,63E-61 | 1,98E-60 | 0,0011106 | 0,0025255 | 0,0016957 | 0,0016957 | TRUE | 0,0167473 | No |
| orf19.7288 | 616 | 507 | 560 | 389 | 5439 | 3979 | 5378 | 5857 | 587 | 536 | 553 | 609 | 4358 | 4524 | 4566 | 4762 | 2562,1 | 571 | 4552 | 7,992 | 2,999 | 54,32 | 0 | 0 | 0,0010226 | 0,0027321 | 0,0018498 | 0,0018498 | TRUE | 0,0193071 | No |
| orf19.729 | 1109 | 1030 | 1042 | 595 | 1716 | 1105 | 1689 | 1604 | 1057 | 1089 | 1029 | 932 | 1375 | 1256 | 1434 | 1304 | 1184,69 | 1027 | 1342 | 1,305 | 0,384 | 5,985 | 2,16E-09 | 3,67E-09 | 0,0027221 | 0,0034772 | 0,0030693 | 0,0030693 | TRUE | 0,0411021 | No |
| orf19.7290 | 1321 | 1161 | 1169 | 720 | 4653 | 2731 | 4415 | 4498 | 1259 | 1228 | 1155 | 1128 | 3728 | 3105 | 3748 | 3657 | 2376,14 | 1192 | 3560 | 2,983 | 1,577 | 22,832 | 2,19E-115 | 1,66E-114 | 0,0048904 | 0,0027823 | 0,003984 | 0,003984 | TRUE | 0,0882998 | No |

| | | | | | | | | | | | | | | | | | | | | | | | | | | | | | | | |
|---|---|---|---|---|---|---|---|---|---|---|---|---|---|---|---|---|---|---|---|---|---|---|---|---|---|---|---|---|---|---|---|
| orf19.7291 | 4096 | 3213 | 3769 | 2474 | 6554 | 4496 | 6433 | 6476 | 3905 | 3397 | 3723 | 3876 | 5252 | 5112 | 5462 | 5266 | 4499,07 | 3725 | 5273 | 1,416 | 0,502 | 9,95 | 2,52E-23 | 6,05E-23 | 0,002017 | 0,0024562 | 0,0022082 | 0,0022082 | TRUE | 0,0430183 | No |
| orf19.729.1 | 1528 | 1315 | 1444 | 1011 | 442 | 327 | 434 | 447 | 1457 | 1390 | 1427 | 1584 | 354 | 372 | 368 | 363 | 914,43 | 1464 | 364 | 0,249 | -2,005 | -31,098 | 2,59E-212 | 3,74E-211 | 1,00E-08 | 0,0038868 | 0,0023842 | 0,0023842 | TRUE | 0,0328894 | No |
| orf19.7292 | 2406 | 1944 | 2107 | 1329 | 4625 | 3010 | 4510 | 4852 | 2294 | 2056 | 2082 | 2082 | 3706 | 3423 | 3829 | 3945 | 2926,94 | 2128 | 3726 | 1,75 | 0,808 | 14,088 | 4,48E-45 | 1,53E-44 | 0,0026846 | 0,0026522 | 0,0027723 | 0,0027723 | TRUE | 0,0356653 | No |
| orf19.7293 | 446 | 413 | 437 | 278 | 860 | 615 | 770 | 908 | 425 | 437 | 432 | 436 | 689 | 699 | 654 | 738 | 563,69 | 432 | 695 | 1,609 | 0,686 | 10,042 | 9,99E-24 | 2,41E-23 | 1,00E-08 | 0,005004 | 0,0025514 | 0,0025514 | TRUE | 0,0219131 | No |
| orf19.7295 | 846 | 719 | 858 | 610 | 1307 | 1024 | 1251 | 1381 | 807 | 760 | 848 | 956 | 1047 | 1164 | 1062 | 1123 | 970,83 | 843 | 1099 | 1,308 | 0,388 | 5,321 | 1,03E-07 | 1,66E-07 | 0,0042524 | 0,0037825 | 0,0040233 | 0,0040233 | TRUE | 0,0875352 | No |
| orf19.7296 | 7941 | 6914 | 7200 | 4954 | 2395 | 1872 | 2314 | 2580 | 7571 | 7311 | 7113 | 7761 | 1919 | 2129 | 1965 | 2098 | 4733,15 | 7439 | 2028 | 0,272 | -1,876 | -38,948 | 0 | 0 | 0,0015679 | 0,0024382 | 0,0019316 | 0,0019316 | TRUE | 0,0158275 | No |
| orf19.7297 | 2203 | 1940 | 2009 | 1207 | 3130 | 2206 | 3024 | 3115 | 2100 | 2051 | 1985 | 1891 | 2508 | 2508 | 2567 | 2533 | 2267,96 | 2007 | 2529 | 1,258 | 0,331 | 7,602 | 2,92E-14 | 5,78E-14 | 0,0003195 | 0,0028152 | 0,0013648 | 0,0013648 | TRUE | 0,0163005 | No |
| orf19.7298 | 1607 | 1624 | 1714 | 1002 | 2893 | 2063 | 2576 | 2862 | 1532 | 1717 | 1693 | 1570 | 2318 | 2346 | 2187 | 2327 | 1961,28 | 1628 | 2294 | 1,408 | 0,494 | 9,491 | 2,28E-21 | 5,24E-21 | 0,0013603 | 0,0029284 | 0,0020572 | 0,0020572 | TRUE | 0,0208273 | No |
| orf19.7299.1 | 6 | 7 | 6 | 5 | 7 | 3 | 3 | 4 | 6 | 7 | 6 | 8 | 6 | 3 | 3 | 3 | 5,21 | 7 | 4 | 0,562 | -0,832 | -1,364 | 0,172542 | 0,1968867 | 1,00E-08 | 0,3170719 | 0,1546889 | 0,1546889 | TRUE | 0,2450077 | No |
| orf19.73 | 5017 | 5057 | 5243 | 3071 | 2813 | 2126 | 2464 | 2457 | 4783 | 5347 | 5180 | 4811 | 2254 | 2417 | 2092 | 1998 | 3610,24 | 5030 | 2190 | 0,435 | -1,201 | -18,495 | 2,27E-76 | 1,18E-75 | 0,0046479 | 0,0025461 | 0,0037342 | 0,0037342 | TRUE | 0,0575456 | No |
| orf19.730 | 2085 | 1697 | 1934 | 1258 | 687 | 490 | 691 | 710 | 1988 | 1794 | 1911 | 1971 | 550 | 557 | 587 | 577 | 1241,89 | 1916 | 568 | 0,297 | -1,753 | -30,951 | 2,47E-210 | 3,53E-209 | 1,00E-08 | 0,0034134 | 0,0020026 | 0,0020026 | TRUE | 0,0222673 | No |
| orf19.7300 | 2112 | 2121 | 2280 | 1400 | 1055 | 860 | 934 | 977 | 2014 | 2243 | 2252 | 2193 | 845 | 978 | 793 | 794 | 1514,06 | 2176 | 852 | 0,391 | -1,355 | -18,848 | 3,03E-79 | 1,64E-78 | 0,0054168 | 0,0031757 | 0,0041747 | 0,0041747 | TRUE | 0,1104678 | No |
| orf19.7301 | 198 | 165 | 157 | 109 | 608 | 415 | 551 | 573 | 189 | 174 | 155 | 171 | 487 | 472 | 468 | 466 | 322,73 | 172 | 473 | 2,744 | 1,456 | 16,151 | 1,12E-58 | 4,65E-58 | 1,00E-08 | 0,0071788 | 0,0036494 | 0,0036494 | TRUE | 0,055738 | No |
| orf19.7303 | 10 | 11 | 9 | 3 | 5 | 1 | 3 | 0 | 10 | 12 | 9 | 5 | 4 | 1 | 3 | 0 | 5,31 | 9 | 2 | 0,222 | -2,169 | -2,953 | 0,0031433 | 0,0041118 | 1,00E-08 | 0,3115467 | 0,229191 | 0,229191 | TRUE | 0,5990437 | No |
| orf19.7304 | 1088 | 1166 | 1251 | 719 | 458 | 318 | 423 | 400 | 1037 | 1233 | 1236 | 1126 | 367 | 362 | 359 | 325 | 755,67 | 1158 | 353 | 0,305 | -1,715 | -21,788 | 3,03E-105 | 2,07E-104 | 0,0041428 | 0,0042641 | 0,0041979 | 0,0041979 | TRUE | 0,0626175 | No |
| orf19.7305 | 504 | 480 | 435 | 230 | 702 | 482 | 673 | 656 | 481 | 508 | 430 | 360 | 562 | 548 | 571 | 533 | 499,18 | 445 | 554 | 1,239 | 0,309 | 3,296 | 0,0009803 | 0,001323 | 0,007542 | 0,0053805 | 0,0063645 | 0,0063645 | TRUE | 0,1625856 | No |
| orf19.7306 | 681 | 622 | 663 | 407 | 584 | 444 | 547 | 548 | 649 | 658 | 655 | 638 | 468 | 505 | 464 | 446 | 560,29 | 650 | 471 | 0,722 | -0,47 | -7,154 | 8,43E-13 | 1,60E-12 | 1,00E-08 | 0,0050217 | 0,0023452 | 0,0023452 | TRUE | 0,0247221 | No |
| orf19.7307 | 2994 | 2678 | 2864 | 2183 | 4680 | 4020 | 4312 | 4605 | 2854 | 2832 | 2829 | 3420 | 3750 | 4571 | 3661 | 3744 | 3457,68 | 2984 | 3932 | 1,318 | 0,399 | 4,787 | 1,70E-06 | 2,60E-06 | 0,0102924 | 0,0025661 | 0,0063613 | 0,0063613 | TRUE | 0,1447552 | No |
| orf19.7308 | 4789 | 3920 | 4311 | 2869 | 6715 | 4645 | 6195 | 6824 | 4566 | 4145 | 4259 | 4494 | 5381 | 5282 | 5260 | 5549 | 4866,8 | 4366 | 5368 | 1,23 | 0,299 | 6,973 | 3,11E-12 | 5,79E-12 | 0,0010447 | 0,0024286 | 0,0015499 | 0,0015499 | TRUE | 0,0142026 | No |
| orf19.731 | 2220 | 1870 | 2056 | 1372 | 3665 | 2686 | 3577 | 3818 | 2117 | 1977 | 2031 | 2149 | 2937 | 3054 | 3037 | 3104 | 2550,8 | 2068 | 3033 | 1,468 | 0,554 | 12,901 | 4,42E-38 | 1,36E-37 | 0,0004495 | 0,002735 | 0,0013483 | 0,0013483 | TRUE | 0,0104939 | No |
| orf19.7310 | 8335 | 7769 | 8074 | 5074 | 2764 | 2048 | 2599 | 2744 | 7947 | 8215 | 7976 | 7949 | 2215 | 2329 | 2207 | 2231 | 5133,43 | 8022 | 2246 | 0,28 | -1,838 | -53,014 | 0 | 0 | 0,0001257 | 0,0024111 | 0,0008857 | 0,0008857 | TRUE | 0,0070001 | No |
| orf19.7311 | 754 | 712 | 757 | 538 | 633 | 538 | 652 | 696 | 719 | 753 | 748 | 843 | 507 | 612 | 554 | 566 | 662,6 | 766 | 560 | 0,732 | -0,451 | -5,934 | 2,96E-09 | 5,00E-09 | 0,0033394 | 0,0045693 | 0,0040051 | 0,0040051 | TRUE | 0,0491019 | No |
| orf19.7312 | 2362 | 2122 | 2121 | 1380 | 4223 | 3053 | 3851 | 4261 | 2252 | 2244 | 2095 | 2162 | 3384 | 3472 | 3269 | 3465 | 2792,78 | 2188 | 3398 | 1,552 | 0,634 | 14,84 | 8,08E-50 | 2,98E-49 | 0,0006015 | 0,0026792 | 0,0013632 | 0,0013632 | TRUE | 0,0106512 | No |
| orf19.7313 | 3293 | 2965 | 3225 | 2156 | 2255 | 1604 | 2123 | 2266 | 3140 | 3135 | 3186 | 3377 | 1807 | 1824 | 1802 | 1842 | 2514,22 | 3210 | 1819 | 0,567 | -0,817 | -20,01 | 4,48E-89 | 2,66E-88 | 0,0002424 | 0,0027443 | 0,0011811 | 0,0011811 | TRUE | 0,0138355 | No |
| orf19.7314 | 4952 | 4213 | 4436 | 3023 | 2168 | 1583 | 2021 | 2280 | 4721 | 4455 | 4382 | 4736 | 1737 | 1800 | 1716 | 1854 | 3175,1 | 4574 | 1777 | 0,389 | -1,364 | -29,633 | 5,64E-193 | 7,50E-192 | 0,0011067 | 0,0026084 | 0,0016589 | 0,0016589 | TRUE | 0,0108423 | No |
| orf19.7316 | 1517 | 1263 | 1532 | 884 | 1539 | 1036 | 1476 | 1618 | 1446 | 1335 | 1513 | 1385 | 1233 | 1178 | 1253 | 1316 | 1332,49 | 1420 | 1245 | 0,877 | -0,19 | -3,295 | 0,0009852 | 0,0013293 | 0,0017645 | 0,0033235 | 0,0024307 | 0,0024307 | TRUE | 0,0241166 | No |
| orf19.7317 | 1787 | 1563 | 1757 | 1004 | 1264 | 973 | 1237 | 1276 | 1704 | 1653 | 1736 | 1573 | 1013 | 1106 | 1050 | 1038 | 1358,98 | 1666 | 1052 | 0,629 | -0,669 | -12,821 | 1,24E-37 | 3,80E-37 | 0,0007056 | 0,0032995 | 0,0018501 | 0,0018501 | TRUE | 0,0154622 | No |
| orf19.7319 | 1590 | 1427 | 1570 | 957 | 1297 | 984 | 1270 | 1316 | 1516 | 1509 | 1551 | 1499 | 1039 | 1119 | 1078 | 1070 | 1297,67 | 1519 | 1076 | 0,708 | -0,499 | -10,621 | 2,39E-26 | 6,09E-26 | 1,00E-08 | 0,0033565 | 0,0013378 | 0,0013378 | TRUE | 0,0076137 | No |
| orf19.732 | 4191 | 3278 | 3569 | 2526 | 3855 | 2688 | 3667 | 3964 | 3996 | 3466 | 3526 | 3957 | 3089 | 3056 | 3113 | 3223 | 3428,31 | 3736 | 3120 | 0,836 | -0,259 | -4,644 | 3,41E-06 | 5,17E-06 | 0,0026624 | 0,0025702 | 0,0026858 | 0,0026858 | TRUE | 0,0287418 | No |
| orf19.7320 | 105 | 81 | 94 | 77 | 152 | 103 | 144 | 152 | 100 | 86 | 93 | 121 | 122 | 117 | 122 | 124 | 110,5 | 100 | 121 | 1,23 | 0,298 | 2,131 | 0,033068 | 0,0403531 | 1,00E-08 | 0,0169501 | 0,0095168 | 0,0095168 | TRUE | 0,143759 | No |
| orf19.7321 | 621 | 553 | 670 | 369 | 880 | 620 | 897 | 880 | 592 | 585 | 662 | 578 | 705 | 705 | 762 | 716 | 662,99 | 604 | 722 | 1,193 | 0,254 | 3,784 | 0,0001543 | 0,0002163 | 0,0010861 | 0,0045678 | 0,0027874 | 0,0027874 | TRUE | 0,0502906 | No |
| orf19.7322 | 5513 | 4619 | 4705 | 3432 | 8368 | 5956 | 7807 | 8494 | 5256 | 4884 | 4648 | 5376 | 6705 | 6773 | 6628 | 6906 | 5897,08 | 5041 | 6753 | 1,341 | 0,423 | 8,477 | 2,31E-17 | 4,90E-17 | 0,0021371 | 0,0023697 | 0,0022107 | 0,0022107 | TRUE | 0,0337398 | No |
| orf19.7323 | 5907 | 5677 | 5986 | 3727 | 5267 | 3593 | 5167 | 5463 | 5632 | 6003 | 5914 | 5839 | 4220 | 4086 | 4387 | 4442 | 5065,14 | 5847 | 4284 | 0,733 | -0,448 | -10,987 | 4,39E-28 | 1,15E-27 | 0,0008136 | 0,0024154 | 0,0013955 | 0,0013955 | TRUE | 0,0113796 | No |
| orf19.7324 | 386 | 317 | 330 | 226 | 373 | 250 | 415 | 418 | 368 | 335 | 326 | 354 | 299 | 284 | 352 | 340 | 332,33 | 346 | 319 | 0,925 | -0,112 | -1,195 | 0,2321542 | 0,2612655 | 0,0038275 | 0,0070319 | 0,0054751 | 0,0054751 | TRUE | 0,0558565 | No |
| orf19.7325 | 4315 | 3563 | 3964 | 2487 | 4711 | 3028 | 4351 | 4698 | 4114 | 3767 | 3916 | 3896 | 3775 | 3443 | 3694 | 3820 | 3803,15 | 3923 | 3683 | 0,939 | -0,091 | -1,935 | 0,0530302 | 0,0635825 | 0,0013566 | 0,002523 | 0,0018414 | 0,0018414 | TRUE | 0,0226591 | No |
| orf19.7326 | 2863 | 2642 | 2776 | 1943 | 3073 | 2333 | 2926 | 3250 | 2730 | 2794 | 2742 | 3044 | 2462 | 2653 | 2484 | 2643 | 2693,9 | 2828 | 2560 | 0,907 | -0,141 | -2,822 | 0,0047753 | 0,0061795 | 0,0016355 | 0,0027008 | 0,0020378 | 0,0020378 | TRUE | 0,0303472 | No |
| orf19.7327 | 5006 | 4125 | 4486 | 3409 | 14992 | 10869 | 14262 | 15114 | 4773 | 4362 | 4432 | 5340 | 12013 | 12359 | 12108 | 12289 | 8459,46 | 4727 | 12192 | 2,582 | 1,369 | 22,847 | 1,58E-115 | 1,20E-114 | 0,0043584 | 0,0022854 | 0,0032904 | 0,0032904 | TRUE | 0,0906726 | No |
| orf19.7328 | 730 | 708 | 680 | 485 | 2652 | 1763 | 2397 | 2714 | 696 | 749 | 672 | 760 | 2125 | 2005 | 2035 | 2207 | 1405,95 | 719 | 2093 | 2,921 | 1,547 | 26,279 | 3,32E-152 | 3,39E-151 | 0,0014187 | 0,0032591 | 0,0023322 | 0,0023322 | TRUE | 0,0230618 | No |
| orf19.7329 | 5413 | 4499 | 4563 | 3312 | 2961 | 1983 | 2814 | 2923 | 5161 | 4757 | 4508 | 5188 | 2373 | 2255 | 2389 | 2377 | 3625,9 | 4904 | 2348 | 0,479 | -1,061 | -19,705 | 1,94E-86 | 1,12E-85 | 0,0021916 | 0,0025441 | 0,0024776 | 0,0024776 | TRUE | 0,0362278 | No |
| orf19.733 | 2122 | 1678 | 1940 | 1338 | 1700 | 1104 | 1661 | 1712 | 2023 | 1774 | 1917 | 2096 | 1362 | 1255 | 1410 | 1392 | 1653,71 | 1952 | 1355 | 0,695 | -0,524 | -8,347 | 6,98E-17 | 1,46E-16 | 0,0030377 | 0,0030841 | 0,0031649 | 0,0031649 | TRUE | 0,0454249 | No |
| orf19.7330 | 32 | 43 | 41 | 28 | 26 | 30 | 18 | 21 | 31 | 45 | 41 | 44 | 21 | 34 | 15 | 17 | 30,96 | 40 | 22 | 0,536 | -0,899 | -3,057 | 0,0022346 | 0,002954 | 0,0375287 | 0,0551316 | 0,0481321 | 0,0481321 | TRUE | 0,6963302 | No |
| orf19.7331 | 198 | 180 | 237 | 141 | 113 | 84 | 133 | 133 | 189 | 190 | 234 | 221 | 91 | 96 | 113 | 108 | 155,15 | 208 | 102 | 0,49 | -1,029 | -7,859 | 3,88E-15 | 7,84E-15 | 0,0055081 | 0,0126737 | 0,0094069 | 0,0094069 | TRUE | 0,0842121 | No |
| orf19.7332 | 35068 | 32443 | 34566 | 20422 | 40374 | 27876 | 37629 | 40342 | 33434 | 34304 | 34148 | 31992 | 32351 | 31698 | 31947 | 32802 | 32834,35 | 33470 | 32200 | 0,962 | -0,056 | -1,716 | 0,0860771 | 0,1010966 | 0,0005692 | 0,0021413 | 0,0009922 | 0,0009922 | TRUE | 0,0106707 | No |
| orf19.7335 | 5784 | 4940 | 5316 | 3115 | 5984 | 4068 | 5963 | 6165 | 5514 | 5223 | 5252 | 4880 | 4795 | 4626 | 5063 | 5013 | 5045,64 | 5217 | 4874 | 0,934 | -0,099 | -2,035 | 0,0418176 | 0,0506431 | 0,0018419 | 0,0024167 | 0,0020682 | 0,0020682 | TRUE | 0,0222776 | No |
| orf19.7336 | 5 | 16 | 13 | 3 | 2 | 0 | 0 | 0 | 5 | 17 | 13 | 5 | 2 | 0 | 0 | 0 | 5,1 | 10 | 0 | 0,045 | -4,49 | -3,809 | 0,0001396 | 0,000196 | 0,350202 | 0,3237973 | 0,3323307 | 0,3323307 | TRUE | 0,5815225 | No |
| orf19.7337 | 2352 | 2429 | 2416 | 1497 | 1793 | 1325 | 1582 | 1656 | 2242 | 2568 | 2387 | 2345 | 1437 | 1507 | 1343 | 1346 | 1896,95 | 2386 | 1408 | 0,59 | -0,762 | -12,993 | 1,35E-38 | 4,18E-38 | 0,0024328 | 0,0029568 | 0,0027501 | 0,0027501 | TRUE | 0,0327102 | No |
| orf19.7339 | 510 | 456 | 472 | 285 | 175 | 122 | 184 | 196 | 486 | 482 | 466 | 446 | 140 | 139 | 156 | 159 | 309,46 | 470 | 148 | 0,316 | -1,661 | -18,186 | 6,64E-74 | 3,36E-73 | 1,00E-08 | 0,007397 | 0,003844 | 0,003844 | TRUE | 0,0255737 | No |
| orf19.734 | 14428 | 12173 | 13318 | 8421 | 1530 | 1037 | 1526 | 1587 | 13756 | 12871 | 13157 | 13192 | 1226 | 1179 | 1296 | 1290 | 7245,86 | 13244 | 1248 | 0,094 | -3,406 | -78,694 | 0 | 0 | 0,0007995 | 0,0023179 | 0,0013996 | 0,0013996 | TRUE | 0,0149293 | No |
| orf19.7340 | 13 | 26 | 19 | 26 | 12 | 11 | 7 | 9 | 12 | 27 | 19 | 41 | 10 | 13 | 6 | 7 | 16,85 | 25 | 9 | 0,36 | -1,476 | -3,371 | 0,0007483 | 0,001019 | 0,1222532 | 0,099555 | 0,1098201 | 0,1098201 | TRUE | 0,1932685 | No |
| orf19.7341 | 4699 | 3927 | 4271 | 2649 | 2196 | 1529 | 2189 | 2264 | 4480 | 4152 | 4219 | 4150 | 1760 | 1739 | 1858 | 1841 | 3024,87 | 4250 | 1800 | 0,423 | -1,24 | -27,848 | 1,13E-170 | 1,29E-169 | 0,0008869 | 0,0026341 | 0,0015233 | 0,0015233 | TRUE | 0,0164266 | No |
| orf19.7341.1 | 30955 | 32246 | 32201 | 17266 | 14602 | 9058 | 13480 | 12636 | 29512 | 34096 | 31812 | 27048 | 11700 | 10300 | 11444 | 10274 | 20773,33 | 30617 | 10930 | 0,357 | -1,486 | -20,445 | 6,62E-93 | 4,06E-92 | 0,0071555 | 0,0021703 | 0,005017 | 0,005017 | TRUE | 0,0751524 | No |
| orf19.7342 | 795 | 813 | 882 | 568 | 357 | 262 | 304 | 241 | 758 | 860 | 871 | 890 | 286 | 298 | 258 | 196 | 552,09 | 845 | 260 | 0,307 | -1,706 | -15,184 | 4,52E-52 | 1,72E-51 | 0,0144887 | 0,0050652 | 0,0097281 | 0,0097281 | TRUE | 0,3152641 | No |
| orf19.7343 | 1816 | 1808 | 1946 | 994 | 1842 | 1237 | 1633 | 1706 | 1731 | 1912 | 1922 | 1557 | 1476 | 1407 | 1386 | 1387 | 1597,35 | 1780 | 1414 | 0,793 | -0,335 | -4,905 | 9,32E-07 | 1,44E-06 | 0,0043156 | 0,0031192 | 0,0038553 | 0,0038553 | TRUE | 0,0807582 | No |
| orf19.7344 | 1231 | 1462 | 1522 | 872 | 676 | 523 | 634 | 575 | 1174 | 1546 | 1504 | 1366 | 542 | 595 | 538 | 468 | 966,41 | 1398 | 536 | 0,383 | -1,386 | -14,327 | 1,49E-46 | 5,25E-46 | 0,0114093 | 0,0037903 | 0,0077487 | 0,0077487 | TRUE | 0,1435895 | No |
| orf19.7345 | 1240 | 1343 | 1306 | 747 | 674 | 468 | 610 | 634 | 1182 | 1420 | 1290 | 1170 | 540 | 532 | 518 | 516 | 896,04 | 1266 | 526 | 0,415 | -1,268 | -17,371 | 1,37E-67 | 6,36E-67 | 0,0037446 | 0,0039237 | 0,0038228 | 0,0038228 | TRUE | 0,0807499 | No |
| orf19.7347 | 794 | 737 | 841 | 587 | 1888 | 1333 | 1858 | 1835 | 757 | 779 | 831 | 920 | 1513 | 1516 | 1577 | 1492 | 1173,08 | 822 | 1524 | 1,864 | 0,898 | 14,364 | 8,72E-47 | 3,08E-46 | 0,0020319 | 0,0034909 | 0,0027777 | 0,0027777 | TRUE | 0,0672622 | No |
| orf19.7348 | 1 | 2 | 2 | 4 | 6 | 2 | 5 | 1 | 1 | 2 | 2 | 6 | 5 | 2 | 4 | 1 | 2,93 | 3 | 3 | 1,166 | 0,222 | 0,249 | 0,8030529 | 0,8251879 | 0,1710705 | 0,5622279 | 0,4013391 | 0,4013391 | TRUE | 0,4981679 | No |
| orf19.7349 | 6439 | 5910 | 6496 | 4123 | 7177 | 5426 | 6688 | 7252 | 6139 | 6249 | 6417 | 6459 | 5751 | 6170 | 5678 | 5897 | 6094,94 | 6316 | 5874 | 0,93 | -0,105 | -2,696 | 0,0070224 | 0,0089925 | 0,0007431 | 0,0023607 | 0,0012879 | 0,0012879 | TRUE | 0,0136616 | No |
| orf19.7350 | 24412 | 18996 | 21408 | 11558 | 55844 | 30659 | 53305 | 53005 | 23274 | 20086 | 21149 | 18106 | 44746 | 34862 | 45256 | 43098 | 31322,18 | 20654 | 41990 | 2,033 | 1,024 | 11,478 | 1,71E-30 | 4,65E-30 | 0,012037 | 0,0021437 | 0,0076032 | 0,0076032 | TRUE | 0,159869 | No |
| orf19.7353 | 746 | 709 | 739 | 472 | 1427 | 1106 | 1385 | 1462 | 711 | 750 | 730 | 739 | 1143 | 1258 | 1176 | 1189 | 962 | 732 | 1192 | 1,625 | 0,701 | 12,606 | 1,96E-36 | 5,85E-36 | 1,00E-08 | 0,003798 | 0,0018288 | 0,0018288 | TRUE | 0,015594 | No |
| orf19.7354 | 1131 | 1243 | 1305 | 672 | 2305 | 1724 | 2152 | 1930 | 1078 | 1314 | 1289 | 1053 | 1847 | 1960 | 1827 | 1569 | 1492,27 | 1184 | 1801 | 1,519 | 0,603 | 6,77 | 1,29E-11 | 2,37E-11 | 0,0099908 | 0,0031916 | 0,0068889 | 0,0068889 | TRUE | 0,0911896 | No |
| orf19.7355 | 651 | 699 | 660 | 453 | 984 | 761 | 901 | 890 | 621 | 739 | 652 | 710 | 788 | 865 | 765 | 724 | 732,98 | 680 | 786 | 1,155 | 0,208 | 2,677 | 0,0074386 | 0,0095084 | 0,0045468 | 0,0043313 | 0,004392 | 0,004392 | TRUE | 0,0524912 | No |
| orf19.7356 | 414 | 418 | 445 | 273 | 707 | 452 | 654 | 656 | 395 | 442 | 440 | 428 | 566 | 514 | 555 | 533 | 484,13 | 426 | 542 | 1,276 | 0,352 | 4,835 | 1,33E-06 | 2,05E-06 | 1,00E-08 | 0,0054827 | 0,0029465 | 0,0029465 | TRUE | 0,0314187 | No |
| orf19.7357 | 1907 | 1816 | 1927 | 1257 | 2140 | 1691 | 2048 | 2011 | 1818 | 1920 | 1904 | 1969 | 1715 | 1923 | 1739 | 1635 | 1827,82 | 1903 | 1753 | 0,921 | -0,119 | -2,084 | 0,0371541 | 0,0451581 | 0,0022245 | 0,0029896 | 0,0025873 | 0,0025873 | TRUE | 0,0493927 | No |
| orf19.7358 | 881 | 1070 | 1104 | 549 | 899 | 696 | 799 | 692 | 840 | 1131 | 1091 | 860 | 720 | 791 | 678 | 563 | 834,35 | 980 | 688 | 0,7 | -0,514 | -4,321 | 1,55E-05 | 2,28E-05 | 0,0200352 | 0,0040592 | 0,0123639 | 0,0123639 | TRUE | 0,1757412 | No |
| orf19.7359 | 2808 | 2569 | 2752 | 1798 | 2049 | 1502 | 1897 | 2173 | 2677 | 2716 | 2719 | 2817 | 1642 | 1708 | 1611 | 1767 | 2207 | 2732 | 1682 | 0,616 | -0,699 | -15,585 | 9,14E-55 | 3,60E-54 | 0,0005723 | 0,0028352 | 0,0014612 | 0,0014612 | TRUE | 0,0148013 | No |
| orf19.736 | 1859 | 1875 | 1984 | 1221 | 968 | 672 | 835 | 827 | 1772 | 1983 | 1960 | 1913 | 776 | 764 | 709 | 672 | 1318,6 | 1907 | 730 | 0,383 | -1,386 | -21,99 | 3,61E-107 | 2,53E-106 | 0,0025602 | 0,0033365 | 0,0029067 | 0,0029067 | TRUE | 0,0346528 | No |
| orf19.7360 | 498 | 588 | 575 | 313 | 513 | 384 | 503 | 513 | 475 | 622 | 568 | 490 | 411 | 437 | 427 | 417 | 480,84 | 539 | 423 | 0,783 | -0,353 | -3,847 | 0,0001194 | 0,0001683 | 0,0064359 | 0,0055059 | 0,0059808 | 0,0059808 | TRUE | 0,1255842 | No |
| orf19.7361 | 3184 | 3080 | 3278 | 1946 | 3168 | 2273 | 3026 | 2966 | 3036 | 3257 | 3238 | 3049 | 2538 | 2585 | 2569 | 2412 | 2835,36 | 3145 | 2526 | 0,802 | -0,318 | -7,172 | 7,37E-13 | 1,40E-12 | 0,000866 | 0,0026704 | 0,0015262 | 0,0015262 | TRUE | 0,0114198 | No |
| orf19.7362 | 1051 | 1066 | 1091 | 726 | 839 | 600 | 798 | 851 | 1002 | 1127 | 1078 | 1137 | 672 | 682 | 677 | 692 | 883,53 | 1086 | 681 | 0,628 | -0,67 | -11,245 | 2,46E-29 | 6,54E-29 | 0,0008768 | 0,0039496 | 0,0022402 | 0,0022402 | TRUE | 0,0335886 | No |
| orf19.7363 | 10888 | 10306 | 11175 | 6791 | 8497 | 6176 | 7903 | 8645 | 10381 | 10897 | 11040 | 10638 | 6808 | 7023 | 6710 | 7029 | 8815,75 | 10739 | 6892 | 0,642 | -0,64 | -18,39 | 1,58E-75 | 8,18E-75 | 0,0004757 | 0,0022775 | 0,0010461 | 0,0010461 | TRUE | 0,0063564 | No |
| orf19.7364 | 36 | 31 | 28 | 20 | 30 | 36 | 30 | 51 | 34 | 33 | 28 | 31 | 24 | 41 | 25 | 41 | 32,25 | 32 | 33 | 1,038 | 0,053 | 0,196 | 0,8446013 | 0,8641015 | 0,0233897 | 0,0530022 | 0,0398233 | 0,0398233 | TRUE | 0,1557185 | No |
| orf19.7365 | 1185 | 1239 | 1310 | 827 | 230 | 239 | 205 | 223 | 1130 | 1310 | 1294 | 1296 | 184 | 272 | 174 | 181 | 730,12 | 1258 | 203 | 0,16 | -2,644 | -22,874 | 8,47E-116 | 6,49E-115 | 0,019003 | 0,0043401 | 0,0101609 | 0,0101609 | TRUE | 0,5273779 | No |

| | | | | | | | | | | | | | | | | | | | | | | | | | | | | | | | |
|---|---|---|---|---|---|---|---|---|---|---|---|---|---|---|---|---|---|---|---|---|---|---|---|---|---|---|---|---|---|---|---|
| orf19.7366 | 2973 | 2787 | 3005 | 1482 | 3321 | 2014 | 3126 | 3105 | 2834 | 2947 | 2969 | 2322 | 2661 | 2290 | 2654 | 2525 | 2650,17 | 2768 | 2532 | 0,914 | -0,129 | -1,628 | 0,1034897 | 0,1206592 | 0,0078397 | 0,0027108 | 0,0056916 | 0,0056916 | TRUE | 0,1384789 | No |
| orf19.7367 | 2879 | 3039 | 3096 | 1890 | 3202 | 2458 | 2900 | 2987 | 2745 | 3213 | 3059 | 2961 | 2566 | 2795 | 2462 | 2429 | 2778,62 | 2994 | 2563 | 0,855 | -0,225 | -3,621 | 0,0002931 | 0,0004064 | 0,0037453 | 0,0026822 | 0,0033604 | 0,0033604 | TRUE | 0,0440074 | No |
| orf19.7368 | 1624 | 1425 | 1577 | 1046 | 3755 | 2624 | 3512 | 3736 | 1548 | 1507 | 1558 | 1639 | 3009 | 2984 | 2982 | 3038 | 2282,94 | 1563 | 3003 | 1,926 | 0,945 | 23,261 | 1,11E-119 | 8,81E-119 | 1,00E-08 | 0,0028105 | 0,0010785 | 0,0010785 | TRUE | 0,0109535 | No |
| orf19.7369 | 613 | 705 | 663 | 415 | 920 | 707 | 875 | 783 | 584 | 745 | 655 | 650 | 737 | 804 | 743 | 637 | 694,45 | 658 | 730 | 1,107 | 0,147 | 1,616 | 0,1061097 | 0,1235108 | 0,0082765 | 0,0044556 | 0,0064602 | 0,0064602 | TRUE | 0,0951558 | No |
| orf19.7370 | 3240 | 3039 | 3196 | 1952 | 2428 | 1783 | 2187 | 2242 | 3089 | 3213 | 3157 | 3058 | 1945 | 2027 | 1857 | 1823 | 2521,28 | 3129 | 1913 | 0,61 | -0,712 | -15,567 | 1,21E-54 | 4,76E-54 | 0,0008441 | 0,0027425 | 0,0015957 | 0,0015957 | TRUE | 0,0184124 | No |
| orf19.7371 | 2356 | 2379 | 2527 | 1546 | 2460 | 1876 | 2433 | 2503 | 2246 | 2515 | 2496 | 2422 | 1971 | 2133 | 2066 | 2035 | 2235,64 | 2420 | 2051 | 0,847 | -0,239 | -4,745 | 2,09E-06 | 3,19E-06 | 0,0014595 | 0,0028257 | 0,0019875 | 0,0019875 | TRUE | 0,0296762 | No |
| orf19.7372 | 1580 | 1420 | 1548 | 1001 | 1872 | 1322 | 1755 | 1840 | 1506 | 1501 | 1529 | 1568 | 1500 | 1503 | 1490 | 1496 | 1511,82 | 1526 | 1497 | 0,982 | -0,026 | -0,612 | 0,5405139 | 0,5763735 | 1,00E-08 | 0,0031773 | 0,0010703 | 0,0010703 | TRUE | 0,0034989 | No |
| orf19.7374 | 8162 | 7571 | 8377 | 5243 | 5091 | 3637 | 4612 | 5184 | 7782 | 8005 | 8276 | 8213 | 4079 | 4136 | 3916 | 4215 | 6077,7 | 8069 | 4086 | 0,507 | -0,981 | -25,536 | 7,86E-144 | 7,61E-143 | 0,00062 | 0,0023614 | 0,001239 | 0,001239 | TRUE | 0,0097454 | No |
| orf19.7375 | 361 | 328 | 307 | 201 | 497 | 348 | 456 | 509 | 344 | 347 | 303 | 315 | 398 | 396 | 387 | 414 | 363,01 | 327 | 399 | 1,216 | 0,283 | 3,507 | 0,0004537 | 0,0006236 | 1,00E-08 | 0,0066143 | 0,003423 | 0,003423 | TRUE | 0,0322051 | No |
| orf19.7376 | 494 | 540 | 518 | 346 | 710 | 501 | 748 | 750 | 471 | 571 | 512 | 542 | 569 | 570 | 635 | 610 | 559,9 | 524 | 596 | 1,141 | 0,19 | 2,438 | 0,0147751 | 0,0185299 | 0,0030417 | 0,0050238 | 0,0040121 | 0,0040121 | TRUE | 0,0571509 | No |
| orf19.7377 | 256 | 241 | 216 | 153 | 252 | 180 | 212 | 224 | 244 | 255 | 213 | 240 | 202 | 205 | 180 | 182 | 215,09 | 238 | 192 | 0,806 | -0,31 | -2,981 | 0,0028733 | 0,0037686 | 1,00E-08 | 0,009725 | 0,0057337 | 0,0057337 | TRUE | 0,058139 | No |
| orf19.7378 | 3691 | 3379 | 3597 | 2264 | 4147 | 2805 | 4058 | 3998 | 3519 | 3573 | 3553 | 3547 | 3323 | 3190 | 3445 | 3251 | 3425,05 | 3548 | 3302 | 0,931 | -0,103 | -2,732 | 0,006302 | 0,0080944 | 0,0002982 | 0,0025707 | 0,001067 | 0,001067 | TRUE | 0,0102962 | No |
| orf19.7379 | 6918 | 6502 | 6870 | 4523 | 9365 | 7116 | 9349 | 10075 | 6596 | 6875 | 6787 | 7086 | 7504 | 8092 | 7937 | 8192 | 7383,46 | 6836 | 7931 | 1,161 | 0,215 | 5,255 | 1,48E-07 | 2,36E-07 | 0,0009928 | 0,0023137 | 0,0014676 | 0,0014676 | TRUE | 0,0162108 | No |
| orf19.738 | 4165 | 3847 | 4302 | 2598 | 3929 | 2754 | 3545 | 3880 | 3971 | 4068 | 4250 | 4070 | 3148 | 3132 | 3010 | 3155 | 3600,34 | 4090 | 3111 | 0,761 | -0,395 | -10,3 | 7,05E-25 | 1,74E-24 | 0,0004236 | 0,0025473 | 0,0011291 | 0,0011291 | TRUE | 0,0088162 | No |
| orf19.7380 | 233 | 193 | 227 | 145 | 400 | 273 | 394 | 427 | 222 | 204 | 224 | 227 | 321 | 310 | 335 | 347 | 273,78 | 219 | 328 | 1,501 | 0,585 | 6,453 | 1,10E-10 | 1,96E-10 | 1,00E-08 | 0,0080884 | 0,0040092 | 0,0040092 | TRUE | 0,0244089 | No |
| orf19.7381 | 10097 | 10064 | 10954 | 6822 | 3231 | 2543 | 2989 | 3304 | 9626 | 10641 | 10822 | 10687 | 2589 | 2892 | 2538 | 2686 | 6560,12 | 10444 | 2676 | 0,256 | -1,965 | -35,94 | 7,32E-283 | 1,50E-281 | 0,0027824 | 0,0023416 | 0,002651 | 0,002651 | TRUE | 0,0346253 | No |
| orf19.7382 | 11169 | 9580 | 10161 | 6894 | 18193 | 13621 | 17127 | 18795 | 10648 | 10130 | 10038 | 10800 | 14578 | 15488 | 14541 | 15282 | 12688,09 | 10404 | 14972 | 1,439 | 0,525 | 13,185 | 1,06E-39 | 3,34E-39 | 0,0010703 | 0,0022207 | 0,001441 | 0,001441 | TRUE | 0,0077939 | No |
| orf19.7383 | 2855 | 2672 | 2733 | 1778 | 3742 | 2555 | 3504 | 3582 | 2722 | 2825 | 2700 | 2785 | 2998 | 2905 | 2975 | 2913 | 2852,94 | 2758 | 2948 | 1,07 | 0,097 | 2,644 | 0,0081928 | 0,0104578 | 1,00E-08 | 0,0026668 | 0,0009397 | 0,0009397 | TRUE | 0,0035135 | No |
| orf19.7384 | 24913 | 22133 | 23988 | 13916 | 34633 | 23758 | 32915 | 35189 | 23752 | 23403 | 23698 | 21800 | 27750 | 27015 | 27945 | 28612 | 25496,85 | 23163 | 27830 | 1,201 | 0,265 | 6,911 | 4,80E-12 | 8,90E-12 | 0,0010174 | 0,0021557 | 0,0013671 | 0,0013671 | TRUE | 0,0189522 | No |
| orf19.7385 | 414 | 316 | 370 | 223 | 663 | 487 | 667 | 744 | 395 | 334 | 366 | 349 | 531 | 554 | 566 | 605 | 462,49 | 361 | 564 | 1,559 | 0,641 | 8,018 | 1,08E-15 | 2,20E-15 | 0,0019582 | 0,0056414 | 0,0038025 | 0,0038025 | TRUE | 0,0482568 | No |
| orf19.7386 | 2722 | 2540 | 2682 | 1853 | 1965 | 1510 | 1926 | 1905 | 2595 | 2686 | 2650 | 2903 | 1574 | 1717 | 1635 | 1549 | 2163,61 | 2708 | 1619 | 0,598 | -0,742 | -14,17 | 1,41E-45 | 4,87E-45 | 0,0016483 | 0,0028502 | 0,0021476 | 0,0021476 | TRUE | 0,0267105 | No |
| orf19.7387 | 10073 | 9012 | 9484 | 5945 | 9013 | 6103 | 8425 | 9163 | 9604 | 9529 | 9369 | 9313 | 7222 | 6940 | 7153 | 7450 | 8322,46 | 9454 | 7191 | 0,761 | -0,394 | -11,833 | 2,64E-32 | 7,42E-32 | 0,0003676 | 0,0022886 | 0,0009453 | 0,0009453 | TRUE | 0,007178 | No |
| orf19.7388 | 10961 | 9606 | 10507 | 6582 | 9322 | 6547 | 8978 | 9747 | 10450 | 10157 | 10380 | 10311 | 7469 | 7445 | 7622 | 7925 | 8969,97 | 10324 | 7615 | 0,738 | -0,439 | -13,394 | 6,56E-41 | 2,11E-40 | 0,0003426 | 0,0022743 | 0,0009177 | 0,0009177 | TRUE | 0,0092368 | No |
| orf19.7389 | 707 | 647 | 700 | 451 | 2023 | 1490 | 1865 | 2136 | 674 | 684 | 692 | 707 | 1621 | 1694 | 1583 | 1737 | 1173,95 | 689 | 1659 | 2,41 | 1,269 | 23,052 | 1,41E-117 | 1,10E-116 | 1,00E-08 | 0,0034899 | 0,0018297 | 0,0018297 | TRUE | 0,0127366 | No |
| orf19.7391 | 4073 | 3494 | 3880 | 2489 | 1957 | 1397 | 1876 | 1979 | 3883 | 3694 | 3833 | 3899 | 1568 | 1589 | 1593 | 1609 | 2708,54 | 3827 | 1590 | 0,415 | -1,267 | -32,79 | 8,09E-236 | 1,28E-234 | 1,00E-08 | 0,0026975 | 0,0010066 | 0,0010066 | TRUE | 0,0066906 | No |
| orf19.7392 | 39678 | 32854 | 36561 | 24303 | 38942 | 27269 | 35991 | 40509 | 37829 | 34739 | 36119 | 38072 | 31203 | 31007 | 30556 | 32938 | 34057,83 | 36690 | 31426 | 0,857 | -0,223 | -5,332 | 9,69E-08 | 1,56E-07 | 0,0014206 | 0,0021395 | 0,0016548 | 0,0016548 | TRUE | 0,0156891 | No |
| orf19.7393 | 3289 | 3002 | 3243 | 2161 | 4255 | 3027 | 4087 | 4272 | 3136 | 3174 | 3204 | 3385 | 3409 | 3442 | 3470 | 3474 | 3336,73 | 3225 | 3449 | 1,071 | 0,099 | 2,649 | 0,008072 | 0,0103056 | 0,0001225 | 0,0025834 | 0,0010384 | 0,0010384 | TRUE | 0,0124431 | No |
| orf19.7394 | 10600 | 9323 | 10321 | 6443 | 16566 | 12189 | 15841 | 17303 | 10106 | 9858 | 10196 | 10093 | 13274 | 13860 | 13449 | 14069 | 11863,14 | 10063 | 13663 | 1,358 | 0,441 | 13,774 | 3,66E-43 | 1,22E-42 | 0,0003458 | 0,0022297 | 0,0008959 | 0,0008959 | TRUE | 0,0049819 | No |
| orf19.7395 | 0 | 2 | 2 | 3 | 1 | 1 | 3 | 4 | 0 | 2 | 2 | 5 | 1 | 1 | 3 | 3 | 2,07 | 2 | 2 | 0,958 | -0,061 | -0,059 | 0,9531171 | 0,957921 | 1,00E-08 | 0,7968196 | 0,5444601 | 0,5444601 | TRUE | 0,2827434 | No |
| orf19.7396 | 674 | 624 | 689 | 464 | 763 | 535 | 686 | 673 | 643 | 660 | 681 | 727 | 611 | 608 | 582 | 547 | 632,41 | 678 | 587 | 0,869 | -0,203 | -2,99 | 0,0027859 | 0,00366 | 0,0010247 | 0,0046875 | 0,0028287 | 0,0028287 | TRUE | 0,0257254 | No |
| orf19.7397 | 1396 | 1312 | 1473 | 859 | 1471 | 1109 | 1407 | 1564 | 1331 | 1387 | 1455 | 1346 | 1179 | 1261 | 1195 | 1272 | 1303,12 | 1380 | 1227 | 0,888 | -0,172 | -3,259 | 0,0011184 | 0,0015037 | 0,0008497 | 0,0033513 | 0,0019018 | 0,0019018 | TRUE | 0,016858 | No |
| orf19.7397.1 | 37 | 37 | 52 | 70 | 31 | 37 | 28 | 26 | 35 | 39 | 51 | 110 | 25 | 42 | 24 | 21 | 43,41 | 59 | 28 | 0,478 | -1,064 | -2,945 | 0,0032311 | 0,0042232 | 0,1666475 | 0,0399175 | 0,0995687 | 0,0995687 | TRUE | 2,2506546 | No |
| orf19.7397.3 | 4 | 4 | 5 | 0 | 9 | 7 | 2 | 1 | 4 | 4 | 5 | 0 | 7 | 8 | 2 | 1 | 3,83 | 3 | 4 | 1,276 | 0,352 | 0,405 | 0,6853415 | 0,7165597 | 0,4963346 | 0,4304408 | 0,4563584 | 0,4563584 | TRUE | 0,1298703 | No |
| orf19.7398 | 0 | 0 | 0 | 0 | 0 | 0 | 0 | 0 | 0 | 0 | 0 | 0 | 0 | 0 | 0 | 0 | 0 | 0 | 0 | NA | NA | NA | NA | NA | NA | NA | NA | NA | NA | NA | NA |
| orf19.7398.1 | 2 | 1 | 0 | 0 | 3 | 4 | 3 | 3 | 2 | 1 | 0 | 0 | 2 | 5 | 3 | 2 | 1,86 | 1 | 3 | 3,645 | 1,866 | 1,595 | 0,1107316 | 0,1286091 | 1,00E-08 | 0,8834962 | 0,4756595 | 0,4756595 | TRUE | 0,4100081 | No |
| orf19.74 | 1188 | 1462 | 1366 | 843 | 1103 | 864 | 882 | 864 | 1133 | 1546 | 1349 | 1321 | 884 | 982 | 749 | 703 | 1083,27 | 1337 | 830 | 0,62 | -0,691 | -6,147 | 7,92E-10 | 1,36E-09 | 0,018564 | 0,003607 | 0,0111684 | 0,0111684 | TRUE | 0,1790474 | No |
| orf19.740 | 45082 | 42743 | 46629 | 28225 | 8261 | 5847 | 8303 | 8262 | 42981 | 45195 | 46065 | 44216 | 6619 | 6649 | 7049 | 6718 | 25686,5 | 44614 | 6759 | 0,152 | -2,723 | -74,508 | 0 | 0 | 0,0007568 | 0,0021552 | 0,0012038 | 0,0012038 | TRUE | 0,0102772 | No |
| orf19.7400 | 1611 | 1342 | 1483 | 1002 | 2192 | 1541 | 2080 | 2406 | 1536 | 1419 | 1465 | 1570 | 1756 | 1752 | 1766 | 1956 | 1652,56 | 1498 | 1808 | 1,209 | 0,274 | 4,879 | 1,07E-06 | 1,65E-06 | 0,0017258 | 0,0030848 | 0,0024067 | 0,0024067 | TRUE | 0,0376457 | No |
| orf19.7401 | 1744 | 1560 | 1758 | 1077 | 2332 | 1707 | 2349 | 2443 | 1663 | 1650 | 1737 | 1687 | 1869 | 1941 | 1994 | 1986 | 1815,8 | 1684 | 1948 | 1,156 | 0,21 | 4,729 | 2,26E-06 | 3,45E-06 | 1,00E-08 | 0,0029955 | 0,0013263 | 0,0013263 | TRUE | 0,0094607 | No |
| orf19.7402 | 1897 | 1649 | 1688 | 1083 | 2542 | 1737 | 2334 | 2557 | 1809 | 1744 | 1668 | 1697 | 2037 | 1975 | 1982 | 2079 | 1873,62 | 1730 | 2018 | 1,167 | 0,223 | 4,925 | 8,46E-07 | 1,31E-06 | 0,0004034 | 0,0029676 | 0,0014166 | 0,0014166 | TRUE | 0,012145 | No |
| orf19.7403 | 2844 | 2535 | 2761 | 1654 | 1971 | 1447 | 1921 | 1991 | 2711 | 2680 | 2728 | 2591 | 1579 | 1645 | 1631 | 1619 | 2148,13 | 2678 | 1618 | 0,603 | -0,729 | -18,177 | 7,79E-74 | 3,93E-73 | 1,00E-08 | 0,0028556 | 0,0010576 | 0,0010576 | TRUE | 0,0051545 | No |
| orf19.7404 | 2683 | 2087 | 2234 | 1589 | 3294 | 2143 | 3130 | 3343 | 2558 | 2207 | 2207 | 2489 | 2639 | 2437 | 2657 | 2718 | 2489,08 | 2365 | 2613 | 1,106 | 0,145 | 2,336 | 0,0194718 | 0,0241919 | 0,0036153 | 0,0027509 | 0,0033075 | 0,0033075 | TRUE | 0,0384002 | No |
| orf19.7405 | 1399 | 1229 | 1264 | 841 | 1177 | 842 | 1166 | 1147 | 1334 | 1300 | 1249 | 1317 | 943 | 957 | 990 | 933 | 1127,82 | 1300 | 956 | 0,735 | -0,443 | -8,788 | 1,52E-18 | 3,30E-18 | 1,00E-08 | 0,0035471 | 0,0015449 | 0,0015449 | TRUE | 0,0088818 | No |
| orf19.7406 | 1254 | 1145 | 1307 | 739 | 1336 | 951 | 1281 | 1418 | 1196 | 1211 | 1291 | 1158 | 1070 | 1081 | 1088 | 1153 | 1155,94 | 1214 | 1098 | 0,903 | -0,146 | -2,696 | 0,0070274 | 0,0089971 | 0,0007683 | 0,0035117 | 0,0019661 | 0,0019661 | TRUE | 0,0223379 | No |
| orf19.7409 | 832 | 698 | 725 | 492 | 1857 | 1275 | 1692 | 1874 | 793 | 738 | 716 | 771 | 1488 | 1450 | 1436 | 1524 | 1114,53 | 754 | 1474 | 1,956 | 0,968 | 17,772 | 1,16E-70 | 5,56E-70 | 0,0003073 | 0,0035645 | 0,0018066 | 0,0018066 | TRUE | 0,0163918 | No |
| orf19.7409.1 | 374 | 340 | 331 | 221 | 546 | 402 | 448 | 510 | 357 | 360 | 327 | 346 | 437 | 457 | 380 | 415 | 384,86 | 348 | 422 | 1,214 | 0,28 | 3,319 | 0,0009035 | 0,0012225 | 0,0018936 | 0,0063575 | 0,0041433 | 0,0041433 | TRUE | 0,0520239 | No |
| orf19.741 | 8 | 11 | 11 | 5 | 7 | 2 | 8 | 7 | 8 | 12 | 11 | 8 | 6 | 2 | 7 | 6 | 7,29 | 10 | 5 | 0,546 | -0,873 | -1,665 | 0,0959418 | 0,1122487 | 1,00E-08 | 0,2273 | 0,1206521 | 0,1206521 | TRUE | 0,2150572 | No |
| orf19.7410 | 4638 | 4449 | 4637 | 2964 | 7503 | 5639 | 7042 | 7738 | 4422 | 4704 | 4581 | 4643 | 6012 | 6412 | 5979 | 6292 | 5380,58 | 4588 | 6174 | 1,346 | 0,428 | 10,84 | 2,23E-27 | 5,77E-27 | 0,0006872 | 0,0023964 | 0,0013047 | 0,0013047 | TRUE | 0,0080354 | No |
| orf19.7411 | 665 | 497 | 579 | 379 | 1435 | 1058 | 1405 | 1478 | 634 | 526 | 572 | 594 | 1150 | 1203 | 1193 | 1202 | 884,09 | 582 | 1187 | 2,041 | 1,029 | 16,008 | 1,13E-57 | 4,61E-57 | 0,0011592 | 0,0039485 | 0,0026327 | 0,0026327 | TRUE | 0,0509298 | No |
| orf19.7412 | 4149 | 3729 | 4087 | 2410 | 4352 | 2952 | 4187 | 4345 | 3956 | 3943 | 4038 | 3775 | 3487 | 3357 | 3555 | 3533 | 3705,37 | 3928 | 3483 | 0,886 | -0,174 | -4,564 | 5,03E-06 | 7,56E-06 | 0,0003552 | 0,0025344 | 0,0011256 | 0,0011256 | TRUE | 0,0076805 | No |
| orf19.7413 | 777 | 758 | 745 | 542 | 1216 | 872 | 1141 | 1170 | 741 | 801 | 736 | 849 | 974 | 992 | 969 | 951 | 876,66 | 782 | 972 | 1,247 | 0,319 | 5,282 | 1,28E-07 | 2,05E-07 | 0,0005988 | 0,0039642 | 0,0023164 | 0,0023164 | TRUE | 0,0345703 | No |
| orf19.7414 | 2323 | 2073 | 2245 | 1470 | 3154 | 2290 | 3027 | 3389 | 2215 | 2192 | 2218 | 2303 | 2527 | 2604 | 2570 | 2756 | 2423 | 2232 | 2614 | 1,173 | 0,23 | 5,264 | 1,41E-07 | 2,25E-07 | 0,000634 | 0,0027689 | 0,0014038 | 0,0014038 | TRUE | 0,0165859 | No |
| orf19.7417 | 3 | 1 | 1 | 1 | 0 | 0 | 2 | 0 | 3 | 1 | 1 | 2 | 0 | 0 | 2 | 0 | 1,02 | 2 | 0 | 0,274 | -1,865 | -1,196 | 0,2316329 | 0,2607709 | 1,00E-08 | 1,6098307 | 1,0546062 | 1,0546062 | TRUE | 0,8858791 | No |
| orf19.7419 | 48 | 37 | 39 | 18 | 146 | 81 | 131 | 122 | 46 | 39 | 39 | 28 | 117 | 92 | 111 | 99 | 71,39 | 38 | 105 | 2,725 | 1,446 | 7,623 | 2,48E-14 | 4,92E-14 | 0,0001848 | 0,0250904 | 0,0159594 | 0,0159594 | TRUE | 0,1391054 | No |
| orf19.742 | 3265 | 2797 | 2989 | 1939 | 2435 | 1869 | 2343 | 2734 | 3113 | 2957 | 2953 | 3038 | 1951 | 2125 | 1989 | 2223 | 2543,65 | 3015 | 2072 | 0,687 | -0,542 | -10,534 | 6,02E-26 | 1,52E-25 | 0,001602 | 0,0027368 | 0,0021352 | 0,0021352 | TRUE | 0,0302536 | No |
| orf19.7420 | 1 | 1 | 1 | 1 | 3 | 2 | 4 | 1 | 1 | 1 | 1 | 2 | 2 | 2 | 3 | 1 | 1,68 | 1 | 2 | 1,996 | 0,997 | 0,905 | 0,3656886 | 0,400525 | 1,00E-08 | 0,9785216 | 0,4666527 | 0,4666527 | TRUE | 0,2342673 | No |
| orf19.7421 | 2425 | 1898 | 2094 | 1500 | 3436 | 2114 | 3186 | 3409 | 2312 | 2007 | 2069 | 2350 | 2753 | 2404 | 2705 | 2772 | 2421,39 | 2184 | 2658 | 1,219 | 0,286 | 4,257 | 2,07E-05 | 3,02E-05 | 0,0046318 | 0,0027694 | 0,003897 | 0,003897 | TRUE | 0,0493258 | No |
| orf19.7422 | 4606 | 3435 | 3945 | 2529 | 6444 | 4102 | 6403 | 6636 | 4391 | 3632 | 3897 | 3962 | 5163 | 4664 | 5436 | 5396 | 4567,76 | 3970 | 5165 | 1,301 | 0,38 | 5,657 | 1,54E-08 | 2,55E-08 | 0,0052012 | 0,0024507 | 0,0040971 | 0,0040971 | TRUE | 0,0628999 | No |
| orf19.7424 | 1 | 0 | 2 | 1 | 1 | 0 | 3 | 0 | 1 | 0 | 2 | 2 | 1 | 0 | 3 | 0 | 0,98 | 1 | 1 | 0,776 | -0,365 | -0,238 | 0,8115736 | 0,8332504 | 1,00E-08 | 1,6766459 | 1,2328898 | 1,2328898 | TRUE | 0,8618635 | No |
| orf19.7425 | 914 | 870 | 929 | 594 | 1127 | 769 | 1127 | 1031 | 871 | 920 | 918 | 931 | 903 | 874 | 957 | 838 | 901,52 | 910 | 893 | 0,983 | -0,025 | -0,412 | 0,6805933 | 0,7121787 | 0,001119 | 0,0039125 | 0,0023542 | 0,0023542 | TRUE | 0,0280122 | No |
| orf19.7426 | 919 | 788 | 911 | 521 | 955 | 698 | 916 | 959 | 876 | 833 | 900 | 816 | 765 | 794 | 778 | 780 | 817,73 | 856 | 779 | 0,907 | -0,141 | -2,483 | 0,0130201 | 0,0164193 | 1,00E-08 | 0,0040991 | 0,0018721 | 0,0018721 | TRUE | 0,0136913 | No |
| orf19.7427 | 6879 | 6008 | 6528 | 4271 | 5482 | 3830 | 5137 | 5350 | 6558 | 6353 | 6449 | 6691 | 4393 | 4355 | 4361 | 4350 | 5438,73 | 6513 | 4365 | 0,671 | -0,577 | -18,32 | 5,71E-75 | 2,92E-74 | 6,52E-05 | 0,0023932 | 0,0007621 | 0,0007621 | TRUE | 0,0038365 | No |
| orf19.7428 | 3791 | 3264 | 3658 | 1990 | 4611 | 3069 | 4386 | 4813 | 3614 | 3451 | 3614 | 3117 | 3695 | 3490 | 3724 | 3913 | 3577,29 | 3449 | 3706 | 1,073 | 0,102 | 1,788 | 0,0737317 | 0,0872394 | 0,0030255 | 0,0025503 | 0,0028468 | 0,0028468 | TRUE | 0,0488376 | No |
| orf19.7433 | 6234 | 5462 | 6141 | 3799 | 6925 | 4835 | 6661 | 7091 | 5943 | 5775 | 6067 | 5951 | 5549 | 5498 | 5655 | 5766 | 5775,54 | 5934 | 5617 | 0,947 | -0,079 | -2,362 | 0,0181957 | 0,0226683 | 0,0002676 | 0,0023756 | 0,0008982 | 0,0008982 | TRUE | 0,0040452 | No |
| orf19.7434 | 1040 | 1065 | 1070 | 627 | 589 | 408 | 525 | 522 | 992 | 1126 | 1057 | 982 | 472 | 464 | 446 | 424 | 745,37 | 1039 | 452 | 0,434 | -1,206 | -17,437 | 4,34E-68 | 2,03E-67 | 0,0019907 | 0,0042941 | 0,0030633 | 0,0030633 | TRUE | 0,037647 | No |
| orf19.7436 | 9072 | 8106 | 9347 | 6214 | 2820 | 2208 | 2635 | 3033 | 8649 | 8571 | 9234 | 9735 | 2260 | 2511 | 2237 | 2466 | 5707,78 | 9047 | 2368 | 0,262 | -1,934 | -33,671 | 1,53E-248 | 2,57E-247 | 0,0032115 | 0,0023789 | 0,002917 | 0,002917 | TRUE | 0,0314567 | No |
| orf19.7436.1 | 607 | 514 | 532 | 393 | 790 | 550 | 735 | 789 | 579 | 543 | 526 | 616 | 633 | 625 | 624 | 642 | 598,42 | 566 | 631 | 1,12 | 0,163 | 2,44 | 0,0147065 | 0,0184511 | 1,00E-08 | 0,004835 | 0,0026136 | 0,0026136 | TRUE | 0,0344006 | No |
| orf19.7437 | 777 | 643 | 708 | 468 | 1744 | 1096 | 1663 | 1666 | 741 | 680 | 699 | 733 | 1397 | 1246 | 1412 | 1355 | 1032,93 | 713 | 1352 | 1,9 | 0,926 | 15,152 | 7,38E-52 | 2,79E-51 | 0,0015046 | 0,0036808 | 0,0024733 | 0,0024733 | TRUE | 0,0316832 | No |
| orf19.7438 | 5535 | 4887 | 5339 | 3549 | 9055 | 6602 | 8536 | 9612 | 5277 | 5167 | 5274 | 5560 | 7256 | 7507 | 7247 | 7815 | 6387,95 | 5320 | 7456 | 1,403 | 0,488 | 12,02 | 2,79E-33 | 7,94E-33 | 0,0009306 | 0,0023483 | 0,0014173 | 0,0014173 | TRUE | 0,0130382 | No |
| orf19.744 | 4888 | 4289 | 4834 | 3059 | 6691 | 5079 | 6319 | 6850 | 4660 | 4535 | 4776 | 4792 | 5361 | 5775 | 5365 | 5570 | 5104,25 | 4691 | 5518 | 1,176 | 0,234 | 5,911 | 3,40E-09 | 5,73E-09 | 0,0006998 | 0,002413 | 0,0013054 | 0,0013054 | TRUE | 0,011698 | No |
| orf19.7440 | 44 | 66 | 58 | 38 | 125 | 82 | 96 | 119 | 42 | 70 | 57 | 60 | 100 | 93 | 82 | 97 | 75,03 | 57 | 93 | 1,636 | 0,71 | 3,884 | 0,0001029 | 0,0001456 | 0,0074237 | 0,0239751 | 0,017528 | 0,017528 | TRUE | 0,3069403 | No |
| orf19.7441 | 1527 | 1532 | 1600 | 1046 | 3117 | 2375 | 2938 | 2804 | 1456 | 1620 | 1581 | 1639 | 2498 | 2701 | 2494 | 2280 | 2033,43 | 1574 | 2493 | 1,585 | 0,664 | 10,71 | 9,18E-27 | 2,35E-26 | 0,0030479 | 0,0028987 | 0,0031568 | 0,0031568 | TRUE | 0,0402937 | No |
| orf19.7442 | 1214 | 1101 | 1176 | 694 | 3509 | 2363 | 3270 | 3435 | 1157 | 1164 | 1162 | 1087 | 2812 | 2687 | 2776 | 2793 | 1954,79 | 1142 | 2767 | 2,417 | 1,273 | 29,058 | 1,20E-185 | 1,51E-184 | 1,00E-08 | 0,0029312 | 0,0011944 | 0,0011944 | TRUE | 0,0106558 | No |

| | | | | | | | | | | | | | | | | | | | | | | | | | | | | | | | |
|---|---|---|---|---|---|---|---|---|---|---|---|---|---|---|---|---|---|---|---|---|---|---|---|---|---|---|---|---|---|---|---|
| orf19.7443 | 477 | 580 | 575 | 360 | 529 | 401 | 443 | 340 | 455 | 613 | 568 | 564 | 424 | 456 | 376 | 276 | 466,56 | 550 | 383 | 0,696 | -0,524 | -3,736 | 0,000187 | 0,0002611 | 0,0285788 | 0,0056105 | 0,0166703 | 0,0166703 | TRUE | 0,4125271 | No |
| orf19.7444 | 784 | 788 | 786 | 483 | 333 | 230 | 319 | 310 | 747 | 833 | 776 | 757 | 267 | 262 | 271 | 252 | 520,63 | 778 | 263 | 0,337 | -1,567 | -21,303 | 1,06E-100 | 6,96E-100 | 1,00E-08 | 0,005245 | 0,0027879 | 0,0027879 | TRUE | 0,0269961 | No |
| orf19.7445 | 8828 | 8407 | 9260 | 5898 | 4932 | 3721 | 4728 | 4849 | 8417 | 8889 | 9148 | 9240 | 3952 | 4231 | 4014 | 3943 | 6479,14 | 8924 | 4035 | 0,452 | -1,145 | -26,484 | 1,46E-154 | 1,52E-153 | 0,0011712 | 0,0023447 | 0,0016219 | 0,0016219 | TRUE | 0,0180196 | No |
| orf19.7446 | 214 | 208 | 211 | 126 | 882 | 615 | 736 | 728 | 204 | 220 | 208 | 197 | 707 | 699 | 625 | 592 | 431,58 | 207 | 656 | 3,15 | 1,655 | 17,95 | 4,80E-72 | 2,37E-71 | 0,0038535 | 0,0058957 | 0,0048058 | 0,0048058 | TRUE | 0,051672 | No |
| orf19.7447 | 39 | 28 | 39 | 18 | 88 | 68 | 96 | 91 | 37 | 30 | 39 | 28 | 71 | 77 | 82 | 74 | 54,61 | 34 | 76 | 2,239 | 1,163 | 5,781 | 7,41E-09 | 1,24E-08 | 1,00E-08 | 0,0321597 | 0,0165942 | 0,0165942 | TRUE | 0,0706088 | No |
| orf19.7448 | 1618 | 1380 | 1466 | 961 | 2492 | 1834 | 2329 | 2527 | 1543 | 1459 | 1448 | 1505 | 1997 | 2085 | 1977 | 2055 | 1758,71 | 1489 | 2028 | 1,362 | 0,446 | 9,976 | 1,94E-23 | 4,66E-23 | 1,00E-08 | 0,0030249 | 0,0013207 | 0,0013207 | TRUE | 0,0079422 | No |
| orf19.7449 | 2298 | 2246 | 2315 | 1452 | 1689 | 1282 | 1591 | 1632 | 2191 | 2375 | 2287 | 2275 | 1353 | 1458 | 1351 | 1327 | 1827,03 | 2282 | 1372 | 0,601 | -0,736 | -15,163 | 6,25E-52 | 2,37E-51 | 0,0006673 | 0,00299 | 0,0016879 | 0,0016879 | TRUE | 0,0197274 | No |
| orf19.745 | 6994 | 6542 | 6985 | 4995 | 4308 | 3550 | 4069 | 4218 | 6668 | 6917 | 6901 | 7825 | 3452 | 4037 | 3455 | 3430 | 5335,44 | 7078 | 3594 | 0,508 | -0,978 | -14,537 | 7,06E-48 | 2,54E-47 | 0,0057164 | 0,002399 | 0,0041451 | 0,0041451 | TRUE | 0,0828197 | No |
| orf19.7450 | 1042 | 1020 | 1028 | 606 | 1381 | 983 | 1335 | 1392 | 993 | 1079 | 1016 | 949 | 1107 | 1118 | 1133 | 1132 | 1065,8 | 1009 | 1122 | 1,11 | 0,15 | 2,808 | 0,0049824 | 0,0064396 | 4,62E-05 | 0,0036318 | 0,0017998 | 0,0017998 | TRUE | 0,0247493 | No |
| orf19.7451 | 6407 | 5970 | 6432 | 4274 | 2508 | 2000 | 2386 | 2684 | 6108 | 6313 | 6354 | 6695 | 2010 | 2274 | 2026 | 2182 | 4245,3 | 6368 | 2123 | 0,333 | -1,585 | -30,904 | 1,05E-209 | 1,49E-208 | 0,0021137 | 0,002478 | 0,002228 | 0,002228 | TRUE | 0,0267131 | No |
| orf19.7452 | 113 | 125 | 120 | 65 | 356 | 240 | 330 | 332 | 108 | 132 | 119 | 102 | 285 | 273 | 280 | 270 | 196,07 | 115 | 277 | 2,394 | 1,26 | 11,294 | 1,41E-29 | 3,78E-29 | 1,00E-08 | 0,0104654 | 0,0055443 | 0,0055443 | TRUE | 0,095633 | No |
| orf19.7453 | 65 | 56 | 70 | 42 | 168 | 121 | 173 | 208 | 62 | 59 | 69 | 66 | 135 | 138 | 147 | 169 | 105,54 | 64 | 147 | 2,304 | 1,204 | 7,886 | 3,12E-15 | 6,33E-15 | 1,00E-08 | 0,0176482 | 0,0106761 | 0,0106761 | TRUE | 0,1129326 | No |
| orf19.7454 | 1238 | 1094 | 1165 | 759 | 1475 | 1082 | 1428 | 1544 | 1180 | 1157 | 1151 | 1189 | 1182 | 1230 | 1212 | 1255 | 1194,62 | 1169 | 1220 | 1,044 | 0,062 | 1,294 | 0,1957792 | 0,2221314 | 1,00E-08 | 0,0034657 | 0,0013566 | 0,0013566 | TRUE | 0,0055827 | No |
| orf19.7455 | 343 | 300 | 351 | 206 | 17 | 11 | 27 | 22 | 327 | 317 | 347 | 323 | 14 | 13 | 23 | 18 | 172,58 | 328 | 17 | 0,052 | -4,277 | -22,513 | 3,06E-112 | 2,26E-111 | 1,00E-08 | 0,0116052 | 0,006906 | 0,006906 | TRUE | 0,3242486 | No |
| orf19.7456 | 56 | 52 | 38 | 31 | 41 | 39 | 42 | 31 | 53 | 55 | 38 | 49 | 33 | 44 | 36 | 25 | 41,57 | 49 | 34 | 0,702 | -0,51 | -2,124 | 0,0336464 | 0,0410198 | 0,0153665 | 0,0415911 | 0,0307694 | 0,0307694 | TRUE | 0,2549756 | No |
| orf19.7457 | 1015 | 852 | 940 | 544 | 880 | 630 | 902 | 925 | 968 | 901 | 929 | 852 | 705 | 716 | 766 | 752 | 823,6 | 912 | 735 | 0,804 | -0,316 | -5,146 | 2,66E-07 | 4,21E-07 | 0,0008704 | 0,0040848 | 0,0023913 | 0,0023913 | TRUE | 0,020489 | No |
| orf19.7458 | 393 | 322 | 328 | 209 | 479 | 311 | 462 | 424 | 375 | 340 | 324 | 327 | 384 | 354 | 392 | 345 | 355,13 | 342 | 369 | 1,077 | 0,108 | 1,248 | 0,2119915 | 0,2398011 | 0,0017246 | 0,0067147 | 0,0042863 | 0,0042863 | TRUE | 0,0513468 | No |
| orf19.7459 | 16260 | 15324 | 16376 | 11299 | 10592 | 8199 | 10179 | 10609 | 15502 | 16203 | 16178 | 17700 | 8487 | 9323 | 8642 | 8626 | 12582,74 | 16396 | 8770 | 0,535 | -0,903 | -18,078 | 4,74E-73 | 2,37E-72 | 0,0024028 | 0,0022218 | 0,0023097 | 0,0023097 | TRUE | 0,0347384 | No |
| orf19.746 | 2492 | 2880 | 2943 | 1722 | 2247 | 1674 | 2069 | 2009 | 2376 | 3045 | 2907 | 2698 | 1800 | 1903 | 1757 | 1634 | 2265,02 | 2756 | 1774 | 0,643 | -0,637 | -8,185 | 2,73E-16 | 5,65E-16 | 0,0069521 | 0,0028162 | 0,0053614 | 0,0053614 | TRUE | 0,1071041 | No |
| orf19.7460 | 2177 | 1798 | 1961 | 1235 | 3274 | 2289 | 3008 | 3072 | 2076 | 1901 | 1937 | 1935 | 2623 | 2603 | 2554 | 2498 | 2265,8 | 1962 | 2570 | 1,308 | 0,388 | 8,773 | 1,74E-18 | 3,78E-18 | 0,0005446 | 0,0028159 | 0,0014147 | 0,0014147 | TRUE | 0,0192286 | No |
| orf19.7463 | 510 | 431 | 518 | 337 | 412 | 261 | 401 | 405 | 486 | 456 | 512 | 528 | 330 | 297 | 340 | 329 | 409,78 | 496 | 324 | 0,658 | -0,604 | -7,525 | 5,28E-14 | 1,04E-13 | 0,0009146 | 0,006098 | 0,0036882 | 0,0036882 | TRUE | 0,0331283 | No |
| orf19.7464 | 16 | 8 | 11 | 3 | 10 | 7 | 11 | 10 | 15 | 8 | 11 | 5 | 8 | 8 | 9 | 8 | 9,09 | 10 | 8 | 0,824 | -0,279 | -0,592 | 0,5540097 | 0,588993 | 1,00E-08 | 0,1827135 | 0,1049981 | 0,1049981 | TRUE | 0,2304395 | No |
| orf19.7466 | 6592 | 5967 | 6412 | 3819 | 99362 | 68486 | 89714 | 101795 | 6285 | 6309 | 6334 | 5983 | 79616 | 77875 | 76167 | 82769 | 42667,21 | 6228 | 79107 | 12,695 | 3,666 | 96,39 | 0 | 0 | 0,0008689 | 0,0021298 | 0,0012934 | 0,0012934 | TRUE | 0,0119133 | No |
| orf19.7468 | 3510 | 3095 | 3249 | 2171 | 1117 | 818 | 1060 | 1065 | 3346 | 3273 | 3210 | 3401 | 895 | 930 | 900 | 866 | 2102,59 | 3308 | 898 | 0,271 | -1,882 | -41,969 | 0 | 0 | 1,00E-08 | 0,0028722 | 0,0012647 | 0,0012647 | TRUE | 0,006762 | No |
| orf19.7469 | 30259 | 25123 | 27164 | 15885 | 16000 | 10679 | 15603 | 17128 | 28849 | 26564 | 26835 | 24885 | 12820 | 12143 | 13247 | 13927 | 19908,77 | 26783 | 13034 | 0,487 | -1,039 | -18,594 | 3,58E-77 | 1,88E-76 | 0,0034142 | 0,0021738 | 0,0029447 | 0,0029447 | TRUE | 0,033009 | No |
| orf19.747 | 3951 | 3509 | 3896 | 2307 | 2117 | 1383 | 2076 | 2185 | 3767 | 3710 | 3849 | 3614 | 1696 | 1573 | 1763 | 1777 | 2718,51 | 3735 | 1702 | 0,456 | -1,133 | -23,539 | 1,65E-122 | 1,34E-121 | 0,0012753 | 0,0026953 | 0,0018108 | 0,0018108 | TRUE | 0,0296641 | No |
| orf19.7472 | 1645 | 1446 | 1651 | 994 | 1062 | 675 | 932 | 1061 | 1568 | 1529 | 1631 | 1557 | 851 | 768 | 791 | 863 | 1194,74 | 1571 | 818 | 0,521 | -0,939 | -16,969 | 1,38E-64 | 6,20E-64 | 0,0007167 | 0,0034656 | 0,0020408 | 0,0020408 | TRUE | 0,0187928 | No |
| orf19.7473 | 1862 | 1861 | 2018 | 1295 | 409 | 320 | 366 | 412 | 1775 | 1968 | 1994 | 2029 | 328 | 364 | 311 | 335 | 1137,82 | 1942 | 334 | 0,172 | -2,541 | -36,558 | 1,32E-292 | 2,85E-291 | 0,0025474 | 0,0035343 | 0,0030127 | 0,0030127 | TRUE | 0,041041 | No |
| orf19.7475 | 940 | 1016 | 1033 | 670 | 787 | 624 | 700 | 705 | 896 | 1074 | 1021 | 1050 | 631 | 710 | 594 | 573 | 818,53 | 1010 | 627 | 0,62 | -0,69 | -8,389 | 4,90E-17 | 1,03E-16 | 0,0059158 | 0,0040972 | 0,0052217 | 0,0052217 | TRUE | 0,0875863 | No |
| orf19.7477 | 5167 | 4813 | 5093 | 2718 | 7242 | 4510 | 6948 | 6908 | 4926 | 5089 | 5031 | 4258 | 5803 | 5128 | 5899 | 5617 | 5218,92 | 4826 | 5612 | 1,162 | 0,217 | 3,331 | 0,0008643 | 0,0011717 | 0,0047883 | 0,0024059 | 0,0038724 | 0,0038724 | TRUE | 0,0746409 | No |
| orf19.7478 | 3094 | 2933 | 3107 | 1952 | 3649 | 2747 | 3536 | 3595 | 2950 | 3101 | 3069 | 3058 | 2924 | 3124 | 3002 | 2923 | 3018,87 | 3044 | 2993 | 0,983 | -0,025 | -0,636 | 0,5247909 | 0,5610776 | 0,0004032 | 0,0026352 | 0,0011926 | 0,0011926 | TRUE | 0,0099518 | No |
| orf19.7479 | 5064 | 4775 | 5134 | 3062 | 5833 | 4293 | 5450 | 5722 | 4828 | 5049 | 5072 | 4797 | 4674 | 4882 | 4627 | 4653 | 4822,57 | 4936 | 4709 | 0,953 | -0,069 | -1,865 | 0,0622216 | 0,0740739 | 0,0004569 | 0,0024317 | 0,0011243 | 0,0011243 | TRUE | 0,0071621 | No |
| orf19.748 | 3843 | 3840 | 4160 | 2340 | 3974 | 2849 | 3493 | 3992 | 3664 | 4060 | 4110 | 3666 | 3184 | 3240 | 2966 | 3246 | 3516,86 | 3875 | 3159 | 0,815 | -0,296 | -5,424 | 5,83E-08 | 9,47E-08 | 0,0024606 | 0,0025581 | 0,0025699 | 0,0025699 | TRUE | 0,0207443 | No |
| orf19.7480 | 6 | 5 | 2 | 4 | 4 | 6 | 4 | 6 | 6 | 5 | 2 | 6 | 3 | 7 | 3 | 5 | 4,69 | 5 | 4 | 0,95 | -0,074 | -0,114 | 0,9095268 | 0,9205309 | 1,00E-08 | 0,3518844 | 0,1945636 | 0,1945636 | TRUE | 0,3364315 | No |
| orf19.7481 | 2810 | 2452 | 2714 | 1891 | 2685 | 2040 | 2405 | 2678 | 2679 | 2593 | 2681 | 2962 | 2151 | 2320 | 2042 | 2177 | 2450,7 | 2729 | 2172 | 0,797 | -0,328 | -5,838 | 5,27E-09 | 8,84E-09 | 0,0025748 | 0,0027613 | 0,0026172 | 0,0026172 | TRUE | 0,0381752 | No |
| orf19.7482 | 38 | 62 | 44 | 20 | 47 | 25 | 27 | 22 | 36 | 66 | 43 | 31 | 38 | 28 | 23 | 18 | 35,44 | 44 | 27 | 0,6 | -0,736 | -2,401 | 0,0163663 | 0,0204691 | 0,0758039 | 0,0484264 | 0,060953 | 0,060953 | TRUE | 0,8071914 | No |
| orf19.7483 | 6166 | 6127 | 6502 | 3869 | 2238 | 1711 | 2106 | 2201 | 5879 | 6479 | 6423 | 6061 | 1793 | 1946 | 1788 | 1790 | 4019,74 | 6210 | 1829 | 0,294 | -1,765 | -36,025 | 3,34E-284 | 6,89E-283 | 0,0015085 | 0,0024997 | 0,0019706 | 0,0019706 | TRUE | 0,0210905 | No |
| orf19.7484 | 1973 | 1606 | 1766 | 1127 | 4123 | 2910 | 3755 | 4333 | 1881 | 1698 | 1745 | 1766 | 3304 | 3309 | 3188 | 3523 | 2551,63 | 1772 | 3331 | 1,879 | 0,91 | 18,174 | 8,30E-74 | 4,19E-73 | 0,0014747 | 0,0027348 | 0,0019554 | 0,0019554 | TRUE | 0,0217136 | No |
| orf19.7485 | 3583 | 3114 | 3342 | 1922 | 4099 | 2591 | 3874 | 4069 | 3416 | 3293 | 3302 | 3011 | 3284 | 2946 | 3289 | 3308 | 3231,16 | 3256 | 3207 | 0,985 | -0,022 | -0,406 | 0,6846583 | 0,7160801 | 0,0022125 | 0,0025994 | 0,0025552 | 0,0025552 | TRUE | 0,0354433 | No |
| orf19.7486 | 947 | 941 | 976 | 602 | 1948 | 1296 | 1822 | 1837 | 903 | 995 | 964 | 943 | 1561 | 1474 | 1547 | 1494 | 1235,02 | 951 | 1519 | 1,598 | 0,676 | 13,045 | 6,83E-39 | 2,13E-38 | 0,0003073 | 0,0034207 | 0,001697 | 0,001697 | TRUE | 0,0155799 | No |
| orf19.7487 | 13 | 16 | 14 | 7 | 38 | 30 | 27 | 36 | 12 | 17 | 14 | 11 | 30 | 34 | 23 | 29 | 21,36 | 14 | 29 | 2,125 | 1,087 | 3,4 | 0,0006749 | 0,0009212 | 1,00E-08 | 0,0789666 | 0,0423589 | 0,0423589 | TRUE | 0,1475163 | No |
| orf19.7488 | 7342 | 6906 | 6739 | 4565 | 7339 | 5254 | 6845 | 7306 | 7000 | 7302 | 6657 | 7151 | 5881 | 5974 | 5811 | 5940 | 6464,69 | 7028 | 5902 | 0,84 | -0,252 | -6,636 | 3,23E-11 | 5,86E-11 | 0,0006368 | 0,0023453 | 0,0012247 | 0,0012247 | TRUE | 0,015574 | No |
| orf19.7489 | 2466 | 2133 | 2424 | 1515 | 4049 | 2794 | 3694 | 4183 | 2351 | 2255 | 2395 | 2373 | 3244 | 3177 | 3136 | 3401 | 2791,65 | 2344 | 3240 | 1,383 | 0,468 | 10,839 | 2,26E-27 | 5,85E-27 | 0,0007153 | 0,0026794 | 0,0014119 | 0,0014119 | TRUE | 0,0138802 | No |
| orf19.7489.3 | 1259 | 1048 | 1087 | 613 | 1181 | 875 | 1134 | 1147 | 1200 | 1108 | 1074 | 960 | 946 | 995 | 963 | 933 | 1022,4 | 1086 | 959 | 0,88 | -0,185 | -2,711 | 0,0066995 | 0,0085895 | 0,0032554 | 0,0036972 | 0,003466 | 0,003466 | TRUE | 0,0654186 | No |
| orf19.7490 | 298 | 279 | 303 | 162 | 548 | 382 | 490 | 549 | 284 | 295 | 299 | 254 | 439 | 434 | 416 | 446 | 358,51 | 283 | 434 | 1,524 | 0,608 | 7,477 | 7,59E-14 | 1,48E-13 | 1,00E-08 | 0,0066711 | 0,00337 | 0,00337 | TRUE | 0,042238 | No |
| orf19.7491 | 72 | 68 | 82 | 47 | 303 | 217 | 272 | 315 | 69 | 72 | 81 | 74 | 243 | 247 | 231 | 256 | 158,97 | 74 | 244 | 3,308 | 1,726 | 13,58 | 5,28E-42 | 1,73E-41 | 1,00E-08 | 0,0124196 | 0,0061849 | 0,0061849 | TRUE | 0,0455374 | No |
| orf19.7492 | 2033 | 1863 | 1958 | 1220 | 2826 | 1984 | 2866 | 2951 | 1938 | 1970 | 1934 | 1911 | 2264 | 2256 | 2433 | 2399 | 2138,34 | 1938 | 2338 | 1,206 | 0,271 | 6,168 | 6,90E-10 | 1,19E-09 | 0,000502 | 0,0028591 | 0,0013697 | 0,0013697 | TRUE | 0,0089946 | No |
| orf19.7494 | 215 | 214 | 240 | 141 | 726 | 506 | 648 | 717 | 205 | 226 | 237 | 221 | 582 | 575 | 550 | 583 | 397,43 | 222 | 572 | 2,576 | 1,365 | 16,813 | 1,97E-63 | 8,67E-63 | 1,00E-08 | 0,0062225 | 0,0030544 | 0,0030544 | TRUE | 0,035561 | No |
| orf19.7495 | 173 | 177 | 180 | 85 | 454 | 353 | 467 | 462 | 165 | 187 | 178 | 133 | 364 | 401 | 396 | 376 | 275,05 | 166 | 384 | 2,291 | 1,196 | 11,627 | 3,00E-31 | 8,26E-31 | 0,0013294 | 0,0080608 | 0,0056711 | 0,0056711 | TRUE | 0,138726 | No |
| orf19.7497 | 1791 | 1629 | 1824 | 1011 | 2698 | 2066 | 2690 | 2917 | 1708 | 1722 | 1802 | 1584 | 2162 | 2349 | 2284 | 2372 | 1997,8 | 1704 | 2292 | 1,342 | 0,424 | 8,042 | 8,83E-16 | 1,81E-15 | 0,0015642 | 0,0029131 | 0,0021466 | 0,0021466 | TRUE | 0,0246499 | No |
| orf19.7498 | 3053 | 2794 | 3039 | 1982 | 9062 | 6184 | 8536 | 9341 | 2911 | 2954 | 3002 | 3105 | 7261 | 7032 | 7247 | 7595 | 5138,4 | 2993 | 7284 | 2,437 | 1,285 | 32,506 | 8,82E-232 | 1,37E-230 | 0,0005841 | 0,0024108 | 0,0012511 | 0,0012511 | TRUE | 0,01013 | No |
| orf19.7499 | 2746 | 2220 | 2476 | 1642 | 3244 | 2335 | 3148 | 3297 | 2618 | 2347 | 2446 | 2572 | 2599 | 2655 | 2673 | 2681 | 2573,94 | 2496 | 2652 | 1,063 | 0,088 | 1,945 | 0,0517262 | 0,062124 | 0,0007889 | 0,0027292 | 0,0015786 | 0,0015786 | TRUE | 0,01962 | No |
| orf19.7499.1 | 564 | 486 | 514 | 338 | 1025 | 676 | 962 | 995 | 538 | 514 | 508 | 529 | 821 | 769 | 817 | 809 | 663,08 | 522 | 804 | 1,542 | 0,625 | 10,217 | 1,66E-24 | 4,05E-24 | 1,00E-08 | 0,0045675 | 0,001973 | 0,001973 | TRUE | 0,0093756 | No |
| orf19.750 | 234 | 160 | 177 | 143 | 353 | 277 | 334 | 376 | 223 | 169 | 175 | 224 | 283 | 315 | 284 | 306 | 247,28 | 198 | 297 | 1,506 | 0,591 | 5,255 | 1,48E-07 | 2,37E-07 | 0,0065125 | 0,0087311 | 0,0077853 | 0,0077853 | TRUE | 0,1003924 | No |
| orf19.7500 | 5575 | 5261 | 5527 | 3518 | 4218 | 3111 | 3819 | 4020 | 5315 | 5563 | 5460 | 5511 | 3380 | 3537 | 3242 | 3269 | 4409,69 | 5462 | 3357 | 0,614 | -0,703 | -17,603 | 2,33E-69 | 1,10E-68 | 0,0006695 | 0,0024636 | 0,0012954 | 0,0012954 | TRUE | 0,0152149 | No |
| orf19.7501 | 28628 | 24688 | 26588 | 15327 | 30874 | 19624 | 30202 | 30714 | 27294 | 26104 | 26266 | 24011 | 24738 | 22314 | 25641 | 24973 | 25167,83 | 25919 | 24416 | 0,942 | -0,086 | -1,581 | 0,1139488 | 0,1321053 | 0,0031285 | 0,0021565 | 0,002817 | 0,002817 | TRUE | 0,0409987 | No |
| orf19.7502 | 9371 | 11307 | 11023 | 6778 | 11905 | 9823 | 10552 | 9336 | 8934 | 11956 | 10890 | 10618 | 9539 | 11170 | 8959 | 7591 | 9957,02 | 10600 | 9315 | 0,879 | -0,187 | -1,306 | 0,19148 | 0,2174082 | 0,0194905 | 0,0022562 | 0,0108865 | 0,0194905 | TRUE | 0,219425 | No |
| orf19.7503 | 1135 | 1070 | 1123 | 789 | 664 | 566 | 630 | 660 | 1082 | 1131 | 1109 | 1236 | 532 | 644 | 535 | 537 | 850,76 | 1140 | 562 | 0,492 | -1,023 | -13,638 | 2,38E-42 | 7,85E-42 | 0,0041908 | 0,0040212 | 0,0041062 | 0,0041062 | TRUE | 0,1049209 | No |
| orf19.7504 | 19456 | 18081 | 19554 | 12525 | 21387 | 14685 | 20272 | 20897 | 18549 | 19118 | 19317 | 19621 | 17137 | 16698 | 17211 | 16991 | 18080,4 | 19151 | 17009 | 0,888 | -0,171 | -5,78 | 7,47E-09 | 1,25E-08 | 0,0002928 | 0,0021821 | 0,0007817 | 0,0007817 | TRUE | 0,0055156 | No |
| orf19.7506 | 2010 | 1888 | 2010 | 1168 | 3061 | 2040 | 2846 | 3104 | 1916 | 1996 | 1986 | 1830 | 2453 | 2320 | 2416 | 2524 | 2180,06 | 1932 | 2428 | 1,256 | 0,329 | 7,033 | 2,02E-12 | 3,77E-12 | 0,0007844 | 0,0028444 | 0,0016216 | 0,0016216 | TRUE | 0,0137337 | No |
| orf19.7507 | 1100 | 893 | 961 | 621 | 1410 | 928 | 1424 | 1470 | 1049 | 944 | 949 | 973 | 1130 | 1055 | 1209 | 1195 | 1063,05 | 979 | 1147 | 1,173 | 0,23 | 3,658 | 0,0002547 | 0,000354 | 0,0022211 | 0,0036358 | 0,0028471 | 0,0028471 | TRUE | 0,0328506 | No |
| orf19.7509.1 | 2937 | 2151 | 2217 | 1565 | 8694 | 5656 | 7960 | 8635 | 2800 | 2274 | 2190 | 2452 | 6966 | 6431 | 6758 | 7021 | 4611,64 | 2429 | 6794 | 2,797 | 1,484 | 20,794 | 4,90E-96 | 3,09E-95 | 0,0065908 | 0,0024473 | 0,0045936 | 0,0045936 | TRUE | 0,1314232 | No |
| orf19.7509.2 | 2736 | 2060 | 2188 | 1100 | 3951 | 2224 | 3805 | 4014 | 2608 | 2178 | 2162 | 1723 | 3166 | 2529 | 3230 | 3264 | 2607,54 | 2168 | 3047 | 1,404 | 0,49 | 4,416 | 1,01E-05 | 1,49E-05 | 0,0198948 | 0,002721 | 0,0114249 | 0,0114249 | TRUE | 0,2265174 | No |
| orf19.751 | 38 | 32 | 40 | 15 | 87 | 66 | 86 | 78 | 36 | 34 | 40 | 23 | 70 | 75 | 73 | 63 | 51,78 | 33 | 70 | 2,064 | 1,046 | 5,042 | 4,61E-07 | 7,24E-07 | 1,00E-08 | 0,033798 | 0,0186089 | 0,0186089 | TRUE | 0,1701742 | No |
| orf19.7510 | 4325 | 3399 | 4002 | 2645 | 5194 | 3542 | 4964 | 5130 | 4123 | 3594 | 3954 | 4144 | 4162 | 4028 | 4214 | 4171 | 4048,69 | 3954 | 4144 | 1,049 | 0,069 | 1,363 | 0,173036 | 0,1973798 | 0,0019313 | 0,0024968 | 0,0021948 | 0,0021948 | TRUE | 0,0458248 | No |
| orf19.7511 | 5344 | 4792 | 5180 | 3184 | 5652 | 4046 | 5187 | 5771 | 5095 | 5067 | 5117 | 4988 | 4529 | 4601 | 4404 | 4692 | 4811,58 | 5067 | 4556 | 0,899 | -0,154 | -4,497 | 6,90E-06 | 1,03E-05 | 0,0002406 | 0,0024325 | 0,000915 | 0,000915 | TRUE | 0,0062714 | No |
| orf19.7512 | 1485 | 1222 | 1324 | 879 | 1285 | 958 | 1224 | 1354 | 1416 | 1292 | 1308 | 1377 | 1030 | 1089 | 1039 | 1101 | 1206,49 | 1348 | 1065 | 0,79 | -0,341 | -6,369 | 1,90E-10 | 3,37E-10 | 0,0007682 | 0,0034522 | 0,0019118 | 0,0019118 | TRUE | 0,0151357 | No |
| orf19.7513 | 1876 | 1443 | 1585 | 970 | 2696 | 1957 | 2608 | 2886 | 1789 | 1526 | 1566 | 1520 | 2160 | 2225 | 2214 | 2347 | 1918,25 | 1600 | 2236 | 1,396 | 0,481 | 7,929 | 2,21E-15 | 4,49E-15 | 0,0030747 | 0,0029472 | 0,0029855 | 0,0029855 | TRUE | 0,0778446 | No |
| orf19.7514 | 40286 | 32257 | 35024 | 23935 | 42466 | 28982 | 40445 | 43976 | 38408 | 34108 | 34600 | 37495 | 34027 | 32955 | 34338 | 35757 | 35211,01 | 36153 | 34269 | 0,948 | -0,077 | -1,596 | 0,1104937 | 0,1283796 | 0,0022494 | 0,0021379 | 0,0022126 | 0,0022126 | TRUE | 0,021725 | No |
| orf19.7516 | 2078 | 1697 | 1914 | 1288 | 2980 | 2043 | 2781 | 3087 | 1981 | 1794 | 1891 | 2018 | 2388 | 2323 | 2361 | 2510 | 2158,25 | 1921 | 2396 | 1,249 | 0,321 | 6,378 | 1,80E-10 | 3,19E-10 | 0,0012954 | 0,002852 | 0,0019483 | 0,0019483 | TRUE | 0,0236615 | No |
| orf19.7517 | 273 | 224 | 209 | 143 | 564 | 377 | 510 | 494 | 260 | 237 | 206 | 224 | 452 | 429 | 433 | 402 | 330,36 | 232 | 429 | 1,845 | 0,884 | 9,523 | 1,68E-21 | 3,88E-21 | 0,0020932 | 0,0070613 | 0,0048266 | 0,0048266 | TRUE | 0,0822339 | No |
| orf19.7518 | 1308 | 1104 | 1149 | 836 | 2807 | 1903 | 2533 | 2723 | 1247 | 1167 | 1135 | 1310 | 2249 | 2164 | 2151 | 2214 | 1704,59 | 1215 | 2194 | 1,812 | 0,857 | 16,287 | 1,21E-59 | 5,10E-59 | 0,0010711 | 0,0030545 | 0,0019941 | 0,0019941 | TRUE | 0,0291845 | No |
| orf19.7519 | 2354 | 1799 | 1963 | 1330 | 1894 | 1256 | 1816 | 1934 | 2244 | 1902 | 1939 | 2084 | 1518 | 1428 | 1542 | 1573 | 1778,67 | 2042 | 1515 | 0,742 | -0,43 | -6,912 | 4,77E-12 | 8,85E-12 | 0,0029801 | 0,0030144 | 0,0031387 | 0,0031387 | TRUE | 0,0560384 | No |

| | | | | | | | | | | | | | | | | | | | | | | | | | | | | | | | |
|---|---|---|---|---|---|---|---|---|---|---|---|---|---|---|---|---|---|---|---|---|---|---|---|---|---|---|---|---|---|---|---|
| orf19.752 | 1849 | 1675 | 1835 | 1136 | 3833 | 2843 | 3542 | 3813 | 1763 | 1771 | 1813 | 1780 | 3071 | 3233 | 3007 | 3100 | 2442,23 | 1782 | 3103 | 1,74 | 0,799 | 19,517 | 7,91E-85 | 4,49E-84 | 1,00E-08 | 0,0027636 | 0,0011523 | 0,0011523 | TRUE | 0,009147 | No |
| orf19.7520 | 7323 | 5773 | 6383 | 4397 | 56393 | 38790 | 53174 | 59398 | 6982 | 6104 | 6306 | 6888 | 45186 | 44108 | 45144 | 48296 | 26126,8 | 6570 | 45684 | 6,956 | 2,798 | 53,445 | 0 | 0 | 0,0028228 | 0,0021541 | 0,0025375 | 0,0025375 | TRUE | 0,0278078 | No |
| orf19.7521 | 1009 | 819 | 980 | 628 | 2052 | 1405 | 1960 | 2107 | 962 | 866 | 968 | 984 | 1644 | 1598 | 1664 | 1713 | 1299,87 | 945 | 1655 | 1,754 | 0,811 | 14,808 | 1,31E-49 | 4,80E-49 | 0,0009101 | 0,0033544 | 0,0020167 | 0,0020167 | TRUE | 0,0381198 | No |
| orf19.7522 | 793 | 669 | 703 | 471 | 1206 | 845 | 1237 | 1321 | 756 | 707 | 694 | 738 | 966 | 961 | 1050 | 1074 | 868,41 | 724 | 1013 | 1,401 | 0,487 | 7,715 | 1,21E-14 | 2,41E-14 | 0,001514 | 0,003982 | 0,0026071 | 0,0026071 | TRUE | 0,0199882 | No |
| orf19.7523 | 3170 | 2637 | 3015 | 1830 | 4491 | 2961 | 4321 | 4556 | 3022 | 2788 | 2979 | 2867 | 3599 | 3367 | 3669 | 3704 | 3249,28 | 2914 | 3585 | 1,23 | 0,299 | 6,436 | 1,23E-10 | 2,19E-10 | 0,0011457 | 0,0025966 | 0,0017547 | 0,0017547 | TRUE | 0,0196382 | No |
| orf19.7527 | 2967 | 2497 | 2714 | 1793 | 2267 | 1507 | 2174 | 2308 | 2829 | 2640 | 2681 | 2809 | 1816 | 1714 | 1846 | 1877 | 2276,42 | 2740 | 1813 | 0,663 | -0,593 | -12,812 | 1,41E-37 | 4,28E-37 | 0,0007277 | 0,0028125 | 0,001608 | 0,001608 | TRUE | 0,0154539 | No |
| orf19.7529 | 18782 | 15228 | 17404 | 10995 | 8762 | 5703 | 7938 | 8771 | 17907 | 16102 | 17194 | 17224 | 7021 | 6485 | 6739 | 7132 | 11975,32 | 17107 | 6844 | 0,4 | -1,321 | -28,744 | 1,09E-181 | 1,32E-180 | 0,001719 | 0,0022284 | 0,0019311 | 0,0019311 | TRUE | 0,0192239 | No |
| orf19.753 | 1363 | 1332 | 1514 | 854 | 2379 | 1751 | 2277 | 2432 | 1299 | 1408 | 1496 | 1338 | 1906 | 1991 | 1933 | 1977 | 1668,66 | 1385 | 1952 | 1,407 | 0,493 | 9,272 | 1,83E-20 | 4,16E-20 | 0,0013194 | 0,0030752 | 0,0020795 | 0,0020795 | TRUE | 0,0358155 | No |
| orf19.7531 | 3731 | 3036 | 3158 | 2298 | 1836 | 1253 | 1749 | 1865 | 3557 | 3210 | 3120 | 3600 | 1471 | 1425 | 1485 | 1516 | 2423,03 | 3372 | 1474 | 0,438 | -1,192 | -20,763 | 9,43E-96 | 5,93E-95 | 0,0026426 | 0,0027689 | 0,0026924 | 0,0026924 | TRUE | 0,0309303 | No |
| orf19.7534 | 18264 | 14646 | 16874 | 11800 | 23832 | 16641 | 22670 | 25151 | 17413 | 15486 | 16670 | 18485 | 19096 | 18922 | 19247 | 20450 | 18221,17 | 17014 | 19429 | 1,142 | 0,192 | 3,471 | 0,0005186 | 0,0007118 | 0,0033139 | 0,0021814 | 0,0028796 | 0,0028796 | TRUE | 0,0446289 | No |
| orf19.7537 | 151 | 98 | 146 | 103 | 730 | 435 | 635 | 658 | 144 | 104 | 144 | 161 | 585 | 495 | 539 | 535 | 338,36 | 138 | 538 | 3,928 | 1,974 | 17,341 | 2,33E-67 | 1,08E-66 | 0,0087044 | 0,0069438 | 0,0075522 | 0,0075522 | TRUE | 0,293091 | No |
| orf19.7538 | 902 | 792 | 925 | 552 | 2212 | 1407 | 2029 | 2136 | 860 | 837 | 914 | 865 | 1772 | 1600 | 1723 | 1737 | 1288,45 | 869 | 1708 | 1,967 | 0,976 | 17,789 | 8,65E-71 | 4,18E-70 | 0,0009424 | 0,0033656 | 0,0019849 | 0,0019849 | TRUE | 0,0206782 | No |
| orf19.7539 | 629 | 442 | 602 | 410 | 1298 | 923 | 1311 | 1317 | 600 | 467 | 595 | 642 | 1040 | 1050 | 1113 | 1071 | 822,19 | 576 | 1068 | 1,86 | 0,895 | 10,575 | 3,88E-26 | 9,83E-26 | 0,0072735 | 0,0040883 | 0,0054872 | 0,0054872 | TRUE | 0,1897113 | No |
| orf19.7539.1 | 1549 | 1253 | 1334 | 913 | 2581 | 1885 | 2575 | 2670 | 1477 | 1325 | 1318 | 1430 | 2068 | 2143 | 2186 | 2171 | 1764,81 | 1388 | 2142 | 1,545 | 0,628 | 12,149 | 5,82E-34 | 1,67E-33 | 0,0011161 | 0,0030216 | 0,0019485 | 0,0019485 | TRUE | 0,0250667 | No |
| orf19.754 | 5302 | 4624 | 4932 | 3056 | 14135 | 9643 | 13861 | 14562 | 5055 | 4889 | 4872 | 4787 | 11326 | 10965 | 11768 | 11840 | 8187,88 | 4901 | 11475 | 2,341 | 1,227 | 31,92 | 1,43E-223 | 2,14E-222 | 0,0007013 | 0,0022918 | 0,001265 | 0,001265 | TRUE | 0,0107632 | No |
| orf19.7544 | 279 | 213 | 260 | 142 | 229 | 115 | 256 | 188 | 266 | 225 | 257 | 222 | 183 | 131 | 217 | 153 | 206,87 | 242 | 171 | 0,706 | -0,501 | -3,425 | 0,0006146 | 0,0008403 | 0,0216855 | 0,0100281 | 0,0156611 | 0,0156611 | TRUE | 0,353518 | No |
| orf19.7545 | 36 | 41 | 41 | 30 | 249 | 170 | 227 | 256 | 34 | 43 | 41 | 47 | 200 | 193 | 193 | 208 | 119,86 | 41 | 198 | 4,875 | 2,285 | 14,5 | 1,22E-47 | 4,35E-47 | 1,00E-08 | 0,0157899 | 0,0080533 | 0,0080533 | TRUE | 0,1055183 | No |
| orf19.7546 | 7478 | 7132 | 7512 | 4854 | 6953 | 4944 | 6355 | 6426 | 7129 | 7541 | 7421 | 7604 | 5571 | 5622 | 5395 | 5225 | 6438,65 | 7424 | 5453 | 0,735 | -0,445 | -11,43 | 2,96E-30 | 8,03E-30 | 0,0007379 | 0,0023463 | 0,0012971 | 0,0012971 | TRUE | 0,0097526 | No |
| orf19.7547 | 3309 | 2993 | 3012 | 1910 | 3363 | 2299 | 3028 | 3239 | 3155 | 3165 | 2976 | 2992 | 2695 | 2614 | 2571 | 2634 | 2850,05 | 3072 | 2628 | 0,855 | -0,226 | -5,589 | 2,29E-08 | 3,76E-08 | 0,0004062 | 0,0026674 | 0,0012126 | 0,0012126 | TRUE | 0,0059204 | No |
| orf19.7548 | 278 | 245 | 246 | 197 | 492 | 341 | 447 | 431 | 265 | 259 | 243 | 309 | 394 | 388 | 380 | 350 | 323,46 | 269 | 378 | 1,415 | 0,501 | 5,455 | 4,90E-08 | 7,97E-08 | 0,0018353 | 0,0071674 | 0,0048326 | 0,0048326 | TRUE | 0,0902352 | No |
| orf19.7549 | 216 | 208 | 211 | 135 | 1280 | 922 | 1134 | 1241 | 206 | 220 | 208 | 211 | 1026 | 1048 | 963 | 1009 | 611,45 | 211 | 1012 | 4,783 | 2,258 | 30,014 | 6,52E-198 | 8,85E-197 | 1,00E-08 | 0,0047765 | 0,0023721 | 0,0023721 | TRUE | 0,012552 | No |
| orf19.755 | 1047 | 939 | 969 | 537 | 1415 | 927 | 1262 | 1325 | 998 | 993 | 957 | 841 | 1134 | 1054 | 1071 | 1077 | 1015,78 | 947 | 1084 | 1,141 | 0,19 | 3,031 | 0,0024359 | 0,0032114 | 0,001847 | 0,0037077 | 0,002763 | 0,002763 | TRUE | 0,0594822 | No |
| orf19.7550 | 652 | 556 | 640 | 397 | 71 | 55 | 74 | 85 | 622 | 588 | 632 | 622 | 57 | 63 | 63 | 69 | 339,38 | 616 | 63 | 0,102 | -3,293 | -29,972 | 2,30E-197 | 3,12E-196 | 1,00E-08 | 0,0069292 | 0,0036419 | 0,0036419 | TRUE | 0,0468128 | No |
| orf19.7551 | 5826 | 5170 | 5529 | 3329 | 5683 | 4090 | 5501 | 5926 | 5554 | 5467 | 5462 | 5215 | 4554 | 4651 | 4670 | 4818 | 5048,91 | 5424 | 4673 | 0,861 | -0,216 | -5,975 | 2,29E-09 | 3,90E-09 | 0,0003706 | 0,0024165 | 0,001057 | 0,001057 | TRUE | 0,0077379 | No |
| orf19.7552 | 16602 | 14540 | 16299 | 8957 | 20729 | 13564 | 19882 | 20847 | 15828 | 15374 | 16102 | 14032 | 16610 | 15423 | 16880 | 16951 | 15899,91 | 15334 | 16466 | 1,074 | 0,102 | 1,977 | 0,0480528 | 0,0578866 | 0,0026409 | 0,0021945 | 0,0025167 | 0,0025167 | TRUE | 0,0395353 | No |
| orf19.7553 | 232 | 245 | 285 | 140 | 295 | 210 | 309 | 298 | 221 | 259 | 282 | 219 | 236 | 239 | 262 | 242 | 245,11 | 245 | 245 | 0,995 | -0,007 | -0,07 | 0,9439541 | 0,949759 | 0,0041805 | 0,0087898 | 0,0066806 | 0,0066806 | TRUE | 0,1122152 | No |
| orf19.7554 | 860 | 834 | 969 | 563 | 1420 | 965 | 1250 | 1340 | 820 | 882 | 957 | 882 | 1138 | 1097 | 1061 | 1090 | 990,86 | 885 | 1096 | 1,239 | 0,309 | 5,115 | 3,14E-07 | 4,96E-07 | 0,0013939 | 0,0037483 | 0,0024619 | 0,0024619 | TRUE | 0,0384522 | No |
| orf19.7555 | 38 | 19 | 34 | 25 | 28 | 13 | 31 | 17 | 36 | 20 | 34 | 39 | 22 | 15 | 26 | 14 | 25,8 | 32 | 19 | 0,608 | -0,718 | -2,25 | 0,0244698 | 0,0301781 | 0,0460775 | 0,0657214 | 0,0575574 | 0,0575574 | TRUE | 0,4400189 | No |
| orf19.7556 | 439 | 428 | 470 | 267 | 584 | 390 | 553 | 454 | 419 | 453 | 464 | 418 | 468 | 443 | 469 | 369 | 437,97 | 438 | 437 | 0,996 | -0,006 | -0,061 | 0,9512457 | 0,9561907 | 0,0058625 | 0,0058402 | 0,0058566 | 0,0058566 | TRUE | 0,1343001 | No |
| orf19.7557 | 20 | 17 | 12 | 6 | 31 | 14 | 19 | 36 | 19 | 18 | 12 | 9 | 25 | 16 | 16 | 29 | 18,06 | 14 | 22 | 1,462 | 0,548 | 1,494 | 0,1352187 | 0,1558299 | 0,0417725 | 0,0930187 | 0,0713225 | 0,0713225 | TRUE | 0,2950484 | No |
| orf19.7558 | 4315 | 3892 | 4308 | 2736 | 4263 | 3142 | 4064 | 4254 | 4114 | 4115 | 4256 | 4286 | 3416 | 3573 | 3450 | 3459 | 3833,62 | 4193 | 3474 | 0,829 | -0,271 | -7,691 | 1,46E-14 | 2,92E-14 | 7,76E-05 | 0,0025196 | 0,00093 | 0,00093 | TRUE | 0,0041739 | No |
| orf19.756 | 10 | 8 | 8 | 8 | 14 | 6 | 3 | 15 | 10 | 8 | 8 | 13 | 11 | 7 | 3 | 12 | 8,9 | 10 | 8 | 0,874 | -0,194 | -0,373 | 0,7094262 | 0,7393187 | 0,0877572 | 0,1865429 | 0,1466294 | 0,1466294 | TRUE | 0,6664887 | No |
| orf19.7561 | 27732 | 22638 | 25991 | 17695 | 7119 | 4908 | 6976 | 7403 | 26439 | 23937 | 25677 | 27720 | 5704 | 5581 | 5923 | 6019 | 15875,01 | 25943 | 5807 | 0,224 | -2,159 | -42,864 | 0 | 0 | 0,0023108 | 0,0021947 | 0,0023386 | 0,0023386 | TRUE | 0,0331638 | No |
| orf19.7563 | 734 | 652 | 723 | 488 | 873 | 612 | 789 | 880 | 700 | 689 | 714 | 764 | 700 | 696 | 670 | 716 | 706,09 | 717 | 696 | 0,973 | -0,04 | -0,658 | 0,5102176 | 0,5463498 | 1,00E-08 | 0,0044166 | 0,002093 | 0,002093 | TRUE | 0,019803 | No |
| orf19.7564 | 2143 | 2059 | 2305 | 1340 | 2796 | 2020 | 2602 | 2778 | 2043 | 2177 | 2277 | 2099 | 2240 | 2297 | 2209 | 2259 | 2200,21 | 2149 | 2251 | 1,047 | 0,066 | 1,43 | 0,1526896 | 0,1750786 | 0,0006415 | 0,0028375 | 0,0015687 | 0,0015687 | TRUE | 0,0201456 | No |
| orf19.7565 | 2321 | 2056 | 2172 | 1353 | 3761 | 2720 | 3649 | 3770 | 2213 | 2174 | 2146 | 2120 | 3014 | 3093 | 3098 | 3065 | 2615,23 | 2163 | 3068 | 1,417 | 0,502 | 13,639 | 2,36E-42 | 7,76E-42 | 1,00E-08 | 0,0027191 | 0,0008985 | 0,0008985 | TRUE | 0,0029549 | No |
| orf19.7566 | 33293 | 28720 | 31850 | 20347 | 72665 | 50852 | 69373 | 73283 | 31741 | 30368 | 31465 | 31875 | 58224 | 57823 | 58897 | 59586 | 44997,45 | 31362 | 58632 | 1,87 | 0,903 | 32,251 | 3,46E-228 | 5,29E-227 | 0,0002929 | 0,0021278 | 0,0007272 | 0,0007272 | TRUE | 0,0055878 | No |
| orf19.7567 | 4165 | 3736 | 4026 | 2721 | 1003 | 745 | 945 | 1093 | 3971 | 3950 | 3977 | 4263 | 804 | 847 | 802 | 889 | 2437,87 | 4040 | 836 | 0,207 | -2,273 | -44,593 | 0 | 0 | 0,0009274 | 0,0027648 | 0,0018218 | 0,0018218 | TRUE | 0,0235803 | No |
| orf19.7568 | 102 | 89 | 96 | 47 | 75 | 40 | 65 | 49 | 97 | 94 | 95 | 74 | 60 | 45 | 55 | 40 | 70,05 | 90 | 50 | 0,555 | -0,85 | -4,567 | 4,94E-06 | 7,44E-06 | 0,0066949 | 0,0255293 | 0,0182494 | 0,0182494 | TRUE | 0,1851314 | No |
| orf19.7569 | 24272 | 21260 | 22155 | 15441 | 49909 | 37755 | 46798 | 50174 | 23141 | 22480 | 21887 | 24189 | 39991 | 42931 | 39731 | 40796 | 31893,21 | 22924 | 40862 | 1,783 | 0,834 | 19,621 | 1,01E-85 | 5,81E-85 | 0,0015118 | 0,0021428 | 0,0017005 | 0,0017005 | TRUE | 0,0167121 | No |
| orf19.757 | 2388 | 2221 | 2423 | 1454 | 2136 | 1681 | 2001 | 2176 | 2277 | 2348 | 2394 | 2278 | 1712 | 1911 | 1699 | 1769 | 2048,46 | 2324 | 1773 | 0,761 | -0,393 | -7,931 | 2,17E-15 | 4,41E-15 | 0,0011497 | 0,0028928 | 0,0018671 | 0,0018671 | TRUE | 0,0317051 | No |
| orf19.7570 | 15452 | 14110 | 15778 | 8905 | 7425 | 5047 | 7032 | 7296 | 14732 | 14920 | 15587 | 13950 | 5949 | 5739 | 5970 | 5932 | 10347,44 | 14797 | 5898 | 0,399 | -1,327 | -32,555 | 1,78E-232 | 2,78E-231 | 0,0010423 | 0,00225 | 0,0014827 | 0,0014827 | TRUE | 0,0178121 | No |
| orf19.7571 | 8293 | 7668 | 8056 | 5354 | 9690 | 7238 | 9190 | 9860 | 7906 | 8108 | 7959 | 8387 | 7764 | 8230 | 7802 | 8017 | 8021,78 | 8090 | 7953 | 0,983 | -0,024 | -0,68 | 0,4967119 | 0,5326027 | 0,0005565 | 0,002296 | 0,0010898 | 0,0010898 | TRUE | 0,0071751 | No |
| orf19.757.1 | 677 | 556 | 584 | 481 | 720 | 545 | 629 | 661 | 645 | 588 | 577 | 754 | 577 | 620 | 534 | 537 | 603,99 | 641 | 567 | 0,887 | -0,173 | -1,894 | 0,0582088 | 0,0694911 | 0,0082446 | 0,0048097 | 0,0063058 | 0,0063058 | TRUE | 0,1498042 | No |
| orf19.7572 | 1742 | 1606 | 1762 | 1049 | 2587 | 1782 | 2536 | 2662 | 1661 | 1698 | 1741 | 1643 | 2073 | 2026 | 2153 | 2164 | 1894,96 | 1686 | 2104 | 1,248 | 0,32 | 7,197 | 6,14E-13 | 1,17E-12 | 0,0003102 | 0,0029577 | 0,0013486 | 0,0013486 | TRUE | 0,007024 | No |
| orf19.7574 | 3034 | 2838 | 3122 | 1975 | 3291 | 2283 | 2849 | 3323 | 2893 | 3001 | 3084 | 3094 | 2637 | 2596 | 2419 | 2702 | 2803,16 | 3018 | 2588 | 0,858 | -0,221 | -4,645 | 3,40E-06 | 5,15E-06 | 0,0012917 | 0,002677 | 0,0018049 | 0,0018049 | TRUE | 0,0240613 | No |
| orf19.7576 | 1261 | 1012 | 1127 | 662 | 875 | 564 | 831 | 854 | 1202 | 1070 | 1113 | 1037 | 701 | 641 | 706 | 694 | 895,63 | 1106 | 686 | 0,62 | -0,69 | -10,733 | 7,10E-27 | 1,82E-26 | 0,0018792 | 0,0039245 | 0,0028166 | 0,0028166 | TRUE | 0,0419844 | No |
| orf19.7577 | 3839 | 3440 | 3641 | 2435 | 3095 | 2429 | 3068 | 3184 | 3660 | 3637 | 3597 | 3815 | 2480 | 2762 | 2605 | 2589 | 3143,06 | 3677 | 2609 | 0,709 | -0,496 | -11,253 | 2,25E-29 | 6,00E-29 | 0,0008896 | 0,0026137 | 0,0015373 | 0,0015373 | TRUE | 0,0181085 | No |
| orf19.7578 | 93 | 75 | 71 | 63 | 151 | 96 | 126 | 167 | 89 | 79 | 70 | 99 | 121 | 109 | 107 | 136 | 101,21 | 84 | 118 | 1,421 | 0,507 | 3,253 | 0,0011424 | 0,001534 | 0,005666 | 0,0183133 | 0,0129004 | 0,0129004 | TRUE | 0,1203261 | No |
| orf19.7579 | 791 | 651 | 717 | 517 | 807 | 603 | 745 | 840 | 754 | 688 | 708 | 810 | 647 | 686 | 633 | 683 | 701,06 | 740 | 662 | 0,897 | -0,158 | -2,316 | 0,0205393 | 0,0254636 | 0,001558 | 0,0044333 | 0,0030092 | 0,0030092 | TRUE | 0,0413224 | No |
| orf19.758 | 1645 | 1355 | 1517 | 957 | 1462 | 1176 | 1453 | 1606 | 1568 | 1433 | 1499 | 1499 | 1171 | 1337 | 1234 | 1306 | 1380,88 | 1500 | 1262 | 0,84 | -0,251 | -4,399 | 1,09E-05 | 1,61E-05 | 0,0017546 | 0,0032803 | 0,0023982 | 0,0023982 | TRUE | 0,0284398 | No |
| orf19.7580 | 660 | 522 | 501 | 290 | 646 | 403 | 619 | 683 | 629 | 552 | 495 | 454 | 518 | 458 | 526 | 555 | 523,4 | 532 | 514 | 0,963 | -0,054 | -0,531 | 0,5951106 | 0,6292296 | 0,0108043 | 0,0052283 | 0,0080039 | 0,0080039 | TRUE | 0,1758452 | No |
| orf19.7581 | 7330 | 6267 | 6980 | 4019 | 3436 | 2279 | 3440 | 3590 | 6988 | 6627 | 6896 | 6296 | 2753 | 2591 | 2921 | 2919 | 4748,83 | 6702 | 2796 | 0,417 | -1,261 | -24,072 | 4,86E-128 | 4,16E-127 | 0,0023229 | 0,002437 | 0,002391 | 0,002391 | TRUE | 0,0284793 | No |
| orf19.7583 | 1847 | 1794 | 1901 | 1146 | 3866 | 2774 | 3527 | 3709 | 1761 | 1897 | 1878 | 1795 | 3098 | 3154 | 2994 | 3016 | 2449,16 | 1833 | 3066 | 1,671 | 0,741 | 17,209 | 2,28E-66 | 1,04E-65 | 0,0003422 | 0,0027617 | 0,0013277 | 0,0013277 | TRUE | 0,0093862 | No |
| orf19.7585 | 1523 | 1271 | 1408 | 864 | 9501 | 6896 | 8885 | 9849 | 1452 | 1344 | 1391 | 1354 | 7613 | 7841 | 7543 | 8008 | 4568,27 | 1385 | 7751 | 5,588 | 2,482 | 58,032 | 0 | 0 | 0,0006311 | 0,0024507 | 0,0012958 | 0,0012958 | TRUE | 0,0132697 | No |
| orf19.7586 | 630 | 563 | 608 | 429 | 261 | 225 | 243 | 260 | 601 | 595 | 601 | 672 | 209 | 256 | 206 | 211 | 418,92 | 617 | 220 | 0,356 | -1,488 | -16,955 | 1,77E-64 | 7,91E-64 | 0,0022552 | 0,0060107 | 0,0044579 | 0,0044579 | TRUE | 0,1116729 | No |
| orf19.7588 | 721 | 721 | 760 | 415 | 1271 | 872 | 1149 | 1109 | 687 | 762 | 751 | 650 | 1018 | 992 | 975 | 902 | 842,23 | 712 | 972 | 1,359 | 0,443 | 6,447 | 1,14E-10 | 2,04E-10 | 0,0024811 | 0,0040407 | 0,0032831 | 0,0032831 | TRUE | 0,035481 | No |
| orf19.7589 | 789 | 712 | 710 | 435 | 960 | 659 | 904 | 870 | 752 | 753 | 701 | 681 | 769 | 749 | 767 | 707 | 735,17 | 722 | 748 | 1,034 | 0,048 | 0,769 | 0,4418127 | 0,4781559 | 0,0005953 | 0,0043246 | 0,0023867 | 0,0023867 | TRUE | 0,0170045 | No |
| orf19.759 | 1321 | 1182 | 1236 | 797 | 3514 | 2579 | 3294 | 3552 | 1259 | 1250 | 1221 | 1249 | 2816 | 2933 | 2797 | 2888 | 2051,47 | 1245 | 2858 | 2,296 | 1,199 | 28,57 | 1,60E-179 | 1,91E-178 | 1,00E-08 | 0,0028916 | 0,0010857 | 0,0010857 | TRUE | 0,0035041 | No |
| orf19.7590 | 4896 | 4438 | 4748 | 2802 | 14102 | 9719 | 12770 | 14031 | 4668 | 4693 | 4691 | 4389 | 11300 | 11051 | 10842 | 11409 | 7880,2 | 4610 | 11150 | 2,416 | 1,273 | 35,097 | 7,56E-270 | 1,42E-268 | 0,0005676 | 0,0022996 | 0,0011009 | 0,0011009 | TRUE | 0,0117635 | No |
| orf19.7592 | 35265 | 33549 | 36495 | 22061 | 58406 | 43223 | 54708 | 57648 | 33621 | 35474 | 36054 | 34560 | 46799 | 49148 | 46447 | 46873 | 41122,02 | 34927 | 47317 | 1,355 | 0,438 | 12,487 | 8,78E-36 | 2,60E-35 | 0,0007731 | 0,0021312 | 0,0011559 | 0,0011559 | TRUE | 0,0082839 | No |
| orf19.7593 | 2854 | 2709 | 2849 | 1744 | 3795 | 2735 | 3514 | 3436 | 2721 | 2864 | 2815 | 2732 | 3041 | 3110 | 2983 | 2794 | 2882,49 | 2783 | 2982 | 1,071 | 0,098 | 2,157 | 0,0309998 | 0,0379528 | 0,0010632 | 0,0026609 | 0,0016438 | 0,0016438 | TRUE | 0,0221351 | No |
| orf19.7594 | 707 | 701 | 683 | 430 | 687 | 482 | 670 | 651 | 674 | 741 | 675 | 674 | 550 | 548 | 569 | 529 | 620,04 | 691 | 549 | 0,794 | -0,332 | -5,138 | 2,77E-07 | 4,39E-07 | 1,00E-08 | 0,0047393 | 0,0023939 | 0,0023939 | TRUE | 0,0289801 | No |
| orf19.7595 | 258 | 245 | 292 | 166 | 247 | 175 | 217 | 240 | 246 | 259 | 288 | 260 | 198 | 199 | 184 | 195 | 228,73 | 263 | 194 | 0,736 | -0,442 | -4,516 | 6,31E-06 | 9,44E-06 | 1,00E-08 | 0,0092697 | 0,004807 | 0,004807 | TRUE | 0,0500267 | No |
| orf19.7596 | 164 | 142 | 151 | 89 | 169 | 117 | 145 | 153 | 156 | 150 | 149 | 139 | 135 | 133 | 123 | 124 | 138,88 | 148 | 129 | 0,863 | -0,213 | -1,791 | 0,0733681 | 0,0868253 | 1,00E-08 | 0,0139135 | 0,0064207 | 0,0064207 | TRUE | 0,0142773 | No |
| orf19.7597 | 4 | 1 | 6 | 4 | 5 | 3 | 5 | 6 | 4 | 1 | 6 | 6 | 4 | 3 | 4 | 5 | 4,2 | 4 | 4 | 0,999 | -0,002 | -0,003 | 0,9979964 | 0,9981528 | 1,00E-08 | 0,3929537 | 0,2179782 | 0,2179782 | TRUE | 0,442509 | No |
| orf19.7598 | 5774 | 4958 | 5529 | 3549 | 5479 | 3711 | 5209 | 5333 | 5505 | 5242 | 5462 | 5560 | 4390 | 4220 | 4422 | 4336 | 4892,22 | 5442 | 4342 | 0,798 | -0,325 | -9,206 | 3,38E-20 | 7,63E-20 | 0,0002606 | 0,0024269 | 0,0009884 | 0,0009884 | TRUE | 0,0074638 | No |
| orf19.7599 | 13512 | 11795 | 12869 | 8855 | 14082 | 10183 | 13700 | 14712 | 12882 | 12472 | 12713 | 13872 | 11284 | 11579 | 11631 | 11962 | 12299,39 | 12985 | 11614 | 0,895 | -0,16 | -3,854 | 0,000116 | 0,0001636 | 0,0012992 | 0,0022248 | 0,0015802 | 0,0015802 | TRUE | 0,0253547 | No |
| orf19.76 | 17714 | 16596 | 17809 | 10622 | 18878 | 13405 | 17190 | 18143 | 16888 | 17548 | 17594 | 16640 | 15126 | 15243 | 14594 | 14752 | 16048,17 | 17168 | 14929 | 0,869 | -0,202 | -6,115 | 9,66E-10 | 1,66E-09 | 0,0005126 | 0,0021936 | 0,0009855 | 0,0009855 | TRUE | 0,0051034 | No |
| orf19.760 | 781 | 619 | 689 | 420 | 934 | 554 | 831 | 936 | 745 | 655 | 681 | 658 | 748 | 630 | 706 | 761 | 697,83 | 685 | 711 | 1,04 | 0,056 | 0,737 | 0,4610865 | 0,4974963 | 0,0039079 | 0,0044442 | 0,0041628 | 0,0041628 | TRUE | 0,065492 | No |
| orf19.7600 | 11623 | 9376 | 10329 | 6532 | 14328 | 9503 | 13859 | 14773 | 11081 | 9914 | 10204 | 10233 | 11481 | 10806 | 11766 | 12012 | 10937,06 | 10358 | 11516 | 1,112 | 0,153 | 3,164 | 0,0015549 | 0,0020727 | 0,0020811 | 0,0022414 | 0,0021522 | 0,0021522 | TRUE | 0,0272761 | No |
| orf19.7601 | 2451 | 2138 | 2331 | 1580 | 2638 | 1936 | 2599 | 2631 | 2337 | 2261 | 2303 | 2475 | 2114 | 2201 | 2207 | 2139 | 2254,54 | 2344 | 2165 | 0,925 | -0,113 | -2,596 | 0,0094198 | 0,0119927 | 0,0004187 | 0,0028195 | 0,0013592 | 0,0013592 | TRUE | 0,015558 | No |
| orf19.7602 | 1674 | 1510 | 1530 | 953 | 932 | 678 | 859 | 964 | 1596 | 1597 | 1511 | 1493 | 747 | 771 | 729 | 784 | 1153,48 | 1549 | 758 | 0,488 | -1,034 | -19,581 | 2,26E-85 | 1,29E-84 | 1,00E-08 | 0,0035147 | 0,001729 | 0,001729 | TRUE | 0,0076764 | No |

| | | | | | | | | | | | | | | | | | | | | | | | | | | | | | | | |
|---|---|---|---|---|---|---|---|---|---|---|---|---|---|---|---|---|---|---|---|---|---|---|---|---|---|---|---|---|---|---|---|
| orf19.7603 | 1382 | 1144 | 1277 | 719 | 2077 | 1297 | 2030 | 2164 | 1318 | 1210 | 1262 | 1126 | 1664 | 1475 | 1723 | 1760 | 1442,15 | 1229 | 1656 | 1,346 | 0,429 | 6,194 | 5,88E-10 | 1,02E-09 | 0,0041717 | 0,0032298 | 0,0038723 | 0,0038723 | TRUE | 0,0628487 | No |
| orf19.7604 | 849 | 730 | 811 | 554 | 1387 | 982 | 1274 | 1490 | 809 | 772 | 801 | 868 | 1111 | 1117 | 1082 | 1212 | 971,44 | 812 | 1130 | 1,395 | 0,48 | 7,924 | 2,31E-15 | 4,68E-15 | 0,0013604 | 0,0037815 | 0,0024429 | 0,0024429 | TRUE | 0,0289755 | No |
| orf19.7605 | 10849 | 8330 | 9551 | 6309 | 8505 | 5442 | 7805 | 8378 | 10343 | 8808 | 9435 | 9883 | 6815 | 6188 | 6626 | 6812 | 8113,94 | 9617 | 6610 | 0,688 | -0,54 | -9,669 | 4,08E-22 | 9,54E-22 | 0,0031431 | 0,0022936 | 0,002875 | 0,002875 | TRUE | 0,0393934 | No |
| orf19.7606 | 25 | 47 | 32 | 29 | 36 | 36 | 28 | 29 | 24 | 50 | 32 | 45 | 29 | 41 | 24 | 24 | 33,46 | 38 | 30 | 0,778 | -0,362 | -1,216 | 0,2239228 | 0,2527152 | 0,059105 | 0,051157 | 0,0546876 | 0,0546876 | TRUE | 0,2265588 | No |
| orf19.7608 | 4 | 3 | 3 | 3 | 8 | 6 | 7 | 7 | 4 | 3 | 3 | 5 | 6 | 7 | 6 | 6 | 4,94 | 4 | 6 | 1,724 | 0,786 | 1,259 | 0,208147 | 0,2357029 | 1,00E-08 | 0,3344906 | 0,1484712 | 0,1484712 | TRUE | 0,0301945 | No |
| orf19.7609 | 20 | 12 | 11 | 11 | 29 | 15 | 24 | 32 | 19 | 13 | 11 | 17 | 23 | 17 | 20 | 26 | 18,32 | 15 | 22 | 1,47 | 0,556 | 1,639 | 0,1011514 | 0,1180534 | 1,00E-08 | 0,0917249 | 0,0527996 | 0,0527996 | TRUE | 0,1988799 | No |
| orf19.761 | 4390 | 3520 | 4209 | 2864 | 2703 | 1901 | 2584 | 2662 | 4185 | 3722 | 4158 | 4487 | 2166 | 2162 | 2194 | 2164 | 3154,72 | 4138 | 2172 | 0,525 | -0,929 | -16,584 | 9,04E-62 | 3,91E-61 | 0,0024493 | 0,0026117 | 0,0026715 | 0,0026715 | TRUE | 0,0555863 | No |
| orf19.7610 | 3386 | 3101 | 3416 | 2127 | 6775 | 5000 | 6758 | 6968 | 3228 | 3279 | 3375 | 3332 | 5429 | 5685 | 5738 | 5666 | 4466,38 | 3304 | 5630 | 1,704 | 0,769 | 21,366 | 2,75E-101 | 1,82E-100 | 0,0003338 | 0,0024589 | 0,0009947 | 0,0009947 | TRUE | 0,0071493 | No |
| orf19.7611 | 6296 | 6450 | 6126 | 4224 | 4593 | 3633 | 3983 | 4062 | 6003 | 6820 | 6052 | 6617 | 3680 | 4131 | 3382 | 3303 | 4998,41 | 6373 | 3624 | 0,568 | -0,815 | -11,105 | 1,18E-28 | 3,12E-28 | 0,0071207 | 0,0024198 | 0,0049606 | 0,0049606 | TRUE | 0,1066983 | No |
| orf19.7612 | 22724 | 20731 | 21695 | 13760 | 3432 | 2460 | 3058 | 3342 | 21665 | 21920 | 21433 | 21556 | 2750 | 2797 | 2596 | 2717 | 12179,31 | 21644 | 2715 | 0,125 | -2,996 | -90,405 | 0 | 0 | 0,0001199 | 0,0022261 | 0,000864 | 0,000864 | TRUE | 0,0105632 | No |
| orf19.7613 | 5032 | 4132 | 4557 | 2812 | 7954 | 5387 | 7876 | 8033 | 4797 | 4369 | 4502 | 4405 | 6373 | 6126 | 6687 | 6532 | 5473,83 | 4518 | 6430 | 1,423 | 0,509 | 11,295 | 1,39E-29 | 3,72E-29 | 0,0013676 | 0,0023912 | 0,0017522 | 0,0017522 | TRUE | 0,0214253 | No |
| orf19.7614 | 4 | 3 | 5 | 2 | 11 | 5 | 7 | 10 | 4 | 3 | 5 | 3 | 9 | 6 | 6 | 8 | 5,45 | 4 | 7 | 1,888 | 0,917 | 1,517 | 0,1293301 | 0,1492864 | 1,00E-08 | 0,3031392 | 0,1439388 | 0,1439388 | TRUE | 0,0790159 | No |
| orf19.7615 | 527 | 441 | 436 | 276 | 980 | 701 | 840 | 888 | 502 | 466 | 431 | 432 | 785 | 797 | 713 | 722 | 606,17 | 458 | 754 | 1,641 | 0,714 | 9,503 | 2,03E-21 | 4,69E-21 | 0,0024717 | 0,0047999 | 0,0036168 | 0,0036168 | TRUE | 0,0518706 | No |
| orf19.7617 | 2015 | 1565 | 1748 | 1112 | 2824 | 1808 | 2712 | 2692 | 1921 | 1655 | 1727 | 1742 | 2263 | 2056 | 2302 | 2189 | 1981,84 | 1761 | 2202 | 1,251 | 0,323 | 5,481 | 4,22E-08 | 6,88E-08 | 0,0025974 | 0,0029198 | 0,0028075 | 0,0028075 | TRUE | 0,0468598 | No |
| orf19.7618 | 8933 | 7456 | 7980 | 5337 | 10685 | 7035 | 10166 | 11068 | 8517 | 7884 | 7883 | 8361 | 8562 | 7999 | 8631 | 8999 | 8354,48 | 8161 | 8548 | 1,048 | 0,067 | 1,431 | 0,1523732 | 0,1747471 | 0,0018089 | 0,0022878 | 0,0020083 | 0,0020083 | TRUE | 0,0224828 | No |
| orf19.7619 | 9827 | 8924 | 9659 | 5677 | 7368 | 5083 | 7131 | 7355 | 9369 | 9436 | 9542 | 8893 | 5904 | 5780 | 6054 | 5980 | 7619,82 | 9310 | 5930 | 0,637 | -0,651 | -18,46 | 4,35E-76 | 2,25E-75 | 0,0004859 | 0,0023068 | 0,0010596 | 0,0010596 | TRUE | 0,0106561 | No |
| orf19.762 | 32 | 30 | 21 | 23 | 45 | 26 | 49 | 38 | 31 | 32 | 21 | 36 | 36 | 30 | 42 | 31 | 32,14 | 30 | 35 | 1,181 | 0,24 | 0,922 | 0,3566425 | 0,3912888 | 1,00E-08 | 0,053176 | 0,0335082 | 0,0335082 | TRUE | 0,288769 | No |
| orf19.7620 | 1062 | 866 | 925 | 550 | 936 | 619 | 846 | 885 | 1013 | 916 | 914 | 862 | 750 | 704 | 718 | 720 | 824,41 | 926 | 723 | 0,779 | -0,36 | -5,671 | 1,42E-08 | 2,35E-08 | 0,0014191 | 0,0040829 | 0,0026496 | 0,0026496 | TRUE | 0,0480679 | No |
| orf19.7621 | 417 | 376 | 392 | 233 | 1113 | 779 | 1089 | 1097 | 398 | 398 | 387 | 365 | 892 | 886 | 925 | 892 | 642,69 | 387 | 899 | 2,315 | 1,211 | 19,001 | 1,66E-80 | 9,11E-80 | 1,00E-08 | 0,004646 | 0,0019729 | 0,0019729 | TRUE | 0,012849 | No |
| orf19.7622 | 1214 | 1059 | 1123 | 669 | 940 | 618 | 922 | 993 | 1157 | 1120 | 1109 | 1048 | 753 | 703 | 783 | 807 | 935,09 | 1108 | 762 | 0,687 | -0,542 | -8,902 | 5,49E-19 | 1,21E-18 | 0,0011093 | 0,0038472 | 0,0024664 | 0,0024664 | TRUE | 0,0292045 | No |
| orf19.7624 | 9807 | 8206 | 9061 | 5570 | 10489 | 7015 | 10273 | 10520 | 9350 | 8677 | 8951 | 8726 | 8405 | 7977 | 8722 | 8554 | 8670,07 | 8926 | 8414 | 0,943 | -0,085 | -2,047 | 0,04063 | 0,0492516 | 0,0011565 | 0,0022807 | 0,0015464 | 0,0015464 | TRUE | 0,0146991 | No |
| orf19.7625 | 1893 | 1733 | 1803 | 1165 | 3376 | 2340 | 3305 | 3217 | 1805 | 1832 | 1781 | 1825 | 2705 | 2661 | 2806 | 2616 | 2253,87 | 1811 | 2697 | 1,49 | 0,575 | 13,872 | 9,33E-44 | 3,14E-43 | 1,00E-08 | 0,0028198 | 0,001177 | 0,001177 | TRUE | 0,0091081 | No |
| orf19.7626 | 3779 | 3203 | 3297 | 2139 | 8884 | 5948 | 8416 | 8965 | 3603 | 3387 | 3257 | 3351 | 7118 | 6763 | 7145 | 7289 | 5239,26 | 3400 | 7079 | 2,082 | 1,058 | 24,188 | 2,98E-129 | 2,56E-128 | 0,0011332 | 0,0024047 | 0,001609 | 0,001609 | TRUE | 0,0204099 | No |
| orf19.7627 | 953 | 885 | 879 | 654 | 860 | 637 | 892 | 931 | 909 | 936 | 868 | 1025 | 689 | 724 | 757 | 757 | 833,12 | 934 | 732 | 0,786 | -0,347 | -5,232 | 1,68E-07 | 2,68E-07 | 0,0018711 | 0,0040621 | 0,0030155 | 0,0030155 | TRUE | 0,0451413 | No |
| orf19.7629 | 530 | 505 | 538 | 353 | 814 | 580 | 771 | 816 | 505 | 534 | 531 | 553 | 652 | 660 | 655 | 663 | 594,2 | 531 | 658 | 1,242 | 0,312 | 5,011 | 5,42E-07 | 8,49E-07 | 1,00E-08 | 0,0048545 | 0,0019955 | 0,0019955 | TRUE | 0,0126859 | No |
| orf19.763 | 3687 | 3331 | 3395 | 1769 | 3220 | 1985 | 3039 | 3406 | 3515 | 3522 | 3354 | 2771 | 2580 | 2257 | 2580 | 2769 | 2918,64 | 3290 | 2546 | 0,774 | -0,37 | -4,483 | 7,37E-06 | 1,10E-05 | 0,0088371 | 0,0026538 | 0,0062117 | 0,0062117 | TRUE | 0,1338316 | No |
| orf19.7631 | 126 | 110 | 144 | 88 | 362 | 262 | 353 | 356 | 120 | 116 | 142 | 138 | 290 | 298 | 300 | 289 | 211,71 | 129 | 294 | 2,287 | 1,193 | 11,213 | 3,53E-29 | 9,37E-29 | 1,00E-08 | 0,0098467 | 0,0050483 | 0,0050483 | TRUE | 0,0593453 | No |
| orf19.7632 | 666 | 550 | 625 | 388 | 554 | 397 | 521 | 626 | 635 | 582 | 617 | 608 | 444 | 451 | 442 | 509 | 536,05 | 610 | 462 | 0,756 | -0,403 | -5,528 | 3,24E-08 | 5,31E-08 | 0,0013615 | 0,0051542 | 0,0032137 | 0,0032137 | TRUE | 0,0581915 | No |
| orf19.7634 | 97 | 88 | 106 | 50 | 67 | 52 | 56 | 56 | 92 | 93 | 105 | 78 | 54 | 59 | 48 | 46 | 71,81 | 92 | 52 | 0,551 | -0,86 | -4,935 | 8,00E-07 | 1,24E-06 | 1,00E-08 | 0,0249565 | 0,0144751 | 0,0144751 | TRUE | 0,0784458 | No |
| orf19.7635 | 13785 | 11738 | 13103 | 8562 | 18370 | 12068 | 17552 | 18900 | 13143 | 12411 | 12945 | 13413 | 14719 | 13722 | 14902 | 15368 | 13827,77 | 12978 | 14678 | 1,131 | 0,178 | 4,034 | 5,48E-05 | 7,85E-05 | 0,0015501 | 0,00221 | 0,0017978 | 0,0017978 | TRUE | 0,023277 | No |
| orf19.7637 | 1879 | 1650 | 1748 | 1033 | 1938 | 1316 | 1925 | 1943 | 1791 | 1745 | 1727 | 1618 | 1553 | 1496 | 1634 | 1580 | 1643,08 | 1720 | 1566 | 0,909 | -0,137 | -2,756 | 0,0058505 | 0,0075297 | 0,0007651 | 0,0030906 | 0,0017733 | 0,0017733 | TRUE | 0,0172736 | No |
| orf19.7638 | 2796 | 2197 | 2387 | 1727 | 2261 | 1513 | 2099 | 2399 | 2666 | 2323 | 2358 | 2705 | 1812 | 1720 | 1782 | 1951 | 2164,63 | 2513 | 1816 | 0,724 | -0,466 | -7,182 | 6,87E-13 | 1,31E-12 | 0,0039817 | 0,0028498 | 0,0035796 | 0,0035796 | TRUE | 0,0311613 | No |
| orf19.764 | 376 | 415 | 439 | 272 | 844 | 663 | 786 | 772 | 358 | 439 | 434 | 426 | 676 | 754 | 667 | 628 | 547,78 | 414 | 681 | 1,645 | 0,718 | 8,297 | 1,06E-16 | 2,22E-16 | 0,0052564 | 0,0050886 | 0,005173 | 0,005173 | TRUE | 0,1018085 | No |
| orf19.7642 | 645 | 577 | 598 | 400 | 1358 | 959 | 1183 | 1253 | 615 | 610 | 591 | 627 | 1088 | 1090 | 1004 | 1019 | 830,52 | 611 | 1050 | 1,721 | 0,784 | 13,186 | 1,05E-39 | 3,32E-39 | 1,00E-08 | 0,0040682 | 0,0020501 | 0,0020501 | TRUE | 0,010376 | No |
| orf19.7643 | 1641 | 1380 | 1514 | 996 | 3856 | 2505 | 3628 | 3750 | 1565 | 1459 | 1496 | 1560 | 3090 | 2848 | 3080 | 3049 | 2268,38 | 1520 | 3017 | 1,988 | 0,991 | 21,105 | 7,14E-99 | 4,62E-98 | 0,0008217 | 0,0028151 | 0,0016003 | 0,0016003 | TRUE | 0,0164402 | No |
| orf19.7644 | 397 | 338 | 350 | 254 | 802 | 479 | 710 | 756 | 378 | 357 | 346 | 398 | 643 | 545 | 603 | 615 | 485,54 | 370 | 602 | 1,635 | 0,709 | 8,933 | 4,15E-19 | 9,15E-19 | 0,002076 | 0,0054729 | 0,003797 | 0,003797 | TRUE | 0,043485 | No |
| orf19.7645 | 1703 | 1423 | 1510 | 971 | 3433 | 2237 | 3361 | 3606 | 1624 | 1505 | 1492 | 1521 | 2751 | 2544 | 2853 | 2932 | 2152,63 | 1536 | 2770 | 1,805 | 0,852 | 15,162 | 6,32E-52 | 2,39E-51 | 0,0022709 | 0,002854 | 0,0025038 | 0,0025038 | TRUE | 0,0355497 | No |
| orf19.7646 | 443 | 348 | 424 | 241 | 463 | 330 | 432 | 429 | 422 | 368 | 419 | 378 | 371 | 375 | 367 | 349 | 381,07 | 397 | 366 | 0,917 | -0,124 | -1,548 | 0,1217364 | 0,1408012 | 1,00E-08 | 0,0064 | 0,0035709 | 0,0035709 | TRUE | 0,0308679 | No |
| orf19.7648 | 575 | 496 | 542 | 392 | 454 | 323 | 370 | 431 | 548 | 524 | 535 | 614 | 364 | 367 | 314 | 350 | 452,23 | 555 | 349 | 0,63 | -0,667 | -8,173 | 3,02E-16 | 6,24E-16 | 0,0023166 | 0,005722 | 0,0041095 | 0,0041095 | TRUE | 0,0524913 | No |
| orf19.7650 | 5963 | 4866 | 5582 | 3385 | 6418 | 4187 | 6215 | 6426 | 5685 | 5145 | 5514 | 5303 | 5143 | 4761 | 5277 | 5225 | 5256,57 | 5412 | 5102 | 0,943 | -0,085 | -1,768 | 0,0769904 | 0,0909433 | 0,0017504 | 0,0024036 | 0,0020337 | 0,0020337 | TRUE | 0,0240392 | No |
| orf19.7652 | 9060 | 7461 | 8625 | 5245 | 6947 | 4598 | 6756 | 7049 | 8638 | 7889 | 8521 | 8217 | 5566 | 5228 | 5736 | 5732 | 6940,77 | 8316 | 5566 | 0,669 | -0,579 | -12,701 | 5,86E-37 | 1,77E-36 | 0,0015296 | 0,0023278 | 0,0018483 | 0,0018483 | TRUE | 0,019839 | No |
| orf19.7654 | 5214 | 4071 | 4576 | 3111 | 4288 | 2927 | 4272 | 4414 | 4971 | 4305 | 4521 | 4874 | 3436 | 3328 | 3627 | 3589 | 4081,22 | 4668 | 3495 | 0,749 | -0,416 | -7,563 | 3,93E-14 | 7,73E-14 | 0,0026536 | 0,0024936 | 0,0026622 | 0,0026622 | TRUE | 0,0334656 | No |
| orf19.7655 | 4447 | 3755 | 4085 | 2527 | 9968 | 6242 | 8864 | 9914 | 4240 | 3970 | 4036 | 3959 | 7987 | 7098 | 7525 | 8061 | 5859,47 | 4051 | 7668 | 1,893 | 0,92 | 18,38 | 1,89E-75 | 9,76E-75 | 0,0019711 | 0,0023715 | 0,0022111 | 0,0022111 | TRUE | 0,0301947 | No |
| orf19.7657 | 3291 | 2509 | 2725 | 1852 | 2981 | 2103 | 2825 | 3009 | 3138 | 2653 | 2692 | 2901 | 2389 | 2391 | 2398 | 2447 | 2626,1 | 2846 | 2406 | 0,846 | -0,242 | -4,24 | 2,24E-05 | 3,26E-05 | 0,0026201 | 0,0027165 | 0,0027436 | 0,0027436 | TRUE | 0,0599639 | No |
| orf19.7658 | 1920 | 1623 | 1793 | 1198 | 3212 | 2139 | 3003 | 3298 | 1831 | 1716 | 1771 | 1877 | 2574 | 2432 | 2550 | 2682 | 2178,96 | 1799 | 2560 | 1,426 | 0,512 | 10,61 | 2,66E-26 | 6,78E-26 | 0,0009237 | 0,0028448 | 0,0017443 | 0,0017443 | TRUE | 0,0129507 | No |
| orf19.7660 | 630 | 556 | 539 | 417 | 907 | 635 | 900 | 954 | 601 | 588 | 532 | 653 | 727 | 722 | 764 | 776 | 670,36 | 594 | 747 | 1,265 | 0,339 | 4,796 | 1,61E-06 | 2,47E-06 | 0,002115 | 0,0045406 | 0,0032559 | 0,0032559 | TRUE | 0,0564979 | No |
| orf19.7661 | 1002 | 890 | 924 | 597 | 2175 | 1509 | 2094 | 2198 | 955 | 941 | 913 | 935 | 1743 | 1716 | 1778 | 1787 | 1346 | 936 | 1756 | 1,877 | 0,908 | 19,425 | 4,75E-84 | 2,68E-83 | 1,00E-08 | 0,0033111 | 0,0012487 | 0,0012487 | TRUE | 0,003723 | No |
| orf19.7662 | 3772 | 3143 | 3445 | 2066 | 4056 | 2531 | 3981 | 4154 | 3596 | 3323 | 3403 | 3236 | 3250 | 2878 | 3380 | 3378 | 3305,59 | 3390 | 3222 | 0,95 | -0,073 | -1,23 | 0,2185279 | 0,2468449 | 0,0033243 | 0,002588 | 0,0031109 | 0,0031109 | TRUE | 0,0611204 | No |
| orf19.7663 | 756 | 588 | 718 | 420 | 858 | 524 | 864 | 767 | 721 | 622 | 709 | 658 | 687 | 596 | 734 | 624 | 668,78 | 678 | 660 | 0,975 | -0,037 | -0,451 | 0,6523433 | 0,6846386 | 0,0053342 | 0,0045463 | 0,0049615 | 0,0049615 | TRUE | 0,065669 | No |
| orf19.7664 | 1677 | 1495 | 1650 | 1137 | 4867 | 3166 | 4584 | 4491 | 1599 | 1581 | 1630 | 1781 | 3900 | 3600 | 3892 | 3652 | 2704,26 | 1648 | 3761 | 2,289 | 1,194 | 23,11 | 3,69E-118 | 2,87E-117 | 0,0018273 | 0,0026984 | 0,0021043 | 0,0021043 | TRUE | 0,032612 | No |
| orf19.7665 | 1224 | 910 | 986 | 677 | 1872 | 1189 | 1772 | 1796 | 1167 | 962 | 974 | 1061 | 1500 | 1352 | 1504 | 1460 | 1247,56 | 1041 | 1454 | 1,398 | 0,484 | 6,869 | 6,48E-12 | 1,20E-11 | 0,0042339 | 0,0034074 | 0,0039102 | 0,0039102 | TRUE | 0,0843748 | No |
| orf19.7666 | 18 | 13 | 16 | 11 | 94 | 72 | 71 | 99 | 17 | 14 | 16 | 17 | 75 | 82 | 60 | 80 | 45,24 | 16 | 74 | 4,668 | 2,223 | 8,633 | 5,97E-18 | 1,28E-17 | 1,00E-08 | 0,0383854 | 0,0229456 | 0,0229456 | TRUE | 0,1588775 | No |
| orf19.7667 | 1528 | 1313 | 1358 | 990 | 1963 | 1462 | 1898 | 2065 | 1457 | 1388 | 1342 | 1551 | 1573 | 1662 | 1611 | 1679 | 1532,92 | 1434 | 1631 | 1,14 | 0,189 | 3,439 | 0,0005838 | 0,0007993 | 0,0015781 | 0,0031624 | 0,0022236 | 0,0022236 | TRUE | 0,032439 | No |
| orf19.7668 | 416 | 306 | 351 | 218 | 479 | 302 | 427 | 444 | 397 | 324 | 347 | 342 | 384 | 343 | 363 | 361 | 357,4 | 352 | 363 | 1,029 | 0,042 | 0,476 | 0,6339926 | 0,6663658 | 0,0021564 | 0,0066853 | 0,0045121 | 0,0045121 | TRUE | 0,0886091 | No |
| orf19.767 | 1189 | 1295 | 935 | 708 | 1900 | 1614 | 1118 | 1152 | 1134 | 1369 | 924 | 1109 | 1522 | 1835 | 949 | 937 | 1222,4 | 1134 | 1311 | 1,156 | 0,209 | 0,773 | 0,4393013 | 0,4755185 | 0,0691725 | 0,0034345 | 0,0317492 | 0,0691725 | TRUE | 0,3320734 | No |
| orf19.7670 | 1873 | 1434 | 1706 | 1074 | 2228 | 1461 | 2121 | 2360 | 1786 | 1516 | 1685 | 1682 | 1785 | 1661 | 1801 | 1919 | 1729,5 | 1667 | 1792 | 1,075 | 0,105 | 1,65 | 0,0990194 | 0,1156797 | 0,0031988 | 0,0030406 | 0,0032694 | 0,0032694 | TRUE | 0,0459672 | No |
| orf19.7672 | 2284 | 1796 | 1983 | 1400 | 3937 | 2624 | 3608 | 4001 | 2178 | 1899 | 1959 | 2193 | 3155 | 2984 | 3063 | 3253 | 2585,43 | 2057 | 3114 | 1,516 | 0,6 | 10,432 | 1,76E-25 | 4,41E-25 | 0,0028227 | 0,0027263 | 0,0027589 | 0,0027589 | TRUE | 0,0321818 | No |
| orf19.7673 | 425 | 307 | 338 | 196 | 1686 | 845 | 1546 | 1584 | 405 | 325 | 334 | 307 | 1351 | 961 | 1313 | 1288 | 785,38 | 343 | 1228 | 3,576 | 1,838 | 15,842 | 1,61E-56 | 6,44E-56 | 0,0179597 | 0,0041819 | 0,0109301 | 0,0109301 | TRUE | 0,2523221 | No |
| orf19.7675 | 751 | 545 | 602 | 453 | 1803 | 1130 | 1651 | 1956 | 716 | 576 | 595 | 710 | 1445 | 1285 | 1402 | 1590 | 1039,79 | 649 | 1430 | 2,209 | 1,144 | 12,887 | 5,30E-38 | 1,63E-37 | 0,008888 | 0,0036703 | 0,006382 | 0,006382 | TRUE | 0,0683652 | No |
| orf19.7676 | 19956 | 15522 | 16642 | 12942 | 4540 | 3070 | 4042 | 4655 | 19026 | 16413 | 16441 | 20274 | 3638 | 3491 | 3432 | 3785 | 10812,34 | 18038 | 3586 | 0,199 | -2,33 | -32,67 | 4,12E-234 | 6,47E-233 | 0,0064726 | 0,0022431 | 0,0047306 | 0,0047306 | TRUE | 0,0847633 | No |
| orf19.7678 | 5979 | 4593 | 4954 | 3449 | 13604 | 8461 | 12648 | 13619 | 5700 | 4857 | 4894 | 5403 | 10900 | 9621 | 10738 | 11074 | 7898,38 | 5214 | 10583 | 2,031 | 1,022 | 15,913 | 5,17E-57 | 2,08E-56 | 0,0048109 | 0,0022992 | 0,0038114 | 0,0038114 | TRUE | 0,0491144 | No |
| orf19.768 | 559 | 546 | 578 | 316 | 1220 | 851 | 1098 | 1156 | 533 | 577 | 571 | 495 | 978 | 968 | 932 | 940 | 749,21 | 544 | 954 | 1,746 | 0,804 | 12,701 | 5,87E-37 | 1,77E-36 | 3,89E-05 | 0,0042828 | 0,0023604 | 0,0023604 | TRUE | 0,0348851 | No |
| orf19.7680 | 523 | 396 | 447 | 293 | 673 | 381 | 681 | 733 | 499 | 419 | 442 | 459 | 539 | 433 | 578 | 596 | 495,57 | 455 | 536 | 1,184 | 0,243 | 2,373 | 0,017658 | 0,0220112 | 0,0106235 | 0,0054044 | 0,0080438 | 0,0080438 | TRUE | 0,18624 | No |
| orf19.769 | 91 | 76 | 69 | 42 | 109 | 77 | 118 | 109 | 87 | 80 | 68 | 66 | 87 | 88 | 100 | 89 | 83,1 | 75 | 91 | 1,2 | 0,262 | 1,645 | 0,0998794 | 0,1166418 | 1,00E-08 | 0,0218499 | 0,0122048 | 0,0122048 | TRUE | 0,1003159 | No |
| orf19.771 | 662 | 641 | 599 | 386 | 362 | 237 | 318 | 346 | 631 | 678 | 592 | 605 | 290 | 269 | 270 | 281 | 452,03 | 626 | 278 | 0,444 | -1,173 | -15,135 | 9,51E-52 | 3,59E-51 | 1,00E-08 | 0,0057236 | 0,0032718 | 0,0032718 | TRUE | 0,0362769 | No |
| orf19.77.1 | 3 | 7 | 10 | 3 | 3 | 4 | 0 | 4 | 3 | 7 | 10 | 5 | 2 | 5 | 0 | 3 | 4,38 | 6 | 2 | 0,396 | -1,337 | -1,726 | 0,0842627 | 0,0991116 | 0,1735439 | 0,376901 | 0,3073128 | 0,3073128 | TRUE | 0,3327637 | No |
| orf19.772 | 2789 | 2453 | 2566 | 1584 | 6601 | 4631 | 6312 | 7052 | 2659 | 2594 | 2535 | 2481 | 5289 | 5266 | 5359 | 5734 | 3989,62 | 2567 | 5412 | 2,107 | 1,075 | 24,515 | 1,02E-132 | 9,12E-132 | 0,0008737 | 0,0025028 | 0,0015446 | 0,0015446 | TRUE | 0,0197294 | No |
| orf19.772.1 | 2162 | 1971 | 1981 | 1374 | 1528 | 1191 | 1427 | 1374 | 2061 | 2084 | 1957 | 2152 | 1224 | 1354 | 1212 | 1117 | 1645,27 | 2064 | 1227 | 0,594 | -0,752 | -12,113 | 8,98E-34 | 2,57E-33 | 0,0030597 | 0,0030892 | 0,0030587 | 0,0030587 | TRUE | 0,0559159 | No |
| orf19.773 | 3965 | 3905 | 3989 | 2551 | 3689 | 2679 | 3493 | 3476 | 3780 | 4129 | 3941 | 3996 | 2956 | 3046 | 2966 | 2826 | 3455,04 | 3962 | 2948 | 0,744 | -0,426 | -9,932 | 3,01E-23 | 7,21E-23 | 0,0007462 | 0,0025665 | 0,0014752 | 0,0014752 | TRUE | 0,0119202 | No |
| orf19.775 | 3400 | 3349 | 3393 | 2121 | 1658 | 1240 | 1526 | 1742 | 3242 | 3541 | 3352 | 3323 | 1329 | 1410 | 1296 | 1416 | 2363,47 | 3364 | 1363 | 0,405 | -1,305 | -26,568 | 1,58E-155 | 1,66E-154 | 0,001195 | 0,002786 | 0,0018214 | 0,0018214 | TRUE | 0,0154566 | No |
| orf19.776 | 117 | 110 | 110 | 65 | 108 | 76 | 78 | 96 | 112 | 116 | 109 | 102 | 87 | 86 | 66 | 78 | 94,45 | 110 | 79 | 0,719 | -0,476 | -3,177 | 0,0014885 | 0,0019858 | 1,00E-08 | 0,0194754 | 0,0108225 | 0,0108225 | TRUE | 0,120718 | No |
| orf19.778 | 10178 | 8599 | 8989 | 6013 | 9308 | 6509 | 8504 | 9716 | 9704 | 9092 | 8880 | 9420 | 7458 | 7401 | 7220 | 7900 | 8384,42 | 9274 | 7495 | 0,808 | -0,307 | -7,07 | 1,55E-12 | 2,91E-12 | 0,0013437 | 0,0022871 | 0,0016908 | 0,0016908 | TRUE | 0,0163025 | No |
| orf19.779 | 849 | 735 | 796 | 451 | 1922 | 1290 | 1746 | 1756 | 809 | 777 | 786 | 707 | 1540 | 1467 | 1482 | 1428 | 1124,56 | 770 | 1479 | 1,914 | 0,937 | 16,583 | 9,17E-62 | 3,97E-61 | 0,0006789 | 0,0035513 | 0,0020373 | 0,0020373 | TRUE | 0,0306252 | No |

| | | | | | | | | | | | | | | | | | | | | | | | | | | | | | | | |
|---|---|---|---|---|---|---|---|---|---|---|---|---|---|---|---|---|---|---|---|---|---|---|---|---|---|---|---|---|---|---|---|
| orf19.780 | 2115 | 1923 | 2251 | 1285 | 10413 | 7439 | 9374 | 10686 | 2016 | 2033 | 2224 | 2013 | 8344 | 8459 | 7958 | 8689 | 5217,03 | 2072 | 8362 | 4,033 | 2,012 | 42,339 | 0 | 0 | 0,0014718 | 0,002406 | 0,0018437 | 0,0018437 | TRUE | 0,0303659 | No |
| orf19.781 | 170 | 158 | 157 | 113 | 454 | 282 | 374 | 437 | 162 | 167 | 155 | 177 | 364 | 321 | 318 | 355 | 252,32 | 165 | 340 | 2,066 | 1,047 | 10,426 | 1,88E-25 | 4,70E-25 | 1,00E-08 | 0,0085985 | 0,0049947 | 0,0049947 | TRUE | 0,0273151 | No |
| orf19.782 | 4160 | 3853 | 4334 | 2745 | 5201 | 3848 | 5122 | 5356 | 3966 | 4074 | 4282 | 4300 | 4167 | 4376 | 4349 | 4355 | 4233,55 | 4156 | 4312 | 1,038 | 0,054 | 1,325 | 0,1852608 | 0,2107218 | 0,0007028 | 0,0024791 | 0,0013541 | 0,0013541 | TRUE | 0,0116396 | No |
| orf19.783 | 3769 | 3518 | 3611 | 2497 | 3354 | 2473 | 3327 | 3422 | 3593 | 3720 | 3567 | 3912 | 2687 | 2812 | 2825 | 2782 | 3237,34 | 3698 | 2776 | 0,752 | -0,412 | -9,638 | 5,55E-22 | 1,29E-21 | 0,000733 | 0,0025985 | 0,0014407 | 0,0014407 | TRUE | 0,0172826 | No |
| orf19.787 | 17 | 24 | 30 | 21 | 26 | 24 | 18 | 14 | 16 | 25 | 30 | 33 | 21 | 27 | 15 | 11 | 22,36 | 26 | 18 | 0,717 | -0,48 | -1,379 | 0,1678681 | 0,1917935 | 0,0636699 | 0,0755102 | 0,0704764 | 0,0704764 | TRUE | 0,380168 | No |
| orf19.787.1 | 6930 | 6727 | 6584 | 4589 | 3394 | 2602 | 3027 | 3145 | 6607 | 7113 | 6504 | 7189 | 2720 | 2959 | 2570 | 2557 | 4777,32 | 6853 | 2702 | 0,394 | -1,344 | -23,345 | 1,55E-120 | 1,24E-119 | 0,0033318 | 0,002435 | 0,0029343 | 0,0029343 | TRUE | 0,0486378 | No |
| orf19.789 | 5776 | 5511 | 5687 | 3568 | 14751 | 10364 | 12902 | 14618 | 5507 | 5827 | 5618 | 5589 | 11820 | 11785 | 10954 | 11886 | 8623,2 | 5635 | 11611 | 2,06 | 1,043 | 26,306 | 1,65E-152 | 1,69E-151 | 0,0008353 | 0,0022817 | 0,0013706 | 0,0013706 | TRUE | 0,0178344 | No |
| orf19.789.1 | 837 | 734 | 771 | 427 | 802 | 534 | 758 | 681 | 798 | 776 | 762 | 669 | 643 | 607 | 644 | 554 | 681,47 | 751 | 612 | 0,812 | -0,301 | -3,958 | 7,56E-05 | 0,0001077 | 0,0036896 | 0,0045006 | 0,0040828 | 0,0040828 | TRUE | 0,0570098 | No |
| orf19.791 | 0 | 0 | 0 | 0 | 0 | 0 | 0 | 0 | 0 | 0 | 0 | 0 | 0 | 0 | 0 | 0 | 0 | 0 | 0 | NA | NA | NA | NA | NA | NA | NA | NA | NA | NA | NA | NA |
| orf19.792 | 2084 | 2296 | 2190 | 1822 | 2661 | 1861 | 2480 | 2547 | 1987 | 2428 | 2164 | 2854 | 2132 | 2116 | 2106 | 2071 | 2232,14 | 2358 | 2106 | 0,895 | -0,161 | -1,759 | 0,0785428 | 0,0927256 | 0,0122659 | 0,0028269 | 0,0075508 | 0,0075508 | TRUE | 0,2376732 | No |
| orf19.793 | 1452 | 1588 | 1708 | 1013 | 1479 | 1154 | 1287 | 1153 | 1384 | 1679 | 1687 | 1587 | 1185 | 1312 | 1093 | 937 | 1358,14 | 1584 | 1132 | 0,714 | -0,487 | -4,973 | 6,59E-07 | 1,03E-06 | 0,0126614 | 0,0033002 | 0,0084481 | 0,0084481 | TRUE | 0,1612954 | No |
| orf19.794 | 1193 | 1182 | 1234 | 828 | 1261 | 851 | 1101 | 1148 | 1137 | 1250 | 1219 | 1297 | 1010 | 968 | 935 | 933 | 1093,7 | 1226 | 962 | 0,786 | -0,347 | -5,965 | 2,45E-09 | 4,16E-09 | 0,0013383 | 0,0035925 | 0,0023291 | 0,0023291 | TRUE | 0,0288534 | No |
| orf19.795 | 2424 | 2347 | 2415 | 1365 | 1885 | 1358 | 1830 | 1818 | 2311 | 2482 | 2386 | 2138 | 1510 | 1544 | 1554 | 1478 | 1925,41 | 2329 | 1522 | 0,652 | -0,617 | -11,648 | 2,36E-31 | 6,50E-31 | 0,0014725 | 0,002944 | 0,0021599 | 0,0021599 | TRUE | 0,0340268 | No |
| orf19.796 | 1238 | 1188 | 1213 | 784 | 1264 | 969 | 1222 | 1279 | 1180 | 1256 | 1198 | 1228 | 1013 | 1102 | 1037 | 1040 | 1131,88 | 1216 | 1048 | 0,861 | -0,215 | -4,192 | 2,77E-05 | 4,02E-05 | 1,00E-08 | 0,0035419 | 0,0016463 | 0,0016463 | TRUE | 0,013111 | No |
| orf19.797 | 2262 | 1902 | 2027 | 1331 | 2621 | 1784 | 2499 | 2700 | 2157 | 2011 | 2002 | 2085 | 2100 | 2029 | 2122 | 2195 | 2087,62 | 2064 | 2112 | 1,024 | 0,034 | 0,746 | 0,4559187 | 0,4925868 | 0,0006811 | 0,0028778 | 0,0015278 | 0,0015278 | TRUE | 0,0119662 | No |
| orf19.798 | 2777 | 2479 | 2548 | 1631 | 4858 | 3465 | 4593 | 4770 | 2648 | 2621 | 2517 | 2555 | 3893 | 3940 | 3899 | 3878 | 3243,94 | 2585 | 3902 | 1,509 | 0,593 | 16,825 | 1,60E-63 | 7,04E-63 | 1,00E-08 | 0,0025974 | 0,0008624 | 0,0008624 | TRUE | 0,0042312 | No |
| orf19.799 | 184 | 131 | 181 | 131 | 312 | 191 | 323 | 307 | 175 | 139 | 179 | 205 | 250 | 217 | 274 | 250 | 211,12 | 174 | 248 | 1,433 | 0,519 | 4,169 | 3,05E-05 | 4,43E-05 | 0,0103113 | 0,0098682 | 0,0098687 | 0,0098687 | TRUE | 0,2018841 | No |
| orf19.8 | 116 | 124 | 117 | 88 | 248 | 204 | 241 | 227 | 111 | 131 | 116 | 138 | 199 | 232 | 205 | 185 | 164,38 | 124 | 205 | 1,662 | 0,733 | 5,878 | 4,15E-09 | 6,97E-09 | 0,0032706 | 0,01208 | 0,0082183 | 0,0082183 | TRUE | 0,078064 | No |
| orf19.801 | 6166 | 5090 | 5890 | 3765 | 5332 | 3699 | 5265 | 5425 | 5879 | 5382 | 5819 | 5898 | 4272 | 4206 | 4470 | 4411 | 5042,12 | 5744 | 4340 | 0,756 | -0,404 | -9,419 | 4,57E-21 | 1,05E-20 | 0,001025 | 0,0024169 | 0,0015635 | 0,0015635 | TRUE | 0,0221297 | No |
| orf19.802 | 1813 | 1467 | 1656 | 1126 | 2047 | 1384 | 2074 | 2124 | 1728 | 1551 | 1636 | 1764 | 1640 | 1574 | 1761 | 1727 | 1672,67 | 1670 | 1676 | 1,005 | 0,008 | 0,131 | 0,8956454 | 0,9101948 | 0,0022475 | 0,0030729 | 0,002597 | 0,002597 | TRUE | 0,0274507 | No |
| orf19.804 | 462 | 410 | 464 | 263 | 1346 | 977 | 1297 | 1419 | 440 | 434 | 458 | 412 | 1079 | 1111 | 1101 | 1154 | 773,59 | 436 | 1111 | 2,539 | 1,344 | 21,757 | 5,94E-105 | 4,04E-104 | 1,00E-08 | 0,0042137 | 0,0019918 | 0,0019918 | TRUE | 0,0131759 | No |
| orf19.804.1 | 6019 | 4147 | 4227 | 2955 | 7757 | 4772 | 6528 | 7599 | 5738 | 4385 | 4176 | 4629 | 6215 | 5426 | 5542 | 6179 | 5286,38 | 4732 | 5840 | 1,234 | 0,304 | 3,356 | 0,0007919 | 0,0010756 | 0,0130806 | 0,0024019 | 0,0076687 | 0,0076687 | TRUE | 0,2516102 | No |
| orf19.805 | 904 | 743 | 840 | 510 | 1676 | 1079 | 1608 | 1641 | 862 | 786 | 830 | 799 | 1343 | 1227 | 1365 | 1334 | 1068,2 | 819 | 1317 | 1,608 | 0,685 | 11,843 | 2,34E-32 | 6,57E-32 | 0,0010187 | 0,0036284 | 0,0021939 | 0,0021939 | TRUE | 0,0239912 | No |
| orf19.806 | 279 | 200 | 252 | 146 | 735 | 479 | 699 | 761 | 266 | 211 | 249 | 229 | 589 | 545 | 593 | 619 | 412,62 | 239 | 586 | 2,45 | 1,293 | 14,624 | 1,97E-48 | 7,10E-48 | 0,0023932 | 0,0060705 | 0,0044193 | 0,0044193 | TRUE | 0,0728195 | No |
| orf19.807 | 2730 | 2289 | 2475 | 1478 | 2591 | 1783 | 2539 | 2707 | 2603 | 2420 | 2445 | 2315 | 2076 | 2027 | 2156 | 2201 | 2280,46 | 2446 | 2115 | 0,864 | -0,211 | -4,244 | 2,20E-05 | 3,20E-05 | 0,00127 | 0,0028113 | 0,0019278 | 0,0019278 | TRUE | 0,0227544 | No |
| orf19.808 | 3821 | 3470 | 3855 | 2349 | 3516 | 2511 | 3432 | 3547 | 3643 | 3669 | 3808 | 3680 | 2817 | 2855 | 2914 | 2884 | 3283,81 | 3700 | 2868 | 0,775 | -0,368 | -10,405 | 2,34E-25 | 5,84E-25 | 1,00E-08 | 0,0025913 | 0,0008928 | 0,0008928 | TRUE | 0,0049336 | No |
| orf19.809 | 10894 | 9526 | 10956 | 6318 | 12702 | 9050 | 12846 | 13275 | 10386 | 10073 | 10823 | 9897 | 10178 | 10291 | 10906 | 10794 | 10418,53 | 10295 | 10542 | 1,024 | 0,034 | 0,81 | 0,4178023 | 0,4538354 | 0,0012303 | 0,0022489 | 0,0015801 | 0,0015801 | TRUE | 0,0146143 | No |
| orf19.81 | 2468 | 3131 | 3019 | 1723 | 1976 | 1320 | 1722 | 1454 | 2353 | 3311 | 2982 | 2699 | 1583 | 1501 | 1462 | 1182 | 2134,22 | 2836 | 1432 | 0,505 | -0,986 | -9,238 | 2,52E-20 | 5,70E-20 | 0,017022 | 0,0028606 | 0,0104398 | 0,0104398 | TRUE | 0,1382755 | No |
| orf19.810 | 713 | 615 | 699 | 426 | 771 | 565 | 743 | 819 | 680 | 650 | 691 | 667 | 618 | 642 | 631 | 666 | 655,61 | 672 | 639 | 0,95 | -0,073 | -1,205 | 0,2282239 | 0,2572054 | 1,00E-08 | 0,0045957 | 0,0020214 | 0,0020214 | TRUE | 0,0100644 | No |
| orf19.811 | 739 | 649 | 725 | 412 | 1131 | 713 | 1035 | 1070 | 705 | 686 | 716 | 645 | 906 | 811 | 879 | 870 | 777,27 | 688 | 866 | 1,257 | 0,33 | 5,349 | 8,84E-08 | 1,43E-07 | 0,0005628 | 0,0042037 | 0,0023404 | 0,0023404 | TRUE | 0,0204734 | No |
| orf19.812 | 42 | 20 | 42 | 18 | 50 | 25 | 34 | 34 | 40 | 21 | 41 | 28 | 40 | 28 | 29 | 28 | 31,99 | 32 | 31 | 0,949 | -0,076 | -0,27 | 0,7875187 | 0,8109016 | 0,0332978 | 0,0534245 | 0,0447634 | 0,0447634 | TRUE | 0,3664003 | No |
| orf19.813 | 37436 | 34497 | 36870 | 21246 | 43147 | 29124 | 40987 | 44084 | 35691 | 36476 | 36424 | 33283 | 34572 | 33117 | 34798 | 35845 | 35025,72 | 35468 | 34583 | 0,975 | -0,037 | -0,879 | 0,3795292 | 0,4145457 | 0,0013812 | 0,0021382 | 0,0016385 | 0,0016385 | TRUE | 0,0209127 | No |
| orf19.814 | 4982 | 4400 | 4903 | 2967 | 2307 | 1674 | 2248 | 2358 | 4750 | 4652 | 4844 | 4648 | 1849 | 1903 | 1909 | 1917 | 3308,97 | 4724 | 1894 | 0,401 | -1,319 | -36,13 | 7,67E-286 | 1,60E-284 | 1,00E-08 | 0,0025875 | 0,0009252 | 0,0009252 | TRUE | 0,0036049 | No |
| orf19.815 | 11540 | 11105 | 11905 | 7181 | 9367 | 6797 | 8085 | 8932 | 11002 | 11742 | 11761 | 11249 | 7506 | 7729 | 6864 | 7263 | 9389,46 | 11438 | 7340 | 0,642 | -0,64 | -14,011 | 1,33E-44 | 4,53E-44 | 0,0016603 | 0,0022661 | 0,0018968 | 0,0018968 | TRUE | 0,0232753 | No |
| orf19.816 | 709 | 634 | 696 | 416 | 534 | 417 | 514 | 528 | 676 | 670 | 688 | 652 | 428 | 474 | 436 | 429 | 556,67 | 672 | 442 | 0,655 | -0,61 | -9,254 | 2,17E-20 | 4,92E-20 | 1,00E-08 | 0,0050408 | 0,0023253 | 0,0023253 | TRUE | 0,024785 | No |
| orf19.819 | 904 | 773 | 838 | 559 | 2714 | 1850 | 2374 | 2578 | 862 | 817 | 828 | 876 | 2175 | 2104 | 2016 | 2096 | 1471,59 | 846 | 2098 | 2,484 | 1,313 | 26,092 | 4,49E-150 | 4,52E-149 | 1,00E-08 | 0,003207 | 0,0015565 | 0,0015565 | TRUE | 0,008585 | No |
| orf19.820 | 1352 | 1142 | 1205 | 742 | 2035 | 1346 | 1856 | 2075 | 1289 | 1208 | 1190 | 1162 | 1631 | 1531 | 1576 | 1687 | 1409,17 | 1212 | 1606 | 1,324 | 0,405 | 7,519 | 5,53E-14 | 1,08E-13 | 0,0011467 | 0,0032564 | 0,0020401 | 0,0020401 | TRUE | 0,0223165 | No |
| orf19.822 | 271052 | 224248 | 228545 | 141921 | 15760 | 10314 | 14306 | 15845 | 258419 | 237114 | 225781 | 222327 | 12628 | 11728 | 12146 | 12883 | 124128,18 | 235910 | 12346 | 0,052 | -4,256 | -78,61 | 0 | 0 | 0,0031889 | 0,0021045 | 0,0027777 | 0,0027777 | TRUE | 0,0505779 | No |
| orf19.823 | 988 | 1262 | 1242 | 1826 | 97 | 156 | 89 | 81 | 942 | 1334 | 1227 | 2861 | 78 | 177 | 76 | 66 | 845,05 | 1591 | 99 | 0,062 | -4,008 | -8,235 | 1,80E-16 | 3,74E-16 | 0,2227195 | 0,0040342 | 0,0849814 | 0,2227195 | TRUE | 0,9244277 | No |
| orf19.825 | 2018 | 1730 | 1857 | 1217 | 3316 | 2301 | 3374 | 3636 | 1924 | 1829 | 1835 | 1906 | 2657 | 2616 | 2865 | 2956 | 2323,58 | 1874 | 2774 | 1,482 | 0,567 | 10,83 | 2,48E-27 | 6,42E-27 | 0,0017893 | 0,0027979 | 0,0021734 | 0,0021734 | TRUE | 0,0239959 | No |
| orf19.826 | 820 | 781 | 852 | 480 | 815 | 594 | 754 | 845 | 782 | 826 | 842 | 752 | 653 | 675 | 640 | 687 | 732,11 | 800 | 664 | 0,827 | -0,274 | -4,432 | 9,32E-06 | 1,38E-05 | 1,00E-08 | 0,004334 | 0,0022904 | 0,0022904 | TRUE | 0,0166242 | No |
| orf19.827 | 1449 | 1207 | 1374 | 942 | 1741 | 1073 | 1606 | 1607 | 1381 | 1276 | 1357 | 1476 | 1395 | 1220 | 1363 | 1307 | 1347 | 1372 | 1321 | 0,965 | -0,051 | -0,824 | 0,4099341 | 0,446076 | 0,0024853 | 0,0033102 | 0,0028873 | 0,0028873 | TRUE | 0,0302913 | No |
| orf19.827.1 | 11957 | 9751 | 9817 | 6475 | 24531 | 15344 | 21844 | 25039 | 11400 | 10310 | 9698 | 10143 | 19656 | 17448 | 18545 | 20359 | 14694,99 | 10388 | 19002 | 1,829 | 0,871 | 14,054 | 7,30E-45 | 2,49E-44 | 0,0045772 | 0,002203 | 0,0036142 | 0,0036142 | TRUE | 0,0528698 | No |
| orf19.8278 | 2106 | 2043 | 2095 | 1289 | 2138 | 1394 | 1863 | 1674 | 2008 | 2160 | 2070 | 2019 | 1713 | 1585 | 1582 | 1361 | 1812,25 | 2064 | 1560 | 0,756 | -0,404 | -5,926 | 3,11E-09 | 5,25E-09 | 0,0042029 | 0,0029973 | 0,0039122 | 0,0039122 | TRUE | 0,0909663 | No |
| orf19.828 | 2450 | 2219 | 2289 | 1472 | 2357 | 1608 | 2243 | 2398 | 2336 | 2346 | 2261 | 2306 | 1889 | 1828 | 1904 | 1950 | 2102,57 | 2312 | 1893 | 0,819 | -0,288 | -7,073 | 1,52E-12 | 2,85E-12 | 1,00E-08 | 0,0028722 | 0,0011104 | 0,0011104 | TRUE | 0,0058416 | No |
| orf19.829 | 5573 | 5088 | 5406 | 3553 | 6506 | 4663 | 6064 | 6546 | 5313 | 5380 | 5341 | 5566 | 5213 | 5302 | 5148 | 5323 | 5323,24 | 5400 | 5246 | 0,972 | -0,041 | -1,256 | 0,2091544 | 0,2367597 | 9,13E-05 | 0,0023997 | 0,0008161 | 0,0008161 | TRUE | 0,0048503 | No |
| orf19.831 | 2854 | 2710 | 2919 | 1813 | 1440 | 1069 | 1322 | 1383 | 2721 | 2865 | 2884 | 2840 | 1154 | 1216 | 1122 | 1125 | 1990,82 | 2828 | 1154 | 0,408 | -1,295 | -28,354 | 7,40E-177 | 8,71E-176 | 0,0003917 | 0,002916 | 0,0014161 | 0,0014161 | TRUE | 0,0140446 | No |
| orf19.832 | 1183 | 1155 | 1277 | 711 | 1107 | 806 | 1083 | 1035 | 1128 | 1221 | 1262 | 1114 | 887 | 916 | 919 | 842 | 1036,13 | 1181 | 891 | 0,753 | -0,41 | -6,722 | 1,79E-11 | 3,28E-11 | 0,001727 | 0,0036759 | 0,0025941 | 0,0025941 | TRUE | 0,0253971 | No |
| orf19.834 | 3889 | 3854 | 4106 | 2434 | 2931 | 2200 | 2779 | 2759 | 3708 | 4075 | 4056 | 3813 | 2349 | 2502 | 2359 | 2243 | 3138,13 | 3913 | 2363 | 0,603 | -0,729 | -14,55 | 5,81E-48 | 2,09E-47 | 0,001632 | 0,0026145 | 0,0020776 | 0,0020776 | TRUE | 0,0181201 | No |
| orf19.835 | 1152 | 1124 | 1186 | 719 | 2478 | 1785 | 2519 | 2607 | 1098 | 1188 | 1172 | 1126 | 1986 | 2030 | 2139 | 2120 | 1607,3 | 1146 | 2069 | 1,804 | 0,851 | 17,034 | 4,56E-65 | 2,05E-64 | 0,0007367 | 0,0031128 | 0,0016944 | 0,0016944 | TRUE | 0,0106518 | No |
| orf19.836 | 209 | 214 | 197 | 133 | 55 | 35 | 57 | 44 | 199 | 226 | 195 | 208 | 44 | 40 | 48 | 36 | 124,57 | 207 | 42 | 0,203 | -2,298 | -15,087 | 1,98E-51 | 7,43E-51 | 1,00E-08 | 0,0152721 | 0,0090391 | 0,0090391 | TRUE | 0,0933232 | No |
| orf19.836.1 | 339 | 249 | 258 | 118 | 735 | 400 | 639 | 730 | 323 | 263 | 255 | 185 | 589 | 455 | 543 | 594 | 400,76 | 256 | 545 | 2,111 | 1,078 | 7,791 | 6,65E-15 | 1,34E-14 | 0,0275371 | 0,0061883 | 0,0153771 | 0,0153771 | TRUE | 0,343039 | No |
| orf19.837 | 1214 | 1133 | 1126 | 611 | 1837 | 1159 | 1608 | 1580 | 1157 | 1198 | 1112 | 957 | 1472 | 1318 | 1365 | 1285 | 1233,08 | 1106 | 1360 | 1,226 | 0,294 | 3,999 | 6,37E-05 | 9,10E-05 | 0,005245 | 0,0034228 | 0,0043618 | 0,0043618 | TRUE | 0,0901526 | No |
| orf19.837.1 | 919 | 856 | 945 | 499 | 2358 | 1434 | 2126 | 2224 | 876 | 905 | 934 | 782 | 1889 | 1631 | 1805 | 1808 | 1328,73 | 874 | 1783 | 2,035 | 1,025 | 15,3 | 7,70E-53 | 2,95E-52 | 0,003577 | 0,003327 | 0,0034211 | 0,0034211 | TRUE | 0,0533121 | No |
| orf19.838.1 | 19928 | 16350 | 17684 | 10478 | 58108 | 36728 | 54638 | 60344 | 18999 | 17288 | 17470 | 16414 | 46560 | 41763 | 46387 | 49065 | 31743,48 | 17543 | 45944 | 2,619 | 1,389 | 23,54 | 1,60E-122 | 1,30E-121 | 0,0040188 | 0,002143 | 0,0033024 | 0,0033024 | TRUE | 0,0459043 | No |
| orf19.839 | 3240 | 2873 | 3104 | 1856 | 4930 | 3599 | 4763 | 5122 | 3089 | 3038 | 3066 | 2908 | 3950 | 4092 | 4044 | 4165 | 3543,99 | 3025 | 4063 | 1,341 | 0,423 | 11,268 | 1,88E-29 | 5,02E-29 | 0,0002209 | 0,0025546 | 0,0010605 | 0,0010605 | TRUE | 0,0075391 | No |
| orf19.84 | 5341 | 4742 | 5175 | 3394 | 1182 | 943 | 1099 | 1160 | 5092 | 5014 | 5112 | 5317 | 947 | 1072 | 933 | 943 | 3053,88 | 5134 | 974 | 0,189 | -2,402 | -47,628 | 0 | 0 | 0,0011665 | 0,0026289 | 0,0018724 | 0,0018724 | TRUE | 0,0507732 | No |
| orf19.840 | 18678 | 14805 | 15522 | 11308 | 46518 | 32415 | 43557 | 48017 | 17807 | 15654 | 15334 | 17715 | 37274 | 36859 | 36980 | 39042 | 27083,14 | 16628 | 37539 | 2,258 | 1,175 | 21,094 | 8,97E-99 | 5,79E-98 | 0,0034402 | 0,0021519 | 0,0029353 | 0,0029353 | TRUE | 0,0336126 | No |
| orf19.841 | 1848 | 1658 | 1763 | 1135 | 2303 | 1555 | 2188 | 2324 | 1762 | 1753 | 1742 | 1778 | 1845 | 1768 | 1858 | 1890 | 1799,43 | 1759 | 1840 | 1,048 | 0,067 | 1,604 | 0,1086186 | 0,1262929 | 1,00E-08 | 0,0030037 | 0,0011344 | 0,0011344 | TRUE | 0,0077319 | No |
| orf19.842 | 16568 | 15340 | 15962 | 9439 | 7661 | 4897 | 7258 | 7583 | 15796 | 16220 | 15769 | 14787 | 6139 | 5568 | 6162 | 6166 | 10825,77 | 15643 | 6009 | 0,384 | -1,38 | -29,627 | 6,73E-193 | 8,93E-192 | 0,0017947 | 0,0022429 | 0,0019739 | 0,0019739 | TRUE | 0,0291213 | No |
| orf19.843 | 1277 | 1212 | 1236 | 723 | 2067 | 1328 | 2039 | 2046 | 1217 | 1282 | 1221 | 1133 | 1656 | 1510 | 1731 | 1664 | 1426,71 | 1213 | 1640 | 1,351 | 0,434 | 7,408 | 1,29E-13 | 2,50E-13 | 0,0019282 | 0,0032421 | 0,0025621 | 0,0025621 | TRUE | 0,0327302 | No |
| orf19.844 | 10225 | 9579 | 10630 | 6246 | 2103 | 1516 | 1993 | 2168 | 9748 | 10129 | 10501 | 9785 | 1685 | 1724 | 1692 | 1763 | 5878,36 | 10041 | 1716 | 0,171 | -2,549 | -62,678 | 0 | 0 | 0,000747 | 0,0023706 | 0,0012724 | 0,0012724 | TRUE | 0,0117519 | No |
| orf19.846 | 6823 | 6156 | 6779 | 3942 | 5120 | 3547 | 5014 | 5282 | 6505 | 6509 | 6697 | 6175 | 4103 | 4033 | 4257 | 4295 | 5321,74 | 6472 | 4172 | 0,644 | -0,634 | -15,954 | 2,68E-57 | 1,08E-56 | 0,0007426 | 0,0023998 | 0,0013217 | 0,0013217 | TRUE | 0,0110704 | No |
| orf19.847 | 4618 | 4469 | 4563 | 2951 | 3973 | 2947 | 3605 | 3896 | 4403 | 4725 | 4508 | 4623 | 3183 | 3351 | 3061 | 3168 | 3877,72 | 4565 | 3191 | 0,699 | -0,517 | -12,214 | 2,61E-34 | 7,56E-34 | 0,0008214 | 0,0025147 | 0,0014573 | 0,0014573 | TRUE | 0,0133094 | No |
| orf19.848 | 0 | 1 | 0 | 0 | 0 | 0 | 0 | 0 | 0 | 1 | 0 | 0 | 0 | 0 | 0 | 0 | 0,13 | 0 | 0 | 0,483 | -1,049 | -0,297 | 0,7665832 | 0,7911318 | 1,00E-08 | 12,424597 | 10 | 10 | TRUE | 0,3159984 | No |
| orf19.849 | 84 | 103 | 88 | 64 | 44 | 40 | 45 | 51 | 80 | 109 | 87 | 100 | 35 | 45 | 38 | 41 | 67,08 | 94 | 40 | 0,426 | -1,232 | -6,561 | 5,34E-11 | 9,62E-11 | 1,00E-08 | 0,0265698 | 0,0167281 | 0,0167281 | TRUE | 0,1207656 | No |
| orf19.85 | 2040 | 2072 | 2058 | 1183 | 1173 | 751 | 1042 | 1033 | 1945 | 2191 | 2033 | 1853 | 940 | 854 | 885 | 840 | 1442,57 | 2006 | 880 | 0,438 | -1,19 | -18,518 | 1,48E-76 | 7,73E-76 | 0,003136 | 0,0032295 | 0,0031759 | 0,0031759 | TRUE | 0,0463456 | No |
| orf19.850 | 1828 | 1711 | 1925 | 1125 | 1751 | 1191 | 1581 | 1666 | 1743 | 1809 | 1902 | 1762 | 1403 | 1354 | 1342 | 1355 | 1583,78 | 1804 | 1364 | 0,755 | -0,405 | -8,544 | 1,29E-17 | 2,76E-17 | 0,0004434 | 0,003128 | 0,0015139 | 0,0015139 | TRUE | 0,0166269 | No |
| orf19.851 | 3812 | 3786 | 4011 | 2491 | 1177 | 943 | 1157 | 1206 | 3634 | 4003 | 3962 | 3902 | 943 | 1072 | 982 | 981 | 2435,07 | 3875 | 994 | 0,256 | -1,965 | -36,644 | 5,80E-294 | 1,26E-292 | 0,0015981 | 0,0027656 | 0,0021658 | 0,0021658 | TRUE | 0,0307124 | No |
| orf19.852 | 875 | 783 | 854 | 640 | 1133 | 907 | 1081 | 1173 | 834 | 828 | 844 | 1003 | 908 | 1031 | 918 | 954 | 914,89 | 877 | 953 | 1,089 | 0,123 | 1,67 | 0,0948267 | 0,1110052 | 0,0043138 | 0,0038859 | 0,0041064 | 0,0041064 | TRUE | 0,1003699 | No |
| orf19.853 | 105 | 86 | 102 | 58 | 218 | 126 | 197 | 236 | 100 | 91 | 101 | 91 | 175 | 143 | 167 | 192 | 132,47 | 96 | 169 | 1,771 | 0,825 | 6,042 | 1,52E-09 | 2,60E-09 | 1,00E-08 | 0,0144858 | 0,0094701 | 0,0094701 | TRUE | 0,1016271 | No |
| orf19.854 | 3364 | 2845 | 3273 | 1984 | 4025 | 2754 | 3810 | 4164 | 3207 | 3008 | 3233 | 3108 | 3225 | 3132 | 3235 | 3386 | 3191,74 | 3139 | 3244 | 1,034 | 0,048 | 1,12 | 0,2626005 | 0,2931514 | 0,0006424 | 0,0026057 | 0,0014171 | 0,0014171 | TRUE | 0,0105141 | No |
| orf19.855 | 42 | 28 | 29 | 16 | 23 | 14 | 29 | 17 | 40 | 30 | 29 | 25 | 18 | 16 | 25 | 14 | 24,52 | 31 | 18 | 0,586 | -0,772 | -2,547 | 0,0108541 | 0,0137529 | 1,00E-08 | 0,0690548 | 0,0458938 | 0,0458938 | TRUE | 0,3245329 | No |

| | | | | | | | | | | | | | | | | | | | | | | | | | | | | | | | |
|---|---|---|---|---|---|---|---|---|---|---|---|---|---|---|---|---|---|---|---|---|---|---|---|---|---|---|---|---|---|---|---|
| orf19.856 | 1225 | 1129 | 1222 | 738 | 1040 | 746 | 1011 | 1059 | 1168 | 1194 | 1207 | 1156 | 833 | 848 | 858 | 861 | 1015,75 | 1181 | 850 | 0,719 | -0,476 | -9,551 | 1,29E-21 | 2,99E-21 | 1,00E-08 | 0,0037077 | 0,001385 | 0,001385 | TRUE | 0,0026251 | No |
| orf19.857 | 1140 | 981 | 1060 | 671 | 2297 | 1637 | 2195 | 2416 | 1087 | 1037 | 1047 | 1051 | 1841 | 1861 | 1864 | 1964 | 1469,05 | 1056 | 1882 | 1,783 | 0,834 | 17,562 | 4,83E-69 | 2,27E-68 | 1,00E-08 | 0,0032089 | 0,0014006 | 0,0014006 | TRUE | 0,0107053 | No |
| orf19.86 | 3072 | 2647 | 2765 | 1596 | 2209 | 1350 | 2090 | 2213 | 2929 | 2799 | 2732 | 2500 | 1770 | 1535 | 1774 | 1799 | 2229,79 | 2740 | 1720 | 0,628 | -0,672 | -10,292 | 7,68E-25 | 1,89E-24 | 0,0041057 | 0,0028276 | 0,0036278 | 0,0036278 | TRUE | 0,0608209 | No |
| orf19.860 | 462 | 383 | 387 | 281 | 1495 | 994 | 1446 | 1538 | 440 | 405 | 382 | 440 | 1198 | 1130 | 1228 | 1251 | 809,29 | 417 | 1202 | 2,892 | 1,532 | 22,537 | 1,82E-112 | 1,34E-111 | 0,0011255 | 0,0041201 | 0,0027265 | 0,0027265 | TRUE | 0,0377349 | No |
| orf19.861 | 751 | 596 | 589 | 434 | 565 | 402 | 538 | 558 | 716 | 630 | 582 | 680 | 453 | 457 | 457 | 454 | 553,53 | 652 | 455 | 0,699 | -0,517 | -6,718 | 1,84E-11 | 3,36E-11 | 0,0027874 | 0,0050575 | 0,0038513 | 0,0038513 | TRUE | 0,0647939 | No |
| orf19.862 | 380 | 337 | 340 | 229 | 514 | 356 | 458 | 532 | 362 | 356 | 336 | 359 | 412 | 405 | 389 | 433 | 381,41 | 353 | 410 | 1,161 | 0,215 | 2,78 | 0,0054309 | 0,007008 | 1,00E-08 | 0,0063961 | 0,0030824 | 0,0030824 | TRUE | 0,0174331 | No |
| orf19.863 | 2798 | 2557 | 2854 | 1599 | 3893 | 2539 | 3553 | 3997 | 2668 | 2704 | 2819 | 2505 | 3119 | 2887 | 3016 | 3250 | 2871,07 | 2674 | 3068 | 1,147 | 0,198 | 3,777 | 0,0001587 | 0,0002224 | 0,001974 | 0,0026632 | 0,0022728 | 0,0022728 | TRUE | 0,0205824 | No |
| orf19.864 | 2896 | 2773 | 3084 | 1874 | 1702 | 1207 | 1659 | 1730 | 2761 | 2932 | 3047 | 2936 | 1364 | 1372 | 1408 | 1407 | 2153,36 | 2919 | 1388 | 0,476 | -1,072 | -23,431 | 2,04E-121 | 1,65E-120 | 0,0007213 | 0,0028538 | 0,0014951 | 0,0014951 | TRUE | 0,0168836 | No |
| orf19.866 | 1467 | 1376 | 1473 | 895 | 14188 | 9989 | 13156 | 14489 | 1399 | 1455 | 1455 | 1402 | 11368 | 11358 | 11169 | 11781 | 6423,5 | 1428 | 11419 | 7,993 | 2,999 | 77,93 | 0 | 0 | 1,00E-08 | 0,0023469 | 0,0009917 | 0,0009917 | TRUE | 0,0056096 | No |
| orf19.867 | 971 | 828 | 917 | 526 | 1826 | 1203 | 1709 | 1782 | 926 | 876 | 906 | 824 | 1463 | 1368 | 1451 | 1449 | 1157,76 | 883 | 1433 | 1,619 | 0,695 | 12,859 | 7,64E-38 | 2,34E-37 | 0,0003473 | 0,0035095 | 0,0018633 | 0,0018633 | TRUE | 0,0203607 | No |
| orf19.868 | 29071 | 25846 | 28612 | 19574 | 35602 | 27306 | 34623 | 36647 | 27716 | 27329 | 28266 | 30664 | 28527 | 31049 | 29395 | 29798 | 29092,89 | 28494 | 29692 | 1,042 | 0,06 | 1,298 | 0,1943322 | 0,2205681 | 0,0019579 | 0,0021477 | 0,001992 | 0,001992 | TRUE | 0,0319523 | No |
| orf19.87 | 212 | 196 | 186 | 112 | 576 | 418 | 550 | 640 | 202 | 207 | 184 | 175 | 462 | 475 | 467 | 520 | 336,59 | 192 | 481 | 2,492 | 1,317 | 14,401 | 5,08E-47 | 1,80E-46 | 1,00E-08 | 0,0069693 | 0,0042259 | 0,0042259 | TRUE | 0,0370085 | No |
| orf19.871 | 5993 | 5496 | 5894 | 3703 | 5362 | 3878 | 5171 | 5532 | 5714 | 5811 | 5823 | 5801 | 4296 | 4410 | 4390 | 4498 | 5092,86 | 5787 | 4398 | 0,76 | -0,396 | -12,624 | 1,56E-36 | 4,69E-36 | 1,00E-08 | 0,0024137 | 0,000745 | 0,000745 | TRUE | 0,0030835 | No |
| orf19.872 | 1356 | 1165 | 1305 | 913 | 1809 | 1183 | 1689 | 1756 | 1293 | 1232 | 1289 | 1430 | 1449 | 1345 | 1434 | 1428 | 1362,57 | 1311 | 1414 | 1,083 | 0,115 | 2,044 | 0,0409055 | 0,0495761 | 0,0013923 | 0,0032963 | 0,0022668 | 0,0022668 | TRUE | 0,0403637 | No |
| orf19.873 | 140 | 131 | 147 | 77 | 118 | 102 | 95 | 138 | 133 | 139 | 145 | 121 | 95 | 116 | 81 | 112 | 117,65 | 134 | 101 | 0,743 | -0,428 | -2,912 | 0,0035939 | 0,0046897 | 0,0069166 | 0,0160466 | 0,0121914 | 0,0121914 | TRUE | 0,1834126 | No |
| orf19.873.1 | 4854 | 4656 | 4764 | 2960 | 9901 | 7407 | 9277 | 9453 | 4628 | 4923 | 4706 | 4637 | 7933 | 8422 | 7876 | 7686 | 6351,55 | 4724 | 7979 | 1,688 | 0,756 | 18,282 | 1,15E-74 | 5,85E-74 | 0,0009718 | 0,0023498 | 0,0014652 | 0,0014652 | TRUE | 0,0167489 | No |
| orf19.874 | 2007 | 1636 | 1886 | 1177 | 1176 | 795 | 1170 | 1251 | 1913 | 1730 | 1863 | 1844 | 942 | 904 | 993 | 1017 | 1400,89 | 1838 | 964 | 0,525 | -0,929 | -16,557 | 1,44E-61 | 6,20E-61 | 0,0014793 | 0,0032633 | 0,0022521 | 0,0022521 | TRUE | 0,0194003 | No |
| orf19.875 | 1649 | 1588 | 1635 | 988 | 2050 | 1400 | 1929 | 2019 | 1572 | 1679 | 1615 | 1548 | 1643 | 1592 | 1638 | 1642 | 1616,01 | 1604 | 1629 | 1,015 | 0,021 | 0,477 | 0,6330734 | 0,6655093 | 1,00E-08 | 0,0031073 | 0,0013177 | 0,0013177 | TRUE | 0,0119548 | No |
| orf19.876 | 1326 | 1025 | 1168 | 975 | 2525 | 2070 | 2402 | 2731 | 1264 | 1084 | 1154 | 1527 | 2023 | 2354 | 2039 | 2221 | 1708,26 | 1257 | 2159 | 1,721 | 0,783 | 8,262 | 1,43E-16 | 2,98E-16 | 0,0138212 | 0,0030524 | 0,0079738 | 0,0079738 | TRUE | 0,2401437 | No |
| orf19.878 | 3098 | 2417 | 2554 | 1727 | 1768 | 1251 | 1792 | 1777 | 2954 | 2556 | 2523 | 2705 | 1417 | 1422 | 1521 | 1445 | 2067,91 | 2684 | 1451 | 0,541 | -0,887 | -14,927 | 2,19E-50 | 8,12E-50 | 0,0028649 | 0,0028853 | 0,0028731 | 0,0028731 | TRUE | 0,0572161 | No |
| orf19.879 | 971 | 879 | 992 | 658 | 2678 | 2011 | 2563 | 2819 | 926 | 929 | 980 | 1031 | 2146 | 2287 | 2176 | 2292 | 1595,82 | 966 | 2225 | 2,308 | 1,207 | 23,04 | 1,85E-117 | 1,44E-116 | 0,0008939 | 0,0031202 | 0,001852 | 0,001852 | TRUE | 0,020592 | No |
| orf19.88 | 6833 | 5704 | 5885 | 3957 | 39335 | 27308 | 37314 | 40945 | 6515 | 6031 | 5814 | 6199 | 31518 | 31052 | 31679 | 33292 | 19012,47 | 6140 | 31885 | 5,194 | 2,377 | 54,568 | 0 | 0 | 0,0014962 | 0,0021776 | 0,0017164 | 0,0017164 | TRUE | 0,0212201 | No |
| orf19.880 | 3606 | 3163 | 3394 | 2377 | 3421 | 2405 | 3214 | 3530 | 3438 | 3344 | 3353 | 3724 | 2741 | 2735 | 2729 | 2870 | 3116,72 | 3465 | 2769 | 0,8 | -0,321 | -6,987 | 2,81E-12 | 5,24E-12 | 0,0011827 | 0,0026181 | 0,0017058 | 0,0017058 | TRUE | 0,0289448 | No |
| orf19.881 | 1011 | 892 | 925 | 640 | 1428 | 1064 | 1428 | 1469 | 964 | 943 | 914 | 1003 | 1144 | 1210 | 1212 | 1194 | 1073,04 | 956 | 1190 | 1,247 | 0,319 | 6,07 | 1,28E-09 | 2,19E-09 | 1,00E-08 | 0,0036214 | 0,0016873 | 0,0016873 | TRUE | 0,0109836 | No |
| orf19.882 | 11494 | 10177 | 11057 | 6781 | 2859 | 1988 | 2705 | 2953 | 10958 | 10761 | 10923 | 10623 | 2291 | 2261 | 2297 | 2401 | 6564,28 | 10816 | 2312 | 0,214 | -2,225 | -65,288 | 0 | 0 | 0,0001041 | 0,0023414 | 0,0008707 | 0,0008707 | TRUE | 0,008246 | No |
| orf19.886 | 6730 | 5711 | 6645 | 4130 | 5914 | 4057 | 5267 | 6248 | 6416 | 6039 | 6565 | 6470 | 4739 | 4613 | 4472 | 5080 | 5549,16 | 6372 | 4726 | 0,742 | -0,431 | -8,767 | 1,83E-18 | 3,97E-18 | 0,0019253 | 0,0023872 | 0,0021357 | 0,0021357 | TRUE | 0,0312967 | No |
| orf19.889 | 428 | 399 | 399 | 295 | 701 | 531 | 647 | 776 | 408 | 422 | 394 | 462 | 562 | 604 | 549 | 631 | 504 | 422 | 586 | 1,397 | 0,482 | 6,151 | 7,69E-10 | 1,32E-09 | 0,0022985 | 0,005349 | 0,0038075 | 0,0038075 | TRUE | 0,0410007 | No |
| orf19.89 | 1715 | 1387 | 1494 | 1020 | 4535 | 3070 | 4246 | 4587 | 1635 | 1467 | 1476 | 1598 | 3634 | 3491 | 3605 | 3730 | 2579,32 | 1544 | 3615 | 2,344 | 1,229 | 24,998 | 6,43E-138 | 5,94E-137 | 0,0012799 | 0,0027278 | 0,0018336 | 0,0018336 | TRUE | 0,021204 | No |
| orf19.891 | 5248 | 4880 | 5190 | 3338 | 9748 | 7258 | 9280 | 10261 | 5003 | 5160 | 5127 | 5229 | 7811 | 8253 | 7879 | 8343 | 6600,68 | 5130 | 8072 | 1,574 | 0,654 | 17,78 | 1,01E-70 | 4,88E-70 | 0,0004991 | 0,00234 | 0,001135 | 0,001135 | TRUE | 0,0064375 | No |
| orf19.893 | 40 | 44 | 40 | 36 | 97 | 76 | 91 | 106 | 38 | 47 | 40 | 56 | 78 | 86 | 77 | 86 | 63,52 | 45 | 82 | 1,845 | 0,884 | 4,731 | 2,23E-06 | 3,40E-06 | 1,00E-08 | 0,0279398 | 0,0155099 | 0,0155099 | TRUE | 0,1536649 | No |
| orf19.894 | 598 | 494 | 522 | 324 | 1323 | 945 | 1209 | 1300 | 570 | 522 | 516 | 508 | 1060 | 1075 | 1026 | 1057 | 791,73 | 529 | 1054 | 1,987 | 0,991 | 16,455 | 7,68E-61 | 3,27E-60 | 1,00E-08 | 0,0041651 | 0,0020073 | 0,0020073 | TRUE | 0,0340029 | No |
| orf19.895 | 5031 | 4450 | 4769 | 3210 | 6946 | 4948 | 6497 | 7086 | 4797 | 4705 | 4711 | 5029 | 5566 | 5626 | 5516 | 5762 | 5213,91 | 4810 | 5618 | 1,169 | 0,225 | 6,247 | 4,19E-10 | 7,33E-10 | 0,0004162 | 0,0024062 | 0,0010502 | 0,0010502 | TRUE | 0,0105949 | No |
| orf19.896 | 4679 | 4308 | 4739 | 2811 | 4197 | 2935 | 3647 | 4021 | 4461 | 4555 | 4682 | 4404 | 3363 | 3337 | 3096 | 3269 | 3895,92 | 4526 | 3266 | 0,721 | -0,471 | -11,392 | 4,59E-30 | 1,24E-29 | 0,0006856 | 0,0025127 | 0,001382 | 0,001382 | TRUE | 0,0150966 | No |
| orf19.897 | 2370 | 1987 | 2142 | 1511 | 654 | 444 | 659 | 618 | 2260 | 2101 | 2116 | 2367 | 524 | 505 | 559 | 502 | 1366,82 | 2211 | 522 | 0,237 | -2,077 | -33,48 | 9,57E-246 | 1,59E-244 | 0,0020243 | 0,0032925 | 0,0025907 | 0,0025907 | TRUE | 0,0265818 | No |
| orf19.898 | 1106 | 860 | 934 | 682 | 2464 | 1823 | 2256 | 2382 | 1054 | 909 | 923 | 1068 | 1974 | 2073 | 1915 | 1937 | 1481,78 | 988 | 1975 | 2,001 | 1 | 15,807 | 2,77E-56 | 1,11E-55 | 0,0032485 | 0,0031993 | 0,0030502 | 0,0030502 | TRUE | 0,0341512 | No |
| orf19.899 | 680 | 620 | 578 | 406 | 997 | 726 | 1017 | 1025 | 648 | 656 | 571 | 636 | 799 | 826 | 863 | 833 | 729,02 | 628 | 830 | 1,324 | 0,405 | 6,239 | 4,40E-10 | 7,68E-10 | 0,0009609 | 0,0043435 | 0,0026101 | 0,0026101 | TRUE | 0,0466667 | No |
| orf19.9 | 128 | 119 | 108 | 73 | 117 | 77 | 99 | 108 | 122 | 126 | 107 | 114 | 94 | 88 | 84 | 88 | 102,76 | 117 | 88 | 0,753 | -0,409 | -2,943 | 0,0032481 | 0,0042446 | 1,00E-08 | 0,0180693 | 0,0087916 | 0,0087916 | TRUE | 0,0419643 | No |
| orf19.90 | 1234 | 1148 | 1274 | 751 | 1567 | 1050 | 1457 | 1575 | 1176 | 1214 | 1259 | 1176 | 1256 | 1194 | 1237 | 1281 | 1224,07 | 1206 | 1242 | 1,029 | 0,042 | 0,839 | 0,4016432 | 0,4373525 | 1,00E-08 | 0,0034326 | 0,0015744 | 0,0015744 | TRUE | 0,0105176 | No |
| orf19.900 | 5177 | 4131 | 4581 | 3143 | 6232 | 4406 | 6089 | 6493 | 4936 | 4368 | 4526 | 4924 | 4994 | 5010 | 5170 | 5279 | 4900,69 | 4688 | 5113 | 1,091 | 0,126 | 2,547 | 0,0108687 | 0,0137687 | 0,0019459 | 0,0024263 | 0,0021439 | 0,0021439 | TRUE | 0,0257376 | No |
| orf19.903 | 13840 | 11224 | 11723 | 8984 | 8707 | 6085 | 8442 | 9026 | 13195 | 11868 | 11581 | 14074 | 6977 | 6919 | 7167 | 7339 | 9890,01 | 12680 | 7100 | 0,56 | -0,836 | -13,516 | 1,27E-41 | 4,11E-41 | 0,00444 | 0,0022573 | 0,0035677 | 0,0035677 | TRUE | 0,0663832 | No |
| orf19.904 | 1222 | 1124 | 1165 | 747 | 633 | 451 | 575 | 646 | 1165 | 1188 | 1151 | 1170 | 507 | 513 | 488 | 525 | 838,52 | 1168 | 508 | 0,435 | -1,201 | -21,509 | 1,29E-102 | 8,61E-102 | 1,00E-08 | 0,0040494 | 0,0016395 | 0,0016395 | TRUE | 0,0086941 | No |
| orf19.905 | 2095 | 1813 | 1982 | 1259 | 4067 | 3003 | 3752 | 3980 | 1997 | 1917 | 1958 | 1972 | 3259 | 3415 | 3185 | 3236 | 2617,46 | 1961 | 3274 | 1,668 | 0,738 | 18,202 | 5,02E-74 | 2,54E-73 | 0,0002888 | 0,0027186 | 0,0011574 | 0,0011574 | TRUE | 0,0097065 | No |
| orf19.906 | 5158 | 4870 | 5354 | 3189 | 4612 | 3220 | 4366 | 4545 | 4918 | 5149 | 5289 | 4996 | 3695 | 3661 | 3707 | 3696 | 4388,89 | 5088 | 3690 | 0,725 | -0,464 | -12,816 | 1,34E-37 | 4,07E-37 | 0,0003638 | 0,0024654 | 0,0010261 | 0,0010261 | TRUE | 0,0088414 | No |
| orf19.908 | 4697 | 4526 | 4748 | 2804 | 6403 | 4488 | 5797 | 6015 | 4478 | 4786 | 4691 | 4393 | 5131 | 5103 | 4922 | 4891 | 4799,14 | 4587 | 5012 | 1,092 | 0,127 | 3,111 | 0,0018627 | 0,0024726 | 0,0008129 | 0,0024334 | 0,0013783 | 0,0013783 | TRUE | 0,0102382 | No |
| orf19.909 | 12697 | 11384 | 12262 | 7703 | 5938 | 4286 | 5627 | 5727 | 12105 | 12037 | 12114 | 12067 | 4758 | 4874 | 4777 | 4657 | 8423,58 | 12081 | 4766 | 0,394 | -1,342 | -47,27 | 0 | 0 | 1,00E-08 | 0,0022862 | 0,0006341 | 0,0006341 | TRUE | 0,0029235 | No |
| orf19.909.1 | 887 | 755 | 739 | 567 | 1318 | 893 | 1057 | 1222 | 846 | 798 | 730 | 888 | 1056 | 1015 | 897 | 994 | 903,1 | 816 | 990 | 1,217 | 0,284 | 3,77 | 0,0001633 | 0,0002286 | 0,0042761 | 0,0039094 | 0,0043008 | 0,0043008 | TRUE | 0,0598179 | No |
| orf19.91 | 1132 | 1030 | 1110 | 664 | 1445 | 1009 | 1354 | 1398 | 1079 | 1089 | 1097 | 1040 | 1158 | 1147 | 1150 | 1137 | 1112,06 | 1076 | 1148 | 1,065 | 0,091 | 1,895 | 0,0580855 | 0,069357 | 1,00E-08 | 0,0035677 | 0,0012852 | 0,0012852 | TRUE | 0,0050846 | No |
| orf19.910 | 4263 | 4033 | 4210 | 2780 | 3799 | 2837 | 3555 | 4005 | 4064 | 4264 | 4159 | 4355 | 3044 | 3226 | 3018 | 3256 | 3673,42 | 4210 | 3136 | 0,745 | -0,425 | -9,878 | 5,16E-23 | 1,23E-22 | 0,0008399 | 0,0025382 | 0,0014972 | 0,0014972 | TRUE | 0,0084922 | No |
| orf19.913.2 | 2776 | 2468 | 2610 | 1578 | 9336 | 5846 | 9222 | 9058 | 2647 | 2610 | 2578 | 2472 | 7481 | 6647 | 7829 | 7365 | 4953,65 | 2577 | 7330 | 2,844 | 1,508 | 27,656 | 2,35E-168 | 2,66E-167 | 0,0023681 | 0,0024227 | 0,0025754 | 0,0025754 | TRUE | 0,0474465 | No |
| orf19.915 | 1236 | 1418 | 1373 | 946 | 486 | 407 | 411 | 428 | 1178 | 1499 | 1356 | 1482 | 389 | 463 | 349 | 348 | 883,16 | 1379 | 387 | 0,28 | -1,836 | -17,962 | 3,89E-72 | 1,93E-71 | 0,0132547 | 0,0039504 | 0,0084629 | 0,0084629 | TRUE | 0,1869429 | No |
| orf19.916 | 3922 | 4065 | 4182 | 2162 | 2586 | 1745 | 2233 | 2307 | 3739 | 4298 | 4131 | 3387 | 2072 | 1984 | 1896 | 1876 | 2922,96 | 3889 | 1957 | 0,503 | -0,992 | -13,607 | 3,62E-42 | 1,19E-41 | 0,0060651 | 0,002653 | 0,0047315 | 0,0047315 | TRUE | 0,0894239 | No |
| orf19.918 | 3911 | 3299 | 3748 | 2269 | 3759 | 2519 | 3284 | 3767 | 3729 | 3488 | 3703 | 3555 | 3012 | 2864 | 2788 | 3063 | 3275,19 | 3619 | 2932 | 0,81 | -0,304 | -6,648 | 2,97E-11 | 5,40E-11 | 0,0010479 | 0,0025926 | 0,001696 | 0,001696 | TRUE | 0,0136492 | No |
| orf19.92 | 2071 | 1863 | 2096 | 1236 | 4076 | 2701 | 3784 | 4166 | 1974 | 1970 | 2071 | 1936 | 3266 | 3071 | 3213 | 3387 | 2611,06 | 1988 | 3234 | 1,627 | 0,702 | 15,404 | 1,54E-53 | 5,95E-53 | 0,0009007 | 0,0027201 | 0,0015738 | 0,0015738 | TRUE | 0,0134514 | No |
| orf19.920 | 2694 | 2665 | 2678 | 1653 | 3807 | 2481 | 3352 | 3361 | 2568 | 2818 | 2646 | 2590 | 3050 | 2821 | 2846 | 2733 | 2758,96 | 2656 | 2862 | 1,078 | 0,108 | 2,156 | 0,0310874 | 0,0380381 | 0,0015248 | 0,0026864 | 0,0020522 | 0,0020522 | TRUE | 0,0241801 | No |
| orf19.921 | 5304 | 5323 | 5801 | 3505 | 1964 | 1478 | 1878 | 1865 | 5057 | 5628 | 5731 | 5491 | 1574 | 1681 | 1594 | 1516 | 3533,99 | 5477 | 1591 | 0,29 | -1,784 | -33,719 | 3,09E-249 | 5,21E-248 | 0,0021496 | 0,0025559 | 0,0023053 | 0,0023053 | TRUE | 0,0331667 | No |
| orf19.922 | 2653 | 2655 | 2782 | 1663 | 4795 | 3701 | 4299 | 4390 | 2529 | 2807 | 2748 | 2605 | 3842 | 4208 | 3650 | 3569 | 3245 | 2672 | 3817 | 1,427 | 0,513 | 8,484 | 2,17E-17 | 4,60E-17 | 0,0034975 | 0,0025973 | 0,0031856 | 0,0031856 | TRUE | 0,056841 | No |
| orf19.923 | 775 | 674 | 758 | 515 | 1125 | 867 | 1039 | 1004 | 739 | 713 | 749 | 807 | 901 | 986 | 882 | 816 | 824,11 | 752 | 896 | 1,193 | 0,255 | 3,601 | 0,0003174 | 0,0004392 | 0,0030954 | 0,0040836 | 0,0035538 | 0,0035538 | TRUE | 0,0507812 | No |
| orf19.925 | 527 | 652 | 670 | 354 | 1550 | 1274 | 1375 | 1315 | 502 | 689 | 662 | 555 | 1242 | 1449 | 1167 | 1069 | 916,94 | 602 | 1232 | 2,042 | 1,03 | 9,179 | 4,34E-20 | 9,76E-20 | 0,0176421 | 0,0038819 | 0,0107944 | 0,0107944 | TRUE | 0,1658028 | No |
| orf19.926 | 1310 | 1151 | 1244 | 787 | 1902 | 1420 | 1866 | 2064 | 1249 | 1217 | 1229 | 1233 | 1524 | 1615 | 1584 | 1678 | 1416,12 | 1232 | 1600 | 1,299 | 0,377 | 7,705 | 1,31E-14 | 2,62E-14 | 1,00E-08 | 0,0032507 | 0,0015646 | 0,0015646 | TRUE | 0,0136856 | No |
| orf19.927 | 215 | 242 | 219 | 144 | 296 | 231 | 255 | 229 | 205 | 256 | 216 | 226 | 237 | 263 | 216 | 186 | 225,67 | 226 | 226 | 0,997 | -0,005 | -0,037 | 0,9705385 | 0,9732841 | 0,0109999 | 0,0093671 | 0,0101557 | 0,0101557 | TRUE | 0,1567164 | No |
| orf19.928 | 1209 | 1051 | 1168 | 814 | 1122 | 815 | 1091 | 1149 | 1153 | 1111 | 1154 | 1275 | 899 | 927 | 926 | 934 | 1047,41 | 1173 | 922 | 0,788 | -0,344 | -6,133 | 8,63E-10 | 1,48E-09 | 0,0006685 | 0,0036589 | 0,0020611 | 0,0020611 | TRUE | 0,0361648 | No |
| orf19.929 | 102 | 102 | 111 | 73 | 92 | 98 | 110 | 97 | 97 | 108 | 110 | 114 | 74 | 111 | 93 | 79 | 98,32 | 107 | 89 | 0,828 | -0,272 | -1,691 | 0,0907432 | 0,1063225 | 0,0088527 | 0,0187916 | 0,0145077 | 0,0145077 | TRUE | 0,2363487 | No |
| orf19.93 | 5490 | 4324 | 4780 | 3957 | 5397 | 3839 | 4959 | 5441 | 5234 | 4572 | 4722 | 6199 | 4324 | 4365 | 4210 | 4424 | 4756,4 | 5182 | 4331 | 0,836 | -0,258 | -3,141 | 0,0016842 | 0,0022426 | 0,0099911 | 0,0024365 | 0,0062524 | 0,0062524 | TRUE | 0,2099603 | No |
| orf19.930 | 64742 | 53137 | 56479 | 37722 | 112130 | 75673 | 103322 | 115327 | 61724 | 56186 | 55796 | 59094 | 89847 | 86047 | 87720 | 93772 | 73773,11 | 58200 | 89346 | 1,535 | 0,618 | 13,846 | 1,35E-43 | 4,54E-43 | 0,0018123 | 0,0021135 | 0,0019023 | 0,0019023 | TRUE | 0,0204309 | No |
| orf19.931 | 1163 | 1141 | 1110 | 719 | 1218 | 876 | 1033 | 1128 | 1109 | 1206 | 1097 | 1126 | 976 | 996 | 877 | 917 | 1038,05 | 1134 | 942 | 0,829 | -0,27 | -4,456 | 8,36E-06 | 1,24E-05 | 0,0016839 | 0,003673 | 0,0025684 | 0,0025684 | TRUE | 0,0253352 | No |
| orf19.932 | 6300 | 5670 | 6128 | 3893 | 2171 | 1610 | 2025 | 2281 | 6006 | 5995 | 6054 | 6099 | 1740 | 1831 | 1719 | 1855 | 3912,29 | 6038 | 1786 | 0,296 | -1,758 | -45,828 | 0 | 0 | 1,95E-05 | 0,002511 | 0,0010699 | 0,0010699 | TRUE | 0,0087914 | No |
| orf19.933 | 4326 | 3750 | 4215 | 2403 | 3891 | 2768 | 3669 | 3806 | 4124 | 3965 | 4164 | 3764 | 3118 | 3147 | 3115 | 3095 | 3561,6 | 4004 | 3119 | 0,778 | -0,363 | -8,748 | 2,17E-18 | 4,71E-18 | 0,000673 | 0,0025523 | 0,0013677 | 0,0013677 | TRUE | 0,0185634 | No |
| orf19.934 | 832 | 764 | 793 | 509 | 2064 | 1182 | 1932 | 1910 | 793 | 808 | 783 | 797 | 1654 | 1344 | 1640 | 1553 | 1171,62 | 795 | 1548 | 1,949 | 0,963 | 13,55 | 7,98E-42 | 2,60E-41 | 0,0041266 | 0,0034927 | 0,0038523 | 0,0038523 | TRUE | 0,0910633 | No |
| orf19.935 | 24 | 12 | 15 | 24 | 129 | 106 | 112 | 140 | 23 | 13 | 15 | 38 | 103 | 121 | 95 | 114 | 65,1 | 22 | 108 | 5,145 | 2,363 | 9,491 | 2,28E-21 | 5,25E-21 | 0,0367132 | 0,0273124 | 0,0289669 | 0,0289669 | TRUE | 0,8381316 | No |
| orf19.936 | 8 | 9 | 7 | 4 | 10 | 3 | 3 | 5 | 8 | 10 | 7 | 6 | 8 | 3 | 3 | 4 | 6,05 | 8 | 4 | 0,599 | -0,738 | -1,266 | 0,2055488 | 0,2329261 | 1,00E-08 | 0,2736964 | 0,1567097 | 0,1567097 | TRUE | 0,6397564 | No |
| orf19.937 | 7 | 11 | 4 | 2 | 1 | 0 | 0 | 1 | 7 | 12 | 4 | 3 | 1 | 0 | 0 | 1 | 3,38 | 6 | 0 | 0,069 | -3,866 | -3,217 | 0,0012935 | 0,0017296 | 1,00E-08 | 0,4885101 | 0,3546916 | 0,3546916 | TRUE | 0,3656389 | No |
| orf19.938 | 358 | 316 | 362 | 228 | 27 | 25 | 25 | 27 | 341 | 334 | 358 | 357 | 22 | 28 | 21 | 22 | 185,44 | 348 | 23 | 0,066 | -3,918 | -23,502 | 3,87E-122 | 3,14E-121 | 1,00E-08 | 0,0109456 | 0,0058437 | 0,0058437 | TRUE | 0,0985337 | No |
| orf19.939 | 7891 | 7226 | 7977 | 4994 | 11561 | 8572 | 10946 | 11633 | 7523 | 7641 | 7881 | 7823 | 9263 | 9747 | 9293 | 9459 | 8578,77 | 7717 | 9440 | 1,223 | 0,291 | 8,802 | 1,35E-18 | 2,94E-18 | 0,0003466 | 0,0022827 | 0,0009281 | 0,0009281 | TRUE | 0,0057001 | No |

| | | | | | | | | | | | | | | | | | | | | | | | | | | | | | | | |
|---|---|---|---|---|---|---|---|---|---|---|---|---|---|---|---|---|---|---|---|---|---|---|---|---|---|---|---|---|---|---|---|
| orf19.94 | 4167 | 3740 | 3905 | 2130 | 3161 | 2130 | 2962 | 3114 | 3973 | 3955 | 3858 | 3337 | 2533 | 2422 | 2515 | 2532 | 3140,43 | 3781 | 2500 | 0,661 | -0,598 | -10,31 | 6,33E-25 | 1,57E-24 | 0,0029314 | 0,0026141 | 0,002902 | 0,002902 | TRUE | 0,0729542 | No |
| orf19.940 | 3815 | 4108 | 4219 | 2433 | 1562 | 1237 | 1352 | 1417 | 3637 | 4344 | 4168 | 3811 | 1252 | 1407 | 1148 | 1152 | 2614,81 | 3990 | 1240 | 0,31 | -1,689 | -21,956 | 7,60E-107 | 5,29E-106 | 0,007356 | 0,0027192 | 0,0051802 | 0,0051802 | TRUE | 0,0956733 | No |
| orf19.941 | 24587 | 23159 | 23541 | 14967 | 23078 | 16265 | 20741 | 22209 | 23441 | 24488 | 23256 | 23447 | 18492 | 18495 | 17609 | 18058 | 20910,65 | 23658 | 18164 | 0,768 | -0,381 | -11,847 | 2,24E-32 | 6,31E-32 | 0,0004863 | 0,0021698 | 0,0009473 | 0,0009473 | TRUE | 0,0068567 | No |
| orf19.942 | 96 | 86 | 107 | 66 | 40 | 43 | 37 | 46 | 92 | 91 | 106 | 103 | 32 | 49 | 31 | 37 | 67,66 | 98 | 37 | 0,378 | -1,404 | -7,408 | 1,28E-13 | 2,49E-13 | 1,00E-08 | 0,0263565 | 0,0167184 | 0,0167184 | TRUE | 0,2589926 | No |
| orf19.943 | 3300 | 3148 | 3234 | 2042 | 4314 | 3419 | 4121 | 4516 | 3146 | 3329 | 3195 | 3199 | 3457 | 3888 | 3499 | 3672 | 3422,95 | 3217 | 3629 | 1,127 | 0,173 | 3,631 | 0,0002819 | 0,0003911 | 0,0013647 | 0,002571 | 0,0018727 | 0,0018727 | TRUE | 0,0272579 | No |
| orf19.944 | 3359 | 3230 | 3471 | 2090 | 4596 | 3093 | 4312 | 4416 | 3202 | 3415 | 3429 | 3274 | 3683 | 3517 | 3661 | 3591 | 3471,5 | 3330 | 3613 | 1,085 | 0,118 | 2,99 | 0,0027916 | 0,0036667 | 0,0003787 | 0,0025642 | 0,0011939 | 0,0011939 | TRUE | 0,0087422 | No |
| orf19.945 | 1239 | 1035 | 1109 | 706 | 1668 | 1108 | 1501 | 1746 | 1181 | 1094 | 1096 | 1106 | 1337 | 1260 | 1274 | 1420 | 1220,95 | 1119 | 1323 | 1,182 | 0,241 | 4,198 | 2,70E-05 | 3,92E-05 | 0,001537 | 0,0034361 | 0,002334 | 0,002334 | TRUE | 0,0296128 | No |
| orf19.946 | 1787 | 1564 | 1554 | 970 | 7055 | 4611 | 6601 | 7023 | 1704 | 1654 | 1535 | 1520 | 5653 | 5243 | 5604 | 5710 | 3577,86 | 1603 | 5552 | 3,46 | 1,791 | 35,201 | 1,92E-271 | 3,66E-270 | 0,0017596 | 0,0025502 | 0,0020541 | 0,0020541 | TRUE | 0,0218681 | No |
| orf19.947 | 1690 | 1376 | 1491 | 875 | 1832 | 1168 | 1738 | 1758 | 1611 | 1455 | 1473 | 1371 | 1468 | 1328 | 1476 | 1429 | 1451,36 | 1478 | 1425 | 0,964 | -0,053 | -0,879 | 0,3791858 | 0,4142415 | 0,0024897 | 0,0032226 | 0,0028463 | 0,0028463 | TRUE | 0,0451015 | No |
| orf19.948 | 1 | 1 | 1 | 4 | 3 | 2 | 5 | 1 | 1 | 1 | 1 | 6 | 2 | 2 | 4 | 1 | 2,38 | 2 | 2 | 1,164 | 0,219 | 0,221 | 0,8254019 | 0,8459508 | 0,2382128 | 0,6933845 | 0,5003144 | 0,5003144 | TRUE | 0,8328848 | No |
| orf19.949 | 918 | 901 | 955 | 556 | 569 | 442 | 579 | 534 | 875 | 953 | 943 | 871 | 456 | 503 | 492 | 434 | 690,83 | 910 | 471 | 0,515 | -0,956 | -13,796 | 2,70E-43 | 9,01E-43 | 0,0016506 | 0,004468 | 0,0030474 | 0,0030474 | TRUE | 0,0326609 | No |
| orf19.95 | 81 | 75 | 100 | 54 | 70 | 57 | 75 | 78 | 77 | 79 | 99 | 85 | 56 | 65 | 64 | 63 | 73,49 | 85 | 62 | 0,728 | -0,459 | -2,738 | 0,0061806 | 0,0079449 | 1,00E-08 | 0,0244335 | 0,0132132 | 0,0132132 | TRUE | 0,1250024 | No |
| orf19.951 | 9358 | 9665 | 10208 | 6020 | 1177 | 831 | 1084 | 1164 | 8922 | 10220 | 10085 | 9431 | 943 | 945 | 920 | 946 | 5301,41 | 9664 | 938 | 0,097 | -3,364 | -62,012 | 0 | 0 | 0,0022304 | 0,002401 | 0,0022928 | 0,0022928 | TRUE | 0,0331949 | No |
| orf19.952 | 635 | 567 | 679 | 442 | 582 | 473 | 586 | 644 | 605 | 600 | 671 | 692 | 466 | 538 | 498 | 524 | 574,18 | 642 | 506 | 0,789 | -0,341 | -4,476 | 7,59E-06 | 1,13E-05 | 0,0027168 | 0,0049508 | 0,003816 | 0,003816 | TRUE | 0,0338529 | No |
| orf19.953.1 | 1617 | 1375 | 1494 | 1010 | 3248 | 2309 | 3031 | 3180 | 1542 | 1454 | 1476 | 1582 | 2603 | 2626 | 2573 | 2586 | 2055,09 | 1514 | 2597 | 1,718 | 0,781 | 18,358 | 2,86E-75 | 1,47E-74 | 1,00E-08 | 0,0028902 | 0,0011959 | 0,0011959 | TRUE | 0,0096422 | No |
| orf19.954 | 2724 | 2284 | 2516 | 1543 | 3437 | 2241 | 3243 | 3529 | 2597 | 2415 | 2486 | 2417 | 2754 | 2548 | 2753 | 2869 | 2604,97 | 2479 | 2731 | 1,102 | 0,14 | 2,861 | 0,0042277 | 0,005491 | 0,0012781 | 0,0027216 | 0,0019137 | 0,0019137 | TRUE | 0,0237248 | No |
| orf19.955 | 2170 | 1918 | 1944 | 1298 | 2095 | 1393 | 2008 | 2099 | 2069 | 2028 | 1920 | 2033 | 1679 | 1584 | 1705 | 1707 | 1840,61 | 2012 | 1669 | 0,83 | -0,269 | -5,778 | 7,54E-09 | 1,26E-08 | 0,0005496 | 0,0029833 | 0,0015275 | 0,0015275 | TRUE | 0,0130683 | No |
| orf19.956 | 991 | 923 | 1028 | 611 | 1126 | 802 | 1063 | 1188 | 945 | 976 | 1016 | 957 | 902 | 912 | 902 | 966 | 947,01 | 974 | 920 | 0,946 | -0,081 | -1,479 | 0,1392018 | 0,1601888 | 1,00E-08 | 0,003825 | 0,0018072 | 0,0018072 | TRUE | 0,0136892 | No |
| orf19.96 | 11272 | 11367 | 11968 | 6858 | 14111 | 9940 | 13410 | 13444 | 10747 | 12019 | 11823 | 10743 | 11307 | 11303 | 11385 | 10931 | 11282,27 | 11333 | 11232 | 0,991 | -0,013 | -0,287 | 0,7740019 | 0,7981426 | 0,0016465 | 0,0022368 | 0,0020025 | 0,0020025 | TRUE | 0,0202494 | No |
| orf19.961.2 | 1284 | 1148 | 1106 | 664 | 1849 | 1022 | 1778 | 1857 | 1224 | 1214 | 1093 | 1040 | 1482 | 1162 | 1510 | 1510 | 1279,24 | 1143 | 1416 | 1,239 | 0,309 | 3,452 | 0,0005571 | 0,0007632 | 0,0092781 | 0,0033748 | 0,0068914 | 0,0068914 | TRUE | 0,1705531 | No |
| orf19.962 | 4907 | 3809 | 4174 | 2530 | 6612 | 4273 | 6439 | 6692 | 4678 | 4028 | 4124 | 3963 | 5298 | 4859 | 5467 | 5441 | 4732,18 | 4198 | 5266 | 1,254 | 0,327 | 5,292 | 1,21E-07 | 1,94E-07 | 0,0041731 | 0,0024382 | 0,0034378 | 0,0034378 | TRUE | 0,0728104 | No |
| orf19.964 | 4202 | 3877 | 4125 | 2474 | 4313 | 2869 | 4039 | 4266 | 4006 | 4099 | 4075 | 3876 | 3456 | 3262 | 3429 | 3469 | 3709,04 | 4014 | 3404 | 0,848 | -0,238 | -6,246 | 4,20E-10 | 7,35E-10 | 0,0003475 | 0,002534 | 0,0011224 | 0,0011224 | TRUE | 0,0090741 | No |
| orf19.966 | 3866 | 2964 | 3206 | 2233 | 3929 | 2532 | 3827 | 3954 | 3686 | 3134 | 3167 | 3498 | 3148 | 2879 | 3249 | 3215 | 3247,07 | 3371 | 3123 | 0,927 | -0,109 | -1,72 | 0,085343 | 0,1002714 | 0,004155 | 0,0025969 | 0,0035713 | 0,0035713 | TRUE | 0,049436 | No |
| orf19.967 | 2165 | 1721 | 1887 | 1308 | 4127 | 2913 | 3801 | 4244 | 2064 | 1820 | 1864 | 2049 | 3307 | 3312 | 3227 | 3451 | 2636,76 | 1949 | 3324 | 1,707 | 0,772 | 14,739 | 3,62E-49 | 1,32E-48 | 0,0019168 | 0,002714 | 0,0022082 | 0,0022082 | TRUE | 0,0240818 | No |
| orf19.968 | 16506 | 12580 | 13337 | 9678 | 25911 | 16691 | 23674 | 26421 | 15737 | 13302 | 13176 | 15161 | 20762 | 18979 | 20099 | 21483 | 17337,26 | 14344 | 20331 | 1,418 | 0,503 | 7,685 | 1,54E-14 | 3,06E-14 | 0,0053867 | 0,002186 | 0,0040629 | 0,0040629 | TRUE | 0,0526976 | No |
| orf19.969 | 12714 | 10275 | 11122 | 7444 | 21345 | 15360 | 19925 | 22034 | 12121 | 10865 | 10987 | 11661 | 17103 | 17466 | 16916 | 17916 | 14379,46 | 11408 | 17350 | 1,521 | 0,605 | 13,773 | 3,71E-43 | 1,24E-42 | 0,0015643 | 0,0022055 | 0,0017778 | 0,0017778 | TRUE | 0,022014 | No |
| orf19.97 | 48269 | 43659 | 46590 | 27529 | 56572 | 39712 | 52513 | 56232 | 46019 | 46164 | 46027 | 43126 | 45330 | 45156 | 44583 | 45722 | 45265,78 | 45334 | 45198 | 0,997 | -0,005 | -0,141 | 0,8875503 | 0,902831 | 0,0004811 | 0,0021276 | 0,0009717 | 0,0009717 | TRUE | 0,0130421 | No |
| orf19.970 | 764 | 664 | 694 | 463 | 1081 | 816 | 1087 | 1118 | 728 | 702 | 686 | 725 | 866 | 928 | 923 | 909 | 808,42 | 710 | 906 | 1,276 | 0,352 | 6,172 | 6,73E-10 | 1,17E-09 | 1,00E-08 | 0,0041223 | 0,0018423 | 0,0018423 | TRUE | 0,0110279 | No |
| orf19.971 | 547 | 485 | 533 | 336 | 789 | 552 | 767 | 909 | 522 | 513 | 527 | 526 | 632 | 628 | 651 | 739 | 592,18 | 522 | 662 | 1,272 | 0,347 | 4,754 | 1,99E-06 | 3,04E-06 | 0,0021332 | 0,0048639 | 0,0033631 | 0,0033631 | TRUE | 0,0733572 | No |
| orf19.972 | 338 | 247 | 270 | 224 | 343 | 239 | 301 | 356 | 322 | 261 | 267 | 351 | 275 | 272 | 256 | 289 | 286,58 | 300 | 273 | 0,914 | -0,129 | -1,198 | 0,2308449 | 0,2600214 | 0,0071737 | 0,0078205 | 0,0076441 | 0,0076441 | TRUE | 0,1240028 | No |
| orf19.974 | 638 | 545 | 659 | 353 | 1444 | 986 | 1350 | 1486 | 608 | 576 | 651 | 553 | 1157 | 1121 | 1146 | 1208 | 877,65 | 597 | 1158 | 1,933 | 0,951 | 15,084 | 2,06E-51 | 7,72E-51 | 0,0009616 | 0,0039621 | 0,0024984 | 0,0024984 | TRUE | 0,0429848 | No |
| orf19.976 | 12047 | 10710 | 11657 | 7107 | 6763 | 4711 | 6703 | 6989 | 11486 | 11324 | 11516 | 11133 | 5419 | 5357 | 5691 | 5683 | 8451,11 | 11365 | 5538 | 0,487 | -1,037 | -30,239 | 7,26E-201 | 9,94E-200 | 0,0004365 | 0,0022856 | 0,0009994 | 0,0009994 | TRUE | 0,0056451 | No |
| orf19.978 | 4798 | 4030 | 4590 | 2919 | 7080 | 4782 | 6862 | 7144 | 4574 | 4261 | 4534 | 4573 | 5673 | 5438 | 5826 | 5809 | 5086 | 4486 | 5686 | 1,268 | 0,343 | 8,436 | 3,28E-17 | 6,92E-17 | 0,0007966 | 0,0024141 | 0,0013832 | 0,0013832 | TRUE | 0,0139572 | No |
| orf19.979 | 6339 | 5623 | 5955 | 4092 | 71014 | 51384 | 65435 | 74587 | 6044 | 5946 | 5883 | 6410 | 56901 | 58428 | 55554 | 60646 | 31976,57 | 6071 | 57882 | 9,541 | 3,254 | 77,407 | 0 | 0 | 0,0013462 | 0,0021426 | 0,0015969 | 0,0015969 | TRUE | 0,0166392 | No |
| orf19.980 | 7867 | 6686 | 7536 | 4950 | 5118 | 3828 | 5012 | 5466 | 7500 | 7070 | 7445 | 7754 | 4101 | 4353 | 4255 | 4444 | 5865,31 | 7442 | 4288 | 0,576 | -0,795 | -18,69 | 5,94E-78 | 3,16E-77 | 0,001063 | 0,0023712 | 0,0015575 | 0,0015575 | TRUE | 0,0138215 | No |
| orf19.984 | 87 | 75 | 87 | 65 | 206 | 159 | 184 | 216 | 83 | 79 | 86 | 102 | 165 | 181 | 156 | 176 | 128,47 | 88 | 170 | 1,953 | 0,966 | 7,224 | 5,06E-13 | 9,68E-13 | 1,00E-08 | 0,0148722 | 0,0081097 | 0,0081097 | TRUE | 0,0832633 | No |
| orf19.985 | 2473 | 2321 | 2626 | 1637 | 2475 | 1879 | 2320 | 2605 | 2358 | 2454 | 2594 | 2564 | 1983 | 2137 | 1970 | 2118 | 2272,26 | 2492 | 2052 | 0,823 | -0,281 | -5,589 | 2,28E-08 | 3,76E-08 | 0,001263 | 0,0028139 | 0,0019762 | 0,0019762 | TRUE | 0,0165407 | No |
| orf19.986 | 38637 | 32894 | 35259 | 22100 | 26597 | 18408 | 25407 | 27008 | 36836 | 34781 | 34833 | 34621 | 21311 | 20932 | 21570 | 21960 | 28355,54 | 35268 | 21443 | 0,608 | -0,718 | -21,612 | 1,39E-103 | 9,38E-103 | 0,0005967 | 0,0021492 | 0,001023 | 0,001023 | TRUE | 0,0110279 | No |
| orf19.988 | 2580 | 2203 | 2371 | 1621 | 1468 | 1101 | 1489 | 1546 | 2460 | 2329 | 2342 | 2539 | 1176 | 1252 | 1264 | 1257 | 1827,53 | 2418 | 1237 | 0,512 | -0,965 | -19,67 | 3,92E-86 | 2,26E-85 | 0,0008549 | 0,0029897 | 0,0017189 | 0,0017189 | TRUE | 0,0138158 | No |
| orf19.989 | 4212 | 3871 | 4153 | 2976 | 4346 | 3403 | 4136 | 4592 | 4016 | 4093 | 4103 | 4662 | 3482 | 3870 | 3511 | 3734 | 3933,83 | 4218 | 3649 | 0,866 | -0,208 | -3,577 | 0,000348 | 0,0004809 | 0,0034429 | 0,0025087 | 0,002995 | 0,002995 | TRUE | 0,0591859 | No |
| orf19.99 | 1124 | 908 | 926 | 690 | 2084 | 1393 | 2019 | 2203 | 1072 | 960 | 915 | 1081 | 1670 | 1584 | 1714 | 1791 | 1348,33 | 1007 | 1690 | 1,683 | 0,751 | 11,225 | 3,06E-29 | 8,15E-29 | 0,0036271 | 0,003309 | 0,0034706 | 0,0034706 | TRUE | 0,045998 | No |
| orf19.991 | 3328 | 2850 | 3137 | 1928 | 3112 | 2177 | 2995 | 3007 | 3173 | 3014 | 3099 | 3020 | 2494 | 2475 | 2543 | 2445 | 2782,81 | 3076 | 2489 | 0,809 | -0,307 | -8,102 | 5,41E-16 | 1,11E-15 | 1,00E-08 | 0,0026813 | 0,0010129 | 0,0010129 | TRUE | 0,0055354 | No |
| orf19.992 | 3853 | 3421 | 3667 | 2197 | 4829 | 3390 | 4675 | 4978 | 3673 | 3617 | 3623 | 3442 | 3869 | 3855 | 3969 | 4048 | 3761,97 | 3589 | 3935 | 1,096 | 0,132 | 3,503 | 0,0004606 | 0,0006328 | 0,0003074 | 0,0025277 | 0,0010879 | 0,0010879 | TRUE | 0,0084869 | No |
| orf19.993 | 241 | 207 | 216 | 131 | 460 | 304 | 414 | 416 | 230 | 219 | 213 | 205 | 369 | 346 | 351 | 338 | 283,91 | 217 | 351 | 1,614 | 0,691 | 7,798 | 6,27E-15 | 1,26E-14 | 1,00E-08 | 0,0078746 | 0,0037061 | 0,0037061 | TRUE | 0,0181131 | No |
| orf19.994 | 3 | 1 | 2 | 3 | 6 | 3 | 4 | 0 | 3 | 1 | 2 | 5 | 5 | 3 | 3 | 0 | 2,78 | 3 | 3 | 1,133 | 0,181 | 0,194 | 0,8460235 | 0,8651403 | 1,00E-08 | 0,5935578 | 0,4577919 | 0,4577919 | TRUE | 0,7344824 | No |
| orf19.996 | 118 | 135 | 112 | 73 | 319 | 245 | 303 | 309 | 113 | 143 | 111 | 114 | 256 | 279 | 257 | 251 | 190,37 | 120 | 261 | 2,165 | 1,114 | 9,625 | 6,27E-22 | 1,46E-21 | 1,00E-08 | 0,0107162 | 0,0065301 | 0,0065301 | TRUE | 0,1689926 | No |
| orf19.997 | 824 | 796 | 695 | 453 | 1329 | 906 | 1145 | 1268 | 786 | 842 | 687 | 710 | 1065 | 1030 | 972 | 1031 | 890,21 | 756 | 1024 | 1,353 | 0,436 | 6,086 | 1,15E-09 | 1,98E-09 | 0,0036271 | 0,0039357 | 0,0037594 | 0,0037594 | TRUE | 0,0689475 | No |
| orf19.998 | 175 | 183 | 157 | 122 | 636 | 438 | 580 | 661 | 167 | 193 | 155 | 191 | 510 | 498 | 492 | 537 | 343,01 | 176 | 509 | 2,905 | 1,539 | 16,503 | 3,47E-61 | 1,49E-60 | 0,0005701 | 0,006878 | 0,0043042 | 0,0043042 | TRUE | 0,0736652 | No |
| orf19.999 | 6022 | 4984 | 5414 | 3877 | 1261 | 958 | 1148 | 1360 | 5741 | 5270 | 5349 | 6074 | 1010 | 1089 | 975 | 1106 | 3326,69 | 5608 | 1045 | 0,186 | -2,424 | -39,221 | 0 | 0 | 0,0032503 | 0,0025848 | 0,0031404 | 0,0031404 | TRUE | 0,0371033 | No |
| RDN18 | 80311 | 73414 | 80379 | 46846 | 42552 | 32637 | 33245 | 30643 | 76568 | 77626 | 79407 | 73387 | 34096 | 37111 | 28225 | 24916 | 53916,86 | 76747 | 31087 | 0,405 | -1,304 | -13,171 | 1,28E-39 | 4,02E-39 | 0,0163134 | 0,0021217 | 0,0093946 | 0,0093946 | TRUE | 0,2188093 | No |
| RDN25 | 73724 | 71932 | 77944 | 51278 | 137754 | 115742 | 93181 | 68353 | 70288 | 76059 | 77001 | 80330 | 110378 | 131609 | 79110 | 55578 | 85044,13 | 75920 | 94169 | 1,24 | 0,311 | 1,194 | 0,2322966 | 0,2613798 | 0,0650348 | 0,0021106 | 0,0299329 | 0,0650348 | TRUE | 0,1810157 | No |
| RDN5 | 276 | 705 | 672 | 479 | 148 | 242 | 172 | 215 | 263 | 745 | 664 | 750 | 119 | 275 | 146 | 175 | 392,18 | 606 | 179 | 0,295 | -1,763 | -4,215 | 2,50E-05 | 3,64E-05 | 0,1646573 | 0,0062779 | 0,0716742 | 0,1646573 | TRUE | 1,7139377 | No |
| RDN58 | 2598 | 2571 | 2991 | 2394 | 723 | 700 | 805 | 1019 | 2477 | 2719 | 2955 | 3750 | 579 | 796 | 683 | 829 | 1848,48 | 2975 | 722 | 0,243 | -2,043 | -11,706 | 1,18E-31 | 3,29E-31 | 0,0284579 | 0,0029795 | 0,0148859 | 0,0284579 | TRUE | 0,370764 | No |
| RPR1 | 169 | 131 | 143 | 95 | 73 | 44 | 86 | 85 | 161 | 139 | 141 | 149 | 58 | 50 | 73 | 69 | 105,05 | 148 | 62 | 0,428 | -1,223 | -8,026 | 1,01E-15 | 2,06E-15 | 0,0010152 | 0,0177213 | 0,011477 | 0,011477 | TRUE | 0,1317269 | No |
| SCR1 | 501 | 410 | 394 | 243 | 195 | 125 | 151 | 176 | 478 | 434 | 389 | 381 | 156 | 142 | 128 | 143 | 281,35 | 420 | 142 | 0,338 | -1,565 | -14,339 | 1,25E-46 | 4,40E-46 | 0,0059833 | 0,0079272 | 0,0069822 | 0,0069822 | TRUE | 0,0998007 | No |
| snR10a | 588 | 563 | 540 | 408 | 276 | 235 | 237 | 239 | 561 | 595 | 533 | 639 | 221 | 267 | 201 | 194 | 401,55 | 582 | 221 | 0,378 | -1,402 | -13,529 | 1,06E-41 | 3,44E-41 | 0,0087457 | 0,0061802 | 0,0073186 | 0,0073186 | TRUE | 0,1983056 | No |
| snR10b | 0 | 0 | 1 | 1 | 3 | 0 | 0 | 0 | 0 | 0 | 1 | 2 | 2 | 0 | 0 | 0 | 0,62 | 1 | 0 | 1,096 | 0,132 | 0,062 | 0,9507497 | 0,9558428 | 1,00E-08 | 2,6512453 | 2,7803441 | 2,7803441 | TRUE | 1,4547666 | No |
| snR10c | 1360 | 1192 | 1128 | 847 | 685 | 436 | 619 | 690 | 1297 | 1260 | 1114 | 1327 | 549 | 496 | 526 | 561 | 891,18 | 1250 | 533 | 0,428 | -1,225 | -17,101 | 1,45E-65 | 6,57E-65 | 0,0033903 | 0,0039337 | 0,0036386 | 0,0036386 | TRUE | 0,0649633 | No |
| snR128 | 2875 | 2539 | 2784 | 1733 | 2932 | 2113 | 2779 | 2600 | 2741 | 2685 | 2750 | 2715 | 2349 | 2403 | 2359 | 2114 | 2514,53 | 2723 | 2306 | 0,846 | -0,241 | -4,937 | 7,95E-07 | 1,23E-06 | 0,0013509 | 0,0027442 | 0,0018791 | 0,0018791 | TRUE | 0,0387637 | No |
| snR13 | 890 | 688 | 701 | 526 | 728 | 430 | 624 | 702 | 849 | 727 | 693 | 824 | 583 | 489 | 530 | 571 | 658,17 | 773 | 543 | 0,705 | -0,504 | -5,881 | 4,08E-09 | 6,85E-09 | 0,0063547 | 0,0045859 | 0,0055096 | 0,0055096 | TRUE | 0,0598383 | No |
| snR161a | 130 | 76 | 88 | 65 | 154 | 72 | 130 | 113 | 124 | 80 | 87 | 102 | 123 | 82 | 110 | 92 | 100,07 | 98 | 102 | 1,041 | 0,059 | 0,32 | 0,7489664 | 0,7770983 | 0,0260957 | 0,0184985 | 0,0220427 | 0,0220427 | TRUE | 0,3330104 | No |
| snR17A | 3673 | 3282 | 3188 | 2409 | 4254 | 3710 | 4040 | 4459 | 3502 | 3470 | 3149 | 3774 | 3409 | 4219 | 3430 | 3626 | 3572,27 | 3474 | 3671 | 1,057 | 0,08 | 1,047 | 0,2951902 | 0,3278152 | 0,0075775 | 0,0025509 | 0,0052692 | 0,0052692 | TRUE | 0,1215373 | No |
| snR18 | 0 | 0 | 0 | 0 | 0 | 0 | 0 | 0 | 0 | 0 | 0 | 0 | 0 | 0 | 0 | 0 | 0 | 0 | 0 | NA | NA | NA | NA | NA | NA | NA | NA | NA | NA | NA | NA |
| snR189a | 3 | 3 | 1 | 1 | 1 | 2 | 2 | 0 | 3 | 3 | 1 | 2 | 1 | 2 | 2 | 0 | 1,67 | 2 | 1 | 0,523 | -0,934 | -0,832 | 0,4053736 | 0,4412639 | 1,00E-08 | 0,9852526 | 0,5628943 | 0,5628943 | TRUE | 0,3208066 | No |
| snR189b | 118 | 99 | 92 | 88 | 44 | 39 | 34 | 40 | 113 | 105 | 91 | 138 | 35 | 44 | 29 | 33 | 73,36 | 112 | 35 | 0,316 | -1,664 | -8,358 | 6,36E-17 | 1,33E-16 | 0,014172 | 0,0244713 | 0,0202332 | 0,0202332 | TRUE | 0,194971 | No |
| snR189c | 183 | 205 | 203 | 144 | 92 | 68 | 83 | 93 | 174 | 217 | 201 | 226 | 74 | 77 | 70 | 76 | 139,31 | 204 | 74 | 0,365 | -1,454 | -10,759 | 5,39E-27 | 1,39E-26 | 1,00E-08 | 0,0138772 | 0,0088013 | 0,0088013 | TRUE | 0,1099816 | No |
| snR189d | 0 | 1 | 1 | 0 | 1 | 0 | 0 | 1 | 0 | 1 | 1 | 0 | 1 | 0 | 0 | 1 | 0,46 | 0 | 0 | 0,785 | -0,349 | -0,17 | 0,8649671 | 0,8826091 | 1,00E-08 | 3,5912989 | 2,1815598 | 2,1815598 | TRUE | 0,1343204 | No |
| snR190 | 1277 | 1862 | 1795 | 872 | 1573 | 1105 | 1414 | 878 | 1217 | 1969 | 1773 | 1366 | 1260 | 1256 | 1200 | 714 | 1344,61 | 1581 | 1108 | 0,7 | -0,514 | -2,079 | 0,0376047 | 0,0456884 | 0,0579489 | 0,0033124 | 0,0272944 | 0,0579489 | TRUE | 0,330622 | No |
| snR24 | 0 | 0 | 0 | 0 | 0 | 0 | 0 | 0 | 0 | 0 | 0 | 0 | 0 | 0 | 0 | 0 | 0 | 0 | 0 | NA | NA | NA | NA | NA | NA | NA | NA | NA | NA | NA | NA |
| snR31a | 44 | 29 | 34 | 25 | 21 | 25 | 21 | 27 | 42 | 31 | 34 | 39 | 17 | 28 | 18 | 22 | 28,8 | 36 | 21 | 0,578 | -0,791 | -2,879 | 0,0039901 | 0,0051951 | 1,00E-08 | 0,0591013 | 0,0359656 | 0,0359656 | TRUE | 0,2397167 | No |
| snR32a | 37 | 39 | 30 | 10 | 62 | 34 | 45 | 53 | 35 | 41 | 30 | 16 | 50 | 39 | 38 | 43 | 36,43 | 30 | 42 | 1,363 | 0,447 | 1,697 | 0,089749 | 0,1052543 | 0,0209073 | 0,0471591 | 0,0383617 | 0,0383617 | TRUE | 0,4605315 | No |
| snR33a | 53991 | 35181 | 38180 | 32840 | 11108 | 7657 | 10297 | 11599 | 51475 | 37199 | 37718 | 51446 | 8901 | 8707 | 8742 | 9431 | 26702,29 | 44460 | 8945 | 0,201 | -2,313 | -17,283 | 6,33E-67 | 2,92E-66 | 0,0171515 | 0,0021528 | 0,009785 | 0,0171515 | TRUE | 0,0480162 | No |
| snR35a | 15 | 19 | 25 | 15 | 32 | 23 | 28 | 27 | 14 | 20 | 25 | 23 | 26 | 26 | 24 | 22 | 22,51 | 20 | 24 | 1,19 | 0,251 | 0,84 | 0,4011694 | 0,4369351 | 1,00E-08 | 0,0750218 | 0,040187 | 0,040187 | TRUE | 0,2677749 | No |

| | | | | | | | | | | | | | | | | | | | | | | | | | | | | | | | |
|---|---|---|---|---|---|---|---|---|---|---|---|---|---|---|---|---|---|---|---|---|---|---|---|---|---|---|---|---|---|---|---|
| snR35c | 36 | 17 | 38 | 19 | 25 | 28 | 23 | 30 | 34 | 18 | 38 | 30 | 20 | 32 | 20 | 24 | 26,92 | 30 | 24 | 0,789 | -0,342 | -1,122 | 0,2616681 | 0,2921617 | 0,0354745 | 0,0630741 | 0,0515206 | 0,0515206 | TRUE | 0,494299 | No |
| snR39b | 0 | 0 | 0 | 0 | 0 | 0 | 0 | 0 | 0 | 0 | 0 | 0 | 0 | 0 | 0 | 0 | 0 | 0 | 0 | NA | NA | NA | NA | NA | NA | NA | NA | NA | NA | NA | NA |
| snR3a | 87 | 83 | 88 | 44 | 102 | 56 | 72 | 80 | 83 | 88 | 87 | 69 | 82 | 64 | 61 | 65 | 74,77 | 82 | 68 | 0,828 | -0,273 | -1,597 | 0,1103458 | 0,128231 | 1,00E-08 | 0,0240509 | 0,0147246 | 0,0147246 | TRUE | 0,1859805 | No |
| snR40 | 30 | 36 | 25 | 23 | 54 | 40 | 34 | 52 | 29 | 38 | 25 | 36 | 43 | 45 | 29 | 42 | 35,91 | 32 | 40 | 1,263 | 0,337 | 1,327 | 0,1846521 | 0,2101418 | 1,00E-08 | 0,0478117 | 0,0332284 | 0,0332284 | TRUE | 0,2841826 | No |
| snR41 | 1747 | 1609 | 1700 | 1006 | 1317 | 816 | 1198 | 1134 | 1666 | 1701 | 1679 | 1576 | 1055 | 928 | 1017 | 922 | 1318,07 | 1656 | 980 | 0,592 | -0,756 | -12,863 | 7,25E-38 | 2,22E-37 | 0,0017446 | 0,003337 | 0,0025188 | 0,0025188 | TRUE | 0,0320125 | No |
| snR41b | 602 | 568 | 520 | 495 | 396 | 263 | 358 | 336 | 574 | 601 | 514 | 775 | 317 | 299 | 304 | 273 | 457,15 | 616 | 298 | 0,487 | -1,039 | -8,897 | 5,75E-19 | 1,26E-18 | 0,0152467 | 0,0056829 | 0,0106759 | 0,0106759 | TRUE | 0,335058 | No |
| snR42a | 157 | 131 | 117 | 125 | 97 | 73 | 84 | 96 | 150 | 139 | 116 | 196 | 78 | 83 | 71 | 78 | 113,71 | 150 | 78 | 0,522 | -0,939 | -5,606 | 2,07E-08 | 3,41E-08 | 0,0183714 | 0,0165303 | 0,017376 | 0,017376 | TRUE | 0,304467 | No |
| snR43a | 3 | 4 | 1 | 3 | 6 | 7 | 5 | 9 | 3 | 4 | 1 | 5 | 5 | 8 | 4 | 7 | 4,64 | 3 | 6 | 1,959 | 0,97 | 1,412 | 0,1578954 | 0,1808202 | 1,00E-08 | 0,3560706 | 0,1989661 | 0,1989661 | TRUE | 0,3209957 | No |
| snR43b | 29 | 25 | 17 | 19 | 50 | 32 | 35 | 48 | 28 | 26 | 17 | 30 | 40 | 36 | 30 | 39 | 30,73 | 25 | 36 | 1,461 | 0,547 | 2,048 | 0,0405189 | 0,0491262 | 1,00E-08 | 0,0555223 | 0,0338631 | 0,0338631 | TRUE | 0,3287617 | No |
| snR44a | 20 | 22 | 24 | 18 | 34 | 32 | 38 | 24 | 19 | 23 | 24 | 28 | 27 | 36 | 32 | 20 | 26,21 | 24 | 29 | 1,228 | 0,296 | 1,01 | 0,3126815 | 0,345912 | 1,00E-08 | 0,0647462 | 0,043156 | 0,043156 | TRUE | 0,3473089 | No |
| snR44b | 62 | 65 | 64 | 42 | 91 | 77 | 90 | 101 | 59 | 69 | 63 | 66 | 73 | 88 | 76 | 82 | 71,98 | 64 | 80 | 1,241 | 0,311 | 1,874 | 0,0609833 | 0,0726947 | 1,00E-08 | 0,0249009 | 0,0121767 | 0,0121767 | TRUE | 0,0348991 | No |
| snR44c | 12 | 18 | 17 | 17 | 27 | 30 | 26 | 22 | 11 | 19 | 17 | 27 | 22 | 34 | 22 | 18 | 21,2 | 18 | 24 | 1,313 | 0,393 | 1,142 | 0,2532475 | 0,2836533 | 0,0412239 | 0,0795358 | 0,063761 | 0,063761 | TRUE | 0,4161617 | No |
| snR44d | 382 | 414 | 339 | 254 | 471 | 279 | 350 | 307 | 364 | 438 | 335 | 398 | 377 | 317 | 297 | 250 | 347,02 | 384 | 310 | 0,81 | -0,304 | -2,379 | 0,0173449 | 0,0216407 | 0,0187707 | 0,0068227 | 0,0127947 | 0,0127947 | TRUE | 0,2521288 | No |
| snR46a | 0 | 0 | 0 | 0 | 1 | 0 | 0 | 0 | 0 | 0 | 0 | 0 | 1 | 0 | 0 | 0 | 0,1 | 0 | 0 | 1,314 | 0,394 | 0,111 | 0,9112982 | 0,9205309 | 1,00E-08 | 16,395074 | 10 | 10 | TRUE | 0,2859086 | No |
| snR47 | 706 | 490 | 535 | 317 | 440 | 252 | 337 | 319 | 673 | 518 | 529 | 497 | 353 | 287 | 286 | 259 | 425,12 | 554 | 296 | 0,534 | -0,906 | -7,586 | 3,30E-14 | 6,51E-14 | 0,016069 | 0,0059535 | 0,0111765 | 0,0111765 | TRUE | 0,2493289 | No |
| snR48 | 1252 | 1381 | 1302 | 581 | 628 | 375 | 521 | 325 | 1194 | 1460 | 1286 | 910 | 503 | 426 | 442 | 264 | 810,81 | 1212 | 409 | 0,337 | -1,569 | -6,57 | 5,02E-11 | 9,04E-11 | 0,0532028 | 0,0041163 | 0,0258017 | 0,0532028 | TRUE | 0,386645 | No |
| snR49a | 0 | 0 | 0 | 0 | 0 | 0 | 0 | 0 | 0 | 0 | 0 | 0 | 0 | 0 | 0 | 0 | 0 | 0 | 0 | NA | NA | NA | NA | NA | NA | NA | NA | NA | NA | NA | NA |
| snR50 | 2 | 5 | 5 | 2 | 3 | 8 | 10 | 5 | 2 | 5 | 5 | 3 | 2 | 9 | 8 | 4 | 4,92 | 4 | 6 | 1,537 | 0,62 | 0,9 | 0,3682731 | 0,4032173 | 1,00E-08 | 0,3361319 | 0,2357059 | 0,2357059 | TRUE | 0,1956763 | No |
| snR52 | 11 | 8 | 11 | 4 | 25 | 21 | 26 | 25 | 10 | 8 | 11 | 6 | 20 | 24 | 22 | 20 | 15,3 | 9 | 22 | 2,324 | 1,217 | 3,268 | 0,0010818 | 0,0014564 | 1,00E-08 | 0,1094128 | 0,0524398 | 0,0524398 | TRUE | 0,0741754 | No |
| snR54 | 0 | 0 | 0 | 0 | 0 | 0 | 0 | 0 | 0 | 0 | 0 | 0 | 0 | 0 | 0 | 0 | 0 | 0 | 0 | NA | NA | NA | NA | NA | NA | NA | NA | NA | NA | NA | NA |
| snR55 | 13 | 7 | 13 | 11 | 5 | 1 | 6 | 5 | 12 | 7 | 13 | 17 | 4 | 1 | 5 | 4 | 8,02 | 12 | 4 | 0,301 | -1,731 | -3,129 | 0,001753 | 0,0023318 | 1,00E-08 | 0,2067745 | 0,1284619 | 0,1284619 | TRUE | 0,1656682 | No |
| snR56 | 570 | 388 | 434 | 344 | 367 | 319 | 307 | 330 | 543 | 410 | 429 | 539 | 294 | 363 | 261 | 268 | 388,39 | 480 | 296 | 0,616 | -0,699 | -5,522 | 3,36E-08 | 5,49E-08 | 0,0194612 | 0,0063188 | 0,0126901 | 0,0126901 | TRUE | 0,2423594 | No |
| snR57 | 99 | 126 | 129 | 35 | 30 | 35 | 34 | 18 | 94 | 133 | 127 | 55 | 24 | 40 | 29 | 15 | 64,65 | 102 | 27 | 0,256 | -1,964 | -6,364 | 1,97E-10 | 3,49E-10 | 0,126192 | 0,027487 | 0,069113 | 0,069113 | TRUE | 0,2621013 | No |
| snR57b | 833 | 853 | 790 | 348 | 444 | 259 | 348 | 251 | 794 | 902 | 780 | 545 | 356 | 295 | 295 | 204 | 521,44 | 755 | 288 | 0,38 | -1,396 | -6,3 | 2,97E-10 | 5,23E-10 | 0,0448378 | 0,0052401 | 0,0232807 | 0,0448378 | TRUE | 0,41021 | No |
| snR58 | 287 | 234 | 196 | 152 | 235 | 194 | 208 | 207 | 274 | 247 | 194 | 238 | 188 | 221 | 177 | 168 | 213,32 | 238 | 188 | 0,788 | -0,344 | -2,667 | 0,0076519 | 0,009777 | 0,0128707 | 0,009788 | 0,0112677 | 0,0112677 | TRUE | 0,1878608 | No |
| snR5a | 362 | 285 | 254 | 229 | 367 | 268 | 231 | 262 | 345 | 301 | 251 | 359 | 294 | 305 | 196 | 213 | 283,01 | 314 | 252 | 0,803 | -0,317 | -2,02 | 0,043376 | 0,0524209 | 0,033437 | 0,0078928 | 0,0201335 | 0,0201335 | TRUE | 0,2207631 | No |
| snR5b | 29 | 9 | 20 | 11 | 31 | 19 | 24 | 32 | 28 | 10 | 20 | 17 | 25 | 22 | 20 | 26 | 20,87 | 19 | 23 | 1,243 | 0,314 | 0,924 | 0,3555131 | 0,3901839 | 0,0311248 | 0,0807482 | 0,061782 | 0,061782 | TRUE | 0,3209306 | No |
| snR5c | 6 | 6 | 7 | 0 | 14 | 14 | 15 | 15 | 6 | 6 | 7 | 0 | 11 | 16 | 13 | 12 | 8,88 | 5 | 13 | 2,544 | 1,347 | 2,613 | 0,0089655 | 0,0114234 | 1,00E-08 | 0,1869686 | 0,1114688 | 0,1114688 | TRUE | 0,4367526 | No |
| snR5d | 215 | 195 | 175 | 171 | 403 | 254 | 324 | 330 | 205 | 206 | 173 | 268 | 323 | 289 | 275 | 268 | 250,88 | 213 | 289 | 1,37 | 0,454 | 3,627 | 0,000287 | 0,000398 | 0,0135596 | 0,0086358 | 0,0108436 | 0,0108436 | TRUE | 0,2815152 | No |
| snR60 | 0 | 0 | 3 | 2 | 2 | 2 | 2 | 0 | 0 | 0 | 3 | 3 | 2 | 2 | 2 | 0 | 1,46 | 2 | 2 | 0,941 | -0,088 | -0,065 | 0,9478903 | 0,9531182 | 1,00E-08 | 1,1274983 | 1,0098581 | 1,0098581 | TRUE | 0,1386839 | No |
| snR61 | 427 | 505 | 463 | 348 | 215 | 176 | 181 | 146 | 407 | 534 | 457 | 545 | 172 | 200 | 154 | 119 | 323,55 | 486 | 161 | 0,331 | -1,596 | -11,03 | 2,73E-28 | 7,17E-28 | 0,0260088 | 0,0071659 | 0,0161601 | 0,0161601 | TRUE | 0,3460792 | No |
| snR62 | 200 | 223 | 176 | 126 | 83 | 63 | 50 | 56 | 191 | 236 | 174 | 197 | 67 | 72 | 42 | 46 | 127,98 | 200 | 57 | 0,281 | -1,831 | -10,343 | 4,48E-25 | 1,11E-24 | 0,0260469 | 0,0149205 | 0,0193157 | 0,0193157 | TRUE | 0,2457697 | No |
| snR63 | 226 | 182 | 156 | 80 | 268 | 165 | 266 | 203 | 215 | 192 | 154 | 125 | 215 | 188 | 226 | 165 | 185,07 | 172 | 198 | 1,146 | 0,196 | 1,208 | 0,2270372 | 0,2559132 | 0,029408 | 0,0109629 | 0,0197858 | 0,0197858 | TRUE | 0,2416104 | No |
| snR64 | 18 | 12 | 7 | 13 | 12 | 9 | 10 | 7 | 17 | 13 | 7 | 20 | 10 | 10 | 8 | 6 | 11,4 | 14 | 8 | 0,603 | -0,73 | -1,615 | 0,106413 | 0,1238412 | 1,00E-08 | 0,1461799 | 0,1038953 | 0,1038953 | TRUE | 0,2696177 | No |
| snR65 | 20 | 7 | 16 | 6 | 8 | 4 | 9 | 12 | 19 | 7 | 16 | 9 | 6 | 5 | 8 | 10 | 10 | 13 | 7 | 0,547 | -0,871 | -1,812 | 0,069967 | 0,0830172 | 1,00E-08 | 0,1662162 | 0,1173001 | 0,1173001 | TRUE | 0,157145 | No |
| snR66 | 82 | 93 | 132 | 113 | 458 | 344 | 337 | 258 | 78 | 98 | 130 | 177 | 367 | 391 | 286 | 210 | 217,25 | 121 | 314 | 2,604 | 1,381 | 4,355 | 1,33E-05 | 1,96E-05 | 0,0904243 | 0,0096491 | 0,0449063 | 0,0904243 | TRUE | 0,3787556 | No |
| snR67 | 2 | 2 | 8 | 5 | 2 | 3 | 1 | 3 | 2 | 2 | 8 | 8 | 2 | 3 | 1 | 2 | 3,51 | 5 | 2 | 0,42 | -1,252 | -1,491 | 0,13603 | 0,1566518 | 1,00E-08 | 0,4702096 | 0,3412124 | 0,3412124 | TRUE | 0,0840951 | No |
| snR68 | 36 | 57 | 40 | 46 | 28 | 29 | 20 | 12 | 34 | 60 | 40 | 72 | 22 | 33 | 17 | 10 | 36,04 | 52 | 20 | 0,396 | -1,336 | -3,808 | 0,00014 | 0,0001966 | 0,1347511 | 0,0476494 | 0,0850715 | 0,0850715 | TRUE | 0,2760307 | No |
| snR69 | 488 | 389 | 422 | 372 | 310 | 188 | 223 | 241 | 465 | 411 | 417 | 583 | 248 | 214 | 189 | 196 | 340,46 | 469 | 212 | 0,454 | -1,14 | -8,949 | 3,58E-19 | 7,90E-19 | 0,017617 | 0,0069139 | 0,012258 | 0,012258 | TRUE | 0,2853998 | No |
| snR70 | 1546 | 1457 | 1509 | 921 | 996 | 687 | 966 | 843 | 1474 | 1541 | 1491 | 1443 | 798 | 781 | 820 | 685 | 1129,11 | 1487 | 771 | 0,518 | -0,949 | -15,014 | 5,92E-51 | 2,21E-50 | 0,0022989 | 0,0035454 | 0,0028775 | 0,0028775 | TRUE | 0,0694475 | No |
| snR71 | 11 | 13 | 14 | 6 | 12 | 7 | 4 | 1 | 10 | 14 | 14 | 9 | 10 | 8 | 3 | 1 | 8,66 | 12 | 6 | 0,449 | -1,156 | -2,025 | 0,0429069 | 0,0518736 | 0,1648831 | 0,1917775 | 0,1831007 | 0,1831007 | TRUE | 0,5860404 | No |
| snR72 | 5 | 11 | 7 | 8 | 5 | 4 | 5 | 5 | 5 | 12 | 7 | 13 | 4 | 5 | 4 | 4 | 6,59 | 9 | 4 | 0,481 | -1,056 | -1,851 | 0,0642384 | 0,0764463 | 1,00E-08 | 0,2512846 | 0,1428032 | 0,1428032 | TRUE | 0,1701729 | No |
| snR73 | 419 | 407 | 397 | 299 | 472 | 327 | 351 | 391 | 399 | 430 | 392 | 468 | 378 | 372 | 298 | 318 | 382,05 | 422 | 342 | 0,81 | -0,304 | -3,059 | 0,0022201 | 0,0029355 | 0,007338 | 0,0063889 | 0,0068574 | 0,0068574 | TRUE | 0,0848995 | No |
| snR74 | 82 | 172 | 112 | 106 | 63 | 72 | 49 | 47 | 78 | 182 | 111 | 166 | 50 | 82 | 42 | 38 | 93,61 | 134 | 53 | 0,393 | -1,346 | -4,738 | 2,16E-06 | 3,30E-06 | 0,1174722 | 0,0196303 | 0,0648204 | 0,0648204 | TRUE | 0,2157977 | No |
| snR75 | 163 | 226 | 207 | 171 | 71 | 45 | 63 | 63 | 155 | 239 | 204 | 268 | 57 | 51 | 53 | 51 | 134,94 | 216 | 53 | 0,248 | -2,014 | -11,202 | 4,00E-29 | 1,06E-28 | 0,0260072 | 0,014259 | 0,0200801 | 0,0200801 | TRUE | 0,1500075 | No |
| snR76 | 409 | 399 | 420 | 228 | 209 | 139 | 163 | 181 | 390 | 422 | 415 | 357 | 167 | 158 | 138 | 147 | 274,38 | 396 | 152 | 0,384 | -1,38 | -13,756 | 4,68E-43 | 1,55E-42 | 0,0017197 | 0,0080754 | 0,0053469 | 0,0053469 | TRUE | 0,0506372 | No |
| snR77 | 259 | 321 | 356 | 231 | 156 | 131 | 122 | 118 | 247 | 339 | 352 | 362 | 125 | 149 | 104 | 96 | 221,67 | 325 | 118 | 0,363 | -1,462 | -9,317 | 1,20E-20 | 2,73E-20 | 0,0281914 | 0,0094981 | 0,0181146 | 0,0181146 | TRUE | 0,3065142 | No |
| snR77b | 103 | 237 | 220 | 127 | 98 | 91 | 74 | 27 | 98 | 251 | 217 | 199 | 79 | 103 | 63 | 22 | 128,98 | 191 | 67 | 0,348 | -1,523 | -3,01 | 0,0026108 | 0,003437 | 0,2363193 | 0,0148209 | 0,1075471 | 0,2363193 | TRUE | 0,9152131 | No |
| snR78 | 51 | 96 | 100 | 32 | 27 | 20 | 29 | 10 | 49 | 102 | 99 | 50 | 22 | 23 | 25 | 8 | 47,02 | 75 | 20 | 0,255 | -1,971 | -5,668 | 1,44E-08 | 2,39E-08 | 0,150276 | 0,0370088 | 0,0853433 | 0,0853433 | TRUE | 0,1410759 | No |
| snR79 | 118 | 170 | 154 | 126 | 101 | 54 | 78 | 55 | 113 | 180 | 152 | 197 | 81 | 61 | 66 | 45 | 111,88 | 160 | 63 | 0,397 | -1,331 | -6,375 | 1,83E-10 | 3,25E-10 | 0,0482946 | 0,0167667 | 0,0313087 | 0,0313087 | TRUE | 0,2394216 | No |
| snR87 | 280 | 271 | 268 | 168 | 182 | 128 | 145 | 143 | 267 | 287 | 265 | 263 | 146 | 146 | 123 | 116 | 201,52 | 270 | 133 | 0,489 | -1,033 | -9,33 | 1,06E-20 | 2,42E-20 | 1,00E-08 | 0,0102387 | 0,0063443 | 0,0063443 | TRUE | 0,0756044 | No |
| snR8a | 235 | 234 | 215 | 133 | 406 | 275 | 350 | 367 | 224 | 247 | 212 | 208 | 325 | 313 | 297 | 298 | 265,72 | 223 | 308 | 1,378 | 0,463 | 4,936 | 7,98E-07 | 1,24E-06 | 1,00E-08 | 0,0082703 | 0,0045046 | 0,0045046 | TRUE | 0,0584479 | No |
| snR8b | 18 | 28 | 23 | 22 | 41 | 30 | 37 | 36 | 17 | 30 | 23 | 34 | 33 | 34 | 31 | 29 | 28,95 | 26 | 32 | 1,254 | 0,327 | 1,189 | 0,2343682 | 0,2636178 | 1,00E-08 | 0,0588058 | 0,0364968 | 0,0364968 | TRUE | 0,2899491 | No |
| snR8c | 101 | 122 | 113 | 87 | 103 | 52 | 64 | 65 | 96 | 129 | 112 | 136 | 83 | 59 | 54 | 53 | 90,26 | 118 | 62 | 0,531 | -0,914 | -4,898 | 9,68E-07 | 1,50E-06 | 0,0230383 | 0,0202826 | 0,0215168 | 0,0215168 | TRUE | 0,4707229 | No |
| SNRNAU1 | 163 | 137 | 121 | 68 | 44 | 29 | 35 | 42 | 155 | 145 | 120 | 107 | 35 | 33 | 30 | 34 | 82,3 | 132 | 33 | 0,249 | -2,004 | -10,555 | 4,80E-26 | 1,22E-25 | 1,00E-08 | 0,0220409 | 0,0169526 | 0,0169526 | TRUE | 0,0791543 | No |
| SNRNAU2 | 39 | 37 | 34 | 22 | 20 | 16 | 10 | 14 | 37 | 39 | 34 | 34 | 16 | 18 | 8 | 11 | 24,81 | 36 | 13 | 0,369 | -1,44 | -4,724 | 2,32E-06 | 3,53E-06 | 1,00E-08 | 0,0682803 | 0,040483 | 0,040483 | TRUE | 0,295598 | No |
| SNRNAU4 | 151 | 162 | 180 | 78 | 96 | 50 | 94 | 89 | 144 | 171 | 178 | 122 | 77 | 57 | 80 | 72 | 112,65 | 154 | 72 | 0,465 | -1,105 | -6,819 | 9,17E-12 | 1,69E-11 | 0,0137304 | 0,0166662 | 0,015421 | 0,015421 | TRUE | 0,1633048 | No |
| SNRNAU5 | 20 | 20 | 19 | 16 | 4 | 3 | 3 | 3 | 19 | 21 | 19 | 25 | 3 | 3 | 3 | 2 | 11,96 | 21 | 3 | 0,138 | -2,857 | -5,574 | 2,49E-08 | 4,09E-08 | 1,00E-08 | 0,1394129 | 0,0713989 | 0,0713989 | TRUE | 0,0628111 | No |
| SNRNAU6 | 31 | 21 | 35 | 17 | 16 | 15 | 18 | 19 | 30 | 22 | 35 | 27 | 13 | 17 | 15 | 15 | 21,7 | 28 | 15 | 0,53 | -0,915 | -2,979 | 0,0028895 | 0,0037891 | 1,00E-08 | 0,0777657 | 0,0414617 | 0,0414617 | TRUE | 0,1558276 | No |
| tA(AGC)1 | 0 | 0 | 0 | 1 | 1 | 0 | 1 | 3 | 0 | 0 | 0 | 2 | 1 | 0 | 1 | 2 | 0,71 | 0 | 1 | 3,105 | 1,635 | 0,912 | 0,3615184 | 0,3962296 | 1,00E-08 | 2,3244004 | 1,6590776 | 1,6590776 | TRUE | 0,6253314 | No |
| tA(AGC)2 | 0 | 0 | 0 | 0 | 0 | 0 | 0 | 0 | 0 | 0 | 0 | 0 | 0 | 0 | 0 | 0 | 0 | 0 | 0 | NA | NA | NA | NA | NA | NA | NA | NA | NA | NA | NA | NA |
| tA(AGC)3 | 0 | 0 | 0 | 0 | 0 | 0 | 0 | 0 | 0 | 0 | 0 | 0 | 0 | 0 | 0 | 0 | 0 | 0 | 0 | NA | NA | NA | NA | NA | NA | NA | NA | NA | NA | NA | NA |
| tA(AGC)4 | 0 | 0 | 1 | 0 | 0 | 0 | 0 | 0 | 0 | 0 | 1 | 0 | 0 | 0 | 0 | 0 | 0,12 | 0 | 0 | 0,483 | -1,049 | -0,297 | 0,7665832 | 0,7911318 | 1,00E-08 | 13,298132 | 10 | 10 | TRUE | 0,2757288 | No |
| tA(AGC)5 | 0 | 0 | 0 | 0 | 5 | 3 | 3 | 4 | 0 | 0 | 0 | 0 | 4 | 3 | 3 | 3 | 1,65 | 0 | 3 | 16,075 | 4,007 | 2,996 | 0,0027344 | 0,0035952 | 1,00E-08 | 0,9959048 | 0,583286 | 0,583286 | TRUE | 0,0475064 | No |
| tA(AGC)7 | 0 | 0 | 0 | 1 | 0 | 0 | 0 | 0 | 0 | 0 | 0 | 2 | 0 | 0 | 0 | 0 | 0,2 | 0 | 0 | 0,483 | -1,049 | -0,297 | 0,7665832 | 0,7911318 | 1,00E-08 | 8,3868857 | 10 | 10 | TRUE | 0,6253243 | No |
| tA(UGC)1 | 1 | 0 | 0 | 0 | 0 | 0 | 0 | 0 | 1 | 0 | 0 | 0 | 0 | 0 | 0 | 0 | 0,12 | 0 | 0 | 0,483 | -1,049 | -0,297 | 0,7665832 | 0,7911318 | 1,00E-08 | 13,779457 | 10 | 10 | TRUE | 0,2560039 | No |
| tA(UGC)2 | 0 | 0 | 0 | 0 | 0 | 0 | 0 | 0 | 0 | 0 | 0 | 0 | 0 | 0 | 0 | 0 | 0 | 0 | 0 | NA | NA | NA | NA | NA | NA | NA | NA | NA | NA | NA | NA |
| tC(GCA)1 | 0 | 0 | 0 | 0 | 0 | 0 | 0 | 0 | 0 | 0 | 0 | 0 | 0 | 0 | 0 | 0 | 0 | 0 | 0 | NA | NA | NA | NA | NA | NA | NA | NA | NA | NA | NA | NA |
| tC(GCA)2 | 0 | 0 | 0 | 0 | 0 | 0 | 0 | 1 | 0 | 0 | 0 | 0 | 0 | 0 | 0 | 1 | 0,1 | 0 | 0 | 1,314 | 0,394 | 0,111 | 0,9112982 | 0,9205309 | 1,00E-08 | 16,156685 | 10 | 10 | TRUE | 0,2944906 | No |
| tD(GUC)2 | 0 | 0 | 1 | 0 | 0 | 0 | 0 | 0 | 0 | 0 | 1 | 0 | 0 | 0 | 0 | 0 | 0,12 | 0 | 0 | 0,483 | -1,049 | -0,297 | 0,7665832 | 0,7911318 | 1,00E-08 | 13,298132 | 10 | 10 | TRUE | 0,2757288 | No |
| tD(GUC)3 | 0 | 0 | 0 | 0 | 0 | 0 | 0 | 1 | 0 | 0 | 0 | 0 | 0 | 0 | 0 | 1 | 0,1 | 0 | 0 | 1,314 | 0,394 | 0,111 | 0,9112982 | 0,9205309 | 1,00E-08 | 16,156685 | 10 | 10 | TRUE | 0,2944906 | No |
| tD(GUC)4 | 0 | 0 | 0 | 1 | 0 | 0 | 0 | 0 | 0 | 0 | 0 | 2 | 0 | 0 | 0 | 0 | 0,2 | 0 | 0 | 0,483 | -1,049 | -0,297 | 0,7665832 | 0,7911318 | 1,00E-08 | 8,3868857 | 10 | 10 | TRUE | 0,6253243 | No |
| tD(GUC)5 | 1 | 1 | 0 | 0 | 0 | 0 | 0 | 0 | 1 | 1 | 0 | 0 | 0 | 0 | 0 | 0 | 0,25 | 0 | 0 | 0,297 | -1,753 | -0,582 | 0,560743 | 0,595457 | 1,00E-08 | 6,5345431 | 6,7949003 | 6,7949003 | TRUE | 0,050013 | No |

| | | | | | | | | | | | | | | | | | | | | | | | | | | | | | | | |
|---|---|---|---|---|---|---|---|---|---|---|---|---|---|---|---|---|---|---|---|---|---|---|---|---|---|---|---|---|---|---|---|
| tD(GUC)6 | 0 | 0 | 0 | 0 | 0 | 1 | 1 | 1 | 0 | 0 | 0 | 0 | 0 | 1 | 1 | 1 | 0,35 | 0 | 1 | 3,35 | 1,744 | 0,745 | 0,4562856 | 0,4928997 | 1,00E-08 | 4,694617 | 3,6184189 | 3,6184189 | TRUE | 0,193549 | No |
| tD(GUC)7 | 0 | 0 | 0 | 0 | 0 | 0 | 1 | 0 | 0 | 0 | 0 | 0 | 0 | 0 | 1 | 0 | 0,11 | 0 | 0 | 1,314 | 0,394 | 0,111 | 0,9112982 | 0,9205309 | 1,00E-08 | 15,473589 | 10 | 10 | TRUE | 0,3207401 | No |
| tE(CUC)1 | 1 | 0 | 0 | 0 | 0 | 0 | 0 | 0 | 1 | 0 | 0 | 0 | 0 | 0 | 0 | 0 | 0,12 | 0 | 0 | 0,483 | -1,049 | -0,297 | 0,7665832 | 0,7911318 | 1,00E-08 | 13,779457 | 10 | 10 | TRUE | 0,2560039 | No |
| TER1 | 3391 | 2952 | 3071 | 2110 | 599 | 413 | 551 | 557 | 3233 | 3121 | 3034 | 3305 | 480 | 470 | 468 | 453 | 1820,48 | 3173 | 468 | 0,147 | -2,762 | -51,504 | 0 | 0 | 1,00E-08 | 0,0029932 | 0,001633 | 0,001633 | TRUE | 0,0106791 | No |
| tE(UUC)1 | 8 | 4 | 6 | 3 | 4 | 6 | 4 | 3 | 8 | 4 | 6 | 5 | 3 | 7 | 3 | 2 | 4,79 | 6 | 4 | 0,673 | -0,571 | -0,887 | 0,3752251 | 0,4102658 | 1,00E-08 | 0,3446307 | 0,1839642 | 0,1839642 | TRUE | 0,3175233 | No |
| tE(UUC)2 | 0 | 0 | 0 | 0 | 0 | 0 | 0 | 2 | 0 | 0 | 0 | 0 | 0 | 0 | 0 | 2 | 0,2 | 0 | 0 | 2,125 | 1,087 | 0,308 | 0,7579036 | 0,7857324 | 1,00E-08 | 8,0793879 | 10 | 10 | TRUE | 0,8724902 | No |
| tE(UUC)3 | 1 | 0 | 0 | 1 | 0 | 0 | 0 | 0 | 1 | 0 | 0 | 2 | 0 | 0 | 0 | 0 | 0,31 | 1 | 0 | 0,291 | -1,783 | -0,632 | 0,5276276 | 0,5637628 | 1,00E-08 | 5,2145966 | 5,7409504 | 5,7409504 | TRUE | 0,1754167 | No |
| tE(UUC)4 | 0 | 0 | 0 | 0 | 1 | 1 | 1 | 0 | 0 | 0 | 0 | 0 | 1 | 1 | 1 | 0 | 0,35 | 0 | 1 | 3,339 | 1,739 | 0,742 | 0,4578918 | 0,4943837 | 1,00E-08 | 4,7145231 | 3,6246089 | 3,6246089 | TRUE | 0,1890691 | No |
| tE(UUC)5 | 0 | 0 | 0 | 0 | 0 | 0 | 1 | 0 | 0 | 0 | 0 | 0 | 0 | 0 | 1 | 0 | 0,11 | 0 | 0 | 1,314 | 0,394 | 0,111 | 0,9112982 | 0,9205309 | 1,00E-08 | 15,473589 | 10 | 10 | TRUE | 0,3207401 | No |
| tE(UUC)6 | 2 | 1 | 0 | 2 | 1 | 1 | 3 | 3 | 2 | 1 | 0 | 3 | 1 | 1 | 3 | 2 | 1,63 | 2 | 2 | 1,225 | 0,293 | 0,259 | 0,7953055 | 0,8177407 | 1,00E-08 | 1,0107943 | 0,5826372 | 0,5826372 | TRUE | 0,2728953 | No |
| tE(UUC)7 | 7 | 7 | 4 | 1 | 10 | 13 | 10 | 8 | 7 | 7 | 4 | 2 | 8 | 15 | 8 | 7 | 7,17 | 5 | 10 | 1,817 | 0,862 | 1,521 | 0,1281827 | 0,1480424 | 0,0228735 | 0,2309953 | 0,1510645 | 0,1510645 | TRUE | 0,4090857 | No |
| tF(GAA)1 | 3 | 3 | 2 | 3 | 15 | 3 | 6 | 12 | 3 | 3 | 2 | 5 | 12 | 3 | 5 | 10 | 5,37 | 3 | 8 | 2,514 | 1,33 | 1,949 | 0,0512697 | 0,0616221 | 1,00E-08 | 0,3076383 | 0,2083566 | 0,2083566 | TRUE | 0,1206188 | No |
| tF(GAA)2 | 0 | 0 | 1 | 0 | 0 | 0 | 0 | 0 | 0 | 0 | 1 | 0 | 0 | 0 | 0 | 0 | 0,12 | 0 | 0 | 0,483 | -1,049 | -0,297 | 0,7665832 | 0,7911318 | 1,00E-08 | 13,298132 | 10 | 10 | TRUE | 0,2757288 | No |
| tF(GAA)3 | 1 | 2 | 0 | 0 | 0 | 0 | 0 | 0 | 1 | 2 | 0 | 0 | 0 | 0 | 0 | 0 | 0,38 | 1 | 0 | 0,209 | -2,258 | -0,857 | 0,3913348 | 0,427148 | 1,00E-08 | 4,2832625 | 4,9460942 | 4,9460942 | TRUE | 0,4068721 | No |
| tF(GAA)4 | 0 | 0 | 0 | 0 | 0 | 1 | 0 | 0 | 0 | 0 | 0 | 0 | 0 | 1 | 0 | 0 | 0,14 | 0 | 0 | 1,314 | 0,394 | 0,111 | 0,9112982 | 0,9205309 | 1,00E-08 | 11,553708 | 10 | 10 | TRUE | 0,5388425 | No |
| tF(GAA)5 | 0 | 0 | 0 | 0 | 1 | 0 | 0 | 0 | 0 | 0 | 0 | 0 | 1 | 0 | 0 | 0 | 0,1 | 0 | 0 | 1,314 | 0,394 | 0,111 | 0,9112982 | 0,9205309 | 1,00E-08 | 16,395074 | 10 | 10 | TRUE | 0,2859086 | No |
| tG(CCC)1 | 2 | 1 | 0 | 0 | 0 | 0 | 3 | 0 | 2 | 1 | 0 | 0 | 0 | 0 | 3 | 0 | 0,69 | 1 | 1 | 0,836 | -0,258 | -0,126 | 0,8999303 | 0,9139671 | 3,5583326 | 2,3854858 | 2,6419241 | 2,6419241 | TRUE | 1,4555722 | No |
| tG(GCC)1 | 0 | 0 | 0 | 0 | 0 | 0 | 0 | 0 | 0 | 0 | 0 | 0 | 0 | 0 | 0 | 0 | 0 | 0 | 0 | NA | NA | NA | NA | NA | NA | NA | NA | NA | NA | NA | NA |
| tG(GCC)2 | 0 | 0 | 0 | 1 | 0 | 0 | 0 | 0 | 0 | 0 | 0 | 2 | 0 | 0 | 0 | 0 | 0,2 | 0 | 0 | 0,483 | -1,049 | -0,297 | 0,7665832 | 0,7911318 | 1,00E-08 | 8,3868857 | 10 | 10 | TRUE | 0,6253243 | No |
| tG(GCC)3 | 0 | 0 | 0 | 0 | 0 | 0 | 0 | 0 | 0 | 0 | 0 | 0 | 0 | 0 | 0 | 0 | 0 | 0 | 0 | NA | NA | NA | NA | NA | NA | NA | NA | NA | NA | NA | NA |
| tG(GCC)4 | 0 | 1 | 2 | 0 | 0 | 0 | 0 | 0 | 0 | 1 | 2 | 0 | 0 | 0 | 0 | 0 | 0,38 | 1 | 0 | 0,211 | -2,247 | -0,851 | 0,3950081 | 0,4307891 | 1,00E-08 | 4,3325994 | 4,9805062 | 4,9805062 | TRUE | 0,2460318 | No |
| tG(GCC)5 | 0 | 0 | 1 | 0 | 0 | 0 | 0 | 0 | 0 | 0 | 1 | 0 | 0 | 0 | 0 | 0 | 0,12 | 0 | 0 | 0,483 | -1,049 | -0,297 | 0,7665832 | 0,7911318 | 1,00E-08 | 13,298132 | 10 | 10 | TRUE | 0,2757288 | No |
| tG(GCC)6 | 3 | 0 | 0 | 0 | 3 | 2 | 1 | 3 | 3 | 0 | 0 | 0 | 2 | 2 | 1 | 2 | 1,35 | 1 | 2 | 2,535 | 1,342 | 0,98 | 0,3271686 | 0,3608128 | 1,00E-08 | 1,2153446 | 0,8684553 | 0,8684553 | TRUE | 1,4521852 | No |
| tG(UCC)1 | 0 | 0 | 0 | 0 | 0 | 0 | 0 | 0 | 0 | 0 | 0 | 0 | 0 | 0 | 0 | 0 | 0 | 0 | 0 | NA | NA | NA | NA | NA | NA | NA | NA | NA | NA | NA | NA |
| tG(UCC)2 | 0 | 2 | 0 | 0 | 0 | 0 | 0 | 0 | 0 | 2 | 0 | 0 | 0 | 0 | 0 | 0 | 0,26 | 0 | 0 | 0,287 | -1,802 | -0,512 | 0,6086527 | 0,6425898 | 10 | 6,2133439 | 10 | 10 | TRUE | 0,8530664 | No |
| tH(GUG)1 | 2 | 0 | 0 | 0 | 1 | 1 | 0 | 1 | 2 | 0 | 0 | 0 | 1 | 1 | 0 | 1 | 0,58 | 0 | 1 | 1,25 | 0,322 | 0,166 | 0,8678668 | 0,8852075 | 1,00E-08 | 2,8218743 | 1,9899716 | 1,9899716 | TRUE | 0,8296956 | No |
| tH(GUG)3 | 10 | 4 | 9 | 2 | 14 | 10 | 9 | 10 | 10 | 4 | 9 | 3 | 11 | 11 | 8 | 8 | 8,02 | 6 | 10 | 1,423 | 0,509 | 0,995 | 0,3198688 | 0,3530674 | 1,00E-08 | 0,2068548 | 0,1220552 | 0,1220552 | TRUE | 0,2056924 | No |
| tI(AAU)1 | 0 | 0 | 0 | 0 | 0 | 0 | 0 | 0 | 0 | 0 | 0 | 0 | 0 | 0 | 0 | 0 | 0 | 0 | 0 | NA | NA | NA | NA | NA | NA | NA | NA | NA | NA | NA | NA |
| tI(AAU)2 | 0 | 0 | 0 | 0 | 1 | 0 | 0 | 0 | 0 | 0 | 0 | 0 | 1 | 0 | 0 | 0 | 0,1 | 0 | 0 | 1,314 | 0,394 | 0,111 | 0,9112982 | 0,9205309 | 1,00E-08 | 16,395074 | 10 | 10 | TRUE | 0,2859086 | No |
| tI(AAU)3 | 0 | 0 | 0 | 0 | 8 | 3 | 2 | 2 | 0 | 0 | 0 | 0 | 6 | 3 | 2 | 2 | 1,64 | 0 | 3 | 16,007 | 4,001 | 2,831 | 0,0046342 | 0,0059981 | 0,3447115 | 1,0013006 | 0,7824184 | 0,7824184 | TRUE | 0,7375252 | No |
| tI(AAU)4 | 0 | 0 | 0 | 0 | 0 | 0 | 0 | 0 | 0 | 0 | 0 | 0 | 0 | 0 | 0 | 0 | 0 | 0 | 0 | NA | NA | NA | NA | NA | NA | NA | NA | NA | NA | NA | NA |
| tI(AAU)5 | 0 | 0 | 0 | 0 | 0 | 0 | 1 | 0 | 0 | 0 | 0 | 0 | 0 | 0 | 1 | 0 | 0,11 | 0 | 0 | 1,314 | 0,394 | 0,111 | 0,9112982 | 0,9205309 | 1,00E-08 | 15,473589 | 10 | 10 | TRUE | 0,3207401 | No |
| tI(UAU)1 | 0 | 0 | 1 | 0 | 0 | 0 | 0 | 1 | 0 | 0 | 1 | 0 | 0 | 0 | 0 | 1 | 0,23 | 0 | 0 | 0,797 | -0,328 | -0,104 | 0,9167894 | 0,924614 | 1,00E-08 | 7,2953851 | 7,451208 | 7,451208 | TRUE | 0,2944896 | No |
| tK(CUU)1 | 3 | 6 | 2 | 3 | 2 | 3 | 2 | 5 | 3 | 6 | 2 | 5 | 2 | 3 | 2 | 4 | 3,33 | 4 | 3 | 0,683 | -0,55 | -0,702 | 0,4828994 | 0,5192755 | 1,00E-08 | 0,4948378 | 0,2803064 | 0,2803064 | TRUE | 0,289692 | No |
| tK(CUU)2 | 1 | 1 | 5 | 3 | 8 | 3 | 2 | 4 | 1 | 1 | 5 | 5 | 6 | 3 | 2 | 3 | 3,3 | 3 | 4 | 1,321 | 0,402 | 0,48 | 0,6309181 | 0,6636809 | 0,1513754 | 0,4992279 | 0,3530322 | 0,3530322 | TRUE | 0,2453545 | No |
| tK(UUU)1 | 0 | 0 | 0 | 0 | 0 | 1 | 0 | 2 | 0 | 0 | 0 | 0 | 0 | 1 | 0 | 2 | 0,35 | 0 | 1 | 3,329 | 1,735 | 0,632 | 0,527086 | 0,5633729 | 1,00E-08 | 4,7555809 | 5,5760801 | 5,5760801 | TRUE | 0,1301645 | No |
| tK(UUU)2 | 0 | 0 | 0 | 0 | 0 | 0 | 0 | 0 | 0 | 0 | 0 | 0 | 0 | 0 | 0 | 0 | 0 | 0 | 0 | NA | NA | NA | NA | NA | NA | NA | NA | NA | NA | NA | NA |
| tK(UUU)3 | 1 | 0 | 0 | 0 | 0 | 0 | 1 | 0 | 1 | 0 | 0 | 0 | 0 | 0 | 1 | 0 | 0,23 | 0 | 0 | 0,797 | -0,328 | -0,105 | 0,9167686 | 0,924614 | 1,00E-08 | 7,2897753 | 7,4464653 | 7,4464653 | TRUE | 0,320739 | No |
| tK(UUU)4 | 0 | 0 | 0 | 0 | 0 | 0 | 1 | 0 | 0 | 0 | 0 | 0 | 0 | 0 | 1 | 0 | 0,11 | 0 | 0 | 1,314 | 0,394 | 0,111 | 0,9112982 | 0,9205309 | 1,00E-08 | 15,473589 | 10 | 10 | TRUE | 0,3207401 | No |
| tK(UUU)5 | 0 | 0 | 0 | 0 | 1 | 0 | 1 | 4 | 0 | 0 | 0 | 0 | 1 | 0 | 1 | 3 | 0,61 | 0 | 1 | 6,079 | 2,604 | 1,294 | 0,195583 | 0,2219483 | 2,0323418 | 2,6813017 | 2,5313523 | 2,5313523 | TRUE | 0,8362634 | No |
| tL(AAG)1 | 3 | 3 | 2 | 4 | 3 | 1 | 3 | 4 | 3 | 3 | 2 | 6 | 2 | 1 | 3 | 3 | 2,95 | 4 | 2 | 0,702 | -0,511 | -0,628 | 0,5299093 | 0,5658218 | 1,00E-08 | 0,5583248 | 0,2853704 | 0,2853704 | TRUE | 0,2299279 | No |
| tL(AAG)2 | 0 | 0 | 1 | 1 | 1 | 0 | 0 | 1 | 0 | 0 | 1 | 2 | 1 | 0 | 0 | 1 | 0,52 | 1 | 0 | 0,773 | -0,371 | -0,184 | 0,853914 | 0,8723703 | 1,00E-08 | 3,1529135 | 2,0361027 | 2,0361027 | TRUE | 0,2291554 | No |
| tL(CAA)1 | 0 | 0 | 2 | 0 | 0 | 0 | 0 | 0 | 0 | 0 | 2 | 0 | 0 | 0 | 0 | 0 | 0,25 | 0 | 0 | 0,3 | -1,735 | -0,492 | 0,6224706 | 0,655876 | 1,00E-08 | 6,6501114 | 10 | 10 | TRUE | 0,8329296 | No |
| tL(CAA)2 | 0 | 0 | 1 | 0 | 0 | 0 | 0 | 0 | 0 | 0 | 1 | 0 | 0 | 0 | 0 | 0 | 0,12 | 0 | 0 | 0,483 | -1,049 | -0,297 | 0,7665832 | 0,7911318 | 1,00E-08 | 13,298132 | 10 | 10 | TRUE | 0,2757288 | No |
| tL(CAA)3 | 0 | 1 | 0 | 0 | 0 | 0 | 1 | 1 | 0 | 1 | 0 | 0 | 0 | 0 | 1 | 1 | 0,34 | 0 | 0 | 1,309 | 0,388 | 0,161 | 0,8723241 | 0,8891853 | 1,00E-08 | 4,8321704 | 3,6556742 | 3,6556742 | TRUE | 0,3160008 | No |
| tL(CAA)4 | 71 | 48 | 74 | 40 | 50 | 40 | 50 | 70 | 68 | 51 | 73 | 63 | 40 | 45 | 42 | 57 | 54,89 | 64 | 46 | 0,726 | -0,462 | -2,251 | 0,0243653 | 0,0300609 | 1,00E-08 | 0,0320035 | 0,0219779 | 0,0219779 | TRUE | 0,2194147 | No |
| tL(CAA)5 | 0 | 0 | 0 | 0 | 0 | 0 | 0 | 0 | 0 | 0 | 0 | 0 | 0 | 0 | 0 | 0 | 0 | 0 | 0 | NA | NA | NA | NA | NA | NA | NA | NA | NA | NA | NA | NA |
| tL(CAA)6 | 2 | 0 | 1 | 0 | 0 | 0 | 0 | 0 | 2 | 0 | 1 | 0 | 0 | 0 | 0 | 0 | 0,36 | 1 | 0 | 0,217 | -2,201 | -0,824 | 0,4102084 | 0,4462059 | 1,00E-08 | 4,5397905 | 5,1233955 | 5,1233955 | TRUE | 0,2800721 | No |
| tL(UAA)1 | 0 | 0 | 0 | 0 | 0 | 0 | 0 | 0 | 0 | 0 | 0 | 0 | 0 | 0 | 0 | 0 | 0 | 0 | 0 | NA | NA | NA | NA | NA | NA | NA | NA | NA | NA | NA | NA |
| tL(UAA)2 | 0 | 0 | 1 | 0 | 0 | 0 | 0 | 0 | 0 | 0 | 1 | 0 | 0 | 0 | 0 | 0 | 0,12 | 0 | 0 | 0,483 | -1,049 | -0,297 | 0,7665832 | 0,7911318 | 1,00E-08 | 13,298132 | 10 | 10 | TRUE | 0,2757288 | No |
| tL(UAA)3 | 1 | 0 | 1 | 0 | 1 | 0 | 0 | 0 | 1 | 0 | 1 | 0 | 1 | 0 | 0 | 0 | 0,34 | 0 | 0 | 0,498 | -1,006 | -0,418 | 0,6759875 | 0,7078234 | 1,00E-08 | 4,7914805 | 3,62201 | 3,62201 | TRUE | 0,2357743 | No |
| tL(UAA)4 | 1 | 0 | 0 | 1 | 1 | 1 | 0 | 0 | 1 | 0 | 0 | 2 | 1 | 1 | 0 | 0 | 0,56 | 1 | 0 | 0,798 | -0,325 | -0,163 | 0,8703823 | 0,8874896 | 1,00E-08 | 2,9483283 | 1,9590969 | 1,9590969 | TRUE | 0,2354105 | No |
| tL(UAA)5 | 0 | 0 | 0 | 0 | 0 | 0 | 0 | 0 | 0 | 0 | 0 | 0 | 0 | 0 | 0 | 0 | 0 | 0 | 0 | NA | NA | NA | NA | NA | NA | NA | NA | NA | NA | NA | NA |
| tM(CAU)1 | 0 | 0 | 0 | 0 | 0 | 0 | 0 | 0 | 0 | 0 | 0 | 0 | 0 | 0 | 0 | 0 | 0 | 0 | 0 | NA | NA | NA | NA | NA | NA | NA | NA | NA | NA | NA | NA |
| tM(CAU)2 | 1 | 1 | 0 | 0 | 0 | 0 | 0 | 0 | 1 | 1 | 0 | 0 | 0 | 0 | 0 | 0 | 0,25 | 0 | 0 | 0,297 | -1,753 | -0,582 | 0,560743 | 0,595457 | 1,00E-08 | 6,5345431 | 6,7949003 | 6,7949003 | TRUE | 0,050013 | No |
| tM(CAU)3 | 0 | 0 | 0 | 1 | 1 | 0 | 1 | 0 | 0 | 0 | 0 | 2 | 1 | 0 | 1 | 0 | 0,4 | 0 | 0 | 1,304 | 0,382 | 0,165 | 0,8686233 | 0,8858375 | 1,00E-08 | 4,0853872 | 3,1912868 | 3,1912868 | TRUE | 0,6253303 | No |
| tM(CAU)4 | 0 | 0 | 0 | 0 | 0 | 0 | 0 | 0 | 0 | 0 | 0 | 0 | 0 | 0 | 0 | 0 | 0 | 0 | 0 | NA | NA | NA | NA | NA | NA | NA | NA | NA | NA | NA | NA |
| tN(GUU)1 | 2 | 0 | 0 | 0 | 0 | 0 | 0 | 1 | 2 | 0 | 0 | 0 | 0 | 0 | 0 | 1 | 0,34 | 0 | 0 | 0,505 | -0,985 | -0,351 | 0,7253273 | 0,7546574 | 1,00E-08 | 4,8314335 | 5,6068172 | 5,6068172 | TRUE | 0,8242672 | No |
| tN(GUU)2 | 0 | 0 | 0 | 0 | 0 | 0 | 1 | 0 | 0 | 0 | 0 | 0 | 0 | 0 | 1 | 0 | 0,11 | 0 | 0 | 1,314 | 0,394 | 0,111 | 0,9112982 | 0,9205309 | 1,00E-08 | 15,473589 | 10 | 10 | TRUE | 0,3207401 | No |
| tN(GUU)3 | 0 | 1 | 0 | 0 | 1 | 1 | 0 | 1 | 0 | 1 | 0 | 0 | 1 | 1 | 0 | 1 | 0,48 | 0 | 1 | 1,995 | 0,996 | 0,502 | 0,6158651 | 0,6495596 | 1,00E-08 | 3,4507163 | 2,130333 | 2,130333 | TRUE | 0,3160017 | No |
| tN(GUU)4 | 0 | 1 | 0 | 0 | 1 | 0 | 0 | 0 | 0 | 1 | 0 | 0 | 1 | 0 | 0 | 0 | 0,23 | 0 | 0 | 0,797 | -0,328 | -0,106 | 0,9159347 | 0,9244826 | 1,00E-08 | 7,0691906 | 7,259296 | 7,259296 | TRUE | 0,3159987 | No |
| tP(AGG)1 | 1 | 0 | 2 | 1 | 3 | 3 | 5 | 6 | 1 | 0 | 2 | 2 | 2 | 3 | 4 | 5 | 2,43 | 1 | 4 | 3,396 | 1,764 | 1,752 | 0,0797502 | 0,0940639 | 1,00E-08 | 0,6779668 | 0,3521866 | 0,3521866 | TRUE | 0,2352574 | No |
| tP(UGG)1 | 0 | 0 | 1 | 0 | 0 | 2 | 0 | 1 | 0 | 0 | 1 | 0 | 0 | 2 | 0 | 1 | 0,51 | 0 | 1 | 2,172 | 1,119 | 0,51 | 0,6100253 | 0,6438259 | 1,00E-08 | 3,2253171 | 3,0180088 | 3,0180088 | TRUE | 0,5414727 | No |
| tP(UGG)2 | 2 | 0 | 1 | 0 | 0 | 0 | 0 | 3 | 2 | 0 | 1 | 0 | 0 | 0 | 0 | 2 | 0,67 | 1 | 0 | 0,824 | -0,28 | -0,136 | 0,8919929 | 0,9070608 | 1,00E-08 | 2,46465 | 2,6472757 | 2,6472757 | TRUE | 1,4549369 | No |
| tP(UGG)3 | 4 | 4 | 2 | 3 | 3 | 5 | 3 | 6 | 4 | 4 | 2 | 5 | 2 | 6 | 3 | 5 | 3,78 | 4 | 4 | 1,055 | 0,077 | 0,107 | 0,9148574 | 0,9236875 | 1,00E-08 | 0,4365527 | 0,2283521 | 0,2283521 | TRUE | 0,1625345 | No |
| tP(UGG)4 | 0 | 0 | 0 | 0 | 0 | 0 | 0 | 0 | 0 | 0 | 0 | 0 | 0 | 0 | 0 | 0 | 0 | 0 | 0 | NA | NA | NA | NA | NA | NA | NA | NA | NA | NA | NA | NA |
| tP(UGG)5 | 0 | 0 | 0 | 1 | 0 | 0 | 0 | 0 | 0 | 0 | 0 | 2 | 0 | 0 | 0 | 0 | 0,2 | 0 | 0 | 0,483 | -1,049 | -0,297 | 0,7665832 | 0,7911318 | 1,00E-08 | 8,3868857 | 10 | 10 | TRUE | 0,6253243 | No |
| tQ(CUG)1 | 0 | 0 | 0 | 0 | 0 | 1 | 0 | 0 | 0 | 0 | 0 | 0 | 0 | 1 | 0 | 0 | 0,14 | 0 | 0 | 1,314 | 0,394 | 0,111 | 0,9112982 | 0,9205309 | 1,00E-08 | 11,553708 | 10 | 10 | TRUE | 0,5388425 | No |
| tQ(UUG)1 | 0 | 0 | 2 | 1 | 0 | 1 | 0 | 1 | 0 | 0 | 2 | 2 | 0 | 1 | 0 | 1 | 0,69 | 1 | 0 | 0,525 | -0,93 | -0,494 | 0,621391 | 0,655063 | 1,00E-08 | 2,3935541 | 1,8162196 | 1,8162196 | TRUE | 0,1960596 | No |
| tQ(UUG)3 | 0 | 2 | 0 | 1 | 1 | 0 | 0 | 0 | 0 | 2 | 0 | 2 | 1 | 0 | 0 | 0 | 0,56 | 1 | 0 | 0,304 | -1,716 | -0,804 | 0,4215334 | 0,4575059 | 1,00E-08 | 2,9323797 | 2,747433 | 2,747433 | TRUE | 0,1784741 | No |
| tQ(UUG)4 | 0 | 0 | 0 | 0 | 0 | 0 | 0 | 0 | 0 | 0 | 0 | 0 | 0 | 0 | 0 | 0 | 0 | 0 | 0 | NA | NA | NA | NA | NA | NA | NA | NA | NA | NA | NA | NA |
| tQ(UUG)5 | 0 | 1 | 1 | 0 | 0 | 0 | 0 | 0 | 0 | 1 | 1 | 0 | 0 | 0 | 0 | 0 | 0,26 | 0 | 0 | 0,293 | -1,769 | -0,59 | 0,555394 | 0,5901697 | 1,00E-08 | 6,4243102 | 6,7246415 | 6,7246415 | TRUE | 0,048134 | No |
| tR(ACG)1 | 0 | 0 | 1 | 0 | 0 | 0 | 1 | 0 | 0 | 0 | 1 | 0 | 0 | 0 | 1 | 0 | 0,23 | 0 | 0 | 0,797 | -0,328 | -0,105 | 0,9162546 | 0,924513 | 1,00E-08 | 7,1528476 | 7,3304376 | 7,3304376 | TRUE | 0,3207391 | No |

| | | | | | | | | | | | | | | | | | | | | | | | | | | | | | | | |
|---|---|---|---|---|---|---|---|---|---|---|---|---|---|---|---|---|---|---|---|---|---|---|---|---|---|---|---|---|---|---|---|
| tR(ACG)2 | 0 | 0 | 0 | 0 | 0 | 0 | 1 | 0 | 0 | 0 | 0 | 0 | 0 | 0 | 1 | 0 | 0,11 | 0 | 0 | 1,314 | 0,394 | 0,111 | 0,9112982 | 0,9205309 | 1,00E-08 | 15,473589 | 10 | 10 | TRUE | 0,3207401 | No |
| tR(CCG)1 | 6 | 6 | 8 | 9 | 22 | 11 | 17 | 25 | 6 | 6 | 8 | 14 | 18 | 13 | 14 | 20 | 12,37 | 8 | 16 | 2,011 | 1,008 | 2,344 | 0,019085 | 0,0237206 | 1,00E-08 | 0,1348198 | 0,0820625 | 0,0820625 | TRUE | 0,3584955 | No |
| tR(CCU)1 | 53 | 78 | 59 | 50 | 11 | 5 | 4 | 13 | 51 | 82 | 58 | 78 | 9 | 6 | 3 | 11 | 37,26 | 67 | 7 | 0,108 | -3,21 | -9,256 | 2,12E-20 | 4,81E-20 | 0,045364 | 0,0461567 | 0,0458939 | 0,0458939 | TRUE | 0,2256155 | No |
| tR(UCU)1 | 3 | 1 | 1 | 0 | 0 | 0 | 0 | 1 | 3 | 1 | 1 | 0 | 0 | 0 | 0 | 1 | 0,71 | 1 | 0 | 0,21 | -2,248 | -1,262 | 0,2067895 | 0,2342904 | 1,00E-08 | 2,2990421 | 1,6100781 | 1,6100781 | TRUE | 0,2946501 | No |
| tR(UCU)2 | 0 | 0 | 0 | 0 | 0 | 0 | 0 | 0 | 0 | 0 | 0 | 0 | 0 | 0 | 0 | 0 | 0 | 0 | 0 | NA | NA | NA | NA | NA | NA | NA | NA | NA | NA | NA | NA |
| tR(UCU)3 | 0 | 0 | 0 | 0 | 0 | 0 | 0 | 0 | 0 | 0 | 0 | 0 | 0 | 0 | 0 | 0 | 0 | 0 | 0 | NA | NA | NA | NA | NA | NA | NA | NA | NA | NA | NA | NA |
| tR(UCU)4 | 4 | 2 | 6 | 3 | 0 | 3 | 5 | 3 | 4 | 2 | 6 | 5 | 0 | 3 | 4 | 2 | 3,33 | 4 | 2 | 0,6 | -0,738 | -0,885 | 0,3759864 | 0,4109215 | 1,00E-08 | 0,494953 | 0,3522269 | 0,3522269 | TRUE | 0,6662333 | No |
| tR(UCU)5 | 0 | 0 | 0 | 0 | 0 | 0 | 0 | 0 | 0 | 0 | 0 | 0 | 0 | 0 | 0 | 0 | 0 | 0 | 0 | NA | NA | NA | NA | NA | NA | NA | NA | NA | NA | NA | NA |
| tS(AGA)1 | 2 | 4 | 2 | 1 | 0 | 4 | 2 | 2 | 2 | 4 | 2 | 2 | 0 | 5 | 2 | 2 | 2,19 | 2 | 2 | 0,76 | -0,396 | -0,392 | 0,69531 | 0,7263116 | 1,00E-08 | 0,7504869 | 0,5134002 | 0,5134002 | TRUE | 0,5485751 | No |
| tS(AGA)2 | 0 | 0 | 0 | 0 | 0 | 0 | 0 | 1 | 0 | 0 | 0 | 0 | 0 | 0 | 0 | 1 | 0,1 | 0 | 0 | 1,314 | 0,394 | 0,111 | 0,9112982 | 0,9205309 | 1,00E-08 | 16,156685 | 10 | 10 | TRUE | 0,2944906 | No |
| tS(AGA)3 | 1 | 0 | 1 | 0 | 0 | 0 | 0 | 1 | 1 | 0 | 1 | 0 | 0 | 0 | 0 | 1 | 0,34 | 0 | 0 | 0,498 | -1,006 | -0,418 | 0,675636 | 0,7075716 | 1,00E-08 | 4,7709204 | 3,6096772 | 3,6096772 | TRUE | 0,2446768 | No |
| tS(AGA)4 | 2 | 0 | 0 | 1 | 2 | 0 | 0 | 0 | 2 | 0 | 0 | 2 | 2 | 0 | 0 | 0 | 0,63 | 1 | 0 | 0,514 | -0,961 | -0,457 | 0,6474857 | 0,6796523 | 1,00E-08 | 2,5898649 | 2,6285235 | 2,6285235 | TRUE | 0,5288446 | No |
| tS(CAG)1 | 0 | 0 | 1 | 0 | 0 | 4 | 2 | 4 | 0 | 0 | 1 | 0 | 0 | 5 | 2 | 3 | 1,31 | 0 | 2 | 6,867 | 2,78 | 1,776 | 0,0757627 | 0,0895429 | 1,00E-08 | 1,2546594 | 1,1621269 | 1,1621269 | TRUE | 0,1695532 | No |
| tS(CGA)1 | 0 | 0 | 1 | 0 | 0 | 2 | 1 | 0 | 0 | 0 | 1 | 0 | 0 | 2 | 1 | 0 | 0,51 | 0 | 1 | 2,192 | 1,132 | 0,517 | 0,6053513 | 0,639474 | 1,00E-08 | 3,1971705 | 3,0131217 | 3,0131217 | TRUE | 0,5319202 | No |
| tS(GCU)1 | 10 | 13 | 8 | 5 | 54 | 22 | 34 | 38 | 10 | 14 | 8 | 8 | 43 | 25 | 29 | 31 | 20,88 | 10 | 32 | 3,273 | 1,711 | 4,735 | 2,19E-06 | 3,34E-06 | 1,00E-08 | 0,0807153 | 0,055241 | 0,055241 | TRUE | 0,4256657 | No |
| tS(GCU)2 | 8 | 6 | 5 | 7 | 27 | 10 | 27 | 35 | 8 | 6 | 5 | 11 | 22 | 11 | 23 | 28 | 14,28 | 8 | 21 | 2,952 | 1,562 | 3,55 | 0,000385 | 0,0005306 | 1,00E-08 | 0,1170472 | 0,0882374 | 0,0882374 | TRUE | 0,2661603 | No |
| tS(UGA)1 | 0 | 0 | 0 | 0 | 1 | 0 | 0 | 0 | 0 | 0 | 0 | 0 | 1 | 0 | 0 | 0 | 0,1 | 0 | 0 | 1,314 | 0,394 | 0,111 | 0,9112982 | 0,9205309 | 1,00E-08 | 16,395074 | 10 | 10 | TRUE | 0,2859086 | No |
| tS(UGA)2 | 0 | 0 | 1 | 0 | 0 | 0 | 0 | 0 | 0 | 0 | 1 | 0 | 0 | 0 | 0 | 0 | 0,12 | 0 | 0 | 0,483 | -1,049 | -0,297 | 0,7665832 | 0,7911318 | 1,00E-08 | 13,298132 | 10 | 10 | TRUE | 0,2757288 | No |
| tS(UGA)3 | 0 | 0 | 0 | 0 | 0 | 0 | 0 | 0 | 0 | 0 | 0 | 0 | 0 | 0 | 0 | 0 | 0 | 0 | 0 | NA | NA | NA | NA | NA | NA | NA | NA | NA | NA | NA | NA |
| tT(AGU)1 | 0 | 0 | 0 | 0 | 0 | 0 | 0 | 0 | 0 | 0 | 0 | 0 | 0 | 0 | 0 | 0 | 0 | 0 | 0 | NA | NA | NA | NA | NA | NA | NA | NA | NA | NA | NA | NA |
| tT(AGU)3 | 0 | 0 | 0 | 0 | 0 | 0 | 1 | 0 | 0 | 0 | 0 | 0 | 0 | 0 | 1 | 0 | 0,11 | 0 | 0 | 1,314 | 0,394 | 0,111 | 0,9112982 | 0,9205309 | 1,00E-08 | 15,473589 | 10 | 10 | TRUE | 0,3207401 | No |
| tT(AGU)4 | 0 | 0 | 0 | 0 | 0 | 0 | 0 | 0 | 0 | 0 | 0 | 0 | 0 | 0 | 0 | 0 | 0 | 0 | 0 | NA | NA | NA | NA | NA | NA | NA | NA | NA | NA | NA | NA |
| tT(AGU)6 | 1 | 0 | 0 | 0 | 0 | 0 | 0 | 0 | 1 | 0 | 0 | 0 | 0 | 0 | 0 | 0 | 0,12 | 0 | 0 | 0,483 | -1,049 | -0,297 | 0,7665832 | 0,7911318 | 1,00E-08 | 13,779457 | 10 | 10 | TRUE | 0,2560039 | No |
| tT(CGU)1 | 0 | 0 | 0 | 0 | 0 | 0 | 0 | 0 | 0 | 0 | 0 | 0 | 0 | 0 | 0 | 0 | 0 | 0 | 0 | NA | NA | NA | NA | NA | NA | NA | NA | NA | NA | NA | NA |
| tT(UGU)1 | 0 | 0 | 0 | 0 | 0 | 0 | 0 | 0 | 0 | 0 | 0 | 0 | 0 | 0 | 0 | 0 | 0 | 0 | 0 | NA | NA | NA | NA | NA | NA | NA | NA | NA | NA | NA | NA |
| tT(UGU)2 | 0 | 0 | 0 | 0 | 0 | 0 | 0 | 1 | 0 | 0 | 0 | 0 | 0 | 0 | 0 | 1 | 0,1 | 0 | 0 | 1,314 | 0,394 | 0,111 | 0,9112982 | 0,9205309 | 1,00E-08 | 16,156685 | 10 | 10 | TRUE | 0,2944906 | No |
| tV(AAC)1 | 1 | 0 | 0 | 0 | 0 | 0 | 0 | 0 | 1 | 0 | 0 | 0 | 0 | 0 | 0 | 0 | 0,12 | 0 | 0 | 0,483 | -1,049 | -0,297 | 0,7665832 | 0,7911318 | 1,00E-08 | 13,779457 | 10 | 10 | TRUE | 0,2560039 | No |
| tV(AAC)2 | 0 | 0 | 0 | 0 | 0 | 0 | 0 | 0 | 0 | 0 | 0 | 0 | 0 | 0 | 0 | 0 | 0 | 0 | 0 | NA | NA | NA | NA | NA | NA | NA | NA | NA | NA | NA | NA |
| tV(AAC)3 | 0 | 0 | 0 | 0 | 0 | 0 | 0 | 1 | 0 | 0 | 0 | 0 | 0 | 0 | 0 | 1 | 0,1 | 0 | 0 | 1,314 | 0,394 | 0,111 | 0,9112982 | 0,9205309 | 1,00E-08 | 16,156685 | 10 | 10 | TRUE | 0,2944906 | No |
| tV(AAC)4 | 0 | 0 | 0 | 0 | 0 | 1 | 0 | 0 | 0 | 0 | 0 | 0 | 0 | 1 | 0 | 0 | 0,14 | 0 | 0 | 1,314 | 0,394 | 0,111 | 0,9112982 | 0,9205309 | 1,00E-08 | 11,553708 | 10 | 10 | TRUE | 0,5388425 | No |
| tV(AAC)5 | 2 | 1 | 0 | 1 | 0 | 1 | 1 | 0 | 2 | 1 | 0 | 2 | 0 | 1 | 1 | 0 | 0,81 | 1 | 0 | 0,406 | -1,301 | -0,78 | 0,4352326 | 0,4712743 | 1,00E-08 | 2,0176873 | 1,2388696 | 1,2388696 | TRUE | 0,2626319 | No |
| tV(AAC)6 | 1 | 2 | 0 | 0 | 0 | 0 | 1 | 0 | 1 | 2 | 0 | 0 | 0 | 0 | 1 | 0 | 0,49 | 1 | 0 | 0,342 | -1,549 | -0,71 | 0,4775331 | 0,5138514 | 1,00E-08 | 3,3553663 | 2,8604469 | 2,8604469 | TRUE | 0,4682425 | No |
| tV(CAC)1 | 1 | 0 | 2 | 1 | 0 | 0 | 0 | 0 | 1 | 0 | 2 | 2 | 0 | 0 | 0 | 0 | 0,56 | 1 | 0 | 0,147 | -2,765 | -1,353 | 0,1761186 | 0,2007167 | 1,00E-08 | 2,9237865 | 2,5087153 | 2,5087153 | TRUE | 0,2313726 | No |
| tV(UAC)1 | 8 | 9 | 9 | 10 | 56 | 40 | 46 | 43 | 8 | 10 | 9 | 16 | 45 | 45 | 39 | 35 | 25,76 | 11 | 41 | 4,1 | 2,035 | 6,184 | 6,25E-10 | 1,08E-09 | 1,00E-08 | 0,0658323 | 0,0378365 | 0,0378365 | TRUE | 0,2386802 | No |
| tW(CCA)1 | 0 | 0 | 1 | 1 | 0 | 0 | 0 | 0 | 0 | 0 | 1 | 2 | 0 | 0 | 0 | 0 | 0,32 | 1 | 0 | 0,286 | -1,805 | -0,642 | 0,5211041 | 0,5575394 | 1,00E-08 | 5,1441702 | 5,7036175 | 5,7036175 | TRUE | 0,1583835 | No |
| tW(CCA)2 | 1 | 0 | 0 | 0 | 1 | 0 | 1 | 1 | 1 | 0 | 0 | 0 | 1 | 0 | 1 | 1 | 0,43 | 0 | 1 | 1,875 | 0,907 | 0,449 | 0,6535285 | 0,6857697 | 1,00E-08 | 3,8464499 | 2,2187566 | 2,2187566 | TRUE | 0,2560066 | No |
| tY(GUA)1 | 0 | 1 | 0 | 0 | 0 | 1 | 1 | 2 | 0 | 1 | 0 | 0 | 0 | 1 | 1 | 2 | 0,58 | 0 | 1 | 2,641 | 1,401 | 0,747 | 0,4550599 | 0,4918254 | 1,00E-08 | 2,8149848 | 1,8823243 | 1,8823243 | TRUE | 0,3160016 | No |
| tY(GUA)2 | 1 | 0 | 0 | 0 | 0 | 0 | 0 | 1 | 1 | 0 | 0 | 0 | 0 | 0 | 0 | 1 | 0,22 | 0 | 0 | 0,797 | -0,328 | -0,104 | 0,9173116 | 0,9248483 | 1,00E-08 | 7,4378804 | 7,5713978 | 7,5713978 | TRUE | 0,2944896 | No |
| tY(GUA)3 | 8 | 4 | 5 | 5 | 30 | 12 | 14 | 12 | 8 | 4 | 5 | 8 | 24 | 14 | 12 | 10 | 10,49 | 6 | 15 | 2,452 | 1,294 | 2,587 | 0,0096785 | 0,0123099 | 1,00E-08 | 0,1585464 | 0,1187347 | 0,1187347 | TRUE | 0,9496109 | No |
| tY(GUA)4 | 0 | 0 | 0 | 0 | 0 | 0 | 0 | 0 | 0 | 0 | 0 | 0 | 0 | 0 | 0 | 0 | 0 | 0 | 0 | NA | NA | NA | NA | NA | NA | NA | NA | NA | NA | NA | NA |

$P_{TET}$-*ZCF15* doxycyclin-treated up-regulated genes

| | WT dox | | | | ZCF15 | | | | norm. WT dox | | | | norm ZCF15 | | | | | | | | | | | | | | | | | | |
|---|---|---|---|---|---|---|---|---|---|---|---|---|---|---|---|---|---|---|---|---|---|---|---|---|---|---|---|---|---|---|---|
| Gene ID | 1 | 2 | 3 | 4 | 1 | 2 | 3 | 4 | 1 | 2 | 3 | 4 | 1 | 2 | 3 | 4 | baseMean | WT | ZCF15 | FC | log2FC | stat | pvalue | padj | dispGeneEst | dispFit | dispMAP | dispersion | betaConv | maxCooks | outlier |
| orf19.935 | 24 | 12 | 15 | 24 | 1836 | 1692 | 1620 | 2299 | 22 | 12 | 14 | 36 | 1633 | 1837 | 1659 | 1754 | 870,89 | 21 | 1721 | 87,11 | 6,445 | 36,304 | 1,38E-288 | 3,41E-287 | 0,0029539 | 0,0035539 | 0,0033735 | 0,0033735 | TRUE | 0,775043 | No |
| orf19.4571 | 86 | 65 | 82 | 42 | 5807 | 5012 | 5341 | 7123 | 78 | 66 | 78 | 63 | 5165 | 5442 | 5470 | 5433 | 2724,36 | 71 | 5378 | 74,571 | 6,221 | 64,481 | 0 | 0 | 1,00E-08 | 0,0025219 | 0,0015517 | 0,0015517 | TRUE | 0,0390507 | No |
| orf19.4569 | 66 | 48 | 54 | 36 | 3048 | 2634 | 2511 | 3506 | 60 | 49 | 51 | 54 | 2711 | 2860 | 2572 | 2674 | 1378,84 | 54 | 2704 | 50,5 | 5,658 | 50,004 | 0 | 0 | 1,00E-08 | 0,0029951 | 0,0022902 | 0,0022902 | TRUE | 0,0730836 | No |
| orf19.7288 | 616 | 507 | 560 | 389 | 29490 | 24463 | 25914 | 36007 | 561 | 513 | 530 | 584 | 26230 | 26560 | 26539 | 27466 | 13623 | 547 | 26699 | 49,006 | 5,615 | 117,856 | 0 | 0 | 0,0003905 | 0,002134 | 0,0011871 | 0,0011871 | TRUE | 0,0193757 | No |
| orf19.5449 | 60 | 51 | 54 | 44 | 2867 | 2275 | 2342 | 3313 | 55 | 52 | 51 | 66 | 2550 | 2470 | 2399 | 2527 | 1271,15 | 56 | 2486 | 45,282 | 5,501 | 50,058 | 0 | 0 | 1,00E-08 | 0,0030762 | 0,0018305 | 0,0018305 | TRUE | 0,0820559 | No |
| orf19.535 | 58 | 38 | 39 | 18 | 1348 | 1096 | 1101 | 1710 | 53 | 38 | 37 | 27 | 1199 | 1190 | 1128 | 1304 | 622,01 | 39 | 1205 | 30,153 | 4,914 | 36,265 | 5,78E-288 | 1,42E-286 | 0,0038782 | 0,0041608 | 0,0040755 | 0,0040755 | TRUE | 0,4504749 | No |
| orf19.3711 | 59 | 50 | 61 | 29 | 1540 | 1306 | 1353 | 1862 | 54 | 51 | 58 | 44 | 1370 | 1418 | 1386 | 1420 | 724,91 | 52 | 1398 | 26,804 | 4,744 | 41,618 | 0 | 0 | 1,00E-08 | 0,0038593 | 0,00206 | 0,00206 | TRUE | 0,0531348 | No |
| orf19.5728 | 269 | 262 | 243 | 173 | 6816 | 5446 | 5756 | 7510 | 245 | 265 | 230 | 260 | 6063 | 5913 | 5895 | 5729 | 3074,85 | 250 | 5900 | 23,701 | 4,567 | 74,484 | 0 | 0 | 1,00E-08 | 0,0024666 | 0,0014057 | 0,0014057 | TRUE | 0,0337691 | No |
| orf19.4504 | 510 | 443 | 489 | 273 | 11643 | 8656 | 10024 | 13371 | 464 | 448 | 463 | 410 | 10356 | 9398 | 10266 | 10200 | 5250,59 | 446 | 10055 | 22,414 | 4,486 | 78,199 | 0 | 0 | 0,0016198 | 0,0022886 | 0,0019272 | 0,0019272 | TRUE | 0,0274543 | No |
| orf19.6445 | 1076 | 850 | 960 | 673 | 22443 | 16400 | 18367 | 26464 | 980 | 860 | 909 | 1011 | 19962 | 17806 | 18810 | 20187 | 10065,51 | 940 | 19191 | 20,468 | 4,355 | 72,188 | 0 | 0 | 0,0035588 | 0,0021683 | 0,0028981 | 0,0028981 | TRUE | 0,0386224 | No |
| orf19.7156 | 1801 | 1614 | 1753 | 1117 | 34981 | 29255 | 31989 | 44016 | 1640 | 1632 | 1659 | 1678 | 31114 | 31763 | 32761 | 33576 | 16977,86 | 1652 | 32304 | 19,564 | 4,29 | 107,739 | 0 | 0 | 0,0007301 | 0,0021148 | 0,0011834 | 0,0011834 | TRUE | 0,0086598 | No |
| orf19.1985 | 643 | 584 | 680 | 414 | 13105 | 11448 | 10233 | 14367 | 585 | 591 | 644 | 622 | 11656 | 12429 | 10480 | 10959 | 5995,81 | 610 | 11381 | 18,659 | 4,222 | 65,29 | 0 | 0 | 0,0037748 | 0,0022573 | 0,0030952 | 0,0030952 | TRUE | 0,0467813 | No |
| orf19.359 | 28 | 18 | 31 | 22 | 570 | 435 | 431 | 644 | 25 | 18 | 29 | 33 | 507 | 472 | 441 | 491 | 252,25 | 26 | 478 | 18,382 | 4,2 | 25,28 | 5,31E-141 | 6,51E-140 | 0,0015631 | 0,0072738 | 0,0053043 | 0,0053043 | TRUE | 0,2709133 | No |
| orf19.5306 | 0 | 0 | 0 | 0 | 2 | 1 | 5 | 8 | 0 | 0 | 0 | 0 | 2 | 1 | 5 | 6 | 1,76 | 0 | 4 | 18,126 | 4,18 | 3,041 | 0,002356 | 0,0032328 | 0,4678609 | 0,7521909 | 0,6840427 | 0,6840427 | TRUE | 0,0568797 | No |
| orf19.3924 | 26 | 27 | 36 | 12 | 504 | 408 | 429 | 581 | 24 | 27 | 34 | 18 | 448 | 443 | 439 | 443 | 234,61 | 26 | 443 | 16,764 | 4,067 | 25,044 | 2,02E-138 | 2,44E-137 | 1,00E-08 | 0,0076676 | 0,0043809 | 0,0043809 | TRUE | 0,2863188 | No |
| orf19.1409.3 | 19 | 14 | 18 | 9 | 290 | 225 | 245 | 323 | 17 | 14 | 17 | 14 | 258 | 244 | 251 | 246 | 132,69 | 16 | 250 | 15,887 | 3,99 | 19,092 | 2,92E-81 | 2,11E-80 | 1,00E-08 | 0,0119923 | 0,0066214 | 0,0066214 | TRUE | 0,0282969 | No |
| orf19.272 | 2394 | 2254 | 2424 | 1579 | 37475 | 32988 | 32808 | 44075 | 2180 | 2280 | 2294 | 2372 | 33333 | 35816 | 33600 | 33621 | 18186,78 | 2282 | 34092 | 14,963 | 3,903 | 94,557 | 0 | 0 | 0,0008628 | 0,0021097 | 0,0013866 | 0,0013866 | TRUE | 0,0141466 | No |
| orf19.4934 | 25 | 21 | 12 | 14 | 327 | 229 | 270 | 392 | 23 | 21 | 11 | 21 | 291 | 249 | 277 | 299 | 148,93 | 19 | 279 | 14,768 | 3,884 | 19,511 | 8,80E-85 | 6,60E-84 | 0,0031718 | 0,0109072 | 0,0085406 | 0,0085406 | TRUE | 0,4626867 | No |
| orf19.1449 | 221 | 238 | 260 | 138 | 3832 | 3034 | 3026 | 4227 | 201 | 241 | 246 | 207 | 3408 | 3294 | 3099 | 3224 | 1740,15 | 224 | 3256 | 14,494 | 3,857 | 54,028 | 0 | 0 | 0,0017569 | 0,0027961 | 0,0023903 | 0,0023903 | TRUE | 0,0653596 | No |
| orf19.6838 | 1739 | 1501 | 1552 | 1057 | 25022 | 20622 | 20373 | 29763 | 1583 | 1518 | 1469 | 1588 | 22256 | 22390 | 20865 | 22703 | 11796,5 | 1540 | 22054 | 14,341 | 3,842 | 86,118 | 0 | 0 | 0,000901 | 0,002149 | 0,001541 | 0,001541 | TRUE | 0,0160973 | No |
| orf19.7585 | 1523 | 1271 | 1408 | 864 | 19585 | 16173 | 16443 | 23270 | 1387 | 1285 | 1333 | 1298 | 17420 | 17559 | 16840 | 17751 | 9359,05 | 1326 | 17392 | 13,103 | 3,712 | 92,183 | 0 | 0 | 0,0004613 | 0,0021782 | 0,001127 | 0,001127 | TRUE | 0,012104 | No |
| orf19.4211 | 1934 | 1835 | 1865 | 1212 | 25508 | 21802 | 22277 | 30863 | 1761 | 1856 | 1765 | 1821 | 22689 | 23671 | 22814 | 23543 | 12489,86 | 1801 | 23179 | 12,883 | 3,687 | 100,011 | 0 | 0 | 0,0003748 | 0,0021428 | 0,0009874 | 0,0009874 | TRUE | 0,0056423 | No |
| orf19.4741 | 18 | 13 | 16 | 7 | 193 | 172 | 142 | 263 | 16 | 13 | 15 | 11 | 172 | 187 | 145 | 201 | 94,96 | 14 | 176 | 12,496 | 3,643 | 15,354 | 3,35E-53 | 1,64E-52 | 1,00E-08 | 0,0159487 | 0,0140938 | 0,0140938 | TRUE | 0,136426 | No |
| orf19.2602 | 994 | 801 | 915 | 609 | 11406 | 10209 | 10707 | 14885 | 905 | 810 | 866 | 915 | 10145 | 11084 | 10965 | 11354 | 5880,63 | 874 | 10887 | 12,475 | 3,641 | 67,028 | 0 | 0 | 0,0017974 | 0,0022617 | 0,0021764 | 0,0021764 | TRUE | 0,0287884 | No |
| orf19.1255 | 61 | 42 | 58 | 43 | 730 | 599 | 627 | 864 | 56 | 42 | 55 | 65 | 649 | 650 | 642 | 659 | 352,29 | 54 | 650 | 12,126 | 3,6 | 30,058 | 1,74E-198 | 2,96E-197 | 1,00E-08 | 0,0057867 | 0,0032315 | 0,0032315 | TRUE | 0,1735978 | No |
| orf19.7520 | 7323 | 5773 | 6383 | 4397 | 83468 | 68154 | 71775 | 1E+05 | 6667 | 5839 | 6041 | 6605 | 74242 | 73996 | 73507 | 78749 | 40705,78 | 6288 | 75124 | 11,951 | 3,579 | 71,507 | 0 | 0 | 0,0025489 | 0,0020695 | 0,0023144 | 0,0023144 | TRUE | 0,028233 | No |
| orf19.7055 | 0 | 5 | 2 | 0 | 27 | 21 | 20 | 24 | 0 | 5 | 2 | 0 | 24 | 23 | 20 | 18 | 11,57 | 2 | 21 | 11,662 | 3,544 | 5,543 | 2,97E-08 | 5,23E-08 | 1,26E-08 | 0,116217 | 0,0828481 | 0,0828481 | TRUE | 1,2382898 | No |
| orf19.133 | 0 | 0 | 0 | 0 | 5 | 1 | 1 | 3 | 0 | 0 | 0 | 0 | 4 | 1 | 1 | 2 | 1,11 | 0 | 2 | 11,397 | 3,511 | 2,338 | 0,0193815 | 0,0248906 | 1,00E-08 | 1,1967514 | 0,9576772 | 0,9576772 | TRUE | 0,5045898 | No |
| orf19.54 | 302 | 250 | 282 | 203 | 3334 | 2988 | 3013 | 3968 | 275 | 253 | 267 | 305 | 2965 | 3244 | 3086 | 3027 | 1677,72 | 275 | 3080 | 11,28 | 3,496 | 53,279 | 0 | 0 | 0,0013161 | 0,0028244 | 0,002038 | 0,002038 | TRUE | 0,0459039 | No |
| orf19.3420 | 0 | 0 | 0 | 0 | 1 | 3 | 4 | 1 | 0 | 0 | 0 | 0 | 1 | 3 | 4 | 1 | 1,13 | 0 | 2 | 11,187 | 3,484 | 2,29 | 0,0220331 | 0,0281542 | 1,00E-08 | 1,1754821 | 1,0116419 | 1,0116419 | TRUE | 0,0843906 | No |
| orf19.3263 | 27 | 26 | 31 | 19 | 333 | 304 | 280 | 364 | 25 | 26 | 29 | 29 | 296 | 330 | 287 | 278 | 162,43 | 27 | 298 | 10,978 | 3,457 | 20,24 | 4,31E-91 | 3,40E-90 | 1,00E-08 | 0,0101698 | 0,0069494 | 0,0069494 | TRUE | 0,0580025 | No |
| orf19.1724 | 0 | 1 | 1 | 1 | 9 | 8 | 7 | 12 | 0 | 1 | 1 | 2 | 8 | 9 | 7 | 9 | 4,56 | 1 | 8 | 10,409 | 3,38 | 3,525 | 0,0004239 | 0,0006126 | 1,00E-08 | 0,2917809 | 0,1641802 | 0,1641802 | TRUE | 0,2342945 | No |
| orf19.5557 | 630 | 685 | 640 | 444 | 7315 | 6352 | 6430 | 8304 | 574 | 693 | 606 | 667 | 6506 | 6897 | 6585 | 6334 | 3607,69 | 635 | 6580 | 10,4 | 3,379 | 56,745 | 0 | 0 | 0,0025903 | 0,0024032 | 0,0024868 | 0,0024868 | TRUE | 0,0519943 | No |
| orf19.1655 | 2404 | 2002 | 2213 | 1558 | 24385 | 20873 | 20954 | 30277 | 2189 | 2025 | 2094 | 2340 | 21690 | 22662 | 21460 | 23096 | 12194,42 | 2162 | 22227 | 10,299 | 3,364 | 68,288 | 0 | 0 | 0,0022699 | 0,0021453 | 0,0020605 | 0,0020605 | TRUE | 0,0344876 | No |
| orf19.1288 | 6296 | 5354 | 5631 | 3891 | 61786 | 51815 | 54425 | 78199 | 5732 | 5415 | 5329 | 5845 | 54957 | 56257 | 55738 | 59651 | 31115,45 | 5580 | 56651 | 10,157 | 3,344 | 76,899 | 0 | 0 | 0,0015184 | 0,0020795 | 0,0017118 | 0,0017118 | TRUE | 0,0156082 | No |
| orf19.1652 | 11107 | 9726 | 10252 | 7484 | 1E+05 | 99219 | 95980 | 1E+05 | 10112 | 9837 | 9703 | 11242 | 1E+05 | 1E+05 | 98296 | 1E+05 | 56985,62 | 10224 | 103748 | 10,153 | 3,344 | 62,941 | 0 | 0 | 0,0031932 | 0,0020602 | 0,0026541 | 0,0026541 | TRUE | 0,0542029 | No |
| orf19.5302 | 127 | 104 | 111 | 80 | 1146 | 1130 | 1064 | 1522 | 116 | 105 | 105 | 120 | 1019 | 1227 | 1090 | 1161 | 617,86 | 112 | 1124 | 10,128 | 3,34 | 33,776 | 4,48E-250 | 9,30E-249 | 1,00E-08 | 0,004175 | 0,0041996 | 0,0041996 | TRUE | 0,0477225 | No |
| orf19.6487 | 16 | 19 | 19 | 19 | 216 | 210 | 167 | 237 | 15 | 19 | 18 | 29 | 192 | 228 | 171 | 181 | 106,53 | 20 | 193 | 9,973 | 3,318 | 15,677 | 2,17E-55 | 1,10E-54 | 0,0107981 | 0,0144371 | 0,0132509 | 0,0132509 | TRUE | 0,2736019 | No |
| orf19.5723 | 3726 | 3173 | 3397 | 2376 | 34673 | 31097 | 30148 | 43630 | 3392 | 3209 | 3215 | 3569 | 30841 | 33763 | 30875 | 33281 | 17768,18 | 3346 | 32190 | 9,629 | 3,267 | 65,922 | 0 | 0 | 0,0022476 | 0,0021114 | 0,0021835 | 0,0021835 | TRUE | 0,0231791 | No |
| orf19.2753 | 2106 | 2040 | 2206 | 1375 | 21217 | 18341 | 19199 | 25774 | 1917 | 2063 | 2088 | 2065 | 18872 | 19913 | 19662 | 19661 | 10780,22 | 2033 | 19527 | 9,61 | 3,265 | 81,574 | 0 | 0 | 0,0006453 | 0,0021596 | 0,0012496 | 0,0012496 | TRUE | 0,0188859 | No |
| orf19.1274 | 12 | 8 | 15 | 13 | 132 | 124 | 113 | 154 | 11 | 8 | 14 | 20 | 117 | 135 | 116 | 117 | 67,25 | 13 | 121 | 9,56 | 3,257 | 13,054 | 6,02E-39 | 2,30E-38 | 1,00E-08 | 0,0216809 | 0,0143089 | 0,0143089 | TRUE | 0,2570811 | No |
| orf19.1704 | 6422 | 5352 | 5903 | 3504 | 60387 | 47565 | 50334 | 70701 | 5847 | 5413 | 5587 | 5264 | 53712 | 51642 | 51548 | 53931 | 29118,06 | 5528 | 52708 | 9,528 | 3,252 | 80,384 | 0 | 0 | 0,0011931 | 0,0020824 | 0,0014658 | 0,0014658 | TRUE | 0,0185318 | No |
| orf19.5217 | 494 | 443 | 447 | 301 | 4637 | 3673 | 3853 | 5328 | 450 | 448 | 423 | 452 | 4124 | 3988 | 3946 | 4064 | 2236,94 | 443 | 4030 | 9,109 | 3,187 | 61,889 | 0 | 0 | 1,00E-08 | 0,0026276 | 0,0012322 | 0,0012322 | TRUE | 0,011682 | No |
| orf19.6958 | 61 | 40 | 57 | 33 | 517 | 437 | 417 | 583 | 56 | 40 | 54 | 50 | 460 | 474 | 427 | 445 | 250,7 | 50 | 452 | 9,011 | 3,172 | 24,437 | 6,89E-132 | 7,82E-131 | 1,00E-08 | 0,0073062 | 0,0046066 | 0,0046066 | TRUE | 0,1449372 | No |
| orf19.360 | 3 | 3 | 0 | 5 | 28 | 27 | 25 | 34 | 3 | 3 | 0 | 8 | 25 | 29 | 26 | 26 | 14,88 | 4 | 26 | 8,979 | 3,167 | 6,095 | 1,09E-09 | 2,04E-09 | 1,00E-08 | 0,0908178 | 0,0621341 | 0,0621341 | TRUE | 0,7183618 | No |
| orf19.4567 | 41 | 41 | 36 | 23 | 360 | 305 | 316 | 432 | 37 | 41 | 34 | 35 | 320 | 331 | 324 | 330 | 181,49 | 37 | 326 | 8,814 | 3,14 | 21,43 | 7,00E-102 | 6,21E-101 | 1,00E-08 | 0,0093156 | 0,0049357 | 0,0049357 | TRUE | 0,051163 | No |
| orf19.7466 | 6592 | 5967 | 6412 | 3819 | 56276 | 46929 | 50893 | 68287 | 6002 | 6035 | 6068 | 5737 | 50056 | 50952 | 52121 | 52090 | 28632,51 | 5960 | 51305 | 8,601 | 3,105 | 95,982 | 0 | 0 | 0,0003865 | 0,0020832 | 0,0009052 | 0,0009052 | TRUE | 0,0072922 | No |
| orf19.2451 | 568 | 590 | 571 | 409 | 5283 | 5080 | 4358 | 6097 | 517 | 597 | 540 | 614 | 4699 | 5515 | 4463 | 4651 | 2699,64 | 567 | 4832 | 8,546 | 3,095 | 39,742 | 0 | 0 | 0,0066281 | 0,0025263 | 0,0047777 | 0,0047777 | TRUE | 0,109782 | No |
| orf19.1679 | 64 | 39 | 51 | 37 | 480 | 394 | 435 | 542 | 58 | 39 | 48 | 56 | 427 | 428 | 445 | 413 | 239,4 | 50 | 428 | 8,534 | 3,093 | 23,768 | 7,21E-125 | 7,67E-124 | 1,00E-08 | 0,0075549 | 0,0046528 | 0,0046528 | TRUE | 0,1789353 | No |
| orf19.6782 | 170 | 162 | 158 | 111 | 1587 | 1320 | 1273 | 1626 | 155 | 164 | 150 | 167 | 1412 | 1433 | 1304 | 1240 | 752,96 | 159 | 1347 | 8,517 | 3,09 | 35,971 | 2,40E-283 | 5,76E-282 | 1,00E-08 | 0,0037914 | 0,0033836 | 0,0033836 | TRUE | 0,0356916 | No |
| orf19.4591 | 2955 | 2653 | 2862 | 1929 | 25281 | 22606 | 21890 | 30630 | 2690 | 2683 | 2709 | 2898 | 22487 | 24544 | 22418 | 23365 | 12974,16 | 2745 | 23204 | 8,463 | 3,081 | 69,413 | 0 | 0 | 0,0013377 | 0,0021388 | 0,0016757 | 0,0016757 | TRUE | 0,0184323 | No |
| orf19.7596 | 164 | 142 | 151 | 89 | 1366 | 1113 | 1120 | 1584 | 149 | 144 | 143 | 134 | 1215 | 1208 | 1147 | 1208 | 668,53 | 142 | 1194 | 8,348 | 3,061 | 38,134 | 0 | 0 | 1,00E-08 | 0,004013 | 0,0021068 | 0,0021068 | TRUE | 0,0123695 | No |
| orf19.6830 | 456 | 436 | 438 | 268 | 4107 | 3154 | 3274 | 4505 | 415 | 441 | 415 | 403 | 3653 | 3424 | 3353 | 3436 | 1942,51 | 418 | 3466 | 8,273 | 3,048 | 53,581 | 0 | 0 | 1,00E-08 | 0,0027171 | 0,0017047 | 0,0017047 | TRUE | 0,0160098 | No |
| orf19.4570 | 140 | 129 | 135 | 91 | 1287 | 926 | 1011 | 1457 | 127 | 130 | 128 | 137 | 1145 | 1005 | 1035 | 1111 | 602,42 | 130 | 1074 | 8,268 | 3,047 | 34,149 | 1,36E-255 | 2,93E-254 | 1,00E-08 | 0,0042299 | 0,0031429 | 0,0031429 | TRUE | 0,0229248 | No |
| orf19.2652 | 95 | 106 | 118 | 62 | 908 | 818 | 734 | 983 | 86 | 107 | 112 | 93 | 808 | 888 | 752 | 750 | 449,48 | 100 | 800 | 7,99 | 2,998 | 28,102 | 9,20E-174 | 1,40E-172 | 0,005133 | 0,004976 | 0,0050384 | 0,0050384 | TRUE | 0,097445 | No |
| orf19.5949 | 6066 | 5619 | 6144 | 3856 | 50021 | 41473 | 43371 | 58689 | 5523 | 5683 | 5815 | 5792 | 44492 | 45028 | 44417 | 44768 | 25189,87 | 5703 | 44676 | 7,837 | 2,97 | 101,501 | 0 | 0 | 5,37E-05 | 0,0020895 | 0,0007175 | 0,0007175 | TRUE | 0,0057006 | No |
| orf19.6254 | 3570 | 3022 | 3291 | 2158 | 26110 | 22164 | 23298 | 32011 | 3250 | 3056 | 3115 | 3242 | 23224 | 24064 | 23860 | 24418 | 13528,67 | 3166 | 23892 | 7,551 | 2,917 | 81,476 | 0 | 0 | 0,0004386 | 0,0021347 | 0,001041 | 0,001041 | TRUE | 0,0065548 | No |
| orf19.6336 | 73 | 46 | 52 | 52 | 433 | 382 | 439 | 605 | 66 | 47 | 49 | 78 | 385 | 415 | 450 | 461 | 243,91 | 60 | 428 | 7,264 | 2,861 | 20,8 | 4,32E-96 | 3,58E-95 | 0,0099256 | 0,0074529 | 0,0080768 | 0,0080768 | TRUE | 0,2425753 | No |

| | | | | | | | | | | | | | | | | | | | | | | | | | | | | | | | |
|---|---|---|---|---|---|---|---|---|---|---|---|---|---|---|---|---|---|---|---|---|---|---|---|---|---|---|---|---|---|---|---|
| orf19.3925 | 8 | 12 | 10 | 5 | 88 | 65 | 61 | 68 | 7 | 12 | 9 | 8 | 78 | 71 | 62 | 52 | 37,45 | 9 | 66 | 7,147 | 2,837 | 9,081 | 1,08E-19 | 2,71E-19 | 1,00E-08 | 0,0373127 | 0,0290833 | 0,0290833 | TRUE | 0,1317936 | No |
| orf19.310 | 41 | 25 | 34 | 21 | 259 | 189 | 217 | 312 | 37 | 25 | 32 | 32 | 230 | 205 | 222 | 238 | 127,77 | 32 | 224 | 7,076 | 2,823 | 16,938 | 2,35E-64 | 1,37E-63 | 1,00E-08 | 0,0123761 | 0,0079516 | 0,0079516 | TRUE | 0,1351781 | No |
| orf19.4128 | 1045 | 955 | 1083 | 594 | 7869 | 6273 | 6630 | 8535 | 951 | 966 | 1025 | 892 | 6999 | 6811 | 6790 | 6511 | 3868,13 | 958 | 6778 | 7,047 | 2,817 | 57,813 | 0 | 0 | 0,0012629 | 0,0023785 | 0,0016541 | 0,0016541 | TRUE | 0,0254812 | No |
| orf19.7545 | 36 | 41 | 41 | 30 | 304 | 259 | 279 | 335 | 33 | 41 | 39 | 45 | 270 | 281 | 286 | 256 | 156,37 | 40 | 273 | 6,999 | 2,807 | 18,708 | 4,22E-78 | 2,93E-77 | 1,00E-08 | 0,0104848 | 0,0063594 | 0,0063594 | TRUE | 0,108516 | No |
| orf19.1743 | 3455 | 2959 | 3085 | 1981 | 23815 | 19054 | 19655 | 28437 | 3146 | 2993 | 2920 | 2976 | 21183 | 20687 | 20129 | 21692 | 11965,62 | 3009 | 20923 | 6,953 | 2,798 | 69,844 | 0 | 0 | 0,0007743 | 0,0021474 | 0,0013394 | 0,0013394 | TRUE | 0,01193 | No |
| orf19.5233 | 0 | 0 | 1 | 1 | 6 | 2 | 5 | 3 | 0 | 0 | 1 | 2 | 5 | 2 | 5 | 2 | 2,17 | 1 | 4 | 6,952 | 2,798 | 2,282 | 0,0224647 | 0,0286654 | 1,00E-08 | 0,6105857 | 0,4008308 | 0,4008308 | TRUE | 0,2014173 | No |
| orf19.5724 | 8 | 5 | 11 | 4 | 57 | 49 | 48 | 62 | 7 | 5 | 10 | 6 | 51 | 53 | 49 | 47 | 28,64 | 7 | 50 | 6,823 | 2,77 | 8,221 | 2,02E-16 | 4,63E-16 | 1,00E-08 | 0,0481629 | 0,0277525 | 0,0277525 | TRUE | 0,2716088 | No |
| orf19.5672 | 42 | 51 | 43 | 40 | 327 | 282 | 296 | 441 | 38 | 52 | 41 | 60 | 291 | 306 | 303 | 336 | 178,4 | 48 | 309 | 6,657 | 2,735 | 18,934 | 5,98E-80 | 4,23E-79 | 0,0023141 | 0,0094419 | 0,0071543 | 0,0071543 | TRUE | 0,1703452 | No |
| orf19.7094 | 12 | 16 | 15 | 10 | 118 | 75 | 85 | 122 | 11 | 16 | 14 | 15 | 105 | 81 | 87 | 93 | 52,85 | 14 | 92 | 6,591 | 2,72 | 10,841 | 2,20E-27 | 6,66E-27 | 1,00E-08 | 0,0270308 | 0,0175332 | 0,0175332 | TRUE | 0,121386 | No |
| orf19.4335 | 278 | 296 | 240 | 175 | 1847 | 1573 | 1633 | 2228 | 253 | 299 | 227 | 263 | 1643 | 1708 | 1672 | 1700 | 970,64 | 260 | 1681 | 6,463 | 2,692 | 37,491 | 1,28E-307 | 3,38E-306 | 1,00E-08 | 0,003398 | 0,0026293 | 0,0026293 | TRUE | 0,1220873 | No |
| orf19.979 | 6339 | 5623 | 5955 | 4092 | 40706 | 35221 | 35633 | 50305 | 5771 | 5687 | 5636 | 6147 | 36207 | 38240 | 36493 | 38373 | 21569,22 | 5810 | 37328 | 6,43 | 2,685 | 67,091 | 0 | 0 | 0,0011232 | 0,0020983 | 0,0014325 | 0,0014325 | TRUE | 0,0177545 | No |
| orf19.2460 | 13 | 9 | 7 | 6 | 64 | 54 | 56 | 81 | 12 | 9 | 7 | 9 | 57 | 59 | 57 | 62 | 33,91 | 9 | 59 | 6,399 | 2,678 | 8,825 | 1,09E-18 | 2,68E-18 | 1,00E-08 | 0,0409945 | 0,023016 | 0,023016 | TRUE | 0,164924 | No |
| orf19.6066 | 1586 | 1433 | 1488 | 980 | 10185 | 8946 | 8537 | 11955 | 1444 | 1449 | 1408 | 1472 | 9059 | 9713 | 8743 | 9119 | 5301,01 | 1443 | 9158 | 6,349 | 2,667 | 58,799 | 0 | 0 | 0,0011229 | 0,0022862 | 0,0015518 | 0,0015518 | TRUE | 0,0200373 | No |
| orf19.5169 | 201 | 179 | 175 | 131 | 1250 | 1079 | 1076 | 1468 | 183 | 181 | 166 | 197 | 1112 | 1171 | 1102 | 1120 | 653,94 | 182 | 1126 | 6,24 | 2,641 | 34,661 | 3,07E-263 | 6,74E-262 | 1,00E-08 | 0,0040571 | 0,002228 | 0,002228 | TRUE | 0,0379613 | No |
| orf19.341 | 376 | 291 | 346 | 213 | 2246 | 1835 | 1863 | 2701 | 342 | 294 | 327 | 320 | 1998 | 1992 | 1908 | 2060 | 1155,3 | 321 | 1990 | 6,191 | 2,63 | 41,438 | 0 | 0 | 0,0005998 | 0,0031804 | 0,0019819 | 0,0019819 | TRUE | 0,0388258 | No |
| orf19.780 | 2115 | 1923 | 2251 | 1285 | 13286 | 11627 | 12052 | 16098 | 1926 | 1945 | 2130 | 1930 | 11817 | 12624 | 12343 | 12280 | 7124,35 | 1983 | 12266 | 6,181 | 2,628 | 59,19 | 0 | 0 | 0,0012203 | 0,0022224 | 0,0015834 | 0,0015834 | TRUE | 0,0313365 | No |
| orf19.3684 | 2405 | 2012 | 2137 | 1482 | 14144 | 12590 | 12154 | 17424 | 2190 | 2035 | 2022 | 2226 | 12581 | 13669 | 12447 | 13291 | 7557,69 | 2118 | 12997 | 6,142 | 2,619 | 53,282 | 0 | 0 | 0,0018552 | 0,0022118 | 0,0020292 | 0,0020292 | TRUE | 0,0147057 | No |
| orf19.4884 | 111 | 120 | 118 | 89 | 763 | 660 | 678 | 971 | 101 | 121 | 112 | 134 | 679 | 717 | 694 | 741 | 412,26 | 117 | 708 | 6,132 | 2,616 | 27,484 | 2,69E-166 | 3,91E-165 | 0,0005364 | 0,0052413 | 0,0034563 | 0,0034563 | TRUE | 0,0867725 | No |
| orf19.3442 | 3612 | 3387 | 3625 | 2371 | 23488 | 20333 | 19424 | 26813 | 3289 | 3426 | 3431 | 3562 | 20892 | 22076 | 19893 | 20453 | 12127,51 | 3427 | 20828 | 6,083 | 2,605 | 59,23 | 0 | 0 | 0,0012929 | 0,0021459 | 0,0016775 | 0,0016775 | TRUE | 0,0198111 | No |
| orf19.2590 | 13 | 4 | 4 | 10 | 51 | 51 | 48 | 66 | 12 | 4 | 4 | 15 | 45 | 55 | 49 | 50 | 29,37 | 9 | 50 | 6,073 | 2,602 | 7,615 | 2,64E-14 | 5,72E-14 | 1,00E-08 | 0,0470212 | 0,0385889 | 0,0385889 | TRUE | 0,483341 | No |
| orf19.4114 | 2387 | 2152 | 2370 | 1423 | 14572 | 11951 | 12870 | 17627 | 2173 | 2177 | 2243 | 2138 | 12961 | 12975 | 13181 | 13446 | 7661,71 | 2183 | 13141 | 6,016 | 2,589 | 75,133 | 0 | 0 | 1,00E-08 | 0,0022094 | 0,0008591 | 0,0008591 | TRUE | 0,0042435 | No |
| orf19.355 | 99 | 88 | 81 | 62 | 577 | 472 | 495 | 703 | 90 | 89 | 77 | 93 | 513 | 512 | 507 | 536 | 302,23 | 87 | 517 | 5,968 | 2,577 | 24,663 | 2,67E-134 | 3,09E-133 | 1,00E-08 | 0,0064079 | 0,0034923 | 0,0034923 | TRUE | 0,0681858 | No |
| orf19.6219 | 134 | 128 | 93 | 75 | 725 | 625 | 628 | 905 | 122 | 129 | 88 | 113 | 645 | 679 | 643 | 690 | 388,63 | 113 | 664 | 5,884 | 2,557 | 25,19 | 5,18E-140 | 6,33E-139 | 0,001218 | 0,0054361 | 0,0044936 | 0,0044936 | TRUE | 0,2546358 | No |
| orf19.3375 | 2 | 3 | 1 | 0 | 10 | 8 | 9 | 12 | 2 | 3 | 1 | 0 | 9 | 9 | 9 | 9 | 5,22 | 2 | 9 | 5,771 | 2,529 | 3,39 | 0,0006993 | 0,0009969 | 1,00E-08 | 0,2551454 | 0,1439851 | 0,1439851 | TRUE | 0,3070413 | No |
| orf19.4481 | 10 | 8 | 10 | 6 | 57 | 43 | 50 | 69 | 9 | 8 | 9 | 9 | 51 | 47 | 51 | 53 | 29,61 | 9 | 50 | 5,649 | 2,498 | 8,014 | 1,11E-15 | 2,51E-15 | 1,00E-08 | 0,0466468 | 0,0249301 | 0,0249301 | TRUE | 0,017629 | No |
| orf19.5215 | 2342 | 1940 | 2077 | 1465 | 12971 | 10281 | 11248 | 15704 | 2132 | 1962 | 1966 | 2201 | 11537 | 11162 | 11519 | 11979 | 6807,36 | 2065 | 11549 | 5,602 | 2,486 | 52,866 | 0 | 0 | 0,001734 | 0,0022311 | 0,0018218 | 0,0018218 | TRUE | 0,0215357 | No |
| orf19.1275 | 282 | 239 | 258 | 185 | 1529 | 1307 | 1407 | 1905 | 257 | 242 | 244 | 278 | 1360 | 1419 | 1441 | 1453 | 836,71 | 255 | 1418 | 5,593 | 2,484 | 36,468 | 3,58E-291 | 8,95E-290 | 1,00E-08 | 0,0036158 | 0,002033 | 0,002033 | TRUE | 0,0301555 | No |
| orf19.3573 | 1537 | 1469 | 1540 | 916 | 8706 | 7457 | 7475 | 10195 | 1399 | 1486 | 1457 | 1376 | 7744 | 8096 | 7655 | 7777 | 4623,83 | 1430 | 7818 | 5,46 | 2,449 | 59,276 | 0 | 0 | 0,0003806 | 0,0023227 | 0,001205 | 0,001205 | TRUE | 0,0081074 | No |
| orf19.4157 | 403 | 331 | 348 | 237 | 2060 | 1795 | 1729 | 2636 | 367 | 335 | 329 | 356 | 1832 | 1949 | 1771 | 2011 | 1118,71 | 347 | 1891 | 5,46 | 2,449 | 35,798 | 1,18E-280 | 2,78E-279 | 0,0022294 | 0,0032178 | 0,0027083 | 0,0027083 | TRUE | 0,0229646 | No |
| orf19.5753 | 5390 | 4940 | 5312 | 3376 | 30012 | 25770 | 25461 | 35691 | 4907 | 4996 | 5027 | 5071 | 26695 | 27979 | 26075 | 27225 | 15997,09 | 5000 | 26994 | 5,4 | 2,433 | 70,719 | 0 | 0 | 0,0003858 | 0,0021196 | 0,0010115 | 0,0010115 | TRUE | 0,0073388 | No |
| orf19.120 | 197 | 233 | 206 | 138 | 1204 | 1083 | 996 | 1300 | 179 | 236 | 195 | 207 | 1071 | 1176 | 1020 | 992 | 634,46 | 204 | 1065 | 5,22 | 2,384 | 26,181 | 4,41E-151 | 5,78E-150 | 0,0058034 | 0,0041191 | 0,0049058 | 0,0049058 | TRUE | 0,1269995 | No |
| orf19.7127.1 | 58 | 47 | 40 | 35 | 249 | 181 | 284 | 358 | 53 | 48 | 38 | 53 | 221 | 197 | 291 | 273 | 146,59 | 48 | 246 | 5,187 | 2,375 | 13,785 | 3,13E-43 | 1,29E-42 | 0,0214916 | 0,0110487 | 0,0153783 | 0,0153783 | TRUE | 0,0815132 | No |
| orf19.3671 | 1 | 1 | 0 | 2 | 4 | 9 | 3 | 8 | 1 | 1 | 0 | 3 | 4 | 10 | 3 | 6 | 3,43 | 1 | 6 | 5,152 | 2,365 | 2,491 | 0,0127486 | 0,0165896 | 0,0956334 | 0,3873026 | 0,2951937 | 0,2951937 | TRUE | 0,411922 | No |
| orf19.7275 | 7 | 6 | 11 | 7 | 46 | 41 | 37 | 58 | 6 | 6 | 10 | 11 | 41 | 45 | 38 | 44 | 25,12 | 8 | 42 | 5,132 | 2,36 | 7,052 | 1,76E-12 | 3,62E-12 | 1,00E-08 | 0,0546316 | 0,0317529 | 0,0317529 | TRUE | 0,1245818 | No |
| orf19.313 | 7 | 8 | 4 | 5 | 31 | 36 | 31 | 41 | 6 | 8 | 4 | 8 | 28 | 39 | 32 | 31 | 19,43 | 6 | 32 | 5,103 | 2,351 | 6,127 | 8,97E-10 | 1,68E-09 | 1,00E-08 | 0,0700231 | 0,0434942 | 0,0434942 | TRUE | 0,2191745 | No |
| orf19.2810 | 387 | 320 | 361 | 241 | 1900 | 1722 | 1627 | 2341 | 352 | 324 | 342 | 362 | 1690 | 1870 | 1666 | 1786 | 1048,91 | 345 | 1753 | 5,094 | 2,349 | 35,263 | 2,14E-272 | 4,90E-271 | 0,0016232 | 0,0032964 | 0,0024455 | 0,0024455 | TRUE | 0,0232906 | No |
| orf19.2587 | 6 | 7 | 5 | 6 | 42 | 23 | 32 | 44 | 5 | 7 | 5 | 9 | 37 | 25 | 33 | 34 | 19,37 | 6 | 32 | 5,091 | 2,348 | 6,112 | 9,81E-10 | 1,84E-09 | 1,00E-08 | 0,0702385 | 0,0440727 | 0,0440727 | TRUE | 0,1409643 | No |
| orf19.1313 | 602 | 517 | 550 | 394 | 2889 | 2609 | 2624 | 3635 | 548 | 523 | 521 | 592 | 2570 | 2833 | 2687 | 2773 | 1630,72 | 546 | 2716 | 4,996 | 2,321 | 39,95 | 0 | 0 | 0,0012563 | 0,0028471 | 0,0020861 | 0,0020861 | TRUE | 0,0309494 | No |
| orf19.849 | 84 | 103 | 88 | 64 | 497 | 436 | 459 | 527 | 76 | 104 | 83 | 96 | 442 | 473 | 470 | 402 | 268,45 | 90 | 447 | 4,992 | 2,32 | 19,868 | 7,67E-88 | 5,94E-87 | 0,0046453 | 0,0069579 | 0,0061017 | 0,0061017 | TRUE | 0,1263717 | No |
| orf19.4833 | 11923 | 9778 | 10533 | 7209 | 57142 | 46905 | 50194 | 69756 | 10855 | 9889 | 9969 | 10829 | 50826 | 50926 | 51405 | 53210 | 30988,7 | 10386 | 51592 | 4,969 | 2,313 | 54,858 | 0 | 0 | 0,0014478 | 0,0020796 | 0,0016466 | 0,0016466 | TRUE | 0,0126123 | No |
| orf19.5005 | 3667 | 3324 | 3587 | 2231 | 18992 | 15313 | 15967 | 21705 | 3339 | 3362 | 3395 | 3351 | 16893 | 16626 | 16352 | 16557 | 9984,25 | 3362 | 16607 | 4,94 | 2,304 | 77,032 | 0 | 0 | 1,00E-08 | 0,0021693 | 0,0006715 | 0,0006715 | TRUE | 0,0016502 | No |
| orf19.7277 | 10 | 0 | 8 | 7 | 39 | 25 | 29 | 50 | 9 | 0 | 8 | 11 | 35 | 27 | 30 | 38 | 19,61 | 7 | 32 | 4,931 | 2,302 | 5,621 | 1,90E-08 | 3,38E-08 | 0,0241141 | 0,0694076 | 0,0665742 | 0,0665742 | TRUE | 1,217546 | No |
| orf19.4789 | 62 | 60 | 76 | 50 | 362 | 301 | 294 | 439 | 56 | 61 | 72 | 75 | 322 | 327 | 301 | 335 | 193,62 | 66 | 321 | 4,923 | 2,3 | 18,349 | 3,37E-75 | 2,24E-74 | 1,00E-08 | 0,0088598 | 0,0055307 | 0,0055307 | TRUE | 0,0915868 | No |
| orf19.4507 | 79 | 91 | 70 | 48 | 442 | 352 | 332 | 488 | 72 | 92 | 66 | 72 | 393 | 382 | 340 | 372 | 223,74 | 76 | 372 | 4,92 | 2,299 | 18,915 | 8,54E-80 | 6,03E-79 | 0,0017346 | 0,0079413 | 0,0059214 | 0,0059214 | TRUE | 0,2074357 | No |
| orf19.5717 | 2 | 11 | 5 | 4 | 30 | 17 | 35 | 42 | 2 | 11 | 5 | 6 | 27 | 18 | 36 | 32 | 17,09 | 6 | 28 | 4,869 | 2,284 | 5,154 | 2,55E-07 | 4,34E-07 | 0,0864629 | 0,0793389 | 0,081076 | 0,081076 | TRUE | 0,7156669 | No |
| orf19.3329 | 861 | 640 | 724 | 491 | 3698 | 3142 | 3358 | 4555 | 784 | 647 | 685 | 738 | 3289 | 3411 | 3439 | 3475 | 2058,52 | 714 | 3404 | 4,773 | 2,255 | 38,612 | 0 | 0 | 0,0020178 | 0,0026787 | 0,0023865 | 0,0023865 | TRUE | 0,0588878 | No |
| orf19.532 | 723 | 605 | 558 | 386 | 3373 | 2640 | 2503 | 3830 | 658 | 612 | 528 | 580 | 3000 | 2866 | 2563 | 2922 | 1716,19 | 594 | 2838 | 4,77 | 2,254 | 30,699 | 5,87E-207 | 1,03E-205 | 0,00515 | 0,0028067 | 0,004109 | 0,004109 | TRUE | 0,0722982 | No |
| orf19.1345 | 47 | 28 | 38 | 15 | 181 | 150 | 142 | 217 | 43 | 28 | 36 | 23 | 161 | 163 | 145 | 166 | 95,55 | 32 | 159 | 4,768 | 2,253 | 12,725 | 4,29E-37 | 1,60E-36 | 1,00E-08 | 0,015862 | 0,0112917 | 0,0112917 | TRUE | 0,3215573 | No |
| orf19.2509.1 | 88 | 89 | 71 | 55 | 446 | 334 | 352 | 491 | 80 | 90 | 67 | 83 | 397 | 363 | 360 | 375 | 226,79 | 80 | 374 | 4,695 | 2,231 | 19,279 | 8,08E-83 | 5,96E-82 | 1,00E-08 | 0,0078618 | 0,004966 | 0,004966 | TRUE | 0,1203113 | No |
| orf19.3051 | 1054 | 947 | 975 | 623 | 4822 | 4056 | 4471 | 5830 | 960 | 958 | 923 | 936 | 4289 | 4404 | 4579 | 4447 | 2686,84 | 944 | 4430 | 4,69 | 2,23 | 50,456 | 0 | 0 | 1,00E-08 | 0,0025287 | 0,0012041 | 0,0012041 | TRUE | 0,0061675 | No |
| orf19.4463 | 9 | 4 | 8 | 3 | 39 | 26 | 25 | 37 | 8 | 4 | 8 | 5 | 35 | 28 | 26 | 28 | 17,63 | 6 | 29 | 4,674 | 2,225 | 5,681 | 1,34E-08 | 2,39E-08 | 1,00E-08 | 0,076954 | 0,0468341 | 0,0468341 | TRUE | 0,145965 | No |
| orf19.7614 | 4 | 3 | 5 | 2 | 25 | 18 | 13 | 17 | 4 | 3 | 5 | 3 | 22 | 20 | 13 | 13 | 10,31 | 4 | 17 | 4,632 | 2,212 | 4,276 | 1,90E-05 | 2,97E-05 | 1,00E-08 | 0,130175 | 0,0849694 | 0,0849694 | TRUE | 0,1531679 | No |
| orf19.866 | 1467 | 1376 | 1473 | 895 | 6844 | 5922 | 6254 | 8271 | 1336 | 1392 | 1394 | 1344 | 6088 | 6430 | 6405 | 6309 | 3837,13 | 1366 | 6308 | 4,613 | 2,206 | 54,509 | 0 | 0 | 1,00E-08 | 0,0023813 | 0,0011066 | 0,0011066 | TRUE | 0,0067536 | No |
| orf19.4697 | 982 | 930 | 1015 | 564 | 4687 | 3812 | 4264 | 5339 | 894 | 941 | 961 | 847 | 4169 | 4139 | 4367 | 4073 | 2548,71 | 911 | 4187 | 4,583 | 2,196 | 44,114 | 0 | 0 | 0,0008726 | 0,0025553 | 0,0016822 | 0,0016822 | TRUE | 0,0219429 | No |
| orf19.1709 | 1862 | 1714 | 1679 | 921 | 9256 | 6170 | 6941 | 9533 | 1695 | 1734 | 1589 | 1383 | 8233 | 6699 | 7108 | 7272 | 4464,18 | 1600 | 7328 | 4,571 | 2,193 | 28,5 | 1,17E-178 | 1,80E-177 | 0,0082311 | 0,0023329 | 0,0052843 | 0,0052843 | TRUE | 0,0950902 | No |
| orf19.7539 | 629 | 442 | 602 | 410 | 2694 | 2223 | 2462 | 3571 | 573 | 447 | 570 | 616 | 2396 | 2414 | 2521 | 2724 | 1532,56 | 552 | 2514 | 4,574 | 2,193 | 26,493 | 1,17E-154 | 1,57E-153 | 0,0090066 | 0,002899 | 0,0054183 | 0,0054183 | TRUE | 0,1907434 | No |
| orf19.4886 | 86 | 82 | 80 | 42 | 360 | 351 | 355 | 416 | 78 | 83 | 76 | 63 | 320 | 381 | 364 | 317 | 210,28 | 75 | 346 | 4,544 | 2,184 | 17,441 | 4,06E-68 | 2,48E-67 | 1,00E-08 | 0,0083192 | 0,0067079 | 0,0067079 | TRUE | 0,0711232 | No |
| orf19.5711 | 958 | 929 | 1002 | 663 | 4739 | 3964 | 3987 | 5657 | 872 | 940 | 948 | 996 | 4215 | 4304 | 4083 | 4315 | 2584,18 | 939 | 4229 | 4,519 | 2,176 | 44,682 | 0 | 0 | 0,0006813 | 0,0025482 | 0,0015953 | 0,0015953 | TRUE | 0,0281036 | No |
| orf19.5556 | 1 | 2 | 1 | 0 | 8 | 5 | 3 | 4 | 1 | 2 | 1 | 0 | 7 | 5 | 3 | 3 | 2,82 | 1 | 4 | 4,486 | 2,165 | 2,232 | 0,0255825 | 0,0325136 | 1,00E-08 | 0,4707332 | 0,2933168 | 0,2933168 | TRUE | 0,2791562 | No |
| orf19.2124 | 2947 | 2637 | 2731 | 1792 | 13954 | 10504 | 11143 | 16099 | 2683 | 2667 | 2585 | 2692 | 12412 | 11404 | 11412 | 12280 | 7266,88 | 2657 | 11877 | 4,473 | 2,161 | 49,759 | 0 | 0 | 0,0012124 | 0,0022188 | 0,0015695 | 0,0015695 | TRUE | 0,0111649 | No |
| orf19.4688 | 1842 | 1626 | 1834 | 1243 | 8312 | 7321 | 7364 | 9676 | 1677 | 1645 | 1736 | 1867 | 7393 | 7949 | 7542 | 7381 | 4648,61 | 1731 | 7566 | 4,38 | 2,131 | 44,156 | 0 | 0 | 0,0017516 | 0,0023212 | 0,0018629 | 0,0018629 | TRUE | 0,0306615 | No |

| | | | | | | | | | | | | | | | | | | | | | | | | | | | | | | | |
|---|---|---|---|---|---|---|---|---|---|---|---|---|---|---|---|---|---|---|---|---|---|---|---|---|---|---|---|---|---|---|---|
| orf19.4316 | 700 | 694 | 722 | 461 | 3293 | 2793 | 2904 | 3740 | 637 | 702 | 683 | 692 | 2929 | 3032 | 2974 | 2853 | 1812,93 | 678 | 2947 | 4,347 | 2,12 | 41,967 | 0 | 0 | 0,0004094 | 0,0027657 | 0,0015049 | 0,0015049 | TRUE | 0,0216682 | No |
| orf19.5469 | 363 | 321 | 300 | 197 | 1357 | 1137 | 1452 | 1897 | 330 | 325 | 284 | 296 | 1207 | 1234 | 1487 | 1447 | 826,32 | 309 | 1344 | 4,346 | 2,12 | 23,711 | 2,75E-124 | 2,92E-123 | 0,0076146 | 0,0036357 | 0,0055946 | 0,0055946 | TRUE | 0,0603981 | No |
| orf19.3059.1 | 0 | 4 | 1 | 2 | 13 | 10 | 5 | 7 | 0 | 4 | 1 | 3 | 12 | 11 | 5 | 5 | 5,11 | 2 | 8 | 4,347 | 2,12 | 2,765 | 0,0056911 | 0,0075949 | 0,1331346 | 0,2605689 | 0,2258807 | 0,2258807 | TRUE | 0,3351669 | No |
| orf19.7024 | 6 | 3 | 4 | 4 | 27 | 21 | 18 | 17 | 5 | 3 | 4 | 6 | 24 | 23 | 18 | 13 | 12,06 | 4 | 20 | 4,331 | 2,115 | 4,426 | 9,58E-06 | 1,52E-05 | 1,00E-08 | 0,1115396 | 0,0771525 | 0,0771525 | TRUE | 0,3497532 | No |
| orf19.756 | 10 | 8 | 8 | 8 | 48 | 33 | 29 | 60 | 9 | 8 | 8 | 12 | 43 | 36 | 30 | 46 | 23,85 | 9 | 39 | 4,318 | 2,11 | 6,249 | 4,13E-10 | 7,83E-10 | 1,00E-08 | 0,0574318 | 0,038584 | 0,038584 | TRUE | 0,1570015 | No |
| orf19.4212 | 370 | 318 | 344 | 206 | 1551 | 1320 | 1329 | 1861 | 337 | 322 | 326 | 309 | 1380 | 1433 | 1361 | 1420 | 860,86 | 324 | 1398 | 4,31 | 2,108 | 33,931 | 2,31E-252 | 4,85E-251 | 1,00E-08 | 0,0035715 | 0,0017351 | 0,0017351 | TRUE | 0,0089366 | No |
| orf19.6501 | 46 | 31 | 38 | 32 | 179 | 144 | 165 | 235 | 42 | 31 | 36 | 48 | 159 | 156 | 169 | 179 | 102,63 | 39 | 166 | 4,293 | 2,102 | 12,772 | 2,35E-37 | 8,80E-37 | 1,00E-08 | 0,0149081 | 0,0095426 | 0,0095426 | TRUE | 0,1292059 | No |
| orf19.3879 | 69 | 69 | 72 | 64 | 333 | 298 | 324 | 388 | 63 | 70 | 68 | 96 | 296 | 324 | 332 | 296 | 193,05 | 74 | 312 | 4,291 | 2,101 | 16,441 | 9,70E-61 | 5,36E-60 | 0,0019503 | 0,0088797 | 0,006877 | 0,006877 | TRUE | 0,2618649 | No |
| orf19.4669 | 485 | 582 | 534 | 334 | 2483 | 2112 | 2067 | 2609 | 442 | 589 | 505 | 502 | 2209 | 2293 | 2117 | 1990 | 1330,74 | 510 | 2152 | 4,226 | 2,079 | 26,124 | 1,92E-150 | 2,51E-149 | 0,0072362 | 0,0030297 | 0,0048106 | 0,0048106 | TRUE | 0,1335996 | No |
| orf19.5777 | 286 | 213 | 300 | 188 | 1225 | 1059 | 1015 | 1441 | 260 | 215 | 284 | 282 | 1090 | 1150 | 1039 | 1099 | 677,53 | 260 | 1094 | 4,215 | 2,076 | 25,647 | 4,61E-145 | 5,85E-144 | 0,0034408 | 0,0039868 | 0,0038085 | 0,0038085 | TRUE | 0,1537936 | No |
| orf19.2599 | 2339 | 1889 | 2144 | 1456 | 9574 | 7940 | 8089 | 11740 | 2130 | 1911 | 2029 | 2187 | 8516 | 8621 | 8284 | 8955 | 5329,03 | 2064 | 8594 | 4,169 | 2,06 | 42,27 | 0 | 0 | 0,0018567 | 0,0022849 | 0,0019647 | 0,0019647 | TRUE | 0,0301448 | No |
| orf19.3733 | 1623 | 1452 | 1525 | 996 | 6759 | 5931 | 5719 | 8082 | 1478 | 1469 | 1443 | 1496 | 6012 | 6439 | 5857 | 6165 | 3794,87 | 1472 | 6118 | 4,16 | 2,057 | 46,624 | 0 | 0 | 0,0008652 | 0,0023851 | 0,0014273 | 0,0014273 | TRUE | 0,01491 | No |
| orf19.6661 | 353 | 307 | 353 | 214 | 1531 | 1217 | 1271 | 1742 | 321 | 310 | 334 | 321 | 1362 | 1321 | 1302 | 1329 | 825,13 | 322 | 1328 | 4,126 | 2,045 | 32,99 | 1,15E-238 | 2,29E-237 | 1,00E-08 | 0,003638 | 0,0016906 | 0,0016906 | TRUE | 0,0082597 | No |
| orf19.7149 | 921 | 776 | 818 | 597 | 3767 | 2964 | 3357 | 4503 | 839 | 785 | 774 | 897 | 3351 | 3218 | 3438 | 3435 | 2091,99 | 824 | 3360 | 4,097 | 2,035 | 37,943 | 0 | 0 | 0,0011968 | 0,0026685 | 0,001969 | 0,001969 | TRUE | 0,0362486 | No |
| orf19.3318 | 313 | 316 | 327 | 187 | 1306 | 1061 | 1233 | 1682 | 285 | 320 | 309 | 281 | 1162 | 1152 | 1263 | 1283 | 756,79 | 299 | 1215 | 4,058 | 2,021 | 27,821 | 2,44E-170 | 3,64E-169 | 0,0019535 | 0,0037825 | 0,0029036 | 0,0029036 | TRUE | 0,0240076 | No |
| orf19.1641 | 38 | 37 | 50 | 32 | 203 | 147 | 157 | 221 | 35 | 37 | 47 | 48 | 181 | 160 | 161 | 169 | 104,62 | 42 | 168 | 4,054 | 2,019 | 12,577 | 2,81E-36 | 1,03E-35 | 1,00E-08 | 0,0146639 | 0,0091711 | 0,0091711 | TRUE | 0,1154556 | No |
| orf19.3668 | 4258 | 3577 | 3770 | 2602 | 16387 | 14344 | 14074 | 20508 | 3877 | 3618 | 3568 | 3909 | 14576 | 15574 | 14414 | 15644 | 9397,2 | 3743 | 15052 | 4,024 | 2,009 | 42,541 | 0 | 0 | 0,0017923 | 0,0021776 | 0,0019659 | 0,0019659 | TRUE | 0,0120808 | No |
| orf19.4505 | 162 | 138 | 129 | 99 | 612 | 496 | 537 | 786 | 147 | 140 | 122 | 149 | 544 | 539 | 550 | 600 | 348,78 | 140 | 558 | 4,026 | 2,009 | 21,282 | 1,65E-100 | 1,44E-99 | 1,00E-08 | 0,0058245 | 0,0039022 | 0,0039022 | TRUE | 0,0785305 | No |
| orf19.2583 | 9 | 7 | 13 | 4 | 38 | 37 | 34 | 36 | 8 | 7 | 12 | 6 | 34 | 40 | 35 | 27 | 21,23 | 8 | 34 | 3,932 | 1,975 | 5,656 | 1,55E-08 | 2,76E-08 | 1,00E-08 | 0,0642607 | 0,0418286 | 0,0418286 | TRUE | 0,3150605 | No |
| orf19.6398 | 410 | 344 | 392 | 256 | 1622 | 1344 | 1405 | 1847 | 373 | 348 | 371 | 385 | 1443 | 1459 | 1439 | 1409 | 903,31 | 369 | 1438 | 3,901 | 1,964 | 33,052 | 1,46E-239 | 2,94E-238 | 1,00E-08 | 0,0034994 | 0,0016231 | 0,0016231 | TRUE | 0,0168709 | No |
| orf19.1258 | 92 | 91 | 70 | 52 | 343 | 261 | 329 | 414 | 84 | 92 | 66 | 78 | 305 | 283 | 337 | 316 | 195,17 | 80 | 310 | 3,878 | 1,955 | 15,784 | 3,99E-56 | 2,06E-55 | 0,0021613 | 0,0088055 | 0,0066061 | 0,0066061 | TRUE | 0,1453049 | No |
| orf19.6191 | 22 | 14 | 18 | 9 | 73 | 53 | 76 | 71 | 20 | 14 | 17 | 14 | 65 | 58 | 78 | 54 | 39,9 | 16 | 64 | 3,855 | 1,947 | 7,508 | 5,98E-14 | 1,28E-13 | 1,00E-08 | 0,0351435 | 0,0250082 | 0,0250082 | TRUE | 0,1931335 | No |
| orf19.7281 | 2115 | 1813 | 1971 | 1252 | 8157 | 6659 | 6973 | 9506 | 1926 | 1834 | 1865 | 1881 | 7255 | 7230 | 7141 | 7251 | 4547,88 | 1876 | 7219 | 3,847 | 1,944 | 55,977 | 0 | 0 | 1,00E-08 | 0,0023275 | 0,000808 | 0,000808 | TRUE | 0,0041783 | No |
| orf19.4011 | 46 | 34 | 53 | 29 | 189 | 158 | 154 | 206 | 42 | 34 | 50 | 44 | 168 | 172 | 158 | 157 | 103,06 | 42 | 164 | 3,845 | 1,943 | 12,247 | 1,75E-34 | 6,20E-34 | 1,00E-08 | 0,0148545 | 0,0088715 | 0,0088715 | TRUE | 0,1346819 | No |
| orf19.2941 | 3 | 7 | 5 | 3 | 18 | 20 | 12 | 29 | 3 | 7 | 5 | 5 | 16 | 22 | 12 | 22 | 11,4 | 5 | 18 | 3,82 | 1,934 | 4,048 | 5,17E-05 | 7,87E-05 | 1,00E-08 | 0,1179325 | 0,0816421 | 0,0816421 | TRUE | 0,2209575 | No |
| orf19.1257 | 12 | 9 | 10 | 9 | 53 | 37 | 38 | 45 | 11 | 9 | 9 | 14 | 47 | 40 | 39 | 34 | 25,45 | 11 | 40 | 3,798 | 1,925 | 6,149 | 7,80E-10 | 1,46E-09 | 1,00E-08 | 0,0539507 | 0,0323717 | 0,0323717 | TRUE | 0,115233 | No |
| orf19.4821 | 15 | 19 | 14 | 8 | 80 | 48 | 38 | 75 | 14 | 19 | 13 | 12 | 71 | 52 | 39 | 57 | 34,69 | 14 | 55 | 3,758 | 1,91 | 6,587 | 4,49E-11 | 8,84E-11 | 1,00E-08 | 0,0401144 | 0,0360966 | 0,0360966 | TRUE | 0,1053154 | No |
| orf19.278 | 1349 | 1157 | 1295 | 908 | 5013 | 4334 | 4504 | 6314 | 1228 | 1170 | 1226 | 1364 | 4459 | 4706 | 4613 | 4816 | 2947,68 | 1247 | 4648 | 3,74 | 1,903 | 36,96 | 5,08E-299 | 1,32E-297 | 0,0015725 | 0,0024852 | 0,0020127 | 0,0020127 | TRUE | 0,0425215 | No |
| orf19.5000 | 4355 | 3924 | 4197 | 2534 | 16458 | 13149 | 14825 | 19103 | 3965 | 3969 | 3972 | 3806 | 14639 | 14276 | 15183 | 14572 | 9297,74 | 3928 | 14668 | 3,731 | 1,9 | 54,735 | 0 | 0 | 0,00033 | 0,0021791 | 0,0009877 | 0,0009877 | TRUE | 0,0067845 | No |
| orf19.2061 | 7 | 4 | 1 | 1 | 13 | 18 | 14 | 7 | 6 | 4 | 1 | 2 | 12 | 20 | 14 | 5 | 7,96 | 3 | 13 | 3,733 | 1,9 | 3,019 | 0,0025396 | 0,0034742 | 0,2168446 | 0,1680675 | 0,184675 | 0,184675 | TRUE | 0,2625376 | No |
| orf19.5307 | 1372 | 1154 | 1221 | 781 | 4781 | 3952 | 4379 | 5937 | 1249 | 1167 | 1156 | 1173 | 4253 | 4291 | 4485 | 4529 | 2787,73 | 1186 | 4390 | 3,698 | 1,887 | 41,057 | 0 | 0 | 0,0007364 | 0,0025109 | 0,0014699 | 0,0014699 | TRUE | 0,0164304 | No |
| orf19.6475 | 32 | 26 | 29 | 20 | 129 | 90 | 91 | 140 | 29 | 26 | 27 | 30 | 115 | 98 | 93 | 107 | 65,67 | 28 | 103 | 3,679 | 1,879 | 9,658 | 4,56E-22 | 1,21E-21 | 1,00E-08 | 0,0221525 | 0,0130286 | 0,0130286 | TRUE | 0,056596 | No |
| orf19.5542 | 67 | 55 | 82 | 58 | 295 | 232 | 238 | 337 | 61 | 56 | 78 | 87 | 262 | 252 | 244 | 257 | 162,06 | 70 | 254 | 3,67 | 1,876 | 14,136 | 2,28E-45 | 9,82E-45 | 1,00E-08 | 0,0101885 | 0,0074082 | 0,0074082 | TRUE | 0,1652704 | No |
| orf19.5874 | 2 | 1 | 1 | 1 | 6 | 5 | 3 | 7 | 2 | 1 | 1 | 2 | 5 | 5 | 3 | 5 | 3,06 | 2 | 4 | 3,668 | 1,875 | 2,149 | 0,0316614 | 0,0399212 | 1,00E-08 | 0,4341181 | 0,2321376 | 0,2321376 | TRUE | 0,1074016 | No |
| orf19.206 | 1439 | 1228 | 1302 | 945 | 5160 | 4436 | 4637 | 6244 | 1310 | 1242 | 1232 | 1420 | 4590 | 4816 | 4749 | 4763 | 3015,21 | 1301 | 4730 | 3,647 | 1,867 | 38,151 | 0 | 0 | 0,0010695 | 0,0024751 | 0,0017856 | 0,0017856 | TRUE | 0,0399493 | No |
| orf19.3653 | 5454 | 4726 | 5023 | 3349 | 19669 | 16366 | 16976 | 24091 | 4966 | 4780 | 4754 | 5031 | 17495 | 17769 | 17386 | 18377 | 11319,53 | 4883 | 17757 | 3,639 | 1,864 | 52,186 | 0 | 0 | 0,0005297 | 0,0021537 | 0,0010878 | 0,0010878 | TRUE | 0,0068354 | No |
| orf19.2452 | 974 | 1041 | 1057 | 629 | 3879 | 3398 | 3517 | 4463 | 887 | 1053 | 1000 | 945 | 3450 | 3689 | 3602 | 3404 | 2253,83 | 971 | 3536 | 3,639 | 1,863 | 32,229 | 6,98E-228 | 1,34E-226 | 0,0025348 | 0,0026231 | 0,0025248 | 0,0025248 | TRUE | 0,0448409 | No |
| orf19.6065 | 1117 | 1020 | 1061 | 615 | 4151 | 3318 | 3417 | 4817 | 1017 | 1032 | 1004 | 924 | 3692 | 3602 | 3499 | 3674 | 2305,63 | 994 | 3617 | 3,627 | 1,859 | 39,711 | 0 | 0 | 0,0003829 | 0,00261 | 0,0014386 | 0,0014386 | TRUE | 0,0226471 | No |
| orf19.7049 | 911 | 902 | 898 | 592 | 3801 | 3051 | 2812 | 3984 | 829 | 912 | 850 | 889 | 3381 | 3313 | 2880 | 3039 | 2011,64 | 870 | 3153 | 3,627 | 1,859 | 29,201 | 1,90E-187 | 3,04E-186 | 0,0034721 | 0,0026937 | 0,0031243 | 0,0031243 | TRUE | 0,0407859 | No |
| orf19.36.1 | 196 | 118 | 120 | 91 | 663 | 418 | 414 | 684 | 178 | 119 | 114 | 137 | 590 | 454 | 424 | 522 | 317,17 | 137 | 498 | 3,628 | 1,859 | 12,997 | 1,27E-38 | 4,85E-38 | 0,0277428 | 0,006202 | 0,0148013 | 0,0148013 | TRUE | 0,4238926 | No |
| orf19.4287 | 808 | 629 | 729 | 432 | 2804 | 2200 | 2451 | 3210 | 736 | 636 | 690 | 649 | 2494 | 2389 | 2510 | 2449 | 1569,01 | 678 | 2460 | 3,62 | 1,856 | 33,952 | 1,15E-252 | 2,43E-251 | 0,0006308 | 0,0028789 | 0,0018931 | 0,0018931 | TRUE | 0,0409908 | No |
| orf19.5468 | 94 | 80 | 77 | 44 | 303 | 276 | 250 | 376 | 86 | 81 | 73 | 66 | 270 | 300 | 256 | 287 | 177,18 | 76 | 278 | 3,603 | 1,849 | 14,905 | 3,08E-50 | 1,42E-49 | 1,00E-08 | 0,0094925 | 0,0062346 | 0,0062346 | TRUE | 0,061086 | No |
| orf19.2693 | 2146 | 1846 | 1969 | 1151 | 7687 | 6036 | 6426 | 8763 | 1954 | 1867 | 1863 | 1729 | 6837 | 6553 | 6581 | 6684 | 4258,7 | 1853 | 6664 | 3,587 | 1,843 | 42,866 | 0 | 0 | 0,0006418 | 0,0023472 | 0,0014139 | 0,0014139 | TRUE | 0,0219865 | No |
| orf19.5934 | 281 | 275 | 281 | 156 | 998 | 847 | 933 | 1243 | 256 | 278 | 266 | 234 | 888 | 920 | 956 | 948 | 593,15 | 258 | 928 | 3,568 | 1,835 | 25,354 | 8,19E-142 | 1,01E-140 | 1,00E-08 | 0,0042641 | 0,0024844 | 0,0024844 | TRUE | 0,0330548 | No |
| orf19.4394 | 17 | 11 | 18 | 19 | 60 | 61 | 61 | 84 | 15 | 11 | 17 | 29 | 53 | 66 | 62 | 64 | 39,79 | 18 | 61 | 3,552 | 1,829 | 7,051 | 1,78E-12 | 3,67E-12 | 1,00E-08 | 0,0352358 | 0,0263917 | 0,0263917 | TRUE | 0,4953001 | No |
| orf19.700 | 2001 | 1942 | 2045 | 1130 | 7471 | 5868 | 6690 | 8474 | 1822 | 1964 | 1935 | 1697 | 6645 | 6371 | 6851 | 6464 | 4218,81 | 1854 | 6583 | 3,542 | 1,825 | 35,928 | 1,13E-282 | 2,68E-281 | 0,0022048 | 0,0023501 | 0,002116 | 0,002116 | TRUE | 0,035819 | No |
| orf19.6315 | 91 | 74 | 75 | 54 | 331 | 257 | 259 | 331 | 83 | 75 | 71 | 81 | 294 | 279 | 265 | 252 | 175,12 | 78 | 272 | 3,528 | 1,819 | 14,774 | 2,17E-49 | 9,87E-49 | 1,00E-08 | 0,0095803 | 0,0059833 | 0,0059833 | TRUE | 0,0341987 | No |
| orf19.7597 | 4 | 1 | 6 | 4 | 16 | 14 | 9 | 22 | 4 | 1 | 6 | 6 | 14 | 15 | 9 | 17 | 8,97 | 4 | 14 | 3,512 | 1,812 | 3,412 | 0,0006458 | 0,0009227 | 1,00E-08 | 0,1492832 | 0,1041065 | 0,1041065 | TRUE | 0,4428141 | No |
| orf19.5572 | 412 | 378 | 398 | 228 | 1460 | 1217 | 1214 | 1699 | 375 | 382 | 377 | 342 | 1299 | 1321 | 1243 | 1296 | 829,48 | 369 | 1290 | 3,478 | 1,798 | 29,263 | 3,11E-188 | 4,99E-187 | 1,00E-08 | 0,0036296 | 0,0018312 | 0,0018312 | TRUE | 0,0207341 | No |
| orf19.3884 | 338 | 270 | 299 | 202 | 1089 | 853 | 1041 | 1407 | 308 | 273 | 283 | 303 | 969 | 926 | 1066 | 1073 | 650,17 | 292 | 1008 | 3,465 | 1,793 | 22,927 | 2,49E-116 | 2,48E-115 | 0,0030603 | 0,0040688 | 0,0035766 | 0,0035766 | TRUE | 0,0349624 | No |
| orf19.7495 | 173 | 177 | 180 | 85 | 630 | 491 | 542 | 743 | 158 | 179 | 170 | 128 | 560 | 533 | 555 | 567 | 356,23 | 159 | 554 | 3,452 | 1,787 | 19,463 | 2,27E-84 | 1,70E-83 | 1,00E-08 | 0,0057453 | 0,0039473 | 0,0039473 | TRUE | 0,1315604 | No |
| orf19.893 | 40 | 44 | 40 | 36 | 171 | 154 | 118 | 188 | 36 | 45 | 38 | 54 | 152 | 167 | 121 | 143 | 94,55 | 43 | 146 | 3,441 | 1,783 | 10,201 | 1,96E-24 | 5,56E-24 | 0,0091733 | 0,0160083 | 0,0135999 | 0,0135999 | TRUE | 0,1546617 | No |
| orf19.2281 | 125 | 99 | 107 | 60 | 397 | 312 | 337 | 473 | 114 | 100 | 101 | 90 | 353 | 339 | 345 | 361 | 225,39 | 101 | 350 | 3,42 | 1,774 | 16,673 | 2,06E-62 | 1,16E-61 | 1,00E-08 | 0,0078979 | 0,0043594 | 0,0043594 | TRUE | 0,0688425 | No |
| orf19.3982 | 234 | 195 | 219 | 124 | 761 | 678 | 661 | 881 | 213 | 197 | 207 | 186 | 677 | 736 | 677 | 672 | 445,72 | 201 | 690 | 3,412 | 1,771 | 22,01 | 2,33E-107 | 2,14E-106 | 1,00E-08 | 0,0050007 | 0,0029122 | 0,0029122 | TRUE | 0,0223051 | No |
| orf19.1628 | 1150 | 1005 | 1053 | 631 | 3894 | 3259 | 3208 | 4473 | 1047 | 1016 | 997 | 948 | 3464 | 3538 | 3285 | 3412 | 2213,41 | 1002 | 3425 | 3,409 | 1,769 | 37,111 | 1,90E-301 | 4,94E-300 | 0,0004857 | 0,0026338 | 0,0015132 | 0,0015132 | TRUE | 0,01339 | No |
| orf19.2496 | 2143 | 1778 | 1878 | 1371 | 7318 | 5940 | 6094 | 8663 | 1951 | 1798 | 1777 | 2059 | 6509 | 6449 | 6241 | 6608 | 4174,22 | 1896 | 6452 | 3,409 | 1,769 | 34,781 | 4,76E-265 | 1,07E-263 | 0,0023078 | 0,0023535 | 0,0021283 | 0,0021283 | TRUE | 0,037135 | No |
| orf19.3931 | 1858 | 1574 | 1686 | 1158 | 6181 | 5290 | 5405 | 7473 | 1692 | 1592 | 1596 | 1740 | 5498 | 5743 | 5535 | 5700 | 3636,98 | 1655 | 5619 | 3,402 | 1,766 | 41,467 | 0 | 0 | 0,0005058 | 0,0024002 | 0,0013341 | 0,0013341 | TRUE | 0,0125706 | No |
| orf19.7136 | 1671 | 1298 | 1486 | 935 | 5068 | 4059 | 4963 | 6627 | 1521 | 1313 | 1406 | 1405 | 4508 | 4407 | 5083 | 5055 | 3087,21 | 1411 | 4763 | 3,374 | 1,755 | 27,28 | 7,32E-164 | 1,05E-162 | 0,0040071 | 0,0024649 | 0,0034928 | 0,0034928 | TRUE | 0,0345497 | No |
| orf19.7537 | 151 | 98 | 146 | 103 | 498 | 375 | 436 | 612 | 137 | 99 | 138 | 155 | 443 | 407 | 447 | 467 | 286,62 | 132 | 441 | 3,362 | 1,749 | 15,695 | 1,65E-55 | 8,39E-55 | 0,0072578 | 0,0066459 | 0,0068115 | 0,0068115 | TRUE | 0,294068 | No |
| orf19.6257 | 4608 | 4220 | 4502 | 2739 | 15666 | 12979 | 13802 | 18703 | 4195 | 4268 | 4261 | 4114 | 13934 | 14092 | 14135 | 14267 | 9158,3 | 4210 | 14107 | 3,349 | 1,744 | 58,774 | 0 | 0 | 1,00E-08 | 0,0021813 | 0,0006844 | 0,0006844 | TRUE | 0,0025314 | No |
| orf19.1631 | 2288 | 2192 | 2248 | 1314 | 8037 | 6630 | 6773 | 8900 | 2083 | 2217 | 2128 | 1974 | 7149 | 7198 | 6936 | 6789 | 4559,23 | 2100 | 7018 | 3,336 | 1,738 | 39,509 | 0 | 0 | 0,001113 | 0,0023268 | 0,0015356 | 0,0015356 | TRUE | 0,0178664 | No |

| | | | | | | | | | | | | | | | | | | | | | | | | | | | | | | | |
|---|---|---|---|---|---|---|---|---|---|---|---|---|---|---|---|---|---|---|---|---|---|---|---|---|---|---|---|---|---|---|---|
| orf19.1064 | 4444 | 4108 | 4375 | 2750 | 14815 | 12917 | 13200 | 18570 | 4046 | 4155 | 4141 | 4131 | 13177 | 14024 | 13518 | 14165 | 8919,74 | 4118 | 13721 | 3,333 | 1,737 | 47,629 | 0 | 0 | 0,0006167 | 0,0021851 | 0,0011117 | 0,0011117 | TRUE | 0,0087726 | No |
| orf19.1239 | 10 | 3 | 6 | 4 | 19 | 25 | 17 | 25 | 9 | 3 | 6 | 6 | 17 | 27 | 17 | 19 | 13,04 | 6 | 20 | 3,33 | 1,735 | 3,976 | 7,01E-05 | 0,0001059 | 1,00E-08 | 0,1033122 | 0,071527 | 0,071527 | TRUE | 0,3688113 | No |
| orf19.4195.1 | 2467 | 2025 | 2161 | 1220 | 8082 | 5566 | 6822 | 9028 | 2246 | 2048 | 2045 | 1833 | 7189 | 6043 | 6987 | 6887 | 4409,63 | 2043 | 6776 | 3,312 | 1,728 | 25,103 | 4,66E-139 | 5,64E-138 | 0,0057638 | 0,0023366 | 0,0042175 | 0,0042175 | TRUE | 0,0644127 | No |
| orf19.2072 | 6 | 9 | 10 | 17 | 41 | 31 | 48 | 44 | 5 | 9 | 9 | 26 | 36 | 34 | 49 | 34 | 25,3 | 12 | 38 | 3,305 | 1,725 | 4,738 | 2,16E-06 | 3,54E-06 | 0,1129647 | 0,0542469 | 0,0688838 | 0,0688838 | TRUE | 1,3782883 | No |
| orf19.3461 | 11 | 10 | 18 | 11 | 50 | 39 | 45 | 54 | 10 | 10 | 17 | 17 | 44 | 42 | 46 | 41 | 28,47 | 14 | 43 | 3,296 | 1,721 | 5,975 | 2,30E-09 | 4,25E-09 | 1,00E-08 | 0,0484322 | 0,0288019 | 0,0288019 | TRUE | 0,1943367 | No |
| orf19.5137 | 12 | 19 | 14 | 9 | 59 | 42 | 33 | 71 | 11 | 19 | 13 | 14 | 52 | 46 | 34 | 54 | 30,37 | 14 | 46 | 3,295 | 1,72 | 5,861 | 4,59E-09 | 8,38E-09 | 1,00E-08 | 0,0455364 | 0,0354014 | 0,0354014 | TRUE | 0,2633815 | No |
| orf19.1847 | 1025 | 872 | 1033 | 685 | 3308 | 2999 | 3081 | 4176 | 933 | 882 | 978 | 1029 | 2942 | 3256 | 3155 | 3185 | 2045,13 | 956 | 3134 | 3,289 | 1,718 | 30,384 | 9,02E-203 | 1,56E-201 | 0,0021718 | 0,0026829 | 0,0023563 | 0,0023563 | TRUE | 0,0310484 | No |
| orf19.2551 | 7293 | 6391 | 6606 | 4724 | 24236 | 19944 | 20097 | 29306 | 6640 | 6464 | 6252 | 7096 | 21557 | 21654 | 20582 | 22355 | 14074,93 | 6613 | 21537 | 3,259 | 1,705 | 36,839 | 4,38E-297 | 1,12E-295 | 0,0019156 | 0,0021309 | 0,0019529 | 0,0019529 | TRUE | 0,0286743 | No |
| orf19.4834 | 26 | 31 | 24 | 16 | 95 | 74 | 87 | 102 | 24 | 31 | 23 | 24 | 84 | 80 | 89 | 78 | 54,19 | 26 | 83 | 3,252 | 1,701 | 8,188 | 2,66E-16 | 6,10E-16 | 1,00E-08 | 0,0264141 | 0,0150499 | 0,0150499 | TRUE | 0,157608 | No |
| orf19.7127 | 26 | 19 | 28 | 16 | 89 | 68 | 69 | 104 | 24 | 19 | 26 | 24 | 79 | 74 | 71 | 79 | 49,55 | 23 | 76 | 3,251 | 1,701 | 7,893 | 2,96E-15 | 6,59E-15 | 1,00E-08 | 0,0286962 | 0,0158014 | 0,0158014 | TRUE | 0,0887459 | No |
| orf19.7395 | 0 | 2 | 2 | 3 | 6 | 3 | 5 | 14 | 0 | 2 | 2 | 5 | 5 | 3 | 5 | 11 | 4,1 | 2 | 6 | 3,251 | 1,701 | 2,111 | 0,0347465 | 0,0436729 | 0,1578397 | 0,3240745 | 0,2710572 | 0,2710572 | TRUE | 0,7867741 | No |
| orf19.4991 | 974 | 937 | 1036 | 692 | 3207 | 3095 | 3052 | 4089 | 887 | 948 | 980 | 1039 | 2853 | 3360 | 3126 | 3119 | 2039 | 964 | 3114 | 3,239 | 1,696 | 26,437 | 5,09E-154 | 6,80E-153 | 0,0035037 | 0,0026849 | 0,0032402 | 0,0032402 | TRUE | 0,0389917 | No |
| orf19.3876 | 79 | 63 | 91 | 48 | 255 | 226 | 240 | 312 | 72 | 64 | 86 | 72 | 227 | 245 | 246 | 238 | 156,23 | 74 | 239 | 3,241 | 1,696 | 13,366 | 9,57E-41 | 3,77E-40 | 1,00E-08 | 0,0104925 | 0,0063206 | 0,0063206 | TRUE | 0,1370624 | No |
| orf19.7209 | 95 | 81 | 89 | 47 | 319 | 255 | 270 | 292 | 86 | 82 | 84 | 71 | 284 | 277 | 277 | 223 | 172,89 | 81 | 265 | 3,24 | 1,696 | 12,945 | 2,51E-38 | 9,52E-38 | 1,00E-08 | 0,0096778 | 0,0081994 | 0,0081994 | TRUE | 0,1405591 | No |
| orf19.5616 | 42 | 40 | 39 | 35 | 143 | 143 | 114 | 176 | 38 | 40 | 37 | 53 | 127 | 155 | 117 | 134 | 87,7 | 42 | 133 | 3,228 | 1,691 | 9,722 | 2,43E-22 | 6,51E-22 | 0,0045633 | 0,0170989 | 0,0126851 | 0,0126851 | TRUE | 0,1489173 | No |
| orf19.118 | 18331 | 17865 | 18334 | 11007 | 62301 | 52894 | 53277 | 71001 | 16690 | 18069 | 17351 | 16534 | 55415 | 57428 | 54562 | 54160 | 36276,16 | 17161 | 55391 | 3,227 | 1,69 | 43,217 | 0 | 0 | 0,0011289 | 0,0020734 | 0,0014296 | 0,0014296 | TRUE | 0,0155011 | No |
| orf19.3908 | 23 | 20 | 26 | 19 | 68 | 72 | 82 | 101 | 21 | 20 | 25 | 29 | 60 | 78 | 84 | 77 | 49,25 | 24 | 75 | 3,22 | 1,687 | 7,497 | 6,52E-14 | 1,40E-13 | 1,00E-08 | 0,0288597 | 0,0195611 | 0,0195611 | TRUE | 0,1611493 | No |
| orf19.3447 | 304 | 383 | 331 | 244 | 1165 | 1130 | 1024 | 1285 | 277 | 387 | 313 | 367 | 1036 | 1227 | 1049 | 980 | 704,49 | 336 | 1073 | 3,204 | 1,68 | 16,327 | 6,37E-60 | 3,48E-59 | 0,0135616 | 0,0039121 | 0,0081242 | 0,0081242 | TRUE | 0,1674587 | No |
| orf19.936 | 8 | 9 | 7 | 4 | 26 | 20 | 24 | 32 | 7 | 9 | 7 | 6 | 23 | 22 | 25 | 24 | 15,36 | 7 | 24 | 3,204 | 1,68 | 4,428 | 9,52E-06 | 1,51E-05 | 1,00E-08 | 0,088061 | 0,0462882 | 0,0462882 | TRUE | 0,0770383 | No |
| orf19.7100 | 190 | 195 | 194 | 133 | 687 | 549 | 593 | 769 | 173 | 197 | 184 | 200 | 611 | 596 | 607 | 587 | 394,33 | 188 | 600 | 3,203 | 1,679 | 20,542 | 9,08E-94 | 7,33E-93 | 1,00E-08 | 0,005387 | 0,0028091 | 0,0028091 | TRUE | 0,0345944 | No |
| orf19.356 | 208 | 195 | 193 | 105 | 677 | 540 | 558 | 766 | 189 | 197 | 183 | 158 | 602 | 586 | 571 | 584 | 383,9 | 182 | 586 | 3,197 | 1,677 | 20,018 | 3,85E-89 | 3,01E-88 | 1,00E-08 | 0,005478 | 0,0030467 | 0,0030467 | TRUE | 0,0618247 | No |
| orf19.7043.1 | 739 | 749 | 662 | 450 | 2743 | 1967 | 2076 | 2638 | 673 | 758 | 627 | 676 | 2440 | 2136 | 2126 | 2012 | 1430,83 | 684 | 2178 | 3,189 | 1,673 | 22,255 | 1,01E-109 | 9,50E-109 | 0,0054246 | 0,0029603 | 0,0044246 | 0,0044246 | TRUE | 0,0798826 | No |
| orf19.2114 | 201 | 157 | 183 | 154 | 616 | 576 | 563 | 788 | 183 | 159 | 173 | 231 | 548 | 625 | 577 | 601 | 387,16 | 186 | 588 | 3,189 | 1,673 | 16,903 | 4,27E-64 | 2,47E-63 | 0,0063021 | 0,005449 | 0,0057178 | 0,0057178 | TRUE | 0,2246305 | No |
| orf19.3152 | 1455 | 1354 | 1331 | 860 | 4647 | 4139 | 3917 | 5272 | 1325 | 1369 | 1260 | 1292 | 4133 | 4494 | 4012 | 4022 | 2738,23 | 1312 | 4165 | 3,174 | 1,666 | 31,848 | 1,38E-222 | 2,60E-221 | 0,0018107 | 0,0025194 | 0,0021061 | 0,0021061 | TRUE | 0,0338939 | No |
| orf19.3250 | 1316 | 1340 | 1391 | 783 | 4638 | 3612 | 4075 | 5013 | 1198 | 1355 | 1316 | 1176 | 4125 | 3922 | 4173 | 3824 | 2636,29 | 1261 | 4011 | 3,174 | 1,666 | 29,671 | 1,84E-193 | 3,02E-192 | 0,0025783 | 0,0025381 | 0,0024858 | 0,0024858 | TRUE | 0,0294373 | No |
| orf19.5641 | 17672 | 13823 | 15433 | 9742 | 52250 | 40579 | 46903 | 64973 | 16090 | 13980 | 14606 | 14634 | 46475 | 44057 | 48035 | 49562 | 30929,85 | 14828 | 47032 | 3,172 | 1,665 | 31,565 | 1,10E-218 | 2,03E-217 | 0,0030059 | 0,0020797 | 0,0026277 | 0,0026277 | TRUE | 0,0404029 | No |
| orf19.3429 | 298 | 318 | 317 | 216 | 1059 | 910 | 927 | 1244 | 271 | 322 | 300 | 324 | 942 | 988 | 949 | 949 | 630,71 | 304 | 957 | 3,161 | 1,66 | 23,552 | 1,19E-122 | 1,24E-121 | 1,00E-08 | 0,0041315 | 0,0025251 | 0,0025251 | TRUE | 0,0645249 | No |
| orf19.7279 | 136 | 120 | 120 | 112 | 487 | 378 | 383 | 525 | 124 | 121 | 114 | 168 | 433 | 410 | 392 | 400 | 270,41 | 132 | 409 | 3,161 | 1,66 | 15,338 | 4,25E-53 | 2,07E-52 | 0,0031728 | 0,0069222 | 0,0060004 | 0,0060004 | TRUE | 0,2766466 | No |
| orf19.6844 | 30395 | 23819 | 25861 | 18126 | 88736 | 73809 | 79342 | 1E+05 | 27673 | 24090 | 24475 | 27228 | 78928 | 80136 | 81256 | 85613 | 53675 | 25866 | 81483 | 3,15 | 1,656 | 31,052 | 1,07E-211 | 1,92E-210 | 0,0031481 | 0,0020616 | 0,0027045 | 0,0027045 | TRUE | 0,0272572 | No |
| orf19.7385 | 414 | 316 | 370 | 223 | 1146 | 973 | 1119 | 1449 | 377 | 320 | 350 | 335 | 1019 | 1056 | 1146 | 1105 | 713,59 | 346 | 1082 | 3,123 | 1,643 | 22,861 | 1,14E-115 | 1,13E-114 | 0,0019151 | 0,0038882 | 0,0029869 | 0,0029869 | TRUE | 0,0462572 | No |
| orf19.2903 | 195 | 270 | 237 | 138 | 778 | 662 | 655 | 845 | 178 | 273 | 224 | 207 | 692 | 719 | 671 | 645 | 451,04 | 220 | 682 | 3,086 | 1,626 | 15,491 | 4,00E-54 | 1,99E-53 | 0,011694 | 0,0049658 | 0,0074654 | 0,0074654 | TRUE | 0,2922038 | No |
| orf19.857 | 1140 | 981 | 1060 | 671 | 3394 | 2834 | 2988 | 4332 | 1038 | 992 | 1003 | 1008 | 3019 | 3077 | 3060 | 3304 | 2062,7 | 1010 | 3115 | 3,084 | 1,625 | 33,292 | 5,09E-243 | 1,03E-241 | 0,0009096 | 0,0026774 | 0,0016072 | 0,0016072 | TRUE | 0,0210635 | No |
| orf19.3306 | 20 | 20 | 21 | 14 | 59 | 59 | 57 | 89 | 18 | 20 | 20 | 21 | 52 | 64 | 58 | 68 | 40,27 | 20 | 60 | 3,082 | 1,624 | 6,8 | 1,05E-11 | 2,11E-11 | 1,00E-08 | 0,0348423 | 0,0202444 | 0,0202444 | TRUE | 0,0785146 | No |
| orf19.1591 | 1521 | 1327 | 1370 | 1009 | 4693 | 3962 | 3950 | 5868 | 1385 | 1342 | 1297 | 1516 | 4174 | 4302 | 4045 | 4476 | 2817,07 | 1385 | 4249 | 3,078 | 1,622 | 29,35 | 2,42E-189 | 3,91E-188 | 0,0026215 | 0,0025059 | 0,0024323 | 0,0024323 | TRUE | 0,044299 | No |
| orf19.3735 | 12 | 11 | 9 | 18 | 44 | 41 | 37 | 58 | 11 | 11 | 9 | 27 | 39 | 45 | 38 | 44 | 27,92 | 14 | 42 | 3,069 | 1,618 | 5,162 | 2,45E-07 | 4,17E-07 | 0,0107955 | 0,0493435 | 0,0437884 | 0,0437884 | TRUE | 1,0168811 | No |
| orf19.6570 | 142 | 130 | 154 | 99 | 446 | 412 | 415 | 554 | 129 | 131 | 146 | 149 | 397 | 447 | 425 | 423 | 280,86 | 139 | 423 | 3,059 | 1,613 | 16,719 | 9,52E-63 | 5,41E-62 | 1,00E-08 | 0,0067405 | 0,0039911 | 0,0039911 | TRUE | 0,0230383 | No |
| orf19.5342.2 | 190 | 123 | 157 | 101 | 594 | 332 | 471 | 596 | 173 | 124 | 149 | 152 | 528 | 360 | 482 | 455 | 302,94 | 150 | 456 | 3,056 | 1,612 | 12,354 | 4,66E-35 | 1,67E-34 | 0,0185962 | 0,0063977 | 0,0117365 | 0,0117365 | TRUE | 0,2253574 | No |
| orf19.804 | 462 | 410 | 464 | 263 | 1403 | 1131 | 1323 | 1666 | 421 | 415 | 439 | 395 | 1248 | 1228 | 1355 | 1271 | 846,39 | 418 | 1276 | 3,045 | 1,607 | 25,518 | 1,26E-143 | 1,57E-142 | 1,00E-08 | 0,0035978 | 0,0021673 | 0,0021673 | TRUE | 0,0204385 | No |
| orf19.5875 | 254 | 198 | 222 | 151 | 719 | 600 | 651 | 892 | 231 | 200 | 210 | 227 | 640 | 651 | 667 | 680 | 438,32 | 217 | 660 | 3,045 | 1,606 | 20,427 | 9,60E-93 | 7,71E-92 | 1,00E-08 | 0,0050508 | 0,0027812 | 0,0027812 | TRUE | 0,0301899 | No |
| orf19.218 | 28 | 20 | 24 | 15 | 84 | 58 | 75 | 82 | 25 | 20 | 23 | 23 | 75 | 63 | 77 | 63 | 46 | 23 | 70 | 3,033 | 1,601 | 7,147 | 8,85E-13 | 1,84E-12 | 1,00E-08 | 0,0307533 | 0,0182078 | 0,0182078 | TRUE | 0,0462458 | No |
| orf19.5784 | 786 | 710 | 707 | 463 | 2310 | 2033 | 2027 | 2808 | 716 | 718 | 669 | 695 | 2055 | 2207 | 2076 | 2142 | 1409,77 | 700 | 2120 | 3,029 | 1,599 | 30,842 | 7,16E-209 | 1,26E-207 | 1,00E-08 | 0,002974 | 0,0015971 | 0,0015971 | TRUE | 0,0113525 | No |
| orf19.7282 | 2788 | 2166 | 2466 | 1625 | 7850 | 6340 | 7203 | 9851 | 2538 | 2191 | 2334 | 2441 | 6982 | 6883 | 7377 | 7514 | 4782,61 | 2376 | 7189 | 3,027 | 1,598 | 29,991 | 1,30E-197 | 2,19E-196 | 0,00249 | 0,0023132 | 0,0024341 | 0,0024341 | TRUE | 0,0338369 | No |
| orf19.6113 | 90 | 64 | 77 | 37 | 242 | 154 | 237 | 284 | 82 | 65 | 73 | 56 | 215 | 167 | 243 | 217 | 139,62 | 69 | 210 | 3,024 | 1,597 | 10,358 | 3,83E-25 | 1,10E-24 | 0,0152822 | 0,0114986 | 0,0129502 | 0,0129502 | TRUE | 0,1969574 | No |
| orf19.7513 | 1876 | 1443 | 1585 | 970 | 5144 | 4182 | 4465 | 6291 | 1708 | 1459 | 1500 | 1457 | 4575 | 4540 | 4573 | 4799 | 3076,51 | 1531 | 4622 | 3,014 | 1,592 | 28,753 | 8,34E-182 | 1,30E-180 | 0,0028001 | 0,0024664 | 0,0024906 | 0,0024906 | TRUE | 0,0750424 | No |
| orf19.5323 | 1022 | 784 | 868 | 609 | 2883 | 2402 | 2424 | 3541 | 930 | 793 | 821 | 915 | 2564 | 2608 | 2482 | 2701 | 1726,94 | 865 | 2589 | 2,999 | 1,584 | 26,541 | 3,29E-155 | 4,45E-154 | 0,0026776 | 0,002802 | 0,0026199 | 0,0026199 | TRUE | 0,0373385 | No |
| orf19.1326 | 267 | 212 | 204 | 136 | 739 | 618 | 600 | 824 | 243 | 214 | 193 | 204 | 657 | 671 | 614 | 629 | 428,27 | 214 | 643 | 2,996 | 1,583 | 18,669 | 8,92E-78 | 6,14E-77 | 0,0011318 | 0,0051215 | 0,0036959 | 0,0036959 | TRUE | 0,1037056 | No |
| orf19.3336 | 6 | 7 | 6 | 2 | 24 | 10 | 11 | 27 | 5 | 7 | 6 | 3 | 21 | 11 | 11 | 21 | 10,66 | 5 | 16 | 2,976 | 1,573 | 3,289 | 0,0010053 | 0,0014188 | 1,00E-08 | 0,1259421 | 0,0938647 | 0,0938647 | TRUE | 0,064698 | No |
| orf19.6518 | 453 | 330 | 400 | 304 | 1227 | 1075 | 1132 | 1649 | 412 | 334 | 379 | 457 | 1091 | 1167 | 1159 | 1258 | 782,14 | 396 | 1169 | 2,974 | 1,572 | 18,601 | 3,15E-77 | 2,16E-76 | 0,0070259 | 0,003726 | 0,0050975 | 0,0050975 | TRUE | 0,1236433 | No |
| orf19.3048 | 161 | 133 | 169 | 117 | 457 | 436 | 452 | 625 | 147 | 135 | 160 | 176 | 406 | 473 | 463 | 477 | 304,54 | 154 | 455 | 2,973 | 1,572 | 15,522 | 2,45E-54 | 1,23E-53 | 0,0037631 | 0,0063747 | 0,0053403 | 0,0053403 | TRUE | 0,072876 | No |
| orf19.3437 | 88 | 89 | 89 | 69 | 310 | 247 | 244 | 335 | 80 | 90 | 84 | 104 | 276 | 268 | 250 | 256 | 175,92 | 90 | 262 | 2,968 | 1,57 | 13,275 | 3,22E-40 | 1,25E-39 | 1,00E-08 | 0,0095462 | 0,0056446 | 0,0056446 | TRUE | 0,0788545 | No |
| orf19.2591 | 51 | 33 | 47 | 29 | 138 | 116 | 118 | 167 | 46 | 33 | 44 | 44 | 123 | 126 | 121 | 127 | 83,1 | 42 | 124 | 2,961 | 1,566 | 9,446 | 3,52E-21 | 9,19E-21 | 1,00E-08 | 0,017934 | 0,010032 | 0,010032 | TRUE | 0,153983 | No |
| orf19.6948 | 1612 | 1383 | 1507 | 982 | 4814 | 3866 | 4152 | 5663 | 1468 | 1399 | 1426 | 1475 | 4282 | 4197 | 4252 | 4320 | 2852,38 | 1442 | 4263 | 2,96 | 1,565 | 40,285 | 0 | 0 | 1,00E-08 | 0,0025001 | 0,0009691 | 0,0009691 | TRUE | 0,0048195 | No |
| orf19.3708 | 15 | 13 | 17 | 5 | 42 | 39 | 41 | 42 | 14 | 13 | 16 | 8 | 37 | 42 | 42 | 32 | 25,52 | 13 | 38 | 2,939 | 1,555 | 5,182 | 2,19E-07 | 3,74E-07 | 1,00E-08 | 0,0538072 | 0,0335287 | 0,0335287 | TRUE | 0,1697446 | No |
| orf19.4716 | 16048 | 13178 | 14286 | 9270 | 45594 | 36142 | 38772 | 54422 | 14611 | 13328 | 13520 | 13925 | 40554 | 39240 | 39707 | 41514 | 27050,01 | 13846 | 40254 | 2,907 | 1,54 | 39,435 | 0 | 0 | 0,0010837 | 0,0020859 | 0,0014138 | 0,0014138 | TRUE | 0,0171631 | No |
| orf19.2929 | 9400 | 9250 | 9451 | 6113 | 27997 | 24986 | 24819 | 35803 | 8558 | 9355 | 8945 | 9183 | 24902 | 27128 | 25418 | 27311 | 17599,97 | 9010 | 26190 | 2,907 | 1,54 | 34,044 | 4,90E-254 | 1,05E-252 | 0,001701 | 0,0021121 | 0,001887 | 0,001887 | TRUE | 0,0140647 | No |
| orf19.5839 | 634 | 493 | 565 | 393 | 1775 | 1422 | 1669 | 2027 | 577 | 499 | 535 | 590 | 1579 | 1544 | 1709 | 1546 | 1072,39 | 550 | 1594 | 2,905 | 1,538 | 23,532 | 1,94E-122 | 2,02E-121 | 0,0024568 | 0,0032688 | 0,0028353 | 0,0028353 | TRUE | 0,0461404 | No |
| orf19.4066 | 1124 | 1439 | 1401 | 873 | 3950 | 3736 | 3737 | 4524 | 1023 | 1455 | 1326 | 1311 | 3513 | 4056 | 3827 | 3451 | 2495,48 | 1279 | 3712 | 2,903 | 1,538 | 16,745 | 6,21E-63 | 3,54E-62 | 0,0123543 | 0,0025664 | 0,0075523 | 0,0075523 | TRUE | 0,2229753 | No |
| orf19.4551 | 2484 | 2417 | 2567 | 1702 | 7890 | 6853 | 6721 | 8842 | 2262 | 2445 | 2429 | 2557 | 7018 | 7440 | 6883 | 6745 | 4722,31 | 2423 | 7022 | 2,901 | 1,537 | 31,245 | 2,60E-214 | 4,70E-213 | 0,0018082 | 0,0023168 | 0,0020332 | 0,0020332 | TRUE | 0,0247108 | No |
| orf19.489 | 553 | 498 | 535 | 345 | 1708 | 1376 | 1406 | 1844 | 503 | 504 | 506 | 518 | 1519 | 1494 | 1440 | 1407 | 986,43 | 508 | 1465 | 2,886 | 1,529 | 26,962 | 4,10E-160 | 5,67E-159 | 1,00E-08 | 0,0033762 | 0,0017198 | 0,0017198 | TRUE | 0,0092508 | No |
| orf19.6229 | 7493 | 6494 | 7055 | 5035 | 21449 | 18944 | 18835 | 27107 | 6822 | 6568 | 6677 | 7563 | 19078 | 20568 | 19289 | 20677 | 13405,41 | 6908 | 19903 | 2,883 | 1,528 | 29,39 | 7,44E-190 | 1,20E-188 | 0,0028694 | 0,0021356 | 0,002494 | 0,002494 | TRUE | 0,0487438 | No |
| orf19.6337 | 25 | 16 | 28 | 19 | 68 | 61 | 64 | 98 | 23 | 16 | 26 | 29 | 60 | 66 | 66 | 75 | 45,12 | 24 | 67 | 2,885 | 1,528 | 6,697 | 2,13E-11 | 4,24E-11 | 1,00E-08 | 0,0313115 | 0,0201938 | 0,0201938 | TRUE | 0,2567313 | No |

| | | | | | | | | | | | | | | | | | | | | | | | | | | | | | | | |
|---|---|---|---|---|---|---|---|---|---|---|---|---|---|---|---|---|---|---|---|---|---|---|---|---|---|---|---|---|---|---|---|
| orf19.4933 | 5888 | 5212 | 5547 | 3822 | 16476 | 15116 | 14966 | 20737 | 5361 | 5271 | 5250 | 5741 | 14655 | 16412 | 15327 | 15818 | 10479,4 | 5406 | 15553 | 2,879 | 1,526 | 32,468 | 3,05E-231 | 5,91E-230 | 0,0018767 | 0,0021631 | 0,0019905 | 0,0019905 | TRUE | 0,020547 | No |
| orf19.5474 | 359 | 275 | 274 | 141 | 838 | 619 | 836 | 1111 | 327 | 278 | 259 | 212 | 745 | 672 | 856 | 847 | 524,65 | 269 | 780 | 2,88 | 1,526 | 13,272 | 3,38E-40 | 1,32E-39 | 0,0186689 | 0,0045549 | 0,0101186 | 0,0101186 | TRUE | 0,2341424 | No |
| orf19.7440 | 44 | 66 | 58 | 38 | 174 | 128 | 167 | 209 | 40 | 67 | 55 | 57 | 155 | 139 | 171 | 159 | 105,37 | 55 | 156 | 2,876 | 1,524 | 9,478 | 2,59E-21 | 6,77E-21 | 0,0057088 | 0,0145734 | 0,0120542 | 0,0120542 | TRUE | 0,3164766 | No |
| orf19.7447 | 39 | 28 | 39 | 18 | 92 | 94 | 90 | 127 | 36 | 28 | 37 | 27 | 82 | 102 | 92 | 97 | 62,59 | 32 | 93 | 2,875 | 1,524 | 7,891 | 2,99E-15 | 6,66E-15 | 1,00E-08 | 0,0231429 | 0,014434 | 0,014434 | TRUE | 0,0715298 | No |
| orf19.5578 | 464 | 430 | 477 | 321 | 1390 | 1169 | 1288 | 1694 | 422 | 435 | 451 | 482 | 1236 | 1269 | 1319 | 1292 | 863,48 | 448 | 1279 | 2,871 | 1,521 | 25,095 | 5,69E-139 | 6,88E-138 | 3,30E-05 | 0,0035669 | 0,0019686 | 0,0019686 | TRUE | 0,0251051 | No |
| orf19.5565 | 298 | 302 | 264 | 199 | 908 | 711 | 773 | 1100 | 271 | 305 | 250 | 299 | 808 | 772 | 792 | 839 | 541,98 | 281 | 803 | 2,869 | 1,521 | 19,82 | 2,00E-87 | 1,54E-86 | 0,001041 | 0,0044744 | 0,0031687 | 0,0031687 | TRUE | 0,0662581 | No |
| orf19.6142 | 13 | 6 | 5 | 4 | 18 | 18 | 22 | 33 | 12 | 6 | 5 | 6 | 16 | 20 | 23 | 25 | 13,99 | 7 | 21 | 2,863 | 1,518 | 3,683 | 0,0002304 | 0,0003392 | 1,00E-08 | 0,0964768 | 0,0679955 | 0,0679955 | TRUE | 0,6232164 | No |
| orf19.1037 | 93 | 97 | 93 | 52 | 284 | 253 | 240 | 305 | 85 | 98 | 88 | 78 | 253 | 275 | 246 | 233 | 169,33 | 87 | 252 | 2,858 | 1,515 | 12,476 | 1,01E-35 | 3,65E-35 | 1,00E-08 | 0,0098383 | 0,0062556 | 0,0062556 | TRUE | 0,0633599 | No |
| orf19.403 | 26 | 23 | 19 | 17 | 89 | 51 | 60 | 78 | 24 | 23 | 18 | 26 | 79 | 55 | 61 | 59 | 43,24 | 23 | 64 | 2,858 | 1,515 | 6,407 | 1,48E-10 | 2,87E-10 | 1,00E-08 | 0,0325862 | 0,0227328 | 0,0227328 | TRUE | 0,2385409 | No |
| orf19.1212 | 804 | 711 | 753 | 557 | 2358 | 2027 | 1997 | 2825 | 732 | 719 | 713 | 837 | 2097 | 2201 | 2045 | 2155 | 1437,34 | 750 | 2124 | 2,847 | 1,509 | 26,223 | 1,44E-151 | 1,90E-150 | 0,0013611 | 0,0029561 | 0,0022435 | 0,0022435 | TRUE | 0,0615302 | No |
| orf19.5353 | 13 | 11 | 14 | 6 | 29 | 35 | 33 | 44 | 12 | 11 | 13 | 9 | 26 | 38 | 34 | 34 | 22,05 | 11 | 33 | 2,843 | 1,507 | 4,741 | 2,13E-06 | 3,49E-06 | 1,00E-08 | 0,0619535 | 0,0367406 | 0,0367406 | TRUE | 0,1516263 | No |
| orf19.3577.1 | 97 | 86 | 84 | 69 | 305 | 224 | 223 | 337 | 88 | 87 | 79 | 104 | 271 | 243 | 228 | 257 | 169,8 | 90 | 250 | 2,826 | 1,499 | 12,232 | 2,09E-34 | 7,38E-34 | 1,00E-08 | 0,0098169 | 0,006567 | 0,006567 | TRUE | 0,0772845 | No |
| orf19.7666 | 18 | 13 | 16 | 11 | 43 | 57 | 29 | 56 | 16 | 13 | 15 | 17 | 38 | 62 | 30 | 43 | 29,22 | 15 | 43 | 2,819 | 1,495 | 4,917 | 8,78E-07 | 1,46E-06 | 1,00E-08 | 0,0472471 | 0,0430607 | 0,0430607 | TRUE | 0,3205208 | No |
| orf19.6287 | 4746 | 4065 | 4354 | 3002 | 13325 | 11273 | 11402 | 16066 | 4321 | 4111 | 4121 | 4509 | 11852 | 12239 | 11677 | 12255 | 8135,79 | 4266 | 12006 | 2,817 | 1,494 | 36,71 | 5,14E-295 | 1,30E-293 | 0,0010588 | 0,0021994 | 0,0014259 | 0,0014259 | TRUE | 0,0169923 | No |
| orf19.1473 | 356 | 357 | 380 | 202 | 1101 | 888 | 883 | 1268 | 324 | 361 | 360 | 303 | 979 | 964 | 904 | 967 | 645,4 | 337 | 954 | 2,816 | 1,494 | 21,152 | 2,63E-99 | 2,28E-98 | 0,000622 | 0,0040838 | 0,0027262 | 0,0027262 | TRUE | 0,0398925 | No |
| orf19.6837 | 279 | 213 | 250 | 183 | 803 | 625 | 655 | 883 | 254 | 215 | 237 | 275 | 714 | 679 | 671 | 674 | 464,77 | 245 | 684 | 2,809 | 1,49 | 18,658 | 1,08E-77 | 7,43E-77 | 6,52E-05 | 0,0048793 | 0,0032574 | 0,0032574 | TRUE | 0,0710103 | No |
| orf19.639 | 164 | 140 | 144 | 108 | 483 | 374 | 418 | 498 | 149 | 142 | 136 | 162 | 430 | 406 | 428 | 380 | 279,13 | 147 | 411 | 2,803 | 1,487 | 15,302 | 7,41E-53 | 3,59E-52 | 1,00E-08 | 0,0067695 | 0,0043045 | 0,0043045 | TRUE | 0,0360347 | No |
| orf19.4055 | 243 | 172 | 178 | 117 | 586 | 432 | 527 | 720 | 221 | 174 | 168 | 176 | 521 | 469 | 540 | 549 | 352,33 | 185 | 520 | 2,804 | 1,487 | 14,833 | 9,02E-50 | 4,14E-49 | 0,0062323 | 0,0057864 | 0,0058835 | 0,0058835 | TRUE | 0,2073318 | No |
| orf19.3107 | 97 | 102 | 94 | 45 | 301 | 227 | 249 | 289 | 88 | 103 | 89 | 68 | 268 | 246 | 255 | 220 | 167,21 | 87 | 247 | 2,802 | 1,487 | 11,471 | 1,84E-30 | 5,99E-30 | 0,0050261 | 0,0099372 | 0,008205 | 0,008205 | TRUE | 0,1499361 | No |
| orf19.3019 | 414 | 359 | 385 | 223 | 1060 | 932 | 984 | 1414 | 377 | 363 | 364 | 335 | 943 | 1012 | 1008 | 1079 | 685,06 | 360 | 1010 | 2,797 | 1,484 | 21,196 | 1,05E-99 | 9,09E-99 | 0,0015906 | 0,0039653 | 0,0027697 | 0,0027697 | TRUE | 0,0265191 | No |
| orf19.860 | 462 | 383 | 387 | 281 | 1173 | 1022 | 1116 | 1499 | 421 | 387 | 366 | 422 | 1043 | 1110 | 1143 | 1143 | 754,46 | 399 | 1110 | 2,789 | 1,48 | 21,809 | 1,90E-105 | 1,72E-104 | 0,0013154 | 0,0037879 | 0,0026582 | 0,0026582 | TRUE | 0,0368014 | No |
| orf19.5365 | 841 | 782 | 861 | 562 | 2318 | 2094 | 2193 | 3094 | 766 | 791 | 815 | 844 | 2062 | 2274 | 2246 | 2360 | 1519,63 | 804 | 2236 | 2,787 | 1,479 | 25,29 | 4,15E-141 | 5,10E-140 | 0,0020032 | 0,0029063 | 0,0024053 | 0,0024053 | TRUE | 0,0336938 | No |
| orf19.1541 | 30 | 40 | 40 | 23 | 109 | 95 | 91 | 127 | 27 | 40 | 38 | 35 | 97 | 103 | 93 | 97 | 66,29 | 35 | 98 | 2,788 | 1,479 | 8,006 | 1,18E-15 | 2,66E-15 | 1,00E-08 | 0,0219637 | 0,012846 | 0,012846 | TRUE | 0,1927509 | No |
| orf19.3105 | 950 | 787 | 890 | 595 | 2518 | 2320 | 2193 | 3177 | 865 | 796 | 842 | 894 | 2240 | 2519 | 2246 | 2423 | 1603,11 | 849 | 2357 | 2,78 | 1,475 | 24,974 | 1,17E-137 | 1,39E-136 | 0,0022257 | 0,002861 | 0,0025179 | 0,0025179 | TRUE | 0,0251107 | No |
| orf19.6329 | 445 | 368 | 439 | 270 | 1191 | 1022 | 1101 | 1484 | 405 | 372 | 415 | 406 | 1059 | 1110 | 1128 | 1132 | 753,37 | 400 | 1107 | 2,771 | 1,471 | 23,318 | 2,93E-120 | 3,00E-119 | 1,00E-08 | 0,0037905 | 0,0020648 | 0,0020648 | TRUE | 0,0256654 | No |
| orf19.1651 | 14 | 22 | 16 | 9 | 49 | 35 | 46 | 63 | 13 | 22 | 15 | 14 | 44 | 38 | 47 | 48 | 30,05 | 16 | 44 | 2,771 | 1,47 | 5,348 | 8,89E-08 | 1,54E-07 | 1,00E-08 | 0,0459952 | 0,028956 | 0,028956 | TRUE | 0,3473984 | No |
| orf19.3935 | 4 | 3 | 2 | 2 | 12 | 5 | 7 | 11 | 4 | 3 | 2 | 3 | 11 | 5 | 7 | 8 | 5,4 | 3 | 8 | 2,767 | 1,468 | 2,329 | 0,0198407 | 0,0254445 | 1,00E-08 | 0,2464674 | 0,1399983 | 0,1399983 | TRUE | 0,1713653 | No |
| orf19.4122 | 1979 | 1839 | 1890 | 1267 | 5772 | 4578 | 4900 | 6728 | 1802 | 1860 | 1789 | 1903 | 5134 | 4970 | 5018 | 5132 | 3451,07 | 1838 | 5064 | 2,76 | 1,465 | 38,337 | 0 | 0 | 5,63E-05 | 0,0024198 | 0,0010169 | 0,0010169 | TRUE | 0,0059812 | No |
| orf19.6214 | 1834 | 1572 | 1705 | 1163 | 4803 | 4272 | 4515 | 6134 | 1670 | 1590 | 1614 | 1747 | 4272 | 4638 | 4624 | 4679 | 3104,21 | 1655 | 4553 | 2,755 | 1,462 | 30,308 | 9,13E-202 | 1,58E-200 | 0,0012019 | 0,0024626 | 0,001807 | 0,001807 | TRUE | 0,0212409 | No |
| orf19.3422 | 824 | 771 | 868 | 502 | 2336 | 2093 | 2083 | 2736 | 750 | 780 | 821 | 754 | 2078 | 2272 | 2133 | 2087 | 1459,51 | 776 | 2142 | 2,754 | 1,462 | 26,937 | 8,05E-160 | 1,11E-158 | 0,0010407 | 0,0029421 | 0,0019206 | 0,0019206 | TRUE | 0,0196745 | No |
| orf19.2492 | 38 | 22 | 24 | 23 | 85 | 76 | 73 | 102 | 35 | 22 | 23 | 35 | 76 | 83 | 75 | 78 | 53,1 | 29 | 78 | 2,753 | 1,461 | 6,985 | 2,85E-12 | 5,83E-12 | 1,00E-08 | 0,0269146 | 0,0172123 | 0,0172123 | TRUE | 0,1894433 | No |
| orf19.2479 | 161 | 155 | 163 | 98 | 418 | 404 | 428 | 548 | 147 | 157 | 154 | 147 | 372 | 439 | 438 | 418 | 283,95 | 151 | 417 | 2,748 | 1,459 | 14,93 | 2,09E-50 | 9,69E-50 | 1,00E-08 | 0,0066892 | 0,0045316 | 0,0045316 | TRUE | 0,0613994 | No |
| orf19.5806 | 60096 | 48214 | 53462 | 36525 | 2E+05 | 1E+05 | 1E+05 | 2E+05 | 54715 | 48763 | 50597 | 54866 | 1E+05 | 1E+05 | 1E+05 | 1E+05 | 97388,59 | 52235 | 142542 | 2,729 | 1,448 | 30,163 | 7,27E-200 | 1,24E-198 | 0,0022959 | 0,0020506 | 0,0022018 | 0,0022018 | TRUE | 0,0245296 | No |
| orf19.95 | 81 | 75 | 100 | 54 | 263 | 173 | 217 | 316 | 74 | 76 | 95 | 81 | 234 | 188 | 222 | 241 | 151,3 | 82 | 221 | 2,729 | 1,448 | 10,717 | 8,46E-27 | 2,53E-26 | 0,0059742 | 0,010768 | 0,0088972 | 0,0088972 | TRUE | 0,1287385 | No |
| orf19.5399 | 31 | 31 | 15 | 18 | 70 | 73 | 57 | 96 | 28 | 31 | 14 | 27 | 62 | 79 | 58 | 73 | 46,74 | 25 | 68 | 2,727 | 1,448 | 6,031 | 1,63E-09 | 3,03E-09 | 0,018762 | 0,0302984 | 0,0272494 | 0,0272494 | TRUE | 0,6027793 | No |
| orf19.7446 | 214 | 208 | 211 | 126 | 610 | 543 | 514 | 676 | 195 | 210 | 200 | 189 | 543 | 590 | 526 | 516 | 371,04 | 198 | 544 | 2,726 | 1,447 | 16,933 | 2,55E-64 | 1,48E-63 | 1,00E-08 | 0,0055972 | 0,0034808 | 0,0034808 | TRUE | 0,036299 | No |
| orf19.3029 | 3221 | 3017 | 3105 | 2089 | 9013 | 7679 | 7963 | 10847 | 2933 | 3051 | 2939 | 3138 | 8017 | 8337 | 8155 | 8274 | 5605,49 | 3015 | 8196 | 2,722 | 1,445 | 39,533 | 0 | 0 | 0,0003793 | 0,0022727 | 0,0010465 | 0,0010465 | TRUE | 0,0083612 | No |
| orf19.1462 | 2175 | 1851 | 2029 | 1372 | 5828 | 4875 | 5269 | 7105 | 1980 | 1872 | 1920 | 2061 | 5184 | 5293 | 5396 | 5420 | 3640,77 | 1958 | 5323 | 2,723 | 1,445 | 34,971 | 6,24E-268 | 1,42E-266 | 0,0004509 | 0,0023999 | 0,001277 | 0,001277 | TRUE | 0,0133082 | No |
| orf19.2752 | 2150 | 2164 | 2305 | 1391 | 6203 | 5419 | 5752 | 7383 | 1957 | 2189 | 2181 | 2090 | 5517 | 5884 | 5891 | 5632 | 3917,57 | 2104 | 5731 | 2,723 | 1,445 | 30,27 | 2,88E-201 | 4,95E-200 | 0,0014075 | 0,0023742 | 0,0018514 | 0,0018514 | TRUE | 0,0283055 | No |
| orf19.7389 | 707 | 647 | 700 | 451 | 1940 | 1721 | 1751 | 2340 | 644 | 654 | 662 | 677 | 1726 | 1869 | 1793 | 1785 | 1226,29 | 659 | 1793 | 2,722 | 1,445 | 27,663 | 1,92E-168 | 2,84E-167 | 1,00E-08 | 0,0031142 | 0,0015475 | 0,0015475 | TRUE | 0,0093431 | No |
| orf19.288 | 2035 | 1854 | 2040 | 1341 | 5757 | 4796 | 5113 | 6719 | 1853 | 1875 | 1931 | 2014 | 5121 | 5207 | 5236 | 5125 | 3545,3 | 1918 | 5172 | 2,701 | 1,434 | 36,94 | 1,05E-298 | 2,70E-297 | 2,19E-05 | 0,0024096 | 0,0010758 | 0,0010758 | TRUE | 0,0121108 | No |
| orf19.6342 | 1003 | 879 | 932 | 428 | 2821 | 1763 | 2380 | 2822 | 913 | 889 | 882 | 643 | 2509 | 1914 | 2437 | 2153 | 1542,57 | 832 | 2253 | 2,701 | 1,433 | 13,458 | 2,78E-41 | 1,11E-40 | 0,0178594 | 0,0028934 | 0,0100407 | 0,0100407 | TRUE | 0,2609707 | No |
| orf19.5141 | 15 | 25 | 20 | 18 | 61 | 54 | 63 | 62 | 14 | 25 | 19 | 27 | 54 | 59 | 65 | 47 | 38,7 | 21 | 56 | 2,698 | 1,432 | 5,663 | 1,48E-08 | 2,65E-08 | 0,007787 | 0,0361703 | 0,0274347 | 0,0274347 | TRUE | 0,3632959 | No |
| orf19.6628 | 617 | 433 | 463 | 321 | 1509 | 1041 | 1369 | 1705 | 562 | 438 | 438 | 482 | 1342 | 1130 | 1402 | 1301 | 886,89 | 480 | 1294 | 2,695 | 1,43 | 15,722 | 1,07E-55 | 5,50E-55 | 0,0095464 | 0,0035265 | 0,0064639 | 0,0064639 | TRUE | 0,1634995 | No |
| orf19.4033 | 587 | 459 | 619 | 335 | 1522 | 1214 | 1417 | 1957 | 534 | 464 | 586 | 503 | 1354 | 1318 | 1451 | 1493 | 962,94 | 522 | 1404 | 2,686 | 1,425 | 18,978 | 2,59E-80 | 1,84E-79 | 0,0047856 | 0,0034088 | 0,0040579 | 0,0040579 | TRUE | 0,0821648 | No |
| orf19.781 | 170 | 158 | 157 | 113 | 463 | 417 | 401 | 539 | 155 | 160 | 149 | 170 | 412 | 453 | 411 | 411 | 289,91 | 158 | 422 | 2,675 | 1,42 | 15,368 | 2,70E-53 | 1,33E-52 | 1,00E-08 | 0,0065935 | 0,0037147 | 0,0037147 | TRUE | 0,0269202 | No |
| orf19.6737 | 1027 | 880 | 939 | 672 | 2798 | 2466 | 2283 | 3176 | 935 | 890 | 889 | 1009 | 2489 | 2677 | 2338 | 2423 | 1706,26 | 931 | 2482 | 2,673 | 1,418 | 23,656 | 1,02E-123 | 1,07E-122 | 0,0025598 | 0,0028112 | 0,0026848 | 0,0026848 | TRUE | 0,0341381 | No |
| orf19.5805 | 773 | 610 | 726 | 475 | 1900 | 1605 | 1898 | 2468 | 704 | 617 | 687 | 714 | 1690 | 1743 | 1944 | 1883 | 1247,54 | 680 | 1815 | 2,671 | 1,418 | 21,292 | 1,35E-100 | 1,18E-99 | 0,0033157 | 0,0030959 | 0,0032084 | 0,0032084 | TRUE | 0,0468667 | No |
| orf19.7608 | 4 | 3 | 3 | 3 | 11 | 9 | 6 | 14 | 4 | 3 | 3 | 5 | 10 | 10 | 6 | 11 | 6,3 | 4 | 9 | 2,667 | 1,415 | 2,445 | 0,0144798 | 0,0187925 | 1,00E-08 | 0,2117132 | 0,1165944 | 0,1165944 | TRUE | 0,1470214 | No |
| orf19.3311 | 189 | 161 | 164 | 92 | 470 | 401 | 414 | 535 | 172 | 163 | 155 | 138 | 418 | 435 | 424 | 408 | 289,23 | 157 | 421 | 2,656 | 1,409 | 15,286 | 9,51E-53 | 4,61E-52 | 1,00E-08 | 0,0066043 | 0,003704 | 0,003704 | TRUE | 0,0506692 | No |
| orf19.1148 | 4 | 3 | 7 | 6 | 14 | 10 | 15 | 23 | 4 | 3 | 7 | 9 | 12 | 11 | 15 | 18 | 9,82 | 6 | 14 | 2,654 | 1,408 | 2,925 | 0,0034417 | 0,0046593 | 1,00E-08 | 0,1366094 | 0,090197 | 0,090197 | TRUE | 0,221424 | No |
| orf19.2360 | 8826 | 8153 | 8552 | 5580 | 23573 | 20792 | 20818 | 28786 | 8036 | 8246 | 8094 | 8382 | 20967 | 22574 | 21320 | 21958 | 14947,19 | 8190 | 21705 | 2,651 | 1,407 | 39,653 | 0 | 0 | 0,0006044 | 0,0021254 | 0,0011212 | 0,0011212 | TRUE | 0,0088328 | No |
| orf19.6910 | 77 | 61 | 62 | 44 | 218 | 146 | 163 | 209 | 70 | 62 | 59 | 66 | 194 | 159 | 167 | 159 | 116,92 | 64 | 170 | 2,65 | 1,406 | 9,764 | 1,61E-22 | 4,32E-22 | 1,00E-08 | 0,0133354 | 0,0088857 | 0,0088857 | TRUE | 0,1020767 | No |
| orf19.3839 | 2254 | 2082 | 2253 | 1386 | 6107 | 4985 | 5481 | 7493 | 2052 | 2106 | 2132 | 2082 | 5432 | 5412 | 5613 | 5716 | 3818,18 | 2093 | 5543 | 2,649 | 1,405 | 36,771 | 5,38E-296 | 1,37E-294 | 0,0003503 | 0,002383 | 0,0010618 | 0,0010618 | TRUE | 0,0055006 | No |
| orf19.2734 | 114 | 103 | 100 | 68 | 274 | 245 | 264 | 380 | 104 | 104 | 95 | 102 | 244 | 266 | 270 | 290 | 184,34 | 101 | 268 | 2,649 | 1,405 | 12,235 | 2,02E-34 | 7,12E-34 | 1,00E-08 | 0,0092032 | 0,0056763 | 0,0056763 | TRUE | 0,0427696 | No |
| orf19.6007 | 2445 | 2961 | 2822 | 1801 | 8011 | 7153 | 6646 | 8212 | 2226 | 2995 | 2671 | 2705 | 7126 | 7766 | 6806 | 6264 | 4819,9 | 2649 | 6990 | 2,639 | 1,4 | 16,393 | 2,15E-60 | 1,18E-59 | 0,0108549 | 0,0023111 | 0,0067363 | 0,0067363 | TRUE | 0,142275 | No |
| orf19.398 | 124 | 113 | 127 | 96 | 323 | 303 | 331 | 430 | 113 | 114 | 120 | 144 | 287 | 329 | 339 | 328 | 221,86 | 123 | 321 | 2,639 | 1,4 | 12,701 | 5,84E-37 | 2,17E-36 | 0,0023022 | 0,0079913 | 0,0058304 | 0,0058304 | TRUE | 0,1047614 | No |
| orf19.1089 | 1271 | 1531 | 1466 | 870 | 3897 | 3679 | 3403 | 4278 | 1157 | 1548 | 1387 | 1307 | 3466 | 3994 | 3485 | 3263 | 2451,12 | 1350 | 3552 | 2,63 | 1,395 | 16,35 | 4,36E-60 | 2,39E-59 | 0,0104596 | 0,002576 | 0,0064625 | 0,0064625 | TRUE | 0,1192783 | No |
| orf19.3618 | 2010 | 1675 | 1842 | 1420 | 5079 | 4863 | 4568 | 6424 | 1830 | 1694 | 1743 | 2133 | 4518 | 5280 | 4678 | 4900 | 3347,05 | 1850 | 4844 | 2,623 | 1,391 | 19,02 | 1,16E-80 | 8,26E-80 | 0,0074849 | 0,0024317 | 0,0047497 | 0,0047497 | TRUE | 0,12254 | No |
| orf19.2060 | 54 | 36 | 41 | 33 | 121 | 104 | 111 | 155 | 49 | 36 | 39 | 50 | 108 | 113 | 114 | 118 | 78,3 | 44 | 113 | 2,623 | 1,391 | 8,232 | 1,84E-16 | 4,24E-16 | 1,00E-08 | 0,0189079 | 0,0110317 | 0,0110317 | TRUE | 0,0902988 | No |
| orf19.5645 | 3024 | 2384 | 2672 | 1892 | 7610 | 6296 | 6571 | 9466 | 2753 | 2411 | 2529 | 2842 | 6769 | 6836 | 6730 | 7221 | 4761,26 | 2634 | 6889 | 2,619 | 1,389 | 25,256 | 9,72E-141 | 1,19E-139 | 0,0030273 | 0,0023145 | 0,0026313 | 0,0026313 | TRUE | 0,0389977 | No |

| | | | | | | | | | | | | | | | | | | | | | | | | | | | | | | | |
|---|---|---|---|---|---|---|---|---|---|---|---|---|---|---|---|---|---|---|---|---|---|---|---|---|---|---|---|---|---|---|---|
| orf19.2673 | 474 | 463 | 457 | 248 | 1313 | 955 | 1165 | 1426 | 432 | 468 | 433 | 373 | 1168 | 1037 | 1193 | 1088 | 773,81 | 426 | 1122 | 2,617 | 1,388 | 18,416 | 9,69E-76 | 6,49E-75 | 0,0042353 | 0,0037442 | 0,003789 | 0,003789 | TRUE | 0,0686826 | No |
| orf19.3358 | 2489 | 1927 | 2081 | 1468 | 6258 | 4844 | 5259 | 7451 | 2266 | 1949 | 1969 | 2205 | 5566 | 5259 | 5386 | 5684 | 3785,6 | 2097 | 5474 | 2,612 | 1,385 | 24,429 | 8,36E-132 | 9,46E-131 | 0,0031255 | 0,002386 | 0,0027453 | 0,0027453 | TRUE | 0,0378467 | No |
| orf19.3065 | 1035 | 935 | 1024 | 648 | 2778 | 2370 | 2360 | 3309 | 942 | 946 | 969 | 973 | 2471 | 2573 | 2417 | 2524 | 1726,96 | 958 | 2496 | 2,609 | 1,383 | 30,278 | 2,21E-201 | 3,80E-200 | 1,00E-08 | 0,0028019 | 0,0012593 | 0,0012593 | TRUE | 0,005362 | No |
| orf19.1208 | 20 | 20 | 27 | 19 | 55 | 53 | 55 | 96 | 18 | 20 | 26 | 29 | 49 | 58 | 56 | 73 | 41,07 | 23 | 59 | 2,605 | 1,382 | 5,652 | 1,59E-08 | 2,83E-08 | 1,00E-08 | 0,0342024 | 0,026165 | 0,026165 | TRUE | 0,2634204 | No |
| orf19.1344 | 1589 | 1830 | 1816 | 1063 | 4867 | 4188 | 4248 | 5250 | 1447 | 1851 | 1719 | 1597 | 4329 | 4547 | 4350 | 4005 | 2980,54 | 1654 | 4308 | 2,605 | 1,381 | 19,357 | 1,79E-83 | 1,33E-82 | 0,0062385 | 0,0024802 | 0,0044537 | 0,0044537 | TRUE | 0,0888075 | No |
| orf19.7380 | 233 | 193 | 227 | 145 | 579 | 499 | 523 | 762 | 212 | 195 | 215 | 218 | 515 | 542 | 536 | 581 | 376,7 | 210 | 544 | 2,595 | 1,376 | 16,433 | 1,10E-60 | 6,10E-60 | 1,00E-08 | 0,0055437 | 0,0033364 | 0,0033364 | TRUE | 0,0280814 | No |
| orf19.6993 | 12993 | 10874 | 11723 | 7632 | 32117 | 26707 | 28136 | 40643 | 11830 | 10998 | 11095 | 11464 | 28567 | 28996 | 28815 | 31003 | 20345,97 | 11347 | 29345 | 2,586 | 1,371 | 33,731 | 2,04E-249 | 4,20E-248 | 0,001205 | 0,0021019 | 0,0015234 | 0,0015234 | TRUE | 0,0177852 | No |
| orf19.2160 | 44 | 14 | 27 | 21 | 68 | 50 | 79 | 119 | 40 | 14 | 26 | 32 | 60 | 54 | 81 | 91 | 49,72 | 28 | 72 | 2,587 | 1,371 | 4,989 | 6,08E-07 | 1,02E-06 | 0,0797469 | 0,0286052 | 0,0468034 | 0,0468034 | TRUE | 0,3558108 | No |
| orf19.2474 | 1256 | 1113 | 1202 | 813 | 3273 | 2765 | 2834 | 4076 | 1144 | 1126 | 1138 | 1221 | 2911 | 3002 | 2902 | 3109 | 2069,11 | 1157 | 2981 | 2,584 | 1,369 | 29,125 | 1,71E-186 | 2,71E-185 | 0,0006286 | 0,0026755 | 0,0015019 | 0,0015019 | TRUE | 0,0144226 | No |
| orf19.539 | 831 | 739 | 780 | 491 | 2182 | 1780 | 1856 | 2519 | 757 | 747 | 738 | 738 | 1941 | 1933 | 1901 | 1922 | 1334,43 | 745 | 1924 | 2,581 | 1,368 | 29,132 | 1,40E-186 | 2,23E-185 | 1,00E-08 | 0,0030269 | 0,0011601 | 0,0011601 | TRUE | 0,0013918 | No |
| orf19.5648 | 531 | 454 | 444 | 279 | 1412 | 948 | 1166 | 1473 | 483 | 459 | 420 | 419 | 1256 | 1029 | 1194 | 1124 | 798,11 | 445 | 1151 | 2,578 | 1,366 | 17,428 | 5,04E-68 | 3,08E-67 | 0,0049129 | 0,0036922 | 0,004293 | 0,004293 | TRUE | 0,0585548 | No |
| orf19.2059 | 1442 | 1292 | 1403 | 957 | 3965 | 3179 | 3345 | 4493 | 1313 | 1307 | 1328 | 1438 | 3527 | 3452 | 3426 | 3427 | 2402,03 | 1346 | 3458 | 2,576 | 1,365 | 31,579 | 7,21E-219 | 1,33E-217 | 1,95E-05 | 0,002587 | 0,0012605 | 0,0012605 | TRUE | 0,0216415 | No |
| orf19.7464 | 16 | 8 | 11 | 3 | 31 | 22 | 20 | 37 | 15 | 8 | 10 | 5 | 28 | 24 | 20 | 28 | 17,22 | 10 | 25 | 2,568 | 1,361 | 3,749 | 0,0001775 | 0,0002627 | 1,00E-08 | 0,0787604 | 0,0538905 | 0,0538905 | TRUE | 0,4687184 | No |
| orf19.1926 | 247 | 234 | 217 | 189 | 665 | 586 | 596 | 747 | 225 | 237 | 205 | 284 | 591 | 636 | 610 | 570 | 419,84 | 238 | 602 | 2,557 | 1,355 | 14,951 | 1,54E-50 | 7,16E-50 | 0,0041709 | 0,0051834 | 0,0048356 | 0,0048356 | TRUE | 0,1553523 | No |
| orf19.3911 | 7965 | 6358 | 6457 | 4885 | 19486 | 15951 | 15855 | 23941 | 7252 | 6430 | 6111 | 7338 | 17332 | 17318 | 16238 | 18262 | 12035,21 | 6783 | 17288 | 2,55 | 1,35 | 21,317 | 7,88E-101 | 6,91E-100 | 0,0050082 | 0,0021468 | 0,0037485 | 0,0037485 | TRUE | 0,0544458 | No |
| orf19.669 | 158 | 157 | 134 | 83 | 380 | 318 | 379 | 458 | 144 | 159 | 127 | 125 | 338 | 345 | 388 | 349 | 246,86 | 139 | 355 | 2,548 | 1,349 | 12,898 | 4,61E-38 | 1,74E-37 | 0,0019782 | 0,0073881 | 0,0053761 | 0,0053761 | TRUE | 0,0970365 | No |
| orf19.4685 | 11 | 10 | 18 | 6 | 45 | 25 | 27 | 32 | 10 | 10 | 17 | 9 | 40 | 27 | 28 | 24 | 20,68 | 12 | 30 | 2,541 | 1,346 | 3,958 | 7,56E-05 | 0,0001139 | 1,00E-08 | 0,0659275 | 0,0500722 | 0,0500722 | TRUE | 0,4264209 | No |
| orf19.4592 | 2153 | 2011 | 2093 | 1331 | 5586 | 4739 | 4925 | 6685 | 1960 | 2034 | 1981 | 1999 | 4969 | 5145 | 5044 | 5099 | 3528,92 | 1994 | 5064 | 2,541 | 1,345 | 37,496 | 1,07E-307 | 2,85E-306 | 1,00E-08 | 0,0024114 | 0,0008749 | 0,0008749 | TRUE | 0,0024058 | No |
| orf19.5700 | 57 | 49 | 59 | 33 | 148 | 110 | 148 | 164 | 52 | 50 | 56 | 50 | 132 | 119 | 152 | 125 | 91,83 | 52 | 132 | 2,54 | 1,345 | 8,505 | 1,82E-17 | 4,32E-17 | 1,00E-08 | 0,0164229 | 0,0102436 | 0,0102436 | TRUE | 0,0981289 | No |
| orf19.5576 | 4774 | 4731 | 4807 | 3119 | 12960 | 11066 | 11282 | 15044 | 4347 | 4785 | 4549 | 4685 | 11528 | 12015 | 11554 | 11476 | 8117,25 | 4592 | 11643 | 2,537 | 1,343 | 33,851 | 3,51E-251 | 7,32E-250 | 0,0008815 | 0,0021998 | 0,001354 | 0,001354 | TRUE | 0,016078 | No |
| orf19.2163 | 1074 | 945 | 1083 | 605 | 2676 | 2064 | 2581 | 3358 | 978 | 956 | 1025 | 909 | 2380 | 2241 | 2643 | 2562 | 1711,66 | 967 | 2456 | 2,537 | 1,343 | 21,067 | 1,60E-98 | 1,37E-97 | 0,0034543 | 0,0028088 | 0,0031564 | 0,0031564 | TRUE | 0,0414539 | No |
| orf19.1906 | 78 | 81 | 68 | 55 | 205 | 172 | 192 | 247 | 71 | 82 | 64 | 83 | 182 | 187 | 197 | 188 | 131,76 | 75 | 188 | 2,536 | 1,343 | 10,243 | 1,27E-24 | 3,62E-24 | 1,00E-08 | 0,0120631 | 0,0068895 | 0,0068895 | TRUE | 0,085964 | No |
| orf19.5369 | 1293 | 1064 | 1173 | 901 | 3293 | 2709 | 2893 | 3999 | 1177 | 1076 | 1110 | 1353 | 2929 | 2941 | 2963 | 3050 | 2075,05 | 1179 | 2971 | 2,529 | 1,339 | 21,089 | 9,96E-99 | 8,57E-98 | 0,0038899 | 0,0026736 | 0,0032526 | 0,0032526 | TRUE | 0,108689 | No |
| orf19.31 | 210 | 157 | 153 | 112 | 437 | 351 | 429 | 606 | 191 | 159 | 145 | 168 | 389 | 381 | 439 | 462 | 291,8 | 166 | 418 | 2,524 | 1,336 | 12,316 | 7,39E-35 | 2,64E-34 | 0,0074948 | 0,006564 | 0,0069595 | 0,0069595 | TRUE | 0,1301528 | No |
| orf19.2812 | 357 | 297 | 351 | 205 | 857 | 761 | 728 | 1130 | 325 | 300 | 332 | 308 | 762 | 826 | 746 | 862 | 557,7 | 316 | 799 | 2,522 | 1,335 | 17,391 | 9,69E-68 | 5,90E-67 | 0,0023474 | 0,0044057 | 0,0033897 | 0,0033897 | TRUE | 0,0356949 | No |
| orf19.1939 | 20 | 18 | 19 | 8 | 46 | 43 | 45 | 50 | 18 | 18 | 18 | 12 | 41 | 47 | 46 | 38 | 29,78 | 16 | 43 | 2,522 | 1,334 | 5,002 | 5,68E-07 | 9,54E-07 | 1,00E-08 | 0,0463954 | 0,026179 | 0,026179 | TRUE | 0,0977659 | No |
| orf19.1224 | 5623 | 5785 | 5913 | 4173 | 15671 | 14463 | 13465 | 18321 | 5120 | 5851 | 5596 | 6269 | 13939 | 15703 | 13790 | 13975 | 10030,24 | 5709 | 14352 | 2,515 | 1,331 | 20,559 | 6,42E-94 | 5,19E-93 | 0,0053681 | 0,0021687 | 0,003898 | 0,003898 | TRUE | 0,0590642 | No |
| orf19.1978 | 955 | 873 | 937 | 575 | 2525 | 1973 | 2060 | 2997 | 869 | 883 | 887 | 864 | 2246 | 2142 | 2110 | 2286 | 1535,85 | 876 | 2196 | 2,508 | 1,327 | 26,779 | 5,65E-158 | 7,74E-157 | 1,00E-08 | 0,0028971 | 0,001534 | 0,001534 | TRUE | 0,0095081 | No |
| orf19.5644 | 274 | 251 | 275 | 187 | 690 | 667 | 635 | 818 | 249 | 254 | 260 | 281 | 614 | 724 | 650 | 624 | 457,09 | 261 | 653 | 2,507 | 1,326 | 16,049 | 5,77E-58 | 3,07E-57 | 0,0024989 | 0,0049271 | 0,0037886 | 0,0037886 | TRUE | 0,0612681 | No |
| orf19.1192 | 745 | 680 | 680 | 471 | 1798 | 1534 | 1750 | 2268 | 678 | 688 | 644 | 708 | 1599 | 1665 | 1792 | 1730 | 1188,02 | 680 | 1696 | 2,503 | 1,324 | 22,874 | 8,50E-116 | 8,40E-115 | 0,0012817 | 0,003149 | 0,0021509 | 0,0021509 | TRUE | 0,0183782 | No |
| orf19.2905 | 16 | 18 | 15 | 4 | 33 | 36 | 35 | 44 | 15 | 18 | 14 | 6 | 29 | 39 | 36 | 34 | 23,85 | 13 | 34 | 2,5 | 1,322 | 4,309 | 1,64E-05 | 2,57E-05 | 1,00E-08 | 0,0574179 | 0,0381736 | 0,0381736 | TRUE | 0,3202702 | No |
| orf19.1632 | 404 | 419 | 435 | 241 | 1084 | 947 | 1008 | 1183 | 368 | 424 | 412 | 362 | 964 | 1028 | 1032 | 902 | 686,55 | 392 | 982 | 2,498 | 1,321 | 17,522 | 9,72E-69 | 5,98E-68 | 0,0032241 | 0,0039611 | 0,0036151 | 0,0036151 | TRUE | 0,0367948 | No |
| orf19.6488 | 8 | 7 | 11 | 5 | 26 | 17 | 19 | 26 | 7 | 7 | 10 | 8 | 23 | 18 | 19 | 20 | 14,14 | 8 | 20 | 2,495 | 1,319 | 3,459 | 0,0005413 | 0,000777 | 1,00E-08 | 0,0954271 | 0,0514609 | 0,0514609 | TRUE | 0,1292553 | No |
| orf19.5517 | 2123 | 1805 | 1901 | 1279 | 5297 | 4339 | 4286 | 6310 | 1933 | 1826 | 1799 | 1921 | 4712 | 4711 | 4389 | 4813 | 3263 | 1870 | 4656 | 2,493 | 1,318 | 28,5 | 1,18E-178 | 1,81E-177 | 0,001147 | 0,0024419 | 0,0016646 | 0,0016646 | TRUE | 0,0180256 | No |
| orf19.7522 | 793 | 669 | 703 | 471 | 1923 | 1563 | 1709 | 2282 | 722 | 677 | 665 | 708 | 1710 | 1697 | 1750 | 1741 | 1208,73 | 693 | 1724 | 2,492 | 1,317 | 25,884 | 1,01E-147 | 1,31E-146 | 1,00E-08 | 0,0031299 | 0,0014451 | 0,0014451 | TRUE | 0,0113382 | No |
| orf19.6294 | 310 | 387 | 370 | 229 | 914 | 832 | 846 | 1074 | 282 | 391 | 350 | 344 | 813 | 903 | 866 | 819 | 596,22 | 342 | 850 | 2,489 | 1,316 | 14,815 | 1,16E-49 | 5,33E-49 | 0,0071874 | 0,0042526 | 0,0054523 | 0,0054523 | TRUE | 0,1722643 | No |
| orf19.5014 | 203 | 172 | 179 | 118 | 544 | 390 | 408 | 558 | 185 | 174 | 169 | 177 | 484 | 423 | 418 | 426 | 307,03 | 176 | 438 | 2,484 | 1,312 | 14,181 | 1,21E-45 | 5,20E-45 | 1,00E-08 | 0,0063395 | 0,004144 | 0,004144 | TRUE | 0,0595329 | No |
| orf19.3926 | 51 | 33 | 29 | 29 | 96 | 80 | 107 | 118 | 46 | 33 | 27 | 44 | 85 | 87 | 110 | 90 | 65,33 | 38 | 93 | 2,482 | 1,311 | 6,627 | 3,44E-11 | 6,79E-11 | 0,0099572 | 0,0222567 | 0,0183652 | 0,0183652 | TRUE | 0,2715124 | No |
| orf19.7163 | 170 | 127 | 126 | 88 | 382 | 275 | 322 | 467 | 155 | 128 | 119 | 132 | 340 | 299 | 330 | 356 | 232,38 | 134 | 331 | 2,478 | 1,309 | 11,948 | 6,66E-33 | 2,27E-32 | 0,0036387 | 0,0077217 | 0,0061504 | 0,0061504 | TRUE | 0,1333182 | No |
| orf19.4807 | 1352 | 1273 | 1380 | 860 | 3647 | 2913 | 3085 | 4060 | 1231 | 1288 | 1306 | 1292 | 3244 | 3163 | 3159 | 3097 | 2222,42 | 1279 | 3166 | 2,476 | 1,308 | 30,978 | 1,06E-210 | 1,88E-209 | 1,00E-08 | 0,0026314 | 0,0011463 | 0,0011463 | TRUE | 0,0083477 | No |
| orf19.3821 | 550 | 476 | 496 | 300 | 1282 | 1036 | 1232 | 1551 | 501 | 481 | 469 | 451 | 1140 | 1125 | 1262 | 1183 | 826,52 | 476 | 1178 | 2,468 | 1,304 | 20,373 | 2,89E-92 | 2,31E-91 | 0,0011754 | 0,0036353 | 0,0024152 | 0,0024152 | TRUE | 0,0268079 | No |
| orf19.6053 | 2990 | 2660 | 3051 | 1801 | 7366 | 6135 | 6824 | 9104 | 2722 | 2690 | 2887 | 2705 | 6552 | 6661 | 6989 | 6945 | 4768,93 | 2751 | 6787 | 2,465 | 1,302 | 31,502 | 8,04E-218 | 1,47E-216 | 0,0007094 | 0,002314 | 0,0013755 | 0,0013755 | TRUE | 0,0138338 | No |
| orf19.6305 | 988 | 1124 | 1084 | 626 | 2887 | 2278 | 2462 | 3042 | 900 | 1137 | 1026 | 940 | 2568 | 2473 | 2521 | 2320 | 1735,7 | 1001 | 2470 | 2,466 | 1,302 | 18,508 | 1,76E-76 | 1,19E-75 | 0,0055075 | 0,0027981 | 0,0040274 | 0,0040274 | TRUE | 0,1011239 | No |
| orf19.5210 | 395 | 315 | 393 | 291 | 974 | 879 | 874 | 1206 | 360 | 319 | 372 | 437 | 866 | 954 | 895 | 920 | 640,38 | 372 | 909 | 2,462 | 1,3 | 15,913 | 5,17E-57 | 2,69E-56 | 0,0050087 | 0,0040999 | 0,004441 | 0,004441 | TRUE | 0,1353766 | No |
| orf19.6141 | 867 | 764 | 847 | 549 | 2209 | 1766 | 1861 | 2647 | 789 | 773 | 802 | 825 | 1965 | 1917 | 1906 | 2019 | 1374,45 | 797 | 1952 | 2,455 | 1,296 | 26,324 | 1,03E-152 | 1,36E-151 | 1,00E-08 | 0,0029981 | 0,001416 | 0,001416 | TRUE | 0,0067841 | No |
| orf19.1544 | 312 | 271 | 294 | 219 | 846 | 626 | 685 | 924 | 284 | 274 | 278 | 329 | 752 | 680 | 702 | 705 | 500,49 | 291 | 710 | 2,456 | 1,296 | 16,973 | 1,30E-64 | 7,57E-64 | 0,0007529 | 0,0046765 | 0,0031008 | 0,0031008 | TRUE | 0,0679399 | No |
| orf19.5873 | 539 | 527 | 518 | 328 | 1328 | 1090 | 1261 | 1660 | 491 | 533 | 490 | 493 | 1181 | 1183 | 1291 | 1266 | 866,13 | 502 | 1230 | 2,453 | 1,294 | 20,9 | 5,41E-97 | 4,57E-96 | 0,0009337 | 0,0035622 | 0,0022402 | 0,0022402 | TRUE | 0,0212912 | No |
| orf19.5649 | 45 | 42 | 30 | 27 | 90 | 105 | 88 | 117 | 41 | 42 | 28 | 41 | 80 | 114 | 90 | 89 | 65,73 | 38 | 93 | 2,453 | 1,294 | 6,589 | 4,43E-11 | 8,73E-11 | 0,01072 | 0,022135 | 0,0180184 | 0,0180184 | TRUE | 0,2425876 | No |
| orf19.2394 | 28 | 31 | 35 | 32 | 93 | 64 | 94 | 105 | 25 | 31 | 33 | 48 | 83 | 69 | 96 | 80 | 58,33 | 34 | 82 | 2,448 | 1,292 | 6,17 | 6,81E-10 | 1,28E-09 | 0,0108437 | 0,0246855 | 0,0205522 | 0,0205522 | TRUE | 0,3443784 | No |
| orf19.4679 | 23209 | 19566 | 21758 | 14503 | 55419 | 46508 | 49834 | 69361 | 21131 | 19789 | 20592 | 21786 | 49293 | 50495 | 51036 | 52909 | 35878,89 | 20824 | 50933 | 2,446 | 1,291 | 32,271 | 1,79E-228 | 3,44E-227 | 0,0012157 | 0,0020738 | 0,0015015 | 0,0015015 | TRUE | 0,0136874 | No |
| orf19.5032 | 33778 | 27708 | 29922 | 20112 | 79699 | 62769 | 70002 | 99986 | 30753 | 28024 | 28318 | 30211 | 70890 | 68150 | 71691 | 76270 | 50538,42 | 29326 | 71750 | 2,447 | 1,291 | 27,365 | 7,22E-165 | 1,04E-163 | 0,0021453 | 0,0020632 | 0,0021131 | 0,0021131 | TRUE | 0,0220628 | No |
| orf19.3828 | 10 | 12 | 12 | 7 | 36 | 22 | 26 | 30 | 9 | 12 | 11 | 11 | 32 | 24 | 27 | 23 | 18,57 | 11 | 26 | 2,446 | 1,291 | 3,835 | 0,0001254 | 0,0001868 | 1,00E-08 | 0,0731867 | 0,0418513 | 0,0418513 | TRUE | 0,1448789 | No |
| orf19.6788 | 906 | 942 | 939 | 563 | 2574 | 1921 | 2048 | 2719 | 825 | 953 | 889 | 846 | 2289 | 2086 | 2097 | 2074 | 1507,33 | 878 | 2136 | 2,432 | 1,282 | 21,514 | 1,14E-102 | 1,02E-101 | 0,0021432 | 0,0029134 | 0,0025802 | 0,0025802 | TRUE | 0,0396258 | No |
| orf19.7237 | 451 | 374 | 378 | 295 | 1062 | 887 | 920 | 1300 | 411 | 378 | 358 | 443 | 945 | 963 | 942 | 992 | 678,91 | 398 | 960 | 2,432 | 1,282 | 18,379 | 1,92E-75 | 1,28E-74 | 0,0008292 | 0,0039828 | 0,0028349 | 0,0028349 | TRUE | 0,0562832 | No |
| orf19.799 | 184 | 131 | 181 | 131 | 459 | 392 | 385 | 503 | 168 | 132 | 171 | 197 | 408 | 426 | 394 | 384 | 284,99 | 167 | 403 | 2,431 | 1,282 | 12,316 | 7,42E-35 | 2,65E-34 | 0,0044908 | 0,0066722 | 0,0060183 | 0,0060183 | TRUE | 0,2015411 | No |
| orf19.4450 | 493 | 397 | 446 | 295 | 1084 | 918 | 1059 | 1459 | 449 | 402 | 422 | 443 | 964 | 997 | 1085 | 1113 | 734,25 | 429 | 1040 | 2,428 | 1,28 | 17,848 | 2,98E-71 | 1,89E-70 | 0,0026584 | 0,0038361 | 0,0032392 | 0,0032392 | TRUE | 0,0296547 | No |
| orf19.6629 | 462 | 369 | 402 | 265 | 1055 | 839 | 949 | 1299 | 421 | 373 | 380 | 398 | 938 | 911 | 972 | 991 | 673,06 | 393 | 953 | 2,426 | 1,279 | 19,262 | 1,12E-82 | 8,22E-82 | 1,00E-08 | 0,0039997 | 0,0023872 | 0,0023872 | TRUE | 0,0295827 | No |
| orf19.540 | 393 | 303 | 385 | 215 | 1000 | 712 | 813 | 1041 | 358 | 306 | 364 | 323 | 889 | 773 | 833 | 794 | 580,1 | 338 | 822 | 2,426 | 1,279 | 16,293 | 1,11E-59 | 6,04E-59 | 0,0031076 | 0,0043142 | 0,003771 | 0,003771 | TRUE | 0,0497744 | No |
| orf19.6724 | 2556 | 1877 | 2189 | 1615 | 5657 | 4995 | 5016 | 7184 | 2327 | 1898 | 2072 | 2426 | 5032 | 5423 | 5137 | 5480 | 3724,39 | 2181 | 5268 | 2,418 | 1,274 | 18,035 | 1,04E-72 | 6,74E-72 | 0,0064901 | 0,0023917 | 0,0044571 | 0,0044571 | TRUE | 0,0917486 | No |
| orf19.6313 | 216 | 277 | 265 | 162 | 639 | 564 | 564 | 768 | 197 | 280 | 251 | 243 | 568 | 612 | 578 | 586 | 414,39 | 243 | 586 | 2,418 | 1,274 | 13,485 | 1,92E-41 | 7,67E-41 | 0,0061546 | 0,0052248 | 0,0055654 | 0,0055654 | TRUE | 0,2021344 | No |
| orf19.5475 | 15 | 14 | 22 | 20 | 40 | 51 | 41 | 67 | 14 | 14 | 21 | 30 | 36 | 55 | 42 | 51 | 32,84 | 20 | 46 | 2,419 | 1,274 | 4,482 | 7,38E-06 | 1,18E-05 | 0,0333456 | 0,042261 | 0,0394458 | 0,0394458 | TRUE | 0,3988709 | No |

| | | | | | | | | | | | | | | | | | | | | | | | | | | | | | | | |
|---|---|---|---|---|---|---|---|---|---|---|---|---|---|---|---|---|---|---|---|---|---|---|---|---|---|---|---|---|---|---|---|
| orf19.6547 | 15 | 5 | 6 | 8 | 26 | 16 | 20 | 33 | 14 | 5 | 6 | 12 | 23 | 17 | 20 | 25 | 15,32 | 9 | 21 | 2,419 | 1,274 | 3,258 | 0,0011236 | 0,0015816 | 0,0032337 | 0,0882623 | 0,0661956 | 0,0661956 | TRUE | 0,5065467 | No |
| orf19.7130 | 41 | 42 | 33 | 30 | 115 | 62 | 94 | 137 | 37 | 42 | 31 | 45 | 102 | 67 | 96 | 105 | 65,81 | 39 | 92 | 2,413 | 1,271 | 6,291 | 3,16E-10 | 6,02E-10 | 0,0175451 | 0,0221099 | 0,0203828 | 0,0203828 | TRUE | 0,3033091 | No |
| orf19.6179 | 7 | 4 | 6 | 6 | 19 | 18 | 12 | 14 | 6 | 4 | 6 | 9 | 17 | 20 | 12 | 11 | 10,57 | 6 | 15 | 2,414 | 1,271 | 2,774 | 0,0055444 | 0,0074099 | 1,00E-08 | 0,1270695 | 0,0831226 | 0,0831226 | TRUE | 0,1493082 | No |
| orf19.4076 | 8342 | 7612 | 8011 | 5007 | 20752 | 16659 | 17834 | 24208 | 7595 | 7699 | 7582 | 7521 | 18458 | 18087 | 18264 | 18466 | 12959,04 | 7599 | 18319 | 2,41 | 1,269 | 47,769 | 0 | 0 | 1,00E-08 | 0,002139 | 0,0005815 | 0,0005815 | TRUE | 0,0009403 | No |
| orf19.2686 | 2340 | 2121 | 2202 | 1516 | 5534 | 5191 | 4886 | 6850 | 2130 | 2145 | 2084 | 2277 | 4922 | 5636 | 5004 | 5225 | 3678,04 | 2159 | 5197 | 2,409 | 1,269 | 23,955 | 8,14E-127 | 8,81E-126 | 0,00219 | 0,0023962 | 0,0023529 | 0,0023529 | TRUE | 0,0389459 | No |
| orf19.1317 | 1246 | 1141 | 1211 | 802 | 2992 | 2611 | 2717 | 3757 | 1134 | 1154 | 1146 | 1205 | 2661 | 2835 | 2783 | 2866 | 1972,97 | 1160 | 2786 | 2,407 | 1,267 | 27,536 | 6,58E-167 | 9,62E-166 | 0,0004915 | 0,0027066 | 0,0014029 | 0,0014029 | TRUE | 0,0112662 | No |
| orf19.348 | 1110 | 934 | 1107 | 631 | 2574 | 2180 | 2379 | 3163 | 1011 | 945 | 1048 | 948 | 2289 | 2367 | 2436 | 2413 | 1682,04 | 988 | 2376 | 2,401 | 1,263 | 24,984 | 9,14E-138 | 1,09E-136 | 0,0007074 | 0,0028224 | 0,0017176 | 0,0017176 | TRUE | 0,0200383 | No |
| orf19.6460 | 1909 | 1876 | 2080 | 1249 | 5070 | 4404 | 4239 | 5607 | 1738 | 1897 | 1969 | 1876 | 4510 | 4782 | 4341 | 4277 | 3173,7 | 1870 | 4478 | 2,394 | 1,259 | 23,565 | 8,76E-123 | 9,15E-122 | 0,0020897 | 0,0024532 | 0,0023503 | 0,0023503 | TRUE | 0,028659 | No |
| orf19.3590 | 2096 | 1812 | 1956 | 1446 | 5327 | 4359 | 4250 | 6081 | 1908 | 1833 | 1851 | 2172 | 4738 | 4733 | 4353 | 4639 | 3278,29 | 1941 | 4616 | 2,383 | 1,253 | 21,695 | 2,29E-104 | 2,07E-103 | 0,0035689 | 0,00244 | 0,0028239 | 0,0028239 | TRUE | 0,0728391 | No |
| orf19.6222 | 106 | 103 | 97 | 67 | 250 | 218 | 232 | 312 | 97 | 104 | 92 | 101 | 222 | 237 | 238 | 238 | 165,97 | 98 | 234 | 2,383 | 1,253 | 10,954 | 6,37E-28 | 1,95E-27 | 1,00E-08 | 0,0099962 | 0,0051578 | 0,0051578 | TRUE | 0,0205459 | No |
| orf19.6385 | 16005 | 12914 | 14236 | 10114 | 35735 | 30692 | 32422 | 46637 | 14572 | 13061 | 13473 | 15193 | 31785 | 33323 | 33204 | 35575 | 23773,29 | 14075 | 33472 | 2,379 | 1,25 | 22,629 | 2,24E-113 | 2,18E-112 | 0,0034566 | 0,0020926 | 0,0028802 | 0,0028802 | TRUE | 0,0346136 | No |
| orf19.1587 | 102 | 110 | 99 | 79 | 262 | 227 | 231 | 344 | 93 | 111 | 94 | 119 | 233 | 246 | 237 | 262 | 174,37 | 104 | 244 | 2,379 | 1,25 | 10,588 | 3,38E-26 | 9,90E-26 | 1,00E-08 | 0,0096128 | 0,0063234 | 0,0063234 | TRUE | 0,0661202 | No |
| orf19.6023 | 1500 | 1423 | 1564 | 922 | 3755 | 3155 | 3219 | 4493 | 1366 | 1439 | 1480 | 1385 | 3340 | 3425 | 3297 | 3427 | 2394,93 | 1418 | 3372 | 2,377 | 1,249 | 28,972 | 1,48E-184 | 2,34E-183 | 0,0001923 | 0,0025886 | 0,0012693 | 0,0012693 | TRUE | 0,0111061 | No |
| orf19.556 | 995 | 1094 | 991 | 651 | 2641 | 2395 | 2054 | 2982 | 906 | 1106 | 938 | 978 | 2349 | 2600 | 2104 | 2275 | 1656,98 | 982 | 2332 | 2,375 | 1,248 | 16,13 | 1,56E-58 | 8,36E-58 | 0,0069677 | 0,0028343 | 0,0049973 | 0,0049973 | TRUE | 0,0893269 | No |
| orf19.4229 | 850 | 731 | 812 | 495 | 2055 | 1634 | 1780 | 2302 | 774 | 739 | 768 | 744 | 1828 | 1774 | 1823 | 1756 | 1275,77 | 756 | 1795 | 2,37 | 1,245 | 25,442 | 8,77E-143 | 1,09E-141 | 1,00E-08 | 0,0030725 | 0,0013347 | 0,0013347 | TRUE | 0,0031717 | No |
| orf19.258 | 428 | 397 | 410 | 283 | 1113 | 905 | 912 | 1161 | 390 | 402 | 388 | 425 | 990 | 983 | 934 | 886 | 674,56 | 401 | 948 | 2,37 | 1,245 | 18,399 | 1,33E-75 | 8,89E-75 | 1,00E-08 | 0,0039953 | 0,0025675 | 0,0025675 | TRUE | 0,0250644 | No |
| orf19.6086 | 486 | 386 | 406 | 278 | 1059 | 876 | 925 | 1349 | 442 | 390 | 384 | 418 | 942 | 951 | 947 | 1029 | 688,01 | 408 | 967 | 2,371 | 1,245 | 18,112 | 2,57E-73 | 1,68E-72 | 0,0013793 | 0,003957 | 0,0027494 | 0,0027494 | TRUE | 0,0414295 | No |
| orf19.3405 | 470 | 424 | 477 | 343 | 1156 | 1016 | 1063 | 1400 | 428 | 429 | 451 | 515 | 1028 | 1103 | 1089 | 1068 | 763,91 | 456 | 1072 | 2,367 | 1,243 | 18,582 | 4,53E-77 | 3,09E-76 | 0,000952 | 0,0037663 | 0,0026794 | 0,0026794 | TRUE | 0,0741802 | No |
| orf19.7414 | 2323 | 2073 | 2245 | 1470 | 5230 | 4723 | 4928 | 6979 | 2115 | 2097 | 2125 | 2208 | 4652 | 5128 | 5047 | 5324 | 3586,85 | 2136 | 5038 | 2,36 | 1,239 | 25,034 | 2,63E-138 | 3,16E-137 | 0,0017531 | 0,0024053 | 0,002007 | 0,002007 | TRUE | 0,0327262 | No |
| orf19.7480 | 6 | 5 | 2 | 4 | 12 | 9 | 11 | 14 | 5 | 5 | 2 | 6 | 11 | 10 | 11 | 11 | 7,6 | 4 | 11 | 2,358 | 1,238 | 2,377 | 0,0174726 | 0,0225345 | 1,00E-08 | 0,1758211 | 0,0987945 | 0,0987945 | TRUE | 0,3404749 | No |
| orf19.3554 | 3454 | 3385 | 3386 | 2089 | 8655 | 7387 | 6922 | 9860 | 3145 | 3424 | 3205 | 3138 | 7698 | 8020 | 7089 | 7521 | 5404,96 | 3228 | 7582 | 2,348 | 1,232 | 25,022 | 3,51E-138 | 4,21E-137 | 0,001878 | 0,0022814 | 0,0020978 | 0,0020978 | TRUE | 0,0231987 | No |
| orf19.2740 | 659 | 605 | 668 | 416 | 1601 | 1349 | 1387 | 1945 | 600 | 612 | 632 | 625 | 1424 | 1465 | 1420 | 1484 | 1032,72 | 617 | 1448 | 2,349 | 1,232 | 23,373 | 7,99E-121 | 8,22E-120 | 1,00E-08 | 0,0033162 | 0,0014814 | 0,0014814 | TRUE | 0,0045452 | No |
| orf19.2593 | 94 | 84 | 93 | 69 | 252 | 175 | 195 | 296 | 86 | 85 | 88 | 104 | 224 | 190 | 200 | 226 | 150,23 | 91 | 210 | 2,349 | 1,232 | 9,755 | 1,75E-22 | 4,71E-22 | 1,00E-08 | 0,0108301 | 0,0071824 | 0,0071824 | TRUE | 0,0663946 | No |
| orf19.3544 | 292 | 317 | 319 | 196 | 767 | 612 | 726 | 896 | 266 | 321 | 302 | 294 | 682 | 664 | 744 | 683 | 494,56 | 296 | 693 | 2,347 | 1,231 | 15,771 | 4,95E-56 | 2,54E-55 | 0,0014837 | 0,0047081 | 0,003368 | 0,003368 | TRUE | 0,0591254 | No |
| orf19.2129 | 706 | 667 | 642 | 477 | 1692 | 1384 | 1558 | 2043 | 643 | 675 | 608 | 717 | 1505 | 1503 | 1596 | 1558 | 1100,39 | 661 | 1540 | 2,343 | 1,229 | 20,897 | 5,67E-97 | 4,79E-96 | 0,000911 | 0,0032375 | 0,0022016 | 0,0022016 | TRUE | 0,0335899 | No |
| orf19.2691 | 1018 | 972 | 979 | 731 | 2554 | 2144 | 2228 | 2996 | 927 | 983 | 927 | 1098 | 2272 | 2328 | 2282 | 2285 | 1637,64 | 984 | 2292 | 2,341 | 1,227 | 22,072 | 5,85E-108 | 5,43E-107 | 0,0014177 | 0,0028437 | 0,0022147 | 0,0022147 | TRUE | 0,0649184 | No |
| orf19.7565 | 2321 | 2056 | 2172 | 1353 | 5254 | 4499 | 4776 | 6424 | 2113 | 2079 | 2056 | 2032 | 4673 | 4885 | 4891 | 4900 | 3453,76 | 2070 | 4837 | 2,335 | 1,223 | 32,575 | 9,20E-233 | 1,80E-231 | 1,00E-08 | 0,0024195 | 0,0009984 | 0,0009984 | TRUE | 0,0064271 | No |
| orf19.5262 | 43 | 21 | 24 | 14 | 59 | 72 | 51 | 84 | 39 | 21 | 23 | 21 | 52 | 78 | 52 | 64 | 43,89 | 26 | 62 | 2,329 | 1,22 | 4,817 | 1,46E-06 | 2,41E-06 | 0,0370099 | 0,0321375 | 0,0338538 | 0,0338538 | TRUE | 0,7923798 | No |
| orf19.7200 | 377 | 426 | 423 | 270 | 1079 | 918 | 861 | 1097 | 343 | 431 | 400 | 406 | 960 | 997 | 882 | 837 | 656,88 | 395 | 919 | 2,328 | 1,219 | 14,459 | 2,20E-47 | 9,70E-47 | 0,0059826 | 0,004048 | 0,0049585 | 0,0049585 | TRUE | 0,096427 | No |
| orf19.577 | 4089 | 3534 | 3854 | 2133 | 9296 | 7165 | 8289 | 11023 | 3723 | 3574 | 3647 | 3204 | 8268 | 7779 | 8489 | 8408 | 5886,72 | 3537 | 8236 | 2,326 | 1,218 | 23,263 | 1,05E-119 | 1,07E-118 | 0,0025936 | 0,0022614 | 0,002422 | 0,002422 | TRUE | 0,0466623 | No |
| orf19.3266 | 188 | 157 | 202 | 164 | 521 | 428 | 393 | 567 | 171 | 159 | 191 | 246 | 463 | 465 | 402 | 433 | 316,32 | 192 | 441 | 2,326 | 1,218 | 10,659 | 1,58E-26 | 4,67E-26 | 0,0130923 | 0,0062131 | 0,0086483 | 0,0086483 | TRUE | 0,3300265 | No |
| orf19.7595 | 258 | 245 | 292 | 166 | 665 | 567 | 591 | 704 | 235 | 248 | 276 | 249 | 591 | 616 | 605 | 537 | 419,72 | 252 | 587 | 2,324 | 1,217 | 14,562 | 4,90E-48 | 2,18E-47 | 0,0021107 | 0,0051843 | 0,0037936 | 0,0037936 | TRUE | 0,0516566 | No |
| orf19.3325 | 1907 | 1657 | 1718 | 1132 | 4175 | 3545 | 4040 | 5187 | 1736 | 1676 | 1626 | 1700 | 3714 | 3849 | 4137 | 3957 | 2799,38 | 1684 | 3914 | 2,324 | 1,216 | 25,647 | 4,60E-145 | 5,84E-144 | 0,0012049 | 0,0025089 | 0,0017214 | 0,0017214 | TRUE | 0,0177039 | No |
| orf19.7357 | 1907 | 1816 | 1927 | 1257 | 4764 | 4148 | 4125 | 5181 | 1736 | 1837 | 1824 | 1888 | 4237 | 4504 | 4225 | 3952 | 3025,31 | 1821 | 4230 | 2,324 | 1,216 | 23,959 | 7,44E-127 | 8,07E-126 | 0,0017936 | 0,0024737 | 0,0020685 | 0,0020685 | TRUE | 0,0241401 | No |
| orf19.1779 | 20322 | 17443 | 19167 | 13593 | 44897 | 42662 | 41468 | 58421 | 18502 | 17642 | 18140 | 20419 | 39934 | 46319 | 42468 | 44564 | 30998,59 | 18676 | 43321 | 2,32 | 1,214 | 20,857 | 1,31E-96 | 1,09E-95 | 0,0040818 | 0,0020796 | 0,0032163 | 0,0032163 | TRUE | 0,0479908 | No |
| orf19.2738 | 3229 | 2865 | 3144 | 2100 | 7291 | 6545 | 6819 | 9359 | 2940 | 2898 | 2976 | 3155 | 6485 | 7106 | 6984 | 7139 | 4960,17 | 2992 | 6928 | 2,318 | 1,213 | 26,411 | 1,03E-153 | 1,37E-152 | 0,0013508 | 0,0023033 | 0,0017777 | 0,0017777 | TRUE | 0,0227901 | No |
| orf19.6169 | 450 | 443 | 490 | 339 | 1184 | 959 | 995 | 1436 | 410 | 448 | 464 | 509 | 1053 | 1041 | 1019 | 1095 | 754,93 | 458 | 1052 | 2,315 | 1,211 | 17,727 | 2,57E-70 | 1,63E-69 | 0,0015074 | 0,0037868 | 0,002862 | 0,002862 | TRUE | 0,058709 | No |
| orf19.1207 | 9 | 3 | 9 | 4 | 16 | 14 | 11 | 25 | 8 | 3 | 9 | 6 | 14 | 15 | 11 | 19 | 10,69 | 6 | 15 | 2,309 | 1,208 | 2,679 | 0,0073803 | 0,0097755 | 1,00E-08 | 0,1256076 | 0,0827516 | 0,0827516 | TRUE | 0,3437343 | No |
| orf19.2417 | 811 | 720 | 810 | 504 | 1875 | 1607 | 1763 | 2209 | 738 | 728 | 767 | 757 | 1668 | 1745 | 1806 | 1685 | 1236,67 | 748 | 1726 | 2,308 | 1,207 | 23,118 | 3,03E-118 | 3,06E-117 | 1,00E-08 | 0,0031052 | 0,00163 | 0,00163 | TRUE | 0,0116463 | No |
| orf19.7274 | 162 | 116 | 135 | 98 | 334 | 270 | 325 | 416 | 147 | 117 | 128 | 147 | 297 | 293 | 333 | 317 | 222,52 | 135 | 310 | 2,308 | 1,207 | 11,285 | 1,56E-29 | 4,94E-29 | 0,0008214 | 0,0079735 | 0,005521 | 0,005521 | TRUE | 0,0792997 | No |
| orf19.889 | 428 | 399 | 399 | 295 | 969 | 849 | 958 | 1224 | 390 | 404 | 378 | 443 | 862 | 922 | 981 | 934 | 664,05 | 404 | 925 | 2,303 | 1,204 | 17,014 | 6,51E-65 | 3,82E-64 | 0,0015233 | 0,0040263 | 0,0029697 | 0,0029697 | TRUE | 0,0418423 | No |
| orf19.5438 | 1702 | 1341 | 1437 | 933 | 3769 | 2805 | 3259 | 4351 | 1550 | 1356 | 1360 | 1402 | 3352 | 3045 | 3338 | 3319 | 2340,23 | 1417 | 3264 | 2,303 | 1,203 | 21,335 | 5,38E-101 | 4,72E-100 | 0,0023826 | 0,0026015 | 0,0025317 | 0,0025317 | TRUE | 0,0503297 | No |
| orf19.362 | 70 | 55 | 57 | 44 | 149 | 110 | 130 | 209 | 64 | 56 | 54 | 66 | 133 | 119 | 133 | 159 | 97,99 | 60 | 136 | 2,3 | 1,201 | 7,642 | 2,14E-14 | 4,66E-14 | 1,00E-08 | 0,0155179 | 0,0113905 | 0,0113905 | TRUE | 0,1504116 | No |

$P_{TET}$-*ZCF15* doxycycline-treated down-regulated genes

| | WT dox | | | | ZCF15 | | | | norm. WT dox | | | | norm. ZCF15 | | | | | | | | | | | | | | | | | | |
|---|---|---|---|---|---|---|---|---|---|---|---|---|---|---|---|---|---|---|---|---|---|---|---|---|---|---|---|---|---|---|---|
| Gene ID | 1 | 2 | 3 | 4 | 1 | 2 | 3 | 4 | 1 | 2 | 3 | 4 | 1 | 2 | 3 | 4 | baseMean | WT | ZCF15 | FC | log2FC | stat | pvalue | padj | dispGeneEst | dispFit | dispMAP | dispersion | betaConv | maxCooks | outlier |
| orf19.3374 | 16501 | 13556 | 14242 | 10402 | 336 | 281 | 235 | 421 | 15023 | 13710 | 13479 | 15625 | 299 | 305 | 241 | 321 | 7375,49 | 14459 | 292 | 0,02 | -5,628 | -72,049 | 0 | 0 | 0,0074802 | 0,0022161 | 0,004229 | 0,004229 | TRUE | 0,1494169 | No |
| orf19.937 | 7 | 11 | 4 | 2 | 0 | 0 | 0 | 0 | 6 | 11 | 4 | 3 | 0 | 0 | 0 | 0 | 3,04 | 6 | 0 | 0,028 | -5,168 | -4,178 | 2,94E-05 | 4,55E-05 | 0,2202387 | 0,4371395 | 0,3827454 | 0,3827454 | TRUE | 0,3398331 | No |
| orf19.3988 | 9452 | 8183 | 8926 | 5710 | 301 | 243 | 268 | 376 | 8606 | 8276 | 8448 | 8577 | 268 | 264 | 274 | 287 | 4374,96 | 8477 | 273 | 0,032 | -4,951 | -90,418 | 0 | 0 | 1,00E-08 | 0,002339 | 0,0011243 | 0,0011243 | TRUE | 0,0147275 | No |
| orf19.3664 | 10653 | 9513 | 9835 | 6309 | 430 | 324 | 339 | 445 | 9699 | 9621 | 9308 | 9477 | 382 | 352 | 347 | 339 | 4940,8 | 9526 | 355 | 0,037 | -4,746 | -91,825 | 0 | 0 | 1,00E-08 | 0,0023044 | 0,0012017 | 0,0012017 | TRUE | 0,0326558 | No |
| orf19.7455 | 343 | 300 | 351 | 206 | 19 | 10 | 13 | 13 | 312 | 303 | 332 | 309 | 17 | 11 | 13 | 10 | 163,54 | 314 | 13 | 0,04 | -4,63 | -21,551 | 5,13E-103 | 4,61E-102 | 1,00E-08 | 0,0101145 | 0,0062501 | 0,0062501 | TRUE | 0,2343678 | No |
| orf19.3621 | 747 | 688 | 755 | 504 | 33 | 38 | 31 | 37 | 680 | 696 | 715 | 757 | 29 | 41 | 32 | 28 | 372,27 | 712 | 32 | 0,045 | -4,466 | -31,837 | 2,00E-222 | 3,75E-221 | 1,00E-08 | 0,0055855 | 0,0037558 | 0,0037558 | TRUE | 0,1923731 | No |
| orf19.4907 | 12038 | 11315 | 11786 | 7531 | 557 | 517 | 535 | 714 | 10960 | 11444 | 11154 | 11313 | 495 | 561 | 548 | 545 | 5877,56 | 11218 | 537 | 0,048 | -4,386 | -92,709 | 0 | 0 | 0,0003137 | 0,0022618 | 0,0012347 | 0,0012347 | TRUE | 0,0327823 | No |
| orf19.4526 | 39009 | 35560 | 36958 | 23720 | 2030 | 1703 | 1735 | 2343 | 35516 | 35965 | 34977 | 35631 | 1806 | 1849 | 1777 | 1787 | 18663,57 | 35522 | 1805 | 0,051 | -4,3 | -134,08 | 0 | 0 | 1,00E-08 | 0,0021078 | 0,0007139 | 0,0007139 | TRUE | 0,0030688 | No |
| orf19.1321 | 13694 | 11416 | 13298 | 8965 | 834 | 762 | 765 | 1065 | 12468 | 11546 | 12585 | 13467 | 742 | 827 | 783 | 812 | 6653,88 | 12516 | 791 | 0,063 | -3,984 | -68,629 | 0 | 0 | 0,0028887 | 0,0022355 | 0,0026028 | 0,0026028 | TRUE | 0,0332436 | No |
| orf19.1562 | 32160 | 29870 | 31589 | 22185 | 2205 | 1994 | 1853 | 2562 | 29280 | 30210 | 29896 | 33325 | 1961 | 2165 | 1898 | 1954 | 16336,3 | 30678 | 1994 | 0,065 | -3,944 | -71,562 | 0 | 0 | 0,0031014 | 0,0021179 | 0,002665 | 0,002665 | TRUE | 0,0411335 | No |
| orf19.3378 | 94 | 70 | 68 | 44 | 2 | 2 | 4 | 13 | 86 | 71 | 64 | 66 | 2 | 2 | 4 | 10 | 38,1 | 72 | 4 | 0,067 | -3,909 | -10,322 | 5,60E-25 | 1,61E-24 | 0,017338 | 0,0367095 | 0,0336797 | 0,0336797 | TRUE | 1,6974797 | No |
| orf19.2048 | 15558 | 12582 | 13136 | 9718 | 1149 | 842 | 799 | 1117 | 14165 | 12725 | 12432 | 14598 | 1022 | 914 | 818 | 852 | 7190,85 | 13480 | 902 | 0,067 | -3,902 | -52,684 | 0 | 0 | 0,0072484 | 0,0022207 | 0,00471 | 0,00471 | TRUE | 0,0979584 | No |
| orf19.265 | 5820 | 5882 | 5786 | 3606 | 468 | 359 | 324 | 478 | 5299 | 5949 | 5476 | 5417 | 416 | 390 | 332 | 365 | 2955,38 | 5535 | 376 | 0,068 | -3,881 | -58,04 | 0 | 0 | 0,0040336 | 0,002484 | 0,0029563 | 0,0029563 | TRUE | 0,0663322 | No |
| orf19.4279 | 21197 | 19906 | 21241 | 12835 | 1556 | 1299 | 1353 | 1802 | 19299 | 20133 | 20103 | 19280 | 1384 | 1410 | 1386 | 1375 | 10546,16 | 19704 | 1389 | 0,07 | -3,828 | -103,51 | 0 | 0 | 1,00E-08 | 0,0021623 | 0,00095 | 0,00095 | TRUE | 0,0026192 | No |
| orf19.3373 | 14416 | 12283 | 13125 | 9620 | 1051 | 892 | 866 | 1226 | 13125 | 12423 | 12422 | 14451 | 935 | 968 | 887 | 935 | 7018,24 | 13105 | 931 | 0,071 | -3,814 | -64,734 | 0 | 0 | 0,0032135 | 0,0022252 | 0,0027918 | 0,0027918 | TRUE | 0,0577991 | No |
| orf19.4664 | 3626 | 3316 | 3423 | 1968 | 273 | 199 | 248 | 288 | 3301 | 3354 | 3240 | 2956 | 243 | 216 | 254 | 220 | 1722,93 | 3213 | 233 | 0,072 | -3,788 | -53,422 | 0 | 0 | 0,0024993 | 0,0028037 | 0,0026641 | 0,0026641 | TRUE | 0,0372901 | No |
| orf19.176 | 2920 | 2632 | 2665 | 1796 | 222 | 161 | 188 | 295 | 2659 | 2662 | 2522 | 2698 | 197 | 175 | 193 | 225 | 1416,3 | 2635 | 198 | 0,076 | -3,726 | -52,926 | 0 | 0 | 0,0008275 | 0,0029697 | 0,00223 | 0,00223 | TRUE | 0,1077848 | No |
| orf19.4975 | 5466 | 4689 | 5034 | 3614 | 401 | 398 | 341 | 505 | 4977 | 4742 | 4764 | 5429 | 357 | 432 | 349 | 385 | 2679,41 | 4978 | 381 | 0,076 | -3,71 | -52,67 | 0 | 0 | 0,0050264 | 0,00253 | 0,0034296 | 0,0034296 | TRUE | 0,0865393 | No |
| orf19.4908 | 117 | 89 | 109 | 74 | 7 | 6 | 12 | 9 | 107 | 90 | 103 | 111 | 6 | 7 | 12 | 7 | 55,34 | 103 | 8 | 0,077 | -3,703 | -12,731 | 3,95E-37 | 1,48E-36 | 1,00E-08 | 0,0259059 | 0,017211 | 0,017211 | TRUE | 0,3575953 | No |
| orf19.7027 | 14477 | 12915 | 14901 | 8440 | 1159 | 902 | 1030 | 1361 | 13181 | 13062 | 14102 | 12678 | 1031 | 979 | 1055 | 1038 | 7140,84 | 13256 | 1026 | 0,077 | -3,691 | -76,935 | 0 | 0 | 0,0013894 | 0,002222 | 0,0017156 | 0,0017156 | TRUE | 0,0226513 | No |
| orf19.5140 | 1597 | 1337 | 1486 | 903 | 115 | 112 | 109 | 135 | 1454 | 1352 | 1406 | 1356 | 102 | 122 | 112 | 103 | 750,94 | 1392 | 110 | 0,078 | -3,678 | -43,119 | 0 | 0 | 1,00E-08 | 0,0037961 | 0,0023463 | 0,0023463 | TRUE | 0,0493289 | No |
| orf19.1354 | 1E+05 | 1E+05 | 1E+05 | 82571 | 12531 | 10954 | 10498 | 14308 | 1E+05 | 1E+05 | 1E+05 | 1E+05 | 11146 | 11893 | 10751 | 10914 | 73608,98 | 136042 | 11176 | 0,082 | -3,606 | -65,323 | 0 | 0 | 0,0033919 | 0,002055 | 0,0028817 | 0,0028817 | TRUE | 0,0432144 | No |
| orf19.6028 | 332 | 342 | 333 | 214 | 31 | 22 | 33 | 30 | 302 | 346 | 315 | 321 | 28 | 24 | 34 | 23 | 174,12 | 321 | 27 | 0,084 | -3,581 | -22,04 | 1,20E-107 | 1,11E-106 | 1,00E-08 | 0,0096239 | 0,0064305 | 0,0064305 | TRUE | 0,1672742 | No |
| orf19.7085 | 13245 | 11224 | 12349 | 8677 | 1133 | 937 | 957 | 1335 | 12059 | 11352 | 11687 | 13034 | 1008 | 1017 | 980 | 1018 | 6519,48 | 12033 | 1006 | 0,084 | -3,58 | -68,339 | 0 | 0 | 0,0020524 | 0,0022396 | 0,0021261 | 0,0021261 | TRUE | 0,0377589 | No |
| orf19.4280 | 8023 | 7385 | 7803 | 4774 | 755 | 653 | 659 | 854 | 7305 | 7469 | 7385 | 7171 | 672 | 709 | 675 | 651 | 4004,59 | 7332 | 677 | 0,092 | -3,441 | -78,556 | 0 | 0 | 1,00E-08 | 0,0023669 | 0,0010805 | 0,0010805 | TRUE | 0,011483 | No |
| orf19.449 | 38067 | 35063 | 37919 | 24321 | 3742 | 2934 | 3286 | 4249 | 34658 | 35462 | 35887 | 36534 | 3328 | 3186 | 3365 | 3241 | 19457,77 | 35635 | 3280 | 0,092 | -3,441 | -100,91 | 0 | 0 | 0,0003321 | 0,0021049 | 0,0009595 | 0,0009595 | TRUE | 0,0043263 | No |
| orf19.2655 | 24787 | 23324 | 24963 | 14754 | 2514 | 2046 | 2213 | 2855 | 22568 | 23590 | 23625 | 22163 | 2236 | 2221 | 2266 | 2178 | 12605,88 | 22986 | 2225 | 0,097 | -3,369 | -89,335 | 0 | 0 | 0,0006434 | 0,0021418 | 0,0011328 | 0,0011328 | TRUE | 0,0070024 | No |
| orf19.4450.1 | 4231 | 3527 | 3514 | 2520 | 438 | 338 | 301 | 447 | 3852 | 3567 | 3326 | 3785 | 390 | 367 | 308 | 341 | 1992,03 | 3632 | 352 | 0,097 | -3,369 | -45,579 | 0 | 0 | 0,0050198 | 0,0027002 | 0,0037709 | 0,0037709 | TRUE | 0,0737259 | No |
| orf19.5288 | 2E+05 | 1E+05 | 2E+05 | 1E+05 | 16814 | 13499 | 13675 | 20038 | 2E+05 | 1E+05 | 1E+05 | 2E+05 | 14956 | 14656 | 14005 | 15285 | 83467,54 | 152210 | 14726 | 0,097 | -3,369 | -77,4 | 0 | 0 | 0,0016082 | 0,0020528 | 0,0017856 | 0,0017856 | TRUE | 0,0132679 | No |
| orf19.5735.3 | 3054 | 2483 | 2673 | 1687 | 281 | 228 | 247 | 335 | 2781 | 2511 | 2530 | 2534 | 250 | 248 | 253 | 256 | 1420,21 | 2589 | 252 | 0,097 | -3,364 | -50,953 | 0 | 0 | 1,00E-08 | 0,0029672 | 0,0021311 | 0,0021311 | TRUE | 0,0309915 | No |
| orf19.3904 | 10 | 19 | 20 | 6 | 0 | 3 | 1 | 2 | 9 | 19 | 19 | 9 | 0 | 3 | 1 | 2 | 7,76 | 14 | 2 | 0,098 | -3,349 | -4,545 | 5,48E-06 | 8,81E-06 | 0,103643 | 0,1722986 | 0,1519162 | 0,1519162 | TRUE | 0,2272579 | No |
| orf19.7227 | 2577 | 2082 | 2284 | 1559 | 254 | 204 | 210 | 294 | 2346 | 2106 | 2162 | 2342 | 226 | 221 | 215 | 224 | 1230,27 | 2239 | 222 | 0,099 | -3,334 | -47,434 | 0 | 0 | 1,00E-08 | 0,0031108 | 0,0024093 | 0,0024093 | TRUE | 0,0193462 | No |
| orf19.822 | 3E+05 | 2E+05 | 2E+05 | 1E+05 | 28221 | 20859 | 21674 | 30402 | 2E+05 | 2E+05 | 2E+05 | 2E+05 | 25102 | 22647 | 22197 | 23191 | 124525,7 | 225767 | 23284 | 0,103 | -3,277 | -58,449 | 0 | 0 | 0,0037329 | 0,0020476 | 0,0029986 | 0,0029986 | TRUE | 0,0481334 | No |
| orf19.5079 | 81762 | 70731 | 78184 | 51925 | 8264 | 7290 | 7675 | 10382 | 74441 | 71537 | 73994 | 78000 | 7351 | 7915 | 7860 | 7919 | 41127,06 | 74493 | 7761 | 0,104 | -3,263 | -80,555 | 0 | 0 | 0,0011728 | 0,0020691 | 0,0015085 | 0,0015085 | TRUE | 0,015583 | No |
| orf19.734 | 14428 | 12173 | 13318 | 8421 | 1448 | 1293 | 1283 | 1802 | 13136 | 12312 | 12604 | 12650 | 1288 | 1404 | 1314 | 1375 | 7010,26 | 12676 | 1345 | 0,106 | -3,237 | -76,834 | 0 | 0 | 0,0006032 | 0,0022255 | 0,0013147 | 0,0013147 | TRUE | 0,0100501 | No |
| orf19.2175 | 21385 | 18393 | 20029 | 12354 | 2267 | 1888 | 1923 | 2686 | 19470 | 18603 | 18956 | 18558 | 2016 | 2050 | 1969 | 2049 | 10458,82 | 18897 | 2021 | 0,107 | -3,225 | -91,872 | 0 | 0 | 0,0003321 | 0,0021633 | 0,0009238 | 0,0009238 | TRUE | 0,0051403 | No |
| orf19.3302 | 23393 | 22188 | 23519 | 14726 | 2609 | 2296 | 2272 | 3024 | 21298 | 22441 | 22259 | 22121 | 2321 | 2493 | 2327 | 2307 | 12195,69 | 22030 | 2362 | 0,107 | -3,223 | -84,195 | 0 | 0 | 0,0006408 | 0,0021453 | 0,0011846 | 0,0011846 | TRUE | 0,016335 | No |
| orf19.7086 | 7339 | 6452 | 6870 | 4375 | 800 | 690 | 700 | 910 | 6682 | 6526 | 6502 | 6572 | 712 | 749 | 717 | 694 | 3644,11 | 6570 | 718 | 0,109 | -3,197 | -75,208 | 0 | 0 | 1,00E-08 | 0,0023995 | 0,001004 | 0,001004 | TRUE | 0,0095601 | No |
| orf19.2770 | 5078 | 4615 | 4754 | 3049 | 613 | 424 | 485 | 652 | 4623 | 4668 | 4499 | 4580 | 545 | 460 | 497 | 497 | 2546,23 | 4592 | 500 | 0,109 | -3,196 | -61,663 | 0 | 0 | 0,0001341 | 0,0025558 | 0,0015345 | 0,0015345 | TRUE | 0,0432724 | No |
| orf19.7365 | 1185 | 1239 | 1310 | 827 | 155 | 109 | 147 | 161 | 1079 | 1253 | 1240 | 1242 | 138 | 118 | 151 | 123 | 667,96 | 1204 | 132 | 0,11 | -3,185 | -34,683 | 1,40E-263 | 3,11E-262 | 0,004311 | 0,0040147 | 0,0041366 | 0,0041366 | TRUE | 0,0810948 | No |
| orf19.5686 | 5857 | 4399 | 4676 | 2874 | 584 | 451 | 486 | 725 | 5333 | 4449 | 4425 | 4317 | 519 | 490 | 498 | 553 | 2573,02 | 4631 | 515 | 0,111 | -3,167 | -41,669 | 0 | 0 | 0,0063091 | 0,0025504 | 0,0045259 | 0,0045259 | TRUE | 0,1275848 | No |
| orf19.2839 | 4738 | 4177 | 4483 | 2617 | 492 | 411 | 490 | 641 | 4314 | 4225 | 4243 | 3931 | 438 | 446 | 502 | 489 | 2323,36 | 4178 | 469 | 0,112 | -3,156 | -54,464 | 0 | 0 | 0,0014965 | 0,0026056 | 0,0021033 | 0,0021033 | TRUE | 0,0246764 | No |
| orf19.1785 | 29607 | 27220 | 29944 | 20314 | 3587 | 3030 | 3115 | 4101 | 26956 | 27530 | 28339 | 30515 | 3191 | 3290 | 3190 | 3128 | 15767,36 | 28335 | 3200 | 0,113 | -3,147 | -68,534 | 0 | 0 | 0,0015017 | 0,0021208 | 0,0018598 | 0,0018598 | TRUE | 0,0326023 | No |
| orf19.1861 | 15241 | 13677 | 14933 | 9177 | 1857 | 1495 | 1560 | 1986 | 13876 | 13833 | 14133 | 13785 | 1652 | 1623 | 1598 | 1515 | 7751,83 | 13907 | 1597 | 0,115 | -3,124 | -82,955 | 0 | 0 | 0,0001377 | 0,0022074 | 0,0010308 | 0,0010308 | TRUE | 0,0156527 | No |
| orf19.6842 | 11970 | 10749 | 12105 | 7345 | 1392 | 1191 | 1237 | 1779 | 10898 | 10871 | 11456 | 11033 | 1238 | 1293 | 1267 | 1357 | 6176,8 | 11064 | 1289 | 0,117 | -3,101 | -73,673 | 0 | 0 | 0,0004961 | 0,0022509 | 0,0012907 | 0,0012907 | TRUE | 0,0166622 | No |
| orf19.4867 | 11634 | 10837 | 11881 | 7471 | 1485 | 1230 | 1195 | 1681 | 10592 | 10960 | 11244 | 11223 | 1321 | 1335 | 1224 | 1282 | 6147,76 | 11005 | 1290 | 0,117 | -3,092 | -73,018 | 0 | 0 | 0,0005498 | 0,0022519 | 0,0013113 | 0,0013113 | TRUE | 0,0137668 | No |
| orf19.6387 | 32317 | 26284 | 29280 | 19494 | 3649 | 3040 | 3288 | 4385 | 29423 | 26583 | 27711 | 29283 | 3246 | 3301 | 3367 | 3345 | 15782,4 | 28250 | 3315 | 0,117 | -3,091 | -72,215 | 0 | 0 | 0,0010575 | 0,0021207 | 0,0015999 | 0,0015999 | TRUE | 0,0193267 | No |
| orf19.5626 | 27896 | 26829 | 28764 | 19222 | 3555 | 3101 | 3131 | 4256 | 25398 | 27135 | 27223 | 28875 | 3162 | 3367 | 3207 | 3247 | 15201,47 | 27158 | 3246 | 0,12 | -3,065 | -66,757 | 0 | 0 | 0,0015039 | 0,0021239 | 0,0018607 | 0,0018607 | TRUE | 0,0233109 | No |
| orf19.6608 | 2937 | 2812 | 3061 | 1752 | 352 | 328 | 314 | 438 | 2674 | 2844 | 2897 | 2632 | 313 | 356 | 322 | 334 | 1546,46 | 2762 | 331 | 0,12 | -3,063 | -48,828 | 0 | 0 | 0,001476 | 0,0028912 | 0,0021772 | 0,0021772 | TRUE | 0,0260229 | No |
| orf19.5348 | 11880 | 9871 | 11245 | 7228 | 1430 | 1129 | 1197 | 1773 | 10816 | 9983 | 10642 | 10858 | 1272 | 1226 | 1226 | 1352 | 5921,97 | 10575 | 1269 | 0,12 | -3,056 | -63,691 | 0 | 0 | 0,0012532 | 0,0022601 | 0,0017933 | 0,0017933 | TRUE | 0,0246981 | No |
| orf19.823 | 988 | 1262 | 1242 | 1826 | 177 | 296 | 112 | 195 | 900 | 1276 | 1175 | 2743 | 157 | 321 | 115 | 149 | 854,57 | 1524 | 186 | 0,122 | -3,038 | -6,339 | 2,31E-10 | 4,42E-10 | 0,2177788 | 0,0035828 | 0,0787746 | 0,2177788 | TRUE | 1,0256197 | No |
| orf19.147 | 13321 | 13905 | 15000 | 9272 | 1875 | 1705 | 1653 | 2193 | 12128 | 14063 | 14196 | 13928 | 1668 | 1851 | 1693 | 1673 | 7650,05 | 13579 | 1721 | 0,127 | -2,981 | -49,854 | 0 | 0 | 0,0034201 | 0,0022097 | 0,003122 | 0,003122 | TRUE | 0,0636232 | No |
| orf19.2414 | 9629 | 8265 | 8602 | 5697 | 1280 | 935 | 1111 | 1359 | 8767 | 8359 | 8141 | 8558 | 1139 | 1015 | 1138 | 1037 | 4769,11 | 8456 | 1082 | 0,128 | -2,966 | -59,778 | 0 | 0 | 0,0016343 | 0,002314 | 0,0018695 | 0,0018695 | TRUE | 0,0190417 | No |
| orf19.2173 | 13142 | 11478 | 12638 | 7630 | 1756 | 1366 | 1525 | 2077 | 11965 | 11609 | 11961 | 11461 | 1562 | 1483 | 1562 | 1584 | 6648,42 | 11749 | 1548 | 0,132 | -2,923 | -76,905 | 0 | 0 | 0,0003295 | 0,0022357 | 0,0010405 | 0,0010405 | TRUE | 0,0090676 | No |
| orf19.6116 | 2015 | 1700 | 1939 | 1119 | 269 | 201 | 240 | 301 | 1835 | 1719 | 1835 | 1681 | 239 | 218 | 246 | 230 | 1000,35 | 1768 | 233 | 0,132 | -2,923 | -42,588 | 0 | 0 | 1,00E-08 | 0,0033575 | 0,0022302 | 0,0022302 | TRUE | 0,0176329 | No |
| orf19.4436 | 412 | 392 | 394 | 271 | 60 | 45 | 44 | 76 | 375 | 396 | 373 | 407 | 53 | 49 | 45 | 58 | 219,6 | 388 | 51 | 0,134 | -2,901 | -23,096 | 5,08E-118 | 5,12E-117 | 1,00E-08 | 0,0080525 | 0,0048261 | 0,0048261 | TRUE | 0,0750598 | No |
| orf19.5848 | 6486 | 5569 | 5956 | 4021 | 825 | 735 | 773 | 1036 | 5905 | 5632 | 5637 | 6040 | 734 | 798 | 792 | 790 | 3291,05 | 5804 | 778 | 0,134 | -2,898 | -60,145 | 0 | 0 | 0,0008561 | 0,0024384 | 0,0015382 | 0,0015382 | TRUE | 0,0180356 | No |
| orf19.7283 | 20935 | 16851 | 18239 | 12063 | 2852 | 2114 | 2504 | 3274 | 19060 | 17043 | 17262 | 18121 | 2537 | 2295 | 2564 | 2497 | 10172,42 | 17872 | 2473 | 0,138 | -2,852 | -56,088 | 0 | 0 | 0,0022312 | 0,0021669 | 0,0022653 | 0,0022653 | TRUE | 0,0277797 | No |
| orf19.5069 | 10880 | 10053 | 10907 | 6945 | 1602 | 1290 | 1429 | 1824 | 9906 | 10168 | 10322 | 10432 | 1425 | 1401 | 1463 | 1391 | 5813,58 | 10207 | 1420 | 0,139 | -2,846 | -74,783 | 0 | 0 | 1,00E-08 | 0,0022642 | 0,0010092 | 0,0010092 | TRUE | 0,0049363 | No |
| orf19.7612 | 22724 | 20731 | 21695 | 13760 | 3277 | 2706 | 2730 | 3778 | 20689 | 20967 | 20532 | 20670 | 2915 | 2938 | 2796 | 2882 | 11798,62 | 20714 | 2883 | 0,139 | -2,845 | -92,349 | 0 | 0 | 3,37E-05 | 0,002149 | 0,0007236 | 0,0007236 | TRUE | 0,0047632 | No |
| orf19.3117 | 318 | 240 | 215 | 165 | 34 | 34 | 35 | 47 | 290 | 243 | 203 | 248 | 30 | 37 | 36 | 36 | 140,31 | 246 | 35 | 0,141 | -2,828 | -16,754 | 5,25E-63 | 3,00E-62 | 1,00E-08 | 0,0114521 | 0,011775 | 0,011775 | TRUE | 0,0943788 | No |

| | | | | | | | | | | | | | | | | | | | | | | | | | | | | | | | |
|---|---|---|---|---|---|---|---|---|---|---|---|---|---|---|---|---|---|---|---|---|---|---|---|---|---|---|---|---|---|---|---|
| orf19.6604 | 5319 | 6054 | 6020 | 3740 | 909 | 807 | 742 | 938 | 4843 | 6123 | 5697 | 5618 | 809 | 876 | 760 | 716 | 3180,16 | 5570 | 790 | 0,142 | -2,82 | -35,814 | 6,61E-281 | 1,56E-279 | 0,0074179 | 0,0024524 | 0,0052638 | 0,0052638 | TRUE | 0,0951692 | No |
| orf19.6881 | 8603 | 7504 | 8042 | 5410 | 1352 | 1049 | 1045 | 1425 | 7833 | 7589 | 7611 | 8127 | 1203 | 1139 | 1070 | 1087 | 4457,32 | 7790 | 1125 | 0,144 | -2,792 | -58,385 | 0 | 0 | 0,0012403 | 0,0023334 | 0,0017112 | 0,0017112 | TRUE | 0,0268804 | No |
| orf19.4972 | 258 | 263 | 275 | 168 | 37 | 28 | 42 | 52 | 235 | 266 | 260 | 252 | 33 | 30 | 43 | 40 | 144,94 | 253 | 36 | 0,145 | -2,789 | -18,486 | 2,65E-76 | 1,79E-75 | 1,00E-08 | 0,0111512 | 0,0070993 | 0,0070993 | TRUE | 0,0866673 | No |
| orf19.3551 | 311 | 243 | 257 | 140 | 43 | 30 | 39 | 45 | 283 | 246 | 243 | 210 | 38 | 33 | 40 | 34 | 140,94 | 246 | 36 | 0,147 | -2,769 | -17,345 | 2,14E-67 | 1,30E-66 | 1,00E-08 | 0,0114096 | 0,0095026 | 0,0095026 | TRUE | 0,1004608 | No |
| orf19.2962 | 916 | 899 | 972 | 572 | 148 | 117 | 144 | 161 | 834 | 909 | 920 | 859 | 132 | 127 | 147 | 123 | 506,42 | 880 | 132 | 0,15 | -2,741 | -31,548 | 1,90E-218 | 3,49E-217 | 1,00E-08 | 0,0046455 | 0,0031184 | 0,0031184 | TRUE | 0,058904 | No |
| orf19.951 | 9358 | 9665 | 10208 | 6020 | 1618 | 1274 | 1366 | 1740 | 8520 | 9775 | 9661 | 9043 | 1439 | 1383 | 1399 | 1327 | 5318,47 | 9250 | 1387 | 0,15 | -2,738 | -50,345 | 0 | 0 | 0,002554 | 0,0022854 | 0,0024452 | 0,0024452 | TRUE | 0,0349813 | No |
| orf19.413 | 4678 | 4447 | 4784 | 2895 | 734 | 640 | 621 | 877 | 4259 | 4498 | 4528 | 4349 | 653 | 695 | 636 | 669 | 2535,73 | 4408 | 663 | 0,15 | -2,733 | -56,769 | 0 | 0 | 1,00E-08 | 0,002558 | 0,0014018 | 0,0014018 | TRUE | 0,0108457 | No |
| orf19.5760 | 5180 | 3905 | 4443 | 3473 | 755 | 647 | 640 | 912 | 4716 | 3949 | 4205 | 5217 | 672 | 702 | 655 | 696 | 2601,59 | 4522 | 681 | 0,151 | -2,729 | -33,647 | 3,43E-248 | 7,05E-247 | 0,0082418 | 0,0025448 | 0,0055146 | 0,0055146 | TRUE | 0,1153981 | No |
| orf19.6621 | 15333 | 14187 | 15491 | 8977 | 2391 | 1903 | 2172 | 2825 | 13960 | 14349 | 14661 | 13485 | 2127 | 2066 | 2224 | 2155 | 8128,32 | 14114 | 2143 | 0,152 | -2,72 | -65,542 | 0 | 0 | 0,0007586 | 0,0021995 | 0,0013973 | 0,0013973 | TRUE | 0,0107696 | No |
| orf19.3537 | 2762 | 2208 | 2140 | 1683 | 410 | 343 | 343 | 458 | 2515 | 2233 | 2025 | 2528 | 365 | 372 | 351 | 349 | 1342,38 | 2325 | 359 | 0,155 | -2,693 | -33,497 | 5,26E-246 | 1,08E-244 | 0,0061627 | 0,0030211 | 0,0046775 | 0,0046775 | TRUE | 0,0922839 | No |
| orf19.2398 | 19972 | 17786 | 19466 | 12883 | 3121 | 2639 | 2804 | 3842 | 18184 | 17989 | 18423 | 19352 | 2776 | 2865 | 2872 | 2931 | 10673,88 | 18487 | 2861 | 0,155 | -2,691 | -70,988 | 0 | 0 | 0,0006931 | 0,0021608 | 0,001188 | 0,001188 | TRUE | 0,0119402 | No |
| orf19.2050 | 7142 | 6226 | 6871 | 4275 | 1115 | 920 | 971 | 1315 | 6502 | 6297 | 6503 | 6422 | 992 | 999 | 994 | 1003 | 3714,01 | 6431 | 997 | 0,155 | -2,689 | -69,538 | 0 | 0 | 1,00E-08 | 0,0023927 | 0,0008856 | 0,0008856 | TRUE | 0,0024703 | No |
| orf19.2768 | 40960 | 40990 | 42868 | 23944 | 6848 | 5857 | 5853 | 7667 | 37292 | 41457 | 40571 | 35968 | 6091 | 6359 | 5994 | 5848 | 22447,57 | 38822 | 6073 | 0,156 | -2,677 | -50,905 | 0 | 0 | 0,0027659 | 0,0020959 | 0,0025655 | 0,0025655 | TRUE | 0,0298634 | No |
| orf19.211 | 1916 | 1627 | 1802 | 1232 | 330 | 223 | 256 | 377 | 1744 | 1646 | 1705 | 1851 | 294 | 242 | 262 | 288 | 1003,93 | 1736 | 272 | 0,157 | -2,669 | -37,549 | 0 | 0 | 0,0022279 | 0,0033528 | 0,0028353 | 0,0028353 | TRUE | 0,0550405 | No |
| orf19.1290 | 4791 | 4396 | 4762 | 2926 | 842 | 696 | 656 | 832 | 4362 | 4446 | 4507 | 4395 | 749 | 756 | 672 | 635 | 2565,16 | 4428 | 703 | 0,158 | -2,659 | -47,278 | 0 | 0 | 0,0018437 | 0,002552 | 0,0022496 | 0,0022496 | TRUE | 0,0549964 | No |
| orf19.938 | 358 | 316 | 362 | 228 | 58 | 41 | 54 | 76 | 326 | 320 | 343 | 342 | 52 | 45 | 55 | 58 | 192,5 | 333 | 52 | 0,159 | -2,655 | -20,891 | 6,53E-97 | 5,51E-96 | 1,00E-08 | 0,0088993 | 0,0051193 | 0,0051193 | TRUE | 0,0729315 | No |
| orf19.1189 | 27688 | 22718 | 25701 | 17458 | 4329 | 3548 | 3812 | 5361 | 25209 | 22977 | 24324 | 26225 | 3851 | 3852 | 3904 | 4089 | 14303,75 | 24684 | 3924 | 0,159 | -2,653 | -55,912 | 0 | 0 | 0,0017754 | 0,0021294 | 0,002021 | 0,002021 | TRUE | 0,0264985 | No |
| orf19.1152 | 55367 | 49905 | 53037 | 32479 | 9128 | 7416 | 8052 | 10179 | 50409 | 50474 | 50195 | 48789 | 8119 | 8052 | 8246 | 7765 | 29005,99 | 49967 | 8046 | 0,161 | -2,635 | -84,238 | 0 | 0 | 0,0003511 | 0,0020826 | 0,0008711 | 0,0008711 | TRUE | 0,0069113 | No |
| orf19.6267 | 537 | 500 | 534 | 309 | 91 | 81 | 71 | 104 | 489 | 506 | 505 | 464 | 81 | 88 | 73 | 79 | 285,64 | 491 | 80 | 0,163 | -2,619 | -24,876 | 1,37E-136 | 1,61E-135 | 1,00E-08 | 0,0066618 | 0,0037686 | 0,0037686 | TRUE | 0,0325343 | No |
| orf19.5118 | 48886 | 42179 | 46387 | 31412 | 7728 | 6805 | 6954 | 10014 | 44509 | 42660 | 43901 | 47186 | 6874 | 7388 | 7122 | 7639 | 25909,72 | 44564 | 7256 | 0,163 | -2,618 | -56,992 | 0 | 0 | 0,0018503 | 0,002088 | 0,0019513 | 0,0019513 | TRUE | 0,019127 | No |
| orf19.836 | 209 | 214 | 197 | 133 | 32 | 34 | 43 | 30 | 190 | 216 | 186 | 200 | 28 | 37 | 44 | 23 | 115,66 | 198 | 33 | 0,163 | -2,613 | -15,086 | 2,00E-51 | 9,50E-51 | 0,0048996 | 0,0134587 | 0,0117679 | 0,0117679 | TRUE | 0,380069 | No |
| orf19.4432 | 18400 | 16196 | 17266 | 10464 | 3030 | 2430 | 2631 | 3501 | 16752 | 16381 | 16341 | 15719 | 2695 | 2638 | 2694 | 2671 | 9486,34 | 16298 | 2674 | 0,164 | -2,608 | -75,784 | 0 | 0 | 0,000367 | 0,0021763 | 0,0009293 | 0,0009293 | TRUE | 0,0068347 | No |
| orf19.7550 | 652 | 556 | 640 | 397 | 114 | 84 | 104 | 120 | 594 | 562 | 606 | 596 | 101 | 91 | 107 | 92 | 343,58 | 590 | 98 | 0,165 | -2,596 | -26,84 | 1,10E-158 | 1,52E-157 | 1,00E-08 | 0,0058818 | 0,0033182 | 0,0033182 | TRUE | 0,0334277 | No |
| orf19.125 | 9690 | 9205 | 9602 | 5680 | 1741 | 1378 | 1427 | 1919 | 8822 | 9310 | 9087 | 8532 | 1549 | 1496 | 1461 | 1464 | 5215,23 | 8938 | 1492 | 0,167 | -2,583 | -59,625 | 0 | 0 | 0,0009055 | 0,0022903 | 0,0014288 | 0,0014288 | TRUE | 0,0110358 | No |
| orf19.2047 | 1643 | 1426 | 1521 | 931 | 267 | 237 | 216 | 328 | 1496 | 1442 | 1439 | 1399 | 237 | 257 | 221 | 250 | 842,79 | 1444 | 241 | 0,167 | -2,58 | -38,448 | 0 | 0 | 1,00E-08 | 0,0036044 | 0,0020321 | 0,0020321 | TRUE | 0,0325862 | No |
| orf19.7473 | 1862 | 1861 | 2018 | 1295 | 318 | 309 | 334 | 385 | 1695 | 1882 | 1910 | 1945 | 283 | 335 | 342 | 294 | 1085,84 | 1858 | 314 | 0,168 | -2,572 | -34,355 | 1,18E-258 | 2,57E-257 | 0,0037804 | 0,0032536 | 0,0035989 | 0,0035989 | TRUE | 0,0475704 | No |
| orf19.2296 | 1E+05 | 82301 | 92018 | 62502 | 16210 | 14304 | 14292 | 20205 | 92267 | 83239 | 87087 | 93888 | 14418 | 15530 | 14637 | 15413 | 52059,73 | 89120 | 15000 | 0,168 | -2,571 | -54,094 | 0 | 0 | 0,0021282 | 0,0020624 | 0,0021328 | 0,0021328 | TRUE | 0,0241838 | No |
| orf19.729.1 | 1528 | 1315 | 1444 | 1011 | 269 | 221 | 232 | 302 | 1391 | 1330 | 1367 | 1519 | 239 | 240 | 238 | 230 | 819,2 | 1402 | 237 | 0,169 | -2,564 | -36,019 | 4,23E-284 | 1,02E-282 | 1,00E-08 | 0,0036496 | 0,0025158 | 0,0025158 | TRUE | 0,0346253 | No |
| orf19.3474 | 1928 | 1942 | 2126 | 1201 | 377 | 270 | 333 | 401 | 1755 | 1964 | 2012 | 1804 | 335 | 293 | 341 | 306 | 1101,38 | 1884 | 319 | 0,169 | -2,564 | -35,453 | 2,57E-275 | 5,97E-274 | 0,0033141 | 0,0032364 | 0,0032742 | 0,0032742 | TRUE | 0,0294575 | No |
| orf19.4673 | 1537 | 1419 | 1564 | 941 | 292 | 238 | 225 | 298 | 1399 | 1435 | 1480 | 1414 | 260 | 258 | 230 | 227 | 838,02 | 1432 | 244 | 0,17 | -2,557 | -37,784 | 0 | 0 | 1,00E-08 | 0,0036134 | 0,0021153 | 0,0021153 | TRUE | 0,0277728 | No |
| orf19.4170 | 1398 | 1276 | 1297 | 824 | 231 | 193 | 227 | 280 | 1273 | 1291 | 1227 | 1238 | 205 | 210 | 232 | 214 | 736,21 | 1257 | 215 | 0,171 | -2,548 | -36,868 | 1,50E-297 | 3,87E-296 | 1,00E-08 | 0,0038313 | 0,0020025 | 0,0020025 | TRUE | 0,0293459 | No |
| orf19.2107.1 | 1E+05 | 78158 | 83372 | 44334 | 18323 | 10213 | 13622 | 18705 | 1E+05 | 79048 | 78904 | 66597 | 16298 | 11088 | 13951 | 14268 | 47611,96 | 81322 | 13901 | 0,171 | -2,548 | -15,168 | 5,77E-52 | 2,77E-51 | 0,0270846 | 0,0020648 | 0,0131677 | 0,0270846 | TRUE | 0,2015577 | No |
| orf19.1153 | 7748 | 6907 | 7682 | 4561 | 1356 | 1129 | 1154 | 1602 | 7054 | 6986 | 7270 | 6851 | 1206 | 1226 | 1182 | 1222 | 4124,67 | 7040 | 1209 | 0,172 | -2,542 | -64,175 | 0 | 0 | 1,00E-08 | 0,0023573 | 0,0010449 | 0,0010449 | TRUE | 0,0059025 | No |
| orf19.3363 | 30621 | 27690 | 29468 | 17871 | 5348 | 4386 | 4611 | 6257 | 27879 | 28005 | 27889 | 26845 | 4757 | 4762 | 4722 | 4773 | 16204,06 | 27654 | 4754 | 0,172 | -2,541 | -86,302 | 0 | 0 | 0,0001901 | 0,0021185 | 0,0007145 | 0,0007145 | TRUE | 0,0046592 | No |
| orf19.5114.1 | 22816 | 16304 | 17523 | 9896 | 3679 | 2444 | 2891 | 4106 | 20773 | 16490 | 16584 | 14865 | 3272 | 2654 | 2961 | 3132 | 10091,34 | 17178 | 3005 | 0,175 | -2,515 | -27,232 | 2,70E-163 | 3,83E-162 | 0,0144613 | 0,0021679 | 0,0080081 | 0,0080081 | TRUE | 0,243099 | No |
| orf19.3038 | 70134 | 61043 | 66769 | 41592 | 12025 | 10147 | 11018 | 14894 | 63854 | 61738 | 63191 | 62478 | 10696 | 11017 | 11284 | 11361 | 36952,36 | 62815 | 11090 | 0,177 | -2,502 | -79,967 | 0 | 0 | 0,000403 | 0,0020728 | 0,0008896 | 0,0008896 | TRUE | 0,0070565 | No |
| orf19.5908 | 10566 | 9139 | 10326 | 6845 | 1842 | 1566 | 1678 | 2368 | 9620 | 9243 | 9773 | 10282 | 1638 | 1700 | 1718 | 1806 | 5722,67 | 9730 | 1716 | 0,177 | -2,502 | -52,63 | 0 | 0 | 0,001339 | 0,0022679 | 0,001843 | 0,001843 | TRUE | 0,0174803 | No |
| orf19.4936.1 | 17231 | 15923 | 16570 | 9585 | 3314 | 2504 | 2731 | 3450 | 15688 | 16104 | 15682 | 14398 | 2948 | 2719 | 2797 | 2632 | 9120,95 | 15468 | 2774 | 0,179 | -2,48 | -49,942 | 0 | 0 | 0,0020316 | 0,0021818 | 0,002165 | 0,002165 | TRUE | 0,026185 | No |
| orf19.2749 | 5668 | 6031 | 6434 | 3841 | 1172 | 1036 | 978 | 1315 | 5160 | 6100 | 6089 | 5770 | 1042 | 1125 | 1002 | 1003 | 3411,39 | 5780 | 1043 | 0,18 | -2,471 | -38,445 | 0 | 0 | 0,0042469 | 0,0024242 | 0,0034269 | 0,0034269 | TRUE | 0,0643743 | No |
| orf19.486 | 592 | 508 | 553 | 320 | 86 | 86 | 105 | 126 | 539 | 514 | 523 | 481 | 76 | 93 | 108 | 96 | 303,79 | 514 | 93 | 0,181 | -2,469 | -23,474 | 7,52E-122 | 7,74E-121 | 0,000274 | 0,0063854 | 0,0045978 | 0,0045978 | TRUE | 0,1501343 | No |
| orf19.3571 | 255 | 224 | 222 | 180 | 59 | 35 | 34 | 56 | 232 | 227 | 210 | 270 | 52 | 38 | 35 | 43 | 138,4 | 235 | 42 | 0,181 | -2,464 | -15,89 | 7,44E-57 | 3,86E-56 | 0,0061167 | 0,0115816 | 0,0098206 | 0,0098206 | TRUE | 0,219955 | No |
| orf19.4911 | 478 | 426 | 472 | 301 | 96 | 69 | 75 | 109 | 435 | 431 | 447 | 452 | 85 | 75 | 77 | 83 | 260,65 | 441 | 80 | 0,182 | -2,454 | -23,266 | 9,75E-120 | 9,94E-119 | 1,00E-08 | 0,0071052 | 0,0037076 | 0,0037076 | TRUE | 0,0174188 | No |
| orf19.6249 | 2885 | 2568 | 2819 | 1622 | 507 | 439 | 477 | 623 | 2627 | 2597 | 2668 | 2437 | 451 | 477 | 489 | 475 | 1527,46 | 2582 | 473 | 0,183 | -2,452 | -44,464 | 0 | 0 | 1,00E-08 | 0,0029018 | 0,0017274 | 0,0017274 | TRUE | 0,0161622 | No |
| orf19.6919 | 126 | 152 | 142 | 80 | 30 | 23 | 18 | 33 | 115 | 154 | 134 | 120 | 27 | 25 | 18 | 25 | 77,28 | 131 | 24 | 0,183 | -2,452 | -12,298 | 9,29E-35 | 3,31E-34 | 1,00E-08 | 0,0191297 | 0,0147086 | 0,0147086 | TRUE | 0,1451303 | No |
| orf19.4943 | 8732 | 7984 | 8528 | 5467 | 1704 | 1321 | 1365 | 2050 | 7950 | 8075 | 8071 | 8212 | 1516 | 1434 | 1398 | 1564 | 4777,49 | 8077 | 1478 | 0,183 | -2,448 | -57,1 | 0 | 0 | 0,0005541 | 0,0023135 | 0,0013822 | 0,0013822 | TRUE | 0,0192056 | No |
| orf19.3669 | 17247 | 15782 | 17267 | 11387 | 3294 | 2887 | 2911 | 3939 | 15703 | 15962 | 16342 | 17105 | 2930 | 3134 | 2981 | 3005 | 9645,19 | 16278 | 3012 | 0,185 | -2,434 | -59,811 | 0 | 0 | 0,0008593 | 0,002174 | 0,001402 | 0,001402 | TRUE | 0,0140855 | No |
| orf19.1331 | 5106 | 5566 | 5505 | 3189 | 1118 | 879 | 913 | 1161 | 4649 | 5629 | 5210 | 4790 | 994 | 954 | 935 | 886 | 3006 | 5070 | 942 | 0,186 | -2,429 | -36,076 | 5,49E-285 | 1,33E-283 | 0,0047978 | 0,0024765 | 0,0037501 | 0,0037501 | TRUE | 0,0674986 | No |
| orf19.2079 | 4523 | 3582 | 3630 | 2955 | 868 | 688 | 650 | 953 | 4118 | 3623 | 3435 | 4439 | 772 | 747 | 666 | 727 | 2315,85 | 3904 | 728 | 0,187 | -2,422 | -29,586 | 2,25E-192 | 3,68E-191 | 0,0077544 | 0,0026074 | 0,0056534 | 0,0056534 | TRUE | 0,0946079 | No |
| orf19.740 | 45082 | 42743 | 46629 | 28225 | 8854 | 7478 | 7930 | 10486 | 41045 | 43230 | 44130 | 42398 | 7875 | 8119 | 8121 | 7999 | 25364,79 | 42701 | 8028 | 0,188 | -2,411 | -71,551 | 0 | 0 | 0,000479 | 0,0020891 | 0,00102 | 0,00102 | TRUE | 0,0084087 | No |
| orf19.7006 | 7566 | 6403 | 7318 | 4580 | 1357 | 1115 | 1368 | 1737 | 6889 | 6476 | 6926 | 6880 | 1207 | 1211 | 1401 | 1325 | 4039,22 | 6793 | 1286 | 0,189 | -2,401 | -44,756 | 0 | 0 | 0,0025591 | 0,0023641 | 0,0023207 | 0,0023207 | TRUE | 0,0418598 | No |
| orf19.6578 | 18536 | 17412 | 18835 | 11809 | 3671 | 3123 | 3278 | 4360 | 16876 | 17610 | 17826 | 17739 | 3265 | 3391 | 3357 | 3326 | 10423,76 | 17513 | 3335 | 0,19 | -2,393 | -70,522 | 0 | 0 | 0,0003884 | 0,0021637 | 0,0009348 | 0,0009348 | TRUE | 0,0074083 | No |
| orf19.7310 | 8335 | 7769 | 8074 | 5074 | 1603 | 1360 | 1443 | 1957 | 7589 | 7858 | 7641 | 7622 | 1426 | 1477 | 1478 | 1493 | 4572,81 | 7678 | 1468 | 0,191 | -2,387 | -65,133 | 0 | 0 | 1,00E-08 | 0,0023259 | 0,0009015 | 0,0009015 | TRUE | 0,0047405 | No |
| orf19.2665 | 39545 | 35601 | 38783 | 23638 | 7568 | 6480 | 6862 | 9046 | 36004 | 36007 | 36705 | 35508 | 6731 | 7035 | 7028 | 6900 | 21489,77 | 36056 | 6924 | 0,192 | -2,381 | -80,993 | 0 | 0 | 0,0001896 | 0,0020985 | 0,0007476 | 0,0007476 | TRUE | 0,0042569 | No |
| orf19.6814 | 1E+05 | 81646 | 89182 | 67659 | 19142 | 16441 | 16840 | 24508 | 95929 | 82576 | 84403 | 1E+05 | 17026 | 17850 | 17246 | 18695 | 54419,97 | 91136 | 17704 | 0,194 | -2,364 | -35,651 | 2,28E-278 | 5,35E-277 | 0,0058969 | 0,0020613 | 0,0041918 | 0,0041918 | TRUE | 0,0736226 | No |
| orf19.3281 | 829 | 745 | 752 | 543 | 164 | 131 | 160 | 186 | 755 | 753 | 712 | 816 | 146 | 142 | 164 | 142 | 453,68 | 759 | 148 | 0,196 | -2,353 | -27,24 | 2,21E-163 | 3,14E-162 | 1,00E-08 | 0,0049487 | 0,0033221 | 0,0033221 | TRUE | 0,0480005 | No |
| orf19.2552 | 943 | 906 | 930 | 601 | 186 | 183 | 162 | 222 | 859 | 916 | 880 | 903 | 165 | 199 | 166 | 169 | 532,15 | 890 | 175 | 0,196 | -2,352 | -29,773 | 8,87E-195 | 1,47E-193 | 2,80E-05 | 0,0045194 | 0,0027205 | 0,0027205 | TRUE | 0,0804854 | No |
| orf19.6763 | 39002 | 34465 | 37600 | 22679 | 7781 | 6224 | 6902 | 9121 | 35510 | 34858 | 35585 | 34067 | 6921 | 6758 | 7069 | 6958 | 20965,55 | 35005 | 6926 | 0,198 | -2,337 | -76,15 | 0 | 0 | 0,0002603 | 0,0021 | 0,0008221 | 0,0008221 | TRUE | 0,0039279 | No |
| orf19.3047 | 5448 | 5084 | 5647 | 3470 | 1158 | 949 | 1019 | 1354 | 4960 | 5142 | 5344 | 5212 | 1030 | 1030 | 1044 | 1033 | 3099,47 | 5164 | 1034 | 0,2 | -2,32 | -54,152 | 0 | 0 | 1,00E-08 | 0,0024632 | 0,0012074 | 0,0012074 | TRUE | 0,0089444 | No |
| orf19.6888 | 15625 | 13967 | 15316 | 10006 | 3237 | 2714 | 2830 | 3783 | 14226 | 14126 | 14495 | 15031 | 2879 | 2947 | 2898 | 2886 | 8685,96 | 14470 | 2902 | 0,201 | -2,318 | -66,742 | 0 | 0 | 0,000394 | 0,0021891 | 0,0009596 | 0,0009596 | TRUE | 0,0081115 | No |
| orf19.2356 | 28074 | 25541 | 28171 | 17897 | 5462 | 5080 | 5245 | 7000 | 25560 | 25832 | 26661 | 26884 | 4858 | 5515 | 5372 | 5340 | 15752,82 | 26234 | 5271 | 0,201 | -2,315 | -51,918 | 0 | 0 | 0,0016129 | 0,0021209 | 0,0018014 | 0,0018014 | TRUE | 0,0340341 | No |
| orf19.3499 | 2073 | 1863 | 1968 | 1327 | 438 | 389 | 363 | 469 | 1887 | 1884 | 1863 | 1993 | 390 | 422 | 372 | 358 | 1146,12 | 1907 | 386 | 0,202 | -2,31 | -37,766 | 0 | 0 | 0,0004987 | 0,0031896 | 0,0020988 | 0,0020988 | TRUE | 0,0448728 | No |
| orf19.508 | 2538 | 2303 | 2476 | 1642 | 514 | 449 | 473 | 627 | 2311 | 2329 | 2343 | 2467 | 457 | 487 | 484 | 478 | 1419,65 | 2362 | 476 | 0,202 | -2,308 | -43,559 | 0 | 0 | 1,00E-08 | 0,0029675 | 0,0014921 | 0,0014921 | TRUE | 0,0097568 | No |
| orf19.730 | 2085 | 1697 | 1934 | 1258 | 421 | 324 | 378 | 481 | 1898 | 1716 | 1830 | 1890 | 374 | 352 | 387 | 367 | 1101,87 | 1834 | 370 | 0,202 | -2,308 | -37,319 | 8,09E-305 | 2,12E-303 | 1,00E-08 | 0,0032359 | 0,0021214 | 0,0021214 | TRUE | 0,0226315 | No |
| orf19.4873 | 12611 | 11711 | 11972 | 7825 | 2746 | 2320 | 2150 | 2982 | 11482 | 11844 | 11330 | 11754 | 2442 | 2519 | 2202 | 2275 | 6981,12 | 11602 | 2360 | 0,203 | -2,298 | -49,011 | 0 | 0 | 0,0017964 | 0,0022262 | 0,0018675 | 0,0018675 | TRUE | 0,0242703 | No |
| orf19.335 | 1305 | 1536 | 1503 | 999 | 316 | 288 | 276 | 368 | 1188 | 1553 | 1422 | 1501 | 281 | 313 | 283 | 281 | 852,74 | 1416 | 290 | 0,204 | -2,292 | -25,428 | 1,24E-142 | 1,53E-141 | 0,0079859 | 0,0035861 | 0,0058051 | 0,0058051 | TRUE | 0,1445579 | No |

| | | | | | | | | | | | | | | | | | | | | | | | | | | | | | | | |
|---|---|---|---|---|---|---|---|---|---|---|---|---|---|---|---|---|---|---|---|---|---|---|---|---|---|---|---|---|---|---|---|
| orf19.1381 | 2848 | 2370 | 2455 | 1564 | 566 | 416 | 482 | 697 | 2593 | 2397 | 2323 | 2349 | 503 | 452 | 494 | 532 | 1455,4 | 2416 | 495 | 0,205 | -2,283 | -36,412 | 2,74E-290 | 6,83E-289 | 0,0022804 | 0,0029447 | 0,0026136 | 0,0026136 | TRUE | 0,0387301 | No |
| orf19.6489 | 10048 | 8935 | 9554 | 6141 | 2171 | 1751 | 1833 | 2380 | 9148 | 9037 | 9042 | 9225 | 1931 | 1901 | 1877 | 1815 | 5497,08 | 9113 | 1881 | 0,206 | -2,277 | -64,859 | 0 | 0 | 1,00E-08 | 0,0022773 | 0,0008761 | 0,0008761 | TRUE | 0,0072839 | No |
| orf19.3448 | 1776 | 2920 | 2753 | 1405 | 538 | 506 | 462 | 546 | 1617 | 2953 | 2605 | 2111 | 479 | 549 | 473 | 416 | 1400,47 | 2322 | 479 | 0,206 | -2,277 | -11,048 | 2,25E-28 | 6,94E-28 | 0,0395929 | 0,0029803 | 0,0187939 | 0,0395929 | TRUE | 0,1044109 | No |
| orf19.2743 | 1197 | 922 | 1006 | 773 | 256 | 179 | 220 | 269 | 1090 | 933 | 952 | 1161 | 228 | 194 | 225 | 205 | 623,52 | 1034 | 213 | 0,207 | -2,275 | -24,615 | 8,72E-134 | 1,00E-132 | 0,0069958 | 0,0041556 | 0,0054914 | 0,0054914 | TRUE | 0,0764878 | No |
| orf19.5045 | 8709 | 9723 | 10201 | 5931 | 2096 | 1863 | 1896 | 2351 | 7929 | 9834 | 9654 | 8909 | 1864 | 2023 | 1942 | 1793 | 5493,59 | 9082 | 1906 | 0,21 | -2,254 | -32,547 | 2,34E-232 | 4,56E-231 | 0,0054724 | 0,0022775 | 0,0043006 | 0,0043006 | TRUE | 0,0899202 | No |
| orf19.554 | 652 | 580 | 610 | 408 | 144 | 119 | 136 | 145 | 594 | 587 | 577 | 613 | 128 | 129 | 139 | 111 | 359,7 | 593 | 127 | 0,213 | -2,232 | -24,484 | 2,19E-132 | 2,50E-131 | 1,00E-08 | 0,0057095 | 0,0033905 | 0,0033905 | TRUE | 0,0899571 | No |
| orf19.2553 | 329 | 348 | 357 | 200 | 64 | 73 | 64 | 97 | 300 | 352 | 338 | 300 | 57 | 79 | 66 | 74 | 195,69 | 322 | 69 | 0,213 | -2,231 | -17,562 | 4,83E-69 | 2,99E-68 | 0,0039513 | 0,0087875 | 0,0070907 | 0,0070907 | TRUE | 0,1313509 | No |
| orf19.1930 | 147 | 141 | 144 | 86 | 32 | 26 | 24 | 44 | 134 | 143 | 136 | 129 | 28 | 28 | 25 | 34 | 82,09 | 136 | 29 | 0,214 | -2,227 | -12,572 | 3,00E-36 | 1,10E-35 | 1,00E-08 | 0,0181284 | 0,0102151 | 0,0102151 | TRUE | 0,1074917 | No |
| orf19.675 | 10778 | 8632 | 9305 | 6927 | 2173 | 1893 | 1950 | 2748 | 9813 | 8730 | 8806 | 10405 | 1933 | 2055 | 1997 | 2096 | 5729,55 | 9438 | 2020 | 0,214 | -2,223 | -35,958 | 3,76E-283 | 8,99E-282 | 0,0039163 | 0,0022676 | 0,0033835 | 0,0033835 | TRUE | 0,0573273 | No |
| orf19.6121 | 8452 | 7340 | 8294 | 5299 | 1852 | 1480 | 1661 | 2207 | 7695 | 7424 | 7850 | 7960 | 1647 | 1607 | 1701 | 1684 | 4695,88 | 7732 | 1660 | 0,215 | -2,219 | -55,15 | 0 | 0 | 0,0005477 | 0,0023183 | 0,0012035 | 0,0012035 | TRUE | 0,0087814 | No |
| orf19.851 | 3812 | 3786 | 4011 | 2491 | 833 | 784 | 803 | 1034 | 3471 | 3829 | 3796 | 3742 | 741 | 851 | 822 | 789 | 2255,12 | 3710 | 801 | 0,216 | -2,214 | -39,677 | 0 | 0 | 0,0017062 | 0,0026228 | 0,0022598 | 0,0022598 | TRUE | 0,029862 | No |
| orf19.6554 | 6990 | 7925 | 7525 | 5061 | 1784 | 1579 | 1508 | 1892 | 6364 | 8015 | 7122 | 7602 | 1587 | 1714 | 1544 | 1443 | 4424,04 | 7276 | 1572 | 0,216 | -2,211 | -29,837 | 1,28E-195 | 2,13E-194 | 0,0067841 | 0,0023356 | 0,0049029 | 0,0049029 | TRUE | 0,0873922 | No |
| orf19.5531 | 2360 | 2050 | 2154 | 1444 | 525 | 426 | 432 | 591 | 2149 | 2073 | 2039 | 2169 | 467 | 463 | 442 | 451 | 1281,56 | 2108 | 456 | 0,216 | -2,208 | -40,89 | 0 | 0 | 1,00E-08 | 0,0030678 | 0,001525 | 0,001525 | TRUE | 0,0059032 | No |
| orf19.7483 | 6166 | 6127 | 6502 | 3869 | 1453 | 1243 | 1234 | 1660 | 5614 | 6197 | 6154 | 5812 | 1292 | 1350 | 1264 | 1266 | 3618,51 | 5944 | 1293 | 0,217 | -2,202 | -45,493 | 0 | 0 | 0,0014034 | 0,0024021 | 0,0017974 | 0,0017974 | TRUE | 0,0176036 | No |
| orf19.6247.1 | 4590 | 3806 | 3990 | 1710 | 1038 | 604 | 738 | 1043 | 4179 | 3849 | 3776 | 2569 | 923 | 656 | 756 | 796 | 2187,96 | 3593 | 783 | 0,218 | -2,199 | -12,663 | 9,48E-37 | 3,52E-36 | 0,0282225 | 0,0026408 | 0,0150998 | 0,0282225 | TRUE | 0,4449436 | No |
| orf19.649 | 8118 | 8269 | 8736 | 4802 | 1908 | 1600 | 1682 | 2217 | 7391 | 8363 | 8268 | 7213 | 1697 | 1737 | 1723 | 1691 | 4760,44 | 7809 | 1712 | 0,219 | -2,19 | -38,955 | 0 | 0 | 0,0029579 | 0,0023145 | 0,0026943 | 0,0026943 | TRUE | 0,0314872 | No |
| orf19.5451 | 681 | 529 | 652 | 574 | 136 | 147 | 134 | 207 | 620 | 535 | 617 | 862 | 121 | 160 | 137 | 158 | 401,26 | 658 | 144 | 0,219 | -2,189 | -15,527 | 2,28E-54 | 1,14E-53 | 0,0273955 | 0,0053292 | 0,01502 | 0,01502 | TRUE | 0,487116 | No |
| orf19.6905 | 4254 | 3622 | 3913 | 2791 | 983 | 750 | 838 | 1121 | 3873 | 3663 | 3703 | 4193 | 874 | 814 | 858 | 855 | 2354,27 | 3858 | 850 | 0,221 | -2,179 | -39,029 | 0 | 0 | 0,0020813 | 0,0025981 | 0,0023047 | 0,0023047 | TRUE | 0,039728 | No |
| orf19.844 | 10225 | 9579 | 10630 | 6246 | 2335 | 1916 | 2144 | 2813 | 9309 | 9688 | 10060 | 9382 | 2077 | 2080 | 2196 | 2146 | 5867,38 | 9610 | 2125 | 0,221 | -2,178 | -52,881 | 0 | 0 | 0,0008376 | 0,0022622 | 0,0013524 | 0,0013524 | TRUE | 0,0122623 | No |
| orf19.6601.1 | 23400 | 22578 | 22798 | 13295 | 5815 | 4422 | 4550 | 5711 | 21305 | 22835 | 21576 | 19971 | 5172 | 4801 | 4660 | 4356 | 13084,61 | 21422 | 4747 | 0,222 | -2,174 | -37,242 | 1,43E-303 | 3,73E-302 | 0,0038754 | 0,002138 | 0,0031508 | 0,0031508 | TRUE | 0,0447029 | No |
| orf19.2838 | 2453 | 2390 | 2564 | 1499 | 596 | 493 | 475 | 678 | 2233 | 2417 | 2427 | 2252 | 530 | 535 | 486 | 517 | 1424,74 | 2332 | 517 | 0,222 | -2,173 | -38,049 | 0 | 0 | 0,0011589 | 0,0029642 | 0,002002 | 0,002002 | TRUE | 0,0175036 | No |
| orf19.450 | 51 | 51 | 59 | 58 | 17 | 14 | 9 | 17 | 46 | 52 | 56 | 87 | 15 | 15 | 9 | 13 | 36,69 | 60 | 13 | 0,222 | -2,172 | -7,384 | 1,54E-13 | 3,28E-13 | 0,0397012 | 0,0380458 | 0,0386766 | 0,0386766 | TRUE | 0,6225518 | No |
| orf19.2094 | 6578 | 6162 | 6594 | 4214 | 1513 | 1281 | 1392 | 1768 | 5989 | 6232 | 6241 | 6330 | 1346 | 1391 | 1426 | 1349 | 3787,84 | 6198 | 1378 | 0,222 | -2,17 | -53,957 | 0 | 0 | 0,0003364 | 0,0023858 | 0,0011276 | 0,0011276 | TRUE | 0,0065645 | No |
| orf19.238 | 7210 | 6274 | 6632 | 3910 | 1586 | 1318 | 1311 | 1823 | 6564 | 6345 | 6277 | 5873 | 1411 | 1431 | 1343 | 1391 | 3829,35 | 6265 | 1394 | 0,222 | -2,169 | -46,328 | 0 | 0 | 0,0012276 | 0,002382 | 0,0016848 | 0,0016848 | TRUE | 0,0208189 | No |
| orf19.2770.1 | 7699 | 6124 | 6291 | 4266 | 1722 | 1294 | 1286 | 1873 | 7010 | 6194 | 5954 | 6408 | 1532 | 1405 | 1317 | 1429 | 3905,98 | 6392 | 1421 | 0,222 | -2,169 | -34,835 | 7,22E-266 | 1,63E-264 | 0,0038578 | 0,0023752 | 0,0033099 | 0,0033099 | TRUE | 0,0524503 | No |
| orf19.716 | 1044 | 1115 | 1084 | 575 | 261 | 203 | 219 | 290 | 951 | 1128 | 1026 | 864 | 232 | 220 | 224 | 221 | 608,24 | 992 | 224 | 0,226 | -2,147 | -23,734 | 1,61E-124 | 1,71E-123 | 0,0062854 | 0,0042089 | 0,0052374 | 0,0052374 | TRUE | 0,0974288 | No |
| orf19.5636 | 23644 | 18043 | 19957 | 15914 | 5169 | 4345 | 4350 | 6587 | 21527 | 18249 | 18887 | 23905 | 4598 | 4717 | 4455 | 5025 | 12670,37 | 20642 | 4699 | 0,228 | -2,135 | -27,311 | 3,16E-164 | 4,53E-163 | 0,0090726 | 0,0021413 | 0,0057449 | 0,0057449 | TRUE | 0,1381384 | No |
| orf19.4323 | 1469 | 1444 | 1402 | 781 | 378 | 293 | 319 | 316 | 1337 | 1460 | 1327 | 1173 | 336 | 318 | 327 | 241 | 815,01 | 1324 | 306 | 0,23 | -2,123 | -21,374 | 2,36E-101 | 2,07E-100 | 0,0126351 | 0,0036579 | 0,0075339 | 0,0075339 | TRUE | 0,2487844 | No |
| orf19.467 | 11616 | 10141 | 11279 | 7755 | 2592 | 2256 | 2504 | 3410 | 10576 | 10257 | 10675 | 11649 | 2306 | 2449 | 2564 | 2601 | 6634,59 | 10789 | 2480 | 0,23 | -2,12 | -39,46 | 0 | 0 | 0,0026601 | 0,0022361 | 0,0025347 | 0,0025347 | TRUE | 0,0347018 | No |
| orf19.2730 | 2418 | 1715 | 2096 | 1483 | 504 | 374 | 495 | 667 | 2201 | 1735 | 1984 | 2228 | 448 | 406 | 507 | 509 | 1252,19 | 2037 | 468 | 0,23 | -2,12 | -22,981 | 7,24E-117 | 7,23E-116 | 0,0102301 | 0,003092 | 0,0069105 | 0,0069105 | TRUE | 0,1210337 | No |
| orf19.1353 | 244 | 173 | 205 | 176 | 48 | 59 | 41 | 63 | 222 | 175 | 194 | 264 | 43 | 64 | 42 | 48 | 131,54 | 214 | 49 | 0,23 | -2,12 | -12,327 | 6,44E-35 | 2,30E-34 | 0,0234528 | 0,0120796 | 0,0163483 | 0,0163483 | TRUE | 0,220111 | No |
| orf19.3538 | 5906 | 5193 | 5902 | 3730 | 1305 | 1190 | 1258 | 1692 | 5377 | 5252 | 5586 | 5603 | 1161 | 1292 | 1288 | 1291 | 3356,23 | 5454 | 1258 | 0,231 | -2,117 | -44,005 | 0 | 0 | 0,001158 | 0,0024306 | 0,0017512 | 0,0017512 | TRUE | 0,0330969 | No |
| orf19.340 | 581 | 551 | 588 | 324 | 148 | 100 | 138 | 147 | 529 | 557 | 556 | 487 | 132 | 109 | 141 | 112 | 327,89 | 532 | 124 | 0,231 | -2,115 | -21,112 | 6,16E-99 | 5,32E-98 | 0,0023578 | 0,0060658 | 0,0048617 | 0,0048617 | TRUE | 0,0901831 | No |
| orf19.6350 | 216 | 165 | 184 | 138 | 41 | 44 | 43 | 58 | 197 | 167 | 174 | 207 | 36 | 48 | 44 | 44 | 114,69 | 186 | 43 | 0,232 | -2,109 | -13,583 | 5,06E-42 | 2,04E-41 | 1,00E-08 | 0,0135553 | 0,0094739 | 0,0094739 | TRUE | 0,0883984 | No |
| orf19.4980 | 61165 | 56134 | 59582 | 36289 | 14496 | 12330 | 12194 | 17364 | 55688 | 56774 | 56389 | 54512 | 12894 | 13387 | 12488 | 13245 | 34422,1 | 55841 | 13004 | 0,233 | -2,102 | -63,084 | 0 | 0 | 0,0005712 | 0,0020754 | 0,0010214 | 0,0010214 | TRUE | 0,0086384 | No |
| orf19.3369 | 22127 | 23586 | 24885 | 15033 | 5990 | 5110 | 5181 | 6396 | 20146 | 23855 | 23551 | 22582 | 5328 | 5548 | 5306 | 4879 | 13899,33 | 22534 | 5265 | 0,234 | -2,098 | -34,859 | 3,13E-266 | 7,10E-265 | 0,0040408 | 0,0021321 | 0,0033668 | 0,0033668 | TRUE | 0,0625331 | No |
| orf19.5749 | 4462 | 4053 | 4311 | 2718 | 1085 | 863 | 941 | 1255 | 4062 | 4099 | 4080 | 4083 | 965 | 937 | 964 | 957 | 2518,44 | 4081 | 956 | 0,234 | -2,094 | -52,754 | 0 | 0 | 1,00E-08 | 0,0025615 | 0,0008937 | 0,0008937 | TRUE | 0,0018962 | No |
| orf19.6391 | 161 | 143 | 158 | 135 | 33 | 27 | 49 | 54 | 147 | 145 | 150 | 203 | 29 | 29 | 50 | 41 | 99,2 | 161 | 37 | 0,235 | -2,087 | -11,01 | 3,42E-28 | 1,05E-27 | 0,0259004 | 0,0153539 | 0,018745 | 0,018745 | TRUE | 0,3399697 | No |
| orf19.347 | 4986 | 4248 | 4594 | 3217 | 1107 | 959 | 1087 | 1451 | 4540 | 4296 | 4348 | 4832 | 985 | 1041 | 1113 | 1107 | 2782,76 | 4504 | 1062 | 0,236 | -2,084 | -36,849 | 3,04E-297 | 7,78E-296 | 0,0023096 | 0,0025117 | 0,0025111 | 0,0025111 | TRUE | 0,0292924 | No |
| orf19.882 | 11494 | 10177 | 11057 | 6781 | 2681 | 2224 | 2458 | 3274 | 10465 | 10293 | 10464 | 10186 | 2385 | 2415 | 2517 | 2497 | 6402,8 | 10352 | 2454 | 0,237 | -2,077 | -60,685 | 0 | 0 | 9,07E-05 | 0,0022433 | 0,0008826 | 0,0008826 | TRUE | 0,0044944 | No |
| orf19.3698 | 338 | 436 | 437 | 232 | 102 | 78 | 96 | 113 | 308 | 441 | 414 | 349 | 91 | 85 | 98 | 86 | 233,84 | 378 | 90 | 0,238 | -2,072 | -15,407 | 1,48E-53 | 7,30E-53 | 0,0143218 | 0,0076862 | 0,0107719 | 0,0107719 | TRUE | 0,1055258 | No |
| orf19.550 | 23973 | 20045 | 22250 | 14692 | 5836 | 4531 | 5121 | 6602 | 21826 | 20273 | 21058 | 22070 | 5191 | 4919 | 5245 | 5036 | 13202,26 | 21307 | 5098 | 0,239 | -2,063 | -51,259 | 0 | 0 | 0,0009879 | 0,0021371 | 0,001439 | 0,001439 | TRUE | 0,0130444 | No |
| orf19.715 | 1894 | 1897 | 1908 | 1257 | 524 | 458 | 373 | 554 | 1724 | 1919 | 1806 | 1888 | 466 | 497 | 382 | 423 | 1138,11 | 1834 | 442 | 0,241 | -2,054 | -27,612 | 7,93E-168 | 1,16E-166 | 0,0050067 | 0,0031977 | 0,0039613 | 0,0039613 | TRUE | 0,0903521 | No |
| orf19.3232 | 438 | 422 | 411 | 290 | 112 | 86 | 82 | 154 | 399 | 427 | 389 | 436 | 100 | 93 | 84 | 117 | 255,58 | 413 | 98 | 0,242 | -2,048 | -18,596 | 3,48E-77 | 2,37E-76 | 0,0022467 | 0,0072057 | 0,0056715 | 0,0056715 | TRUE | 0,1853269 | No |
| orf19.5961 | 4536 | 3928 | 4210 | 2650 | 1145 | 869 | 926 | 1300 | 4130 | 3973 | 3984 | 3981 | 1018 | 943 | 948 | 992 | 2496,2 | 4017 | 975 | 0,243 | -2,04 | -46,649 | 0 | 0 | 0,0001731 | 0,0025662 | 0,0012256 | 0,0012256 | TRUE | 0,0101846 | No |
| orf19.6594 | 24334 | 23595 | 24880 | 14880 | 6168 | 5320 | 5536 | 7167 | 22155 | 23864 | 23547 | 22352 | 5486 | 5776 | 5670 | 5467 | 14289,57 | 22980 | 5600 | 0,244 | -2,037 | -52,069 | 0 | 0 | 0,0008922 | 0,0021295 | 0,0013638 | 0,0013638 | TRUE | 0,0082151 | No |
| orf19.4722 | 2758 | 2324 | 2631 | 1528 | 656 | 514 | 610 | 781 | 2511 | 2350 | 2490 | 2295 | 583 | 558 | 625 | 596 | 1501,11 | 2412 | 590 | 0,245 | -2,031 | -36,35 | 2,65E-289 | 6,58E-288 | 0,0011331 | 0,002917 | 0,0019863 | 0,0019863 | TRUE | 0,0163892 | No |
| orf19.3888 | 4151 | 3525 | 3802 | 2528 | 1025 | 804 | 859 | 1229 | 3779 | 3565 | 3598 | 3797 | 912 | 873 | 880 | 937 | 2292,75 | 3685 | 900 | 0,245 | -2,03 | -43,043 | 0 | 0 | 0,0006197 | 0,0026132 | 0,0014724 | 0,0014724 | TRUE | 0,0096811 | No |
| orf19.5053 | 1292 | 1213 | 1340 | 844 | 318 | 329 | 283 | 376 | 1176 | 1227 | 1268 | 1268 | 283 | 357 | 290 | 287 | 769,48 | 1235 | 304 | 0,245 | -2,027 | -27,034 | 5,85E-161 | 8,13E-160 | 0,0028273 | 0,0037538 | 0,0034215 | 0,0034215 | TRUE | 0,1452991 | No |
| orf19.3651 | 20804 | 16377 | 17502 | 11579 | 4940 | 3836 | 3991 | 5785 | 18941 | 16564 | 16564 | 17394 | 4394 | 4165 | 4087 | 4413 | 10815,16 | 17366 | 4265 | 0,246 | -2,025 | -38,686 | 0 | 0 | 0,0026223 | 0,0021592 | 0,0024921 | 0,0024921 | TRUE | 0,045952 | No |
| orf19.6245 | 14100 | 12871 | 13791 | 8222 | 3594 | 2873 | 3132 | 4033 | 12837 | 13018 | 13052 | 12351 | 3197 | 3119 | 3208 | 3076 | 7982,22 | 12814 | 3150 | 0,246 | -2,025 | -57,8 | 0 | 0 | 0,0003862 | 0,0022025 | 0,0009885 | 0,0009885 | TRUE | 0,0070321 | No |
| orf19.7567 | 4165 | 3736 | 4026 | 2721 | 1028 | 955 | 961 | 1212 | 3792 | 3779 | 3810 | 4087 | 914 | 1037 | 984 | 925 | 2416,02 | 3867 | 965 | 0,249 | -2,004 | -38,162 | 0 | 0 | 0,0016059 | 0,0025838 | 0,0020244 | 0,0020244 | TRUE | 0,029146 | No |
| orf19.1944 | 20649 | 17719 | 20142 | 12906 | 4990 | 4386 | 4708 | 6215 | 18800 | 17921 | 19063 | 19387 | 4438 | 4762 | 4822 | 4741 | 11741,65 | 18793 | 4691 | 0,25 | -2,002 | -48,873 | 0 | 0 | 0,0010514 | 0,0021495 | 0,0014839 | 0,0014839 | TRUE | 0,0160648 | No |
| orf19.2049 | 681 | 658 | 623 | 377 | 174 | 148 | 153 | 195 | 620 | 665 | 590 | 566 | 155 | 161 | 157 | 149 | 382,79 | 610 | 156 | 0,253 | -1,982 | -22,28 | 5,78E-110 | 5,45E-109 | 1,00E-08 | 0,005488 | 0,0037216 | 0,0037216 | TRUE | 0,042492 | No |
| orf19.7084 | 19530 | 16783 | 18836 | 11159 | 4754 | 3835 | 4561 | 5936 | 17781 | 16974 | 17827 | 16763 | 4229 | 4164 | 4671 | 4528 | 10866,99 | 17336 | 4398 | 0,254 | -1,979 | -42,725 | 0 | 0 | 0,0018167 | 0,0021586 | 0,0019233 | 0,0019233 | TRUE | 0,0208866 | No |
| orf19.4377 | 2574 | 2458 | 2626 | 1636 | 695 | 582 | 631 | 779 | 2344 | 2486 | 2485 | 2458 | 618 | 632 | 646 | 594 | 1532,86 | 2443 | 622 | 0,254 | -1,975 | -38,706 | 0 | 0 | 1,00E-08 | 0,0028988 | 0,0015293 | 0,0015293 | TRUE | 0,0121557 | No |
| orf19.5131 | 5690 | 5102 | 5669 | 3449 | 1460 | 1271 | 1263 | 1790 | 5181 | 5160 | 5365 | 5181 | 1299 | 1380 | 1293 | 1365 | 3278,03 | 5222 | 1334 | 0,256 | -1,968 | -48,478 | 0 | 0 | 0,0002005 | 0,00244 | 0,0011305 | 0,0011305 | TRUE | 0,0057257 | No |
| orf19.34 | 383 | 334 | 354 | 206 | 92 | 67 | 94 | 120 | 349 | 338 | 335 | 309 | 82 | 73 | 96 | 92 | 209,17 | 333 | 86 | 0,257 | -1,958 | -17,288 | 5,79E-67 | 3,48E-66 | 1,00E-08 | 0,0083524 | 0,0053087 | 0,0053087 | TRUE | 0,0835098 | No |
| orf19.542 | 4983 | 4098 | 4554 | 2720 | 1263 | 939 | 1087 | 1513 | 4537 | 4145 | 4310 | 4086 | 1123 | 1019 | 1113 | 1154 | 2685,95 | 4270 | 1102 | 0,258 | -1,952 | -36,491 | 1,52E-291 | 3,81E-290 | 0,001744 | 0,0025288 | 0,0021986 | 0,0021986 | TRUE | 0,0296667 | No |
| orf19.6211 | 5670 | 4992 | 5503 | 3288 | 1460 | 1202 | 1299 | 1789 | 5162 | 5049 | 5208 | 4939 | 1299 | 1305 | 1330 | 1365 | 3207,13 | 5090 | 1325 | 0,26 | -1,942 | -48,248 | 0 | 0 | 1,00E-08 | 0,0024489 | 0,0010978 | 0,0010978 | TRUE | 0,0054096 | No |
| orf19.4372 | 5333 | 5067 | 5410 | 3454 | 1394 | 1308 | 1293 | 1713 | 4855 | 5125 | 5120 | 5188 | 1240 | 1420 | 1324 | 1307 | 3197,46 | 5072 | 1323 | 0,261 | -1,94 | -40,017 | 0 | 0 | 0,0013707 | 0,0024502 | 0,0017977 | 0,0017977 | TRUE | 0,0283998 | No |
| orf19.84 | 5341 | 4742 | 5175 | 3394 | 1357 | 1273 | 1191 | 1744 | 4863 | 4796 | 4898 | 5098 | 1207 | 1382 | 1220 | 1330 | 3099,25 | 4914 | 1285 | 0,262 | -1,935 | -37,957 | 0 | 0 | 0,0019002 | 0,0024632 | 0,002022 | 0,002022 | TRUE | 0,0293262 | No |
| orf19.1352 | 4902 | 4264 | 4794 | 2897 | 1368 | 992 | 1163 | 1496 | 4463 | 4313 | 4537 | 4352 | 1217 | 1077 | 1191 | 1141 | 2786,32 | 4416 | 1156 | 0,262 | -1,932 | -41,168 | 0 | 0 | 0,0007529 | 0,0025111 | 0,0015906 | 0,0015906 | TRUE | 0,0239147 | No |
| orf19.4887 | 13227 | 12112 | 12871 | 8651 | 3452 | 3137 | 3146 | 4298 | 12043 | 12250 | 12181 | 12995 | 3070 | 3406 | 3222 | 3279 | 7805,73 | 12367 | 3244 | 0,262 | -1,931 | -44,484 | 0 | 0 | 0,0012565 | 0,0022063 | 0,0016211 | 0,0016211 | TRUE | 0,0158702 | No |
| orf19.4631 | 3360 | 3463 | 3468 | 2520 | 900 | 1031 | 770 | 1144 | 3059 | 3502 | 3282 | 3785 | 801 | 1119 | 789 | 873 | 2151,29 | 3407 | 896 | 0,263 | -1,929 | -19,215 | 2,78E-82 | 2,04E-81 | 0,0179911 | 0,0026511 | 0,0089984 | 0,0089984 | TRUE | 0,3321476 | No |

| | | | | | | | | | | | | | | | | | | | | | | | | | | | | | | | |
|---|---|---|---|---|---|---|---|---|---|---|---|---|---|---|---|---|---|---|---|---|---|---|---|---|---|---|---|---|---|---|---|
| orf19.3444 | 509 | 349 | 373 | 289 | 114 | 115 | 92 | 134 | 463 | 353 | 353 | 434 | 101 | 125 | 94 | 102 | 253,28 | 401 | 106 | 0,263 | -1,926 | -15,085 | 2,04E-51 | 9,69E-51 | 0,0133059 | 0,0072526 | 0,0098856 | 0,0098856 | TRUE | 0,1271922 | No |
| orf19.2948 | 1707 | 1436 | 1615 | 1078 | 472 | 353 | 404 | 528 | 1554 | 1452 | 1528 | 1619 | 420 | 383 | 414 | 403 | 971,74 | 1538 | 405 | 0,264 | -1,923 | -31,471 | 2,18E-217 | 3,98E-216 | 1,00E-08 | 0,0033964 | 0,0020859 | 0,0020859 | TRUE | 0,0168352 | No |
| orf19.6640 | 9078 | 8090 | 8555 | 5240 | 2457 | 1982 | 2049 | 2783 | 8265 | 8182 | 8097 | 7871 | 2185 | 2152 | 2098 | 2123 | 5121,73 | 8104 | 2140 | 0,264 | -1,922 | -54,36 | 0 | 0 | 1,00E-08 | 0,0022949 | 0,0009155 | 0,0009155 | TRUE | 0,004338 | No |
| orf19.6882 | 38975 | 32439 | 36365 | 21310 | 10111 | 7881 | 8753 | 11886 | 35485 | 32809 | 34416 | 32011 | 8993 | 8557 | 8964 | 9067 | 21287,72 | 33680 | 8895 | 0,264 | -1,921 | -45,693 | 0 | 0 | 0,0013183 | 0,0020991 | 0,001629 | 0,001629 | TRUE | 0,0159424 | No |
| orf19.6590 | 123 | 128 | 101 | 68 | 37 | 25 | 27 | 37 | 112 | 129 | 96 | 102 | 33 | 27 | 28 | 28 | 69,39 | 110 | 29 | 0,264 | -1,921 | -9,985 | 1,77E-23 | 4,89E-23 | 1,00E-08 | 0,0210747 | 0,0147064 | 0,0147064 | TRUE | 0,1464037 | No |
| orf19.3923 | 420 | 374 | 371 | 297 | 124 | 81 | 110 | 132 | 382 | 378 | 351 | 446 | 110 | 88 | 113 | 101 | 246,19 | 389 | 103 | 0,266 | -1,911 | -16,749 | 5,79E-63 | 3,30E-62 | 0,0053197 | 0,0074029 | 0,0065975 | 0,0065975 | TRUE | 0,0969927 | No |
| orf19.3940.1 | 74206 | 42907 | 49512 | 40372 | 17793 | 12679 | 12480 | 20786 | 67561 | 43396 | 46859 | 60645 | 15826 | 13766 | 12781 | 15856 | 34586,26 | 54615 | 14557 | 0,267 | -1,908 | -11,308 | 1,20E-29 | 3,80E-29 | 0,0273 | 0,0020752 | 0,0133377 | 0,0273 | TRUE | 0,1038138 | No |
| orf19.6200 | 27 | 17 | 22 | 16 | 6 | 4 | 8 | 7 | 25 | 17 | 21 | 24 | 5 | 4 | 8 | 5 | 13,73 | 22 | 6 | 0,268 | -1,9 | -4,612 | 3,99E-06 | 6,47E-06 | 1,00E-08 | 0,0982461 | 0,0580792 | 0,0580792 | TRUE | 0,1581242 | No |
| orf19.3084 | 2281 | 1752 | 1942 | 1114 | 539 | 413 | 518 | 677 | 2077 | 1772 | 1838 | 1673 | 479 | 448 | 530 | 516 | 1166,85 | 1840 | 493 | 0,268 | -1,899 | -24,294 | 2,29E-130 | 2,55E-129 | 0,0060949 | 0,0031691 | 0,0046276 | 0,0046276 | TRUE | 0,0913238 | No |
| orf19.6349 | 1797 | 2101 | 2138 | 1150 | 579 | 479 | 550 | 565 | 1636 | 2125 | 2023 | 1727 | 515 | 520 | 563 | 431 | 1192,66 | 1878 | 507 | 0,269 | -1,892 | -19,439 | 3,58E-84 | 2,67E-83 | 0,0130778 | 0,0031446 | 0,0078848 | 0,0078848 | TRUE | 0,0996576 | No |
| orf19.897 | 2370 | 1987 | 2142 | 1511 | 615 | 564 | 546 | 753 | 2158 | 2010 | 2027 | 2270 | 547 | 612 | 559 | 574 | 1344,67 | 2116 | 573 | 0,271 | -1,884 | -31,012 | 3,77E-211 | 6,73E-210 | 0,0019648 | 0,0030194 | 0,0024734 | 0,0024734 | TRUE | 0,0269703 | No |
| orf19.4753 | 8483 | 7877 | 8677 | 4862 | 2422 | 1973 | 2150 | 2700 | 7723 | 7967 | 8212 | 7303 | 2154 | 2142 | 2202 | 2060 | 4970,44 | 7801 | 2140 | 0,274 | -1,868 | -40,076 | 0 | 0 | 0,001484 | 0,0023028 | 0,0017987 | 0,0017987 | TRUE | 0,0218862 | No |
| orf19.4914.1 | 42186 | 39346 | 39854 | 21332 | 12767 | 8487 | 10179 | 12537 | 38409 | 39794 | 37718 | 32044 | 11356 | 9215 | 10425 | 9563 | 23565,42 | 36991 | 10140 | 0,274 | -1,867 | -24,331 | 9,24E-131 | 1,03E-129 | 0,0086186 | 0,0020931 | 0,0055984 | 0,0055984 | TRUE | 0,0990521 | No |
| orf19.1862 | 7477 | 6116 | 6479 | 3788 | 2077 | 1496 | 1578 | 2266 | 6807 | 6186 | 6132 | 5690 | 1847 | 1624 | 1616 | 1729 | 3953,92 | 6204 | 1704 | 0,275 | -1,864 | -29,461 | 9,08E-191 | 1,47E-189 | 0,0042699 | 0,0023711 | 0,0034837 | 0,0034837 | TRUE | 0,0524708 | No |
| orf19.1800 | 10424 | 9833 | 10849 | 6225 | 2973 | 2491 | 2755 | 3447 | 9491 | 9945 | 10268 | 9351 | 2644 | 2705 | 2821 | 2629 | 6231,75 | 9764 | 2700 | 0,276 | -1,856 | -41,987 | 0 | 0 | 0,0011369 | 0,002249 | 0,0016473 | 0,0016473 | TRUE | 0,0147938 | No |
| orf19.3861 | 7648 | 6665 | 7311 | 4313 | 2043 | 1680 | 1918 | 2503 | 6963 | 6741 | 6919 | 6479 | 1817 | 1824 | 1964 | 1909 | 4327,11 | 6776 | 1878 | 0,277 | -1,851 | -42,989 | 0 | 0 | 0,0007835 | 0,0023423 | 0,0014526 | 0,0014526 | TRUE | 0,0108777 | No |
| orf19.7381 | 10097 | 10064 | 10954 | 6822 | 3023 | 2765 | 2657 | 3519 | 9193 | 10179 | 10367 | 10248 | 2689 | 3002 | 2721 | 2684 | 6385,32 | 9997 | 2774 | 0,277 | -1,85 | -34,427 | 1,01E-259 | 2,20E-258 | 0,0026774 | 0,0022439 | 0,0025509 | 0,0025509 | TRUE | 0,0365647 | No |
| orf19.5275 | 488 | 517 | 510 | 333 | 153 | 136 | 132 | 163 | 444 | 523 | 483 | 500 | 136 | 148 | 135 | 124 | 311,67 | 488 | 136 | 0,278 | -1,848 | -19,064 | 4,99E-81 | 3,57E-80 | 1,00E-08 | 0,0062755 | 0,0044821 | 0,0044821 | TRUE | 0,0440009 | No |
| orf19.4035 | 9465 | 7588 | 8482 | 5375 | 2652 | 1938 | 2161 | 3046 | 8617 | 7674 | 8027 | 8074 | 2359 | 2104 | 2213 | 2324 | 5174,14 | 8098 | 2250 | 0,278 | -1,847 | -36,138 | 5,82E-286 | 1,42E-284 | 0,0020907 | 0,0022923 | 0,0022339 | 0,0022339 | TRUE | 0,0230487 | No |
| orf19.1502 | 483 | 561 | 566 | 346 | 161 | 150 | 130 | 180 | 440 | 567 | 536 | 520 | 143 | 163 | 133 | 137 | 329,88 | 516 | 144 | 0,279 | -1,842 | -17,313 | 3,74E-67 | 2,26E-66 | 0,0072388 | 0,0060415 | 0,0065818 | 0,0065818 | TRUE | 0,1216932 | No |
| orf19.4828 | 12077 | 10530 | 11767 | 6704 | 3417 | 2643 | 2942 | 4012 | 10996 | 10650 | 11136 | 10070 | 3039 | 2870 | 3013 | 3060 | 6854,34 | 10713 | 2996 | 0,28 | -1,839 | -42,108 | 0 | 0 | 0,0011109 | 0,0022297 | 0,0016246 | 0,0016246 | TRUE | 0,019485 | No |
| orf19.7078 | 565 | 443 | 464 | 275 | 165 | 106 | 117 | 170 | 514 | 448 | 439 | 413 | 147 | 115 | 120 | 130 | 290,75 | 454 | 128 | 0,282 | -1,826 | -17,002 | 7,90E-65 | 4,63E-64 | 0,0058322 | 0,0065804 | 0,0062725 | 0,0062725 | TRUE | 0,1013324 | No |
| orf19.3083 | 2265 | 1788 | 2188 | 1480 | 620 | 503 | 602 | 803 | 2062 | 1808 | 2071 | 2223 | 551 | 546 | 617 | 613 | 1311,39 | 2041 | 582 | 0,286 | -1,808 | -24,803 | 8,32E-136 | 9,71E-135 | 0,0049928 | 0,0030443 | 0,0040377 | 0,0040377 | TRUE | 0,0711196 | No |
| orf19.3555 | 10230 | 8880 | 9872 | 5584 | 2805 | 2324 | 2621 | 3430 | 9314 | 8981 | 9343 | 8388 | 2495 | 2523 | 2684 | 2616 | 5793,13 | 9006 | 2580 | 0,286 | -1,805 | -38,471 | 0 | 0 | 0,0014487 | 0,002265 | 0,001872 | 0,001872 | TRUE | 0,025497 | No |
| orf19.4127 | 10413 | 8902 | 9841 | 5774 | 2911 | 2423 | 2551 | 3480 | 9481 | 9003 | 9314 | 8673 | 2589 | 2631 | 2613 | 2655 | 5869,77 | 9118 | 2622 | 0,287 | -1,799 | -45,341 | 0 | 0 | 0,0007545 | 0,0022621 | 0,0012742 | 0,0012742 | TRUE | 0,0127375 | No |
| orf19.6022 | 96 | 83 | 110 | 70 | 27 | 28 | 29 | 34 | 87 | 84 | 104 | 105 | 24 | 30 | 30 | 26 | 61,33 | 95 | 28 | 0,288 | -1,793 | -9,054 | 1,38E-19 | 3,45E-19 | 1,00E-08 | 0,0235755 | 0,0150456 | 0,0150456 | TRUE | 0,0589165 | No |
| orf19.6981 | 931 | 974 | 974 | 565 | 284 | 259 | 243 | 341 | 848 | 985 | 922 | 849 | 253 | 281 | 249 | 260 | 580,76 | 901 | 261 | 0,289 | -1,792 | -22,792 | 5,56E-115 | 5,46E-114 | 0,0027263 | 0,0043116 | 0,0035545 | 0,0035545 | TRUE | 0,0471883 | No |
| orf19.410.3 | 1153 | 931 | 1045 | 717 | 328 | 266 | 267 | 411 | 1050 | 942 | 989 | 1077 | 292 | 289 | 273 | 314 | 653,11 | 1014 | 292 | 0,289 | -1,791 | -24,407 | 1,43E-131 | 1,61E-130 | 0,0018925 | 0,0040596 | 0,0030479 | 0,0030479 | TRUE | 0,0311243 | No |
| orf19.2196 | 983 | 939 | 1045 | 589 | 293 | 275 | 246 | 351 | 895 | 950 | 989 | 885 | 261 | 299 | 252 | 268 | 599,66 | 930 | 270 | 0,289 | -1,79 | -23,944 | 1,07E-126 | 1,16E-125 | 0,0015257 | 0,0042399 | 0,0030593 | 0,0030593 | TRUE | 0,052182 | No |
| orf19.6540 | 26616 | 22236 | 24327 | 15514 | 7214 | 6149 | 6632 | 9268 | 24233 | 22489 | 23023 | 23305 | 6417 | 6676 | 6792 | 7070 | 15000,54 | 23262 | 6739 | 0,29 | -1,787 | -43,516 | 0 | 0 | 0,0011723 | 0,0021251 | 0,0015278 | 0,0015278 | TRUE | 0,0136406 | No |
| orf19.5552 | 4808 | 4416 | 4769 | 2861 | 1542 | 1170 | 1208 | 1627 | 4377 | 4466 | 4513 | 4298 | 1372 | 1270 | 1237 | 1241 | 2846,87 | 4414 | 1280 | 0,29 | -1,786 | -39,2 | 0 | 0 | 0,0007665 | 0,002501 | 0,0015071 | 0,0015071 | TRUE | 0,0285577 | No |
| orf19.1433.1 | 4 | 4 | 5 | 9 | 1 | 1 | 1 | 5 | 4 | 4 | 5 | 14 | 1 | 1 | 1 | 4 | 4,09 | 7 | 2 | 0,29 | -1,786 | -2,197 | 0,027992 | 0,0354629 | 0,2254498 | 0,3247014 | 0,2882503 | 0,2882503 | TRUE | 0,8852627 | No |
| orf19.6544 | 15501 | 13228 | 14532 | 8970 | 4243 | 3648 | 4067 | 5225 | 14113 | 13379 | 13753 | 13474 | 3774 | 3961 | 4165 | 3986 | 8825,6 | 13680 | 3972 | 0,29 | -1,785 | -45,005 | 0 | 0 | 0,0008853 | 0,0021867 | 0,0013534 | 0,0013534 | TRUE | 0,0137061 | No |
| orf19.285 | 6130 | 5272 | 5735 | 3081 | 1834 | 1319 | 1528 | 1929 | 5581 | 5332 | 5428 | 4628 | 1631 | 1432 | 1565 | 1471 | 3383,58 | 5242 | 1525 | 0,291 | -1,782 | -27,215 | 4,30E-163 | 6,06E-162 | 0,0044246 | 0,0024274 | 0,0037102 | 0,0037102 | TRUE | 0,0741238 | No |
| orf19.5177 | 2857 | 2534 | 2720 | 1662 | 848 | 693 | 718 | 976 | 2601 | 2563 | 2574 | 2497 | 754 | 752 | 735 | 745 | 1652,67 | 2559 | 746 | 0,292 | -1,778 | -39,334 | 0 | 0 | 1,00E-08 | 0,0028363 | 0,0011291 | 0,0011291 | TRUE | 0,0029237 | No |
| orf19.1868 | 7428 | 6458 | 7075 | 4361 | 2256 | 1790 | 1891 | 2442 | 6763 | 6532 | 6696 | 6551 | 2007 | 1943 | 1937 | 1863 | 4286,34 | 6636 | 1938 | 0,292 | -1,777 | -46,828 | 0 | 0 | 0,0002498 | 0,0023452 | 0,0010607 | 0,0010607 | TRUE | 0,0088019 | No |
| orf19.4646 | 315 | 251 | 295 | 260 | 82 | 92 | 74 | 135 | 287 | 254 | 279 | 391 | 73 | 100 | 76 | 103 | 195,25 | 303 | 88 | 0,293 | -1,772 | -11,308 | 1,19E-29 | 3,80E-29 | 0,0276026 | 0,0088027 | 0,0164988 | 0,0164988 | TRUE | 0,3943722 | No |
| orf19.4309 | 3683 | 3306 | 3350 | 2226 | 1117 | 911 | 915 | 1260 | 3353 | 3344 | 3170 | 3344 | 994 | 989 | 937 | 961 | 2136,5 | 3303 | 970 | 0,294 | -1,767 | -39,355 | 0 | 0 | 1,00E-08 | 0,0026553 | 0,001292 | 0,001292 | TRUE | 0,0091037 | No |
| orf19.4771 | 5948 | 5715 | 6135 | 3878 | 1869 | 1666 | 1642 | 2046 | 5415 | 5780 | 5806 | 5825 | 1662 | 1809 | 1682 | 1561 | 3692,58 | 5706 | 1678 | 0,294 | -1,767 | -34,672 | 2,11E-263 | 4,64E-262 | 0,0019121 | 0,0023948 | 0,0021214 | 0,0021214 | TRUE | 0,0324805 | No |
| orf19.2870 | 2828 | 2341 | 2453 | 1299 | 861 | 582 | 659 | 838 | 2575 | 2368 | 2322 | 1951 | 766 | 632 | 675 | 639 | 1490,89 | 2304 | 678 | 0,294 | -1,766 | -20,365 | 3,39E-92 | 2,70E-91 | 0,0091894 | 0,0029231 | 0,0063009 | 0,0063009 | TRUE | 0,1244536 | No |
| orf19.5551 | 6529 | 5747 | 6293 | 3782 | 1917 | 1537 | 1704 | 2321 | 5944 | 5812 | 5956 | 5681 | 1705 | 1669 | 1745 | 1770 | 3785,4 | 5848 | 1722 | 0,295 | -1,763 | -45,708 | 0 | 0 | 0,0002308 | 0,002386 | 0,0010664 | 0,0010664 | TRUE | 0,0050764 | No |
| orf19.7060 | 1566 | 1322 | 1503 | 908 | 451 | 348 | 451 | 521 | 1426 | 1337 | 1422 | 1364 | 401 | 378 | 462 | 397 | 898,44 | 1387 | 410 | 0,295 | -1,762 | -26,663 | 1,27E-156 | 1,73E-155 | 0,0014116 | 0,0035073 | 0,0026683 | 0,0026683 | TRUE | 0,080441 | No |
| orf19.3315 | 2353 | 2155 | 2460 | 1559 | 725 | 584 | 681 | 884 | 2142 | 2180 | 2328 | 2342 | 645 | 634 | 697 | 674 | 1455,32 | 2248 | 662 | 0,295 | -1,76 | -31,696 | 1,77E-220 | 3,28E-219 | 0,0012108 | 0,0029447 | 0,0020185 | 0,0020185 | TRUE | 0,0135789 | No |
| orf19.3475 | 1546 | 1442 | 1516 | 1079 | 460 | 407 | 430 | 599 | 1408 | 1458 | 1435 | 1621 | 409 | 442 | 440 | 457 | 958,74 | 1480 | 437 | 0,296 | -1,756 | -26,566 | 1,66E-155 | 2,24E-154 | 0,0021091 | 0,0034149 | 0,0027642 | 0,0027642 | TRUE | 0,0453912 | No |
| orf19.5755 | 3292 | 3064 | 3368 | 1981 | 1076 | 848 | 890 | 1142 | 2997 | 3099 | 3188 | 2976 | 957 | 921 | 911 | 871 | 1989,97 | 3065 | 915 | 0,298 | -1,746 | -35,955 | 4,19E-283 | 9,98E-282 | 0,00068 | 0,0027008 | 0,0015781 | 0,0015781 | TRUE | 0,0138312 | No |
| orf19.6288 | 158 | 127 | 130 | 92 | 42 | 33 | 45 | 52 | 144 | 128 | 123 | 138 | 37 | 36 | 46 | 40 | 86,56 | 133 | 40 | 0,298 | -1,746 | -10,628 | 2,21E-26 | 6,50E-26 | 1,00E-08 | 0,0172983 | 0,0102243 | 0,0102243 | TRUE | 0,0777311 | No |
| orf19.5960 | 38765 | 35479 | 38516 | 28931 | 12406 | 11230 | 10545 | 14476 | 35294 | 35883 | 36452 | 43459 | 11035 | 12193 | 10799 | 11042 | 24519,65 | 37772 | 11267 | 0,298 | -1,745 | -25,652 | 3,98E-145 | 5,06E-144 | 0,0065879 | 0,0020909 | 0,0043908 | 0,0043908 | TRUE | 0,1255355 | No |
| orf19.6598 | 3418 | 2989 | 3315 | 2202 | 1078 | 825 | 890 | 1291 | 3112 | 3023 | 3137 | 3308 | 959 | 896 | 911 | 985 | 2041,37 | 3145 | 938 | 0,299 | -1,742 | -34,689 | 1,16E-263 | 2,57E-262 | 0,0009629 | 0,0026841 | 0,0017528 | 0,0017528 | TRUE | 0,0142588 | No |
| orf19.1277 | 15782 | 13278 | 14984 | 10449 | 4495 | 4260 | 4142 | 5730 | 14369 | 13429 | 14181 | 15696 | 3998 | 4625 | 4242 | 4371 | 9363,92 | 14419 | 4309 | 0,299 | -1,742 | -29,965 | 2,78E-197 | 4,68E-196 | 0,0037785 | 0,0021781 | 0,0031021 | 0,0031021 | TRUE | 0,0430975 | No |
| orf19.5528 | 1002 | 886 | 972 | 624 | 314 | 232 | 254 | 395 | 912 | 896 | 920 | 937 | 279 | 252 | 260 | 301 | 594,78 | 916 | 273 | 0,3 | -1,737 | -24,406 | 1,47E-131 | 1,66E-130 | 1,00E-08 | 0,004258 | 0,0025865 | 0,0025865 | TRUE | 0,0590193 | No |
| orf19.2381 | 802 | 635 | 690 | 500 | 264 | 197 | 184 | 255 | 730 | 642 | 653 | 751 | 235 | 214 | 188 | 195 | 451,02 | 694 | 208 | 0,3 | -1,737 | -19,089 | 3,09E-81 | 2,23E-80 | 0,0049025 | 0,0049659 | 0,0049306 | 0,0049306 | TRUE | 0,0876426 | No |
| orf19.4665 | 2799 | 2139 | 2429 | 1335 | 790 | 561 | 704 | 890 | 2548 | 2163 | 2299 | 2005 | 703 | 609 | 721 | 679 | 1465,95 | 2254 | 678 | 0,301 | -1,734 | -22,019 | 1,89E-107 | 1,74E-106 | 0,00654 | 0,0029381 | 0,0050298 | 0,0050298 | TRUE | 0,0938641 | No |
| orf19.4593 | 10418 | 9199 | 10176 | 6298 | 3070 | 2538 | 2936 | 3801 | 9485 | 9304 | 9631 | 9461 | 2731 | 2756 | 3007 | 2899 | 6159,09 | 9470 | 2848 | 0,301 | -1,733 | -43,319 | 0 | 0 | 0,0008265 | 0,0022515 | 0,0013165 | 0,0013165 | TRUE | 0,016544 | No |
| orf19.1513 | 6521 | 6281 | 6545 | 3767 | 2053 | 1700 | 1762 | 2364 | 5937 | 6353 | 6194 | 5659 | 1826 | 1846 | 1805 | 1803 | 3927,77 | 6036 | 1820 | 0,301 | -1,731 | -37,866 | 0 | 0 | 0,0012251 | 0,0023733 | 0,0016603 | 0,0016603 | TRUE | 0,0206877 | No |
| orf19.5671 | 5360 | 4573 | 4938 | 3464 | 1643 | 1342 | 1431 | 1917 | 4880 | 4625 | 4673 | 5203 | 1461 | 1457 | 1466 | 1462 | 3153,53 | 4845 | 1462 | 0,302 | -1,728 | -36,142 | 5,01E-286 | 1,23E-284 | 0,0013186 | 0,0024559 | 0,0017634 | 0,0017634 | TRUE | 0,0288478 | No |
| orf19.2519 | 522 | 446 | 466 | 328 | 179 | 131 | 137 | 158 | 475 | 451 | 441 | 493 | 159 | 142 | 140 | 121 | 302,79 | 465 | 140 | 0,302 | -1,728 | -17,566 | 4,52E-69 | 2,80E-68 | 0,0017364 | 0,0063997 | 0,0048181 | 0,0048181 | TRUE | 0,1151558 | No |
| orf19.7676 | 19956 | 15522 | 16642 | 12942 | 5834 | 4735 | 4881 | 7243 | 18169 | 15699 | 15750 | 19441 | 5189 | 5141 | 4999 | 5525 | 11239,11 | 17265 | 5214 | 0,302 | -1,727 | -24,757 | 2,62E-135 | 3,06E-134 | 0,0065064 | 0,0021546 | 0,0045543 | 0,0045543 | TRUE | 0,0876302 | No |
| orf19.7184 | 2812 | 2509 | 2696 | 1952 | 980 | 775 | 720 | 975 | 2560 | 2538 | 2552 | 2932 | 872 | 841 | 737 | 744 | 1721,97 | 2646 | 798 | 0,302 | -1,727 | -24,452 | 4,82E-132 | 5,49E-131 | 0,0051738 | 0,0028042 | 0,0040009 | 0,0040009 | TRUE | 0,0623315 | No |
| orf19.4645 | 5027 | 5704 | 5826 | 3310 | 1783 | 1492 | 1535 | 2004 | 4577 | 5769 | 5514 | 4972 | 1586 | 1620 | 1572 | 1529 | 3392,29 | 5208 | 1577 | 0,303 | -1,725 | -24,935 | 3,08E-137 | 3,65E-136 | 0,005034 | 0,0024264 | 0,0041936 | 0,0041936 | TRUE | 0,0824685 | No |
| orf19.2030 | 11158 | 8570 | 9276 | 6447 | 3318 | 2590 | 2577 | 3789 | 10159 | 8668 | 8779 | 9684 | 2951 | 2812 | 2639 | 2890 | 6072,82 | 9322 | 2823 | 0,303 | -1,723 | -28,614 | 4,53E-180 | 7,00E-179 | 0,003822 | 0,0022545 | 0,0032586 | 0,0032586 | TRUE | 0,045161 | No |
| orf19.173 | 24307 | 22932 | 25099 | 15373 | 7519 | 6429 | 7034 | 9267 | 22130 | 23193 | 23754 | 23093 | 6688 | 6980 | 7204 | 7069 | 15013,88 | 23042 | 6985 | 0,303 | -1,722 | -45,823 | 0 | 0 | 0,0007925 | 0,002125 | 0,0012661 | 0,0012661 | TRUE | 0,0100848 | No |
| orf19.2769 | 16305 | 13916 | 14030 | 8185 | 5242 | 3440 | 3936 | 5366 | 14845 | 14075 | 13278 | 12295 | 4663 | 3735 | 4031 | 4093 | 8876,82 | 13623 | 4130 | 0,303 | -1,722 | -23,689 | 4,69E-124 | 4,95E-123 | 0,0074092 | 0,0021858 | 0,0049217 | 0,0049217 | TRUE | 0,0917014 | No |
| orf19.2247 | 102 | 96 | 90 | 81 | 28 | 25 | 35 | 41 | 93 | 97 | 85 | 122 | 25 | 27 | 36 | 31 | 64,5 | 99 | 30 | 0,303 | -1,721 | -8,636 | 5,80E-18 | 1,39E-17 | 1,00E-08 | 0,0225184 | 0,0170804 | 0,0170804 | TRUE | 0,1826615 | No |
| orf19.2772 | 6301 | 5902 | 6326 | 3990 | 1961 | 1749 | 1730 | 2346 | 5737 | 5969 | 5987 | 5994 | 1744 | 1899 | 1772 | 1790 | 3861,39 | 5922 | 1801 | 0,304 | -1,718 | -42,455 | 0 | 0 | 0,0004443 | 0,0023791 | 0,0012216 | 0,0012216 | TRUE | 0,0153647 | No |
| orf19.551 | 18926 | 17083 | 18210 | 11867 | 6014 | 4910 | 5101 | 6877 | 17231 | 17278 | 17234 | 17826 | 5349 | 5331 | 5224 | 5246 | 11339,91 | 17392 | 5288 | 0,304 | -1,718 | -58,743 | 0 | 0 | 0,0001163 | 0,0021535 | 0,0007015 | 0,0007015 | TRUE | 0,0033474 | No |

| | | | | | | | | | | | | | | | | | | | | | | | | | | | | | | | |
|---|---|---|---|---|---|---|---|---|---|---|---|---|---|---|---|---|---|---|---|---|---|---|---|---|---|---|---|---|---|---|---|
| orf19.1481 | 7286 | 6788 | 7489 | 4549 | 2140 | 1962 | 2050 | 2882 | 6634 | 6865 | 7088 | 6833 | 1903 | 2130 | 2099 | 2198 | 4468,93 | 6855 | 2082 | 0,304 | -1,718 | -34,775 | 5,81E-265 | 1,30E-263 | 0,0017965 | 0,0023326 | 0,0020421 | 0,0020421 | TRUE | 0,0415056 | No |
| orf19.713 | 3530 | 3025 | 3351 | 2260 | 1190 | 881 | 903 | 1259 | 3214 | 3059 | 3171 | 3395 | 1058 | 957 | 925 | 960 | 2092,48 | 3210 | 975 | 0,304 | -1,717 | -31,353 | 8,81E-216 | 1,60E-214 | 0,0018131 | 0,0026683 | 0,0022325 | 0,0022325 | TRUE | 0,0400769 | No |
| orf19.5042 | 59 | 87 | 77 | 45 | 23 | 22 | 19 | 29 | 54 | 88 | 73 | 68 | 20 | 24 | 19 | 22 | 46,01 | 71 | 21 | 0,304 | -1,716 | -7,249 | 4,19E-13 | 8,81E-13 | 1,00E-08 | 0,0307464 | 0,0245529 | 0,0245529 | TRUE | 0,122815 | No |
| orf19.5618 | 1455 | 1489 | 1539 | 980 | 516 | 430 | 404 | 549 | 1325 | 1506 | 1457 | 1472 | 459 | 467 | 414 | 419 | 939,71 | 1440 | 440 | 0,305 | -1,712 | -25,618 | 9,53E-145 | 1,20E-143 | 0,0022541 | 0,0034428 | 0,0028531 | 0,0028531 | TRUE | 0,0362583 | No |
| orf19.7054 | 54 | 43 | 43 | 38 | 10 | 12 | 14 | 27 | 49 | 43 | 41 | 57 | 9 | 13 | 14 | 21 | 30,91 | 48 | 14 | 0,305 | -1,712 | -5,978 | 2,26E-09 | 4,18E-09 | 0,0098261 | 0,0447723 | 0,035155 | 0,035155 | TRUE | 0,5445115 | No |
| orf19.6117 | 8545 | 6833 | 7600 | 5667 | 2451 | 2154 | 2243 | 3236 | 7780 | 6911 | 7193 | 8513 | 2180 | 2339 | 2297 | 2468 | 4960,06 | 7599 | 2321 | 0,306 | -1,71 | -25,56 | 4,29E-144 | 5,38E-143 | 0,0052866 | 0,0023033 | 0,0040272 | 0,0040272 | TRUE | 0,0789149 | No |
| orf19.7304 | 1088 | 1166 | 1251 | 719 | 374 | 338 | 338 | 411 | 991 | 1179 | 1184 | 1080 | 333 | 367 | 346 | 314 | 724,15 | 1108 | 340 | 0,306 | -1,71 | -21,419 | 8,88E-102 | 7,85E-101 | 0,0046657 | 0,0038612 | 0,0042601 | 0,0042601 | TRUE | 0,0650283 | No |
| orf19.2917.1 | 715 | 550 | 585 | 455 | 218 | 161 | 169 | 268 | 651 | 556 | 554 | 683 | 194 | 175 | 173 | 204 | 398,82 | 611 | 186 | 0,307 | -1,702 | -17,164 | 4,93E-66 | 2,95E-65 | 0,0069354 | 0,0053493 | 0,006068 | 0,006068 | TRUE | 0,0674353 | No |
| orf19.4255 | 434 | 339 | 381 | 292 | 126 | 123 | 102 | 160 | 395 | 343 | 361 | 439 | 112 | 134 | 104 | 122 | 251,17 | 384 | 118 | 0,308 | -1,698 | -15 | 7,33E-51 | 3,44E-50 | 0,006433 | 0,0072964 | 0,0069516 | 0,0069516 | TRUE | 0,0910016 | No |
| orf19.3234.1 | 749 | 644 | 629 | 363 | 253 | 167 | 178 | 231 | 682 | 651 | 595 | 545 | 225 | 181 | 182 | 176 | 404,84 | 618 | 191 | 0,308 | -1,697 | -16,785 | 3,13E-63 | 1,80E-62 | 0,0081521 | 0,0053 | 0,0065119 | 0,0065119 | TRUE | 0,1599216 | No |
| orf19.4843 | 6186 | 5757 | 6168 | 4012 | 1979 | 1755 | 1688 | 2374 | 5632 | 5823 | 5837 | 6027 | 1760 | 1905 | 1729 | 1811 | 3815,52 | 5830 | 1801 | 0,309 | -1,694 | -39,195 | 0 | 0 | 0,0008167 | 0,0023832 | 0,001442 | 0,001442 | TRUE | 0,0172925 | No |
| orf19.395 | 14978 | 11641 | 12454 | 8741 | 4495 | 3535 | 3555 | 5348 | 13637 | 11774 | 11787 | 13130 | 3998 | 3838 | 3641 | 4079 | 8235,48 | 12582 | 3889 | 0,309 | -1,693 | -28,493 | 1,43E-178 | 2,19E-177 | 0,0038566 | 0,0021974 | 0,0032293 | 0,0032293 | TRUE | 0,039375 | No |
| orf19.842 | 16568 | 15340 | 15962 | 9439 | 5709 | 4174 | 4466 | 5759 | 15084 | 15515 | 15107 | 14179 | 5078 | 4532 | 4574 | 4393 | 9807,65 | 14971 | 4644 | 0,31 | -1,689 | -32,869 | 6,06E-237 | 1,20E-235 | 0,0025992 | 0,0021717 | 0,0023996 | 0,0023996 | TRUE | 0,0485898 | No |
| orf19.3433 | 568 | 570 | 554 | 365 | 182 | 174 | 145 | 226 | 517 | 576 | 524 | 548 | 162 | 189 | 148 | 172 | 354,74 | 541 | 168 | 0,31 | -1,689 | -18,738 | 2,43E-78 | 1,69E-77 | 0,0011948 | 0,0057609 | 0,004038 | 0,004038 | TRUE | 0,0650597 | No |
| orf19.1877 | 632 | 583 | 644 | 402 | 200 | 166 | 183 | 253 | 575 | 590 | 609 | 604 | 178 | 180 | 187 | 193 | 389,62 | 594 | 184 | 0,311 | -1,684 | -21,072 | 1,44E-98 | 1,24E-97 | 1,00E-08 | 0,0054275 | 0,0027196 | 0,0027196 | TRUE | 0,0118621 | No |
| orf19.3932 | 11765 | 9706 | 10734 | 6896 | 3723 | 2892 | 2998 | 4267 | 10712 | 9817 | 10159 | 10359 | 3311 | 3140 | 3070 | 3255 | 6727,8 | 10262 | 3194 | 0,311 | -1,683 | -39,935 | 0 | 0 | 0,0009757 | 0,0022334 | 0,0015069 | 0,0015069 | TRUE | 0,0108658 | No |
| orf19.5094 | 1565 | 1401 | 1487 | 890 | 466 | 415 | 463 | 539 | 1425 | 1417 | 1407 | 1337 | 414 | 451 | 474 | 411 | 917,06 | 1396 | 438 | 0,312 | -1,681 | -27,13 | 4,33E-162 | 6,05E-161 | 0,0004048 | 0,0034775 | 0,0022333 | 0,0022333 | TRUE | 0,0368857 | No |
| orf19.1770 | 8255 | 6835 | 6983 | 4786 | 2683 | 2057 | 2084 | 2743 | 7516 | 6913 | 6609 | 7189 | 2386 | 2233 | 2134 | 2092 | 4634,16 | 7057 | 2211 | 0,313 | -1,674 | -30,009 | 7,56E-198 | 1,28E-196 | 0,002855 | 0,0023221 | 0,0027008 | 0,0027008 | TRUE | 0,0349296 | No |
| orf19.3997 | 75794 | 57465 | 64327 | 42099 | 22165 | 17219 | 18973 | 27419 | 69007 | 58120 | 60880 | 63239 | 19715 | 18695 | 19431 | 20915 | 41250,29 | 62812 | 19689 | 0,313 | -1,674 | -29,521 | 1,54E-191 | 2,51E-190 | 0,0037729 | 0,002069 | 0,0030554 | 0,0030554 | TRUE | 0,0541128 | No |
| orf19.2623 | 15021 | 13342 | 14429 | 9492 | 4723 | 4107 | 4199 | 5653 | 13676 | 13494 | 13656 | 14258 | 4201 | 4459 | 4300 | 4312 | 9044,59 | 13771 | 4318 | 0,314 | -1,673 | -48,165 | 0 | 0 | 0,0003965 | 0,0021831 | 0,0010114 | 0,0010114 | TRUE | 0,006755 | No |
| orf19.2832 | 4022 | 3903 | 4244 | 2561 | 1386 | 1125 | 1206 | 1547 | 3662 | 3947 | 4017 | 3847 | 1233 | 1221 | 1235 | 1180 | 2542,79 | 3868 | 1217 | 0,315 | -1,669 | -36,108 | 1,73E-285 | 4,20E-284 | 0,0008754 | 0,0025565 | 0,0015281 | 0,0015281 | TRUE | 0,0163902 | No |
| orf19.1285 | 5777 | 5533 | 5973 | 3978 | 1945 | 1665 | 1722 | 2314 | 5260 | 5596 | 5653 | 5976 | 1730 | 1808 | 1764 | 1765 | 3693,83 | 5621 | 1767 | 0,314 | -1,669 | -35,367 | 5,58E-274 | 1,29E-272 | 0,0013907 | 0,0023946 | 0,0017781 | 0,0017781 | TRUE | 0,0229335 | No |
| orf19.4531 | 26415 | 24471 | 26708 | 15803 | 8489 | 7168 | 7627 | 10017 | 24050 | 24750 | 25277 | 23739 | 7551 | 7782 | 7811 | 7641 | 16075,01 | 24454 | 7696 | 0,315 | -1,668 | -50,387 | 0 | 0 | 0,0004135 | 0,0021192 | 0,0009701 | 0,0009701 | TRUE | 0,0062902 | No |
| orf19.7022 | 1326 | 1250 | 1287 | 738 | 451 | 378 | 349 | 456 | 1207 | 1264 | 1218 | 1109 | 401 | 410 | 357 | 348 | 789,37 | 1200 | 379 | 0,315 | -1,667 | -23,245 | 1,58E-119 | 1,61E-118 | 0,0025522 | 0,0037105 | 0,0032601 | 0,0032601 | TRUE | 0,0395554 | No |
| orf19.4617 | 1383 | 1431 | 1331 | 836 | 504 | 382 | 368 | 532 | 1259 | 1447 | 1260 | 1256 | 448 | 415 | 377 | 406 | 858,46 | 1306 | 412 | 0,315 | -1,665 | -22,63 | 2,19E-113 | 2,13E-112 | 0,0037382 | 0,0035758 | 0,0036553 | 0,0036553 | TRUE | 0,0649991 | No |
| orf19.5610 | 3599 | 3539 | 3692 | 2163 | 1229 | 1044 | 976 | 1408 | 3277 | 3579 | 3494 | 3249 | 1093 | 1133 | 1000 | 1074 | 2237,45 | 3400 | 1075 | 0,316 | -1,662 | -30,825 | 1,23E-208 | 2,16E-207 | 0,0016844 | 0,0026274 | 0,0021985 | 0,0021985 | TRUE | 0,0255664 | No |
| CaalfMp05 | 23 | 40 | 27 | 32 | 11 | 14 | 8 | 12 | 21 | 40 | 26 | 48 | 10 | 15 | 8 | 9 | 22,17 | 34 | 10 | 0,316 | -1,662 | -4,571 | 4,86E-06 | 7,85E-06 | 0,073509 | 0,0616255 | 0,0662982 | 0,0662982 | TRUE | 0,0975802 | No |
| orf19.6420 | 8235 | 6767 | 7567 | 5932 | 2416 | 2257 | 2358 | 3406 | 7498 | 6844 | 7161 | 8911 | 2149 | 2450 | 2415 | 2598 | 5003,31 | 7604 | 2403 | 0,316 | -1,661 | -20,398 | 1,73E-92 | 1,38E-91 | 0,0098505 | 0,002301 | 0,0061036 | 0,0061036 | TRUE | 0,1621312 | No |
| orf19.4174 | 2658 | 2222 | 2304 | 1676 | 812 | 677 | 697 | 1029 | 2420 | 2247 | 2181 | 2518 | 722 | 735 | 714 | 785 | 1540,19 | 2342 | 739 | 0,316 | -1,66 | -27,102 | 9,45E-162 | 1,32E-160 | 0,0026129 | 0,0028947 | 0,002742 | 0,002742 | TRUE | 0,029342 | No |
| orf19.4802 | 5052 | 4436 | 4921 | 2801 | 1581 | 1350 | 1432 | 1781 | 4600 | 4487 | 4657 | 4208 | 1406 | 1466 | 1467 | 1359 | 2956,01 | 4488 | 1424 | 0,317 | -1,659 | -34,29 | 1,12E-257 | 2,41E-256 | 0,0013071 | 0,0024839 | 0,0017987 | 0,0017987 | TRUE | 0,0205352 | No |
| orf19.3841 | 7339 | 6369 | 7080 | 4522 | 2377 | 1973 | 2066 | 2732 | 6682 | 6442 | 6701 | 6793 | 2114 | 2142 | 2116 | 2084 | 4384,12 | 6654 | 2114 | 0,318 | -1,654 | -46,416 | 0 | 0 | 1,00E-08 | 0,0023383 | 0,000918 | 0,000918 | TRUE | 0,005642 | No |
| orf19.428 | 1758 | 1659 | 1708 | 1038 | 571 | 494 | 491 | 677 | 1601 | 1678 | 1616 | 1559 | 508 | 536 | 503 | 516 | 1064,71 | 1614 | 516 | 0,319 | -1,647 | -30,257 | 4,21E-201 | 7,22E-200 | 1,00E-08 | 0,0032777 | 0,0016112 | 0,0016112 | TRUE | 0,0085218 | No |
| orf19.4618 | 10564 | 8608 | 9382 | 6692 | 3194 | 2864 | 2741 | 4104 | 9618 | 8706 | 8879 | 10052 | 2841 | 3110 | 2807 | 3131 | 6143 | 9314 | 2972 | 0,319 | -1,647 | -27,988 | 2,27E-172 | 3,39E-171 | 0,0036707 | 0,0022521 | 0,0031122 | 0,0031122 | TRUE | 0,0343051 | No |
| orf19.2529.1 | 17392 | 12949 | 13002 | 7510 | 5280 | 3176 | 4278 | 5559 | 15835 | 13097 | 12305 | 11281 | 4696 | 3448 | 4381 | 4240 | 8660,49 | 13130 | 4191 | 0,319 | -1,647 | -11,713 | 1,09E-31 | 3,64E-31 | 0,0188491 | 0,0021895 | 0,010008 | 0,0188491 | TRUE | 0,2355037 | No |
| orf19.1309 | 747 | 542 | 604 | 380 | 246 | 166 | 157 | 258 | 680 | 548 | 572 | 571 | 219 | 180 | 161 | 197 | 390,92 | 593 | 189 | 0,32 | -1,644 | -15,703 | 1,45E-55 | 7,40E-55 | 0,0095124 | 0,0054162 | 0,0071662 | 0,0071662 | TRUE | 0,1203226 | No |
| orf19.3360 | 9339 | 9626 | 9941 | 6252 | 3391 | 2805 | 2954 | 3636 | 8503 | 9736 | 9408 | 9392 | 3016 | 3045 | 3025 | 2774 | 6112,33 | 9260 | 2965 | 0,32 | -1,643 | -31,398 | 2,14E-216 | 3,90E-215 | 0,0022699 | 0,0022531 | 0,0024151 | 0,0024151 | TRUE | 0,0373705 | No |
| orf19.5092 | 19 | 19 | 16 | 9 | 7 | 4 | 4 | 8 | 17 | 19 | 15 | 14 | 6 | 4 | 4 | 6 | 10,74 | 16 | 5 | 0,321 | -1,639 | -3,689 | 0,000225 | 0,0003319 | 1,00E-08 | 0,1249988 | 0,0691756 | 0,0691756 | TRUE | 0,0658099 | No |
| orf19.3902 | 36 | 48 | 44 | 38 | 17 | 18 | 7 | 20 | 33 | 49 | 42 | 57 | 15 | 20 | 7 | 15 | 29,64 | 45 | 14 | 0,322 | -1,637 | -5,414 | 6,18E-08 | 1,08E-07 | 0,0355018 | 0,046602 | 0,0432029 | 0,0432029 | TRUE | 0,2472006 | No |
| orf19.7561 | 27732 | 22638 | 25991 | 17695 | 8428 | 7211 | 8014 | 11029 | 25249 | 22896 | 24598 | 26581 | 7496 | 7829 | 8207 | 8413 | 16408,69 | 24831 | 7986 | 0,322 | -1,636 | -30,216 | 1,48E-200 | 2,53E-199 | 0,0030863 | 0,0021175 | 0,0027369 | 0,0027369 | TRUE | 0,0336707 | No |
| orf19.6884 | 355 | 297 | 358 | 225 | 126 | 75 | 112 | 141 | 323 | 300 | 339 | 338 | 112 | 81 | 115 | 108 | 214,52 | 325 | 104 | 0,322 | -1,636 | -14,466 | 1,99E-47 | 8,78E-47 | 0,0014808 | 0,008195 | 0,0061829 | 0,0061829 | TRUE | 0,1839196 | No |
| orf19.4772 | 1200 | 1128 | 1179 | 813 | 403 | 349 | 335 | 514 | 1093 | 1141 | 1116 | 1221 | 358 | 379 | 343 | 392 | 755,38 | 1143 | 368 | 0,323 | -1,628 | -24,278 | 3,35E-130 | 3,71E-129 | 0,0011287 | 0,0037858 | 0,0025848 | 0,0025848 | TRUE | 0,0248279 | No |
| orf19.1979 | 22144 | 20492 | 21615 | 12990 | 7416 | 5952 | 6352 | 8674 | 20161 | 20725 | 20457 | 19513 | 6596 | 6462 | 6505 | 6617 | 13379,59 | 20214 | 6545 | 0,324 | -1,627 | -51,304 | 0 | 0 | 0,0002604 | 0,0021357 | 0,0008677 | 0,0008677 | TRUE | 0,0065148 | No |
| orf19.419 | 7407 | 6750 | 6900 | 4020 | 2489 | 1942 | 2057 | 2700 | 6744 | 6827 | 6530 | 6039 | 2214 | 2108 | 2107 | 2060 | 4328,52 | 6535 | 2122 | 0,324 | -1,624 | -33,176 | 2,37E-241 | 4,77E-240 | 0,0015201 | 0,0023422 | 0,0019977 | 0,0019977 | TRUE | 0,0308869 | No |
| orf19.7602 | 1674 | 1510 | 1530 | 953 | 549 | 450 | 481 | 608 | 1524 | 1527 | 1448 | 1432 | 488 | 489 | 493 | 464 | 983,02 | 1483 | 484 | 0,325 | -1,621 | -28,66 | 1,19E-180 | 1,85E-179 | 1,00E-08 | 0,0033808 | 0,0017454 | 0,0017454 | TRUE | 0,0096571 | No |
| orf19.693 | 169 | 176 | 216 | 119 | 70 | 61 | 56 | 64 | 154 | 178 | 204 | 179 | 62 | 66 | 57 | 49 | 118,71 | 179 | 58 | 0,326 | -1,617 | -10,704 | 9,78E-27 | 2,91E-26 | 1,00E-08 | 0,0131645 | 0,0109409 | 0,0109409 | TRUE | 0,1330856 | No |
| orf19.1484 | 1039 | 1252 | 1224 | 773 | 410 | 369 | 387 | 421 | 946 | 1266 | 1158 | 1161 | 365 | 401 | 396 | 321 | 751,82 | 1133 | 371 | 0,326 | -1,615 | -16,525 | 2,42E-61 | 1,35E-60 | 0,0113766 | 0,0037941 | 0,0074303 | 0,0074303 | TRUE | 0,1519585 | No |
| orf19.7252 | 9 | 3 | 7 | 5 | 3 | 2 | 1 | 3 | 8 | 3 | 7 | 8 | 3 | 2 | 1 | 2 | 4,19 | 6 | 2 | 0,327 | -1,612 | -2,229 | 0,025793 | 0,0327677 | 1,00E-08 | 0,3173452 | 0,1926029 | 0,1926029 | TRUE | 0,3126284 | No |
| orf19.4777 | 3414 | 3062 | 3315 | 1916 | 1061 | 908 | 1041 | 1335 | 3108 | 3097 | 3137 | 2878 | 944 | 986 | 1066 | 1018 | 2029,34 | 3055 | 1004 | 0,328 | -1,608 | -31,018 | 3,04E-211 | 5,45E-210 | 0,0012075 | 0,002688 | 0,0019395 | 0,0019395 | TRUE | 0,0202253 | No |
| orf19.2608 | 2E+05 | 1E+05 | 1E+05 | 82223 | 45928 | 36647 | 41368 | 58390 | 1E+05 | 1E+05 | 1E+05 | 1E+05 | 40852 | 39788 | 42366 | 44540 | 84664,81 | 127443 | 41886 | 0,329 | -1,605 | -32,469 | 2,89E-231 | 5,63E-230 | 0,002537 | 0,0020526 | 0,0023332 | 0,0023332 | TRUE | 0,0293409 | No |
| orf19.5159 | 3966 | 3631 | 3816 | 2338 | 1410 | 1043 | 1133 | 1568 | 3611 | 3672 | 3611 | 3512 | 1254 | 1132 | 1160 | 1196 | 2393,72 | 3602 | 1186 | 0,329 | -1,602 | -35,988 | 1,30E-283 | 3,14E-282 | 0,0002127 | 0,0025889 | 0,0013599 | 0,0013599 | TRUE | 0,0178665 | No |
| orf19.7042 | 1119 | 1105 | 1079 | 712 | 404 | 347 | 350 | 398 | 1019 | 1118 | 1021 | 1070 | 359 | 377 | 358 | 304 | 703,15 | 1057 | 350 | 0,33 | -1,601 | -21,477 | 2,54E-102 | 2,27E-101 | 0,0029315 | 0,0039157 | 0,0034899 | 0,0034899 | TRUE | 0,0989289 | No |
| orf19.1464 | 5196 | 4681 | 5211 | 2968 | 1834 | 1376 | 1582 | 2004 | 4731 | 4734 | 4932 | 4458 | 1631 | 1494 | 1620 | 1529 | 3141,16 | 4714 | 1568 | 0,333 | -1,588 | -33,023 | 3,74E-239 | 7,50E-238 | 0,0012162 | 0,0024576 | 0,0018096 | 0,0018096 | TRUE | 0,0154921 | No |
| orf19.2361 | 3823 | 3544 | 3918 | 2484 | 1329 | 1161 | 1181 | 1550 | 3481 | 3584 | 3708 | 3731 | 1182 | 1261 | 1209 | 1182 | 2417,37 | 3626 | 1208 | 0,333 | -1,586 | -35,329 | 2,11E-273 | 4,86E-272 | 0,0005664 | 0,0025835 | 0,0013991 | 0,0013991 | TRUE | 0,0096204 | No |
| orf19.4082 | 7723 | 6095 | 6396 | 4586 | 2508 | 1987 | 2055 | 2939 | 7031 | 6164 | 6053 | 6889 | 2231 | 2157 | 2105 | 2242 | 4359,08 | 6534 | 2184 | 0,334 | -1,58 | -27,527 | 8,44E-167 | 1,23E-165 | 0,0029245 | 0,0023401 | 0,0028692 | 0,0028692 | TRUE | 0,0327798 | No |
| orf19.6690 | 2460 | 2267 | 2436 | 1374 | 871 | 663 | 737 | 965 | 2240 | 2293 | 2305 | 2064 | 775 | 720 | 755 | 736 | 1485,93 | 2226 | 746 | 0,335 | -1,578 | -29,098 | 3,78E-186 | 5,99E-185 | 0,0011147 | 0,002926 | 0,001958 | 0,001958 | TRUE | 0,0265975 | No |
| orf19.3679 | 1802 | 1721 | 1785 | 1095 | 638 | 574 | 515 | 708 | 1641 | 1741 | 1689 | 1645 | 567 | 623 | 527 | 540 | 1121,7 | 1679 | 564 | 0,336 | -1,576 | -26,377 | 2,52E-153 | 3,34E-152 | 0,0010521 | 0,0032147 | 0,0022784 | 0,0022784 | TRUE | 0,0534209 | No |
| orf19.5245 | 1781 | 1677 | 1871 | 1222 | 630 | 568 | 546 | 766 | 1622 | 1696 | 1771 | 1836 | 560 | 617 | 559 | 584 | 1155,57 | 1731 | 580 | 0,335 | -1,576 | -26,214 | 1,82E-151 | 2,39E-150 | 0,0015799 | 0,0031802 | 0,0023538 | 0,0023538 | TRUE | 0,0221297 | No |
| orf19.3969 | 6451 | 5738 | 6325 | 3964 | 2119 | 1814 | 1988 | 2681 | 5873 | 5803 | 5986 | 5955 | 1885 | 1969 | 2036 | 2045 | 3944,08 | 5904 | 1984 | 0,336 | -1,573 | -39,969 | 0 | 0 | 0,0003465 | 0,0023719 | 0,0011607 | 0,0011607 | TRUE | 0,0141145 | No |
| orf19.5058 | 21958 | 19774 | 21530 | 12778 | 7462 | 6266 | 6634 | 8608 | 19992 | 19999 | 20376 | 19195 | 6637 | 6803 | 6794 | 6566 | 13295,32 | 19890 | 6700 | 0,337 | -1,571 | -48,294 | 0 | 0 | 0,0003331 | 0,0021364 | 0,0009188 | 0,0009188 | TRUE | 0,0066432 | No |
| orf19.6084 | 11794 | 10222 | 10864 | 7148 | 3937 | 3315 | 3471 | 4668 | 10738 | 10338 | 10282 | 10737 | 3502 | 3599 | 3555 | 3561 | 7039,02 | 10524 | 3554 | 0,338 | -1,566 | -47,278 | 0 | 0 | 0,0002628 | 0,0022247 | 0,0008708 | 0,0008708 | TRUE | 0,0029464 | No |
| orf19.4003 | 778 | 748 | 744 | 483 | 296 | 225 | 243 | 292 | 708 | 757 | 704 | 726 | 263 | 244 | 249 | 223 | 484,21 | 724 | 245 | 0,338 | -1,566 | -20,614 | 2,04E-94 | 1,66E-93 | 1,00E-08 | 0,0047652 | 0,0028983 | 0,0028983 | TRUE | 0,0480849 | No |
| orf19.3748 | 59 | 51 | 76 | 34 | 23 | 20 | 16 | 25 | 54 | 52 | 72 | 51 | 20 | 22 | 16 | 19 | 38,24 | 57 | 19 | 0,338 | -1,566 | -6,38 | 1,77E-10 | 3,40E-10 | 1,00E-08 | 0,0365811 | 0,0245548 | 0,0245548 | TRUE | 0,2750113 | No |
| orf19.5442 | 2757 | 2331 | 2642 | 1564 | 927 | 729 | 808 | 1101 | 2510 | 2358 | 2500 | 2349 | 825 | 791 | 827 | 840 | 1625,11 | 2429 | 821 | 0,338 | -1,565 | -31,846 | 1,50E-222 | 2,81E-221 | 1,00E-08 | 0,0028499 | 0,0015318 | 0,0015318 | TRUE | 0,0065003 | No |
| orf19.5266 | 64 | 45 | 47 | 22 | 19 | 18 | 13 | 17 | 58 | 46 | 44 | 33 | 17 | 20 | 13 | 13 | 30,5 | 45 | 16 | 0,338 | -1,564 | -5,566 | 2,61E-08 | 4,61E-08 | 1,00E-08 | 0,0453422 | 0,0342798 | 0,0342798 | TRUE | 0,1365796 | No |
| orf19.4842 | 1438 | 1104 | 1183 | 940 | 474 | 377 | 416 | 546 | 1309 | 1117 | 1120 | 1412 | 422 | 409 | 426 | 416 | 828,86 | 1240 | 418 | 0,338 | -1,563 | -18,76 | 1,61E-78 | 1,13E-77 | 0,0065396 | 0,0036308 | 0,0051173 | 0,0051173 | TRUE | 0,0991751 | No |

| | | | | | | | | | | | | | | | | | | | | | | | | | | | | | | | |
|---|---|---|---|---|---|---|---|---|---|---|---|---|---|---|---|---|---|---|---|---|---|---|---|---|---|---|---|---|---|---|---|
| orf19.5041 | 225 | 202 | 176 | 125 | 70 | 71 | 55 | 85 | 205 | 204 | 167 | 188 | 62 | 77 | 56 | 65 | 128 | 191 | 65 | 0,34 | -1,556 | -11,022 | 2,98E-28 | 9,18E-28 | 1,00E-08 | 0,0123572 | 0,0091764 | 0,0091764 | TRUE | 0,1168016 | No |
| orf19.2947 | 4150 | 3605 | 3792 | 2438 | 1416 | 1136 | 1252 | 1603 | 3778 | 3646 | 3589 | 3662 | 1259 | 1233 | 1282 | 1223 | 2459,17 | 3669 | 1249 | 0,34 | -1,555 | -37,761 | 0 | 0 | 1,00E-08 | 0,0025742 | 0,0011079 | 0,0011079 | TRUE | 0,0051648 | No |
| orf19.2722 | 723 | 668 | 696 | 398 | 249 | 224 | 193 | 291 | 658 | 676 | 659 | 598 | 221 | 243 | 198 | 222 | 434,34 | 648 | 221 | 0,34 | -1,555 | -19,071 | 4,41E-81 | 3,17E-80 | 0,0006518 | 0,0050784 | 0,0034559 | 0,0034559 | TRUE | 0,0506773 | No |
| orf19.7345 | 1240 | 1343 | 1306 | 747 | 499 | 388 | 373 | 544 | 1129 | 1358 | 1236 | 1122 | 444 | 421 | 382 | 415 | 813,43 | 1211 | 416 | 0,343 | -1,544 | -19,716 | 1,56E-86 | 1,19E-85 | 0,0049893 | 0,003661 | 0,0043235 | 0,0043235 | TRUE | 0,0799591 | No |
| orf19.2841 | 8829 | 9009 | 9147 | 5089 | 3355 | 2649 | 2798 | 3672 | 8038 | 9112 | 8657 | 7644 | 2984 | 2876 | 2866 | 2801 | 5622,27 | 8363 | 2882 | 0,344 | -1,538 | -27,14 | 3,32E-162 | 4,67E-161 | 0,0030589 | 0,002272 | 0,0028568 | 0,0028568 | TRUE | 0,0442079 | No |
| orf19.4094 | 60 | 65 | 59 | 55 | 22 | 26 | 19 | 28 | 55 | 66 | 56 | 83 | 20 | 28 | 19 | 21 | 43,43 | 65 | 22 | 0,345 | -1,536 | -6,459 | 1,05E-10 | 2,04E-10 | 1,00E-08 | 0,0324538 | 0,024804 | 0,024804 | TRUE | 0,2387998 | No |
| orf19.2988 | 2356 | 2005 | 2120 | 1461 | 787 | 683 | 717 | 956 | 2145 | 2028 | 2006 | 2195 | 700 | 742 | 734 | 729 | 1409,88 | 2094 | 726 | 0,347 | -1,526 | -29,075 | 7,43E-186 | 1,17E-184 | 0,0007425 | 0,002974 | 0,0017469 | 0,0017469 | TRUE | 0,0116685 | No |
| orf19.5785 | 1876 | 1758 | 1884 | 1175 | 645 | 623 | 584 | 791 | 1708 | 1778 | 1783 | 1765 | 574 | 676 | 598 | 603 | 1185,71 | 1758 | 613 | 0,348 | -1,523 | -26,585 | 1,01E-155 | 1,37E-154 | 0,0006485 | 0,0031511 | 0,0020846 | 0,0020846 | TRUE | 0,0535062 | No |
| orf19.1999 | 1911 | 1601 | 1723 | 1055 | 660 | 496 | 606 | 715 | 1740 | 1619 | 1631 | 1585 | 587 | 539 | 621 | 545 | 1108,27 | 1644 | 573 | 0,348 | -1,523 | -25,007 | 5,17E-138 | 6,18E-137 | 0,0015052 | 0,003229 | 0,0024206 | 0,0024206 | TRUE | 0,0352696 | No |
| orf19.6478 | 13153 | 13445 | 13848 | 7578 | 5034 | 4035 | 4357 | 5406 | 11975 | 13598 | 13106 | 11383 | 4478 | 4381 | 4462 | 4124 | 8438,37 | 12516 | 4361 | 0,348 | -1,522 | -25,73 | 5,44E-146 | 6,94E-145 | 0,0037917 | 0,0021936 | 0,0032093 | 0,0032093 | TRUE | 0,0448279 | No |
| orf19.4435 | 4059 | 3940 | 4004 | 2733 | 1500 | 1275 | 1304 | 1810 | 3696 | 3985 | 3789 | 4105 | 1334 | 1384 | 1335 | 1381 | 2626,24 | 3894 | 1358 | 0,349 | -1,517 | -31,969 | 2,90E-224 | 5,47E-223 | 0,0011094 | 0,00254 | 0,0016808 | 0,0016808 | TRUE | 0,0155114 | No |
| orf19.4800 | 2528 | 2253 | 2426 | 1470 | 911 | 700 | 811 | 1020 | 2302 | 2279 | 2296 | 2208 | 810 | 760 | 831 | 778 | 1532,93 | 2271 | 795 | 0,35 | -1,516 | -31,631 | 1,40E-219 | 2,60E-218 | 1,00E-08 | 0,0028988 | 0,0013836 | 0,0013836 | TRUE | 0,0102191 | No |
| orf19.33 | 4209 | 3677 | 3783 | 2480 | 1604 | 1120 | 1205 | 1719 | 3832 | 3719 | 3580 | 3725 | 1427 | 1216 | 1234 | 1311 | 2505,59 | 3714 | 1297 | 0,35 | -1,516 | -27,622 | 6,13E-168 | 9,03E-167 | 0,0024222 | 0,0025642 | 0,0023878 | 0,0023878 | TRUE | 0,0541995 | No |
| orf19.6875 | 1793 | 1700 | 1745 | 1047 | 638 | 533 | 550 | 777 | 1632 | 1719 | 1651 | 1573 | 567 | 579 | 563 | 593 | 1109,78 | 1644 | 576 | 0,35 | -1,515 | -28,031 | 6,79E-173 | 1,02E-171 | 1,00E-08 | 0,0032273 | 0,0016697 | 0,0016697 | TRUE | 0,0111788 | No |
| orf19.7124 | 16015 | 12375 | 13844 | 8237 | 5251 | 3935 | 4585 | 6253 | 14581 | 12516 | 13102 | 12373 | 4671 | 4272 | 4696 | 4770 | 8872,59 | 13143 | 4602 | 0,35 | -1,514 | -25,618 | 9,65E-145 | 1,22E-143 | 0,0039351 | 0,0021859 | 0,0032122 | 0,0032122 | TRUE | 0,0665364 | No |
| orf19.1735 | 485 | 479 | 473 | 269 | 202 | 135 | 149 | 190 | 442 | 484 | 448 | 404 | 180 | 147 | 153 | 145 | 300,19 | 444 | 156 | 0,35 | -1,514 | -15,313 | 6,28E-53 | 3,05E-52 | 0,0032081 | 0,0064375 | 0,005206 | 0,005206 | TRUE | 0,1163313 | No |
| orf19.1375 | 27705 | 23957 | 25968 | 16330 | 9534 | 8040 | 8220 | 11705 | 25224 | 24230 | 24576 | 24530 | 8480 | 8729 | 8418 | 8929 | 16639,65 | 24640 | 8639 | 0,351 | -1,512 | -46,563 | 0 | 0 | 0,0004107 | 0,0021164 | 0,0009366 | 0,0009366 | TRUE | 0,0063737 | No |
| orf19.2748 | 1392 | 1582 | 1567 | 949 | 548 | 512 | 488 | 637 | 1267 | 1600 | 1483 | 1426 | 487 | 556 | 500 | 486 | 975,62 | 1444 | 507 | 0,351 | -1,511 | -19,102 | 2,41E-81 | 1,74E-80 | 0,0059869 | 0,003391 | 0,0047159 | 0,0047159 | TRUE | 0,0847824 | No |
| orf19.3296 | 197 | 164 | 171 | 100 | 74 | 37 | 62 | 79 | 179 | 166 | 162 | 150 | 66 | 40 | 63 | 60 | 110,88 | 164 | 57 | 0,351 | -1,511 | -9,934 | 2,97E-23 | 8,15E-23 | 0,0021636 | 0,0139509 | 0,0109428 | 0,0109428 | TRUE | 0,3035444 | No |
| orf19.7451 | 6407 | 5970 | 6432 | 4274 | 2374 | 1999 | 2093 | 2834 | 5833 | 6038 | 6087 | 6420 | 2112 | 2170 | 2143 | 2162 | 4120,76 | 6094 | 2147 | 0,353 | -1,504 | -36,632 | 8,96E-294 | 2,26E-292 | 0,0007165 | 0,0023576 | 0,0013131 | 0,0013131 | TRUE | 0,015109 | No |
| orf19.1034 | 2002 | 1872 | 1942 | 1091 | 764 | 558 | 616 | 825 | 1823 | 1893 | 1838 | 1639 | 680 | 606 | 631 | 629 | 1217,3 | 1798 | 636 | 0,354 | -1,5 | -24,302 | 1,88E-130 | 2,10E-129 | 0,0021471 | 0,0031222 | 0,0026272 | 0,0026272 | TRUE | 0,0396347 | No |
| orf19.4947 | 7612 | 6886 | 6472 | 4453 | 2793 | 2421 | 1956 | 3049 | 6930 | 6964 | 6125 | 6689 | 2484 | 2629 | 2003 | 2326 | 4518,87 | 6677 | 2360 | 0,353 | -1,5 | -19,825 | 1,82E-87 | 1,40E-86 | 0,0077678 | 0,0023293 | 0,0052224 | 0,0052224 | TRUE | 0,1244751 | No |
| orf19.4557 | 576 | 539 | 535 | 320 | 224 | 161 | 176 | 231 | 524 | 545 | 506 | 481 | 199 | 175 | 180 | 176 | 348,39 | 514 | 182 | 0,355 | -1,496 | -17,362 | 1,61E-67 | 9,76E-67 | 1,00E-08 | 0,0058288 | 0,0035459 | 0,0035459 | TRUE | 0,0421372 | No |
| orf19.635 | 10 | 7 | 11 | 13 | 2 | 4 | 6 | 5 | 9 | 7 | 10 | 20 | 2 | 4 | 6 | 4 | 7,78 | 12 | 4 | 0,355 | -1,496 | -2,678 | 0,007416 | 0,0098211 | 1,00E-08 | 0,1719306 | 0,1344415 | 0,1344415 | TRUE | 0,558177 | No |
| orf19.4347 | 8411 | 7558 | 8546 | 5481 | 3075 | 2514 | 2863 | 3725 | 7658 | 7644 | 8088 | 8233 | 2735 | 2730 | 2932 | 2841 | 5357,68 | 7906 | 2810 | 0,356 | -1,492 | -34,77 | 6,93E-265 | 1,55E-263 | 0,0009903 | 0,0022836 | 0,0015334 | 0,0015334 | TRUE | 0,0101647 | No |
| orf19.2108 | 4061 | 3383 | 3667 | 2304 | 1440 | 1160 | 1173 | 1642 | 3697 | 3422 | 3470 | 3461 | 1281 | 1259 | 1201 | 1253 | 2380,56 | 3512 | 1248 | 0,355 | -1,492 | -32,908 | 1,71E-237 | 3,41E-236 | 0,000682 | 0,0025919 | 0,0014481 | 0,0014481 | TRUE | 0,0157238 | No |
| orf19.2685 | 321 | 307 | 299 | 238 | 114 | 113 | 93 | 157 | 292 | 310 | 283 | 358 | 101 | 123 | 95 | 120 | 210,29 | 311 | 110 | 0,356 | -1,491 | -12,557 | 3,66E-36 | 1,34E-35 | 0,0063387 | 0,0083187 | 0,0075738 | 0,0075738 | TRUE | 0,1009928 | No |
| orf19.5259 | 664 | 619 | 667 | 433 | 240 | 227 | 210 | 290 | 605 | 626 | 631 | 650 | 213 | 246 | 215 | 221 | 426,06 | 628 | 224 | 0,356 | -1,488 | -19,196 | 4,02E-82 | 2,93E-81 | 1,00E-08 | 0,0051375 | 0,0028369 | 0,0028369 | TRUE | 0,0447381 | No |
| orf19.3264 | 340 | 381 | 377 | 237 | 145 | 132 | 122 | 141 | 310 | 385 | 357 | 356 | 129 | 143 | 125 | 108 | 239,06 | 352 | 126 | 0,357 | -1,487 | -13,16 | 1,48E-39 | 5,74E-39 | 0,006223 | 0,0075628 | 0,007017 | 0,007017 | TRUE | 0,1156288 | No |
| orf19.4317 | 2151 | 2085 | 2080 | 1282 | 903 | 655 | 667 | 857 | 1958 | 2109 | 1969 | 1926 | 803 | 711 | 683 | 654 | 1351,58 | 1990 | 713 | 0,358 | -1,483 | -22,384 | 5,67E-111 | 5,38E-110 | 0,0036734 | 0,0030144 | 0,0032868 | 0,0032868 | TRUE | 0,0893769 | No |
| orf19.6641 | 1179 | 1032 | 1030 | 733 | 431 | 367 | 337 | 487 | 1073 | 1044 | 975 | 1101 | 383 | 398 | 345 | 371 | 711,44 | 1048 | 374 | 0,358 | -1,483 | -21,662 | 4,68E-104 | 4,23E-103 | 0,0014012 | 0,0038938 | 0,0027417 | 0,0027417 | TRUE | 0,0296272 | No |
| orf19.6072.1 | 852 | 823 | 833 | 627 | 378 | 286 | 286 | 336 | 776 | 832 | 788 | 942 | 336 | 311 | 293 | 256 | 566,78 | 834 | 299 | 0,358 | -1,48 | -15,998 | 1,32E-57 | 6,94E-57 | 0,0080269 | 0,0043677 | 0,0060056 | 0,0060056 | TRUE | 0,1156934 | No |
| orf19.3707 | 29758 | 26245 | 28293 | 17328 | 11098 | 8824 | 9101 | 12336 | 27093 | 26544 | 26777 | 26029 | 9871 | 9580 | 9321 | 9410 | 18078,24 | 26611 | 9546 | 0,359 | -1,479 | -46,809 | 0 | 0 | 0,0003695 | 0,0021101 | 0,00089 | 0,00089 | TRUE | 0,006505 | No |
| orf19.193 | 433 | 509 | 497 | 287 | 196 | 154 | 158 | 195 | 394 | 515 | 470 | 431 | 174 | 167 | 162 | 149 | 307,83 | 452 | 163 | 0,359 | -1,477 | -13,979 | 2,09E-44 | 8,86E-44 | 0,0070641 | 0,0063284 | 0,0066696 | 0,0066696 | TRUE | 0,1010311 | No |
| orf19.2914 | 688 | 542 | 637 | 376 | 244 | 191 | 204 | 277 | 626 | 548 | 603 | 565 | 217 | 207 | 209 | 211 | 398,36 | 586 | 211 | 0,36 | -1,473 | -18,067 | 5,81E-73 | 3,78E-72 | 1,00E-08 | 0,0053531 | 0,0032731 | 0,0032731 | TRUE | 0,0268047 | No |
| orf19.1417 | 1139 | 988 | 1012 | 600 | 434 | 309 | 338 | 446 | 1037 | 999 | 958 | 901 | 386 | 335 | 346 | 340 | 662,9 | 974 | 352 | 0,361 | -1,471 | -20,359 | 3,89E-92 | 3,10E-91 | 0,0020712 | 0,0040298 | 0,0031365 | 0,0031365 | TRUE | 0,0498545 | No |
| orf19.6689 | 4526 | 4341 | 4373 | 2606 | 1708 | 1383 | 1514 | 1861 | 4121 | 4390 | 4139 | 3915 | 1519 | 1502 | 1551 | 1420 | 2819,42 | 4141 | 1498 | 0,361 | -1,469 | -29,783 | 6,54E-195 | 1,09E-193 | 0,00146 | 0,0025056 | 0,0018952 | 0,0018952 | TRUE | 0,0198 | No |
| orf19.1136 | 197 | 202 | 173 | 113 | 91 | 49 | 61 | 82 | 179 | 204 | 164 | 170 | 81 | 53 | 62 | 63 | 122,04 | 179 | 65 | 0,363 | -1,463 | -9,863 | 6,04E-23 | 1,64E-22 | 0,0070667 | 0,0128616 | 0,0110093 | 0,0110093 | TRUE | 0,2554803 | No |
| orf19.2341 | 5624 | 4982 | 5407 | 3320 | 2102 | 1701 | 1844 | 2322 | 5120 | 5039 | 5117 | 4987 | 1870 | 1847 | 1888 | 1771 | 3454,97 | 5066 | 1844 | 0,364 | -1,46 | -38,808 | 0 | 0 | 1,00E-08 | 0,0024194 | 0,0009987 | 0,0009987 | TRUE | 0,009186 | No |
| orf19.2067 | 4618 | 4025 | 4257 | 2472 | 1761 | 1202 | 1462 | 1922 | 4204 | 4071 | 4029 | 3713 | 1566 | 1305 | 1497 | 1466 | 2731,54 | 4004 | 1458 | 0,364 | -1,457 | -23,897 | 3,27E-126 | 3,52E-125 | 0,0036034 | 0,0025206 | 0,0031141 | 0,0031141 | TRUE | 0,0586952 | No |
| orf19.3758 | 4932 | 4944 | 5002 | 3324 | 1888 | 1752 | 1709 | 2205 | 4490 | 5000 | 4734 | 4993 | 1679 | 1902 | 1750 | 1682 | 3278,94 | 4804 | 1753 | 0,365 | -1,455 | -26,424 | 7,19E-154 | 9,57E-153 | 0,0025379 | 0,0024399 | 0,0025321 | 0,0025321 | TRUE | 0,0386108 | No |
| orf19.2203.1 | 59 | 56 | 55 | 43 | 20 | 24 | 16 | 29 | 54 | 57 | 52 | 65 | 18 | 26 | 16 | 22 | 38,67 | 57 | 20 | 0,365 | -1,455 | -6,128 | 8,87E-10 | 1,66E-09 | 1,00E-08 | 0,0361984 | 0,0220197 | 0,0220197 | TRUE | 0,144518 | No |
| orf19.6373 | 8204 | 6875 | 7796 | 4820 | 2847 | 2379 | 2644 | 3637 | 7469 | 6953 | 7378 | 7240 | 2532 | 2583 | 2708 | 2774 | 4954,84 | 7260 | 2649 | 0,365 | -1,454 | -33,389 | 2,00E-244 | 4,08E-243 | 0,0010199 | 0,0023036 | 0,0015704 | 0,0015704 | TRUE | 0,0127892 | No |
| orf19.2638.1 | 2999 | 2620 | 2603 | 1476 | 1205 | 759 | 856 | 1208 | 2730 | 2650 | 2464 | 2217 | 1072 | 824 | 877 | 921 | 1719,37 | 2515 | 924 | 0,367 | -1,445 | -17,047 | 3,69E-65 | 2,18E-64 | 0,009782 | 0,0028053 | 0,0061849 | 0,0061849 | TRUE | 0,1395536 | No |
| orf19.1367 | 162 | 186 | 177 | 96 | 79 | 50 | 67 | 61 | 147 | 188 | 168 | 144 | 70 | 54 | 69 | 47 | 110,88 | 162 | 60 | 0,367 | -1,445 | -8,935 | 4,07E-19 | 1,01E-18 | 0,0140837 | 0,0139508 | 0,0139905 | 0,0139905 | TRUE | 0,2415641 | No |
| orf19.7007 | 23 | 17 | 21 | 12 | 7 | 7 | 11 | 5 | 21 | 17 | 20 | 18 | 6 | 8 | 11 | 4 | 13,12 | 19 | 7 | 0,368 | -1,441 | -3,524 | 0,000426 | 0,0006149 | 1,00E-08 | 0,1027415 | 0,0664841 | 0,0664841 | TRUE | 0,3900042 | No |
| orf19.3670 | 6097 | 5715 | 6019 | 3658 | 2338 | 1936 | 2039 | 2672 | 5551 | 5780 | 5696 | 5495 | 2080 | 2102 | 2088 | 2038 | 3853,81 | 5630 | 2077 | 0,369 | -1,44 | -39,515 | 0 | 0 | 1,00E-08 | 0,0023798 | 0,0009548 | 0,0009548 | TRUE | 0,0038484 | No |
| orf19.4370 | 2691 | 2490 | 2641 | 1615 | 1019 | 859 | 899 | 1179 | 2450 | 2518 | 2499 | 2426 | 906 | 933 | 921 | 899 | 1694,11 | 2473 | 915 | 0,369 | -1,437 | -32,749 | 3,11E-235 | 6,14E-234 | 1,00E-08 | 0,0028168 | 0,0011223 | 0,0011223 | TRUE | 0,001992 | No |
| orf19.2006.1 | 2255 | 1972 | 1955 | 1188 | 880 | 650 | 619 | 946 | 2053 | 1994 | 1850 | 1785 | 783 | 706 | 634 | 722 | 1315,79 | 1920 | 711 | 0,37 | -1,433 | -20,324 | 7,96E-92 | 6,31E-91 | 0,0045156 | 0,003041 | 0,0038405 | 0,0038405 | TRUE | 0,0618009 | No |
| orf19.5894 | 30 | 25 | 27 | 22 | 5 | 9 | 16 | 14 | 27 | 25 | 26 | 33 | 4 | 10 | 16 | 11 | 19,06 | 28 | 10 | 0,371 | -1,432 | -4,045 | 5,22E-05 | 7,95E-05 | 1,00E-08 | 0,0713429 | 0,0554324 | 0,0554324 | TRUE | 0,5837237 | No |
| orf19.100 | 2501 | 2105 | 2433 | 1417 | 942 | 742 | 854 | 1006 | 2277 | 2129 | 2303 | 2129 | 838 | 806 | 875 | 767 | 1515,34 | 2210 | 822 | 0,371 | -1,431 | -25,443 | 8,48E-143 | 1,05E-141 | 0,0016037 | 0,0029088 | 0,0022237 | 0,0022237 | TRUE | 0,0252233 | No |
| orf19.4779 | 7221 | 6279 | 6822 | 4356 | 2643 | 2165 | 2381 | 3243 | 6574 | 6351 | 6456 | 6543 | 2351 | 2351 | 2438 | 2474 | 4442,31 | 6481 | 2404 | 0,371 | -1,43 | -39,855 | 0 | 0 | 0,0001544 | 0,0023344 | 0,0009593 | 0,0009593 | TRUE | 0,0050102 | No |
| orf19.6134 | 805 | 720 | 793 | 506 | 313 | 281 | 253 | 347 | 733 | 728 | 751 | 760 | 278 | 305 | 259 | 265 | 509,88 | 743 | 277 | 0,372 | -1,427 | -19,769 | 5,48E-87 | 4,21E-86 | 1,00E-08 | 0,0046278 | 0,0025941 | 0,0025941 | TRUE | 0,0481022 | No |
| orf19.3021 | 2105 | 1788 | 1903 | 1197 | 760 | 658 | 634 | 907 | 1917 | 1808 | 1801 | 1798 | 676 | 714 | 649 | 692 | 1256,94 | 1831 | 683 | 0,373 | -1,424 | -27,094 | 1,15E-161 | 1,60E-160 | 1,00E-08 | 0,003088 | 0,0016765 | 0,0016765 | TRUE | 0,0124268 | No |
| orf19.592 | 2635 | 2036 | 2442 | 1536 | 965 | 737 | 827 | 1150 | 2399 | 2059 | 2311 | 2307 | 858 | 800 | 847 | 877 | 1557,42 | 2269 | 846 | 0,373 | -1,422 | -23,715 | 2,52E-124 | 2,68E-123 | 0,0024863 | 0,0028852 | 0,0026646 | 0,0026646 | TRUE | 0,0478467 | No |
| orf19.6440 | 6923 | 5801 | 6557 | 3893 | 2422 | 1993 | 2284 | 3134 | 6303 | 5867 | 6206 | 5848 | 2154 | 2164 | 2339 | 2391 | 4158,95 | 6056 | 2262 | 0,374 | -1,421 | -28,811 | 1,54E-182 | 2,41E-181 | 0,0017578 | 0,0023546 | 0,0020392 | 0,0020392 | TRUE | 0,0184579 | No |
| orf19.1336.2 | 1645 | 1464 | 1435 | 728 | 624 | 448 | 474 | 661 | 1498 | 1481 | 1358 | 1094 | 555 | 486 | 485 | 504 | 932,64 | 1358 | 508 | 0,373 | -1,421 | -15,099 | 1,64E-51 | 7,83E-51 | 0,0112028 | 0,0034534 | 0,0071905 | 0,0071905 | TRUE | 0,1959215 | No |
| orf19.409 | 53612 | 43511 | 47000 | 30771 | 19595 | 15689 | 16259 | 22959 | 48811 | 44007 | 44481 | 46223 | 17429 | 17034 | 16651 | 17513 | 31518,75 | 45880 | 17157 | 0,374 | -1,419 | -34,321 | 3,76E-258 | 8,12E-257 | 0,0013363 | 0,0020789 | 0,0016032 | 0,0016032 | TRUE | 0,0227492 | No |
| orf19.1691 | 406 | 373 | 376 | 268 | 164 | 140 | 126 | 177 | 370 | 377 | 356 | 403 | 146 | 152 | 129 | 135 | 258,41 | 376 | 140 | 0,374 | -1,419 | -14,698 | 6,62E-49 | 2,99E-48 | 1,00E-08 | 0,0071491 | 0,004211 | 0,004211 | TRUE | 0,0278902 | No |
| orf19.3192 | 6510 | 6062 | 6419 | 4334 | 2554 | 2235 | 2200 | 2973 | 5927 | 6131 | 6075 | 6510 | 2272 | 2427 | 2253 | 2268 | 4232,84 | 6161 | 2305 | 0,374 | -1,418 | -32,02 | 5,67E-225 | 1,07E-223 | 0,0010417 | 0,0023491 | 0,0015931 | 0,0015931 | TRUE | 0,0171646 | No |
| orf19.3406 | 1231 | 1091 | 1167 | 754 | 472 | 365 | 415 | 559 | 1121 | 1103 | 1104 | 1133 | 420 | 396 | 425 | 426 | 766,1 | 1115 | 417 | 0,375 | -1,417 | -24,257 | 5,54E-130 | 6,13E-129 | 1,00E-08 | 0,0037613 | 0,0016808 | 0,0016808 | TRUE | 0,0116315 | No |
| orf19.1488 | 460 | 504 | 478 | 317 | 210 | 165 | 160 | 217 | 419 | 510 | 452 | 476 | 187 | 179 | 164 | 166 | 319,06 | 464 | 174 | 0,375 | -1,416 | -14,872 | 5,03E-50 | 2,32E-49 | 0,0032606 | 0,0061774 | 0,0048651 | 0,0048651 | TRUE | 0,0543288 | No |
| orf19.4691 | 183 | 170 | 192 | 107 | 75 | 60 | 65 | 77 | 167 | 172 | 182 | 161 | 67 | 65 | 67 | 59 | 117,27 | 170 | 64 | 0,375 | -1,414 | -10,453 | 1,42E-25 | 4,13E-25 | 1,00E-08 | 0,0133017 | 0,0072021 | 0,0072021 | TRUE | 0,0368536 | No |
| orf19.5526 | 130 | 115 | 119 | 90 | 56 | 48 | 42 | 48 | 118 | 116 | 113 | 135 | 50 | 52 | 43 | 37 | 83 | 120 | 46 | 0,375 | -1,414 | -8,496 | 1,97E-17 | 4,67E-17 | 1,00E-08 | 0,0179517 | 0,0118085 | 0,0118085 | TRUE | 0,1673476 | No |
| orf19.6639 | 3986 | 3582 | 3768 | 2397 | 1527 | 1266 | 1324 | 1767 | 3629 | 3623 | 3566 | 3601 | 1358 | 1375 | 1356 | 1348 | 2481,9 | 3605 | 1359 | 0,377 | -1,408 | -37,411 | 2,63E-306 | 6,94E-305 | 1,00E-08 | 0,0025693 | 0,0008674 | 0,0008674 | TRUE | 0,0006714 | No |

| | | | | | | | | | | | | | | | | | | | | | | | | | | | | | | | |
|---|---|---|---|---|---|---|---|---|---|---|---|---|---|---|---|---|---|---|---|---|---|---|---|---|---|---|---|---|---|---|---|
| orf19.2620 | 262 | 203 | 255 | 184 | 99 | 75 | 81 | 139 | 239 | 205 | 241 | 276 | 88 | 81 | 83 | 106 | 165,01 | 240 | 90 | 0,377 | -1,406 | -10,64 | 1,95E-26 | 5,76E-26 | 0,0082819 | 0,0100427 | 0,0093596 | 0,0093596 | TRUE | 0,1655333 | No |
| orf19.904 | 1222 | 1124 | 1165 | 747 | 496 | 371 | 415 | 548 | 1113 | 1137 | 1103 | 1122 | 441 | 403 | 425 | 418 | 770,13 | 1119 | 422 | 0,377 | -1,405 | -23,96 | 7,32E-127 | 7,95E-126 | 1,00E-08 | 0,0037523 | 0,001723 | 0,001723 | TRUE | 0,0112073 | No |
| orf19.4888 | 3555 | 3353 | 3399 | 2097 | 1432 | 1072 | 1200 | 1629 | 3237 | 3391 | 3217 | 3150 | 1274 | 1164 | 1229 | 1243 | 2237,99 | 3249 | 1228 | 0,378 | -1,404 | -30,518 | 1,51E-204 | 2,64E-203 | 0,0005914 | 0,0026273 | 0,0014858 | 0,0014858 | TRUE | 0,0137943 | No |
| orf19.1112 | 1208 | 1156 | 1244 | 727 | 477 | 451 | 394 | 529 | 1100 | 1169 | 1177 | 1092 | 424 | 490 | 404 | 404 | 782,42 | 1134 | 430 | 0,378 | -1,404 | -19,943 | 1,71E-88 | 1,33E-87 | 0,0024824 | 0,0037254 | 0,0032005 | 0,0032005 | TRUE | 0,0940468 | No |
| orf19.430 | 2152 | 1795 | 1993 | 1257 | 772 | 634 | 751 | 934 | 1959 | 1815 | 1886 | 1888 | 687 | 688 | 769 | 712 | 1300,72 | 1887 | 714 | 0,378 | -1,403 | -25,837 | 3,40E-147 | 4,39E-146 | 0,0006502 | 0,0030526 | 0,0018928 | 0,0018928 | TRUE | 0,0301427 | No |
| orf19.48 | 4019 | 4635 | 4748 | 3005 | 1737 | 1661 | 1636 | 2022 | 3659 | 4688 | 4494 | 4514 | 1545 | 1803 | 1675 | 1542 | 2990,09 | 4339 | 1641 | 0,378 | -1,403 | -17,658 | 8,87E-70 | 5,54E-69 | 0,0079682 | 0,0024788 | 0,0056522 | 0,0056522 | TRUE | 0,1366825 | No |
| orf19.2478 | 94 | 75 | 77 | 34 | 31 | 28 | 23 | 37 | 86 | 76 | 73 | 51 | 28 | 30 | 24 | 28 | 49,39 | 72 | 28 | 0,38 | -1,398 | -6,372 | 1,87E-10 | 3,59E-10 | 1,00E-08 | 0,0287821 | 0,0218795 | 0,0218795 | TRUE | 0,2483254 | No |
| orf19.1670 | 1433 | 1482 | 1608 | 860 | 616 | 545 | 518 | 617 | 1305 | 1499 | 1522 | 1292 | 548 | 592 | 530 | 471 | 969,75 | 1404 | 535 | 0,38 | -1,397 | -17,031 | 4,84E-65 | 2,84E-64 | 0,0070884 | 0,0033992 | 0,0051995 | 0,0051995 | TRUE | 0,0816683 | No |
| orf19.3439 | 3312 | 3561 | 3734 | 2208 | 1468 | 1209 | 1233 | 1627 | 3015 | 3602 | 3534 | 3317 | 1306 | 1313 | 1263 | 1241 | 2323,74 | 3367 | 1281 | 0,38 | -1,395 | -22,864 | 1,07E-115 | 1,05E-114 | 0,0033835 | 0,0026055 | 0,0030504 | 0,0030504 | TRUE | 0,0617316 | No |
| orf19.4841 | 3360 | 2828 | 3135 | 1863 | 1243 | 975 | 1121 | 1498 | 3059 | 2860 | 2967 | 2799 | 1106 | 1059 | 1148 | 1143 | 2017,47 | 2921 | 1114 | 0,381 | -1,392 | -28,484 | 1,85E-178 | 2,81E-177 | 0,0009077 | 0,0026918 | 0,0016888 | 0,0016888 | TRUE | 0,0125775 | No |
| orf19.3071 | 7979 | 6977 | 7846 | 5231 | 2888 | 2697 | 2888 | 3741 | 7265 | 7056 | 7426 | 7858 | 2569 | 2928 | 2958 | 2854 | 5114,08 | 7401 | 2827 | 0,382 | -1,388 | -25,681 | 1,91E-145 | 2,43E-144 | 0,0027035 | 0,0022953 | 0,0025688 | 0,0025688 | TRUE | 0,0461418 | No |
| orf19.999 | 6022 | 4984 | 5414 | 3877 | 2170 | 2003 | 1877 | 2848 | 5483 | 5041 | 5124 | 5824 | 1930 | 2175 | 1922 | 2172 | 3708,86 | 5368 | 2050 | 0,382 | -1,388 | -22,184 | 4,90E-109 | 4,59E-108 | 0,0042214 | 0,0023932 | 0,0034302 | 0,0034302 | TRUE | 0,0389925 | No |
| orf19.668 | 978 | 964 | 976 | 762 | 403 | 349 | 382 | 485 | 890 | 975 | 924 | 1145 | 358 | 379 | 391 | 370 | 679,04 | 984 | 374 | 0,382 | -1,388 | -16,56 | 1,36E-61 | 7,60E-61 | 0,005945 | 0,0039824 | 0,0049501 | 0,0049501 | TRUE | 0,135631 | No |
| orf19.1210 | 4505 | 4400 | 4617 | 2773 | 1838 | 1504 | 1575 | 2188 | 4102 | 4450 | 4370 | 4165 | 1635 | 1633 | 1613 | 1669 | 2954,57 | 4272 | 1638 | 0,383 | -1,383 | -32,164 | 5,56E-227 | 1,06E-225 | 0,0006946 | 0,0024841 | 0,0013647 | 0,0013647 | TRUE | 0,0096506 | No |
| orf19.1611 | 328 | 358 | 337 | 215 | 146 | 131 | 120 | 141 | 299 | 362 | 319 | 323 | 130 | 142 | 123 | 108 | 225,64 | 326 | 126 | 0,384 | -1,382 | -12,352 | 4,73E-35 | 1,69E-34 | 0,0047434 | 0,0078913 | 0,006645 | 0,006645 | TRUE | 0,1102604 | No |
| orf19.2408 | 1691 | 1617 | 1604 | 1030 | 689 | 594 | 564 | 744 | 1540 | 1635 | 1518 | 1547 | 613 | 645 | 578 | 568 | 1080,4 | 1560 | 601 | 0,384 | -1,38 | -23,786 | 4,62E-125 | 4,94E-124 | 0,0007383 | 0,0032597 | 0,0021078 | 0,0021078 | TRUE | 0,027203 | No |
| orf19.7468 | 3510 | 3095 | 3249 | 2171 | 1314 | 1166 | 1153 | 1637 | 3196 | 3130 | 3075 | 3261 | 1169 | 1266 | 1181 | 1249 | 2190,79 | 3166 | 1216 | 0,384 | -1,379 | -30,421 | 2,89E-203 | 5,02E-202 | 0,0003587 | 0,00264 | 0,0014199 | 0,0014199 | TRUE | 0,0084857 | No |
| orf19.6706 | 12393 | 11890 | 12588 | 6971 | 5002 | 4017 | 4441 | 5536 | 11283 | 12025 | 11913 | 10472 | 4449 | 4361 | 4548 | 4223 | 7909,4 | 11423 | 4395 | 0,385 | -1,379 | -27,22 | 3,76E-163 | 5,31E-162 | 0,0022347 | 0,002204 | 0,0023108 | 0,0023108 | TRUE | 0,0378264 | No |
| orf19.2023 | 27155 | 21870 | 24432 | 15567 | 10013 | 8183 | 8589 | 12220 | 24723 | 22119 | 23123 | 23384 | 8906 | 8884 | 8796 | 9322 | 16157,23 | 23337 | 8977 | 0,385 | -1,378 | -32,814 | 3,77E-236 | 7,46E-235 | 0,0012951 | 0,0021188 | 0,0016193 | 0,0016193 | TRUE | 0,0197121 | No |
| orf19.4270 | 1055 | 1112 | 1134 | 710 | 457 | 406 | 401 | 487 | 961 | 1125 | 1073 | 1067 | 406 | 441 | 411 | 371 | 731,8 | 1056 | 407 | 0,385 | -1,378 | -18,666 | 9,40E-78 | 6,46E-77 | 0,0032925 | 0,0038421 | 0,0035746 | 0,0035746 | TRUE | 0,0467932 | No |
| orf19.1482 | 1737 | 1310 | 1469 | 792 | 661 | 442 | 525 | 667 | 1581 | 1325 | 1390 | 1190 | 588 | 480 | 538 | 509 | 950,08 | 1372 | 529 | 0,385 | -1,378 | -15,348 | 3,67E-53 | 1,80E-52 | 0,0091085 | 0,0034274 | 0,0064617 | 0,0064617 | TRUE | 0,1279055 | No |
| orf19.6874 | 3680 | 3316 | 3663 | 2503 | 1459 | 1219 | 1342 | 1783 | 3350 | 3354 | 3467 | 3760 | 1298 | 1323 | 1374 | 1360 | 2410,82 | 3483 | 1339 | 0,385 | -1,377 | -27,304 | 3,78E-164 | 5,41E-163 | 0,0014698 | 0,002585 | 0,0019388 | 0,0019388 | TRUE | 0,0331378 | No |
| orf19.3529 | 231 | 221 | 249 | 152 | 95 | 89 | 85 | 104 | 210 | 224 | 236 | 228 | 84 | 97 | 87 | 79 | 155,67 | 224 | 87 | 0,385 | -1,376 | -11,489 | 1,50E-30 | 4,91E-30 | 1,00E-08 | 0,0105231 | 0,006016 | 0,006016 | TRUE | 0,0504482 | No |
| orf19.420 | 32 | 23 | 12 | 13 | 4 | 8 | 9 | 14 | 29 | 23 | 11 | 20 | 4 | 9 | 9 | 11 | 14,43 | 21 | 8 | 0,385 | -1,375 | -3,213 | 0,001314 | 0,0018415 | 0,0903168 | 0,0935967 | 0,0925558 | 0,0925558 | TRUE | 0,2225741 | No |
| orf19.3158 | 2542 | 2333 | 2520 | 1614 | 1030 | 902 | 877 | 1146 | 2314 | 2360 | 2385 | 2424 | 916 | 979 | 898 | 874 | 1643,9 | 2371 | 917 | 0,386 | -1,372 | -28,118 | 5,87E-174 | 8,92E-173 | 0,0002365 | 0,0028406 | 0,0015498 | 0,0015498 | TRUE | 0,0237923 | No |
| orf19.4784 | 1763 | 1507 | 1579 | 1077 | 686 | 555 | 591 | 779 | 1605 | 1524 | 1494 | 1618 | 610 | 603 | 605 | 594 | 1081,72 | 1560 | 603 | 0,387 | -1,37 | -25,412 | 1,85E-142 | 2,28E-141 | 1,00E-08 | 0,0032582 | 0,0016752 | 0,0016752 | TRUE | 0,0098927 | No |
| orf19.3672 | 6920 | 6218 | 6519 | 4217 | 2670 | 2371 | 2307 | 3160 | 6300 | 6289 | 6170 | 6335 | 2375 | 2574 | 2363 | 2410 | 4351,96 | 6274 | 2430 | 0,387 | -1,369 | -35,396 | 1,98E-274 | 4,59E-273 | 0,0003103 | 0,0023406 | 0,0011572 | 0,0011572 | TRUE | 0,0183764 | No |
| orf19.3414 | 9053 | 8538 | 9260 | 5360 | 3772 | 2981 | 3111 | 4284 | 8242 | 8635 | 8764 | 8052 | 3355 | 3237 | 3186 | 3268 | 5842,31 | 8423 | 3262 | 0,387 | -1,369 | -33,762 | 7,08E-250 | 1,47E-248 | 0,0007325 | 0,0022631 | 0,001372 | 0,001372 | TRUE | 0,0103995 | No |
| orf19.6586 | 3178 | 3100 | 3298 | 2015 | 1402 | 1138 | 1155 | 1385 | 2893 | 3135 | 3121 | 3027 | 1247 | 1236 | 1183 | 1056 | 2112,35 | 3044 | 1180 | 0,387 | -1,369 | -23,243 | 1,68E-119 | 1,70E-118 | 0,0026184 | 0,0026624 | 0,0027561 | 0,0027561 | TRUE | 0,063268 | No |
| orf19.3984 | 11600 | 12418 | 12609 | 7504 | 5112 | 4335 | 4367 | 5604 | 10561 | 12559 | 11933 | 11272 | 4547 | 4707 | 4472 | 4275 | 8040,87 | 11581 | 4500 | 0,388 | -1,364 | -23,97 | 5,78E-127 | 6,28E-126 | 0,0033418 | 0,0022013 | 0,0029611 | 0,0029611 | TRUE | 0,0434361 | No |
| orf19.6534 | 7 | 6 | 9 | 9 | 5 | 4 | 2 | 3 | 6 | 6 | 9 | 14 | 4 | 4 | 2 | 2 | 5,95 | 9 | 3 | 0,389 | -1,364 | -2,257 | 0,024039 | 0,0306134 | 1,00E-08 | 0,2240274 | 0,144074 | 0,144074 | TRUE | 0,2913987 | No |
| orf19.864 | 2896 | 2773 | 3084 | 1874 | 1188 | 1000 | 1092 | 1426 | 2637 | 2805 | 2919 | 2815 | 1057 | 1086 | 1118 | 1088 | 1940,45 | 2794 | 1087 | 0,389 | -1,362 | -28,096 | 1,08E-173 | 1,64E-172 | 0,0008858 | 0,0027178 | 0,0016333 | 0,0016333 | TRUE | 0,0181775 | No |
| orf19.6820 | 4509 | 3916 | 4405 | 2884 | 1717 | 1512 | 1615 | 2147 | 4105 | 3961 | 4169 | 4332 | 1527 | 1642 | 1654 | 1638 | 2878,45 | 4142 | 1615 | 0,39 | -1,358 | -29,675 | 1,63E-193 | 2,68E-192 | 0,0009488 | 0,0024959 | 0,0015914 | 0,0015914 | TRUE | 0,016423 | No |
| orf19.1881 | 1402 | 1276 | 1230 | 835 | 499 | 439 | 515 | 664 | 1276 | 1291 | 1164 | 1254 | 444 | 477 | 527 | 507 | 867,47 | 1246 | 489 | 0,392 | -1,351 | -20,143 | 3,09E-90 | 2,42E-89 | 0,0022202 | 0,0035598 | 0,0029312 | 0,0029312 | TRUE | 0,0459471 | No |
| orf19.1826 | 20168 | 20208 | 21038 | 13363 | 8410 | 7798 | 7413 | 9735 | 18362 | 20438 | 19911 | 20073 | 7480 | 8466 | 7592 | 7426 | 13718,62 | 19696 | 7741 | 0,393 | -1,348 | -25,467 | 4,58E-143 | 5,71E-142 | 0,0029526 | 0,0021333 | 0,0026019 | 0,0026019 | TRUE | 0,048322 | No |
| orf19.5030 | 1773 | 1568 | 1688 | 949 | 701 | 539 | 610 | 809 | 1614 | 1586 | 1598 | 1426 | 624 | 585 | 625 | 617 | 1084,22 | 1556 | 613 | 0,393 | -1,347 | -22,977 | 8,01E-117 | 7,98E-116 | 0,0011617 | 0,0032554 | 0,0021947 | 0,0021947 | TRUE | 0,0346412 | No |
| orf19.6913 | 1222 | 1221 | 1248 | 692 | 490 | 447 | 449 | 555 | 1113 | 1235 | 1181 | 1039 | 436 | 485 | 460 | 423 | 796,56 | 1142 | 451 | 0,393 | -1,347 | -18,695 | 5,42E-78 | 3,75E-77 | 0,0032397 | 0,0036954 | 0,0034752 | 0,0034752 | TRUE | 0,0396308 | No |
| orf19.524 | 1362 | 1123 | 1324 | 757 | 545 | 356 | 504 | 636 | 1240 | 1136 | 1253 | 1137 | 485 | 387 | 516 | 485 | 829,83 | 1192 | 468 | 0,393 | -1,347 | -16,621 | 4,94E-62 | 2,78E-61 | 0,0066992 | 0,0036289 | 0,0048549 | 0,0048549 | TRUE | 0,1507804 | No |
| orf19.1240 | 9978 | 9008 | 9729 | 6093 | 4028 | 3348 | 3596 | 4582 | 9085 | 9111 | 9208 | 9153 | 3583 | 3635 | 3683 | 3495 | 6368,9 | 9139 | 3599 | 0,394 | -1,345 | -42,439 | 0 | 0 | 8,11E-06 | 0,0022444 | 0,000776 | 0,000776 | TRUE | 0,0047787 | No |
| orf19.2351 | 1290 | 1159 | 1164 | 827 | 529 | 387 | 432 | 661 | 1174 | 1172 | 1102 | 1242 | 471 | 420 | 442 | 504 | 815,99 | 1172 | 459 | 0,394 | -1,345 | -19,462 | 2,30E-84 | 1,72E-83 | 0,0023073 | 0,0036559 | 0,0031115 | 0,0031115 | TRUE | 0,0532412 | No |
| orf19.5827 | 153 | 154 | 159 | 126 | 66 | 62 | 58 | 82 | 139 | 156 | 150 | 189 | 59 | 67 | 59 | 63 | 110,35 | 158 | 62 | 0,394 | -1,345 | -9,107 | 8,49E-20 | 2,14E-19 | 1,00E-08 | 0,0140083 | 0,0100178 | 0,0100178 | TRUE | 0,1466972 | No |
| orf19.5846 | 3361 | 2888 | 3216 | 2093 | 1342 | 1135 | 1134 | 1576 | 3060 | 2921 | 3044 | 3144 | 1194 | 1232 | 1161 | 1202 | 2119,77 | 3042 | 1197 | 0,394 | -1,344 | -30,57 | 3,03E-205 | 5,31E-204 | 1,00E-08 | 0,0026602 | 0,00129 | 0,00129 | TRUE | 0,0086477 | No |
| orf19.3027 | 5752 | 5138 | 5548 | 3635 | 2306 | 2008 | 2017 | 2692 | 5237 | 5197 | 5251 | 5460 | 2051 | 2180 | 2066 | 2053 | 3686,86 | 5286 | 2088 | 0,395 | -1,341 | -34,72 | 3,98E-264 | 8,88E-263 | 0,0002442 | 0,0023953 | 0,0011055 | 0,0011055 | TRUE | 0,0103406 | No |
| orf19.6992 | 11882 | 10421 | 11239 | 7027 | 4468 | 4065 | 4083 | 5564 | 10818 | 10540 | 10637 | 10556 | 3974 | 4413 | 4182 | 4244 | 7420,44 | 10638 | 4203 | 0,395 | -1,34 | -34,629 | 9,23E-263 | 2,02E-261 | 0,000783 | 0,002215 | 0,0012763 | 0,0012763 | TRUE | 0,0164694 | No |
| orf19.6408 | 6033 | 5022 | 5788 | 3760 | 2244 | 1937 | 2214 | 2887 | 5493 | 5079 | 5478 | 5648 | 1996 | 2103 | 2267 | 2202 | 3783,32 | 5424 | 2142 | 0,395 | -1,34 | -25,761 | 2,44E-146 | 3,13E-145 | 0,0020996 | 0,0023862 | 0,0022806 | 0,0022806 | TRUE | 0,0259508 | No |
| orf19.5258 | 1249 | 1257 | 1330 | 791 | 545 | 459 | 466 | 623 | 1137 | 1271 | 1259 | 1188 | 485 | 498 | 477 | 475 | 848,87 | 1214 | 484 | 0,398 | -1,328 | -21,57 | 3,43E-103 | 3,09E-102 | 1,00E-08 | 0,0035932 | 0,0022362 | 0,0022362 | TRUE | 0,0237392 | No |
| orf19.2241 | 5701 | 4703 | 5235 | 3228 | 2257 | 1831 | 1971 | 2471 | 5191 | 4757 | 4954 | 4849 | 2008 | 1988 | 2019 | 1885 | 3456,18 | 4938 | 1975 | 0,4 | -1,324 | -29,925 | 9,30E-197 | 1,56E-195 | 0,0009813 | 0,0024192 | 0,0015329 | 0,0015329 | TRUE | 0,0147647 | No |
| orf19.932 | 6300 | 5670 | 6128 | 3893 | 2500 | 2188 | 2330 | 2966 | 5736 | 5735 | 5800 | 5848 | 2224 | 2376 | 2386 | 2262 | 4045,74 | 5780 | 2312 | 0,4 | -1,323 | -35,167 | 6,47E-271 | 1,48E-269 | 0,0001417 | 0,0023635 | 0,0010632 | 0,0010632 | TRUE | 0,0078414 | No |
| orf19.3505 | 8757 | 7587 | 8261 | 5476 | 3435 | 2915 | 3117 | 4273 | 7973 | 7673 | 7818 | 8226 | 3055 | 3165 | 3192 | 3259 | 5545,29 | 7922 | 3168 | 0,4 | -1,322 | -34,377 | 5,65E-259 | 1,23E-257 | 0,0005097 | 0,0022752 | 0,0012036 | 0,0012036 | TRUE | 0,0077831 | No |
| orf19.6512 | 1283 | 1249 | 1285 | 731 | 514 | 474 | 447 | 624 | 1168 | 1263 | 1216 | 1098 | 457 | 515 | 458 | 476 | 831,39 | 1186 | 476 | 0,4 | -1,321 | -19,963 | 1,15E-88 | 8,94E-88 | 0,0018202 | 0,0036259 | 0,0027705 | 0,0027705 | TRUE | 0,0313287 | No |
| orf19.215 | 10232 | 8792 | 9481 | 6411 | 4048 | 3352 | 3621 | 4984 | 9316 | 8892 | 8973 | 9630 | 3601 | 3639 | 3708 | 3802 | 6445,17 | 9203 | 3688 | 0,401 | -1,318 | -33,293 | 4,88E-243 | 9,94E-242 | 0,0007121 | 0,002242 | 0,0013207 | 0,0013207 | TRUE | 0,0115689 | No |
| orf19.6062 | 5416 | 4374 | 4552 | 2898 | 2237 | 1555 | 1682 | 2387 | 4931 | 4424 | 4308 | 4353 | 1990 | 1688 | 1723 | 1821 | 3154,7 | 4504 | 1806 | 0,401 | -1,318 | -20,814 | 3,20E-96 | 2,66E-95 | 0,0043376 | 0,0024558 | 0,0034748 | 0,0034748 | TRUE | 0,0570417 | No |
| orf19.4690 | 181 | 158 | 171 | 96 | 62 | 66 | 64 | 83 | 165 | 160 | 162 | 144 | 55 | 72 | 66 | 63 | 110,79 | 158 | 64 | 0,402 | -1,316 | -9,414 | 4,77E-21 | 1,24E-20 | 1,00E-08 | 0,0139607 | 0,0080882 | 0,0080882 | TRUE | 0,0765404 | No |
| orf19.3376 | 135 | 149 | 134 | 89 | 51 | 48 | 60 | 73 | 123 | 151 | 127 | 134 | 45 | 52 | 61 | 56 | 93,59 | 134 | 54 | 0,402 | -1,316 | -8,428 | 3,52E-17 | 8,27E-17 | 1,00E-08 | 0,0161516 | 0,0107153 | 0,0107153 | TRUE | 0,0966735 | No |
| orf19.6277 | 50 | 46 | 31 | 26 | 13 | 11 | 22 | 24 | 46 | 47 | 29 | 39 | 12 | 12 | 23 | 18 | 28,1 | 40 | 16 | 0,402 | -1,316 | -4,462 | 8,12E-06 | 1,30E-05 | 0,0258317 | 0,0490505 | 0,0413289 | 0,0413289 | TRUE | 0,3087043 | No |
| orf19.5524 | 49 | 58 | 56 | 37 | 22 | 19 | 16 | 36 | 45 | 59 | 53 | 56 | 20 | 21 | 16 | 27 | 36,99 | 53 | 21 | 0,404 | -1,309 | -5,366 | 8,05E-08 | 1,40E-07 | 1,00E-08 | 0,0377522 | 0,0252066 | 0,0252066 | TRUE | 0,3036038 | No |
| orf19.3742 | 808 | 655 | 702 | 485 | 325 | 252 | 285 | 358 | 736 | 662 | 664 | 729 | 289 | 274 | 292 | 273 | 489,83 | 698 | 282 | 0,404 | -1,306 | -17,493 | 1,63E-68 | 1,00E-67 | 1,00E-08 | 0,0047338 | 0,0029338 | 0,0029338 | TRUE | 0,018319 | No |
| orf19.29 | 5432 | 5613 | 5711 | 3754 | 2476 | 2185 | 2161 | 2611 | 4946 | 5677 | 5405 | 5639 | 2202 | 2372 | 2213 | 1992 | 3805,76 | 5417 | 2195 | 0,405 | -1,304 | -20,924 | 3,27E-97 | 2,77E-96 | 0,0040312 | 0,0023841 | 0,0034174 | 0,0034174 | TRUE | 0,048073 | No |
| orf19.2959.1 | 2842 | 2460 | 2334 | 1436 | 1155 | 811 | 930 | 1267 | 2588 | 2488 | 2209 | 2157 | 1027 | 881 | 952 | 966 | 1658,54 | 2360 | 956 | 0,405 | -1,303 | -18,291 | 9,85E-75 | 6,51E-74 | 0,0049599 | 0,0028335 | 0,0041583 | 0,0041583 | TRUE | 0,0510965 | No |
| orf19.7594 | 707 | 701 | 683 | 430 | 283 | 254 | 272 | 351 | 644 | 709 | 646 | 646 | 252 | 276 | 279 | 268 | 464,85 | 661 | 269 | 0,405 | -1,303 | -17,38 | 1,18E-67 | 7,15E-67 | 1,00E-08 | 0,0048788 | 0,0028553 | 0,0028553 | TRUE | 0,028506 | No |
| orf19.212 | 1767 | 1562 | 1724 | 1180 | 764 | 643 | 622 | 863 | 1609 | 1580 | 1632 | 1773 | 680 | 698 | 637 | 658 | 1158,21 | 1648 | 668 | 0,406 | -1,3 | -22,72 | 2,83E-114 | 2,77E-113 | 0,0010871 | 0,0031776 | 0,0021189 | 0,0021189 | TRUE | 0,0283826 | No |
| orf19.4939 | 382 | 436 | 419 | 275 | 171 | 158 | 174 | 195 | 348 | 441 | 397 | 413 | 152 | 172 | 178 | 149 | 281,12 | 400 | 163 | 0,406 | -1,3 | -12,39 | 2,97E-35 | 1,07E-34 | 0,0058642 | 0,006736 | 0,0063529 | 0,0063529 | TRUE | 0,0930816 | No |
| orf19.1286 | 144 | 88 | 116 | 67 | 48 | 50 | 35 | 57 | 131 | 89 | 110 | 101 | 43 | 54 | 36 | 43 | 75,86 | 108 | 44 | 0,406 | -1,299 | -7,001 | 2,53E-12 | 5,20E-12 | 0,0144143 | 0,0194517 | 0,0175037 | 0,0175037 | TRUE | 0,2284314 | No |
| orf19.903 | 13840 | 11224 | 11723 | 8984 | 5766 | 4393 | 4576 | 6741 | 12601 | 11352 | 11095 | 13495 | 5129 | 4770 | 4686 | 5142 | 8533,69 | 12136 | 4932 | 0,407 | -1,298 | -19,992 | 6,51E-89 | 5,07E-88 | 0,00523 | 0,0021918 | 0,0039127 | 0,0039127 | TRUE | 0,0689544 | No |
| orf19.4837.1 | 666 | 577 | 612 | 319 | 270 | 176 | 257 | 292 | 606 | 584 | 579 | 479 | 240 | 191 | 263 | 223 | 395,69 | 562 | 229 | 0,407 | -1,298 | -12,597 | 2,19E-36 | 8,06E-36 | 0,0097461 | 0,0053755 | 0,0072063 | 0,0072063 | TRUE | 0,126622 | No |

| | | | | | | | | | | | | | | | | | | | | | | | | | | | | | | | |
|---|---|---|---|---|---|---|---|---|---|---|---|---|---|---|---|---|---|---|---|---|---|---|---|---|---|---|---|---|---|---|---|
| orf19.940 | 3815 | 4108 | 4219 | 2433 | 1796 | 1537 | 1482 | 1885 | 3473 | 4155 | 3993 | 3655 | 1597 | 1669 | 1518 | 1438 | 2687,22 | 3819 | 1556 | 0,407 | -1,297 | -19,281 | 7,79E-83 | 5,76E-82 | 0,0047602 | 0,0025286 | 0,0039053 | 0,0039053 | TRUE | 0,0462839 | No |
| orf19.7215.3 | 2742 | 2200 | 2368 | 1428 | 1114 | 795 | 857 | 1286 | 2496 | 2225 | 2241 | 2145 | 991 | 863 | 878 | 981 | 1602,55 | 2277 | 928 | 0,408 | -1,294 | -19,539 | 5,07E-85 | 3,83E-84 | 0,0038361 | 0,0028613 | 0,0034699 | 0,0034699 | TRUE | 0,0516544 | No |
| orf19.5751 | 4615 | 4492 | 4752 | 3051 | 1973 | 1812 | 1738 | 2327 | 4202 | 4543 | 4497 | 4583 | 1755 | 1967 | 1780 | 1775 | 3137,82 | 4456 | 1819 | 0,408 | -1,293 | -25,743 | 3,85E-146 | 4,93E-145 | 0,0017197 | 0,002458 | 0,0020445 | 0,0020445 | TRUE | 0,0351896 | No |
| orf19.7434 | 1040 | 1065 | 1070 | 627 | 446 | 361 | 430 | 522 | 947 | 1077 | 1013 | 942 | 397 | 392 | 440 | 398 | 700,72 | 995 | 407 | 0,408 | -1,293 | -18,403 | 1,25E-75 | 8,34E-75 | 0,002117 | 0,0039222 | 0,0030519 | 0,0030519 | TRUE | 0,0371239 | No |
| orf19.7342 | 795 | 813 | 882 | 568 | 359 | 353 | 295 | 415 | 724 | 822 | 835 | 853 | 319 | 383 | 302 | 317 | 569,41 | 808 | 330 | 0,408 | -1,292 | -15,148 | 7,86E-52 | 3,75E-51 | 0,0056894 | 0,0043569 | 0,0049076 | 0,0049076 | TRUE | 0,1242072 | No |
| orf19.2498 | 11072 | 9143 | 10296 | 6861 | 4408 | 3516 | 4079 | 5490 | 10081 | 9247 | 9744 | 10306 | 3921 | 3817 | 4177 | 4188 | 6935,21 | 9844 | 4026 | 0,409 | -1,289 | -26,59 | 8,80E-156 | 1,20E-154 | 0,0019265 | 0,0022275 | 0,0020872 | 0,0020872 | TRUE | 0,0203772 | No |
| orf19.7482 | 38 | 62 | 44 | 20 | 19 | 14 | 17 | 26 | 35 | 63 | 42 | 30 | 17 | 15 | 17 | 20 | 29,79 | 42 | 17 | 0,409 | -1,289 | -4,379 | 1,19E-05 | 1,89E-05 | 0,04043 | 0,0463784 | 0,0439473 | 0,0439473 | TRUE | 0,715212 | No |
| orf19.4941 | 28783 | 25073 | 28101 | 17479 | 11923 | 9743 | 10670 | 13972 | 26206 | 25359 | 26595 | 26256 | 10605 | 10578 | 10927 | 10658 | 18398,04 | 26104 | 10692 | 0,41 | -1,288 | -43,57 | 0 | 0 | 0,0002275 | 0,0021088 | 0,0007746 | 0,0007746 | TRUE | 0,0045475 | No |
| orf19.1714 | 96 | 125 | 114 | 111 | 54 | 52 | 55 | 51 | 87 | 126 | 108 | 167 | 48 | 56 | 56 | 39 | 86,02 | 122 | 50 | 0,41 | -1,285 | -6,173 | 6,70E-10 | 1,26E-09 | 0,0420204 | 0,0173935 | 0,0277289 | 0,0277289 | TRUE | 0,2268522 | No |
| orf19.711 | 23492 | 19694 | 21152 | 14136 | 9379 | 7811 | 8161 | 11439 | 21388 | 19918 | 20018 | 21235 | 8342 | 8481 | 8358 | 8726 | 14558,3 | 20640 | 8477 | 0,411 | -1,284 | -33,976 | 5,06E-253 | 1,07E-251 | 0,0008199 | 0,0021278 | 0,0012896 | 0,0012896 | TRUE | 0,0074251 | No |
| orf19.4413 | 9297 | 7643 | 8344 | 5335 | 3939 | 2860 | 3153 | 4390 | 8465 | 7730 | 7897 | 8014 | 3504 | 3105 | 3229 | 3349 | 5661,5 | 8026 | 3297 | 0,411 | -1,283 | -26,687 | 6,67E-157 | 9,12E-156 | 0,0018132 | 0,0022703 | 0,0020111 | 0,0020111 | TRUE | 0,0215956 | No |
| orf19.703 | 1682 | 1396 | 1479 | 1009 | 702 | 556 | 588 | 756 | 1531 | 1412 | 1400 | 1516 | 624 | 604 | 602 | 577 | 1033,21 | 1465 | 602 | 0,411 | -1,283 | -22,183 | 5,05E-109 | 4,73E-108 | 0,0009624 | 0,0033156 | 0,0020708 | 0,0020708 | TRUE | 0,0126596 | No |
| orf19.916 | 3922 | 4065 | 4182 | 2162 | 1789 | 1358 | 1513 | 1979 | 3571 | 4111 | 3958 | 3248 | 1591 | 1474 | 1550 | 1510 | 2626,56 | 3722 | 1531 | 0,411 | -1,283 | -18,206 | 4,66E-74 | 3,06E-73 | 0,0054031 | 0,00254 | 0,0043159 | 0,0043159 | TRUE | 0,0869071 | No |
| orf19.5902 | 5768 | 5316 | 5793 | 3986 | 2447 | 2133 | 2283 | 2971 | 5252 | 5377 | 5483 | 5988 | 2177 | 2316 | 2338 | 2266 | 3899,37 | 5525 | 2274 | 0,412 | -1,28 | -25,558 | 4,53E-144 | 5,67E-143 | 0,0017902 | 0,0023758 | 0,0021043 | 0,0021043 | TRUE | 0,0376241 | No |
| orf19.1236 | 2785 | 2663 | 2654 | 1989 | 1298 | 1111 | 981 | 1378 | 2536 | 2693 | 2512 | 2988 | 1155 | 1206 | 1005 | 1051 | 1893,14 | 2682 | 1104 | 0,412 | -1,279 | -17,681 | 5,90E-70 | 3,71E-69 | 0,0059936 | 0,0027348 | 0,0044034 | 0,0044034 | TRUE | 0,069192 | No |
| orf19.1135 | 411 | 507 | 497 | 265 | 194 | 175 | 194 | 217 | 374 | 513 | 470 | 398 | 173 | 190 | 199 | 166 | 310,27 | 439 | 182 | 0,412 | -1,279 | -11,102 | 1,23E-28 | 3,82E-28 | 0,0120323 | 0,0062946 | 0,0089488 | 0,0089488 | TRUE | 0,1357955 | No |
| orf19.2897 | 112 | 80 | 105 | 69 | 57 | 27 | 43 | 45 | 102 | 81 | 99 | 104 | 51 | 29 | 44 | 34 | 68,04 | 96 | 40 | 0,412 | -1,279 | -6,656 | 2,81E-11 | 5,58E-11 | 0,0108318 | 0,0214534 | 0,0181564 | 0,0181564 | TRUE | 0,2867167 | No |
| orf19.3273 | 317 | 272 | 260 | 190 | 141 | 108 | 105 | 134 | 289 | 275 | 246 | 285 | 125 | 117 | 108 | 102 | 193,45 | 274 | 113 | 0,413 | -1,276 | -11,34 | 8,28E-30 | 2,65E-29 | 1,00E-08 | 0,0088656 | 0,0060789 | 0,0060789 | TRUE | 0,0600247 | No |
| orf19.699 | 805 | 1362 | 1277 | 641 | 485 | 476 | 447 | 478 | 733 | 1378 | 1209 | 963 | 431 | 517 | 458 | 365 | 756,56 | 1071 | 443 | 0,413 | -1,275 | -5,719 | 1,07E-08 | 1,93E-08 | 0,0462018 | 0,0037831 | 0,022049 | 0,0462018 | TRUE | 0,1369651 | No |
| orf19.2969 | 22894 | 19524 | 21950 | 13394 | 9201 | 7482 | 8628 | 11204 | 20844 | 19746 | 20774 | 20120 | 8184 | 8123 | 8836 | 8547 | 14396,76 | 20371 | 8422 | 0,413 | -1,274 | -32,566 | 1,26E-232 | 2,46E-231 | 0,0010071 | 0,0021288 | 0,0013887 | 0,0013887 | TRUE | 0,0131871 | No |
| orf19.2850 | 6961 | 6192 | 6785 | 4059 | 2944 | 2274 | 2693 | 3336 | 6338 | 6263 | 6421 | 6097 | 2619 | 2469 | 2758 | 2545 | 4438,64 | 6280 | 2598 | 0,413 | -1,274 | -29,813 | 2,66E-195 | 4,43E-194 | 0,0010023 | 0,0023346 | 0,0014884 | 0,0014884 | TRUE | 0,0204187 | No |
| orf19.4358 | 1458 | 1217 | 1269 | 766 | 595 | 437 | 470 | 712 | 1327 | 1231 | 1201 | 1151 | 529 | 474 | 481 | 543 | 867,26 | 1228 | 507 | 0,413 | -1,274 | -18,58 | 4,65E-77 | 3,17E-76 | 0,002738 | 0,0035602 | 0,0031598 | 0,0031598 | TRUE | 0,0365534 | No |
| orf19.929 | 102 | 102 | 111 | 73 | 43 | 36 | 51 | 53 | 93 | 103 | 105 | 110 | 38 | 39 | 52 | 40 | 72,59 | 103 | 42 | 0,414 | -1,272 | -7,381 | 1,57E-13 | 3,34E-13 | 1,00E-08 | 0,0202348 | 0,0123435 | 0,0123435 | TRUE | 0,1705665 | No |
| orf19.3749 | 15278 | 14480 | 15053 | 9386 | 6720 | 5463 | 5767 | 7584 | 13910 | 14645 | 14246 | 14099 | 5977 | 5931 | 5906 | 5785 | 10062,54 | 14225 | 5900 | 0,415 | -1,27 | -40,454 | 0 | 0 | 0,0002751 | 0,0021683 | 0,0008292 | 0,0008292 | TRUE | 0,0048658 | No |
| orf19.2876 | 3371 | 2850 | 3096 | 1994 | 1355 | 1044 | 1197 | 1772 | 3069 | 2882 | 2930 | 2995 | 1205 | 1133 | 1226 | 1352 | 2099,16 | 2969 | 1229 | 0,415 | -1,27 | -22,624 | 2,53E-113 | 2,44E-112 | 0,0023366 | 0,0026663 | 0,0024642 | 0,0024642 | TRUE | 0,0564196 | No |
| orf19.1535 | 144 | 113 | 136 | 66 | 55 | 36 | 55 | 69 | 131 | 114 | 129 | 99 | 49 | 39 | 56 | 53 | 83,78 | 118 | 49 | 0,415 | -1,27 | -7,588 | 3,25E-14 | 7,03E-14 | 1,00E-08 | 0,0178051 | 0,013018 | 0,013018 | TRUE | 0,1356154 | No |
| orf19.6536 | 432 | 436 | 466 | 263 | 207 | 156 | 164 | 226 | 393 | 441 | 441 | 395 | 184 | 169 | 168 | 172 | 295,53 | 418 | 173 | 0,415 | -1,269 | -13,95 | 3,14E-44 | 1,32E-43 | 1,00E-08 | 0,006507 | 0,0039831 | 0,0039831 | TRUE | 0,0210013 | No |
| orf19.3675 | 7745 | 7553 | 7704 | 5487 | 3580 | 3107 | 2929 | 3917 | 7051 | 7639 | 7291 | 8242 | 3184 | 3373 | 3000 | 2988 | 5346,16 | 7556 | 3136 | 0,415 | -1,268 | -21,548 | 5,59E-103 | 5,00E-102 | 0,0037227 | 0,0022841 | 0,0031071 | 0,0031071 | TRUE | 0,0449473 | No |
| orf19.7053 | 41900 | 40319 | 43666 | 27996 | 18160 | 16271 | 16287 | 22392 | 38148 | 40778 | 41326 | 42054 | 16153 | 17666 | 16680 | 17081 | 28735,77 | 40576 | 16895 | 0,416 | -1,264 | -29,248 | 4,83E-188 | 7,73E-187 | 0,0015389 | 0,002083 | 0,0017535 | 0,0017535 | TRUE | 0,0199239 | No |
| orf19.3967 | 33248 | 29380 | 31722 | 19913 | 13681 | 11449 | 12394 | 16618 | 30271 | 29715 | 30022 | 29913 | 12169 | 12430 | 12693 | 12676 | 21236,09 | 29980 | 12492 | 0,417 | -1,263 | -45,845 | 0 | 0 | 0,0001494 | 0,0020992 | 0,0006735 | 0,0006735 | TRUE | 0,0037536 | No |
| orf19.6588 | 2563 | 2467 | 2531 | 1472 | 1080 | 910 | 952 | 1327 | 2334 | 2495 | 2395 | 2211 | 961 | 988 | 975 | 1012 | 1671,37 | 2359 | 984 | 0,417 | -1,263 | -24,583 | 1,93E-133 | 2,22E-132 | 0,0010501 | 0,0028274 | 0,0018313 | 0,0018313 | TRUE | 0,0197148 | No |
| orf19.3245 | 1839 | 1731 | 1774 | 1123 | 805 | 688 | 638 | 936 | 1674 | 1751 | 1679 | 1687 | 716 | 747 | 653 | 714 | 1202,66 | 1698 | 708 | 0,417 | -1,263 | -23,589 | 4,95E-123 | 5,19E-122 | 0,0001878 | 0,0031354 | 0,0017751 | 0,0017751 | TRUE | 0,0294983 | No |
| orf19.2853 | 1457 | 1446 | 1509 | 1029 | 662 | 583 | 568 | 780 | 1327 | 1462 | 1428 | 1546 | 589 | 633 | 582 | 595 | 1020,17 | 1441 | 600 | 0,417 | -1,263 | -20,256 | 3,16E-91 | 2,50E-90 | 0,00188 | 0,0033319 | 0,0025776 | 0,0025776 | TRUE | 0,0346586 | No |
| orf19.4792 | 17765 | 15331 | 17092 | 11488 | 7290 | 6461 | 6735 | 9069 | 16174 | 15506 | 16176 | 17257 | 6484 | 7015 | 6897 | 6918 | 11553,41 | 16278 | 6828 | 0,42 | -1,253 | -28,72 | 2,17E-181 | 3,37E-180 | 0,0014458 | 0,0021514 | 0,0017269 | 0,0017269 | TRUE | 0,0197514 | No |
| orf19.5016 | 1137 | 1023 | 1016 | 765 | 533 | 412 | 416 | 531 | 1035 | 1035 | 962 | 1149 | 474 | 447 | 426 | 405 | 741,63 | 1045 | 438 | 0,42 | -1,252 | -16,903 | 4,26E-64 | 2,47E-63 | 0,0035407 | 0,0038182 | 0,0036854 | 0,0036854 | TRUE | 0,0492247 | No |
| orf19.21 | 630 | 532 | 573 | 428 | 272 | 226 | 221 | 322 | 574 | 538 | 542 | 643 | 242 | 245 | 226 | 246 | 407,02 | 574 | 240 | 0,42 | -1,252 | -14,96 | 1,35E-50 | 6,29E-50 | 0,0020235 | 0,0052826 | 0,0038489 | 0,0038489 | TRUE | 0,0663795 | No |
| orf19.6985 | 1824 | 1919 | 2034 | 1231 | 927 | 744 | 741 | 943 | 1661 | 1941 | 1925 | 1849 | 825 | 808 | 759 | 719 | 1310,78 | 1844 | 778 | 0,421 | -1,247 | -18,519 | 1,45E-76 | 9,82E-76 | 0,0038217 | 0,0030448 | 0,0034574 | 0,0034574 | TRUE | 0,0560978 | No |
| orf19.1446 | 3728 | 3390 | 3747 | 2246 | 1490 | 1360 | 1507 | 1907 | 3394 | 3429 | 3546 | 3374 | 1325 | 1477 | 1543 | 1455 | 2442,85 | 3436 | 1450 | 0,422 | -1,246 | -24,486 | 2,07E-132 | 2,37E-131 | 0,0015788 | 0,0025778 | 0,0020074 | 0,0020074 | TRUE | 0,0406281 | No |
| orf19.411 | 19940 | 17605 | 18706 | 10985 | 8349 | 6677 | 7423 | 9610 | 18155 | 17806 | 17706 | 16501 | 7426 | 7249 | 7602 | 7331 | 12471,63 | 17542 | 7402 | 0,422 | -1,245 | -31,806 | 5,32E-222 | 9,92E-221 | 0,0009236 | 0,0021429 | 0,0013784 | 0,0013784 | TRUE | 0,0191674 | No |
| orf19.708 | 399 | 360 | 426 | 226 | 175 | 143 | 153 | 204 | 363 | 364 | 403 | 339 | 156 | 155 | 157 | 156 | 261,66 | 367 | 156 | 0,422 | -1,243 | -13,115 | 2,71E-39 | 1,04E-38 | 1,00E-08 | 0,0070856 | 0,0041964 | 0,0041964 | TRUE | 0,0491192 | No |
| orf19.543 | 774 | 689 | 713 | 465 | 332 | 283 | 275 | 380 | 705 | 697 | 675 | 699 | 295 | 307 | 282 | 290 | 493,61 | 694 | 294 | 0,423 | -1,242 | -17,967 | 3,56E-72 | 2,28E-71 | 1,00E-08 | 0,0047132 | 0,0022292 | 0,0022292 | TRUE | 0,0099243 | No |
| orf19.5246 | 327 | 285 | 328 | 220 | 145 | 130 | 123 | 162 | 298 | 288 | 310 | 330 | 129 | 141 | 126 | 124 | 218,32 | 306 | 130 | 0,424 | -1,239 | -12,154 | 5,43E-34 | 1,89E-33 | 1,00E-08 | 0,0080879 | 0,0046347 | 0,0046347 | TRUE | 0,0316542 | No |
| orf19.7377 | 256 | 241 | 216 | 153 | 85 | 90 | 104 | 139 | 233 | 244 | 204 | 230 | 76 | 98 | 107 | 106 | 162,12 | 228 | 97 | 0,424 | -1,239 | -9,663 | 4,32E-22 | 1,15E-21 | 0,0056305 | 0,0101854 | 0,008602 | 0,008602 | TRUE | 0,2194405 | No |
| orf19.4353 | 2075 | 2111 | 2270 | 1345 | 991 | 852 | 839 | 1091 | 1889 | 2135 | 2148 | 2020 | 881 | 925 | 859 | 832 | 1461,37 | 2048 | 874 | 0,426 | -1,23 | -20,79 | 5,29E-96 | 4,36E-95 | 0,0022504 | 0,002941 | 0,0025615 | 0,0025615 | TRUE | 0,0347612 | No |
| orf19.789.1 | 837 | 734 | 771 | 427 | 363 | 265 | 309 | 398 | 762 | 742 | 730 | 641 | 323 | 288 | 316 | 304 | 513,27 | 719 | 308 | 0,427 | -1,229 | -15,945 | 3,07E-57 | 1,60E-56 | 0,001961 | 0,0046107 | 0,003446 | 0,003446 | TRUE | 0,0544386 | No |
| orf19.1848.1 | 503 | 388 | 342 | 267 | 188 | 155 | 144 | 245 | 458 | 392 | 324 | 401 | 167 | 168 | 147 | 187 | 280,62 | 394 | 167 | 0,427 | -1,229 | -10,379 | 3,08E-25 | 8,88E-25 | 0,0123327 | 0,0067444 | 0,0093148 | 0,0093148 | TRUE | 0,1704398 | No |
| orf19.1228 | 3775 | 3579 | 3831 | 2404 | 1654 | 1410 | 1600 | 1922 | 3437 | 3620 | 3626 | 3611 | 1471 | 1531 | 1639 | 1466 | 2550,05 | 3574 | 1527 | 0,427 | -1,228 | -25,767 | 2,07E-146 | 2,66E-145 | 0,0010816 | 0,002555 | 0,001724 | 0,001724 | TRUE | 0,0291101 | No |
| orf19.6982 | 1135 | 1267 | 1175 | 728 | 557 | 461 | 484 | 581 | 1033 | 1281 | 1112 | 1094 | 495 | 501 | 496 | 443 | 806,9 | 1130 | 484 | 0,427 | -1,227 | -15,698 | 1,55E-55 | 7,91E-55 | 0,0051387 | 0,0036741 | 0,0044228 | 0,0044228 | TRUE | 0,0990667 | No |
| orf19.6030 | 2492 | 2098 | 2276 | 1358 | 1048 | 779 | 924 | 1242 | 2269 | 2122 | 2154 | 2040 | 932 | 846 | 946 | 947 | 1532,05 | 2146 | 918 | 0,428 | -1,225 | -22,23 | 1,76E-109 | 1,65E-108 | 0,0013406 | 0,0028993 | 0,0021587 | 0,0021587 | TRUE | 0,0316195 | No |
| orf19.5320 | 271 | 227 | 274 | 184 | 121 | 101 | 101 | 145 | 247 | 230 | 259 | 276 | 108 | 110 | 103 | 111 | 180,42 | 253 | 108 | 0,428 | -1,223 | -10,976 | 4,99E-28 | 1,53E-27 | 1,00E-08 | 0,0093588 | 0,0054955 | 0,0054955 | TRUE | 0,0415189 | No |
| orf19.5602 | 2277 | 2055 | 2197 | 1428 | 984 | 845 | 850 | 1209 | 2073 | 2078 | 2079 | 2145 | 875 | 917 | 871 | 922 | 1495,16 | 2094 | 896 | 0,429 | -1,222 | -26,468 | 2,24E-154 | 3,01E-153 | 1,00E-08 | 0,0029205 | 0,0012713 | 0,0012713 | TRUE | 0,0050933 | No |
| orf19.2792 | 978 | 818 | 955 | 599 | 423 | 367 | 350 | 493 | 890 | 827 | 904 | 900 | 376 | 398 | 358 | 376 | 628,82 | 880 | 377 | 0,429 | -1,222 | -18,548 | 8,49E-77 | 5,76E-76 | 1,00E-08 | 0,0041378 | 0,002327 | 0,002327 | TRUE | 0,0199491 | No |
| orf19.4086 | 716 | 613 | 557 | 411 | 311 | 228 | 239 | 346 | 652 | 620 | 527 | 617 | 277 | 248 | 245 | 264 | 431,16 | 604 | 258 | 0,429 | -1,222 | -13,924 | 4,53E-44 | 1,90E-43 | 0,0042759 | 0,0051008 | 0,004708 | 0,004708 | TRUE | 0,0910999 | No |
| orf19.3212 | 1980 | 2007 | 2057 | 1261 | 940 | 771 | 810 | 1046 | 1803 | 2030 | 1947 | 1894 | 836 | 837 | 830 | 798 | 1371,77 | 1918 | 825 | 0,43 | -1,218 | -22,59 | 5,45E-113 | 5,25E-112 | 0,0011385 | 0,003 | 0,0019465 | 0,0019465 | TRUE | 0,0212298 | No |
| orf19.6302 | 31 | 23 | 11 | 7 | 11 | 2 | 9 | 13 | 28 | 23 | 10 | 11 | 10 | 2 | 9 | 10 | 12,94 | 18 | 8 | 0,431 | -1,215 | -2,538 | 0,011157 | 0,0145867 | 0,1837585 | 0,104142 | 0,1321474 | 0,1321474 | TRUE | 0,1997813 | No |
| orf19.2480.1 | 17088 | 14277 | 15811 | 9393 | 7331 | 5533 | 6383 | 8384 | 15558 | 14440 | 14964 | 14110 | 6521 | 6007 | 6537 | 6395 | 10566,42 | 14768 | 6365 | 0,431 | -1,214 | -27,363 | 7,64E-165 | 1,10E-163 | 0,0015213 | 0,002162 | 0,0017818 | 0,0017818 | TRUE | 0,0172321 | No |
| orf19.3446 | 667 | 692 | 703 | 348 | 321 | 254 | 270 | 321 | 607 | 700 | 665 | 523 | 286 | 276 | 277 | 245 | 447,24 | 624 | 271 | 0,431 | -1,213 | -12,491 | 8,35E-36 | 3,03E-35 | 0,008196 | 0,0049907 | 0,0064613 | 0,0064613 | TRUE | 0,1256265 | No |
| orf19.1777 | 11727 | 12362 | 12818 | 7314 | 5478 | 4851 | 4958 | 6267 | 10677 | 12503 | 12131 | 10987 | 4873 | 5267 | 5078 | 4781 | 8286,89 | 11574 | 5000 | 0,432 | -1,212 | -20,878 | 8,51E-97 | 7,15E-96 | 0,0036432 | 0,0021964 | 0,0030956 | 0,0030956 | TRUE | 0,0356668 | No |
| orf19.7581 | 7330 | 6267 | 6980 | 4019 | 3165 | 2435 | 2823 | 3586 | 6674 | 6338 | 6606 | 6037 | 2815 | 2644 | 2891 | 2735 | 4592,58 | 6414 | 2771 | 0,432 | -1,211 | -26,06 | 1,04E-149 | 1,34E-148 | 0,0014107 | 0,0023247 | 0,0018226 | 0,0018226 | TRUE | 0,0184566 | No |
| orf19.317 | 1105 | 914 | 1027 | 634 | 482 | 348 | 420 | 565 | 1006 | 924 | 972 | 952 | 429 | 378 | 430 | 431 | 690,31 | 964 | 417 | 0,433 | -1,207 | -18,42 | 9,08E-76 | 6,09E-75 | 0,0005411 | 0,0039506 | 0,002449 | 0,002449 | TRUE | 0,0423586 | No |
| orf19.7150 | 12441 | 10435 | 12253 | 8258 | 5269 | 4605 | 4941 | 6744 | 11327 | 10554 | 11596 | 12405 | 4687 | 5000 | 5060 | 5144 | 8221,63 | 11470 | 4973 | 0,434 | -1,205 | -22,432 | 1,91E-111 | 1,82E-110 | 0,0028309 | 0,0021977 | 0,0026323 | 0,0026323 | TRUE | 0,036257 | No |
| orf19.4900 | 1156 | 1224 | 1241 | 755 | 583 | 482 | 486 | 602 | 1052 | 1238 | 1174 | 1134 | 519 | 523 | 498 | 459 | 824,73 | 1150 | 500 | 0,434 | -1,205 | -17,089 | 1,78E-65 | 1,06E-64 | 0,0031072 | 0,0036387 | 0,0033688 | 0,0033688 | TRUE | 0,0413136 | No |
| orf19.2483 | 2182 | 1678 | 1694 | 1469 | 949 | 743 | 738 | 1101 | 1987 | 1697 | 1603 | 2207 | 844 | 807 | 756 | 840 | 1342,51 | 1874 | 812 | 0,434 | -1,204 | -13,128 | 2,29E-39 | 8,79E-39 | 0,0109113 | 0,003021 | 0,0072115 | 0,0072115 | TRUE | 0,1678934 | No |
| orf19.2436 | 8296 | 7833 | 8404 | 5187 | 3783 | 3168 | 3281 | 4461 | 7553 | 7922 | 7954 | 7792 | 3365 | 3440 | 3360 | 3403 | 5598,52 | 7805 | 3392 | 0,435 | -1,202 | -35,635 | 3,99E-278 | 9,32E-277 | 0,0002448 | 0,002273 | 0,0008864 | 0,0008864 | TRUE | 0,0059794 | No |
| orf19.5241 | 7932 | 6839 | 7505 | 4650 | 3406 | 2800 | 3112 | 3964 | 7222 | 6917 | 7103 | 6985 | 3030 | 3040 | 3187 | 3024 | 5063,36 | 7057 | 3070 | 0,435 | -1,202 | -33,899 | 6,97E-252 | 1,46E-250 | 0,0001943 | 0,0022979 | 0,0009777 | 0,0009777 | TRUE | 0,0078529 | No |

| | | | | | | | | | | | | | | | | | | | | | | | | | | | | | | | |
|---|---|---|---|---|---|---|---|---|---|---|---|---|---|---|---|---|---|---|---|---|---|---|---|---|---|---|---|---|---|---|---|
| orf19.7056 | 316 | 257 | 295 | 199 | 130 | 116 | 140 | 140 | 288 | 260 | 279 | 299 | 116 | 126 | 143 | 107 | 202,19 | 282 | 123 | 0,435 | -1,201 | -10,696 | 1,06E-26 | 3,15E-26 | 0,0020891 | 0,0085706 | 0,0063941 | 0,0063941 | TRUE | 0,1275211 | No |

P$_{TET}$-*ZCF26* doxycycline-treated up-regulated genes

| | WT dox | | | | ZCF26 | | | | norm. WT dox | | | | norm. ZCF26 | | | | | | | | | | | | | | | | | | |
|---|---|---|---|---|---|---|---|---|---|---|---|---|---|---|---|---|---|---|---|---|---|---|---|---|---|---|---|---|---|---|
| Gene ID | 1 | 2 | 3 | 4 | 1 | 2 | 3 | 4 | 1 | 2 | 3 | 4 | 1 | 2 | 3 | 4 | baseMean | WT | ZCF26 | FC | log2FC | stat | pvalue | padj | dispGeneEst | dispFit | dispMAP | dispersion | betaConv | maxCooks | outlier |
| orf19.4573 | 239 | 168 | 220 | 165 | 21821 | 15509 | 20814 | 22198 | 228 | 178 | 217 | 258 | 17485 | 17635 | 17671 | 18049 | 8965,13 | 220 | 17710 | 81,307 | 6,345 | 96,513 | 0 | 0 | 0,0001649 | 0,0022744 | 0,0015891 | 0,0015891 | TRUE | 0,178534 | No |
| orf19.4571 | 86 | 65 | 82 | 42 | 2085 | 1561 | 2032 | 2182 | 82 | 69 | 81 | 66 | 1671 | 1775 | 1725 | 1774 | 905,31 | 74 | 1736 | 23,042 | 4,526 | 44,923 | 0 | 0 | 1,00E-08 | 0,0039049 | 0,0021813 | 0,0021813 | TRUE | 0,0410372 | No |
| orf19.2451 | 568 | 590 | 571 | 409 | 12254 | 10390 | 10969 | 11047 | 542 | 624 | 564 | 641 | 9819 | 11814 | 9313 | 8982 | 5287,28 | 593 | 9982 | 16,885 | 4,078 | 45,734 | 0 | 0 | 0,0100767 | 0,0024018 | 0,0066382 | 0,0066382 | TRUE | 0,1860867 | No |
| orf19.7466 | 6592 | 5967 | 6412 | 3819 | 99362 | 68486 | 89714 | 1E+05 | 6285 | 6309 | 6334 | 5983 | 79616 | 77875 | 76167 | 82769 | 42667,21 | 6228 | 79107 | 12,695 | 3,666 | 96,39 | 0 | 0 | 0,0008689 | 0,0021298 | 0,0012934 | 0,0012934 | TRUE | 0,0119133 | No |
| orf19.4569 | 66 | 48 | 54 | 36 | 845 | 682 | 812 | 836 | 63 | 51 | 53 | 56 | 677 | 775 | 689 | 680 | 380,64 | 56 | 705 | 12,587 | 3,654 | 29,184 | 3,09E-187 | 3,91E-186 | 1,00E-08 | 0,0064048 | 0,0045575 | 0,0045575 | TRUE | 0,0744146 | No |
| orf19.535 | 58 | 38 | 39 | 18 | 669 | 425 | 543 | 655 | 55 | 40 | 39 | 28 | 536 | 483 | 461 | 533 | 271,89 | 40 | 503 | 12,077 | 3,594 | 23,994 | 3,22E-127 | 2,74E-126 | 0,005861 | 0,0081302 | 0,0074204 | 0,0074204 | TRUE | 0,4565635 | No |
| orf19.7156 | 1801 | 1614 | 1753 | 1117 | 22784 | 15881 | 21868 | 23447 | 1717 | 1707 | 1732 | 1750 | 18256 | 18058 | 18566 | 19065 | 10106,26 | 1726 | 18486 | 10,716 | 3,422 | 94,706 | 0 | 0 | 1,00E-08 | 0,0022537 | 0,0009053 | 0,0009053 | TRUE | 0,0054418 | No |
| orf19.4591 | 2955 | 2653 | 2862 | 1929 | 35487 | 27771 | 33973 | 37294 | 2817 | 2805 | 2827 | 3022 | 28435 | 31578 | 28843 | 30324 | 16331,39 | 2868 | 29795 | 10,4 | 3,378 | 72,889 | 0 | 0 | 0,0015574 | 0,0021918 | 0,0018517 | 0,0018517 | TRUE | 0,0198018 | No |
| orf19.1655 | 2404 | 2002 | 2213 | 1558 | 27977 | 20060 | 26261 | 29787 | 2292 | 2117 | 2186 | 2441 | 22417 | 22810 | 22295 | 24220 | 12597,26 | 2259 | 22936 | 10,17 | 3,346 | 66,549 | 0 | 0 | 0,0023647 | 0,0022216 | 0,0021591 | 0,0021591 | TRUE | 0,0329333 | No |
| orf19.5949 | 6066 | 5619 | 6144 | 3856 | 71785 | 49681 | 63731 | 74569 | 5783 | 5941 | 6070 | 6041 | 57519 | 56492 | 54107 | 60632 | 31573,13 | 5959 | 57188 | 9,599 | 3,263 | 77,195 | 0 | 0 | 0,00123 | 0,0021433 | 0,0016136 | 0,0016136 | TRUE | 0,0201641 | No |
| orf19.979 | 6339 | 5623 | 5955 | 4092 | 71014 | 51384 | 65435 | 74587 | 6044 | 5946 | 5883 | 6410 | 56901 | 58428 | 55554 | 60646 | 31976,57 | 6071 | 57882 | 9,541 | 3,254 | 77,407 | 0 | 0 | 0,0013462 | 0,0021426 | 0,0015969 | 0,0015969 | TRUE | 0,0166392 | No |
| orf19.1288 | 6296 | 5354 | 5631 | 3891 | 64810 | 44689 | 59613 | 68681 | 6003 | 5661 | 5563 | 6095 | 51930 | 50815 | 50611 | 55844 | 29065,41 | 5830 | 52300 | 8,974 | 3,166 | 67,416 | 0 | 0 | 0,0019357 | 0,0021478 | 0,0020128 | 0,0020128 | TRUE | 0,0255141 | No |
| orf19.4335 | 278 | 296 | 240 | 175 | 3036 | 2140 | 2821 | 2971 | 265 | 313 | 237 | 274 | 2433 | 2433 | 2395 | 2416 | 1345,75 | 272 | 2419 | 8,904 | 3,154 | 45,957 | 0 | 0 | 1,00E-08 | 0,0033113 | 0,0022972 | 0,0022972 | TRUE | 0,123259 | No |
| orf19.5449 | 60 | 51 | 54 | 44 | 633 | 417 | 575 | 598 | 57 | 54 | 53 | 69 | 507 | 474 | 488 | 486 | 273,65 | 58 | 489 | 8,514 | 3,09 | 25,037 | 2,44E-138 | 2,26E-137 | 1,00E-08 | 0,0080913 | 0,0041203 | 0,0041203 | TRUE | 0,0827094 | No |
| orf19.4934 | 25 | 21 | 12 | 14 | 253 | 134 | 204 | 166 | 24 | 22 | 12 | 22 | 203 | 152 | 173 | 135 | 92,89 | 20 | 166 | 8,386 | 3,068 | 13,002 | 1,19E-38 | 3,69E-38 | 0,0274711 | 0,0197679 | 0,0228211 | 0,0228211 | TRUE | 0,3057674 | No |
| orf19.1985 | 643 | 584 | 680 | 414 | 6355 | 4963 | 5804 | 6541 | 613 | 618 | 672 | 649 | 5092 | 5643 | 4928 | 5318 | 2941,54 | 638 | 5245 | 8,228 | 3,041 | 50,532 | 0 | 0 | 0,0024187 | 0,0026495 | 0,0025139 | 0,0025139 | TRUE | 0,0312908 | No |
| orf19.1641 | 38 | 37 | 50 | 32 | 460 | 300 | 428 | 427 | 36 | 39 | 49 | 50 | 369 | 341 | 363 | 347 | 199,39 | 44 | 355 | 8,233 | 3,041 | 21,2 | 9,56E-100 | 6,25E-99 | 1,00E-08 | 0,0103258 | 0,0057338 | 0,0057338 | TRUE | 0,113359 | No |
| orf19.7288 | 616 | 507 | 560 | 389 | 5439 | 3979 | 5378 | 5857 | 587 | 536 | 553 | 609 | 4358 | 4524 | 4566 | 4762 | 2562,1 | 571 | 4552 | 7,992 | 2,999 | 54,32 | 0 | 0 | 0,0010226 | 0,0027321 | 0,0018498 | 0,0018498 | TRUE | 0,0193071 | No |
| orf19.866 | 1467 | 1376 | 1473 | 895 | 14188 | 9989 | 13156 | 14489 | 1399 | 1455 | 1455 | 1402 | 11368 | 11358 | 11169 | 11781 | 6423,5 | 1428 | 11419 | 7,993 | 2,999 | 77,93 | 0 | 0 | 1,00E-08 | 0,0023469 | 0,0009917 | 0,0009917 | TRUE | 0,0056096 | No |
| orf19.360 | 3 | 3 | 0 | 5 | 35 | 14 | 36 | 28 | 3 | 3 | 0 | 8 | 28 | 16 | 31 | 23 | 13,89 | 4 | 24 | 7,896 | 2,981 | 5,316 | 1,06E-07 | 1,71E-07 | 0,0588922 | 0,1202571 | 0,1057343 | 0,1057343 | TRUE | 0,6302373 | No |
| orf19.2551 | 7293 | 6391 | 6606 | 4724 | 64331 | 46073 | 60007 | 68724 | 6953 | 6758 | 6526 | 7400 | 51547 | 52389 | 50946 | 55879 | 29799,73 | 6909 | 52690 | 7,631 | 2,932 | 60,876 | 0 | 0 | 0,0022097 | 0,0021464 | 0,0021378 | 0,0021378 | TRUE | 0,0271955 | No |
| orf19.6229 | 7493 | 6494 | 7055 | 5035 | 65953 | 47343 | 62459 | 69811 | 7144 | 6867 | 6970 | 7888 | 52846 | 53833 | 53027 | 56763 | 30667,19 | 7217 | 54117 | 7,504 | 2,908 | 58,928 | 0 | 0 | 0,0025026 | 0,0021448 | 0,0022523 | 0,0022523 | TRUE | 0,046612 | No |
| orf19.1255 | 61 | 42 | 58 | 43 | 515 | 355 | 413 | 518 | 58 | 44 | 57 | 67 | 413 | 404 | 351 | 421 | 226,92 | 56 | 397 | 7,072 | 2,822 | 20,417 | 1,19E-92 | 7,30E-92 | 0,0046609 | 0,0093269 | 0,00737 | 0,00737 | TRUE | 0,174391 | No |
| orf19.7520 | 7323 | 5773 | 6383 | 4397 | 56393 | 38790 | 53174 | 59398 | 6982 | 6104 | 6306 | 6888 | 45186 | 44108 | 45144 | 48296 | 26126,8 | 6570 | 45684 | 6,956 | 2,798 | 53,445 | 0 | 0 | 0,0028228 | 0,0021541 | 0,0025375 | 0,0025375 | TRUE | 0,0278078 | No |
| orf19.7093 | 333 | 282 | 316 | 190 | 2622 | 1726 | 2555 | 2680 | 317 | 298 | 312 | 298 | 2101 | 1963 | 2169 | 2179 | 1204,66 | 306 | 2103 | 6,853 | 2,777 | 41,781 | 0 | 0 | 1,00E-08 | 0,0034542 | 0,0022214 | 0,0022214 | TRUE | 0,0233419 | No |
| orf19.7127.1 | 58 | 47 | 40 | 35 | 377 | 246 | 450 | 455 | 55 | 50 | 40 | 55 | 302 | 280 | 382 | 370 | 191,64 | 50 | 334 | 6,748 | 2,754 | 16,8 | 2,44E-63 | 1,07E-62 | 0,0163936 | 0,0106588 | 0,0132667 | 0,0132667 | TRUE | 0,1051163 | No |
| orf19.272 | 2394 | 2254 | 2424 | 1579 | 19622 | 14677 | 18548 | 19385 | 2282 | 2383 | 2395 | 2474 | 15723 | 16689 | 15747 | 15762 | 9181,83 | 2384 | 15980 | 6,712 | 2,747 | 69,242 | 0 | 0 | 0,0006579 | 0,0022701 | 0,0012472 | 0,0012472 | TRUE | 0,0107943 | No |
| orf19.313 | 7 | 8 | 4 | 5 | 57 | 33 | 53 | 56 | 7 | 8 | 4 | 8 | 46 | 38 | 45 | 46 | 25,08 | 7 | 44 | 6,601 | 2,723 | 7,405 | 1,31E-13 | 2,54E-13 | 1,00E-08 | 0,0675567 | 0,0361194 | 0,0361194 | TRUE | 0,220939 | No |
| orf19.2590 | 13 | 4 | 4 | 10 | 65 | 55 | 59 | 76 | 12 | 4 | 4 | 16 | 52 | 63 | 50 | 62 | 32,84 | 9 | 57 | 6,561 | 2,714 | 7,819 | 5,33E-15 | 1,07E-14 | 0,0040683 | 0,0520828 | 0,0436021 | 0,0436021 | TRUE | 0,4850802 | No |
| orf19.4481 | 10 | 8 | 10 | 6 | 78 | 45 | 66 | 90 | 10 | 8 | 10 | 9 | 62 | 51 | 56 | 73 | 35,02 | 9 | 60 | 6,543 | 2,71 | 8,56 | 1,13E-17 | 2,41E-17 | 1,00E-08 | 0,0489774 | 0,0297491 | 0,0297491 | TRUE | 0,1744311 | No |
| orf19.5728 | 269 | 262 | 243 | 173 | 2044 | 1451 | 1945 | 2060 | 256 | 277 | 240 | 271 | 1638 | 1650 | 1651 | 1675 | 957,32 | 261 | 1654 | 6,356 | 2,668 | 40,697 | 0 | 0 | 1,00E-08 | 0,0038064 | 0,00173 | 0,00173 | TRUE | 0,0346743 | No |
| orf19.4211 | 1934 | 1835 | 1865 | 1212 | 14532 | 10655 | 13706 | 14937 | 1844 | 1940 | 1842 | 1899 | 11644 | 12116 | 11636 | 12145 | 6883,32 | 1881 | 11885 | 6,321 | 2,66 | 70,369 | 0 | 0 | 0,0004185 | 0,0023298 | 0,0010368 | 0,0010368 | TRUE | 0,0058215 | No |
| orf19.2509.1 | 88 | 89 | 71 | 55 | 706 | 474 | 588 | 598 | 84 | 94 | 70 | 86 | 566 | 539 | 499 | 486 | 303,05 | 84 | 522 | 6,274 | 2,649 | 22,616 | 3,04E-113 | 2,28E-112 | 0,0033728 | 0,0075092 | 0,0056721 | 0,0056721 | TRUE | 0,121614 | No |
| orf19.1652 | 11107 | 9726 | 10252 | 7484 | 80882 | 60408 | 75601 | 85131 | 10589 | 10284 | 10128 | 11724 | 64808 | 68689 | 64185 | 69220 | 38703,46 | 10681 | 66726 | 6,25 | 2,644 | 50,6 | 0 | 0 | 0,0029627 | 0,0021337 | 0,0025631 | 0,0025631 | TRUE | 0,0520094 | No |
| orf19.6254 | 3570 | 3022 | 3291 | 2158 | 25234 | 17210 | 24120 | 25751 | 3404 | 3195 | 3251 | 3381 | 20219 | 19569 | 20478 | 20938 | 11804,4 | 3308 | 20301 | 6,141 | 2,618 | 69,075 | 0 | 0 | 0,0006235 | 0,0022304 | 0,0011878 | 0,0011878 | TRUE | 0,0071387 | No |
| orf19.3573 | 1537 | 1469 | 1540 | 916 | 11432 | 8200 | 10530 | 10992 | 1465 | 1553 | 1521 | 1435 | 9160 | 9324 | 8940 | 8938 | 5292,09 | 1494 | 9090 | 6,075 | 2,603 | 65,447 | 0 | 0 | 0,0003332 | 0,0024015 | 0,0010959 | 0,0010959 | TRUE | 0,0083141 | No |
| orf19.6086 | 486 | 386 | 406 | 278 | 3199 | 2181 | 2987 | 3413 | 463 | 408 | 401 | 436 | 2563 | 2480 | 2536 | 2775 | 1507,8 | 427 | 2588 | 6,067 | 2,601 | 39,722 | 0 | 0 | 0,0021574 | 0,0031802 | 0,0026383 | 0,0026383 | TRUE | 0,0437416 | No |
| orf19.5809 | 66 | 55 | 67 | 57 | 495 | 340 | 498 | 513 | 63 | 58 | 66 | 89 | 397 | 387 | 423 | 417 | 237,47 | 69 | 406 | 6,007 | 2,587 | 20,707 | 2,98E-95 | 1,86E-94 | 1,00E-08 | 0,0090056 | 0,0057084 | 0,0057084 | TRUE | 0,2376182 | No |
| orf19.1631 | 2288 | 2192 | 2248 | 1314 | 16512 | 12058 | 15406 | 15722 | 2181 | 2318 | 2221 | 2058 | 13231 | 13711 | 13080 | 12783 | 7697,89 | 2194 | 13201 | 6,005 | 2,586 | 59,001 | 0 | 0 | 0,0012724 | 0,0023046 | 0,0015553 | 0,0015553 | TRUE | 0,0189816 | No |
| orf19.7055 | 0 | 5 | 2 | 0 | 14 | 10 | 10 | 18 | 0 | 5 | 2 | 0 | 11 | 11 | 8 | 15 | 6,62 | 2 | 11 | 6,006 | 2,586 | 3,515 | 0,0004405 | 0,0006059 | 1,58E-08 | 0,2500331 | 0,1908359 | 0,1908359 | TRUE | 1,2466369 | No |
| orf19.6958 | 61 | 40 | 57 | 33 | 388 | 281 | 344 | 399 | 58 | 42 | 56 | 52 | 311 | 320 | 292 | 324 | 181,92 | 52 | 312 | 5,954 | 2,574 | 18,642 | 1,48E-77 | 7,81E-77 | 1,00E-08 | 0,0111167 | 0,0063197 | 0,0063197 | TRUE | 0,1437255 | No |
| orf19.5672 | 42 | 51 | 43 | 40 | 343 | 257 | 332 | 346 | 40 | 54 | 42 | 63 | 275 | 292 | 282 | 281 | 166,17 | 50 | 282 | 5,81 | 2,539 | 17,618 | 1,81E-69 | 8,58E-69 | 1,00E-08 | 0,011972 | 0,0069877 | 0,0069877 | TRUE | 0,1676927 | No |
| orf19.3577.1 | 97 | 86 | 84 | 69 | 720 | 466 | 606 | 640 | 92 | 91 | 83 | 108 | 577 | 530 | 514 | 520 | 314,52 | 94 | 535 | 5,79 | 2,534 | 23,314 | 3,22E-120 | 2,57E-119 | 1,00E-08 | 0,0073116 | 0,004528 | 0,004528 | TRUE | 0,073439 | No |
| orf19.1274 | 12 | 8 | 15 | 13 | 95 | 79 | 83 | 83 | 11 | 8 | 15 | 20 | 76 | 90 | 70 | 67 | 44,87 | 14 | 76 | 5,693 | 2,509 | 9,046 | 1,48E-19 | 3,30E-19 | 0,0036271 | 0,0386809 | 0,0263878 | 0,0263878 | TRUE | 0,2604642 | No |
| orf19.7149 | 921 | 776 | 818 | 597 | 5988 | 4187 | 5700 | 6110 | 878 | 821 | 808 | 935 | 4798 | 4761 | 4839 | 4968 | 2851,03 | 860 | 4842 | 5,65 | 2,498 | 50,231 | 0 | 0 | 0,0004518 | 0,0026672 | 0,0016313 | 0,0016313 | TRUE | 0,034078 | No |
| orf19.359 | 28 | 18 | 31 | 22 | 181 | 151 | 161 | 198 | 27 | 19 | 31 | 34 | 145 | 172 | 137 | 161 | 90,65 | 28 | 154 | 5,618 | 2,49 | 12,603 | 2,04E-36 | 6,10E-36 | 0,0045127 | 0,0202031 | 0,0142972 | 0,0142972 | TRUE | 0,2722486 | No |
| orf19.7585 | 1523 | 1271 | 1408 | 864 | 9501 | 6896 | 8885 | 9849 | 1452 | 1344 | 1391 | 1354 | 7613 | 7841 | 7543 | 8008 | 4568,27 | 1385 | 7751 | 5,588 | 2,482 | 58,032 | 0 | 0 | 0,0006311 | 0,0024507 | 0,0012958 | 0,0012958 | TRUE | 0,0132697 | No |
| orf19.3924 | 26 | 27 | 36 | 12 | 184 | 143 | 170 | 196 | 25 | 29 | 36 | 19 | 147 | 163 | 144 | 159 | 90,18 | 27 | 153 | 5,559 | 2,475 | 12,938 | 2,75E-38 | 8,49E-38 | 1,00E-08 | 0,0202984 | 0,0121796 | 0,0121796 | TRUE | 0,2858677 | No |
| orf19.6219 | 134 | 128 | 93 | 75 | 813 | 554 | 752 | 863 | 128 | 135 | 92 | 117 | 651 | 630 | 638 | 702 | 386,75 | 118 | 655 | 5,557 | 2,474 | 23,239 | 1,83E-119 | 1,45E-118 | 0,0035746 | 0,0063367 | 0,0055017 | 0,0055017 | TRUE | 0,2552174 | No |
| orf19.5645 | 3024 | 2384 | 2672 | 1892 | 18607 | 13028 | 17699 | 19329 | 2883 | 2521 | 2640 | 2964 | 14909 | 14814 | 15026 | 15716 | 8934,17 | 2752 | 15116 | 5,5 | 2,459 | 46,169 | 0 | 0 | 0,0029008 | 0,0022751 | 0,0024905 | 0,0024905 | TRUE | 0,038482 | No |
| orf19.54 | 302 | 250 | 282 | 203 | 1873 | 1547 | 1789 | 1835 | 288 | 264 | 279 | 318 | 1501 | 1759 | 1519 | 1492 | 927,45 | 287 | 1568 | 5,48 | 2,454 | 30,034 | 3,53E-198 | 4,81E-197 | 0,0044699 | 0,0038616 | 0,0041738 | 0,0041738 | TRUE | 0,0790089 | No |
| orf19.3684 | 2405 | 2012 | 2137 | 1482 | 14831 | 11082 | 13966 | 14853 | 2293 | 2127 | 2111 | 2322 | 11884 | 12601 | 11857 | 12077 | 7159 | 2213 | 12105 | 5,475 | 2,453 | 55,429 | 0 | 0 | 0,0012711 | 0,0023206 | 0,0015892 | 0,0015892 | TRUE | 0,0118926 | No |
| orf19.7049 | 911 | 902 | 898 | 592 | 6455 | 4492 | 5608 | 5798 | 869 | 954 | 887 | 927 | 5172 | 5108 | 4761 | 4714 | 2924,04 | 909 | 4939 | 5,438 | 2,443 | 45,467 | 0 | 0 | 0,0016814 | 0,0026528 | 0,0020654 | 0,0020654 | TRUE | 0,0142191 | No |
| orf19.4212 | 370 | 318 | 344 | 206 | 2282 | 1610 | 2145 | 2301 | 353 | 336 | 340 | 323 | 1829 | 1831 | 1821 | 1871 | 1087,85 | 338 | 1838 | 5,419 | 2,438 | 41,32 | 0 | 0 | 1,00E-08 | 0,0036006 | 0,0014673 | 0,0014673 | TRUE | 0,0099414 | No |
| orf19.1326 | 267 | 212 | 204 | 136 | 1459 | 1035 | 1422 | 1493 | 255 | 224 | 202 | 213 | 1169 | 1177 | 1207 | 1214 | 707,56 | 224 | 1192 | 5,316 | 2,41 | 31,742 | 4,13E-221 | 6,14E-220 | 1,00E-08 | 0,0044118 | 0,0026886 | 0,0026886 | TRUE | 0,1082133 | No |
| orf19.310 | 41 | 25 | 34 | 21 | 186 | 165 | 188 | 250 | 39 | 26 | 34 | 33 | 149 | 188 | 160 | 203 | 103,94 | 33 | 175 | 5,272 | 2,398 | 12,557 | 3,66E-36 | 1,09E-35 | 1,00E-08 | 0,0178873 | 0,0157467 | 0,0157467 | TRUE | 0,1050308 | No |
| orf19.1704 | 6422 | 5352 | 5903 | 3504 | 37594 | 25599 | 36092 | 38823 | 6123 | 5659 | 5832 | 5489 | 30123 | 29108 | 30642 | 31567 | 18067,83 | 5776 | 30360 | 5,253 | 2,393 | 54,659 | 0 | 0 | 0,001529 | 0,0021822 | 0,001729 | 0,001729 | TRUE | 0,0200264 | No |

| | | | | | | | | | | | | | | | | | | | | | | | | | | | | | | | |
|---|---|---|---|---|---|---|---|---|---|---|---|---|---|---|---|---|---|---|---|---|---|---|---|---|---|---|---|---|---|---|---|
| orf19.7136 | 1671 | 1298 | 1486 | 935 | 9871 | 6295 | 9292 | 9589 | 1593 | 1372 | 1468 | 1465 | 7909 | 7158 | 7889 | 7797 | 4581,41 | 1474 | 7688 | 5,212 | 2,382 | 43,175 | 0 | 0 | 0,0024255 | 0,0024497 | 0,0024834 | 0,0024834 | TRUE | 0,0370078 | No |
| orf19.2496 | 2143 | 1778 | 1878 | 1371 | 12815 | 9165 | 11885 | 12826 | 2043 | 1880 | 1855 | 2148 | 10268 | 10421 | 10090 | 10429 | 6141,87 | 1982 | 10302 | 5,21 | 2,381 | 48,822 | 0 | 0 | 0,001635 | 0,0023586 | 0,0019565 | 0,0019565 | TRUE | 0,0352317 | No |
| orf19.88 | 6833 | 5704 | 5885 | 3957 | 39335 | 27308 | 37314 | 40945 | 6515 | 6031 | 5814 | 6199 | 31518 | 31052 | 31679 | 33292 | 19012,47 | 6140 | 31885 | 5,194 | 2,377 | 54,568 | 0 | 0 | 0,0014962 | 0,0021776 | 0,0017164 | 0,0017164 | TRUE | 0,0212201 | No |
| orf19.1589.1 | 172 | 139 | 155 | 75 | 916 | 543 | 950 | 1078 | 164 | 147 | 153 | 117 | 734 | 617 | 807 | 877 | 452,01 | 145 | 759 | 5,176 | 2,372 | 18,834 | 4,00E-79 | 2,16E-78 | 0,0166947 | 0,0057238 | 0,0108313 | 0,0108313 | TRUE | 0,1318666 | No |
| orf19.935 | 24 | 12 | 15 | 24 | 129 | 106 | 112 | 140 | 23 | 13 | 15 | 38 | 103 | 121 | 95 | 114 | 65,1 | 22 | 108 | 5,145 | 2,363 | 9,491 | 2,28E-21 | 5,25E-21 | 0,0367132 | 0,0273124 | 0,0289669 | 0,0289669 | TRUE | 0,8381316 | No |
| orf19.4697 | 982 | 930 | 1015 | 564 | 6330 | 4294 | 5612 | 5883 | 936 | 983 | 1003 | 884 | 5072 | 4883 | 4765 | 4783 | 2913,57 | 952 | 4876 | 5,108 | 2,353 | 47,802 | 0 | 0 | 0,0008535 | 0,0026548 | 0,0016489 | 0,0016489 | TRUE | 0,023059 | No |
| orf19.5874 | 2 | 1 | 1 | 1 | 11 | 5 | 10 | 6 | 2 | 1 | 1 | 2 | 9 | 6 | 8 | 5 | 4,17 | 2 | 7 | 5,105 | 2,352 | 2,814 | 0,0048948 | 0,0063302 | 1,00E-08 | 0,3955197 | 0,2050411 | 0,2050411 | TRUE | 0,139093 | No |
| orf19.4041 | 793 | 618 | 691 | 460 | 4596 | 2965 | 4166 | 4570 | 756 | 653 | 683 | 721 | 3683 | 3371 | 3537 | 3716 | 2139,95 | 703 | 3577 | 5,091 | 2,348 | 40,06 | 0 | 0 | 0,0018531 | 0,0028585 | 0,0023779 | 0,0023779 | TRUE | 0,0341343 | No |
| orf19.2244 | 805 | 711 | 788 | 549 | 4855 | 3598 | 4565 | 5038 | 767 | 752 | 778 | 860 | 3890 | 4091 | 3876 | 4096 | 2388,91 | 789 | 3988 | 5,073 | 2,343 | 45,37 | 0 | 0 | 0,000854 | 0,0027786 | 0,001738 | 0,001738 | TRUE | 0,0360651 | No |
| orf19.2599 | 2339 | 1889 | 2144 | 1456 | 13318 | 9603 | 12470 | 14128 | 2230 | 1997 | 2118 | 2281 | 10671 | 10919 | 10587 | 11487 | 6536,44 | 2156 | 10916 | 5,068 | 2,341 | 46,922 | 0 | 0 | 0,00203 | 0,0023425 | 0,0020888 | 0,0020888 | TRUE | 0,029721 | No |
| orf19.7043.1 | 739 | 749 | 662 | 450 | 4991 | 3301 | 4293 | 3738 | 705 | 792 | 654 | 705 | 3999 | 3754 | 3645 | 3039 | 2161,53 | 714 | 3609 | 5,056 | 2,338 | 26,709 | 3,67E-157 | 3,91E-156 | 0,0096022 | 0,0028509 | 0,0064451 | 0,0064451 | TRUE | 0,1383544 | No |
| orf19.3329 | 861 | 640 | 724 | 491 | 4573 | 3246 | 4423 | 4727 | 821 | 677 | 715 | 769 | 3664 | 3691 | 3755 | 3844 | 2241,98 | 746 | 3738 | 5,018 | 2,327 | 39,85 | 0 | 0 | 0,0019289 | 0,0028236 | 0,0024035 | 0,0024035 | TRUE | 0,0615496 | No |
| orf19.3590 | 2096 | 1812 | 1956 | 1446 | 12312 | 9106 | 11480 | 13066 | 1998 | 1916 | 1932 | 2265 | 9865 | 10354 | 9746 | 10624 | 6087,73 | 2028 | 10147 | 5,015 | 2,326 | 40,79 | 0 | 0 | 0,0036275 | 0,002361 | 0,002801 | 0,002801 | TRUE | 0,0703616 | No |
| orf19.5557 | 630 | 685 | 640 | 444 | 3945 | 3007 | 3697 | 3964 | 601 | 724 | 632 | 696 | 3161 | 3419 | 3139 | 3223 | 1949,35 | 663 | 3236 | 4,891 | 2,29 | 36,294 | 1,99E-288 | 4,24E-287 | 0,0027389 | 0,0029336 | 0,0028385 | 0,0028385 | TRUE | 0,0505041 | No |
| orf19.7545 | 36 | 41 | 41 | 30 | 249 | 170 | 227 | 256 | 34 | 43 | 41 | 47 | 200 | 193 | 193 | 208 | 119,86 | 41 | 198 | 4,875 | 2,285 | 14,5 | 1,22E-47 | 4,35E-47 | 1,00E-08 | 0,0157899 | 0,0080533 | 0,0080533 | TRUE | 0,1055183 | No |
| orf19.3051 | 1054 | 947 | 975 | 623 | 6013 | 4295 | 5657 | 5674 | 1005 | 1001 | 963 | 976 | 4818 | 4884 | 4803 | 4614 | 2882,94 | 986 | 4780 | 4,842 | 2,276 | 52,621 | 0 | 0 | 1,00E-08 | 0,0026608 | 0,0011371 | 0,0011371 | TRUE | 0,0067219 | No |
| orf19.3671 | 1 | 1 | 0 | 2 | 6 | 4 | 12 | 3 | 1 | 1 | 0 | 3 | 5 | 5 | 10 | 2 | 3,39 | 1 | 6 | 4,827 | 2,271 | 2,306 | 0,0211203 | 0,0261686 | 0,1958775 | 0,486302 | 0,365504 | 0,365504 | TRUE | 0,6204155 | No |
| orf19.3461 | 11 | 10 | 18 | 11 | 83 | 61 | 75 | 81 | 10 | 11 | 18 | 17 | 67 | 69 | 64 | 66 | 40,18 | 14 | 66 | 4,812 | 2,267 | 8,406 | 4,23E-17 | 8,91E-17 | 1,00E-08 | 0,04295 | 0,0229871 | 0,0229871 | TRUE | 0,195509 | No |
| orf19.2587 | 6 | 7 | 5 | 6 | 46 | 22 | 36 | 42 | 6 | 7 | 5 | 9 | 37 | 25 | 31 | 34 | 19,26 | 7 | 32 | 4,8 | 2,263 | 5,792 | 6,95E-09 | 1,16E-08 | 1,00E-08 | 0,0873578 | 0,0493866 | 0,0493866 | TRUE | 0,1162672 | No |
| orf19.7549 | 216 | 208 | 211 | 135 | 1280 | 922 | 1134 | 1241 | 206 | 220 | 208 | 211 | 1026 | 1048 | 963 | 1009 | 611,45 | 211 | 1012 | 4,783 | 2,258 | 30,014 | 6,52E-198 | 8,85E-197 | 1,00E-08 | 0,0047765 | 0,0023721 | 0,0023721 | TRUE | 0,012552 | No |
| orf19.5164 | 358 | 341 | 351 | 271 | 2179 | 1580 | 1999 | 2104 | 341 | 361 | 347 | 425 | 1746 | 1797 | 1697 | 1711 | 1052,95 | 368 | 1738 | 4,762 | 2,252 | 33,786 | 3,17E-250 | 5,41E-249 | 0,000421 | 0,0036506 | 0,0024814 | 0,0024814 | TRUE | 0,0979634 | No |
| orf19.1275 | 282 | 239 | 258 | 185 | 1511 | 1131 | 1474 | 1583 | 269 | 253 | 255 | 290 | 1211 | 1286 | 1251 | 1287 | 762,7 | 267 | 1259 | 4,747 | 2,247 | 32,438 | 8,06E-231 | 1,24E-229 | 1,00E-08 | 0,004244 | 0,0021613 | 0,0021613 | TRUE | 0,028985 | No |
| orf19.6445 | 1076 | 850 | 960 | 673 | 5860 | 3705 | 5561 | 6065 | 1026 | 899 | 948 | 1054 | 4695 | 4213 | 4721 | 4931 | 2811,04 | 982 | 4640 | 4,737 | 2,244 | 34,194 | 3,02E-256 | 5,29E-255 | 0,0039505 | 0,0026754 | 0,0034654 | 0,0034654 | TRUE | 0,0461496 | No |
| orf19.4583 | 167 | 159 | 161 | 98 | 937 | 635 | 936 | 940 | 159 | 168 | 159 | 154 | 751 | 722 | 795 | 764 | 458,97 | 160 | 758 | 4,731 | 2,242 | 26,439 | 4,82E-154 | 4,97E-153 | 1,00E-08 | 0,0056687 | 0,0028662 | 0,0028662 | TRUE | 0,0124198 | No |
| orf19.2452 | 974 | 1041 | 1057 | 629 | 6112 | 4426 | 5517 | 5449 | 929 | 1101 | 1044 | 985 | 4897 | 5033 | 4684 | 4431 | 2887,94 | 1015 | 4761 | 4,689 | 2,229 | 35,504 | 4,28E-276 | 8,35E-275 | 0,0034073 | 0,0026598 | 0,0031385 | 0,0031385 | TRUE | 0,0422534 | No |
| orf19.4570 | 140 | 129 | 135 | 91 | 795 | 547 | 761 | 781 | 133 | 136 | 133 | 143 | 637 | 622 | 646 | 635 | 385,74 | 136 | 635 | 4,674 | 2,225 | 25,144 | 1,64E-139 | 1,53E-138 | 1,00E-08 | 0,0063478 | 0,0027488 | 0,0027488 | TRUE | 0,0067067 | No |
| orf19.7666 | 18 | 13 | 16 | 11 | 94 | 72 | 71 | 99 | 17 | 14 | 16 | 17 | 75 | 82 | 60 | 80 | 45,24 | 16 | 74 | 4,668 | 2,223 | 8,633 | 5,97E-18 | 1,28E-17 | 1,00E-08 | 0,0383854 | 0,0229456 | 0,0229456 | TRUE | 0,1588775 | No |
| orf19.3306 | 20 | 20 | 21 | 14 | 118 | 101 | 100 | 107 | 19 | 21 | 21 | 22 | 95 | 115 | 85 | 87 | 58,02 | 21 | 96 | 4,6 | 2,202 | 9,612 | 7,09E-22 | 1,65E-21 | 1,00E-08 | 0,0303884 | 0,0188426 | 0,0188426 | TRUE | 0,1657867 | No |
| orf19.5753 | 5390 | 4940 | 5312 | 3376 | 29878 | 21700 | 27096 | 29438 | 5139 | 5223 | 5248 | 5289 | 23940 | 24675 | 23004 | 23936 | 14556,77 | 5225 | 23889 | 4,573 | 2,193 | 65,693 | 0 | 0 | 0,0004765 | 0,0022041 | 0,0009439 | 0,0009439 | TRUE | 0,0075977 | No |
| orf19.4688 | 1842 | 1626 | 1834 | 1243 | 10161 | 7736 | 9579 | 9318 | 1756 | 1719 | 1812 | 1947 | 8142 | 8797 | 8133 | 7576 | 4985,21 | 1808 | 8162 | 4,519 | 2,176 | 38,124 | 0 | 0 | 0,0029444 | 0,0024206 | 0,0027619 | 0,0027619 | TRUE | 0,0331689 | No |
| orf19.5648 | 531 | 454 | 444 | 279 | 2770 | 1608 | 2528 | 2718 | 506 | 480 | 439 | 437 | 2220 | 1828 | 2146 | 2210 | 1283,28 | 466 | 2101 | 4,506 | 2,172 | 27,265 | 1,12E-163 | 1,24E-162 | 0,0058313 | 0,0033707 | 0,0046802 | 0,0046802 | TRUE | 0,0901324 | No |
| orf19.6191 | 22 | 14 | 18 | 9 | 112 | 56 | 99 | 86 | 21 | 15 | 18 | 14 | 90 | 64 | 84 | 70 | 46,88 | 17 | 77 | 4,491 | 2,167 | 8,461 | 2,65E-17 | 5,62E-17 | 1,00E-08 | 0,0371134 | 0,0250245 | 0,0250245 | TRUE | 0,1040885 | No |
| orf19.3311 | 189 | 161 | 164 | 92 | 929 | 579 | 885 | 995 | 180 | 170 | 162 | 144 | 744 | 658 | 751 | 809 | 452,46 | 164 | 740 | 4,486 | 2,165 | 22,183 | 5,01E-109 | 3,59E-108 | 0,0046059 | 0,0057201 | 0,0051851 | 0,0051851 | TRUE | 0,0628191 | No |
| orf19.6487 | 16 | 19 | 19 | 19 | 99 | 86 | 104 | 115 | 15 | 20 | 19 | 30 | 79 | 98 | 88 | 94 | 55,35 | 21 | 90 | 4,422 | 2,145 | 9,212 | 3,19E-20 | 7,20E-20 | 1,00E-08 | 0,0317556 | 0,0197332 | 0,0197332 | TRUE | 0,2765485 | No |
| orf19.5005 | 3667 | 3324 | 3587 | 2231 | 19411 | 13649 | 17846 | 19451 | 3496 | 3515 | 3544 | 3495 | 15553 | 15520 | 15151 | 15816 | 9511,21 | 3512 | 15510 | 4,415 | 2,142 | 70,305 | 0 | 0 | 1,00E-08 | 0,0022639 | 0,0007032 | 0,0007032 | TRUE | 0,0029953 | No |
| orf19.6782 | 170 | 162 | 158 | 111 | 891 | 700 | 861 | 835 | 162 | 171 | 156 | 174 | 714 | 796 | 731 | 679 | 447,9 | 166 | 730 | 4,41 | 2,141 | 23,693 | 4,29E-124 | 3,55E-123 | 1,00E-08 | 0,0057571 | 0,0038686 | 0,0038686 | TRUE | 0,0412725 | No |
| orf19.4040 | 3025 | 2327 | 2603 | 1809 | 14508 | 9882 | 13641 | 14845 | 2884 | 2461 | 2572 | 2834 | 11625 | 11237 | 11581 | 12070 | 7157,88 | 2688 | 11628 | 4,33 | 2,114 | 38,616 | 0 | 0 | 0,0030123 | 0,0023207 | 0,0026311 | 0,0026311 | TRUE | 0,0392878 | No |
| orf19.4833 | 11923 | 9778 | 10533 | 7209 | 57227 | 39412 | 54828 | 61427 | 11367 | 10339 | 10406 | 11293 | 45854 | 44815 | 46549 | 49946 | 28821,16 | 10851 | 46791 | 4,313 | 2,109 | 42,764 | 0 | 0 | 0,0023566 | 0,0021482 | 0,0022743 | 0,0022743 | TRUE | 0,0252559 | No |
| orf19.5748 | 17 | 18 | 20 | 12 | 102 | 66 | 89 | 95 | 16 | 19 | 20 | 19 | 82 | 75 | 76 | 77 | 47,92 | 18 | 78 | 4,212 | 2,075 | 8,832 | 1,03E-18 | 2,25E-18 | 1,00E-08 | 0,036353 | 0,0172217 | 0,0172217 | TRUE | 0,0415784 | No |
| orf19.1064 | 4444 | 4108 | 4375 | 2750 | 21852 | 16347 | 20583 | 23127 | 4237 | 4344 | 4322 | 4308 | 17509 | 18588 | 17475 | 18804 | 11198,42 | 4303 | 18094 | 4,205 | 2,072 | 54,289 | 0 | 0 | 0,0008288 | 0,0022379 | 0,0012436 | 0,0012436 | TRUE | 0,0086426 | No |
| orf19.2163 | 1074 | 945 | 1083 | 605 | 5446 | 3385 | 5054 | 5297 | 1024 | 999 | 1070 | 948 | 4364 | 3849 | 4291 | 4307 | 2606,42 | 1010 | 4203 | 4,153 | 2,054 | 35,658 | 1,78E-278 | 3,56E-277 | 0,0023865 | 0,0027212 | 0,002526 | 0,002526 | TRUE | 0,0382331 | No |
| orf19.3931 | 1858 | 1574 | 1686 | 1158 | 8578 | 6407 | 8468 | 8955 | 1771 | 1664 | 1666 | 1814 | 6873 | 7285 | 7189 | 7281 | 4443,07 | 1729 | 7157 | 4,146 | 2,052 | 47,135 | 0 | 0 | 0,0008516 | 0,0024608 | 0,0014318 | 0,0014318 | TRUE | 0,0116965 | No |
| orf19.1208 | 20 | 20 | 27 | 19 | 135 | 72 | 118 | 121 | 19 | 21 | 27 | 30 | 108 | 82 | 100 | 98 | 60,66 | 24 | 97 | 4,113 | 2,04 | 9,3 | 1,40E-20 | 3,18E-20 | 1,00E-08 | 0,0291596 | 0,0183177 | 0,0183177 | TRUE | 0,1281201 | No |
| orf19.2360 | 8826 | 8153 | 8552 | 5580 | 43611 | 32677 | 38543 | 44000 | 8415 | 8621 | 8449 | 8741 | 34944 | 37157 | 32723 | 35776 | 21853,16 | 8556 | 35150 | 4,109 | 2,039 | 46,523 | 0 | 0 | 0,0014592 | 0,0021664 | 0,0017659 | 0,0017659 | TRUE | 0,0264638 | No |
| orf19.6910 | 77 | 61 | 62 | 44 | 348 | 257 | 326 | 312 | 73 | 64 | 61 | 69 | 279 | 292 | 277 | 254 | 171,2 | 67 | 276 | 4,106 | 2,038 | 15,656 | 3,04E-55 | 1,20E-54 | 1,00E-08 | 0,0116817 | 0,0063825 | 0,0063825 | TRUE | 0,0458015 | No |
| orf19.5211 | 628 | 489 | 505 | 377 | 2781 | 1936 | 2622 | 2911 | 599 | 517 | 499 | 591 | 2228 | 2201 | 2226 | 2367 | 1403,5 | 552 | 2256 | 4,104 | 2,037 | 31,155 | 4,33E-213 | 6,25E-212 | 0,0024572 | 0,0032611 | 0,0028893 | 0,0028893 | TRUE | 0,0496753 | No |
| orf19.2469 | 161 | 146 | 149 | 94 | 713 | 522 | 759 | 815 | 153 | 154 | 147 | 147 | 571 | 594 | 644 | 663 | 384,28 | 150 | 618 | 4,101 | 2,036 | 21,538 | 6,88E-103 | 4,61E-102 | 1,00E-08 | 0,0063639 | 0,0041795 | 0,0041795 | TRUE | 0,0326879 | No |
| orf19.3065 | 1035 | 935 | 1024 | 648 | 4905 | 3564 | 4842 | 5130 | 987 | 989 | 1012 | 1015 | 3930 | 4053 | 4111 | 4171 | 2533,37 | 1001 | 4066 | 4,067 | 2,024 | 46,523 | 0 | 0 | 1,00E-08 | 0,0027394 | 0,0011499 | 0,0011499 | TRUE | 0,0063676 | No |
| orf19.780 | 2115 | 1923 | 2251 | 1285 | 10413 | 7439 | 9374 | 10686 | 2016 | 2033 | 2224 | 2013 | 8344 | 8459 | 7958 | 8689 | 5217,03 | 2072 | 8362 | 4,033 | 2,012 | 42,339 | 0 | 0 | 0,0014718 | 0,002406 | 0,0018437 | 0,0018437 | TRUE | 0,0303659 | No |
| orf19.2479 | 161 | 155 | 163 | 98 | 728 | 587 | 760 | 807 | 153 | 164 | 161 | 154 | 583 | 667 | 645 | 656 | 398,02 | 158 | 638 | 4,028 | 2,01 | 22,103 | 2,99E-108 | 2,13E-107 | 1,00E-08 | 0,0062165 | 0,0037362 | 0,0037362 | TRUE | 0,0405329 | No |
| orf19.5479 | 4 | 3 | 1 | 0 | 10 | 7 | 13 | 9 | 4 | 3 | 1 | 0 | 8 | 8 | 11 | 7 | 5,29 | 2 | 8 | 3,994 | 1,998 | 2,802 | 0,0050799 | 0,006563 | 1,00E-08 | 0,3126114 | 0,1792199 | 0,1792199 | TRUE | 0,3308939 | No |
| orf19.1429 | 85 | 76 | 85 | 46 | 412 | 250 | 400 | 384 | 81 | 80 | 84 | 72 | 330 | 284 | 340 | 312 | 197,96 | 79 | 316 | 3,971 | 1,99 | 16,154 | 1,06E-58 | 4,40E-58 | 1,00E-08 | 0,0103855 | 0,0062309 | 0,0062309 | TRUE | 0,0485325 | No |
| orf19.4157 | 403 | 331 | 348 | 237 | 1791 | 1300 | 1620 | 1790 | 384 | 350 | 344 | 371 | 1435 | 1478 | 1375 | 1455 | 899,17 | 362 | 1436 | 3,965 | 1,987 | 31,194 | 1,30E-213 | 1,89E-212 | 1,00E-08 | 0,0039173 | 0,0020519 | 0,0020519 | TRUE | 0,0229652 | No |
| orf19.5369 | 1293 | 1064 | 1173 | 901 | 5982 | 4102 | 5694 | 6246 | 1233 | 1125 | 1159 | 1411 | 4793 | 4664 | 4834 | 5079 | 3037,3 | 1232 | 4842 | 3,945 | 1,98 | 30,372 | 1,29E-202 | 1,79E-201 | 0,0043816 | 0,0026319 | 0,0035301 | 0,0035301 | TRUE | 0,1062025 | No |
| orf19.4356 | 188 | 219 | 196 | 136 | 1035 | 727 | 904 | 964 | 179 | 232 | 194 | 213 | 829 | 827 | 767 | 784 | 503,1 | 204 | 802 | 3,94 | 1,978 | 22,948 | 1,56E-116 | 1,20E-115 | 0,0014177 | 0,0053549 | 0,0038467 | 0,0038467 | TRUE | 0,0991999 | No |
| orf19.7537 | 151 | 98 | 146 | 103 | 730 | 435 | 635 | 658 | 144 | 104 | 144 | 161 | 585 | 495 | 539 | 535 | 338,36 | 138 | 538 | 3,928 | 1,974 | 17,341 | 2,33E-67 | 1,08E-66 | 0,0087044 | 0,0069438 | 0,0075522 | 0,0075522 | TRUE | 0,293091 | No |
| orf19.1632 | 404 | 419 | 435 | 241 | 2036 | 1485 | 1871 | 1883 | 385 | 443 | 430 | 378 | 1631 | 1689 | 1588 | 1531 | 1009,37 | 409 | 1610 | 3,921 | 1,971 | 29,409 | 4,19E-190 | 5,43E-189 | 0,001543 | 0,0037179 | 0,0026776 | 0,0026776 | TRUE | 0,0348274 | No |
| orf19.2344 | 2352 | 1820 | 1963 | 1502 | 10098 | 7042 | 9493 | 10359 | 2242 | 1924 | 1939 | 2353 | 8091 | 8007 | 8060 | 8423 | 5130 | 2114 | 8145 | 3,858 | 1,948 | 30,287 | 1,72E-201 | 2,38E-200 | 0,0050896 | 0,0024113 | 0,0036504 | 0,0036504 | TRUE | 0,0659492 | No |
| orf19.1212 | 804 | 711 | 753 | 557 | 3662 | 2754 | 3463 | 3599 | 767 | 752 | 744 | 873 | 2934 | 3132 | 2940 | 2926 | 1883,37 | 784 | 2983 | 3,824 | 1,935 | 34,298 | 8,49E-258 | 1,51E-256 | 0,0013833 | 0,0029631 | 0,0021888 | 0,0021888 | TRUE | 0,0593073 | No |
| orf19.3318 | 313 | 316 | 327 | 187 | 1555 | 1071 | 1372 | 1415 | 298 | 334 | 323 | 293 | 1246 | 1218 | 1165 | 1151 | 753,46 | 312 | 1195 | 3,815 | 1,932 | 27,911 | 1,96E-171 | 2,25E-170 | 1,00E-08 | 0,0042704 | 0,0024501 | 0,0024501 | TRUE | 0,024284 | No |

| | | | | | | | | | | | | | | | | | | | | | | | | | | | | | | | |
|---|---|---|---|---|---|---|---|---|---|---|---|---|---|---|---|---|---|---|---|---|---|---|---|---|---|---|---|---|---|---|---|
| orf19.3911 | 7965 | 6358 | 6457 | 4885 | 32960 | 23350 | 30264 | 35115 | 7594 | 6723 | 6379 | 7653 | 26410 | 26551 | 25694 | 28552 | 16944,36 | 7087 | 26802 | 3,783 | 1,92 | 30,226 | 1,09E-200 | 1,48E-199 | 0,0049444 | 0,0021882 | 0,0037785 | 0,0037785 | TRUE | 0,0554176 | No |
| orf19.5469 | 363 | 321 | 300 | 197 | 1646 | 1037 | 1452 | 1414 | 346 | 339 | 296 | 309 | 1319 | 1179 | 1233 | 1150 | 771,37 | 322 | 1220 | 3,775 | 1,917 | 25,024 | 3,37E-138 | 3,13E-137 | 0,002904 | 0,0042198 | 0,003542 | 0,003542 | TRUE | 0,039759 | No |
| orf19.639 | 164 | 140 | 144 | 108 | 727 | 509 | 685 | 694 | 156 | 148 | 142 | 169 | 583 | 579 | 582 | 564 | 365,37 | 154 | 577 | 3,773 | 1,916 | 21,74 | 8,60E-105 | 5,83E-104 | 1,00E-08 | 0,006585 | 0,0030604 | 0,0030604 | TRUE | 0,0337568 | No |
| orf19.3177 | 509 | 444 | 477 | 258 | 2206 | 1379 | 2072 | 2234 | 485 | 469 | 471 | 404 | 1768 | 1568 | 1759 | 1816 | 1092,67 | 457 | 1728 | 3,759 | 1,91 | 26,701 | 4,54E-157 | 4,84E-156 | 0,0033067 | 0,0035939 | 0,0034414 | 0,0034414 | TRUE | 0,0592835 | No |
| orf19.3879 | 69 | 69 | 72 | 64 | 356 | 245 | 311 | 384 | 66 | 73 | 71 | 100 | 285 | 279 | 264 | 312 | 181,28 | 78 | 285 | 3,757 | 1,91 | 14,185 | 1,13E-45 | 3,92E-45 | 0,0039484 | 0,0111486 | 0,0085276 | 0,0085276 | TRUE | 0,2621173 | No |
| orf19.3884 | 338 | 270 | 299 | 202 | 1461 | 938 | 1406 | 1391 | 322 | 285 | 295 | 316 | 1171 | 1067 | 1194 | 1131 | 722,69 | 304 | 1141 | 3,751 | 1,907 | 26,524 | 5,17E-155 | 5,41E-154 | 0,0011685 | 0,0043632 | 0,0027466 | 0,0027466 | TRUE | 0,0213863 | No |
| orf19.5723 | 3726 | 3173 | 3397 | 2376 | 15721 | 11837 | 14802 | 16496 | 3552 | 3355 | 3356 | 3722 | 12597 | 13460 | 12567 | 13413 | 8252,7 | 3496 | 13009 | 3,724 | 1,897 | 40,515 | 0 | 0 | 0,0017575 | 0,0022902 | 0,0019097 | 0,0019097 | TRUE | 0,0216928 | No |
| orf19.6636 | 76 | 86 | 84 | 48 | 354 | 243 | 370 | 398 | 72 | 91 | 83 | 75 | 284 | 276 | 314 | 324 | 189,91 | 80 | 300 | 3,721 | 1,896 | 15,006 | 6,67E-51 | 2,49E-50 | 1,00E-08 | 0,010737 | 0,0069617 | 0,0069617 | TRUE | 0,0725203 | No |
| orf19.2921 | 105 | 121 | 114 | 80 | 504 | 377 | 522 | 537 | 100 | 128 | 113 | 125 | 404 | 429 | 443 | 437 | 272,29 | 116 | 428 | 3,702 | 1,888 | 17,953 | 4,56E-72 | 2,25E-71 | 1,00E-08 | 0,0081212 | 0,0047862 | 0,0047862 | TRUE | 0,0981508 | No |
| orf19.5217 | 494 | 443 | 447 | 301 | 2158 | 1450 | 2039 | 2076 | 471 | 468 | 442 | 472 | 1729 | 1649 | 1731 | 1688 | 1081,19 | 463 | 1699 | 3,676 | 1,878 | 33,424 | 6,15E-245 | 1,02E-243 | 1,00E-08 | 0,0036099 | 0,0015683 | 0,0015683 | TRUE | 0,0121938 | No |
| orf19.2114 | 201 | 157 | 183 | 154 | 832 | 646 | 856 | 864 | 192 | 166 | 181 | 241 | 667 | 735 | 727 | 703 | 451,27 | 195 | 708 | 3,675 | 1,878 | 19,562 | 3,27E-85 | 1,86E-84 | 0,0044354 | 0,0057297 | 0,0053204 | 0,0053204 | TRUE | 0,2194925 | No |
| orf19.4716 | 16048 | 13178 | 14286 | 9270 | 66175 | 44415 | 62823 | 67619 | 15300 | 13934 | 14113 | 14522 | 53024 | 50504 | 53336 | 54981 | 33714,3 | 14467 | 52961 | 3,661 | 1,872 | 44,365 | 0 | 0 | 0,0014319 | 0,00214 | 0,0016633 | 0,0016633 | TRUE | 0,0185867 | No |
| orf19.398 | 124 | 113 | 127 | 96 | 615 | 403 | 520 | 559 | 118 | 119 | 125 | 150 | 493 | 458 | 441 | 455 | 295,07 | 128 | 462 | 3,642 | 1,865 | 18,331 | 4,71E-75 | 2,41E-74 | 0,0006899 | 0,0076557 | 0,0045996 | 0,0045996 | TRUE | 0,100602 | No |
| orf19.5341 | 7571 | 5535 | 5844 | 4185 | 28414 | 19218 | 26619 | 31039 | 7218 | 5853 | 5773 | 6556 | 22767 | 21853 | 22599 | 25238 | 14732,13 | 6350 | 23114 | 3,64 | 1,864 | 25,34 | 1,15E-141 | 1,10E-140 | 0,0075774 | 0,0022027 | 0,0050907 | 0,0050907 | TRUE | 0,1045216 | No |
| orf19.5805 | 773 | 610 | 726 | 475 | 3157 | 2211 | 3100 | 3250 | 737 | 645 | 717 | 744 | 2530 | 2514 | 2632 | 2643 | 1645,18 | 711 | 2580 | 3,636 | 1,862 | 34,004 | 1,97E-253 | 3,41E-252 | 0,0007345 | 0,0030893 | 0,00192 | 0,00192 | TRUE | 0,0464261 | No |
| orf19.2812 | 357 | 297 | 351 | 205 | 1493 | 1036 | 1422 | 1502 | 340 | 314 | 347 | 321 | 1196 | 1178 | 1207 | 1221 | 765,65 | 330 | 1200 | 3,624 | 1,857 | 29,152 | 7,78E-187 | 9,81E-186 | 1,00E-08 | 0,0042358 | 0,0018535 | 0,0018535 | TRUE | 0,0150337 | No |
| orf19.4128 | 1045 | 955 | 1083 | 594 | 4492 | 3354 | 4173 | 4365 | 996 | 1010 | 1070 | 931 | 3599 | 3814 | 3543 | 3549 | 2313,96 | 1002 | 3626 | 3,608 | 1,851 | 35,768 | 3,53E-280 | 7,17E-279 | 0,0011561 | 0,0028008 | 0,001891 | 0,001891 | TRUE | 0,0246882 | No |
| orf19.3437 | 88 | 89 | 89 | 69 | 437 | 319 | 383 | 364 | 84 | 94 | 88 | 108 | 350 | 363 | 325 | 296 | 213,5 | 94 | 334 | 3,601 | 1,848 | 15,001 | 7,19E-51 | 2,68E-50 | 0,0042374 | 0,0097815 | 0,0072331 | 0,0072331 | TRUE | 0,0782646 | No |
| orf19.6838 | 1739 | 1501 | 1552 | 1057 | 7112 | 5112 | 6443 | 7482 | 1658 | 1587 | 1533 | 1656 | 5699 | 5813 | 5470 | 6084 | 3687,41 | 1608 | 5766 | 3,588 | 1,843 | 37,904 | 0 | 0 | 0,0014373 | 0,0025366 | 0,0018434 | 0,0018434 | TRUE | 0,0169849 | No |
| orf19.4076 | 8342 | 7612 | 8011 | 5007 | 34971 | 24856 | 32575 | 36455 | 7953 | 8049 | 7914 | 7844 | 28021 | 28264 | 27656 | 29641 | 18167,74 | 7940 | 28396 | 3,576 | 1,838 | 55,747 | 0 | 0 | 0,000417 | 0,0021817 | 0,0009576 | 0,0009576 | TRUE | 0,0107315 | No |
| orf19.7673 | 425 | 307 | 338 | 196 | 1686 | 845 | 1546 | 1584 | 405 | 325 | 334 | 307 | 1351 | 961 | 1313 | 1288 | 785,38 | 343 | 1228 | 3,576 | 1,838 | 15,842 | 1,61E-56 | 6,44E-56 | 0,0179597 | 0,0041819 | 0,0109301 | 0,0109301 | TRUE | 0,2523221 | No |
| orf19.1544 | 312 | 271 | 294 | 219 | 1355 | 910 | 1310 | 1327 | 297 | 287 | 290 | 343 | 1086 | 1035 | 1112 | 1079 | 691,15 | 304 | 1078 | 3,573 | 1,837 | 26,092 | 4,49E-150 | 4,52E-149 | 1,00E-08 | 0,0044669 | 0,0025056 | 0,0025056 | TRUE | 0,0647223 | No |
| orf19.5934 | 281 | 275 | 281 | 156 | 1164 | 797 | 1216 | 1230 | 268 | 291 | 278 | 244 | 933 | 906 | 1032 | 1000 | 619,01 | 270 | 968 | 3,568 | 1,835 | 23,326 | 2,42E-120 | 1,94E-119 | 0,0021378 | 0,0047437 | 0,0034315 | 0,0034315 | TRUE | 0,0354282 | No |
| orf19.3411 | 204 | 166 | 179 | 102 | 782 | 581 | 693 | 817 | 194 | 176 | 177 | 160 | 627 | 661 | 588 | 664 | 405,82 | 177 | 635 | 3,566 | 1,834 | 20,416 | 1,21E-92 | 7,42E-92 | 1,00E-08 | 0,0061372 | 0,0039274 | 0,0039274 | TRUE | 0,0496585 | No |
| orf19.1639 | 104 | 66 | 82 | 46 | 339 | 282 | 340 | 344 | 99 | 70 | 81 | 72 | 272 | 321 | 289 | 280 | 185,33 | 80 | 290 | 3,56 | 1,832 | 13,911 | 5,43E-44 | 1,83E-43 | 0,0037201 | 0,0109505 | 0,0082792 | 0,0082792 | TRUE | 0,2424799 | No |
| orf19.5564 | 437 | 422 | 427 | 267 | 1812 | 1357 | 1836 | 1782 | 417 | 446 | 422 | 418 | 1452 | 1543 | 1559 | 1449 | 963,2 | 426 | 1501 | 3,521 | 1,816 | 29,587 | 2,21E-192 | 2,92E-191 | 1,00E-08 | 0,0037959 | 0,002013 | 0,002013 | TRUE | 0,0124066 | No |
| orf19.7281 | 2115 | 1813 | 1971 | 1252 | 8512 | 5984 | 8196 | 8630 | 2016 | 1917 | 1947 | 1961 | 6820 | 6804 | 6958 | 7017 | 4430,26 | 1960 | 6900 | 3,519 | 1,815 | 50,936 | 0 | 0 | 1,00E-08 | 0,0024619 | 0,0008695 | 0,0008695 | TRUE | 0,004879 | No |
| orf19.5711 | 958 | 929 | 1002 | 663 | 4292 | 3032 | 3938 | 4348 | 913 | 982 | 990 | 1039 | 3439 | 3448 | 3343 | 3535 | 2211,19 | 981 | 3441 | 3,519 | 1,815 | 37,327 | 5,94E-305 | 1,34E-303 | 0,000524 | 0,0028338 | 0,0015691 | 0,0015691 | TRUE | 0,0264261 | No |
| orf19.446.1 | 533 | 467 | 462 | 294 | 2168 | 1537 | 1863 | 2078 | 508 | 494 | 456 | 461 | 1737 | 1748 | 1582 | 1690 | 1084,39 | 480 | 1689 | 3,511 | 1,812 | 28,818 | 1,29E-182 | 1,58E-181 | 0,0013449 | 0,0036054 | 0,002372 | 0,002372 | TRUE | 0,0222522 | No |
| orf19.4287 | 808 | 629 | 729 | 432 | 3094 | 2045 | 3049 | 3154 | 770 | 665 | 720 | 677 | 2479 | 2325 | 2589 | 2565 | 1598,74 | 708 | 2490 | 3,509 | 1,811 | 29,414 | 3,59E-190 | 4,65E-189 | 0,0022115 | 0,0031183 | 0,0026716 | 0,0026716 | TRUE | 0,0429253 | No |
| orf19.288 | 2035 | 1854 | 2040 | 1341 | 8499 | 6276 | 8146 | 8760 | 1940 | 1960 | 2015 | 2101 | 6810 | 7136 | 6916 | 7123 | 4500,2 | 2004 | 6996 | 3,497 | 1,806 | 45,478 | 0 | 0 | 0,0004753 | 0,0024561 | 0,00117 | 0,00117 | TRUE | 0,0112744 | No |
| orf19.1591 | 1521 | 1327 | 1370 | 1009 | 6100 | 4549 | 5720 | 6486 | 1450 | 1403 | 1353 | 1581 | 4888 | 5173 | 4856 | 5274 | 3247,21 | 1447 | 5048 | 3,497 | 1,806 | 33,074 | 6,98E-240 | 1,13E-238 | 0,0025238 | 0,0025969 | 0,002386 | 0,002386 | TRUE | 0,0423913 | No |
| orf19.3019 | 414 | 359 | 385 | 223 | 1604 | 1152 | 1580 | 1621 | 395 | 380 | 380 | 349 | 1285 | 1310 | 1341 | 1318 | 844,82 | 376 | 1314 | 3,476 | 1,797 | 29,369 | 1,38E-189 | 1,78E-188 | 1,00E-08 | 0,0040348 | 0,0017881 | 0,0017881 | TRUE | 0,020121 | No |
| orf19.946 | 1787 | 1564 | 1554 | 970 | 7055 | 4611 | 6601 | 7023 | 1704 | 1654 | 1535 | 1520 | 5653 | 5243 | 5604 | 5710 | 3577,86 | 1603 | 5552 | 3,46 | 1,791 | 35,201 | 1,92E-271 | 3,66E-270 | 0,0017596 | 0,0025502 | 0,0020541 | 0,0020541 | TRUE | 0,0218681 | No |
| orf19.7279.1 | 2912 | 2468 | 2654 | 1710 | 11694 | 8092 | 10880 | 11248 | 2776 | 2610 | 2622 | 2679 | 9370 | 9201 | 9237 | 9146 | 5955,09 | 2672 | 9238 | 3,458 | 1,79 | 52,085 | 0 | 0 | 0,0001361 | 0,002367 | 0,0008756 | 0,0008756 | TRUE | 0,0091402 | No |
| orf19.7279 | 136 | 120 | 120 | 112 | 558 | 419 | 528 | 606 | 130 | 127 | 119 | 175 | 447 | 476 | 448 | 493 | 301,89 | 138 | 466 | 3,444 | 1,784 | 16,333 | 5,71E-60 | 2,40E-59 | 0,0038345 | 0,0075301 | 0,0063864 | 0,0063864 | TRUE | 0,2738706 | No |
| orf19.7130 | 41 | 42 | 33 | 30 | 193 | 109 | 147 | 181 | 39 | 44 | 33 | 47 | 155 | 124 | 125 | 147 | 89,21 | 41 | 138 | 3,429 | 1,778 | 9,891 | 4,54E-23 | 1,08E-22 | 1,00E-08 | 0,0204968 | 0,014022 | 0,014022 | TRUE | 0,1419439 | No |
| orf19.3925 | 8 | 12 | 10 | 5 | 42 | 30 | 36 | 41 | 8 | 13 | 10 | 8 | 34 | 34 | 31 | 33 | 21,21 | 10 | 33 | 3,43 | 1,778 | 5,248 | 1,54E-07 | 2,46E-07 | 1,00E-08 | 0,0794965 | 0,0390709 | 0,0390709 | TRUE | 0,1637722 | No |
| orf19.5806 | 60096 | 48214 | 53462 | 36525 | 2E+05 | 2E+05 | 2E+05 | 2E+05 | 57295 | 50980 | 52815 | 57218 | 2E+05 | 2E+05 | 2E+05 | 2E+05 | 120837,7 | 54577 | 187098 | 3,428 | 1,777 | 37,649 | 0 | 0 | 0,0021452 | 0,0021049 | 0,002129 | 0,002129 | TRUE | 0,0241175 | No |
| orf19.7276.1 | 136 | 132 | 112 | 78 | 520 | 329 | 564 | 549 | 130 | 140 | 111 | 122 | 417 | 374 | 479 | 446 | 277,26 | 126 | 429 | 3,423 | 1,775 | 15,346 | 3,76E-53 | 1,45E-52 | 0,0066698 | 0,0080132 | 0,0073775 | 0,0073775 | TRUE | 0,0799198 | No |
| orf19.4831 | 3253 | 2724 | 2989 | 1867 | 12437 | 8573 | 11875 | 13261 | 3101 | 2880 | 2953 | 2925 | 9965 | 9748 | 10082 | 10782 | 6554,65 | 2965 | 10144 | 3,421 | 1,774 | 39,569 | 0 | 0 | 0,0012034 | 0,0023418 | 0,001697 | 0,001697 | TRUE | 0,0219734 | No |
| orf19.489 | 553 | 498 | 535 | 345 | 2243 | 1575 | 2217 | 2171 | 527 | 527 | 529 | 540 | 1797 | 1791 | 1882 | 1765 | 1169,8 | 531 | 1809 | 3,413 | 1,771 | 32,736 | 4,86E-235 | 7,66E-234 | 1,00E-08 | 0,0034949 | 0,0015147 | 0,0015147 | TRUE | 0,0091665 | No |
| orf19.5323 | 1022 | 784 | 868 | 609 | 3847 | 2557 | 3644 | 3936 | 974 | 829 | 858 | 954 | 3082 | 2908 | 3094 | 3200 | 1987,37 | 904 | 3071 | 3,405 | 1,767 | 29,054 | 1,39E-185 | 1,73E-184 | 0,0029354 | 0,0029174 | 0,0027864 | 0,0027864 | TRUE | 0,03733 | No |
| orf19.3996 | 103 | 83 | 102 | 58 | 403 | 315 | 338 | 397 | 98 | 88 | 101 | 91 | 323 | 358 | 287 | 323 | 208,56 | 94 | 323 | 3,398 | 1,765 | 14,637 | 1,62E-48 | 5,88E-48 | 1,00E-08 | 0,009964 | 0,0067138 | 0,0067138 | TRUE | 0,063046 | No |
| orf19.5646 | 407 | 311 | 342 | 233 | 1531 | 1018 | 1412 | 1510 | 388 | 329 | 338 | 365 | 1227 | 1158 | 1199 | 1228 | 778,83 | 355 | 1203 | 3,393 | 1,763 | 25,974 | 9,75E-149 | 9,67E-148 | 0,0001731 | 0,0041995 | 0,0024827 | 0,0024827 | TRUE | 0,0530021 | No |
| orf19.7209 | 95 | 81 | 89 | 47 | 348 | 235 | 351 | 383 | 91 | 86 | 88 | 74 | 279 | 267 | 298 | 311 | 186,66 | 85 | 289 | 3,394 | 1,763 | 14,447 | 2,63E-47 | 9,36E-47 | 1,00E-08 | 0,0108877 | 0,0062633 | 0,0062633 | TRUE | 0,0482781 | No |
| orf19.7015 | 13605 | 10792 | 11269 | 8168 | 51082 | 34646 | 46025 | 54617 | 12971 | 11411 | 11133 | 12796 | 40931 | 39396 | 39075 | 44409 | 26515,05 | 12078 | 40953 | 3,391 | 1,762 | 28,312 | 2,43E-176 | 2,84E-175 | 0,0047353 | 0,0021532 | 0,0036629 | 0,0036629 | TRUE | 0,0395678 | No |
| orf19.206 | 1439 | 1228 | 1302 | 945 | 5630 | 4054 | 5341 | 5801 | 1372 | 1298 | 1286 | 1480 | 4511 | 4610 | 4534 | 4717 | 2976,15 | 1359 | 4593 | 3,39 | 1,761 | 35,965 | 3,00E-283 | 6,17E-282 | 0,0010284 | 0,002643 | 0,0017911 | 0,0017911 | TRUE | 0,0382314 | No |
| orf19.3391 | 1260 | 1086 | 1165 | 810 | 4870 | 3501 | 4691 | 5160 | 1201 | 1148 | 1151 | 1269 | 3902 | 3981 | 3983 | 4196 | 2603,84 | 1192 | 4016 | 3,376 | 1,756 | 36,79 | 2,69E-296 | 5,91E-295 | 0,0008914 | 0,0027218 | 0,0016042 | 0,0016042 | TRUE | 0,0193318 | No |
| orf19.6337 | 25 | 16 | 28 | 19 | 111 | 64 | 109 | 89 | 24 | 17 | 28 | 30 | 89 | 73 | 93 | 72 | 53,1 | 25 | 82 | 3,377 | 1,756 | 7,647 | 2,05E-14 | 4,07E-14 | 1,00E-08 | 0,0330123 | 0,0223154 | 0,0223154 | TRUE | 0,2561024 | No |
| orf19.2299 | 905 | 682 | 779 | 492 | 3490 | 2066 | 3138 | 3369 | 863 | 721 | 770 | 771 | 2796 | 2349 | 2664 | 2739 | 1709,18 | 781 | 2637 | 3,376 | 1,755 | 24,19 | 2,87E-129 | 2,47E-128 | 0,0046283 | 0,0030519 | 0,0041677 | 0,0041677 | TRUE | 0,0642173 | No |
| orf19.5302 | 127 | 104 | 111 | 80 | 505 | 347 | 467 | 451 | 121 | 110 | 110 | 125 | 405 | 395 | 396 | 367 | 253,55 | 116 | 391 | 3,368 | 1,752 | 16,903 | 4,29E-64 | 1,91E-63 | 1,00E-08 | 0,0085668 | 0,0043942 | 0,0043942 | TRUE | 0,0206272 | No |
| orf19.4532 | 2466 | 2173 | 2272 | 1497 | 9402 | 6923 | 8909 | 9966 | 2351 | 2298 | 2245 | 2345 | 7534 | 7872 | 7564 | 8103 | 5038,88 | 2310 | 7768 | 3,365 | 1,75 | 43,055 | 0 | 0 | 0,0007492 | 0,0024171 | 0,0012863 | 0,0012863 | TRUE | 0,0104934 | No |
| orf19.5000 | 4355 | 3924 | 4197 | 2534 | 17330 | 11804 | 16367 | 17245 | 4152 | 4149 | 4146 | 3970 | 13886 | 13422 | 13896 | 14022 | 8955,33 | 4104 | 13806 | 3,361 | 1,749 | 53,954 | 0 | 0 | 0,0002106 | 0,0022746 | 0,0008399 | 0,0008399 | TRUE | 0,0053933 | No |
| orf19.7237 | 451 | 374 | 378 | 295 | 1730 | 1167 | 1625 | 1766 | 430 | 395 | 373 | 462 | 1386 | 1327 | 1380 | 1436 | 898,72 | 415 | 1382 | 3,35 | 1,744 | 25,473 | 3,89E-143 | 3,75E-142 | 0,0013009 | 0,0039182 | 0,0028279 | 0,0028279 | TRUE | 0,0543211 | No |
| orf19.5365 | 841 | 782 | 861 | 562 | 3346 | 2482 | 3206 | 3672 | 802 | 827 | 851 | 880 | 2681 | 2822 | 2722 | 2986 | 1821,32 | 840 | 2803 | 3,344 | 1,742 | 31,534 | 3,00E-218 | 4,43E-217 | 0,0015104 | 0,0029928 | 0,0021019 | 0,0021019 | TRUE | 0,0240507 | No |
| orf19.362 | 70 | 55 | 57 | 44 | 280 | 140 | 261 | 274 | 67 | 58 | 56 | 69 | 224 | 159 | 222 | 223 | 134,76 | 62 | 207 | 3,344 | 1,742 | 10,869 | 1,62E-27 | 4,19E-27 | 0,0127323 | 0,0142755 | 0,013563 | 0,013563 | TRUE | 0,242362 | No |
| orf19.1155 | 274 | 256 | 256 | 177 | 1100 | 740 | 1051 | 1124 | 261 | 271 | 253 | 277 | 881 | 841 | 892 | 914 | 573,89 | 266 | 882 | 3,338 | 1,739 | 24,125 | 1,36E-128 | 1,17E-127 | 1,00E-08 | 0,0049523 | 0,0023873 | 0,0023873 | TRUE | 0,0112654 | No |
| orf19.1874 | 4 | 2 | 3 | 0 | 9 | 4 | 13 | 11 | 4 | 2 | 3 | 0 | 7 | 5 | 11 | 9 | 5,08 | 2 | 8 | 3,337 | 1,739 | 2,454 | 0,0141471 | 0,0177842 | 1,00E-08 | 0,3253577 | 0,1964403 | 0,1964403 | TRUE | 0,187086 | No |
| orf19.4443 | 1474 | 1230 | 1377 | 787 | 5753 | 3433 | 5358 | 5592 | 1405 | 1301 | 1360 | 1233 | 4610 | 3904 | 4549 | 4547 | 2863,52 | 1325 | 4402 | 3,319 | 1,731 | 26,531 | 4,21E-155 | 4,42E-154 | 0,0038119 | 0,0026647 | 0,0035607 | 0,0035607 | TRUE | 0,0697478 | No |

| | | | | | | | | | | | | | | | | | | | | | | | | | | | | | | | |
|---|---|---|---|---|---|---|---|---|---|---|---|---|---|---|---|---|---|---|---|---|---|---|---|---|---|---|---|---|---|---|---|
| orf19.7188 | 2064 | 1674 | 1782 | 1568 | 7898 | 6282 | 7022 | 8489 | 1968 | 1770 | 1760 | 2456 | 6328 | 7143 | 5962 | 6902 | 4286,29 | 1988 | 6584 | 3,316 | 1,729 | 17,76 | 1,45E-70 | 6,98E-70 | 0,0163416 | 0,0024743 | 0,0087576 | 0,0087576 | TRUE | 0,2953737 | No |
| orf19.1181 | 93 | 69 | 91 | 50 | 336 | 245 | 310 | 355 | 89 | 73 | 90 | 78 | 269 | 279 | 263 | 289 | 178,69 | 82 | 275 | 3,314 | 1,729 | 14,204 | 8,63E-46 | 3,00E-45 | 1,00E-08 | 0,0112799 | 0,0059212 | 0,0059212 | TRUE | 0,0643711 | No |
| orf19.7491 | 72 | 68 | 82 | 47 | 303 | 217 | 272 | 315 | 69 | 72 | 81 | 74 | 243 | 247 | 231 | 256 | 158,97 | 74 | 244 | 3,308 | 1,726 | 13,58 | 5,28E-42 | 1,73E-41 | 1,00E-08 | 0,0124196 | 0,0061849 | 0,0061849 | TRUE | 0,0455374 | No |
| orf19.2468 | 8 | 8 | 3 | 3 | 20 | 18 | 22 | 30 | 8 | 8 | 3 | 5 | 16 | 20 | 19 | 24 | 12,91 | 6 | 20 | 3,305 | 1,725 | 3,86 | 0,0001135 | 0,0001602 | 1,00E-08 | 0,129231 | 0,077232 | 0,077232 | TRUE | 0,320359 | No |
| orf19.4507 | 79 | 91 | 70 | 48 | 312 | 217 | 316 | 341 | 75 | 96 | 69 | 75 | 250 | 247 | 268 | 277 | 169,77 | 79 | 260 | 3,302 | 1,723 | 13,241 | 5,07E-40 | 1,61E-39 | 0,0005151 | 0,0117625 | 0,007534 | 0,007534 | TRUE | 0,2078301 | No |
| orf19.2396 | 643 | 568 | 628 | 396 | 2543 | 1844 | 2304 | 2450 | 613 | 601 | 620 | 620 | 2038 | 2097 | 1956 | 1992 | 1317,12 | 614 | 2021 | 3,293 | 1,719 | 33,024 | 3,73E-239 | 6,00E-238 | 1,00E-08 | 0,0033379 | 0,0014739 | 0,0014739 | TRUE | 0,0073082 | No |
| orf19.3266 | 188 | 157 | 202 | 164 | 768 | 615 | 698 | 841 | 179 | 166 | 200 | 257 | 615 | 699 | 593 | 684 | 424,1 | 200 | 648 | 3,268 | 1,708 | 14,833 | 8,90E-50 | 3,28E-49 | 0,0141175 | 0,0059628 | 0,0092197 | 0,0092197 | TRUE | 0,3268646 | No |
| orf19.5572 | 412 | 378 | 398 | 228 | 1575 | 1122 | 1452 | 1551 | 393 | 400 | 393 | 357 | 1262 | 1276 | 1233 | 1261 | 821,81 | 386 | 1258 | 3,245 | 1,698 | 27,992 | 2,05E-172 | 2,36E-171 | 1,00E-08 | 0,0040892 | 0,0017464 | 0,0017464 | TRUE | 0,0217194 | No |
| orf19.5342.2 | 190 | 123 | 157 | 101 | 652 | 348 | 677 | 653 | 181 | 130 | 155 | 158 | 522 | 396 | 575 | 531 | 331,05 | 156 | 506 | 3,245 | 1,698 | 12,775 | 2,27E-37 | 6,87E-37 | 0,0184893 | 0,007051 | 0,0125015 | 0,0125015 | TRUE | 0,2405204 | No |
| orf19.4336 | 12170 | 10109 | 10141 | 7408 | 43509 | 31396 | 40771 | 45534 | 11603 | 10689 | 10018 | 11605 | 34863 | 35700 | 34614 | 37023 | 23264,45 | 10979 | 35550 | 3,239 | 1,695 | 32,209 | 1,34E-227 | 2,04E-226 | 0,0028599 | 0,0021619 | 0,0025982 | 0,0025982 | TRUE | 0,0425678 | No |
| orf19.6403.1 | 3499 | 2828 | 2957 | 2316 | 13482 | 8799 | 11998 | 12969 | 3336 | 2990 | 2921 | 3628 | 10803 | 10005 | 10186 | 10545 | 6801,85 | 3219 | 10385 | 3,23 | 1,692 | 25,728 | 5,66E-146 | 5,57E-145 | 0,0055023 | 0,0023327 | 0,0039342 | 0,0039342 | TRUE | 0,0861307 | No |
| orf19.2150 | 1079 | 842 | 907 | 639 | 3872 | 2464 | 3634 | 4055 | 1029 | 890 | 896 | 1001 | 3103 | 2802 | 3085 | 3297 | 2012,84 | 954 | 3072 | 3,225 | 1,689 | 24,949 | 2,21E-137 | 2,04E-136 | 0,0040738 | 0,002907 | 0,003667 | 0,003667 | TRUE | 0,0416404 | No |
| orf19.2991 | 123 | 93 | 104 | 90 | 395 | 345 | 453 | 454 | 117 | 98 | 103 | 141 | 317 | 392 | 385 | 369 | 240,23 | 115 | 366 | 3,218 | 1,686 | 13,518 | 1,23E-41 | 4,00E-41 | 0,008625 | 0,0089259 | 0,00881 | 0,00881 | TRUE | 0,1794198 | No |
| orf19.5875 | 254 | 198 | 222 | 151 | 913 | 568 | 883 | 942 | 242 | 209 | 219 | 237 | 732 | 646 | 750 | 766 | 475,05 | 227 | 724 | 3,2 | 1,678 | 19,159 | 8,13E-82 | 4,52E-81 | 0,0030053 | 0,0055475 | 0,0042903 | 0,0042903 | TRUE | 0,0574718 | No |
| orf19.4669 | 485 | 582 | 534 | 334 | 2135 | 1648 | 2048 | 1829 | 462 | 615 | 528 | 523 | 1711 | 1874 | 1739 | 1487 | 1117,39 | 532 | 1703 | 3,199 | 1,677 | 18,143 | 1,46E-73 | 7,34E-73 | 0,0097921 | 0,0035607 | 0,0068891 | 0,0068891 | TRUE | 0,1335766 | No |
| orf19.5694 | 153 | 129 | 171 | 118 | 603 | 421 | 582 | 684 | 146 | 136 | 169 | 185 | 483 | 479 | 494 | 556 | 331,03 | 159 | 503 | 3,195 | 1,676 | 15,94 | 3,37E-57 | 1,36E-56 | 0,0050072 | 0,0070513 | 0,0061937 | 0,0061937 | TRUE | 0,0976441 | No |
| orf19.2738 | 3229 | 2865 | 3144 | 2100 | 12203 | 8894 | 11305 | 12730 | 3079 | 3029 | 3106 | 3290 | 9778 | 10113 | 9598 | 10351 | 6542,93 | 3126 | 9960 | 3,19 | 1,674 | 40,214 | 0 | 0 | 0,0009537 | 0,0023422 | 0,0014379 | 0,0014379 | TRUE | 0,0140013 | No |
| orf19.6375 | 8818 | 7232 | 7436 | 4467 | 31359 | 19089 | 29326 | 31109 | 8407 | 7647 | 7346 | 6998 | 25127 | 21706 | 24898 | 25295 | 15927,87 | 7600 | 24256 | 3,191 | 1,674 | 25,233 | 1,75E-140 | 1,66E-139 | 0,0054825 | 0,0021944 | 0,0041356 | 0,0041356 | TRUE | 0,0626502 | No |
| orf19.5475 | 15 | 14 | 22 | 20 | 74 | 45 | 94 | 76 | 14 | 15 | 22 | 31 | 59 | 51 | 80 | 62 | 41,78 | 20 | 63 | 3,187 | 1,672 | 6,065 | 1,32E-09 | 2,25E-09 | 0,0318673 | 0,0413909 | 0,0379217 | 0,0379217 | TRUE | 0,4435191 | No |
| orf19.3334 | 20239 | 15571 | 16619 | 10620 | 68789 | 44639 | 64487 | 71619 | 19296 | 16464 | 16418 | 16637 | 55119 | 50759 | 54749 | 58233 | 35959,29 | 17204 | 54715 | 3,18 | 1,669 | 26,925 | 1,13E-159 | 1,23E-158 | 0,0048196 | 0,0021369 | 0,0036514 | 0,0036514 | TRUE | 0,0822808 | No |
| orf19.3876 | 79 | 63 | 91 | 48 | 311 | 231 | 270 | 295 | 75 | 67 | 90 | 75 | 249 | 263 | 229 | 240 | 161 | 77 | 245 | 3,18 | 1,669 | 12,811 | 1,42E-37 | 4,33E-37 | 1,00E-08 | 0,0122896 | 0,0072761 | 0,0072761 | TRUE | 0,1348004 | No |
| orf19.7106 | 590 | 467 | 561 | 342 | 2043 | 1484 | 2063 | 2154 | 563 | 494 | 554 | 536 | 1637 | 1687 | 1751 | 1751 | 1121,7 | 537 | 1706 | 3,178 | 1,668 | 28,01 | 1,22E-172 | 1,41E-171 | 0,000784 | 0,003555 | 0,0021062 | 0,0021062 | TRUE | 0,0359936 | No |
| orf19.1873 | 1242 | 1056 | 1152 | 807 | 4570 | 3437 | 4367 | 4412 | 1184 | 1117 | 1138 | 1264 | 3662 | 3908 | 3708 | 3587 | 2445,99 | 1176 | 3716 | 3,168 | 1,663 | 32,556 | 1,72E-232 | 2,70E-231 | 0,0014803 | 0,0027625 | 0,0019082 | 0,0019082 | TRUE | 0,0268841 | No |
| orf19.6948 | 1612 | 1383 | 1507 | 982 | 5886 | 4195 | 5573 | 5936 | 1537 | 1462 | 1489 | 1538 | 4716 | 4770 | 4731 | 4827 | 3133,84 | 1506 | 4761 | 3,163 | 1,661 | 43,968 | 0 | 0 | 1,00E-08 | 0,0026152 | 0,0009057 | 0,0009057 | TRUE | 0,0046545 | No |
| orf19.7282 | 2788 | 2166 | 2466 | 1625 | 10037 | 6395 | 9500 | 9810 | 2658 | 2290 | 2436 | 2546 | 8042 | 7272 | 8065 | 7976 | 5160,77 | 2482 | 7839 | 3,159 | 1,66 | 29,768 | 1,00E-194 | 1,35E-193 | 0,0028205 | 0,0024094 | 0,0027005 | 0,0027005 | TRUE | 0,0333597 | No |
| orf19.1866 | 436 | 317 | 340 | 236 | 1487 | 914 | 1419 | 1432 | 416 | 335 | 336 | 370 | 1191 | 1039 | 1205 | 1164 | 757,04 | 364 | 1150 | 3,161 | 1,66 | 19,924 | 2,52E-88 | 1,48E-87 | 0,0051895 | 0,0042601 | 0,0047437 | 0,0047437 | TRUE | 0,1153888 | No |
| orf19.7446 | 214 | 208 | 211 | 126 | 882 | 615 | 736 | 728 | 204 | 220 | 208 | 197 | 707 | 699 | 625 | 592 | 431,58 | 207 | 656 | 3,15 | 1,655 | 17,95 | 4,80E-72 | 2,37E-71 | 0,0038535 | 0,0058957 | 0,0048058 | 0,0048058 | TRUE | 0,051672 | No |
| orf19.5328 | 1885 | 1605 | 1722 | 1045 | 6946 | 4701 | 6193 | 6612 | 1797 | 1697 | 1701 | 1637 | 5566 | 5345 | 5258 | 5376 | 3547,19 | 1708 | 5386 | 3,148 | 1,654 | 39,257 | 0 | 0 | 0,0005266 | 0,0025542 | 0,0012947 | 0,0012947 | TRUE | 0,0150826 | No |
| orf19.5517 | 2123 | 1805 | 1901 | 1279 | 7481 | 5690 | 7091 | 7474 | 2024 | 1909 | 1878 | 2004 | 5994 | 6470 | 6020 | 6077 | 4046,99 | 1954 | 6140 | 3,144 | 1,653 | 37,118 | 1,43E-301 | 3,17E-300 | 0,0008706 | 0,002497 | 0,0015429 | 0,0015429 | TRUE | 0,0155898 | No |
| orf19.5700 | 57 | 49 | 59 | 33 | 217 | 117 | 235 | 212 | 54 | 52 | 58 | 52 | 174 | 133 | 200 | 172 | 111,87 | 54 | 170 | 3,144 | 1,653 | 9,826 | 8,70E-23 | 2,06E-22 | 1,00E-08 | 0,0167684 | 0,0142989 | 0,0142989 | TRUE | 0,1928919 | No |
| orf19.5466 | 12074 | 9100 | 9664 | 6308 | 42722 | 24497 | 37677 | 40867 | 11511 | 9622 | 9547 | 9882 | 34232 | 27855 | 31988 | 33229 | 20983,23 | 10140 | 31826 | 3,138 | 1,65 | 21,907 | 2,20E-106 | 1,53E-105 | 0,0079483 | 0,0021695 | 0,0053805 | 0,0053805 | TRUE | 0,1016572 | No |
| orf19.6385 | 16005 | 12914 | 14236 | 10114 | 56724 | 39613 | 52725 | 60513 | 15259 | 13655 | 14064 | 15844 | 45451 | 45044 | 44763 | 49203 | 30410,36 | 14706 | 46115 | 3,137 | 1,649 | 29,959 | 3,33E-197 | 4,50E-196 | 0,0033585 | 0,0021453 | 0,0028634 | 0,0028634 | TRUE | 0,0328829 | No |
| orf19.6626 | 237 | 186 | 220 | 147 | 880 | 557 | 815 | 837 | 226 | 197 | 217 | 230 | 705 | 633 | 692 | 681 | 447,65 | 218 | 678 | 3,129 | 1,646 | 20 | 5,55E-89 | 3,29E-88 | 1,00E-08 | 0,0057591 | 0,0032838 | 0,0032838 | TRUE | 0,0455907 | No |
| orf19.5468 | 94 | 80 | 77 | 44 | 374 | 188 | 295 | 295 | 90 | 85 | 76 | 69 | 300 | 214 | 250 | 240 | 165,37 | 80 | 251 | 3,128 | 1,645 | 11,272 | 1,80E-29 | 4,82E-29 | 0,0114911 | 0,0120199 | 0,011766 | 0,011766 | TRUE | 0,1945592 | No |
| orf19.1154 | 2071 | 1698 | 1779 | 1223 | 7174 | 5153 | 6660 | 7354 | 1974 | 1795 | 1757 | 1916 | 5748 | 5859 | 5654 | 5980 | 3835,6 | 1860 | 5810 | 3,124 | 1,644 | 35,067 | 2,16E-269 | 4,01E-268 | 0,0013069 | 0,0025193 | 0,0017303 | 0,0017303 | TRUE | 0,0224231 | No |
| orf19.4311 | 2458 | 2068 | 2212 | 1327 | 8884 | 5741 | 8144 | 8554 | 2343 | 2187 | 2185 | 2079 | 7118 | 6528 | 6914 | 6955 | 4538,76 | 2198 | 6879 | 3,125 | 1,644 | 34,678 | 1,71E-263 | 3,10E-262 | 0,0014897 | 0,002453 | 0,0018373 | 0,0018373 | TRUE | 0,0240355 | No |
| orf19.5142 | 81 | 86 | 82 | 52 | 320 | 209 | 299 | 344 | 77 | 91 | 81 | 81 | 256 | 238 | 254 | 280 | 169,78 | 82 | 257 | 3,117 | 1,64 | 13,137 | 2,01E-39 | 6,30E-39 | 1,00E-08 | 0,011762 | 0,0065253 | 0,0065253 | TRUE | 0,0450549 | No |
| orf19.1345 | 47 | 28 | 38 | 15 | 127 | 105 | 126 | 130 | 45 | 30 | 38 | 23 | 102 | 119 | 107 | 106 | 71,16 | 34 | 108 | 3,114 | 1,639 | 8,362 | 6,19E-17 | 1,30E-16 | 1,00E-08 | 0,0251644 | 0,0167917 | 0,0167917 | TRUE | 0,3277344 | No |
| orf19.4684.2 | 4860 | 3794 | 3905 | 2553 | 17591 | 9741 | 15494 | 15954 | 4633 | 4012 | 3858 | 3999 | 14095 | 11076 | 13154 | 12972 | 8475,04 | 4126 | 12824 | 3,108 | 1,636 | 21,1 | 7,97E-99 | 5,15E-98 | 0,0081629 | 0,002285 | 0,0056044 | 0,0056044 | TRUE | 0,1026376 | No |
| orf19.6026 | 166 | 188 | 183 | 112 | 685 | 546 | 619 | 644 | 158 | 199 | 181 | 175 | 549 | 621 | 526 | 524 | 366,52 | 178 | 555 | 3,106 | 1,635 | 16,51 | 3,12E-61 | 1,34E-60 | 0,0042462 | 0,006571 | 0,0054845 | 0,0054845 | TRUE | 0,0710055 | No |
| orf19.2644 | 2837 | 2685 | 2665 | 1956 | 10356 | 8345 | 9894 | 10600 | 2705 | 2839 | 2633 | 3064 | 8298 | 9489 | 8400 | 8619 | 5755,81 | 2810 | 8702 | 3,1 | 1,632 | 27,237 | 2,35E-163 | 2,62E-162 | 0,0039029 | 0,0023765 | 0,0031963 | 0,0031963 | TRUE | 0,0450145 | No |
| orf19.4033 | 587 | 459 | 619 | 335 | 2092 | 1416 | 1983 | 2215 | 560 | 485 | 612 | 525 | 1676 | 1610 | 1684 | 1801 | 1119,03 | 546 | 1693 | 3,099 | 1,632 | 22,553 | 1,26E-112 | 9,37E-112 | 0,0043066 | 0,0035585 | 0,0037369 | 0,0037369 | TRUE | 0,0807766 | No |
| orf19.2158 | 272 | 212 | 237 | 166 | 958 | 664 | 891 | 912 | 259 | 224 | 234 | 260 | 768 | 755 | 756 | 742 | 499,79 | 244 | 755 | 3,1 | 1,632 | 21,576 | 3,01E-103 | 2,02E-102 | 1,00E-08 | 0,0053765 | 0,0026264 | 0,0026264 | TRUE | 0,0337011 | No |
| orf19.3789 | 7622 | 6689 | 6565 | 4638 | 27057 | 19878 | 23669 | 27681 | 7267 | 7073 | 6486 | 7266 | 21680 | 22603 | 20095 | 22507 | 14372 | 7023 | 21721 | 3,094 | 1,629 | 30,843 | 7,04E-209 | 9,93E-208 | 0,0026865 | 0,0022055 | 0,0025798 | 0,0025798 | TRUE | 0,0326133 | No |
| orf19.3354 | 9353 | 7345 | 7884 | 4287 | 30962 | 18825 | 28656 | 31809 | 8917 | 7766 | 7789 | 6716 | 24809 | 21406 | 24329 | 25864 | 15949,4 | 7797 | 24102 | 3,09 | 1,628 | 20,009 | 4,56E-89 | 2,70E-88 | 0,0096683 | 0,0021942 | 0,0062663 | 0,0062663 | TRUE | 0,1142404 | No |
| orf19.3453 | 164 | 166 | 169 | 112 | 664 | 469 | 593 | 625 | 156 | 176 | 167 | 175 | 532 | 533 | 503 | 508 | 343,91 | 168 | 519 | 3,09 | 1,628 | 18,505 | 1,89E-76 | 9,83E-76 | 1,00E-08 | 0,0068655 | 0,0032702 | 0,0032702 | TRUE | 0,0272822 | No |
| orf19.1421 | 162 | 120 | 143 | 109 | 583 | 374 | 538 | 574 | 154 | 127 | 141 | 171 | 467 | 425 | 457 | 467 | 301,16 | 148 | 454 | 3,092 | 1,628 | 16,334 | 5,61E-60 | 2,36E-59 | 1,19E-08 | 0,0075433 | 0,004786 | 0,004786 | TRUE | 0,0933686 | No |
| orf19.2202 | 128 | 122 | 125 | 86 | 463 | 317 | 505 | 489 | 122 | 129 | 123 | 135 | 371 | 360 | 429 | 398 | 258,38 | 127 | 390 | 3,077 | 1,622 | 15,276 | 1,11E-52 | 4,24E-52 | 1,00E-08 | 0,0084459 | 0,0052988 | 0,0052988 | TRUE | 0,0531711 | No |
| orf19.2521 | 430 | 392 | 445 | 257 | 1560 | 1078 | 1521 | 1673 | 410 | 414 | 440 | 403 | 1250 | 1226 | 1291 | 1360 | 849,26 | 417 | 1282 | 3,073 | 1,62 | 25,078 | 8,70E-139 | 8,12E-138 | 1,00E-08 | 0,0040246 | 0,002323 | 0,002323 | TRUE | 0,0206247 | No |
| orf19.6096 | 768 | 604 | 663 | 410 | 2659 | 1643 | 2559 | 2505 | 732 | 639 | 655 | 642 | 2131 | 1868 | 2173 | 2037 | 1359,54 | 667 | 2052 | 3,073 | 1,62 | 23,78 | 5,33E-125 | 4,47E-124 | 0,0034787 | 0,003299 | 0,0033975 | 0,0033975 | TRUE | 0,0534581 | No |
| orf19.3735 | 12 | 11 | 9 | 18 | 48 | 48 | 47 | 52 | 11 | 12 | 9 | 28 | 38 | 55 | 40 | 42 | 29,42 | 15 | 44 | 3,07 | 1,618 | 4,905 | 9,33E-07 | 1,45E-06 | 0,048215 | 0,0578942 | 0,0550936 | 0,0550936 | TRUE | 1,0261386 | No |
| orf19.6938 | 2529 | 1977 | 2112 | 1392 | 8328 | 5615 | 8029 | 8526 | 2411 | 2090 | 2086 | 2181 | 6673 | 6385 | 6817 | 6932 | 4446,93 | 2192 | 6702 | 3,057 | 1,612 | 29,696 | 8,54E-194 | 1,14E-192 | 0,0026327 | 0,0024605 | 0,0025059 | 0,0025059 | TRUE | 0,0569415 | No |
| orf19.7258 | 1715 | 1278 | 1428 | 927 | 5799 | 3470 | 5469 | 5702 | 1635 | 1351 | 1411 | 1452 | 4647 | 3946 | 4643 | 4636 | 2965,13 | 1462 | 4468 | 3,055 | 1,611 | 22,138 | 1,35E-108 | 9,66E-108 | 0,0059328 | 0,002645 | 0,0046024 | 0,0046024 | TRUE | 0,078658 | No |
| orf19.2249 | 361 | 273 | 328 | 202 | 1258 | 835 | 1089 | 1240 | 344 | 289 | 324 | 316 | 1008 | 949 | 925 | 1008 | 645,45 | 318 | 972 | 3,052 | 1,61 | 21,821 | 1,48E-105 | 1,01E-104 | 0,0013096 | 0,0046351 | 0,0030214 | 0,0030214 | TRUE | 0,0481856 | No |
| orf19.3087 | 17946 | 13740 | 14644 | 10516 | 58710 | 39926 | 56940 | 61398 | 17110 | 14528 | 14467 | 16474 | 47043 | 45399 | 48342 | 49922 | 31660,62 | 15645 | 47676 | 3,048 | 1,608 | 26,38 | 2,31E-153 | 2,37E-152 | 0,0044781 | 0,0021431 | 0,0035234 | 0,0035234 | TRUE | 0,0490197 | No |
| orf19.3325.3 | 8301 | 6301 | 6284 | 3776 | 26507 | 15517 | 24394 | 26759 | 7914 | 6663 | 6208 | 5915 | 21239 | 17644 | 20710 | 21758 | 13506,43 | 6675 | 20338 | 3,046 | 1,607 | 17,973 | 3,15E-72 | 1,56E-71 | 0,0127051 | 0,0022128 | 0,0075734 | 0,0075734 | TRUE | 0,1910722 | No |
| orf19.6065 | 1117 | 1020 | 1061 | 615 | 3974 | 2728 | 3730 | 3943 | 1065 | 1079 | 1048 | 963 | 3184 | 3102 | 3167 | 3206 | 2101,76 | 1039 | 3165 | 3,037 | 1,603 | 35,153 | 1,04E-270 | 1,97E-269 | 1,00E-08 | 0,0028725 | 0,0013186 | 0,0013186 | TRUE | 0,0236441 | No |
| orf19.727 | 46 | 41 | 46 | 39 | 176 | 141 | 165 | 167 | 44 | 43 | 45 | 61 | 141 | 160 | 140 | 136 | 96,37 | 48 | 144 | 3,026 | 1,598 | 9,74 | 2,04E-22 | 4,79E-22 | 1,00E-08 | 0,0191285 | 0,0110773 | 0,0110773 | TRUE | 0,168694 | No |
| orf19.3358 | 2489 | 1927 | 2081 | 1468 | 8373 | 5453 | 7661 | 8653 | 2373 | 2038 | 2056 | 2300 | 6709 | 6201 | 6504 | 7036 | 4401,94 | 2192 | 6612 | 3,02 | 1,594 | 25,776 | 1,66E-146 | 1,65E-145 | 0,0040153 | 0,0024643 | 0,0033498 | 0,0033498 | TRUE | 0,0396679 | No |
| orf19.2111.2 | 6678 | 5169 | 5233 | 3337 | 20674 | 14553 | 19265 | 21537 | 6367 | 5466 | 5170 | 5228 | 16565 | 16548 | 16356 | 17512 | 11151,34 | 5558 | 16745 | 3,012 | 1,591 | 24,761 | 2,35E-135 | 2,14E-134 | 0,0053003 | 0,0022385 | 0,0038361 | 0,0038361 | TRUE | 0,118125 | No |

| | | | | | | | | | | | | | | | | | | | | | | | | | | | | | | | |
|---|---|---|---|---|---|---|---|---|---|---|---|---|---|---|---|---|---|---|---|---|---|---|---|---|---|---|---|---|---|---|---|
| orf19.31 | 210 | 157 | 153 | 112 | 627 | 460 | 585 | 692 | 200 | 166 | 151 | 175 | 502 | 523 | 497 | 563 | 347,2 | 173 | 521 | 3,008 | 1,589 | 15,978 | 1,83E-57 | 7,42E-57 | 0,0035856 | 0,0068203 | 0,0054258 | 0,0054258 | TRUE | 0,1358275 | No |
| orf19.6005 | 348 | 350 | 361 | 229 | 1273 | 1016 | 1244 | 1257 | 332 | 370 | 357 | 359 | 1020 | 1155 | 1056 | 1022 | 708,84 | 354 | 1063 | 3 | 1,585 | 22,058 | 8,02E-108 | 5,66E-107 | 0,0016906 | 0,0044076 | 0,0029635 | 0,0029635 | TRUE | 0,0383178 | No |
| orf19.2673 | 474 | 463 | 457 | 248 | 1717 | 1043 | 1660 | 1702 | 452 | 490 | 451 | 389 | 1376 | 1186 | 1409 | 1384 | 892,05 | 446 | 1339 | 2,995 | 1,583 | 19,691 | 2,60E-86 | 1,50E-85 | 0,0053811 | 0,0039319 | 0,0046181 | 0,0046181 | TRUE | 0,0719649 | No |
| orf19.2072 | 6 | 9 | 10 | 17 | 42 | 37 | 33 | 51 | 6 | 10 | 10 | 27 | 34 | 42 | 28 | 41 | 24,62 | 13 | 36 | 2,989 | 1,58 | 4,119 | 3,80E-05 | 5,48E-05 | 0,1136463 | 0,0687819 | 0,0828691 | 0,0828691 | TRUE | 1,3875125 | No |
| orf19.7290 | 1321 | 1161 | 1169 | 720 | 4653 | 2731 | 4415 | 4498 | 1259 | 1228 | 1155 | 1128 | 3728 | 3105 | 3748 | 3657 | 2376,14 | 1192 | 3560 | 2,983 | 1,577 | 22,832 | 2,19E-115 | 1,66E-114 | 0,0048904 | 0,0027823 | 0,003984 | 0,003984 | TRUE | 0,0882998 | No |
| orf19.1409.3 | 19 | 14 | 18 | 9 | 70 | 34 | 73 | 46 | 18 | 15 | 18 | 14 | 56 | 39 | 62 | 37 | 32,37 | 16 | 48 | 2,978 | 1,574 | 5,318 | 1,05E-07 | 1,68E-07 | 1,00E-08 | 0,0528205 | 0,0410995 | 0,0410995 | TRUE | 0,1178663 | No |
| orf19.6265 | 8804 | 6569 | 6532 | 4459 | 27383 | 17490 | 24851 | 27756 | 8394 | 6946 | 6453 | 6985 | 21941 | 19888 | 21098 | 22568 | 14284,17 | 7194 | 21374 | 2,971 | 1,571 | 20,969 | 1,26E-97 | 8,08E-97 | 0,0080882 | 0,0022062 | 0,0052916 | 0,0052916 | TRUE | 0,1547292 | No |
| orf19.236 | 10652 | 9144 | 9387 | 5525 | 35477 | 22979 | 33379 | 35711 | 10156 | 9669 | 9273 | 8655 | 28427 | 26129 | 28339 | 29036 | 18710,47 | 9438 | 27983 | 2,964 | 1,567 | 28,58 | 1,17E-179 | 1,40E-178 | 0,003202 | 0,002179 | 0,0028137 | 0,0028137 | TRUE | 0,0374822 | No |
| orf19.3690.2 | 6008 | 5918 | 5344 | 3462 | 21713 | 15792 | 19253 | 19037 | 5728 | 6258 | 5279 | 5423 | 17398 | 17957 | 16346 | 15479 | 11233,48 | 5672 | 16795 | 2,96 | 1,566 | 24,528 | 7,48E-133 | 6,70E-132 | 0,0049092 | 0,0022374 | 0,0037882 | 0,0037882 | TRUE | 0,0590157 | No |
| orf19.3711 | 59 | 50 | 61 | 29 | 197 | 138 | 192 | 204 | 56 | 53 | 60 | 45 | 158 | 157 | 163 | 166 | 107,31 | 54 | 161 | 2,962 | 1,566 | 10,509 | 7,83E-26 | 1,97E-25 | 1,00E-08 | 0,0173922 | 0,0083879 | 0,0083879 | TRUE | 0,058382 | No |
| orf19.5231.2 | 1311 | 1117 | 1193 | 685 | 4534 | 2699 | 4310 | 4318 | 1250 | 1181 | 1179 | 1073 | 3633 | 3069 | 3659 | 3511 | 2319,34 | 1171 | 3468 | 2,958 | 1,565 | 22,945 | 1,67E-116 | 1,28E-115 | 0,0042956 | 0,0027992 | 0,0038569 | 0,0038569 | TRUE | 0,071649 | No |
| orf19.6630 | 364 | 246 | 252 | 142 | 1007 | 646 | 963 | 1048 | 347 | 260 | 249 | 222 | 807 | 735 | 818 | 852 | 536,21 | 270 | 803 | 2,958 | 1,565 | 13,844 | 1,38E-43 | 4,64E-43 | 0,0154963 | 0,0051533 | 0,0096443 | 0,0096443 | TRUE | 0,4270511 | No |
| orf19.3268 | 11881 | 9287 | 9628 | 7778 | 38936 | 28030 | 36325 | 39932 | 11327 | 9820 | 9512 | 12185 | 31198 | 31873 | 30840 | 32469 | 21152,81 | 10711 | 31595 | 2,951 | 1,561 | 21,755 | 6,20E-105 | 4,21E-104 | 0,007129 | 0,0021689 | 0,0048804 | 0,0048804 | TRUE | 0,1040031 | No |
| orf19.4933 | 5888 | 5212 | 5547 | 3822 | 19866 | 15336 | 19299 | 20505 | 5614 | 5511 | 5480 | 5987 | 15918 | 17438 | 16385 | 16673 | 11125,7 | 5648 | 16604 | 2,942 | 1,557 | 35,864 | 1,11E-281 | 2,27E-280 | 0,0014327 | 0,0022389 | 0,0016824 | 0,0016824 | TRUE | 0,0191644 | No |
| orf19.6214 | 1834 | 1572 | 1705 | 1163 | 6342 | 4464 | 5761 | 6481 | 1749 | 1662 | 1684 | 1822 | 5082 | 5076 | 4891 | 5270 | 3404,42 | 1729 | 5080 | 2,943 | 1,557 | 35,082 | 1,26E-269 | 2,36E-268 | 0,0008083 | 0,0025736 | 0,0014786 | 0,0014786 | TRUE | 0,0138848 | No |
| orf19.2343.1 | 369 | 312 | 345 | 241 | 1257 | 861 | 1302 | 1239 | 352 | 330 | 341 | 378 | 1007 | 979 | 1105 | 1007 | 687,39 | 350 | 1024 | 2,942 | 1,557 | 21,709 | 1,70E-104 | 1,15E-103 | 0,0014822 | 0,0044799 | 0,0029046 | 0,0029046 | TRUE | 0,0337294 | No |
| orf19.657 | 1246 | 1058 | 1103 | 803 | 4410 | 3167 | 3646 | 4223 | 1188 | 1119 | 1090 | 1258 | 3534 | 3601 | 3095 | 3434 | 2289,77 | 1164 | 3416 | 2,941 | 1,556 | 24,354 | 5,32E-131 | 4,66E-130 | 0,0034811 | 0,0028083 | 0,0033063 | 0,0033063 | TRUE | 0,048685 | No |
| orf19.6906 | 16576 | 13650 | 14234 | 9265 | 54518 | 35732 | 50848 | 55803 | 15803 | 14433 | 14062 | 14514 | 43684 | 40630 | 43170 | 45373 | 28958,7 | 14703 | 43214 | 2,939 | 1,555 | 31,835 | 2,14E-222 | 3,20E-221 | 0,002305 | 0,002148 | 0,0022445 | 0,0022445 | TRUE | 0,0312717 | No |
| orf19.5693 | 1036 | 792 | 922 | 560 | 3371 | 2121 | 3309 | 3325 | 988 | 837 | 911 | 877 | 2701 | 2412 | 2809 | 2704 | 1779,87 | 903 | 2656 | 2,939 | 1,555 | 23,325 | 2,45E-120 | 1,96E-119 | 0,0035679 | 0,0030138 | 0,0034788 | 0,0034788 | TRUE | 0,0490057 | No |
| orf19.1462 | 2175 | 1851 | 2029 | 1372 | 7548 | 5040 | 6952 | 7678 | 2074 | 1957 | 2004 | 2149 | 6048 | 5731 | 5902 | 6243 | 4013,59 | 2046 | 5981 | 2,928 | 1,55 | 34,239 | 6,39E-257 | 1,13E-255 | 0,0010615 | 0,0025004 | 0,0016179 | 0,0016179 | TRUE | 0,0125596 | No |
| orf19.2992 | 6796 | 5684 | 5906 | 4283 | 23336 | 17149 | 20313 | 21801 | 6479 | 6010 | 5835 | 6710 | 18698 | 19500 | 17246 | 17726 | 12275,48 | 6258 | 18292 | 2,924 | 1,548 | 27,309 | 3,35E-164 | 3,74E-163 | 0,0034712 | 0,002225 | 0,0029719 | 0,0029719 | TRUE | 0,0280795 | No |
| orf19.4555 | 4196 | 3375 | 3868 | 2814 | 14223 | 10457 | 12813 | 14750 | 4000 | 3569 | 3821 | 4408 | 11396 | 11891 | 10878 | 11993 | 7744,62 | 3950 | 11540 | 2,924 | 1,548 | 24,639 | 4,83E-134 | 4,38E-133 | 0,0047604 | 0,0023033 | 0,0036104 | 0,0036104 | TRUE | 0,072183 | No |
| orf19.2107 | 869 | 683 | 739 | 520 | 2782 | 1809 | 2814 | 2900 | 828 | 722 | 730 | 815 | 2229 | 2057 | 2389 | 2358 | 1516,07 | 774 | 2258 | 2,924 | 1,548 | 22,547 | 1,45E-112 | 1,07E-111 | 0,003649 | 0,0031743 | 0,0036026 | 0,0036026 | TRUE | 0,042369 | No |
| orf19.7328 | 730 | 708 | 680 | 485 | 2652 | 1763 | 2397 | 2714 | 696 | 749 | 672 | 760 | 2125 | 2005 | 2035 | 2207 | 1405,95 | 719 | 2093 | 2,921 | 1,547 | 26,279 | 3,32E-152 | 3,39E-151 | 0,0014187 | 0,0032591 | 0,0023322 | 0,0023322 | TRUE | 0,0230618 | No |
| orf19.5952 | 272 | 249 | 295 | 159 | 889 | 692 | 903 | 1042 | 259 | 263 | 291 | 249 | 712 | 787 | 767 | 847 | 522,03 | 266 | 778 | 2,917 | 1,545 | 18,273 | 1,36E-74 | 6,91E-74 | 0,0032507 | 0,0052365 | 0,0041962 | 0,0041962 | TRUE | 0,0482374 | No |
| orf19.5097 | 740 | 683 | 731 | 424 | 2537 | 1760 | 2381 | 2641 | 706 | 722 | 722 | 664 | 2033 | 2001 | 2021 | 2147 | 1377,13 | 704 | 2050 | 2,908 | 1,54 | 29,317 | 6,23E-189 | 7,99E-188 | 1,00E-08 | 0,0032835 | 0,0016411 | 0,0016411 | TRUE | 0,0135737 | No |
| orf19.998 | 175 | 183 | 157 | 122 | 636 | 438 | 580 | 661 | 167 | 193 | 155 | 191 | 510 | 498 | 492 | 537 | 343,01 | 176 | 509 | 2,905 | 1,539 | 16,503 | 3,47E-61 | 1,49E-60 | 0,0005701 | 0,006878 | 0,0043042 | 0,0043042 | TRUE | 0,0736652 | No |
| orf19.4909.1 | 10814 | 8262 | 8737 | 5351 | 33443 | 20874 | 31289 | 33725 | 10310 | 8736 | 8631 | 8383 | 26797 | 23736 | 26564 | 27422 | 17572,29 | 9015 | 26130 | 2,898 | 1,535 | 22,022 | 1,76E-107 | 1,24E-106 | 0,0065826 | 0,0021847 | 0,0045879 | 0,0045879 | TRUE | 0,1146667 | No |
| orf19.7048.1 | 5817 | 4405 | 4654 | 3228 | 18430 | 11842 | 16948 | 18228 | 5546 | 4658 | 4598 | 5057 | 14767 | 13465 | 14389 | 14821 | 9662,61 | 4965 | 14360 | 2,893 | 1,533 | 24,486 | 2,09E-132 | 1,86E-131 | 0,0046621 | 0,0022612 | 0,0036184 | 0,0036184 | TRUE | 0,077097 | No |
| orf19.860 | 462 | 383 | 387 | 281 | 1495 | 994 | 1446 | 1538 | 440 | 405 | 382 | 440 | 1198 | 1130 | 1228 | 1251 | 809,29 | 417 | 1202 | 2,892 | 1,532 | 22,537 | 1,82E-112 | 1,34E-111 | 0,0011255 | 0,0041201 | 0,0027265 | 0,0027265 | TRUE | 0,0377349 | No |
| orf19.5270 | 182 | 170 | 196 | 113 | 697 | 470 | 600 | 609 | 174 | 180 | 194 | 177 | 558 | 534 | 509 | 495 | 352,68 | 181 | 524 | 2,892 | 1,532 | 17,116 | 1,13E-65 | 5,13E-65 | 1,00E-08 | 0,0067468 | 0,0037815 | 0,0037815 | TRUE | 0,0258666 | No |
| orf19.6724 | 2556 | 1877 | 2189 | 1615 | 7884 | 6090 | 7535 | 8131 | 2437 | 1985 | 2163 | 2530 | 6317 | 6925 | 6397 | 6611 | 4420,58 | 2279 | 6562 | 2,883 | 1,528 | 21,414 | 9,91E-102 | 6,57E-101 | 0,0065003 | 0,0024627 | 0,0045717 | 0,0045717 | TRUE | 0,0910063 | No |
| orf19.1926 | 247 | 234 | 217 | 189 | 918 | 715 | 760 | 795 | 235 | 247 | 214 | 296 | 736 | 813 | 645 | 646 | 479,2 | 248 | 710 | 2,878 | 1,525 | 14,006 | 1,44E-44 | 4,89E-44 | 0,012549 | 0,0055176 | 0,0084802 | 0,0084802 | TRUE | 0,1586946 | No |
| orf19.1709 | 1862 | 1714 | 1679 | 921 | 6192 | 3955 | 5687 | 6130 | 1775 | 1812 | 1659 | 1443 | 4961 | 4497 | 4828 | 4984 | 3245,03 | 1672 | 4818 | 2,875 | 1,524 | 22,377 | 6,63E-111 | 4,85E-110 | 0,0054993 | 0,0025973 | 0,0040235 | 0,0040235 | TRUE | 0,0963519 | No |
| orf19.364 | 1595 | 1317 | 1392 | 913 | 4999 | 3305 | 4976 | 5469 | 1521 | 1393 | 1375 | 1430 | 4006 | 3758 | 4225 | 4447 | 2769,21 | 1430 | 4109 | 2,875 | 1,523 | 24,853 | 2,38E-136 | 2,17E-135 | 0,0029811 | 0,0026842 | 0,0031058 | 0,0031058 | TRUE | 0,0394547 | No |
| orf19.2245 | 642 | 597 | 603 | 365 | 2003 | 1612 | 2097 | 2119 | 612 | 631 | 596 | 572 | 1605 | 1833 | 1780 | 1723 | 1169,01 | 603 | 1735 | 2,87 | 1,521 | 24,426 | 9,10E-132 | 8,07E-131 | 0,0018303 | 0,0034958 | 0,0025405 | 0,0025405 | TRUE | 0,031227 | No |
| orf19.532 | 723 | 605 | 558 | 386 | 2265 | 1653 | 1951 | 2206 | 689 | 640 | 551 | 605 | 1815 | 1880 | 1656 | 1794 | 1203,69 | 621 | 1786 | 2,871 | 1,521 | 20,942 | 2,24E-97 | 1,42E-96 | 0,0043172 | 0,0034553 | 0,0039153 | 0,0039153 | TRUE | 0,0737668 | No |
| orf19.342 | 35 | 33 | 24 | 23 | 101 | 84 | 109 | 117 | 33 | 35 | 24 | 36 | 81 | 96 | 93 | 95 | 61,51 | 32 | 91 | 2,869 | 1,521 | 7,57 | 3,73E-14 | 7,35E-14 | 1,00E-08 | 0,0287824 | 0,0163443 | 0,0163443 | TRUE | 0,225131 | No |
| orf19.5467 | 12 | 12 | 17 | 9 | 49 | 32 | 54 | 44 | 11 | 13 | 17 | 14 | 39 | 36 | 46 | 36 | 26,54 | 14 | 39 | 2,87 | 1,521 | 5,109 | 3,23E-07 | 5,10E-07 | 1,00E-08 | 0,0639642 | 0,033446 | 0,033446 | TRUE | 0,113547 | No |
| orf19.6257 | 4608 | 4220 | 4502 | 2739 | 15896 | 11018 | 14460 | 15936 | 4393 | 4462 | 4448 | 4291 | 12737 | 12528 | 12276 | 12957 | 8511,64 | 4398 | 12624 | 2,869 | 1,52 | 46,161 | 0 | 0 | 0,000326 | 0,0022842 | 0,0008785 | 0,0008785 | TRUE | 0,0042749 | No |
| orf19.2740 | 659 | 605 | 668 | 416 | 2256 | 1639 | 2127 | 2337 | 628 | 640 | 660 | 652 | 1808 | 1864 | 1806 | 1900 | 1244,62 | 645 | 1844 | 2,861 | 1,517 | 29,427 | 2,50E-190 | 3,24E-189 | 1,00E-08 | 0,0034105 | 0,001445 | 0,001445 | TRUE | 0,0053099 | No |
| orf19.4114 | 2387 | 2152 | 2370 | 1423 | 7976 | 5739 | 7633 | 8259 | 2276 | 2275 | 2341 | 2229 | 6391 | 6526 | 6480 | 6715 | 4404,27 | 2280 | 6528 | 2,86 | 1,516 | 41,932 | 0 | 0 | 0,0002399 | 0,0024641 | 0,0009411 | 0,0009411 | TRUE | 0,0046478 | No |
| orf19.2364 | 1588 | 1419 | 1502 | 1014 | 5410 | 3758 | 4985 | 5568 | 1514 | 1500 | 1484 | 1588 | 4335 | 4273 | 4232 | 4527 | 2931,79 | 1522 | 4342 | 2,859 | 1,516 | 35,299 | 6,17E-273 | 1,18E-271 | 0,0005359 | 0,0026513 | 0,0012985 | 0,0012985 | TRUE | 0,0101909 | No |
| orf19.4650 | 1129 | 969 | 1088 | 649 | 3739 | 2664 | 3466 | 3739 | 1076 | 1025 | 1075 | 1017 | 2996 | 3029 | 2943 | 3040 | 2025,06 | 1048 | 3002 | 2,858 | 1,515 | 34,927 | 2,91E-267 | 5,36E-266 | 1,00E-08 | 0,0029021 | 0,0011269 | 0,0011269 | TRUE | 0,0041013 | No |
| orf19.4036 | 762 | 643 | 732 | 470 | 2523 | 1669 | 2547 | 2574 | 726 | 680 | 723 | 736 | 2022 | 1898 | 2162 | 2093 | 1380,06 | 716 | 2044 | 2,858 | 1,515 | 25,708 | 9,63E-146 | 9,45E-145 | 0,0016548 | 0,003281 | 0,002334 | 0,002334 | TRUE | 0,0266797 | No |
| orf19.2953 | 2016 | 1753 | 1780 | 1150 | 6498 | 4452 | 6245 | 6603 | 1922 | 1854 | 1758 | 1802 | 5207 | 5062 | 5302 | 5369 | 3534,43 | 1834 | 5235 | 2,853 | 1,513 | 35,51 | 3,40E-276 | 6,65E-275 | 0,0005696 | 0,0025558 | 0,0013513 | 0,0013513 | TRUE | 0,0133242 | No |
| orf19.4991 | 974 | 937 | 1036 | 692 | 3553 | 2688 | 3199 | 3501 | 929 | 991 | 1023 | 1084 | 2847 | 3056 | 2716 | 2847 | 1936,61 | 1007 | 2866 | 2,854 | 1,513 | 25,751 | 3,12E-146 | 3,09E-145 | 0,0023881 | 0,0029391 | 0,0025994 | 0,0025994 | TRUE | 0,0330027 | No |
| orf19.5750 | 4209 | 3815 | 4109 | 2438 | 14031 | 9990 | 13366 | 14087 | 4013 | 4034 | 4059 | 3819 | 11243 | 11360 | 11348 | 11454 | 7666,15 | 3981 | 11351 | 2,847 | 1,51 | 47,708 | 0 | 0 | 1,38E-05 | 0,0023055 | 0,0007812 | 0,0007812 | TRUE | 0,0082356 | No |
| orf19.6169 | 450 | 443 | 490 | 339 | 1667 | 1196 | 1599 | 1667 | 429 | 468 | 484 | 531 | 1336 | 1360 | 1358 | 1355 | 915,15 | 478 | 1352 | 2,846 | 1,509 | 23,694 | 4,15E-124 | 3,44E-123 | 1,00E-08 | 0,0038854 | 0,0023908 | 0,0023908 | TRUE | 0,0563548 | No |
| orf19.913.2 | 2776 | 2468 | 2610 | 1578 | 9336 | 5846 | 9222 | 9058 | 2647 | 2610 | 2578 | 2472 | 7481 | 6647 | 7829 | 7365 | 4953,65 | 2577 | 7330 | 2,844 | 1,508 | 27,656 | 2,35E-168 | 2,66E-167 | 0,0023681 | 0,0024227 | 0,0025754 | 0,0025754 | TRUE | 0,0474465 | No |
| orf19.5563 | 1649 | 1344 | 1471 | 896 | 5230 | 3437 | 5067 | 5172 | 1572 | 1421 | 1453 | 1404 | 4191 | 3908 | 4302 | 4205 | 2807,01 | 1462 | 4152 | 2,835 | 1,504 | 29,227 | 8,74E-188 | 1,11E-186 | 0,0015645 | 0,0026762 | 0,00205 | 0,00205 | TRUE | 0,0315131 | No |
| orf19.3920 | 639 | 529 | 570 | 368 | 1967 | 1398 | 1889 | 2166 | 609 | 559 | 563 | 576 | 1576 | 1590 | 1604 | 1761 | 1104,85 | 577 | 1633 | 2,829 | 1,5 | 24,226 | 1,17E-129 | 1,01E-128 | 0,0016163 | 0,0035774 | 0,0024418 | 0,0024418 | TRUE | 0,0345664 | No |
| orf19.5685 | 4357 | 3552 | 3982 | 2430 | 14052 | 8755 | 13288 | 14441 | 4154 | 3756 | 3934 | 3807 | 11259 | 9955 | 11281 | 11742 | 7486,04 | 3913 | 11059 | 2,826 | 1,499 | 26,086 | 5,24E-150 | 5,26E-149 | 0,0032159 | 0,0023106 | 0,0029865 | 0,0029865 | TRUE | 0,0548395 | No |
| orf19.405 | 974 | 818 | 911 | 595 | 3121 | 2195 | 2989 | 3291 | 929 | 865 | 900 | 932 | 2501 | 2496 | 2538 | 2676 | 1729,48 | 906 | 2553 | 2,82 | 1,496 | 30,142 | 1,37E-199 | 1,86E-198 | 0,0005944 | 0,0030406 | 0,0015723 | 0,0015723 | TRUE | 0,0130408 | No |
| orf19.341 | 376 | 291 | 346 | 213 | 1138 | 872 | 1086 | 1190 | 358 | 308 | 342 | 334 | 912 | 992 | 922 | 968 | 641,83 | 336 | 948 | 2,82 | 1,496 | 21,052 | 2,18E-98 | 1,40E-97 | 0,0006481 | 0,0046494 | 0,0027195 | 0,0027195 | TRUE | 0,0380979 | No |
| orf19.6312 | 13314 | 11277 | 11986 | 8021 | 42161 | 31335 | 38697 | 44063 | 12693 | 11924 | 11841 | 12565 | 33782 | 35631 | 32854 | 35827 | 23389,74 | 12256 | 34524 | 2,817 | 1,494 | 34,987 | 3,56E-268 | 6,58E-267 | 0,0014341 | 0,0021615 | 0,0016932 | 0,0016932 | TRUE | 0,0129904 | No |
| orf19.30 | 2565 | 2389 | 2494 | 1575 | 8693 | 6445 | 7724 | 8667 | 2445 | 2526 | 2464 | 2467 | 6965 | 7329 | 6558 | 7047 | 4725,18 | 2476 | 6975 | 2,816 | 1,494 | 34,147 | 1,46E-255 | 2,56E-254 | 0,0011037 | 0,0024388 | 0,0015465 | 0,0015465 | TRUE | 0,019755 | No |
| orf19.5322 | 776 | 604 | 696 | 388 | 2372 | 1584 | 2264 | 2374 | 740 | 639 | 688 | 608 | 1901 | 1801 | 1922 | 1930 | 1278,51 | 669 | 1888 | 2,814 | 1,492 | 23,628 | 1,98E-123 | 1,62E-122 | 0,0020902 | 0,0033755 | 0,0027609 | 0,0027609 | TRUE | 0,0611901 | No |
| orf19.6317 | 3590 | 3502 | 3677 | 2263 | 12628 | 9379 | 11295 | 12094 | 3423 | 3703 | 3633 | 3545 | 10118 | 10665 | 9589 | 9834 | 6813,68 | 3576 | 10052 | 2,81 | 1,491 | 32,747 | 3,30E-235 | 5,21E-234 | 0,0013761 | 0,0023323 | 0,0017879 | 0,0017879 | TRUE | 0,0203606 | No |
| orf19.5191 | 41 | 32 | 25 | 23 | 114 | 86 | 110 | 113 | 39 | 34 | 25 | 36 | 91 | 98 | 93 | 92 | 63,51 | 34 | 94 | 2,811 | 1,491 | 7,654 | 1,94E-14 | 3,86E-14 | 1,00E-08 | 0,0279451 | 0,0150163 | 0,0150163 | TRUE | 0,2407941 | No |

| | | | | | | | | | | | | | | | | | | | | | | | | | | | | | | | |
|---|---|---|---|---|---|---|---|---|---|---|---|---|---|---|---|---|---|---|---|---|---|---|---|---|---|---|---|---|---|---|---|
| orf19.3447 | 304 | 383 | 331 | 244 | 1264 | 974 | 1131 | 1041 | 290 | 405 | 327 | 382 | 1013 | 1108 | 960 | 846 | 666,38 | 351 | 982 | 2,805 | 1,488 | 13,465 | 2,52E-41 | 8,15E-41 | 0,0146722 | 0,0045552 | 0,00966 | 0,00966 | TRUE | 0,163067 | No |
| orf19.6253 | 16846 | 13493 | 14563 | 9070 | 51024 | 34584 | 48525 | 53817 | 16061 | 14267 | 14387 | 14209 | 40884 | 39325 | 41197 | 43758 | 28011,07 | 14731 | 41291 | 2,803 | 1,487 | 28,945 | 3,25E-184 | 4,02E-183 | 0,0027526 | 0,0021499 | 0,0024858 | 0,0024858 | TRUE | 0,0453993 | No |
| orf19.1234 | 39 | 42 | 53 | 33 | 163 | 103 | 161 | 159 | 37 | 44 | 52 | 52 | 131 | 117 | 137 | 129 | 87,42 | 46 | 128 | 2,8 | 1,485 | 8,811 | 1,24E-18 | 2,69E-18 | 1,00E-08 | 0,0208735 | 0,0117729 | 0,0117729 | TRUE | 0,1590151 | No |
| orf19.7509.1 | 2937 | 2151 | 2217 | 1565 | 8694 | 5656 | 7960 | 8635 | 2800 | 2274 | 2190 | 2452 | 6966 | 6431 | 6758 | 7021 | 4611,64 | 2429 | 6794 | 2,797 | 1,484 | 20,794 | 4,90E-96 | 3,09E-95 | 0,0065908 | 0,0024473 | 0,0045936 | 0,0045936 | TRUE | 0,1314232 | No |
| orf19.446.2 | 470 | 431 | 397 | 312 | 1566 | 1141 | 1439 | 1467 | 448 | 456 | 392 | 489 | 1255 | 1297 | 1222 | 1193 | 843,94 | 446 | 1242 | 2,795 | 1,483 | 21,302 | 1,09E-100 | 7,18E-100 | 0,0017851 | 0,0040368 | 0,0030355 | 0,0030355 | TRUE | 0,0782958 | No |
| orf19.3507 | 2570 | 2105 | 2141 | 1403 | 7741 | 5206 | 7385 | 8239 | 2450 | 2226 | 2115 | 2198 | 6203 | 5920 | 6270 | 6699 | 4260,03 | 2247 | 6273 | 2,79 | 1,481 | 26,07 | 7,91E-150 | 7,93E-149 | 0,0029474 | 0,0024767 | 0,0027753 | 0,0027753 | TRUE | 0,046127 | No |
| orf19.3107 | 97 | 102 | 94 | 45 | 309 | 245 | 304 | 304 | 92 | 108 | 93 | 70 | 248 | 279 | 258 | 247 | 174,39 | 91 | 258 | 2,792 | 1,481 | 11,737 | 8,21E-32 | 2,29E-31 | 1,00E-08 | 0,0115062 | 0,0075345 | 0,0075345 | TRUE | 0,1511219 | No |
| orf19.5017 | 70 | 63 | 61 | 42 | 217 | 173 | 201 | 226 | 67 | 67 | 60 | 66 | 174 | 197 | 171 | 184 | 123,05 | 65 | 182 | 2,791 | 1,481 | 10,591 | 3,27E-26 | 8,32E-26 | 1,00E-08 | 0,0154345 | 0,0077536 | 0,0077536 | TRUE | 0,0316767 | No |
| orf19.4651 | 1459 | 1492 | 1505 | 992 | 5131 | 3778 | 4898 | 5112 | 1391 | 1578 | 1487 | 1554 | 4111 | 4296 | 4158 | 4157 | 2841,45 | 1502 | 4180 | 2,787 | 1,479 | 31,183 | 1,83E-213 | 2,65E-212 | 0,0009128 | 0,0026691 | 0,0016777 | 0,0016777 | TRUE | 0,031224 | No |
| orf19.2382 | 453 | 438 | 455 | 278 | 1512 | 1104 | 1457 | 1556 | 432 | 463 | 449 | 436 | 1212 | 1255 | 1237 | 1265 | 843,63 | 445 | 1242 | 2,788 | 1,479 | 25,239 | 1,51E-140 | 1,43E-139 | 1,00E-08 | 0,0040375 | 0,0016926 | 0,0016926 | TRUE | 0,0087602 | No |
| orf19.6673 | 531 | 431 | 510 | 344 | 1726 | 1175 | 1640 | 1781 | 506 | 456 | 504 | 539 | 1383 | 1336 | 1392 | 1448 | 945,53 | 501 | 1390 | 2,785 | 1,478 | 23,676 | 6,43E-124 | 5,31E-123 | 0,0006561 | 0,0038278 | 0,0023025 | 0,0023025 | TRUE | 0,0422853 | No |
| orf19.5996.1 | 17196 | 13051 | 13830 | 8931 | 52049 | 33173 | 46618 | 51714 | 16395 | 13800 | 13663 | 13991 | 41705 | 37721 | 39578 | 42048 | 27362,61 | 14462 | 40263 | 2,784 | 1,477 | 23,221 | 2,77E-119 | 2,19E-118 | 0,0052031 | 0,0021513 | 0,0038373 | 0,0038373 | TRUE | 0,0993904 | No |
| orf19.1449 | 221 | 238 | 260 | 138 | 837 | 536 | 780 | 816 | 211 | 252 | 257 | 216 | 671 | 609 | 662 | 663 | 442,66 | 234 | 651 | 2,78 | 1,475 | 17,235 | 1,46E-66 | 6,70E-66 | 0,0012414 | 0,0058005 | 0,0039782 | 0,0039782 | TRUE | 0,060484 | No |
| orf19.1743 | 3455 | 2959 | 3085 | 1981 | 10744 | 7672 | 9926 | 11292 | 3294 | 3129 | 3048 | 3103 | 8609 | 8724 | 8427 | 9181 | 5939,37 | 3144 | 8735 | 2,778 | 1,474 | 34,548 | 1,50E-261 | 2,70E-260 | 0,0009566 | 0,0023677 | 0,0015178 | 0,0015178 | TRUE | 0,0145909 | No |
| orf19.5684.1 | 110 | 95 | 90 | 78 | 373 | 227 | 318 | 385 | 105 | 100 | 89 | 122 | 299 | 258 | 270 | 313 | 194,56 | 104 | 285 | 2,775 | 1,472 | 11,818 | 3,14E-32 | 8,80E-32 | 0,0045307 | 0,0105305 | 0,007941 | 0,007941 | TRUE | 0,1004303 | No |
| orf19.6072 | 370 | 333 | 346 | 232 | 1242 | 825 | 1143 | 1218 | 353 | 352 | 342 | 363 | 995 | 938 | 970 | 990 | 663,02 | 352 | 973 | 2,77 | 1,47 | 22,77 | 9,03E-115 | 6,82E-114 | 1,00E-08 | 0,0045677 | 0,0019658 | 0,0019658 | TRUE | 0,0065803 | No |
| orf19.7131 | 582 | 443 | 543 | 407 | 1800 | 1279 | 1842 | 1964 | 555 | 468 | 536 | 638 | 1442 | 1454 | 1564 | 1597 | 1031,84 | 549 | 1514 | 2,771 | 1,47 | 18,141 | 1,50E-73 | 7,53E-73 | 0,0064782 | 0,0036825 | 0,0049868 | 0,0049868 | TRUE | 0,1204099 | No |
| orf19.4149.1 | 14018 | 11936 | 12490 | 8860 | 44696 | 31907 | 41929 | 45241 | 13365 | 12621 | 12339 | 13880 | 35814 | 36281 | 35598 | 36785 | 24585,2 | 13051 | 36120 | 2,769 | 1,469 | 34,704 | 6,75E-264 | 1,23E-262 | 0,0014826 | 0,0021581 | 0,0016659 | 0,0016659 | TRUE | 0,0219172 | No |
| orf19.6975 | 18719 | 14823 | 16028 | 11352 | 57017 | 39729 | 53914 | 60130 | 17847 | 15673 | 15834 | 17784 | 45686 | 45175 | 45773 | 48891 | 31582,92 | 16784 | 46381 | 2,764 | 1,467 | 27,224 | 3,34E-163 | 3,71E-162 | 0,0031352 | 0,0021433 | 0,0027452 | 0,0027452 | TRUE | 0,0242727 | No |
| orf19.3076 | 859 | 680 | 728 | 410 | 2769 | 1386 | 2599 | 2493 | 819 | 719 | 719 | 642 | 2219 | 1576 | 2207 | 2027 | 1365,97 | 725 | 2007 | 2,765 | 1,467 | 13,764 | 4,20E-43 | 1,40E-42 | 0,0164721 | 0,0032933 | 0,0099186 | 0,0099186 | TRUE | 0,2494228 | No |
| orf19.6325.1 | 153 | 104 | 130 | 113 | 496 | 267 | 490 | 511 | 146 | 110 | 128 | 177 | 397 | 304 | 416 | 415 | 261,73 | 140 | 383 | 2,763 | 1,466 | 10,177 | 2,51E-24 | 6,12E-24 | 0,0239958 | 0,0083646 | 0,0147402 | 0,0147402 | TRUE | 0,2629703 | No |
| orf19.6744 | 146 | 103 | 150 | 75 | 419 | 327 | 425 | 446 | 139 | 109 | 148 | 117 | 336 | 372 | 361 | 363 | 243,1 | 128 | 358 | 2,761 | 1,465 | 13,261 | 3,90E-40 | 1,24E-39 | 0,0012847 | 0,0088453 | 0,0061821 | 0,0061821 | TRUE | 0,121733 | No |
| orf19.6398 | 410 | 344 | 392 | 256 | 1318 | 981 | 1179 | 1325 | 391 | 364 | 387 | 401 | 1056 | 1115 | 1001 | 1077 | 724,1 | 386 | 1062 | 2,757 | 1,463 | 21,927 | 1,45E-106 | 1,00E-105 | 1,00E-08 | 0,0043588 | 0,0024087 | 0,0024087 | TRUE | 0,0180794 | No |
| orf19.198 | 5538 | 4894 | 5401 | 3618 | 17617 | 13185 | 17164 | 18944 | 5280 | 5175 | 5336 | 5668 | 14116 | 14993 | 14572 | 15403 | 10067,76 | 5365 | 14771 | 2,755 | 1,462 | 34,118 | 4,02E-255 | 7,01E-254 | 0,0013188 | 0,0022544 | 0,0016288 | 0,0016288 | TRUE | 0,0168928 | No |
| orf19.687 | 219 | 183 | 185 | 140 | 679 | 493 | 656 | 668 | 209 | 193 | 183 | 219 | 544 | 561 | 557 | 543 | 376,14 | 201 | 551 | 2,755 | 1,462 | 17,448 | 3,58E-68 | 1,67E-67 | 1,00E-08 | 0,0064564 | 0,0031597 | 0,0031597 | TRUE | 0,0413039 | No |
| orf19.1440.1 | 9 | 23 | 9 | 9 | 58 | 37 | 44 | 33 | 9 | 24 | 9 | 14 | 46 | 42 | 37 | 27 | 26,08 | 14 | 38 | 2,752 | 1,461 | 4,073 | 4,64E-05 | 6,66E-05 | 0,0819873 | 0,0650525 | 0,0715364 | 0,0715364 | TRUE | 1,1453259 | No |
| orf19.7024 | 6 | 3 | 4 | 4 | 16 | 9 | 15 | 19 | 6 | 3 | 4 | 6 | 13 | 10 | 13 | 15 | 8,79 | 5 | 13 | 2,752 | 1,461 | 2,879 | 0,0039948 | 0,0052001 | 1,00E-08 | 0,1888086 | 0,0946478 | 0,0946478 | TRUE | 0,0925018 | No |
| orf19.1635 | 17038 | 14160 | 14512 | 10374 | 51754 | 37111 | 49271 | 54678 | 16244 | 14972 | 14336 | 16251 | 41469 | 42199 | 41831 | 44458 | 28970,13 | 15451 | 42489 | 2,75 | 1,46 | 29,679 | 1,45E-193 | 1,94E-192 | 0,0023563 | 0,002148 | 0,0022768 | 0,0022768 | TRUE | 0,0289321 | No |
| orf19.3002 | 14698 | 12055 | 12543 | 8012 | 43372 | 30666 | 40743 | 46769 | 14013 | 12747 | 12391 | 12551 | 34753 | 34870 | 34591 | 38028 | 24242,89 | 12926 | 35560 | 2,751 | 1,46 | 28,74 | 1,21E-181 | 1,46E-180 | 0,0026408 | 0,002159 | 0,0024225 | 0,0024225 | TRUE | 0,0394663 | No |
| orf19.7127 | 26 | 19 | 28 | 16 | 72 | 69 | 83 | 78 | 25 | 20 | 28 | 25 | 58 | 78 | 70 | 63 | 45,96 | 24 | 67 | 2,751 | 1,46 | 6,319 | 2,63E-10 | 4,63E-10 | 1,00E-08 | 0,0378195 | 0,0219025 | 0,0219025 | TRUE | 0,0946698 | No |
| orf19.6100 | 689 | 486 | 517 | 380 | 1951 | 1277 | 1881 | 2017 | 657 | 514 | 511 | 595 | 1563 | 1452 | 1597 | 1640 | 1066,14 | 569 | 1563 | 2,75 | 1,459 | 17,692 | 4,86E-70 | 2,33E-69 | 0,0073623 | 0,0036313 | 0,0052597 | 0,0052597 | TRUE | 0,1375684 | No |
| orf19.5014 | 203 | 172 | 179 | 118 | 687 | 440 | 584 | 589 | 194 | 182 | 177 | 185 | 550 | 500 | 496 | 479 | 345,33 | 184 | 506 | 2,749 | 1,459 | 16,112 | 2,10E-58 | 8,64E-58 | 1,00E-08 | 0,0068459 | 0,0039767 | 0,0039767 | TRUE | 0,042279 | No |
| orf19.1700 | 10363 | 9784 | 9502 | 6754 | 33930 | 26423 | 30795 | 33275 | 9880 | 10345 | 9387 | 10581 | 27187 | 30045 | 26145 | 27056 | 18828,24 | 10048 | 27608 | 2,748 | 1,458 | 26,77 | 7,20E-158 | 7,78E-157 | 0,0031637 | 0,0021785 | 0,0027791 | 0,0027791 | TRUE | 0,0431178 | No |
| orf19.7301 | 198 | 165 | 157 | 109 | 608 | 415 | 551 | 573 | 189 | 174 | 155 | 171 | 487 | 472 | 468 | 466 | 322,73 | 172 | 473 | 2,744 | 1,456 | 16,151 | 1,12E-58 | 4,65E-58 | 1,00E-08 | 0,0071788 | 0,0036494 | 0,0036494 | TRUE | 0,055738 | No |
| orf19.3846 | 1735 | 1571 | 1630 | 1012 | 5536 | 3861 | 5248 | 5621 | 1654 | 1661 | 1610 | 1585 | 4436 | 4390 | 4456 | 4570 | 3045,37 | 1628 | 4463 | 2,739 | 1,454 | 38,074 | 0 | 0 | 1,00E-08 | 0,0026304 | 0,0009568 | 0,0009568 | TRUE | 0,0032216 | No |
| orf19.4836 | 2272 | 1798 | 1930 | 1347 | 6865 | 4733 | 6592 | 6892 | 2166 | 1901 | 1907 | 2110 | 5501 | 5382 | 5597 | 5604 | 3770,88 | 2021 | 5521 | 2,734 | 1,451 | 28,585 | 1,04E-179 | 1,25E-178 | 0,0020293 | 0,0025267 | 0,0021153 | 0,0021153 | TRUE | 0,0306514 | No |
| orf19.396 | 314 | 286 | 310 | 185 | 1006 | 657 | 1061 | 1010 | 299 | 302 | 306 | 290 | 806 | 747 | 901 | 821 | 559,12 | 299 | 819 | 2,733 | 1,451 | 18,202 | 4,97E-74 | 2,52E-73 | 0,0025866 | 0,0050278 | 0,0036967 | 0,0036967 | TRUE | 0,0536026 | No |
| orf19.4463 | 9 | 4 | 8 | 3 | 23 | 15 | 22 | 21 | 9 | 4 | 8 | 5 | 18 | 17 | 19 | 17 | 12,08 | 6 | 18 | 2,734 | 1,451 | 3,366 | 0,0007618 | 0,0010365 | 1,00E-08 | 0,1379944 | 0,0685758 | 0,0685758 | TRUE | 0,1437501 | No |
| orf19.5660.1 | 2592 | 2101 | 2156 | 1572 | 8033 | 5263 | 7563 | 7995 | 2471 | 2222 | 2130 | 2463 | 6437 | 5985 | 6421 | 6501 | 4328,51 | 2322 | 6336 | 2,733 | 1,45 | 25,602 | 1,44E-144 | 1,41E-143 | 0,0030584 | 0,0024706 | 0,002768 | 0,002768 | TRUE | 0,0377992 | No |
| orf19.2068 | 544 | 494 | 531 | 320 | 1691 | 1322 | 1706 | 1662 | 519 | 522 | 525 | 501 | 1355 | 1503 | 1448 | 1351 | 965,6 | 517 | 1414 | 2,73 | 1,449 | 23,488 | 5,45E-122 | 4,42E-121 | 1,00E-08 | 0,0037917 | 0,002259 | 0,002259 | TRUE | 0,0203703 | No |
| orf19.3029 | 3221 | 3017 | 3105 | 2089 | 10259 | 7738 | 10009 | 10857 | 3071 | 3190 | 3067 | 3273 | 8220 | 8799 | 8498 | 8828 | 5868,17 | 3150 | 8586 | 2,728 | 1,448 | 35,56 | 5,80E-277 | 1,14E-275 | 0,0007641 | 0,0023711 | 0,001361 | 0,001361 | TRUE | 0,0101217 | No |
| orf19.7419 | 48 | 37 | 39 | 18 | 146 | 81 | 131 | 122 | 46 | 39 | 39 | 28 | 117 | 92 | 111 | 99 | 71,39 | 38 | 105 | 2,725 | 1,446 | 7,623 | 2,48E-14 | 4,92E-14 | 0,0001848 | 0,0250904 | 0,0159594 | 0,0159594 | TRUE | 0,1391054 | No |
| orf19.5658 | 1039 | 805 | 850 | 584 | 2991 | 2045 | 2887 | 3205 | 991 | 851 | 840 | 915 | 2397 | 2325 | 2451 | 2606 | 1671,91 | 899 | 2445 | 2,721 | 1,444 | 22,563 | 9,94E-113 | 7,39E-112 | 0,0031507 | 0,0030733 | 0,0031251 | 0,0031251 | TRUE | 0,060607 | No |
| orf19.3980 | 985 | 924 | 1079 | 635 | 3311 | 2349 | 3130 | 3472 | 939 | 977 | 1066 | 995 | 2653 | 2671 | 2657 | 2823 | 1847,66 | 994 | 2701 | 2,718 | 1,442 | 27,658 | 2,25E-168 | 2,55E-167 | 0,0009696 | 0,0029799 | 0,0018826 | 0,0018826 | TRUE | 0,0305946 | No |
| orf19.7270 | 3741 | 2872 | 3242 | 1766 | 11668 | 6100 | 10612 | 10910 | 3567 | 3037 | 3203 | 2767 | 9349 | 6936 | 9010 | 8871 | 5842,33 | 3144 | 8542 | 2,716 | 1,442 | 15,161 | 6,45E-52 | 2,44E-51 | 0,0135044 | 0,0023723 | 0,0084542 | 0,0084542 | TRUE | 0,1948572 | No |
| orf19.3465 | 20221 | 17666 | 17905 | 11984 | 61806 | 45188 | 57751 | 64083 | 19279 | 18680 | 17688 | 18774 | 49523 | 51383 | 49030 | 52106 | 34557,78 | 18605 | 50510 | 2,715 | 1,441 | 37,7 | 0 | 0 | 0,0010129 | 0,0021388 | 0,0013642 | 0,0013642 | TRUE | 0,0135742 | No |
| orf19.2184 | 234 | 182 | 201 | 135 | 721 | 470 | 668 | 686 | 223 | 192 | 199 | 211 | 578 | 534 | 567 | 558 | 382,83 | 206 | 559 | 2,714 | 1,441 | 17,194 | 2,93E-66 | 1,34E-65 | 1,00E-08 | 0,0063801 | 0,0032537 | 0,0032537 | TRUE | 0,0387368 | No |
| orf19.5851 | 771 | 703 | 747 | 513 | 2475 | 1739 | 2416 | 2646 | 735 | 743 | 738 | 804 | 1983 | 1977 | 2051 | 2151 | 1397,9 | 755 | 2040 | 2,713 | 1,44 | 26,47 | 2,14E-154 | 2,21E-153 | 0,0008714 | 0,0032658 | 0,0018806 | 0,0018806 | TRUE | 0,0187893 | No |
| orf19.3818 | 295 | 245 | 306 | 191 | 949 | 664 | 897 | 1000 | 281 | 259 | 302 | 299 | 760 | 755 | 762 | 813 | 528,99 | 285 | 772 | 2,714 | 1,44 | 19,179 | 5,55E-82 | 3,09E-81 | 1,00E-08 | 0,0051951 | 0,0028868 | 0,0028868 | TRUE | 0,0436539 | No |
| orf19.1032 | 4068 | 3561 | 3902 | 2678 | 13151 | 9225 | 12490 | 13352 | 3878 | 3765 | 3855 | 4195 | 10538 | 10490 | 10604 | 10856 | 7272,66 | 3923 | 10622 | 2,711 | 1,439 | 36,086 | 3,72E-285 | 7,71E-284 | 0,0007671 | 0,002317 | 0,0013411 | 0,0013411 | TRUE | 0,0247398 | No |
| orf19.1002 | 38 | 55 | 61 | 34 | 150 | 140 | 161 | 181 | 36 | 58 | 60 | 53 | 120 | 159 | 137 | 147 | 96,39 | 52 | 141 | 2,712 | 1,439 | 8,008 | 1,17E-15 | 2,39E-15 | 0,013882 | 0,0191245 | 0,0170954 | 0,0170954 | TRUE | 0,3951631 | No |
| orf19.6574 | 29 | 33 | 28 | 18 | 92 | 90 | 85 | 93 | 28 | 35 | 28 | 28 | 74 | 102 | 72 | 76 | 55,28 | 30 | 81 | 2,712 | 1,439 | 6,589 | 4,43E-11 | 7,99E-11 | 1,00E-08 | 0,031793 | 0,0215508 | 0,0215508 | TRUE | 0,2656279 | No |
| orf19.5666 | 899 | 729 | 768 | 436 | 2714 | 1538 | 2707 | 2590 | 857 | 771 | 759 | 683 | 2175 | 1749 | 2298 | 2106 | 1424,66 | 768 | 2082 | 2,708 | 1,437 | 15,928 | 4,05E-57 | 1,63E-56 | 0,0095458 | 0,0032438 | 0,0068733 | 0,0068733 | TRUE | 0,1378404 | No |
| orf19.5665 | 546 | 452 | 498 | 322 | 1701 | 1122 | 1652 | 1659 | 521 | 478 | 492 | 504 | 1363 | 1276 | 1403 | 1349 | 923,14 | 499 | 1348 | 2,705 | 1,436 | 23,837 | 1,37E-125 | 1,15E-124 | 1,00E-08 | 0,0038699 | 0,0020351 | 0,0020351 | TRUE | 0,0143974 | No |
| orf19.3267 | 83 | 74 | 89 | 65 | 262 | 233 | 266 | 282 | 79 | 78 | 88 | 102 | 210 | 265 | 226 | 229 | 159,64 | 87 | 232 | 2,699 | 1,432 | 10,713 | 8,89E-27 | 2,28E-26 | 0,004091 | 0,0123763 | 0,00877 | 0,00877 | TRUE | 0,0940257 | No |
| orf19.6217 | 1285 | 1010 | 1223 | 832 | 3957 | 2846 | 3675 | 4170 | 1225 | 1068 | 1208 | 1303 | 3171 | 3236 | 3120 | 3391 | 2215,26 | 1201 | 3230 | 2,694 | 1,43 | 23,732 | 1,69E-124 | 1,40E-123 | 0,0032296 | 0,0028325 | 0,002878 | 0,002878 | TRUE | 0,066391 | No |
| orf19.3429 | 298 | 318 | 317 | 216 | 1069 | 811 | 945 | 1025 | 284 | 336 | 313 | 338 | 857 | 922 | 802 | 833 | 585,79 | 318 | 854 | 2,693 | 1,429 | 18,042 | 9,17E-73 | 4,57E-72 | 0,0024982 | 0,0048941 | 0,0037392 | 0,0037392 | TRUE | 0,0615062 | No |
| orf19.3954.1 | 39 | 38 | 52 | 32 | 158 | 110 | 145 | 127 | 37 | 40 | 51 | 50 | 127 | 125 | 123 | 103 | 82,11 | 44 | 120 | 2,692 | 1,429 | 8,068 | 7,15E-16 | 1,47E-15 | 1,00E-08 | 0,0220867 | 0,013858 | 0,013858 | TRUE | 0,1107796 | No |
| orf19.5438 | 1702 | 1341 | 1437 | 933 | 5159 | 3148 | 4930 | 4968 | 1623 | 1418 | 1420 | 1462 | 4134 | 3580 | 4186 | 4039 | 2732,52 | 1481 | 3985 | 2,691 | 1,428 | 21,98 | 4,50E-107 | 3,14E-106 | 0,0039373 | 0,0026922 | 0,0035631 | 0,0035631 | TRUE | 0,0560414 | No |
| orf19.5893 | 6832 | 5490 | 5945 | 3696 | 20413 | 12958 | 19489 | 20547 | 6514 | 5805 | 5873 | 5790 | 16356 | 14734 | 16546 | 16707 | 11040,64 | 5996 | 16086 | 2,682 | 1,424 | 25,743 | 3,85E-146 | 3,79E-145 | 0,0031337 | 0,00224 | 0,0028156 | 0,0028156 | TRUE | 0,041688 | No |

| | | | | | | | | | | | | | | | | | | | | | | | | | | | | | | |
|---|---|---|---|---|---|---|---|---|---|---|---|---|---|---|---|---|---|---|---|---|---|---|---|---|---|---|---|---|---|---|
| orf19.4472 | 41 | 34 | 34 | 29 | 108 | 94 | 133 | 124 | 39 | 36 | 34 | 45 | 87 | 107 | 113 | 101 | 70,15 | 38 | 102 | 2,67 | 1,417 | 7,488 | 6,97E-14 | 1,36E-13 | 1,00E-08 | 0,0254959 | 0,0154023 | 0,0154023 | TRUE | 0,1083676 | No |
| orf19.6392 | 452 | 375 | 459 | 270 | 1420 | 901 | 1374 | 1489 | 431 | 397 | 453 | 423 | 1138 | 1025 | 1167 | 1211 | 780,43 | 426 | 1135 | 2,666 | 1,415 | 19,174 | 6,16E-82 | 3,42E-81 | 0,0029346 | 0,0041951 | 0,0035228 | 0,0035228 | TRUE | 0,0487025 | No |
| orf19.4919 | 376 | 295 | 320 | 226 | 1110 | 757 | 1014 | 1155 | 358 | 312 | 316 | 354 | 889 | 861 | 861 | 939 | 611,35 | 335 | 888 | 2,657 | 1,41 | 19,233 | 1,94E-82 | 1,09E-81 | 0,0010488 | 0,004777 | 0,0029892 | 0,0029892 | TRUE | 0,0309961 | No |
| orf19.4624 | 422 | 378 | 351 | 207 | 1222 | 950 | 1097 | 1166 | 402 | 400 | 347 | 324 | 979 | 1080 | 931 | 948 | 676,48 | 368 | 984 | 2,656 | 1,409 | 16,979 | 1,17E-64 | 5,23E-64 | 0,0048012 | 0,0045184 | 0,0046507 | 0,0046507 | TRUE | 0,0612101 | No |
| orf19.2346 | 989 | 891 | 962 | 692 | 3284 | 2186 | 3076 | 3226 | 943 | 942 | 950 | 1084 | 2631 | 2486 | 2612 | 2623 | 1783,88 | 980 | 2588 | 2,654 | 1,408 | 26,205 | 2,34E-151 | 2,37E-150 | 0,0010253 | 0,0030117 | 0,0020262 | 0,0020262 | TRUE | 0,053294 | No |
| orf19.6830 | 456 | 436 | 438 | 268 | 1525 | 1044 | 1266 | 1434 | 435 | 461 | 433 | 420 | 1222 | 1187 | 1075 | 1166 | 799,77 | 437 | 1162 | 2,655 | 1,408 | 20,998 | 6,79E-98 | 4,35E-97 | 0,0014929 | 0,0041442 | 0,0026618 | 0,0026618 | TRUE | 0,0314694 | No |
| orf19.1470 | 10651 | 8522 | 9081 | 8039 | 32529 | 25283 | 30347 | 33772 | 10155 | 9011 | 8971 | 12594 | 26065 | 28749 | 25764 | 27460 | 18596,02 | 10183 | 27010 | 2,653 | 1,408 | 15,088 | 1,95E-51 | 7,32E-51 | 0,015116 | 0,0021796 | 0,0082943 | 0,0082943 | TRUE | 0,3089196 | No |
| orf19.2059 | 1442 | 1292 | 1403 | 957 | 4565 | 3254 | 4460 | 4588 | 1375 | 1366 | 1386 | 1499 | 3658 | 3700 | 3787 | 3730 | 2562,63 | 1406 | 3719 | 2,652 | 1,407 | 33,08 | 5,72E-240 | 9,31E-239 | 5,81E-05 | 0,002732 | 0,0012178 | 0,0012178 | TRUE | 0,0203607 | No |
| orf19.6541 | 19339 | 15612 | 16947 | 10505 | 56023 | 37500 | 54360 | 57762 | 18438 | 16508 | 16742 | 16457 | 44890 | 42641 | 46151 | 46966 | 31098,99 | 17036 | 45162 | 2,651 | 1,406 | 28,757 | 7,30E-182 | 8,88E-181 | 0,0023483 | 0,0021441 | 0,0022547 | 0,0022547 | TRUE | 0,0376966 | No |
| orf19.7265 | 978 | 768 | 834 | 563 | 2946 | 1746 | 2737 | 3013 | 932 | 812 | 824 | 882 | 2361 | 1985 | 2324 | 2450 | 1571,23 | 862 | 2280 | 2,646 | 1,404 | 18,699 | 5,05E-78 | 2,69E-77 | 0,0057142 | 0,0031363 | 0,0045644 | 0,0045644 | TRUE | 0,0895331 | No |
| orf19.5262 | 43 | 21 | 24 | 14 | 90 | 61 | 93 | 89 | 41 | 22 | 24 | 22 | 72 | 69 | 79 | 72 | 50,21 | 27 | 73 | 2,646 | 1,404 | 6,058 | 1,38E-09 | 2,35E-09 | 1,00E-08 | 0,0347952 | 0,0254072 | 0,0254072 | TRUE | 0,8041147 | No |
| orf19.1340 | 142 | 118 | 153 | 76 | 422 | 331 | 421 | 423 | 135 | 125 | 151 | 119 | 338 | 376 | 357 | 344 | 243,28 | 132 | 354 | 2,644 | 1,403 | 13,376 | 8,38E-41 | 2,69E-40 | 1,00E-08 | 0,0088403 | 0,0051437 | 0,0051437 | TRUE | 0,0910831 | No |
| orf19.2352 | 437 | 357 | 383 | 208 | 1240 | 790 | 1246 | 1267 | 417 | 377 | 378 | 326 | 994 | 898 | 1058 | 1030 | 684,78 | 374 | 995 | 2,643 | 1,402 | 17,057 | 3,09E-65 | 1,39E-64 | 0,0049473 | 0,004489 | 0,0045554 | 0,0045554 | TRUE | 0,0732306 | No |
| orf19.5858 | 2808 | 2122 | 2282 | 1372 | 7811 | 4871 | 7311 | 8057 | 2677 | 2244 | 2254 | 2149 | 6259 | 5539 | 6207 | 6551 | 4235,02 | 2331 | 6139 | 2,631 | 1,396 | 18,87 | 2,00E-79 | 1,08E-78 | 0,0071339 | 0,002479 | 0,0049407 | 0,0049407 | TRUE | 0,12216 | No |
| orf19.4406 | 426 | 397 | 366 | 332 | 1404 | 1043 | 1249 | 1332 | 406 | 420 | 362 | 520 | 1125 | 1186 | 1060 | 1083 | 770,25 | 427 | 1114 | 2,627 | 1,394 | 15,216 | 2,77E-52 | 1,05E-51 | 0,0089945 | 0,0042229 | 0,0063282 | 0,0063282 | TRUE | 0,2210673 | No |
| orf19.3572.3 | 11419 | 9967 | 10117 | 6395 | 33985 | 22752 | 32402 | 34387 | 10887 | 10539 | 9995 | 10018 | 27231 | 25871 | 27509 | 27960 | 18751,21 | 10360 | 27143 | 2,62 | 1,389 | 33,213 | 7,01E-242 | 1,15E-240 | 0,0013296 | 0,0021788 | 0,00161 | 0,00161 | TRUE | 0,0144427 | No |
| orf19.838.1 | 19928 | 16350 | 17684 | 10478 | 58108 | 36728 | 54638 | 60344 | 18999 | 17288 | 17470 | 16414 | 46560 | 41763 | 46387 | 49065 | 31743,48 | 17543 | 45944 | 2,619 | 1,389 | 23,54 | 1,60E-122 | 1,30E-121 | 0,0040188 | 0,002143 | 0,0033024 | 0,0033024 | TRUE | 0,0459043 | No |
| orf19.4308 | 397 | 353 | 373 | 261 | 1219 | 841 | 1126 | 1351 | 378 | 373 | 368 | 409 | 977 | 956 | 956 | 1098 | 689,58 | 382 | 997 | 2,619 | 1,389 | 18,774 | 1,24E-78 | 6,66E-78 | 0,0024306 | 0,0044723 | 0,0033447 | 0,0033447 | TRUE | 0,057435 | No |
| orf19.2257 | 1604 | 1403 | 1507 | 898 | 4682 | 3464 | 4598 | 4796 | 1529 | 1483 | 1489 | 1407 | 3752 | 3939 | 3904 | 3900 | 2675,25 | 1477 | 3874 | 2,616 | 1,387 | 33,078 | 6,25E-240 | 1,01E-238 | 0,000206 | 0,002705 | 0,0011954 | 0,0011954 | TRUE | 0,0106386 | No |
| orf19.1249 | 738 | 587 | 640 | 414 | 2101 | 1458 | 2021 | 2165 | 704 | 621 | 632 | 649 | 1683 | 1658 | 1716 | 1760 | 1177,83 | 652 | 1704 | 2,616 | 1,387 | 24,546 | 4,74E-133 | 4,27E-132 | 0,0005104 | 0,0034853 | 0,0019489 | 0,0019489 | TRUE | 0,038109 | No |
| orf19.4016 | 6671 | 5375 | 5913 | 4318 | 19085 | 14441 | 18762 | 20526 | 6360 | 5683 | 5841 | 6764 | 15292 | 16421 | 15929 | 16690 | 11122,59 | 6162 | 16083 | 2,611 | 1,385 | 23,886 | 4,27E-126 | 3,61E-125 | 0,0038002 | 0,0022389 | 0,0031093 | 0,0031093 | TRUE | 0,051595 | No |
| orf19.2149 | 854 | 646 | 750 | 490 | 2488 | 1578 | 2360 | 2507 | 814 | 683 | 741 | 768 | 1994 | 1794 | 2004 | 2038 | 1354,47 | 752 | 1958 | 2,608 | 1,383 | 20,784 | 6,02E-96 | 3,79E-95 | 0,0030165 | 0,0033035 | 0,0032748 | 0,0032748 | TRUE | 0,0457896 | No |
| orf19.2510 | 113 | 172 | 192 | 83 | 520 | 426 | 457 | 377 | 108 | 182 | 190 | 130 | 417 | 484 | 388 | 307 | 275,61 | 152 | 399 | 2,608 | 1,383 | 7,761 | 8,42E-15 | 1,69E-14 | 0,0474194 | 0,0080486 | 0,0256754 | 0,0256754 | TRUE | 0,2338531 | No |
| orf19.2966 | 268 | 265 | 241 | 135 | 764 | 605 | 737 | 811 | 256 | 280 | 238 | 211 | 612 | 688 | 626 | 659 | 446,32 | 246 | 646 | 2,602 | 1,38 | 15,642 | 3,74E-55 | 1,48E-54 | 0,0027605 | 0,0057701 | 0,0045304 | 0,0045304 | TRUE | 0,0881859 | No |
| orf19.6625 | 961 | 874 | 921 | 513 | 2922 | 1858 | 2835 | 2934 | 916 | 924 | 910 | 804 | 2341 | 2113 | 2407 | 2386 | 1600,05 | 888 | 2312 | 2,595 | 1,376 | 22,029 | 1,52E-107 | 1,07E-106 | 0,0027238 | 0,0031174 | 0,0029235 | 0,0029235 | TRUE | 0,0430601 | No |
| orf19.5488 | 321 | 276 | 298 | 200 | 1041 | 594 | 952 | 976 | 306 | 292 | 294 | 313 | 834 | 675 | 808 | 794 | 539,62 | 301 | 778 | 2,591 | 1,374 | 16,212 | 4,18E-59 | 1,74E-58 | 0,0038765 | 0,005134 | 0,0044658 | 0,0044658 | TRUE | 0,0876987 | No |
| orf19.6113 | 90 | 64 | 77 | 37 | 239 | 127 | 238 | 260 | 86 | 68 | 76 | 58 | 192 | 144 | 202 | 211 | 129,61 | 72 | 187 | 2,586 | 1,371 | 8,352 | 6,69E-17 | 1,40E-16 | 0,0172077 | 0,0147592 | 0,0158302 | 0,0158302 | TRUE | 0,2364007 | No |
| orf19.2994 | 21751 | 17635 | 19237 | 12741 | 63953 | 42106 | 60997 | 63242 | 20737 | 18647 | 19004 | 19959 | 51244 | 47878 | 51786 | 51422 | 35084,72 | 19587 | 50582 | 2,583 | 1,369 | 30,504 | 2,32E-204 | 3,22E-203 | 0,0017644 | 0,0021381 | 0,001897 | 0,001897 | TRUE | 0,0193184 | No |
| orf19.7327 | 5006 | 4125 | 4486 | 3409 | 14992 | 10869 | 14262 | 15114 | 4773 | 4362 | 4432 | 5340 | 12013 | 12359 | 12108 | 12289 | 8459,46 | 4727 | 12192 | 2,582 | 1,369 | 22,847 | 1,58E-115 | 1,20E-114 | 0,0043584 | 0,0022854 | 0,0032904 | 0,0032904 | TRUE | 0,0906726 | No |
| orf19.2383 | 133 | 130 | 127 | 112 | 450 | 295 | 419 | 471 | 127 | 137 | 125 | 175 | 361 | 335 | 356 | 383 | 249,99 | 141 | 359 | 2,582 | 1,368 | 12,29 | 1,03E-34 | 3,00E-34 | 0,0029204 | 0,0086593 | 0,006661 | 0,006661 | TRUE | 0,2149613 | No |
| orf19.2115 | 2784 | 2207 | 2510 | 1548 | 7946 | 5038 | 7900 | 8265 | 2654 | 2334 | 2480 | 2425 | 6367 | 5729 | 6707 | 6720 | 4426,92 | 2473 | 6381 | 2,58 | 1,367 | 22,088 | 4,09E-108 | 2,91E-107 | 0,0037635 | 0,0024622 | 0,0033817 | 0,0033817 | TRUE | 0,0571032 | No |
| orf19.5351 | 5765 | 4410 | 4902 | 2748 | 16297 | 9425 | 15563 | 15798 | 5496 | 4663 | 4843 | 4305 | 13058 | 10717 | 13213 | 12845 | 8642,56 | 4827 | 12458 | 2,58 | 1,367 | 16,689 | 1,58E-62 | 6,90E-62 | 0,0095719 | 0,0022813 | 0,006297 | 0,006297 | TRUE | 0,1079314 | No |
| orf19.7494 | 215 | 214 | 240 | 141 | 726 | 506 | 648 | 717 | 205 | 226 | 237 | 221 | 582 | 575 | 550 | 583 | 397,43 | 222 | 572 | 2,576 | 1,365 | 16,813 | 1,97E-63 | 8,67E-63 | 1,00E-08 | 0,0062225 | 0,0030544 | 0,0030544 | TRUE | 0,035561 | No |
| orf19.114 | 153 | 187 | 188 | 107 | 531 | 385 | 556 | 566 | 146 | 198 | 186 | 168 | 425 | 438 | 472 | 460 | 311,56 | 174 | 449 | 2,576 | 1,365 | 13,42 | 4,59E-41 | 1,48E-40 | 0,0032484 | 0,0073613 | 0,0057489 | 0,0057489 | TRUE | 0,1488546 | No |
| orf19.3490 | 39 | 28 | 29 | 18 | 105 | 52 | 111 | 100 | 37 | 30 | 29 | 28 | 84 | 59 | 94 | 81 | 55,31 | 31 | 80 | 2,576 | 1,365 | 6,165 | 7,04E-10 | 1,22E-09 | 1,00E-08 | 0,031779 | 0,023689 | 0,023689 | TRUE | 0,2474981 | No |
| orf19.3726 | 895 | 816 | 917 | 523 | 2808 | 1857 | 2665 | 2749 | 853 | 863 | 906 | 819 | 2250 | 2112 | 2263 | 2235 | 1537,58 | 860 | 2215 | 2,57 | 1,362 | 26,812 | 2,35E-158 | 2,54E-157 | 0,0004218 | 0,0031591 | 0,0016281 | 0,0016281 | TRUE | 0,0150549 | No |
| orf19.6662 | 391 | 306 | 334 | 196 | 1077 | 707 | 1024 | 1112 | 373 | 324 | 330 | 307 | 863 | 804 | 869 | 904 | 596,72 | 334 | 860 | 2,57 | 1,362 | 17,846 | 3,10E-71 | 1,51E-70 | 0,0019252 | 0,0048428 | 0,0034094 | 0,0034094 | TRUE | 0,0746153 | No |
| orf19.278 | 1349 | 1157 | 1295 | 908 | 4056 | 3083 | 3825 | 4106 | 1286 | 1223 | 1279 | 1422 | 3250 | 3506 | 3247 | 3339 | 2319,11 | 1302 | 3336 | 2,566 | 1,36 | 25,543 | 6,49E-144 | 6,30E-143 | 0,0016315 | 0,0027993 | 0,0021545 | 0,0021545 | TRUE | 0,0409555 | No |
| orf19.7176 | 797 | 752 | 815 | 511 | 2393 | 1771 | 2436 | 2585 | 760 | 795 | 805 | 801 | 1917 | 2014 | 2068 | 2102 | 1407,74 | 790 | 2025 | 2,565 | 1,359 | 25,623 | 8,49E-145 | 8,28E-144 | 1,00E-08 | 0,0032576 | 0,0017738 | 0,0017738 | TRUE | 0,0161833 | No |
| orf19.4395 | 1295 | 1164 | 1246 | 823 | 3869 | 2729 | 3832 | 4052 | 1235 | 1231 | 1231 | 1289 | 3100 | 3103 | 3253 | 3295 | 2217,11 | 1246 | 3188 | 2,563 | 1,358 | 30,45 | 1,21E-203 | 1,68E-202 | 0,0004523 | 0,0028318 | 0,0013191 | 0,0013191 | TRUE | 0,0062365 | No |
| orf19.1480 | 1974 | 1675 | 1770 | 1287 | 5896 | 4129 | 5530 | 5974 | 1882 | 1771 | 1749 | 2016 | 4724 | 4695 | 4695 | 4857 | 3298,69 | 1854 | 4743 | 2,564 | 1,358 | 27,887 | 3,85E-171 | 4,41E-170 | 0,0013167 | 0,002589 | 0,0018801 | 0,0018801 | TRUE | 0,0378222 | No |
| orf19.5982 | 17187 | 13356 | 14325 | 8991 | 46968 | 31052 | 44260 | 49143 | 16386 | 14122 | 14152 | 14085 | 37634 | 35309 | 37577 | 39958 | 26152,8 | 14686 | 37620 | 2,561 | 1,357 | 22,965 | 1,04E-116 | 8,01E-116 | 0,0041883 | 0,0021541 | 0,0033035 | 0,0033035 | TRUE | 0,0745448 | No |
| orf19.4478 | 282 | 233 | 277 | 176 | 911 | 578 | 771 | 835 | 269 | 246 | 274 | 276 | 730 | 657 | 655 | 679 | 473,16 | 266 | 680 | 2,562 | 1,357 | 17,251 | 1,11E-66 | 5,08E-66 | 1,00E-08 | 0,0055613 | 0,0031968 | 0,0031968 | TRUE | 0,0295243 | No |
| orf19.6085 | 18708 | 15845 | 16961 | 10695 | 53749 | 36911 | 52373 | 55067 | 17836 | 16754 | 16756 | 16754 | 43068 | 41971 | 44464 | 44775 | 30297,26 | 17025 | 43570 | 2,559 | 1,356 | 36,71 | 5,02E-295 | 1,10E-293 | 0,0008941 | 0,0021455 | 0,0012664 | 0,0012664 | TRUE | 0,0126884 | No |
| orf19.3225 | 815 | 814 | 888 | 500 | 2621 | 1987 | 2445 | 2490 | 777 | 861 | 877 | 783 | 2100 | 2259 | 2076 | 2025 | 1469,77 | 824 | 2115 | 2,559 | 1,355 | 22,554 | 1,23E-112 | 9,16E-112 | 0,001996 | 0,0032084 | 0,0025777 | 0,0025777 | TRUE | 0,0246671 | No |
| orf19.2051 | 406 | 387 | 437 | 292 | 1337 | 911 | 1245 | 1373 | 387 | 409 | 432 | 457 | 1071 | 1036 | 1057 | 1116 | 745,75 | 421 | 1070 | 2,555 | 1,353 | 20,352 | 4,45E-92 | 2,71E-91 | 0,0004625 | 0,004293 | 0,0024988 | 0,0024988 | TRUE | 0,0352332 | No |
| orf19.2281 | 125 | 99 | 107 | 60 | 339 | 227 | 335 | 338 | 119 | 105 | 106 | 94 | 272 | 258 | 284 | 275 | 189,07 | 106 | 272 | 2,552 | 1,352 | 11,901 | 1,16E-32 | 3,28E-32 | 1,00E-08 | 0,0107755 | 0,0055587 | 0,0055587 | TRUE | 0,0714323 | No |
| orf19.5156 | 708 | 652 | 604 | 442 | 2056 | 1548 | 1928 | 2095 | 675 | 689 | 597 | 692 | 1647 | 1760 | 1637 | 1703 | 1175,18 | 663 | 1687 | 2,547 | 1,349 | 22,313 | 2,76E-110 | 2,00E-109 | 0,0014798 | 0,0034884 | 0,0023995 | 0,0023995 | TRUE | 0,0564916 | No |
| orf19.115 | 129 | 149 | 110 | 59 | 399 | 286 | 368 | 347 | 123 | 158 | 109 | 92 | 320 | 325 | 312 | 282 | 215,14 | 120 | 310 | 2,546 | 1,348 | 10,034 | 1,08E-23 | 2,61E-23 | 0,0141054 | 0,0097231 | 0,0113035 | 0,0113035 | TRUE | 0,4162414 | No |
| orf19.5025 | 14259 | 13157 | 13528 | 8291 | 43165 | 29696 | 39724 | 42978 | 13594 | 13912 | 13364 | 12988 | 34587 | 33767 | 33725 | 34945 | 23860,45 | 13464 | 34256 | 2,543 | 1,347 | 41,753 | 0 | 0 | 0,0005052 | 0,0021601 | 0,0009444 | 0,0009444 | TRUE | 0,0067121 | No |
| orf19.6286 | 474 | 369 | 431 | 242 | 1327 | 857 | 1306 | 1303 | 452 | 390 | 426 | 379 | 1063 | 974 | 1109 | 1059 | 731,62 | 412 | 1051 | 2,544 | 1,347 | 18,375 | 2,07E-75 | 1,06E-74 | 0,0024506 | 0,0043355 | 0,0033881 | 0,0033881 | TRUE | 0,0501917 | No |
| orf19.3250 | 1316 | 1340 | 1391 | 783 | 4238 | 3046 | 3886 | 4025 | 1255 | 1417 | 1374 | 1227 | 3396 | 3464 | 3299 | 3273 | 2337,95 | 1318 | 3358 | 2,542 | 1,346 | 25,161 | 1,07E-139 | 1,00E-138 | 0,0016848 | 0,0027936 | 0,0021893 | 0,0021893 | TRUE | 0,0298362 | No |
| orf19.1959 | 1320 | 1259 | 1266 | 894 | 4092 | 2844 | 4031 | 4104 | 1258 | 1331 | 1251 | 1400 | 3279 | 3234 | 3422 | 3337 | 2314,1 | 1310 | 3318 | 2,541 | 1,345 | 28,123 | 5,16E-174 | 6,01E-173 | 0,0007061 | 0,0028008 | 0,0016333 | 0,0016333 | TRUE | 0,02285 | No |
| orf19.4447 | 1600 | 1246 | 1444 | 880 | 4521 | 2919 | 4250 | 4682 | 1525 | 1317 | 1427 | 1379 | 3623 | 3319 | 3608 | 3807 | 2500,61 | 1412 | 3589 | 2,541 | 1,345 | 22,514 | 3,00E-112 | 2,22E-111 | 0,0028511 | 0,0027479 | 0,0029057 | 0,0029057 | TRUE | 0,0363588 | No |
| orf19.6374 | 630 | 563 | 623 | 340 | 1845 | 1184 | 1876 | 1908 | 601 | 595 | 615 | 533 | 1478 | 1346 | 1593 | 1551 | 1039,1 | 586 | 1492 | 2,54 | 1,345 | 19,008 | 1,46E-80 | 8,02E-80 | 0,0034514 | 0,0036714 | 0,0035535 | 0,0035535 | TRUE | 0,0499758 | No |
| orf19.7102 | 377 | 304 | 310 | 235 | 1110 | 679 | 1054 | 1067 | 359 | 321 | 306 | 368 | 889 | 772 | 895 | 868 | 597,4 | 338 | 856 | 2,54 | 1,345 | 16,462 | 6,86E-61 | 2,93E-60 | 0,0033262 | 0,0048397 | 0,0042325 | 0,0042325 | TRUE | 0,0488592 | No |
| orf19.804 | 462 | 410 | 464 | 263 | 1346 | 977 | 1297 | 1419 | 440 | 434 | 458 | 412 | 1079 | 1111 | 1101 | 1154 | 773,59 | 436 | 1111 | 2,539 | 1,344 | 21,757 | 5,94E-105 | 4,04E-104 | 1,00E-08 | 0,0042137 | 0,0019918 | 0,0019918 | TRUE | 0,0131759 | No |
| orf19.2798 | 187 | 155 | 168 | 116 | 511 | 387 | 537 | 537 | 178 | 164 | 166 | 182 | 409 | 440 | 456 | 437 | 303,99 | 172 | 436 | 2,531 | 1,34 | 14,632 | 1,77E-48 | 6,38E-48 | 1,00E-08 | 0,0074925 | 0,0038013 | 0,0038013 | TRUE | 0,0198769 | No |
| orf19.2707.1 | 2067 | 1418 | 1569 | 1108 | 5648 | 3141 | 5486 | 5336 | 1971 | 1499 | 1550 | 1736 | 4526 | 3572 | 4658 | 4339 | 2981,15 | 1689 | 4274 | 2,531 | 1,34 | 13,836 | 1,55E-43 | 5,19E-43 | 0,0139035 | 0,002642 | 0,0085682 | 0,0085682 | TRUE | 0,1555376 | No |
| orf19.4055 | 243 | 172 | 178 | 117 | 655 | 409 | 561 | 607 | 232 | 182 | 176 | 183 | 525 | 465 | 476 | 494 | 341,55 | 193 | 490 | 2,529 | 1,338 | 13,314 | 1,93E-40 | 6,13E-40 | 0,0045652 | 0,0068985 | 0,0059171 | 0,0059171 | TRUE | 0,2121408 | No |

| | | | | | | | | | | | | | | | | | | | | | | | | | | | | | | | |
|---|---|---|---|---|---|---|---|---|---|---|---|---|---|---|---|---|---|---|---|---|---|---|---|---|---|---|---|---|---|---|---|
| orf19.3983 | 992 | 951 | 974 | 642 | 3049 | 2329 | 2872 | 2894 | 946 | 1006 | 962 | 1006 | 2443 | 2648 | 2438 | 2353 | 1725,26 | 980 | 2470 | 2,523 | 1,335 | 24,844 | 2,98E-136 | 2,72E-135 | 0,001338 | 0,003043 | 0,0020188 | 0,0020188 | TRUE | 0,0273223 | No |
| orf19.5852 | 259 | 281 | 271 | 168 | 825 | 646 | 790 | 797 | 247 | 297 | 268 | 263 | 661 | 735 | 671 | 648 | 473,66 | 269 | 679 | 2,521 | 1,334 | 16,207 | 4,49E-59 | 1,87E-58 | 0,0017322 | 0,0055577 | 0,0037712 | 0,0037712 | TRUE | 0,0592299 | No |
| orf19.6663 | 12813 | 10175 | 10635 | 8442 | 37674 | 26088 | 34140 | 35430 | 12216 | 10759 | 10506 | 13225 | 30187 | 29664 | 28985 | 28808 | 20543,75 | 11676 | 29411 | 2,52 | 1,333 | 19,634 | 7,98E-86 | 4,59E-85 | 0,0061433 | 0,0021712 | 0,0043668 | 0,0043668 | TRUE | 0,0966027 | No |
| orf19.3929 | 778 | 667 | 694 | 480 | 2165 | 1591 | 2184 | 2261 | 742 | 705 | 686 | 752 | 1735 | 1809 | 1854 | 1838 | 1265,13 | 721 | 1809 | 2,514 | 1,33 | 24,856 | 2,21E-136 | 2,02E-135 | 0,0004289 | 0,0033891 | 0,0017258 | 0,0017258 | TRUE | 0,013248 | No |
| orf19.5419 | 4880 | 4043 | 4373 | 2986 | 13873 | 9681 | 13539 | 13976 | 4653 | 4275 | 4320 | 4678 | 11116 | 11008 | 11495 | 11364 | 7863,49 | 4482 | 11246 | 2,511 | 1,328 | 32,094 | 5,28E-226 | 8,01E-225 | 0,001121 | 0,0023001 | 0,0014797 | 0,0014797 | TRUE | 0,0115952 | No |
| orf19.6126 | 5981 | 5076 | 5667 | 3410 | 17187 | 11402 | 15910 | 18413 | 5702 | 5367 | 5598 | 5342 | 13771 | 12965 | 13508 | 14972 | 9653,18 | 5502 | 13804 | 2,508 | 1,327 | 26,41 | 1,06E-153 | 1,09E-152 | 0,0022291 | 0,0022614 | 0,0022888 | 0,0022888 | TRUE | 0,0397082 | No |
| orf19.2951 | 699 | 637 | 691 | 468 | 2053 | 1587 | 1970 | 2174 | 666 | 674 | 683 | 733 | 1645 | 1805 | 1673 | 1768 | 1205,69 | 689 | 1723 | 2,506 | 1,326 | 23,179 | 7,38E-119 | 5,80E-118 | 0,0010066 | 0,0034531 | 0,0020669 | 0,0020669 | TRUE | 0,0185403 | No |
| orf19.6415.1 | 4106 | 2937 | 3079 | 2163 | 10793 | 6973 | 9436 | 11217 | 3915 | 3106 | 3042 | 3388 | 8648 | 7929 | 8011 | 9120 | 5894,88 | 3363 | 8427 | 2,506 | 1,326 | 16,766 | 4,31E-63 | 1,89E-62 | 0,0089474 | 0,0023698 | 0,0057833 | 0,0057833 | TRUE | 0,1507643 | No |
| orf19.2784 | 22 | 31 | 26 | 33 | 97 | 90 | 91 | 77 | 21 | 33 | 26 | 52 | 78 | 102 | 77 | 63 | 56,38 | 33 | 80 | 2,507 | 1,326 | 5,088 | 3,63E-07 | 5,71E-07 | 0,0590708 | 0,0312119 | 0,0420211 | 0,0420211 | TRUE | 0,6967671 | No |
| orf19.4931.1 | 11232 | 9633 | 9728 | 6462 | 32027 | 21838 | 29736 | 32008 | 10708 | 10186 | 9610 | 10123 | 25662 | 24832 | 25246 | 26026 | 17799,12 | 10157 | 25442 | 2,505 | 1,325 | 33,447 | 2,82E-245 | 4,69E-244 | 0,0010915 | 0,0021835 | 0,0014336 | 0,0014336 | TRUE | 0,0166353 | No |
| orf19.5928 | 7115 | 5470 | 5372 | 4482 | 19574 | 14028 | 16817 | 20209 | 6783 | 5784 | 5307 | 7021 | 15684 | 15951 | 14278 | 16432 | 10905,01 | 6224 | 15586 | 2,505 | 1,325 | 15,931 | 3,85E-57 | 1,55E-56 | 0,0101463 | 0,0022418 | 0,0065257 | 0,0065257 | TRUE | 0,1202682 | No |
| orf19.4099 | 14727 | 12743 | 13838 | 8735 | 42608 | 29627 | 39945 | 43764 | 14041 | 13474 | 13671 | 13684 | 34141 | 33689 | 33913 | 35584 | 24024,47 | 13718 | 34332 | 2,503 | 1,324 | 42,661 | 0 | 0 | 0,0003902 | 0,0021596 | 0,0008703 | 0,0008703 | TRUE | 0,0074047 | No |
| orf19.4115 | 16 | 8 | 8 | 4 | 25 | 24 | 31 | 30 | 15 | 8 | 8 | 6 | 20 | 27 | 26 | 24 | 16,99 | 9 | 24 | 2,503 | 1,324 | 3,534 | 0,0004087 | 0,0005624 | 1,00E-08 | 0,0987333 | 0,0595592 | 0,0595592 | TRUE | 0,6072513 | No |
| orf19.6699 | 339 | 260 | 314 | 200 | 950 | 694 | 879 | 933 | 323 | 275 | 310 | 313 | 761 | 789 | 746 | 759 | 534,61 | 305 | 764 | 2,5 | 1,322 | 18,181 | 7,27E-74 | 3,68E-73 | 1,00E-08 | 0,0051625 | 0,0026701 | 0,0026701 | TRUE | 0,0531804 | No |
| orf19.1914 | 543 | 417 | 458 | 335 | 1546 | 1047 | 1379 | 1499 | 518 | 441 | 452 | 525 | 1239 | 1191 | 1171 | 1219 | 844,35 | 484 | 1205 | 2,499 | 1,321 | 19,545 | 4,59E-85 | 2,61E-84 | 0,0013206 | 0,0040359 | 0,0028584 | 0,0028584 | TRUE | 0,040438 | No |
| orf19.3173 | 613 | 494 | 531 | 295 | 1655 | 996 | 1619 | 1735 | 584 | 522 | 525 | 462 | 1326 | 1133 | 1375 | 1411 | 917,17 | 523 | 1311 | 2,498 | 1,321 | 15,23 | 2,22E-52 | 8,47E-52 | 0,0072356 | 0,0038815 | 0,0058101 | 0,0058101 | TRUE | 0,0977235 | No |
| orf19.6785 | 9560 | 8163 | 8326 | 4930 | 26740 | 17784 | 23964 | 27282 | 9114 | 8631 | 8225 | 7723 | 21426 | 20222 | 20345 | 22183 | 14733,8 | 8423 | 21044 | 2,497 | 1,32 | 23,771 | 6,71E-125 | 5,61E-124 | 0,0033375 | 0,0022027 | 0,0028755 | 0,0028755 | TRUE | 0,0375838 | No |
| orf19.3312 | 545 | 517 | 524 | 403 | 1633 | 1212 | 1598 | 1786 | 520 | 547 | 518 | 631 | 1308 | 1378 | 1357 | 1452 | 963,84 | 554 | 1374 | 2,497 | 1,32 | 18,943 | 5,06E-80 | 2,76E-79 | 0,0028004 | 0,0037948 | 0,0033194 | 0,0033194 | TRUE | 0,0897463 | No |
| orf19.1836 | 152 | 152 | 139 | 85 | 508 | 297 | 424 | 414 | 145 | 161 | 137 | 133 | 407 | 338 | 360 | 337 | 252,18 | 144 | 360 | 2,497 | 1,32 | 12,027 | 2,56E-33 | 7,31E-33 | 0,0042893 | 0,008602 | 0,0064818 | 0,0064818 | TRUE | 0,0902707 | No |
| orf19.2994.1 | 12694 | 10396 | 10736 | 7298 | 35555 | 23475 | 32585 | 36454 | 12102 | 10992 | 10606 | 11433 | 28489 | 26693 | 27665 | 29641 | 19702,64 | 11283 | 28122 | 2,493 | 1,318 | 26,017 | 3,19E-149 | 3,17E-148 | 0,0025244 | 0,0021746 | 0,0023981 | 0,0023981 | TRUE | 0,0295525 | No |
| orf19.87 | 212 | 196 | 186 | 112 | 576 | 418 | 550 | 640 | 202 | 207 | 184 | 175 | 462 | 475 | 467 | 520 | 336,59 | 192 | 481 | 2,492 | 1,317 | 14,401 | 5,08E-47 | 1,80E-46 | 1,00E-08 | 0,0069693 | 0,0042259 | 0,0042259 | TRUE | 0,0370085 | No |
| orf19.1180 | 296 | 209 | 281 | 190 | 865 | 518 | 817 | 852 | 282 | 221 | 278 | 298 | 693 | 589 | 694 | 693 | 468,37 | 270 | 667 | 2,487 | 1,315 | 13,491 | 1,76E-41 | 5,70E-41 | 0,0072231 | 0,0055969 | 0,0063655 | 0,0063655 | TRUE | 0,1655541 | No |
| orf19.819 | 904 | 773 | 838 | 559 | 2714 | 1850 | 2374 | 2578 | 862 | 817 | 828 | 876 | 2175 | 2104 | 2016 | 2096 | 1471,59 | 846 | 2098 | 2,484 | 1,313 | 26,092 | 4,49E-150 | 4,52E-149 | 1,00E-08 | 0,003207 | 0,0015565 | 0,0015565 | TRUE | 0,008585 | No |
| orf19.1440.2 | 240 | 212 | 213 | 101 | 703 | 363 | 634 | 663 | 229 | 224 | 210 | 158 | 563 | 413 | 538 | 539 | 359,38 | 205 | 513 | 2,484 | 1,312 | 10,477 | 1,11E-25 | 2,77E-25 | 0,0166917 | 0,00666 | 0,0114913 | 0,0114913 | TRUE | 0,2152931 | No |
| orf19.4316 | 700 | 694 | 722 | 461 | 2175 | 1720 | 1983 | 2037 | 667 | 734 | 713 | 722 | 1743 | 1956 | 1684 | 1656 | 1234,38 | 709 | 1760 | 2,481 | 1,311 | 19,461 | 2,36E-84 | 1,33E-83 | 0,0032264 | 0,0034214 | 0,0033099 | 0,0033099 | TRUE | 0,0657677 | No |
| orf19.2761 | 253 | 291 | 290 | 185 | 829 | 671 | 817 | 818 | 241 | 308 | 286 | 290 | 664 | 763 | 694 | 665 | 488,9 | 281 | 696 | 2,479 | 1,31 | 14,762 | 2,56E-49 | 9,35E-49 | 0,0043144 | 0,0054497 | 0,0049223 | 0,0049223 | TRUE | 0,1129232 | No |
| orf19.2445 | 92 | 79 | 84 | 68 | 301 | 208 | 246 | 245 | 88 | 84 | 83 | 107 | 241 | 237 | 209 | 199 | 155,81 | 90 | 222 | 2,48 | 1,31 | 9,873 | 5,46E-23 | 1,30E-22 | 0,0038286 | 0,0126289 | 0,0086592 | 0,0086592 | TRUE | 0,1041226 | No |
| orf19.6475 | 32 | 26 | 29 | 20 | 93 | 77 | 95 | 62 | 31 | 27 | 29 | 31 | 75 | 88 | 81 | 50 | 51,39 | 30 | 74 | 2,479 | 1,31 | 5,548 | 2,89E-08 | 4,74E-08 | 1,00E-08 | 0,034041 | 0,0284938 | 0,0284938 | TRUE | 0,4328333 | No |
| orf19.1565 | 984 | 875 | 921 | 589 | 2759 | 1972 | 2709 | 2956 | 938 | 925 | 910 | 923 | 2211 | 2242 | 2300 | 2404 | 1606,55 | 924 | 2289 | 2,478 | 1,309 | 26,632 | 2,90E-156 | 3,06E-155 | 1,00E-08 | 0,0031133 | 0,0015208 | 0,0015208 | TRUE | 0,0140006 | No |
| orf19.3223.1 | 1620 | 1469 | 1384 | 934 | 4700 | 3038 | 4271 | 4717 | 1544 | 1553 | 1367 | 1463 | 3766 | 3454 | 3626 | 3835 | 2576,26 | 1482 | 3670 | 2,477 | 1,309 | 23,56 | 9,88E-123 | 8,08E-122 | 0,0021271 | 0,0027286 | 0,0024617 | 0,0024617 | TRUE | 0,0346918 | No |
| orf19.4026 | 1179 | 863 | 981 | 521 | 3042 | 1900 | 2807 | 3083 | 1124 | 913 | 969 | 816 | 2437 | 2160 | 2383 | 2507 | 1663,71 | 956 | 2372 | 2,476 | 1,308 | 15,183 | 4,58E-52 | 1,74E-51 | 0,010117 | 0,0030782 | 0,0063501 | 0,0063501 | TRUE | 0,1687586 | No |
| orf19.4880 | 263 | 207 | 252 | 126 | 704 | 485 | 718 | 692 | 251 | 219 | 249 | 197 | 564 | 551 | 610 | 563 | 400,47 | 229 | 572 | 2,477 | 1,308 | 14,81 | 1,27E-49 | 4,65E-49 | 0,0015429 | 0,0061912 | 0,0043046 | 0,0043046 | TRUE | 0,0742574 | No |
| orf19.4839 | 1240 | 933 | 1082 | 738 | 3331 | 2348 | 3319 | 3326 | 1182 | 987 | 1069 | 1156 | 2669 | 2670 | 2818 | 2704 | 1906,86 | 1098 | 2715 | 2,475 | 1,307 | 22,07 | 6,09E-108 | 4,31E-107 | 0,002762 | 0,0029523 | 0,002693 | 0,002693 | TRUE | 0,0568315 | No |
| orf19.3782.2 | 1261 | 1009 | 1120 | 559 | 3447 | 1994 | 3225 | 3409 | 1202 | 1067 | 1106 | 876 | 2762 | 2267 | 2738 | 2772 | 1848,81 | 1063 | 2635 | 2,473 | 1,307 | 14,09 | 4,37E-45 | 1,50E-44 | 0,0117729 | 0,0029794 | 0,007559 | 0,007559 | TRUE | 0,1580666 | No |
| orf19.2216 | 238 | 229 | 225 | 145 | 693 | 501 | 649 | 721 | 227 | 242 | 222 | 227 | 555 | 570 | 551 | 586 | 397,58 | 230 | 566 | 2,462 | 1,3 | 16,417 | 1,44E-60 | 6,10E-60 | 1,00E-08 | 0,006221 | 0,0028167 | 0,0028167 | TRUE | 0,0155527 | No |
| orf19.5212 | 11 | 17 | 17 | 13 | 43 | 47 | 35 | 52 | 10 | 18 | 17 | 20 | 34 | 53 | 30 | 42 | 28,19 | 16 | 40 | 2,462 | 1,3 | 4,155 | 3,26E-05 | 4,72E-05 | 0,0319452 | 0,0603365 | 0,0482337 | 0,0482337 | TRUE | 0,2426374 | No |
| orf19.3442 | 3612 | 3387 | 3625 | 2371 | 10927 | 7941 | 10144 | 10779 | 3444 | 3581 | 3581 | 3714 | 8755 | 9030 | 8612 | 8764 | 6185,26 | 3580 | 8790 | 2,458 | 1,297 | 36,112 | 1,49E-285 | 3,11E-284 | 0,0004103 | 0,0023567 | 0,0010311 | 0,0010311 | TRUE | 0,0080518 | No |
| orf19.3405 | 470 | 424 | 477 | 343 | 1374 | 1095 | 1315 | 1468 | 448 | 448 | 471 | 537 | 1101 | 1245 | 1116 | 1194 | 820,14 | 476 | 1164 | 2,456 | 1,296 | 17,722 | 2,84E-70 | 1,36E-69 | 0,0030311 | 0,0040933 | 0,0035711 | 0,0035711 | TRUE | 0,0736721 | No |
| orf19.3819 | 98 | 77 | 93 | 61 | 259 | 196 | 255 | 294 | 93 | 81 | 92 | 96 | 208 | 223 | 216 | 239 | 156,03 | 90 | 222 | 2,451 | 1,294 | 10,424 | 1,93E-25 | 4,81E-25 | 1,00E-08 | 0,0126144 | 0,0066365 | 0,0066365 | TRUE | 0,0441437 | No |
| orf19.806 | 279 | 200 | 252 | 146 | 735 | 479 | 699 | 761 | 266 | 211 | 249 | 229 | 589 | 545 | 593 | 619 | 412,62 | 239 | 586 | 2,45 | 1,293 | 14,624 | 1,97E-48 | 7,10E-48 | 0,0023932 | 0,0060705 | 0,0044193 | 0,0044193 | TRUE | 0,0728195 | No |
| orf19.2871 | 7784 | 6696 | 7059 | 4763 | 21785 | 14932 | 20704 | 23039 | 7421 | 7080 | 6974 | 7461 | 17456 | 16979 | 17578 | 18733 | 12460,22 | 7234 | 17686 | 2,446 | 1,29 | 30,269 | 2,92E-201 | 4,01E-200 | 0,0013227 | 0,0022231 | 0,0016421 | 0,0016421 | TRUE | 0,0194503 | No |
| orf19.5032 | 33778 | 27708 | 29922 | 20112 | 93652 | 61655 | 89032 | 97063 | 32204 | 29298 | 29560 | 31507 | 75041 | 70107 | 75588 | 78921 | 52778,15 | 30642 | 74914 | 2,445 | 1,29 | 26,737 | 1,76E-157 | 1,88E-156 | 0,0022405 | 0,0021224 | 0,0022117 | 0,0022117 | TRUE | 0,022832 | No |
| orf19.7266 | 630 | 509 | 576 | 316 | 1808 | 942 | 1718 | 1737 | 601 | 538 | 569 | 495 | 1449 | 1071 | 1459 | 1412 | 949,21 | 551 | 1348 | 2,444 | 1,289 | 12,753 | 2,98E-37 | 9,04E-37 | 0,012865 | 0,003821 | 0,0084613 | 0,0084613 | TRUE | 0,2241596 | No |
| orf19.4490 | 14942 | 12863 | 12830 | 9295 | 41787 | 30159 | 38159 | 42142 | 14246 | 13601 | 12675 | 14561 | 33483 | 34293 | 32397 | 34265 | 23690,13 | 13771 | 33610 | 2,441 | 1,288 | 26,964 | 3,86E-160 | 4,21E-159 | 0,0021117 | 0,0021606 | 0,0021364 | 0,0021364 | TRUE | 0,035202 | No |
| orf19.5873 | 539 | 527 | 518 | 328 | 1549 | 1118 | 1488 | 1657 | 514 | 557 | 512 | 514 | 1241 | 1271 | 1263 | 1347 | 902,46 | 524 | 1280 | 2,443 | 1,288 | 21,438 | 5,95E-102 | 3,96E-101 | 1,00E-08 | 0,0039106 | 0,002057 | 0,002057 | TRUE | 0,0216555 | No |
| orf19.5591 | 2676 | 2392 | 2503 | 1782 | 7606 | 6264 | 6962 | 7503 | 2551 | 2529 | 2473 | 2792 | 6094 | 7123 | 5911 | 6101 | 4446,68 | 2586 | 6307 | 2,44 | 1,287 | 19,622 | 1,00E-85 | 5,73E-85 | 0,0049334 | 0,0024605 | 0,0038426 | 0,0038426 | TRUE | 0,0916754 | No |
| orf19.7498 | 3053 | 2794 | 3039 | 1982 | 9062 | 6184 | 8536 | 9341 | 2911 | 2954 | 3002 | 3105 | 7261 | 7032 | 7247 | 7595 | 5138,4 | 2993 | 7284 | 2,437 | 1,285 | 32,506 | 8,82E-232 | 1,37E-230 | 0,0005841 | 0,0024108 | 0,0012511 | 0,0012511 | TRUE | 0,01013 | No |
| orf19.7100 | 190 | 195 | 194 | 133 | 565 | 457 | 566 | 567 | 181 | 206 | 192 | 208 | 453 | 520 | 481 | 461 | 337,66 | 197 | 479 | 2,437 | 1,285 | 14,108 | 3,37E-45 | 1,16E-44 | 1,00E-08 | 0,0069539 | 0,0041909 | 0,0041909 | TRUE | 0,0360863 | No |
| orf19.4660 | 20674 | 17816 | 18599 | 11675 | 56123 | 39348 | 53229 | 59032 | 19710 | 18838 | 18374 | 18290 | 44970 | 44742 | 45191 | 47999 | 32264,24 | 18803 | 45726 | 2,432 | 1,282 | 32,92 | 1,15E-237 | 1,84E-236 | 0,001107 | 0,0021422 | 0,0014168 | 0,0014168 | TRUE | 0,0137389 | No |
| orf19.5941 | 642 | 546 | 572 | 333 | 1758 | 1126 | 1629 | 1806 | 612 | 577 | 565 | 522 | 1409 | 1280 | 1383 | 1468 | 977,08 | 569 | 1385 | 2,427 | 1,279 | 19,133 | 1,34E-81 | 7,46E-81 | 0,0021042 | 0,0037717 | 0,0029916 | 0,0029916 | TRUE | 0,0310692 | No |
| orf19.6057 | 876 | 638 | 783 | 499 | 2262 | 1561 | 2329 | 2303 | 835 | 675 | 774 | 782 | 1812 | 1775 | 1977 | 1873 | 1312,79 | 766 | 1859 | 2,427 | 1,279 | 18,667 | 9,26E-78 | 4,90E-77 | 0,0036271 | 0,003342 | 0,0035374 | 0,0035374 | TRUE | 0,079294 | No |
| orf19.3871 | 31 | 29 | 43 | 21 | 107 | 73 | 98 | 96 | 30 | 31 | 42 | 33 | 86 | 83 | 83 | 78 | 58,2 | 34 | 82 | 2,427 | 1,279 | 6,461 | 1,04E-10 | 1,86E-10 | 1,00E-08 | 0,0303028 | 0,0159356 | 0,0159356 | TRUE | 0,2290623 | No |
| orf19.6460 | 1909 | 1876 | 2080 | 1249 | 5802 | 4546 | 5353 | 5631 | 1820 | 1984 | 2055 | 1957 | 4649 | 5169 | 4545 | 4579 | 3344,57 | 1954 | 4736 | 2,423 | 1,277 | 22,377 | 6,58E-111 | 4,81E-110 | 0,0026919 | 0,0025822 | 0,0027432 | 0,0027432 | TRUE | 0,0456128 | No |
| orf19.7590 | 4896 | 4438 | 4748 | 2802 | 14102 | 9719 | 12770 | 14031 | 4668 | 4693 | 4691 | 4389 | 11300 | 11051 | 10842 | 11409 | 7880,2 | 4610 | 11150 | 2,416 | 1,273 | 35,097 | 7,56E-270 | 1,42E-268 | 0,0005676 | 0,0022996 | 0,0011009 | 0,0011009 | TRUE | 0,0117635 | No |
| orf19.7442 | 1214 | 1101 | 1176 | 694 | 3509 | 2363 | 3270 | 3435 | 1157 | 1164 | 1162 | 1087 | 2812 | 2687 | 2776 | 2793 | 1954,79 | 1142 | 2767 | 2,417 | 1,273 | 29,058 | 1,20E-185 | 1,51E-184 | 1,00E-08 | 0,0029312 | 0,0011944 | 0,0011944 | TRUE | 0,0106558 | No |
| orf19.300 | 1720 | 1360 | 1463 | 998 | 4478 | 3092 | 4473 | 4666 | 1640 | 1438 | 1445 | 1563 | 3588 | 3516 | 3798 | 3794 | 2597,75 | 1522 | 3674 | 2,416 | 1,273 | 23,113 | 3,45E-118 | 2,70E-117 | 0,0021636 | 0,0027233 | 0,0024203 | 0,0024203 | TRUE | 0,0355463 | No |
| orf19.3982 | 234 | 195 | 219 | 124 | 621 | 511 | 557 | 604 | 223 | 206 | 216 | 194 | 498 | 581 | 473 | 491 | 360,32 | 210 | 511 | 2,414 | 1,271 | 13,308 | 2,07E-40 | 6,60E-40 | 0,003842 | 0,0066481 | 0,0052427 | 0,0052427 | TRUE | 0,0940344 | No |
| orf19.4495 | 5836 | 5236 | 5610 | 3384 | 16352 | 11568 | 15380 | 16782 | 5564 | 5536 | 5542 | 5301 | 13102 | 13154 | 13058 | 13645 | 9362,87 | 5486 | 13240 | 2,412 | 1,27 | 39,034 | 0 | 0 | 0,0002954 | 0,0022666 | 0,0008803 | 0,0008803 | TRUE | 0,0058238 | No |
| orf19.7389 | 707 | 647 | 700 | 451 | 2023 | 1490 | 1865 | 2136 | 674 | 684 | 692 | 707 | 1621 | 1694 | 1583 | 1737 | 1173,95 | 689 | 1659 | 2,41 | 1,269 | 23,052 | 1,41E-117 | 1,10E-116 | 1,00E-08 | 0,0034899 | 0,0018297 | 0,0018297 | TRUE | 0,0127366 | No |
| orf19.3527 | 4427 | 3683 | 3989 | 2647 | 12214 | 8136 | 11798 | 12242 | 4221 | 3894 | 3941 | 4147 | 9787 | 9251 | 10016 | 9954 | 6901,36 | 4051 | 9752 | 2,409 | 1,268 | 29,431 | 2,18E-190 | 2,84E-189 | 0,0011606 | 0,0023292 | 0,0015985 | 0,0015985 | TRUE | 0,0143861 | No |

| | | | | | | | | | | | | | | | | | | | | | | | | | | | | | | | |
|---|---|---|---|---|---|---|---|---|---|---|---|---|---|---|---|---|---|---|---|---|---|---|---|---|---|---|---|---|---|---|---|
| orf19.5112 | 5727 | 4746 | 5123 | 3670 | 15657 | 11252 | 14417 | 16653 | 5460 | 5018 | 5061 | 5749 | 12546 | 12795 | 12240 | 13540 | 9051,14 | 5322 | 12780 | 2,403 | 1,265 | 23,65 | 1,18E-123 | 9,68E-123 | 0,0028963 | 0,0022727 | 0,0026062 | 0,0026062 | TRUE | 0,0345154 | No |
| orf19.5081 | 10991 | 9044 | 9989 | 6339 | 30103 | 19741 | 28715 | 30332 | 10479 | 9563 | 9868 | 9930 | 24121 | 22447 | 24379 | 24663 | 16931,23 | 9960 | 23902 | 2,4 | 1,263 | 28,878 | 2,23E-183 | 2,74E-182 | 0,0015043 | 0,0021883 | 0,0017619 | 0,0017619 | TRUE | 0,0204471 | No |
| orf19.6844 | 30395 | 23819 | 25861 | 18126 | 79640 | 54492 | 76936 | 84029 | 28978 | 25186 | 25548 | 28395 | 63813 | 61962 | 65318 | 68324 | 45940,62 | 27027 | 64854 | 2,4 | 1,263 | 22,852 | 1,41E-115 | 1,08E-114 | 0,0033935 | 0,002127 | 0,002907 | 0,002907 | TRUE | 0,0291118 | No |
| orf19.6569 | 449 | 463 | 493 | 244 | 1410 | 1031 | 1191 | 1227 | 428 | 490 | 487 | 382 | 1130 | 1172 | 1011 | 998 | 762,23 | 447 | 1078 | 2,399 | 1,263 | 14,341 | 1,21E-46 | 4,27E-46 | 0,0077714 | 0,0042453 | 0,0057771 | 0,0057771 | TRUE | 0,0937413 | No |
| orf19.5367 | 559 | 493 | 517 | 341 | 1575 | 1060 | 1530 | 1537 | 533 | 521 | 511 | 534 | 1262 | 1205 | 1299 | 1250 | 889,4 | 525 | 1254 | 2,394 | 1,259 | 21,999 | 2,93E-107 | 2,05E-106 | 1,00E-08 | 0,0039374 | 0,0017299 | 0,0017299 | TRUE | 0,0075453 | No |
| orf19.4550 | 1022 | 926 | 974 | 634 | 2867 | 2120 | 2666 | 2918 | 974 | 979 | 962 | 993 | 2297 | 2411 | 2263 | 2373 | 1656,6 | 977 | 2336 | 2,391 | 1,258 | 27,188 | 8,94E-163 | 9,84E-162 | 1,00E-08 | 0,0030824 | 0,001292 | 0,001292 | TRUE | 0,0052715 | No |
| orf19.1613 | 5363 | 4999 | 5488 | 3636 | 15711 | 12109 | 14805 | 15397 | 5113 | 5286 | 5422 | 5696 | 12589 | 13769 | 12569 | 12519 | 9120,36 | 5379 | 12862 | 2,392 | 1,258 | 26,273 | 3,86E-152 | 3,93E-151 | 0,0019797 | 0,0022713 | 0,0020635 | 0,0020635 | TRUE | 0,0273669 | No |
| orf19.4019 | 53 | 48 | 60 | 30 | 153 | 100 | 144 | 170 | 51 | 51 | 59 | 47 | 123 | 114 | 122 | 138 | 88,04 | 52 | 124 | 2,384 | 1,253 | 7,709 | 1,27E-14 | 2,54E-14 | 1,00E-08 | 0,0207403 | 0,0111918 | 0,0111918 | TRUE | 0,0775035 | No |
| orf19.4504 | 510 | 443 | 489 | 273 | 1418 | 898 | 1308 | 1459 | 486 | 468 | 483 | 428 | 1136 | 1021 | 1110 | 1186 | 789,94 | 466 | 1113 | 2,381 | 1,252 | 17,899 | 1,19E-71 | 5,85E-71 | 0,0021364 | 0,0041698 | 0,0031013 | 0,0031013 | TRUE | 0,0353021 | No |
| orf19.2731 | 147 | 122 | 117 | 57 | 390 | 200 | 370 | 350 | 140 | 129 | 116 | 89 | 312 | 227 | 314 | 285 | 201,58 | 118 | 284 | 2,382 | 1,252 | 8,587 | 8,92E-18 | 1,91E-17 | 0,0192463 | 0,0102364 | 0,0141847 | 0,0141847 | TRUE | 0,2153236 | No |
| orf19.1037 | 93 | 97 | 93 | 52 | 269 | 201 | 248 | 270 | 89 | 103 | 92 | 81 | 216 | 229 | 211 | 220 | 154,84 | 91 | 219 | 2,38 | 1,251 | 10,242 | 1,28E-24 | 3,15E-24 | 1,00E-08 | 0,0126949 | 0,0062364 | 0,0062364 | TRUE | 0,0638539 | No |
| orf19.4632 | 11607 | 10271 | 10175 | 7688 | 32173 | 25230 | 28369 | 32128 | 11066 | 10860 | 10052 | 12044 | 25779 | 28689 | 24085 | 26123 | 18587,28 | 11006 | 26169 | 2,378 | 1,25 | 19,232 | 2,01E-82 | 1,12E-81 | 0,0053218 | 0,0021796 | 0,0039903 | 0,0039903 | TRUE | 0,051335 | No |
| orf19.5870 | 484 | 449 | 450 | 266 | 1334 | 897 | 1289 | 1351 | 461 | 475 | 445 | 417 | 1069 | 1020 | 1094 | 1098 | 759,9 | 450 | 1070 | 2,375 | 1,248 | 19,698 | 2,26E-86 | 1,30E-85 | 1,44E-05 | 0,004252 | 0,0022035 | 0,0022035 | TRUE | 0,0219149 | No |
| orf19.2128 | 755 | 563 | 595 | 444 | 1987 | 1253 | 1825 | 1954 | 720 | 595 | 588 | 696 | 1592 | 1425 | 1549 | 1589 | 1094,2 | 650 | 1539 | 2,375 | 1,248 | 16,504 | 3,42E-61 | 1,47E-60 | 0,0052931 | 0,0035918 | 0,0043326 | 0,0043326 | TRUE | 0,0699913 | No |
| orf19.2232 | 12829 | 10735 | 10913 | 6754 | 33848 | 21696 | 30838 | 35340 | 12231 | 11351 | 10781 | 10581 | 27121 | 24670 | 26181 | 28735 | 18956,42 | 11236 | 26677 | 2,374 | 1,247 | 20,962 | 1,47E-97 | 9,36E-97 | 0,0041194 | 0,0021779 | 0,0033344 | 0,0033344 | TRUE | 0,0435946 | No |
| orf19.1679 | 64 | 39 | 51 | 37 | 172 | 99 | 132 | 165 | 61 | 41 | 50 | 58 | 138 | 113 | 112 | 134 | 88,4 | 52 | 124 | 2,374 | 1,247 | 7,201 | 6,00E-13 | 1,14E-12 | 0,0060426 | 0,0206643 | 0,0146405 | 0,0146405 | TRUE | 0,1777541 | No |
| orf19.493 | 25663 | 20052 | 20963 | 12759 | 65196 | 39935 | 62356 | 66569 | 24467 | 21202 | 20709 | 19988 | 52240 | 45410 | 52940 | 54127 | 36385,34 | 21592 | 51179 | 2,37 | 1,245 | 17,327 | 2,96E-67 | 1,37E-66 | 0,0071036 | 0,0021364 | 0,0049258 | 0,0049258 | TRUE | 0,0985062 | No |
| orf19.4195.1 | 2467 | 2025 | 2161 | 1220 | 6652 | 3778 | 6377 | 6409 | 2352 | 2141 | 2135 | 1911 | 5330 | 4296 | 5414 | 5211 | 3598,8 | 2135 | 5063 | 2,37 | 1,245 | 15,387 | 2,01E-53 | 7,77E-53 | 0,0083452 | 0,0025475 | 0,0059337 | 0,0059337 | TRUE | 0,1256816 | No |
| orf19.3653 | 5454 | 4726 | 5023 | 3349 | 14744 | 10680 | 13999 | 15312 | 5200 | 4997 | 4962 | 5246 | 11814 | 12144 | 11885 | 12450 | 8587,36 | 5101 | 12073 | 2,368 | 1,244 | 35,108 | 5,13E-270 | 9,66E-269 | 0,0004899 | 0,0022825 | 0,0010574 | 0,0010574 | TRUE | 0,0054696 | No |
| orf19.6538 | 843 | 687 | 732 | 476 | 2312 | 1528 | 2056 | 2177 | 804 | 726 | 723 | 746 | 1853 | 1737 | 1746 | 1770 | 1263,08 | 750 | 1776 | 2,369 | 1,244 | 22,707 | 3,81E-114 | 2,87E-113 | 0,000638 | 0,0033912 | 0,0018863 | 0,0018863 | TRUE | 0,0305794 | No |
| orf19.4503 | 967 | 720 | 864 | 581 | 2546 | 1689 | 2525 | 2518 | 922 | 761 | 854 | 910 | 2040 | 1921 | 2144 | 2047 | 1449,83 | 862 | 2038 | 2,369 | 1,244 | 18,822 | 5,02E-79 | 2,71E-78 | 0,0035233 | 0,0032238 | 0,0033248 | 0,0033248 | TRUE | 0,074025 | No |
| orf19.2310.1 | 9479 | 7163 | 7316 | 5824 | 25297 | 16499 | 22053 | 24897 | 9037 | 7574 | 7228 | 9124 | 20270 | 18761 | 18723 | 20244 | 13869,93 | 8241 | 19500 | 2,367 | 1,243 | 16,445 | 9,10E-61 | 3,87E-60 | 0,007972 | 0,0022097 | 0,0053981 | 0,0053981 | TRUE | 0,0838321 | No |
| orf19.5231 | 38 | 49 | 39 | 18 | 129 | 80 | 109 | 104 | 36 | 52 | 39 | 28 | 103 | 91 | 93 | 85 | 65,78 | 39 | 93 | 2,367 | 1,243 | 6,322 | 2,58E-10 | 4,54E-10 | 0,0021636 | 0,0270537 | 0,0181856 | 0,0181856 | TRUE | 0,3575836 | No |
| orf19.1224 | 5623 | 5785 | 5913 | 4173 | 17520 | 13486 | 16378 | 16143 | 5361 | 6117 | 5841 | 6537 | 14038 | 15335 | 13905 | 13126 | 10032,53 | 5964 | 14101 | 2,365 | 1,242 | 18,847 | 3,08E-79 | 1,67E-78 | 0,0053732 | 0,0022549 | 0,0040459 | 0,0040459 | TRUE | 0,0566971 | No |
| orf19.4674.1 | 310 | 276 | 255 | 149 | 814 | 579 | 727 | 773 | 296 | 292 | 252 | 233 | 652 | 658 | 617 | 629 | 453,63 | 268 | 639 | 2,365 | 1,242 | 14,852 | 6,75E-50 | 2,49E-49 | 0,000949 | 0,0057107 | 0,003948 | 0,003948 | TRUE | 0,0664983 | No |
| orf19.4367 | 617 | 594 | 603 | 395 | 1796 | 1291 | 1615 | 1793 | 588 | 628 | 596 | 619 | 1439 | 1468 | 1371 | 1458 | 1020,86 | 608 | 1434 | 2,362 | 1,24 | 22,373 | 7,25E-111 | 5,29E-110 | 1,00E-08 | 0,0036996 | 0,001721 | 0,001721 | TRUE | 0,0105768 | No |
| orf19.1872 | 3379 | 2693 | 2876 | 2111 | 8928 | 6160 | 8575 | 9090 | 3222 | 2848 | 2841 | 3307 | 7154 | 7004 | 7280 | 7391 | 5130,83 | 3054 | 7207 | 2,362 | 1,24 | 22,206 | 3,03E-109 | 2,18E-108 | 0,0031713 | 0,0024113 | 0,0027494 | 0,0027494 | TRUE | 0,0358603 | No |
| orf19.6007 | 2445 | 2961 | 2822 | 1801 | 8433 | 6381 | 7669 | 6909 | 2331 | 3131 | 2788 | 2821 | 6757 | 7256 | 6511 | 5618 | 4651,58 | 2768 | 6536 | 2,361 | 1,24 | 13,515 | 1,28E-41 | 4,17E-41 | 0,0122522 | 0,0024443 | 0,007811 | 0,007811 | TRUE | 0,1386146 | No |
| orf19.5064.1 | 725 | 577 | 568 | 340 | 1857 | 1212 | 1554 | 1828 | 691 | 610 | 561 | 533 | 1488 | 1378 | 1319 | 1486 | 1008,36 | 599 | 1418 | 2,36 | 1,239 | 15,289 | 9,08E-53 | 3,48E-52 | 0,0068722 | 0,0037196 | 0,0050536 | 0,0050536 | TRUE | 0,129126 | No |
| orf19.5281 | 1603 | 1477 | 1600 | 1002 | 4456 | 3333 | 4196 | 4669 | 1528 | 1562 | 1581 | 1570 | 3570 | 3790 | 3562 | 3796 | 2619,93 | 1560 | 3680 | 2,359 | 1,238 | 28,537 | 4,06E-179 | 4,84E-178 | 0,0005943 | 0,002718 | 0,0013261 | 0,0013261 | TRUE | 0,0058206 | No |
| orf19.6092 | 1430 | 1173 | 1358 | 933 | 3972 | 2697 | 3609 | 4174 | 1363 | 1240 | 1342 | 1462 | 3183 | 3067 | 3064 | 3394 | 2264,26 | 1352 | 3177 | 2,356 | 1,236 | 21,195 | 1,07E-99 | 6,99E-99 | 0,0027411 | 0,0028164 | 0,0027089 | 0,0027089 | TRUE | 0,0362927 | No |
| orf19.6328 | 207 | 143 | 172 | 121 | 571 | 330 | 494 | 502 | 197 | 151 | 170 | 190 | 458 | 375 | 419 | 408 | 296,05 | 177 | 415 | 2,355 | 1,236 | 11,598 | 4,24E-31 | 1,16E-30 | 0,0054849 | 0,0076374 | 0,0066723 | 0,0066723 | TRUE | 0,1045194 | No |
| orf19.1089 | 1271 | 1531 | 1466 | 870 | 4297 | 3770 | 3597 | 3081 | 1212 | 1619 | 1448 | 1363 | 3443 | 4287 | 3054 | 2505 | 2366,33 | 1410 | 3322 | 2,355 | 1,236 | 6,713 | 1,91E-11 | 3,49E-11 | 0,0320215 | 0,0027851 | 0,0165083 | 0,0320215 | TRUE | 0,1908753 | No |
| orf19.667.1 | 13216 | 10336 | 10359 | 8168 | 35750 | 23660 | 30807 | 34301 | 12600 | 10929 | 10234 | 12796 | 28645 | 26904 | 26155 | 27890 | 19519,04 | 11640 | 27398 | 2,354 | 1,235 | 17,58 | 3,48E-69 | 1,64E-68 | 0,0065463 | 0,0021754 | 0,0046797 | 0,0046797 | TRUE | 0,0809821 | No |
| orf19.6264.4 | 7119 | 5447 | 5871 | 3676 | 18913 | 10771 | 17196 | 18128 | 6787 | 5760 | 5800 | 5759 | 15154 | 12248 | 14599 | 14740 | 10105,81 | 6026 | 14185 | 2,354 | 1,235 | 16,364 | 3,47E-60 | 1,47E-59 | 0,0076743 | 0,0022538 | 0,0053461 | 0,0053461 | TRUE | 0,103111 | No |
| orf19.6524 | 5095 | 4206 | 4504 | 3204 | 13478 | 9731 | 12900 | 13888 | 4858 | 4447 | 4450 | 5019 | 10800 | 11065 | 10952 | 11292 | 7860,31 | 4694 | 11027 | 2,351 | 1,234 | 26,259 | 5,71E-152 | 5,79E-151 | 0,0018899 | 0,0023002 | 0,0019587 | 0,0019587 | TRUE | 0,0254591 | No |
| orf19.437 | 4873 | 4353 | 4774 | 2780 | 13352 | 8874 | 12771 | 13929 | 4646 | 4603 | 4716 | 4355 | 10699 | 10091 | 10843 | 11326 | 7659,64 | 4580 | 10740 | 2,344 | 1,229 | 26,957 | 4,70E-160 | 5,12E-159 | 0,0015034 | 0,0023056 | 0,0018308 | 0,0018308 | TRUE | 0,0199817 | No |
| orf19.89 | 1715 | 1387 | 1494 | 1020 | 4535 | 3070 | 4246 | 4587 | 1635 | 1467 | 1476 | 1598 | 3634 | 3491 | 3605 | 3730 | 2579,32 | 1544 | 3615 | 2,344 | 1,229 | 24,998 | 6,43E-138 | 5,94E-137 | 0,0012799 | 0,0027278 | 0,0018336 | 0,0018336 | TRUE | 0,021204 | No |
| orf19.2036 | 487 | 421 | 459 | 309 | 1276 | 957 | 1295 | 1363 | 464 | 445 | 453 | 484 | 1022 | 1088 | 1099 | 1108 | 770,66 | 462 | 1079 | 2,343 | 1,229 | 19,758 | 6,83E-87 | 3,98E-86 | 1,00E-08 | 0,0042218 | 0,0020904 | 0,0020904 | TRUE | 0,015905 | No |
| orf19.754 | 5302 | 4624 | 4932 | 3056 | 14135 | 9643 | 13861 | 14562 | 5055 | 4889 | 4872 | 4787 | 11326 | 10965 | 11768 | 11840 | 8187,88 | 4901 | 11475 | 2,341 | 1,227 | 31,92 | 1,43E-223 | 2,14E-222 | 0,0007013 | 0,0022918 | 0,001265 | 0,001265 | TRUE | 0,0107632 | No |
| orf19.2329.1 | 15691 | 13976 | 13706 | 8625 | 41666 | 28892 | 38258 | 42150 | 14960 | 14778 | 13540 | 13512 | 33386 | 32853 | 32481 | 34272 | 23722,59 | 14198 | 33248 | 2,341 | 1,227 | 27,597 | 1,21E-167 | 1,36E-166 | 0,0017114 | 0,0021605 | 0,0018469 | 0,0018469 | TRUE | 0,0160203 | No |
| orf19.4007 | 259 | 200 | 231 | 164 | 655 | 504 | 642 | 687 | 247 | 211 | 228 | 257 | 525 | 573 | 545 | 559 | 393,14 | 236 | 550 | 2,341 | 1,227 | 14,639 | 1,59E-48 | 5,76E-48 | 1,00E-08 | 0,0062677 | 0,0035693 | 0,0035693 | TRUE | 0,0518657 | No |
| orf19.5289 | 5 | 3 | 11 | 4 | 13 | 20 | 16 | 16 | 5 | 3 | 11 | 6 | 10 | 23 | 14 | 13 | 10,6 | 6 | 15 | 2,339 | 1,226 | 2,451 | 0,0142286 | 0,0178761 | 0,0687025 | 0,156944 | 0,1214253 | 0,1214253 | TRUE | 0,670497 | No |
| orf19.2129 | 706 | 667 | 642 | 477 | 2037 | 1338 | 1933 | 1999 | 673 | 705 | 634 | 747 | 1632 | 1521 | 1641 | 1625 | 1147,49 | 690 | 1605 | 2,338 | 1,225 | 20,4 | 1,68E-92 | 1,03E-91 | 0,0012007 | 0,0035221 | 0,0023717 | 0,0023717 | TRUE | 0,0344876 | No |
| orf19.3307 | 261 | 242 | 276 | 177 | 748 | 565 | 736 | 730 | 249 | 256 | 273 | 277 | 599 | 642 | 625 | 594 | 439,36 | 264 | 615 | 2,338 | 1,225 | 15,861 | 1,18E-56 | 4,73E-56 | 1,00E-08 | 0,0058283 | 0,0028872 | 0,0028872 | TRUE | 0,0166483 | No |
| orf19.4253 | 524 | 555 | 555 | 307 | 1616 | 1063 | 1469 | 1480 | 500 | 587 | 548 | 481 | 1295 | 1209 | 1247 | 1203 | 883,72 | 529 | 1238 | 2,335 | 1,223 | 17,905 | 1,07E-71 | 5,26E-71 | 0,0020555 | 0,0039492 | 0,0030672 | 0,0030672 | TRUE | 0,061605 | No |
| orf19.4712 | 25 | 17 | 15 | 15 | 47 | 32 | 57 | 76 | 24 | 18 | 15 | 23 | 38 | 36 | 48 | 62 | 33,05 | 20 | 46 | 2,333 | 1,222 | 4,219 | 2,45E-05 | 3,57E-05 | 0,0322587 | 0,0517779 | 0,0428898 | 0,0428898 | TRUE | 0,4535647 | No |
| orf19.4600.1 | 662 | 530 | 533 | 372 | 1712 | 1062 | 1598 | 1748 | 631 | 560 | 527 | 583 | 1372 | 1208 | 1357 | 1421 | 957,28 | 575 | 1340 | 2,332 | 1,221 | 16,456 | 7,63E-61 | 3,25E-60 | 0,0041346 | 0,0038065 | 0,0039806 | 0,0039806 | TRUE | 0,0551437 | No |
| orf19.1409.1 | 9708 | 8118 | 8666 | 5476 | 25745 | 18106 | 23248 | 25284 | 9256 | 8584 | 8561 | 8578 | 20629 | 20588 | 19737 | 20558 | 14561,44 | 8745 | 20378 | 2,33 | 1,22 | 32,448 | 5,78E-231 | 8,96E-230 | 0,0008706 | 0,002204 | 0,0012716 | 0,0012716 | TRUE | 0,0191838 | No |
| orf19.2965 | 2193 | 2016 | 2040 | 1416 | 6026 | 4238 | 5789 | 6250 | 2091 | 2132 | 2015 | 2218 | 4828 | 4819 | 4915 | 5082 | 3512,52 | 2114 | 4911 | 2,328 | 1,219 | 28,993 | 7,99E-185 | 9,94E-184 | 0,0006564 | 0,0025587 | 0,0013398 | 0,0013398 | TRUE | 0,0120228 | No |
| orf19.1113 | 535 | 444 | 511 | 279 | 1413 | 971 | 1329 | 1393 | 510 | 469 | 505 | 437 | 1132 | 1104 | 1128 | 1133 | 802,34 | 480 | 1124 | 2,328 | 1,219 | 19,72 | 1,44E-86 | 8,36E-86 | 1,00E-08 | 0,0041377 | 0,0021196 | 0,0021196 | TRUE | 0,0345167 | No |
| orf19.4228 | 1165 | 982 | 936 | 665 | 3136 | 1923 | 2819 | 3038 | 1111 | 1038 | 925 | 1042 | 2513 | 2187 | 2393 | 2470 | 1709,8 | 1029 | 2391 | 2,326 | 1,218 | 18,168 | 9,33E-74 | 4,70E-73 | 0,0037104 | 0,0030516 | 0,0035809 | 0,0035809 | TRUE | 0,0582458 | No |
| orf19.2711.1 | 197 | 179 | 177 | 116 | 551 | 363 | 485 | 540 | 188 | 189 | 175 | 182 | 442 | 413 | 412 | 439 | 304,85 | 184 | 426 | 2,326 | 1,218 | 13,702 | 9,90E-43 | 3,28E-42 | 1,00E-08 | 0,0074773 | 0,0035161 | 0,0035161 | TRUE | 0,0122813 | No |
| orf19.3940 | 80 | 74 | 70 | 53 | 196 | 159 | 209 | 239 | 76 | 78 | 69 | 83 | 157 | 181 | 177 | 194 | 127,04 | 76 | 177 | 2,326 | 1,218 | 8,75 | 2,13E-18 | 4,61E-18 | 1,00E-08 | 0,0150157 | 0,0088028 | 0,0088028 | TRUE | 0,0692566 | No |
| orf19.7018 | 16107 | 13885 | 14334 | 10241 | 43339 | 29852 | 41543 | 44229 | 15356 | 14682 | 14161 | 16043 | 34726 | 33944 | 35270 | 35962 | 25018,07 | 15060 | 34976 | 2,323 | 1,216 | 27,447 | 7,52E-166 | 8,40E-165 | 0,0017087 | 0,0021569 | 0,0018356 | 0,0018356 | TRUE | 0,0232491 | No |
| orf19.4229 | 850 | 731 | 812 | 495 | 2295 | 1590 | 2213 | 2239 | 810 | 773 | 802 | 775 | 1839 | 1808 | 1879 | 1821 | 1313,4 | 790 | 1837 | 2,321 | 1,215 | 25,214 | 2,78E-140 | 2,62E-139 | 1,00E-08 | 0,0033414 | 0,0012828 | 0,0012828 | TRUE | 0,0035646 | No |
| orf19.3680 | 388 | 428 | 425 | 261 | 1178 | 898 | 1058 | 1194 | 370 | 453 | 420 | 409 | 944 | 1021 | 898 | 971 | 685,66 | 413 | 958 | 2,321 | 1,215 | 16,082 | 3,42E-58 | 1,41E-57 | 0,002534 | 0,0044859 | 0,0036569 | 0,0036569 | TRUE | 0,0626981 | No |
| orf19.6002 | 10673 | 10072 | 10401 | 7337 | 30460 | 23306 | 28075 | 29571 | 10176 | 10650 | 10275 | 11494 | 24407 | 26501 | 23836 | 24044 | 17672,72 | 10649 | 24697 | 2,32 | 1,214 | 23,608 | 3,19E-123 | 2,62E-122 | 0,0027258 | 0,0021842 | 0,0024699 | 0,0024699 | TRUE | 0,0343847 | No |
| orf19.3422 | 824 | 771 | 868 | 502 | 2330 | 1727 | 2107 | 2367 | 786 | 815 | 858 | 786 | 1867 | 1964 | 1789 | 1925 | 1348,61 | 811 | 1886 | 2,321 | 1,214 | 22,18 | 5,41E-109 | 3,88E-108 | 0,0009412 | 0,0033088 | 0,0019534 | 0,0019534 | TRUE | 0,0182091 | No |
| orf19.6021 | 283 | 247 | 251 | 169 | 768 | 465 | 730 | 798 | 270 | 261 | 248 | 265 | 615 | 529 | 620 | 649 | 432,05 | 261 | 603 | 2,319 | 1,214 | 13,879 | 8,50E-44 | 2,86E-43 | 0,0031236 | 0,0058915 | 0,0044662 | 0,0044662 | TRUE | 0,0753409 | No |

| | | | | | | | | | | | | | | | | | | | | | | | | | | | | | | | |
|---|---|---|---|---|---|---|---|---|---|---|---|---|---|---|---|---|---|---|---|---|---|---|---|---|---|---|---|---|---|---|---|
| orf19.1415 | 16264 | 14770 | 15800 | 10398 | 44578 | 32851 | 41373 | 46438 | 15506 | 15617 | 15609 | 16289 | 35719 | 37355 | 35125 | 37759 | 26122,37 | 15755 | 36490 | 2,316 | 1,212 | 33,525 | 2,06E-246 | 3,45E-245 | 0,000802 | 0,0021541 | 0,0012073 | 0,0012073 | TRUE | 0,0077575 | No |
| orf19.6367 | 43685 | 34842 | 38478 | 27585 | 1E+05 | 79964 | 1E+05 | 1E+05 | 41649 | 36841 | 38013 | 43213 | 89791 | 90926 | 91459 | 97709 | 66200,15 | 39929 | 92471 | 2,316 | 1,212 | 21,698 | 2,14E-104 | 1,45E-103 | 0,0035349 | 0,0021161 | 0,0029774 | 0,0029774 | TRUE | 0,0374146 | No |
| orf19.4450 | 493 | 397 | 446 | 295 | 1257 | 898 | 1206 | 1347 | 470 | 420 | 441 | 462 | 1007 | 1021 | 1024 | 1095 | 742,5 | 448 | 1037 | 2,317 | 1,212 | 18,675 | 7,90E-78 | 4,19E-77 | 1,00E-08 | 0,0043026 | 0,0023709 | 0,0023709 | TRUE | 0,0213137 | No |
| orf19.292 | 106 | 77 | 70 | 60 | 282 | 156 | 233 | 239 | 101 | 81 | 69 | 94 | 226 | 177 | 198 | 194 | 142,64 | 86 | 199 | 2,317 | 1,212 | 8,352 | 6,69E-17 | 1,40E-16 | 0,0086192 | 0,0136022 | 0,0115215 | 0,0115215 | TRUE | 0,1852827 | No |
| orf19.7621 | 417 | 376 | 392 | 233 | 1113 | 779 | 1089 | 1097 | 398 | 398 | 387 | 365 | 892 | 886 | 925 | 892 | 642,69 | 387 | 899 | 2,315 | 1,211 | 19,001 | 1,66E-80 | 9,11E-80 | 1,00E-08 | 0,004646 | 0,0019729 | 0,0019729 | TRUE | 0,012849 | No |
| orf19.7080 | 788 | 682 | 644 | 579 | 2104 | 1541 | 2031 | 2194 | 751 | 721 | 636 | 907 | 1686 | 1752 | 1724 | 1784 | 1245,25 | 754 | 1736 | 2,314 | 1,211 | 14,173 | 1,35E-45 | 4,66E-45 | 0,0088257 | 0,0034098 | 0,0059994 | 0,0059994 | TRUE | 0,2040825 | No |
| orf19.2179.2 | 15850 | 12553 | 12815 | 8071 | 39558 | 25423 | 37049 | 39599 | 15111 | 13273 | 12660 | 12644 | 31697 | 28908 | 31454 | 32198 | 22243,16 | 13422 | 31064 | 2,314 | 1,21 | 19,475 | 1,79E-84 | 1,02E-83 | 0,0048071 | 0,0021651 | 0,0036552 | 0,0036552 | TRUE | 0,0880857 | No |
| orf19.6094 | 1082 | 863 | 927 | 545 | 2699 | 1650 | 2720 | 2769 | 1032 | 913 | 916 | 854 | 2163 | 1876 | 2309 | 2251 | 1539,15 | 929 | 2150 | 2,313 | 1,21 | 15,553 | 1,51E-54 | 5,93E-54 | 0,0061813 | 0,003158 | 0,0049945 | 0,0049945 | TRUE | 0,0868894 | No |
| orf19.6634 | 1720 | 1502 | 1685 | 1017 | 4605 | 3263 | 4370 | 4788 | 1640 | 1588 | 1665 | 1593 | 3690 | 3710 | 3710 | 3893 | 2686,15 | 1622 | 3751 | 2,312 | 1,209 | 29,52 | 1,61E-191 | 2,12E-190 | 1,00E-08 | 0,0027025 | 0,0011461 | 0,0011461 | TRUE | 0,008089 | No |
| orf19.4051 | 4386 | 3332 | 3890 | 2249 | 10915 | 7061 | 9941 | 11866 | 4182 | 3523 | 3843 | 3523 | 8746 | 8029 | 8440 | 9648 | 6241,72 | 3768 | 8716 | 2,312 | 1,209 | 17,17 | 4,42E-66 | 2,01E-65 | 0,0063395 | 0,0023543 | 0,0045636 | 0,0045636 | TRUE | 0,0669006 | No |
| orf19.879 | 971 | 879 | 992 | 658 | 2678 | 2011 | 2563 | 2819 | 926 | 929 | 980 | 1031 | 2146 | 2287 | 2176 | 2292 | 1595,82 | 966 | 2225 | 2,308 | 1,207 | 23,04 | 1,85E-117 | 1,44E-116 | 0,0008939 | 0,0031202 | 0,001852 | 0,001852 | TRUE | 0,020592 | No |
| orf19.386 | 1943 | 1654 | 1725 | 1236 | 5187 | 3807 | 4814 | 5054 | 1852 | 1749 | 1704 | 1936 | 4156 | 4329 | 4087 | 4109 | 2990,41 | 1810 | 4170 | 2,307 | 1,206 | 25,019 | 3,80E-138 | 3,52E-137 | 0,0012104 | 0,0026403 | 0,0018135 | 0,0018135 | TRUE | 0,0238869 | No |
| orf19.2506 | 782 | 710 | 778 | 478 | 2120 | 1431 | 2178 | 2178 | 746 | 751 | 769 | 749 | 1699 | 1627 | 1849 | 1771 | 1244,95 | 754 | 1736 | 2,306 | 1,206 | 21,078 | 1,27E-98 | 8,20E-98 | 0,0012843 | 0,0034101 | 0,002142 | 0,002142 | TRUE | 0,0227849 | No |
| orf19.1552 | 3135 | 2407 | 2418 | 1645 | 7591 | 5028 | 7045 | 7897 | 2989 | 2545 | 2389 | 2577 | 6082 | 5717 | 5981 | 6421 | 4337,71 | 2625 | 6050 | 2,305 | 1,205 | 17,775 | 1,10E-70 | 5,28E-70 | 0,0055559 | 0,0024698 | 0,0041215 | 0,0041215 | TRUE | 0,1079602 | No |
| orf19.5225.2 | 14223 | 11965 | 12432 | 9059 | 37149 | 27179 | 35361 | 37617 | 13560 | 12651 | 12282 | 14191 | 29766 | 30905 | 30021 | 30586 | 21745,44 | 13171 | 30320 | 2,303 | 1,203 | 24,965 | 1,47E-137 | 1,35E-136 | 0,002239 | 0,0021668 | 0,0021744 | 0,0021744 | TRUE | 0,0327675 | No |

$P_{TET}$-*ZCF26* doxycycline-treated down-regulated genes

| | WT dox | | | | ZCF26 | | | | norm. WT dox | | | | norm. ZCF26 | | | | | | | | | | | | | | | | | | |
|---|---|---|---|---|---|---|---|---|---|---|---|---|---|---|---|---|---|---|---|---|---|---|---|---|---|---|---|---|---|---|---|
| Gene ID | 1 | 2 | 3 | 4 | 1 | 2 | 3 | 4 | 1 | 2 | 3 | 4 | 1 | 2 | 3 | 4 | baseMean | WT | ZCF26 | FC | log2FC | stat | pvalue | padj | dispGeneEst | dispFit | dispMAP | dispersion | betaConv | maxCooks | outlier |
| orf19.3904 | 10 | 19 | 20 | 6 | 0 | 0 | 1 | 0 | 10 | 20 | 20 | 9 | 0 | 0 | 1 | 0 | 7,45 | 15 | 0 | 0,018 | -5,78 | -5,132 | 2,86E-07 | 4,52E-07 | 1,00E-08 | 0,2223685 | 0,180924 | 0,180924 | TRUE | 0,2613074 | No |
| orf19.3374 | 16501 | 13556 | 14242 | 10402 | 536 | 371 | 527 | 550 | 15732 | 14334 | 14070 | 16295 | 429 | 422 | 447 | 447 | 7772,08 | 15108 | 436 | 0,029 | -5,111 | -80,094 | 0 | 0 | 0,0032301 | 0,0023025 | 0,0028551 | 0,0028551 | TRUE | 0,0339104 | No |
| orf19.1562 | 32160 | 29870 | 31589 | 22185 | 1367 | 1010 | 1297 | 1301 | 30661 | 31584 | 31207 | 34754 | 1095 | 1148 | 1101 | 1058 | 16576,07 | 32052 | 1100 | 0,034 | -4,865 | -93,92 | 0 | 0 | 0,0015938 | 0,0021903 | 0,0021539 | 0,0021539 | TRUE | 0,0392306 | No |
| orf19.7085 | 13245 | 11224 | 12349 | 8677 | 569 | 437 | 578 | 622 | 12628 | 11868 | 12200 | 13593 | 456 | 497 | 491 | 506 | 6529,69 | 12572 | 488 | 0,039 | -4,69 | -78,583 | 0 | 0 | 0,0025457 | 0,0023427 | 0,0024617 | 0,0024617 | TRUE | 0,0360437 | No |
| orf19.3664 | 10653 | 9513 | 9835 | 6309 | 498 | 362 | 490 | 485 | 10156 | 10059 | 9716 | 9883 | 399 | 412 | 416 | 394 | 5179,47 | 9954 | 405 | 0,041 | -4,62 | -96,54 | 0 | 0 | 1,00E-08 | 0,0024083 | 0,0010478 | 0,0010478 | TRUE | 0,0042464 | No |
| orf19.7336 | 5 | 16 | 13 | 3 | 2 | 0 | 0 | 0 | 5 | 17 | 13 | 5 | 2 | 0 | 0 | 0 | 5,1 | 10 | 0 | 0,045 | -4,49 | -3,809 | 0,0001396 | 0,000196 | 0,350202 | 0,3237973 | 0,3323307 | 0,3323307 | TRUE | 0,5815225 | No |
| orf19.1354 | 147029 | 144734 | 147807 | 82571 | 8425 | 6453 | 7890 | 7868 | 140176 | 153038 | 146019 | 129352 | 6751 | 7338 | 6699 | 6397 | 74471,21 | 142146 | 6796 | 0,048 | -4,387 | -73,912 | 0 | 0 | 0,0040553 | 0,0021133 | 0,003315 | 0,003315 | TRUE | 0,0449514 | No |
| orf19.3988 | 9452 | 8183 | 8926 | 5710 | 527 | 351 | 481 | 575 | 9011 | 8652 | 8818 | 8945 | 422 | 399 | 408 | 468 | 4640,54 | 8856 | 424 | 0,048 | -4,379 | -85,088 | 0 | 0 | 0,0002935 | 0,0024451 | 0,0014369 | 0,0014369 | TRUE | 0,0560185 | No |
| orf19.6200 | 27 | 17 | 22 | 16 | 0 | 1 | 1 | 3 | 26 | 18 | 22 | 25 | 0 | 1 | 1 | 2 | 11,87 | 23 | 1 | 0,049 | -4,355 | -5,971 | 2,36E-09 | 4,00E-09 | 1,00E-08 | 0,1404427 | 0,0856191 | 0,0856191 | TRUE | 0,4767147 | No |
| orf19.449 | 38067 | 35063 | 37919 | 24321 | 2225 | 1645 | 2192 | 2299 | 36293 | 37075 | 37460 | 38100 | 1783 | 1871 | 1861 | 1869 | 19538,95 | 37232 | 1846 | 0,05 | -4,335 | -125,584 | 0 | 0 | 0,0003301 | 0,0021753 | 0,0008875 | 0,0008875 | TRUE | 0,0066481 | No |
| orf19.3621 | 747 | 688 | 755 | 504 | 52 | 31 | 38 | 48 | 712 | 727 | 746 | 790 | 42 | 35 | 32 | 39 | 390,41 | 744 | 37 | 0,05 | -4,317 | -33,066 | 9,24E-240 | 1,49E-238 | 1,00E-08 | 0,0062969 | 0,0037484 | 0,0037484 | TRUE | 0,0669834 | No |
| orf19.5140 | 1597 | 1337 | 1486 | 903 | 70 | 77 | 93 | 95 | 1523 | 1414 | 1468 | 1415 | 56 | 88 | 79 | 77 | 764,84 | 1455 | 75 | 0,051 | -4,297 | -43,039 | 0 | 0 | 0,0009136 | 0,004238 | 0,003213 | 0,003213 | TRUE | 0,2943515 | No |
| orf19.7455 | 343 | 300 | 351 | 206 | 17 | 11 | 27 | 22 | 327 | 317 | 347 | 323 | 14 | 13 | 23 | 18 | 172,58 | 328 | 17 | 0,052 | -4,277 | -22,513 | 3,06E-112 | 2,26E-111 | 1,00E-08 | 0,0116052 | 0,006906 | 0,006906 | TRUE | 0,3242486 | No |
| orf19.822 | 271052 | 224248 | 228545 | 141921 | 15760 | 10314 | 14306 | 15845 | 258419 | 237114 | 225781 | 222327 | 12628 | 11728 | 12146 | 12883 | 124128,2 | 235910 | 12346 | 0,052 | -4,256 | -78,61 | 0 | 0 | 0,0031889 | 0,0021045 | 0,0027777 | 0,0027777 | TRUE | 0,0505779 | No |
| orf19.3378 | 94 | 70 | 68 | 44 | 8 | 5 | 2 | 4 | 90 | 74 | 67 | 69 | 6 | 6 | 2 | 3 | 39,6 | 75 | 4 | 0,056 | -4,164 | -10,784 | 4,10E-27 | 1,06E-26 | 1,00E-08 | 0,0435552 | 0,0304541 | 0,0304541 | TRUE | 0,3448418 | No |
| orf19.1321 | 13694 | 11416 | 13298 | 8965 | 888 | 615 | 901 | 911 | 13056 | 12071 | 13137 | 14044 | 712 | 699 | 765 | 741 | 6903,07 | 13077 | 729 | 0,056 | -4,163 | -72,717 | 0 | 0 | 0,0026024 | 0,0023291 | 0,0024936 | 0,0024936 | TRUE | 0,0327304 | No |
| orf19.2048 | 15558 | 12582 | 13136 | 9718 | 980 | 769 | 924 | 1130 | 14833 | 13304 | 12977 | 15224 | 785 | 874 | 784 | 919 | 7462,57 | 14084 | 840 | 0,06 | -4,067 | -57,618 | 0 | 0 | 0,0054399 | 0,0023113 | 0,0042138 | 0,0042138 | TRUE | 0,048966 | No |
| orf19.823 | 988 | 1262 | 1242 | 1826 | 97 | 156 | 89 | 81 | 942 | 1334 | 1227 | 2861 | 78 | 177 | 76 | 66 | 845,05 | 1591 | 99 | 0,062 | -4,008 | -8,235 | 1,80E-16 | 3,74E-16 | 0,2227195 | 0,0040342 | 0,0849814 | 0,2227195 | TRUE | 0,9244277 | No |
| orf19.3302 | 23393 | 22188 | 23519 | 14726 | 1741 | 1322 | 1642 | 1788 | 22303 | 23461 | 23235 | 23069 | 1395 | 1503 | 1394 | 1454 | 12226,68 | 23017 | 1436 | 0,062 | -4,004 | -103,685 | 0 | 0 | 0,0004137 | 0,0022256 | 0,001097 | 0,001097 | TRUE | 0,0107006 | No |
| orf19.4907 | 12038 | 11315 | 11786 | 7531 | 936 | 666 | 817 | 982 | 11477 | 11964 | 11643 | 11798 | 750 | 757 | 694 | 798 | 6235,21 | 11720 | 750 | 0,064 | -3,966 | -86,122 | 0 | 0 | 0,0004568 | 0,0023546 | 0,0013953 | 0,0013953 | TRUE | 0,0314925 | No |
| orf19.5288 | 175079 | 144775 | 154847 | 104159 | 12876 | 9093 | 12136 | 13434 | 166919 | 153081 | 152974 | 163171 | 10317 | 10340 | 10303 | 10923 | 84753,52 | 159036 | 10471 | 0,066 | -3,925 | -94,213 | 0 | 0 | 0,0013457 | 0,0021107 | 0,0016211 | 0,0016211 | TRUE | 0,0136688 | No |
| orf19.938 | 358 | 316 | 362 | 228 | 27 | 25 | 25 | 27 | 341 | 334 | 358 | 357 | 22 | 28 | 21 | 22 | 185,44 | 348 | 23 | 0,066 | -3,918 | -23,502 | 3,87E-122 | 3,14E-121 | 1,00E-08 | 0,0109456 | 0,0058437 | 0,0058437 | TRUE | 0,0985337 | No |
| orf19.3373 | 14416 | 12283 | 13125 | 9620 | 1140 | 894 | 1031 | 1136 | 13744 | 12988 | 12966 | 15070 | 913 | 1017 | 875 | 924 | 7312,16 | 13692 | 932 | 0,068 | -3,878 | -61,275 | 0 | 0 | 0,0039032 | 0,0023158 | 0,003327 | 0,003327 | TRUE | 0,0556332 | No |
| orf19.937 | 7 | 11 | 4 | 2 | 1 | 0 | 0 | 1 | 7 | 12 | 4 | 3 | 1 | 0 | 0 | 1 | 3,38 | 6 | 0 | 0,069 | -3,866 | -3,217 | 0,0012935 | 0,0017296 | 1,00E-08 | 0,4885101 | 0,3546916 | 0,3546916 | TRUE | 0,3656389 | No |
| orf19.5079 | 81762 | 70731 | 78184 | 51925 | 6652 | 4702 | 6334 | 6538 | 77951 | 74789 | 77238 | 81343 | 5330 | 5347 | 5378 | 5316 | 41586,52 | 77830 | 5343 | 0,069 | -3,865 | -112,058 | 0 | 0 | 0,000483 | 0,0021308 | 0,0010516 | 0,0010516 | TRUE | 0,0112518 | No |
| orf19.265 | 5820 | 5882 | 5786 | 3606 | 520 | 395 | 476 | 425 | 5549 | 6219 | 5716 | 5649 | 417 | 449 | 404 | 346 | 3093,59 | 5783 | 404 | 0,07 | -3,845 | -54,387 | 0 | 0 | 0,0053043 | 0,002622 | 0,0035959 | 0,0035959 | TRUE | 0,1118794 | No |
| orf19.4975 | 5466 | 4689 | 5034 | 3614 | 441 | 310 | 431 | 460 | 5211 | 4958 | 4973 | 5662 | 353 | 352 | 366 | 374 | 2781,21 | 5201 | 361 | 0,07 | -3,844 | -60,744 | 0 | 0 | 0,0023871 | 0,0026816 | 0,0025089 | 0,0025089 | TRUE | 0,0419955 | No |
| orf19.2962 | 916 | 899 | 972 | 572 | 80 | 62 | 81 | 72 | 873 | 951 | 960 | 896 | 64 | 70 | 69 | 59 | 492,76 | 920 | 66 | 0,071 | -3,821 | -36,135 | 6,31E-286 | 1,32E-284 | 1,00E-08 | 0,0054233 | 0,0033378 | 0,0033378 | TRUE | 0,0484793 | No |
| orf19.6575 | 2 | 1 | 5 | 5 | 1 | 0 | 0 | 0 | 2 | 1 | 5 | 8 | 1 | 0 | 0 | 0 | 2,07 | 4 | 0 | 0,071 | -3,815 | -2,778 | 0,0054655 | 0,0070512 | 1,00E-08 | 0,7963499 | 0,6633922 | 0,6633922 | TRUE | 0,2046112 | No |
| orf19.2655 | 24787 | 23324 | 24963 | 14754 | 2120 | 1478 | 1993 | 2169 | 23632 | 24662 | 24661 | 23113 | 1699 | 1681 | 1692 | 1764 | 12862,86 | 24017 | 1709 | 0,071 | -3,813 | -97,651 | 0 | 0 | 0,0007035 | 0,0022189 | 0,0011796 | 0,0011796 | TRUE | 0,0077372 | No |
| orf19.4867 | 11634 | 10837 | 11881 | 7471 | 966 | 746 | 937 | 1088 | 11092 | 11459 | 11737 | 11704 | 774 | 848 | 796 | 885 | 6161,75 | 11498 | 826 | 0,072 | -3,801 | -77,927 | 0 | 0 | 0,00132 | 0,0023577 | 0,0016955 | 0,0016955 | TRUE | 0,0301625 | No |
| orf19.7027 | 14477 | 12915 | 14901 | 8440 | 1295 | 841 | 1177 | 1289 | 13802 | 13656 | 14721 | 13222 | 1038 | 956 | 999 | 1048 | 7430,25 | 13850 | 1010 | 0,073 | -3,775 | -76,59 | 0 | 0 | 0,0015682 | 0,0023123 | 0,0018519 | 0,0018519 | TRUE | 0,0219326 | No |
| orf19.5626 | 27896 | 26829 | 28764 | 19222 | 2625 | 1925 | 2383 | 2462 | 26596 | 28368 | 28416 | 30112 | 2103 | 2189 | 2023 | 2002 | 15226,21 | 28373 | 2079 | 0,073 | -3,771 | -77,302 | 0 | 0 | 0,0017746 | 0,0021991 | 0,0020511 | 0,0020511 | TRUE | 0,0217922 | No |
| orf19.1785 | 29607 | 27220 | 29944 | 20314 | 2807 | 2153 | 2667 | 2648 | 28227 | 28782 | 29582 | 31823 | 2249 | 2448 | 2264 | 2153 | 15941,03 | 29604 | 2278 | 0,077 | -3,701 | -70,442 | 0 | 0 | 0,0025735 | 0,0021943 | 0,0024356 | 0,0024356 | TRUE | 0,0310197 | No |
| orf19.6387 | 32317 | 26284 | 29280 | 19494 | 2973 | 2046 | 2869 | 3090 | 30811 | 27792 | 28926 | 30538 | 2382 | 2326 | 2436 | 2512 | 15965,49 | 29517 | 2414 | 0,082 | -3,611 | -78,501 | 0 | 0 | 0,0013959 | 0,0021941 | 0,0018284 | 0,0018284 | TRUE | 0,0189598 | No |
| orf19.2839 | 4738 | 4177 | 4483 | 2617 | 459 | 293 | 440 | 454 | 4517 | 4417 | 4429 | 4100 | 368 | 333 | 374 | 369 | 2363,24 | 4366 | 361 | 0,083 | -3,593 | -61,758 | 0 | 0 | 0,0011694 | 0,002786 | 0,0018947 | 0,0018947 | TRUE | 0,0260794 | No |
| orf19.4450.1 | 4231 | 3527 | 3514 | 2520 | 391 | 283 | 356 | 396 | 4034 | 3729 | 3471 | 3948 | 313 | 322 | 302 | 322 | 2055,21 | 3796 | 315 | 0,083 | -3,592 | -53,093 | 0 | 0 | 0,0027934 | 0,0028902 | 0,0028339 | 0,0028339 | TRUE | 0,0407421 | No |
| orf19.2768 | 40960 | 40990 | 42868 | 23944 | 4385 | 3108 | 4106 | 3939 | 39051 | 43342 | 42350 | 37510 | 3514 | 3534 | 3486 | 3203 | 21998,52 | 40563 | 3434 | 0,085 | -3,563 | -63,698 | 0 | 0 | 0,0030957 | 0,0021659 | 0,0028609 | 0,0028609 | TRUE | 0,0313213 | No |
| orf19.2553 | 329 | 348 | 357 | 200 | 41 | 30 | 26 | 35 | 314 | 368 | 353 | 313 | 33 | 34 | 22 | 28 | 183,14 | 337 | 29 | 0,086 | -3,533 | -21,905 | 2,32E-106 | 1,60E-105 | 1,00E-08 | 0,0110566 | 0,0081474 | 0,0081474 | TRUE | 0,2002543 | No |
| orf19.2398 | 19972 | 17786 | 19466 | 12883 | 2044 | 1518 | 1990 | 2009 | 19041 | 18806 | 19231 | 20182 | 1638 | 1726 | 1690 | 1634 | 10493,37 | 19315 | 1672 | 0,086 | -3,531 | -89,839 | 0 | 0 | 0,0006704 | 0,0022478 | 0,0011876 | 0,0011876 | TRUE | 0,0109747 | No |
| orf19.2107.1 | 110649 | 78158 | 83372 | 44334 | 10052 | 5055 | 9164 | 9961 | 105492 | 82642 | 82364 | 69452 | 8054 | 5748 | 7780 | 8099 | 46203,88 | 84988 | 7420 | 0,087 | -3,518 | -20,986 | 8,75E-98 | 5,60E-97 | 0,0269281 | 0,0021268 | 0,0140254 | 0,0269281 | TRUE | 0,1743244 | No |
| orf19.4526 | 39009 | 35560 | 36958 | 23720 | 4238 | 2906 | 3692 | 3970 | 37191 | 37600 | 36511 | 37159 | 3396 | 3304 | 3134 | 3228 | 20190,42 | 37115 | 3266 | 0,088 | -3,507 | -103,342 | 0 | 0 | 0,0004306 | 0,0021726 | 0,0009539 | 0,0009539 | TRUE | 0,0091749 | No |
| orf19.553 | 11 | 16 | 18 | 9 | 1 | 1 | 2 | 2 | 10 | 17 | 18 | 14 | 1 | 1 | 2 | 2 | 8,07 | 15 | 2 | 0,089 | -3,487 | -4,884 | 1,04E-06 | 1,61E-06 | 1,00E-08 | 0,2055833 | 0,1176853 | 0,1176853 | TRUE | 0,2021995 | No |
| orf19.2770 | 5078 | 4615 | 4754 | 3049 | 527 | 329 | 536 | 566 | 4841 | 4880 | 4697 | 4776 | 422 | 374 | 455 | 460 | 2613,21 | 4798 | 428 | 0,09 | -3,48 | -61,2 | 0 | 0 | 0,0003402 | 0,0027196 | 0,001954 | 0,001954 | TRUE | 0,0711451 | No |
| orf19.6028 | 332 | 342 | 333 | 214 | 38 | 29 | 36 | 35 | 317 | 362 | 329 | 335 | 30 | 33 | 31 | 28 | 183,1 | 336 | 30 | 0,091 | -3,461 | -22,634 | 2,00E-113 | 1,50E-112 | 1,00E-08 | 0,0110585 | 0,0062677 | 0,0062677 | TRUE | 0,0325932 | No |
| orf19.6501 | 46 | 31 | 38 | 32 | 3 | 2 | 4 | 8 | 44 | 33 | 38 | 50 | 2 | 2 | 3 | 7 | 22,36 | 41 | 4 | 0,092 | -3,449 | -7,887 | 3,08E-15 | 6,25E-15 | 1,00E-08 | 0,0755203 | 0,0514118 | 0,0514118 | TRUE | 0,4469592 | No |
| orf19.6078.1 | 0 | 4 | 3 | 3 | 0 | 0 | 0 | 1 | 0 | 4 | 3 | 5 | 0 | 0 | 0 | 1 | 1,59 | 3 | 0 | 0,093 | -3,425 | -2,285 | 0,0223134 | 0,0275934 | 1,00E-08 | 1,0358756 | 0,9648712 | 0,9648712 | TRUE | 0,6505023 | No |
| orf19.734 | 14428 | 12173 | 13318 | 8421 | 1530 | 1037 | 1526 | 1587 | 13756 | 12871 | 13157 | 13192 | 1226 | 1179 | 1296 | 1290 | 7245,86 | 13244 | 1248 | 0,094 | -3,406 | -78,694 | 0 | 0 | 0,0007995 | 0,0023179 | 0,0013996 | 0,0013996 | TRUE | 0,0149293 | No |
| orf19.147 | 13321 | 13905 | 15000 | 9272 | 1697 | 1353 | 1606 | 1450 | 12700 | 14703 | 14819 | 14525 | 1360 | 1538 | 1363 | 1179 | 7773,41 | 14187 | 1360 | 0,096 | -3,384 | -44,336 | 0 | 0 | 0,0077615 | 0,0023025 | 0,0052287 | 0,0052287 | TRUE | 0,0971987 | No |
| orf19.7227 | 2577 | 2082 | 2284 | 1559 | 275 | 186 | 262 | 300 | 2457 | 2201 | 2256 | 2442 | 220 | 211 | 222 | 244 | 1281,9 | 2339 | 224 | 0,096 | -3,376 | -47,049 | 0 | 0 | 0,0021594 | 0,0033721 | 0,0027317 | 0,0027317 | TRUE | 0,0384831 | No |
| orf19.6842 | 11970 | 10749 | 12105 | 7345 | 1307 | 973 | 1306 | 1503 | 11412 | 11366 | 11959 | 11506 | 1047 | 1106 | 1109 | 1222 | 6340,91 | 11561 | 1121 | 0,097 | -3,366 | -68,368 | 0 | 0 | 0,0017477 | 0,0023502 | 0,0018811 | 0,0018811 | TRUE | 0,0459826 | No |
| orf19.951 | 9358 | 9665 | 10208 | 6020 | 1177 | 831 | 1084 | 1164 | 8922 | 10220 | 10085 | 9431 | 943 | 945 | 920 | 946 | 5301,41 | 9664 | 938 | 0,097 | -3,364 | -62,012 | 0 | 0 | 0,0022304 | 0,002401 | 0,0022928 | 0,0022928 | TRUE | 0,0331949 | No |
| orf19.4280 | 8023 | 7385 | 7803 | 4774 | 895 | 690 | 920 | 892 | 7649 | 7809 | 7709 | 7479 | 717 | 785 | 781 | 725 | 4206,65 | 7662 | 752 | 0,098 | -3,352 | -75,424 | 0 | 0 | 0,0001946 | 0,0024816 | 0,0012307 | 0,0012307 | TRUE | 0,0116227 | No |
| orf19.4936.1 | 17231 | 15923 | 16570 | 9585 | 2122 | 1398 | 1898 | 1915 | 16428 | 16837 | 16370 | 15015 | 1700 | 1590 | 1611 | 1557 | 8888,48 | 16162 | 1614 | 0,1 | -3,323 | -68,095 | 0 | 0 | 0,0014691 | 0,002276 | 0,0019762 | 0,0019762 | TRUE | 0,0275293 | No |
| orf19.2552 | 943 | 906 | 930 | 601 | 112 | 84 | 114 | 114 | 899 | 958 | 919 | 941 | 90 | 96 | 97 | 93 | 511,5 | 929 | 94 | 0,101 | -3,312 | -36,428 | 1,51E-290 | 3,25E-289 | 1,00E-08 | 0,0053012 | 0,0026019 | 0,0026019 | TRUE | 0,0085621 | No |
| orf19.7550 | 652 | 556 | 640 | 397 | 71 | 55 | 74 | 85 | 622 | 588 | 632 | 622 | 57 | 63 | 63 | 69 | 339,38 | 616 | 63 | 0,102 | -3,293 | -29,972 | 2,30E-197 | 3,12E-196 | 1,00E-08 | 0,0069292 | 0,0036419 | 0,0036419 | TRUE | 0,0468128 | No |
| orf19.2175 | 21385 | 18393 | 20029 | 12354 | 2541 | 1810 | 2451 | 2563 | 20388 | 19448 | 19787 | 19353 | 2036 | 2058 | 2081 | 2084 | 10904,44 | 19744 | 2065 | 0,105 | -3,258 | -95,755 | 0 | 0 | 1,00E-08 | 0,0022419 | 0,000867 | 0,000867 | TRUE | 0,0059374 | No |
| orf19.5735.3 | 3054 | 2483 | 2673 | 1687 | 346 | 244 | 341 | 360 | 2912 | 2625 | 2641 | 2643 | 277 | 277 | 290 | 293 | 1494,68 | 2705 | 284 | 0,105 | -3,25 | -50,326 | 0 | 0 | 1,00E-08 | 0,0031898 | 0,0022362 | 0,0022362 | TRUE | 0,0329104 | No |
| orf19.5348 | 11880 | 9871 | 11245 | 7228 | 1484 | 1006 | 1395 | 1533 | 11326 | 10437 | 11109 | 11323 | 1189 | 1144 | 1184 | 1246 | 6119,94 | 11049 | 1191 | 0,108 | -3,212 | -70,631 | 0 | 0 | 0,0011044 | 0,0023596 | 0,0015609 | 0,0015609 | TRUE | 0,0170316 | No |
| orf19.6608 | 2937 | 2812 | 3061 | 1752 | 373 | 292 | 360 | 390 | 2800 | 2973 | 3024 | 2745 | 299 | 332 | 306 | 317 | 1599,46 | 2886 | 314 | 0,108 | -3,208 | -51,37 | 0 | 0 | 1,00E-08 | 0,0031178 | 0,0021286 | 0,0021286 | TRUE | 0,0155948 | No |
| orf19.4908 | 117 | 89 | 109 | 74 | 14 | 8 | 12 | 19 | 112 | 94 | 108 | 116 | 11 | 9 | 10 | 15 | 59,4 | 108 | 11 | 0,109 | -3,199 | -12,706 | 5,49E-37 | 1,66E-36 | 1,00E-08 | 0,0297321 | 0,0177061 | 0,0177061 | TRUE | 0,2411154 | No |
| orf19.467 | 11616 | 10141 | 11279 | 7755 | 1484 | 1118 | 1423 | 1544 | 11075 | 10723 | 11143 | 12149 | 1189 | 1271 | 1208 | 1255 | 6251,57 | 11272 | 1231 | 0,109 | -3,195 | -63,489 | 0 | 0 | 0,0018238 | 0,0023539 | 0,0020198 | 0,0020198 | TRUE | 0,0328948 | No |
| orf19.2838 | 2453 | 2390 | 2564 | 1499 | 342 | 286 | 318 | 302 | 2339 | 2527 | 2533 | 2348 | 274 | 325 | 270 | 246 | 1357,73 | 2437 | 279 | 0,114 | -3,138 | -42,369 | 0 | 0 | 0,0036321 | 0,0033006 | 0,003431 | 0,003431 | TRUE | 0,1255042 | No |
| orf19.4432 | 18400 | 16196 | 17266 | 10464 | 2373 | 1732 | 2290 | 2481 | 17542 | 17125 | 17057 | 16392 | 1901 | 1969 | 1944 | 2017 | 9493,7 | 17029 | 1958 | 0,115 | -3,121 | -82,777 | 0 | 0 | 0,0005595 | 0,0022642 | 0,0011043 | 0,0011043 | TRUE | 0,0075913 | No |
| orf19.6267 | 537 | 500 | 534 | 309 | 78 | 47 | 71 | 73 | 512 | 529 | 528 | 484 | 62 | 53 | 60 | 59 | 285,98 | 513 | 58 | 0,115 | -3,118 | -27,191 | 8,23E-163 | 9,07E-162 | 1,00E-08 | 0,0078326 | 0,0040666 | 0,0040666 | TRUE | 0,0260658 | No |
| orf19.2173 | 13142 | 11478 | 12638 | 7630 | 1741 | 1248 | 1621 | 1860 | 12529 | 12137 | 12485 | 11953 | 1395 | 1419 | 1376 | 1512 | 6850,83 | 12276 | 1426 | 0,116 | -3,106 | -74,967 | 0 | 0 | 0,0006481 | 0,0023309 | 0,00129 | 0,00129 | TRUE | 0,0211278 | No |
| orf19.4943 | 8732 | 7984 | 8528 | 5467 | 1203 | 909 | 1126 | 1246 | 8325 | 8442 | 8425 | 8564 | 964 | 1034 | 956 | 1013 | 4715,37 | 8439 | 992 | 0,117 | -3,091 | -77,95 | 0 | 0 | 1,00E-08 | 0,0024395 | 0,0009939 | 0,0009939 | TRUE | 0,0085374 | No |

| | | | | | | | | | | | | | | | | | | | | | | | | | | | | | | | |
|---|---|---|---|---|---|---|---|---|---|---|---|---|---|---|---|---|---|---|---|---|---|---|---|---|---|---|---|---|---|---|---|
| orf19.3038 | 70134 | 61043 | 66769 | 41592 | 9817 | 6738 | 8973 | 9906 | 66865 | 64545 | 65961 | 65156 | 7866 | 7662 | 7618 | 8055 | 36716,04 | 65632 | 7800 | 0,119 | -3,073 | -100,527 | 0 | 0 | 0,0003544 | 0,002136 | 0,0008314 | 0,0008314 | TRUE | 0,005926 | No |
| orf19.2749 | 5668 | 6031 | 6434 | 3841 | 868 | 672 | 894 | 840 | 5404 | 6377 | 6356 | 6017 | 696 | 764 | 759 | 683 | 3381,97 | 6038 | 726 | 0,12 | -3,059 | -45,858 | 0 | 0 | 0,0043063 | 0,0025768 | 0,0035643 | 0,0035643 | TRUE | 0,0619166 | No |
| orf19.176 | 2920 | 2632 | 2665 | 1796 | 389 | 305 | 385 | 423 | 2784 | 2783 | 2633 | 2814 | 312 | 347 | 327 | 344 | 1542,81 | 2754 | 332 | 0,12 | -3,053 | -52,464 | 0 | 0 | 1,00E-08 | 0,0031555 | 0,0017087 | 0,0017087 | TRUE | 0,020942 | No |
| orf19.4664 | 3626 | 3316 | 3423 | 1968 | 530 | 337 | 490 | 525 | 3457 | 3506 | 3382 | 3083 | 425 | 383 | 416 | 427 | 1884,82 | 3357 | 413 | 0,123 | -3,022 | -48,666 | 0 | 0 | 0,002054 | 0,0029624 | 0,0024592 | 0,0024592 | TRUE | 0,0348671 | No |
| orf19.3363 | 30621 | 27690 | 29468 | 17871 | 4361 | 3128 | 4032 | 4653 | 29194 | 29279 | 29112 | 27996 | 3494 | 3557 | 3423 | 3783 | 16229,7 | 28895 | 3564 | 0,123 | -3,019 | -76,46 | 0 | 0 | 0,0009635 | 0,0021924 | 0,0013527 | 0,0013527 | TRUE | 0,0212984 | No |
| orf19.6116 | 2015 | 1700 | 1939 | 1119 | 305 | 171 | 279 | 300 | 1921 | 1798 | 1916 | 1753 | 244 | 194 | 237 | 244 | 1038,35 | 1847 | 230 | 0,125 | -2,997 | -40,609 | 0 | 0 | 0,0020642 | 0,0036725 | 0,0030087 | 0,0030087 | TRUE | 0,100584 | No |
| orf19.7612 | 22724 | 20731 | 21695 | 13760 | 3432 | 2460 | 3058 | 3342 | 21665 | 21920 | 21433 | 21556 | 2750 | 2797 | 2596 | 2717 | 12179,31 | 21644 | 2715 | 0,125 | -2,996 | -90,405 | 0 | 0 | 0,0001199 | 0,0022261 | 0,000864 | 0,000864 | TRUE | 0,0105632 | No |
| orf19.3117 | 318 | 240 | 215 | 165 | 38 | 37 | 32 | 39 | 303 | 254 | 212 | 258 | 30 | 42 | 27 | 32 | 144,9 | 257 | 33 | 0,126 | -2,988 | -16,841 | 1,23E-63 | 5,41E-63 | 0,0155745 | 0,0134223 | 0,0143555 | 0,0143555 | TRUE | 0,1338644 | No |
| orf19.7283 | 20935 | 16851 | 18239 | 12063 | 2965 | 1938 | 2889 | 2959 | 19959 | 17818 | 18018 | 18897 | 2376 | 2204 | 2453 | 2406 | 10516,37 | 18673 | 2360 | 0,126 | -2,983 | -58,966 | 0 | 0 | 0,0021107 | 0,0022474 | 0,0022391 | 0,0022391 | TRUE | 0,0265164 | No |
| orf19.4873 | 12611 | 11711 | 11972 | 7825 | 1916 | 1394 | 1800 | 1910 | 12023 | 12383 | 11827 | 12258 | 1535 | 1585 | 1528 | 1553 | 6836,64 | 12123 | 1550 | 0,128 | -2,968 | -83,776 | 0 | 0 | 1,00E-08 | 0,0023314 | 0,0008711 | 0,0008711 | TRUE | 0,0033231 | No |
| orf19.4972 | 258 | 263 | 275 | 168 | 36 | 27 | 39 | 52 | 246 | 278 | 272 | 263 | 29 | 31 | 33 | 42 | 149,23 | 265 | 34 | 0,128 | -2,964 | -19,057 | 5,76E-81 | 3,18E-80 | 1,00E-08 | 0,0130936 | 0,0080672 | 0,0080672 | TRUE | 0,236124 | No |
| orf19.5611 | 236 | 236 | 240 | 118 | 42 | 28 | 28 | 33 | 225 | 250 | 237 | 185 | 34 | 32 | 24 | 27 | 126,57 | 224 | 29 | 0,128 | -2,962 | -16,785 | 3,14E-63 | 1,38E-62 | 1,00E-08 | 0,0150632 | 0,0121034 | 0,0121034 | TRUE | 0,1284051 | No |
| orf19.2665 | 39545 | 35601 | 38783 | 23638 | 6125 | 4103 | 5802 | 6017 | 37702 | 37644 | 38314 | 37030 | 4908 | 4665 | 4926 | 4892 | 21260,14 | 37672 | 4848 | 0,129 | -2,957 | -97,001 | 0 | 0 | 0,0002603 | 0,0021685 | 0,0007857 | 0,0007857 | TRUE | 0,0074266 | No |
| orf19.1930 | 147 | 141 | 144 | 86 | 17 | 26 | 15 | 24 | 140 | 149 | 142 | 135 | 14 | 30 | 13 | 20 | 80,21 | 142 | 19 | 0,129 | -2,957 | -13,678 | 1,38E-42 | 4,55E-42 | 1,00E-08 | 0,0225621 | 0,0166822 | 0,0166822 | TRUE | 0,6153218 | No |
| orf19.3902 | 36 | 48 | 44 | 38 | 8 | 7 | 6 | 6 | 34 | 51 | 43 | 60 | 6 | 8 | 5 | 5 | 26,55 | 47 | 6 | 0,129 | -2,957 | -7,988 | 1,38E-15 | 2,80E-15 | 1,00E-08 | 0,0639288 | 0,0455633 | 0,0455633 | TRUE | 0,0814141 | No |
| orf19.4127 | 10413 | 8902 | 9841 | 5774 | 1636 | 981 | 1578 | 1493 | 9928 | 9413 | 9722 | 9045 | 1311 | 1115 | 1340 | 1214 | 5385,96 | 9527 | 1245 | 0,131 | -2,934 | -49,947 | 0 | 0 | 0,0035478 | 0,0023961 | 0,0028963 | 0,0028963 | TRUE | 0,0555397 | No |
| orf19.6888 | 15625 | 13967 | 15316 | 10006 | 2517 | 1809 | 2288 | 2431 | 14897 | 14768 | 15131 | 15675 | 2017 | 2057 | 1943 | 1977 | 8557,96 | 15118 | 1998 | 0,132 | -2,92 | -79,059 | 0 | 0 | 0,0004652 | 0,0022831 | 0,0010493 | 0,0010493 | TRUE | 0,0073437 | No |
| orf19.4887 | 13227 | 12112 | 12871 | 8651 | 2128 | 1678 | 1928 | 2115 | 12611 | 12807 | 12715 | 13552 | 1705 | 1908 | 1637 | 1720 | 7331,84 | 12921 | 1742 | 0,135 | -2,892 | -57,2 | 0 | 0 | 0,0023372 | 0,0023152 | 0,0021554 | 0,0021554 | TRUE | 0,0470249 | No |
| orf19.6621 | 15333 | 14187 | 15491 | 8977 | 2454 | 1700 | 2288 | 2603 | 14618 | 15001 | 15304 | 14063 | 1966 | 1933 | 1943 | 2116 | 8368,03 | 14746 | 1990 | 0,135 | -2,889 | -64,617 | 0 | 0 | 0,0012067 | 0,0022875 | 0,0016577 | 0,0016577 | TRUE | 0,0226901 | No |
| orf19.5686 | 5857 | 4399 | 4676 | 2874 | 831 | 560 | 819 | 787 | 5584 | 4651 | 4619 | 4502 | 666 | 637 | 695 | 640 | 2749,37 | 4839 | 660 | 0,136 | -2,875 | -38,647 | 0 | 0 | 0,0059161 | 0,0026885 | 0,004523 | 0,004523 | TRUE | 0,1316747 | No |
| orf19.4279 | 21197 | 19906 | 21241 | 12835 | 3395 | 2525 | 3268 | 3578 | 20209 | 21048 | 20984 | 20107 | 2720 | 2871 | 2775 | 2909 | 11702,9 | 20587 | 2819 | 0,137 | -2,869 | -78,236 | 0 | 0 | 0,0004973 | 0,0022316 | 0,0011059 | 0,0011059 | TRUE | 0,0068736 | No |
| orf19.6881 | 8603 | 7504 | 8042 | 5410 | 1356 | 1002 | 1335 | 1376 | 8202 | 7935 | 7945 | 8475 | 1087 | 1139 | 1133 | 1119 | 4629,31 | 8139 | 1120 | 0,138 | -2,862 | -68,611 | 0 | 0 | 0,0005512 | 0,002446 | 0,0012037 | 0,0012037 | TRUE | 0,0090567 | No |
| orf19.1861 | 15241 | 13677 | 14933 | 9177 | 2444 | 1765 | 2373 | 2495 | 14531 | 14462 | 14752 | 14376 | 1958 | 2007 | 2015 | 2029 | 8266,2 | 14530 | 2002 | 0,138 | -2,86 | -90,807 | 0 | 0 | 1,00E-08 | 0,0022899 | 0,0006911 | 0,0006911 | TRUE | 0,0027913 | No |
| orf19.4673 | 1537 | 1419 | 1564 | 941 | 237 | 207 | 248 | 254 | 1465 | 1500 | 1545 | 1474 | 190 | 235 | 211 | 207 | 853,42 | 1496 | 211 | 0,14 | -2,839 | -40,177 | 0 | 0 | 2,19E-05 | 0,0040152 | 0,0022989 | 0,0022989 | TRUE | 0,0584701 | No |
| orf19.3369 | 22127 | 23586 | 24885 | 15033 | 4248 | 3231 | 3761 | 3620 | 21096 | 24939 | 24584 | 23550 | 3404 | 3674 | 3193 | 2943 | 13422,89 | 23542 | 3304 | 0,14 | -2,834 | -39,112 | 0 | 0 | 0,0068988 | 0,0022136 | 0,0048842 | 0,0048842 | TRUE | 0,0683406 | No |
| orf19.5114.1 | 22816 | 16304 | 17523 | 9896 | 3346 | 1943 | 3037 | 3272 | 21753 | 17239 | 17311 | 15503 | 2681 | 2209 | 2578 | 2660 | 10241,87 | 17952 | 2532 | 0,141 | -2,825 | -29,818 | 2,26E-195 | 3,04E-194 | 0,0145157 | 0,0022516 | 0,0084163 | 0,0084163 | TRUE | 0,2488598 | No |
| orf19.7167 | 51 | 30 | 31 | 29 | 8 | 4 | 4 | 9 | 49 | 32 | 31 | 45 | 6 | 5 | 3 | 7 | 22,26 | 39 | 5 | 0,141 | -2,825 | -7,139 | 9,42E-13 | 1,78E-12 | 1,00E-08 | 0,075855 | 0,0553767 | 0,0553767 | TRUE | 0,1076923 | No |
| orf19.413 | 4678 | 4447 | 4784 | 2895 | 805 | 603 | 735 | 817 | 4460 | 4702 | 4726 | 4535 | 645 | 686 | 624 | 664 | 2630,3 | 4606 | 655 | 0,142 | -2,817 | -58,631 | 0 | 0 | 0,0005205 | 0,0027155 | 0,0014143 | 0,0014143 | TRUE | 0,0114827 | No |
| orf19.4309 | 3683 | 3306 | 3350 | 2226 | 601 | 449 | 599 | 576 | 3511 | 3496 | 3309 | 3487 | 482 | 511 | 509 | 468 | 1971,58 | 3451 | 492 | 0,142 | -2,812 | -54,254 | 0 | 0 | 1,00E-08 | 0,0029241 | 0,001512 | 0,001512 | TRUE | 0,0126377 | No |
| orf19.1370 | 5 | 14 | 5 | 4 | 1 | 1 | 1 | 2 | 5 | 15 | 5 | 6 | 1 | 1 | 1 | 2 | 4,4 | 8 | 1 | 0,143 | -2,805 | -3,192 | 0,0014137 | 0,001888 | 1,00E-08 | 0,3753628 | 0,2671873 | 0,2671873 | TRUE | 1,1030784 | No |
| orf19.5908 | 10566 | 9139 | 10326 | 6845 | 1786 | 1280 | 1692 | 1857 | 10074 | 9663 | 10201 | 10723 | 1431 | 1455 | 1436 | 1510 | 5811,75 | 10165 | 1458 | 0,144 | -2,801 | -62,663 | 0 | 0 | 0,0011244 | 0,0023738 | 0,0015568 | 0,0015568 | TRUE | 0,0162444 | No |
| orf19.419 | 7407 | 6750 | 6900 | 4020 | 1248 | 924 | 1118 | 1216 | 7062 | 7137 | 6817 | 6298 | 1000 | 1051 | 949 | 989 | 3912,71 | 6828 | 997 | 0,146 | -2,778 | -51,421 | 0 | 0 | 0,0021156 | 0,0025109 | 0,0022728 | 0,0022728 | TRUE | 0,0324804 | No |
| orf19.4914.1 | 42186 | 39346 | 39854 | 21332 | 7777 | 4379 | 7118 | 7238 | 40220 | 41603 | 39372 | 33418 | 6231 | 4979 | 6043 | 5885 | 22219 | 38653 | 5784 | 0,15 | -2,74 | -34,274 | 1,93E-257 | 3,42E-256 | 0,0088352 | 0,0021652 | 0,0060484 | 0,0060484 | TRUE | 0,1064163 | No |
| orf19.4531 | 26415 | 24471 | 26708 | 15803 | 4756 | 3355 | 4380 | 4914 | 25184 | 25875 | 26385 | 24756 | 3811 | 3815 | 3719 | 3996 | 14692,5 | 25550 | 3835 | 0,15 | -2,736 | -73,593 | 0 | 0 | 0,0006569 | 0,002203 | 0,001189 | 0,001189 | TRUE | 0,0098806 | No |
| orf19.211 | 1916 | 1627 | 1802 | 1232 | 327 | 226 | 337 | 352 | 1827 | 1720 | 1780 | 1930 | 262 | 257 | 286 | 286 | 1043,57 | 1814 | 273 | 0,151 | -2,728 | -40,388 | 0 | 0 | 2,91E-07 | 0,0036646 | 0,0024463 | 0,0024463 | TRUE | 0,0204681 | No |
| orf19.5848 | 6486 | 5569 | 5956 | 4021 | 1108 | 818 | 1056 | 1171 | 6184 | 5889 | 5884 | 6299 | 888 | 930 | 897 | 952 | 3490,24 | 6064 | 917 | 0,151 | -2,726 | -59,135 | 0 | 0 | 0,000791 | 0,0025617 | 0,001461 | 0,001461 | TRUE | 0,0088625 | No |
| orf19.740 | 45082 | 42743 | 46629 | 28225 | 8261 | 5847 | 8303 | 8262 | 42981 | 45195 | 46065 | 44216 | 6619 | 6649 | 7049 | 6718 | 25686,5 | 44614 | 6759 | 0,152 | -2,723 | -74,508 | 0 | 0 | 0,0007568 | 0,0021552 | 0,0012038 | 0,0012038 | TRUE | 0,0102772 | No |
| orf19.6919 | 126 | 152 | 142 | 80 | 26 | 19 | 24 | 25 | 120 | 161 | 140 | 125 | 21 | 22 | 20 | 20 | 78,7 | 136 | 21 | 0,152 | -2,723 | -13,219 | 6,78E-40 | 2,14E-39 | 1,00E-08 | 0,0229542 | 0,0152891 | 0,0152891 | TRUE | 0,1461004 | No |
| orf19.4980 | 61165 | 56134 | 59582 | 36289 | 10693 | 7994 | 10141 | 11418 | 58314 | 59355 | 58861 | 56849 | 8568 | 9090 | 8610 | 9284 | 33616,29 | 58345 | 8888 | 0,152 | -2,715 | -73,432 | 0 | 0 | 0,0008824 | 0,0021401 | 0,0012532 | 0,0012532 | TRUE | 0,0112097 | No |
| orf19.6578 | 18536 | 17412 | 18835 | 11809 | 3425 | 2429 | 3242 | 3573 | 17672 | 18411 | 18607 | 18499 | 2744 | 2762 | 2752 | 2905 | 10544,21 | 18297 | 2791 | 0,153 | -2,712 | -75,985 | 0 | 0 | 0,0003692 | 0,002247 | 0,0010329 | 0,0010329 | TRUE | 0,0094662 | No |
| orf19.5118 | 48886 | 42179 | 46387 | 31412 | 8722 | 6184 | 8307 | 9059 | 46608 | 44599 | 45826 | 49209 | 6989 | 7032 | 7053 | 7366 | 26834,99 | 46560 | 7110 | 0,153 | -2,711 | -68,124 | 0 | 0 | 0,0010639 | 0,0021525 | 0,0014463 | 0,0014463 | TRUE | 0,0178462 | No |
| orf19.6489 | 10048 | 8935 | 9554 | 6141 | 1879 | 1252 | 1675 | 1876 | 9580 | 9448 | 9438 | 9620 | 1506 | 1424 | 1422 | 1525 | 5495,33 | 9522 | 1469 | 0,154 | -2,694 | -71,902 | 0 | 0 | 3,11E-06 | 0,0023901 | 0,0009862 | 0,0009862 | TRUE | 0,0079689 | No |
| orf19.7086 | 7339 | 6452 | 6870 | 4375 | 1334 | 919 | 1171 | 1393 | 6997 | 6822 | 6787 | 6854 | 1069 | 1045 | 994 | 1133 | 3962,55 | 6865 | 1060 | 0,155 | -2,694 | -59,978 | 0 | 0 | 0,0004697 | 0,0025056 | 0,0014351 | 0,0014351 | TRUE | 0,026356 | No |
| orf19.6554 | 6990 | 7925 | 7525 | 5061 | 1417 | 1258 | 1314 | 1276 | 6664 | 8380 | 7434 | 7928 | 1135 | 1430 | 1116 | 1038 | 4390,64 | 7602 | 1180 | 0,155 | -2,69 | -28,34 | 1,12E-176 | 1,31E-175 | 0,0146952 | 0,0024652 | 0,0082004 | 0,0082004 | TRUE | 0,2403567 | No |
| orf19.6601.1 | 23400 | 22578 | 22798 | 13295 | 4502 | 3027 | 4217 | 4107 | 22309 | 23873 | 22522 | 20827 | 3607 | 3442 | 3580 | 3339 | 12937,65 | 22383 | 3492 | 0,156 | -2,68 | -54,548 | 0 | 0 | 0,0019722 | 0,0022182 | 0,002166 | 0,002166 | TRUE | 0,0265396 | No |
| orf19.6882 | 38975 | 32439 | 36365 | 21310 | 6790 | 4528 | 6578 | 7138 | 37158 | 34300 | 35925 | 33383 | 5441 | 5149 | 5585 | 5804 | 20343,12 | 35192 | 5495 | 0,156 | -2,679 | -54,324 | 0 | 0 | 0,0022491 | 0,002172 | 0,0022386 | 0,0022386 | TRUE | 0,021478 | No |
| orf19.1944 | 20649 | 17719 | 20142 | 12906 | 3790 | 2716 | 3668 | 3896 | 19687 | 18736 | 19898 | 20218 | 3037 | 3088 | 3114 | 3168 | 11368,2 | 19635 | 3102 | 0,158 | -2,662 | -72,486 | 0 | 0 | 0,0006538 | 0,0022357 | 0,0011225 | 0,0011225 | TRUE | 0,0116282 | No |
| orf19.6282 | 798 | 773 | 761 | 529 | 131 | 112 | 162 | 162 | 761 | 817 | 752 | 829 | 105 | 127 | 138 | 132 | 457,53 | 790 | 126 | 0,159 | -2,655 | -28,305 | 3,01E-176 | 3,51E-175 | 0,001781 | 0,0056799 | 0,0041909 | 0,0041909 | TRUE | 0,1388366 | No |
| orf19.7365 | 1185 | 1239 | 1310 | 827 | 230 | 239 | 205 | 223 | 1130 | 1310 | 1294 | 1296 | 184 | 272 | 174 | 181 | 730,12 | 1258 | 203 | 0,16 | -2,644 | -22,874 | 8,47E-116 | 6,49E-115 | 0,019003 | 0,0043401 | 0,0101609 | 0,0101609 | TRUE | 0,5273779 | No |
| orf19.3364 | 834 | 790 | 864 | 488 | 159 | 114 | 159 | 159 | 795 | 835 | 854 | 764 | 127 | 130 | 135 | 129 | 471,22 | 812 | 130 | 0,16 | -2,642 | -30,647 | 2,91E-206 | 4,07E-205 | 1,00E-08 | 0,0055756 | 0,0030492 | 0,0030492 | TRUE | 0,0161087 | No |
| orf19.3537 | 2762 | 2208 | 2140 | 1683 | 518 | 324 | 466 | 479 | 2633 | 2335 | 2114 | 2637 | 415 | 368 | 396 | 389 | 1410,89 | 2430 | 392 | 0,162 | -2,628 | -32,048 | 2,36E-225 | 3,55E-224 | 0,0065895 | 0,003255 | 0,0050885 | 0,0050885 | TRUE | 0,0934915 | No |
| orf19.1152 | 55367 | 49905 | 53037 | 32479 | 10852 | 7477 | 9985 | 10224 | 52786 | 52768 | 52396 | 50880 | 8695 | 8502 | 8477 | 8313 | 30352,25 | 52208 | 8497 | 0,163 | -2,619 | -90,892 | 0 | 0 | 0,0002283 | 0,0021454 | 0,0007343 | 0,0007343 | TRUE | 0,00355 | No |
| orf19.4737 | 1674 | 1540 | 1589 | 1033 | 308 | 238 | 301 | 336 | 1596 | 1628 | 1570 | 1618 | 247 | 271 | 256 | 273 | 932,32 | 1603 | 262 | 0,163 | -2,618 | -41,64 | 0 | 0 | 1,00E-08 | 0,0038524 | 0,0017466 | 0,0017466 | TRUE | 0,0172486 | No |
| orf19.5843 | 2802 | 2790 | 2926 | 1856 | 574 | 439 | 513 | 584 | 2671 | 2950 | 2891 | 2908 | 460 | 499 | 436 | 475 | 1661,14 | 2855 | 468 | 0,163 | -2,613 | -44,085 | 0 | 0 | 0,0015206 | 0,0030797 | 0,0022229 | 0,0022229 | TRUE | 0,0239777 | No |
| orf19.1189 | 27688 | 22718 | 25701 | 17458 | 5125 | 3611 | 5131 | 5427 | 26398 | 24021 | 25390 | 27349 | 4107 | 4106 | 4356 | 4413 | 15017,42 | 25790 | 4246 | 0,165 | -2,602 | -52,885 | 0 | 0 | 0,0020727 | 0,0022006 | 0,0021988 | 0,0021988 | TRUE | 0,0260584 | No |
| orf19.675 | 10778 | 8632 | 9305 | 6927 | 2005 | 1413 | 1925 | 2023 | 10276 | 9127 | 9192 | 10852 | 1607 | 1607 | 1634 | 1645 | 5742,42 | 9862 | 1623 | 0,165 | -2,602 | -42,825 | 0 | 0 | 0,0032586 | 0,0023772 | 0,0032142 | 0,0032142 | TRUE | 0,0550241 | No |
| orf19.1290 | 4791 | 4396 | 4762 | 2926 | 971 | 693 | 895 | 902 | 4568 | 4648 | 4704 | 4584 | 778 | 788 | 760 | 733 | 2695,42 | 4626 | 765 | 0,165 | -2,599 | -60,654 | 0 | 0 | 1,00E-08 | 0,0027004 | 0,001059 | 0,001059 | TRUE | 0,0093012 | No |
| orf19.6594 | 24334 | 23595 | 24880 | 14880 | 4942 | 3524 | 4666 | 5094 | 23200 | 24949 | 24579 | 23310 | 3960 | 4007 | 3961 | 4142 | 14013,53 | 24010 | 4018 | 0,167 | -2,579 | -67,294 | 0 | 0 | 0,0007137 | 0,0022084 | 0,0012763 | 0,0012763 | TRUE | 0,0084836 | No |
| orf19.1433.1 | 4 | 4 | 5 | 9 | 1 | 0 | 2 | 2 | 4 | 4 | 5 | 14 | 1 | 0 | 2 | 2 | 3,9 | 7 | 1 | 0,168 | -2,576 | -2,797 | 0,0051584 | 0,0066618 | 1,00E-08 | 0,4229975 | 0,3199374 | 0,3199374 | TRUE | 0,8874377 | No |
| orf19.3669 | 17247 | 15782 | 17267 | 11387 | 3508 | 2616 | 3247 | 3552 | 16443 | 16687 | 17058 | 17838 | 2811 | 2975 | 2757 | 2888 | 9932,18 | 17006 | 2858 | 0,168 | -2,574 | -62,802 | 0 | 0 | 0,0009203 | 0,0022566 | 0,0014234 | 0,0014234 | TRUE | 0,0130412 | No |
| orf19.2608 | 150148 | 120586 | 134822 | 82223 | 28649 | 18183 | 26996 | 28427 | 143150 | 127504 | 133191 | 128807 | 22956 | 20676 | 22919 | 23114 | 77789,63 | 133163 | 22416 | 0,168 | -2,57 | -49,981 | 0 | 0 | 0,0027288 | 0,0021124 | 0,0025171 | 0,0025171 | TRUE | 0,0333044 | No |
| orf19.7054 | 54 | 43 | 43 | 38 | 8 | 4 | 10 | 16 | 51 | 45 | 42 | 60 | 6 | 5 | 8 | 13 | 28,93 | 50 | 8 | 0,168 | -2,57 | -7,744 | 9,61E-15 | 1,92E-14 | 0,0008154 | 0,0588512 | 0,0411046 | 0,0411046 | TRUE | 0,5708054 | No |
| orf19.844 | 10225 | 9579 | 10630 | 6246 | 2103 | 1516 | 1993 | 2168 | 9748 | 10129 | 10501 | 9785 | 1685 | 1724 | 1692 | 1763 | 5878,36 | 10041 | 1716 | 0,171 | -2,549 | -62,678 | 0 | 0 | 0,000747 | 0,0023706 | 0,0012724 | 0,0012724 | TRUE | 0,0117519 | No |
| orf19.7473 | 1862 | 1861 | 2018 | 1295 | 409 | 320 | 366 | 412 | 1775 | 1968 | 1994 | 2029 | 328 | 364 | 311 | 335 | 1137,82 | 1942 | 334 | 0,172 | -2,541 | -36,558 | 1,32E-292 | 2,85E-291 | 0,0025474 | 0,0035343 | 0,0030127 | 0,0030127 | TRUE | 0,041041 | No |
| orf19.550 | 23973 | 20045 | 22250 | 14692 | 4761 | 3271 | 4601 | 4756 | 22856 | 21195 | 21981 | 23016 | 3815 | 3719 | 3906 | 3867 | 13044,37 | 22262 | 3827 | 0,172 | -2,54 | -65,444 | 0 | 0 | 0,000738 | 0,0022172 | 0,0013043 | 0,0013043 | TRUE | 0,0127367 | No |
| orf19.4094 | 60 | 65 | 59 | 55 | 11 | 11 | 16 | 14 | 57 | 69 | 58 | 86 | 9 | 13 | 14 | 11 | 39,58 | 68 | 12 | 0,172 | -2,538 | -8,941 | 3,86E-19 | 8,52E-19 | 1,00E-08 | 0,0435707 | 0,0304806 | 0,0304806 | TRUE | 0,2387387 | No |
| orf19.5755 | 3292 | 3064 | 3368 | 1981 | 709 | 490 | 684 | 641 | 3139 | 3240 | 3327 | 3103 | 568 | 557 | 581 | 521 | 1879,52 | 3202 | 557 | 0,174 | -2,525 | -47,945 | 0 | 0 | 0,0006631 | 0,0029649 | 0,0016886 | 0,0016886 | TRUE | 0,0234206 | No |
| orf19.4941 | 28783 | 25073 | 28101 | 17479 | 6017 | 4067 | 5807 | 5886 | 27441 | 26512 | 27761 | 27382 | 4821 | 4625 | 4930 | 4786 | 16032,21 | 27274 | 4790 | 0,176 | -2,509 | -76,521 | 0 | 0 | 0,0003773 | 0,0021937 | 0,0009183 | 0,0009183 | TRUE | 0,0063183 | No |
| orf19.2766 | 9 | 12 | 12 | 4 | 0 | 1 | 4 | 3 | 9 | 13 | 12 | 6 | 0 | 1 | 3 | 2 | 5,8 | 10 | 2 | 0,176 | -2,503 | -3,461 | 0,0005381 | 0,0007382 | 1,00E-08 | 0,2854074 | 0,1946521 | 0,1946521 | TRUE | 0,4774382 | No |
| orf19.4792 | 17765 | 15331 | 17092 | 11488 | 3669 | 2675 | 3539 | 3806 | 16937 | 16211 | 16885 | 17997 | 2940 | 3042 | 3005 | 3095 | 10013,78 | 17008 | 3020 | 0,178 | -2,493 | -59,933 | 0 | 0 | 0,0009132 | 0,0022552 | 0,0014809 | 0,0014809 | TRUE | 0,0184472 | No |

| | | | | | | | | | | | | | | | | | | | | | | | | | | | | | | | |
|---|---|---|---|---|---|---|---|---|---|---|---|---|---|---|---|---|---|---|---|---|---|---|---|---|---|---|---|---|---|---|---|
| orf19.2414 | 9629 | 8265 | 8602 | 5697 | 1928 | 1399 | 1804 | 2004 | 9180 | 8739 | 8498 | 8925 | 1545 | 1591 | 1532 | 1629 | 5204,84 | 8836 | 1574 | 0,178 | -2,489 | -59,697 | 0 | 0 | 0,000764 | 0,0024067 | 0,0013219 | 0,0013219 | TRUE | 0,0086708 | No |
| orf19.508 | 2538 | 2303 | 2476 | 1642 | 540 | 411 | 521 | 517 | 2420 | 2435 | 2446 | 2572 | 433 | 467 | 442 | 420 | 1454,49 | 2468 | 440 | 0,178 | -2,488 | -45,811 | 0 | 0 | 1,00E-08 | 0,0032201 | 0,0015939 | 0,0015939 | TRUE | 0,0166934 | No |
| orf19.649 | 8118 | 8269 | 8736 | 4802 | 1883 | 1327 | 1721 | 1655 | 7740 | 8743 | 8630 | 7523 | 1509 | 1509 | 1461 | 1346 | 4807,56 | 8159 | 1456 | 0,178 | -2,487 | -39,59 | 0 | 0 | 0,0037242 | 0,0024328 | 0,0034153 | 0,0034153 | TRUE | 0,0330863 | No |
| orf19.1353 | 244 | 173 | 205 | 176 | 76 | 36 | 37 | 33 | 233 | 183 | 203 | 276 | 61 | 41 | 31 | 27 | 131,73 | 224 | 40 | 0,18 | -2,475 | -11,286 | 1,54E-29 | 4,11E-29 | 0,0613713 | 0,0145552 | 0,0325233 | 0,0325233 | TRUE | 0,726712 | No |
| orf19.4783 | 115 | 103 | 134 | 77 | 25 | 12 | 28 | 32 | 110 | 109 | 132 | 121 | 20 | 14 | 24 | 26 | 69,38 | 118 | 21 | 0,18 | -2,47 | -11,724 | 9,62E-32 | 2,68E-31 | 0,0017197 | 0,0257575 | 0,017041 | 0,017041 | TRUE | 0,2277738 | No |
| orf19.2772 | 6301 | 5902 | 6326 | 3990 | 1379 | 1008 | 1241 | 1456 | 6007 | 6241 | 6249 | 6251 | 1105 | 1146 | 1054 | 1184 | 3654,57 | 6187 | 1122 | 0,181 | -2,463 | -54,868 | 0 | 0 | 0,0007275 | 0,0025406 | 0,001447 | 0,001447 | TRUE | 0,0206744 | No |
| orf19.3707 | 29758 | 26245 | 28293 | 17328 | 6234 | 4460 | 5857 | 6310 | 28371 | 27751 | 27951 | 27145 | 4995 | 5071 | 4973 | 5131 | 16423,45 | 27804 | 5042 | 0,181 | -2,463 | -83,96 | 0 | 0 | 0,0002113 | 0,0021913 | 0,0007179 | 0,0007179 | TRUE | 0,0030589 | No |
| orf19.3360 | 9339 | 9626 | 9941 | 6252 | 2212 | 1639 | 2127 | 1985 | 8904 | 10178 | 9821 | 9794 | 1772 | 1864 | 1806 | 1614 | 5719,1 | 9674 | 1764 | 0,182 | -2,456 | -42,741 | 0 | 0 | 0,0029616 | 0,0023784 | 0,0028602 | 0,0028602 | TRUE | 0,0399761 | No |
| orf19.1331 | 5106 | 5566 | 5505 | 3189 | 1204 | 975 | 1114 | 1069 | 4868 | 5885 | 5438 | 4996 | 965 | 1109 | 946 | 869 | 3134,49 | 5297 | 972 | 0,183 | -2,449 | -30,723 | 2,78E-207 | 3,90E-206 | 0,0082842 | 0,0026151 | 0,0055345 | 0,0055345 | TRUE | 0,1029754 | No |
| orf19.1277 | 15782 | 13278 | 14984 | 10449 | 3351 | 2573 | 3148 | 3403 | 15046 | 14040 | 14803 | 16369 | 2685 | 2926 | 2673 | 2767 | 8913,54 | 15064 | 2763 | 0,183 | -2,447 | -45,442 | 0 | 0 | 0,0026753 | 0,0022755 | 0,002587 | 0,002587 | TRUE | 0,0411104 | No |
| orf19.6488 | 8 | 7 | 11 | 5 | 1 | 2 | 3 | 1 | 8 | 7 | 11 | 8 | 1 | 2 | 3 | 1 | 5,02 | 8 | 2 | 0,184 | -2,442 | -3,299 | 0,0009687 | 0,0013082 | 1,00E-08 | 0,3291311 | 0,1760321 | 0,1760321 | TRUE | 0,1616939 | No |
| orf19.1868 | 7428 | 6458 | 7075 | 4361 | 1608 | 1062 | 1505 | 1650 | 7082 | 6829 | 6989 | 6832 | 1288 | 1208 | 1278 | 1342 | 4105,86 | 6933 | 1279 | 0,185 | -2,436 | -58,833 | 0 | 0 | 0,0003103 | 0,0024912 | 0,0012177 | 0,0012177 | TRUE | 0,0153239 | No |
| orf19.3932.1 | 1946 | 1361 | 1406 | 1212 | 409 | 299 | 314 | 357 | 1855 | 1439 | 1389 | 1899 | 328 | 340 | 267 | 290 | 975,83 | 1646 | 306 | 0,186 | -2,427 | -20,954 | 1,74E-97 | 1,11E-96 | 0,0186268 | 0,0037739 | 0,0110829 | 0,0110829 | TRUE | 0,0522427 | No |
| orf19.999 | 6022 | 4984 | 5414 | 3877 | 1261 | 958 | 1148 | 1360 | 5741 | 5270 | 5349 | 6074 | 1010 | 1089 | 975 | 1106 | 3326,69 | 5608 | 1045 | 0,186 | -2,424 | -39,221 | 0 | 0 | 0,0032503 | 0,0025848 | 0,0031404 | 0,0031404 | TRUE | 0,0371033 | No |
| orf19.3968 | 6 | 14 | 9 | 8 | 5 | 0 | 3 | 1 | 6 | 15 | 9 | 13 | 4 | 0 | 3 | 1 | 6,16 | 11 | 2 | 0,187 | -2,415 | -3,358 | 0,0007858 | 0,0010676 | 0,1052108 | 0,2684525 | 0,2147782 | 0,2147782 | TRUE | 0,3792608 | No |
| orf19.2047 | 1643 | 1426 | 1521 | 931 | 374 | 241 | 319 | 357 | 1566 | 1508 | 1503 | 1458 | 300 | 274 | 271 | 290 | 896,27 | 1509 | 284 | 0,188 | -2,41 | -37,795 | 0 | 0 | 1,00E-08 | 0,0039232 | 0,0019705 | 0,0019705 | TRUE | 0,0168278 | No |
| orf19.3984 | 11600 | 12418 | 12609 | 7504 | 2893 | 2217 | 2572 | 2601 | 11059 | 13130 | 12456 | 11755 | 2318 | 2521 | 2184 | 2115 | 7192,4 | 12100 | 2284 | 0,189 | -2,406 | -35,764 | 4,05E-280 | 8,21E-279 | 0,0054398 | 0,0023196 | 0,0041063 | 0,0041063 | TRUE | 0,0575681 | No |
| orf19.2787 | 9 | 11 | 11 | 7 | 1 | 1 | 4 | 3 | 9 | 12 | 11 | 11 | 1 | 1 | 3 | 2 | 6,23 | 11 | 2 | 0,189 | -2,405 | -3,636 | 0,0002773 | 0,000385 | 1,00E-08 | 0,2657551 | 0,1431152 | 0,1431152 | TRUE | 0,2574342 | No |
| orf19.84 | 5341 | 4742 | 5175 | 3394 | 1182 | 943 | 1099 | 1160 | 5092 | 5014 | 5112 | 5317 | 947 | 1072 | 933 | 943 | 3053,88 | 5134 | 974 | 0,189 | -2,402 | -47,628 | 0 | 0 | 0,0011665 | 0,0026289 | 0,0018724 | 0,0018724 | TRUE | 0,0507732 | No |
| orf19.3997 | 75794 | 57465 | 64327 | 42099 | 15638 | 10124 | 14904 | 16331 | 72261 | 60762 | 63549 | 65950 | 12530 | 11512 | 12653 | 13279 | 39062,12 | 65630 | 12494 | 0,19 | -2,393 | -39,298 | 0 | 0 | 0,0044424 | 0,0021333 | 0,0035183 | 0,0035183 | TRUE | 0,0567549 | No |
| orf19.5045 | 8709 | 9723 | 10201 | 5931 | 2326 | 1790 | 2035 | 1982 | 8303 | 10281 | 10078 | 9291 | 1864 | 2035 | 1728 | 1612 | 5648,9 | 9488 | 1810 | 0,191 | -2,392 | -29,303 | 9,59E-189 | 1,23E-187 | 0,0091212 | 0,0023819 | 0,0060925 | 0,0060925 | TRUE | 0,0869223 | No |
| orf19.6249 | 2885 | 2568 | 2819 | 1622 | 618 | 451 | 617 | 660 | 2751 | 2715 | 2785 | 2541 | 495 | 513 | 524 | 537 | 1607,53 | 2698 | 517 | 0,191 | -2,385 | -43,809 | 0 | 0 | 1,00E-08 | 0,0031127 | 0,0017776 | 0,0017776 | TRUE | 0,0172492 | No |
| orf19.6763 | 39002 | 34465 | 37600 | 22679 | 8934 | 5773 | 8412 | 8788 | 37184 | 36442 | 37145 | 35528 | 7159 | 6564 | 7142 | 7145 | 21788,73 | 36575 | 7002 | 0,192 | -2,384 | -62,014 | 0 | 0 | 0,0009826 | 0,0021666 | 0,0013408 | 0,0013408 | TRUE | 0,02115 | No |
| orf19.2050 | 7142 | 6226 | 6871 | 4275 | 1605 | 1124 | 1525 | 1672 | 6809 | 6583 | 6788 | 6697 | 1286 | 1278 | 1295 | 1359 | 4011,95 | 6719 | 1304 | 0,194 | -2,364 | -61,89 | 0 | 0 | 1,00E-08 | 0,0025005 | 0,0009759 | 0,0009759 | TRUE | 0,0099509 | No |
| orf19.3551 | 311 | 243 | 257 | 140 | 64 | 35 | 69 | 61 | 297 | 257 | 254 | 219 | 51 | 40 | 59 | 50 | 153,24 | 257 | 50 | 0,195 | -2,36 | -15,579 | 1,02E-54 | 4,00E-54 | 0,0085301 | 0,0128059 | 0,0110303 | 0,0110303 | TRUE | 0,1101494 | No |
| orf19.2023 | 27155 | 21870 | 24432 | 15567 | 5800 | 3992 | 5624 | 6237 | 25889 | 23125 | 24137 | 24387 | 4647 | 4539 | 4775 | 5071 | 14571,23 | 24384 | 4758 | 0,195 | -2,357 | -48,486 | 0 | 0 | 0,0020962 | 0,002204 | 0,0021531 | 0,0021531 | TRUE | 0,024013 | No |
| orf19.3474 | 1928 | 1942 | 2126 | 1201 | 508 | 327 | 445 | 466 | 1838 | 2053 | 2100 | 1881 | 407 | 372 | 378 | 379 | 1176,11 | 1968 | 384 | 0,195 | -2,357 | -34,942 | 1,69E-267 | 3,12E-266 | 0,0024967 | 0,0034873 | 0,0029261 | 0,0029261 | TRUE | 0,0255874 | No |
| orf19.4372 | 5333 | 5067 | 5410 | 3454 | 1311 | 949 | 1178 | 1253 | 5084 | 5358 | 5345 | 5411 | 1050 | 1079 | 1000 | 1019 | 3168,26 | 5300 | 1037 | 0,196 | -2,354 | -53,905 | 0 | 0 | 0,0004977 | 0,0026095 | 0,0012954 | 0,0012954 | TRUE | 0,0092601 | No |
| orf19.5094 | 1565 | 1401 | 1487 | 890 | 353 | 263 | 367 | 311 | 1492 | 1481 | 1469 | 1394 | 283 | 299 | 312 | 253 | 872,88 | 1459 | 287 | 0,196 | -2,353 | -33,742 | 1,41E-249 | 2,40E-248 | 0,0007237 | 0,0039723 | 0,0027315 | 0,0027315 | TRUE | 0,074871 | No |
| orf19.6920 | 1485 | 2036 | 1871 | 1111 | 474 | 313 | 423 | 378 | 1416 | 2153 | 1848 | 1740 | 380 | 356 | 359 | 307 | 1069,95 | 1789 | 350 | 0,196 | -2,352 | -20,468 | 4,19E-93 | 2,58E-92 | 0,0184703 | 0,0036258 | 0,0110921 | 0,0110921 | TRUE | 0,0924089 | No |
| orf19.2094 | 6578 | 6162 | 6594 | 4214 | 1502 | 1214 | 1468 | 1563 | 6271 | 6516 | 6514 | 6601 | 1204 | 1380 | 1246 | 1271 | 3875,47 | 6476 | 1275 | 0,197 | -2,347 | -50,041 | 0 | 0 | 0,0010047 | 0,0025149 | 0,0016744 | 0,0016744 | TRUE | 0,0343731 | No |
| orf19.5451 | 681 | 529 | 652 | 574 | 171 | 126 | 145 | 171 | 649 | 559 | 644 | 899 | 137 | 143 | 123 | 139 | 411,8 | 688 | 136 | 0,198 | -2,339 | -16,968 | 1,41E-64 | 6,31E-64 | 0,0223131 | 0,0060785 | 0,0141353 | 0,0141353 | TRUE | 0,4794284 | No |
| orf19.6678 | 367 | 332 | 352 | 246 | 90 | 67 | 85 | 79 | 350 | 351 | 348 | 385 | 72 | 76 | 72 | 64 | 214,84 | 358 | 71 | 0,198 | -2,334 | -20,144 | 3,03E-90 | 1,82E-89 | 1,00E-08 | 0,0097336 | 0,005073 | 0,005073 | TRUE | 0,0420847 | No |
| orf19.7676 | 19956 | 15522 | 16642 | 12942 | 4540 | 3070 | 4042 | 4655 | 19026 | 16413 | 16441 | 20274 | 3638 | 3491 | 3432 | 3785 | 10812,34 | 18038 | 3586 | 0,199 | -2,33 | -32,67 | 4,12E-234 | 6,47E-233 | 0,0064726 | 0,0022431 | 0,0047306 | 0,0047306 | TRUE | 0,0847633 | No |
| orf19.6420 | 8235 | 6767 | 7567 | 5932 | 1913 | 1441 | 1827 | 1960 | 7851 | 7155 | 7475 | 9293 | 1533 | 1639 | 1551 | 1594 | 4761,36 | 7944 | 1579 | 0,199 | -2,33 | -31,043 | 1,43E-211 | 2,05E-210 | 0,0069905 | 0,0024361 | 0,0050593 | 0,0050593 | TRUE | 0,1579514 | No |
| orf19.693 | 169 | 176 | 216 | 119 | 46 | 34 | 46 | 43 | 161 | 186 | 213 | 186 | 37 | 39 | 39 | 35 | 112,07 | 186 | 38 | 0,2 | -2,323 | -14,077 | 5,26E-45 | 1,80E-44 | 1,00E-08 | 0,0167419 | 0,0112758 | 0,0112758 | TRUE | 0,1077961 | No |
| orf19.3315 | 2353 | 2155 | 2460 | 1559 | 578 | 425 | 553 | 575 | 2243 | 2279 | 2430 | 2442 | 463 | 483 | 469 | 468 | 1409,74 | 2348 | 471 | 0,2 | -2,319 | -41,384 | 0 | 0 | 1,00E-08 | 0,003256 | 0,0018292 | 0,0018292 | TRUE | 0,0112003 | No |
| orf19.3448 | 1776 | 2920 | 2753 | 1405 | 651 | 516 | 620 | 400 | 1693 | 3088 | 2720 | 2201 | 522 | 587 | 526 | 325 | 1457,68 | 2426 | 490 | 0,202 | -2,308 | -8,914 | 4,94E-19 | 1,09E-18 | 0,063278 | 0,0032177 | 0,0288808 | 0,063278 | TRUE | 0,1243341 | No |
| orf19.836 | 209 | 214 | 197 | 133 | 55 | 35 | 57 | 44 | 199 | 226 | 195 | 208 | 44 | 40 | 48 | 36 | 124,57 | 207 | 42 | 0,203 | -2,298 | -15,087 | 1,98E-51 | 7,43E-51 | 1,00E-08 | 0,0152721 | 0,0090391 | 0,0090391 | TRUE | 0,0933232 | No |
| orf19.4843 | 6186 | 5757 | 6168 | 4012 | 1476 | 1149 | 1398 | 1582 | 5898 | 6087 | 6093 | 6285 | 1183 | 1307 | 1187 | 1286 | 3665,72 | 6091 | 1241 | 0,204 | -2,297 | -49,438 | 0 | 0 | 0,0010627 | 0,0025392 | 0,0016201 | 0,0016201 | TRUE | 0,0141562 | No |
| orf19.2623 | 15021 | 13342 | 14429 | 9492 | 3611 | 2615 | 3401 | 3671 | 14321 | 14107 | 14254 | 14870 | 2893 | 2973 | 2887 | 2985 | 8661,47 | 14388 | 2934 | 0,204 | -2,293 | -68,689 | 0 | 0 | 0,0002957 | 0,0022808 | 0,0008805 | 0,0008805 | TRUE | 0,006018 | No |
| orf19.5552 | 4808 | 4416 | 4769 | 2861 | 1165 | 817 | 1080 | 1211 | 4584 | 4669 | 4711 | 4482 | 933 | 929 | 917 | 985 | 2776,32 | 4612 | 941 | 0,204 | -2,293 | -53,27 | 0 | 0 | 1,00E-08 | 0,0026827 | 0,0011843 | 0,0011843 | TRUE | 0,0121801 | No |
| orf19.2356 | 28074 | 25541 | 28171 | 17897 | 6962 | 5075 | 6619 | 6743 | 26766 | 27006 | 27830 | 28037 | 5578 | 5771 | 5619 | 5483 | 16511,27 | 27410 | 5613 | 0,205 | -2,288 | -71,32 | 0 | 0 | 0,0003394 | 0,0021907 | 0,0008884 | 0,0008884 | TRUE | 0,0042097 | No |
| orf19.5499 | 160 | 182 | 179 | 98 | 52 | 34 | 37 | 34 | 153 | 192 | 177 | 154 | 42 | 39 | 31 | 28 | 101,84 | 169 | 35 | 0,205 | -2,287 | -12,917 | 3,61E-38 | 1,11E-37 | 0,0073399 | 0,0182135 | 0,0139963 | 0,0139963 | TRUE | 0,1592563 | No |
| orf19.3893 | 156 | 129 | 162 | 112 | 42 | 24 | 33 | 45 | 149 | 136 | 160 | 175 | 34 | 27 | 28 | 37 | 93,27 | 155 | 32 | 0,205 | -2,286 | -12,88 | 5,81E-38 | 1,78E-37 | 1,00E-08 | 0,0196947 | 0,0126239 | 0,0126239 | TRUE | 0,0911475 | No |
| orf19.4771 | 5948 | 5715 | 6135 | 3878 | 1518 | 1167 | 1362 | 1511 | 5671 | 6043 | 6061 | 6075 | 1216 | 1327 | 1156 | 1229 | 3597,22 | 5962 | 1232 | 0,206 | -2,277 | -46,032 | 0 | 0 | 0,0015286 | 0,0025477 | 0,001892 | 0,001892 | TRUE | 0,0299888 | No |
| orf19.7567 | 4165 | 3736 | 4026 | 2721 | 1003 | 745 | 945 | 1093 | 3971 | 3950 | 3977 | 4263 | 804 | 847 | 802 | 889 | 2437,87 | 4040 | 836 | 0,207 | -2,273 | -44,593 | 0 | 0 | 0,0009274 | 0,0027648 | 0,0018218 | 0,0018218 | TRUE | 0,0235803 | No |
| orf19.6604 | 5319 | 6054 | 6020 | 3740 | 1556 | 1210 | 1401 | 1299 | 5071 | 6401 | 5947 | 5859 | 1247 | 1376 | 1189 | 1056 | 3518,35 | 5820 | 1217 | 0,209 | -2,259 | -26,516 | 6,29E-155 | 6,57E-154 | 0,0097821 | 0,0025579 | 0,006507 | 0,006507 | TRUE | 0,095559 | No |
| orf19.6211 | 5670 | 4992 | 5503 | 3288 | 1393 | 932 | 1279 | 1459 | 5406 | 5278 | 5436 | 5151 | 1116 | 1060 | 1086 | 1186 | 3214,94 | 5318 | 1112 | 0,209 | -2,256 | -49,291 | 0 | 0 | 0,0007834 | 0,002602 | 0,0015058 | 0,0015058 | TRUE | 0,0244694 | No |
| orf19.3277 | 26 | 17 | 24 | 20 | 8 | 4 | 9 | 2 | 25 | 18 | 24 | 31 | 6 | 5 | 8 | 2 | 14,75 | 24 | 5 | 0,209 | -2,255 | -5,035 | 4,78E-07 | 7,49E-07 | 1,00E-08 | 0,1133785 | 0,0820489 | 0,0820489 | TRUE | 0,4319131 | No |
| orf19.1153 | 7748 | 6907 | 7682 | 4561 | 1935 | 1314 | 1794 | 1979 | 7387 | 7303 | 7589 | 7145 | 1550 | 1494 | 1523 | 1609 | 4450,14 | 7356 | 1544 | 0,21 | -2,251 | -56,316 | 0 | 0 | 0,0004401 | 0,0024602 | 0,0011695 | 0,0011695 | TRUE | 0,0097097 | No |
| orf19.3047 | 5448 | 5084 | 5647 | 3470 | 1344 | 1055 | 1338 | 1398 | 5194 | 5376 | 5579 | 5436 | 1077 | 1200 | 1136 | 1137 | 3266,7 | 5396 | 1138 | 0,21 | -2,248 | -49,054 | 0 | 0 | 0,000697 | 0,0025939 | 0,0015203 | 0,0015203 | TRUE | 0,0153935 | No |
| orf19.6950 | 1693 | 1497 | 1688 | 1152 | 452 | 294 | 418 | 427 | 1614 | 1583 | 1668 | 1805 | 362 | 334 | 355 | 347 | 1008,47 | 1668 | 350 | 0,211 | -2,248 | -34,343 | 1,77E-258 | 3,16E-257 | 0,0015332 | 0,0037194 | 0,0025068 | 0,0025068 | TRUE | 0,0341186 | No |
| orf19.3071 | 7979 | 6977 | 7846 | 5231 | 1940 | 1469 | 1867 | 2198 | 7607 | 7377 | 7751 | 8195 | 1554 | 1670 | 1585 | 1787 | 4690,91 | 7732 | 1649 | 0,213 | -2,229 | -41,017 | 0 | 0 | 0,0024973 | 0,0024413 | 0,0024918 | 0,0024918 | TRUE | 0,0396319 | No |
| orf19.882 | 11494 | 10177 | 11057 | 6781 | 2859 | 1988 | 2705 | 2953 | 10958 | 10761 | 10923 | 10623 | 2291 | 2261 | 2297 | 2401 | 6564,28 | 10816 | 2312 | 0,214 | -2,225 | -65,288 | 0 | 0 | 0,0001041 | 0,0023414 | 0,0008707 | 0,0008707 | TRUE | 0,008246 | No |
| orf19.3264 | 340 | 381 | 377 | 237 | 102 | 94 | 84 | 72 | 324 | 403 | 372 | 371 | 82 | 107 | 71 | 59 | 223,65 | 368 | 80 | 0,214 | -2,224 | -15,096 | 1,72E-51 | 6,46E-51 | 0,0213246 | 0,0094327 | 0,0136576 | 0,0136576 | TRUE | 0,4357498 | No |
| orf19.6288 | 158 | 127 | 130 | 92 | 37 | 30 | 37 | 31 | 151 | 134 | 128 | 144 | 30 | 34 | 31 | 25 | 84,73 | 139 | 30 | 0,214 | -2,221 | -12,427 | 1,86E-35 | 5,48E-35 | 1,00E-08 | 0,021469 | 0,0118298 | 0,0118298 | TRUE | 0,0887483 | No |
| orf19.3670 | 6097 | 5715 | 6019 | 3658 | 1618 | 1112 | 1456 | 1545 | 5813 | 6043 | 5946 | 5730 | 1296 | 1264 | 1236 | 1256 | 3573,21 | 5883 | 1263 | 0,215 | -2,22 | -56,449 | 0 | 0 | 1,00E-08 | 0,0025508 | 0,0010366 | 0,0010366 | TRUE | 0,0040228 | No |
| orf19.2841 | 8829 | 9009 | 9147 | 5089 | 2417 | 1669 | 2249 | 2202 | 8417 | 9526 | 9036 | 7972 | 1937 | 1898 | 1909 | 1790 | 5310,78 | 8738 | 1884 | 0,215 | -2,215 | -37,186 | 1,16E-302 | 2,60E-301 | 0,0032043 | 0,0024004 | 0,0031045 | 0,0031045 | TRUE | 0,0448397 | No |
| orf19.3555 | 10230 | 8880 | 9872 | 5584 | 2503 | 1705 | 2474 | 2555 | 9753 | 9389 | 9753 | 8748 | 2006 | 1939 | 2100 | 2077 | 5720,64 | 9411 | 2030 | 0,216 | -2,212 | -45,665 | 0 | 0 | 0,001522 | 0,0023783 | 0,0019742 | 0,0019742 | TRUE | 0,0268777 | No |
| orf19.5610 | 3599 | 3539 | 3692 | 2163 | 958 | 741 | 906 | 856 | 3431 | 3742 | 3647 | 3388 | 768 | 843 | 769 | 696 | 2160,56 | 3552 | 769 | 0,216 | -2,212 | -35,756 | 5,26E-280 | 1,06E-278 | 0,0031174 | 0,0028512 | 0,0029347 | 0,0029347 | TRUE | 0,0484933 | No |
| orf19.1048 | 564 | 522 | 546 | 326 | 131 | 114 | 126 | 152 | 538 | 552 | 539 | 511 | 105 | 130 | 107 | 124 | 325,61 | 535 | 116 | 0,216 | -2,212 | -22,983 | 6,84E-117 | 5,28E-116 | 1,00E-08 | 0,0071338 | 0,0040172 | 0,0040172 | TRUE | 0,049975 | No |
| orf19.5760 | 5180 | 3905 | 4443 | 3473 | 1236 | 929 | 1127 | 1327 | 4939 | 4129 | 4389 | 5441 | 990 | 1056 | 957 | 1079 | 2872,5 | 4724 | 1020 | 0,216 | -2,21 | -26,592 | 8,49E-156 | 8,94E-155 | 0,0084113 | 0,0026629 | 0,0060778 | 0,0060778 | TRUE | 0,123676 | No |
| orf19.486 | 592 | 508 | 553 | 320 | 160 | 98 | 132 | 139 | 564 | 537 | 546 | 501 | 128 | 111 | 112 | 113 | 326,74 | 537 | 116 | 0,216 | -2,21 | -23,085 | 6,56E-118 | 5,11E-117 | 1,00E-08 | 0,0071164 | 0,0039511 | 0,0039511 | TRUE | 0,0535636 | No |
| orf19.6814 | 105363 | 81646 | 89182 | 67659 | 25071 | 18395 | 24052 | 26323 | 100452 | 86330 | 88103 | 105991 | 20089 | 20917 | 20420 | 21403 | 57963,24 | 95219 | 20707 | 0,217 | -2,201 | -33,844 | 4,39E-251 | 7,53E-250 | 0,0053666 | 0,0021196 | 0,0040365 | 0,0040365 | TRUE | 0,0709933 | No |
| orf19.3337 | 28155 | 24424 | 27176 | 18157 | 7176 | 5248 | 7001 | 7413 | 26843 | 25825 | 26847 | 28444 | 5750 | 5967 | 5944 | 6027 | 16455,99 | 26990 | 5922 | 0,219 | -2,188 | -56,638 | 0 | 0 | 0,000872 | 0,0021911 | 0,0013372 | 0,0013372 | TRUE | 0,0159169 | No |
| orf19.5131 | 5690 | 5102 | 5669 | 3449 | 1463 | 1097 | 1377 | 1495 | 5425 | 5395 | 5600 | 5403 | 1172 | 1247 | 1169 | 1216 | 3328,41 | 5456 | 1201 | 0,22 | -2,186 | -54,681 | 0 | 0 | 1,00E-08 | 0,0025846 | 0,0010596 | 0,0010596 | TRUE | 0,0072504 | No |
| orf19.3083 | 2265 | 1788 | 2188 | 1480 | 570 | 427 | 563 | 570 | 2159 | 1891 | 2162 | 2318 | 457 | 486 | 478 | 463 | 1301,72 | 2132 | 471 | 0,221 | -2,179 | -30,937 | 3,84E-210 | 5,46E-209 | 0,0036997 | 0,0033526 | 0,0035505 | 0,0035505 | TRUE | 0,0703884 | No |
| orf19.6540 | 26616 | 22236 | 24327 | 15514 | 6699 | 4616 | 6192 | 6900 | 25375 | 23512 | 24033 | 24304 | 5368 | 5249 | 5257 | 5610 | 14838,42 | 24306 | 5371 | 0,221 | -2,178 | -56,603 | 0 | 0 | 0,0008626 | 0,0022019 | 0,0013154 | 0,0013154 | TRUE | 0,0110622 | No |
| orf19.4436 | 412 | 392 | 394 | 271 | 106 | 87 | 109 | 104 | 393 | 414 | 389 | 425 | 85 | 99 | 93 | 85 | 247,75 | 405 | 90 | 0,222 | -2,172 | -20,444 | 6,75E-93 | 4,14E-92 | 1,00E-08 | 0,0087185 | 0,0045132 | 0,0045132 | TRUE | 0,0334562 | No |

| | | | | | | | | | | | | | | | | | | | | | | | | | | | | | | | |
|---|---|---|---|---|---|---|---|---|---|---|---|---|---|---|---|---|---|---|---|---|---|---|---|---|---|---|---|---|---|---|---|
| orf19.347 | 4986 | 4248 | 4594 | 3217 | 1253 | 946 | 1249 | 1278 | 4754 | 4492 | 4538 | 5040 | 1004 | 1076 | 1060 | 1039 | 2875,32 | 4706 | 1045 | 0,222 | -2,171 | -42,256 | 0 | 0 | 0,0015667 | 0,0026623 | 0,0019879 | 0,0019879 | TRUE | 0,026618 | No |
| orf19.7303 | 10 | 11 | 9 | 3 | 5 | 1 | 3 | 0 | 10 | 12 | 9 | 5 | 4 | 1 | 3 | 0 | 5,31 | 9 | 2 | 0,222 | -2,169 | -2,953 | 0,0031433 | 0,0041118 | 1,00E-08 | 0,3115467 | 0,229191 | 0,229191 | TRUE | 0,5990437 | No |
| orf19.7006 | 7566 | 6403 | 7318 | 4580 | 2010 | 1339 | 1820 | 2006 | 7213 | 6770 | 7229 | 7175 | 1611 | 1523 | 1545 | 1631 | 4337,17 | 7097 | 1578 | 0,222 | -2,168 | -51,448 | 0 | 0 | 0,0007199 | 0,0024698 | 0,0013435 | 0,0013435 | TRUE | 0,0118432 | No |
| orf19.4631 | 3360 | 3463 | 3468 | 2520 | 971 | 833 | 896 | 848 | 3203 | 3662 | 3426 | 3948 | 778 | 947 | 761 | 690 | 2176,79 | 3560 | 794 | 0,223 | -2,167 | -23,651 | 1,16E-123 | 9,57E-123 | 0,0124497 | 0,0028456 | 0,0073414 | 0,0073414 | TRUE | 0,1940748 | No |
| orf19.5053 | 1292 | 1213 | 1340 | 844 | 367 | 265 | 353 | 315 | 1232 | 1283 | 1324 | 1322 | 294 | 301 | 300 | 256 | 788,94 | 1290 | 288 | 0,223 | -2,166 | -31,303 | 4,19E-215 | 6,12E-214 | 0,0007445 | 0,0041724 | 0,0026159 | 0,0026159 | TRUE | 0,0656023 | No |
| orf19.7561 | 27732 | 22638 | 25991 | 17695 | 7119 | 4908 | 6976 | 7403 | 26439 | 23937 | 25677 | 27720 | 5704 | 5581 | 5923 | 6019 | 15875,01 | 25943 | 5807 | 0,224 | -2,159 | -42,864 | 0 | 0 | 0,0023108 | 0,0021947 | 0,0023386 | 0,0023386 | TRUE | 0,0331638 | No |
| orf19.3841 | 7339 | 6369 | 7080 | 4522 | 1874 | 1437 | 1777 | 1996 | 6997 | 6734 | 6994 | 7084 | 1502 | 1634 | 1509 | 1623 | 4259,61 | 6952 | 1567 | 0,225 | -2,151 | -50,185 | 0 | 0 | 0,0007538 | 0,0024767 | 0,0013977 | 0,0013977 | TRUE | 0,0095477 | No |
| orf19.5069 | 10880 | 10053 | 10907 | 6945 | 2917 | 2091 | 2922 | 2955 | 10373 | 10630 | 10775 | 10880 | 2337 | 2378 | 2481 | 2403 | 6531,99 | 10664 | 2400 | 0,225 | -2,151 | -60,8 | 0 | 0 | 0,0003161 | 0,0023426 | 0,0009635 | 0,0009635 | TRUE | 0,0064134 | No |
| orf19.2638 | 467 | 443 | 460 | 300 | 152 | 94 | 119 | 106 | 445 | 468 | 454 | 470 | 122 | 107 | 101 | 86 | 281,74 | 459 | 104 | 0,226 | -2,144 | -20,119 | 5,06E-90 | 3,03E-89 | 1,00E-08 | 0,0079189 | 0,0054073 | 0,0054073 | TRUE | 0,153645 | No |
| orf19.4170 | 1398 | 1276 | 1297 | 824 | 384 | 243 | 326 | 400 | 1333 | 1349 | 1281 | 1291 | 308 | 276 | 277 | 325 | 805,03 | 1314 | 296 | 0,226 | -2,143 | -31,55 | 1,79E-218 | 2,66E-217 | 0,0001941 | 0,0041308 | 0,0025089 | 0,0025089 | TRUE | 0,0488086 | No |
| orf19.3672 | 6920 | 6218 | 6519 | 4217 | 1850 | 1322 | 1720 | 1854 | 6597 | 6575 | 6440 | 6606 | 1482 | 1503 | 1460 | 1507 | 4021,48 | 6554 | 1488 | 0,227 | -2,139 | -61,095 | 0 | 0 | 1,00E-08 | 0,0024996 | 0,0007919 | 0,0007919 | TRUE | 0,0019572 | No |
| orf19.4828 | 12077 | 10530 | 11767 | 6704 | 3194 | 2135 | 3009 | 3276 | 11514 | 11134 | 11625 | 10502 | 2559 | 2428 | 2555 | 2664 | 6872,55 | 11194 | 2552 | 0,228 | -2,133 | -46,104 | 0 | 0 | 0,0014202 | 0,0023302 | 0,0018303 | 0,0018303 | TRUE | 0,0207222 | No |
| orf19.2481 | 541 | 462 | 499 | 379 | 140 | 96 | 145 | 163 | 516 | 489 | 493 | 594 | 112 | 109 | 123 | 133 | 320,99 | 523 | 119 | 0,23 | -2,121 | -20,17 | 1,81E-90 | 1,09E-89 | 0,0041353 | 0,0072063 | 0,0058276 | 0,0058276 | TRUE | 0,0866873 | No |
| orf19.635 | 10 | 7 | 11 | 13 | 2 | 4 | 4 | 2 | 10 | 7 | 11 | 20 | 2 | 5 | 3 | 2 | 7,42 | 12 | 3 | 0,23 | -2,121 | -3,402 | 0,0006686 | 0,0009128 | 1,00E-08 | 0,2234438 | 0,16039 | 0,16039 | TRUE | 0,5619064 | No |
| orf19.3278 | 26585 | 22972 | 24811 | 16191 | 7067 | 5020 | 6596 | 7332 | 25346 | 24290 | 24511 | 25364 | 5663 | 5708 | 5600 | 5962 | 15305,41 | 24878 | 5733 | 0,23 | -2,117 | -62,175 | 0 | 0 | 0,0004919 | 0,0021986 | 0,0010131 | 0,0010131 | TRUE | 0,0089476 | No |
| orf19.4035 | 9465 | 7588 | 8482 | 5375 | 2572 | 1534 | 2175 | 2656 | 9024 | 8023 | 8379 | 8420 | 2061 | 1744 | 1847 | 2160 | 5207,27 | 8462 | 1953 | 0,231 | -2,114 | -31,245 | 2,61E-214 | 3,80E-213 | 0,0055548 | 0,0024066 | 0,0041017 | 0,0041017 | TRUE | 0,0612899 | No |
| orf19.6121 | 8452 | 7340 | 8294 | 5299 | 2296 | 1658 | 2151 | 2358 | 8058 | 7761 | 8194 | 8301 | 1840 | 1885 | 1826 | 1917 | 4972,82 | 8078 | 1867 | 0,231 | -2,113 | -54,554 | 0 | 0 | 0,0004982 | 0,0024215 | 0,0011319 | 0,0011319 | TRUE | 0,0085252 | No |
| orf19.3439 | 3312 | 3561 | 3734 | 2208 | 1048 | 770 | 1040 | 833 | 3158 | 3765 | 3689 | 3459 | 840 | 876 | 883 | 677 | 2168,29 | 3518 | 819 | 0,233 | -2,104 | -24,601 | 1,23E-133 | 1,11E-132 | 0,0089177 | 0,0028485 | 0,0063208 | 0,0063208 | TRUE | 0,1642667 | No |
| orf19.5058 | 21958 | 19774 | 21530 | 12778 | 5956 | 4276 | 5829 | 5928 | 20935 | 20909 | 21270 | 20017 | 4772 | 4862 | 4949 | 4820 | 12816,69 | 20783 | 4851 | 0,233 | -2,1 | -64,623 | 0 | 0 | 0,0003896 | 0,0022194 | 0,0008945 | 0,0008945 | TRUE | 0,0073532 | No |
| orf19.2020 | 11044 | 9528 | 10493 | 6820 | 3061 | 2158 | 2578 | 3240 | 10529 | 10075 | 10366 | 10684 | 2453 | 2454 | 2189 | 2634 | 6422,95 | 10414 | 2432 | 0,234 | -2,098 | -38,461 | 0 | 0 | 0,0028219 | 0,0023469 | 0,0026195 | 0,0026195 | TRUE | 0,0559419 | No |
| orf19.6601 | 11864 | 11490 | 11880 | 6829 | 3510 | 2308 | 3321 | 3112 | 11311 | 12149 | 11736 | 10698 | 2812 | 2624 | 2820 | 2530 | 7085,16 | 11474 | 2696 | 0,235 | -2,089 | -38,967 | 0 | 0 | 0,0025618 | 0,002323 | 0,002546 | 0,002546 | TRUE | 0,0249149 | No |
| orf19.335 | 1305 | 1536 | 1503 | 999 | 473 | 380 | 367 | 334 | 1244 | 1624 | 1485 | 1565 | 379 | 432 | 312 | 272 | 914,05 | 1480 | 349 | 0,235 | -2,089 | -16,623 | 4,72E-62 | 2,04E-61 | 0,0248539 | 0,0038876 | 0,0135054 | 0,0135054 | TRUE | 0,2842496 | No |
| orf19.4646 | 315 | 251 | 295 | 260 | 96 | 73 | 78 | 87 | 300 | 265 | 291 | 407 | 77 | 83 | 66 | 71 | 195,17 | 316 | 74 | 0,235 | -2,087 | -13,682 | 1,30E-42 | 4,30E-42 | 0,0189359 | 0,010504 | 0,0145239 | 0,0145239 | TRUE | 0,3861615 | No |
| orf19.4593 | 10418 | 9199 | 10176 | 6298 | 2923 | 1983 | 2752 | 2977 | 9932 | 9727 | 10053 | 9866 | 2342 | 2255 | 2336 | 2421 | 6116,53 | 9894 | 2338 | 0,237 | -2,08 | -60,074 | 0 | 0 | 0,0001611 | 0,0023597 | 0,0009034 | 0,0009034 | TRUE | 0,0067252 | No |
| orf19.897 | 2370 | 1987 | 2142 | 1511 | 654 | 444 | 659 | 618 | 2260 | 2101 | 2116 | 2367 | 524 | 505 | 559 | 502 | 1366,82 | 2211 | 522 | 0,237 | -2,077 | -33,48 | 9,57E-246 | 1,59E-244 | 0,0020243 | 0,0032925 | 0,0025907 | 0,0025907 | TRUE | 0,0265818 | No |
| orf19.2296 | 101341 | 82301 | 92018 | 62502 | 27901 | 19967 | 24885 | 27399 | 96618 | 87023 | 90905 | 97913 | 22356 | 22704 | 21127 | 22278 | 57615,52 | 93115 | 22116 | 0,238 | -2,074 | -44,914 | 0 | 0 | 0,001928 | 0,0021198 | 0,0020223 | 0,0020223 | TRUE | 0,0237669 | No |
| orf19.1539 | 277 | 318 | 302 | 212 | 103 | 62 | 87 | 79 | 264 | 336 | 298 | 332 | 83 | 70 | 74 | 64 | 190,24 | 308 | 73 | 0,238 | -2,072 | -15,612 | 6,01E-55 | 2,37E-54 | 0,0069775 | 0,010722 | 0,0089967 | 0,0089967 | TRUE | 0,1070689 | No |
| orf19.1800 | 10424 | 9833 | 10849 | 6225 | 3056 | 2214 | 2925 | 2801 | 9938 | 10397 | 10718 | 9752 | 2449 | 2518 | 2483 | 2277 | 6316,49 | 10201 | 2432 | 0,238 | -2,07 | -43,657 | 0 | 0 | 0,0015448 | 0,0023512 | 0,0019197 | 0,0019197 | TRUE | 0,0223873 | No |
| orf19.5524 | 49 | 58 | 56 | 37 | 18 | 9 | 14 | 19 | 47 | 61 | 55 | 58 | 14 | 10 | 12 | 15 | 34,17 | 55 | 13 | 0,239 | -2,066 | -7,568 | 3,78E-14 | 7,44E-14 | 1,00E-08 | 0,0501493 | 0,0277555 | 0,0277555 | TRUE | 0,1008634 | No |
| orf19.1826 | 20168 | 20208 | 21038 | 13363 | 6125 | 4798 | 5447 | 5778 | 19228 | 21367 | 20784 | 20934 | 4908 | 5456 | 4624 | 4698 | 12749,86 | 20578 | 4922 | 0,239 | -2,065 | -35,728 | 1,44E-279 | 2,89E-278 | 0,0037771 | 0,0022201 | 0,0030896 | 0,0030896 | TRUE | 0,0641796 | No |
| orf19.6350 | 216 | 165 | 184 | 138 | 63 | 42 | 57 | 49 | 206 | 174 | 182 | 216 | 50 | 48 | 48 | 40 | 120,6 | 194 | 46 | 0,24 | -2,058 | -13,443 | 3,41E-41 | 1,10E-40 | 1,00E-08 | 0,0157053 | 0,0100862 | 0,0100862 | TRUE | 0,088453 | No |
| orf19.3444 | 509 | 349 | 373 | 289 | 112 | 89 | 130 | 126 | 485 | 369 | 368 | 453 | 90 | 101 | 110 | 102 | 259,91 | 419 | 101 | 0,241 | -2,052 | -16,009 | 1,10E-57 | 4,49E-57 | 0,0116316 | 0,0084085 | 0,010021 | 0,010021 | TRUE | 0,1275693 | No |
| orf19.3084 | 2281 | 1752 | 1942 | 1114 | 592 | 342 | 588 | 604 | 2175 | 1853 | 1919 | 1745 | 474 | 389 | 499 | 491 | 1193,05 | 1923 | 463 | 0,241 | -2,05 | -23,286 | 6,13E-120 | 4,88E-119 | 0,0087156 | 0,0034675 | 0,0061887 | 0,0061887 | TRUE | 0,1262603 | No |
| orf19.554 | 652 | 580 | 610 | 408 | 203 | 143 | 158 | 173 | 622 | 613 | 603 | 639 | 163 | 163 | 134 | 141 | 384,59 | 619 | 150 | 0,242 | -2,048 | -23,21 | 3,58E-119 | 2,83E-118 | 1,00E-08 | 0,0063605 | 0,0036043 | 0,0036043 | TRUE | 0,0538823 | No |
| orf19.1445 | 85 | 74 | 78 | 44 | 36 | 10 | 19 | 20 | 81 | 78 | 77 | 69 | 29 | 11 | 16 | 16 | 47,23 | 76 | 18 | 0,242 | -2,047 | -8,118 | 4,75E-16 | 9,78E-16 | 0,0021636 | 0,0368517 | 0,0298188 | 0,0298188 | TRUE | 0,9185599 | No |
| orf19.2274 | 49 | 53 | 57 | 36 | 18 | 13 | 15 | 13 | 47 | 56 | 56 | 56 | 14 | 15 | 13 | 11 | 33,5 | 54 | 13 | 0,243 | -2,042 | -7,514 | 5,74E-14 | 1,12E-13 | 1,00E-08 | 0,0511081 | 0,0265484 | 0,0265484 | TRUE | 0,0863104 | No |
| orf19.4347 | 8411 | 7558 | 8546 | 5481 | 2525 | 1772 | 2349 | 2500 | 8019 | 7992 | 8443 | 8586 | 2023 | 2015 | 1994 | 2033 | 5138,09 | 8260 | 2016 | 0,244 | -2,034 | -51,739 | 0 | 0 | 0,0006342 | 0,0024108 | 0,0011939 | 0,0011939 | TRUE | 0,008309 | No |
| orf19.6274 | 201 | 164 | 156 | 131 | 53 | 44 | 55 | 47 | 192 | 173 | 154 | 205 | 42 | 50 | 47 | 38 | 112,72 | 181 | 44 | 0,244 | -2,033 | -12,564 | 3,35E-36 | 9,96E-36 | 1,00E-08 | 0,0166571 | 0,0119504 | 0,0119504 | TRUE | 0,1098978 | No |
| orf19.5992 | 3279 | 2860 | 3159 | 2280 | 940 | 705 | 929 | 982 | 3126 | 3024 | 3121 | 3572 | 753 | 802 | 789 | 798 | 1998,1 | 3211 | 786 | 0,245 | -2,03 | -32,904 | 1,91E-237 | 3,04E-236 | 0,0029103 | 0,002913 | 0,0029103 | 0,0029103 | TRUE | 0,0667973 | No |
| orf19.7022 | 1326 | 1250 | 1287 | 738 | 378 | 289 | 371 | 354 | 1264 | 1322 | 1271 | 1156 | 303 | 329 | 315 | 288 | 780,97 | 1253 | 309 | 0,245 | -2,029 | -28,984 | 1,05E-184 | 1,31E-183 | 0,0014075 | 0,0041937 | 0,0028143 | 0,0028143 | TRUE | 0,0294213 | No |
| orf19.6349 | 1797 | 2101 | 2138 | 1150 | 633 | 468 | 574 | 494 | 1713 | 2222 | 2112 | 1802 | 507 | 532 | 487 | 402 | 1222,1 | 1962 | 482 | 0,245 | -2,028 | -19,971 | 9,77E-89 | 5,77E-88 | 0,0128706 | 0,0034348 | 0,0086871 | 0,0086871 | TRUE | 0,1184084 | No |
| orf19.3906 | 5 | 9 | 15 | 9 | 7 | 2 | 1 | 2 | 5 | 10 | 15 | 14 | 6 | 2 | 1 | 2 | 6,69 | 11 | 3 | 0,246 | -2,023 | -2,984 | 0,0028474 | 0,0037355 | 0,1754657 | 0,2473384 | 0,2196808 | 0,2196808 | TRUE | 0,7045511 | No |
| orf19.1513 | 6521 | 6281 | 6545 | 3767 | 1980 | 1390 | 1749 | 1941 | 6217 | 6641 | 6466 | 5901 | 1587 | 1581 | 1485 | 1578 | 3931,96 | 6306 | 1558 | 0,247 | -2,019 | -41,203 | 0 | 0 | 0,0015866 | 0,0025089 | 0,0019289 | 0,0019289 | TRUE | 0,0220055 | No |
| orf19.173 | 24307 | 22932 | 25099 | 15373 | 7517 | 5153 | 7046 | 7315 | 23174 | 24248 | 24795 | 24083 | 6023 | 5859 | 5982 | 5948 | 15014,03 | 24075 | 5953 | 0,247 | -2,016 | -62,215 | 0 | 0 | 0,0004453 | 0,0022006 | 0,0009095 | 0,0009095 | TRUE | 0,0078679 | No |
| orf19.3538 | 5906 | 5193 | 5902 | 3730 | 1720 | 1213 | 1726 | 1757 | 5631 | 5491 | 5831 | 5843 | 1378 | 1379 | 1465 | 1429 | 3555,87 | 5699 | 1413 | 0,248 | -2,011 | -47,838 | 0 | 0 | 0,000563 | 0,002553 | 0,0012822 | 0,0012822 | TRUE | 0,0075493 | No |
| orf19.3770 | 7402 | 7204 | 7481 | 4199 | 2272 | 1565 | 2062 | 2179 | 7057 | 7617 | 7391 | 6578 | 1820 | 1780 | 1751 | 1772 | 4470,65 | 7161 | 1781 | 0,249 | -2,009 | -39,078 | 0 | 0 | 0,0020622 | 0,0024585 | 0,0022074 | 0,0022074 | TRUE | 0,0355954 | No |
| orf19.409 | 53612 | 43511 | 47000 | 30771 | 14642 | 10263 | 13905 | 15442 | 51113 | 46007 | 46432 | 48204 | 11732 | 11670 | 11805 | 12556 | 29939,98 | 47939 | 11941 | 0,249 | -2,005 | -45,022 | 0 | 0 | 0,0017017 | 0,0021461 | 0,0018565 | 0,0018565 | TRUE | 0,0244233 | No |
| orf19.729.1 | 1528 | 1315 | 1444 | 1011 | 442 | 327 | 434 | 447 | 1457 | 1390 | 1427 | 1584 | 354 | 372 | 368 | 363 | 914,43 | 1464 | 364 | 0,249 | -2,005 | -31,098 | 2,59E-212 | 3,74E-211 | 1,00E-08 | 0,0038868 | 0,0023842 | 0,0023842 | TRUE | 0,0328894 | No |
| orf19.699 | 805 | 1362 | 1277 | 641 | 395 | 319 | 324 | 202 | 767 | 1440 | 1262 | 1004 | 317 | 363 | 275 | 164 | 698,99 | 1118 | 280 | 0,25 | -2,001 | -6,559 | 5,43E-11 | 9,77E-11 | 0,0873398 | 0,0044403 | 0,039269 | 0,0873398 | TRUE | 0,1784166 | No |
| orf19.3120 | 40 | 29 | 41 | 26 | 10 | 4 | 9 | 20 | 38 | 31 | 41 | 41 | 8 | 5 | 8 | 16 | 23,31 | 38 | 9 | 0,25 | -2 | -5,664 | 1,48E-08 | 2,45E-08 | 0,0171177 | 0,0725222 | 0,0572996 | 0,0572996 | TRUE | 1,0450777 | No |
| orf19.1979 | 22144 | 20492 | 21615 | 12990 | 6723 | 4619 | 6204 | 6462 | 21112 | 21668 | 21354 | 20350 | 5387 | 5252 | 5267 | 5254 | 13205,41 | 21121 | 5290 | 0,25 | -1,997 | -62,28 | 0 | 0 | 0,0003802 | 0,0022156 | 0,0008767 | 0,0008767 | TRUE | 0,0072242 | No |
| orf19.4698 | 2517 | 2627 | 2843 | 1456 | 851 | 575 | 780 | 709 | 2400 | 2778 | 2809 | 2281 | 682 | 654 | 662 | 576 | 1605,17 | 2567 | 644 | 0,25 | -1,997 | -24,427 | 8,87E-132 | 7,87E-131 | 0,0075529 | 0,0031142 | 0,0055076 | 0,0055076 | TRUE | 0,0657569 | No |
| orf19.3368 | 319 | 333 | 307 | 241 | 107 | 77 | 107 | 87 | 304 | 352 | 303 | 378 | 86 | 88 | 91 | 71 | 208,99 | 334 | 84 | 0,251 | -1,994 | -15,716 | 1,17E-55 | 4,64E-55 | 0,0065973 | 0,0099476 | 0,0084495 | 0,0084495 | TRUE | 0,1133744 | No |
| orf19.6640 | 9078 | 8090 | 8555 | 5240 | 2618 | 1817 | 2553 | 2677 | 8655 | 8554 | 8452 | 8209 | 2098 | 2066 | 2167 | 2177 | 5297,16 | 8468 | 2127 | 0,251 | -1,993 | -54,62 | 0 | 0 | 0,0002944 | 0,0024012 | 0,0010021 | 0,0010021 | TRUE | 0,0049354 | No |
| orf19.7017 | 86 | 70 | 92 | 52 | 26 | 15 | 29 | 24 | 82 | 74 | 91 | 81 | 21 | 17 | 25 | 20 | 51,3 | 82 | 21 | 0,252 | -1,991 | -9,038 | 1,60E-19 | 3,56E-19 | 1,00E-08 | 0,0340987 | 0,0183753 | 0,0183753 | TRUE | 0,1053309 | No |
| orf19.6391 | 161 | 143 | 158 | 135 | 51 | 26 | 53 | 62 | 153 | 151 | 156 | 211 | 41 | 30 | 45 | 50 | 104,76 | 168 | 42 | 0,252 | -1,99 | -10,9 | 1,15E-27 | 3,00E-27 | 0,0185108 | 0,0177637 | 0,0180404 | 0,0180404 | TRUE | 0,2757519 | No |
| orf19.4784 | 1763 | 1507 | 1579 | 1077 | 506 | 384 | 444 | 528 | 1681 | 1593 | 1560 | 1687 | 405 | 437 | 377 | 429 | 1021,22 | 1630 | 412 | 0,252 | -1,986 | -31,368 | 5,58E-216 | 8,17E-215 | 0,0011317 | 0,0036991 | 0,0024214 | 0,0024214 | TRUE | 0,0382155 | No |
| orf19.668 | 978 | 964 | 976 | 762 | 340 | 203 | 299 | 341 | 932 | 1019 | 964 | 1194 | 272 | 231 | 254 | 277 | 643 | 1027 | 258 | 0,253 | -1,981 | -21,122 | 4,95E-99 | 3,21E-98 | 0,0076921 | 0,0046448 | 0,0061766 | 0,0061766 | TRUE | 0,1335913 | No |
| orf19.3923 | 420 | 374 | 371 | 297 | 127 | 74 | 127 | 141 | 400 | 395 | 367 | 465 | 102 | 84 | 108 | 115 | 254,5 | 407 | 102 | 0,254 | -1,978 | -16,799 | 2,49E-63 | 1,09E-62 | 0,0062945 | 0,0085427 | 0,0075995 | 0,0075995 | TRUE | 0,1153295 | No |
| orf19.1920 | 32 | 12 | 18 | 10 | 4 | 6 | 9 | 3 | 31 | 13 | 18 | 16 | 3 | 7 | 8 | 2 | 12,09 | 20 | 5 | 0,254 | -1,976 | -3,918 | 8,93E-05 | 0,0001268 | 0,1073056 | 0,1378521 | 0,1251259 | 0,1251259 | TRUE | 0,8986598 | No |
| orf19.4645 | 5027 | 5704 | 5826 | 3310 | 1814 | 1353 | 1564 | 1518 | 4793 | 6031 | 5756 | 5185 | 1454 | 1538 | 1328 | 1234 | 3414,86 | 5441 | 1388 | 0,255 | -1,972 | -23,51 | 3,19E-122 | 2,60E-121 | 0,0093338 | 0,0025721 | 0,0063302 | 0,0063302 | TRUE | 0,0793863 | No |
| orf19.6544 | 15501 | 13228 | 14532 | 8970 | 4502 | 3140 | 4265 | 4677 | 14779 | 13987 | 14356 | 14052 | 3607 | 3570 | 3621 | 3803 | 8971,91 | 14294 | 3650 | 0,255 | -1,969 | -54,44 | 0 | 0 | 0,0004948 | 0,0022743 | 0,0010945 | 0,0010945 | TRUE | 0,0097396 | No |
| orf19.851 | 3812 | 3786 | 4011 | 2491 | 1177 | 943 | 1157 | 1206 | 3634 | 4003 | 3962 | 3902 | 943 | 1072 | 982 | 981 | 2435,07 | 3875 | 994 | 0,256 | -1,965 | -36,644 | 5,80E-294 | 1,26E-292 | 0,0015981 | 0,0027656 | 0,0021658 | 0,0021658 | TRUE | 0,0307124 | No |
| orf19.7381 | 10097 | 10064 | 10954 | 6822 | 3231 | 2543 | 2989 | 3304 | 9626 | 10641 | 10822 | 10687 | 2589 | 2892 | 2538 | 2686 | 6560,12 | 10444 | 2676 | 0,256 | -1,965 | -35,94 | 7,32E-283 | 1,50E-281 | 0,0027824 | 0,0023416 | 0,002651 | 0,002651 | TRUE | 0,0346253 | No |
| orf19.551 | 18926 | 17083 | 18210 | 11867 | 5807 | 4175 | 5383 | 5783 | 18044 | 18063 | 17990 | 18590 | 4653 | 4747 | 4570 | 4702 | 11419,96 | 18172 | 4668 | 0,257 | -1,961 | -67,016 | 0 | 0 | 0,0001211 | 0,0022351 | 0,0006954 | 0,0006954 | TRUE | 0,0028535 | No |
| orf19.7071 | 11 | 10 | 5 | 8 | 3 | 3 | 3 | 2 | 10 | 11 | 5 | 13 | 2 | 3 | 3 | 2 | 6,07 | 10 | 2 | 0,258 | -1,956 | -3,04 | 0,002367 | 0,0031238 | 1,00E-08 | 0,2728021 | 0,1574226 | 0,1574226 | TRUE | 0,386521 | No |
| orf19.1481 | 7286 | 6788 | 7489 | 4549 | 2260 | 1616 | 2218 | 2292 | 6946 | 7177 | 7398 | 7126 | 1811 | 1838 | 1883 | 1864 | 4505,45 | 7162 | 1849 | 0,258 | -1,954 | -52,218 | 0 | 0 | 0,0003684 | 0,0024557 | 0,0010243 | 0,0010243 | TRUE | 0,0061864 | No |
| orf19.3713 | 4025 | 5711 | 4514 | 3380 | 1737 | 1398 | 1223 | 1290 | 3837 | 6039 | 4459 | 5295 | 1392 | 1590 | 1038 | 1049 | 3087,39 | 4908 | 1267 | 0,258 | -1,954 | -9,367 | 7,45E-21 | 1,70E-20 | 0,0413329 | 0,0026231 | 0,0198214 | 0,0413329 | TRUE | 0,1164692 | No |
| orf19.3712 | 15372 | 16687 | 16874 | 9718 | 5409 | 3987 | 4821 | 4476 | 14656 | 17644 | 16670 | 15224 | 4334 | 4534 | 4093 | 3639 | 10099,21 | 16048 | 4150 | 0,258 | -1,952 | -25,815 | 5,96E-147 | 5,91E-146 | 0,0076976 | 0,0022539 | 0,0053487 | 0,0053487 | TRUE | 0,0838405 | No |
| orf19.5551 | 6529 | 5747 | 6293 | 3782 | 1975 | 1298 | 1894 | 2027 | 6225 | 6077 | 6217 | 5925 | 1583 | 1476 | 1608 | 1648 | 3844,7 | 6111 | 1579 | 0,259 | -1,951 | -45,036 | 0 | 0 | 0,0008235 | 0,0025183 | 0,0014278 | 0,0014278 | TRUE | 0,0213758 | No |

| | | | | | | | | | | | | | | | | | | | | | | | | | | | | | | | |
|---|---|---|---|---|---|---|---|---|---|---|---|---|---|---|---|---|---|---|---|---|---|---|---|---|---|---|---|---|---|---|---|
| orf19.7436 | 9072 | 8106 | 9347 | 6214 | 2820 | 2208 | 2635 | 3033 | 8649 | 8571 | 9234 | 9735 | 2260 | 2511 | 2237 | 2466 | 5707,78 | 9047 | 2368 | 0,262 | -1,934 | -33,671 | 1,53E-248 | 2,57E-247 | 0,0032115 | 0,0023789 | 0,002917 | 0,002917 | TRUE | 0,0314567 | No |
| orf19.6690 | 2460 | 2267 | 2436 | 1374 | 747 | 577 | 709 | 727 | 2345 | 2397 | 2407 | 2152 | 599 | 656 | 602 | 591 | 1468,64 | 2325 | 612 | 0,262 | -1,932 | -33,294 | 4,74E-243 | 7,84E-242 | 0,0014733 | 0,0032093 | 0,0022607 | 0,0022607 | TRUE | 0,0281699 | No |
| orf19.4789 | 62 | 60 | 76 | 50 | 25 | 12 | 19 | 26 | 59 | 63 | 75 | 78 | 20 | 14 | 16 | 21 | 43,36 | 69 | 18 | 0,262 | -1,931 | -7,92 | 2,38E-15 | 4,82E-15 | 1,00E-08 | 0,0399548 | 0,0242663 | 0,0242663 | TRUE | 0,0948645 | No |
| orf19.203 | 3379 | 2955 | 3321 | 2300 | 1104 | 728 | 1043 | 1066 | 3222 | 3125 | 3281 | 3603 | 885 | 828 | 886 | 867 | 2086,83 | 3308 | 866 | 0,263 | -1,93 | -34,294 | 9,61E-258 | 1,71E-256 | 0,0019957 | 0,0028781 | 0,0023546 | 0,0023546 | TRUE | 0,0418834 | No |
| orf19.2030 | 11158 | 8570 | 9276 | 6447 | 3251 | 2143 | 2978 | 3303 | 10638 | 9062 | 9164 | 10100 | 2605 | 2437 | 2528 | 2686 | 6152,34 | 9741 | 2564 | 0,263 | -1,925 | -32,129 | 1,72E-226 | 2,62E-225 | 0,0035667 | 0,0023582 | 0,0032154 | 0,0032154 | TRUE | 0,0475832 | No |
| orf19.215 | 10232 | 8792 | 9481 | 6411 | 3066 | 2160 | 3099 | 3181 | 9755 | 9296 | 9366 | 10043 | 2457 | 2456 | 2631 | 2586 | 6073,92 | 9615 | 2532 | 0,264 | -1,924 | -45,116 | 0 | 0 | 0,0009566 | 0,0023616 | 0,0015109 | 0,0015109 | TRUE | 0,0106563 | No |
| orf19.3932 | 11765 | 9706 | 10734 | 6896 | 3459 | 2430 | 3287 | 3655 | 11217 | 10263 | 10604 | 10803 | 2772 | 2763 | 2791 | 2972 | 6772,99 | 10722 | 2824 | 0,264 | -1,924 | -45,03 | 0 | 0 | 0,0010066 | 0,0023337 | 0,0015426 | 0,0015426 | TRUE | 0,0152571 | No |
| orf19.1580 | 293 | 308 | 303 | 201 | 107 | 74 | 81 | 102 | 279 | 326 | 299 | 315 | 86 | 84 | 69 | 83 | 192,6 | 305 | 80 | 0,264 | -1,921 | -15,98 | 1,76E-57 | 7,14E-57 | 1,00E-08 | 0,0106162 | 0,0065174 | 0,0065174 | TRUE | 0,0949283 | No |
| orf19.2685 | 321 | 307 | 299 | 238 | 97 | 91 | 108 | 89 | 306 | 325 | 295 | 373 | 78 | 103 | 92 | 72 | 205,52 | 325 | 86 | 0,265 | -1,918 | -14,634 | 1,71E-48 | 6,18E-48 | 0,009034 | 0,0100804 | 0,0095548 | 0,0095548 | TRUE | 0,1477788 | No |
| orf19.5894 | 30 | 25 | 27 | 22 | 5 | 12 | 7 | 10 | 29 | 26 | 27 | 34 | 4 | 14 | 6 | 8 | 18,49 | 29 | 8 | 0,265 | -1,917 | -5,025 | 5,04E-07 | 7,89E-07 | 1,26E-08 | 0,0909034 | 0,0621689 | 0,0621689 | TRUE | 0,557821 | No |
| orf19.2108 | 4061 | 3383 | 3667 | 2304 | 1244 | 791 | 1127 | 1271 | 3872 | 3577 | 3623 | 3609 | 997 | 899 | 957 | 1033 | 2320,91 | 3670 | 972 | 0,265 | -1,915 | -36,087 | 3,59E-285 | 7,46E-284 | 0,0015609 | 0,0027987 | 0,0020923 | 0,0020923 | TRUE | 0,0274915 | No |
| orf19.6592 | 581 | 552 | 598 | 358 | 181 | 144 | 183 | 180 | 554 | 584 | 591 | 561 | 145 | 164 | 155 | 146 | 362,46 | 572 | 152 | 0,265 | -1,914 | -22,229 | 1,81E-109 | 1,31E-108 | 1,00E-08 | 0,0066212 | 0,0032214 | 0,0032214 | TRUE | 0,0214001 | No |
| orf19.4435 | 4059 | 3940 | 4004 | 2733 | 1338 | 1005 | 1243 | 1305 | 3870 | 4166 | 3956 | 4281 | 1072 | 1143 | 1055 | 1061 | 2575,51 | 4068 | 1083 | 0,266 | -1,91 | -38,008 | 0 | 0 | 0,0013216 | 0,0027288 | 0,0018744 | 0,0018744 | TRUE | 0,0153477 | No |
| orf19.4898 | 2257 | 1904 | 2154 | 1244 | 725 | 465 | 626 | 679 | 2152 | 2013 | 2128 | 1949 | 581 | 529 | 531 | 552 | 1304,38 | 2060 | 548 | 0,266 | -1,91 | -32,881 | 4,09E-237 | 6,48E-236 | 0,0012008 | 0,00335 | 0,0021559 | 0,0021559 | TRUE | 0,0193468 | No |
| orf19.1486 | 2013 | 2099 | 2271 | 1619 | 704 | 571 | 734 | 663 | 1919 | 2219 | 2244 | 2536 | 564 | 649 | 623 | 539 | 1411,75 | 2230 | 594 | 0,266 | -1,909 | -21,646 | 6,65E-104 | 4,49E-103 | 0,0086652 | 0,0032543 | 0,0064624 | 0,0064624 | TRUE | 0,1002104 | No |
| orf19.3281 | 829 | 745 | 752 | 543 | 255 | 183 | 251 | 268 | 790 | 788 | 743 | 851 | 204 | 208 | 213 | 218 | 501,88 | 793 | 211 | 0,267 | -1,906 | -24,54 | 5,49E-133 | 4,94E-132 | 1,00E-08 | 0,0053628 | 0,0029798 | 0,0029798 | TRUE | 0,0250606 | No |
| orf19.34 | 383 | 334 | 354 | 206 | 135 | 82 | 102 | 104 | 365 | 353 | 350 | 323 | 108 | 93 | 87 | 85 | 220,41 | 348 | 93 | 0,267 | -1,904 | -16,864 | 8,33E-64 | 3,69E-63 | 1,00E-08 | 0,0095405 | 0,0058684 | 0,0058684 | TRUE | 0,1308389 | No |
| orf19.4691 | 183 | 170 | 192 | 107 | 69 | 43 | 56 | 48 | 174 | 180 | 190 | 168 | 55 | 49 | 48 | 39 | 112,78 | 178 | 48 | 0,267 | -1,904 | -12,587 | 2,49E-36 | 7,44E-36 | 1,00E-08 | 0,0166491 | 0,0095349 | 0,0095349 | TRUE | 0,1402323 | No |
| orf19.3567 | 2209 | 2099 | 2057 | 1385 | 697 | 506 | 689 | 689 | 2106 | 2219 | 2032 | 2170 | 558 | 575 | 585 | 560 | 1350,79 | 2132 | 570 | 0,267 | -1,903 | -35,624 | 5,89E-278 | 1,18E-276 | 1,00E-08 | 0,0033068 | 0,0016963 | 0,0016963 | TRUE | 0,0122819 | No |
| orf19.3734 | 3928 | 3761 | 3996 | 2289 | 1322 | 855 | 1210 | 1264 | 3745 | 3977 | 3948 | 3586 | 1059 | 972 | 1027 | 1028 | 2417,72 | 3814 | 1022 | 0,268 | -1,9 | -37,255 | 8,87E-304 | 1,99E-302 | 0,0013456 | 0,0027704 | 0,0019132 | 0,0019132 | TRUE | 0,0186024 | No |
| orf19.5749 | 4462 | 4053 | 4311 | 2718 | 1422 | 1009 | 1324 | 1452 | 4254 | 4286 | 4259 | 4258 | 1139 | 1147 | 1124 | 1181 | 2705,97 | 4264 | 1148 | 0,269 | -1,893 | -49,522 | 0 | 0 | 1,00E-08 | 0,0026981 | 0,0008838 | 0,0008838 | TRUE | 0,0047328 | No |
| orf19.3969 | 6451 | 5738 | 6325 | 3964 | 1977 | 1497 | 1861 | 2197 | 6150 | 6067 | 6249 | 6210 | 1584 | 1702 | 1580 | 1786 | 3916,07 | 6169 | 1663 | 0,27 | -1,892 | -40,26 | 0 | 0 | 0,001373 | 0,0025106 | 0,0017598 | 0,0017598 | TRUE | 0,0315934 | No |
| orf19.7084 | 19530 | 16783 | 18836 | 11159 | 6019 | 4138 | 5987 | 6039 | 18620 | 17746 | 18608 | 17481 | 4823 | 4705 | 5083 | 4910 | 11497,04 | 18114 | 4880 | 0,269 | -1,892 | -48,439 | 0 | 0 | 0,0008839 | 0,0022341 | 0,0013425 | 0,0013425 | TRUE | 0,0094802 | No |
| orf19.7468 | 3510 | 3095 | 3249 | 2171 | 1117 | 818 | 1060 | 1065 | 3346 | 3273 | 3210 | 3401 | 895 | 930 | 900 | 866 | 2102,59 | 3308 | 898 | 0,271 | -1,882 | -41,969 | 0 | 0 | 1,00E-08 | 0,0028722 | 0,0012647 | 0,0012647 | TRUE | 0,006762 | No |
| orf19.1450 | 13 | 12 | 19 | 9 | 3 | 2 | 6 | 7 | 12 | 13 | 19 | 14 | 2 | 2 | 5 | 6 | 9,18 | 14 | 4 | 0,272 | -1,881 | -3,647 | 0,0002657 | 0,0003691 | 1,00E-08 | 0,1810078 | 0,1044161 | 0,1044161 | TRUE | 0,1965874 | No |
| orf19.2770.1 | 7699 | 6124 | 6291 | 4266 | 2385 | 1581 | 2014 | 2268 | 7340 | 6475 | 6215 | 6683 | 1911 | 1798 | 1710 | 1844 | 4247,01 | 6678 | 1816 | 0,272 | -1,879 | -31,5 | 8,56E-218 | 1,26E-216 | 0,0032091 | 0,0024779 | 0,0030858 | 0,0030858 | TRUE | 0,0550753 | No |
| orf19.2853 | 1457 | 1446 | 1509 | 1029 | 488 | 410 | 467 | 476 | 1389 | 1529 | 1491 | 1612 | 391 | 466 | 396 | 387 | 957,69 | 1505 | 410 | 0,272 | -1,879 | -25,444 | 8,17E-143 | 7,84E-142 | 0,0040089 | 0,0038057 | 0,0037727 | 0,0037727 | TRUE | 0,0888524 | No |
| orf19.7296 | 7941 | 6914 | 7200 | 4954 | 2395 | 1872 | 2314 | 2580 | 7571 | 7311 | 7113 | 7761 | 1919 | 2129 | 1965 | 2098 | 4733,15 | 7439 | 2028 | 0,272 | -1,876 | -38,948 | 0 | 0 | 0,0015679 | 0,0024382 | 0,0019316 | 0,0019316 | TRUE | 0,0158275 | No |
| orf19.3550 | 1176 | 1035 | 1117 | 694 | 362 | 251 | 359 | 395 | 1121 | 1094 | 1103 | 1087 | 290 | 285 | 305 | 321 | 700,96 | 1101 | 300 | 0,273 | -1,872 | -28,945 | 3,23E-184 | 4,00E-183 | 1,00E-08 | 0,0044336 | 0,0020349 | 0,0020349 | TRUE | 0,0254121 | No |
| orf19.6084 | 11794 | 10222 | 10864 | 7148 | 3780 | 2658 | 3475 | 3712 | 11244 | 10808 | 10733 | 11198 | 3029 | 3022 | 2950 | 3018 | 7000,35 | 10996 | 3005 | 0,273 | -1,871 | -56,122 | 0 | 0 | 0,0002884 | 0,0023258 | 0,0008678 | 0,0008678 | TRUE | 0,0031986 | No |
| orf19.4802 | 5052 | 4436 | 4921 | 2801 | 1675 | 1067 | 1553 | 1550 | 4817 | 4691 | 4861 | 4388 | 1342 | 1213 | 1318 | 1260 | 2986,33 | 4689 | 1283 | 0,274 | -1,869 | -36,852 | 2,67E-297 | 5,90E-296 | 0,0013993 | 0,0026411 | 0,0020017 | 0,0020017 | TRUE | 0,0218266 | No |
| orf19.2079 | 4523 | 3582 | 3630 | 2955 | 1397 | 1000 | 1191 | 1484 | 4312 | 3788 | 3586 | 4629 | 1119 | 1137 | 1011 | 1207 | 2598,65 | 4079 | 1118 | 0,274 | -1,866 | -22,254 | 1,03E-109 | 7,49E-109 | 0,0087187 | 0,0027231 | 0,0062121 | 0,0062121 | TRUE | 0,0986744 | No |
| orf19.2832 | 4022 | 3903 | 4244 | 2561 | 1332 | 994 | 1300 | 1407 | 3835 | 4127 | 4193 | 4012 | 1067 | 1130 | 1104 | 1144 | 2576,42 | 4042 | 1111 | 0,275 | -1,863 | -39,331 | 0 | 0 | 0,0009572 | 0,0027286 | 0,0016145 | 0,0016145 | TRUE | 0,0150889 | No |
| orf19.450 | 51 | 51 | 59 | 58 | 29 | 7 | 22 | 21 | 49 | 54 | 58 | 91 | 23 | 8 | 19 | 17 | 39,83 | 63 | 17 | 0,276 | -1,86 | -6,138 | 8,36E-10 | 1,44E-09 | 0,0686277 | 0,0433134 | 0,0530974 | 0,0530974 | TRUE | 0,6282744 | No |
| orf19.2049 | 681 | 658 | 623 | 377 | 251 | 147 | 195 | 210 | 649 | 696 | 615 | 591 | 201 | 167 | 166 | 171 | 406,96 | 638 | 176 | 0,276 | -1,855 | -19,951 | 1,48E-88 | 8,73E-88 | 0,0034472 | 0,0061259 | 0,0049067 | 0,0049067 | TRUE | 0,1047345 | No |
| orf19.6478 | 13153 | 13445 | 13848 | 7578 | 4637 | 3335 | 4077 | 4317 | 12540 | 14216 | 13681 | 11871 | 3715 | 3792 | 3461 | 3510 | 8348,42 | 13077 | 3620 | 0,277 | -1,854 | -30,281 | 2,03E-201 | 2,80E-200 | 0,0040497 | 0,002288 | 0,0034335 | 0,0034335 | TRUE | 0,0465959 | No |
| orf19.2730 | 2418 | 1715 | 2096 | 1483 | 736 | 494 | 707 | 740 | 2305 | 1813 | 2071 | 2323 | 590 | 562 | 600 | 602 | 1358,24 | 2128 | 588 | 0,277 | -1,853 | -22,901 | 4,50E-116 | 3,46E-115 | 0,0068323 | 0,0033001 | 0,0052599 | 0,0052599 | TRUE | 0,1202315 | No |
| orf19.6302 | 31 | 23 | 11 | 7 | 10 | 6 | 5 | 3 | 30 | 24 | 11 | 11 | 8 | 7 | 4 | 2 | 12,15 | 19 | 5 | 0,278 | -1,849 | -3,544 | 0,0003945 | 0,0005433 | 0,1640077 | 0,1371897 | 0,1488014 | 0,1488014 | TRUE | 0,1298465 | No |
| orf19.7060 | 1566 | 1322 | 1503 | 908 | 482 | 330 | 485 | 538 | 1493 | 1398 | 1485 | 1422 | 386 | 375 | 412 | 437 | 926,1 | 1450 | 402 | 0,278 | -1,845 | -29,026 | 3,09E-185 | 3,85E-184 | 0,0007833 | 0,0038642 | 0,0023916 | 0,0023916 | TRUE | 0,0397365 | No |
| orf19.4227 | 94 | 76 | 98 | 67 | 32 | 25 | 20 | 40 | 90 | 80 | 97 | 105 | 26 | 28 | 17 | 33 | 59,42 | 93 | 26 | 0,279 | -1,841 | -8,487 | 2,12E-17 | 4,49E-17 | 0,0057145 | 0,0297255 | 0,0219229 | 0,0219229 | TRUE | 0,374561 | No |
| orf19.7310 | 8335 | 7769 | 8074 | 5074 | 2764 | 2048 | 2599 | 2744 | 7947 | 8215 | 7976 | 7949 | 2215 | 2329 | 2207 | 2231 | 5133,43 | 8022 | 2246 | 0,28 | -1,838 | -53,014 | 0 | 0 | 0,0001257 | 0,0024111 | 0,0008857 | 0,0008857 | TRUE | 0,0070001 | No |
| orf19.2125 | 5095 | 3726 | 3907 | 3136 | 1529 | 1040 | 1496 | 1515 | 4858 | 3940 | 3860 | 4913 | 1225 | 1183 | 1270 | 1232 | 2809,93 | 4393 | 1228 | 0,28 | -1,838 | -22,275 | 6,44E-110 | 4,67E-109 | 0,00816 | 0,0026756 | 0,0060484 | 0,0060484 | TRUE | 0,0586742 | No |
| orf19.915 | 1236 | 1418 | 1373 | 946 | 486 | 407 | 411 | 428 | 1178 | 1499 | 1356 | 1482 | 389 | 463 | 349 | 348 | 883,16 | 1379 | 387 | 0,28 | -1,836 | -17,962 | 3,89E-72 | 1,93E-71 | 0,0132547 | 0,0039504 | 0,0084629 | 0,0084629 | TRUE | 0,1869429 | No |
| orf19.1430 | 76 | 69 | 93 | 53 | 27 | 25 | 28 | 21 | 72 | 73 | 92 | 83 | 22 | 28 | 24 | 17 | 51,4 | 80 | 23 | 0,28 | -1,835 | -8,191 | 2,60E-16 | 5,38E-16 | 1,00E-08 | 0,0340328 | 0,0214097 | 0,0214097 | TRUE | 0,1818148 | No |
| orf19.1464 | 5196 | 4681 | 5211 | 2968 | 1702 | 1212 | 1699 | 1657 | 4954 | 4950 | 5148 | 4650 | 1364 | 1378 | 1442 | 1347 | 3154,07 | 4926 | 1383 | 0,281 | -1,834 | -39,673 | 0 | 0 | 0,0010209 | 0,0026118 | 0,0016146 | 0,0016146 | TRUE | 0,0164669 | No |
| orf19.3675 | 7745 | 7553 | 7704 | 5487 | 2715 | 2177 | 2497 | 2594 | 7384 | 7986 | 7611 | 8596 | 2175 | 2475 | 2120 | 2109 | 5057,11 | 7894 | 2220 | 0,281 | -1,831 | -28,325 | 1,69E-176 | 1,98E-175 | 0,0048926 | 0,002416 | 0,0037405 | 0,0037405 | TRUE | 0,070904 | No |
| orf19.5636 | 23644 | 18043 | 19957 | 15914 | 7458 | 5365 | 6943 | 7741 | 22542 | 19078 | 19716 | 24930 | 5976 | 6100 | 5895 | 6294 | 13816,38 | 21566 | 6066 | 0,281 | -1,83 | -24,008 | 2,29E-127 | 1,95E-126 | 0,0081117 | 0,0022101 | 0,0054804 | 0,0054804 | TRUE | 0,1344342 | No |
| orf19.285 | 6130 | 5272 | 5735 | 3081 | 2044 | 1208 | 1884 | 1913 | 5844 | 5574 | 5666 | 4827 | 1638 | 1374 | 1600 | 1555 | 3509,66 | 5478 | 1542 | 0,282 | -1,829 | -25,7 | 1,17E-145 | 1,14E-144 | 0,0056725 | 0,0025591 | 0,0044697 | 0,0044697 | TRUE | 0,0766041 | No |
| orf19.2529.1 | 17392 | 12949 | 13002 | 7510 | 5219 | 2710 | 4750 | 5116 | 16581 | 13692 | 12845 | 11765 | 4182 | 3082 | 4033 | 4160 | 8792,34 | 13721 | 3864 | 0,282 | -1,828 | -12,514 | 6,29E-36 | 1,86E-35 | 0,0203478 | 0,002278 | 0,0112774 | 0,0203478 | TRUE | 0,2411228 | No |
| orf19.5614 | 7951 | 6518 | 7279 | 4557 | 2452 | 1745 | 2464 | 2582 | 7580 | 6892 | 7191 | 7139 | 1965 | 1984 | 2092 | 2099 | 4617,8 | 7200 | 2035 | 0,283 | -1,823 | -40,769 | 0 | 0 | 0,0011727 | 0,0024468 | 0,0016218 | 0,0016218 | TRUE | 0,0156686 | No |
| orf19.3651 | 20804 | 16377 | 17502 | 11579 | 6231 | 4519 | 5899 | 6674 | 19834 | 17317 | 17290 | 18139 | 4993 | 5139 | 5008 | 5427 | 11643,31 | 18145 | 5142 | 0,283 | -1,819 | -34,236 | 7,02E-257 | 1,23E-255 | 0,0027595 | 0,0022323 | 0,0025948 | 0,0025948 | TRUE | 0,0483607 | No |
| orf19.2519 | 522 | 446 | 466 | 328 | 195 | 106 | 154 | 172 | 498 | 472 | 460 | 514 | 156 | 121 | 131 | 140 | 311,35 | 486 | 137 | 0,284 | -1,816 | -18,548 | 8,51E-77 | 4,45E-76 | 0,0008898 | 0,0073647 | 0,0048254 | 0,0048254 | TRUE | 0,0971651 | No |
| orf19.5245 | 1781 | 1677 | 1871 | 1222 | 607 | 513 | 570 | 626 | 1698 | 1773 | 1848 | 1914 | 486 | 583 | 484 | 509 | 1162,07 | 1808 | 516 | 0,284 | -1,814 | -26,261 | 5,39E-152 | 5,48E-151 | 0,0032974 | 0,0035042 | 0,0034016 | 0,0034016 | TRUE | 0,0835505 | No |
| orf19.6905 | 4254 | 3622 | 3913 | 2791 | 1429 | 1012 | 1296 | 1462 | 4056 | 3830 | 3866 | 4372 | 1145 | 1151 | 1100 | 1189 | 2588,53 | 4031 | 1146 | 0,285 | -1,813 | -33,272 | 9,66E-243 | 1,59E-241 | 0,0020566 | 0,0027256 | 0,0023206 | 0,0023206 | TRUE | 0,037928 | No |
| orf19.1502 | 483 | 561 | 566 | 346 | 189 | 164 | 186 | 151 | 460 | 593 | 559 | 542 | 151 | 186 | 158 | 123 | 346,68 | 538 | 154 | 0,285 | -1,809 | -14,82 | 1,08E-49 | 3,98E-49 | 0,0142024 | 0,0068273 | 0,0103582 | 0,0103582 | TRUE | 0,209381 | No |
| orf19.301 | 2355 | 1843 | 2078 | 1646 | 760 | 555 | 732 | 805 | 2245 | 1949 | 2053 | 2579 | 609 | 631 | 621 | 655 | 1417,68 | 2206 | 629 | 0,286 | -1,808 | -21,669 | 4,04E-104 | 2,73E-103 | 0,0077645 | 0,0032494 | 0,0057215 | 0,0057215 | TRUE | 0,1508827 | No |
| orf19.7053 | 41900 | 40319 | 43666 | 27996 | 14764 | 11188 | 13785 | 15066 | 39947 | 42632 | 43138 | 43857 | 11830 | 12722 | 11703 | 12250 | 27259,96 | 42394 | 12126 | 0,286 | -1,806 | -42,247 | 0 | 0 | 0,0014611 | 0,0021515 | 0,0017052 | 0,0017052 | TRUE | 0,0185243 | No |
| orf19.6689 | 4526 | 4341 | 4373 | 2606 | 1553 | 1078 | 1495 | 1507 | 4315 | 4590 | 4320 | 4082 | 1244 | 1226 | 1269 | 1225 | 2784,05 | 4327 | 1241 | 0,287 | -1,802 | -38,054 | 0 | 0 | 0,0010754 | 0,002681 | 0,0016648 | 0,0016648 | TRUE | 0,0201556 | No |
| orf19.7042 | 1119 | 1105 | 1079 | 712 | 431 | 278 | 365 | 363 | 1067 | 1168 | 1066 | 1115 | 345 | 316 | 310 | 295 | 710,38 | 1104 | 316 | 0,287 | -1,801 | -25,478 | 3,46E-143 | 3,34E-142 | 0,0013395 | 0,0044026 | 0,002883 | 0,002883 | TRUE | 0,0464629 | No |
| orf19.4899 | 1082 | 863 | 936 | 660 | 357 | 228 | 303 | 387 | 1032 | 913 | 925 | 1034 | 286 | 259 | 257 | 315 | 627,49 | 976 | 279 | 0,287 | -1,799 | -21,495 | 1,72E-102 | 1,15E-101 | 0,0043924 | 0,0047079 | 0,0045561 | 0,0045561 | TRUE | 0,0839067 | No |
| orf19.6520 | 61 | 51 | 49 | 41 | 23 | 14 | 19 | 17 | 58 | 54 | 48 | 64 | 18 | 16 | 16 | 14 | 36,13 | 56 | 16 | 0,288 | -1,794 | -7,016 | 2,28E-12 | 4,27E-12 | 1,00E-08 | 0,0475386 | 0,0252226 | 0,0252226 | TRUE | 0,0738292 | No |
| orf19.2769 | 16305 | 13916 | 14030 | 8185 | 5413 | 3265 | 4971 | 5153 | 15545 | 14714 | 13860 | 12822 | 4337 | 3713 | 4220 | 4190 | 9175,27 | 14235 | 4115 | 0,289 | -1,79 | -26,719 | 2,81E-157 | 3,01E-156 | 0,0053931 | 0,0022702 | 0,0041645 | 0,0041645 | TRUE | 0,054206 | No |
| orf19.921 | 5304 | 5323 | 5801 | 3505 | 1964 | 1478 | 1878 | 1865 | 5057 | 5628 | 5731 | 5491 | 1574 | 1681 | 1594 | 1516 | 3533,99 | 5477 | 1591 | 0,29 | -1,784 | -33,719 | 3,09E-249 | 5,21E-248 | 0,0021496 | 0,0025559 | 0,0023053 | 0,0023053 | TRUE | 0,0331667 | No |
| orf19.1446 | 3728 | 3390 | 3747 | 2246 | 1257 | 983 | 1262 | 1221 | 3554 | 3584 | 3702 | 3518 | 1007 | 1118 | 1071 | 993 | 2318,51 | 3590 | 1047 | 0,291 | -1,781 | -36,749 | 1,19E-295 | 2,61E-294 | 0,0007392 | 0,0027995 | 0,0016711 | 0,0016711 | TRUE | 0,0226253 | No |
| orf19.3376 | 135 | 149 | 134 | 89 | 46 | 30 | 54 | 55 | 129 | 158 | 132 | 139 | 37 | 34 | 46 | 45 | 89,95 | 140 | 40 | 0,292 | -1,777 | -10,525 | 6,60E-26 | 1,66E-25 | 1,00E-08 | 0,0203449 | 0,0124487 | 0,0124487 | TRUE | 0,082592 | No |
| orf19.2458 | 21292 | 19620 | 21289 | 14171 | 7723 | 5642 | 7075 | 7482 | 20300 | 20746 | 21032 | 22200 | 6188 | 6415 | 6007 | 6084 | 13621,29 | 21070 | 6174 | 0,293 | -1,771 | -44,603 | 0 | 0 | 0,0010164 | 0,0022118 | 0,0014152 | 0,0014152 | TRUE | 0,01576 | No |
| orf19.4318 | 17510 | 16362 | 18446 | 11314 | 6289 | 4569 | 6091 | 6265 | 16694 | 17301 | 18223 | 17724 | 5039 | 5195 | 5171 | 5094 | 11305,17 | 17486 | 5125 | 0,293 | -1,771 | -47,881 | 0 | 0 | 0,0006172 | 0,0022365 | 0,0011939 | 0,0011939 | TRUE | 0,0114816 | No |
| orf19.7483 | 6166 | 6127 | 6502 | 3869 | 2238 | 1711 | 2106 | 2201 | 5879 | 6479 | 6423 | 6061 | 1793 | 1946 | 1788 | 1790 | 4019,74 | 6210 | 1829 | 0,294 | -1,765 | -36,025 | 3,34E-284 | 6,89E-283 | 0,0015085 | 0,0024997 | 0,0019706 | 0,0019706 | TRUE | 0,0210905 | No |
| orf19.2876 | 3371 | 2850 | 3096 | 1994 | 1119 | 772 | 1114 | 1145 | 3214 | 3014 | 3059 | 3124 | 897 | 878 | 946 | 931 | 2007,61 | 3103 | 913 | 0,295 | -1,763 | -37,908 | 0 | 0 | 0,000411 | 0,0029091 | 0,0014089 | 0,0014089 | TRUE | 0,0075278 | No |
| orf19.1477 | 1899 | 1910 | 1981 | 1264 | 784 | 528 | 666 | 615 | 1810 | 2020 | 1957 | 1980 | 628 | 600 | 565 | 500 | 1257,66 | 1942 | 573 | 0,295 | -1,76 | -24,375 | 3,15E-131 | 2,78E-130 | 0,0042475 | 0,0033968 | 0,0039386 | 0,0039386 | TRUE | 0,0907123 | No |

| | | | | | | | | | | | | | | | | | | | | | | | | | | | | | | | |
|---|---|---|---|---|---|---|---|---|---|---|---|---|---|---|---|---|---|---|---|---|---|---|---|---|---|---|---|---|---|---|---|
| orf19.932 | 6300 | 5670 | 6128 | 3893 | 2171 | 1610 | 2025 | 2281 | 6006 | 5995 | 6054 | 6099 | 1740 | 1831 | 1719 | 1855 | 3912,29 | 6038 | 1786 | 0,296 | -1,758 | -45,828 | 0 | 0 | 1,95E-05 | 0,002511 | 0,0010699 | 0,0010699 | TRUE | 0,0087914 | No |
| orf19.4722 | 2758 | 2324 | 2631 | 1528 | 935 | 626 | 879 | 952 | 2629 | 2457 | 2599 | 2394 | 749 | 712 | 746 | 774 | 1632,63 | 2520 | 745 | 0,296 | -1,757 | -33,202 | 1,00E-241 | 1,64E-240 | 0,0009956 | 0,003097 | 0,0018695 | 0,0018695 | TRUE | 0,0128257 | No |
| orf19.1334 | 783 | 731 | 882 | 484 | 292 | 223 | 258 | 282 | 747 | 773 | 871 | 758 | 234 | 254 | 219 | 229 | 510,61 | 787 | 234 | 0,296 | -1,757 | -21,194 | 1,09E-99 | 7,10E-99 | 0,0027252 | 0,0053069 | 0,0039767 | 0,0039767 | TRUE | 0,0634758 | No |
| orf19.6861 | 1176 | 1418 | 1325 | 793 | 507 | 376 | 433 | 412 | 1121 | 1499 | 1309 | 1242 | 406 | 428 | 368 | 335 | 838,52 | 1293 | 384 | 0,296 | -1,754 | -17,525 | 9,17E-69 | 4,32E-68 | 0,0118445 | 0,0040494 | 0,008021 | 0,008021 | TRUE | 0,1400754 | No |
| orf19.730 | 2085 | 1697 | 1934 | 1258 | 687 | 490 | 691 | 710 | 1988 | 1794 | 1911 | 1971 | 550 | 557 | 587 | 577 | 1241,89 | 1916 | 568 | 0,297 | -1,753 | -30,951 | 2,47E-210 | 3,53E-209 | 1,00E-08 | 0,0034134 | 0,0020026 | 0,0020026 | TRUE | 0,0222673 | No |
| orf19.6577 | 12358 | 11112 | 11530 | 7942 | 4371 | 3261 | 4015 | 4335 | 11782 | 11750 | 11391 | 12442 | 3502 | 3708 | 3409 | 3525 | 7688,44 | 11841 | 3536 | 0,299 | -1,744 | -41,7 | 0 | 0 | 0,0010665 | 0,0023048 | 0,0015055 | 0,0015055 | TRUE | 0,0139456 | No |
| orf19.2397.3 | 6566 | 6653 | 6954 | 3814 | 2508 | 1692 | 2386 | 2300 | 6260 | 7035 | 6870 | 5975 | 2010 | 1924 | 2026 | 1870 | 4246,1 | 6535 | 1958 | 0,299 | -1,74 | -29,221 | 1,06E-187 | 1,35E-186 | 0,003191 | 0,002478 | 0,00309 | 0,00309 | TRUE | 0,039657 | No |
| orf19.5172 | 13 | 7 | 8 | 4 | 4 | 0 | 3 | 5 | 12 | 7 | 8 | 6 | 3 | 0 | 3 | 4 | 5,47 | 8 | 2 | 0,3 | -1,737 | -2,608 | 0,0091196 | 0,0116176 | 1,00E-08 | 0,3021007 | 0,1898014 | 0,1898014 | TRUE | 0,3549311 | No |
| orf19.5902 | 5768 | 5316 | 5793 | 3986 | 2106 | 1552 | 2077 | 2125 | 5499 | 5621 | 5723 | 6244 | 1687 | 1765 | 1763 | 1728 | 3753,85 | 5772 | 1736 | 0,301 | -1,733 | -35,385 | 2,91E-274 | 5,60E-273 | 0,0016119 | 0,0025287 | 0,001947 | 0,001947 | TRUE | 0,0358074 | No |
| orf19.2989 | 4404 | 3676 | 4081 | 2654 | 1495 | 1048 | 1505 | 1512 | 4199 | 3887 | 4032 | 4158 | 1198 | 1192 | 1278 | 1229 | 2646,45 | 4069 | 1224 | 0,301 | -1,732 | -38,088 | 0 | 0 | 0,0007389 | 0,0027117 | 0,0014819 | 0,0014819 | TRUE | 0,0110899 | No |
| orf19.6798 | 27587 | 23611 | 26920 | 17883 | 9844 | 7003 | 9384 | 9937 | 26301 | 24966 | 26594 | 28015 | 7888 | 7963 | 7967 | 8080 | 17221,68 | 26469 | 7974 | 0,301 | -1,731 | -43,224 | 0 | 0 | 0,0010569 | 0,0021866 | 0,0014624 | 0,0014624 | TRUE | 0,0187113 | No |
| orf19.5266 | 64 | 45 | 47 | 22 | 25 | 11 | 12 | 18 | 61 | 48 | 46 | 34 | 20 | 13 | 10 | 15 | 30,86 | 47 | 14 | 0,302 | -1,729 | -5,769 | 7,96E-09 | 1,33E-08 | 0,0259839 | 0,0553009 | 0,0438188 | 0,0438188 | TRUE | 0,2053285 | No |
| orf19.6588 | 2563 | 2467 | 2531 | 1472 | 943 | 645 | 909 | 886 | 2444 | 2609 | 2500 | 2306 | 756 | 733 | 772 | 720 | 1604,95 | 2465 | 745 | 0,302 | -1,727 | -31,689 | 2,20E-220 | 3,27E-219 | 0,0012462 | 0,0031143 | 0,0020252 | 0,0020252 | TRUE | 0,0210059 | No |
| orf19.4317 | 2151 | 2085 | 2080 | 1282 | 836 | 603 | 679 | 723 | 2051 | 2205 | 2055 | 2008 | 670 | 686 | 576 | 588 | 1354,8 | 2080 | 630 | 0,302 | -1,727 | -26,046 | 1,50E-149 | 1,50E-148 | 0,0033927 | 0,0033032 | 0,0032389 | 0,0032389 | TRUE | 0,0382856 | No |
| orf19.6641 | 1179 | 1032 | 1030 | 733 | 383 | 309 | 398 | 412 | 1124 | 1091 | 1018 | 1148 | 307 | 351 | 338 | 335 | 714,03 | 1095 | 333 | 0,304 | -1,72 | -24,533 | 6,58E-133 | 5,90E-132 | 0,0014096 | 0,0043908 | 0,0028667 | 0,0028667 | TRUE | 0,0325124 | No |
| orf19.3571 | 255 | 224 | 222 | 180 | 91 | 75 | 89 | 80 | 243 | 237 | 219 | 282 | 73 | 85 | 76 | 65 | 160,01 | 245 | 75 | 0,304 | -1,717 | -12,865 | 7,08E-38 | 2,17E-37 | 0,0038521 | 0,0123526 | 0,0088168 | 0,0088168 | TRUE | 0,0952758 | No |
| orf19.2172 | 11777 | 11357 | 11125 | 6229 | 4273 | 3000 | 3837 | 4064 | 11228 | 12009 | 10990 | 9758 | 3424 | 3411 | 3258 | 3304 | 7172,79 | 10996 | 3349 | 0,304 | -1,716 | -28,609 | 5,21E-180 | 6,26E-179 | 0,0036346 | 0,0023202 | 0,0032704 | 0,0032704 | TRUE | 0,0693043 | No |
| orf19.2948 | 1707 | 1436 | 1615 | 1078 | 604 | 415 | 562 | 640 | 1627 | 1518 | 1595 | 1689 | 484 | 472 | 477 | 520 | 1047,93 | 1607 | 488 | 0,305 | -1,715 | -28,682 | 6,49E-181 | 7,83E-180 | 0,00088 | 0,0036581 | 0,0021732 | 0,0021732 | TRUE | 0,0234916 | No |
| orf19.7304 | 1088 | 1166 | 1251 | 719 | 458 | 318 | 423 | 400 | 1037 | 1233 | 1236 | 1126 | 367 | 362 | 359 | 325 | 755,67 | 1158 | 353 | 0,305 | -1,715 | -21,788 | 3,03E-105 | 2,07E-104 | 0,0041428 | 0,0042641 | 0,0041979 | 0,0041979 | TRUE | 0,0626175 | No |
| orf19.6555 | 523 | 572 | 524 | 338 | 205 | 148 | 196 | 193 | 499 | 605 | 518 | 529 | 164 | 168 | 166 | 157 | 350,81 | 538 | 164 | 0,305 | -1,715 | -18,146 | 1,39E-73 | 6,97E-73 | 1,00E-08 | 0,0067716 | 0,0048161 | 0,0048161 | TRUE | 0,0856488 | No |
| orf19.7184 | 2812 | 2509 | 2696 | 1952 | 1061 | 736 | 972 | 1051 | 2681 | 2653 | 2663 | 3058 | 850 | 837 | 825 | 855 | 1802,75 | 2764 | 842 | 0,305 | -1,712 | -29,39 | 7,29E-190 | 9,41E-189 | 0,0021978 | 0,0030021 | 0,0025253 | 0,0025253 | TRUE | 0,0590811 | No |
| orf19.5785 | 1876 | 1758 | 1884 | 1175 | 705 | 534 | 658 | 646 | 1789 | 1859 | 1861 | 1841 | 565 | 607 | 559 | 525 | 1200,67 | 1838 | 564 | 0,306 | -1,708 | -30,946 | 2,86E-210 | 4,08E-209 | 7,32E-05 | 0,0034588 | 0,0018274 | 0,0018274 | TRUE | 0,0276914 | No |
| orf19.7021 | 33380 | 30412 | 31617 | 19542 | 12110 | 8599 | 10805 | 12181 | 31824 | 32157 | 31235 | 30614 | 9703 | 9778 | 9173 | 9904 | 20548,52 | 31458 | 9640 | 0,306 | -1,707 | -48,145 | 0 | 0 | 0,0007001 | 0,0021712 | 0,0011426 | 0,0011426 | TRUE | 0,0130342 | No |
| orf19.6645 | 18657 | 15509 | 16704 | 11931 | 6630 | 4606 | 6203 | 6698 | 17787 | 16399 | 16502 | 18691 | 5312 | 5237 | 5266 | 5446 | 11330,13 | 17345 | 5315 | 0,307 | -1,706 | -35,196 | 2,31E-271 | 4,38E-270 | 0,0019882 | 0,0022362 | 0,0021394 | 0,0021394 | TRUE | 0,0330022 | No |
| orf19.7342 | 795 | 813 | 882 | 568 | 357 | 262 | 304 | 241 | 758 | 860 | 871 | 890 | 286 | 298 | 258 | 196 | 552,09 | 845 | 260 | 0,307 | -1,706 | -15,184 | 4,52E-52 | 1,72E-51 | 0,0144887 | 0,0050652 | 0,0097281 | 0,0097281 | TRUE | 0,3152641 | No |
| orf19.6245 | 14100 | 12871 | 13791 | 8222 | 5073 | 3598 | 4814 | 5195 | 13443 | 13609 | 13624 | 12880 | 4065 | 4091 | 4087 | 4224 | 8752,98 | 13389 | 4117 | 0,307 | -1,702 | -50,564 | 0 | 0 | 0,0003828 | 0,0022789 | 0,0009373 | 0,0009373 | TRUE | 0,0077509 | No |
| orf19.1777 | 11727 | 12362 | 12818 | 7314 | 4708 | 3540 | 4171 | 4368 | 11180 | 13071 | 12663 | 11458 | 3772 | 4025 | 3541 | 3552 | 7907,86 | 12093 | 3722 | 0,308 | -1,701 | -27,024 | 7,62E-161 | 8,33E-160 | 0,004515 | 0,0022989 | 0,0036371 | 0,0036371 | TRUE | 0,0362599 | No |
| orf19.6408 | 6033 | 5022 | 5788 | 3760 | 2098 | 1503 | 2182 | 2129 | 5752 | 5310 | 5718 | 5890 | 1681 | 1709 | 1853 | 1731 | 3705,49 | 5668 | 1744 | 0,308 | -1,7 | -34,664 | 2,80E-263 | 5,05E-262 | 0,001409 | 0,0025344 | 0,001954 | 0,001954 | TRUE | 0,0219988 | No |
| orf19.4630 | 11614 | 10251 | 10895 | 7145 | 4069 | 3110 | 3919 | 4175 | 11073 | 10839 | 10763 | 11193 | 3260 | 3536 | 3327 | 3395 | 7173,34 | 10967 | 3380 | 0,308 | -1,699 | -46,674 | 0 | 0 | 0,0005177 | 0,0023202 | 0,0010888 | 0,0010888 | TRUE | 0,0112743 | No |
| orf19.5960 | 38765 | 35479 | 38516 | 28931 | 15264 | 11634 | 13766 | 14285 | 36958 | 37515 | 38050 | 45322 | 12231 | 13229 | 11687 | 11615 | 25825,84 | 39461 | 12190 | 0,309 | -1,695 | -24,315 | 1,34E-130 | 1,17E-129 | 0,0067605 | 0,0021549 | 0,0046162 | 0,0046162 | TRUE | 0,1221676 | No |
| orf19.2947 | 4150 | 3605 | 3792 | 2438 | 1419 | 992 | 1414 | 1562 | 3957 | 3812 | 3746 | 3819 | 1137 | 1128 | 1200 | 1270 | 2508,67 | 3834 | 1184 | 0,309 | -1,694 | -34,807 | 1,88E-265 | 3,44E-264 | 0,0010775 | 0,0027458 | 0,00175 | 0,00175 | TRUE | 0,0293051 | No |
| orf19.2917.1 | 715 | 550 | 585 | 455 | 234 | 172 | 243 | 243 | 682 | 582 | 578 | 713 | 187 | 196 | 206 | 198 | 417,61 | 639 | 197 | 0,309 | -1,694 | -17,68 | 6,02E-70 | 2,88E-69 | 0,0053472 | 0,0060229 | 0,0056731 | 0,0056731 | TRUE | 0,0648189 | No |
| orf19.940 | 3815 | 4108 | 4219 | 2433 | 1562 | 1237 | 1352 | 1417 | 3637 | 4344 | 4168 | 3811 | 1252 | 1407 | 1148 | 1152 | 2614,81 | 3990 | 1240 | 0,31 | -1,689 | -21,956 | 7,60E-107 | 5,29E-106 | 0,007356 | 0,0027192 | 0,0051802 | 0,0051802 | TRUE | 0,0956733 | No |
| orf19.6781 | 3003 | 3587 | 3714 | 2160 | 1329 | 1055 | 1297 | 1105 | 2863 | 3793 | 3669 | 3384 | 1065 | 1200 | 1101 | 898 | 2246,6 | 3427 | 1066 | 0,311 | -1,686 | -17,187 | 3,34E-66 | 1,52E-65 | 0,0133664 | 0,0028221 | 0,0086609 | 0,0086609 | TRUE | 0,1509322 | No |
| orf19.5267 | 138 | 120 | 122 | 95 | 47 | 43 | 55 | 39 | 132 | 127 | 121 | 149 | 38 | 49 | 47 | 32 | 86,59 | 132 | 42 | 0,311 | -1,685 | -9,626 | 6,22E-22 | 1,45E-21 | 0,001439 | 0,021052 | 0,0142828 | 0,0142828 | TRUE | 0,201717 | No |
| orf19.4599 | 238 | 242 | 234 | 179 | 93 | 70 | 92 | 95 | 227 | 256 | 231 | 280 | 75 | 80 | 78 | 77 | 162,98 | 248 | 78 | 0,313 | -1,677 | -13,269 | 3,49E-40 | 1,11E-39 | 1,00E-08 | 0,0121655 | 0,0073322 | 0,0073322 | TRUE | 0,0697088 | No |
| orf19.4772 | 1200 | 1128 | 1179 | 813 | 457 | 389 | 365 | 468 | 1144 | 1193 | 1165 | 1274 | 366 | 442 | 310 | 381 | 784,26 | 1194 | 375 | 0,313 | -1,676 | -18,636 | 1,64E-77 | 8,64E-77 | 0,0083931 | 0,0041849 | 0,0060944 | 0,0060944 | TRUE | 0,1566361 | No |
| orf19.3698 | 338 | 436 | 437 | 232 | 174 | 112 | 153 | 121 | 322 | 461 | 432 | 363 | 139 | 127 | 130 | 98 | 259,18 | 394 | 124 | 0,313 | -1,675 | -11,852 | 2,09E-32 | 5,89E-32 | 0,0203768 | 0,0084262 | 0,0141547 | 0,0141547 | TRUE | 0,131103 | No |
| orf19.4056 | 26344 | 22994 | 25417 | 17877 | 9952 | 7186 | 9438 | 9792 | 25116 | 24313 | 25110 | 28005 | 7974 | 8171 | 8013 | 7962 | 16833,04 | 25636 | 8030 | 0,313 | -1,674 | -35,577 | 3,21E-277 | 6,34E-276 | 0,0019693 | 0,0021888 | 0,0020505 | 0,0020505 | TRUE | 0,0470173 | No |
| orf19.216 | 5434 | 4831 | 5353 | 3051 | 1965 | 1336 | 1979 | 1974 | 5181 | 5108 | 5288 | 4780 | 1574 | 1519 | 1680 | 1605 | 3341,95 | 5089 | 1594 | 0,313 | -1,674 | -34,63 | 8,84E-263 | 1,59E-261 | 0,0012656 | 0,0025826 | 0,0018537 | 0,0018537 | TRUE | 0,0195922 | No |
| orf19.6817 | 2736 | 2468 | 2695 | 1732 | 991 | 728 | 1006 | 1041 | 2608 | 2610 | 2662 | 2713 | 794 | 828 | 854 | 846 | 1739,52 | 2648 | 830 | 0,314 | -1,672 | -36,389 | 6,39E-290 | 1,37E-288 | 1,00E-08 | 0,0030352 | 0,0012791 | 0,0012791 | TRUE | 0,0113241 | No |
| orf19.2745 | 20 | 17 | 21 | 11 | 7 | 5 | 3 | 12 | 19 | 18 | 21 | 17 | 6 | 6 | 3 | 10 | 12,33 | 19 | 6 | 0,315 | -1,666 | -3,771 | 0,0001626 | 0,0002277 | 1,00E-08 | 0,1352819 | 0,0830931 | 0,0830931 | TRUE | 0,5725479 | No |
| orf19.4323 | 1469 | 1444 | 1402 | 781 | 564 | 401 | 527 | 487 | 1401 | 1527 | 1385 | 1223 | 452 | 456 | 447 | 396 | 910,9 | 1384 | 438 | 0,315 | -1,665 | -21,154 | 2,56E-99 | 1,67E-98 | 0,005086 | 0,0038938 | 0,0045216 | 0,0045216 | TRUE | 0,0681598 | No |
| orf19.6874 | 3680 | 3316 | 3663 | 2503 | 1405 | 1023 | 1296 | 1480 | 3508 | 3506 | 3619 | 3921 | 1126 | 1163 | 1100 | 1203 | 2393,4 | 3638 | 1148 | 0,316 | -1,663 | -31,389 | 2,90E-216 | 4,26E-215 | 0,0017305 | 0,0027773 | 0,0021503 | 0,0021503 | TRUE | 0,0316614 | No |
| orf19.7339 | 510 | 456 | 472 | 285 | 175 | 122 | 184 | 196 | 486 | 482 | 466 | 446 | 140 | 139 | 156 | 159 | 309,46 | 470 | 148 | 0,316 | -1,661 | -18,186 | 6,64E-74 | 3,36E-73 | 1,00E-08 | 0,007397 | 0,003844 | 0,003844 | TRUE | 0,0255737 | No |
| orf19.2850 | 6961 | 6192 | 6785 | 4059 | 2633 | 1763 | 2436 | 2630 | 6637 | 6547 | 6703 | 6359 | 2110 | 2005 | 2068 | 2138 | 4320,8 | 6562 | 2080 | 0,317 | -1,657 | -44,043 | 0 | 0 | 0,0002267 | 0,0024713 | 0,0010585 | 0,0010585 | TRUE | 0,0067144 | No |
| orf19.1363 | 34139 | 32582 | 33618 | 22204 | 13542 | 10044 | 12358 | 12362 | 32548 | 34451 | 33211 | 34784 | 10851 | 11421 | 10492 | 10051 | 22226,18 | 33748 | 10704 | 0,317 | -1,657 | -36,171 | 1,72E-286 | 3,64E-285 | 0,0018641 | 0,0021652 | 0,0019574 | 0,0019574 | TRUE | 0,0246137 | No |
| orf19.4255 | 434 | 339 | 381 | 292 | 131 | 136 | 154 | 149 | 414 | 358 | 376 | 457 | 105 | 155 | 131 | 121 | 264,69 | 401 | 128 | 0,317 | -1,657 | -13,358 | 1,07E-40 | 3,41E-40 | 0,0122813 | 0,0082943 | 0,0098618 | 0,0098618 | TRUE | 0,1822013 | No |
| orf19.6247.1 | 4590 | 3806 | 3990 | 1710 | 1652 | 886 | 1368 | 1565 | 4376 | 4024 | 3942 | 2679 | 1324 | 1007 | 1161 | 1272 | 2473,26 | 3755 | 1191 | 0,317 | -1,656 | -9,468 | 2,85E-21 | 6,54E-21 | 0,0288802 | 0,0027551 | 0,0150245 | 0,0288802 | TRUE | 0,4497184 | No |
| orf19.5177 | 2857 | 2534 | 2720 | 1662 | 1009 | 766 | 971 | 1107 | 2724 | 2679 | 2687 | 2604 | 808 | 871 | 824 | 900 | 1762,24 | 2674 | 851 | 0,318 | -1,654 | -33,698 | 6,14E-249 | 1,03E-247 | 0,0003714 | 0,003023 | 0,0015779 | 0,0015779 | TRUE | 0,0197178 | No |
| orf19.3433 | 568 | 570 | 554 | 365 | 219 | 161 | 214 | 220 | 542 | 603 | 547 | 572 | 175 | 183 | 182 | 179 | 372,8 | 566 | 180 | 0,318 | -1,654 | -19,738 | 1,01E-86 | 5,87E-86 | 1,00E-08 | 0,0064955 | 0,0032822 | 0,0032822 | TRUE | 0,0241686 | No |
| orf19.6983 | 5226 | 5546 | 5779 | 3226 | 2219 | 1607 | 2009 | 1947 | 4982 | 5864 | 5709 | 5054 | 1778 | 1827 | 1706 | 1583 | 3562,93 | 5402 | 1724 | 0,319 | -1,65 | -24,402 | 1,63E-131 | 1,44E-130 | 0,0048417 | 0,0025521 | 0,004025 | 0,004025 | TRUE | 0,0403271 | No |
| orf19.6440 | 6923 | 5801 | 6557 | 3893 | 2532 | 1714 | 2500 | 2453 | 6600 | 6134 | 6478 | 6099 | 2029 | 1949 | 2122 | 1995 | 4175,66 | 6328 | 2024 | 0,32 | -1,645 | -36,568 | 9,35E-293 | 2,03E-291 | 0,0010034 | 0,0024845 | 0,0016327 | 0,0016327 | TRUE | 0,0130175 | No |
| orf19.5030 | 1773 | 1568 | 1688 | 949 | 657 | 406 | 617 | 705 | 1690 | 1658 | 1668 | 1487 | 526 | 462 | 524 | 573 | 1073,47 | 1626 | 521 | 0,321 | -1,639 | -23,02 | 2,93E-117 | 2,27E-116 | 0,0038261 | 0,0036208 | 0,0036673 | 0,0036673 | TRUE | 0,0631455 | No |
| orf19.238 | 7210 | 6274 | 6632 | 3910 | 2644 | 1755 | 2570 | 2609 | 6874 | 6634 | 6552 | 6125 | 2119 | 1996 | 2182 | 2121 | 4325,3 | 6546 | 2104 | 0,321 | -1,637 | -34,27 | 2,17E-257 | 3,84E-256 | 0,0014086 | 0,0024709 | 0,0018934 | 0,0018934 | TRUE | 0,0221289 | No |
| orf19.2722 | 723 | 668 | 696 | 398 | 263 | 202 | 268 | 257 | 689 | 706 | 688 | 623 | 211 | 230 | 228 | 209 | 447,95 | 676 | 220 | 0,322 | -1,634 | -20,587 | 3,57E-94 | 2,21E-93 | 1,00E-08 | 0,0057566 | 0,0031916 | 0,0031916 | TRUE | 0,0283482 | No |
| orf19.2743 | 1197 | 922 | 1006 | 773 | 455 | 288 | 411 | 430 | 1141 | 975 | 994 | 1211 | 365 | 327 | 349 | 350 | 713,94 | 1080 | 348 | 0,323 | -1,63 | -19,287 | 6,88E-83 | 3,86E-82 | 0,0056315 | 0,0043911 | 0,0050546 | 0,0050546 | TRUE | 0,0736276 | No |
| orf19.3751 | 2854 | 2557 | 2802 | 1745 | 1094 | 767 | 1024 | 1121 | 2721 | 2704 | 2768 | 2734 | 877 | 872 | 869 | 911 | 1807 | 2732 | 882 | 0,323 | -1,629 | -37,842 | 0 | 0 | 1,00E-08 | 0,0029999 | 0,0010711 | 0,0010711 | TRUE | 0,0061322 | No |
| orf19.4361 | 75 | 49 | 50 | 34 | 25 | 12 | 24 | 23 | 72 | 52 | 49 | 53 | 20 | 14 | 20 | 19 | 37,34 | 56 | 18 | 0,325 | -1,624 | -6,31 | 2,78E-10 | 4,90E-10 | 1,00E-08 | 0,0460619 | 0,0296053 | 0,0296053 | TRUE | 0,2947739 | No |
| orf19.2480.1 | 17088 | 14277 | 15811 | 9393 | 6448 | 4137 | 6046 | 6228 | 16292 | 15096 | 15620 | 14715 | 5167 | 4704 | 5133 | 5064 | 10223,73 | 15431 | 5017 | 0,325 | -1,621 | -35,086 | 1,12E-269 | 2,09E-268 | 0,001705 | 0,0022519 | 0,0019237 | 0,0019237 | TRUE | 0,0209795 | No |
| orf19.5634 | 4214 | 3498 | 3830 | 2331 | 1540 | 1036 | 1487 | 1541 | 4018 | 3699 | 3784 | 3652 | 1234 | 1178 | 1262 | 1253 | 2509,88 | 3788 | 1232 | 0,325 | -1,62 | -33,963 | 7,96E-253 | 1,37E-251 | 0,0010246 | 0,0027455 | 0,0016749 | 0,0016749 | TRUE | 0,0205597 | No |
| orf19.3758 | 4932 | 4944 | 5002 | 3324 | 2050 | 1506 | 1924 | 1899 | 4702 | 5228 | 4942 | 5207 | 1643 | 1712 | 1633 | 1544 | 3326,39 | 5020 | 1633 | 0,325 | -1,62 | -31,753 | 2,87E-221 | 4,27E-220 | 0,0015797 | 0,0025849 | 0,0021145 | 0,0021145 | TRUE | 0,0223343 | No |
| CaalfMp05 | 23 | 40 | 27 | 32 | 14 | 11 | 14 | 12 | 22 | 42 | 27 | 50 | 11 | 13 | 12 | 10 | 23,3 | 35 | 12 | 0,326 | -1,616 | -4,56 | 5,10E-06 | 7,67E-06 | 1,00E-08 | 0,0725611 | 0,0649908 | 0,0649908 | TRUE | 0,0974586 | No |
| orf19.3190 | 7022 | 6486 | 6792 | 4490 | 2759 | 2043 | 2559 | 2748 | 6695 | 6858 | 6710 | 7034 | 2211 | 2323 | 2173 | 2234 | 4529,66 | 6824 | 2235 | 0,327 | -1,611 | -43,019 | 0 | 0 | 0,0002807 | 0,0024538 | 0,0010637 | 0,0010637 | TRUE | 0,0078918 | No |
| orf19.2024 | 39 | 32 | 26 | 17 | 13 | 11 | 11 | 11 | 37 | 34 | 26 | 27 | 10 | 13 | 9 | 9 | 20,57 | 31 | 10 | 0,328 | -1,607 | -4,828 | 1,38E-06 | 2,12E-06 | 1,00E-08 | 0,08192 | 0,0449691 | 0,0449691 | TRUE | 0,1375224 | No |
| orf19.2724 | 58125 | 58917 | 61853 | 41939 | 24543 | 19747 | 23112 | 23099 | 55416 | 62297 | 61105 | 65700 | 19666 | 22454 | 19622 | 18782 | 40630,14 | 61130 | 20131 | 0,329 | -1,603 | -24,434 | 7,52E-132 | 6,69E-131 | 0,0055684 | 0,0021317 | 0,0041019 | 0,0041019 | TRUE | 0,0736985 | No |
| orf19.3694 | 411 | 408 | 451 | 227 | 186 | 127 | 165 | 128 | 392 | 431 | 446 | 356 | 149 | 144 | 140 | 104 | 270,25 | 406 | 134 | 0,329 | -1,603 | -13,074 | 4,67E-39 | 1,46E-38 | 0,011814 | 0,0081667 | 0,0097403 | 0,0097403 | TRUE | 0,2565675 | No |
| orf19.5954 | 100 | 71 | 63 | 41 | 22 | 28 | 22 | 39 | 95 | 75 | 62 | 64 | 18 | 32 | 19 | 32 | 49,59 | 74 | 25 | 0,331 | -1,595 | -6,231 | 4,64E-10 | 8,09E-10 | 0,0405234 | 0,0351996 | 0,0373748 | 0,0373748 | TRUE | 0,3507843 | No |
| orf19.715 | 1894 | 1897 | 1908 | 1257 | 838 | 600 | 681 | 749 | 1806 | 2006 | 1885 | 1969 | 671 | 682 | 578 | 609 | 1275,82 | 1916 | 635 | 0,331 | -1,594 | -24,364 | 4,14E-131 | 3,64E-130 | 0,0027775 | 0,0033782 | 0,0031145 | 0,0031145 | TRUE | 0,043072 | No |
| orf19.1482 | 1737 | 1310 | 1469 | 792 | 607 | 406 | 576 | 574 | 1656 | 1385 | 1451 | 1241 | 486 | 462 | 489 | 467 | 954,61 | 1433 | 476 | 0,332 | -1,593 | -18,774 | 1,24E-78 | 6,63E-78 | 0,0070595 | 0,0038112 | 0,0055828 | 0,0055828 | TRUE | 0,1319274 | No |

| | | | | | | | | | | | | | | | | | | | | | | | | | | | | | | | |
|---|---|---|---|---|---|---|---|---|---|---|---|---|---|---|---|---|---|---|---|---|---|---|---|---|---|---|---|---|---|---|---|
| orf19.1352 | 4902 | 4264 | 4794 | 2897 | 1962 | 1220 | 1819 | 1988 | 4674 | 4509 | 4736 | 4538 | 1572 | 1387 | 1544 | 1616 | 3072,07 | 4614 | 1530 | 0,332 | -1,59 | -31,377 | 4,22E-216 | 6,19E-215 | 0,0018193 | 0,0026257 | 0,0020533 | 0,0020533 | TRUE | 0,0447018 | No |
| orf19.3815 | 7932 | 6947 | 7849 | 5236 | 3150 | 2242 | 2960 | 3280 | 7562 | 7346 | 7754 | 8202 | 2524 | 2549 | 2513 | 2667 | 5139,72 | 7716 | 2563 | 0,332 | -1,589 | -35,336 | 1,66E-273 | 3,18E-272 | 0,001151 | 0,0024107 | 0,0016942 | 0,0016942 | TRUE | 0,0213187 | No |
| orf19.4174 | 2658 | 2222 | 2304 | 1676 | 984 | 687 | 975 | 1045 | 2534 | 2349 | 2276 | 2626 | 788 | 781 | 828 | 850 | 1629,05 | 2446 | 812 | 0,333 | -1,589 | -26,259 | 5,70E-152 | 5,79E-151 | 0,0024715 | 0,0030992 | 0,0027344 | 0,0027344 | TRUE | 0,0277962 | No |
| orf19.2620 | 262 | 203 | 255 | 184 | 90 | 75 | 103 | 109 | 250 | 215 | 252 | 288 | 72 | 85 | 87 | 89 | 167,26 | 251 | 83 | 0,333 | -1,586 | -11,992 | 3,91E-33 | 1,11E-32 | 0,0058089 | 0,0119078 | 0,0092141 | 0,0092141 | TRUE | 0,1033475 | No |
| orf19.4753 | 8483 | 7877 | 8677 | 4862 | 3380 | 2363 | 3238 | 3358 | 8088 | 8329 | 8572 | 7617 | 2708 | 2687 | 2749 | 2730 | 5434,98 | 8152 | 2718 | 0,333 | -1,585 | -36,21 | 4,20E-287 | 8,89E-286 | 0,0012034 | 0,0023934 | 0,0016061 | 0,0016061 | TRUE | 0,0230542 | No |
| orf19.7451 | 6407 | 5970 | 6432 | 4274 | 2508 | 2000 | 2386 | 2684 | 6108 | 6313 | 6354 | 6695 | 2010 | 2274 | 2026 | 2182 | 4245,3 | 6368 | 2123 | 0,333 | -1,585 | -30,904 | 1,05E-209 | 1,49E-208 | 0,0021137 | 0,002478 | 0,002228 | 0,002228 | TRUE | 0,0267131 | No |
| orf19.2308 | 30929 | 28640 | 30843 | 19118 | 12502 | 9033 | 12117 | 11756 | 29487 | 30283 | 30470 | 29949 | 10017 | 10271 | 10287 | 9559 | 20040,6 | 30047 | 10034 | 0,334 | -1,583 | -46,588 | 0 | 0 | 0,0005881 | 0,0021732 | 0,0010452 | 0,0010452 | TRUE | 0,0124862 | No |
| orf19.1187 | 36280 | 33438 | 37018 | 23905 | 14300 | 11034 | 13843 | 15232 | 34589 | 35356 | 36570 | 37448 | 11458 | 12547 | 11753 | 12385 | 24013,35 | 35991 | 12036 | 0,334 | -1,58 | -37,005 | 9,67E-300 | 2,14E-298 | 0,001464 | 0,0021597 | 0,0016995 | 0,0016995 | TRUE | 0,0127859 | No |
| orf19.1509 | 9211 | 9071 | 9649 | 5932 | 3834 | 2871 | 3723 | 3719 | 8782 | 9591 | 9532 | 9293 | 3072 | 3265 | 3161 | 3024 | 6214,95 | 9300 | 3130 | 0,336 | -1,572 | -36,281 | 3,19E-288 | 6,79E-287 | 0,0010736 | 0,0023555 | 0,0015985 | 0,0015985 | TRUE | 0,0174847 | No |
| orf19.2833 | 88 | 84 | 100 | 75 | 39 | 37 | 36 | 34 | 84 | 89 | 99 | 117 | 31 | 42 | 31 | 28 | 65,07 | 97 | 33 | 0,337 | -1,57 | -7,803 | 6,04E-15 | 1,22E-14 | 0,0070695 | 0,0273256 | 0,019396 | 0,019396 | TRUE | 0,2041744 | No |
| orf19.6981 | 931 | 974 | 974 | 565 | 407 | 305 | 359 | 366 | 888 | 1030 | 962 | 885 | 326 | 347 | 305 | 298 | 630,02 | 941 | 319 | 0,337 | -1,568 | -19,885 | 5,53E-88 | 3,25E-87 | 0,0032701 | 0,0046974 | 0,00397 | 0,00397 | TRUE | 0,047806 | No |
| orf19.7444 | 784 | 788 | 786 | 483 | 333 | 230 | 319 | 310 | 747 | 833 | 776 | 757 | 267 | 262 | 271 | 252 | 520,63 | 778 | 263 | 0,337 | -1,567 | -21,303 | 1,06E-100 | 6,96E-100 | 1,00E-08 | 0,005245 | 0,0027879 | 0,0027879 | TRUE | 0,0269961 | No |
| orf19.4534 | 11102 | 9086 | 10092 | 6261 | 4115 | 2827 | 4131 | 4328 | 10585 | 9607 | 9970 | 9808 | 3297 | 3215 | 3507 | 3519 | 6688,51 | 9992 | 3384 | 0,339 | -1,561 | -33,282 | 7,03E-243 | 1,16E-241 | 0,0016402 | 0,0023368 | 0,0019256 | 0,0019256 | TRUE | 0,0196232 | No |
| orf19.5933 | 72 | 71 | 86 | 45 | 33 | 25 | 30 | 27 | 69 | 75 | 85 | 70 | 26 | 28 | 25 | 22 | 50,18 | 75 | 25 | 0,339 | -1,561 | -7,357 | 1,88E-13 | 3,64E-13 | 1,00E-08 | 0,0348095 | 0,01832 | 0,01832 | TRUE | 0,0809403 | No |
| orf19.1309 | 747 | 542 | 604 | 380 | 267 | 162 | 244 | 287 | 712 | 573 | 597 | 595 | 214 | 184 | 207 | 233 | 414,49 | 619 | 210 | 0,339 | -1,559 | -15,638 | 4,00E-55 | 1,58E-54 | 0,0069916 | 0,0060525 | 0,0065185 | 0,0065185 | TRUE | 0,1245145 | No |
| orf19.35 | 18923 | 20636 | 21636 | 11880 | 8622 | 6287 | 8024 | 7691 | 18041 | 21820 | 21374 | 18611 | 6909 | 7149 | 6812 | 6254 | 13371,16 | 19962 | 6781 | 0,34 | -1,558 | -22,59 | 5,47E-113 | 4,08E-112 | 0,0060155 | 0,0022141 | 0,004476 | 0,004476 | TRUE | 0,0516168 | No |
| orf19.1653 | 2277 | 2200 | 2223 | 1227 | 977 | 608 | 913 | 860 | 2171 | 2326 | 2196 | 1922 | 783 | 691 | 775 | 699 | 1445,49 | 2154 | 737 | 0,342 | -1,547 | -21,971 | 5,41E-107 | 3,77E-106 | 0,0040232 | 0,0032272 | 0,0038955 | 0,0038955 | TRUE | 0,0601078 | No |
| orf19.5866 | 27 | 24 | 20 | 14 | 11 | 9 | 8 | 8 | 26 | 25 | 20 | 22 | 9 | 10 | 7 | 7 | 15,64 | 23 | 8 | 0,343 | -1,544 | -4,133 | 3,59E-05 | 5,19E-05 | 1,00E-08 | 0,1070447 | 0,0544325 | 0,0544325 | TRUE | 0,073658 | No |
| orf19.4214 | 42 | 38 | 44 | 26 | 17 | 14 | 12 | 21 | 40 | 40 | 43 | 41 | 14 | 16 | 10 | 17 | 27,65 | 41 | 14 | 0,344 | -1,541 | -5,46 | 4,75E-08 | 7,74E-08 | 1,00E-08 | 0,0614663 | 0,0315209 | 0,0315209 | TRUE | 0,1819455 | No |
| orf19.3861 | 7648 | 6665 | 7311 | 4313 | 3015 | 2087 | 2925 | 3057 | 7292 | 7047 | 7223 | 6757 | 2416 | 2373 | 2483 | 2486 | 4759,49 | 7080 | 2440 | 0,345 | -1,537 | -39,121 | 0 | 0 | 0,000606 | 0,0024363 | 0,0012204 | 0,0012204 | TRUE | 0,0110212 | No |
| orf19.2361 | 3823 | 3544 | 3918 | 2484 | 1572 | 1172 | 1557 | 1610 | 3645 | 3747 | 3871 | 3891 | 1260 | 1333 | 1322 | 1309 | 2547,16 | 3788 | 1306 | 0,345 | -1,537 | -35,762 | 4,32E-280 | 8,71E-279 | 0,0004413 | 0,0027359 | 0,0012828 | 0,0012828 | TRUE | 0,0080826 | No |
| orf19.100 | 2501 | 2105 | 2433 | 1417 | 1022 | 680 | 872 | 1044 | 2384 | 2226 | 2404 | 2220 | 819 | 773 | 740 | 849 | 1551,86 | 2308 | 795 | 0,345 | -1,537 | -26,467 | 2,31E-154 | 2,39E-153 | 0,0018644 | 0,0031493 | 0,0024347 | 0,0024347 | TRUE | 0,0273133 | No |
| orf19.6512 | 1283 | 1249 | 1285 | 731 | 518 | 373 | 528 | 523 | 1223 | 1321 | 1269 | 1145 | 415 | 424 | 448 | 425 | 833,9 | 1240 | 428 | 0,345 | -1,536 | -23,519 | 2,63E-122 | 2,14E-121 | 0,00134 | 0,0040602 | 0,0026064 | 0,0026064 | TRUE | 0,0281257 | No |
| orf19.1366 | 94 | 64 | 81 | 50 | 31 | 30 | 31 | 31 | 90 | 68 | 80 | 78 | 25 | 34 | 26 | 25 | 53,26 | 79 | 28 | 0,345 | -1,535 | -7,293 | 3,02E-13 | 5,82E-13 | 1,00E-08 | 0,0329168 | 0,0191884 | 0,0191884 | TRUE | 0,1312589 | No |
| orf19.1285 | 5777 | 5533 | 5973 | 3978 | 2413 | 1877 | 2384 | 2501 | 5508 | 5850 | 5901 | 6232 | 1933 | 2134 | 2024 | 2034 | 3952,01 | 5873 | 2031 | 0,346 | -1,532 | -30,751 | 1,20E-207 | 1,68E-206 | 0,0016487 | 0,0025067 | 0,0020665 | 0,0020665 | TRUE | 0,0214158 | No |
| orf19.6556 | 11712 | 11539 | 11786 | 7553 | 5321 | 3441 | 4844 | 4816 | 11166 | 12201 | 11643 | 11832 | 4264 | 3913 | 4113 | 3916 | 7880,93 | 11710 | 4052 | 0,346 | -1,531 | -34,743 | 1,74E-264 | 3,18E-263 | 0,0013314 | 0,0022996 | 0,0017059 | 0,0017059 | TRUE | 0,0152967 | No |
| orf19.6447 | 9477 | 7534 | 8049 | 5073 | 3732 | 2264 | 3474 | 3568 | 9035 | 7966 | 7952 | 7947 | 2990 | 2574 | 2949 | 2901 | 5539,45 | 8225 | 2854 | 0,347 | -1,526 | -24,86 | 2,00E-136 | 1,84E-135 | 0,0040544 | 0,0023877 | 0,003396 | 0,003396 | TRUE | 0,0540697 | No |
| orf19.149 | 17 | 12 | 11 | 15 | 7 | 8 | 3 | 6 | 16 | 13 | 11 | 23 | 6 | 9 | 3 | 5 | 10,67 | 16 | 6 | 0,347 | -1,526 | -3,11 | 0,0018697 | 0,0024814 | 0,0379079 | 0,1559129 | 0,112402 | 0,112402 | TRUE | 0,3342698 | No |
| orf19.5241 | 7932 | 6839 | 7505 | 4650 | 3188 | 2190 | 3084 | 3177 | 7562 | 7231 | 7414 | 7284 | 2554 | 2490 | 2618 | 2583 | 4967,32 | 7373 | 2561 | 0,348 | -1,525 | -43,904 | 0 | 0 | 0,0002055 | 0,0024218 | 0,0009077 | 0,0009077 | TRUE | 0,0039427 | No |
| orf19.6321 | 4179 | 3036 | 3266 | 2859 | 1508 | 1247 | 1490 | 1580 | 3984 | 3210 | 3226 | 4479 | 1208 | 1418 | 1265 | 1285 | 2509,46 | 3725 | 1294 | 0,347 | -1,525 | -15,12 | 1,19E-51 | 4,48E-51 | 0,0154589 | 0,0027456 | 0,0092765 | 0,0092765 | TRUE | 0,2032696 | No |
| orf19.4800 | 2528 | 2253 | 2426 | 1470 | 1007 | 740 | 981 | 1023 | 2410 | 2382 | 2397 | 2303 | 807 | 841 | 833 | 832 | 1600,61 | 2373 | 828 | 0,348 | -1,521 | -33,742 | 1,42E-249 | 2,40E-248 | 1,00E-08 | 0,0031171 | 0,0011776 | 0,0011776 | TRUE | 0,0042975 | No |
| orf19.5503 | 51 | 43 | 45 | 27 | 23 | 20 | 14 | 14 | 49 | 45 | 44 | 42 | 18 | 23 | 12 | 11 | 30,66 | 45 | 16 | 0,348 | -1,521 | -5,433 | 5,56E-08 | 9,03E-08 | 1,00E-08 | 0,0556426 | 0,0348586 | 0,0348586 | TRUE | 0,2958369 | No |
| orf19.6598 | 3418 | 2989 | 3315 | 2202 | 1409 | 963 | 1391 | 1441 | 3259 | 3160 | 3275 | 3450 | 1129 | 1095 | 1181 | 1172 | 2215,03 | 3286 | 1144 | 0,349 | -1,519 | -32,412 | 1,83E-230 | 2,82E-229 | 0,0006692 | 0,0028325 | 0,0015479 | 0,0015479 | TRUE | 0,0127139 | No |
| orf19.1854 | 3489 | 3062 | 3215 | 2048 | 1482 | 993 | 1269 | 1390 | 3326 | 3238 | 3176 | 3208 | 1187 | 1129 | 1077 | 1130 | 2184,08 | 3237 | 1131 | 0,349 | -1,517 | -33,753 | 9,69E-250 | 1,65E-248 | 0,000336 | 0,002843 | 0,0013718 | 0,0013718 | TRUE | 0,0143103 | No |
| orf19.251 | 11165 | 9856 | 9932 | 5974 | 4459 | 3060 | 4051 | 4397 | 10645 | 10421 | 9812 | 9359 | 3573 | 3479 | 3439 | 3575 | 6787,92 | 10059 | 3516 | 0,349 | -1,517 | -31,91 | 1,92E-223 | 2,88E-222 | 0,0016221 | 0,0023332 | 0,0019866 | 0,0019866 | TRUE | 0,0262863 | No |
| orf19.465 | 36 | 18 | 28 | 32 | 12 | 10 | 13 | 16 | 34 | 19 | 28 | 50 | 10 | 11 | 11 | 13 | 22,02 | 33 | 11 | 0,35 | -1,516 | -4,201 | 2,65E-05 | 3,86E-05 | 0,0575173 | 0,0766476 | 0,0680606 | 0,0680606 | TRUE | 0,1892995 | No |
| orf19.5528 | 1002 | 886 | 972 | 624 | 422 | 291 | 401 | 404 | 955 | 937 | 960 | 978 | 338 | 331 | 340 | 328 | 645,98 | 958 | 334 | 0,35 | -1,515 | -24,197 | 2,39E-129 | 2,06E-128 | 1,00E-08 | 0,004633 | 0,0018549 | 0,0018549 | TRUE | 0,0024191 | No |
| orf19.271 | 71 | 63 | 84 | 49 | 37 | 20 | 22 | 38 | 68 | 67 | 83 | 77 | 30 | 23 | 19 | 31 | 49,5 | 74 | 26 | 0,35 | -1,515 | -6,774 | 1,25E-11 | 2,30E-11 | 1,00E-08 | 0,0352598 | 0,0231138 | 0,0231138 | TRUE | 0,2366737 | No |
| orf19.1691 | 406 | 373 | 376 | 268 | 183 | 128 | 144 | 167 | 387 | 394 | 371 | 420 | 147 | 146 | 122 | 136 | 265,37 | 393 | 138 | 0,35 | -1,514 | -15,336 | 4,41E-53 | 1,69E-52 | 1,00E-08 | 0,0082784 | 0,0046908 | 0,0046908 | TRUE | 0,0603123 | No |
| orf19.1484 | 1039 | 1252 | 1224 | 773 | 551 | 428 | 450 | 433 | 991 | 1324 | 1209 | 1211 | 442 | 487 | 382 | 352 | 799,61 | 1184 | 416 | 0,35 | -1,513 | -13,806 | 2,33E-43 | 7,79E-43 | 0,0152752 | 0,0041447 | 0,0099765 | 0,0099765 | TRUE | 0,1475841 | No |
| orf19.6595 | 65 | 55 | 57 | 35 | 26 | 19 | 28 | 19 | 62 | 58 | 56 | 55 | 21 | 22 | 24 | 15 | 39,12 | 58 | 20 | 0,351 | -1,512 | -6,362 | 1,99E-10 | 3,52E-10 | 1,00E-08 | 0,044067 | 0,0228169 | 0,0228169 | TRUE | 0,1761228 | No |
| orf19.1830 | 3333 | 3492 | 3543 | 2149 | 1553 | 994 | 1488 | 1453 | 3178 | 3692 | 3500 | 3367 | 1244 | 1130 | 1263 | 1181 | 2319,51 | 3434 | 1204 | 0,351 | -1,51 | -25,476 | 3,65E-143 | 3,52E-142 | 0,0028663 | 0,0027991 | 0,0028378 | 0,0028378 | TRUE | 0,0317222 | No |
| orf19.6639 | 3986 | 3582 | 3768 | 2397 | 1676 | 1148 | 1550 | 1634 | 3800 | 3788 | 3722 | 3755 | 1343 | 1305 | 1316 | 1329 | 2544,75 | 3766 | 1323 | 0,351 | -1,508 | -40,322 | 0 | 0 | 1,00E-08 | 0,0027365 | 0,0008584 | 0,0008584 | TRUE | 0,001216 | No |
| orf19.2436 | 8296 | 7833 | 8404 | 5187 | 3535 | 2559 | 3360 | 3552 | 7909 | 8282 | 8302 | 8126 | 2832 | 2910 | 2853 | 2888 | 5512,86 | 8155 | 2871 | 0,352 | -1,507 | -44,927 | 0 | 0 | 0,0002291 | 0,0023891 | 0,0008553 | 0,0008553 | TRUE | 0,0052162 | No |
| orf19.3127 | 2096 | 1679 | 2029 | 1234 | 821 | 561 | 809 | 901 | 1998 | 1775 | 2004 | 1933 | 658 | 638 | 687 | 733 | 1303,3 | 1928 | 679 | 0,353 | -1,504 | -23,815 | 2,34E-125 | 1,96E-124 | 0,0025056 | 0,0033511 | 0,0028813 | 0,0028813 | TRUE | 0,0351038 | No |
| orf19.1240 | 9978 | 9008 | 9729 | 6093 | 4221 | 3056 | 3782 | 4197 | 9513 | 9525 | 9611 | 9545 | 3382 | 3475 | 3211 | 3413 | 6459,33 | 9548 | 3370 | 0,353 | -1,503 | -43,633 | 0 | 0 | 0,0003048 | 0,0023455 | 0,000948 | 0,000948 | TRUE | 0,0123207 | No |
| orf19.3888 | 4151 | 3525 | 3802 | 2528 | 1663 | 1240 | 1566 | 1678 | 3958 | 3727 | 3756 | 3960 | 1333 | 1410 | 1330 | 1364 | 2604,68 | 3850 | 1359 | 0,353 | -1,503 | -34,521 | 3,82E-261 | 6,84E-260 | 0,0005319 | 0,0027216 | 0,0013454 | 0,0013454 | TRUE | 0,0070077 | No |
| orf19.1207 | 9 | 3 | 9 | 4 | 5 | 1 | 3 | 2 | 9 | 3 | 9 | 6 | 4 | 1 | 3 | 2 | 4,53 | 7 | 2 | 0,353 | -1,503 | -2,107 | 0,0351355 | 0,0427699 | 1,00E-08 | 0,3646761 | 0,2210239 | 0,2210239 | TRUE | 0,3347922 | No |
| orf19.6824 | 1104 | 1085 | 1100 | 675 | 447 | 346 | 449 | 497 | 1053 | 1147 | 1087 | 1057 | 358 | 393 | 381 | 404 | 735,1 | 1086 | 384 | 0,353 | -1,501 | -22,901 | 4,53E-116 | 3,47E-115 | 1,00E-08 | 0,0043248 | 0,0024514 | 0,0024514 | TRUE | 0,0256657 | No |
| orf19.6706 | 12393 | 11890 | 12588 | 6971 | 5401 | 3562 | 5028 | 5212 | 11815 | 12572 | 12436 | 10920 | 4328 | 4050 | 4269 | 4238 | 8078,54 | 11936 | 4221 | 0,354 | -1,5 | -29,588 | 2,14E-192 | 2,83E-191 | 0,0021783 | 0,0022945 | 0,0023147 | 0,0023147 | TRUE | 0,0394398 | No |
| orf19.3940.1 | 74206 | 42907 | 49512 | 40372 | 25582 | 18020 | 21495 | 26394 | 70747 | 45369 | 48913 | 63245 | 20498 | 20490 | 18249 | 21461 | 38621,58 | 57068 | 20174 | 0,354 | -1,5 | -9,452 | 3,31E-21 | 7,61E-21 | 0,0241708 | 0,0021338 | 0,0126833 | 0,0241708 | TRUE | 0,1233602 | No |
| orf19.6164 | 17 | 10 | 6 | 2 | 6 | 2 | 2 | 5 | 16 | 11 | 6 | 3 | 5 | 2 | 2 | 4 | 6,09 | 9 | 3 | 0,354 | -1,496 | -2,249 | 0,0244974 | 0,0302005 | 0,1695054 | 0,2718793 | 0,2280098 | 0,2280098 | TRUE | 0,1917827 | No |
| orf19.4959 | 6207 | 5421 | 5885 | 3898 | 2520 | 1853 | 2459 | 2641 | 5918 | 5732 | 5814 | 6106 | 2019 | 2107 | 2088 | 2147 | 3991,41 | 5892 | 2090 | 0,355 | -1,495 | -38,455 | 0 | 0 | 0,0004195 | 0,0025026 | 0,0011413 | 0,0011413 | TRUE | 0,0068827 | No |
| orf19.5671 | 5360 | 4573 | 4938 | 3464 | 2232 | 1532 | 2017 | 2380 | 5110 | 4835 | 4878 | 5427 | 1788 | 1742 | 1712 | 1935 | 3428,55 | 5062 | 1794 | 0,355 | -1,495 | -27,196 | 7,25E-163 | 8,02E-162 | 0,0024467 | 0,0025702 | 0,0025401 | 0,0025401 | TRUE | 0,0341598 | No |
| orf19.6536 | 432 | 436 | 466 | 263 | 155 | 154 | 202 | 189 | 412 | 461 | 460 | 412 | 124 | 175 | 171 | 154 | 296,22 | 436 | 156 | 0,355 | -1,492 | -13,444 | 3,32E-41 | 1,07E-40 | 0,0073757 | 0,0076342 | 0,0076742 | 0,0076742 | TRUE | 0,2128656 | No |
| orf19.5031 | 10198 | 8895 | 9985 | 6676 | 4153 | 3182 | 4103 | 4449 | 9723 | 9405 | 9864 | 10458 | 3328 | 3618 | 3483 | 3617 | 6687,17 | 9862 | 3512 | 0,356 | -1,49 | -32,339 | 1,96E-229 | 3,02E-228 | 0,0015154 | 0,0023368 | 0,0018534 | 0,0018534 | TRUE | 0,019748 | No |
| orf19.510 | 44 | 41 | 42 | 19 | 19 | 19 | 12 | 13 | 42 | 43 | 41 | 30 | 15 | 22 | 10 | 11 | 26,77 | 39 | 14 | 0,356 | -1,49 | -4,867 | 1,13E-06 | 1,75E-06 | 1,19E-08 | 0,0634292 | 0,0443612 | 0,0443612 | TRUE | 0,4123632 | No |
| orf19.7586 | 630 | 563 | 608 | 429 | 261 | 225 | 243 | 260 | 601 | 595 | 601 | 672 | 209 | 256 | 206 | 211 | 418,92 | 617 | 220 | 0,356 | -1,488 | -16,955 | 1,77E-64 | 7,91E-64 | 0,0022552 | 0,0060107 | 0,0044579 | 0,0044579 | TRUE | 0,1116729 | No |
| orf19.1227 | 133 | 120 | 129 | 76 | 44 | 35 | 60 | 64 | 127 | 127 | 127 | 119 | 35 | 40 | 51 | 52 | 84,78 | 125 | 44 | 0,357 | -1,488 | -8,813 | 1,22E-18 | 2,65E-18 | 1,00E-08 | 0,0214586 | 0,0129769 | 0,0129769 | TRUE | 0,1892621 | No |
| orf19.1778 | 93 | 97 | 84 | 58 | 46 | 33 | 36 | 32 | 89 | 103 | 83 | 91 | 37 | 38 | 31 | 26 | 62,01 | 92 | 33 | 0,357 | -1,488 | -7,612 | 2,69E-14 | 5,32E-14 | 1,00E-08 | 0,0285715 | 0,0168949 | 0,0168949 | TRUE | 0,1497231 | No |
| orf19.7341.1 | 30955 | 32246 | 32201 | 17266 | 14602 | 9058 | 13480 | 12636 | 29512 | 34096 | 31812 | 27048 | 11700 | 10300 | 11444 | 10274 | 20773,33 | 30617 | 10930 | 0,357 | -1,486 | -20,445 | 6,62E-93 | 4,06E-92 | 0,0071555 | 0,0021703 | 0,005017 | 0,005017 | TRUE | 0,0751524 | No |
| orf19.1611 | 328 | 358 | 337 | 215 | 171 | 112 | 155 | 114 | 313 | 379 | 333 | 337 | 137 | 127 | 132 | 93 | 231,21 | 340 | 122 | 0,359 | -1,48 | -11,723 | 9,69E-32 | 2,70E-31 | 0,0110817 | 0,0091928 | 0,0099943 | 0,0099943 | TRUE | 0,2923186 | No |
| orf19.5159 | 3966 | 3631 | 3816 | 2338 | 1704 | 1193 | 1589 | 1639 | 3781 | 3839 | 3770 | 3663 | 1365 | 1357 | 1349 | 1333 | 2557,07 | 3763 | 1351 | 0,359 | -1,479 | -38,432 | 0 | 0 | 1,00E-08 | 0,0027334 | 0,0009435 | 0,0009435 | TRUE | 0,0035767 | No |
| orf19.29 | 5432 | 5613 | 5711 | 3754 | 2591 | 1960 | 2382 | 2224 | 5179 | 5935 | 5642 | 5881 | 2076 | 2229 | 2022 | 1808 | 3846,51 | 5659 | 2034 | 0,359 | -1,477 | -21,901 | 2,56E-106 | 1,77E-105 | 0,0050647 | 0,0025181 | 0,0040496 | 0,0040496 | TRUE | 0,0677959 | No |
| orf19.542 | 4983 | 4098 | 4554 | 2720 | 1946 | 1393 | 1844 | 2102 | 4751 | 4333 | 4499 | 4261 | 1559 | 1584 | 1566 | 1709 | 3032,72 | 4461 | 1604 | 0,36 | -1,476 | -28,793 | 2,66E-182 | 3,24E-181 | 0,0015888 | 0,0026327 | 0,0021185 | 0,0021185 | TRUE | 0,0239026 | No |
| orf19.7340 | 13 | 26 | 19 | 26 | 12 | 11 | 7 | 9 | 12 | 27 | 19 | 41 | 10 | 13 | 6 | 7 | 16,85 | 25 | 9 | 0,36 | -1,476 | -3,371 | 0,0007483 | 0,001019 | 0,1222532 | 0,099555 | 0,1098201 | 0,1098201 | TRUE | 0,1932685 | No |
| orf19.6484 | 229 | 171 | 164 | 134 | 97 | 53 | 79 | 86 | 218 | 181 | 162 | 210 | 78 | 60 | 67 | 70 | 130,76 | 193 | 69 | 0,36 | -1,475 | -9,919 | 3,45E-23 | 8,24E-23 | 0,0087163 | 0,0146482 | 0,0118958 | 0,0118958 | TRUE | 0,1303163 | No |
| orf19.3499 | 2073 | 1863 | 1968 | 1327 | 895 | 660 | 802 | 886 | 1976 | 1970 | 1944 | 2079 | 717 | 750 | 681 | 720 | 1354,77 | 1992 | 717 | 0,36 | -1,474 | -28,85 | 5,09E-183 | 6,26E-182 | 1,00E-08 | 0,0033032 | 0,001603 | 0,001603 | TRUE | 0,0135625 | No |
| orf19.6001 | 612 | 631 | 672 | 378 | 272 | 197 | 281 | 275 | 583 | 667 | 664 | 592 | 218 | 224 | 239 | 224 | 426,35 | 626 | 226 | 0,36 | -1,473 | -17,6 | 2,48E-69 | 1,17E-68 | 1,00E-08 | 0,0059423 | 0,0038693 | 0,0038693 | TRUE | 0,0288303 | No |
| orf19.2988 | 2356 | 2005 | 2120 | 1461 | 986 | 692 | 921 | 973 | 2246 | 2120 | 2094 | 2289 | 790 | 787 | 782 | 791 | 1487,41 | 2187 | 788 | 0,36 | -1,472 | -29,153 | 7,67E-187 | 9,69E-186 | 1,00E-08 | 0,0031951 | 0,0016253 | 0,0016253 | TRUE | 0,0106876 | No |

| | | | | | | | | | | | | | | | | | | | | | | | | | | | | | | | |
|---|---|---|---|---|---|---|---|---|---|---|---|---|---|---|---|---|---|---|---|---|---|---|---|---|---|---|---|---|---|---|---|
| orf19.5429 | 2080 | 1836 | 2032 | 1303 | 863 | 645 | 819 | 928 | 1983 | 1941 | 2007 | 2041 | 691 | 733 | 695 | 755 | 1355,98 | 1993 | 718 | 0,361 | -1,472 | -29,057 | 1,26E-185 | 1,57E-184 | 1,00E-08 | 0,0033021 | 0,0015613 | 0,0015613 | TRUE | 0,0147828 | No |
| orf19.6124 | 1746 | 1630 | 1700 | 1077 | 756 | 546 | 701 | 757 | 1665 | 1724 | 1679 | 1687 | 606 | 621 | 595 | 616 | 1149 | 1689 | 610 | 0,361 | -1,471 | -29,531 | 1,17E-191 | 1,53E-190 | 1,00E-08 | 0,0035203 | 0,0013222 | 0,0013222 | TRUE | 0,0028488 | No |
| orf19.5259 | 664 | 619 | 667 | 433 | 307 | 223 | 235 | 306 | 633 | 655 | 659 | 678 | 246 | 254 | 200 | 249 | 446,59 | 656 | 237 | 0,361 | -1,47 | -17,397 | 8,74E-68 | 4,07E-67 | 0,0015168 | 0,0057678 | 0,0041224 | 0,0041224 | TRUE | 0,1295911 | No |
| orf19.3678 | 2926 | 2855 | 3033 | 1925 | 1256 | 926 | 1324 | 1343 | 2790 | 3019 | 2996 | 3016 | 1006 | 1053 | 1124 | 1092 | 2011,97 | 2955 | 1069 | 0,362 | -1,466 | -28,399 | 2,07E-177 | 2,44E-176 | 0,0012903 | 0,0029073 | 0,0019511 | 0,0019511 | TRUE | 0,0197366 | No |
| orf19.644 | 217 | 195 | 198 | 131 | 103 | 60 | 83 | 89 | 207 | 206 | 196 | 205 | 83 | 68 | 70 | 72 | 138,44 | 204 | 73 | 0,362 | -1,464 | -11,498 | 1,35E-30 | 3,68E-30 | 1,00E-08 | 0,0139517 | 0,006809 | 0,006809 | TRUE | 0,0697786 | No |
| orf19.5635 | 3366 | 2570 | 2830 | 2270 | 1360 | 1014 | 1258 | 1407 | 3209 | 2717 | 2796 | 3556 | 1090 | 1153 | 1068 | 1144 | 2091,65 | 3070 | 1114 | 0,363 | -1,462 | -17,875 | 1,86E-71 | 9,11E-71 | 0,007923 | 0,0028763 | 0,0058366 | 0,0058366 | TRUE | 0,1348542 | No |
| orf19.2341 | 5624 | 4982 | 5407 | 3320 | 2418 | 1711 | 2275 | 2314 | 5362 | 5268 | 5342 | 5201 | 1937 | 1946 | 1931 | 1882 | 3608,53 | 5293 | 1924 | 0,363 | -1,461 | -42,191 | 0 | 0 | 1,00E-08 | 0,0025463 | 0,0008137 | 0,0008137 | TRUE | 0,0027471 | No |
| orf19.3440 | 118 | 133 | 119 | 81 | 64 | 31 | 50 | 61 | 113 | 141 | 118 | 127 | 51 | 35 | 42 | 50 | 84,52 | 125 | 44 | 0,364 | -1,46 | -8,501 | 1,87E-17 | 3,98E-17 | 1,00E-08 | 0,0215174 | 0,0139689 | 0,0139689 | TRUE | 0,1280625 | No |
| orf19.3475 | 1546 | 1442 | 1516 | 1079 | 676 | 534 | 631 | 696 | 1474 | 1525 | 1498 | 1690 | 542 | 607 | 536 | 566 | 1054,64 | 1547 | 563 | 0,364 | -1,459 | -22,372 | 7,40E-111 | 5,40E-110 | 0,0020557 | 0,0036481 | 0,0029241 | 0,0029241 | TRUE | 0,0436597 | No |
| orf19.6117 | 8545 | 6833 | 7600 | 5667 | 3469 | 2681 | 3381 | 3520 | 8147 | 7225 | 7508 | 8878 | 2780 | 3049 | 2870 | 2862 | 5414,78 | 7940 | 2890 | 0,364 | -1,458 | -22,477 | 6,90E-112 | 5,08E-111 | 0,0047357 | 0,0023945 | 0,0038141 | 0,0038141 | TRUE | 0,0761896 | No |
| orf19.5588 | 86 | 72 | 63 | 43 | 31 | 32 | 24 | 31 | 82 | 76 | 62 | 67 | 25 | 36 | 20 | 25 | 49,32 | 72 | 26 | 0,364 | -1,457 | -6,451 | 1,11E-10 | 1,98E-10 | 0,0059069 | 0,0353846 | 0,024342 | 0,024342 | TRUE | 0,3105278 | No |
| orf19.1307 | 764 | 711 | 698 | 501 | 337 | 247 | 300 | 333 | 728 | 752 | 690 | 785 | 270 | 281 | 255 | 271 | 503,87 | 739 | 269 | 0,365 | -1,455 | -19,433 | 4,09E-84 | 2,31E-83 | 1,00E-08 | 0,0053499 | 0,0029702 | 0,0029702 | TRUE | 0,0236143 | No |
| orf19.428 | 1758 | 1659 | 1708 | 1038 | 742 | 545 | 708 | 800 | 1676 | 1754 | 1687 | 1626 | 595 | 620 | 601 | 650 | 1151,19 | 1686 | 616 | 0,365 | -1,453 | -26,762 | 8,99E-158 | 9,66E-157 | 1,00E-08 | 0,0035176 | 0,0017758 | 0,0017758 | TRUE | 0,0176191 | No |
| orf19.4850 | 10183 | 9749 | 10635 | 6082 | 4595 | 3357 | 4257 | 4368 | 9708 | 10308 | 10506 | 9528 | 3682 | 3817 | 3614 | 3552 | 6839,46 | 10012 | 3666 | 0,366 | -1,451 | -32,466 | 3,19E-231 | 4,95E-230 | 0,0013291 | 0,0023313 | 0,0017389 | 0,0017389 | TRUE | 0,0134641 | No |
| orf19.2870 | 2828 | 2341 | 2453 | 1299 | 1213 | 704 | 997 | 1117 | 2696 | 2475 | 2423 | 2035 | 972 | 801 | 846 | 908 | 1644,61 | 2407 | 882 | 0,366 | -1,449 | -16,65 | 3,02E-62 | 1,31E-61 | 0,00907 | 0,0030896 | 0,0065355 | 0,0065355 | TRUE | 0,1273913 | No |
| orf19.69.2 | 17 | 32 | 15 | 13 | 12 | 9 | 9 | 5 | 16 | 34 | 15 | 20 | 10 | 10 | 8 | 4 | 14,6 | 21 | 8 | 0,366 | -1,448 | -3,307 | 0,0009434 | 0,0012748 | 0,0835059 | 0,1145669 | 0,100793 | 0,100793 | TRUE | 0,8970614 | No |
| orf19.1381 | 2848 | 2370 | 2455 | 1564 | 1134 | 768 | 1118 | 1197 | 2715 | 2506 | 2425 | 2450 | 909 | 873 | 949 | 973 | 1725,13 | 2524 | 926 | 0,367 | -1,446 | -25,587 | 2,12E-144 | 2,06E-143 | 0,0018766 | 0,003043 | 0,0023609 | 0,0023609 | TRUE | 0,0322902 | No |
| orf19.4982 | 1212 | 1257 | 1341 | 732 | 551 | 423 | 541 | 546 | 1156 | 1329 | 1325 | 1147 | 442 | 481 | 459 | 444 | 847,73 | 1239 | 456 | 0,367 | -1,445 | -19,884 | 5,64E-88 | 3,31E-87 | 0,0033241 | 0,0040281 | 0,0036411 | 0,0036411 | TRUE | 0,0281287 | No |
| orf19.487 | 1847 | 1627 | 1827 | 1228 | 782 | 600 | 819 | 792 | 1761 | 1720 | 1805 | 1924 | 627 | 682 | 695 | 644 | 1232,25 | 1802 | 662 | 0,367 | -1,444 | -24,713 | 7,85E-135 | 7,14E-134 | 0,0013488 | 0,0034237 | 0,0022936 | 0,0022936 | TRUE | 0,0227427 | No |
| orf19.48 | 4019 | 4635 | 4748 | 3005 | 2105 | 1641 | 1888 | 1859 | 3832 | 4901 | 4691 | 4707 | 1687 | 1866 | 1603 | 1512 | 3099,72 | 4533 | 1667 | 0,368 | -1,444 | -17,213 | 2,13E-66 | 9,72E-66 | 0,0090839 | 0,002621 | 0,0063676 | 0,0063676 | TRUE | 0,1331535 | No |
| orf19.5442 | 2757 | 2331 | 2642 | 1564 | 1159 | 741 | 1110 | 1248 | 2629 | 2465 | 2610 | 2450 | 929 | 843 | 942 | 1015 | 1735,22 | 2538 | 932 | 0,368 | -1,443 | -24,237 | 9,18E-130 | 7,94E-129 | 0,002577 | 0,0030375 | 0,0027054 | 0,0027054 | TRUE | 0,0462597 | No |
| orf19.4910 | 178 | 164 | 153 | 101 | 86 | 52 | 73 | 62 | 170 | 173 | 151 | 158 | 69 | 59 | 62 | 50 | 111,61 | 163 | 60 | 0,368 | -1,441 | -9,815 | 9,66E-23 | 2,29E-22 | 1,00E-08 | 0,0168018 | 0,0098845 | 0,0098845 | TRUE | 0,1191335 | No |
| orf19.5282 | 9071 | 7863 | 8643 | 5760 | 3867 | 2846 | 3669 | 4049 | 8648 | 8314 | 8538 | 9023 | 3099 | 3236 | 3115 | 3292 | 5908,25 | 8631 | 3186 | 0,369 | -1,438 | -35,831 | 3,64E-281 | 7,42E-280 | 0,0007376 | 0,0023692 | 0,0013404 | 0,0013404 | TRUE | 0,0110415 | No |
| orf19.524 | 1362 | 1123 | 1324 | 757 | 612 | 366 | 534 | 584 | 1299 | 1187 | 1308 | 1186 | 490 | 416 | 453 | 475 | 851,82 | 1245 | 458 | 0,369 | -1,438 | -20,944 | 2,15E-97 | 1,37E-96 | 0,0022305 | 0,0040188 | 0,0031083 | 0,0031083 | TRUE | 0,0406039 | No |
| orf19.3021 | 2105 | 1788 | 1903 | 1197 | 857 | 615 | 846 | 897 | 2007 | 1891 | 1880 | 1875 | 687 | 699 | 718 | 729 | 1310,78 | 1913 | 708 | 0,37 | -1,434 | -28,037 | 5,71E-173 | 6,63E-172 | 1,00E-08 | 0,0033439 | 0,0015902 | 0,0015902 | TRUE | 0,0136696 | No |
| orf19.2758 | 414 | 335 | 360 | 269 | 179 | 131 | 151 | 177 | 395 | 354 | 356 | 421 | 143 | 149 | 128 | 144 | 261,31 | 382 | 141 | 0,371 | -1,432 | -13,962 | 2,66E-44 | 9,02E-44 | 1,00E-08 | 0,0083746 | 0,0054706 | 0,0054706 | TRUE | 0,0484209 | No |
| orf19.4911 | 478 | 426 | 472 | 301 | 227 | 121 | 207 | 227 | 456 | 450 | 466 | 472 | 182 | 138 | 176 | 185 | 315,47 | 461 | 170 | 0,372 | -1,427 | -15,005 | 6,81E-51 | 2,54E-50 | 1,00E-08 | 0,0072959 | 0,0048706 | 0,0048706 | TRUE | 0,1489473 | No |
| orf19.1008 | 57 | 54 | 43 | 41 | 19 | 19 | 23 | 30 | 54 | 57 | 42 | 64 | 15 | 22 | 20 | 24 | 37,36 | 54 | 20 | 0,372 | -1,426 | -5,6 | 2,14E-08 | 3,53E-08 | 1,00E-08 | 0,0460367 | 0,0298844 | 0,0298844 | TRUE | 0,1902421 | No |
| orf19.2408 | 1691 | 1617 | 1604 | 1030 | 678 | 559 | 734 | 776 | 1612 | 1710 | 1585 | 1614 | 543 | 636 | 623 | 631 | 1119,14 | 1630 | 608 | 0,373 | -1,425 | -22,828 | 2,41E-115 | 1,83E-114 | 0,0016333 | 0,0035584 | 0,00266 | 0,00266 | TRUE | 0,0636426 | No |
| orf19.4779 | 7221 | 6279 | 6822 | 4356 | 3186 | 2206 | 2985 | 3087 | 6884 | 6639 | 6739 | 6824 | 2553 | 2508 | 2534 | 2510 | 4649,08 | 6772 | 2526 | 0,373 | -1,422 | -44,004 | 0 | 0 | 1,00E-08 | 0,0024444 | 0,0007431 | 0,0007431 | TRUE | 0,0021432 | No |
| orf19.4245 | 1241 | 1045 | 1139 | 769 | 538 | 348 | 513 | 558 | 1183 | 1105 | 1125 | 1205 | 431 | 396 | 436 | 454 | 791,76 | 1154 | 429 | 0,373 | -1,422 | -22,246 | 1,24E-109 | 9,00E-109 | 0,0006044 | 0,004165 | 0,0024033 | 0,0024033 | TRUE | 0,0276891 | No |
| orf19.3505 | 8757 | 7587 | 8261 | 5476 | 3904 | 2649 | 3540 | 3942 | 8349 | 8022 | 8161 | 8578 | 3128 | 3012 | 3005 | 3205 | 5682,71 | 8278 | 3088 | 0,373 | -1,421 | -36,447 | 7,77E-291 | 1,68E-289 | 0,0006157 | 0,0023802 | 0,0012479 | 0,0012479 | TRUE | 0,0080218 | No |
| orf19.5585 | 920 | 792 | 833 | 591 | 367 | 305 | 342 | 447 | 877 | 837 | 823 | 926 | 294 | 347 | 290 | 363 | 594,75 | 866 | 324 | 0,374 | -1,42 | -16,653 | 2,90E-62 | 1,26E-61 | 0,0051173 | 0,0048519 | 0,0049508 | 0,0049508 | TRUE | 0,086343 | No |
| orf19.3428 | 9425 | 8804 | 9674 | 5936 | 4300 | 3068 | 4092 | 4297 | 8986 | 9309 | 9557 | 9299 | 3445 | 3489 | 3474 | 3494 | 6381,61 | 9288 | 3476 | 0,374 | -1,418 | -42,879 | 0 | 0 | 0,0002879 | 0,0023486 | 0,0008612 | 0,0008612 | TRUE | 0,0060394 | No |
| orf19.395 | 14978 | 11641 | 12454 | 8741 | 6045 | 4258 | 5635 | 6396 | 14280 | 12309 | 12303 | 13693 | 4844 | 4842 | 4784 | 5201 | 9031,93 | 13146 | 4918 | 0,374 | -1,418 | -24,618 | 8,14E-134 | 7,36E-133 | 0,0034549 | 0,0022731 | 0,0030551 | 0,0030551 | TRUE | 0,0416469 | No |
| orf19.430 | 2152 | 1795 | 1993 | 1257 | 872 | 647 | 875 | 957 | 2052 | 1898 | 1969 | 1969 | 699 | 736 | 743 | 778 | 1355,39 | 1972 | 739 | 0,375 | -1,417 | -26,636 | 2,58E-156 | 2,73E-155 | 0,000687 | 0,0033027 | 0,0018278 | 0,0018278 | TRUE | 0,0171468 | No |
| orf19.2684 | 1866 | 1854 | 1941 | 1128 | 830 | 634 | 800 | 886 | 1779 | 1960 | 1918 | 1767 | 665 | 721 | 679 | 720 | 1276,2 | 1856 | 696 | 0,374 | -1,417 | -24,332 | 9,03E-131 | 7,89E-130 | 0,0015068 | 0,0033778 | 0,0023133 | 0,0023133 | TRUE | 0,0170891 | No |
| orf19.638 | 14229 | 12308 | 12655 | 8815 | 6176 | 4479 | 5570 | 6240 | 13566 | 13014 | 12502 | 13809 | 4949 | 5093 | 4729 | 5074 | 9091,92 | 13223 | 4961 | 0,375 | -1,414 | -32,124 | 2,02E-226 | 3,07E-225 | 0,0013618 | 0,0022719 | 0,0017287 | 0,0017287 | TRUE | 0,0165133 | No |
| orf19.4366 | 44 | 33 | 42 | 34 | 25 | 9 | 21 | 18 | 42 | 35 | 41 | 53 | 20 | 10 | 18 | 15 | 29,29 | 43 | 16 | 0,376 | -1,413 | -4,945 | 7,61E-07 | 1,18E-06 | 1,00E-08 | 0,0581463 | 0,0373411 | 0,0373411 | TRUE | 0,1998982 | No |
| orf19.3027 | 5752 | 5138 | 5548 | 3635 | 2488 | 1857 | 2375 | 2677 | 5484 | 5433 | 5481 | 5694 | 1994 | 2112 | 2016 | 2177 | 3798,77 | 5523 | 2075 | 0,376 | -1,412 | -34 | 2,19E-253 | 3,79E-252 | 0,000623 | 0,0025235 | 0,0013397 | 0,0013397 | TRUE | 0,0140463 | No |
| orf19.2748 | 1392 | 1582 | 1567 | 949 | 737 | 552 | 624 | 648 | 1327 | 1673 | 1548 | 1487 | 591 | 628 | 530 | 527 | 1038,68 | 1509 | 569 | 0,376 | -1,41 | -16,811 | 2,02E-63 | 8,90E-63 | 0,0072856 | 0,003672 | 0,005594 | 0,005594 | TRUE | 0,0816441 | No |
| orf19.2825 | 7112 | 6561 | 6923 | 4081 | 3318 | 2297 | 3003 | 2875 | 6781 | 6937 | 6839 | 6393 | 2659 | 2612 | 2550 | 2338 | 4638,5 | 6738 | 2540 | 0,377 | -1,409 | -28,35 | 8,36E-177 | 9,82E-176 | 0,0018234 | 0,0024453 | 0,002112 | 0,002112 | TRUE | 0,0353087 | No |
| orf19.2885 | 54 | 63 | 52 | 33 | 23 | 23 | 24 | 24 | 51 | 67 | 51 | 52 | 18 | 26 | 20 | 20 | 38,2 | 55 | 21 | 0,377 | -1,409 | -5,791 | 7,00E-09 | 1,17E-08 | 1,00E-08 | 0,0450678 | 0,0254603 | 0,0254603 | TRUE | 0,1614495 | No |
| orf19.1368 | 2545 | 2250 | 2518 | 1694 | 1143 | 822 | 1092 | 1189 | 2426 | 2379 | 2488 | 2654 | 916 | 935 | 927 | 967 | 1711,4 | 2487 | 936 | 0,377 | -1,407 | -27,681 | 1,19E-168 | 1,34E-167 | 0,000905 | 0,0030507 | 0,0017784 | 0,0017784 | TRUE | 0,0229127 | No |
| orf19.6707 | 4011 | 3772 | 3939 | 2328 | 1836 | 1272 | 1685 | 1786 | 3824 | 3988 | 3891 | 3647 | 1471 | 1446 | 1431 | 1452 | 2643,88 | 3838 | 1450 | 0,378 | -1,405 | -32,957 | 3,37E-238 | 5,41E-237 | 0,0004775 | 0,0027123 | 0,001292 | 0,001292 | TRUE | 0,0126084 | No |
| orf19.3529 | 231 | 221 | 249 | 152 | 110 | 90 | 97 | 103 | 220 | 234 | 246 | 238 | 88 | 102 | 82 | 84 | 161,82 | 234 | 89 | 0,378 | -1,404 | -11,653 | 2,23E-31 | 6,15E-31 | 1,00E-08 | 0,0122375 | 0,0065409 | 0,0065409 | TRUE | 0,0795882 | No |
| orf19.942 | 96 | 86 | 107 | 66 | 40 | 43 | 37 | 46 | 92 | 91 | 106 | 103 | 32 | 49 | 31 | 37 | 67,66 | 98 | 37 | 0,378 | -1,404 | -7,408 | 1,28E-13 | 2,49E-13 | 1,00E-08 | 0,0263565 | 0,0167184 | 0,0167184 | TRUE | 0,2589926 | No |
| orf19.6492 | 1970 | 2270 | 2457 | 1481 | 1128 | 823 | 987 | 911 | 1878 | 2400 | 2427 | 2320 | 904 | 936 | 838 | 741 | 1555,51 | 2256 | 855 | 0,378 | -1,402 | -15,057 | 3,08E-51 | 1,15E-50 | 0,0115676 | 0,0031468 | 0,0075483 | 0,0075483 | TRUE | 0,1565798 | No |
| orf19.3852 | 181 | 186 | 200 | 122 | 90 | 67 | 74 | 94 | 173 | 197 | 198 | 191 | 72 | 76 | 63 | 76 | 130,69 | 190 | 72 | 0,379 | -1,4 | -10,495 | 9,06E-26 | 2,27E-25 | 1,00E-08 | 0,014655 | 0,0079428 | 0,0079428 | TRUE | 0,06924 | No |
| orf19.3967 | 33248 | 29380 | 31722 | 19913 | 14690 | 10464 | 13621 | 15083 | 31698 | 31066 | 31338 | 31195 | 11771 | 11899 | 11564 | 12264 | 21599,3 | 31324 | 11874 | 0,379 | -1,399 | -48,321 | 0 | 0 | 0,000274 | 0,0021673 | 0,00075 | 0,00075 | TRUE | 0,0060562 | No |
| orf19.6277 | 50 | 46 | 31 | 26 | 16 | 15 | 20 | 21 | 48 | 49 | 31 | 41 | 13 | 17 | 17 | 17 | 28,95 | 42 | 16 | 0,379 | -1,398 | -4,903 | 9,43E-07 | 1,46E-06 | 1,00E-08 | 0,0588074 | 0,0368646 | 0,0368646 | TRUE | 0,277686 | No |
| orf19.2381 | 802 | 635 | 690 | 500 | 313 | 242 | 325 | 366 | 765 | 671 | 682 | 783 | 251 | 275 | 276 | 298 | 500,06 | 725 | 275 | 0,38 | -1,397 | -16,498 | 3,81E-61 | 1,63E-60 | 0,0036078 | 0,0053747 | 0,0044884 | 0,0044884 | TRUE | 0,0425793 | No |
| orf19.3693 | 33 | 32 | 27 | 13 | 15 | 10 | 10 | 14 | 31 | 34 | 27 | 20 | 12 | 11 | 8 | 11 | 19,45 | 28 | 10 | 0,38 | -1,397 | -4,154 | 3,27E-05 | 4,74E-05 | 1,00E-08 | 0,086508 | 0,0479303 | 0,0479303 | TRUE | 0,1450043 | No |
| CaalfMp06 | 5 | 19 | 13 | 13 | 9 | 2 | 7 | 7 | 5 | 20 | 13 | 20 | 7 | 2 | 6 | 6 | 9,9 | 14 | 5 | 0,38 | -1,397 | -2,547 | 0,010872 | 0,0137702 | 0,1665639 | 0,167971 | 0,167441 | 0,167441 | TRUE | 0,4556861 | No |
| orf19.3760 | 111 | 95 | 88 | 52 | 49 | 28 | 46 | 40 | 106 | 100 | 87 | 81 | 39 | 32 | 39 | 33 | 64,67 | 94 | 36 | 0,38 | -1,395 | -7,378 | 1,61E-13 | 3,12E-13 | 1,00E-08 | 0,0274806 | 0,0160211 | 0,0160211 | TRUE | 0,0744855 | No |
| orf19.5877 | 13681 | 12497 | 13790 | 8256 | 6264 | 4318 | 6016 | 6281 | 13043 | 13214 | 13623 | 12933 | 5019 | 4910 | 5108 | 5107 | 9119,72 | 13203 | 5036 | 0,381 | -1,39 | -43,006 | 0 | 0 | 0,000253 | 0,0022713 | 0,0008724 | 0,0008724 | TRUE | 0,0056339 | No |
| orf19.6992 | 11882 | 10421 | 11239 | 7027 | 5198 | 3835 | 4995 | 5235 | 11328 | 11019 | 11103 | 11008 | 4165 | 4361 | 4241 | 4257 | 7685,17 | 11114 | 4256 | 0,383 | -1,386 | -45,644 | 0 | 0 | 9,73E-05 | 0,0023049 | 0,0007296 | 0,0007296 | TRUE | 0,0031753 | No |
| orf19.736 | 1859 | 1875 | 1984 | 1221 | 968 | 672 | 835 | 827 | 1772 | 1983 | 1960 | 1913 | 776 | 764 | 709 | 672 | 1318,6 | 1907 | 730 | 0,383 | -1,386 | -21,99 | 3,61E-107 | 2,53E-106 | 0,0025602 | 0,0033365 | 0,0029067 | 0,0029067 | TRUE | 0,0346528 | No |
| orf19.7344 | 1231 | 1462 | 1522 | 872 | 676 | 523 | 634 | 575 | 1174 | 1546 | 1504 | 1366 | 542 | 595 | 538 | 468 | 966,41 | 1398 | 536 | 0,383 | -1,386 | -14,327 | 1,49E-46 | 5,25E-46 | 0,0114093 | 0,0037903 | 0,0077487 | 0,0077487 | TRUE | 0,1435895 | No |
| orf19.2203.1 | 59 | 56 | 55 | 43 | 26 | 20 | 31 | 25 | 56 | 59 | 54 | 67 | 21 | 23 | 26 | 20 | 40,92 | 59 | 22 | 0,383 | -1,386 | -6,034 | 1,60E-09 | 2,73E-09 | 1,00E-08 | 0,0422135 | 0,0214813 | 0,0214813 | TRUE | 0,0856771 | No |
| orf19.3897 | 24 | 15 | 31 | 10 | 12 | 3 | 12 | 11 | 23 | 16 | 31 | 16 | 10 | 3 | 10 | 9 | 14,65 | 22 | 8 | 0,383 | -1,386 | -3,323 | 0,0008907 | 0,001206 | 0,0451479 | 0,1141751 | 0,0871948 | 0,0871948 | TRUE | 0,4715857 | No |
| orf19.1193 | 124718 | 104779 | 116396 | 73853 | 54443 | 38048 | 51742 | 56075 | 118905 | 110791 | 114988 | 115695 | 43624 | 43264 | 43929 | 45594 | 79598,65 | 115095 | 44103 | 0,383 | -1,384 | -41,287 | 0 | 0 | 0,0006727 | 0,0021119 | 0,0010643 | 0,0010643 | TRUE | 0,0077769 | No |
| orf19.2317 | 101 | 77 | 67 | 54 | 39 | 24 | 47 | 33 | 96 | 81 | 66 | 85 | 31 | 27 | 40 | 27 | 56,72 | 82 | 31 | 0,383 | -1,384 | -6,478 | 9,32E-11 | 1,67E-10 | 0,0121088 | 0,0310383 | 0,0228633 | 0,0228633 | TRUE | 0,2499423 | No |
| orf19.3748 | 59 | 51 | 76 | 34 | 35 | 23 | 18 | 28 | 56 | 54 | 75 | 53 | 28 | 26 | 15 | 23 | 41,35 | 60 | 23 | 0,383 | -1,383 | -5,519 | 3,40E-08 | 5,56E-08 | 0,0163402 | 0,0418028 | 0,031623 | 0,031623 | TRUE | 0,337979 | No |
| orf19.1417 | 1139 | 988 | 1012 | 600 | 523 | 332 | 465 | 458 | 1086 | 1045 | 1000 | 940 | 419 | 378 | 395 | 372 | 704,26 | 1018 | 391 | 0,384 | -1,382 | -19,556 | 3,67E-85 | 2,09E-84 | 0,001866 | 0,0044227 | 0,0031054 | 0,0031054 | TRUE | 0,0283319 | No |
| orf19.7124 | 16015 | 12375 | 13844 | 8237 | 6729 | 4232 | 6466 | 6635 | 15269 | 13085 | 13677 | 12904 | 5392 | 4812 | 5490 | 5395 | 9502,78 | 13734 | 5272 | 0,384 | -1,381 | -21,988 | 3,73E-107 | 2,61E-106 | 0,0046187 | 0,0022641 | 0,003664 | 0,003664 | TRUE | 0,0694332 | No |
| orf19.842 | 16568 | 15340 | 15962 | 9439 | 7661 | 4897 | 7258 | 7583 | 15796 | 16220 | 15769 | 14787 | 6139 | 5568 | 6162 | 6166 | 10825,77 | 15643 | 6009 | 0,384 | -1,38 | -29,627 | 6,73E-193 | 8,93E-192 | 0,0017947 | 0,0022429 | 0,0019739 | 0,0019739 | TRUE | 0,0291213 | No |
| orf19.7007 | 23 | 17 | 21 | 12 | 8 | 4 | 13 | 10 | 22 | 18 | 21 | 19 | 6 | 5 | 11 | 8 | 13,7 | 20 | 8 | 0,384 | -1,38 | -3,473 | 0,0005154 | 0,0007075 | 1,00E-08 | 0,1219662 | 0,0658227 | 0,0658227 | TRUE | 0,3003006 | No |
| orf19.610 | 7007 | 5678 | 6610 | 4667 | 3179 | 2158 | 3017 | 3250 | 6680 | 6004 | 6530 | 7311 | 2547 | 2454 | 2561 | 2643 | 4591,3 | 6631 | 2551 | 0,385 | -1,377 | -23,201 | 4,42E-119 | 3,48E-118 | 0,0033897 | 0,0024489 | 0,0031217 | 0,0031217 | TRUE | 0,0569141 | No |
| orf19.6820 | 4509 | 3916 | 4405 | 2884 | 2047 | 1471 | 2004 | 2034 | 4299 | 4141 | 4352 | 4518 | 1640 | 1673 | 1701 | 1654 | 2997,16 | 4328 | 1667 | 0,385 | -1,375 | -33,079 | 5,91E-240 | 9,60E-239 | 0,0005058 | 0,0026391 | 0,0012617 | 0,0012617 | TRUE | 0,0100907 | No |

| | | | | | | | | | | | | | | | | | | | | | | | | | | | | | | | |
|---|---|---|---|---|---|---|---|---|---|---|---|---|---|---|---|---|---|---|---|---|---|---|---|---|---|---|---|---|---|---|---|
| orf19.2515 | 11265 | 11190 | 11752 | 6926 | 5246 | 3936 | 5078 | 5407 | 10740 | 11832 | 11610 | 10850 | 4203 | 4476 | 4311 | 4396 | 7802,31 | 11258 | 4346 | 0,386 | -1,374 | -31,065 | 7,08E-212 | 1,02E-210 | 0,0013158 | 0,0023017 | 0,0017249 | 0,0017249 | TRUE | 0,0143944 | No |
| orf19.260 | 6578 | 6064 | 6574 | 3882 | 3067 | 2188 | 2726 | 3091 | 6271 | 6412 | 6494 | 6081 | 2457 | 2488 | 2314 | 2513 | 4379,03 | 6314 | 2443 | 0,387 | -1,371 | -33,112 | 2,00E-240 | 3,27E-239 | 0,0006937 | 0,0024662 | 0,001374 | 0,001374 | TRUE | 0,0154972 | No |
| orf19.5754 | 10340 | 10691 | 11027 | 6505 | 5160 | 3880 | 4624 | 4767 | 9858 | 11304 | 10894 | 10190 | 4135 | 4412 | 3926 | 3876 | 7324,35 | 10562 | 4087 | 0,387 | -1,37 | -23,733 | 1,64E-124 | 1,36E-123 | 0,0034876 | 0,0023155 | 0,0030402 | 0,0030402 | TRUE | 0,0342567 | No |
| orf19.584 | 7 | 10 | 11 | 7 | 4 | 3 | 7 | 3 | 7 | 11 | 11 | 11 | 3 | 3 | 6 | 2 | 6,76 | 10 | 4 | 0,387 | -1,369 | -2,417 | 0,0156604 | 0,0196209 | 1,00E-08 | 0,2449813 | 0,1322168 | 0,1322168 | TRUE | 0,2990106 | No |
| orf19.5961 | 4536 | 3928 | 4210 | 2650 | 2034 | 1499 | 1831 | 2000 | 4325 | 4153 | 4159 | 4151 | 1630 | 1704 | 1555 | 1626 | 2912,92 | 4197 | 1629 | 0,388 | -1,368 | -32,92 | 1,13E-237 | 1,80E-236 | 0,0003444 | 0,0026549 | 0,0012489 | 0,0012489 | TRUE | 0,0111702 | No |
| orf19.6078 | 3114 | 2668 | 2897 | 1646 | 1377 | 883 | 1252 | 1452 | 2969 | 2821 | 2862 | 2579 | 1103 | 1004 | 1063 | 1181 | 1947,68 | 2808 | 1088 | 0,388 | -1,368 | -21,888 | 3,41E-106 | 2,35E-105 | 0,0031977 | 0,0029343 | 0,0031385 | 0,0031385 | TRUE | 0,0395697 | No |
| orf19.4670 | 5949 | 5515 | 6067 | 3678 | 2785 | 1993 | 2672 | 2775 | 5672 | 5831 | 5994 | 5762 | 2232 | 2266 | 2269 | 2256 | 4035,15 | 5815 | 2256 | 0,388 | -1,367 | -38,839 | 0 | 0 | 1,00E-08 | 0,0024982 | 0,0008941 | 0,0008941 | TRUE | 0,0053914 | No |
| orf19.4415 | 1534 | 1327 | 1496 | 1008 | 696 | 485 | 719 | 702 | 1463 | 1403 | 1478 | 1579 | 558 | 551 | 610 | 571 | 1026,63 | 1481 | 572 | 0,388 | -1,367 | -22,754 | 1,32E-114 | 9,93E-114 | 0,0010882 | 0,0036906 | 0,0023066 | 0,0023066 | TRUE | 0,0234395 | No |
| orf19.7314 | 4952 | 4213 | 4436 | 3023 | 2168 | 1583 | 2021 | 2280 | 4721 | 4455 | 4382 | 4736 | 1737 | 1800 | 1716 | 1854 | 3175,1 | 4574 | 1777 | 0,389 | -1,364 | -29,633 | 5,64E-193 | 7,50E-192 | 0,0011067 | 0,0026084 | 0,0016589 | 0,0016589 | TRUE | 0,0108423 | No |
| orf19.3156 | 1810 | 1628 | 1779 | 1091 | 831 | 566 | 845 | 810 | 1726 | 1721 | 1757 | 1709 | 666 | 644 | 717 | 659 | 1199,89 | 1728 | 672 | 0,389 | -1,363 | -26,304 | 1,73E-152 | 1,77E-151 | 1,00E-08 | 0,0034597 | 0,0015918 | 0,0015918 | TRUE | 0,0252636 | No |
| orf19.6973 | 12171 | 12133 | 12924 | 7151 | 5995 | 4297 | 5570 | 5539 | 11604 | 12829 | 12768 | 11202 | 4804 | 4886 | 4729 | 4504 | 8415,66 | 12101 | 4731 | 0,391 | -1,356 | -25,244 | 1,30E-140 | 1,24E-139 | 0,0027169 | 0,0022864 | 0,0026291 | 0,0026291 | TRUE | 0,0300817 | No |
| orf19.7300 | 2112 | 2121 | 2280 | 1400 | 1055 | 860 | 934 | 977 | 2014 | 2243 | 2252 | 2193 | 845 | 978 | 793 | 794 | 1514,06 | 2176 | 852 | 0,391 | -1,355 | -18,848 | 3,03E-79 | 1,64E-78 | 0,0054168 | 0,0031757 | 0,0041747 | 0,0041747 | TRUE | 0,1104678 | No |
| orf19.3900 | 9397 | 7973 | 9040 | 5511 | 4248 | 2893 | 4072 | 4327 | 8959 | 8430 | 8931 | 8633 | 3404 | 3290 | 3457 | 3518 | 6077,77 | 8738 | 3417 | 0,391 | -1,354 | -35,613 | 8,71E-278 | 1,73E-276 | 0,0005501 | 0,0023614 | 0,001193 | 0,001193 | TRUE | 0,0073154 | No |
| orf19.6770 | 8877 | 7978 | 8670 | 6032 | 4060 | 3161 | 4009 | 4184 | 8463 | 8436 | 8565 | 9449 | 3253 | 3594 | 3404 | 3402 | 6070,84 | 8728 | 3413 | 0,391 | -1,354 | -27,1 | 9,82E-162 | 1,08E-160 | 0,0021041 | 0,0023617 | 0,0022036 | 0,0022036 | TRUE | 0,0370051 | No |
| orf19.5258 | 1249 | 1257 | 1330 | 791 | 584 | 470 | 586 | 603 | 1191 | 1329 | 1314 | 1239 | 468 | 534 | 498 | 490 | 882,89 | 1268 | 498 | 0,391 | -1,354 | -20,887 | 7,01E-97 | 4,44E-96 | 0,0016077 | 0,003951 | 0,0026956 | 0,0026956 | TRUE | 0,0262639 | No |
| orf19.7150 | 12441 | 10435 | 12253 | 8258 | 5722 | 4122 | 5451 | 5984 | 11861 | 11034 | 12105 | 12937 | 4585 | 4687 | 4628 | 4866 | 8337,7 | 11984 | 4692 | 0,392 | -1,353 | -26,526 | 4,81E-155 | 5,04E-154 | 0,0022841 | 0,0022882 | 0,0023548 | 0,0023548 | TRUE | 0,034745 | No |
| orf19.2367 | 1423 | 1439 | 1467 | 672 | 699 | 425 | 670 | 612 | 1357 | 1522 | 1449 | 1053 | 560 | 483 | 569 | 498 | 936,25 | 1345 | 528 | 0,392 | -1,352 | -12,805 | 1,53E-37 | 4,66E-37 | 0,0149728 | 0,003845 | 0,0094413 | 0,0094413 | TRUE | 0,2460679 | No |
| orf19.6999 | 3 | 14 | 13 | 5 | 5 | 4 | 5 | 3 | 3 | 15 | 13 | 8 | 4 | 5 | 4 | 2 | 6,7 | 10 | 4 | 0,393 | -1,348 | -2,151 | 0,0314597 | 0,0384567 | 1,00E-08 | 0,2472525 | 0,2009853 | 0,2009853 | TRUE | 0,2446471 | No |
| orf19.2432 | 6562 | 6629 | 7026 | 4325 | 3332 | 2368 | 3143 | 3198 | 6256 | 7009 | 6941 | 6775 | 2670 | 2693 | 2668 | 2600 | 4701,62 | 6745 | 2658 | 0,394 | -1,344 | -29,666 | 2,12E-193 | 2,82E-192 | 0,0013483 | 0,0024405 | 0,0017185 | 0,0017185 | TRUE | 0,0297271 | No |
| orf19.787.1 | 6930 | 6727 | 6584 | 4589 | 3394 | 2602 | 3027 | 3145 | 6607 | 7113 | 6504 | 7189 | 2720 | 2959 | 2570 | 2557 | 4777,32 | 6853 | 2702 | 0,394 | -1,344 | -23,345 | 1,55E-120 | 1,24E-119 | 0,0033318 | 0,002435 | 0,0029343 | 0,0029343 | TRUE | 0,0486378 | No |
| orf19.909 | 12697 | 11384 | 12262 | 7703 | 5938 | 4286 | 5627 | 5727 | 12105 | 12037 | 12114 | 12067 | 4758 | 4874 | 4777 | 4657 | 8423,58 | 12081 | 4766 | 0,394 | -1,342 | -47,27 | 0 | 0 | 1,00E-08 | 0,0022862 | 0,0006341 | 0,0006341 | TRUE | 0,0029235 | No |
| orf19.4286 | 4476 | 4169 | 4432 | 2616 | 2104 | 1470 | 2069 | 2040 | 4267 | 4408 | 4378 | 4098 | 1686 | 1672 | 1757 | 1659 | 2990,59 | 4288 | 1694 | 0,395 | -1,342 | -32,02 | 5,74E-225 | 8,63E-224 | 0,0005041 | 0,0026403 | 0,0012907 | 0,0012907 | TRUE | 0,0100944 | No |
| orf19.4837 | 514 | 383 | 466 | 349 | 226 | 154 | 231 | 238 | 490 | 405 | 460 | 547 | 181 | 175 | 196 | 194 | 330,99 | 476 | 186 | 0,394 | -1,342 | -12,844 | 9,35E-38 | 2,85E-37 | 0,0067807 | 0,0070518 | 0,0069176 | 0,0069176 | TRUE | 0,1117729 | No |
| orf19.2262 | 2957 | 2610 | 2879 | 1698 | 1356 | 906 | 1312 | 1406 | 2819 | 2760 | 2844 | 2660 | 1087 | 1030 | 1114 | 1143 | 1932,12 | 2771 | 1094 | 0,395 | -1,341 | -28,003 | 1,47E-172 | 1,70E-171 | 0,0005542 | 0,0029411 | 0,0015905 | 0,0015905 | TRUE | 0,0165161 | No |
| orf19.2074 | 254 | 194 | 204 | 122 | 105 | 69 | 101 | 104 | 242 | 205 | 202 | 191 | 84 | 78 | 86 | 85 | 146,61 | 210 | 83 | 0,395 | -1,339 | -10,336 | 4,87E-25 | 1,21E-24 | 1,00E-08 | 0,0132907 | 0,0081302 | 0,0081302 | TRUE | 0,1207534 | No |
| orf19.713 | 3530 | 3025 | 3351 | 2260 | 1710 | 1093 | 1534 | 1716 | 3365 | 3199 | 3310 | 3540 | 1370 | 1243 | 1302 | 1395 | 2340,69 | 3354 | 1328 | 0,397 | -1,334 | -25,665 | 2,87E-145 | 2,81E-144 | 0,001546 | 0,0027927 | 0,0020886 | 0,0020886 | TRUE | 0,0208405 | No |
| orf19.1727 | 14135 | 13832 | 14788 | 9072 | 6936 | 5256 | 6438 | 6916 | 13476 | 14626 | 14609 | 14212 | 5558 | 5977 | 5466 | 5623 | 9943,26 | 14231 | 5656 | 0,397 | -1,332 | -30,921 | 6,26E-210 | 8,87E-209 | 0,00133 | 0,0022564 | 0,001663 | 0,001663 | TRUE | 0,0173767 | No |
| orf19.2282 | 1179 | 1030 | 1047 | 618 | 541 | 361 | 506 | 499 | 1124 | 1089 | 1034 | 968 | 433 | 410 | 430 | 406 | 736,86 | 1054 | 420 | 0,398 | -1,331 | -19,617 | 1,10E-85 | 6,31E-85 | 0,001508 | 0,0043195 | 0,0028267 | 0,0028267 | TRUE | 0,0319356 | No |
| orf19.7570 | 15452 | 14110 | 15778 | 8905 | 7425 | 5047 | 7032 | 7296 | 14732 | 14920 | 15587 | 13950 | 5949 | 5739 | 5970 | 5932 | 10347,44 | 14797 | 5898 | 0,399 | -1,327 | -32,555 | 1,78E-232 | 2,78E-231 | 0,0010423 | 0,00225 | 0,0014827 | 0,0014827 | TRUE | 0,0178121 | No |
| orf19.5305 | 3624 | 2964 | 3130 | 2256 | 1735 | 1120 | 1523 | 1605 | 3455 | 3134 | 3092 | 3534 | 1390 | 1274 | 1293 | 1305 | 2309,65 | 3304 | 1316 | 0,399 | -1,326 | -22,824 | 2,65E-115 | 2,01E-114 | 0,0026882 | 0,0028022 | 0,0027333 | 0,0027333 | TRUE | 0,0255517 | No |
| orf19.4553 | 6165 | 5595 | 6243 | 4119 | 2901 | 2198 | 2848 | 3089 | 5878 | 5916 | 6167 | 6453 | 2324 | 2499 | 2418 | 2512 | 4270,9 | 6104 | 2438 | 0,4 | -1,323 | -28,911 | 8,64E-184 | 1,07E-182 | 0,0011965 | 0,0024757 | 0,0017361 | 0,0017361 | TRUE | 0,0174493 | No |
| orf19.7529 | 18782 | 15228 | 17404 | 10995 | 8762 | 5703 | 7938 | 8771 | 17907 | 16102 | 17194 | 17224 | 7021 | 6485 | 6739 | 7132 | 11975,32 | 17107 | 6844 | 0,4 | -1,321 | -28,744 | 1,09E-181 | 1,32E-180 | 0,001719 | 0,0022284 | 0,0019311 | 0,0019311 | TRUE | 0,0192239 | No |
| orf19.3981 | 412 | 344 | 390 | 260 | 199 | 126 | 187 | 191 | 393 | 364 | 385 | 407 | 159 | 143 | 159 | 155 | 270,74 | 387 | 154 | 0,4 | -1,321 | -14,196 | 9,68E-46 | 3,36E-45 | 1,00E-08 | 0,0081558 | 0,0039968 | 0,0039968 | TRUE | 0,02008 | No |
| orf19.814 | 4982 | 4400 | 4903 | 2967 | 2307 | 1674 | 2248 | 2358 | 4750 | 4652 | 4844 | 4648 | 1849 | 1903 | 1909 | 1917 | 3308,97 | 4724 | 1894 | 0,401 | -1,319 | -36,13 | 7,67E-286 | 1,60E-284 | 1,00E-08 | 0,0025875 | 0,0009252 | 0,0009252 | TRUE | 0,0036049 | No |
| orf19.212 | 1767 | 1562 | 1724 | 1180 | 851 | 596 | 820 | 857 | 1685 | 1652 | 1703 | 1849 | 682 | 678 | 696 | 697 | 1205,07 | 1722 | 688 | 0,401 | -1,319 | -24,283 | 2,96E-130 | 2,57E-129 | 0,0006481 | 0,0034538 | 0,0018611 | 0,0018611 | TRUE | 0,0266977 | No |
| orf19.6276 | 1667 | 1613 | 1804 | 1009 | 848 | 618 | 770 | 792 | 1589 | 1706 | 1782 | 1581 | 679 | 703 | 654 | 644 | 1167,2 | 1664 | 670 | 0,402 | -1,316 | -21,809 | 1,89E-105 | 1,30E-104 | 0,0017332 | 0,003498 | 0,0024945 | 0,0024945 | TRUE | 0,0277047 | No |
| orf19.6928 | 19667 | 17258 | 18386 | 12115 | 9044 | 6440 | 8657 | 9699 | 18750 | 18248 | 18164 | 18979 | 7247 | 7323 | 7350 | 7886 | 12993,31 | 18535 | 7452 | 0,402 | -1,314 | -34,7 | 7,88E-264 | 1,43E-262 | 0,0008733 | 0,0022176 | 0,0012876 | 0,0012876 | TRUE | 0,0190327 | No |
| orf19.5536 | 521 | 451 | 504 | 334 | 231 | 196 | 226 | 252 | 497 | 477 | 498 | 523 | 185 | 223 | 192 | 205 | 349,93 | 499 | 201 | 0,402 | -1,314 | -15,266 | 1,28E-52 | 4,91E-52 | 1,00E-08 | 0,0067833 | 0,0037665 | 0,0037665 | TRUE | 0,0496175 | No |
| orf19.2638.1 | 2999 | 2620 | 2603 | 1476 | 1417 | 856 | 1260 | 1297 | 2859 | 2770 | 2572 | 2312 | 1135 | 973 | 1070 | 1055 | 1843,3 | 2628 | 1058 | 0,403 | -1,313 | -17,934 | 6,42E-72 | 3,16E-71 | 0,0053577 | 0,002982 | 0,0045089 | 0,0045089 | TRUE | 0,0764577 | No |
| orf19.11 | 480 | 348 | 412 | 230 | 215 | 114 | 206 | 200 | 458 | 368 | 407 | 360 | 172 | 130 | 175 | 163 | 279,04 | 398 | 160 | 0,402 | -1,313 | -11,294 | 1,40E-29 | 3,75E-29 | 0,0098017 | 0,0079754 | 0,0088031 | 0,0088031 | TRUE | 0,1506414 | No |
| orf19.3753 | 9456 | 8391 | 9476 | 6216 | 4374 | 3466 | 4251 | 4753 | 9015 | 8872 | 9361 | 9738 | 3505 | 3941 | 3609 | 3865 | 6488,3 | 9246 | 3730 | 0,403 | -1,31 | -26,115 | 2,45E-150 | 2,48E-149 | 0,0021565 | 0,0023443 | 0,0022359 | 0,0022359 | TRUE | 0,0201893 | No |
| orf19.6022 | 96 | 83 | 110 | 70 | 52 | 35 | 43 | 51 | 92 | 88 | 109 | 110 | 42 | 40 | 37 | 41 | 69,63 | 100 | 40 | 0,403 | -1,31 | -7,309 | 2,69E-13 | 5,19E-13 | 1,00E-08 | 0,0256711 | 0,0140962 | 0,0140962 | TRUE | 0,058274 | No |
| orf19.3331 | 23800 | 21332 | 23257 | 15506 | 11327 | 8403 | 10985 | 11635 | 22691 | 22556 | 22976 | 24291 | 9076 | 9555 | 9326 | 9460 | 16241,36 | 23128 | 9354 | 0,404 | -1,306 | -36,169 | 1,85E-286 | 3,90E-285 | 0,0007372 | 0,0021924 | 0,0011798 | 0,0011798 | TRUE | 0,013812 | No |
| orf19.1760 | 11030 | 10671 | 11070 | 7002 | 5429 | 4064 | 5054 | 5433 | 10516 | 11283 | 10936 | 10969 | 4350 | 4621 | 4291 | 4418 | 7672,98 | 10926 | 4420 | 0,404 | -1,306 | -34,033 | 7,19E-254 | 1,25E-252 | 0,0007331 | 0,0023053 | 0,0012621 | 0,0012621 | TRUE | 0,0110356 | No |
| orf19.775 | 3400 | 3349 | 3393 | 2121 | 1658 | 1240 | 1526 | 1742 | 3242 | 3541 | 3352 | 3323 | 1329 | 1410 | 1296 | 1416 | 2363,47 | 3364 | 1363 | 0,405 | -1,305 | -26,568 | 1,58E-155 | 1,66E-154 | 0,001195 | 0,002786 | 0,0018214 | 0,0018214 | TRUE | 0,0154566 | No |
| orf19.4192 | 3207 | 2828 | 3000 | 2033 | 1455 | 1148 | 1396 | 1576 | 3058 | 2990 | 2964 | 3185 | 1166 | 1305 | 1185 | 1281 | 2141,77 | 3049 | 1234 | 0,405 | -1,305 | -25,471 | 4,10E-143 | 3,94E-142 | 0,0013372 | 0,0028579 | 0,0019741 | 0,0019741 | TRUE | 0,0169559 | No |
| orf19.1841 | 51 | 87 | 80 | 40 | 30 | 29 | 34 | 36 | 49 | 92 | 79 | 63 | 24 | 33 | 29 | 29 | 49,68 | 71 | 29 | 0,405 | -1,305 | -5,309 | 1,10E-07 | 1,77E-07 | 0,0338646 | 0,0351392 | 0,034541 | 0,034541 | TRUE | 0,1259707 | No |
| orf19.3301 | 29617 | 26467 | 29114 | 17610 | 13963 | 10164 | 13626 | 14040 | 28237 | 27986 | 28762 | 27587 | 11188 | 11557 | 11568 | 11416 | 19787,6 | 28143 | 11432 | 0,406 | -1,3 | -46,435 | 0 | 0 | 0,0001906 | 0,0021743 | 0,0006937 | 0,0006937 | TRUE | 0,0026835 | No |
| orf19.5343 | 323 | 288 | 345 | 231 | 156 | 119 | 156 | 172 | 308 | 305 | 341 | 362 | 125 | 135 | 132 | 140 | 230,97 | 329 | 133 | 0,406 | -1,299 | -12,403 | 2,53E-35 | 7,43E-35 | 1,00E-08 | 0,0091999 | 0,0054865 | 0,0054865 | TRUE | 0,0438339 | No |
| orf19.1228 | 3775 | 3579 | 3831 | 2404 | 1928 | 1324 | 1834 | 1798 | 3599 | 3784 | 3785 | 3766 | 1545 | 1506 | 1557 | 1462 | 2625,43 | 3734 | 1518 | 0,407 | -1,298 | -31,265 | 1,41E-214 | 2,06E-213 | 0,0003392 | 0,0027167 | 0,0012112 | 0,0012112 | TRUE | 0,0077437 | No |
| orf19.6530 | 3802 | 3413 | 3695 | 2511 | 1833 | 1364 | 1758 | 1861 | 3625 | 3609 | 3650 | 3934 | 1469 | 1551 | 1493 | 1513 | 2605,37 | 3704 | 1506 | 0,407 | -1,297 | -29,065 | 9,98E-186 | 1,25E-184 | 0,0006955 | 0,0027215 | 0,0014646 | 0,0014646 | TRUE | 0,0197841 | No |
| orf19.5443 | 6086 | 5043 | 5542 | 3193 | 2743 | 1801 | 2638 | 2839 | 5802 | 5332 | 5475 | 5002 | 2198 | 2048 | 2240 | 2308 | 3800,68 | 5403 | 2198 | 0,407 | -1,297 | -23,14 | 1,82E-118 | 1,43E-117 | 0,0027169 | 0,0025233 | 0,0027105 | 0,0027105 | TRUE | 0,0301842 | No |
| orf19.831 | 2854 | 2710 | 2919 | 1813 | 1440 | 1069 | 1322 | 1383 | 2721 | 2865 | 2884 | 2840 | 1154 | 1216 | 1122 | 1125 | 1990,82 | 2828 | 1154 | 0,408 | -1,295 | -28,354 | 7,40E-177 | 8,71E-176 | 0,0003917 | 0,002916 | 0,0014161 | 0,0014161 | TRUE | 0,0140446 | No |
| orf19.3455 | 2098 | 2156 | 2258 | 1292 | 1038 | 864 | 1065 | 951 | 2000 | 2280 | 2231 | 2024 | 832 | 982 | 904 | 773 | 1503,27 | 2134 | 873 | 0,408 | -1,293 | -16,729 | 8,02E-63 | 3,51E-62 | 0,0065233 | 0,0031835 | 0,0049592 | 0,0049592 | TRUE | 0,0814989 | No |
| orf19.1210 | 4505 | 4400 | 4617 | 2773 | 2356 | 1641 | 2138 | 2125 | 4295 | 4652 | 4561 | 4344 | 1888 | 1866 | 1815 | 1728 | 3143,68 | 4463 | 1824 | 0,408 | -1,292 | -27,739 | 2,37E-169 | 2,70E-168 | 0,0011777 | 0,0026136 | 0,0017115 | 0,0017115 | TRUE | 0,0156416 | No |
| orf19.1169 | 6683 | 5818 | 6486 | 3468 | 3040 | 2098 | 3024 | 3169 | 6372 | 6152 | 6408 | 5433 | 2436 | 2386 | 2567 | 2577 | 4291,15 | 6091 | 2492 | 0,409 | -1,29 | -22,065 | 6,78E-108 | 4,79E-107 | 0,0031231 | 0,0024739 | 0,0030125 | 0,0030125 | TRUE | 0,0629899 | No |
| orf19.1364 | 2693 | 2579 | 2780 | 1850 | 1389 | 974 | 1294 | 1417 | 2567 | 2727 | 2746 | 2898 | 1113 | 1108 | 1099 | 1152 | 1926,27 | 2734 | 1118 | 0,409 | -1,288 | -25,751 | 3,15E-146 | 3,11E-145 | 0,0010678 | 0,0029437 | 0,0017978 | 0,0017978 | TRUE | 0,0207073 | No |
| orf19.4665 | 2799 | 2139 | 2429 | 1335 | 1232 | 743 | 1210 | 1229 | 2669 | 2262 | 2400 | 2091 | 987 | 845 | 1027 | 999 | 1659,98 | 2356 | 964 | 0,41 | -1,288 | -15,598 | 7,50E-55 | 2,96E-54 | 0,0079676 | 0,0030804 | 0,0058414 | 0,0058414 | TRUE | 0,0973083 | No |
| orf19.7105 | 848 | 766 | 907 | 628 | 432 | 341 | 402 | 439 | 808 | 810 | 896 | 984 | 346 | 388 | 341 | 357 | 616,3 | 874 | 358 | 0,41 | -1,287 | -15,651 | 3,28E-55 | 1,30E-54 | 0,0044574 | 0,0047554 | 0,004602 | 0,004602 | TRUE | 0,0768779 | No |
| orf19.4191.1 | 1438 | 1479 | 1591 | 929 | 753 | 529 | 752 | 752 | 1371 | 1564 | 1572 | 1455 | 603 | 602 | 638 | 611 | 1052,09 | 1490 | 614 | 0,412 | -1,28 | -20,367 | 3,25E-92 | 1,98E-91 | 0,0019896 | 0,0036519 | 0,0026891 | 0,0026891 | TRUE | 0,0375478 | No |
| orf19.2006.1 | 2255 | 1972 | 1955 | 1188 | 1057 | 760 | 943 | 998 | 2150 | 2085 | 1931 | 1861 | 847 | 864 | 801 | 811 | 1418,83 | 2007 | 831 | 0,413 | -1,276 | -21,138 | 3,54E-99 | 2,30E-98 | 0,0022633 | 0,0032485 | 0,0026802 | 0,0026802 | TRUE | 0,0278907 | No |
| orf19.1620 | 7 | 13 | 13 | 6 | 7 | 4 | 6 | 3 | 7 | 14 | 13 | 9 | 6 | 5 | 5 | 2 | 7,54 | 11 | 4 | 0,413 | -1,276 | -2,361 | 0,0182093 | 0,0226807 | 1,00E-08 | 0,2197342 | 0,1277532 | 0,1277532 | TRUE | 0,2732377 | No |
| orf19.4951 | 2724 | 2847 | 2993 | 1878 | 1475 | 1109 | 1373 | 1413 | 2597 | 3010 | 2957 | 2942 | 1182 | 1261 | 1166 | 1149 | 2032,96 | 2876 | 1190 | 0,413 | -1,275 | -21,541 | 6,37E-103 | 4,28E-102 | 0,0027199 | 0,0028989 | 0,0027916 | 0,0027916 | TRUE | 0,0532603 | No |
| orf19.5042 | 59 | 87 | 77 | 45 | 39 | 33 | 36 | 29 | 56 | 92 | 76 | 70 | 31 | 38 | 31 | 24 | 52,22 | 74 | 31 | 0,413 | -1,275 | -5,586 | 2,32E-08 | 3,82E-08 | 0,0206142 | 0,0335362 | 0,0278255 | 0,0278255 | TRUE | 0,146969 | No |
| orf19.5087 | 3086 | 2535 | 2895 | 1682 | 1410 | 938 | 1397 | 1505 | 2942 | 2680 | 2860 | 2635 | 1130 | 1067 | 1186 | 1224 | 1965,46 | 2779 | 1152 | 0,414 | -1,271 | -21,852 | 7,41E-106 | 5,10E-105 | 0,0023252 | 0,0029267 | 0,0026569 | 0,0026569 | TRUE | 0,0276958 | No |
| orf19.4961 | 39259 | 35803 | 38734 | 25804 | 19785 | 14539 | 18564 | 19301 | 37429 | 37857 | 38266 | 40423 | 15853 | 16532 | 15761 | 15694 | 27226,85 | 38494 | 15960 | 0,415 | -1,27 | -35,135 | 1,94E-270 | 3,67E-269 | 0,0008327 | 0,0021516 | 0,0012128 | 0,0012128 | TRUE | 0,0138267 | No |
| orf19.4557 | 576 | 539 | 535 | 320 | 270 | 188 | 278 | 278 | 549 | 570 | 529 | 501 | 216 | 214 | 236 | 226 | 380,14 | 537 | 223 | 0,415 | -1,27 | -15,517 | 2,67E-54 | 1,05E-53 | 1,00E-08 | 0,0064105 | 0,0034034 | 0,0034034 | TRUE | 0,019891 | No |
| orf19.2878 | 81 | 72 | 95 | 48 | 41 | 22 | 49 | 41 | 77 | 76 | 94 | 75 | 33 | 25 | 42 | 33 | 56,9 | 80 | 33 | 0,415 | -1,27 | -6,229 | 4,69E-10 | 8,17E-10 | 1,00E-08 | 0,0309468 | 0,0197618 | 0,0197618 | TRUE | 0,2056004 | No |
| orf19.5713 | 6403 | 5970 | 6462 | 4157 | 3290 | 2336 | 3029 | 3238 | 6105 | 6313 | 6384 | 6512 | 2636 | 2656 | 2572 | 2633 | 4476,24 | 6328 | 2624 | 0,415 | -1,269 | -35,408 | 1,28E-274 | 2,47E-273 | 0,0002949 | 0,0024581 | 0,0009749 | 0,0009749 | TRUE | 0,0069805 | No |
| orf19.6248 | 45 | 66 | 60 | 32 | 29 | 18 | 31 | 27 | 43 | 70 | 59 | 50 | 23 | 20 | 26 | 22 | 39,26 | 56 | 23 | 0,415 | -1,269 | -5,122 | 3,02E-07 | 4,77E-07 | 1,00E-08 | 0,0439139 | 0,0296219 | 0,0296219 | TRUE | 0,1657401 | No |

| | | | | | | | | | | | | | | | | | | | | | | | | | | | | | | | |
|---|---|---|---|---|---|---|---|---|---|---|---|---|---|---|---|---|---|---|---|---|---|---|---|---|---|---|---|---|---|---|---|
| orf19.3749 | 15278 | 14480 | 15053 | 9386 | 7727 | 5589 | 6925 | 7703 | 14566 | 15311 | 14871 | 14704 | 6191 | 6355 | 5879 | 6263 | 10517,56 | 14863 | 6172 | 0,415 | -1,268 | -35,019 | 1,15E-268 | 2,14E-267 | 0,0006439 | 0,0022474 | 0,0011493 | 0,0011493 | TRUE | 0,0124838 | No |
| orf19.7345 | 1240 | 1343 | 1306 | 747 | 674 | 468 | 610 | 634 | 1182 | 1420 | 1290 | 1170 | 540 | 532 | 518 | 516 | 896,04 | 1266 | 526 | 0,415 | -1,268 | -17,371 | 1,37E-67 | 6,36E-67 | 0,0037446 | 0,0039237 | 0,0038228 | 0,0038228 | TRUE | 0,0807499 | No |
| orf19.7391 | 4073 | 3494 | 3880 | 2489 | 1957 | 1397 | 1876 | 1979 | 3883 | 3694 | 3833 | 3899 | 1568 | 1589 | 1593 | 1609 | 2708,54 | 3827 | 1590 | 0,415 | -1,267 | -32,79 | 8,09E-236 | 1,28E-234 | 1,00E-08 | 0,0026975 | 0,0010066 | 0,0010066 | TRUE | 0,0066906 | No |
| orf19.263.1 | 7503 | 5863 | 6582 | 3739 | 3572 | 1989 | 3246 | 3446 | 7153 | 6199 | 6502 | 5857 | 2862 | 2262 | 2756 | 2802 | 4549,25 | 6428 | 2670 | 0,416 | -1,267 | -15,762 | 5,65E-56 | 2,25E-55 | 0,0086539 | 0,0024522 | 0,00595 | 0,00595 | TRUE | 0,1268161 | No |
| orf19.2565 | 2550 | 2343 | 2507 | 1446 | 1266 | 854 | 1191 | 1261 | 2431 | 2477 | 2477 | 2265 | 1014 | 971 | 1011 | 1025 | 1709,06 | 2412 | 1005 | 0,417 | -1,264 | -26,092 | 4,52E-150 | 4,55E-149 | 0,0005139 | 0,003052 | 0,0015762 | 0,0015762 | TRUE | 0,018655 | No |
| orf19.1106 | 277 | 302 | 300 | 195 | 140 | 118 | 153 | 146 | 264 | 319 | 296 | 305 | 112 | 134 | 130 | 119 | 210,03 | 296 | 124 | 0,417 | -1,263 | -11,333 | 9,01E-30 | 2,42E-29 | 1,00E-08 | 0,0099088 | 0,0064202 | 0,0064202 | TRUE | 0,0646752 | No |
| orf19.7581 | 7330 | 6267 | 6980 | 4019 | 3436 | 2279 | 3440 | 3590 | 6988 | 6627 | 6896 | 6296 | 2753 | 2591 | 2921 | 2919 | 4748,83 | 6702 | 2796 | 0,417 | -1,261 | -24,072 | 4,86E-128 | 4,16E-127 | 0,0023229 | 0,002437 | 0,002391 | 0,002391 | TRUE | 0,0284793 | No |
| orf19.1999 | 1911 | 1601 | 1723 | 1055 | 877 | 585 | 847 | 958 | 1822 | 1693 | 1702 | 1653 | 703 | 665 | 719 | 779 | 1216,95 | 1718 | 716 | 0,418 | -1,26 | -20,755 | 1,12E-95 | 7,00E-95 | 0,0017668 | 0,0034405 | 0,0025901 | 0,0025901 | TRUE | 0,0410715 | No |
| orf19.2005 | 18724 | 16446 | 17945 | 11023 | 9214 | 6314 | 8496 | 9305 | 17851 | 17390 | 17728 | 17268 | 7383 | 7180 | 7213 | 7566 | 12447,3 | 17559 | 7336 | 0,418 | -1,259 | -40,636 | 0 | 0 | 0,0002871 | 0,0022232 | 0,0008291 | 0,0008291 | TRUE | 0,0054858 | No |
| orf19.5607 | 10441 | 9627 | 10041 | 6245 | 5209 | 3500 | 4909 | 5305 | 9954 | 10179 | 9920 | 9783 | 4174 | 3980 | 4168 | 4313 | 7058,9 | 9959 | 4159 | 0,418 | -1,259 | -35,591 | 1,93E-277 | 3,83E-276 | 0,0004512 | 0,0023239 | 0,0010381 | 0,0010381 | TRUE | 0,0097544 | No |
| orf19.2241 | 5701 | 4703 | 5235 | 3228 | 2576 | 1803 | 2646 | 2788 | 5435 | 4973 | 5172 | 5057 | 2064 | 2050 | 2246 | 2267 | 3658,03 | 5159 | 2157 | 0,418 | -1,258 | -25,056 | 1,49E-138 | 1,39E-137 | 0,0017606 | 0,0025401 | 0,0021034 | 0,0021034 | TRUE | 0,0160457 | No |
| orf19.4803 | 14 | 11 | 14 | 9 | 6 | 5 | 6 | 8 | 13 | 12 | 14 | 14 | 5 | 6 | 5 | 7 | 9,38 | 13 | 6 | 0,418 | -1,258 | -2,728 | 0,0063652 | 0,0081723 | 1,00E-08 | 0,1772275 | 0,0815041 | 0,0815041 | TRUE | 0,0420209 | No |
| orf19.2927 | 1507 | 1968 | 1961 | 1166 | 1051 | 789 | 803 | 773 | 1437 | 2081 | 1937 | 1827 | 842 | 897 | 682 | 629 | 1291,39 | 1820 | 762 | 0,418 | -1,257 | -10,08 | 6,75E-24 | 1,64E-23 | 0,0242994 | 0,0033627 | 0,0140387 | 0,0140387 | TRUE | 0,2460589 | No |
| orf19.664 | 85 | 93 | 87 | 63 | 65 | 24 | 53 | 32 | 81 | 98 | 86 | 99 | 52 | 27 | 45 | 26 | 64,3 | 91 | 38 | 0,419 | -1,256 | -5,548 | 2,89E-08 | 4,74E-08 | 0,0371981 | 0,0276262 | 0,0313215 | 0,0313215 | TRUE | 0,5426258 | No |
| orf19.5275 | 488 | 517 | 510 | 333 | 301 | 155 | 255 | 263 | 465 | 547 | 504 | 522 | 241 | 176 | 216 | 214 | 360,65 | 510 | 212 | 0,419 | -1,255 | -12,653 | 1,08E-36 | 3,24E-36 | 0,0057957 | 0,0066439 | 0,0062492 | 0,0062492 | TRUE | 0,1241661 | No |
| orf19.3150 | 2578 | 2455 | 2396 | 1612 | 1262 | 962 | 1245 | 1247 | 2458 | 2596 | 2367 | 2525 | 1011 | 1094 | 1057 | 1014 | 1765,25 | 2486 | 1044 | 0,42 | -1,253 | -24,975 | 1,13E-137 | 1,04E-136 | 0,0009072 | 0,0030214 | 0,0017635 | 0,0017635 | TRUE | 0,0129837 | No |
| orf19.3406 | 1231 | 1091 | 1167 | 754 | 562 | 391 | 653 | 617 | 1174 | 1154 | 1153 | 1181 | 450 | 445 | 554 | 502 | 826,53 | 1166 | 488 | 0,419 | -1,253 | -17,299 | 4,80E-67 | 2,22E-66 | 0,0031664 | 0,0040778 | 0,0036443 | 0,0036443 | TRUE | 0,0989342 | No |
| orf19.609 | 48 | 45 | 45 | 37 | 27 | 21 | 19 | 25 | 46 | 48 | 44 | 58 | 22 | 24 | 16 | 20 | 34,72 | 49 | 20 | 0,42 | -1,251 | -5,019 | 5,20E-07 | 8,14E-07 | 1,00E-08 | 0,0493855 | 0,0263832 | 0,0263832 | TRUE | 0,1226019 | No |
| orf19.1122 | 60 | 77 | 79 | 53 | 44 | 29 | 33 | 36 | 57 | 81 | 78 | 83 | 35 | 33 | 28 | 29 | 53,15 | 75 | 31 | 0,421 | -1,248 | -5,887 | 3,93E-09 | 6,62E-09 | 1,00E-08 | 0,0329822 | 0,0214731 | 0,0214731 | TRUE | 0,2509684 | No |
| orf19.5416 | 7673 | 6941 | 7685 | 4787 | 3859 | 2722 | 3687 | 3966 | 7315 | 7339 | 7592 | 7499 | 3092 | 3095 | 3130 | 3225 | 5286 | 7436 | 3136 | 0,422 | -1,245 | -37,305 | 1,38E-304 | 3,10E-303 | 0,0001712 | 0,0024019 | 0,0008519 | 0,0008519 | TRUE | 0,0045508 | No |
| orf19.4540 | 6306 | 5452 | 5987 | 3765 | 3042 | 2206 | 2885 | 3148 | 6012 | 5765 | 5915 | 5898 | 2437 | 2508 | 2449 | 2560 | 4193,05 | 5898 | 2488 | 0,422 | -1,245 | -35,261 | 2,31E-272 | 4,41E-271 | 0,0001642 | 0,0024829 | 0,0009223 | 0,0009223 | TRUE | 0,0047277 | No |
| orf19.6644 | 182 | 192 | 245 | 167 | 101 | 93 | 122 | 100 | 174 | 203 | 242 | 262 | 81 | 106 | 104 | 81 | 156,47 | 220 | 93 | 0,422 | -1,243 | -7,877 | 3,35E-15 | 6,79E-15 | 0,0209945 | 0,0125848 | 0,0165324 | 0,0165324 | TRUE | 0,1440593 | No |
| orf19.7341 | 4699 | 3927 | 4271 | 2649 | 2196 | 1529 | 2189 | 2264 | 4480 | 4152 | 4219 | 4150 | 1760 | 1739 | 1858 | 1841 | 3024,87 | 4250 | 1800 | 0,423 | -1,24 | -27,848 | 1,13E-170 | 1,29E-169 | 0,0008869 | 0,0026341 | 0,0015233 | 0,0015233 | TRUE | 0,0164266 | No |
| orf19.1835 | 4228 | 4317 | 4578 | 2829 | 2266 | 1751 | 2163 | 2203 | 4031 | 4565 | 4523 | 4432 | 1816 | 1991 | 1836 | 1791 | 3123,05 | 4388 | 1858 | 0,423 | -1,24 | -22,716 | 3,15E-114 | 2,37E-113 | 0,0022149 | 0,002617 | 0,0024938 | 0,0024938 | TRUE | 0,0372377 | No |
| orf19.2237 | 76 | 56 | 67 | 41 | 40 | 26 | 28 | 32 | 72 | 59 | 66 | 64 | 32 | 30 | 24 | 26 | 46,69 | 65 | 28 | 0,423 | -1,24 | -5,796 | 6,80E-09 | 1,14E-08 | 1,00E-08 | 0,0372597 | 0,0194992 | 0,0194992 | TRUE | 0,0838774 | No |
| orf19.5558 | 1625 | 1262 | 1508 | 1082 | 794 | 541 | 726 | 857 | 1549 | 1334 | 1490 | 1695 | 636 | 615 | 616 | 697 | 1079,13 | 1517 | 641 | 0,424 | -1,239 | -15,881 | 8,60E-57 | 3,45E-56 | 0,0057585 | 0,0036128 | 0,0047759 | 0,0047759 | TRUE | 0,0784529 | No |
| orf19.5113 | 4356 | 3074 | 3294 | 2418 | 1954 | 1273 | 1709 | 2037 | 4153 | 3250 | 3254 | 3788 | 1566 | 1448 | 1451 | 1656 | 2570,73 | 3611 | 1530 | 0,424 | -1,238 | -14,822 | 1,06E-49 | 3,89E-49 | 0,0089779 | 0,00273 | 0,0062527 | 0,0062527 | TRUE | 0,1260983 | No |
| orf19.5620 | 795 | 777 | 751 | 485 | 449 | 290 | 383 | 362 | 758 | 822 | 742 | 760 | 360 | 330 | 325 | 294 | 548,78 | 770 | 327 | 0,425 | -1,235 | -15,963 | 2,31E-57 | 9,35E-57 | 0,0021373 | 0,0050832 | 0,0036606 | 0,0036606 | TRUE | 0,0563548 | No |
| orf19.2067 | 4618 | 4025 | 4257 | 2472 | 2251 | 1469 | 2199 | 2181 | 4403 | 4256 | 4206 | 3873 | 1804 | 1670 | 1867 | 1773 | 2981,38 | 4184 | 1778 | 0,425 | -1,234 | -23,324 | 2,55E-120 | 2,03E-119 | 0,0019614 | 0,002642 | 0,0023041 | 0,0023041 | TRUE | 0,0293337 | No |
| orf19.849 | 84 | 103 | 88 | 64 | 44 | 40 | 45 | 51 | 80 | 109 | 87 | 100 | 35 | 45 | 38 | 41 | 67,08 | 94 | 40 | 0,426 | -1,232 | -6,561 | 5,34E-11 | 9,62E-11 | 1,00E-08 | 0,0265698 | 0,0167281 | 0,0167281 | TRUE | 0,1207656 | No |
| orf19.4530.1 | 8342 | 6990 | 7902 | 5093 | 4084 | 2810 | 4108 | 4059 | 7953 | 7391 | 7806 | 7978 | 3272 | 3195 | 3488 | 3300 | 5548,1 | 7782 | 3314 | 0,426 | -1,231 | -28,85 | 5,14E-183 | 6,31E-182 | 0,0010233 | 0,0023872 | 0,0015407 | 0,0015407 | TRUE | 0,0151454 | No |
| orf19.4653 | 12 | 8 | 14 | 14 | 4 | 6 | 8 | 8 | 11 | 8 | 14 | 22 | 3 | 7 | 7 | 7 | 9,87 | 14 | 6 | 0,426 | -1,231 | -2,505 | 0,0122387 | 0,0154675 | 1,00E-08 | 0,1683885 | 0,1137902 | 0,1137902 | TRUE | 0,3409633 | No |
| orf19.3759 | 6728 | 6236 | 6900 | 4342 | 3451 | 2647 | 3352 | 3364 | 6414 | 6594 | 6817 | 6802 | 2765 | 3010 | 2846 | 2735 | 4747,86 | 6657 | 2839 | 0,426 | -1,23 | -28,7 | 3,82E-181 | 4,62E-180 | 0,0010161 | 0,0024371 | 0,0015225 | 0,0015225 | TRUE | 0,0191016 | No |
| orf19.3563 | 3398 | 3019 | 3348 | 2390 | 1739 | 1320 | 1704 | 1751 | 3240 | 3192 | 3308 | 3744 | 1393 | 1501 | 1447 | 1424 | 2406,02 | 3371 | 1441 | 0,428 | -1,225 | -21,028 | 3,67E-98 | 2,36E-97 | 0,0026589 | 0,0027737 | 0,0027801 | 0,0027801 | TRUE | 0,0648186 | No |
| orf19.3456 | 866 | 870 | 900 | 591 | 487 | 347 | 453 | 435 | 826 | 920 | 889 | 926 | 390 | 395 | 385 | 354 | 635,45 | 890 | 381 | 0,428 | -1,224 | -17,591 | 2,87E-69 | 1,36E-68 | 0,0010865 | 0,0046751 | 0,0028501 | 0,0028501 | TRUE | 0,0293815 | No |
| orf19.420 | 32 | 23 | 12 | 13 | 13 | 11 | 8 | 10 | 31 | 24 | 12 | 20 | 10 | 13 | 7 | 8 | 15,61 | 22 | 10 | 0,428 | -1,224 | -3,009 | 0,0026197 | 0,0034481 | 1,00E-08 | 0,1072609 | 0,0857583 | 0,0857583 | TRUE | 0,2002529 | No |
| orf19.4617 | 1383 | 1431 | 1331 | 836 | 745 | 507 | 727 | 680 | 1319 | 1513 | 1315 | 1310 | 597 | 577 | 617 | 553 | 974,97 | 1364 | 586 | 0,429 | -1,22 | -17,902 | 1,14E-71 | 5,61E-71 | 0,002893 | 0,0037753 | 0,0032832 | 0,0032832 | TRUE | 0,0657571 | No |
| orf19.5299.1 | 1261 | 1184 | 1348 | 588 | 651 | 431 | 621 | 598 | 1202 | 1252 | 1332 | 921 | 522 | 490 | 527 | 486 | 841,52 | 1177 | 506 | 0,429 | -1,22 | -12,206 | 2,90E-34 | 8,37E-34 | 0,0122375 | 0,0040424 | 0,0082349 | 0,0082349 | TRUE | 0,2426381 | No |
| orf19.4299 | 1788 | 1584 | 1745 | 1145 | 916 | 619 | 910 | 924 | 1705 | 1675 | 1724 | 1794 | 734 | 704 | 773 | 751 | 1232,36 | 1724 | 740 | 0,431 | -1,215 | -23,563 | 9,27E-123 | 7,59E-122 | 1,00E-08 | 0,0034236 | 0,0016286 | 0,0016286 | TRUE | 0,0115755 | No |
| orf19.113 | 2155 | 2036 | 2074 | 1309 | 1154 | 843 | 1000 | 1050 | 2055 | 2153 | 2049 | 2051 | 925 | 959 | 849 | 854 | 1486,61 | 2077 | 897 | 0,431 | -1,215 | -22,686 | 6,12E-114 | 4,59E-113 | 0,0010446 | 0,0031957 | 0,0019839 | 0,0019839 | TRUE | 0,0236236 | No |
| orf19.2710 | 4553 | 4036 | 4442 | 2258 | 2200 | 1451 | 2265 | 2202 | 4341 | 4268 | 4388 | 3537 | 1763 | 1650 | 1923 | 1790 | 2957,5 | 4134 | 1782 | 0,431 | -1,215 | -16,496 | 3,91E-61 | 1,67E-60 | 0,0061426 | 0,0026464 | 0,004822 | 0,004822 | TRUE | 0,1120209 | No |
| orf19.703 | 1682 | 1396 | 1479 | 1009 | 818 | 566 | 782 | 826 | 1604 | 1476 | 1461 | 1581 | 655 | 644 | 664 | 672 | 1094,5 | 1530 | 659 | 0,431 | -1,214 | -21,801 | 2,29E-105 | 1,56E-104 | 1,00E-08 | 0,0035914 | 0,0019348 | 0,0019348 | TRUE | 0,0139703 | No |
| orf19.3483 | 10181 | 11277 | 11144 | 6422 | 6063 | 4499 | 5225 | 4919 | 9706 | 11924 | 11009 | 10060 | 4858 | 5116 | 4436 | 4000 | 7638,7 | 10675 | 4602 | 0,431 | -1,214 | -14,703 | 6,18E-49 | 2,24E-48 | 0,009827 | 0,0023062 | 0,0064032 | 0,0064032 | TRUE | 0,094969 | No |
| orf19.5228 | 6173 | 5548 | 6109 | 3945 | 3147 | 2410 | 3026 | 3091 | 5885 | 5866 | 6035 | 6180 | 2522 | 2740 | 2569 | 2513 | 4288,89 | 5992 | 2586 | 0,431 | -1,213 | -29,554 | 5,86E-192 | 7,73E-191 | 0,0007663 | 0,0024741 | 0,0013508 | 0,0013508 | TRUE | 0,018603 | No |
| orf19.4892 | 12078 | 11737 | 12118 | 7205 | 6236 | 4651 | 6072 | 6064 | 11515 | 12410 | 11971 | 11287 | 4997 | 5289 | 5155 | 4931 | 8444,37 | 11796 | 5093 | 0,431 | -1,213 | -28,528 | 5,33E-179 | 6,32E-178 | 0,0012061 | 0,0022857 | 0,0015999 | 0,0015999 | TRUE | 0,0149873 | No |
| orf19.5041 | 225 | 202 | 176 | 125 | 95 | 78 | 102 | 115 | 215 | 214 | 174 | 196 | 76 | 89 | 87 | 94 | 142,84 | 200 | 86 | 0,431 | -1,213 | -9,173 | 4,60E-20 | 1,03E-19 | 1,00E-08 | 0,0135861 | 0,0088228 | 0,0088228 | TRUE | 0,088848 | No |
| orf19.431 | 7604 | 6593 | 7260 | 4616 | 3758 | 2723 | 3588 | 3960 | 7250 | 6971 | 7172 | 7231 | 3011 | 3096 | 3046 | 3220 | 5124,72 | 7156 | 3093 | 0,432 | -1,21 | -33,54 | 1,24E-246 | 2,08E-245 | 0,0002854 | 0,0024117 | 0,0010262 | 0,0010262 | TRUE | 0,0095504 | No |
| orf19.5389 | 1226 | 1116 | 1192 | 866 | 609 | 515 | 628 | 623 | 1169 | 1180 | 1178 | 1357 | 488 | 586 | 533 | 507 | 874,55 | 1221 | 528 | 0,433 | -1,209 | -16,195 | 5,44E-59 | 2,26E-58 | 0,0041799 | 0,0039687 | 0,0040401 | 0,0040401 | TRUE | 0,0625263 | No |
| orf19.1405 | 722 | 655 | 633 | 435 | 354 | 248 | 376 | 342 | 688 | 693 | 625 | 681 | 284 | 282 | 319 | 278 | 481,33 | 672 | 291 | 0,433 | -1,207 | -15,653 | 3,15E-55 | 1,25E-54 | 1,00E-08 | 0,0055024 | 0,0033422 | 0,0033422 | TRUE | 0,0508672 | No |
| orf19.4842 | 1438 | 1104 | 1183 | 940 | 682 | 466 | 684 | 713 | 1371 | 1167 | 1169 | 1473 | 546 | 530 | 581 | 580 | 927,05 | 1295 | 559 | 0,433 | -1,207 | -14,351 | 1,05E-46 | 3,71E-46 | 0,007019 | 0,0038624 | 0,00556 | 0,00556 | TRUE | 0,0965747 | No |
| orf19.193 | 433 | 509 | 497 | 287 | 280 | 206 | 233 | 204 | 413 | 538 | 491 | 450 | 224 | 234 | 198 | 166 | 339,24 | 473 | 206 | 0,433 | -1,207 | -10,047 | 9,45E-24 | 2,28E-23 | 0,0144051 | 0,0069313 | 0,0105001 | 0,0105001 | TRUE | 0,1917761 | No |
| orf19.7434 | 1040 | 1065 | 1070 | 627 | 589 | 408 | 525 | 522 | 992 | 1126 | 1057 | 982 | 472 | 464 | 446 | 424 | 745,37 | 1039 | 452 | 0,434 | -1,206 | -17,437 | 4,34E-68 | 2,03E-67 | 0,0019907 | 0,0042941 | 0,0030633 | 0,0030633 | TRUE | 0,037647 | No |
| orf19.4888 | 3555 | 3353 | 3399 | 2097 | 1953 | 1250 | 1776 | 1714 | 3389 | 3545 | 3358 | 3285 | 1565 | 1421 | 1508 | 1394 | 2433,17 | 3394 | 1472 | 0,434 | -1,205 | -24,044 | 9,61E-128 | 8,20E-127 | 0,0013048 | 0,0027661 | 0,0019432 | 0,0019432 | TRUE | 0,0222726 | No |
| orf19.5569 | 10795 | 9605 | 10421 | 6263 | 5510 | 3739 | 5302 | 5453 | 10292 | 10156 | 10295 | 9811 | 4415 | 4252 | 4501 | 4434 | 7269,5 | 10138 | 4400 | 0,434 | -1,204 | -35,436 | 4,84E-275 | 9,39E-274 | 0,0003245 | 0,0023171 | 0,0009516 | 0,0009516 | TRUE | 0,0060167 | No |
| orf19.6843 | 7817 | 6687 | 7302 | 3982 | 3923 | 2401 | 3754 | 3812 | 7453 | 7071 | 7214 | 6238 | 3143 | 2730 | 3187 | 3100 | 5016,9 | 6994 | 3040 | 0,435 | -1,202 | -18,158 | 1,12E-73 | 5,62E-73 | 0,0049124 | 0,0024186 | 0,003981 | 0,003981 | TRUE | 0,0635316 | No |
| orf19.2834 | 3465 | 3230 | 3535 | 2148 | 1883 | 1284 | 1748 | 1787 | 3304 | 3415 | 3492 | 3365 | 1509 | 1460 | 1484 | 1453 | 2435,24 | 3394 | 1476 | 0,435 | -1,201 | -29,955 | 3,76E-197 | 5,07E-196 | 1,00E-08 | 0,0027655 | 0,001075 | 0,001075 | TRUE | 0,0048015 | No |
| orf19.6672 | 8119 | 7010 | 7630 | 4685 | 4068 | 2727 | 3838 | 4224 | 7741 | 7412 | 7538 | 7339 | 3260 | 3101 | 3258 | 3435 | 5385,4 | 7508 | 3264 | 0,435 | -1,201 | -29,46 | 9,31E-191 | 1,22E-189 | 0,0008532 | 0,0023962 | 0,0013851 | 0,0013851 | TRUE | 0,0151405 | No |
| orf19.904 | 1222 | 1124 | 1165 | 747 | 633 | 451 | 575 | 646 | 1165 | 1188 | 1151 | 1170 | 507 | 513 | 488 | 525 | 838,52 | 1168 | 508 | 0,435 | -1,201 | -21,509 | 1,29E-102 | 8,61E-102 | 1,00E-08 | 0,0040494 | 0,0016395 | 0,0016395 | TRUE | 0,0086941 | No |
| orf19.73 | 5017 | 5057 | 5243 | 3071 | 2813 | 2126 | 2464 | 2457 | 4783 | 5347 | 5180 | 4811 | 2254 | 2417 | 2092 | 1998 | 3610,24 | 5030 | 2190 | 0,435 | -1,201 | -18,495 | 2,27E-76 | 1,18E-75 | 0,0046479 | 0,0025461 | 0,0037342 | 0,0037342 | TRUE | 0,0575456 | No |
| orf19.1877 | 632 | 583 | 644 | 402 | 294 | 249 | 338 | 341 | 603 | 616 | 636 | 630 | 236 | 283 | 287 | 277 | 445,99 | 621 | 271 | 0,435 | -1,201 | -14,956 | 1,42E-50 | 5,29E-50 | 0,0006099 | 0,0057728 | 0,0036401 | 0,0036401 | TRUE | 0,0917714 | No |

| Chromosome (start-end) | score | signal Value | p-value (-log10) | q-value (-log10) | ORF_left | ORF_right |
|---|---|---|---|---|---|---|
| chr4:707875-708489 | 181992.0 | 49,2993 | 18203.7 | 18199.3 | orf19.3364 | |
| chr2:660301-663071 | 166411.0 | 36,935 | 16645.5 | 16641.1 | orf19.909 | |
| chr4:623628-625018 | 131964.0 | 36,8241 | 13200.6 | 13196.4 | orf19.2693 | |
| chr5:420121-420753 | 141568.0 | 36,4483 | 14161.1 | 14156.8 | orf19.3188 | |
| chrR:523374-524356 | 101989.0 | 31,1712 | 10203.1 | 10199.0 | orf19.3734 | orf19.3733 |
| chrR:1669286-1671350 | 74421.0 | 25,0399 | 7446.08 | 7442.19 | | orf19.1985 |
| chr2:642987-643647 | 85602.0 | 25,0131 | 8564.26 | 8560.28 | orf19.5779 | orf19.5777 |
| chr3:431963-433080 | 67008.0 | 22,3347 | 6704.63 | 6700.82 | orf19.1652 | |
| chr2:1313742-1315290 | 64562.0 | 21,9286 | 6459.99 | 6456.21 | orf19.22.1 | orf19.23 |
| chrR:51541-52559 | 73454.0 | 21,1574 | 7349.32 | 7345.45 | | orf19.7513 |
| chr2:1465213-1466140 | 57214.0 | 20,3256 | 5725.11 | 5721.41 | orf19.2272 | orf19.2274 |
| chr1:1080222-1082319 | 48541.0 | 18,226 | 4857.73 | 4854.12 | | orf19.4056 |
| chr1:3060027-3061295 | 44593.0 | 18,0735 | 4462.87 | 4459.32 | orf19.5026 | |
| chr2:1218155-1222263 | 51162.0 | 17,8595 | 5119.89 | 5116.25 | orf19.5217 | |
| chr2:1218155-1222263 | 51162.0 | 17,8595 | 5119.89 | 5116.25 | orf19.4122 | |
| chr1:918488-921560 | 54864.0 | 17,5817 | 5490.16 | 5486.48 | orf19.6838 | |
| chr4:1495039-1496117 | 47685.0 | 17,205 | 4772.19 | 4768.59 | | orf19.3139 |
| chr7:284035-284550 | 39933.0 | 16,7848 | 3996.85 | 3993.38 | | orf19.6920 |
| chr2:936958-937975 | 39762.0 | 16,7367 | 3979.69 | 3976.22 | | orf19.4504 |
| chr1:435307-441844 | 47492.0 | 16,7095 | 4752.86 | 4749.27 | orf19.3668 | |
| chr1:3005862-3006560 | 37922.0 | 16,2177 | 3795.71 | 3792.27 | | orf19.5005 |
| chr6:35410-37507 | 50554.0 | 16,1287 | 5059.1 | 5055.47 | | orf19.1189 |
| chr2:1215452-1216381 | 40505.0 | 15,9197 | 4054.07 | 4050.58 | orf19.5215 | |
| chr2:1132118-1134566 | 39079.0 | 15,3802 | 3911.44 | 3907.98 | orf19.6869 | orf19.6871 |
| chr5:686543-687480 | 34012.0 | 15,0939 | 3404.57 | 3401.22 | orf19.4337 | orf19.4339 |
| chr3:865944-866949 | 37880.0 | 13,9412 | 3791.46 | 3788.03 | | orf19.5859 |
| chr2:131527-133650 | 34913.0 | 13,826 | 3494.69 | 3491.31 | orf19.2047 | orf19.2047 |
| chr2:131527-133650 | 34913.0 | 13,826 | 3494.69 | 3491.31 | | orf19.2046 |
| chr2:827297-828948 | 31010.0 | 13,8256 | 3104.31 | 3101.03 | orf19.829 | orf19.828 |
| chr4:787820-788565 | 29754.0 | 13,411 | 2978.69 | 2975.45 | orf19.1309 | orf19.1308 |
| chrR:443946-444747 | 35714.0 | 13,2439 | 3574.79 | 3571.4 | orf19.2602 | |
| chr1:2270114-2271413 | 31226.0 | 13,2168 | 3125.91 | 3122.62 | orf19.4898 | orf19.4899 |
| chr1:65805-67624 | 30565.0 | 12,8029 | 3059.86 | 3056.59 | orf19.6066 | orf19.6065 |
| chr3:312341-314205 | 26295.0 | 12,7776 | 2632.72 | 2629.55 | orf19.1704 | orf19.1702 |
| chr6:111825-112431 | 30424.0 | 12,6069 | 3045.73 | 3042.46 | orf19.4196 | orf19.4195.1 |
| chr3:734618-735828 | 29639.0 | 12,3663 | 2967.2 | 2963.95 | orf19.344 | orf19.345 |
| chr1:2996467-2997094 | 24920.0 | 12,348 | 2495.14 | 2492.01 | | orf19.5000 |
| chr3:1414306-1414896 | 29089.0 | 12,1882 | 2912.17 | 2908.94 | orf19.7414 | |
| chr3:607665-609196 | 26337.0 | 12,0136 | 2636.96 | 2633.79 | orf19.272 | |
| chr6:615542-617704 | 26025.0 | 11,929 | 2605.72 | 2602.56 | orf19.5576 | orf19.5578 |
| chr5:111444-113904 | 33381.0 | 11,8511 | 3341.51 | 3338.17 | orf19.935 | |
| chr5:177976-178823 | 23808.0 | 11,7765 | 2383.95 | 2380.86 | orf19.577 | orf19.578 |
| chrR:1698594-1700220 | 21402.0 | 11,2208 | 2143.29 | 2140.26 | | orf19.3711 |
| chr2:258174-259422 | 23390.0 | 11,1169 | 2342.1 | 2339.02 | orf19.1449 | |
| chr1:2044955-2045971 | 22135.0 | 10,9745 | 2216.57 | 2213.53 | orf19.4789 | |
| chr5:415559-416930 | 20441.0 | 10,9046 | 2047.11 | 2044.12 | orf19.3190 | |
| chrR:589755-591285 | 25089.0 | 10,8362 | 2512.06 | 2508.92 | orf19.2822 | orf19.2823 |
| chr7:297080-298585 | 19573.0 | 10,6159 | 1960.3 | 1957.34 | orf19.6586 | orf19.6585 |
| chr1:1572859-1573633 | 21250.0 | 10,5862 | 2128.03 | 2125.01 | | orf19.4438 |
| chr6:166457-166984 | 19623.0 | 10,5661 | 1965.3 | 1962.33 | | orf19.86 |
| chr3:152559-153870 | 22746.0 | 10,5579 | 2277.69 | 2274.62 | | orf19.3071 |
| chr1:2563457-2564331 | 20305.0 | 10,4995 | 2033.57 | 2030.58 | orf19.1149 | |
| chr3:160874-161702 | 21484.0 | 10,3814 | 2151.44 | 2148.41 | orf19.1288 | orf19.1289 |
| chr2:613465-614209 | 23403.0 | 10,1867 | 2343.43 | 2340.34 | | orf19.5797 |
| chr4:853079-853825 | 18290.0 | 10,1829 | 1831.97 | 1829.04 | orf19.1208 | orf19.5312 |

| | | | | | | |
|---|---|---|---|---|---|---|
| chr7:909075-913741 | 25509.0 | 10,1782 | 2554.12 | 2550.98 | orf19.7152 | orf19.7151 |
| chr3:796529-798193 | 22587.0 | 9,89276 | 2261.77 | 2258.71 | | orf19.6937 |
| chrR:990163-991506 | 19629.0 | 9,8831 | 1965.9 | 1962.93 | orf19.539 | |
| chrR:208821-209780 | 17298.0 | 9,84267 | 1732.76 | 1729.87 | | orf19.3263 |
| chr3:667172-667965 | 19553.0 | 9,74601 | 1958.36 | 1955.4 | orf19.310 | |
| chr7:899209-902096 | 20756.0 | 9,73205 | 2078.65 | 2075.64 | | orf19.7156 |
| chrR:1911285-1913112 | 21546.0 | 9,64615 | 2157.71 | 2154.68 | orf19.7286 | orf19.7288 |
| chr2:104781-105495 | 20165.0 | 9,61973 | 2019.58 | 2016.59 | | orf19.2063 |
| chr1:2291374-2292606 | 16184.0 | 9,45433 | 1621.34 | 1618.48 | orf19.4906 | |
| chr4:709395-710265 | 16057.0 | 9,40965 | 1608.64 | 1605.79 | orf19.3365 | orf19.3366 |
| chr6:668408-669334 | 16785.0 | 9,3715 | 1681.39 | 1678.51 | orf19.5604 | |
| chrR:1376755-1377673 | 15737.0 | 9,29624 | 1576.55 | 1573.71 | orf19.723 | orf19.721 |
| chr5:238058-239563 | 15640.0 | 9,26187 | 1566.86 | 1564.02 | orf19.1961 | |
| chr6:985330-985977 | 19115.0 | 9,24539 | 1914.51 | 1911.57 | orf19.2124 | orf19.2125 |
| chrR:2210729-2211222 | 15387.0 | 9,13045 | 1541.56 | 1538.73 | orf19.7611 | |
| chrR:1233181-1234207 | 18304.0 | 9,06139 | 1833.4 | 1830.48 | | orf19.6627 |
| chr6:739795-740317 | 14499.0 | 8,85291 | 1452.79 | 1449.99 | | orf19.5713 |
| chr2:1478310-1478812 | 16587.0 | 8,72218 | 1661.58 | 1658.71 | orf19.2281 | |
| chr1:444445-448224 | 14945.0 | 8,67466 | 1497.38 | 1494.57 | orf19.3669 | |
| chr1:2852632-2853815 | 18871.0 | 8,65146 | 1890.04 | 1887.11 | | orf19.4936 |
| chr1:1544943-1545539 | 15199.0 | 8,64218 | 1522.78 | 1519.96 | | orf19.4447 |
| chr5:497248-497795 | 13858.0 | 8,61921 | 1388.67 | 1385.9 | | orf19.4231 |
| chr4:920869-921799 | 15874.0 | 8,60414 | 1590.29 | 1587.44 | | orf19.1415 |
| chr2:1067519-1069590 | 15115.0 | 8,53388 | 1514.37 | 1511.55 | | orf19.3548.1 |
| chr5:910536-911096 | 16257.0 | 8,49922 | 1628.6 | 1625.74 | orf19.1282 | |
| chr4:625281-626302 | 13858.0 | 8,44137 | 1388.66 | 1385.89 | orf19.2691 | |
| chr4:635777-636653 | 13417.0 | 8,33495 | 1344.48 | 1341.72 | orf19.2686 | |
| chr2:1362314-1363495 | 13041.0 | 8,31678 | 1306.89 | 1304.15 | orf19.1223 | orf19.1224 |
| chr4:238664-239946 | 12986.0 | 8,21092 | 1301.4 | 1298.67 | orf19.4673 | orf19.4672 |
| chrR:1295814-1296389 | 13333.0 | 8,18849 | 1336.1 | 1333.35 | | orf19.3859 |
| chr2:1310277-1311031 | 12657.0 | 8,12154 | 1268.48 | 1265.75 | orf19.21 | |
| chr2:1079394-1080154 | 13106.0 | 8,09417 | 1313.41 | 1310.66 | orf19.3554 | orf19.3555 |
| chr6:790827-794076 | 16212.0 | 8,02185 | 1624.14 | 1621.29 | | orf19.5741 |
| chr1:3112036-3112607 | 13425.0 | 7,97951 | 1345.33 | 1342.58 | orf19.7218 | orf19.7219 |
| chr3:174431-175139 | 14489.0 | 7,97849 | 1451.72 | 1448.92 | | orf19.6165 |
| chr1:2335820-2337219 | 13047.0 | 7,89974 | 1307.52 | 1304.77 | orf19.999 | |
| chr5:681252-682188 | 11680.0 | 7,80128 | 1170.72 | 1168.03 | orf19.4335 | orf19.4336 |
| chr7:453361-454370 | 14627.0 | 7,79011 | 1465.53 | 1462.72 | orf19.6514 | |
| chr5:696126-697188 | 12423.0 | 7,77418 | 1245.07 | 1242.36 | orf19.4341 | orf19.4342 |
| chrR:1704069-1706092 | 12332.0 | 7,76178 | 1235.93 | 1233.22 | orf19.3713 | orf19.3714 |
| chrR:670221-671440 | 13289.0 | 7,75442 | 1331.71 | 1328.96 | orf19.2849 | orf19.2850 |
| chr1:1946936-1947442 | 10998.0 | 7,53666 | 1102.47 | 1099.82 | orf19.4752 | orf19.4753 |
| chr3:1194761-1195210 | 10857.0 | 7,48167 | 1088.43 | 1085.79 | orf19.6975 | orf19.6976 |
| chr4:574482-576198 | 11184.0 | 7,47563 | 1121.1 | 1118.44 | | orf19.2723 |
| chr4:1306932-1309222 | 10155.0 | 7,2033 | 1018.17 | 1015.56 | orf19.1277 | |
| chr2:765447-766873 | 10708.0 | 7,19614 | 1073.44 | 1070.81 | | orf19.856 |
| chr1:2225772-2227642 | 11895.0 | 7,0988 | 1192.21 | 1189.52 | orf19.4883 | orf19.4884 |
| chr7:144709-145468 | 9890.0 | 7,09676 | 991.635 | 989.039 | | orf19.7042 |
| chr2:167837-170091 | 11535.0 | 7,06391 | 1156.19 | 1153.51 | | orf19.2020 |
| chr2:829296-829904 | 14049.0 | 7,0521 | 1407.74 | 1404.96 | orf19.829 | orf19.828 |
| chr3:903665-905038 | 11772.0 | 7,04893 | 1179.95 | 1177.26 | orf19.5879 | orf19.5880 |
| chr5:168589-170675 | 10006.0 | 7,03814 | 1003.27 | 1000.67 | | orf19.575 |
| chr1:1207644-1208273 | 12142.0 | 6,97557 | 1216.98 | 1214.28 | orf19.2474 | orf19.2473 |
| chr5:123167-124416 | 10879.0 | 6,96824 | 1090.62 | 1087.97 | | orf19.932 |
| chr3:970791-971991 | 13398.0 | 6,96107 | 1342.58 | 1339.82 | orf19.5917 | orf19.5917.3 |
| chr2:790178-790978 | 10975.0 | 6,93557 | 1100.19 | 1097.54 | | orf19.847 |

| | | | | | | |
|---|---|---|---|---|---|---|
| chr1:2485716-2486709 | 11780.0 | 6,93419 | 1180.76 | 1178.07 | orf19.670.2 | |
| chr6:291454-292422 | 9307.0 | 6,85963 | 933.305 | 930.738 | orf19.3439 | |
| chr5:44644-46163 | 10251.0 | 6,80128 | 1027.73 | 1025.11 | | orf19.968 |
| chr7:888437-889220 | 10779.0 | 6,76297 | 1080.6 | 1077.96 | orf19.7166 | |
| chr4:862273-864479 | 8965.0 | 6,71872 | 899.134 | 896.585 | orf19.5308 | orf19.5307 |
| chr5:653154-655149 | 8941.0 | 6,70841 | 896.648 | 894.101 | orf19.4318 | |
| chrR:1373755-1374418 | 9605.0 | 6,68629 | 963.103 | 960.521 | orf19.723 | orf19.721 |
| chrR:1802054-1803691 | 10541.0 | 6,62444 | 1056.77 | 1054.14 | orf19.6398 | orf19.6399 |
| chr6:837286-838121 | 9512.0 | 6,61831 | 953.787 | 951.209 | orf19.5759 | |
| chr5:1136173-1137103 | 9660.0 | 6,59514 | 968.647 | 966.062 | orf19.4010 | orf19.4011 |
| chr4:129370-131331 | 10896.0 | 6,57597 | 1092.29 | 1089.64 | orf19.4167 | orf19.4166 |
| chr3:1224979-1225844 | 8489.0 | 6,5194 | 851.433 | 848.91 | | orf19.6984 |
| chr3:760457-760836 | 9971.0 | 6,49887 | 999.795 | 997.194 | orf19.356 | orf19.357 |
| chr4:1540616-1541088 | 10047.0 | 6,48108 | 1007.31 | 1004.71 | orf19.3117 | |
| chr3:853701-854524 | 9057.0 | 6,45103 | 908.258 | 905.704 | orf19.918 | orf19.913.2 |
| chrR:1943295-1943904 | 9729.0 | 6,45079 | 975.526 | 972.938 | orf19.7306 | |
| chr4:521506-522846 | 8172.0 | 6,38537 | 819.79 | 817.285 | | orf19.2747 |
| chr2:2165887-2166848 | 8671.0 | 6,37229 | 869.666 | 867.134 | | orf19.5323 |
| chr2:1484119-1484833 | 9802.0 | 6,32604 | 982.862 | 980.27 | | orf19.2286 |
| chr2:518176-518737 | 8982.0 | 6,27747 | 900.777 | 898.227 | | orf19.1582 |
| chr1:31443-31871 | 8813.0 | 6,26652 | 883.895 | 881.354 | | orf19.6085 |
| chr5:825210-825670 | 9017.0 | 6,25643 | 904.27 | 901.719 | | orf19.1121 |
| chr1:2870325-2871779 | 8800.0 | 6,23738 | 882.571 | 880.032 | orf19.4941 | orf19.4942 |
| chr2:199826-201239 | 7772.0 | 6,21353 | 779.742 | 777.262 | orf19.2005 | |
| chr2:17666-18460 | 8977.0 | 6,15174 | 900.333 | 897.784 | orf19.1907 | orf19.1906 |
| chrR:730730-732043 | 7614.0 | 6,1448 | 763.891 | 761.419 | orf19.2398 | orf19.2397.3 |
| chr1:1047276-1047700 | 7590.0 | 6,13449 | 761.521 | 759.052 | orf19.53 | orf19.54 |
| chrR:1875716-1876622 | 9773.0 | 6,13339 | 979.928 | 977.338 | orf19.6445 | |
| chr4:1254831-1255907 | 8371.0 | 6,12786 | 839.681 | 837.165 | | orf19.1255 |
| chr1:1988882-1989898 | 7551.0 | 6,1173 | 757.577 | 755.11 | | orf19.4769 |
| chr1:231905-232585 | 8988.0 | 6,10336 | 901.449 | 898.899 | orf19.3302 | |
| chrR:39014-42875 | 8613.0 | 5,98492 | 863.877 | 861.347 | orf19.7520 | |
| chrR:2111004-2111852 | 7486.0 | 5,97585 | 751.132 | 748.669 | | orf19.7563 |
| chr3:197137-198003 | 7206.0 | 5,96609 | 723.133 | 720.687 | | orf19.2501 |
| chr3:1239352-1240098 | 7160.0 | 5,94547 | 718.473 | 716.03 | orf19.6987 | orf19.6988 |
| chr1:2844899-2845820 | 7220.0 | 5,92432 | 724.473 | 722.026 | orf19.4933 | orf19.4934 |
| chrR:87047-87767 | 8510.0 | 5,90356 | 853.606 | 851.082 | orf19.7499.1 | orf19.7499 |
| chr2:315624-316101 | 7013.0 | 5,88017 | 703.778 | 701.344 | orf19.1486 | |
| chr1:2003392-2004646 | 8090.0 | 5,83165 | 811.541 | 809.041 | orf19.4772 | orf19.4773 |
| chr5:428958-429425 | 7495.0 | 5,80501 | 752.038 | 749.574 | | orf19.3182 |
| chr4:1455463-1456349 | 7601.0 | 5,79896 | 762.655 | 760.185 | orf19.2882 | orf19.2881 |
| chr4:442875-444337 | 7538.0 | 5,79621 | 756.276 | 753.809 | orf19.4567 | |
| chr2:1424277-1425247 | 7327.0 | 5,75989 | 735.185 | 732.732 | orf19.2248 | |
| chr3:69331-69821 | 8008.0 | 5,75248 | 803.341 | 800.846 | orf19.5417 | orf19.5416 |
| chrR:12196-13822 | 8174.0 | 5,72539 | 819.929 | 817.424 | orf19.7539.1 | orf19.7539 |
| chr1:2314324-2314988 | 7303.0 | 5,70754 | 732.786 | 730.334 | | orf19.986 |
| chr6:284148-284737 | 6525.0 | 5,66022 | 654.956 | 652.553 | orf19.3442 | |
| chr1:2024160-2025092 | 7039.0 | 5,59105 | 706.342 | 703.907 | orf19.4779 | orf19.4780 |
| chr1:404584-406904 | 6752.0 | 5,58676 | 677.716 | 675.299 | | orf19.4527 |
| chr1:1303013-1303454 | 6352.0 | 5,58118 | 637.672 | 635.281 | orf19.1826 | |
| chr2:350003-350442 | 6285.0 | 5,55025 | 630.947 | 628.561 | orf19.1497 | orf19.1499 |
| chr4:113244-113913 | 8173.0 | 5,53404 | 819.829 | 817.324 | | orf19.4175 |
| chr2:1394203-1395205 | 6011.0 | 5,42309 | 603.53 | 601.163 | orf19.2231 | |
| chr1:534496-536667 | 6027.0 | 5,41235 | 605.073 | 602.705 | orf19.2942 | |
| chrR:1607304-1609036 | 6572.0 | 5,39504 | 659.639 | 657.233 | | orf19.6126 |
| chr2:2105108-2105773 | 5945.0 | 5,39216 | 596.918 | 594.555 | | orf19.1756 |

| | | | | | | |
|---|---|---|---|---|---|---|
| chr1:2440571-2441735 | 5983.0 | 5,35244 | 600.685 | 598.32 | | orf19.2308 |
| chr1:1085832-1086879 | 5997.0 | 5,3483 | 602.125 | 599.759 | orf19.450 | |
| chr6:217220-217731 | 6618.0 | 5,34687 | 664.25 | 661.842 | | orf19.118 |
| chr1:666209-667597 | 6289.0 | 5,33065 | 631.306 | 628.919 | orf19.3010.1 | |
| chr6:250438-251733 | 5815.0 | 5,32111 | 583.932 | 581.58 | orf19.136 | orf19.137 |
| chr7:39751-41319 | 6707.0 | 5,2895 | 673.122 | 670.708 | | orf19.7094 |
| chr5:789981-790781 | 7561.0 | 5,28471 | 758.62 | 756.152 | orf19.6660 | orf19.6661 |
| chr7:319813-320678 | 5510.0 | 5,18596 | 553.415 | 551.085 | | orf19.6577 |
| chr2:1149379-1149921 | 5528.0 | 5,1675 | 555.152 | 552.821 | | orf19.6874 |
| chr4:1497581-1497958 | 6303.0 | 5,16617 | 632.712 | 630.325 | | orf19.3138 |
| chr5:529261-529665 | 5410.0 | 5,13785 | 543.412 | 541.089 | orf19.4245 | orf19.4246 |
| chr2:1456582-1457327 | 6321.0 | 5,12665 | 634.525 | 632.136 | orf19.2268 | orf19.2269 |
| chr6:520348-520778 | 5482.0 | 5,11276 | 550.547 | 548.219 | orf19.5520 | orf19.5521 |
| chr3:1289179-1289488 | 5339.0 | 5,10348 | 536.301 | 533.984 | orf19.7356 | orf19.7357 |
| chr4:279124-279598 | 5357.0 | 5,09629 | 538.075 | 535.757 | | orf19.4650 |
| chr2:1951111-1951615 | 5317.0 | 5,07764 | 534.016 | 531.7 | | orf19.1401 |
| chrR:1219300-1219953 | 5248.0 | 5,03822 | 527.128 | 524.818 | orf19.6638 | orf19.6637 |
| chr5:628148-628682 | 6776.0 | 5,03461 | 680.065 | 677.646 | orf19.4304 | |
| chr6:755438-756707 | 5410.0 | 5,03432 | 543.368 | 541.045 | orf19.5722 | orf19.5723 |
| chr6:524078-524616 | 5915.0 | 5,02424 | 593.889 | 591.529 | | orf19.5525 |
| chr3:728623-729005 | 5343.0 | 5,00058 | 536.621 | 534.303 | orf19.341 | |
| chr3:926732-927191 | 5786.0 | 4,96141 | 580.974 | 578.623 | orf19.5896 | orf19.5897 |
| chr5:815038-816069 | 6546.0 | 4,959 | 657.08 | 654.676 | orf19.6679 | orf19.6680 |
| chr6:171203-171991 | 5343.0 | 4,92907 | 536.643 | 534.326 | orf19.88 | orf19.89 |
| chr2:1415287-1415767 | 5169.0 | 4,92512 | 519.239 | 516.936 | orf19.2242 | |
| chr1:1250184-1250612 | 4975.0 | 4,92477 | 499.803 | 497.517 | orf19.2452 | orf19.2451 |
| chr2:346966-347606 | 4864.0 | 4,86979 | 488.736 | 486.459 | orf19.1497 | orf19.1499 |
| chr3:710576-711326 | 4837.0 | 4,85604 | 485.982 | 483.707 | orf19.334 | orf19.335 |
| chr2:596935-597703 | 5414.0 | 4,85318 | 543.724 | 541.401 | orf19.5805 | |
| chr2:1877785-1879789 | 6527.0 | 4,84453 | 655.187 | 652.784 | orf19.4076 | orf19.4077 |
| chr3:962139-962486 | 5677.0 | 4,79415 | 570.068 | 567.725 | orf19.5911 | orf19.5912 |
| chr6:819570-820851 | 4625.0 | 4,7495 | 464.802 | 462.545 | orf19.5753 | |
| chr1:3050531-3050944 | 4675.0 | 4,74942 | 469.803 | 467.542 | | orf19.5025 |
| chr3:1475248-1475852 | 5311.0 | 4,73366 | 533.431 | 531.116 | | orf19.7436 |
| chr3:1721665-1723174 | 6405.0 | 4,72918 | 642.952 | 640.558 | orf19.6736 | orf19.6734 |
| chr6:613339-614086 | 4578.0 | 4,72545 | 460.061 | 457.809 | orf19.5575 | |
| chr6:56406-57545 | 5582.0 | 4,71588 | 560.542 | 558.207 | orf19.1193 | |
| chrR:1156949-1157884 | 2613.0 | 4,69789 | 263.438 | 261.397 | orf19.5286 | orf19.5285 |
| chr4:441532-441981 | 4941.0 | 4,66621 | 496.403 | 494.12 | orf19.4568 | |
| chr1:2395358-2395963 | 5158.0 | 4,65318 | 518.174 | 515.872 | orf19.2337 | orf19.2336 |
| chr5:312180-313065 | 4540.0 | 4,64617 | 456.321 | 454.072 | orf19.1927 | orf19.1926 |
| chr1:2923278-2923805 | 4390.0 | 4,62922 | 441.252 | 439.017 | orf19.4966 | orf19.4967 |
| chr6:513544-514187 | 4784.0 | 4,6245 | 480.737 | 478.466 | | orf19.5517 |
| chr1:2541670-2542211 | 5257.0 | 4,60783 | 528.11 | 525.799 | | orf19.1161 |
| chr1:2519435-2520397 | 4290.0 | 4,57767 | 431.28 | 429.054 | orf19.656 | orf19.655 |
| chr7:197728-198226 | 4290.0 | 4,57767 | 431.28 | 429.054 | | orf19.7020 |
| chrR:1577382-1579062 | 4733.0 | 4,57495 | 475.653 | 473.387 | orf19.6146 | |
| chr2:2047652-2048317 | 4270.0 | 4,56736 | 429.294 | 427.07 | orf19.1783 | orf19.1782.1 |
| chr2:2022258-2023764 | 4264.0 | 4,56392 | 428.633 | 426.409 | orf19.1363 | orf19.1362 |
| chr1:172622-173665 | 5611.0 | 4,53977 | 563.526 | 561.188 | | orf19.6017 |
| chr6:392636-393164 | 5261.0 | 4,51919 | 528.478 | 526.168 | orf19.3392 | orf19.3391 |
| chr4:1044702-1045676 | 4446.0 | 4,51242 | 446.928 | 444.688 | orf19.3794 | orf19.3793 |
| chr5:228632-228998 | 4165.0 | 4,51237 | 418.754 | 416.54 | orf19.1964 | |
| chr1:281172-282378 | 4895.0 | 4,45872 | 491.859 | 489.579 | orf19.3330.3 | orf19.3331 |
| chr1:844000-844419 | 4054.0 | 4,45395 | 407.647 | 405.443 | orf19.4476 | |
| chr4:907667-908281 | 4397.0 | 4,43033 | 441.936 | 439.7 | orf19.1426 | orf19.1424 |

| | | | | | | |
|---|---|---|---|---|---|---|
| chr3:1406862-1407539 | 7145.0 | 4,35995 | 716.961 | 714.519 | orf19.7417 | |
| chrR:1314064-1315171 | 5069.0 | 4,3596 | 509.277 | 506.982 | | orf19.3868 |
| chr1:2776166-2776510 | 3848.0 | 4,34397 | 387.004 | 384.819 | | orf19.6348 |
| chr4:226430-227092 | 3816.0 | 4,32679 | 383.81 | 381.628 | orf19.4679 | |
| chrR:815675-816185 | 4184.0 | 4,27888 | 420.633 | 418.417 | orf19.4365 | orf19.4366 |
| chr3:277142-277506 | 4072.0 | 4,27509 | 409.476 | 407.271 | orf19.1721 | orf19.1720 |
| chr1:305630-306503 | 3840.0 | 4,27209 | 386.198 | 384.014 | orf19.3340 | |
| chr7:709267-709944 | 3724.0 | 4,2699 | 374.654 | 372.481 | orf19.5129 | orf19.5126 |
| chrR:1380442-1380790 | 3695.0 | 4,26149 | 371.751 | 369.582 | | orf19.720 |
| chr6:706959-707305 | 3683.0 | 4,25462 | 370.489 | 368.321 | orf19.5625 | orf19.5626 |
| chrR:2090225-2090929 | 3862.0 | 4,25261 | 388.39 | 386.204 | | orf19.7551 |
| chr1:3010445-3010903 | 4985.0 | 4,24412 | 500.879 | 498.592 | orf19.5006.1 | orf19.5007 |
| chr3:946165-946901 | 4088.0 | 4,24246 | 411.023 | 408.816 | orf19.5906 | |
| chr5:1076829-1077713 | 4349.0 | 4,23572 | 437.165 | 434.934 | orf19.3981 | orf19.3982 |
| chr3:1566666-1567587 | 3864.0 | 4,2287 | 388.601 | 386.415 | | orf19.6817 |
| chr2:166482-167211 | 4045.0 | 4,19248 | 406.753 | 404.55 | | orf19.2021 |
| chr4:667106-667889 | 4773.0 | 4,19129 | 479.576 | 477.307 | | orf19.2672 |
| chr1:3143720-3144530 | 4323.0 | 4,1837 | 434.621 | 432.391 | orf19.7235 | |
| chr2:1339548-1340821 | 3508.0 | 4,15839 | 352.97 | 350.819 | orf19.34 | orf19.35 |
| chr1:520692-521040 | 3495.0 | 4,15152 | 351.729 | 349.58 | orf19.2934 | orf19.2935 |
| chr2:1230811-1232235 | 3692.0 | 4,11341 | 371.468 | 369.299 | | orf19.4118 |
| chr1:1214857-1215614 | 3722.0 | 4,10571 | 374.376 | 372.204 | | orf19.2467 |
| chr6:765993-766616 | 3650.0 | 4,09727 | 367.236 | 365.071 | orf19.5728 | orf19.5729 |
| chr1:944922-946228 | 3814.0 | 4,09103 | 383.623 | 381.441 | orf19.6852 | orf19.6852.1 |
| chr5:506244-507208 | 4356.0 | 4,07761 | 437.877 | 435.645 | orf19.4234 | orf19.4235 |
| chr3:566180-566629 | 3317.0 | 4,05186 | 333.902 | 331.771 | orf19.251 | |
| chr4:1300669-1301361 | 4120.0 | 4,01311 | 414.287 | 412.077 | orf19.1274 | orf19.1275 |
| chr2:1934958-1936176 | 3624.0 | 3,9977 | 364.62 | 362.458 | orf19.1604 | |
| chr3:1627294-1628194 | 3584.0 | 3,96588 | 360.656 | 358.498 | orf19.6790 | orf19.6789 |
| chr7:301126-301535 | 3148.0 | 3,95563 | 316.982 | 314.871 | orf19.6586 | orf19.6585 |
| chr4:1099749-1101609 | 3485.0 | 3,94626 | 350.652 | 348.503 | orf19.740 | |
| chr4:1505504-1506618 | 3713.0 | 3,94115 | 373.477 | 371.306 | orf19.3134 | orf19.3133 |
| chr2:576485-577199 | 3909.0 | 3,93731 | 393.126 | 390.935 | orf19.5816 | orf19.5815 |
| chr1:1260325-1260756 | 3077.0 | 3,91439 | 309.821 | 307.717 | orf19.2446 | orf19.2445 |
| chrR:914169-914575 | 3556.0 | 3,90709 | 357.8 | 355.644 | orf19.489 | orf19.490 |
| chr1:1806713-1807877 | 3512.0 | 3,89615 | 353.384 | 351.232 | orf19.5102 | |
| chrR:105407-105774 | 3442.0 | 3,88308 | 346.355 | 344.211 | | orf19.7490 |
| chrR:1262734-1263132 | 3131.0 | 3,86053 | 315.307 | 313.197 | orf19.454 | orf19.451 |
| chr1:1096101-1096854 | 2982.0 | 3,8594 | 300.359 | 298.265 | orf19.448 | |
| chr6:633559-634231 | 3246.0 | 3,82248 | 326.724 | 324.602 | | orf19.5585 |
| chr4:987317-987639 | 3538.0 | 3,79206 | 355.954 | 353.8 | | orf19.3822 |
| chr3:953486-954653 | 3061.0 | 3,78928 | 308.228 | 306.126 | orf19.5908 | |
| chr2:1656039-1656954 | 3306.0 | 3,78362 | 332.743 | 330.614 | orf19.2168.3 | orf19.2168 |
| chrR:1526038-1529492 | 2848.0 | 3,78036 | 286.93 | 284.851 | orf19.2356 | |
| chrR:1093477-1094221 | 2907.0 | 3,75617 | 292.865 | 290.78 | | orf19.646 |
| chr1:801101-801692 | 2785.0 | 3,74255 | 280.581 | 278.509 | | orf19.1027 |
| chr2:2135838-2136263 | 2767.0 | 3,73224 | 278.857 | 276.788 | orf19.1743 | |
| chr1:298663-299249 | 2750.0 | 3,72193 | 277.138 | 275.07 | | orf19.3337 |
| chr2:1010262-1010567 | 2750.0 | 3,72193 | 277.138 | 275.07 | | orf19.4128 |
| chr4:120457-120883 | 3014.0 | 3,6959 | 303.592 | 301.495 | orf19.4172 | orf19.4171 |
| chr3:901330-901979 | 3315.0 | 3,68024 | 333.694 | 331.564 | orf19.5876 | orf19.5877 |
| chr2:1870767-1871139 | 2974.0 | 3,67841 | 299.539 | 297.446 | orf19.4072 | |
| chr4:516447-516924 | 2614.0 | 3,63945 | 263.511 | 261.461 | orf19.2749 | orf19.2748 |
| chr2:934484-935123 | 2810.0 | 3,63558 | 283.074 | 281.0 | | orf19.4505 |
| chr1:1221018-1221420 | 2597.0 | 3,62914 | 261.824 | 259.776 | | orf19.2461 |
| chr5:132993-134452 | 2586.0 | 3,62227 | 260.702 | 258.655 | | orf19.930 |

| | | | | | | |
|---|---|---|---|---|---|---|
| chr5:1106657-1107257 | 2624.0 | 3,60542 | 264.503 | 262.452 | | orf19.3997 |
| chr1:2954470-2955098 | 2713.0 | 3,60211 | 273.426 | 271.363 | | orf19.4980 |
| chr1:889847-890629 | 2553.0 | 3,60165 | 257.344 | 255.302 | orf19.1064 | |
| chr5:709620-710110 | 2754.0 | 3,59908 | 277.501 | 275.433 | | orf19.4347 |
| chr7:613589-614514 | 2946.0 | 3,59733 | 296.735 | 294.645 | orf19.5178 | orf19.5177 |
| chr2:627086-627629 | 3699.0 | 3,57711 | 372.124 | 369.954 | | orf19.5785 |
| chr4:461784-462212 | 2992.0 | 3,55128 | 301.37 | 299.276 | orf19.2770.1 | |
| chr1:1610656-1611267 | 2464.0 | 3,54666 | 248.464 | 246.434 | | orf19.4423 |
| chr2:2056429-2057037 | 2409.0 | 3,5123 | 242.968 | 240.945 | | orf19.1777 |
| chr3:1760556-1761003 | 2695.0 | 3,5098 | 271.637 | 269.576 | orf19.6715 | orf19.6713 |
| chr5:450262-451473 | 2533.0 | 3,48468 | 255.345 | 253.306 | orf19.3171 | |
| chr7:628370-628931 | 2400.0 | 3,47167 | 242.042 | 240.021 | orf19.5171 | orf19.5170.1 |
| chr1:895908-897592 | 2473.0 | 3,44725 | 249.41 | 247.378 | | orf19.1069 |
| chr1:2811122-2811840 | 2377.0 | 3,43986 | 239.775 | 237.757 | | orf19.4921.1 |
| chr1:2080561-2080920 | 2690.0 | 3,40545 | 271.07 | 269.01 | | orf19.4807 |
| chr1:2179852-2180284 | 2236.0 | 3,40232 | 225.669 | 223.671 | orf19.4864 | orf19.4865 |
| chr3:763429-764319 | 2646.0 | 3,38996 | 266.71 | 264.655 | orf19.359 | orf19.360 |
| chr1:1836152-1836699 | 2371.0 | 3,37842 | 239.143 | 237.126 | orf19.5117 | orf19.5118 |
| chr6:560230-561017 | 2703.0 | 3,3664 | 272.403 | 270.341 | orf19.5542 | |
| chr1:1078376-1078860 | 2167.0 | 3,35764 | 218.768 | 216.781 | | orf19.4056 |
| chr4:289866-290631 | 2167.0 | 3,35764 | 218.768 | 216.781 | | orf19.4646 |
| chr3:646754-648107 | 2339.0 | 3,35239 | 235.965 | 233.952 | orf19.301 | orf19.302 |
| chr1:2119787-2120198 | 2398.0 | 3,33814 | 241.831 | 239.81 | orf19.4825 | orf19.4826 |
| chr1:3039810-3041115 | 2344.0 | 3,3038 | 236.445 | 234.432 | orf19.5021 | orf19.5022 |
| chr1:1944920-1945271 | 2056.0 | 3,27377 | 207.599 | 205.63 | orf19.4752 | orf19.4753 |
| chr1:1326203-1326548 | 2021.0 | 3,26142 | 204.161 | 202.198 | orf19.1821 | |
| chr2:373002-373287 | 2011.0 | 3,25454 | 203.132 | 201.17 | orf19.1509 | |
| chrR:174790-175169 | 2269.0 | 3,2313 | 228.971 | 226.969 | orf19.3282 | |
| chr6:430763-431378 | 2537.0 | 3,18887 | 255.745 | 253.705 | orf19.3501 | orf19.3499 |
| chr3:1144801-1145513 | 2184.0 | 3,18135 | 220.433 | 218.443 | orf19.5999 | orf19.6000 |
| chr1:1914008-1914775 | 2186.0 | 3,14335 | 220.65 | 218.66 | | orf19.4737 |
| chr1:1318325-1318757 | 1809.0 | 3,11708 | 182.927 | 180.998 | orf19.1822 | |
| chr2:1743966-1744256 | 1790.0 | 3,10333 | 180.948 | 179.022 | | orf19.3618 |
| chr1:1469856-1470887 | 2135.0 | 3,09703 | 215.488 | 213.507 | orf19.6220.4 | |
| chr4:509777-510205 | 1765.0 | 3,08615 | 178.485 | 176.563 | orf19.2752 | orf19.2751 |
| chr1:625618-626549 | 1968.0 | 3,07284 | 198.769 | 196.814 | orf19.2990 | orf19.2991 |
| chr6:273224-274044 | 1741.0 | 3,06896 | 176.034 | 174.117 | | orf19.3447 |
| chr1:1548075-1548852 | 2002.0 | 3,01705 | 202.257 | 200.297 | orf19.4446 | orf19.4445 |

TAP-Zcf26 ChIP sequencing data

| Chromosome (start-end) | score | signal Value | p-value (-log10) | q-value (-log10) | ORF_left | ORF_right |
|---|---|---|---|---|---|---|
| chr4:707893-708488 | 127330.0 | 62,9095 | 12737.4 | 12733.1 | orf19.3364 | |
| chr5:420140-420726 | 91163.0 | 41,3254 | 9120.53 | 9116.33 | orf19.3188 | |
| chr4:623644-624818 | 63315.0 | 35,4729 | 6335.47 | 6331.55 | orf19.2693 | |
| chr2:660365-663080 | 73620.0 | 34,5094 | 7366.13 | 7362.1 | orf19.909 | |
| chrR:523320-524358 | 55297.0 | 32,1814 | 5533.54 | 5529.7 | orf19.3734 | orf19.3733 |
| chrR:1669924-1671286 | 55753.0 | 31,3453 | 5579.16 | 5575.31 | | orf19.1985 |
| chr1:3060057-3061222 | 45342.0 | 29,313 | 4538.0 | 4534.3 | orf19.5026 | |
| chr2:1465224-1466129 | 43474.0 | 26,3074 | 4351.14 | 4347.47 | orf19.2272 | orf19.2274 |
| chrR:51584-52427 | 52800.0 | 25,3771 | 5283.85 | 5280.06 | | orf19.7513 |
| chr3:866371-866964 | 51543.0 | 24,9443 | 5158.13 | 5154.36 | | orf19.5859 |
| chr1:3005885-3006519 | 33060.0 | 23,4544 | 3309.55 | 3306.04 | | orf19.5005 |
| chr2:1313753-1315181 | 34192.0 | 23,4347 | 3422.83 | 3419.29 | orf19.22.1 | orf19.23 |
| chrR:990071-991725 | 35020.0 | 22,5176 | 3505.66 | 3502.1 | orf19.539 | |
| chr3:431945-433088 | 34146.0 | 22,3485 | 3418.24 | 3414.7 | orf19.1652 | |
| chr4:1494983-1496133 | 34980.0 | 21,4988 | 3501.63 | 3498.08 | | orf19.3139 |
| chr6:35420-36087 | 43800.0 | 21,4938 | 4383.72 | 4380.04 | | orf19.1189 |
| chr2:643041-643564 | 35293.0 | 20,839 | 3532.91 | 3529.34 | orf19.5779 | orf19.5777 |
| chrR:2210673-2211240 | 28309.0 | 20,6729 | 2834.32 | 2830.92 | orf19.7611 | |
| chr2:131505-133610 | 28817.0 | 19,9693 | 2885.14 | 2881.73 | orf19.2047 | orf19.2047 |
| chr2:131505-133610 | 28817.0 | 19,9693 | 2885.14 | 2881.73 | | orf19.2046 |
| chr1:435357-441859 | 29948.0 | 19,5881 | 2998.28 | 2994.84 | orf19.3668 | |
| chr2:1219375-1222282 | 32026.0 | 19,5444 | 3206.09 | 3202.61 | orf19.4122 | |
| chr6:706880-707366 | 25334.0 | 19,5239 | 2536.79 | 2533.47 | orf19.5625 | orf19.5626 |
| chr4:787872-788561 | 26157.0 | 19,03 | 2619.08 | 2615.73 | orf19.1309 | orf19.1308 |
| chr1:2996458-2997139 | 24112.0 | 18,879 | 2414.59 | 2411.3 | | orf19.5000 |
| chr2:1132109-1134569 | 27373.0 | 18,6828 | 2740.72 | 2737.34 | orf19.6869 | orf19.6871 |
| chr1:1080225-1082306 | 26581.0 | 18,5937 | 2661.47 | 2658.11 | | orf19.4056 |
| chr2:936947-937838 | 22438.0 | 17,9829 | 2247.14 | 2243.89 | | orf19.4504 |
| chr6:111854-112364 | 27461.0 | 17,4721 | 2749.57 | 2746.19 | orf19.4196 | orf19.4195.1 |
| chr6:615470-617700 | 20917.0 | 17,1547 | 2094.91 | 2091.71 | orf19.5576 | orf19.5578 |
| chr3:796532-798220 | 29074.0 | 17,1108 | 2910.9 | 2907.48 | | orf19.6937 |
| chr3:734610-735863 | 23125.0 | 16,8706 | 2315.76 | 2312.5 | orf19.344 | orf19.345 |
| chr5:111372-114011 | 21069.0 | 16,7909 | 2110.18 | 2106.98 | orf19.935 | |
| chr3:607699-609218 | 20139.0 | 16,5371 | 2017.08 | 2013.91 | orf19.272 | |
| chr6:668408-669334 | 22030.0 | 16,5063 | 2206.28 | 2203.05 | orf19.5604 | |
| chrR:443871-444721 | 25395.0 | 16,5034 | 2542.91 | 2539.58 | orf19.2602 | |
| chr1:2044944-2045983 | 23506.0 | 16,3428 | 2353.91 | 2350.63 | orf19.4789 | |
| chr6:739783-740330 | 18815.0 | 15,987 | 1884.67 | 1881.55 | | orf19.5713 |
| chr1:2270132-2271713 | 21690.0 | 15,7607 | 2172.25 | 2169.02 | orf19.4898 | orf19.4899 |
| chr1:918470-921435 | 23747.0 | 15,5121 | 2378.02 | 2374.74 | orf19.6838 | |
| chrR:1295793-1296377 | 19095.0 | 15,3879 | 1912.67 | 1909.54 | | orf19.3859 |
| chr6:837268-838154 | 20135.0 | 15,3675 | 2016.74 | 2013.57 | orf19.5759 | |
| chrR:589760-591270 | 20830.0 | 15,3459 | 2086.25 | 2083.05 | orf19.2822 | orf19.2823 |
| chr1:65758-67669 | 21006.0 | 15,1478 | 2103.88 | 2100.68 | orf19.6066 | orf19.6065 |
| chr1:1545018-1545857 | 19587.0 | 14,953 | 1961.86 | 1958.7 | | orf19.4447 |
| chr3:312914-314211 | 16662.0 | 14,7583 | 1669.34 | 1666.3 | orf19.1704 | orf19.1702 |
| chr2:827984-828922 | 16791.0 | 14,6928 | 1682.17 | 1679.12 | orf19.829 | orf19.828 |
| chr3:174444-175167 | 22067.0 | 14,3057 | 2210.0 | 2206.77 | | orf19.6165 |
| chr7:284058-284518 | 15813.0 | 14,2627 | 1584.31 | 1581.31 | | orf19.6920 |
| chr1:1085836-1087019 | 16031.0 | 14,2033 | 1606.2 | 1603.19 | orf19.450 | |
| chrR:87018-87792 | 20028.0 | 14,1731 | 2005.97 | 2002.8 | orf19.7499.1 | orf19.7499 |
| chr4:625273-626179 | 16067.0 | 13,9951 | 1609.73 | 1606.72 | orf19.2691 | |
| chr5:177979-178801 | 15410.0 | 13,7341 | 1543.99 | 1541.02 | orf19.577 | orf19.578 |
| chr3:160869-161667 | 16346.0 | 13,2198 | 1637.67 | 1634.65 | orf19.1288 | orf19.1289 |

| | | | | | | |
|---|---|---|---|---|---|---|
| chr5:123092-124516 | 16690.0 | 13,1979 | 1672.12 | 1669.08 | | orf19.932 |
| chr3:197120-198029 | 13955.0 | 13,1562 | 1398.49 | 1395.59 | | orf19.2501 |
| chr7:297097-298699 | 13944.0 | 13,1494 | 1397.37 | 1394.47 | orf19.6586 | orf19.6585 |
| chr1:2485688-2486753 | 17078.0 | 13,0313 | 1710.95 | 1707.89 | orf19.670.2 | |
| chr6:166433-166975 | 13642.0 | 12,9661 | 1367.16 | 1364.27 | | orf19.86 |
| chr5:686501-687476 | 13631.0 | 12,9593 | 1366.05 | 1363.16 | orf19.4337 | orf19.4339 |
| chr5:1136153-1137112 | 15036.0 | 12,9539 | 1506.65 | 1503.69 | orf19.4010 | orf19.4011 |
| chr1:2563419-2564355 | 14144.0 | 12,7336 | 1417.36 | 1414.45 | orf19.1149 | |
| chr7:899542-900575 | 17356.0 | 12,5978 | 1738.68 | 1735.61 | | orf19.7156 |
| chrR:1233184-1233927 | 16032.0 | 12,4269 | 1606.26 | 1603.24 | | orf19.6627 |
| chr3:667235-667941 | 15814.0 | 12,2458 | 1584.42 | 1581.42 | orf19.310 | |
| chr2:315599-316101 | 12429.0 | 12,2194 | 1245.81 | 1242.99 | orf19.1486 | |
| chr1:1988856-1989887 | 12408.0 | 12,2058 | 1243.63 | 1240.81 | | orf19.4769 |
| chr5:653254-655141 | 12288.0 | 12,1311 | 1231.66 | 1228.84 | orf19.4318 | |
| chr2:1215520-1216129 | 13675.0 | 12,0837 | 1370.43 | 1367.54 | orf19.5215 | |
| chrR:1801495-1803290 | 16478.0 | 12,0413 | 1650.88 | 1647.85 | orf19.6398 | orf19.6399 |
| chr1:1207637-1208216 | 16068.0 | 11,8513 | 1609.84 | 1606.83 | orf19.2474 | orf19.2473 |
| chr5:696125-697197 | 12543.0 | 11,8347 | 1257.18 | 1254.34 | orf19.4341 | orf19.4342 |
| chr3:1289169-1290022 | 11780.0 | 11,8121 | 1180.84 | 1178.05 | orf19.7356 | orf19.7357 |
| chrR:1875771-1876605 | 17131.0 | 11,8058 | 1716.17 | 1713.11 | orf19.6445 | |
| chr3:970764-972020 | 17373.0 | 11,7591 | 1740.43 | 1737.36 | orf19.5917 | orf19.5917.3 |
| chr4:709445-710276 | 11555.0 | 11,6695 | 1158.32 | 1155.53 | orf19.3365 | orf19.3366 |
| chr3:1414355-1414859 | 13905.0 | 11,5656 | 1393.45 | 1390.55 | orf19.7414 | |
| chr2:104790-105591 | 14595.0 | 11,5621 | 1462.51 | 1459.58 | | orf19.2063 |
| chr1:1572881-1573625 | 11256.0 | 11,4794 | 1128.45 | 1125.69 | | orf19.4438 |
| chr2:2165909-2167038 | 12748.0 | 11,4594 | 1277.74 | 1274.9 | | orf19.5323 |
| chr2:613479-614230 | 14835.0 | 11,3746 | 1486.53 | 1483.59 | | orf19.5797 |
| chr1:2852619-2853789 | 14483.0 | 11,3708 | 1451.27 | 1448.34 | | orf19.4936 |
| chr3:69093-69854 | 13913.0 | 11,3496 | 1394.28 | 1391.38 | orf19.5417 | orf19.5416 |
| chr4:1254752-1255934 | 12240.0 | 11,3196 | 1226.9 | 1224.08 | | orf19.1255 |
| chr2:258734-259229 | 12546.0 | 11,3149 | 1257.52 | 1254.68 | orf19.1449 | |
| chr7:888399-889155 | 13492.0 | 11,2957 | 1352.18 | 1349.3 | orf19.7166 | |
| chr3:853701-854606 | 12760.0 | 11,2724 | 1278.95 | 1276.1 | orf19.918 | orf19.913.2 |
| chr2:1655971-1656986 | 10940.0 | 11,1786 | 1096.82 | 1094.08 | orf19.2168.3 | orf19.2168 |
| chr3:1194827-1195213 | 10477.0 | 10,9771 | 1050.5 | 1047.78 | orf19.6975 | orf19.6976 |
| chrR:1698564-1699873 | 10373.0 | 10,9092 | 1040.08 | 1037.37 | | orf19.3711 |
| chr1:444447-448221 | 11061.0 | 10,9085 | 1108.88 | 1106.13 | orf19.3669 | |
| chr2:518201-518743 | 12368.0 | 10,8924 | 1239.65 | 1236.83 | | orf19.1582 |
| chr5:44641-45641 | 12089.0 | 10,8514 | 1211.77 | 1208.96 | | orf19.968 |
| chrR:1373731-1374446 | 11032.0 | 10,6121 | 1105.96 | 1103.21 | orf19.723 | orf19.721 |
| chr4:920837-921805 | 11546.0 | 10,5599 | 1157.43 | 1154.65 | | orf19.1415 |
| chr2:1079337-1080118 | 10513.0 | 10,3727 | 1054.1 | 1051.38 | orf19.3554 | orf19.3555 |
| chr7:144648-145612 | 9468.0 | 10,3118 | 949.542 | 946.889 | | orf19.7042 |
| chr2:1484217-1484824 | 12073.0 | 10,1669 | 1210.13 | 1207.32 | | orf19.2286 |
| chrR:12206-13899 | 12645.0 | 10,1357 | 1267.36 | 1264.53 | orf19.7539.1 | orf19.7539 |
| chr4:461749-462223 | 11092.0 | 10,1023 | 1112.01 | 1109.25 | orf19.2770.1 | |
| chr5:238040-239554 | 9404.0 | 10,0676 | 943.131 | 940.482 | orf19.1961 | |
| chr1:231537-232697 | 10639.0 | 10,0674 | 1066.72 | 1063.99 | orf19.3302 | |
| chrR:1943279-1943900 | 10630.0 | 10,0542 | 1065.82 | 1063.1 | orf19.7306 | |
| chrR:39841-40971 | 11808.0 | 9,89211 | 1183.61 | 1180.81 | orf19.7520 | |
| chr3:1239326-1240108 | 8754.0 | 9,82982 | 878.105 | 875.499 | orf19.6987 | orf19.6988 |
| chr2:1310376-1310993 | 8655.0 | 9,76193 | 868.163 | 865.564 | orf19.21 | |
| chr4:1306956-1309222 | 8635.0 | 9,74836 | 866.178 | 863.581 | orf19.1277 | |
| chr3:152633-153896 | 9973.0 | 9,70693 | 1000.04 | 997.36 | | orf19.3071 |
| chr1:1303004-1303464 | 8467.0 | 9,63295 | 849.357 | 846.77 | orf19.1826 | |
| chr1:2923241-2923801 | 8597.0 | 9,63116 | 862.335 | 859.74 | orf19.4966 | orf19.4967 |

| | | | | | | |
|---|---|---|---|---|---|---|
| chrR:1376753-1377705 | 10311.0 | 9,62693 | 1033.88 | 1031.17 | orf19.723 | orf19.721 |
| chr5:910553-911086 | 9346.0 | 9,55852 | 937.342 | 934.697 | orf19.1282 | |
| chr3:1225006-1225790 | 8339.0 | 9,5447 | 836.553 | 833.976 | | orf19.6984 |
| chr1:1946927-1947475 | 8280.0 | 9,50397 | 830.661 | 828.088 | orf19.4752 | orf19.4753 |
| chr6:755781-756723 | 8761.0 | 9,42362 | 878.762 | 876.156 | orf19.5722 | orf19.5723 |
| chr5:415544-416896 | 8163.0 | 9,42251 | 818.911 | 816.346 | orf19.3190 | |
| chr4:853067-853814 | 8143.0 | 9,40893 | 816.957 | 814.393 | orf19.1208 | orf19.5312 |
| chrR:1704062-1706610 | 8131.0 | 9,38675 | 815.677 | 813.115 | orf19.3713 | orf19.3714 |
| chr7:453160-454388 | 9452.0 | 9,37068 | 947.891 | 945.24 | orf19.6514 | |
| chr2:168829-170084 | 9644.0 | 9,34411 | 967.084 | 964.421 | | orf19.2020 |
| chr4:239262-239894 | 7939.0 | 9,26637 | 796.516 | 793.968 | orf19.4673 | orf19.4672 |
| chr1:2225790-2227667 | 8932.0 | 9,22552 | 895.874 | 893.257 | orf19.4883 | orf19.4884 |
| chr6:613335-614091 | 7640.0 | 9,05593 | 766.597 | 764.072 | orf19.5575 | |
| chr2:829267-829890 | 11372.0 | 9,05148 | 1140.04 | 1137.27 | orf19.829 | orf19.828 |
| chr1:31466-31870 | 9075.0 | 8,97753 | 910.17 | 907.544 | | orf19.6085 |
| chr1:2870284-2871623 | 8374.0 | 8,96308 | 840.029 | 837.449 | orf19.4941 | orf19.4942 |
| chrR:730808-732044 | 7478.0 | 8,94052 | 750.323 | 747.811 | orf19.2398 | orf19.2397.3 |
| chr1:3112057-3112590 | 8411.0 | 8,91441 | 843.69 | 841.107 | orf19.7218 | orf19.7219 |
| chr3:760441-760847 | 9370.0 | 8,89175 | 939.736 | 937.09 | orf19.356 | orf19.357 |
| chr7:907733-915424 | 11635.0 | 8,82931 | 1166.31 | 1163.53 | orf19.7152 | orf19.7151 |
| chrR:1219314-1219965 | 7277.0 | 8,73852 | 730.286 | 727.789 | orf19.6638 | orf19.6637 |
| chr1:2314349-2314996 | 7970.0 | 8,69846 | 799.592 | 797.042 | | orf19.986 |
| chrR:1380411-1380825 | 7080.0 | 8,6554 | 710.527 | 708.044 | | orf19.720 |
| chr6:171141-172032 | 8393.0 | 8,64254 | 841.899 | 839.317 | orf19.88 | orf19.89 |
| chr2:1155664-1156422 | 7730.0 | 8,62427 | 775.533 | 773.001 | | orf19.6877 |
| chr4:635741-636419 | 7084.0 | 8,50027 | 710.9 | 708.417 | orf19.2686 | |
| chr5:506207-507231 | 8924.0 | 8,43541 | 895.11 | 892.493 | orf19.4234 | orf19.4235 |
| chrR:1608587-1609542 | 8042.0 | 8,39947 | 806.814 | 804.259 | | orf19.6126 |
| chrR:2110978-2111853 | 7223.0 | 8,35311 | 724.865 | 722.372 | | orf19.7563 |
| chr3:566146-566669 | 6578.0 | 8,28882 | 660.245 | 657.804 | orf19.251 | |
| chr4:1540190-1541074 | 7998.0 | 8,26503 | 802.448 | 799.896 | orf19.3117 | |
| chr4:862510-864522 | 6492.0 | 8,19487 | 651.698 | 649.264 | orf19.5308 | orf19.5307 |
| chr4:521454-522858 | 6412.0 | 8,16663 | 643.713 | 641.285 | | orf19.2747 |
| chr2:765328-766928 | 6883.0 | 8,14653 | 690.819 | 688.351 | | orf19.856 |
| chr6:985357-985953 | 8488.0 | 8,14594 | 851.404 | 848.816 | orf19.2124 | orf19.2125 |
| chr1:2291360-2292557 | 6376.0 | 8,13947 | 640.055 | 637.63 | orf19.4906 | |
| chr1:2024178-2025094 | 7272.0 | 8,10081 | 729.737 | 727.24 | orf19.4779 | orf19.4780 |
| chr4:1497309-1498013 | 7429.0 | 8,09149 | 745.485 | 742.977 | | orf19.3138 |
| chr5:681256-682152 | 6294.0 | 8,07837 | 631.846 | 629.427 | orf19.4335 | orf19.4336 |
| chr2:17660-18421 | 8513.0 | 8,05425 | 853.946 | 851.356 | orf19.1907 | orf19.1906 |
| chr2:1478004-1478810 | 7756.0 | 8,02303 | 778.144 | 775.61 | orf19.2281 | |
| chr2:1218217-1219023 | 6844.0 | 7,81013 | 686.908 | 684.444 | orf19.5217 | |
| chr2:1068428-1069556 | 6430.0 | 7,70571 | 645.49 | 643.06 | | orf19.3548.1 |
| chr2:1361339-1363536 | 5781.0 | 7,69143 | 580.548 | 578.17 | orf19.1223 | orf19.1224 |
| chr4:279150-279613 | 5763.0 | 7,67785 | 578.771 | 576.394 | | orf19.4650 |
| chr7:301092-301535 | 5763.0 | 7,67785 | 578.771 | 576.394 | orf19.6586 | orf19.6585 |
| chr4:574487-577284 | 6816.0 | 7,67329 | 684.065 | 681.603 | | orf19.2723 |
| chr1:2776126-2776521 | 5728.0 | 7,6507 | 575.22 | 572.846 | | orf19.6348 |
| chr2:199529-201237 | 6014.0 | 7,64752 | 603.895 | 601.499 | orf19.2005 | |
| chr1:404545-406660 | 6109.0 | 7,64432 | 613.321 | 610.918 | | orf19.4527 |
| chrR:815670-816201 | 6803.0 | 7,63113 | 682.777 | 680.316 | orf19.4365 | orf19.4366 |
| chrR:1314227-1315201 | 8052.0 | 7,5763 | 807.775 | 805.219 | | orf19.3868 |
| chr3:961434-962495 | 7230.0 | 7,57226 | 725.558 | 723.064 | orf19.5911 | orf19.5912 |
| chr1:2003379-2004668 | 7018.0 | 7,57056 | 704.337 | 701.859 | orf19.4772 | orf19.4773 |
| chr1:2844921-2845819 | 5858.0 | 7,56146 | 588.262 | 585.878 | orf19.4933 | orf19.4934 |
| chr1:1096058-1096723 | 5604.0 | 7,55566 | 562.841 | 560.479 | orf19.448 | |

| | | | | | | |
|---|---|---|---|---|---|---|
| chr3:709536-712124 | 5569.0 | 7,5285 | 559.318 | 556.959 | orf19.334 | orf19.335 |
| chr4:129389-131323 | 6741.0 | 7,42972 | 676.598 | 674.142 | orf19.4167 | orf19.4166 |
| chr1:3047105-3048064 | 5403.0 | 7,39952 | 542.671 | 540.326 | orf19.5021 | orf19.5022 |
| chrR:105359-105767 | 6282.0 | 7,38621 | 630.686 | 628.268 | | orf19.7490 |
| chr4:667099-667930 | 7371.0 | 7,33506 | 739.669 | 737.165 | | orf19.2672 |
| chr3:904548-905046 | 6692.0 | 7,31714 | 671.707 | 669.257 | orf19.5879 | orf19.5880 |
| chrR:208821-209790 | 5221.0 | 7,25696 | 524.438 | 522.11 | | orf19.3263 |
| chr4:509710-510256 | 5481.0 | 7,25352 | 550.508 | 548.156 | orf19.2752 | orf19.2751 |
| chrR:1911400-1912638 | 7001.0 | 7,19374 | 702.593 | 700.116 | orf19.7286 | orf19.7288 |
| chr1:2335667-2337174 | 5680.0 | 7,17413 | 570.441 | 568.072 | orf19.999 | |
| chr1:1326178-1326631 | 5083.0 | 7,14834 | 510.667 | 508.352 | orf19.1821 | |
| chrR:670256-671397 | 5476.0 | 7,14617 | 549.967 | 547.616 | orf19.2849 | orf19.2850 |
| chr1:843986-844403 | 5040.0 | 7,1144 | 506.385 | 504.074 | orf19.4476 | |
| chr5:228618-229187 | 5015.0 | 7,09403 | 503.821 | 501.512 | orf19.1964 | |
| chr5:789964-790746 | 6467.0 | 7,08114 | 649.164 | 646.732 | orf19.6660 | orf19.6661 |
| chr6:524162-524603 | 5852.0 | 7,05316 | 587.679 | 585.295 | | orf19.5525 |
| chr6:560246-561053 | 6506.0 | 7,04899 | 653.04 | 650.604 | orf19.5542 | |
| chr2:2047656-2048323 | 4870.0 | 6,97863 | 489.361 | 487.067 | orf19.1783 | orf19.1782.1 |
| chr5:497265-497789 | 4836.0 | 6,95147 | 485.976 | 483.685 | | orf19.4231 |
| chr2:1149387-1149929 | 4917.0 | 6,94519 | 494.091 | 491.793 | | orf19.6874 |
| chr2:790174-790966 | 5106.0 | 6,93073 | 512.99 | 510.673 | | orf19.847 |
| chr1:801084-801710 | 4744.0 | 6,8768 | 476.703 | 474.421 | | orf19.1027 |
| chr5:815067-816058 | 5982.0 | 6,87494 | 600.644 | 598.25 | orf19.6679 | orf19.6680 |
| chr6:633534-634189 | 5687.0 | 6,83753 | 571.11 | 568.74 | | orf19.5585 |
| chr1:2811093-2811863 | 4905.0 | 6,80019 | 492.802 | 490.504 | | orf19.4921.1 |
| chr3:1144840-1145579 | 5407.0 | 6,74825 | 543.074 | 540.728 | orf19.5999 | orf19.6000 |
| chr1:2440566-2441715 | 4635.0 | 6,73068 | 465.827 | 463.555 | | orf19.2308 |
| chr3:1406799-1407551 | 8334.0 | 6,71934 | 836.027 | 833.45 | orf19.7417 | |
| chr6:250449-251737 | 4543.0 | 6,71388 | 456.653 | 454.39 | orf19.136 | orf19.137 |
| chr6:520369-520789 | 4519.0 | 6,69351 | 454.164 | 451.903 | orf19.5520 | orf19.5521 |
| chr2:596924-598395 | 5407.0 | 6,65835 | 543.129 | 540.784 | orf19.5805 | |
| chr1:2110351-2110796 | 4790.0 | 6,61235 | 481.325 | 479.039 | orf19.4821 | |
| chr2:1394222-1395131 | 4395.0 | 6,59168 | 441.78 | 439.532 | orf19.2231 | |
| chr2:1424231-1425218 | 4687.0 | 6,59071 | 471.025 | 468.748 | orf19.2248 | |
| chr6:291459-292341 | 4337.0 | 6,54416 | 436.035 | 433.792 | orf19.3439 | |
| chr1:3131767-3132217 | 5080.0 | 6,47476 | 510.405 | 508.091 | orf19.7229 | |
| chr1:534493-536676 | 4352.0 | 6,44927 | 437.46 | 435.216 | orf19.2942 | |
| chrR:1525996-1529486 | 4505.0 | 6,44607 | 452.771 | 450.512 | orf19.2356 | |
| chr1:3143686-3144816 | 5415.0 | 6,44566 | 543.91 | 541.564 | orf19.7235 | |
| chr5:709597-710141 | 4659.0 | 6,43139 | 468.261 | 465.986 | | orf19.4347 |
| chr5:529259-529665 | 4102.0 | 6,34729 | 412.472 | 410.254 | orf19.4245 | orf19.4246 |
| chr1:1610653-1611279 | 4094.0 | 6,34051 | 411.666 | 409.449 | | orf19.4423 |
| chr1:172612-173687 | 5910.0 | 6,32652 | 593.465 | 591.077 | | orf19.6017 |
| chr4:1099714-1101651 | 5348.0 | 6,31368 | 537.217 | 534.877 | orf19.740 | |
| chr2:1415321-1415761 | 4024.0 | 6,25922 | 404.636 | 402.426 | orf19.2242 | |
| chr5:825221-825652 | 4793.0 | 6,2546 | 481.63 | 479.344 | | orf19.1121 |
| chr4:443132-444212 | 4343.0 | 6,23749 | 436.602 | 434.358 | orf19.4567 | |
| chr3:1721712-1723201 | 5504.0 | 6,14715 | 552.832 | 550.478 | orf19.6736 | orf19.6734 |
| chr6:284148-284719 | 3855.0 | 6,13685 | 387.716 | 385.525 | orf19.3442 | |
| chr1:349299-349996 | 4175.0 | 6,11927 | 419.806 | 417.58 | | orf19.3360 |
| chr3:1566715-1567602 | 3825.0 | 6,10505 | 384.727 | 382.539 | | orf19.6817 |
| chr4:1505769-1506610 | 4445.0 | 6,01777 | 446.786 | 444.533 | orf19.3134 | orf19.3133 |
| chrR:1577899-1579077 | 4320.0 | 6,01166 | 434.296 | 432.055 | orf19.6146 | |
| chr1:3010470-3010910 | 4653.0 | 6,00455 | 467.604 | 465.33 | orf19.5006.1 | orf19.5007 |
| chr2:709703-710069 | 4971.0 | 5,98352 | 499.415 | 497.111 | orf19.882 | |
| chr7:197848-198226 | 3659.0 | 5,96713 | 368.086 | 365.916 | | orf19.7020 |

| | | | | | | |
|---|---|---|---|---|---|---|
| chr1:2179858-2180308 | 3620.0 | 5,93319 | 364.197 | 362.031 | orf19.4864 | orf19.4865 |
| chr1:305614-306491 | 3734.0 | 5,86502 | 375.629 | 373.451 | orf19.3340 | |
| chr3:901348-901985 | 4758.0 | 5,82111 | 478.174 | 475.891 | orf19.5876 | orf19.5877 |
| chr5:628147-628617 | 4824.0 | 5,80529 | 484.708 | 482.418 | orf19.4304 | |
| chr1:1914023-1914802 | 4132.0 | 5,79848 | 415.505 | 413.283 | | orf19.4737 |
| chr1:1047276-1047681 | 3427.0 | 5,76348 | 344.939 | 342.795 | orf19.53 | orf19.54 |
| chr6:790832-791666 | 4376.0 | 5,74193 | 439.936 | 437.689 | | orf19.5741 |
| chr4:987296-987642 | 4414.0 | 5,66998 | 443.698 | 441.448 | | orf19.3822 |
| chr4:113255-113652 | 4275.0 | 5,65777 | 429.791 | 427.555 | | orf19.4175 |
| chr1:281187-282366 | 3838.0 | 5,61533 | 386.039 | 383.85 | orf19.3330.3 | orf19.3331 |
| chr1:889828-890598 | 3246.0 | 5,60055 | 326.751 | 324.629 | orf19.1064 | |
| chr3:728629-729002 | 3397.0 | 5,59078 | 341.877 | 339.737 | orf19.341 | |
| chr2:2105123-2105790 | 3223.0 | 5,58019 | 324.499 | 322.379 | | orf19.1756 |
| chr6:392648-393682 | 4220.0 | 5,5775 | 424.256 | 422.025 | orf19.3392 | orf19.3391 |
| chr1:2910708-2911551 | 5140.0 | 5,53624 | 516.335 | 514.015 | | orf19.4961 |
| chr1:2395336-2395956 | 3829.0 | 5,53375 | 385.184 | 382.996 | orf19.2337 | orf19.2336 |
| chr5:1076826-1077721 | 3577.0 | 5,48144 | 359.944 | 357.783 | orf19.3981 | orf19.3982 |
| chr4:1044698-1045682 | 3338.0 | 5,47208 | 335.986 | 333.852 | orf19.3794 | orf19.3793 |
| chr1:455936-456701 | 3100.0 | 5,4137 | 312.161 | 310.055 | orf19.3674 | orf19.3675 |
| chr5:428970-429408 | 3235.0 | 5,33693 | 325.628 | 323.507 | | orf19.3182 |
| chrR:2090239-2090713 | 3098.0 | 5,33283 | 311.916 | 309.811 | | orf19.7551 |
| chr2:1339490-1340818 | 2906.0 | 5,28828 | 292.746 | 290.666 | orf19.34 | orf19.35 |
| chr2:2022246-2023761 | 2906.0 | 5,28828 | 292.746 | 290.666 | orf19.1363 | orf19.1362 |
| chr3:1475263-1475837 | 3338.0 | 5,28536 | 336.024 | 333.89 | | orf19.7436 |
| chr4:1300693-1301352 | 3918.0 | 5,26643 | 394.068 | 391.87 | orf19.1274 | orf19.1275 |
| chr2:627093-627608 | 4592.0 | 5,25861 | 461.472 | 459.205 | | orf19.5785 |
| chr2:1230770-1231582 | 3076.0 | 5,25319 | 309.727 | 307.624 | | orf19.4118 |
| chr3:926745-927132 | 3438.0 | 5,24886 | 345.991 | 343.846 | orf19.5896 | orf19.5897 |
| chr4:226431-227106 | 2863.0 | 5,24755 | 288.396 | 286.321 | orf19.4679 | |
| chr1:895956-897613 | 2791.0 | 5,17966 | 281.191 | 279.126 | | orf19.1069 |
| chr1:1955602-1956364 | 2762.0 | 5,15251 | 278.325 | 276.263 | orf19.4754 | |
| chr2:1456575-1457327 | 3090.0 | 5,11864 | 311.115 | 309.011 | orf19.2268 | orf19.2269 |
| chr1:2541657-2542208 | 3527.0 | 5,07989 | 354.918 | 352.763 | | orf19.1161 |
| chr5:1106668-1107163 | 2774.0 | 5,07354 | 279.547 | 277.484 | | orf19.3997 |
| chr2:934484-935210 | 2999.0 | 5,05623 | 302.092 | 300.0 | | orf19.4505 |
| chr1:3050519-3050946 | 2722.0 | 5,05601 | 274.337 | 272.281 | | orf19.5025 |
| chr5:133727-134476 | 2656.0 | 5,05068 | 267.658 | 265.612 | | orf19.930 |
| chrR:914164-914553 | 3215.0 | 5,04885 | 323.681 | 321.562 | orf19.489 | orf19.490 |
| chr5:169688-170087 | 2919.0 | 5,04022 | 294.065 | 291.983 | | orf19.575 |
| chr5:450220-451464 | 2870.0 | 5,02117 | 289.09 | 287.015 | orf19.3171 | |
| chr4:120445-120868 | 3020.0 | 5,01828 | 304.109 | 302.013 | orf19.4172 | orf19.4171 |
| chr1:520689-521041 | 2613.0 | 5,00995 | 263.429 | 261.388 | orf19.2934 | orf19.2935 |
| chr2:1877433-1879801 | 3616.0 | 4,99904 | 363.804 | 361.64 | orf19.4076 | orf19.4077 |
| chr7:319865-320481 | 2578.0 | 4,97601 | 259.92 | 257.884 | | orf19.6577 |
| chr2:350002-350429 | 2571.0 | 4,96922 | 259.22 | 257.186 | orf19.1497 | orf19.1499 |
| chrR:776301-779260 | 2954.0 | 4,96682 | 297.54 | 295.454 | | orf19.4393 |
| chr1:1250182-1250581 | 2557.0 | 4,95564 | 257.822 | 255.79 | orf19.2452 | orf19.2451 |
| chr7:39811-41292 | 3288.0 | 4,93938 | 331.012 | 328.885 | | orf19.7094 |
| chr7:613838-614526 | 3062.0 | 4,92109 | 308.325 | 306.224 | orf19.5178 | orf19.5177 |
| chr4:1455489-1456319 | 2621.0 | 4,90455 | 264.152 | 262.111 | orf19.2882 | orf19.2881 |
| chr6:765991-766666 | 2677.0 | 4,88175 | 269.795 | 267.745 | orf19.5728 | orf19.5729 |
| chr2:346967-347594 | 2481.0 | 4,88097 | 250.174 | 248.155 | orf19.1497 | orf19.1499 |
| chr1:1806745-1807539 | 3074.0 | 4,86932 | 309.535 | 307.433 | orf19.5102 | |
| chr1:2519914-2520390 | 2467.0 | 4,86739 | 248.792 | 246.774 | orf19.656 | orf19.655 |
| chr1:2119782-2120147 | 2886.0 | 4,86416 | 290.725 | 288.647 | orf19.4825 | orf19.4826 |
| chr2:1829295-1830344 | 3029.0 | 4,86404 | 305.091 | 302.995 | orf19.207 | |

| | | | | | | |
|---|---|---|---|---|---|---|
| chr3:843601-844041 | 2433.0 | 4,83345 | 245.346 | 243.333 | orf19.2805 | orf19.2804 |
| chr6:513608-514195 | 2557.0 | 4,8 | 257.812 | 255.781 | | orf19.5517 |
| chr1:1214867-1215288 | 2598.0 | 4,78165 | 261.858 | 259.82 | | orf19.2467 |
| chr6:56418-56934 | 3107.0 | 4,77434 | 312.833 | 310.726 | orf19.1193 | |
| chr6:217241-217707 | 2688.0 | 4,72963 | 270.923 | 268.872 | | orf19.118 |
| chr4:441550-441981 | 2517.0 | 4,72624 | 253.772 | 251.747 | orf19.4568 | |
| chr2:1951130-1951645 | 2855.0 | 4,68408 | 287.666 | 285.592 | | orf19.1401 |
| chr4:907656-908563 | 2529.0 | 4,67674 | 254.98 | 252.953 | orf19.1426 | orf19.1424 |
| chr1:2954470-2955113 | 2250.0 | 4,65016 | 227.003 | 225.02 | | orf19.4980 |
| chr1:666180-667596 | 2485.0 | 4,63858 | 250.54 | 248.52 | orf19.3010.1 | |
| chr3:1627278-1628195 | 2464.0 | 4,63694 | 248.493 | 246.476 | orf19.6790 | orf19.6789 |
| chr3:1124610-1126012 | 3113.0 | 4,61122 | 313.466 | 311.358 | | orf19.5994 |
| chr1:1221036-1221449 | 2190.0 | 4,58906 | 220.991 | 219.018 | | orf19.2461 |
| chr1:2080564-2080924 | 2637.0 | 4,56725 | 265.752 | 263.708 | | orf19.4807 |
| chr5:312358-313055 | 2130.0 | 4,52796 | 215.03 | 213.068 | orf19.1927 | orf19.1926 |
| chr2:1010258-1010570 | 2078.0 | 4,47365 | 209.776 | 207.823 | | orf19.4128 |
| chr3:646939-648054 | 2307.0 | 4,4362 | 232.766 | 230.773 | orf19.301 | orf19.302 |
| chr6:430929-431374 | 2686.0 | 4,40336 | 270.686 | 268.636 | orf19.3501 | orf19.3499 |
| chr3:946172-946883 | 2165.0 | 4,3901 | 218.47 | 216.503 | orf19.5906 | |
| chr2:166736-167204 | 2275.0 | 4,37416 | 229.549 | 227.562 | | orf19.2021 |
| chr2:2098024-2098373 | 1955.0 | 4,34467 | 197.467 | 195.535 | orf19.1760 | |
| chr6:819585-820860 | 1948.0 | 4,33788 | 196.826 | 194.895 | orf19.5753 | |
| chr1:2095803-2096459 | 2308.0 | 4,31637 | 232.869 | 230.877 | orf19.4816 | |
| chr5:322465-322831 | 2319.0 | 4,27479 | 233.974 | 231.98 | orf19.4134 | orf19.4135 |
| chr7:709278-709664 | 1885.0 | 4,27 | 190.453 | 188.533 | orf19.5129 | orf19.5126 |
| chr2:576514-577225 | 2377.0 | 4,24833 | 239.775 | 237.771 | orf19.5816 | orf19.5815 |
| chr1:101122-101904 | 2480.0 | 4,22089 | 250.109 | 248.09 | | orf19.6045 |
| chr1:1382672-1382975 | 2026.0 | 4,19134 | 204.641 | 202.697 | orf19.6263 | |
| chr2:2135811-2136256 | 1784.0 | 4,16138 | 180.398 | 178.498 | orf19.1743 | |
| chr3:277159-277469 | 1698.0 | 4,06634 | 171.75 | 169.865 | orf19.1721 | orf19.1720 |
| chr1:298659-299230 | 1668.0 | 4,0324 | 168.695 | 166.816 | | orf19.3337 |
| chrR:1506801-1507301 | 1924.0 | 3,98781 | 194.364 | 192.437 | orf19.2365 | orf19.2364 |
| chrR:1093471-1094161 | 1752.0 | 3,94431 | 177.174 | 175.28 | | orf19.646 |
| chr1:1318313-1318873 | 1589.0 | 3,94415 | 160.838 | 158.975 | orf19.1822 | |
| chr4:289826-290624 | 1583.0 | 3,93736 | 160.239 | 158.377 | | orf19.4646 |
| chr1:1757081-1757869 | 2198.0 | 3,917 | 221.859 | 219.885 | | orf19.5079 |
| chrR:1156956-1157892 | 3308.0 | 3,88839 | 333.023 | 330.894 | orf19.5286 | orf19.5285 |
| chr2:2056550-2057014 | 1536.0 | 3,88305 | 155.473 | 153.62 | | orf19.1777 |
| chr1:1260388-1260746 | 1512.0 | 3,8559 | 153.108 | 151.26 | orf19.2446 | orf19.2445 |
| chr1:35671-37381 | 1905.0 | 3,84585 | 192.484 | 190.561 | orf19.6082 | orf19.6081 |
| chr1:1836097-1836696 | 1593.0 | 3,80183 | 161.18 | 159.317 | orf19.5117 | orf19.5118 |
| chr2:1935138-1936160 | 1707.0 | 3,78875 | 172.683 | 170.798 | orf19.1604 | |
| chr3:953481-954484 | 1540.0 | 3,76868 | 155.855 | 154.002 | orf19.5908 | |
| chr4:913515-914039 | 1742.0 | 3,75358 | 176.094 | 174.201 | orf19.1422 | orf19.1421 |
| chr3:106360-106854 | 1512.0 | 3,73981 | 153.108 | 151.26 | | orf19.5392 |
| chr1:972432-973024 | 1830.0 | 3,73874 | 184.919 | 183.01 | | orf19.778 |
| chr1:248360-248687 | 1637.0 | 3,73161 | 165.663 | 163.791 | | orf19.3311 |
| chr1:625677-626574 | 1672.0 | 3,72991 | 169.128 | 167.249 | orf19.2990 | orf19.2991 |
| chrR:1262777-1263129 | 1376.0 | 3,69284 | 139.422 | 137.604 | orf19.454 | orf19.451 |
| chr1:2694858-2695166 | 1388.0 | 3,69276 | 140.674 | 138.853 | orf19.5528 | orf19.5527 |
| chr2:1834561-1836011 | 1422.0 | 3,6645 | 144.047 | 142.219 | orf19.206 | |
| chr2:883629-883936 | 1593.0 | 3,65538 | 161.189 | 159.325 | | orf19.799 |
| chr6:273231-274039 | 1339.0 | 3,65224 | 135.765 | 133.955 | | orf19.3447 |
| chr1:3003062-3003588 | 1351.0 | 3,62007 | 136.944 | 135.131 | orf19.5003 | orf19.5004 |
| chr4:516443-516916 | 1402.0 | 3,58158 | 142.085 | 140.261 | orf19.2749 | orf19.2748 |
| chr2:1870810-1871131 | 1457.0 | 3,56675 | 147.571 | 145.734 | orf19.4072 | |

| | | | | | | |
|---|---|---|---|---|---|---|
| chr3:763445-764341 | 1595.0 | 3,55782 | 161.45 | 159.586 | orf19.359 | orf19.360 |
| chr1:993603-993970 | 1200.0 | 3,48253 | 121.865 | 120.087 | orf19.767 | |
| chr1:2207782-2208203 | 1787.0 | 3,46839 | 180.678 | 178.778 | orf19.4874 | orf19.4875 |
| chr1:1717257-1717766 | 1173.0 | 3,44858 | 119.148 | 117.376 | orf19.5059 | |
| chr2:1316721-1317437 | 1263.0 | 3,39912 | 128.129 | 126.336 | | orf19.24 |
| chrR:1428398-1429287 | 1120.0 | 3,3807 | 113.777 | 112.019 | orf19.699 | orf19.700 |
| chr1:1944923-1945240 | 1114.0 | 3,37391 | 113.245 | 111.488 | orf19.4752 | orf19.4753 |
| chr1:2014371-2014746 | 1162.0 | 3,33892 | 118.022 | 116.254 | | orf19.4776 |
| chr1:244658-245169 | 1317.0 | 3,29426 | 133.536 | 131.731 | | orf19.3309 |
| chr1:2717924-2718276 | 1000.0 | 3,22456 | 101.757 | 100.031 | orf19.5219 | |
| chr1:1470245-1470852 | 1228.0 | 3,21071 | 124.639 | 122.854 | orf19.6220.4 | |
| chr3:1760594-1760985 | 1171.0 | 3,20405 | 118.881 | 117.11 | orf19.6715 | orf19.6713 |
| chr1:1909924-1910308 | 1241.0 | 3,19766 | 125.984 | 124.196 | | orf19.4735 |
| chrR:174793-175179 | 1117.0 | 3,16818 | 113.51 | 111.752 | orf19.3282 | |
| chr1:1548060-1548575 | 1103.0 | 3,06685 | 112.121 | 110.367 | orf19.4446 | orf19.4445 |
| chr3:774281-774541 | 1077.0 | 3,05856 | 109.54 | 107.793 | | orf19.6952 |
| chr1:1219263-1219719 | 870.0 | 3,04806 | 88.7515 | 87.0616 | | orf19.2462 |
| chrR:1930314-1930601 | 945.0 | 3,03635 | 96.2865 | 94.5751 | orf19.7298 | |
| chr1:2505288-2505662 | 923.0 | 3,03386 | 94.0635 | 92.3587 | | orf19.660 |
| chr1:1654501-1654839 | 836.0 | 3,00054 | 85.3598 | 83.6798 | orf19.2781 | |
| chr7:668371-668782 | 1066.0 | 2,97162 | 108.378 | 106.634 | orf19.5148 | orf19.5147 |
| chr1:2117307-2117559 | 900.0 | 2,95227 | 91.7056 | 90.0071 | orf19.4823 | orf19.4824 |
| chr1:2169065-2169498 | 797.0 | 2,8536 | 81.373 | 79.7054 | orf19.4857 | orf19.4858 |
| chr6:349479-350089 | 836.0 | 2,85214 | 85.2989 | 83.6195 | orf19.3420 | orf19.3419 |
| chr1:3086870-3087190 | 1133.0 | 2,84569 | 115.072 | 113.31 | orf19.7207 | orf19.7208 |
| chr1:730686-731155 | 819.0 | 2,83088 | 83.5936 | 81.919 | orf19.3048 | |
| chr1:344051-345066 | 702.0 | 2,76286 | 71.9337 | 70.2969 | orf19.3358 | orf19.3359 |
| chr1:1973168-1973409 | 661.0 | 2,74257 | 67.8045 | 66.1821 | orf19.4763 | orf19.4764 |
| chr1:2073496-2074039 | 748.0 | 2,71709 | 76.5214 | 74.869 | orf19.4802 | orf19.4803 |

TAP-Zcf15 and TAP-Zcf26 bound and expression level altered genes

| TAP-Zcf15 bound and upregulated | TAP-Zcf15 bound and downregulated | TAP-Zcf26 bound and upregulated | TAP-Zcf26 bound and downregulated |
|---|---|---|---|
| ORF19.2693 | ORF19.2047 | ORF19.1652 | ORF19.3364 |
| ORF19.1652 | ORF19.1309 | ORF19.935 | ORF19.909 |
| ORF19.5217 | ORF19.6586 | ORF19.272 | ORF19.3734 |
| ORF19.4122 | ORF19.3669 | ORF19.6838 | ORF19.2047 |
| ORF19.6838 | ORF19.4673 | ORF19.1704 | ORF19.1309 |
| ORF19.3668 | ORF19.21 | ORF19.1288 | ORF19.4789 |
| ORF19.5215 | ORF19.999 | ORF19.310 | ORF19.4898 |
| ORF19.2602 | ORF19.1277 | ORF19.6398 | ORF19.450 |
| ORF19.6066 | ORF19.3439 | ORF19.6445 | ORF19.1486 |
| ORF19.1704 | ORF19.3117 | ORF19.1449 | ORF19.4318 |
| ORF19.7414 | ORF19.4941 | ORF19.6975 | ORF19.3669 |
| ORF19.272 | ORF19.2398 | ORF19.7520 | ORF19.2770.1 |
| ORF19.5576 | ORF19.3302 | ORF19.1208 | ORF19.3302 |
| ORF19.935 | ORF19.4772 | ORF19.88 | ORF19.1277 |
| ORF19.577 | ORF19.4779 | ORF19.4335 | ORF19.1826 |
| ORF19.1449 | ORF19.1826 | ORF19.2281 | ORF19.3190 |
| ORF19.4789 | ORF19.450 | ORF19.5217 | ORF19.3713 |
| ORF19.1288 | ORF19.34 | ORF19.4933 | ORF19.4673 |
| ORF19.1208 | ORF19.740 | ORF19.5805 | ORF19.4941 |
| ORF19.539 | ORF19.5908 | ORF19.3442 | ORF19.2398 |
| ORF19.310 | ORF19.2356 | ORF19.1064 | ORF19.251 |
| ORF19.2124 | ORF19.2749 | ORF19.341 | ORF19.3117 |
| ORF19.2281 | ORF19.2770.1 | ORF19.1274 | ORF19.4779 |
| ORF19.2691 | ORF19.1189 | ORF19.489 | ORF19.2005 |
| ORF19.2686 | ORF19.3071 | ORF19.4076 | ORF19.4772 |
| ORF19.3554 | ORF19.3555 | ORF19.2452 | ORF19.999 |
| ORF19.4335 | ORF19.2850 | ORF19.5728 | ORF19.3439 |
| ORF19.2474 | ORF19.4753 | ORF19.5753 | ORF19.2356 |
| ORF19.6398 | ORF19.7042 | ORF19.1743 | ORF19.4245 |
| ORF19.356 | ORF19.932 | ORF19.206 | ORF19.740 |
| ORF19.6445 | ORF19.6874 | ORF19.359 | ORF19.882 |
| ORF19.7520 | ORF19.335 | ORF19.3358 | ORF19.3981 |
| ORF19.4933 | ORF19.5626 | ORF19.1985 | ORF19.34 |
| ORF19.4567 | ORF19.2748 | ORF19.5005 | ORF19.1363 |
| ORF19.3442 | ORF19.3997 | ORF19.5000 | ORF19.1193 |
| ORF19.341 | ORF19.4980 | ORF19.4504 | ORF19.301 |
| ORF19.2452 | ORF19.4347 | ORF19.4195.1 | ORF19.1760 |
| ORF19.5805 | ORF19.5177 | ORF19.6065 | ORF19.5908 |
| ORF19.4076 | ORF19.5785 | ORF19.4447 | ORF19.5528 |
| ORF19.5753 | ORF19.1777 | ORF19.7156 | ORF19.2749 |
| ORF19.4679 | ORF19.5118 | ORF19.5323 | ORF19.699 |
| ORF19.5728 | ORF19.4646 | ORF19.1255 | ORF19.4802 |
| ORF19.1274 | ORF19.3499 | ORF19.913.2 | ORF19.2274 |
| ORF19.489 | | ORF19.3711 | ORF19.1189 |
| ORF19.1743 | | ORF19.1415 | ORF19.5626 |
| ORF19.1064 | | ORF19.5723 | ORF19.4056 |
| ORF19.359 | | ORF19.6085 | ORF19.5713 |
| ORF19.5542 | | ORF19.89 | ORF19.4899 |
| ORF19.2752 | | ORF19.6126 | ORF19.6920 |
| ORF19.3733 | | ORF19.4336 | ORF19.932 |
| ORF19.1985 | | ORF19.1224 | ORF19.5416 |
| ORF19.5777 | | ORF19.4650 | ORF19.3555 |
| ORF19.7513 | | ORF19.4934 | ORF19.7042 |
| ORF19.4504 | | ORF19.7288 | ORF19.3071 |
| ORF19.5005 | | ORF19.54 | ORF19.4753 |
| ORF19.6065 | | ORF19.3391 | ORF19.2020 |
| ORF19.4195.1 | | ORF19.3982 | ORF19.2397.3 |
| ORF19.5000 | | ORF19.1275 | ORF19.2125 |
| ORF19.5578 | | ORF19.5025 | ORF19.4366 |
| ORF19.3711 | | ORF19.2451 | ORF19.335 |
| ORF19.3263 | | ORF19.5517 | ORF19.2850 |
| ORF19.7156 | | ORF19.1926 | ORF19.6874 |
| ORF19.7288 | | ORF19.4128 | ORF19.5585 |
| ORF19.1224 | | ORF19.2364 | ORF19.2308 |
| ORF19.4884 | | ORF19.2445 | ORF19.4347 |
| ORF19.5307 | | ORF19.1421 | ORF19.3360 |
| ORF19.4011 | | ORF19.3311 | ORF19.6817 |
| ORF19.5323 | | ORF19.2991 | ORF19.5877 |
| ORF19.1906 | | ORF19.3447 | ORF19.4737 |
| ORF19.54 | | ORF19.360 | ORF19.3331 |
| ORF19.1255 | | | ORF19.4961 |
| ORF19.4934 | | | ORF19.3675 |
| ORF19.7539 | | | ORF19.35 |
| ORF19.118 | | | ORF19.7436 |
| ORF19.7094 | | | ORF19.5785 |
| ORF19.6661 | | | ORF19.3997 |
| ORF19.7357 | | | ORF19.6577 |
| ORF19.5723 | | | ORF19.5177 |
| ORF19.2451 | | | ORF19.4980 |
| ORF19.1926 | | | ORF19.3499 |
| ORF19.5517 | | | ORF19.3337 |
| ORF19.3982 | | | ORF19.4646 |
| ORF19.1275 | | | ORF19.5079 |
| ORF19.4128 | | | ORF19.1777 |
| ORF19.4505 | | | ORF19.5118 |
| ORF19.4807 | | | ORF19.2748 |
| ORF19.360 | | | ORF19.4803 |
| ORF19.3618 | | | |
| ORF19.3447 | | | |
